# Immunoinformatic identification of B cell and T cell epitopes in the SARS-CoV-2 proteome

**DOI:** 10.1101/2020.05.14.093757

**Authors:** Stephen N. Crooke, Inna G. Ovsyannikova, Richard B. Kennedy, Gregory A. Poland

## Abstract

A novel coronavirus (SARS-CoV-2) emerged from China in late 2019 and rapidly spread across the globe, infecting millions of people and generating societal disruption on a level not seen since the 1918 influenza pandemic. A safe and effective vaccine is desperately needed to prevent the continued spread of SARS-CoV-2; yet, rational vaccine design efforts are currently hampered by the lack of knowledge regarding viral epitopes targeted during an immune response, and the need for more in-depth knowledge on betacoronavirus immunology. To that end, we developed a computational workflow using a series of open-source algorithms and webtools to analyze the proteome of SARS-CoV-2 and identify putative T cell and B cell epitopes. Using increasingly stringent selection criteria to select peptides with significant HLA promiscuity and predicted antigenicity, we identified 41 potential T cell epitopes (5 HLA class I, 36 HLA class II) and 6 potential B cell epitopes, respectively. Docking analysis and binding predictions demonstrated enrichment for peptide binding to HLA-B (class I) and HLA-DRB1 (class II) molecules. Overlays of predicted B cell epitopes with the structure of the viral spike (S) glycoprotein revealed that 4 of 6 epitopes were located in the receptor-binding domain of the S protein. To our knowledge, this is the first study to comprehensively analyze all 10 (structural, non-structural and accessory) proteins from SARS-CoV-2 using predictive algorithms to identify potential targets for vaccine development.

**Significance Statement:** The novel coronavirus SARS-CoV-2 recently emerged from China, rapidly spreading and ushering in a global pandemic. Despite intensive research efforts, our knowledge of SARS-CoV-2 immunology and the proteins targeted by the immune response remains relatively limited, making it difficult to rationally design candidate vaccines. We employed a suite of bioinformatic tools, computational algorithms, and structural modeling to comprehensively analyze the entire SARS-CoV-2 proteome for potential T cell and B cell epitopes. Utilizing a set of stringent selection criteria to filter peptide epitopes, we identified 41 T cell epitopes (5 HLA class I, 36 HLA class II) and 6 B cell epitopes that could serve as promising targets for peptide-based vaccine development against this emerging global pathogen.

## Introduction

In December 2019, public health officials in Wuhan, China, reported the first case of severe respiratory disease attributed to infection with the novel coronavirus SARS-CoV-2 (1). Since its emergence, SARS-CoV-2 has spread rapidly via human-to-human transmission (2), threatening to overwhelm healthcare systems around the world and resulting in the declaration of a pandemic by the World Health Organization (3). The disease caused by the virus (COVID-19) is characterized by fever, pneumonia, and other respiratory and inflammatory symptoms that can result in severe inflammation of lung tissue and ultimately death—particularly among older adults or individuals with underlying comorbidities (4–6). As of this writing, the SARS-CoV-2 pandemic has resulted in 4 million confirmed cases of COVID-19 and over 280,000 deaths worldwide (7).

SARS-CoV-2 is the third pathogenic coronavirus to cross the species barrier into humans in the past two decades, preceded by severe acute respiratory syndrome coronavirus (SARS-CoV) (8, 9) and Middle-East respiratory syndrome coronavirus (MERS-CoV) (10). All three of these viruses belong to the ß-coronavirus genus and have either been confirmed (SARS-CoV) or suggested (MERS-CoV, SARS-CoV-2) to originate in bats, with transmission to humans occurring through intermediary animal hosts (11–14). While previous zoonotic spillovers of coronaviruses have been marked by high case fatality rates (~10% for SARS-CoV; ~34% for MERS-CoV), widespread transmission of disease has been relatively limited (8,098 cases of SARS; 2,494 cases of MERS) (15). In contrast, SARS-CoV-2 is estimated to have a lower case fatality rate (~2-4%) but is far more infectious and has achieved world-wide spread in a matter of months (16).

As the number of COVID-19 cases continues to grow, there is an urgent need for a safe and effective vaccine to combat the spread of SARS-CoV-2 and reduce the burden on hospitals and healthcare systems. No licensed vaccine or therapeutic is currently available for SARS-CoV-2, although there are over 100 vaccine candidates reportedly in development worldwide. Seven vaccine candidates have rapidly progressed into Phase I/II clinical trials: adenoviral vector-based vaccines (CanSino Biologics, ChiCTR2000030906; University of Oxford, NCT04324606), nucleic-acid based vaccines encoding for the viral spike (S) protein (Moderna, NCT04283461; Inovio Pharmaceuticals, NCT04336410; BioNTech/Pfizer, 2020-001038-36), and inactivated virus formulations (Sinopharm, ChiCTR2000031809; Sinovac (NCT04352608) (17). While the advancement of these vaccine candidates into clinical testing is promising, it is imperative they meet stringent endpoints for safety (18). Preclinical studies of multiple experimental SARS-CoV vaccines have reported a Th2-type immunopathology in the lungs of vaccinated mice following viral challenge, suggesting hypersensitization of the immune response against certain viral proteins (19–22). Similarly, a modified vaccinia virus Ankara vector expressing the SARS-CoV S protein induced significant hepatitis in immunized ferrets (23). These data suggest that candidate coronavirus vaccines that limit the inclusion of whole viral proteins may have more beneficial safety profiles.

The SARS-CoV-2 genome encodes for 10 unique protein products: 4 structural proteins (surface glycoprotein (S), envelope (E), membrane (M), nucleocapsid (N)); 5 non-structural proteins (open reading frame (ORF)3a, ORF6, ORF7a, ORF8, ORF10); and 1 non-structural polyprotein (ORF1ab) (**Figure 1A, B**) (24). It is currently unknown which epitopes in the SARS-CoV-2 proteome are recognized by the human immune system, although studies of SARS-CoV immune responses suggest that both cellular and humoral responses against structural proteins mediate protection against disease (19, 22, 25–27). It is likely that cellular immune responses against non-structural viral proteins also play a key role in orchestrating protective antiviral immunity (28–30). In lieu of biological data, immunoinformatic algorithms can be employed to predict peptide epitopes based on amino acid properties and known human leukocyte antigen (HLA) binding profiles (31–33). These computational approaches represent a validated methodology for rapidly identifying potential T cell and B cell epitopes for exploratory peptide-based vaccine development and have been recently used to identify target epitopes for MERS-CoV (34) and SARS-CoV-2, although many of these reports focus solely on structural proteins (35–38).

**Figure 1.**
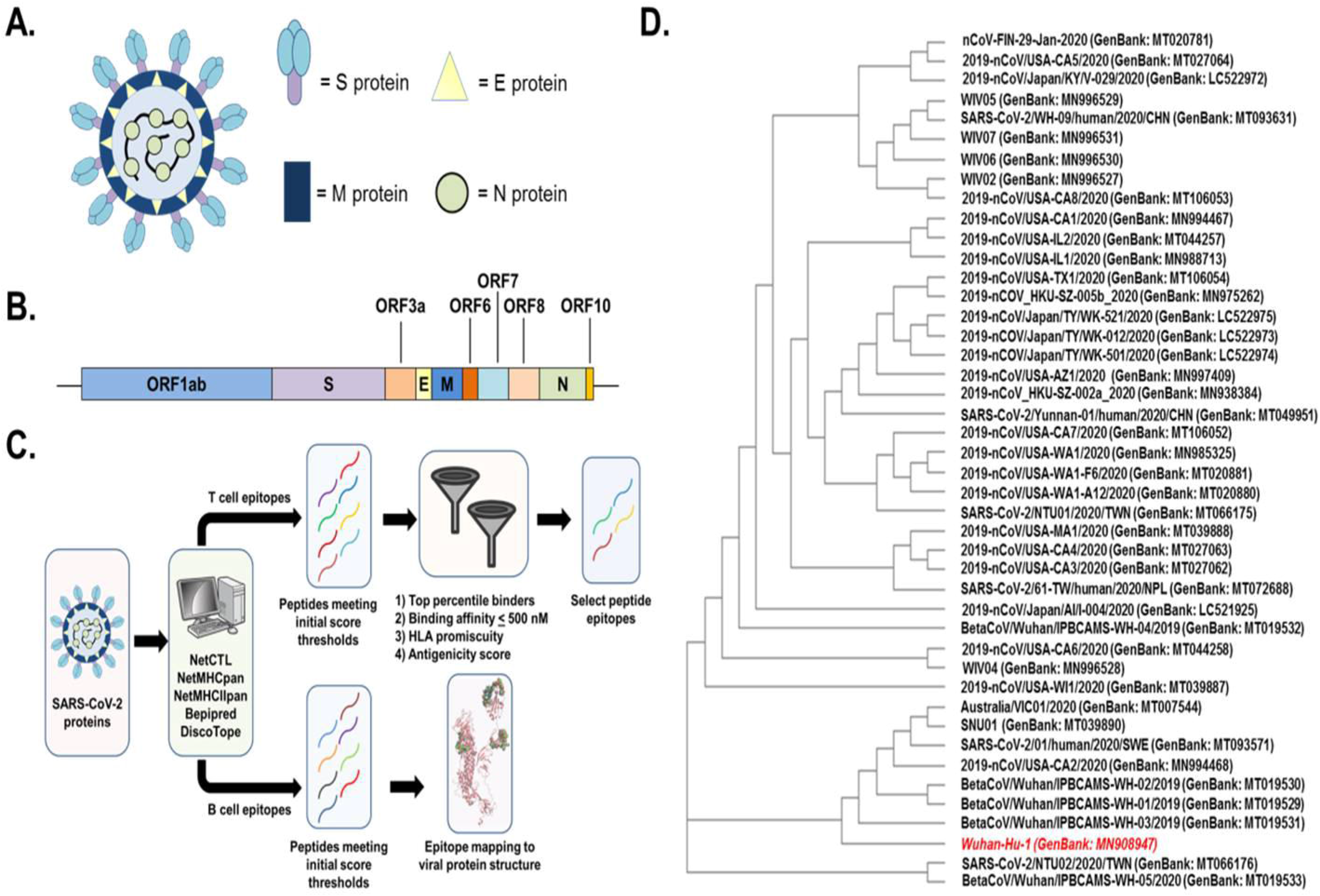
(A) Diagram of SARS-CoV-2 virion structure with the major structural proteins (S, M, N, and E) highlighted. (B) Cartoon representation of the SARS-CoV-2 genome with the 10 major protein-coding regions annotated. The box diagrams are proportional to the protein size. (C) Diagram of peptide identification workflow illustrating the algorithms used (33, 40–43, 45–47, 51, 53) and filtering criterion applied to refine peptide selection. (D) Cladogram illustrating the genetic relationship of SARS-CoV-2 isolates. The original viral isolate and consensus sequence (Wuhan-Hu-1) is highlighted in red.

Herein, we employed a comprehensive immunoinformatics approach to identify putative T cell and B cell epitopes across the entire SARS-CoV-2 proteome (**Figure 1C**). We independently identified peptides from each viral protein that were restricted to either HLA class I or HLA class II molecules across a subset of the most common HLA alleles in the global population. By filtering this list of peptides on the basis of predicted binding affinity, antigenicity, and promiscuity, we produced 5 HLA class I-restricted and 36 HLA class II-restricted peptides as leading candidates for further study. We also evaluated linear and structural B cell epitopes in the SARS-CoV-2 spike protein, with six antigenic regions identified as potential sites for antibody binding. These selected peptides may serve as initial candidates in the rational and accelerated design of a peptide-based vaccine against SARS-CoV-2.

## Methods

### Comparison of genome sequences from SARS-CoV-2 isolates

Genomic sequences for reported SARS-CoV-2 isolates were identified and retrieved from the Virus Pathogen Resource (ViPR) database on February 27, 2020 (https://www.viprbrc.org/brc/home.spg?decorator=corona_ncov). Sequences that did not cover the complete viral genome (~29,900 nucleotides) were excluded from further analysis. Remaining sequences were aligned using the Clustal Omega program (version 1.2.4) from the European Bioinformatics Institute (39) and compared against the first reported genome sequence for SARS-CoV-2 (Wuhan-Hu-1; taxonomy ID: 2697049) (1). Sequences from Wuhan-Hu-1 viral proteins were determined to be representative of those from all viral isolates and were subsequently used for epitope prediction analyses.

### Prediction of SARS-CoV-2 T cell epitopes

Prediction of HLA class I and class II peptide epitopes was carried out with the 10 protein sequences reported for the Wuhan-Hu-1 isolate: E (GenBank accession: QHD43418), M (QHD43419), N (QHD43423), S (QHD43416), ORF3a (QHD43417), ORF6 (QHD43420), ORF7a (QHD43421), ORF8 (QHD43422), ORF10 (QHI42199), ORF1ab (QHD43415).

For CD8^+^ T cell epitope prediction, NetCTL 1.2 (Immune Epitope Database) was initially used to evaluate the binding of nonameric peptides derived from each viral protein to the most common HLA class I supertypes present among the human population (40, 41). HLA class I molecules preferentially bind 9-mer peptides, and most algorithm training datasets have been based on peptides of this length. The weight placed on C-terminal cleavage and antigen transport efficiency was 0.15 and 0.05, respectively. The antigenic score threshold was 0.75. Peptides with scores above this threshold were subsequently analyzed on the NetMHCpan 4.0 server (Technical University of Denmark) to predict binding affinity and percentile rank across representative alleles of each major HLA class I supertype (HLA-A*01:01, HLA-A*02:01, HLA-A*03:01, HLA-A*24:02, HLA-B*07:02, HLA-B*08:01, HLA-B*27:05, HLA-B*40:01, HLA-B*58:01, HLA-B* 15:01), which collectively cover the majority of class I alleles present in the human population (42–44). Thresholds for defining binding strength were set at 0.5% and 2.0% for strong and weak binders, respectively.

For CD4^+^ T cell epitope prediction, NetMHCIIpan 3.2 server (Technical University of Denmark) was used for predicting the binding affinity and percentile rank of 15-mer peptides derived from each viral protein across a reference panel of 27 HLA class II molecules (33, 45). Thresholds for defining binding strength were set at 2% and 10% for strong and weak binders, respectively.

HLA class I and class II peptides with high predicted binding affinities (≤ 500 nM), high percentile ranks (≤ 0.5% for class I; ≤ 2% for class II), and broad HLA coverage (> 3 alleles) were independently analyzed on the VaxiJen 2.0 server (Edward Jenner Institute) (46, 47) using a conservative score threshold (0.7) to predict antigenicity.

### Molecular docking of HLA class I peptides

Docking simulations of 5 HLA class I-restricted SARS-CoV-2 peptides with high antigenicity scores and a commonly shared predicted HLA molecule (HLA-DRB1*15:01) were performed using the GalaxyPepDock server (Seoul National University Laboratory of Computational Biology) (48). The structure of HLA-DRB1*15:01 was accessed from the Protein Data Bank as a co-crystallized structure of the HLA molecule with a nonameric SARS-CoV peptide (PDB ID: 3C9N) (49). The bound nonamer peptide was removed from the structure using Chimera 1.14 (University of California-San Francisco) (50) prior to running simulations. Ten models of each peptide-HLA complex were generated on the basis of minimized energy scores, and the top model for each complex was selected for comparative analysis.

### Prediction and structural modeling of SARS-CoV-2 B cell epitopes

Linear B cell epitope predictions were performed on the three exposed SARS-CoV-2 structural proteins: S (GenBank accession: QHD43416), M (QHD43419), and E (QHD43418) using the BepiPred 1.0 algorithm (51). Epitope probability scores were calculated for each amino acid residue using a threshold of 0.35 (corresponding to > 0.75 specificity and sensitivity below 0.5), and only epitopes ≥ 5 amino acid residues in length were further analyzed. The structure of the SARS-CoV-2 S protein was accessed from the Protein Data Bank (PDB ID: 6VSB) (52). Discontinuous (i.e., structural) B cell epitope predictions for the S protein structure were carried out using DiscoTope 1.1 (53) with a score threshold greater than – 7.7 (corresponding to > 0.75 specificity and sensitivity below 0.5). The main protein structure was modeled in PyMOL (Schrödinger, LLC), with predicted B cell epitopes identified by both BepiPred 1.0 and DiscoTope 1.1 highlighted as spheres.

## Results

### Genetic similarity of SARS-CoV-2 isolates

The primary goal of our study was to identify peptide epitopes that would be broadly applicable in vaccine development efforts against SARS-CoV-2. We identified 64 point mutations and 4 deletions across the genomes of 44 clinical isolates, with all deletions and the majority of mutations (n=45) occurring in the ORF1ab polyprotein (**Supp. Figure S1**). Single-point mutations were also found in the S protein (n=5), N protein (n=5), ORF8 protein (n=3), ORF3a protein (n=2), ORF10 protein (n=2), E protein (n=1), and M protein (n=1). Despite the genetic diversity introduced by these events (**Figure 1D**), matrix analysis determined that > 99% sequence identity was maintained across all viral genomes. Based on these findings and for study feasibility, the genome from the original virus isolate (Wuhan-Hu-1; GenBank: MN908947) was selected as the consensus sequence for all further analyses.

### Prediction of CD8^+^ T cell epitopes in the SARS-CoV-2proteome

We next identified potential CD8^+^ T cell epitopes from all proteins in the SARS-CoV-2 proteome. Using the NetCTL 1.2 predictive algorithm, we analyzed the complete amino acid sequence of each viral protein to generate sets of 9-mer peptides predicted to be recognized across at least one of the major HLA class I supertypes (**Figure 2A, Supp. Figure S2**). This approach yielded a significant number of potential epitopes from each viral protein (ORF10: 9, ORF6: 17, ORF8: 23, E: 25, ORF7: 39, N: 80, M: 87, ORF3a: 87, S: 321, ORF1ab: 2814), with the number directly related to the size of the parent protein. We used the NetMHCpan 4.0 server to further refine the list of potential CD8^+^ T cell epitopes by predicting binding affinity across representative HLA class I alleles (see Methods) and assigning percentile scores to quantify binding propensity. Peptides with percentile rank scores ≤ 0.5% (i.e., strong binders) were filtered using a 500 nM threshold for binding affinity to further delineate 740 candidate HLA class I epitopes from the viral proteome (54). For feasibility reasons, we refined our selection to 83 candidate epitopes by excluding peptides predicted to bind only one HLA molecule (**Supp. Table S1**). The resultant peptides were enriched for predicted binders to HLA-B molecules (HLA-B* 15:01=50; HLA-B*58:01=32; HLA-B*08:01=31) (**Figure 2B**). A final round of selection on the basis of HLA promiscuity (i.e., predicted binding to ≥ 3 HLA molecules) and predicted antigenicity scoring using the VaxiJen 2.0 server produced a subset of five candidate peptides (four ORF1ab, one S protein) as potential targets for vaccine development (**Table 1**) with the hypothesis that increased HLA binding promiscuity meant broader population base coverage by those peptides. These peptides were predicted to provide 74% global population coverage and had higher predicted binding affinities for HLA-B molecules (B*08:01=42.6 nM; B*15:01=67.7 nM; B*58:01=110.3 nM) compared to HLA-A molecules (A*01:01=238.6 nM; A*24:02=142.9 nM), with the exception of one ORF1ab-derived peptide (MMISAGFSL) that was predicted to bind HLA-A*02:01 with high affinity (IC_50_= 6.9 nM) (**Figure 2C**).

**Figure 2.**
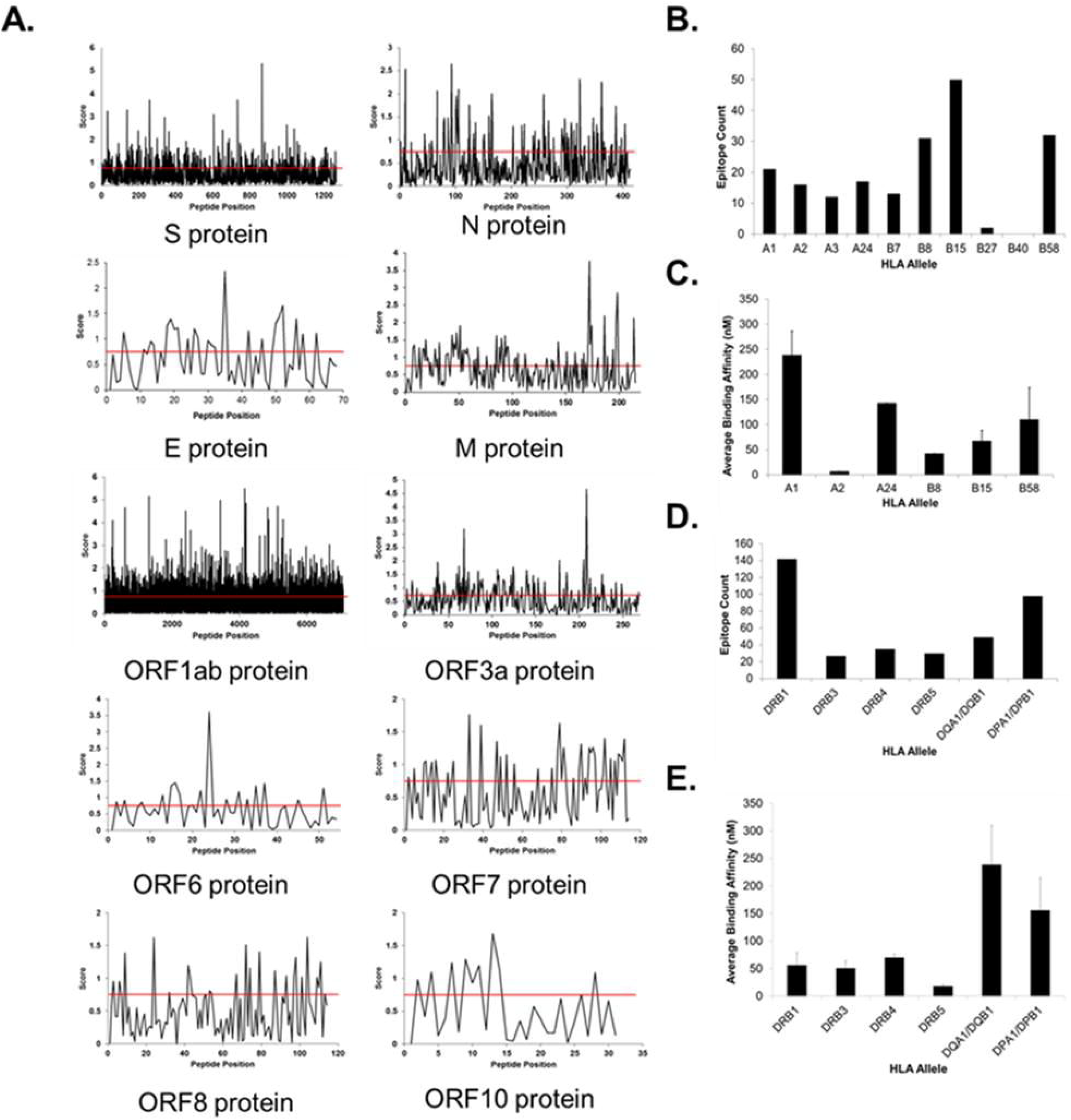
Immunogenicity scoring of peptides in the SARS-CoV-2 proteome with predicted HLA class I and II coverage and binding affinities. (A) Plots illustrating the NetCTL score for each sequential peptide across the entire amino acid sequence for each SARS-CoV-2 protein. Scores presented are the highest score identified across all HLA class I supertypes for each peptide. (B) Total number of predicted peptide epitopes distributed across HLA class I alleles. (C) Average predicted binding affinities by HLA allele for the top candidate class I peptides listed in Table 1. (D) Total number of predicted peptide epitopes distributed across HLA class II alleles. (E) Average predicted binding affinities by HLA allele for the top candidate class II peptides listed in Table 1.

**Table 1.**
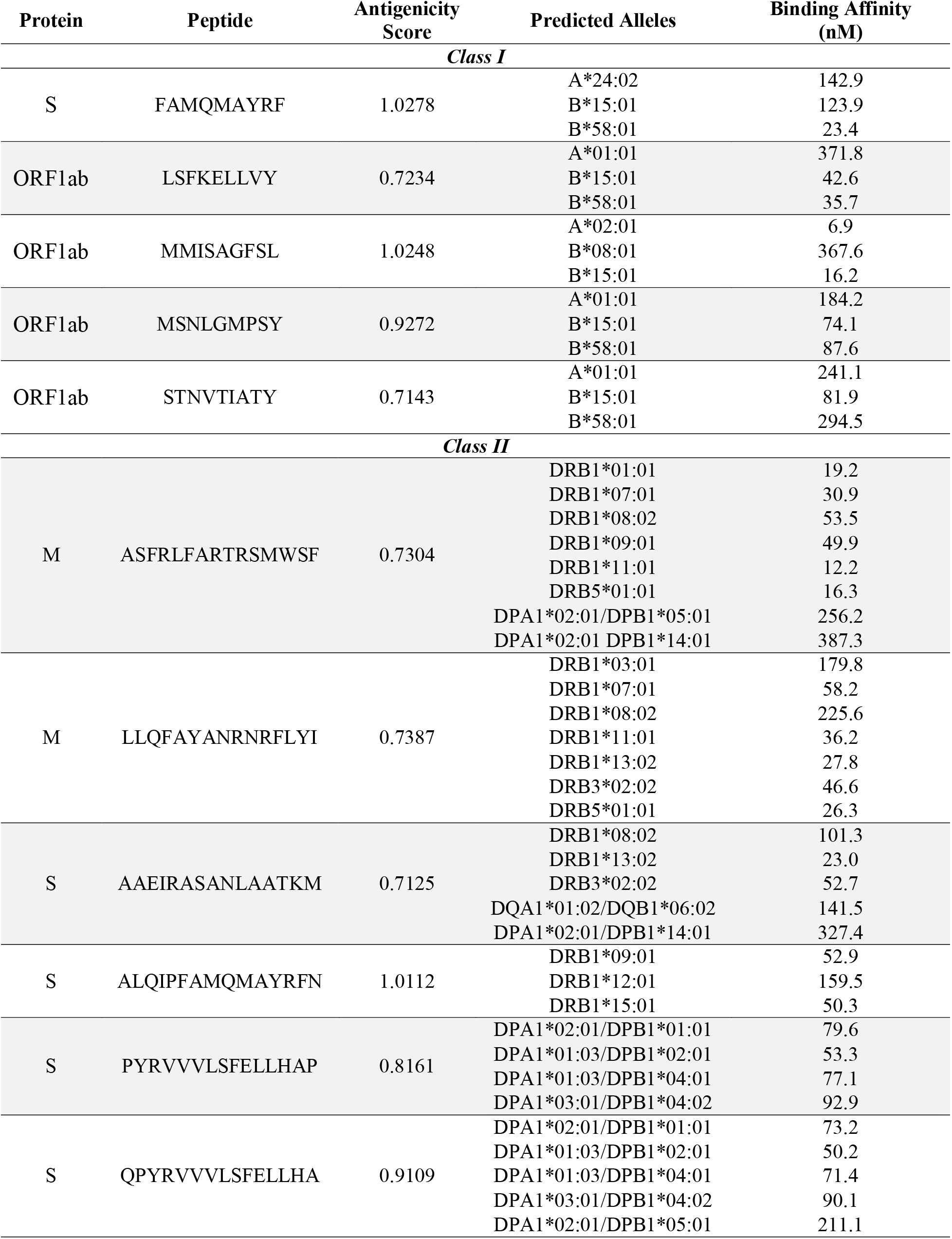
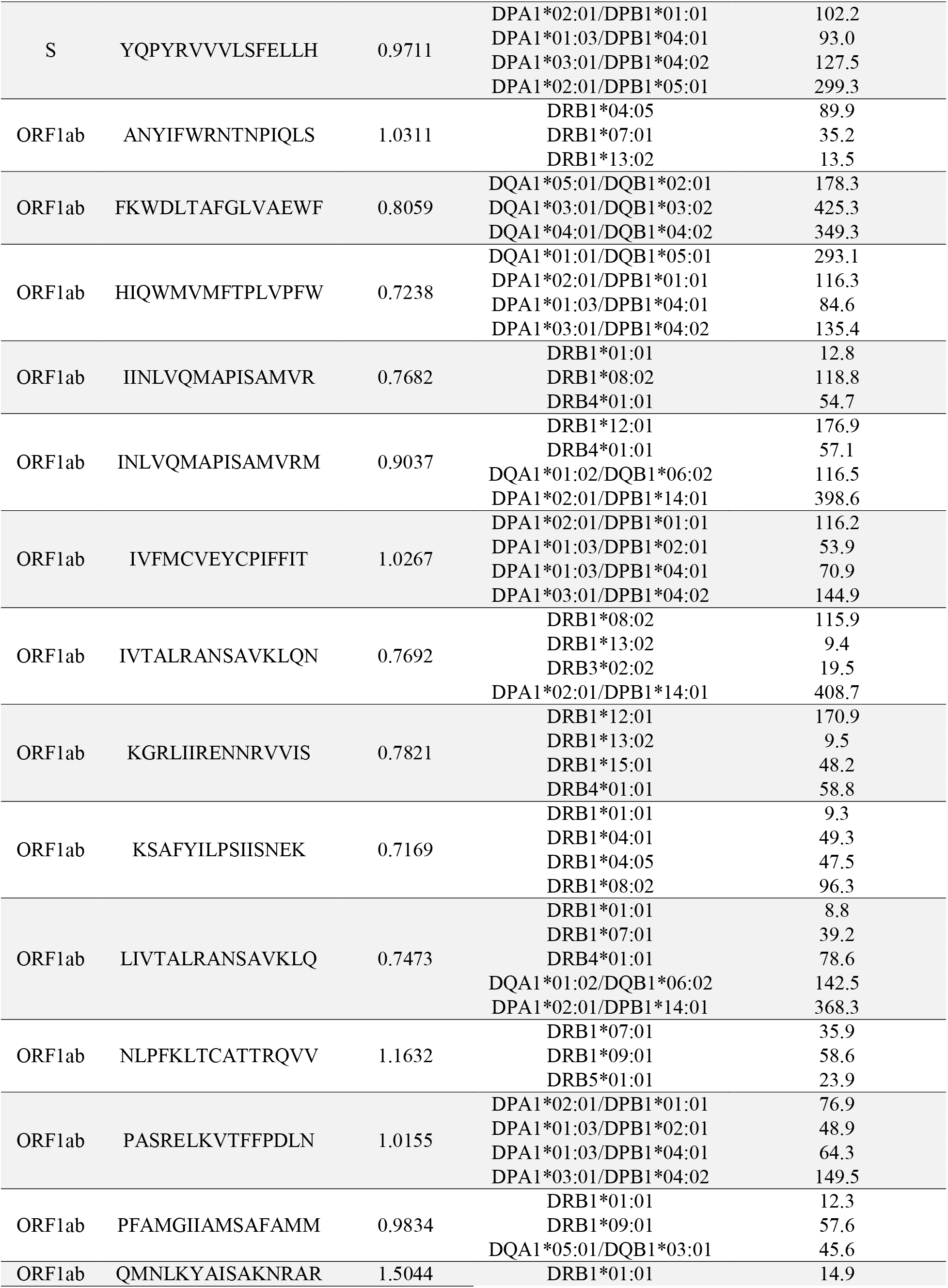
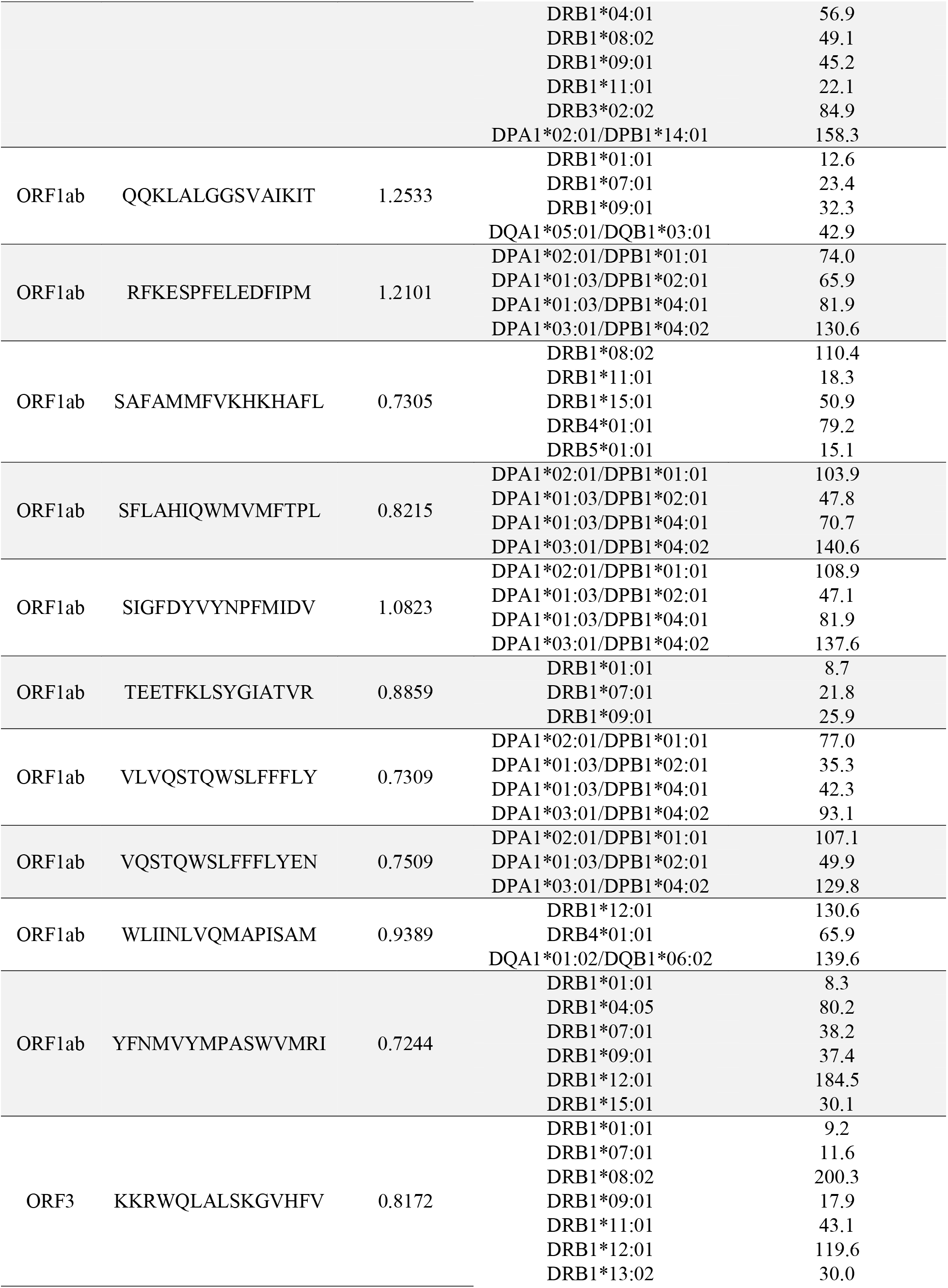
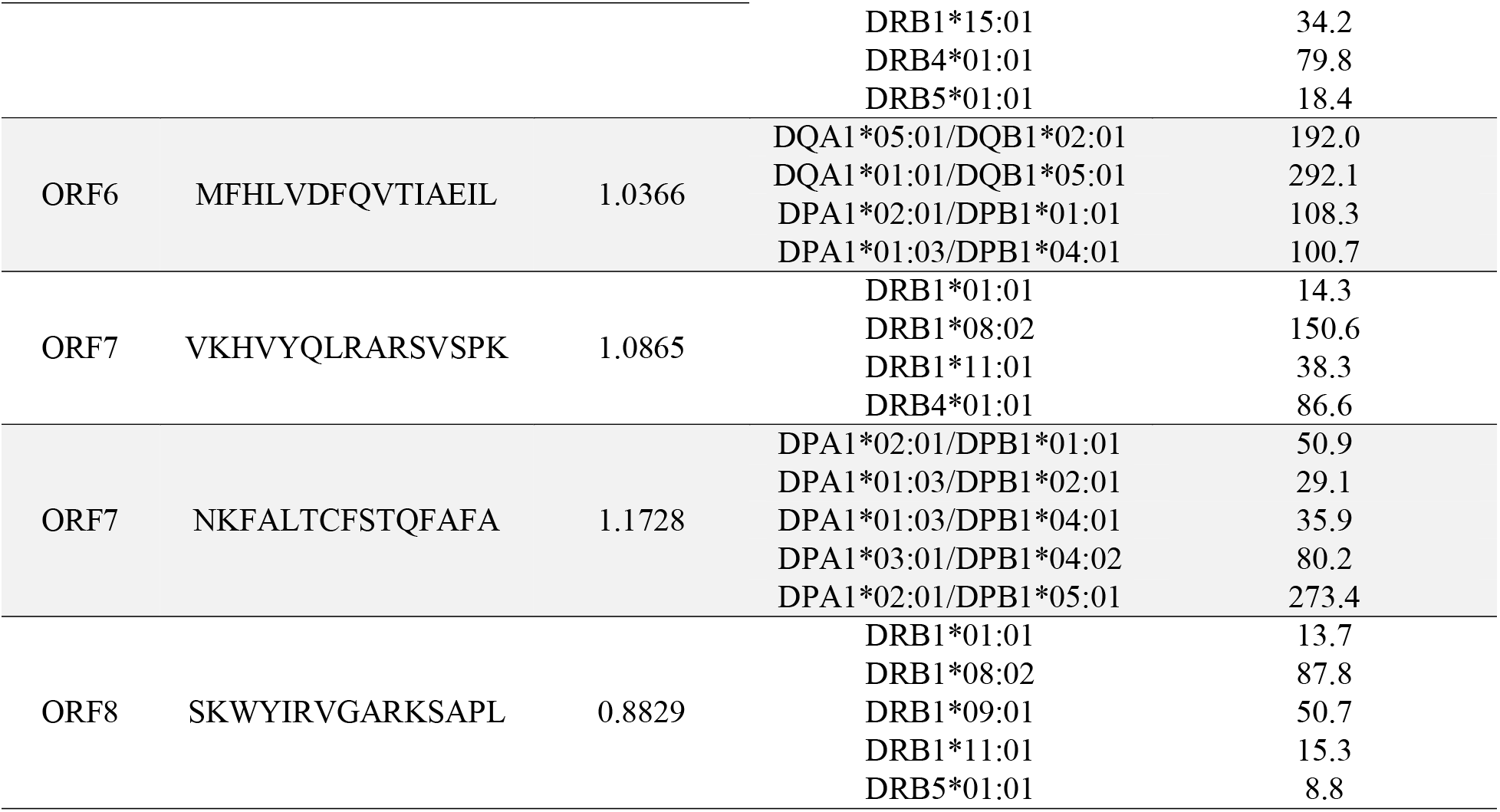
Top predicted HLA class I and class II T cell epitopes.

### Prediction of CD4^+^ T cell epitopes in the SARS-CoV-2proteome

We also sought to identify potential HLA class II peptides from SARS-CoV-2, as the stimulation of CD4^+^ T-helper cells is critical for robust vaccine-induced adaptive immune responses. Using the NetMHCIIpan 3.2 server, we identified 801 candidate HLA class II peptides from the viral proteome predicted to have high binding affinity (≤ 500 nM) and percentile rank scores ≤ 2% across a reference panel of HLA molecules covering > 97% of the population (33, 45). Similar to HLA class I epitope predictions, the number of class II epitopes identified for each viral protein (ORF10: 4, E protein: 7, ORF7: 8, ORF8: 10, ORF6: 14, N: 15, M: 29, ORF3a: 31, S: 96, ORF1ab: 587) was largely proportional to protein size. After excluding peptides predicted to bind to only a single HLA molecule in our panel, we refined our selection to 211 peptides (**Supp. Table S2**), which were enriched for binding to HLA-DRB1 molecules (n=142) (**Figure 2D**). Filtering on HLA promiscuity and predicted antigenicity scores yielded a subset of 36 peptides (24 ORF1ab, 5 S protein, 2 M protein, 2 ORF7, 1 ORF3a, 1 ORF6, 1 ORF8) as CD4^+^ T cell epitopes for further study (**Table 1**). These peptides were predicted to collectively provide 99% population coverage and have significantly higher average binding affinities for HLA-DR alleles (DRB1=56.4 nM; DRB3=50.9 nM; DRB4=70.1 nM; DRB5=18 nM) compared to HLA-DP (155.9 nM) or HLA-DQ (238.6 nM) molecules (**Figure 2E**).

### Characterization of HLA class I peptide docking with HLA-B*15:01

The five candidate HLA class I peptides identified by our computational approach were predicted to provide coverage across six HLA alleles (A*01:01, A*02:01, A*24:02, B*08:01, B*15:01, B*58:01). The peptide FAMQMAYRF was the only candidate predicted to bind to A*24:02 molecules, whereas MMISAGFSL was predicted to uniquely bind A*02:01 and B*08:01 molecules. Four of the five peptides were predicted to bind A*01:01 and B*58:01 molecules, but all were predicted to bind with relatively high affinity (average IC_50_ = 67.7 nM) to HLA-B* 15:01. Therefore, we performed molecular docking studies of each peptide with the molecular structure of HLA-B* 15:01 (PDB: 3C9N).

All peptides were predicted to bind within the peptide binding groove, forming hydrogen bond contacts with numerous amino acid side chains (**Figure 3A**). The binding motif for HLA-B*15:01 is highly selective for residues at the P2 and P9 anchor positions, with a preference for bulky hydrophobic amino acids at the C-terminus (**Figure 3B**) (55). All candidate peptides possessed terminal residues (Phe, Tyr, Leu) that fit into the hydrophobic binding pocket of the HLA groove, further supporting that these peptides should be strong binders of HLA-B*15:01 and promising candidates for vaccine development studies.

**Figure 3.**
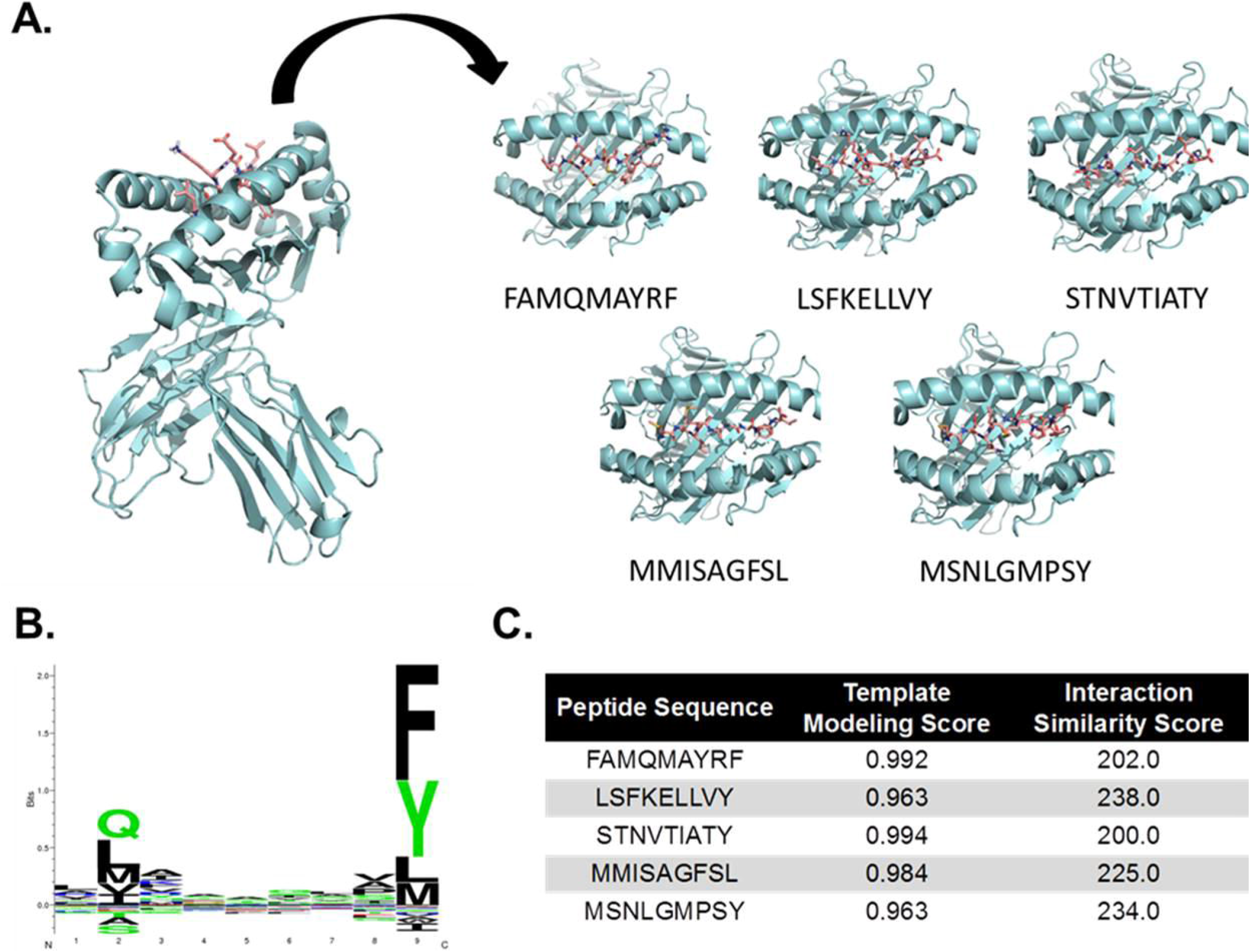
Docking of top predicted HLA class I peptides with a shared HLA molecule. (A) Structural docking model for each indicated peptide with the molecular structure of HLA-B*15:01 (PDB: 3C9N). Individual panels represent top-down views of the peptide binding groove. (B) Binding motif for HLA-B*15:01. (C) Template Modeling and Interaction Similarity scores for the selected peptide docking models shown in panel A. (74, 75)

### Prediction of B cell epitopes in SARS-CoV-2proteins

An effective vaccine should stimulate both cellular and humoral immune responses against the target pathogen; therefore, we also sought to identify potential B cell epitopes from SARS-CoV-2 proteins. We limited our analysis to the primary structural proteins exposed on the virus capsid (S, N, M, and E), as these are the most accessible antigens for engaging B cell receptors. Using the Bepipred 1.0 algorithm, we identified 26 potential linear B cell epitopes in the S protein, 14 potential epitopes in the N protein, and 3 potential epitopes in the M protein (**Table 2**). No epitopes were identified in the E protein. Studies have previously shown the S protein to be the predominant target of neutralizing antibodies against coronaviruses (56, 57), and, as our findings indicate this to likely be the case for SARS-CoV-2, we focused all subsequent analyses on the S protein. While the N protein is also a major target of the antibody response (58), it is unlikely these antibodies have any neutralizing activity based on the viral structure. As epitope conformation can significantly influence recognition by antibodies, we also employed DiscoTope 1.1 to identify discontinuous B cell epitopes in the protein structure. Our analysis identified 14 potential structural epitopes in the S protein (7 in the S1 domain, 7 in the S2 domain), with six regions having significant overlap with our predicted linear epitopes (**Table 2**). Antigenic regions identified in both analyses were modeled using the recently published structure of the SARS-CoV-2 S protein (52) to examine their accessibility for antibody binding. Epitopes in the S2 domain (P792-D796; Y1138-D1146) were clustered near the base of the spike protein, whereas regions in the S1 domain (D405-D428; N440-N450; G496-P507; D568-T573) were exposed on the protein surface (**Figure 4**).

**Figure 4.**
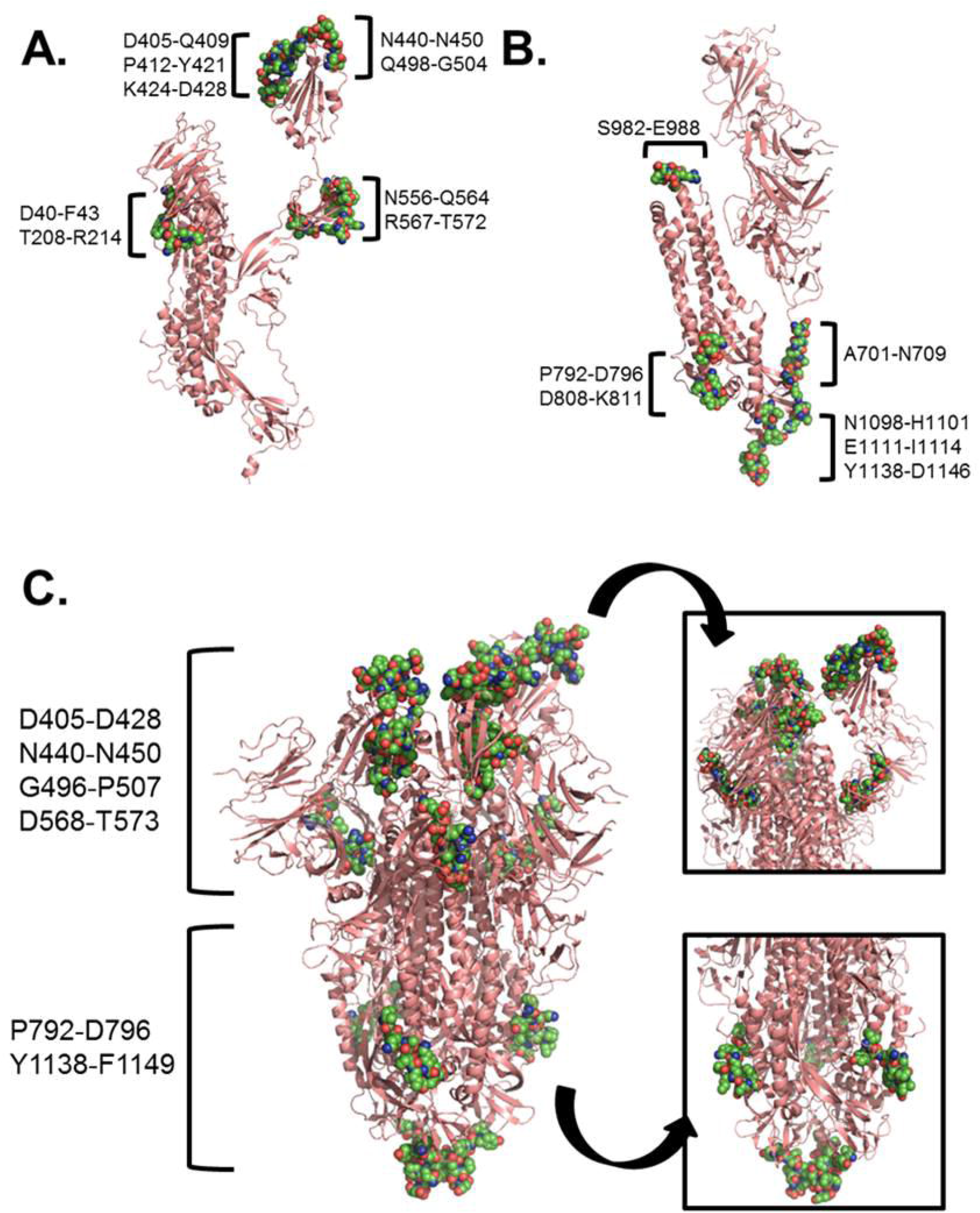
Modeling of predicted B cell epitopes on the crystal structure of the S glycoprotein. Predicted structural epitopes in the S1 domain (A) and S2 domain (B) highlighted on the structure of the S glycoprotein monomer (PDB: 6VSB). (C) Top predicted B cell epitopes identified by both Bepipred and DiscoTope prediction algorithms highlighted on the trimeric structure of the S glycoprotein. Inset panels show the S1 domain (upper) and S2 domain (lower). Predicted epitopes are highlighted as colored atoms (green, blue, red) on the surface of the S protein (salmon).

**Table 2.**
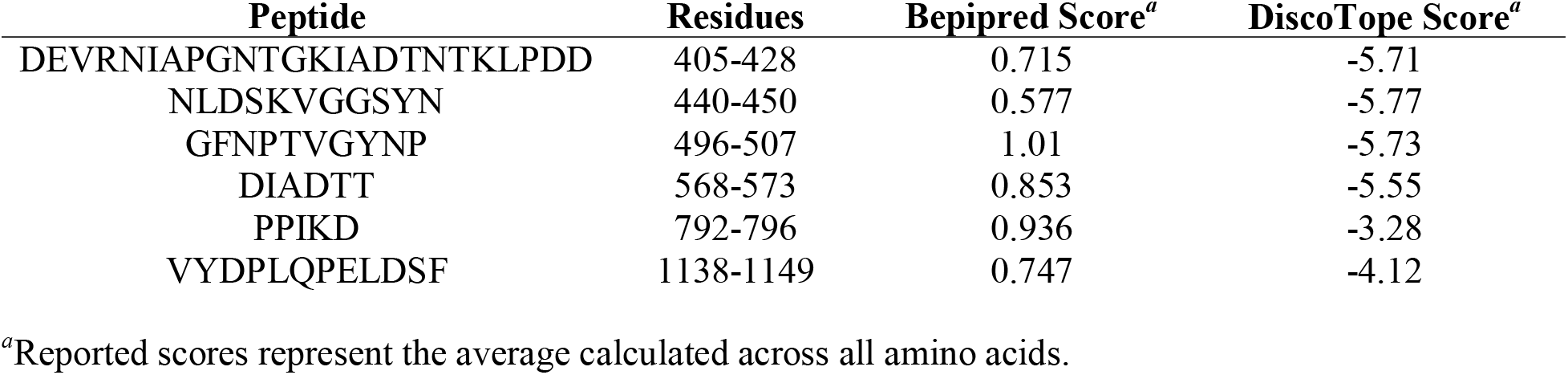
Top predicted B cell epitopes.

## Discussion

In the face of the COVID-19 pandemic, it is imperative that safe and effective vaccines be rapidly developed in order to induce widespread herd immunity in the population and prevent the continued spread of SARS-CoV-2. Our study identified probable peptide targets of both cellular and humoral immune responses against SARS-CoV-2 using computational methodologies to investigate the entire viral proteome *a priori*. Studies such as these are paramount during the early stages of pandemic vaccine development given the relative scarcity of biological data available on the viral immune response, and we employed an approach that allowed us to systematically refine our predictions using increasingly stringent criteria to select a subset of the most promising epitopes for further study. The data we have curated could inform the design of a candidate peptide-based vaccine or diagnostic against SARS-CoV-2.

As selective pressures are known to introduce viral mutations that promote fitness and can lead to evasion of immune responses (59, 60), we first sought to investigate the genetic similarity of all reported SARS-CoV-2 clinical isolates and identify a consensus sequence for use in our epitope prediction studies. We identified 68 mutations/deletions across the 44 genomes of clinical isolates reported as of 27 February 2020. Despite these variations, the viral genomic identity was > 99% conserved across all isolates. As the protein coding sequences were largely conserved, the genome of the original virus isolate (Wuhan-Hu-1) was deemed a representative consensus sequence for analysis of the SARS-CoV-2 proteome.

CD4^+^ and CD8^+^ T cell responses will likely be directed against both structural and non-structural proteins during antiviral immune responses, as all viral proteins are accessible for processing and presentation on the HLA molecules of infected cells. Therefore, we sought to identify T cell epitopes across the entire viral proteome. Our analysis identified 83 potential CD8^+^ T cell epitopes (**Supp. Table S1**) and 211 potential CD4^+^ T cell epitopes (**Supp. Table S2**), with stringent filtering for more promiscuous peptides with high predicted antigenicity yielding a subset of 5 CD8^+^ T cell epitopes and 36 CD4^+^ T cell epitopes (**Table 1**) as potential targets for vaccine development. A single study by Grifoni and colleagues has recently reported the computational identification of 241 CD4^+^ T cell epitopes from SARS-CoV-2 (35), and 22 peptides from our analysis shared sequence homology or were nested within peptides identified in their study. Moreover, seven peptides from this initial report were replicated in our final subset of HLA class II epitopes, supporting that these peptides may be promising vaccine targets.

An increasing number of studies have employed predictive algorithms to identify potential HLA class I epitopes for SARS-CoV-2, although relatively few have comprehensively analyzed the entire viral proteome. A report from Feng *et al*. recently outlined the identification of 499 potential class I epitopes in the main structural proteins from SARS-CoV-2 but did not consider any non-structural proteins (38). Grifoni and colleagues conducted a more rigorous analysis, identifying 628 unique CD8^+^ T cell epitopes across all SARS-CoV-2 proteins but focusing their analyses solely on peptides with sequence homology to known SARS-CoV epitopes (35). Our approach initially identified ~ 3,500 potential CD8^+^ T cell epitopes across all viral proteins, which we refined to a subset of 5 peptides (**Table 1**). One peptide derived from ORF1ab (MMISAGFSL) was predicted to bind HLA-A*02:01 with high affinity (IC_50_= 6.9 nM) (**Figure 2C**). Given the prevalence of this allele in the American and European populations (25-60% frequency) (61), MMISAGFSL may represent a promising epitope capable of providing broad vaccine population coverage.

We also observed a notable enrichment of epitopes predicted to bind HLA-B molecules– particularly HLA-B* 15:01–as we imposed more stringent selection criteria (**Figure 2B**). All five peptides identified by our approach were predicted to be relatively strong binders for this allele (IC_50_ = 67.7 nM), with molecular docking simulations illustrating strong contacts with amino acid residues in the peptide binding groove (**Figure 3 A, B**). A recent computational study identified another HLA-B allele (B*15:03) as having a high capacity for presenting epitopes from SARS-CoV-2 that were conserved among other pathogenic coronaviruses (62). These data collectively suggest the HLA-B locus may be significantly associated with the immune response to SARS-CoV-2 (and potentially other coronaviruses), with further biological studies warranted to determine the true role of host genetics in SARS-CoV-2 immunology.

Lastly, we analyzed the primary structural proteins of SARS-CoV-2 (S, N, M, E proteins) for potential B cell epitopes, as an ideal vaccine would be designed to stimulate both cellular and humoral immunity. Our analysis identified potential linear B cell epitopes in all proteins except for the E protein (**Table 2**). The greatest number of epitopes were predicted in the surface-exposed S protein (n=26), but a significant number of epitopes were also predicted for the N protein (n=14). This is not surprising, as previous reports identified the N protein as a significant target of the humoral response to SARS-CoV (63, 64). As the S protein is the predominant surface protein and has been the primary target of neutralizing antibody responses against other coronaviruses (56, 57), we elected to focus our subsequent analyses solely on antigenic regions in the S protein. We identified 14 potential structural epitopes in the S protein structure and referenced against our linear epitope predictions to identify six regions that were independently identified by both analyses (**Table 2, Figure 4**). Feng *et al*. recently reported the computational identification of 19 surface epitopes in the S protein using Bepipred and the Kolaskar method (38), four of which had significant sequence overlap with the regions identified by our analyses.

To further evaluate the potential of these six antigenic regions as targets for antibody binding, we modeled their surface accessibility on the crystal structure of the SARS-Cov-2 spike protein (52). Four regions in the S1 domain (D405-D428; N440-N450; G496-P507; D568-T573) were solvent exposed (**Figure 4 A, B**), with minimal steric hindrance for antibody accessibility. The S1 domain contains the residues (N331-V524) important for virus binding to angiotensin converting enzyme 2 (ACE2) on the cell surface (65), and studies have shown that antibodies with potent neutralizing activity against SARS-CoV target this domain (66–68). Indeed, three of the four S1 epitopes identified in our analyses are located in the ACE2-binding region, supporting their potential utility in vaccine development against SARS-CoV-2. Two regions were identified in the S2 “stalk” domain of the S protein (**Figure 4 A, C**). While Y1138-D1146 is located at the base of the S protein and likely inaccessible to antibodies, P792-D796 is on the outer face of the protein and has been previously identified as part of a larger B cell epitope that is conserved with SARS-CoV (35). As SARS-CoV S2-specific antibodies have previously been shown to possess antiviral activity (66), it is interesting to speculate whether a strategy similar to targeting the influenza hemagglutinin protein stalk could be employed for developing a broadly reactive coronavirus vaccine.

Our study possessed several strengths and limitations. Rather than restricting our analyses of HLA class I and class II epitopes to specific proteins based on prior studies of SARS-CoV immunology, we investigated the complete proteome of SARS-CoV-2 using an unbiased approach. Furthermore, we employed a multi-tiered strategy for identifying putative B cell and T cell epitopes from all viral proteins studied. Our initial analyses were performed with liberal thresholds for epitope identification, and at each additional step, we imposed more stringent selection criteria to filter these peptides to a subset of B cell and T cell epitopes for further study. Nevertheless, the results of this study are derived purely from computational methods, and it should be noted that computational algorithms can fail to capture a significant number of antigenic peptides (69). Experimental validation with biological samples will ultimately be needed.

During the early stages of a pandemic, access to sufficient biological samples may be extremely limited, so we must continue to utilize methodologies—such as computational predictive algorithms— that allow us to explore the epitope landscape for experimental vaccine development. Our approach in this study allowed us to identify and refine a manageable subset of T cell and B cell epitopes for further testing as components of a SARS-CoV-2 vaccine. Based on our results, our proposed SARS-CoV-2 vaccine formulation could contain the following: 1) one or more B cell peptide epitopes from the S protein to generate protective neutralizing antibodies; and 2) multiple HLA class I and class II-derived peptides from other viral proteins to stimulate robust CD8^+^ and CD4^+^ T cell responses. Based on global allele frequencies, these class I and class II peptides would be expected to collectively provide 74% and 99% population coverage, respectively. While such a vaccine could be readily formulated as a synthetic polypeptide or an adjuvanted peptide mixture, these strategies may not retain the epitope structural features necessary to induce a robust antibody response. Recombinant nanoparticles and assembly into VLPs represent promising alternative vaccine platforms, as they have been extensively used for the controlled display and delivery of peptide-based vaccine components (70–73). By omitting whole viral proteins from the vaccine formulation, a peptide-based SARS-CoV-2 vaccine should have a well-tolerated safety profile and avoid the adverse events previously observed with experimental SARS-CoV vaccines (19–22).

In summary, we have identified 41 potential T cell epitopes (5 HLA class I, 36 HLA class II) and 6 potential B cell epitopes from across the SARS-CoV-2 proteome that are predicted to have broad population coverage and could serve as the basis for designing investigational peptide-based vaccines. Further study on the biological relevance and immunogenicity of these peptides is warranted in an effort to develop a safe and effective vaccine to combat the SARS-CoV-2 pandemic.

## Acknowledgments

The authors would like to thank Caroline L. Vitse for editorial assistance with this manuscript. The research presented here was not supported by any specific funding source.

