## Supplemental Tables 1 and 2 for "Immunoinformatic identification of B cell and T cell epitopes in the SARS-CoV-2 proteome"

**Supplementary Table S1.** Partially refined list of HLA class I epitopes from SARS-CoV-2 proteome.

| **Protein** | **Peptide** | **Predicted Alleles** | **Predicted Binding Affinity (nM)** |
| --- | --- | --- | --- |
| E | LTALRLCAY | HLA-A*01:01, HLA-B*15:01 | 139.4, 60.2 |
| M | ATSRTLSYY | HLA-A*01:01, HLA-A*03:01, HLA-B*15:01, HLA-B*58:01 | 62.4, 184.4, 138.8, 238.6 |
| M | RLFARTRSM | HLA-B*07:02, HLA-B*08:01, HLA-B*15:01 | 90.7, 27.6, 31.6 |
| M | YANRNRFLY | HLA-A*01:01, HLA-B*15:01, HLA-B*58:01 | 119.2, 245.3, 65.7 |
| M | YFIASFRLF | HLA-A*24:02, HLA-B*15:01 | 7.9, 415.6 |
| N | FPRGQGVPI | HLA-B*07:02, HLA-B*08:01 | 4.7, 368.3 |
| N | KAYNVTQAF | HLA-B*15:01, HLA-B*58:01 | 18.9, 17.7 |
| N | KPRQKRTAT | HLA-B*07:02, HLA-B*08:01 | 18.8, 432.6 |
| N | LLLDRLNQL | HLA-A*02:01, HLA-B*08:01 | 11.3, 136.8 |
| N | LSPRWYFYY | HLA-A*01:01, HLA-B*58:01 | 76.9, 430.6 |
| N | SPRWYFYYL | HLA-B*07:02, HLA-B*08:01 | 15.3, 42.1 |
| S | FAMQMAYRF | HLA-A*24:02, HLA-B*15:01, HLA-B*58:01 | 142.9, 123.9, 23.4 |
| S | MIAQYTSAL | HLA-A*02:01, HLA-B*07:02, HLA-B*08:01, HLA-B*15:01 | 82.1, 63.6, 80.4, 71.1 |
| S | SIIAYTMSL | HLA-A*02:01, HLA-B*08:01, HLA-B*15:01 | 16.9, 239.2, 202.9 |
| S | WTAGAAAYY | HLA-A*01:01, HLA-B*15:01, HLA-B*58:01 | 31.1, 65.7, 281.5 |
| S | YLQPRTFLL | HLA-A*02:01, HLA-A*24:02, HLA-B*08:01 | 4.1, 201.3, 23.9 |
| S | YRLFRKSNL | HLA-B*08:01, HLA-B*27:05 | 202.3, 68.3 |
| S | YSSANNCTF | HLA-B*15:01, HLA-B*58:01 | 88.0, 33.1 |
| ORF1ab | AMSAFAMMF | HLA-A*24:02, HLA-B*15:01 | 294.5, 19.1 |
| ORF1ab | ASHMYCSFY | HLA-A*01:01, HLA-B*15:01 | 328.9, 86.2 |
| ORF1ab | CTNYMPYFF | HLA-A*01:01, HLA-B*58:01 | 397.9, 30.1 |
| ORF1ab | FLGRYMSAL | HLA-A*02:01, HLA-B*08:01 | 9.8, 55.1 |
| ORF1ab | FLKTNCCRF | HLA-B*08:01, HLA-B*15:01 | 300.4, 51.4 |
| ORF1ab | FLLNKEMYL | HLA-A*02:01, HLA-B*08:01 | 2.6, 91.4 |
| ORF1ab | FLNRFTTTL | HLA-A*02:01, HLA-B*08:01 | 5.4, 83.7 |
| ORF1ab | FTDGVCLFW | HLA-A*01:01, HLA-B*58:01 | 384.2, 28.6 |
| ORF1ab | FVNLKQLPF | HLA-B*08:01, HLA-B*15:01 | 241.3, 68.6 |
| ORF1ab | HSIGFDYVY | HLA-A*01:01, HLA-B*58:01 | 102.9, 26.2 |
| ORF1ab | ILFTRFFYV | HLA-A*02:01, HLA-B*08:01 | 2.2, 130.8 |
| ORF1ab | IMQLFFSYF | HLA-A*24:02, HLA-B*15:01 | 211.4, 85.8 |
| ORF1ab | IPRRNVATL | HLA-B*07:02, HLA-B*08:01 | 5.3, 101.8 |
| ORF1ab | IQLSSYSLF | HLA-A*24:02, HLA-B*15:01 | 50.2, 11.9 |
| ORF1ab | ISTKHFYWF | HLA-A*24:02, HLA-B*58:01 | 172.1, 44.2 |
| ORF1ab | KLAKKFDTF | HLA-A*24:02, HLA-B*15:01 | 195.7, 58.6 |
| ORF1ab | KLMGHFAWW | HLA-A*24:02, HLA-B*58:01 | 294.3, 49.9 |
| ORF1ab | KLVNKFLAL | HLA-A*02:01, HLA-B*08:01, HLA-B*15:01 | 30.4, 73.8, 236.5 |
| ORF1ab | KMNYQVNGY | HLA-A*03:01, HLA-B*15:01 | 132.3, 49.1 |
| ORF1ab | KSYELQTPF | HLA-B*15:01, HLA-B*58:01 | 28.5, 12.3 |
| ORF1ab | LASHMYCSF | HLA-B*15:01, HLA-B*58:01 | 48.1, 66.1 |
| ORF1ab | LLADKFPVL | HLA-A*02:01, HLA-B*08:01 | 11.1, 106.3 |
| ORF1ab | LLTNMFTPL | HLA-A*02:01, HLA-B*08:01, HLA-B*15:01 | 27.2, 230.1, 200.1 |
| ORF1ab | LMIERFVSL | HLA-A*02:01, HLA-B*08:01, HLA-B*15:01 | 6.8, 6.6, 43.2 |
| ORF1ab | LMNVLTLVY | HLA-A*01:01, HLA-A*03:01, HLA-B*15:01 | 238.7, 115.2, 11.6 |
| ORF1ab | LSFKELLVY | HLA-A*01:01, HLA-B*15:01, HLA-B*58:01 | 371.8, 42.6, 35.7 |
| ORF1ab | LTNIFGTVY | HLA-A*01:01, HLA-B*15:01, HLA-B*58:01 | 64.6, 73.7, 479.9 |
| ORF1ab | LVAEWFLAY | HLA-A*01:01, HLA-A*03:01, HLA-B*15:01 | 191.6, 378.8, 32.1 |
| ORF1ab | MFTPLVPFW | HLA-A*24:02, HLA-B*58:01 | 141.6, 80.6 |
| ORF1ab | MMISAGFSL | HLA-A*02:01, HLA-B*08:01, HLA-B*15:01 | 6.9, 367.6, 16.2 |
| ORF1ab | MMSAPPAQY | HLA-A*01:01, HLA-B*15:01, HLA-B*58:01 | 157.3, 14.4, 447.6 |
| ORF1ab | MPYFFTLLL | B58 | 103.1, 271.0 |
| ORF1ab | MSMTYGQQF | HLA-B*15:01, HLA-B*58:01 | 28.1, 11.4 |
| ORF1ab | MSNLGMPSY | HLA-A*01:01, HLA-B*15:01, HLA-B*58:01 | 184.2, 74.1, 87.6 |
| ORF1ab | MVMCGGSLY | HLA-A*03:01, HLA-B*15:01 | 143.8, 14.1 |
| ORF1ab | NPAWRKAVF | HLA-B*07:02, HLA-B*08:01 | 23.5, 158.4 |
| ORF1ab | QLYLGGMSY | HLA-A*03:01, HLA-B*15:01 | 68.1, 25.2 |
| ORF1ab | RLYYDSMSY | HLA-A*03:01, HLA-B*15:01 | 10.9, 7.9 |
| ORF1ab | RMYIFFASF | HLA-A*24:02, HLA-B*15:01 | 196.9, 13.9 |
| ORF1ab | RQFHQKLLK | HLA-A*03:01, HLA-B*27:05 | 16.9, 104.5 |
| ORF1ab | RTAPHGHVM | HLA-B*07:02, HLA-B*15:01, HLA-B*58:01 | 56.4, 62.2, 189.1 |
| ORF1ab | RTNVYLAVF | HLA-B*15:01, HLA-B*58:01 | 36.4, 24.4 |
| ORF1ab | SMMGFKMNY | HLA-A*03:01, HLA-B*15:01 | 20.3, 23.4 |
| ORF1ab | SSAKSASVY | HLA-A*01:01, HLA-B*15:01 | 444.5, 48.4 |
| ORF1ab | SSLPSYAAF | HLA-B*15:01, HLA-B*58:01 | 49.2, 101.1 |
| ORF1ab | STNVTIATY | HLA-A*01:01, HLA-B*15:01, HLA-B*58:01 | 241.1, 81.9, 294.5 |
| ORF1ab | TQYNRYLAL | HLA-B*08:01, HLA-B*15:01 | 39.5, 73.2 |
| ORF1ab | TVAYFNMVY | HLA-A*01:01, HLA-B*15:01 | 456.8, 57.8 |
| ORF1ab | VAKSHNIAL | HLA-B*07:02, HLA-B*08:01 | 102.0, 98.0 |
| ORF1ab | VMHANYIFW | HLA-A*24:02, HLA-B*58:01 | 188.0, 69.6 |
| ORF1ab | VMYMGTLSY | HLA-A*03:01, HLA-B*15:01 | 9.2, 5.3 |
| ORF1ab | VPMEKLKTL | HLA-B*07:02, HLA-B*08:01 | 17.3, 90.8 |
| ORF1ab | VQSTQWSLF | HLA-A*24:02, HLA-B*15:01 | 258.7, 31.3 |
| ORF1ab | VVYRGTTTY | HLA-A*03:01, HLA-B*15:01 | 70.9, 23.3 |
| ORF1ab | WSMATYYLF | HLA-A*24:02, HLA-B*15:01, HLA-B*58:01 | 15.6, 42.7, 8.6 |
| ORF1ab | YFMRFRRAF | HLA-A*24:02, HLA-B*08:01 | 124.5, 91.6 |
| ORF1ab | YIFFASFYY | HLA-A*01:01, HLA-A*03:01 | 89.8, 195.5 |
| ORF1ab | YLDAYNMMI | HLA-A*01:01, HLA-A*02:01 | 236.2, 2.9 |
| ORF1ab | YLITPVHVM | HLA-A*02:01, HLA-B*15:01 | 26.5, 49.2 |
| ORF1ab | YLRKHFSMM | HLA-B*07:02, HLA-B*08:01, HLA-B*15:01 | 168.3, 5.2, 37.8 |
| ORF1ab | YMPYFFTLL | HLA-A*02:01, HLA-A*24:02 | 11.2, 177.5 |
| ORF1ab | YTEISFMLW | HLA-A*01:01, HLA-B*58:01 | 305.4, 16.9 |
| ORF1ab | YVMHANYIF | HLA-A*24:02, HLA-B*15:01, HLA-B*58:01 | 79.2, 28.8, 151.4 |
| ORF3a | APFLYLYAL | HLA-B*07:02, HLA-B*08:01 | 50.1, 156.7 |
| ORF7 | ITLATCELY | HLA-A*01:01, HLA-B*58:01 | 285.7, 84.4 |

**Supplementary Table S2.** Partially refined list of HLA class II epitopes from SARS-CoV-2 proteome.

| **Protein** | **Peptide** | **Predicted Alleles** | **Predicted Binding Affinity (nM)** |
| --- | --- | --- | --- |
| E | FYVYSRVKNLNSSRV | HLA-DRB1*04:01, HLA-DRB1*04:05, HLA-DRB1*08:02, HLA-DRB1*11:01, HLA-DRB3*02:02, HLA-DRB5*01:01 | 50.1, 69.7, 148.1, 47.2, 130.8, 67.6 |
| E | SFYVYSRVKNLNSSR | HLA-DRB1*11:01, HLA-DRB3*02:02, HLA-DRB5*01:01 | 42.3, 164.0, 62.1 |
| E | LVTLAILTALRLCAY | HLA-DRB1*11:01, HLA-DRB1*12:01, HLA-DRB1*15:01, HLA-DRB4*01:01 | 82.2, 121.5, 75.5, 97.4 |
| M | TLSYYKLGASQRVAG | HLA-DRB1*01:01, HLA-DRB5*01:01 | 7.3, 16.7 |
| M | RTLSYYKLGASQRVA | HLA-DRB1*01:01, HLA-DRB1*04:01, HLA-DRB1*04:05, HLA-DRB1*07:01, HLA-DRB1*09:01 | 7.5, 71.3, 118.8, 31.5, 25.6 |
| M | ASFRLFARTRSMWSF | HLA-DRB1*01:01, HLA-DRB1*07:01, HLA-DRB1*08:02, HLA-DRB1*09:01, HLA-DRB1*11:01, HLA-DRB5*01:01, HLA-DPA1*02:01/DPB1*05:01, HLA-DPA1*02:01/DPB1*14:01 | 19.2, 30.9, 53.5, 49.9, 12.2, 16.3, 256.2, 387.3 |
| M | IASFRLFARTRSMWS | HLA-DRB1*01:01, HLA-DRB1*04:05 | 23.9, 77.8 |
| M | LLQFAYANRNRFLYI | HLA-DRB1*03:01, HLA-DRB1*07:01, HLA-DRB1*08:02, HLA-DRB1*11:01, HLA-DRB1*13:02, HLA-DRB3*02:02, HLA-DRB5*01:01 | 179.8, 58.2, 225.6, 36.2, 27.8, 46.6, 26.3 |
| M | SFRLFARTRSMWSFN | HLA-DRB1*04:01, HLA-DRB1*04:05 | 47.5, 53.7 |
| M | ETNILLNVPLHGTIL | HLA-DRB1*08:02, HLA-DRB1*13:02, HLA-DRB3*02:02 | 228.4, 24.3, 66.5 |
| M | LSYFIASFRLFARTR | HLA-DRB1*11:01, HLA-DRB1*15:01, HLA-DPA1*02:01/DPB1*01:01,HLA-DPA1*03:01/DPB1*04:02, HLA-DPA1*02:01/DPB1*05:01 | 27.4, 36.3, 75.5, 94.8, 112.8 |
| M | ILRGHLRIAGHHLGR | HLA-DRB1*11:01, HLA-DRB4*01:01, HLA-DRB5*01:01 | 37.6, 87.1, 34.2 |
| M | MWLSYFIASFRLFAR | HLA-DRB1*15:01, HLA-DPA1*02:01/DPB1*01:01,HLA-DPA1*01:03/DPB1*02:01, HLA-DPA1*01:03/DPB1*04:01, HLA-DPA1*03:01/DPB1*04:02, HLA-DPA1*02:01/DPB1*05:01 | 31.8, 63.4, 29.8, 42.5, 79.9, 133.8 |
| M | YRINWITGGIAIAMA | HLA-DQA1*05:01/DQB1*03:01, HLA-DQA1*01:02/DQB1*06:02 | 20.2, 56.4 |
| M | GLMWLSYFIASFRLF | HLA-DQA1*01:01/DQB1*05:01, HLA-DPA1*01:03/DPB1*02:01, HLA-DPA1*01:03/DPB1*04:01 | 141.4, 38.2, 54.4 |
| M | SYFIASFRLFARTRS | HLA-DPA1*01:03/DPB1*02:01, HLA-DPA1*01:03/DPB1*04:01 | 52.6, 73.7 |
| N | QIAQFAPSASAFFGM | HLA-DRB1*07:01, HLA-DRB1*09:01, HLA-DQA1*05:01/DQB1*03:01 | 32.2, 30.1, 57.8 |
| N | QIGYYRRATRRIRGG | HLA-DRB1*08:02, HLA-DRB1*11:01 | 111.2, 14.9 |
| N | QQTVTLLPAADLDDF | HLA-DQA1*05:01/DQB1*02:01, HLA-DQA1*04:01/DQB1*04:02 | 188.6, 453.6 |
| S | AAEIRASANLAATKM | HLA-DRB1*08:02, HLA-DRB1*13:02, HLA-DRB3*02:02, HLA-DQA1*01:02/DQB1*06:02, HLA-DPA1*02:01/DPB1*14:01 | 101.3, 23.0, 52.7, 141.5, 327.4 |
| S | ALQIPFAMQMAYRFN | HLA-DRB1*09:01, HLA-DRB1*12:01, HLA-DRB1*15:01 | 52.9, 159.5, 50.3 |
| S | AQKFNGLTVLPPLLT | HLA-DRB1*04:01, HLA-DRB1*04:05 | 63.1, 46.9 |
| S | DLFLPFFSNVTWFHA | HLA-DRB1*04:01, HLA-DRB1*04:05, HLA-DPA1*01:03/DPB1*02:01, HLA-DPA1*01:03/DPB1*04:01 | 74.8, 60.5, 64.7, 100.0 |
| S | DSSSGWTAGAAAYYV | HLA-DQA1*05:01/DQB1*03:01, HLA-DQA1*04:01/DQB1*04:02 | 22.9, 455.2 |
| S | FGEVFNATRFASVYA | HLA-DRB1*07:01, HLA-DPA1*02:01/DPB1*01:01, HLA-DPA1*01:03/DPB1*02:01, HLA-DPA1*01:03/DPB1*04:01, HLA-DPA1*03:01/DPB1*04:02, HLA-DPA1*02:01/DPB1*05:01, HLA-DPA1*02:01/DPB1*14:01 | 25.5, 35.9, 19.6, 24.6, 55.0, 160.5, 278.3 |
| S | GINITRFQTLLALHR | HLA-DRB1*04:01, HLA-DRB1*04:05, HLA-DRB1*12:01, HLA-DRB1*15:01, HLA-DPA1*02:01/DPB1*01:01, HLA-DPA1*03:01/DPB1*04:02, HLA-DPA1*02:01/DPB1*14:01 | 64.3, 45.1, 145.1, 27.6, 106.0, 167.6, 433.6 |
| S | GNYNYLYRLFRKSNL | HLA-DRB1*15:01, HLA-DRB5*01:01 | 57.5, 31.5 |
| S | GWTFGAGAALQIPFA | HLA-DRB1*01:01, HLA-DRB1*09:01, HLA-DQA1*05:01/DQB1*03:01 | 16.2, 24.8, 20.8 |
| S | IIAYTMSLGAENSVA | HLA-DRB1*01:01, HLA-DRB1*04:01, HLA-DRB1*09:01, HLA-DQA1*01:02/DQB1*06:02 | 8.9, 55.0, 24.1, 171.3 |
| S | ITRFQTLLALHRSYL | HLA-DRB1*01:01, HLA-DRB1*12:01, HLA-DRB1*15:01, HLA-DRB5*01:01, HLA-DPA1*02:01/DPB1*05:01, HLA-DPA1*02:01/DPB1*14:01 | 10.7, 108.3, 34.9, 10.1, 156.2, 318.0 |
| S | ITSGWTFGAGAALQI | HLA-DRB1*09:01, HLA-DQA1*05:01/DQB1*03:01 | 36.2, 24.4 |
| S | KRSFIEDLLFNKVTL | HLA-DPA1*02:01/DPB1*01:01, HLA-DPA1*01:03/DPB1*02:01, HLA-DPA1*01:03/DPB1*04:01, HLA-DPA1*03:01/DPB1*04:02, HLA-DPA1*02:01/DPB1*05:01 | 51.5, 36.1, 50.3, 68.5, 158.4 |
| S | LFLPFFSNVTWFHAI | HLA-DRB1*04:05, HLA-DPA1*01:03/DPB1*02:01, HLA-DPA1*01:03/DPB1*04:01 | 62.9, 53.2, 83.1 |
| S | LQIPFAMQMAYRFNG | HLA-DRB4*01:01, HLA-DRB5*01:01 | 68.8, 20.9 |
| S | NCTFEYVSQPFLMDL | HLA-DPA1*02:01/DPB1*01:01, HLA-DPA1*01:03/DPB1*02:01, HLA-DPA1*01:03/DPB1*04:01, HLA-DPA1*03:01/DPB1*04:02 | 53.3, 30.9, 41.3, 70.1 |
| S | NITRFQTLLALHRSY | HLA-DRB1*04:01, HLA-DRB1*04:05, HLA-DRB1*08:02, HLA-DRB4*01:01 | 57.3, 41.9, 125.1, 54.1 |
| S | NYNYLYRLFRKSNLK | HLA-DRB1*08:02, HLA-DRB1*11:01, HLA-DRB1*15:01, HLA-DRB5*01:01, HLA-DPA1*02:01/DPB1*05:01 | 148.6, 16.6, 52.9, 168.8 |
| S | PINLVRDLPQGFSAL | HLA-DRB1*03:01, HLA-DRB3*01:01 | 51.7, 56.8 |
| S | PYRVVVLSFELLHAP | HLA-DPA1*02:01/DPB1*01:01, HLA-DPA1*01:03/DPB1*02:01, HLA-DPA1*01:03/DPB1*04:01, HLA-DPA1*03:01/DPB1*04:02 | 79.6, 53.3, 77.1, 92.9 |
| S | QDLFLPFFSNVTWFH | HLA-DRB1*04:01, HLA-DRB1*15:01 | 88.4, 44.6 |
| S | QPYRVVVLSFELLHA | HLA-DPA1*02:01/DPB1*01:01, HLA-DPA1*01:03/DPB1*02:01, HLA-DPA1*01:03/DPB1*04:01, HLA-DPA1*03:01/DPB1*04:02, HLA-DPA1*02:01/DPB1*05:01 | 73.2, 50.2, 71.4, 90.1, 211.1 |
| S | QQLIRAAEIRASANL | HLA-DRB1*08:02, HLA-DRB4*01:01, HLA-DQA1*01:02/DQB1*06:02, HLA-DPA1*02:01/DPB1*14:01 | 76.1, 42.9, 53.9, 229.4 |
| S | QSLLIVNNATNVVIK | HLA-DRB1*04:01, HLA-DRB1*13:02, HLA-DRB3*02:02 | 77.2, 5.9, 13.1 |
| S | REGVFVSNGTHWFVT | HLA-DRB1*13:02, HLA-DRB3*02:02 | 21.4, 25.8 |
| S | RSFIEDLLFNKVTLA | HLA-DPA1*01:03/DPB1*02:01, HLA-DPA1*01:03/DPB1*04:01 | 45.4, 63.1 |
| S | SIIAYTMSLGAENSV | HLA-DRB1*04:05, HLA-DRB1*07:01 | 68.4, 35.7 |
| S | TQLNRALTGIAVEQD | HLA-DQA1*03:01/DQB1*03:02, HLA-DQA1*04:01/DQB1*04:02 | 445.8, 373.3 |
| S | TRFQTLLALHRSYLT | HLA-DRB1*11:01, HLA-DRB1*12:01 | 20.9, 117.1 |
| S | TSNFRVQPTESIVRF | HLA-DRB1*04:01, HLA-DRB1*04:05 | 52.4, 83.9 |
| S | VFNATRFASVYAWNR | HLA-DRB1*09:01, HLA-DQA1*01:02/DQB1*06:02 | 48.7, 141.4 |
| S | VNFNFNGLTGTGVLT | HLA-DRB1*01:01, HLA-DRB1*09:01 | 14.5, 47.2 |
| S | YQPYRVVVLSFELLH | HLA-DPA1*02:01/DPB1*01:01, HLA-DPA1*01:03/DPB1*04:01, HLA-DPA1*03:01/DPB1*04:02, HLA-DPA1*02:01/DPB1*05:01 | 102.2, 93.0, 127.5, 299.3 |
| ORF1ab | AAIFYLITPVHVMSK | HLA-DRB1*01:01, HLA-DRB1*04:01, HLA-DRB1*04:05, HLA-DRB1*07:01, HLA-DRB1*08:02, HLA-DRB1*09:01, HLA-DRB1*15:01 | 9.9, 77.9, 59.6, 21.3, 116.2, 49.2, 49.4 |
| ORF1ab | AARYMRSLKVPATVS | HLA-DRB1*01:01, HLA-DRB1*04:01, HLA-DRB1*04:05, HLA-DRB1*08:02, HLA-DRB1*11:01, HLA-DRB4*01:01, HLA-DPA1*02:01/DPB1*14:01 | 10.9, 64.7, 82.9, 54.4, 23.9, 59.1, 249.3 |
| ORF1ab | ACRKVQHMVVKAALL | HLA-DRB4*01:01, HLA-DPA1*02:01/DPB1*14:01 | 77.5, 486.3 |
| ORF1ab | AFGLVAEWFLAYILF | HLA-DPA1*01:03/DPB1*02:01, HLA-DPA1*01:03/DPB1*04:01 | 40.2, 66.6 |
| ORF1ab | AHIQWMVMFTPLVPF | HLA-DQA1*01:01/DQB1*05:01, HLA-DPA1*03:01/DPB1*04:02 | 285.1, 141.4 |
| ORF1ab | AIILASFSASTSAFV | HLA-DRB1*01:01, HLA-DRB1*04:01, HLA-DRB1*04:05, HLA-DQA1*05:01/DQB1*03:01, HLA-DQA1*01:02/DQB1*06:02 | 13.8, 76.6, 87.5, 49.3, 152.2 |
| ORF1ab | AKRFKESPFELEDFI | HLA-DPA1*02:01/DPB1*01:01, HLA-DPA1*03:01/DPB1*04:02 | 73.2, 146.7 |
| ORF1ab | ALDISASIVAGGIVA | HLA-DQA1*05:01/DQB1*03:01, HLA-DQA1*01:02/DQB1*06:02 | 32.9, 73.9 |
| ORF1ab | AMMFTSDLATNNLVV | HLA-DRB1*04:01, HLA-DRB3*01:01 | 77.5, 63.1 |
| ORF1ab | ANYIFWRNTNPIQLS | HLA-DRB1*04:05, HLA-DRB1*07:01, HLA-DRB1*13:02 | 89.9, 35.2, 13.5 |
| ORF1ab | ARVVRSIFSRTLETA | HLA-DRB1*15:01, HLA-DPA1*02:01/DPB1*05:01 | 55.9, 282.2 |
| ORF1ab | ARYMRSLKVPATVSV | HLA-DRB1*04:01, HLA-DRB1*04:05, HLA-DRB1*08:02 | 64.7, 87.3, 54.4 |
| ORF1ab | ASFNYLKSPNFSKLI | HLA-DRB1*01:01, HLA-DRB1*04:01, HLA-DRB1*07:01, HLA-DRB5*01:01 | 12.1, 66.5, 36.4, 28.8 |
| ORF1ab | AVNLLTNMFTPLIQP | HLA-DPA1*01:03/DPB1*02:01, HLA-DPA1*01:03/DPB1*04:01, HLA-DPA1*03:01/DPB1*04:02 | 55.8, 95.0, 142.2 |
| ORF1ab | AVVLLILMTARTVYD | HLA-DRB1*01:01, HLA-DRB1*08:02, HLA-DRB4*01:01 | 14.5, 140.1, 77.3 |
| ORF1ab | CTNYMPYFFTLLLQL | HLA-DPA1*02:01/DPB1*01:01, HLA-DPA1*01:03/DPB1*02:01, HLA-DPA1*01:03/DPB1*04:01, HLA-DPA1*03:01/DPB1*04:02 | 108.2, 40.8, 54.1, 112.4 |
| ORF1ab | DMILSLLSKGRLIIR | HLA-DRB1*12:01, HLA-DRB1*15:01, HLA-DRB4*01:01 | 85.4, 38.9, 52.9 |
| ORF1ab | EAARYMRSLKVPATV | HLA-DRB1*01:01, HLA-DRB1*15:01, HLA-DRB4*01:01, HLA-DRB5*01:01, HLA-DPA1*02:01/DPB1*14:01 | 11.5, 50.4, 64.5, 31.0, 288.8 |
| ORF1ab | EASFNYLKSPNFSKL | HLA-DRB1*04:05, HLA-DRB1*09:01, HLA-DRB5*01:01 | 69.6, 47.8, 33.6 |
| ORF1ab | EKLKTLVATAEAELA | HLA-DQA1*05:01/DQB1*02:01, HLA-DQA1*03:01/DQB1*03:02, HLA-DQA1*04:01/DQB1*04:02, HLA-DPA1*02:01/DPB1*14:01 | 117.7, 309.6, 233.5, 476.4 |
| ORF1ab | ELLVYAADPAMHAAS | HLA-DRB1*01:01, HLA-DRB1*03:01, HLA-DRB1*04:01, HLA-DRB1*04:05, HLA-DRB3*01:01, HLA-DRB3*02:02, HLA-DQA1*05:01/DQB1*02:01 | 14.9, 99.3, 37.3, 83.0, 29.6, 66.6, 240.7 |
| ORF1ab | EMYLKLRSDVLLPLT | HLA-DRB3*01:01, HLA-DPA1*02:01/DPB1*01:01, HLA-DPA1*03:01/DPB1*04:02 | 56.9, 93.5, 148.9 |
| ORF1ab | ERLKLFAAETLKATE | HLA-DRB1*01:01, HLA-DRB1*04:01, HLA-DRB1*04:05, HLA-DPA1*03:01/DPB1*04:02, HLA-DPA1*02:01/DPB1*05:01 | 11.9, 47.4, 61.4, 164.7, 160.2 |
| ORF1ab | ERYKLEGYAFEHIVY | HLA-DPA1*02:01/DPB1*01:01, HLA-DPA1*01:03/DPB1*02:01, HLA-DPA1*01:03/DPB1*04:01 | 101.7, 61.4, 83.1 |
| ORF1ab | EWFLAYILFTRFFYV | HLA-DPA1*02:01/DPB1*01:01, HLA-DPA1*01:03/DPB1*02:01, HLA-DPA1*01:03/DPB1*04:01, HLA-DPA1*03:01/DPB1*04:02, HLA-DPA1*02:01/DPB1*05:01 | 79.6, 26.3, 33.1, 73.4, 287.7 |
| ORF1ab | EYSHVVAFNTLLFLM | HLA-DPA1*01:03/DPB1*02:01, HLA-DPA1*01:03/DPB1*04:01, HLA-DPA1*03:01/DPB1*04:02 | 59.2, 98.5, 147.3 |
| ORF1ab | FAMGIIAMSAFAMMF | HLA-DRB1*12:01, HLA-DRB1*15:01, HLA-DQA1*05:01/DQB1*03:01 | 141.0, 38.6, 60.1 |
| ORF1ab | FGEYSHVVAFNTLLF | HLA-DRB1*04:05, HLA-DRB1*07:01, HLA-DRB1*09:01 | 85.9, 29.7, 48.3 |
| ORF1ab | FKWDLTAFGLVAEWF | HLA-DQA1*05:01/DQB1*02:01, HLA-DQA1*03:01/DQB1*03:02, HLA-DQA1*04:01/DQB1*04:02 | 178.3, 425.3, 349.3 |
| ORF1ab | FLELAMDEFIERYKL | HLA-DRB1*03:01, HLA-DPA1*02:01/DPB1*01:01, HLA-DPA1*03:01/DPB1*04:02, HLA-DPA1*02:01/DPB1*05:01 | 98.0, 86.9, 147.1, 288.8 |
| ORF1ab | FYAYLRKHFSMMILS | HLA-DRB1*07:01, HLA-DRB1*09:01, HLA-DRB1*15:01, HLA-DRB4*01:01, HLA-DPA1*02:01/DPB1*05:01, HLA-DPA1*02:01/DPB1*14:01 | 27.8, 49.7, 24.7, 87.6, 202.6, 420.7 |
| ORF1ab | GDQFKHLIPLMYKGL | HLA-DRB1*04:05, HLA-DRB1*12:01, HLA-DRB5*01:01 | 80.6, 157.9, 31.4 |
| ORF1ab | GDYFVLTSHTVMPLS | HLA-DRB1*01:01, HLA-DRB1*04:01, HLA-DRB1*04:05, HLA-DRB1*07:01, HLA-DRB1*09:01 | 10.0, 77.4, 83.9, 19.0, 42.9 |
| ORF1ab | HIQWMVMFTPLVPFW | HLA-DQA1*01:01/DQB1*05:01, HLA-DPA1*02:01/DPB1*01:01, HLA-DPA1*01:03/DPB1*04:01, HLA-DPA1*03:01/DPB1*04:02 | 293.1, 116.3, 84.6, 135.4 |
| ORF1ab | HQKLLKSIAATRGAT | HLA-DRB1*01:01, HLA-DRB1*08:02, HLA-DRB5*01:01 | 9.9, 55.9, 16.7 |
| ORF1ab | IASEFSSLPSYAAFA | HLA-DRB1*04:01, HLA-DRB1*04:05, HLA-DRB1*09:01 | 40.8, 47.9, 36.8 |
| ORF1ab | IDFLELAMDEFIERY | HLA-DQA1*01:01/DQB1*05:01, HLA-DPA1*02:01/DPB1*01:01, HLA-DPA1*03:01/DPB1*04:02 | 114.7, 83.4, 151.0 |
| ORF1ab | IINLVQMAPISAMVR | HLA-DRB1*01:01, HLA-DRB1*08:02, HLA-DRB4*01:01 | 12.8, 118.8, 54.7 |
| ORF1ab | IKNFKSVLYYQNNVF | HLA-DPA1*02:01/DPB1*01:01, HLA-DPA1*01:03/DPB1*04:01, HLA-DPA1*02:01/DPB1*05:01 | 103.0, 83.0, 251.4 |
| ORF1ab | INLVQMAPISAMVRM | HLA-DRB1*12:01, HLA-DRB4*01:01, HLA-DQA1*01:02/DQB1*06:02, HLA-DPA1*02:01/DPB1*14:01 | 176.9, 57.1, 116.5, 398.6 |
| ORF1ab | IVFMCVEYCPIFFIT | HLA-DPA1*02:01/DPB1*01:01, HLA-DPA1*01:03/DPB1*02:01, HLA-DPA1*01:03/DPB1*04:01, HLA-DPA1*03:01/DPB1*04:02 | 116.2, 53.9, 70.9, 144.9 |
| ORF1ab | IVTALRANSAVKLQN | HLA-DRB1*08:02, HLA-DRB1*13:02, HLA-DRB3*02:02, HLA-DPA1*02:01/DPB1*14:01 | 115.9, 9.4, 19.5, 408.7 |
| ORF1ab | KEIKESVQTFFKLVN | HLA-DPA1*02:01/DPB1*01:01, HLA-DPA1*01:03/DPB1*02:01, HLA-DPA1*01:03/DPB1*04:01 | 107.1, 64.3, 94.0 |
| ORF1ab | KEMYLKLRSDVLLPL | HLA-DRB1*01:01, HLA-DRB1*04:05, HLA-DPA1*02:01/DPB1*05:01 | 11.0, 68.0, 285.6 |
| ORF1ab | KGRLIIRENNRVVIS | HLA-DRB1*12:01, HLA-DRB1*13:02, HLA-DRB1*15:01, HLA-DRB4*01:01 | 170.9, 9.5, 48.2, 58.8 |
| ORF1ab | KHFYWFFSNYLKRRV | HLA-DRB1*15:01, HLA-DPA1*02:01/DPB1*01:01, HLA-DPA1*01:03/DPB1*02:01, HLA-DPA1*01:03/DPB1*04:01, HLA-DPA1*03:01/DPB1*04:02, HLA-DPA1*02:01/DPB1*05:01 | 38.8, 57.3, 27.1, 38.3, 88.8, 115.3 |
| ORF1ab | KSAFYILPSIISNEK | HLA-DRB1*01:01, HLA-DRB1*04:01, HLA-DRB1*04:05, HLA-DRB1*08:02 | 9.3, 49.3, 47.5, 96.3 |
| ORF1ab | KTPKYKFVRIQPGQT | HLA-DRB1*04:01, HLA-DRB1*04:05, HLA-DRB1*08:02 | 77.4, 90.3, 112.3 |
| ORF1ab | KYKFVRIQPGQTFSV | HLA-DRB1*01:01, HLA-DRB1*07:01, HLA-DRB1*08:02, HLA-DRB1*09:01, HLA-DRB1*11:01, HLA-DRB1*15:01, HLA-DRB4*01:01, HLA-DRB5*01:01, HLA-DPA1*02:01/DPB1*14:01 | 7.2, 38.8, 67.3, 31.4, 30.6, 51.5, 58.3, 21.5, 421.1 |
| ORF1ab | KYLYFIKGLNNLNRG | HLA-DRB1*08:02, HLA-DRB1*11:01, HLA-DRB1*15:01, HLA-DRB5*01:01 | 139.3, 38.1, 40.6, 32.8 |
| ORF1ab | LASFSASTSAFVETV | HLA-DRB1*07:01, HLA-DRB1*09:01, HLA-DRB3*02:02, HLA-DQA1*05:01/DQB1*03:01, HLA-DQA1*03:01/DQB1*03:02, HLA-DQA1*04:01/DQB1*04:02, HLA-DQA1*01:02/DQB1*06:02 | 13.0, 20.8, 84.8, 62.8, 418.9, 302.9, 147.1 |
| ORF1ab | LCTFLLNKEMYLKLR | HLA-DRB3*02:02, HLA-DPA1*02:01/DPB1*01:01, HLA-DPA1*01:03/DPB1*02:01, HLA-DPA1*01:03/DPB1*04:01, HLA-DPA1*03:01/DPB1*04:02 | 43.0, 96.4, 63.1, 92.6, 130.6 |
| ORF1ab | LEETKFLTENLLLYI | HLA-DPA1*02:01/DPB1*01:01, HLA-DPA1*01:03/DPB1*02:01, HLA-DPA1*01:03/DPB1*04:01, HLA-DPA1*03:01/DPB1*04:02, HLA-DPA1*02:01/DPB1*05:01 | 36.9, 27.9, 37.3, 41.1, 145.1 |
| ORF1ab | LFFFLYENAFLPFAM | HLA-DRB1*04:05, HLA-DQA1*05:01/DQB1*02:01, HLA-DQA1*01:01/DQB1*05:01, HLA-DPA1*02:01/DPB1*01:01, HLA-DPA1*01:03/DPB1*02:01, HLA-DPA1*01:03/DPB1*04:01, HLA-DPA1*03:01/DPB1*04:02, HLA-DPA1*02:01/DPB1*05:01 | 72.4, 188.7, 136.1, 31.2, 14.9, 21.5, 37.1, 155.7 |
| ORF1ab | LGSLIYSTAALGVLM | HLA-DRB1*01:01, HLA-DRB1*07:01, HLA-DRB1*09:01, HLA-DQA1*01:02/DQB1*06:02 | 14.0, 18.3, 36.1, 156.5 |
| ORF1ab | LIVTALRANSAVKLQ | HLA-DRB1*01:01, HLA-DRB1*07:01, HLA-DRB4*01:01, HLA-DQA1*01:02/DQB1*06:02, HLA-DPA1*02:01/DPB1*14:01 | 8.8, 39.2, 78.6, 142.5, 368.3 |
| ORF1ab | LKHFFFAQDGNAAIS | HLA-DRB1*01:01, HLA-DRB1*04:01, HLA-DRB1*04:05, HLA-DRB3*01:01, HLA-DRB3*02:02 | 12.2, 33.6, 66.2, 32.8, 67.0 |
| ORF1ab | LLKSIAATRGATVVI | HLA-DRB1*01:01, HLA-DRB1*07:01, HLA-DRB1*09:01, HLA-DRB1*13:02, HLA-DQA1*01:02/DQB1*06:02, HLA-DPA1*02:01/DPB1*14:01 | 10.9, 14.9, 26.5, 25.2, 161.4, 346.8 |
| ORF1ab | LNSIIKTIQPRVEKK | HLA-DRB1*08:02, HLA-DRB1*12:01, HLA-DRB4*01:01 | 143.2, 170.3, 52.0 |
| ORF1ab | LPSYAAFATAQEAYE | HLA-DQA1*05:01/DQB1*02:01, HLA-DQA1*03:01/DQB1*03:02, HLA-DQA1*04:01/DQB1*04:02 | 163.1, 306.4, 191.8 |
| ORF1ab | LSRVLGLKTLATHGL | HLA-DRB1*08:02, HLA-DRB1*12:01, HLA-DRB4*01:01 | 101.6, 184.9, 81.0 |
| ORF1ab | LTFYLTNDVSFLAHI | HLA-DRB1*03:01, HLA-DRB1*04:05, HLA-DRB3*01:01, HLA-DRB3*02:02 | 89.2, 70.5, 30.5, 40.4 |
| ORF1ab | LVNKFLALCADSIII | HLA-DRB1*01:01, HLA-DRB1*04:01, HLA-DRB1*04:05, HLA-DRB1*07:01, HLA-DRB1*09:01, HLA-DRB1*15:01, HLA-DQA1*05:01/DQB1*02:01, HLA-DQA1*01:01/DQB1*05:01 | 7.4, 56.7, 44.8, 29.9, 33.8, 59.3, 158.9, 234.9 |
| ORF1ab | LVRKIFVDGVPFVVS | HLA-DRB1*03:01, HLA-DRB1*13:02, HLA-DRB3*01:01 | 46.5, 25.6, 22.8 |
| ORF1ab | LYAFASEAARVVRSI | HLA-DRB1*07:01, HLA-DQA1*05:01/DQB1*03:01, HLA-DPA1*02:01/DPB1*14:01 | 28.5, 31.7, 313.2 |
| ORF1ab | MIERFVSLAIDAYPL | HLA-DRB1*04:05, HLA-DQA1*05:01/DQB1*02:01, HLA-DQA1*04:01/DQB1*04:02, HLA-DPA1*02:01/DPB1*14:01 | 83.0, 96.3, 428.4, 392.1 |
| ORF1ab | MLRIMASLVLARKHT | HLA-DRB1*11:01, HLA-DRB1*12:01, HLA-DRB1*15:01, HLA-DRB5*01:01 | 34.2, 97.2, 33.9, 23.7 |
| ORF1ab | MPNMLRIMASLVLAR | HLA-DRB1*08:02, HLA-DRB4*01:01, HLA-DQA1*01:02/DQB1*06:02 | 98.1, 38.6, 93.7 |
| ORF1ab | MQLFFSYFAVHFISN | HLA-DQA1*01:01/DQB1*05:01, HLA-DPA1*01:03/DPB1*02:01, HLA-DPA1*01:03/DPB1*04:01, HLA-DPA1*03:01/DPB1*04:02 | 232.8, 30.3, 45.4, 103.5 |
| ORF1ab | MTYRRLISMMGFKMN | HLA-DRB1*01:01, HLA-DRB1*04:01, HLA-DRB1*04:05, HLA-DRB1*08:02, HLA-DRB1*09:01, HLA-DRB1*11:01, HLA-DRB5*01:01, HLA-DPA1*02:01/DPB1*14:01 | 6.4, 26.7, 27.7, 60.9, 42.4, 12.2, 13.5, 294.7 |
| ORF1ab | NEYRLYLDAYNMMIS | HLA-DRB1*04:05, HLA-DRB3*01:01, HLA-DQA1*01:01/DQB1*05:01 | 85.5, 44.4, 160.5 |
| ORF1ab | NFNVLFSTVFPPTSF | HLA-DPA1*02:01/DPB1*01:01, HLA-DPA1*01:03/DPB1*02:01, HLA-DPA1*01:03/DPB1*04:01, HLA-DPA1*03:01/DPB1*04:02 | 88.8, 34.5, 51.0, 127.9 |
| ORF1ab | NGDFLHFLPRVFSAV | HLA-DRB1*11:01, HLA-DRB1*12:01, HLA-DRB1*15:01 | 30.2, 198.0, 50.0 |
| ORF1ab | NLPFKLTCATTRQVV | HLA-DRB1*07:01, HLA-DRB1*09:01, HLA-DRB5*01:01 | 35.9, 58.6, 23.9 |
| ORF1ab | NLYDKLVSSFLEMKS | HLA-DPA1*01:03/DPB1*02:01, HLA-DPA1*01:03/DPB1*04:01, HLA-DPA1*03:01/DPB1*04:02, HLA-DPA1*02:01/DPB1*05:01 | 36.7, 45.5, 64.9, 140.4 |
| ORF1ab | NMLRIMASLVLARKH | HLA-DRB1*03:01, HLA-DQA1*01:02/DQB1*06:02, HLA-DPA1*02:01/DPB1*05:01, HLA-DPA1*02:01/DPB1*14:01 | 80.3, 164.2, 190.9, 210.1 |
| ORF1ab | NNCYLATALLTLQQI | HLA-DPA1*02:01/DPB1*01:01, HLA-DPA1*01:03/DPB1*02:01, HLA-DPA1*01:03/DPB1*04:01, HLA-DPA1*03:01/DPB1*04:02 | 111.3, 58.7, 79.7, 112.8 |
| ORF1ab | PASRELKVTFFPDLN | HLA-DPA1*02:01/DPB1*01:01, HLA-DPA1*01:03/DPB1*02:01, HLA-DPA1*01:03/DPB1*04:01, HLA-DPA1*03:01/DPB1*04:02 | 76.9, 48.9, 64.3, 149.5 |
| ORF1ab | PFAMGIIAMSAFAMM | HLA-DRB1*01:01, HLA-DRB1*09:01, HLA-DQA1*05:01/DQB1*03:01 | 12.3, 57.6, 45.6 |
| ORF1ab | PNMLRIMASLVLARK | HLA-DRB1*01:01, HLA-DRB1*07:01, HLA-DRB1*08:02, HLA-DRB1*09:01, HLA-DRB1*12:01, HLA-DRB1*15:01, HLA-DRB4*01:01 | 8.9, 30.9, 82.4, 56.9, 69.9, 22.2, 34.2 |
| ORF1ab | QESPFVMMSAPPAQY | HLA-DRB1*01:01, HLA-DRB1*04:05, HLA-DRB3*02:02 | 5.0, 36.2, 53.3 |
| ORF1ab | QGLVASIKNFKSVLY | HLA-DRB1*04:01, HLA-DRB1*04:05, HLA-DRB1*08:02, HLA-DRB1*11:01, HLA-DRB1*12:01, HLA-DRB4*01:01, HLA-DRB5*01:01 | 18.3, 62.8, 86.5, 37.7, 115.4, 86.6, 24.5 |
| ORF1ab | QKLLKSIAATRGATV | HLA-DRB1*01:01, HLA-DRB1*08:02, HLA-DRB4*01:01, HLA-DPA1*02:01/DPB1*14:01 | 9.1, 55.2, 73.1, 198.6 |
| ORF1ab | QLCQYLNTLTLAVPY | HLA-DRB1*01:01, HLA-DRB1*04:01, HLA-DRB1*04:05, HLA-DRB1*07:01 | 16.1, 66.8, 65.9, 35.8 |
| ORF1ab | QMEIDFLELAMDEFI | HLA-DQA1*05:01/DQB1*02:01, HLA-DQA1*03:01/DQB1*03:02, HLA-DQA1*01:01/DQB1*05:01 | 57.2, 450.7, 125.8 |
| ORF1ab | QMNLKYAISAKNRAR | HLA-DRB1*01:01, HLA-DRB1*04:01, HLA-DRB1*08:02, HLA-DRB1*09:01, HLA-DRB1*11:01, HLA-DRB3*02:02, HLA-DPA1*02:01/DPB1*14:01 | 14.9, 56.9, 49.1, 45.2, 22.1, 84.9, 158.3 |
| ORF1ab | QQKLALGGSVAIKIT | HLA-DRB1*01:01, HLA-DRB1*07:01, HLA-DRB1*09:01, HLA-DQA1*05:01/DQB1*03:01 | 12.6, 23.4, 32.3, 42.9 |
| ORF1ab | QTFFKLVNKFLALCA | HLA-DRB1*08:02, HLA-DRB1*12:01, HLA-DRB1*15:01, HLA-DPA1*02:01/DPB1*01:01, HLA-DPA1*01:03/DPB1*02:01 | 133.7, 197.3, 48.8, 81.5, 53.4 |
| ORF1ab | QWLTNIFGTVYEKLK | HLA-DPA1*02:01/DPB1*01:01, HLA-DPA1*01:03/DPB1*02:01, HLA-DPA1*01:03/DPB1*04:01, HLA-DPA1*03:01/DPB1*04:02 | 60.0, 36.2, 50.9, 97.1 |
| ORF1ab | QYNRYLALYNKYKYF | HLA-DRB1*15:01, HLA-DRB5*01:01, HLA-DPA1*02:01/DPB1*05:01 | 50.0, 27.1, 242.9 |
| ORF1ab | RFKESPFELEDFIPM | HLA-DPA1*02:01/DPB1*01:01, HLA-DPA1*01:03/DPB1*02:01, HLA-DPA1*01:03/DPB1*04:01, HLA-DPA1*03:01/DPB1*04:02 | 74.0, 65.9, 81.9, 130.6 |
| ORF1ab | RFYFYTSKTTVASLI | HLA-DRB1*01:01, HLA-DRB1*04:01, HLA-DRB1*04:05, HLA-DRB1*07:01, HLA-DRB1*08:02, HLA-DRB1*09:01, HLA-DRB3*02:02, HLA-DPA1*02:01/DPB1*14:01 | 10.1, 44.5, 75.7, 17.5, 98.6, 30.5, 46.1, 466.8 |
| ORF1ab | RIKIVQMLSDTLKNL | HLA-DRB1*04:01, HLA-DRB1*04:05, HLA-DRB1*15:01, HLA-DRB4*01:01 | 78.1, 58.8, 55.3, 43.4 |
| ORF1ab | RLKLFAAETLKATEE | HLA-DRB1*08:02, HLA-DRB1*15:01, HLA-DPA1*02:01/DPB1*01:01 | 100.9, 58.6, 109.5 |
| ORF1ab | RRLISMMGFKMNYQV | HLA-DRB1*08:02, HLA-DRB1*11:01, HLA-DPA1*02:01/DPB1*05:01 | 134.2, 30.5, 251.6 |
| ORF1ab | SAFAMMFVKHKHAFL | HLA-DRB1*08:02, HLA-DRB1*11:01, HLA-DRB1*15:01, HLA-DRB4*01:01, HLA-DRB5*01:01 | 110.4, 18.3, 50.9, 79.2, 15.1 |
| ORF1ab | SFLAHIQWMVMFTPL | HLA-DPA1*02:01/DPB1*01:01, HLA-DPA1*01:03/DPB1*02:01, HLA-DPA1*01:03/DPB1*04:01, HLA-DPA1*03:01/DPB1*04:02 | 103.9, 47.8, 70.7, 140.6 |
| ORF1ab | SHKLVLSVNPYVCNA | HLA-DRB1*04:01, HLA-DRB1*04:05, HLA-DRB1*08:02, HLA-DRB1*13:02, HLA-DRB1*15:01, HLA-DRB3*02:02 | 49.7, 72.9, 138.5, 11.4, 41.4, 32.5 |
| ORF1ab | SHRFYRLANECAQVL | HLA-DRB1*01:01, HLA-DRB1*04:01, HLA-DRB1*04:05 | 14.3, 50.4, 56.9 |
| ORF1ab | SIGFDYVYNPFMIDV | HLA-DPA1*02:01/DPB1*01:01, HLA-DPA1*01:03/DPB1*02:01, HLA-DPA1*01:03/DPB1*04:01, HLA-DPA1*03:01/DPB1*04:02 | 108.9, 47.1, 81.9, 137.6 |
| ORF1ab | SLLMPILTLTRALTA | HLA-DRB1*11:01, HLA-DRB1*12:01, HLA-DRB4*01:01 | 37.3, 122.2, 59.9 |
| ORF1ab | SPFVMMSAPPAQYEL | HLA-DRB1*08:02, HLA-DRB1*09:01, HLA-DQA1*05:01/DQB1*03:01 | 58.3, 33.8, 65.6 |
| ORF1ab | SPLYAFASEAARVVR | HLA-DRB1*09:01, HLA-DQA1*01:02/DQB1*06:02, HLA-DPA1*02:01/DPB1*14:01 | 28.9, 173.6, 397.7 |
| ORF1ab | SVQTFFKLVNKFLAL | HLA-DRB1*11:01, HLA-DPA1*02:01/DPB1*01:01, HLA-DPA1*01:03/DPB1*04:01, HLA-DPA1*03:01/DPB1*04:02, HLA-DPA1*02:01/DPB1*05:01 | 23.5, 76.9, 70.7, 111.5, 113.3 |
| ORF1ab | TCLAYYFMRFRRAFG | HLA-DRB1*08:02, HLA-DRB1*11:01, HLA-DRB1*15:01 | 143.9, 16.4, 44.7 |
| ORF1ab | TDFVNEFYAYLRKHF | HLA-DPA1*02:01/DPB1*01:01, HLA-DPA1*01:03/DPB1*02:01, HLA-DPA1*01:03/DPB1*04:01, HLA-DPA1*03:01/DPB1*04:02, HLA-DPA1*02:01/DPB1*05:01 | 90.9, 46.9, 67.2, 144.4, 239.2 |
| ORF1ab | TEETFKLSYGIATVR | HLA-DRB1*01:01, HLA-DRB1*07:01, HLA-DRB1*09:01 | 8.7, 21.8, 25.9 |
| ORF1ab | TEKYCALAPNMMVTN | HLA-DRB1*01:01, HLA-DRB1*04:01, HLA-DRB1*04:05, HLA-DRB1*09:01 | 8.5, 55.1, 71.1, 27.8 |
| ORF1ab | TERLKLFAAETLKAT | HLA-DRB1*09:01, HLA-DRB1*15:01, HLA-DPA1*02:01/DPB1*01:01, HLA-DPA1*02:01/DPB1*05:01, HLA-DPA1*02:01/DPB1*14:01 | 51.9, 46.6, 99.4, 161.5, 214.9 |
| ORF1ab | TEVNEFACVVADAVI | HLA-DQA1*05:01/DQB1*02:01, HLA-DQA1*03:01/DQB1*03:02, HLA-DQA1*04:01/DQB1*04:02 | 136.8, 424.7, 351.1 |
| ORF1ab | TNSRIKASMPTTIAK | HLA-DRB1*07:01, HLA-DRB1*09:01, HLA-DRB1*13:02 | 38.1, 51.7, 20.6 |
| ORF1ab | TRYVLMDGSIIQFPN | HLA-DRB1*01:01, HLA-DRB1*03:01, HLA-DRB1*04:01, HLA-DRB1*04:05 | 10.7, 46.6, 58.2, 85.6 |
| ORF1ab | TSAMQTMLFTMLRKL | HLA-DPA1*02:01/DPB1*01:01, HLA-DPA1*01:03/DPB1*04:01, HLA-DPA1*03:01/DPB1*04:02, HLA-DPA1*02:01/DPB1*05:01 | 98.6, 87.4, 113.4, 162.8 |
| ORF1ab | TYRRLISMMGFKMNY | HLA-DRB1*12:01, HLA-DRB1*15:01, HLA-DRB4*01:01, HLA-DRB5*01:01, HLA-DPA1*02:01/DPB1*05:01 | 66.9, 15.3, 34.3, 13.9, 166.5 |
| ORF1ab | VFTGYRVTKNSKVQI | HLA-DRB1*07:01, HLA-DRB1*09:01, HLA-DRB3*02:02, HLA-DRB5*01:01 | 19.1, 50.7, 53.1, 31.4 |
| ORF1ab | VLSFCAFAVDAAKAY | HLA-DQA1*05:01/DQB1*02:01, HLA-DQA1*04:01/DQB1*04:02, HLA-DQA1*01:02/DQB1*06:02 | 203.9, 360.2, 143.4 |
| ORF1ab | VLVQSTQWSLFFFLY | HLA-DPA1*02:01/DPB1*01:01, HLA-DPA1*01:03/DPB1*02:01, HLA-DPA1*01:03/DPB1*04:01, HLA-DPA1*03:01/DPB1*04:02 | 77.0, 35.3, 42.3, 93.1 |
| ORF1ab | VNRFNVAITRAKVGI | HLA-DRB1*08:02, HLA-DRB1*11:01, HLA-DRB5*01:01, HLA-DPA1*02:01/DPB1*14:01 | 78.4, 31.7, 12.1, 389.7 |
| ORF1ab | VNTFSSTFNVPMEKL | HLA-DRB1*04:01, HLA-DRB1*04:05, HLA-DRB1*07:01 | 77.6, 84.5, 38.7 |
| ORF1ab | VQSTQWSLFFFLYEN | HLA-DPA1*02:01/DPB1*01:01, HLA-DPA1*01:03/DPB1*02:01, HLA-DPA1*03:01/DPB1*04:02 | 107.1, 49.9, 129.8 |
| ORF1ab | VTCLAYYFMRFRRAF | HLA-DPA1*02:01/DPB1*01:01, HLA-DPA1*01:03/DPB1*02:01, HLA-DPA1*01:03/DPB1*04:01, HLA-DPA1*03:01/DPB1*04:02, HLA-DPA1*02:01/DPB1*05:01 | 94.2, 49.1, 62.2, 121.3, 132.7 |
| ORF1ab | VVDSYYSLLMPILTL | HLA-DPA1*02:01/DPB1*01:01, HLA-DPA1*01:03/DPB1*02:01, HLA-DPA1*01:03/DPB1*04:01, HLA-DPA1*03:01/DPB1*04:02 | 102.1, 58.0, 84.6, 114.6 |
| ORF1ab | VVLKKLKKSLNVAKS | HLA-DRB1*08:02, HLA-DRB4*01:01, HLA-DRB5*01:01 | 112.7, 76.3, 31.4 |
| ORF1ab | WLIINLVQMAPISAM | HLA-DRB1*12:01, HLA-DRB4*01:01, HLA-DQA1*01:02/DQB1*06:02 | 130.6, 65.9, 139.6 |
| ORF1ab | YFAVHFISNSWLMWL | HLA-DRB1*07:01, HLA-DPA1*02:01/DPB1*01:01, HLA-DPA1*01:03/DPB1*02:01, HLA-DPA1*01:03/DPB1*04:01 | 36.2, 84.0, 33.4, 55.3 |
| ORF1ab | YFMRFRRAFGEYSHV | HLA-DRB1*04:01, HLA-DRB1*04:05, HLA-DRB1*15:01 | 53.9, 50.7, 53.5 |
| ORF1ab | YFNMVYMPASWVMRI | HLA-DRB1*01:01, HLA-DRB1*04:05, HLA-DRB1*07:01, HLA-DRB1*09:01, HLA-DRB1*12:01, HLA-DRB1*15:01 | 8.3, 80.2, 38.2, 37.4, 184.5, 30.1 |
| ORF1ab | YLYFIKGLNNLNRGM | HLA-DRB1*01:01, HLA-DRB1*04:01, HLA-DRB1*04:05, HLA-DRB1*11:01, HLA-DRB3*02:02 | 9.7, 27.4, 32.3, 44.4, 70.3 |
| ORF3 | KKRWQLALSKGVHFV | HLA-DRB1*01:01, HLA-DRB1*07:01, HLA-DRB1*08:02, HLA-DRB1*09:01, HLA-DRB1*11:01, HLA-DRB1*12:01, HLA-DRB1*13:02, HLA-DRB1*15:01, HLA-DRB4*01:01, HLA-DRB5*01:01 | 9.2, 11.6, 200.3, 17.9, 43.1, 119.6, 30.0, 34.2, 79.8, 18.4 |
| ORF3 | SDFVRATATIPIQAS | HLA-DRB1*01:01, HLA-DRB1*07:01, HLA-DRB1*08:02, HLA-DRB1*09:01, HLA-DRB1*13:02 | 12.5, 16.0, 88.5, 27.8, 27.6 |
| ORF3 | PSDFVRATATIPIQA | HLA-DRB1*04:01, HLA-DRB1*04:05, HLA-DRB1*08:02 | 54.4, 90.0, 100.9 |
| ORF3 | YCIPYNSVTSSIVIT | HLA-DRB1*07:01, HLA-DRB1*09:01 | 24.3, 44.7 |
| ORF3 | ASKIITLKKRWQLAL | HLA-DRB1*08:02, HLA-DRB1*11:01 | 121.0, 20.5 |
| ORF3 | FVRATATIPIQASLP | HLA-DRB1*09:01, HLA-DQA1*01:02/DQB1*06:02, HLA-DPA1*02:01/DPB1*14:01 | 45.7, 47.8, 196.9 |
| ORF3 | VFQSASKIITLKKRW | HLA-DRB1*11:01,HLA-DRB5*01:01 | 32.2, 26.3 |
| ORF3 | YFLQSINFVRIIMRL | HLA-DRB1*12:01, HLA-DRB1*15:01, HLA-DRB4*01:01, HLA-DPA1*02:01/DPB1*05:01 | 124.4, 52.2, 81.9, 175.5 |
| ORF3 | VRATATIPIQASLPF | HLA-DQA1*04:01/DQB1*04:02, HLA-DQA1*01:02/DQB1*06:02 | 492.8, 51.5 |
| ORF3 | VYFLQSINFVRIIMR | HLA-DPA1*02:01/DPB1*01:01, HLA-DPA1*03:01/DPB1*04:02 | 58.7, 54.9 |
| ORF3 | VLHSYFTSDYYQLYS | HLA-DPA1*02:01/DPB1*01:01, HLA-DPA1*01:03/DPB1*02:01, HLA-DPA1*01:03/DPB1*04:01, HLA-DPA1*03:01/DPB1*04:02 | 65.6, 33.1, 40.3, 93.4 |
| ORF3 | LVYFLQSINFVRIIM | HLA-DPA1*01:03/DPB1*02:01, HLA-DPA1*01:03/DPB1*04:01 | 24.8, 32.9 |
| ORF6 | HLVDFQVTIAEILLI | HLA-DRB1*07:01, HLA-DQA1*05:01/DQB1*02:01 | 36.7, 198.5 |
| ORF6 | YIINLIIKNLSKSLT | HLA-DRB1*08:02, HLA-DRB1*11:01, HLA-DRB1*12:01, HLA-DRB1*13:02, HLA-DRB3*02:02, HLA-DRB4*01:01 | 113.5, 32.4, 135.4, 15.4, 43.6, 68.3 |
| ORF6 | IINLIIKNLSKSLTE | HLA-DRB1*08:02, HLA-DRB1*11:01 | 118.6, 31.7 |
| ORF6 | EILLIIMRTFKVSIW | HLA-DRB1*08:02, HLA-DRB1*12:01, HLA-DRB1*15:01, HLA-DRB4*01:01 | 130.2, 90.4, 20.2, 62.6 |
| ORF6 | DYIINLIIKNLSKSL | HLA-DRB1*11:01,HLA-DRB1*12:01, HLA-DRB3*02:02, HLA-DRB4*01:01 | 41.9, 166.1, 42.9, 77.1 |
| ORF6 | FKVSIWNLDYIINLI | HLA-DRB3*01:01, HLA-DQA1*01:01/DQB1*05:01 | 66.1, 90.1 |
| ORF6 | MFHLVDFQVTIAEIL | HLA-DQA1*05:01/DQB1*02:01, HLA-DQA1*01:01/DQB1*05:01, HLA-DPA1*02:01/DPB1*01:01, HLA-DPA1*01:03/DPB1*04:01 | 192.0, 292.1, 108.3, 100.7 |
| ORF6 | MRTFKVSIWNLDYII | HLA-DPA1*02:01/DPB1*01:01, HLA-DPA1*01:03/DPB1*02:01, HLA-DPA1*01:03/DPB1*04:01, HLA-DPA1*03:01/DPB1*04:02, HLA-DPA1*02:01/DPB1*05:01 | 79.0, 44.3, 60.7, 110.1, 304.0 |
| ORF7 | VKHVYQLRARSVSPK | HLA-DRB1*01:01, HLA-DRB1*08:02, HLA-DRB1*11:01, HLA-DRB4*01:01 | 14.3, 150.6, 38.3, 86.6 |
| ORF7 | GVKHVYQLRARSVSP | HLA-DRB1*11:01, HLA-DRB4*01:01 | 39.6, 82.1 |
| ORF7 | NKFALTCFSTQFAFA | HLA-DPA1*02:01/DPB1*01:01, HLA-DPA1*01:03/DPB1*02:01, HLA-DPA1*01:03/DPB1*04:01, HLA-DPA1*03:01/DPB1*04:02, HLA-DPA1*02:01/DPB1*05:01 | 50.9, 29.1, 35.9, 80.2, 273.4 |
| ORF7 | EEVQELYSPIFLIVA | HLA-DPA1*02:01/DPB1*01:01, HLA-DPA1*01:03/DPB1*02:01 | 59.7, 30.7 |
| ORF7 | FALTCFSTQFAFACP | HLA-DPA1*01:03/DPB1*02:01, HLA-DPA1*01:03/DPB1*04:01 | 61.9, 76.1 |
| ORF7 | EVQELYSPIFLIVAA | HLA-DPA1*01:03/DPB1*04:01, HLA-DPA1*03:01/DPB1*04:02 | 44.2, 74.6 |
| ORF8 | SKWYIRVGARKSAPL | HLA-DRB1*01:01, HLA-DRB1*08:02, HLA-DRB1*09:01, HLA-DRB1*11:01, HLA-DRB5*01:01 | 13.7, 87.8, 50.7, 15.3, 8.8 |
| ORF8 | LVVRCSFYEDFLEYH | HLA-DQA1*01:01/DQB1*05:01, HLA-DPA1*02:01/DPB1*01:01, HLA-DPA1*01:03/DPB1*04:01, HLA-DPA1*03:01/DPB1*04:02 | 177.0, 87.4, 89.3, 166.8 |
| ORF10 | FAFPFTIYSLLLCRM | HLA-DPA1*02:01/DPB1*01:01, HLA-DPA1*01:03/DPB1*02:01, HLA-DPA1*01:03/DPB1*04:01, HLA-DPA1*03:01/DPB1*04:02 | 97.3, 37.2, 56.9, 104.8 |
| ORF10 | INVFAFPFTIYSLLL | HLA-DPA1*02:01/DPB1*01:01, HLA-DPA1*01:03/DPB1*02:01, HLA-DPA1*01:03/DPB1*04:01, HLA-DPA1*03:01/DPB1*04:02 | 102.1, 36.8, 57.3, 126.7 |
| ORF10 | YINVFAFPFTIYSLL | HLA-DPA1*01:03/DPB1*02:01, HLA-DPA1*03:01/DPB1*04:02 | 42.9, 142.3 |

**Allele reference panels used for all binding predictions in this study:**

Class I: HLA-A*01:01, HLA-A*02:01, HLA-A*03:01, HLA-A*24:02, HLA-B*07:02, HLA-B*08:01, HLA-B*27:05, HLA-B*40:01, HLA-B*58:01, HLA-B*15:01

Class II: HLA-DRB1*01:01, HLA-DRB1*03:01, HLA-DRB1*04:01, HLA-DRB1*04:05, HLA-DRB1*07:01, HLA-DRB1*08:02, HLA-DRB1*09:01, HLA-DRB1*11:01, HLA-DRB1*12:01, HLA-DRB1*13:02, HLA-DRB1*15:01, HLA-DRB3*01:01, HLA-DRB3*02:02, HLA-DRB4*01:01, HLA-DRB5*01:01, HLA-DQA1*05:01/DQB1*02:01, HLA-DQA1*05:01/DQB1*03:01, HLA-DQA1*03:01/DQB1*03:02, HLA-DQA1*04:01/DQB1*04:02, HLA-DQA1*01:01/DQB1*05:01, HLA-DQA1*01:02/DQB1*06:02, HLA-DPA1*02:01/DPB1*01:01, HLA-DPA1*01:03/DPB1*02:01, HLA-DPA1*01:03/DPB1*04:01, HLA-DPA1*03:01/DPB1*04:02, HLA-DPA1*02:01/DPB1*05:01, HLA-DPA1*02:01/DPB1*14:01
