## Supplemental Figure 1 for "Immunoinformatic identification of B cell and T cell epitopes in the SARS-CoV-2 proteome"

```
CLUSTAL O(1.2.4) multiple sequence alignment


gb:MT020781|Organism:Severe      --------------------------------------------------TACCTTCCCA	10
gb:MT007544|Organism:Severe      --------------------------------------ATTAAAGGTTTATACCTTCCCA	22
gb:MN994467|Organism:Severe      --------------------------------------ATTAAAGGTTTATACCTTCCCA	22
gb:MT044257|Organism:Severe      --------------------------------------ATTAAAGGTTTATACCTTCCCA	22
gb:MT106054|Organism:Severe      --------------------------------------ATTAAAGGTTTATACCTTCCCA	22
gb:MT049951|Organism:Severe      --------------------------------------ATTAAAGGTTTATACCTTCCCA	22
gb:MN975262|Organism:Severe      --------------------------------------ATTAAAGGTTTATACCTTCCCA	22
gb:MT106052|Organism:Severe      --------------------------------------ATTAAAGGTTTATACCTTCCCA	22
gb:LC522975|Organism:Severe      -----------------------------------------AAAGGTTTATACCTTCCCA	19
gb:LC522973|Organism:Severe      -----------------------------------------AAAGGTTTATACCTTCCCA	19
gb:LC522974|Organism:Severe      -----------------------------------------AAAGGTTTATACCTTCCCA	19
gb:MN985325|Organism:Severe      --------------------------------------ATTAAAGGTTTATACCTTCCCA	22
gb:MT020881|Organism:Severe      --------------------------------------ATTAAAGGTTTATACCTTCCCA	22
gb:MT020880|Organism:Severe      --------------------------------------ATTAAAGGTTTATACCTTCCCA	22
gb:MT066175|Organism:Severe      --------------------------------------ATTAAAGGTTTATACCTTCCCA	22
gb:MN997409|Organism:Severe      --------------------------------------ATTAAAGGTTTATACCTTCCCA	22
gb:MN938384|Organism:Severe      ------------------------------------------------------------	0
gb:MT044258|Organism:Severe      --------------------------------------ATTAAAGGTTTATACCTTCCCA	22
gb:MT039890|Organism:Severe      --------------------------------------ATTAAAGGTTTATACCTTCCCA	22
gb:MN988713|Organism:Severe      --------------------------------------ATTAAAGGTTTATACCTTCCCA	22
gb:LC521925|Organism:Severe      -----------------------------------------AAAGGTTTATACCTTCCCA	19
gb:MT093571|Organism:Severe      --------------------------------------ATTAAAGGTTTATACCTTCCCA	22
gb:MT039887|Organism:Severe      --------------------------------------ATTAAAGGTTTATACCTTCCCA	22
gb:MT019530|Organism:Severe      --------------------------------------ATTAAAGGTTTATACCTTCCCA	22
gb:MT039888|Organism:Severe      --------------------------------------ATTAAAGGTTTATACCTTCCCA	22
gb:LC522972|Organism:Severe      -----------------------------------------AAAGGTTTATACCTTCCCA	19
gb:MT027063|Organism:Severe      --------------------------------------ATTAAAGGTTTATACCTTCCCA	22
gb:MT027062|Organism:Severe      --------------------------------------ATTAAAGGTTTATACCTTCCCA	22
gb:MT019529|Organism:Severe      --------------------------------------ATTAAAGGTTTATACCTTCCCA	22
gb:MN996529|Organism:Severe      --------------------------------------------------TACCTTCCCA	10
gb:MN996531|Organism:Severe      ---------------------------------------------------ACCTTCCCA	9
gb:MT066176|Organism:Severe      --------------------------------------ATTAAAGGTTTATACCTTCCCA	22
gb:MT027064|Organism:Severe      --------------------------------------ATTAAAGGTTTATACCTTCCCA	22
gb:MN994468|Organism:Severe      --------------------------------------ATTAAAGGTTTATACCTTCCCA	22
gb:MT072688|Organism:Severe      -----------------------------------------------------CTTCCCA	7
gb:MN996527|Organism:Severe      ------------------------------------------------------------	0
gb:MT093631|Organism:Severe      AGAGTGCTTATGAAAATTTTAATCAGCACGAAGTTCTACTTGCACCATTATACCTTCCCA	60
gb:MT106053|Organism:Severe      --------------------------------------ATTAAAGGTTTATACCTTCCCA	22
gb:MT019533|Organism:Severe      --------------------------------------ATTAAAGGTTTATACCTTCCCA	22
gb:MT019531|Organism:Severe      --------------------------------------ATTAAAGGTTTATACCTTCCCA	22
gb:MN996528|Organism:Severe      --------------------------------------ATTAAAGGTTTATACCTTCCCA	22
gb:MN996530|Organism:Severe      ----------------------------------------------------CCTTCCCA	8
gb:MN908947|Organism:Severe      --------------------------------------ATTAAAGGTTTATACCTTCCCA	22
gb:MT019532|Organism:Severe      --------------------------------------ATTAAAGGTTTATACCTTCCCA	22


gb:MT020781|Organism:Severe      GGTAACAAACCAACCAACTTTCGATCTCTTGTAGATCTGTTCTCTAAACGAACTTTAAAA	70
gb:MT007544|Organism:Severe      GGTAACAAACCAACCAACTTTCGATCTCTTGTAGATCTGTTCTCTAAACGAACTTTAAAA	82
gb:MN994467|Organism:Severe      GGTAACAAACCAACCAACTTTCGATCTCTTGTAGATCTGTTCTCTAAACGAACTTTAAAA	82
gb:MT044257|Organism:Severe      GGTAACAAACCAACCAACTTTCGATCTCTTGTAGATCTGTTCTCTAAACGAACTTTAAAA	82
gb:MT106054|Organism:Severe      GGTAACAAACCAACCAACTTTCGATCTCTTGTAGATCTGTTCTCTAAACGAACTTTAAAA	82
gb:MT049951|Organism:Severe      GGTAACAAACCAACCAACTTTCGATCTCTTGTAGATCTGTTCTCTAAACGAAATTTAAAA	82
gb:MN975262|Organism:Severe      GGTAACAAACCAACCAACTTTCGATCTCTTGTAGATCTGTTCTCTAAACGAACTTTAAAA	82
gb:MT106052|Organism:Severe      GGTAACAAACCAACCAACTTTCGATCTCTTGTAGATCTGTTCTCTAAACGAACTTTAAAA	82
gb:LC522975|Organism:Severe      GGTAACAAACCAACCAACTTTCGATCTCTTGTAGATCTGTTCTCTAAACGAACTTTAAAA	79
gb:LC522973|Organism:Severe      GGTAACAAACCAACCAACTTTCGATCTCTTGTAGATCTGTTCTCTAAACGAACTTTAAAA	79
gb:LC522974|Organism:Severe      GGTAACAAACCAACCAACTTTCGATCTCTTGTAGATCTGTTCTCTAAACGAACTTTAAAA	79
gb:MN985325|Organism:Severe      GGTAACAAACCAACCAACTTTCGATCTCTTGTAGATCTGTTCTCTAAACGAACTTTAAAA	82
gb:MT020881|Organism:Severe      GGTAACAAACCAACCAACTTTCGATCTCTTGTAGATCTGTTCTCTAAACGAACTTTAAAA	82
gb:MT020880|Organism:Severe      GGTAACAAACCAACCAACTTTCGATCTCTTGTAGATCTGTTCTCTAAACGAACTTTAAAA	82
gb:MT066175|Organism:Severe      GGTAACAAACCAACCAACTTTCGATCTCTTGTAGATCTGTTCTCTAAACGAACTTTAAAA	82
```

```
gb:MN997409|Organism:Severe      GGTAACAAACCAACCAACTTTCGATCTCTTGTAGATCTGTTCTCTAAACGAACTTTAAAA   82
gb:MN938384|Organism:Severe      ----------CAACCAACTTTCGATCTCTTGTAGATCTGTTCTCTAAACGAACTTTAAAA   50
gb:MT044258|Organism:Severe      GGTAACAAACCAACCAACTTTCGATCTCTTGTAGATCTGTTCTCTAAACGAACTTTAAAA   82
gb:MT039890|Organism:Severe      GGTAACAAACCAACCAACTTTCGATCTCTTGTAGATCTGTTCTCTAAACGAACTTTAAAA   82
gb:MN988713|Organism:Severe      GGTAACAAACCAACCAACTTTCGATCTCTTGTAGATCTGTTCTCTAAACGAACTTTAAAA   82
gb:LC521925|Organism:Severe      GGTAACAAACCAACCAACTTTCGATCTCTTGTAGATCTGTTCTCTAAACGAACTTTAAAA   79
gb:MT093571|Organism:Severe      GGTAACAAACCAACCAACTTTCGATCTCTTGTAGATCTGTTCTCTAAACGAACTTTAAAA   82
gb:MT039887|Organism:Severe      GGTAACAAACCAACCAACTTTCGATCTCTTGTAGATCTGTTCTCTAAACGAACTTTAAAA   82
gb:MT019530|Organism:Severe      GGTAACAAACCAACCAACTTTCGATCTCTTGTAGATCTGTTCTCTAAACGAACTTTAAAA   82
gb:MT039888|Organism:Severe      GGTAACAAACCAACCAACTTTCGATCTCTTGTAGATCTGTTCTCTAAACGAACTTTAAAA   82
gb:LC522972|Organism:Severe      GGTAACAAACCAACCAACTTTCGATCTCTTGTAGATCTGTTCTCTAAACGAACTTTAAAA   79
gb:MT027063|Organism:Severe      GGTAACAAACCAACCAACTTTCGATCTCTTGTAGATCTGTTCTCTAAACGAACTTTAAAA   82
gb:MT027062|Organism:Severe      GGTAACAAACCAACCAACTTTCGATCTCTTGTAGATCTGTTCTCTAAACGAACTTTAAAA   82
gb:MT019529|Organism:Severe      GGTAACAAACCAACCAACTTTCGATCTCTTGTAGATCTGTTCTCTAAACGAACTTTAAAA   82
gb:MN996529|Organism:Severe      GGTAACAAACCAACCAACTTTCGATCTCTTGTAGATCTGTTCTCTAAACGAACTTTAAAA   70
gb:MN996531|Organism:Severe      GGTAACAAACCAACCAACTTTCGATCTCTTGTAGATCTGTTCTCTAAACGAACTTTAAAA   69
gb:MT066176|Organism:Severe      GGTAACAAACCAACCAACTTTCGATCTCTTGTAGATCTGTTCTCTAAACGAACTTTAAAA   82
gb:MT027064|Organism:Severe      GGTAACAAACCAACCAACTTTCGATCTCTTGTAGATCTGTTCTCTAAACGAACTTTAAAA   82
gb:MN994468|Organism:Severe      GGTAACAAACCAACCAACTTTCGATCTCTTGTAGATCTGTTCTCTAAACGAACTTTAAAA   82
gb:MT072688|Organism:Severe      GGTAACAAACCAACCAACTTTCGATCTCTTGTAGATCTGTTCTCTAAACGAACTTTAAAA   67
gb:MN996527|Organism:Severe      -----------AACCAACTTTCGATCTCTTGTAGATCTGTTCTCTAAACGAACTTTAAAA   49
gb:MT093631|Organism:Severe      GGTAACAAACCAACCAACTTTCGATCTCTTGTAGATCTGTTCTCTAAACGAACTTTAAAA   120
gb:MT106053|Organism:Severe      GGTAACAAACCAACCAACTTTCGATCTCTTGTAGATCTGTTCTCTAAACGAACTTTAAAA   82
gb:MT019533|Organism:Severe      GGTAACAAACCAACCAACTTTCGATCTCTTGTAGATCTGTTCTCTAAACGAACTTTAAAA   82
gb:MT019531|Organism:Severe      GGTAACAAACCAACCAACTTTCGATCTCTTGTAGATCTGTTCTCTAAACGAACTTTAAAA   82
gb:MN996528|Organism:Severe      GGTAACAAACCAACCAACTTTCGATCTCTTGTAGATCTGTTCTCTAAACGAACTTTAAAA   82
gb:MN996530|Organism:Severe      GGTAACAAACCAACCAACTTTCGATCTCTTGTAGATCTGTTCTCTAAACGAACTTTAAAA   68
gb:MN908947|Organism:Severe      GGTAACAAACCAACCAACTTTCGATCTCTTGTAGATCTGTTCTCTAAACGAACTTTAAAA   82
gb:MT019532|Organism:Severe      GGTAACAAACCAACCAACTTTCGATCTCTTGTAGATCTGTTCTCTAAACGAACTTTAAAA   82
                                            ***************************************** *******

gb:MT020781|Organism:Severe      TCTGTGTGGCTGTCACTCGGCTGCATGCTTAGTGCACTCACGCAGTATAATTAATAACTA   130
gb:MT007544|Organism:Severe      TCTGTGTGGCTGTCACTCGGCTGCATGCTTAGTGCACTCACGCAGTATAATTAATAACTA   142
gb:MN994467|Organism:Severe      TCTGTGTGGCTGTCACTCGGCTGCATGCTTAGTGCACTCACGCAGTATAATTAATAACTA   142
gb:MT044257|Organism:Severe      TCTGTGTGGCTGTCACTCGGCTGCATGCTTAGTGCACTCACGCAGTATAATTAATAACTA   142
gb:MT106054|Organism:Severe      TCTGTGTGGCTGTCACTCGGCTGCATGCTTAGTGCACTCACGCAGTATAATTAATAACTA   142
gb:MT049951|Organism:Severe      TCTGTGTGGCTGTCACTCGGCTGCATGCTTAGTGCACTCACGCAGTATAATTAATAACTA   142
gb:MN975262|Organism:Severe      TCTGTGTGGCTGTCACTCGGCTGCATGCTTAGTGCACTCACGCAGTATAATTAATAACTA   142
gb:MT106052|Organism:Severe      TCTGTGTGGCTGTCACTCGGCTGCATGCTTAGTGCACTCACGCAGTATAATTAATAACTA   142
gb:LC522975|Organism:Severe      TCTGTGTGGCTGTCACTCGGCTGCATGCTTAGTGCACTCACGCAGTATAATTAATAACTA   139
gb:LC522973|Organism:Severe      TCTGTGTGGCTGTCACTCGGCTGCATGCTTAGTGCACTCACGCAGTATAATTAATAACTA   139
gb:LC522974|Organism:Severe      TCTGTGTGGCTGTCACTCGGCTGCATGCTTAGTGCACTCACGCAGTATAATTAATAACTA   139
gb:MN985325|Organism:Severe      TCTGTGTGGCTGTCACTCGGCTGCATGCTTAGTGCACTCACGCAGTATAATTAATAACTA   142
gb:MT020881|Organism:Severe      TCTGTGTGGCTGTCACTCGGCTGCATGCTTAGTGCACTCACGCAGTATAATTAATAACTA   142
gb:MT020880|Organism:Severe      TCTGTGTGGCTGTCACTCGGCTGCATGCTTAGTGCACTCACGCAGTATAATTAATAACTA   142
gb:MT066175|Organism:Severe      TCTGTGTGGCTGTCACTCGGCTGCATGCTTAGTGCACTCACGCAGTATAATTAATAACTA   142
gb:MN997409|Organism:Severe      TCTGTGTGGCTGTCACTCGGCTGCATGCTTAGTGCACTCACGCAGTATAATTAATAACTA   142
gb:MN938384|Organism:Severe      TCTGTGTGGCTGTCACTCGGCTGCATGCTTAGTGCACTCACGCAGTATAATTAATAACTA   110
gb:MT044258|Organism:Severe      TCTGTGTGGCTGTCACTCGGCTGCATGCTTAGTGCACTCACGCAGTATAATTAATAACTA   142
gb:MT039890|Organism:Severe      TCTGTGTGGCTGTCACTCGGCTGCATGCTTAGTGCACTCACGCAGTATAATTAATAACTA   142
gb:MN988713|Organism:Severe      TCTGTGTGGCTGTCACTCGGCTGCATGCTTAGTGCACTCACGCAGTATAATTAATAACTA   142
gb:LC521925|Organism:Severe      TCTGTGTGGCTGTCACTCGGCTGCATGCTTAGTGCACTCACGCAGTATAATTAATAACTA   139
gb:MT093571|Organism:Severe      TCTGTGTGGCTGTCACTCGGCTGCATGCTTAGTGCACTCACGCAGTATAATTAATAACTA   142
gb:MT039887|Organism:Severe      TCTGTGTGGCTGTCACTCGGCTGCATGCTTAGTGCACTCACGCAGTATAATTAATAACTA   142
gb:MT019530|Organism:Severe      TCTGTGTGGCTGTCACTCGGCAGCATGCCGAGTGCAGCCACACAGTATAATTAATAACTA   142
gb:MT039888|Organism:Severe      TCTGTGTGGCTGTCACTCGGCTGCATGCTTAGTGCACTCACGCAGTATAATTAATAACTA   142
gb:LC522972|Organism:Severe      TCTGTGTGGCTGTCACTCGGCTGCATGCTTAGTGCACTCACGCAGTATAATTAATAACTA   139
gb:MT027063|Organism:Severe      TCTGTGTGGCTGTCACTCGGCTGCATGCTTAGTGCACTCACGCAGTATAATTAATAACTA   142
gb:MT027062|Organism:Severe      TCTGTGTGGCTGTCACTCGGCTGCATGCTTAGTGCACTCACGCAGTATAATTAATAACTA   142
gb:MT019529|Organism:Severe      TCTGTGTGGCTGTCACTCGGCTGCATGCTTAGTGCACTCACGCAGTATAATTAATAACTA   142
gb:MN996529|Organism:Severe      TCTGTGTGGCTGTCACTCGGCTGCATGCTTAGTGCACTCACGCAGTATAATTAATAACTA   130
gb:MN996531|Organism:Severe      TCTGTGTGGCTGTCACTCGGCTGCATGCTTAGTGCACTCACGCAGTATAATTAATAACTA   129
gb:MT066176|Organism:Severe      TCTGTGTGGCTGTCACTCGGCTGCATGCTTAGTGCACTCACGCAGTATAATTAATAACTA   142
gb:MT027064|Organism:Severe      TCTGTGTGGCTGTCACTCGGCTGCATGCTTAGTGCACTCACGCAGTATAATTAATAACTA   142
gb:MN994468|Organism:Severe      TCTGTGTGGCTGTCACTCGGCTGCATGCTTAGTGCACTCACGCAGTATAATTAATAACTA   142
```

```
gb:MT072688|Organism:Severe      TCTGTGTGGCTGTCACTCGGCTGCATGCTTAGTGCACTCACGCAGTATAATTAATAACTA   127
gb:MN996527|Organism:Severe      TCTGTGTGGCTGTCACTCGGCTGCATGCTTAGTGCACTCACGCAGTATAATTAATAACTA   109
gb:MT093631|Organism:Severe      TCTGTGTGGCTGTCACTCGGCTGCATGCTTAGTGCACTCACGCAGTATAATTAATAACTA   180
gb:MT106053|Organism:Severe      TCTGTGTGGCTGTCACTCGGCTGCATGCTTAGTGCACTCACGCAGTATAATTAATAACTA   142
gb:MT019533|Organism:Severe      TCTGTGTGGCTGTCACTCGGCTGCATGCTTAGTGCACTCACGCAGTATAATTAATAACTA   142
gb:MT019531|Organism:Severe      TCTGTGTGGCTGTCACTCGGCTGCATGCTTAGTGCACTCACGCAGTATAATTAATAACTA   142
gb:MN996528|Organism:Severe      TCTGTGTGGCTGTCACTCGGCTGCATGCTTAGTGCACTCACGCAGTATAATTAATAACTA   142
gb:MN996530|Organism:Severe      TCTGTGTGGCTGTCACTCGGCTGCATGCTTAGTGCACTCACGCAGTATAATTAATAACTA   128
gb:MN908947|Organism:Severe      TCTGTGTGGCTGTCACTCGGCTGCATGCTTAGTGCACTCACGCAGTATAATTAATAACTA   142
gb:MT019532|Organism:Severe      TCTGTGTGGCTGTCACTCGGCTGCATGCTTAGTGCACTCACGCAGTATAATTAATAACTA   142
                                 ********************* ******  ******  *** ******************

gb:MT020781|Organism:Severe      ATTACTGTCGTTGACAGGACACGAGTAACTCGTCTATCTTCTGCAGGCTGCTTACGGTTT   190
gb:MT007544|Organism:Severe      ATTACTGTCGTTGACAGGACACGAGTAACTCGTCTATCTTCTGCAGGCTGCTTACGGTTT   202
gb:MN994467|Organism:Severe      ATTACTGTCGTTGACAGGACACGAGTAACTCGTCTATCTTCTGCAGGCTGCTTACGGTTT   202
gb:MT044257|Organism:Severe      ATTACTGTCGTTGACAGGACACGAGTAACTCGTCTATCTTCTGCAGGCTGCTTACGGTTT   202
gb:MT106054|Organism:Severe      ATTACTGTCGTTGACAGGACACGAGTAACTCGTCTATCTTCTGCAGGCTGCTTACGGTTT   202
gb:MT049951|Organism:Severe      ATTACTGTCGTTGACAGGACACGAGTAACTCGTCTATCTTCTGCAGGCTGCTTACGGTTT   202
gb:MN975262|Organism:Severe      ATTACTGTCGTTGACAGGACACGAGTAACTCGTCTATCTTCTGCAGGCTGCTTACGGTTT   202
gb:MT106052|Organism:Severe      ATTACTGTCGTTGACAGGACACGAGTAACTCGTCTATCTTCTGCAGGCTGCTTACGGTTT   202
gb:LC522975|Organism:Severe      ATTACTGTCGTTGACAGGACACGAGTAACTCGTCTATCTTCTGCAGGCTGCTTACGGTTT   199
gb:LC522973|Organism:Severe      ATTACTGTCGTTGACAGGACACGAGTAACTCGTCTATCTTCTGCAGGCTGCTTACGGTTT   199
gb:LC522974|Organism:Severe      ATTACTGTCGTTGACAGGACACGAGTAACTCGTCTATCTTCTGCAGGCTGCTTACGGTTT   199
gb:MN985325|Organism:Severe      ATTACTGTCGTTGACAGGACACGAGTAACTCGTCTATCTTCTGCAGGCTGCTTACGGTTT   202
gb:MT020881|Organism:Severe      ATTACTGTCGTTGACAGGACACGAGTAACTCGTCTATCTTCTGCAGGCTGCTTACGGTTT   202
gb:MT020880|Organism:Severe      ATTACTGTCGTTGACAGGACACGAGTAACTCGTCTATCTTCTGCAGGCTGCTTACGGTTT   202
gb:MT066175|Organism:Severe      ATTACTGTCGTTGACAGGACACGAGTAACTCGTCTATCTTCTGCAGGCTGCTTACGGTTT   202
gb:MN997409|Organism:Severe      ATTACTGTCGTTGACAGGACACGAGTAACTCGTCTATCTTCTGCAGGCTGCTTACGGTTT   202
gb:MN938384|Organism:Severe      ATTACTGTCGTTGACAGGACACGAGTAACTCGTCTATCTTCTGCAGGCTGCTTACGGTTT   170
gb:MT044258|Organism:Severe      ATTACTGTCGTTGACAGGACACGAGTAACTCGTCTATCTTCTGCAGGCTGCTTACGGTTT   202
gb:MT039890|Organism:Severe      ATTACTGTCGTTGACAGGACACGAGTAACTCGTCTATCTTCTGCAGGCTGCTTACGGTTT   202
gb:MN988713|Organism:Severe      ATTACTGTCGTTGACAGGACACGAGTAACTCGTCTATCTTCTGCAGGCTGCTTACGGTTT   202
gb:LC521925|Organism:Severe      ATTACTGTCGTTGACAGGACACGAGTAACTCGTCTATCTTCTGCAGGCTGCTTACGGTTT   199
gb:MT093571|Organism:Severe      ATTACTGTCGTTGACAGGACACGAGTAACTCGTCTATCTTCTGCAGGCTGCTTACGGTTT   202
gb:MT039887|Organism:Severe      ATTACTGTCGTTGACAGGACACGAGTAACTCGTCTATCTTCTGCAGGCTGCTTACGGTTT   202
gb:MT019530|Organism:Severe      ATTACTGTCGTTGACAGGACACGAGTAACTCGTCTATCTTCTGCAGGCTGCTTACGGTTT   202
gb:MT039888|Organism:Severe      ATTACTGTCGTTGACAGGACACGAGTAACTCGTCTATCTTCTGCAGGCTGCTTACGGTTT   202
gb:LC522972|Organism:Severe      ATTACTGTCGTTGACAGGACACGAGTAACTCGTCTATCTTCTGCAGGCTGCTTACGGTTT   199
gb:MT027063|Organism:Severe      ATTACTGTCGTTGACAGGACACGAGTAACTCGTCTATCTTCTGCAGGCTGCTTACGGTTT   202
gb:MT027062|Organism:Severe      ATTACTGTCGTTGACAGGACACGAGTAACTCGTCTATCTTCTGCAGGCTGCTTACGGTTT   202
gb:MT019529|Organism:Severe      ATTACTGTCGTTGACAGGACACGAGTAACTCGTCTATCTTCTGCAGGCTGCTTACGGTTT   202
gb:MN996529|Organism:Severe      ATTACTGTCGTTGACAGGACACGAGTAACTCGTCTATCTTCTGCAGGCTGCTTACGGTTT   190
gb:MN996531|Organism:Severe      ATTACTGTCGTTGACAGGACACGAGTAACTCGTCTATCTTCTGCAGGCTGCTTACGGTTT   189
gb:MT066176|Organism:Severe      ATTACTGTCGTTGACAGGACACGAGTAACTCGTCTATCTTCTGCAGGCTGCTTACGGTTT   202
gb:MT027064|Organism:Severe      ATTACTGTCGTTGACAGGACACGAGTAACTCGTCTATCTTCTGCAGGCTGCTTACGGTTT   202
gb:MN994468|Organism:Severe      ATTACTGTCGTTGACAGGACACGAGTAACTCGTCTATCTTCTGCAGGCTGCTTACGGTTT   202
gb:MT072688|Organism:Severe      ATTACTGTCGTTGACAGGACACGAGTAACTCGTCTATCTTCTGCAGGCTGCTTACGGTTT   187
gb:MN996527|Organism:Severe      ATTACTGTCGTTGACAGGACACGAGTAACTCGTCTATCTTCTGCAGGCTGCTTACGGTTT   169
gb:MT093631|Organism:Severe      ATTACTGTCGTTGACAGGACACGAGTAACTCGTCTATCTTCTGTAGGCTGCTTACGGTTT   240
gb:MT106053|Organism:Severe      ATTACTGTCGTTGACAGGACACGAGTAACTCGTCTATCTTCTGCAGGCTGCTTACGGTTT   202
gb:MT019533|Organism:Severe      ATTACTGTCGTTGACAGGACACGAGTAACTCGTCTATCTTCTGCAGGCTGCTTACGGTTT   202
gb:MT019531|Organism:Severe      ATTACTGTCGTTGACAGGACACGAGTAACTCGTCTATCTTCTGCAGGCTGCTTACGGTTT   202
gb:MN996528|Organism:Severe      ATTACTGTCGTTGACAGGACACGAGTAACTCGTCTATCTTCTGCAGGCTGCTTACGGTTT   202
gb:MN996530|Organism:Severe      ATTACTGTCGTTGACAGGACACGAGTAACTCGTCTATCTTCTGCAGGCTGCTTACGGTTT   188
gb:MN908947|Organism:Severe      ATTACTGTCGTTGACAGGACACGAGTAACTCGTCTATCTTCTGCAGGCTGCTTACGGTTT   202
gb:MT019532|Organism:Severe      ATTACTGTCGTTGACAGGACACGAGTAACTCGTCTATCTTCTGCAGGCTGCTTACGGTTT   202
                                 ******************************************* ****************

gb:MT020781|Organism:Severe      CGTCCGTGTTGCAGCCGATCATCAGCACATCTAGGTTTCGTCCGGGTGTGACCGAAAGGT   250
gb:MT007544|Organism:Severe      CGTCCGTGTTGCAGCCGATCATCAGCACATCTAGGTTTCGTCCGGGTGTGACCGAAAGGT   262
gb:MN994467|Organism:Severe      CGTCCGTGTTGCAGCCGATCATCAGCACATCTAGGTTTCGTCCGGGTGTGACCGAAAGGT   262
gb:MT044257|Organism:Severe      CGTCCGTGTTGCAGCCGATCATCAGCACATCTAGGTTTCGTCCGGGTGTGACCGAAAGGT   262
gb:MT106054|Organism:Severe      CGTCCGTGTTGCAGCCGATCATCAGCACATCTAGGTTTCGTCCGGGTGTGACCGAAAGGT   262
gb:MT049951|Organism:Severe      CGTCCGTGTTGCAGCCGATCATCAGCACATCTAGGTTTCGTCCGGGTGTGACCGAAAGGT   262
gb:MN975262|Organism:Severe      CGTCCGTGTTGCAGCCGATCATCAGCACATCTAGGTTTCGTCCGGGTGTGACCGAAAGGT   262
```

```
gb:MT106052|Organism:Severe      CGTCCGTGTTGCAGCCGATCATCAGCACATCTAGGTTTCGTCCGGGTGTGACCGAAAGGT    262
gb:LC522975|Organism:Severe      CGTCCGTGTTGCAGCCGATCATCAGCACATCTAGGTTTCGTCCGGGTGTGACCGAAAGGT    259
gb:LC522973|Organism:Severe      CGTCCGTGTTGCAGCCGATCATCAGCACATCTAGGTTTCGTCCGGGTGTGACCGAAAGGT    259
gb:LC522974|Organism:Severe      CGTCCGTGTTGCAGCCGATCATCAGCACATCTAGGTTTCGTCCGGGTGTGACCGAAAGGT    259
gb:MN985325|Organism:Severe      CGTCCGTGTTGCAGCCGATCATCAGCACATCTAGGTTTCGTCCGGGTGTGACCGAAAGGT    262
gb:MT020881|Organism:Severe      CGTCCGTGTTGCAGCCGATCATCAGCACATCTAGGTTTCGTCCGGGTGTGACCGAAAGGT    262
gb:MT020880|Organism:Severe      CGTCCGTGTTGCAGCCGATCATCAGCACATCTAGGTTTCGTCCGGGTGTGACCGAAAGGT    262
gb:MT066175|Organism:Severe      CGTCCGTGTTGCAGCCGATCATCAGCACATCTAGGTTTCGTCCGGGTGTGACCGAAAGGT    262
gb:MN997409|Organism:Severe      CGTCCGTGTTGCAGCCGATCATCAGCACATCTAGGTTTCGTCCGGGTGTGACCGAAAGGT    262
gb:MN938384|Organism:Severe      CGTCCGTGTTGCAGCCGATCATCAGCACATCTAGGTTTCGTCCGGGTGTGACCGAAAGGT    230
gb:MT044258|Organism:Severe      CGTCCGTGTTGCAGCCGATCATCAGCACATCTAGGTTTCGTCCGGGTGTGACCGAAAGGT    262
gb:MT039890|Organism:Severe      CGTCCGTGTTGCAGCCGATCATCAGCACATCTAGGTTTCGTCCGGGTGTGACCGAAAGGT    262
gb:MN988713|Organism:Severe      CGTCCGTGTTGCAGCCGATCATCAGCACATCTAGGTTTCGTCCGGGTGTGACCGAAAGGT    262
gb:LC521925|Organism:Severe      CGTCCGTGTTGCAGCCGATCATCAGCACATCTAGGTTTCGTCCGGGTGTGACCGAAAGGT    259
gb:MT093571|Organism:Severe      CGTCCGTGTTGCAGCCGATCATCAGCACATCTAGGTTTCGTCCGGGTGTGACCGAAAGGT    262
gb:MT039887|Organism:Severe      CGTCCGTGTTGCAGCCGATCATCAGCACATCTAGGTTTCGTCCGGGTGTGACCGAAAGGT    262
gb:MT019530|Organism:Severe      CGTCCGTGTTGCAGCCGATCATCAGCACATCTAGGTTTCGTCCGGGTGTGACCGAAAGGT    262
gb:MT039888|Organism:Severe      CGTCCGTGTTGCAGCCGATCATCAGCACATCTAGGTTTCGTCCGGGTGTGACCGAAAGGT    262
gb:LC522972|Organism:Severe      CGTCCGTGTTGCAGCCGATCATCAGCACATCTAGGTTTCGTCCGGGTGTGACCGAAAGGT    259
gb:MT027063|Organism:Severe      CGTCCGTGTTGCAGCCGATCATCAGCACATCTAGGTTTCGTCCGGGTGTGACCGAAAGGT    262
gb:MT027062|Organism:Severe      CGTCCGTGTTGCAGCCGATCATCAGCACATCTAGGTTTCGTCCGGGTGTGACCGAAAGGT    262
gb:MT019529|Organism:Severe      CGTCCGTGTTGCAGCCGATCATCAGCACATCTAGGTTTCGTCCGGGTGTGACCGAAAGGT    262
gb:MN996529|Organism:Severe      CGTCCGTGTTGCAGCCGATCATCAGCACATCTAGGTTTCGTCCGGGTGTGACCGAAAGGT    250
gb:MN996531|Organism:Severe      CGTCCGTGTTGCAGCCGATCATCAGCACATCTAGGTTTCGTCCGGGTGTGACCGAAAGGT    249
gb:MT066176|Organism:Severe      CGTCCGTGTTGCAGCCGATCATCAGCACATCTAGGTTTCGTCCGGGTGTGACCGAAAGGT    262
gb:MT027064|Organism:Severe      CGTCCGTGTTGCAGCCGATCATCAGCACATCTAGGTTTCGTCCGGGTGTGACCGAAAGGT    262
gb:MN994468|Organism:Severe      CGTCCGTGTTGCAGCCGATCATCAGCACATCTAGGTTTCGTCCGGGTGTGACCGAAAGGT    262
gb:MT072688|Organism:Severe      CGTCCGTGTTGCAGCCGATCATCAGCACATCTAGGTTTCGTCCGGGTGTGACCGAAAGGT    247
gb:MN996527|Organism:Severe      CGTCCGTGTTGCAGCCGATCATCAGCACATCTAGGTTTCGTCCGGGTGTGACCGAAAGGT    229
gb:MT093631|Organism:Severe      CGTCCGTGTTGCAGCCGATCATCAGCACATCTAGGTTTCGTCCGGGTGTGACCGAAAGGT    300
gb:MT106053|Organism:Severe      CGTCCGTGTTGCAGCCGATCATCAGCACATCTAGGTTTCGTCCGGGTGTGACCGAAAGGT    262
gb:MT019533|Organism:Severe      CGTCCGTGTTGCAGCCGATCATCAGCACATCTAGGTTTCGTCCGGGTGTGACCGAAAGGT    262
gb:MT019531|Organism:Severe      CGTCCGTGTTGCAGCCGATCATCAGCACATCTAGGTTTCGTCCGGGTGTGACCGAAAGGT    262
gb:MN996528|Organism:Severe      CGTCCGTGTTGCAGCCGATCATCAGCACATCTAGGTTTCGTCCGGGTGTGACCGAAAGGT    262
gb:MN996530|Organism:Severe      CGTCCGTGTTGCAGCCGATCATCAGCACATCTAGGTTTCGTCCGGGTGTGACCGAAAGGT    248
gb:MN908947|Organism:Severe      CGTCCGTGTTGCAGCCGATCATCAGCACATCTAGGTTTCGTCCGGGTGTGACCGAAAGGT    262
gb:MT019532|Organism:Severe      CGTCCGTGTTGCAGCCGATCATCAGCACATCTAGGTTTCGTCCGGGTGTGACCGAAAGGT    262
                                 ************************************************************

gb:MT020781|Organism:Severe      AAGATGGAGAGCCTTGTCCCTGGTTTCAACGAGAAAACACACGTCCAACTCAGTTTGCCT    310
gb:MT007544|Organism:Severe      AAGATGGAGAGCCTTGTCCCTGGTTTCAACGAGAAAACACACGTCCAACTCAGTTTGCCT    322
gb:MN994467|Organism:Severe      AAGATGGAGAGCCTTGTCCCTGGTTTCAACGAGAAAACACACGTCCAACTCAGTTTGCCT    322
gb:MT044257|Organism:Severe      AAGATGGAGAGCCTTGTCCCTGGTTTCAACGAGAAAACACACGTCCAACTCAGTTTGCCT    322
gb:MT106054|Organism:Severe      AAGATGGAGAGCCTTGTCCCTGGTTTCAACGAGAAAACACACGTCCAACTCAGTTTGCCT    322
gb:MT049951|Organism:Severe      AAGATGGAGAGCCTTGTCCCTGGTTTCAACGAGAAAACACACGTCCAACTCAGTTTGCCT    322
gb:MN975262|Organism:Severe      AAGATGGAGAGCCTTGTCCCTGGTTTCAACGAGAAAACACACGTCCAACTCAGTTTGCCT    322
gb:MT106052|Organism:Severe      AAGATGGAGAGCCTTGTCCCTGGTTTCAACGAGAAAACACACGTCCAACTCAGTTTGCCT    322
gb:LC522975|Organism:Severe      AAGATGGAGAGCCTTGTCCCTGGTTTCAACGAGAAAACACACGTCCAACTCAGTTTGCCT    319
gb:LC522973|Organism:Severe      AAGATGGAGAGCCTTGTCCCTGGTTTCAACGAGAAAACACACGTCCAACTCAGTTTGCCT    319
gb:LC522974|Organism:Severe      AAGATGGAGAGCCTTGTCCCTGGTTTCAACGAGAAAACACACGTCCAACTCAGTTTGCCT    319
gb:MN985325|Organism:Severe      AAGATGGAGAGCCTTGTCCCTGGTTTCAACGAGAAAACACACGTCCAACTCAGTTTGCCT    322
gb:MT020881|Organism:Severe      AAGATGGAGAGCCTTGTCCCTGGTTTCAACGAGAAAACACACGTCCAACTCAGTTTGCCT    322
gb:MT020880|Organism:Severe      AAGATGGAGAGCCTTGTCCCTGGTTTCAACGAGAAAACACACGTCCAACTCAGTTTGCCT    322
gb:MT066175|Organism:Severe      AAGATGGAGAGCCTTGTCCCTGGTTTCAACGAGAAAACACACGTCCAACTCAGTTTGCCT    322
gb:MN997409|Organism:Severe      AAGATGGAGAGCCTTGTCCCTGGTTTCAACGAGAAAACACACGTCCAACTCAGTTTGCCT    322
gb:MN938384|Organism:Severe      AAGATGGAGAGCCTTGTCCCTGGTTTCAACGAGAAAACACACGTCCAACTCAGTTTGCCT    290
gb:MT044258|Organism:Severe      AAGATGGAGAGCCTTGTCCCTGGTTTCAACGAGAAAACACACGTCCAACTCAGTTTGCCT    322
gb:MT039890|Organism:Severe      AAGATGGAGAGCCTTGTCCCTGGTTTCAACGAGAAAACACACGTCCAACTCAGTTTGCCT    322
gb:MN988713|Organism:Severe      AAGATGGAGAGCCTTGTCCCTGGTTTCAACGAGAAAACACACGTCCAACTCAGTTTGCCT    322
gb:LC521925|Organism:Severe      AAGATGGAGAGCCTTGTCCCTGGTTTCAACGAGAAAACACACGTCCAACTCAGTTTGCCT    319
gb:MT093571|Organism:Severe      AAGATGGAGAGCCTTGTCCCTGGTTTCAACGAGAAAACACACGTCCAACTCAGTTTGCCT    322
gb:MT039887|Organism:Severe      AAGATGGAGAGCCTTGTCCCTGGTTTCAACGAGAAAACACACGTCCAACTCAGTTTGCCT    322
gb:MT019530|Organism:Severe      AAGATGGAGAGCCTTGTCCCTGGTTTCAACGAGAAAACACACGTCCAACTCAGTTTGCCT    322
gb:MT039888|Organism:Severe      AAGATGGAGAGCCTTGTCCCTGGTTTCAACGAGAAAACACACGTCCAACTCAGTTTGCCT    322
gb:LC522972|Organism:Severe      AAGATGGAGAGCCTTGTCCCTGGTTTCAACGAGAAAACACACGTCCAACTCAGTTTGCCT    319
```

```
gb:MT027063|Organism:Severe      AAGATGGAGAGCCTTGTCCCTGGTTTCAACGAGAAAACACACGTCCAACTCAGTTTGCCT   322
gb:MT027062|Organism:Severe      AAGATGGAGAGCCTTGTCCCTGGTTTCAACGAGAAAACACACGTCCAACTCAGTTTGCCT   322
gb:MT019529|Organism:Severe      AAGATGGAGAGCCTTGTCCCTGGTTTCAACGAGAAAACACACGTCCAACTCAGTTTGCCT   322
gb:MN996529|Organism:Severe      AAGATGGAGAGCCTTGTCCCTGGTTTCAACGAGAAAACACACGTCCAACTCAGTTTGCCT   310
gb:MN996531|Organism:Severe      AAGATGGAGAGCCTTGTCCCTGGTTTCAACGAGAAAACACACGTCCAACTCAGTTTGCCT   309
gb:MT066176|Organism:Severe      AAGATGGAGAGCCTTGTCCCTGGTTTCAACGAGAAAACACACGTCCAACTCAGTTTGCCT   322
gb:MT027064|Organism:Severe      AAGATGGAGAGCCTTGTCCCTGGTTTCAACGAGAAAACACACGTCCAACTCAGTTTGCCT   322
gb:MN994468|Organism:Severe      AAGATGGAGAGCCTTGTCCCTGGTTTCAACGAGAAAACACACGTCCAACTCAGTTTGCCT   322
gb:MT072688|Organism:Severe      AAGATGGAGAGCCTTGTCCCTGGTTTCAACGAGAAAACACACGTCCAACTCAGTTTGCCT   307
gb:MN996527|Organism:Severe      AAGATGGAGAGCCTTGTCCCTGGTTTCAACGAGAAAACACACGTCCAACTCAGTTTGCCT   289
gb:MT093631|Organism:Severe      AAGATGGAGAGCCTTGTCCCTGGTTTCAACGAGAAAACACACGTCCAACTCAGTTTGCCT   360
gb:MT106053|Organism:Severe      AAGATGGAGAGCCTTGTCCCTGGTTTCAACGAGAAAACACACGTCCAACTCAGTTTGCCT   322
gb:MT019533|Organism:Severe      AAGATGGAGAGCCTTGTCCCTGGTTTCAACGAGAAAACACACGTCCAACTCAGTTTGCCT   322
gb:MT019531|Organism:Severe      AAGATGGAGAGCCTTGTCCCTGGTTTCAACGAGAAAACACACGTCCAACTCAGTTTGCCT   322
gb:MN996528|Organism:Severe      AAGATGGAGAGCCTTGTCCCTGGTTTCAACGAGAAAACACACGTCCAACTCAGTTTGCCT   322
gb:MN996530|Organism:Severe      AAGATGGAGAGCCTTGTCCCTGGTTTCAACGAGAAAACACACGTCCAACTCAGTTTGCCT   308
gb:MN908947|Organism:Severe      AAGATGGAGAGCCTTGTCCCTGGTTTCAACGAGAAAACACACGTCCAACTCAGTTTGCCT   322
gb:MT019532|Organism:Severe      AAGATGGAGAGCCTTGTCCCTGGTTTCAACGAGAAAACACACGTCCAACTCAGTTTGCCT   322
                                 ************************************************************

gb:MT020781|Organism:Severe      GTTTTACAGGTTCGCGACGTGCTCGTACGTGGCTTTGGAGACTCCGTGGAGGAGGTCTTA   370
gb:MT007544|Organism:Severe      GTTTTACAGGTTCGCGACGTGCTCGTACGTGGCTTTGGAGACTCCGTGGAGGAGGTCTTA   382
gb:MN994467|Organism:Severe      GTTTTACAGGTTCGCGACGTGCTCGTACGTGGCTTTGGAGACTCCGTGGAGGAGGTCTTA   382
gb:MT044257|Organism:Severe      GTTTTACAGGTTCGCGACGTGCTCGTACGTGGCTTTGGAGACTCCGTGGAGGAGGTCTTA   382
gb:MT106054|Organism:Severe      GTTTTACAGGTTCGCGACGTGCTCGTACGTGGCTTTGGAGACTCCGTGGAGGAGGTCTTA   382
gb:MT049951|Organism:Severe      GTTTTACAGGTTCGCGACGTGCTCGTACGTGGCTTTGGAGACTCCGTGGAGGAGGTCTTA   382
gb:MN975262|Organism:Severe      GTTTTACAGGTTCGCGACGTGCTCGTACGTGGCTTTGGAGACTCCGTGGAGGAGGTCTTA   382
gb:MT106052|Organism:Severe      GTTTTACAGGTTCGCGACGTGCTCGTACGTGGCTTTGGAGACTCCGTGGAGGAGGTCTTA   382
gb:LC522975|Organism:Severe      GTTTTACAGGTTCGCGACGTGCTCGTACGTGGCTTTGGAGACTCCGTGGAGGAGGTCTTA   379
gb:LC522973|Organism:Severe      GTTTTACAGGTTCGCGACGTGCTCGTACGTGGCTTTGGAGACTCCGTGGAGGAGGTCTTA   379
gb:LC522974|Organism:Severe      GTTTTACAGGTTCGCGACGTGCTCGTACGTGGCTTTGGAGACTCCGTGGAGGAGGTCTTA   379
gb:MN985325|Organism:Severe      GTTTTACAGGTTCGCGACGTGCTCGTACGTGGCTTTGGAGACTCCGTGGAGGAGGTCTTA   382
gb:MT020881|Organism:Severe      GTTTTACAGGTTCGCGACGTGCTCGTACGTGGCTTTGGAGACTCCGTGGAGGAGGTCTTA   382
gb:MT020880|Organism:Severe      GTTTTACAGGTTCGCGACGTGCTCGTACGTGGCTTTGGAGACTCCGTGGAGGAGGTCTTA   382
gb:MT066175|Organism:Severe      GTTTTACAGGTTCGCGACGTGCTCGTACGTGGCTTTGGAGACTCCGTGGAGGAGGTCTTA   382
gb:MN997409|Organism:Severe      GTTTTACAGGTTCGCGACGTGCTCGTACGTGGCTTTGGAGACTCCGTGGAGGAGGTCTTA   382
gb:MN938384|Organism:Severe      GTTTTACAGGTTCGCGACGTGCTCGTACGTGGCTTTGGAGACTCCGTGGAGGAGGTCTTA   350
gb:MT044258|Organism:Severe      GTTTTACAGGTTCGCGACGTGCTCGTACGTGGCTTTGGAGACTCCGTGGAGGAGGTCTTA   382
gb:MT039890|Organism:Severe      GTTTTACAGGTTCGCGACGTGCTCGTACGTGGCTTTGGAGACTCCGTGGAGGAGGTCTTA   382
gb:MN988713|Organism:Severe      GTTTTACAGGTTCGCGACGTGCTCGTACGTGGCTTTGGAGACTCCGTGGAGGAGGTCTTA   382
gb:LC521925|Organism:Severe      GTTTTACAGGTTCGCGACGTGCTCGTACGTGGCTTT------------------------   355
gb:MT093571|Organism:Severe      GTTTTACAGGTTCGCGACGTGCTCGTACGTGGCTTTGGAGACTCCGTGGAGGAGGTCTTA   382
gb:MT039887|Organism:Severe      GTTTTACAGGTTCGCGACGTGCTCGTACGTGGCTTTGGAGACTCCGTGGAGGAGGTCTTA   382
gb:MT019530|Organism:Severe      GTTTTACAGGTTCGCGACGTGCTCGTACGTGGCTTTGGAGACTCCGTGGAGGAGGTCTTA   382
gb:MT039888|Organism:Severe      GTTTTACAGGTTCGCGACGTGCTCGTACGTGGCTTTGGAGACTCCGTGGAGGAGGTCTTA   382
gb:LC522972|Organism:Severe      GTTTTACAGGTTCGCGACGTGCTCGTACGTGGCTTTGGAGACTCCGTGGAGGAGGTCTTA   379
gb:MT027063|Organism:Severe      GTTTTACAGGTTCGCGACGTGCTCGTACGTGGCTTTGGAGACTCCGTGGAGGAGGTCTTA   382
gb:MT027062|Organism:Severe      GTTTTACAGGTTCGCGACGTGCTCGTACGTGGCTTTGGAGACTCCGTGGAGGAGGTCTTA   382
gb:MT019529|Organism:Severe      GTTTTACAGGTTCGCGACGTGCTCGTACGTGGCTTTGGAGACTCCGTGGAGGAGGTCTTA   382
gb:MN996529|Organism:Severe      GTTTTACAGGTTCGCGACGTGCTCGTACGTGGCTTTGGAGACTCCGTGGAGGAGGTCTTA   370
gb:MN996531|Organism:Severe      GTTTTACAGGTTCGCGACGTGCTCGTACGTGGCTTTGGAGACTCCGTGGAGGAGGTCTTA   369
gb:MT066176|Organism:Severe      GTTTTACAGGTTCGCGACGTGCTCGTACGTGGCTTTGGAGACTCCGTGGAGGAGGTCTTA   382
gb:MT027064|Organism:Severe      GTTTTACAGGTTCGCGACGTGCTCGTACGTGGCTTTGGAGACTCCGTGGAGGAGGTCTTA   382
gb:MN994468|Organism:Severe      GTTTTACAGGTTCGCGACGTGCTCGTACGTGGCTTTGGAGACTCCGTGGAGGAGGTCTTA   382
gb:MT072688|Organism:Severe      GTTTTACAGGTTCGCGACGTGCTCGTACGTGGCTTTGGAGACTCCGTGGAGGAGGTCTTA   367
gb:MN996527|Organism:Severe      GTTTTACAGGTTCGCGACGTGCTCGTACGTGGCTTTGGAGACTCCGTGGAGGAGGTCTTA   349
gb:MT093631|Organism:Severe      GTTTTACAGGTTCGCGACGTGCTCGTACGTGGCTTTGGAGACTCCGTGGAGGAGGTCTTA   420
gb:MT106053|Organism:Severe      GTTTTACAGGTTCGCGACGTGCTCGTACGTGGCTTTGGAGACTCCGTGGAGGAGGTCTTA   382
gb:MT019533|Organism:Severe      GTTTTACAGGTTCGCGACGTGCTCGTACGTGGCTTTGGAGACTCCGTGGAGGAGGTCTTA   382
gb:MT019531|Organism:Severe      GTTTTACAGGTTCGCGACGTGCTCGTACGTGGCTTTGGAGACTCCGTGGAGGAGGTCTTA   382
gb:MN996528|Organism:Severe      GTTTTACAGGTTCGCGACGTGCTCGTACGTGGCTTTGGAGACTCCGTGGAGGAGGTCTTA   382
gb:MN996530|Organism:Severe      GTTTTACAGGTTCGCGACGTGCTCGTACGTGGCTTTGGAGACTCCGTGGAGGAGGTCTTA   368
gb:MN908947|Organism:Severe      GTTTTACAGGTTCGCGACGTGCTCGTACGTGGCTTTGGAGACTCCGTGGAGGAGGTCTTA   382
gb:MT019532|Organism:Severe      GTTTTACAGGTTCGCGACGTGCTCGTACGTGGCTTTGGAGACTCCGTGGAGGAGGTCTTA   382
                                 ************************************
```

```
gb:MT020781|Organism:Severe      TCAGAGGCACGTCAACATCTTAAAGATGGCACTTGTGGCTTAGTAGAAGTTGAAAAAGGC   430
gb:MT007544|Organism:Severe      TCAGAGGCACGTCAACATCTTAAAGATGGCACTTGTGGCTTAGTAGAAGTTGAAAAAGGC   442
gb:MN994467|Organism:Severe      TCAGAGGCACGTCAACATCTTAAAGATGGCACTTGTGGCTTAGTAGAAGTTGAAAAAGGC   442
gb:MT044257|Organism:Severe      TCAGAGGCACGTCAACATCTTAAAGATGGCACTTGTGGCTTAGTAGAAGTTGAAAAAGGC   442
gb:MT106054|Organism:Severe      TCAGAGGCACGTCAACATCTTAAAGATGGCACTTGTGGCTTAGTAGAAGTTGAAAAAGGC   442
gb:MT049951|Organism:Severe      TCAGAGGCACGTCAACATCTTAAAGATGGCACTTGTGGCTTAGTAGAAGTTGAAAAAGGC   442
gb:MN975262|Organism:Severe      TCAGAGGCACGTCAACATCTTAAAGATGGCACTTGTGGCTTAGTAGAAGTTGAAAAAGGC   442
gb:MT106052|Organism:Severe      TCAGAGGCACGTCAACATCTTAAAGATGGCACTTGTGGCTTAGTAGAAGTTGAAAAAGGC   442
gb:LC522975|Organism:Severe      TCAGAGGCACGTCAACATCTTAAAGATGGCACTTGTGGCTTAGTAGAAGTTGAAAAAGGC   439
gb:LC522973|Organism:Severe      TCAGAGGCACGTCAACATCTTAAAGATGGCACTTGTGGCTTAGTAGAAGTTGAAAAAGGC   439
gb:LC522974|Organism:Severe      TCAGAGGCACGTCAACATCTTAAAGATGGCACTTGTGGCTTAGTAGAAGTTGAAAAAGGC   439
gb:MN985325|Organism:Severe      TCAGAGGCACGTCAACATCTTAAAGATGGCACTTGTGGCTTAGTAGAAGTTGAAAAAGGC   442
gb:MT020881|Organism:Severe      TCAGAGGCACGTCAACATCTTAAAGATGGCACTTGTGGCTTAGTAGAAGTTGAAAAAGGC   442
gb:MT020880|Organism:Severe      TCAGAGGCACGTCAACATCTTAAAGATGGCACTTGTGGCTTAGTAGAAGTTGAAAAAGGC   442
gb:MT066175|Organism:Severe      TCAGAGGCACGTCAACATCTTAAAGATGGCACTTGTGGCTTAGTAGAAGTTGAAAAAGGC   442
gb:MN997409|Organism:Severe      TCAGAGGCACGTCAACATCTTAAAGATGGCACTTGTGGCTTAGTAGAAGTTGAAAAAGGC   442
gb:MN938384|Organism:Severe      TCAGAGGCACGTCAACATCTTAAAGATGGCACTTGTGGCTTAGTAGAAGTTGAAAAAGGC   410
gb:MT044258|Organism:Severe      TCAGAGGCACGTCAACATCTTAAAGATGGCACTTGTGGCTTAGTAGAAGTTGAAAAAGGC   442
gb:MT039890|Organism:Severe      TCAGAGGCACGTCAACATCTTAAAGATGGCACTTGTGGCTTAGTAGAAGTTGAAAAAGGC   442
gb:MN988713|Organism:Severe      TCAGAGGCACGTCAACATCTTAAAGATGGCACTTGTGGCTTAGTAGAAGTTGAAAAAGGC   442
gb:LC521925|Organism:Severe      TCAGAGGCACGTCAACATCTTAAAGATGGCACTTGTGGCTTAGTAGAAGTTGAAAAAGGC   415
gb:MT093571|Organism:Severe      TCAGAGGCACGTCAACATCTTAAAGATGGCACTTGTGGCTTAGTAGAAGTTGAAAAAGGC   442
gb:MT039887|Organism:Severe      TCAGAGGCACGTCAACATCTTAAAGATGGCACTTGTGGCTTAGTAGAAGTTGAAAAAGGC   442
gb:MT019530|Organism:Severe      TCAGAGGCACGTCAACATCTTAAAGATGGCACTTGTGGCTTAGTAGAAGTTGAAAAAGGC   442
gb:MT039888|Organism:Severe      TCAGAGGCACGTCAACATCTTAAAGATGGCACTTGTGGCTTAGTAGAAGTTGAAAAAGGC   442
gb:LC522972|Organism:Severe      TCAGAGGCACGTCAACATCTTAAAGATGGCACTTGTGGCTTAGTAGAAGTTGAAAAAGGC   439
gb:MT027063|Organism:Severe      TCAGAGGCACGTCAACATCTTAAAGATGGCACTTGTGGCTTAGTAGAAGTTGAAAAAGGC   442
gb:MT027062|Organism:Severe      TCAGAGGCACGTCAACATCTTAAAGATGGCACTTGTGGCTTAGTAGAAGTTGAAAAAGGC   442
gb:MT019529|Organism:Severe      TCAGAGGCACGTCAACATCTTAAAGATGGCACTTGTGGCTTAGTAGAAGTTGAAAAAGGC   442
gb:MN996529|Organism:Severe      TCAGAGGCACGTCAACATCTTAAAGATGGCACTTGTGGCTTAGTAGAAGTTGAAAAAGGC   430
gb:MN996531|Organism:Severe      TCAGAGGCACGTCAACATCTTAAAGATGGCACTTGTGGCTTAGTAGAAGTTGAAAAAGGC   429
gb:MT066176|Organism:Severe      TCAGAGGCACGTCAACATCTTAAAGATGGCACTTGTGGCTTAGTAGAAGTTGAAAAAGGC   442
gb:MT027064|Organism:Severe      TCAGAGGCACGTCAACATCTTAAAGATGGCACTTGTGGCTTAGTAGAAGTTGAAAAAGGC   442
gb:MN994468|Organism:Severe      TCAGAGGCACGTCAACATCTTAAAGATGGCACTTGTGGCTTAGTAGAAGTTGAAAAAGGC   442
gb:MT072688|Organism:Severe      TCAGAGGCACGTCAACATCTTAAAGATGGCACTTGTGGCTTAGTAGAAGTTGAAAAAGGC   427
gb:MN996527|Organism:Severe      TCAGAGGCACGTCAACATCTTAAAGATGGCACTTGTGGCTTAGTAGAAGTTGAAAAAGGC   409
gb:MT093631|Organism:Severe      TCAGAGGCACGTCAACATCTTAAAGATGGCACTTGTGGCTTAGTAGAAGTTGAAAAAGGC   480
gb:MT106053|Organism:Severe      TCAGAGGCACGTCAACATCTTAAAGATGGCACTTGTGGCTTAGTAGAAGTTGAAAAAGGC   442
gb:MT019533|Organism:Severe      TCAGAGGCACGTCAACATCTTAAAGATGGCACTTGTGGCTTAGTAGAAGTTGAAAAAGGC   442
gb:MT019531|Organism:Severe      TCAGAGGCACGTCAACATCTTAAAGATGGCACTTGTGGCTTAGTAGAAGTTGAAAAAGGC   442
gb:MN996528|Organism:Severe      TCAGAGGCACGTCAACATCTTAAAGATGGCACTTGTGGCTTAGTAGAAGTTGAAAAAGGC   442
gb:MN996530|Organism:Severe      TCAGAGGCACGTCAACATCTTAAAGATGGCACTTGTGGCTTAGTAGAAGTTGAAAAAGGC   428
gb:MN908947|Organism:Severe      TCAGAGGCACGTCAACATCTTAAAGATGGCACTTGTGGCTTAGTAGAAGTTGAAAAAGGC   442
gb:MT019532|Organism:Severe      TCAGAGGCACGTCAACATCTTAAAGATGGCACTTGTGGCTTAGTAGAAGTTGAAAAAGGC   442
                                 ************************************************************

gb:MT020781|Organism:Severe      GTTTTGCCTCAACTTGAACAGCCCTATGTGTTCATCAAACGTTCGGATGCTCGAACTGCA   490
gb:MT007544|Organism:Severe      GTTTTGCCTCAACTTGAACAGCCCTATGTGTTCATCAAACGTTCGGATGCTCGAACTGCA   502
gb:MN994467|Organism:Severe      GTTTTGCCTCAACTTGAACAGCCCTATGTGTTCATCAAACGTTCGGATGCTCGAACTGCA   502
gb:MT044257|Organism:Severe      GTTTTGCCTCAACTTGAACAGCCCTATGTGTTCATCAAACGTTCGGAAGCTCGAACTGCA   502
gb:MT106054|Organism:Severe      GTTTTGCCTCAACTTGAACAGCCCTATGTGTTCATCAAACGTTCGGATGCTCGAACTGCA   502
gb:MT049951|Organism:Severe      GTTTTGCCTCAACTTGAACAGCCCTATGTGTTCATCAAACGTTCGGATGCTCGAACTGCA   502
gb:MN975262|Organism:Severe      GTTTTGCCTCAACTTGAACAGCCCTATGTGTTCATCAAACGTTCGGATGCTCGAACTGCA   502
gb:MT106052|Organism:Severe      GTTTTGCCTCAACTTGAACAGCCCTATGTGTTCATCAAACGTTCGGATGCTCGAACTGCA   502
gb:LC522975|Organism:Severe      GTTTTGCCTCAACTTGAACAGCCCTATGTGTTCATCAAACGTTCGGATGCTCGAACTGCA   499
gb:LC522973|Organism:Severe      GTTTTGCCTCAACTTGAACAGCCCTATGTGTTCATCAAACGTTCGGATGCTCGAACTGCA   499
gb:LC522974|Organism:Severe      GTTTTGCCTCAACTTGAACAGCCCTATGTGTTCATCAAACGTTCGGATGCTCGAACTGCA   499
gb:MN985325|Organism:Severe      GTTTTGCCTCAACTTGAACAGCCCTATGTGTTCATCAAACGTTCGGATGCTCGAACTGCA   502
gb:MT020881|Organism:Severe      GTTTTGCCTCAACTTGAACAGCCCTATGTGTTCATCAAACGTTCGGATGCTCGAACTGCA   502
gb:MT020880|Organism:Severe      GTTTTGCCTCAACTTGAACAGCCCTATGTGTTCATCAAACGTTCGGATGCTCGAACTGCA   502
gb:MT066175|Organism:Severe      GTTTTGCCTCAACTTGAACAGCCCTATGTGTTCATCAAACGTTCGGATGCTCGAACTGCA   502
gb:MN997409|Organism:Severe      GTTTTGCCTCAACTTGAACAGCCCTATGTGTTCATCAAACGTTCGGATGCTCGAACTGCA   502
gb:MN938384|Organism:Severe      GTTTTGCCTCAACTTGAACAGCCCTATGTGTTCATCAAACGTTCGGATGCTCGAACTGCA   470
gb:MT044258|Organism:Severe      GTTTTGCCTCAACTTGAACAGCCCTATGTGTTCATCAAACGTTCGGATGCTCGAACTGCA   502
```

```
gb:MT039890|Organism:Severe      GTTTTGCCTCAACTTGAACAGCCCTATGTGTTCATCAAACGTTCGGATGCTCGAACTGCA   502
gb:MN988713|Organism:Severe      GTTTTGCCTCAACTTGAACAGCCCTATGTGTTCATCAAACGTTCGGAWGCTCGAACTGCA   502
gb:LC521925|Organism:Severe      GTTTTGCCTCAACTTGAACAGCCCTATGTGTTCATCAAACGTTCGGATGCTCGAACTGCA   475
gb:MT093571|Organism:Severe      GTTTTGCCTCAACTTGAACAGCCCTATGTGTTCATCAAACGTTCGGATGCTCGAACTGCA   502
gb:MT039887|Organism:Severe      GTTTTGCCTCAACTTGAACAGCCCTATGTGTTCATCAAACGTTCGGATGCTCGAACTGCA   502
gb:MT019530|Organism:Severe      GTTTTGCCTCAACTTGAACAGCCCTATGTGTTCATCAAACGTTCGGATGCTCGAACTGCA   502
gb:MT039888|Organism:Severe      GTTTTGCCTCAACTTGAACAGCCCTATGTGTTCATCAAACGTTCGGATGCTCGAACTGCA   502
gb:LC522972|Organism:Severe      GTTTTGCCTCAACTTGAACAGCCCTATGTGTTCATCAAACGTTCGGATGCTCGAACTGCA   499
gb:MT027063|Organism:Severe      GTTTTGCCTCAACTTGAACAGCCCTATGTGTTCATCAAACGTTCGGATGCTCGAACTGCA   502
gb:MT027062|Organism:Severe      GTTTTGCCTCAACTTGAACAGCCCTATGTGTTCATCAAACGTTCGGATGCTCGAACTGCA   502
gb:MT019529|Organism:Severe      GTTTTGCCTCAACTTGAACAGCCCTATGTGTTCATCAAACGTTCGGATGCTCGAACTGCA   502
gb:MN996529|Organism:Severe      GTTTTGCCTCAACTTGAACAGCCCTATGTGTTCATCAAACGTTCGGATGCTCGAACTGCA   490
gb:MN996531|Organism:Severe      GTTTTGCCTCAACTTGAACAGCCCTATGTGTTCATCAAACGTTCGGATGCTCGAACTGCA   489
gb:MT066176|Organism:Severe      GTTTTGCCTCAACTTGAACAGCCCTATGTGTTCATCAAACGTTCGGATGCTCGAACTGCA   502
gb:MT027064|Organism:Severe      GTTTTGCCTCAACTTGAACAGCCCTATGTGTTCATCAAACGTTCGGATGCTCGAACTGCA   502
gb:MN994468|Organism:Severe      GTTTTGCCTCAACTTGAACAGCCCTATGTGTTCATCAAACGTTCGGATGCTCGAACTGCA   502
gb:MT072688|Organism:Severe      GTTTTGCCTCAACTTGAACAGCCCTATGTGTTCATCAAACGTTCGGATGCTCGAACTGCA   487
gb:MN996527|Organism:Severe      GTTTTGCCTCAACTTGAACAGCCCTATGTGTTCATCAAACGTTCGGATGCTCGAACTGCA   469
gb:MT093631|Organism:Severe      GTTTTGCCTCAACTTGAACAGCCCTATGTGTTCATCAAACGTTCGGATGCTCGAACTGCA   540
gb:MT106053|Organism:Severe      GTTTTGCCTCAACTTGAACAGCCCTATGTGTTCATCAAACGTTCGGATGCTCGAACTGCA   502
gb:MT019533|Organism:Severe      GTTTTGCCTCAACTTGAACAGCCCTATGTGTTCATCAAACGTTCGGATGCTCGAACTGCA   502
gb:MT019531|Organism:Severe      GTTTTGCCTCAACTTGAACAGCCCTATGTGTTCATCAAACGTTCGGATGCTCGAACTGCA   502
gb:MN996528|Organism:Severe      GTTTTGCCTCAACTTGAACAGCCCTATGTGTTCATCAAACGTTCGGATGCTCGAACTGCA   502
gb:MN996530|Organism:Severe      GTTTTGCCTCAACTTGAACAGCCCTATGTGTTCATCAAACGTTCGGATGCTCGAACTGCA   488
gb:MN908947|Organism:Severe      GTTTTGCCTCAACTTGAACAGCCCTATGTGTTCATCAAACGTTCGGATGCTCGAACTGCA   502
gb:MT019532|Organism:Severe      GTTTTGCCTCAACTTGAACAGCCCTATGTGTTCATCAAACGTTCGGATGCTCGAACTGCA   502
                                 *********************************************** ************

gb:MT020781|Organism:Severe      CCTCATGGTCATGTTATGGTTGAGCTGGTAGCAGAACTCGAAGGCATTCAGTACGGTCGT   550
gb:MT007544|Organism:Severe      CCTCATGGTCATGTTATGGTTGAGCTGGTAGCAGAACTCGAAGGCATTCAGTACGGTCGT   562
gb:MN994467|Organism:Severe      CCTCATGGTCATGTTATGGTTGAGCTGGTAGCAGAACTCGAAGGCATTCAGTACGGTCGT   562
gb:MT044257|Organism:Severe      CCTCATGGTCATGTTATGGTTGAGCTGGTAGCAGAACTCGAAGGCATTCAGTACGGTCGT   562
gb:MT106054|Organism:Severe      CCTCATGGTCATGTTATGGTTGAGCTGGTAGCAGAACTCGAAGGCATTCAGTACGGTCGT   562
gb:MT049951|Organism:Severe      CCTCATGGTCATGTTATGGTTGAGCTGGTAGCAGAACTCGAAGGCATTCAGTACGGTCGT   562
gb:MN975262|Organism:Severe      CCTCATGGTCATGTTATGGTTGAGCTGGTAGCAGAACTCGAAGGCATTCAGTACGGTCGT   562
gb:MT106052|Organism:Severe      CCTCATGGTCATGTTATGGTTGAGCTGGTAGCAGAACTCGAAGGCATTCAGTACGGTCGT   562
gb:LC522975|Organism:Severe      CCTCATGGTCATGTTATGGTTGAGCTGGTAGCAGAACTCGAAGGCATTCAGTACGGTCGT   559
gb:LC522973|Organism:Severe      CCTCATGGTCATGTTATGGTTGAGCTGGTAGCAGAACTCGAAGGCATTCAGTACGGTCGT   559
gb:LC522974|Organism:Severe      CCTCATGGTCATGTTATGGTTGAGCTGGTAGCAGAACTCGAAGGCATTCAGTACGGTCGT   559
gb:MN985325|Organism:Severe      CCTCATGGTCATGTTATGGTTGAGCTGGTAGCAGAACTCGAAGGCATTCAGTACGGTCGT   562
gb:MT020881|Organism:Severe      CCTCATGGTCATGTTATGGTTGAGCTGGTAGCAGAACTCGAAGGCATTCAGTACGGTCGT   562
gb:MT020880|Organism:Severe      CCTCATGGTCATGTTATGGTTGAGCTGGTAGCAGAACTCGAAGGCATTCAGTACGGTCGT   562
gb:MT066175|Organism:Severe      CCTCATGGTCATGTTATGGTTGAGCTGGTAGCAGAACTCGAAGGCATTCAGTACGGTCGT   562
gb:MN997409|Organism:Severe      CCTCATGGTCATGTTATGGTTGAGCTGGTAGCAGAACTCGAAGGCATTCAGTACGGTCGT   562
gb:MN938384|Organism:Severe      CCTCATGGTCATGTTATGGTTGAGCTGGTAGCAGAACTCGAAGGCATTCAGTACGGTCGT   530
gb:MT044258|Organism:Severe      CCTCA---------------TGAGCTGGTAGCAGAACTCGAAGGCATTCAGTACGGTCGT   547
gb:MT039890|Organism:Severe      CCTCATGGTCATGTTATGGTTGAGCTGGTAGCAGAACTCGAAGGCATTCAGTACGGTCGT   562
gb:MN988713|Organism:Severe      CCTCATGGTCATGTTATGGTTGAGCTGGTAGCAGAACTCGAAGGCATTCAGTACGGTCGT   562
gb:LC521925|Organism:Severe      CCTCATGGTCATGTTATGGTTGAGCTGGTAGCAGAACTCGAAGGCATTCAGTACGGTCGT   535
gb:MT093571|Organism:Severe      CCTCATGGTCATGTTATGGTTGAGCTGGTAGCAGAACTCGAAGGCATTCAGTACGGTCGT   562
gb:MT039887|Organism:Severe      CCTCATGGTCATGTTATGGTTGAGCTGGTAGCAGAACTCGAAGGCATTCAGTACGGTCGT   562
gb:MT019530|Organism:Severe      CCTCATGGTCATGTTATGGTTGAGCTGGTAGCAGAACTCGAAGGCATTCAGTACGGTCGT   562
gb:MT039888|Organism:Severe      CCTCATGGTCATGTTATGGTTGAGCTGGTAGCAGAACTCGAAGGCATTCAGTACGGTCGT   562
gb:LC522972|Organism:Severe      CCTCATGGTCATGTTATGGTTGAGCTGGTAGCAGAACTCGAAGGCATTCAGTACGGTCGT   559
gb:MT027063|Organism:Severe      CCTCATGGTCATGTTATGGTTGAGCTGGTAGCAGAACTCGAAGGCATTCAGTACGGTCGT   562
gb:MT027062|Organism:Severe      CCTCATGGTCATGTTATGGTTGAGCTGGTAGCAGAACTCGAAGGCATTCAGTACGGTCGT   562
gb:MT019529|Organism:Severe      CCTCATGGTCATGTTATGGTTGAGCTGGTAGCAGAACTCGAAGGCATTCAGTACGGTCGT   562
gb:MN996529|Organism:Severe      CCTCATGGTCATGTTATGGTTGAGCTGGTAGCAGAACTCGAAGGCATTCAGTACGGTCGT   550
gb:MN996531|Organism:Severe      CCTCATGGTCATGTTATGGTTGAGCTGGTAGCAGAACTCGAAGGCATTCAGTACGGTCGT   549
gb:MT066176|Organism:Severe      CCTCATGGTCATGTTATGGTTGAGCTGGTAGCAGAACTCGAAGGCATTCAGTACGGTCGT   562
gb:MT027064|Organism:Severe      CCTCATGGTCATGTTATGGTTGAGCTGGTAGCAGAACTCGAAGGCATTCAGTACGGTCGT   562
gb:MN994468|Organism:Severe      CCTCATGGTCATGTTATGGTTGAGCTGGTAGCAGAACTCGAAGGCATTCAGTACGGTCGT   562
gb:MT072688|Organism:Severe      CCTCATGGTCATGTTATGGTTGAGCTGGTAGCAGAACTCGAAGGCATTCAGTACGGTCGT   547
gb:MN996527|Organism:Severe      CCTCATGGTCATGTTATGGTTGAGCTGGTAGCAGAACTCGAAGGCATTCAGTACGGTCGT   529
gb:MT093631|Organism:Severe      CCTCATGGTCATGTTATGGTTGAGCTGGTAGCAGAACTCGAAGGCATTCAGTACGGTCGT   600
```

```
gb:MT106053|Organism:Severe      CCTCATGGTCATGTTATGGTTGAGCTGGTAGCAGAACTCGAAGGCATTCAGTACGGTCGT   562
gb:MT019533|Organism:Severe      CCTCATGGTCATGTTATGGTTGAGCTGGTAGCAGAACTCGAAGGCATTCAGTACGGTCGT   562
gb:MT019531|Organism:Severe      CCTCATGGTCATGTTATGGTTGAGCTGGTAGCAGAACTCGAAGGCATTCAGTACGGTCGT   562
gb:MN996528|Organism:Severe      CCTCATGGTCATGTTATGGTTGAGCTGGTAGCAGAACTCGAAGGCATTCAGTACGGTCGT   562
gb:MN996530|Organism:Severe      CCTCATGGTCATGTTATGGTTGAGCTGGTAGCAGAACTCGAAGGCATTCAGTACGGTCGT   548
gb:MN908947|Organism:Severe      CCTCATGGTCATGTTATGGTTGAGCTGGTAGCAGAACTCGAAGGCATTCAGTACGGTCGT   562
gb:MT019532|Organism:Severe      CCTCATGGTCATGTTATGGTTGAGCTGGTAGCAGAACTCGAAGGCATTCAGTACGGTCGT   562
                                 *****               ****************************************

gb:MT020781|Organism:Severe      AGTGGTGAGACACTTGGTGTCCTTGTCCCTCATGTGGGCGAAATACCAGTGGCTTACCGC   610
gb:MT007544|Organism:Severe      AGTGGTGAGACACTTGGTGTCCTTGTCCCTCATGTGGGCGAAATACCAGTGGCTTACCGC   622
gb:MN994467|Organism:Severe      AGTGGTGAGACACTTGGTGTCCTTGTCCCTCATGTGGGCGAAATACCAGTGGCTTACCGC   622
gb:MT044257|Organism:Severe      AGTGGTGAGACACTTGGTGTCCTTGTCCCTCATGTGGGCGAAATACCAGTGGCTTACCGC   622
gb:MT106054|Organism:Severe      AGTGGTGAGACACTTGGTGTCCTTGTCCCTCATGTGGGCGAAATACCAGTGGCTTACCGC   622
gb:MT049951|Organism:Severe      AGTGGTGAGACACTTGGTGTCCTTGTCCCTCATGTGGGCGAAATACCAGTGGCTTACCGC   622
gb:MN975262|Organism:Severe      AGTGGTGAGACACTTGGTGTCCTTGTCCCTCATGTGGGCGAAATACCAGTGGCTTACCGC   622
gb:MT106052|Organism:Severe      AGTGGTGAGACACTTGGTGTCCTTGTCCCTCATGTGGGCGAAATACCAGTGGCTTACCGC   622
gb:LC522975|Organism:Severe      AGTGGTGAGACACTTGGTGTCCTTGTCCCTCATGTGGGCGAAATACCAGTGGCTTACCGC   619
gb:LC522973|Organism:Severe      AGTGGTGAGACACTTGGTGTCCTTGTCCCTCATGTGGGCGAAATACCAGTGGCTTACCGC   619
gb:LC522974|Organism:Severe      AGTGGTGAGACACTTGGTGTCCTTGTCCCTCATGTGGGCGAAATACCAGTGGCTTACCGC   619
gb:MN985325|Organism:Severe      AGTGGTGAGACACTTGGTGTCCTTGTCCCTCATGTGGGCGAAATACCAGTGGCTTACCGC   622
gb:MT020881|Organism:Severe      AGTGGTGAGACACTTGGTGTCCTTGTCCCTCATGTGGGCGAAATACCAGTGGCTTACCGC   622
gb:MT020880|Organism:Severe      AGTGGTGAGACACTTGGTGTCCTTGTCCCTCATGTGGGCGAAATACCAGTGGCTTACCGC   622
gb:MT066175|Organism:Severe      AGTGGTGAGACACTTGGTGTCCTTGTCCCTCATGTGGGCGAAATACCAGTGGCTTACCGC   622
gb:MN997409|Organism:Severe      AGTGGTGAGACACTTGGTGTCCTTGTCCCTCATGTGGGCGAAATACCAGTGGCTTACCGC   622
gb:MN938384|Organism:Severe      AGTGGTGAGACACTTGGTGTCCTTGTCCCTCATGTGGGCGAAATACCAGTGGCTTACCGC   590
gb:MT044258|Organism:Severe      AGTGGTGAGACACTTGGTGTCCTTGTCCCTCATGTGGGCGAAATACCAGTGGCTTACCGC   607
gb:MT039890|Organism:Severe      AGTGGTGAGACACTTGGTGTCCTTGTCCCTCATGTGGGCGAAATACCAGTGGCTTACCGC   622
gb:MN988713|Organism:Severe      AGTGGTGAGACACTTGGTGTCCTTGTCCCTCATGTGGGCGAAATACCAGTGGCTTACCGC   622
gb:LC521925|Organism:Severe      AGTGGTGAGACACTTGGTGTCCTTGTCCCTCATGTGGGCGAAATACCAGTGGCTTACCGC   595
gb:MT093571|Organism:Severe      AGTGGTGAGACACTTGGTGTCCTTGTCCCTCATGTGGGCGAAATACCAGTGGCTTACCGC   622
gb:MT039887|Organism:Severe      AGTGGTGAGACACTTGGTGTCCTTGTCCCTCATGTGGGCGAAATACCAGTGGCTTACCGC   622
gb:MT019530|Organism:Severe      AGTGGTGAGACACTTGGTGTCCTTGTCCCTCATGTGGGCGAAATACCAGTGGCTTACCGC   622
gb:MT039888|Organism:Severe      AGTGGTGAGACACTTGGTGTCCTTGTCCCTCATGTGGGCGAAATACCAGTGGCTTACCGC   622
gb:LC522972|Organism:Severe      AGTGGTGAGACACTTGGTGTCCTTGTCCCTCATGTGGGCGAAATACCAGTGGCTTACCGC   619
gb:MT027063|Organism:Severe      AGTGGTGAGACACTTGGTGTCCTTGTCCCTCATGTGGGCGAAATACCAGTGACTTACCGC   622
gb:MT027062|Organism:Severe      AGTGGTGAGACACTTGGTGTCCTTGTCCCTCATGTGGGCGAAATACCAGTGACTTACCGC   622
gb:MT019529|Organism:Severe      AGTGGTGAGACACTTGGTGTCCTTGTCCCTCATGTGGGCGAAATACCAGTGGCTTACCGC   622
gb:MN996529|Organism:Severe      AGTGGTGAGACACTTGGTGTCCTTGTCCCTCATGTGGGCGAAATACCAGTGGCTTACCGC   610
gb:MN996531|Organism:Severe      AGTGGTGAGACACTTGGTGTCCTTGTCCCTCATGTGGGCGAAATACCAGTGGCTTACCGC   609
gb:MT066176|Organism:Severe      AGTGGTGAGACACTTGGTGTCCTTGTCCCTCATGTGGGCGAAATACCAGTGGCTTACCGC   622
gb:MT027064|Organism:Severe      AGTGGTGAGACACTTGGTGTCCTTGTCCCTCATGTGGGCGAAATACCAGTGGCTTACCGC   622
gb:MN994468|Organism:Severe      AGTGGTGAGACACTTGGTGTCCTTGTCCCTCATGTGGGCGAAATACCAGTGGCTTACCGC   622
gb:MT072688|Organism:Severe      AGTGGTGAGACACTTGGTGTCCTTGTCCCTCATGTGGGCGAAATACCAGTGGCTTACCGC   607
gb:MN996527|Organism:Severe      AGTGGTGAGACACTTGGTGTCCTTGTCCCTCATGTGGGCGAAATACCAGTGGCTTACCGC   589
gb:MT093631|Organism:Severe      AGTGGTGAGACACTTGGTGTCCTTGTCCCTCATGTGGGCGAAATACCAGTGGCTTACCGC   660
gb:MT106053|Organism:Severe      AGTGGTGAGACACTTGGTGTCCTTGTCCCTCATGTGGGCGAAATACCAGTGGCTTACCGC   622
gb:MT019533|Organism:Severe      AGTGGTGAGACACTTGGTGTCCTTGTCCCTCATGTGGGCGAAATACCAGTGGCTTACCGC   622
gb:MT019531|Organism:Severe      AGTGGTGAGACACTTGGTGTCCTTGTCCCTCATGTGGGCGAAATACCAGTGGCTTACCGC   622
gb:MN996528|Organism:Severe      AGTGGTGAGACACTTGGTGTCCTTGTCCCTCATGTGGGCGAAATACCAGTGGCTTACCGC   622
gb:MN996530|Organism:Severe      AGTGGTGAGACACTTGGTGTCCTTGTCCCTCATGTGGGCGAAATACCAGTGGCTTACCGC   608
gb:MN908947|Organism:Severe      AGTGGTGAGACACTTGGTGTCCTTGTCCCTCATGTGGGCGAAATACCAGTGGCTTACCGC   622
gb:MT019532|Organism:Severe      AGTGGTGAGACACTTGGTGTCCTTGTCCCTCATGTGGGCGAAATACCAGTGGCTTACCGC   622
                                 *************************************************** ********

gb:MT020781|Organism:Severe      AAGGTTCTTCTTCGTAAGAACGGTAATAAAGGAGCTGGTGGCCATAGTTACGGCGCCGAT   670
gb:MT007544|Organism:Severe      AAGGTTCTTCTTCGTAAGAACGGTAATAAAGGAGCTGGTGGCCATAGTTACGGCGCCGAT   682
gb:MN994467|Organism:Severe      AAGGTTCTTCTTCGTAAGAACGGTAATAAAGGAGCTGGTGGCCATAGTTACGGCGCCGAT   682
gb:MT044257|Organism:Severe      AAGGTTCTTCTTCGTAAGAACGGTAATAAAGGAGCTGGTGGCCATAGTTACGGCGCCGAT   682
gb:MT106054|Organism:Severe      AAGGTTCTTCTTCGTAAGAACGGTAATAAAGGAGCTGGTGGCCATAGTTACGGCGCCGAT   682
gb:MT049951|Organism:Severe      AAGGTTCTTCTTCGTAAGAACGGTAATAAAGGAGCTGGTGGCCATAGTTACGGCGCCGAT   682
gb:MN975262|Organism:Severe      AAGGTTCTTCTTCGTAAGAACGGTAATAAAGGAGCTGGTGGCCATAGTTACGGCGCCGAT   682
gb:MT106052|Organism:Severe      AAGGTTCTTCTTCGTAAGAACGGTAATAAAGGAGCTGGTGGCCATAGTTACGGCGCCGAT   682
gb:LC522975|Organism:Severe      AAGGTTCTTCTTCGTAAGAACGGTAATAAAGGAGCTGGTGGCCATAGTTACGGCGCCGAT   679
gb:LC522973|Organism:Severe      AAGGTTCTTCTTCGTAAGAACGGTAATAAAGGAGCTGGTGGCCATAGTTACGGCGCCGAT   679
```

```
gb:LC522974|Organism:Severe      AAGGTTCTTCTTCGTAAGAACGGTAATAAAGGAGCTGGTGGCCATAGTTACGGCGCCGAT   679
gb:MN985325|Organism:Severe      AAGGTTCTTCTTCGTAAGAACGGTAATAAAGGAGCTGGTGGCCATAGTTACGGCGCCGAT   682
gb:MT020881|Organism:Severe      AAGGTTCTTCTTCGTAAGAACGGTAATAAAGGAGCTGGTGGCCATAGTTACGGCGCCGAT   682
gb:MT020880|Organism:Severe      AAGGTTCTTCTTCGTAAGAACGGTAATAAAGGAGCTGGTGGCCATAGTTACGGCGCCGAT   682
gb:MT066175|Organism:Severe      AAGGTTCTTCTTCGTAAGAACGGTAATAAAGGAGCTGGTGGCCATAGTTACGGCGCCGAT   682
gb:MN997409|Organism:Severe      AAGGTTCTTCTTCGTAAGAACGGTAATAAAGGAGCTGGTGGCCATAGTTACGGCGCCGAT   682
gb:MN938384|Organism:Severe      AAGGTTCTTCTTCGTAAGAACGGTAATAAAGGAGCTGGTGGCCATAGTTACGGCGCCGAT   650
gb:MT044258|Organism:Severe      AAGGTTCTTCTTCGTAAGAACGGTAATAAAGGAGCTGGTGGCCATAGTTACGGCGCCGAT   667
gb:MT039890|Organism:Severe      AAGGTTCTTCTTCGTAAGAACGGTAATAAAGGAGCTGGTGGCCATAGTTACGGCGCCGAT   682
gb:MN988713|Organism:Severe      AAGGTTCTTCTTCGTAAGAACGGTAATAAAGGAGCTGGTGGCCATAGTTACGGCGCCGAT   682
gb:LC521925|Organism:Severe      AAGGTTCTTCTTCGTAAGAACGGTAATAAAGGAGCTGGTGGCCATAGTTACGGCGCCGAT   655
gb:MT093571|Organism:Severe      AAGGTTCTTCTTCGTAAGAACGGTAATAAAGGAGCTGGTGGCCATAGTTACGGCGCCGAT   682
gb:MT039887|Organism:Severe      AAGGTTCTTCTTCGTAAGAACGGTAATAAAGGAGCTGGTGGCCATAGTTACGGCGCCGAT   682
gb:MT019530|Organism:Severe      AAGGTTCTTCTTCGTAAGAACGGTAATAAAGGAGCTGGTGGCCATAGTTACGGCGCCGAT   682
gb:MT039888|Organism:Severe      AAGGTTCTTCTTCGTAAGAACGGTAATAAAGGAGCTGGTGGCCATAGTTACGGCGCCGAT   682
gb:LC522972|Organism:Severe      AAGGTTCTTCTTCGTAAGAACGGTAATAAAGGAGCTGGTGGCCATAGTTACGGCGCCGAT   679
gb:MT027063|Organism:Severe      AAGGTTCTTCTTCGTAAGAACGGTAATAAAGGAGCTGGTGGCCATAGTTACGGCGCCGAT   682
gb:MT027062|Organism:Severe      AAGGTTCTTCTTCGTAAGAACGGTAATAAAGGAGCTGGTGGCCATAGTTACGGCGCCGAT   682
gb:MT019529|Organism:Severe      AAGGTTCTTCTTCGTAAGAACGGTAATAAAGGAGCTGGTGGCCATAGTTACGGCGCCGAT   682
gb:MN996529|Organism:Severe      AAGGTTCTTCTTCGTAAGAACGGTAATAAAGGAGCTGGTGGCCATAGTTACGGCGCCGAT   670
gb:MN996531|Organism:Severe      AAGGTTCTTCTTCGTAAGAACGGTAATAAAGGAGCTGGTGGCCATAGTTACGGCGCCGAT   669
gb:MT066176|Organism:Severe      AAGGTTCTTCTTCGTAAGAACGGTAATAAAGGAGCTGGTGGCCATAGTTACGGCGCCGAT   682
gb:MT027064|Organism:Severe      AAGGTTCTTCTTCGTAAGAACGGTAATAAAGGAGCTGGTGGCCATAGTTACGGCGCCGAT   682
gb:MN994468|Organism:Severe      AAGGTTCTTCTTCGTAAGAACGGTAATAAAGGAGCTGGTGGCCATAGTTACGGCGCCGAT   682
gb:MT072688|Organism:Severe      AAGGTTCTTCTTCGTAAGAACGGTAATAAAGGAGCTGGTGGCCATAGTTACGGCGCCGAT   667
gb:MN996527|Organism:Severe      AAGGTTCTTCTTCGTAAGAACGGTAATAAAGGAGCTGGTGGCCATAGTTACGGCGCCGAT   649
gb:MT093631|Organism:Severe      AAGGTTCTTCTTCGTAAGAACGGTAATAAAGGAGCTGGTGGCCATAGTTACGGCGCCGAT   720
gb:MT106053|Organism:Severe      AAGGTTCTTCTTCGTAAGAACGGTAATAAAGGAGCTGGTGGCCATAGTTACGGCGCCGAT   682
gb:MT019533|Organism:Severe      AAGGTTCTTCTTCGTAAGAACGGTAATAAAGGAGCTGGTGGCCATAGTTACGGCGCCGAT   682
gb:MT019531|Organism:Severe      AAGGTTCTTCTTCGTAAGAACGGTAATAAAGGAGCTGGTGGCCATAGTTACGGCGCCGAT   682
gb:MN996528|Organism:Severe      AAGGTTCTTCTTCGTAAGAACGGTAATAAAGGAGCTGGTGGCCATAGTTACGGCGCCGAT   682
gb:MN996530|Organism:Severe      AAGGTTCTTCTTCGTAAGAACGGTAATAAAGGAGCTGGTGGCCATAGTTACGGCGCCGAT   668
gb:MN908947|Organism:Severe      AAGGTTCTTCTTCGTAAGAACGGTAATAAAGGAGCTGGTGGCCATAGTTACGGCGCCGAT   682
gb:MT019532|Organism:Severe      AAGGTTCTTCTTCGTAAGAACGGTAATAAAGGAGCTGGTGGCCATAGTTACGGCGCCGAT   682
                                 ************************************************************

gb:MT020781|Organism:Severe      CTAAAGTCATTTGACTTAGGCGACGAGCTTGGCACTGATCCTTATGAAGATTTTCAAGAA   730
gb:MT007544|Organism:Severe      CTAAAGTCATTTGACTTAGGCGACGAGCTTGGCACTGATCCTTATGAAGATTTTCAAGAA   742
gb:MN994467|Organism:Severe      CTAAAGTCATTTGACTTAGGCGACGAGCTTGGCACTGATCCTTATGAAGATTTTCAAGAA   742
gb:MT044257|Organism:Severe      CTAAAGTCATTTGACTTAGGCGACGAGCTTGGCACTGATCCTTATGAAGATTTTCAAGAA   742
gb:MT106054|Organism:Severe      CTAAAGTCATTTGACTTAGGCGACGAGCTTGGCACTGATCCTTATGAAGATTTTCAAGAA   742
gb:MT049951|Organism:Severe      CTAAAGTCATTTGACTTAGGCGACGAGCTTGGCACTGATCCTTATGAAGATTTTCAAGAA   742
gb:MN975262|Organism:Severe      CTAAAGTCATTTGACTTAGGCGACGAGCTTGGCACTGATCCTTATGAAGATTTTCAAGAA   742
gb:MT106052|Organism:Severe      CTAAAGTCATTTGACTTAGGCGACGAGCTTGGCACTGATCCTTATGAAGATTTTCAAGAA   742
gb:LC522975|Organism:Severe      CTAAAGTCATTTGACTTAGGCGACGAGCTTGGCACTGATCCTTATGAAGATTTTCAAGAA   739
gb:LC522973|Organism:Severe      CTAAAGTCATTTGACTTAGGCGACGAGCTTGGCACTGATCCTTATGAAGATTTTCAAGAA   739
gb:LC522974|Organism:Severe      CTAAAGTCATTTGACTTAGGCGACGAGCTTGGCACTGATCCTTATGAAGATTTTCAAGAA   739
gb:MN985325|Organism:Severe      CTAAAGTCATTTGACTTAGGCGACGAGCTTGGCACTGATCCTTATGAAGATTTTCAAGAA   742
gb:MT020881|Organism:Severe      CTAAAGTCATTTGACTTAGGCGACGAGCTTGGCACTGATCCTTATGAAGATTTTCAAGAA   742
gb:MT020880|Organism:Severe      CTAAAGTCATTTGACTTAGGCGACGAGCTTGGCACTGATCCTTATGAAGATTTTCAAGAA   742
gb:MT066175|Organism:Severe      CTAAAGTCATTTGACTTAGGCGACGAGCTTGGCACTGATCCTTATGAAGATTTTCAAGAA   742
gb:MN997409|Organism:Severe      CTAAAGTCATTTGACTTAGGCGACGAGCTTGGCACTGATCCTTATGAAGATTTTCAAGAA   742
gb:MN938384|Organism:Severe      CTAAAGTCATTTGACTTAGGCGACGAGCTTGGCACTGATCCTTATGAAGATTTTCAAGAA   710
gb:MT044258|Organism:Severe      CTA---------GACTTAGGCGACGAGCTTGGCACTGATCCTTATGAAGATTTTCAAGAA   718
gb:MT039890|Organism:Severe      CTAAAGTCATTTGACTTAGGCGACGAGCTTGGCACTGATCCTTATGAAGATTTTCAAGAA   742
gb:MN988713|Organism:Severe      CTAAAGTCATTTGACTTAGGCGACGAGCTTGGCACTGATCCTTATGAAGATTTTCAAGAA   742
gb:LC521925|Organism:Severe      CTAAAGTCATTTGACTTAGGCGACGAGCTTGGCACTGATCCTTATGAAGATTTTCAAGAA   715
gb:MT093571|Organism:Severe      CTAAAGTCATTTGACTTAGGCGACGAGCTTGGCACTGATCCTTATGAAGATTTTCAAGAA   742
gb:MT039887|Organism:Severe      CTAAAGTCATTTGACTTAGGCGACGAGCTTGGCACTGATCCTTATGAAGATTTTCAAGAA   742
gb:MT019530|Organism:Severe      CTAAAGTCATTTGACTTAGGCGACGAGCTTGGCACTGATCCTTATGAAGATTTTCAAGAA   742
gb:MT039888|Organism:Severe      CTAAAGTCATTTGACTTAGGCGACGAGCTTGGCACTGATCCTTATGAAGATTTTCAAGAA   742
gb:LC522972|Organism:Severe      CTAAAGTCATTTGACTTAGGCGACGAGCTTGGCACTGATCCTTATGAAGATTTTCAAGAA   739
gb:MT027063|Organism:Severe      CTAAAGTCATTTGACTTAGGCGACGAGCTTGGCACTGATCCTTATGAAGATTTTCAAGAA   742
gb:MT027062|Organism:Severe      CTAAAGTCATTTGACTTAGGCGACGAGCTTGGCACTGATCCTTATGAAGATTTTCAAGAA   742
gb:MT019529|Organism:Severe      CTAAAGTCATTTGACTTAGGCGACGAGCTTGGCACTGATCCTTATGAAGATTTTCAAGAA   742
```

```
gb:MN996529|Organism:Severe      CTAAAGTCATTTGACTTAGGCGACGAGCTTGGCACTGATCCTTATGAAGATTTTCAAGAA   730
gb:MN996531|Organism:Severe      CTAAAGTCATTTGACTTAGGCGACGAGCTTGGCACTGATCCTTATGAAGATTTTCAAGAA   729
gb:MT066176|Organism:Severe      CTAAAGTCATTTGACTTAGGCGACGAGCTTGGCACTGATCCTTATGAAGATTTTCAAGAA   742
gb:MT027064|Organism:Severe      CTAAAGTCATTTGACTTAGGCGACGAGCTTGGCACTGATCCTTATGAAGATTTTCAAGAA   742
gb:MN994468|Organism:Severe      CTAAAGTCATTTGACTTAGGCGACGAGCTTGGCACTGATCCTTATGAAGATTTTCAAGAA   742
gb:MT072688|Organism:Severe      CTAAAGTCATTTGACTTAGGCGACGAGCTTGGCACTGATCCTTATGAAGATTTTCAAGAA   727
gb:MN996527|Organism:Severe      CTAAAGTCATTTGACTTAGGCGACGAGCTTGGCACTGATCCTTATGAAGATTTTCAAGAA   709
gb:MT093631|Organism:Severe      CTAAAGTCATTTGACTTAGGCGACGAGCTTGGCACTGATCCTTATGAAGATTTTCAAGAA   780
gb:MT106053|Organism:Severe      CTAAAGTCATTTGACTTAGGCGACGAGCTTGGCACTGATCCTTATGAAGATTTTCAAGAA   742
gb:MT019533|Organism:Severe      CTAAAGTCATTTGACTTAGGCGACGAGCTTGGCACTGATCCTTATGAAGATTTTCAAGAA   742
gb:MT019531|Organism:Severe      CTAAAGTCATTTGACTTAGGCGACGAGCTTGGCACTGATCCTTATGAAGATTTTCAAGAA   742
gb:MN996528|Organism:Severe      CTAAAGTCATTTGACTTAGGCGACGAGCTTGGCACTGATCCTTATGAAGATTTTCAAGAA   742
gb:MN996530|Organism:Severe      CTAAAGTCATTTGACTTAGGCGACGAGCTTGGCACTGATCCTTATGAAGATTTTCAAGAA   728
gb:MN908947|Organism:Severe      CTAAAGTCATTTGACTTAGGCGACGAGCTTGGCACTGATCCTTATGAAGATTTTCAAGAA   742
gb:MT019532|Organism:Severe      CTAAAGTCATTTGACTTAGGCGACGAGCTTGGCACTGATCCTTATGAAGATTTTCAAGAA   742
                                 ***         ************************************************

gb:MT020781|Organism:Severe      AACTGGAACACTAAACATAGCAGTGGTGTTACCCGTGAACTCATGCGTGAGCTTAACGGA   790
gb:MT007544|Organism:Severe      AACTGGAACACTAAACATAGCAGTGGTGTTACCCGTGAACTCATGCGTGAGCTTAACGGA   802
gb:MN994467|Organism:Severe      AACTGGAACACTAAACATAGCAGTGGTGTTACCCGTGAACTCATGCGTGAGCTTAACGGA   802
gb:MT044257|Organism:Severe      AACTGGAACACTAAACATAGCAGTGGTGTTACCCGTGAACTCATGCGTGAGCTTAACGGA   802
gb:MT106054|Organism:Severe      AACTGGAACACTAAACATAGCAGTGGTGTTACCCGTGAACTCATGCGTGAGCTTAACGGA   802
gb:MT049951|Organism:Severe      AACTGGAACACTAAACATAGCAGTGGTGTTACCCGTGAACTCATGCGTGAGCTTAACGGA   802
gb:MN975262|Organism:Severe      AACTGGAACACTAAACATAGCAGTGGTGTTACCCGTGAACTCATGCGTGAGCTTAACGGA   802
gb:MT106052|Organism:Severe      AACTGGAACACTAAACATAGCAGTGGTGTTACCCGTGAACTCATGCGTGAGCTTAACGGA   802
gb:LC522975|Organism:Severe      AACTGGAACACTAAACATAGCAGTGGTGTTACCCGTGAACTCATGCGTGAGCTTAACGGA   799
gb:LC522973|Organism:Severe      AACTGGAACACTAAACATAGCAGTGGTGTTACCCGTGAACTCATGCGTGAGCTTAACGGA   799
gb:LC522974|Organism:Severe      AACTGGAACACTAAACATAGCAGTGGTGTTACCCGTGAACTCATGCGTGAGCTTAACGGA   799
gb:MN985325|Organism:Severe      AACTGGAACACTAAACATAGCAGTGGTGTTACCCGTGAACTCATGCGTGAGCTTAACGGA   802
gb:MT020881|Organism:Severe      AACTGGAACACTAAACATAGCAGTGGTGTTACCCGTGAACTCATGCGTGAGCTTAACGGA   802
gb:MT020880|Organism:Severe      AACTGGAACACTAAACATAGCAGTGGTGTTACCCGTGAACTCATGCGTGAGCTTAACGGA   802
gb:MT066175|Organism:Severe      AACTGGAACACTAAACATAGCAGTGGTGTTACCCGTGAACTCATGCGTGAGCTTAACGGA   802
gb:MN997409|Organism:Severe      AACTGGAACACTAAACATAGCAGTGGTGTTACCCGTGAACTCATGCGTGAGCTTAACGGA   802
gb:MN938384|Organism:Severe      AACTGGAACACTAAACATAGCAGTGGTGTTACCCGTGAACTCATGCGTGAGCTTAACGGA   770
gb:MT044258|Organism:Severe      AACTGGAACACTAAACATAGCAGTGGTGTTACCCGTGAACTCATGCGTGAGCTTAACGGA   778
gb:MT039890|Organism:Severe      AACTGGAACACTAAACATAGCAGTGGTGTTACCCGTGAACTCATGCGTGAGCTTAACGGA   802
gb:MN988713|Organism:Severe      AACTGGAACACTAAACATAGCAGTGGTGTTACCCGTGAACTCATGCGTGAGCTTAACGGA   802
gb:LC521925|Organism:Severe      AACTGGAACACTAAACATAGCAGTGGTGTTACCCGTGAACTCATGCGTGAGCTTAACGGA   775
gb:MT093571|Organism:Severe      AACTGGAACACTAAACATAGCAGTGGTGTTACCCGTGAACTCATGCGTGAGCTTAACGGA   802
gb:MT039887|Organism:Severe      AACTGGAACACTAAACATAGCAGTGGTGTTACCCGTGAACTCATGCGTGAGCTTAACGGA   802
gb:MT019530|Organism:Severe      AACTGGAACACTAAACATAGCAGTGGTGTTACCCGTGAACTCATGCGTGAGCTTAACGGA   802
gb:MT039888|Organism:Severe      AACTGGAACACTAAACATAGCAGTGGTGTTACCCGTGAACTCATGCGTGAGCTTAACGGA   802
gb:LC522972|Organism:Severe      AACTGGAACACTAAACATAGCAGTGGTGTTACCCGTGAACTCATGCGTGAGCTTAACGGA   799
gb:MT027063|Organism:Severe      AACTGGAACACTAAACATAGCAGTGGTGTTACCCGTGAACTCATGCGTGAGCTTAACGGA   802
gb:MT027062|Organism:Severe      AACTGGAACACTAAACATAGCAGTGGTGTTACCCGTGAACTCATGCGTGAGCTTAACGGA   802
gb:MT019529|Organism:Severe      AACTGGAACACTAAACATAGCAGTGGTGTTACCCGTGAACTCATGCGTGAGCTTAACGGA   802
gb:MN996529|Organism:Severe      AACTGGAACACTAAACATAGCAGTGGTGTTACCCGTGAACTCATGCGTGAGCTTAACGGA   790
gb:MN996531|Organism:Severe      AACTGGAACACTAAACATAGCAGTGGTGTTACCCGTGAACTCATGCGTGAGCTTAACGGA   789
gb:MT066176|Organism:Severe      AACTGGAACACTAAACATAGCAGTGGTGTTACCCGTGAACTCATGCGTGAGCTTAACGGA   802
gb:MT027064|Organism:Severe      AACTGGAACACTAAACATAGCAGTGGTGTTACCCGTGAACTCATGCGTGAGCTTAACGGA   802
gb:MN994468|Organism:Severe      AACTGGAACACTAAACATAGCAGTGGTGTTACCCGTGAACTCATGCGTGAGCTTAACGGA   802
gb:MT072688|Organism:Severe      AACTGGAACACTAAACATAGCAGTGGTGTTACCCGTGAACTCATGCGTGAGCTTAACGGA   787
gb:MN996527|Organism:Severe      AACTGGAACACTAAACATAGCAGTGGTGTTACCCGTGAACTCATGCGTGAGCTTAACGGA   769
gb:MT093631|Organism:Severe      AACTGGAACACTAAACATAGCAGTGGTGTTACCCGTGAACTCATGCGTGAGCTTAACGGA   840
gb:MT106053|Organism:Severe      AACTGGAACACTAAACATAGCAGTGGTGTTACCCGTGAACTCATGCGTGAGCTTAACGGA   802
gb:MT019533|Organism:Severe      AACTGGAACACTAAACATAGCAGTGGTGTTACCCGTGAACTCATGCGTGAGCTTAACGGA   802
gb:MT019531|Organism:Severe      AACTGGAACACTAAACATAGCAGTGGTGTTACCCGTGAACTCATGCGTGAGCTTAACGGA   802
gb:MN996528|Organism:Severe      AACTGGAACACTAAACATAGCAGTGGTGTTACCCGTGAACTCATGCGTGAGCTTAACGGA   802
gb:MN996530|Organism:Severe      AACTGGAACACTAAACATAGCAGTGGTGTTACCCGTGAACTCATGCGTGAGCTTAACGGA   788
gb:MN908947|Organism:Severe      AACTGGAACACTAAACATAGCAGTGGTGTTACCCGTGAACTCATGCGTGAGCTTAACGGA   802
gb:MT019532|Organism:Severe      AACTGGAACACTAAACATAGCAGTGGTGTTACCCGTGAACTCATGCGTGAGCTTAACGGA   802
                                 ************************************************************

gb:MT020781|Organism:Severe      GGGGCATACACTCGCTATGTCGATAACAACTTCTGTGGCCCTGATGGCTACCCTCTTGAG   850
gb:MT007544|Organism:Severe      GGGGCATACACTCGCTATGTCGATAACAACTTCTGTGGCCCTGATGGCTACCCTCTTGAG   862
```

```
gb:MN994467|Organism:Severe      GGGGCATACACTCGCTATGTCGATAACAACTTCTGTGGCCCTGATGGCTACCCTCTTGAG	862
gb:MT044257|Organism:Severe      GGGGCATACACTCGCTATGTCGATAACAACTTCTGTGGCCCTGATGGCTACCCTCTTGAG	862
gb:MT106054|Organism:Severe      GGGGCATACACTCGCTATGTCGATAACAACTTCTGTGGCCCTGATGGCTACCCTCTTGAG	862
gb:MT049951|Organism:Severe      GGGGCATACACTCGCTATGTCGATAACAACTTCTGTGGCCCTGATGGCTACCCTCTTGAG	862
gb:MN975262|Organism:Severe      GGGGCATACACTCGCTATGTCGATAACAACTTCTGTGGCCCTGATGGCTACCCTCTTGAG	862
gb:MT106052|Organism:Severe      GGGGCATACACTCGCTATGTCGATAACAACTTCTGTGGCCCTGATGGCTACCCTCTTGAG	862
gb:LC522975|Organism:Severe      GGGGCATACACTCGCTATGTCGATAACAACTTCTGTGGCCCTGATGGCTACCCTCTTGAG	859
gb:LC522973|Organism:Severe      GGGGCATACACTCGCTATGTCGATAACAACTTCTGTGGCCCTGATGGCTACCCTCTTGAG	859
gb:LC522974|Organism:Severe      GGGGCATACACTCGCTATGTCGATAACAACTTCTGTGGCCCTGATGGCTACCCTCTTGAG	859
gb:MN985325|Organism:Severe      GGGGCATACACTCGCTATGTCGATAACAACTTCTGTGGCCCTGATGGCTACCCTCTTGAG	862
gb:MT020881|Organism:Severe      GGGGCATACACTCGCTATGTCGATAACAACTTCTGTGGCCCTGATGGCTACCCTCTTGAG	862
gb:MT020880|Organism:Severe      GGGGCATACACTCGCTATGTCGATAACAACTTCTGTGGCCCTGATGGCTACCCTCTTGAG	862
gb:MT066175|Organism:Severe      GGGGCATACACTCGCTATGTCGATAACAACTTCTGTGGCCCTGATGGCTACCCTCTTGAG	862
gb:MN997409|Organism:Severe      GGGGCATACACTCGCTATGTCGATAACAACTTCTGTGGCCCTGATGGCTACCCTCTTGAG	862
gb:MN938384|Organism:Severe      GGGGCATACACTCGCTATGTCGATAACAACTTCTGTGGCCCTGATGGCTACCCTCTTGAG	830
gb:MT044258|Organism:Severe      GGGGCATACACTCGCTATGTCGATAACAACTTCTGTGGCCCTGATGGCTACCCTCTTGAG	838
gb:MT039890|Organism:Severe      GGGGCATACACTCGCTATGTCGATAACAACTTCTGTGGCCCTGATGGCTACCCTCTTGAG	862
gb:MN988713|Organism:Severe      GGGGCATACACTCGCTATGTCGATAACAACTTCTGTGGCCCTGATGGCTACCCTCTTGAG	862
gb:LC521925|Organism:Severe      GGGGCATACACTCGCTATGTCGATAACAACTTCTGTGGCCCTGATGGCTACCCTCTTGAG	835
gb:MT093571|Organism:Severe      GGGGCATACACTCGCTATGTCGATAACAACTTCTGTGGCCCTGATGGCTACCCTCTTGAG	862
gb:MT039887|Organism:Severe      GGGGCATACACTCGCTATGTCGATAACAACTTCTGTGGCCCTGATGGCTACCCTCTTGAG	862
gb:MT019530|Organism:Severe      GGGGCATACACTCGCTATGTCGATAACAACTTCTGTGGCCCTGATGGCTACCCTCTTGAG	862
gb:MT039888|Organism:Severe      GGGGCATACACTCGCTATGTCGATAACAACTTCTGTGGCCCTGATGGCTACCCTCTTGAG	862
gb:LC522972|Organism:Severe      GGGGCATACACTCGCTATGTCGATAACAACTTCTGTGGCCCTGATGGCTACCCTCTTGAG	859
gb:MT027063|Organism:Severe      GGGGCATACACTCGCTATGTCGATAACAACTTCTGTGGCCCTGATGGCTACCCTCTTGAG	862
gb:MT027062|Organism:Severe      GGGGCATACACTCGCTATGTCGATAACAACTTCTGTGGCCCTGATGGCTACCCTCTTGAG	862
gb:MT019529|Organism:Severe      GGGGCATACACTCGCTATGTCGATAACAACTTCTGTGGCCCTGATGGCTACCCTCTTGAG	862
gb:MN996529|Organism:Severe      GGGGCATACACTCGCTATGTCGATAACAACTTCTGTGGCCCTGATGGCTACCCTCTTGAG	850
gb:MN996531|Organism:Severe      GGGGCATACACTCGCTATGTCGATAACAACTTCTGTGGCCCTGATGGCTACCCTCTTGAG	849
gb:MT066176|Organism:Severe      GGGGCATACACTCGCTATGTCGATAACAACTTCTGTGGCCCTGATGGCTACCCTCTTGAG	862
gb:MT027064|Organism:Severe      GGGGCATACACTCGCTATGTCGATAACAACTTCTGTGGCCCTGATGGCTACCCTCTTGAG	862
gb:MN994468|Organism:Severe      GGGGCATACACTCGCTATGTCGATAACAACTTCTGTGGCCCTGATGGCTACCCTCTTGAG	862
gb:MT072688|Organism:Severe      GGGGCATACACTCGCTATGTCGATAACAACTTCTGTGGCCCTGATGGCTACCCTCTTGAG	847
gb:MN996527|Organism:Severe      GGGGCATACACTCGCTATGTCGATAACAACTTCTGTGGCCCTGATGGCTACCCTCTTGAG	829
gb:MT093631|Organism:Severe      GGGGCATACACTCGCTATGTCGATAACAACTTCTGTGGCCCTGATGGCTACCCTCTTGAG	900
gb:MT106053|Organism:Severe      GGGGCATACACTCGCTATGTCGATAACAACTTCTGTGGCCCTGATGGCTACCCTCTTGAG	862
gb:MT019533|Organism:Severe      GGGGCATACACTCGCTATGTCGATAACAACTTCTGTGGCCCTGATGGCTACCCTCTTGAG	862
gb:MT019531|Organism:Severe      GGGGCATACACTCGCTATGTCGATAACAACTTCTGTGGCCCTGATGGCTACCCTCTTGAG	862
gb:MN996528|Organism:Severe      GGGGCATACACTCGCTATGTCGATAACAACTTCTGTGGCCCTGATGGCTACCCTCTTGAG	862
gb:MN996530|Organism:Severe      GGGGCATACACTCGCTATGTCGATAACAACTTCTGTGGCCCTGATGGCTACCCTCTTGAG	848
gb:MN908947|Organism:Severe      GGGGCATACACTCGCTATGTCGATAACAACTTCTGTGGCCCTGATGGCTACCCTCTTGAG	862
gb:MT019532|Organism:Severe      GGGGCATACACTCGCTATGTCGATAACAACTTCTGTGGCCCTGATGGCTACCCTCTTGAG	862
                                 ************************************************************

gb:MT020781|Organism:Severe      TGCATTAAAGACCTTCTAGCACGTGCTGGTAAAGCTTCATGCACTTTGTCCGAACAACTG	910
gb:MT007544|Organism:Severe      TGCATTAAAGACCTTCTAGCACGTGCTGGTAAAGCTTCATGCACTTTGTCCGAACAACTG	922
gb:MN994467|Organism:Severe      TGCATTAAAGACCTTCTAGCACGTGCTGGTAAAGCTTCATGCACTTTGTCCGAACAACTG	922
gb:MT044257|Organism:Severe      TGCATTAAAGACCTTCTAGCACGTGCTGGTAAAGCTTCATGCACTTTGTCCGAACAACTG	922
gb:MT106054|Organism:Severe      TGCATTAAAGACCTTCTAGCACGTGCTGGTAAAGCTTCATGCACTTTGTCCGAACAACTG	922
gb:MT049951|Organism:Severe      TGCATTAAAGACCTTCTAGCACGTGCTGGTAAAGCTTCATGCACTTTGTCCGAACAACTG	922
gb:MN975262|Organism:Severe      TGCATTAAAGACCTTCTAGCACGTGCTGGTAAAGCTTCATGCACTTTGTCCGAACAACTG	922
gb:MT106052|Organism:Severe      TGCATTAAAGACCTTCTAGCACGTGCTGGTAAAGCTTCATGCACTTTGTCCGAACAACTG	922
gb:LC522975|Organism:Severe      TGCATTAAAGACCTTCTAGCACGTGCTGGTAAAGCTTCATGCACTTTGTCCGAACAACTG	919
gb:LC522973|Organism:Severe      TGCATTAAAGACCTTCTAGCACGTGCTGGTAAAGCTTCATGCACTTTGTCCGAACAACTG	919
gb:LC522974|Organism:Severe      TGCATTAAAGACCTTCTAGCACGTGCTGGTAAAGCTTCATGCACTTTGTCCGAACAACTG	919
gb:MN985325|Organism:Severe      TGCATTAAAGACCTTCTAGCACGTGCTGGTAAAGCTTCATGCACTTTGTCCGAACAACTG	922
gb:MT020881|Organism:Severe      TGCATTAAAGACCTTCTAGCACGTGCTGGTAAAGCTTCATGCACTTTGTCCGAACAACTG	922
gb:MT020880|Organism:Severe      TGCATTAAAGACCTTCTAGCACGTGCTGGTAAAGCTTCATGCACTTTGTCCGAACAACTG	922
gb:MT066175|Organism:Severe      TGCATTAAAGACCTTCTAGCACGTGCTGGTAAAGCTTCATGCACTTTGTCCGAACAACTG	922
gb:MN997409|Organism:Severe      TGCATTAAAGACCTTCTAGCACGTGCTGGTAAAGCTTCATGCACTTTGTCCGAACAACTG	922
gb:MN938384|Organism:Severe      TGCATTAAAGACCTTCTAGCACGTGCTGGTAAAGCTTCATGCACTTTGTCCGAACAACTG	890
gb:MT044258|Organism:Severe      TGCATTAAAGACCTTCTAGCACGTGCTGGTAAAGCTTCATGCACTTTGTCCGAACAACTG	898
gb:MT039890|Organism:Severe      TGCATTAAAGACCTTCTAGCACGTGCTGGTAAAGCTTCATGCACTTTGTCCGAACAACTG	922
gb:MN988713|Organism:Severe      TGCATTAAAGACCTTCTAGCACGTGCTGGTAAAGCTTCATGCACTTTGTCCGAACAACTG	922
gb:LC521925|Organism:Severe      TGCATTAAAGACCTTCTAGCACGTGCTGGTAAAGCTTCATGCACTTTGTCCGAACAACTG	895
```

```
gb:MT093571|Organism:Severe      TGCATTAAAGACCTTCTAGCACGTGCTGGTAAAGCTTCATGCACTTTGTCCGAACAACTG    922
gb:MT039887|Organism:Severe      TGCATTAAAGACCTTCTAGCACGTGCTGGTAAAGCTTCATGCACTTTGTCCGAACAACTG    922
gb:MT019530|Organism:Severe      TGCATTAAAGACCTTCTAGCACGTGCTGGTAAAGCTTCATGCACTTTGTCCGAACAACTG    922
gb:MT039888|Organism:Severe      TGCATTAAAGACCTTCTAGCACGTGCTGGTAAAGCTTCATGCACTTTGTCCGAACAACTG    922
gb:LC522972|Organism:Severe      TGCATTAAAGACCTTCTAGCACGTGCTGGTAAAGCTTCATGCACTTTGTCCGAACAACTG    919
gb:MT027063|Organism:Severe      TGCATTAAAGACCTTCTAGCACGTGCTGGTAAAGCTTCATGCACTTTGTCCGAACAACTG    922
gb:MT027062|Organism:Severe      TGCATTAAAGACCTTCTAGCACGTGCTGGTAAAGCTTCATGCACTTTGTCCGAACAACTG    922
gb:MT019529|Organism:Severe      TGCATTAAAGACCTTCTAGCACGTGCTGGTAAAGCTTCATGCACTTTGTCCGAACAACTG    922
gb:MN996529|Organism:Severe      TGCATTAAAGACCTTCTAGCACGTGCTGGTAAAGCTTCATGCACTTTGTCCGAACAACTG    910
gb:MN996531|Organism:Severe      TGCATTAAAGACCTTCTAGCACGTGCTGGTAAAGCTTCATGCACTTTGTCCGAACAACTG    909
gb:MT066176|Organism:Severe      TGCATTAAAGACCTTCTAGCACGTGCTGGTAAAGCTTCATGCACTTTGTCCGAACAACTG    922
gb:MT027064|Organism:Severe      TGCATTAAAGACCTTCTAGCACGTGCTGGTAAAGCTTCATGCACTTTGTCCGAACAACTG    922
gb:MN994468|Organism:Severe      TGCATTAAAGACCTTCTAGCACGTGCTGGTAAAGCTTCATGCACTTTGTCCGAACAACTG    922
gb:MT072688|Organism:Severe      TGCATTAAAGACCTTCTAGCACGTGCTGGTAAAGCTTCATGCACTTTGTCCGAACAACTG    907
gb:MN996527|Organism:Severe      TGCATTAAAGACCTTCTAGCACGTGCTGGTAAAGCTTCATGCACTTTGTCCGAACAACTG    889
gb:MT093631|Organism:Severe      TGCATTAAAGACCTTCTAGCACGTGCTGGTAAAGCTTCATGCACTTTGTCCGAACAACTG    960
gb:MT106053|Organism:Severe      TGCATTAAAGACCTTCTAGCACGTGCTGGTAAAGCTTCATGCACTTTGTCCGAACAACTG    922
gb:MT019533|Organism:Severe      TGCATTAAAGACCTTCTAGCACGTGCTGGTAAAGCTTCATGCACTTTGTCCGAACAACTG    922
gb:MT019531|Organism:Severe      TGCATTAAAGACCTTCTAGCACGTGCTGGTAAAGCTTCATGCACTTTGTCCGAACAACTG    922
gb:MN996528|Organism:Severe      TGCATTAAAGACCTTCTAGCACGTGCTGGTAAAGCTTCATGCACTTTGTCCGAACAACTG    922
gb:MN996530|Organism:Severe      TGCATTAAAGACCTTCTAGCACGTGCTGGTAAAGCTTCATGCACTTTGTCCGAACAACTG    908
gb:MN908947|Organism:Severe      TGCATTAAAGACCTTCTAGCACGTGCTGGTAAAGCTTCATGCACTTTGTCCGAACAACTG    922
gb:MT019532|Organism:Severe      TGCATTAAAGACCTTCTAGCACGTGCTGGTAAAGCTTCATGCACTTTGTCCGAACAACTG    922
                                 ************************************************************

gb:MT020781|Organism:Severe      GACTTTATTGACACTAAGAGGGGTGTATACTGCTGCCGTGAACATGAGCATGAAATTGCT    970
gb:MT007544|Organism:Severe      GACTTTATTGACACTAAGAGGGGTGTATACTGCTGCCGTGAACATGAGCATGAAATTGCT    982
gb:MN994467|Organism:Severe      GACTTTATTGACACTAAGAGGGGTGTATACTGCTGCCGTGAACATGAGCATGAAATTGCT    982
gb:MT044257|Organism:Severe      GACTTTATTGACACTAAGAGGGGTGTATACTGCTGCCGTGAACATGAGCATGAAATTGCT    982
gb:MT106054|Organism:Severe      GACTTTATTGACACTAAGAGGGGTGTATACTGCTGCCGTGAACATGAGCATGAAATTGCT    982
gb:MT049951|Organism:Severe      GACTTTATTGACACTAAGAGGGGTGTATACTGCTGCCGTGAACATGAGCATGAAATTGCT    982
gb:MN975262|Organism:Severe      GACTTTATTGACACTAAGAGGGGTGTATACTGCTGCCGTGAACATGAGCATGAAATTGCT    982
gb:MT106052|Organism:Severe      GACTTTATTGACACTAAGAGGGGTGTATACTGCTGCCGTGAACATGAGCATGAAATTGCT    982
gb:LC522975|Organism:Severe      GACTTTATTGACACTAAGAGGGGTGTATACTGCTGCCGTGAACATGAGCATGAAATTGCT    979
gb:LC522973|Organism:Severe      GACTTTATTGACACTAAGAGGGGTGTATACTGCTGCCGTGAACATGAGCATGAAATTGCT    979
gb:LC522974|Organism:Severe      GACTTTATTGACACTAAGAGGGGTGTATACTGCTGCCGTGAACATGAGCATGAAATTGCT    979
gb:MN985325|Organism:Severe      GACTTTATTGACACTAAGAGGGGTGTATACTGCTGCCGTGAACATGAGCATGAAATTGCT    982
gb:MT020881|Organism:Severe      GACTTTATTGACACTAAGAGGGGTGTATACTGCTGCCGTGAACATGAGCATGAAATTGCT    982
gb:MT020880|Organism:Severe      GACTTTATTGACACTAAGAGGGGTGTATACTGCTGCCGTGAACATGAGCATGAAATTGCT    982
gb:MT066175|Organism:Severe      GACTTTATTGACACTAAGAGGGGTGTATACTGCTGCCGTGAACATGAGCATGAAATTGCT    982
gb:MN997409|Organism:Severe      GACTTTATTGACACTAAGAGGGGTGTATACTGCTGCCGTGAACATGAGCATGAAATTGCT    982
gb:MN938384|Organism:Severe      GACTTTATTGACACTAAGAGGGGTGTATACTGCTGCCGTGAACATGAGCATGAAATTGCT    950
gb:MT044258|Organism:Severe      GACTTTATTGACACTAAGAGGGGTGTATACTGCTGCCGTGAACATGAGCATGAAATTGCT    958
gb:MT039890|Organism:Severe      GACTTTATTGACACTAAGAGGGGTGTATACTGCTGCCGTGAACATGAGCATGAAATTGCT    982
gb:MN988713|Organism:Severe      GACTTTATTGACACTAAGAGGGGTGTATACTGCTGCCGTGAACATGAGCATGAAATTGCT    982
gb:LC521925|Organism:Severe      GACTTTATTGACACTAAGAGGGGTGTATACTGCTGCCGTGAACATGAGCATGAAATTGCT    955
gb:MT093571|Organism:Severe      GACTTTATTGACACTAAGAGGGGTGTATACTGCTGCCGTGAACATGAGCATGAAATTGCT    982
gb:MT039887|Organism:Severe      GACTTTATTGACACTAAGAGGGGTGTATACTGCTGCCGTGAACATGAGCATGAAATTGCT    982
gb:MT019530|Organism:Severe      GACTTTATTGACACTAAGAGGGGTGTATACTGCTGCCGTGAACATGAGCATGAAATTGCT    982
gb:MT039888|Organism:Severe      GACTTTATTGACACTAAGAGGGGTGTATACTGCTGCCGTGAACATGAGCATGAAATTGCT    982
gb:LC522972|Organism:Severe      GACTTTATTGACACTAAGAGGGGTGTATACTGCTGCCGTGAACATGAGCATGAAATTGCT    979
gb:MT027063|Organism:Severe      GACTTTATTGACACTAAGAGGGGTGTATACTGCTGCCGTGAACATGAGCATGAAATTGCT    982
gb:MT027062|Organism:Severe      GACTTTATTGACACTAAGAGGGGTGTATACTGCTGCCGTGAACATGAGCATGAAATTGCT    982
gb:MT019529|Organism:Severe      GACTTTATTGACACTAAGAGGGGTGTATACTGCTGCCGTGAACATGAGCATGAAATTGCT    982
gb:MN996529|Organism:Severe      GACTTTATTGACACTAAGAGGGGTGTATACTGCTGCCGTGAACATGAGCATGAAATTGCT    970
gb:MN996531|Organism:Severe      GACTTTATTGACACTAAGAGGGGTGTATACTGCTGCCGTGAACATGAGCATGAAATTGCT    969
gb:MT066176|Organism:Severe      GACTTTATTGACACTAAGAGGGGTGTATACTGCTGCCGTGAACATGAGCATGAAATTGCT    982
gb:MT027064|Organism:Severe      GACTTTATTGACACTAAGAGGGGTGTATACTGCTGCCGTGAACATGAGCATGAAATTGCT    982
gb:MN994468|Organism:Severe      GACTTTATTGACACTAAGAGGGGTGTATACTGCTGCCGTGAACATGAGCATGAAATTGCT    982
gb:MT072688|Organism:Severe      GACTTTATTGACACTAAGAGGGGTGTATACTGCTGCCGTGAACATGAGCATGAAATTGCT    967
gb:MN996527|Organism:Severe      GACTTTATTGACACTAAGAGGGGTGTATACTGCTGCCGTGAACATGAGCATGAAATTGCT    949
gb:MT093631|Organism:Severe      GACTTTATTGACACTAAGAGGGGTGTATACTGCTGCCGTGAACATGAGCATGAAATTGCT    1020
gb:MT106053|Organism:Severe      GACTTTATTGACACTAAGAGGGGTGTATACTGCTGCCGTGAACATGAGCATGAAATTGCT    982
gb:MT019533|Organism:Severe      GACTTTATTGACACTAAGAGGGGTGTATACTGCTGCCGTGAACATGAGCATGAAATTGCT    982
gb:MT019531|Organism:Severe      GACTTTATTGACACTAAGAGGGGTGTATACTGCTGCCGTGAACATGAGCATGAAATTGCT    982
```

```
gb:MN996528|Organism:Severe      GACTTTATTGACACTAAGAGGGGTGTATACTGCTGCCGTGAACATGAGCATGAAATTGCT    982
gb:MN996530|Organism:Severe      GACTTTATTGACACTAAGAGGGGTGTATACTGCTGCCGTGAACATGAGCATGAAATTGCT    968
gb:MN908947|Organism:Severe      GACTTTATTGACACTAAGAGGGGTGTATACTGCTGCCGTGAACATGAGCATGAAATTGCT    982
gb:MT019532|Organism:Severe      GACTTTATTGACACTAAGAGGGGTGTATACTGCTGCCGTGAACATGAGCATGAAATTGCT    982
                                 ************************************************************

gb:MT020781|Organism:Severe      TGGTACACGGAACGTTCTGAAAAGAGCTATGAATTGCAGACACCTTTTGAAATTAAATTG    1030
gb:MT007544|Organism:Severe      TGGTACACGGAACGTTCTGAAAAGAGCTATGAATTGCAGACACCTTTTGAAATTAAATTG    1042
gb:MN994467|Organism:Severe      TGGTACACGGAACGTTCTGAAAAGAGCTATGAATTGCAGACACCTTTTGAAATTAAATTG    1042
gb:MT044257|Organism:Severe      TGGTACACGGAACGTTCTGAAAAGAGCTATGAATTGCAGACACCTTTTGAAATTAAATTG    1042
gb:MT106054|Organism:Severe      TGGTACACGGAACGTTCTGAAAAGAGCTATGAATTGCAGACACCTTTTGAAATTAAATTG    1042
gb:MT049951|Organism:Severe      TGGTACACGGAACGTTCTGAAAAGAGCTATGAATTGCAGACACCTTTTGAAATTAAATTG    1042
gb:MN975262|Organism:Severe      TGGTACACGGAACGTTCTGAAAAGAGCTATGAATTGCAGACACCTTTTGAAATTAAATTG    1042
gb:MT106052|Organism:Severe      TGGTACACGGAACGTTCTGAAAAGAGCTATGAATTGCAGACACCTTTTGAAATTAAATTG    1042
gb:LC522975|Organism:Severe      TGGTACACGGAACGTTCTGAAAAGAGCTATGAATTGCAGACACCTTTTGAAATTAAATTG    1039
gb:LC522973|Organism:Severe      TGGTACACGGAACGTTCTGAAAAGAGCTATGAATTGCAGACACCTTTTGAAATTAAATTG    1039
gb:LC522974|Organism:Severe      TGGTACACGGAACGTTCTGAAAAGAGCTATGAATTGCAGACACCTTTTGAAATTAAATTG    1039
gb:MN985325|Organism:Severe      TGGTACACGGAACGTTCTGAAAAGAGCTATGAATTGCAGACACCTTTTGAAATTAAATTG    1042
gb:MT020881|Organism:Severe      TGGTACACGGAACGTTCTGAAAAGAGCTATGAATTGCAGACACCTTTTGAAATTAAATTG    1042
gb:MT020880|Organism:Severe      TGGTACACGGAACGTTCTGAAAAGAGCTATGAATTGCAGACACCTTTTGAAATTAAATTG    1042
gb:MT066175|Organism:Severe      TGGTACACGGAACGTTCTGAAAAGAGCTATGAATTGCAGACACCTTTTGAAATTAAATTG    1042
gb:MN997409|Organism:Severe      TGGTACACGGAACGTTCTGAAAAGAGCTATGAATTGCAGACACCTTTTGAAATTAAATTG    1042
gb:MN938384|Organism:Severe      TGGTACACGGAACGTTCTGAAAAGAGCTATGAATTGCAGACACCTTTTGAAATTAAATTG    1010
gb:MT044258|Organism:Severe      TGGTACACGGAACGTTCTGAAAAGAGCTATGAATTGCAGACACCTTTTGAAATTAAATTG    1018
gb:MT039890|Organism:Severe      TGGTACACGGAACGTTCTGAAAAGAGCTATGAATTGCAGACACCTTTTGAAATTAAATTG    1042
gb:MN988713|Organism:Severe      TGGTACACGGAACGTTCTGAAAAGAGCTATGAATTGCAGACACCTTTTGAAATTAAATTG    1042
gb:LC521925|Organism:Severe      TGGTACACGGAACGTTCTGAAAAGAGCTATGAATTGCAGACACCTTTTGAAATTAAATTG    1015
gb:MT093571|Organism:Severe      TGGTACACGGAACGTTCTGAAAAGAGCTATGAATTGCAGACACCTTTTGAAATTAAATTG    1042
gb:MT039887|Organism:Severe      TGGTACACGGAACGTTCTGAAAAGAGCTATGAATTGCAGACACCTTTTGAAATTAAATTG    1042
gb:MT019530|Organism:Severe      TGGTACACGGAACGTTCTGAAAAGAGCTATGAATTGCAGACACCTTTTGAAATTAAATTG    1042
gb:MT039888|Organism:Severe      TGGTACACGGAACGTTCTGAAAAGAGCTATGAATTGCAGACACCTTTTGAAATTAAATTG    1042
gb:LC522972|Organism:Severe      TGGTACACGGAACGTTCTGAAAAGAGCTATGAATTGCAGACACCTTTTGAAATTAAATTG    1039
gb:MT027063|Organism:Severe      TGGTACACGGAACGTTCTGAAAAGAGCTATGAATTGCAGACACCTTTTGAAATTAAATTG    1042
gb:MT027062|Organism:Severe      TGGTACACGGAACGTTCTGAAAAGAGCTATGAATTGCAGACACCTTTTGAAATTAAATTG    1042
gb:MT019529|Organism:Severe      TGGTACACGGAACGTTCTGAAAAGAGCTATGAATTGCAGACACCTTTTGAAATTAAATTG    1042
gb:MN996529|Organism:Severe      TGGTACACGGAACGTTCTGAAAAGAGCTATGAATTGCAGACACCTTTTGAAATTAAATTG    1030
gb:MN996531|Organism:Severe      TGGTACACGGAACGTTCTGAAAAGAGCTATGAATTGCAGACACCTTTTGAAATTAAATTG    1029
gb:MT066176|Organism:Severe      TGGTACACGGAACGTTCTGAAAAGAGCTATGAATTGCAGACACCTTTTGAAATTAAATTG    1042
gb:MT027064|Organism:Severe      TGGTACACGGAACGTTCTGAAAAGAGCTATGAATTGCAGACACCTTTTGAAATTAAATTG    1042
gb:MN994468|Organism:Severe      TGGTACACGGAACGTTCTGAAAAGAGCTATGAATTGCAGACACCTTTTGAAATTAAATTG    1042
gb:MT072688|Organism:Severe      TGGTACACGGAACGTTCTGAAAAGAGCTATGAATTGCAGACACCTTTTGAAATTAAATTG    1027
gb:MN996527|Organism:Severe      TGGTACACGGAACGTTCTGAAAAGAGCTATGAATTGCAGACACCTTTTGAAATTAAATTG    1009
gb:MT093631|Organism:Severe      TGGTACACGGAACGTTCTGAAAAGAGCTATGAATTGCAGACACCTTTTGAAATTAAATTG    1080
gb:MT106053|Organism:Severe      TGGTACACGGAACGTTCTGAAAAGAGCTATGAATTGCAGACACCTTTTGAAATTAAATTG    1042
gb:MT019533|Organism:Severe      TGGTACACGGAACGTTCTGAAAAGAGCTATGAATTGCAGACACCTTTTGAAATTAAATTG    1042
gb:MT019531|Organism:Severe      TGGTACACGGAACGTTCTGAAAAGAGCTATGAATTGCAGACACCTTTTGAAATTAAATTG    1042
gb:MN996528|Organism:Severe      TGGTACACGGAACGTTCTGAAAAGAGCTATGAATTGCAGACACCTTTTGAAATTAAATTG    1042
gb:MN996530|Organism:Severe      TGGTACACGGAACGTTCTGAAAAGAGCTATGAATTGCAGACACCTTTTGAAATTAAATTG    1028
gb:MN908947|Organism:Severe      TGGTACACGGAACGTTCTGAAAAGAGCTATGAATTGCAGACACCTTTTGAAATTAAATTG    1042
gb:MT019532|Organism:Severe      TGGTACACGGAACGTTCTGAAAAGAGCTATGAATTGCAGACACCTTTTGAAATTAAATTG    1042
                                 ************************************************************

gb:MT020781|Organism:Severe      GCAAAGAAATTTGACACCTTCAATGGGGAATGTCCAAATTTTGTATTTCCCTTAAATTCC    1090
gb:MT007544|Organism:Severe      GCAAAGAAATTTGACACCTTCAATGGGGAATGTCCAAATTTTGTATTTCCCTTAAATTCC    1102
gb:MN994467|Organism:Severe      GCAAAGAAATTTGACACCTTCAATGGGGAATGTCCAAATTTTGTATTTCCCTTAAATTCC    1102
gb:MT044257|Organism:Severe      GCAAAGAAATTTGACACCTTCAATGGGGAATGTCCAAATTTTGTATTTCCCTTAAATTCC    1102
gb:MT106054|Organism:Severe      GCAAAGAAATTTGACACCTTCAATGGGGAATGTCCAAATTTTGTATTTCCCTTAAATTCC    1102
gb:MT049951|Organism:Severe      GCAAAGAAATTTGACACCTTCAATGGGGAATGTCCAAATTTTGTATTTCCCTTAAATTCC    1102
gb:MN975262|Organism:Severe      GCAAAGAAATTTGACACCTTCAATGGGGAATGTCCAAATTTTGTATTTCCCTTAAATTCC    1102
gb:MT106052|Organism:Severe      GCAAAGAAATTTGACACCTTCAATGGGGAATGTCCAAATTTTGTATTTCCCTTAAATTCC    1102
gb:LC522975|Organism:Severe      GCAAAGAAATTTGACACCTTCAATGGGGAATGTCCAAATTTTGTATTTCCCTTAAATTCC    1099
gb:LC522973|Organism:Severe      GCAAAGAAATTTGACACCTTCAATGGGGAATGTCCAAATTTTGTATTTCCCTTAAATTCC    1099
gb:LC522974|Organism:Severe      GCAAAGAAATTTGACACCTTCAATGGGGAATGTCCAAATTTTGTATTTCCCTTAAATTCC    1099
gb:MN985325|Organism:Severe      GCAAAGAAATTTGACACCTTCAATGGGGAATGTCCAAATTTTGTATTTCCCTTAAATTCC    1102
gb:MT020881|Organism:Severe      GCAAAGAAATTTGACACCTTCAATGGGGAATGTCCAAATTTTGTATTTCCCTTAAATTCC    1102
```

```
gb:MT020880|Organism:Severe      GCAAAGAAATTTGACACCTTCAATGGGGAATGTCCAAATTTTGTATTTCCCTTAAATTCC   1102
gb:MT066175|Organism:Severe      GCAAAGAAATTTGACACCTTCAATGGGGAATGTCCAAATTTTGTATTTCCCTTAAATTCC   1102
gb:MN997409|Organism:Severe      GCAAAGAAATTTGACACCTTCAATGGGGAATGTCCAAATTTTGTATTTCCCTTAAATTCC   1102
gb:MN938384|Organism:Severe      GCAAAGAAATTTGACACCTTCAATGGGGAATGTCCAAATTTTGTATTTCCCTTAAATTCC   1070
gb:MT044258|Organism:Severe      GCAAAGAAATTTGACACCTTCAATGGGGAATGTCCAAATTTTGTATTTCCCTTAAATTCC   1078
gb:MT039890|Organism:Severe      GCAAAGAAATTTGACACCTTCAATGGGGAATGTCCAAATTTTGTATTTCCCTTAAATTCC   1102
gb:MN988713|Organism:Severe      GCAAAGAAATTTGACACCTTCAATGGGGAATGTCCAAATTTTGTATTTCCCTTAAATTCC   1102
gb:LC521925|Organism:Severe      GCAAAGAAATTTGACACCTTCAATGGGGAATGTCCAAATTTTGTATTTCCCTTAAATTCC   1075
gb:MT093571|Organism:Severe      GCAAAGAAATTTGACACCTTCAATGGGGAATGTCCAAATTTTGTATTTCCCTTAAATTCC   1102
gb:MT039887|Organism:Severe      GCAAAGAAATTTGACACCTTCAATGGGGAATGTCCAAATTTTGTATTTCCCTTAAATTCC   1102
gb:MT019530|Organism:Severe      GCAAAGAAATTTGACACCTTCAATGGGGAATGTCCAAATTTTGTATTTCCCTTAAATTCC   1102
gb:MT039888|Organism:Severe      GCAAAGAAATTTGACACCTTCAATGGGGAATGTCCAAATTTTGTATTTCCCTTAAATTCC   1102
gb:LC522972|Organism:Severe      GCAAAGAAATTTGACACCTTCAATGGGGAATGTCCAAATTTTGTATTTCCCTTAAATTCC   1099
gb:MT027063|Organism:Severe      GCAAAGAAATTTGACACCTTCAATGGGGAATGTCCAAATTTTGTATTTCCCTTAAATTCC   1102
gb:MT027062|Organism:Severe      GCAAAGAAATTTGACACCTTCAATGGGGAATGTCCAAATTTTGTATTTCCCTTAAATTCC   1102
gb:MT019529|Organism:Severe      GCAAAGAAATTTGACACCTTCAATGGGGAATGTCCAAATTTTGTATTTCCCTTAAATTCC   1102
gb:MN996529|Organism:Severe      GCAAAGAAATTTGACACCTTCAATGGGGAATGTCCAAATTTTGTATTTCCCTTAAATTCC   1090
gb:MN996531|Organism:Severe      GCAAAGAAATTTGACACCTTCAATGGGGAATGTCCAAATTTTGTATTTCCCTTAAATTCC   1089
gb:MT066176|Organism:Severe      GCAAAGAAATTTGACACCTTCAATGGGGAATGTCCAAATTTTGTATTTCCCTTAAATTCC   1102
gb:MT027064|Organism:Severe      GCAAAGAAATTTGACACCTTCAATGGGGAATGTCCAAATTTTGTATTTCCCTTAAATTCC   1102
gb:MN994468|Organism:Severe      GCAAAGAAATTTGACACCTTCAATGGGGAATGTCCAAATTTTGTATTTCCCTTAAATTCC   1102
gb:MT072688|Organism:Severe      GCAAAGAAATTTGACACCTTCAATGGGGAATGTCCAAATTTTGTATTTCCCTTAAATTCC   1087
gb:MN996527|Organism:Severe      GCAAAGAAATTTGACACCTTCAATGGGGAATGTCCAAATTTTGTATTTCCCTTAAATTCC   1069
gb:MT093631|Organism:Severe      GCAAAGAAATTTGACACCTTCAATGGGGAATGTCCAAATTTTGTATTTCCCTTAAATTCC   1140
gb:MT106053|Organism:Severe      GCAAAGAAATTTGACACCTTCAATGGGGAATGTCCAAATTTTGTATTTCCCTTAAATTCC   1102
gb:MT019533|Organism:Severe      GCAAAGAAATTTGACACCTTCAATGGGGAATGTCCAAATTTTGTATTTCCCTTAAATTCC   1102
gb:MT019531|Organism:Severe      GCAAAGAAATTTGACACCTTCAATGGGGAATGTCCAAATTTTGTATTTCCCTTAAATTCC   1102
gb:MN996528|Organism:Severe      GCAAAGAAATTTGACACCTTCAATGGGGAATGTCCAAATTTTGTATTTCCCTTAAATTCC   1102
gb:MN996530|Organism:Severe      GCAAAGAAATTTGACACCTTCAATGGGGAATGTCCAAATTTTGTATTTCCCTTAAATTCC   1088
gb:MN908947|Organism:Severe      GCAAAGAAATTTGACACCTTCAATGGGGAATGTCCAAATTTTGTATTTCCCTTAAATTCC   1102
gb:MT019532|Organism:Severe      GCAAAGAAATTTGACACCTTCAATGGGGAATGTCCAAATTTTGTATTTCCCTTAAATTCC   1102
                                 ************************************************************

gb:MT020781|Organism:Severe      ATAATCAAGACTATTCAACCAAGGGTTGAAAAGAAAAAGCTTGATGGCTTTATGGGTAGA   1150
gb:MT007544|Organism:Severe      ATAATCAAGACTATTCAACCAAGGGTTGAAAAGAAAAAGCTTGATGGCTTTATGGGTAGA   1162
gb:MN994467|Organism:Severe      ATAATCAAGACTATTCAACCAAGGGTTGAAAAGAAAAAGCTTGATGGCTTTATGGGTAGA   1162
gb:MT044257|Organism:Severe      ATAATCAAGACTATTCAACCAAGGGTTGAAAAGAAAAAGCTTGATGGCTTTATGGGTAGA   1162
gb:MT106054|Organism:Severe      ATAATCAAGACTATTCAACCAAGGGTTGAAAAGAAAAAGCTTGATGGCTTTATGGGTAGA   1162
gb:MT049951|Organism:Severe      ATAATCAAGACTATTCAACCAAGGGTTGAAAAGAAAAAGCTTGATGGCTTTATGGGTAGA   1162
gb:MN975262|Organism:Severe      ATAATCAAGACTATTCAACCAAGGGTTGAAAAGAAAAAGCTTGATGGCTTTATGGGTAGA   1162
gb:MT106052|Organism:Severe      ATAATCAAGACTATTCAACCAAGGGTTGAAAAGAAAAAGCTTGATGGCTTTATGGGTAGA   1162
gb:LC522975|Organism:Severe      ATAATCAAGACTATTCAACCAAGGGTTGAAAAGAAAAAGCTTGATGGCTTTATGGGTAGA   1159
gb:LC522973|Organism:Severe      ATAATCAAGACTATTCAACCAAGGGTTGAAAAGAAAAAGCTTGATGGCTTTATGGGTAGA   1159
gb:LC522974|Organism:Severe      ATAATCAAGACTATTCAACCAAGGGTTGAAAAGAAAAAGCTTGATGGCTTTATGGGTAGA   1159
gb:MN985325|Organism:Severe      ATAATCAAGACTATTCAACCAAGGGTTGAAAAGAAAAAGCTTGATGGCTTTATGGGTAGA   1162
gb:MT020881|Organism:Severe      ATAATCAAGACTATTCAACCAAGGGTTGAAAAGAAAAAGCTTGATGGCTTTATGGGTAGA   1162
gb:MT020880|Organism:Severe      ATAATCAAGACTATTCAACCAAGGGTTGAAAAGAAAAAGCTTGATGGCTTTATGGGTAGA   1162
gb:MT066175|Organism:Severe      ATAATCAAGACTATTCAACCAAGGGTTGAAAAGAAAAAGCTTGATGGCTTTATGGGTAGA   1162
gb:MN997409|Organism:Severe      ATAATCAAGACTATTCAACCAAGGGTTGAAAAGAAAAAGCTTGATGGCTTTATGGGTAGA   1162
gb:MN938384|Organism:Severe      ATAATCAAGACTATTCAACCAAGGGTTGAAAAGAAAAAGCTTGATGGCTTTATGGGTAGA   1130
gb:MT044258|Organism:Severe      ATAATCAAGACTATTCAACCAAGGGTTGAAAAGAAAAAGCTTGATGGCTTTATGGGTAGA   1138
gb:MT039890|Organism:Severe      ATAATCAAGACTATTCAACCAAGGGTTGAAAAGAAAAAGCTTGATGGCTTTATGGGTAGA   1162
gb:MN988713|Organism:Severe      ATAATCAAGACTATTCAACCAAGGGTTGAAAAGAAAAAGCTTGATGGCTTTATGGGTAGA   1162
gb:LC521925|Organism:Severe      ATAATCAAGACTATTCAACCAAGGGTTGAAAAGAAAAAGCTTGATGGCTTTATGGGTAGA   1135
gb:MT093571|Organism:Severe      ATAATCAAGACTATTCAACCAAGGGTTGAAAAGAAAAAGCTTGATGGCTTTATGGGTAGA   1162
gb:MT039887|Organism:Severe      ATAATCAAGACTATTCAACCAAGGGTTGAAAAGAAAAAGCTTGATGGCTTTATGGGTAGA   1162
gb:MT019530|Organism:Severe      ATAATCAAGACTATTCAACCAAGGGTTGAAAAGAAAAAGCTTGATGGCTTTATGGGTAGA   1162
gb:MT039888|Organism:Severe      ATAATCAAGACTATTCAACCAAGGGTTGAAAAGAAAAAGCTTGATGGCTTTATGGGTAGA   1162
gb:LC522972|Organism:Severe      ATAATCAAGACTATTCAACCAAGGGTTGAAAAGAAAAAGCTTGATGGCTTTATGGGTAGA   1159
gb:MT027063|Organism:Severe      ATAATCAAGACTATTCAACCAAGGGTTGAAAAGAAAAAGCTTGATGGCTTTATGGGTAGA   1162
gb:MT027062|Organism:Severe      ATAATCAAGACTATTCAACCAAGGGTTGAAAAGAAAAAGCTTGATGGCTTTATGGGTAGA   1162
gb:MT019529|Organism:Severe      ATAATCAAGACTATTCAACCAAGGGTTGAAAAGAAAAAGCTTGATGGCTTTATGGGTAGA   1162
gb:MN996529|Organism:Severe      ATAATCAAGACTATTCAACCAAGGGTTGAAAAGAAAAAGCTTGATGGCTTTATGGGTAGA   1150
gb:MN996531|Organism:Severe      ATAATCAAGACTATTCAACCAAGGGTTGAAAAGAAAAAGCTTGATGGCTTTATGGGTAGA   1149
gb:MT066176|Organism:Severe      ATAATCAAGACTATTCAACCAAGGGTTGAAAAGAAAAAGCTTGATGGCTTTATGGGTAGA   1162
```

```
gb:MT027064|Organism:Severe      ATAATCAAGACTATTCAACCAAGGGTTGAAAAGAAAAAGCTTGATGGCTTTATGGGTAGA   1162
gb:MN994468|Organism:Severe      ATAATCAAGACTATTCAACCAAGGGTTGAAAAGAAAAAGCTTGATGGCTTTATGGGTAGA   1162
gb:MT072688|Organism:Severe      ATAATCAAGACTATTCAACCAAGGGTTGAAAAGAAAAAGCTTGATGGCTTTATGGGTAGA   1147
gb:MN996527|Organism:Severe      ATAATCAAGACTATTCAACCAAGGGTTGAAAAGAAAAAGCTTGATGGCTTTATGGGTAGA   1129
gb:MT093631|Organism:Severe      ATAATCAAGACTATTCAACCAAGGGTTGAAAAGAAAAAGCTTGATGGCTTTATGGGTAGA   1200
gb:MT106053|Organism:Severe      ATAATCAAGACTATTCAACCAAGGGTTGAAAAGAAAAAGCTTGATGGCTTTATGGGTAGA   1162
gb:MT019533|Organism:Severe      ATAATCAAGACTATTCAACCAAGGGTTGAAAAGAAAAAGCTTGATGGCTTTATGGGTAGA   1162
gb:MT019531|Organism:Severe      ATAATCAAGACTATTCAACCAAGGGTTGAAAAGAAAAAGCTTGATGGCTTTATGGGTAGA   1162
gb:MN996528|Organism:Severe      ATAATCAAGACTATTCAACCAAGGGTTGAAAAGAAAAAGCTTGATGGCTTTATGGGTAGA   1162
gb:MN996530|Organism:Severe      ATAATCAAGACTATTCAACCAAGGGTTGAAAAGAAAAAGCTTGATGGCTTTATGGGTAGA   1148
gb:MN908947|Organism:Severe      ATAATCAAGACTATTCAACCAAGGGTTGAAAAGAAAAAGCTTGATGGCTTTATGGGTAGA   1162
gb:MT019532|Organism:Severe      ATAATCAAGACTATTCAACCAAGGGTTGAAAAGAAAAAGCTTGATGGCTTTATGGGTAGA   1162
                                 ************************************************************

gb:MT020781|Organism:Severe      ATTCGATCTGTCTATCCAGTTGCGTCACCAAATGAATGCAACCAAATGTGCCTTTCAACT   1210
gb:MT007544|Organism:Severe      ATTCGATCTGTCTATCCAGTTGCGTCACCAAATGAATGCAACCAAATGTGCCTTTCAACT   1222
gb:MN994467|Organism:Severe      ATTCGATCTGTCTATCCAGTTGCGTCACCAAATGAATGCAACCAAATGTGCCTTTCAACT   1222
gb:MT044257|Organism:Severe      ATTCGATCTGTCTATCCAGTTGCGTCACCAAATGAATGCAACCAAATGTGCCTTTCAACT   1222
gb:MT106054|Organism:Severe      ATTCGATCTGTCTATCCAGTTGCGTCACCAAATGAATGCAACCAAATGTGCCTTTCAACT   1222
gb:MT049951|Organism:Severe      ATTCGATCTGTCTATCCAGTTGCGTCACCAAATGAATGCAACCAAATGTGCCTTTCAACT   1222
gb:MN975262|Organism:Severe      ATTCGATCTGTCTATCCAGTTGCGTCACCAAATGAATGCAACCAAATGTGCCTTTCAACT   1222
gb:MT106052|Organism:Severe      ATTCGATCTGTCTATCCAGTTGCGTCACCAAATGAATGCAACCAAATGTGCCTTTCAACT   1222
gb:LC522975|Organism:Severe      ATTCGATCTGTCTATCCAGTTGCGTCACCAAATGAATGCAACCAAATGTGCCTTTCAACT   1219
gb:LC522973|Organism:Severe      ATTCGATCTGTCTATCCAGTTGCGTCACCAAATGAATGCAACCAAATGTGCCTTTCAACT   1219
gb:LC522974|Organism:Severe      ATTCGATCTGTCTATCCAGTTGCGTCACCAAATGAATGCAACCAAATGTGCCTTTCAACT   1219
gb:MN985325|Organism:Severe      ATTCGATCTGTCTATCCAGTTGCGTCACCAAATGAATGCAACCAAATGTGCCTTTCAACT   1222
gb:MT020881|Organism:Severe      ATTCGATCTGTCTATCCAGTTGCGTCACCAAATGAATGCAACCAAATGTGCCTTTCAACT   1222
gb:MT020880|Organism:Severe      ATTCGATCTGTCTATCCAGTTGCGTCACCAAATGAATGCAACCAAATGTGCCTTTCAACT   1222
gb:MT066175|Organism:Severe      ATTCGATCTGTCTATCCAGTTGCGTCACCAAATGAATGCAACCAAATGTGCCTTTCAACT   1222
gb:MN997409|Organism:Severe      ATTCGATCTGTCTATCCAGTTGCGTCACCAAATGAATGCAACCAAATGTGCCTTTCAACT   1222
gb:MN938384|Organism:Severe      ATTCGATCTGTCTATCCAGTTGCGTCACCAAATGAATGCAACCAAATGTGCCTTTCAACT   1190
gb:MT044258|Organism:Severe      ATTCGATCTGTCTATCCAGTTGCGTCACCAAATGAATGCAACCAAATGTGCCTTTCAACT   1198
gb:MT039890|Organism:Severe      ATTCGATCTGTCTATCCAGTTGCGTCACCAAATGAATGCAACCAAATGTGCCTTTCAACT   1222
gb:MN988713|Organism:Severe      ATTCGATCTGTCTATCCAGTTGCGTCACCAAATGAATGCAACCAAATGTGCCTTTCAACT   1222
gb:LC521925|Organism:Severe      ATTCGATCTGTCTATCCAGTTGCGTCACCAAATGAATGCAACCAAATGTGCCTTTCAACT   1195
gb:MT093571|Organism:Severe      ATTCGATCTGTCTATCCAGTTGCGTCACCAAATGAATGCAACCAAATGTGCCTTTCAACT   1222
gb:MT039887|Organism:Severe      ATTCGATCTGTCTATCCAGTTGCGTCACCAAATGAATGCAACCAAATGTGCCTTTCAACT   1222
gb:MT019530|Organism:Severe      ATTCGATCTGTCTATCCAGTTGCGTCACCAAATGAATGCAACCAAATGTGCCTTTCAACT   1222
gb:MT039888|Organism:Severe      ATTCGATCTGTCTATCCAGTTGCGTCACCAAATGAATGCAACCAAATGTGCCTTTCAACT   1222
gb:LC522972|Organism:Severe      ATTCGATCTGTCTATCCAGTTGCGTCACCAAATGAATGCAACCAAATGTGCCTTTCAACT   1219
gb:MT027063|Organism:Severe      ATTCGATCTGTCTATCCAGTTGCGTCACCAAATGAATGCAACCAAATGTGCCTTTCAACT   1222
gb:MT027062|Organism:Severe      ATTCGATCTGTCTATCCAGTTGCGTCACCAAATGAATGCAACCAAATGTGCCTTTCAACT   1222
gb:MT019529|Organism:Severe      ATTCGATCTGTCTATCCAGTTGCGTCACCAAATGAATGCAACCAAATGTGCCTTTCAACT   1222
gb:MN996529|Organism:Severe      ATTCGATCTGTCTATCCAGTTGCGTCACCAAATGAATGCAACCAAATGTGCCTTTCAACT   1210
gb:MN996531|Organism:Severe      ATTCGATCTGTCTATCCAGTTGCGTCACCAAATGAATGCAACCAAATGTGCCTTTCAACT   1209
gb:MT066176|Organism:Severe      ATTCGATCTGTCTATCCAGTTGCGTCACCAAATGAATGCAACCAAATGTGCCTTTCAACT   1222
gb:MT027064|Organism:Severe      ATTCGATCTGTCTATCCAGTTGCGTCACCAAATGAATGCAACCAAATGTGCCTTTCAACT   1222
gb:MN994468|Organism:Severe      ATTCGATCTGTCTATCCAGTTGCGTCACCAAATGAATGCAACCAAATGTGCCTTTCAACT   1222
gb:MT072688|Organism:Severe      ATTCGATCTGTCTATCCAGTTGCGTCACCAAATGAATGCAACCAAATGTGCCTTTCAACT   1207
gb:MN996527|Organism:Severe      ATTCGATCTGTCTATCCAGTTGCGTCACCAAATGAATGCAACCAAATGTGCCTTTCAACT   1189
gb:MT093631|Organism:Severe      ATTCGATCTGTCTATCCAGTTGCGTCACCAAATGAATGCAACCAAATGTGCCTTTCAACT   1260
gb:MT106053|Organism:Severe      ATTCGATCTGTCTATCCAGTTGCGTCACCAAATGAATGCAACCAAATGTGCCTTTCAACT   1222
gb:MT019533|Organism:Severe      ATTCGATCTGTCTATCCAGTTGCGTCACCAAATGAATGCAACCAAATGTGCCTTTCAACT   1222
gb:MT019531|Organism:Severe      ATTCGATCTGTCTATCCAGTTGCGTCACCAAATGAATGCAACCAAATGTGCCTTTCAACT   1222
gb:MN996528|Organism:Severe      ATTCGATCTGTCTATCCAGTTGCGTCACCAAATGAATGCAACCAAATGTGCCTTTCAACT   1222
gb:MN996530|Organism:Severe      ATTCGATCTGTCTATCCAGTTGCGTCACCAAATGAATGCAACCAAATGTGCCTTTCAACT   1208
gb:MN908947|Organism:Severe      ATTCGATCTGTCTATCCAGTTGCGTCACCAAATGAATGCAACCAAATGTGCCTTTCAACT   1222
gb:MT019532|Organism:Severe      ATTCGATCTGTCTATCCAGTTGCGTCACCAAATGAATGCAACCAAATGTGCCTTTCAACT   1222
                                 ************************************************************

gb:MT020781|Organism:Severe      CTCATGAAGTGTGATCATTGTGGTGAAACTTCATGGCAGACGGGCGATTTTGTTAAAGCC   1270
gb:MT007544|Organism:Severe      CTCATGAAGTGTGATCATTGTGGTGAAACTTCATGGCAGACGGGCGATTTTGTTAAAGCC   1282
gb:MN994467|Organism:Severe      CTCATGAAGTGTGATCATTGTGGTGAAACTTCATGGCAGACGGGCGATTTTGTTAAAGCC   1282
gb:MT044257|Organism:Severe      CTCATGAAGTGTGATCATTGTGGTGAAACTTCATGGCAGACGGGCGATTTTGTTAAAGCC   1282
gb:MT106054|Organism:Severe      CTCATGAAGTGTGATCATTGTGGTGAAACTTCATGGCAGACGGGCGATTTTGTTAAAGCC   1282
```

```
gb:MT049951|Organism:Severe      CTCATGAAGTGTGATCATTGTGGTGAAACTTCATGGCAGACGGGCGATTTTGTTAAAGCC    1282
gb:MN975262|Organism:Severe      CTCATGAAGTGTGATCATTGTGGTGAAACTTCATGGCAGACGGGCGATTTTGTTAAAGCC    1282
gb:MT106052|Organism:Severe      CTCATGAAGTGTGATCATTGTGGTGAAACTTCATGGCAGACGGGCGATTTTGTTAAAGCC    1282
gb:LC522975|Organism:Severe      CTCATGAAGTGTGATCATTGTGGTGAAACTTCATGGCAGACGGGCGATTTTGTTAAAGCC    1279
gb:LC522973|Organism:Severe      CTCATGAAGTGTGATCATTGTGGTGAAACTTCATGGCAGACGGGCGATTTTGTTAAAGCC    1279
gb:LC522974|Organism:Severe      CTCATGAAGTGTGATCATTGTGGTGAAACTTCATGGCAGACGGGCGATTTTGTTAAAGCC    1279
gb:MN985325|Organism:Severe      CTCATGAAGTGTGATCATTGTGGTGAAACTTCATGGCAGACGGGCGATTTTGTTAAAGCC    1282
gb:MT020881|Organism:Severe      CTCATGAAGTGTGATCATTGTGGTGAAACTTCATGGCAGACGGGCGATTTTGTTAAAGCC    1282
gb:MT020880|Organism:Severe      CTCATGAAGTGTGATCATTGTGGTGAAACTTCATGGCAGACGGGCGATTTTGTTAAAGCC    1282
gb:MT066175|Organism:Severe      CTCATGAAGTGTGATCATTGTGGTGAAACTTCATGGCAGACGGGCGATTTTGTTAAAGCC    1282
gb:MN997409|Organism:Severe      CTCATGAAGTGTGATCATTGTGGTGAAACTTCATGGCAGACGGGCGATTTTGTTAAAGCC    1282
gb:MN938384|Organism:Severe      CTCATGAAGTGTGATCATTGTGGTGAAACTTCATGGCAGACGGGCGATTTTGTTAAAGCC    1250
gb:MT044258|Organism:Severe      CTCATGAAGTGTGATCATTGTGGTGAAACTTCATGGCAGACGGGCGATTTTGTTAAAGCC    1258
gb:MT039890|Organism:Severe      CTCATGAAGTGTGATCATTGTGGTGAAACTTCATGGCAGACGGGCGATTTTGTTAAAGCC    1282
gb:MN988713|Organism:Severe      CTCATGAAGTGTGATCATTGTGGTGAAACTTCATGGCAGACGGGCGATTTTGTTAAAGCC    1282
gb:LC521925|Organism:Severe      CTCATGAAGTGTGATCATTGTGGTGAAACTTCATGGCAGACGGGCGATTTTGTTAAAGCC    1255
gb:MT093571|Organism:Severe      CTCATGAAGTGTGATCATTGTGGTGAAACTTCATGGCAGACGGGCGATTTTGTTAAAGCC    1282
gb:MT039887|Organism:Severe      CTCATGAAGTGTGATCATTGTGGTGAAACTTCATGGCAGACGGGCGATTTTGTTAAAGCC    1282
gb:MT019530|Organism:Severe      CTCATGAAGTGTGATCATTGTGGTGAAACTTCATGGCAGACGGGCGATTTTGTTAAAGCC    1282
gb:MT039888|Organism:Severe      CTCATGAAGTGTGATCATTGTGGTGAAACTTCATGGCAGACGGGCGATTTTGTTAAAGCC    1282
gb:LC522972|Organism:Severe      CTCATGAAGTGTGATCATTGTGGTGAAACTTCATGGCAGACGGGCGATTTTGTTAAAGCC    1279
gb:MT027063|Organism:Severe      CTCATGAAGTGTGATCATTGTGGTGAAACTTCATGGCAGACGGGCGATTTTGTTAAAGCC    1282
gb:MT027062|Organism:Severe      CTCATGAAGTGTGATCATTGTGGTGAAACTTCATGGCAGACGGGCGATTTTGTTAAAGCC    1282
gb:MT019529|Organism:Severe      CTCATGAAGTGTGATCATTGTGGTGAAACTTCATGGCAGACGGGCGATTTTGTTAAAGCC    1282
gb:MN996529|Organism:Severe      CTCATGAAGTGTGATCATTGTGGTGAAACTTCATGGCAGACGGGCGATTTTGTTAAAGCC    1270
gb:MN996531|Organism:Severe      CTCATGAAGTGTGATCATTGTGGTGAAACTTCATGGCAGACGGGCGATTTTGTTAAAGCC    1269
gb:MT066176|Organism:Severe      CTCATGAAGTGTGATCATTGTGGTGAAACTTCATGGCAGACGGGCGATTTTGTTAAAGCC    1282
gb:MT027064|Organism:Severe      CTCATGAAGTGTGATCATTGTGGTGAAACTTCATGGCAGACGGGCGATTTTGTTAAAGCC    1282
gb:MN994468|Organism:Severe      CTCATGAAGTGTGATCATTGTGGTGAAACTTCATGGCAGACGGGCGATTTTGTTAAAGCC    1282
gb:MT072688|Organism:Severe      CTCATGAAGTGTGATCATTGTGGTGAAACTTCATGGCAGACGGGCGATTTTGTTAAAGCC    1267
gb:MN996527|Organism:Severe      CTCATGAAGTGTGATCATTGTGGTGAAACTTCATGGCAGACGGGCGATTTTGTTAAAGCC    1249
gb:MT093631|Organism:Severe      CTCATGAAGTGTGATCATTGTGGTGAAACTTCATGGCAGACGGGCGATTTTGTTAAAGCC    1320
gb:MT106053|Organism:Severe      CTCATGAAGTGTGATCATTGTGGTGAAACTTCATGGCAGACGGGCGATTTTGTTAAAGCC    1282
gb:MT019533|Organism:Severe      CTCATGAAGTGTGATCATTGTGGTGAAACTTCATGGCAGACGGGCGATTTTGTTAAAGCC    1282
gb:MT019531|Organism:Severe      CTCATGAAGTGTGATCATTGTGGTGAAACTTCATGGCAGACGGGCGATTTTGTTAAAGCC    1282
gb:MN996528|Organism:Severe      CTCATGAAGTGTGATCATTGTGGTGAAACTTCATGGCAGACGGGCGATTTTGTTAAAGCC    1282
gb:MN996530|Organism:Severe      CTCATGAAGTGTGATCATTGTGGTGAAACTTCATGGCAGACGGGCGATTTTGTTAAAGCC    1268
gb:MN908947|Organism:Severe      CTCATGAAGTGTGATCATTGTGGTGAAACTTCATGGCAGACGGGCGATTTTGTTAAAGCC    1282
gb:MT019532|Organism:Severe      CTCATGAAGTGTGATCATTGTGGTGAAACTTCATGGCAGACGGGCGATTTTGTTAAAGCC    1282
                                 ************************************************************

gb:MT020781|Organism:Severe      ACTTGCGAATTTTGTGGCACTGAGAATTTGACTAAAGAAGGTGCCACTACTTGTGGTTAC    1330
gb:MT007544|Organism:Severe      ACTTGCGAATTTTGTGGCACTGAGAATTTGACTAAAGAAGGTGCCACTACTTGTGGTTAC    1342
gb:MN994467|Organism:Severe      ACTTGCGAATTTTGTGGCACTGAGAATTTGACTAAAGAAGGTGCCACTACTTGTGGTTAC    1342
gb:MT044257|Organism:Severe      ACTTGCGAATTTTGTGGCACTGAGAATTTGACTAAAGAAGGTGCCACTACTTGTGGTTAC    1342
gb:MT106054|Organism:Severe      ACTTGCGAATTTTGTGGCACTGAGAATTTGACTAAAGAAGGTGCCACTACTTGTGGTTAC    1342
gb:MT049951|Organism:Severe      ACTTGCGAATTTTGTGGCACTGAGAATTTGACTAAAGAAGGTGCCACTACTTGTGGTTAC    1342
gb:MN975262|Organism:Severe      ACTTGCGAATTTTGTGGCACTGAGAATTTGACTAAAGAAGGTGCCACTACTTGTGGTTAC    1342
gb:MT106052|Organism:Severe      ACTTGCGAATTTTGTGGCACTGAGAATTTGACTAAAGAAGGTGCCACTACTTGTGGTTAC    1342
gb:LC522975|Organism:Severe      ACTTGCGAATTTTGTGGCACTGAGAATTTGACTAAAGAAGGTGCCACTACTTGTGGTTAC    1339
gb:LC522973|Organism:Severe      ACTTGCGAATTTTGTGGCACTGAGAATTTGACTAAAGAAGGTGCCACTACTTGTGGTTAC    1339
gb:LC522974|Organism:Severe      ACTTGCGAATTTTGTGGCACTGAGAATTTGACTAAAGAAGGTGCCACTACTTGTGGTTAC    1339
gb:MN985325|Organism:Severe      ACTTGCGAATTTTGTGGCACTGAGAATTTGACTAAAGAAGGTGCCACTACTTGTGGTTAC    1342
gb:MT020881|Organism:Severe      ACTTGCGAATTTTGTGGCACTGAGAATTTGACTAAAGAAGGTGCCACTACTTGTGGTTAC    1342
gb:MT020880|Organism:Severe      ACTTGCGAATTTTGTGGCACTGAGAATTTGACTAAAGAAGGTGCCACTACTTGTGGTTAC    1342
gb:MT066175|Organism:Severe      ACTTGCGAATTTTGTGGCACTGAGAATTTGACTAAAGAAGGTGCCACTACTTGTGGTTAC    1342
gb:MN997409|Organism:Severe      ACTTGCGAATTTTGTGGCACTGAGAATTTGACTAAAGAAGGTGCCACTACTTGTGGTTAC    1342
gb:MN938384|Organism:Severe      ACTTGCGAATTTTGTGGCACTGAGAATTTGACTAAAGAAGGTGCCACTACTTGTGGTTAC    1310
gb:MT044258|Organism:Severe      ACTTGCGAATTTTGTGGCACTGAGAATTTGACTAAAGAAGGTGCCACTACTTGTGGTTAC    1318
gb:MT039890|Organism:Severe      ACTTGCGAATTTTGTGGCACTGAGAATTTGACTAAAGAAGGTGCCACTACTTGTGGTTAC    1342
gb:MN988713|Organism:Severe      ACTTGCGAATTTTGTGGCACTGAGAATTTGACTAAAGAAGGTGCCACTACTTGTGGTTAC    1342
gb:LC521925|Organism:Severe      ACTTGCGAATTTTGTGGCACTGAGAATTTGACTAAAGAAGGTGCCACTACTTGTGGTTAC    1315
gb:MT093571|Organism:Severe      ACTTGCGAATTTTGTGGCACTGAGAATTTGACTAAAGAAGGTGCCACTACTTGTGGTTAC    1342
gb:MT039887|Organism:Severe      ACTTGCGAATTTTGTGGCACTGAGAATTTGACTAAAGAAGGTGCCACTACTTGTGGTTAC    1342
gb:MT019530|Organism:Severe      ACTTGCGAATTTTGTGGCACTGAGAATTTGACTAAAGAAGGTGCCACTACTTGTGGTTAC    1342
```

```
gb:MT039888|Organism:Severe      ACTTGCGAATTTTGTGGCACTGAGAATTTGACTAAAGAAGGTGCCACTACTTGTGGTTAC    1342
gb:LC522972|Organism:Severe      ACTTGCGAATTTTGTGGCACTGAGAATTTGACTAAAGAAGGTGCCACTACTTGTGGTTAC    1339
gb:MT027063|Organism:Severe      ACTTGCGAATTTTGTGGCACTGAGAATTTGACTAAAGAAGGTGCCACTACTTGTGGTTAC    1342
gb:MT027062|Organism:Severe      ACTTGCGAATTTTGTGGCACTGAGAATTTGACTAAAGAAGGTGCCACTACTTGTGGTTAC    1342
gb:MT019529|Organism:Severe      ACTTGCGAATTTTGTGGCACTGAGAATTTGACTAAAGAAGGTGCCACTACTTGTGGTTAC    1342
gb:MN996529|Organism:Severe      ACTTGCGAATTTTGTGGCACTGAGAATTTGACTAAAGAAGGTGCCACTACTTGTGGTTAC    1330
gb:MN996531|Organism:Severe      ACTTGCGAATTTTGTGGCACTGAGAATTTGACTAAAGAAGGTGCCACTACTTGTGGTTAC    1329
gb:MT066176|Organism:Severe      ACTTGCGAATTTTGTGGCACTGAGAATTTGACTAAAGAAGGTGCCACTACTTGTGGTTAC    1342
gb:MT027064|Organism:Severe      ACTTGCGAATTTTGTGGCACTGAGAATTTGACTAAAGAAGGTGCCACTACTTGTGGTTAC    1342
gb:MN994468|Organism:Severe      ACTTGCGAATTTTGTGGCACTGAGAATTTGACTAAAGAAGGTGCCACTACTTGTGGTTAC    1342
gb:MT072688|Organism:Severe      ACTTGCGAATTTTGTGGCACTGAGAATTTGACTAAAGAAGGTGCCACTACTTGTGGTTAC    1327
gb:MN996527|Organism:Severe      ACTTGCGAATTTTGTGGCACTGAGAATTTGACTAAAGAAGGTGCCACTACTTGTGGTTAC    1309
gb:MT093631|Organism:Severe      ACTTGCGAATTTTGTGGCACTGAGAATTTGACTAAAGAAGGTGCCACTACTTGTGGTTAC    1380
gb:MT106053|Organism:Severe      ACTTGCGAATTTTGTGGCACTGAGAATTTGACTAAAGAAGGTGCCACTACTTGTGGTTAC    1342
gb:MT019533|Organism:Severe      ACTTGCGAATTTTGTGGCACTGAGAATTTGACTAAAGAAGGTGCCACTACTTGTGGTTAC    1342
gb:MT019531|Organism:Severe      ACTTGCGAATTTTGTGGCACTGAGAATTTGACTAAAGAAGGTGCCACTACTTGTGGTTAC    1342
gb:MN996528|Organism:Severe      ACTTGCGAATTTTGTGGCACTGAGAATTTGACTAAAGAAGGTGCCACTACTTGTGGTTAC    1342
gb:MN996530|Organism:Severe      ACTTGCGAATTTTGTGGCACTGAGAATTTGACTAAAGAAGGTGCCACTACTTGTGGTTAC    1328
gb:MN908947|Organism:Severe      ACTTGCGAATTTTGTGGCACTGAGAATTTGACTAAAGAAGGTGCCACTACTTGTGGTTAC    1342
gb:MT019532|Organism:Severe      ACTTGCGAATTTTGTGGCACTGAGAATTTGACTAAAGAAGGTGCCACTACTTGTGGTTAC    1342
                                 ************************************************************

gb:MT020781|Organism:Severe      TTACCCCAAAATGCTGTTGTTAAAATTTATTGTCCAGCATGTCACAATTCAGAAGTAGGA    1390
gb:MT007544|Organism:Severe      TTACCCCAAAATGCTGTTGTTAAAATTTATTGTCCAGCATGTCACAATTCAGAAGTAGGA    1402
gb:MN994467|Organism:Severe      TTACCCCAAAATGCTGTTGTTAAAATTTATTGTCCAGCATGTCACAATTCAGAAGTAGGA    1402
gb:MT044257|Organism:Severe      TTACCCCAAAATGCTGTTGTTAAAATTTATTGTCCAGCATGTCACAATTCAGAAGTAGGA    1402
gb:MT106054|Organism:Severe      TTACCCCAAAATGCTGTTGTTAAAATTTATTGTCCAGCATGTCACAATTCAGAAGTAGGA    1402
gb:MT049951|Organism:Severe      TTACCCCAAAATGCTGTTGTTAAAATTTATTGTCCAGCATGTCACAATTCAGAAGTAGGA    1402
gb:MN975262|Organism:Severe      TTACCCCAAAATGCTGTTGTTAAAATTTATTGTCCAGCATGTCACAATTCAGAAGTAGGA    1402
gb:MT106052|Organism:Severe      TTACCCCAAAATGCTGTTGTTAAAATTTATTGTCCAGCATGTCACAATTCAGAAGTAGGA    1402
gb:LC522975|Organism:Severe      TTACCCCAAAATGCTGTTGTTAAAATTTATTGTCCAGCATGTCACAATTCAGAAGTAGGA    1399
gb:LC522973|Organism:Severe      TTACCCCAAAATGCTGTTGTTAAAATTTATTGTCCAGCATGTCACAATTCAGAAGTAGGA    1399
gb:LC522974|Organism:Severe      TTACCCCAAAATGCTGTTGTTAAAATTTATTGTCCAGCATGTCACAATTCAGAAGTAGGA    1399
gb:MN985325|Organism:Severe      TTACCCCAAAATGCTGTTGTTAAAATTTATTGTCCAGCATGTCACAATTCAGAAGTAGGA    1402
gb:MT020881|Organism:Severe      TTACCCCAAAATGCTGTTGTTAAAATTTATTGTCCAGCATGTCACAATTCAGAAGTAGGA    1402
gb:MT020880|Organism:Severe      TTACCCCAAAATGCTGTTGTTAAAATTTATTGTCCAGCATGTCACAATTCAGAAGTAGGA    1402
gb:MT066175|Organism:Severe      TTACCCCAAAATGCTGTTGTTAAAATTTATTGTCCAGCATGTCACAATTCAGAAGTAGGA    1402
gb:MN997409|Organism:Severe      TTACCCCAAAATGCTGTTGTTAAAATTTATTGTCCAGCATGTCACAATTCAGAAGTAGGA    1402
gb:MN938384|Organism:Severe      TTACCCCAAAATGCTGTTGTTAAAATTTATTGTCCAGCATGTCACAATTCAGAAGTAGGA    1370
gb:MT044258|Organism:Severe      TTACCCCAAAATGCTGTTGTTAAAATTTATTGTCCAGCATGTCACAATTCAGAAGTAGGA    1378
gb:MT039890|Organism:Severe      TTACCCCAAAATGCTGTTGTTAAAATTTATTGTCCAGCATGTCACAATTCAGAAGTAGGA    1402
gb:MN988713|Organism:Severe      TTACCCCAAAATGCTGTTGTTAAAATTTATTGTCCAGCATGTCACAATTCAGAAGTAGGA    1402
gb:LC521925|Organism:Severe      TTACCCCAAAATGCTGTTGTTAAAATTTATTGTCCAGCATGTCACAATTCAGAAGTAGGA    1375
gb:MT093571|Organism:Severe      TTACCCCAAAATGCTGTTGTTAAAATTTATTGTCCAGCATGTCACAATTCAGAAGTAGGA    1402
gb:MT039887|Organism:Severe      TTACCCCAAAATGCTGTTGTTAAAATTTATTGTCCAGCATGTCACAATTCAGAAGTAGGA    1402
gb:MT019530|Organism:Severe      TTACCCCAAAATGCTGTTGTTAAAATTTATTGTCCAGCATGTCACAATTCAGAAGTAGGA    1402
gb:MT039888|Organism:Severe      TTACCCCAAAATGCTGTTGTTAAAATTTATTGTCCAGCATGTCACAATTCAGAAGTAGGA    1402
gb:LC522972|Organism:Severe      TTACCCCAAAATGCTGTTGTTAAAATTTATTGTCCAGCATGTCACAATTCAGAAGTAGGA    1399
gb:MT027063|Organism:Severe      TTACCCCAAAATGCTGTTGTTAAAATTTATTGTCCAGCATGTCACAATTCAGAAGTAGGA    1402
gb:MT027062|Organism:Severe      TTACCCCAAAATGCTGTTGTTAAAATTTATTGTCCAGCATGTCACAATTCAGAAGTAGGA    1402
gb:MT019529|Organism:Severe      TTACCCCAAAATGCTGTTGTTAAAATTTATTGTCCAGCATGTCACAATTCAGAAGTAGGA    1402
gb:MN996529|Organism:Severe      TTACCCCAAAATGCTGTTGTTAAAATTTATTGTCCAGCATGTCACAATTCAGAAGTAGGA    1390
gb:MN996531|Organism:Severe      TTACCCCAAAATGCTGTTGTTAAAATTTATTGTCCAGCATGTCACAATTCAGAAGTAGGA    1389
gb:MT066176|Organism:Severe      TTACCCCAAAATGCTGTTGTTAAAATTTATTGTCCAGCATGTCACAATTCAGAAGTAGGA    1402
gb:MT027064|Organism:Severe      TTACCCCAAAATGCTGTTGTTAAAATTTATTGTCCAGCATGTCACAATTCAGAAGTAGGA    1402
gb:MN994468|Organism:Severe      TTACCCCAAAATGCTGTTGTTAAAATTTATTGTCCAGCATGTCACAATTCAGAAGTAGGA    1402
gb:MT072688|Organism:Severe      TTACCCCAAAATGCTGTTGTTAAAATTTATTGTCCAGCATGTCACAATTCAGAAGTAGGA    1387
gb:MN996527|Organism:Severe      TTACCCCAAAATGCTGTTGTTAAAATTTATTGTCCAGCATGTCACAATTCAGAAGTAGGA    1369
gb:MT093631|Organism:Severe      TTACCCCAAAATGCTGTTGTTAAAATTTATTGTCCAGCATGTCACAATTCAGAAGTAGGA    1440
gb:MT106053|Organism:Severe      TTACCCCAAAATGCTGTTGTTAAAATTTATTGTCCAGCATGTCACAATTCAGAAGTAGGA    1402
gb:MT019533|Organism:Severe      TTACCCCAAAATGCTGTTGTTAAAATTTATTGTCCAGCATGTCACAATTCAGAAGTAGGA    1402
gb:MT019531|Organism:Severe      TTACCCCAAAATGCTGTTGTTAAAATTTATTGTCCAGCATGTCACAATTCAGAAGTAGGA    1402
gb:MN996528|Organism:Severe      TTACCCCAAAATGCTGTTGTTAAAATTTATTGTCCAGCATGTCACAATTCAGAAGTAGGA    1402
gb:MN996530|Organism:Severe      TTACCCCAAAATGCTGTTGTTAAAATTTATTGTCCAGCATGTCACAATTCAGAAGTAGGA    1388
gb:MN908947|Organism:Severe      TTACCCCAAAATGCTGTTGTTAAAATTTATTGTCCAGCATGTCACAATTCAGAAGTAGGA    1402
```

```
gb:MT019532|Organism:Severe      TTACCCCAAAATGCTGTTGTTAAAATTTATTGTCCAGCATGTCACAATTCAGAAGTAGGA   1402
                                 ************************************************************

gb:MT020781|Organism:Severe      CCTGAGCATAGTCTTGCCGAATACCATAATGAATCTGGCTTGAAAACCATTCTTCGTAAG   1450
gb:MT007544|Organism:Severe      CCTGAGCATAGTCTTGCCGAATACCATAATGAATCTGGCTTGAAAACCATTCTTCGTAAG   1462
gb:MN994467|Organism:Severe      CCTGAGCATAGTCTTGCCGAATACCATAATGAATCTGGCTTGAAAACCATTCTTCGTAAG   1462
gb:MT044257|Organism:Severe      CCTGAGCATAGTCTTGCCGAATACCATAATGAATCTGGCTTGAAAACCATTCTTCGTAAG   1462
gb:MT106054|Organism:Severe      CCTGAGCATAGTCTTGCCGAATACCATAATGAATCTGGCTTGAAAACCATTCTTCGTAAG   1462
gb:MT049951|Organism:Severe      CCTGAGCATAGTCTTGCCGAATACCATAATGAATCTGGCTTGAAAACCATTCTTCGTAAG   1462
gb:MN975262|Organism:Severe      CCTGAGCATAGTCTTGCCGAATACCATAATGAATCTGGCTTGAAAACCATTCTTCGTAAG   1462
gb:MT106052|Organism:Severe      CCTGAGCATAGTCTTGCCGAATACCATAATGAATCTGGCTTGAAAACCATTCTTCGTAAG   1462
gb:LC522975|Organism:Severe      CCTGAGCATAGTCTTGCCGAATACCATAATGAATCTGGCTTGAAAACCATTCTTCGTAAG   1459
gb:LC522973|Organism:Severe      CCTGAGCATAGTCTTGCCGAATACCATAATGAATCTGGCTTGAAAACCATTCTTCGTAAG   1459
gb:LC522974|Organism:Severe      CCTGAGCATAGTCTTGCCGAATACCATAATGAATCTGGCTTGAAAACCATTCTTCGTAAG   1459
gb:MN985325|Organism:Severe      CCTGAGCATAGTCTTGCCGAATACCATAATGAATCTGGCTTGAAAACCATTCTTCGTAAG   1462
gb:MT020881|Organism:Severe      CCTGAGCATAGTCTTGCCGAATACCATAATGAATCTGGCTTGAAAACCATTCTTCGTAAG   1462
gb:MT020880|Organism:Severe      CCTGAGCATAGTCTTGCCGAATACCATAATGAATCTGGCTTGAAAACCATTCTTCGTAAG   1462
gb:MT066175|Organism:Severe      CCTGAGCATAGTCTTGCCGAATACCATAATGAATCTGGCTTGAAAACCATTCTTCGTAAG   1462
gb:MN997409|Organism:Severe      CCTGAGCATAGTCTTGCCGAATACCATAATGAATCTGGCTTGAAAACCATTCTTCGTAAG   1462
gb:MN938384|Organism:Severe      CCTGAGCATAGTCTTGCCGAATACCATAATGAATCTGGCTTGAAAACCATTCTTCGTAAG   1430
gb:MT044258|Organism:Severe      CCTGAGCATAGTCTTGCCGAATACCATAATGAATCTGGCTTGAAAACCATTCTTCGTAAG   1438
gb:MT039890|Organism:Severe      CCTGAGCATAGTCTTGCCGAATACCATAATGAATCTGGCTTGAAAACCATTCTTCGTAAG   1462
gb:MN988713|Organism:Severe      CCTGAGCATAGTCTTGCCGAATACCATAATGAATCTGGCTTGAAAACCATTCTTCGTAAG   1462
gb:LC521925|Organism:Severe      CCTGAGCATAGTCTTGCCGAATACCATAATGAATCTGGCTTGAAAACCATTCTTCGTAAG   1435
gb:MT093571|Organism:Severe      CCTGAGCATAGTCTTGCCGAATACCATAATGAATCTGGCTTGAAAACCATTCTTCGTAAG   1462
gb:MT039887|Organism:Severe      CCTGAGCATAGTCTTGCCGAATACCATAATGAATCTGGCTTGAAAACCATTCTTCGTAAG   1462
gb:MT019530|Organism:Severe      CCTGAGCATAGTCTTGCCGAATACCATAATGAATCTGGCTTGAAAACCATTCTTCGTAAG   1462
gb:MT039888|Organism:Severe      CCTGAGCATAGTCTTGCCGAATACCATAATGAATCTGGCTTGAAAACCATTCTTCGTAAG   1462
gb:LC522972|Organism:Severe      CCTGAGCATAGTCTTGCCGAATACCATAATGAATCTGGCTTGAAAACCATTCTTCGTAAG   1459
gb:MT027063|Organism:Severe      CCTGAGCATAGTCTTGCCGAATACCATAATGAATCTGGCTTGAAAACCATTCTTCGTAAG   1462
gb:MT027062|Organism:Severe      CCTGAGCATAGTCTTGCCGAATACCATAATGAATCTGGCTTGAAAACCATTCTTCGTAAG   1462
gb:MT019529|Organism:Severe      CCTGAGCATAGTCTTGCCGAATACCATAATGAATCTGGCTTGAAAACCATTCTTCGTAAG   1462
gb:MN996529|Organism:Severe      CCTGAGCATAGTCTTGCCGAATACCATAATGAATCTGGCTTGAAAACCATTCTTCGTAAG   1450
gb:MN996531|Organism:Severe      CCTGAGCATAGTCTTGCCGAATACCATAATGAATCTGGCTTGAAAACCATTCTTCGTAAG   1449
gb:MT066176|Organism:Severe      CCTGAGCATAGTCTTGCCGAATACCATAATGAATCTGGCTTGAAAACCATTCTTCGTAAG   1462
gb:MT027064|Organism:Severe      CCTGAGCATAGTCTTGCCGAATACCATAATGAATCTGGCTTGAAAACCATTCTTCGTAAG   1462
gb:MN994468|Organism:Severe      CCTGAGCATAGTCTTGCCGAATACCATAATGAATCTGGCTTGAAAACCATTCTTCGTAAG   1462
gb:MT072688|Organism:Severe      CCTGAGCATAGTCTTGCCGAATACCATAATGAATCTGGCTTGAAAACCATTCTTCGTAAG   1447
gb:MN996527|Organism:Severe      CCTGAGCATAGTCTTGCCGAATACCATAATGAATCTGGCTTGAAAACCATTCTTCGTAAG   1429
gb:MT093631|Organism:Severe      CCTGAGCATAGTCTTGCCGAATACCATAATGAATCTGGCTTGAAAACCATTCTTCGTAAG   1500
gb:MT106053|Organism:Severe      CCTGAGCATAGTCTTGCCGAATACCATAATGAATCTGGCTTGAAAACCATTCTTCGTAAG   1462
gb:MT019533|Organism:Severe      CCTGAGCATAGTCTTGCCGAATACCATAATGAATCTGGCTTGAAAACCATTCTTCGTAAG   1462
gb:MT019531|Organism:Severe      CCTGAGCATAGTCTTGCCGAATACCATAATGAATCTGGCTTGAAAACCATTCTTCGTAAG   1462
gb:MN996528|Organism:Severe      CCTGAGCATAGTCTTGCCGAATACCATAATGAATCTGGCTTGAAAACCATTCTTCGTAAG   1462
gb:MN996530|Organism:Severe      CCTGAGCATAGTCTTGCCGAATACCATAATGAATCTGGCTTGAAAACCATTCTTCGTAAG   1448
gb:MN908947|Organism:Severe      CCTGAGCATAGTCTTGCCGAATACCATAATGAATCTGGCTTGAAAACCATTCTTCGTAAG   1462
gb:MT019532|Organism:Severe      CCTGAGCATAGTCTTGCCGAATACCATAATGAATCTGGCTTGAAAACCATTCTTCGTAAG   1462
                                 ************************************************************

gb:MT020781|Organism:Severe      GGTGGTCGCACTATTGCCTTTGGAGGCTGTGTGTTCTCTTATGTTGGTTGCCATAACAAG   1510
gb:MT007544|Organism:Severe      GGTGGTCGCACTATTGCCTTTGGAGGCTGTGTGTTCTCTTATGTTGGTTGCCATAACAAG   1522
gb:MN994467|Organism:Severe      GGTGGTCGCACTATTGCCTTTGGAGGCTGTGTGTTCTCTTATGTTGGTTGCCATAACAAG   1522
gb:MT044257|Organism:Severe      GGTGGTCGCACTATTGCCTTTGGAGGCTGTGTGTTCTCTTATGTTGGTTGCCATAACAAG   1522
gb:MT106054|Organism:Severe      GGTGGTCGCACTATTGCCTTTGGAGGCTGTGTGTTCTCTTATGTTGGTTGCCATAACAAG   1522
gb:MT049951|Organism:Severe      GGTGGTCGCACTATTGCCTTTGGAGGCTGTGTGTTCTCTTATGTTGGTTGCCATAACAAG   1522
gb:MN975262|Organism:Severe      GGTGGTCGCACTATTGCCTTTGGAGGCTGTGTGTTCTCTTATGTTGGTTGCCATAACAAG   1522
gb:MT106052|Organism:Severe      GGTGGTCGCACTATTGCCTTTGGAGGCTGTGTGTTCTCTTATGTTGGTTGCCATAACAAG   1522
gb:LC522975|Organism:Severe      GGTGGTCGCACTATTGCCTTTGGAGGCTGTGTGTTCTCTTATGTTGGTTGCCATAACAAG   1519
gb:LC522973|Organism:Severe      GGTGGTCGCACTATTGCCTTTGGAGGCTGTGTGTTCTCTTATGTTGGTTGCCATAACAAG   1519
gb:LC522974|Organism:Severe      GGTGGTCGCACTATTGCCTTTGGAGGCTGTGTGTTCTCTTATGTTGGTTGCCATAACAAG   1519
gb:MN985325|Organism:Severe      GGTGGTCGCACTATTGCCTTTGGAGGCTGTGTGTTCTCTTATGTTGGTTGCCATAACAAG   1522
gb:MT020881|Organism:Severe      GGTGGTCGCACTATTGCCTTTGGAGGCTGTGTGTTCTCTTATGTTGGTTGCCATAACAAG   1522
gb:MT020880|Organism:Severe      GGTGGTCGCACTATTGCCTTTGGAGGCTGTGTGTTCTCTTATGTTGGTTGCCATAACAAG   1522
gb:MT066175|Organism:Severe      GGTGGTCGCACTATTGCCTTTGGAGGCTGTGTGTTCTCTTATGTTGGTTGCCATAACAAG   1522
gb:MN997409|Organism:Severe      GGTGGTCGCACTATTGCCTTTGGAGGCTGTGTGTTCTCTTATGTTGGTTGCCATAACAAG   1522
```

```
gb:MN938384|Organism:Severe      GGTGGTCGCACTATTGCCTTTGGAGGCTGTGTGTTCTCTTATGTTGGTTGCCATAACAAG   1490
gb:MT044258|Organism:Severe      GGTGGTCGCACTATTGCCTTTGGAGGCTGTGTGTTCTCTTATGTTGGTTGCCATAACAAG   1498
gb:MT039890|Organism:Severe      GGTGGTCGCACTATTGCCTTTGGAGGCTGTGTGTTCTCTTATGTTGGTTGCCATAACAAG   1522
gb:MN988713|Organism:Severe      GGTGGTCGCACTATTGCCTTTGGAGGCTGTGTGTTCTCTTATGTTGGTTGCCATAACAAG   1522
gb:LC521925|Organism:Severe      GGTGGTCGCACTATTGCCTTTGGAGGCTGTGTGTTCTCTTATGTTGGTTGCCATAACAAG   1495
gb:MT093571|Organism:Severe      GGTGGTCGCACTATTGCCTTTGGAGGCTGTGTGTTCTCTTATGTTGGTTGCCATAACAAG   1522
gb:MT039887|Organism:Severe      GGTGGTCGCACTATTGCCTTTGGAGGCTGTGTGTTCTCTTATGTTGGTTGCCATAACAAG   1522
gb:MT019530|Organism:Severe      GGTGGTCGCACTATTGCCTTTGGAGGCTGTGTGTTCTCTTATGTTGGTTGCCATAACAAG   1522
gb:MT039888|Organism:Severe      GGTGGTCGCACTATTGCCTTTGGAGGCTGTGTGTTCTCTTATGTTGGTTGCCATAACAAG   1522
gb:LC522972|Organism:Severe      GGTGGTCGCACTATTGCCTTTGGAGGCTGTGTGTTCTCTTATGTTGGTTGCCATAACAAG   1519
gb:MT027063|Organism:Severe      GGTGGTCGCACTATTGCCTTTGGAGGCTGTGTGTTCTCTTATGTTGGTTGCCATAACAAG   1522
gb:MT027062|Organism:Severe      GGTGGTCGCACTATTGCCTTTGGAGGCTGTGTGTTCTCTTATGTTGGTTGCCATAACAAG   1522
gb:MT019529|Organism:Severe      GGTGGTCGCACTATTGCCTTTGGAGGCTGTGTGTTCTCTTATGTTGGTTGCCATAACAAG   1522
gb:MN996529|Organism:Severe      GGTGGTCGCACTATTGCCTTTGGAGGCTGTGTGTTCTCTTATGTTGGTTGCCATAACAAG   1510
gb:MN996531|Organism:Severe      GGTGGTCGCACTATTGCCTTTGGAGGCTGTGTGTTCTCTTATGTTGGTTGCCATAACAAG   1509
gb:MT066176|Organism:Severe      GGTGGTCGCACTATTGCCTTTGGAGGCTGTGTGTTCTCTTATGTTGGTTGCCATAACAAG   1522
gb:MT027064|Organism:Severe      GGTGGTCGCACTATTGCCTTTGGAGGCTGTGTGTTCTCTTATGTTGGTTGCCATAACAAG   1522
gb:MN994468|Organism:Severe      GGTGGTCGCACTATTGCCTTTGGAGGCTGTGTGTTCTCTTATGTTGGTTGCCATAACAAG   1522
gb:MT072688|Organism:Severe      GGTGGTCGCACTATTGCCTTTGGAGGCTGTGTGTTCTCTTATGTTGGTTGCCATAACAAG   1507
gb:MN996527|Organism:Severe      GGTGGTCGCACTATTGCCTTTGGAGGCTGTGTGTTCTCTTATGTTGGTTGCCATAACAAG   1489
gb:MT093631|Organism:Severe      GGTGGTCGCACTATTGCCTTTGGAGGCTGTGTGTTCTCTTATGTTGGTTGCCATAACAAG   1560
gb:MT106053|Organism:Severe      GGTGGTCGCACTATTGCCTTTGGAGGCTGTGTGTTCTCTTATGTTGGTTGCCATAACAAG   1522
gb:MT019533|Organism:Severe      GGTGGTCGCACTATTGCCTTTGGAGGCTGTGTGTTCTCTTATGTTGGTTGCCATAACAAG   1522
gb:MT019531|Organism:Severe      GGTGGTCGCACTATTGCCTTTGGAGGCTGTGTGTTCTCTTATGTTGGTTGCCATAACAAG   1522
gb:MN996528|Organism:Severe      GGTGGTCGCACTATTGCCTTTGGAGGCTGTGTGTTCTCTTATGTTGGTTGCCATAACAAG   1522
gb:MN996530|Organism:Severe      GGTGGTCGCACTATTGCCTTTGGAGGCTGTGTGTTCTCTTATGTTGGTTGCCATAACAAG   1508
gb:MN908947|Organism:Severe      GGTGGTCGCACTATTGCCTTTGGAGGCTGTGTGTTCTCTTATGTTGGTTGCCATAACAAG   1522
gb:MT019532|Organism:Severe      GGTGGTCGCACTATTGCCTTTGGAGGCTGTGTGTTCTCTTATGTTGGTTGCCATAACAAG   1522
                                 ************************************************************

gb:MT020781|Organism:Severe      TGTGCCTATTGGGTTCCACGTGCTAGCGCTAACATAGGTTGTAACCATACAGGTGTTGTT   1570
gb:MT007544|Organism:Severe      TGTGCCTATTGGGTTCCACGTGCTAGCGCTAACATAGGTTGTAACCATACAGGTGTTGTT   1582
gb:MN994467|Organism:Severe      TGTGCCTATTGGGTTCCACGTGCTAACGCTAACATAGGTTGTAACCATACAGGTGTTGTT   1582
gb:MT044257|Organism:Severe      TGTGCCTATTGGGTTCCACGTGCTAGCGCTAACATAGGTTGTAACCATACAGGTGTTGTT   1582
gb:MT106054|Organism:Severe      TGTGCCTATTGGGTTCCACGTGCTAGCGCTAACATAGGTTGTAACCATACAGGTGTTGTT   1582
gb:MT049951|Organism:Severe      TGTGCCTATTGGGTTCCACGTGCTAGCGCTAACATAGGTTGTAACCATACAGGTGTTGTT   1582
gb:MN975262|Organism:Severe      TGTGCCTATTGGGTTCCACGTGCTAGCGCTAACATAGGTTGTAACCATACAGGTGTTGTT   1582
gb:MT106052|Organism:Severe      TGTGCCTATTGGGTTCCACGTGCTAGCGCTAACATAGGTTGTAACCATACAGGTGTTGTT   1582
gb:LC522975|Organism:Severe      TGTGCCTATTGGGTTCCACGTGCTAGCGCTAACATAGGTTGTAACCATACAGGTGTTGTT   1579
gb:LC522973|Organism:Severe      TGTGCCTATTGGGTTCCACGTGCTAGCGCTAACATAGGTTGTAACCATACAGGTGTTGTT   1579
gb:LC522974|Organism:Severe      TGTGCCTATTGGGTTCCACGTGCTAGCGCTAACATAGGTTGTAACCATACAGGTGTTGTT   1579
gb:MN985325|Organism:Severe      TGTGCCTATTGGGTTCCACGTGCTAGCGCTAACATAGGTTGTAACCATACAGGTGTTGTT   1582
gb:MT020881|Organism:Severe      TGTGCCTATTGGGTTCCACGTGCTAGCGCTAACATAGGTTGTAACCATACAGGTGTTGTT   1582
gb:MT020880|Organism:Severe      TGTGCCTATTGGGTTCCACGTGCTAGCGCTAACATAGGTTGTAACCATACAGGTGTTGTT   1582
gb:MT066175|Organism:Severe      TGTGCCTATTGGGTTCCACGTGCTAGCGCTAACATAGGTTGTAACCATACAGGTGTTGTT   1582
gb:MN997409|Organism:Severe      TGTGCCTATTGGGTTCCACGTGCTAGCGCTAACATAGGTTGTAACCATACAGGTGTTGTT   1582
gb:MN938384|Organism:Severe      TGTGCCTATTGGGTTCCACGTGCTAGCGCTAACATAGGTTGTAACCATACAGGTGTTGTT   1550
gb:MT044258|Organism:Severe      TGTGCCTATTGGGTTCCACGTGCTAGCGCTAACATAGGTTGTAACCATACAGGTGTTGTT   1558
gb:MT039890|Organism:Severe      TGTGCCTATTGGGTTCCACGTGCTAGCGCTAACATAGGTTGTAACCATACAGGTGTTGTT   1582
gb:MN988713|Organism:Severe      TGTGCCTATTGGGTTCCACGTGCTAGCGCTAACATAGGTTGTAACCATACAGGTGTTGTT   1582
gb:LC521925|Organism:Severe      TGTGCCTATTGGGTTCCACGTGCTAGCGCTAACATAGGTTGTAACCATACAGGTGTTGTT   1555
gb:MT093571|Organism:Severe      TGTGCCTATTGGGTTCCACGTGCTAGCGCTAACATAGGTTGTAACCATACAGGTGTTGTT   1582
gb:MT039887|Organism:Severe      TGTGCCTATTGGGTTCCACGTGCTAGCGCTAACATAGGTTGTAACCATACAGGTGTTGTT   1582
gb:MT019530|Organism:Severe      TGTGCCTATTGGGTTCCACGTGCTAGCGCTAACATAGGTTGTAACCATACAGGTGTTGTT   1582
gb:MT039888|Organism:Severe      TGTGCCTATTGGGTTCCACGTGCTAGCGCTAACATAGGTTGTAACCATACAGGTGTTGTT   1582
gb:LC522972|Organism:Severe      TGTGCCTATTGGGTTCCACGTGCTAGCGCTAACATAGGTTGTAACCATACAGGTGTTGTT   1579
gb:MT027063|Organism:Severe      TGTGCCTATTGGGTTCCACGTGCTAGCGCTAACATAGGTTGTAACCATACAGGTGTTGTT   1582
gb:MT027062|Organism:Severe      TGTGCCTATTGGGTTCCACGTGCTAGCGCTAACATAGGTTGTAACCATACAGGTGTTGTT   1582
gb:MT019529|Organism:Severe      TGTGCCTATTGGGTTCCACGTGCTAGCGCTAACATAGGTTGTAACCATACAGGTGTTGTT   1582
gb:MN996529|Organism:Severe      TGTGCCTATTGGGTTCCACGTGCTAGCGCTAACATAGGTTGTAACCATACAGGTGTTGTT   1570
gb:MN996531|Organism:Severe      TGTGCCTATTGGGTTCCACGTGCTAGCGCTAACATAGGTTGTAACCATACAGGTGTTGTT   1569
gb:MT066176|Organism:Severe      TGTGCCTATTGGGTTCCACGTGCTAGCGCTAACATAGGTTGTAACCATACAGGTGTTGTT   1582
gb:MT027064|Organism:Severe      TGTGCCTATTGGGTTCCACGTGCTAGCGCTAACATAGGTTGTAACCATACAGGTGTTGTT   1582
gb:MN994468|Organism:Severe      TGTGCCTATTGGGTTCCACGTGCTAGCGCTAACATAGGTTGTAACCATACAGGTGTTGTT   1582
gb:MT072688|Organism:Severe      TGTGCCTATTGGGTTCCACGTGCTAGCGCTAACATAGGTTGTAACCATACAGGTGTTGTT   1567
```

```
gb:MN996527|Organism:Severe      TGTGCCTATTGGGTTCCACGTGCTAGCGCTAACATAGGTTGTAACCATACAGGTGTTGTT   1549
gb:MT093631|Organism:Severe      TGTGCCTATTGGGTTCCACGTGCTAGCGCTAACATAGGTTGTAACCATACAGGTGTTGTT   1620
gb:MT106053|Organism:Severe      TGTGCCTATTGGGTTCCACGTGCTAGCGCTAACATAGGTTGTAACCATACAGGTGTTGTT   1582
gb:MT019533|Organism:Severe      TGTGCCTATTGGGTTCCACGTGCTAGCGCTAACATAGGTTGTAACCATACAGGTGTTGTT   1582
gb:MT019531|Organism:Severe      TGTGCCTATTGGGTTCCACGTGCTAGCGCTAACATAGGTTGTAACCATACAGGTGTTGTT   1582
gb:MN996528|Organism:Severe      TGTGCCTATTGGGTTCCACGTGCTAGCGCTAACATAGGTTGTAACCATACAGGTGTTGTT   1582
gb:MN996530|Organism:Severe      TGTGCCTATTGGGTTCCACGTGCTAGCGCTAACATAGGTTGTAACCATACAGGTGTTGTT   1568
gb:MN908947|Organism:Severe      TGTGCCTATTGGGTTCCACGTGCTAGCGCTAACATAGGTTGTAACCATACAGGTGTTGTT   1582
gb:MT019532|Organism:Severe      TGTGCCTATTGGGTTCCACGTGCTAGCGCTAACATAGGTTGTAACCATACAGGTGTTGTT   1582
                                 ************************* **********************************

gb:MT020781|Organism:Severe      GGAGAAGGTTCCGAAGGTCTTAATGACAACCTTCTTGAAATACTCCAAAAAGAGAAAGTC   1630
gb:MT007544|Organism:Severe      GGAGAAGGTTCCGAAGGTCTTAATGACAACCTTCTTGAAATACTCCAAAAAGAGAAAGTC   1642
gb:MN994467|Organism:Severe      GGAGAAGGTTCCGAAGGTCTTAATGACAACCTTCTTGAAATACTCCAAAAAGAGAAAGTC   1642
gb:MT044257|Organism:Severe      GGAGAAGGTTCCGAAGGTCTTAATGACAACCTTCTTGAAATACTCCAAAAAGAGAAAGTC   1642
gb:MT106054|Organism:Severe      GGAGAAGGTTCCGAAGGTCTTAATGACAACCTTCTTGAAATACTCCAAAAAGAGAAAGTC   1642
gb:MT049951|Organism:Severe      GGAGAAGGTTCCGAAGGTCTTAATGACAACCTTCTTGAAATACTCCAAAAAGAGAAAGTC   1642
gb:MN975262|Organism:Severe      GGAGAAGGTTCCGAAGGTCTTAATGACAACCTTCTTGAAATACTCCAAAAAGAGAAAGTC   1642
gb:MT106052|Organism:Severe      GGAGAAGGTTCCGAAGGTCTTAATGACAACCTTCTTGAAATACTCCAAAAAGAGAAAGTC   1642
gb:LC522975|Organism:Severe      GGAGAAGGTTCCGAAGGTCTTAATGACAACCTTCTTGAAATACTCCAAAAAGAGAAAGTC   1639
gb:LC522973|Organism:Severe      GGAGAAGGTTCCGAAGGTCTTAATGACAACCTTCTTGAAATACTCCAAAAAGAGAAAGTC   1639
gb:LC522974|Organism:Severe      GGAGAAGGTTCCGAAGGTCTTAATGACAACCTTCTTGAAATACTCCAAAAAGAGAAAGTC   1639
gb:MN985325|Organism:Severe      GGAGAAGGTTCCGAAGGTCTTAATGACAACCTTCTTGAAATACTCCAAAAAGAGAAAGTC   1642
gb:MT020881|Organism:Severe      GGAGAAGGTTCCGAAGGTCTTAATGACAACCTTCTTGAAATACTCCAAAAAGAGAAAGTC   1642
gb:MT020880|Organism:Severe      GGAGAAGGTTCCGAAGGTCTTAATGACAACCTTCTTGAAATACTCCAAAAAGAGAAAGTC   1642
gb:MT066175|Organism:Severe      GGAGAAGGTTCCGAAGGTCTTAATGACAACCTTCTTGAAATACTCCAAAAAGAGAAAGTC   1642
gb:MN997409|Organism:Severe      GGAGAAGGTTCCGAAGGTCTTAATGACAACCTTCTTGAAATACTCCAAAAAGAGAAAGTC   1642
gb:MN938384|Organism:Severe      GGAGAAGGTTCCGAAGGTCTTAATGACAACCTTCTTGAAATACTCCAAAAAGAGAAAGTC   1610
gb:MT044258|Organism:Severe      GGAGAAGGTTCCGAAGGTCTTAATGACAACCTTCTTGAAATACTCCAAAAAGAGAAAGTC   1618
gb:MT039890|Organism:Severe      GGAGAAGGTTCCGAAGGTCTTAATGACAACCTTCTTGAAATACTCCAAAAAGAGAAAGTC   1642
gb:MN988713|Organism:Severe      GGAGAAGGTTCCGAAGGTCTTAATGACAACCTTCTTGAAATACTCCAAAAAGAGAAAGTC   1642
gb:LC521925|Organism:Severe      GGAGAAGGTTCCGAAGGTCTTAATGACAACCTTCTTGAAATACTCCAAAAAGAGAAAGTC   1615
gb:MT093571|Organism:Severe      GGAGAAGGTTCCGAAGGTCTTAATGACAACCTTCTTGAAATACTCCAAAAAGAGAAAGTC   1642
gb:MT039887|Organism:Severe      GGAGAAGGTTCCGAAGGTCTTAATGACAACCTTCTTGAAATACTCCAAAAAGAGAAAGTC   1642
gb:MT019530|Organism:Severe      GGAGAAGGTTCCGAAGGTCTTAATGACAACCTTCTTGAAATACTCCAAAAAGAGAAAGTC   1642
gb:MT039888|Organism:Severe      GGAGAAGGTTCCGAAGGTCTTAATGACAACCTTCTTGAAATACTCCAAAAAGAGAAAGTC   1642
gb:LC522972|Organism:Severe      GGAGAAGGTTCCGAAGGTCTTAATGACAACCTTCTTGAAATACTCCAAAAAGAGAAAGTC   1639
gb:MT027063|Organism:Severe      GGAGAAGGTTCCGAAGGTCTTAATGACAACCTTCTTGAAATACTCCAAAAAGAGAAAGTC   1642
gb:MT027062|Organism:Severe      GGAGAAGGTTCCGAAGGTCTTAATGACAACCTTCTTGAAATACTCCAAAAAGAGAAAGTC   1642
gb:MT019529|Organism:Severe      GGAGAAGGTTCCGAAGGTCTTAATGACAACCTTCTTGAAATACTCCAAAAAGAGAAAGTC   1642
gb:MN996529|Organism:Severe      GGAGAAGGTTCCGAAGGTCTTAATGACAACCTTCTTGAAATACTCCAAAAAGAGAAAGTC   1630
gb:MN996531|Organism:Severe      GGAGAAGGTTCCGAAGGTCTTAATGACAACCTTCTTGAAATACTCCAAAAAGAGAAAGTC   1629
gb:MT066176|Organism:Severe      GGAGAAGGTTCCGAAGGTCTTAATGACAACCTTCTTGAAATACTCCAAAAAGAGAAAGTC   1642
gb:MT027064|Organism:Severe      GGAGAAGGTTCCGAAGGTCTTAATGACAACCTTCTTGAAATACTCCAAAAAGAGAAAGTC   1642
gb:MN994468|Organism:Severe      GGAGAAGGTTCCGAAGGTCTTAATGACAACCTTCTTGAAATACTCCAAAAAGAGAAAGTC   1642
gb:MT072688|Organism:Severe      GGAGAAGGTTCCGAAGGTCTTAATGACAACCTTCTTGAAATACTCCAAAAAGAGAAAGTC   1627
gb:MN996527|Organism:Severe      GGAGAAGGTTCCGAAGGTCTTAATGACAACCTTCTTGAAATACTCCAAAAAGAGAAAGTC   1609
gb:MT093631|Organism:Severe      GGAGAAGGTTCCGAAGGTCTTAATGACAACCTTCTTGAAATACTCCAAAAAGAGAAAGTC   1680
gb:MT106053|Organism:Severe      GGAGAAGGTTCCGAAGGTCTTAATGACAACCTTCTTGAAATACTCCAAAAAGAGAAAGTC   1642
gb:MT019533|Organism:Severe      GGAGAAGGTTCCGAAGGTCTTAATGACAACCTTCTTGAAATACTCCAAAAAGAGAAAGTC   1642
gb:MT019531|Organism:Severe      GGAGAAGGTTCCGAAGGTCTTAATGACAACCTTCTTGAAATACTCCAAAAAGAGAAAGTC   1642
gb:MN996528|Organism:Severe      GGAGAAGGTTCCGAAGGTCTTAATGACAACCTTCTTGAAATACTCCAAAAAGAGAAAGTC   1642
gb:MN996530|Organism:Severe      GGAGAAGGTTCCGAAGGTCTTAATGACAACCTTCTTGAAATACTCCAAAAAGAGAAAGTC   1628
gb:MN908947|Organism:Severe      GGAGAAGGTTCCGAAGGTCTTAATGACAACCTTCTTGAAATACTCCAAAAAGAGAAAGTC   1642
gb:MT019532|Organism:Severe      GGAGAAGGTTCCGAAGGTCTTAATGACAACCTTCTTGAAATACTCCAAAAAGAGAAAGTC   1642
                                 ************************************************************

gb:MT020781|Organism:Severe      AACATCAATATTGTTGGTGACTTTAAACTTAATGAAGAGATCGCCATTATTTTGGCATCT   1690
gb:MT007544|Organism:Severe      AACATCAATATTGTTGGTGACTTTAAACTTAATGAAGAGATCGCCATTATTTTGGCATCT   1702
gb:MN994467|Organism:Severe      AACATCAATATTGTTGGTGACTTTAAACTTAATGAAGAGATCGCCATTATTTTGGCATCT   1702
gb:MT044257|Organism:Severe      AACATCAATATTGTTGGTGACTTTAAACTTAATGAAGAGATCGCCATTATTTTGGCATCT   1702
gb:MT106054|Organism:Severe      AACATCAATATTGTTGGTGACTTTAAACTTAATGAAGAGATCGCCATTATTTTGGCATCT   1702
gb:MT049951|Organism:Severe      AACATCAATATTGTTGGTGACTTTAAACTTAATGAAGAGATCGCCATTATTTTGGCATCT   1702
gb:MN975262|Organism:Severe      AACATCAATATTGTTGGTGACTTTAAACTTAATGAAGAGATCGCCATTATTTTGGCATCT   1702
gb:MT106052|Organism:Severe      AACATCAATATTGTTGGTGACTTTAAACTTAATGAAGAGATCGCCATTATTTTGGCATCT   1702
```

```
gb:LC522975|Organism:Severe      AACATCAATATTGTTGGTGACTTTAAACTTAATGAAGAGATCGCCATTATTTTGGCATCT    1699
gb:LC522973|Organism:Severe      AACATCAATATTGTTGGTGACTTTAAACTTAATGAAGAGATCGCCATTATTTTGGCATCT    1699
gb:LC522974|Organism:Severe      AACATCAATATTGTTGGTGACTTTAAACTTAATGAAGAGATCGCCATTATTTTGGCATCT    1699
gb:MN985325|Organism:Severe      AACATCAATATTGTTGGTGACTTTAAACTTAATGAAGAGATCGCCATTATTTTGGCATCT    1702
gb:MT020881|Organism:Severe      AACATCAATATTGTTGGTGACTTTAAACTTAATGAAGAGATCGCCATTATTTTGGCATCT    1702
gb:MT020880|Organism:Severe      AACATCAATATTGTTGGTGACTTTAAACTTAATGAAGAGATCGCCATTATTTTGGCATCT    1702
gb:MT066175|Organism:Severe      AACATCAATATTGTTGGTGACTTTAAACTTAATGAAGAGATCGCCATTATTTTGGCATCT    1702
gb:MN997409|Organism:Severe      AACATCAATATTGTTGGTGACTTTAAACTTAATGAAGAGATCGCCATTATTTTGGCATCT    1702
gb:MN938384|Organism:Severe      AACATCAATATTGTTGGTGACTTTAAACTTAATGAAGAGATCGCCATTATTTTGGCATCT    1670
gb:MT044258|Organism:Severe      AACATCAATATTGTTGGTGACTTTAAACTTAATGAAGAGATCGCCATTATTTTGGCATCT    1678
gb:MT039890|Organism:Severe      AACATCAATATTGTTGGTGACTTTAAACTTAATGAAGAGATCGCCATTATTTTGGCATCT    1702
gb:MN988713|Organism:Severe      AACATCAATATTGTTGGTGACTTTAAACTTAATGAAGAGATCGCCATTATTTTGGCATCT    1702
gb:LC521925|Organism:Severe      AACATCAATATTGTTGGTGACTTTAAACTTAATGAAGAGATCGCCATTATTTTGGCATCT    1675
gb:MT093571|Organism:Severe      AACATCAATATTGTTGGTGACTTTAAACTTAATGAAGAGATCGCCATTATTTTGGCATCT    1702
gb:MT039887|Organism:Severe      AACATCAATATTGTTGGTGACTTTAAACTTAATGAAGAGATCGCCATTATTTTGGCATCT    1702
gb:MT019530|Organism:Severe      AACATCAATATTGTTGGTGACTTTAAACTTAATGAAGAGATCGCCATTATTTTGGCATCT    1702
gb:MT039888|Organism:Severe      AACATCAATATTGTTGGTGACTTTAAACTTAATGAAGAGATCGCCATTATTTTGGCATCT    1702
gb:LC522972|Organism:Severe      AACATCAATATTGTTGGTGACTTTAAACTTAATGAAGAGATCGCCATTATTTTGGCATCT    1699
gb:MT027063|Organism:Severe      AACATCAATATTGTTGGTGACTTTAAACTTAATGAAGAGATCGCCATTATTTTGGCATCT    1702
gb:MT027062|Organism:Severe      AACATCAATATTGTTGGTGACTTTAAACTTAATGAAGAGATCGCCATTATTTTGGCATCT    1702
gb:MT019529|Organism:Severe      AACATCAATATTGTTGGTGACTTTAAACTTAATGAAGAGATCGCCATTATTTTGGCATCT    1702
gb:MN996529|Organism:Severe      AACATCAATATTGTTGGTGACTTTAAACTTAATGAAGAGATCGCCATTATTTTGGCATCT    1690
gb:MN996531|Organism:Severe      AACATCAATATTGTTGGTGACTTTAAACTTAATGAAGAGATCGCCATTATTTTGGCATCT    1689
gb:MT066176|Organism:Severe      AACATCAATATTGTTGGTGACTTTAAACTTAATGAAGAGATCGCCATTATTTTGGCATCT    1702
gb:MT027064|Organism:Severe      AACATCAATATTGTTGGTGACTTTAAACTTAATGAAGAGATCGCCATTATTTTGGCATCT    1702
gb:MN994468|Organism:Severe      AACATCAATATTGTTGGTGACTTTAAACTTAATGAAGAGATCGCCATTATTTTGGCATCT    1702
gb:MT072688|Organism:Severe      AACATCAATATTGTTGGTGACTTTAAACTTAATGAAGAGATCGCCATTATTTTGGCATCT    1687
gb:MN996527|Organism:Severe      AACATCAATATTGTTGGTGACTTTAAACTTAATGAAGAGATCGCCATTATTTTGGCATCT    1669
gb:MT093631|Organism:Severe      AACATCAATATTGTTGGTGACTTTAAACTTAATGAAGAGATCGCCATTATTTTGGCATCT    1740
gb:MT106053|Organism:Severe      AACATCAATATTGTTGGTGACTTTAAACTTAATGAAGAGATCGCCATTATTTTGGCATCT    1702
gb:MT019533|Organism:Severe      AACATCAATATTGTTGGTGACTTTAAACTTAATGAAGAGATCGCCATTATTTTGGCATCT    1702
gb:MT019531|Organism:Severe      AACATCAATATTGTTGGTGACTTTAAACTTAATGAAGAGATCGCCATTATTTTGGCATCT    1702
gb:MN996528|Organism:Severe      AACATCAATATTGTTGGTGACTTTAAACTTAATGAAGAGATCGCCATTATTTTGGCATCT    1702
gb:MN996530|Organism:Severe      AACATCAATATTGTTGGTGACTTTAAACTTAATGAAGAGATCGCCATTATTTTGGCATCT    1688
gb:MN908947|Organism:Severe      AACATCAATATTGTTGGTGACTTTAAACTTAATGAAGAGATCGCCATTATTTTGGCATCT    1702
gb:MT019532|Organism:Severe      AACATCAATATTGTTGGTGACTTTAAACTTAATGAAGAGATCGCCATTATTTTGGCATCT    1702
                                 ************************************************************

gb:MT020781|Organism:Severe      TTTTCTGCTTCCACAAGTGCTTTTGTGGAAACTGTGAAAGGTTTGGATTATAAAGCATTC    1750
gb:MT007544|Organism:Severe      TTTTCTGCTTCCACAAGTGCTTTTGTGGAAACTGTGAAAGGTTTGGATTATAAAGCATTC    1762
gb:MN994467|Organism:Severe      TTTTCTGCTTCCACAAGTGCTTTTGTGGAAACTGTGAAAGGTTTGGATTATAAAGCATTC    1762
gb:MT044257|Organism:Severe      TTTTCTGCTTCCACAAGTGCTTTTGTGGAAACTGTGAAAGGTTTGGATTATAAAGCATTC    1762
gb:MT106054|Organism:Severe      TTTTCTGCTTCCACAAGTGCTTTTGTGGAAACTGTGAAAGGTTTGGATTATAAAGCATTC    1762
gb:MT049951|Organism:Severe      TTTTCTGCTTCCACAAGTGCTTTTGTGGAAACTGTGAAAGGTTTGGATTATAAAGCATTC    1762
gb:MN975262|Organism:Severe      TTTTCTGCTTCCACAAGTGCTTTTGTGGAAACTGTGAAAGGTTTGGATTATAAAGCATTC    1762
gb:MT106052|Organism:Severe      TTTTCTGCTTCCACAAGTGCTTTTGTGGAAACTGTGAAAGGTTTGGATTATAAAGCATTC    1762
gb:LC522975|Organism:Severe      TTTTCTGCTTCCACAAGTGCTTTTGTGGAAACTGTGAAAGGTTTGGATTATAAAGCATTC    1759
gb:LC522973|Organism:Severe      TTTTCTGCTTCCACAAGTGCTTTTGTGGAAACTGTGAAAGGTTTGGATTATAAAGCATTC    1759
gb:LC522974|Organism:Severe      TTTTCTGCTTCCACAAGTGCTTTTGTGGAAACTGTGAAAGGTTTGGATTATAAAGCATTC    1759
gb:MN985325|Organism:Severe      TTTTCTGCTTCCACAAGTGCTTTTGTGGAAACTGTGAAAGGTTTGGATTATAAAGCATTC    1762
gb:MT020881|Organism:Severe      TTTTCTGCTTCCACAAGTGCTTTTGTGGAAACTGTGAAAGGTTTGGATTATAAAGCATTC    1762
gb:MT020880|Organism:Severe      TTTTCTGCTTCCACAAGTGCTTTTGTGGAAACTGTGAAAGGTTTGGATTATAAAGCATTC    1762
gb:MT066175|Organism:Severe      TTTTCTGCTTCCACAAGTGCTTTTGTGGAAACTGTGAAAGGTTTGGATTATAAAGCATTC    1762
gb:MN997409|Organism:Severe      TTTTCTGCTTCCACAAGTGCTTTTGTGGAAACTGTGAAAGGTTTGGATTATAAAGCATTC    1762
gb:MN938384|Organism:Severe      TTTTCTGCTTCCACAAGTGCTTTTGTGGAAACTGTGAAAGGTTTGGATTATAAAGCATTC    1730
gb:MT044258|Organism:Severe      TTTTCTGCTTCCACAAGTGCTTTTGTGGAAACTGTGAAAGGTTTGGATTATAAAGCATTC    1738
gb:MT039890|Organism:Severe      TTTTCTGCTTCCACAAGTGCTTTTGTGGAAACTGTGAAAGGTTTGGATTATAAAGCATTC    1762
gb:MN988713|Organism:Severe      TTTTCTGCTTCCACAAGTGCTTTTGTGGAAACTGTGAAAGGTTTGGATTATAAAGCATTC    1762
gb:LC521925|Organism:Severe      TTTTCTGCTTCCACAAGTGCTTTTGTGGAAACTGTGAAAGGTTTGGATTATAAAGCATTC    1735
gb:MT093571|Organism:Severe      TTTTCTGCTTCCACAAGTGCTTTTGTGGAAACTGTGAAAGGTTTGGATTATAAAGCATTC    1762
gb:MT039887|Organism:Severe      TTTTCTGCTTCCACAAGTGCTTTTGTGGAAACTGTGAAAGGTTTGGATTATAAAGCATTC    1762
gb:MT019530|Organism:Severe      TTTTCTGCTTCCACAAGTGCTTTTGTGGAAACTGTGAAAGGTTTGGATTATAAAGCATTC    1762
gb:MT039888|Organism:Severe      TTTTCTGCTTCCACAAGTGCTTTTGTGGAAACTGTGAAAGGTTTGGATTATAAAGCATTC    1762
gb:LC522972|Organism:Severe      TTTTCTGCTTCCACAAGTGCTTTTGTGGAAACTGTGAAAGGTTTGGATTATAAAGCATTC    1759
gb:MT027063|Organism:Severe      TTTTCTGCTTCCACAAGTGCTTTTGTGGAAACTGTGAAAGGTTTGGATTATAAAGCATTC    1762
```

```
gb:MT027062|Organism:Severe      TTTTCTGCTTCCACAAGTGCTTTTGTGGAAACTGTGAAAGGTTTGGATTATAAAGCATTC    1762
gb:MT019529|Organism:Severe      TTTTCTGCTTCCACAAGTGCTTTTGTGGAAACTGTGAAAGGTTTGGATTATAAAGCATTC    1762
gb:MN996529|Organism:Severe      TTTTCTGCTTCCACAAGTGCTTTTGTGGAAACTGTGAAAGGTTTGGATTATAAAGCATTC    1750
gb:MN996531|Organism:Severe      TTTTCTGCTTCCACAAGTGCTTTTGTGGAAACTGTGAAAGGTTTGGATTATAAAGCATTC    1749
gb:MT066176|Organism:Severe      TTTTCTGCTTCCACAAGTGCTTTTGTGGAAACTGTGAAAGGTTTGGATTATAAAGCATTC    1762
gb:MT027064|Organism:Severe      TTTTCTGCTTCCACAAGTGCTTTTGTGGAAACTGTGAAAGGTTTGGATTATAAAGCATTC    1762
gb:MN994468|Organism:Severe      TTTTCTGCTTCCACAAGTGCTTTTGTGGAAACTGTGAAAGGTTTGGATTATAAAGCATTC    1762
gb:MT072688|Organism:Severe      TTTTCTGCTTCCACAAGTGCTTTTGTGGAAACTGTGAAAGGTTTGGATTATAAAGCATTC    1747
gb:MN996527|Organism:Severe      TTTTCTGCTTCCACAAGTGCTTTTGTGGAAACTGTGAAAGGTTTGGATTATAAAGCATTC    1729
gb:MT093631|Organism:Severe      TTTTCTGCTTCCACAAGTGCTTTTGTGGAAACTGTGAAAGGTTTGGATTATAAAGCATTC    1800
gb:MT106053|Organism:Severe      TTTTCTGCTTCCACAAGTGCTTTTGTGGAAACTGTGAAAGGTTTGGATTATAAAGCATTC    1762
gb:MT019533|Organism:Severe      TTTTCTGCTTCCACAAGTGCTTTTGTGGAAACTGTGAAAGGTTTGGATTATAAAGCATTC    1762
gb:MT019531|Organism:Severe      TTTTCTGCTTCCACAAGTGCTTTTGTGGAAACTGTGAAAGGTTTGGATTATAAAGCATTC    1762
gb:MN996528|Organism:Severe      TTTTCTGCTTCCACAAGTGCTTTTGTGGAAACTGTGAAAGGTTTGGATTATAAAGCATTC    1762
gb:MN996530|Organism:Severe      TTTTCTGCTTCCACAAGTGCTTTTGTGGAAACTGTGAAAGGTTTGGATTATAAAGCATTC    1748
gb:MN908947|Organism:Severe      TTTTCTGCTTCCACAAGTGCTTTTGTGGAAACTGTGAAAGGTTTGGATTATAAAGCATTC    1762
gb:MT019532|Organism:Severe      TTTTCTGCTTCCACAAGTGCTTTTGTGGAAACTGTGAAAGGTTTGGATTATAAAGCATTC    1762
                                 ************************************************************

gb:MT020781|Organism:Severe      AAACAAATTGTTGAATCCTGTGGTAATTTTAAAGTTACAAAAGGAAAAGCTAAAAAAGGT    1810
gb:MT007544|Organism:Severe      AAACAAATTGTTGAATCCTGTGGTAATTTTAAAGTTACAAAAGGAAAAGCTAAAAAAGGT    1822
gb:MN994467|Organism:Severe      AAACAAATTGTTGAATCCTGTGGTAATTTTAAAGTTACAAAAGGAAAAGCTAAAAAAGGT    1822
gb:MT044257|Organism:Severe      AAACAAATTGTTGAATCCTGTGGTAATTTTAAAGTTACAAAAGGAAAAGCTAAAAAAGGT    1822
gb:MT106054|Organism:Severe      AAACAAATTGTTGAATCCTGTGGTAATTTTAAAGTTACAAAAGGAAAAGCTAAAAAAGGT    1822
gb:MT049951|Organism:Severe      AAACAAATTGTTGAATCCTGTGGTAATTTTAAAGTTACAAAAGGAAAAGCTAAAAAAGGT    1822
gb:MN975262|Organism:Severe      AAACAAATTGTTGAATCCTGTGGTAATTTTAAAGTTACAAAAGGAAAAGCTAAAAAAGGT    1822
gb:MT106052|Organism:Severe      AAACAAATTGTTGAATCCTGTGGTAATTTTAAAGTTACAAAAGGAAAAGCTAAAAAAGGT    1822
gb:LC522975|Organism:Severe      AAACAAATTGTTGAATCCTGTGGTAATTTTAAAGTTACAAAAGGAAAAGCTAAAAAAGGT    1819
gb:LC522973|Organism:Severe      AAACAAATTGTTGAATCCTGTGGTAATTTTAAAGTTACAAAAGGAAAAGCTAAAAAAGGT    1819
gb:LC522974|Organism:Severe      AAACAAATTGTTGAATCCTGTGGTAATTTTAAAGTTACAAAAGGAAAAGCTAAAAAAGGT    1819
gb:MN985325|Organism:Severe      AAACAAATTGTTGAATCCTGTGGTAATTTTAAAGTTACAAAAGGAAAAGCTAAAAAAGGT    1822
gb:MT020881|Organism:Severe      AAACAAATTGTTGAATCCTGTGGTAATTTTAAAGTTACAAAAGGAAAAGCTAAAAAAGGT    1822
gb:MT020880|Organism:Severe      AAACAAATTGTTGAATCCTGTGGTAATTTTAAAGTTACAAAAGGAAAAGCTAAAAAAGGT    1822
gb:MT066175|Organism:Severe      AAACAAATTGTTGAATCCTGTGGTAATTTTAAAGTTACAAAAGGAAAAGCTAAAAAAGGT    1822
gb:MN997409|Organism:Severe      AAACAAATTGTTGAATCCTGTGGTAATTTTAAAGTTACAAAAGGAAAAGCTAAAAAAGGT    1822
gb:MN938384|Organism:Severe      AAACAAATTGTTGAATCCTGTGGTAATTTTAAAGTTACAAAAGGAAAAGCTAAAAAAGGT    1790
gb:MT044258|Organism:Severe      AAACAAATTGTTGAATCCTGTGGTAATTTTAAAGTTACAAAAGGAAAAGCTAAAAAAGGT    1798
gb:MT039890|Organism:Severe      AAACAAATTGTTGAATCCTGTGGTAATTTTAAAGTTACAAAAGGAAAAGCTAAAAAAGGT    1822
gb:MN988713|Organism:Severe      AAACAAATTGTTGAATCCTGTGGTAATTTTAAAGTTACAAAAGGAAAAGCTAAAAAAGGT    1822
gb:LC521925|Organism:Severe      AAACAAATTGTTGAATCCTGTGGTAATTTTAAAGTTACAAAAGGAAAAGCTAAAAAAGGT    1795
gb:MT093571|Organism:Severe      AAACAAATTGTTGAATCCTGTGGTAATTTTAAAGTTACAAAAGGAAAAGCTAAAAAAGGT    1822
gb:MT039887|Organism:Severe      AAACAAATTGTTGAATCCTGTGGTAATTTTAAAGTTACAAAAGGAAAAGCTAAAAAAGGT    1822
gb:MT019530|Organism:Severe      AAACAAATTGTTGAATCCTGTGGTAATTTTAAAGTTACAAAAGGAAAAGCTAAAAAAGGT    1822
gb:MT039888|Organism:Severe      AAACAAATTGTTGAATCCTGTGGTAATTTTAAAGTTACAAAAGGAAAAGCTAAAAAAGGT    1822
gb:LC522972|Organism:Severe      AAACAAATTGTTGAATCCTGTGGTAATTTTAAAGTTACAAAAGGAAAAGCTAAAAAAGGT    1819
gb:MT027063|Organism:Severe      AAACAAATTGTTGAATCCTGTGGTAATTTTAAAGTTACAAAAGGAAAAGCTAAAAAAGGT    1822
gb:MT027062|Organism:Severe      AAACAAATTGTTGAATCCTGTGGTAATTTTAAAGTTACAAAAGGAAAAGCTAAAAAAGGT    1822
gb:MT019529|Organism:Severe      AAACAAATTGTTGAATCCTGTGGTAATTTTAAAGTTACAAAAGGAAAAGCTAAAAAAGGT    1822
gb:MN996529|Organism:Severe      AAACAAATTGTTGAATCCTGTGGTAATTTTAAAGTTACAAAAGGAAAAGCTAAAAAAGGT    1810
gb:MN996531|Organism:Severe      AAACAAATTGTTGAATCCTGTGGTAATTTTAAAGTTACAAAAGGAAAAGCTAAAAAAGGT    1809
gb:MT066176|Organism:Severe      AAACAAATTGTTGAATCCTGTGGTAATTTTAAAGTTACAAAAGGAAAAGCTAAAAAAGGT    1822
gb:MT027064|Organism:Severe      AAACAAATTGTTGAATCCTGTGGTAATTTTAAAGTTACAAAAGGAAAAGCTAAAAAAGGT    1822
gb:MN994468|Organism:Severe      AAACAAATTGTTGAATCCTGTGGTAATTTTAAAGTTACAAAAGGAAAAGCTAAAAAAGGT    1822
gb:MT072688|Organism:Severe      AAACAAATTGTTGAATCCTGTGGTAATTTTAAAGTTACAAAAGGAAAAGCTAAAAAAGGT    1807
gb:MN996527|Organism:Severe      AAACAAATTGTTGAATCCTGTGGTAATTTTAAAGTTACAAAAGGAAAAGCTAAAAAAGGT    1789
gb:MT093631|Organism:Severe      AAACAAATTGTTGAATCCTGTGGTAATTTTAAAGTTACAAAAGGAAAAGCTAAAAAAGGT    1860
gb:MT106053|Organism:Severe      AAACAAATTGTTGAATCCTGTGGTAATTTTAAAGTTACAAAAGGAAAAGCTAAAAAAGGT    1822
gb:MT019533|Organism:Severe      AAACAAATTGTTGAATCCTGTGGTAATTTTAAAGTTACAAAAGGAAAAGCTAAAAAAGGT    1822
gb:MT019531|Organism:Severe      AAACAAATTGTTGAATCCTGTGGTAATTTTAAAGTTACAAAAGGAAAAGCTAAAAAAGGT    1822
gb:MN996528|Organism:Severe      AAACAAATTGTTGAATCCTGTGGTAATTTTAAAGTTACAAAAGGAAAAGCTAAAAAAGGT    1822
gb:MN996530|Organism:Severe      AAACAAATTGTTGAATCCTGTGGTAATTTTAAAGTTACAAAAGGAAAAGCTAAAAAAGGT    1808
gb:MN908947|Organism:Severe      AAACAAATTGTTGAATCCTGTGGTAATTTTAAAGTTACAAAAGGAAAAGCTAAAAAAGGT    1822
gb:MT019532|Organism:Severe      AAACAAATTGTTGAATCCTGTGGTAATTTTAAAGTTACAAAAGGAAAAGCTAAAAAAGGT    1822
                                 ************************************************************
```

```
gb:MT020781|Organism:Severe      GCCTGGAATATTGGTGAACAGAAATCAATACTGAGTCCTCTTTATGCATTTGCATCAGAG   1870
gb:MT007544|Organism:Severe      GCCTGGAATATTGGTGAACAGAAATCAATACTGAGTCCTCTTTATGCATTTGCATCAGAG   1882
gb:MN994467|Organism:Severe      GCCTGGAATATTGGTGAACAGAAATCAATACTGAGTCCTCTTTATGCATTTGCATCAGAG   1882
gb:MT044257|Organism:Severe      GCCTGGAATATTGGTGAACAGAAATCAATACTGAGTCCTCTTTATGCATTTGCATCAGAG   1882
gb:MT106054|Organism:Severe      GCCTGGAATATTGGTGAACAGAAATCAATACTGAGTCCTCTTTATGCATTTGCATCAGAG   1882
gb:MT049951|Organism:Severe      GCCTGGAATATTGGTGAACAGAAATCAATACTGAGTCCTCTTTATGCATTTGCATCAGAG   1882
gb:MN975262|Organism:Severe      GCCTGGAATATTGGTGAACAGAAATCAATACTGAGTCCTCTTTATGCATTTGCATCAGAG   1882
gb:MT106052|Organism:Severe      GCCTGGAATATTGGTGAACAGAAATCAATACTGAGTCCTCTTTATGCATTTGCATCAGAG   1882
gb:LC522975|Organism:Severe      GCCTGGAATATTGGTGAACAGAAATCAATACTGAGTCCTCTTTATGCATTTGCATCAGAG   1879
gb:LC522973|Organism:Severe      GCCTGGAATATTGGTGAACAGAAATCAATACTGAGTCCTCTTTATGCATTTGCATCAGAG   1879
gb:LC522974|Organism:Severe      GCCTGGAATATTGGTGAACAGAAATCAATACTGAGTCCTCTTTATGCATTTGCATCAGAG   1879
gb:MN985325|Organism:Severe      GCCTGGAATATTGGTGAACAGAAATCAATACTGAGTCCTCTTTATGCATTTGCATCAGAG   1882
gb:MT020881|Organism:Severe      GCCTGGAATATTGGTGAACAGAAATCAATACTGAGTCCTCTTTATGCATTTGCATCAGAG   1882
gb:MT020880|Organism:Severe      GCCTGGAATATTGGTGAACAGAAATCAATACTGAGTCCTCTTTATGCATTTGCATCAGAG   1882
gb:MT066175|Organism:Severe      GCCTGGAATATTGGTGAACAGAAATCAATACTGAGTCCTCTTTATGCATTTGCATCAGAG   1882
gb:MN997409|Organism:Severe      GCCTGGAATATTGGTGAACAGAAATCAATACTGAGTCCTCTTTATGCATTTGCATCAGAG   1882
gb:MN938384|Organism:Severe      GCCTGGAATATTGGTGAACAGAAATCAATACTGAGTCCTCTTTATGCATTTGCATCAGAG   1850
gb:MT044258|Organism:Severe      GCCTGGAATATTGGTGAACAGAAATCAATACTGAGTCCTCTTTATGCATTTGCATCAGAG   1858
gb:MT039890|Organism:Severe      GCCTGGAATATTGGTGAACAGAAATCAATACTGAGTCCTCTTTATGCATTTGCATCAGAG   1882
gb:MN988713|Organism:Severe      GCCTGGAATATTGGTGAACAGAAATCAATACTGAGTCCTCTTTATGCATTTGCATCAGAG   1882
gb:LC521925|Organism:Severe      GCCTGGAATATTGGTGAACAGAAATCAATACTGAGTCCTCTTTATGCATTTGCATCAGAG   1855
gb:MT093571|Organism:Severe      GCCTGGAATATTGGTGAACAGAAATCAATACTGAGTCCTCTTTATGCATTTGCATCAGAG   1882
gb:MT039887|Organism:Severe      GCCTGGAATATTGGTGAACAGAAATCAATACTGAGTCCTCTTTATGCATTTGCATCAGAG   1882
gb:MT019530|Organism:Severe      GCCTGGAATATTGGTGAACAGAAATCAATACTGAGTCCTCTTTATGCATTTGCATCAGAG   1882
gb:MT039888|Organism:Severe      GCCTGGAATATTGGTGAACAGAAATCAATACTGAGTCCTCTTTATGCATTTGCATCAGAG   1882
gb:LC522972|Organism:Severe      GCCTGGAATATTGGTGAACAGAAATCAATACTGAGTCCTCTTTATGCATTTGCATCAGAG   1879
gb:MT027063|Organism:Severe      GCCTGGAATATTGGTGAACAGAAATCAATACTGAGTCCTCTTTATGCATTTGCATCAGAG   1882
gb:MT027062|Organism:Severe      GCCTGGAATATTGGTGAACAGAAATCAATACTGAGTCCTCTTTATGCATTTGCATCAGAG   1882
gb:MT019529|Organism:Severe      GCCTGGAATATTGGTGAACAGAAATCAATACTGAGTCCTCTTTATGCATTTGCATCAGAG   1882
gb:MN996529|Organism:Severe      GCCTGGAATATTGGTGAACAGAAATCAATACTGAGTCCTCTTTATGCATTTGCATCAGAG   1870
gb:MN996531|Organism:Severe      GCCTGGAATATTGGTGAACAGAAATCAATACTGAGTCCTCTTTATGCATTTGCATCAGAG   1869
gb:MT066176|Organism:Severe      GCCTGGAATATTGGTGAACAGAAATCAATACTGAGTCCTCTTTATGCATTTGCATCAGAG   1882
gb:MT027064|Organism:Severe      GCCTGGAATATTGGTGAACAGAAATCAATACTGAGTCCTCTTTATGCATTTGCATCAGAG   1882
gb:MN994468|Organism:Severe      GCCTGGAATATTGGTGAACAGAAATCAATACTGAGTCCTCTTTATGCATTTGCATCAGAG   1882
gb:MT072688|Organism:Severe      GCCTGGAATATTGGTGAACAGAAATCAATACTGAGTCCTCTTTATGCATTTGCATCAGAG   1867
gb:MN996527|Organism:Severe      GCCTGGAATATTGGTGAACAGAAATCAATACTGAGTCCTCTTTATGCATTTGCATCAGAG   1849
gb:MT093631|Organism:Severe      GCCTGGAATATTGGTGAACAGAAATCAATACTGAGTCCTCTTTATGCATTTGCATCAGAG   1920
gb:MT106053|Organism:Severe      GCCTGGAATATTGGTGAACAGAAATCAATACTGAGTCCTCTTTATGCATTTGCATCAGAG   1882
gb:MT019533|Organism:Severe      GCCTGGAATATTGGTGAACAGAAATCAATACTGAGTCCTCTTTATGCATTTGCATCAGAG   1882
gb:MT019531|Organism:Severe      GCCTGGAATATTGGTGAACAGAAATCAATACTGAGTCCTCTTTATGCATTTGCATCAGAG   1882
gb:MN996528|Organism:Severe      GCCTGGAATATTGGTGAACAGAAATCAATACTGAGTCCTCTTTATGCATTTGCATCAGAG   1882
gb:MN996530|Organism:Severe      GCCTGGAATATTGGTGAACAGAAATCAATACTGAGTCCTCTTTATGCATTTGCATCAGAG   1868
gb:MN908947|Organism:Severe      GCCTGGAATATTGGTGAACAGAAATCAATACTGAGTCCTCTTTATGCATTTGCATCAGAG   1882
gb:MT019532|Organism:Severe      GCCTGGAATATTGGTGAACAGAAATCAATACTGAGTCCTCTTTATGCATTTGCATCAGAG   1882
                                 ************************************************************

gb:MT020781|Organism:Severe      GCTGCTCGTGTTGTACGATCAATTTTCTCCCGCACTCTTGAAACTGCTCAAAATTCTGTG   1930
gb:MT007544|Organism:Severe      GCTGCTCGTGTTGTACGATCAATTTTCTCCCGCACTCTTGAAACTGCTCAAAATTCTGTG   1942
gb:MN994467|Organism:Severe      GCTGCTCGTGTTGTACGATCAATTTTCTCCCGCACTCTTGAAACTGCTCAAAATTCTGTG   1942
gb:MT044257|Organism:Severe      GCTGCTCGTGTTGTACGATCAATTTTCTCCCGCACTCTTGAAACTGCTCAAAATTCTGTG   1942
gb:MT106054|Organism:Severe      GCTGCTCGTGTTGTACGATCAATTTTCTCCCGCACTCTTGAAACTGCTCAAAATTCTGTG   1942
gb:MT049951|Organism:Severe      GCTGCTCGTGTTGTACGATCAATTTTCTCCCGCACTCTTGAAACTGCTCAAAATTCTGTG   1942
gb:MN975262|Organism:Severe      GCTGCTCGTGTTGTACGATCAATTTTCTCCCGCACTCTTGAAACTGCTCAAAATTCTGTG   1942
gb:MT106052|Organism:Severe      GCTGCTCGTGTTGTACGATCAATTTTCTCCCGCACTCTTGAAACTGCTCAAAATTCTGTG   1942
gb:LC522975|Organism:Severe      GCTGCTCGTGTTGTACGATCAATTTTCTCCCGCACTCTTGAAACTGCTCAAAATTCTGTG   1939
gb:LC522973|Organism:Severe      GCTGCTCGTGTTGTACGATCAATTTTCTCCCGCACTCTTGAAACTGCTCAAAATTCTGTG   1939
gb:LC522974|Organism:Severe      GCTGCTCGTGTTGTACGATCAATTTTCTCCCGCACTCTTGAAACTGCTCAAAATTCTGTG   1939
gb:MN985325|Organism:Severe      GCTGCTCGTGTTGTACGATCAATTTTCTCCCGCACTCTTGAAACTGCTCAAAATTCTGTG   1942
gb:MT020881|Organism:Severe      GCTGCTCGTGTTGTACGATCAATTTTCTCCCGCACTCTTGAAACTGCTCAAAATTCTGTG   1942
gb:MT020880|Organism:Severe      GCTGCTCGTGTTGTACGATCAATTTTCTCCCGCACTCTTGAAACTGCTCAAAATTCTGTG   1942
gb:MT066175|Organism:Severe      GCTGCTCGTGTTGTACGATCAATTTTCTCCCGCACTCTTGAAACTGCTCAAAATTCTGTG   1942
gb:MN997409|Organism:Severe      GCTGCTCGTGTTGTACGATCAATTTTCTCCCGCACTCTTGAAACTGCTCAAAATTCTGTG   1942
gb:MN938384|Organism:Severe      GCTGCTCGTGTTGTACGATCAATTTTCTCCCGCACTCTTGAAACTGCTCAAAATTCTGTG   1910
gb:MT044258|Organism:Severe      GCTGCTCGTGTTGTACGATCAATTTTCTCCCGCACTCTTGAAACTGCTCAAAATTCTGTG   1918
gb:MT039890|Organism:Severe      GCTGCTCGTGTTGTACGATCAATTTTCTCCCGCACTCTTGAAACTGCTCAAAATTCTGTG   1942
```

```
gb:MN988713|Organism:Severe      GCTGCTCGTGTTGTACGATCAATTTTCTCCCGCACTCTTGAAACTGCTCAAAATTCTGTG   1942
gb:LC521925|Organism:Severe      GCTGCTCGTGTTGTACGATCAATTTTCTCTCGCACTCTTGAAACTGCTCAAAATTCTGTG   1915
gb:MT093571|Organism:Severe      GCTGCTCGTGTTGTACGATCAATTTTCTCCCGCACTCTTGAAACTGCTCAAAATTCTGTG   1942
gb:MT039887|Organism:Severe      GCTGCTCGTGTTGTACGATCAATTTTCTCCCGCACTCTTGAAACTGCTCAAAATTCTGTG   1942
gb:MT019530|Organism:Severe      GCTGCTCGTGTTGTACGATCAATTTTCTCCCGCACTCTTGAAACTGCTCAAAATTCTGTG   1942
gb:MT039888|Organism:Severe      GCTGCTCGTGTTGTACGATCAATTTTCTCCCGCACTCTTGAAACTGCTCAAAATTCTGTG   1942
gb:LC522972|Organism:Severe      GCTGCTCGTGTTGTACGATCAATTTTCTCCCGCACTCTTGAAACTGCTCAAAATTCTGTG   1939
gb:MT027063|Organism:Severe      GCTGCTCGTGTTGTACGATCAATTTTCTCCCGCACTCTTGAAACTGCTCAAAATTCTGTG   1942
gb:MT027062|Organism:Severe      GCTGCTCGTGTTGTACGATCAATTTTCTCCCGCACTCTTGAAACTGCTCAAAATTCTGTG   1942
gb:MT019529|Organism:Severe      GCTGCTCGTGTTGTACGATCAATTTTCTCCCGCACTCTTGAAACTGCTCAAAATTCTGTG   1942
gb:MN996529|Organism:Severe      GCTGCTCGTGTTGTACGATCAATTTTCTCCCGCACTCTTGAAACTGCTCAAAATTCTGTG   1930
gb:MN996531|Organism:Severe      GCTGCTCGTGTTGTACGATCAATTTTCTCCCGCACTCTTGAAACTGCTCAAAATTCTGTG   1929
gb:MT066176|Organism:Severe      GCTGCTCGTGTTGTACGATCAATTTTCTCCCGCACTCTTGAAACTGCTCAAAATTCTGTG   1942
gb:MT027064|Organism:Severe      GCTGCTCGTGTTGTACGATCAATTTTCTCCCGCACTCTTGAAACTGCTCAAAATTCTGTG   1942
gb:MN994468|Organism:Severe      GCTGCTCGTGTTGTACGATCAATTTTCTCCCGCACTCTTGAAACTGCTCAAAATTCTGTG   1942
gb:MT072688|Organism:Severe      GCTGCTCGTGTTGTACGATCAATTTTCTCCCGCACTCTTGAAACTGCTCAAAATTCTGTG   1927
gb:MN996527|Organism:Severe      GCTGCTCGTGTTGTACGATCAATTTTCTCCCGCACTCTTGAAACTGCTCAAAATTCTGTG   1909
gb:MT093631|Organism:Severe      GCTGCTCGTGTTGTACGATCAATTTTCTCCCGCACTCTTGAAACTGCTCAAAATTCTGTG   1980
gb:MT106053|Organism:Severe      GCTGCTCGTGTTGTACGATCAATTTTCTCCCGCACTCTTGAAACTGCTCAAAATTCTGTG   1942
gb:MT019533|Organism:Severe      GCTGCTCGTGTTGTACGATCAATTTTCTCCCGCACTCTTGAAACTGCTCAAAATTCTGTG   1942
gb:MT019531|Organism:Severe      GCTGCTCGTGTTGTACGATCAATTTTCTCCCGCACTCTTGAAACTGCTCAAAATTCTGTG   1942
gb:MN996528|Organism:Severe      GCTGCTCGTGTTGTACGATCAATTTTCTCCCGCACTCTTGAAACTGCTCAAAATTCTGTG   1942
gb:MN996530|Organism:Severe      GCTGCTCGTGTTGTACGATCAATTTTCTCCCGCACTCTTGAAACTGCTCAAAATTCTGTG   1928
gb:MN908947|Organism:Severe      GCTGCTCGTGTTGTACGATCAATTTTCTCCCGCACTCTTGAAACTGCTCAAAATTCTGTG   1942
gb:MT019532|Organism:Severe      GCTGCTCGTGTTGTACGATCAATTTTCTCCCGCACTCTTGAAACTGCTCAAAATTCTGTG   1942
                                 ***************************** ******************************

gb:MT020781|Organism:Severe      CGTGTTTTACAGAAGGCCGCTATAACAATACTAGATGGAATTTCACAGTATTCACTGAGA   1990
gb:MT007544|Organism:Severe      CGTGTTTTACAGAAGGCCGCTATAACAATACTAGATGGAATTTCACAGTATTCACTGAGA   2002
gb:MN994467|Organism:Severe      CGTGTTTTACAGAAGGCCGCTATAACAATACTAGATGGAATTTCACAGTATTCACTGAGA   2002
gb:MT044257|Organism:Severe      CGTGTTTTACAGAAGGCCGCTATAACAATACTAGATGGAATTTCACAGTATTCACTGAGA   2002
gb:MT106054|Organism:Severe      CGTGTTTTACAGAAGGCCGCTATAACAATACTAGATGGAATTTCACAGTATTCACTGAGA   2002
gb:MT049951|Organism:Severe      CGTGTTTTACAGAAGGCCGCTATAACAATACTAGATGGAATTTCACAGTATTCACTGAGA   2002
gb:MN975262|Organism:Severe      CGTGTTTTACAGAAGGCCGCTATAACAATACTAGATGGAATTTCACAGTATTCACTGAGA   2002
gb:MT106052|Organism:Severe      CGTGTTTTACAGAAGGCCGCTATAACAATACTAGATGGAATTTCACAGTATTCACTGAGA   2002
gb:LC522975|Organism:Severe      CGTGTTTTACAGAAGGCCGCTATAACAATACTAGATGGAATTTCACAGTATTCACTGAGA   1999
gb:LC522973|Organism:Severe      CGTGTTTTACAGAAGGCCGCTATAACAATACTAGATGGAATTTCACAGTATTCACTGAGA   1999
gb:LC522974|Organism:Severe      CGTGTTTTACAGAAGGCCGCTATAACAATACTAGATGGAATTTCACAGTATTCACTGAGA   1999
gb:MN985325|Organism:Severe      CGTGTTTTACAGAAGGCCGCTATAACAATACTAGATGGAATTTCACAGTATTCACTGAGA   2002
gb:MT020881|Organism:Severe      CGTGTTTTACAGAAGGCCGCTATAACAATACTAGATGGAATTTCACAGTATTCACTGAGA   2002
gb:MT020880|Organism:Severe      CGTGTTTTACAGAAGGCCGCTATAACAATACTAGATGGAATTTCACAGTATTCACTGAGA   2002
gb:MT066175|Organism:Severe      CGTGTTTTACAGAAGGCCGCTATAACAATACTAGATGGAATTTCACAGTATTCACTGAGA   2002
gb:MN997409|Organism:Severe      CGTGTTTTACAGAAGGCCGCTATAACAATACTAGATGGAATTTCACAGTATTCACTGAGA   2002
gb:MN938384|Organism:Severe      CGTGTTTTACAGAAGGCCGCTATAACAATACTAGATGGAATTTCACAGTATTCACTGAGA   1970
gb:MT044258|Organism:Severe      CGTGTTTTACAGAAGGCCGCTATAACAATACTAGATGGAATTTCACAGTATTCACTGAGA   1978
gb:MT039890|Organism:Severe      CGTGTTTTACAGAAGGCCGCTATAACAATACTAGATGGAATTTCACAGTATTCACTGAGA   2002
gb:MN988713|Organism:Severe      CGTGTTTTACAGAAGGCCGCTATAACAATACTAGATGGAATTTCACAGTATTCACTGAGA   2002
gb:LC521925|Organism:Severe      CGTGTTTTACAGAAGGCCGCTATAACAATACTAGATGGAATTTCACAGTATTCACTGAGA   1975
gb:MT093571|Organism:Severe      CGTGTTTTACAGAAGGCCGCTATAACAATACTAGATGGAATTTCACAGTATTCACTGAGA   2002
gb:MT039887|Organism:Severe      CGTGTTTTACAGAAGGCCGCTATAACAATACTAGATGGAATTTCACAGTATTCACTGAGA   2002
gb:MT019530|Organism:Severe      CGTGTTTTACAGAAGGCCGCTATAACAATACTAGATGGAATTTCACAGTATTCACTGAGA   2002
gb:MT039888|Organism:Severe      CGTGTTTTACAGAAGGCCGCTATAACAATACTAGATGGAATTTCACAGTATTCACTGAGA   2002
gb:LC522972|Organism:Severe      CGTGTTTTACAGAAGGCCGCTATAACAATACTAGATGGAATTTCACAGTATTCACTGAGA   1999
gb:MT027063|Organism:Severe      CGTGTTTTACAGAAGGCCGCTATAACAATACTAGATGGAATTTCACAGTATTCACTGAGA   2002
gb:MT027062|Organism:Severe      CGTGTTTTACAGAAGGCCGCTATAACAATACTAGATGGAATTTCACAGTATTCACTGAGA   2002
gb:MT019529|Organism:Severe      CGTGTTTTACAGAAGGCCGCTATAACAATACTAGATGGAATTTCACAGTATTCACTGAGA   2002
gb:MN996529|Organism:Severe      CGTGTTTTACAGAAGGCCGCTATAACAATACTAGATGGAATTTCACAGTATTCACTGAGA   1990
gb:MN996531|Organism:Severe      CGTGTTTTACAGAAGGCCGCTATAACAATACTAGATGGAATTTCACAGTATTCACTGAGA   1989
gb:MT066176|Organism:Severe      CGTGTTTTACAGAAGGCCGCTATAACAATACTAGATGGAATTTCACAGTATTCACTGAGA   2002
gb:MT027064|Organism:Severe      CGTGTTTTACAGAAGGCCGCTATAACAATACTAGATGGAATTTCACAGTATTCACTGAGA   2002
gb:MN994468|Organism:Severe      CGTGTTTTACAGAAGGCCGCTATAACAATACTAGATGGAATTTCACAGTATTCACTGAGA   2002
gb:MT072688|Organism:Severe      CGTGTTTTACAGAAGGCCGCTATAACAATACTAGATGGAATTTCACAGTATTCACTGAGA   1987
gb:MN996527|Organism:Severe      CGTGTTTTACAGAAGGCCGCTATAACAATACTAGATGGAATTTCACAGTATTCACTGAGA   1969
gb:MT093631|Organism:Severe      CGTGTTTTACAGAAGGCCGCTATAACAATACTAGATGGAATTTCACAGTATTCACTGAGA   2040
gb:MT106053|Organism:Severe      CGTGTTTTACAGAAGGCCGCTATAACAATACTAGATGGAATTTCACAGTATTCACTGAGA   2002
```

```
gb:MT019533|Organism:Severe      CGTGTTTTACAGAAGGCCGCTATAACAATACTAGATGGAATTTCACAGTATTCACTGAGA   2002
gb:MT019531|Organism:Severe      CGTGTTTTACAGAAGGCCGCTATAACAATACTAGATGGAATTTCACAGTATTCACTGAGA   2002
gb:MN996528|Organism:Severe      CGTGTTTTACAGAAGGCCGCTATAACAATACTAGATGGAATTTCACAGTATTCACTGAGA   2002
gb:MN996530|Organism:Severe      CGTGTTTTACAGAAGGCCGCTATAACAATACTAGATGGAATTTCACAGTATTCACTGAGA   1988
gb:MN908947|Organism:Severe      CGTGTTTTACAGAAGGCCGCTATAACAATACTAGATGGAATTTCACAGTATTCACTGAGA   2002
gb:MT019532|Organism:Severe      CGTGTTTTACAGAAGGCCGCTATAACAATACTAGATGGAATTTCACAGTATTCACTGAGA   2002
                                 ************************************************************

gb:MT020781|Organism:Severe      CTCATTGATGCTATGATGTTCACATCTGATTTGGCTACTAACAATCTAGTTGTAATGGCC   2050
gb:MT007544|Organism:Severe      CTCATTGATGCTATGATGTTCACATCTGATTTGGCTACTAACAATCTAGTTGTAATGGCC   2062
gb:MN994467|Organism:Severe      CTCATTGATGCTATGATGTTCACATCTGATTTGGCTACTAACAATCTAGTTGTAATGGCC   2062
gb:MT044257|Organism:Severe      CTCATTGATGCTATGATGTTCACATCTGATTTGGCTACTAACAATCTAGTTGTAATGGCC   2062
gb:MT106054|Organism:Severe      CTCATTGATGCTATGATGTTCACATCTGATTTGGCTACTAACAATCTAGTTGTAATGGCC   2062
gb:MT049951|Organism:Severe      CTCATTGATGCTATGATGTTCACATCTGATTTGGCTACTAACAATCTAGTTGTAATGGCC   2062
gb:MN975262|Organism:Severe      CTCATTGATGCTATGATGTTCACATCTGATTTGGCTACTAACAATCTAGTTGTAATGGCC   2062
gb:MT106052|Organism:Severe      CTCATTGATGCTATGATGTTCACATCTGATTTGGCTACTAACAATCTAGTTGTAATGGCC   2062
gb:LC522975|Organism:Severe      CTCATTGATGCTATGATGTTCACATCTGATTTGGCTACTAACAATCTAGTTGTAATGGCC   2059
gb:LC522973|Organism:Severe      CTCATTGATGCTATGATGTTCACATCTGATTTGGCTACTAACAATCTAGTTGTAATGGCC   2059
gb:LC522974|Organism:Severe      CTCATTGATGCTATGATGTTCACATCTGATTTGGCTACTAACAATCTAGTTGTAATGGCC   2059
gb:MN985325|Organism:Severe      CTCATTGATGCTATGATGTTCACATCTGATTTGGCTACTAACAATCTAGTTGTAATGGCC   2062
gb:MT020881|Organism:Severe      CTCATTGATGCTATGATGTTCACATCTGATTTGGCTACTAACAATCTAGTTGTAATGGCC   2062
gb:MT020880|Organism:Severe      CTCATTGATGCTATGATGTTCACATCTGATTTGGCTACTAACAATCTAGTTGTAATGGCC   2062
gb:MT066175|Organism:Severe      CTCATTGATGCTATGATGTTCACATCTGATTTGGCTACTAACAATCTAGTTGTAATGGCC   2062
gb:MN997409|Organism:Severe      CTCATTGATGCTATGATGTTCACATCTGATTTGGCTACTAACAATCTAGTTGTAATGGCC   2062
gb:MN938384|Organism:Severe      CTCATTGATGCTATGATGTTCACATCTGATTTGGCTACTAACAATCTAGTTGTAATGGCC   2030
gb:MT044258|Organism:Severe      CTCATTGATGCTATGATGTTCACATCTGATTTGGCTACTAACAATCTAGTTGTAATGGCC   2038
gb:MT039890|Organism:Severe      CTCATTGATGCTATGATGTTCACATCTGATTTGGCTACTAACAATCTAGTTGTAATGGCC   2062
gb:MN988713|Organism:Severe      CTCATTGATGCTATGATGTTCACATCTGATTTGGCTACTAACAATCTAGTTGTAATGGCC   2062
gb:LC521925|Organism:Severe      CTCATTGATGCTATGATGTTCACATCTGATTTGGCTACTAACAATCTAGTTGTAATGGCC   2035
gb:MT093571|Organism:Severe      CTCATTGATGCTATGATGTTCACATCTGATTTGGCTACTAACAATCTAGTTGTAATGGCC   2062
gb:MT039887|Organism:Severe      CTCATTGATGCTATGATGTTCACATCTGATTTGGCTACTAACAATCTAGTTGTAATGGCC   2062
gb:MT019530|Organism:Severe      CTCATTGATGCTATGATGTTCACATCTGATTTGGCTACTAACAATCTAGTTGTAATGGCC   2062
gb:MT039888|Organism:Severe      CTCATTGATGCTATGATGTTCACATCTGATTTGGCTACTAACAATCTAGTTGTAATGGCC   2062
gb:LC522972|Organism:Severe      CTCATTGATGCTATGATGTTCACATCTGATTTGGCTACTAACAATCTAGTTGTAATGGCC   2059
gb:MT027063|Organism:Severe      CTCATTGATGCTATGATGTTCACATCTGATTTGGCTACTAACAATCTAGTTGTAATGGCC   2062
gb:MT027062|Organism:Severe      CTCATTGATGCTATGATGTTCACATCTGATTTGGCTACTAACAATCTAGTTGTAATGGCC   2062
gb:MT019529|Organism:Severe      CTCATTGATGCTATGATGTTCACATCTGATTTGGCTACTAACAATCTAGTTGTAATGGCC   2062
gb:MN996529|Organism:Severe      CTCATTGATGCTATGATGTTCACATCTGATTTGGCTACTAACAATCTAGTTGTAATGGCC   2050
gb:MN996531|Organism:Severe      CTCATTGATGCTATGATGTTCACATCTGATTTGGCTACTAACAATCTAGTTGTAATGGCC   2049
gb:MT066176|Organism:Severe      CTCATTGATGCTATGATGTTCACATCTGATTTGGCTACTAACAATCTAGTTGTAATGGCC   2062
gb:MT027064|Organism:Severe      CTCATTGATGCTATGATGTTCACATCTGATTTGGCTACTAACAATCTAGTTGTAATGGCC   2062
gb:MN994468|Organism:Severe      CTCATTGATGCTATGATGTTCACATCTGATTTGGCTACTAACAATCTAGTTGTAATGGCC   2062
gb:MT072688|Organism:Severe      CTCATTGATGCTATGATGTTCACATCTGATTTGGCTACTAACAATCTAGTTGTAATGGCC   2047
gb:MN996527|Organism:Severe      CTCATTGATGCTATGATGTTCACATCTGATTTGGCTACTAACAATCTAGTTGTAATGGCC   2029
gb:MT093631|Organism:Severe      CTCATTGATGCTATGATGTTCACATCTGATTTGGCTACTAACAATCTAGTTGTAATGGCC   2100
gb:MT106053|Organism:Severe      CTCATTGATGCTATGATGTTCACATCTGATTTGGCTACTAACAATCTAGTTGTAATGGCC   2062
gb:MT019533|Organism:Severe      CTCATTGATGCTATGATGTTCACATCTGATTTGGCTACTAACAATCTAGTTGTAATGGCC   2062
gb:MT019531|Organism:Severe      CTCATTGATGCTATGATGTTCACATCTGATTTGGCTACTAACAATCTAGTTGTAATGGCC   2062
gb:MN996528|Organism:Severe      CTCATTGATGCTATGATGTTCACATCTGATTTGGCTACTAACAATCTAGTTGTAATGGCC   2062
gb:MN996530|Organism:Severe      CTCATTGATGCTATGATGTTCACATCTGATTTGGCTACTAACAATCTAGTTGTAATGGCC   2048
gb:MN908947|Organism:Severe      CTCATTGATGCTATGATGTTCACATCTGATTTGGCTACTAACAATCTAGTTGTAATGGCC   2062
gb:MT019532|Organism:Severe      CTCATTGATGCTATGATGTTCACATCTGATTTGGCTACTAACAATCTAGTTGTAATGGCC   2062
                                 ************************************************************

gb:MT020781|Organism:Severe      TACATTACAGGTGGTGTTGTTCAGTTGACTTCGCAGTGGCTAACTAACATCTTTGGCACT   2110
gb:MT007544|Organism:Severe      TACATTACAGGTGGTGTTGTTCAGTTGACTTCGCAGTGGCTAACTAACATCTTTGGCACT   2122
gb:MN994467|Organism:Severe      TACATTACAGGTGGTGTTGTTCAGTTGACTTCGCAGTGGCTAACTAACATCTTTGGCACT   2122
gb:MT044257|Organism:Severe      TACATTACAGGTGGTGTTGTTCAGTTGACTTCGCAGTGGCTAACTAACATCTTTGGCACT   2122
gb:MT106054|Organism:Severe      TACATTACAGGTGGTGTTGTTCAGTTGACTTCGCAGTGGCTAACTAACATCTTTGGCACT   2122
gb:MT049951|Organism:Severe      TACATTACAGGTGGTGTTGTTCAGTTGACTTCGCAGTGGCTAACTAACATCTTTGGCACT   2122
gb:MN975262|Organism:Severe      TACATTACAGGTGGTGTTGTTCAGTTGACTTCGCAGTGGCTAACTAACATCTTTGGCACT   2122
gb:MT106052|Organism:Severe      TACATTACAGGTGGTGTTGTTCAGTTGACTTCGCAGTGGCTAACTAACATCTTTGGCACT   2122
gb:LC522975|Organism:Severe      TACATTACAGGTGGTGTTGTTCAGTTGACTTCGCAGTGGCTAACTAACATCTTTGGCACT   2119
gb:LC522973|Organism:Severe      TACATTACAGGTGGTGTTGTTCAGTTGACTTCGCAGTGGCTAACTAACATCTTTGGCACT   2119
gb:LC522974|Organism:Severe      TACATTACAGGTGGTGTTGTTCAGTTGACTTCGCAGTGGCTAACTAACATCTTTGGCACT   2119
```

```
gb:MN985325|Organism:Severe      TACATTACAGGTGGTGTTGTTCAGTTGACTTCGCAGTGGCTAACTAACATCTTTGGCACT   2122
gb:MT020881|Organism:Severe      TACATTACAGGTGGTGTTGTTCAGTTGACTTCGCAGTGGCTAACTAACATCTTTGGCACT   2122
gb:MT020880|Organism:Severe      TACATTACAGGTGGTGTTGTTCAGTTGACTTCGCAGTGGCTAACTAACATCTTTGGCACT   2122
gb:MT066175|Organism:Severe      TACATTACAGGTGGTGTTGTTCAGTTGACTTCGCAGTGGCTAACTAACATCTTTGGCACT   2122
gb:MN997409|Organism:Severe      TACATTACAGGTGGTGTTGTTCAGTTGACTTCGCAGTGGCTAACTAACATCTTTGGCACT   2122
gb:MN938384|Organism:Severe      TACATTACAGGTGGTGTTGTTCAGTTGACTTCGCAGTGGCTAACTAACATCTTTGGCACT   2090
gb:MT044258|Organism:Severe      TACATTACAGGTGGTGTTGTTCAGTTGACTTCGCAGTGGCTAACTAACATCTTTGGCACT   2098
gb:MT039890|Organism:Severe      TACATTACAGGTGGTGTTGTTCAGTTGACTTCGCAGTGGCTAACTAACATCTTTGGCACT   2122
gb:MN988713|Organism:Severe      TACATTACAGGTGGTGTTGTTCAGTTGACTTCGCAGTGGCTAACTAACATCTTTGGCACT   2122
gb:LC521925|Organism:Severe      TACATTACAGGTGGTGTTGTTCAGTTGACTTCGCAGTGGCTAACTAACATCTTTGGCACT   2095
gb:MT093571|Organism:Severe      TACATTACAGGTGGTGTTGTTCAGTTGACTTCGCAGTGGCTAACTAACATCTTTGGCACT   2122
gb:MT039887|Organism:Severe      TACATTACAGGTGGTGTTGTTCAGTTGACTTCGCAGTGGCTAACTAACATCTTTGGCACT   2122
gb:MT019530|Organism:Severe      TACATTACAGGTGGTGTTGTTCAGTTGACTTCGCAGTGGCTAACTAACATCTTTGGCACT   2122
gb:MT039888|Organism:Severe      TACATTACAGGTGGTGTTGTTCAGTTGACTTCGCAGTGGCTAACTAACATCTTTGGCACT   2122
gb:LC522972|Organism:Severe      TACATTACAGGTGGTGTTGTTCAGTTGACTTCGCAGTGGCTAACTAACATCTTTGGCACT   2119
gb:MT027063|Organism:Severe      TACATTACAGGTGGTGTTGTTCAGTTGACTTCGCAGTGGCTAACTAACATCTTTGGCACT   2122
gb:MT027062|Organism:Severe      TACATTACAGGTGGTGTTGTTCAGTTGACTTCGCAGTGGCTAACTAACATCTTTGGCACT   2122
gb:MT019529|Organism:Severe      TACATTACAGGTGGTGTTGTTCAGTTGACTTCGCAGTGGCTAACTAACATCTTTGGCACT   2122
gb:MN996529|Organism:Severe      TACATTACAGGTGGTGTTGTTCAGTTGACTTCGCAGTGGCTAACTAACATCTTTGGCACT   2110
gb:MN996531|Organism:Severe      TACATTACAGGTGGTGTTGTTCAGTTGACTTCGCAGTGGCTAACTAACATCTTTGGCACT   2109
gb:MT066176|Organism:Severe      TACATTACAGGTGGTGTTGTTCAGTTGACTTCGCAGTGGCTAACTAACATCTTTGGCACT   2122
gb:MT027064|Organism:Severe      TACATTACAGGTGGTGTTGTTCAGTTGATTTCGCAGTGGCTAACTAACATCTTTGGCACT   2122
gb:MN994468|Organism:Severe      TACATTACAGGTGGTGTTGTTCAGTTGACTTCGCAGTGGCTAACTAACATCTTTGGCACT   2122
gb:MT072688|Organism:Severe      TACATTACAGGTGGTGTTGTTCAGTTGACTTCGCAGTGGCTAACTAACATCTTTGGCACT   2107
gb:MN996527|Organism:Severe      TACATTACAGGTGGTGTTGTTCAGTTGACTTCGCAGTGGCTAACTAACATCTTTGGCACT   2089
gb:MT093631|Organism:Severe      TACATTACAGGTGGTGTTGTTCAGTTGACTTCGCAGTGGCTAACTAACATCTTTGGCACT   2160
gb:MT106053|Organism:Severe      TACATTACAGGTGGTGTTGTTCAGTTGACTTCGCAGTGGCTAACTAACATCTTTGGCACT   2122
gb:MT019533|Organism:Severe      TACATTACAGGTGGTGTTGTTCAGTTGACTTCGCAGTGGCTAACTAACATCTTTGGCACT   2122
gb:MT019531|Organism:Severe      TACATTACAGGTGGTGTTGTTCAGTTGACTTCGCAGTGGCTAACTAACATCTTTGGCACT   2122
gb:MN996528|Organism:Severe      TACATTACAGGTGGTGTTGTTCAGTTGACTTCGCAGTGGCTAACTAACATCTTTGGCACT   2122
gb:MN996530|Organism:Severe      TACATTACAGGTGGTGTTGTTCAGTTGACTTCGCAGTGGCTAACTAACATCTTTGGCACT   2108
gb:MN908947|Organism:Severe      TACATTACAGGTGGTGTTGTTCAGTTGACTTCGCAGTGGCTAACTAACATCTTTGGCACT   2122
gb:MT019532|Organism:Severe      TACATTACAGGTGGTGTTGTTCAGTTGACTTCGCAGTGGCTAACTAACATCTTTGGCACT   2122
                                 **************************** *******************************

gb:MT020781|Organism:Severe      GTTTATGAAAAACTCAAACCCGTCCTTGATTGGCTTGAAGAGAAGTTTAAGGAAGGTGTA   2170
gb:MT007544|Organism:Severe      GTTTATGAAAAACTCAAACCCGTCCTTGATTGGCTTGAAGAGAAGTTTAAGGAAGGTGTA   2182
gb:MN994467|Organism:Severe      GTTTATGAAAAACTCAAACCCGTCCTTGATTGGCTTGAAGAGAAGTTTAAGGAAGGTGTA   2182
gb:MT044257|Organism:Severe      GTTTATGAAAAACTCAAACCCGTCCTTGATTGGCTTGAAGAGAAGTTTAAGGAAGGTGTA   2182
gb:MT106054|Organism:Severe      GTTTATGAAAAACTCAAACCCGTCCTTGATTGGCTTGAAGAGAAGTTTAAGGAAGGTGTA   2182
gb:MT049951|Organism:Severe      GTTTATGAAAAACTCAAACCCGTCCTTGATTGGCTTGAAGAGAAGTTTAAGGAAGGTGTA   2182
gb:MN975262|Organism:Severe      GTTTATGAAAAACTCAAACCCGTCCTTGATTGGCTTGAAGAGAAGTTTAAGGAAGGTGTA   2182
gb:MT106052|Organism:Severe      GTTTATGAAAAACTCAAACCCGTCCTTGATTGGCTTGAAGAGAAGTTTAAGGAAGGTGTA   2182
gb:LC522975|Organism:Severe      GTTTATGAAAAACTCAAACCCGTCCTTGATTGGCTTGAAGAGAAGTTTAAGGAAGGTGTA   2179
gb:LC522973|Organism:Severe      GTTTATGAAAAACTCAAACCCGTCCTTGATTGGCTTGAAGAGAAGTTTAAGGAAGGTGTA   2179
gb:LC522974|Organism:Severe      GTTTATGAAAAACTCAAACCCGTCCTTGATTGGCTTGAAGAGAAGTTTAAGGAAGGTGTA   2179
gb:MN985325|Organism:Severe      GTTTATGAAAAACTCAAACCCGTCCTTGATTGGCTTGAAGAGAAGTTTAAGGAAGGTGTA   2182
gb:MT020881|Organism:Severe      GTTTATGAAAAACTCAAACCCGTCCTTGATTGGCTTGAAGAGAAGTTTAAGGAAGGTGTA   2182
gb:MT020880|Organism:Severe      GTTTATGAAAAACTCAAACCCGTCCTTGATTGGCTTGAAGAGAAGTTTAAGGAAGGTGTA   2182
gb:MT066175|Organism:Severe      GTTTATGAAAAACTCAAACCCGTCCTTGATTGGCTTGAAGAGAAGTTTAAGGAAGGTGTA   2182
gb:MN997409|Organism:Severe      GTTTATGAAAAACTCAAACCCGTCCTTGATTGGCTTGAAGAGAAGTTTAAGGAAGGTGTA   2182
gb:MN938384|Organism:Severe      GTTTATGAAAAACTCAAACCCGTCCTTGATTGGCTTGAAGAGAAGTTTAAGGAAGGTGTA   2150
gb:MT044258|Organism:Severe      GTTTATGAAAAACTCAAACCCGTCCTTGATTGGCTTGAAGAGAAGTTTAAGGAAGGTGTA   2158
gb:MT039890|Organism:Severe      GTTTATGAAAAACTCAAACCCGTCCTTGATTGGCTTGAAGAGAAGTTTAAGGAAGGTGTA   2182
gb:MN988713|Organism:Severe      GTTTATGAAAAACTCAAACCCGTCCTTGATTGGCTTGAAGAGAAGTTTAAGGAAGGTGTA   2182
gb:LC521925|Organism:Severe      GTTTATGAAAAACTCAAACCCGTCCTTGATTGGCTTGAAGAGAAGTTTAAGGAAGGTGTA   2155
gb:MT093571|Organism:Severe      GTTTATGAAAAACTCAAACCCGTCCTTGATTGGCTTGAAGAGAAGTTTAAGGAAGGTGTA   2182
gb:MT039887|Organism:Severe      GTTTATGAAAAACTCAAACCCGTCCTTGATTGGCTTGAAGAGAAGTTTAAGGAAGGTGTA   2182
gb:MT019530|Organism:Severe      GTTTATGAAAAACTCAAACCCGTCCTTGATTGGCTTGAAGAGAAGTTTAAGGAAGGTGTA   2182
gb:MT039888|Organism:Severe      GTTTATGAAAAACTCAAACCCGTCCTTGATTGGCTTGAAGAGAAGTTTAAGGAAGGTGTA   2182
gb:LC522972|Organism:Severe      GTTTATGAAAAACTCAAACCCGTCCTTGATTGGCTTGAAGAGAAGTTTAAGGAAGGTGTA   2179
gb:MT027063|Organism:Severe      GTTTATGAAAAACTCAAACCCGTCCTTGATTGGCTTGAAGAGAAGTTTAAGGAAGGTGTA   2182
gb:MT027062|Organism:Severe      GTTTATGAAAAACTCAAACCCGTCCTTGATTGGCTTGAAGAGAAGTTTAAGGAAGGTGTA   2182
gb:MT019529|Organism:Severe      GTTTATGAAAAACTCAAACCCGTCCTTGATTGGCTTGAAGAGAAGTTTAAGGAAGGTGTA   2182
gb:MN996529|Organism:Severe      GTTTATGAAAAACTCAAACCCGTCCTTGATTGGCTTGAAGAGAAGTTTAAGGAAGGTGTA   2170
```

```
gb:MN996531|Organism:Severe      GTTTATGAAAAACTCAAACCCGTCCTTGATTGGCTTGAAGAGAAGTTTAAGGAAGGTGTA   2169
gb:MT066176|Organism:Severe      GTTTATGAAAAACTCAAACCCGTCCTTGATTGGCTTGAAGAGAAGTTTAAGGAAGGTGTA   2182
gb:MT027064|Organism:Severe      GTTTATGAAAAACTCAAACCCGTCCTTGATTGGCTTGAAGAGAAGTTTAAGGAAGGTGTA   2182
gb:MN994468|Organism:Severe      GTTTATGAAAAACTCAAACCCGTCCTTGATTGGCTTGAAGAGAAGTTTAAGGAAGGTGTA   2182
gb:MT072688|Organism:Severe      GTTTATGAAAAACTCAAACCCGTCCTTGATTGGCTTGAAGAGAAGTTTAAGGAAGGTGTA   2167
gb:MN996527|Organism:Severe      GTTTATGAAAAACTCAAACCCGTCCTTGATTGGCTTGAAGAGAAGTTTAAGGAAGGTGTA   2149
gb:MT093631|Organism:Severe      GTTTATGAAAAACTCAAACCCGTCCTTGATTGGCTTGAAGAGAAGTTTAAGGAAGGTGTA   2220
gb:MT106053|Organism:Severe      GTTTATGAAAAACTCAAACCCGTCCTTGATTGGCTTGAAGAGAAGTTTAAGGAAGGTGTA   2182
gb:MT019533|Organism:Severe      GTTTATGAAAAACTCAAACCCGTCCTTGATTGGCTTGAAGAGAAGTTTAAGGAAGGTGTA   2182
gb:MT019531|Organism:Severe      GTTTATGAAAAACTCAAACCCGTCCTTGATTGGCTTGAAGAGAAGTTTAAGGAAGGTGTA   2182
gb:MN996528|Organism:Severe      GTTTATGAAAAACTCAAACCCGTCCTTGATTGGCTTGAAGAGAAGTTTAAGGAAGGTGTA   2182
gb:MN996530|Organism:Severe      GTTTATGAAAAACTCAAACCCGTCCTTGATTGGCTTGAAGAGAAGTTTAAGGAAGGTGTA   2168
gb:MN908947|Organism:Severe      GTTTATGAAAAACTCAAACCCGTCCTTGATTGGCTTGAAGAGAAGTTTAAGGAAGGTGTA   2182
gb:MT019532|Organism:Severe      GTTTATGAAAAACTCAAACCCGTCCTTGATTGGCTTGAAGAGAAGTTTAAGGAAGGTGTA   2182
                                 ************************************************************

gb:MT020781|Organism:Severe      GAGTTTCTTAGAGACGGTTGGGAAATTGTTAAATTTATCTCAACCTGTGCTTGTGAAATT   2230
gb:MT007544|Organism:Severe      GAGTTTCTTAGAGACGGTTGGGAAATTGTTAAATTTATCTCAACCTGTGCTTGTGAAATT   2242
gb:MN994467|Organism:Severe      GAGTTTCTTAGAGACGGTTGGGAAATTGTTAAATTTATCTCAACCTGTGCTTGTGAAATT   2242
gb:MT044257|Organism:Severe      GAGTTTCTTAGAGACGGTTGGGAAATTGTTAAATTTATCTCAACCTGTGCTTGTGAAATT   2242
gb:MT106054|Organism:Severe      GAGTTTCTTAGAGACGGTTGGGAAATTGTTAAATTTATCTCAACCTGTGCTTGTGAAATT   2242
gb:MT049951|Organism:Severe      GAGTTTCTTAGAGACGGTTGGGAAATTGTTAAATTTATCTCAACCTGTGCTTGTGAAATT   2242
gb:MN975262|Organism:Severe      GAGTTTCTTAGAGACGGTTGGGAAATTGTTAAATTTATCTCAACCTGTGCTTGTGAAATT   2242
gb:MT106052|Organism:Severe      GAGTTTCTTAGAGACGGTTGGGAAATTGTTAAATTTATCTCAACCTGTGCTTGTGAAATT   2242
gb:LC522975|Organism:Severe      GAGTTTCTTAGAGACGGTTGGGAAATTGTTAAATTTATCTCAACCTGTGCTTGTGAAATT   2239
gb:LC522973|Organism:Severe      GAGTTTCTTAGAGACGGTTGGGAAATTGTTAAATTTATCTCAACCTGTGCTTGTGAAATT   2239
gb:LC522974|Organism:Severe      GAGTTTCTTAGAGACGGTTGGGAAATTGTTAAATTTATCTCAACCTGTGCTTGTGAAATT   2239
gb:MN985325|Organism:Severe      GAGTTTCTTAGAGACGGTTGGGAAATTGTTAAATTTATCTCAACCTGTGCTTGTGAAATT   2242
gb:MT020881|Organism:Severe      GAGTTTCTTAGAGACGGTTGGGAAATTGTTAAATTTATCTCAACCTGTGCTTGTGAAATT   2242
gb:MT020880|Organism:Severe      GAGTTTCTTAGAGACGGTTGGGAAATTGTTAAATTTATCTCAACCTGTGCTTGTGAAATT   2242
gb:MT066175|Organism:Severe      GAGTTTCTTAGAGACGGTTGGGAAATTGTTAAATTTATCTCAACCTGTGCTTGTGAAATT   2242
gb:MN997409|Organism:Severe      GAGTTTCTTAGAGACGGTTGGGAAATTGTTAAATTTATCTCAACCTGTGCTTGTGAAATT   2242
gb:MN938384|Organism:Severe      GAGTTTCTTAGAGACGGTTGGGAAATTGTTAAATTTATCTCAACCTGTGCTTGTGAAATT   2210
gb:MT044258|Organism:Severe      GAGTTTCTTAGAGACGGTTGGGAAATTGTTAAATTTATCTCAACCTGTGCTTGTGAAATT   2218
gb:MT039890|Organism:Severe      GAGTTTCTTAGAGACGGTTGGGAAATTGTTAAATTTATCTCAACCTGTGCTTGTGAAATT   2242
gb:MN988713|Organism:Severe      GAGTTTCTTAGAGACGGTTGGGAAATTGTTAAATTTATCTCAACCTGTGCTTGTGAAATT   2242
gb:LC521925|Organism:Severe      GAGTTTCTTAGAGACGGTTGGGAAATTGTTAAATTTATCTCAACCTGTGCTTGTGAAATT   2215
gb:MT093571|Organism:Severe      GAGTTTCTTAGAGACGGTTGGGAAATTGTTAAATTTATCTCAACCTGTGCTTGTGAAATT   2242
gb:MT039887|Organism:Severe      GAGTTTCTTAGAGACGGTTGGGAAATTGTTAAATTTATCTCAACCTGTGCTTGTGAAATT   2242
gb:MT019530|Organism:Severe      GAGTTTCTTAGAGACGGTTGGGAAATTGTTAAATTTATCTCAACCTGTGCTTGTGAAATT   2242
gb:MT039888|Organism:Severe      GAGTTTCTTAGAGACGGTTGGGAAATTGTTAAATTTATCTCAACCTGTGCTTGTGAAATT   2242
gb:LC522972|Organism:Severe      GAGTTTCTTAGAGACGGTTGGGAAATTGTTAAATTTATCTCAACCTGTGCTTGTGAAATT   2239
gb:MT027063|Organism:Severe      GAGTTTCTTAGAGACGGTTGGGAAATTGTTAAATTTATCTCAACCTGTGCTTGTGAAATT   2242
gb:MT027062|Organism:Severe      GAGTTTCTTAGAGACGGTTGGGAAATTGTTAAATTTATCTCAACCTGTGCTTGTGAAATT   2242
gb:MT019529|Organism:Severe      GAGTTTCTTAGAGACGGTTGGGAAATTGTTAAATTTATCTCAACCTGTGCTTGTGAAATT   2242
gb:MN996529|Organism:Severe      GAGTTTCTTAGAGACGGTTGGGAAATTGTTAAATTTATCTCAACCTGTGCTTGTGAAATT   2230
gb:MN996531|Organism:Severe      GAGTTTCTTAGAGACGGTTGGGAAATTGTTAAATTTATCTCAACCTGTGCTTGTGAAATT   2229
gb:MT066176|Organism:Severe      GAGTTTCTTAGAGACGGTTGGGAAATTGTTAAATTTATCTCAACCTGTGCTTGTGAAATT   2242
gb:MT027064|Organism:Severe      GAGTTTCTTAGAGACGGTTGGGAAATTGTTAAATTTATCTCAACCTGTGCTTGTGAAATT   2242
gb:MN994468|Organism:Severe      GAGTTTCTTAGAGACGGTTGGGAAATTGTTAAATTTATCTCAACCTGTGCTTGTGAAATT   2242
gb:MT072688|Organism:Severe      GAGTTTCTTAGAGACGGTTGGGAAATTGTTAAATTTATCTCAACCTGTGCTTGTGAAATT   2227
gb:MN996527|Organism:Severe      GAGTTTCTTAGAGACGGTTGGGAAATTGTTAAATTTATCTCAACCTGTGCTTGTGAAATT   2209
gb:MT093631|Organism:Severe      GAGTTTCTTAGAGACGGTTGGGAAATTGTTAAATTTATCTCAACCTGTGCTTGTGAAATT   2280
gb:MT106053|Organism:Severe      GAGTTTCTTAGAGACGGTTGGGAAATTGTTAAATTTATCTCAACCTGTGCTTGTGAAATT   2242
gb:MT019533|Organism:Severe      GAGTTTCTTAGAGACGGTTGGGAAATTGTTAAATTTATCTCAACCTGTGCTTGTGAAATT   2242
gb:MT019531|Organism:Severe      GAGTTTCTTAGAGACGGTTGGGAAATTGTTAAATTTATCTCAACCTGTGCTTGTGAAATT   2242
gb:MN996528|Organism:Severe      GAGTTTCTTAGAGACGGTTGGGAAATTGTTAAATTTATCTCAACCTGTGCTTGTGAAATT   2242
gb:MN996530|Organism:Severe      GAGTTTCTTAGAGACGGTTGGGAAATTGTTAAATTTATCTCAACCTGTGCTTGTGAAATT   2228
gb:MN908947|Organism:Severe      GAGTTTCTTAGAGACGGTTGGGAAATTGTTAAATTTATCTCAACCTGTGCTTGTGAAATT   2242
gb:MT019532|Organism:Severe      GAGTTTCTTAGAGACGGTTGGGAAATTGTTAAATTTATCTCAACCTGTGCTTGTGAAATT   2242
                                 ************************************************************

gb:MT020781|Organism:Severe      GTCGGTGGACAAATTGTCACCTGTGCAAAGGAAATTAAGGAGAGTGTTCAGACATTCTTT   2290
gb:MT007544|Organism:Severe      GTCGGTGGACAAATTGTCACCTGTGCAAAGGAAATTAAGGAGAGTGTTCAGACATTCTTT   2302
gb:MN994467|Organism:Severe      GTCGGTGGACAAATTGTCACCTGTGCAAAGGAAATTAAGGAGAGTGTTCAGACATTCTTT   2302
```

```
gb:MT044257|Organism:Severe      GTCGGTGGACAAATTGTCACCTGTGCAAAGGAAATTAAGGAGAGTGTTCAGACATTCTTT   2302
gb:MT106054|Organism:Severe      GTCGGTGGACAAATTGTCACCTGTGCAAAGGAAATTAAGGAGAGTGTTCAGACATTCTTT   2302
gb:MT049951|Organism:Severe      GTCGGTGGACAAATTGTCACCTGTGCAAAGGAAATTAAGGAGAGTGTTCAGACATTCTTT   2302
gb:MN975262|Organism:Severe      GTCGGTGGACAAATTGTCACCTGTGCAAAGGAAATTAAGGAGAGTGTTCAGACATTCTTT   2302
gb:MT106052|Organism:Severe      GTCGGTGGACAAATTGTCACCTGTGCAAAGGAAATTAAGGAGAGTGTTCAGACATTCTTT   2302
gb:LC522975|Organism:Severe      GTCGGTGGACAAATTGTCACCTGTGCAAAGGAAATTAAGGAGAGTGTTCAGACATTCTTT   2299
gb:LC522973|Organism:Severe      GTCGGTGGACAAATTGTCACCTGTGCAAAGGAAATTAAGGAGAGTGTTCAGACATTCTTT   2299
gb:LC522974|Organism:Severe      GTCGGTGGACAAATTGTCACCTGTGCAAAGGAAATTAAGGAGAGTGTTCAGACATTCTTT   2299
gb:MN985325|Organism:Severe      GTCGGTGGACAAATTGTCACCTGTGCAAAGGAAATTAAGGAGAGTGTTCAGACATTCTTT   2302
gb:MT020881|Organism:Severe      GTCGGTGGACAAATTGTCACCTGTGCAAAGGAAATTAAGGAGAGTGTTCAGACATTCTTT   2302
gb:MT020880|Organism:Severe      GTCGGTGGACAAATTGTCACCTGTGCAAAGGAAATTAAGGAGAGTGTTCAGACATTCTTT   2302
gb:MT066175|Organism:Severe      GTCGGTGGACAAATTGTCACCTGTGCAAAGGAAATTAAGGAGAGTGTTCAGACATTCTTT   2302
gb:MN997409|Organism:Severe      GTCGGTGGACAAATTGTCACCTGTGCAAAGGAAATTAAGGAGAGTGTTCAGACATTCTTT   2302
gb:MN938384|Organism:Severe      GTCGGTGGACAAATTGTCACCTGTGCAAAGGAAATTAAGGAGAGTGTTCAGACATTCTTT   2270
gb:MT044258|Organism:Severe      GTCGGTGGACAAATTGTCACCTGTGCAAAGGAAATTAAGGAGAGTGTTCAGACATTCTTT   2278
gb:MT039890|Organism:Severe      GTCGGTGGACAAATTGTCACCTGTGCAAAGGAAATTAAGGAGAGTGTTCAGACATTCTTT   2302
gb:MN988713|Organism:Severe      GTCGGTGGACAAATTGTCACCTGTGCAAAGGAAATTAAGGAGAGTGTTCAGACATTCTTT   2302
gb:LC521925|Organism:Severe      GTCGGTGGACAAATTGTCACCTGTGCAAAGGAAATTAAGGAGAGTGTTCAGACATTCTTT   2275
gb:MT093571|Organism:Severe      GTCGGTGGACAAATTGTCACCTGTGCAAAGGAAATTAAGGAGAGTGTTCAGACATTCTTT   2302
gb:MT039887|Organism:Severe      GTCGGTGGACAAATTGTCACCTGTGCAAAGGAAATTAAGGAGAGTGTTCAGACATTCTTT   2302
gb:MT019530|Organism:Severe      GTCGGTGGACAAATTGTCACCTGTGCAAAGGAAATTAAGGAGAGTGTTCAGACATTCTTT   2302
gb:MT039888|Organism:Severe      GTCGGTGGACAAATTGTCACCTGTGCAAAGGAAATTAAGGAGAGTGTTCAGACATTCTTT   2302
gb:LC522972|Organism:Severe      GTCGGTGGACAAATTGTCACCTGTGCAAAGGAAATTAAGGAGAGTGTTCAGACATTCTTT   2299
gb:MT027063|Organism:Severe      GTCGGTGGACAAATTGTCACCTGTGCAAAGGAAATTAAGGAGAGTGTTCAGACATTCTTT   2302
gb:MT027062|Organism:Severe      GTCGGTGGACAAATTGTCACCTGTGCAAAGGAAATTAAGGAGAGTGTTCAGACATTCTTT   2302
gb:MT019529|Organism:Severe      GTCGGTGGACAAATTGTCACCTGTGCAAAGGAAATTAAGGAGAGTGTTCAGACATTCTTT   2302
gb:MN996529|Organism:Severe      GTCGGTGGACAAATTGTCACCTGTGCAAAGGAAATTAAGGAGAGTGTTCAGACATTCTTT   2290
gb:MN996531|Organism:Severe      GTCGGTGGACAAATTGTCACCTGTGCAAAGGAAATTAAGGAGAGTGTTCAGACATTCTTT   2289
gb:MT066176|Organism:Severe      GTCGGTGGACAAATTGTCACCTGTGCAAAGGAAATTAAGGAGAGTGTTCAGACATTCTTT   2302
gb:MT027064|Organism:Severe      GTCGGTGGACAAATTGTCACCTGTGCAAAGGAAATTAAGGAGAGTGTTCAGACATTCTTT   2302
gb:MN994468|Organism:Severe      GTCGGTGGACAAATTGTCACCTGTGCAAAGGAAATTAAGGAGAGTGTTCAGACATTCTTT   2302
gb:MT072688|Organism:Severe      GTCGGTGGACAAATTGTCACCTGTGCAAAGGAAATTAAGGAGAGTGTTCAGACATTCTTT   2287
gb:MN996527|Organism:Severe      GTCGGTGGACAAATTGTCACCTGTGCAAAGGAAATTAAGGAGAGTGTTCAGACATTCTTT   2269
gb:MT093631|Organism:Severe      GTCGGTGGACAAATTGTCACCTGTGCAAAGGAAATTAAGGAGAGTGTTCAGACATTCTTT   2340
gb:MT106053|Organism:Severe      GTCGGTGGACAAATTGTCACCTGTGCAAAGGAAATTAAGGAGAGTGTTCAGACATTCTTT   2302
gb:MT019533|Organism:Severe      GTCGGTGGACAAATTGTCACCTGTGCAAAGGAAATTAAGGAGAGTGTTCAGACATTCTTT   2302
gb:MT019531|Organism:Severe      GTCGGTGGACAAATTGTCACCTGTGCAAAGGAAATTAAGGAGAGTGTTCAGACATTCTTT   2302
gb:MN996528|Organism:Severe      GTCGGTGGACAAATTGTCACCTGTGCAAAGGAAATTAAGGAGAGTGTTCAGACATTCTTT   2302
gb:MN996530|Organism:Severe      GTCGGTGGACAAATTGTCACCTGTGCAAAGGAAATTAAGGAGAGTGTTCAGACATTCTTT   2288
gb:MN908947|Organism:Severe      GTCGGTGGACAAATTGTCACCTGTGCAAAGGAAATTAAGGAGAGTGTTCAGACATTCTTT   2302
gb:MT019532|Organism:Severe      GTCGGTGGACAAATTGTCACCTGTGCAAAGGAAATTAAGGAGAGTGTTCAGACATTCTTT   2302
                                 ************************************************************

gb:MT020781|Organism:Severe      AAGCTTGTAAATAAATTTTTGGCTTTGTGTGCTGACTCTATCATTATTGGTGGAGCTAAA   2350
gb:MT007544|Organism:Severe      AAGCTTGTAAATAAATTTTTGGCTTTGTGTGCTGACTCTATCATTATTGGTGGAGCTAAA   2362
gb:MN994467|Organism:Severe      AAGCTTGTAAATAAATTTTTGGCTTTGTGTGCTGACTCTATCATTATTGGTGGAGCTAAA   2362
gb:MT044257|Organism:Severe      AAGCTTGTAAATAAATTTTTGGCTTTGTGTGCTGACTCTATCATTATTGGTGGAGCTAAA   2362
gb:MT106054|Organism:Severe      AAGCTTGTAAATAAATTTTTGGCTTTGTGTGCTGACTCTATCATTATTGGTGGAGCTAAA   2362
gb:MT049951|Organism:Severe      AAGCTTGTAAATAAATTTTTGGCTTTGTGTGCTGACTCTATCATTATTGGTGGAGCTAAA   2362
gb:MN975262|Organism:Severe      AAGCTTGTAAATAAATTTTTGGCTTTGTGTGCTGACTCTATCATTATTGGTGGAGCTAAA   2362
gb:MT106052|Organism:Severe      AAGCTTGTAAATAAATTTTTGGCTTTGTGTGCTGACTCTATCATTATTGGTGGAGCTAAA   2362
gb:LC522975|Organism:Severe      AAGCTTGTAAATAAATTTTTGGCTTTGTGTGCTGACTCTATCATTATTGGTGGAGCTAAA   2359
gb:LC522973|Organism:Severe      AAGCTTGTAAATAAATTTTTGGCTTTGTGTGCTGACTCTATCATTATTGGTGGAGCTAAA   2359
gb:LC522974|Organism:Severe      AAGCTTGTAAATAAATTTTTGGCTTTGTGTGCTGACTCTATCATTATTGGTGGAGCTAAA   2359
gb:MN985325|Organism:Severe      AAGCTTGTAAATAAATTTTTGGCTTTGTGTGCTGACTCTATCATTATTGGTGGAGCTAAA   2362
gb:MT020881|Organism:Severe      AAGCTTGTAAATAAATTTTTGGCTTTGTGTGCTGACTCTATCATTATTGGTGGAGCTAAA   2362
gb:MT020880|Organism:Severe      AAGCTTGTAAATAAATTTTTGGCTTTGTGTGCTGACTCTATCATTATTGGTGGAGCTAAA   2362
gb:MT066175|Organism:Severe      AAGCTTGTAAATAAATTTTTGGCTTTGTGTGCTGACTCTATCATTATTGGTGGAGCTAAA   2362
gb:MN997409|Organism:Severe      AAGCTTGTAAATAAATTTTTGGCTTTGTGTGCTGACTCTATCATTATTGGTGGAGCTAAA   2362
gb:MN938384|Organism:Severe      AAGCTTGTAAATAAATTTTTGGCTTTGTGTGCTGACTCTATCATTATTGGTGGAGCTAAA   2330
gb:MT044258|Organism:Severe      AAGCTTGTAAATAAATTTTTGGCTTTGTGTGCTGACTCTATCATTATTGGTGGAGCTAAA   2338
gb:MT039890|Organism:Severe      AAGCTTGTAAATAAATTTTTGGCTTTGTGTGCTGACTCTATCATTATTGGTGGAGCTAAA   2362
gb:MN988713|Organism:Severe      AAGCTTGTAAATAAATTTTTGGCTTTGTGTGCTGACTCTATCATTATTGGTGGAGCTAAA   2362
gb:LC521925|Organism:Severe      AAGCTTGTAAATAAATTTTTGGCTTTGTGTGCTGACTCTATCATTATTGGTGGAGCTAAA   2335
gb:MT093571|Organism:Severe      AAGCTTGTAAATAAATTTTTGGCTTTGTGTGCTGACTCTATCATTATTGGTGGAGCTAAA   2362
```

```
gb:MT039887|Organism:Severe      AAGCTTGTAAATAAATTTTTGGCTTTGTGTGCTGACTCTATCATTATTGGTGGAGCTAAA    2362
gb:MT019530|Organism:Severe      AAGCTTGTAAATAAATTTTTGGCTTTGTGTGCTGACTCTATCATTATTGGTGGAGCTAAA    2362
gb:MT039888|Organism:Severe      AAGCTTGTAAATAAATTTTTGGCTTTGTGTGCTGACTCTATCATTATTGGTGGAGCTAAA    2362
gb:LC522972|Organism:Severe      AAGCTTGTAAATAAATTTTTGGCTTTGTGTGCTGACTCTATCATTATTGGTGGAGCTAAA    2359
gb:MT027063|Organism:Severe      AAGCTTGTAAATAAATTTTTGGCTTTGTGTGCTGACTCTATCATTATTGGTGGAGCTAAA    2362
gb:MT027062|Organism:Severe      AAGCTTGTAAATAAATTTTTGGCTTTGTGTGCTGACTCTATCATTATTGGTGGAGCTAAA    2362
gb:MT019529|Organism:Severe      AAGCTTGTAAATAAATTTTTGGCTTTGTGTGCTGACTCTATCATTATTGGTGGAGCTAAA    2362
gb:MN996529|Organism:Severe      AAGCTTGTAAATAAATTTTTGGCTTTGTGTGCTGACTCTATCATTATTGGTGGAGCTAAA    2350
gb:MN996531|Organism:Severe      AAGCTTGTAAATAAATTTTTGGCTTTGTGTGCTGACTCTATCATTATTGGTGGAGCTAAA    2349
gb:MT066176|Organism:Severe      AAGCTTGTAAATAAATTTTTGGCTTTGTGTGCTGACTCTATCATTATTGGTGGAGCTAAA    2362
gb:MT027064|Organism:Severe      AAGCTTGTAAATAAATTTTTGGCTTTGTGTGCTGACTCTATCATTATTGGTGGAGCTAAA    2362
gb:MN994468|Organism:Severe      AAGCTTGTAAATAAATTTTTGGCTTTGTGTGCTGACTCTATCATTATTGGTGGAGCTAAA    2362
gb:MT072688|Organism:Severe      AAGCTTGTAAATAAATTTTTGGCTTTGTGTGCTGACTCTATCATTATTGGTGGAGCTAAA    2347
gb:MN996527|Organism:Severe      AAGCTTGTAAATAAATTTTTGGCTTTGTGTGCTGACTCTATCATTATTGGTGGAGCTAAA    2329
gb:MT093631|Organism:Severe      AAGCTTGTAAATAAATTTTTGGCTTTGTGTGCTGACTCTATCATTATTGGTGGAGCTAAA    2400
gb:MT106053|Organism:Severe      AAGCTTGTAAATAAATTTTTGGCTTTGTGTGCTGACTCTATCATTATTGGTGGAGCTAAA    2362
gb:MT019533|Organism:Severe      AAGCTTGTAAATAAATTTTTGGCTTTGTGTGCTGACTCTATCATTATTGGTGGAGCTAAA    2362
gb:MT019531|Organism:Severe      AAGCTTGTAAATAAATTTTTGGCTTTGTGTGCTGACTCTATCATTATTGGTGGAGCTAAA    2362
gb:MN996528|Organism:Severe      AAGCTTGTAAATAAATTTTTGGCTTTGTGTGCTGACTCTATCATTATTGGTGGAGCTAAA    2362
gb:MN996530|Organism:Severe      AAGCTTGTAAATAAATTTTTGGCTTTGTGTGCTGACTCTATCATTATTGGTGGAGCTAAA    2348
gb:MN908947|Organism:Severe      AAGCTTGTAAATAAATTTTTGGCTTTGTGTGCTGACTCTATCATTATTGGTGGAGCTAAA    2362
gb:MT019532|Organism:Severe      AAGCTTGTAAATAAATTTTTGGCTTTGTGTGCTGACTCTATCATTATTGGTGGAGCTAAA    2362
                                 ************************************************************

gb:MT020781|Organism:Severe      CTTAAAGCCTTGAATTTAGGTGAAACATTTGTCACGCACTCAAAGGGATTGTACAGAAAG    2410
gb:MT007544|Organism:Severe      CTTAAAGCCTTGAATTTAGGTGAAACATTTGTCACGCACTCAAAGGGATTGTACAGAAAG    2422
gb:MN994467|Organism:Severe      CTTAAAGCCTTGAATTTAGGTGAAACATTTGTCACGCACTCAAAGGGATTGTACAGAAAG    2422
gb:MT044257|Organism:Severe      CTTAAAGCCTTGAATTTAGGTGAAACATTTGTCACGCACTCAAAGGGATTGTACAGAAAG    2422
gb:MT106054|Organism:Severe      CTTAAAGCCTTGAATTTAGGTGAAACATTTGTCACGCACTCAAAGGGATTGTACAGAAAG    2422
gb:MT049951|Organism:Severe      CTTAAAGCCTTGAATTTAGGTGAAACATTTGTCACGCACTCAAAGGGATTGTACAGAAAG    2422
gb:MN975262|Organism:Severe      CTTAAAGCCTTGAATTTAGGTGAAACATTTGTCACGCACTCAAAGGGATTGTACAGAAAG    2422
gb:MT106052|Organism:Severe      CTTAAAGCCTTGAATTTAGGTGAAACATTTGTCACGCACTCAAAGGGATTGTACAGAAAG    2422
gb:LC522975|Organism:Severe      CTTAAAGCCTTGAATTTAGGTGAAACATTTGTCACGCACTCAAAGGGATTGTACAGAAAG    2419
gb:LC522973|Organism:Severe      CTTAAAGCCTTGAATTTAGGTGAAACATTTGTCACGCACTCAAAGGGATTGTACAGAAAG    2419
gb:LC522974|Organism:Severe      CTTAAAGCCTTGAATTTAGGTGAAACATTTGTCACGCACTCAAAGGGATTGTACAGAAAG    2419
gb:MN985325|Organism:Severe      CTTAAAGCCTTGAATTTAGGTGAAACATTTGTCACGCACTCAAAGGGATTGTACAGAAAG    2422
gb:MT020881|Organism:Severe      CTTAAAGCCTTGAATTTAGGTGAAACATTTGTCACGCACTCAAAGGGATTGTACAGAAAG    2422
gb:MT020880|Organism:Severe      CTTAAAGCCTTGAATTTAGGTGAAACATTTGTCACGCACTCAAAGGGATTGTACAGAAAG    2422
gb:MT066175|Organism:Severe      CTTAAAGCCTTGAATTTAGGTGAAACATTTGTCACGCACTCAAAGGGATTGTACAGAAAG    2422
gb:MN997409|Organism:Severe      CTTAAAGCCTTGAATTTAGGTGAAACATTTGTCACGCACTCAAAGGGATTGTACAGAAAG    2422
gb:MN938384|Organism:Severe      CTTAAAGCCTTGAATTTAGGTGAAACATTTGTCACGCACTCAAAGGGATTGTACAGAAAG    2390
gb:MT044258|Organism:Severe      CTTAAAGCCTTGAATTTAGGTGAAACATTTGTCACGCACTCAAAGGGATTGTACAGAAAG    2398
gb:MT039890|Organism:Severe      CTTAAAGCCTTGAATTTAGGTGAAACATTTGTCACGCACTCAAAGGGATTGTACAGAAAG    2422
gb:MN988713|Organism:Severe      CTTAAAGCCTTGAATTTAGGTGAAACATTTGTCACGCACTCAAAGGGATTGTACAGAAAG    2422
gb:LC521925|Organism:Severe      CTTAAAGCCTTGAATTTAGGTGAAACATTTGTCACGCACTCAAAGGGATTGTACAGAAAG    2395
gb:MT093571|Organism:Severe      CTTAAAGCCTTGAATTTAGGTGAAACATTTGTCACGCACTCAAAGGGATTGTACAGAAAG    2422
gb:MT039887|Organism:Severe      CTTAAAGCCTTGAATTTAGGTGAAACATTTGTCACGCACTCAAAGGGATTGTACAGAAAG    2422
gb:MT019530|Organism:Severe      CTTAAAGCCTTGAATTTAGGTGAAACATTTGTCACGCACTCAAAGGGATTGTACAGAAAG    2422
gb:MT039888|Organism:Severe      CTTAAAGCCTTGAATTTAGGTGAAACATTTGTCACGCACTCAAAGGGATTGTACAGAAAG    2422
gb:LC522972|Organism:Severe      CTTAAAGCCTTGAATTTAGGTGAAACATTTGTCACGCACTCAAAGGGATTGTACAGAAAG    2419
gb:MT027063|Organism:Severe      CTTAAAGCCTTGAATTTAGGTGAAACATTTGTCACGCACTCAAAGGGATTGTACAGAAAG    2422
gb:MT027062|Organism:Severe      CTTAAAGCCTTGAATTTAGGTGAAACATTTGTCACGCACTCAAAGGGATTGTACAGAAAG    2422
gb:MT019529|Organism:Severe      CTTAAAGCCTTGAATTTAGGTGAAACATTTGTCACGCACTCAAAGGGATTGTACAGAAAG    2422
gb:MN996529|Organism:Severe      CTTAAAGCCTTGAATTTAGGTGAAACATTTGTCACGCACTCAAAGGGATTGTACAGAAAG    2410
gb:MN996531|Organism:Severe      CTTAAAGCCTTGAATTTAGGTGAAACATTTGTCACGCACTCAAAGGGATTGTACAGAAAG    2409
gb:MT066176|Organism:Severe      CTTAAAGCCTTGAATTTAGGTGAAACATTTGTCACGCACTCAAAGGGATTGTACAGAAAG    2422
gb:MT027064|Organism:Severe      CTTAAAGCCTTGAATTTAGGTGAAACATTTGTCACGCACTCAAAGGGATTGTACAGAAAG    2422
gb:MN994468|Organism:Severe      CTTAAAGCCTTGAATTTAGGTGAAACATTTGTCACGCACTCAAAGGGATTGTACAGAAAG    2422
gb:MT072688|Organism:Severe      CTTAAAGCCTTGAATTTAGGTGAAACATTTGTCACGCACTCAAAGGGATTGTACAGAAAG    2407
gb:MN996527|Organism:Severe      CTTAAAGCCTTGAATTTAGGTGAAACATTTGTCACGCACTCAAAGGGATTGTACAGAAAG    2389
gb:MT093631|Organism:Severe      CTTAAAGCCTTGAATTTAGGTGAAACATTTGTCACGCACTCAAAGGGATTGTACAGAAAG    2460
gb:MT106053|Organism:Severe      CTTAAAGCCTTGAATTTAGGTGAAACATTTGTCACGCACTCAAAGGGATTGTACAGAAAG    2422
gb:MT019533|Organism:Severe      CTTAAAGCCTTGAATTTAGGTGAAACATTTGTCACGCACTCAAAGGGATTGTACAGAAAG    2422
gb:MT019531|Organism:Severe      CTTAAAGCCTTGAATTTAGGTGAAACATTTGTCACGCACTCAAAGGGATTGTACAGAAAG    2422
gb:MN996528|Organism:Severe      CTTAAAGCCTTGAATTTAGGTGAAACATTTGTCACGCACTCAAAGGGATTGTACAGAAAG    2422
```

```
gb:MN996530|Organism:Severe      CTTAAAGCCTTGAATTTAGGTGAAACATTTGTCACGCACTCAAAGGGATTGTACAGAAAG    2408
gb:MN908947|Organism:Severe      CTTAAAGCCTTGAATTTAGGTGAAACATTTGTCACGCACTCAAAGGGATTGTACAGAAAG    2422
gb:MT019532|Organism:Severe      CTTAAAGCCTTGAATTTAGGTGAAACATTTGTCACGCACTCAAAGGGATTGTACAGAAAG    2422
                                 ************************************************************

gb:MT020781|Organism:Severe      TGTGTTAAATCCAGAGAAGAAACTGGCCTACTCATGCCTCTAAAAGCCCCAAAAGAAATT    2470
gb:MT007544|Organism:Severe      TGTGTTAAATCCAGAGAAGAAACTGGCCTACTCATGCCTCTAAAAGCCCCAAAAGAAATT    2482
gb:MN994467|Organism:Severe      TGTGTTAAATCCAGAGAAGAAACTGGCCTACTCATGCCTCTAAAAGCCCCAAAAGAAATT    2482
gb:MT044257|Organism:Severe      TGTGTTAAATCCAGAGAAGAAACTGGCCTACTCATGCCTCTAAAAGCCCCAAAAGAAATT    2482
gb:MT106054|Organism:Severe      TGTGTTAAATCCAGAGAAGAAACTGGCCTACTCATGCCTCTAAAAGCCCCAAAAGAAATT    2482
gb:MT049951|Organism:Severe      TGTGTTAAATCCAGAGAAGAAACTGGCCTACTCATGCCTCTAAAAGCCCCAAAAGAAATT    2482
gb:MN975262|Organism:Severe      TGTGTTAAATCCAGAGAAGAAACTGGCCTACTCATGCCTCTAAAAGCCCCAAAAGAAATT    2482
gb:MT106052|Organism:Severe      TGTGTTAAATCCAGAGAAGAAACTGGCCTACTCATGCCTCTAAAAGCCCCAAAAGAAATT    2482
gb:LC522975|Organism:Severe      TGTGTTAAATCCAGAGAAGAAACTGGCCTACTCATGCCTCTAAAAGCCCCAAAAGAAATT    2479
gb:LC522973|Organism:Severe      TGTGTTAAATCCAGAGAAGAAACTGGCCTACTCATGCCTCTAAAAGCCCCAAAAGAAATT    2479
gb:LC522974|Organism:Severe      TGTGTTAAATCCAGAGAAGAAACTGGCCTACTCATGCCTCTAAAAGCCCCAAAAGAAATT    2479
gb:MN985325|Organism:Severe      TGTGTTAAATCCAGAGAAGAAACTGGCCTACTCATGCCTCTAAAAGCCCCAAAAGAAATT    2482
gb:MT020881|Organism:Severe      TGTGTTAAATCCAGAGAAGAAACTGGCCTACTCATGCCTCTAAAAGCCCCAAAAGAAATT    2482
gb:MT020880|Organism:Severe      TGTGTTAAATCCAGAGAAGAAACTGGCCTACTCATGCCTCTAAAAGCCCCAAAAGAAATT    2482
gb:MT066175|Organism:Severe      TGTGTTAAATCCAGAGAAGAAACTGGCCTACTCATGCCTCTAAAAGCCCCAAAAGAAATT    2482
gb:MN997409|Organism:Severe      TGTGTTAAATCCAGAGAAGAAACTGGCCTACTCATGCCTCTAAAAGCCCCAAAAGAAATT    2482
gb:MN938384|Organism:Severe      TGTGTTAAATCCAGAGAAGAAACTGGCCTACTCATGCCTCTAAAAGCCCCAAAAGAAATT    2450
gb:MT044258|Organism:Severe      TGTGTTAAATCCAGAGAAGAAACTGGCCTACTCATGCCTCTAAAAGCCCCAAAAGAAATT    2458
gb:MT039890|Organism:Severe      TGTGTTAAATCCAGAGAAGAAACTGGCCTACTCATGCCTCTAAAAGCCCCAAAAGAAATT    2482
gb:MN988713|Organism:Severe      TGTGTTAAATCCAGAGAAGAAACTGGCCTACTCATGCCTCTAAAAGCCCCAAAAGAAATT    2482
gb:LC521925|Organism:Severe      TGTGTTAAATCCAGAGAAGAAACTGGCCTACTCATGCCTCTAAAAGCCCCAAAAGAAATT    2455
gb:MT093571|Organism:Severe      TGTGTTAAATCCAGAGAAGAAACTGGCCTACTCATGCCTCTAAAAGCCCCAAAAGAAATT    2482
gb:MT039887|Organism:Severe      TGTGTTAAATCCAGAGAAGAAACTGGCCTACTCATGCCTCTAAAAGCCCCAAAAGAAATT    2482
gb:MT019530|Organism:Severe      TGTGTTAAATCCAGAGAAGAAACTGGCCTACTCATGCCTCTAAAAGCCCCAAAAGAAATT    2482
gb:MT039888|Organism:Severe      TGTGTTAAATCCAGAGAAGAAACTGGCCTACTCATGCCTCTAAAAGCCCCAAAAGAAATT    2482
gb:LC522972|Organism:Severe      TGTGTTAAATCCAGAGAAGAAACTGGCCTACTCATGCCTCTAAAAGCCCCAAAAGAAATT    2479
gb:MT027063|Organism:Severe      TGTGTTAAATCCAGAGAAGAAACTGGCCTACTCATGCCTCTAAAAGCCCCAAAAGAAATT    2482
gb:MT027062|Organism:Severe      TGTGTTAAATCCAGAGAAGAAACTGGCCTACTCATGCCTCTAAAAGCCCCAAAAGAAATT    2482
gb:MT019529|Organism:Severe      TGTGTTAAATCCAGAGAAGAAACTGGCCTACTCATGCCTCTAAAAGCCCCAAAAGAAATT    2482
gb:MN996529|Organism:Severe      TGTGTTAAATCCAGAGAAGAAACTGGCCTACTCATGCCTCTAAAAGCCCCAAAAGAAATT    2470
gb:MN996531|Organism:Severe      TGTGTTAAATCCAGAGAAGAAACTGGCCTACTCATGCCTCTAAAAGCCCCAAAAGAAATT    2469
gb:MT066176|Organism:Severe      TGTGTTAAATCCAGAGAAGAAACTGGCCTACTCATGCCTCTAAAAGCCCCAAAAGAAATT    2482
gb:MT027064|Organism:Severe      TGTGTTAAATCCAGAGAAGAAACTGGCCTACTCATGCCTCTAAAAGCCCCAAAAGAAATT    2482
gb:MN994468|Organism:Severe      TGTGTTAAATCCAGAGAAGAAACTGGCCTACTCATGCCTCTAAAAGCCCCAAAAGAAATT    2482
gb:MT072688|Organism:Severe      TGTGTTAAATCCAGAGAAGAAACTGGCCTACTCATGCCTCTAAAAGCCCCAAAAGAAATT    2467
gb:MN996527|Organism:Severe      TGTGTTAAATCCAGAGAAGAAACTGGCCTACTCATGCCTCTAAAAGCCCCAAAAGAAATT    2449
gb:MT093631|Organism:Severe      TGTGTTAAATCCAGAGAAGAAACTGGCCTACTCATGCCTCTAAAAGCCCCAAAAGAAATT    2520
gb:MT106053|Organism:Severe      TGTGTTAAATCCAGAGAAGAAACTGGCCTACTCATGCCTCTAAAAGCCCCAAAAGAAATT    2482
gb:MT019533|Organism:Severe      TGTGTTAAATCCAGAGAAGAAACTGGCCTACTCATGCCTCTAAAAGCCCCAAAAGAAATT    2482
gb:MT019531|Organism:Severe      TGTGTTAAATCCAGAGAAGAAACTGGCCTACTCATGCCTCTAAAAGCCCCAAAAGAAATT    2482
gb:MN996528|Organism:Severe      TGTGTTAAATCCAGAGAAGAAACTGGCCTACTCATGCCTCTAAAAGCCCCAAAAGAAATT    2482
gb:MN996530|Organism:Severe      TGTGTTAAATCCAGAGAAGAAACTGGCCTACTCATGCCTCTAAAAGCCCCAAAAGAAATT    2468
gb:MN908947|Organism:Severe      TGTGTTAAATCCAGAGAAGAAACTGGCCTACTCATGCCTCTAAAAGCCCCAAAAGAAATT    2482
gb:MT019532|Organism:Severe      TGTGTTAAATCCAGAGAAGAAACTGGCCTACTCATGCCTCTAAAAGCCCCAAAAGAAATT    2482
                                 ************************************************************

gb:MT020781|Organism:Severe      ATCTTCTTAGAGGGAGAAACACTTCCCACAGAAGTGTTAACAGAGGAAGTTGTCTTGAAA    2530
gb:MT007544|Organism:Severe      ATCTTCTTAGAGGGAGAAACACTTCCCACAGAAGTGTTAACAGAGGAAGTTGTCTTGAAA    2542
gb:MN994467|Organism:Severe      ATCTTCTTAGAGGGAGAAACACTTCCCACAGAAGTGTTAACAGAGGAAGTTGTCTTGAAA    2542
gb:MT044257|Organism:Severe      ATCTTCTTAGAGGGAGAAACACTTCCCACAGAAGTGTTAACAGAGGAAGTTGTCTTGAAA    2542
gb:MT106054|Organism:Severe      ATCTTCTTAGAGGGAGAAACACTTCCCACAGAAGTGTTAACAGAGGAAGTTGTCTTGAAA    2542
gb:MT049951|Organism:Severe      ATCTTCTTAGAGGGAGAAACACTTCCCACAGAAGTGTTAACAGAGGAAGTTGTCTTGAAA    2542
gb:MN975262|Organism:Severe      ATCTTCTTAGAGGGAGAAACACTTCCCACAGAAGTGTTAACAGAGGAAGTTGTCTTGAAA    2542
gb:MT106052|Organism:Severe      ATCTTCTTAGAGGGAGAAACACTTCCCACAGAAGTGTTAACAGAGGAAGTTGTCTTGAAA    2542
gb:LC522975|Organism:Severe      ATCTTCTTAGAGGGAGAAACACTTCCCACAGAAGTGTTAACAGAGGAAGTTGTCTTGAAA    2539
gb:LC522973|Organism:Severe      ATCTTCTTAGAGGGAGAAACACTTCCCACAGAAGTGTTAACAGAGGAAGTTGTCTTGAAA    2539
gb:LC522974|Organism:Severe      ATCTTCTTAGAGGGAGAAACACTTCCCACAGAAGTGTTAACAGAGGAAGTTGTCTTGAAA    2539
gb:MN985325|Organism:Severe      ATCTTCTTAGAGGGAGAAACACTTCCCACAGAAGTGTTAACAGAGGAAGTTGTCTTGAAA    2542
gb:MT020881|Organism:Severe      ATCTTCTTAGAGGGAGAAACACTTCCCACAGAAGTGTTAACAGAGGAAGTTGTCTTGAAA    2542
gb:MT020880|Organism:Severe      ATCTTCTTAGAGGGAGAAACACTTCCCACAGAAGTGTTAACAGAGGAAGTTGTCTTGAAA    2542
```

```
gb:MT066175|Organism:Severe      ATCTTCTTAGAGGGAGAAACACTTCCCACAGAAGTGTTAACAGAGGAAGTTGTCTTGAAA   2542
gb:MN997409|Organism:Severe      ATCTTCTTAGAGGGAGAAACACTTCCCACAGAAGTGTTAACAGAGGAAGTTGTCTTGAAA   2542
gb:MN938384|Organism:Severe      ATCTTCTTAGAGGGAGAAACACTTCCCACAGAAGTGTTAACAGAGGAAGTTGTCTTGAAA   2510
gb:MT044258|Organism:Severe      ATCTTCTTAGAGGGAGAAACACTTCCCACAGAAGTGTTAACAGAGGAAGTTGTCTTGAAA   2518
gb:MT039890|Organism:Severe      ATCTTCTTAGAGGGAGAAACACTTCCCACAGAAGTGTTAACAGAGGAAGTTGTCTTGAAA   2542
gb:MN988713|Organism:Severe      ATCTTCTTAGAGGGAGAAACACTTCCCACAGAAGTGTTAACAGAGGAAGTTGTCTTGAAA   2542
gb:LC521925|Organism:Severe      ATCTTCTTAGAGGGAGAAACACTTCCCACAGAAGTGTTAACAGAGGAAGTTGTCTTGAAA   2515
gb:MT093571|Organism:Severe      ATCTTCTTAGAGGGAGAAACACTTCCCACAGAAGTGTTAACAGAGGAAGTTGTCTTGAAA   2542
gb:MT039887|Organism:Severe      ATCTTCTTAGAGGGAGAAACACTTCCCACAGAAGTGTTAACAGAGGAAGTTGTCTTGAAA   2542
gb:MT019530|Organism:Severe      ATCTTCTTAGAGGGAGAAACACTTCCCACAGAAGTGTTAACAGAGGAAGTTGTCTTGAAA   2542
gb:MT039888|Organism:Severe      ATCTTCTTAGAGGGAGAAACACTTCCCACAGAAGTGTTAACAGAGGAAGTTGTCTTGAAA   2542
gb:LC522972|Organism:Severe      ATCTTCTTAGAGGGAGAAACACTTCCCACAGAAGTGTTAACAGAGGAAGTTGTCTTGAAA   2539
gb:MT027063|Organism:Severe      ATCTTCTTAGAGGGAGAAACACTTCCCACAGAAGTGTTAACAGAGGAAGTTGTCTTGAAA   2542
gb:MT027062|Organism:Severe      ATCTTCTTAGAGGGAGAAACACTTCCCACAGAAGTGTTAACAGAGGAAGTTGTCTTGAAA   2542
gb:MT019529|Organism:Severe      ATCTTCTTAGAGGGAGAAACACTTCCCACAGAAGTGTTAACAGAGGAAGTTGTCTTGAAA   2542
gb:MN996529|Organism:Severe      ATCTTCTTAGAGGGAGAAACACTTCCCACAGAAGTGTTAACAGAGGAAGTTGTCTTGAAA   2530
gb:MN996531|Organism:Severe      ATCTTCTTAGAGGGAGAAACACTTCCCACAGAAGTGTTAACAGAGGAAGTTGTCTTGAAA   2529
gb:MT066176|Organism:Severe      ATCTTCTTAGAGGGAGAAACACTTCCCACAGAAGTGTTAACAGAGGAAGTTGTCTTGAAA   2542
gb:MT027064|Organism:Severe      ATCTTCTTAGAGGGAGAAACACTTCCCACAGAAGTGTTAACAGAGGAAGTTGTCTTGAAA   2542
gb:MN994468|Organism:Severe      ATCTTCTTAGAGGGAGAAACACTTCCCACAGAAGTGTTAACAGAGGAAGTTGTCTTGAAA   2542
gb:MT072688|Organism:Severe      ATCTTCTTAGAGGGAGAAACACTTCCCACAGAAGTGTTAACAGAGGAAGTTGTCTTGAAA   2527
gb:MN996527|Organism:Severe      ATCTTCTTAGAGGGAGAAACACTTCCCACAGAAGTGTTAACAGAGGAAGTTGTCTTGAAA   2509
gb:MT093631|Organism:Severe      ATCTTCTTAGAGGGAGAAACACTTCCCACAGAAGTGTTAACAGAGGAAGTTGTCTTGAAA   2580
gb:MT106053|Organism:Severe      ATCTTCTTAGAGGGAGAAACACTTCCCACAGAAGTGTTAACAGAGGAAGTTGTCTTGAAA   2542
gb:MT019533|Organism:Severe      ATCTTCTTAGAGGGAGAAACACTTCCCACAGAAGTGTTAACAGAGGAAGTTGTCTTGAAA   2542
gb:MT019531|Organism:Severe      ATCTTCTTAGAGGGAGAAACACTTCCCACAGAAGTGTTAACAGAGGAAGTTGTCTTGAAA   2542
gb:MN996528|Organism:Severe      ATCTTCTTAGAGGGAGAAACACTTCCCACAGAAGTGTTAACAGAGGAAGTTGTCTTGAAA   2542
gb:MN996530|Organism:Severe      ATCTTCTTAGAGGGAGAAACACTTCCCACAGAAGTGTTAACAGAGGAAGTTGTCTTGAAA   2528
gb:MN908947|Organism:Severe      ATCTTCTTAGAGGGAGAAACACTTCCCACAGAAGTGTTAACAGAGGAAGTTGTCTTGAAA   2542
gb:MT019532|Organism:Severe      ATCTTCTTAGAGGGAGAAACACTTCCCACAGAAGTGTTAACAGAGGAAGTTGTCTTGAAA   2542
                                 ************************************************************

gb:MT020781|Organism:Severe      ACTGGTGATTTACAACCATTAGAACAACCTACTAGTGAAGCTGTTGAAGCTCCATTGGTT   2590
gb:MT007544|Organism:Severe      ACTGGTGATTTACAACCATTAGAACAACCTACTAGTGAAGCTGTTGAAGCTCCATTGGTT   2602
gb:MN994467|Organism:Severe      ACTGGTGATTTACAACCATTAGAACAACCTACTAGTGAAGCTGTTGAAGCTCCATTGGTT   2602
gb:MT044257|Organism:Severe      ACTGGTGATTTACAACCATTAGAACAACCTACTAGTGAAGCTGTTGAAGCTCCATTGGTT   2602
gb:MT106054|Organism:Severe      ACTGGTGATTTACAACCATTAGAACAACCTACTAGTGAAGCTGTTGAAGCTCCATTGGTT   2602
gb:MT049951|Organism:Severe      ACTGGTGATTTACAACCATTAGAACAACCTACTAGTGAAGCTGTTGAAGCTCCATTGGTT   2602
gb:MN975262|Organism:Severe      ACTGGTGATTTACAACCATTAGAACAACCTACTAGTGAAGCTGTTGAAGCTCCATTGGTT   2602
gb:MT106052|Organism:Severe      ACTGGTGATTTACAACCATTAGAACAACCTACTAGTGAAGCTGTTGAAGCTCCATTGGTT   2602
gb:LC522975|Organism:Severe      ACTGGTGATTTACAACCATTAGAACAACCTACTAGTGAAGCTGTTGAAGCTCCATTGGTT   2599
gb:LC522973|Organism:Severe      ACTGGTGATTTACAACCATTAGAACAACCTACTAGTGAAGCTGTTGAAGCTCCATTGGTT   2599
gb:LC522974|Organism:Severe      ACTGGTGATTTACAACCATTAGAACAACCTACTAGTGAAGCTGTTGAAGCTCCATTGGTT   2599
gb:MN985325|Organism:Severe      ACTGGTGATTTACAACCATTAGAACAACCTACTAGTGAAGCTGTTGAAGCTCCATTGGTT   2602
gb:MT020881|Organism:Severe      ACTGGTGATTTACAACCATTAGAACAACCTACTAGTGAAGCTGTTGAAGCTCCATTGGTT   2602
gb:MT020880|Organism:Severe      ACTGGTGATTTACAACCATTAGAACAACCTACTAGTGAAGCTGTTGAAGCTCCATTGGTT   2602
gb:MT066175|Organism:Severe      ACTGGTGATTTACAACCATTAGAACAACCTACTAGTGAAGCTGTTGAAGCTCCATTGGTT   2602
gb:MN997409|Organism:Severe      ACTGGTGATTTACAACCATTAGAACAACCTACTAGTGAAGCTGTTGAAGCTCCATTGGTT   2602
gb:MN938384|Organism:Severe      ACTGGTGATTTACAACCATTAGAACAACCTACTAGTGAAGCTGTTGAAGCTCCATTGGTT   2570
gb:MT044258|Organism:Severe      ACTGGTGATTTACAACCATTAGAACAACCTACTAGTGAAGCTGTTGAAGCTCCATTGGTT   2578
gb:MT039890|Organism:Severe      ACTGGTGATTTACAACCATTAGAACAACCTACTAGTGAAGCTGTTGAAGCTCCATTGGTT   2602
gb:MN988713|Organism:Severe      ACTGGTGATTTACAACCATTAGAACAACCTACTAGTGAAGCTGTTGAAGCTCCATTGGTT   2602
gb:LC521925|Organism:Severe      ACTGGTGATTTACAACCATTAGAACAACCTACTAGTGAAGCTGTTGAAGCTCCATTGGTT   2575
gb:MT093571|Organism:Severe      ACTGGTGATTTACAACCATTAGAACAACCTACTAGTGAAGCTGTTGAAGCTCCATTGGTT   2602
gb:MT039887|Organism:Severe      ACTGGTGATTTACAACCATTAGAACAACCTACTAGTGAAGCTGTTGAAGCTCCATTGGTT   2602
gb:MT019530|Organism:Severe      ACTGGTGATTTACAACCATTAGAACAACCTACTAGTGAAGCTGTTGAAGCTCCATTGGTT   2602
gb:MT039888|Organism:Severe      ACTGGTGATTTACAACCATTAGAACAACCTACTAGTGAAGCTGTTGAAGCTCCATTGGTT   2602
gb:LC522972|Organism:Severe      ACTGGTGATTTACAACCATTAGAACAACCTACTAGTGAAGCTGTTGAAGCTCCATTGGTT   2599
gb:MT027063|Organism:Severe      ACTGGTGATTTACAACCATTAGAACAACCTACTAGTGAAGCTGTTGAAGCTCCATTGGTT   2602
gb:MT027062|Organism:Severe      ACTGGTGATTTACAACCATTAGAACAACCTACTAGTGAAGCTGTTGAAGCTCCATTGGTT   2602
gb:MT019529|Organism:Severe      ACTGGTGATTTACAACCATTAGAACAACCTACTAGTGAAGCTGTTGAAGCTCCATTGGTT   2602
gb:MN996529|Organism:Severe      ACTGGTGATTTACAACCATTAGAACAACCTACTAGTGAAGCTGTTGAAGCTCCATTGGTT   2590
gb:MN996531|Organism:Severe      ACTGGTGATTTACAACCATTAGAACAACCTACTAGTGAAGCTGTTGAAGCTCCATTGGTT   2589
gb:MT066176|Organism:Severe      ACTGGTGATTTACAACCATTAGAACAACCTACTAGTGAAGCTGTTGAAGCTCCATTGGTT   2602
gb:MT027064|Organism:Severe      ACTGGTGATTTACAACCATTAGAACAACCTACTAGTGAAGCTGTTGAAGCTCCATTGGTT   2602
```

```
gb:MN994468|Organism:Severe      ACTGGTGATTTACAACCATTAGAACAACCTACTAGTGAAGCTGTTGAAGCTCCATTGGTT   2602
gb:MT072688|Organism:Severe      ACTGGTGATTTACAACCATTAGAACAACCTACTAGTGAAGCTGTTGAAGCTCCATTGGTT   2587
gb:MN996527|Organism:Severe      ACTGGTGATTTACAACCATTAGAACAACCTACTAGTGAAGCTGTTGAAGCTCCATTGGTT   2569
gb:MT093631|Organism:Severe      ACTGGTGATTTACAACCATTAGAACAACCTACTAGTGAAGCTGTTGAAGCTCCATTGGTT   2640
gb:MT106053|Organism:Severe      ACTGGTGATTTACAACCATTAGAACAACCTACTAGTGAAGCTGTTGAAGCTCCATTGGTT   2602
gb:MT019533|Organism:Severe      ACTGGTGATTTACAACCATTAGAACAACCTACTAGTGAAGCTGTTGAAGCTCCATTGGTT   2602
gb:MT019531|Organism:Severe      ACTGGTGATTTACAACCATTAGAACAACCTACTAGTGAAGCTGTTGAAGCTCCATTGGTT   2602
gb:MN996528|Organism:Severe      ACTGGTGATTTACAACCATTAGAACAACCTACTAGTGAAGCTGTTGAAGCTCCATTGGTT   2602
gb:MN996530|Organism:Severe      ACTGGTGATTTACAACCATTAGAACAACCTACTAGTGAAGCTGTTGAAGCTCCATTGGTT   2588
gb:MN908947|Organism:Severe      ACTGGTGATTTACAACCATTAGAACAACCTACTAGTGAAGCTGTTGAAGCTCCATTGGTT   2602
gb:MT019532|Organism:Severe      ACTGGTGATTTACAACCATTAGAACAACCTACTAGTGAAGCTGTTGAAGCTCCATTGGTT   2602
                                 ************************************************************

gb:MT020781|Organism:Severe      GGTACACCAGTTTGTATTAACGGGCTTATGTTGCTCGAAATCAAAGACACAGAAAAGTAC   2650
gb:MT007544|Organism:Severe      GGTACACCAGTTTGTATTAACGGGCTTATGTTGCTCGAAATCAAAGACACAGAAAAGTAC   2662
gb:MN994467|Organism:Severe      GGTACACCAGTTTGTATTAACGGGCTTATGTTGCTCGAAATCAAAGACACAGAAAAGTAC   2662
gb:MT044257|Organism:Severe      GGTACACCAGTTTGTATTAACGGGCTTATGTTGCTCGAAATCAAAGACACAGAAAAGTAC   2662
gb:MT106054|Organism:Severe      GGTACACCAGTTTGTATTAACGGGCTTATGTTGCTCGAAATCAAAGACACAGAAAAGTAC   2662
gb:MT049951|Organism:Severe      GGTACACCAGTTTGTATTAACGGGCTTATGTTGCTCGAAATCAAAGACACAGAAAAGTAC   2662
gb:MN975262|Organism:Severe      GGTACACCAGTTTGTATTAACGGGCTTATGTTGCTCGAAATCAAAGACACAGAAAAGTAC   2662
gb:MT106052|Organism:Severe      GGTACACCAGTTTGTATTAACGGGCTTATGTTGCTCGAAATCAAAGACACAGAAAAGTAC   2662
gb:LC522975|Organism:Severe      GGTACACCAGTTTGTATTAACGGGCTTATGTTGCTCGAAATCAAAGACACAGAAAAGTAT   2659
gb:LC522973|Organism:Severe      GGTACACCAGTTTGTATTAACGGGCTTATGTTGCTCGAAATCAAAGACACAGAAAAGTAT   2659
gb:LC522974|Organism:Severe      GGTACACCAGTTTGTATTAACGGGCTTATGTTGCTCGAAATCAAAGACACAGAAAAGTAT   2659
gb:MN985325|Organism:Severe      GGTACACCAGTTTGTATTAACGGGCTTATGTTGCTCGAAATCAAAGACACAGAAAAGTAC   2662
gb:MT020881|Organism:Severe      GGTACACCAGTTTGTATTAACGGGCTTATGTTGCTCGAAATCAAAGACACAGAAAAGTAC   2662
gb:MT020880|Organism:Severe      GGTACACCAGTTTGTATTAACGGGCTTATGTTGCTCGAAATCAAAGACACAGAAAAGTAC   2662
gb:MT066175|Organism:Severe      GGTACACCAGTTTGTATTAACGGGCTTATGTTGCTCGAAATCAAAGACACAGAAAAGTAC   2662
gb:MN997409|Organism:Severe      GGTACACCAGTTTGTATTAACGGGCTTATGTTGCTCGAAATCAAAGACACAGAAAAGTAC   2662
gb:MN938384|Organism:Severe      GGTACACCAGTTTGTATTAACGGGCTTATGTTGCTCGAAATCAAAGACACAGAAAAGTAC   2630
gb:MT044258|Organism:Severe      GGTACACCAGTTTGTATTAACGGGCTTATGTTGCTCGAAATCAAAGACACAGAAAAGTAC   2638
gb:MT039890|Organism:Severe      GGTACACCAGTTTGTATTAACGGGCTTATGTTGCTCGAAATCAAAGACACAGAAAAGTAC   2662
gb:MN988713|Organism:Severe      GGTACACCAGTTTGTATTAACGGGCTTATGTTGCTCGAAATCAAAGACACAGAAAAGTAC   2662
gb:LC521925|Organism:Severe      GGTACACCAGTTTGTATTAACGGGCTTATGTTGCTCGAAATCAAAGACACAGAAAAGTAC   2635
gb:MT093571|Organism:Severe      GGTACACCAGTTTGTATTAACGGGCTTATGTTGCTCGAAATCAAAGACACAGAAAAGTAC   2662
gb:MT039887|Organism:Severe      GGTACACCAGTTTGTATTAACGGGCTTATGTTGCTCGAAATCAAAGACACAGAAAAGTAC   2662
gb:MT019530|Organism:Severe      GGTACACCAGTTTGTATTAACGGGCTTATGTTGCTCGAAATCAAAGACACAGAAAAGTAC   2662
gb:MT039888|Organism:Severe      GGTACACCAGTTTGTATTAACGGGCTTATGTTGCTCGAAATCAAAGACACAGAAAAGTAC   2662
gb:LC522972|Organism:Severe      GGTACACCAGTTTGTATTAACGGGCTTATGTTGCTCGAAATCAAAGACACAGAAAAGTAC   2659
gb:MT027063|Organism:Severe      GGTACACCAGTTTGTATTAACGGGCTTATGTTGCTCGAAATCAAAGACACAGAAAAGTAC   2662
gb:MT027062|Organism:Severe      GGTACACCAGTTTGTATTAACGGGCTTATGTTGCTCGAAATCAAAGACACAGAAAAGTAC   2662
gb:MT019529|Organism:Severe      GGTACACCAGTTTGTATTAACGGGCTTATGTTGCTCGAAATCAAAGACACAGAAAAGTAC   2662
gb:MN996529|Organism:Severe      GGTACACCAGTTTGTATTAACGGGCTTATGTTGCTCGAAATCAAAGACACAGAAAAGTAC   2650
gb:MN996531|Organism:Severe      GGTACACCAGTTTGTATTAACGGGCTTATGTTGCTCGAAATCAAAGACACAGAAAAGTAC   2649
gb:MT066176|Organism:Severe      GGTACACCAGTTTGTATTAACGGGCTTATGTTGCTCGAAATCAAAGACACAGAAAAGTAC   2662
gb:MT027064|Organism:Severe      GGTACACCAGTTTGTATTAACGGGCTTATGTTGCTCGAAATCAAAGACACAGAAAAGTAC   2662
gb:MN994468|Organism:Severe      GGTACACCAGTTTGTATTAACGGGCTTATGTTGCTCGAAATCAAAGACACAGAAAAGTAC   2662
gb:MT072688|Organism:Severe      GGTACACCAGTTTGTATTAACGGGCTTATGTTGCTCGAAATCAAAGACACAGAAAAGTAC   2647
gb:MN996527|Organism:Severe      GGTACACCAGTTTGTATTAACGGGCTTATGTTGCTCGAAATCAAAGACACAGAAAAGTAC   2629
gb:MT093631|Organism:Severe      GGTACACCAGTTTGTATTAACGGGCTTATGTTGCTCGAAATCAAAGACACAGAAAAGTAC   2700
gb:MT106053|Organism:Severe      GGTACACCAGTTTGTATTAACGGGCTTATGTTGCTCGAAATCAAAGACACAGAAAAGTAC   2662
gb:MT019533|Organism:Severe      GGTACACCAGTTTGTATTAACGGGCTTATGTTGCTCGAAATCAAAGACACAGAAAAGTAC   2662
gb:MT019531|Organism:Severe      GGTACACCAGTTTGTATTAACGGGCTTATGTTGCTCGAAATCAAAGACACAGAAAAGTAC   2662
gb:MN996528|Organism:Severe      GGTACACCAGTTTGTATTAACGGGCTTATGTTGCTCGAAATCAAAGACACAGAAAAGTAC   2662
gb:MN996530|Organism:Severe      GGTACACCAGTTTGTATTAACGGGCTTATGTTGCTCGAAATCAAAGACACAGAAAAGTAC   2648
gb:MN908947|Organism:Severe      GGTACACCAGTTTGTATTAACGGGCTTATGTTGCTCGAAATCAAAGACACAGAAAAGTAC   2662
gb:MT019532|Organism:Severe      GGTACACCAGTTTGTATTAACGGGCTTATGTTGCTCGAAATCAAAGACACAGAAAAGTAC   2662
                                 ***********************************************************

gb:MT020781|Organism:Severe      TGTGCCCTTGCACCTAATATGATGGTAACAAACAATACCTTCACACTCAAAGGCGGTGCA   2710
gb:MT007544|Organism:Severe      TGTGCCCTTGCACCTAATATGATGGTAACAAACAATACCTTCACACTCAAAGGCGGTGCA   2722
gb:MN994467|Organism:Severe      TGTGCCCTTGCACCTAATATGATGGTAACAAACAATACCTTCACACTCAAAGGCGGTGCA   2722
gb:MT044257|Organism:Severe      TGTGCCCTTGCACCTAATATGATGGTAACAAACAATACCTTCACACTCAAAGGCGGTGCA   2722
gb:MT106054|Organism:Severe      TGTGCCCTTGCACCTAATATGATGGTAACAAACAATACCTTCACACTCAAAGGCGGTGCA   2722
gb:MT049951|Organism:Severe      TGTGCCCTTGCACCTAATATGATGGTAACAAACAATACCTTCACACTCAAAGGCGGTGCA   2722
```

```
gb:MN975262|Organism:Severe      TGTGCCCTTGCACCTAATATGATGGTAACAAACAATACCTTCACACTCAAAGGCGGTGCA   2722
gb:MT106052|Organism:Severe      TGTGCCCTTGCACCTAATATGATGGTAACAAACAATACCTTCACACTCAAAGGCGGTGCA   2722
gb:LC522975|Organism:Severe      TGTGCCCTTGCACCTAATATGATGGTAACAAACAATACCTTCACACTCAAAGGCGGTGCA   2719
gb:LC522973|Organism:Severe      TGTGCCCTTGCACCTAATATGATGGTAACAAACAATACCTTCACACTCAAAGGCGGTGCA   2719
gb:LC522974|Organism:Severe      TGTGCCCTTGCACCTAATATGATGGTAACAAACAATACCTTCACACTCAAAGGCGGTGCA   2719
gb:MN985325|Organism:Severe      TGTGCCCTTGCACCTAATATGATGGTAACAAACAATACCTTCACACTCAAAGGCGGTGCA   2722
gb:MT020881|Organism:Severe      TGTGCCCTTGCACCTAATATGATGGTAACAAACAATACCTTCACACTCAAAGGCGGTGCA   2722
gb:MT020880|Organism:Severe      TGTGCCCTTGCACCTAATATGATGGTAACAAACAATACCTTCACACTCAAAGGCGGTGCA   2722
gb:MT066175|Organism:Severe      TGTGCCCTTGCACCTAATATGATGGTAACAAACAATACCTTCACACTCAAAGGCGGTGCA   2722
gb:MN997409|Organism:Severe      TGTGCCCTTGCACCTAATATGATGGTAACAAACAATACCTTCACACTCAAAGGCGGTGCA   2722
gb:MN938384|Organism:Severe      TGTGCCCTTGCACCTAATATGATGGTAACAAACAATACCTTCACACTCAAAGGCGGTGCA   2690
gb:MT044258|Organism:Severe      TGTGCCCTTGCACCTAATATGATGGTAACAAACAATACCTTCACACTCAAAGGCGGTGCA   2698
gb:MT039890|Organism:Severe      TGTGCCCTTGCACCTAATATGATGGTAACAAACAATACCTTCACACTCAAAGGCGGTGCA   2722
gb:MN988713|Organism:Severe      TGTGCCCTTGCACCTAATATGATGGTAACAAACAATACCTTCACACTCAAAGGCGGTGCA   2722
gb:LC521925|Organism:Severe      TGTGCCCTTGCACCTAATATGATGGTAACAAACAATACCTTCACACTCAAAGGCGGTGCA   2695
gb:MT093571|Organism:Severe      TGTGCCCTTGCACCTAATATGATGGTAACAAACAATACCTTCACACTCAAAGGCAGTGCA   2722
gb:MT039887|Organism:Severe      TGTGCCCTTGCACCTAATATGATGGTAACAAACAATACCTTCACACTCAAAGGCGGTGCA   2722
gb:MT019530|Organism:Severe      TGTGCCCTTGCACCTAATATGATGGTAACAAACAATACCTTCACACTCAAAGGCGGTGCA   2722
gb:MT039888|Organism:Severe      TGTGCCCTTGCACCTAATATGATGGTAACAAACAATACCTTCACACTCAAAGGCGGTGCA   2722
gb:LC522972|Organism:Severe      TGTGCCCTTGCACCTAATATGATGGTAACAAACAATACCTTCACACTCAAAGGCGGTGCA   2719
gb:MT027063|Organism:Severe      TGTGCCCTTGCACCTAATATGATGGTAACAAACAATACCTTCACACTCAAAGGCGGTGCA   2722
gb:MT027062|Organism:Severe      TGTGCCCTTGCACCTAATATGATGGTAACAAACAATACCTTCACACTCAAAGGCGGTGCA   2722
gb:MT019529|Organism:Severe      TGTGCCCTTGCACCTAATATGATGGTAACAAACAATACCTTCACACTCAAAGGCGGTGCA   2722
gb:MN996529|Organism:Severe      TGTGCCCTTGCACCTAATATGATGGTAACAAACAATACCTTCACACTCAAAGGCGGTGCA   2710
gb:MN996531|Organism:Severe      TGTGCCCTTGCACCTAATATGATGGTAACAAACAATACCTTCACACTCAAAGGCGGTGCA   2709
gb:MT066176|Organism:Severe      TGTGCCCTTGCACCTAATATGATGGTAACAAACAATACCTTCACACTCAAAGGCGGTGCA   2722
gb:MT027064|Organism:Severe      TGTGCCCTTGCACCTAATATGATGGTAACAAACAATACCTTCACACTCAAAGGCGGTGCA   2722
gb:MN994468|Organism:Severe      TGTGCCCTTGCACCTAATATGATGGTAACAAACAATACCTTCACACTCAAAGGCGGTGCA   2722
gb:MT072688|Organism:Severe      TGTGCCCTTGCACCTAATATGATGGTAACAAACAATACCTTCACACTCAAAGGCGGTGCA   2707
gb:MN996527|Organism:Severe      TGTGCCCTTGCACCTAATATGATGGTAACAAACAATACCTTCACACTCAAAGGCGGTGCA   2689
gb:MT093631|Organism:Severe      TGTGCCCTTGCACCTAATATGATGGTAACAAACAATACCTTCACACTCAAAGGCGGTGCA   2760
gb:MT106053|Organism:Severe      TGTGCCCTTGCACCTAATATGATGGTAACAAACAATACCTTCACACTCAAAGGCGGTGCA   2722
gb:MT019533|Organism:Severe      TGTGCCCTTGCACCTAATATGATGGTAACAAACAATACCTTCACACTCAAAGGCGGTGCA   2722
gb:MT019531|Organism:Severe      TGTGCCCTTGCACCTAATATGATGGTAACAAACAATACCTTCACACTCAAAGGCGGTGCA   2722
gb:MN996528|Organism:Severe      TGTGCCCTTGCACCTAATATGATGGTAACAAACAATACCTTCACACTCAAAGGCGGTGCA   2722
gb:MN996530|Organism:Severe      TGTGCCCTTGCACCTAATATGATGGTAACAAACAATACCTTCACACTCAAAGGCGGTGCA   2708
gb:MN908947|Organism:Severe      TGTGCCCTTGCACCTAATATGATGGTAACAAACAATACCTTCACACTCAAAGGCGGTGCA   2722
gb:MT019532|Organism:Severe      TGTGCCCTTGCACCTAATATGATGGTAACAAACAATACCTTCACACTCAAAGGCGGTGCA   2722
                                 ****************************************************** *****

gb:MT020781|Organism:Severe      CCAACAAAGGTTACTTTTGGTGATGACACTGTGATAGAAGTGCAAGGTTACAAGAGTGTG   2770
gb:MT007544|Organism:Severe      CCAACAAAGGTTACTTTTGGTGATGACACTGTGATAGAAGTGCAAGGTTACAAGAGTGTG   2782
gb:MN994467|Organism:Severe      CCAACAAAGGTTACTTTTGGTGATGACACTGTGATAGAAGTGCAAGGTTACAAGAGTGTG   2782
gb:MT044257|Organism:Severe      CCAACAAAGGTTACTTTTGGTGATGACACTGTGATAGAAGTGCAAGGTTACAAGAGTGTG   2782
gb:MT106054|Organism:Severe      CCAACAAAGGTTACTTTTGGTGATGACACTGTGATAGAAGTGCAAGGTTACAAGAGTGTG   2782
gb:MT049951|Organism:Severe      CCAACAAAGGTTACTTTTGGTGATGACACTGTGATAGAAGTGCAAGGTTACAAGAGTGTG   2782
gb:MN975262|Organism:Severe      CCAACAAAGGTTACTTTTGGTGATGACACTGTGATAGAAGTGCAAGGTTACAAGAGTGTG   2782
gb:MT106052|Organism:Severe      CCAACAAAGGTTACTTTTGGTGATGACACTGTGATAGAAGTGCAAGGTTACAAGAGTGTG   2782
gb:LC522975|Organism:Severe      CCAACAAAGGTTACTTTTGGTGATGACACTGTGATAGAAGTGCAAGGTTACAAGAGTGTG   2779
gb:LC522973|Organism:Severe      CCAACAAAGGTTACTTTTGGTGATGACACTGTGATAGAAGTGCAAGGTTACAAGAGTGTG   2779
gb:LC522974|Organism:Severe      CCAACAAAGGTTACTTTTGGTGATGACACTGTGATAGAAGTGCAAGGTTACAAGAGTGTG   2779
gb:MN985325|Organism:Severe      CCAACAAAGGTTACTTTTGGTGATGACACTGTGATAGAAGTGCAAGGTTACAAGAGTGTG   2782
gb:MT020881|Organism:Severe      CCAACAAAGGTTACTTTTGGTGATGACACTGTGATAGAAGTGCAAGGTTACAAGAGTGTG   2782
gb:MT020880|Organism:Severe      CCAACAAAGGTTACTTTTGGTGATGACACTGTGATAGAAGTGCAAGGTTACAAGAGTGTG   2782
gb:MT066175|Organism:Severe      CCAACAAAGGTTACTTTTGGTGATGACACTGTGATAGAAGTGCAAGGTTACAAGAGTGTG   2782
gb:MN997409|Organism:Severe      CCAACAAAGGTTACTTTTGGTGATGACACTGTGATAGAAGTGCAAGGTTACAAGAGTGTG   2782
gb:MN938384|Organism:Severe      CCAACAAAGGTTACTTTTGGTGATGACACTGTGATAGAAGTGCAAGGTTACAAGAGTGTG   2750
gb:MT044258|Organism:Severe      CCAACAAAGGTTACTTTTGGTGATGACACTGTGATAGAAGTGCAAGGTTACAAGAGTGTG   2758
gb:MT039890|Organism:Severe      CCAACAAAGGTTACTTTTGGTGATGACACTGTGATAGAAGTGCAAGGTTACAAGAGTGTG   2782
gb:MN988713|Organism:Severe      CCAACAAAGGTTACTTTTGGTGATGACACTGTGATAGAAGTGCAAGGTTACAAGAGTGTG   2782
gb:LC521925|Organism:Severe      CCAACAAAGGTTACTTTTGGTGATGACACTGTGATAGAAGTGCAAGGTTACAAGAGTGTG   2755
gb:MT093571|Organism:Severe      CCAACAAAGGTTACTTTTGGTGATGACACTGTGATAGAAGTGCAAGGTTACAAGAGTGTG   2782
gb:MT039887|Organism:Severe      CCAACAAAGGTTACTTTTGGTGATGACACTGTGATAGAAGTGCAAGGTTACAAGAGTGTG   2782
gb:MT019530|Organism:Severe      CCAACAAAGGTTACTTTTGGTGATGACACTGTGATAGAAGTGCAAGGTTACAAGAGTGTG   2782
gb:MT039888|Organism:Severe      CCAACAAAGGTTACTTTTGGTGATGACACTGTGATAGAAGTGCAAGGTTACAAGAGTGTG   2782
```

```
gb:LC522972|Organism:Severe      CCAACAAAGGTTACTTTTGGTGATGACACTGTGATAGAAGTGCAAGGTTACAAGAGTGTG	2779
gb:MT027063|Organism:Severe      CCAACAAAGGTTACTTTTGGTGATGACACTGTGATAGAAGTGCAAGGTTACAAGAGTGTG	2782
gb:MT027062|Organism:Severe      CCAACAAAGGTTACTTTTGGTGATGACACTGTGATAGAAGTGCAAGGTTACAAGAGTGTG	2782
gb:MT019529|Organism:Severe      CCAACAAAGGTTACTTTTGGTGATGACACTGTGATAGAAGTGCAAGGTTACAAGAGTGTG	2782
gb:MN996529|Organism:Severe      CCAACAAAGGTTACTTTTGGTGATGACACTGTGATAGAAGTGCAAGGTTACAAGAGTGTG	2770
gb:MN996531|Organism:Severe      CCAACAAAGGTTACTTTTGGTGATGACACTGTGATAGAAGTGCAAGGTTACAAGAGTGTG	2769
gb:MT066176|Organism:Severe      CCAACAAAGGTTACTTTTGGTGATGACACTGTGATAGAAGTGCAAGGTTACAAGAGTGTG	2782
gb:MT027064|Organism:Severe      CCAACAAAGGTTACTTTTGGTGATGACACTGTGATAGAAGTGCAAGGTTACAAGAGTGTG	2782
gb:MN994468|Organism:Severe      CCAACAAAGGTTACTTTTGGTGATGACACTGTGATAGAAGTGCAAGGTTACAAGAGTGTG	2782
gb:MT072688|Organism:Severe      CCAACAAAGGTTACTTTTGGTGATGACACTGTGATAGAAGTGCAAGGTTACAAGAGTGTG	2767
gb:MN996527|Organism:Severe      CCAACAAAGGTTACTTTTGGTGATGACACTGTGATAGAAGTGCAAGGTTACAAGAGTGTG	2749
gb:MT093631|Organism:Severe      CCAACAAAGGTTACTTTTGGTGATGACACTGTGATAGAAGTGCAAGGTTACAAGAGTGTG	2820
gb:MT106053|Organism:Severe      CCAACAAAGGTTACTTTTGGTGATGACACTGTGATAGAAGTGCAAGGTTACAAGAGTGTG	2782
gb:MT019533|Organism:Severe      CCAACAAAGGTTACTTTTGGTGATGACACTGTGATAGAAGTGCAAGGTTACAAGAGTGTG	2782
gb:MT019531|Organism:Severe      CCAACAAAGGTTACTTTTGGTGATGACACTGTGATAGAAGTGCAAGGTTACAAGAGTGTG	2782
gb:MN996528|Organism:Severe      CCAACAAAGGTTACTTTTGGTGATGACACTGTGATAGAAGTGCAAGGTTACAAGAGTGTG	2782
gb:MN996530|Organism:Severe      CCAACAAAGGTTACTTTTGGTGATGACACTGTGATAGAAGTGCAAGGTTACAAGAGTGTG	2768
gb:MN908947|Organism:Severe      CCAACAAAGGTTACTTTTGGTGATGACACTGTGATAGAAGTGCAAGGTTACAAGAGTGTG	2782
gb:MT019532|Organism:Severe      CCAACAAAGGTTACTTTTGGTGATGACACTGTGATAGAAGTGCAAGGTTACAAGAGTGTG	2782
                                 ************************************************************

gb:MT020781|Organism:Severe      AATATCACTTTTGAACTTGATGAAAGGATTGATAAAGTACTTAATGAGAAGTGCTCTGCC	2830
gb:MT007544|Organism:Severe      AATATCACTTTTGAACTTGATGAAAGGATTGATAAAGTACTTAATGAGAAGTGCTCTGCC	2842
gb:MN994467|Organism:Severe      AATATCACTTTTGAACTTGATGAAAGGATTGATAAAGTACTTAATGAGAAGTGCTCTGCC	2842
gb:MT044257|Organism:Severe      AATATCACTTTTGAACTTGATGAAAGGATTGATAAAGTACTTAATGAGAAGTGCTCTGCC	2842
gb:MT106054|Organism:Severe      AATATCACTTTTGAACTTGATGAAAGGATTGATAAAGTACTTAATGAGAAGTGCTCTGCC	2842
gb:MT049951|Organism:Severe      AATATCACTTTTGAACTTGATGAAAGGATTGATAAAGTACTTAATGAGAAGTGCTCTGCC	2842
gb:MN975262|Organism:Severe      AATATCACTTTTGAACTTGATGAAAGGATTGATAAAGTACTTAATGAGAAGTGCTCTGCC	2842
gb:MT106052|Organism:Severe      AATATCACTTTTGAACTTGATGAAAGGATTGATAAAGTACTTAATGAGAAGTGCTCTGCC	2842
gb:LC522975|Organism:Severe      AATATCACTTTTGAACTTGATGAAAGGATTGATAAAGTACTTAATGAGAAGTGCTCTGCC	2839
gb:LC522973|Organism:Severe      AATATCACTTTTGAACTTGATGAAAGGATTGATAAAGTACTTAATGAGAAGTGCTCTGCC	2839
gb:LC522974|Organism:Severe      AATATCACTTTTGAACTTGATGAAAGGATTGATAAAGTACTTAATGAGAAGTGCTCTGCC	2839
gb:MN985325|Organism:Severe      AATATCACTTTTGAACTTGATGAAAGGATTGATAAAGTACTTAATGAGAAGTGCTCTGCC	2842
gb:MT020881|Organism:Severe      AATATCACTTTTGAACTTGATGAAAGGATTGATAAAGTACTTAATGAGAAGTGCTCTGCC	2842
gb:MT020880|Organism:Severe      AATATCACTTTTGAACTTGATGAAAGGATTGATAAAGTACTTAATGAGAAGTGCTCTGCC	2842
gb:MT066175|Organism:Severe      AATATCACTTTTGAACTTGATGAAAGGATTGATAAAGTACTTAATGAGAAGTGCTCTGCC	2842
gb:MN997409|Organism:Severe      AATATCACTTTTGAACTTGATGAAAGGATTGATAAAGTACTTAATGAGAAGTGCTCTGCC	2842
gb:MN938384|Organism:Severe      AATATCACTTTTGAACTTGATGAAAGGATTGATAAAGTACTTAATGAGAAGTGCTCTGCC	2810
gb:MT044258|Organism:Severe      AATATCACTTTTGAACTTGATGAAAGGATTGATAAAGTACTTAATGAGAAGTGCTCTGCC	2818
gb:MT039890|Organism:Severe      AATATCACTTTTGAACTTGATGAAAGGATTGATAAAGTACTTAATGAGAAGTGCTCTGCC	2842
gb:MN988713|Organism:Severe      AATATCACTTTTGAACTTGATGAAAGGATTGATAAAGTACTTAATGAGAAGTGCTCTGCC	2842
gb:LC521925|Organism:Severe      AATATCACTTTTGAACTTGATGAAAGGATTGATAAAGTACTTAATGAGAAGTGCTCTGCC	2815
gb:MT093571|Organism:Severe      AATATCACTTTTGAACTTGATGAAAGGATTGATAAAGTACTTAATGAGAAGTGCTCTGCC	2842
gb:MT039887|Organism:Severe      AATATCACTTTTGAACTTGATGAAAGGATTGATAAAGTACTTAATGAGAAGTGCTCTGCC	2842
gb:MT019530|Organism:Severe      AATATCACTTTTGAACTTGATGAAAGGATTGATAAAGTACTTAATGAGAAGTGCTCTGCC	2842
gb:MT039888|Organism:Severe      AATATCACTTTTGAACTTGATGAAAGGATTGATAAAGTACTTAATGAGAAGTGCTCTGCC	2842
gb:LC522972|Organism:Severe      AATATCACTTTTGAACTTGATGAAAGGATTGATAAAGTACTTAATGAGAAGTGCTCTGCC	2839
gb:MT027063|Organism:Severe      AATATCACTTTTGAACTTGATGAAAGGATTGATAAAGTACTTAATGAGAAGTGCTCTGCC	2842
gb:MT027062|Organism:Severe      AATATCACTTTTGAACTTGATGAAAGGATTGATAAAGTACTTAATGAGAAGTGCTCTGCC	2842
gb:MT019529|Organism:Severe      AATATCACTTTTGAACTTGATGAAAGGATTGATAAAGTACTTAATGAGAAGTGCTCTGCC	2842
gb:MN996529|Organism:Severe      AATATCACTTTTGAACTTGATGAAAGGATTGATAAAGTACTTAATGAGAAGTGCTCTGCC	2830
gb:MN996531|Organism:Severe      AATATCACTTTTGAACTTGATGAAAGGATTGATAAAGTACTTAATGAGAAGTGCTCTGCC	2829
gb:MT066176|Organism:Severe      AATATCACTTTTGAACTTGATGAAAGGATTGATAAAGTACTTAATGAGAAGTGCTCTGCC	2842
gb:MT027064|Organism:Severe      AATATCACTTTTGAACTTGATGAAAGGATTGATAAAGTACTTAATGAGAAGTGCTCTGCC	2842
gb:MN994468|Organism:Severe      AATATCACTTTTGAACTTGATGAAAGGATTGATAAAGTACTTAATGAGAAGTGCTCTGCC	2842
gb:MT072688|Organism:Severe      AATATCACTTTTGAACTTGATGAAAGGATTGATAAAGTACTTAATGAGAAGTGCTCTGCC	2827
gb:MN996527|Organism:Severe      AATATCACTTTTGAACTTGATGAAAGGATTGATAAAGTACTTAATGAGAAGTGCTCTGCC	2809
gb:MT093631|Organism:Severe      AATATCACTTTTGAACTTGATGAAAGGATTGATAAAGTACTTAATGAGAAGTGCTCTGCC	2880
gb:MT106053|Organism:Severe      AATATCACTTTTGAACTTGATGAAAGGATTGATAAAGTACTTAATGAGAAGTGCTCTGCC	2842
gb:MT019533|Organism:Severe      AATATCACTTTTGAACTTGATGAAAGGATTGATAAAGTACTTAATGAGAAGTGCTCTGCC	2842
gb:MT019531|Organism:Severe      AATATCACTTTTGAACTTGATGAAAGGATTGATAAAGTACTTAATGAGAAGTGCTCTGCC	2842
gb:MN996528|Organism:Severe      AATATCACTTTTGAACTTGATGAAAGGATTGATAAAGTACTTAATGAGAAGTGCTCTGCC	2842
gb:MN996530|Organism:Severe      AATATCACTTTTGAACTTGATGAAAGGATTGATAAAGTACTTAATGAGAAGTGCTCTGCC	2828
gb:MN908947|Organism:Severe      AATATCACTTTTGAACTTGATGAAAGGATTGATAAAGTACTTAATGAGAAGTGCTCTGCC	2842
gb:MT019532|Organism:Severe      AATATCACTTTTGAACTTGATGAAAGGATTGATAAAGTACTTAATGAGAAGTGCTCTGCC	2842
```

```
                                   ************************************************************

gb:MT020781|Organism:Severe        TATACAGTTGAACTCGGTACAGAAGTAAATGAGTTCGCCTGTGTTGTGGCAGATGCTGTC    2890
gb:MT007544|Organism:Severe        TATACAGTTGAACTCGGTACAGAAGTAAATGAGTTCGCCTGTGTTGTGGCAGATGCTGTC    2902
gb:MN994467|Organism:Severe        TATACAGTTGAACTCGGTACAGAAGTAAATGAGTTCGCCTGTGTTGTGGCAGATGCTGTC    2902
gb:MT044257|Organism:Severe        TATACAGTTGAACTCGGTACAGAAGTAAATGAGTTCGCCTGTGTTGTGGCAGATGCTGTC    2902
gb:MT106054|Organism:Severe        TATACAGTTGAACTCGGTACAGAAGTAAATGAGTTCGCCTGTGTTGTGGCAGATGCTGTC    2902
gb:MT049951|Organism:Severe        TATACAGTTGAACTCGGTACAGAAGTAAATGAGTTCGCCTGTGTTGTGGCAGATGCTGTC    2902
gb:MN975262|Organism:Severe        TATACAGTTGAACTCGGTACAGAAGTAAATGAGTTCGCCTGTGTTGTGGCAGATGCTGTC    2902
gb:MT106052|Organism:Severe        TATACAGTTGAACTCGGTACAGAAGTAAATGAGTTCGCCTGTGTTGTGGCAGATGCTGTC    2902
gb:LC522975|Organism:Severe        TATACAGTTGAACTCGGTACAGAAGTAAATGAGTTCGCCTGTGTTGTGGCAGATGCTGTC    2899
gb:LC522973|Organism:Severe        TATACAGTTGAACTCGGTACAGAAGTAAATGAGTTCGCCTGTGTTGTGGCAGATGCTGTC    2899
gb:LC522974|Organism:Severe        TATACAGTTGAACTCGGTACAGAAGTAAATGAGTTCGCCTGTGTTGTGGCAGATGCTGTC    2899
gb:MN985325|Organism:Severe        TATACAGTTGAACTCGGTACAGAAGTAAATGAGTTCGCCTGTGTTGTGGCAGATGCTGTC    2902
gb:MT020881|Organism:Severe        TATACAGTTGAACTCGGTACAGAAGTAAATGAGTTCGCCTGTGTTGTGGCAGATGCTGTC    2902
gb:MT020880|Organism:Severe        TATACAGTTGAACTCGGTACAGAAGTAAATGAGTTCGCCTGTGTTGTGGCAGATGCTGTC    2902
gb:MT066175|Organism:Severe        TATACAGTTGAACTCGGTACAGAAGTAAATGAGTTCGCCTGTGTTGTGGCAGATGCTGTC    2902
gb:MN997409|Organism:Severe        TATACAGTTGAACTCGGTACAGAAGTAAATGAGTTCGCCTGTGTTGTGGCAGATGCTGTC    2902
gb:MN938384|Organism:Severe        TATACAGTTGAACTCGGTACAGAAGTAAATGAGTTCGCCTGTGTTGTGGCAGATGCTGTC    2870
gb:MT044258|Organism:Severe        TATACAGTTGAACTCGGTACAGAAGTAAATGAGTTCGCCTGTGTTGTGGCAGATGCTGTC    2878
gb:MT039890|Organism:Severe        TATACAGTTGAACTCGGTACAGAAGTAAATGAGTTCGCCTGTGTTGTGGCAGATGCTGTC    2902
gb:MN988713|Organism:Severe        TATACAGTTGAACTCGGTACAGAAGTAAATGAGTTCGCCTGTGTTGTGGCAGATGCTGTC    2902
gb:LC521925|Organism:Severe        TATACAGTTGAACTCGGTACAGAAGTAAATGAGTTCGCCTGTGTTGTGGCAGATGCTGTC    2875
gb:MT093571|Organism:Severe        TATACAGTTGAACTCGGTACAGAAGTAAATGAGTTCGCCTGTGTTGTGGCAGATGCTGTC    2902
gb:MT039887|Organism:Severe        TATACAGTTGAACTCGGTACAGAAGTAAATGAGTTCGCCTGTGTTGTGGCAGATGCTGTC    2902
gb:MT019530|Organism:Severe        TATACAGTTGAACTCGGTACAGAAGTAAATGAGTTCGCCTGTGTTGTGGCAGATGCTGTC    2902
gb:MT039888|Organism:Severe        TATACAGTTGAACTCGGTACAGAAGTAAATGAGTTCGCCTGTGTTGTGGCAGATGCTGTC    2902
gb:LC522972|Organism:Severe        TATACAGTTGAACTCGGTACAGAAGTAAATGAGTTCGCCTGTGTTGTGGCAGATGCTGTC    2899
gb:MT027063|Organism:Severe        TATACAGTTGAACTCGGTACAGAAGTAAATGAGTTCGCCTGTGTTGTGGCAGATGCTGTC    2902
gb:MT027062|Organism:Severe        TATACAGTTGAACTCGGTACAGAAGTAAATGAGTTCGCCTGTGTTGTGGCAGATGCTGTC    2902
gb:MT019529|Organism:Severe        TATACAGTTGAACTCGGTACAGAAGTAAATGAGTTCGCCTGTGTTGTGGCAGATGCTGTC    2902
gb:MN996529|Organism:Severe        TATACAGTTGAACTCGGTACAGAAGTAAATGAGTTCGCCTGTGTTGTGGCAGATGCTGTC    2890
gb:MN996531|Organism:Severe        TATACAGTTGAACTCGGTACAGAAGTAAATGAGTTCGCCTGTGTTGTGGCAGATGCTGTC    2889
gb:MT066176|Organism:Severe        TATACAGTTGAACTCGGTACAGAAGTAAATGAGTTCGCCTGTGTTGTGGCAGATGCTGTC    2902
gb:MT027064|Organism:Severe        TATACAGTTGAACTCGGTACAGAAGTAAATGAGTTCGCCTGTGTTGTGGCAGATGCTGTC    2902
gb:MN994468|Organism:Severe        TATACAGTTGAACTCGGTACAGAAGTAAATGAGTTCGCCTGTGTTGTGGCAGATGCTGTC    2902
gb:MT072688|Organism:Severe        TATACAGTTGAACTCGGTACAGAAGTAAATGAGTTCGCCTGTGTTGTGGCAGATGCTGTC    2887
gb:MN996527|Organism:Severe        TATACAGTTGAACTCGGTACAGAAGTAAATGAGTTCGCCTGTGTTGTGGCAGATGCTGTC    2869
gb:MT093631|Organism:Severe        TATACAGTTGAACTCGGTACAGAAGTAAATGAGTTCGCCTGTGTTGTGGCAGATGCTGTC    2940
gb:MT106053|Organism:Severe        TATACAGTTGAACTCGGTACAGAAGTAAATGAGTTCGCCTGTGTTGTGGCAGATGCTGTC    2902
gb:MT019533|Organism:Severe        TATACAGTTGAACTCGGTACAGAAGTAAATGAGTTCGCCTGTGTTGTGGCAGATGCTGTC    2902
gb:MT019531|Organism:Severe        TATACAGTTGAACTCGGTACAGAAGTAAATGAGTTCGCCTGTGTTGTGGCAGATGCTGTC    2902
gb:MN996528|Organism:Severe        TATACAGTTGAACTCGGTACAGAAGTAAATGAGTTCGCCTGTGTTGTGGCAGATGCTGTC    2902
gb:MN996530|Organism:Severe        TATACAGTTGAACTCGGTACAGAAGTAAATGAGTTCGCCTGTGTTGTGGCAGATGCTGTC    2888
gb:MN908947|Organism:Severe        TATACAGTTGAACTCGGTACAGAAGTAAATGAGTTCGCCTGTGTTGTGGCAGATGCTGTC    2902
gb:MT019532|Organism:Severe        TATACAGTTGAACTCGGTACAGAAGTAAATGAGTTCGCCTGTGTTGTGGCAGATGCTGTC    2902
                                   ************************************************************

gb:MT020781|Organism:Severe        ATAAAAACTTTGCAACCAGTATCTGAATTACTTACACCACTGGGCATTGATTTAGATGAG    2950
gb:MT007544|Organism:Severe        ATAAAAACTTTGCAACCAGTATCTGAATTACTTACACCACTGGGCATTGATTTAGATGAG    2962
gb:MN994467|Organism:Severe        ATAAAAACTTTGCAACCAGTATCTGAATTACTTACACCACTGGGCATTGATTTAGATGAG    2962
gb:MT044257|Organism:Severe        ATAAAAACTTTGCAACCAGTATCTGAATTACTTACACCACTGGGCATTGATTTAGATGAG    2962
gb:MT106054|Organism:Severe        ATAAAAACTTTGCAACCAGTATCTGAATTACTTACACCACTGGGCATTGATTTAGATGAG    2962
gb:MT049951|Organism:Severe        ATAAAAACTTTGCAACCAGTATCTGAATTACTTACACCACTGGGCATTGATTTAGATGAG    2962
gb:MN975262|Organism:Severe        ATAAAAACTTTGCAACCAGTATCTGAATTACTTACACCACTGGGCATTGATTTAGATGAG    2962
gb:MT106052|Organism:Severe        ATAAAAACTTTGCAACCAGTATCTGAATTACTTACACCACTGGGCATTGATTTAGATGAG    2962
gb:LC522975|Organism:Severe        ATAAAAACTTTGCAACCAGTATCTGAATTACTTACACCACTGGGCATTGATTTAGATGAG    2959
gb:LC522973|Organism:Severe        ATAAAAACTTTGCAACCAGTATCTGAATTACTTACACCACTGGGCATTGATTTAGATGAG    2959
gb:LC522974|Organism:Severe        ATAAAAACTTTGCAACCAGTATCTGAATTACTTACACCACTGGGCATTGATTTAGATGAG    2959
gb:MN985325|Organism:Severe        ATAAAAACTTTGCAACCAGTATCTGAATTACTTACACCACTGGGCATTGATTTAGATGAG    2962
gb:MT020881|Organism:Severe        ATAAAAACTTTGCAACCAGTATCTGAATTACTTACACCACTGGGCATTGATTTAGATGAG    2962
gb:MT020880|Organism:Severe        ATAAAAACTTTGCAACCAGTATCTGAATTACTTACACCACTGGGCATTGATTTAGATGAG    2962
gb:MT066175|Organism:Severe        ATAAAAACTTTGCAACCAGTATCTGAATTACTTACACCACTGGGCATTGATTTAGATGAG    2962
gb:MN997409|Organism:Severe        ATAAAAACTTTGCAACCAGTATCTGAATTACTTACACCACTGGGCATTGATTTAGATGAG    2962
gb:MN938384|Organism:Severe        ATAAAAACTTTGCAACCAGTATCTGAATTACTTACACCACTGGGCATTGATTTAGATGAG    2930
```

```
gb:MT044258|Organism:Severe      ATAAAAACTTTGCAACCAGTATCTGAATTACTTACACCACTGGGCATTGATTTAGATGAG    2938
gb:MT039890|Organism:Severe      ATAAAAACTTTGCAACCAGTATCTGAATTACTTACACCACTGGGCATTGATTTAGATGAG    2962
gb:MN988713|Organism:Severe      ATAAAAACTTTGCAACCAGTATCTGAATTACTTACACCACTGGGCATTGATTTAGATGAG    2962
gb:LC521925|Organism:Severe      ATAAAAACTTTGCAACCAGTATCTGAATTACTTACACCACTGGGCATTGATTTAGATGAG    2935
gb:MT093571|Organism:Severe      ATAAAAACTTTGCAACCAGTATCTGAATTACTTACACCACTGGGCATTGATTTAGATGAG    2962
gb:MT039887|Organism:Severe      ATAAAAACTTTGCAACCAGTATCTGAATTACTTACACCACTGGGCATTGATTTAGATGAG    2962
gb:MT019530|Organism:Severe      ATAAAAACTTTGCAACCAGTATCTGAATTACTTACACCACTGGGCATTGATTTAGATGAG    2962
gb:MT039888|Organism:Severe      ATAAAAACTTTGCAACCAGTATCTGAATTACTTACACCACTGGGCATTGATTTAGATGAG    2962
gb:LC522972|Organism:Severe      ATAAAAACTTTGCAACCAGTATCTGAATTACTTACACCACTGGGCATTGATTTAGATGAG    2959
gb:MT027063|Organism:Severe      ATAAAAACTTTGCAACCAGTATCTGAATTACTTACACCACTGGGCATTGATTTAGATGAG    2962
gb:MT027062|Organism:Severe      ATAAAAACTTTGCAACCAGTATCTGAATTACTTACACCACTGGGCATTGATTTAGATGAG    2962
gb:MT019529|Organism:Severe      ATAAAAACTTTGCAACCAGTATCTGAATTACTTACACCACTGGGCATTGATTTAGATGAG    2962
gb:MN996529|Organism:Severe      ATAAAAACTTTGCAACCAGTATCTGAATTACTTACACCACTGGGCATTGATTTAGATGAG    2950
gb:MN996531|Organism:Severe      ATAAAAACTTTGCAACCAGTATCTGAATTACTTACACCACTGGGCATTGATTTAGATGAG    2949
gb:MT066176|Organism:Severe      ATAAAAACTTTGCAACCAGTATCTGAATTACTTACACCACTGGGCATTGATTTAGATGAG    2962
gb:MT027064|Organism:Severe      ATAAAAACTTTGCAACCAGTATCTGAATTACTTACACCACTGGGCATTGATTTAGATGAG    2962
gb:MN994468|Organism:Severe      ATAAAAACTTTGCAACCAGTATCTGAATTACTTACACCACTGGGCATTGATTTAGATGAG    2962
gb:MT072688|Organism:Severe      ATAAAAACTTTGCAACCAGTATCTGAATTACTTACACCACTGGGCATTGATTTAGATGAG    2947
gb:MN996527|Organism:Severe      ATAAAAACTTTGCAACCAGTATCTGAATTACTTACACCACTGGGCATTGATTTAGATGAG    2929
gb:MT093631|Organism:Severe      ATAAAAACTTTGCAACCAGTATCTGAATTACTTACACCACTGGGCATTGATTTAGATGAG    3000
gb:MT106053|Organism:Severe      ATAAAAACTTTGCAACCAGTATCTGAATTACTTACACCACTGGGCATTGATTTAGATGAG    2962
gb:MT019533|Organism:Severe      ATAAAAACTTTGCAACCAGTATCTGAATTACTTACACCACTGGGCATTGATTTAGATGAG    2962
gb:MT019531|Organism:Severe      ATAAAAACTTTGCAACCAGTATCTGAATTACTTACACCACTGGGCATTGATTTAGATGAG    2962
gb:MN996528|Organism:Severe      ATAAAAACTTTGCAACCAGTATCTGAATTACTTACACCACTGGGCATTGATTTAGATGAG    2962
gb:MN996530|Organism:Severe      ATAAAAACTTTGCAACCAGTATCTGAATTACTTACACCACTGGGCATTGATTTAGATGAG    2948
gb:MN908947|Organism:Severe      ATAAAAACTTTGCAACCAGTATCTGAATTACTTACACCACTGGGCATTGATTTAGATGAG    2962
gb:MT019532|Organism:Severe      ATAAAAACTTTGCAACCAGTATCTGAATTACTTACACCACTGGGCATTGATTTAGATGAG    2962
                                 ************************************************************

gb:MT020781|Organism:Severe      TGGAGTATGGCTACATACTACTTATTTGATGAGTCTGGTGAGTTTAAATTGGCTTCACAT    3010
gb:MT007544|Organism:Severe      TGGAGTATGGCTACATACTACTTATTTGATGAGTCTGGTGAGTTTAAATTGGCTTCACAT    3022
gb:MN994467|Organism:Severe      TGGAGTATGGCTACATACTACTTATTTGATGAGTCTGGTGAGTTTAAATTGGCTTCACAT    3022
gb:MT044257|Organism:Severe      TGGAGTATGGCTACATACTACTTATTTGATGAGTCTGGTGAGTTTAAATTGGCTTCACAT    3022
gb:MT106054|Organism:Severe      TGGAGTATGGCTACATACTACTTATTTGATGAGTCTGGTGAGTTTAAATTGGCTTCACAT    3022
gb:MT049951|Organism:Severe      TGGAGTATGGCTACATACTACTTATTTGATGAGTCTGGTGAGTTTAAATTGGCTTCACAT    3022
gb:MN975262|Organism:Severe      TGGAGTATGGCTACATACTACTTATTTGATGAGTCTGGTGAGTTTAAATTGGCTTCACAT    3022
gb:MT106052|Organism:Severe      TGGAGTATGGCTACATACTACTTATTTGATGAGTCTGGTGAGTTTAAATTGGCTTCACAT    3022
gb:LC522975|Organism:Severe      TGGAGTATGGCTACATACTACTTATTTGATGAGTCTGGTGAGTTTAAATTGGCTTCACAT    3019
gb:LC522973|Organism:Severe      TGGAGTATGGCTACATACTACTTATTTGATGAGTCTGGTGAGTTTAAATTGGCTTCACAT    3019
gb:LC522974|Organism:Severe      TGGAGTATGGCTACATACTACTTATTTGATGAGTCTGGTGAGTTTAAATTGGCTTCACAT    3019
gb:MN985325|Organism:Severe      TGGAGTATGGCTACATACTACTTATTTGATGAGTCTGGTGAGTTTAAATTGGCTTCACAT    3022
gb:MT020881|Organism:Severe      TGGAGTATGGCTACATACTACTTATTTGATGAGTCTGGTGAGTTTAAATTGGCTTCACAT    3022
gb:MT020880|Organism:Severe      TGGAGTATGGCTACATACTACTTATTTGATGAGTCTGGTGAGTTTAAATTGGCTTCACAT    3022
gb:MT066175|Organism:Severe      TGGAGTATGGCTACATACTACTTATTTGATGAGTCTGGTGAGTTTAAATTGGCTTCACAT    3022
gb:MN997409|Organism:Severe      TGGAGTATGGCTACATACTACTTATTTGATGAGTCTGGTGAGTTTAAATTGGCTTCACAT    3022
gb:MN938384|Organism:Severe      TGGAGTATGGCTACATACTACTTATTTGATGAGTCTGGTGAGTTTAAATTGGCTTCACAT    2990
gb:MT044258|Organism:Severe      TGGAGTATGGCTACATACTACTTATTTGATGAGTCTGGTGAGTTTAAATTGGCTTCACAT    2998
gb:MT039890|Organism:Severe      TGGAGTATTGCTACATACTACTTATTTGATGAGTCTGGTGAGTTTAAATTGGCTTCACAT    3022
gb:MN988713|Organism:Severe      TGGAGTATGGCTACATACTACTTATTTGATGAGTCTGGTGAGTTTAAATTGGCTTCACAT    3022
gb:LC521925|Organism:Severe      TGGAGTATGGCTACATACTACTTATTTGATGAGTCTGGTGAGTTTAAATTGGCTTCACAT    2995
gb:MT093571|Organism:Severe      TGGAGTATGGCTACATACTACTTATTTGATGAGTCTGGTGAGTTTAAATTGGCTTCACAT    3022
gb:MT039887|Organism:Severe      TGGAGTATGGCTACATACTACTTATTTGATGAGTCTGGTGAGTTTAAATTGGCTTCACAT    3022
gb:MT019530|Organism:Severe      TGGAGTATGGCTACATACTACTTATTTGATGAGTCTGGTGAGTTTAAATTGGCTTCACAT    3022
gb:MT039888|Organism:Severe      TGGAGTATGGCTACATACTACTTATTTGATGAGTCTGGTGAGTTTAAATTGGCTTCACAT    3022
gb:LC522972|Organism:Severe      TGGAGTATGGCTACATACTACTTATTTGATGAGTCTGGTGAGTTTAAATTGGCTTCACAT    3019
gb:MT027063|Organism:Severe      TGGAGTATGGCTACATACTACTTATTTGATGAGTCTGGTGAGTTTAAATTGGCTTCACAT    3022
gb:MT027062|Organism:Severe      TGGAGTATGGCTACATACTACTTATTTGATGAGTCTGGTGAGTTTAAATTGGCTTCACAT    3022
gb:MT019529|Organism:Severe      TGGAGTATGGCTACATACTACTTATTTGATGAGTCTGGTGAGTTTAAATTGGCTTCACAT    3022
gb:MN996529|Organism:Severe      TGGAGTATGGCTACATACTACTTATTTGATGAGTCTGGTGAGTTTAAATTGGCTTCACAT    3010
gb:MN996531|Organism:Severe      TGGAGTATGGCTACATACTACTTATTTGATGAGTCTGGTGAGTTTAAATTGGCTTCACAT    3009
gb:MT066176|Organism:Severe      TGGAGTATGGCTACATACTACTTATTTGATGAGTCTGGTGAGTTTAAATTGGCTTCACAT    3022
gb:MT027064|Organism:Severe      TGGAGTATGGCTACATACTACTTATTTGATGAGTCTGGTGAGTTTAAATTGGCTTCACAT    3022
gb:MN994468|Organism:Severe      TGGAGTATGGCTACATACTACTTATTTGATGAGTCTGGTGAGTTTAAATTGGCTTCACAT    3022
gb:MT072688|Organism:Severe      TGGAGTATGGCTACATACTACTTATTTGATGAGTCTGGTGAGTTTAAATTGGCTTCACAT    3007
gb:MN996527|Organism:Severe      TGGAGTATGGCTACATACTACTTATTTGATGAGTCTGGTGAGTTTAAATTGGCTTCACAT    2989
```

```
gb:MT093631|Organism:Severe      TGGAGTATGGCTACATACTACTTATTTGATGAGTCTGGTGAGTTTAAATTGGCTTCACAT   3060
gb:MT106053|Organism:Severe      TGGAGTATGGCTACATACTACTTATTTGATGAGTCTGGTGAGTTTAAATTGGCTTCACAT   3022
gb:MT019533|Organism:Severe      TGGAGTATGGCTACATACTACTTATTTGATGAGTCTGGTGAGTTTAAATTGGCTTCACAT   3022
gb:MT019531|Organism:Severe      TGGAGTATGGCTACATACTACTTATTTGATGAGTCTGGTGAGTTTAAATTGGCTTCACAT   3022
gb:MN996528|Organism:Severe      TGGAGTATGGCTACATACTACTTATTTGATGAGTCTGGTGAGTTTAAATTGGCTTCACAT   3022
gb:MN996530|Organism:Severe      TGGAGTATGGCTACATACTACTTATTTGATGAGTCTGGTGAGTTTAAATTGGCTTCACAT   3008
gb:MN908947|Organism:Severe      TGGAGTATGGCTACATACTACTTATTTGATGAGTCTGGTGAGTTTAAATTGGCTTCACAT   3022
gb:MT019532|Organism:Severe      TGGAGTATGGCTACATACTACTTATTTGATGAGTCTGGTGAGTTTAAATTGGCTTCACAT   3022
                                 ******** ***************************************************

gb:MT020781|Organism:Severe      ATGTATTGTTCTTTCTACCCTCCAGATGAGGATGAAGAAGAAGGTGATTGTGAAGAAGAA   3070
gb:MT007544|Organism:Severe      ATGTATTGTTCTTTCTACCCTCCAGATGAGGATGAAGAAGAAGGTGATTGTGAAGAAGAA   3082
gb:MN994467|Organism:Severe      ATGTATTGTTCTTTCTACCCTCCAGATGAGGATGAAGAAGAAGGTGATTGTGAAGAAGAA   3082
gb:MT044257|Organism:Severe      ATGTATTGTTCTTTCTACCCTCCAGATGAGGATGAAGAAGAAGGTGATTGTGAAGAAGAA   3082
gb:MT106054|Organism:Severe      ATGTATTGTTCTTTCTACCCTCCAGATGAGGATGAAGAAGAAGGTGATTGTGAAGAAGAA   3082
gb:MT049951|Organism:Severe      ATGTATTGTTCTTTCTACCCTCCAGATGAGGATGAAGAAGAAGGTGATTGTGAAGAAGAA   3082
gb:MN975262|Organism:Severe      ATGTATTGTTCTTTCTACCCTCCAGATGAGGATGAAGAAGAAGGTGATTGTGAAGAAGAA   3082
gb:MT106052|Organism:Severe      ATGTATTGTTCTTTCTACCCTCCAGATGAGGATGAAGAAGAAGGTGATTGTGAAGAAGAA   3082
gb:LC522975|Organism:Severe      ATGTATTGTTCTTTCTACCCTCCAGATGAGGATGAAGAAGAAGGTGATTGTGAAGAAGAA   3079
gb:LC522973|Organism:Severe      ATGTATTGTTCTTTCTACCCTCCAGATGAGGATGAAGAAGAAGGTGATTGTGAAGAAGAA   3079
gb:LC522974|Organism:Severe      ATGTATTGTTCTTTCTACCCTCCAGATGAGGATGAAGAAGAAGGTGATTGTGAAGAAGAA   3079
gb:MN985325|Organism:Severe      ATGTATTGTTCTTTCTACCCTCCAGATGAGGATGAAGAAGAAGGTGATTGTGAAGAAGAA   3082
gb:MT020881|Organism:Severe      ATGTATTGTTCTTTCTACCCTCCAGATGAGGATGAAGAAGAAGGTGATTGTGAAGAAGAA   3082
gb:MT020880|Organism:Severe      ATGTATTGTTCTTTCTACCCTCCAGATGAGGATGAAGAAGAAGGTGATTGTGAAGAAGAA   3082
gb:MT066175|Organism:Severe      ATGTATTGTTCTTTCTACCCTCCAGATGAGGATGAAGAAGAAGGTGATTGTGAAGAAGAA   3082
gb:MN997409|Organism:Severe      ATGTATTGTTCTTTCTACCCTCCAGATGAGGATGAAGAAGAAGGTGATTGTGAAGAAGAA   3082
gb:MN938384|Organism:Severe      ATGTATTGTTCTTTCTACCCTCCAGATGAGGATGAAGAAGAAGGTGATTGTGAAGAAGAA   3050
gb:MT044258|Organism:Severe      ATGTATTGTTCTTTCTACCCTCCAGATGAGGATGAAGAAGAAGGTGATTGTGAAGAAGAA   3058
gb:MT039890|Organism:Severe      ATGTATTGTTCTTTCTACCCTCCAGATGAGGATGAAGAAGAAGGTGATTGTGAAGAAGAA   3082
gb:MN988713|Organism:Severe      ATGTATTGTTCTTTCTACCCTCCAGATGAGGATGAAGAAGAAGGTGATTGTGAAGAAGAA   3082
gb:LC521925|Organism:Severe      ATGTATTGTTCTTTCTACCCTCCAGATGAGGATGAAGAAGAAGGTGATTGTGAAGAAGAA   3055
gb:MT093571|Organism:Severe      ATGTATTGTTCTTTCTACCCTCCAGATGAGGATGAAGAAGAAGGTGATTGTGAAGAAGAA   3082
gb:MT039887|Organism:Severe      ATGTATTGTTCTTTCTACCCTCCAGATGAGGATGAAGAAGAAGGTGATTGTGAAGAAGAA   3082
gb:MT019530|Organism:Severe      ATGTATTGTTCTTTCTACCCTCCAGATGAGGATGAAGAAGAAGGTGATTGTGAAGAAGAA   3082
gb:MT039888|Organism:Severe      ATGTATTGTTCTTTCTACCCTCCAGATGAGGATGAAGAAGAAGGTGATTGTGAAGAAGAA   3082
gb:LC522972|Organism:Severe      ATGTATTGTTCTTTCTACCCTCCAGATGAGGATGAAGAAGAAGGTGATTGTGAAGAAGAA   3079
gb:MT027063|Organism:Severe      ATGTATTGTTCTTTCTACCCTCCAGATGAGGATGAAGAAGAAGGTGATTGTGAAGAAGAA   3082
gb:MT027062|Organism:Severe      ATGTATTGTTCTTTCTACCCTCCAGATGAGGATGAAGAAGAAGGTGATTGTGAAGAAGAA   3082
gb:MT019529|Organism:Severe      ATGTATTGTTCTTTCTACCCTCCAGATGAGGATGAAGAAGAAGGTGATTGTGAAGAAGAA   3082
gb:MN996529|Organism:Severe      ATGTATTGTTCTTTCTACCCTCCAGATGAGGATGAAGAAGAAGGTGATTGTGAAGAAGAA   3070
gb:MN996531|Organism:Severe      ATGTATTGTTCTTTCTACCCTCCAGATGAGGATGAAGAAGAAGGTGATTGTGAAGAAGAA   3069
gb:MT066176|Organism:Severe      ATGTATTGTTCTTTCTACCCTCCAGATGAGGATGAAGAAGAAGGTGATTGTGAAGAAGAA   3082
gb:MT027064|Organism:Severe      ATGTATTGTTCTTTCTACCCTCCAGATGAGGATGAAGAAGAAGGTGATTGTGAAGAAGAA   3082
gb:MN994468|Organism:Severe      ATGTATTGTTCTTTCTACCCTCCAGATGAGGATGAAGAAGAAGGTGATTGTGAAGAAGAA   3082
gb:MT072688|Organism:Severe      ATGTATTGTTCTTTCTACCCTCCAGATGAGGATGAAGAAGAAGGTGATTGTGAAGAAGAA   3067
gb:MN996527|Organism:Severe      ATGTATTGTTCTTTCTACCCTCCAGATGAGGATGAAGAAGAAGGTGATTGTGAAGAAGAA   3049
gb:MT093631|Organism:Severe      ATGTATTGTTCTTTCTACCCTCCAGATGAGGATGAAGAAGAAGGTGATTGTGAAGAAGAA   3120
gb:MT106053|Organism:Severe      ATGTATTGTTCTTTCTACCCTCCAGATGAGGATGAAGAAGAAGGTGATTGTGAAGAAGAA   3082
gb:MT019533|Organism:Severe      ATGTATTGTTCTTTCTACCCTCCAGATGAGGATGAAGAAGAAGGTGATTGTGAAGAAGAA   3082
gb:MT019531|Organism:Severe      ATGTATTGTTCTTTCTACCCTCCAGATGAGGATGAAGAAGAAGGTGATTGTGAAGAAGAA   3082
gb:MN996528|Organism:Severe      ATGTATTGTTCTTTCTACCCTCCAGATGAGGATGAAGAAGAAGGTGATTGTGAAGAAGAA   3082
gb:MN996530|Organism:Severe      ATGTATTGTTCTTTCTACCCTCCAGATGAGGATGAAGAAGAAGGTGATTGTGAAGAAGAA   3068
gb:MN908947|Organism:Severe      ATGTATTGTTCTTTCTACCCTCCAGATGAGGATGAAGAAGAAGGTGATTGTGAAGAAGAA   3082
gb:MT019532|Organism:Severe      ATGTATTGTTCTTTCTACCCTCCAGATGAGGATGAAGAAGAAGGTGATTGTGAAGAAGAA   3082
                                 ************************************************************

gb:MT020781|Organism:Severe      GAGTTTGAGCCATCAACTCAATATGAGTATGGTACTGAAGATGATTACCAAGGTAAACCT   3130
gb:MT007544|Organism:Severe      GAGTTTGAGCCATCAACTCAATATGAGTATGGTACTGAAGATGATTACCAAGGTAAACCT   3142
gb:MN994467|Organism:Severe      GAGTTTGAGCCATCAACTCAATATGAGTATGGTACTGAAGATGATTACCAAGGTAAACCT   3142
gb:MT044257|Organism:Severe      GAGTTTGAGCCATCAACTCAATATGAGTATGGTACTGAAGATGATTACCAAGGTAAACCT   3142
gb:MT106054|Organism:Severe      GAGTTTGAGCCATCAACTCAATATGAGTATGGTACTGAAGATGATTACCAAGGTAAACCT   3142
gb:MT049951|Organism:Severe      GAGTTTGAGCCATCAACTCAATATGAGTATGGTACTGAAGATGATTACCAAGGTAAACCT   3142
gb:MN975262|Organism:Severe      GAGTTTGAGCCATCAACTCAATATGAGTATGGTACTGAAGATGATTACCAAGGTAAACCT   3142
gb:MT106052|Organism:Severe      GAGTTTGAGCCATCAACTCAATATGAGTATGGTACTGAAGATGATTACCAAGGTAAACCT   3142
gb:LC522975|Organism:Severe      GAGTTTGAGCCATCAACTCAATATGAGTATGGTACTGAAGATGATTACCAAGGTAAACCT   3139
```

```
gb:LC522973|Organism:Severe      GAGTTTGAGCCATCAACTCAATATGAGTATGGTACTGAAGATGATTACCAAGGTAAACCT   3139
gb:LC522974|Organism:Severe      GAGTTTGAGCCATCAACTCAATATGAGTATGGTACTGAAGATGATTACCAAGGTAAACCT   3139
gb:MN985325|Organism:Severe      GAGTTTGAGCCATCAACTCAATATGAGTATGGTACTGAAGATGATTACCAAGGTAAACCT   3142
gb:MT020881|Organism:Severe      GAGTTTGAGCCATCAACTCAATATGAGTATGGTACTGAAGATGATTACCAAGGTAAACCT   3142
gb:MT020880|Organism:Severe      GAGTTTGAGCCATCAACTCAATATGAGTATGGTACTGAAGATGATTACCAAGGTAAACCT   3142
gb:MT066175|Organism:Severe      GAGTTTGAGCCATCAACTCAATATGAGTATGGTACTGAAGATGATTACCAAGGTAAACCT   3142
gb:MN997409|Organism:Severe      GAGTTTGAGCCATCAACTCAATATGAGTATGGTACTGAAGATGATTACCAAGGTAAACCT   3142
gb:MN938384|Organism:Severe      GAGTTTGAGCCATCAACTCAATATGAGTATGGTACTGAAGATGATTACCAAGGTAAACCT   3110
gb:MT044258|Organism:Severe      GAGTTTGAGCCATCAACTCAATATGAGTATGGTACTGAAGATGATTACCAAGGTAAACCT   3118
gb:MT039890|Organism:Severe      GAGTTTGAGCCATCAACTCAATATGAGTATGGTACTGAAGATGATTACCAAGGTAAACCT   3142
gb:MN988713|Organism:Severe      GAGTTTGAGCCATCAACTCAATATGAGTATGGTACTGAAGATGATTACCAAGGTAAACCT   3142
gb:LC521925|Organism:Severe      GAGTTTGAGCCATCAACTCAATATGAGTATGGTACTGAAGATGATTACCAAGGTAAACCT   3115
gb:MT093571|Organism:Severe      GAGTTTGAGCCATCAACTCAATATGAGTATGGTACTGAAGATGATTACCAAGGTAAACCT   3142
gb:MT039887|Organism:Severe      GAGTTTGAGCCATCAACTCAATATGAGTATGGTACTGAAGATGATTACCAAGGTAAACCT   3142
gb:MT019530|Organism:Severe      GAGTTTGAGCCATCAACTCAATATGAGTATGGTACTGAAGATGATTACCAAGGTAAACCT   3142
gb:MT039888|Organism:Severe      GAGTTTGAGCCATCAACTCAATATGAGTATGGTACTGAAGATGATTACCAAGGTAAACCT   3142
gb:LC522972|Organism:Severe      GAGTTTGAGCCATCAACTCAATATGAGTATGGTACTGAAGATGATTACCAAGGTAAACCT   3139
gb:MT027063|Organism:Severe      GAGTTTGAGCCATCAACTCAATATGAGTATGGTACTGAAGATGATTACCAAGGTAAACCT   3142
gb:MT027062|Organism:Severe      GAGTTTGAGCCATCAACTCAATATGAGTATGGTACTGAAGATGATTACCAAGGTAAACCT   3142
gb:MT019529|Organism:Severe      GAGTTTGAGCCATCAACTCAATATGAGTATGGTACTGAAGATGATTACCAAGGTAAACCT   3142
gb:MN996529|Organism:Severe      GAGTTTGAGCCATCAACTCAATATGAGTATGGTACTGAAGATGATTACCAAGGTAAACCT   3130
gb:MN996531|Organism:Severe      GAGTTTGAGCCATCAACTCAATATGAGTATGGTACTGAAGATGATTACCAAGGTAAACCT   3129
gb:MT066176|Organism:Severe      GAGTTTGAGCCATCAACTCAATATGAGTATGGTACTGAAGATGATTACCAAGGTAAACCT   3142
gb:MT027064|Organism:Severe      GAGTTTGAGCCATCAACTCAATATGAGTATGGTACTGAAGATGATTACCAAGGTAAACCT   3142
gb:MN994468|Organism:Severe      GAGTTTGAGCCATCAACTCAATATGAGTATGGTACTGAAGATGATTACCAAGGTAAACCT   3142
gb:MT072688|Organism:Severe      GAGTTTGAGCCATCAACTCAATATGAGTATGGTACTGAAGATGATTACCAAGGTAAACCT   3127
gb:MN996527|Organism:Severe      GAGTTTGAGCCATCAACTCAATATGAGTATGGTACTGAAGATGATTACCAAGGTAAACCT   3109
gb:MT093631|Organism:Severe      GAGTTTGAGCCATCAACTCAATATGAGTATGGTACTGAAGATGATTACCAAGGTAAACCT   3180
gb:MT106053|Organism:Severe      GAGTTTGAGCCATCAACTCAATATGAGTATGGTACTGAAGATGATTACCAAGGTAAACCT   3142
gb:MT019533|Organism:Severe      GAGTTTGAGCCATCAACTCAATATGAGTATGGTACTGAAGATGATTACCAAGGTAAACCT   3142
gb:MT019531|Organism:Severe      GAGTTTGAGCCATCAACTCAATATGAGTATGGTACTGAAGATGATTACCAAGGTAAACCT   3142
gb:MN996528|Organism:Severe      GAGTTTGAGCCATCAACTCAATATGAGTATGGTACTGAAGATGATTACCAAGGTAAACCT   3142
gb:MN996530|Organism:Severe      GAGTTTGAGCCATCAACTCAATATGAGTATGGTACTGAAGATGATTACCAAGGTAAACCT   3128
gb:MN908947|Organism:Severe      GAGTTTGAGCCATCAACTCAATATGAGTATGGTACTGAAGATGATTACCAAGGTAAACCT   3142
gb:MT019532|Organism:Severe      GAGTTTGAGCCATCAACTCAATATGAGTATGGTACTGAAGATGATTACCAAGGTAAACCT   3142
                                 ************************************************************

gb:MT020781|Organism:Severe      TTGGAATTTGGTGCCACTTCTGCTGCTCTTCAACCTGAAGAAGAGCAAGAAGAAGATTGG   3190
gb:MT007544|Organism:Severe      TTGGAATTTGGTGCCACTTCTGCTGCTCTTCAACCTGAAGAAGAGCAAGAAGAAGATTGG   3202
gb:MN994467|Organism:Severe      TTGGAATTTGGTGCCACTTCTGCTGCTCTTCAACCTGAAGAAGAGCAAGAAGAAGATTGG   3202
gb:MT044257|Organism:Severe      TTGGAATTTGGTGCCACTTCTGCTGCTCTTCAACTTGAAGAAGAGCAAGAAGAAGATTGG   3202
gb:MT106054|Organism:Severe      TTGGAATTTGGTGCCACTTCTGCTGCTCTTCAACCTGAAGAAGAGCAAGAAGAAGATTGG   3202
gb:MT049951|Organism:Severe      TTGGAATTTGGTGCCACTTCTGCTGCTCTTCAACCTGAAGAAGAGCAAGAAGAAGATTGG   3202
gb:MN975262|Organism:Severe      TTGGAATTTGGTGCCACTTCTGCTGCTCTTCAACCTGAAGAAGAGCAAGAAGAAGATTGG   3202
gb:MT106052|Organism:Severe      TTGGAATTTGGTGCCACTTCTGCTGCTCTTCAACCTGAAGAAGAGCAAGAAGAAGATTGG   3202
gb:LC522975|Organism:Severe      TTGGAATTTGGTGCCACTTCTGCTGCTCTTCAACCTGAAGAAGAGCAAGAAGAAGATTGG   3199
gb:LC522973|Organism:Severe      TTGGAATTTGGTGCCACTTCTGCTGCTCTTCAACCTGAAGAAGAGCAAGAAGAAGATTGG   3199
gb:LC522974|Organism:Severe      TTGGAATTTGGTGCCACTTCTGCTGCTCTTCAACCTGAAGAAGAGCAAGAAGAAGATTGG   3199
gb:MN985325|Organism:Severe      TTGGAATTTGGTGCCACTTCTGCTGCTCTTCAACCTGAAGAAGAGCAAGAAGAAGATTGG   3202
gb:MT020881|Organism:Severe      TTGGAATTTGGTGCCACTTCTGCTGCTCTTCAACCTGAAGAAGAGCAAGAAGAAGATTGG   3202
gb:MT020880|Organism:Severe      TTGGAATTTGGTGCCACTTCTGCTGCTCTTCAACCTGAAGAAGAGCAAGAAGAAGATTGG   3202
gb:MT066175|Organism:Severe      TTGGAATTTGGTGCCACTTCTGCTGCTCTTCAACCTGAAGAAGAGCAAGAAGAAGATTGG   3202
gb:MN997409|Organism:Severe      TTGGAATTTGGTGCCACTTCTGCTGCTCTTCAACCTGAAGAAGAGCAAGAAGAAGATTGG   3202
gb:MN938384|Organism:Severe      TTGGAATTTGGTGCCACTTCTGCTGCTCTTCAACCTGAAGAAGAGCAAGAAGAAGATTGG   3170
gb:MT044258|Organism:Severe      TTGGAATTTGGTGCCACTTCTGCTGCTCTTCAACCTGAAGAAGAGCAAGAAGAAGATTGG   3178
gb:MT039890|Organism:Severe      TTGGAATTTGGTGCCACTTCTGCTGCTCTTCAACCTGAAGAAGAGCAAGAAGAAGATTGG   3202
gb:MN988713|Organism:Severe      TTGGAATTTGGTGCCACTTCTGCTGCTCTTCAACYTGAAGAAGAGCAAGAAGAAGATTGG   3202
gb:LC521925|Organism:Severe      TTGGAATTTGGTGCCACTTCTGCTGCTCTTCAACCTGAAGAAGAGCAAGAAGAAGATTGG   3175
gb:MT093571|Organism:Severe      TTGGAATTTGGTGCCACTTCTGCTGCTCTTCAACCTGAAGAAGAGCAAGAAGAAGATTGG   3202
gb:MT039887|Organism:Severe      TTGGAATTTGGTGCCACTTCTGCTGCTCTTCAACCTGAAGAAGAGCAAGAAGAAGATTGG   3202
gb:MT019530|Organism:Severe      TTGGAATTTGGTGCCACTTCTGCTGCTCTTCAACCTGAAGAAGAGCAAGAAGAAGATTGG   3202
gb:MT039888|Organism:Severe      TTGGAATTTGGTGCCACTTCTGCTGCTCTTCAACCTGAAGAAGAGCAAGAAGAAGATTGG   3202
gb:LC522972|Organism:Severe      TTGGAATTTGGTGCCACTTCTGCTGCTCTTCAACCTGAAGAAGAGCAAGAAGAAGATTGG   3199
gb:MT027063|Organism:Severe      TTGGAATTTGGTGCCACTTCTGCTGCTCTTCAACCTGAAGAAGAGCAAGAAGAAGATTGG   3202
gb:MT027062|Organism:Severe      TTGGAATTTGGTGCCACTTCTGCTGCTCTTCAACCTGAAGAAGAGCAAGAAGAAGATTGG   3202
```

```
gb:MT019529|Organism:Severe      TTGGAATTTGGTGCCACTTCTGCTGCTCTTCAACCTGAAGAAGAGCAAGAAGAAGATTGG   3202
gb:MN996529|Organism:Severe      TTGGAATTTGGTGCCACTTCTGCTGCTCTTCAACCTGAAGAAGAGCAAGAAGAAGATTGG   3190
gb:MN996531|Organism:Severe      TTGGAATTTGGTGCCACTTCTGCTGCTCTTCAACCTGAAGAAGAGCAAGAAGAAGATTGG   3189
gb:MT066176|Organism:Severe      TTGGAATTTGGTGCCACTTCTGCTGCTCTTCAACCTGAAGAAGAGCAAGAAGAAGATTGG   3202
gb:MT027064|Organism:Severe      TTGGAATTTGGTGCCACTTCTGCTGCTCTTCAACCTGAAGAAGAGCAAGAAGAAGATTGG   3202
gb:MN994468|Organism:Severe      TTGGAATTTGGTGCCACTTCTGCTGCTCTTCAACCTGAAGAAGAGCAAGAAGAAGATTGG   3202
gb:MT072688|Organism:Severe      TTGGAATTTGGTGCCACTTCTGCTGCTCTTCAACCTGAAGAAGAGCAAGAAGAAGATTGG   3187
gb:MN996527|Organism:Severe      TTGGAATTTGGTGCCACTTCTGCTGCTCTTCAACCTGAAGAAGAGCAAGAAGAAGATTGG   3169
gb:MT093631|Organism:Severe      TTGGAATTTGGTGCCACTTCTGCTGCTCTTCAACCTGAAGAAGAGCAAGAAGAAGATTGG   3240
gb:MT106053|Organism:Severe      TTGGAATTTGGTGCCACTTCTGCTGCTCTTCAACCTGAAGAAGAGCAAGAAGAAGATTGG   3202
gb:MT019533|Organism:Severe      TTGGAATTTGGTGCCACTTCTGCTGCTCTTCAACCTGAAGAAGAGCAAGAAGAAGATTGG   3202
gb:MT019531|Organism:Severe      TTGGAATTTGGTGCCACTTCTGCTGCTCTTCAACCTGAAGAAGAGCAAGAAGAAGATTGG   3202
gb:MN996528|Organism:Severe      TTGGAATTTGGTGCCACTTCTGCTGCTCTTCAACCTGAAGAAGAGCAAGAAGAAGATTGG   3202
gb:MN996530|Organism:Severe      TTGGAATTTGGTGCCACTTCTGCTGCTCTTCAACCTGAAGAAGAGCAAGAAGAAGATTGG   3188
gb:MN908947|Organism:Severe      TTGGAATTTGGTGCCACTTCTGCTGCTCTTCAACCTGAAGAAGAGCAAGAAGAAGATTGG   3202
gb:MT019532|Organism:Severe      TTGGAATTTGGTGCCACTTCTGCTGCTCTTCAACCTGAAGAAGAGCAAGAAGAAGATTGG   3202
                                 ********************************** *************************

gb:MT020781|Organism:Severe      TTAGATGATGATAGTCAACAAACTGTTGGTCAACAAGACGGCAGTGAGGACAATCAGACA   3250
gb:MT007544|Organism:Severe      TTAGATGATGATAGTCAACAAACTGTTGGTCAACAAGACGGCAGTGAGGACAATCAGACA   3262
gb:MN994467|Organism:Severe      TTAGATGATGATAGTCAACAAACTGTTGGTCAACAAGACGGCAGTGAGGACAATCAGACA   3262
gb:MT044257|Organism:Severe      TTAGATGATGATAGTCAACAAACTGTTGGTCAACAAGACGGCAGTGAGGACAATCAGACA   3262
gb:MT106054|Organism:Severe      TTAGATGATGATAGTCAACAAACTGTTGGTCAACAAGACGGCAGTGAGGACAATCAGACA   3262
gb:MT049951|Organism:Severe      TTAGATGATGATAGTCAACAAACTGTTGGTCAACAAGACGGCAGTGAGGACAATCAGACA   3262
gb:MN975262|Organism:Severe      TTAGATGATGATAGTCAACAAACTGTTGGTCAACAAGACGGCAGTGAGGACAATCAGACA   3262
gb:MT106052|Organism:Severe      TTAGATGATGATAGTCAACAAACTGTTGGTCAACAAGACGGCAGTGAGGACAATCAGACA   3262
gb:LC522975|Organism:Severe      TTAGATGATGATAGTCAACAAACTGTTGGTCAACAAGACGGCAGTGAGGACAATCAGACA   3259
gb:LC522973|Organism:Severe      TTAGATGATGATAGTCAACAAACTGTTGGTCAACAAGACGGCAGTGAGGACAATCAGACA   3259
gb:LC522974|Organism:Severe      TTAGATGATGATAGTCAACAAACTGTTGGTCAACAAGACGGCAGTGAGGACAATCAGACA   3259
gb:MN985325|Organism:Severe      TTAGATGATGATAGTCAACAAACTGTTGGTCAACAAGACGGCAGTGAGGACAATCAGACA   3262
gb:MT020881|Organism:Severe      TTAGATGATGATAGTCAACAAACTGTTGGTCAACAAGACGGCAGTGAGGACAATCAGACA   3262
gb:MT020880|Organism:Severe      TTAGATGATGATAGTCAACAAACTGTTGGTCAACAAGACGGCAGTGAGGACAATCAGACA   3262
gb:MT066175|Organism:Severe      TTAGATGATGATAGTCAACAAACTGTTGGTCAACAAGACGGCAGTGAGGACAATCAGACA   3262
gb:MN997409|Organism:Severe      TTAGATGATGATAGTCAACAAACTGTTGGTCAACAAGACGGCAGTGAGGACAATCAGACA   3262
gb:MN938384|Organism:Severe      TTAGATGATGATAGTCAACAAACTGTTGGTCAACAAGACGGCAGTGAGGACAATCAGACA   3230
gb:MT044258|Organism:Severe      TTAGATGATGATAGTCAACAAACTGTTGGTCAACAAGACGGCAGTGAGGACAATCAGACA   3238
gb:MT039890|Organism:Severe      TTAGATGATGATAGTCAACAAACTGTTGGTCAACAAGACGGCAGTGAGGACAATCAGACA   3262
gb:MN988713|Organism:Severe      TTAGATGATGATAGTCAACAAACTGTTGGTCAACAAGACGGCAGTGAGGACAATCAGACA   3262
gb:LC521925|Organism:Severe      TTAGATGATGATAGTCAACAAACTGTTGGTCAACAAGACGGCAGTGAGGACAATCAGACA   3235
gb:MT093571|Organism:Severe      TTAGATGATGATAGTCAACAAACTGTTGGTCAACAAGACGGCAGTGAGGACAATCAGACA   3262
gb:MT039887|Organism:Severe      TTAGATGATGATAGTCAACAAACTGTTGGTCAACAAGACGGCAGTGAGGACAATCAGACA   3262
gb:MT019530|Organism:Severe      TTAGATGATGATAGTCAACAAACTGTTGGTCAACAAGACGGCAGTGAGGACAATCAGACA   3262
gb:MT039888|Organism:Severe      TTAGATGATGATAGTCAACAAACTGTTGGTCAACAAGACGGCAGTGAGGACAATCAGACA   3262
gb:LC522972|Organism:Severe      TTAGATGATGATAGTCAACAAACTGTTGGTCAACAAGACGGCAGTGAGGACAATCAGACA   3259
gb:MT027063|Organism:Severe      TTAGATGATGATAGTCAACAAACTGTTGGTCAACAAGACGGCAGTGAGGACAATCAGACA   3262
gb:MT027062|Organism:Severe      TTAGATGATGATAGTCAACAAACTGTTGGTCAACAAGACGGCAGTGAGGACAATCAGACA   3262
gb:MT019529|Organism:Severe      TTAGATGATGATAGTCAACAAACTGTTGGTCAACAAGACGGCAGTGAGGACAATCAGACA   3262
gb:MN996529|Organism:Severe      TTAGATGATGATAGTCAACAAACTGTTGGTCAACAAGACGGCAGTGAGGACAATCAGACA   3250
gb:MN996531|Organism:Severe      TTAGATGATGATAGTCAACAAACTGTTGGTCAACAAGACGGCAGTGAGGACAATCAGACA   3249
gb:MT066176|Organism:Severe      TTAGATGATGATAGTCAACAAACTGTTGGTCAACAAGACGGCAGTGAGGACAATCAGACA   3262
gb:MT027064|Organism:Severe      TTAGATGATGATAGTCAACAAACTGTTGGTCAACAAGACGGCAGTGAGGACAATCAGACA   3262
gb:MN994468|Organism:Severe      TTAGATGATGATAGTCAACAAACTGTTGGTCAACAAGACGGCAGTGAGGACAATCAGACA   3262
gb:MT072688|Organism:Severe      TTAGATGATGATAGTCAACAAACTGTTGGTCAACAAGACGGCAGTGAGGACAATCAGACA   3247
gb:MN996527|Organism:Severe      TTAGATGATGATAGTCAACAAACTGTTGGTCAACAAGACGGCAGTGAGGACAATCAGACA   3229
gb:MT093631|Organism:Severe      TTAGATGATGATAGTCAACAAACTGTTGGTCAACAAGACGGCAGTGAGGACAATCAGACA   3300
gb:MT106053|Organism:Severe      TTAGATGATGATAGTCAACAAACTGTTGGTCAACAAGACGGCAGTGAGGACAATCAGACA   3262
gb:MT019533|Organism:Severe      TTAGATGATGATAGTCAACAAACTGTTGGTCAACAAGACGGCAGTGAGGACAATCAGACA   3262
gb:MT019531|Organism:Severe      TTAGATGATGATAGTCAACAAACTGTTGGTCAACAAGACGGCAGTGAGGACAATCAGACA   3262
gb:MN996528|Organism:Severe      TTAGATGATGATAGTCAACAAACTGTTGGTCAACAAGACGGCAGTGAGGACAATCAGACA   3262
gb:MN996530|Organism:Severe      TTAGATGATGATAGTCAACAAACTGTTGGTCAACAAGACGGCAGTGAGGACAATCAGACA   3248
gb:MN908947|Organism:Severe      TTAGATGATGATAGTCAACAAACTGTTGGTCAACAAGACGGCAGTGAGGACAATCAGACA   3262
gb:MT019532|Organism:Severe      TTAGATGATGATAGTCAACAAACTGTTGGTCAACAAGACGGCAGTGAGGACAATCAGACA   3262
                                 ************************************************************

gb:MT020781|Organism:Severe      ACTACTATTCAAACAATTGTTGAGGTTCAACCTCAATTAGAGATGGAACTTACACCAGTT   3310
```

```
gb:MT007544|Organism:Severe      ACTACTATTCAAACAATTGTTGAGGTTCAACCTCAATTAGAGATGGAACTTACACCAGTT   3322
gb:MN994467|Organism:Severe      ACTACTATTCAAACAATTGTTGAGGTTCAACCTCAATTAGAGATGGAACTTACACCAGTT   3322
gb:MT044257|Organism:Severe      ACTACTATTCAAACAATTGTTGAGGTTCAACCTCAATTAGAGATGGAACTTACACCAGTT   3322
gb:MT106054|Organism:Severe      ACTACTATTCAAACAATTGTTGAGGTTCAACCTCAATTAGAGATGGAACTTACACCAGTT   3322
gb:MT049951|Organism:Severe      ACTACTATTCAAACAATTGTTGAGGTTCAACCTCAATTAGAGATGGAACTTACACCAGTT   3322
gb:MN975262|Organism:Severe      ACTACTATTCAAACAATTGTTGAGGTTCAACCTCAATTAGAGATGGAACTTACACCAGTT   3322
gb:MT106052|Organism:Severe      ACTACTATTCAAACAATTGTTGAGGTTCAACCTCAATTAGAGATGGAACTTACACCAGTT   3322
gb:LC522975|Organism:Severe      ACTACTATTCAAACAATTGTTGAGGTTCAACCTCAATTAGAGATGGAACTTACACCAGTT   3319
gb:LC522973|Organism:Severe      ACTACTATTCAAACAATTGTTGAGGTTCAACCTCAATTAGAGATGGAACTTACACCAGTT   3319
gb:LC522974|Organism:Severe      ACTACTATTCAAACAATTGTTGAGGTTCAACCTCAATTAGAGATGGAACTTACACCAGTT   3319
gb:MN985325|Organism:Severe      ACTACTATTCAAACAATTGTTGAGGTTCAACCTCAATTAGAGATGGAACTTACACCAGTT   3322
gb:MT020881|Organism:Severe      ACTACTATTCAAACAATTGTTGAGGTTCAACCTCAATTAGAGATGGAACTTACACCAGTT   3322
gb:MT020880|Organism:Severe      ACTACTATTCAAACAATTGTTGAGGTTCAACCTCAATTAGAGATGGAACTTACACCAGTT   3322
gb:MT066175|Organism:Severe      ACTACTATTCAAACAATTGTTGAGGTTCAACCTCAATTAGAGATGGAACTTACACCAGTT   3322
gb:MN997409|Organism:Severe      ACTACTATTCAAACAATTGTTGAGGTTCAACCTCAATTAGAGATGGAACTTACACCAGTT   3322
gb:MN938384|Organism:Severe      ACTACTATTCAAACAATTGTTGAGGTTCAACCTCAATTAGAGATGGAACTTACACCAGTT   3290
gb:MT044258|Organism:Severe      ACTACTATTCAAACAATTGTTGAGGTTCAACCTCAATTAGAGATGGAACTTACACCAGTT   3298
gb:MT039890|Organism:Severe      ACTACTATTCAAACAATTGTTGAGGTTCAACCTCAATTAGAGATGGAACTTACACCAGTT   3322
gb:MN988713|Organism:Severe      ACTACTATTCAAACAATTGTTGAGGTTCAACCTCAATTAGAGATGGAACTTACACCAGTT   3322
gb:LC521925|Organism:Severe      ACTACTATTCAAACAATTGTTGAGGTTCAACCTCAATTAGAGATGGAACTTACACCAGTT   3295
gb:MT093571|Organism:Severe      ACTACTATTCAAACAATTGTTGAGGTTCAACCTCAATTAGAGATGGAACTTACACCAGTT   3322
gb:MT039887|Organism:Severe      ACTACTATTCAAACAATTGTTGAGGTTCAACCTCAATTAGAGATGGAACTTACACCAGTT   3322
gb:MT019530|Organism:Severe      ACTACTATTCAAACAATTGTTGAGGTTCAACCTCAATTAGAGATGGAACTTACACCAGTT   3322
gb:MT039888|Organism:Severe      ACTACTATTCAAACAATTGTTGAGGTTCAACCTCAATTAGAGATGGAACTTACACCAGTT   3322
gb:LC522972|Organism:Severe      ACTACTATTCAAACAATTGTTGAGGTTCAACCTCAATTAGAGATGGAACTTACACCAGTT   3319
gb:MT027063|Organism:Severe      ACTACTATTCAAACAATTGTTGAGGTTCAACCTCAATTAGAGATGGAACTTACACCAGTT   3322
gb:MT027062|Organism:Severe      ACTACTATTCAAACAATTGTTGAGGTTCAACCTCAATTAGAGATGGAACTTACACCAGTT   3322
gb:MT019529|Organism:Severe      ACTACTATTCAAACAATTGTTGAGGTTCAACCTCAATTAGAGATGGAACTTACACCAGTT   3322
gb:MN996529|Organism:Severe      ACTACTATTCAAACAATTGTTGAGGTTCAACCTCAATTAGAGATGGAACTTACACCAGTT   3310
gb:MN996531|Organism:Severe      ACTACTATTCAAACAATTGTTGAGGTTCAACCTCAATTAGAGATGGAACTTACACCAGTT   3309
gb:MT066176|Organism:Severe      ACTACTATTCAAACAATTGTTGAGGTTCAACCTCAATTAGAGATGGAACTTACACCAGTT   3322
gb:MT027064|Organism:Severe      ACTACTATTCAAACAATTGTTGAGGTTCAACCTCAATTAGAGATGGAACTTACACCAGTT   3322
gb:MN994468|Organism:Severe      ACTACTATTCAAACAATTGTTGAGGTTCAACCTCAATTAGAGATGGAACTTACACCAGTT   3322
gb:MT072688|Organism:Severe      ACTACTATTCAAACAATTGTTGAGGTTCAACCTCAATTAGAGATGGAACTTACACCAGTT   3307
gb:MN996527|Organism:Severe      ACTACTATTCAAACAATTGTTGAGGTTCAACCTCAATTAGAGATGGAACTTACACCAGTT   3289
gb:MT093631|Organism:Severe      ACTACTATTCAAACAATTGTTGAGGTTCAACCTCAATTAGAGATGGAACTTACACCAGTT   3360
gb:MT106053|Organism:Severe      ACTACTATTCAAACAATTGTTGAGGTTCAACCTCAATTAGAGATGGAACTTACACCAGTT   3322
gb:MT019533|Organism:Severe      ACTACTATTCAAACAATTGTTGAGGTTCAACCTCAATTAGAGATGGAACTTACACCAGTT   3322
gb:MT019531|Organism:Severe      ACTACTATTCAAACAATTGTTGAGGTTCAACCTCAATTAGAGATGGAACTTACACCAGTT   3322
gb:MN996528|Organism:Severe      ACTACTATTCAAACAATTGTTGAGGTTCAACCTCAATTAGAGATGGAACTTACACCAGTT   3322
gb:MN996530|Organism:Severe      ACTACTATTCAAACAATTGTTGAGGTTCAACCTCAATTAGAGATGGAACTTACACCAGTT   3308
gb:MN908947|Organism:Severe      ACTACTATTCAAACAATTGTTGAGGTTCAACCTCAATTAGAGATGGAACTTACACCAGTT   3322
gb:MT019532|Organism:Severe      ACTACTATTCAAACAATTGTTGAGGTTCAACCTCAATTAGAGATGGAACTTACACCAGTT   3322
                                 ************************************************************

gb:MT020781|Organism:Severe      GTTCAGACTATTGAAGTGAATAGTTTAGTGGTTATTTAAAACTTACTGACAATGTATAC   3370
gb:MT007544|Organism:Severe      GTTCAGACTATTGAAGTGAATAGTTTAGTGGTTATTTAAAACTTACTGACAATGTATAC   3382
gb:MN994467|Organism:Severe      GTTCAGACTATTGAAGTGAATAGTTTAGTGGTTATTTAAAACTTACTGACAATGTATAC   3382
gb:MT044257|Organism:Severe      GTTCAGACTATTGAAGTGAATAGTTTAGTGGTTATTTAAAACTTACTGACAATGTATAC   3382
gb:MT106054|Organism:Severe      GTTCAGACTATTGAAGTGAATAGTTTAGTGGTTATTTAAAACTTACTGACAATGTATAC   3382
gb:MT049951|Organism:Severe      GTTCAGACTATTGAAGTGAATAGTTTAGTGGTTATTTAAAACTTACTGACAATGTATAC   3382
gb:MN975262|Organism:Severe      GTTCAGACTATTGAAGTGAATAGTTTAGTGGTTATTTAAAACTTACTGACAATGTATAC   3382
gb:MT106052|Organism:Severe      GTTCAGACTATTGAAGTGAATAGTTTAGTGGTTATTTAAAACTTACTGACAATGTATAC   3382
gb:LC522975|Organism:Severe      GTTCAGACTATTGAAGTGAATAGTTTAGTGGTTATTTAAAACTTACTGACAATGTATAC   3379
gb:LC522973|Organism:Severe      GTTCAGACTATTGAAGTGAATAGTTTAGTGGTTATTTAAAACTTACTGACAATGTATAC   3379
gb:LC522974|Organism:Severe      GTTCAGACTATTGAAGTGAATAGTTTAGTGGTTATTTAAAACTTACTGACAATGTATAC   3379
gb:MN985325|Organism:Severe      GTTCAGACTATTGAAGTGAATAGTTTAGTGGTTATTTAAAACTTACTGACAATGTATAC   3382
gb:MT020881|Organism:Severe      GTTCAGACTATTGAAGTGAATAGTTTAGTGGTTATTTAAAACTTACTGACAATGTATAC   3382
gb:MT020880|Organism:Severe      GTTCAGACTATTGAAGTGAATAGTTTAGTGGTTATTTAAAACTTACTGACAATGTATAC   3382
gb:MT066175|Organism:Severe      GTTCAGACTATTGAAGTGAATAGTTTAGTGGTTATTTAAAACTTACTGACAATGTATAC   3382
gb:MN997409|Organism:Severe      GTTCAGACTATTGAAGTGAATAGTTTAGTGGTTATTTAAAACTTACTGACAATGTATAC   3382
gb:MN938384|Organism:Severe      GTTCAGACTATTGAAGTGAATAGTTTAGTGGTTATTTAAAACTTACTGACAATGTATAC   3350
gb:MT044258|Organism:Severe      GTTCAGACTATTGAAGTGAATAGTTTAGTGGTTATTTAAAACTTACTGACAATGTATAC   3358
gb:MT039890|Organism:Severe      GTTCAGACTATTGAAGTGAATAGTTTAGTGGTTATTTAAAACTTACTGACAATGTATAC   3382
gb:MN988713|Organism:Severe      GTTCAGACTATTGAAGTGAATAGTTTAGTGGTTATTTAAAACTTACTGACAATGTATAC   3382
```

```
gb:LC521925|Organism:Severe      GTTCAGACTATTGAAGTGAATAGTTTTAGTGGTTATTTAAAACTTACTGACAATGTATAC   3355
gb:MT093571|Organism:Severe      GTTCAGACTATTGAAGTGAATAGTTTTAGTGGTTATTTAAAACTTACTGACAATGTATAC   3382
gb:MT039887|Organism:Severe      GTTCAGACTATTGAAGTGAATAGTTTTAGTGGTTATTTAAAACTTACTGACAATGTATAC   3382
gb:MT019530|Organism:Severe      GTTCAGACTATTGAAGTGAATAGTTTTAGTGGTTATTTAAAACTTACTGACAATGTATAC   3382
gb:MT039888|Organism:Severe      GTTCAGACTATTGAAGTGAATAGTTTTAGTGGTTATTTAAAACTTACTGACAATGTATAC   3382
gb:LC522972|Organism:Severe      GTTCAGACTATTGAAGTGAATAGTTTTAGTGGTTATTTAAAACTTACTGACAATGTATAC   3379
gb:MT027063|Organism:Severe      GTTCAGACTATTGAAGTGAATAGTTTTAGTGGTTATTTAAAACTTACTGACAATGTATAC   3382
gb:MT027062|Organism:Severe      GTTCAGACTATTGAAGTGAATAGTTTTAGTGGTTATTTAAAACTTACTGACAATGTATAC   3382
gb:MT019529|Organism:Severe      GTTCAGACTATTGAAGTGAATAGTTTTAGTGGTTATTTAAAACTTACTGACAATGTATAC   3382
gb:MN996529|Organism:Severe      GTTCAGACTATTGAAGTGAATAGTTTTAGTGGTTATTTAAAACTTACTGACAATGTATAC   3370
gb:MN996531|Organism:Severe      GTTCAGACTATTGAAGTGAATAGTTTTAGTGGTTATTTAAAACTTACTGACAATGTATAC   3369
gb:MT066176|Organism:Severe      GTTCAGACTATTGAAGTGAATAGTTTTAGTGGTTATTTAAAACTTACTGACAATGTATAC   3382
gb:MT027064|Organism:Severe      GTTCAGACTATTGAAGTGAATAGTTTTAGTGGTTATTTAAAACTTACTGACAATGTATAC   3382
gb:MN994468|Organism:Severe      GTTCAGACTATTGAAGTGAATAGTTTTAGTGGTTATTTAAAACTTACTGACAATGTATAC   3382
gb:MT072688|Organism:Severe      GTTCAGACTATTGAAGTGAATAGTTTTAGTGGTTATTTAAAACTTACTGACAATGTATAC   3367
gb:MN996527|Organism:Severe      GTTCAGACTATTGAAGTGAATAGTTTTAGTGGTTATTTAAAACTTACTGACAATGTATAC   3349
gb:MT093631|Organism:Severe      GTTCAGACTATTGAAGTGAATAGTTTTAGTGGTTATTTAAAACTTACTGACAATGTATAC   3420
gb:MT106053|Organism:Severe      GTTCAGACTATTGAAGTGAATAGTTTTAGTGGTTATTTAAAACTTACTGACAATGTATAC   3382
gb:MT019533|Organism:Severe      GTTCAGACTATTGAAGTGAATAGTTTTAGTGGTTATTTAAAACTTACTGACAATGTATAC   3382
gb:MT019531|Organism:Severe      GTTCAGACTATTGAAGTGAATAGTTTTAGTGGTTATTTAAAACTTACTGACAATGTATAC   3382
gb:MN996528|Organism:Severe      GTTCAGACTATTGAAGTGAATAGTTTTAGTGGTTATTTAAAACTTACTGACAATGTATAC   3382
gb:MN996530|Organism:Severe      GTTCAGACTATTGAAGTGAATAGTTTTAGTGGTTATTTAAAACTTACTGACAATGTATAC   3368
gb:MN908947|Organism:Severe      GTTCAGACTATTGAAGTGAATAGTTTTAGTGGTTATTTAAAACTTACTGACAATGTATAC   3382
gb:MT019532|Organism:Severe      GTTCAGACTATTGAAGTGAATAGTTTTAGTGGTTATTTAAAACTTACTGACAATGTATAC   3382
                                 ************************************************************

gb:MT020781|Organism:Severe      ATTAAAAATGCAGACATTGTGGAAGAAGCTAAAAAGGTAAAACCAACAGTGGTTGTTAAT   3430
gb:MT007544|Organism:Severe      ATTAAAAATGCAGACATTGTGGAAGAAGCTAAAAAGGTAAAACCAACAGTGGTTGTTAAT   3442
gb:MN994467|Organism:Severe      ATTAAAAATGCAGACATTGTGGAAGAAGCTAAAAAGGTAAAACCAACAGTGGTTGTTAAT   3442
gb:MT044257|Organism:Severe      ATTAAAAATGCAGACATTGTGGAAGAAGCTAAAAAGGTAAAACCAACAGTGGTTGTTAAT   3442
gb:MT106054|Organism:Severe      ATTAAAAATGCAGACATTGTGGAAGAAGCTAAAAAGGTAAAACCAACAGTGGTTGTTAAT   3442
gb:MT049951|Organism:Severe      ATTAAAAATGCAGACATTGTGGAAGAAGCTAAAAAGGTAAAACCAACAGTGGTTGTTAAT   3442
gb:MN975262|Organism:Severe      ATTAAAAATGCAGACATTGTGGAAGAAGCTAAAAAGGTAAAACCAACAGTGGTTGTTAAT   3442
gb:MT106052|Organism:Severe      ATTAAAAATGCAGACATTGTGGAAGAAGCTAAAAAGGTAAAACCAACAGTGGTTGTTAAT   3442
gb:LC522975|Organism:Severe      ATTAAAAATGCAGACATTGTGGAAGAAGCTAAAAAGGTAAAACCAACAGTGGTTGTTAAT   3439
gb:LC522973|Organism:Severe      ATTAAAAATGCAGACATTGTGGAAGAAGCTAAAAAGGTAAAACCAACAGTGGTTGTTAAT   3439
gb:LC522974|Organism:Severe      ATTAAAAATGCAGACATTGTGGAAGAAGCTAAAAAGGTAAAACCAACAGTGGTTGTTAAT   3439
gb:MN985325|Organism:Severe      ATTAAAAATGCAGACATTGTGGAAGAAGCTAAAAAGGTAAAACCAACAGTGGTTGTTAAT   3442
gb:MT020881|Organism:Severe      ATTAAAAATGCAGACATTGTGGAAGAAGCTAAAAAGGTAAAACCAACAGTGGTTGTTAAT   3442
gb:MT020880|Organism:Severe      ATTAAAAATGCAGACATTGTGGAAGAAGCTAAAAAGGTAAAACCAACAGTGGTTGTTAAT   3442
gb:MT066175|Organism:Severe      ATTAAAAATGCAGACATTGTGGAAGAAGCTAAAAAGGTAAAACCAACAGTGGTTGTTAAT   3442
gb:MN997409|Organism:Severe      ATTAAAAATGCAGACATTGTGGAAGAAGCTAAAAAGGTAAAACCAACAGTGGTTGTTAAT   3442
gb:MN938384|Organism:Severe      ATTAAAAATGCAGACATTGTGGAAGAAGCTAAAAAGGTAAAACCAACAGTGGTTGTTAAT   3410
gb:MT044258|Organism:Severe      ATTAAAAATGCAGACATTGTGGAAGAAGCTAAAAAGGTAAAACCAACAGTGGTTGTTAAT   3418
gb:MT039890|Organism:Severe      ATTAAAAATGCAGACATTGTGGAAGAAGCTAAAAAGGTAAAACCAACAGTGGTTGTTAAT   3442
gb:MN988713|Organism:Severe      ATTAAAAATGCAGACATTGTGGAAGAAGCTAAAAAGGTAAAACCAACAGTGGTTGTTAAT   3442
gb:LC521925|Organism:Severe      ATTAAAAATGCAGACATTGTGGAAGAAGCTAAAAAGGTAAAACCAACAGTGGTTGTTAAT   3415
gb:MT093571|Organism:Severe      ATTAAAAATGCAGACATTGTGGAAGAAGCTAAAAAGGTAAAACCAACAGTGGTTGTTAAT   3442
gb:MT039887|Organism:Severe      ATTAAAAATGCAGACATTGTGGAAGAAGCTAAAAAGGTAAAACCAACAGTGGTTGTTAAT   3442
gb:MT019530|Organism:Severe      ATTAAAAATGCAGACATTGTGGAAGAAGCTAAAAAGGTAAAACCAACAGTGGTTGTTAAT   3442
gb:MT039888|Organism:Severe      ATTAAAAATGCAGACATTGTGGAAGAAGCTAAAAAGGTAAAACCAACAGTGGTTGTTAAT   3442
gb:LC522972|Organism:Severe      ATTAAAAATGCAGACATTGTGGAAGAAGCTAAAAAGGTAAAACCAACAGTGGTTGTTAAT   3439
gb:MT027063|Organism:Severe      ATTAAAAATGCAGACATTGTGGAAGAAGCTAAAAAGGTAAAACCAACAGTGGTTGTTAAT   3442
gb:MT027062|Organism:Severe      ATTAAAAATGCAGACATTGTGGAAGAAGCTAAAAAGGTAAAACCAACAGTGGTTGTTAAT   3442
gb:MT019529|Organism:Severe      ATTAAAAATGCAGACATTGTGGAAGAAGCTAAAAAGGTAAAACCAACAGTGGTTGTTAAT   3442
gb:MN996529|Organism:Severe      ATTAAAAATGCAGACATTGTGGAAGAAGCTAAAAAGGTAAAACCAACAGTGGTTGTTAAT   3430
gb:MN996531|Organism:Severe      ATTAAAAATGCAGACATTGTGGAAGAAGCTAAAAAGGTAAAACCAACAGTGGTTGTTAAT   3429
gb:MT066176|Organism:Severe      ATTAAAAATGCAGACATTGTGGAAGAAGCTAAAAAGGTAAAACCAACAGTGGTTGTTAAT   3442
gb:MT027064|Organism:Severe      ATTAAAAATGCAGACATTGTGGAAGAAGCTAAAAAGGTAAAACCAACAGTGGTTGTTAAT   3442
gb:MN994468|Organism:Severe      ATTAAAAATGCAGACATTGTGGAAGAAGCTAAAAAGGTAAAACCAACAGTGGTTGTTAAT   3442
gb:MT072688|Organism:Severe      ATTAAAAATGCAGACATTGTGGAAGAAGCTAAAAAGGTAAAACCAACAGTGGTTGTTAAT   3427
gb:MN996527|Organism:Severe      ATTAAAAATGCAGACATTGTGGAAGAAGCTAAAAAGGTAAAACCAACAGTGGTTGTTAAT   3409
gb:MT093631|Organism:Severe      ATTAAAAATGCAGACATTGTGGAAGAAGCTAAAAAGGTAAAACCAACAGTGGTTGTTAAT   3480
gb:MT106053|Organism:Severe      ATTAAAAATGCAGACATTGTGGAAGAAGCTAAAAAGGTAAAACCAACAGTGGTTGTTAAT   3442
gb:MT019533|Organism:Severe      ATTAAAAATGCAGACATTGTGGAAGAAGCTAAAAAGGTAAAACCAACAGTGGTTGTTAAT   3442
```

```
gb:MT019531|Organism:Severe      ATTAAAAATGCAGACATTGTGGAAGAAGCTAAAAAGGTAAAACCAACAGTGGTTGTTAAT    3442
gb:MN996528|Organism:Severe      ATTAAAAATGCAGACATTGTGGAAGAAGCTAAAAAGGTAAAACCAACAGTGGTTGTTAAT    3442
gb:MN996530|Organism:Severe      ATTAAAAATGCAGACATTGTGGAAGAAGCTAAAAAGGTAAAACCAACAGTGGTTGTTAAT    3428
gb:MN908947|Organism:Severe      ATTAAAAATGCAGACATTGTGGAAGAAGCTAAAAAGGTAAAACCAACAGTGGTTGTTAAT    3442
gb:MT019532|Organism:Severe      ATTAAAAATGCAGACATTGTGGAAGAAGCTAAAAAGGTAAAACCAACAGTGGTTGTTAAT    3442
                                 ************************************************************

gb:MT020781|Organism:Severe      GCAGCCAATGTTTACCTTAAACATGGAGGAGGTGTTGCAGGAGCCTTAAATAAGGCTACT    3490
gb:MT007544|Organism:Severe      GCAGCCAATGTTTACCTTAAACATGGAGGAGGTGTTGCAGGAGCCTTAAATAAGGCTACT    3502
gb:MN994467|Organism:Severe      GCAGCCAATGTTTACCTTAAACATGGAGGAGGTGTTGCAGGAGCCTTAAATAAGGCTACT    3502
gb:MT044257|Organism:Severe      GCAGCCAATGTTTACCTTAAACATGGAGGAGGTGTTGCAGGAGCCTTAAATAAGGCTACT    3502
gb:MT106054|Organism:Severe      GCAGCCAATGTTTACCTTAAACATGGAGGAGGTGTTGCAGGAGCCTTAAATAAGGCTACT    3502
gb:MT049951|Organism:Severe      GCAGCCAATGTTTACCTTAAACATGGAGGAGGTGTTGCAGGAGCCTTAAATAAGGCTACT    3502
gb:MN975262|Organism:Severe      GCAGCCAATGTTTACCTTAAACATGGAGGAGGTGTTGCAGGAGCCTTAAATAAGGCTACT    3502
gb:MT106052|Organism:Severe      GCAGCCAATGTTTACCTTAAACATGGAGGAGGTGTTGCAGGAGCCTTAAATAAGGCTACT    3502
gb:LC522975|Organism:Severe      GCAGCCAATGTTTACCTTAAACATGGAGGAGGTGTTGCAGGAGCCTTAAATAAGGCTACT    3499
gb:LC522973|Organism:Severe      GCAGCCAATGTTTACCTTAAACATGGAGGAGGTGTTGCAGGAGCCTTAAATAAGGCTACT    3499
gb:LC522974|Organism:Severe      GCAGCCAATGTTTACCTTAAACATGGAGGAGGTGTTGCAGGAGCCTTAAATAAGGCTACT    3499
gb:MN985325|Organism:Severe      GCAGCCAATGTTTACCTTAAACATGGAGGAGGTGTTGCAGGAGCCTTAAATAAGGCTACT    3502
gb:MT020881|Organism:Severe      GCAGCCAATGTTTACCTTAAACATGGAGGAGGTGTTGCAGGAGCCTTAAATAAGGCTACT    3502
gb:MT020880|Organism:Severe      GCAGCCAATGTTTACCTTAAACATGGAGGAGGTGTTGCAGGAGCCTTAAATAAGGCTACT    3502
gb:MT066175|Organism:Severe      GCAGCCAATGTTTACCTTAAACATGGAGGAGGTGTTGCAGGAGCCTTAAATAAGGCTACT    3502
gb:MN997409|Organism:Severe      GCAGCCAATGTTTACCTTAAACATGGAGGAGGTGTTGCAGGAGCCTTAAATAAGGCTACT    3502
gb:MN938384|Organism:Severe      GCAGCCAATGTTTACCTTAAACATGGAGGAGGTGTTGCAGGAGCCTTAAATAAGGCTACT    3470
gb:MT044258|Organism:Severe      GCAGCCAATGTTTACCTTAAACATGGAGGAGGTGTTGCAGGAGCCTTAAATAAGGCTACT    3478
gb:MT039890|Organism:Severe      GCAGCCAATGTTTACCTTAAACATGGAGGAGGTGTTGCAGGAGCCTTAAATAAGGCTACT    3502
gb:MN988713|Organism:Severe      GCAGCCAATGTTTACCTTAAACATGGAGGAGGTGTTGCAGGAGCCTTAAATAAGGCTACT    3502
gb:LC521925|Organism:Severe      GCAGCCAATGTTTACCTTAAACATGGAGGAGGTGTTGCAGGAGCCTTAAATAAGGCTACT    3475
gb:MT093571|Organism:Severe      GCAGCCAATGTTTACCTTAAACATGGAGGAGGTGTTGCAGGAGCCTTAAATAAGGCTACT    3502
gb:MT039887|Organism:Severe      GCAGCCAATGTTTACCTTAAACATGGAGGAGGTGTTGCAGGAGCCTTAAATAAGGCTACT    3502
gb:MT019530|Organism:Severe      GCAGCCAATGTTTACCTTAAACATGGAGGAGGTGTTGCAGGAGCCTTAAATAAGGCTACT    3502
gb:MT039888|Organism:Severe      GCAGCCAATGTTTACCTTAAACATGGAGGAGGTGTTGCAGGAGCCTTAAATAAGGCTACT    3502
gb:LC522972|Organism:Severe      GCAGCCAATGTTTACCTTAAACATGGAGGAGGTGTTGCAGGAGCCTTAAATAAGGCTACT    3499
gb:MT027063|Organism:Severe      GCAGCCAATGTTTACCTTAAACATGGAGGAGGTGTTGCAGGAGCCTTAAATAAGGCTACT    3502
gb:MT027062|Organism:Severe      GCAGCCAATGTTTACCTTAAACATGGAGGAGGTGTTGCAGGAGCCTTAAATAAGGCTACT    3502
gb:MT019529|Organism:Severe      GCAGCCAATGTTTACCTTAAACATGGAGGAGGTGTTGCAGGAGCCTTAAATAAGGCTACT    3502
gb:MN996529|Organism:Severe      GCAGCCAATGTTTACCTTAAACATGGAGGAGGTGTTGCAGGAGCCTTAAATAAGGCTACT    3490
gb:MN996531|Organism:Severe      GCAGCCAATGTTTACCTTAAACATGGAGGAGGTGTTGCAGGAGCCTTAAATAAGGCTACT    3489
gb:MT066176|Organism:Severe      GCAGCCAATGTTTACCTTAAACATGGAGGAGGTGTTGCAGGAGCCTTAAATAAGGCTACT    3502
gb:MT027064|Organism:Severe      GCAGCCAATGTTTACCTTAAACATGGAGGAGGTGTTGCAGGAGCCTTAAATAAGGCTACT    3502
gb:MN994468|Organism:Severe      GCAGCCAATGTTTACCTTAAACATGGAGGAGGTGTTGCAGGAGCCTTAAATAAGGCTACT    3502
gb:MT072688|Organism:Severe      GCAGCCAATGTTTACCTTAAACATGGAGGAGGTGTTGCAGGAGCCTTAAATAAGGCTACT    3487
gb:MN996527|Organism:Severe      GCAGCCAATGTTTACCTTAAACATGGAGGAGGTGTTGCAGGAGCCTTAAATAAGGCTACT    3469
gb:MT093631|Organism:Severe      GCAGCCAATGTTTACCTTAAACATGGAGGAGGTGTTGCAGGAGCCTTAAATAAGGCTACT    3540
gb:MT106053|Organism:Severe      GCAGCCAATGTTTACCTTAAACATGGAGGAGGTGTTGCAGGAGCCTTAAATAAGGCTACT    3502
gb:MT019533|Organism:Severe      GCAGCCAATGTTTACCTTAAACATGGAGGAGGTGTTGCAGGAGCCTTAAATAAGGCTACT    3502
gb:MT019531|Organism:Severe      GCAGCCAATGTTTACCTTAAACATGGAGGAGGTGTTGCAGGAGCCTTAAATAAGGCTACT    3502
gb:MN996528|Organism:Severe      GCAGCCAATGTTTACCTTAAACATGGAGGAGGTGTTGCAGGAGCCTTAAATAAGGCTACT    3502
gb:MN996530|Organism:Severe      GCAGCCAATGTTTACCTTAAACATGGAGGAGGTGTTGCAGGAGCCTTAAATAAGGCTACT    3488
gb:MN908947|Organism:Severe      GCAGCCAATGTTTACCTTAAACATGGAGGAGGTGTTGCAGGAGCCTTAAATAAGGCTACT    3502
gb:MT019532|Organism:Severe      GCAGCCAATGTTTACCTTAAACATGGAGGAGGTGTTGCAGGAGCCTTAAATAAGGCTACT    3502
                                 ************************************************************

gb:MT020781|Organism:Severe      AACAATGCCATGCAAGTTGAATCTGATGATTACATAGCTACTAATGGACCACTTAAAGTG    3550
gb:MT007544|Organism:Severe      AACAATGCCATGCAAGTTGAATCTGATGATTACATAGCTACTAATGGACCACTTAAAGTG    3562
gb:MN994467|Organism:Severe      AACAATGCCATGCAAGTTGAATCTGATGATTACATAGCTACTAATGGACCACTTAAAGTG    3562
gb:MT044257|Organism:Severe      AACAATGCCATGCAAGTTGAATCTGATGATTACATAGCTACTAATGGACCACTTAAAGTG    3562
gb:MT106054|Organism:Severe      AACAATGCCATGCAAGTTGAATCTGATGATTACATAGCTACTAATGGACCACTTAAAGTG    3562
gb:MT049951|Organism:Severe      AACAATGCCATGCAAGTTGAATCTGATGATTACATAGCTACTAATGGACCACTTAAAGTG    3562
gb:MN975262|Organism:Severe      AACAATGCCATGCAAGTTGAATCTGATGATTACATAGCTACTAATGGACCACTTAAAGTG    3562
gb:MT106052|Organism:Severe      AACAATGCCATGCAAGTTGAATCTGATGATTACATAGCTACTAATGGACCACTTAAAGTG    3562
gb:LC522975|Organism:Severe      AACAATGCCATGCAAGTTGAATCTGATGATTACATAGCTACTAATGGACCACTTAAAGTG    3559
gb:LC522973|Organism:Severe      AACAATGCCATGCAAGTTGAATCTGATGATTACATAGCTACTAATGGACCACTTAAAGTG    3559
gb:LC522974|Organism:Severe      AACAATGCCATGCAAGTTGAATCTGATGATTACATAGCTACTAATGGACCACTTAAAGTG    3559
gb:MN985325|Organism:Severe      AACAATGCCATGCAAGTTGAATCTGATGATTACATAGCTACTAATGGACCACTTAAAGTG    3562
```

```
gb:MT020881|Organism:Severe      AACAATGCCATGCAAGTTGAATCTGATGATTACATAGCTACTAATGGACCACTTAAAGTG   3562
gb:MT020880|Organism:Severe      AACAATGCCATGCAAGTTGAATCTGATGATTACATAGCTACTAATGGACCACTTAAAGTG   3562
gb:MT066175|Organism:Severe      AACAATGCCATGCAAGTTGAATCTGATGATTACATAGCTACTAATGGACCACTTAAAGTG   3562
gb:MN997409|Organism:Severe      AACAATGCCATGCAAGTTGAATCTGATGATTACATAGCTACTAATGGACCACTTAAAGTG   3562
gb:MN938384|Organism:Severe      AACAATGCCATGCAAGTTGAATCTGATGATTACATAGCTACTAATGGACCACTTAAAGTG   3530
gb:MT044258|Organism:Severe      AACAATGCCATGCAAGTTGAATCTGATGATTACATAGCTACTAATGGACCACTTAAAGTG   3538
gb:MT039890|Organism:Severe      AACAATGCCATGCAAGTTGAATCTGATGATTACATAGCTACTAATGGACCACTTAAAGTG   3562
gb:MN988713|Organism:Severe      AACAATGCCATGCAAGTTGAATCTGATGATTACATAGCTACTAATGGACCACTTAAAGTG   3562
gb:LC521925|Organism:Severe      AACAATGCCATGCAAGTTGAATCTGATGATTACATAGCTACTAATGGACCACTTAAAGTG   3535
gb:MT093571|Organism:Severe      AACAATGCCATGCAAGTTGAATCTGATGATTACATAGCTACTAATGGACCACTTAAAGTG   3562
gb:MT039887|Organism:Severe      AACAATGCCATGCAAGTTGAATCTGATGATTACATAGCTACTAATGGACCACTTAAAGTG   3562
gb:MT019530|Organism:Severe      AACAATGCCATGCAAGTTGAATCTGATGATTACATAGCTACTAATGGACCACTTAAAGTG   3562
gb:MT039888|Organism:Severe      AACAATGCCATGCAATTTGAATCTGATGATTACATAGCTACTAATGGACCACTTAAAGTG   3562
gb:LC522972|Organism:Severe      AACAATGCCATGCAAGTTGAATCTGATGATTACATAGCTACTAATGGACCACTTAAAGTG   3559
gb:MT027063|Organism:Severe      AACAATGCCATGCAAGTTGAATCTGATGATTACATAGCTACTAATGGACCACTTAAAGTG   3562
gb:MT027062|Organism:Severe      AACAATGCCATGCAAGTTGAATCTGATGATTACATAGCTACTAATGGACCACTTAAAGTG   3562
gb:MT019529|Organism:Severe      AACAATGCCATGCAAGTTGAATCTGATGATTACATAGCTACTAATGGACCACTTAAAGTG   3562
gb:MN996529|Organism:Severe      AACAATGCCATGCAAGTTGAATCTGATGATTACATAGCTACTAATGGACCACTTAAAGTG   3550
gb:MN996531|Organism:Severe      AACAATGCCATGCAAGTTGAATCTGATGATTACATAGCTACTAATGGACCACTTAAAGTG   3549
gb:MT066176|Organism:Severe      AACAATGCCATGCAAGTTGAATCTGATGATTACATAGCTACTAATGGACCACTTAAAGTG   3562
gb:MT027064|Organism:Severe      AACAATGCCATGCAAGTTGAATCTGATGATTACATAGCTACTAATGGACCACTTAAAGTG   3562
gb:MN994468|Organism:Severe      AACAATGCCATGCAAGTTGAATCTGATGATTACATAGCTACTAATGGACCACTTAAAGTG   3562
gb:MT072688|Organism:Severe      AACAATGCCATGCAAGTTGAATCTGATGATTACATAGCTACTAATGGACCACTTAAAGTG   3547
gb:MN996527|Organism:Severe      AACAATGCCATGCAAGTTGAATCTGATGATTACATAGCTACTAATGGACCACTTAAAGTG   3529
gb:MT093631|Organism:Severe      AACAATGCCATGCAAGTTGAATCTGATGATTACATAGCTACTAATGGACCACTTAAAGTG   3600
gb:MT106053|Organism:Severe      AACAATGCCATGCAAGTTGAATCTGATGATTACATAGCTACTAATGGACCACTTAAAGTG   3562
gb:MT019533|Organism:Severe      AACAATGCCATGCAAGTTGAATCTGATGATTACATAGCTACTAATGGACCACTTAAAGTG   3562
gb:MT019531|Organism:Severe      AACAATGCCATGCAAGTTGAATCTGATGATTACATAGCTACTAATGGACCACTTAAAGTG   3562
gb:MN996528|Organism:Severe      AACAATGCCATGCAAGTTGAATCTGATGATTACATAGCTACTAATGGACCACTTAAAGTG   3562
gb:MN996530|Organism:Severe      AACAATGCCATGCAAGTTGAATCTGATGATTACATAGCTACTAATGGACCACTTAAAGTG   3548
gb:MN908947|Organism:Severe      AACAATGCCATGCAAGTTGAATCTGATGATTACATAGCTACTAATGGACCACTTAAAGTG   3562
gb:MT019532|Organism:Severe      AACAATGCCATGCAAGTTGAATCTGATGATTACATAGCTACTAATGGACCACTTAAAGTG   3562
                                 *************** ********************************************

gb:MT020781|Organism:Severe      GGTGGTAGTTGTGTTTTAAGCGGACACAATCTTGCTAAACACTGTCTTCATGTTGTCGGC   3610
gb:MT007544|Organism:Severe      GGTGGTAGTTGTGTTTTAAGCGGACACAATCTTGCTAAACACTGTCTTCATGTTGTCGGC   3622
gb:MN994467|Organism:Severe      GGTGGTAGTTGTGTTTTAAGCGGACACAATCTTGCTAAACACTGTCTTCATGTTGTCGGC   3622
gb:MT044257|Organism:Severe      GGTGGTAGTTGTGTTTTAAGCGGACACAATCTTGCTAAACACTGTCTTCATGTTGTCGGC   3622
gb:MT106054|Organism:Severe      GGTGGTAGTTGTGTTTTAAGCGGACACAATCTTGCTAAACACTGTCTTCATGTTGTCGGC   3622
gb:MT049951|Organism:Severe      GGTGGTAGTTGTGTTTTAAGCGGACACAATCTTGCTAAACACTGTCTTCATGTTGTCGGC   3622
gb:MN975262|Organism:Severe      GGTGGTAGTTGTGTTTTAAGCGGACACAATCTTGCTAAACACTGTCTTCATGTTGTCGGC   3622
gb:MT106052|Organism:Severe      GGTGGTAGTTGTGTTTTAAGCGGACACAATCTTGCTAAACACTGTCTTCATGTTGTCGGC   3622
gb:LC522975|Organism:Severe      GGTGGTAGTTGTGTTTTAAGCGGACACAATCTTGCTAAACACTGTCTTCATGTTGTCGGC   3619
gb:LC522973|Organism:Severe      GGTGGTAGTTGTGTTTTAAGCGGACACAATCTTGCTAAACACTGTCTTCATGTTGTCGGC   3619
gb:LC522974|Organism:Severe      GGTGGTAGTTGTGTTTTAAGCGGACACAATCTTGCTAAACACTGTCTTCATGTTGTCGGC   3619
gb:MN985325|Organism:Severe      GGTGGTAGTTGTGTTTTAAGCGGACACAATCTTGCTAAACACTGTCTTCATGTTGTCGGC   3622
gb:MT020881|Organism:Severe      GGTGGTAGTTGTGTTTTAAGCGGACACAATCTTGCTAAACACTGTCTTCATGTTGTCGGC   3622
gb:MT020880|Organism:Severe      GGTGGTAGTTGTGTTTTAAGCGGACACAATCTTGCTAAACACTGTCTTCATGTTGTCGGC   3622
gb:MT066175|Organism:Severe      GGTGGTAGTTGTGTTTTAAGCGGACACAATCTTGCTAAACACTGTCTTCATGTTGTCGGC   3622
gb:MN997409|Organism:Severe      GGTGGTAGTTGTGTTTTAAGCGGACACAATCTTGCTAAACACTGTCTTCATGTTGTCGGC   3622
gb:MN938384|Organism:Severe      GGTGGTAGTTGTGTTTTAAGCGGACACAATCTTGCTAAACACTGTCTTCATGTTGTCGGC   3590
gb:MT044258|Organism:Severe      GGTGGTAGTTGTGTTTTAAGCGGACACAATCTTGCTAAACACTGTCTTCATGTTGTCGGC   3598
gb:MT039890|Organism:Severe      GGTGGTAGTTGTGTTTTAAGCGGACACAATCTTGCTAAACACTGTCTTCATGTTGTCGGC   3622
gb:MN988713|Organism:Severe      GGTGGTAGTTGTGTTTTAAGCGGACACAATCTTGCTAAACACTGTCTTCATGTTGTCGGC   3622
gb:LC521925|Organism:Severe      GGTGGTAGTTGTGTTTTAAGCGGACACAATCTTGCTAAACACTGTCTTCATGTTGTCGGC   3595
gb:MT093571|Organism:Severe      GGTGGTAGTTGTGTTTTAAGCGGACACAATCTTGCTAAACACTGTCTTCATGTTGTCGGC   3622
gb:MT039887|Organism:Severe      GGTGGTAGTTGTGTTTTAAGCGGACACAATCTTGCTAAACACTGTCTTCATGTTGTCGGC   3622
gb:MT019530|Organism:Severe      GGTGGTAGTTGTGTTTTAAGCGGACACAATCTTGCTAAACACTGTCTTCATGTTGTCGGC   3622
gb:MT039888|Organism:Severe      GGTGGTAGTTGTGTTTTAAGCGGACACAATCTTGCTAAACACTGTCTTCATGTTGTCGGC   3622
gb:LC522972|Organism:Severe      GGTGGTAGTTGTGTTTTAAGCGGACACAATCTTGCTAAACACTGTCTTCATGTTGTCGGC   3619
gb:MT027063|Organism:Severe      GGTGGTAGTTGTGTTTTAAGCGGACACAATCTTGCTAAACACTGTCTTCATGTTGTCGGC   3622
gb:MT027062|Organism:Severe      GGTGGTAGTTGTGTTTTAAGCGGACACAATCTTGCTAAACACTGTCTTCATGTTGTCGGC   3622
gb:MT019529|Organism:Severe      GGTGGTAGTTGTGTTTTAAGCGGACACAATCTTGCTAAACACTGTCTTCATGTTGTCGGC   3622
gb:MN996529|Organism:Severe      GGTGGTAGTTGTGTTTTAAGCGGACACAATCTTGCTAAACACTGTCTTCATGTTGTCGGC   3610
gb:MN996531|Organism:Severe      GGTGGTAGTTGTGTTTTAAGCGGACACAATCTTGCTAAACACTGTCTTCATGTTGTCGGC   3609
```

```
gb:MT066176|Organism:Severe      GGTGGTAGTTGTGTTTTAAGCGGACACAATCTTGCTAAACACTGTCTTCATGTTGTCGGC   3622
gb:MT027064|Organism:Severe      GGTGGTAGTTGTGTTTTAAGCGGACACAATCTTGCTAAACACTGTCTTCATGTTGTCGGC   3622
gb:MN994468|Organism:Severe      GGTGGTAGTTGTGTTTTAAGCGGACACAATCTTGCTAAACACTGTCTTCATGTTGTCGGC   3622
gb:MT072688|Organism:Severe      GGTGGTAGTTGTGTTTTAAGCGGACACAATCTTGCTAAACACTGTCTTCATGTTGTCGGC   3607
gb:MN996527|Organism:Severe      GGTGGTAGTTGTGTTTTAAGCGGACACAATCTTGCTAAACACTGTCTTCATGTTGTCGGC   3589
gb:MT093631|Organism:Severe      GGTGGTAGTTGTGTTTTAAGCGGACACAATCTTGCTAAACACTGTCTTCATGTTGTCGGC   3660
gb:MT106053|Organism:Severe      GGTGGTAGTTGTGTTTTAAGCGGACACAATCTTGCTAAACACTGTCTTCATGTTGTCGGC   3622
gb:MT019533|Organism:Severe      GGTGGTAGTTGTGTTTTAAGCGGACACAATCTTGCTAAACACTGTCTTCATGTTGTCGGC   3622
gb:MT019531|Organism:Severe      GGTGGTAGTTGTGTTTTAAGCGGACACAATCTTGCTAAACACTGTCTTCATGTTGTCGGC   3622
gb:MN996528|Organism:Severe      GGTGGTAGTTGTGTTTTAAGCGGACACAATCTTGCTAAACACTGTCTTCATGTTGTCGGC   3622
gb:MN996530|Organism:Severe      GGTGGTAGTTGTGTTTTAAGCGGACACAATCTTGCTAAACACTGTCTTCATGTTGTCGGC   3608
gb:MN908947|Organism:Severe      GGTGGTAGTTGTGTTTTAAGCGGACACAATCTTGCTAAACACTGTCTTCATGTTGTCGGC   3622
gb:MT019532|Organism:Severe      GGTGGTAGTTGTGTTTTAAGCGGACACAATCTTGCTAAACACTGTCTTCATGTTGTCGGC   3622
                                 ************************************************************

gb:MT020781|Organism:Severe      CCAAATGTTAACAAAGGTGAAGACATTCAACTTCTTAAGAGTGCTTATGAAAATTTTAAT   3670
gb:MT007544|Organism:Severe      CCAAATGTTAACAAAGGTGAAGACATTCAACTTCTTAAGAGTGCTTATGAAAATTTTAAT   3682
gb:MN994467|Organism:Severe      CCAAATGTTAACAAAGGTGAAGACATTCAACTTCTTAAGAGTGCTTATGAAAATTTTAAT   3682
gb:MT044257|Organism:Severe      CCAAATGTTAACAAAGGTGAAGACATTCAACTTCTTAAGAGTGCTTATGAAAATTTTAAT   3682
gb:MT106054|Organism:Severe      CCAAATGTTAACAAAGGTGAAGACATTCAACTTCTTAAGAGTGCTTATGAAAATTTTAAT   3682
gb:MT049951|Organism:Severe      CCAAATGTTAACAAAGGTGAAGACATTCAACTTCTTAAGAGTGCTTATGAAAATTTTAAT   3682
gb:MN975262|Organism:Severe      CCAAATGTTAACAAAGGTGAAGACATTCAACTTCTTAAGAGTGCTTATGAAAATTTTAAT   3682
gb:MT106052|Organism:Severe      CCAAATGTTAACAAAGGTGAAGACATTCAACTTCTTAAGAGTGCTTATGAAAATTTTAAT   3682
gb:LC522975|Organism:Severe      CCAAATGTTAACAAAGGTGAAGACATTCAACTTCTTAAGAGTGCTTATGAAAATTTTAAT   3679
gb:LC522973|Organism:Severe      CCAAATGTTAACAAAGGTGAAGACATTCAACTTCTTAAGAGTGCTTATGAAAATTTTAAT   3679
gb:LC522974|Organism:Severe      CCAAATGTTAACAAAGGTGAAGACATTCAACTTCTTAAGAGTGCTTATGAAAATTTTAAT   3679
gb:MN985325|Organism:Severe      CCAAATGTTAACAAAGGTGAAGACATTCAACTTCTTAAGAGTGCTTATGAAAATTTTAAT   3682
gb:MT020881|Organism:Severe      CCAAATGTTAACAAAGGTGAAGACATTCAACTTCTTAAGAGTGCTTATGAAAATTTTAAT   3682
gb:MT020880|Organism:Severe      CCAAATGTTAACAAAGGTGAAGACATTCAACTTCTTAAGAGTGCTTATGAAAATTTTAAT   3682
gb:MT066175|Organism:Severe      CCAAATGTTAACAAAGGTGAAGACATTCAACTTCTTAAGAGTGCTTATGAAAATTTTAAT   3682
gb:MN997409|Organism:Severe      CCAAATGTTAACAAAGGTGAAGACATTCAACTTCTTAAGAGTGCTTATGAAAATTTTAAT   3682
gb:MN938384|Organism:Severe      CCAAATGTTAACAAAGGTGAAGACATTCAACTTCTTAAGAGTGCTTATGAAAATTTTAAT   3650
gb:MT044258|Organism:Severe      CCAAATGTTAACAAAGGTGAAGACATTCAACTTCTTAAGAGTGCTTATGAAAATTTTAAT   3658
gb:MT039890|Organism:Severe      CCAAATGTTAACAAAGGTGAAGACATTCAACTTCTTAAGAGTGCTTATGAAAATTTTAAT   3682
gb:MN988713|Organism:Severe      CCAAATGTTAACAAAGGTGAAGACATTCAACTTCTTAAGAGTGCTTATGAAAATTTTAAT   3682
gb:LC521925|Organism:Severe      CCAAATGTTAACAAAGGTGAAGACATTCAACTTCTTAAGAGTGCTTATGAAAATTTTAAT   3655
gb:MT093571|Organism:Severe      CCAAATGTTAACAAAGGTGAAGACATTCAACTTCTTAAGAGTGCTTATGAAAATTTTAAT   3682
gb:MT039887|Organism:Severe      CCAAATGTTAACAAAGGTGAAGACATTCAACTTCTTAAGAGTGCTTATGAAAATTTTAAT   3682
gb:MT019530|Organism:Severe      CCAAATGTTAACAAAGGTGAAGACATTCAACTTCTTAAGAGTGCTTATGAAAATTTTAAT   3682
gb:MT039888|Organism:Severe      CCAAATGTTAACAAAGGTGAAGACATTCAACTTCTTAAGAGTGCTTATGAAAATTTTAAT   3682
gb:LC522972|Organism:Severe      CCAAATGTTAACAAAGGTGAAGACATTCAACTTCTTAAGAGTGCTTATGAAAATTTTAAT   3679
gb:MT027063|Organism:Severe      CCAAATGTTAACAAAGGTGAAGACATTCAACTTCTTAAGAGTGCTTATGAAAATTTTAAT   3682
gb:MT027062|Organism:Severe      CCAAATGTTAACAAAGGTGAAGACATTCAACTTCTTAAGAGTGCTTATGAAAATTTTAAT   3682
gb:MT019529|Organism:Severe      CCAAATGTTAACAAAGGTGAAGACATTCAACTTCTTAAGAGTGCTTATGAAAATTTTAAT   3682
gb:MN996529|Organism:Severe      CCAAATGTTAACAAAGGTGAAGACATTCAACTTCTTAAGAGTGCTTATGAAAATTTTAAT   3670
gb:MN996531|Organism:Severe      CCAAATGTTAACAAAGGTGAAGACATTCAACTTCTTAAGAGTGCTTATGAAAATTTTAAT   3669
gb:MT066176|Organism:Severe      CCAAATGTTAACAAAGGTGAAGACATTCAACTTCTTAAGAGTGCTTATGAAAATTTTAAT   3682
gb:MT027064|Organism:Severe      CCAAATGTTAACAAAGGTGAAGACATTCAACTTCTTAAGAGTGCTTATGAAAATTTTAAT   3682
gb:MN994468|Organism:Severe      CCAAATGTTAACAAAGGTGAAGACATTCAACTTCTTAAGAGTGCTTATGAAAATTTTAAT   3682
gb:MT072688|Organism:Severe      CCAAATGTTAACAAAGGTGAAGACATTCAACTTCTTAAGAGTGCTTATGAAAATTTTAAT   3667
gb:MN996527|Organism:Severe      CCAAATGTTAACAAAGGTGAAGACATTCAACTTCTTAAGAGTGCTTATGAAAATTTTAAT   3649
gb:MT093631|Organism:Severe      CCAAATGTTAACAAAGGTGAAGACATTCAACTTCTTAAGAGTGCTTATGAAAATTTTAAT   3720
gb:MT106053|Organism:Severe      CCAAATGTTAACAAAGGTGAAGACATTCAACTTCTTAAGAGTGCTTATGAAAATTTTAAT   3682
gb:MT019533|Organism:Severe      CCAAATGTTAACAAAGGTGAAGACATTCAACTTCTTAAGAGTGCTTATGAAAATTTTAAT   3682
gb:MT019531|Organism:Severe      CCAAATGTTAACAAAGGTGAAGACATTCAACTTCTTAAGAGTGCTTATGAAAATTTTAAT   3682
gb:MN996528|Organism:Severe      CCAAATGTTAACAAAGGTGAAGACATTCAACTTCTTAAGAGTGCTTATGAAAATTTTAAT   3682
gb:MN996530|Organism:Severe      CCAAATGTTAACAAAGGTGAAGACATTCAACTTCTTAAGAGTGCTTATGAAAATTTTAAT   3668
gb:MN908947|Organism:Severe      CCAAATGTTAACAAAGGTGAAGACATTCAACTTCTTAAGAGTGCTTATGAAAATTTTAAT   3682
gb:MT019532|Organism:Severe      CCAAATGTTAACAAAGGTGAAGACATTCAACTTCTTAAGAGTGCTTATGAAAATTTTAAT   3682
                                 ************************************************************

gb:MT020781|Organism:Severe      CAGCACGAAGTTCTACTTGCACCATTATTATCAGCTGGTATTTTTGGTGCTGACCCTATA   3730
gb:MT007544|Organism:Severe      CAGCACGAAGTTCTACTTGCACCATTATTATCAGCTGGTATTTTTGGTGCTGACCCTATA   3742
gb:MN994467|Organism:Severe      CAGCACGAAGTTCTACTTGCACCATTATTATCAGCTGGTATTTTTGGTGCTGACCCTATA   3742
gb:MT044257|Organism:Severe      CAGCACGAAGTTCTACTTGCACCATTATTATCAGCTGGTATTTTTGGTGCTGACCCTATA   3742
```

```
gb:MT106054|Organism:Severe      CAGCACGAAGTTCTACTTGCACCATTATTATCAGCTGGTATTTTTGGTGCTGACCCTATA   3742
gb:MT049951|Organism:Severe      CAGCACGAAGTTCTACTTGCACCATTATTATCAGCTGGTATTTTTGGTGCTGACCCTATA   3742
gb:MN975262|Organism:Severe      CAGCACGAAGTTCTACTTGCACCATTATTATCAGCTGGTATTTTTGGTGCTGACCCTATA   3742
gb:MT106052|Organism:Severe      CAGCACGAAGTTCTACTTGCACCATTATTATCAGCTGGTATTTTTGGTGCTGACCCTATA   3742
gb:LC522975|Organism:Severe      CAGCACGAAGTTCTACTTGCACCATTATTATCAGCTGGTATTTTTGGTGCTGACCCTATA   3739
gb:LC522973|Organism:Severe      CAGCACGAAGTTCTACTTGCACCATTATTATCAGCTGGTATTTTTGGTGCTGACCCTATA   3739
gb:LC522974|Organism:Severe      CAGCACGAAGTTCTACTTGCACCATTATTATCAGCTGGTATTTTTGGTGCTGACCCTATA   3739
gb:MN985325|Organism:Severe      CAGCACGAAGTTCTACTTGCACCATTATTATCAGCTGGTATTTTTGGTGCTGACCCTATA   3742
gb:MT020881|Organism:Severe      CAGCACGAAGTTCTACTTGCACCATTATTATCAGCTGGTATTTTTGGTGCTGACCCTATA   3742
gb:MT020880|Organism:Severe      CAGCACGAAGTTCTACTTGCACCATTATTATCAGCTGGTATTTTTGGTGCTGACCCTATA   3742
gb:MT066175|Organism:Severe      CAGCACGAAGTTCTACTTGCACCATTATTATCAGCTGGTATTTTTGGTGCTGACCCTATA   3742
gb:MN997409|Organism:Severe      CAGCACGAAGTTCTACTTGCACCATTATTATCAGCTGGTATTTTTGGTGCTGACCCTATA   3742
gb:MN938384|Organism:Severe      CAGCACGAAGTTCTACTTGCACCATTATTATCAGCTGGTATTTTTGGTGCTGACCCTATA   3710
gb:MT044258|Organism:Severe      CAGCACGAAGTTCTACTTGCACCATTATTATCAGCTGGTATTTTTGGTGCTGACCCTATA   3718
gb:MT039890|Organism:Severe      CAGCACGAAGTTCTACTTGCACCATTATTATCAGCTGGTATTTTTGGTGCTGACCCTATA   3742
gb:MN988713|Organism:Severe      CAGCACGAAGTTCTACTTGCACCATTATTATCAGCTGGTATTTTTGGTGCTGACCCTATA   3742
gb:LC521925|Organism:Severe      CAGCACGAAGTTCTACTTGCACCATTATTATCAGCTGGTATTTTTGGTGCTGACCCTATA   3715
gb:MT093571|Organism:Severe      CAGCACGAAGTTCTACTTGCACCATTATTATCAGCTGGTATTTTTGGTGCTGACCCTATA   3742
gb:MT039887|Organism:Severe      CAGCACGAAGTTCTACTTGCACCATTATTATCAGCTGGTATTTTTGGTGCTGACCCTATA   3742
gb:MT019530|Organism:Severe      CAGCACGAAGTTCTACTTGCACCATTATTATCAGCTGGTATTTTTGGTGCTGACCCTATA   3742
gb:MT039888|Organism:Severe      CAGCACGAAGTTCTACTTGCACCATTATTATCAGCTGGTATTTTTGGTGCTGACCCTATA   3742
gb:LC522972|Organism:Severe      CAGCACGAAGTTCTACTTGCACCATTATTATCAGCTGGTATTTTTGGTGCTGACCCTATA   3739
gb:MT027063|Organism:Severe      CAGCACGAAGTTCTACTTGCACCATTATTATCAGCTGGTATTTTTGGTGCTGACCCTATA   3742
gb:MT027062|Organism:Severe      CAGCACGAAGTTCTACTTGCACCATTATTATCAGCTGGTATTTTTGGTGCTGACCCTATA   3742
gb:MT019529|Organism:Severe      CAGCACGAAGTTCTACTTGCACCATTATTATCAGCTGGTATTTTTGGTGCTGACCCTATA   3742
gb:MN996529|Organism:Severe      CAGCACGAAGTTCTACTTGCACCATTATTATCAGCTGGTATTTTTGGTGCTGACCCTATA   3730
gb:MN996531|Organism:Severe      CAGCACGAAGTTCTACTTGCACCATTATTATCAGCTGGTATTTTTGGTGCTGACCCTATA   3729
gb:MT066176|Organism:Severe      CAGCACGAAGTTCTACTTGCACCATTATTATCAGCTGGTATTTTTGGTGCTGACCCTATA   3742
gb:MT027064|Organism:Severe      CAGCACGAAGTTCTACTTGCACCATTATTATCAGCTGGTATTTTTGGTGCTGACCCTATA   3742
gb:MN994468|Organism:Severe      CAGCACGAAGTTCTACTTGCACCATTATTATCAGCTGGTATTTTTGGTGCTGACCCTATA   3742
gb:MT072688|Organism:Severe      CAGCACGAAGTTCTACTTGCACCATTATTATCAGCTGGTATTTTTGGTGCTGACCCTATA   3727
gb:MN996527|Organism:Severe      CAGCACGAAGTTCTACTTGCACCATTATTATCAGCTGGTATTTTTGGTGCTGACCCTATA   3709
gb:MT093631|Organism:Severe      CAGCACGAAGTTCTACTTGCACCATTATTATCAGCTGGTATTTTTGGTGCTGACCCTATA   3780
gb:MT106053|Organism:Severe      CAGCACGAAGTTCTACTTGCACCATTATTATCAGCTGGTATTTTTGGTGCTGACCCTATA   3742
gb:MT019533|Organism:Severe      CAGCACGAAGTTCTACTTGCACCATTATTATCAGCTGGTATTTTTGGTGCTGACCCTATA   3742
gb:MT019531|Organism:Severe      CAGCACGAAGTTCTACTTGCACCATTATTATCAGCTGGTATTTTTGGTGCTGACCCTATA   3742
gb:MN996528|Organism:Severe      CAGCACGAAGTTCTACTTGCACCATTATTATCAGCTGGTATTTTTGGTGCTGACCCTATA   3742
gb:MN996530|Organism:Severe      CAGCACGAAGTTCTACTTGCACCATTATTATCAGCTGGTATTTTTGGTGCTGACCCTATA   3728
gb:MN908947|Organism:Severe      CAGCACGAAGTTCTACTTGCACCATTATTATCAGCTGGTATTTTTGGTGCTGACCCTATA   3742
gb:MT019532|Organism:Severe      CAGCACGAAGTTCTACTTGCACCATTATTATCAGCTGGTATTTTTGGTGCTGACCCTATA   3742
                                 ************************************************************

gb:MT020781|Organism:Severe      CATTCTTTAAGAGTTTGTGTAGATACTGTTCGCACAAATGTCTACTTAGCTGTCTTTGAT   3790
gb:MT007544|Organism:Severe      CATTCTTTAAGAGTTTGTGTAGATACTGTTCGCACAAATGTCTACTTAGCTGTCTTTGAT   3802
gb:MN994467|Organism:Severe      CATTCTTTAAGAGTTTGTGTAGATACTGTTCGCACAAATGTCTACTTAGCTGTCTTTGAT   3802
gb:MT044257|Organism:Severe      CATTCTTTAAGAGTTTGTGTAGATACTGTTCGCACAAATGTCTACTTAGCTGTCTTTGAT   3802
gb:MT106054|Organism:Severe      CATTCTTTAAGAGTTTGTGTAGATACTGTTCGCACAAATGTCTACTTAGCTGTCTTTGAT   3802
gb:MT049951|Organism:Severe      CATTCTTTAAGAGTTTGTGTAGATACTGTTCGCACAAATGTCTACTTAGCTGTCTTTGAT   3802
gb:MN975262|Organism:Severe      CATTCTTTAAGAGTTTGTGTAGATACTGTTCGCACAAATGTCTACTTAGCTGTCTTTGAT   3802
gb:MT106052|Organism:Severe      CATTCTTTAAGAGTTTGTGTAGATACTGTTCGCACAAATGTCTACTTAGCTGTCTTTGAT   3802
gb:LC522975|Organism:Severe      CATTCTTTAAGAGTTTGTGTAGATACTGTTCGCACAAATGTCTACTTAGCTGTCTTTGAT   3799
gb:LC522973|Organism:Severe      CATTCTTTAAGAGTTTGTGTAGATACTGTTCGCACAAATGTCTACTTAGTTGTCTTTGAT   3799
gb:LC522974|Organism:Severe      CATTCTTTAAGAGTTTGTGTAGATACTGTTCGCACAAATGTCTACTTAGCTGTCTTTGAT   3799
gb:MN985325|Organism:Severe      CATTCTTTAAGAGTTTGTGTAGATACTGTTCGCACAAATGTCTACTTAGCTGTCTTTGAT   3802
gb:MT020881|Organism:Severe      CATTCTTTAAGAGTTTGTGTAGATACTGTTCGCACAAATGTCTACTTAGCTGTCTTTGAT   3802
gb:MT020880|Organism:Severe      CATTCTTTAAGAGTTTGTGTAGATACTGTTCGCACAAATGTCTACTTAGCTGTCTTTGAT   3802
gb:MT066175|Organism:Severe      CATTCTTTAAGAGTTTGTGTAGATACTGTTCGCACAAATGTCTACTTAGCTGTCTTTGAT   3802
gb:MN997409|Organism:Severe      CATTCTTTAAGAGTTTGTGTAGATACTGTTCGCACAAATGTCTACTTAGCTGTCTTTGAT   3802
gb:MN938384|Organism:Severe      CATTCTTTAAGAGTTTGTGTAGATACTGTTCGCACAAATGTCTACTTAGCTGTCTTTGAT   3770
gb:MT044258|Organism:Severe      CATTCTTTAAGAGTTTGTGTAGATACTGTTCGCACAAATGTCTACTTAGCTGTCTTTGAT   3778
gb:MT039890|Organism:Severe      CATTCTTTAAGAGTTTGTGTAGATACTGTTCGCACAAATGTCTACTTAGCTGTCTTTGAT   3802
gb:MN988713|Organism:Severe      CATTCTTTAAGAGTTTGTGTAGATACTGTTCGCACAAATGTCTACTTAGCTGTCTTTGAT   3802
gb:LC521925|Organism:Severe      CATTCTTTAAGAGTTTGTGTAGATACTGTTCGCACAAATGTCTACTTAGCTGTCTTTGAT   3775
gb:MT093571|Organism:Severe      CATTCTTTAAGAGTTTGTGTAGATACTGTTCGCACAAATGTCTACTTAGCTGTCTTTGAT   3802
gb:MT039887|Organism:Severe      CATTCTTTAAGAGTTTGTGTAGATACTGTTCGCACAAATGTCTACTTAGCTGTCTTTGAT   3802
```

```
gb:MT019530|Organism:Severe      CATTCTTTAAGAGTTTGTGTAGATACTGTTCGCACAAATGTCTACTTAGCTGTCTTTGAT	3802
gb:MT039888|Organism:Severe      CATTCTTTAAGAGTTTGTGTAGATACTGTTCGCACAAATGTCTACTTAGCTGTCTTTGAT	3802
gb:LC522972|Organism:Severe      CATTCTTTAAGAGTTTGTGTAGATACTGTTCGCACAAATGTCTACTTAGCTGTCTTTGAT	3799
gb:MT027063|Organism:Severe      CATTCTTTAAGAGTTTGTGTAGATACTGTTCGCACAAATGTCTACTTAGCTGTCTTTGAT	3802
gb:MT027062|Organism:Severe      CATTCTTTAAGAGTTTGTGTAGATACTGTTCGCACAAATGTCTACTTAGCTGTCTTTGAT	3802
gb:MT019529|Organism:Severe      CATTCTTTAAGAGTTTGTGTAGATACTGTTCGCACGAATGTCTACTTAGCTGTCTTTGAT	3802
gb:MN996529|Organism:Severe      CATTCTTTAAGAGTTTGTGTAGATACTGTTCGCACAAATGTCTACTTAGCTGTCTTTGAT	3790
gb:MN996531|Organism:Severe      CATTCTTTAAGAGTTTGTGTAGATACTGTTCGCACAAATGTCTACTTAGCTGTCTTTGAT	3789
gb:MT066176|Organism:Severe      CATTCTTTAAGAGTTTGTGTAGATACTGTTCGCACAAATGTCTACTTAGCTGTCTTTGAT	3802
gb:MT027064|Organism:Severe      CATTCTTTAAGAGTTTGTGTAGATACTGTTCGCACAAATGTCTACTTAGCTGTCTTTGAT	3802
gb:MN994468|Organism:Severe      CATTCTTTAAGAGTTTGTGTAGATACTGTTCGCACAAATGTCTACTTAGCTGTCTTTGAT	3802
gb:MT072688|Organism:Severe      CATTCTTTAAGAGTTTGTGTAGATACTGTTCGCACAAATGTCTACTTAGCTGTCTTTGAT	3787
gb:MN996527|Organism:Severe      CATTCTTTAAGAGTTTGTGTAGATACTGTTCGCACAAATGTCTACTTAGCTGTCTTTGAT	3769
gb:MT093631|Organism:Severe      CATTCTTTAAGAGTTTGTGTAGATACTGTTCGCACAAATGTCTACTTAGCTGTCTTTGAT	3840
gb:MT106053|Organism:Severe      CATTCTTTAAGAGTTTGTGTAGATACTGTTCGCACAAATGTCTACTTAGCTGTCTTTGAT	3802
gb:MT019533|Organism:Severe      CATTCTTTAAGAGTTTGTGTAGATACTGTTCGCACAAATGTCTACTTAGCTGTCTTTGAT	3802
gb:MT019531|Organism:Severe      CATTCTTTAAGAGTTTGTGTAGATACTGTTCGCACAAATGTCTACTTAGCTGTCTTTGAT	3802
gb:MN996528|Organism:Severe      CATTCTTTAAGAGTTTGTGTAGATACTGTTCGCACAAATGTCTACTTAGCTGTCTTTGAT	3802
gb:MN996530|Organism:Severe      CATTCTTTAAGAGTTTGTGTAGATACTGTTCGCACAAATGTCTACTTAGCTGTCTTTGAT	3788
gb:MN908947|Organism:Severe      CATTCTTTAAGAGTTTGTGTAGATACTGTTCGCACAAATGTCTACTTAGCTGTCTTTGAT	3802
gb:MT019532|Organism:Severe      CATTCTTTAAGAGTTTGTGTAGATACTGTTCGCACAAATGTCTACTTAGCTGTCTTTGAT	3802
                                 *********************************** ************* **********

gb:MT020781|Organism:Severe      AAAAATCTCTATGACAAACTTGTTTCAAGCTTTTTGGAAATGAAGAGTGAAAAGCAAGTT	3850
gb:MT007544|Organism:Severe      AAAAATCTCTATGACAAACTTGTTTCAAGCTTTTTGGAAATGAAGAGTGAAAAGCAAGTT	3862
gb:MN994467|Organism:Severe      AAAAATCTCTATGACAAACTTGTTTCAAGCTTTTTGGAAATGAAGAGTGAAAAGCAAGTT	3862
gb:MT044257|Organism:Severe      AAAAATCTCTATGACAAACTTGTTTCAAGCTTTTTGGAAATGAAGAGTGAAAAGCAAGTT	3862
gb:MT106054|Organism:Severe      AAAAATCTCTATGACAAACTTGTTTCAAGCTTTTTGGAAATGAAGAGTGAAAAGCAAGTT	3862
gb:MT049951|Organism:Severe      AAAAATCTCTATGACAAACTTGTTTCAAGCTTTTTGGAAATGAAGAGTGAAAAGCAAGTT	3862
gb:MN975262|Organism:Severe      AAAAATCTCTATGACAAACTTGTTTCAAGCTTTTTGGAAATGAAGAGTGAAAAGCAAGTT	3862
gb:MT106052|Organism:Severe      AAAAATCTCTATGACAAACTTGTTTCAAGCTTTTTGGAAATGAAGAGTGAAAAGCAAGTT	3862
gb:LC522975|Organism:Severe      AAAAATCTCTATGACAAACTTGTTTCAAGCTTTTTGGAAATGAAGAGTGAAAAGCAAGTT	3859
gb:LC522973|Organism:Severe      AAAAATCTCTATGACAAACTTGTTTCAAGCTTTTTGGAAATGAAGAGTGAAAAGCAAGTT	3859
gb:LC522974|Organism:Severe      AAAAATCTCTATGACAAACTTGTTTCAAGCTTTTTGGAAATGAAGAGTGAAAAGCAAGTT	3859
gb:MN985325|Organism:Severe      AAAAATCTCTATGACAAACTTGTTTCAAGCTTTTTGGAAATGAAGAGTGAAAAGCAAGTT	3862
gb:MT020881|Organism:Severe      AAAAATCTCTATGACAAACTTGTTTCAAGCTTTTTGGAAATGAAGAGTGAAAAGCAAGTT	3862
gb:MT020880|Organism:Severe      AAAAATCTCTATGACAAACTTGTTTCAAGCTTTTTGGAAATGAAGAGTGAAAAGCAAGTT	3862
gb:MT066175|Organism:Severe      AAAAATCTCTATGACAAACTTGTTTCAAGCTTTTTGGAAATGAAGAGTGAAAAGCAAGTT	3862
gb:MN997409|Organism:Severe      AAAAATCTCTATGACAAACTTGTTTCAAGCTTTTTGGAAATGAAGAGTGAAAAGCAAGTT	3862
gb:MN938384|Organism:Severe      AAAAATCTCTATGACAAACTTGTTTCAAGCTTTTTGGAAATGAAGAGTGAAAAGCAAGTT	3830
gb:MT044258|Organism:Severe      AAAAATCTCTATGACAAACTTGTTTCAAGCTTTTTGGAAATGAAGAGTGAAAAGCAAGTT	3838
gb:MT039890|Organism:Severe      AAAAATCTCTATGACAAACTTGTTTCAAGCTTTTTGGAAATGAAGAGTGAAAAGCAAGTT	3862
gb:MN988713|Organism:Severe      AAAAATCTCTATGACAAACTTGTTTCAAGCTTTTTGGAAATGAAGAGTGAAAAGCAAGTT	3862
gb:LC521925|Organism:Severe      AAAAATCTCTATGACAAACTTGTTTCAAGCTTTTTGGAAATGAAGAGTGAAAAGCAAGTT	3835
gb:MT093571|Organism:Severe      AAAAATCTCTATGACAAACTTGTTTCAAGCTTTTTGGAAATGAAGAGTGAAAAGCAAGTT	3862
gb:MT039887|Organism:Severe      AAAAATCTCTATGACAAACTTGTTTCAAGCTTTTTGGAAATGAAGAGTGAAAAGCAAGTT	3862
gb:MT019530|Organism:Severe      AAAAATCTCTATGACAAACTTGTTTCAAGCTTTTTGGAAATGAAGAGTGAAAAGCAAGTT	3862
gb:MT039888|Organism:Severe      AAAAATCTCTATGACAAACTTGTTTCAAGCTTTTTGGAAATGAAGAGTGAAAAGCAAGTT	3862
gb:LC522972|Organism:Severe      AAAAATCTCTATGACAAACTTGTTTCAAGCTTTTTGGAAATGAAGAGTGAAAAGCAAGTT	3859
gb:MT027063|Organism:Severe      AAAAATCTCTATGACAAACTTGTTTCAAGCTTTTTGGAAATGAAGAGTGAAAAGCAAGTT	3862
gb:MT027062|Organism:Severe      AAAAATCTCTATGACAAACTTGTTTCAAGCTTTTTGGAAATGAAGAGTGAAAAGCAAGTT	3862
gb:MT019529|Organism:Severe      AAAAATCTCTATGACAAACTTGTTTCAAGCTTTTTGGAAATGAAGAGTGAAAAGCAAGTT	3862
gb:MN996529|Organism:Severe      AAAAATCTCTATGACAAACTTGTTTCAAGCTTTTTGGAAATGAAGAGTGAAAAGCAAGTT	3850
gb:MN996531|Organism:Severe      AAAAATCTCTATGACAAACTTGTTTCAAGCTTTTTGGAAATGAAGAGTGAAAAGCAAGTT	3849
gb:MT066176|Organism:Severe      AAAAATCTCTATGACAAACTTGTTTCAAGCTTTTTGGAAATGAAGAGTGAAAAGCAAGTT	3862
gb:MT027064|Organism:Severe      AAAAATCTCTATGACAAACTTGTTTCAAGCTTTTTGGAAATGAAGAGTGAAAAGCAAGTT	3862
gb:MN994468|Organism:Severe      AAAAATCTCTATGACAAACTTGTTTCAAGCTTTTTGGAAATGAAGAGTGAAAAGCAAGTT	3862
gb:MT072688|Organism:Severe      AAAAATCTCTATGACAAACTTGTTTCAAGCTTTTTGGAAATGAAGAGTGAAAAGCAAGTT	3847
gb:MN996527|Organism:Severe      AAAAATCTCTATGACAAACTTGTTTCAAGCTTTTTGGAAATGAAGAGTGAAAAGCAAGTT	3829
gb:MT093631|Organism:Severe      AAAAATCTCTATGACAAACTTGTTTCAAGCTTTTTGGAAATGAAGAGTGAAAAGCAAGTT	3900
gb:MT106053|Organism:Severe      AAAAATCTCTATGACAAACTTGTTTCAAGCTTTTTGGAAATGAAGAGTGAAAAGCAAGTT	3862
gb:MT019533|Organism:Severe      AAAAATCTCTATGACAAACTTGTTTCAAGCTTTTTGGAAATGAAGAGTGAAAAGCAAGTT	3862
gb:MT019531|Organism:Severe      AAAAATCTCTATGACAAACTTGTTTCAAGCTTTTTGGAAATGAAGAGTGAAAAGCAAGTT	3862
gb:MN996528|Organism:Severe      AAAAATCTCTATGACAAACTTGTTTCAAGCTTTTTGGAAATGAAGAGTGAAAAGCAAGTT	3862
gb:MN996530|Organism:Severe      AAAAATCTCTATGACAAACTTGTTTCAAGCTTTTTGGAAATGAAGAGTGAAAAGCAAGTT	3848
```

```
gb:MN908947|Organism:Severe      AAAAATCTCTATGACAAACTTGTTTCAAGCTTTTTGGAAATGAAGAGTGAAAAGCAAGTT    3862
gb:MT019532|Organism:Severe      AAAAATCTCTATGACAAACTTGTTTCAAGCTTTTTGGAAATGAAGAGTGAAAAGCAAGTT    3862
                                 ************************************************************

gb:MT020781|Organism:Severe      GAACAAAAGATCGCTGAGATTCCTAAAGAGGAAGTTAAGCCATTTATAACTGAAAGTAAA    3910
gb:MT007544|Organism:Severe      GAACAAAAGATCGCTGAGATTCCTAAAGAGGAAGTTAAGCCATTTATAACTGAAAGTAAA    3922
gb:MN994467|Organism:Severe      GAACAAAAGATCGCTGAGATTCCTAAAGAGGAAGTTAAGCCATTTATAACTGAAAGTAAA    3922
gb:MT044257|Organism:Severe      GAACAAAAGATCGCTGAGATTCCTAAAGAGGAAGTTAAGCCATTTATAACTGAAAGTAAA    3922
gb:MT106054|Organism:Severe      GAACAAAAGATCGCTGAGATTCCTAAAGAGGAAGTTAAGCCATTTATAACTGAAAGTAAA    3922
gb:MT049951|Organism:Severe      GAACAAAAGATCGCTGAGATTCCTAAAGAGGAAGTTAAGCCATTTATAACTGAAAGTAAA    3922
gb:MN975262|Organism:Severe      GAACAAAAGATCGCTGAGATTCCTAAAGAGGAAGTTAAGCCATTTATAACTGAAAGTAAA    3922
gb:MT106052|Organism:Severe      GAACAAAAGATCGCTGAGATTCCTAAAGAGGAAGTTAAGCCATTTATAACTGAAAGTAAA    3922
gb:LC522975|Organism:Severe      GAACAAAAGATCGCTGAGATTCCTAAAGAGGAAGTTAAGCCATTTATAACTGAAAGTAAA    3919
gb:LC522973|Organism:Severe      GAACAAAAGATCGCTGAGATTCCTAAAGAGGAAGTTAAGCCATTTATAACTGAAAGTAAA    3919
gb:LC522974|Organism:Severe      GAACAAAAGATCGCTGAGATTCCTAAAGAGGAAGTTAAGCCATTTATAACTGAAAGTAAA    3919
gb:MN985325|Organism:Severe      GAACAAAAGATCGCTGAGATTCCTAAAGAGGAAGTTAAGCCATTTATAACTGAAAGTAAA    3922
gb:MT020881|Organism:Severe      GAACAAAAGATCGCTGAGATTCCTAAAGAGGAAGTTAAGCCATTTATAACTGAAAGTAAA    3922
gb:MT020880|Organism:Severe      GAACAAAAGATCGCTGAGATTCCTAAAGAGGAAGTTAAGCCATTTATAACTGAAAGTAAA    3922
gb:MT066175|Organism:Severe      GAACAAAAGATCGCTGAGATTCCTAAAGAGGAAGTTAAGCCATTTATAACTGAAAGTAAA    3922
gb:MN997409|Organism:Severe      GAACAAAAGATCGCTGAGATTCCTAAAGAGGAAGTTAAGCCATTTATAACTGAAAGTAAA    3922
gb:MN938384|Organism:Severe      GAACAAAAGATCGCTGAGATTCCTAAAGAGGAAGTTAAGCCATTTATAACTGAAAGTAAA    3890
gb:MT044258|Organism:Severe      GAACAAAAGATCGCTGAGATTCCTAAAGAGGAAGTTAAGCCATTTATAACTGAAAGTAAA    3898
gb:MT039890|Organism:Severe      GAACAAAAGATCGCTGAGATTCCTAAAGAGGAAGTTAAGCCATTTATAACTGAAAGTAAA    3922
gb:MN988713|Organism:Severe      GAACAAAAGATCGCTGAGATTCCTAAAGAGGAAGTTAAGCCATTTATAACTGAAAGTAAA    3922
gb:LC521925|Organism:Severe      GAACAAAAGATCGCTGAGATTCCTAAAGAGGAAGTTAAGCCATTTATAACTGAAAGTAAA    3895
gb:MT093571|Organism:Severe      GAACAAAAGATCGCTGAGATTCCTAAAGAGGAAGTTAAGCCATTTATAACTGAAAGTAAA    3922
gb:MT039887|Organism:Severe      GAACAAAAGATCGCTGAGATTCCTAAAGAGGAAGTTAAGCCATTTATAACTGAAAGTAAA    3922
gb:MT019530|Organism:Severe      GAACAAAAGATCGCTGAGATTCCTAAAGAGGAAGTTAAGCCATTTATAACTGAAAGTAAA    3922
gb:MT039888|Organism:Severe      GAACAAAAGATCGCTGAGATTCCTAAAGAGGAAGTTAAGCCATTTATAACTGAAAGTAAA    3922
gb:LC522972|Organism:Severe      GAACAAAAGATCGCTGAGATTCCTAAAGAGGAAGTTAAGCCATTTATAACTGAAAGTAAA    3919
gb:MT027063|Organism:Severe      GAACAAAAGATCGCTGAGATTCCTAAAGAGGAAGTTAAGCCATTTATAACTGAAAGTAAA    3922
gb:MT027062|Organism:Severe      GAACAAAAGATCGCTGAGATTCCTAAAGAGGAAGTTAAGCCATTTATAACTGAAAGTAAA    3922
gb:MT019529|Organism:Severe      GAACAAAAGATCGCTGAGATTCCTAAAGAGGAAGTTAAGCCATTTATAACTGAAAGTAAA    3922
gb:MN996529|Organism:Severe      GAACAAAAGATCGCTGAGATTCCTAAAGAGGAAGTTAAGCCATTTATAACTGAAAGTAAA    3910
gb:MN996531|Organism:Severe      GAACAAAAGATCGCTGAGATTCCTAAAGAGGAAGTTAAGCCATTTATAACTGAAAGTAAA    3909
gb:MT066176|Organism:Severe      GAACAAAAGATCGCTGAGATTCCTAAAGAGGAAGTTAAGCCATTTATAACTGAAAGTAAA    3922
gb:MT027064|Organism:Severe      GAACAAAAGATCGCTGAGATTCCTAAAGAGGAAGTTAAGCCATTTATAACTGAAAGTAAA    3922
gb:MN994468|Organism:Severe      GAACAAAAGATCGCTGAGATTCCTAAAGAGGAAGTTAAGCCATTTATAACTGAAAGTAAA    3922
gb:MT072688|Organism:Severe      GAACAAAAGATCGCTGAGATTCCTAAAGAGGAAGTTAAGCCATTTATAACTGAAAGTAAA    3907
gb:MN996527|Organism:Severe      GAACAAAAGATCGCTGAGATTCCTAAAGAGGAAGTTAAGCCATTTATAACTGAAAGTAAA    3889
gb:MT093631|Organism:Severe      GAACAAAAGATCGCTGAGATTCCTAAAGAGGAAGTTAAGCCATTTATAACTGAAAGTAAA    3960
gb:MT106053|Organism:Severe      GAACAAAAGATCGCTGAGATTCCTAAAGAGGAAGTTAAGCCATTTATAACTGAAAGTAAA    3922
gb:MT019533|Organism:Severe      GAACAAAAGATCGCTGAGATTCCTAAAGAGGAAGTTAAGCCATTTATAACTGAAAGTAAA    3922
gb:MT019531|Organism:Severe      GAACAAAAGATCGCTGAGATTCCTAAAGAGGAAGTTAAGCCATTTATAACTGAAAGTAAA    3922
gb:MN996528|Organism:Severe      GAACAAAAGATCGCTGAGATTCCTAAAGAGGAAGTTAAGCCATTTATAACTGAAAGTAAA    3922
gb:MN996530|Organism:Severe      GAACAAAAGATCGCTGAGATTCCTAAAGAGGAAGTTAAGCCATTTATAACTGAAAGTAAA    3908
gb:MN908947|Organism:Severe      GAACAAAAGATCGCTGAGATTCCTAAAGAGGAAGTTAAGCCATTTATAACTGAAAGTAAA    3922
gb:MT019532|Organism:Severe      GAACAAAAGATCGCTGAGATTCCTAAAGAGGAAGTTAAGCCATTTATAACTGAAAGTAAA    3922
                                 ************************************************************

gb:MT020781|Organism:Severe      CCTTCAGTTGAACAGAGAAAACAAGATGATAAGAAAATCAAAGCTTGTGTTGAAGAAGTT    3970
gb:MT007544|Organism:Severe      CCTTCAGTTGAACAGAGAAAACAAGATGATAAGAAAATCAAAGCTTGTGTTGAAGAAGTT    3982
gb:MN994467|Organism:Severe      CCTTCAGTTGAACAGAGAAAACAAGATGATAAGAAAATCAAAGCTTGTGTTGAAGAAGTT    3982
gb:MT044257|Organism:Severe      CCTTCAGTTGAACAGAGAAAACAAGATGATAAGAAAATCAAAGCTTGTGTTGAAGAAGTT    3982
gb:MT106054|Organism:Severe      CCTTCAGTTGAACAGAGAAAACAAGATGATAAGAAAATCAAAGCTTGTGTTGAAGAAGTT    3982
gb:MT049951|Organism:Severe      CCTTCAGTTGAACAGAGAAAACAAGATGATAAGAAAATCAAAGCTTGTGTTGAAGAAGTT    3982
gb:MN975262|Organism:Severe      CCTTCAGTTGAACAGAGAAAACAAGATGATAAGAAAATCAAAGCTTGTGTTGAAGAAGTT    3982
gb:MT106052|Organism:Severe      CCTTCAGTTGAACAGAGAAAACAAGATGATAAGAAAATCAAAGCTTGTGTTGAAGAAGTT    3982
gb:LC522975|Organism:Severe      CCTTCAGTTGAACAGAGAAAACAAGATGATAAGAAAATCAAAGCTTGTGTTGAAGAAGTT    3979
gb:LC522973|Organism:Severe      CCTTCAGTTGAACAGAGAAAACAAGATGATAAGAAAATCAAAGCTTGTGTTGAAGAAGTT    3979
gb:LC522974|Organism:Severe      CCTTCAGTTGAACAGAGAAAACAAGATGATAAGAAAATCAAAGCTTGTGTTGAAGAAGTT    3979
gb:MN985325|Organism:Severe      CCTTCAGTTGAACAGAGAAAACAAGATGATAAGAAAATCAAAGCTTGTGTTGAAGAAGTT    3982
gb:MT020881|Organism:Severe      CCTTCAGTTGAACAGAGAAAACAAGATGATAAGAAAATCAAAGCTTGTGTTGAAGAAGTT    3982
gb:MT020880|Organism:Severe      CCTTCAGTTGAACAGAGAAAACAAGATGATAAGAAAATCAAAGCTTGTGTTGAAGAAGTT    3982
gb:MT066175|Organism:Severe      CCTTCAGTTGAACAGAGAAAACAAGATGATAAGAAAATCAAAGCTTGTGTTGAAGAAGTT    3982
```

```
gb:MN997409|Organism:Severe      CCTTCAGTTGAACAGAGAAAACAAGATGATAAGAAAATCAAAGCTTGTGTTGAAGAAGTT   3982
gb:MN938384|Organism:Severe      CCTTCAGTTGAACAGAGAAAACAAGATGATAAGAAAATCAAAGCTTGTGTTGAAGAAGTT   3950
gb:MT044258|Organism:Severe      CCTTCAGTTGAACAGAGAAAACAAGATGATAAGAAAATCAAAGCTTGTGTTGAAGAAGTT   3958
gb:MT039890|Organism:Severe      CCTTCAGTTGAACAGAGAAAACAAGATGATAAGAAAATCAAAGCTTGTGTTGAAGAAGTT   3982
gb:MN988713|Organism:Severe      CCTTCAGTTGAACAGAGAAAACAAGATGATAAGAAAATCAAAGCTTGTGTTGAAGAAGTT   3982
gb:LC521925|Organism:Severe      CCTTCAGTTGAACAGAGAAAACAAGATGATAAGAAAATCAAAGCTTGTGTTGAAGAAGTT   3955
gb:MT093571|Organism:Severe      CCTTCAGTTGAACAGAGAAAACAAGATGATAAGAAAATCAAAGCTTGTGTTGAAGAAGTT   3982
gb:MT039887|Organism:Severe      CCTTCAGTTGAACAGAGAAAACAAGATGATAAGAAAATCAAAGCTTGTGTTGAAGAAGTT   3982
gb:MT019530|Organism:Severe      CCTTCAGTTGAACAGAGAAAACAAGATGATAAGAAAATCAAAGCTTGTGTTGAAGAAGTT   3982
gb:MT039888|Organism:Severe      CCTTCAGTTGAACAGAGAAAACAAGATGATAAGAAAATCAAAGCTTGTGTTGAAGAAGTT   3982
gb:LC522972|Organism:Severe      CCTTCAGTTGAACAGAGAAAACAAGATGATAAGAAAATCAAAGCTTGTGTTGAAGAAGTT   3979
gb:MT027063|Organism:Severe      CCTTCAGTTGAACAGAGAAAACAAGATGATAAGAAAATCAAAGCTTGTGTTGAAGAAGTT   3982
gb:MT027062|Organism:Severe      CCTTCAGTTGAACAGAGAAAACAAGATGATAAGAAAATCAAAGCTTGTGTTGAAGAAGTT   3982
gb:MT019529|Organism:Severe      CCTTCAGTTGAACAGAGAAAACAAGATGATAAGAAAATCAAAGCTTGTGTTGAAGAAGTT   3982
gb:MN996529|Organism:Severe      CCTTCAGTTGAACAGAGAAAACAAGATGATAAGAAAATCAAAGCTTGTGTTGAAGAAGTT   3970
gb:MN996531|Organism:Severe      CCTTCAGTTGAACAGAGAAAACAAGATGATAAGAAAATCAAAGCTTGTGTTGAAGAAGTT   3969
gb:MT066176|Organism:Severe      CCTTCAGTTGAACAGAGAAAACAAGATGATAAGAAAATCAAAGCTTGTGTTGAAGAAGTT   3982
gb:MT027064|Organism:Severe      CCTTCAGTTGAACAGAGAAAACAAGATGATAAGAAAATCAAAGCTTGTGTTGAAGAAGTT   3982
gb:MN994468|Organism:Severe      CCTTCAGTTGAACAGAGAAAACAAGATGATAAGAAAATCAAAGCTTGTGTTGAAGAAGTT   3982
gb:MT072688|Organism:Severe      CCTTCAGTTGAACAGAGAAAACAAGATGATAAGAAAATCAAAGCTTGTGTTGAAGAAGTT   3967
gb:MN996527|Organism:Severe      CCTTCAGTTGAACAGAGAAAACAAGATGATAAGAAAATCAAAGCTTGTGTTGAAGAAGTT   3949
gb:MT093631|Organism:Severe      CCTTCAGTTGAACAGAGAAAACAAGATGATAAGAAAATCAAAGCTTGTGTTGAAGAAGTT   4020
gb:MT106053|Organism:Severe      CCTTCAGTTGAACAGAGAAAACAAGATGATAAGAAAATCAAAGCTTGTGTTGAAGAAGTT   3982
gb:MT019533|Organism:Severe      CCTTCAGTTGAACAGAGAAAACAAGATGATAAGAAAATCAAAGCTTGTGTTGAAGAAGTT   3982
gb:MT019531|Organism:Severe      CCTTCAGTTGAACAGAGAAAACAAGATGATAAGAAAATCAAAGCTTGTGTTGAAGAAGTT   3982
gb:MN996528|Organism:Severe      CCTTCAGTTGAACAGAGAAAACAAGATGATAAGAAAATCAAAGCTTGTGTTGAAGAAGTT   3982
gb:MN996530|Organism:Severe      CCTTCAGTTGAACAGAGAAAACAAGATGATAAGAAAATCAAAGCTTGTGTTGAAGAAGTT   3968
gb:MN908947|Organism:Severe      CCTTCAGTTGAACAGAGAAAACAAGATGATAAGAAAATCAAAGCTTGTGTTGAAGAAGTT   3982
gb:MT019532|Organism:Severe      CCTTCAGTTGAACAGAGAAAACAAGATGATAAGAAAATCAAAGCTTGTGTTGAAGAAGTT   3982
                                 ************************************************************

gb:MT020781|Organism:Severe      ACAACAACTCTGGAAGAAACTAAGTTCCTCACAGAAAACTTGTTACTTTATATTGACATT   4030
gb:MT007544|Organism:Severe      ACAACAACTCTGGAAGAAACTAAGTTCCTCACAGAAAACTTGTTACTTTATATTGACATT   4042
gb:MN994467|Organism:Severe      ACAACAACTCTGGAAGAAACTAAGTTCCTCACAGAAAACTTGTTACTTTATATTGACATT   4042
gb:MT044257|Organism:Severe      ACAACAACTCTGGAAGAAACTAAGTTCCTCACAGAAAACTTGTTACTTTATATTGACATT   4042
gb:MT106054|Organism:Severe      ACAACAACTCTGGAAGAAACTAAGTTCCTCACAGAAAACTTGTTACTTTATATTGACATT   4042
gb:MT049951|Organism:Severe      ACAACAACTCTGGAAGAAACTAAGTTCCTCACAGAAAACTTGTTACTTTATATTGACATT   4042
gb:MN975262|Organism:Severe      ACAACAACTCTGGAAGAAACTAAGTTCCTCACAGAAAACTTGTTACTTTATATTGACATT   4042
gb:MT106052|Organism:Severe      ACAACAACTCTGGAAGAAACTAAGTTCCTCACAGAAAACTTGTTACTTTATATTGACATT   4042
gb:LC522975|Organism:Severe      ACAACAACTCTGGAAGAAACTAAGTTCCTCACAGAAAACTTGTTACTTTATATTGACATT   4039
gb:LC522973|Organism:Severe      ACAACAACTCTGGAAGAAACTAAGTTCCTCACAGAAAACTTGTTACTTTATATTGACATT   4039
gb:LC522974|Organism:Severe      ACAACAACTCTGGAAGAAACTAAGTTCCTCACAGAAAACTTGTTACTTTATATTGACATT   4039
gb:MN985325|Organism:Severe      ACAACAACTCTGGAAGAAACTAAGTTCCTCACAGAAAACTTGTTACTTTATATTGACATT   4042
gb:MT020881|Organism:Severe      ACAACAACTCTGGAAGAAACTAAGTTCCTCACAGAAAACTTGTTACTTTATATTGACATT   4042
gb:MT020880|Organism:Severe      ACAACAACTCTGGAAGAAACTAAGTTCCTCACAGAAAACTTGTTACTTTATATTGACATT   4042
gb:MT066175|Organism:Severe      ACAACAACTCTGGAAGAAACTAAGTTCCTCACAGAAAACTTGTTACTTTATATTGACATT   4042
gb:MN997409|Organism:Severe      ACAACAACTCTGGAAGAAACTAAGTTCCTCACAGAAAACTTGTTACTTTATATTGACATT   4042
gb:MN938384|Organism:Severe      ACAACAACTCTGGAAGAAACTAAGTTCCTCACAGAAAACTTGTTACTTTATATTGACATT   4010
gb:MT044258|Organism:Severe      ACAACAACTCTGGAAGAAACTAAGTTCCTCACAGAAAACTTGTTACTTTATATTGACATT   4018
gb:MT039890|Organism:Severe      ACAACAACTCTGGAAGAAACTAAGTTCCTCACAGAAAACTTGTTACTTTATATTGACATT   4042
gb:MN988713|Organism:Severe      ACAACAACTCTGGAAGAAACTAAGTTCCTCACAGAAAACTTGTTACTTTATATTGACATT   4042
gb:LC521925|Organism:Severe      ACAACAACTCTGGAAGAAACTAAGTTCCTCACAGAAAACTTGTTACTTTATATTGACATT   4015
gb:MT093571|Organism:Severe      ACAACAACTCTGGAAGAAACTAAGTTCCTCACAGAAAACTTGTTACTTTATATTGACATT   4042
gb:MT039887|Organism:Severe      ACAACAACTCTGGAAGAAACTAAGTTCCTCACAGAAAACTTGTTACTTTATATTGACATT   4042
gb:MT019530|Organism:Severe      ACAACAACTCTGGAAGAAACTAAGTTCCTCACAGAAAACTTGTTACTTTATATTGACATT   4042
gb:MT039888|Organism:Severe      ACAACAACTCTGGAAGAAACTAAGTTCCTCACAGAAAACTTGTTACTTTATATTGACATT   4042
gb:LC522972|Organism:Severe      ACAACAACTCTGGAAGAAACTAAGTTCCTCACAGAAAACTTGTTACTTTATATTGACATT   4039
gb:MT027063|Organism:Severe      ACAACAACTCTGGAAGAAACTAAGTTCCTCACAGAAAACTTGTTACTTTATATTGACATT   4042
gb:MT027062|Organism:Severe      ACAACAACTCTGGAAGAAACTAAGTTCCTCACAGAAAACTTGTTACTTTATATTGACATT   4042
gb:MT019529|Organism:Severe      ACAACAACTCTGGAAGAAACTAAGTTCCTCACAGAAAACTTGTTACTTTATATTGACATT   4042
gb:MN996529|Organism:Severe      ACAACAACTCTGGAAGAAACTAAGTTCCTCACAGAAAACTTGTTACTTTATATTGACATT   4030
gb:MN996531|Organism:Severe      ACAACAACTCTGGAAGAAACTAAGTTCCTCACAGAAAACTTGTTACTTTATATTGACATT   4029
gb:MT066176|Organism:Severe      ACAACAACTCTGGAAGAAACTAAGTTCCTCACAGAAAACTTGTTACTTTATATTGACATT   4042
gb:MT027064|Organism:Severe      ACAACAACTCTGGAAGAAACTAAGTTCCTCACAGAAAACTTGTTACTTTATATTGACATT   4042
gb:MN994468|Organism:Severe      ACAACAACTCTGGAAGAAACTAAGTTCCTCACAGAAAACTTGTTACTTTATATTGACATT   4042
```

```
gb:MT072688|Organism:Severe      ACAACAACTCTGGAAGAAACTAAGTTCCTCACAGAAAACTTGTTACTTTATATTGACATT   4027
gb:MN996527|Organism:Severe      ACAACAACTCTGGAAGAAACTAAGTTCCTCACAGAAAACTTGTTACTTTATATTGACATT   4009
gb:MT093631|Organism:Severe      ACAACAACTCTGGAAGAAACTAAGTTCCTCACAGAAAACTTGTTACTTTATATTGACATT   4080
gb:MT106053|Organism:Severe      ACAACAACTCTGGAAGAAACTAAGTTCCTCACAGAAAACTTGTTACTTTATATTGACATT   4042
gb:MT019533|Organism:Severe      ACAACAACTCTGGAAGAAACTAAGTTCCTCACAGAAAACTTGTTACTTTATATTGACATT   4042
gb:MT019531|Organism:Severe      ACAACAACTCTGGAAGAAACTAAGTTCCTCACAGAAAACTTGTTACTTTATATTGACATT   4042
gb:MN996528|Organism:Severe      ACAACAACTCTGGAAGAAACTAAGTTCCTCACAGAAAACTTGTTACTTTATATTGACATT   4042
gb:MN996530|Organism:Severe      ACAACAACTCTGGAAGAAACTAAGTTCCTCACAGAAAACTTGTTACTTTATATTGACATT   4028
gb:MN908947|Organism:Severe      ACAACAACTCTGGAAGAAACTAAGTTCCTCACAGAAAACTTGTTACTTTATATTGACATT   4042
gb:MT019532|Organism:Severe      ACAACAACTCTGGAAGAAACTAAGTTCCTCACAGAAAACTTGTTACTTTATATTGACATT   4042
                                 ************************************************************

gb:MT020781|Organism:Severe      AATGGCAATCTTCATCCAGATTCTGCCACTCTTGTTAGTGACATTGACATCACTTTCTTA   4090
gb:MT007544|Organism:Severe      AATGGCAATCTTCATCCAGATTCTGCCACTCTTGTTAGTGACATTGACATCACTTTCTTA   4102
gb:MN994467|Organism:Severe      AATGGCAATCTTCATCCAGATTCTGCCACTCTTGTTAGTGACATTGACATCACTTTCTTA   4102
gb:MT044257|Organism:Severe      AATGGCAATCTTCATCCAGATTCTGCCACTCTTGTTAGTGACATTGACATCACTTTCTTA   4102
gb:MT106054|Organism:Severe      AATGGCAATCTTCATCCAGATTCTGCCACTCTTGTTAGTGACATTGACATCACTTTCTTA   4102
gb:MT049951|Organism:Severe      AATGGCAATCTTCATCCAGATTCTGCCACTCTTGTTAGTGACATTGACATCACTTTCTTA   4102
gb:MN975262|Organism:Severe      AATGGCAATCTTCATCCAGATTCTGCCACTCTTGTTAGTGACATTGACATCACTTTCTTA   4102
gb:MT106052|Organism:Severe      AATGGCAATCTTCATCCAGATTCTGCCACTCTTGTTAGTGACATTGACATCACTTTCTTA   4102
gb:LC522975|Organism:Severe      AATGGCAATCTTCATCCAGATTCTGCCACTCTTGTTAGTGACATTGACATCACTTTCTTA   4099
gb:LC522973|Organism:Severe      AATGGCAATCTTCATCCAGATTCTGCCACTCTTGTTAGTGACATTGACATCACTTTCTTA   4099
gb:LC522974|Organism:Severe      AATGGCAATCTTCATCCAGATTCTGCCACTCTTGTTAGTGACATTGACATCACTTTCTTA   4099
gb:MN985325|Organism:Severe      AATGGCAATCTTCATCCAGATTCTGCCACTCTTGTTAGTGACATTGACATCACTTTCTTA   4102
gb:MT020881|Organism:Severe      AATGGCAATCTTCATCCAGATTCTGCCACTCTTGTTAGTGACATTGACATCACTTTCTTA   4102
gb:MT020880|Organism:Severe      AATGGCAATCTTCATCCAGATTCTGCCACTCTTGTTAGTGACATTGACATCACTTTCTTA   4102
gb:MT066175|Organism:Severe      AATGGCAATCTTCATCCAGATTCTGCCACTCTTGTTAGTGACATTGACATCACTTTCTTA   4102
gb:MN997409|Organism:Severe      AATGGCAATCTTCATCCAGATTCTGCCACTCTTGTTAGTGACATTGACATCACTTTCTTA   4102
gb:MN938384|Organism:Severe      AATGGCAATCTTCATCCAGATTCTGCCACTCTTGTTAGTGACATTGACATCACTTTCTTA   4070
gb:MT044258|Organism:Severe      AATGGCAATCTTCATCCAGATTCTGCCACTCTTGTTAGTGACATTGACATCACTTTCTTA   4078
gb:MT039890|Organism:Severe      AATGGCAATCTTCATCCAGATTCTGCCACTCTTGTTAGTGACATTGACATCACTTTCTTA   4102
gb:MN988713|Organism:Severe      AATGGCAATCTTCATCCAGATTCTGCCACTCTTGTTAGTGACATTGACATCACTTTCTTA   4102
gb:LC521925|Organism:Severe      AATGGCAATCTTCATCCAGATTCTGCCACTCTTGTTAGTGACATTGACATCACTTTCTTA   4075
gb:MT093571|Organism:Severe      AATGGCAATCTTCATCCAGATTCTGCCACTCTTGTTAGTGACATTGACATCACTTTCTTA   4102
gb:MT039887|Organism:Severe      AATGGCAATCTTCATCCAGATTCTGCCACTCTTGTTAGTGACATTGACATCACTTTCTTA   4102
gb:MT019530|Organism:Severe      AATGGCAATCTTCATCCAGATTCTGCCACTCTTGTTAGTGACATTGACATCACTTTCTTA   4102
gb:MT039888|Organism:Severe      AATGGCAATCTTCATCCAGATTCTGCCACTCTTGTTAGTGACATTGACATCACTTTCTTA   4102
gb:LC522972|Organism:Severe      AATGGCAATCTTCATCCAGATTCTGCCACTCTTGTTAGTGACATTGACATCACTTTCTTA   4099
gb:MT027063|Organism:Severe      AATGGCAATCTTCATCCAGATTCTGCCACTCTTGTTAGTGACATTGACATCACTTTCTTA   4102
gb:MT027062|Organism:Severe      AATGGCAATCTTCATCCAGATTCTGCCACTCTTGTTAGTGACATTGACATCACTTTCTTA   4102
gb:MT019529|Organism:Severe      AATGGCAATCTTCATCCAGATTCTGCCACTCTTGTTAGTGACATTGACATCACTTTCTTA   4102
gb:MN996529|Organism:Severe      AATGGCAATCTTCATCCAGATTCTGCCACTCTTGTTAGTGACATTGACATCACTTTCTTA   4090
gb:MN996531|Organism:Severe      AATGGCAATCTTCATCCAGATTCTGCCACTCTTGTTAGTGACATTGACATCACTTTCTTA   4089
gb:MT066176|Organism:Severe      AATGGCAATCTTCATCCAGATTCTGCCACTCTTGTTAGTGACATTGACATCACTTTCTTA   4102
gb:MT027064|Organism:Severe      AATGGCAATCTTCATCCAGATTCTGCCACTCTTGTTAGTGACATTGACATCACTTTCTTA   4102
gb:MN994468|Organism:Severe      AATGGCAATCTTCATCCAGATTCTGCCACTCTTGTTAGTGACATTGACATCACTTTCTTA   4102
gb:MT072688|Organism:Severe      AATGGCAATCTTCATCCAGATTCTGCCACTCTTGTTAGTGACATTGACATCACTTTCTTA   4087
gb:MN996527|Organism:Severe      AATGGCAATCTTCATCCAGATTCTGCCACTCTTGTTAGTGACATTGACATCACTTTCTTA   4069
gb:MT093631|Organism:Severe      AATGGCAATCTTCATCCAGATTCTGCCACTCTTGTTAGTGACATTGACATCACTTTCTTA   4140
gb:MT106053|Organism:Severe      AATGGCAATCTTCATCCAGATTCTGCCACTCTTGTTAGTGACATTGACATCACTTTCTTA   4102
gb:MT019533|Organism:Severe      AATGGCAATCTTCATCCAGATTCTGCCACTCTTGTTAGTGACATTGACATCACTTTCTTA   4102
gb:MT019531|Organism:Severe      AATGGCAATCTTCATCCAGATTCTGCCACTCTTGTTAGTGACATTGACATCACTTTCTTA   4102
gb:MN996528|Organism:Severe      AATGGCAATCTTCATCCAGATTCTGCCACTCTTGTTAGTGACATTGACATCACTTTCTTA   4102
gb:MN996530|Organism:Severe      AATGGCAATCTTCATCCAGATTCTGCCACTCTTGTTAGTGACATTGACATCACTTTCTTA   4088
gb:MN908947|Organism:Severe      AATGGCAATCTTCATCCAGATTCTGCCACTCTTGTTAGTGACATTGACATCACTTTCTTA   4102
gb:MT019532|Organism:Severe      AATGGCAATCTTCATCCAGATTCTGCCACTCTTGTTAGTGACATTGACATCACTTTCTTA   4102
                                 ************************************************************

gb:MT020781|Organism:Severe      AAGAAAGATGCTCCATATATAGTGGGTGATGTTGTTCAAGAGGGTGTTTTAACTGCTGTG   4150
gb:MT007544|Organism:Severe      AAGAAAGATGCTCCATATATAGTGGGTGATGTTGTTCAAGAGGGTGTTTTAACTGCTGTG   4162
gb:MN994467|Organism:Severe      AAGAAAGATGCTCCATATATAGTGGGTGATGTTGTTCAAGAGGGTGTTTTAACTGCTGTG   4162
gb:MT044257|Organism:Severe      AAGAAAGATGCTCCATATATAGTGGGTGATGTTGTTCAAGAGGGTGTTTTAACTGCTGTG   4162
gb:MT106054|Organism:Severe      AAGAAAGATGCTCCATATATAGTGGGTGATGTTGTTCAAGAGGGTGTTTTAACTGCTGTG   4162
gb:MT049951|Organism:Severe      AAGAAAGATGCTCCATATATAGTGGGTGATGTTGTTCAAGAGGGTGTTTTAACTGCTGTG   4162
gb:MN975262|Organism:Severe      AAGAAAGATGCTCCATATATAGTGGGTGATGTTGTTCAAGAGGGTGTTTTAACTGCTGTG   4162
```

```
gb:MT106052|Organism:Severe      AAGAAAGATGCTCCATATATAGTGGGTGATGTTGTTCAAGAGGGTGTTTTAACTGCTGTG   4162
gb:LC522975|Organism:Severe      AAGAAAGATGCTCCATATATAGTGGGTGATGTTGTTCAAGAGGGTGTTTTAACTGCTGTG   4159
gb:LC522973|Organism:Severe      AAGAAAGATGCTCCATATATAGTGGGTGATGTTGTTCAAGAGGGTGTTTTAACTGCTGTG   4159
gb:LC522974|Organism:Severe      AAGAAAGATGCTCCATATATAGTGGGTGATGTTGTTCAAGAGGGTGTTTTAACTGCTGTG   4159
gb:MN985325|Organism:Severe      AAGAAAGATGCTCCATATATAGTGGGTGATGTTGTTCAAGAGGGTGTTTTAACTGCTGTG   4162
gb:MT020881|Organism:Severe      AAGAAAGATGCTCCATATATAGTGGGTGATGTTGTTCAAGAGGGTGTTTTAACTGCTGTG   4162
gb:MT020880|Organism:Severe      AAGAAAGATGCTCCATATATAGTGGGTGATGTTGTTCAAGAGGGTGTTTTAACTGCTGTG   4162
gb:MT066175|Organism:Severe      AAGAAAGATGCTCCATATATAGTGGGTGATGTTGTTCAAGAGGGTGTTTTAACTGCTGTG   4162
gb:MN997409|Organism:Severe      AAGAAAGATGCTCCATATATAGTGGGTGATGTTGTTCAAGAGGGTGTTTTAACTGCTGTG   4162
gb:MN938384|Organism:Severe      AAGAAAGATGCTCCATATATAGTGGGTGATGTTGTTCAAGAGGGTGTTTTAACTGCTGTG   4130
gb:MT044258|Organism:Severe      AAGAAAGATGCTCCATATATAGTGGGTGATGTTGTTCAAGAGGGTGTTTTAACTGCTGTG   4138
gb:MT039890|Organism:Severe      AAGAAAGATGCTCCATATATAGTGGGTGATGTTGTTCAAGAGGGTGTTTTAACTGCTGTG   4162
gb:MN988713|Organism:Severe      AAGAAAGATGCTCCATATATAGTGGGTGATGTTGTTCAAGAGGGTGTTTTAACTGCTGTG   4162
gb:LC521925|Organism:Severe      AAGAAAGATGCTCCATATATAGTGGGTGATGTTGTTCAAGAGGGTGTTTTAACTGCTGTG   4135
gb:MT093571|Organism:Severe      AAGAAAGATGCTCCATATATAGTGGGTGATGTTGTTCAAGAGGGTGTTTTAACTGCTGTG   4162
gb:MT039887|Organism:Severe      AAGAAAGATGCTCCATATATAGTGGGTGATGTTGTTCAAGAGGGTGTTTTAACTGCTGTG   4162
gb:MT019530|Organism:Severe      AAGAAAGATGCTCCATATATAGTGGGTGATGTTGTTCAAGAGGGTGTTTTAACTGCTGTG   4162
gb:MT039888|Organism:Severe      AAGAAAGATGCTCCATATATAGTGGGTGATGTTGTTCAAGAGGGTGTTTTAACTGCTGTG   4162
gb:LC522972|Organism:Severe      AAGAAAGATGCTCCATATATAGTGGGTGATGTTGTTCAAGAGGGTGTTTTAACTGCTGTG   4159
gb:MT027063|Organism:Severe      AAGAAAGATGCTCCATATATAGTGGGTGATGTTGTTCAAGAGGGTGTTTTAACTGCTGTG   4162
gb:MT027062|Organism:Severe      AAGAAAGATGCTCCATATATAGTGGGTGATGTTGTTCAAGAGGGTGTTTTAACTGCTGTG   4162
gb:MT019529|Organism:Severe      AAGAAAGATGCTCCATATATAGTGGGTGATGTTGTTCAAGAGGGTGTTTTAACTGCTGTG   4162
gb:MN996529|Organism:Severe      AAGAAAGATGCTCCATATATAGTGGGTGATGTTGTTCAAGAGGGTGTTTTAACTGCTGTG   4150
gb:MN996531|Organism:Severe      AAGAAAGATGCTCCATATATAGTGGGTGATGTTGTTCAAGAGGGTGTTTTAACTGCTGTG   4149
gb:MT066176|Organism:Severe      AAGAAAGATGCTCCATATATAGTGGGTGATGTTGTTCAAGAGGGTGTTTTAACTGCTGTG   4162
gb:MT027064|Organism:Severe      AAGAAAGATGCTCCATATATAGTGGGTGATGTTGTTCAAGAGGGTGTTTTAACTGCTGTG   4162
gb:MN994468|Organism:Severe      AAGAAAGATGCTCCATATATAGTGGGTGATGTTGTTCAAGAGGGTGTTTTAACTGCTGTG   4162
gb:MT072688|Organism:Severe      AAGAAAGATGCTCCATATATAGTGGGTGATGTTGTTCAAGAGGGTGTTTTAACTGCTGTG   4147
gb:MN996527|Organism:Severe      AAGAAAGATGCTCCATATATAGTGGGTGATGTTGTTCAAGAGGGTGTTTTAACTGCTGTG   4129
gb:MT093631|Organism:Severe      AAGAAAGATGCTCCATATATAGTGGGTGATGTTGTTCAAGAGGGTGTTTTAACTGCTGTG   4200
gb:MT106053|Organism:Severe      AAGAAAGATGCTCCATATATAGTGGGTGATGTTGTTCAAGAGGGTGTTTTAACTGCTGTG   4162
gb:MT019533|Organism:Severe      AAGAAAGATGCTCCATATATAGTGGGTGATGTTGTTCAAGAGGGTGTTTTAACTGCTGTG   4162
gb:MT019531|Organism:Severe      AAGAAAGATGCTCCATATATAGTGGGTGATGTTGTTCAAGAGGGTGTTTTAACTGCTGTG   4162
gb:MN996528|Organism:Severe      AAGAAAGATGCTCCATATATAGTGGGTGATGTTGTTCAAGAGGGTGTTTTAACTGCTGTG   4162
gb:MN996530|Organism:Severe      AAGAAAGATGCTCCATATATAGTGGGTGATGTTGTTCAAGAGGGTGTTTTAACTGCTGTG   4148
gb:MN908947|Organism:Severe      AAGAAAGATGCTCCATATATAGTGGGTGATGTTGTTCAAGAGGGTGTTTTAACTGCTGTG   4162
gb:MT019532|Organism:Severe      AAGAAAGATGCTCCATATATAGTGGGTGATGTTGTTCAAGAGGGTGTTTTAACTGCTGTG   4162
                                 ************************************************************

gb:MT020781|Organism:Severe      GTTATACCTACTAAAAAGGCTGGTGGCACTACTGAAATGCTAGCGAAAGCTTTGAGAAAA   4210
gb:MT007544|Organism:Severe      GTTATACCTACTAAAAAGGCTGGTGGCACTACTGAAATGCTAGCGAAAGCTTTGAGAAAA   4222
gb:MN994467|Organism:Severe      GTTATACCTACTAAAAAGGCTGGTGGCACTACTGAAATGCTAGCGAAAGCTTTGAGAAAA   4222
gb:MT044257|Organism:Severe      GTTATACCTACTAAAAAGGCTGGTGGCACTACTGAAATGCTAGCGAAAGCTTTGAGAAAA   4222
gb:MT106054|Organism:Severe      GTTATACCTACTAAAAAGGCTGGTGGCACTACTGAAATGCTAGCGAAAGCTTTGAGAAAA   4222
gb:MT049951|Organism:Severe      GTTATACCTACTAAAAAGGCTGGTGGCACTACTGAAATGCTAGCGAAAGCTTTGAGAAAA   4222
gb:MN975262|Organism:Severe      GTTATACCTACTAAAAAGGCTGGTGGCACTACTGAAATGCTAGCGAAAGCTTTGAGAAAA   4222
gb:MT106052|Organism:Severe      GTTATACCTACTAAAAAGGCTGGTGGCACTACTGAAATGCTAGCGAAAGCTTTGAGAAAA   4222
gb:LC522975|Organism:Severe      GTTATACCTACTAAAAAGGCTGGTGGCACTACTGAAATGCTAGCGAAAGCTTTGAGAAAA   4219
gb:LC522973|Organism:Severe      GTTATACCTACTAAAAAGGCTGGTGGCACTACTGAAATGCTAGCGAAAGCTTTGAGAAAA   4219
gb:LC522974|Organism:Severe      GTTATACCTACTAAAAAGGCTGGTGGCACTACTGAAATGCTAGCGAAAGCTTTGAGAAAA   4219
gb:MN985325|Organism:Severe      GTTATACCTACTAAAAAGGCTGGTGGCACTACTGAAATGCTAGCGAAAGCTTTGAGAAAA   4222
gb:MT020881|Organism:Severe      GTTATACCTACTAAAAAGGCTGGTGGCACTACTGAAATGCTAGCGAAAGCTTTGAGAAAA   4222
gb:MT020880|Organism:Severe      GTTATACCTACTAAAAAGGCTGGTGGCACTACTGAAATGCTAGCGAAAGCTTTGAGAAAA   4222
gb:MT066175|Organism:Severe      GTTATACCTACTAAAAAGGCTGGTGGCACTACTGAAATGCTAGCGAAAGCTTTGAGAAAA   4222
gb:MN997409|Organism:Severe      GTTATACCTACTAAAAAGGCTGGTGGCACTACTGAAATGCTAGCGAAAGCTTTGAGAAAA   4222
gb:MN938384|Organism:Severe      GTTATACCTACTAAAAAGGCTGGTGGCACTACTGAAATGCTAGCGAAAGCTTTGAGAAAA   4190
gb:MT044258|Organism:Severe      GTTATACCTACTAAAAAGGCTGGTGGCACTACTGAAATGCTAGCGAAAGCTTTGAGAAAA   4198
gb:MT039890|Organism:Severe      GTTATACCTACTAAAAAGGCTGGTGGCACTACTGAAATGCTAGCGAAAGCTTTGAGAAAA   4222
gb:MN988713|Organism:Severe      GTTATACCTACTAAAAAGGCTGGTGGCACTACTGAAATGCTAGCGAAAGCTTTGAGAAAA   4222
gb:LC521925|Organism:Severe      GTTATACCTACTAAAAAGGCTGGTGGCACTACTGAAATGCTAGCGAAAGCTTTGAGAAAA   4195
gb:MT093571|Organism:Severe      GTTATACCTACTAAAAAGGCTGGTGGCACTACTGAAATGCTAGCGAAAGCTTTGAGAAAA   4222
gb:MT039887|Organism:Severe      GTTATACCTACTAAAAAGGCTGGTGGCACTACTGAAATGCTAGCGAAAGCTTTGAGAAAA   4222
gb:MT019530|Organism:Severe      GTTATACCTACTAAAAAGGCTGGTGGCACTACTGAAATGCTAGCGAAAGCTTTGAGAAAA   4222
gb:MT039888|Organism:Severe      GTTATACCTACTAAAAAGGCTGGTGGCACTACTGAAATGCTAGCGAAAGCTTTGAGAAAA   4222
gb:LC522972|Organism:Severe      GTTATACCTACTAAAAAGGCTGGTGGCACTACTGAAATGCTAGCGAAAGCTTTGAGAAAA   4219
```

```
gb:MT027063|Organism:Severe      GTTATACCTACTAAAAAGGCTGGTGGCACTACTGAAATGCTAGCGAAAGCTTTGAGAAAA    4222
gb:MT027062|Organism:Severe      GTTATACCTACTAAAAAGGCTGGTGGCACTACTGAAATGCTAGCGAAAGCTTTGAGAAAA    4222
gb:MT019529|Organism:Severe      GTTATACCTACTAAAAAGGCTGGTGGCACTACTGAAATGCTAGCGAAAGCTTTGAGAAAA    4222
gb:MN996529|Organism:Severe      GTTATACCTACTAAAAAGGCTGGTGGCACTACTGAAATGCTAGCGAAAGCTTTGAGAAAA    4210
gb:MN996531|Organism:Severe      GTTATACCTACTAAAAAGGCTGGTGGCACTACTGAAATGCTAGCGAAAGCTTTGAGAAAA    4209
gb:MT066176|Organism:Severe      GTTATACCTACTAAAAAGGCTGGTGGCACTACTGAAATGCTAGCGAAAGCTTTGAGAAAA    4222
gb:MT027064|Organism:Severe      GTTATACCTACTAAAAAGGCTGGTGGCACTACTGAAATGCTAGCGAAAGCTTTGAGAAAA    4222
gb:MN994468|Organism:Severe      GTTATACCTACTAAAAAGGCTGGTGGCACTACTGAAATGCTAGCGAAAGCTTTGAGAAAA    4222
gb:MT072688|Organism:Severe      GTTATACCTACTAAAAAGGCTGGTGGCACTACTGAAATGCTAGCGAAAGCTTTGAGAAAA    4207
gb:MN996527|Organism:Severe      GTTATACCTACTAAAAAGGCTGGTGGCACTACTGAAATGCTAGCGAAAGCTTTGAGAAAA    4189
gb:MT093631|Organism:Severe      GTTATACCTACTAAAAAGGCTGGTGGCACTACTGAAATGCTAGCGAAAGCTTTGAGAAAA    4260
gb:MT106053|Organism:Severe      GTTATACCTACTAAAAAGGCTGGTGGCACTACTGAAATGCTAGCGAAAGCTTTGAGAAAA    4222
gb:MT019533|Organism:Severe      GTTATACCTACTAAAAAGGCTGGTGGCACTACTGAAATGCTAGCGAAAGCTTTGAGAAAA    4222
gb:MT019531|Organism:Severe      GTTATACCTACTAAAAAGGCTGGTGGCACTACTGAAATGCTAGCGAAAGCTTTGAGAAAA    4222
gb:MN996528|Organism:Severe      GTTATACCTACTAAAAAGGCTGGTGGCACTACTGAAATGCTAGCGAAAGCTTTGAGAAAA    4222
gb:MN996530|Organism:Severe      GTTATACCTACTAAAAAGGCTGGTGGCACTACTGAAATGCTAGCGAAAGCTTTGAGAAAA    4208
gb:MN908947|Organism:Severe      GTTATACCTACTAAAAAGGCTGGTGGCACTACTGAAATGCTAGCGAAAGCTTTGAGAAAA    4222
gb:MT019532|Organism:Severe      GTTATACCTACTAAAAAGGCTGGTGGCACTACTGAAATGCTAGCGAAAGCTTTGAGAAAA    4222
                                 ************************************************************

gb:MT020781|Organism:Severe      GTGCCAACAGACAATTATATAACCACTTACCCGGGTCAGGGTTTAAATGGTTACACTGTA    4270
gb:MT007544|Organism:Severe      GTGCCAACAGACAATTATATAACCACTTACCCGGGTCAGGGTTTAAATGGTTACACTGTA    4282
gb:MN994467|Organism:Severe      GTGCCAACAGACAATTATATAACCACTTACCCGGGTCAGGGTTTAAATGGTTACACTGTA    4282
gb:MT044257|Organism:Severe      GTGCCAACAGACAATTATATAACCACTTACCCGGGTCAGGGTTTAAATGGTTACACTGTA    4282
gb:MT106054|Organism:Severe      GTGCCAACAGACAATTATATAACCACTTACCCGGGTCAGGGTTTAAATGGTTACACTGTA    4282
gb:MT049951|Organism:Severe      GTGCCAACAGACAATTATATAACCACTTACCCGGGTCAGGGTTTAAATGGTTACACTGTA    4282
gb:MN975262|Organism:Severe      GTGCCAACAGACAATTATATAACCACTTACCCGGGTCAGGGTTTAAATGGTTACACTGTA    4282
gb:MT106052|Organism:Severe      GTGCCAACAGACAATTATATAACCACTTACCCGGGTCAGGGTTTAAATGGTTACACTGTA    4282
gb:LC522975|Organism:Severe      GTGCCAACAGACAATTATATAACCACTTACCCGGGTCAGGGTTTAAATGGTTACACTGTA    4279
gb:LC522973|Organism:Severe      GTGCCAACAGACAATTATATAACCACTTACCCGGGTCAGGGTTTAAATGGTTACACTGTA    4279
gb:LC522974|Organism:Severe      GTGCCAACAGACAATTATATAACCACTTACCCGGGTCAGGGTTTAAATGGTTACACTGTA    4279
gb:MN985325|Organism:Severe      GTGCCAACAGACAATTATATAACCACTTACCCGGGTCAGGGTTTAAATGGTTACACTGTA    4282
gb:MT020881|Organism:Severe      GTGCCAACAGACAATTATATAACCACTTACCCGGGTCAGGGTTTAAATGGTTACACTGTA    4282
gb:MT020880|Organism:Severe      GTGCCAACAGACAATTATATAACCACTTACCCGGGTCAGGGTTTAAATGGTTACACTGTA    4282
gb:MT066175|Organism:Severe      GTGCCAACAGACAATTATATAACCACTTACCCGGGTCAGGGTTTAAATGGTTACACTGTA    4282
gb:MN997409|Organism:Severe      GTGCCAACAGACAATTATATAACCACTTACCCGGGTCAGGGTTTAAATGGTTACACTGTA    4282
gb:MN938384|Organism:Severe      GTGCCAACAGACAATTATATAACCACTTACCCGGGTCAGGGTTTAAATGGTTACACTGTA    4250
gb:MT044258|Organism:Severe      GTGCCAACAGACAATTATATAACCACTTACCCGGGTCAGGGTTTAAATGGTTACACTGTA    4258
gb:MT039890|Organism:Severe      GTGCCAACAGACAATTATATAACCACTTACCCGGGTCAGGGTTTAAATGGTTACACTGTA    4282
gb:MN988713|Organism:Severe      GTGCCAACAGACAATTATATAACCACTTACCCGGGTCAGGGTTTAAATGGTTACACTGTA    4282
gb:LC521925|Organism:Severe      GTGCCAACAGACAATTATATAACCACTTACCCGGGTCAGGGTTTAAATGGTTACACTGTA    4255
gb:MT093571|Organism:Severe      GTGCCAACAGACAATTATATAACCACTTACCCGGGTCAGGGTTTAAATGGTTACACTGTA    4282
gb:MT039887|Organism:Severe      GTGCCAACAGACAATTATATAACCACTTACCCGGGTCAGGGTTTAAATGGTTACACTGTA    4282
gb:MT019530|Organism:Severe      GTGCCAACAGACAATTATATAACCACTTACCCGGGTCAGGGTTTAAATGGTTACACTGTA    4282
gb:MT039888|Organism:Severe      GTGCCAACAGACAATTATATAACCACTTACCCGGGTCAGGGTTTAAATGGTTACACTGTA    4282
gb:LC522972|Organism:Severe      GTGCCAACAGACAATTATATAACCACTTACCCGGGTCAGGGTTTAAATGGTTACACTGTA    4279
gb:MT027063|Organism:Severe      GTGCCAACAGACAATTATATAACCACTTACCCGGGTCAGGGTTTAAATGGTTACACTGTA    4282
gb:MT027062|Organism:Severe      GTGCCAACAGACAATTATATAACCACTTACCCGGGTCAGGGTTTAAATGGTTACACTGTA    4282
gb:MT019529|Organism:Severe      GTGCCAACAGACAATTATATAACCACTTACCCGGGTCAGGGTTTAAATGGTTACACTGTA    4282
gb:MN996529|Organism:Severe      GTGCCAACAGACAATTATATAACCACTTACCCGGGTCAGGGTTTAAATGGTTACACTGTA    4270
gb:MN996531|Organism:Severe      GTGCCAACAGACAATTATATAACCACTTACCCGGGTCAGGGTTTAAATGGTTACACTGTA    4269
gb:MT066176|Organism:Severe      GTGCCAACAGACAATTATATAACCACTTACCCGGGTCAGGGTTTAAATGGTTACACTGTA    4282
gb:MT027064|Organism:Severe      GTGCCAACAGACAATTATATAACCACTTACCCGGGTCAGGGTTTAAATGGTTACACTGTA    4282
gb:MN994468|Organism:Severe      GTGCCAACAGACAATTATATAACCACTTACCCGGGTCAGGGTTTAAATGGTTACACTGTA    4282
gb:MT072688|Organism:Severe      GTGCCAACAGACAATTATATAACCACTTACCCGGGTCAGGGTTTAAATGGTTACACTGTA    4267
gb:MN996527|Organism:Severe      GTGCCAACAGACAATTATATAACCACTTACCCGGGTCAGGGTTTAAATGGTTACACTGTA    4249
gb:MT093631|Organism:Severe      GTGCCAACAGACAATTATATAACCACTTACCCGGGTCAGGGTTTAAATGGTTACACTGTA    4320
gb:MT106053|Organism:Severe      GTGCCAACAGACAATTATATAACCACTTACCCGGGTCAGGGTTTAAATGGTTACACTGTA    4282
gb:MT019533|Organism:Severe      GTGCCAACAGACAATTATATAACCACTTACCCGGGTCAGGGTTTAAATGGTTACACTGTA    4282
gb:MT019531|Organism:Severe      GTGCCAACAGACAATTATATAACCACTTACCCGGGTCAGGGTTTAAATGGTTACACTGTA    4282
gb:MN996528|Organism:Severe      GTGCCAACAGACAATTATATAACCACTTACCCGGGTCAGGGTTTAAATGGTTACACTGTA    4282
gb:MN996530|Organism:Severe      GTGCCAACAGACAATTATATAACCACTTACCCGGGTCAGGGTTTAAATGGTTACACTGTA    4268
gb:MN908947|Organism:Severe      GTGCCAACAGACAATTATATAACCACTTACCCGGGTCAGGGTTTAAATGGTTACACTGTA    4282
gb:MT019532|Organism:Severe      GTGCCAACAGACAATTATATAACCACTTACCCGGGTCAGGGTTTAAATGGTTACACTGTA    4282
                                 ************************************************************
```

```
gb:MT020781|Organism:Severe      GAGGAGGCAAAGACAGTGCTTAAAAAGTGTAAAAGTGCCTTTTACATTCTACCATCTATT   4330
gb:MT007544|Organism:Severe      GAGGAGGCAAAGACAGTGCTTAAAAAGTGTAAAAGTGCCTTTTACATTCTACCATCTATT   4342
gb:MN994467|Organism:Severe      GAGGAGGCAAAGACAGTGCTTAAAAAGTGTAAAAGTGCCTTTTACATTCTACCATCTATT   4342
gb:MT044257|Organism:Severe      GAGGAGGCAAAGACAGTGCTTAAAAAGTGTAAAAGTGCCTTTTACATTCTACCATCTATT   4342
gb:MT106054|Organism:Severe      GAGGAGGCAAAGACAGTGCTTAAAAAGTGTAAAAGTGCCTTTTACATTCTACCATCTATT   4342
gb:MT049951|Organism:Severe      GAGGAGGCAAAGACAGTGCTTAAAAAGTGTAAAAGTGCCTTTTACATTCTACCATCTATT   4342
gb:MN975262|Organism:Severe      GAGGAGGCAAAGACAGTGCTTAAAAAGTGTAAAAGTGCCTTTTACATTCTACCATCTATT   4342
gb:MT106052|Organism:Severe      GAGGAGGCAAAGACAGTGCTTAAAAAGTGTAAAAGTGCCTTTTACATTCTACCATCTATT   4342
gb:LC522975|Organism:Severe      GAGGAGGCAAAGACAGTGCTTAAAAAGTGTAAAAGTGCCTTTTACATTCTACCATCTATT   4339
gb:LC522973|Organism:Severe      GAGGAGGCAAAGACAGTGCTTAAAAAGTGTAAAAGTGCCTTTTACATTCTACCATCTATT   4339
gb:LC522974|Organism:Severe      GAGGAGGCAAAGACAGTGCTTAAAAAGTGTAAAAGTGCCTTTTACATTCTACCATCTATT   4339
gb:MN985325|Organism:Severe      GAGGAGGCAAAGACAGTGCTTAAAAAGTGTAAAAGTGCCTTTTACATTCTACCATCTATT   4342
gb:MT020881|Organism:Severe      GAGGAGGCAAAGACAGTGCTTAAAAAGTGTAAAAGTGCCTTTTACATTCTACCATCTATT   4342
gb:MT020880|Organism:Severe      GAGGAGGCAAAGACAGTGCTTAAAAAGTGTAAAAGTGCCTTTTACATTCTACCATCTATT   4342
gb:MT066175|Organism:Severe      GAGGAGGCAAAGACAGTGCTTAAAAAGTGTAAAAGTGCCTTTTACATTCTACCATCTATT   4342
gb:MN997409|Organism:Severe      GAGGAGGCAAAGACAGTGCTTAAAAAGTGTAAAAGTGCCTTTTACATTCTACCATCTATT   4342
gb:MN938384|Organism:Severe      GAGGAGGCAAAGACAGTGCTTAAAAAGTGTAAAAGTGCCTTTTACATTCTACCATCTATT   4310
gb:MT044258|Organism:Severe      GAGGAGGCAAAGACAGTGCTTAAAAAGTGTAAAAGTGCCTTTTACATTCTACCATCTATT   4318
gb:MT039890|Organism:Severe      GAGGAGGCAAAGACAGTGCTTAAAAAGTGTAAAAGTGCCTTTTACATTCTACCATCTATT   4342
gb:MN988713|Organism:Severe      GAGGAGGCAAAGACAGTGCTTAAAAAGTGTAAAAGTGCCTTTTACATTCTACCATCTATT   4342
gb:LC521925|Organism:Severe      GAGGAGGCAAAGACAGTGCTTAAAAAGTGTAAAAGTGCCTTTTACATTCTACCATCTATT   4315
gb:MT093571|Organism:Severe      GAGGAGGCAAAGACAGTGCTTAAAAAGTGTAAAAGTGCCTTTTACATTCTACCATCTATT   4342
gb:MT039887|Organism:Severe      GAGGAGGCAAAGACAGTGCTTAAAAAGTGTAAAAGTGCCTTTTACATTCTACCATCTATT   4342
gb:MT019530|Organism:Severe      GAGGAGGCAAAGACAGTGCTTAAAAAGTGTAAAAGTGCCTTTTACATTCTACCATCTATT   4342
gb:MT039888|Organism:Severe      GAGGAGGCAAAGACAGTGCTTAAAAAGTGTAAAAGTGCCTTTTACATTCTACCATCTATT   4342
gb:LC522972|Organism:Severe      GAGGAGGCAAAGACAGTGCTTAAAAAGTGTAAAAGTGCCTTTTACATTCTACCATCTATT   4339
gb:MT027063|Organism:Severe      GAGGAGGCAAAGACAGTGCTTAAAAAGTGTAAAAGTGCCTTTTACATTCTACCATCTATT   4342
gb:MT027062|Organism:Severe      GAGGAGGCAAAGACAGTGCTTAAAAAGTGTAAAAGTGCCTTTTACATTCTACCATCTATT   4342
gb:MT019529|Organism:Severe      GAGGAGGCAAAGACAGTGCTTAAAAAGTGTAAAAGTGCCTTTTACATTCTACCATCTATT   4342
gb:MN996529|Organism:Severe      GAGGAGGCAAAGACAGTGCTTAAAAAGTGTAAAAGTGCCTTTTACATTCTACCATCTATT   4330
gb:MN996531|Organism:Severe      GAGGAGGCAAAGACAGTGCTTAAAAAGTGTAAAAGTGCCTTTTACATTCTACCATCTATT   4329
gb:MT066176|Organism:Severe      GAGGAGGCAAAGACAGTGCTTAAAAAGTGTAAAAGTGCCTTTTACATTCTACCATCTATT   4342
gb:MT027064|Organism:Severe      GAGGAGGCAAAGACAGTGCTTAAAAAGTGTAAAAGTGCCTTTTACATTCTACCATCTATT   4342
gb:MN994468|Organism:Severe      GAGGAGGCAAAGACAGTGCTTAAAAAGTGTAAAAGTGCCTTTTACATTCTACCATCTATT   4342
gb:MT072688|Organism:Severe      GAGGAGGCAAAGACAGTGCTTAAAAAGTGTAAAAGTGCCTTTTACATTCTACCATCTATT   4327
gb:MN996527|Organism:Severe      GAGGAGGCAAAGACAGTGCTTAAAAAGTGTAAAAGTGCCTTTTACATTCTACCATCTATT   4309
gb:MT093631|Organism:Severe      GAGGAGGCAAAGACAGTGCTTAAAAAGTGTAAAAGTGCCTTTTACATTCTACCATCTATT   4380
gb:MT106053|Organism:Severe      GAGGAGGCAAAGACAGTGCTTAAAAAGTGTAAAAGTGCCTTTTACATTCTACCATCTATT   4342
gb:MT019533|Organism:Severe      GAGGAGGCAAAGACAGTGCTTAAAAAGTGTAAAAGTGCCTTTTACATTCTACCATCTATT   4342
gb:MT019531|Organism:Severe      GAGGAGGCAAAGACAGTGCTTAAAAAGTGTAAAAGTGCCTTTTACATTCTACCATCTATT   4342
gb:MN996528|Organism:Severe      GAGGAGGCAAAGACAGTGCTTAAAAAGTGTAAAAGTGCCTTTTACATTCTACCATCTATT   4342
gb:MN996530|Organism:Severe      GAGGAGGCAAAGACAGTGCTTAAAAAGTGTAAAAGTGCCTTTTACATTCTACCATCTATT   4328
gb:MN908947|Organism:Severe      GAGGAGGCAAAGACAGTGCTTAAAAAGTGTAAAAGTGCCTTTTACATTCTACCATCTATT   4342
gb:MT019532|Organism:Severe      GAGGAGGCAAAGACAGTGCTTAAAAAGTGTAAAAGTGCCTTTTACATTCTACCATCTATT   4342
                                 ************************************************************

gb:MT020781|Organism:Severe      ATCTCTAATGAGAAGCAAGAAATTCTTGGAACTGTTTCTTGGAATTTGCGAGAAATGCTT   4390
gb:MT007544|Organism:Severe      ATCTCTAATGAGAAGCAAGAAATTCTTGGAACTGTTTCTTGGAATTTGCGAGAAATGCTT   4402
gb:MN994467|Organism:Severe      ATCTCTAATGAGAAGCAAGAAATTCTTGGAACTGTTTCTTGGAATTTGCGAGAAATGCTT   4402
gb:MT044257|Organism:Severe      ATCTCTAATGAGAAGCAAGAAATTCTTGGAACTGTTTCTTGGAATTTGCGAGAAATGCTT   4402
gb:MT106054|Organism:Severe      ATCTCTAATGAGAAGCAAGAAATTCTTGGAACTGTTTCTTGGAATTTGCGAGAAATGCTT   4402
gb:MT049951|Organism:Severe      ATCTCTAATGAGAAGCAAGAAATTCTTGGAACTGTTTCTTGGAATTTGCGAGAAATGCTT   4402
gb:MN975262|Organism:Severe      ATCTCTAATGAGAAGCAAGAAATTCTTGGAACTGTTTCTTGGAATTTGCGAGAAATGCTT   4402
gb:MT106052|Organism:Severe      ATCTCTAATGAGAAGCAAGAAATTCTTGGAACTGTTTCTTGGAATTTGCGAGAAATGCTT   4402
gb:LC522975|Organism:Severe      ATCTCTAATGAGAAGCAAGAAATTCTTGGAACTGTTTCTTGGAATTTGCGAGAAATGCTT   4399
gb:LC522973|Organism:Severe      ATCTCTAATGAGAAGCAAGAAATTCTTGGAACTGTTTCTTGGAATTTGCGAGAAATGCTT   4399
gb:LC522974|Organism:Severe      ATCTCTAATGAGAAGCAAGAAATTCTTGGAACTGTTTCTTGGAATTTGCGAGAAATGCTT   4399
gb:MN985325|Organism:Severe      ATCTCTAATGAGAAGCAAGAAATTCTTGGAACTGTTTCTTGGAATTTGCGAGAAATGCTT   4402
gb:MT020881|Organism:Severe      ATCTCTAATGAGAAGCAAGAAATTCTTGGAACTGTTTCTTGGAATTTGCGAGAAATGCTT   4402
gb:MT020880|Organism:Severe      ATCTCTAATGAGAAGCAAGAAATTCTTGGAACTGTTTCTTGGAATTTGCGAGAAATGCTT   4402
gb:MT066175|Organism:Severe      ATCTCTAATGAGAAGCAAGAAATTCTTGGAACTGTTTCTTGGAATTTGCGAGAAATGCTT   4402
gb:MN997409|Organism:Severe      ATCTCTAATGAGAAGCAAGAAATTCTTGGAACTGTTTCTTGGAATTTGCGAGAAATGCTT   4402
gb:MN938384|Organism:Severe      ATCTCTAATGAGAAGCAAGAAATTCTTGGAACTGTTTCTTGGAATTTGCGAGAAATGCTT   4370
gb:MT044258|Organism:Severe      ATCTCTAATGAGAAGCAAGAAATTCTTGGAACTGTTTCTTGGAATTTGCGAGAAATGCTT   4378
```

```
gb:MT039890|Organism:Severe      ATCTCTAATGAGAAGCAAGAAATTCTTGGAACTGTTTCTTGGAATTTGCGAGAAATGCTT   4402
gb:MN988713|Organism:Severe      ATCTCTAATGAGAAGCAAGAAATTCTTGGAACTGTTTCTTGGAATTTGCGAGAAATGCTT   4402
gb:LC521925|Organism:Severe      ATCTCTAATGAGAAGCAAGAAATTCTTGGAACTGTTTCTTGGAATTTGCGAGAAATGCTT   4375
gb:MT093571|Organism:Severe      ATCTCTAATGAGAAGCAAGAAATTCTTGGAACTGTTTCTTGGAATTTGCGAGAAATGCTT   4402
gb:MT039887|Organism:Severe      ATCTCTAATGAGAAGCAAGAAATTCTTGGAACTGTTTCTTGGAATTTGCGAGAAATGCTT   4402
gb:MT019530|Organism:Severe      ATCTCTAATGAGAAGCAAGAAATTCTTGGAACTGTTTCTTGGAATTTGCGAGAAATGCTT   4402
gb:MT039888|Organism:Severe      ATCTCTAATGAGAAGCAAGAAATTCTTGGAACTGTTTCTTGGAATTTGCGAGAAATGCTT   4402
gb:LC522972|Organism:Severe      ATCTCTAATGAGAAGCAAGAAATTCTTGGAACTGTTTCTTGGAATTTGCGAGAAATGCTT   4399
gb:MT027063|Organism:Severe      ATCTCTAATGAGAAGCAAGAAATTCTTGGAACTGTTTCTTGGAATTTGCGAGAAATGCTT   4402
gb:MT027062|Organism:Severe      ATCTCTAATGAGAAGCAAGAAATTCTTGGAACTGTTTCTTGGAATTTGCGAGAAATGCTT   4402
gb:MT019529|Organism:Severe      ATCTCTAATGAGAAGCAAGAAATTCTTGGAACTGTTTCTTGGAATTTGCGAGAAATGCTT   4402
gb:MN996529|Organism:Severe      ATCTCTAATGAGAAGCAAGAAATTCTTGGAACTGTTTCTTGGAATTTGCGAGAAATGCTT   4390
gb:MN996531|Organism:Severe      ATCTCTAATGAGAAGCAAGAAATTCTTGGAACTGTTTCTTGGAATTTGCGAGAAATGCTT   4389
gb:MT066176|Organism:Severe      ATCTCTAATGAGAAGCAAGAAATTCTTGGAACTGTTTCTTGGAATTTGCGAGAAATGCTT   4402
gb:MT027064|Organism:Severe      ATCTCTAATGAGAAGCAAGAAATTCTTGGAACTGTTTCTTGGAATTTGCGAGAAATGCTT   4402
gb:MN994468|Organism:Severe      ATCTCTAATGAGAAGCAAGAAATTCTTGGAACTGTTTCTTGGAATTTGCGAGAAATGCTT   4402
gb:MT072688|Organism:Severe      ATCTCTAATGAGAAGCAAGAAATTCTTGGAACTGTTTCTTGGAATTTGCGAGAAATGCTT   4387
gb:MN996527|Organism:Severe      ATCTCTAATGAGAAGCAAGAAATTCTTGGAACTGTTTCTTGGAATTTGCGAGAAATGCTT   4369
gb:MT093631|Organism:Severe      ATCTCTAATGAGAAGCAAGAAATTCTTGGAACTGTTTCTTGGAATTTGCGAGAAATGCTT   4440
gb:MT106053|Organism:Severe      ATCTCTAATGAGAAGCAAGAAATTCTTGGAACTGTTTCTTGGAATTTGCGAGAAATGCTT   4402
gb:MT019533|Organism:Severe      ATCTCTAATGAGAAGCAAGAAATTCTTGGAACTGTTTCTTGGAATTTGCGAGAAATGCTT   4402
gb:MT019531|Organism:Severe      ATCTCTAATGAGAAGCAAGAAATTCTTGGAACTGTTTCTTGGAATTTGCGAGAAATGCTT   4402
gb:MN996528|Organism:Severe      ATCTCTAATGAGAAGCAAGAAATTCTTGGAACTGTTTCTTGGAATTTGCGAGAAATGCTT   4402
gb:MN996530|Organism:Severe      ATCTCTAATGAGAAGCAAGAAATTCTTGGAACTGTTTCTTGGAATTTGCGAGAAATGCTT   4388
gb:MN908947|Organism:Severe      ATCTCTAATGAGAAGCAAGAAATTCTTGGAACTGTTTCTTGGAATTTGCGAGAAATGCTT   4402
gb:MT019532|Organism:Severe      ATCTCTAATGAGAAGCAAGAAATTCTTGGAACTGTTTCTTGGAATTTGCGAGAAATGCTT   4402
                                 ************************************************************

gb:MT020781|Organism:Severe      GCACATGCAGAAGAAACACGCAAATTAATGCCTGTCTGTGTGGAAACTAAAGCCATAGTT   4450
gb:MT007544|Organism:Severe      GCACATGCAGAAGAAACACGCAAATTAATGCCTGTCTGTGTGGAAACTAAAGCCATAGTT   4462
gb:MN994467|Organism:Severe      GCACATGCAGAAGAAACACGCAAATTAATGCCTGTCTGTGTGGAAACTAAAGCCATAGTT   4462
gb:MT044257|Organism:Severe      GCACATGCAGAAGAAACACGCAAATTAATGCCTGTCTGTGTGGAAACTAAAGCCATAGTT   4462
gb:MT106054|Organism:Severe      GCACATGCAGAAGAAACACGCAAATTAATGCCTGTCTGTGTGGAAACTAAAGCCATAGTT   4462
gb:MT049951|Organism:Severe      GCACATGCAGAAGAAACACGCAAATTAATGCCTGTCTGTGTGGAAACTAAAGCCATAGTT   4462
gb:MN975262|Organism:Severe      GCACATGCAGAAGAAACACGCAAATTAATGCCTGTCTGTGTGGAAACTAAAGCCATAGTT   4462
gb:MT106052|Organism:Severe      GCACATGCAGAAGAAACACGCAAATTAATGCCTGTCTGTGTGGAAACTAAAGCCATAGTT   4462
gb:LC522975|Organism:Severe      GCACATGCAGAAGAAACACGCAAATTAATGCCTGTCTGTGTGGAAACTAAAGCCATAGTT   4459
gb:LC522973|Organism:Severe      GCACATGCAGAAGAAACACGCAAATTAATGCCTGTCTGTGTGGAAACTAAAGCCATAGTT   4459
gb:LC522974|Organism:Severe      GCACATGCAGAAGAAACACGCAAATTAATGCCTGTCTGTGTGGAAACTAAAGCCATAGTT   4459
gb:MN985325|Organism:Severe      GCACATGCAGAAGAAACACGCAAATTAATGCCTGTCTGTGTGGAAACTAAAGCCATAGTT   4462
gb:MT020881|Organism:Severe      GCACATGCAGAAGAAACACGCAAATTAATGCCTGTCTGTGTGGAAACTAAAGCCATAGTT   4462
gb:MT020880|Organism:Severe      GCACATGCAGAAGAAACACGCAAATTAATGCCTGTCTGTGTGGAAACTAAAGCCATAGTT   4462
gb:MT066175|Organism:Severe      GCACATGCAGAAGAAACACGCAAATTAATGCCTGTCTGTGTGGAAACTAAAGCCATAGTT   4462
gb:MN997409|Organism:Severe      GCACATGCAGAAGAAACACGCAAATTAATGCCTGTCTGTGTGGAAACTAAAGCCATAGTT   4462
gb:MN938384|Organism:Severe      GCACATGCAGAAGAAACACGCAAATTAATGCCTGTCTGTGTGGAAACTAAAGCCATAGTT   4430
gb:MT044258|Organism:Severe      GCACATGCAGAAGAAACACGCAAATTAATGCCTGTCTGTGTGGAAACTAAAGCCATAGTT   4438
gb:MT039890|Organism:Severe      GCACATGCAGAAGAAACACGCAAATTAATGCCTGTCTGTGTGGAAACTAAAGCCATAGTT   4462
gb:MN988713|Organism:Severe      GCACATGCAGAAGAAACACGCAAATTAATGCCTGTCTGTGTGGAAACTAAAGCCATAGTT   4462
gb:LC521925|Organism:Severe      GCACATGCAGAAGAAACACGCAAATTAATGCCTGTCTGTGTGGAAACTAAAGCCATAGTT   4435
gb:MT093571|Organism:Severe      GCACATGCAGAAGAAACACGCAAATTAATGCCTGTCTGTGTGGAAACTAAAGCCATAGTT   4462
gb:MT039887|Organism:Severe      GCACATGCAGAAGAAACACGCAAATTAATGCCTGTCTGTGTGGAAACTAAAGCCATAGTT   4462
gb:MT019530|Organism:Severe      GCACATGCAGAAGAAACACGCAAATTAATGCCTGTCTGTGTGGAAACTAAAGCCATAGTT   4462
gb:MT039888|Organism:Severe      GCACATGCAGAAGAAACACGCAAATTAATGCCTGTCTGTGTGGAAACTAAAGCCATAGTT   4462
gb:LC522972|Organism:Severe      GCACATGCAGAAGAAACACGCAAATTAATGCCTGTCTGTGTGGAAACTAAAGCCATAGTT   4459
gb:MT027063|Organism:Severe      GCACATGCAGAAGAAACACGCAAATTAATGCCTGTCTGTGTGGAAACTAAAGCCATAGTT   4462
gb:MT027062|Organism:Severe      GCACATGCAGAAGAAACACGCAAATTAATGCCTGTCTGTGTGGAAACTAAAGCCATAGTT   4462
gb:MT019529|Organism:Severe      GCACATGCAGAAGAAACACGCAAATTAATGCCTGTCTGTGTGGAAACTAAAGCCATAGTT   4462
gb:MN996529|Organism:Severe      GCACATGCAGAAGAAACACGCAAATTAATGCCTGTCTGTGTGGAAACTAAAGCCATAGTT   4450
gb:MN996531|Organism:Severe      GCACATGCAGAAGAAACACGCAAATTAATGCCTGTCTGTGTGGAAACTAAAGCCATAGTT   4449
gb:MT066176|Organism:Severe      GCACATGCAGAAGAAACACGCAAATTAATGCCTGTCTGTGTGGAAACTAAAGCCATAGTT   4462
gb:MT027064|Organism:Severe      GCACATGCAGAAGAAACACGCAAATTAATGCCTGTCTGTGTGGAAACTAAAGCCATAGTT   4462
gb:MN994468|Organism:Severe      GCACATGCAGAAGAAACACGCAAATTAATGCCTGTCTGTGTGGAAACTAAAGCCATAGTT   4462
gb:MT072688|Organism:Severe      GCACATGCAGAAGAAACACGCAAATTAATGCCTGTCTGTGTGGAAACTAAAGCCATAGTT   4447
gb:MN996527|Organism:Severe      GCACATGCAGAAGAAACACGCAAATTAATGCCTGTCTGTGTGGAAACTAAAGCCATAGTT   4429
gb:MT093631|Organism:Severe      GCACATGCAGAAGAAACACGCAAATTAATGCCTGTCTGTGTGGAAACTAAAGCCATAGTT   4500
```

```
gb:MT106053|Organism:Severe      GCACATGCAGAAGAAACACGCAAATTAATGCCTGTCTGTGTGGAAACTAAAGCCATAGTT    4462
gb:MT019533|Organism:Severe      GCACATGCAGAAGAAACACGCAAATTAATGCCTGTCTGTGTGGAAACTAAAGCCATAGTT    4462
gb:MT019531|Organism:Severe      GCACATGCAGAAGAAACACGCAAATTAATGCCTGTCTGTGTGGAAACTAAAGCCATAGTT    4462
gb:MN996528|Organism:Severe      GCACATGCAGAAGAAACACGCAAATTAATGCCTGTCTGTGTGGAAACTAAAGCCATAGTT    4462
gb:MN996530|Organism:Severe      GCACATGCAGAAGAAACACGCAAATTAATGCCTGTCTGTGTGGAAACTAAAGCCATAGTT    4448
gb:MN908947|Organism:Severe      GCACATGCAGAAGAAACACGCAAATTAATGCCTGTCTGTGTGGAAACTAAAGCCATAGTT    4462
gb:MT019532|Organism:Severe      GCACATGCAGAAGAAACACGCAAATTAATGCCTGTCTGTGTGGAAACTAAAGCCATAGTT    4462
                                 ************************************************************

gb:MT020781|Organism:Severe      TCAACTATACAGCGTAAATATAAGGGTATTAAAATACAAGAGGGTGTGGTTGATTATGGT    4510
gb:MT007544|Organism:Severe      TCAACTATACAGCGTAAATATAAGGGTATTAAAATACAAGAGGGTGTGGTTGATTATGGT    4522
gb:MN994467|Organism:Severe      TCAACTATACAGCGTAAATATAAGGGTATTAAAATACAAGAGGGTGTGGTTGATTATGGT    4522
gb:MT044257|Organism:Severe      TCAACTATACAGCGTAAATATAAGGGTATTAAAATACAAGAGGGTGTGGTTGATTATGGT    4522
gb:MT106054|Organism:Severe      TCAACTATACAGCGTAAATATAAGGGTATTAAAATACAAGAGGGTGTGGTTGATTATGGT    4522
gb:MT049951|Organism:Severe      TCAACTATACAGCGTAAATATAAGGGTATTAAAATACAAGAGGGTGTGGTTGATTATGGT    4522
gb:MN975262|Organism:Severe      TCAACTATACAGCGTAAATATAAGGGTATTAAAATACAAGAGGGTGTGGTTGATTATGGT    4522
gb:MT106052|Organism:Severe      TCAACTATACAGCGTAAATATAAGGGTATTAAAATACAAGAGGGTGTGGTTGATTATGGT    4522
gb:LC522975|Organism:Severe      TCAACTATACAGCGTAAATATAAGGGTATTAAAATACAAGAGGGTGTGGTTGATTATGGT    4519
gb:LC522973|Organism:Severe      TCAACTATACAGCGTAAATATAAGGGTATTAAAATACAAGAGGGTGTGGTTGATTATGGT    4519
gb:LC522974|Organism:Severe      TCAACTATACAGCGTAAATATAAGGGTATTAAAATACAAGAGGGTGTGGTTGATTATGGT    4519
gb:MN985325|Organism:Severe      TCAACTATACAGCGTAAATATAAGGGTATTAAAATACAAGAGGGTGTGGTTGATTATGGT    4522
gb:MT020881|Organism:Severe      TCAACTATACAGCGTAAATATAAGGGTATTAAAATACAAGAGGGTGTGGTTGATTATGGT    4522
gb:MT020880|Organism:Severe      TCAACTATACAGCGTAAATATAAGGGTATTAAAATACAAGAGGGTGTGGTTGATTATGGT    4522
gb:MT066175|Organism:Severe      TCAACTATACAGCGTAAATATAAGGGTATTAAAATACAAGAGGGTGTGGTTGATTATGGT    4522
gb:MN997409|Organism:Severe      TCAACTATACAGCGTAAATATAAGGGTATTAAAATACAAGAGGGTGTGGTTGATTATGGT    4522
gb:MN938384|Organism:Severe      TCAACTATACAGCGTAAATATAAGGGTATTAAAATACAAGAGGGTGTGGTTGATTATGGT    4490
gb:MT044258|Organism:Severe      TCAACTATACAGCGTAAATATAAGGGTATTAAAATACAAGAGGGTGTGGTTGATTATGGT    4498
gb:MT039890|Organism:Severe      TCAACTATACAGCGTAAATATAAGGGTATTAAAATACAAGAGGGTGTGGTTGATTATGGT    4522
gb:MN988713|Organism:Severe      TCAACTATACAGCGTAAATATAAGGGTATTAAAATACAAGAGGGTGTGGTTGATTATGGT    4522
gb:LC521925|Organism:Severe      TCAACTATACAGCGTAAATATAAGGGTATTAAAATACAAGAGGGTGTGGTTGATTATGGT    4495
gb:MT093571|Organism:Severe      TCAACTATACAGCGTAAATATAAGGGTATTAAAATACAAGAGGGTGTGGTTGATTATGGT    4522
gb:MT039887|Organism:Severe      TCAACTATACAGCGTAAATATAAGGGTATTAAAATACAAGAGGGTGTGGTTGATTATGGT    4522
gb:MT019530|Organism:Severe      TCAACTATACAGCGTAAATATAAGGGTATTAAAATACAAGAGGGTGTGGTTGATTATGGT    4522
gb:MT039888|Organism:Severe      TCAACTATACAGCGTAAATATAAGGGTATTAAAATACAAGAGGGTGTGGTTGATTATGGT    4522
gb:LC522972|Organism:Severe      TCAACTATACAGCGTAAATATAAGGGTATTAAAATACAAGAGGGTGTGGTTGATTATGGT    4519
gb:MT027063|Organism:Severe      TCAACTATACAGCGTAAATATAAGGGTATTAAAATACAAGAGGGTGTGGTTGATTATGGT    4522
gb:MT027062|Organism:Severe      TCAACTATACAGCGTAAATATAAGGGTATTAAAATACAAGAGGGTGTGGTTGATTATGGT    4522
gb:MT019529|Organism:Severe      TCAACTATACAGCGTAAATATAAGGGTATTAAAATACAAGAGGGTGTGGTTGATTATGGT    4522
gb:MN996529|Organism:Severe      TCAACTATACAGCGTAAATATAAGGGTATTAAAATACAAGAGGGTGTGGTTGATTATGGT    4510
gb:MN996531|Organism:Severe      TCAACTATACAGCGTAAATATAAGGGTATTAAAATACAAGAGGGTGTGGTTGATTATGGT    4509
gb:MT066176|Organism:Severe      TCAACTATACAGCGTAAATATAAGGGTATTAAAATACAAGAGGGTGTGGTTGATTATGGT    4522
gb:MT027064|Organism:Severe      TCAACTATACAGCGTAAATATAAGGGTATTAAAATACAAGAGGGTGTGGTTGATTATGGT    4522
gb:MN994468|Organism:Severe      TCAACTATACAGCGTAAATATAAGGGTATTAAAATACAAGAGGGTGTGGTTGATTATGGT    4522
gb:MT072688|Organism:Severe      TCAACTATACAGCGTAAATATAAGGGTATTAAAATACAAGAGGGTGTGGTTGATTATGGT    4507
gb:MN996527|Organism:Severe      TCAACTATACAGCGTAAATATAAGGGTATTAAAATACAAGAGGGTGTGGTTGATTATGGT    4489
gb:MT093631|Organism:Severe      TCAACTATACAGCGTAAATATAAGGGTATTAAAATACAAGAGGGTGTGGTTGATTATGGT    4560
gb:MT106053|Organism:Severe      TCAACTATACAGCGTAAATATAAGGGTATTAAAATACAAGAGGGTGTGGTTGATTATGGT    4522
gb:MT019533|Organism:Severe      TCAACTATACAGCGTAAATATAAGGGTATTAAAATACAAGAGGGTGTGGTTGATTATGGT    4522
gb:MT019531|Organism:Severe      TCAACTATACAGCGTAAATATAAGGGTATTAAAATACAAGAGGGTGTGGTTGATTATGGT    4522
gb:MN996528|Organism:Severe      TCAACTATACAGCGTAAATATAAGGGTATTAAAATACAAGAGGGTGTGGTTGATTATGGT    4522
gb:MN996530|Organism:Severe      TCAACTATACAGCGTAAATATAAGGGTATTAAAATACAAGAGGGTGTGGTTGATTATGGT    4508
gb:MN908947|Organism:Severe      TCAACTATACAGCGTAAATATAAGGGTATTAAAATACAAGAGGGTGTGGTTGATTATGGT    4522
gb:MT019532|Organism:Severe      TCAACTATACAGCGTAAATATAAGGGTATTAAAATACAAGAGGGTGTGGTTGATTATGGT    4522
                                 ************************************************************

gb:MT020781|Organism:Severe      GCTAGATTTTACTTTTACACCAGTAAAACAACTGTAGCGTCACTTATCAACACACTTAAC    4570
gb:MT007544|Organism:Severe      GCTAGATTTTACTTTTACACCAGTAAAACAACTGTAGCGTCACTTATCAACACACTTAAC    4582
gb:MN994467|Organism:Severe      GCTAGATTTTACTTTTACACCAGTAAAACAACTGTAGCGTCACTTATCAACACACTTAAC    4582
gb:MT044257|Organism:Severe      GCTAGATTTTACTTTTACACCAGTAAAACAACTGTAGCGTCACTTATCAACACACTTAAC    4582
gb:MT106054|Organism:Severe      GCTAGATTTTACTTTTACACCAGTAAAACAACTGTAGCGTCACTTATCAACACACTTAAC    4582
gb:MT049951|Organism:Severe      GCTAGATTTTACTTTTACACCAGTAAAACAACTGTAGCGTCACTTATCAACACACTTAAC    4582
gb:MN975262|Organism:Severe      GCTAGATTTTACTTTTACACCAGTAAAACAACTGTAGCGTCACTTATCAACACACTTAAC    4582
gb:MT106052|Organism:Severe      GCTAGATTTTACTTTTACACCAGTAAAACAACTGTAGCGTCACTTATCAACACACTTAAC    4582
gb:LC522975|Organism:Severe      GCTAGATTTTACTTTTACACCAGTAAAACAACTGTAGCGTCACTTATCAACACACTTAAC    4579
gb:LC522973|Organism:Severe      GCTAGATTTTACTTTTACACCAGTAAAACAACTGTAGCGTCACTTATCAACACACTTAAC    4579
```

```
gb:LC522974|Organism:Severe      GCTAGATTTTACTTTTACACCAGTAAAACAACTGTAGCGTCACTTATCAACACACTTAAC   4579
gb:MN985325|Organism:Severe      GCTAGATTTTACTTTTACACCAGTAAAACAACTGTAGCGTCACTTATCAACACACTTAAC   4582
gb:MT020881|Organism:Severe      GCTAGATTTTACTTTTACACCAGTAAAACAACTGTAGCGTCACTTATCAACACACTTAAC   4582
gb:MT020880|Organism:Severe      GCTAGATTTTACTTTTACACCAGTAAAACAACTGTAGCGTCACTTATCAACACACTTAAC   4582
gb:MT066175|Organism:Severe      GCTAGATTTTACTTTTACACCAGTAAAACAACTGTAGCGTCACTTATCAACACACTTAAC   4582
gb:MN997409|Organism:Severe      GCTAGATTTTACTTTTACACCAGTAAAACAACTGTAGCGTCACTTATCAACACACTTAAC   4582
gb:MN938384|Organism:Severe      GCTAGATTTTACTTTTACACCAGTAAAACAACTGTAGCGTCACTTATCAACACACTTAAC   4550
gb:MT044258|Organism:Severe      GCTAGATTTTACTTTTACACCAGTAAAACAACTGTAGCGTCACTTATCAACACACTTAAC   4558
gb:MT039890|Organism:Severe      GCTAGATTTTACTTTTACACCAGTAAAACAACTGTAGCGTCACTTATCAACACACTTAAC   4582
gb:MN988713|Organism:Severe      GCTAGATTTTACTTTTACACCAGTAAAACAACTGTAGCGTCACTTATCAACACACTTAAC   4582
gb:LC521925|Organism:Severe      GCTAGATTTTACTTTTACACCAGTAAAACAACTGTAGCGTCACTTATCAACACACTTAAC   4555
gb:MT093571|Organism:Severe      GCTAGATTTTACTTTTACACCAGTAAAACAACTGTAGCGTCACTTATCAACACACTTAAC   4582
gb:MT039887|Organism:Severe      GCTAGATTTTACTTTTACACCAGTAAAACAACTGTAGCGTCACTTATCAACACACTTAAC   4582
gb:MT019530|Organism:Severe      GCTAGATTTTACTTTTACACCAGTAAAACAACTGTAGCGTCACTTATCAACACACTTAAC   4582
gb:MT039888|Organism:Severe      GCTAGATTTTACTTTTACACCAGTAAAACAACTGTAGCGTCACTTATCAACACACTTAAC   4582
gb:LC522972|Organism:Severe      GCTAGATTTTACTTTTACACCAGTAAAACAACTGTAGCGTCACTTATCAACACACTTAAC   4579
gb:MT027063|Organism:Severe      GCTAGATTTTACTTTTACACCAGTAAAACAACTGTAGCGTCACTTATCAACACACTTAAC   4582
gb:MT027062|Organism:Severe      GCTAGATTTTACTTTTACACCAGTAAAACAACTGTAGCGTCACTTATCAACACACTTAAC   4582
gb:MT019529|Organism:Severe      GCTAGATTTTACTTTTACACCAGTAAAACAACTGTAGCGTCACTTATCAACACACTTAAC   4582
gb:MN996529|Organism:Severe      GCTAGATTTTACTTTTACACCAGTAAAACAACTGTAGCGTCACTTATCAACACACTTAAC   4570
gb:MN996531|Organism:Severe      GCTAGATTTTACTTTTACACCAGTAAAACAACTGTAGCGTCACTTATCAACACACTTAAC   4569
gb:MT066176|Organism:Severe      GCTAGATTTTACTTTTACACCAGTAAAACAACTGTAGCGTCACTTATCAACACACTTAAC   4582
gb:MT027064|Organism:Severe      GCTAGATTTTACTTTTACACCAGTAAAACAACTGTAGCGTCACTTATCAACACACTTAAC   4582
gb:MN994468|Organism:Severe      GCTAGATTTTACTTTTACACCAGTAAAACAACTGTAGCGTCACTTATCAACACACTTAAC   4582
gb:MT072688|Organism:Severe      GCTAGATTTTACTTTTACACCAGTAAAACAACTGTAGCGTCACTTATCAACACACTTAAC   4567
gb:MN996527|Organism:Severe      GCTAGATTTTACTTTTACACCAGTAAAACAACTGTAGCGTCACTTATCAACACACTTAAC   4549
gb:MT093631|Organism:Severe      GCTAGATTTTACTTTTACACCAGTAAAACAACTGTAGCGTCACTTATCAACACACTTAAC   4620
gb:MT106053|Organism:Severe      GCTAGATTTTACTTTTACACCAGTAAAACAACTGTAGCGTCACTTATCAACACACTTAAC   4582
gb:MT019533|Organism:Severe      GCTAGATTTTACTTTTACACCAGTAAAACAACTGTAGCGTCACTTATCAACACACTTAAC   4582
gb:MT019531|Organism:Severe      GCTAGATTTTACTTTTACACCAGTAAAACAACTGTAGCGTCACTTATCAACACACTTAAC   4582
gb:MN996528|Organism:Severe      GCTAGATTTTACTTTTACACCAGTAAAACAACTGTAGCGTCACTTATCAACACACTTAAC   4582
gb:MN996530|Organism:Severe      GCTAGATTTTACTTTTACACCAGTAAAACAACTGTAGCGTCACTTATCAACACACTTAAC   4568
gb:MN908947|Organism:Severe      GCTAGATTTTACTTTTACACCAGTAAAACAACTGTAGCGTCACTTATCAACACACTTAAC   4582
gb:MT019532|Organism:Severe      GCTAGATTTTACTTTTACACCAGTAAAACAACTGTAGCGTCACTTATCAACACACTTAAC   4582
                                 ************************************************************

gb:MT020781|Organism:Severe      GATCTAAATGAAACTCTTGTTACAATGCCACTTGGCTATGTAACACATGGCTTAAATTTG   4630
gb:MT007544|Organism:Severe      GATCTAAATGAAACTCTTGTTACAATGCCACTTGGCTATGTAACACATGGCTTAAATTTG   4642
gb:MN994467|Organism:Severe      GATCTAAATGAAACTCTTGTTACAATGCCACTTGGCTATGTAACACATGGCTTAAATTTG   4642
gb:MT044257|Organism:Severe      GATCTAAATGAAACTCTTGTTACAATGCCACTTGGCTATGTAACACATGGCTTAAATTTG   4642
gb:MT106054|Organism:Severe      GATCTAAATGAAACTCTTGTTACAATGCCACTTGGCTATGTAACACATGGCTTAAATTTG   4642
gb:MT049951|Organism:Severe      GATCTAAATGAAACTCTTGTTACAATGCCACTTGGCTATGTAACACATGGCTTAAATTTG   4642
gb:MN975262|Organism:Severe      GATCTAAATGAAACTCTTGTTACAATGCCACTTGGCTATGTAACACATGGCTTAAATTTG   4642
gb:MT106052|Organism:Severe      GATCTAAATGAAACTCTTGTTACAATGCCACTTGGCTATGTAACACATGGCTTAAATTTG   4642
gb:LC522975|Organism:Severe      GATCTAAATGAAACTCTTGTTACAATGCCACTTGGCTATGTAACACATGGCTTAAATTTG   4639
gb:LC522973|Organism:Severe      GATCTAAATGAAACTCTTGTTACAATGCCACTTGGCTATGTAACACATGGCTTAAATTTG   4639
gb:LC522974|Organism:Severe      GATCTAAATGAAACTCTTGTTACAATGCCACTTGGCTATGTAACACATGGCTTAAATTTG   4639
gb:MN985325|Organism:Severe      GATCTAAATGAAACTCTTGTTACAATGCCACTTGGCTATGTAACACATGGCTTAAATTTG   4642
gb:MT020881|Organism:Severe      GATCTAAATGAAACTCTTGTTACAATGCCACTTGGCTATGTAACACATGGCTTAAATTTG   4642
gb:MT020880|Organism:Severe      GATCTAAATGAAACTCTTGTTACAATGCCACTTGGCTATGTAACACATGGCTTAAATTTG   4642
gb:MT066175|Organism:Severe      GATCTAAATGAAACTCTTGTTACAATGCCACTTGGCTATGTAACACATGGCTTAAATTTG   4642
gb:MN997409|Organism:Severe      GATCTAAATGAAACTCTTGTTACAATGCCACTTGGCTATGTAACACATGGCTTAAATTTG   4642
gb:MN938384|Organism:Severe      GATCTAAATGAAACTCTTGTTACAATGCCACTTGGCTATGTAACACATGGCTTAAATTTG   4610
gb:MT044258|Organism:Severe      GATCTAAATGAAACTCTTGTTACAATGCCACTTGGCTATGTAACACATGGCTTAAATTTG   4618
gb:MT039890|Organism:Severe      GATCTAAATGAAACTCTTGTTACAATGCCACTTGGCTATGTAACACATGGCTTAAATTTG   4642
gb:MN988713|Organism:Severe      GATCTAAATGAAACTCTTGTTACAATGCCACTTGGCTATGTAACACATGGCTTAAATTTG   4642
gb:LC521925|Organism:Severe      GATCTAAATGAAACTCTTGTTACAATGCCACTTGGCTATGTAACACATGGCTTAAATTTG   4615
gb:MT093571|Organism:Severe      GATCTAAATGAAACTCTTGTTACAATGCCACTTGGCTATGTAACACATGGCTTAAATTTG   4642
gb:MT039887|Organism:Severe      GATCTAAATGAAACTCTTGTTACAATGCCACTTGGCTATGTAACACATGGCTTAAATTTG   4642
gb:MT019530|Organism:Severe      GATCTAAATGAAACTCTTGTTACAATGCCACTTGGCTATGTAACACATGGCTTAAATTTG   4642
gb:MT039888|Organism:Severe      GATCTAAATGAAACTCTTGTTACAATGCCACTTGGCTATGTAACACATGGCTTAAATTTG   4642
gb:LC522972|Organism:Severe      GATCTAAATGAAACTCTTGTTACAATGCCACTTGGCTATGTAACACATGGCTTAAATTTG   4639
gb:MT027063|Organism:Severe      GATCTAAATGAAACTCTTGTTACAATGCCACTTGGCTATGTAACACATGGCTTAAATTTG   4642
gb:MT027062|Organism:Severe      GATCTAAATGAAACTCTTGTTACAATGCCACTTGGCTATGTAACACATGGCTTAAATTTG   4642
gb:MT019529|Organism:Severe      GATCTAAATGAAACTCTTGTTACAATGCCACTTGGCTATGTAACACATGGCTTAAATTTG   4642
```

```
gb:MN996529|Organism:Severe      GATCTAAATGAAACTCTTGTTACAATGCCACTTGGCTATGTAACACATGGCTTAAATTTG   4630
gb:MN996531|Organism:Severe      GATCTAAATGAAACTCTTGTTACAATGCCACTTGGCTATGTAACACATGGCTTAAATTTG   4629
gb:MT066176|Organism:Severe      GATCTAAATGAAACTCTTGTTACAATGCCACTTGGCTATGTAACACATGGCTTAAATTTG   4642
gb:MT027064|Organism:Severe      GATCTAAATGAAACTCTTGTTACAATGCCACTTGGCTATGTAACACATGGCTTAAATTTG   4642
gb:MN994468|Organism:Severe      GATCTAAATGAAACTCTTGTTACAATGCCACTTGGCTATGTAACACATGGCTTAAATTTG   4642
gb:MT072688|Organism:Severe      GATCTAAATGAAACTCTTGTTACAATGCCACTTGGCTATGTAACACATGGCTTAAATTTG   4627
gb:MN996527|Organism:Severe      GATCTAAATGAAACTCTTGTTACAATGCCACTTGGCTATGTAACACATGGCTTAAATTTG   4609
gb:MT093631|Organism:Severe      GATCTAAATGAAACTCTTGTTACAATGCCACTTGGCTATGTAACACATGGCTTAAATTTG   4680
gb:MT106053|Organism:Severe      GATCTAAATGAAACTCTTGTTACAATGCCACTTGGCTATGTAACACATGGCTTAAATTTG   4642
gb:MT019533|Organism:Severe      GATCTAAATGAAACTCTTGTTACAATGCCACTTGGCTATGTAACACATGGCTTAAATTTG   4642
gb:MT019531|Organism:Severe      GATCTAAATGAAACTCTTGTTACAATGCCACTTGGCTATGTAACACATGGCTTAAATTTG   4642
gb:MN996528|Organism:Severe      GATCTAAATGAAACTCTTGTTACAATGCCACTTGGCTATGTAACACATGGCTTAAATTTG   4642
gb:MN996530|Organism:Severe      GATCTAAATGAAACTCTTGTTACAATGCCACTTGGCTATGTAACACATGGCTTAAATTTG   4628
gb:MN908947|Organism:Severe      GATCTAAATGAAACTCTTGTTACAATGCCACTTGGCTATGTAACACATGGCTTAAATTTG   4642
gb:MT019532|Organism:Severe      GATCTAAATGAAACTCTTGTTACAATGCCACTTGGCTATGTAACACATGGCTTAAATTTG   4642
                                 ************************************************************

gb:MT020781|Organism:Severe      GAAGAAGCTGCTCGGTATATGAGATCTCTCAAAGTGCCAGCTACAGTTTCTGTTTCTTCA   4690
gb:MT007544|Organism:Severe      GAAGAAGCTGCTCGGTATATGAGATCTCTCAAAGTGCCAGCTACAGTTTCTGTTTCTTCA   4702
gb:MN994467|Organism:Severe      GAAGAAGCTGCTCGGTATATGAGATCTCTCAAAGTGCCAGCTACAGTTTCTGTTTCTTCA   4702
gb:MT044257|Organism:Severe      GAAGAAGCTGCTCGGTATATGAGATCTCTCAAAGTGCCAGCTACAGTTTCTGTTTCTTCA   4702
gb:MT106054|Organism:Severe      GAAGAAGCTGCTCGGTATATGAGATCTCTCAAAGTGCCAGCTACAGTTTCTGTTTCTTCA   4702
gb:MT049951|Organism:Severe      GAAGAAGCTGCTCGGTATATGAGATCTCTCAAAGTGCCAGCTACAGTTTCTGTTTCTTCA   4702
gb:MN975262|Organism:Severe      GAAGAAGCTGCTCGGTATATGAGATCTCTCAAAGTGCCAGCTACAGTTTCTGTTTCTTCA   4702
gb:MT106052|Organism:Severe      GAAGAAGCTGCTCGGTATATGAGATCTCTCAAAGTGCCAGCTACAGTTTCTGTTTCTTCA   4702
gb:LC522975|Organism:Severe      GAAGAAGCTGCTCGGTATATGAGATCTCTCAAAGTGCCAGCTACAGTTTCTGTTTCTTCA   4699
gb:LC522973|Organism:Severe      GAAGAAGCTGCTCGGTATATGAGATCTCTCAAAGTGCCAGCTACAGTTTCTGTTTCTTCA   4699
gb:LC522974|Organism:Severe      GAAGAAGCTGCTCGGTATATGAGATCTCTCAAAGTGCCAGCTACAGTTTCTGTTTCTTCA   4699
gb:MN985325|Organism:Severe      GAAGAAGCTGCTCGGTATATGAGATCTCTCAAAGTGCCAGCTACAGTTTCTGTTTCTTCA   4702
gb:MT020881|Organism:Severe      GAAGAAGCTGCTCGGTATATGAGATCTCTCAAAGTGCCAGCTACAGTTTCTGTTTCTTCA   4702
gb:MT020880|Organism:Severe      GAAGAAGCTGCTCGGTATATGAGATCTCTCAAAGTGCCAGCTACAGTTTCTGTTTCTTCA   4702
gb:MT066175|Organism:Severe      GAAGAAGCTGCTCGGTATATGAGATCTCTCAAAGTGCCAGCTACAGTTTCTGTTTCTTCA   4702
gb:MN997409|Organism:Severe      GAAGAAGCTGCTCGGTATATGAGATCTCTCAAAGTGCCAGCTACAGTTTCTGTTTCTTCA   4702
gb:MN938384|Organism:Severe      GAAGAAGCTGCTCGGTATATGAGATCTCTCAAAGTGCCAGCTACAGTTTCTGTTTCTTCA   4670
gb:MT044258|Organism:Severe      GAAGAAGCTGCTCGGTATATGAGATCTCTCAAAGTGCCAGCTACAGTTTCTGTTTCTTCA   4678
gb:MT039890|Organism:Severe      GAAGAAGCTGCTCGGTATATGAGATCTCTCAAAGTGCCAGCTACAGTTTCTGTTTCTTCA   4702
gb:MN988713|Organism:Severe      GAAGAAGCTGCTCGGTATATGAGATCTCTCAAAGTGCCAGCTACAGTTTCTGTTTCTTCA   4702
gb:LC521925|Organism:Severe      GAAGAAGCTGCTCGGTATATGAGATCTCTCAAAGTGCCAGCTACAGTTTCTGTTTCTTCA   4675
gb:MT093571|Organism:Severe      GAAGAAGCTGCTCGGTATATGAGATCTCTCAAAGTGCCAGCTACAGTTTCTGTTTCTTCA   4702
gb:MT039887|Organism:Severe      GAAGAAGCTGCTCGGTATATGAGATCTCTCAAAGTGCCAGCTACAGTTTCTGTTTCTTCA   4702
gb:MT019530|Organism:Severe      GAAGAAGCTGCTCGGTATATGAGATCTCTCAAAGTGCCAGCTACAGTTTCTGTTTCTTCA   4702
gb:MT039888|Organism:Severe      GAAGAAGCTGCTCGGTATATGAGATCTCTCAAAGTGCCAGCTACAGTTTCTGTTTCTTCA   4702
gb:LC522972|Organism:Severe      GAAGAAGCTGCTCGGTATATGAGATCTCTCAAAGTGCCAGCTACAGTTTCTGTTTCTTCA   4699
gb:MT027063|Organism:Severe      GAAGAAGCTGCTCGGTATATGAGATCTCTCAAAGTGCCAGCTACAGTTTCTGTTTCTTCA   4702
gb:MT027062|Organism:Severe      GAAGAAGCTGCTCGGTATATGAGATCTCTCAAAGTGCCAGCTACAGTTTCTGTTTCTTCA   4702
gb:MT019529|Organism:Severe      GAAGAAGCTGCTCGGTATATGAGATCTCTCAAAGTGCCAGCTACAGTTTCTGTTTCTTCA   4702
gb:MN996529|Organism:Severe      GAAGAAGCTGCTCGGTATATGAGATCTCTCAAAGTGCCAGCTACAGTTTCTGTTTCTTCA   4690
gb:MN996531|Organism:Severe      GAAGAAGCTGCTCGGTATATGAGATCTCTCAAAGTGCCAGCTACAGTTTCTGTTTCTTCA   4689
gb:MT066176|Organism:Severe      GAAGAAGCTGCTCGGTATATGAGATCTCTCAAAGTGCCAGCTACAGTTTCTGTTTCTTCA   4702
gb:MT027064|Organism:Severe      GAAGAAGCTGCTCGGTATATGAGATCTCTCAAAGTGCCAGCTACAGTTTCTGTTTCTTCA   4702
gb:MN994468|Organism:Severe      GAAGAAGCTGCTCGGTATATGAGATCTCTCAAAGTGCCAGCTACAGTTTCTGTTTCTTCA   4702
gb:MT072688|Organism:Severe      GAAGAAGCTGCTCGGTATATGAGATCTCTCAAAGTGCCAGCTACAGTTTCTGTTTCTTCA   4687
gb:MN996527|Organism:Severe      GAAGAAGCTGCTCGGTATATGAGATCTCTCAAAGTGCCAGCTACAGTTTCTGTTTCTTCA   4669
gb:MT093631|Organism:Severe      GAAGAAGCTGCTCGGTATATGAGATCTCTCAAAGTGCCAGCTACAGTTTCTGTTTCTTCA   4740
gb:MT106053|Organism:Severe      GAAGAAGCTGCTCGGTATATGAGATCTCTCAAAGTGCCAGCTACAGTTTCTGTTTCTTCA   4702
gb:MT019533|Organism:Severe      GAAGAAGCTGCTCGGTATATGAGATCTCTCAAAGTGCCAGCTACAGTTTCTGTTTCTTCA   4702
gb:MT019531|Organism:Severe      GAAGAAGCTGCTCGGTATATGAGATCTCTCAAAGTGCCAGCTACAGTTTCTGTTTCTTCA   4702
gb:MN996528|Organism:Severe      GAAGAAGCTGCTCGGTATATGAGATCTCTCAAAGTGCCAGCTACAGTTTCTGTTTCTTCA   4702
gb:MN996530|Organism:Severe      GAAGAAGCTGCTCGGTATATGAGATCTCTCAAAGTGCCAGCTACAGTTTCTGTTTCTTCA   4688
gb:MN908947|Organism:Severe      GAAGAAGCTGCTCGGTATATGAGATCTCTCAAAGTGCCAGCTACAGTTTCTGTTTCTTCA   4702
gb:MT019532|Organism:Severe      GAAGAAGCTGCTCGGTATATGAGATCTCTCAAAGTGCCAGCTACAGTTTCTGTTTCTTCA   4702
                                 ************************************************************

gb:MT020781|Organism:Severe      CCTGATGCTGTTACAGCGTATAATGGTTATCTTACTTCTTCTTCTAAAACACCTGAAGAA   4750
gb:MT007544|Organism:Severe      CCTGATGCTGTTACAGCGTATAATGGTTATCTTACTTCTTCTTCTAAAACACCTGAAGAA   4762
```

```
gb:MN994467|Organism:Severe      CCTGATGCTGTTACAGCGTATAATGGTTATCTTACTTCTTCTTCTAAAACACCTGAAGAA    4762
gb:MT044257|Organism:Severe      CCTGATGCTGTTACAGCGTATAATGGTTATCTTACTTCTTCTTCTAAAACACCTGAAGAA    4762
gb:MT106054|Organism:Severe      CCTGATGCTGTTACAGCGTATAATGGTTATCTTACTTCTTCTTCTAAAACACCTGAAGAA    4762
gb:MT049951|Organism:Severe      CCTGATGCTGTTACAGCGTATAATGGTTATCTTACTTCTTCTTCTAAAACACCTGAAGAA    4762
gb:MN975262|Organism:Severe      CCTGATGCTGTTACAGCGTATAATGGTTATCTTACTTCTTCTTCTAAAACACCTGAAGAA    4762
gb:MT106052|Organism:Severe      CCTGATGCTGTTACAGCGTATAATGGTTATCTTACTTCTTCTTCTAAAACACCTGAAGAA    4762
gb:LC522975|Organism:Severe      CCTGATGCTGTTACAGCGTATAATGGTTATCTTACTTCTTCTTCTAAAACACCTGAAGAA    4759
gb:LC522973|Organism:Severe      CCTGATGCTGTTACAGCGTATAATGGTTATCTTACTTCTTCTTCTAAAACACCTGAAGAA    4759
gb:LC522974|Organism:Severe      CCTGATGCTGTTACAGCGTATAATGGTTATCTTACTTCTTCTTCTAAAACACCTGAAGAA    4759
gb:MN985325|Organism:Severe      CCTGATGCTGTTACAGCGTATAATGGTTATCTTACTTCTTCTTCTAAAACACCTGAAGAA    4762
gb:MT020881|Organism:Severe      CCTGATGCTGTTACAGCGTATAATGGTTATCTTACTTCTTCTTCTAAAACACCTGAAGAA    4762
gb:MT020880|Organism:Severe      CCTGATGCTGTTACAGCGTATAATGGTTATCTTACTTCTTCTTCTAAAACACCTGAAGAA    4762
gb:MT066175|Organism:Severe      CCTGATGCTGTTACAGCGTATAATGGTTATCTTACTTCTTCTTCTAAAACACCTGAAGAA    4762
gb:MN997409|Organism:Severe      CCTGATGCTGTTACAGCGTATAATGGTTATCTTACTTCTTCTTCTAAAACACCTGAAGAA    4762
gb:MN938384|Organism:Severe      CCTGATGCTGTTACAGCGTATAATGGTTATCTTACTTCTTCTTCTAAAACACCTGAAGAA    4730
gb:MT044258|Organism:Severe      CCTGATGCTGTTACAGCGTATAATGGTTATCTTACTTCTTCTTCTAAAACACCTGAAGAA    4738
gb:MT039890|Organism:Severe      CCTGATGCTGTTACAGCGTATAATGGTTATCTTACTTCTTCTTCTAAAACACCTGAAGAA    4762
gb:MN988713|Organism:Severe      CCTGATGCTGTTACAGCGTATAATGGTTATCTTACTTCTTCTTCTAAAACACCTGAAGAA    4762
gb:LC521925|Organism:Severe      CCTGATGCTGTTACAGCGTATAATGGTTATCTTACTTCTTCTTCTAAAACACCTGAAGAA    4735
gb:MT093571|Organism:Severe      CCTGATGCTGTTACAGCGTATAATGGTTATCTTACTTCTTCTTCTAAAACACCTGAAGAA    4762
gb:MT039887|Organism:Severe      CCTGATGCTGTTACAGCGTATAATGGTTATCTTACTTCTTCTTCTAAAACACCTGAAGAA    4762
gb:MT019530|Organism:Severe      CCTGATGCTGTTACAGCGTATAATGGTTATCTTACTTCTTCTTCTAAAACACCTGAAGAA    4762
gb:MT039888|Organism:Severe      CCTGATGCTGTTACAGCGTATAATGGTTATCTTACTTCTTCTTCTAAAACACCTGAAGAA    4762
gb:LC522972|Organism:Severe      CCTGATGCTGTTACAGCGTATAATGGTTATCTTACTTCTTCTTCTAAAACACCTGAAGAA    4759
gb:MT027063|Organism:Severe      CCTGATGCTGTTACAGCGTATAATGGTTATCTTACTTCTTCTTCTAAAACACCTGAAGAA    4762
gb:MT027062|Organism:Severe      CCTGATGCTGTTACAGCGTATAATGGTTATCTTACTTCTTCTTCTAAAACACCTGAAGAA    4762
gb:MT019529|Organism:Severe      CCTGATGCTGTTACAGCGTATAATGGTTATCTTACTTCTTCTTCTAAAACACCTGAAGAA    4762
gb:MN996529|Organism:Severe      CCTGATGCTGTTACAGCGTATAATGGTTATCTTACTTCTTCTTCTAAAACACCTGAAGAA    4750
gb:MN996531|Organism:Severe      CCTGATGCTGTTACAGCGTATAATGGTTATCTTACTTCTTCTTCTAAAACACCTGAAGAA    4749
gb:MT066176|Organism:Severe      CCTGATGCTGTTACAGCGTATAATGGTTATCTTACTTCTTCTTCTAAAACACCTGAAGAA    4762
gb:MT027064|Organism:Severe      CCTGATGCTGTTACAGCGTATAATGGTTATCTTACTTCTTCTTCTAAAACACCTGAAGAA    4762
gb:MN994468|Organism:Severe      CCTGATGCTGTTACAGCGTATAATGGTTATCTTACTTCTTCTTCTAAAACACCTGAAGAA    4762
gb:MT072688|Organism:Severe      CCTGATGCTGTTACAGCGTATAATGGTTATCTTACTTCTTCTTCTAAAACACCTGAAGAA    4747
gb:MN996527|Organism:Severe      CCTGATGCTGTTACAGCGTATAATGGTTATCTTACTTCTTCTTCTAAAACACCTGAAGAA    4729
gb:MT093631|Organism:Severe      CCTGATGCTGTTACAGCGTATAATGGTTATCTTACTTCTTCTTCTAAAACACCTGAAGAA    4800
gb:MT106053|Organism:Severe      CCTGATGCTGTTACAGCGTATAATGGTTATCTTACTTCTTCTTCTAAAACACCTGAAGAA    4762
gb:MT019533|Organism:Severe      CCTGATGCTGTTACAGCGTATAATGGTTATCTTACTTCTTCTTCTAAAACACCTGAAGAA    4762
gb:MT019531|Organism:Severe      CCTGATGCTGTTACAGCGTATAATGGTTATCTTACTTCTTCTTCTAAAACACCTGAAGAA    4762
gb:MN996528|Organism:Severe      CCTGATGCTGTTACAGCGTATAATGGTTATCTTACTTCTTCTTCTAAAACACCTGAAGAA    4762
gb:MN996530|Organism:Severe      CCTGATGCTGTTACAGCGTATAATGGTTATCTTACTTCTTCTTCTAAAACACCTGAAGAA    4748
gb:MN908947|Organism:Severe      CCTGATGCTGTTACAGCGTATAATGGTTATCTTACTTCTTCTTCTAAAACACCTGAAGAA    4762
gb:MT019532|Organism:Severe      CCTGATGCTGTTACAGCGTATAATGGTTATCTTACTTCTTCTTCTAAAACACCTGAAGAA    4762
                                 ************************************************************

gb:MT020781|Organism:Severe      CATTTTATTGAAACCATCTCACTTGCTGGTTCCTATAAAGATTGGTCCTATTCTGGACAA    4810
gb:MT007544|Organism:Severe      CATTTTATTGAAACCATCTCACTTGCTGGTTCCTATAAAGATTGGTCCTATTCTGGACAA    4822
gb:MN994467|Organism:Severe      CATTTTATTGAAACCATCTCACTTGCTGGTTCCTATAAAGATTGGTCCTATTCTGGACAA    4822
gb:MT044257|Organism:Severe      CATTTTATTGAAACCATCTCACTTGCTGGTTCCTATAAAGATTGGTCCTATTCTGGACAA    4822
gb:MT106054|Organism:Severe      CATTTTATTGAAACCATCTCACTTGCTGGTTCCTATAAAGATTGGTCCTATTCTGGACAA    4822
gb:MT049951|Organism:Severe      CATTTTATTGAAACCATCTCACTTGCTGGTTCCTATAAAGATTGGTCCTATTCTGGACAA    4822
gb:MN975262|Organism:Severe      CATTTTATTGAAACCATCTCACTTGCTGGTTCCTATAAAGATTGGTCCTATTCTGGACAA    4822
gb:MT106052|Organism:Severe      CATTTTATTGAAACCATCTCACTTGCTGGTTCCTATAAAGATTGGTCCTATTCTGGACAA    4822
gb:LC522975|Organism:Severe      CATTTTATTGAAACCATCTCACTTGCTGGTTCCTATAAAGATTGGTCCTATTCTGGACAA    4819
gb:LC522973|Organism:Severe      CATTTTATTGAAACCATCTCACTTGCTGGTTCCTATAAAGATTGGTCCTATTCTGGACAA    4819
gb:LC522974|Organism:Severe      CATTTTATTGAAACCATCTCACTTGCTGGTTCCTATAAAGATTGGTCCTATTCTGGACAA    4819
gb:MN985325|Organism:Severe      CATTTTATTGAAACCATCTCACTTGCTGGTTCCTATAAAGATTGGTCCTATTCTGGACAA    4822
gb:MT020881|Organism:Severe      CATTTTATTGAAACCATCTCACTTGCTGGTTCCTATAAAGATTGGTCCTATTCTGGACAA    4822
gb:MT020880|Organism:Severe      CATTTTATTGAAACCATCTCACTTGCTGGTTCCTATAAAGATTGGTCCTATTCTGGACAA    4822
gb:MT066175|Organism:Severe      CATTTTATTGAAACCATCTCACTTGCTGGTTCCTATAAAGATTGGTCCTATTCTGGACAA    4822
gb:MN997409|Organism:Severe      CATTTTATTGAAACCATCTCACTTGCTGGTTCCTATAAAGATTGGTCCTATTCTGGACAA    4822
gb:MN938384|Organism:Severe      CATTTTATTGAAACCATCTCACTTGCTGGTTCCTATAAAGATTGGTCCTATTCTGGACAA    4790
gb:MT044258|Organism:Severe      CATTTTATTGAAACCATCTCACTTGCTGGTTCCTATAAAGATTGGTCCTATTCTGGACAA    4798
gb:MT039890|Organism:Severe      CATTTTATTGAAACCATCTCACTTGCTGGTTCCTATAAAGATTGGTCCTATTCTGGACAA    4822
gb:MN988713|Organism:Severe      CATTTTATTGAAACCATCTCACTTGCTGGTTCCTATAAAGATTGGTCCTATTCTGGACAA    4822
gb:LC521925|Organism:Severe      CATTTTATTGAAACCATCTCACTTGCTGGTTCCTATAAAGATTGGTCCTATTCTGGACAA    4795
```

```
gb:MT093571|Organism:Severe      CATTTTATTGAAACCATCTCACTTGCTGGTTCCTATAAAGATTGGTCCTATTCTGGACAA   4822
gb:MT039887|Organism:Severe      CATTTTATTGAAACCATCTCACTTGCTGGTTCCTATAAAGATTGGTCCTATTCTGGACAA   4822
gb:MT019530|Organism:Severe      CATTTTATTGAAACCATCTCACTTGCTGGTTCCTATAAAGATTGGTCCTATTCTGGACAA   4822
gb:MT039888|Organism:Severe      CATTTTATTGAAACCATCTCACTTGCTGGTTCCTATAAAGATTGGTCCTATTCTGGACAA   4822
gb:LC522972|Organism:Severe      CATTTTATTGAAACCATCTCACTTGCTGGTTCCTATAAAGATTGGTCCTATTCTGGACAA   4819
gb:MT027063|Organism:Severe      CATTTTATTGAAACCATCTCACTTGCTGGTTCCTATAAAGATTGGTCCTATTCTGGACAA   4822
gb:MT027062|Organism:Severe      CATTTTATTGAAACCATCTCACTTGCTGGTTCCTATAAAGATTGGTCCTATTCTGGACAA   4822
gb:MT019529|Organism:Severe      CATTTTATTGAAACCATCTCACTTGCTGGTTCCTATAAAGATTGGTCCTATTCTGGACAA   4822
gb:MN996529|Organism:Severe      CATTTTATTGAAACCATCTCACTTGCTGGTTCCTATAAAGATTGGTCCTATTCTGGACAA   4810
gb:MN996531|Organism:Severe      CATTTTATTGAAACCATCTCACTTGCTGGTTCCTATAAAGATTGGTCCTATTCTGGACAA   4809
gb:MT066176|Organism:Severe      CATTTTATTGAAACCATCTCACTTGCTGGTTCCTATAAAGATTGGTCCTATTCTGGACAA   4822
gb:MT027064|Organism:Severe      CATTTTATTGAAACCATCTCACTTGCTGGTTCCTATAAAGATTGGTCCTATTCTGGACAA   4822
gb:MN994468|Organism:Severe      CATTTTATTGAAACCATCTCACTTGCTGGTTCCTATAAAGATTGGTCCTATTCTGGACAA   4822
gb:MT072688|Organism:Severe      CATTTTATTGAAACCATCTCACTTGCTGGTTCCTATAAAGATTGGTCCTATTCTGGACAA   4807
gb:MN996527|Organism:Severe      CATTTTATTGAAACCATCTCACTTGCTGGTTCCTATAAAGATTGGTCCTATTCTGGACAA   4789
gb:MT093631|Organism:Severe      CATTTTATTGAAACCATCTCACTTGCTGGTTCCTATAAAGATTGGTCCTATTCTGGACAA   4860
gb:MT106053|Organism:Severe      CATTTTATTGAAACCATCTCACTTGCTGGTTCCTATAAAGATTGGTCCTATTCTGGACAA   4822
gb:MT019533|Organism:Severe      CATTTTATTGAAACCATCTCACTTGCTGGTTCCTATAAAGATTGGTCCTATTCTGGACAA   4822
gb:MT019531|Organism:Severe      CATTTTATTGAAACCATCTCACTTGCTGGTTCCTATAAAGATTGGTCCTATTCTGGACAA   4822
gb:MN996528|Organism:Severe      CATTTTATTGAAACCATCTCACTTGCTGGTTCCTATAAAGATTGGTCCTATTCTGGACAA   4822
gb:MN996530|Organism:Severe      CATTTTATTGAAACCATCTCACTTGCTGGTTCCTATAAAGATTGGTCCTATTCTGGACAA   4808
gb:MN908947|Organism:Severe      CATTTTATTGAAACCATCTCACTTGCTGGTTCCTATAAAGATTGGTCCTATTCTGGACAA   4822
gb:MT019532|Organism:Severe      CATTTTATTGAAACCATCTCACTTGCTGGTTCCTATAAAGATTGGTCCTATTCTGGACAA   4822
                                 ************************************************************

gb:MT020781|Organism:Severe      TCTACACAACTAGGTATAGAATTTCTTAAGAGAGGTGATAAAAGTGTATATTACACTAGT   4870
gb:MT007544|Organism:Severe      TCTACACAACTAGGTATAGAATTTCTTAAGAGAGGTGATAAAAGTGTATATTACACTAGT   4882
gb:MN994467|Organism:Severe      TCTACACAACTAGGTATAGAATTTCTTAAGAGAGGTGATAAAAGTGTATATTACACTAGT   4882
gb:MT044257|Organism:Severe      TCTACACAACTAGGTATAGAATTTCTTAAGAGAGGTGATAAAAGTGTATATTACACTAGT   4882
gb:MT106054|Organism:Severe      TCTACACAACTAGGTATAGAATTTCTTAAGAGAGGTGATAAAAGTGTATATTACACTAGT   4882
gb:MT049951|Organism:Severe      TCTACACAACTAGGTATAGAATTTCTTAAGAGAGGTGATAAAAGTGTATATTACACTAGT   4882
gb:MN975262|Organism:Severe      TCTACACAACTAGGTATAGAATTTCTTAAGAGAGGTGATAAAAGTGTATATTACACTAGT   4882
gb:MT106052|Organism:Severe      TCTACACAACTAGGTATAGAATTTCTTAAGAGAGGTGATAAAAGTGTATATTACACTAGT   4882
gb:LC522975|Organism:Severe      TCTACACAACTAGGTATAGAATTTCTTAAGAGAGGTGATAAAAGTGTATATTACACTAGT   4879
gb:LC522973|Organism:Severe      TCTACACAACTAGGTATAGAATTTCTTAAGAGAGGTGATAAAAGTGTATATTACACTAGT   4879
gb:LC522974|Organism:Severe      TCTACACAACTAGGTATAGAATTTCTTAAGAGAGGTGATAAAAGTGTATATTACACTAGT   4879
gb:MN985325|Organism:Severe      TCTACACAACTAGGTATAGAATTTCTTAAGAGAGGTGATAAAAGTGTATATTACACTAGT   4882
gb:MT020881|Organism:Severe      TCTACACAACTAGGTATAGAATTTCTTAAGAGAGGTGATAAAAGTGTATATTACACTAGT   4882
gb:MT020880|Organism:Severe      TCTACACAACTAGGTATAGAATTTCTTAAGAGAGGTGATAAAAGTGTATATTACACTAGT   4882
gb:MT066175|Organism:Severe      TCTACACAACTAGGTATAGAATTTCTTAAGAGAGGTGATAAAAGTGTATATTACACTAGT   4882
gb:MN997409|Organism:Severe      TCTACACAACTAGGTATAGAATTTCTTAAGAGAGGTGATAAAAGTGTATATTACACTAGT   4882
gb:MN938384|Organism:Severe      TCTACACAACTAGGTATAGAATTTCTTAAGAGAGGTGATAAAAGTGTATATTACACTAGT   4850
gb:MT044258|Organism:Severe      TCTACACAACTAGGTATAGAATTTCTTAAGAGAGGTGATAAAAGTGTATATTACACTAGT   4858
gb:MT039890|Organism:Severe      TCTACACAACTAGGTATAGAATTTCTTAAGAGAGGTGATAAAAGTGTATATTACACTAGT   4882
gb:MN988713|Organism:Severe      TCTACACAACTAGGTATAGAATTTCTTAAGAGAGGTGATAAAAGTGTATATTACACTAGT   4882
gb:LC521925|Organism:Severe      TCTACACAACTAGGTATAGAATTTCTTAAGAGAGGTGATAAAAGTGTATATTACACTAGT   4855
gb:MT093571|Organism:Severe      TCTACACAACTAGGTATAGAATTTCTTAAGAGAGGTGATAAAAGTGTATATTACACTAGT   4882
gb:MT039887|Organism:Severe      TCTACACAACTAGGTATAGAATTTCTTAAGAGAGGTGATAAAAGTGTATATTACACTAGT   4882
gb:MT019530|Organism:Severe      TCTACACAACTAGGTATAGAATTTCTTAAGAGAGGTGATAAAAGTGTATATTACACTAGT   4882
gb:MT039888|Organism:Severe      TCTACACAACTAGGTATAGAATTTCTTAAGAGAGGTGATAAAAGTGTATATTACACTAGT   4882
gb:LC522972|Organism:Severe      TCTACACAACTAGGTATAGAATTTCTTAAGAGAGGTGATAAAAGTGTATATTACACTAGT   4879
gb:MT027063|Organism:Severe      TCTACACAACTAGGTATAGAATTTCTTAAGAGAGGTGATAAAAGTGTATATTACACTAGT   4882
gb:MT027062|Organism:Severe      TCTACACAACTAGGTATAGAATTTCTTAAGAGAGGTGATAAAAGTGTATATTACACTAGT   4882
gb:MT019529|Organism:Severe      TCTACACAACTAGGTATAGAATTTCTTAAGAGAGGTGATAAAAGTGTATATTACACTAGT   4882
gb:MN996529|Organism:Severe      TCTACACAACTAGGTATAGAATTTCTTAAGAGAGGTGATAAAAGTGTATATTACACTAGT   4870
gb:MN996531|Organism:Severe      TCTACACAACTAGGTATAGAATTTCTTAAGAGAGGTGATAAAAGTGTATATTACACTAGT   4869
gb:MT066176|Organism:Severe      TCTACACAACTAGGTATAGAATTTCTTAAGAGAGGTGATAAAAGTGTATATTACACTAGT   4882
gb:MT027064|Organism:Severe      TCTACACAACTAGGTATAGAATTTCTTAAGAGAGGTGATAAAAGTGTATATTACACTAGT   4882
gb:MN994468|Organism:Severe      TCTACACAACTAGGTATAGAATTTCTTAAGAGAGGTGATAAAAGTGTATATTACACTAGT   4882
gb:MT072688|Organism:Severe      TCTACACAACTAGGTATAGAATTTCTTAAGAGAGGTGATAAAAGTGTATATTACACTAGT   4867
gb:MN996527|Organism:Severe      TCTACACAACTAGGTATAGAATTTCTTAAGAGAGGTGATAAAAGTGTATATTACACTAGT   4849
gb:MT093631|Organism:Severe      TCTACACAACTAGGTATAGAATTTCTTAAGAGAGGTGATAAAAGTGTATATTACACTAGT   4920
gb:MT106053|Organism:Severe      TCTACACAACTAGGTATAGAATTTCTTAAGAGAGGTGATAAAAGTGTATATTACACTAGT   4882
gb:MT019533|Organism:Severe      TCTACACAACTAGGTATAGAATTTCTTAAGAGAGGTGATAAAAGTGTATATTACACTAGT   4882
gb:MT019531|Organism:Severe      TCTACACAACTAGGTATAGAATTTCTTAAGAGAGGTGATAAAAGTGTATATTACACTAGT   4882
```

```
gb:MN996528|Organism:Severe      TCTACACAACTAGGTATAGAATTTCTTAAGAGAGGTGATAAAAGTGTATATTACACTAGT   4882
gb:MN996530|Organism:Severe      TCTACACAACTAGGTATAGAATTTCTTAAGAGAGGTGATAAAAGTGTATATTACACTAGT   4868
gb:MN908947|Organism:Severe      TCTACACAACTAGGTATAGAATTTCTTAAGAGAGGTGATAAAAGTGTATATTACACTAGT   4882
gb:MT019532|Organism:Severe      TCTACACAACTAGGTATAGAATTTCTTAAGAGAGGTGATAAAAGTGTATATTACACTAGT   4882
                                 ************************************************************

gb:MT020781|Organism:Severe      AATCCTACCACATTCCACCTAGATGGTGAAGTTATCACCTTTGACAATCTTAAGACACTT   4930
gb:MT007544|Organism:Severe      AATCCTACCACATTCCACCTAGATGGTGAAGTTATCACCTTTGACAATCTTAAGACACTT   4942
gb:MN994467|Organism:Severe      AATCCTACCACATTCCACCTAGATGGTGAAGTTATCACCTTTGACAATCTTAAGACACTT   4942
gb:MT044257|Organism:Severe      AATCCTACCACATTCCACCTAGATGGTGAAGTTATCACCTTTGACAATCTTAAGACACTT   4942
gb:MT106054|Organism:Severe      AATCCTACCACATTCCACCTAGATGGTGAAGTTATCACCTTTGACAATCTTAAGACACTT   4942
gb:MT049951|Organism:Severe      AATCCTACCACATTCCACCTAGATGGTGAAGTTATCACCTTTGACAATCTTAAGACACTT   4942
gb:MN975262|Organism:Severe      AATCCTACCACATTCCACCTAGATGGTGAAGTTATCACCTTTGACAATCTTAAGACACTT   4942
gb:MT106052|Organism:Severe      AATCCTACCACATTCCACCTAGATGGTGAAGTTATCACCTTTGACAATCTTAAGACACTT   4942
gb:LC522975|Organism:Severe      AATCCTACCACATTCCACCTAGATGGTGAAGTTATCACCTTTGACAATCTTAAGACACTT   4939
gb:LC522973|Organism:Severe      AATCCTACCACATTCCACCTAGATGGTGAAGTTATCACCTTTGACAATCTTAAGACACTT   4939
gb:LC522974|Organism:Severe      AATCCTACCACATTCCACCTAGATGGTGAAGTTATCACCTTTGACAATCTTAAGACACTT   4939
gb:MN985325|Organism:Severe      AATCCTACCACATTCCACCTAGATGGTGAAGTTATCACCTTTGACAATCTTAAGACACTT   4942
gb:MT020881|Organism:Severe      AATCCTACCACATTCCACCTAGATGGTGAAGTTATCACCTTTGACAATCTTAAGACACTT   4942
gb:MT020880|Organism:Severe      AATCCTACCACATTCCACCTAGATGGTGAAGTTATCACCTTTGACAATCTTAAGACACTT   4942
gb:MT066175|Organism:Severe      AATCCTACCACATTCCACCTAGATGGTGAAGTTATCACCTTTGACAATCTTAAGACACTT   4942
gb:MN997409|Organism:Severe      AATCCTACCACATTCCACCTAGATGGTGAAGTTATCACCTTTGACAATCTTAAGACACTT   4942
gb:MN938384|Organism:Severe      AATCCTACCACATTCCACCTAGATGGTGAAGTTATCACCTTTGACAATCTTAAGACACTT   4910
gb:MT044258|Organism:Severe      AATCCTACCACATTCCACCTAGATGGTGAAGTTATCACCTTTGACAATCTTAAGACACTT   4918
gb:MT039890|Organism:Severe      AATCCTACCACATTCCACCTAGATGGTGAAGTTATCACCTTTGACAATCTTAAGACACTT   4942
gb:MN988713|Organism:Severe      AATCCTACCACATTCCACCTAGATGGTGAAGTTATCACCTTTGACAATCTTAAGACACTT   4942
gb:LC521925|Organism:Severe      AATCCTACCACATTCCACCTAGATGGTGAAGTTATCACCTTTGACAATCTTAAGACACTT   4915
gb:MT093571|Organism:Severe      AATCCTACCACATTCCACCTAGATGGTGAAGTTATCACCTTTGACAATCTTAAGACACTT   4942
gb:MT039887|Organism:Severe      AATCCTACCACATTCCACCTAGATGGTGAAGTTATCACCTTTGACAATCTTAAGACACTT   4942
gb:MT019530|Organism:Severe      AATCCTACCACATTCCACCTAGATGGTGAAGTTATCACCTTTGACAATCTTAAGACACTT   4942
gb:MT039888|Organism:Severe      AATCCTACCACATTCCACCTAGATGGTGAAGTTATCACCTTTGACAATCTTAAGACACTT   4942
gb:LC522972|Organism:Severe      AATCCTACCACATTCCACCTAGATGGTGAAGTTATCACCTTTGACAATCTTAAGACACTT   4939
gb:MT027063|Organism:Severe      AATCCTACCACATTCCACCTAGATGGTGAAGTTATCACCTTTGACAATCTTAAGACACTT   4942
gb:MT027062|Organism:Severe      AATCCTACCACATTCCACCTAGATGGTGAAGTTATCACCTTTGACAATCTTAAGACACTT   4942
gb:MT019529|Organism:Severe      AATCCTACCACATTCCACCTAGATGGTGAAGTTATCACCTTTGACAATCTTAAGACACTT   4942
gb:MN996529|Organism:Severe      AATCCTACCACATTCCACCTAGATGGTGAAGTTATCACCTTTGACAATCTTAAGACACTT   4930
gb:MN996531|Organism:Severe      AATCCTACCACATTCCACCTAGATGGTGAAGTTATCACCTTTGACAATCTTAAGACACTT   4929
gb:MT066176|Organism:Severe      AATCCTACCACATTCCACCTAGATGGTGAAGTTATCACCTTTGACAATCTTAAGACACTT   4942
gb:MT027064|Organism:Severe      AATCCTACCACATTCCACCTAGATGGTGAAGTTATCACCTTTGACAATCTTAAGACACTT   4942
gb:MN994468|Organism:Severe      AATCCTACCACATTCCACCTAGATGGTGAAGTTATCACCTTTGACAATCTTAAGACACTT   4942
gb:MT072688|Organism:Severe      AATCCTACCACATTCCACCTAGATGGTGAAGTTATCACCTTTGACAATCTTAAGACACTT   4927
gb:MN996527|Organism:Severe      AATCCTACCACATTCCACCTAGATGGTGAAGTTATCACCTTTGACAATCTTAAGACACTT   4909
gb:MT093631|Organism:Severe      AATCCTACCACATTCCACCTAGATGGTGAAGTTATCACCTTTGACAATCTTAAGACACTT   4980
gb:MT106053|Organism:Severe      AATCCTACCACATTCCACCTAGATGGTGAAGTTATCACCTTTGACAATCTTAAGACACTT   4942
gb:MT019533|Organism:Severe      AATCCTACCACATTCCACCTAGATGGTGAAGTTATCACCTTTGACAATCTTAAGACACTT   4942
gb:MT019531|Organism:Severe      AATCCTACCACATTCCACCTAGATGGTGAAGTTATCACCTTTGACAATCTTAAGACACTT   4942
gb:MN996528|Organism:Severe      AATCCTACCACATTCCACCTAGATGGTGAAGTTATCACCTTTGACAATCTTAAGACACTT   4942
gb:MN996530|Organism:Severe      AATCCTACCACATTCCACCTAGATGGTGAAGTTATCACCTTTGACAATCTTAAGACACTT   4928
gb:MN908947|Organism:Severe      AATCCTACCACATTCCACCTAGATGGTGAAGTTATCACCTTTGACAATCTTAAGACACTT   4942
gb:MT019532|Organism:Severe      AATCCTACCACATTCCACCTAGATGGTGAAGTTATCACCTTTGACAATCTTAAGACACTT   4942
                                 ************************************************************

gb:MT020781|Organism:Severe      CTTTCTTTGAGAGAAGTGAGGACTATTAAGGTGTTTACAACAGTAGACAACATTAACCTC   4990
gb:MT007544|Organism:Severe      CTTTCTTTGAGAGAAGTGAGGACTATTAAGGTGTTTACAACAGTAGACAACATTAACCTC   5002
gb:MN994467|Organism:Severe      CTTTCTTTGAGAGAAGTGAGGACTATTAAGGTGTTTACAACAGTAGACAACATTAACCTC   5002
gb:MT044257|Organism:Severe      CTTTCTTTGAGAGAAGTGAGGACTATTAAGGTGTTTACAACAGTAGACAACATTAACCTC   5002
gb:MT106054|Organism:Severe      CTTTCTTTGAGAGAAGTGAGGACTATTAAGGTGTTTACAACAGTAGACAACATTAACCTC   5002
gb:MT049951|Organism:Severe      CTTTCTTTGAGAGAAGTGAGGACTATTAAGGTGTTTACAACAGTAGACAACATTAACCTC   5002
gb:MN975262|Organism:Severe      CTTTCTTTGAGAGAAGTGAGGACTATTAAGGTGTTTACAACAGTAGACAACATTAACCTC   5002
gb:MT106052|Organism:Severe      CTTTCTTTGAGAGAAGTGAGGACTATTAAGGTGTTTACAACAGTAGACAACATTAACCTC   5002
gb:LC522975|Organism:Severe      CTTTCTTTGAGAGAAGTGAGGACTATTAAGGTGTTTACAACAGTAGACAACATTAACCTC   4999
gb:LC522973|Organism:Severe      CTTTCTTTGAGAGAAGTGAGGACTATTAAGGTGTTTACAACAGTAGACAACATTAACCTC   4999
gb:LC522974|Organism:Severe      CTTTCTTTGAGAGAAGTGAGGACTATTAAGGTGTTTACAACAGTAGACAACATTAACCTC   4999
gb:MN985325|Organism:Severe      CTTTCTTTGAGAGAAGTGAGGACTATTAAGGTGTTTACAACAGTAGACAACATTAACCTC   5002
gb:MT020881|Organism:Severe      CTTTCTTTGAGAGAAGTGAGGACTATTAAGGTGTTTACAACAGTAGACAACATTAACCTC   5002
```

```
gb:MT020880|Organism:Severe      CTTTCTTTGAGAGAAGTGAGGACTATTAAGGTGTTTACAACAGTAGACAACATTAACCTC    5002
gb:MT066175|Organism:Severe      CTTTCTTTGAGAGAAGTGAGGACTATTAAGGTGTTTACAACAGTAGACAACATTAACCTC    5002
gb:MN997409|Organism:Severe      CTTTCTTTGAGAGAAGTGAGGACTATTAAGGTGTTTACAACAGTAGACAACATTAACCTC    5002
gb:MN938384|Organism:Severe      CTTTCTTTGAGAGAAGTGAGGACTATTAAGGTGTTTACAACAGTAGACAACATTAACCTC    4970
gb:MT044258|Organism:Severe      CTTTCTTTGAGAGAAGTGAGGACTATTAAGGTGTTTACAACAGTAGACAACATTAACCTC    4978
gb:MT039890|Organism:Severe      CTTTCTTTGAGAGAAGTGAGGACTATTAAGGTGTTTACAACAGTAGACAACATTAACCTC    5002
gb:MN988713|Organism:Severe      CTTTCTTTGAGAGAAGTGAGGACTATTAAGGTGTTTACAACAGTAGACAACATTAACCTC    5002
gb:LC521925|Organism:Severe      CTTTCTTTGAGAGAAGTGAGGACTATTAAGGTGTTTACAACAGTAGACAACATTAACCTC    4975
gb:MT093571|Organism:Severe      CTTTCTTTGAGAGAAGTGAGGACTATTAAGGTGTTTACAACAGTAGACAACATTAACCTC    5002
gb:MT039887|Organism:Severe      CTTTCTTTGAGAGAAGTGAGGACTATTAAGGTGTTTACAACAGTAGACAACATTAACCTC    5002
gb:MT019530|Organism:Severe      CTTTCTTTGAGAGAAGTGAGGACTATTAAGGTGTTTACAACAGTAGACAACATTAACCTC    5002
gb:MT039888|Organism:Severe      CTTTCTTTGAGAGAAGTGAGGACTATTAAGGTGTTTACAACAGTAGACAACATTAACCTC    5002
gb:LC522972|Organism:Severe      CTTTCTTTGAGAGAAGTGAGGACTATTAAGGTGTTTACAACAGTAGACAACATTAACCTC    4999
gb:MT027063|Organism:Severe      CTTTCTTTGAGAGAAGTGAGGACTATTAAGGTGTTTACAACAGTAGACAACATTAACCTC    5002
gb:MT027062|Organism:Severe      CTTTCTTTGAGAGAAGTGAGGACTATTAAGGTGTTTACAACAGTAGACAACATTAACCTC    5002
gb:MT019529|Organism:Severe      CTTTCTTTGAGAGAAGTGAGGACTATTAAGGTGTTTACAACAGTAGACAACATTAACCTC    5002
gb:MN996529|Organism:Severe      CTTTCTTTGAGAGAAGTGAGGACTATTAAGGTGTTTACAACAGTAGACAACATTAACCTC    4990
gb:MN996531|Organism:Severe      CTTTCTTTGAGAGAAGTGAGGACTATTAAGGTGTTTACAACAGTAGACAACATTAACCTC    4989
gb:MT066176|Organism:Severe      CTTTCTTTGAGAGAAGTGAGGACTATTAAGGTGTTTACAACAGTAGACAACATTAACCTC    5002
gb:MT027064|Organism:Severe      CTTTCTTTGAGAGAAGTGAGGACTATTAAGGTGTTTACAACAGTAGACAACATTAACCTC    5002
gb:MN994468|Organism:Severe      CTTTCTTTGAGAGAAGTGAGGACTATTAAGGTGTTTACAACAGTAGACAACATTAACCTC    5002
gb:MT072688|Organism:Severe      CTTTCTTTGAGAGAAGTGAGGACTATTAAGGTGTTTACAACAGTAGACAACATTAACCTC    4987
gb:MN996527|Organism:Severe      CTTTCTTTGAGAGAAGTGAGGACTATTAAGGTGTTTACAACAGTAGACAACATTAACCTC    4969
gb:MT093631|Organism:Severe      CTTTCTTTGAGAGAAGTGAGGACTATTAAGGTGTTTACAACAGTAGACAACATTAACCTC    5040
gb:MT106053|Organism:Severe      CTTTCTTTGAGAGAAGTGAGGACTATTAAGGTGTTTACAACAGTAGACAACATTAACCTC    5002
gb:MT019533|Organism:Severe      CTTTCTTTGAGAGAAGTGAGGACTATTAAGGTGTTTACAACAGTAGACAACATTAACCTC    5002
gb:MT019531|Organism:Severe      CTTTCTTTGAGAGAAGTGAGGACTATTAAGGTGTTTACAACAGTAGACAACATTAACCTC    5002
gb:MN996528|Organism:Severe      CTTTCTTTGAGAGAAGTGAGGACTATTAAGGTGTTTACAACAGTAGACAACATTAACCTC    5002
gb:MN996530|Organism:Severe      CTTTCTTTGAGAGAAGTGAGGACTATTAAGGTGTTTACAACAGTAGACAACATTAACCTC    4988
gb:MN908947|Organism:Severe      CTTTCTTTGAGAGAAGTGAGGACTATTAAGGTGTTTACAACAGTAGACAACATTAACCTC    5002
gb:MT019532|Organism:Severe      CTTTCTTTGAGAGAAGTGAGGACTATTAAGGTGTTTACAACAGTAGACAACATTAACCTC    5002
                                 ************************************************************

gb:MT020781|Organism:Severe      CACACGCAAGTTGTGGACATGTCAATGACATATGGACAACAGTTTGGTCCAACTTATTTG    5050
gb:MT007544|Organism:Severe      CACACGCAAGTTGTGGACATGTCAATGACATATGGACAACAGTTTGGTCCAACTTATTTG    5062
gb:MN994467|Organism:Severe      CACACGCAAGTTGTGGACATGTCAATGACATATGGACAACAGTTTGGTCCAACTTATTTG    5062
gb:MT044257|Organism:Severe      CACACGCAAGTTGTGGACATGTCAATGACATATGGACAACAGTTTGGTCCAACTTATTTG    5062
gb:MT106054|Organism:Severe      CACACGCAAGTTGTGGACATGTCAATGACATATGGACAACAGTTTGGTCCAACTTATTTG    5062
gb:MT049951|Organism:Severe      CACACGCAAGTTGTGGACATGTCAATGACATATGGACAACAGTTTGGTCCAACTTATTTG    5062
gb:MN975262|Organism:Severe      CACACGCAAGTTGTGGACATGTCAATGACATATGGACAACAGTTTGGTCCAACTTATTTG    5062
gb:MT106052|Organism:Severe      CACACGCAAGTTGTGGACATGTCAATGACATATGGACAACAGTTTGGTCCAACTTATTTG    5062
gb:LC522975|Organism:Severe      CACACGCAAGTTGTGGACATGTCAATGACATATGGACAACAGTTTGGTCCAACTTATTTG    5059
gb:LC522973|Organism:Severe      CACACGCAAGTTGTGGACATGTCAATGACATATGGACAACAGTTTGGTCCAACTTATTTG    5059
gb:LC522974|Organism:Severe      CACACGCAAGTTGTGGACATGTCAATGACATATGGACAACAGTTTGGTCCAACTTATTTG    5059
gb:MN985325|Organism:Severe      CACACGCAAGTTGTGGACATGTCAATGACATATGGACAACAGTTTGGTCCAACTTATTTG    5062
gb:MT020881|Organism:Severe      CACACGCAAGTTGTGGACATGTCAATGACATATGGACAACAGTTTGGTCCAACTTATTTG    5062
gb:MT020880|Organism:Severe      CACACGCAAGTTGTGGACATGTCAATGACATATGGACAACAGTTTGGTCCAACTTATTTG    5062
gb:MT066175|Organism:Severe      CACACGCAAGTTGTGGACATGTCAATGACATATGGACAACAGTTTGGTCCAACTTATTTG    5062
gb:MN997409|Organism:Severe      CACACGCAAGTTGTGGACATGTCAATGACATATGGACAACAGTTTGGTCCAACTTATTTG    5062
gb:MN938384|Organism:Severe      CACACGCAAGTTGTGGACATGTCAATGACATATGGACAACAGTTTGGTCCAACTTATTTG    5030
gb:MT044258|Organism:Severe      CACACGCAAGTTGTGGACATGTCAATGACATATGGACAACAGTTTGGTCCAACTTATTTG    5038
gb:MT039890|Organism:Severe      CACACGCAAGTTGTGGACATGTCAATGACATATGGACAACAGTTTGGTCCAACTTATTTG    5062
gb:MN988713|Organism:Severe      CACACGCAAGTTGTGGACATGTCAATGACATATGGACAACAGTTTGGTCCAACTTATTTG    5062
gb:LC521925|Organism:Severe      CACACGCAAGTTGTGGACATGTCAATGACATATGGACAACAGTTTGGTCCAACTTATTTG    5035
gb:MT093571|Organism:Severe      CACACGCAAGTTGTGGACATGTCAATGACATATGGACAACAGTTTGGTCCAACTTATTTG    5062
gb:MT039887|Organism:Severe      CACACGCAAGTTGTGGACATGTCAATGACATATGGACAACAGTTTGGTCCAACTTATTTG    5062
gb:MT019530|Organism:Severe      CACACGCAAGTTGTGGACATGTCAATGACATATGGACAACAGTTTGGTCCAACTTATTTG    5062
gb:MT039888|Organism:Severe      CACACGCAAGTTGTGGACATGTCAATGACATATGGACAACAGTTTGGTCCAACTTATTTG    5062
gb:LC522972|Organism:Severe      CACACGCAAGTTGTGGACATGTCAATGACATATGGACAACAGTTTGGTCCAACTTATTTG    5059
gb:MT027063|Organism:Severe      CACACGCAAGTTGTGGACATGTCAATGACATATGGACAACAGTTTGGTCCAACTTATTTG    5062
gb:MT027062|Organism:Severe      CACACGCAAGTTGTGGACATGTCAATGACATATGGACAACAGTTTGGTCCAACTTATTTG    5062
gb:MT019529|Organism:Severe      CACACGCAAGTTGTGGACATGTCAATGACATATGGACAACAGTTTGGTCCAACTTATTTG    5062
gb:MN996529|Organism:Severe      CACACGCAAGTTGTGGACATGTCAATGACATATGGACAACAGTTTGGTCCAACTTATTTG    5050
gb:MN996531|Organism:Severe      CACACGCAAGTTGTGGACATGTCAATGACATATGGACAACAGTTTGGTCCAACTTATTTG    5049
gb:MT066176|Organism:Severe      CACACGCAAGTTGTGGACATGTCAATGACATATGGACAACAGTTTGGTCCAACTTATTTG    5062
```

```
gb:MT027064|Organism:Severe      CACACGCAAGTTGTGGACATGTCAATGACATATGGACAACAGTTTGGTCCAACTTATTTG    5062
gb:MN994468|Organism:Severe      CACACGCAAGTTGTGGACATGTCAATGACATATGGACAACAGTTTGGTCCAACTTATTTG    5062
gb:MT072688|Organism:Severe      CACACGCAAGTTGTGGACATGTCAATGACATATGGACAACAGTTTGGTCCAACTTATTTG    5047
gb:MN996527|Organism:Severe      CACACGCAAGTTGTGGACATGTCAATGACATATGGACAACAGTTTGGTCCAACTTATTTG    5029
gb:MT093631|Organism:Severe      CACACGCAAGTTGTGGACATGTCAATGACATATGGACAACAGTTTGGTCCAACTTATTTG    5100
gb:MT106053|Organism:Severe      CACACGCAAGTTGTGGACATGTCAATGACATATGGACAACAGTTTGGTCCAACTTATTTG    5062
gb:MT019533|Organism:Severe      CACACGCAAGTTGTGGACATGTCAATGACATATGGACAACAGTTTGGTCCAACTTATTTG    5062
gb:MT019531|Organism:Severe      CACACGCAAGTTGTGGACATGTCAATGACATATGGACAACAGTTTGGTCCAACTTATTTG    5062
gb:MN996528|Organism:Severe      CACACGCAAGTTGTGGACATGTCAATGACATATGGACAACAGTTTGGTCCAACTTATTTG    5062
gb:MN996530|Organism:Severe      CACACGCAAGTTGTGGACATGTCAATGACATATGGACAACAGTTTGGTCCAACTTATTTG    5048
gb:MN908947|Organism:Severe      CACACGCAAGTTGTGGACATGTCAATGACATATGGACAACAGTTTGGTCCAACTTATTTG    5062
gb:MT019532|Organism:Severe      CACACGCAAGTTGTGGACATGTCAATGACATATGGACAACAGTTTGGTCCAACTTATTTG    5062
                                 ************************************************************

gb:MT020781|Organism:Severe      GATGGAGCTGATGTTACTAAAATAAAACCTCATAATTCACATGAAGGTAAAACATTTTAT    5110
gb:MT007544|Organism:Severe      GATGGAGCTGATGTTACTAAAATAAAACCTCATAATTCACATGAAGGTAAAACATTTTAT    5122
gb:MN994467|Organism:Severe      GATGGAGCTGATGTTACTAAAATAAAACCTCATAATTCACATGAAGGTAAAACATTTTAT    5122
gb:MT044257|Organism:Severe      GATGGAGCTGATGTTACTAAAATAAAACCTCATAATTCACATGAAGGTAAAACATTTTAT    5122
gb:MT106054|Organism:Severe      GATGGAGCTGATGTTACTAAAATAAAACCTCATAATTCACATGAAGGTAAAACATTTTAT    5122
gb:MT049951|Organism:Severe      GATGGAGCTGATGTTACTAAAATAAAACCTCATAATTCACATGAAGGTAAAACATTTTAT    5122
gb:MN975262|Organism:Severe      GATGGAGCTGATGTTACTAAAATAAAACCTCATAATTCACATGAAGGTAAAACATTTTAT    5122
gb:MT106052|Organism:Severe      GATGGAGCTGATGTTACTAAAATAAAACCTCATAATTCACATGAAGGTAAAACATTTTAT    5122
gb:LC522975|Organism:Severe      GATGGAGCTGATGTTACTAAAATAAAACCTCATAATTCACATGAAGGTAAAACATTTTAT    5119
gb:LC522973|Organism:Severe      GATGGAGCTGATGTTACTAAAATAAAACCTCATAATTCACATGAAGGTAAAACATTTTAT    5119
gb:LC522974|Organism:Severe      GATGGAGCTGATGTTACTAAAATAAAACCTCATAATTCACATGAAGGTAAAACATTTTAT    5119
gb:MN985325|Organism:Severe      GATGGAGCTGATGTTACTAAAATAAAACCTCATAATTCACATGAAGGTAAAACATTTTAT    5122
gb:MT020881|Organism:Severe      GATGGAGCTGATGTTACTAAAATAAAACCTCATAATTCACATGAAGGTAAAACATTTTAT    5122
gb:MT020880|Organism:Severe      GATGGAGCTGATGTTACTAAAATAAAACCTCATAATTCACATGAAGGTAAAACATTTTAT    5122
gb:MT066175|Organism:Severe      GATGGAGCTGATGTTACTAAAATAAAACCTCATAATTCACATGAAGGTAAAACATTTTAT    5122
gb:MN997409|Organism:Severe      GATGGAGCTGATGTTACTAAAATAAAACCTCATAATTCACATGAAGGTAAAACATTTTAT    5122
gb:MN938384|Organism:Severe      GATGGAGCTGATGTTACTAAAATAAAACCTCATAATTCACATGAAGGTAAAACATTTTAT    5090
gb:MT044258|Organism:Severe      GATGGAGCTGATGTTACTAAAATAAAACCTCATAATTCACATGAAGGTAAAACATTTTAT    5098
gb:MT039890|Organism:Severe      GATGGAGCTGATGTTACTAAAATAAAACCTCATAATTCACATGAAGGTAAAACATTTTAT    5122
gb:MN988713|Organism:Severe      GATGGAGCTGATGTTACTAAAATAAAACCTCATAATTCACATGAAGGTAAAACATTTTAT    5122
gb:LC521925|Organism:Severe      GATGGAGCTGATGTTACTAAAATAAAACCTCATAATTCACATGAAGGTAAAACATTTTAT    5095
gb:MT093571|Organism:Severe      GATGGAGCTGATGTTACTAAAATAAAACCTCATAATTCACATGAAGGTAAAACATTTTAT    5122
gb:MT039887|Organism:Severe      GATGGAGCTGATGTTACTAAAATAAAACCTCATAATTCACATGAAGGTAAAACATTTTAT    5122
gb:MT019530|Organism:Severe      GATGGAGCTGATGTTACTAAAATAAAACCTCATAATTCACATGAAGGTAAAACATTTTAT    5122
gb:MT039888|Organism:Severe      GATGGAGCTGATGTTACTAAAATAAAACCTCATAATTCACATGAAGGTAAAACATTTTAT    5122
gb:LC522972|Organism:Severe      GATGGAGCTGATGTTACTAAAATAAAACCTCATAATTCACATGAAGGTAAAACATTTTAT    5119
gb:MT027063|Organism:Severe      GATGGAGCTGATGTTACTAAAGTAAAACCTCATAATTCACATGAAGGTAAAACATTTTAT    5122
gb:MT027062|Organism:Severe      GATGGAGCTGATGTTACTAAAGTAAAACCTCATAATTCACATGAAGGTAAAACATTTTAT    5122
gb:MT019529|Organism:Severe      GATGGAGCTGATGTTACTAAAATAAAACCTCATAATTCACATGAAGGTAAAACATTTTAT    5122
gb:MN996529|Organism:Severe      GATGGAGCTGATGTTACTAAAATAAAACCTCATAATTCACATGAAGGTAAAACATTTTAT    5110
gb:MN996531|Organism:Severe      GATGGAGCTGATGTTACTAAAATAAAACCTCATAATTCACATGAAGGTAAAACATTTTAT    5109
gb:MT066176|Organism:Severe      GATGGAGCTGATGTTACTAAAATAAAACCTCATAATTCACATGAAGGTAAAACATTTTAT    5122
gb:MT027064|Organism:Severe      GATGGAGCTGATGTTACTAAAATAAAACCTCATAATTCACATGAAGGTAAAACATTTTAT    5122
gb:MN994468|Organism:Severe      GATGGAGCTGATGTTACTAAAATAAAACCTCATAATTCACATGAAGGTAAAACATTTTAT    5122
gb:MT072688|Organism:Severe      GATGGAGCTGATGTTACTAAAATAAAACCTCATAATTCACATGAAGGTAAAACATTTTAT    5107
gb:MN996527|Organism:Severe      GATGGAGCTGATGTTACTAAAATAAAACCTCATAATTCACATGAAGGTAAAACATTTTAT    5089
gb:MT093631|Organism:Severe      GATGGAGCTGATGTTACTAAAATAAAACCTCATAATTCACATGAAGGTAAAACATTTTAT    5160
gb:MT106053|Organism:Severe      GATGGAGCTGATGTTACTAAAATAAAACCTCATAATTCACATGAAGGTAAAACATTTTAT    5122
gb:MT019533|Organism:Severe      GATGGAGCTGATGTTACTAAAATAAAACCTCATAATTCACATGAAGGTAAAACATTTTAT    5122
gb:MT019531|Organism:Severe      GATGGAGCTGATGTTACTAAAATAAAACCTCATAATTCACATGAAGGTAAAACATTTTAT    5122
gb:MN996528|Organism:Severe      GATGGAGCTGATGTTACTAAAATAAAACCTCATAATTCACATGAAGGTAAAACATTTTAT    5122
gb:MN996530|Organism:Severe      GATGGAGCTGATGTTACTAAAATAAAACCTCATAATTCACATGAAGGTAAAACATTTTAT    5108
gb:MN908947|Organism:Severe      GATGGAGCTGATGTTACTAAAATAAAACCTCATAATTCACATGAAGGTAAAACATTTTAT    5122
gb:MT019532|Organism:Severe      GATGGAGCTGATGTTACTAAAATAAAACCTCATAATTCACATGAAGGTAAAACATTTTAT    5122
                                 ********************* **************************************

gb:MT020781|Organism:Severe      GTTTTACCTAATGATGACACTCTACGTGTTGAGGCTTTTGAGTACTACCACACAACTGAT    5170
gb:MT007544|Organism:Severe      GTTTTACCTAATGATGACACTCTACGTGTTGAGGCTTTTGAGTACTACCACACAACTGAT    5182
gb:MN994467|Organism:Severe      GTTTTACCTAATGATGACACTCTACGTGTTGAGGCTTTTGAGTACTACCACACAACTGAT    5182
gb:MT044257|Organism:Severe      GTTTTACCTAATGATGACACTCTACGTGTTGAGGCTTTTGAGTACTACCACACAACTGAT    5182
gb:MT106054|Organism:Severe      GTTTTACCTAATGATGACACTCTACGTGTTGAGGCTTTTGAGTACTACCACACAACTGAT    5182
```

```
gb:MT049951|Organism:Severe      GTTTTACCTAATGATGACACTCTACGTGTTGAGGCTTTTGAGTACTACCACACAACTGAT    5182
gb:MN975262|Organism:Severe      GTTTTACCTAATGATGACACTCTACGTGTTGAGGCTTTTGAGTACTACCACACAACTGAT    5182
gb:MT106052|Organism:Severe      GTTTTACCTAATGATGACACTCTACGTGTTGAGGCTTTTGAGTACTACCACACAACTGAT    5182
gb:LC522975|Organism:Severe      GTTTTACCTAATGATGACACTCTACGTGTTGAGGCTTTTGAGTACTACCACACAACTGAT    5179
gb:LC522973|Organism:Severe      GTTTTACCTAATGATGACACTCTACGTGTTGAGGCTTTTGAGTACTACCACACAACTGAT    5179
gb:LC522974|Organism:Severe      GTTTTACCTAATGATGACACTCTACGTGTTGAGGCTTTTGAGTACTACCACACAACTGAT    5179
gb:MN985325|Organism:Severe      GTTTTACCTAATGATGACACTCTACGTGTTGAGGCTTTTGAGTACTACCACACAACTGAT    5182
gb:MT020881|Organism:Severe      GTTTTACCTAATGATGACACTCTACGTGTTGAGGCTTTTGAGTACTACCACACAACTGAT    5182
gb:MT020880|Organism:Severe      GTTTTACCTAATGATGACACTCTACGTGTTGAGGCTTTTGAGTACTACCACACAACTGAT    5182
gb:MT066175|Organism:Severe      GTTTTACCTAATGATGACACTCTACGTGTTGAGGCTTTTGAGTACTACCACACAACTGAT    5182
gb:MN997409|Organism:Severe      GTTTTACCTAATGATGACACTCTACGTGTTGAGGCTTTTGAGTACTACCACACAACTGAT    5182
gb:MN938384|Organism:Severe      GTTTTACCTAATGATGACACTCTACGTGTTGAGGCTTTTGAGTACTACCACACAACTGAT    5150
gb:MT044258|Organism:Severe      GTTTTACCTAATGATGACACTCTACGTGTTGAGGCTTTTGAGTACTACCACACAACTGAT    5158
gb:MT039890|Organism:Severe      GTTTTACCTAATGATGACACTCTACGTGTTGAGGCTTTTGAGTACTACCACACAACTGAT    5182
gb:MN988713|Organism:Severe      GTTTTACCTAATGATGACACTCTACGTGTTGAGGCTTTTGAGTACTACCACACAACTGAT    5182
gb:LC521925|Organism:Severe      GTTTTACCTAATGATGACACTCTACGTGTTGAGGCTTTTGAGTACTACCACACAACTGAT    5155
gb:MT093571|Organism:Severe      GTTTTACCTAATGATGACACTCTACGTGTTGAGGCTTTTGAGTACTACCACACAACTGAT    5182
gb:MT039887|Organism:Severe      GTTTTACCTAATGATGACACTCTACGTGTTGAGGCTTTTGAGTACTACCACACAACTGAT    5182
gb:MT019530|Organism:Severe      GTTTTACCTAATGATGACACTCTACGTGTTGAGGCTTTTGAGTACTACCACACAACTGAT    5182
gb:MT039888|Organism:Severe      GTTTTACCTAATGATGACACTCTACGTGTTGAGGCTTTTGAGTACTACCACACAACTGAT    5182
gb:LC522972|Organism:Severe      GTTTTACCTAATGATGACACTCTACGTGTTGAGGCTTTTGAGTACTACCACACAACTGAT    5179
gb:MT027063|Organism:Severe      GTTTTACCTAATGATGACACTCTACGTGTTGAGGCTTTTGAGTACTACCACACAACTGAT    5182
gb:MT027062|Organism:Severe      GTTTTACCTAATGATGACACTCTACGTGTTGAGGCTTTTGAGTACTACCACACAACTGAT    5182
gb:MT019529|Organism:Severe      GTTTTACCTAATGATGACACTCTACGTGTTGAGGCTTTTGAGTACTACCACACAACTGAT    5182
gb:MN996529|Organism:Severe      GTTTTACCTAATGATGACACTCTACGTGTTGAGGCTTTTGAGTACTACCACACAACTGAT    5170
gb:MN996531|Organism:Severe      GTTTTACCTAATGATGACACTCTACGTGTTGAGGCTTTTGAGTACTACCACACAACTGAT    5169
gb:MT066176|Organism:Severe      GTTTTACCTAATGATGACACTCTACGTGTTGAGGCTTTTGAGTACTACCACACAACTGAT    5182
gb:MT027064|Organism:Severe      GTTTTACCTAATGATGACACTCTACGTGTTGAGGCTTTTGAGTACTACCACACAACTGAT    5182
gb:MN994468|Organism:Severe      GTTTTACCTAATGATGACACTCTACGTGTTGAGGCTTTTGAGTACTACCACACAACTGAT    5182
gb:MT072688|Organism:Severe      GTTTTACCTAATGATGACACTCTACGTGTTGAGGCTTTTGAGTACTACCACACAACTGAT    5167
gb:MN996527|Organism:Severe      GTTTTACCTAATGATGACACTCTACGTGTTGAGGCTTTTGAGTACTACCACACAACTGAT    5149
gb:MT093631|Organism:Severe      GTTTTACCTAATGATGACACTCTACGTGTTGAGGCTTTTGAGTACTACCACACAACTGAT    5220
gb:MT106053|Organism:Severe      GTTTTACCTAATGATGACACTCTACGTGTTGAGGCTTTTGAGTACTACCACACAACTGAT    5182
gb:MT019533|Organism:Severe      GTTTTACCTAATGATGACACTCTACGTGTTGAGGCTTTTGAGTACTACCACACAACTGAT    5182
gb:MT019531|Organism:Severe      GTTTTACCTAATGATGACACTCTACGTGTTGAGGCTTTTGAGTACTACCACACAACTGAT    5182
gb:MN996528|Organism:Severe      GTTTTACCTAATGATGACACTCTACGTGTTGAGGCTTTTGAGTACTACCACACAACTGAT    5182
gb:MN996530|Organism:Severe      GTTTTACCTAATGATGACACTCTACGTGTTGAGGCTTTTGAGTACTACCACACAACTGAT    5168
gb:MN908947|Organism:Severe      GTTTTACCTAATGATGACACTCTACGTGTTGAGGCTTTTGAGTACTACCACACAACTGAT    5182
gb:MT019532|Organism:Severe      GTTTTACCTAATGATGACACTCTACGTGTTGAGGCTTTTGAGTACTACCACACAACTGAT    5182
                                 ************************************************************

gb:MT020781|Organism:Severe      CCTAGTTTTCTGGGTAGGTACATGTCAGCATTAAATCACACTAAAAAGTGGAAATACCCA    5230
gb:MT007544|Organism:Severe      CCTAGTTTTCTGGGTAGGTACATGTCAGCATTAAATCACACTAAAAAGTGGAAATACCCA    5242
gb:MN994467|Organism:Severe      CCTAGTTTTCTGGGTAGGTACATGTCAGCATTAAATCACACTAAAAAGTGGAAATACCCA    5242
gb:MT044257|Organism:Severe      CCTAGTTTTCTGGGTAGGTACATGTCAGCATTAAATCACACTAAAAAGTGGAAATACCCA    5242
gb:MT106054|Organism:Severe      CCTAGTTTTCTGGGTAGGTACATGTCAGCATTAAATCACACTAAAAAGTGGAAATACCCA    5242
gb:MT049951|Organism:Severe      CCTAGTTTTCTGGGTAGGTACATGTCAGCATTAAATCACACTAAAAAGTGGAAATACCCA    5242
gb:MN975262|Organism:Severe      CCTAGTTTTCTGGGTAGGTACATGTCAGCATTAAATCACACTAAAAAGTGGAAATACCCA    5242
gb:MT106052|Organism:Severe      CCTAGTTTTCTGGGTAGGTACATGTCAGCATTAAATCACACTAAAAAGTGGAAATACCCA    5242
gb:LC522975|Organism:Severe      CCTAGTTTTCTGGGTAGGTACATGTCAGCATTAAATCACACTAAAAAGTGGAAATACCCA    5239
gb:LC522973|Organism:Severe      CCTAGTTTTCTGGGTAGGTACATGTCAGCATTAAATCACACTAAAAAGTGGAAATACCCA    5239
gb:LC522974|Organism:Severe      CCTAGTTTTCTGGGTAGGTACATGTCAGCATTAAATCACACTAAAAAGTGGAAATACCCA    5239
gb:MN985325|Organism:Severe      CCTAGTTTTCTGGGTAGGTACATGTCAGCATTAAATCACACTAAAAAGTGGAAATACCCA    5242
gb:MT020881|Organism:Severe      CCTAGTTTTCTGGGTAGGTACATGTCAGCATTAAATCACACTAAAAAGTGGAAATACCCA    5242
gb:MT020880|Organism:Severe      CCTAGTTTTCTGGGTAGGTACATGTCAGCATTAAATCACACTAAAAAGTGGAAATACCCA    5242
gb:MT066175|Organism:Severe      CCTAGTTTTCTGGGTAGGTACATGTCAGCATTAAATCACACTAAAAAGTGGAAATACCCA    5242
gb:MN997409|Organism:Severe      CCTAGTTTTCTGGGTAGGTACATGTCAGCATTAAATCACACTAAAAAGTGGAAATACCCA    5242
gb:MN938384|Organism:Severe      CCTAGTTTTCTGGGTAGGTACATGTCAGCATTAAATCACACTAAAAAGTGGAAATACCCA    5210
gb:MT044258|Organism:Severe      CCTAGTTTTCTGGGTAGGTACATGTCAGCATTAAATCACACTAAAAAGTGGAAATACCCA    5218
gb:MT039890|Organism:Severe      CCTAGTTTTCTGGGTAGGTACATGTCAGCATTAAATCACACTAAAAAGTGGAAATACCCA    5242
gb:MN988713|Organism:Severe      CCTAGTTTTCTGGGTAGGTACATGTCAGCATTAAATCACACTAAAAAGTGGAAATACCCA    5242
gb:LC521925|Organism:Severe      CCTAGTTTTCTGGGTAGGTACATGTCAGCATTAAATCACACTAAAAAGTGGAAATACCCA    5215
gb:MT093571|Organism:Severe      CCTAGTTTTCTGGGTAGGTACATGTCAGCATTAAATCACACTAAAAAGTGGAAATACCCA    5242
gb:MT039887|Organism:Severe      CCTAGTTTTCTGGGTAGGTACATGTCAGCATTAAATCACACTAAAAAGTGGAAATACCCA    5242
gb:MT019530|Organism:Severe      CCTAGTTTTCTGGGTAGGTACATGTCAGCATTAAATCACACTAAAAAGTGGAAATACCCA    5242
```

```
gb:MT039888|Organism:Severe      CCTAGTTTTCTGGGTAGGTACATGTCAGCATTAAATCACACTAAAAAGTGGAAATACCCA   5242
gb:LC522972|Organism:Severe      CCTAGTTTTCTGGGTAGGTACATGTCAGCATTAAATCACACTAAAAAGTGGAAATACCCA   5239
gb:MT027063|Organism:Severe      CCTAGTTTTCTGGGTAGGTACATGTCAGCATTAAATCACACTAAAAAGTGGAAATACCCA   5242
gb:MT027062|Organism:Severe      CCTAGTTTTCTGGGTAGGTACATGTCAGCATTAAATCACACTAAAAAGTGGAAATACCCA   5242
gb:MT019529|Organism:Severe      CCTAGTTTTCTGGGTAGGTACATGTCAGCATTAAATCACACTAAAAAGTGGAAATACCCA   5242
gb:MN996529|Organism:Severe      CCTAGTTTTCTGGGTAGGTACATGTCAGCATTAAATCACACTAAAAAGTGGAAATACCCA   5230
gb:MN996531|Organism:Severe      CCTAGTTTTCTGGGTAGGTACATGTCAGCATTAAATCACACTAAAAAGTGGAAATACCCA   5229
gb:MT066176|Organism:Severe      CCTAGTTTTCTGGGTAGGTACATGTCAGCATTAAATCACACTAAAAAGTGGAAATACCCA   5242
gb:MT027064|Organism:Severe      CCTAGTTTTCTGGGTAGGTACATGTCAGCATTAAATCACACTAAAAAGTGGAAATACCCA   5242
gb:MN994468|Organism:Severe      CCTAGTTTTCTGGGTAGGTACATGTCAGCATTAAATCACACTAAAAAGTGGAAATACCCA   5242
gb:MT072688|Organism:Severe      CCTAGTTTTCTGGGTAGGTACATGTCAGCATTAAATCACACTAAAAAGTGGAAATACCCA   5227
gb:MN996527|Organism:Severe      CCTAGTTTTCTGGGTAGGTACATGTCAGCATTAAATCACACTAAAAAGTGGAAATACCCA   5209
gb:MT093631|Organism:Severe      CCTAGTTTTCTGGGTAGGTACATGTCAGCATTAAATCACACTAAAAAGTGGAAATACCCA   5280
gb:MT106053|Organism:Severe      CCTAGTTTTCTGGGTAGGTACATGTCAGCATTAAATCACACTAAAAAGTGGAAATACCCA   5242
gb:MT019533|Organism:Severe      CCTAGTTTTCTGGGTAGGTACATGTCAGCATTAAATCACACTAAAAAGTGGAAATACCCA   5242
gb:MT019531|Organism:Severe      CCTAGTTTTCTGGGTAGGTACATGTCAGCATTAAATCACACTAAAAAGTGGAAATACCCA   5242
gb:MN996528|Organism:Severe      CCTAGTTTTCTGGGTAGGTACATGTCAGCATTAAATCACACTAAAAAGTGGAAATACCCA   5242
gb:MN996530|Organism:Severe      CCTAGTTTTCTGGGTAGGTACATGTCAGCATTAAATCACACTAAAAAGTGGAAATACCCA   5228
gb:MN908947|Organism:Severe      CCTAGTTTTCTGGGTAGGTACATGTCAGCATTAAATCACACTAAAAAGTGGAAATACCCA   5242
gb:MT019532|Organism:Severe      CCTAGTTTTCTGGGTAGGTACATGTCAGCATTAAATCACACTAAAAAGTGGAAATACCCA   5242
                                 ************************************************************

gb:MT020781|Organism:Severe      CAAGTTAATGGTTTAACTTCTATTAAATGGGCAGATAACAACTGTTATCTTGCCACTGCA   5290
gb:MT007544|Organism:Severe      CAAGTTAATGGTTTAACTTCTATTAAATGGGCAGATAACAACTGTTATCTTGCCACTGCA   5302
gb:MN994467|Organism:Severe      CAAGTTAATGGTTTAACTTCTATTAAATGGGCAGATAACAACTGTTATCTTGCCACTGCA   5302
gb:MT044257|Organism:Severe      CAAGTTAATGGTTTAACTTCTATTAAATGGGCAGATAACAACTGTTATCTTGCCACTGCA   5302
gb:MT106054|Organism:Severe      CAAGTTAATGGTTTAACTTCTATTAAATGGGCAGATAACAACTGTTATCTTGCCACTGCA   5302
gb:MT049951|Organism:Severe      CAAGTTAATGGTTTAACTTCTATTAAATGGGCAGATAACAACTGTTATCTTGCCACTGCA   5302
gb:MN975262|Organism:Severe      CAAGTTAATGGTTTAACTTCTATTAAATGGGCAGATAACAACTGTTATCTTGCCACTGCA   5302
gb:MT106052|Organism:Severe      CAAGTTAATGGTTTAACTTCTATTAAATGGGCAGATAACAACTGTTATCTTGCCACTGCA   5302
gb:LC522975|Organism:Severe      CAAGTTAATGGTTTAACTTCTATTAAATGGGCAGATAACAACTGTTATCTTGCCACTGCA   5299
gb:LC522973|Organism:Severe      CAAGTTAATGGTTTAACTTCTATTAAATGGGCAGATAACAACTGTTATCTTGCCACTGCA   5299
gb:LC522974|Organism:Severe      CAAGTTAATGGTTTAACTTCTATTAAATGGGCAGATAACAACTGTTATCTTGCCACTGCA   5299
gb:MN985325|Organism:Severe      CAAGTTAATGGTTTAACTTCTATTAAATGGGCAGATAACAACTGTTATCTTGCCACTGCA   5302
gb:MT020881|Organism:Severe      CAAGTTAATGGTTTAACTTCTATTAAATGGGCAGATAACAACTGTTATCTTGCCACTGCA   5302
gb:MT020880|Organism:Severe      CAAGTTAATGGTTTAACTTCTATTAAATGGGCAGATAACAACTGTTATCTTGCCACTGCA   5302
gb:MT066175|Organism:Severe      CAAGTTAATGGTTTAACTTCTATTAAATGGGCAGATAACAACTGTTATCTTGCCACTGCA   5302
gb:MN997409|Organism:Severe      CAAGTTAATGGTTTAACTTCTATTAAATGGGCAGATAACAACTGTTATCTTGCCACTGCA   5302
gb:MN938384|Organism:Severe      CAAGTTAATGGTTTAACTTCTATTAAATGGGCAGATAACAACTGTTATCTTGCCACTGCA   5270
gb:MT044258|Organism:Severe      CAAGTTAATGGTTTAACTTCTATTAAATGGGCAGATAACAACTGTTATCTTGCCACTGCA   5278
gb:MT039890|Organism:Severe      CAAGTTAATGGTTTAACTTCTATTAAATGGGCAGATAACAACTGTTATCTTGCCACTGCA   5302
gb:MN988713|Organism:Severe      CAAGTTAATGGTTTAACTTCTATTAAATGGGCAGATAACAACTGTTATCTTGCCACTGCA   5302
gb:LC521925|Organism:Severe      CAAGTTAATGGTTTAACTTCTATTAAATGGGCAGATAACAACTGTTATCTTGCCACTGCA   5275
gb:MT093571|Organism:Severe      CAAGTTAATGGTTTAACTTCTATTAAATGGGCAGATAACAACTGTTATCTTGCCACTGCA   5302
gb:MT039887|Organism:Severe      CAAGTTAATGGTTTAACTTCTATTAAATGGGCAGATAACAACTGTTATCTTGCCACTGCA   5302
gb:MT019530|Organism:Severe      CAAGTTAATGGTTTAACTTCTATTAAATGGGCAGATAACAACTGTTATCTTGCCACTGCA   5302
gb:MT039888|Organism:Severe      CAAGTTAATGGTTTAACTTCTATTAAATGGGCAGATAACAACTGTTATCTTGCCACTGCA   5302
gb:LC522972|Organism:Severe      CAAGTTAATGGTTTAACTTCTATTAAATGGGCAGATAACAACTGTTATCTTGCCACTGCA   5299
gb:MT027063|Organism:Severe      CAAGTTAATGGTTTAACTTCTATTAAATGGGCAGATAACAACTGTTATCTTGCCACTGCA   5302
gb:MT027062|Organism:Severe      CAAGTTAATGGTTTAACTTCTATTAAATGGGCAGATAACAACTGTTATCTTGCCACTGCA   5302
gb:MT019529|Organism:Severe      CAAGTTAATGGTTTAACTTCTATTAAATGGGCAGATAACAACTGTTATCTTGCCACTGCA   5302
gb:MN996529|Organism:Severe      CAAGTTAATGGTTTAACTTCTATTAAATGGGCAGATAACAACTGTTATCTTGCCACTGCA   5290
gb:MN996531|Organism:Severe      CAAGTTAATGGTTTAACTTCTATTAAATGGGCAGATAACAACTGTTATCTTGCCACTGCA   5289
gb:MT066176|Organism:Severe      CAAGTTAATGGTTTAACTTCTATTAAATGGGCAGATAACAACTGTTATCTTGCCACTGCA   5302
gb:MT027064|Organism:Severe      CAAGTTAATGGTTTAACTTCTATTAAATGGGCAGATAACAACTGTTATCTTGCCACTGCA   5302
gb:MN994468|Organism:Severe      CAAGTTAATGGTTTAACTTCTATTAAATGGGCAGATAACAACTGTTATCTTGCCACTGCA   5302
gb:MT072688|Organism:Severe      CAAGTTAATGGTTTAACTTCTATTAAATGGGCAGATAACAACTGTTATCTTGCCACTGCA   5287
gb:MN996527|Organism:Severe      CAAGTTAATGGTTTAACTTCTATTAAATGGGCAGATAACAACTGTTATCTTGCCACTGCA   5269
gb:MT093631|Organism:Severe      CAAGTTAATGGTTTAACTTCTATTAAATGGGCAGATAACAACTGTTATCTTGCCACTGCA   5340
gb:MT106053|Organism:Severe      CAAGTTAATGGTTTAACTTCTATTAAATGGGCAGATAACAACTGTTATCTTGCCACTGCA   5302
gb:MT019533|Organism:Severe      CAAGTTAATGGTTTAACTTCTATTAAATGGGCAGATAACAACTGTTATCTTGCCACTGCA   5302
gb:MT019531|Organism:Severe      CAAGTTAATGGTTTAACTTCTATTAAATGGGCAGATAACAACTGTTATCTTGCCACTGCA   5302
gb:MN996528|Organism:Severe      CAAGTTAATGGTTTAACTTCTATTAAATGGGCAGATAACAACTGTTATCTTGCCACTGCA   5302
gb:MN996530|Organism:Severe      CAAGTTAATGGTTTAACTTCTATTAAATGGGCAGATAACAACTGTTATCTTGCCACTGCA   5288
gb:MN908947|Organism:Severe      CAAGTTAATGGTTTAACTTCTATTAAATGGGCAGATAACAACTGTTATCTTGCCACTGCA   5302
```

```
gb:MT019532|Organism:Severe      CAAGTTAATGGTTTAACTTCTATTAAATGGGCAGATAACAACTGTTATCTTGCCACTGCA   5302
                                 ************************************************************

gb:MT020781|Organism:Severe      TTGTTAACACTCCAACAAATAGAGTTGAAGTTTAATCCACCTGCTCTACAAGATGCTTAT   5350
gb:MT007544|Organism:Severe      TTGTTAACACTCCAACAAATAGAGTTGAAGTTTAATCCACCTGCTCTACAAGATGCTTAT   5362
gb:MN994467|Organism:Severe      TTGTTAACACTCCAACAAATAGAGTTGAAGTTTAATCCACCTGCTCTACAAGATGCTTAT   5362
gb:MT044257|Organism:Severe      TTGTTAACACTCCAACAAATAGAGTTGAAGTTTAATCCACCTGCTCTACAAGATGCTTAT   5362
gb:MT106054|Organism:Severe      TTGTTAACACTCCAACAAATAGAGTTGAAGTTTAATCCACCTGCTCTACAAGATGCTTAT   5362
gb:MT049951|Organism:Severe      TTGTTAACACTCCAACAAATAGAGTTGAAGTTTAATCCACCTGCTCTACAAGATGCTTAT   5362
gb:MN975262|Organism:Severe      TTGTTAACACTCCAACAAATAGAGTTGAAGTTTAATCCACCTGCTCTACAAGATGCTTAT   5362
gb:MT106052|Organism:Severe      TTGTTAACACTCCAACAAATAGAGTTGAAGTTTAATCCACCTGCTCTACAAGATGCTTAT   5362
gb:LC522975|Organism:Severe      TTGTTAACACTCCAACAAATAGAGTTGAAGTTTAATCCACCTGCTCTACAAGATGCTTAT   5359
gb:LC522973|Organism:Severe      TTGTTAACACTCCAACAAATAGAGTTGAAGTTTAATCCACCTGCTCTACAAGATGCTTAT   5359
gb:LC522974|Organism:Severe      TTGTTAACACTCCAACAAATAGAGTTGAAGTTTAATCCACCTGCTCTACAAGATGCTTAT   5359
gb:MN985325|Organism:Severe      TTGTTAACACTCCAACAAATAGAGTTGAAGTTTAATCCACCTGCTCTACAAGATGCTTAT   5362
gb:MT020881|Organism:Severe      TTGTTAACACTCCAACAAATAGAGTTGAAGTTTAATCCACCTGCTCTACAAGATGCTTAT   5362
gb:MT020880|Organism:Severe      TTGTTAACACTCCAACAAATAGAGTTGAAGTTTAATCCACCTGCTCTACAAGATGCTTAT   5362
gb:MT066175|Organism:Severe      TTGTTAACACTCCAACAAATAGAGTTGAAGTTTAATCCACCTGCTCTACAAGATGCTTAT   5362
gb:MN997409|Organism:Severe      TTGTTAACACTCCAACAAATAGAGTTGAAGTTTAATCCACCTGCTCTACAAGATGCTTAT   5362
gb:MN938384|Organism:Severe      TTGTTAACACTCCAACAAATAGAGTTGAAGTTTAATCCACCTGCTCTACAAGATGCTTAT   5330
gb:MT044258|Organism:Severe      TTGTTAACACTCCAACAAATAGAGTTGAAGTTTAATCCACCTGCTCTACAAGATGCTTAT   5338
gb:MT039890|Organism:Severe      TTGTTAACACTCCAACAAATAGAGTTGAAGTTTAATCCACCTGCTCTACAAGATGCTTAT   5362
gb:MN988713|Organism:Severe      TTGTTAACACTCCAACAAATAGAGTTGAAGTTTAATCCACCTGCTCTACAAGATGCTTAT   5362
gb:LC521925|Organism:Severe      TTGTTAACACTCCAACAAATAGAGTTGAAGTTTAATCCACCTGCTCTACAAGATGCTTAT   5335
gb:MT093571|Organism:Severe      TTGTTAACACTCCAACAAATAGAGTTGAAGTTTAATCCACCTGCTCTACAAGATGCTTAT   5362
gb:MT039887|Organism:Severe      TTGTTAACACTCCAACAAATAGAGTTGAAGTTTAATCCACCTGCTCTACAAGATGCTTAT   5362
gb:MT019530|Organism:Severe      TTGTTAACACTCCAACAAATAGAGTTGAAGTTTAATCCACCTGCTCTACAAGATGCTTAT   5362
gb:MT039888|Organism:Severe      TTGTTAACACTCCAACAAATAGAGTTGAAGTTTAATCCACCTGCTCTACAAGATGCTTAT   5362
gb:LC522972|Organism:Severe      TTGTTAACACTCCAACAAATAGAGTTGAAGTTTAATCCACCTGCTCTACAAGATGCTTAT   5359
gb:MT027063|Organism:Severe      TTGTTAACACTCCAACAAATAGAGTTGAAGTTTAATCCACCTGCTCTACAAGATGCTTAT   5362
gb:MT027062|Organism:Severe      TTGTTAACACTCCAACAAATAGAGTTGAAGTTTAATCCACCTGCTCTACAAGATGCTTAT   5362
gb:MT019529|Organism:Severe      TTGTTAACACTCCAACAAATAGAGTTGAAGTTTAATCCACCTGCTCTACAAGATGCTTAT   5362
gb:MN996529|Organism:Severe      TTGTTAACACTCCAACAAATAGAGTTGAAGTTTAATCCACCTGCTCTACAAGATGCTTAT   5350
gb:MN996531|Organism:Severe      TTGTTAACACTCCAACAAATAGAGTTGAAGTTTAATCCACCTGCTCTACAAGATGCTTAT   5349
gb:MT066176|Organism:Severe      TTGTTAACACTCCAACAAATAGAGTTGAAGTTTAATCCACCTGCTCTACAAGATGCTTAT   5362
gb:MT027064|Organism:Severe      TTGTTAACACTCCAACAAATAGAGTTGAAGTTTAATCCACCTGCTCTACAAGATGCTTAT   5362
gb:MN994468|Organism:Severe      TTGTTAACACTCCAACAAATAGAGTTGAAGTTTAATCCACCTGCTCTACAAGATGCTTAT   5362
gb:MT072688|Organism:Severe      TTGTTAACACTCCAACAAATAGAGTTGAAGTTTAATCCACCTGCTCTACAAGATGCTTAT   5347
gb:MN996527|Organism:Severe      TTGTTAACACTCCAACAAATAGAGTTGAAGTTTAATCCACCTGCTCTACAAGATGCTTAT   5329
gb:MT093631|Organism:Severe      TTGTTAACACTCCAACAAATAGAGTTGAAGTTTAATCCACCTGCTCTACAAGATGCTTAT   5400
gb:MT106053|Organism:Severe      TTGTTAACACTCCAACAAATAGAGTTGAAGTTTAATCCACCTGCTCTACAAGATGCTTAT   5362
gb:MT019533|Organism:Severe      TTGTTAACACTCCAACAAATAGAGTTGAAGTTTAATCCACCTGCTCTACAAGATGCTTAT   5362
gb:MT019531|Organism:Severe      TTGTTAACACTCCAACAAATAGAGTTGAAGTTTAATCCACCTGCTCTACAAGATGCTTAT   5362
gb:MN996528|Organism:Severe      TTGTTAACACTCCAACAAATAGAGTTGAAGTTTAATCCACCTGCTCTACAAGATGCTTAT   5362
gb:MN996530|Organism:Severe      TTGTTAACACTCCAACAAATAGAGTTGAAGTTTAATCCACCTGCTCTACAAGATGCTTAT   5348
gb:MN908947|Organism:Severe      TTGTTAACACTCCAACAAATAGAGTTGAAGTTTAATCCACCTGCTCTACAAGATGCTTAT   5362
gb:MT019532|Organism:Severe      TTGTTAACACTCCAACAAATAGAGTTGAAGTTTAATCCACCTGCTCTACAAGATGCTTAT   5362
                                 ************************************************************

gb:MT020781|Organism:Severe      TACAGAGCAAGGGCTGGTGAAGCTGCTAACTTTTGTGCACTTATCTTAGCCTACTGTAAT   5410
gb:MT007544|Organism:Severe      TACAGAGCAAGGGCTGGTGAAGCTGCTAACTTTTGTGCACTTATCTTAGCCTACTGTAAT   5422
gb:MN994467|Organism:Severe      TACAGAGCAAGGGCTGGTGAAGCTGCTAACTTTTGTGCACTTATCTTAGCCTACTGTAAT   5422
gb:MT044257|Organism:Severe      TACAGAGCAAGGGCTGGTGAAGCTGCTAACTTTTGTGCACTTATCTTAGCCTACTGTAAT   5422
gb:MT106054|Organism:Severe      TACAGAGCAAGGGCTGGTGAAGCTGCTAACTTTTGTGCACTTATCTTAGCCTACTGTAAT   5422
gb:MT049951|Organism:Severe      TACAGAGCAAGGGCTGGTGAAGCTGCTAACTTTTGTGCACTTATCTTAGCCTACTGTAAT   5422
gb:MN975262|Organism:Severe      TACAGAGCAAGGGCTGGTGAAGCTGCTAACTTTTGTGCACTTATCTTAGCCTACTGTAAT   5422
gb:MT106052|Organism:Severe      TACAGAGCAAGGGCTGGTGAAGCTGCTAACTTTTGTGCACTTATCTTAGCCTACTGTAAT   5422
gb:LC522975|Organism:Severe      TACAGAGCAAGGGCTGGTGAAGCTGCTAACTTTTGTGCACTTATCTTAGCCTACTGTAAT   5419
gb:LC522973|Organism:Severe      TACAGAGCAAGGGCTGGTGAAGCTGCTAACTTTTGTGCACTTATCTTAGCCTACTGTAAT   5419
gb:LC522974|Organism:Severe      TACAGAGCAAGGGCTGGTGAAGCTGCTAACTTTTGTGCACTTATCTTAGCCTACTGTAAT   5419
gb:MN985325|Organism:Severe      TACAGAGCAAGGGCTGGTGAAGCTGCTAACTTTTGTGCACTTATCTTAGCCTACTGTAAT   5422
gb:MT020881|Organism:Severe      TACAGAGCAAGGGCTGGTGAAGCTGCTAACTTTTGTGCACTTATCTTAGCCTACTGTAAT   5422
gb:MT020880|Organism:Severe      TACAGAGCAAGGGCTGGTGAAGCTGCTAACTTTTGTGCACTTATCTTAGCCTACTGTAAT   5422
gb:MT066175|Organism:Severe      TACAGAGCAAGGGCTGGTGAAGCTGCTAACTTTTGTGCACTTATCTTAGCCTACTGTAAT   5422
gb:MN997409|Organism:Severe      TACAGAGCAAGGGCTGGTGAAGCTGCTAACTTTTGTGCACTTATCTTAGCCTACTGTAAT   5422
```

```
gb:MN938384|Organism:Severe      TACAGAGCAAGGGCTGGTGAAGCTGCTAACTTTTGTGCACTTATCTTAGCCTACTGTAAT   5390
gb:MT044258|Organism:Severe      TACAGAGCAAGGGCTGGTGAAGCTGCTAACTTTTGTGCACTTATCTTAGCCTACTGTAAT   5398
gb:MT039890|Organism:Severe      TACAGAGCAAGGGCTGGTGAAGCTGCTAACTTTTGTGCACTTATCTTAGCCTACTGTAAT   5422
gb:MN988713|Organism:Severe      TACAGAGCAAGGGCTGGTGAAGCTGCTAACTTTTGTGCACTTATCTTAGCCTACTGTAAT   5422
gb:LC521925|Organism:Severe      TACAGAGCAAGGGCTGGTGAAGCTGCTAACTTTTGTGCACTTATCTTAGCCTACTGTAAT   5395
gb:MT093571|Organism:Severe      TACAGAGCAAGGGCTGGTGAAGCTGCTAACTTTTGTGCACTTATCTTAGCCTACTGTAAT   5422
gb:MT039887|Organism:Severe      TACAGAGCAAGGGCTGGTGAAGCTGCTAACTTTTGTGCACTTATCTTAGCCTACTGTAAT   5422
gb:MT019530|Organism:Severe      TACAGAGCAAGGGCTGGTGAAGCTGCTAACTTTTGTGCACTTATCTTAGCCTACTGTAAT   5422
gb:MT039888|Organism:Severe      TACAGAGCAAGGGCTGGTGAAGCTGCTAACTTTTGTGCACTTATCTTAGCCTACTGTAAT   5422
gb:LC522972|Organism:Severe      TACAGAGCAAGGGCTGGTGAAGCTGCTAACTTTTGTGCACTTATCTTAGCCTACTGTAAT   5419
gb:MT027063|Organism:Severe      TACAGAGCAAGGGCTGGTGAAGCTGCTAACTTTTGTGCACTTATCTTAGCCTACTGTAAT   5422
gb:MT027062|Organism:Severe      TACAGAGCAAGGGCTGGTGAAGCTGCTAACTTTTGTGCACTTATCTTAGCCTACTGTAAT   5422
gb:MT019529|Organism:Severe      TACAGAGCAAGGGCTGGTGAAGCTGCTAACTTTTGTGCACTTATCTTAGCCTACTGTAAT   5422
gb:MN996529|Organism:Severe      TACAGAGCAAGGGCTGGTGAAGCTGCTAACTTTTGTGCACTTATCTTAGCCTACTGTAAT   5410
gb:MN996531|Organism:Severe      TACAGAGCAAGGGCTGGTGAAGCTGCTAACTTTTGTGCACTTATCTTAGCCTACTGTAAT   5409
gb:MT066176|Organism:Severe      TACAGAGCAAGGGCTGGTGAAGCTGCTAACTTTTGTGCACTTATCTTAGCCTACTGTAAT   5422
gb:MT027064|Organism:Severe      TACAGAGCAAGGGCTGGTGAAGCTGCTAACTTTTGTGCACTTATCTTAGCCTACTGTAAT   5422
gb:MN994468|Organism:Severe      TACAGAGCAAGGGCTGGTGAAGCTGCTAACTTTTGTGCACTTATCTTAGCCTACTGTAAT   5422
gb:MT072688|Organism:Severe      TACAGAGCAAGGGCTGGTGAAGCTGCTAACTTTTGTGCACTTATCTTAGCCTACTGTAAT   5407
gb:MN996527|Organism:Severe      TACAGAGCAAGGGCTGGTGAAGCTGCTAACTTTTGTGCACTTATCTTAGCCTACTGTAAT   5389
gb:MT093631|Organism:Severe      TACAGAGCAAGGGCTGGTGAAGCTGCTAACTTTTGTGCACTTATCTTAGCCTACTGTAAT   5460
gb:MT106053|Organism:Severe      TACAGAGCAAGGGCTGGTGAAGCTGCTAACTTTTGTGCACTTATCTTAGCCTACTGTAAT   5422
gb:MT019533|Organism:Severe      TACAGAGCAAGGGCTGGTGAAGCTGCTAACTTTTGTGCACTTATCTTAGCCTACTGTAAT   5422
gb:MT019531|Organism:Severe      TACAGAGCAAGGGCTGGTGAAGCTGCTAACTTTTGTGCACTTATCTTAGCCTACTGTAAT   5422
gb:MN996528|Organism:Severe      TACAGAGCAAGGGCTGGTGAAGCTGCTAACTTTTGTGCACTTATCTTAGCCTACTGTAAT   5422
gb:MN996530|Organism:Severe      TACAGAGCAAGGGCTGGTGAAGCTGCTAACTTTTGTGCACTTATCTTAGCCTACTGTAAT   5408
gb:MN908947|Organism:Severe      TACAGAGCAAGGGCTGGTGAAGCTGCTAACTTTTGTGCACTTATCTTAGCCTACTGTAAT   5422
gb:MT019532|Organism:Severe      TACAGAGCAAGGGCTGGTGAAGCTGCTAACTTTTGTGCACTTATCTTAGCCTACTGTAAT   5422
                                 ************************************************************

gb:MT020781|Organism:Severe      AAGACAGTAGGTGAGTTAGGTGATGTTAGAGAAACAATGAGTTACTTGTTTCAACATGCC   5470
gb:MT007544|Organism:Severe      AAGACAGTAGGTGAGTTAGGTGATGTTAGAGAAACAATGAGTTACTTGTTTCAACATGCC   5482
gb:MN994467|Organism:Severe      AAGACAGTAGGTGAGTTAGGTGATGTTAGAGAAACAATGAGTTACTTGTTTCAACATGCC   5482
gb:MT044257|Organism:Severe      AAGACAGTAGGTGAGTTAGGTGATGTTAGAGAAACAATGAGTTACTTGTTTCAACATGCC   5482
gb:MT106054|Organism:Severe      AAGACAGTAGGTGAGTTAGGTGATGTTAGAGAAACAATGAGTTACTTGTTTCAACATGCC   5482
gb:MT049951|Organism:Severe      AAGACAGTAGGTGAGTTAGGTGATGTTAGAGAAACAATGAGTTACTTGTTTCAACATGCC   5482
gb:MN975262|Organism:Severe      AAGACAGTAGGTGAGTTAGGTGATGTTAGAGAAACAATGAGTTACTTGTTTCAACATGCC   5482
gb:MT106052|Organism:Severe      AAGACAGTAGGTGAGTTAGGTGATGTTAGAGAAACAATGAGTTACTTGTTTCAACATGCC   5482
gb:LC522975|Organism:Severe      AAGACAGTAGGTGAGTTAGGTGATGTTAGAGAAACAATGAGTTACTTGTTTCAACATGCC   5479
gb:LC522973|Organism:Severe      AAGACAGTAGGTGAGTTAGGTGATGTTAGAGAAACAATGAGTTACTTGTTTCAACATGCC   5479
gb:LC522974|Organism:Severe      AAGACAGTAGGTGAGTTAGGTGATGTTAGAGAAACAATGAGTTACTTGTTTCAACATGCC   5479
gb:MN985325|Organism:Severe      AAGACAGTAGGTGAGTTAGGTGATGTTAGAGAAACAATGAGTTACTTGTTTCAACATGCC   5482
gb:MT020881|Organism:Severe      AAGACAGTAGGTGAGTTAGGTGATGTTAGAGAAACAATGAGTTACTTGTTTCAACATGCC   5482
gb:MT020880|Organism:Severe      AAGACAGTAGGTGAGTTAGGTGATGTTAGAGAAACAATGAGTTACTTGTTTCAACATGCC   5482
gb:MT066175|Organism:Severe      AAGACAGTAGGTGAGTTAGGTGATGTTAGAGAAACAATGAGTTACTTGTTTCAACATGCC   5482
gb:MN997409|Organism:Severe      AAGACAGTAGGTGAGTTAGGTGATGTTAGAGAAACAATGAGTTACTTGTTTCAACATGCC   5482
gb:MN938384|Organism:Severe      AAGACAGTAGGTGAGTTAGGTGATGTTAGAGAAACAATGAGTTACTTGTTTCAACATGCC   5450
gb:MT044258|Organism:Severe      AAGACAGTAGGTGAGTTAGGTGATGTTAGAGAAACAATGAGTTACTTGTTTCAACATGCC   5458
gb:MT039890|Organism:Severe      AAGACAGTAGGTGAGTTAGGTGATGTTAGAGAAACAATGAGTTACTTGTTTCAACATGCC   5482
gb:MN988713|Organism:Severe      AAGACAGTAGGTGAGTTAGGTGATGTTAGAGAAACAATGAGTTACTTGTTTCAACATGCC   5482
gb:LC521925|Organism:Severe      AAGACAGTAGGTGAGTTAGGTGATGTTAGAGAAACAATGAGTTACTTGTTTCAACATGCC   5455
gb:MT093571|Organism:Severe      AAGACAGTAGGTGAGTTAGGTGATGTTAGAGAAACAATGAGTTACTTGTTTCAACATGCC   5482
gb:MT039887|Organism:Severe      AAGACAGTAGGTGAGTTAGGTGATGTTAGAGAAACAATGAGTTACTTGTTTCAACATGCC   5482
gb:MT019530|Organism:Severe      AAGACAGTAGGTGAGTTAGGTGATGTTAGAGAAACAATGAGTTACTTGTTTCAACATGCC   5482
gb:MT039888|Organism:Severe      AAGACAGTAGGTGAGTTAGGTGATGTTAGAGAAACAATGAGTTACTTGTTTCAACATRCC   5482
gb:LC522972|Organism:Severe      AAGACAGTAGGTGAGTTAGGTGATGTTAGAGAAACAATGAGTTACTTGTTTCAACATGCC   5479
gb:MT027063|Organism:Severe      AAGACAGTAGGTGAGTTAGGTGATGTTAGAGAAACAATGAGTTACTTGTTTCAACATGCC   5482
gb:MT027062|Organism:Severe      AAGACAGTAGGTGAGTTAGGTGATGTTAGAGAAACAATGAGTTACTTGTTTCAACATGCC   5482
gb:MT019529|Organism:Severe      AAGACAGTAGGTGAGTTAGGTGATGTTAGAGAAACAATGAGTTACTTGTTTCAACATGCC   5482
gb:MN996529|Organism:Severe      AAGACAGTAGGTGAGTTAGGTGATGTTAGAGAAACAATGAGTTACTTGTTTCAACATGCC   5470
gb:MN996531|Organism:Severe      AAGACAGTAGGTGAGTTAGGTGATGTTAGAGAAACAATGAGTTACTTGTTTCAACATGCC   5469
gb:MT066176|Organism:Severe      AAGACAGTAGGTGAGTTAGGTGATGTTAGAGAAACAATGAGTTACTTGTTTCAACATGCC   5482
gb:MT027064|Organism:Severe      AAGACAGTAGGTGAGTTAGGTGATGTTAGAGAAACAATGAGTTACTTGTTTCAACATGCC   5482
gb:MN994468|Organism:Severe      AAGACAGTAGGTGAGTTAGGTGATGTTAGAGAAACAATGAGTTACTTGTTTCAACATGCC   5482
gb:MT072688|Organism:Severe      AAGACAGTAGGTGAGTTAGGTGATGTTAGAGAAACAATGAGTTACTTGTTTCAACATGCC   5467
```

```
gb:MN996527|Organism:Severe      AAGACAGTAGGTGAGTTAGGTGATGTTAGAGAAACAATGAGTTACTTGTTTCAACATGCC   5449
gb:MT093631|Organism:Severe      AAGACAGTAGGTGAGTTAGGTGATGTTAGAGAAACAATGAGTTACTTGTTTCAACATGCC   5520
gb:MT106053|Organism:Severe      AAGACAGTAGGTGAGTTAGGTGATGTTAGAGAAACAATGAGTTACTTGTTTCAACATGCC   5482
gb:MT019533|Organism:Severe      AAGACAGTAGGTGAGTTAGGTGATGTTAGAGAAACAATGAGTTACTTGTTTCAACATGCC   5482
gb:MT019531|Organism:Severe      AAGACAGTAGGTGAGTTAGGTGATGTTAGAGAAACAATGAGTTACTTGTTTCAACATGCC   5482
gb:MN996528|Organism:Severe      AAGACAGTAGGTGAGTTAGGTGATGTTAGAGAAACAATGAGTTACTTGTTTCAACATGCC   5482
gb:MN996530|Organism:Severe      AAGACAGTAGGTGAGTTAGGTGATGTTAGAGAAACAATGAGTTACTTGTTTCAACATGCC   5468
gb:MN908947|Organism:Severe      AAGACAGTAGGTGAGTTAGGTGATGTTAGAGAAACAATGAGTTACTTGTTTCAACATGCC   5482
gb:MT019532|Organism:Severe      AAGACAGTAGGTGAGTTAGGTGATGTTAGAGAAACAATGAGTTACTTGTTTCAACATGCC   5482
                                 ********************************************************* **

gb:MT020781|Organism:Severe      AATTTAGATTCTTGCAAAAGAGTCTTGAACGTGGTGTGTAAAACTTGTGGACAACAGCAG   5530
gb:MT007544|Organism:Severe      AATTTAGATTCTTGCAAAAGAGTCTTGAACGTGGTGTGTAAAACTTGTGGACAACAGCAG   5542
gb:MN994467|Organism:Severe      AATTTAGATTCTTGCAAAAGAGTCTTGAACGTGGTGTGTAAAACTTGTGGACAACAGCAG   5542
gb:MT044257|Organism:Severe      AATTTAGATTCTTGCAAAAGAGTCTTGAACGTGGTGTGTAAAACTTGTGGACAACAGCAG   5542
gb:MT106054|Organism:Severe      AATTTAGATTCTTGCAAAAGAGTCTTGAACGTGGTGTGTAAAACTTGTGGACAACAGCAG   5542
gb:MT049951|Organism:Severe      AATTTAGATTCTTGCAAAAGAGTCTTGAACGTGGTGTGTAAAACTTGTGGACAACAGCAG   5542
gb:MN975262|Organism:Severe      AATTTAGATTCTTGCAAAAGAGTCTTGAACGTGGTGTGTAAAACTTGTGGACAACAGCAG   5542
gb:MT106052|Organism:Severe      AATTTAGATTCTTGCAAAAGAGTCTTGAACGTGGTGTGTAAAACTTGTGGACAACAGCAG   5542
gb:LC522975|Organism:Severe      AATTTAGATTCTTGCAAAAGAGTCTTGAACGTGGTGTGTAAAACTTGTGGACAACAGCAG   5539
gb:LC522973|Organism:Severe      AATTTAGATTCTTGCAAAAGAGTCTTGAACGTGGTGTGTAAAACTTGTGGACAACAGCAG   5539
gb:LC522974|Organism:Severe      AATTTAGATTCTTGCAAAAGAGTCTTGAACGTGGTGTGTAAAACTTGTGGACAACAGCAG   5539
gb:MN985325|Organism:Severe      AATTTAGATTCTTGCAAAAGAGTCTTGAACGTGGTGTGTAAAACTTGTGGACAACAGCAG   5542
gb:MT020881|Organism:Severe      AATTTAGATTCTTGCAAAAGAGTCTTGAACGTGGTGTGTAAAACTTGTGGACAACAGCAG   5542
gb:MT020880|Organism:Severe      AATTTAGATTCTTGCAAAAGAGTCTTGAACGTGGTGTGTAAAACTTGTGGACAACAGCAG   5542
gb:MT066175|Organism:Severe      AATTTAGATTCTTGCAAAAGAGTCTTGAACGTGGTGTGTAAAACTTGTGGACAACAGCAG   5542
gb:MN997409|Organism:Severe      AATTTAGATTCTTGCAAAAGAGTCTTGAACGTGGTGTGTAAAACTTGTGGACAACAGCAG   5542
gb:MN938384|Organism:Severe      AATTTAGATTCTTGCAAAAGAGTCTTGAACGTGGTGTGTAAAACTTGTGGACAACAGCAG   5510
gb:MT044258|Organism:Severe      AATTTAGATTCTTGCAAAAGAGTCTTGAACGTGGTGTGTAAAACTTGTGGACAACAGCAG   5518
gb:MT039890|Organism:Severe      AATTTAGATTCTTGCAAAAGAGTCTTGAACGTGGTGTGTAAAACTTGTGGACAACAGCAG   5542
gb:MN988713|Organism:Severe      AATTTAGATTCTTGCAAAAGAGTCTTGAACGTGGTGTGTAAAACTTGTGGACAACAGCAG   5542
gb:LC521925|Organism:Severe      AATTTAGATTCTTGCAAAAGAGTCTTGAACGTGGTGTGTAAAACTTGTGGACAACAGCAG   5515
gb:MT093571|Organism:Severe      AATTTAGATTCTTGCAAAAGAGTCTTGAACGTGGTGTGTAAAACTTGTGGACAACAGCAG   5542
gb:MT039887|Organism:Severe      AATTTAGATTCTTGCAAAAGAGTCTTGAACGTGGTGTGTAAAACTTGTGGACAACAGCAG   5542
gb:MT019530|Organism:Severe      AATTTAGATTCTTGCAAAAGAGTCTTGAACGTGGTGTGTAAAACTTGTGGACAACAGCAG   5542
gb:MT039888|Organism:Severe      AATTTAGATTCTTGCAAAAGAGTCTTGAACGTGGTGTGTAAAACTTGTGGACAACAGCAG   5542
gb:LC522972|Organism:Severe      AATTTAGATTCTTGCAAAAGAGTCTTGAACGTGGTGTGTAAAACTTGTGGACAACAGCAG   5539
gb:MT027063|Organism:Severe      AATTTAGATTCTTGCAAAAGAGTCTTGAACGTGGTGTGTAAAACTTGTGGACAACAGCAG   5542
gb:MT027062|Organism:Severe      AATTTAGATTCTTGCAAAAGAGTCTTGAACGTGGTGTGTAAAACTTGTGGACAACAGCAG   5542
gb:MT019529|Organism:Severe      AATTTAGATTCTTGCAAAAGAGTCTTGAACGTGGTGTGTAAAACTTGTGGACAACAGCAG   5542
gb:MN996529|Organism:Severe      AATTTAGATTCTTGCAAAAGAGTCTTGAACGTGGTGTGTAAAACTTGTGGACAACAGCAG   5530
gb:MN996531|Organism:Severe      AATTTAGATTCTTGCAAAAGAGTCTTGAACGTGGTGTGTAAAACTTGTGGACAACAGCAG   5529
gb:MT066176|Organism:Severe      AATTTAGATTCTTGCAAAAGAGTCTTGAACGTGGTGTGTAAAACTTGTGGACAACAGCAG   5542
gb:MT027064|Organism:Severe      AATTTAGATTCTTGCAAAAGAGTCTTGAACGTGGTGTGTAAAACTTGTGGACAACAGCAG   5542
gb:MN994468|Organism:Severe      AATTTAGATTCTTGCAAAAGAGTCTTGAACGTGGTGTGTAAAACTTGTGGACAACAGCAG   5542
gb:MT072688|Organism:Severe      AATTTAGATTCTTGCAAAAGAGTCTTGAACGTGGTGTGTAAAACTTGTGGACAACAGCAG   5527
gb:MN996527|Organism:Severe      AATTTAGATTCTTGCAAAAGAGTCTTGAACGTGGTGTGTAAAACTTGTGGACAACAGCAG   5509
gb:MT093631|Organism:Severe      AATTTAGATTCTTGCAAAAGAGTCTTGAACGTGGTGTGTAAAACTTGTGGACAACAGCAG   5580
gb:MT106053|Organism:Severe      AATTTAGATTCTTGCAAAAGAGTCTTGAACGTGGTGTGTAAAACTTGTGGACAACAGCAG   5542
gb:MT019533|Organism:Severe      AATTTAGATTCTTGCAAAAGAGTCTTGAACGTGGTGTGTAAAACTTGTGGACAACAGCAG   5542
gb:MT019531|Organism:Severe      AATTTAGATTCTTGCAAAAGAGTCTTGAACGTGGTGTGTAAAACTTGTGGACAACAGCAG   5542
gb:MN996528|Organism:Severe      AATTTAGATTCTTGCAAAAGAGTCTTGAACGTGGTGTGTAAAACTTGTGGACAACAGCAG   5542
gb:MN996530|Organism:Severe      AATTTAGATTCTTGCAAAAGAGTCTTGAACGTGGTGTGTAAAACTTGTGGACAACAGCAG   5528
gb:MN908947|Organism:Severe      AATTTAGATTCTTGCAAAAGAGTCTTGAACGTGGTGTGTAAAACTTGTGGACAACAGCAG   5542
gb:MT019532|Organism:Severe      AATTTAGATTCTTGCAAAAGAGTCTTGAACGTGGTGTGTAAAACTTGTGGACAACAGCAG   5542
                                 ************************************************************

gb:MT020781|Organism:Severe      ACAACCCTTAAGGGTGTAGAAGCTGTTATGTACATGGGCACACTTTCTTATGAACAATTT   5590
gb:MT007544|Organism:Severe      ACAACCCTTAAGGGTGTAGAAGCTGTTATGTACATGGGCACACTTTCTTATGAACAATTT   5602
gb:MN994467|Organism:Severe      ACAACCCTTAAGGGTGTAGAAGCTGTTATGTACATGGGCACACTTTCTTATGAACAATTT   5602
gb:MT044257|Organism:Severe      ACAACCCTTAAGGGTGTAGAAGCTGTTATGTACATGGGCACACTTTCTTATGAACAATTT   5602
gb:MT106054|Organism:Severe      ACAACCCTTAAGGGTGTAGAAGCTGTTATGTACATGGGCACACTTTCTTATGAACAATTT   5602
gb:MT049951|Organism:Severe      ACAACCCTTAAGGGTGTAGAAGCTGTTATGTACATGGGCACACTTTCTTATGAACAATTT   5602
gb:MN975262|Organism:Severe      ACAACCCTTAAGGGTGTAGAAGCTGTTATGTACATGGGCACACTTTCTTATGAACAATTT   5602
gb:MT106052|Organism:Severe      ACAACCCTTAAGGGTGTAGAAGCTGTTATGTACATGGGCACACTTTCTTATGAACAATTT   5602
```

```
gb:LC522975|Organism:Severe      ACAACCCTTAAGGGTGTAGAAGCTGTTATGTACATGGGCACACTTTCTTATGAACAATTT    5599
gb:LC522973|Organism:Severe      ACAACCCTTAAGGGTGTAGAAGCTGTTATGTACATGGGCACACTTTCTTATGAACAATTT    5599
gb:LC522974|Organism:Severe      ACAACCCTTAAGGGTGTAGAAGCTGTTATGTACATGGGCACACTTTCTTATGAACAATTT    5599
gb:MN985325|Organism:Severe      ACAACCCTTAAGGGTGTAGAAGCTGTTATGTACATGGGCACACTTTCTTATGAACAATTT    5602
gb:MT020881|Organism:Severe      ACAACCCTTAAGGGTGTAGAAGCTGTTATGTACATGGGCACACTTTCTTATGAACAATTT    5602
gb:MT020880|Organism:Severe      ACAACCCTTAAGGGTGTAGAAGCTGTTATGTACATGGGCACACTTTCTTATGAACAATTT    5602
gb:MT066175|Organism:Severe      ACAACCCTTAAGGGTGTAGAAGCTGTTATGTACATGGGCACACTTTCTTATGAACAATTT    5602
gb:MN997409|Organism:Severe      ACAACCCTTAAGGGTGTAGAAGCTGTTATGTACATGGGCACACTTTCTTATGAACAATTT    5602
gb:MN938384|Organism:Severe      ACAACCCTTAAGGGTGTAGAAGCTGTTATGTACATGGGCACACTTTCTTATGAACAATTT    5570
gb:MT044258|Organism:Severe      ACAACCCTTAAGGGTGTAGAAGCTGTTATGTACATGGGCACACTTTCTTATGAACAATTT    5578
gb:MT039890|Organism:Severe      ACAACCCTTAAGGGTGTAGAAGCTGTTATGTACATGGGCACACTTTCTTATGAACAATTT    5602
gb:MN988713|Organism:Severe      ACAACCCTTAAGGGTGTAGAAGCTGTTATGTACATGGGCACACTTTCTTATGAACAATTT    5602
gb:LC521925|Organism:Severe      ACAACCCTTAAGGGTGTAGAAGCTGTTATGTACATGGGCACACTTTCTTATGAACAATTT    5575
gb:MT093571|Organism:Severe      ACAACCCTTAAGGGTGTAGAAGCTGTTATGTACATGGGCACACTTTCTTATGAACAATTT    5602
gb:MT039887|Organism:Severe      ACAACCCTTAAGGGTGTAGAAGCTGTTATGTACATGGGCACACTTTCTTATGAACAATTT    5602
gb:MT019530|Organism:Severe      ACAACCCTTAAGGGTGTAGAAGCTGTTATGTACATGGGCACACTTTCTTATGAACAATTT    5602
gb:MT039888|Organism:Severe      ACAACCCTTAAGGGTGTAGAAGCTGTTATGTACATGGGCACACTTTCTTATGAACAATTT    5602
gb:LC522972|Organism:Severe      ACAACCCTTAAGGGTGTAGAAGCTGTTATGTACATGGGCACACTTTCTTATGAACAATTT    5599
gb:MT027063|Organism:Severe      ACAACCCTTAAGGGTGTAGAAGCTGTTATGTACATGGGCACACTTTCTTATGAACAATTT    5602
gb:MT027062|Organism:Severe      ACAACCCTTAAGGGTGTAGAAGCTGTTATGTACATGGGCACACTTTCTTATGAACAATTT    5602
gb:MT019529|Organism:Severe      ACAACCCTTAAGGGTGTAGAAGCTGTTATGTACATGGGCACACTTTCTTATGAACAATTT    5602
gb:MN996529|Organism:Severe      ACAACCCTTAAGGGTGTAGAAGCTGTTATGTACATGGGCACACTTTCTTATGAACAATTT    5590
gb:MN996531|Organism:Severe      ACAACCCTTAAGGGTGTAGAAGCTGTTATGTACATGGGCACACTTTCTTATGAACAATTT    5589
gb:MT066176|Organism:Severe      ACAACCCTTAAGGGTGTAGAAGCTGTTATGTACATGGGCACACTTTCTTATGAACAATTT    5602
gb:MT027064|Organism:Severe      ACAACCCTTAAGGGTGTAGAAGCTGTTATGTACATGGGCACACTTTCTTATGAACAATTT    5602
gb:MN994468|Organism:Severe      ACAACCCTTAAGGGTGTAGAAGCTGTTATGTACATGGGCACACTTTCTTATGAACAATTT    5602
gb:MT072688|Organism:Severe      ACAACCCTTAAGGGTGTAGAAGCTGTTATGTACATGGGCACACTTTCTTATGAACAATTT    5587
gb:MN996527|Organism:Severe      ACAACCCTTAAGGGTGTAGAAGCTGTTATGTACATGGGCACACTTTCTTATGAACAATTT    5569
gb:MT093631|Organism:Severe      ACAACCCTTAAGGGTGTAGAAGCTGTTATGTACATGGGCACACTTTCTTATGAACAATTT    5640
gb:MT106053|Organism:Severe      ACAACCCTTAAGGGTGTAGAAGCTGTTATGTACATGGGCACACTTTCTTATGAACAATTT    5602
gb:MT019533|Organism:Severe      ACAACCCTTAAGGGTGTAGAAGCTGTTATGTACATGGGCACACTTTCTTATGAACAATTT    5602
gb:MT019531|Organism:Severe      ACAACCCTTAAGGGTGTAGAAGCTGTTATGTACATGGGCACACTTTCTTATGAACAATTT    5602
gb:MN996528|Organism:Severe      ACAACCCTTAAGGGTGTAGAAGCTGTTATGTACATGGGCACACTTTCTTATGAACAATTT    5602
gb:MN996530|Organism:Severe      ACAACCCTTAAGGGTGTAGAAGCTGTTATGTACATGGGCACACTTTCTTATGAACAATTT    5588
gb:MN908947|Organism:Severe      ACAACCCTTAAGGGTGTAGAAGCTGTTATGTACATGGGCACACTTTCTTATGAACAATTT    5602
gb:MT019532|Organism:Severe      ACAACCCTTAAGGGTGTAGAAGCTGTTATGTACATGGGCACACTTTCTTATGAACAATTT    5602
                                 ************************************************************

gb:MT020781|Organism:Severe      AAGAAAGGTGTTCAGATACCTTGTACGTGTGGTAAACAAGCTACAAAATATCTAGTACAA    5650
gb:MT007544|Organism:Severe      AAGAAAGGTGTTCAGATACCTTGTACGTGTGGTAAACAAGCTACAAAATATCTAGTACAA    5662
gb:MN994467|Organism:Severe      AAGAAAGGTGTTCAGATACCTTGTACGTGTGGTAAACAAGCTACAAAATATCTAGTACAA    5662
gb:MT044257|Organism:Severe      AAGAAAGGTGTTCAGATACCTTGTACGTGTGGTAAACAAGCTACAAAATATCTAGTACAA    5662
gb:MT106054|Organism:Severe      AAGAAAGGTGTTCAGATACCTTGTACGTGTGGTAAACAAGCTACAAAATATCTAGTACAA    5662
gb:MT049951|Organism:Severe      AAGAAAGGTGTTCAGATACCTTGTACGTGTGGTAAACAAGCTACAAAATATCTAGTACAA    5662
gb:MN975262|Organism:Severe      AAGAAAGGTGTTCAGATACCTTGTACGTGTGGTAAACAAGCTACAAAATATCTAGTACAA    5662
gb:MT106052|Organism:Severe      AAGAAAGGTGTTCAGATACCTTGTACGTGTGGTAAACAAGCTACAAAATATCTAGTACAA    5662
gb:LC522975|Organism:Severe      AAGAAAGGTGTTCAGATACCTTGTACGTGTGGTAAACAAGCTACAAAATATCTAGTACAA    5659
gb:LC522973|Organism:Severe      AAGAAAGGTGTTCAGATACCTTGTACGTGTGGTAAACAAGCTACAAAATATCTAGTACAA    5659
gb:LC522974|Organism:Severe      AAGAAAGGTGTTCAGATACCTTGTACGTGTGGTAAACAAGCTACAAAATATCTAGTACAA    5659
gb:MN985325|Organism:Severe      AAGAAAGGTGTTCAGATACCTTGTACGTGTGGTAAACAAGCTACAAAATATCTAGTACAA    5662
gb:MT020881|Organism:Severe      AAGAAAGGTGTTCAGATACCTTGTACGTGTGGTAAACAAGCTACAAAATATCTAGTACAA    5662
gb:MT020880|Organism:Severe      AAGAAAGGTGTTCAGATACCTTGTACGTGTGGTAAACAAGCTACAAAATATCTAGTACAA    5662
gb:MT066175|Organism:Severe      AAGAAAGGTGTTCAGATACCTTGTACGTGTGGTAAACAAGCTACAAAATATCTAGTACAA    5662
gb:MN997409|Organism:Severe      AAGAAAGGTGTTCAGATACCTTGTACGTGTGGTAAACAAGCTACAAAATATCTAGTACAA    5662
gb:MN938384|Organism:Severe      AAGAAAGGTGTTCAGATACCTTGTACGTGTGGTAAACAAGCTACAAAATATCTAGTACAA    5630
gb:MT044258|Organism:Severe      AAGAAAGGTGTTCAGATACCTTGTACGTGTGGTAAACAAGCTACAAAATATCTAGTACAA    5638
gb:MT039890|Organism:Severe      AAGAAAGGTGTTCAGATACCTTGTACGTGTGGTAAACAAGCTACAAAATATCTAGTACAA    5662
gb:MN988713|Organism:Severe      AAGAAAGGTGTTCAGATACCTTGTACGTGTGGTAAACAAGCTACAAAATATCTAGTACAA    5662
gb:LC521925|Organism:Severe      AAGAAAGGTGTTCAGATACCTTGTACGTGTGGTAAACAAGCTACAAAATATCTAGTACAA    5635
gb:MT093571|Organism:Severe      AAGAAAGGTGTTCAGATACCTTGTACGTGTGGTAAACAAGCTACAAAATATCTAGTACAA    5662
gb:MT039887|Organism:Severe      AAGAAAGGTGTTCAGATACCTTGTACGTGTGGTAAACAAGCTACAAAATATCTAGTACAA    5662
gb:MT019530|Organism:Severe      AAGAAAGGTGTTCAGATACCTTGTACGTGTGGTAAACAAGCTACAAAATATCTAGTACAA    5662
gb:MT039888|Organism:Severe      AAGAAAGGTGTTCAGATACCTTGTACGTGTGGTAAACAAGCTACAAAATATCTAGTACAA    5662
gb:LC522972|Organism:Severe      AAGAAAGGTGTTCAGATACCTTGTACGTGTGGTAAACAAGCTACAAAATATCTAGTACAA    5659
gb:MT027063|Organism:Severe      AAGAAAGGTGTTCAGATACCTTGTACGTGTGGTAAACAAGCTACAAAATATCTAGTACAA    5662
```

```
gb:MT027062|Organism:Severe      AAGAAAGGTGTTCAGATACCTTGTACGTGTGGTAAACAAGCTACAAAATATCTAGTACAA    5662
gb:MT019529|Organism:Severe      AAGAAAGGTGTTCAGATACCTTGTACGTGTGGTAAACAAGCTACAAAATATCTAGTACAA    5662
gb:MN996529|Organism:Severe      AAGAAAGGTGTTCAGATACCTTGTACGTGTGGTAAACAAGCTACAAAATATCTAGTACAA    5650
gb:MN996531|Organism:Severe      AAGAAAGGTGTTCAGATACCTTGTACGTGTGGTAAACAAGCTACAAAATATCTAGTACAA    5649
gb:MT066176|Organism:Severe      AAGAAAGGTGTTCAGATACCTTGTACGTGTGGTAAACAAGCTACAAAATATCTAGTACAA    5662
gb:MT027064|Organism:Severe      AAGAAAGGTGTTCAGATACCTTGTACGTGTGGTAAACAAGCTACAAAATATCTAGTACAA    5662
gb:MN994468|Organism:Severe      AAGAAAGGTGTTCAGATACCTTGTACGTGTGGTAAACAAGCTACAAAATATCTAGTACAA    5662
gb:MT072688|Organism:Severe      AAGAAAGGTGTTCAGATACCTTGTACGTGTGGTAAACAAGCTACAAAATATCTAGTACAA    5647
gb:MN996527|Organism:Severe      AAGAAAGGTGTTCAGATACCTTGTACGTGTGGTAAACAAGCTACAAAATATCTAGTACAA    5629
gb:MT093631|Organism:Severe      AAGAAAGGTGTTCAGATACCTTGTACGTGTGGTAAACAAGCTACAAAATATCTAGTACAA    5700
gb:MT106053|Organism:Severe      AAGAAAGGTGTTCAGATACCTTGTACGTGTGGTAAACAAGCTACAAAATATCTAGTACAA    5662
gb:MT019533|Organism:Severe      AAGAAAGGTGTTCAGATACCTTGTACGTGTGGTAAACAAGCTACAAAATATCTAGTACAA    5662
gb:MT019531|Organism:Severe      AAGAAAGGTGTTCAGATACCTTGTACGTGTGGTAAACAAGCTACAAAATATCTAGTACAA    5662
gb:MN996528|Organism:Severe      AAGAAAGGTGTTCAGATACCTTGTACGTGTGGTAAACAAGCTACAAAATATCTAGTACAA    5662
gb:MN996530|Organism:Severe      AAGAAAGGTGTTCAGATACCTTGTACGTGTGGTAAACAAGCTACAAAATATCTAGTACAA    5648
gb:MN908947|Organism:Severe      AAGAAAGGTGTTCAGATACCTTGTACGTGTGGTAAACAAGCTACAAAATATCTAGTACAA    5662
gb:MT019532|Organism:Severe      AAGAAAGGTGTTCAGATACCTTGTACGTGTGGTAAACAAGCTACAAAATATCTAGTACAA    5662
                                 ************************************************************

gb:MT020781|Organism:Severe      CAGGAGTCACCTTTTGTTATGATGTCAGCACCACCTGCTCAGTATGAACTTAAGCATGGT    5710
gb:MT007544|Organism:Severe      CAGGAGTCACCTTTTGTTATGATGTCAGCACCACCTGCTCAGTATGAACTTAAGCATGGT    5722
gb:MN994467|Organism:Severe      CAGGAGTCACCTTTTGTTATGATGTCAGCACCACCTGCTCAGTATGAACTTAAGCATGGT    5722
gb:MT044257|Organism:Severe      CAGGAGTCACCTTTTGTTATGATGTCAGCACCACCTGCTCAGTATGAACTTAAGCATGGT    5722
gb:MT106054|Organism:Severe      CAGGAGTCACCTTTTGTTATGATGTCAGCACCACCTGCTCAGTATGAACTTAAGCATGGT    5722
gb:MT049951|Organism:Severe      CAGGAGTCACCTTTTGTTATGATGTCAGCACCACCTGCTCAGTATGAACTTAAGCATGGT    5722
gb:MN975262|Organism:Severe      CAGGAGTCACCTTTTGTTATGATGTCAGCACCACCTGCTCAGTATGAACTTAAGCATGGT    5722
gb:MT106052|Organism:Severe      CAGGAGTCACCTTTTGTTATGATGTCAGCACCACCTGCTCAGTATGAACTTAAGCATGGT    5722
gb:LC522975|Organism:Severe      CAGGAGTCACCTTTTGTTATGATGTCAGCACCACCTGCTCAGTATGAACTTAAGCATGGT    5719
gb:LC522973|Organism:Severe      CAGGAGTCACCTTTTGTTATGATGTCAGCACCACCTGCTCAGTATGAACTTAAGCATGGT    5719
gb:LC522974|Organism:Severe      CAGGAGTCACCTTTTGTTATGATGTCAGCACCACCTGCTCAGTATGAACTTAAGCATGGT    5719
gb:MN985325|Organism:Severe      CAGGAGTCACCTTTTGTTATGATGTCAGCACCACCTGCTCAGTATGAACTTAAGCATGGT    5722
gb:MT020881|Organism:Severe      CAGGAGTCACCTTTTGTTATGATGTCAGCACCACCTGCTCAGTATGAACTTAAGCATGGT    5722
gb:MT020880|Organism:Severe      CAGGAGTCACCTTTTGTTATGATGTCAGCACCACCTGCTCAGTATGAACTTAAGCATGGT    5722
gb:MT066175|Organism:Severe      CAGGAGTCACCTTTTGTTATGATGTCAGCACCACCTGCTCAGTATGAACTTAAGCATGGT    5722
gb:MN997409|Organism:Severe      CAGGAGTCACCTTTTGTTATGATGTCAGCACCACCTGCTCAGTATGAACTTAAGCATGGT    5722
gb:MN938384|Organism:Severe      CAGGAGTCACCTTTTGTTATGATGTCAGCACCACCTGCTCAGTATGAACTTAAGCATGGT    5690
gb:MT044258|Organism:Severe      CAGGAGTCACCTTTTGTTATGATGTCAGCACCACCTGCTCAGTATGAACTTAAGCATGGT    5698
gb:MT039890|Organism:Severe      CAGGAGTCACCTTTTGTTATGATGTCAGCACCACCTGCTCAGTATGAACTTAAGCATGGT    5722
gb:MN988713|Organism:Severe      CAGGAGTCACCTTTTGTTATGATGTCAGCACCACCTGCTCAGTATGAACTTAAGCATGGT    5722
gb:LC521925|Organism:Severe      CAGGAGTCACCTTTTGTTATGATGTCAGCACCACCTGCTCAGTATGAACTTAAGCATGGT    5695
gb:MT093571|Organism:Severe      CAGGAGTCACCTTTTGTTATGATGTCAGCACCACCTGCTCAGTATGAACTTAAGCATGGT    5722
gb:MT039887|Organism:Severe      CAGGAGTCACCTTTTGTTATGATGTCAGCACCACCTGCTCAGTATGAACTTAAGCATGGT    5722
gb:MT019530|Organism:Severe      CAGGAGTCACCTTTTGTTATGATGTCAGCACCACCTGCTCAGTATGAACTTAAGCATGGT    5722
gb:MT039888|Organism:Severe      CAGGAGTCACCTTTTGTTATGATGTCAGCACCACCTGCTCAGTATGAACTTAAGCATGGT    5722
gb:LC522972|Organism:Severe      CAGGAGTCACCTTTTGTTATGATGTCAGCACCACCTGCTCAGTATGAACTTAAGCATGGT    5719
gb:MT027063|Organism:Severe      CAGGAGTCACCTTTTGTTATGATGTCAGCACCACCTGCTCAGTATGAACTTAAGCATGGT    5722
gb:MT027062|Organism:Severe      CAGGAGTCACCTTTTGTTATGATGTCAGCACCACCTGCTCAGTATGAACTTAAGCATGGT    5722
gb:MT019529|Organism:Severe      CAGGAGTCACCTTTTGTTATGATGTCAGCACCACCTGCTCAGTATGAACTTAAGCATGGT    5722
gb:MN996529|Organism:Severe      CAGGAGTCACCTTTTGTTATGATGTCAGCACCACCTGCTCAGTATGAACTTAAGCATGGT    5710
gb:MN996531|Organism:Severe      CAGGAGTCACCTTTTGTTATGATGTCAGCACCACCTGCTCAGTATGAACTTAAGCATGGT    5709
gb:MT066176|Organism:Severe      CAGGAGTCACCTTTTGTTATGATGTCAGCACCACCTGCTCAGTATGAACTTAAGCATGGT    5722
gb:MT027064|Organism:Severe      CAGGAGTCACCTTTTGTTATGATGTCAGCACCACCTGCTCAGTATGAACTTAAGCATGGT    5722
gb:MN994468|Organism:Severe      CAGGAGTCACCTTTTGTTATGATGTCAGCACCACCTGCTCAGTATGAACTTAAGCATGGT    5722
gb:MT072688|Organism:Severe      CAGGAGTCACCTTTTGTTATGATGTCAGCACCACCTGCTCAGTATGAACTTAAGCATGGT    5707
gb:MN996527|Organism:Severe      CAGGAGTCACCTTTTGTTATGATGTCAGCACCACCTGCTCAGTATGAACTTAAGCATGGT    5689
gb:MT093631|Organism:Severe      CAGGAGTCACCTTTTGTTATGATGTCAGCACCACCTGCTCAGTATGAACTTAAGCATGGT    5760
gb:MT106053|Organism:Severe      CAGGAGTCACCTTTTGTTATGATGTCAGCACCACCTGCTCAGTATGAACTTAAGCATGGT    5722
gb:MT019533|Organism:Severe      CAGGAGTCACCTTTTGTTATGATGTCAGCACCACCTGCTCAGTATGAACTTAAGCATGGT    5722
gb:MT019531|Organism:Severe      CAGGAGTCACCTTTTGTTATGATGTCAGCACCACCTGCTCAGTATGAACTTAAGCATGGT    5722
gb:MN996528|Organism:Severe      CAGGAGTCACCTTTTGTTATGATGTCAGCACCACCTGCTCAGTATGAACTTAAGCATGGT    5722
gb:MN996530|Organism:Severe      CAGGAGTCACCTTTTGTTATGATGTCAGCACCACCTGCTCAGTATGAACTTAAGCATGGT    5708
gb:MN908947|Organism:Severe      CAGGAGTCACCTTTTGTTATGATGTCAGCACCACCTGCTCAGTATGAACTTAAGCATGGT    5722
gb:MT019532|Organism:Severe      CAGGAGTCACCTTTTGTTATGATGTCAGCACCACCTGCTCAGTATGAACTTAAGCATGGT    5722
                                 ************************************************************
```

```
gb:MT020781|Organism:Severe      ACATTTACTTGTGCTAGTGAGTACACTGGTAATTACCAGTGTGGTCACTATAAACATATA   5770
gb:MT007544|Organism:Severe      ACATTTACTTGTGCTAGTGAGTACACTGGTAATTACCAGTGTGGTCACTATAAACATATA   5782
gb:MN994467|Organism:Severe      ACATTTACTTGTGCTAGTGAGTACACTGGTAATTACCAGTGTGGTCACTATAAACATATA   5782
gb:MT044257|Organism:Severe      ACATTTACTTGTGCTAGTGAGTACACTGGTAATTACCAGTGTGGTCACTATAAACATATA   5782
gb:MT106054|Organism:Severe      ACATTTACTTGTGCTAGTGAGTACACTGGTAATTACCAGTGTGGTCACTATAAACATATA   5782
gb:MT049951|Organism:Severe      ACATTTACTTGTGCTAGTGAGTACACTGGTAATTACCAGTGTGGTCACTATAAACATATA   5782
gb:MN975262|Organism:Severe      ACATTTACTTGTGCTAGTGAGTACACTGGTAATTACCAGTGTGGTCACTATAAACATATA   5782
gb:MT106052|Organism:Severe      ACATTTACTTGTGCTAGTGAGTACACTGGTAATTACCAGTGTGGTCACTATAAACATATA   5782
gb:LC522975|Organism:Severe      ACATTTACTTGTGCTAGTGAGTACACTGGTAATTACCAGTGTGGTCACTATAAACATATA   5779
gb:LC522973|Organism:Severe      ACATTTACTTGTGCTAGTGAGTACACTGGTAATTACCAGTGTGGTCACTATAAACATATA   5779
gb:LC522974|Organism:Severe      ACATTTACTTGTGCTAGTGAGTACACTGGTAATTACCAGTGTGGTCACTATAAACATATA   5779
gb:MN985325|Organism:Severe      ACATTTACTTGTGCTAGTGAGTACACTGGTAATTACCAGTGTGGTCACTATAAACATATA   5782
gb:MT020881|Organism:Severe      ACATTTACTTGTGCTAGTGAGTACACTGGTAATTACCAGTGTGGTCACTATAAACATATA   5782
gb:MT020880|Organism:Severe      ACATTTACTTGTGCTAGTGAGTACACTGGTAATTACCAGTGTGGTCACTATAAACATATA   5782
gb:MT066175|Organism:Severe      ACATTTACTTGTGCTAGTGAGTACACTGGTAATTACCAGTGTGGTCACTATAAACATATA   5782
gb:MN997409|Organism:Severe      ACATTTACTTGTGCTAGTGAGTACACTGGTAATTACCAGTGTGGTCACTATAAACATATA   5782
gb:MN938384|Organism:Severe      ACATTTACTTGTGCTAGTGAGTACACTGGTAATTACCAGTGTGGTCACTATAAACATATA   5750
gb:MT044258|Organism:Severe      ACATTTACTTGTGCTAGTGAGTACACTGGTAATTACCAGTGTGGTCACTATAAACATATA   5758
gb:MT039890|Organism:Severe      ACATTTACTTGTGCTAGTGAGTACACTGGTAATTACCAGTGTGGTCACTATAAACATATA   5782
gb:MN988713|Organism:Severe      ACATTTACTTGTGCTAGTGAGTACACTGGTAATTACCAGTGTGGTCACTATAAACATATA   5782
gb:LC521925|Organism:Severe      ACATTTACTTGTGCTAGTGAGTACACTGGTAATTACCAGTGTGGTCACTATAAACATATA   5755
gb:MT093571|Organism:Severe      ACATTTACTTGTGCTAGTGAGTACACTGGTAATTACCAGTGTGGTCACTATAAACATATA   5782
gb:MT039887|Organism:Severe      ACATTTACTTGTGCTAGTGAGTACACTGGTAATTACCAGTGTGGTCACTATAAACATATA   5782
gb:MT019530|Organism:Severe      ACATTTACTTGTGCTAGTGAGTACACTGGTAATTACCAGTGTGGTCACTATAAACATATA   5782
gb:MT039888|Organism:Severe      ACATTTACTTGTGCTAGTGAGTACACTGGTAATTACCAGTGTGGTCACTATAAACATATA   5782
gb:LC522972|Organism:Severe      ACATTTACTTGTGCTAGTGAGTACACTGGTAATTACCAGTGTGGTCACTATAAACATATA   5779
gb:MT027063|Organism:Severe      ACATTTACTTGTGCTAGTGAGTACACTGGTAATTACCAGTGTGGTCACTATAAACATATA   5782
gb:MT027062|Organism:Severe      ACATTTACTTGTGCTAGTGAGTACACTGGTAATTACCAGTGTGGTCACTATAAACATATA   5782
gb:MT019529|Organism:Severe      ACATTTACTTGTGCTAGTGAGTACACTGGTAATTACCAGTGTGGTCACTATAAACATATA   5782
gb:MN996529|Organism:Severe      ACATTTACTTGTGCTAGTGAGTACACTGGTAATTACCAGTGTGGTCACTATAAACATATA   5770
gb:MN996531|Organism:Severe      ACATTTACTTGTGCTAGTGAGTACACTGGTAATTACCAGTGTGGTCACTATAAACATATA   5769
gb:MT066176|Organism:Severe      ACATTTACTTGTGCTAGTGAGTACACTGGTAATTACCAGTGTGGTCACTATAAACATATA   5782
gb:MT027064|Organism:Severe      ACATTTACTTGTGCTAGTGAGTACACTGGTAATTACCAGTGTGGTCACTATAAACATATA   5782
gb:MN994468|Organism:Severe      ACATTTACTTGTGCTAGTGAGTACACTGGTAATTACCAGTGTGGTCACTATAAACATATA   5782
gb:MT072688|Organism:Severe      ACATTTACTTGTGCTAGTGAGTACACTGGTAATTACCAGTGTGGTCACTATAAACATATA   5767
gb:MN996527|Organism:Severe      ACATTTACTTGTGCTAGTGAGTACACTGGTAATTACCAGTGTGGTCACTATAAACATATA   5749
gb:MT093631|Organism:Severe      ACATTTACTTGTGCTAGTGAGTACACTGGTAATTACCAGTGTGGTCACTATAAACATATA   5820
gb:MT106053|Organism:Severe      ACATTTACTTGTGCTAGTGAGTACACTGGTAATTACCAGTGTGGTCACTATAAACATATA   5782
gb:MT019533|Organism:Severe      ACATTTACTTGTGCTAGTGAGTACACTGGTAATTACCAGTGTGGTCACTATAAACATATA   5782
gb:MT019531|Organism:Severe      ACATTTACTTGTGCTAGTGAGTACACTGGTAATTACCAGTGTGGTCACTATAAACATATA   5782
gb:MN996528|Organism:Severe      ACATTTACTTGTGCTAGTGAGTACACTGGTAATTACCAGTGTGGTCACTATAAACATATA   5782
gb:MN996530|Organism:Severe      ACATTTACTTGTGCTAGTGAGTACACTGGTAATTACCAGTGTGGTCACTATAAACATATA   5768
gb:MN908947|Organism:Severe      ACATTTACTTGTGCTAGTGAGTACACTGGTAATTACCAGTGTGGTCACTATAAACATATA   5782
gb:MT019532|Organism:Severe      ACATTTACTTGTGCTAGTGAGTACACTGGTAATTACCAGTGTGGTCACTATAAACATATA   5782
                                 ************************************************************

gb:MT020781|Organism:Severe      ACTTCTAAAGAAACTTTGTATTGCATAGACGGTGCTTTACTTACAAAGTCCTCAGAATAC   5830
gb:MT007544|Organism:Severe      ACTTCTAAAGAAACTTTGTATTGCATAGACGGTGCTTTACTTACAAAGTCCTCAGAATAC   5842
gb:MN994467|Organism:Severe      ACTTCTAAAGAAACTTTGTATTGCATAGACGGTGCTTTACTTACAAAGTCCTCAGAATAC   5842
gb:MT044257|Organism:Severe      ACTTCTAAAGAAACTTTGTATTGCATAGACGGTGCTTTACTTACAAAGTCCTCAGAATAC   5842
gb:MT106054|Organism:Severe      ACTTCTAAAGAAACTTTGTATTGCATAGACGGTGCTTTACTTACAAAGTCCTCAGAATAC   5842
gb:MT049951|Organism:Severe      ACTTCTAAAGAAACTTTGTATTGCATAGACGGTGCTTTACTTACAAAGTCCTCAGAATAC   5842
gb:MN975262|Organism:Severe      ACTTCTAAAGAAACTTTGTATTGCATAGACGGTGCTTTACTTACAAAGTCCTCAGAATAC   5842
gb:MT106052|Organism:Severe      ACTTCTAAAGAAACTTTGTATTGCATAGACGGTGCTTTACTTACAAAGTCCTCAGAATAC   5842
gb:LC522975|Organism:Severe      ACTTCTAAAGAAACTTTGTATTGCATAGACGGTGCTTTACTTACAAAGTCCTCAGAATAC   5839
gb:LC522973|Organism:Severe      ACTTCTAAAGAAACTTTGTATTGCATAGACGGTGCTTTACTTACAAAGTCCTCAGAATAC   5839
gb:LC522974|Organism:Severe      ACTTCTAAAGAAACTTTGTATTGCATAGACGGTGCTTTACTTACAAAGTCCTCAGAATAC   5839
gb:MN985325|Organism:Severe      ACTTCTAAAGAAACTTTGTATTGCATAGACGGTGCTTTACTTACAAAGTCCTCAGAATAC   5842
gb:MT020881|Organism:Severe      ACTTCTAAAGAAACTTTGTATTGCATAGACGGTGCTTTACTTACAAAGTCCTCAGAATAC   5842
gb:MT020880|Organism:Severe      ACTTCTAAAGAAACTTTGTATTGCATAGACGGTGCTTTACTTACAAAGTCCTCAGAATAC   5842
gb:MT066175|Organism:Severe      ACTTCTAAAGAAACTTTGTATTGCATAGACGGTGCTTTACTTACAAAGTCCTCAGAATAC   5842
gb:MN997409|Organism:Severe      ACTTCTAAAGAAACTTTGTATTGCATAGACGGTGCTTTACTTACAAAGTCCTCAGAATAC   5842
gb:MN938384|Organism:Severe      ACTTCTAAAGAAACTTTGTATTGCATAGACGGTGCTTTACTTACAAAGTCCTCAGAATAC   5810
gb:MT044258|Organism:Severe      ACTTCTAAAGAAACTTTGTATTGCATAGACGGTGCTTTACTTACAAAGTCCTCAGAATAC   5818
gb:MT039890|Organism:Severe      ACTTCTAAAGAAACTTTGTATTGCATAGACGGTGCTTTACTTACAAAGTCCTCAGAATAC   5842
```

```
gb:MN988713|Organism:Severe      ACTTCTAAAGAAACTTTGTATTGCATAGACGGTGCTTTACTTACAAAGTCCTCAGAATAC   5842
gb:LC521925|Organism:Severe      ACTTCTAAAGAAACTTTGTATTGCATAGACGGTGCTTTACTTACAAAGTCCTCAGAATAC   5815
gb:MT093571|Organism:Severe      ACTTCTAAAGAAACTTTGTATTGCATAGACGGTGCTTTACTTACAAAGTCCTCAGAATAC   5842
gb:MT039887|Organism:Severe      ACTTCTAAAGAAACTTTGTATTGCATAGACGGTGCTTTACTTACAAAGTCCTCAGAATAC   5842
gb:MT019530|Organism:Severe      ACTTCTAAAGAAACTTTGTATTGCATAGACGGTGCTTTACTTACAAAGTCCTCAGAATAC   5842
gb:MT039888|Organism:Severe      ACTTCTAAAGAAACTTTGTATTGCATAGACGGTGCTTTACTTACAAAGTCCTCAGAATAC   5842
gb:LC522972|Organism:Severe      ACTTCTAAAGAAACTTTGTATTGCATAGACGGTGCTTTACTTACAAAGTCCTCAGAATAC   5839
gb:MT027063|Organism:Severe      ACTTCTAAAGAAACTTTGTATTGCATAGACGGTGCTTTACTTACAAAGTCCTCAGAATAC   5842
gb:MT027062|Organism:Severe      ACTTCTAAAGAAACTTTGTATTGCATAGACGGTGCTTTACTTACAAAGTCCTCAGAATAC   5842
gb:MT019529|Organism:Severe      ACTTCTAAAGAAACTTTGTATTGCATAGACGGTGCTTTACTTACAAAGTCCTCAGAATAC   5842
gb:MN996529|Organism:Severe      ACTTCTAAAGAAACTTTGTATTGCATAGACGGTGCTTTACTTACAAAGTCCTCAGAATAC   5830
gb:MN996531|Organism:Severe      ACTTCTAAAGAAACTTTGTATTGCATAGACGGTGCTTTACTTACAAAGTCCTCAGAATAC   5829
gb:MT066176|Organism:Severe      ACTTCTAAAGAAACTTTGTATTGCATAGACGGTGCTTTACTTACAAAGTCCTCAGAATAC   5842
gb:MT027064|Organism:Severe      ACTTCTAAAGAAACTTTGTATTGCATAGACGGTGCTTTACTTACAAAGTCCTCAGAATAC   5842
gb:MN994468|Organism:Severe      ACTTCTAAAGAAACTTTGTATTGCATAGACGGTGCTTTACTTACAAAGTCCTCAGAATAC   5842
gb:MT072688|Organism:Severe      ACTTCTAAAGAAACTTTGTATTGCATAGACGGTGCTTTACTTACAAAGTCCTCAGAATAC   5827
gb:MN996527|Organism:Severe      ACTTCTAAAGAAACTTTGTATTGCATAGACGGTGCTTTACTTACAAAGTCCTCAGAATAC   5809
gb:MT093631|Organism:Severe      ACTTCTAAAGAAACTTTGTATTGCATAGACGGTGCTTTACTTACAAAGTCCTCAGAATAC   5880
gb:MT106053|Organism:Severe      ACTTCTAAAGAAACTTTGTATTGCATAGACGGTGCTTTACTTACAAAGTCCTCAGAATAC   5842
gb:MT019533|Organism:Severe      ACTTCTAAAGAAACTTTGTATTGCATAGACGGTGCTTTACTTACAAAGTCCTCAGAATAC   5842
gb:MT019531|Organism:Severe      ACTTCTAAAGAAACTTTGTATTGCATAGACGGTGCTTTACTTACAAAGTCCTCAGAATAC   5842
gb:MN996528|Organism:Severe      ACTTCTAAAGAAACTTTGTATTGCATAGACGGTGCTTTACTTACAAAGTCCTCAGAATAC   5842
gb:MN996530|Organism:Severe      ACTTCTAAAGAAACTTTGTATTGCATAGACGGTGCTTTACTTACAAAGTCCTCAGAATAC   5828
gb:MN908947|Organism:Severe      ACTTCTAAAGAAACTTTGTATTGCATAGACGGTGCTTTACTTACAAAGTCCTCAGAATAC   5842
gb:MT019532|Organism:Severe      ACTTCTAAAGAAACTTTGTATTGCATAGACGGTGCTTTACTTACAAAGTCCTCAGAATAC   5842
                                 ************************************************************

gb:MT020781|Organism:Severe      AAAGGTCCTATTACGGATGTTTTCTACAAAGAAAACAGTTACACAACAACCATAAAACCA   5890
gb:MT007544|Organism:Severe      AAAGGTCCTATTACGGATGTTTTCTACAAAGAAAACAGTTACACAACAACCATAAAACCA   5902
gb:MN994467|Organism:Severe      AAAGGTCCTATTACGGATGTTTTCTACAAAGAAAACAGTTACACAACAACCATAAAACCA   5902
gb:MT044257|Organism:Severe      AAAGGTCCTATTACGGATGTTTTCTACAAAGAAAACAGTTACACAACAACCATAAAACCA   5902
gb:MT106054|Organism:Severe      AAAGGTCCTATTACGGATGTTTTCTACAAAGAAAACAGTTACACAACAACCATAAAACCA   5902
gb:MT049951|Organism:Severe      AAAGGTCCTATTACGGATGTTTTCTACAAAGAAAACAGTTACACAACAACCATAAAACCA   5902
gb:MN975262|Organism:Severe      AAAGGTCCTATTACGGATGTTTTCTACAAAGAAAACAGTTACACAACAACCATAAAACCA   5902
gb:MT106052|Organism:Severe      AAAGGTCCTATTACGGATGTTTTCTACAAAGAAAACAGTTACACAACAACCATAAAACCA   5902
gb:LC522975|Organism:Severe      AAAGGTCCTATTACGGATGTTTTCTACAAAGAAAACAGTTACACAACAACCATAAAACCA   5899
gb:LC522973|Organism:Severe      AAAGGTCCTATTACGGATGTTTTCTACAAAGAAAACAGTTACACAACAACCATAAAACCA   5899
gb:LC522974|Organism:Severe      AAAGGTCCTATTACGGATGTTTTCTACAAAGAAAACAGTTACACAACAACCATAAAACCA   5899
gb:MN985325|Organism:Severe      AAAGGTCCTATTACGGATGTTTTCTACAAAGAAAACAGTTACACAACAACCATAAAACCA   5902
gb:MT020881|Organism:Severe      AAAGGTCCTATTACGGATGTTTTCTACAAAGAAAACAGTTACACAACAACCATAAAACCA   5902
gb:MT020880|Organism:Severe      AAAGGTCCTATTACGGATGTTTTCTACAAAGAAAACAGTTACACAACAACCATAAAACCA   5902
gb:MT066175|Organism:Severe      AAAGGTCCTATTACGGATGTTTTCTACAAAGAAAACAGTTACACAACAACCATAAAACCA   5902
gb:MN997409|Organism:Severe      AAAGGTCCTATTACGGATGTTTTCTACAAAGAAAACAGTTACACAACAACCATAAAACCA   5902
gb:MN938384|Organism:Severe      AAAGGTCCTATTACGGATGTTTTCTACAAAGAAAACAGTTACACAACAACCATAAAACCA   5870
gb:MT044258|Organism:Severe      AAAGGTCCTATTACGGATGTTTTCTACAAAGAAAACAGTTACACAACAACCATAAAACCA   5878
gb:MT039890|Organism:Severe      AAAGGTCCTATTACGGATGTTTTCTACAAAGAAAACAGTTACACAACAACCATAAAACCA   5902
gb:MN988713|Organism:Severe      AAAGGTCCTATTACGGATGTTTTCTACAAAGAAAACAGTTACACAACAACCATAAAACCA   5902
gb:LC521925|Organism:Severe      AAAGGTCCTATTACGGATGTTTTCTACAAAGAAAACAGTTACACAACAACCATAAAACCA   5875
gb:MT093571|Organism:Severe      AAAGGTCCTATTACGGATGTTTTCTACAAAGAAAACAGTTACACAACAACCATAAAACCA   5902
gb:MT039887|Organism:Severe      AAAGGTCCTATTACGGATGTTTTCTACAAAGAAAACAGTTACACAACAACCATAAAACCA   5902
gb:MT019530|Organism:Severe      AAAGGTCCTATTACGGATGTTTTCTACAAAGAAAACAGTTACACAACAACCATAAAACCA   5902
gb:MT039888|Organism:Severe      AAAGGTCCTATTACGGATGTTTTCTACAAAGAAAACAGTTACACAACAACCATAAAACCA   5902
gb:LC522972|Organism:Severe      AAAGGTCCTATTACGGATGTTTTCTACAAAGAAAACAGTTACACAACAACCATAAAACCA   5899
gb:MT027063|Organism:Severe      AAAGGTCCTATTACGGATGTTTTCTACAAAGAAAACAGTTACACAACAACCATAAAACCA   5902
gb:MT027062|Organism:Severe      AAAGGTCCTATTACGGATGTTTTCTACAAAGAAAACAGTTACACAACAACCATAAAACCA   5902
gb:MT019529|Organism:Severe      AAAGGTCCTATTACGGATGTTTTCTACAAAGAAAACAGTTACACAACAACCATAAAACCA   5902
gb:MN996529|Organism:Severe      AAAGGTCCTATTACGGATGTTTTCTACAAAGAAAACAGTTACACAACAACCATAAAACCA   5890
gb:MN996531|Organism:Severe      AAAGGTCCTATTACGGATGTTTTCTACAAAGAAAACAGTTACACAACAACCATAAAACCA   5889
gb:MT066176|Organism:Severe      AAAGGTCCTATTACGGATGTTTTCTACAAAGAAAACAGTTACACAACAACCATAAAACCA   5902
gb:MT027064|Organism:Severe      AAAGGTCCTATTACGGATGTTTTCTACAAAGAAAACAGTTACACAACAACCATAAAACCA   5902
gb:MN994468|Organism:Severe      AAAGGTCCTATTACGGATGTTTTCTACAAAGAAAACAGTTACACAACAACCATAAAACCA   5902
gb:MT072688|Organism:Severe      AAAGGTCCTATTACGGATGTTTTCTACAAAGAAAACAGTTACACAACAACCATAAAACCA   5887
gb:MN996527|Organism:Severe      AAAGGTCCTATTACGGATGTTTTCTACAAAGAAAACAGTTACACAACAACCATAAAACCA   5869
gb:MT093631|Organism:Severe      AAAGGTCCTATTACGGATGTTTTCTACAAAGAAAACAGTTACACAACAACCATAAAACCA   5940
gb:MT106053|Organism:Severe      AAAGGTCCTATTACGGATGTTTTCTACAAAGAAAACAGTTACACAACAACCATAAAACCA   5902
```

```
gb:MT019533|Organism:Severe      AAAGGTCCTATTACGGATGTTTTCTACAAAGAAAACAGTTACACAACAACCATAAAACCA   5902
gb:MT019531|Organism:Severe      AAAGGTCCTATTACGGATGTTTTCTACAAAGAAAACAGTTACACAACAACCATAAAACCA   5902
gb:MN996528|Organism:Severe      AAAGGTCCTATTACGGATGTTTTCTACAAAGAAAACAGTTACACAACAACCATAAAACCA   5902
gb:MN996530|Organism:Severe      AAAGGTCCTATTACGGATGTTTTCTACAAAGAAAACAGTTACACAACAACCATAAAACCA   5888
gb:MN908947|Organism:Severe      AAAGGTCCTATTACGGATGTTTTCTACAAAGAAAACAGTTACACAACAACCATAAAACCA   5902
gb:MT019532|Organism:Severe      AAAGGTCCTATTACGGATGTTTTCTACAAAGAAAACAGTTACACAACAACCATAAAACCA   5902
                                 ************************************************************

gb:MT020781|Organism:Severe      GTTACTTATAAATTGGATGGTGTTGTTTGTACAGAAATTGACCCTAAGTTGGACAATTAT   5950
gb:MT007544|Organism:Severe      GTTACTTATAAATTGGATGGTGTTGTTTGTACAGAAATTGACCCTAAGTTGGACAATTAT   5962
gb:MN994467|Organism:Severe      GTTACTTATAAATTGGATGGTGTTGTTTGTACAGAAATTGACCCTAAGTTGGACAATTAT   5962
gb:MT044257|Organism:Severe      GTTACTTATAAATTGGATGGTGTTGTTTGTACAGAAATTGACCCTAAGTTGGACAATTAT   5962
gb:MT106054|Organism:Severe      GTTACTTATAAATTGGATGGTGTTGTTTGTACAGAAATTGACCCTAAGTTGGACAATTAT   5962
gb:MT049951|Organism:Severe      GTTACTTATAAATTGGATGGTGTTGTTTGTACAGAAATTGACCCTAAGTTGGACAATTAT   5962
gb:MN975262|Organism:Severe      GTTACTTATAAATTGGATGGTGTTGTTTGTACAGAAATTGACCCTAAGTTGGACAATTAT   5962
gb:MT106052|Organism:Severe      GTTACTTATAAATTGGATGGTGTTGTTTGTACAGAAATTGACCCTAAGTTGGACAATTAT   5962
gb:LC522975|Organism:Severe      GTTACTTATAAATTGGATGGTGTTGTTTGTACAGAAATTGACCCTAAGTTGGACAATTAT   5959
gb:LC522973|Organism:Severe      GTTACTTATAAATTGGATGGTGTTGTTTGTACAGAAATTGACCCTAAGTTGGACAATTAT   5959
gb:LC522974|Organism:Severe      GTTACTTATAAATTGGATGGTGTTGTTTGTACAGAAATTGACCCTAAGTTGGACAATTAT   5959
gb:MN985325|Organism:Severe      GTTACTTATAAATTGGATGGTGTTGTTTGTACAGAAATTGACCCTAAGTTGGACAATTAT   5962
gb:MT020881|Organism:Severe      GTTACTTATAAATTGGATGGTGTTGTTTGTACAGAAATTGACCCTAAGTTGGACAATTAT   5962
gb:MT020880|Organism:Severe      GTTACTTATAAATTGGATGGTGTTGTTTGTACAGAAATTGACCCTAAGTTGGACAATTAT   5962
gb:MT066175|Organism:Severe      GTTACTTATAAATTGGATGGTGTTGTTTGTACAGAAATTGACCCTAAGTTGGACAATTAT   5962
gb:MN997409|Organism:Severe      GTTACTTATAAATTGGATGGTGTTGTTTGTACAGAAATTGACCCTAAGTTGGACAATTAT   5962
gb:MN938384|Organism:Severe      GTTACTTATAAATTGGATGGTGTTGTTTGTACAGAAATTGACCCTAAGTTGGACAATTAT   5930
gb:MT044258|Organism:Severe      GTTACTTATAAATTGGATGGTGTTGTTTGTACAGAAATTGACCCTAAGTTGGACAATTAT   5938
gb:MT039890|Organism:Severe      GTTACTTATAAATTGGATGGTGTTGTTTGTACAGAAATTGACCCTAAGTTGGACAATTAT   5962
gb:MN988713|Organism:Severe      GTTACTTATAAATTGGATGGTGTTGTTTGTACAGAAATTGACCCTAAGTTGGACAATTAT   5962
gb:LC521925|Organism:Severe      GTTACTTATAAATTGGATGGTGTTGTTTGTACAGAAATTGACCCTAAGTTGGACAATTAT   5935
gb:MT093571|Organism:Severe      GTTACTTATAAATTGGATGGTGTTGTTTGTACAGAAATTGACCCTAAGTTGGACAATTAT   5962
gb:MT039887|Organism:Severe      GTTACTTATAAATTGGATGGTGTTGTTTGTACAGAAATTGACCCTAAGTTGGACAATTAT   5962
gb:MT019530|Organism:Severe      GTTACTTATAAATTGGATGGTGTTGTTTGTACAGAAATTGACCCTAAGTTGGACAATTAT   5962
gb:MT039888|Organism:Severe      GTTACTTATAAATTGGATGGTGTTGTTTGTACAGAAATTGACCCTAAGTTGGACAATTAT   5962
gb:LC522972|Organism:Severe      GTTACTTATAAATTGGATGGTGTTGTTTGTACAGAAATTGACCCTAAGTTGGACAATTAT   5959
gb:MT027063|Organism:Severe      GTTACTTATAAATTGGATGGTGTTGTTTGTACAGAAATTGACCCTAAGTTGGACAATTAT   5962
gb:MT027062|Organism:Severe      GTTACTTATAAATTGGATGGTGTTGTTTGTACAGAAATTGACCCTAAGTTGGACAATTAT   5962
gb:MT019529|Organism:Severe      GTTACTTATAAATTGGATGGTGTTGTTTGTACAGAAATTGACCCTAAGTTGGACAATTAT   5962
gb:MN996529|Organism:Severe      GTTACTTATAAATTGGATGGTGTTGTTTGTACAGAAATTGACCCTAAGTTGGACAATTAT   5950
gb:MN996531|Organism:Severe      GTTACTTATAAATTGGATGGTGTTGTTTGTACAGAAATTGACCCTAAGTTGGACAATTAT   5949
gb:MT066176|Organism:Severe      GTTACTTATAAATTGGATGGTGTTGTTTGTACAGAAATTGACCCTAAGTTGGACAATTAT   5962
gb:MT027064|Organism:Severe      GTTACTTATAAATTGGATGGTGTTGTTTGTACAGAAATTGACCCTAAGTTGGACAATTAT   5962
gb:MN994468|Organism:Severe      GTTACTTATAAATTGGATGGTGTTGTTTGTACAGAAATTGACCCTAAGTTGGACAATTAT   5962
gb:MT072688|Organism:Severe      GTTACTTATAAATTGGATGGTGTTGTTTGTACAGAAATTGACCCTAAGTTGGACAATTAT   5947
gb:MN996527|Organism:Severe      GTTACTTATAAATTGGATGGTGTTGTTTGTACAGAAATTGACCCTAAGTTGGACAATTAT   5929
gb:MT093631|Organism:Severe      GTTACTTATAAATTGGATGGTGTTGTTTGTACAGAAATTGACCCTAAGTTGGACAATTAT   6000
gb:MT106053|Organism:Severe      GTTACTTATAAATTGGATGGTGTTGTTTGTACAGAAATTGACCCTAAGTTGGACAATTAT   5962
gb:MT019533|Organism:Severe      GTTACTTATAAATTGGATGGTGTTGTTTGTACAGAAATTGACCCTAAGTTGGACAATTAT   5962
gb:MT019531|Organism:Severe      GTTACTTATAAATTGGATGGTGTTGTTTGTACAGAAATTGACCCTAAGTTGGACAATTAT   5962
gb:MN996528|Organism:Severe      GTTACTTATAAATTGGATGGTGTTGTTTGTACAGAAATTGACCCTAAGTTGGACAATTAT   5962
gb:MN996530|Organism:Severe      GTTACTTATAAATTGGATGGTGTTGTTTGTACAGAAATTGACCCTAAGTTGGACAATTAT   5948
gb:MN908947|Organism:Severe      GTTACTTATAAATTGGATGGTGTTGTTTGTACAGAAATTGACCCTAAGTTGGACAATTAT   5962
gb:MT019532|Organism:Severe      GTTACTTATAAATTGGATGGTGTTGTTTGTACAGAAATTGACCCTAAGTTGGACAATTAT   5962
                                 ************************************************************

gb:MT020781|Organism:Severe      TATAAGAAAGACAATTCTTATTTCACAGAGCAACCAATTGATCTTGTACCAAACCAACCA   6010
gb:MT007544|Organism:Severe      TATAAGAAAGACAATTCTTATTTCACAGAGCAACCAATTGATCTTGTACCAAACCAACCA   6022
gb:MN994467|Organism:Severe      TATAAGAAAGACAATTCTTATTTCACAGAGCAACCAATTGATCTTGTACCAAACCAACCA   6022
gb:MT044257|Organism:Severe      TATAAGAAAGACAATTCTTATTTCACAGAGCAACCAATTGATCTTGTACCAAACCAACCA   6022
gb:MT106054|Organism:Severe      TATAAGAAAGACAATTCTTATTTCACAGAGCAACCAATTGATCTTGTACCAAACCAACCA   6022
gb:MT049951|Organism:Severe      TATAAGAAAGACAATTCTTATTTCACAGAGCAACCAATTGATCTTGTACCAAACCAACCA   6022
gb:MN975262|Organism:Severe      TATAAGAAAGACAATTCTTATTTCACAGAGCAACCAATTGATCTTGTACCAAACCAACCA   6022
gb:MT106052|Organism:Severe      TATAAGAAAGACAATTCTTATTTCACAGAGCAACCAATTGATCTTGTACCAAACCAACCA   6022
gb:LC522975|Organism:Severe      TATAAGAAAGACAATTCTTATTTCACAGAGCAACCAATTGATCTTGTACCAAACCAACCA   6019
gb:LC522973|Organism:Severe      TATAAGAAAGACAATTCTTATTTCACAGAGCAACCAATTGATCTTGTACCAAACCAACCA   6019
gb:LC522974|Organism:Severe      TATAAGAAAGACAATTCTTATTTCACAGAGCAACCAATTGATCTTGTACCAAACCAACCA   6019
```

```
gb:MN985325|Organism:Severe      TATAAGAAAGACAATTCTTATTTCACAGAGCAACCAATTGATCTTGTACCAAACCAACCA   6022
gb:MT020881|Organism:Severe      TATAAGAAAGACAATTCTTATTTCACAGAGCAACCAATTGATCTTGTACCAAACCAACCA   6022
gb:MT020880|Organism:Severe      TATAAGAAAGACAATTCTTATTTCACAGAGCAACCAATTGATCTTGTACCAAACCAACCA   6022
gb:MT066175|Organism:Severe      TATAAGAAAGACAATTCTTATTTCACAGAGCAACCAATTGATCTTGTACCAAACCAACCA   6022
gb:MN997409|Organism:Severe      TATAAGAAAGACAATTCTTATTTCACAGAGCAACCAATTGATCTTGTACCAAACCAACCA   6022
gb:MN938384|Organism:Severe      TATAAGAAAGACAATTCTTATTTCACAGAGCAACCAATTGATCTTGTACCAAACCAACCA   5990
gb:MT044258|Organism:Severe      TATAAGAAAGACAATTCTTATTTCACAGAGCAACCAATTGATCTTGTACCAAACCAACCA   5998
gb:MT039890|Organism:Severe      TATAAGAAAGACAATTCTTATTTCACAGAGCAACCAATTGATCTTGTACCAAACCAACCA   6022
gb:MN988713|Organism:Severe      TATAAGAAAGACAATTCTTATTTCACAGAGCAACCAATTGATCTTGTACCAAACCAACCA   6022
gb:LC521925|Organism:Severe      TATAAGAAAGACAATTCTTATTTCACAGAGCAACCAATTGATCTTGTACCAAACCAACCA   5995
gb:MT093571|Organism:Severe      TATAAGAAAGACAATTCTTATTTCACAGAGCAACCAATTGATCTTGTACCAAACCAACCA   6022
gb:MT039887|Organism:Severe      TATAAGAAAGACAATTCTTATTTCACAGAGCAACCAATTGATCTTGTACCAAACCAACCA   6022
gb:MT019530|Organism:Severe      TATAAGAAAGACAATTCTTATTTCACAGAGCAACCAATTGATCTTGTACCAAACCAACCA   6022
gb:MT039888|Organism:Severe      TATAAGAAAGACAATTCTTATTTCACAGAGCAACCAATTGATCTTGTACCAAACCAACCA   6022
gb:LC522972|Organism:Severe      TATAAGAAAGACAATTCTTATTTCACAGAGCAACCAATTGATCTTGTACCAAACCAACCA   6019
gb:MT027063|Organism:Severe      TATAAGAAAGACAATTCTTATTTCACAGAGCAACCAATTGATCTTGTACCAAACCAACCA   6022
gb:MT027062|Organism:Severe      TATAAGAAAGACAATTCTTATTTCACAGAGCAACCAATTGATCTTGTACCAAACCAACCA   6022
gb:MT019529|Organism:Severe      TATAAGAAAGACAATTCTTATTTCACAGAGCAACCAATTGATCTTGTACCAAACCAACCA   6022
gb:MN996529|Organism:Severe      TATAAGAAAGACAATTCTTATTTCACAGAGCAACCAATTGATCTTGTACCAAACCAACCA   6010
gb:MN996531|Organism:Severe      TATAAGAAAGACAATTCTTATTTCACAGAGCAACCAATTGATCTTGTACCAAACCAACCA   6009
gb:MT066176|Organism:Severe      TATAAGAAAGACAATTCTTATTTCACAGAGCAACCAATTGATCTTGTACCAAACCAACCA   6022
gb:MT027064|Organism:Severe      TATAAGAAAGACAATTCTTATTTCACAGAGCAACCAATTGATCTTGTACCAAACCAACCA   6022
gb:MN994468|Organism:Severe      TATAAGAAAGACAATTCTTATTTCACAGAGCAACCAATTGATCTTGTACCAAACCAACCA   6022
gb:MT072688|Organism:Severe      TATAAGAAAGACAATTCTTATTTCACAGAGCAACCAATTGATCTTGTACCAAACCAACCA   6007
gb:MN996527|Organism:Severe      TATAAGAAAGACAATTCTTATTTCACAGAGCAACCAATTGATCTTGTACCAAACCAACCA   5989
gb:MT093631|Organism:Severe      TATAAGAAAGACAATTCTTATTTCACAGAGCAACCAATTGATCTTGTACCAAACCAACCA   6060
gb:MT106053|Organism:Severe      TATAAGAAAGACAATTCTTATTTCACAGAGCAACCAATTGATCTTGTACCAAACCAACCA   6022
gb:MT019533|Organism:Severe      TATAAGAAAGACAATTCTTATTTCACAGAGCAACCAATTGATCTTGTACCAAACCAACCA   6022
gb:MT019531|Organism:Severe      TATAAGAAAGACAATTCTTATTTCACAGAGCAACCAATTGATCTTGTACCAAACCAACCA   6022
gb:MN996528|Organism:Severe      TATAAGAAAGACAATTCTTATTTCACAGAGCAACCAATTGATCTTGTACCAAACCAACCA   6022
gb:MN996530|Organism:Severe      TATAAGAAAGACAATTCTTATTTCACAGAGCAACCAATTGATCTTGTACCAAACCAACCA   6008
gb:MN908947|Organism:Severe      TATAAGAAAGACAATTCTTATTTCACAGAGCAACCAATTGATCTTGTACCAAACCAACCA   6022
gb:MT019532|Organism:Severe      TATAAGAAAGACAATTCTTATTTCACAGAGCAACCAATTGATCTTGTACCAAACCAACCA   6022
                                 ************************************************************

gb:MT020781|Organism:Severe      TATCCAAACGCAAGCTTCGATAATTTTAAGTTTGTATGTGATAATATCAAATTTGCTGAT   6070
gb:MT007544|Organism:Severe      TATCCAAACGCAAGCTTCGATAATTTTAAGTTTGTATGTGATAATATCAAATTTGCTGAT   6082
gb:MN994467|Organism:Severe      TATCCAAACGCAAGCTTCGATAATTTTAAGTTTGTATGTGATAATATCAAATTTGCTGAT   6082
gb:MT044257|Organism:Severe      TATCCAAACGCAAGCTTCGATAATTTTAAGTTTGTATGTGATAATATCAAATTTGCTGAT   6082
gb:MT106054|Organism:Severe      TATCCAAACGCAAGCTTCGATAATTTTAAGTTTGTATGTGATAATATCAAATTTGCTGAT   6082
gb:MT049951|Organism:Severe      TATCCAAACGCAAGCTTCGATAATTTTAAGTTTGTATGTGATAATATCAAATTTGCTGAT   6082
gb:MN975262|Organism:Severe      TATCCAAACGCAAGCTTCGATAATTTTAAGTTTGTATGTGATAATATCAAATTTGCTGAT   6082
gb:MT106052|Organism:Severe      TATCCAAACGCAAGCTTCGATAATTTTAAGTTTGTATGTGATAATATCAAATTTGCTGAT   6082
gb:LC522975|Organism:Severe      TATCCAAACGCAAGCTTCGATAATTTTAAGTTTGTATGTGATAATATCAAATTTGCTGAT   6079
gb:LC522973|Organism:Severe      TATCCAAACGCAAGCTTCGATAATTTTAAGTTTGTATGTGATAATATCAAATTTGCTGAT   6079
gb:LC522974|Organism:Severe      TATCCAAACGCAAGCTTCGATAATTTTAAGTTTGTATGTGATAATATCAAATTTGCTGAT   6079
gb:MN985325|Organism:Severe      TATCCAAACGCAAGCTTCGATAATTTTAAGTTTGTATGTGATAATATCAAATTTGCTGAT   6082
gb:MT020881|Organism:Severe      TATCCAAACGCAAGCTTCGATAATTTTAAGTTTGTATGTGATAATATCAAATTTGCTGAT   6082
gb:MT020880|Organism:Severe      TATCCAAACGCAAGCTTCGATAATTTTAAGTTTGTATGTGATAATATCAAATTTGCTGAT   6082
gb:MT066175|Organism:Severe      TATCCAAACGCAAGCTTCGATAATTTTAAGTTTGTATGTGATAATATCAAATTTGCTGAT   6082
gb:MN997409|Organism:Severe      TATCCAAACGCAAGCTTCGATAATTTTAAGTTTGTATGTGATAATATCAAATTTGCTGAT   6082
gb:MN938384|Organism:Severe      TATCCAAACGCAAGCTTCGATAATTTTAAGTTTGTATGTGATAATATCAAATTTGCTGAT   6050
gb:MT044258|Organism:Severe      TATCCAAACGCAAGCTTCGATAATTTTAAGTTTGTATGTGATAATATCAAATTTGCTGAT   6058
gb:MT039890|Organism:Severe      TATCCAAATGCAAGCTTCGATAATTTTAAGTTTGTATGTGATAATATCAAATTTGCTGAT   6082
gb:MN988713|Organism:Severe      TATCCAAACGCAAGCTTCGATAATTTTAAGTTTGTATGTGATAATATCAAATTTGCTGAT   6082
gb:LC521925|Organism:Severe      TATCCAAACGCAAGCTTCGATAATTTTAAGTTTGTATGTGATAATATCAAATTTGCTGAT   6055
gb:MT093571|Organism:Severe      TATCCAAACGCAAGCTTCGATAATTTTAAGTTTGTATGTGATAATATCAAATTTGCTGAT   6082
gb:MT039887|Organism:Severe      TATCCAAACGCAAGCTTCGATAATTTTAAGTTTGTATGTGATAATATCAAATTTGCTGAT   6082
gb:MT019530|Organism:Severe      TATCCAAACGCAAGCTTCGATAATTTTAAGTTTGTATGTGATAATATCAAATTTGCTGAT   6082
gb:MT039888|Organism:Severe      TATCCAAACGCAAGCTTCGATAATTTTAAGTTTGTATGTGATAATATCAAATTTGCTGAT   6082
gb:LC522972|Organism:Severe      TATCCAAACGCAAGCTTCGATAATTTTAAGTTTGTATGTGATAATATCAAATTTGCTGAT   6079
gb:MT027063|Organism:Severe      TATCCAAACGCAAGCTTCGATAATTTTAAGTTTGTATGTGATAATATCAAATTTGCTGAT   6082
gb:MT027062|Organism:Severe      TATCCAAACGCAAGCTTCGATAATTTTAAGTTTGTATGTGATAATATCAAATTTGCTGAT   6082
gb:MT019529|Organism:Severe      TATCCAAACGCAAGCTTCGATAATTTTAAGTTTGTATGTGATAATATCAAATTTGCTGAT   6082
gb:MN996529|Organism:Severe      TATCCAAACGCAAGCTTCGATAATTTTAAGTTTGTATGTGATAATATCAAATTTGCTGAT   6070
```

```
gb:MN996531|Organism:Severe      TATCCAAACGCAAGCTTCGATAATTTTAAGTTTGTATGTGATAATATCAAATTTGCTGAT   6069
gb:MT066176|Organism:Severe      TATCCAAACGCAAGCTTCGATAATTTTAAGTTTGTATGTGATAATATCAAATTTGCTGAT   6082
gb:MT027064|Organism:Severe      TATCCAAACGCAAGCTTCGATAATTTTAAGTTTGTATGTGATAATATCAAATTTGCTGAT   6082
gb:MN994468|Organism:Severe      TATCCAAACGCAAGCTTCGATAATTTTAAGTTTGTATGTGATAATATCAAATTTGCTGAT   6082
gb:MT072688|Organism:Severe      TATCCAAACGCAAGCTTCGATAATTTTAAGTTTGTATGTGATAATATCAAATTTGCTGAT   6067
gb:MN996527|Organism:Severe      TATCCAAACGCAAGCTTCGATAATTTTAAGTTTGTATGTGATAATATCAAATTTGCTGAT   6049
gb:MT093631|Organism:Severe      TATCCAAACGCAAGCTTCGATAATTTTAAGTTTGTATGTGATAATATCAAATTTGCTGAT   6120
gb:MT106053|Organism:Severe      TATCCAAACGCAAGCTTCGATAATTTTAAGTTTGTATGTGATAATATCAAATTTGCTGAT   6082
gb:MT019533|Organism:Severe      TATCCAAACGCAAGCTTCGATAATTTTAAGTTTGTATGTGATAATATCAAATTTGCTGAT   6082
gb:MT019531|Organism:Severe      TATCCAAACGCAAGCTTCGATAATTTTAAGTTTGTATGTGATAATATCAAATTTGCTGAT   6082
gb:MN996528|Organism:Severe      TATCCAAACGCAAGCTTCGATAATTTTAAGTTTGTATGTGATAATATCAAATTTGCTGAT   6082
gb:MN996530|Organism:Severe      TATCCAAACGCAAGCTTCGATAATTTTAAGTTTGTATGTGATAATATCAAATTTGCTGAT   6068
gb:MN908947|Organism:Severe      TATCCAAACGCAAGCTTCGATAATTTTAAGTTTGTATGTGATAATATCAAATTTGCTGAT   6082
gb:MT019532|Organism:Severe      TATCCAAACGCAAGCTTCGATAATTTTAAGTTTGTATGTGATAATATCAAATTTGCTGAT   6082
                                 ******** ***************************************************

gb:MT020781|Organism:Severe      GATTTAAACCAGTTAACTGGTTATAAGAAACCTGCTTCAAGAGAGCTTAAAGTTACATTT   6130
gb:MT007544|Organism:Severe      GATTTAAACCAGTTAACTGGTTATAAGAAACCTGCTTCAAGAGAGCTTAAAGTTACATTT   6142
gb:MN994467|Organism:Severe      GATTTAAACCAGTTAACTGGTTATAAGAAACCTGCTTCAAGAGAGCTTAAAGTTACATTT   6142
gb:MT044257|Organism:Severe      GATTTAAACCAGTTAACTGGTTATAAGAAACCTGCTTCAAGAGAGCTTAAAGTTACATTT   6142
gb:MT106054|Organism:Severe      GATTTAAACCAGTTAACTGGTTATAAGAAACCTGCTTCAAGAGAGCTTAAAGTTACATTT   6142
gb:MT049951|Organism:Severe      GATTTAAACCAGTTAACTGGTTATAAGAAACCTGCTTCAAGAGAGCTTAAAGTTACATTT   6142
gb:MN975262|Organism:Severe      GATTTAAACCAGTTAACTGGTTATAAGAAACCTGCTTCAAGAGAGCTTAAAGTTACATTT   6142
gb:MT106052|Organism:Severe      GATTTAAACCAGTTAACTGGTTATAAGAAACCTGCTTCAAGAGAGCTTAAAGTTACATTT   6142
gb:LC522975|Organism:Severe      GATTTAAACCAGTTAACTGGTTATAAGAAACCTGCTTCAAGAGAGCTTAAAGTTACATTT   6139
gb:LC522973|Organism:Severe      GATTTAAACCAGTTAACTGGTTATAAGAAACCTGCTTCAAGAGAGCTTAAAGTTACATTT   6139
gb:LC522974|Organism:Severe      GATTTAAACCAGTTAACTGGTTATAAGAAACCTGCTTCAAGAGAGCTTAAAGTTACATTT   6139
gb:MN985325|Organism:Severe      GATTTAAACCAGTTAACTGGTTATAAGAAACCTGCTTCAAGAGAGCTTAAAGTTACATTT   6142
gb:MT020881|Organism:Severe      GATTTAAACCAGTTAACTGGTTATAAGAAACCTGCTTCAAGAGAGCTTAAAGTTACATTT   6142
gb:MT020880|Organism:Severe      GATTTAAACCAGTTAACTGGTTATAAGAAACCTGCTTCAAGAGAGCTTAAAGTTACATTT   6142
gb:MT066175|Organism:Severe      GATTTAAACCAGTTAACTGGTTATAAGAAACCTGCTTCAAGAGAGCTTAAAGTTACATTT   6142
gb:MN997409|Organism:Severe      GATTTAAACCAGTTAACTGGTTATAAGAAACCTGCTTCAAGAGAGCTTAAAGTTACATTT   6142
gb:MN938384|Organism:Severe      GATTTAAACCAGTTAACTGGTTATAAGAAACCTGCTTCAAGAGAGCTTAAAGTTACATTT   6110
gb:MT044258|Organism:Severe      GATTTAAACCAGTTAACTGGTTATAAGAAACCTGCTTCAAGAGAGCTTAAAGTTACATTT   6118
gb:MT039890|Organism:Severe      GATTTAAACCAGTTAACTGGTTATAAGAAACCTGCTTCAAGAGAGCTTAAAGTTACATTT   6142
gb:MN988713|Organism:Severe      GATTTAAACCAGTTAACTGGTTATAAGAAACCTGCTTCAAGAGAGCTTAAAGTTACATTT   6142
gb:LC521925|Organism:Severe      GATTTAAACCAGTTAACTGGTTATAAGAAACCTGCTTCAAGAGAGCTTAAAGTTACATTT   6115
gb:MT093571|Organism:Severe      GATTTAAACCAGTTAACTGGTTATAAGAAACCTGCTTCAAGAGAGCTTAAAGTTACATTT   6142
gb:MT039887|Organism:Severe      GATTTAAACCAGTTAACTGGTTATAAGAAACCTGCTTCAAGAGAGCTTAAAGTTACATTT   6142
gb:MT019530|Organism:Severe      GATTTAAACCAGTTAACTGGTTATAAGAAACCTGCTTCAAGAGAGCTTAAAGTTACATTT   6142
gb:MT039888|Organism:Severe      GATTTAAACCAGTTAACTGGTTATAAGAAACCTGCTTCAAGAGAGCTTAAAGTTACATTT   6142
gb:LC522972|Organism:Severe      GATTTAAACCAGTTAACTGGTTATAAGAAACCTGCTTCAAGAGAGCTTAAAGTTACATTT   6139
gb:MT027063|Organism:Severe      GATTTAAACCAGTTAACTGGTTATAAGAAACCTGCTTCAAGAGAGCTTAAAGTTACATTT   6142
gb:MT027062|Organism:Severe      GATTTAAACCAGTTAACTGGTTATAAGAAACCTGCTTCAAGAGAGCTTAAAGTTACATTT   6142
gb:MT019529|Organism:Severe      GATTTAAACCAGTTAACTGGTTATAAGAAACCTGCTTCAAGAGAGCTTAAAGTTACATTT   6142
gb:MN996529|Organism:Severe      GATTTAAACCAGTTAACTGGTTATAAGAAACCTGCTTCAAGAGAGCTTAAAGTTACATTT   6130
gb:MN996531|Organism:Severe      GATTTAAACCAGTTAACTGGTTATAAGAAACCTGCTTCAAGAGAGCTTAAAGTTACATTT   6129
gb:MT066176|Organism:Severe      GATTTAAACCAGTTAACTGGTTATAAGAAACCTGCTTCAAGAGAGCTTAAAGTTACATTT   6142
gb:MT027064|Organism:Severe      GATTTAAACCAGTTAACTGGTTATAAGAAACCTGCTTCAAGAGAGCTTAAAGTTACATTT   6142
gb:MN994468|Organism:Severe      GATTTAAACCAGTTAACTGGTTATAAGAAACCTGCTTCAAGAGAGCTTAAAGTTACATTT   6142
gb:MT072688|Organism:Severe      GATTTAAACCAGTTAACTGGTTATAAGAAACCTGCTTCAAGAGAGCTTAAAGTTACATTT   6127
gb:MN996527|Organism:Severe      GATTTAAACCAGTTAACTGGTTATAAGAAACCTGCTTCAAGAGAGCTTAAAGTTACATTT   6109
gb:MT093631|Organism:Severe      GATTTAAACCAGTTAACTGGTTATAAGAAACCTGCTTCAAGAGAGCTTAAAGTTACATTT   6180
gb:MT106053|Organism:Severe      GATTTAAACCAGTTAACTGGTTATAAGAAACCTGCTTCAAGAGAGCTTAAAGTTACATTT   6142
gb:MT019533|Organism:Severe      GATTTAAACCAGTTAACTGGTTATAAGAAACCTGCTTCAAGAGAGCTTAAAGTTACATTT   6142
gb:MT019531|Organism:Severe      GATTTAAACCAGTTAACTGGTTATAAGAAACCTGCTTCAAGAGAGCTTAAAGTTACATTT   6142
gb:MN996528|Organism:Severe      GATTTAAACCAGTTAACTGGTTATAAGAAACCTGCTTCAAGAGAGCTTAAAGTTACATTT   6142
gb:MN996530|Organism:Severe      GATTTAAACCAGTTAACTGGTTATAAGAAACCTGCTTCAAGAGAGCTTAAAGTTACATTT   6128
gb:MN908947|Organism:Severe      GATTTAAACCAGTTAACTGGTTATAAGAAACCTGCTTCAAGAGAGCTTAAAGTTACATTT   6142
gb:MT019532|Organism:Severe      GATTTAAACCAGTTAACTGGTTATAAGAAACCTGCTTCAAGAGAGCTTAAAGTTACATTT   6142
                                 ************************************************************

gb:MT020781|Organism:Severe      TTCCCTGACTTAAATGGTGATGTGGTGGCTATTGATTATAAACACTACACACCCTCTTTT   6190
gb:MT007544|Organism:Severe      TTCCCTGACTTAAATGGTGATGTGGTGGCTATTGATTATAAACACTACACACCCTCTTTT   6202
gb:MN994467|Organism:Severe      TTCCCTGACTTAAATGGTGATGTGGTGGCTATTGATTATAAACACTACACACCCTCTTTT   6202
```

```
gb:MT044257|Organism:Severe      TTCCCTGACTTAAATGGTGATGTGGTGGCTATTGATTATAAACACTACACACCCTCTTTT   6202
gb:MT106054|Organism:Severe      TTCCCTGACTTAAATGGTGATGTGGTGGCTATTGATTATAAACACTACACACCCTCTTTT   6202
gb:MT049951|Organism:Severe      TTCCCTGACTTAAATGGTGATGTGGTGGCTATTGATTATAAACACTACACACCCTCTTTT   6202
gb:MN975262|Organism:Severe      TTCCCTGACTTAAATGGTGATGTGGTGGCTATTGATTATAAACACTACACACCCTCTTTT   6202
gb:MT106052|Organism:Severe      TTCCCTGACTTAAATGGTGATGTGGTGGCTATTGATTATAAACACTACACACCCTCTTTT   6202
gb:LC522975|Organism:Severe      TTCCCTGACTTAAATGGTGATGTGGTGGCTATTGATTATAAACACTACACACCCTCTTTT   6199
gb:LC522973|Organism:Severe      TTCCCTGACTTAAATGGTGATGTGGTGGCTATTGATTATAAACACTACACACCCTCTTTT   6199
gb:LC522974|Organism:Severe      TTCCCTGACTTAAATGGTGATGTGGTGGCTATTGATTATAAACACTACACACCCTCTTTT   6199
gb:MN985325|Organism:Severe      TTCCCTGACTTAAATGGTGATGTGGTGGCTATTGATTATAAACACTACACACCCTCTTTT   6202
gb:MT020881|Organism:Severe      TTCCCTGACTTAAATGGTGATGTGGTGGCTATTGATTATAAACACTACACACCCTCTTTT   6202
gb:MT020880|Organism:Severe      TTCCCTGACTTAAATGGTGATGTGGTGGCTATTGATTATAAACACTACACACCCTCTTTT   6202
gb:MT066175|Organism:Severe      TTCCCTGACTTAAATGGTGATGTGGTGGCTATTGATTATAAACACTACACACCCTCTTTT   6202
gb:MN997409|Organism:Severe      TTCCCTGACTTAAATGGTGATGTGGTGGCTATTGATTATAAACACTACACACCCTCTTTT   6202
gb:MN938384|Organism:Severe      TTCCCTGACTTAAATGGTGATGTGGTGGCTATTGATTATAAACACTACACACCCTCTTTT   6170
gb:MT044258|Organism:Severe      TTCCCTGACTTAAATGGTGATGTGGTGGCTATTGATTATAAACACTACACACCCTCTTTT   6178
gb:MT039890|Organism:Severe      TTCCCTGACTTAAATGGTGATGTGGTGGCTATTGATTATAAACACTACACACCCTCTTTT   6202
gb:MN988713|Organism:Severe      TTCCCTGACTTAAATGGTGATGTGGTGGCTATTGATTATAAACACTACACACCCTCTTTT   6202
gb:LC521925|Organism:Severe      TTCCCTGACTTAAATGGTGATGTGGTGGCTATTGATTATAAACACTACACACCCTCTTTT   6175
gb:MT093571|Organism:Severe      TTCCCTGACTTAAATGGTGATGTGGTGGCTATTGATTATAAACACTACACACCCTCTTTT   6202
gb:MT039887|Organism:Severe      TTCCCTGACTTAAATGGTGATGTGGTGGCTATTGATTATAAACACTACACACCCTCTTTT   6202
gb:MT019530|Organism:Severe      TTCCCTGACTTAAATGGTGATGTGGTGGCTATTGATTATAAACACTACACACCCTCTTTT   6202
gb:MT039888|Organism:Severe      TTCCCTGACTTAAATGGTGATGTGGTGGCTATTGATTATAAACACTACACACCCTCTTTT   6202
gb:LC522972|Organism:Severe      TTCCCTGACTTAAATGGTGATGTGGTGGCTATTGATTATAAACACTACACACCCTCTTTT   6199
gb:MT027063|Organism:Severe      TTCCCTGACTTAAATGGTGATGTGGTGGCTATTGATTATAAACACTACACACCCTCTTTT   6202
gb:MT027062|Organism:Severe      TTCCCTGACTTAAATGGTGATGTGGTGGCTATTGATTATAAACACTACACACCCTCTTTT   6202
gb:MT019529|Organism:Severe      TTCCCTGACTTAAATGGTGATGTGGTGGCTATTGATTATAAACACTACACACCCTCTTTT   6202
gb:MN996529|Organism:Severe      TTCCCTGACTTAAATGGTGATGTGGTGGCTATTGATTATAAACACTACACACCCTCTTTT   6190
gb:MN996531|Organism:Severe      TTCCCTGACTTAAATGGTGATGTGGTGGCTATTGATTATAAACACTACACACCCTCTTTT   6189
gb:MT066176|Organism:Severe      TTCCCTGACTTAAATGGTGATGTGGTGGCTATTGATTATAAACACTACACACCCTCTTTT   6202
gb:MT027064|Organism:Severe      TTCCCTGACTTAAATGGTGATGTGGTGGCTATTGATTATAAACACTACACACCCTCTTTT   6202
gb:MN994468|Organism:Severe      TTCCCTGACTTAAATGGTGATGTGGTGGCTATTGATTATAAACACTACACACCCTCTTTT   6202
gb:MT072688|Organism:Severe      TTCCCTGACTTAAATGGTGATGTGGTGGCTATTGATTATAAACACTACACACCCTCTTTT   6187
gb:MN996527|Organism:Severe      TTCCCTGACTTAAATGGTGATGTGGTGGCTATTGATTATAAACACTACACACCCTCTTTT   6169
gb:MT093631|Organism:Severe      TTCCCTGACTTAAATGGTGATGTGGTGGCTATTGATTATAAACACTACACACCCTCTTTT   6240
gb:MT106053|Organism:Severe      TTCCCTGACTTAAATGGTGATGTGGTGGCTATTGATTATAAACACTACACACCCTCTTTT   6202
gb:MT019533|Organism:Severe      TTCCCTGACTTAAATGGTGATGTGGTGGCTATTGATTATAAACACTACACACCCTCTTTT   6202
gb:MT019531|Organism:Severe      TTCCCTGACTTAAATGGTGATGTGGTGGCTATTGATTATAAACACTACACACCCTCTTTT   6202
gb:MN996528|Organism:Severe      TTCCCTGACTTAAATGGTGATGTGGTGGCTATTGATTATAAACACTACACACCCTCTTTT   6202
gb:MN996530|Organism:Severe      TTCCCTGACTTAAATGGTGATGTGGTGGCTATTGATTATAAACACTACACACCCTCTTTT   6188
gb:MN908947|Organism:Severe      TTCCCTGACTTAAATGGTGATGTGGTGGCTATTGATTATAAACACTACACACCCTCTTTT   6202
gb:MT019532|Organism:Severe      TTCCCTGACTTAAATGGTGATGTGGTGGCTATTGATTATAAACACTACACACCCTCTTTT   6202
                                 ************************************************************

gb:MT020781|Organism:Severe      AAGAAAGGAGCTAAATTGTTACATAAACCTATTGTTTGGCATGTTAACAATGCAACTAAT   6250
gb:MT007544|Organism:Severe      AAGAAAGGAGCTAAATTGTTACATAAACCTATTGTTTGGCATGTTAACAATGCAACTAAT   6262
gb:MN994467|Organism:Severe      AAGAAAGGAGCTAAATTGTTACATAAACCTATTGTTTGGCATGTTAACAATGCAACTAAT   6262
gb:MT044257|Organism:Severe      AAGAAAGGAGCTAAATTGTTACATAAACCTATTGTTTGGCATGTTAACAATGCAACTAAT   6262
gb:MT106054|Organism:Severe      AAGAAAGGAGCTAAATTGTTACATAAACCTATTGTTTGGCATGTTAACAATGCAACTAAT   6262
gb:MT049951|Organism:Severe      AAGAAAGGAGCTAAATTGTTACATAAACCTATTGTTTGGCATGTTAACAATGCAACTAAT   6262
gb:MN975262|Organism:Severe      AAGAAAGGAGCTAAATTGTTACATAAACCTATTGTTTGGCATGTTAACAATGCAACTAAT   6262
gb:MT106052|Organism:Severe      AAGAAAGGAGCTAAATTGTTACATAAACCTATTGTTTGGCATGTTAACAATGCAACTAAT   6262
gb:LC522975|Organism:Severe      AAGAAAGGAGCTAAATTGTTACATAAACCTATTGTTTGGCATGTTAACAATGCAACTAAT   6259
gb:LC522973|Organism:Severe      AAGAAAGGAGCTAAATTGTTACATAAACCTATTGTTTGGCATGTTAACAATGCAACTAAT   6259
gb:LC522974|Organism:Severe      AAGAAAGGAGCTAAATTGTTACATAAACCTATTGTTTGGCATGTTAACAATGCAACTAAT   6259
gb:MN985325|Organism:Severe      AAGAAAGGAGCTAAATTGTTACATAAACCTATTGTTTGGCATGTTAACAATGCAACTAAT   6262
gb:MT020881|Organism:Severe      AAGAAAGGAGCTAAATTGTTACATAAACCTATTGTTTGGCATGTTAACAATGCAACTAAT   6262
gb:MT020880|Organism:Severe      AAGAAAGGAGCTAAATTGTTACATAAACCTATTGTTTGGCATGTTAACAATGCAACTAAT   6262
gb:MT066175|Organism:Severe      AAGAAAGGAGCTAAATTGTTACATAAACCTATTGTTTGGCATGTTAACAATGCAACTAAT   6262
gb:MN997409|Organism:Severe      AAGAAAGGAGCTAAATTGTTACATAAACCTATTGTTTGGCATGTTAACAATGCAACTAAT   6262
gb:MN938384|Organism:Severe      AAGAAAGGAGCTAAATTGTTACATAAACCTATTGTTTGGCATGTTAACAATGCAACTAAT   6230
gb:MT044258|Organism:Severe      AAGAAAGGAGCTAAATTGTTACATAAACCTATTGTTTGGCATGTTAACAATGCAACTAAT   6238
gb:MT039890|Organism:Severe      AAGAAAGGAGCTAAATTGTTACATAAACCTATTGTTTGGCATGTTAACAATGCAACTAAT   6262
gb:MN988713|Organism:Severe      AAGAAAGGAGCTAAATTGTTACATAAACCTATTGTTTGGCATGTTAACAATGCAACTAAT   6262
gb:LC521925|Organism:Severe      AAGAAAGGAGCTAAATTGTTACATAAACCTATTGTTTGGCATGTTAACAATGCAACTAAT   6235
gb:MT093571|Organism:Severe      AAGAAAGGAGCTAAATTGTTACATAAACCTATTGTTTGGCATGTTAACAATGCAACTAAT   6262
```

```
gb:MT039887|Organism:Severe      AAGAAAGGAGCTAAATTGTTACATAAACCTATTGTTTGGCATGTTAACAATGCAACTAAT    6262
gb:MT019530|Organism:Severe      AAGAAAGGAGCTAAATTGTTACATAAACCTATTGTTTGGCATGTTAACAATGCAACTAAT    6262
gb:MT039888|Organism:Severe      AAGAAAGGAGCTAAATTGTTACATAAACCTATTGTTTGGCATGTTAACAATGCAACTAAT    6262
gb:LC522972|Organism:Severe      AAGAAAGGAGCTAAATTGTTACATAAACCTATTGTTTGGCATGTTAACAATGCAACTAAT    6259
gb:MT027063|Organism:Severe      AAGAAAGGAGCTAAATTGTTACATAAACCTATTGTTTGGCATGTTAACAATGCAACTAAT    6262
gb:MT027062|Organism:Severe      AAGAAAGGAGCTAAATTGTTACATAAACCTATTGTTTGGCATGTTAACAATGCAACTAAT    6262
gb:MT019529|Organism:Severe      AAGAAAGGAGCTAAATTGTTACATAAACCTATTGTTTGGCATGTTAACAATGCAACTAAT    6262
gb:MN996529|Organism:Severe      AAGAAAGGAGCTAAATTGTTACATAAACCTATTGTTTGGCATGTTAACAATGCAACTAAT    6250
gb:MN996531|Organism:Severe      AAGAAAGGAGCTAAATTGTTACATAAACCTATTGTTTGGCATGTTAACAATGCAACTAAT    6249
gb:MT066176|Organism:Severe      AAGAAAGGAGCTAAATTGTTACATAAACCTATTGTTTGGCATGTTAACAATGCAACTAAT    6262
gb:MT027064|Organism:Severe      AAGAAAGGAGCTAAATTGTTACATAAACCTATTGTTTGGCATGTTAACAATGCAACTAAT    6262
gb:MN994468|Organism:Severe      AAGAAAGGAGCTAAATTGTTACATAAACCTATTGTTTGGCATGTTAACAATGCAACTAAT    6262
gb:MT072688|Organism:Severe      AAGAAAGGAGCTAAATTGTTACATAAACCTATTGTTTGGCATGTTAACAATGCAACTAAT    6247
gb:MN996527|Organism:Severe      AAGAAAGGAGCTAAATTGTTACATAAACCTATTGTTTGGCATGTTAACAATGCAACTAAT    6229
gb:MT093631|Organism:Severe      AAGAAAGGAGCTAAATTGTTACATAAACCTATTGTTTGGCATGTTAACAATGCAACTAAT    6300
gb:MT106053|Organism:Severe      AAGAAAGGAGCTAAATTGTTACATAAACCTATTGTTTGGCATGTTAACAATGCAACTAAT    6262
gb:MT019533|Organism:Severe      AAGAAAGGAGCTAAATTGTTACATAAACCTATTGTTTGGCATGTTAACAATGCAACTAAT    6262
gb:MT019531|Organism:Severe      AAGAAAGGAGCTAAATTGTTACATAAACCTATTGTTTGGCATGTTAACAATGCAACTAAT    6262
gb:MN996528|Organism:Severe      AAGAAAGGAGCTAAATTGTTACATAAACCTATTGTTTGGCATGTTAACAATGCAACTAAT    6262
gb:MN996530|Organism:Severe      AAGAAAGGAGCTAAATTGTTACATAAACCTATTGTTTGGCATGTTAACAATGCAACTAAT    6248
gb:MN908947|Organism:Severe      AAGAAAGGAGCTAAATTGTTACATAAACCTATTGTTTGGCATGTTAACAATGCAACTAAT    6262
gb:MT019532|Organism:Severe      AAGAAAGGAGCTAAATTGTTACATAAACCTATTGTTTGGCATGTTAACAATGCAACTAAT    6262
                                 ************************************************************

gb:MT020781|Organism:Severe      AAAGCCACGTATAAACCAAATACCTGGTGTATACGTTGTCTTTGGAGCACAAAACCAGTT    6310
gb:MT007544|Organism:Severe      AAAGCCACGTATAAACCAAATACCTGGTGTATACGTTGTCTTTGGAGCACAAAACCAGTT    6322
gb:MN994467|Organism:Severe      AAAGCCACGTATAAACCAAATACCTGGTGTATACGTTGTCTTTGGAGCACAAAACCAGTT    6322
gb:MT044257|Organism:Severe      AAAGCCACGTATAAACCAAATACCTGGTGTATACGTTGTCTTTGGAGCACAAAACCAGTT    6322
gb:MT106054|Organism:Severe      AAAGCCACGTATAAACCAAATACCTGGTGTATACGTTGTCTTTGGAGCACAAAACCAGTT    6322
gb:MT049951|Organism:Severe      AAAGCCACGTATAAACCAAATACCTGGTGTATACGTTGTCTTTGGAGCACAAAACCAGTT    6322
gb:MN975262|Organism:Severe      AAAGCCACGTATAAACCAAATACCTGGTGTATACGTTGTCTTTGGAGCACAAAACCAGTT    6322
gb:MT106052|Organism:Severe      AAAGCCACGTATAAACCAAATACCTGGTGTATACGTTGTCTTTGGAGCACAAAACCAGTT    6322
gb:LC522975|Organism:Severe      AAAGCCACGTATAAACCAAATACCTGGTGTATACGTTGTCTTTGGAGCACAAAACCAGTT    6319
gb:LC522973|Organism:Severe      AAAGCCACGTATAAACCAAATACCTGGTGTATACGTTGTCTTTGGAGCACAAAACCAGTT    6319
gb:LC522974|Organism:Severe      AAAGCCACGTATAAACCAAATACCTGGTGTATACGTTGTCTTTGGAGCACAAAACCAGTT    6319
gb:MN985325|Organism:Severe      AAAGCCACGTATAAACCAAATACCTGGTGTATACGTTGTCTTTGGAGCACAAAACCAGTT    6322
gb:MT020881|Organism:Severe      AAAGCCACGTATAAACCAAATACCTGGTGTATACGTTGTCTTTGGAGCACAAAACCAGTT    6322
gb:MT020880|Organism:Severe      AAAGCCACGTATAAACCAAATACCTGGTGTATACGTTGTCTTTGGAGCACAAAACCAGTT    6322
gb:MT066175|Organism:Severe      AAAGCCACGTATAAACCAAATACCTGGTGTATACGTTGTCTTTGGAGCACAAAACCAGTT    6322
gb:MN997409|Organism:Severe      AAAGCCACGTATAAACCAAATACCTGGTGTATACGTTGTCTTTGGAGCACAAAACCAGTT    6322
gb:MN938384|Organism:Severe      AAAGCCACGTATAAACCAAATACCTGGTGTATACGTTGTCTTTGGAGCACAAAACCAGTT    6290
gb:MT044258|Organism:Severe      AAAGCCACGTATAAACCAAATACCTGGTGTATACGTTGTCTTTGGAGCACAAAACCAGTT    6298
gb:MT039890|Organism:Severe      AAAGCCACGTATAAACCAAATACCTGGTGTATACGTTGTCTTTGGAGCACAAAACCAGTT    6322
gb:MN988713|Organism:Severe      AAAGCCACGTATAAACCAAATACCTGGTGTATACGTTGTCTTTGGAGCACAAAACCAGTT    6322
gb:LC521925|Organism:Severe      AAAGCCACGTATAAACCAAATACCTGGTGTATACGTTGTCTTTGGAGCACAAAACCAGTT    6295
gb:MT093571|Organism:Severe      AAAGCCACGTATAAACCAAATACCTGGTGTATACGTTGTCTTTGGAGCACAAAACCAGTT    6322
gb:MT039887|Organism:Severe      AAAGCCACGTATAAACCAAATACCTGGTGTATACGTTGTCTTTGGAGCACAAAACCAGTT    6322
gb:MT019530|Organism:Severe      AAAGCCACGTATAAACCAAATACCTGGTGTATACGTTGTCTTTGGAGCACAAAACCAGTT    6322
gb:MT039888|Organism:Severe      AAAGCCACGTATAAACCAAATACCTGGTGTATACGTTGTCTTTGGAGCACAAAACCAGTT    6322
gb:LC522972|Organism:Severe      AAAGCCACGTATAAACCAAATACCTGGTGTATACGTTGTCTTTGGAGCACAAAACCAGTT    6319
gb:MT027063|Organism:Severe      AAAGCCACGTATAAACCAAATACCTGGTGTATACGTTGTCTTTGGAGCACAAAACCAGTT    6322
gb:MT027062|Organism:Severe      AAAGCCACGTATAAACCAAATACCTGGTGTATACGTTGTCTTTGGAGCACAAAACCAGTT    6322
gb:MT019529|Organism:Severe      AAAGCCACGTATAAACCAAATACCTGGTGTATACGTTGTCTTTGGAGCACAAAACCAGTT    6322
gb:MN996529|Organism:Severe      AAAGCCACGTATAAACCAAATACCTGGTGTATACGTTGTCTTTGGAGCACAAAACCAGTT    6310
gb:MN996531|Organism:Severe      AAAGCCACGTATAAACCAAATACCTGGTGTATACGTTGTCTTTGGAGCACAAAACCAGTT    6309
gb:MT066176|Organism:Severe      AAAGCCACGTATAAACCAAATACCTGGTGTATACGTTGTCTTTGGAGCACAAAACCAGTT    6322
gb:MT027064|Organism:Severe      AAAGCCACGTATAAACCAAATACCTGGTGTATACGTTGTCTTTGGAGCACAAAACCAGTT    6322
gb:MN994468|Organism:Severe      AAAGCCACGTATAAACCAAATACCTGGTGTATACGTTGTCTTTGGAGCACAAAACCAGTT    6322
gb:MT072688|Organism:Severe      AAAGCCACGTATAAACCAAATACCTGGTGTATACGTTGTCTTTGGAGCACAAAACCAGTT    6307
gb:MN996527|Organism:Severe      AAAGCCACGTATAAACCAAATACCTGGTGTATACGTTGTCTTTGGAGCACAAAACCAGTT    6289
gb:MT093631|Organism:Severe      AAAGCCACGTATAAACCAAATACCTGGTGTATACGTTGTCTTTGGAGCACAAAACCAGTT    6360
gb:MT106053|Organism:Severe      AAAGCCACGTATAAACCAAATACCTGGTGTATACGTTGTCTTTGGAGCACAAAACCAGTT    6322
gb:MT019533|Organism:Severe      AAAGCCACGTATAAACCAAATACCTGGTGTATACGTTGTCTTTGGAGCACAAAACCAGTT    6322
gb:MT019531|Organism:Severe      AAAGCCACGTATAAACCAAATACCTGGTGTATACGTTGTCTTTGGAGCACAAAACCAGTT    6322
gb:MN996528|Organism:Severe      AAAGCCACGTATAAACCAAATACCTGGTGTATACGTTGTCTTTGGAGCACAAAACCAGTT    6322
```

```
gb:MN996530|Organism:Severe      AAAGCCACGTATAAACCAAATACCTGGTGTATACGTTGTCTTTGGAGCACAAAACCAGTT   6308
gb:MN908947|Organism:Severe      AAAGCCACGTATAAACCAAATACCTGGTGTATACGTTGTCTTTGGAGCACAAAACCAGTT   6322
gb:MT019532|Organism:Severe      AAAGCCACGTATAAACCAAATACCTGGTGTATACGTTGTCTTTGGAGCACAAAACCAGTT   6322
                                 ************************************************************

gb:MT020781|Organism:Severe      GAAACATCAAATTCGTTTGATGTACTGAAGTCAGAGGACGCGCAGGGAATGGATAATCTT   6370
gb:MT007544|Organism:Severe      GAAACATCAAATTCGTTTGATGTACTGAAGTCAGAGGACGCGCAGGGAATGGATAATCTT   6382
gb:MN994467|Organism:Severe      GAAACATCAAATTCGTTTGATGTACTGAAGTCAGAGGACGCGCAGGGAATGGATAATCTT   6382
gb:MT044257|Organism:Severe      GAAACATCAAATTCGTTTGATGTACTGAAGTCAGAGGACGCGCAGGGAATGGATAATCTT   6382
gb:MT106054|Organism:Severe      GAAACATCAAATTCGTTTGATGTACTGAAGTCAGAGGACGCGCAGGGAATGGATAATCTT   6382
gb:MT049951|Organism:Severe      GAAACATCAAATTCGTTTGATGTACTGAAGTCAGAGGACGCGCAGGGAATGGATAATCTT   6382
gb:MN975262|Organism:Severe      GAAACATCAAATTCGTTTGATGTACTGAAGTCAGAGGACGCGCAGGGAATGGATAATCTT   6382
gb:MT106052|Organism:Severe      GAAACATCAAATTCGTTTGATGTACTGAAGTCAGAGGACGCGCAGGGAATGGATAATCTT   6382
gb:LC522975|Organism:Severe      GAAACATCAAATTCGTTTGATGTACTGAAGTCAGAGGACGCGCAGGGAATGGATAATCTT   6379
gb:LC522973|Organism:Severe      GAAACATCAAATTCGTTTGATGTACTGAAGTCAGAGGACGCGCAGGGAATGGATAATCTT   6379
gb:LC522974|Organism:Severe      GAAACATCAAATTCGTTTGATGTACTGAAGTCAGAGGACGCGCAGGGAATGGATAATCTT   6379
gb:MN985325|Organism:Severe      GAAACATCAAATTCGTTTGATGTACTGAAGTCAGAGGACGCGCAGGGAATGGATAATCTT   6382
gb:MT020881|Organism:Severe      GAAACATCAAATTCGTTTGATGTACTGAAGTCAGAGGACGCGCAGGGAATGGATAATCTT   6382
gb:MT020880|Organism:Severe      GAAACATCAAATTCGTTTGATGTACTGAAGTCAGAGGACGCGCAGGGAATGGATAATCTT   6382
gb:MT066175|Organism:Severe      GAAACATCAAATTCGTTTGATGTACTGAAGTCAGAGGACGCGCAGGGAATGGATAATCTT   6382
gb:MN997409|Organism:Severe      GAAACATCAAATTCGTTTGATGTACTGAAGTCAGAGGACGCGCAGGGAATGGATAATCTT   6382
gb:MN938384|Organism:Severe      GAAACATCAAATTCGTTTGATGTACTGAAGTCAGAGGACGCGCAGGGAATGGATAATCTT   6350
gb:MT044258|Organism:Severe      GAAACATCAAATTCGTTTGATGTACTGAAGTCAGAGGACGCGCAGGGAATGGATAATCTT   6358
gb:MT039890|Organism:Severe      GAAACATCAAATTCGTTTGATGTACTGAAGTCAGAGGACGCGCAGGGAATGGATAATCTT   6382
gb:MN988713|Organism:Severe      GAAACATCAAATTCGTTTGATGTACTGAAGTCAGAGGACGCGCAGGGAATGGATAATCTT   6382
gb:LC521925|Organism:Severe      GAAACATCAAATTCGTTTGATGTACTGAAGTCAGAGGACGCGCAGGGAATGGATAATCTT   6355
gb:MT093571|Organism:Severe      GAAACATCAAATTCGTTTGATGTACTGAAGTCAGAGGACGCGCAGGGAATGGATAATCTT   6382
gb:MT039887|Organism:Severe      GAAACATCAAATTCGTTTGATGTACTGAAGTCAGAGGACGCGCAGGGAATGGATAATCTT   6382
gb:MT019530|Organism:Severe      GAAACATCAAATTCGTTTGATGTACTGAAGTCAGAGGACGCGCAGGGAATGGATAATCTT   6382
gb:MT039888|Organism:Severe      GAAACATCAAATTCGTTTGATGTACTGAAGTCAGAGGACGCGCAGGGAATGGATAATCTT   6382
gb:LC522972|Organism:Severe      GAAACATCAAATTCGTTTGATGTACTGAAGTCAGAGGACGCGCAGGGAATGGATAATCTT   6379
gb:MT027063|Organism:Severe      GAAACATCAAATTCGTTTGATGTACTGAAGTCAGAGGACGCGCAGGGAATGGATAATCTT   6382
gb:MT027062|Organism:Severe      GAAACATCAAATTCGTTTGATGTACTGAAGTCAGAGGACGCGCAGGGAATGGATAATCTT   6382
gb:MT019529|Organism:Severe      GAAACATCAAATTCGTTTGATGTACTGAAGTCAGAGGACGCGCAGGGAATGGATAATCTT   6382
gb:MN996529|Organism:Severe      GAAACATCAAATTCGTTTGATGTACTGAAGTCAGAGGACGCGCAGGGAATGGATAATCTT   6370
gb:MN996531|Organism:Severe      GAAACATCAAATTCGTTTGATGTACTGAAGTCAGAGGACGCGCAGGGAATGGATAATCTT   6369
gb:MT066176|Organism:Severe      GAAACATCAAATTCGTTTGATGTACTGAAGTCAGAGGACGCGCAGGGAATGGATAATCTT   6382
gb:MT027064|Organism:Severe      GAAACATCAAATTCGTTTGATGTACTGAAGTCAGAGGACGCGCAGGGAATGGATAATCTT   6382
gb:MN994468|Organism:Severe      GAAACATCAAATTCGTTTGATGTACTGAAGTCAGAGGACGCGCAGGGAATGGATAATCTT   6382
gb:MT072688|Organism:Severe      GAAACATCAAATTCGTTTGATGTACTGAAGTCAGAGGACGCGCAGGGAATGGATAATCTT   6367
gb:MN996527|Organism:Severe      GAAACATCAAATTCGTTTGATGTACTGAAGTCAGAGGACGCGCAGGGAATGGATAATCTT   6349
gb:MT093631|Organism:Severe      GAAACATCAAATTCGTTTGATGTACTGAAGTCAGAGGACGCGCAGGGAATGGATAATCTT   6420
gb:MT106053|Organism:Severe      GAAACATCAAATTCGTTTGATGTACTGAAGTCAGAGGACGCGCAGGGAATGGATAATCTT   6382
gb:MT019533|Organism:Severe      GAAACATCAAATTCGTTTGATGTACTGAAGTCAGAGGACGCGCAGGGAATGGATAATCTT   6382
gb:MT019531|Organism:Severe      GAAACATCAAATTCGTTTGATGTACTGAAGTCAGAGGACGCGCAGGGAATGGATAATCTT   6382
gb:MN996528|Organism:Severe      GAAACATCAAATTCGTTTGATGTACTGAAGTCAGAGGACGCGCAGGGAATGGATAATCTT   6382
gb:MN996530|Organism:Severe      GAAACATCAAATTCGTTTGATGTACTGAAGTCAGAGGACGCGCAGGGAATGGATAATCTT   6368
gb:MN908947|Organism:Severe      GAAACATCAAATTCGTTTGATGTACTGAAGTCAGAGGACGCGCAGGGAATGGATAATCTT   6382
gb:MT019532|Organism:Severe      GAAACATCAAATTCGTTTGATGTACTGAAGTCAGAGGACGCGCAGGGAATGGATAATCTT   6382
                                 ************************************************************

gb:MT020781|Organism:Severe      GCCTGCGAAGATCTAAAACCAGTCTCTGAAGAAGTAGTGGAAAATCCTACCATACAGAAA   6430
gb:MT007544|Organism:Severe      GCCTGCGAAGATCTAAAACCAGTCTCTGAAGAAGTAGTGGAAAATCCTACCATACAGAAA   6442
gb:MN994467|Organism:Severe      GCCTGCGAAGATCTAAAACCAGTCTCTGAAGAAGTAGTGGAAAATCCTACCATACAGAAA   6442
gb:MT044257|Organism:Severe      GCCTGCGAAGATCTAAAACCAGTCTCTGAAGAAGTAGTGGAAAATCCTACCATACAGAAA   6442
gb:MT106054|Organism:Severe      GCCTGCGAAGATCTAAAACCAGTCTCTGAAGAAGTAGTGGAAAATCCTACCATACAGAAA   6442
gb:MT049951|Organism:Severe      GCCTGCGAAGATCTAAAACCAGTCTCTGAAGAAGTAGTGGAAAATCCTACCATACAGAAA   6442
gb:MN975262|Organism:Severe      GCCTGCGAAGATCTAAAACCAGTCTCTGAAGAAGTAGTGGAAAATCCTACCATACAGAAA   6442
gb:MT106052|Organism:Severe      GCCTGCGAAGATCTAAAACCAGTCTCTGAAGAAGTAGTGGAAAATCCTACCATACAGAAA   6442
gb:LC522975|Organism:Severe      GCCTGCGAAGATCTAAAACCAGTCTCTGAAGAAGTAGTGGAAAATCCTACCATACAGAAA   6439
gb:LC522973|Organism:Severe      GCCTGCGAAGATCTAAAACCAGTCTCTGAAGAAGTAGTGGAAAATCCTACCATACAGAAA   6439
gb:LC522974|Organism:Severe      GCCTGCGAAGATCTAAAACCAGTCTCTGAAGAAGTAGTGGAAAATCCTACCATACAGAAA   6439
gb:MN985325|Organism:Severe      GCCTGCGAAGATCTAAAACCAGTCTCTGAAGAAGTAGTGGAAAATCCTACCATACAGAAA   6442
gb:MT020881|Organism:Severe      GCCTGCGAAGATCTAAAACCAGTCTCTGAAGAAGTAGTGGAAAATCCTACCATACAGAAA   6442
gb:MT020880|Organism:Severe      GCCTGCGAAGATCTAAAACCAGTCTCTGAAGAAGTAGTGGAAAATCCTACCATACAGAAA   6442
```

```
gb:MT066175|Organism:Severe      GCCTGCGAAGATCTAAAACCAGTCTCTGAAGAAGTAGTGGAAAATCCTACCATACAGAAA   6442
gb:MN997409|Organism:Severe      GCCTGCGAAGATCTAAAACCAGTCTCTGAAGAAGTAGTGGAAAATCCTACCATACAGAAA   6442
gb:MN938384|Organism:Severe      GCCTGCGAAGATCTAAAACCAGTCTCTGAAGAAGTAGTGGAAAATCCTACCATACAGAAA   6410
gb:MT044258|Organism:Severe      GCCTGCGAAGATCTAAAACCAGTCTCTGAAGAAGTAGTGGAAAATCCTACCATACAGAAA   6418
gb:MT039890|Organism:Severe      GCCTGCGAAGATCTAAAACCAGTCTCTGAAGAAGTAGTGGAAAATCCTACCATACAGAAA   6442
gb:MN988713|Organism:Severe      GCCTGCGAAGATCTAAAACCAGTCTCTGAAGAAGTAGTGGAAAATCCTACCATACAGAAA   6442
gb:LC521925|Organism:Severe      GCCTGCGAAGATCTAAAACCAGTCTCTGAAGAAGTAGTGGAAAATCCTACCATACAGAAA   6415
gb:MT093571|Organism:Severe      GCCTGCGAAGATCTAAAACCAGTCTCTGAAGAAGTAGTGGAAAATCCTACCATACAGAAA   6442
gb:MT039887|Organism:Severe      GCCTGCGAAGATCTAAAACCAGTCTCTGAAGAAGTAGTGGAAAATCCTACCATACAGAAA   6442
gb:MT019530|Organism:Severe      GCCTGCGAAGATCTAAAACCAGTCTCTGAAGAAGTAGTGGAAAATCCTACCATACAGAAA   6442
gb:MT039888|Organism:Severe      GCCTGCGAAGATCTAAAACCAGTCTCTGAAGAAGTAGTGGAAAATCCTACCATACAGAAA   6442
gb:LC522972|Organism:Severe      GCCTGCGAAGATCTAAAACCAGTCTCTGAAGAAGTAGTGGAAAATCCTACCATACAGAAA   6439
gb:MT027063|Organism:Severe      GCCTGCGAAGATCTAAAACCAGTCTCTGAAGAAGTAGTGGAAAATCCTACCATACAGAAA   6442
gb:MT027062|Organism:Severe      GCCTGCGAAGATCTAAAACCAGTCTCTGAAGAAGTAGTGGAAAATCCTACCATACAGAAA   6442
gb:MT019529|Organism:Severe      GCCTGCGAAGATCTAAAACCAGTCTCTGAAGAAGTAGTGGAAAATCCTACCATACAGAAA   6442
gb:MN996529|Organism:Severe      GCCTGCGAAGATCTAAAACCAGTCTCTGAAGAAGTAGTGGAAAATCCTACCATACAGAAA   6430
gb:MN996531|Organism:Severe      GCCTGCGAAGATCTAAAACCAGTCTCTGAAGAAGTAGTGGAAAATCCTACCATACAGAAA   6429
gb:MT066176|Organism:Severe      GCCTGCGAAGATCTAAAACCAGTCTCTGAAGAAGTAGTGGAAAATCCTACCATACAGAAA   6442
gb:MT027064|Organism:Severe      GCCTGCGAAGATCTAAAACCAGTCTCTGAAGAAGTAGTGGAAAATCCTACCATACAGAAA   6442
gb:MN994468|Organism:Severe      GCCTGCGAAGATCTAAAACCAGTCTCTGAAGAAGTAGTGGAAAATCCTACCATACAGAAA   6442
gb:MT072688|Organism:Severe      GCCTGCGAAGATCTAAAACCAGTCTCTGAAGAAGTAGTGGAAAATCCTACCATACAGAAA   6427
gb:MN996527|Organism:Severe      GCCTGCGAAGATCTAAAACCAGTCTCTGAAGAAGTAGTGGAAAATCCTACCATACAGAAA   6409
gb:MT093631|Organism:Severe      GCCTGCGAAGATCTAAAACCAGTCTCTGAAGAAGTAGTGGAAAATCCTACCATACAGAAA   6480
gb:MT106053|Organism:Severe      GCCTGCGAAGATCTAAAACCAGTCTCTGAAGAAGTAGTGGAAAATCCTACCATACAGAAA   6442
gb:MT019533|Organism:Severe      GCCTGCGAAGATCTAAAACCAGTCTCTGAAGAAGTAGTGGAAAATCCTACCATACAGAAA   6442
gb:MT019531|Organism:Severe      GCCTGCGAAGATCTAAAACCAGTCTCTGAAGAAGTAGTGGAAAATCCTACCATACAGAAA   6442
gb:MN996528|Organism:Severe      GCCTGCGAAGATCTAAAACCAGTCTCTGAAGAAGTAGTGGAAAATCCTACCATACAGAAA   6442
gb:MN996530|Organism:Severe      GCCTGCGAAGATCTAAAACCAGTCTCTGAAGAAGTAGTGGAAAATCCTACCATACAGAAA   6428
gb:MN908947|Organism:Severe      GCCTGCGAAGATCTAAAACCAGTCTCTGAAGAAGTAGTGGAAAATCCTACCATACAGAAA   6442
gb:MT019532|Organism:Severe      GCCTGCGAAGATCTAAAACCAGTCTCTGAAGAAGTAGTGGAAAATCCTACCATACAGAAA   6442
                                 ************************************************************

gb:MT020781|Organism:Severe      GACGTTCTTGAGTGTAATGTGAAAACTACCGAAGTTGTAGGAGACATTATACTTAAACCA   6490
gb:MT007544|Organism:Severe      GACGTTCTTGAGTGTAATGTGAAAACTACCGAAGTTGTAGGAGACATTATACTTAAACCA   6502
gb:MN994467|Organism:Severe      GACGTTCTTGAGTGTAATGTGAAAACTACCGAAGTTGTAGGAGACATTATACTTAAACCA   6502
gb:MT044257|Organism:Severe      GACGTTCTTGAGTGTAATGTGAAAACTACCGAAGTTGTAGGAGACATTATACTTAAACCA   6502
gb:MT106054|Organism:Severe      GACGTTCTTGAGTGTAATGTGAAAACTACCGAAGTTGTAGGAGACATTATACTTAAACCA   6502
gb:MT049951|Organism:Severe      GACGTTCTTGAGTGTAATGTGAAAACTACCGAAGTTGTAGGAGACATTATACTTAAACCA   6502
gb:MN975262|Organism:Severe      GACGTTCTTGAGTGTAATGTGAAAACTACCGAAGTTGTAGGAGACATTATACTTAAACCA   6502
gb:MT106052|Organism:Severe      GACGTTCTTGAGTGTAATGTGAAAACTACCGAAGTTGTAGGAGACATTATACTTAAACCA   6502
gb:LC522975|Organism:Severe      GACGTTCTTGAGTGTAATGTGAAAACTACCGAAGTTGTAGGAGACATTATACTTAAACCA   6499
gb:LC522973|Organism:Severe      GACGTTCTTGAGTGTAATGTGAAAACTACCGAAGTTGTAGGAGACATTATACTTAAACCA   6499
gb:LC522974|Organism:Severe      GACGTTCTTGAGTGTAATGTGAAAACTACCGAAGTTGTAGGAGACATTATACTTAAACCA   6499
gb:MN985325|Organism:Severe      GACGTTCTTGAGTGTAATGTGAAAACTACCGAAGTTGTAGGAGACATTATACTTAAACCA   6502
gb:MT020881|Organism:Severe      GACGTTCTTGAGTGTAATGTGAAAACTACCGAAGTTGTAGGAGACATTATACTTAAACCA   6502
gb:MT020880|Organism:Severe      GACGTTCTTGAGTGTAATGTGAAAACTACCGAAGTTGTAGGAGACATTATACTTAAACCA   6502
gb:MT066175|Organism:Severe      GACGTTCTTGAGTGTAATGTGAAAACTACCGAAGTTGTAGGAGACATTATACTTAAACCA   6502
gb:MN997409|Organism:Severe      GACGTTCTTGAGTGTAATGTGAAAACTACCGAAGTTGTAGGAGACATTATACTTAAACCA   6502
gb:MN938384|Organism:Severe      GACGTTCTTGAGTGTAATGTGAAAACTACCGAAGTTGTAGGAGACATTATACTTAAACCA   6470
gb:MT044258|Organism:Severe      GACGTTCTTGAGTGTAATGTGAAAACTACCGAAGTTGTAGGAGACATTATACTTAAACCA   6478
gb:MT039890|Organism:Severe      GACGTTCTTGAGTGTAATGTGAAAACTACCGAAGTTGTAGGAGACATTATACTTAAACCA   6502
gb:MN988713|Organism:Severe      GACGTTCTTGAGTGTAATGTGAAAACTACCGAAGTTGTAGGAGACATTATACTTAAACCA   6502
gb:LC521925|Organism:Severe      GACGTTCTTGAGTGTAATGTGAAAACTACCGAAGTTGTAGGAGACATTATACTTAAACCA   6475
gb:MT093571|Organism:Severe      GACGTTCTTGAGTGTAATGTGAAAACTACCGAAGTTGTAGGAGACATTATACTTAAACCA   6502
gb:MT039887|Organism:Severe      GACGTTCTTGAGTGTAATGTGAAAACTACCGAAGTTGTAGGAGACATTATACTTAAACCA   6502
gb:MT019530|Organism:Severe      GACGTTCTTGAGTGTAATGTGAAAACTACCGAAGTTGTAGGAGACATTATACTTAAACCA   6502
gb:MT039888|Organism:Severe      GACGTTCTTGAGTGTAATGTGAAAACTACCGAAGTTGTAGGAGACATTATACTTAAACCA   6502
gb:LC522972|Organism:Severe      GACGTTCTTGAGTGTAATGTGAAAACTACCGAAGTTGTAGGAGACATTATACTTAAACCA   6499
gb:MT027063|Organism:Severe      GACGTTCTTGAGTGTAATGTGAAAACTACCGAAGTTGTAGGAGACATTATACTTAAACCA   6502
gb:MT027062|Organism:Severe      GACGTTCTTGAGTGTAATGTGAAAACTACCGAAGTTGTAGGAGACATTATACTTAAACCA   6502
gb:MT019529|Organism:Severe      GACGTTCTTGAGTGTAATGTGAAAACTACCGAAGTTGTAGGAGACATTATACTTAAACCA   6502
gb:MN996529|Organism:Severe      GACGTTCTTGAGTGTAATGTGAAAACTACCGAAGTTGTAGGAGACATTATACTTAAACCA   6490
gb:MN996531|Organism:Severe      GACGTTCTTGAGTGTAATGTGAAAACTACCGAAGTTGTAGGAGACATTATACTTAAACCA   6489
gb:MT066176|Organism:Severe      GACGTTCTTGAGTGTAATGTGAAAACTACCGAAGTTGTAGGAGACATTATACTTAAACCA   6502
gb:MT027064|Organism:Severe      GACGTTCTTGAGTGTAATGTGAAAACTACCGAAGTTGTAGGAGACATTATACTTAAACCA   6502
```

```
gb:MN994468|Organism:Severe      GACGTTCTTGAGTGTAATGTGAAAACTACCGAAGTTGTAGGAGACATTATACTTAAACCA    6502
gb:MT072688|Organism:Severe      GACGTTCTTGAGTGTAATGTGAAAACTACCGAAGTTGTAGGAGACATTATACTTAAACCA    6487
gb:MN996527|Organism:Severe      GACGTTCTTGAGTGTAATGTGAAAACTACCGAAGTTGTAGGAGACATTATACTTAAACCA    6469
gb:MT093631|Organism:Severe      GACGTTCTTGAGTGTAATGTGAAAACTACCGAAGTTGTAGGAGACATTATACTTAAACCA    6540
gb:MT106053|Organism:Severe      GACGTTCTTGAGTGTAATGTGAAAACTACCGAAGTTGTAGGAGACATTATACTTAAACCA    6502
gb:MT019533|Organism:Severe      GACGTTCTTGAGTGTAATGTGAAAACTACCGAAGTTGTAGGAGACATTATACTTAAACCA    6502
gb:MT019531|Organism:Severe      GACGTTCTTGAGTGTAATGTGAAAACTACCGAAGTTGTAGGAGACATTATACTTAAACCA    6502
gb:MN996528|Organism:Severe      GACGTTCTTGAGTGTAATGTGAAAACTACCGAAGTTGTAGGAGACATTATACTTAAACCA    6502
gb:MN996530|Organism:Severe      GACGTTCTTGAGTGTAATGTGAAAACTACCGAAGTTGTAGGAGACATTATACTTAAACCA    6488
gb:MN908947|Organism:Severe      GACGTTCTTGAGTGTAATGTGAAAACTACCGAAGTTGTAGGAGACATTATACTTAAACCA    6502
gb:MT019532|Organism:Severe      GACGTTCTTGAGTGTAATGTGAAAACTACCGAAGTTGTAGGAGACATTATACTTAAACCA    6502
                                 ************************************************************

gb:MT020781|Organism:Severe      GCAAATAATAGTTTAAAAATTACAGAAGAGGTTGGCCACACAGATCTAATGGCTGCTTAT    6550
gb:MT007544|Organism:Severe      GCAAATAATAGTTTAAAAATTACAGAAGAGGTTGGCCACACAGATCTAATGGCTGCTTAT    6562
gb:MN994467|Organism:Severe      GCAAATAATAGTTTAAAAATTACAGAAGAGGTTGGCCACACAGATCTAATGGCTGCTTAT    6562
gb:MT044257|Organism:Severe      GCAAATAATAGTTTAAAAATTACAGAAGAGGTTGGCCACACAGATCTAATGGCTGCTTAT    6562
gb:MT106054|Organism:Severe      GCAAATAATAGTTTAAAAATTACAGAAGAGGTTGGCCACACAGATCTAATGGCTGCTTAT    6562
gb:MT049951|Organism:Severe      GCAAATAATAGTTTAAAAATTACAGAAGAGGTTGGCCACACAGATCTAATGGCTGCTTAT    6562
gb:MN975262|Organism:Severe      GCAAATAATAGTTTAAAAATTACAGAAGAGGTTGGCCACACAGATCTAATGGCTGCTTAT    6562
gb:MT106052|Organism:Severe      GCAAATAATAGTTTAAAAATTACAGAAGAGGTTGGCCACACAGATCTAATGGCTGCTTAT    6562
gb:LC522975|Organism:Severe      GCAAATAATAGTTTAAAAATTACAGAAGAGGTTGGCCACACAGATCTAATGGCTGCTTAT    6559
gb:LC522973|Organism:Severe      GCAAATAATAGTTTAAAAATTACAGAAGAGGTTGGCCACACAGATCTAATGGCTGCTTAT    6559
gb:LC522974|Organism:Severe      GCAAATAATAGTTTAAAAATTACAGAAGAGGTTGGCCACACAGATCTAATGGCTGCTTAT    6559
gb:MN985325|Organism:Severe      GCAAATAATAGTTTAAAAATTACAGAAGAGGTTGGCCACACAGATCTAATGGCTGCTTAT    6562
gb:MT020881|Organism:Severe      GCAAATAATAGTTTAAAAATTACAGAAGAGGTTGGCCACACAGATCTAATGGCTGCTTAT    6562
gb:MT020880|Organism:Severe      GCAAATAATAGTTTAAAAATTACAGAAGAGGTTGGCCACACAGATCTAATGGCTGCTTAT    6562
gb:MT066175|Organism:Severe      GCAAATAATAGTTTAAAAATTACAGAAGAGGTTGGCCACACAGATCTAATGGCTGCTTAT    6562
gb:MN997409|Organism:Severe      GCAAATAATAGTTTAAAAATTACAGAAGAGGTTGGCCACACAGATCTAATGGCTGCTTAT    6562
gb:MN938384|Organism:Severe      GCAAATAATAGTTTAAAAATTACAGAAGAGGTTGGCCACACAGATCTAATGGCTGCTTAT    6530
gb:MT044258|Organism:Severe      GCAAATAATAGTTTAAAAATTACAGAAGAGGTTGGCCACACAGATCTAATGGCTGCTTAT    6538
gb:MT039890|Organism:Severe      GCAAATAATAGTTTAAAAATTACAGAAGAGGTTGGCCACACAGATCTAATGGCTGCTTAT    6562
gb:MN988713|Organism:Severe      GCAAATAATAGTTTAAAAATTACAGAAGAGGTTGGCCACACAGATCTAATGGCTGCTTAT    6562
gb:LC521925|Organism:Severe      GCAAATAATAGTTTAAAAATTACAGAAGAGGTTGGCCACACAGATCTAATGGCTGCTTAT    6535
gb:MT093571|Organism:Severe      GCAAATAATAGTTTAAAAATTACAGAAGAGGTTGGCCACACAGATCTAATGGCTGCTTAT    6562
gb:MT039887|Organism:Severe      GCAAATAATAGTTTAAAAATTACAGAAGAGGTTGGCCACACAGATCTAATGGCTGCTTAT    6562
gb:MT019530|Organism:Severe      GCAAATAATAGTTTAAAAATTACAGAAGAGGTTGGCCACACAGATCTAATGGCTGCTTAT    6562
gb:MT039888|Organism:Severe      GCAAATAATAGTTTAAAAATTACAGAAGAGGTTGGCCACACAGATCTAATGGCTGCTTAT    6562
gb:LC522972|Organism:Severe      GCAAATAATAGTTTAAAAATTACAGAAGAGGTTGGCCACACAGATCTAATGGCTGCTTAT    6559
gb:MT027063|Organism:Severe      GCAAATAATAGTTTAAAAATTACAGAAGAGGTTGGCCACACAGATCTAATGGCTGCTTAT    6562
gb:MT027062|Organism:Severe      GCAAATAATAGTTTAAAAATTACAGAAGAGGTTGGCCACACAGATCTAATGGCTGCTTAT    6562
gb:MT019529|Organism:Severe      GCAAATAATAGTTTAAAAATTACAGAAGAGGTTGGCCACACAGATCTAATGGCTGCTTAT    6562
gb:MN996529|Organism:Severe      GCAAATAATAGTTTAAAAATTACAGAAGAGGTTGGCCACACAGATCTAATGGCTGCTTAT    6550
gb:MN996531|Organism:Severe      GCAAATAATAGTTTAAAAATTACAGAAGAGGTTGGCCACACAGATCTAATGGCTGCTTAT    6549
gb:MT066176|Organism:Severe      GCAAATAATAGTTTAAAAATTACAGAAGAGGTTGGCCACACAGATCTAATGGCTGCTTAT    6562
gb:MT027064|Organism:Severe      GCAAATAATAGTTTAAAAATTACAGAAGAGGTTGGCCACACAGATCTAATGGCTGCTTAT    6562
gb:MN994468|Organism:Severe      GCAAATAATAGTTTAAAAATTACAGAAGAGGTTGGCCACACAGATCTAATGGCTGCTTAT    6562
gb:MT072688|Organism:Severe      GCAAATAATAGTTTAAAAATTACAGAAGAGGTTGGCCACACAGATCTAATGGCTGCTTAT    6547
gb:MN996527|Organism:Severe      GCAAATAATAGTTTAAAAATTACAGAAGAGGTTGGCCACACAGATCTAATGGCTGCTTAT    6529
gb:MT093631|Organism:Severe      GCAAATAATAGTTTAAAAATTACAGAAGAGGTTGGCCACACAGATCTAATGGCTGCTTAT    6600
gb:MT106053|Organism:Severe      GCAAATAATAGTTTAAAAATTACAGAAGAGGTTGGCCACACAGATCTAATGGCTGCTTAT    6562
gb:MT019533|Organism:Severe      GCAAATAATAGTTTAAAAATTACAGAAGAGGTTGGCCACACAGATCTAATGGCTGCTTAT    6562
gb:MT019531|Organism:Severe      GCAAATAATAGTTTAAAAATTACAGAAGAGGTTGGCCACACAGATCTAATGGCTGCTTAT    6562
gb:MN996528|Organism:Severe      GCAAATAATAGTTTAAAAATTACAGAAGAGGTTGGCCACACAGATCTAATGGCTGCTTAT    6562
gb:MN996530|Organism:Severe      GCAAATAATAGTTTAAAAATTACAGAAGAGGTTGGCCACACAGATCTAATGGCTGCTTAT    6548
gb:MN908947|Organism:Severe      GCAAATAATAGTTTAAAAATTACAGAAGAGGTTGGCCACACAGATCTAATGGCTGCTTAT    6562
gb:MT019532|Organism:Severe      GCAAATAATAGTTTAAAAATTACAGAAGAGGTTGGCCACACAGATCTAATGGCTGCTTAT    6562
                                 ************************************************************

gb:MT020781|Organism:Severe      GTAGACAATTCTAGTCTTACTATTAAGAAACCTAATGAATTATCTAGAGTATTAGGTTTG    6610
gb:MT007544|Organism:Severe      GTAGACAATTCTAGTCTTACTATTAAGAAACCTAATGAATTATCTAGAGTATTAGGTTTG    6622
gb:MN994467|Organism:Severe      GTAGACAATTCTAGTCTTACTATTAAGAAACCTAATGAATTATCTAGAGTATTAGGTTTG    6622
gb:MT044257|Organism:Severe      GTAGACAATTCTAGTCTTACTATTAAGAAACCTAATGAATTATCTAGAGTATTAGGTTTG    6622
gb:MT106054|Organism:Severe      GTAGACAATTCTAGTCTTACTATTAAGAAACCTAATGAATTATCTAGAGTATTAGGTTTG    6622
gb:MT049951|Organism:Severe      GTAGACAATTCTAGTCTTACTATTAAGAAACCTAATGAATTATCTAGAGTATTAGGTTTG    6622
```

```
gb:MN975262|Organism:Severe      GTAGACAATTCTAGTCTTACTATTAAGAAACCTAATGAATTATCTAGAGTATTAGGTTTG    6622
gb:MT106052|Organism:Severe      GTAGACAATTCTAGTCTTACTATTAAGAAACCTAATGAATTATCTAGAGTATTAGGTTTG    6622
gb:LC522975|Organism:Severe      GTAGACAATTCTAGTCTTACTATTAAGAAACCTAATGAATTATCTAGAGTATTAGGTTTG    6619
gb:LC522973|Organism:Severe      GTAGACAATTCTAGTCTTACTATTAAGAAACCTAATGAATTATCTAGAGTATTAGGTTTG    6619
gb:LC522974|Organism:Severe      GTAGACAATTCTAGTCTTACTATTAAGAAACCTAATGAATTATCTAGAGTATTAGGTTTG    6619
gb:MN985325|Organism:Severe      GTAGACAATTCTAGTCTTACTATTAAGAAACCTAATGAATTATCTAGAGTATTAGGTTTG    6622
gb:MT020881|Organism:Severe      GTAGACAATTCTAGTCTTACTATTAAGAAACCTAATGAATTATCTAGAGTATTAGGTTTG    6622
gb:MT020880|Organism:Severe      GTAGACAATTCTAGTCTTACTATTAAGAAACCTAATGAATTATCTAGAGTATTAGGTTTG    6622
gb:MT066175|Organism:Severe      GTAGACAATTCTAGTCTTACTATTAAGAAACCTAATGAATTATCTAGAGTATTAGGTTTG    6622
gb:MN997409|Organism:Severe      GTAGACAATTCTAGTCTTACTATTAAGAAACCTAATGAATTATCTAGAGTATTAGGTTTG    6622
gb:MN938384|Organism:Severe      GTAGACAATTCTAGTCTTACTATTAAGAAACCTAATGAATTATCTAGAGTATTAGGTTTG    6590
gb:MT044258|Organism:Severe      GTAGACAATTCTAGTCTTACTATTAAGAAACCTAATGAATTATCTAGAGTATTAGGTTTG    6598
gb:MT039890|Organism:Severe      GTAGACAATTCTAGTCTTACTATTAAGAAACCTAATGAATTATCTAGAGTATTAGGTTTG    6622
gb:MN988713|Organism:Severe      GTAGACAATTCTAGTCTTACTATTAAGAAACCTAATGAATTATCTAGAGTATTAGGTTTG    6622
gb:LC521925|Organism:Severe      GTAGACAATTCTAGTCTTACTATTAAGAAACCTAATGAATTATCTAGAGTATTAGGTTTG    6595
gb:MT093571|Organism:Severe      GTAGACAATTCTAGTCTTACTATTAAGAAACCTAATGAATTATCTAGAGTATTAGGTTTG    6622
gb:MT039887|Organism:Severe      GTAGACAATTCTAGTCTTACTATTAAGAAACCTAATGAATTATCTAGAGTATTAGGTTTG    6622
gb:MT019530|Organism:Severe      GTAGACAATTCTAGTCTTACTATTAAGAAACCTAATGAATTATCTAGAGTATTAGGTTTG    6622
gb:MT039888|Organism:Severe      GTAGACAATTCTAGTCTTACTATTAAGAAACCTAATGAATTATCTAGAGTATTAGGTTTG    6622
gb:LC522972|Organism:Severe      GTAGACAATTCTAGTCTTACTATTAAGAAACCTAATGAATTATCTAGAGTATTAGGTTTG    6619
gb:MT027063|Organism:Severe      GTAGACAATTCTAGTCTTACTATTAAGAAACCTAATGAATTATCTAGAGTATTAGGTTTG    6622
gb:MT027062|Organism:Severe      GTAGACAATTCTAGTCTTACTATTAAGAAACCTAATGAATTATCTAGAGTATTAGGTTTG    6622
gb:MT019529|Organism:Severe      GTAGACAATTCTAGTCTTACTATTAAGAAACCTAATGAATTATCTAGAGTATTAGGTTTG    6622
gb:MN996529|Organism:Severe      GTAGACAATTCTAGTCTTACTATTAAGAAACCTAATGAATTATCTAGAGTATTAGGTTTG    6610
gb:MN996531|Organism:Severe      GTAGACAATTCTAGTCTTACTATTAAGAAACCTAATGAATTATCTAGAGTATTAGGTTTG    6609
gb:MT066176|Organism:Severe      GTAGACAATTCTAGTCTTACTATTAAGAAACCTAATGAATTATCTAGAGTATTAGGTTTG    6622
gb:MT027064|Organism:Severe      GTAGACAATTCTAGTCTTACTATTAAGAAACCTAATGAATTATCTAGAGTATTAGGTTTG    6622
gb:MN994468|Organism:Severe      GTAGACAATTCTAGTCTTACTATTAAGAAACCTAATGAATTATCTAGAGTATTAGGTTTG    6622
gb:MT072688|Organism:Severe      GTAGACAATTCTAGTCTTACTATTAAGAAACCTAATGAATTATCTAGAGTATTAGGTTTG    6607
gb:MN996527|Organism:Severe      GTAGACAATTCTAGTCTTACTATTAAGAAACCTAATGAATTATCTAGAGTATTAGGTTTG    6589
gb:MT093631|Organism:Severe      GTAGACAATTCTAGTCTTACTATTAAGAAACCTAATGAATTATCTAGAGTATTAGGTTTG    6660
gb:MT106053|Organism:Severe      GTAGACAATTCTAGTCTTACTATTAAGAAACCTAATGAATTATCTAGAGTATTAGGTTTG    6622
gb:MT019533|Organism:Severe      GTAGACAATTCTAGTCTTACTATTAAGAAACCTAATGAATTATCTAGAGTATTAGGTTTG    6622
gb:MT019531|Organism:Severe      GTAGACAATTCTAGTCTTACTATTAAGAAACCTAATGAATTATCTAGAGTATTAGGTTTG    6622
gb:MN996528|Organism:Severe      GTAGACAATTCTAGTCTTACTATTAAGAAACCTAATGAATTATCTAGAGTATTAGGTTTG    6622
gb:MN996530|Organism:Severe      GTAGACAATTCTAGTCTTACTATTAAGAAACCTAATGAATTATCTAGAGTATTAGGTTTG    6608
gb:MN908947|Organism:Severe      GTAGACAATTCTAGTCTTACTATTAAGAAACCTAATGAATTATCTAGAGTATTAGGTTTG    6622
gb:MT019532|Organism:Severe      GTAGACAATTCTAGTCTTACTATTAAGAAACCTAATGAATTATCTAGAGTATTAGGTTTG    6622
                                 ************************************************************

gb:MT020781|Organism:Severe      AAAACCCTTGCTACTCATGGTTTAGCTGCTGTTAATAGTGTCCCTTGGGATACTATAGCT    6670
gb:MT007544|Organism:Severe      AAAACCCTTGCTACTCATGGTTTAGCTGCTGTTAATAGTGTCCCTTGGGATACTATAGCT    6682
gb:MN994467|Organism:Severe      AAAACCCTTGCTACTCATGGTTTAGCTGCTGTTAATAGTGTCCCTTGGGATACTATAGCT    6682
gb:MT044257|Organism:Severe      AAAACCCTTGCTACTCATGGTTTAGCTGCTGTTAATAGTGTCCCTTGGGATACTATAGCT    6682
gb:MT106054|Organism:Severe      AAAACCCTTGCTACTCATGGTTTAGCTGCTGTTAATAGTGTCCCTTGGGATACTATAGCT    6682
gb:MT049951|Organism:Severe      AAAACCCTTGCTACTCATGGTTTAGCTGCTGTTAATAGTGTCCCTTGGGATACTATAGCT    6682
gb:MN975262|Organism:Severe      AAAACCCTTGCTACTCATGGTTTAGCTGCTGTTAATAGTGTCCCTTGGGATACTATAGCT    6682
gb:MT106052|Organism:Severe      AAAACCCTTGCTACTCATGGTTTAGCTGCTGTTAATAGTGTCCCTTGGGATACTATAGCT    6682
gb:LC522975|Organism:Severe      AAAACCCTTGCTACTCATGGTTTAGCTGCTGTTAATAGTGTCCCTTGGGATACTATAGCT    6679
gb:LC522973|Organism:Severe      AAAACCCTTGCTACTCATGGTTTAGCTGCTGTTAATAGTGTCCCTTGGGATACTATAGCT    6679
gb:LC522974|Organism:Severe      AAAACCCTTGCTACTCATGGTTTAGCTGCTGTTAATAGTGTCCCTTGGGATACTATAGCT    6679
gb:MN985325|Organism:Severe      AAAACCCTTGCTACTCATGGTTTAGCTGCTGTTAATAGTGTCCCTTGGGATACTATAGCT    6682
gb:MT020881|Organism:Severe      AAAACCCTTGCTACTCATGGTTTAGCTGCTGTTAATAGTGTCCCTTGGGATACTATAGCT    6682
gb:MT020880|Organism:Severe      AAAACCCTTGCTACTCATGGTTTAGCTGCTGTTAATAGTGTCCCTTGGGATACTATAGCT    6682
gb:MT066175|Organism:Severe      AAAACCCTTGCTACTCATGGTTTAGCTGCTGTTAATAGTGTCCCTTGGGATACTATAGCT    6682
gb:MN997409|Organism:Severe      AAAACCCTTGCTACTCATGGTTTAGCTGCTGTTAATAGTGTCCCTTGGGATACTATAGCT    6682
gb:MN938384|Organism:Severe      AAAACCCTTGCTACTCATGGTTTAGCTGCTGTTAATAGTGTCCCTTGGGATACTATAGCT    6650
gb:MT044258|Organism:Severe      AAAACCCTTGCTACTCATGGTTTAGCTGCTGTTAATAGTGTCCCTTGGGATACTATAGCT    6658
gb:MT039890|Organism:Severe      AAAACCCTTGCTACTCATGGTTTAGCTGCTGTTAATAGTGTCCCTTGGGATACTATAGCT    6682
gb:MN988713|Organism:Severe      AAAACCCTTGCTACTCATGGTTTAGCTGCTGTTAATAGTGTCCCTTGGGATACTATAGCT    6682
gb:LC521925|Organism:Severe      AAAACCCTTGCTACTCATGGTTTAGCTGCTGTTAATAGTGTCCCTTGGGATACTATAGCT    6655
gb:MT093571|Organism:Severe      AAAACCCTTGCTACTCATGGTTTAGCTGCTGTTAATAGTGTCCCTTGGGATACTATAGCT    6682
gb:MT039887|Organism:Severe      AAAACCCTTGCTACTCATGGTTTAGCTGCTGTTAATAGTGTCCCTTGGGATACTATAGCT    6682
gb:MT019530|Organism:Severe      AAAACCCTTGCTACTCATGGTTTAGCTGCTGTTAATAGTGTCCCTTGGGATACTATAGCT    6682
gb:MT039888|Organism:Severe      AAAACCCTTGCTACTCATGGTTTAGCTGCTGTTAATAGTGTCCCTTGGGATACTATAGCT    6682
```

```
gb:LC522972|Organism:Severe      AAAACCCTTGCTACTCATGGTTTAGCTGCTGTTAATAGTGTCCCTTGGGATACTATAGCT   6679
gb:MT027063|Organism:Severe      AAAACCCTTGCTACTCATGGTTTAGCTGCTGTTAATAGTGTCCCTTGGGATACTATAGCT   6682
gb:MT027062|Organism:Severe      AAAACCCTTGCTACTCATGGTTTAGCTGCTGTTAATAGTGTCCCTTGGGATACTATAGCT   6682
gb:MT019529|Organism:Severe      AAAACCCTTGCTACTCATGGTTTAGCTGCTGTTAATAGTGTCCCTTGGGATACTATAGCT   6682
gb:MN996529|Organism:Severe      AAAACCCTTGCTACTCATGGTTTAGCTGCTGTTAATAGTGTCCCTTGGGATACTATAGCT   6670
gb:MN996531|Organism:Severe      AAAACCCTTGCTACTCATGGTTTAGCTGCTGTTAATAGTGTCCCTTGGGATACTATAGCT   6669
gb:MT066176|Organism:Severe      AAAACCCTTGCTACTCATGGTTTAGCTGCTGTTAATAGTGTCCCTTGGGATACTATAGCT   6682
gb:MT027064|Organism:Severe      AAAACCCTTGCTACTCATGGTTTAGCTGCTGTTAATAGTGTCCCTTGGGATACTATAGCT   6682
gb:MN994468|Organism:Severe      AAAACCCTTGCTACTCATGGTTTAGCTGCTGTTAATAGTGTCCCTTGGGATACTATAGCT   6682
gb:MT072688|Organism:Severe      AAAACCCTTGCTACTCATGGTTTAGCTGCTGTTAATAGTGTCCCTTGGGATACTATAGCT   6667
gb:MN996527|Organism:Severe      AAAACCCTTGCTACTCATGGTTTAGCTGCTGTTAATAGTGTCCCTTGGGATACTATAGCT   6649
gb:MT093631|Organism:Severe      AAAACCCTTGCTACTCATGGTTTAGCTGCTGTTAATAGTGTCCCTTGGGATACTATAGCT   6720
gb:MT106053|Organism:Severe      AAAACCCTTGCTACTCATGGTTTAGCTGCTGTTAATAGTGTCCCTTGGGATACTATAGCT   6682
gb:MT019533|Organism:Severe      AAAACCCTTGCTACTCATGGTTTAGCTGCTGTTAATAGTGTCCCTTGGGATACTATAGCT   6682
gb:MT019531|Organism:Severe      AAAACCCTTGCTACTCATGGTTTAGCTGCTGTTAATAGTGTCCCTTGGGATACTATAGCT   6682
gb:MN996528|Organism:Severe      AAAACCCTTGCTACTCATGGTTTAGCTGCTGTTAATAGTGTCCCTTGGGATACTATAGCT   6682
gb:MN996530|Organism:Severe      AAAACCCTTGCTACTCATGGTTTAGCTGCTGTTAATAGTGTCCCTTGGGATACTATAGCT   6668
gb:MN908947|Organism:Severe      AAAACCCTTGCTACTCATGGTTTAGCTGCTGTTAATAGTGTCCCTTGGGATACTATAGCT   6682
gb:MT019532|Organism:Severe      AAAACCCTTGCTACTCATGGTTTAGCTGCTGTTAATAGTGTCCCTTGGGATACTATAGCT   6682
                                 ************************************************************

gb:MT020781|Organism:Severe      AATTATGCTAAGCCTTTTCTTAACAAAGTTGTTAGTACAACTACTAACATAGTTACACGG   6730
gb:MT007544|Organism:Severe      AATTATGCTAAGCCTTTTCTTAACAAAGTTGTTAGTACAACTACTAACATAGTTACACGG   6742
gb:MN994467|Organism:Severe      AATTATGCTAAGCCTTTTCTTAACAAAGTTGTTAGTACAACTACTAACATAGTTACACGG   6742
gb:MT044257|Organism:Severe      AATTATGCTAAGCCTTTTCTTAACAAAGTTGTTAGTACAACTACTAACATAGTTACACGG   6742
gb:MT106054|Organism:Severe      AATTATGCTAAGCCTTTTCTTAACAAAGTTGTTAGTACAACTACTAACATAGTTACACGG   6742
gb:MT049951|Organism:Severe      AATTATGCTAAGCCTTTTCTTAACAAAGTTGTTAGTACAACTACTAACATAGTTACACGG   6742
gb:MN975262|Organism:Severe      AATTATGCTAAGCCTTTTCTTAACAAAGTTGTTAGTACAACTACTAACATAGTTACACGG   6742
gb:MT106052|Organism:Severe      AATTATGCTAAGCCTTTTCTTAACAAAGTTGTTAGTACAACTACTAACATAGTTACACGG   6742
gb:LC522975|Organism:Severe      AATTATGCTAAGCCTTTTCTTAACAAAGTTGTTAGTACAACTACTAACATAGTTACACGG   6739
gb:LC522973|Organism:Severe      AATTATGCTAAGCCTTTTCTTAACAAAGTTGTTAGTACAACTACTAACATAGTTACACGG   6739
gb:LC522974|Organism:Severe      AATTATGCTAAGCCTTTTCTTAACAAAGTTGTTAGTACAACTACTAACATAGTTACACGG   6739
gb:MN985325|Organism:Severe      AATTATGCTAAGCCTTTTCTTAACAAAGTTGTTAGTACAACTACTAACATAGTTACACGG   6742
gb:MT020881|Organism:Severe      AATTATGCTAAGCCTTTTCTTAACAAAGTTGTTAGTACAACTACTAACATAGTTACACGG   6742
gb:MT020880|Organism:Severe      AATTATGCTAAGCCTTTTCTTAACAAAGTTGTTAGTACAACTACTAACATAGTTACACGG   6742
gb:MT066175|Organism:Severe      AATTATGCTAAGCCTTTTCTTAACAAAGTTGTTAGTACAACTACTAACATAGTTACACGG   6742
gb:MN997409|Organism:Severe      AATTATGCTAAGCCTTTTCTTAACAAAGTTGTTAGTACAACTACTAACATAGTTACACGG   6742
gb:MN938384|Organism:Severe      AATTATGCTAAGCCTTTTCTTAACAAAGTTGTTAGTACAACTACTAACATAGTTACACGG   6710
gb:MT044258|Organism:Severe      AATTATGCTAAGCCTTTTCTTAACAAAGTTGTTAGTACAACTACTAACATAGTTACACGG   6718
gb:MT039890|Organism:Severe      AATTATGCTAAGCCTTTTCTTAACAAAGTTGTTAGTACAACTACTAACATAGTTACACGG   6742
gb:MN988713|Organism:Severe      AATTATGCTAAGCCTTTTCTTAACAAAGTTGTTAGTACAACTACTAACATAGTTACACGG   6742
gb:LC521925|Organism:Severe      AATTATGCTAAGCCTTTTCTTAACAAAGTTGTTAGTACAACTACTAACATAGTTACACGG   6715
gb:MT093571|Organism:Severe      AATTATGCTAAGCCTTTTCTTAACAAAGTTGTTAGTACAACTACTAACATAGTTACACGG   6742
gb:MT039887|Organism:Severe      AATTATGCTAAGCCTTTTCTTAACAAAGTTGTTAGTACAACTACTAACATAGTTACACGG   6742
gb:MT019530|Organism:Severe      AATTATGCTAAGCCTTTTCTTAACAAAGTTGTTAGTACAACTACTAACATAGTTACACGG   6742
gb:MT039888|Organism:Severe      AATTATGCTAAGCCTTTTCTTAACAAAGTTGTTAGTACAACTACTAACATAGTTACACGG   6742
gb:LC522972|Organism:Severe      AATTATGCTAAGCCTTTTCTTAACAAAGTTGTTAGTACAACTACTAACATAGTTACACGG   6739
gb:MT027063|Organism:Severe      AATTATGCTAAGCCTTTTCTTAACAAAGTTGTTAGTACAACTACTAACATAGTTACACGG   6742
gb:MT027062|Organism:Severe      AATTATGCTAAGCCTTTTCTTAACAAAGTTGTTAGTACAACTACTAACATAGTTACACGG   6742
gb:MT019529|Organism:Severe      AATTATGCTAAGCCTTTTCTTAACAAAGTTGTTAGTACAACTACTAACATAGTTACACGG   6742
gb:MN996529|Organism:Severe      AATTATGCTAAGCCTTTTCTTAACAAAGTTGTTAGTACAACTACTAACATAGTTACACGG   6730
gb:MN996531|Organism:Severe      AATTATGCTAAGCCTTTTCTTAACAAAGTTGTTAGTACAACTACTAACATAGTTACACGG   6729
gb:MT066176|Organism:Severe      AATTATGCTAAGCCTTTTCTTAACAAAGTTGTTAGTACAACTACTAACATAGTTACACGG   6742
gb:MT027064|Organism:Severe      AATTATGCTAAGCCTTTTCTTAACAAAGTTGTTAGTACAACTACTAACATAGTTACACGG   6742
gb:MN994468|Organism:Severe      AATTATGCTAAGCCTTTTCTTAACAAAGTTGTTAGTACAACTACTAACATAGTTACACGG   6742
gb:MT072688|Organism:Severe      AATTATGCTAAGCCTTTTCTTAACAAAGTTGTTAGTACAACTACTAACATAGTTACACGG   6727
gb:MN996527|Organism:Severe      AATTATGCTAAGCCTTTTCTTAACAAAGTTGTTAGTACAACTACTAACATAGTTACACGG   6709
gb:MT093631|Organism:Severe      AATTATGCTAAGCCTTTTCTTAACAAAGTTGTTAGTACAACTACTAACATAGTTACACGG   6780
gb:MT106053|Organism:Severe      AATTATGCTAAGCCTTTTCTTAACAAAGTTGTTAGTACAACTACTAACATAGTTACACGG   6742
gb:MT019533|Organism:Severe      AATTATGCTAAGCCTTTTCTTAACAAAGTTGTTAGTACAACTACTAACATAGTTACACGG   6742
gb:MT019531|Organism:Severe      AATTATGCTAAGCCTTTTCTTAACAAAGTTGTTAGTACAACTACTAACATAGTTACACGG   6742
gb:MN996528|Organism:Severe      AATTATGCTAAGCCTTTTCTTAACAAAGTTGTTAGTACAACTACTAACATAGTTACACGG   6742
gb:MN996530|Organism:Severe      AATTATGCTAAGCCTTTTCTTAACAAAGTTGTTAGTACAACTACTAACATAGTTACACGG   6728
gb:MN908947|Organism:Severe      AATTATGCTAAGCCTTTTCTTAACAAAGTTGTTAGTACAACTACTAACATAGTTACACGG   6742
gb:MT019532|Organism:Severe      AATTATGCTAAGCCTTTTCTTAACAAAGTTGTTAGTACAACTACTAACATAGTTACACGG   6742
```

```
                                  ************************************************************

gb:MT020781|Organism:Severe      TGTTTAAACCGTGTTTGTANNNNNNNNNNNNNNNNNNNNTTTACTTTATTGCTACAATTG   6790
gb:MT007544|Organism:Severe      TGTTTAAACCGTGTTTGTACTAATTATATGCCTTATTTCTTTACTTTATTGCTACAATTG   6802
gb:MN994467|Organism:Severe      TGTTTAAACCGTGTTTGTACTAATTATATGCCTTATTTCTTTACTTTATTGCTACAATTG   6802
gb:MT044257|Organism:Severe      TGTTTAAACCGTGTTTGTACTAATTATATGCCTTATTTCTTTACTTTATTGCTACAATTG   6802
gb:MT106054|Organism:Severe      TGTTTAAACCGTGTTTGTACTAATTATATGCCTTATTTCTTTACTTTATTGCTACAATTG   6802
gb:MT049951|Organism:Severe      TGTTTAAACCGTGTTTGTACTAATTATATGCCTTATTTCTTTACTTTATTGCTACAATTG   6802
gb:MN975262|Organism:Severe      TGTTTAAACCGTGTTTGTACTAATTATATGCCTTATTTCTTTACTTTATTGCTACAATTG   6802
gb:MT106052|Organism:Severe      TGTTTAAACCGTGTTTGTACTAATTATATGCCTTATTTCTTTACTTTATTGCTACAATTG   6802
gb:LC522975|Organism:Severe      TGTTTAAACCGTGTTTGTACTAATTATATGCCTTATTTCTTTACTTTATTGCTACAATTG   6799
gb:LC522973|Organism:Severe      TGTTTAAACCGTGTTTGTACTAATTATATGCCTTATTTCTTTACTTTATTGCTACAATTG   6799
gb:LC522974|Organism:Severe      TGTTTAAACCGTGTTTGTACTAATTATATGCCTTATTTCTTTACTTTATTGCTACAATTG   6799
gb:MN985325|Organism:Severe      TGTTTAAACCGTGTTTGTACTAATTATATGCCTTATTTCTTTACTTTATTGCTACAATTG   6802
gb:MT020881|Organism:Severe      TGTTTAAACCGTGTTTGTACTAATTATATGCCTTATTTCTTTACTTTATTGCTACAATTG   6802
gb:MT020880|Organism:Severe      TGTTTAAACCGTGTTTGTACTAATTATATGCCTTATTTCTTTACTTTATTGCTACAATTG   6802
gb:MT066175|Organism:Severe      TGTTTAAACCGTGTTTGTACTAATTATATGCCTTATTTCTTTACTTTATTGCTACAATTG   6802
gb:MN997409|Organism:Severe      TGTTTAAACCGTGTTTGTACTAATTATATGCCTTATTTCTTTACTTTATTGCTACAATTG   6802
gb:MN938384|Organism:Severe      TGTTTAAACCGTGTTTGTACTAATTATATGCCTTATTTCTTTACTTTATTGCTACAATTG   6770
gb:MT044258|Organism:Severe      TGTTTAAACCGTGTTTGTACTAATTATATGCCTTATTTCTTTACTTTATTGCTACAATTG   6778
gb:MT039890|Organism:Severe      TGTTTAAACCGTGTTTGTACTAATTATATGCCTTATTTCTTTACTTTATTGCTACAATTG   6802
gb:MN988713|Organism:Severe      TGTTTAAACCGTGTTTGTACTAATTATATGCCTTATTTCTTTACTTTATTGCTACAATTG   6802
gb:LC521925|Organism:Severe      TGTTTAAACCGTGTTTGTACTAATTATATGCCTTATTTCTTTACTTTATTGCTACAATTG   6775
gb:MT093571|Organism:Severe      TGTTTAAACCGTGTTTGTACTAATTATATGCCTTATTTCTTTACTTTATTGCTACAATTG   6802
gb:MT039887|Organism:Severe      TGTTTAAACCGTGTTTGTACTAATTATATGCCTTATTTCTTTACTTTATTGCTACAATTG   6802
gb:MT019530|Organism:Severe      TGTTTAAACCGTGTTTGTACTAATTATATGCCTTATTTCTTTACTTTATTGCTACAATTG   6802
gb:MT039888|Organism:Severe      TGTTTAAACCGTGTTTGTACTAATTATATGCCTTATTTCTTTACTTTATTGCTACAATTG   6802
gb:LC522972|Organism:Severe      TGTTTAAACCGTGTTTGTACTAATTATATGCCTTATTTCTTTACTTTATTGCTACAATTG   6799
gb:MT027063|Organism:Severe      TGTTTAAACCGTGTTTGTACTAATTATATGCCTTATTTCTTTACTTTATTGCTACAATTG   6802
gb:MT027062|Organism:Severe      TGTTTAAACCGTGTTTGTACTAATTATATGCCTTATTTCTTTACTTTATTGCTACAATTG   6802
gb:MT019529|Organism:Severe      TGTTTAAACCGTGTTTGTACTAATTATATGCCTTATTTCTTTACTTTATTGCTACAATTG   6802
gb:MN996529|Organism:Severe      TGTTTAAACCGTGTTTGTACTAATTATATGCCTTATTTCTTTACTTTATTGCTACAATTG   6790
gb:MN996531|Organism:Severe      TGTTTAAACCGTGTTTGTACTAATTATATGCCTTATTTCTTTACTTTATTGCTACAATTG   6789
gb:MT066176|Organism:Severe      TGTTTAAACCGTGTTTGTACTAATTATATGCCTTATTTCTTTACTTTATTGCTACAATTG   6802
gb:MT027064|Organism:Severe      TGTTTAAACCGTGTTTGTACTAATTATATGCCTTATTTCTTTACTTTATTGCTACAATTG   6802
gb:MN994468|Organism:Severe      TGTTTAAACCGTGTTTGTACTAATTATATGCCTTATTTCTTTACTTTATTGCTACAATTG   6802
gb:MT072688|Organism:Severe      TGTTTAAACCGTGTTTGTACTAATTATATGCCTTATTTCTTTACTTTATTGCTACAATTG   6787
gb:MN996527|Organism:Severe      TGTTTAAACCGTGTTTGTACTAATTATATGCCTTATTTCTTTACTTTATTGCTACAATTG   6769
gb:MT093631|Organism:Severe      TGTTTAAACCGTGTTTGTACTAATTATATGCCTTATTTCTTTACTTTATTGCTACAATTG   6840
gb:MT106053|Organism:Severe      TGTTTAAACCGTGTTTGTACTAATTATATGCCTTATTTCTTTACTTTATTGCTACAATTG   6802
gb:MT019533|Organism:Severe      TGTTTAAACCGTGTTTGTACTAATTATATGCCTTATTTCTTTACTTTATTGCTACAATTG   6802
gb:MT019531|Organism:Severe      TGTTTAAACCGTGTTTGTACTAATTATATGCCTTATTTCTTTACTTTATTGCTACAATTG   6802
gb:MN996528|Organism:Severe      TGTTTAAACCGTGTTTGTACTAATTATATGCCTTATTTCTTTACTTTATTGCTACAATTG   6802
gb:MN996530|Organism:Severe      TGTTTAAACCGTGTTTGTACTAATTATATGCCTTATTTCTTTACTTTATTGCTACAATTG   6788
gb:MN908947|Organism:Severe      TGTTTAAACCGTGTTTGTACTAATTATATGCCTTATTTCTTTACTTTATTGCTACAATTG   6802
gb:MT019532|Organism:Severe      TGTTTAAACCGTGTTTGTACTAATTATATGCCTTATTTCTTTACTTTATTGCTACAATTG   6802
                                 *******************                    *********************

gb:MT020781|Organism:Severe      TGTACTTTTACTAGAAGTACAAATTCTAGAATTAAAGCATCTATGCCGACTACTATAGCA   6850
gb:MT007544|Organism:Severe      TGTACTTTTACTAGAAGTACAAATTCTAGAATTAAAGCATCTATGCCGACTACTATAGCA   6862
gb:MN994467|Organism:Severe      TGTACTTTTACTAGAAGTACAAATTCTAGAATTAAAGCATCTATGCCGACTACTATAGCA   6862
gb:MT044257|Organism:Severe      TGTACTTTTACTAGAAGTACAAATTCTAGAATTAAAGCATCTATGCCGACTACTATAGCA   6862
gb:MT106054|Organism:Severe      TGTACTTTTACTAGAAGTACAAATTCTAGAATTAAAGCATCTATGCCGACTACTATAGCA   6862
gb:MT049951|Organism:Severe      TGTACTTTTACTAGAAGTACAAATTCTAGAATTAAAGCATCTATGCCGACTACTATAGCA   6862
gb:MN975262|Organism:Severe      TGTACTTTTACTAGAAGTACAAATTCTAGAATTAAAGCATCTATGCCGACTACTATAGCA   6862
gb:MT106052|Organism:Severe      TGTACTTTTACTAGAAGTACAAATTCTAGAATTAAAGCATCTATGCCGACTACTATAGCA   6862
gb:LC522975|Organism:Severe      TGTACTTTTACTAGAAGTACAAATTCTAGAATTAAAGCATCTATGCCGACTACTATAGCA   6859
gb:LC522973|Organism:Severe      TGTACTTTTACTAGAAGTACAAATTCTAGAATTAAAGCATCTATGCCGACTACTATAGCA   6859
gb:LC522974|Organism:Severe      TGTACTTTTACTAGAAGTACAAATTCTAGAATTAAAGCATCTATGCCGACTACTATAGCA   6859
gb:MN985325|Organism:Severe      TGTACTTTTACTAGAAGTACAAATTCTAGAATTAAAGCATCTATGCCGACTACTATAGCA   6862
gb:MT020881|Organism:Severe      TGTACTTTTACTAGAAGTACAAATTCTAGAATTAAAGCATCTATGCCGACTACTATAGCA   6862
gb:MT020880|Organism:Severe      TGTACTTTTACTAGAAGTACAAATTCTAGAATTAAAGCATCTATGCCGACTACTATAGCA   6862
gb:MT066175|Organism:Severe      TGTACTTTTACTAGAAGTACAAATTCTAGAATTAAAGCATCTATGCCGACTACTATAGCA   6862
gb:MN997409|Organism:Severe      TGTACTTTTACTAGAAGTACAAATTCTAGAATTAAAGCATCTATGCCGACTACTATAGCA   6862
gb:MN938384|Organism:Severe      TGTACTTTTACTAGAAGTACAAATTCTAGAATTAAAGCATCTATGCCGACTACTATAGCA   6830
```

```
gb:MT044258|Organism:Severe      TGTACTTTTACTAGAAGTACAAATTCTAGAATTAAAGCATCTATGCCGACTACTATAGCA   6838
gb:MT039890|Organism:Severe      TGTACTTTTACTAGAAGTACAAATTCTAGAATTAAAGCATCTATGCCGACTACTATAGCA   6862
gb:MN988713|Organism:Severe      TGTACTTTTACTAGAAGTACAAATTCTAGAATTAAAGCATCTATGCCGACTACTATAGCA   6862
gb:LC521925|Organism:Severe      TGTACTTTTACTAGAAGTACAAATTCTAGAATTAAAGCATCTATGCCGACTACTATAGCA   6835
gb:MT093571|Organism:Severe      TGTACTTTTACTAGAAGTACAAATTCTAGAATTAAAGCATCTATGCCGACTACTATAGCA   6862
gb:MT039887|Organism:Severe      TGTACTTTTACTAGAAGTACAAATTCTAGAATTAAAGCATCTATGCCGACTACTATAGCA   6862
gb:MT019530|Organism:Severe      TGTACTTTTACTAGAAGTACAAATTCTAGAATTAAAGCATCTATGCCGACTACTATAGCA   6862
gb:MT039888|Organism:Severe      TGTACTTTTACTAGAAGTACAAATTCTAGAATTAAAGCATCTATGCCGACTACTATAGCA   6862
gb:LC522972|Organism:Severe      TGTACTTTTACTAGAAGTACAAATTCTAGAATTAAAGCATCTATGCCGACTACTATAGCA   6859
gb:MT027063|Organism:Severe      TGTACTTTTACTAGAAGTACAAATTCTAGAATTAAAGCATCTATGCCGACTACTATAGCA   6862
gb:MT027062|Organism:Severe      TGTACTTTTACTAGAAGTACAAATTCTAGAATTAAAGCATCTATGCCGACTACTATAGCA   6862
gb:MT019529|Organism:Severe      TGTACTTTTACTAGAAGTACAAATTCTAGAATTAAAGCATCTATGCCGACTACTATAGCA   6862
gb:MN996529|Organism:Severe      TGTACTTTTACTAGAAGTACAAATTCTAGAATTAAAGCATCTATGCCGACTACTATAGCA   6850
gb:MN996531|Organism:Severe      TGTACTTTTACTAGAAGTACAAATTCTAGAATTAAAGCATCTATGCCGACTACTATAGCA   6849
gb:MT066176|Organism:Severe      TGTACTTTTACTAGAAGTACAAATTCTAGAATTAAAGCATCTATGCCGACTACTATAGCA   6862
gb:MT027064|Organism:Severe      TGTACTTTTACTAGAAGTACAAATTCTAGAATTAAAGCATCTATGCCGACTACTATAGCA   6862
gb:MN994468|Organism:Severe      TGTACTTTTACTAGAAGTACAAATTCTAGAATTAAAGCATCTATGCCGACTACTATAGCA   6862
gb:MT072688|Organism:Severe      TGTACTTTTACTAGAAGTACAAATTCTAGAATTAAAGCATCTATGCCGACTACTATAGCA   6847
gb:MN996527|Organism:Severe      TGTACTTTTACTAGAAGTACAAATTCTAGAATTAAAGCATCTATGCCGACTACTATAGCA   6829
gb:MT093631|Organism:Severe      TGTACTTTTACTAGAAGTACAAATTCTAGAATTAAAGCATCTATGCCGACTACTATAGCA   6900
gb:MT106053|Organism:Severe      TGTACTTTTACTAGAAGTACAAATTCTAGAATTAAAGCATCTATGCCGACTACTATAGCA   6862
gb:MT019533|Organism:Severe      TGTACTTTTACTAGAAGTACAAATTCTAGAATTAAAGCATCTATGCCGACTACTATAGCA   6862
gb:MT019531|Organism:Severe      TGTACTTTTACTAGAAGTACAAATTCTAGAATTAAAGCATCTATGCCGACTACTATAGCA   6862
gb:MN996528|Organism:Severe      TGTACTTTTACTAGAAGTACAAATTCTAGAATTAAAGCATCTATGCCGACTACTATAGCA   6862
gb:MN996530|Organism:Severe      TGTACTTTTACTAGAAGTACAAATTCTAGAATTAAAGCATCTATGCCGACTACTATAGCA   6848
gb:MN908947|Organism:Severe      TGTACTTTTACTAGAAGTACAAATTCTAGAATTAAAGCATCTATGCCGACTACTATAGCA   6862
gb:MT019532|Organism:Severe      TGTACTTTTACTAGAAGTACAAATTCTAGAATTAAAGCATCTATGCCGACTACTATAGCA   6862
                                 ************************************************************

gb:MT020781|Organism:Severe      AAGAATACTGTTAAGAGTGTCGGTAAATTTTGTCTAGAGGCTTCATTTAATTATTTGAAG   6910
gb:MT007544|Organism:Severe      AAGAATACTGTTAAGAGTGTCGGTAAATTTTGTCTAGAGGCTTCATTTAATTATTTGAAG   6922
gb:MN994467|Organism:Severe      AAGAATACTGTTAAGAGTGTCGGTAAATTTTGTCTAGAGGCTTCATTTAATTATTTGAAG   6922
gb:MT044257|Organism:Severe      AAGAATACTGTTAAGAGTGTCGGTAAATTTTGTCTAGAGGCTTCATTTAATTATTTGAAG   6922
gb:MT106054|Organism:Severe      AAGAATACTGTTAAGAGTGTCGGTAAATTTTGTCTAGAGGCTTCATTTAATTATTTGAAG   6922
gb:MT049951|Organism:Severe      AAGAATACTGTTAAGAGTGTCGGTAAATTTTGTCTAGAGGCTTCATTTAATTATTTGAAG   6922
gb:MN975262|Organism:Severe      AAGAATACTGTTAAGAGTGTCGGTAAATTTTGTCTAGAGGCTTCATTTAATTATTTGAAG   6922
gb:MT106052|Organism:Severe      AAGAATACTGTTAAGAGTGTCGGTAAATTTTGTCTAGAGGCTTCATTTAATTATTTGAAG   6922
gb:LC522975|Organism:Severe      AAGAATACTGTTAAGAGTGTCGGTAAATTTTGTCTAGAGGCTTCATTTAATTATTTGAAG   6919
gb:LC522973|Organism:Severe      AAGAATACTGTTAAGAGTGTCGGTAAATTTTGTCTAGAGGCTTCATTTAATTATTTGAAG   6919
gb:LC522974|Organism:Severe      AAGAATACTGTTAAGAGTGTCGGTAAATTTTGTCTAGAGGCTTCATTTAATTATTTGAAG   6919
gb:MN985325|Organism:Severe      AAGAATACTGTTAAGAGTGTCGGTAAATTTTGTCTAGAGGCTTCATTTAATTATTTGAAG   6922
gb:MT020881|Organism:Severe      AAGAATACTGTTAAGAGTGTCGGTAAATTTTGTCTAGAGGCTTCATTTAATTATTTGAAG   6922
gb:MT020880|Organism:Severe      AAGAATACTGTTAAGAGTGTCGGTAAATTTTGTCTAGAGGCTTCATTTAATTATTTGAAG   6922
gb:MT066175|Organism:Severe      AAGAATACTGTTAAGAGTGTCGGTAAATTTTGTCTAGAGGCTTCATTTAATTATTTGAAG   6922
gb:MN997409|Organism:Severe      AAGAATACTGTTAAGAGTGTCGGTAAATTTTGTCTAGAGGCTTCATTTAATTATTTGAAG   6922
gb:MN938384|Organism:Severe      AAGAATACTGTTAAGAGTGTCGGTAAATTTTGTCTAGAGGCTTCATTTAATTATTTGAAG   6890
gb:MT044258|Organism:Severe      AAGAATACTGTTAAGAGTGTCGGTAAATTTTGTCTAGAGGCTTCATTTAATTATTTGAAG   6898
gb:MT039890|Organism:Severe      AAGAATACTGTTAAGAGTGTCGGTAAATTTTGTCTAGAGGCTTCATTTAATTATTTGAAG   6922
gb:MN988713|Organism:Severe      AAGAATACTGTTAAGAGTGTCGGTAAATTTTGTCTAGAGGCTTCATTTAATTATTTGAAG   6922
gb:LC521925|Organism:Severe      AAGAATACTGTTAAGAGTGTCGGTAAATTTTGTCTAGAGGCTTCATTTAATTATTTGAAG   6895
gb:MT093571|Organism:Severe      AAGAATACTGTTAAGAGTGTCGGTAAATTTTGTCTAGAGGCTTCATTTAATTATTTGAAG   6922
gb:MT039887|Organism:Severe      AAGAATACTGTTAAGAGTGTCGGTAAATTTTGTCTAGAGGCTTCATTTAATTATTTGAAG   6922
gb:MT019530|Organism:Severe      AAGAATACTGTTAAGAGTGTCGGTAAATTTTGTCTAGAGGCTTCATTTAATTATTTGAAG   6922
gb:MT039888|Organism:Severe      AAGAATACTGTTAAGAGTGTCGGTAAATTTTGTCTAGAGGCTTCATTTAATTATTTGAAG   6922
gb:LC522972|Organism:Severe      AAGAATACTGTTAAGAGTGTCGGTAAATTTTGTCTAGAGGCTTCATTTAATTATTTGAAG   6919
gb:MT027063|Organism:Severe      AAGAATACTGTTAAGAGTGTCGGTAAATTTTGTCTAGAGGCTTCATTTAATTATTTGAAG   6922
gb:MT027062|Organism:Severe      AAGAATACTGTTAAGAGTGTCGGTAAATTTTGTCTAGAGGCTTCATTTAATTATTTGAAG   6922
gb:MT019529|Organism:Severe      AAGAATACTGTTAAGAGTGTCGGTAAATTTTGTCTAGAGGCTTCATTTAATTATTTGAAG   6922
gb:MN996529|Organism:Severe      AAGAATACTGTTAAGAGTGTCGGTAAATTTTGTCTAGAGGCTTCATTTAATTATTTGAAG   6910
gb:MN996531|Organism:Severe      AAGAATACTGTTAAGAGTGTCGGTAAATTTTGTCTAGAGGCTTCATTTAATTATTTGAAG   6909
gb:MT066176|Organism:Severe      AAGAATACTGTTAAGAGTGTCGGTAAATTTTGTCTAGAGGCTTCATTTAATTATTTGAAG   6922
gb:MT027064|Organism:Severe      AAGAATACTGTTAAGAGTGTCGGTAAATTTTGTCTAGAGGCTTCATTTAATTATTTGAAG   6922
gb:MN994468|Organism:Severe      AAGAATACTGTTAAGAGTGTCGGTAAATTTTGTCTAGAGGCTTCATTTAATTATTTGAAG   6922
gb:MT072688|Organism:Severe      AAGAATACTGTTAAGAGTGTCGGTAAATTTTGTCTAGAGGCTTCATTTAATTATTTGAAG   6907
gb:MN996527|Organism:Severe      AAGAATACTGTTAAGAGTGTCGGTAAATTTTGTCTAGAGGCTTCATTTAATTATTTGAAG   6889
```

```
gb:MT093631|Organism:Severe      AAGAATACTGTTAAGAGTGTCGGTAAATTTTGTCTAGAGGCTTCATTTAATTATTTGAAG   6960
gb:MT106053|Organism:Severe      AAGAATACTGTTAAGAGTGTCGGTAAATTTTGTCTAGAGGCTTCATTTAATTATTTGAAG   6922
gb:MT019533|Organism:Severe      AAGAATACTGTTAAGAGTGTCGGTAAATTTTGTCTAGAGGCTTCATTTAATTATTTGAAG   6922
gb:MT019531|Organism:Severe      AAGAATACTGTTAAGAGTGTCGGTAAATTTTGTCTAGAGGCTTCATTTAATTATTTGAAG   6922
gb:MN996528|Organism:Severe      AAGAATACTGTTAAGAGTGTCGGTAAATTTTGTCTAGAGGCTTCATTTAATTATTTGAAG   6922
gb:MN996530|Organism:Severe      AAGAATACTGTTAAGAGTGTCGGTAAATTTTGTCTAGAGGCTTCATTTAATTATTTGAAG   6908
gb:MN908947|Organism:Severe      AAGAATACTGTTAAGAGTGTCGGTAAATTTTGTCTAGAGGCTTCATTTAATTATTTGAAG   6922
gb:MT019532|Organism:Severe      AAGAATACTGTTAAGAGTGTCGGTAAATTTTGTCTAGAGGCTTCATTTAATTATTTGAAG   6922
                                 ************************************************************

gb:MT020781|Organism:Severe      TCACCTAATTTTTCTAAACTGATAAATATTATAATTTGGTTTTTACTATTAAGTGTTTGC   6970
gb:MT007544|Organism:Severe      TCACCTAATTTTTCTAAACTGATAAATATTATAATTTGGTTTTTACTATTAAGTGTTTGC   6982
gb:MN994467|Organism:Severe      TCACCTAATTTTTCTAAACTGATAAATATTATAATTTGGTTTTTACTATTAAGTGTTTGC   6982
gb:MT044257|Organism:Severe      TCACCTAATTTTTCTAAACTGATAAATATTATAATTTGGTTTTTACTATTAAGTGTTTGC   6982
gb:MT106054|Organism:Severe      TCACCTAATTTTTCTAAACTGATAAATATTATAATTTGGTTTTTACTATTAAGTGTTTGC   6982
gb:MT049951|Organism:Severe      TCACCTAATTTTTCTAAACTGATAAATATTATAATTTGGTTTTTACTATTAAGTGTTTGC   6982
gb:MN975262|Organism:Severe      TCACCTAATTTTTCTAAACTGATAAATATTATAATTTGGTTTTTACTATTAAGTGTTTGC   6982
gb:MT106052|Organism:Severe      TCACCTAATTTTTCTAAACTGATAAATATTATAATTTGGTTTTTACTATTAAGTGTTTGC   6982
gb:LC522975|Organism:Severe      TCACCTAATTTTTCTAAACTGATAAATATTATAATTTGGTTTTTACTATTAAGTGTTTGC   6979
gb:LC522973|Organism:Severe      TCACCTAATTTTTCTAAACTGATAAATATTATAATTTGGTTTTTACTATTAAGTGTTTGC   6979
gb:LC522974|Organism:Severe      TCACCTAATTTTTCTAAACTGATAAATATTATAATTTGGTTTTTACTATTAAGTGTTTGC   6979
gb:MN985325|Organism:Severe      TCACCTAATTTTTCTAAACTGATAAATATTATAATTTGGTTTTTACTATTAAGTGTTTGC   6982
gb:MT020881|Organism:Severe      TCACCTAATTTTTCTAAACTGATAAATATTATAATTTGGTTTTTACTATTAAGTGTTTGC   6982
gb:MT020880|Organism:Severe      TCACCTAATTTTTCTAAACTGATAAATATTATAATTTGGTTTTTACTATTAAGTGTTTGC   6982
gb:MT066175|Organism:Severe      TCACCTAATTTTTCTAAACTGATAAATATTATAATTTGGTTTTTACTATTAAGTGTTTGC   6982
gb:MN997409|Organism:Severe      TCACCTAATTTTTCTAAACTGATAAATATTATAATTTGGTTTTTACTATTAAGTGTTTGC   6982
gb:MN938384|Organism:Severe      TCACCTAATTTTTCTAAACTGATAAATATTATAATTTGGTTTTTACTATTAAGTGTTTGC   6950
gb:MT044258|Organism:Severe      TCACCTAATTTTTCTAAACTGATAAATATTATAATTTGGTTTTTACTATTAAGTGTTTGC   6958
gb:MT039890|Organism:Severe      TCACCTAATTTTTCTAAACTGATAAATATTATAATTTGGTTTTTACTATTAAGTGTTTGC   6982
gb:MN988713|Organism:Severe      TCACCTAATTTTTCTAAACTGATAAATATTATAATTTGGTTTTTACTATTAAGTGTTTGC   6982
gb:LC521925|Organism:Severe      TCACCTAATTTTTCTAAACTGATAAATATTATAATTTGGTTTTTACTATTAAGTGTTTGC   6955
gb:MT093571|Organism:Severe      TCACCTAATTTTTCTAAACTGATAAATATTATAATTTGGTTTTTACTATTAAGTGTTTGC   6982
gb:MT039887|Organism:Severe      TCACCTAATTTTTCTAAACTGATAAATATTATAATTTGGTTTTTACTATTAAGTGTTTGC   6982
gb:MT019530|Organism:Severe      TCACCTAATTTTTCTAAACTGATAAATATTATAATTTGGTTTTTACTATTAAGTGTTTGC   6982
gb:MT039888|Organism:Severe      TCACCTAATTTTTCTAAACTGATAAATATTATAATTTGGTTTTTACTATTAAGTGTTTGC   6982
gb:LC522972|Organism:Severe      TCACCTAATTTTTCTAAACTGATAAATATTATAATTTGGTTTTTACTATTAAGTGTTTGC   6979
gb:MT027063|Organism:Severe      TCACCTAATTTTTCTAAACTGATAAATATTATAATTTGGTTTTTACTATTAAGTGTTTGC   6982
gb:MT027062|Organism:Severe      TCACCTAATTTTTCTAAACTGATAAATATTATAATTTGGTTTTTACTATTAAGTGTTTGC   6982
gb:MT019529|Organism:Severe      TCACCTAATTTTTCTAAACTGATAAATATTATAATTTGGTTTTTACTATTAAGTGTTTGC   6982
gb:MN996529|Organism:Severe      TCACCTAATTTTTCTAAACTGATAAATATTATAATTTGGTTTTTACTATTAAGTGTTTGC   6970
gb:MN996531|Organism:Severe      TCACCTAATTTTTCTAAACTGATAAATATTATAATTTGGTTTTTACTATTAAGTGTTTGC   6969
gb:MT066176|Organism:Severe      TCACCTAATTTTTCTAAACTGATAAATATTATAATTTGGTTTTTACTATTAAGTGTTTGC   6982
gb:MT027064|Organism:Severe      TCACCTAATTTTTCTAAACTGATAAATATTATAATTTGGTTTTTACTATTAAGTGTTTGC   6982
gb:MN994468|Organism:Severe      TCACCTAATTTTTCTAAACTGATAAATATTATAATTTGGTTTTTACTATTAAGTGTTTGC   6982
gb:MT072688|Organism:Severe      TCACCTAATTTTTCTAAACTGATAAATATTATAATTTGGTTTTTACTATTAAGTGTTTGC   6967
gb:MN996527|Organism:Severe      TCACCTAATTTTTCTAAACTGATAAATATTATAATTTGGTTTTTACTATTAAGTGTTTGC   6949
gb:MT093631|Organism:Severe      TCACCTAATTTTTCTAAACTGATAAATATTATAATTTGGTTTTTACTATTAAGTGTTTGC   7020
gb:MT106053|Organism:Severe      TCACCTAATTTTTCTAAACTGATAAATATTATAATTTGGTTTTTACTATTAAGTGTTTGC   6982
gb:MT019533|Organism:Severe      TCACCTAATTTTTCTAAACTGATAAATATTATAATTTGGTTTTTACTATTAAGTGTTTGC   6982
gb:MT019531|Organism:Severe      TCACCTAATTTTTCTAAACTGATAAATATTATAATTTGGTTTTTACTATTAAGTGTTTGC   6982
gb:MN996528|Organism:Severe      TCACCTAATTTTTCTAAACTGATAAATATTATAATTTGGTTTTTACTATTAAGTGTTTGC   6982
gb:MN996530|Organism:Severe      TCACCTAATTTTTCTAAACTGATAAATATTATAATTTGGTTTTTACTATTAAGTGTTTGC   6968
gb:MN908947|Organism:Severe      TCACCTAATTTTTCTAAACTGATAAATATTATAATTTGGTTTTTACTATTAAGTGTTTGC   6982
gb:MT019532|Organism:Severe      TCACCTAATTTTTCTAAACTGATAAATATTATAATTTGGTTTTTACTATTAAGTGTTTGC   6982
                                 ************************************************************

gb:MT020781|Organism:Severe      CTAGGTTCTTTAATCTACTCAACCGCTGCTTTAGGTGTTTTAATGTCTAATTTAGGCATG   7030
gb:MT007544|Organism:Severe      CTAGGTTCTTTAATCTACTCAACCGCTGCTTTAGGTGTTTTAATGTCTAATTTAGGCATG   7042
gb:MN994467|Organism:Severe      CTAGGTTCTTTAATCTACTCAACCGCTGCTTTAGGTGTTTTAATGTCTAATTTAGGCATG   7042
gb:MT044257|Organism:Severe      CTAGGTTCTTTAATCTACTCAACCGCTGCTTTAGGTGTTTTAATGTCTAATTTAGGCATG   7042
gb:MT106054|Organism:Severe      CTAGGTTCTTTAATCTACTCAACCGCTGCTTTAGGTGTTTTAATGTCTAATTTAGGCATG   7042
gb:MT049951|Organism:Severe      CTAGGTTCTTTAATCTACTCAACCGCTGCTTTAGGTGTTTTAATGTCTAATTTAGGCATG   7042
gb:MN975262|Organism:Severe      CTAGGTTCTTTAATCTACTCAACCGCTGCTTTAGGTGTTTTAATGTCTAATTTAGGCATG   7042
gb:MT106052|Organism:Severe      CTAGGTTCTTTAATCTACTCAACCGCTGCTTTAGGTGTTTTAATGTCTAATTTAGGCATG   7042
gb:LC522975|Organism:Severe      CTAGGTTCTTTAATCTACTCAACCGCTGCTTTAGGTGTTTTAATGTCTAATTTAGGCATG   7039
```

```
gb:LC522973|Organism:Severe      CTAGGTTCTTTAATCTACTCAACCGCTGCTTTAGGTGTTTTAATGTCTAATTTAGGCATG   7039
gb:LC522974|Organism:Severe      CTAGGTTCTTTAATCTACTCAACCGCTGCTTTAGGTGTTTTAATGTCTAATTTAGGCATG   7039
gb:MN985325|Organism:Severe      CTAGGTTCTTTAATCTACTCAACCGCTGCTTTAGGTGTTTTAATGTCTAATTTAGGCATG   7042
gb:MT020881|Organism:Severe      CTAGGTTCTTTAATCTACTCAACCGCTGCTTTAGGTGTTTTAATGTCTAATTTAGGCATG   7042
gb:MT020880|Organism:Severe      CTAGGTTCTTTAATCTACTCAACCGCTGCTTTAGGTGTTTTAATGTCTAATTTAGGCATG   7042
gb:MT066175|Organism:Severe      CTAGGTTCTTTAATCTACTCAACCGCTGCTTTAGGTGTTTTAATGTCTAATTTAGGCATG   7042
gb:MN997409|Organism:Severe      CTAGGTTCTTTAATCTACTCAACCGCTGCTTTAGGTGTTTTAATGTCTAATTTAGGCATG   7042
gb:MN938384|Organism:Severe      CTAGGTTCTTTAATCTACTCAACCGCTGCTTTAGGTGTTTTAATGTCTAATTTAGGCATG   7010
gb:MT044258|Organism:Severe      CTAGGTTCTTTAATCTACTCAACCGCTGCTTTAGGTGTTTTAATGTCTAATTTAGGCATG   7018
gb:MT039890|Organism:Severe      CTAGGTTCTTTAATCTACTCAACCGCTGCTTTAGGTGTTTTAATGTCTAATTTAGGCATG   7042
gb:MN988713|Organism:Severe      CTAGGTTCTTTAATCTACTCAACCGCTGCTTTAGGTGTTTTAATGTCTAATTTAGGCATG   7042
gb:LC521925|Organism:Severe      CTAGGTTCTTTAATCTACTCAACCGCTGCTTTAGGTGTTTTAATGTCTAATTTAGGCATG   7015
gb:MT093571|Organism:Severe      CTAGGTTCTTTAATCTACTCAACCGCTGCTTTAGGTGTTTTAATGTCTAATTTAGGCATG   7042
gb:MT039887|Organism:Severe      CTAGGTTCTTTAATCTACTCAACCGCTGCTTTAGGTGTTTTAATGTCTAATTTAGGCATG   7042
gb:MT019530|Organism:Severe      CTAGGTTCTTTAATCTACTCAACCGCTGCTTTAGGTGTTTTAATGTCTAATTTAGGCATG   7042
gb:MT039888|Organism:Severe      CTAGGTTCTTTAATCTACTCAACCGCTGCTTTAGGTGTTTTAATGTCTAATTTAGGCATG   7042
gb:LC522972|Organism:Severe      CTAGGTTCTTTAATCTACTCAACCGCTGCTTTAGGTGTTTTAATGTCTAATTTAGGCATG   7039
gb:MT027063|Organism:Severe      CTAGGTTCTTTAATCTACTCAACCGCTGCTTTAGGTGTTTTAATGTCTAATTTAGGCATG   7042
gb:MT027062|Organism:Severe      CTAGGTTCTTTAATCTACTCAACCGCTGCTTTAGGTGTTTTAATGTCTAATTTAGGCATG   7042
gb:MT019529|Organism:Severe      CTAGGTTCTTTAATCTACTCAACCGCTGCTTTAGGTGTTTTAATGTCTAATTTAGGCATG   7042
gb:MN996529|Organism:Severe      CTAGGTTCTTTAATCTACTCAACCGCTGCTTTAAGTGTTTTAATGTCTAATTTAGGCATG   7030
gb:MN996531|Organism:Severe      CTAGGTTCTTTAATCTACTCAACCGCTGCTTTAGGTGTTTTAATGTCTAATTTAGGCATG   7029
gb:MT066176|Organism:Severe      CTAGGTTCTTTAATCTACTCAACCGCTGCTTTAGGTGTTTTAATGTCTAATTTAGGCATG   7042
gb:MT027064|Organism:Severe      CTAGGTTCTTTAATCTACTCAACCGCTGCTTTAGGTGTTTTAATGTCTAATTTAGGCATG   7042
gb:MN994468|Organism:Severe      CTAGGTTCTTTAATCTACTCAACCGCTGCTTTAGGTGTTTTAATGTCTAATTTAGGCATG   7042
gb:MT072688|Organism:Severe      CTAGGTTCTTTAATCTACTCAACCGCTGCTTTAGGTGTTTTAATGTCTAATTTAGGCATG   7027
gb:MN996527|Organism:Severe      CTAGGTTCTTTAATCTACTCAACCGCTGCTTTAGGTGTTTTAATGTCTAATTTAGGCATG   7009
gb:MT093631|Organism:Severe      CTAGGTTCTTTAATCTACTCAACCGCTGCTTTAGGTGTTTTAATGTCTAATTTAGGCATG   7080
gb:MT106053|Organism:Severe      CTAGGTTCTTTAATCTACTCAACCGCTGCTTTAGGTGTTTTAATGTCTAATTTAGGCATG   7042
gb:MT019533|Organism:Severe      CTAGGTTCTTTAATCTACTCAACCGCTGCTTTAGGTGTTTTAATGTCTAATTTAGGCATG   7042
gb:MT019531|Organism:Severe      CTAGGTTCTTTAACCTACTCAACCGCTGCTTTAGGTGTTTTAATGTCTAATTTAGGCATG   7042
gb:MN996528|Organism:Severe      CTAGGTTCTTTAATCTACTCAACCGCTGCTTTAGGTGTTTTAATGTCTAATTTAGGCATG   7042
gb:MN996530|Organism:Severe      CTAGGTTCTTTAATCTACTCAACCGCTGCTTTAGGTGTTTTAATGTCTAATTTAGGCATG   7028
gb:MN908947|Organism:Severe      CTAGGTTCTTTAATCTACTCAACCGCTGCTTTAGGTGTTTTAATGTCTAATTTAGGCATG   7042
gb:MT019532|Organism:Severe      CTAGGTTCTTTAATCTACTCAACCGCTGCTTTAGGTGTTTTAATGTCTAATTTAGGCATG   7042
                                 ************* ******************* **************************

gb:MT020781|Organism:Severe      CCTTCTTACTGTACTGGTTACAGAGAAGGCTATTTGAACTCTACTAATGTCACTATTGCA   7090
gb:MT007544|Organism:Severe      CCTTCTTACTGTACTGGTTACAGAGAAGGCTATTTGAACTCTACTAATGTCACTATTGCA   7102
gb:MN994467|Organism:Severe      CCTTCTTACTGTACTGGTTACAGAGAAGGCTATTTGAACTCTACTAATGTCACTATTGCA   7102
gb:MT044257|Organism:Severe      CCTTCTTACTGTACTGGTTACAGAGAAGGCTATTTGAACTCTACTAATGTCACTATTGCA   7102
gb:MT106054|Organism:Severe      CCTTCTTACTGTACTGGTTACAGAGAAGGCTATTTGAACTCTACTAATGTCACTATTGCA   7102
gb:MT049951|Organism:Severe      CCTTCTTACTGTACTGGTTACAGAGAAGGCTATTTGAACTCTACTAATGTCACTATTGCA   7102
gb:MN975262|Organism:Severe      CCTTCTTACTGTACTGGTTACAGAGAAGGCTATTTGAACTCTACTAATGTCACTATTGCA   7102
gb:MT106052|Organism:Severe      CCTTCTTACTGTACTGGTTACAGAGAAGGCTATTTGAACTCTACTAATGTCACTATTGCA   7102
gb:LC522975|Organism:Severe      CCTTCTTACTGTACTGGTTACAGAGAAGGCTATTTGAACTCTACTAATGTCACTATTGCA   7099
gb:LC522973|Organism:Severe      CCTTCTTACTGTACTGGTTACAGAGAAGGCTATTTGAACTCTACTAATGTCACTATTGCA   7099
gb:LC522974|Organism:Severe      CCTTCTTACTGTACTGGTTACAGAGAAGGCTATTTGAACTCTACTAATGTCACTATTGCA   7099
gb:MN985325|Organism:Severe      CCTTCTTACTGTACTGGTTACAGAGAAGGCTATTTGAACTCTACTAATGTCACTATTGCA   7102
gb:MT020881|Organism:Severe      CCTTCTTACTGTACTGGTTACAGAGAAGGCTATTTGAACTCTACTAATGTCACTATTGCA   7102
gb:MT020880|Organism:Severe      CCTTCTTACTGTACTGGTTACAGAGAAGGCTATTTGAACTCTACTAATGTCACTATTGCA   7102
gb:MT066175|Organism:Severe      CCTTCTTACTGTACTGGTTACAGAGAAGGCTATTTGAACTCTACTAATGTCACTATTGCA   7102
gb:MN997409|Organism:Severe      CCTTCTTACTGTACTGGTTACAGAGAAGGCTATTTGAACTCTACTAATGTCACTATTGCA   7102
gb:MN938384|Organism:Severe      CCTTCTTACTGTACTGGTTACAGAGAAGGCTATTTGAACTCTACTAATGTCACTATTGCA   7070
gb:MT044258|Organism:Severe      CCTTCTTACTGTACTGGTTACAGAGAAGGCTATTTGAACTCTACTAATGTCACTATTGCA   7078
gb:MT039890|Organism:Severe      CCTTCTTACTGTACTGGTTACAGAGAAGGCTATTTGAACTCTACTAATGTCACTATTGCA   7102
gb:MN988713|Organism:Severe      CCTTCTTACTGTACTGGTTACAGAGAAGGCTATTTGAACTCTACTAATGTCACTATTGCA   7102
gb:LC521925|Organism:Severe      CCTTCTTACTGTACTGGTTACAGAGAAGGCTATTTGAACTCTACTAATGTCACTATTGCA   7075
gb:MT093571|Organism:Severe      CCTTCTTACTGTACTGGTTACAGAGAAGGCTATTTGAACTCTACTAATGTCACTATTGCA   7102
gb:MT039887|Organism:Severe      CCTTCTTACTGTACTGGTTACAGAGAAGGCTATTTGAACTCTACTAATGTCACTATTGCA   7102
gb:MT019530|Organism:Severe      CCTTCTTACTGTACTGGTTACAGAGAAGGCTATTTGAACTCTACTAATGTCACTATTGCA   7102
gb:MT039888|Organism:Severe      CCTTCTTACTGTACTGGTTACAGAGAAGGCTATTTGAACTCTACTAATGTCACTATTGCA   7102
gb:LC522972|Organism:Severe      CCTTCTTACTGTACTGGTTACAGAGAAGGCTATTTGAACTCTACTAATGTCACTATTGCA   7099
gb:MT027063|Organism:Severe      CCTTCTTACTGTACTGGTTACAGAGAAGGCTATTTGAACTCTACTAATGTCACTATTGCA   7102
gb:MT027062|Organism:Severe      CCTTCTTACTGTACTGGTTACAGAGAAGGCTATTTGAACTCTACTAATGTCACTATTGCA   7102
```

```
gb:MT019529|Organism:Severe      CCTTCTTACTGTACTGGTTACAGAGAAGGCTATTTGAACTCTACTAATGTCACTATTGCA   7102
gb:MN996529|Organism:Severe      CCTTCTTACTGTACTGGTTACAGAGAAGGCTATTTGAACTCTACTAATGTCACTATTGCA   7090
gb:MN996531|Organism:Severe      CCTTCTTACTGTACTGGTTACAGAGAAGGCTATTTGAACTCTACTAATGTCACTATTGCA   7089
gb:MT066176|Organism:Severe      CCTTCTTACTGTACTGGTTACAGAGAAGGCTATTTGAACTCTACTAATGTCACTATTGCA   7102
gb:MT027064|Organism:Severe      CCTTCTTACTGTACTGGTTACAGAGAAGGCTATTTGAACTCTACTAATGTCACTATTGCA   7102
gb:MN994468|Organism:Severe      CCTTCTTACTGTACTGGTTACAGAGAAGGCTATTTGAACTCTACTAATGTCACTATTGCA   7102
gb:MT072688|Organism:Severe      CCTTCTTACTGTACTGGTTACAGAGAAGGCTATTTGAACTCTACTAATGTCACTATTGCA   7087
gb:MN996527|Organism:Severe      CCTTCTTACTGTACTGGTTACAGAGAAGGCTATTTGAACTCTACTAATGTCACTATTGCA   7069
gb:MT093631|Organism:Severe      CCTTCTTACTGTACTGGTTACAGAGAAGGCTATTTGAACTCTACTAATGTCACTATTGCA   7140
gb:MT106053|Organism:Severe      CCTTCTTACTGTACTGGTTACAGAGAAGGCTATTTGAACTCTACTAATGTCACTATTGCA   7102
gb:MT019533|Organism:Severe      CCTTCTTACTGTACTGGTTACAGAGAAGGCTATTTGAACTCTACTAATGTCACTATTGCA   7102
gb:MT019531|Organism:Severe      CCTTCTTACTGTACTGGTTACAGAGAAGGCTATTTGAACTCTACTAATGTCACTATTGCA   7102
gb:MN996528|Organism:Severe      CCTTCTTACTGTACTGGTTACAGAGAAGGCTATTTGAACTCTACTAATGTCACTATTGCA   7102
gb:MN996530|Organism:Severe      CCTTCTTACTGTACTGGTTACAGAGAAGGCTATTTGAACTCTACTAATGTCACTATTGCA   7088
gb:MN908947|Organism:Severe      CCTTCTTACTGTACTGGTTACAGAGAAGGCTATTTGAACTCTACTAATGTCACTATTGCA   7102
gb:MT019532|Organism:Severe      CCTTCTTACTGTACTGGTTACAGAGAAGGCTATTTGAACTCTACTAATGTCACTATTGCA   7102
                                 ************************************************************

gb:MT020781|Organism:Severe      ACCTACTGTACTGGTTCTATACCTTGTAGTGTTTGTCTTAGTGGTTTAGATTCTTTAGAC   7150
gb:MT007544|Organism:Severe      ACCTACTGTACTGGTTCTATACCTTGTAGTGTTTGTCTTAGTGGTTTAGATTCTTTAGAC   7162
gb:MN994467|Organism:Severe      ACCTACTGTACTGGTTCTATACCTTGTAGTGTTTGTCTTAGTGGTTTAGATTCTTTAGAC   7162
gb:MT044257|Organism:Severe      ACCTACTGTACTGGTTCTATACCTTGTAGTGTTTGTCTTAGTGGTTTAGATTCTTTAGAC   7162
gb:MT106054|Organism:Severe      ACCTACTGTACTGGTTCTATACCTTGTAGTGTTTGTCTTAGTGGTTTAGATTCTTTAGAC   7162
gb:MT049951|Organism:Severe      ACCTACTGTACTGGTTCTATACCTTGTAGTGTTTGTCTTAGTGGTTTAGATTCTTTAGAC   7162
gb:MN975262|Organism:Severe      ACCTACTGTACTGGTTCTATACCTTGTAGTGTTTGTCTTAGTGGTTTAGATTCTTTAGAC   7162
gb:MT106052|Organism:Severe      ACCTACTGTACTGGTTCTATACCTTGTAGTGTTTGTCTTAGTGGTTTAGATTCTTTAGAC   7162
gb:LC522975|Organism:Severe      ACCTACTGTACTGGTTCTATACCTTGTAGTGTTTGTCTTAGTGGTTTAGATTCTTTAGAC   7159
gb:LC522973|Organism:Severe      ACCTACTGTACTGGTTCTATACCTTGTAGTGTTTGTCTTAGTGGTTTAGATTCTTTAGAC   7159
gb:LC522974|Organism:Severe      ACCTACTGTACTGGTTCTATACCTTGTAGTGTTTGTCTTAGTGGTTTAGATTCTTTAGAC   7159
gb:MN985325|Organism:Severe      ACCTACTGTACTGGTTCTATACCTTGTAGTGTTTGTCTTAGTGGTTTAGATTCTTTAGAC   7162
gb:MT020881|Organism:Severe      ACCTACTGTACTGGTTCTATACCTTGTAGTGTTTGTCTTAGTGGTTTAGATTCTTTAGAC   7162
gb:MT020880|Organism:Severe      ACCTACTGTACTGGTTCTATACCTTGTAGTGTTTGTCTTAGTGGTTTAGATTCTTTAGAC   7162
gb:MT066175|Organism:Severe      ACCTACTGTACTGGTTCTATACCTTGTAGTGTTTGTCTTAGTGGTTTAGATTCTTTAGAC   7162
gb:MN997409|Organism:Severe      ACCTACTGTACTGGTTCTATACCTTGTAGTGTTTGTCTTAGTGGTTTAGATTCTTTAGAC   7162
gb:MN938384|Organism:Severe      ACCTACTGTACTGGTTCTATACCTTGTAGTGTTTGTCTTAGTGGTTTAGATTCTTTAGAC   7130
gb:MT044258|Organism:Severe      ACCTACTGTACTGGTTCTATACCTTGTAGTGTTTGTCTTAGTGGTTTAGATTCTTTAGAC   7138
gb:MT039890|Organism:Severe      ACCTACTGTACTGGTTCTATACCTTGTAGTGTTTGTCTTAGTGGTTTAGATTCTTTAGAC   7162
gb:MN988713|Organism:Severe      ACCTACTGTACTGGTTCTATACCTTGTAGTGTTTGTCTTAGTGGTTTAGATTCTTTAGAC   7162
gb:LC521925|Organism:Severe      ACCTACTGTACTGGTTCTATACCTTGTAGTGTTTGTCTTAGTGGTTTAGATTCTTTAGAC   7135
gb:MT093571|Organism:Severe      ACCTACTGTACTGGTTCTATACCTTGTAGTGTTTGTCTTAGTGGTTTAGATTCTTTAGAC   7162
gb:MT039887|Organism:Severe      ACCTACTGTACTGGTTCTATACCTTGTAGTGTTTGTCTTAGTGGTTTAGATTCTTTAGAC   7162
gb:MT019530|Organism:Severe      ACCTACTGTACTGGTTCTATACCTTGTAGTGTTTGTCTTAGTGGTTTAGATTCTTTAGAC   7162
gb:MT039888|Organism:Severe      ACCTACTGTACTGGTTCTATACCTTGTAGTGTTTGTCTTAGTGGTTTAGATTCTTTAGAC   7162
gb:LC522972|Organism:Severe      ACCTACTGTACTGGTTCTATACCTTGTAGTGTTTGTCTTAGTGGTTTAGATTCTTTAGAC   7159
gb:MT027063|Organism:Severe      ACCTACTGTACTGGTTCTATACCTTGTAGTGTTTGTCTTAGTGGTTTAGATTCTTTAGAC   7162
gb:MT027062|Organism:Severe      ACCTACTGTACTGGTTCTATACCTTGTAGTGTTTGTCTTAGTGGTTTAGATTCTTTAGAC   7162
gb:MT019529|Organism:Severe      ACCTACTGTACTGGTTCTATACCTTGTAGTGTTTGTCTTAGTGGTTTAGATTCTTTAGAC   7162
gb:MN996529|Organism:Severe      ACCTACTGTACTGGTTCTATACCTTGTAGTGTTTGTCTTAGTGGTTTAGATTCTTTAGAC   7150
gb:MN996531|Organism:Severe      ACCTACTGTACTGGTTCTATACCTTGTAGTGTTTGTCTTAGTGGTTTAGATTCTTTAGAC   7149
gb:MT066176|Organism:Severe      ACCTACTGTACTGGTTCTATACCTTGTAGTGTTTGTCTTAGTGGTTTAGATTCTTTAGAC   7162
gb:MT027064|Organism:Severe      ACCTACTGTACTGGTTCTATACCTTGTAGTGTTTGTCTTAGTGGTTTAGATTCTTTAGAC   7162
gb:MN994468|Organism:Severe      ACCTACTGTACTGGTTCTATACCTTGTAGTGTTTGTCTTAGTGGTTTAGATTCTTTAGAC   7162
gb:MT072688|Organism:Severe      ACCTACTGTACTGGTTCTATACCTTGTAGTGTTTGTCTTAGTGGTTTAGATTCTTTAGAC   7147
gb:MN996527|Organism:Severe      ACCTACTGTACTGGTTCTATACCTTGTAGTGTTTGTCTTAGTGGTTTAGATTCTTTAGAC   7129
gb:MT093631|Organism:Severe      ACCTACTGTACTGGTTCTATACCTTGTAGTGTTTGTCTTAGTGGTTTAGATTCTTTAGAC   7200
gb:MT106053|Organism:Severe      ACCTACTGTACTGGTTCTATACCTTGTAGTGTTTGTCTTAGTGGTTTAGATTCTTTAGAC   7162
gb:MT019533|Organism:Severe      ACCTACTGTACTGGTTCTATACCTTGTAGTGTTTGTCTTAGTGGTTTAGATTCTTTAGAC   7162
gb:MT019531|Organism:Severe      ACCTACTGTACTGGTTCTATACCTTGTAGTGTTTGTCTTAGTGGTTTAGATTCTTTAGAC   7162
gb:MN996528|Organism:Severe      ACCTACTGTACTGGTTCTATACCTTGTAGTGTTTGTCTTAGTGGTTTAGATTCTTTAGAC   7162
gb:MN996530|Organism:Severe      ACCTACTGTACTGGTTCTATACCTTGTAGTGTTTGTCTTAGTGGTTTAGATTCTTTAGAC   7148
gb:MN908947|Organism:Severe      ACCTACTGTACTGGTTCTATACCTTGTAGTGTTTGTCTTAGTGGTTTAGATTCTTTAGAC   7162
gb:MT019532|Organism:Severe      ACCTACTGTACTGGTTCTATACCTTGTAGTGTTTGTCTTAGTGGTTTAGATTCTTTAGAC   7162
                                 ************************************************************

gb:MT020781|Organism:Severe      ACCTATCCTTCTTTAGAAACTATACAAATTACCATTTCATCTTTTAAATGGGATTTAACT   7210
```

```
gb:MT007544|Organism:Severe      ACCTATCCTTCTTTAGAAACTATACAAATTACCATTTCATCTTTTAAATGGGATTTAACT   7222
gb:MN994467|Organism:Severe      ACCTATCCTTCTTTAGAAACTATACAAATTACCATTTCATCTTTTAAATGGGATTTAACT   7222
gb:MT044257|Organism:Severe      ACCTATCCTTCTTTAGAAACTATACAAATTACCATTTCATCTTTTAAATGGGATTTAACT   7222
gb:MT106054|Organism:Severe      ACCTATCCTTCTTTAGAAACTATACAAATTACCATTTCATCTTTTAAATGGGATTTAACT   7222
gb:MT049951|Organism:Severe      ACCTATCCTTCTTTAGAAACTATACAAATTACCATTTCATCTTTTAAATGGGATTTAACT   7222
gb:MN975262|Organism:Severe      ACCTATCCTTCTTTAGAAACTATACAAATTACCATTTCATCTTTTAAATGGGATTTAACT   7222
gb:MT106052|Organism:Severe      ACCTATCCTTCTTTAGAAACTATACAAATTACCATTTCATCTTTTAAATGGGATTTAACT   7222
gb:LC522975|Organism:Severe      ACCTATCCTTCTTTAGAAACTATACAAATTACCATTTCATCTTTTAAATGGGATTTAACT   7219
gb:LC522973|Organism:Severe      ACCTATCCTTCTTTAGAAACTATACAAATTACCATTTCATCTTTTAAATGGGATTTAACT   7219
gb:LC522974|Organism:Severe      ACCTATCCTTCTTTAGAAACTATACAAATTACCATTTCATCTTTTAAATGGGATTTAACT   7219
gb:MN985325|Organism:Severe      ACCTATCCTTCTTTAGAAACTATACAAATTACCATTTCATCTTTTAAATGGGATTTAACT   7222
gb:MT020881|Organism:Severe      ACCTATCCTTCTTTAGAAACTATACAAATTACCATTTCATCTTTTAAATGGGATTTAACT   7222
gb:MT020880|Organism:Severe      ACCTATCCTTCTTTAGAAACTATACAAATTACCATTTCATCTTTTAAATGGGATTTAACT   7222
gb:MT066175|Organism:Severe      ACCTATCCTTCTTTAGAAACTATACAAATTACCATTTCATCTTTTAAATGGGATTTAACT   7222
gb:MN997409|Organism:Severe      ACCTATCCTTCTTTAGAAACTATACAAATTACCATTTCATCTTTTAAATGGGATTTAACT   7222
gb:MN938384|Organism:Severe      ACCTATCCTTCTTTAGAAACTATACAAATTACCATTTCATCTTTTAAATGGGATTTAACT   7190
gb:MT044258|Organism:Severe      ACCTATCCTTCTTTAGAAACTATACAAATTACCATTTCATCTTTTAAATGGGATTTAACT   7198
gb:MT039890|Organism:Severe      ACCTATCCTTCTTTAGAAACTATACAAATTACCATTTCATCTTTTAAATGGGATTTAACT   7222
gb:MN988713|Organism:Severe      ACCTATCCTTCTTTAGAAACTATACAAATTACCATTTCATCTTTTAAATGGGATTTAACT   7222
gb:LC521925|Organism:Severe      ACCTATCCTTCTTTAGAAACTATACAAATTACCATTTCATCTTTTAAATGGGATTTAACT   7195
gb:MT093571|Organism:Severe      ACCTATCCTTCTTTAGAAACTATACAAATTACCATTTCATCTTTTAAATGGGATTTAACT   7222
gb:MT039887|Organism:Severe      ACCTATCCTTCTTTAGAAACTATACAAATTACCATTTCATCTTTTAAATGGGATTTAACT   7222
gb:MT019530|Organism:Severe      ACCTATCCTTCTTTAGAAACTATACAAATTACCATTTCATCTTTTAAATGGGATTTAACT   7222
gb:MT039888|Organism:Severe      ACCTATCCTTCTTTAGAAACTATACAAATTACCATTTCATCTTTTAAATGGGATTTAACT   7222
gb:LC522972|Organism:Severe      ACCTATCCTTCTTTAGAAACTATACAAATTACCATTTCATCTTTTAAATGGGATTTAACT   7219
gb:MT027063|Organism:Severe      ACCTATCCTTCTTTAGAAACTATACAAATTACCATTTCATCTTTTAAATGGGATTTAACT   7222
gb:MT027062|Organism:Severe      ACCTATCCTTCTTTAGAAACTATACAAATTACCATTTCATCTTTTAAATGGGATTTAACT   7222
gb:MT019529|Organism:Severe      ACCTATCCTTCTTTAGAAACTATACAAATTACCATTTCATCTTTTAAATGGGATTTAACT   7222
gb:MN996529|Organism:Severe      ACCTATCCTTCTTTAGAAACTATACAAATTACCATTTCATCTTTTAAATGGGATTTAACT   7210
gb:MN996531|Organism:Severe      ACCTATCCTTCTTTAGAAACTATACAAATTACCATTTCATCTTTTAAATGGGATTTAACT   7209
gb:MT066176|Organism:Severe      ACCTATCCTTCTTTAGAAACTATACAAATTACCATTTCATCTTTTAAATGGGATTTAACT   7222
gb:MT027064|Organism:Severe      ACCTATCCTTCTTTAGAAACTATACAAATTACCATTTCATCTTTTAAATGGGATTTAACT   7222
gb:MN994468|Organism:Severe      ACCTATCCTTCTTTAGAAACTATACAAATTACCATTTCATCTTTTAAATGGGATTTAACT   7222
gb:MT072688|Organism:Severe      ACCTATCCTTCTTTAGAAACTATACAAATTACCATTTCATCTTTTAAATGGGATTTAACT   7207
gb:MN996527|Organism:Severe      ACCTATCCTTCTTTAGAAACTATACAAATTACCATTTCATCTTTTAAATGGGATTTAACT   7189
gb:MT093631|Organism:Severe      ACCTATCCTTCTTTAGAAACTATACAAATTACCATTTCATCTTTTAAATGGGATTTAACT   7260
gb:MT106053|Organism:Severe      ACCTATCCTTCTTTAGAAACTATACAAATTACCATTTCATCTTTTAAATGGGATTTAACT   7222
gb:MT019533|Organism:Severe      ACCTATCCTTCTTTAGAAACTATACAAATTACCATTTCATCTTTTAAATGGGATTTAACT   7222
gb:MT019531|Organism:Severe      ACCTATCCTTCTTTAGAAACTATACAAATTACCATTTCATCTTTTAAATGGGATTTAACT   7222
gb:MN996528|Organism:Severe      ACCTATCCTTCTTTAGAAACTATACAAATTACCATTTCATCTTTTAAATGGGATTTAACT   7222
gb:MN996530|Organism:Severe      ACCTATCCTTCTTTAGAAACTATACAAATTACCATTTCATCTTTTAAATGGGATTTAACT   7208
gb:MN908947|Organism:Severe      ACCTATCCTTCTTTAGAAACTATACAAATTACCATTTCATCTTTTAAATGGGATTTAACT   7222
gb:MT019532|Organism:Severe      ACCTATCCTTCTTTAGAAACTATACAAATTACCATTTCATCTTTTAAATGGGATTTAACT   7222
                                 ************************************************************

gb:MT020781|Organism:Severe      GCTTTTGGCTTAGTTGCAGAGTGGTTTTTGGCATATATTCTTTTCACTAGGTTTTTCTAT   7270
gb:MT007544|Organism:Severe      GCTTTTGGCTTAGTTGCAGAGTGGTTTTTGGCATATATTCTTTTCACTAGGTTTTTCTAT   7282
gb:MN994467|Organism:Severe      GCTTTTGGCTTAGTTGCAGAGTGGTTTTTGGCATATATTCTTTTCACTAGGTTTTTCTAT   7282
gb:MT044257|Organism:Severe      GCTTTTGGCTTAGTTGCAGAGTGGTTTTTGGCATATATTCTTTTCACTAGGTTTTTCTAT   7282
gb:MT106054|Organism:Severe      GCTTTTGGCTTAGTTGCAGAGTGGTTTTTGGCATATATTCTTTTCACTAGGTTTTTCTAT   7282
gb:MT049951|Organism:Severe      GCTTTTGGCTTAGTTGCAGAGTGGTTTTTGGCATATATTCTTTTCACTAGGTTTTTCTAT   7282
gb:MN975262|Organism:Severe      GCTTTTGGCTTAGTTGCAGAGTGGTTTTTGGCATATATTCTTTTCACTAGGTTTTTCTAT   7282
gb:MT106052|Organism:Severe      GCTTTTGGCTTAGTTGCAGAGTGGTTTTTGGCATATATTCTTTTCACTAGGTTTTTCTAT   7282
gb:LC522975|Organism:Severe      GCTTTTGGCTTAGTTGCAGAGTGGTTTTTGGCATATATTCTTTTCACTAGGTTTTTCTAT   7279
gb:LC522973|Organism:Severe      GCTTTTGGCTTAGTTGCAGAGTGGTTTTTGGCATATATTCTTTTCACTAGGTTTTTCTAT   7279
gb:LC522974|Organism:Severe      GCTTTTGGCTTAGTTGCAGAGTGGTTTTTGGCATATATTCTTTTCACTAGGTTTTTCTAT   7279
gb:MN985325|Organism:Severe      GCTTTTGGCTTAGTTGCAGAGTGGTTTTTGGCATATATTCTTTTCACTAGGTTTTTCTAT   7282
gb:MT020881|Organism:Severe      GCTTTTGGCTTAGTTGCAGAGTGGTTTTTGGCATATATTCTTTTCACTAGGTTTTTCTAT   7282
gb:MT020880|Organism:Severe      GCTTTTGGCTTAGTTGCAGAGTGGTTTTTGGCATATATTCTTTTCACTAGGTTTTTCTAT   7282
gb:MT066175|Organism:Severe      GCTTTTGGCTTAGTTGCAGAGTGGTTTTTGGCATATATTCTTTTCACTAGGTTTTTCTAT   7282
gb:MN997409|Organism:Severe      GCTTTTGGCTTAGTTGCAGAGTGGTTTTTGGCATATATTCTTTTCACTAGGTTTTTCTAT   7282
gb:MN938384|Organism:Severe      GCTTTTGGCTTAGTTGCAGAGTGGTTTTTGGCATATATTCTTTTCACTAGGTTTTTCTAT   7250
gb:MT044258|Organism:Severe      GCTTTTGGCTTAGTTGCAGAGTGGTTTTTGGCATATATTCTTTTCACTAGGTTTTTCTAT   7258
gb:MT039890|Organism:Severe      GCTTTTGGCTTAGTTGCAGAGTGGTTTTTGGCATATATTCTTTTCACTAGGTTTTTCTAT   7282
gb:MN988713|Organism:Severe      GCTTTTGGCTTAGTTGCAGAGTGGTTTTTGGCATATATTCTTTTCACTAGGTTTTTCTAT   7282
```

```
gb:LC521925|Organism:Severe      GCTTTTGGCTTAGTTGCAGAGTGGTTTTTGGCATATATTCTTTTCACTAGGTTTTTCTAT   7255
gb:MT093571|Organism:Severe      GCTTTTGGCTTAGTTGCAGAGTGGTTTTTGGCATATATTCTTTTCACTAGGTTTTTCTAT   7282
gb:MT039887|Organism:Severe      GCTTTTGGCTTAGTTGCAGAGTGGTTTTTGGCATATATTCTTTTCACTAGGTTTTTCTAT   7282
gb:MT019530|Organism:Severe      GCTTTTGGCTTAGTTGCAGAGTGGTTTTTGGCATATATTCTTTTCACTAGGTTTTTCTAT   7282
gb:MT039888|Organism:Severe      GCTTTTGGCTTAGTTGCAGAGTGGTTTTTGGCATATATTCTTTTCACTAGGTTTTTCTAT   7282
gb:LC522972|Organism:Severe      GCTTTTGGCTTAGTTGCAGAGTGGTTTTTGGCATATATTCTTTTCACTAGGTTTTTCTAT   7279
gb:MT027063|Organism:Severe      GCTTTTGGCTTAGTTGCAGAGTGGTTTTTGGCATATATTCTTTTCACTAGGTTTTTCTAT   7282
gb:MT027062|Organism:Severe      GCTTTTGGCTTAGTTGCAGAGTGGTTTTTGGCATATATTCTTTTCACTAGGTTTTTCTAT   7282
gb:MT019529|Organism:Severe      GCTTTTGGCTTAGTTGCAGAGTGGTTTTTGGCATATATTCTTTTCACTAGGTTTTTCTAT   7282
gb:MN996529|Organism:Severe      GCTTTTGGCTTAGTTGCAGAGTGGTTTTTGGCATATATTCTTTTCACTAGGTTTTTCTAT   7270
gb:MN996531|Organism:Severe      GCTTTTGGCTTAGTTGCAGAGTGGTTTTTGGCATATATTCTTTTCACTAGGTTTTTCTAT   7269
gb:MT066176|Organism:Severe      GCTTTTGGCTTAGTTGCAGAGTGGTTTTTGGCATATATTCTTTTCACTAGGTTTTTCTAT   7282
gb:MT027064|Organism:Severe      GCTTTTGGCTTAGTTGCAGAGTGGTTTTTGGCATATATTCTTTTCACTAGGTTTTTCTAT   7282
gb:MN994468|Organism:Severe      GCTTTTGGCTTAGTTGCAGAGTGGTTTTTGGCATATATTCTTTTCACTAGGTTTTTCTAT   7282
gb:MT072688|Organism:Severe      GCTTTTGGCTTAGTTGCAGAGTGGTTTTTGGCATATATTCTTTTCACTAGGTTTTTCTAT   7267
gb:MN996527|Organism:Severe      GCTTTTGGCTTAGTTGCAGAGTGGTTTTTGGCATATATTCTTTTCACTAGGTTTTTCTAT   7249
gb:MT093631|Organism:Severe      GCTTTTGGCTTAGTTGCAGAGTGGTTTTTGGCATATATTCTTTTCACTAGGTTTTTCTAT   7320
gb:MT106053|Organism:Severe      GCTTTTGGCTTAGTTGCAGAGTGGTTTTTGGCATATATTCTTTTCACTAGGTTTTTCTAT   7282
gb:MT019533|Organism:Severe      GCTTTTGGCTTAGTTGCAGAGTGGTTTTTGGCATATATTCTTTTCACTAGGTTTTTCTAT   7282
gb:MT019531|Organism:Severe      GCTTTTGGCTTAGTTGCAGAGTGGTTTTTGGCATATATTCTTTTCACTAGGTTTTTCTAT   7282
gb:MN996528|Organism:Severe      GCTTTTGGCTTAGTTGCAGAGTGGTTTTTGGCATATATTCTTTTCACTAGGTTTTTCTAT   7282
gb:MN996530|Organism:Severe      GCTTTTGGCTTAGTTGCAGAGTGGTTTTTGGCATATATTCTTTTCACTAGGTTTTTCTAT   7268
gb:MN908947|Organism:Severe      GCTTTTGGCTTAGTTGCAGAGTGGTTTTTGGCATATATTCTTTTCACTAGGTTTTTCTAT   7282
gb:MT019532|Organism:Severe      GCTTTTGGCTTAGTTGCAGAGTGGTTTTTGGCATATATTCTTTTCACTAGGTTTTTCTAT   7282
                                 ************************************************************

gb:MT020781|Organism:Severe      GTACTTGGATTGGCTGCAATCATGCAATTGTTTTTCAGCTATTTTGCAGTACATTTTATT   7330
gb:MT007544|Organism:Severe      GTACTTGGATTGGCTGCAATCATGCAATTGTTTTTCAGCTATTTTGCAGTACATTTTATT   7342
gb:MN994467|Organism:Severe      GTACTTGGATTGGCTGCAATCATGCAATTGTTTTTCAGCTATTTTGCAGTACATTTTATT   7342
gb:MT044257|Organism:Severe      GTACTTGGATTGGCTGCAATCATGCAATTGTTTTTCAGCTATTTTGCAGTACATTTTATT   7342
gb:MT106054|Organism:Severe      GTACTTGGATTGGCTGCAATCATGCAATTGTTTTTCAGCTATTTTGCAGTACATTTTATT   7342
gb:MT049951|Organism:Severe      GTACTTGGATTGGCTGCAATCATGCAATTGTTTTTCAGCTATTTTGCAGTACATTTTATT   7342
gb:MN975262|Organism:Severe      GTACTTGGATTGGCTGCAATCATGCAATTGTTTTTCAGCTATTTTGCAGTACATTTTATT   7342
gb:MT106052|Organism:Severe      GTACTTGGATTGGCTGCAATCATGCAATTGTTTTTCAGCTATTTTGCAGTACATTTTATT   7342
gb:LC522975|Organism:Severe      GTACTTGGATTGGCTGCAATCATGCAATTGTTTTTCAGCTATTTTGCAGTACATTTTATT   7339
gb:LC522973|Organism:Severe      GTACTTGGATTGGCTGCAATCATGCAATTGTTTTTCAGCTATTTTGCAGTACATTTTATT   7339
gb:LC522974|Organism:Severe      GTACTTGGATTGGCTGCAATCATGCAATTGTTTTTCAGCTATTTTGCAGTACATTTTATT   7339
gb:MN985325|Organism:Severe      GTACTTGGATTGGCTGCAATCATGCAATTGTTTTTCAGCTATTTTGCAGTACATTTTATT   7342
gb:MT020881|Organism:Severe      GTACTTGGATTGGCTGCAATCATGCAATTGTTTTTCAGCTATTTTGCAGTACATTTTATT   7342
gb:MT020880|Organism:Severe      GTACTTGGATTGGCTGCAATCATGCAATTGTTTTTCAGCTATTTTGCAGTACATTTTATT   7342
gb:MT066175|Organism:Severe      GTACTTGGATTGGCTGCAATCATGCAATTGTTTTTCAGCTATTTTGCAGTACATTTTATT   7342
gb:MN997409|Organism:Severe      GTACTTGGATTGGCTGCAATCATGCAATTGTTTTTCAGCTATTTTGCAGTACATTTTATT   7342
gb:MN938384|Organism:Severe      GTACTTGGATTGGCTGCAATCATGCAATTGTTTTTCAGCTATTTTGCAGTACATTTTATT   7310
gb:MT044258|Organism:Severe      GTACTTGGATTGGCTGCAATCATGCAATTGTTTTTCAGCTATTTTGCAGTACATTTTATT   7318
gb:MT039890|Organism:Severe      GTACTTGGATTGGCTGCAATCATGCAATTGTTTTTCAGCTATTTTGCAGTACATTTTATT   7342
gb:MN988713|Organism:Severe      GTACTTGGATTGGCTGCAATCATGCAATTGTTTTTCAGCTATTTTGCAGTACATTTTATT   7342
gb:LC521925|Organism:Severe      GTACTTGGATTGGCTGCAATCATGCAATTGTTTTTCAGCTATTTTGCAGTACATTTTATT   7315
gb:MT093571|Organism:Severe      GTACTTGGATTGGCTGCAATCATGCAATTGTTTTTCAGCTATTTTGCAGTACATTTTATT   7342
gb:MT039887|Organism:Severe      GTACTTGGATTGGCTGCAATCATGCAATTGTTTTTCAGCTATTTTGCAGTACATTTTATT   7342
gb:MT019530|Organism:Severe      GTACTTGGATTGGCTGCAATCATGCAATTGTTTTTCAGCTATTTTGCAGTACATTTTATT   7342
gb:MT039888|Organism:Severe      GTACTTGGATTGGCTGCAATCATGCAATTGTTTTTCAGCTATTTTGCAGTACATTTTATT   7342
gb:LC522972|Organism:Severe      GTACTTGGATTGGCTGCAATCATGCAATTGTTTTTCAGCTATTTTGCAGTACATTTTATT   7339
gb:MT027063|Organism:Severe      GTACTTGGATTGGCTGCAATCATGCAATTGTTTTTCAGCTATTTTGCAGTACATTTTATT   7342
gb:MT027062|Organism:Severe      GTACTTGGATTGGCTGCAATCATGCAATTGTTTTTCAGCTATTTTGCAGTACATTTTATT   7342
gb:MT019529|Organism:Severe      GTACTTGGATTGGCTGCAATCATGCAATTGTTTTTCAGCTATTTTGCAGTACATTTTATT   7342
gb:MN996529|Organism:Severe      GTACTTGGATTGGCTGCAATCATGCAATTGTTTTTCAGCTATTTTGCAGTACATTTTATT   7330
gb:MN996531|Organism:Severe      GTACTTGGATTGGCTGCAATCATGCAATTGTTTTTCAGCTATTTTGCAGTACATTTTATT   7329
gb:MT066176|Organism:Severe      GTACTTGGATTGGCTGCAATCATGCAATTGTTTTTCAGCTATTTTGCAGTACATTTTATT   7342
gb:MT027064|Organism:Severe      GTACTTGGATTGGCTGCAATCATGCAATTGTTTTTCAGCTATTTTGCAGTACATTTTATT   7342
gb:MN994468|Organism:Severe      GTACTTGGATTGGCTGCAATCATGCAATTGTTTTTCAGCTATTTTGCAGTACATTTTATT   7342
gb:MT072688|Organism:Severe      GTACTTGGATTGGCTGCAATCATGCAATTGTTTTTCAGCTATTTTGCAGTACATTTTATT   7327
gb:MN996527|Organism:Severe      GTACTTGGATTGGCTGCAATCATGCAATTGTTTTTCAGCTATTTTGCAGTACATTTTATT   7309
gb:MT093631|Organism:Severe      GTACTTGGATTGGCTGCAATCATGCAATTGTTTTTCAGCTATTTTGCAGTACATTTTATT   7380
gb:MT106053|Organism:Severe      GTACTTGGATTGGCTGCAATCATGCAATTGTTTTTCAGCTATTTTGCAGTACATTTTATT   7342
gb:MT019533|Organism:Severe      GTACTTGGATTGGCTGCAATCATGCAATTGTTTTTCAGCTATTTTGCAGTACATTTTATT   7342
```

```
gb:MT019531|Organism:Severe      GTACTTGGATTGGCTGCAATCATGCAATTGTTTTTCAGCTATTTTGCAGTACATTTTATT    7342
gb:MN996528|Organism:Severe      GTACTTGGATTGGCTGCAATCATGCAATTGTTTTTCAGCTATTTTGCAGTACATTTTATT    7342
gb:MN996530|Organism:Severe      GTACTTGGATTGGCTGCAATCATGCAATTGTTTTTCAGCTATTTTGCAGTACATTTTATT    7328
gb:MN908947|Organism:Severe      GTACTTGGATTGGCTGCAATCATGCAATTGTTTTTCAGCTATTTTGCAGTACATTTTATT    7342
gb:MT019532|Organism:Severe      GTACTTGGATTGGCTGCAATCATGCAATTGTTTTTCAGCTATTTTGCAGTACATTTTATT    7342
                                 ************************************************************

gb:MT020781|Organism:Severe      AGTAATTCTTGGCTTATGTGGTTAATANNNNNNNNNNNNNAAATGGCCCCGATTTCAGCT    7390
gb:MT007544|Organism:Severe      AGTAATTCTTGGCTTATGTGGTTAATAATTAATCTTGTACAAATGGCCCCGATTTCAGCT    7402
gb:MN994467|Organism:Severe      AGTAATTCTTGGCTTATGTGGTTAATAATTAATCTTGTACAAATGGCCCCGATTTCAGCT    7402
gb:MT044257|Organism:Severe      AGTAATTCTTGGCTTATGTGGTTAATAATTAATCTTGTACAAATGGCCCCGATTTCAGCT    7402
gb:MT106054|Organism:Severe      AGTAATTCTTGGCTTATGTGGTTAATAATTAATCTTGTACAAATGGCCCCGATTTCAGCT    7402
gb:MT049951|Organism:Severe      AGTAATTCTTGGCTTATGTGGTTAATAATTAATCTTGTACAAATGGCCCCGATTTCAGCT    7402
gb:MN975262|Organism:Severe      AGTAATTCTTGGCTTATGTGGTTAATAATTAATCTTGTACAAATGGCCCCGATTTCAGCT    7402
gb:MT106052|Organism:Severe      AGTAATTCTTGGCTTATGTGGTTAATAATTAATCTTGTACAAATGGCCCCGATTTCAGCT    7402
gb:LC522975|Organism:Severe      AGTAATTCTTGGCTTATGTGGTTAATAATTAATCTTGTACAAATGGCCCCGATTTCAGCT    7399
gb:LC522973|Organism:Severe      AGTAATTCTTGGCTTATGTGGTTAATAATTAATCTTGTACAAATGGCCCCGATTTCAGCT    7399
gb:LC522974|Organism:Severe      AGTAATTCTTGGCTTATGTGGTTAATAATTAATCTTGTACAAATGGCCCCGATTTCAGCT    7399
gb:MN985325|Organism:Severe      AGTAATTCTTGGCTTATGTGGTTAATAATTAATCTTGTACAAATGGCCCCGATTTCAGCT    7402
gb:MT020881|Organism:Severe      AGTAATTCTTGGCTTATGTGGTTAATAATTAATCTTGTACAAATGGCCCCGATTTCAGCT    7402
gb:MT020880|Organism:Severe      AGTAATTCTTGGCTTATGTGGTTAATAATTAATCTTGTACAAATGGCCCCGATTTCAGCT    7402
gb:MT066175|Organism:Severe      AGTAATTCTTGGCTTATGTGGTTAATAATTAATCTTGTACAAATGGCCCCGATTTCAGCT    7402
gb:MN997409|Organism:Severe      AGTAATTCTTGGCTTATGTGGTTAATAATTAATCTTGTACAAATGGCCCCGATTTCAGCT    7402
gb:MN938384|Organism:Severe      AGTAATTCTTGGCTTATGTGGTTAATAATTAATCTTGTACAAATGGCCCCGATTTCAGCT    7370
gb:MT044258|Organism:Severe      AGTAATTCTTGGCTTATGTGGTTAATAATTAATCTTGTACAAATGGCCCCGATTTCAGCT    7378
gb:MT039890|Organism:Severe      AGTAATTCTTGGCTTATGTGGTTAATAATTAATCTTGTACAAATGGCCCCGATTTCAGCT    7402
gb:MN988713|Organism:Severe      AGTAATTCTTGGCTTATGTGGTTAATAATTAATCTTGTACAAATGGCCCCGATTTCAGCT    7402
gb:LC521925|Organism:Severe      AGTAATTCTTGGCTTATGTGGTTAATAATTAATCTTGTACAAATGGCCCCGATTTCAGCT    7375
gb:MT093571|Organism:Severe      AGTAATTCTTGGCTTATGTGGTTAATAATTAATCTTGTACAAATGGCCCCGATTTCAGCT    7402
gb:MT039887|Organism:Severe      AGTAATTCTTGGCTTATGTGGTTAATAATTAATCTTGTACAAATGGCCCCGATTTCAGCT    7402
gb:MT019530|Organism:Severe      AGTAATTCTTGGCTTATGTGGTTAATAATTAATCTTGTACAAATGGCCCCGATTTCAGCT    7402
gb:MT039888|Organism:Severe      AGTAATTCTTGGCTTATGTGGTTAATAATTAATCTTGTACAAATGGCCCCGATTTCAGCT    7402
gb:LC522972|Organism:Severe      AGTAATTCTTGGCTTATGTGGTTAATAATTAATCTTGTACAAATGGCCCCGATTTCAGCT    7399
gb:MT027063|Organism:Severe      AGTAATTCTTGGCTTATGTGGTTAATAATTAATCTTGTACAAATGGCCCCGATTTCAGCT    7402
gb:MT027062|Organism:Severe      AGTAATTCTTGGCTTATGTGGTTAATAATTAATCTTGTACAAATGGCCCCGATTTCAGCT    7402
gb:MT019529|Organism:Severe      AGTAATTCTTGGCTTATGTGGTTAATAATTAATCTTGTACAAATGGCCCCGATTTCAGCT    7402
gb:MN996529|Organism:Severe      AGTAATTCTTGGCTTATGTGGTTAATAATTAATCTTGTACAAATGGCCCCGATTTCAGCT    7390
gb:MN996531|Organism:Severe      AGTAATTCTTGGCTTATGTGGTTAATAATTAATCTTGTACAAATGGCCCCGATTTCAGCT    7389
gb:MT066176|Organism:Severe      AGTAATTCTTGGCTTATGTGGTTAATAATTAATCTTGTACAAATGGCCCCGATTTCAGCT    7402
gb:MT027064|Organism:Severe      AGTAATTCTTGGCTTATGTGGTTAATAATTAATCTTGTACAAATGGCCCCGATTTCAGCT    7402
gb:MN994468|Organism:Severe      AGTAATTCTTGGCTTATGTGGTTAATAATTAATCTTGTACAAATGGCCCCGATTTCAGCT    7402
gb:MT072688|Organism:Severe      AGTAATTCTTGGCTTATGTGGTTAATAATTAATCTTGTACAAATGGCCCCGATTTCAGCT    7387
gb:MN996527|Organism:Severe      AGTAATTCTTGGCTTATGTGGTTAATAATTAATCTTGTACAAATGGCCCCGATTTCAGCT    7369
gb:MT093631|Organism:Severe      AGTAATTCTTGGCTTATGTGGTTAATAATTAATCTTGTACAAATGGCCCCGATTTCAGCT    7440
gb:MT106053|Organism:Severe      AGTAATTCTTGGCTTATGTGGTTAATAATTAATCTTGTACAAATGGCCCCGATTTCAGCT    7402
gb:MT019533|Organism:Severe      AGTAATTCTTGGCTTATGTGGTTAATAATTAATCTTGTACAAATGGCCCCGATTTCAGCT    7402
gb:MT019531|Organism:Severe      AGTAATTCTTGGCTTATGTGGTTAATAATTAATCTTGTACAAATGGCCCCGATTTCAGCT    7402
gb:MN996528|Organism:Severe      AGTAATTCTTGGCTTATGTGGTTAATAATTAATCTTGTACAAATGGCCCCGATTTCAGCT    7402
gb:MN996530|Organism:Severe      AGTAATTCTTGGCTTATGTGGTTAATAATTAATCTTGTACAAATGGCCCCGATTTCAGCT    7388
gb:MN908947|Organism:Severe      AGTAATTCTTGGCTTATGTGGTTAATAATTAATCTTGTACAAATGGCCCCGATTTCAGCT    7402
gb:MT019532|Organism:Severe      AGTAATTCTTGGCTTATGTGGTTAATAATTAATCTTGTACAAATGGCCCCGATTTCAGCT    7402
                                 ***************************             ********************

gb:MT020781|Organism:Severe      ATGGTTAGAATGTACATCTTCTTTGCATCATTTTATTATGTATGGAAAAGTTATGTGCAT    7450
gb:MT007544|Organism:Severe      ATGGTTAGAATGTACATCTTCTTTGCATCATTTTATTATGTATGGAAAAGTTATGTGCAT    7462
gb:MN994467|Organism:Severe      ATGGTTAGAATGTACATCTTCTTTGCATCATTTTATTATGTATGGAAAAGTTATGTGCAT    7462
gb:MT044257|Organism:Severe      ATGGTTAGAATGTACATCTTCTTTGCATCATTTTATTATGTATGGAAAAGTTATGTGCAT    7462
gb:MT106054|Organism:Severe      ATGGTTAGAATGTACATCTTCTTTGCATCATTTTATTATGTATGGAAAAGTTATGTGCAT    7462
gb:MT049951|Organism:Severe      ATGGTTAGAATGTACATCTTCTTTGCATCATTTTATTATGTATGGAAAAGTTATGTGCAT    7462
gb:MN975262|Organism:Severe      ATGGTTAGAATGTACATCTTCTTTGCATCATTTTATTATGTATGGAAAAGTTATGTGCAT    7462
gb:MT106052|Organism:Severe      ATGGTTAGAATGTACATCTTCTTTGCATCATTTTATTATGTATGGAAAAGTTATGTGCAT    7462
gb:LC522975|Organism:Severe      ATGGTTAGAATGTACATCTTCTTTGCATCATTTTATTATGTATGGAAAAGTTATGTGCAT    7459
gb:LC522973|Organism:Severe      ATGGTTAGAATGTACATCTTCTTTGCATCATTTTATTATGTATGGAAAAGTTATGTGCAT    7459
gb:LC522974|Organism:Severe      ATGGTTAGAATGTACATCTTCTTTGCATCATTTTATTATGTATGGAAAAGTTATGTGCAT    7459
gb:MN985325|Organism:Severe      ATGGTTAGAATGTACATCTTCTTTGCATCATTTTATTATGTATGGAAAAGTTATGTGCAT    7462
```

```
gb:MT020881|Organism:Severe      ATGGTTAGAATGTACATCTTCTTTGCATCATTTTATTATGTATGGAAAAGTTATGTGCAT    7462
gb:MT020880|Organism:Severe      ATGGTTAGAATGTACATCTTCTTTGCATCATTTTATTATGTATGGAAAAGTTATGTGCAT    7462
gb:MT066175|Organism:Severe      ATGGTTAGAATGTACATCTTCTTTGCATCATTTTATTATGTATGGAAAAGTTATGTGCAT    7462
gb:MN997409|Organism:Severe      ATGGTTAGAATGTACATCTTCTTTGCATCATTTTATTATGTATGGAAAAGTTATGTGCAT    7462
gb:MN938384|Organism:Severe      ATGGTTAGAATGTACATCTTCTTTGCATCATTTTATTATGTATGGAAAAGTTATGTGCAT    7430
gb:MT044258|Organism:Severe      ATGGTTAGAATGTACATCTTCTTTGCATCATTTTATTATGTATGGAAAAGTTATGTGCAT    7438
gb:MT039890|Organism:Severe      ATGGTTAGAATGTACATCTTCTTTGCATCATTTTATTATGTATGGAAAAGTTATGTGCAT    7462
gb:MN988713|Organism:Severe      ATGGTTAGAATGTACATCTTCTTTGCATCATTTTATTATGTATGGAAAAGTTATGTGCAT    7462
gb:LC521925|Organism:Severe      ATGGTTAGAATGTACATCTTCTTTGCATCATTTTATTATGTATGGAAAAGTTATGTGCAT    7435
gb:MT093571|Organism:Severe      ATGGTTAGAATGTACATCTTCTTTGCATCATTTTATTATGTATGGAAAAGTTATGTGCAT    7462
gb:MT039887|Organism:Severe      ATGGTTAGAATGTACATCTTCTTTGCATCATTTTATTATGTATGGAAAAGTTATGTGCAT    7462
gb:MT019530|Organism:Severe      ATGGTTAGAATGTACATCTTCTTTGCATCATTTTATTATGTATGGAAAAGTTATGTGCAT    7462
gb:MT039888|Organism:Severe      ATGGTTAGAATGTACATCTTCTTTGCATCATTTTATTATGTATGGAAAAGTTATGTGCAT    7462
gb:LC522972|Organism:Severe      ATGGTTAGAATGTACATCTTCTTTGCATCATTTTATTATGTATGGAAAAGTTATGTGCAT    7459
gb:MT027063|Organism:Severe      ATGGTTAGAATGTACATCTTCTTTGCATCATTTTATTATGTATGGAAAAGTTATGTGCAT    7462
gb:MT027062|Organism:Severe      ATGGTTAGAATGTACATCTTCTTTGCATCATTTTATTATGTATGGAAAAGTTATGTGCAT    7462
gb:MT019529|Organism:Severe      ATGGTTAGAATGTACATCTTCTTTGCATCATTTTATTATGTATGGAAAAGTTATGTGCAT    7462
gb:MN996529|Organism:Severe      ATGGTTAGAATGTACATCTTCTTTGCATCATTTTATTATGTATGGAAAAGTTATGTGCAT    7450
gb:MN996531|Organism:Severe      ATGGTTAGAATGTACATCTTCTTTGCATCATTTTATTATGTATGGAAAAGTTATGTGCAT    7449
gb:MT066176|Organism:Severe      ATGGTTAGAATGTACATCTTCTTTGCATCATTTTATTATGTATGGAAAAGTTATGTGCAT    7462
gb:MT027064|Organism:Severe      ATGGTTAGAATGTACATCTTCTTTGCATCATTTTATTATGTATGGAAAAGTTATGTGCAT    7462
gb:MN994468|Organism:Severe      ATGGTTAGAATGTACATCTTCTTTGCATCATTTTATTATGTATGGAAAAGTTATGTGCAT    7462
gb:MT072688|Organism:Severe      ATGGTTAGAATGTACATCTTCTTTGCATCATTTTATTATGTATGGAAAAGTTATGTGCAT    7447
gb:MN996527|Organism:Severe      ATGGTTAGAATGTACATCTTCTTTGCATCATTTTATTATGTATGGAAAAGTTATGTGCAT    7429
gb:MT093631|Organism:Severe      ATGGTTAGAATGTACATCTTCTTTGCATCATTTTATTATGTATGGAAAAGTTATGTGCAT    7500
gb:MT106053|Organism:Severe      ATGGTTAGAATGTACATCTTCTTTGCATCATTTTATTATGTATGGAAAAGTTATGTGCAT    7462
gb:MT019533|Organism:Severe      ATGGTTAGAATGTACATCTTCTTTGCATCATTTTATTATGTATGGAAAAGTTATGTGCAT    7462
gb:MT019531|Organism:Severe      ATGGTTAGAATGTACATCTTCTTTGCATCATTTTATTATGTATGGAAAAGTTATGTGCAT    7462
gb:MN996528|Organism:Severe      ATGGTTAGAATGTACATCTTCTTTGCATCATTTTATTATGTATGGAAAAGTTATGTGCAT    7462
gb:MN996530|Organism:Severe      ATGGTTAGAATGTACATCTTCTTTGCATCATTTTATTATGTATGGAAAAGTTATGTGCAT    7448
gb:MN908947|Organism:Severe      ATGGTTAGAATGTACATCTTCTTTGCATCATTTTATTATGTATGGAAAAGTTATGTGCAT    7462
gb:MT019532|Organism:Severe      ATGGTTAGAATGTACATCTTCTTTGCATCATTTTATTATGTATGGAAAAGTTATGTGCAT    7462
                                 ************************************************************

gb:MT020781|Organism:Severe      GTTGTAGACGGTTGTAATTCATCAACTTGTATGATGTGTTACAAACGTAATAGAGCAACA    7510
gb:MT007544|Organism:Severe      GTTGTAGACGGTTGTAATTCATCAACTTGTATGATGTGTTACAAACGTAATAGAGCAACA    7522
gb:MN994467|Organism:Severe      GTTGTAGACGGTTGTAATTCATCAACTTGTATGATGTGTTACAAACGTAATAGAGCAACA    7522
gb:MT044257|Organism:Severe      GTTGTAGACGGTTGTAATTCATCAACTTGTATGATGTGTTACAAACGTAATAGAGCAACA    7522
gb:MT106054|Organism:Severe      GTTGTAGACGGTTGTAATTCATCAACTTGTATGATGTGTTACAAACGTAATAGAGCAACA    7522
gb:MT049951|Organism:Severe      GTTGTAGACGGTTGTAATTCATCAACTTGTATGATGTGTTACAAACGTAATAGAGCAACA    7522
gb:MN975262|Organism:Severe      GTTGTAGACGGTTGTAATTCATCAACTTGTATGATGTGTTACAAACGTAATAGAGCAACA    7522
gb:MT106052|Organism:Severe      GTTGTAGACGGTTGTAATTCATCAACTTGTATGATGTGTTACAAACGTAATAGAGCAACA    7522
gb:LC522975|Organism:Severe      GTTGTAGACGGTTGTAATTCATCAACTTGTATGATGTGTTACAAACGTAATAGAGCAACA    7519
gb:LC522973|Organism:Severe      GTTGTAGACGGTTGTAATTCATCAACTTGTATGATGTGTTACAAACGTAATAGAGCAACA    7519
gb:LC522974|Organism:Severe      GTTGTAGACGGTTGTAATTCATCAACTTGTATGATGTGTTACAAACGTAATAGAGCAACA    7519
gb:MN985325|Organism:Severe      GTTGTAGACGGTTGTAATTCATCAACTTGTATGATGTGTTACAAACGTAATAGAGCAACA    7522
gb:MT020881|Organism:Severe      GTTGTAGACGGTTGTAATTCATCAACTTGTATGATGTGTTACAAACGTAATAGAGCAACA    7522
gb:MT020880|Organism:Severe      GTTGTAGACGGTTGTAATTCATCAACTTGTATGATGTGTTACAAACGTAATAGAGCAACA    7522
gb:MT066175|Organism:Severe      GTTGTAGACGGTTGTAATTCATCAACTTGTATGATGTGTTACAAACGTAATAGAGCAACA    7522
gb:MN997409|Organism:Severe      GTTGTAGACGGTTGTAATTCATCAACTTGTATGATGTGTTACAAACGTAATAGAGCAACA    7522
gb:MN938384|Organism:Severe      GTTGTAGACGGTTGTAATTCATCAACTTGTATGATGTGTTACAAACGTAATAGAGCAACA    7490
gb:MT044258|Organism:Severe      GTTGTAGACGGTTGTAATTCATCAACTTGTATGATGTGTTACAAACGTAATAGAGCAACA    7498
gb:MT039890|Organism:Severe      GTTGTAGACGGTTGTAATTCATCAACTTGTATGATGTGTTACAAACGTAATAGAGCAACA    7522
gb:MN988713|Organism:Severe      GTTGTAGACGGTTGTAATTCATCAACTTGTATGATGTGTTACAAACGTAATAGAGCAACA    7522
gb:LC521925|Organism:Severe      GTTGTAGACGGTTGTAATTCATCAACTTGTATGATGTGTTACAAACGTAATAGAGCAACA    7495
gb:MT093571|Organism:Severe      GTTGTAGACGGTTGTAATTCATCAACTTGTATGATGTGTTACAAACGTAATAGAGCAACA    7522
gb:MT039887|Organism:Severe      GTTGTAGACGGTTGTAATTCATCAACTTGTATGATGTGTTACAAACGTAATAGAGCAACA    7522
gb:MT019530|Organism:Severe      GTTGTAGACGGTTGTAATTCATCAACTTGTATGATGTGTTACAAACGTAATAGAGCAACA    7522
gb:MT039888|Organism:Severe      GTTGTAGACGGTTGTAATTCATCAACTTGTATGATGTGTTACAAACGTAATAGAGCAACA    7522
gb:LC522972|Organism:Severe      GTTGTAGACGGTTGTAATTCATCAACTTGTATGATGTGTTACAAACGTAATAGAGCAACA    7519
gb:MT027063|Organism:Severe      GTTGTAGACGGTTGTAATTCATCAACTTGTATGATGTGTTACAAACGTAATAGAGCAACA    7522
gb:MT027062|Organism:Severe      GTTGTAGACGGTTGTAATTCATCAACTTGTATGATGTGTTACAAACGTAATAGAGCAACA    7522
gb:MT019529|Organism:Severe      GTTGTAGACGGTTGTAATTCATCAACTTGTATGATGTGTTACAAACGTAATAGAGCAACA    7522
gb:MN996529|Organism:Severe      GTTGTAGACGGTTGTAATTCATCAACTTGTATGATGTGTTACAAACGTAATAGAGCAACA    7510
gb:MN996531|Organism:Severe      GTTGTAGACGGTTGTAATTCATCAACTTGTATGATGTGTTACAAACGTAATAGAGCAACA    7509
```

```
gb:MT066176|Organism:Severe      GTTGTAGACGGTTGTAATTCATCAACTTGTATGATGTGTTACAAACGTAATAGAGCAACA	7522
gb:MT027064|Organism:Severe      GTTGTAGACGGTTGTAATTCATCAACTTGTATGATGTGTTACAAACGTAATAGAGCAACA	7522
gb:MN994468|Organism:Severe      GTTGTAGACGGTTGTAATTCATCAACTTGTATGATGTGTTACAAACGTAATAGAGCAACA	7522
gb:MT072688|Organism:Severe      GTTGTAGACGGTTGTAATTCATCAACTTGTATGATGTGTTACAAACGTAATAGAGCAACA	7507
gb:MN996527|Organism:Severe      GTTGTAGACGGTTGTAATTCATCAACTTGTATGATGTGTTACAAACGTAATAGAGCAACA	7489
gb:MT093631|Organism:Severe      GTTGTAGACGGTTGTAATTCATCAACTTGTATGATGTGTTACAAACGTAATAGAGCAACA	7560
gb:MT106053|Organism:Severe      GTTGTAGACGGTTGTAATTCATCAACTTGTATGATGTGTTACAAACGTAATAGAGCAACA	7522
gb:MT019533|Organism:Severe      GTTGTAGACGGTTGTAATTCATCAACTTGTATGATGTGTTACAAACGTAATAGAGCAACA	7522
gb:MT019531|Organism:Severe      GTTGTAGACGGTTGTAATTCATCAACTTGTATGATGTGTTACAAACGTAATAGAGCAACA	7522
gb:MN996528|Organism:Severe      GTTGTAGACGGTTGTAATTCATCAACTTGTATGATGTGTTACAAACGTAATAGAGCAACA	7522
gb:MN996530|Organism:Severe      GTTGTAGACGGTTGTAATTCATCAACTTGTATGATGTGTTACAAACGTAATAGAGCAACA	7508
gb:MN908947|Organism:Severe      GTTGTAGACGGTTGTAATTCATCAACTTGTATGATGTGTTACAAACGTAATAGAGCAACA	7522
gb:MT019532|Organism:Severe      GTTGTAGACGGTTGTAATTCATCAACTTGTATGATGTGTTACAAACGTAATAGAGCAACA	7522
                                 ************************************************************

gb:MT020781|Organism:Severe      AGAGTCGAATGTACAACTATTGTTAATGGTGTTAGAAGGTCCTTTTATGTCTATGCTAAT	7570
gb:MT007544|Organism:Severe      AGAGTCGAATGTACAACTATTGTTAATGGTGTTAGAAGGTCCTTTTATGTCTATGCTAAT	7582
gb:MN994467|Organism:Severe      AGAGTCGAATGTACAACTATTGTTAATGGTGTTAGAAGGTCCTTTTATGTCTATGCTAAT	7582
gb:MT044257|Organism:Severe      AGAGTCGAATGTACAACTATTGTTAATGGTGTTAGAAGGTCCTTTTATGTCTATGCTAAT	7582
gb:MT106054|Organism:Severe      AGAGTCGAATGTACAACTATTGTTAATGGTGTTAGAAGGTCCTTTTATGTCTATGCTAAT	7582
gb:MT049951|Organism:Severe      AGAGTCGAATGTACAACTATTGTTAATGGTGTTAGAAGGTCCTTTTATGTCTATGCTAAT	7582
gb:MN975262|Organism:Severe      AGAGTCGAATGTACAACTATTGTTAATGGTGTTAGAAGGTCCTTTTATGTCTATGCTAAT	7582
gb:MT106052|Organism:Severe      AGAGTCGAATGTACAACTATTGTTAATGGTGTTAGAAGGTCCTTTTATGTCTATGCTAAT	7582
gb:LC522975|Organism:Severe      AGAGTCGAATGTACAACTATTGTTAATGGTGTTAGAAGGTCCTTTTATGTCTATGCTAAT	7579
gb:LC522973|Organism:Severe      AGAGTCGAATGTACAACTATTGTTAATGGTGTTAGAAGGTCCTTTTATGTCTATGCTAAT	7579
gb:LC522974|Organism:Severe      AGAGTCGAATGTACAACTATTGTTAATGGTGTTAGAAGGTCCTTTTATGTCTATGCTAAT	7579
gb:MN985325|Organism:Severe      AGAGTCGAATGTACAACTATTGTTAATGGTGTTAGAAGGTCCTTTTATGTCTATGCTAAT	7582
gb:MT020881|Organism:Severe      AGAGTCGAATGTACAACTATTGTTAATGGTGTTAGAAGGTCCTTTTATGTCTATGCTAAT	7582
gb:MT020880|Organism:Severe      AGAGTCGAATGTACAACTATTGTTAATGGTGTTAGAAGGTCCTTTTATGTCTATGCTAAT	7582
gb:MT066175|Organism:Severe      AGAGTCGAATGTACAACTATTGTTAATGGTGTTAGAAGGTCCTTTTATGTCTATGCTAAT	7582
gb:MN997409|Organism:Severe      AGAGTCGAATGTACAACTATTGTTAATGGTGTTAGAAGGTCCTTTTATGTCTATGCTAAT	7582
gb:MN938384|Organism:Severe      AGAGTCGAATGTACAACTATTGTTAATGGTGTTAGAAGGTCCTTTTATGTCTATGCTAAT	7550
gb:MT044258|Organism:Severe      AGAGTCGAATGTACAACTATTGTTAATGGTGTTAGAAGGTCCTTTTATGTCTATGCTAAT	7558
gb:MT039890|Organism:Severe      AGAGTCGAATGTACAACTATTGTTAATGGTGTTAGAAGGTCCTTTTATGTCTATGCTAAT	7582
gb:MN988713|Organism:Severe      AGAGTCGAATGTACAACTATTGTTAATGGTGTTAGAAGGTCCTTTTATGTCTATGCTAAT	7582
gb:LC521925|Organism:Severe      AGAGTCGAATGTACAACTATTGTTAATGGTGTTAGAAGGTCCTTTTATGTCTATGCTAAT	7555
gb:MT093571|Organism:Severe      AGAGTCGAATGTACAACTATTGTTAATGGTGTTAGAAGGTCCTTTTATGTCTATGCTAAT	7582
gb:MT039887|Organism:Severe      AGAGTCGAATGTACAACTATTGTTAATGGTGTTAGAAGGTCCTTTTATGTCTATGCTAAT	7582
gb:MT019530|Organism:Severe      AGAGTCGAATGTACAACTATTGTTAATGGTGTTAGAAGGTCCTTTTATGTCTATGCTAAT	7582
gb:MT039888|Organism:Severe      AGAGTCGAATGTACAACTATTGTTAATGGTGTTAGAAGGTCCTTTTATGTCTATGCTAAT	7582
gb:LC522972|Organism:Severe      AGAGTCGAATGTACAACTATTGTTAATGGTGTTAGAAGGTCCTTTTATGTCTATGCTAAT	7579
gb:MT027063|Organism:Severe      AGAGTCGAATGTACAACTATTGTTAATGGTGTTAGAAGGTCCTTTTATGTCTATGCTAAT	7582
gb:MT027062|Organism:Severe      AGAGTCGAATGTACAACTATTGTTAATGGTGTTAGAAGGTCCTTTTATGTCTATGCTAAT	7582
gb:MT019529|Organism:Severe      AGAGTCGAATGTACAACTATTGTTAATGGTGTTAGAAGGTCCTTTTATGTCTATGCTAAT	7582
gb:MN996529|Organism:Severe      AGAGTCGAATGTACAACTATTGTTAATGGTGTTAGAAGGTCCTTTTATGTCTATGCTAAT	7570
gb:MN996531|Organism:Severe      AGAGTCGAATGTACAACTATTGTTAATGGTGTTAGAAGGTCCTTTTATGTCTATGCTAAT	7569
gb:MT066176|Organism:Severe      AGAGTCGAATGTACAACTATTGTTAATGGTGTTAGAAGGTCCTTTTATGTCTATGCTAAT	7582
gb:MT027064|Organism:Severe      AGAGTCGAATGTACAACTATTGTTAATGGTGTTAGAAGGTCCTTTTATGTCTATGCTAAT	7582
gb:MN994468|Organism:Severe      AGAGTCGAATGTACAACTATTGTTAATGGTGTTAGAAGGTCCTTTTATGTCTATGCTAAT	7582
gb:MT072688|Organism:Severe      AGAGTCGAATGTACAACTATTGTTAATGGTGTTAGAAGGTCCTTTTATGTCTATGCTAAT	7567
gb:MN996527|Organism:Severe      AGAGTCGAATGTACAACTATTGTTAATGGTGTTAGAAGGTCCTTTTATGTCTATGCTAAT	7549
gb:MT093631|Organism:Severe      AGAGTCGAATGTACAACTATTGTTAATGGTGTTAGAAGGTCCTTTTATGTCTATGCTAAT	7620
gb:MT106053|Organism:Severe      AGAGTCGAATGTACAACTATTGTTAATGGTGTTAGAAGGTCCTTTTATGTCTATGCTAAT	7582
gb:MT019533|Organism:Severe      AGAGTCGAATGTACAACTATTGTTAATGGTGTTAGAAGGTCCTTTTATGTCTATGCTAAT	7582
gb:MT019531|Organism:Severe      AGAGTCGAATGTACAACTATTGTTAATGGTGTTAGAAGGTCCTTTTATGTCTATGCTAAT	7582
gb:MN996528|Organism:Severe      AGAGTCGAATGTACAACTATTGTTAATGGTGTTAGAAGGTCCTTTTATGTCTATGCTAAT	7582
gb:MN996530|Organism:Severe      AGAGTCGAATGTACAACTATTGTTAATGGTGTTAGAAGGTCCTTTTATGTCTATGCTAAT	7568
gb:MN908947|Organism:Severe      AGAGTCGAATGTACAACTATTGTTAATGGTGTTAGAAGGTCCTTTTATGTCTATGCTAAT	7582
gb:MT019532|Organism:Severe      AGAGTCGAATGTACAACTATTGTTAATGGTGTTAGAAGGTCCTTTTATGTCTATGCTAAT	7582
                                 ************************************************************

gb:MT020781|Organism:Severe      GGAGGTAAAGGCTTTTGCAAACTACACAATTGGAATTGTGTTAATTGTGATACATTCTGT	7630
gb:MT007544|Organism:Severe      GGAGGTAAAGGCTTTTGCAAACTACACAATTGGAATTGTGTTAATTGTGATACATTCTGT	7642
gb:MN994467|Organism:Severe      GGAGGTAAAGGCTTTTGCAAACTACACAATTGGAATTGTGTTAATTGTGATACATTCTGT	7642
gb:MT044257|Organism:Severe      GGAGGTAAAGGCTTTTGCAAACTACACAATTGGAATTGTGTTAATTGTGATACATTCTGT	7642
```

```
gb:MT106054|Organism:Severe      GGAGGTAAAGGCTTTTGCAAACTACACAATTGGAATTGTGTTAATTGTGATACATTCTGT   7642
gb:MT049951|Organism:Severe      GGAGGTAAAGGCTTTTGCAAACTACACAATTGGAATTGTGTTAATTGTGATACATTCTGT   7642
gb:MN975262|Organism:Severe      GGAGGTAAAGGCTTTTGCAAACTACACAATTGGAATTGTGTTAATTGTGATACATTCTGT   7642
gb:MT106052|Organism:Severe      GGAGGTAAAGGCTTTTGCAAACTACACAATTGGAATTGTGTTAATTGTGATACATTCTGT   7642
gb:LC522975|Organism:Severe      GGAGGTAAAGGCTTTTGCAAACTACACAATTGGAATTGTGTTAATTGTGATACATTCTGT   7639
gb:LC522973|Organism:Severe      GGAGGTAAAGGCTTTTGCAAACTACACAATTGGAATTGTGTTAATTGTGATACATTCTGT   7639
gb:LC522974|Organism:Severe      GGAGGTAAAGGCTTTTGCAAACTACACAATTGGAATTGTGTTAATTGTGATACATTCTGT   7639
gb:MN985325|Organism:Severe      GGAGGTAAAGGCTTTTGCAAACTACACAATTGGAATTGTGTTAATTGTGATACATTCTGT   7642
gb:MT020881|Organism:Severe      GGAGGTAAAGGCTTTTGCAAACTACACAATTGGAATTGTGTTAATTGTGATACATTCTGT   7642
gb:MT020880|Organism:Severe      GGAGGTAAAGGCTTTTGCAAACTACACAATTGGAATTGTGTTAATTGTGATACATTCTGT   7642
gb:MT066175|Organism:Severe      GGAGGTAAAGGCTTTTGCAAACTACACAATTGGAATTGTGTTAATTGTGATACATTCTGT   7642
gb:MN997409|Organism:Severe      GGAGGTAAAGGCTTTTGCAAACTACACAATTGGAATTGTGTTAATTGTGATACATTCTGT   7642
gb:MN938384|Organism:Severe      GGAGGTAAAGGCTTTTGCAAACTACACAATTGGAATTGTGTTAATTGTGATACATTCTGT   7610
gb:MT044258|Organism:Severe      GGAGGTAAAGGCTTTTGCAAACTACACAATTGGAATTGTGTTAATTGTGATACATTCTGT   7618
gb:MT039890|Organism:Severe      GGAGGTAAAGGCTTTTGCAAACTACACAATTGGAATTGTGTTAATTGTGATACATTCTGT   7642
gb:MN988713|Organism:Severe      GGAGGTAAAGGCTTTTGCAAACTACACAATTGGAATTGTGTTAATTGTGATACATTCTGT   7642
gb:LC521925|Organism:Severe      GGAGGTAAAGGCTTTTGCAAACTACACAATTGGAATTGTGTTAATTGTGATACATTCTGT   7615
gb:MT093571|Organism:Severe      GGAGGTAAAGGCTTTTGCAAACTACACAATTGGAATTGTGTTAATTGTGATACATTCTGT   7642
gb:MT039887|Organism:Severe      GGAGGTAAAGGCTTTTGCAAACTACACAATTGGAATTGTGTTAATTGTGATACATTCTGT   7642
gb:MT019530|Organism:Severe      GGAGGTAAAGGCTTTTGCAAACTACACAATTGGAATTGTGTTAATTGTGATACATTCTGT   7642
gb:MT039888|Organism:Severe      GGAGGTAAAGGCTTTTGCAAACTACACAATTGGAATTGTGTTAATTGTGATACATTCTGT   7642
gb:LC522972|Organism:Severe      GGAGGTAAAGGCTTTTGCAAACTACACAATTGGAATTGTGTTAATTGTGATACATTCTGT   7639
gb:MT027063|Organism:Severe      GGAGGTAAAGGCTTTTGCAAACTACACAATTGGAATTGTGTTAATTGTGATACATTCTGT   7642
gb:MT027062|Organism:Severe      GGAGGTAAAGGCTTTTGCAAACTACACAATTGGAATTGTGTTAATTGTGATACATTCTGT   7642
gb:MT019529|Organism:Severe      GGAGGTAAAGGCTTTTGCAAACTACACAATTGGAATTGTGTTAATTGTGATACATTCTGT   7642
gb:MN996529|Organism:Severe      GGAGGTAAAGGCTTTTGCAAACTACACAATTGGAATTGTGTTAATTGTGATACATTCTGT   7630
gb:MN996531|Organism:Severe      GGAGGTAAAGGCTTTTGCAAACTACACAATTGGAATTGTGTTAATTGTGATACATTCTGT   7629
gb:MT066176|Organism:Severe      GGAGGTAAAGGCTTTTGCAAACTACACAATTGGAATTGTGTTAATTGTGATACATTCTGT   7642
gb:MT027064|Organism:Severe      GGAGGTAAAGGCTTTTGCAAACTACACAATTGGAATTGTGTTAATTGTGATACATTCTGT   7642
gb:MN994468|Organism:Severe      GGAGGTAAAGGCTTTTGCAAACTACACAATTGGAATTGTGTTAATTGTGATACATTCTGT   7642
gb:MT072688|Organism:Severe      GGAGGTAAAGGCTTTTGCAAACTACACAATTGGAATTGTGTTAATTGTGATACATTCTGT   7627
gb:MN996527|Organism:Severe      GGAGGTAAAGGCTTTTGCAAACTACACAATTGGAATTGTGTTAATTGTGATACATTCTGT   7609
gb:MT093631|Organism:Severe      GGAGGTAAAGGCTTTTGCAAACTACACAATTGGAATTGTGTTAATTGTGATACATTCTGT   7680
gb:MT106053|Organism:Severe      GGAGGTAAAGGCTTTTGCAAACTACACAATTGGAATTGTGTTAATTGTGATACATTCTGT   7642
gb:MT019533|Organism:Severe      GGAGGTAAAGGCTTTTGCAAACTACACAATTGGAATTGTGTTAATTGTGATACATTCTGT   7642
gb:MT019531|Organism:Severe      GGAGGTAAAGGCTTTTGCAAACTACACAATTGGAATTGTGTTAATTGTGATACATTCTGT   7642
gb:MN996528|Organism:Severe      GGAGGTAAAGGCTTTTGCAAACTACACAATTGGAATTGTGTTAATTGTGATACATTCTGT   7642
gb:MN996530|Organism:Severe      GGAGGTAAAGGCTTTTGCAAACTACACAATTGGAATTGTGTTAATTGTGATACATTCTGT   7628
gb:MN908947|Organism:Severe      GGAGGTAAAGGCTTTTGCAAACTACACAATTGGAATTGTGTTAATTGTGATACATTCTGT   7642
gb:MT019532|Organism:Severe      GGAGGTAAAGGCTTTTGCAAACTACACAATTGGAATTGTGTTAATTGTGATACATTCTGT   7642
                                 ************************************************************

gb:MT020781|Organism:Severe      GCTGGTAGTACATTTATTAGTGATGAAGTTGCGAGAGACTTGTCACTACAGTTTAAAAGA   7690
gb:MT007544|Organism:Severe      GCTGGTAGTACATTTATTAGTGATGAAGTTGCGAGAGACTTGTCACTACAGTTTAAAAGA   7702
gb:MN994467|Organism:Severe      GCTGGTAGTACATTTATTAGTGATGAAGTTGCGAGAGACTTGTCACTACAGTTTAAAAGA   7702
gb:MT044257|Organism:Severe      GCTGGTAGTACATTTATTAGTGATGAAGTTGCGAGAGACTTGTCACTACAGTTTAAAAGA   7702
gb:MT106054|Organism:Severe      GCTGGTAGTACATTTATTAGTGATGAAGTTGCGAGAGACTTGTCACTACAGTTTAAAAGA   7702
gb:MT049951|Organism:Severe      GCTGGTAGTACATTTATTAGTGATGAAGTTGCGAGAGACTTGTCACTACAGTTTAAAAGA   7702
gb:MN975262|Organism:Severe      GCTGGTAGTACATTTATTAGTGATGAAGTTGCGAGAGACTTGTCACTACAGTTTAAAAGA   7702
gb:MT106052|Organism:Severe      GCTGGTAGTACATTTATTAGTGATGAAGTTGCGAGAGACTTGTCACTACAGTTTAAAAGA   7702
gb:LC522975|Organism:Severe      GCTGGTAGTACATTTATTAGTGATGAAGTTGCGAGAGACTTGTCACTACAGTTTAAAAGA   7699
gb:LC522973|Organism:Severe      GCTGGTAGTACATTTATTAGTGATGAAGTTGCGAGAGACTTGTCACTACAGTTTAAAAGA   7699
gb:LC522974|Organism:Severe      GCTGGTAGTACATTTATTAGTGATGAAGTTGCGAGAGACTTGTCACTACAGTTTAAAAGA   7699
gb:MN985325|Organism:Severe      GCTGGTAGTACATTTATTAGTGATGAAGTTGCGAGAGACTTGTCACTACAGTTTAAAAGA   7702
gb:MT020881|Organism:Severe      GCTGGTAGTACATTTATTAGTGATGAAGTTGCGAGAGACTTGTCACTACAGTTTAAAAGA   7702
gb:MT020880|Organism:Severe      GCTGGTAGTACATTTATTAGTGATGAAGTTGCGAGAGACTTGTCACTACAGTTTAAAAGA   7702
gb:MT066175|Organism:Severe      GCTGGTAGTACATTTATTAGTGATGAAGTTGCGAGAGACTTGTCACTACAGTTTAAAAGA   7702
gb:MN997409|Organism:Severe      GCTGGTAGTACATTTATTAGTGATGAAGTTGCGAGAGACTTGTCACTACAGTTTAAAAGA   7702
gb:MN938384|Organism:Severe      GCTGGTAGTACATTTATTAGTGATGAAGTTGCGAGAGACTTGTCACTACAGTTTAAAAGA   7670
gb:MT044258|Organism:Severe      GCTGGTAGTACATTTATTAGTGATGAAGTTGCGAGAGACTTGTCACTACAGTTTAAAAGA   7678
gb:MT039890|Organism:Severe      GCTGGTAGTACATTTATTAGTGATGAAGTTGCGAGAGACTTGTCACTACAGTTTAAAAGA   7702
gb:MN988713|Organism:Severe      GCTGGTAGTACATTTATTAGTGATGAAGTTGCGAGAGACTTGTCACTACAGTTTAAAAGA   7702
gb:LC521925|Organism:Severe      GCTGGTAGTACATTTATTAGTGATGAAGTTGCGAGAGACTTGTCACTACAGTTTAAAAGA   7675
gb:MT093571|Organism:Severe      GCTGGTAGTACATTTATTAGTGATGAAGTTGCGAGAGACTTGTCACTACAGTTTAAAAGA   7702
gb:MT039887|Organism:Severe      GCTGGTAGTACATTTATTAGTGATGAAGTTGCGAGAGACTTGTCACTACAGTTTAAAAGA   7702
```

```
gb:MT019530|Organism:Severe      GCTGGTAGTACATTTATTAGTGATGAAGTTGCGAGAGACTTGTCACTACAGTTTAAAAGA   7702
gb:MT039888|Organism:Severe      GCTGGTAGTACATTTATTAGTGATGAAGTTGCGAGAGACTTGTCACTACAGTTTAAAAGA   7702
gb:LC522972|Organism:Severe      GCTGGTAGTACATTTATTAGTGATGAAGTTGCGAGAGACTTGTCACTACAGTTTAAAAGA   7699
gb:MT027063|Organism:Severe      GCTGGTAGTACATTTATTAGTGATGAAGTTGCGAGAGACTTGTCACTACAGTTTAAAAGA   7702
gb:MT027062|Organism:Severe      GCTGGTAGTACATTTATTAGTGATGAAGTTGCGAGAGACTTGTCACTACAGTTTAAAAGA   7702
gb:MT019529|Organism:Severe      GCTGGTAGTACATTTATTAGTGATGAAGTTGCGAGAGACTTGTCACTACAGTTTAAAAGA   7702
gb:MN996529|Organism:Severe      GCTGGTAGTACATTTATTAGTGATGAAGTTGCGAGAGACTTGTCACTACAGTTTAAAAGA   7690
gb:MN996531|Organism:Severe      GCTGGTAGTACATTTATTAGTGATGAAGTTGCGAGAGACTTGTCACTACAGTTTAAAAGA   7689
gb:MT066176|Organism:Severe      GCTGGTAGTACATTTATTAGTGATGAAGTTGCGAGAGACTTGTCACTACAGTTTAAAAGA   7702
gb:MT027064|Organism:Severe      GCTGGTAGTACATTTATTAGTGATGAAGTTGCGAGAGACTTGTCACTACAGTTTAAAAGA   7702
gb:MN994468|Organism:Severe      GCTGGTAGTACATTTATTAGTGATGAAGTTGCGAGAGACTTGTCACTACAGTTTAAAAGA   7702
gb:MT072688|Organism:Severe      GCTGGTAGTACATTTATTAGTGATGAAGTTGCGAGAGACTTGTCACTACAGTTTAAAAGA   7687
gb:MN996527|Organism:Severe      GCTGGTAGTACATTTATTAGTGATGAAGTTGCGAGAGACTTGTCACTACAGTTTAAAAGA   7669
gb:MT093631|Organism:Severe      GCTGGTAGTACATTTATTAGTGATGAAGTTGCGAGAGACTTGTCACTACAGTTTAAAAGA   7740
gb:MT106053|Organism:Severe      GCTGGTAGTACATTTATTAGTGATGAAGTTGCGAGAGACTTGTCACTACAGTTTAAAAGA   7702
gb:MT019533|Organism:Severe      GCTGGTAGTACATTTATTAGTGATGAAGTTGCGAGAGACTTGTCACTACAGTTTAAAAGA   7702
gb:MT019531|Organism:Severe      GCTGGTAGTACATTTATTAGTGATGAAGTTGCGAGAGACTTGTCACTACAGTTTAAAAGA   7702
gb:MN996528|Organism:Severe      GCTGGTAGTACATTTATTAGTGATGAAGTTGCGAGAGACTTGTCACTACAGTTTAAAAGA   7702
gb:MN996530|Organism:Severe      GCTGGTAGTACATTTATTAGTGATGAAGTTGCGAGAGACTTGTCACTACAGTTTAAAAGA   7688
gb:MN908947|Organism:Severe      GCTGGTAGTACATTTATTAGTGATGAAGTTGCGAGAGACTTGTCACTACAGTTTAAAAGA   7702
gb:MT019532|Organism:Severe      GCTGGTAGTACATTTATTAGTGATGAAGTTGCGAGAGACTTGTCACTACAGTTTAAAAGA   7702
                                 ************************************************************

gb:MT020781|Organism:Severe      CCAATAAATCCTACTGACCAGTCTTCTTACATCGTTGATAGTGTTACAGTGAAGAATGGT   7750
gb:MT007544|Organism:Severe      CCAATAAATCCTACTGACCAGTCTTCTTACATCGTTGATAGTGTTACAGTGAAGAATGGT   7762
gb:MN994467|Organism:Severe      CCAATAAATCCTACTGACCAGTCTTCTTACATCGTTGATAGTGTTACAGTGAAGAATGGT   7762
gb:MT044257|Organism:Severe      CCAATAAATCCTACTGACCAGTCTTCTTACATCGTTGATAGTGTTACAGTGAAGAATGGT   7762
gb:MT106054|Organism:Severe      CCAATAAATCCTACTGACCAGTCTTCTTACATCGTTGATAGTGTTACAGTGAAGAATGGT   7762
gb:MT049951|Organism:Severe      CCAATAAATCCTACTGACCAGTCTTCTTACATCGTTGATAGTGTTACAGTGAAGAATGGT   7762
gb:MN975262|Organism:Severe      CCAATAAATCCTACTGACCAGTCTTCTTACATCGTTGATAGTGTTACAGTGAAGAATGGT   7762
gb:MT106052|Organism:Severe      CCAATAAATCCTACTGACCAGTCTTCTTACATCGTTGATAGTGTTACAGTGAAGAATGGT   7762
gb:LC522975|Organism:Severe      CCAATAAATCCTACTGACCAGTCTTCTTACATCGTTGATAGTGTTACAGTGAAGAATGGT   7759
gb:LC522973|Organism:Severe      CCAATAAATCCTACTGACCAGTCTTCTTACATCGTTGATAGTGTTACAGTGAAGAATGGT   7759
gb:LC522974|Organism:Severe      CCAATAAATCCTACTGACCAGTCTTCTTACATCGTTGATAGTGTTACAGTGAAGAATGGT   7759
gb:MN985325|Organism:Severe      CCAATAAATCCTACTGACCAGTCTTCTTACATCGTTGATAGTGTTACAGTGAAGAATGGT   7762
gb:MT020881|Organism:Severe      CCAATAAATCCTACTGACCAGTCTTCTTACATCGTTGATAGTGTTACAGTGAAGAATGGT   7762
gb:MT020880|Organism:Severe      CCAATAAATCCTACTGACCAGTCTTCTTACATCGTTGATAGTGTTACAGTGAAGAATGGT   7762
gb:MT066175|Organism:Severe      CCAATAAATCCTACTGACCAGTCTTCTTACATCGTTGATAGTGTTACAGTGAAGAATGGT   7762
gb:MN997409|Organism:Severe      CCAATAAATCCTACTGACCAGTCTTCTTACATCGTTGATAGTGTTACAGTGAAGAATGGT   7762
gb:MN938384|Organism:Severe      CCAATAAATCCTACTGACCAGTCTTCTTACATCGTTGATAGTGTTACAGTGAAGAATGGT   7730
gb:MT044258|Organism:Severe      CCAATAAATCCTACTGACCAGTCTTCTTACATCGTTGATAGTGTTACAGTGAAGAATGGT   7738
gb:MT039890|Organism:Severe      CCAATAAATCCTACTGACCAGTCTTCTTACATCGTTGATAGTGTTACAGTGAAGAATGGT   7762
gb:MN988713|Organism:Severe      CCAATAAATCCTACTGACCAGTCTTCTTACATCGTTGATAGTGTTACAGTGAAGAATGGT   7762
gb:LC521925|Organism:Severe      CCAATAAATCCTACTGACCAGTCTTCTTACATCGTTGATAGTGTTACAGTGAAGAATGGT   7735
gb:MT093571|Organism:Severe      CCAATAAATCCTACTGACCAGTCTTCTTACATCGTTGATAGTGTTACAGTGAAGAATGGT   7762
gb:MT039887|Organism:Severe      CCAATAAATCCTACTGACCAGTCTTCTTACATCGTTGATAGTGTTACAGTGAAGAATGGT   7762
gb:MT019530|Organism:Severe      CCAATAAATCCTACTGACCAGTCTTCTTACATCGTTGATAGTGTTACAGTGAAGAATGGT   7762
gb:MT039888|Organism:Severe      CCAATAAATCCTACTGACCAGTCTTCTTACATCGTTGATAGTGTTACAGTGAAGAATGGT   7762
gb:LC522972|Organism:Severe      CCAATAAATCCTACTGACCAGTCTTCTTACATCGTTGATAGTGTTACAGTGAAGAATGGT   7759
gb:MT027063|Organism:Severe      CCAATAAATCCTACTGACCAGTCTTCTTACATCGTTGATAGTGTTACAGTGAAGAATGGT   7762
gb:MT027062|Organism:Severe      CCAATAAATCCTACTGACCAGTCTTCTTACATCGTTGATAGTGTTACAGTGAAGAATGGT   7762
gb:MT019529|Organism:Severe      CCAATAAATCCTACTGACCAGTCTTCTTACATCGTTGATAGTGTTACAGTGAAGAATGGT   7762
gb:MN996529|Organism:Severe      CCAATAAATCCTACTGACCAGTCTTCTTACATCGTTGATAGTGTTACAGTGAAGAATGGT   7750
gb:MN996531|Organism:Severe      CCAATAAATCCTACTGACCAGTCTTCTTACATCGTTGATAGTGTTACAGTGAAGAATGGT   7749
gb:MT066176|Organism:Severe      CCAATAAATCCTACTGACCAGTCTTCTTACATCGTTGATAGTGTTACAGTGAAGAATGGT   7762
gb:MT027064|Organism:Severe      CCAATAAATCCTACTGACCAGTCTTCTTACATCGTTGATAGTGTTACAGTGAAGAATGGT   7762
gb:MN994468|Organism:Severe      CCAATAAATCCTACTGACCAGTCTTCTTACATCGTTGATAGTGTTACAGTGAAGAATGGT   7762
gb:MT072688|Organism:Severe      CCAATAAATCCTACTGACCAGTCTTCTTACATCGTTGATAGTGTTACAGTGAAGAATGGT   7747
gb:MN996527|Organism:Severe      CCAATAAATCCTACTGACCAGTCTTCTTACATCGTTGATAGTGTTACAGTGAAGAATGGT   7729
gb:MT093631|Organism:Severe      CCAATAAATCCTACTGACCAGTCTTCTTACATCGTTGATAGTGTTACAGTGAAGAATGGT   7800
gb:MT106053|Organism:Severe      CCAATAAATCCTACTGACCAGTCTTCTTACATCGTTGATAGTGTTACAGTGAAGAATGGT   7762
gb:MT019533|Organism:Severe      CCAATAAATCCTACTGACCAGTCTTCTTACATCGTTGATAGTGTTACAGTGAAGAATGGT   7762
gb:MT019531|Organism:Severe      CCAATAAATCCTACTGACCAGTCTTCTTACATCGTTGATAGTGTTACAGTGAAGAATGGT   7762
gb:MN996528|Organism:Severe      CCAATAAATCCTACTGACCAGTCTTCTTACATCGTTGATAGTGTTACAGTGAAGAATGGT   7762
gb:MN996530|Organism:Severe      CCAATAAATCCTACTGACCAGTCTTCTTACATCGTTGATAGTGTTACAGTGAAGAATGGT   7748
```

```
gb:MN908947|Organism:Severe      CCAATAAATCCTACTGACCAGTCTTCTTACATCGTTGATAGTGTTACAGTGAAGAATGGT    7762
gb:MT019532|Organism:Severe      CCAATAAATCCTACTGACCAGTCTTCTTACATCGTTGATAGTGTTACAGTGAAGAATGGT    7762
                                 ************************************************************

gb:MT020781|Organism:Severe      TCCATCCATCTTTACTTTGATAAAGCTGGTCAAAAGACTTATGAAAGACATTCTCTCTCT    7810
gb:MT007544|Organism:Severe      TCCATCCATCTTTACTTTGATAAAGCTGGTCAAAAGACTTATGAAAGACATTCTCTCTCT    7822
gb:MN994467|Organism:Severe      TCCATCCATCTTTACTTTGATAAAGCTGGTCAAAAGACTTATGAAAGACATTCTCTCTCT    7822
gb:MT044257|Organism:Severe      TCCATCCATCTTTACTTTGATAAAGCTGGTCAAAAGACTTATGAAAGACATTCTCTCTCT    7822
gb:MT106054|Organism:Severe      TCCATCCATCTTTACTTTGATAAAGCTGGTCAAAAGACTTATGAAAGACATTCTCTCTCT    7822
gb:MT049951|Organism:Severe      TCCATCCATCTTTACTTTGATAAAGCTGGTCAAAAGACTTATGAAAGACATTCTCTCTCT    7822
gb:MN975262|Organism:Severe      TCCATCCATCTTTACTTTGATAAAGCTGGTCAAAAGACTTATGAAAGACATTCTCTCTCT    7822
gb:MT106052|Organism:Severe      TCCATCCATCTTTACTTTGATAAAGCTGGTCAAAAGACTTATGAAAGACATTCTCTCTCT    7822
gb:LC522975|Organism:Severe      TCCATCCATCTTTACTTTGATAAAGCTGGTCAAAAGACTTATGAAAGACATTCTCTCTCT    7819
gb:LC522973|Organism:Severe      TCCATCCATCTTTACTTTGATAAAGCTGGTCAAAAGACTTATGAAAGACATTCTCTCTCT    7819
gb:LC522974|Organism:Severe      TCCATCCATCTTTACTTTGATAAAGCTGGTCAAAAGACTTATGAAAGACATTCTCTCTCT    7819
gb:MN985325|Organism:Severe      TCCATCCATCTTTACTTTGATAAAGCTGGTCAAAAGACTTATGAAAGACATTCTCTCTCT    7822
gb:MT020881|Organism:Severe      TCCATCCATCTTTACTTTGATAAAGCTGGTCAAAAGACTTATGAAAGACATTCTCTCTCT    7822
gb:MT020880|Organism:Severe      TCCATCCATCTTTACTTTGATAAAGCTGGTCAAAAGACTTATGAAAGACATTCTCTCTCT    7822
gb:MT066175|Organism:Severe      TCCATCCATCTTTACTTTGATAAAGCTGGTCAAAAGACTTATGAAAGACATTCTCTCTCT    7822
gb:MN997409|Organism:Severe      TCCATCCATCTTTACTTTGATAAAGCTGGTCAAAAGACTTATGAAAGACATTCTCTCTCT    7822
gb:MN938384|Organism:Severe      TCCATCCATCTTTACTTTGATAAAGCTGGTCAAAAGACTTATGAAAGACATTCTCTCTCT    7790
gb:MT044258|Organism:Severe      TCCATCCATCTTTACTTTGATAAAGCTGGTCAAAAGACTTATGAAAGACATTCTCTCTCT    7798
gb:MT039890|Organism:Severe      TCCATCCATCTTTACTTTGATAAAGCTGGTCAAAAGACTTATGAAAGACATTCTCTCTCT    7822
gb:MN988713|Organism:Severe      TCCATCCATCTTTACTTTGATAAAGCTGGTCAAAAGACTTATGAAAGACATTCTCTCTCT    7822
gb:LC521925|Organism:Severe      TCCATCCATCTTTACTTTGATAAAGCTGGTCAAAAGACTTATGAAAGACATTCTCTCTCT    7795
gb:MT093571|Organism:Severe      TCCATCCATCTTTACTTTGATAAAGCTGGTCAAAAGACTTATGAAAGACATTCTCTCTCT    7822
gb:MT039887|Organism:Severe      TCCATCCATCTTTACTTTGATAAAGCTGGTCAAAAGACTTATGAAAGACATTCTCTCTCT    7822
gb:MT019530|Organism:Severe      TCCATCCATCTTTACTTTGATAAAGCTGGTCAAAAGACTTATGAAAGACATTCTCTCTCT    7822
gb:MT039888|Organism:Severe      TCCATCCATCTTTACTTTGATAAAGCTGGTCAAAAGACTTATGAAAGACATTCTCTCTCT    7822
gb:LC522972|Organism:Severe      TCCATCCATCTTTACTTTGATAAAGCTGGTCAAAAGACTTATGAAAGACATTCTCTCTCT    7819
gb:MT027063|Organism:Severe      TCCATCCATCTTTACTTTGATAAAGCTGGTCAAAAGACTTATGAAAGACATTCTCTCTCT    7822
gb:MT027062|Organism:Severe      TCCATCCATCTTTACTTTGATAAAGCTGGTCAAAAGACTTATGAAAGACATTCTCTCTCT    7822
gb:MT019529|Organism:Severe      TCCATCCATCTTTACTTTGATAAAGCTGGTCAAAAGACTTATGAAAGACATTCTCTCTCT    7822
gb:MN996529|Organism:Severe      TCCATCCATCTTTACTTTGATAAAGCTGGTCAAAAGACTTATGAAAGACATTCTCTCTCT    7810
gb:MN996531|Organism:Severe      TCCATCCATCTTTACTTTGATAAAGCTGGTCAAAAGACTTATGAAAGACATTCTCTCTCT    7809
gb:MT066176|Organism:Severe      TCCATCCATCTTTACTTTGATAAAGCTGGTCAAAAGACTTATGAAAGACATTCTCTCTCT    7822
gb:MT027064|Organism:Severe      TCCATCCATCTTTACTTTGATAAAGCTGGTCAAAAGACTTATGAAAGACATTCTCTCTCT    7822
gb:MN994468|Organism:Severe      TCCATCCATCTTTACTTTGATAAAGCTGGTCAAAAGACTTATGAAAGACATTCTCTCTCT    7822
gb:MT072688|Organism:Severe      TCCATCCATCTTTACTTTGATAAAGCTGGTCAAAAGACTTATGAAAGACATTCTCTCTCT    7807
gb:MN996527|Organism:Severe      TCCATCCATCTTTACTTTGATAAAGCTGGTCAAAAGACTTATGAAAGACATTCTCTCTCT    7789
gb:MT093631|Organism:Severe      TCCATCCATCTTTACTTTGATAAAGCTGGTCAAAAGACTTATGAAAGACATTCTCTCTCT    7860
gb:MT106053|Organism:Severe      TCCATCCATCTTTACTTTGATAAAGCTGGTCAAAAGACTTATGAAAGACATTCTCTCTCT    7822
gb:MT019533|Organism:Severe      TCCATCCATCTTTACTTTGATAAAGCTGGTCAAAAGACTTATGAAAGACATTCTCTCTCT    7822
gb:MT019531|Organism:Severe      TCCATCCATCTTTACTTTGATAAAGCTGGTCAAAAGACTTATGAAAGACATTCTCTCTCT    7822
gb:MN996528|Organism:Severe      TCCATCCATCTTTACTTTGATAAAGCTGGTCAAAAGACTTATGAAAGACATTCTCTCTCT    7822
gb:MN996530|Organism:Severe      TCCATCCATCTTTACTTTGATAAAGCTGGTCAAAAGACTTATGAAAGACATTCTCTCTCT    7808
gb:MN908947|Organism:Severe      TCCATCCATCTTTACTTTGATAAAGCTGGTCAAAAGACTTATGAAAGACATTCTCTCTCT    7822
gb:MT019532|Organism:Severe      TCCATCCATCTTTACTTTGATAAAGCTGGTCAAAAGACTTATGAAAGACATTCTCTCTCT    7822
                                 ************************************************************

gb:MT020781|Organism:Severe      CATTTTGTTAACTTAGACAACCTGAGAGCTAATAACACTAAAGGTTCATTGCCTATTAAT    7870
gb:MT007544|Organism:Severe      CATTTTGTTAACTTAGACAACCTGAGAGCTAATAACACTAAAGGTTCATTGCCTATTAAT    7882
gb:MN994467|Organism:Severe      CATTTTGTTAACTTAGACAACCTGAGAGCTAATAACACTAAAGGTTCATTGCCTATTAAT    7882
gb:MT044257|Organism:Severe      CATTTTGTTAACTTAGACAACCTGAGAGCTAATAACACTAAAGGTTCATTGCCTATTAAT    7882
gb:MT106054|Organism:Severe      CATTTTGTTAACTTAGACAACCTGAGAGCTAATAACACTAAAGGTTCATTGCCTATTAAT    7882
gb:MT049951|Organism:Severe      CATTTTGTTAACTTAGACAACCTGAGAGCTAATAACACTAAAGGTTCATTGCCTATTAAT    7882
gb:MN975262|Organism:Severe      CATTTTGTTAACTTAGACAACCTGAGAGCTAATAACACTAAAGGTTCATTGCCTATTAAT    7882
gb:MT106052|Organism:Severe      CATTTTGTTAACTTAGACAACCTGAGAGCTAATAACACTAAAGGTTCATTGCCTATTAAT    7882
gb:LC522975|Organism:Severe      CATTTTGTTAACTTAGACAACCTGAGAGCTAATAACACTAAAGGTTCATTGCCTATTAAT    7879
gb:LC522973|Organism:Severe      CATTTTGTTAACTTAGACAACCTGAGAGCTAATAACACTAAAGGTTCATTGCCTATTAAT    7879
gb:LC522974|Organism:Severe      CATTTTGTTAACTTAGACAACCTGAGAGCTAATAACACTAAAGGTTCATTGCCTATTAAT    7879
gb:MN985325|Organism:Severe      CATTTTGTTAACTTAGACAACCTGAGAGCTAATAACACTAAAGGTTCATTGCCTATTAAT    7882
gb:MT020881|Organism:Severe      CATTTTGTTAACTTAGACAACCTGAGAGCTAATAACACTAAAGGTTCATTGCCTATTAAT    7882
gb:MT020880|Organism:Severe      CATTTTGTTAACTTAGACAACCTGAGAGCTAATAACACTAAAGGTTCATTGCCTATTAAT    7882
gb:MT066175|Organism:Severe      CATTTTGTTAACTTAGACAACCTGAGAGCTAATAACACTAAAGGTTCATTGCCTATTAAT    7882
```

```
gb:MN997409|Organism:Severe      CATTTTGTTAACTTAGACAACCTGAGAGCTAATAACACTAAAGGTTCATTGCCTATTAAT   7882
gb:MN938384|Organism:Severe      CATTTTGTTAACTTAGACAACCTGAGAGCTAATAACACTAAAGGTTCATTGCCTATTAAT   7850
gb:MT044258|Organism:Severe      CATTTTGTTAACTTAGACAACCTGAGAGCTAATAACACTAAAGGTTCATTGCCTATTAAT   7858
gb:MT039890|Organism:Severe      CATTTTGTTAACTTAGACAACCTGAGAGCTAATAACACTAAAGGTTCATTGCCTATTAAT   7882
gb:MN988713|Organism:Severe      CATTTTGTTAACTTAGACAACCTGAGAGCTAATAACACTAAAGGTTCATTGCCTATTAAT   7882
gb:LC521925|Organism:Severe      CATTTTGTTAACTTAGACAACCTGAGAGCTAATAACACTAAAGGTTCATTGCCTATTAAT   7855
gb:MT093571|Organism:Severe      CATTTTGTTAACTTAGACAACCTGAGAGCTAATAACACTAAAGGTTCATTGCCTATTAAT   7882
gb:MT039887|Organism:Severe      CATTTTGTTAACTTAGACAACCTGAGAGCTAATAACACTAAAGGTTCATTGCCTATTAAT   7882
gb:MT019530|Organism:Severe      CATTTTGTTAACTTAGACAACCTGAGAGCTAATAACACTAAAGGTTCATTGCCTATTAAT   7882
gb:MT039888|Organism:Severe      CATTTTGTTAACTTAGACAACCTGAGAGCTAATAACACTAAAGGTTCATTGCCTATTAAT   7882
gb:LC522972|Organism:Severe      CATTTTGTTAACTTAGACAACCTGAGAGCTAATAACACTAAAGGTTCATTGCCTATTAAT   7879
gb:MT027063|Organism:Severe      CATTTTGTTAACTTAGACAACCTGAGAGCTAATAACACTAAAGGTTCATTGCCTATTAAT   7882
gb:MT027062|Organism:Severe      CATTTTGTTAACTTAGACAACCTGAGAGCTAATAACACTAAAGGTTCATTGCCTATTAAT   7882
gb:MT019529|Organism:Severe      CATTTTGTTAACTTAGACAACCTGAGAGCTAATAACACTAAAGGTTCATTGCCTATTAAT   7882
gb:MN996529|Organism:Severe      CATTTTGTTAACTTAGACAACCTGAGAGCTAATAACACTAAAGGTTCATTGCCTATTAAT   7870
gb:MN996531|Organism:Severe      CATTTTGTTAACTTAGACAACCTGAGAGCTAATAACACTAAAGGTTCATTGCCTATTAAT   7869
gb:MT066176|Organism:Severe      CATTTTGTTAACTTAGACAACCTGAGAGCTAATAACACTAAAGGTTCATTGCCTATTAAT   7882
gb:MT027064|Organism:Severe      CATTTTGTTAACTTAGACAACCTGAGAGCTAATAACACTAAAGGTTCATTGCCTATTAAT   7882
gb:MN994468|Organism:Severe      CATTTTGTTAACTTAGACAACCTGAGAGCTAATAACACTAAAGGTTCATTGCCTATTAAT   7882
gb:MT072688|Organism:Severe      CATTTTGTTAACTTAGACAACCTGAGAGCTAATAACACTAAAGGTTCATTGCCTATTAAT   7867
gb:MN996527|Organism:Severe      CATTTTGTTAACTTAGACAACCTGAGAGCTAATAACACTAAAGGTTCATTGCCTATTAAT   7849
gb:MT093631|Organism:Severe      CATTTTGTTAACTTAGACAACCTGAGAGCTAATAACACTAAAGGTTCATTGCCTATTAAT   7920
gb:MT106053|Organism:Severe      CATTTTGTTAACTTAGACAACCTGAGAGCTAATAACACTAAAGGTTCATTGCCTATTAAT   7882
gb:MT019533|Organism:Severe      CATTTTGTTAACTTAGACAACCTGAGAGCTAATAACACTAAAGTTTCATTGCCTATTAAT   7882
gb:MT019531|Organism:Severe      CATTTTGTTAACTTAGACAACCTGAGAGCTAATAACACTAAAGGTTCATTGCCTATTAAT   7882
gb:MN996528|Organism:Severe      CATTTTGTTAACTTAGACAACCTGAGAGCTAATAACACTAAAGGTTCATTGCCTATTAAT   7882
gb:MN996530|Organism:Severe      CATTTTGTTAACTTAGACAACCTGAGAGCTAATAACACTAAAGGTTCATTGCCTATTAAT   7868
gb:MN908947|Organism:Severe      CATTTTGTTAACTTAGACAACCTGAGAGCTAATAACACTAAAGGTTCATTGCCTATTAAT   7882
gb:MT019532|Organism:Severe      CATTTTGTTAACTTAGACAACCTGAGAGCTAATAACACTAAAGGTTCATTGCCTATTAAT   7882
                                 ******************************************* ****************

gb:MT020781|Organism:Severe      GTTATAGTTTTTGATGGTAAATCAAAATGTGAAGAATCATCTGCAAAATCAGCGTCTGTT   7930
gb:MT007544|Organism:Severe      GTTATAGTTTTTGATGGTAAATCAAAATGTGAAGAATCATCTGCAAAATCAGCGTCTGTT   7942
gb:MN994467|Organism:Severe      GTTATAGTTTTTGATGGTAAATCAAAATGTGAAGAATCATCTGCAAAATCAGCGTCTGTT   7942
gb:MT044257|Organism:Severe      GTTATAGTTTTTGATGGTAAATCAAAATGTGAAGAATCATCTGCAAAATCAGCGTCTGTT   7942
gb:MT106054|Organism:Severe      GTTATAGTTTTTGATGGTAAATCAAAATGTGAAGAATCATCTGCAAAATCAGCGTCTGTT   7942
gb:MT049951|Organism:Severe      GTTATAGTTTTTGATGGTAAATCAAAATGTGAAGAATCATCTGCAAAATCAGCGTCTGTT   7942
gb:MN975262|Organism:Severe      GTTATAGTTTTTGATGGTAAATCAAAATGTGAAGAATCATCTGCAAAATCAGCGTCTGTT   7942
gb:MT106052|Organism:Severe      GTTATAGTTTTTGATGGTAAATCAAAATGTGAAGAATCATCTGCAAAATCAGCGTCTGTT   7942
gb:LC522975|Organism:Severe      GTTATAGTTTTTGATGGTAAATCAAAATGTGAAGAATCATCTGCAAAATCAGCGTCTGTT   7939
gb:LC522973|Organism:Severe      GTTATAGTTTTTGATGGTAAATCAAAATGTGAAGAATCATCTGCAAAATCAGCGTCTGTT   7939
gb:LC522974|Organism:Severe      GTTATAGTTTTTGATGGTAAATCAAAATGTGAAGAATCATCTGCAAAATCAGCGTCTGTT   7939
gb:MN985325|Organism:Severe      GTTATAGTTTTTGATGGTAAATCAAAATGTGAAGAATCATCTGCAAAATCAGCGTCTGTT   7942
gb:MT020881|Organism:Severe      GTTATAGTTTTTGATGGTAAATCAAAATGTGAAGAATCATCTGCAAAATCAGCGTCTGTT   7942
gb:MT020880|Organism:Severe      GTTATAGTTTTTGATGGTAAATCAAAATGTGAAGAATCATCTGCAAAATCAGCGTCTGTT   7942
gb:MT066175|Organism:Severe      GTTATAGTTTTTGATGGTAAATCAAAATGTGAAGAATCATCTGCAAAATCAGCGTCTGTT   7942
gb:MN997409|Organism:Severe      GTTATAGTTTTTGATGGTAAATCAAAATGTGAAGAATCATCTGCAAAATCAGCGTCTGTT   7942
gb:MN938384|Organism:Severe      GTTATAGTTTTTGATGGTAAATCAAAATGTGAAGAATCATCTGCAAAATCAGCGTCTGTT   7910
gb:MT044258|Organism:Severe      GTTATAGTTTTTGATGGTAAATCAAAATGTGAAGAATCATCTGCAAAATCAGCGTCTGTT   7918
gb:MT039890|Organism:Severe      GTTATAGTTTTTGATGGTAAATCAAAATGTGAAGAATCATCTGCAAAATCAGCGTCTGTT   7942
gb:MN988713|Organism:Severe      GTTATAGTTTTTGATGGTAAATCAAAATGTGAAGAATCATCTGCAAAATCAGCGTCTGTT   7942
gb:LC521925|Organism:Severe      GTTATAGTTTTTGATGGTAAATCAAAATGTGAAGAATCATCTGCAAAATCAGCGTCTGTT   7915
gb:MT093571|Organism:Severe      GTTATAGTTTTTGATGGTAAATCAAAATGTGAAGAATCATCTGCAAAATCAGCGTCTGTT   7942
gb:MT039887|Organism:Severe      GTTATAGTTTTTGATGGTAAATCAAAATGTGAAGAATCATCTGCAAAATCAGCGTCTGTT   7942
gb:MT019530|Organism:Severe      GTTATAGTTTTTGATGGTAAATCAAAATGTGAAGAATCATCTGCAAAATCAGCGTCTGTT   7942
gb:MT039888|Organism:Severe      GTTATAGTTTTTGATGGTAAATCAAAATGTGAAGAATCATCTGCAAAATCAGCGTCTGTT   7942
gb:LC522972|Organism:Severe      GTTATAGTTTTTGATGGTAAATCAAAATGTGAAGAATCATCTGCAAAATCAGCGTCTGTT   7939
gb:MT027063|Organism:Severe      GTTATAGTTTTTGATGGTAAATCAAAATGTGAAGAATCATCTGCAAAATCAGCGTCTGTT   7942
gb:MT027062|Organism:Severe      GTTATAGTTTTTGATGGTAAATCAAAATGTGAAGAATCATCTGCAAAATCAGCGTCTGTT   7942
gb:MT019529|Organism:Severe      GTTATAGTTTTTGATGGTAAATCAAAATGTGAAGAATCATCTGCAAAATCAGCGTCTGTT   7942
gb:MN996529|Organism:Severe      GTTATAGTTTTTGATGGTAAATCAAAATGTGAAGAATCATCTGCAAAATCAGCGTCTGTT   7930
gb:MN996531|Organism:Severe      GTTATAGTTTTTGATGGTAAATCAAAATGTGAAGAATCATCTGCAAAATCAGCGTCTGTT   7929
gb:MT066176|Organism:Severe      GTTATAGTTTTTGATGGTAAATCAAAATGTGAAGAATCATCTGCAAAATCAGCGTCTGTT   7942
gb:MT027064|Organism:Severe      GTTATAGTTTTTGATGGTAAATCAAAATGTGAAGAATCATCTGCAAAATCAGCGTCTGTT   7942
gb:MN994468|Organism:Severe      GTTATAGTTTTTGATGGTAAATCAAAATGTGAAGAATCATCTGCAAAATCAGCGTCTGTT   7942
```

```
gb:MT072688|Organism:Severe      GTTATAGTTTTTGATGGTAAATCAAAATGTGAAGAATCATCTGCAAAATCAGCGTCTGTT   7927
gb:MN996527|Organism:Severe      GTTATAGTTTTTGATGGTAAATCAAAATGTGAAGAATCATCTGCAAAATCAGCGTCTGTT   7909
gb:MT093631|Organism:Severe      GTTATAGTTTTTGATGGTAAATCAAAATGTGAAGAATCATCTGCAAAATCAGCGTCTGTT   7980
gb:MT106053|Organism:Severe      GTTATAGTTTTTGATGGTAAATCAAAATGTGAAGAATCATCTGCAAAATCAGCGTCTGTT   7942
gb:MT019533|Organism:Severe      GTTATAGTTTTTGATGGTAAATCAAAATGTGAAGAATCATCTGCAAAATCAGCGTCTGTT   7942
gb:MT019531|Organism:Severe      GTTATAGTTTTTGATGGTAAATCAAAATGTGAAGAATCATCTGCAAAATCAGCGTCTGTT   7942
gb:MN996528|Organism:Severe      GTTATAGTTTTTGATGGTAAATCAAAATGTGAAGAATCATCTGCAAAATCAGCGTCTGTT   7942
gb:MN996530|Organism:Severe      GTTATAGTTTTTGATGGTAAATCAAAATGTGAAGAATCATCTGCAAAATCAGCGTCTGTT   7928
gb:MN908947|Organism:Severe      GTTATAGTTTTTGATGGTAAATCAAAATGTGAAGAATCATCTGCAAAATCAGCGTCTGTT   7942
gb:MT019532|Organism:Severe      GTTATAGTTTTTGATGGTAAATCAAAATGTGAAGAATCATCTGCAAAATCAGCGTCTGTT   7942
                                 ************************************************************

gb:MT020781|Organism:Severe      TACTACAGTCAGCTTATGTGTCAACCTATACTGTTACTAGATCAGGCATTAGTGTCTGAT   7990
gb:MT007544|Organism:Severe      TACTACAGTCAGCTTATGTGTCAACCTATACTGTTACTAGATCAGGCATTAGTGTCTGAT   8002
gb:MN994467|Organism:Severe      TACTACAGTCAGCTTATGTGTCAACCTATACTGTTACTAGATCAGGCATTAGTGTCTGAT   8002
gb:MT044257|Organism:Severe      TACTACAGTCAGCTTATGTGTCAACCTATACTGTTACTAGATCAGGCATTAGTGTCTGAT   8002
gb:MT106054|Organism:Severe      TACTACAGTCAGCTTATGTGTCAACCTATACTGTTACTAGATCAGGCATTAGTGTCTGAT   8002
gb:MT049951|Organism:Severe      TACTACAGTCAGCTTATGTGTCAACCTATACTGTTACTAGATCAGGCATTAGTGTCTGAT   8002
gb:MN975262|Organism:Severe      TACTACAGTCAGCTTATGTGTCAACCTATACTGTTACTAGATCAGGCATTAGTGTCTGAT   8002
gb:MT106052|Organism:Severe      TACTACAGTCAGCTTATGTGTCAACCTATACTGTTACTAGATCAGGCATTAGTGTCTGAT   8002
gb:LC522975|Organism:Severe      TACTACAGTCAGCTTATGTGTCAACCTATACTGTTACTAGATCAGGCATTAGTGTCTGAT   7999
gb:LC522973|Organism:Severe      TACTACAGTCAGCTTATGTGTCAACCTATACTGTTACTAGATCAGGCATTAGTGTCTGAT   7999
gb:LC522974|Organism:Severe      TACTACAGTCAGCTTATGTGTCAACCTATACTGTTACTAGATCAGGCATTAGTGTCTGAT   7999
gb:MN985325|Organism:Severe      TACTACAGTCAGCTTATGTGTCAACCTATACTGTTACTAGATCAGGCATTAGTGTCTGAT   8002
gb:MT020881|Organism:Severe      TACTACAGTCAGCTTATGTGTCAACCTATACTGTTACTAGATCAGGCATTAGTGTCTGAT   8002
gb:MT020880|Organism:Severe      TACTACAGTCAGCTTATGTGTCAACCTATACTGTTACTAGATCAGGCATTAGTGTCTGAT   8002
gb:MT066175|Organism:Severe      TACTACAGTCAGCTTATGTGTCAACCTATACTGTTACTAGATCAGGCATTAGTGTCTGAT   8002
gb:MN997409|Organism:Severe      TACTACAGTCAGCTTATGTGTCAACCTATACTGTTACTAGATCAGGCATTAGTGTCTGAT   8002
gb:MN938384|Organism:Severe      TACTACAGTCAGCTTATGTGTCAACCTATACTGTTACTAGATCAGGCATTAGTGTCTGAT   7970
gb:MT044258|Organism:Severe      TACTACAGTCAGCTTATGTGTCAACCTATACTGTTACTAGATCAGGCATTAGTGTCTGAT   7978
gb:MT039890|Organism:Severe      TACTACAGTCAGCTTATGTGTCAACCTATACTGTTACTAGATCAGGCATTAGTGTCTGAT   8002
gb:MN988713|Organism:Severe      TACTACAGTCAGCTTATGTGTCAACCTATACTGTTACTAGATCAGGCATTAGTGTCTGAT   8002
gb:LC521925|Organism:Severe      TACTACAGTCAGCTTATGTGTCAACCTATACTGTTACTAGATCAGGCATTAGTGTCTGAT   7975
gb:MT093571|Organism:Severe      TACTACAGTCAGCTTATGTGTCAACCTATACTGTTACTAGATCAGGCATTAGTGTCTGAT   8002
gb:MT039887|Organism:Severe      TACTACAGTCAGCTTATGTGTCAACCTATACTGTTACTAGATCAGGCATTAGTGTCTGAT   8002
gb:MT019530|Organism:Severe      TACTACAGTCAGCTTATGTGTCAACCTATACTGTTACTAGATCAGGCATTAGTGTCTGAT   8002
gb:MT039888|Organism:Severe      TACTACAGTCAGCTTATGTGTCAACCTATACTGTTACTAGATCAGGCATTAGTGTCTGAT   8002
gb:LC522972|Organism:Severe      TACTACAGTCAGCTTATGTGTCAACCTATACTGTTACTAGATCAGGCATTAGTGTCTGAT   7999
gb:MT027063|Organism:Severe      TACTACAGTCAGCTTATGTGTCAACCTATACTGTTACTAGATCAGGCATTAGTGTCTGAT   8002
gb:MT027062|Organism:Severe      TACTACAGTCAGCTTATGTGTCAACCTATACTGTTACTAGATCAGGCATTAGTGTCTGAT   8002
gb:MT019529|Organism:Severe      TACTACAGTCAGCTTATGTGTCAACCTATACTGTTACTAGATCAGGCATTAGTGTCTGAT   8002
gb:MN996529|Organism:Severe      TACTACAGTCAGCTTATGTGTCAACCTATACTGTTACTAGATCAGGCATTAGTGTCTGAT   7990
gb:MN996531|Organism:Severe      TACTACAGTCAGCTTATGTGTCAACCTATACTGTTACTAGATCAGGCATTAGTGTCTGCT   7989
gb:MT066176|Organism:Severe      TACTACAGTCAGCTTATGTGTCAACCTATACTGTTACTAGATCAGGCATTAGTGTCTGAT   8002
gb:MT027064|Organism:Severe      TACTACAGTCAGCTTATGTGTCAACCTATACTGTTACTAGATCAGGCATTAGTGTCTGAT   8002
gb:MN994468|Organism:Severe      TACTACAGTCAGCTTATGTGTCAACCTATACTGTTACTAGATCAGGCATTAGTGTCTGAT   8002
gb:MT072688|Organism:Severe      TACTACAGTCAGCTTATGTGTCAACCTATACTGTTACTAGATCAGGCATTAGTGTCTGAT   7987
gb:MN996527|Organism:Severe      TACTACAGTCAGCTTATGTGTCAACCTATACTGTTACTAGATCAGGCATTAGTGTCTGAT   7969
gb:MT093631|Organism:Severe      TACTACAGTCAGCTTATGTGTCAACCTATACTGTTACTAGATCAGGCATTAGTGTCTGAT   8040
gb:MT106053|Organism:Severe      TACTACAGTCAGCTTATGTGTCAACCTATACTGTTACTAGATCAGGCATTAGTGTCTGAT   8002
gb:MT019533|Organism:Severe      TACTACAGTCAGCTTATGTGTCAACCTATACTGTTACTAGATCAGGCATTAGTGTCTGAT   8002
gb:MT019531|Organism:Severe      TACTACAGTCAGCTTATGTGTCAACCTATACTGTTACTAGATCAGGCATTAGTGTCTGAT   8002
gb:MN996528|Organism:Severe      TACTACAGTCAGCTTATGTGTCAACCTATACTGTTACTAGATCAGGCATTAGTGTCTGAT   8002
gb:MN996530|Organism:Severe      TACTACAGTCAGCTTATGTGTCAACCTATACTGTTACTAGATCAGGCATTAGTGTCTGAT   7988
gb:MN908947|Organism:Severe      TACTACAGTCAGCTTATGTGTCAACCTATACTGTTACTAGATCAGGCATTAGTGTCTGAT   8002
gb:MT019532|Organism:Severe      TACTACAGTCAGCTTATGTGTCAACCTATACTGTTACTAGATCAGGCATTAGTGTCTGAT   8002
                                 ********************************************************** *

gb:MT020781|Organism:Severe      GTTGGTGATAGTGCGGAAGTTGCAGTTAAAATGTTTGATGCTTACGTTAATACGTTTTCA   8050
gb:MT007544|Organism:Severe      GTTGGTGATAGTGCGGAAGTTGCAGTTAAAATGTTTGATGCTTACGTTAATACGTTTTCA   8062
gb:MN994467|Organism:Severe      GTTGGTGATAGTGCGGAAGTTGCAGTTAAAATGTTTGATGCTTACGTTAATACGTTTTCA   8062
gb:MT044257|Organism:Severe      GTTGGTGATAGTGCGGAAGTTGCAGTTAAAATGTTTGATGCTTACGTTAATACGTTTTCA   8062
gb:MT106054|Organism:Severe      GTTGGTGATAGTGCGGAAGTTGCAGTTAAAATGTTTGATGCTTACGTTAATACGTTTTCA   8062
gb:MT049951|Organism:Severe      GTTGGTGATAGTGCGGAAGTTGCAGTTAAAATGTTTGATGCTTACGTTAATACGTTTTCA   8062
gb:MN975262|Organism:Severe      GTTGGTGATAGTGCGGAAGTTGCAGTTAAAATGTTTGATGCTTACGTTAATACGTTTTCA   8062
```

```
gb:MT106052|Organism:Severe      GTTGGTGATAGTGCGGAAGTTGCAGTTAAAATGTTTGATGCTTACGTTAATACGTTTTCA   8062
gb:LC522975|Organism:Severe      GTTGGTGATAGTGCGGAAGTTGCAGTTAAAATGTTTGATGCTTACGTTAATACGTTTTCA   8059
gb:LC522973|Organism:Severe      GTTGGTGATAGTGCGGAAGTTGCAGTTAAAATGTTTGATGCTTACGTTAATACGTTTTCA   8059
gb:LC522974|Organism:Severe      GTTGGTGATAGTGCGGAAGTTGCAGTTAAAATGTTTGATGCTTACGTTAATACGTTTTCA   8059
gb:MN985325|Organism:Severe      GTTGGTGATAGTGCGGAAGTTGCAGTTAAAATGTTTGATGCTTACGTTAATACGTTTTCA   8062
gb:MT020881|Organism:Severe      GTTGGTGATAGTGCGGAAGTTGCAGTTAAAATGTTTGATGCTTACGTTAATACGTTTTCA   8062
gb:MT020880|Organism:Severe      GTTGGTGATAGTGCGGAAGTTGCAGTTAAAATGTTTGATGCTTACGTTAATACGTTTTCA   8062
gb:MT066175|Organism:Severe      GTTGGTGATAGTGCGGAAGTTGCAGTTAAAATGTTTGATGCTTACGTTAATACGTTTTCA   8062
gb:MN997409|Organism:Severe      GTTGGTGATAGTGCGGAAGTTGCAGTTAAAATGTTTGATGCTTACGTTAATACGTTTTCA   8062
gb:MN938384|Organism:Severe      GTTGGTGATAGTGCGGAAGTTGCAGTTAAAATGTTTGATGCTTACGTTAATACGTTTTCA   8030
gb:MT044258|Organism:Severe      GTTGGTGATAGTGCGGAAGTTGCAGTTAAAATGTTTGATGCTTACGTTAATACGTTTTCA   8038
gb:MT039890|Organism:Severe      GTTGGTGATAGTGCGGAAGTTGCAGTTAAAATGTTTGATGCTTACGTTAATACGTTTTCA   8062
gb:MN988713|Organism:Severe      GTTGGTGATAGTGCGGAAGTTGCAGTTAAAATGTTTGATGCTTACGTTAATACGTTTTCA   8062
gb:LC521925|Organism:Severe      GTTGGTGATAGTGCGGAAGTTGCAGTTAAAATGTTTGATGCTTACGTTAATACGTTTTCA   8035
gb:MT093571|Organism:Severe      GTTGGTGATAGTGCGGAAGTTGCAGTTAAAATGTTTGATGCTTACGTTAATACGTTTTCA   8062
gb:MT039887|Organism:Severe      GTTGGTGATAGTGCGGAAGTTGCAGTTAAAATGTTTGATGCTTACGTTAATACGTTTTCA   8062
gb:MT019530|Organism:Severe      GTTGGTGATAGTGCGGAAGTTGCAGTTAAAATGTTTGATGCTTACGTTAATACGTTTTCA   8062
gb:MT039888|Organism:Severe      GTTGGTGATAGTGCGGAAGTTGCAGTTAAAATGTTTGATGCTTACGTTAATACGTTTTCA   8062
gb:LC522972|Organism:Severe      GTTGGTGATAGTGCGGAAGTTGCAGTTAAAATGTTTGATGCTTACGTTAATACGTTTTCA   8059
gb:MT027063|Organism:Severe      GTTGGTGATAGTGCGGAAGTTGCAGTTAAAATGTTTGATGCTTACGTTAATACGTTTTCA   8062
gb:MT027062|Organism:Severe      GTTGGTGATAGTGCGGAAGTTGCAGTTAAAATGTTTGATGCTTACGTTAATACGTTTTCA   8062
gb:MT019529|Organism:Severe      GTTGGTGATAGTGCGGAAGTTGCAGTTAAAATGTTTGATGCTTACGTTAATACGTTTTCA   8062
gb:MN996529|Organism:Severe      GTTGGTGATAGTGCGGAAGTTGCAGTTAAAATGTTTGATGCTTACGTTAATACGTTTTCA   8050
gb:MN996531|Organism:Severe      GTTGGTGATAGTGCGGAAGTTGCAGTTAAAATGTTTGATGCTTACGTTAATACGTTTTCA   8049
gb:MT066176|Organism:Severe      GTTGGTGATAGTGCGGAAGTTGCAGTTAAAATGTTTGATGCTTACGTTAATACGTTTTCA   8062
gb:MT027064|Organism:Severe      GTTGGTGATAGTGCGGAAGTTGCAGTTAAAATGTTTGATGCTTACGTTAATACGTTTTCA   8062
gb:MN994468|Organism:Severe      GTTGGTGATAGTGCGGAAGTTGCAGTTAAAATGTTTGATGCTTACGTTAATACGTTTTCA   8062
gb:MT072688|Organism:Severe      GTTGGTGATAGTGCGGAAGTTGCAGTTAAAATGTTTGATGCTTACGTTAATACGTTTTCA   8047
gb:MN996527|Organism:Severe      GTTGGTGATAGTGCGGAAGTTGCAGTTAAAATGTTTGATGCTTACGTTAATACGTTTTCA   8029
gb:MT093631|Organism:Severe      GTTGGTGATAGTGCGGAAGTTGCAGTTAAAATGTTTGATGCTTACGTTAATACGTTTTCA   8100
gb:MT106053|Organism:Severe      GTTGGTGATAGTGCGGAAGTTGCAGTTAAAATGTTTGATGCTTACGTTAATACGTTTTCA   8062
gb:MT019533|Organism:Severe      GTTGGTGATAGTGCGGAAGTTGCAGTTAAAATGTTTGATGCTTACGTTAATACGTTTTCA   8062
gb:MT019531|Organism:Severe      GTTGGTGATAGTGCGGAAGTTGCAGTTAAAATGTTTGATGCTTACGTTAATACGTTTTCA   8062
gb:MN996528|Organism:Severe      GTTGGTGATAGTGCGGAAGTTGCAGTTAAAATGTTTGATGCTTACGTTAATACGTTTTCA   8062
gb:MN996530|Organism:Severe      GTTGGTGATAGTGCGGAAGTTGCAGTTAAAATGTTTGATGCTTACGTTAATACGTTTTCA   8048
gb:MN908947|Organism:Severe      GTTGGTGATAGTGCGGAAGTTGCAGTTAAAATGTTTGATGCTTACGTTAATACGTTTTCA   8062
gb:MT019532|Organism:Severe      GTTGGTGATAGTGCGGAAGTTGCAGTTAAAATGTTTGATGCTTACGTTAATACGTTTTCA   8062
                                 ************************************************************

gb:MT020781|Organism:Severe      TCAACTTTTAACGTACCAATGGAAAAACTCAAAACACTAGTTGCAACTGCAGAAGCTGAA   8110
gb:MT007544|Organism:Severe      TCAACTTTTAACGTACCAATGGAAAAACTCAAAACACTAGTTGCAACTGCAGAAGCTGAA   8122
gb:MN994467|Organism:Severe      TCAACTTTTAACGTACCAATGGAAAAACTCAAAACACTAGTTGCAACTGCAGAAGCTGAA   8122
gb:MT044257|Organism:Severe      TCAACTTTTAACGTACCAATGGAAAAACTCAAAACACTAGTTGCAACTGCAGAAGCTGAA   8122
gb:MT106054|Organism:Severe      TCAACTTTTAACGTACCAATGGAAAAACTCAAAACACTAGTTGCAACTGCAGAAGCTGAA   8122
gb:MT049951|Organism:Severe      TCAACTTTTAACGTACCAATGGAAAAACTCAAAACACTAGTTGCAACTGCAGAAGCTGAA   8122
gb:MN975262|Organism:Severe      TCAACTTTTAACGTACCAATGGAAAAACTCAAAACACTAGTTGCAACTGCAGAAGCTGAA   8122
gb:MT106052|Organism:Severe      TCAACTTTTAACGTACCAATGGAAAAACTCAAAACACTAGTTGCAACTGCAGAAGCTGAA   8122
gb:LC522975|Organism:Severe      TCAACTTTTAACGTACCAATGGAAAAACTCAAAACACTAGTTGCAACTGCAGAAGCTGAA   8119
gb:LC522973|Organism:Severe      TCAACTTTTAACGTACCAATGGAAAAACTCAAAACACTAGTTGCAACTGCAGAAGCTGAA   8119
gb:LC522974|Organism:Severe      TCAACTTTTAACGTACCAATGGAAAAACTCAAAACACTAGTTGCAACTGCAGAAGCTGAA   8119
gb:MN985325|Organism:Severe      TCAACTTTTAACGTACCAATGGAAAAACTCAAAACACTAGTTGCAACTGCAGAAGCTGAA   8122
gb:MT020881|Organism:Severe      TCAACTTTTAACGTACCAATGGAAAAACTCAAAACACTAGTTGCAACTGCAGAAGCTGAA   8122
gb:MT020880|Organism:Severe      TCAACTTTTAACGTACCAATGGAAAAACTCAAAACACTAGTTGCAACTGCAGAAGCTGAA   8122
gb:MT066175|Organism:Severe      TCAACTTTTAACGTACCAATGGAAAAACTCAAAACACTAGTTGCAACTGCAGAAGCTGAA   8122
gb:MN997409|Organism:Severe      TCAACTTTTAACGTACCAATGGAAAAACTCAAAACACTAGTTGCAACTGCAGAAGCTGAA   8122
gb:MN938384|Organism:Severe      TCAACTTTTAACGTACCAATGGAAAAACTCAAAACACTAGTTGCAACTGCAGAAGCTGAA   8090
gb:MT044258|Organism:Severe      TCAACTTTTAACGTACCAATGGAAAAACTCAAAACACTAGTTGCAACTGCAGAAGCTGAA   8098
gb:MT039890|Organism:Severe      TCAACTTTTAACGTACCAATGGAAAAACTCAAAACACTAGTTGCAACTGCAGAAGCTGAA   8122
gb:MN988713|Organism:Severe      TCAACTTTTAACGTACCAATGGAAAAACTCAAAACACTAGTTGCAACTGCAGAAGCTGAA   8122
gb:LC521925|Organism:Severe      TCAACTTTTAACGTACCAATGGAAAAACTCAAAACACTAGTTGCAACTGCAGAAGCTGAA   8095
gb:MT093571|Organism:Severe      TCAACTTTTAACGTACCAATGGAAAAACTCAAAACACTAGTTGCAACTGCAGAAGCTGAA   8122
gb:MT039887|Organism:Severe      TCAACTTTTAACGTACCAATGGAAAAACTCAAAACACTAGTTGCAACTGCAGAAGCTGAA   8122
gb:MT019530|Organism:Severe      TCAACTTTTAACGTACCAATGGAAAAACTCAAAACACTAGTTGCAACTGCAGAAGCTGAA   8122
gb:MT039888|Organism:Severe      TCAACTTTTAACGTACCAATGGAAAAACTCAAAACACTAGTTGCAACTGCAGAAGCTGAA   8122
gb:LC522972|Organism:Severe      TCAACTTTTAACGTACCAATGGAAAAACTCAAAACACTAGTTGCAACTGCAGAAGCTGAA   8119
```

```
gb:MT027063|Organism:Severe      TCAACTTTTAACGTACCAATGGAAAAACTCAAAACACTAGTTGCAACTGCAGAAGCTGAA   8122
gb:MT027062|Organism:Severe      TCAACTTTTAACGTACCAATGGAAAAACTCAAAACACTAGTTGCAACTGCAGAAGCTGAA   8122
gb:MT019529|Organism:Severe      TCAACTTTTAACGTACCAATGGAAAAACTCAAAACACTAGTTGCAACTGCAGAAGCTGAA   8122
gb:MN996529|Organism:Severe      TCAACTTTTAACGTACCAATGGAAAAACTCAAAACACTAGTTGCAACTGCAGAAGCTGAA   8110
gb:MN996531|Organism:Severe      TCAACTTTTAACGTACCAATGGAAAAACTCAAAACACTAGTTGCAACTGCAGAAGCTGAA   8109
gb:MT066176|Organism:Severe      TCAACTTTTAACGTACCAATGGAAAAACTCAAAACACTAGTTGCAACTGCAGAAGCTGAA   8122
gb:MT027064|Organism:Severe      TCAACTTTTAACGTACCAATGGAAAAACTCAAAACACTAGTTGCAACTGCAGAAGCTGAA   8122
gb:MN994468|Organism:Severe      TCAACTTTTAACGTACCAATGGAAAAACTCAAAACACTAGTTGCAACTGCAGAAGCTGAA   8122
gb:MT072688|Organism:Severe      TCAACTTTTAACGTACCAATGGAAAAACTCAAAACACTAGTTGCAACTGCAGAAGCTGAA   8107
gb:MN996527|Organism:Severe      TCAACTTTTAACGTACCAATGGAAAAACTCAAAACACTAGTTGCAACTGCAGAAGCTGAA   8089
gb:MT093631|Organism:Severe      TCAACTTTTAACGTACCAATGGAAAAACTCAAAACACTAGTTGCAACTGCAGAAGCTGAA   8160
gb:MT106053|Organism:Severe      TCAACTTTTAACGTACCAATGGAAAAACTCAAAACACTAGTTGCAACTGCAGAAGCTGAA   8122
gb:MT019533|Organism:Severe      TCAACTTTTAACGTACCAATGGAAAAACTCAAAACACTAGTTGCAACTGCAGAAGCTGAA   8122
gb:MT019531|Organism:Severe      TCAACTTTTAACGTACCAATGGAAAAACTCAAAACACTAGTTGCAACTGCAGAAGCTGAA   8122
gb:MN996528|Organism:Severe      TCAACTTTTAACGTACCAATGGAAAAACTCAAAACACTAGTTGCAACTGCAGAAGCTGAA   8122
gb:MN996530|Organism:Severe      TCAACTTTTAACGTACCAATGGAAAAACTCAAAACACTAGTTGCAACTGCAGAAGCTGAA   8108
gb:MN908947|Organism:Severe      TCAACTTTTAACGTACCAATGGAAAAACTCAAAACACTAGTTGCAACTGCAGAAGCTGAA   8122
gb:MT019532|Organism:Severe      TCAACTTTTAACGTACCAATGGAAAAACTCAAAACACTAGTTGCAACTGCAGAAGCTGAA   8122
                                 ************************************************************

gb:MT020781|Organism:Severe      CTTGCAAAGAATGTGTCCTTAGACAATGTCTTATCTACTTTTATTTCAGCAGCTCGGCAA   8170
gb:MT007544|Organism:Severe      CTTGCAAAGAATGTGTCCTTAGACAATGTCTTATCTACTTTTATTTCAGCAGCTCGGCAA   8182
gb:MN994467|Organism:Severe      CTTGCAAAGAATGTGTCCTTAGACAATGTCTTATCTACTTTTATTTCAGCAGCTCGGCAA   8182
gb:MT044257|Organism:Severe      CTTGCAAAGAATGTGTCCTTAGACAATGTCTTATCTACTTTTATTTCAGCAGCTCGGCAA   8182
gb:MT106054|Organism:Severe      CTTGCAAAGAATGTGTCCTTAGACAATGTCTTATCTACTTTTATTTCAGCAGCTCGGCAA   8182
gb:MT049951|Organism:Severe      CTTGCAAAGAATGTGTCCTTAGACAATGTCTTATCTACTTTTATTTCAGCAGCTCGGCAA   8182
gb:MN975262|Organism:Severe      CTTGCAAAGAATGTGTCCTTAGACAATGTCTTATCTACTTTTATTTCAGCAGCTCGGCAA   8182
gb:MT106052|Organism:Severe      CTTGCAAAGAATGTGTCCTTAGACAATGTCTTATCTACTTTTATTTCAGCAGCTCGGCAA   8182
gb:LC522975|Organism:Severe      CTTGCAAAGAATGTGTCCTTAGACAATGTCTTATCTACTTTTATTTCAGCAGCTCGGCAA   8179
gb:LC522973|Organism:Severe      CTTGCAAAGAATGTGTCCTTAGACAATGTCTTATCTACTTTTATTTCAGCAGCTCGGCAA   8179
gb:LC522974|Organism:Severe      CTTGCAAAGAATGTGTCCTTAGACAATGTCTTATCTACTTTTATTTCAGCAGCTCGGCAA   8179
gb:MN985325|Organism:Severe      CTTGCAAAGAATGTGTCCTTAGACAATGTCTTATCTACTTTTATTTCAGCAGCTCGGCAA   8182
gb:MT020881|Organism:Severe      CTTGCAAAGAATGTGTCCTTAGACAATGTCTTATCTACTTTTATTTCAGCAGCTCGGCAA   8182
gb:MT020880|Organism:Severe      CTTGCAAAGAATGTGTCCTTAGACAATGTCTTATCTACTTTTATTTCAGCAGCTCGGCAA   8182
gb:MT066175|Organism:Severe      CTTGCAAAGAATGTGTCCTTAGACAATGTCTTATCTACTTTTATTTCAGCAGCTCGGCAA   8182
gb:MN997409|Organism:Severe      CTTGCAAAGAATGTGTCCTTAGACAATGTCTTATCTACTTTTATTTCAGCAGCTCGGCAA   8182
gb:MN938384|Organism:Severe      CTTGCAAAGAATGTGTCCTTAGACAATGTCTTATCTACTTTTATTTCAGCAGCTCGGCAA   8150
gb:MT044258|Organism:Severe      CTTGCAAAGAATGTGTCCTTAGACAATGTCTTATCTACTTTTATTTCAGCAGCTCGGCAA   8158
gb:MT039890|Organism:Severe      CTTGCAAAGAATGTGTCCTTAGACAATGTCTTATCTACTTTTATTTCAGCAGCTCGGCAA   8182
gb:MN988713|Organism:Severe      CTTGCAAAGAATGTGTCCTTAGACAATGTCTTATCTACTTTTATTTCAGCAGCTCGGCAA   8182
gb:LC521925|Organism:Severe      CTTGCAAAGAATGTGTCCTTAGACAATGTCTTATCTACTTTTATTTCAGCAGCTCGGCAA   8155
gb:MT093571|Organism:Severe      CTTGCAAAGAATGTGTCCTTAGACAATGTCTTATCTACTTTTATTTCAGCAGCTCGGCAA   8182
gb:MT039887|Organism:Severe      CTTGCAAAGAATGTGTCCTTAGACAATGTCTTATCTACTTTTATTTCAGCAGCTCGGCAA   8182
gb:MT019530|Organism:Severe      CTTGCAAAGAATGTGTCCTTAGACAATGTCTTATCTACTTTTATTTCAGCAGCTCGGCAA   8182
gb:MT039888|Organism:Severe      CTTGCAAAGAATGTGTCCTTAGACAATGTCTTATCTACTTTTATTTCAGCAGCTCGGCAA   8182
gb:LC522972|Organism:Severe      CTTGCAAAGAATGTGTCCTTAGACAATGTCTTATCTACTTTTATTTCAGCAGCTCGGCAA   8179
gb:MT027063|Organism:Severe      CTTGCAAAGAATGTGTCCTTAGACAATGTCTTATCTACTTTTATTTCAGCAGCTCGGCAA   8182
gb:MT027062|Organism:Severe      CTTGCAAAGAATGTGTCCTTAGACAATGTCTTATCTACTTTTATTTCAGCAGCTCGGCAA   8182
gb:MT019529|Organism:Severe      CTTGCAAAGAATGTGTCCTTAGACAATGTCTTATCTACTTTTATTTCAGCAGCTCGGCAA   8182
gb:MN996529|Organism:Severe      CTTGCAAAGAATGTGTCCTTAGACAATGTCTTATCTACTTTTATTTCAGCAGCTCGGCAA   8170
gb:MN996531|Organism:Severe      CTTGCAAAGAATGTGTCCTTAGACAATGTCTTATCTACTTTTATTTCAGCAGCTCGGCAA   8169
gb:MT066176|Organism:Severe      CTTGCAAAGAATGTGTCCTTAGACAATGTCTTATCTACTTTTATTTCAGCAGCTCGGCAA   8182
gb:MT027064|Organism:Severe      CTTGCAAAGAATGTGTCCTTAGACAATGTCTTATCTACTTTTATTTCAGCAGCTCGGCAA   8182
gb:MN994468|Organism:Severe      CTTGCAAAGAATGTGTCCTTAGACAATGTCTTATCTACTTTTATTTCAGCAGCTCGGCAA   8182
gb:MT072688|Organism:Severe      CTTGCAAAGAATGTGTCCTTAGACAATGTCTTATCTACTTTTATTTCAGCAGCTCGGCAA   8167
gb:MN996527|Organism:Severe      CTTGCAAAGAATGTGTCCTTAGACAATGTCTTATCTACTTTTATTTCAGCAGCTCGGCAA   8149
gb:MT093631|Organism:Severe      CTTGCAAAGAATGTGTCCTTAGACAATGTCTTATCTACTTTTATTTCAGCAGCTCGGCAA   8220
gb:MT106053|Organism:Severe      CTTGCAAAGAATGTGTCCTTAGACAATGTCTTATCTACTTTTATTTCAGCAGCTCGGCAA   8182
gb:MT019533|Organism:Severe      CTTGCAAAGAATGTGTCCTTAGACAATGTCTTATCTACTTTTATTTCAGCAGCTCGGCAA   8182
gb:MT019531|Organism:Severe      CTTGCAAAGAATGTGTCCTTAGACAATGTCTTATCTACTTTTATTTCAGCAGCTCGGCAA   8182
gb:MN996528|Organism:Severe      CTTGCAAAGAATGTGTCCTTAGACAATGTCTTATCTACTTTTATTTCAGCAGCTCGGCAA   8182
gb:MN996530|Organism:Severe      CTTGCAAAGAATGTGTCCTTAGACAATGTCTTATCTACTTTTATTTCAGCAGCTCGGCAA   8168
gb:MN908947|Organism:Severe      CTTGCAAAGAATGTGTCCTTAGACAATGTCTTATCTACTTTTATTTCAGCAGCTCGGCAA   8182
gb:MT019532|Organism:Severe      CTTGCAAAGAATGTGTCCTTAGACAATGTCTTATCTACTTTTATTTCAGCAGCTCGGCAA   8182
                                 ************************************************************
```

```
gb:MT020781|Organism:Severe      GGGTTTGTTGATTCAGATGTAGAAACTAAAGATGTTGTTGAATGTCTTAAATTGTCACAT   8230
gb:MT007544|Organism:Severe      GGGTTTGTTGATTCAGATGTAGAAACTAAAGATGTTGTTGAATGTCTTAAATTGTCACAT   8242
gb:MN994467|Organism:Severe      GGGTTTGTTGATTCAGATGTAGAAACTAAAGATGTTGTTGAATGTCTTAAATTGTCACAT   8242
gb:MT044257|Organism:Severe      GGGTTTGTTGATTCAGATGTAGAAACTAAAGATGTTGTTGAATGTCTTAAATTGTCACAT   8242
gb:MT106054|Organism:Severe      GGGTTTGTTGATTCAGATGTAGAAACTAAAGATGTTGTTGAATGTCTTAAATTGTCACAT   8242
gb:MT049951|Organism:Severe      GGGTTTGTTGATTCAGATGTAGAAACTAAAGATGTTGTTGAATGTCTTAAATTGTCACAT   8242
gb:MN975262|Organism:Severe      GGGTTTGTTGATTCAGATGTAGAAACTAAAGATGTTGTTGAATGTCTTAAATTGTCACAT   8242
gb:MT106052|Organism:Severe      GGGTTTGTTGATTCAGATGTAGAAACTAAAGATGTTGTTGAATGTCTTAAATTGTCACAT   8242
gb:LC522975|Organism:Severe      GGGTTTGTTGATTCAGATGTAGAAACTAAAGATGTTGTTGAATGTCTTAAATTGTCACAT   8239
gb:LC522973|Organism:Severe      GGGTTTGTTGATTCAGATGTAGAAACTAAAGATGTTGTTGAATGTCTTAAATTGTCACAT   8239
gb:LC522974|Organism:Severe      GGGTTTGTTGATTCAGATGTAGAAACTAAAGATGTTGTTGAATGTCTTAAATTGTCACAT   8239
gb:MN985325|Organism:Severe      GGGTTTGTTGATTCAGATGTAGAAACTAAAGATGTTGTTGAATGTCTTAAATTGTCACAT   8242
gb:MT020881|Organism:Severe      GGGTTTGTTGATTCAGATGTAGAAACTAAAGATGTTGTTGAATGTCTTAAATTGTCACAT   8242
gb:MT020880|Organism:Severe      GGGTTTGTTGATTCAGATGTAGAAACTAAAGATGTTGTTGAATGTCTTAAATTGTCACAT   8242
gb:MT066175|Organism:Severe      GGGTTTGTTGATTCAGATGTAGAAACTAAAGATGTTGTTGAATGTCTTAAATTGTCACAT   8242
gb:MN997409|Organism:Severe      GGGTTTGTTGATTCAGATGTAGAAACTAAAGATGTTGTTGAATGTCTTAAATTGTCACAT   8242
gb:MN938384|Organism:Severe      GGGTTTGTTGATTCAGATGTAGAAACTAAAGATGTTGTTGAATGTCTTAAATTGTCACAT   8210
gb:MT044258|Organism:Severe      GGGTTTGTTGATTCAGATGTAGAAACTAAAGATGTTGTTGAATGTCTTAAATTGTCACAT   8218
gb:MT039890|Organism:Severe      GGGTTTGTTGATTCAGATGTAGAAACTAAAGATGTTGTTGAATGTCTTAAATTGTCACAT   8242
gb:MN988713|Organism:Severe      GGGTTTGTTGATTCAGATGTAGAAACTAAAGATGTTGTTGAATGTCTTAAATTGTCACAT   8242
gb:LC521925|Organism:Severe      GGGTTTGTTGATTCAGATGTAGAAACTAAAGATGTTGTTGAATGTCTTAAATTGTCACAT   8215
gb:MT093571|Organism:Severe      GGGTTTGTTGATTCAGATGTAGAAACTAAAGATGTTGTTGAATGTCTTAAATTGTCACAT   8242
gb:MT039887|Organism:Severe      GGGTTTGTTGATTCAGATGTAGAAACTAAAGATGTTGTTGAATGTCTTAAATTGTCACAT   8242
gb:MT019530|Organism:Severe      GGGTTTGTTGATTCAGATGTAGAAACTAAAGATGTTGTTGAATGTCTTAAATTGTCACAT   8242
gb:MT039888|Organism:Severe      GGGTTTGTTGATTCAGATGTAGAAACTAAAGATGTTGTTGAATGTCTTAAATTGTCACAT   8242
gb:LC522972|Organism:Severe      GGGTTTGTTGATTCAGATGTAGAAACTAAAGATGTTGTTGAATGTCTTAAATTGTCACAT   8239
gb:MT027063|Organism:Severe      GGGTTTGTTGATTCAGATGTAGAAACTAAAGATGTTGTTGAATGTCTTAAATTGTCACAT   8242
gb:MT027062|Organism:Severe      GGGTTTGTTGATTCAGATGTAGAAACTAAAGATGTTGTTGAATGTCTTAAATTGTCACAT   8242
gb:MT019529|Organism:Severe      GGGTTTGTTGATTCAGATGTAGAAACTAAAGATGTTGTTGAATGTCTTAAATTGTCACAT   8242
gb:MN996529|Organism:Severe      GGGTTTGTTGATTCAGATGTAGAAACTAAAGATGTTGTTGAATGTCTTAAATTGTCACAT   8230
gb:MN996531|Organism:Severe      GGGTTTGTTGATTCAGATGTAGAAACTAAAGATGTTGTTGAATGTCTTAAATTGTCACAT   8229
gb:MT066176|Organism:Severe      GGGTTTGTTGATTCAGATGTAGAAACTAAAGATGTTGTTGAATGTCTTAAATTGTCACAT   8242
gb:MT027064|Organism:Severe      GGGTTTGTTGATTCAGATGTAGAAACTAAAGATGTTGTTGAATGTCTTAAATTGTCACAT   8242
gb:MN994468|Organism:Severe      GGGTTTGTTGATTCAGATGTAGAAACTAAAGATGTTGTTGAATGTCTTAAATTGTCACAT   8242
gb:MT072688|Organism:Severe      GGGTTTGTTGATTCAGATGTAGAAACTAAAGATGTTGTTGAATGTCTTAAATTGTCACAT   8227
gb:MN996527|Organism:Severe      GGGTTTGTTGATTCAGATGTAGAAACTAAAGATGTTGTTGAATGTCTTAAATTGTCACAT   8209
gb:MT093631|Organism:Severe      GGGTTTGTTGATTCAGATGTAGAAACTAAAGATGTTGTTGAATGTCTTAAATTGTCACAT   8280
gb:MT106053|Organism:Severe      GGGTTTGTTGATTCAGATGTAGAAACTAAAGATGTTGTTGAATGTCTTAAATTGTCACAT   8242
gb:MT019533|Organism:Severe      GGGTTTGTTGATTCAGATGTAGAAACTAAAGATGTTGTTGAATGTCTTAAATTGTCACAT   8242
gb:MT019531|Organism:Severe      GGGTTTGTTGATTCAGATGTAGAAACTAAAGATGTTGTTGAATGTCTTAAATTGTCACAT   8242
gb:MN996528|Organism:Severe      GGGTTTGTTGATTCAGATGTAGAAACTAAAGATGTTGTTGAATGTCTTAAATTGTCACAT   8242
gb:MN996530|Organism:Severe      GGGTTTGTTGATTCAGATGTAGAAACTAAAGATGTTGTTGAATGTCTTAAATTGTCACAT   8228
gb:MN908947|Organism:Severe      GGGTTTGTTGATTCAGATGTAGAAACTAAAGATGTTGTTGAATGTCTTAAATTGTCACAT   8242
gb:MT019532|Organism:Severe      GGGTTTGTTGATTCAGATGTAGAAACTAAAGATGTTGTTGAATGTCTTAAATTGTCACAT   8242
                                 ************************************************************

gb:MT020781|Organism:Severe      CAATCTGACATAGAAGTTACTGGCGATAGTTGTAATAACTATATGCTCACCTATAACAAA   8290
gb:MT007544|Organism:Severe      CAATCTGACATAGAAGTTACTGGCGATAGTTGTAATAACTATATGCTCACCTATAACAAA   8302
gb:MN994467|Organism:Severe      CAATCTGACATAGAAGTTACTGGCGATAGTTGTAATAACTATATGCTCACCTATAACAAA   8302
gb:MT044257|Organism:Severe      CAATCTGACATAGAAGTTACTGGCGATAGTTGTAATAACTATATGCTCACCTATAACAAA   8302
gb:MT106054|Organism:Severe      CAATCTGACATAGAAGTTACTGGCGATAGTTGTAATAACTATATGCTCACCTATAACAAA   8302
gb:MT049951|Organism:Severe      CAATCTGACATAGAAGTTACTGGCGATAGTTGTAATAACTATATGCTCACCTATAACAAA   8302
gb:MN975262|Organism:Severe      CAATCTGACATAGAAGTTACTGGCGATAGTTGTAATAACTATATGCTCACCTATAACAAA   8302
gb:MT106052|Organism:Severe      CAATCTGACATAGAAGTTACTGGCGATAGTTGTAATAACTATATGCTCACCTATAACAAA   8302
gb:LC522975|Organism:Severe      CAATCTGACATAGAAGTTACTGGCGATAGTTGTAATAACTATATGCTCACCTATAACAAA   8299
gb:LC522973|Organism:Severe      CAATCTGACATAGAAGTTACTGGCGATAGTTGTAATAACTATATGCTCACCTATAACAAA   8299
gb:LC522974|Organism:Severe      CAATCTGACATAGAAGTTACTGGCGATAGTTGTAATAACTATATGCTCACCTATAACAAA   8299
gb:MN985325|Organism:Severe      CAATCTGACATAGAAGTTACTGGCGATAGTTGTAATAACTATATGCTCACCTATAACAAA   8302
gb:MT020881|Organism:Severe      CAATCTGACATAGAAGTTACTGGCGATAGTTGTAATAACTATATGCTCACCTATAACAAA   8302
gb:MT020880|Organism:Severe      CAATCTGACATAGAAGTTACTGGCGATAGTTGTAATAACTATATGCTCACCTATAACAAA   8302
gb:MT066175|Organism:Severe      CAATCTGACATAGAAGTTACTGGCGATAGTTGTAATAACTATATGCTCACCTATAACAAA   8302
gb:MN997409|Organism:Severe      CAATCTGACATAGAAGTTACTGGCGATAGTTGTAATAACTATATGCTCACCTATAACAAA   8302
gb:MN938384|Organism:Severe      CAATCTGACATAGAAGTTACTGGCGATAGTTGTAATAACTATATGCTCACCTATAACAAA   8270
gb:MT044258|Organism:Severe      CAATCTGACATAGAAGTTACTGGCGATAGTTGTAATAACTATATGCTCACCTATAACAAA   8278
```

```
gb:MT039890|Organism:Severe      CAATCTGACATAGAAGTTACTGGCGATAGTTGTAATAACTATATGCTCACCTATAACAAA   8302
gb:MN988713|Organism:Severe      CAATCTGACATAGAAGTTACTGGCGATAGTTGTAATAACTATATGCTCACCTATAACAAA   8302
gb:LC521925|Organism:Severe      CAATCTGACATAGAAGTTACTGGCGATAGTTGTAATAACTATATGCTCACCTATAACAAA   8275
gb:MT093571|Organism:Severe      CAATCTGACATAGAAGTTACTGGCGATAGTTGTAATAACTATATGCTCACCTATAACAAA   8302
gb:MT039887|Organism:Severe      CAATCTGACATAGAAGTTACTGGCGATAGTTGTAATAACTATATGCTCACCTATAACAAA   8302
gb:MT019530|Organism:Severe      CAATCTGACATAGAAGTTACTGGCGATAGTTGTAATAACTATATGCTCACCTATAACAAA   8302
gb:MT039888|Organism:Severe      CAATCTGACATAGAAGTTACTGGCGATAGTTGTAATAACTATATGCTCACCTATAACAAA   8302
gb:LC522972|Organism:Severe      CAATCTGACATAGAAGTTACTGGCGATAGTTGTAATAACTATATGCTCACCTATAACAAA   8299
gb:MT027063|Organism:Severe      CAATCTGACATAGAAGTTACTGGCGATAGTTGTAATAACTATATGCTCACCTATAACAAA   8302
gb:MT027062|Organism:Severe      CAATCTGACATAGAAGTTACTGGCGATAGTTGTAATAACTATATGCTCACCTATAACAAA   8302
gb:MT019529|Organism:Severe      CAATCTGACATAGAAGTTACTGGCGATAGTTGTAATAACTATATGCTCACCTATAACAAA   8302
gb:MN996529|Organism:Severe      CAATCTGACATAGAAGTTACTGGCGATAGTTGTAATAACTATATGCTCACCTATAACAAA   8290
gb:MN996531|Organism:Severe      CAATCTGACATAGAAGTTACTGGCGATAGTTGTAATAACTATATGCTCACCTATAACAAA   8289
gb:MT066176|Organism:Severe      CAATCTGACATAGAAGTTACTGGCGATAGTTGTAATAACTATATGCTCACCTATAACAAA   8302
gb:MT027064|Organism:Severe      CAATCTGACATAGAAGTTACTGGCGATAGTTGTAATAACTATATGCTCACCTATAACAAA   8302
gb:MN994468|Organism:Severe      CAATCTGACATAGAAGTTACTGGCGATAGTTGTAATAACTATATGCTCACCTATAACAAA   8302
gb:MT072688|Organism:Severe      CAATCTGACATAGAAGTTACTGGCGATAGTTGTAATAACTATATGCTCACCTATAACAAA   8287
gb:MN996527|Organism:Severe      CAATCTGACATAGAAGTTACTGGCGATAGTTGTAATAACTATATGCTCACCTATAACAAA   8269
gb:MT093631|Organism:Severe      CAATCTGACATAGAAGTTACTGGCGATAGTTGTAATAACTATATGCTCACCTATAACAAA   8340
gb:MT106053|Organism:Severe      CAATCTGACATAGAAGTTACTGGCGATAGTTGTAATAACTATATGCTCACCTATAACAAA   8302
gb:MT019533|Organism:Severe      CAATCTGACATAGAAGTTACTGGCGATAGTTGTAATAACTATATGCTCACCTATAACAAA   8302
gb:MT019531|Organism:Severe      CAATCTGACATAGAAGTTACTGGCGATAGTTGTAATAACTATATGCTCACCTATAACAAA   8302
gb:MN996528|Organism:Severe      CAATCTGACATAGAAGTTACTGGCGATAGTTGTAATAACTATATGCTCACCTATAACAAA   8302
gb:MN996530|Organism:Severe      CAATCTGACATAGAAGTTACTGGCGATAGTTGTAATAACTATATGCTCACCTATAACAAA   8288
gb:MN908947|Organism:Severe      CAATCTGACATAGAAGTTACTGGCGATAGTTGTAATAACTATATGCTCACCTATAACAAA   8302
gb:MT019532|Organism:Severe      CAATCTGACATAGAAGTTACTGGCGATAGTTGTAATAACTATATGCTCACCTATAACAAA   8302
                                 ************************************************************

gb:MT020781|Organism:Severe      GTTGAAAACATGACACCCCGTGACCTTGGTGCTTGTATTGACTGTAGTGCGCGTCATATT   8350
gb:MT007544|Organism:Severe      GTTGAAAACATGACACCCCGTGACCTTGGTGCTTGTATTGACTGTAGTGCGCGTCATATT   8362
gb:MN994467|Organism:Severe      GTTGAAAACATGACACCCCGTGACCTTGGTGCTTGTATTGACTGTAGTGCGCGTCATATT   8362
gb:MT044257|Organism:Severe      GTTGAAAACATGACACCCCGTGACCTTGGTGCTTGTATTGACTGTAGTGCGCGTCATATT   8362
gb:MT106054|Organism:Severe      GTTGAAAACATGACACCCCGTGACCTTGGTGCTTGTATTGACTGTAGTGCGCGTCATATT   8362
gb:MT049951|Organism:Severe      GTTGAAAACATGACACCCCGTGACCTTGGTGCTTGTATTGACTGTAGTGCGCGTCATATT   8362
gb:MN975262|Organism:Severe      GTTGAAAACATGACACCCCGTGACCTTGGTGCTTGTATTGACTGTAGTGCGCGTCATATT   8362
gb:MT106052|Organism:Severe      GTTGAAAACATGACACCCCGTGACCTTGGTGCTTGTATTGACTGTAGTGCGCGTCATATT   8362
gb:LC522975|Organism:Severe      GTTGAAAACATGACACCCCGTGACCTTGGTGCTTGTATTGACTGTAGTGCGCGTCATATT   8359
gb:LC522973|Organism:Severe      GTTGAAAACATGACACCCCGTGACCTTGGTGCTTGTATTGACTGTAGTGCGCGTCATATT   8359
gb:LC522974|Organism:Severe      GTTGAAAACATGACACCCCGTGACCTTGGTGCTTGTATTGACTGTAGTGCGCGTCATATT   8359
gb:MN985325|Organism:Severe      GTTGAAAACATGACACCCCGTGACCTTGGTGCTTGTATTGACTGTAGTGCGCGTCATATT   8362
gb:MT020881|Organism:Severe      GTTGAAAACATGACACCCCGTGACCTTGGTGCTTGTATTGACTGTAGTGCGCGTCATATT   8362
gb:MT020880|Organism:Severe      GTTGAAAACATGACACCCCGTGACCTTGGTGCTTGTATTGACTGTAGTGCGCGTCATATT   8362
gb:MT066175|Organism:Severe      GTTGAAAACATGACACCCCGTGACCTTGGTGCTTGTATTGACTGTAGTGCGCGTCATATT   8362
gb:MN997409|Organism:Severe      GTTGAAAACATGACACCCCGTGACCTTGGTGCTTGTATTGACTGTAGTGCGCGTCATATT   8362
gb:MN938384|Organism:Severe      GTTGAAAACATGACACCCCGTGACCTTGGTGCTTGTATTGACTGTAGTGCGCGTCATATT   8330
gb:MT044258|Organism:Severe      GTTGAAAACATGACACCCCGTGACCTTGGTGCTTGTATTGACTGTAGTGCGCGTCATATT   8338
gb:MT039890|Organism:Severe      GTTGAAAACATGACACCCCGTGACCTTGGTGCTTGTATTGACTGTAGTGCGCGTCATATT   8362
gb:MN988713|Organism:Severe      GTTGAAAACATGACACCCCGTGACCTTGGTGCTTGTATTGACTGTAGTGCGCGTCATATT   8362
gb:LC521925|Organism:Severe      GTTGAAAACATGACACCCCGTGACCTTGGTGCTTGTATTGACTGTAGTGCGCGTCATATT   8335
gb:MT093571|Organism:Severe      GTTGAAAACATGACACCCCGTGACCTTGGTGCTTGTATTGACTGTAGTGCGCGTCATATT   8362
gb:MT039887|Organism:Severe      GTTGAAAACATGACACCCCGTGACCTTGGTGCTTGTATTGACTGTAGTGCGCGTCATATT   8362
gb:MT019530|Organism:Severe      GTTGAAAACATGACACCCCGTGACCTTGGTGCTTGTATTGACTGTAGTGCGCGTCATATT   8362
gb:MT039888|Organism:Severe      GTTGAAAACATGACACCCCGTGACCTTGGTGCTTGTATTGACTGTAGTGCGCGTCATATT   8362
gb:LC522972|Organism:Severe      GTTGAAAACATGACACCCCGTGACCTTGGTGCTTGTATTGACTGTAGTGCGCGTCATATT   8359
gb:MT027063|Organism:Severe      GTTGAAAACATGACACCCCGTGACCTTGGTGCTTGTATTGACTGTAGTGCGCGTCATATT   8362
gb:MT027062|Organism:Severe      GTTGAAAACATGACACCCCGTGACCTTGGTGCTTGTATTGACTGTAGTGCGCGTCATATT   8362
gb:MT019529|Organism:Severe      GTTGAAAACATGACACCCCGTGACCTTGGTGCTTGTATTGACTGTAGTGCGCGTCATATT   8362
gb:MN996529|Organism:Severe      GTTGAAAACATGACACCCCGTGACCTTGGTGCTTGTATTGACTGTAGTGCGCGTCATATT   8350
gb:MN996531|Organism:Severe      GTTGAAAACATGACACCCCGTGACCTTGGTGCTTGTATTGACTGTAGTGCGCGTCATATT   8349
gb:MT066176|Organism:Severe      GTTGAAAACATGACACCCCGTGACCTTGGTGCTTGTATTGACTGTAGTGCGCGTCATATT   8362
gb:MT027064|Organism:Severe      GTTGAAAACATGACACCCCGTGACCTTGGTGCTTGTATTGACTGTAGTGCGCGTCATATT   8362
gb:MN994468|Organism:Severe      GTTGAAAACATGACACCCCGTGACCTTGGTGCTTGTATTGACTGTAGTGCGCGTCATATT   8362
gb:MT072688|Organism:Severe      GTTGAAAACATGACACCCCGTGACCTTGGTGCTTGTATTGACTGTAGTGCGCGTCATATT   8347
gb:MN996527|Organism:Severe      GTTGAAAACATGACACCCCGTGACCTTGGTGCTTGTATTGACTGTAGTGCGCGTCATATT   8329
gb:MT093631|Organism:Severe      GTTGAAAACATGACACCCCGTGACCTTGGTGCTTGTATTGACTGTAGTGCGCGTCATATT   8400
```

```
gb:MT106053|Organism:Severe      GTTGAAAACATGACACCCCGTGACCTTGGTGCTTGTATTGACTGTAGTGCGCGTCATATT   8362
gb:MT019533|Organism:Severe      GTTGAAAACATGACACCCCGTGACCTTGGTGCTTGTATTGACTGTAGTGCGCGTCATATT   8362
gb:MT019531|Organism:Severe      GTTGAAAACATGACACCCCGTGACCTTGGTGCTTGTATTGACTGTAGTGCGCGTCATATT   8362
gb:MN996528|Organism:Severe      GTTGAAAACATGACACCCCGTGACCTTGGTGCTTGTATTGACTGTAGTGCGCGTCATATT   8362
gb:MN996530|Organism:Severe      GTTGAAAACATGACACCCCGTGACCTTGGTGCTTGTATTGACTGTAGTGCGCGTCATATT   8348
gb:MN908947|Organism:Severe      GTTGAAAACATGACACCCCGTGACCTTGGTGCTTGTATTGACTGTAGTGCGCGTCATATT   8362
gb:MT019532|Organism:Severe      GTTGAAAACATGACACCCCGTGACCTTGGTGCTTGTATTGACTGTAGTGCGCGTCATATT   8362
                                 ************************************************************

gb:MT020781|Organism:Severe      AATGCGCAGGTAGCAAAAAGTCACAACATTGCTTTGATATGGAACGTTAAAGATTTCATG   8410
gb:MT007544|Organism:Severe      AATGCGCAGGTAGCAAAAAGTCACAACATTGCTTTGATATGGAACGTTAAAGATTTCATG   8422
gb:MN994467|Organism:Severe      AATGCGCAGGTAGCAAAAAGTCACAACATTGCTTTGATATGGAACGTTAAAGATTTCATG   8422
gb:MT044257|Organism:Severe      AATGCGCAGGTAGCAAAAAGTCACAACATTGCTTTGATATGGAACGTTAAAGATTTCATG   8422
gb:MT106054|Organism:Severe      AATGCGCAGGTAGCAAAAAGTCACAACATTGCTTTGATATGGAACGTTAAAGATTTCATG   8422
gb:MT049951|Organism:Severe      AATGCGCAGGTAGCAAAAAGTCACAACATTGCTTTGATATGGAACGTTAAAGATTTCATG   8422
gb:MN975262|Organism:Severe      AATGCGCAGGTAGCAAAAAGTCACAACATTGCTTTGATATGGAACGTTAAAGATTTCATG   8422
gb:MT106052|Organism:Severe      AATGCGCAGGTAGCAAAAAGTCACAACATTGCTTTGATATGGAACGTTAAAGATTTCATG   8422
gb:LC522975|Organism:Severe      AATGCGCAGGTAGCAAAAAGTCACAACATTGCTTTGATATGGAACGTTAAAGATTTCATG   8419
gb:LC522973|Organism:Severe      AATGCGCAGGTAGCAAAAAGTCACAACATTGCTTTGATATGGAACGTTAAAGATTTCATG   8419
gb:LC522974|Organism:Severe      AATGCGCAGGTAGCAAAAAGTCACAACATTGCTTTGATATGGAACGTTAAAGATTTCATG   8419
gb:MN985325|Organism:Severe      AATGCGCAGGTAGCAAAAAGTCACAACATTGCTTTGATATGGAACGTTAAAGATTTCATG   8422
gb:MT020881|Organism:Severe      AATGCGCAGGTAGCAAAAAGTCACAACATTGCTTTGATATGGAACGTTAAAGATTTCATG   8422
gb:MT020880|Organism:Severe      AATGCGCAGGTAGCAAAAAGTCACAACATTGCTTTGATATGGAACGTTAAAGATTTCATG   8422
gb:MT066175|Organism:Severe      AATGCGCAGGTAGCAAAAAGTCACAACATTGCTTTGATATGGAACGTTAAAGATTTCATG   8422
gb:MN997409|Organism:Severe      AATGCGCAGGTAGCAAAAAGTCACAACATTGCTTTGATATGGAACGTTAAAGATTTCATG   8422
gb:MN938384|Organism:Severe      AATGCGCAGGTAGCAAAAAGTCACAACATTGCTTTGATATGGAACGTTAAAGATTTCATG   8390
gb:MT044258|Organism:Severe      AATGCGCAGGTAGCAAAAAGTCACAACATTGCTTTGATATGGAACGTTAAAGATTTCATG   8398
gb:MT039890|Organism:Severe      AATGCGCAGGTAGCAAAAAGTCACAACATTGCTTTGATATGGAACGTTAAAGATTTCATG   8422
gb:MN988713|Organism:Severe      AATGCGCAGGTAGCAAAAAGTCACAACATTGCTTTGATATGGAACGTTAAAGATTTCATG   8422
gb:LC521925|Organism:Severe      AATGCGCAGGTAGCAAAAAGTCACAACATTGCTTTGATATGGAACGTTAAAGATTTCATG   8395
gb:MT093571|Organism:Severe      AATGCGCAGGTAGCAAAAAGTCACAACATTGCTTTGATATGGAACGTTAAAGATTTCATG   8422
gb:MT039887|Organism:Severe      AATGCGCAGGTAGCAAAAAGTCACAACATTGCTTTGATATGGAACGTTAAAGATTTCATG   8422
gb:MT019530|Organism:Severe      AATGCGCAGGTAGCAAAAAGTCACAACATTGCTTTGATATGGAACGTTAAAGATTTCATG   8422
gb:MT039888|Organism:Severe      AATGCGCAGGTAGCAAAAAGTCACAACATTGCTTTGATATGGAACGTTAAAGATTTCATG   8422
gb:LC522972|Organism:Severe      AATGCGCAGGTAGCAAAAAGTCACAACATTGCTTTGATATGGAACGTTAAAGATTTCATG   8419
gb:MT027063|Organism:Severe      AATGCGCAGGTAGCAAAAAGTCACAACATTGCTTTGATATGGAACGTTAAAGATTTCATG   8422
gb:MT027062|Organism:Severe      AATGCGCAGGTAGCAAAAAGTCACAACATTGCTTTGATATGGAACGTTAAAGATTTCATG   8422
gb:MT019529|Organism:Severe      AATGCGCAGGTAGCAAAAAGTCACAGCATTGCTTTGATATGGAACGTTAAAGATTTCATG   8422
gb:MN996529|Organism:Severe      AATGCGCAGGTAGCAAAAAGTCACAACATTGCTTTGATATGGAACGTTAAAGATTTCATG   8410
gb:MN996531|Organism:Severe      AATGCGCAGGTAGCAAAAAGTCACAACATTGCTTTGATATGGAACGTTAAAGATTTCATG   8409
gb:MT066176|Organism:Severe      AATGCGCAGGTAGCAAAAAGTCACAACATTGCTTTGATATGGAACGTTAAAGATTTCATG   8422
gb:MT027064|Organism:Severe      AATGCGCAGGTAGCAAAAAGTCACAACATTGCTTTGATATGGAACGTTAAAGATTTCATG   8422
gb:MN994468|Organism:Severe      AATGCGCAGGTAGCAAAAAGTCACAACATTGCTTTGATATGGAACGTTAAAGATTTCATG   8422
gb:MT072688|Organism:Severe      AATGCGCAGGTAGCAAAAAGTCACAACATTGCTTTGATATGGAACGTTAAAGATTTCATG   8407
gb:MN996527|Organism:Severe      AATGCGCAGGTAGCAAAAAGTCACAACATTGCTTTGATATGGAACGTTAAAGATTTCATG   8389
gb:MT093631|Organism:Severe      AATGCGCAGGTAGCAAAAAGTCACAACATTGCTTTGATATGGAACGTTAAAGATTTCATG   8460
gb:MT106053|Organism:Severe      AATGCGCAGGTAGCAAAAAGTCACAACATTGCTTTGATATGGAACGTTAAAGATTTCATG   8422
gb:MT019533|Organism:Severe      AATGCGCAGGTAGCAAAAAGTCACAACATTGCTTTGATATGGAACGTTAAAGATTTCATG   8422
gb:MT019531|Organism:Severe      AATGCGCAGGTAGCAAAAAGTCACAACATTGCTTTGATATGGAACGTTAAAGATTTCATG   8422
gb:MN996528|Organism:Severe      AATGCGCAGGTAGCAAAAAGTCACAACATTGCTTTGATATGGAACGTTAAAGATTTCATG   8422
gb:MN996530|Organism:Severe      AATGCGCAGGTAGCAAAAAGTCACAACATTGCTTTGATATGGAACGTTAAAGATTTCATG   8408
gb:MN908947|Organism:Severe      AATGCGCAGGTAGCAAAAAGTCACAACATTGCTTTGATATGGAACGTTAAAGATTTCATG   8422
gb:MT019532|Organism:Severe      AATGCGCAGGTAGCAAAAAGTCACAACATTGCTTTGATATGGAACGTTAAAGATTTCATG   8422
                                 ************************* **********************************

gb:MT020781|Organism:Severe      TCATTGTCTGAACAACTACGAAAACAAATACGTAGTGCTGCTAAAAAGAATAACTTACCT   8470
gb:MT007544|Organism:Severe      TCATTGTCTGAACAACTACGAAAACAAATACGTAGTGCTGCTAAAAAGAATAACTTACCT   8482
gb:MN994467|Organism:Severe      TCATTGTCTGAACAACTACGAAAACAAATACGTAGTGCTGCTAAAAAGAATAACTTACCT   8482
gb:MT044257|Organism:Severe      TCATTGTCTGAACAACTACGAAAACAAATACGTAGTGCTGCTAAAAAGAATAACTTACCT   8482
gb:MT106054|Organism:Severe      TCATTGTCTGAACAACTACGAAAACAAATACGTAGTGCTGCTAAAAAGAATAACTTACCT   8482
gb:MT049951|Organism:Severe      TCATTGTCTGAACAACTACGAAAACAAATACGTAGTGCTGCTAAAAAGAATAACTTACCT   8482
gb:MN975262|Organism:Severe      TCATTGTCTGAACAACTACGAAAACAAATACGTAGTGCTGCTAAAAAGAATAACTTACCT   8482
gb:MT106052|Organism:Severe      TCATTGTCTGAACAACTACGAAAACAAATACGTAGTGCTGCTAAAAAGAATAACTTACCT   8482
gb:LC522975|Organism:Severe      TCATTGTCTGAACAACTACGAAAACAAATACGTAGTGCTGCTAAAAAGAATAACTTACCT   8479
gb:LC522973|Organism:Severe      TCATTGTCTGAACAACTACGAAAACAAATACGTAGTGCTGCTAAAAAGAATAACTTACCT   8479
```

```
gb:LC522974|Organism:Severe      TCATTGTCTGAACAACTACGAAAACAAATACGTAGTGCTGCTAAAAAGAATAACTTACCT   8479
gb:MN985325|Organism:Severe      TCATTGTCTGAACAACTACGAAAACAAATACGTAGTGCTGCTAAAAAGAATAACTTACCT   8482
gb:MT020881|Organism:Severe      TCATTGTCTGAACAACTACGAAAACAAATACGTAGTGCTGCTAAAAAGAATAACTTACCT   8482
gb:MT020880|Organism:Severe      TCATTGTCTGAACAACTACGAAAACAAATACGTAGTGCTGCTAAAAAGAATAACTTACCT   8482
gb:MT066175|Organism:Severe      TCATTGTCTGAACAACTACGAAAACAAATACGTAGTGCTGCTAAAAAGAATAACTTACCT   8482
gb:MN997409|Organism:Severe      TCATTGTCTGAACAACTACGAAAACAAATACGTAGTGCTGCTAAAAAGAATAACTTACCT   8482
gb:MN938384|Organism:Severe      TCATTGTCTGAACAACTACGAAAACAAATACGTAGTGCTGCTAAAAAGAATAACTTACCT   8450
gb:MT044258|Organism:Severe      TCATTGTCTGAACAACTACGAAAACAAATACGTAGTGCTGCTAAAAAGAATAACTTACCT   8458
gb:MT039890|Organism:Severe      TCATTGTCTGAACAACTACGAAAACAAATACGTAGTGCTGCTAAAAAGAATAACTTACCT   8482
gb:MN988713|Organism:Severe      TCATTGTCTGAACAACTACGAAAACAAATACGTAGTGCTGCTAAAAAGAATAACTTACCT   8482
gb:LC521925|Organism:Severe      TCATTGTCTGAACAACTACGAAAACAAATACGTAGTGCTGCTAAAAAGAATAACTTACCT   8455
gb:MT093571|Organism:Severe      TCATTGTCTGAACAACTACGAAAACAAATACGTAGTGCTGCTAAAAAGAATAACTTACCT   8482
gb:MT039887|Organism:Severe      TCATTGTCTGAACAACTACGAAAACAAATACGTAGTGCTGCTAAAAAGAATAACTTACCT   8482
gb:MT019530|Organism:Severe      TCATTGTCTGAACAACTACGAAAACAAATACGTAGTGCTGCTAAAAAGAATAACTTACCT   8482
gb:MT039888|Organism:Severe      TCATTGTCTGAACAACTACGAAAACAAATACGTAGTGCTGCTAAAAAGAATAACTTACCT   8482
gb:LC522972|Organism:Severe      TCATTGTCTGAACAACTACGAAAACAAATACGTAGTGCTGCTAAAAAGAATAACTTACCT   8479
gb:MT027063|Organism:Severe      TCATTGTCTGAACAACTACGAAAACAAATACGTAGTGCTGCTAAAAAGAATAACTTACCT   8482
gb:MT027062|Organism:Severe      TCATTGTCTGAACAACTACGAAAACAAATACGTAGTGCTGCTAAAAAGAATAACTTACCT   8482
gb:MT019529|Organism:Severe      TCATTGTCTGAACAACTACGAAAACAAATACGTAGTGCTGCTAAAAAGAATAACTTACCT   8482
gb:MN996529|Organism:Severe      TCATTGTCTGAACAACTACGAAAACAAATACGTAGTGCTGCTAAAAAGAATAACTTACCT   8470
gb:MN996531|Organism:Severe      TCATTGTCTGAACAACTACGAAAACAAATACGTAGTGCTGCTAAAAAGAATAACTTACCT   8469
gb:MT066176|Organism:Severe      TCATTGTCTGAACAACTACGAAAACAAATACGTAGTGCTGCTAAAAAGAATAACTTACCT   8482
gb:MT027064|Organism:Severe      TCATTGTCTGAACAACTACGAAAACAAATACGTAGTGCTGCTAAAAAGAATAACTTACCT   8482
gb:MN994468|Organism:Severe      TCATTGTCTGAACAACTACGAAAACAAATACGTAGTGCTGCTAAAAAGAATAACTTACCT   8482
gb:MT072688|Organism:Severe      TCATTGTCTGAACAACTACGAAAACAAATACGTAGTGCTGCTAAAAAGAATAACTTACCT   8467
gb:MN996527|Organism:Severe      TCATTGTCTGAACAACTACGAAAACAAATACGTAGTGCTGCTAAAAAGAATAACTTACCT   8449
gb:MT093631|Organism:Severe      TCATTGTCTGAACAACTACGAAAACAAATACGTAGTGCTGCTAAAAAGAATAACTTACCT   8520
gb:MT106053|Organism:Severe      TCATTGTCTGAACAACTACGAAAACAAATACGTAGTGCTGCTAAAAAGAATAACTTACCT   8482
gb:MT019533|Organism:Severe      TCATTGTCTGAACAACTACGAAAACAAATACGTAGTGCTGCTAAAAAGAATAACTTACCT   8482
gb:MT019531|Organism:Severe      TCATTGTCTGAACAACTACGAAAACAAATACGTAGTGCTGCTAAAAAGAATAACTTACCT   8482
gb:MN996528|Organism:Severe      TCATTGTCTGAACAACTACGAAAACAAATACGTAGTGCTGCTAAAAAGAATAACTTACCT   8482
gb:MN996530|Organism:Severe      TCATTGTCTGAACAACTACGAAAACAAATACGTAGTGCTGCTAAAAAGAATAACTTACCT   8468
gb:MN908947|Organism:Severe      TCATTGTCTGAACAACTACGAAAACAAATACGTAGTGCTGCTAAAAAGAATAACTTACCT   8482
gb:MT019532|Organism:Severe      TCATTGTCTGAACAACTACGAAAACAAATACGTAGTGCTGCTAAAAAGAATAACTTACCT   8482
                                 ************************************************************

gb:MT020781|Organism:Severe      TTTAAGTTGACATGTGCAACTACTAGACAAGTTGTTAATGTTGTAACAACAAAGATAGCA   8530
gb:MT007544|Organism:Severe      TTTAAGTTGACATGTGCAACTACTAGACAAGTTGTTAATGTTGTAACAACAAAGATAGCA   8542
gb:MN994467|Organism:Severe      TTTAAGTTGACATGTGCAACTACTAGACAAGTTGTTAATGTTGTAACAACAAAGATAGCA   8542
gb:MT044257|Organism:Severe      TTTAAGTTGACATGTGCAACTACTAGACAAGTTGTTAATGTTGTAACAACAAAGATAGCA   8542
gb:MT106054|Organism:Severe      TTTAAGTTGACATGTGCAACTACTAGACAAGTTGTTAATGTTGTAACAACAAAGATAGCA   8542
gb:MT049951|Organism:Severe      TTTAAGTTGACATGTGCAACTACTAGACAAGTTGTTAATGTTGTAACAACAAAGATAGCA   8542
gb:MN975262|Organism:Severe      TTTAAGTTGACATGTGCAACTACTAGACAAGTTGTTAATGTTGTAACAACAAAGATAGCA   8542
gb:MT106052|Organism:Severe      TTTAAGTTGACATGTGCAACTACTAGACAAGTTGTTAATGTTGTAACAACAAAGATAGCA   8542
gb:LC522975|Organism:Severe      TTTAAGTTGACATGTGCAACTACTAGACAAGTTGTTAATGTTGTAACAACAAAGATAGCA   8539
gb:LC522973|Organism:Severe      TTTAAGTTGACATGTGCAACTACTAGACAAGTTGTTAATGTTGTAACAACAAAGATAGCA   8539
gb:LC522974|Organism:Severe      TTTAAGTTGACATGTGCAACTACTAGACAAGTTGTTAATGTTGTAACAACAAAGATAGCA   8539
gb:MN985325|Organism:Severe      TTTAAGTTGACATGTGCAACTACTAGACAAGTTGTTAATGTTGTAACAACAAAGATAGCA   8542
gb:MT020881|Organism:Severe      TTTAAGTTGACATGTGCAACTACTAGACAAGTTGTTAATGTTGTAACAACAAAGATAGCA   8542
gb:MT020880|Organism:Severe      TTTAAGTTGACATGTGCAACTACTAGACAAGTTGTTAATGTTGTAACAACAAAGATAGCA   8542
gb:MT066175|Organism:Severe      TTTAAGTTGACATGTGCAACTACTAGACAAGTTGTTAATGTTGTAACAACAAAGATAGCA   8542
gb:MN997409|Organism:Severe      TTTAAGTTGACATGTGCAACTACTAGACAAGTTGTTAATGTTGTAACAACAAAGATAGCA   8542
gb:MN938384|Organism:Severe      TTTAAGTTGACATGTGCAACTACTAGACAAGTTGTTAATGTTGTAACAACAAAGATAGCA   8510
gb:MT044258|Organism:Severe      TTTAAGTTGACATGTGCAACTACTAGACAAGTTGTTAATGTTGTAACAACAAAGATAGCA   8518
gb:MT039890|Organism:Severe      TTTAAGTTGACATGTGCAACTACTAGACAAGTTGTTAATGTTGTAACAACAAAGATAGCA   8542
gb:MN988713|Organism:Severe      TTTAAGTTGACATGTGCAACTACTAGACAAGTTGTTAATGTTGTAACAACAAAGATAGCA   8542
gb:LC521925|Organism:Severe      TTTAAGTTGACATGTGCAACTACTAGACAAGTTGTTAATGTTGTAACAACAAAGATAGCA   8515
gb:MT093571|Organism:Severe      TTTAAGTTGACATGTGCAACTACTAGACAAGTTGTTAATGTTGTAACAACAAAGATAGCA   8542
gb:MT039887|Organism:Severe      TTTAAGTTGACATGTGCAACTACTAGACAAGTTGTTAATGTTGTAACAACAAAGATAGCA   8542
gb:MT019530|Organism:Severe      TTTAAGTTGACATGTGCAACTACTAGACAAGTTGTTAATGTTGTAACAACAAAGATAGCA   8542
gb:MT039888|Organism:Severe      TTTAAGTTGACATGTGCAACTACTAGACAAGTTGTTAATGTTGTAACAACAAAGATAGCA   8542
gb:LC522972|Organism:Severe      TTTAAGTTGACATGTGCAACTACTAGACAAGTTGTTAATGTTGTAACAACAAAGATAGCA   8539
gb:MT027063|Organism:Severe      TTTAAGTTGACATGTGCAACTACTAGACAAGTTGTTAATGTTGTAACAACAAAGATAGCA   8542
gb:MT027062|Organism:Severe      TTTAAGTTGACATGTGCAACTACTAGACAAGTTGTTAATGTTGTAACAACAAAGATAGCA   8542
gb:MT019529|Organism:Severe      TTTAAGTTGACATGTGCAACTACTAGACAAGTTGTTAATGTTGTAACAACAAAGATAGCA   8542
```

```
gb:MN996529|Organism:Severe      TTTAAGTTGACATGTGCAACTACTAGACAAGTTGTTAATGTTGTAACAACAAAGATAGCA   8530
gb:MN996531|Organism:Severe      TTTAAGTTGACATGTGCAACTACTAGACAAGTTGTTAATGTTGTAACAACAAAGATAGCA   8529
gb:MT066176|Organism:Severe      TTTAAGTTGACATGTGCAACTACTAGACAAGTTGTTAATGTTGTAACAACAAAGATAGCA   8542
gb:MT027064|Organism:Severe      TTTAAGTTGACATGTGCAACTACTAGACAAGTTGTTAATGTTGTAACAACAAAGATAGCA   8542
gb:MN994468|Organism:Severe      TTTAAGTTGACATGTGCAACTACTAGACAAGTTGTTAATGTTGTAACAACAAAGATAGCA   8542
gb:MT072688|Organism:Severe      TTTAAGTTGACATGTGCAACTACTAGACAAGTTGTTAATGTTGTAACAACAAAGATAGCA   8527
gb:MN996527|Organism:Severe      TTTAAGTTGACATGTGCAACTACTAGACAAGTTGTTAATGTTGTAACAACAAAGATAGCA   8509
gb:MT093631|Organism:Severe      TTTAAGTTGACATGTGCAACTACTAGACAAGTTGTTAATGTTGTAACAACAAAGATAGCA   8580
gb:MT106053|Organism:Severe      TTTAAGTTGACATGTGCAACTACTAGACAAGTTGTTAATGTTGTAACAACAAAGATAGCA   8542
gb:MT019533|Organism:Severe      TTTAAGTTGACATGTGCAACTACTAGACAAGTTGTTAATGTTGTAACAACAAAGATAGCA   8542
gb:MT019531|Organism:Severe      TTTAAGTTGACATGTGCAACTACTAGACAAGTTGTTAATGTTGTAACAACAAAGATAGCA   8542
gb:MN996528|Organism:Severe      TTTAAGTTGACATGTGCAACTACTAGACAAGTTGTTAATGTTGTAACAACAAAGATAGCA   8542
gb:MN996530|Organism:Severe      TTTAAGTTGACATGTGCAACTACTAGACAAGTTGTTAATGTTGTAACAACAAAGATAGCA   8528
gb:MN908947|Organism:Severe      TTTAAGTTGACATGTGCAACTACTAGACAAGTTGTTAATGTTGTAACAACAAAGATAGCA   8542
gb:MT019532|Organism:Severe      TTTAAGTTGACATGTGCAACTACTAGACAAGTTGTTAATGTTGTAACAACAAAGATAGCA   8542
                                 ************************************************************

gb:MT020781|Organism:Severe      CTTAAGGGTGGTAAAATTGTTAATAATTGGTTGAAGCAGTTAATTAAAGTTACACTTGTG   8590
gb:MT007544|Organism:Severe      CTTAAGGGTGGTAAAATTGTTAATAATTGGTTGAAGCAGTTAATTAAAGTTACACTTGTG   8602
gb:MN994467|Organism:Severe      CTTAAGGGTGGTAAAATTGTTAATAATTGGTTGAAGCAGTTAATTAAAGTTACACTTGTG   8602
gb:MT044257|Organism:Severe      CTTAAGGGTGGTAAAATTGTTAATAATTGGTTGAAGCAGTTAATTAAAGTTACACTTGTG   8602
gb:MT106054|Organism:Severe      CTTAAGGGTGGTAAAATTGTTAATAATTGGTTGAAGCAGTTAATTAAAGTTACACTTGTG   8602
gb:MT049951|Organism:Severe      CTTAAGGGTGGTAAAATTGTTAATAATTGGTTGAAGCAGTTAATTAAAGTTACACTTGTG   8602
gb:MN975262|Organism:Severe      CTTAAGGGTGGTAAAATTGTTAATAATTGGTTGAAGCAGTTAATTAAAGTTACACTTGTG   8602
gb:MT106052|Organism:Severe      CTTAAGGGTGGTAAAATTGTTAATAATTGGTTGAAGCAGTTAATTAAAGTTACACTTGTG   8602
gb:LC522975|Organism:Severe      CTTAAGGGTGGTAAAATTGTTAATAATTGGTTGAAGCAGTTAATTAAAGTTACACTTGTG   8599
gb:LC522973|Organism:Severe      CTTAAGGGTGGTAAAATTGTTAATAATTGGTTGAAGCAGTTAATTAAAGTTACACTTGTG   8599
gb:LC522974|Organism:Severe      CTTAAGGGTGGTAAAATTGTTAATAATTGGTTGAAGCAGTTAATTAAAGTTACACTTGTG   8599
gb:MN985325|Organism:Severe      CTTAAGGGTGGTAAAATTGTTAATAATTGGTTGAAGCAGTTAATTAAAGTTACACTTGTG   8602
gb:MT020881|Organism:Severe      CTTAAGGGTGGTAAAATTGTTAATAATTGGTTGAAGCAGTTAATTAAAGTTACACTTGTG   8602
gb:MT020880|Organism:Severe      CTTAAGGGTGGTAAAATTGTTAATAATTGGTTGAAGCAGTTAATTAAAGTTACACTTGTG   8602
gb:MT066175|Organism:Severe      CTTAAGGGTGGTAAAATTGTTAATAATTGGTTGAAGCAGTTAATTAAAGTTACACTTGTG   8602
gb:MN997409|Organism:Severe      CTTAAGGGTGGTAAAATTGTTAATAATTGGTTGAAGCAGTTAATTAAAGTTACACTTGTG   8602
gb:MN938384|Organism:Severe      CTTAAGGGTGGTAAAATTGTTAATAATTGGTTGAAGCAGTTAATTAAAGTTACACTTGTG   8570
gb:MT044258|Organism:Severe      CTTAAGGGTGGTAAAATTGTTAATAATTGGTTGAAGCAGTTAATTAAAGTTACACTTGTG   8578
gb:MT039890|Organism:Severe      CTTAAGGGTGGTAAAATTGTTAATAATTGGTTGAAGCAGTTAATTAAAGTTACACTTGTG   8602
gb:MN988713|Organism:Severe      CTTAAGGGTGGTAAAATTGTTAATAATTGGTTGAAGCAGTTAATTAAAGTTACACTTGTG   8602
gb:LC521925|Organism:Severe      CTTAAGGGTGGTAAAATTGTTAATAATTGGTTGAAGCAGTTAATTAAAGTTACACTTGTG   8575
gb:MT093571|Organism:Severe      CTTAAGGGTGGTAAAATTGTTAATAATTGGTTGAAGCAGTTAATTAAAGTTACACTTGTG   8602
gb:MT039887|Organism:Severe      CTTAAGGGTGGTAAAATTGTTAATAATTGGTTGAAGCAGTTAATTAAAGTTACACTTGTG   8602
gb:MT019530|Organism:Severe      CTTAAGGGTGGTAAAATTGTTAATAATTGGTTGAAGCAGTTAATTAAAGTTACACTTGTG   8602
gb:MT039888|Organism:Severe      CTTAAGGGTGGTAAAATTGTTAATAATTGGTTGAAGCAGTTAATTAAAGTTACACTTGTG   8602
gb:LC522972|Organism:Severe      CTTAAGGGTGGTAAAATTGTTAATAATTGGTTGAAGCAGTTAATTAAAGTTACACTTGTG   8599
gb:MT027063|Organism:Severe      CTTAAGGGTGGTAAAATTGTTAATAATTGGTTGAAGCAGTTAATTAAAGTTACACTTGTG   8602
gb:MT027062|Organism:Severe      CTTAAGGGTGGTAAAATTGTTAATAATTGGTTGAAGCAGTTAATTAAAGTTACACTTGTG   8602
gb:MT019529|Organism:Severe      CTTAAGGGTGGTAAAATTGTTAATAATTGGTTGAAGCAGTTAATTAAAGTTACACTTGTG   8602
gb:MN996529|Organism:Severe      CTTAAGGGTGGTAAAATTGTTAATAATTGGTTGAAGCAGTTAATTAAAGTTACACTTGTG   8590
gb:MN996531|Organism:Severe      CTTAAGGGTGGTAAAATTGTTAATAATTGGTTGAAGCAGTTAATTAAAGTTACACTTGTG   8589
gb:MT066176|Organism:Severe      CTTAAGGGTGGTAAAATTGTTAATAATTGGTTGAAGCAGTTAATTAAAGTTACACTTGTG   8602
gb:MT027064|Organism:Severe      CTTAAGGGTGGTAAAATTGTTAATAATTGGTTGAAGCAGTTAATTAAAGTTACACTTGTG   8602
gb:MN994468|Organism:Severe      CTTAAGGGTGGTAAAATTGTTAATAATTGGTTGAAGCAGTTAATTAAAGTTACACTTGTG   8602
gb:MT072688|Organism:Severe      CTTAAGGGTGGTAAAATTGTTAATAATTGGTTGAAGCAGTTAATTAAAGTTACACTTGTG   8587
gb:MN996527|Organism:Severe      CTTAAGGGTGGTAAAATTGTTAATAATTGGTTGAAGCAGTTAATTAAAGTTACACTTGTG   8569
gb:MT093631|Organism:Severe      CTTAAGGGTGGTAAAATTGTTAATAATTGGTTGAAGCAGTTAATTAAAGTTACACTTGTG   8640
gb:MT106053|Organism:Severe      CTTAAGGGTGGTAAAATTGTTAATAATTGGTTGAAGCAGTTAATTAAAGTTACACTTGTG   8602
gb:MT019533|Organism:Severe      CTTAAGGGTGGTAAAATTGTTAATAATTGGTTGAAGCAGTTAATTAAAGTTACACTTGTG   8602
gb:MT019531|Organism:Severe      CTTAAGGGTGGTAAAATTGTTAATAATTGGTTGAAGCAGTTAATTAAAGTTACACTTGTG   8602
gb:MN996528|Organism:Severe      CTTAAGGGTGGTAAAATTGTTAATAATTGGTTGAAGCAGTTAATTAAAGTTACACTTGTG   8602
gb:MN996530|Organism:Severe      CTTAAGGGTGGTAAAATTGTTAATAATTGGTTGAAGCAGTTAATTAAAGTTACACTTGTG   8588
gb:MN908947|Organism:Severe      CTTAAGGGTGGTAAAATTGTTAATAATTGGTTGAAGCAGTTAATTAAAGTTACACTTGTG   8602
gb:MT019532|Organism:Severe      CTTAAGGGTGGTAAAATTGTTAATAATTGGTTGAAGCAGTTAATTAAAGTTACACTTGTG   8602
                                 ************************************************************

gb:MT020781|Organism:Severe      TTCCTTTTTGTTGCTGCTATTTTCTATTTAATAACACCTGTTCATGTCATGTCTAAACAT   8650
gb:MT007544|Organism:Severe      TTCCTTTTTGTTGCTGCTATTTTCTATTTAATAACACCTGTTCATGTCATGTCTAAACAT   8662
```

```
gb:MN994467|Organism:Severe      TTCCTTTTTGTTGCTGCTATTTTCTATTTAATAACACCTGTTCATGTCATGTCTAAACAT    8662
gb:MT044257|Organism:Severe      TTCCTTTTTGTTGCTGCTATTTTCTATTTAATAACACCTGTTCATGTCATGTCTAAACAT    8662
gb:MT106054|Organism:Severe      TTCCTTTTTGTTGCTGCTATTTTCTATTTAATAACACCTGTTCATGTCATGTCTAAACAT    8662
gb:MT049951|Organism:Severe      TTCCTTTTTGTTGCTGCTATTTTCTATTTAATAACACCTGTTCATGTCATGTCTAAACAT    8662
gb:MN975262|Organism:Severe      TTCCTTTTTGTTGCTGCTATTTTCTATTTAATAACACCTGTTCATGTCATGTCTAAACAT    8662
gb:MT106052|Organism:Severe      TTCCTTTTTGTTGCTGCTATTTTCTATTTAATAACACCTGTTCATGTCATGTCTAAACAT    8662
gb:LC522975|Organism:Severe      TTCCTTTTTGTTGCTGCTATTTTCTATTTAATAACACCTGTTCATGTCATGTCTAAACAT    8659
gb:LC522973|Organism:Severe      TTCCTTTTTGTTGCTGCTATTTTCTATTTAATAACACCTGTTCATGTCATGTCTAAACAT    8659
gb:LC522974|Organism:Severe      TTCCTTTTTGTTGCTGCTATTTTCTATTTAATAACACCTGTTCATGTCATGTCTAAACAT    8659
gb:MN985325|Organism:Severe      TTCCTTTTTGTTGCTGCTATTTTCTATTTAATAACACCTGTTCATGTCATGTCTAAACAT    8662
gb:MT020881|Organism:Severe      TTCCTTTTTGTTGCTGCTATTTTCTATTTAATAACACCTGTTCATGTCATGTCTAAACAT    8662
gb:MT020880|Organism:Severe      TTCCTTTTTGTTGCTGCTATTTTCTATTTAATAACACCTGTTCATGTCATGTCTAAACAT    8662
gb:MT066175|Organism:Severe      TTCCTTTTTGTTGCTGCTATTTTCTATTTAATAACACCTGTTCATGTCATGTCTAAACAT    8662
gb:MN997409|Organism:Severe      TTCCTTTTTGTTGCTGCTATTTTCTATTTAATAACACCTGTTCATGTCATGTCTAAACAT    8662
gb:MN938384|Organism:Severe      TTCCTTTTTGTTGCTGCTATTTTCTATTTAATAACACCTGTTCATGTCATGTCTAAACAT    8630
gb:MT044258|Organism:Severe      TTCCTTTTTGTTGCTGCTATTTTCTATTTAATAACACCTGTTCATGTCATGTCTAAACAT    8638
gb:MT039890|Organism:Severe      TTCCTTTTTGTTGCTGCTATTTTCTATTTAATAACACCTGTTCATGTCATGTCTAAACAT    8662
gb:MN988713|Organism:Severe      TTCCTTTTTGTTGCTGCTATTTTCTATTTAATAACACCTGTTCATGTCATGTCTAAACAT    8662
gb:LC521925|Organism:Severe      TTCCTTTTTGTTGCTGCTATTTTCTATTTAATAACACCTGTTCATGTCATGTCTAAACAT    8635
gb:MT093571|Organism:Severe      TTCCTTTTTGTTGCTGCTATTTTCTATTTAATAACACCTGTTCATGTCATGTCTAAACAT    8662
gb:MT039887|Organism:Severe      TTCCTTTTTGTTGCTGCTATTTTCTATTTAATAACACCTGTTCATGTCATGTCTAAACAT    8662
gb:MT019530|Organism:Severe      TTCCTTTTTGTTGCTGCTATTTTCTATTTAATAACACCTGTTCATGTCATGTCTAAACAT    8662
gb:MT039888|Organism:Severe      TTCCTTTTTGTTGCTGCTATTTTCTATTTAATAACACCTGTTCATGTCATGTCTAAACAT    8662
gb:LC522972|Organism:Severe      TTCCTTTTTGTTGCTGCTATTTTCTATTTAATAACACCTGTTCATGTCATGTCTAAACAT    8659
gb:MT027063|Organism:Severe      TTCCTTTTTGTTGCTGCTATTTTCTATTTAATAACACCTGTTCATGTCATGTCTAAACAT    8662
gb:MT027062|Organism:Severe      TTCCTTTTTGTTGCTGCTATTTTCTATTTAATAACACCTGTTCATGTCATGTCTAAACAT    8662
gb:MT019529|Organism:Severe      TTCCTTTTTGTTGCTGCTATTTTCTATTTAATAACACCTGTTCATGTCATGTCTAAACAT    8662
gb:MN996529|Organism:Severe      TTCCTTTTTGTTGCTGCTATTTTCTATTTAATAACACCTGTTCATGTCATGTCTAAACAT    8650
gb:MN996531|Organism:Severe      TTCCTTTTTGTTGCTGCTATTTTCTATTTAATAACACCTGTTCATGTCATGTCTAAACAT    8649
gb:MT066176|Organism:Severe      TTCCTTTTTGTTGCTGCTATTTTCTATTTAATAACACCTGTTCATGTCATGTCTAAACAT    8662
gb:MT027064|Organism:Severe      TTCCTTTTTGTTGCTGCTATTTTCTATTTAATAACACCTGTTCATGTCATGTCTAAACAT    8662
gb:MN994468|Organism:Severe      TTCCTTTTTGTTGCTGCTATTTTCTATTTAATAACACCTGTTCATGTCATGTCTAAACAT    8662
gb:MT072688|Organism:Severe      TTCCTTTTTGTTGCTGCTATTTTCTATTTAATAACACCTGTTCATGTCATGTCTAAACAT    8647
gb:MN996527|Organism:Severe      TTCCTTTTTGTTGCTGCTATTTTCTATTTAATAACACCTGTTCATGTCATGTCTAAACAT    8629
gb:MT093631|Organism:Severe      TTCCTTTTTGTTGCTGCTATTTTCTATTTAATAACACCTGTTCATGTCATGTCTAAACAT    8700
gb:MT106053|Organism:Severe      TTCCTTTTTGTTGCTGCTATTTTCTATTTAATAACACCTGTTCATGTCATGTCTAAACAT    8662
gb:MT019533|Organism:Severe      TTCCTTTTTGTTGCTGCTATTTTCTATTTAATAACACCTGTTCATGTCATGTCTAAACAT    8662
gb:MT019531|Organism:Severe      TTCCTTTTTGTTGCTGCTATTTTCTATTTAATAACACCTGTTCATGTCATGTCTAAACAT    8662
gb:MN996528|Organism:Severe      TTCCTTTTTGTTGCTGCTATTTTCTATTTAATAACACCTGTTCATGTCATGTCTAAACAT    8662
gb:MN996530|Organism:Severe      TTCCTTTTTGTTGCTGCTATTTTCTATTTAATAACACCTGTTCATGTCATGTCTAAACAT    8648
gb:MN908947|Organism:Severe      TTCCTTTTTGTTGCTGCTATTTTCTATTTAATAACACCTGTTCATGTCATGTCTAAACAT    8662
gb:MT019532|Organism:Severe      TTCCTTTTTGTTGCTGCTATTTTCTATTTAATAACACCTGTTCATGTCATGTCTAAACAT    8662
                                 ************************************************************

gb:MT020781|Organism:Severe      ACTGACTTTTCAAGTGAAATCATAGGATACAAGGCTATTGATGGTGGTGTCACTCGTGAC    8710
gb:MT007544|Organism:Severe      ACTGACTTTTCAAGTGAAATCATAGGATACAAGGCTATTGATGGTGGTGTCACTCGTGAC    8722
gb:MN994467|Organism:Severe      ACTGACTTTTCAAGTGAAATCATAGGATACAAGGCTATTGATGGTGGTGTCACTCGTGAC    8722
gb:MT044257|Organism:Severe      ACTGACTTTTCAAGTGAAATCATAGGATACAAGGCTATTGATGGTGGTGTCACTCGTGAC    8722
gb:MT106054|Organism:Severe      ACTGACTTTTCAAGTGAAATCATAGGATACAAGGCTATTGATGGTGGTGTCACTCGTGAC    8722
gb:MT049951|Organism:Severe      ACTGACTTTTCAAGTGAAATCATAGGATACAAGGCTATTGATGGTGGTGTCACTCGTGAC    8722
gb:MN975262|Organism:Severe      ACTGACTTTTCAAGTGAAATCATAGGATACAAGGCTATTGATGGTGGTGTCACTCGTGAC    8722
gb:MT106052|Organism:Severe      ACTGACTTTTCAAGTGAAATCATAGGATACAAGGCTATTGATGGTGGTGTCACTCGTGAC    8722
gb:LC522975|Organism:Severe      ACTGACTTTTCAAGTGAAATCATAGGATACAAGGCTATTGATGGTGGTGTCACTCGTGAC    8719
gb:LC522973|Organism:Severe      ACTGACTTTTCAAGTGAAATCATAGGATACAAGGCTATTGATGGTGGTGTCACTCGTGAC    8719
gb:LC522974|Organism:Severe      ACTGACTTTTCAAGTGAAATCATAGGATACAAGGCTATTGATGGTGGTGTCACTCGTGAC    8719
gb:MN985325|Organism:Severe      ACTGACTTTTCAAGTGAAATCATAGGATACAAGGCTATTGATGGTGGTGTCACTCGTGAC    8722
gb:MT020881|Organism:Severe      ACTGACTTTTCAAGTGAAATCATAGGATACAAGGCTATTGATGGTGGTGTCACTCGTGAC    8722
gb:MT020880|Organism:Severe      ACTGACTTTTCAAGTGAAATCATAGGATACAAGGCTATTGATGGTGGTGTCACTCGTGAC    8722
gb:MT066175|Organism:Severe      ACTGACTTTTCAAGTGAAATCATAGGATACAAGGCTATTGATGGTGGTGTCACTCGTGAC    8722
gb:MN997409|Organism:Severe      ACTGACTTTTCAAGTGAAATCATAGGATACAAGGCTATTGATGGTGGTGTCACTCGTGAC    8722
gb:MN938384|Organism:Severe      ACTGACTTTTCAAGTGAAATCATAGGATACAAGGCTATTGATGGTGGTGTCACTCGTGAC    8690
gb:MT044258|Organism:Severe      ACTGACTTTTCAAGTGAAATCATAGGATACAAGGCTATTGATGGTGGTGTCACTCGTGAC    8698
gb:MT039890|Organism:Severe      ACTGACTTTTCAAGTGAAATCATAGGATACAAGGCTATTGATGGTGGTGTCACTCGTGAC    8722
gb:MN988713|Organism:Severe      ACTGACTTTTCAAGTGAAATCATAGGATACAAGGCTATTGATGGTGGTGTCACTCGTGAC    8722
gb:LC521925|Organism:Severe      ACTGACTTTTCAAGTGAAATCATAGGATACAAGGCTATTGATGGTGGTGTCACTCGTGAC    8695
```

```
gb:MT093571|Organism:Severe      ACTGACTTTTCAAGTGAAATCATAGGATACAAGGCTATTGATGGTGGTGTCACTCGTGAC    8722
gb:MT039887|Organism:Severe      ACTGACTTTTCAAGTGAAATCATAGGATACAAGGCTATTGATGGTGGTGTCACTCGTGAC    8722
gb:MT019530|Organism:Severe      ACTGACTTTTCAAGTGAAATCATAGGATACAAGGCTATTGATGGTGGTGTCACTCGTGAC    8722
gb:MT039888|Organism:Severe      ACTGACTTTTCAAGTGAAATCATAGGATACAAGGCTATTGATGGTGGTGTCACTCGTGAC    8722
gb:LC522972|Organism:Severe      ACTGACTTTTCAAGTGAAATCATAGGATACAAGGCTATTGATGGTGGTGTCACTCGTGAC    8719
gb:MT027063|Organism:Severe      ACTGACTTTTCAAGTGAAATCATAGGATACAAGGCTATTGATGGTGGTGTCACTCGTGAC    8722
gb:MT027062|Organism:Severe      ACTGACTTTTCAAGTGAAATCATAGGATACAAGGCTATTGATGGTGGTGTCACTCGTGAC    8722
gb:MT019529|Organism:Severe      ACTGACTTTTCAAGTGAAATCATAGGATACAAGGCTATTGATGGTGGTGTCACTCGTGAC    8722
gb:MN996529|Organism:Severe      ACTGACTTTTCAAGTGAAATCATAGGATACAAGGCTATTGATGGTGGTGTCACTCGTGAC    8710
gb:MN996531|Organism:Severe      ACTGACTTTTCAAGTGAAATCATAGGATACAAGGCTATTGATGGTGGTGTCACTCGTGAC    8709
gb:MT066176|Organism:Severe      ACTGACTTTTCAAGTGAAATCATAGGATACAAGGCTATTGATGGTGGTGTCACTCGTGAC    8722
gb:MT027064|Organism:Severe      ACTGACTTTTCAAGTGAAATCATAGGATACAAGGCTATTGATGGTGGTGTCACTCGTGAC    8722
gb:MN994468|Organism:Severe      ACTGACTTTTCAAGTGAAATCATAGGATACAAGGCTATTGATGGTGGTGTCACTCGTGAC    8722
gb:MT072688|Organism:Severe      ACTGACTTTTCAAGTGAAATCATAGGATACAAGGCTATTGATGGTGGTGTCACTCGTGAC    8707
gb:MN996527|Organism:Severe      ACTGACTTTTCAAGTGAAATCATAGGATACAAGGCTATTGATGGTGGTGTCACTCGTGAC    8689
gb:MT093631|Organism:Severe      ACTGACTTTTCAAGTGAAATCATAGGATACAAGGCTATTGATGGTGGTGTCACTCGTGAC    8760
gb:MT106053|Organism:Severe      ACTGACTTTTCAAGTGAAATCATAGGATACAAGGCTATTGATGGTGGTGTCACTCGTGAC    8722
gb:MT019533|Organism:Severe      ACTGACTTTTCAAGTGAAATCATAGGATACAAGGCTATTGATGGTGGTGTCACTCGTGAC    8722
gb:MT019531|Organism:Severe      ACTGACTTTTCAAGTGAAATCATAGGATACAAGGCTATTGATGGTGGTGTCACTCGTGAC    8722
gb:MN996528|Organism:Severe      ACTGACTTTTCAAGTGAAATCATAGGATACAAGGCTATTGATGGTGGTGTCACTCGTGAC    8722
gb:MN996530|Organism:Severe      ACTGACTTTTCAAGTGAAATCATAGGATACAAGGCTATTGATGGTGGTGTCACTCGTGAC    8708
gb:MN908947|Organism:Severe      ACTGACTTTTCAAGTGAAATCATAGGATACAAGGCTATTGATGGTGGTGTCACTCGTGAC    8722
gb:MT019532|Organism:Severe      ACTGACTTTTCAAGTGAAATCATAGGATACAAGGCTATTGATGGTGGTGTCACTCGTGAC    8722
                                 ************************************************************

gb:MT020781|Organism:Severe      ATAGCATCTACAGATACTTGTTTTGCTAACAAACATGCTGATTTTGACACATGGTTTAGC    8770
gb:MT007544|Organism:Severe      ATAGCATCTACAGATACTTGTTTTGCTAACAAACATGCTGATTTTGACACATGGTTTAGC    8782
gb:MN994467|Organism:Severe      ATAGCATCTACAGATACTTGTTTTGCTAACAAACATGCTGATTTTGACACATGGTTTAGT    8782
gb:MT044257|Organism:Severe      ATAGCATCTACAGATACTTGTTTTGCTAACAAACATGCTGATTTTGACACATGGTTTAGT    8782
gb:MT106054|Organism:Severe      ATAGCATCTACAGATACTTGTTTTGCTAACAAACATGCTGATTTTGACACATGGTTTAGT    8782
gb:MT049951|Organism:Severe      ATAGCATCTACAGATACTTGTTTTGCTAACAAACATGCTGATTTTGACACATGGTTTAGT    8782
gb:MN975262|Organism:Severe      ATAGCATCTACAGATACTTGTTTTGCTAACAAACATGCTGATTTTGACACATGGTTTAGT    8782
gb:MT106052|Organism:Severe      ATAGCATCTACAGATACTTGTTTTGCTAACAAACATGCTGATTTTGACACATGGTTTAGT    8782
gb:LC522975|Organism:Severe      ATAGCATCTACAGATACTTGTTTTGCTAACAAACATGCTGATTTTGACACATGGTTTAGT    8779
gb:LC522973|Organism:Severe      ATAGCATCTACAGATACTTGTTTTGCTAACAAACATGCTGATTTTGACACATGGTTTAGT    8779
gb:LC522974|Organism:Severe      ATAGCATCTACAGATACTTGTTTTGCTAACAAACATGCTGATTTTGACACATGGTTTAGT    8779
gb:MN985325|Organism:Severe      ATAGCATCTACAGATACTTGTTTTGCTAACAAACATGCTGATTTTGACACATGGTTTAGT    8782
gb:MT020881|Organism:Severe      ATAGCATCTACAGATACTTGTTTTGCTAACAAACATGCTGATTTTGACACATGGTTTAGT    8782
gb:MT020880|Organism:Severe      ATAGCATCTACAGATACTTGTTTTGCTAACAAACATGCTGATTTTGACACATGGTTTAGT    8782
gb:MT066175|Organism:Severe      ATAGCATCTACAGATACTTGTTTTGCTAACAAACATGCTGATTTTGACACATGGTTTAGT    8782
gb:MN997409|Organism:Severe      ATAGCATCTACAGATACTTGTTTTGCTAACAAACATGCTGATTTTGACACATGGTTTAGT    8782
gb:MN938384|Organism:Severe      ATAGCATCTACAGATACTTGTTTTGCTAACAAACATGCTGATTTTGACACATGGTTTAGT    8750
gb:MT044258|Organism:Severe      ATAGCATCTACAGATACTTGTTTTGCTAACAAACATGCTGATTTTGACACATGGTTTAGC    8758
gb:MT039890|Organism:Severe      ATAGCATCTACAGATACTTGTTTTGCTAACAAACATGCTGATTTTGACACATGGTTTAGC    8782
gb:MN988713|Organism:Severe      ATAGCATCTACAGATACTTGTTTTGCTAACAAACATGCTGATTTTGACACATGGTTTAGY    8782
gb:LC521925|Organism:Severe      ATAGCATCTACAGATACTTGTTTTGCTAACAAACATGCTGATTTTGACACATGGTTTAGC    8755
gb:MT093571|Organism:Severe      ATAGCATCTACAGATACTTGTTTTGCTAACAAACATGCTGATTTTGACACATGGTTTAGC    8782
gb:MT039887|Organism:Severe      ATAGCATCTACAGATACTTGTTTTGCTAACAAACATGCTGATTTTGACACATGGTTTAGC    8782
gb:MT019530|Organism:Severe      ATAGCATCTACAGATACTTGTTTTGCTAACAAACATGCTGATTTTGACACATGGTTTAGC    8782
gb:MT039888|Organism:Severe      ATAGCATCTACAGATACTTGTTTTGCTAACAAACATGCTGATTTTGACACATGGTTTAGC    8782
gb:LC522972|Organism:Severe      ATAGCATCTACAGATACTTGTTTTGCTAACAAACATGCTGATTTTGACACATGGTTTAGC    8779
gb:MT027063|Organism:Severe      ATAGCATCTACAGATACTTGTTTTGCTAACAAACATGCTGATTTTGACACATGGTTTAGC    8782
gb:MT027062|Organism:Severe      ATAGCATCTACAGATACTTGTTTTGCTAACAAACATGCTGATTTTGACACATGGTTTAGC    8782
gb:MT019529|Organism:Severe      ATAGCATCTACAGATACTTGTTTTGCTAACAAACATGCTGATTTTGACACATGGTTTAGC    8782
gb:MN996529|Organism:Severe      ATAGCATCTACAGATACTTGTTTTGCTAACAAACATGCTGATTTTGACACATGGTTTAGC    8770
gb:MN996531|Organism:Severe      ATAGCATCTACAGATACTTGTTTTGCTAACAAACATGCTGATTTTGACACATGGTTTAGC    8769
gb:MT066176|Organism:Severe      ATAGCATCTACAGATACTTGTTTTGCTAACAAACATGCTGATTTTGACACATGGTTTAGC    8782
gb:MT027064|Organism:Severe      ATAGCATCTACAGATACTTGTTTTGCTAACAAACATGCTGATTTTGACACATGGTTTAGC    8782
gb:MN994468|Organism:Severe      ATAGCATCTACAGATACTTGTTTTGCTAACAAACATGCTGATTTTGACACATGGTTTAGC    8782
gb:MT072688|Organism:Severe      ATAGCATCTACAGATACTTGTTTTGCTAACAAACATGCTGATTTTGACACATGGTTTAGC    8767
gb:MN996527|Organism:Severe      ATAGCATCTACAGATACTTGTTTTGCTAACAAACATGCTGATTTTGACACATGGTTTAGC    8749
gb:MT093631|Organism:Severe      ATAGCATCTACAGATACTTGTTTTGCTAACAAACATGCTGATTTTGACACATGGTTTAGC    8820
gb:MT106053|Organism:Severe      ATAGCATCTACAGATACTTGTTTTGCTAACAAACATGCTGATTTTGACACATGGTTTAGC    8782
gb:MT019533|Organism:Severe      ATAGCATCTACAGATACTTGTTTTGCTAACAAACATGCTGATTTTGACACATGGTTTAGC    8782
gb:MT019531|Organism:Severe      ATAGCATCTACAGATACTTGTTTTGCTAACAAACATGCTGATTTTGACACATGGTTTAGC    8782
```

```
gb:MN996528|Organism:Severe      ATAGCATCTACAGATACTTGTTTTGCTAACAAACATGCTGATTTTGACACATGGTTTAGC   8782
gb:MN996530|Organism:Severe      ATAGCATCTACAGATACTTGTTTTGCTAACAAACATGCTGATTTTGACACATGGTTTAGC   8768
gb:MN908947|Organism:Severe      ATAGCATCTACAGATACTTGTTTTGCTAACAAACATGCTGATTTTGACACATGGTTTAGC   8782
gb:MT019532|Organism:Severe      ATAGCATCTACAGATACTTGTTTTGCTAACAAACATGCTGATTTTGACACATGGTTTAGC   8782
                                 ************************************************************

gb:MT020781|Organism:Severe      CAGCGTGGTGGTAGTTATACTAATGACAAAGCTTGCCCATTGATTGCTGCAGTCATAACA   8830
gb:MT007544|Organism:Severe      CAGCGTGGTGGTAGTTATACTAATGACAAAGCTTGCCCATTGATTGCTGCAGTCATAACA   8842
gb:MN994467|Organism:Severe      CAGCGTGGTGGTAGTTATACTAATGACAAAGCTTGCCCATTGATTGCTGCAGTCATAACA   8842
gb:MT044257|Organism:Severe      CAGCGTGGTGGTAGTTATACTAATGACAAAGCTTGCCCATTGATTGCTGCAGTCATAACA   8842
gb:MT106054|Organism:Severe      CAGCGTGGTGGTAGTTATACTAATGACAAAGCTTGCCCATTGATTGCTGCAGTCATAACA   8842
gb:MT049951|Organism:Severe      CAGCGTGGTGGTAGTTATACTAATGACAAAGCTTGCCCATTGATTGCTGCAGTCATAACA   8842
gb:MN975262|Organism:Severe      CAGCGTGGTGGTAGTTATACTAATGACAAAGCTTGCCCATTGATTGCTGCAGTCATAACA   8842
gb:MT106052|Organism:Severe      CAGCGTGGTGGTAGTTATACTAATGACAAAGCTTGCCCATTGATTGCTGCAGTCATAACA   8842
gb:LC522975|Organism:Severe      CAGCGTGGTGGTAGTTATACTAATGACAAAGCTTGCCCATTGATTGCTGCAGTCATAACA   8839
gb:LC522973|Organism:Severe      CAGCGTGGTGGTAGTTATACTAATGACAAAGCTTGCCCATTGATTGCTGCAGTCATAACA   8839
gb:LC522974|Organism:Severe      CAGCGTGGTGGTAGTTATACTAATGACAAAGCTTGCCCATTGATTGCTGCAGTCATAACA   8839
gb:MN985325|Organism:Severe      CAGCGTGGTGGTAGTTATACTAATGACAAAGCTTGCCCATTGATTGCTGCAGTCATAACA   8842
gb:MT020881|Organism:Severe      CAGCGTGGTGGTAGTTATACTAATGACAAAGCTTGCCCATTGATTGCTGCAGTCATAACA   8842
gb:MT020880|Organism:Severe      CAGCGTGGTGGTAGTTATACTAATGACAAAGCTTGCCCATTGATTGCTGCAGTCATAACA   8842
gb:MT066175|Organism:Severe      CAGCGTGGTGGTAGTTATACTAATGACAAAGCTTGCCCATTGATTGCTGCAGTCATAACA   8842
gb:MN997409|Organism:Severe      CAGCGTGGTGGTAGTTATACTAATGACAAAGCTTGCCCATTGATTGCTGCAGTCATAACA   8842
gb:MN938384|Organism:Severe      CAGCGTGGTGGTAGTTATACTAATGACAAAGCTTGCCCATTGATTGCTGCAGTCATAACA   8810
gb:MT044258|Organism:Severe      CAGCGTGGTGGTAGTTATACTAATGACAAAGCTTGCCCATTGATTGCTGCAGTCATAACA   8818
gb:MT039890|Organism:Severe      CAGCGTGGTGGTAGTTATACTAATGACAAAGCTTGCCCATTGATTGCTGCAGTCATAACA   8842
gb:MN988713|Organism:Severe      CAGCGTGGTGGTAGTTATACTAATGACAAAGCTTGCCCATTGATTGCTGCAGTCATAACA   8842
gb:LC521925|Organism:Severe      CAGCGTGGTGGTAGTTATACTAATGACAAAGCTTGCCCATTGATTGCTGCAGTCATAACA   8815
gb:MT093571|Organism:Severe      CAGCGTGGTGGTAGTTATACTAATGACAAAGCTTGCCCATTGATTGCTGCAGTCATAACA   8842
gb:MT039887|Organism:Severe      CAGCGTGGTGGTAGTTATACTAATGACAAAGCTTGCCCATTGATTGCTGCAGTCATAACA   8842
gb:MT019530|Organism:Severe      CAGCGTGGTGGTAGTTATACTAATGACAAAGCTTGCCCATTGATTGCTGCAGTCATAACA   8842
gb:MT039888|Organism:Severe      CAGCGTGGTGGTAGTTATACTAATGACAAAGCTTGCCCATTGATTGCTGCAGTCATAACA   8842
gb:LC522972|Organism:Severe      CAGCGTGGTGGTAGTTATACTAATGACAAAGCTTGCCCATTGATTGCTGCAGTCATAACA   8839
gb:MT027063|Organism:Severe      CAGCGTGGTGGTAGTTATACTAATGACAAAGCTTGCCCATTGATTGCTGCAGTCATAACA   8842
gb:MT027062|Organism:Severe      CAGCGTGGTGGTAGTTATACTAATGACAAAGCTTGCCCATTGATTGCTGCAGTCATAACA   8842
gb:MT019529|Organism:Severe      CAGCGTGGTGGTAGTTATACTAATGACAAAGCTTGCCCATTGATTGCTGCAGTCATAACA   8842
gb:MN996529|Organism:Severe      CAGCGTGGTGGTAGTTATACTAATGACAAAGCTTGCCCATTGATTGCTGCAGTCATAACA   8830
gb:MN996531|Organism:Severe      CAGCGTGGTGGTAGTTATACTAATGACAAAGCTTGCCCATTGATTGCTGCAGTCATAACA   8829
gb:MT066176|Organism:Severe      CAGCGTGGTGGTAGTTATACTAATGACAAAGCTTGCCCATTGATTGCTGCAGTCATAACA   8842
gb:MT027064|Organism:Severe      CAGCGTGGTGGTAGTTATACTAATGACAAAGCTTGCCCATTGATTGCTGCAGTCATAACA   8842
gb:MN994468|Organism:Severe      CAGCGTGGTGGTAGTTATACTAATGACAAAGCTTGCCCATTGATTGCTGCAGTCATAACA   8842
gb:MT072688|Organism:Severe      CAGCGTGGTGGTAGTTATACTAATGACAAAGCTTGCCCATTGATTGCTGCAGTCATAACA   8827
gb:MN996527|Organism:Severe      CAGCGTGGTGGTAGTTATACTAATGACAAAGCTTGCCCATTGATTGCTGCAGTCATAACA   8809
gb:MT093631|Organism:Severe      CAGCGTGGTGGTAGTTATACTAATGACAAAGCTTGCCCATTGATTGCTGCAGTCATAACA   8880
gb:MT106053|Organism:Severe      CAGCGTGGTGGTAGTTATACTAATGACAAAGCTTGCCCATTGATTGCTGCAGTCATAACA   8842
gb:MT019533|Organism:Severe      CAGCGTGGTGGTAGTTATACTAATGACAAAGCTTGCCCATTGATTGCTGCAGTCATAACA   8842
gb:MT019531|Organism:Severe      CAGCGTGGTGGTAGTTATACTAATGACAAAGCTTGCCCATTGATTGCTGCAGTCATAACA   8842
gb:MN996528|Organism:Severe      CAGCGTGGTGGTAGTTATACTAATGACAAAGCTTGCCCATTGATTGCTGCAGTCATAACA   8842
gb:MN996530|Organism:Severe      CAGCGTGGTGGTAGTTATACTAATGACAAAGCTTGCCCATTGATTGCTGCAGTCATAACA   8828
gb:MN908947|Organism:Severe      CAGCGTGGTGGTAGTTATACTAATGACAAAGCTTGCCCATTGATTGCTGCAGTCATAACA   8842
gb:MT019532|Organism:Severe      CAGCGTGGTGGTAGTTATACTAATGACAAAGCTTGCCCATTGATTGCTGCAGTCATAACA   8842
                                 ************************************************************

gb:MT020781|Organism:Severe      AGAGAAGTGGGTTTTGTCGTGCCTGGTTTGCCTGGCACGATATTACGCACAACTAATGGT   8890
gb:MT007544|Organism:Severe      AGAGAAGTGGGTTTTGTCGTGCCTGGTTTGCCTGGCACGATATTACGCACAACTAATGGT   8902
gb:MN994467|Organism:Severe      AGAGAAGTGGGTTTTGTCGTGCCTGGTTTGCCTGGCACGATATTACGCACAACTAATGGT   8902
gb:MT044257|Organism:Severe      AGAGAAGTGGGTTTTGTCGTGCCTGGTTTGCCTGGCACGATATTACGCACAACTAATGGT   8902
gb:MT106054|Organism:Severe      AGAGAAGTGGGTTTTGTCGTGCCTGGTTTGCCTGGCACGATATTACGCACAACTAATGGT   8902
gb:MT049951|Organism:Severe      AGAGAAGTGGGTTTTGTCGTGCCTGGTTTGCCTGGCACGATATTACGCACAACTAATGGT   8902
gb:MN975262|Organism:Severe      AGAGAAGTGGGTTTTGTCGTGCCTGGTTTGCCTGGCACGATATTACGCACAACTAATGGT   8902
gb:MT106052|Organism:Severe      AGAGAAGTGGGTTTTGTCGTGCCTGGTTTGCCTGGCACGATATTACGCACAACTAATGGT   8902
gb:LC522975|Organism:Severe      AGAGAAGTGGGTTTTGTCGTGCCTGGTTTGCCTGGCACGATATTACGCACAACTAATGGT   8899
gb:LC522973|Organism:Severe      AGAGAAGTGGGTTTTGTCGTGCCTGGTTTGCCTGGCACGATATTACGCACAACTAATGGT   8899
gb:LC522974|Organism:Severe      AGAGAAGTGGGTTTTGTCGTGCCTGGTTTGCCTGGCACGATATTACGCACAACTAATGGT   8899
gb:MN985325|Organism:Severe      AGAGAAGTGGGTTTTGTCGTGCCTGGTTTGCCTGGCACGATATTACGCACAACTAATGGT   8902
gb:MT020881|Organism:Severe      AGAGAAGTGGGTTTTGTCGTGCCTGGTTTGCCTGGCACGATATTACGCACAACTAATGGT   8902
```

```
gb:MT020880|Organism:Severe      AGAGAAGTGGGTTTTGTCGTGCCTGGTTTGCCTGGCACGATATTACGCACAACTAATGGT   8902
gb:MT066175|Organism:Severe      AGAGAAGTGGGTTTTGTCGTGCCTGGTTTGCCTGGCACGATATTACGCACAACTAATGGT   8902
gb:MN997409|Organism:Severe      AGAGAAGTGGGTTTTGTCGTGCCTGGTTTGCCTGGCACGATATTACGCACAACTAATGGT   8902
gb:MN938384|Organism:Severe      AGAGAAGTGGGTTTTGTCGTGCCTGGTTTGCCTGGCACGATATTACGCACAACTAATGGT   8870
gb:MT044258|Organism:Severe      AGAGAAGTGGGTTTTGTCGTGCCTGGTTTGCCTGGCACGATATTACGCACAACTAATGGT   8878
gb:MT039890|Organism:Severe      AGAGAAGTGGGTTTTGTCGTGCCTGGTTTGCCTGGCACGATATTACGCACAACTAATGGT   8902
gb:MN988713|Organism:Severe      AGAGAAGTGGGTTTTGTCGTGCCTGGTTTGCCTGGCACGATATTACGCACAACTAATGGT   8902
gb:LC521925|Organism:Severe      AGAGAAGTGGGTTTTGTCGTGCCTGGTTTGCCTGGCACGATATTACGCACAACTAATGGT   8875
gb:MT093571|Organism:Severe      AGAGAAGTGGGTTTTGTCGTGCCTGGTTTGCCTGGCACGATATTACGCACAACTAATGGT   8902
gb:MT039887|Organism:Severe      AGAGAAGTGGGTTTTGTCGTGCCTGGTTTGCCTGGCACGATATTACGCACAACTAATGGT   8902
gb:MT019530|Organism:Severe      AGAGAAGTGGGTTTTGTCGTGCCTGGTTTGCCTGGCACGATATTACGCACAACTAATGGT   8902
gb:MT039888|Organism:Severe      AGAGAAGTGGGTTTTGTCGTGCCTGGTTTGCCTGGCACGATATTACGCACAACTAATGGT   8902
gb:LC522972|Organism:Severe      AGAGAAGTGGGTTTTGTCGTGCCTGGTTTGCCTGGCACGATATTACGCACAACTAATGGT   8899
gb:MT027063|Organism:Severe      AGAGAAGTGGGTTTTGTCGTGCCTGGTTTGCCTGGCACGATATTACGCACAACTAATGGT   8902
gb:MT027062|Organism:Severe      AGAGAAGTGGGTTTTGTCGTGCCTGGTTTGCCTGGCACGATATTACGCACAACTAATGGT   8902
gb:MT019529|Organism:Severe      AGAGAAGTGGGTTTTGTCGTGCCTGGTTTGCCTGGCACGATATTACGCACAACTAATGGT   8902
gb:MN996529|Organism:Severe      AGAGAAGTGGGTTTTGTCGTGCCTGGTTTGCCTGGCACGATATTACGCACAACTAATGGT   8890
gb:MN996531|Organism:Severe      AGAGAAGTGGGTTTTGTCGTGCCTGGTTTGCCTGGCACGATATTACGCACAACTAATGGT   8889
gb:MT066176|Organism:Severe      AGAGAAGTGGGTTTTGTCGTGCCTGGTTTGCCTGGCACGATATTACGCACAACTAATGGT   8902
gb:MT027064|Organism:Severe      AGAGAAGTGGGTTTTGTCGTGCCTGGTTTGCCTGGCACGATATTACGCACAACTAATGGT   8902
gb:MN994468|Organism:Severe      AGAGAAGTGGGTTTTGTCGTGCCTGGTTTGCCTGGCACGATATTACGCACAACTAATGGT   8902
gb:MT072688|Organism:Severe      AGAGAAGTGGGTTTTGTCGTGCCTGGTTTGCCTGGCACGATATTACGCACAACTAATGGT   8887
gb:MN996527|Organism:Severe      AGAGAAGTGGGTTTTGTCGTGCCTGGTTTGCCTGGCACGATATTACGCACAACTAATGGT   8869
gb:MT093631|Organism:Severe      AGAGAAGTGGGTTTTGTCGTGCCTGGTTTGCCTGGCACGATATTACGCACAACTAATGGT   8940
gb:MT106053|Organism:Severe      AGAGAAGTGGGTTTTGTCGTGCCTGGTTTGCCTGGCACGATATTACGCACAACTAATGGT   8902
gb:MT019533|Organism:Severe      AGAGAAGTGGGTTTTGTCGTGCCTGGTTTGCCTGGCACGATATTACGCACAACTAATGGT   8902
gb:MT019531|Organism:Severe      AGAGAAGTGGGTTTTGTCGTGCCTGGTTTGCCTGGCACGATATTACGCACAACTAATGGT   8902
gb:MN996528|Organism:Severe      AGAGAAGTGGGTTTTGTCGTGCCTGGTTTGCCTGGCACGATATTACGCACAACTAATGGT   8902
gb:MN996530|Organism:Severe      AGAGAAGTGGGTTTTGTCGTGCCTGGTTTGCCTGGCACGATATTACGCACAACTAATGGT   8888
gb:MN908947|Organism:Severe      AGAGAAGTGGGTTTTGTCGTGCCTGGTTTGCCTGGCACGATATTACGCACAACTAATGGT   8902
gb:MT019532|Organism:Severe      AGAGAAGTGGGTTTTGTCGTGCCTGGTTTGCCTGGCACGATATTACGCACAACTAATGGT   8902
                                 ************************************************************

gb:MT020781|Organism:Severe      GACTTTTTGCATTTCTTACCTAGAGTTTTTAGTGCAGTTGGTAACATCTGTTACACACCA   8950
gb:MT007544|Organism:Severe      GACTTTTTGCATTTCTTACCTAGAGTTTTTAGTGCAGTTGGTAACATCTGTTACACACCA   8962
gb:MN994467|Organism:Severe      GACTTTTTGCATTTCTTACCTAGAGTTTTTAGTGCAGTTGGTAACATCTGTTACACACCA   8962
gb:MT044257|Organism:Severe      GACTTTTTGCATTTCTTACCTAGAGTTTTTAGTGCAGTTGGTAACATCTGTTACACACCA   8962
gb:MT106054|Organism:Severe      GACTTTTTGCATTTCTTACCTAGAGTTTTTAGTGCAGTTGGTAACATCTGTTACACACCA   8962
gb:MT049951|Organism:Severe      GACTTTTTGCATTTCTTACCTAGAGTTTTTAGTGCAGTTGGTAACATCTGTTACACACCA   8962
gb:MN975262|Organism:Severe      GACTTTTTGCATTTCTTACCTAGAGTTTTTAGTGCAGTTGGTAACATCTGTTACACACCA   8962
gb:MT106052|Organism:Severe      GACTTTTTGCATTTCTTACCTAGAGTTTTTAGTGCAGTTGGTAACATCTGTTACACACCA   8962
gb:LC522975|Organism:Severe      GACTTTTTGCATTTCTTACCTAGAGTTTTTAGTGCAGTTGGTAACATCTGTTACACACCA   8959
gb:LC522973|Organism:Severe      GACTTTTTGCATTTCTTACCTAGAGTTTTTAGTGCAGTTGGTAACATCTGTTACACACCA   8959
gb:LC522974|Organism:Severe      GACTTTTTGCATTTCTTACCTAGAGTTTTTAGTGCAGTTGGTAACATCTGTTACACACCA   8959
gb:MN985325|Organism:Severe      GACTTTTTGCATTTCTTACCTAGAGTTTTTAGTGCAGTTGGTAACATCTGTTACACACCA   8962
gb:MT020881|Organism:Severe      GACTTTTTGCATTTCTTACCTAGAGTTTTTAGTGCAGTTGGTAACATCTGTTACACACCA   8962
gb:MT020880|Organism:Severe      GACTTTTTGCATTTCTTACCTAGAGTTTTTAGTGCAGTTGGTAACATCTGTTACACACCA   8962
gb:MT066175|Organism:Severe      GACTTTTTGCATTTCTTACCTAGAGTTTTTAGTGCAGTTGGTAACATCTGTTACACACCA   8962
gb:MN997409|Organism:Severe      GACTTTTTGCATTTCTTACCTAGAGTTTTTAGTGCAGTTGGTAACATCTGTTACACACCA   8962
gb:MN938384|Organism:Severe      GACTTTTTGCATTTCTTACCTAGAGTTTTTAGTGCAGTTGGTAACATCTGTTACACACCA   8930
gb:MT044258|Organism:Severe      GACTTTTTGCATTTCTTACCTAGAGTTTTTAGTGCAGTTGGTAACATCTGTTACACACCA   8938
gb:MT039890|Organism:Severe      GACTTTTTGCATTTCTTACCTAGAGTTTTTAGTGCAGTTGGTAACATCTGTTACACACCA   8962
gb:MN988713|Organism:Severe      GACTTTTTGCATTTCTTACCTAGAGTTTTTAGTGCAGTTGGTAACATCTGTTACACACCA   8962
gb:LC521925|Organism:Severe      GACTTTTTGCATTTCTTACCTAGAGTTTTTAGTGCAGTTGGTAACATCTGTTACACACCA   8935
gb:MT093571|Organism:Severe      GACTTTTTGCATTTCTTACCTAGAGTTTTTAGTGCAGTTGGTAACATCTGTTACACACCA   8962
gb:MT039887|Organism:Severe      GACTTTTTGCATTTCTTACCTAGAGTTTTTAGTGCAGTTGGTAACATCTGTTACACACCA   8962
gb:MT019530|Organism:Severe      GACTTTTTGCATTTCTTACCTAGAGTTTTTAGTGCAGTTGGTAACATCTGTTACACACCA   8962
gb:MT039888|Organism:Severe      GACTTTTTGCATTTCTTACCTAGAGTTTTTAGTGCAGTTGGTAACATCTGTTACACACCA   8962
gb:LC522972|Organism:Severe      GACTTTTTGCATTTCTTACCTAGAGTTTTTAGTGCAGTTGGTAACATCTGTTACACACCA   8959
gb:MT027063|Organism:Severe      GACTTTTTGCATTTCTTACCTAGAGTTTTTAGTGCAGTTGGTAACATCTGTTACACACCA   8962
gb:MT027062|Organism:Severe      GACTTTTTGCATTTCTTACCTAGAGTTTTTAGTGCAGTTGGTAACATCTGTTACACACCA   8962
gb:MT019529|Organism:Severe      GACTTTTTGCATTTCTTACCTAGAGTTTTTAGTGCAGTTGGTAACATCTGTTACACACCA   8962
gb:MN996529|Organism:Severe      GACTTTTTGCATTTCTTACCTAGAGTTTTTAGTGCAGTTGGTAACATCTGTTACACACCA   8950
gb:MN996531|Organism:Severe      GACTTTTTGCATTTCTTACCTAGAGTTTTTAGTGCAGTTGGTAACATCTGTTACACACCA   8949
gb:MT066176|Organism:Severe      GACTTTTTGCATTTCTTACCTAGAGTTTTTAGTGCAGTTGGTAACATCTGTTACACACCA   8962
```

```
gb:MT027064|Organism:Severe      GACTTTTTGCATTTCTTACCTAGAGTTTTTAGTGCAGTTGGTAACATCTGTTACACACCA   8962
gb:MN994468|Organism:Severe      GACTTTTTGCATTTCTTACCTAGAGTTTTTAGTGCAGTTGGTAACATCTGTTACACACCA   8962
gb:MT072688|Organism:Severe      GACTTTTTGCATTTCTTACCTAGAGTTTTTAGTGCAGTTGGTAACATCTGTTACACACCA   8947
gb:MN996527|Organism:Severe      GACTTTTTGCATTTCTTACCTAGAGTTTTTAGTGCAGTTGGTAACATCTGTTACACACCA   8929
gb:MT093631|Organism:Severe      GACTTTTTGCATTTCTTACCTAGAGTTTTTAGTGCAGTTGGTAACATCTGTTACACACCA   9000
gb:MT106053|Organism:Severe      GACTTTTTGCATTTCTTACCTAGAGTTTTTAGTGCAGTTGGTAACATCTGTTACACACCA   8962
gb:MT019533|Organism:Severe      GACTTTTTGCATTTCTTACCTAGAGTTTTTAGTGCAGTTGGTAACATCTGTTACACACCA   8962
gb:MT019531|Organism:Severe      GACTTTTTGCATTTCTTACCTAGAGTTTTTAGTGCAGTTGGTAACATCTGTTACACACCA   8962
gb:MN996528|Organism:Severe      GACTTTTTGCATTTCTTACCTAGAGTTTTTAGTGCAGTTGGTAACATCTGTTACACACCA   8962
gb:MN996530|Organism:Severe      GACTTTTTGCATTTCTTACCTAGAGTTTTTAGTGCAGTTGGTAACATCTGTTACACACCA   8948
gb:MN908947|Organism:Severe      GACTTTTTGCATTTCTTACCTAGAGTTTTTAGTGCAGTTGGTAACATCTGTTACACACCA   8962
gb:MT019532|Organism:Severe      GACTTTTTGCATTTCTTACCTAGAGTTTTTAGTGCAGTTGGTAACATCTGTTACACACCA   8962
                                 ************************************************************

gb:MT020781|Organism:Severe      TCAAAACTTATAGAGTACACTGACTTTGCAACATCAGCTTGTGTTTTGGCTGCTGAATGT   9010
gb:MT007544|Organism:Severe      TCAAAACTTATAGAGTACACTGACTTTGCAACATCAGCTTGTGTTTTGGCTGCTGAATGT   9022
gb:MN994467|Organism:Severe      TCAAAACTTATAGAGTACACTGACTTTGCAACATCAGCTTGTGTTTTGGCTGCTGAATGT   9022
gb:MT044257|Organism:Severe      TCAAAACTTATAGAGTACACTGACTTTGCAACATCAGCTTGTGTTTTGGCTGCTGAATGT   9022
gb:MT106054|Organism:Severe      TCAAAACTTATAGAGTACACTGACTTTGCAACATCAGCTTGTGTTTTGGCTGCTGAATGT   9022
gb:MT049951|Organism:Severe      TCAAAACTTATAGAGTACACTGACTTTGCAACATCAGCTTGTGTTTTGGCTGCTGAATGT   9022
gb:MN975262|Organism:Severe      TCAAAACTTATAGAGTACACTGACTTTGCAACATCAGCTTGTGTTTTGGCTGCTGAATGT   9022
gb:MT106052|Organism:Severe      TCAAAACTTATAGAGTACACTGACTTTGCAACATCAGCTTGTGTTTTGGCTGCTGAATGT   9022
gb:LC522975|Organism:Severe      TCAAAACTTATAGAGTACACTGACTTTGCAACATCAGCTTGTGTTTTGGCTGCTGAATGT   9019
gb:LC522973|Organism:Severe      TCAAAACTTATAGAGTACACTGACTTTGCAACATCAGCTTGTGTTTTGGCTGCTGAATGT   9019
gb:LC522974|Organism:Severe      TCAAAACTTATAGAGTACACTGACTTTGCAACATCAGCTTGTGTTTTGGCTGCTGAATGT   9019
gb:MN985325|Organism:Severe      TCAAAACTTATAGAGTACACTGACTTTGCAACATCAGCTTGTGTTTTGGCTGCTGAATGT   9022
gb:MT020881|Organism:Severe      TCAAAACTTATAGAGTACACTGACTTTGCAACATCAGCTTGTGTTTTGGCTGCTGAATGT   9022
gb:MT020880|Organism:Severe      TCAAAACTTATAGAGTACACTGACTTTGCAACATCAGCTTGTGTTTTGGCTGCTGAATGT   9022
gb:MT066175|Organism:Severe      TCAAAACTTATAGAGTACACTGACTTTGCAACATCAGCTTGTGTTTTGGCTGCTGAATGT   9022
gb:MN997409|Organism:Severe      TCAAAACTTATAGAGTACACTGACTTTGCAACATCAGCTTGTGTTTTGGCTGCTGAATGT   9022
gb:MN938384|Organism:Severe      TCAAAACTTATAGAGTACACTGACTTTGCAACATCAGCTTGTGTTTTGGCTGCTGAATGT   8990
gb:MT044258|Organism:Severe      TCAAAACTTATAGAGTACACTGACTTTGCAACATCAGCTTGTGTTTTGGCTGCTGAATGT   8998
gb:MT039890|Organism:Severe      TCAAAACTTATAGAGTACACTGACTTTGCAACATCAGCTTGTGTTTTGGCTGCTGAATGT   9022
gb:MN988713|Organism:Severe      TCAAAACTTATAGAGTACACTGACTTTGCAACATCAGCTTGTGTTTTGGCTGCTGAATGT   9022
gb:LC521925|Organism:Severe      TCAAAACTTATAGAGTACACTGACTTTGCAACATCAGCTTGTGTTTTGGCTGCTGAATGT   8995
gb:MT093571|Organism:Severe      TCAAAACTTATAGAGTACACTGACTTTGCAACATCAGCTTGTGTTTTGGCTGCTGAATGT   9022
gb:MT039887|Organism:Severe      TCAAAACTTATAGAGTACACTGACTTTGCAACATCAGCTTGTGTTTTGGCTGCTGAATGT   9022
gb:MT019530|Organism:Severe      TCAAAACTTATAGAGTACACTGACTTTGCAACATCAGCTTGTGTTTTGGCTGCTGAATGT   9022
gb:MT039888|Organism:Severe      TCAAAACTTATAGAGTACACTGACTTTGCAACATCAGCTTGTGTTTTGGCTGCTGAATGT   9022
gb:LC522972|Organism:Severe      TCAAAACTTATAGAGTACACTGACTTTGCAACATCAGCTTGTGTTTTGGCTGCTGAATGT   9019
gb:MT027063|Organism:Severe      TCAAAACTTATAGAGTACACTGACTTTGCAACATCAGCTTGTGTTTTGGCTGCTGAATGT   9022
gb:MT027062|Organism:Severe      TCAAAACTTATAGAGTACACTGACTTTGCAACATCAGCTTGTGTTTTGGCTGCTGAATGT   9022
gb:MT019529|Organism:Severe      TCAAAACTTATAGAGTACACTGACATTGCAACATCAGCTTGTGTTTTGGCTGCTGAATGT   9022
gb:MN996529|Organism:Severe      TCAAAACTTATAGAGTACACTGACTTTGCAACATCAGCTTGTGTTTTGGCTGCTGAATGT   9010
gb:MN996531|Organism:Severe      TCAAAACTTATAGAGTACACTGACTTTGCAACATCAGCTTGTGTTTTGGCTGCTGAATGT   9009
gb:MT066176|Organism:Severe      TCAAAACTTATAGAGTACACTGACTTTGCAACATCAGCTTGTGTTTTGGCTGCTGAATGT   9022
gb:MT027064|Organism:Severe      TCAAAACTTATAGAGTACACTGACTTTGCAACATCAGCTTGTGTTTTGGCTGCTGAATGT   9022
gb:MN994468|Organism:Severe      TCAAAACTTATAGAGTACACTGACTTTGCAACATCAGCTTGTGTTTTGGCTGCTGAATGT   9022
gb:MT072688|Organism:Severe      TCAAAACTTATAGAGTACACTGACTTTGCAACATCAGCTTGTGTTTTGGCTGCTGAATGT   9007
gb:MN996527|Organism:Severe      TCAAAACTTATAGAGTACACTGACTTTGCAACATCAGCTTGTGTTTTGGCTGCTGAATGT   8989
gb:MT093631|Organism:Severe      TCAAAACTTATAGAGTACACTGACTTTGCAACATCAGCTTGTGTTTTGGCTGCTGAATGT   9060
gb:MT106053|Organism:Severe      TCAAAACTTATAGAGTACACTGACTTTGCAACATCAGCTTGTGTTTTGGCTGCTGAATGT   9022
gb:MT019533|Organism:Severe      TCAAAACTTATAGAGTACACTGACTTTGCAACATCAGCTTGTGTTTTGGCTGCTGAATGT   9022
gb:MT019531|Organism:Severe      TCAAAACTTATAGAGTACACTGACTTTGCAACATCAGCTTGTGTTTTGGCTGCTGAATGT   9022
gb:MN996528|Organism:Severe      TCAAAACTTATAGAGTACACTGACTTTGCAACATCAGCTTGTGTTTTGGCTGCTGAATGT   9022
gb:MN996530|Organism:Severe      TCAAAACTTATAGAGTACACTGACTTTGCAACATCAGCTTGTGTTTTGGCTGCTGAATGT   9008
gb:MN908947|Organism:Severe      TCAAAACTTATAGAGTACACTGACTTTGCAACATCAGCTTGTGTTTTGGCTGCTGAATGT   9022
gb:MT019532|Organism:Severe      TCAAAACTTATAGAGTACACTGACTTTGCAACATCAGCTTGTGTTTTGGCTGCTGAATGT   9022
                                 ************************ ***********************************

gb:MT020781|Organism:Severe      ACAATTTTTAAAGATGCTTCTGGTAAGCCAGTACCATATTGTTATGATACCAATGTACTA   9070
gb:MT007544|Organism:Severe      ACAATTTTTAAAGATGCTTCTGGTAAGCCAGTACCATATTGTTATGATACCAATGTACTA   9082
gb:MN994467|Organism:Severe      ACAATTTTTAAAGATGCTTCTGGTAAGCCAGTACCATATTGTTATGATACCAATGTACTA   9082
gb:MT044257|Organism:Severe      ACAATTTTTAAAGATGCTTCTGGTAAGCCAGTACCATATTGTTATGATACCAATGTACTA   9082
gb:MT106054|Organism:Severe      ACAATTTTTAAAGATGCTTCTGGTAAGCCAGTACCATATTGTTATGATACCAATGTACTA   9082
```

```
gb:MT049951|Organism:Severe      ACAATTTTTAAAGATGCTTCTGGTAAGCCAGTACCATATTGTTATGATACCAATGTACTA   9082
gb:MN975262|Organism:Severe      ACAATTTTTAAAGATGCTTCTGGTAAGCCAGTACCATATTGTTATGATACCAATGTACTA   9082
gb:MT106052|Organism:Severe      ACAATTTTTAAAGATGCTTCTGGTAAGCCAGTACCATATTGTTATGATACCAATGTACTA   9082
gb:LC522975|Organism:Severe      ACAATTTTTAAAGATGCTTCTGGTAAGCCAGTACCATATTGTTATGATACCAATGTACTA   9079
gb:LC522973|Organism:Severe      ACAATTTTTAAAGATGCTTCTGGTAAGCCAGTACCATATTGTTATGATACCAATGTACTA   9079
gb:LC522974|Organism:Severe      ACAATTTTTAAAGATGCTTCTGGTAAGCCAGTACCATATTGTTATGATACCAATGTACTA   9079
gb:MN985325|Organism:Severe      ACAATTTTTAAAGATGCTTCTGGTAAGCCAGTACCATATTGTTATGATACCAATGTACTA   9082
gb:MT020881|Organism:Severe      ACAATTTTTAAAGATGCTTCTGGTAAGCCAGTACCATATTGTTATGATACCAATGTACTA   9082
gb:MT020880|Organism:Severe      ACAATTTTTAAAGATGCTTCTGGTAAGCCAGTACCATATTGTTATGATACCAATGTACTA   9082
gb:MT066175|Organism:Severe      ACAATTTTTAAAGATGCTTCTGGTAAGCCAGTACCATATTGTTATGATACCAATGTACTA   9082
gb:MN997409|Organism:Severe      ACAATTTTTAAAGATGCTTCTGGTAAGCCAGTACCATATTGTTATGATACCAATGTACTA   9082
gb:MN938384|Organism:Severe      ACAATTTTTAAAGATGCTTCTGGTAAGCCAGTACCATATTGTTATGATACCAATGTACTA   9050
gb:MT044258|Organism:Severe      ACAATTTTTAAAGATGCTTCTGGTAAGCCAGTACCATATTGTTATGATACCAATGTACTA   9058
gb:MT039890|Organism:Severe      ACAATTTTTAAAGATGCTTCTGGTAAGCCAGTACCATATTGTTATGATACCAATGTACTA   9082
gb:MN988713|Organism:Severe      ACAATTTTTAAAGATGCTTCTGGTAAGCCAGTACCATATTGTTATGATACCAATGTACTA   9082
gb:LC521925|Organism:Severe      ACAATTTTTAAAGATGCTTCTGGTAAGCCAGTACCATATTGTTATGATACCAATGTACTA   9055
gb:MT093571|Organism:Severe      ACAATTTTTAAAGATGCTTCTGGTAAGCCAGTACCATATTGTTATGATACCAATGTACTA   9082
gb:MT039887|Organism:Severe      ACAATTTTTAAAGATGCTTCTGGTAAGCCAGTACCATATTGTTATGATACCAATGTACTA   9082
gb:MT019530|Organism:Severe      ACAATTTTTAAAGATGCTTCTGGTAAGCCAGTACCATATTGTTATGATACCAATGTACTA   9082
gb:MT039888|Organism:Severe      ACAATTTTTAAAGATGCTTCTGGTAAGCCAGTACCATATTGTTATGATACCAATGTACTA   9082
gb:LC522972|Organism:Severe      ACAATTTTTAAAGATGCTTCTGGTAAGCCAGTACCATATTGTTATGATACCAATGTACTA   9079
gb:MT027063|Organism:Severe      ACAATTTTTAAAGATGCTTCTGGTAAGCCAGTACCATATTGTTATGATACCAATGTACTA   9082
gb:MT027062|Organism:Severe      ACAATTTTTAAAGATGCTTCTGGTAAGCCAGTACCATATTGTTATGATACCAATGTACTA   9082
gb:MT019529|Organism:Severe      ACAATTTTTAAAGATGCTTCTGGTAAGCCAGTACCATATTGTTATGATACCAATGTACTA   9082
gb:MN996529|Organism:Severe      ACAATTTTTAAAGATGCTTCTGGTAAGCCAGTACCATATTGTTATGATACCAATGTACTA   9070
gb:MN996531|Organism:Severe      ACAATTTTTAAAGATGCTTCTGGTAAGCCAGTACCATATTGTTATGATACCAATGTACTA   9069
gb:MT066176|Organism:Severe      ACAATTTTTAAGGATGCTTCTGGTAAGCCAGTACCATATTGTTATGATACCAATGTACTA   9082
gb:MT027064|Organism:Severe      ACAATTTTTAAAGATGCTTCTGGTAAGCCAGTACCATATTGTTATGATACCAATGTACTA   9082
gb:MN994468|Organism:Severe      ACAATTTTTAAAGATGCTTCTGGTAAGCCAGTACCATATTGTTATGATACCAATGTACTA   9082
gb:MT072688|Organism:Severe      ACAATTTTTAAAGATGCTTCTGGTAAGCCAGTACCATATTGTTATGATACCAATGTACTA   9067
gb:MN996527|Organism:Severe      ACAATTTTTAAAGATGCTTCTGGTAAGCCAGTACCATATTGTTATGATACCAATGTACTA   9049
gb:MT093631|Organism:Severe      ACAATTTTTAAAGATGCTTCTGGTAAGCCAGTACCATATTGTTATGATACCAATGTACTA   9120
gb:MT106053|Organism:Severe      ACAATTTTTAAAGATGCTTCTGGTAAGCCAGTACCATATTGTTATGATACCAATGTACTA   9082
gb:MT019533|Organism:Severe      ACAATTTTTAAAGATGCTTCTGGTAAGCCAGTACCATATTGTTATGATACCAATGTACTA   9082
gb:MT019531|Organism:Severe      ACAATTTTTAAAGATGCTTCTGGTAAGCCAGTACCATATTGTTATGATACCAATGTACTA   9082
gb:MN996528|Organism:Severe      ACAATTTTTAAAGATGCTTCTGGTAAGCCAGTACCATATTGTTATGATACCAATGTACTA   9082
gb:MN996530|Organism:Severe      ACAATTTTTAAAGATGCTTCTGGTAAGCCAGTACCATATTGTTATGATACCAATGTACTA   9068
gb:MN908947|Organism:Severe      ACAATTTTTAAAGATGCTTCTGGTAAGCCAGTACCATATTGTTATGATACCAATGTACTA   9082
gb:MT019532|Organism:Severe      ACAATTTTTAAAGATGCTTCTGGTAAGCCAGTACCATATTGTTATGATACCAATGTACTA   9082
                                 *********** ************************************************

gb:MT020781|Organism:Severe      GAAGGTTCTGTTGCTTATGAAAGTTTACGCCCTGACACACGTTATGTGCTCATGGATGGC   9130
gb:MT007544|Organism:Severe      GAAGGTTCTGTTGCTTATGAAAGTTTACGCCCTGACACACGTTATGTGCTCATGGATGGC   9142
gb:MN994467|Organism:Severe      GAAGGTTCTGTTGCTTATGAAAGTTTACGCCCTGACACACGTTATGTGCTCATGGATGGC   9142
gb:MT044257|Organism:Severe      GAAGGTTCTGTTGCTTATGAAAGTTTACGCCCTGACACACGTTATGTGCTCATGGATGGC   9142
gb:MT106054|Organism:Severe      GAAGGTTCTGTTGCTTATGAAAGTTTACGCCCTGACACACGTTATGTGCTCATGGATGGC   9142
gb:MT049951|Organism:Severe      GAAGGTTCTGTTGCTTATGAAAGTTTACGCCCTGACACACGTTATGTGCTCATGGATGGC   9142
gb:MN975262|Organism:Severe      GAAGGTTCTGTTGCTTATGAAAGTTTACGCCCTGACACACGTTATGTGCTCATGGATGGC   9142
gb:MT106052|Organism:Severe      GAAGGTTCTGTTGCTTATGAAAGTTTACGCCCTGACACACGTTATGTGCTCATGGATGGC   9142
gb:LC522975|Organism:Severe      GAAGGTTCTGTTGCTTATGAAAGTTTACGCCCTGACACACGTTATGTGCTCATGGATGGC   9139
gb:LC522973|Organism:Severe      GAAGGTTCTGTTGCTTATGAAAGTTTACGCCCTGACACACGTTATGTGCTCATGGATGGC   9139
gb:LC522974|Organism:Severe      GAAGGTTCTGTTGCTTATGAAAGTTTACGCCCTGACACACGTTATGTGCTCATGGATGGC   9139
gb:MN985325|Organism:Severe      GAAGGTTCTGTTGCTTATGAAAGTTTACGCCCTGACACACGTTATGTGCTCATGGATGGC   9142
gb:MT020881|Organism:Severe      GAAGGTTCTGTTGCTTATGAAAGTTTACGCCCTGACACACGTTATGTGCTCATGGATGGC   9142
gb:MT020880|Organism:Severe      GAAGGTTCTGTTGCTTATGAAAGTTTACGCCCTGACACACGTTATGTGCTCATGGATGGC   9142
gb:MT066175|Organism:Severe      GAAGGTTCTGTTGCTTATGAAAGTTTACGCCCTGACACACGTTATGTGCTCATGGATGGC   9142
gb:MN997409|Organism:Severe      GAAGGTTCTGTTGCTTATGAAAGTTTACGCCCTGACACACGTTATGTGCTCATGGATGGC   9142
gb:MN938384|Organism:Severe      GAAGGTTCTGTTGCTTATGAAAGTTTACGCCCTGACACACGTTATGTGCTCATGGATGGC   9110
gb:MT044258|Organism:Severe      GAAGGTTCTGTTGCTTATGAAAGTTTACGCCCTGACACACGTTATGTGCTCATGGATGGC   9118
gb:MT039890|Organism:Severe      GAAGGTTCTGTTGCTTATGAAAGTTTACGCCCTGACACACGTTATGTGCTCATGGATGGC   9142
gb:MN988713|Organism:Severe      GAAGGTTCTGTTGCTTATGAAAGTTTACGCCCTGACACACGTTATGTGCTCATGGATGGC   9142
gb:LC521925|Organism:Severe      GAAGGTTCTGTTGCTTATGAAAGTTTACGCCCTGACACACGTTATGTGCTCATGGATGGC   9115
gb:MT093571|Organism:Severe      GAAGGTTCTGTTGCTTATGAAAGTTTACGCCCTGACACACGTTATGTGCTCATGGATGGC   9142
gb:MT039887|Organism:Severe      GAAGGTTCTGTTGCTTATGAAAGTTTACGCCCTGACACACGTTATGTGCTCATGGATGGC   9142
gb:MT019530|Organism:Severe      GAAGGTTCTGTTGCTTATGAAAGTTTACGCCCTGACACACGTTATGTGCTCATGGATGGC   9142
```

```
gb:MT039888|Organism:Severe      GAAGGTTCTGTTGCTTATGAAAGTTTACGCCCTGACACACGTTATGTGCTCATGGATGGC   9142
gb:LC522972|Organism:Severe      GAAGGTTCTGTTGCTTATGAAAGTTTACGCCCTGACACACGTTATGTGCTCATGGATGGC   9139
gb:MT027063|Organism:Severe      GAAGGTTCTGTTGCTTATGAAAGTTTACGCCCTGACACACGTTATGTGCTCATGGATGGC   9142
gb:MT027062|Organism:Severe      GAAGGTTCTGTTGCTTATGAAAGTTTACGCCCTGACACACGTTATGTGCTCATGGATGGC   9142
gb:MT019529|Organism:Severe      GAAGGTTCTGTTGCTTATGAAAGTTTACGCCCTGACACACGTTATGTGCTCATGGATGGC   9142
gb:MN996529|Organism:Severe      GAAGGTTCTGTTGCTTATGAAAGTTTACGCCCTGACACACGTTATGTGCTCATGGATGGC   9130
gb:MN996531|Organism:Severe      GAAGGTTCTGTTGCTTATGAAAGTTTACGCCCTGACACACGTTATGTGCTCATGGATGGC   9129
gb:MT066176|Organism:Severe      GAAGGTTCTGTTGCTTATGAAAGTTTACGCCCTGACACACGTTATGTGCTCATGGATGGC   9142
gb:MT027064|Organism:Severe      GAAGGTTCTGTTGCTTATGAAAGTTTACGCCCTGACACACGTTATGTGCTCATGGATGGC   9142
gb:MN994468|Organism:Severe      GAAGGTTCTGTTGCTTATGAAAGTTTACGCCCTGACACACGTTATGTGCTCATGGATGGC   9142
gb:MT072688|Organism:Severe      GAAGGTTCTGTTGCTTATGAAAGTTTACGCCCTGACACACGTTATGTGCTCATGGATGGC   9127
gb:MN996527|Organism:Severe      GAAGGTTCTGTTGCTTATGAAAGTTTACGCCCTGACACACGTTATGTGCTCATGGATGGC   9109
gb:MT093631|Organism:Severe      GAAGGTTCTGTTGCTTATGAAAGTTTACGCCCTGACACACGTTATGTGCTCATGGATGGC   9180
gb:MT106053|Organism:Severe      GAAGGTTCTGTTGCTTATGAAAGTTTACGCCCTGACACACGTTATGTGCTCATGGATGGC   9142
gb:MT019533|Organism:Severe      GAAGGTTCTGTTGCTTATGAAAGTTTACGCCCTGACACACGTTATGTGCTCATGGATGGC   9142
gb:MT019531|Organism:Severe      GAAGGTTCTGTTGCTTATGAAAGTTTACGCCCTGACACACGTTATGTGCTCATGGATGGC   9142
gb:MN996528|Organism:Severe      GAAGGTTCTGTTGCTTATGAAAGTTTACGCCCTGACACACGTTATGTGCTCATGGATGGC   9142
gb:MN996530|Organism:Severe      GAAGGTTCTGTTGCTTATGAAAGTTTACGCCCTGACACACGTTATGTGCTCATGGATGGC   9128
gb:MN908947|Organism:Severe      GAAGGTTCTGTTGCTTATGAAAGTTTACGCCCTGACACACGTTATGTGCTCATGGATGGC   9142
gb:MT019532|Organism:Severe      GAAGGTTCTGTTGCTTATGAAAGTTTACGCCCTGACACACGTTATGTGCTCATGGATGGC   9142
                                 ************************************************************

gb:MT020781|Organism:Severe      TCTATTATTCAATTTCCTAACACCTACCTTGAAGGTTCTGTTAGAGTGGTAACAACTTTT   9190
gb:MT007544|Organism:Severe      TCTATTATTCAATTTCCTAACACCTACCTTGAAGGTTCTGTTAGAGTGGTAACAACTTTT   9202
gb:MN994467|Organism:Severe      TCTATTATTCAATTTCCTAACACCTACCTTGAAGGTTCTGTTAGAGTGGTAACAACTTTT   9202
gb:MT044257|Organism:Severe      TCTATTATTCAATTTCCTAACACCTACCTTGAAGGTTCTGTTAGAGTGGTAACAACTTTT   9202
gb:MT106054|Organism:Severe      TCTATTATTCAATTTCCTAACACCTACCTTGAAGGTTCTGTTAGAGTGGTAACAACTTTT   9202
gb:MT049951|Organism:Severe      TCTATTATTCAATTTCCTAACACCTACCTTGAAGGTTCTGTTAGAGTGGTAACAACTTTT   9202
gb:MN975262|Organism:Severe      TCTATTATTCAATTTCCTAACACCTACCTTGAAGGTTCTGTTAGAGTGGTAACAACTTTT   9202
gb:MT106052|Organism:Severe      TCTATTATTCAATTTCCTAACACCTACCTTGAAGGTTCTGTTAGAGTGGTAACAACTTTT   9202
gb:LC522975|Organism:Severe      TCTATTATTCAATTTCCTAACACCTACCTTGAAGGTTCTGTTAGAGTGGTAACAACTTTT   9199
gb:LC522973|Organism:Severe      TCTATTATTCAATTTCCTAACACCTACCTTGAAGGTTCTGTTAGAGTGGTAACAACTTTT   9199
gb:LC522974|Organism:Severe      TCTATTATTCAATTTCCTAACACCTACCTTGAAGGTTCTGTTAGAGTGGTAACAACTTTT   9199
gb:MN985325|Organism:Severe      TCTATTATTCAATTTCCTAACACCTACCTTGAAGGTTCTGTTAGAGTGGTAACAACTTTT   9202
gb:MT020881|Organism:Severe      TCTATTATTCAATTTCCTAACACCTACCTTGAAGGTTCTGTTAGAGTGGTAACAACTTTT   9202
gb:MT020880|Organism:Severe      TCTATTATTCAATTTCCTAACACCTACCTTGAAGGTTCTGTTAGAGTGGTAACAACTTTT   9202
gb:MT066175|Organism:Severe      TCTATTATTCAATTTCCTAACACCTACCTTGAAGGTTCTGTTAGAGTGGTAACAACTTTT   9202
gb:MN997409|Organism:Severe      TCTATTATTCAATTTCCTAACACCTACCTTGAAGGTTCTGTTAGAGTGGTAACAACTTTT   9202
gb:MN938384|Organism:Severe      TCTATTATTCAATTTCCTAACACCTACCTTGAAGGTTCTGTTAGAGTGGTAACAACTTTT   9170
gb:MT044258|Organism:Severe      TCTATTATTCAATTTCCTAACACCTACCTTGAAGGTTCTGTTAGAGTGGTAACAACTTTT   9178
gb:MT039890|Organism:Severe      TCTATTATTCAATTTCCTAACACCTACCTTGAAGGTTCTGTTAGAGTGGTAACAACTTTT   9202
gb:MN988713|Organism:Severe      TCTATTATTCAATTTCCTAACACCTACCTTGAAGGTTCTGTTAGAGTGGTAACAACTTTT   9202
gb:LC521925|Organism:Severe      TCTATTATTCAATTTCCTAACACCTACCTTGAAGGTTCTGTTAGAGTGGTAACAACTTTT   9175
gb:MT093571|Organism:Severe      TCTATTATTCAATTTCCTAACACCTACCTTGAAGGTTCTGTTAGAGTGGTAACAACTTTT   9202
gb:MT039887|Organism:Severe      TCTATTATTCAATTTCCTAACACCTACCTTGAAGGTTCTGTTAGAGTGGTAACAACTTTT   9202
gb:MT019530|Organism:Severe      TCTATTATTCAATTTCCTAACACCTACCTTGAAGGTTCTGTTAGAGTGGTAACAACTTTT   9202
gb:MT039888|Organism:Severe      TCTATTATTCAATTTCCTAACACCTACCTTGAAGGTTCTGTTAGAGTGGTAACAACTTTT   9202
gb:LC522972|Organism:Severe      TCTATTATTCAATTTCCTAACACCTACCTTGAAGGTTCTGTTAGAGTGGTAACAACTTTT   9199
gb:MT027063|Organism:Severe      TCTATTATTCAATTTCCTAACACCTACCTTGAAGGTTCTGTTAGAGTGGTAACAACTTTT   9202
gb:MT027062|Organism:Severe      TCTATTATTCAATTTCCTAACACCTACCTTGAAGGTTCTGTTAGAGTGGTAACAACTTTT   9202
gb:MT019529|Organism:Severe      TCTATTATTCAATTTCCTAACACCTACCTTGAAGGTTCTGTTAGAGTGGTAACAACTTTT   9202
gb:MN996529|Organism:Severe      TCTATTATTCAATTTCCTAACACCTACCTTGAAGGTTCTGTTAGAGTGGTAACAACTTTT   9190
gb:MN996531|Organism:Severe      TCTATTATTCAATTTCCTAACACCTACCTTGAAGGTTCTGTTAGAGTGGTAACAACTTTT   9189
gb:MT066176|Organism:Severe      TCTATTATTCAATTTCCTAACACCTACCTTGAAGGTTCTGTTAGAGTGGTAACAACTTTT   9202
gb:MT027064|Organism:Severe      TCTATTATTCAATTTCCTAACACCTACCTTGAAGGTTCTGTTAGAGTGGTAACAACTTTT   9202
gb:MN994468|Organism:Severe      TCTATTATTCAATTTCCTAACACCTACCTTGAAGGTTCTGTTAGAGTGGTAACAACTTTT   9202
gb:MT072688|Organism:Severe      TCTATTATTCAATTTCCTAACACCTACCTTGAAGGTTCTGTTAGAGTGGTAACAACTTTT   9187
gb:MN996527|Organism:Severe      TCTATTATTCAATTTCCTAACACCTACCTTGAAGGTTCTGTTAGAGTGGTAACAACTTTT   9169
gb:MT093631|Organism:Severe      TCTATTATTCAATTTCCTAACACCTACCTTGAAGGTTCTGTTAGAGTGGTAACAACTTTT   9240
gb:MT106053|Organism:Severe      TCTATTATTCAATTTCCTAACACCTACCTTGAAGGTTCTGTTAGAGTGGTAACAACTTTT   9202
gb:MT019533|Organism:Severe      TCTATTATTCAATTTCCTAACACCTACCTTGAAGGTTCTGTTAGAGTGGTAACAACTTTT   9202
gb:MT019531|Organism:Severe      TCTATTATTCAATTTCCTAACACCTACCTTGAAGGTTCTGTTAGAGTGGTAACAACTTTT   9202
gb:MN996528|Organism:Severe      TCTATTATTCAATTTCCTAACACCTACCTTGAAGGTTCTGTTAGAGTGGTAACAACTTTT   9202
gb:MN996530|Organism:Severe      TCTATTATTCAATTTCCTAACACCTACCTTGAAGGTTCTGTTAGAGTGGTAACAACTTTT   9188
gb:MN908947|Organism:Severe      TCTATTATTCAATTTCCTAACACCTACCTTGAAGGTTCTGTTAGAGTGGTAACAACTTTT   9202
```

```
gb:MT019532|Organism:Severe      TCTATTATTCAATTTCCTAACACCTACCTTGAAGGTTCTGTTAGAGTGGTAACAACTTTT   9202
                                 ************************************************************

gb:MT020781|Organism:Severe      GATTCTGAGTACTGTAGGCACGGCACTTGTGAAAGATCAGAAGCTGGTGTTTGTGTATCT   9250
gb:MT007544|Organism:Severe      GATTCTGAGTACTGTAGGCACGGCACTTGTGAAAGATCAGAAGCTGGTGTTTGTGTATCT   9262
gb:MN994467|Organism:Severe      GATTCTGAGTACTGTAGGCACGGCACTTGTGAAAGATCAGAAGCTGGTGTTTGTGTATCT   9262
gb:MT044257|Organism:Severe      GATTCTGAGTACTGTAGGCACGGCACTTGTGAAAGATCAGAAGCTGGTGTTTGTGTATCT   9262
gb:MT106054|Organism:Severe      GATTCTGAGTACTGTAGGCACGGCACTTGTGAAAGATCAGAAGCTGGTGTTTGTGTATCT   9262
gb:MT049951|Organism:Severe      GATTCTGAGTACTGTAGGCACGGCACTTGTGAAAGATCAGAAGCTGGTGTTTGTGTATCT   9262
gb:MN975262|Organism:Severe      GATTCTGAGTACTGTAGGCACGGCACTTGTGAAAGATCAGAAGCTGGTGTTTGTGTATCT   9262
gb:MT106052|Organism:Severe      GATTCTGAGTACTGTAGGCACGGCACTTGTGAAAGATCAGAAGCTGGTGTTTGTGTATCT   9262
gb:LC522975|Organism:Severe      GATTCTGAGTACTGTAGGCACGGCACTTGTGAAAGATCAGAAGCTGGTGTTTGTGTATCT   9259
gb:LC522973|Organism:Severe      GATTCTGAGTACTGTAGGCACGGCACTTGTGAAAGATCAGAAGCTGGTGTTTGTGTATCT   9259
gb:LC522974|Organism:Severe      GATTCTGAGTACTGTAGGCACGGCACTTGTGAAAGATCAGAAGCTGGTGTTTGTGTATCT   9259
gb:MN985325|Organism:Severe      GATTCTGAGTACTGTAGGCACGGCACTTGTGAAAGATCAGAAGCTGGTGTTTGTGTATCT   9262
gb:MT020881|Organism:Severe      GATTCTGAGTACTGTAGGCACGGCACTTGTGAAAGATCAGAAGCTGGTGTTTGTGTATCT   9262
gb:MT020880|Organism:Severe      GATTCTGAGTACTGTAGGCACGGCACTTGTGAAAGATCAGAAGCTGGTGTTTGTGTATCT   9262
gb:MT066175|Organism:Severe      GATTCTGAGTACTGTAGGCACGGCACTTGTGAAAGATCAGAAGCTGGTGTTTGTGTATCT   9262
gb:MN997409|Organism:Severe      GATTCTGAGTACTGTAGGCACGGCACTTGTGAAAGATCAGAAGCTGGTGTTTGTGTATCT   9262
gb:MN938384|Organism:Severe      GATTCTGAGTACTGTAGGCACGGCACTTGTGAAAGATCAGAAGCTGGTGTTTGTGTATCT   9230
gb:MT044258|Organism:Severe      GATTCTGAGTACTGTAGGCACGGCACTTGTGAAAGATCAGAAGCTGGTGTTTGTGTATCT   9238
gb:MT039890|Organism:Severe      GATTCTGAGTACTGTAGGCACGGCACTTGTGAAAGATCAGAAGCTGGTGTTTGTGTATCT   9262
gb:MN988713|Organism:Severe      GATTCTGAGTACTGTAGGCACGGCACTTGTGAAAGATCAGAAGCTGGTGTTTGTGTATCT   9262
gb:LC521925|Organism:Severe      GATTCTGAGTACTGTAGGCACGGCACTTGTGAAAGATCAGAAGCTGGTGTTTGTGTATCT   9235
gb:MT093571|Organism:Severe      GATTCTGAGTACTGTAGGCACGGCACTTGTGAAAGATCAGAAGCTGGTGTTTGTGTATCT   9262
gb:MT039887|Organism:Severe      GATTCTGAGTACTGTAGGCACGGCACTTGTGAAAGATCAGAAGCTGGTGTTTGTGTATCT   9262
gb:MT019530|Organism:Severe      GATTCTGAGTACTGTAGGCACGGCACTTGTGAAAGATCAGAAGCTGGTGTTTGTGTATCT   9262
gb:MT039888|Organism:Severe      GATTCTGAGTACTGTAGGCACGGCACTTGTGAAAGATCAGAAGCTGGTGTTTGTGTATCT   9262
gb:LC522972|Organism:Severe      GATTCTGAGTACTGTAGGCACGGCACTTGTGAAAGATCAGAAGCTGGTGTTTGTGTATCT   9259
gb:MT027063|Organism:Severe      GATTCTGAGTACTGTAGGCACGGCACTTGTGAAAGATCAGAAGCTGGTGTTTGTGTATCT   9262
gb:MT027062|Organism:Severe      GATTCTGAGTACTGTAGGCACGGCACTTGTGAAAGATCAGAAGCTGGTGTTTGTGTATCT   9262
gb:MT019529|Organism:Severe      GATTCTGAGTACTGTAGGCACGGCACTTGTGAAAGATCAGAAGCTGGTGTTTGTGTATCT   9262
gb:MN996529|Organism:Severe      GATTCTGAGTACTGTAGGCACGGCACTTGTGAAAGATCAGAAGCTGGTGTTTGTGTATCT   9250
gb:MN996531|Organism:Severe      GATTCTGAGTACTGTAGGCACGGCACTTGTGAAAGATCAGAAGCTGGTGTTTGTGTATCT   9249
gb:MT066176|Organism:Severe      GATTCTGAGTACTGTAGGCACGGCACTTGTGAAAGATCAGAAGCTGGTGTTTGTGTATCT   9262
gb:MT027064|Organism:Severe      GATTCTGAGTACTGTAGGCACGGCACTTGTGAAAGATCAGAAGCTGGTGTTTGTGTATCT   9262
gb:MN994468|Organism:Severe      GATTCTGAGTACTGTAGGCACGGCACTTGTGAAAGATCAGAAGCTGGTGTTTGTGTATCT   9262
gb:MT072688|Organism:Severe      GATTCTGAGTACTGTAGGCACGGCACTTGTGAAAGATCAGAAGCTGGTGTTTGTGTATCT   9247
gb:MN996527|Organism:Severe      GATTCTGAGTACTGTAGGCACGGCACTTGTGAAAGATCAGAAGCTGGTGTTTGTGTATCT   9229
gb:MT093631|Organism:Severe      GATTCTGAGTACTGTAGGCACGGCACTTGTGAAAGATCAGAAGCTGGTGTTTGTGTATCT   9300
gb:MT106053|Organism:Severe      GATTCTGAGTACTGTAGGCACGGCACTTGTGAAAGATCAGAAGCTGGTGTTTGTGTATCT   9262
gb:MT019533|Organism:Severe      GATTCTGAGTACTGTAGGCACGGCACTTGTGAAAGATCAGAAGCTGGTGTTTGTGTATCT   9262
gb:MT019531|Organism:Severe      GATTCTGAGTACTGTAGGCACGGCACTTGTGAAAGATCAGAAGCTGGTGTTTGTGTATCT   9262
gb:MN996528|Organism:Severe      GATTCTGAGTACTGTAGGCACGGCACTTGTGAAAGATCAGAAGCTGGTGTTTGTGTATCT   9262
gb:MN996530|Organism:Severe      GATTCTGAGTACTGTAGGCACGGCACTTGTGAAAGATCAGAAGCTGGTGTTTGTGTATCT   9248
gb:MN908947|Organism:Severe      GATTCTGAGTACTGTAGGCACGGCACTTGTGAAAGATCAGAAGCTGGTGTTTGTGTATCT   9262
gb:MT019532|Organism:Severe      GATTCTGAGTACTGTAGGCACGGCACTTGTGAAAGATCAGAAGCTGGTGTTTGTGTATCT   9262
                                 ************************************************************

gb:MT020781|Organism:Severe      ACTAGTGGTAGATGGGTACTTAACAATGATTATTACAGATCTTTACCAGGAGTTTTCTGT   9310
gb:MT007544|Organism:Severe      ACTAGTGGTAGATGGGTACTTAACAATGATTATTACAGATCTTTACCAGGAGTTTTCTGT   9322
gb:MN994467|Organism:Severe      ACTAGTGGTAGATGGGTACTTAACAATGATTATTACAGATCTTTACCAGGAGTTTTCTGT   9322
gb:MT044257|Organism:Severe      ACTAGTGGTAGATGGGTACTTAACAATGATTATTACAGATCTTTACCAGGAGTTTTCTGT   9322
gb:MT106054|Organism:Severe      ACTAGTGGTAGATGGGTACTTAACAATGATTATTACAGATCTTTACCAGGAGTTTTCTGT   9322
gb:MT049951|Organism:Severe      ACTAGTGGTAGATGGGTACTTAACAATGATTATTACAGATCTTTACCAGGAGTTTTCTGT   9322
gb:MN975262|Organism:Severe      ACTAGTGGTAGATGGGTACTTAACAATGATTATTACAGATCTTTACCAGGAGTTTTCTGT   9322
gb:MT106052|Organism:Severe      ACTAGTGGTAGATGGGTACTTAACAATGATTATTACAGATCTTTACCAGGAGTTTTCTGT   9322
gb:LC522975|Organism:Severe      ACTAGTGGTAGATGGGTACTTAACAATGATTATTACAGATCTTTACCAGGAGTTTTCTGT   9319
gb:LC522973|Organism:Severe      ACTAGTGGTAGATGGGTACTTAACAATGATTATTACAGATCTTTACCAGGAGTTTTCTGT   9319
gb:LC522974|Organism:Severe      ACTAGTGGTAGATGGGTACTTAACAATGATTATTACAGATCTTTACCAGGAGTTTTCTGT   9319
gb:MN985325|Organism:Severe      ACTAGTGGTAGATGGGTACTTAACAATGATTATTACAGATCTTTACCAGGAGTTTTCTGT   9322
gb:MT020881|Organism:Severe      ACTAGTGGTAGATGGGTACTTAACAATGATTATTACAGATCTTTACCAGGAGTTTTCTGT   9322
gb:MT020880|Organism:Severe      ACTAGTGGTAGATGGGTACTTAACAATGATTATTACAGATCTTTACCAGGAGTTTTCTGT   9322
gb:MT066175|Organism:Severe      ACTAGTGGTAGATGGGTACTTAACAATGATTATTACAGATCTTTACCAGGAGTTTTCTGT   9322
gb:MN997409|Organism:Severe      ACTAGTGGTAGATGGGTACTTAACAATGATTATTACAGATCTTTACCAGGAGTTTTCTGT   9322
```

```
gb:MN938384|Organism:Severe      ACTAGTGGTAGATGGGTACTTAACAATGATTATTACAGATCTTTACCAGGAGTTTTCTGT   9290
gb:MT044258|Organism:Severe      ACTAGTGGTAGATGGGTACTTAACAATGATTATTACAGATCTTTACCAGGAGTTTTCTGT   9298
gb:MT039890|Organism:Severe      ACTAGTGGTAGATGGGTACTTAACAATGATTATTACAGATCTTTACCAGGAGTTTTCTGT   9322
gb:MN988713|Organism:Severe      ACTAGTGGTAGATGGGTACTTAACAATGATTATTACAGATCTTTACCAGGAGTTTTCTGT   9322
gb:LC521925|Organism:Severe      ACTAGTGGTAGATGGGTACTTAACAATGATTATTACAGATCTTTACCAGGAGTTTTCTGT   9295
gb:MT093571|Organism:Severe      ACTAGTGGTAGGTGGGTACTTAACAATGATTATTACAGATCTTTACCAGGAGTTTTCTGT   9322
gb:MT039887|Organism:Severe      ACTAGTGGTAGATGGGTACTTAACAATGATTATTACAGATCTTTACCAGGAGTTTTCTGT   9322
gb:MT019530|Organism:Severe      ACTAGTGGTAGATGGGTACTTAACAATGATTATTACAGATCTTTACCAGGAGTTTTCTGT   9322
gb:MT039888|Organism:Severe      ACTAGTGGTAGATGGGTACTTAACAATGATTATTACAGATCTTTACCAGGAGTTTTCTGT   9322
gb:LC522972|Organism:Severe      ACTAGTGGTAGATGGGTACTTAACAATGATTATTACAGATCTTTACCAGGAGTTTTCTGT   9319
gb:MT027063|Organism:Severe      ACTAGTGGTAGATGGGTACTTAACAATGATTATTACAGATCTTTACCAGGAGTTTTCTGT   9322
gb:MT027062|Organism:Severe      ACTAGTGGTAGATGGGTACTTAACAATGATTATTACAGATCTTTACCAGGAGTTTTCTGT   9322
gb:MT019529|Organism:Severe      ACTAGTGGTAGATGGGTACTTAACAATGATTATTACAGATCTTTACCAGGAGTTTTCTGT   9322
gb:MN996529|Organism:Severe      ACTAGTGGTAGATGGGTACTTAACAATGATTATTACAGATCTTTACCAGGAGTTTTCTGT   9310
gb:MN996531|Organism:Severe      ACTAGTGGTAGATGGGTACTTAACAATGATTATTACAGATCTTTACCAGGAGTTTTCTGT   9309
gb:MT066176|Organism:Severe      ACTAGTGGTAGATGGGTACTTAACAATGATTATTACAGATCTTTACCAGGAGTTTTCTGT   9322
gb:MT027064|Organism:Severe      ACTAGTGGTAGATGGGTACTTAACAATGATTATTACAGATCTTTACCAGGAGTTTTCTGT   9322
gb:MN994468|Organism:Severe      ACTAGTGGTAGATGGGTACTTAACAATGATTATTACAGATCTTTACCAGGAGTTTTCTGT   9322
gb:MT072688|Organism:Severe      ACTAGTGGTAGATGGGTACTTAACAATGATTATTACAGATCTTTACCAGGAGTTTTCTGT   9307
gb:MN996527|Organism:Severe      ACTAGTGGTAGATGGGTACTTAACAATGATTATTACAGATCTTTACCAGGAGTTTTCTGT   9289
gb:MT093631|Organism:Severe      ACTAGTGGTAGATGGGTACTTAACAATGATTATTACAGATCTTTACCAGGAGTTTTCTGT   9360
gb:MT106053|Organism:Severe      ACTAGTGGTAGATGGGTACTTAACAATGATTATTACAGATCTTTACCAGGAGTTTTCTGT   9322
gb:MT019533|Organism:Severe      ACTAGTGGTAGATGGGTACTTAACAATGATTATTACAGATCTTTACCAGGAGTTTTCTGT   9322
gb:MT019531|Organism:Severe      ACTAGTGGTAGATGGGTACTTAACAATGATTATTACAGATCTTTACCAGGAGTTTTCTGT   9322
gb:MN996528|Organism:Severe      ACTAGTGGTAGATGGGTACTTAACAATGATTATTACAGATCTTTACCAGGAGTTTTCTGT   9322
gb:MN996530|Organism:Severe      ACTAGTGGTAGATGGGTACTTAACAATGATTATTACAGATCTTTACCAGGAGTTTTCTGT   9308
gb:MN908947|Organism:Severe      ACTAGTGGTAGATGGGTACTTAACAATGATTATTACAGATCTTTACCAGGAGTTTTCTGT   9322
gb:MT019532|Organism:Severe      ACTAGTGGTAGATGGGTACTTAACAATGATTATTACAGATCTTTACCAGGAGTTTTCTGT   9322
                                 *********** ************************************************

gb:MT020781|Organism:Severe      GGTGTAGATGCTGTAAATTTACTTACTAATATGTTTACACCACTAATTCAACCTATTGGT   9370
gb:MT007544|Organism:Severe      GGTGTAGATGCTGTAAATTTACTTACTAATATGTTTACACCACTAATTCAACCTATTGGT   9382
gb:MN994467|Organism:Severe      GGTGTAGATGCTGTAAATTTACTTACTAATATGTTTACACCACTAATTCAACCTATTGGT   9382
gb:MT044257|Organism:Severe      GGTGTAGATGCTGTAAATTTACTTACTAATATGTTTACACCACTAATTCAACCTATTGGT   9382
gb:MT106054|Organism:Severe      GGTGTAGATGCTGTAAATTTACTTACTAATATGTTTACACCACTAATTCAACCTATTGGT   9382
gb:MT049951|Organism:Severe      GGTGTAGATGCTGTAAATTTACTTACTAATATGTTTACACCACTAATTCAACCTATTGGT   9382
gb:MN975262|Organism:Severe      GGTGTAGATGCTGTAAATTTACTTACTAATATGTTTACACCACTAATTCAACCTATTGGT   9382
gb:MT106052|Organism:Severe      GGTGTAGATGCTGTAAATTTACTTACTAATATGTTTACACCACTAATTCAACCTATTGGT   9382
gb:LC522975|Organism:Severe      GGTGTAGATGCTGTAAATTTACTTACTAATATGTTTACACCACTAATTCAACCTATTGGT   9379
gb:LC522973|Organism:Severe      GGTGTAGATGCTGTAAATTTACTTACTAATATGTTTACACCACTAATTCAACCTATTGGT   9379
gb:LC522974|Organism:Severe      GGTGTAGATGCTGTAAATTTACTTACTAATATGTTTACACCACTAATTCAACCTATTGGT   9379
gb:MN985325|Organism:Severe      GGTGTAGATGCTGTAAATTTACTTACTAATATGTTTACACCACTAATTCAACCTATTGGT   9382
gb:MT020881|Organism:Severe      GGTGTAGATGCTGTAAATTTACTTACTAATATGTTTACACCACTAATTCAACCTATTGGT   9382
gb:MT020880|Organism:Severe      GGTGTAGATGCTGTAAATTTACTTACTAATATGTTTACACCACTAATTCAACCTATTGGT   9382
gb:MT066175|Organism:Severe      GGTGTAGATGCTGTAAATTTACTTACTAATATGTTTACACCACTAATTCAACCTATTGGT   9382
gb:MN997409|Organism:Severe      GGTGTAGATGCTGTAAATTTACTTACTAATATGTTTACACCACTAATTCAACCTATTGGT   9382
gb:MN938384|Organism:Severe      GGTGTAGATGCTGTAAATTTACTTACTAATATGTTTACACCACTAATTCAACCTATTGGT   9350
gb:MT044258|Organism:Severe      GGTGTAGATGCTGTAAATTTACTTACTAATATGTTTACACCACTAATTCAACCTATTGGT   9358
gb:MT039890|Organism:Severe      GGTGTAGATGCTGTAAATTTACTTACTAATATGTTTACACCACTAATTCAACCTATTGGT   9382
gb:MN988713|Organism:Severe      GGTGTAGATGCTGTAAATTTACTTACTAATATGTTTACACCACTAATTCAACCTATTGGT   9382
gb:LC521925|Organism:Severe      GGTGTAGATGCTGTAAATTTACTTACTAATATGTTTACACCACTAATTCAACCTATTGGT   9355
gb:MT093571|Organism:Severe      GGTGTAGATGCTGTAAATTTACTTACTAATATGTTTACACCACTAATTCAACCTATTGGT   9382
gb:MT039887|Organism:Severe      GGTGTAGATGCTGTAAATTTACTTACTAATATGTTTACACCACTAATTCAACCTATTGGT   9382
gb:MT019530|Organism:Severe      GGTGTAGATGCTGTAAATTTACTTACTAATATGTTTACACCACTAATTCAACCTATTGGT   9382
gb:MT039888|Organism:Severe      GGTGTAGATGCTGTAAATTTACTTACTAATATGTTTACACCACTAATTCAACCTATTGGT   9382
gb:LC522972|Organism:Severe      GGTGTAGATGCTGTAAATTTACTTACTAATATGTTTACACCACTAATTCAACCTATTGGT   9379
gb:MT027063|Organism:Severe      GGTGTAGATGCTGTAAATTTACTTACTAATATGTTTACACCACTAATTCAACCTATTGGT   9382
gb:MT027062|Organism:Severe      GGTGTAGATGCTGTAAATTTACTTACTAATATGTTTACACCACTAATTCAACCTATTGGT   9382
gb:MT019529|Organism:Severe      GGTGTAGATGCTGTAAATTTACTTACTAATATGTTTACACCACTAATTCAACCTATTGGT   9382
gb:MN996529|Organism:Severe      GGTGTAGATGCTGTAAATTTACTTACTAATATGTTTACACCACTAATTCAACCTATTGGT   9370
gb:MN996531|Organism:Severe      GGTGTAGATGCTGTAAATTTACTTACTAATATGTTTACACCACTAATTCAACCTATTGGT   9369
gb:MT066176|Organism:Severe      GGTGTAGATGCTGTAAATTTACTTACTAATATGTTTACACCACTAATTCAACCTATTGGT   9382
gb:MT027064|Organism:Severe      GGTGTAGATGCTGTAAATTTACTTACTAATATGTTTACACCACTAATTCAACCTATTGGT   9382
gb:MN994468|Organism:Severe      GGTGTAGATGCTGTAAATTTACTTACTAATATGTTTACACCACTAATTCAACCTATTGGT   9382
gb:MT072688|Organism:Severe      GGTGTAGATGCTGTAAATTTACTTACTAATATGTTTACACCACTAATTCAACCTATTGGT   9367
```

```
gb:MN996527|Organism:Severe      GGTGTAGATGCTGTAAATTTACTTACTAATATGTTTACACCACTAATTCAACCTATTGGT   9349
gb:MT093631|Organism:Severe      GGTGTAGATGCTGTAAATTTACTTACTAATATGTTTACACCACTAATTCAACCTATTGGT   9420
gb:MT106053|Organism:Severe      GGTGTAGATGCTGTAAATTTACTTACTAATATGTTTACACCACTAATTCAACCTATTGGT   9382
gb:MT019533|Organism:Severe      GGTGTAGATGCTGTAAATTTACTTACTAATATGTTTACACCACTAATTCAACCTATTGGT   9382
gb:MT019531|Organism:Severe      GGTGTAGATGCTGTAAATTTACTTACTAATATGTTTACACCACTAATTCAACCTATTGGT   9382
gb:MN996528|Organism:Severe      GGTGTAGATGCTGTAAATTTACTTACTAATATGTTTACACCACTAATTCAACCTATTGGT   9382
gb:MN996530|Organism:Severe      GGTGTAGATGCTGTAAATTTACTTACTAATATGTTTACACCACTAATTCAACCTATTGGT   9368
gb:MN908947|Organism:Severe      GGTGTAGATGCTGTAAATTTACTTACTAATATGTTTACACCACTAATTCAACCTATTGGT   9382
gb:MT019532|Organism:Severe      GGTGTAGATGCTGTAAATTTACTTACTAATATGTTTACACCACTAATTCAACCTATTGGT   9382
                                 ************************************************************

gb:MT020781|Organism:Severe      GCTTTGGACATATCAGCATCTATAGTAGCTGGTGGTATTGTAGCTATCGTAGTAACATGC   9430
gb:MT007544|Organism:Severe      GCTTTGGACATATCAGCATCTATAGTAGCTGGTGGTATTGTAGCTATCGTAGTAACATGC   9442
gb:MN994467|Organism:Severe      GCTTTGGACATATCAGCATCTATAGTAGCTGGTGGTATTGTAGCTATCGTAGTAACATGC   9442
gb:MT044257|Organism:Severe      GCTTTGGACATATCAGCATCTATAGTAGCTGGTGGTATTGTAGCTATCGTAGTAACATGC   9442
gb:MT106054|Organism:Severe      GCTTTGGACATATCAGCATCTATAGTAGCTGGTGGTATTGTAGCTATCGTAGTAACATGC   9442
gb:MT049951|Organism:Severe      GCTTTGGACATATCAGCATCTATAGTAGCTGGTGGTATTGTAGCTATCGTAGTAACATGC   9442
gb:MN975262|Organism:Severe      GCTTTGGACATATCAGCATCTATAGTAGCTGGTGGTATTGTAGCTATCGTAGTAACATGC   9442
gb:MT106052|Organism:Severe      GCTTTGGACATATCAGCATCTATAGTAGCTGGTGGTATTGTAGCTATCGTAGTAACATGC   9442
gb:LC522975|Organism:Severe      GCTTTGGACATATCAGCATCTATAGTAGCTGGTGGTATTGTAGCTATCGTAGTAACATGC   9439
gb:LC522973|Organism:Severe      GCTTTGGACATATCAGCATCTATAGTAGCTGGTGGTATTGTAGCTATCGTAGTAACATGC   9439
gb:LC522974|Organism:Severe      GCTTTGGACATATCAGCATCTATAGTAGCTGGTGGTATTGTAGCTATCGTAGTAACATGC   9439
gb:MN985325|Organism:Severe      GCTTTGGACATATCAGCATCTATAGTAGCTGGTGGTATTGTAGCTATCGTAGTAACATGC   9442
gb:MT020881|Organism:Severe      GCTTTGGACATATCAGCATCTATAGTAGCTGGTGGTATTGTAGCTATCGTAGTAACATGC   9442
gb:MT020880|Organism:Severe      GCTTTGGACATATCAGCATCTATAGTAGCTGGTGGTATTGTAGCTATCGTAGTAACATGC   9442
gb:MT066175|Organism:Severe      GCTTTGGACATATCAGCATCTATAGTAGCTGGTGGTATTGTAGCTATCGTAGTAACATGC   9442
gb:MN997409|Organism:Severe      GCTTTGGACATATCAGCATCTATAGTAGCTGGTGGTATTGTAGCTATCGTAGTAACATGC   9442
gb:MN938384|Organism:Severe      GCTTTGGACATATCAGCATCTATAGTAGCTGGTGGTATTGTAGCTATCGTAGTAACATGC   9410
gb:MT044258|Organism:Severe      GCTTTGGACATATCAGCATCTATAGTAGCTGGTGGTATTGTAGCTATCGTAGTAACATGC   9418
gb:MT039890|Organism:Severe      GCTTTGGACATATCAGCATCTATAGTAGCTGGTGGTATTGTAGCTATCGTAGTAACATGC   9442
gb:MN988713|Organism:Severe      GCTTTGGACATATCAGCATCTATAGTAGCTGGTGGTATTGTAGCTATCGTAGTAACATGC   9442
gb:LC521925|Organism:Severe      GCTTTGGACATATCAGCATCTATAGTAGCTGGTGGTATTGTAGCTATCGTAGTAACATGC   9415
gb:MT093571|Organism:Severe      GCTTTGGACATATCAGCATCTATAGTAGCTGGTGGTATTGTAGCTATCGTAGTAACATGC   9442
gb:MT039887|Organism:Severe      GCTTTGGACATATCAGCATCTATAGTAGCTGGTGGTATTGTAGCTATCGTAGTAACATGC   9442
gb:MT019530|Organism:Severe      GCTTTGGACATATCAGCATCTATAGTAGCTGGTGGTATTGTAGCTATCGTAGTAACATGC   9442
gb:MT039888|Organism:Severe      GCTTTGGACATATCAGCATCTATAGTAGCTGGTGGTATTGTAGCTATCGTAGTAACATGC   9442
gb:LC522972|Organism:Severe      GCTTTGGACATATCAGCATCTATAGTAGCTGGTGGTATTGTAGCTATCGTAGTAACATGC   9439
gb:MT027063|Organism:Severe      GCTTTGGACATATCAGCATCTATAGTAGCTGGTGGTATTGTAGCTATCGTAGTAACATGC   9442
gb:MT027062|Organism:Severe      GCTTTGGACATATCAGCATCTATAGTAGCTGGTGGTATTGTAGCTATCGTAGTAACATGC   9442
gb:MT019529|Organism:Severe      GCTTTGGACATATCAGCATCTATAGTAGCTGGTGGTATTGTAGCTATCGTAGTAACATGC   9442
gb:MN996529|Organism:Severe      GCTTTGGACATATCAGCATCTATAGTAGCTGGTGGTATTGTAGCTATCGTAGTAACATGC   9430
gb:MN996531|Organism:Severe      GCTTTGGACATATCAGCATCTATAGTAGCTGGTGGTATTGTAGCTATCGTAGTAACATGC   9429
gb:MT066176|Organism:Severe      GCTTTGGACATATCAGCATCTATAGTAGCTGGTGGTATTGTAGCTATCGTAGTAACATGC   9442
gb:MT027064|Organism:Severe      GCTTTGGACATATCAGCATCTATAGTAGCTGGTGGTATTGTAGCTATCGTAGTAACATGC   9442
gb:MN994468|Organism:Severe      GCTTTGGACATATCAGCATCTATAGTAGCTGGTGGTATTGTAGCTATCGTAGTAACATGC   9442
gb:MT072688|Organism:Severe      GCTTTGGACATATCAGCATCTATAGTAGCTGGTGGTATTGTAGCTATCGTAGTAACATGC   9427
gb:MN996527|Organism:Severe      GCTTTGGACATATCAGCATCTATAGTAGCTGGTGGTATTGTAGCTATCGTAGTAACATGC   9409
gb:MT093631|Organism:Severe      GCTTTGGACATATCAGCATCTATAGTAGCTGGTGGTATTGTAGCTATCGTAGTAACATGC   9480
gb:MT106053|Organism:Severe      GCTTTGGACATATCAGCATCTATAGTAGCTGGTGGTATTGTAGCTATCGTAGTAACATGC   9442
gb:MT019533|Organism:Severe      GCTTTGGACATATCAGCATCTATAGTAGCTGGTGGTATTGTAGCTATCGTAGTAACATGC   9442
gb:MT019531|Organism:Severe      GCTTTGGACATATCAGCATCTATAGTAGCTGGTGGTATTGTAGCTATCGTAGTAACATGC   9442
gb:MN996528|Organism:Severe      GCTTTGGACATATCAGCATCTATAGTAGCTGGTGGTATTGTAGCTATCGTAGTAACATGC   9442
gb:MN996530|Organism:Severe      GCTTTGGACATATCAGCATCTATAGTAGCTGGTGGTATTGTAGCTATCGTAGTAACATGC   9428
gb:MN908947|Organism:Severe      GCTTTGGACATATCAGCATCTATAGTAGCTGGTGGTATTGTAGCTATCGTAGTAACATGC   9442
gb:MT019532|Organism:Severe      GCTTTGGACATATCAGCATCTATAGTAGCTGGTGGTATTGTAGCTATCGTAGTAACATGC   9442
                                 ************************************************************

gb:MT020781|Organism:Severe      CTTGCCTACTATTTTATGAGGTTTAGAAGAGCTTTTGGTGAATACAGTCATGTAGTTGCC   9490
gb:MT007544|Organism:Severe      CTTGCCTACTATTTTATGAGGTTTAGAAGAGCTTTTGGTGAATACAGTCATGTAGTTGCC   9502
gb:MN994467|Organism:Severe      CTTGCCTACTATTTTATGAGGTTTAGAAGAGCTTTTGGTGAATACAGTCATGTAGTTGCC   9502
gb:MT044257|Organism:Severe      CTTGCCTACTATTTTATGAGGTTTAGAAGAGCTTTTGGTGAATACAGTCATGTAGTTGCC   9502
gb:MT106054|Organism:Severe      CTTGCCTACTATTTTATGAGGTTTAGAAGAGCTTTTGGTGAATACAGTCATGTAGTTGCC   9502
gb:MT049951|Organism:Severe      CTTGCCTACTATTTTATGAGGTTTAGAAGAGCTTTTGGTGAATACAGTCATGTAGTTGCC   9502
gb:MN975262|Organism:Severe      CTTGCCTACTATTTTATGAGGTTTAGAAGAGCTTTTGGTGAATACAGTCATGTAGTTGCC   9502
gb:MT106052|Organism:Severe      CTTGCCTACTATTTTATGAGGTTTAGAAGAGCTTTTGGTGAATACAGTCATGTAGTTGCC   9502
```

```
gb:LC522975|Organism:Severe      CTTGCCTACTATTTTATGAGGTTTAGAAGAGCTTTTGGTGAATACAGTCATGTAGTTGCC   9499
gb:LC522973|Organism:Severe      CTTGCCTACTATTTTATGAGGTTTAGAAGAGCTTTTGGTGAATACAGTCATGTAGTTGCC   9499
gb:LC522974|Organism:Severe      CTTGCCTACTATTTTATGAGGTTTAGAAGAGCTTTTGGTGAATACAGTCATGTAGTTGCC   9499
gb:MN985325|Organism:Severe      CTTGCCTACTATTTTATGAGGTTTAGAAGAGCTTTTGGTGAATACAGTCATGTAGTTGCC   9502
gb:MT020881|Organism:Severe      CTTGCCTACTATTTTATGAGGTTTAGAAGAGCTTTTGGTGAATACAGTCATGTAGTTGCC   9502
gb:MT020880|Organism:Severe      CTTGCCTACTATTTTATGAGGTTTAGAAGAGCTTTTGGTGAATACAGTCATGTAGTTGCC   9502
gb:MT066175|Organism:Severe      CTTGCCTACTATTTTATGAGGTTTAGAAGAGCTTTTGGTGAATACAGTCATGTAGTTGCC   9502
gb:MN997409|Organism:Severe      CTTGCCTACTATTTTATGAGGTTTAGAAGAGCTTTTGGTGAATACAGTCATGTAGTTGCC   9502
gb:MN938384|Organism:Severe      CTTGCCTACTATTTTATGAGGTTTAGAAGAGCTTTTGGTGAATACAGTCATGTAGTTGCC   9470
gb:MT044258|Organism:Severe      CTTGCCTACTATTTTATGAGGTTTAGAAGAGCTTTTGGTGAATACAGTCATGTAGTTGCC   9478
gb:MT039890|Organism:Severe      CTTGCCTACTATTTTATGAGGTTTAGAAGAGCTTTTGGTGAATACAGTCATGTAGTTGCC   9502
gb:MN988713|Organism:Severe      CTTGCCTACTATTTTATGAGGTTTAGAAGAGCTTTTGGTGAATACAGTCATGTAGTTGCC   9502
gb:LC521925|Organism:Severe      CTTGCCTACTATTTTATGAGGTTTAGAAGAGCTTTTGGTGAATACAGTCATGTAGTTGCC   9475
gb:MT093571|Organism:Severe      CTTGCCTACTATTTTATGAGGTTTAGAAGAGCTTTTGGTGAATACAGTCATGTAGTTGCC   9502
gb:MT039887|Organism:Severe      CTTGCCTACTATTTTATGAGGTTTAGAAGAGCTTTTGGTGAATACAGTCATGTAGTTGCC   9502
gb:MT019530|Organism:Severe      CTTGCCTACTATTTTATGAGGTTTAGAAGAGCTTTTGGTGAATACAGTCATGTAGTTGCC   9502
gb:MT039888|Organism:Severe      CTTGCCTACTATTTTATGAGGTTTAGAAGAGCTTTTGGTGAATACAGTCATGTAGTTGCC   9502
gb:LC522972|Organism:Severe      CTTGCCTACTATTTTATGAGGTTTAGAAGAGCTTTTGGTGAATACAGTCATGTAGTTGCC   9499
gb:MT027063|Organism:Severe      CTTGCCTACTATTTTATGAGGTTTAGAAGAGCTTTTGGTGAATACAGTCATGTAGTTGCC   9502
gb:MT027062|Organism:Severe      CTTGCCTACTATTTTATGAGGTTTAGAAGAGCTTTTGGTGAATACAGTCATGTAGTTGCC   9502
gb:MT019529|Organism:Severe      CTTGCCTACTATTTTATGAGGTTTAGAAGAGCTTTTGGTGAATACAGTCATGTAGTTGCC   9502
gb:MN996529|Organism:Severe      CTTGCCTACTATTTTATGAGGTTTAGAAGAGCTTTTGGTGAATACAGTCATGTAGTTGCC   9490
gb:MN996531|Organism:Severe      CTTGCCTACTATTTTATGAGGTTTAGAAGAGCTTTTGGTGAATACAGTCATGTAGTTGCC   9489
gb:MT066176|Organism:Severe      CTTGCCTACTATTTTATGAGGTTTAGAAGAGCTTTTGGTGAATACAGTTATGTAGTTGCC   9502
gb:MT027064|Organism:Severe      CTTGCCTACTATTTTATGAGGTTTAGAAGAGCTTTTGGTGAATACAGTCATGTAGTTGCC   9502
gb:MN994468|Organism:Severe      CTTGCCTACTATTTTATGAGGTTTAGAAGAGCTTTTGGTGAATACAGTCATGTAGTTGCC   9502
gb:MT072688|Organism:Severe      CTTGCCTACTATTTTATGAGGTTTAGAAGAGCTTTTGGTGAATACAGTCATGTAGTTGCC   9487
gb:MN996527|Organism:Severe      CTTGCCTACTATTTTATGAGGTTTAGAAGAGCTTTTGGTGAATACAGTCATGTAGTTGCC   9469
gb:MT093631|Organism:Severe      CTTGCCTACTATTTTATGAGGTTTAGAAGAGCTTTTGGTGAATACAGTCATGTAGTTGCC   9540
gb:MT106053|Organism:Severe      CTTGCCTACTATTTTATGAGGTTTAGAAGAGCTTTTGGTGAATACAGTCATGTAGTTGCC   9502
gb:MT019533|Organism:Severe      CTTGCCTACTATTTTATGAGGTTTAGAAGAGCTTTTGGTGAATACAGTCATGTAGTTGCC   9502
gb:MT019531|Organism:Severe      CTTGCCTACTATTTTATGAGGTTTAGAAGAGCTTTTGGTGAATACAGTCATGTAGTTGCC   9502
gb:MN996528|Organism:Severe      CTTGCCTACTATTTTATGAGGTTTAGAAGAGCTTTTGGTGAATACAGTCATGTAGTTGCC   9502
gb:MN996530|Organism:Severe      CTTGCCTACTATTTTATGAGGTTTAGAAGAGCTTTTGGTGAATACAGTCATGTAGTTGCC   9488
gb:MN908947|Organism:Severe      CTTGCCTACTATTTTATGAGGTTTAGAAGAGCTTTTGGTGAATACAGTCATGTAGTTGCC   9502
gb:MT019532|Organism:Severe      CTTGCCTACTATTTTATGAGGTTTAGAAGAGCTTTTGGTGAATACAGTCATGTAGTTGCC   9502
                                 ************************************************ ***********

gb:MT020781|Organism:Severe      TTTAATACTTTACTATTCCTTATGTCATTCACTGTACTCTGTTTAACACCAGTTTACTCA   9550
gb:MT007544|Organism:Severe      TTTAATACTTTACTATTCCTTATGTCATTCACTGTACTCTGTTTAACACCAGTTTACTCA   9562
gb:MN994467|Organism:Severe      TTTAATACTTTACTATTCCTTATGTCATTCACTGTACTCTGTTTAACACCAGTTTACTCA   9562
gb:MT044257|Organism:Severe      TTTAATACTTTACTATTCCTTATGTCATTCACTGTACTCTGTTTAACACCAGTTTACTCA   9562
gb:MT106054|Organism:Severe      TTTAATACTTTACTATTCCTTATGTCATTCACTGTACTCTGTTTAACACCAGTTTACTCA   9562
gb:MT049951|Organism:Severe      TTTAATACTTTACTATTCCTTATGTCATTCACTGTACTCTGTTTAACACCAGTTTACTCA   9562
gb:MN975262|Organism:Severe      TTTAATACTTTACTATTCCTTATGTCATTCACTGTACTCTGTTTAACACCAGTTTACTTA   9562
gb:MT106052|Organism:Severe      TTTAATACTTTACTATTCCTTATGTCATTCACTGTACTCTGTTTAACACCAGTTTACTCA   9562
gb:LC522975|Organism:Severe      TTTAATACTTTACTATTCCTTATGTCATTCACTGTACTCTGTTTAACACCAGTTTACTCA   9559
gb:LC522973|Organism:Severe      TTTAATACTTTACTATTCCTTATGTCATTCACTGTACTCTGTTTAACACCAGTTTACTCA   9559
gb:LC522974|Organism:Severe      TTTAATACTTTACTATTCCTTATGTCATTCACTGTACTCTGTTTAACACCAGTTTACTCA   9559
gb:MN985325|Organism:Severe      TTTAATACTTTACTATTCCTTATGTCATTCACTGTACTCTGTTTAACACCAGTTTACTCA   9562
gb:MT020881|Organism:Severe      TTTAATACTTTACTATTCCTTATGTCATTCACTGTACTCTGTTTAACACCAGTTTACTCA   9562
gb:MT020880|Organism:Severe      TTTAATACTTTACTATTCCTTATGTCATTCACTGTACTCTGTTTAACACCAGTTTACTCA   9562
gb:MT066175|Organism:Severe      TTTAATACTTTACTATTCCTTATGTCATTCACTGTACTCTGTTTAACACCAGTTTACTCA   9562
gb:MN997409|Organism:Severe      TTTAATACTTTACTATTCCTTATGTCATTCACTGTACTCTGTTTAACACCAGTTTACTCA   9562
gb:MN938384|Organism:Severe      TTTAATACTTTACTATTCCTTATGTCATTCACTGTACTCTGTTTAACACCAGTTTACTCA   9530
gb:MT044258|Organism:Severe      TTTAATACTTTACTATTCCTTATGTCATTCACTGTACTCTGTTTAACACCAGTTTACTCA   9538
gb:MT039890|Organism:Severe      TTTAATACTTTACTATTCCTTATGTCATTCACTGTACTCTGTTTAACACCAGTTTACTCA   9562
gb:MN988713|Organism:Severe      TTTAATACTTTACTATTCCTTATGTCATTCACTGTACTCTGTTTAACACCAGTTTACTCA   9562
gb:LC521925|Organism:Severe      TTTAATACTTTACTATTCCTTATGTCATTCACTGTACTCTGTTTAACACCAGTTTACTCA   9535
gb:MT093571|Organism:Severe      TTTAATACTTTACTATTCCTTATGTCATTCACTGTACTCTGTTTAACACCAGTTTACTCA   9562
gb:MT039887|Organism:Severe      TTTAATACTTTACTATTCCTTATGTCATTCACTGTACTCTGTTTAACACCAGTTTACTCA   9562
gb:MT019530|Organism:Severe      TTTAATACTTTACTATTCCTTATGTCATTCACTGTACTCTGTTTAACACCAGTTTACTCA   9562
gb:MT039888|Organism:Severe      TTTAATACTTTACTATTCCTTATGTCATTCACTGTACTCTGTTTAACACCAGTTTACTCA   9562
gb:LC522972|Organism:Severe      TTTAATACTTTACTATTCCTTATGTCATTCACTGTACTCTGTTTAACACCAGTTTACTCA   9559
gb:MT027063|Organism:Severe      TTTAATACTTTACTATTCCTTATGTCATTCACTGTACTCTGTTTAACACCAGTTTACTCA   9562
```

```
gb:MT027062|Organism:Severe      TTTAATACTTTACTATTCCTTATGTCATTCACTGTACTCTGTTTAACACCAGTTTACTCA	9562
gb:MT019529|Organism:Severe      TTTAATACTTTACTATTCCTTATGTCATTCACTGTACTCTGTTTAACACCAGTTTACTCA	9562
gb:MN996529|Organism:Severe      TTTAATACTTTACTATTCCTTATGTCATTCACTGTACTCTGTTTAACACCAGTTTACTCA	9550
gb:MN996531|Organism:Severe      TTTAATACTTTACTATTCCTTATGTCATTCATTGTACTCTGTTTAACACCAGTTTACTCA	9549
gb:MT066176|Organism:Severe      TTTAATACTTTACTATTCCTTATGTCATTCACTGTACTCTGTTTAACACCAGTTTACTCA	9562
gb:MT027064|Organism:Severe      TTTAATACTTTACTATTCCTTATGTCATTCACTGTACTCTGTTTAACACCAGTTTACTCA	9562
gb:MN994468|Organism:Severe      TTTAATACTTTACTATTCCTTATGTCATTCACTGTACTCTGTTTAACACCAGTTTACTCA	9562
gb:MT072688|Organism:Severe      TTTAATACTTTACTATTCCTTATGTCATTCACTGTACTCTGTTTAACACCAGTTTACTCA	9547
gb:MN996527|Organism:Severe      TTTAATACTTTACTATTCCTTATGTCATTCACTGTACTCTGTTTAACACCAGTTTACTCA	9529
gb:MT093631|Organism:Severe      TTTAATACTTTACTATTCCTTATGTCATTCACTGTACTCTGTTTAACACCAGTTTACTCA	9600
gb:MT106053|Organism:Severe      TTTAATACTTTACTATTCCTTATGTCATTCACTGTACTCTGTTTAACACCAGTTTACTCA	9562
gb:MT019533|Organism:Severe      TTTAATACTTTACTATTCCTTATGTCATTCACTGTACTCTGTTTAACACCAGTTTACTCA	9562
gb:MT019531|Organism:Severe      TTTAATACTTTACTATTCCTTATGTCATTCACTGTACTCTGTTTAACACCAGTTTACTCA	9562
gb:MN996528|Organism:Severe      TTTAATACTTTACTATTCCTTATGTCATTCACTGTACTCTGTTTAACACCAGTTTACTCA	9562
gb:MN996530|Organism:Severe      TTTAATACTTTACTATTCCTTATGTCATTCACTGTACTCTGTTTAACACCAGTTTACTCA	9548
gb:MN908947|Organism:Severe      TTTAATACTTTACTATTCCTTATGTCATTCACTGTACTCTGTTTAACACCAGTTTACTCA	9562
gb:MT019532|Organism:Severe      TTTAATACTTTACTATTCCTTATGTCATTCACTGTACTCTGTTTAACACCAGTTTACTCA	9562
                                 ******************************* ************************** *

gb:MT020781|Organism:Severe      TTCTTACCTGGTGTTTATTCTGTTATTTACTTGTACTTGACATTTTATCTTACTAATGAT	9610
gb:MT007544|Organism:Severe      TTCTTACCTGGTGTTTATTCTGTTATTTACTTGTACTTGACATTTTATCTTACTAATGAT	9622
gb:MN994467|Organism:Severe      TTCTTACCTGGTGTTTATTCTGTTATTTACTTGTACTTGACATTTTATCTTACTAATGAT	9622
gb:MT044257|Organism:Severe      TTCTTACCTGGTGTTTATTCTGTTATTTACTTGTACTTGACATTTTATCTTACTAATGAT	9622
gb:MT106054|Organism:Severe      TTCTTACCTGGTGTTTATTCTGTTATTTACTTGTACTTGACATTTTATCTTACTAATGAT	9622
gb:MT049951|Organism:Severe      TTCTTACCTGGTGTTTATTCTGTTATTTACTTGTACTTGACATTTTATCTTACTAATGAT	9622
gb:MN975262|Organism:Severe      TTCTTACCTGGTGTTTATTCTGTTATTTACTTGTACTTGACATTTTATCTTACTAATGAT	9622
gb:MT106052|Organism:Severe      TTCTTACCTGGTGTTTATTCTGTTATTTACTTGTACTTGACATTTTATCTTACTAATGAT	9622
gb:LC522975|Organism:Severe      TTCTTACCTGGTGTTTATTCTGTTATTTACTTGTACTTGACATTTTATCTTACTAATGAT	9619
gb:LC522973|Organism:Severe      TTCTTACCTGGTGTTTATTCTGTTATTTACTTGTACTTGACATTTTATCTTACTAATGAT	9619
gb:LC522974|Organism:Severe      TTCTTACCTGGTGTTTATTCTGTTATTTACTTGTACTTGACATTTTATCTTACTAATGAT	9619
gb:MN985325|Organism:Severe      TTCTTACCTGGTGTTTATTCTGTTATTTACTTGTACTTGACATTTTATCTTACTAATGAT	9622
gb:MT020881|Organism:Severe      TTCTTACCTGGTGTTTATTCTGTTATTTACTTGTACTTGACATTTTATCTTACTAATGAT	9622
gb:MT020880|Organism:Severe      TTCTTACCTGGTGTTTATTCTGTTATTTACTTGTACTTGACATTTTATCTTACTAATGAT	9622
gb:MT066175|Organism:Severe      TTCTTACCTGGTGTTTATTCTGTTATTTACTTGTACTTGACATTTTATCTTACTAATGAT	9622
gb:MN997409|Organism:Severe      TTCTTACCTGGTGTTTATTCTGTTATTTACTTGTACTTGACATTTTATCTTACTAATGAT	9622
gb:MN938384|Organism:Severe      TTCTTACCTGGTGTTTATTCTGTTATTTACTTGTACTTGACATTTTATCTTACTAATGAT	9590
gb:MT044258|Organism:Severe      TTCTTACCTGGTGTTTATTCTGTTATTTACTTGTACTTGACATTTTATCTTACTAATGAT	9598
gb:MT039890|Organism:Severe      TTCTTACCTGGTGTTTATTCTGTTATTTACTTGTACTTGACATTTTATCTTACTAATGAT	9622
gb:MN988713|Organism:Severe      TTCTTACCTGGTGTTTATTCTGTTATTTACTTGTACTTGACATTTTATCTTACTAATGAT	9622
gb:LC521925|Organism:Severe      TTCTTACCTGGTGTTTATTCTGTTATTTACTTGTACTTGACATTTTATCTTACTAATGAT	9595
gb:MT093571|Organism:Severe      TTCTTACCTGGTGTTTATTCTGTTATTTACTTGTACTTGACATTTTATCTTACTAATGAT	9622
gb:MT039887|Organism:Severe      TTCTTACCTGGTGTTTATTCTGTTATTTACTTGTACTTGACATTTTATCTTACTAATGAT	9622
gb:MT019530|Organism:Severe      TTCTTACCTGGTGTTTATTCTGTTATTTACTTGTACTTGACATTTTATCTTACTAATGAT	9622
gb:MT039888|Organism:Severe      TTCTTACCTGGTGTTTATTCTGTTATTTACTTGTACTTGACATTTTATCTTACTAATGAT	9622
gb:LC522972|Organism:Severe      TTCTTACCTGGTGTTTATTCTGTTATTTACTTGTACTTGACATTTTATCTTACTAATGAT	9619
gb:MT027063|Organism:Severe      TTCTTACCTGGTGTTTATTCTGTTATTTACTTGTACTTGACATTTTATCTTACTAATGAT	9622
gb:MT027062|Organism:Severe      TTCTTACCTGGTGTTTATTCTGTTATTTACTTGTACTTGACATTTTATCTTACTAATGAT	9622
gb:MT019529|Organism:Severe      TTCTTACCTGGTGTTTATTCTGTTATTTACTTGTACTTGACATTTTATCTTACTAATGAT	9622
gb:MN996529|Organism:Severe      TTCTTACCTGGTGTTTATTCTGTTATTTACTTGTACTTGACATTTTATCTTACTAATGAT	9610
gb:MN996531|Organism:Severe      TTCTTACCTGGTGTTTATTCTGTTATTTACTTGTACTTGACATTTTATCTTACTAATGAT	9609
gb:MT066176|Organism:Severe      TTCTTACCTGGTGTTTATTCTGTTATTTACTTGTACTTGACATTTTATCTTACTAATGAT	9622
gb:MT027064|Organism:Severe      TTCTTACCTGGTGTTTATTCTGTTATTTACTTGTACTTGACATTTTATCTTACTAATGAT	9622
gb:MN994468|Organism:Severe      TTCTTACCTGGTGTTTATTCTGTTATTTACTTGTACTTGACATTTTATCTTACTAATGAT	9622
gb:MT072688|Organism:Severe      TTCTTACCTGGTGTTTATTCTGTTATTTACTTGTACTTGACATTTTATCTTACTAATGAT	9607
gb:MN996527|Organism:Severe      TTCTTACCTGGTGTTTATTCTGTTATTTACTTGTACTTGACATTTTATCTTACTAATGAT	9589
gb:MT093631|Organism:Severe      TTCTTACCTGGTGTTTATTCTGTTATTTACTTGTACTTGACATTTTATCTTACTAATGAT	9660
gb:MT106053|Organism:Severe      TTCTTACCTGGTGTTTATTCTGTTATTTACTTGTACTTGACATTTTATCTTACTAATGAT	9622
gb:MT019533|Organism:Severe      TTCTTACCTGGTGTTTATTCTGTTATTTACTTGTACTTGACATTTTATCTTACTAATGAT	9622
gb:MT019531|Organism:Severe      TTCTTACCTGGTGTTTATTCTGTTATTTACTTGTACTTGACATTTTATCTTACTAATGAT	9622
gb:MN996528|Organism:Severe      TTCTTACCTGGTGTTTATTCTGTTATTTACTTGTACTTGACATTTTATCTTACTAATGAT	9622
gb:MN996530|Organism:Severe      TTCTTACCTGGTGTTTATTCTGTTATTTACTTGTACTTGACATTTTATCTTACTAATGAT	9608
gb:MN908947|Organism:Severe      TTCTTACCTGGTGTTTATTCTGTTATTTACTTGTACTTGACATTTTATCTTACTAATGAT	9622
gb:MT019532|Organism:Severe      TTCTTACCTGGTGTTTATTCTGTTATTTACTTGTACTTGACATTTTATCTTACTAATGAT	9622
                                 ************************************************************
```

```
gb:MT020781|Organism:Severe      GTTTCTTTTTTAGCACATATTCAGTGGATGGTTATGTTCACACCTTTAGTACCTTTCTGG   9670
gb:MT007544|Organism:Severe      GTTTCTTTTTTAGCACATATTCAGTGGATGGTTATGTTCACACCTTTAGTACCTTTCTGG   9682
gb:MN994467|Organism:Severe      GTTTCTTTTTTAGCACATATTCAGTGGATGGTTATGTTCACACCTTTAGTACCTTTCTGG   9682
gb:MT044257|Organism:Severe      GTTTCTTTTTTAGCACATATTCAGTGGATGGTTATGTTCACACCTTTAGTACCTTTCTGG   9682
gb:MT106054|Organism:Severe      GTTTCTTTTTTAGCACATATTCAGTGGATGGTTATGTTCACACCTTTAGTACCTTTCTGG   9682
gb:MT049951|Organism:Severe      GTTTCTTTTTTAGCACATATTCAGTGGATGGTTATGTTCACACCTTTAGTACCTTTCTGG   9682
gb:MN975262|Organism:Severe      GTTTCTTTTTTAGCACATATTCAGTGGATGGTTATGTTCACACCTTTAGTACCTTTCTGG   9682
gb:MT106052|Organism:Severe      GTTTCTTTTTTAGCACATATTCAGTGGATGGTTATGTTCACACCTTTAGTACCTTTCTGG   9682
gb:LC522975|Organism:Severe      GTTTCTTTTTTAGCACATATTCAGTGGATGGTTATGTTCACACCTTTAGTACCTTTCTGG   9679
gb:LC522973|Organism:Severe      GTTTCTTTTTTAGCACATATTCAGTGGATGGTTATGTTCACACCTTTAGTACCTTTCTGG   9679
gb:LC522974|Organism:Severe      GTTTCTTTTTTAGCACATATTCAGTGGATGGTTATGTTCACACCTTTAGTACCTTTCTGG   9679
gb:MN985325|Organism:Severe      GTTTCTTTTTTAGCACATATTCAGTGGATGGTTATGTTCACACCTTTAGTACCTTTCTGG   9682
gb:MT020881|Organism:Severe      GTTTCTTTTTTAGCACATATTCAGTGGATGGTTATGTTCACACCTTTAGTACCTTTCTGG   9682
gb:MT020880|Organism:Severe      GTTTCTTTTTTAGCACATATTCAGTGGATGGTTATGTTCACACCTTTAGTACCTTTCTGG   9682
gb:MT066175|Organism:Severe      GTTTCTTTTTTAGCACATATTCAGTGGATGGTTATGTTCACACCTTTAGTACCTTTCTGG   9682
gb:MN997409|Organism:Severe      GTTTCTTTTTTAGCACATATTCAGTGGATGGTTATGTTCACACCTTTAGTACCTTTCTGG   9682
gb:MN938384|Organism:Severe      GTTTCTTTTTTAGCACATATTCAGTGGATGGTTATGTTCACACCTTTAGTACCTTTCTGG   9650
gb:MT044258|Organism:Severe      GTTTCTTTTTTAGCACATATTCAGTGGATGGTTATGTTCACACCTTTAGTACCTTTCTGG   9658
gb:MT039890|Organism:Severe      GTTTCTTTTTTAGCACATATTCAGTGGATGGTTATGTTCACACCTTTAGTACCTTTCTGG   9682
gb:MN988713|Organism:Severe      GTTTCTTTTTTAGCACATATTCAGTGGATGGTTATGTTCACACCTTTAGTACCTTTCTGG   9682
gb:LC521925|Organism:Severe      GTTTCTTTTTTAGCACATATTCAGTGGATGGTTATGTTCACACCTTTAGTACCTTTCTGG   9655
gb:MT093571|Organism:Severe      GTTTCTTTTTTAGCACATATTCAGTGGATGGTTATGTTCACACCTTTAGTACCTTTCTGG   9682
gb:MT039887|Organism:Severe      GTTTCTTTTTTAGCACATATTCAGTGGATGGTTATGTTCACACCTTTAGTACCTTTCTGG   9682
gb:MT019530|Organism:Severe      GTTTCTTTTTTAGCACATATTCAGTGGATGGTTATGTTCACACCTTTAGTACCTTTCTGG   9682
gb:MT039888|Organism:Severe      GTTTCTTTTTTAGCACATATTCAGTGGATGGTTATGTTCACACCTTTAGTACCTTTCTGG   9682
gb:LC522972|Organism:Severe      GTTTCTTTTTTAGCACATATTCAGTGGATGGTTATGTTCACACCTTTAGTACCTTTCTGG   9679
gb:MT027063|Organism:Severe      GTTTCTTTTTTAGCACATATTCAGTGGATGGTTATGTTCACACCTTTAGTACCTTTCTGG   9682
gb:MT027062|Organism:Severe      GTTTCTTTTTTAGCACATATTCAGTGGATGGTTATGTTCACACCTTTAGTACCTTTCTGG   9682
gb:MT019529|Organism:Severe      GTTTCTTTTTTAGCACATATTCAGTGGATGGTTATGTTCACACCTTTAGTACCTTTCTGG   9682
gb:MN996529|Organism:Severe      GTTTCTTTTTTAGCACATATTCAGTGGATGGTTATGTTCACACCTTTAGTACCTTTCTGG   9670
gb:MN996531|Organism:Severe      GTTTCTTTTTTAGCACATATTCAGTGGATGGTTATGTTCACACCTTTAGTACCTTTCTGG   9669
gb:MT066176|Organism:Severe      GTTTCTTTTTTAGCACATATTCAGTGGATGGTTATGTTCACACCTTTAGTACCTTTCTGG   9682
gb:MT027064|Organism:Severe      GTTTCTTTTTTAGCACATATTCAGTGGATGGTTATGTTCACACCTTTAGTACCTTTCTGG   9682
gb:MN994468|Organism:Severe      GTTTCTTTTTTAGCACATATTCAGTGGATGGTTATGTTCACACCTTTAGTACCTTTCTGG   9682
gb:MT072688|Organism:Severe      GTTTCTTTTTTAGCACATATTCAGTGGATGGTTATGTTCACACCTTTAGTACCTTTCTGG   9667
gb:MN996527|Organism:Severe      GTTTCTTTTTTAGCACATATTCAGTGGATGGTTATGTTCACACCTTTAGTACCTTTCTGG   9649
gb:MT093631|Organism:Severe      GTTTCTTTTTTAGCACATATTCAGTGGATGGTTATGTTCACACCTTTAGTACCTTTCTGG   9720
gb:MT106053|Organism:Severe      GTTTCTTTTTTAGCACATATTCAGTGGATGGTTATGTTCACACCTTTAGTACCTTTCTGG   9682
gb:MT019533|Organism:Severe      GTTTCTTTTTTAGCACATATTCAGTGGATGGTTATGTTCACACCTTTAGTACCTTTCTGG   9682
gb:MT019531|Organism:Severe      GTTTCTTTTTTAGCACATATTCAGTGGATGGTTATGTTCACACCTTTAGTACCTTTCTGG   9682
gb:MN996528|Organism:Severe      GTTTCTTTTTTAGCACATATTCAGTGGATGGTTATGTTCACACCTTTAGTACCTTTCTGG   9682
gb:MN996530|Organism:Severe      GTTTCTTTTTTAGCACATATTCAGTGGATGGTTATGTTCACACCTTTAGTACCTTTCTGG   9668
gb:MN908947|Organism:Severe      GTTTCTTTTTTAGCACATATTCAGTGGATGGTTATGTTCACACCTTTAGTACCTTTCTGG   9682
gb:MT019532|Organism:Severe      GTTTCTTTTTTAGCACATATTCAGTGGATGGTTATGTTCACACCTTTAGTACCTTTCTGG   9682
                                 ************************************************************

gb:MT020781|Organism:Severe      ATAACAATTGCTTATATCATTTGTATTTCCACAAAGCATTTCTATTGGTTCTTTAGTAAT   9730
gb:MT007544|Organism:Severe      ATAACAATTGCTTATATCATTTGTATTTCCACAAAGCATTTCTATTGGTTCTTTAGTAAT   9742
gb:MN994467|Organism:Severe      ATAACAATTGCTTATATCATTTGTATTTCCACAAAGCATTTCTATTGGTTCTTTAGTAAT   9742
gb:MT044257|Organism:Severe      ATAACAATTGCTTATATCATTTGTATTTCCACAAAGCATTTCTATTGGTTCTTTAGTAAT   9742
gb:MT106054|Organism:Severe      ATAACAATTGCTTATATCATTTGTATTTCCACAAAGCATTTCTATTGGTTCTTTAGTAAT   9742
gb:MT049951|Organism:Severe      ATAACAATTGCTTATATCATTTGTATTTCCACAAAGCATTTCTATTGGTTCTTTAGTAAT   9742
gb:MN975262|Organism:Severe      ATAACAATTGCTTATATCATTTGTATTTCCACAAAGCATTTCTATTGGTTCTTTAGTAAT   9742
gb:MT106052|Organism:Severe      ATAACAATTGCTTATATCATTTGTATTTCCACAAAGCATTTCTATTGGTTCTTTAGTAAT   9742
gb:LC522975|Organism:Severe      ATAACAATTGCTTATATCATTTGTATTTCCACAAAGCATTTCTATTGGTTCTTTAGTAAT   9739
gb:LC522973|Organism:Severe      ATAACAATTGCTTATATCATTTGTATTTCCACAAAGCATTTCTATTGGTTCTTTAGTAAT   9739
gb:LC522974|Organism:Severe      ATAACAATTGCTTATATCATTTGTATTTCCACAAAGCATTTCTATTGGTTCTTTAGTAAT   9739
gb:MN985325|Organism:Severe      ATAACAATTGCTTATATCATTTGTATTTCCACAAAGCATTTCTATTGGTTCTTTAGTAAT   9742
gb:MT020881|Organism:Severe      ATAACAATTGCTTATATCATTTGTATTTCCACAAAGCATTTCTATTGGTTCTTTAGTAAT   9742
gb:MT020880|Organism:Severe      ATAACAATTGCTTATATCATTTGTATTTCCACAAAGCATTTCTATTGGTTCTTTAGTAAT   9742
gb:MT066175|Organism:Severe      ATAACAATTGCTTATATCATTTGTATTTCCACAAAGCATTTCTATTGGTTCTTTAGTAAT   9742
gb:MN997409|Organism:Severe      ATAACAATTGCTTATATCATTTGTATTTCCACAAAGCATTTCTATTGGTTCTTTAGTAAT   9742
gb:MN938384|Organism:Severe      ATAACAATTGCTTATATCATTTGTATTTCCACAAAGCATTTCTATTGGTTCTTTAGTAAT   9710
gb:MT044258|Organism:Severe      ATAACAATTGCTTATATCATTTGTATTTCCACAAAGCATTTCTATTGGTTCTTTAGTAAT   9718
gb:MT039890|Organism:Severe      ATAACAATTGCTTATATCATTTGTATTTCCACAAAGCATTTCTATTGGTTCTTTAGTAAT   9742
```

```
gb:MN988713|Organism:Severe      ATAACAATTGCTTATATCATTTGTATTTCCACAAAGCATTTCTATTGGTTCTTTAGTAAT   9742
gb:LC521925|Organism:Severe      ATAACAATTGCTTATATCATTTGTATTTCCACAAAGCATTTCTATTGGTTCTTTAGTAAT   9715
gb:MT093571|Organism:Severe      ATAACAATTGCTTATATCATTTGTATTTCCACAAAGCATTTCTATTGGTTCTTTAGTAAT   9742
gb:MT039887|Organism:Severe      ATAACAATTGCTTATATCATTTGTATTTCCACAAAGCATTTCTATTGGTTCTTTAGTAAT   9742
gb:MT019530|Organism:Severe      ATAACAATTGCTTATATCATTTGTATTTCCACAAAGCATTTCTATTGGTTCTTTAGTAAT   9742
gb:MT039888|Organism:Severe      ATAACAATTGCTTATATCATTTGTATTTCCACAAAGCATTTCTATTGGTTCTTTAGTAAT   9742
gb:LC522972|Organism:Severe      ATAACAATTGCTTATATCATTTGTATTTCCACAAAGCATTTCTATTGGTTCTTTAGTAAT   9739
gb:MT027063|Organism:Severe      ATAACAATTGCTTATATCATTTGTATTTCCACAAAGCATTTCTATTGGTTCTTTAGTAAT   9742
gb:MT027062|Organism:Severe      ATAACAATTGCTTATATCATTTGTATTTCCACAAAGCATTTCTATTGGTTCTTTAGTAAT   9742
gb:MT019529|Organism:Severe      ATAACAATTGCTTATATCATTTGTATTTCCACAAAGCATTTCTATTGGTTCTTTAGTAAT   9742
gb:MN996529|Organism:Severe      ATAACAATTGCTTATATCATTTGTATTTCCACAAAGCATTTCTATTGGTTCTTTAGTAAT   9730
gb:MN996531|Organism:Severe      ATAACAATTGCTTATATCATTTGTATTTCCACAAAGCATTTCTATTGGTTCTTTAGTAAT   9729
gb:MT066176|Organism:Severe      ATAACAATTGCTTATATCATTTGTATTTCCACAAAGCATTTCTATTGGTTCTTTAGTAAT   9742
gb:MT027064|Organism:Severe      ATAACAATTGCTTATATCATTTGTATTTCCACAAAGCATTTCTATTGGTTCTTTAGTAAT   9742
gb:MN994468|Organism:Severe      ATAACAATTGCTTATATCATTTGTATTTCCACAAAGCATTTCTATTGGTTCTTTAGTAAT   9742
gb:MT072688|Organism:Severe      ATAACAATTGCTTATATCATTTGTATTTCCACAAAGCATTTCTATTGGTTCTTTAGTAAT   9727
gb:MN996527|Organism:Severe      ATAACAATTGCTTATATCATTTGTATTTCCACAAAGCATTTCTATTGGTTCTTTAGTAAT   9709
gb:MT093631|Organism:Severe      ATAACAATTGCTTATATCATTTGTATTTCCACAAAGCATTTCTATTGGTTCTTTAGTAAT   9780
gb:MT106053|Organism:Severe      ATAACAATTGCTTATATCATTTGTATTTCCACAAAGCATTTCTATTGGTTCTTTAGTAAT   9742
gb:MT019533|Organism:Severe      ATAACAATTGCTTATATCATTTGTATTTCCACAAAGCATTTCTATTGGTTCTTTAGTAAT   9742
gb:MT019531|Organism:Severe      ATAACAATTGCTTATATCATTTGTATTTCCACAAAGCATTTCTATTGGTTCTTTAGTAAT   9742
gb:MN996528|Organism:Severe      ATAACAATTGCTTATATCATTTGTATTTCCACAAAGCATTTCTATTGGTTCTTTAGTAAT   9742
gb:MN996530|Organism:Severe      ATAACAATTGCTTATATCATTTGTATTTCCACAAAGCATTTCTATTGGTTCTTTAGTAAT   9728
gb:MN908947|Organism:Severe      ATAACAATTGCTTATATCATTTGTATTTCCACAAAGCATTTCTATTGGTTCTTTAGTAAT   9742
gb:MT019532|Organism:Severe      ATAACAATTGCTTATATCATTTGTATTTCCACAAAGCATTTCTATTGGTTCTTTAGTAAT   9742
                                 ************************************************************

gb:MT020781|Organism:Severe      TACCTAAAGAGACGTGTAGTCTTTAATGGTGTTTCCTTTAGTACTTTTGAAGAAGCTGCG   9790
gb:MT007544|Organism:Severe      TACCTAAAGAGACGTGTAGTCTTTAATGGTGTTTCCTTTAGTACTTTTGAAGAAGCTGCG   9802
gb:MN994467|Organism:Severe      TACCTAAAGAGACGTGTAGTCTTTAATGGTGTTTCCTTTAGTACTTTTGAAGAAGCTGCG   9802
gb:MT044257|Organism:Severe      TACCTAAAGAGACGTGTAGTCTTTAATGGTGTTTCCTTTAGTACTTTTGAAGAAGCTGCG   9802
gb:MT106054|Organism:Severe      TACCTAAAGAGACGTGTAGTCTTTAATGGTGTTTCCTTTAGTACTTTTGAAGAAGCTGCG   9802
gb:MT049951|Organism:Severe      TACCTAAAGAGACGTGTAGTCTTTAATGGTGTTTCCTTTAGTACTTTTGAAGAAGCTGCG   9802
gb:MN975262|Organism:Severe      TACCTAAAGAGACGTGTAGTCTTTAATGGTGTTTCCTTTAGTACTTTTGAAGAAGCTGCG   9802
gb:MT106052|Organism:Severe      TACCTAAAGAGACGTGTAGTCTTTAATGGTGTTTCCTTTAGTACTTTTGAAGAAGCTGCG   9802
gb:LC522975|Organism:Severe      TACCTAAAGAGACGTGTAGTCTTTAATGGTGTTTCCTTTAGTACTTTTGAAGAAGCTGCG   9799
gb:LC522973|Organism:Severe      TACCTAAAGAGACGTGTAGTCTTTAATGGTGTTTCCTTTAGTACTTTTGAAGAAGCTGCG   9799
gb:LC522974|Organism:Severe      TACCTAAAGAGACGTGTAGTCTTTAATGGTGTTTCCTTTAGTACTTTTGAAGAAGCTGCG   9799
gb:MN985325|Organism:Severe      TACCTAAAGAGACGTGTAGTCTTTAATGGTGTTTCCTTTAGTACTTTTGAAGAAGCTGCG   9802
gb:MT020881|Organism:Severe      TACCTAAAGAGACGTGTAGTCTTTAATGGTGTTTCCTTTAGTACTTTTGAAGAAGCTGCG   9802
gb:MT020880|Organism:Severe      TACCTAAAGAGACGTGTAGTCTTTAATGGTGTTTCCTTTAGTACTTTTGAAGAAGCTGCG   9802
gb:MT066175|Organism:Severe      TACCTAAAGAGACGTGTAGTCTTTAATGGTGTTTCCTTTAGTACTTTTGAAGAAGCTGCG   9802
gb:MN997409|Organism:Severe      TACCTAAAGAGACGTGTAGTCTTTAATGGTGTTTCCTTTAGTACTTTTGAAGAAGCTGCG   9802
gb:MN938384|Organism:Severe      TACCTAAAGAGACGTGTAGTCTTTAATGGTGTTTCCTTTAGTACTTTTGAAGAAGCTGCG   9770
gb:MT044258|Organism:Severe      TACCTAAAGAGACGTGTAGTCTTTAATGGTGTTTCCTTTAGTACTTTTGAAGAAGCTGCG   9778
gb:MT039890|Organism:Severe      TACCTAAAGAGACGTGTAGTCTTTAATGGTGTTTCCTTTAGTACTTTTGAAGAAGCTGCG   9802
gb:MN988713|Organism:Severe      TACCTAAAGAGACGTGTAGTCTTTAATGGTGTTTCCTTTAGTACTTTTGAAGAAGCTGCG   9802
gb:LC521925|Organism:Severe      TACCTAAAGAGACGTGTAGTCTTTAATGGTGTTTCCTTTAGTACTTTTGAAGAAGCTGCG   9775
gb:MT093571|Organism:Severe      TACCTAAAGAGACGTGTAGTCTTTAATGGTGTTTCCTTTAGTACTTTTGAAGAAGCTGCG   9802
gb:MT039887|Organism:Severe      TACCTAAAGAGACGTGTAGTCTTTAATGGTGTTTCCTTTAGTACTTTTGAAGAAGCTGCG   9802
gb:MT019530|Organism:Severe      TACCTAAAGAGACGTGTAGTCTTTAATGGTGTTTCCTTTAGTACTTTTGAAGAAGCTGCG   9802
gb:MT039888|Organism:Severe      TACCTAAAGAGACGTGTAGTCTTTAATGGTGTTTCCTTTAGTACTTTTGAAGAAGCTGCG   9802
gb:LC522972|Organism:Severe      TACCTAAAGAGACGTGTAGTCTTTAATGGTGTTTCCTTTAGTACTTTTGAAGAAGCTGCG   9799
gb:MT027063|Organism:Severe      TACCTAAAGAGACGTGTAGTCTTTAATGGTGTTTCCTTTAGTACTTTTGAAGAAGCTGCG   9802
gb:MT027062|Organism:Severe      TACCTAAAGAGACGTGTAGTCTTTAATGGTGTTTCCTTTAGTACTTTTGAAGAAGCTGCG   9802
gb:MT019529|Organism:Severe      TACCTAAAGAGACGTGTAGTCTTTAATGGTGTTTCCTTTAGTACTTTTGAAGAAGCTGCG   9802
gb:MN996529|Organism:Severe      TACCTAAAGAGACGTGTAGTCTTTAATGGTGTTTCCTTTAGTACTTTTGAAGAAGCTGCG   9790
gb:MN996531|Organism:Severe      TACCTAAAGAGACGTGTAGTCTTTAATGGTGTTTCCTTTAGTACTTTTGAAGAAGCTGCG   9789
gb:MT066176|Organism:Severe      TACCTAAAGAGACGTGTAGTCTTTAATGGTGTTTCCTTTAGTACTTTTGAAGAAGCTGCG   9802
gb:MT027064|Organism:Severe      TACCTAAAGAGACGTGTAGTCTTTAATGGTGTTTCCTTTAGTACTTTTGAAGAAGCTGCG   9802
gb:MN994468|Organism:Severe      TACCTAAAGAGACGTGTAGTCTTTAATGGTGTTTCCTTTAGTACTTTTGAAGAAGCTGCG   9802
gb:MT072688|Organism:Severe      TACCTAAAGAGACGTGTAGTCTTTAATGGTGTTTCCTTTAGTACTTTTGAAGAAGCTGCG   9787
gb:MN996527|Organism:Severe      TACCTAAAGAGACGTGTAGTCTTTAATGGTGTTTCCTTTAGTACTTTTGAAGAAGCTGCG   9769
gb:MT093631|Organism:Severe      TACCTAAAGAGACGTGTAGTCTTTAATGGTGTTTCCTTTAGTACTTTTGAAGAAGCTGCG   9840
gb:MT106053|Organism:Severe      TACCTAAAGAGACGTGTAGTCTTTAATGGTGTTTCCTTTAGTACTTTTGAAGAAGCTGCG   9802
```

```
gb:MT019533|Organism:Severe      TACCTAAAGAGACGTGTAGTCTTTAATGGTGTTTCCTTTAGTACTTTTGAAGAAGCTGCG   9802
gb:MT019531|Organism:Severe      TACCTAAAGAGACGTGTAGTCTTTAATGGTGTTTCCTTTAGTACTTTTGAAGAAGCTGCG   9802
gb:MN996528|Organism:Severe      TACCTAAAGAGACGTGTAGTCTTTAATGGTGTTTCCTTTAGTACTTTTGAAGAAGCTGCG   9802
gb:MN996530|Organism:Severe      TACCTAAAGAGACGTGTAGTCTTTAATGGTGTTTCCTTTAGTACTTTTGAAGAAGCTGCG   9788
gb:MN908947|Organism:Severe      TACCTAAAGAGACGTGTAGTCTTTAATGGTGTTTCCTTTAGTACTTTTGAAGAAGCTGCG   9802
gb:MT019532|Organism:Severe      TACCTAAAGAGACGTGTAGTCTTTAATGGTGTTTCCTTTAGTACTTTTGAAGAAGCTGCG   9802
                                 ************************************************************

gb:MT020781|Organism:Severe      CTGTGCACCTTTTTGTTAAATAAAGAAATGTATCTAAAGTTGCGTAGTGATGTGCTATTA   9850
gb:MT007544|Organism:Severe      CTGTGCACCTTTTTGTTAAATAAAGAAATGTATCTAAAGTTGCGTAGTGATGTGCTATTA   9862
gb:MN994467|Organism:Severe      CTGTGCACCTTTTTGTTAAATAAAGAAATGTATCTAAAGTTGCGTAGTGATGTGCTATTA   9862
gb:MT044257|Organism:Severe      CTGTGCACCTTTTTGTTAAATAAAGAAATGTATCTAAAGTTGCGTAGTGATGTGCTATTA   9862
gb:MT106054|Organism:Severe      CTGTGCACCTTTTTGTTAAATAAAGAAATGTATCTAAAGTTGCGTAGTGATGTGCTATTA   9862
gb:MT049951|Organism:Severe      CTGTGCACCTTTTTGTTAAATAAAGAAATGTATCTAAAGTTGCGTAGTGATGTGCTATTA   9862
gb:MN975262|Organism:Severe      CTGTGCACCTTTTTGTTAAATAAAGAAATGTATCTAAAGTTGCGTAGTGATGTGCTATTA   9862
gb:MT106052|Organism:Severe      CTGTGCACCTTTTTGTTAAATAAAGAAATGTATCTAAAGTTGCGTAGTGATGTGCTATTA   9862
gb:LC522975|Organism:Severe      CTGTGCACCTTTTTGTTAAATAAAGAAATGTATCTAAAGTTGCGTAGTGATGTGCTATTA   9859
gb:LC522973|Organism:Severe      CTGTGCACCTTTTTGTTAAATAAAGAAATGTATCTAAAGTTGCGTAGTGATGTGCTATTA   9859
gb:LC522974|Organism:Severe      CTGTGCACCTTTTTGTTAAATAAAGAAATGTATCTAAAGTTGCGTAGTGATGTGCTATTA   9859
gb:MN985325|Organism:Severe      CTGTGCACCTTTTTGTTAAATAAAGAAATGTATCTAAAGTTGCGTAGTGATGTGCTATTA   9862
gb:MT020881|Organism:Severe      CTGTGCACCTTTTTGTTAAATAAAGAAATGTATCTAAAGTTGCGTAGTGATGTGCTATTA   9862
gb:MT020880|Organism:Severe      CTGTGCACCTTTTTGTTAAATAAAGAAATGTATCTAAAGTTGCGTAGTGATGTGCTATTA   9862
gb:MT066175|Organism:Severe      CTGTGCACCTTTTTGTTAAATAAAGAAATGTATCTAAAGTTGCGTAGTGATGTGCTATTA   9862
gb:MN997409|Organism:Severe      CTGTGCACCTTTTTGTTAAATAAAGAAATGTATCTAAAGTTGCGTAGTGATGTGCTATTA   9862
gb:MN938384|Organism:Severe      CTGTGCACCTTTTTGTTAAATAAAGAAATGTATCTAAAGTTGCGTAGTGATGTGCTATTA   9830
gb:MT044258|Organism:Severe      CTGTGCACCTTTTTGTTAAATAAAGAAATGTATCTAAAGTTGCGTAGTGATGTGCTATTA   9838
gb:MT039890|Organism:Severe      CTGTGCACCTTTTTGTTAAATAAAGAAATGTATCTAAAGTTGCGTAGTGATGTGCTATTA   9862
gb:MN988713|Organism:Severe      CTGTGCACCTTTTTGTTAAATAAAGAAATGTATCTAAAGTTGCGTAGTGATGTGCTATTA   9862
gb:LC521925|Organism:Severe      CTGTGCACCTTTTTGTTAAATAAAGAAATGTATCTAAAGTTGCGTAGTGATGTGCTATTA   9835
gb:MT093571|Organism:Severe      CTGTGCACCTTTTTGTTAAATAAAGAAATGTATCTAAAGTTGCGTAGTGATGTGCTATTA   9862
gb:MT039887|Organism:Severe      CTGTGCACCTTTTTGTTAAATAAAGAAATGTATCTAAAGTTGCGTAGTGATGTGCTATTA   9862
gb:MT019530|Organism:Severe      CTGTGCACCTTTTTGTTAAATAAAGAAATGTATCTAAAGTTGCGTAGTGATGTGCTATTA   9862
gb:MT039888|Organism:Severe      CTGTGCACCTTTTTGTTAAATAAAGAAATGTATCTAAAGTTGCGTAGTGATGTGCTATTA   9862
gb:LC522972|Organism:Severe      CTGTGCACCTTTTTGTTAAATAAAGAAATGTATCTAAAGTTGCGTAGTGATGTGCTATTA   9859
gb:MT027063|Organism:Severe      CTGTGCACCTTTTTGTTAAATAAAGAAATGTATCTAAAGTTGCGTAGTGATGTGCTATTA   9862
gb:MT027062|Organism:Severe      CTGTGCACCTTTTTGTTAAATAAAGAAATGTATCTAAAGTTGCGTAGTGATGTGCTATTA   9862
gb:MT019529|Organism:Severe      CTGTGCACCTTTTTGTTAAATAAAGAAATGTATCTAAAGTTGCGTAGTGATGTGCTATTA   9862
gb:MN996529|Organism:Severe      CTGTGCACCTTTTTGTTAAATAAAGAAATGTATCTAAAGTTGCGTAGTGATGTGCTATTA   9850
gb:MN996531|Organism:Severe      CTGTGCACCTTTTTGTTAAATAAAGAAATGTATCTAAAGTTGCGTAGTGATGTGCTATTA   9849
gb:MT066176|Organism:Severe      CTGTGCACCTTTTTGTTAAATAAAGAAATGTATCTAAAGTTGCGTAGTGATGTGCTATTA   9862
gb:MT027064|Organism:Severe      CTGTGCACCTTTTTGTTAAATAAAGAAATGTATCTAAAGTTGCGTAGTGATGTGCTATTA   9862
gb:MN994468|Organism:Severe      CTGTGCACCTTTTTGTTAAATAAAGAAATGTATCTAAAGTTGCGTAGTGATGTGCTATTA   9862
gb:MT072688|Organism:Severe      CTGTGCACCTTTTTGTTAAATAAAGAAATGTATCTAAAGTTGCGTAGTGATGTGCTATTA   9847
gb:MN996527|Organism:Severe      CTGTGCACCTTTTTGTTAAATAAAGAAATGTATCTAAAGTTGCGTAGTGATGTGCTATTA   9829
gb:MT093631|Organism:Severe      CTGTGCACCTTTTTGTTAAATAAAGAAATGTATCTAAAGTTGCGTAGTGATGTGCTATTA   9900
gb:MT106053|Organism:Severe      CTGTGCACCTTTTTGTTAAATAAAGAAATGTATCTAAAGTTGCGTAGTGATGTGCTATTA   9862
gb:MT019533|Organism:Severe      CTGTGCACCTTTTTGTTAAATAAAGAAATGTATCTAAAGTTGCGTAGTGATGTGCTATTA   9862
gb:MT019531|Organism:Severe      CTGTGCACCTTTTTGTTAAATAAAGAAATGTATCTAAAGTTGCGTAGTGATGTGCTATTA   9862
gb:MN996528|Organism:Severe      CTGTGCACCTTTTTGTTAAATAAAGAAATGTATCTAAAGTTGCGTAGTGATGTGCTATTA   9862
gb:MN996530|Organism:Severe      CTGTGCACCTTTTTGTTAAATAAAGAAATGTATCTAAAGTTGCGTAGTGATGTGCTATTA   9848
gb:MN908947|Organism:Severe      CTGTGCACCTTTTTGTTAAATAAAGAAATGTATCTAAAGTTGCGTAGTGATGTGCTATTA   9862
gb:MT019532|Organism:Severe      CTGTGCACCTTTTTGTTAAATAAAGAAATGTATCTAAAGTTGCGTAGTGATGTGCTATTA   9862
                                 ************************************************************

gb:MT020781|Organism:Severe      CCTCTTACGCAATATAATAGATACTTAGCTCTTTATAATAAGTACAAGTATTTTAGTGGA   9910
gb:MT007544|Organism:Severe      CCTCTTACGCAATATAATAGATACTTAGCTCTTTATAATAAGTACAAGTATTTTAGTGGA   9922
gb:MN994467|Organism:Severe      CCTCTTACGCAATATAATAGATACTTAGCTCTTTATAATAAGTACAAGTATTTTAGTGGA   9922
gb:MT044257|Organism:Severe      CCTCTTACGCAATATAATAGATACTTAGCTCTTTATAATAAGTACAAGTATTTTAGTGGA   9922
gb:MT106054|Organism:Severe      CCTCTTACGCAATATAATAGATACTTAGCTCTTTATAATAAGTACAAGTATTTTAGTGGA   9922
gb:MT049951|Organism:Severe      CCTCTTACGCAATATAATAGATACTTAGCTCTTTATAATAAGTACAAGTATTTTAGTGGA   9922
gb:MN975262|Organism:Severe      CCTCTTACGCAATATAATAGATACTTAGCTCTTTATAATAAGTACAAGTATTTTAGTGGA   9922
gb:MT106052|Organism:Severe      CCTCTTACGCAATATAATAGATACTTAGCTCTTTATAATAAGTACAAGTATTTTAGTGGA   9922
gb:LC522975|Organism:Severe      CCTCTTACGCAATATAATAGATACTTAGCTCTTTATAATAAGTACAAGTATTTTAGTGGA   9919
gb:LC522973|Organism:Severe      CCTCTTACGCAATATAATAGATACTTAGCTCTTTATAATAAGTACAAGTATTTTAGTGGA   9919
gb:LC522974|Organism:Severe      CCTCTTACGCAATATAATAGATACTTAGCTCTTTATAATAAGTACAAGTATTTTAGTGGA   9919
```

```
gb:MN985325|Organism:Severe      CCTCTTACGCAATATAATAGATACTTAGCTCTTTATAATAAGTACAAGTATTTTAGTGGA   9922
gb:MT020881|Organism:Severe      CCTCTTACGCAATATAATAGATACTTAGCTCTTTATAATAAGTACAAGTATTTTAGTGGA   9922
gb:MT020880|Organism:Severe      CCTCTTACGCAATATAATAGATACTTAGCTCTTTATAATAAGTACAAGTATTTTAGTGGA   9922
gb:MT066175|Organism:Severe      CCTCTTACGCAATATAATAGATACTTAGCTCTTTATAATAAGTACAAGTATTTTAGTGGA   9922
gb:MN997409|Organism:Severe      CCTCTTACGCAATATAATAGATACTTAGCTCTTTATAATAAGTACAAGTATTTTAGTGGA   9922
gb:MN938384|Organism:Severe      CCTCTTACGCAATATAATAGATACTTAGCTCTTTATAATAAGTACAAGTATTTTAGTGGA   9890
gb:MT044258|Organism:Severe      CCTCTTACGCAATATAATAGATACTTAGCTCTTTATAATAAGTACAAGTATTTTAGTGGA   9898
gb:MT039890|Organism:Severe      CCTCTTACGCAATATAATAGATACTTAGCTCTTTATAATAAGTACAAGTATTTTAGTGGA   9922
gb:MN988713|Organism:Severe      CCTCTTACGCAATATAATAGATACTTAGCTCTTTATAATAAGTACAAGTATTTTAGTGGA   9922
gb:LC521925|Organism:Severe      CCTCTTACGCAATATAATAGATACTTAGCTCTTTATAATAAGTACAAGTATTTTAGTGGA   9895
gb:MT093571|Organism:Severe      CCTCTTACGCAATATAATAGATACTTAGCTCTTTATAATAAGTACAAGTATTTTAGTGGA   9922
gb:MT039887|Organism:Severe      CCTCTTACGCAATATAATAGATACTTAGCTCTTTATAATAAGTACAAGTATTTTAGTGGA   9922
gb:MT019530|Organism:Severe      CCTCTTACGCAATATAATAGATACTTAGCTCTTTATAATAAGTACAAGTATTTTAGTGGA   9922
gb:MT039888|Organism:Severe      CCTCTTACGCAATATAATAGATACTTAGCTCTTTATAATAAGTACAAGTATTTTAGTGGA   9922
gb:LC522972|Organism:Severe      CCTCTTACGCAATATAATAGATACTTAGCTCTTTATAATAAGTACAAGTATTTTAGTGGA   9919
gb:MT027063|Organism:Severe      CCTCTTACGCAATATAATAGATACTTAGCTCTTTATAATAAGTACAAGTATTTTAGTGGA   9922
gb:MT027062|Organism:Severe      CCTCTTACGCAATATAATAGATACTTAGCTCTTTATAATAAGTACAAGTATTTTAGTGGA   9922
gb:MT019529|Organism:Severe      CCTCTTACGCAATATAATAGATACTTAGCTCTTTATAATAAGTACAAGTATTTTAGTGGA   9922
gb:MN996529|Organism:Severe      CCTCTTACGCAATATAATAGATACTTAGCTCTTTATAATAAGTACAAGTATTTTAGTGGA   9910
gb:MN996531|Organism:Severe      CCTCTTACGCAATATAATAGATACTTAGCTCTTTATAATAAGTACAAGTATTTTAGTGGA   9909
gb:MT066176|Organism:Severe      CCTCTTACGCAATATAATAGATACTTAGCTCTTTATAATAAGTACAAGTATTTTAGTGGA   9922
gb:MT027064|Organism:Severe      CCTCTTACGCAATATAATAGATACTTAGCTCTTTATAATAAGTACAAGTATTTTAGTGGA   9922
gb:MN994468|Organism:Severe      CCTCTTACGCAATATAATAGATACTTAGCTCTTTATAATAAGTACAAGTATTTTAGTGGA   9922
gb:MT072688|Organism:Severe      CCTCTTACGCAATATAATAGATACTTAGCTCTTTATAATAAGTACAAGTATTTTAGTGGA   9907
gb:MN996527|Organism:Severe      CCTCTTACGCAATATAATAGATACTTAGCTCTTTATAATAAGTACAAGTATTTTAGTGGA   9889
gb:MT093631|Organism:Severe      CCTCTTACGCAATATAATAGATACTTAGCTCTTTATAATAAGTACAAGTATTTTAGTGGA   9960
gb:MT106053|Organism:Severe      CCTCTTACGCAATATAATAGATACTTAGCTCTTTATAATAAGTACAAGTATTTTAGTGGA   9922
gb:MT019533|Organism:Severe      CCTCTTACGCAATATAATAGATACTTAGCTCTTTATAATAAGTACAAGTATTTTAGTGGA   9922
gb:MT019531|Organism:Severe      CCTCTTACGCAATATAATAGATACTTAGCTCTTTATAATAAGTACAAGTATTTTAGTGGA   9922
gb:MN996528|Organism:Severe      CCTCTTACGCAATATAATAGATACTTAGCTCTTTATAATAAGTACAAGTATTTTAGTGGA   9922
gb:MN996530|Organism:Severe      CCTCTTACGCAATATAATAGATACTTAGCTCTTTATAATAAGTACAAGTATTTTAGTGGA   9908
gb:MN908947|Organism:Severe      CCTCTTACGCAATATAATAGATACTTAGCTCTTTATAATAAGTACAAGTATTTTAGTGGA   9922
gb:MT019532|Organism:Severe      CCTCTTACGCAATATAATAGATACTTAGCTCTTTATAATAAGTACAAGTATTTTAGTGGA   9922
                                 ************************************************************

gb:MT020781|Organism:Severe      GCAATGGATACAACTAGCTACAGAGAAGCTGCTTGTTGTCATCTCGCAAAGGCTCTCAAT   9970
gb:MT007544|Organism:Severe      GCAATGGATACAACTAGCTACAGAGAAGCTGCTTGTTGTCATCTCGCAAAGGCTCTCAAT   9982
gb:MN994467|Organism:Severe      GCAATGGATACAACTAGCTACAGAGAAGCTGCTTGTTGTCATCTCGCAAAGGCTCTCAAT   9982
gb:MT044257|Organism:Severe      GCAATGGATACAACTAGCTACAGAGAAGCTGCTTGTTGTCATCTCGCAAAGGCTCTCAAT   9982
gb:MT106054|Organism:Severe      GCAATGGATACAACTAGCTACAGAGAAGCTGCTTGTTGTCATCTCGCAAAGGCTCTCAAT   9982
gb:MT049951|Organism:Severe      GCAATGGATACAACTAGCTACAGAGAAGCTGCTTGTTGTCATCTCGCAAAGGCTCTCAAT   9982
gb:MN975262|Organism:Severe      GCAATGGATACAACTAGCTACAGAGAAGCTGCTTGTTGTCATCTCGCAAAGGCTCTCAAT   9982
gb:MT106052|Organism:Severe      GCAATGGATACAACTAGCTACAGAGAAGCTGCTTGTTGTCATCTCGCAAAGGCTCTCAAT   9982
gb:LC522975|Organism:Severe      GCAATGGATACAACTAGCTACAGAGAAGCTGCTTGTTGTCATCTCGCAAAGGCTCTCAAT   9979
gb:LC522973|Organism:Severe      GCAATGGATACAACTAGCTACAGAGAAGCTGCTTGTTGTCATCTCGCAAAGGCTCTCAAT   9979
gb:LC522974|Organism:Severe      GCAATGGATACAACTAGCTACAGAGAAGCTGCTTGTTGTCATCTCGCAAAGGCTCTCAAT   9979
gb:MN985325|Organism:Severe      GCAATGGATACAACTAGCTACAGAGAAGCTGCTTGTTGTCATCTCGCAAAGGCTCTCAAT   9982
gb:MT020881|Organism:Severe      GCAATGGATACAACTAGCTACAGAGAAGCTGCTTGTTGTCATCTCGCAAAGGCTCTCAAT   9982
gb:MT020880|Organism:Severe      GCAATGGATACAACTAGCTACAGAGAAGCTGCTTGTTGTCATCTCGCAAAGGCTCTCAAT   9982
gb:MT066175|Organism:Severe      GCAATGGATACAACTAGCTACAGAGAAGCTGCTTGTTGTCATCTCGCAAAGGCTCTCAAT   9982
gb:MN997409|Organism:Severe      GCAATGGATACAACTAGCTACAGAGAAGCTGCTTGTTGTCATCTCGCAAAGGCTCTCAAT   9982
gb:MN938384|Organism:Severe      GCAATGGATACAACTAGCTACAGAGAAGCTGCTTGTTGTCATCTCGCAAAGGCTCTCAAT   9950
gb:MT044258|Organism:Severe      GCAATGGATACAACTAGCTACAGAGAAGCTGCTTGTTGTCATCTCGCAAAGGCTCTCAAT   9958
gb:MT039890|Organism:Severe      GCAATGGATACAACTAGCTACAGAGAAGCTGCTTGTTGTCATCTCGCAAAGGCTCTCAAT   9982
gb:MN988713|Organism:Severe      GCAATGGATACAACTAGCTACAGAGAAGCTGCTTGTTGTCATCTCGCAAAGGCTCTCAAT   9982
gb:LC521925|Organism:Severe      GCAATGGATACAACTAGCTACAGAGAAGCTGCTTGTTGTCATCTCGCAAAGGCTCTCAAT   9955
gb:MT093571|Organism:Severe      GCAATGGATACAACTAGCTACAGAGAAGCTGCTTGTTGTCATCTCGCAAAGGCTCTCAAT   9982
gb:MT039887|Organism:Severe      GCAATGGATACAACTAGCTACAGAGAAGCTGCTTGTTGTCATCTCGCAAAGGCTCTCAAT   9982
gb:MT019530|Organism:Severe      GCAATGGATACAACTAGCTACAGAGAAGCTGCTTGTTGTCATCTCGCAAAGGCTCTCAAT   9982
gb:MT039888|Organism:Severe      GCAATGGATACAACTAGCTACAGAGAAGCTGCTTGTTGTCATCTCGCAAAGGCTCTCAAT   9982
gb:LC522972|Organism:Severe      GCAATGGATACAACTAGCTACAGAGAAGCTGCTTGTTGTCATCTCGCAAAGGCTCTCAAT   9979
gb:MT027063|Organism:Severe      GCAATGGATACAACTAGCTACAGAGAAGCTGCTTGTTGTCATCTCGCAAAGGCTCTCAAT   9982
gb:MT027062|Organism:Severe      GCAATGGATACAACTAGCTACAGAGAAGCTGCTTGTTGTCATCTCGCAAAGGCTCTCAAT   9982
gb:MT019529|Organism:Severe      GCAATGGATACAACTAGCTACAGAGAAGCTGCTTGTTGTCATCTCGCAAAGGCTCTCAAT   9982
gb:MN996529|Organism:Severe      GCAATGGATACAACTAGCTACAGAGAAGCTGCTTGTTGTCATCTCGCAAAGGCTCTCAAT   9970
```

```
gb:MN996531|Organism:Severe      GCAATGGATACAACTAGCTACAGAGAAGCTGCTTGTTGTCATCTCGCAAAGGCTCTCAAT   9969
gb:MT066176|Organism:Severe      GCAATGGATACAACTAGCTACAGAGAAGCTGCTTGTTGTCATCTCGCAAAGGCTCTCAAT   9982
gb:MT027064|Organism:Severe      GCAATGGATACAACTAGCTACAGAGAAGCTGCTTGTTGTCATCTCGCAAAGGCTCTCAAT   9982
gb:MN994468|Organism:Severe      GCAATGGATACAACTAGCTACAGAGAAGCTGCTTGTTGTCATCTCGCAAAGGCTCTCAAT   9982
gb:MT072688|Organism:Severe      GCAATGGATACAACTAGCTACAGAGAAGCTGCTTGTTGTCATCTCGCAAAGGCTCTCAAT   9967
gb:MN996527|Organism:Severe      GCAATGGATACAACTAGCTACAGAGAAGCTGCTTGTTGTCATCTCGCAAAGGCTCTCAAT   9949
gb:MT093631|Organism:Severe      GCAATGGATACAACTAGCTACAGAGAAGCTGCTTGTTGTCATCTCGCAAAGGCTCTCAAT   10020
gb:MT106053|Organism:Severe      GCAATGGATACAACTAGCTACAGAGAAGCTGCTTGTTGTCATCTCGCAAAGGCTCTCAAT   9982
gb:MT019533|Organism:Severe      GCAATGGATACAACTAGCTACAGAGAAGCTGCTTGTTGTCATCTCGCAAAGGCTCTCAAT   9982
gb:MT019531|Organism:Severe      GCAATGGATACAACTAGCTACAGAGAAGCTGCTTGTTGTCATCTCGCAAAGGCTCTCAAT   9982
gb:MN996528|Organism:Severe      GCAATGGATACAACTAGCTACAGAGAAGCTGCTTGTTGTCATCTCGCAAAGGCTCTCAAT   9982
gb:MN996530|Organism:Severe      GCAATGGATACAACTAGCTACAGAGAAGCTGCTTGTTGTCATCTCGCAAAGGCTCTCAAT   9968
gb:MN908947|Organism:Severe      GCAATGGATACAACTAGCTACAGAGAAGCTGCTTGTTGTCATCTCGCAAAGGCTCTCAAT   9982
gb:MT019532|Organism:Severe      GCAATGGATACAACTAGCTACAGAGAAGCTGCTTGTTGTCATCTCGCAAAGGCTCTCAAT   9982
                                 ************************************************************

gb:MT020781|Organism:Severe      GACTTCAGTAACTCAGGTTCTGATGTTCTTTACCAACCACCACAAACCTCTATCACCTCA   10030
gb:MT007544|Organism:Severe      GACTTCAGTAACTCAGGTTCTGATGTTCTTTACCAACCACCACAAACCTCTATCACCTCA   10042
gb:MN994467|Organism:Severe      GACTTCAGTAACTCAGGTTCTGATGTTCTTTACCAACCACCACAAACCTCTATCACCTCA   10042
gb:MT044257|Organism:Severe      GACTTCAGTAACTCAGGTTCTGATGTTCTTTACCAACCACCACAAACCTCTATCACCTCA   10042
gb:MT106054|Organism:Severe      GACTTCAGTAACTCAGGTTCTGATGTTCTTTACCAACCACCACAAACCTCTATCACCTCA   10042
gb:MT049951|Organism:Severe      GACTTCAGTAACTCAGGTTCTGATGTTCTTTACCAACCACCACAAACCTCTATCACCTCA   10042
gb:MN975262|Organism:Severe      GACTTCAGTAACTCAGGTTCTGATGTTCTTTACCAACCACCACAAACCTCTATCACCTCA   10042
gb:MT106052|Organism:Severe      GACTTCAGTAACTCAGGTTCTGATGTTCTTTACCAACCACCACAAACCTCTATCACCTCA   10042
gb:LC522975|Organism:Severe      GACTTCAGTAACTCAGGTTCTGATGTTCTTTACCAACCACCACAAACCTCTATCACCTCA   10039
gb:LC522973|Organism:Severe      GACTTCAGTAACTCAGGTTCTGATGTTCTTTACCAACCACCACAAACCTCTATCACCTCA   10039
gb:LC522974|Organism:Severe      GACTTCAGTAACTCAGGTTCTGATGTTCTTTACCAACCACCACAAACCTCTATCACCTCA   10039
gb:MN985325|Organism:Severe      GACTTCAGTAACTCAGGTTCTGATGTTCTTTACCAACCACCACAAACCTCTATCACCTCA   10042
gb:MT020881|Organism:Severe      GACTTCAGTAACTCAGGTTCTGATGTTCTTTACCAACCACCACAAACCTCTATCACCTCA   10042
gb:MT020880|Organism:Severe      GACTTCAGTAACTCAGGTTCTGATGTTCTTTACCAACCACCACAAACCTCTATCACCTCA   10042
gb:MT066175|Organism:Severe      GACTTCAGTAACTCAGGTTCTGATGTTCTTTACCAACCACCACAAACCTCTATCACCTCA   10042
gb:MN997409|Organism:Severe      GACTTCAGTAACTCAGGTTCTGATGTTCTTTACCAACCACCACAAACCTCTATCACCTCA   10042
gb:MN938384|Organism:Severe      GACTTCAGTAACTCAGGTTCTGATGTTCTTTACCAACCACCACAAACCTCTATCACCTCA   10010
gb:MT044258|Organism:Severe      GACTTCAGTAACTCAGGTTCTGATGTTCTTTACCAACCACCACAAACCTCTATCACCTCA   10018
gb:MT039890|Organism:Severe      GACTTCAGTAACTCAGGTTCTGATGTTCTTTACCAACCACCACAAACCTCTATCACCTCA   10042
gb:MN988713|Organism:Severe      GACTTCAGTAACTCAGGTTCTGATGTTCTTTACCAACCACCACAAACCTCTATCACCTCA   10042
gb:LC521925|Organism:Severe      GACTTCAGTAACTCAGGTTCTGATGTTCTTTACCAACCACCACAAACCTCTATCACCTCA   10015
gb:MT093571|Organism:Severe      GACTTCAGTAACTCAGGTTCTGATGTTCTTTACCAACCACCACAAACCTCTATCACCTCA   10042
gb:MT039887|Organism:Severe      GACTTCAGTAACTCAGGTTCTGATGTTCTTTACCAACCACCACAAACCTCTATCACCTCA   10042
gb:MT019530|Organism:Severe      GACTTCAGTAACTCAGGTTCTGATGTTCTTTACCAACCACCACAAACCTCTATCACCTCA   10042
gb:MT039888|Organism:Severe      GACTTCAGTAACTCAGGTTCTGATGTTCTTTACCAACCACCACAAACCTCTATCACCTCA   10042
gb:LC522972|Organism:Severe      GACTTCAGTAACTCAGGTTCTGATGTTCTTTACCAACCACCACAAACCTCTATCACCTCA   10039
gb:MT027063|Organism:Severe      GACTTCAGTAACTCAGGTTCTGATGTTCTTTACCAACCACCACAAACCTCTATCACCTCA   10042
gb:MT027062|Organism:Severe      GACTTCAGTAACTCAGGTTCTGATGTTCTTTACCAACCACCACAAACCTCTATCACCTCA   10042
gb:MT019529|Organism:Severe      GACTTCAGTAACTCAGGTTCTGATGTTCTTTACCAACCACCACAAACCTCTATCACCTCA   10042
gb:MN996529|Organism:Severe      GACTTCAGTAACTCAGGTTCTGATGTTCTTTACCAACCACCACAAACCTCTATCACCTCA   10030
gb:MN996531|Organism:Severe      GACTTCAGTAACTCAGGTTCTGATGTTCTTTACCAACCACCACAAACCTCTATCACCTCA   10029
gb:MT066176|Organism:Severe      GACTTCAGTAACTCAGGTTCTGATGTTCTTTACCAACCACCACAAACCTCTATCACCTCA   10042
gb:MT027064|Organism:Severe      GACTTCAGTAACTCAGGTTCTGATGTTCTTTACCAACCACCACAAACCTCTATCACCTCA   10042
gb:MN994468|Organism:Severe      GACTTCAGTAACTCAGGTTCTGATGTTCTTTACCAACCACCACAAACCTCTATCACCTCA   10042
gb:MT072688|Organism:Severe      GACTTCAGTAACTCAGGTTCTGATGTTCTTTACCAACCACCACAAACCTCTATCACCTCA   10027
gb:MN996527|Organism:Severe      GACTTCAGTAACTCAGGTTCTGATGTTCTTTACCAACCACCACAAACCTCTATCACCTCA   10009
gb:MT093631|Organism:Severe      GACTTCAGTAACTCAGGTTCTGATGTTCTTTACCAACCACCACAAACCTCTATCACCTCA   10080
gb:MT106053|Organism:Severe      GACTTCAGTAACTCAGGTTCTGATGTTCTTTACCAACCACCACAAACCTCTATCACCTCA   10042
gb:MT019533|Organism:Severe      GACTTCAGTAACTCAGGTTCTGATGTTCTTTACCAACCACCACAAACCTCTATCACCTCA   10042
gb:MT019531|Organism:Severe      GACTTCAGTAACTCAGGTTCTGATGTTCTTTACCAACCACCACAAACCTCTATCACCTCA   10042
gb:MN996528|Organism:Severe      GACTTCAGTAACTCAGGTTCTGATGTTCTTTACCAACCACCACAAACCTCTATCACCTCA   10042
gb:MN996530|Organism:Severe      GACTTCAGTAACTCAGGTTCTGATGTTCTTTACCAACCACCACAAACCTCTATCACCTCA   10028
gb:MN908947|Organism:Severe      GACTTCAGTAACTCAGGTTCTGATGTTCTTTACCAACCACCACAAACCTCTATCACCTCA   10042
gb:MT019532|Organism:Severe      GACTTCAGTAACTCAGGTTCTGATGTTCTTTACCAACCACCACAAACCTCTATCACCTCA   10042
                                 ************************************************************

gb:MT020781|Organism:Severe      GCTGTTTTGCAGAGTGGTTTTAGAAAAATGGCATTCCCATCTGGTAAAGTTGAGGGTTGT   10090
gb:MT007544|Organism:Severe      GCTGTTTTGCAGAGTGGTTTTAGAAAAATGGCATTCCCATCTGGTAAAGTTGAGGGTTGT   10102
gb:MN994467|Organism:Severe      GCTGTTTTGCAGAGTGGTTTTAGAAAAATGGCATTCCCATCTGGTAAAGTTGAGGGTTGT   10102
```

```
gb:MT044257|Organism:Severe      GCTGTTTTGCAGAGTGGTTTTAGAAAAATGGCATTCCCATCTGGTAAAGTTGAGGGTTGT   10102
gb:MT106054|Organism:Severe      GCTGTTTTGCAGAGTGGTTTTAGAAAAATGGCATTCCCATCTGGTAAAGTTGAGGGTTGT   10102
gb:MT049951|Organism:Severe      GCTGTTTTGCAGAGTGGTTTTAGAAAAATGGCATTCCCATCTGGTAAAGTTGAGGGTTGT   10102
gb:MN975262|Organism:Severe      GCTGTTTTGCAGAGTGGTTTTAGAAAAATGGCATTCCCATCTGGTAAAGTTGAGGGTTGT   10102
gb:MT106052|Organism:Severe      GCTGTTTTGCAGAGTGGTTTTAGAAAAATGGCATTCCCATCTGGTAAAGTTGAGGGTTGT   10102
gb:LC522975|Organism:Severe      GCTGTTTTGCAGAGTGGTTTTAGAAAAATGGCATTCCCATCTGGTAAAGTTGAGGGTTGT   10099
gb:LC522973|Organism:Severe      GCTGTTTTGCAGAGTGGTTTTAGAAAAATGGCATTCCCATCTGGTAAAGTTGAGGGTTGT   10099
gb:LC522974|Organism:Severe      GCTGTTTTGCAGAGTGGTTTTAGAAAAATGGCATTCCCATCTGGTAAAGTTGAGGGTTGT   10099
gb:MN985325|Organism:Severe      GCTGTTTTGCAGAGTGGTTTTAGAAAAATGGCATTCCCATCTGGTAAAGTTGAGGGTTGT   10102
gb:MT020881|Organism:Severe      GCTGTTTTGCAGAGTGGTTTTAGAAAAATGGCATTCCCATCTGGTAAAGTTGAGGGTTGT   10102
gb:MT020880|Organism:Severe      GCTGTTTTGCAGAGTGGTTTTAGAAAAATGGCATTCCCATCTGGTAAAGTTGAGGGTTGT   10102
gb:MT066175|Organism:Severe      GCTGTTTTGCAGAGTGGTTTTAGAAAAATGGCATTCCCATCTGGTAAAGTTGAGGGTTGT   10102
gb:MN997409|Organism:Severe      GCTGTTTTGCAGAGTGGTTTTAGAAAAATGGCATTCCCATCTGGTAAAGTTGAGGGTTGT   10102
gb:MN938384|Organism:Severe      GCTGTTTTGCAGAGTGGTTTTAGAAAAATGGCATTCCCATCTGGTAAAGTTGAGGGTTGT   10070
gb:MT044258|Organism:Severe      GCTGTTTTGCAGAGTGGTTTTAGAAAAATGGCATTCCCATCTGGTAAAGTTGAGGGTTGT   10078
gb:MT039890|Organism:Severe      GCTGTTTTGCAGAGTGGTTTTAGAAAAATGGCATTCCCATCTGGTAAAGTTGAGGGTTGT   10102
gb:MN988713|Organism:Severe      GCTGTTTTGCAGAGTGGTTTTAGAAAAATGGCATTCCCATCTGGTAAAGTTGAGGGTTGT   10102
gb:LC521925|Organism:Severe      GCTGTTTTGCAGAGTGGTTTTAGAAAAATGGCATTCCCATCTGGTAAAGTTGAGGGTTGT   10075
gb:MT093571|Organism:Severe      GCTGTTTTGCAGAGTGGTTTTAGAAAAATGGCATTCCCATCTGGTAAAGTTGAGGGTTGT   10102
gb:MT039887|Organism:Severe      GCTGTTTTGCAGAGTGGTTTTAGAAAAATGGCATTCCCATCTGGTAAAGTTGAGGGTTGT   10102
gb:MT019530|Organism:Severe      GCTGTTTTGCAGAGTGGTTTTAGAAAAATGGCATTCCCATCTGGTAAAGTTGAGGGTTGT   10102
gb:MT039888|Organism:Severe      GCTGTTTTGCAGAGTGGTTTTAGAAAAATGGCATTCCCATCTGGTAAAGTTGAGGGTTGT   10102
gb:LC522972|Organism:Severe      GCTGTTTTGCAGAGTGGTTTTAGAAAAATGGCATTCCCATCTGGTAAAGTTGAGGGTTGT   10099
gb:MT027063|Organism:Severe      GCTGTTTTGCAGAGTGGTTTTAGAAAAATGGCATTCCCATCTGGTAAAGTTGAGGGTTGT   10102
gb:MT027062|Organism:Severe      GCTGTTTTGCAGAGTGGTTTTAGAAAAATGGCATTCCCATCTGGTAAAGTTGAGGGTTGT   10102
gb:MT019529|Organism:Severe      GCTGTTTTGCAGAGTGGTTTTAGAAAAATGGCATTCCCATCTGGTAAAGTTGAGGGTTGT   10102
gb:MN996529|Organism:Severe      GCTGTTTTGCAGAGTGGTTTTAGAAAAATGGCATTCCCATCTGGTAAAGTTGAGGGTTGT   10090
gb:MN996531|Organism:Severe      GCTGTTTTGCAGAGTGGTTTTAGAAAAATGGCATTCCCATCTGGTAAAGTTGAGGGTTGT   10089
gb:MT066176|Organism:Severe      GCTGTTTTGCAGAGTGGTTTTAGAAAAATGGCATTCCCATCTGGTAAAGTTGAGGGTTGT   10102
gb:MT027064|Organism:Severe      GCTGTTTTGCAGAGTGGTTTTAGAAAAATGGCATTCCCATCTGGTAAAGTTGAGGGTTGT   10102
gb:MN994468|Organism:Severe      GCTGTTTTGCAGAGTGGTTTTAGAAAAATGGCATTCCCATCTGGTAAAGTTGAGGGTTGT   10102
gb:MT072688|Organism:Severe      GCTGTTTTGCAGAGTGGTTTTAGAAAAATGGCATTCCCATCTGGTAAAGTTGAGGGTTGT   10087
gb:MN996527|Organism:Severe      GCTGTTTTGCAGAGTGGTTTTAGAAAAATGGCATTCCCATCTGGTAAAGTTGAGGGTTGT   10069
gb:MT093631|Organism:Severe      GCTGTTTTGCAGAGTGGTTTTAGAAAAATGGCATTCCCATCTGGTAAAGTTGAGGGTTGT   10140
gb:MT106053|Organism:Severe      GCTGTTTTGCAGAGTGGTTTTAGAAAAATGGCATTCCCATCTGGTAAAGTTGAGGGTTGT   10102
gb:MT019533|Organism:Severe      GCTGTTTTGCAGAGTGGTTTTAGAAAAATGGCATTCCCATCTGGTAAAGTTGAGGGTTGT   10102
gb:MT019531|Organism:Severe      GCTGTTTTGCAGAGTGGTTTTAGAAAAATGGCATTCCCATCTGGTAAAGTTGAGGGTTGT   10102
gb:MN996528|Organism:Severe      GCTGTTTTGCAGAGTGGTTTTAGAAAAATGGCATTCCCATCTGGTAAAGTTGAGGGTTGT   10102
gb:MN996530|Organism:Severe      GCTGTTTTGCAGAGTGGTTTTAGAAAAATGGCATTCCCATCTGGTAAAGTTGAGGGTTGT   10088
gb:MN908947|Organism:Severe      GCTGTTTTGCAGAGTGGTTTTAGAAAAATGGCATTCCCATCTGGTAAAGTTGAGGGTTGT   10102
gb:MT019532|Organism:Severe      GCTGTTTTGCAGAGTGGTTTTAGAAAAATGGCATTCCCATCTGGTAAAGTTGAGGGTTGT   10102
                                 ************************************************************

gb:MT020781|Organism:Severe      ATGGTACAAGTAACTTGTGGTACAACTACACTTAACGGTCTTTGGCTTGATGACGTAGTT   10150
gb:MT007544|Organism:Severe      ATGGTACAAGTAACTTGTGGTACAACTACACTTAACGGTCTTTGGCTTGATGACGTAGTT   10162
gb:MN994467|Organism:Severe      ATGGTACAAGTAACTTGTGGTACAACTACACTTAACGGTCTTTGGCTTGATGACGTAGTT   10162
gb:MT044257|Organism:Severe      ATGGTACAAGTAACTTGTGGTACAACTACACTTAACGGTCTTTGGCTTGATGACGTAGTT   10162
gb:MT106054|Organism:Severe      ATGGTACAAGTAACTTGTGGTACAACTACACTTAACGGTCTTTGGCTTGATGACGTAGTT   10162
gb:MT049951|Organism:Severe      ATGGTACAAGTAACTTGTGGTACAACTACACTTAACGGTCTTTGGCTTGATGACGTAGTT   10162
gb:MN975262|Organism:Severe      ATGGTACAAGTAACTTGTGGTACAACTACACTTAACGGTCTTTGGCTTGATGACGTAGTT   10162
gb:MT106052|Organism:Severe      ATGGTACAAGTAACTTGTGGTACAACTACACTTAACGGTCTTTGGCTTGATGACGTAGTT   10162
gb:LC522975|Organism:Severe      ATGGTACAAGTAACTTGTGGTACAACTACACTTAACGGTCTTTGGCTTGATGACGTAGTT   10159
gb:LC522973|Organism:Severe      ATGGTACAAGTAACTTGTGGTACAACTACACTTAACGGTCTTTGGCTTGATGACGTAGTT   10159
gb:LC522974|Organism:Severe      ATGGTACAAGTAACTTGTGGTACAACTACACTTAACGGTCTTTGGCTTGATGACGTAGTT   10159
gb:MN985325|Organism:Severe      ATGGTACAAGTAACTTGTGGTACAACTACACTTAACGGTCTTTGGCTTGATGACGTAGTT   10162
gb:MT020881|Organism:Severe      ATGGTACAAGTAACTTGTGGTACAACTACACTTAACGGTCTTTGGCTTGATGACGTAGTT   10162
gb:MT020880|Organism:Severe      ATGGTACAAGTAACTTGTGGTACAACTACACTTAACGGTCTTTGGCTTGATGACGTAGTT   10162
gb:MT066175|Organism:Severe      ATGGTACAAGTAACTTGTGGTACAACTACACTTAACGGTCTTTGGCTTGATGACGTAGTT   10162
gb:MN997409|Organism:Severe      ATGGTACAAGTAACTTGTGGTACAACTACACTTAACGGTCTTTGGCTTGATGACGTAGTT   10162
gb:MN938384|Organism:Severe      ATGGTACAAGTAACTTGTGGTACAACTACACTTAACGGTCTTTGGCTTGATGACGTAGTT   10130
gb:MT044258|Organism:Severe      ATGGTACAAGTAACTTGTGGTACAACTACACTTAACGGTCTTTGGCTTGATGACGTAGTT   10138
gb:MT039890|Organism:Severe      ATGGTACAAGTAACTTGTGGTACAACTACACTTAACGGTCTTTGGCTTGATGACGTAGTT   10162
gb:MN988713|Organism:Severe      ATGGTACAAGTAACTTGTGGTACAACTACACTTAACGGTCTTTGGCTTGATGACGTAGTT   10162
gb:LC521925|Organism:Severe      ATGGTACAAGTAACTTGTGGTACAACTACACTTAACGGTCTTTGGCTTGATGACGTAGTT   10135
gb:MT093571|Organism:Severe      ATGGTACAAGTAACTTGTGGTACAACTACACTTAACGGTCTTTGGCTTGATGACGTAGTT   10162
```

```
gb:MT039887|Organism:Severe      ATGGTACAAGTAACTTGTGGTACAACTACACTTAACGGTCTTTGGCTTGATGACGTAGTT   10162
gb:MT019530|Organism:Severe      ATGGTACAAGTAACTTGTGGTACAACTACACTTAACGGTCTTTGGCTTGATGACGTAGTT   10162
gb:MT039888|Organism:Severe      ATGGTACAAGTAACTTGTGGTACAACTACACTTAACGGTCTTTGGCTTGATGACGTAGTT   10162
gb:LC522972|Organism:Severe      ATGGTACAAGTAACTTGTGGTACAACTACACTTAACGGTCTTTGGCTTGATGACGTAGTT   10159
gb:MT027063|Organism:Severe      ATGGTACAAGTAACTTGTGGTACAACTACACTTAACGGTCTTTGGCTTGATGACGTAGTT   10162
gb:MT027062|Organism:Severe      ATGGTACAAGTAACTTGTGGTACAACTACACTTAACGGTCTTTGGCTTGATGACGTAGTT   10162
gb:MT019529|Organism:Severe      ATGGTACAAGTAACTTGTGGTACAACTACACTTAACGGTCTTTGGCTTGATGACGTAGTT   10162
gb:MN996529|Organism:Severe      ATGGTACAAGTAACTTGTGGTACAACTACACTTAACGGTCTTTGGCTTGATGACGTAGTT   10150
gb:MN996531|Organism:Severe      ATGGTACAAGTAACTTGTGGTACAACTACACTTAACGGTCTTTGGCTTGATGACGTAGTT   10149
gb:MT066176|Organism:Severe      ATGGTACAAGTAACTTGTGGTACAACTACACTTAACGGTCTTTGGCTTGATGACGTAGTT   10162
gb:MT027064|Organism:Severe      ATGGTACAAGTAACTTGTGGTACAACTACACTTAACGGTCTTTGGCTTGATGACGTAGTT   10162
gb:MN994468|Organism:Severe      ATGGTACAAGTAACTTGTGGTACAACTACACTTAACGGTCTTTGGCTTGATGACGTAGTT   10162
gb:MT072688|Organism:Severe      ATGGTACAAGTAACTTGTGGTACAACTACACTTAACGGTCTTTGGCTTGATGACGTAGTT   10147
gb:MN996527|Organism:Severe      ATGGTACAAGTAACTTGTGGTACAACTACACTTAACGGTCTTTGGCTTGATGACGTAGTT   10129
gb:MT093631|Organism:Severe      ATGGTACAAGTAACTTGTGGTACAACTACACTTAACGGTCTTTGGCTTGATGACGTAGTT   10200
gb:MT106053|Organism:Severe      ATGGTACAAGTAACTTGTGGTACAACTACACTTAACGGTCTTTGGCTTGATGACGTAGTT   10162
gb:MT019533|Organism:Severe      ATGGTACAAGTAACTTGTGGTACAACTACACTTAACGGTCTTTGGCTTGATGACGTAGTT   10162
gb:MT019531|Organism:Severe      ATGGTACAAGTAACTTGTGGTACAACTACACTTAACGGTCTTTGGCTTGATGACGTAGTT   10162
gb:MN996528|Organism:Severe      ATGGTACAAGTAACTTGTGGTACAACTACACTTAACGGTCTTTGGCTTGATGACGTAGTT   10162
gb:MN996530|Organism:Severe      ATGGTACAAGTAACTTGTGGTACAACTACACTTAACGGTCTTTGGCTTGATGACGTAGTT   10148
gb:MN908947|Organism:Severe      ATGGTACAAGTAACTTGTGGTACAACTACACTTAACGGTCTTTGGCTTGATGACGTAGTT   10162
gb:MT019532|Organism:Severe      ATGGTACAAGTAACTTGTGGTACAACTACACTTAACGGTCTTTGGCTTGATGACGTAGTT   10162
                                 ************************************************************

gb:MT020781|Organism:Severe      TACTGTCCAAGACATGTGATCTGCACCTCTGAAGACATGCTTAACCCTAATTATGAAGAT   10210
gb:MT007544|Organism:Severe      TACTGTCCAAGACATGTGATCTGCACCTCTGAAGACATGCTTAACCCTAATTATGAAGAT   10222
gb:MN994467|Organism:Severe      TACTGTCCAAGACATGTGATCTGCACCTCTGAAGACATGCTTAACCCTAATTATGAAGAT   10222
gb:MT044257|Organism:Severe      TACTGTCCAAGACATGTGATCTGCACCTCTGAAGACATGCTTAACCCTAATTATGAAGAT   10222
gb:MT106054|Organism:Severe      TACTGTCCAAGACATGTGATCTGCACCTCTGAAGACATGCTTAACCCTAATTATGAAGAT   10222
gb:MT049951|Organism:Severe      TACTGTCCAAGACATGTGATCTGCACCTCTGAAGACATGCTTAACCCTAATTATGAAGAT   10222
gb:MN975262|Organism:Severe      TACTGTCCAAGACATGTGATCTGCACCTCTGAAGACATGCTTAACCCTAATTATGAAGAT   10222
gb:MT106052|Organism:Severe      TACTGTCCAAGACATGTGATCTGCACCTCTGAAGACATGCTTAACCCTAATTATGAAGAT   10222
gb:LC522975|Organism:Severe      TACTGTCCAAGACATGTGATCTGCACCTCTGAAGACATGCTTAACCCTAATTATGAAGAT   10219
gb:LC522973|Organism:Severe      TACTGTCCAAGACATGTGATCTGCACCTCTGAAGACATGCTTAACCCTAATTATGAAGAT   10219
gb:LC522974|Organism:Severe      TACTGTCCAAGACATGTGATCTGCACCTCTGAAGACATGCTTAACCCTAATTATGAAGAT   10219
gb:MN985325|Organism:Severe      TACTGTCCAAGACATGTGATCTGCACCTCTGAAGACATGCTTAACCCTAATTATGAAGAT   10222
gb:MT020881|Organism:Severe      TACTGTCCAAGACATGTGATCTGCACCTCTGAAGACATGCTTAACCCTAATTATGAAGAT   10222
gb:MT020880|Organism:Severe      TACTGTCCAAGACATGTGATCTGCACCTCTGAAGACATGCTTAACCCTAATTATGAAGAT   10222
gb:MT066175|Organism:Severe      TACTGTCCAAGACATGTGATCTGCACCTCTGAAGACATGCTTAACCCTAATTATGAAGAT   10222
gb:MN997409|Organism:Severe      TACTGTCCAAGACATGTGATCTGCACCTCTGAAGACATGCTTAACCCTAATTATGAAGAT   10222
gb:MN938384|Organism:Severe      TACTGTCCAAGACATGTGATCTGCACCTCTGAAGACATGCTTAACCCTAATTATGAAGAT   10190
gb:MT044258|Organism:Severe      TACTGTCCAAGACATGTGATCTGCACCTCTGAAGACATGCTTAACCCTAATTATGAAGAT   10198
gb:MT039890|Organism:Severe      TACTGTCCAAGACATGTGATCTGCACCTCTGAAGACATGCTTAACCCTAATTATGAAGAT   10222
gb:MN988713|Organism:Severe      TACTGTCCAAGACATGTGATCTGCACCTCTGAAGACATGCTTAACCCTAATTATGAAGAT   10222
gb:LC521925|Organism:Severe      TACTGTCCAAGACATGTGATCTGCACCTCTGAAGACATGCTTAACCCTAATTATGAAGAT   10195
gb:MT093571|Organism:Severe      TACTGTCCAAGACATGTGATCTGCACCTCTGAAGACATGCTTAACCCTAATTATGAAGAT   10222
gb:MT039887|Organism:Severe      TACTGTCCAAGACATGTGATCTGCACCTCTGAAGACATGCTTAACCCTAATTATGAAGAT   10222
gb:MT019530|Organism:Severe      TACTGTCCAAGACATGTGATCTGCACCTCTGAAGACATGCTTAACCCTAATTATGAAGAT   10222
gb:MT039888|Organism:Severe      TACTGTCCAAGACATGTGATCTGCACCTCTGAAGACATGCTTAACCCTAATTATGAAGAT   10222
gb:LC522972|Organism:Severe      TACTGTCCAAGACATGTGATCTGCACCTCTGAAGACATGCTTAACCCTAATTATGAAGAT   10219
gb:MT027063|Organism:Severe      TACTGTCCAAGACATGTGATCTGCACCTCTGAAGACATGCTTAACCCTAATTATGAAGAT   10222
gb:MT027062|Organism:Severe      TACTGTCCAAGACATGTGATCTGCACCTCTGAAGACATGCTTAACCCTAATTATGAAGAT   10222
gb:MT019529|Organism:Severe      TACTGTCCAAGACATGTGATCTGCACCTCTGAAGACATGCTTAACCCTAATTATGAAGAT   10222
gb:MN996529|Organism:Severe      TACTGTCCAAGACATGTGATCTGCACCTCTGAAGACATGCTTAACCCTAATTATGAAGAT   10210
gb:MN996531|Organism:Severe      TACTGTCCAAGACATGTGATCTGCACCTCTGAAGACATGCTTAACCCTAATTATGAAGAT   10209
gb:MT066176|Organism:Severe      TACTGTCCAAGACATGTGATCTGCACCTCTGAAGACATGCTTAACCCTAATTATGAAGAT   10222
gb:MT027064|Organism:Severe      TACTGTCCAAGACATGTGATCTGCACCTCTGAAGACATGCTTAACCCTAATTATGAAGAT   10222
gb:MN994468|Organism:Severe      TACTGTCCAAGACATGTGATCTGCACCTCTGAAGACATGCTTAACCCTAATTATGAAGAT   10222
gb:MT072688|Organism:Severe      TACTGTCCAAGACATGTGATCTGCACCTCTGAAGACATGCTTAACCCTAATTATGAAGAT   10207
gb:MN996527|Organism:Severe      TACTGTCCAAGACATGTGATCTGCACCTCTGAAGACATGCTTAACCCTAATTATGAAGAT   10189
gb:MT093631|Organism:Severe      TACTGTCCAAGACATGTGATCTGCACCTCTGAAGACATGCTTAACCCTAATTATGAAGAT   10260
gb:MT106053|Organism:Severe      TACTGTCCAAGACATGTGATCTGCACCTCTGAAGACATGCTTAACCCTAATTATGAAGAT   10222
gb:MT019533|Organism:Severe      TACTGTCCAAGACATGTGATCTGCACCTCTGAAGACATGCTTAACCCTAATTATGAAGAT   10222
gb:MT019531|Organism:Severe      TACTGTCCAAGACATGTGATCTGCACCTCTGAAGACATGCTTAACCCTAATTATGAAGAT   10222
gb:MN996528|Organism:Severe      TACTGTCCAAGACATGTGATCTGCACCTCTGAAGACATGCTTAACCCTAATTATGAAGAT   10222
```

```
gb:MN996530|Organism:Severe      TACTGTCCAAGACATGTGATCTGCACCTCTGAAGACATGCTTAACCCTAATTATGAAGAT   10208
gb:MN908947|Organism:Severe      TACTGTCCAAGACATGTGATCTGCACCTCTGAAGACATGCTTAACCCTAATTATGAAGAT   10222
gb:MT019532|Organism:Severe      TACTGTCCAAGACATGTGATCTGCACCTCTGAAGACATGCTTAACCCTAATTATGAAGAT   10222
                                 ************************************************************

gb:MT020781|Organism:Severe      TTACTCATTCGTAAGTCTAATCATAATTTCTTGGTACAGGCTGGTAATGTTCAACTCAGG   10270
gb:MT007544|Organism:Severe      TTACTCATTCGTAAGTCTAATCATAATTTCTTGGTACAGGCTGGTAATGTTCAACTCAGG   10282
gb:MN994467|Organism:Severe      TTACTCATTCGTAAGTCTAATCATAATTTCTTGGTACAGGCTGGTAATGTTCAACTCAGG   10282
gb:MT044257|Organism:Severe      TTACTCATTCGTAAGTCTAATCATAATTTCTTGGTACAGGCTGGTAATGTTCAACTCAGG   10282
gb:MT106054|Organism:Severe      TTACTCATTCGTAAGTCTAATCATAATTTCTTGGTACAGGCTGGTAATGTTCAACTCAGG   10282
gb:MT049951|Organism:Severe      TTACTCATTCGTAAGTCTAATCATAATTTCTTGGTACAGGCTGGTAATGTTCAACTCAGG   10282
gb:MN975262|Organism:Severe      TTACTCATTCGTAAGTCTAATCATAATTTCTTGGTACAGGCTGGTAATGTTCAACTCAGG   10282
gb:MT106052|Organism:Severe      TTACTCATTCGTAAGTCTAATCATAATTTCTTGGTACAGGCTGGTAATGTTCAACTCAGG   10282
gb:LC522975|Organism:Severe      TTACTCATTCGTAAGTCTAATCATAATTTCTTGGTACAGGCTGGTAATGTTCAACTCAGG   10279
gb:LC522973|Organism:Severe      TTACTCATTCGTAAGTCTAATCATAATTTCTTGGTACAGGCTGGTAATGTTCAACTCAGG   10279
gb:LC522974|Organism:Severe      TTACTCATTCGTAAGTCTAATCATAATTTCTTGGTACAGGCTGGTAATGTTCAACTCAGG   10279
gb:MN985325|Organism:Severe      TTACTCATTCGTAAGTCTAATCATAATTTCTTGGTACAGGCTGGTAATGTTCAACTCAGG   10282
gb:MT020881|Organism:Severe      TTACTCATTCGTAAGTCTAATCATAATTTCTTGGTACAGGCTGGTAATGTTCAACTCAGG   10282
gb:MT020880|Organism:Severe      TTACTCATTCGTAAGTCTAATCATAATTTCTTGGTACAGGCTGGTAATGTTCAACTCAGG   10282
gb:MT066175|Organism:Severe      TTACTCATTCGTAAGTCTAATCATAATTTCTTGGTACAGGCTGGTAATGTTCAACTCAGG   10282
gb:MN997409|Organism:Severe      TTACTCATTCGTAAGTCTAATCATAATTTCTTGGTACAGGCTGGTAATGTTCAACTCAGG   10282
gb:MN938384|Organism:Severe      TTACTCATTCGTAAGTCTAATCATAATTTCTTGGTACAGGCTGGTAATGTTCAACTCAGG   10250
gb:MT044258|Organism:Severe      TTACTCATTCGTAAGTCTAATCATAATTTCTTGGTACAGGCTGGTAATGTTCAACTCAGG   10258
gb:MT039890|Organism:Severe      TTACTCATTCGTAAGTCTAATCATAATTTCTTGGTACAGGCTGGTAATGTTCAACTCAGG   10282
gb:MN988713|Organism:Severe      TTACTCATTCGTAAGTCTAATCATAATTTCTTGGTACAGGCTGGTAATGTTCAACTCAGG   10282
gb:LC521925|Organism:Severe      TTACTCATTCGTAAGTCTAATCATAATTTCTTGGTACAGGCTGGTAATGTTCAACTCAGG   10255
gb:MT093571|Organism:Severe      TTACTCATTCGTAAGTCTAATCATAATTTCTTGGTACAGGCTGGTAATGTTCAACTCAGG   10282
gb:MT039887|Organism:Severe      TTACTCATTCGTAAGTCTAATCATAATTTCTTGGTACAGGCTGGTAATGTTCAACTCAGG   10282
gb:MT019530|Organism:Severe      TTACTCATTCGTAAGTCTAATCATAATTTCTTGGTACAGGCTGGTAATGTTCAACTCAGG   10282
gb:MT039888|Organism:Severe      TTACTCATTCGTAAGTCTAATCATAATTTCTTGGTACAGGCTGGTAATGTTCAACTCAGG   10282
gb:LC522972|Organism:Severe      TTACTCATTCGTAAGTCTAATCATAATTTCTTGGTACAGGCTGGTAATGTTCAACTCAGG   10279
gb:MT027063|Organism:Severe      TTACTCATTCGTAAGTCTAATCATAATTTCTTGGTACAGGCTGGTAATGTTCAACTCAGG   10282
gb:MT027062|Organism:Severe      TTACTCATTCGTAAGTCTAATCATAATTTCTTGGTACAGGCTGGTAATGTTCAACTCAGG   10282
gb:MT019529|Organism:Severe      TTACTCATTCGTAAGTCTAATCATAATTTCTTGGTACAGGCTGGTAATGTTCAACTCAGG   10282
gb:MN996529|Organism:Severe      TTACTCATTCGTAAGTCTAATCATAATTTCTTGGTACAGGCTGGTAATGTTCAACTCAGG   10270
gb:MN996531|Organism:Severe      TTACTCATTCGTAAGTCTAATCATAATTTCTTGGTACAGGCTGGTAATGTTCAACTCAGG   10269
gb:MT066176|Organism:Severe      TTACTCATTCGTAAGTCTAATCATAATTTCTTGGTACAGGCTGGTAATGTTCAACTCAGG   10282
gb:MT027064|Organism:Severe      TTACTCATTCGTAAGTCTAATCATAATTTCTTGGTACAGGCTGGTAATGTTCAACTCAGG   10282
gb:MN994468|Organism:Severe      TTACTCATTCGTAAGTCTAATCATAATTTCTTGGTACAGGCTGGTAATGTTCAACTCAGG   10282
gb:MT072688|Organism:Severe      TTACTCATTCGTAAGTCTAATCATAATTTCTTGGTACAGGCTGGTAATGTTCAACTCAGG   10267
gb:MN996527|Organism:Severe      TTACTCATTCGTAAGTCTAATCATAATTTCTTGGTACAGGCTGGTAATGTTCAACTCAGG   10249
gb:MT093631|Organism:Severe      TTACTCATTCGTAAGTCTAATCATAATTTCTTGGTACAGGCTGGTAATGTTCAACTCAGG   10320
gb:MT106053|Organism:Severe      TTACTCATTCGTAAGTCTAATCATAATTTCTTGGTACAGGCTGGTAATGTTCAACTCAGG   10282
gb:MT019533|Organism:Severe      TTACTCATTCGTAAGTCTAATCATAATTTCTTGGTACAGGCTGGTAATGTTCAACTCAGG   10282
gb:MT019531|Organism:Severe      TTACTCATTCGTAAGTCTAATCATAATTTCTTGGTACAGGCTGGTAATGTTCAACTCAGG   10282
gb:MN996528|Organism:Severe      TTACTCATTCGTAAGTCTAATCATAATTTCTTGGTACAGGCTGGTAATGTTCAACTCAGG   10282
gb:MN996530|Organism:Severe      TTACTCATTCGTAAGTCTAATCATAATTTCTTGGTACAGGCTGGTAATGTTCAACTCAGG   10268
gb:MN908947|Organism:Severe      TTACTCATTCGTAAGTCTAATCATAATTTCTTGGTACAGGCTGGTAATGTTCAACTCAGG   10282
gb:MT019532|Organism:Severe      TTACTCATTCGTAAGTCTAATCATAATTTCTTGGTACAGGCTGGTAATGTTCAACTCAGG   10282
                                 ************************************************************

gb:MT020781|Organism:Severe      GTTATTGGACATTCTATGCAAAATTGTGTACTTAAGCTTAAGGTTGATACAGCCAATCCT   10330
gb:MT007544|Organism:Severe      GTTATTGGACATTCTATGCAAAATTGTGTACTTAAGCTTAAGGTTGATACAGCCAATCCT   10342
gb:MN994467|Organism:Severe      GTTATTGGACATTCTATGCAAAATTGTGTACTTAAGCTTAAGGTTGATACAGCCAATCCT   10342
gb:MT044257|Organism:Severe      GTTATTGGACATTCTATGCAAAATTGTGTACTTAAGCTTAAGGTTGATACAGCCAATCCT   10342
gb:MT106054|Organism:Severe      GTTATTGGACATTCTATGCAAAATTGTGTACTTAAGCTTAAGGTTGATACAGCCAATCCT   10342
gb:MT049951|Organism:Severe      GTTATTGGACATTCTATGCAAAATTGTGTACTTAAGCTTAAGGTTGATACAGCCAATCCT   10342
gb:MN975262|Organism:Severe      GTTATTGGACATTCTATGCAAAATTGTGTACTTAAGCTTAAGGTTGATACAGCCAATCCT   10342
gb:MT106052|Organism:Severe      GTTATTGGACATTCTATGCAAAATTGTGTACTTAAGCTTAAGGTTGATACAGCCAATCCT   10342
gb:LC522975|Organism:Severe      GTTATTGGACATTCTATGCAAAATTGTGTACTTAAGCTTAAGGTTGATACAGCCAATCCT   10339
gb:LC522973|Organism:Severe      GTTATTGGACATTCTATGCAAAATTGTGTACTTAAGCTTAAGGTTGATACAGCCAATCCT   10339
gb:LC522974|Organism:Severe      GTTATTGGACATTCTATGCAAAATTGTGTACTTAAGCTTAAGGTTGATACAGCCAATCCT   10339
gb:MN985325|Organism:Severe      GTTATTGGACATTCTATGCAAAATTGTGTACTTAAGCTTAAGGTTGATACAGCCAATCCT   10342
gb:MT020881|Organism:Severe      GTTATTGGACATTCTATGCAAAATTGTGTACTTAAGCTTAAGGTTGATACAGCCAATCCT   10342
gb:MT020880|Organism:Severe      GTTATTGGACATTCTATGCAAAATTGTGTACTTAAGCTTAAGGTTGATACAGCCAATCCT   10342
```

```
gb:MT066175|Organism:Severe      GTTATTGGACATTCTATGCAAAATTGTGTACTTAAGCTTAAGGTTGATACAGCCAATCCT   10342
gb:MN997409|Organism:Severe      GTTATTGGACATTCTATGCAAAATTGTGTACTTAAGCTTAAGGTTGATACAGCCAATCCT   10342
gb:MN938384|Organism:Severe      GTTATTGGACATTCTATGCAAAATTGTGTACTTAAGCTTAAGGTTGATACAGCCAATCCT   10310
gb:MT044258|Organism:Severe      GTTATTGGACATTCTATGCAAAATTGTGTACTTAAGCTTAAGGTTGATACAGCCAATCCT   10318
gb:MT039890|Organism:Severe      GTTATTGGACATTCTATGCAAAATTGTGTACTTAAGCTTAAGGTTGATACAGCCAATCCT   10342
gb:MN988713|Organism:Severe      GTTATTGGACATTCTATGCAAAATTGTGTACTTAAGCTTAAGGTTGATACAGCCAATCCT   10342
gb:LC521925|Organism:Severe      GTTATTGGACATTCTATGCAAAATTGTGTACTTAAGCTTAAGGTTGATACAGCCAATCCT   10315
gb:MT093571|Organism:Severe      GTTATTGGACATTCTATGCAAAATTGTGTACTTAAGCTTAAGGTTGATACAGCCAATCCT   10342
gb:MT039887|Organism:Severe      GTTATTGGACATTCTATGCAAAATTGTGTACTTAAGCTTAAGGTTGATACAGCCAATCCT   10342
gb:MT019530|Organism:Severe      GTTATTGGACATTCTATGCAAAATTGTGTACTTAAGCTTAAGGTTGATACAGCCAATCCT   10342
gb:MT039888|Organism:Severe      GTTATTGGACATTCTATGCAAAATTGTGTACTTAAGCTTAAGGTTGATACAGCCAATCCT   10342
gb:LC522972|Organism:Severe      GTTATTGGACATTCTATGCAAAATTGTGTACTTAAGCTTAAGGTTGATACAGCCAATCCT   10339
gb:MT027063|Organism:Severe      GTTATTGGACATTCTATGCAAAATTGTGTACTTAAGCTTAAGGTTGATACAGCCAATCCT   10342
gb:MT027062|Organism:Severe      GTTATTGGACATTCTATGCAAAATTGTGTACTTAAGCTTAAGGTTGATACAGCCAATCCT   10342
gb:MT019529|Organism:Severe      GTTATTGGACATTCTATGCAAAATTGTGTACTTAAGCTTAAGGTTGATACAGCCAATCCT   10342
gb:MN996529|Organism:Severe      GTTATTGGACATTCTATGCAAAATTGTGTACTTAAGCTTAAGGTTGATACAGCCAATCCT   10330
gb:MN996531|Organism:Severe      GTTATTGGACATTCTATGCAAAATTGTGTACTTAAGCTTAAGGTTGATACAGCCAATCCT   10329
gb:MT066176|Organism:Severe      GTTATTGGACATTCTATGCAAAATTGTGTACTTAAGCTTAAGGTTGATACAGCCAATCCT   10342
gb:MT027064|Organism:Severe      GTTATTGGACATTCTATGCAAAATTGTGTACTTAAGCTTAAGGTTGATACAGCCAATCCT   10342
gb:MN994468|Organism:Severe      GTTATTGGACATTCTATGCAAAATTGTGTACTTAAGCTTAAGGTTGATACAGCCAATCCT   10342
gb:MT072688|Organism:Severe      GTTATTGGACATTCTATGCAAAATTGTGTACTTAAGCTTAAGGTTGATACAGCCAATCCT   10327
gb:MN996527|Organism:Severe      GTTATTGGACATTCTATGCAAAATTGTGTACTTAAGCTTAAGGTTGATACAGCCAATCCT   10309
gb:MT093631|Organism:Severe      GTTATTGGACATTCTATGCAAAATTGTGTACTTAAGCTTAAGGTTGATACAGCCAATCCT   10380
gb:MT106053|Organism:Severe      GTTATTGGACATTCTATGCAAAATTGTGTACTTAAGCTTAAGGTTGATACAGCCAATCCT   10342
gb:MT019533|Organism:Severe      GTTATTGGACATTCTATGCAAAATTGTGTACTTAAGCTTAAGGTTGATACAGCCAATCCT   10342
gb:MT019531|Organism:Severe      GTTATTGGACATTCTATGCAAAATTGTGTACTTAAGCTTAAGGTTGATACAGCCAATCCT   10342
gb:MN996528|Organism:Severe      GTTATTGGACATTCTATGCAAAATTGTGTACTTAAGCTTAAGGTTGATACAGCCAATCCT   10342
gb:MN996530|Organism:Severe      GTTATTGGACATTCTATGCAAAATTGTGTACTTAAGCTTAAGGTTGATACAGCCAATCCT   10328
gb:MN908947|Organism:Severe      GTTATTGGACATTCTATGCAAAATTGTGTACTTAAGCTTAAGGTTGATACAGCCAATCCT   10342
gb:MT019532|Organism:Severe      GTTATTGGACATTCTATGCAAAATTGTGTACTTAAGCTTAAGGTTGATACAGCCAATCCT   10342
                                 ************************************************************

gb:MT020781|Organism:Severe      AAGACACCTAAGTATAAGTTTGTTCGCATTCAACCAGGACAGACTTTTTCAGTGTTAGCT   10390
gb:MT007544|Organism:Severe      AAGACACCTAAGTATAAGTTTGTTCGCATTCAACCAGGACAGACTTTTTCAGTGTTAGCT   10402
gb:MN994467|Organism:Severe      AAGACACCTAAGTATAAGTTTGTTCGCATTCAACCAGGACAGACTTTTTCAGTGTTAGCT   10402
gb:MT044257|Organism:Severe      AAGACACCTAAGTATAAGTTTGTTCGCATTCAACCAGGACAGACTTTTTCAGTGTTAGCT   10402
gb:MT106054|Organism:Severe      AAGACACCTAAGTATAAGTTTGTTCGCATTCAACCAGGACAGACTTTTTCAGTGTTAGCT   10402
gb:MT049951|Organism:Severe      AAGACACCTAAGTATAAGTTTGTTCGCATTCAACCAGGACAGACTTTTTCAGTGTTAGCT   10402
gb:MN975262|Organism:Severe      AAGACACCTAAGTATAAGTTTGTTCGCATTCAACCAGGACAGACTTTTTCAGTGTTAGCT   10402
gb:MT106052|Organism:Severe      AAGACACCTAAGTATAAGTTTGTTCGCATTCAACCAGGACAGACTTTTTCAGTGTTAGCT   10402
gb:LC522975|Organism:Severe      AAGACACCTAAGTATAAGTTTGTTCGCATTCAACCAGGACAGACTTTTTCAGTGTTAGCT   10399
gb:LC522973|Organism:Severe      AAGACACCTAAGTATAAGTTTGTTCGCATTCAACCAGGACAGACTTTTTCAGTGTTAGCT   10399
gb:LC522974|Organism:Severe      AAGACACCTAAGTATAAGTTTGTTCGCATTCAACCAGGACAGACTTTTTCAGTGTTAGCT   10399
gb:MN985325|Organism:Severe      AAGACACCTAAGTATAAGTTTGTTCGCATTCAACCAGGACAGACTTTTTCAGTGTTAGCT   10402
gb:MT020881|Organism:Severe      AAGACACCTAAGTATAAGTTTGTTCGCATTCAACCAGGACAGACTTTTTCAGTGTTAGCT   10402
gb:MT020880|Organism:Severe      AAGACACCTAAGTATAAGTTTGTTCGCATTCAACCAGGACAGACTTTTTCAGTGTTAGCT   10402
gb:MT066175|Organism:Severe      AAGACACCTAAGTATAAGTTTGTTCGCATTCAACCAGGACAGACTTTTTCAGTGTTAGCT   10402
gb:MN997409|Organism:Severe      AAGACACCTAAGTATAAGTTTGTTCGCATTCAACCAGGACAGACTTTTTCAGTGTTAGCT   10402
gb:MN938384|Organism:Severe      AAGACACCTAAGTATAAGTTTGTTCGCATTCAACCAGGACAGACTTTTTCAGTGTTAGCT   10370
gb:MT044258|Organism:Severe      AAGACACCTAAGTATAAGTTTGTTCGCATTCAACCAGGACAGACTTTTTCAGTGTTAGCT   10378
gb:MT039890|Organism:Severe      AAGACACCTAAGTATAAGTTTGTTCGCATTCAACCAGGACAGACTTTTTCAGTGTTAGCT   10402
gb:MN988713|Organism:Severe      AAGACACCTAAGTATAAGTTTGTTCGCATTCAACCAGGACAGACTTTTTCAGTGTTAGCT   10402
gb:LC521925|Organism:Severe      AAGACACCTAAGTATAAGTTTGTTCGCATTCAACCAGGACAGACTTTTTCAGTGTTAGCT   10375
gb:MT093571|Organism:Severe      AAGACACCTAAGTATAAGTTTGTTCGCATTCAACCAGGACAGACTTTTTCAGTGTTAGCT   10402
gb:MT039887|Organism:Severe      AAGACACCTAAGTATAAGTTTGTTCGCATTCAACCAGGACAGACTTTTTCAGTGTTAGCT   10402
gb:MT019530|Organism:Severe      AAGACACCTAAGTATAAGTTTGTTCGCATTCAACCAGGACAGACTTTTTCAGTGTTAGCT   10402
gb:MT039888|Organism:Severe      AAGACACCTAAGTATAAGTTTGTTCGCATTCAACCAGGACAGACTTTTTCAGTGTTAGCT   10402
gb:LC522972|Organism:Severe      AAGACACCTAAGTATAAGTTTGTTCGCATTCAACCAGGACAGACTTTTTCAGTGTTAGCT   10399
gb:MT027063|Organism:Severe      AAGACACCTAAGTATAAGTTTGTTCGCATTCAACCAGGACAGACTTTTTCAGTGTTAGCT   10402
gb:MT027062|Organism:Severe      AAGACACCTAAGTATAAGTTTGTTCGCATTCAACCAGGACAGACTTTTTCAGTGTTAGCT   10402
gb:MT019529|Organism:Severe      AAGACACCTAAGTATAAGTTTGTTCGCATTCAACCAGGACAGACTTTTTCAGTGTTAGCT   10402
gb:MN996529|Organism:Severe      AAGACACCTAAGTATAAGTTTGTTCGCATTCAACCAGGACAGACTTTTTCAGTGTTAGCT   10390
gb:MN996531|Organism:Severe      AAGACACCTAAGTATAAGTTTGTTCGCATTCAACCAGGACAGACTTTTTCAGTGTTAGCT   10389
gb:MT066176|Organism:Severe      AAGACACCTAAGTATAAGTTTGTTCGCATTCAACCAGGACAGACTTTTTCAGTGTTAGCT   10402
gb:MT027064|Organism:Severe      AAGACACCTAAGTATAAGTTTGTTCGCATTCAACCAGGACAGACTTTTTCAGTGTTAGCT   10402
```

```
gb:MN994468|Organism:Severe      AAGACACCTAAGTATAAGTTTGTTCGCATTCAACCAGGACAGACTTTTTCAGTGTTAGCT   10402
gb:MT072688|Organism:Severe      AAGACACCTAAGTATAAGTTTGTTCGCATTCAACCAGGACAGACTTTTTCAGTGTTAGCT   10387
gb:MN996527|Organism:Severe      AAGACACCTAAGTATAAGTTTGTTCGCATTCAACCAGGACAGACTTTTTCAGTGTTAGCT   10369
gb:MT093631|Organism:Severe      AAGACACCTAAGTATAAGTTTGTTCGCATTCAACCAGGACAGACTTTTTCAGTGTTAGCT   10440
gb:MT106053|Organism:Severe      AAGACACCTAAGTATAAGTTTGTTCGCATTCAACCAGGACAGACTTTTTCAGTGTTAGCT   10402
gb:MT019533|Organism:Severe      AAGACACCTAAGTATAAGTTTGTTCGCATTCAACCAGGACAGACTTTTTCAGTGTTAGCT   10402
gb:MT019531|Organism:Severe      AAGACACCTAAGTATAAGTTTGTTCGCATTCAACCAGGACAGACTTTTTCAGTGTTAGCT   10402
gb:MN996528|Organism:Severe      AAGACACCTAAGTATAAGTTTGTTCGCATTCAACCAGGACAGACTTTTTCAGTGTTAGCT   10402
gb:MN996530|Organism:Severe      AAGACACCTAAGTATAAGTTTGTTCGCATTCAACCAGGACAGACTTTTTCAGTGTTAGCT   10388
gb:MN908947|Organism:Severe      AAGACACCTAAGTATAAGTTTGTTCGCATTCAACCAGGACAGACTTTTTCAGTGTTAGCT   10402
gb:MT019532|Organism:Severe      AAGACACCTAAGTATAAGTTTGTTCGCATTCAACCAGGACAGACTTTTTCAGTGTTAGCT   10402
                                 ************************************************************

gb:MT020781|Organism:Severe      TGTTACAATGGTTCACCATCTGGTGTTTACCAATGTGCTATGAGGCCCAATTTCACTATT   10450
gb:MT007544|Organism:Severe      TGTTACAATGGTTCACCATCTGGTGTTTACCAATGTGCTATGAGGCCCAATTTCACTATT   10462
gb:MN994467|Organism:Severe      TGTTACAATGGTTCACCATCTGGTGTTTACCAATGTGCTATGAGGCCCAATTTCACTATT   10462
gb:MT044257|Organism:Severe      TGTTACAATGGTTCACCATCTGGTGTTTACCAATGTGCTATGAGGCCCAATTTCACTATT   10462
gb:MT106054|Organism:Severe      TGTTACAATGGTTCACCATCTGGTGTTTACCAATGTGCTATGAGGCCCAATTTCACTATT   10462
gb:MT049951|Organism:Severe      TGTTACAATGGTTCACCATCTGGTGTTTACCAATGTGCTATGAGGCCCAATTTCACTATT   10462
gb:MN975262|Organism:Severe      TGTTACAATGGTTCACCATCTGGTGTTTACCAATGTGCTATGAGGCCCAATTTCACTATT   10462
gb:MT106052|Organism:Severe      TGTTACAATGGTTCACCATCTGGTGTTTACCAATGTGCTATGAGGCCCAATTTCACTATT   10462
gb:LC522975|Organism:Severe      TGTTACAATGGTTCACCATCTGGTGTTTACCAATGTGCTATGAGGCCCAATTTCACTATT   10459
gb:LC522973|Organism:Severe      TGTTACAATGGTTCACCATCTGGTGTTTACCAATGTGCTATGAGGCCCAATTTCACTATT   10459
gb:LC522974|Organism:Severe      TGTTACAATGGTTCACCATCTGGTGTTTACCAATGTGCTATGAGGCCCAATTTCACTATT   10459
gb:MN985325|Organism:Severe      TGTTACAATGGTTCACCATCTGGTGTTTACCAATGTGCTATGAGGCCCAATTTCACTATT   10462
gb:MT020881|Organism:Severe      TGTTACAATGGTTCACCATCTGGTGTTTACCAATGTGCTATGAGGCCCAATTTCACTATT   10462
gb:MT020880|Organism:Severe      TGTTACAATGGTTCACCATCTGGTGTTTACCAATGTGCTATGAGGCCCAATTTCACTATT   10462
gb:MT066175|Organism:Severe      TGTTACAATGGTTCACCATCTGGTGTTTACCAATGTGCTATGAGGCCCAATTTCACTATT   10462
gb:MN997409|Organism:Severe      TGTTACAATGGTTCACCATCTGGTGTTTACCAATGTGCTATGAGGCCCAATTTCACTATT   10462
gb:MN938384|Organism:Severe      TGTTACAATGGTTCACCATCTGGTGTTTACCAATGTGCTATGAGGCCCAATTTCACTATT   10430
gb:MT044258|Organism:Severe      TGTTACAATGGTTCACCATCTGGTGTTTACCAATGTGCTATGAGGCCCAATTTCACTATT   10438
gb:MT039890|Organism:Severe      TGTTACAATGGTTCACCATCTGGTGTTTACCAATGTGCTATGAGGCCCAATTTCACTATT   10462
gb:MN988713|Organism:Severe      TGTTACAATGGTTCACCATCTGGTGTTTACCAATGTGCTATGAGGCCCAATTTCACTATT   10462
gb:LC521925|Organism:Severe      TGTTACAATGGTTCACCATCTGGTGTTTACCAATGTGCTATGAGGCCCAATTTCACTATT   10435
gb:MT093571|Organism:Severe      TGTTACAATGGTTCACCATCTGGTGTTTACCAATGTGCTATGAGGCCCAATTTCACTATT   10462
gb:MT039887|Organism:Severe      TGTTACAATGGTTCACCATCTGGTGTTTACCAATGTGCTATGAGGCCCAATTTCACTATT   10462
gb:MT019530|Organism:Severe      TGTTACAATGGTTCACCATCTGGTGTTTACCAATGTGCTATGAGGCCCAATTTCACTATT   10462
gb:MT039888|Organism:Severe      TGTTACAATGGTTCACCATCTGGTGTTTACCAATGTGCTATGAGGCCCAATTTCACTATT   10462
gb:LC522972|Organism:Severe      TGTTACAATGGTTCACCATCTGGTGTTTACCAATGTGCTATGAGGCCCAATTTCACTATT   10459
gb:MT027063|Organism:Severe      TGTTACAATGGTTCACCATCTGGTGTTTACCAATGTGCTATGAGGCCCAATTTCACTATT   10462
gb:MT027062|Organism:Severe      TGTTACAATGGTTCACCATCTGGTGTTTACCAATGTGCTATGAGGCCCAATTTCACTATT   10462
gb:MT019529|Organism:Severe      TGTTACAATGGTTCACCATCTGGTGTTTACCAATGTGCTATGAGGCCCAATTTCACTATT   10462
gb:MN996529|Organism:Severe      TGTTACAATGGTTCACCATCTGGTGTTTACCAATGTGCTATGAGGCCCAATTTCACTATT   10450
gb:MN996531|Organism:Severe      TGTTACAATGGTTCACCATCTGGTGTTTACCAATGTGCTATGAGGCCCAATTTCACTATT   10449
gb:MT066176|Organism:Severe      TGTTACAATGGTTCACCATCTGGTGTTTACCAATGTGCTATGAGGCCCAATTTCACTATT   10462
gb:MT027064|Organism:Severe      TGTTACAATGGTTCACCATCTGGTGTTTACCAATGTGCTATGAGGCCCAATTTCACTATT   10462
gb:MN994468|Organism:Severe      TGTTACAATGGTTCACCATCTGGTGTTTACCAATGTGCTATGAGGCCCAATTTCACTATT   10462
gb:MT072688|Organism:Severe      TGTTACAATGGTTCACCATCTGGTGTTTACCAATGTGCTATGAGGCCCAATTTCACTATT   10447
gb:MN996527|Organism:Severe      TGTTACAATGGTTCACCATCTGGTGTTTACCAATGTGCTATGAGGCCCAATTTCACTATT   10429
gb:MT093631|Organism:Severe      TGTTACAATGGTTCACCATCTGGTGTTTACCAATGTGCTATGAGGCCCAATTTCACTATT   10500
gb:MT106053|Organism:Severe      TGTTACAATGGTTCACCATCTGGTGTTTACCAATGTGCTATGAGGCCCAATTTCACTATT   10462
gb:MT019533|Organism:Severe      TGTTACAATGGTTCACCATCTGGTGTTTACCAATGTGCTATGAGGCCCAATTTCACTATT   10462
gb:MT019531|Organism:Severe      TGTTACAATGGTTCACCATCTGGTGTTTACCAATGTGCTATGAGGCCCAATTTCACTATT   10462
gb:MN996528|Organism:Severe      TGTTACAATGGTTCACCATCTGGTGTTTACCAATGTGCTATGAGGCCCAATTTCACTATT   10462
gb:MN996530|Organism:Severe      TGTTACAATGGTTCACCATCTGGTGTTTACCAATGTGCTATGAGGCCCAATTTCACTATT   10448
gb:MN908947|Organism:Severe      TGTTACAATGGTTCACCATCTGGTGTTTACCAATGTGCTATGAGGCCCAATTTCACTATT   10462
gb:MT019532|Organism:Severe      TGTTACAATGGTTCACCATCTGGTGTTTACCAATGTGCTATGAGGCCCAATTTCACTATT   10462
                                 ************************************************************

gb:MT020781|Organism:Severe      AAGGGTTCATTCCTTAATGGTTCATGTGGTAGTGTTGGTTTTAACATAGATTATGACTGT   10510
gb:MT007544|Organism:Severe      AAGGGTTCATTCCTTAATGGTTCATGTGGTAGTGTTGGTTTTAACATAGATTATGACTGT   10522
gb:MN994467|Organism:Severe      AAGGGTTCATTCCTTAATGGTTCATGTGGTAGTGTTGGTTTTAACATAGATTATGACTGT   10522
gb:MT044257|Organism:Severe      AAGGGTTCATTCCTTAATGGTTCATGTGGTAGTGTTGGTTTTAACATAGATTATGACTGT   10522
gb:MT106054|Organism:Severe      AAGGGTTCATTCCTTAATGGTTCATGTGGTAGTGTTGGTTTTAACATAGATTATGACTGT   10522
gb:MT049951|Organism:Severe      AAGGGTTCATTCCTTAATGGTTCATGTGGTAGTGTTGGTTTTAACATAGATTATGACTGT   10522
```

```
gb:MN975262|Organism:Severe      AAGGGTTCATTCCTTAATGGTTCATGTGGTAGTGTTGGTTTTAACATAGATTATGACTGT   10522
gb:MT106052|Organism:Severe      AAGGGTTCATTCCTTAATGGTTCATGTGGTAGTGTTGGTTTTAACATAGATTATGACTGT   10522
gb:LC522975|Organism:Severe      AAGGGTTCATTCCTTAATGGTTCATGTGGTAGTGTTGGTTTTAACATAGATTATGACTGT   10519
gb:LC522973|Organism:Severe      AAGGGTTCATTCCTTAATGGTTCATGTGGTAGTGTTGGTTTTAACATAGATTATGACTGT   10519
gb:LC522974|Organism:Severe      AAGGGTTCATTCCTTAATGGTTCATGTGGTAGTGTTGGTTTTAACATAGATTATGACTGT   10519
gb:MN985325|Organism:Severe      AAGGGTTCATTCCTTAATGGTTCATGTGGTAGTGTTGGTTTTAACATAGATTATGACTGT   10522
gb:MT020881|Organism:Severe      AAGGGTTCATTCCTTAATGGTTCATGTGGTAGTGTTGGTTTTAACATAGATTATGACTGT   10522
gb:MT020880|Organism:Severe      AAGGGTTCATTCCTTAATGGTTCATGTGGTAGTGTTGGTTTTAACATAGATTATGACTGT   10522
gb:MT066175|Organism:Severe      AAGGGTTCATTCCTTAATGGTTCATGTGGTAGTGTTGGTTTTAACATAGATTATGACTGT   10522
gb:MN997409|Organism:Severe      AAGGGTTCATTCCTTAATGGTTCATGTGGTAGTGTTGGTTTTAACATAGATTATGACTGT   10522
gb:MN938384|Organism:Severe      AAGGGTTCATTCCTTAATGGTTCATGTGGTAGTGTTGGTTTTAACATAGATTATGACTGT   10490
gb:MT044258|Organism:Severe      AAGGGTTCATTCCTTAATGGTTCATGTGGTAGTGTTGGTTTTAACATAGATTATGACTGT   10498
gb:MT039890|Organism:Severe      AAGGGTTCATTCCTTAATGGTTCATGTGGTAGTGTTGGTTTTAACATAGATTATGACTGT   10522
gb:MN988713|Organism:Severe      AAGGGTTCATTCCTTAATGGTTCATGTGGTAGTGTTGGTTTTAACATAGATTATGACTGT   10522
gb:LC521925|Organism:Severe      AAGGGTTCATTCCTTAATGGTTCATGTGGTAGTGTTGGTTTTAACATAGATTATGACTGT   10495
gb:MT093571|Organism:Severe      AAGGGTTCATTCCTTAATGGTTCATGTGGTAGTGTTGGTTTTAACATAGATTATGACTGT   10522
gb:MT039887|Organism:Severe      AAGGGTTCATTCCTTAATGGTTCATGTGGTAGTGTTGGTTTTAACATAGATTATGACTGT   10522
gb:MT019530|Organism:Severe      AAGGGTTCATTCCTTAATGGTTCATGTGGTAGTGTTGGTTTTAACATAGATTATGACTGT   10522
gb:MT039888|Organism:Severe      AAGGGTTCATTCCTTAATGGTTCATGTGGTAGTGTTGGTTTTAACATAGATTATGACTGT   10522
gb:LC522972|Organism:Severe      AAGGGTTCATTCCTTAATGGTTCATGTGGTAGTGTTGGTTTTAACATAGATTATGACTGT   10519
gb:MT027063|Organism:Severe      AAGGGTTCATTCCTTAATGGTTCATGTGGTAGTGTTGGTTTTAACATAGATTATGACTGT   10522
gb:MT027062|Organism:Severe      AAGGGTTCATTCCTTAATGGTTCATGTGGTAGTGTTGGTTTTAACATAGATTATGACTGT   10522
gb:MT019529|Organism:Severe      AAGGGTTCATTCCTTAATGGTTCATGTGGTAGTGTTGGTTTTAACATAGATTATGACTGT   10522
gb:MN996529|Organism:Severe      AAGGGTTCATTCCTTAATGGTTCATGTGGTAGTGTTGGTTTTAACATAGATTATGACTGT   10510
gb:MN996531|Organism:Severe      AAGGGTTCATTCCTTAATGGTTCATGTGGTAGTGTTGGTTTTAACATAGATTATGACTGT   10509
gb:MT066176|Organism:Severe      AAGGGTTCATTCCTTAATGGTTCATGTGGTAGTGTTGGTTTTAACATAGATTATGACTGT   10522
gb:MT027064|Organism:Severe      AAGGGTTCATTCCTTAATGGTTCATGTGGTAGTGTTGGTTTTAACATAGATTATGACTGT   10522
gb:MN994468|Organism:Severe      AAGGGTTCATTCCTTAATGGTTCATGTGGTAGTGTTGGTTTTAACATAGATTATGACTGT   10522
gb:MT072688|Organism:Severe      AAGGGTTCATTCCTTAATGGTTCATGTGGTAGTGTTGGTTTTAACATAGATTATGACTGT   10507
gb:MN996527|Organism:Severe      AAGGGTTCATTCCTTAATGGTTCATGTGGTAGTGTTGGTTTTAACATAGATTATGACTGT   10489
gb:MT093631|Organism:Severe      AAGGGTTCATTCCTTAATGGTTCATGTGGTAGTGTTGGTTTTAACATAGATTATGACTGT   10560
gb:MT106053|Organism:Severe      AAGGGTTCATTCCTTAATGGTTCATGTGGTAGTGTTGGTTTTAACATAGATTATGACTGT   10522
gb:MT019533|Organism:Severe      AAGGGTTCATTCCTTAATGGTTCATGTGGTAGTGTTGGTTTTAACATAGATTATGACTGT   10522
gb:MT019531|Organism:Severe      AAGGGTTCATTCCTTAATGGTTCATGTGGTAGTGTTGGTTTTAACATAGATTATGACTGT   10522
gb:MN996528|Organism:Severe      AAGGGTTCATTCCTTAATGGTTCATGTGGTAGTGTTGGTTTTAACATAGATTATGACTGT   10522
gb:MN996530|Organism:Severe      AAGGGTTCATTCCTTAATGGTTCATGTGGTAGTGTTGGTTTTAACATAGATTATGACTGT   10508
gb:MN908947|Organism:Severe      AAGGGTTCATTCCTTAATGGTTCATGTGGTAGTGTTGGTTTTAACATAGATTATGACTGT   10522
gb:MT019532|Organism:Severe      AAGGGTTCATTCCTTAATGGTTCATGTGGTAGTGTTGGTTTTAACATAGATTATGACTGT   10522
                                 ************************************************************

gb:MT020781|Organism:Severe      GTCTCTTTTTGTTACATGCACCATATGGAATTACCAACTGGAGTTCATGCTGGCACAGAC   10570
gb:MT007544|Organism:Severe      GTCTCTTTTTGTTACATGCACCATATGGAATTACCAACTGGAGTTCATGCTGGCACAGAC   10582
gb:MN994467|Organism:Severe      GTCTCTTTTTGTTACATGCACCATATGGAATTACCAACTGGAGTTCATGCTGGCACAGAC   10582
gb:MT044257|Organism:Severe      GTCTCTTTTTGTTACATGCACCATATGGAATTACCAACTGGAGTTCATGCTGGCACAGAC   10582
gb:MT106054|Organism:Severe      GTCTCTTTTTGTTACATGCACCATATGGAATTACCAACTGGAGTTCATGCTGGCACAGAC   10582
gb:MT049951|Organism:Severe      GTCTCTTTTTGTTACATGCACCATATGGAATTACCAACTGGAGTTCATGCTGGCACAGAC   10582
gb:MN975262|Organism:Severe      GTCTCTTTTTGTTACATGCACCATATGGAATTACCAACTGGAGTTCATGCTGGCACAGAC   10582
gb:MT106052|Organism:Severe      GTCTCTTTTTGTTACATGCACCATATGGAATTACCAACTGGAGTTCATGCTGGCACAGAC   10582
gb:LC522975|Organism:Severe      GTCTCTTTTTGTTACATGCACCATATGGAATTACCAACTGGAGTTCATGCTGGCACAGAC   10579
gb:LC522973|Organism:Severe      GTCTCTTTTTGTTACATGCACCATATGGAATTACCAACTGGAGTTCATGCTGGCACAGAC   10579
gb:LC522974|Organism:Severe      GTCTCTTTTTGTTACATGCACCATATGGAATTACCAACTGGAGTTCATGCTGGCACAGAC   10579
gb:MN985325|Organism:Severe      GTCTCTTTTTGTTACATGCACCATATGGAATTACCAACTGGAGTTCATGCTGGCACAGAC   10582
gb:MT020881|Organism:Severe      GTCTCTTTTTGTTACATGCACCATATGGAATTACCAACTGGAGTTCATGCTGGCACAGAC   10582
gb:MT020880|Organism:Severe      GTCTCTTTTTGTTACATGCACCATATGGAATTACCAACTGGAGTTCATGCTGGCACAGAC   10582
gb:MT066175|Organism:Severe      GTCTCTTTTTGTTACATGCACCATATGGAATTACCAACTGGAGTTCATGCTGGCACAGAC   10582
gb:MN997409|Organism:Severe      GTCTCTTTTTGTTACATGCACCATATGGAATTACCAACTGGAGTTCATGCTGGCACAGAC   10582
gb:MN938384|Organism:Severe      GTCTCTTTTTGTTACATGCACCATATGGAATTACCAACTGGAGTTCATGCTGGCACAGAC   10550
gb:MT044258|Organism:Severe      GTCTCTTTTTGTTACATGCACCATATGGAATTACCAACTGGAGTTCATGCTGGCACAGAC   10558
gb:MT039890|Organism:Severe      GTCTCTTTTTGTTACATGCACCATATGGAATTACCAACTGGAGTTCATGCTGGCACAGAC   10582
gb:MN988713|Organism:Severe      GTCTCTTTTTGTTACATGCACCATATGGAATTACCAACTGGAGTTCATGCTGGCACAGAC   10582
gb:LC521925|Organism:Severe      GTCTCTTTTTGTTACATGCACCATATGGAATTACCAACTGGAGTTCATGCTGGCACAGAC   10555
gb:MT093571|Organism:Severe      GTCTCTTTTTGTTACATGCACCATATGGAATTACCAACTGGAGTTCATGCTGGCACAGAC   10582
gb:MT039887|Organism:Severe      GTCTCTTTTTGTTACATGCACCATATGGAATTACCAACTGGAGTTCATGCTGGCACAGAC   10582
gb:MT019530|Organism:Severe      GTCTCTTTTTGTTACATGCACCATATGGAATTACCAACTGGAGTTCATGCTGGCACAGAC   10582
gb:MT039888|Organism:Severe      GTCTCTTTTTGTTACATGCACCATATGGAATTACCAACTGGAGTTCATGCTGGCACAGAC   10582
```

```
gb:LC522972|Organism:Severe      GTCTCTTTTTGTTACATGCACCATATGGAATTACCAACTGGAGTTCATGCTGGCACAGAC   10579
gb:MT027063|Organism:Severe      GTCTCTTTTTGTTACATGCACCATATGGAATTACCAACTGGAGTTCATGCTGGCACAGAC   10582
gb:MT027062|Organism:Severe      GTCTCTTTTTGTTACATGCACCATATGGAATTACCAACTGGAGTTCATGCTGGCACAGAC   10582
gb:MT019529|Organism:Severe      GTCTCTTTTTGTTACATGCACCATATGGAATTACCAACTGGAGTTCATGCTGGCACAGAC   10582
gb:MN996529|Organism:Severe      GTCTCTTTTTGTTACATGCACCATATGGAATTACCAACTGGAGTTCATGCTGGCACAGAC   10570
gb:MN996531|Organism:Severe      GTCTCTTTTTGTTACATGCACCATATGGAATTACCAACTGGAGTTCATGCTGGCACAGAC   10569
gb:MT066176|Organism:Severe      GTCTCTTTTTGTTACATGCACCATATGGAATTACCAACTGGAGTTCATGCTGGCACAGAC   10582
gb:MT027064|Organism:Severe      GTCTCTTTTTGTTACATGCACCATATGGAATTACCAACTGGAGTTCATGCTGGCACAGAC   10582
gb:MN994468|Organism:Severe      GTCTCTTTTTGTTACATGCACCATATGGAATTACCAACTGGAGTTCATGCTGGCACAGAC   10582
gb:MT072688|Organism:Severe      GTCTCTTTTTGTTACATGCACCATATGGAATTACCAACTGGAGTTCATGCTGGCACAGAC   10567
gb:MN996527|Organism:Severe      GTCTCTTTTTGTTACATGCACCATATGGAATTACCAACTGGAGTTCATGCTGGCACAGAC   10549
gb:MT093631|Organism:Severe      GTCTCTTTTTGTTACATGCACCATATGGAATTACCAACTGGAGTTCATGCTGGCACAGAC   10620
gb:MT106053|Organism:Severe      GTCTCTTTTTGTTACATGCACCATATGGAATTACCAACTGGAGTTCATGCTGGCACAGAC   10582
gb:MT019533|Organism:Severe      GTCTCTTTTTGTTACATGCACCATATGGAATTACCAACTGGAGTTCATGCTGGCACAGAC   10582
gb:MT019531|Organism:Severe      GTCTCTTTTTGTTACATGCACCATATGGAATTACCAACTGGAGTTCATGCTGGCACAGAC   10582
gb:MN996528|Organism:Severe      GTCTCTTTTTGTTACATGCACCATATGGAATTACCAACTGGAGTTCATGCTGGCACAGAC   10582
gb:MN996530|Organism:Severe      GTCTCTTTTTGTTACATGCACCATATGGAATTACCAACTGGAGTTCATGCTGGCACAGAC   10568
gb:MN908947|Organism:Severe      GTCTCTTTTTGTTACATGCACCATATGGAATTACCAACTGGAGTTCATGCTGGCACAGAC   10582
gb:MT019532|Organism:Severe      GTCTCTTTTTGTTACATGCACCATATGGAATTACCAACTGGAGTTCATGCTGGCACAGAC   10582
                                 ************************************************************

gb:MT020781|Organism:Severe      TTAGAAGGTAACTTTTATGGACCTTTTGTTGACAGGCAAACAGCACAAGCAGCTGGTACG   10630
gb:MT007544|Organism:Severe      TTAGAAGGTAACTTTTATGGACCTTTTGTTGACAGGCAAACAGCACAAGCAGCTGGTACG   10642
gb:MN994467|Organism:Severe      TTAGAAGGTAACTTTTATGGACCTTTTGTTGACAGGCAAACAGCACAAGCAGCTGGTACG   10642
gb:MT044257|Organism:Severe      TTAGAAGGTAACTTTTATGGACCTTTTGTTGACAGGCAAACAGCACAAGCAGCTGGTACG   10642
gb:MT106054|Organism:Severe      TTAGAAGGTAACTTTTATGGACCTTTTGTTGACAGGCAAACAGCACAAGCAGCTGGTACG   10642
gb:MT049951|Organism:Severe      TTAGAAGGTAACTTTTATGGACCTTTTGTTGACAGGCAAACAGCACAAGCAGCTGGTACG   10642
gb:MN975262|Organism:Severe      TTAGAAGGTAACTTTTATGGACCTTTTGTTGACAGGCAAACAGCACAAGCAGCTGGTACG   10642
gb:MT106052|Organism:Severe      TTAGAAGGTAACTTTTATGGACCTTTTGTTGACAGGCAAACAGCACAAGCAGCTGGTACG   10642
gb:LC522975|Organism:Severe      TTAGAAGGTAACTTTTATGGACCTTTTGTTGACAGGCAAACAGCACAAGCAGCTGGTACG   10639
gb:LC522973|Organism:Severe      TTAGAAGGTAACTTTTATGGACCTTTTGTTGACAGGCAAACAGCACAAGCAGCTGGTACG   10639
gb:LC522974|Organism:Severe      TTAGAAGGTAACTTTTATGGACCTTTTGTTGACAGGCAAACAGCACAAGCAGCTGGTACG   10639
gb:MN985325|Organism:Severe      TTAGAAGGTAACTTTTATGGACCTTTTGTTGACAGGCAAACAGCACAAGCAGCTGGTACG   10642
gb:MT020881|Organism:Severe      TTAGAAGGTAACTTTTATGGACCTTTTGTTGACAGGCAAACAGCACAAGCAGCTGGTACG   10642
gb:MT020880|Organism:Severe      TTAGAAGGTAACTTTTATGGACCTTTTGTTGACAGGCAAACAGCACAAGCAGCTGGTACG   10642
gb:MT066175|Organism:Severe      TTAGAAGGTAACTTTTATGGACCTTTTGTTGACAGGCAAACAGCACAAGCAGCTGGTACG   10642
gb:MN997409|Organism:Severe      TTAGAAGGTAACTTTTATGGACCTTTTGTTGACAGGCAAACAGCACAAGCAGCTGGTACG   10642
gb:MN938384|Organism:Severe      TTAGAAGGTAACTTTTATGGACCTTTTGTTGACAGGCAAACAGCACAAGCAGCTGGTACG   10610
gb:MT044258|Organism:Severe      TTAGAAGGTAACTTTTATGGACCTTTTGTTGACAGGCAAACAGCACAAGCAGCTGGTACG   10618
gb:MT039890|Organism:Severe      TTAGAAGGTAACTTTTATGGACCTTTTGTTGACAGGCAAACAGCACAAGCAGCTGGTACG   10642
gb:MN988713|Organism:Severe      TTAGAAGGTAACTTTTATGGACCTTTTGTTGACAGGCAAACAGCACAAGCAGCTGGTACG   10642
gb:LC521925|Organism:Severe      TTAGAAGGTAACTTTTATGGACCTTTTGTTGACAGGCAAACAGCACAAGCAGCTGGTACG   10615
gb:MT093571|Organism:Severe      TTAGAAGGTAACTTTTATGGACCTTTTGTTGACAGGCAAACAGCACAAGCAGCTGGTACG   10642
gb:MT039887|Organism:Severe      TTAGAAGGTAACTTTTATGGACCTTTTGTTGACAGGCAAACAGCACAAGCAGCTGGTACG   10642
gb:MT019530|Organism:Severe      TTAGAAGGTAACTTTTATGGACCTTTTGTTGACAGGCAAACAGCACAAGCAGCTGGTACG   10642
gb:MT039888|Organism:Severe      TTAGAAGGTAACTTTTATGGACCTTTTGTTGACAGGCAAACAGCACAAGCAGCTGGTACG   10642
gb:LC522972|Organism:Severe      TTAGAAGGTAACTTTTATGGACCTTTTGTTGACAGGCAAACAGCACAAGCAGCTGGTACG   10639
gb:MT027063|Organism:Severe      TTAGAAGGTAACTTTTATGGACCTTTTGTTGACAGGCAAACAGCACAAGCAGCTGGTACG   10642
gb:MT027062|Organism:Severe      TTAGAAGGTAACTTTTATGGACCTTTTGTTGACAGGCAAACAGCACAAGCAGCTGGTACG   10642
gb:MT019529|Organism:Severe      TTAGAAGGTAACTTTTATGGACCTTTTGTTGACAGGCAAACAGCACAAGCAGCTGGTACG   10642
gb:MN996529|Organism:Severe      TTAGAAGGTAACTTTTATGGACCTTTTGTTGACAGGCAAACAGCACAAGCAGCTGGTACG   10630
gb:MN996531|Organism:Severe      TTAGAAGGTAACTTTTATGGACCTTTTGTTGACAGGCAAACAGCACAAGCAGCTGGTACG   10629
gb:MT066176|Organism:Severe      TTAGAAGGTAACTTTTATGGACCTTTTGTTGACAGGCAAACAGCACAAGCAGCTGGTACG   10642
gb:MT027064|Organism:Severe      TTAGAAGGTAACTTTTATGGACCTTTTGTTGACAGGCAAACAGCACAAGCAGCTGGTACG   10642
gb:MN994468|Organism:Severe      TTAGAAGGTAACTTTTATGGACCTTTTGTTGACAGGCAAACAGCACAAGCAGCTGGTACG   10642
gb:MT072688|Organism:Severe      TTAGAAGGTAACTTTTATGGACCTTTTGTTGACAGGCAAACAGCACAAGCAGCTGGTACG   10627
gb:MN996527|Organism:Severe      TTAGAAGGTAACTTTTATGGACCTTTTGTTGACAGGCAAACAGCACAAGCAGCTGGTACG   10609
gb:MT093631|Organism:Severe      TTAGAAGGTAACTTTTATGGACCTTTTGTTGACAGGCAAACAGCACAAGCAGCTGGTACG   10680
gb:MT106053|Organism:Severe      TTAGAAGGTAACTTTTATGGACCTTTTGTTGACAGGCAAACAGCACAAGCAGCTGGTACG   10642
gb:MT019533|Organism:Severe      TTAGAAGGTAACTTTTATGGACCTTTTGTTGACAGGCAAACAGCACAAGCAGCTGGTACG   10642
gb:MT019531|Organism:Severe      TTAGAAGGTAACTTTTATGGACCTTTTGTTGACAGGCAAACAGCACAAGCAGCTGGTACG   10642
gb:MN996528|Organism:Severe      TTAGAAGGTAACTTTTATGGACCTTTTGTTGACAGGCAAACAGCACAAGCAGCTGGTACG   10642
gb:MN996530|Organism:Severe      TTAGAAGGTAACTTTTATGGACCTTTTGTTGACAGGCAAACAGCACAAGCAGCTGGTACG   10628
gb:MN908947|Organism:Severe      TTAGAAGGTAACTTTTATGGACCTTTTGTTGACAGGCAAACAGCACAAGCAGCTGGTACG   10642
gb:MT019532|Organism:Severe      TTAGAAGGTAACTTTTATGGACCTTTTGTTGACAGGCAAACAGCACAAGCAGCTGGTACG   10642
```

```
                                 ************************************************************

gb:MT020781|Organism:Severe      GACACAACTATTACAGTTAATGTTTTAGCTTGGTTGTACGCTGCTGTTATAAATGGAGAC   10690
gb:MT007544|Organism:Severe      GACACAACTATTACAGTTAATGTTTTAGCTTGGTTGTACGCTGCTGTTATAAATGGAGAC   10702
gb:MN994467|Organism:Severe      GACACAACTATTACAGTTAATGTTTTAGCTTGGTTGTACGCTGCTGTTATAAATGGAGAC   10702
gb:MT044257|Organism:Severe      GACACAACTATTACAGTTAATGTTTTAGCTTGGTTGTACGCTGCTGTTATAAATGGAGAC   10702
gb:MT106054|Organism:Severe      GACACAACTATTACAGTTAATGTTTTAGCTTGGTTGTACGCTGCTGTTATAAATGGAGAC   10702
gb:MT049951|Organism:Severe      GACACAACTATTACAGTTAATGTTTTAGCTTGGTTGTACGCTGCTGTTATAAATGGAGAC   10702
gb:MN975262|Organism:Severe      GACACAACTATTACAGTTAATGTTTTAGCTTGGTTGTACGCTGCTGTTATAAATGGAGAC   10702
gb:MT106052|Organism:Severe      GACACAACTATTACAGTTAATGTTTTAGCTTGGTTGTACGCTGCTGTTATAAATGGAGAC   10702
gb:LC522975|Organism:Severe      GACACAACTATTACAGTTAATGTTTTAGCTTGGTTGTACGCTGCTGTTATAAATGGAGAC   10699
gb:LC522973|Organism:Severe      GACACAACTATTACAGTTAATGTTTTAGCTTGGTTGTACGCTGCTGTTATAAATGGAGAC   10699
gb:LC522974|Organism:Severe      GACACAACTATTACAGTTAATGTTTTAGCTTGGTTGTACGCTGCTGTTATAAATGGAGAC   10699
gb:MN985325|Organism:Severe      GACACAACTATTACAGTTAATGTTTTAGCTTGGTTGTACGCTGCTGTTATAAATGGAGAC   10702
gb:MT020881|Organism:Severe      GACACAACTATTACAGTTAATGTTTTAGCTTGGTTGTACGCTGCTGTTATAAATGGAGAC   10702
gb:MT020880|Organism:Severe      GACACAACTATTACAGTTAATGTTTTAGCTTGGTTGTACGCTGCTGTTATAAATGGAGAC   10702
gb:MT066175|Organism:Severe      GACACAACTATTACAGTTAATGTTTTAGCTTGGTTGTACGCTGCTGTTATAAATGGAGAC   10702
gb:MN997409|Organism:Severe      GACACAACTATTACAGTTAATGTTTTAGCTTGGTTGTACGCTGCTGTTATAAATGGAGAC   10702
gb:MN938384|Organism:Severe      GACACAACTATTACAGTTAATGTTTTAGCTTGGTTGTACGCTGCTGTTATAAATGGAGAC   10670
gb:MT044258|Organism:Severe      GACACAACTATTACAGTTAATGTTTTAGCTTGGTTGTACGCTGCTGTTATAAATGGAGAC   10678
gb:MT039890|Organism:Severe      GACACAACTATTACAGTTAATGTTTTAGCTTGGTTGTACGCTGCTGTTATAAATGGAGAC   10702
gb:MN988713|Organism:Severe      GACACAACTATTACAGTTAATGTTTTAGCTTGGTTGTACGCTGCTGTTATAAATGGAGAC   10702
gb:LC521925|Organism:Severe      GACACAACTATTACAGTTAATGTTTTAGCTTGGTTGTACGCTGCTGTTATAAATGGAGAC   10675
gb:MT093571|Organism:Severe      GACACAACTATTACAGTTAATGTTTTAGCTTGGTTGTACGCTGCTGTTATAAATGGAGAC   10702
gb:MT039887|Organism:Severe      GACACAACTATTACAGTTAATGTTTTAGCTTGGTTGTACGCTGCTGTTATAAATGGAGAC   10702
gb:MT019530|Organism:Severe      GACACAACTATTACAGTTAATGTTTTAGCTTGGTTGTACGCTGCTGTTATAAATGGAGAC   10702
gb:MT039888|Organism:Severe      GACACAACTATTACAGTTAATGTTTTAGCTTGGTTGTACGCTGCTGTTATAAATGGAGAC   10702
gb:LC522972|Organism:Severe      GACACAACTATTACAGTTAATGTTTTAGCTTGGTTGTACGCTGCTGTTATAAATGGAGAC   10699
gb:MT027063|Organism:Severe      GACACAACTATTACAGTTAATGTTTTAGCTTGGTTGTACGCTGCTGTTATAAATGGAGAC   10702
gb:MT027062|Organism:Severe      GACACAACTATTACAGTTAATGTTTTAGCTTGGTTGTACGCTGCTGTTATAAATGGAGAC   10702
gb:MT019529|Organism:Severe      GACACAACTATTACAGTTAATGTTTTAGCTTGGTTGTACGCTGCTGTTATAAATGGAGAC   10702
gb:MN996529|Organism:Severe      GACACAACTATTACAGTTAATGTTTTAGCTTGGTTGTACGCTGCTGTTATAAATGGAGAC   10690
gb:MN996531|Organism:Severe      GACACAACTATTACAGTTAATGTTTTAGCTTGGTTGTACGCTGCTGTTATAAATGGAGAC   10689
gb:MT066176|Organism:Severe      GACACAACTATTACAGTTAATGTTTTAGCTTGGTTGTACGCTGCTGTTATAAATGGAGAC   10702
gb:MT027064|Organism:Severe      GACACAACTATTACAGTTAATGTTTTAGCTTGGTTGTACGCTGCTGTTATAAATGGAGAC   10702
gb:MN994468|Organism:Severe      GACACAACTATTACAGTTAATGTTTTAGCTTGGTTGTACGCTGCTGTTATAAATGGAGAC   10702
gb:MT072688|Organism:Severe      GACACAACTATTACAGTTAATGTTTTAGCTTGGTTGTACGCTGCTGTTATAAATGGAGAC   10687
gb:MN996527|Organism:Severe      GACACAACTATTACAGTTAATGTTTTAGCTTGGTTGTACGCTGCTGTTATAAATGGAGAC   10669
gb:MT093631|Organism:Severe      GACACAACTATTACAGTTAATGTTTTAGCTTGGTTGTACGCTGCTGTTATAAATGGAGAC   10740
gb:MT106053|Organism:Severe      GACACAACTATTACAGTTAATGTTTTAGCTTGGTTGTACGCTGCTGTTATAAATGGAGAC   10702
gb:MT019533|Organism:Severe      GACACAACTATTACAGTTAATGTTTTAGCTTGGTTGTACGCTGCTGTTATAAATGGAGAC   10702
gb:MT019531|Organism:Severe      GACACAACTATTACAGTTAATGTTTTAGCTTGGTTGTACGCTGCTGTTATAAATGGAGAC   10702
gb:MN996528|Organism:Severe      GACACAACTATTACAGTTAATGTTTTAGCTTGGTTGTACGCTGCTGTTATAAATGGAGAC   10702
gb:MN996530|Organism:Severe      GACACAACTATTACAGTTAATGTTTTAGCTTGGTTGTACGCTGCTGTTATAAATGGAGAC   10688
gb:MN908947|Organism:Severe      GACACAACTATTACAGTTAATGTTTTAGCTTGGTTGTACGCTGCTGTTATAAATGGAGAC   10702
gb:MT019532|Organism:Severe      GACACAACTATTACAGTTAATGTTTTAGCTTGGTTGTACGCTGCTGTTATAAATGGAGAC   10702
                                 ************************************************************

gb:MT020781|Organism:Severe      AGGTGGTTTCTCAATCGATTTACCACAACTCTTAATGACTTTAACCTTGTGGCTATGAAG   10750
gb:MT007544|Organism:Severe      AGGTGGTTTCTCAATCGATTTACCACAACTCTTAATGACTTTAACCTTGTGGCTATGAAG   10762
gb:MN994467|Organism:Severe      AGGTGGTTTCTCAATCGATTTACCACAACTCTTAATGACTTTAACCTTGTGGCTATGAAG   10762
gb:MT044257|Organism:Severe      AGGTGGTTTCTCAATCGATTTACCACAACTCTTAATGACTTTAACCTTGTGGCTATGAAG   10762
gb:MT106054|Organism:Severe      AGGTGGTTTCTCAATCGATTTACCACAACTCTTAATGACTTTAACCTTGTGGCTATGAAG   10762
gb:MT049951|Organism:Severe      AGGTGGTTTCTCAATCGATTTACCACAACTCTTAATGACTTTAACCTTGTGGCTATGAAG   10762
gb:MN975262|Organism:Severe      AGGTGGTTTCTCAATCGATTTACCACAACTCTTAATGACTTTAACCTTGTGGCTATGAAG   10762
gb:MT106052|Organism:Severe      AGGTGGTTTCTCAATCGATTTACCACAACTCTTAATGACTTTAACCTTGTGGCTATGAAG   10762
gb:LC522975|Organism:Severe      AGGTGGTTTCTCAATCGATTTACCACAACTCTTAATGACTTTAACCTTGTGGCTATGAAG   10759
gb:LC522973|Organism:Severe      AGGTGGTTTCTCAATCGATTTACCACAACTCTTAATGACTTTAACCTTGTGGCTATGAAG   10759
gb:LC522974|Organism:Severe      AGGTGGTTTCTCAATCGATTTACCACAACTCTTAATGACTTTAACCTTGTGGCTATGAAG   10759
gb:MN985325|Organism:Severe      AGGTGGTTTCTCAATCGATTTACCACAACTCTTAATGACTTTAACCTTGTGGCTATGAAG   10762
gb:MT020881|Organism:Severe      AGGTGGTTTCTCAATCGATTTACCACAACTCTTAATGACTTTAACCTTGTGGCTATGAAG   10762
gb:MT020880|Organism:Severe      AGGTGGTTTCTCAATCGATTTACCACAACTCTTAATGACTTTAACCTTGTGGCTATGAAG   10762
gb:MT066175|Organism:Severe      AGGTGGTTTCTCAATCGATTTACCACAACTCTTAATGACTTTAACCTTGTGGCTATGAAG   10762
gb:MN997409|Organism:Severe      AGGTGGTTTCTCAATCGATTTACCACAACTCTTAATGACTTTAACCTTGTGGCTATGAAG   10762
gb:MN938384|Organism:Severe      AGGTGGTTTCTCAATCGATTTACCACAACTCTTAATGACTTTAACCTTGTGGCTATGAAG   10730
```

```
gb:MT044258|Organism:Severe      AGGTGGTTTCTCAATCGATTTACCACAACTCTTAATGACTTTAACCTTGTGGCTATGAAG   10738
gb:MT039890|Organism:Severe      AGGTGGTTTCTCAATCGATTTACCACAACTCTTAATGACTTTAACCTTGTGGCTATGAAG   10762
gb:MN988713|Organism:Severe      AGGTGGTTTCTCAATCGATTTACCACAACTCTTAATGACTTTAACCTTGTGGCTATGAAG   10762
gb:LC521925|Organism:Severe      AGGTGGTTTCTCAATCGATTTACCACAACTCTTAATGACTTTAACCTTGTGGCTATGAAG   10735
gb:MT093571|Organism:Severe      AGGTGGTTTCTCAATCGATTTACCACAACTCTTAATGACTTTAACCTTGTGGCTATGAAG   10762
gb:MT039887|Organism:Severe      AGGTGGTTTCTCAATCGATTTACCACAACTCTTAATGACTTTAACCTTGTGGCTATGAAG   10762
gb:MT019530|Organism:Severe      AGGTGGTTTCTCAATCGATTTACCACAACTCTTAATGACTTTAACCTTGTGGCTATGAAG   10762
gb:MT039888|Organism:Severe      AGGTGGTTTCTCAATCGATTTACCACAACTCTTAATGACTTTAACCTTGTGGCTATGAAG   10762
gb:LC522972|Organism:Severe      AGGTGGTTTCTCAATCGATTTACCACAACTCTTAATGACTTTAACCTTGTGGCTATGAAG   10759
gb:MT027063|Organism:Severe      AGGTGGTTTCTCAATCGATTTACCACAACTCTTAATGACTTTAACCTTGTGGCTATGAAG   10762
gb:MT027062|Organism:Severe      AGGTGGTTTCTCAATCGATTTACCACAACTCTTAATGACTTTAACCTTGTGGCTATGAAG   10762
gb:MT019529|Organism:Severe      AGGTGGTTTCTCAATCGATTTACCACAACTCTTAATGACTTTAACCTTGTGGCTATGAAG   10762
gb:MN996529|Organism:Severe      AGGTGGTTTCTCAATCGATTTACCACAACTCTTAATGACTTTAACCTTGTGGCTATGAAG   10750
gb:MN996531|Organism:Severe      AGGTGGTTTCTCAATCGATTTACCACAACTCTTAATGACTTTAACCTTGTGGCTATGAAG   10749
gb:MT066176|Organism:Severe      AGGTGGTTTCTCAATCGATTTACCACAACTCTTAATGACTTTAACCTTGTGGCTATGAAG   10762
gb:MT027064|Organism:Severe      AGGTGGTTTCTCAATCGATTTACCACAACTCTTAATGACTTTAACCTTGTGGCTATGAAG   10762
gb:MN994468|Organism:Severe      AGGTGGTTTCTCAATCGATTTACCACAACTCTTAATGACTTTAACCTTGTGGCTATGAAG   10762
gb:MT072688|Organism:Severe      AGGTGGTTTCTCAATCGATTTACCACAACTCTTAATGACTTTAACCTTGTGGCTATGAAG   10747
gb:MN996527|Organism:Severe      AGGTGGTTTCTCAATCGATTTACCACAACTCTTAATGACTTTAACCTTGTGGCTATGAAG   10729
gb:MT093631|Organism:Severe      AGGTGGTTTCTCAATCGATTTACCACAACTCTTAATGACTTTAACCTTGTGGCTATGAAG   10800
gb:MT106053|Organism:Severe      AGGTGGTTTCTCAATCGATTTACCACAACTCTTAATGACTTTAACCTTGTGGCTATGAAG   10762
gb:MT019533|Organism:Severe      AGGTGGTTTCTCAATCGATTTACCACAACTCTTAATGACTTTAACCTTGTGGCTATGAAG   10762
gb:MT019531|Organism:Severe      AGGTGGTTTCTCAATCGATTTACCACAACTCTTAATGACTTTAACCTTGTGGCTATGAAG   10762
gb:MN996528|Organism:Severe      AGGTGGTTTCTCAATCGATTTACCACAACTCTTAATGACTTTAACCTTGTGGCTATGAAG   10762
gb:MN996530|Organism:Severe      AGGTGGTTTCTCAATCGATTTACCACAACTCTTAATGACTTTAACCTTGTGGCTATGAAG   10748
gb:MN908947|Organism:Severe      AGGTGGTTTCTCAATCGATTTACCACAACTCTTAATGACTTTAACCTTGTGGCTATGAAG   10762
gb:MT019532|Organism:Severe      AGGTGGTTTCTCAATCGATTTACCACAACTCTTAATGACTTTAACCTTGTGGCTATGAAG   10762
                                 ************************************************************

gb:MT020781|Organism:Severe      TACAATTATGAACCTCTAACACAAGACCATGTTGACATACTAGGACCTCTTTCTGCTCAA   10810
gb:MT007544|Organism:Severe      TACAATTATGAACCTCTAACACAAGACCATGTTGACATACTAGGACCTCTTTCTGCTCAA   10822
gb:MN994467|Organism:Severe      TACAATTATGAACCTCTAACACAAGACCATGTTGACATACTAGGACCTCTTTCTGCTCAA   10822
gb:MT044257|Organism:Severe      TACAATTATGAACCTCTAACACAAGACCATGTTGACATACTAGGACCTCTTTCTGCTCAA   10822
gb:MT106054|Organism:Severe      TACAATTATGAACCTCTAACACAAGACCATGTTGACATACTAGGACCTCTTTCTGCTCAA   10822
gb:MT049951|Organism:Severe      TACAATTATGAACCTCTAACACAAGACCATGTTGACATACTAGGACCTCTTTCTGCTCAA   10822
gb:MN975262|Organism:Severe      TACAATTATGAACCTCTAACACAAGACCATGTTGACATACTAGGACCTCTTTCTGCTCAA   10822
gb:MT106052|Organism:Severe      TACAATTATGAACCTCTAACACAAGACCATGTTGACATACTAGGACCTCTTTCTGCTCAA   10822
gb:LC522975|Organism:Severe      TACAATTATGAACCTCTAACACAAGACCATGTTGACATACTAGGACCTCTTTCTGCTCAA   10819
gb:LC522973|Organism:Severe      TACAATTATGAACCTCTAACACAAGACCATGTTGACATACTAGGACCTCTTTCTGCTCAA   10819
gb:LC522974|Organism:Severe      TACAATTATGAACCTCTAACACAAGACCATGTTGACATACTAGGACCTCTTTCTGCTCAA   10819
gb:MN985325|Organism:Severe      TACAATTATGAACCTCTAACACAAGACCATGTTGACATACTAGGACCTCTTTCTGCTCAA   10822
gb:MT020881|Organism:Severe      TACAATTATGAACCTCTAACACAAGACCATGTTGACATACTAGGACCTCTTTCTGCTCAA   10822
gb:MT020880|Organism:Severe      TACAATTATGAACCTCTAACACAAGACCATGTTGACATACTAGGACCTCTTTCTGCTCAA   10822
gb:MT066175|Organism:Severe      TACAATTATGAACCTCTAACACAAGACCATGTTGACATACTAGGACCTCTTTCTGCTCAA   10822
gb:MN997409|Organism:Severe      TACAATTATGAACCTCTAACACAAGACCATGTTGACATACTAGGACCTCTTTCTGCTCAA   10822
gb:MN938384|Organism:Severe      TACAATTATGAACCTCTAACACAAGACCATGTTGACATACTAGGACCTCTTTCTGCTCAA   10790
gb:MT044258|Organism:Severe      TACAATTATGAACCTCTAACACAAGACCATGTTGACATACTAGGACCTCTTTCTGCTCAA   10798
gb:MT039890|Organism:Severe      TACAATTATGAACCTCTAACACAAGACCATGTTGACATACTAGGACCTCTTTCTGCTCAA   10822
gb:MN988713|Organism:Severe      TACAATTATGAACCTCTAACACAAGACCATGTTGACATACTAGGACCTCTTTCTGCTCAA   10822
gb:LC521925|Organism:Severe      TACAATTATGAACCTCTAACACAAGACCATGTTGACATACTAGGACCTCTTTCTGCTCAA   10795
gb:MT093571|Organism:Severe      TACAATTATGAACCTCTAACACAAGACCATGTTGACATACTAGGACCTCTTTCTGCTCAA   10822
gb:MT039887|Organism:Severe      TACAATTATGAACCTCTAACACAAGACCATGTTGACATACTAGGACCTCTTTCTGCTCAA   10822
gb:MT019530|Organism:Severe      TACAATTATGAACCTCTAACACAAGACCATGTTGACATACTAGGACCTCTTTCTGCTCAA   10822
gb:MT039888|Organism:Severe      TACAATTATGAACCTCTAACACAAGACCATGTTGACATACTAGGACCTCTTTCTGCTCAA   10822
gb:LC522972|Organism:Severe      TACAATTATGAACCTCTAACACAAGACCATGTTGACATACTAGGACCTCTTTCTGCTCAA   10819
gb:MT027063|Organism:Severe      TACAATTATGAACCTCTAACACAAGACCATGTTGACATACTAGGACCTCTTTCTGCTCAA   10822
gb:MT027062|Organism:Severe      TACAATTATGAACCTCTAACACAAGACCATGTTGACATACTAGGACCTCTTTCTGCTCAA   10822
gb:MT019529|Organism:Severe      TACAATTATGAACCTCTAACACAAGACCATGTTGACATACTAGGACCTCTTTCTGCTCAA   10822
gb:MN996529|Organism:Severe      TACAATTATGAACCTCTAACACAAGACCATGTTGACATACTAGGACCTCTTTCTGCTCAA   10810
gb:MN996531|Organism:Severe      TACAATTATGAACCTCTAACACAAGACCATGTTGACATACTAGGACCTCTTTCTGCTCAA   10809
gb:MT066176|Organism:Severe      TACAATTATGAACCTCTAACACAAGACCATGTTGACATACTAGGACCTCTTTCTGCTCAA   10822
gb:MT027064|Organism:Severe      TACAATTATGAACCTCTAACACAAGACCATGTTGACATACTAGGACCTCTTTCTGCTCAA   10822
gb:MN994468|Organism:Severe      TACAATTATGAACCTCTAACACAAGACCATGTTGACATACTAGGACCTCTTTCTGCTCAA   10822
gb:MT072688|Organism:Severe      TACAATTATGAACCTCTAACACAAGACCATGTTGACATACTAGGACCTCTTTCTGCTCAA   10807
gb:MN996527|Organism:Severe      TACAATTATGAACCTCTAACACAAGACCATGTTGACATACTAGGACCTCTTTCTGCTCAA   10789
```

```
gb:MT093631|Organism:Severe      TACAATTATGAACCTCTAACACAAGACCATGTTGACATACTAGGACCTCTTTCTGCTCAA   10860
gb:MT106053|Organism:Severe      TACAATTATGAACCTCTAACACAAGACCATGTTGACATACTAGGACCTCTTTCTGCTCAA   10822
gb:MT019533|Organism:Severe      TACAATTATGAACCTCTAACACAAGACCATGTTGACATACTAGGACCTCTTTCTGCTCAA   10822
gb:MT019531|Organism:Severe      TACAATTATGAACCTCTAACACAAGACCATGTTGACATACTAGGACCTCTTTCTGCTCAA   10822
gb:MN996528|Organism:Severe      TACAATTATGAACCTCTAACACAAGACCATGTTGACATACTAGGACCTCTTTCTGCTCAA   10822
gb:MN996530|Organism:Severe      TACAATTATGAACCTCTAACACAAGACCATGTTGACATACTAGGACCTCTTTCTGCTCAA   10808
gb:MN908947|Organism:Severe      TACAATTATGAACCTCTAACACAAGACCATGTTGACATACTAGGACCTCTTTCTGCTCAA   10822
gb:MT019532|Organism:Severe      TACAATTATGAACCTCTAACACAAGACCATGTTGACATACTAGGACCTCTTTCTGCTCAA   10822
                                 ************************************************************

gb:MT020781|Organism:Severe      ACTGGAATTGCCGTTTTAGATATGTGTGCTTCATTAAAAGAATTACTGCAAAATGGTATG   10870
gb:MT007544|Organism:Severe      ACTGGAATTGCCGTTTTAGATATGTGTGCTTCATTAAAAGAATTACTGCAAAATGGTATG   10882
gb:MN994467|Organism:Severe      ACTGGAATTGCCGTTTTAGATATGTGTGCTTCATTAAAAGAATTACTGCAAAATGGTATG   10882
gb:MT044257|Organism:Severe      ACTGGAATTGCCGTTTTAGATATGTGTGCTTCATTAAAAGAATTACTGCAAAATGGTATG   10882
gb:MT106054|Organism:Severe      ACTGGAATTGCCGTTTTAGATATGTGTGCTTCATTAAAAGAATTACTGCAAAATGGTATG   10882
gb:MT049951|Organism:Severe      ACTGGAATTGCCGTTTTAGATATGTGTGCTTCATTAAAAGAATTACTGCAAAATGGTATG   10882
gb:MN975262|Organism:Severe      ACTGGAATTGCCGTTTTAGATATGTGTGCTTCATTAAAAGAATTACTGCAAAATGGTATG   10882
gb:MT106052|Organism:Severe      ACTGGAATTGCCGTTTTAGATATGTGTGCTTCATTAAAAGAATTACTGCAAAATGGTATG   10882
gb:LC522975|Organism:Severe      ACTGGAATTGCCGTTTTAGATATGTGTGCTTCATTAAAAGAATTACTGCAAAATGGTATG   10879
gb:LC522973|Organism:Severe      ACTGGAATTGCCGTTTTAGATATGTGTGCTTCATTAAAAGAATTACTGCAAAATGGTATG   10879
gb:LC522974|Organism:Severe      ACTGGAATTGCCGTTTTAGATATGTGTGCTTCATTAAAAGAATTACTGCAAAATGGTATG   10879
gb:MN985325|Organism:Severe      ACTGGAATTGCCGTTTTAGATATGTGTGCTTCATTAAAAGAATTACTGCAAAATGGTATG   10882
gb:MT020881|Organism:Severe      ACTGGAATTGCCGTTTTAGATATGTGTGCTTCATTAAAAGAATTACTGCAAAATGGTATG   10882
gb:MT020880|Organism:Severe      ACTGGAATTGCCGTTTTAGATATGTGTGCTTCATTAAAAGAATTACTGCAAAATGGTATG   10882
gb:MT066175|Organism:Severe      ACTGGAATTGCCGTTTTAGATATGTGTGCTTCATTAAAAGAATTACTGCAAAATGGTATG   10882
gb:MN997409|Organism:Severe      ACTGGAATTGCCGTTTTAGATATGTGTGCTTCATTAAAAGAATTACTGCAAAATGGTATG   10882
gb:MN938384|Organism:Severe      ACTGGAATTGCCGTTTTAGATATGTGTGCTTCATTAAAAGAATTACTGCAAAATGGTATG   10850
gb:MT044258|Organism:Severe      ACTGGAATTGCCGTTTTAGATATGTGTGCTTCATTAAAAGAATTACTGCAAAATGGTATG   10858
gb:MT039890|Organism:Severe      ACTGGAATTGCCGTTTTAGATATGTGTGCTTCATTAAAAGAATTACTGCAAAATGGTATG   10882
gb:MN988713|Organism:Severe      ACTGGAATTGCCGTTTTAGATATGTGTGCTTCATTAAAAGAATTACTGCAAAATGGTATG   10882
gb:LC521925|Organism:Severe      ACTGGAATTGCCGTTTTAGATATGTGTGCTTCATTAAAAGAATTACTGCAAAATGGTATG   10855
gb:MT093571|Organism:Severe      ACTGGAATTGCCGTTTTAGATATGTGTGCTTCATTAAAAGAATTACTGCAAAATGGTATG   10882
gb:MT039887|Organism:Severe      ACTGGAATTGCCGTTTTAGATATGTGTGCTTCATTAAAAGAATTACTGCAAAATGGTATG   10882
gb:MT019530|Organism:Severe      ACTGGAATTGCCGTTTTAGATATGTGTGCTTCATTAAAAGAATTACTGCAAAATGGTATG   10882
gb:MT039888|Organism:Severe      ACTGGAATTGCCGTTTTAGATATGTGTGCTTCATTAAAAGAATTACTGCAAAATGGTATG   10882
gb:LC522972|Organism:Severe      ACTGGAATTGCCGTTTTAGATATGTGTGCTTCATTAAAAGAATTACTGCAAAATGGTATG   10879
gb:MT027063|Organism:Severe      ACTGGAATTGCCGTTTTAGATATGTGTGCTTCATTAAAAGAATTACTGCAAAATGGTATG   10882
gb:MT027062|Organism:Severe      ACTGGAATTGCCGTTTTAGATATGTGTGCTTCATTAAAAGAATTACTGCAAAATGGTATG   10882
gb:MT019529|Organism:Severe      ACTGGAATTGCCGTTTTAGATATGTGTGCTTCATTAAAAGAATTACTGCAAAATGGTATG   10882
gb:MN996529|Organism:Severe      ACTGGAATTGCCGTTTTAGATATGTGTGCTTCATTAAAAGAATTACTGCAAAATGGTATG   10870
gb:MN996531|Organism:Severe      ACTGGAATTGCCGTTTTAGATATGTGTGCTTCATTAAAAGAATTACTGCAAAATGGTATG   10869
gb:MT066176|Organism:Severe      ACTGGAATTGCCGTTTTAGATATGTGTGCTTCATTAAAAGAATTACTGCAAAATGGTATG   10882
gb:MT027064|Organism:Severe      ACTGGAATTGCCGTTTTAGATATGTGTGCTTCATTAAAAGAATTACTGCAAAATGGTATG   10882
gb:MN994468|Organism:Severe      ACTGGAATTGCCGTTTTAGATATGTGTGCTTCATTAAAAGAATTACTGCAAAATGGTATG   10882
gb:MT072688|Organism:Severe      ACTGGAATTGCCGTTTTAGATATGTGTGCTTCATTAAAAGAATTACTGCAAAATGGTATG   10867
gb:MN996527|Organism:Severe      ACTGGAATTGCCGTTTTAGATATGTGTGCTTCATTAAAAGAATTACTGCAAAATGGTATG   10849
gb:MT093631|Organism:Severe      ACTGGAATTGCCGTTTTAGATATGTGTGCTTCATTAAAAGAATTACTGCAAAATGGTATG   10920
gb:MT106053|Organism:Severe      ACTGGAATTGCCGTTTTAGATATGTGTGCTTCATTAAAAGAATTACTGCAAAATGGTATG   10882
gb:MT019533|Organism:Severe      ACTGGAATTGCCGTTTTAGATATGTGTGCTTCATTAAAAGAATTACTGCAAAATGGTATG   10882
gb:MT019531|Organism:Severe      ACTGGAATTGCCGTTTTAGATATGTGTGCTTCATTAAAAGAATTACTGCAAAATGGTATG   10882
gb:MN996528|Organism:Severe      ACTGGAATTGCCGTTTTAGATATGTGTGCTTCATTAAAAGAATTACTGCAAAATGGTATG   10882
gb:MN996530|Organism:Severe      ACTGGAATTGCCGTTTTAGATATGTGTGCTTCATTAAAAGAATTACTGCAAAATGGTATG   10868
gb:MN908947|Organism:Severe      ACTGGAATTGCCGTTTTAGATATGTGTGCTTCATTAAAAGAATTACTGCAAAATGGTATG   10882
gb:MT019532|Organism:Severe      ACTGGAATTGCCGTTTTAGATATGTGTGCTTCATTAAAAGAATTACTGCAAAATGGTATG   10882
                                 ************************************************************

gb:MT020781|Organism:Severe      AATGGACGTACCATATTGGGTAGTGCTTTATTAGAAGATGAATTTACACCTTTTGATGTT   10930
gb:MT007544|Organism:Severe      AATGGACGTACCATATTGGGTAGTGCTTTATTAGAAGATGAATTTACACCTTTTGATGTT   10942
gb:MN994467|Organism:Severe      AATGGACGTACCATATTGGGTAGTGCTTTATTAGAAGATGAATTTACACCTTTTGATGTT   10942
gb:MT044257|Organism:Severe      AATGGACGTACCATATTGGGTAGTGCTTTATTAGAAGATGAATTTACACCTTTTGATGTT   10942
gb:MT106054|Organism:Severe      AATGGACGTACCATATTGGGTAGTGCTTTATTAGAAGATGAATTTACACCTTTTGATGTT   10942
gb:MT049951|Organism:Severe      AATGGACGTACCATATTGGGTAGTGCTTTATTAGAAGATGAATTTACACCTTTTGATGTT   10942
gb:MN975262|Organism:Severe      AATGGACGTACCATATTGGGTAGTGCTTTATTAGAAGATGAATTTACACCTTTTGATGTT   10942
gb:MT106052|Organism:Severe      AATGGACGTACCATATTGGGTAGTGCTTTATTAGAAGATGAATTTACACCTTTTGATGTT   10942
gb:LC522975|Organism:Severe      AATGGACGTACCATATTGGGTAGTGCTTTATTAGAAGATGAATTTACACCTTTTGATGTT   10939
```

```
gb:LC522973|Organism:Severe      AATGGACGTACCATATTGGGTAGTGCTTTATTAGAAGATGAATTTACACCTTTTGATGTT   10939
gb:LC522974|Organism:Severe      AATGGACGTACCATATTGGGTAGTGCTTTATTAGAAGATGAATTTACACCTTTTGATGTT   10939
gb:MN985325|Organism:Severe      AATGGACGTACCATATTGGGTAGTGCTTTATTAGAAGATGAATTTACACCTTTTGATGTT   10942
gb:MT020881|Organism:Severe      AATGGACGTACCATATTGGGTAGTGCTTTATTAGAAGATGAATTTACACCTTTTGATGTT   10942
gb:MT020880|Organism:Severe      AATGGACGTACCATATTGGGTAGTGCTTTATTAGAAGATGAATTTACACCTTTTGATGTT   10942
gb:MT066175|Organism:Severe      AATGGACGTACCATATTGGGTAGTGCTTTATTAGAAGATGAATTTACACCTTTTGATGTT   10942
gb:MN997409|Organism:Severe      AATGGACGTACCATATTGGGTAGTGCTTTATTAGAAGATGAATTTACACCTTTTGATGTT   10942
gb:MN938384|Organism:Severe      AATGGACGTACCATATTGGGTAGTGCTTTATTAGAAGATGAATTTACACCTTTTGATGTT   10910
gb:MT044258|Organism:Severe      AATGGACGTACCATATTGGGTAGTGCTTTATTAGAAGATGAATTTACACCTTTTGATGTT   10918
gb:MT039890|Organism:Severe      AATGGACGTACCATATTGGGTAGTGCTTTATTAGAAGATGAATTTACACCTTTTGATGTT   10942
gb:MN988713|Organism:Severe      AATGGACGTACCATATTGGGTAGTGCTTTATTAGAAGATGAATTTACACCTTTTGATGTT   10942
gb:LC521925|Organism:Severe      AATGGACGTACCATATTGGGTAGTGCTTTATTAGAAGATGAATTTACACCTTTTGATGTT   10915
gb:MT093571|Organism:Severe      AATGGACGTACCATATTGGGTAGTGCTTTATTAGAAGATGAATTTACACCTTTTGATGTT   10942
gb:MT039887|Organism:Severe      AATGGACGTACCATATTGGGTAGTGCTTTATTAGAAGATGAATTTACACCTTTTGATGTT   10942
gb:MT019530|Organism:Severe      AATGGACGTACCATATTGGGTAGTGCTTTATTAGAAGATGAATTTACACCTTTTGATGTT   10942
gb:MT039888|Organism:Severe      AATGGACGTACCATATTGGGTAGTGCTTTATTAGAAGATGAATTTACACCTTTTGATGTT   10942
gb:LC522972|Organism:Severe      AATGGACGTACCATATTGGGTAGTGCTTTATTAGAAGATGAATTTACACCTTTTGATGTT   10939
gb:MT027063|Organism:Severe      AATGGACGTACCATATTGGGTAGTGCTTTATTAGAAGATGAATTTACACCTTTTGATGTT   10942
gb:MT027062|Organism:Severe      AATGGACGTACCATATTGGGTAGTGCTTTATTAGAAGATGAATTTACACCTTTTGATGTT   10942
gb:MT019529|Organism:Severe      AATGGACGTACCATATTGGGTAGTGCTTTATTAGAAGATGAATTTACACCTTTTGATGTT   10942
gb:MN996529|Organism:Severe      AATGGACGTACCATATTGGGTAGTGCTTTATTAGAAGATGAATTTACACCTTTTGATGTT   10930
gb:MN996531|Organism:Severe      AATGGACGTACCATATTGGGTAGTGCTTTATTAGAAGATGAATTTACACCTTTTGATGTT   10929
gb:MT066176|Organism:Severe      AATGGACGTACCATATTGGGTAGTGCTTTATTAGAAGATGAATTTACACCTTTTGATGTT   10942
gb:MT027064|Organism:Severe      AATGGACGTACCATATTGGGTAGTGCTTTATTAGAAGATGAATTTACACCTTTTGATGTT   10942
gb:MN994468|Organism:Severe      AATGGACGTACCATATTGGGTAGTGCTTTATTAGAAGATGAATTTACACCTTTTGATGTT   10942
gb:MT072688|Organism:Severe      AATGGACGTACCATATTGGGTAGTGCTTTATTAGAAGATGAATTTACACCTTTTGATGTT   10927
gb:MN996527|Organism:Severe      AATGGACGTACCATATTGGGTAGTGCTTTATTAGAAGATGAATTTACACCTTTTGATGTT   10909
gb:MT093631|Organism:Severe      AATGGACGTACCATATTGGGTAGTGCTTTATTAGAAGATGAATTTACACCTTTTGATGTT   10980
gb:MT106053|Organism:Severe      AATGGACGTACCATATTGGGTAGTGCTTTATTAGAAGATGAATTTACACCTTTTGATGTT   10942
gb:MT019533|Organism:Severe      AATGGACGTACCATATTGGGTAGTGCTTTATTAGAAGATGAATTTACACCTTTTGATGTT   10942
gb:MT019531|Organism:Severe      AATGGACGTACCATATTGGGTAGTGCTTTATTAGAAGATGAATTTACACCTTTTGATGTT   10942
gb:MN996528|Organism:Severe      AATGGACGTACCATATTGGGTAGTGCTTTATTAGAAGATGAATTTACACCTTTTGATGTT   10942
gb:MN996530|Organism:Severe      AATGGACGTACCATATTGGGTAGTGCTTTATTAGAAGATGAATTTACACCTTTTGATGTT   10928
gb:MN908947|Organism:Severe      AATGGACGTACCATATTGGGTAGTGCTTTATTAGAAGATGAATTTACACCTTTTGATGTT   10942
gb:MT019532|Organism:Severe      AATGGACGTACCATATTGGGTAGTGCTTTATTAGAAGATGAATTTACACCTTTTGATGTT   10942
                                 ************************************************************

gb:MT020781|Organism:Severe      GTTAGACAATGCTCAGGTGTTACTTTCCAAAGTGCAGTGAAAAGAACAATCAAGGGTACA   10990
gb:MT007544|Organism:Severe      GTTAGACAATGCTCAGGTGTTACTTTCCAAAGTGCAGTGAAAAGAACAATCAAGGGTACA   11002
gb:MN994467|Organism:Severe      GTTAGACAATGCTCAGGTGTTACTTTCCAAAGTGCAGTGAAAAGAACAATCAAGGGTACA   11002
gb:MT044257|Organism:Severe      GTTAGACAATGCTCAGGTGTTACTTTCCAAAGTGCAGTGAAAAGAACAATCAAGGGTACA   11002
gb:MT106054|Organism:Severe      GTTAGACAATGCTCAGGTGTTACTTTCCAAAGTGCAGTGAAAAGAACAATCAAGGGTACA   11002
gb:MT049951|Organism:Severe      GTTAGACAATGCTCAGGTGTTACTTTCCAAAGTGCAGTGAAAAGAACAATCAAGGGTACA   11002
gb:MN975262|Organism:Severe      GTTAGACAATGCTCAGGTGTTACTTTCCAAAGTGCAGTGAAAAGAACAATCAAGGGTACA   11002
gb:MT106052|Organism:Severe      GTTAGACAATGCTCAGGTGTTACTTTCCAAAGTGCAGTGAAAAGAACAATCAAGGGTACA   11002
gb:LC522975|Organism:Severe      GTTAGACAATGCTCAGGTGTTACTTTCCAAAGTGCAGTGAAAAGAACAATCAAGGGTACA   10999
gb:LC522973|Organism:Severe      GTTAGACAATGCTCAGGTGTTACTTTCCAAAGTGCAGTGAAAAGAACAATCAAGGGTACA   10999
gb:LC522974|Organism:Severe      GTTAGACAATGCTCAGGTGTTACTTTCCAAAGTGCAGTGAAAAGAACAATCAAGGGTACA   10999
gb:MN985325|Organism:Severe      GTTAGACAATGCTCAGGTGTTACTTTCCAAAGTGCAGTGAAAAGAACAATCAAGGGTACA   11002
gb:MT020881|Organism:Severe      GTTAGACAATGCTCAGGTGTTACTTTCCAAAGTGCAGTGAAAAGAACAATCAAGGGTACA   11002
gb:MT020880|Organism:Severe      GTTAGACAATGCTCAGGTGTTACTTTCCAAAGTGCAGTGAAAAGAACAATCAAGGGTACA   11002
gb:MT066175|Organism:Severe      GTTAGACAATGCTCAGGTGTTACTTTCCAAAGTGCAGTGAAAAGAACAATCAAGGGTACA   11002
gb:MN997409|Organism:Severe      GTTAGACAATGCTCAGGTGTTACTTTCCAAAGTGCAGTGAAAAGAACAATCAAGGGTACA   11002
gb:MN938384|Organism:Severe      GTTAGACAATGCTCAGGTGTTACTTTCCAAAGTGCAGTGAAAAGAACAATCAAGGGTACA   10970
gb:MT044258|Organism:Severe      GTTAGACAATGCTCAGGTGTTACTTTCCAAAGTGCAGTGAAAAGAACAATCAAGGGTACA   10978
gb:MT039890|Organism:Severe      GTTAGACAATGCTCAGGTGTTACTTTCCAAAGTGCAGTGAAAAGAACAATCAAGGGTACA   11002
gb:MN988713|Organism:Severe      GTTAGACAATGCTCAGGTGTTACTTTCCAAAGTGCAGTGAAAAGAACAATCAAGGGTACA   11002
gb:LC521925|Organism:Severe      GTTAGACAATGCTCAGGTGTTACTTTCCAAAGTGCAGTGAAAAGAACAATCAAGGGTACA   10975
gb:MT093571|Organism:Severe      GTTAGACAATGCTCAGGTGTTACTTTCCAAAGTGCAGTGAAAAGAACAATCAAGGGTACA   11002
gb:MT039887|Organism:Severe      GTTAGACAATGCTCAGGTGTTACTTTCCAAAGTGCAGTGAAAAGAACAATCAAGGGTACA   11002
gb:MT019530|Organism:Severe      GTTAGACAATGCTCAGGTGTTACTTTCCAAAGTGCAGTGAAAAGAACAATCAAGGGTACA   11002
gb:MT039888|Organism:Severe      GTTAGACAATGCTCAGGTGTTACTTTCCAAAGTGCAGTGAAAAGAACAATCAAGGGTACA   11002
gb:LC522972|Organism:Severe      GTTAGACAATGCTCAGGTGTTACTTTCCAAAGTGCAGTGAAAAGAACAATCAAGGGTACA   10999
gb:MT027063|Organism:Severe      GTTAGACAATGCTCAGGTGTTACTTTCCAAAGTGCAGTGAAAAGAACAATCAAGGGTACA   11002
gb:MT027062|Organism:Severe      GTTAGACAATGCTCAGGTGTTACTTTCCAAAGTGCAGTGAAAAGAACAATCAAGGGTACA   11002
```

```
gb:MT019529|Organism:Severe      GTTAGACAATGCTCAGGTGTTACTTTCCAAAGTGCAGTGAAAAGAACAATCAAGGGTACA    11002
gb:MN996529|Organism:Severe      GTTAGACAATGCTCAGGTGTTACTTTCCAAAGTGCAGTGAAAAGAACAATCAAGGGTACA    10990
gb:MN996531|Organism:Severe      GTTAGACAATGCTCAGGTGTTACTTTCCAAAGTGCAGTGAAAAGAACAATCAAGGGTACA    10989
gb:MT066176|Organism:Severe      GTTAGACAATGCTCAGGTGTTACTTTCCAAAGTGCAGTGAAAAGAACAATCAAGGGTACA    11002
gb:MT027064|Organism:Severe      GTTAGACAATGCTCAGGTGTTACTTTCCAAAGTGCAGTGAAAAGAACAATCAAGGGTACA    11002
gb:MN994468|Organism:Severe      GTTAGACAATGCTCAGGTGTTACTTTCCAAAGTGCAGTGAAAAGAACAATCAAGGGTACA    11002
gb:MT072688|Organism:Severe      GTTAGACAATGCTCAGGTGTTACTTTCCAAAGTGCAGTGAAAAGAACAATCAAGGGTACA    10987
gb:MN996527|Organism:Severe      GTTAGACAATGCTCAGGTGTTACTTTCCAAAGTGCAGTGAAAAGAACAATCAAGGGTACA    10969
gb:MT093631|Organism:Severe      GTTAGACAATGCTCAGGTGTTACTTTCCAAAGTGCAGTGAAAAGAACAATCAAGGGTACA    11040
gb:MT106053|Organism:Severe      GTTAGACAATGCTCAGGTGTTACTTTCCAAAGTGCAGTGAAAAGAACAATCAAGGGTACA    11002
gb:MT019533|Organism:Severe      GTTAGACAATGCTCAGGTGTTACTTTCCAAAGTGCAGTGAAAAGAACAATCAAGGGTACA    11002
gb:MT019531|Organism:Severe      GTTAGACAATGCTCAGGTGTTACTTTCCAAAGTGCAGTGAAAAGAACAATCAAGGGTACA    11002
gb:MN996528|Organism:Severe      GTTAGACAATGCTCAGGTGTTACTTTCCAAAGTGCAGTGAAAAGAACAATCAAGGGTACA    11002
gb:MN996530|Organism:Severe      GTTAGACAATGCTCAGGTGTTACTTTCCAAAGTGCAGTGAAAAGAACAATCAAGGGTACA    10988
gb:MN908947|Organism:Severe      GTTAGACAATGCTCAGGTGTTACTTTCCAAAGTGCAGTGAAAAGAACAATCAAGGGTACA    11002
gb:MT019532|Organism:Severe      GTTAGACAATGCTCAGGTGTTACTTTCCAAAGTGCAGTGAAAAGAACAATCAAGGGTACA    11002
                                 ************************************************************

gb:MT020781|Organism:Severe      CACCACTGGTTGTTACTCACAATTTTGACTTCACTTTTAGTTTTAGTCCAGAGTACTCAA    11050
gb:MT007544|Organism:Severe      CACCACTGGTTGTTACTCACAATTTTGACTTCACTTTTAGTTTTAGTCCAGAGTACTCAA    11062
gb:MN994467|Organism:Severe      CACCACTGGTTGTTACTCACAATTTTGACTTCACTTTTAGTTTTAGTCCAGAGTACTCAA    11062
gb:MT044257|Organism:Severe      CACCACTGGTTGTTACTCACAATTTTGACTTCACTTTTAGTTTTAGTCCAGAGTACTCAA    11062
gb:MT106054|Organism:Severe      CACCACTGGTTGTTACTCACAATTTTGACTTCACTTTTAGTTTTAGTCCAGAGTACTCAA    11062
gb:MT049951|Organism:Severe      CACCACTGGTTGTTACTCACAATTTTGACTTCACTTTTAGTTTTAGTCCAGAGTACTCAA    11062
gb:MN975262|Organism:Severe      CACCACTGGTTGTTACTCACAATTTTGACTTCACTTTTAGTTTTAGTCCAGAGTACTCAA    11062
gb:MT106052|Organism:Severe      CACCACTGGTTGTTACTCACAATTTTGACTTCACTTTTAGTTTTAGTCCAGAGTACTCAA    11062
gb:LC522975|Organism:Severe      CACCACTGGTTGTTACTCACAATTTTGACTTCACTTTTAGTTTTAGTCCAGAGTACTCAA    11059
gb:LC522973|Organism:Severe      CACCACTGGTTGTTACTCACAATTTTGACTTCACTTTTAGTTTTAGTCCAGAGTACTCAA    11059
gb:LC522974|Organism:Severe      CACCACTGGTTGTTACTCACAATTTTGACTTCACTTTTAGTTTTAGTCCAGAGTACTCAA    11059
gb:MN985325|Organism:Severe      CACCACTGGTTGTTACTCACAATTTTGACTTCACTTTTAGTTTTAGTCCAGAGTACTCAA    11062
gb:MT020881|Organism:Severe      CACCACTGGTTGTTACTCACAATTTTGACTTCACTTTTAGTTTTAGTCCAGAGTACTCAA    11062
gb:MT020880|Organism:Severe      CACCACTGGTTGTTACTCACAATTTTGACTTCACTTTTAGTTTTAGTCCAGAGTACTCAA    11062
gb:MT066175|Organism:Severe      CACCACTGGTTGTTACTCACAATTTTGACTTCACTTTTAGTTTTAGTCCAGAGTACTCAA    11062
gb:MN997409|Organism:Severe      CACCACTGGTTGTTACTCACAATTTTGACTTCACTTTTAGTTTTAGTCCAGAGTACTCAA    11062
gb:MN938384|Organism:Severe      CACCACTGGTTGTTACTCACAATTTTGACTTCACTTTTAGTTTTAGTCCAGAGTACTCAA    11030
gb:MT044258|Organism:Severe      CACCACTGGTTGTTACTCACAATTTTGACTTCACTTTTAGTTTTAGTCCAGAGTACTCAA    11038
gb:MT039890|Organism:Severe      CACCACTGGTTGTTACTCACAATTTTGACTTCACTTTTAGTTTTAGTCCAGAGTACTCAA    11062
gb:MN988713|Organism:Severe      CACCACTGGTTGTTACTCACAATTTTGACTTCACTTTTAGTTTTAGTCCAGAGTACTCAA    11062
gb:LC521925|Organism:Severe      CACCACTGGTTGTTACTCACAATTTTGACTTCACTTTTAGTTTTAGTCCAGAGTACTCAA    11035
gb:MT093571|Organism:Severe      CACCACTGGTTGTTACTCACAATTTTGACTTCACTTTTAGTTTTAGTCCAGAGTACTCAA    11062
gb:MT039887|Organism:Severe      CACCACTGGTTGTTACTCACAATTTTGACTTCACTTTTAGTTTTAGTCCAGAGTACTCAA    11062
gb:MT019530|Organism:Severe      CACCACTGGTTGTTACTCACAATTTTGACTTCACTTTTAGTTTTAGTCCAGAGTACTCAA    11062
gb:MT039888|Organism:Severe      CACCACTGGTTGTTACTCACAATTTTGACTTCACTTTTAGTTTTAGTCCAGAGTACTCAA    11062
gb:LC522972|Organism:Severe      CACCACTGGTTGTTACTCACAATTTTGACTTCACTTTTAGTTTTAGTCCAGAGTACTCAA    11059
gb:MT027063|Organism:Severe      CACCACTGGTTGTTACTCACAATTTTGACTTCACTTTTAGTTTTAGTCCAGAGTACTCAA    11062
gb:MT027062|Organism:Severe      CACCACTGGTTGTTACTCACAATTTTGACTTCACTTTTAGTTTTAGTCCAGAGTACTCAA    11062
gb:MT019529|Organism:Severe      CACCACTGGTTGTTACTCACAATTTTGACTTCACTTTTAGTTTTAGTCCAGAGTACTCAA    11062
gb:MN996529|Organism:Severe      CACCACTGGTTGTTACTCACAATTTTGACTTCACTTTTAGTTTTAGTCCAGAGTACTCAA    11050
gb:MN996531|Organism:Severe      CACCACTGGTTGTTACTCACAATTTTGACTTCACTTTTAGTTTTAGTCCAGAGTACTCAA    11049
gb:MT066176|Organism:Severe      CACCACTGGTTGTTACTCACAATTTTGACTTCACTTTTAGTTTTAGTCCAGAGTACTCAA    11062
gb:MT027064|Organism:Severe      CACCACTGGTTGTTACTCACAATTTTGACTTCACTTTTAGTTTTAGTCCAGAGTACTCAA    11062
gb:MN994468|Organism:Severe      CACCACTGGTTGTTACTCACAATTTTGACTTCACTTTTAGTTTTAGTCCAGAGTACTCAA    11062
gb:MT072688|Organism:Severe      CACCACTGGTTGTTACTCACAATTTTGACTTCACTTTTAGTTTTAGTCCAGAGTACTCAA    11047
gb:MN996527|Organism:Severe      CACCACTGGTTGTTACTCACAATTTTGACTTCACTTTTAGTTTTAGTCCAGAGTACTCAA    11029
gb:MT093631|Organism:Severe      CACCACTGGTTGTTACTCACAATTTTGACTTCACTTTTAGTTTTAGTCCAGAGTACTCAA    11100
gb:MT106053|Organism:Severe      CACCACTGGTTGTTACTCACAATTTTGACTTCACTTTTAGTTTTAGTCCAGAGTACTCAA    11062
gb:MT019533|Organism:Severe      CACCACTGGTTGTTACTCACAATTTTGACTTCACTTTTAGTTTTAGTCCAGAGTACTCAA    11062
gb:MT019531|Organism:Severe      CACCACTGGTTGTTACTCACAATTTTGACTTCACTTTTAGTTTTAGTCCAGAGTACTCAA    11062
gb:MN996528|Organism:Severe      CACCACTGGTTGTTACTCACAATTTTGACTTCACTTTTAGTTTTAGTCCAGAGTACTCAA    11062
gb:MN996530|Organism:Severe      CACCACTGGTTGTTACTCACAATTTTGACTTCACTTTTAGTTTTAGTCCAGAGTACTCAA    11048
gb:MN908947|Organism:Severe      CACCACTGGTTGTTACTCACAATTTTGACTTCACTTTTAGTTTTAGTCCAGAGTACTCAA    11062
gb:MT019532|Organism:Severe      CACCACTGGTTGTTACTCACAATTTTGACTTCACTTTTAGTTTTAGTCCAGAGTACTCAA    11062
                                 ************************************************************

gb:MT020781|Organism:Severe      TGGTCTTTGTTCTTTTTTTTGTATGAAAATGCCTTTTTACCTTTTGCTATGGGTATTATT    11110
```

```
gb:MT007544|Organism:Severe      TGGTCTTTGTTCTTTTTTTTGTATGAAAATGCCTTTTTACCTTTTGCTATGGGTATTATT   11122
gb:MN994467|Organism:Severe      TGGTCTTTGTTCTTTTTTTTGTATGAAAATGCCTTTTTACCTTTTGCTATGGGTATTATT   11122
gb:MT044257|Organism:Severe      TGGTCTTTGTTCTTTTTTTTGTATGAAAATGCCTTTTTACCTTTTGCTATGGGTATTATT   11122
gb:MT106054|Organism:Severe      TGGTCTTTGTTCTTTTTTTTGTATGAAAATGCCTTTTTACCTTTTGCTATGGGTATTATT   11122
gb:MT049951|Organism:Severe      TGGTCTTTGTTCTTTTTTTTCTATGAAAATGCCTTTTTACCTTTTGCTATGGGTATTATT   11122
gb:MN975262|Organism:Severe      TGGTCTTTGTTCTTTTTTTTGTATGAAAATGCCTTTTTACCTTTTGCTATGGGTATTATT   11122
gb:MT106052|Organism:Severe      TGGTCTTTGTTCTTTTTTTTGTATGAAAATGCCTTTTTACCTTTTGCTATGGGTATTATT   11122
gb:LC522975|Organism:Severe      TGGTCTTTGTTCTTTTTTTTGTATGAAAATGCCTTTTTACCTTTTGCTATGGGTATTATT   11119
gb:LC522973|Organism:Severe      TGGTCTTTGTTCTTTTTTTTGTATGAAAATGCCTTTTTACCTTTTGCTATGGGTATTATT   11119
gb:LC522974|Organism:Severe      TGGTCTTTGTTCTTTTTTTTGTATGAAAATGCCTTTTTACCTTTTGCTATGGGTATTATT   11119
gb:MN985325|Organism:Severe      TGGTCTTTGTTCTTTTTTTTGTATGAAAATGCCTTTTTACCTTTTGCTATGGGTATTATT   11122
gb:MT020881|Organism:Severe      TGGTCTTTGTTCTTTTTTTTGTATGAAAATGCCTTTTTACCTTTTGCTATGGGTATTATT   11122
gb:MT020880|Organism:Severe      TGGTCTTTGTTCTTTTTTTTGTATGAAAATGCCTTTTTACCTTTTGCTATGGGTATTATT   11122
gb:MT066175|Organism:Severe      TGGTCTTTGTTCTTTTTTTTGTATGAAAATGCCTTTTTACCTTTTGCTATGGGTATTATT   11122
gb:MN997409|Organism:Severe      TGGTCTTTGTTCTTTTTTTTTTATGAAAATGCCTTTTTACCTTTTGCTATGGGTATTATT   11122
gb:MN938384|Organism:Severe      TGGTCTTTGTTCTTTTTTTTGTATGAAAATGCCTTTTTACCTTTTGCTATGGGTATTATT   11090
gb:MT044258|Organism:Severe      TGGTCTTTGTTCTTTTTTTTGTATGAAAATGCCTTTTTACCTTTTGCTATGGGTATTATT   11098
gb:MT039890|Organism:Severe      TGGTCTTTGTTCTTTTTTTTGTATGAAAATGCCTTTTTACCTTTTGCTATGGGTATTATT   11122
gb:MN988713|Organism:Severe      TGGTCTTTGTTCTTTTTTTTGTATGAAAATGCCTTTTTACCTTTTGCTATGGGTATTATT   11122
gb:LC521925|Organism:Severe      TGGTCTTTGTTCTTTTTTTTGTATGAAAATGCCTTTTTACCTTTTGCTATGGGTATTATT   11095
gb:MT093571|Organism:Severe      TGGTCTTTGTTCTTTTTTTTGTATGAAAATGCCTTTTTACCTTTTGCTATGGGTATTATT   11122
gb:MT039887|Organism:Severe      TGGTCTTTGTTCTTTTTTTTGTATGAAAATGCCTTTTTACCTTTTGCTATGGGTATTATT   11122
gb:MT019530|Organism:Severe      TGGTCTTTGTTCTTTTTTTTGTATGAAAATGCCTTTTTACCTTTTGCTATGGGTATTATT   11122
gb:MT039888|Organism:Severe      TGGTCTTTGTTCTTTTTTTTGTATGAAAATGCCTTTTTACCTTTTGCTATGGGTATTATT   11122
gb:LC522972|Organism:Severe      TGGTCTTTGTTCTTTTTTTTGTATGAAAATGCCTTTTTACCTTTTGCTATGGGTATTATT   11119
gb:MT027063|Organism:Severe      TGGTCTTTGTTCTTTTTTTTGTATGAAAATGCCTTTTTACCTTTTGCTATGGGTATTATT   11122
gb:MT027062|Organism:Severe      TGGTCTTTGTTCTTTTTTTTGTATGAAAATGCCTTTTTACCTTTTGCTATGGGTATTATT   11122
gb:MT019529|Organism:Severe      TGGTCTTTGTTCTTTTTTTTGTATGAAAATGCCTTTTTACCTTTTGCTATGGGTATTATT   11122
gb:MN996529|Organism:Severe      TGGTCTTTGTTCTTTTTTTTGTATGAAAATGCCTTTTTACCTTTTGCTATGGGTATTATT   11110
gb:MN996531|Organism:Severe      TGGTCTTTGTTCTTTTTTTTGTATGAAAATGCCTTTTTACCTTTTGCTATGGGTATTATT   11109
gb:MT066176|Organism:Severe      TGGTCTTTGTTCTTTTTTTTGTATGAAAATGCCTTTTTACCTTTTGCTATGGGTATTATT   11122
gb:MT027064|Organism:Severe      TGGTCTTTGTTCTTTTTTTTGTATGAAAATGCCTTTTTACCTTTTGCTATGGGTATTATT   11122
gb:MN994468|Organism:Severe      TGGTCTTTGTTCTTTTTTTTGTATGAAAATGCCTTTTTACCTTTTGCTATGGGTATTATT   11122
gb:MT072688|Organism:Severe      TGGTCTTTGTTCTTTTTTTTGTATGAAAATGCCTTTTTACCTTTTGCTATGGGTATTATT   11107
gb:MN996527|Organism:Severe      TGGTCTTTGTTCTTTTTTTTGTATGAAAATGCCTTTTTACCTTTTGCTATGGGTATTATT   11089
gb:MT093631|Organism:Severe      TGGTCTTTGTTCTTTTTTTTGTATGAAAATGCCTTTTTACCTTTTGCTATGGGTATTATT   11160
gb:MT106053|Organism:Severe      TGGTCTTTGTTCTTTTTTTTGTATGAAAATGCCTTTTTACCTTTTGCTATGGGTATTATT   11122
gb:MT019533|Organism:Severe      TGGTCTTTGTTCTTTTTTTTGTATGAAAATGCCTTTTTACCTTTTGCTATGGGTATTATT   11122
gb:MT019531|Organism:Severe      TGGTCTTTGTTCTTTTTTTTGTATGAAAATGCCTTTTTACCTTTTGCTATGGGTATTATT   11122
gb:MN996528|Organism:Severe      TGGTCTTTGTTCTTTTTTTTGTATGAAAATGCCTTTTTACCTTTTGCTATGGGTATTATT   11122
gb:MN996530|Organism:Severe      TGGTCTTTGTTCTTTTTTTTGTATGAAAATGCCTTTTTACCTTTTGCTATGGGTATTATT   11108
gb:MN908947|Organism:Severe      TGGTCTTTGTTCTTTTTTTTGTATGAAAATGCCTTTTTACCTTTTGCTATGGGTATTATT   11122
gb:MT019532|Organism:Severe      TGGTCTTTGTTCTTTTTTTTGTATGAAAATGCCTTTTTACCTTTTGCTATGGGTATTATT   11122
                                 ******************** ***************************************

gb:MT020781|Organism:Severe      GCTATGTCTGCTTTTGCAATGATGTTTGTCAAACATAAGCATGCATTTCTCTGTTTGTTT   11170
gb:MT007544|Organism:Severe      GCTATGTCTGCTTTTGCAATGATGTTTGTCAAACATAAGCATGCATTTCTCTGTTTGTTT   11182
gb:MN994467|Organism:Severe      GCTATGTCTGCTTTTGCAATGATGTTTGTCAAACATAAGCATGCATTTCTCTGTTTGTTT   11182
gb:MT044257|Organism:Severe      GCTATGTCTGCTTTTGCAATGATGTTTGTCAAACATAAGCATGCATTTCTCTGTTTGTTT   11182
gb:MT106054|Organism:Severe      GCTATGTCTGCTTTTGCAATGATGTTTGTCAAACATAAGCATGCATTTCTCTGTTTGTTT   11182
gb:MT049951|Organism:Severe      GCTATGTCTGCTTTTGCAATGATGTTTGTCAAACATAAGCATGCATTTCTCTGTTTGTTT   11182
gb:MN975262|Organism:Severe      GCTATGTCTGCTTTTGCAATGATGTTTGTCAAACATAAGCATGCATTTCTCTGTTTGTTT   11182
gb:MT106052|Organism:Severe      GCTATGTCTGCTTTTGCAATGATGTTTGTCAAACATAAGCATGCATTTCTCTGTTTGTTT   11182
gb:LC522975|Organism:Severe      GCTATGTCTGCTTTTGCAATGATGTTTGTCAAACATAAGCATGCATTTCTCTGTTTGTTT   11179
gb:LC522973|Organism:Severe      GCTATGTCTGCTTTTGCAATGATGTTTGTCAAACATAAGCATGCATTTCTCTGTTTGTTT   11179
gb:LC522974|Organism:Severe      GCTATGTCTGCTTTTGCAATGATGTTTGTCAAACATAAGCATGCATTTCTCTGTTTGTTT   11179
gb:MN985325|Organism:Severe      GCTATGTCTGCTTTTGCAATGATGTTTGTCAAACATAAGCATGCATTTCTCTGTTTGTTT   11182
gb:MT020881|Organism:Severe      GCTATGTCTGCTTTTGCAATGATGTTTGTCAAACATAAGCATGCATTTCTCTGTTTGTTT   11182
gb:MT020880|Organism:Severe      GCTATGTCTGCTTTTGCAATGATGTTTGTCAAACATAAGCATGCATTTCTCTGTTTGTTT   11182
gb:MT066175|Organism:Severe      GCTATGTCTGCTTTTGCAATGATGTTTGTCAAACATAAGCATGCATTTCTCTGTTTGTTT   11182
gb:MN997409|Organism:Severe      GCTATGTCTGCTTTTGCAATGATGTTTGTCAAACATAAGCATGCATTTCTCTGTTTGTTT   11182
gb:MN938384|Organism:Severe      GCTATGTCTGCTTTTGCAATGATGTTTGTCAAACATAAGCATGCATTTCTCTGTTTGTTT   11150
gb:MT044258|Organism:Severe      GCTATGTCTGCTTTTGCAATGATGTTTGTCAAACATAAGCATGCATTTCTCTGTTTGTTT   11158
gb:MT039890|Organism:Severe      GCTATGTCTGCTTTTGCAATGATGTTTGTCAAACATAAGCATGCATTTCTCTGTTTGTTT   11182
gb:MN988713|Organism:Severe      GCTATGTCTGCTTTTGCAATGATGTTTGTCAAACATAAGCATGCATTTCTCTGTTTGTTT   11182
```

```
gb:LC521925|Organism:Severe      GCTATGTCTGCTTTTGCAATGATGTTTGTCAAACATAAGCATGCATTTCTCTGTTTGTTT   11155
gb:MT093571|Organism:Severe      GCTATGTCTGCTTTTGCAATGATGTTTGTCAAACATAAGCATGCATTTCTCTGTTTGTTT   11182
gb:MT039887|Organism:Severe      GCTATGTCTGCTTTTGCAATGATGTTTGTCAAACATAAGCATGCATTTCTCTGTTTGTTT   11182
gb:MT019530|Organism:Severe      GCTATGTCTGCTTTTGCAATGATGTTTGTCAAACATAAGCATGCATTTCTCTGTTTGTTT   11182
gb:MT039888|Organism:Severe      GCTATGTCTGCTTTTGCAATGATGTTTGTCAAACATAAGCATGCATTTCTCTGTTTGTTT   11182
gb:LC522972|Organism:Severe      GCTATGTCTGCTTTTGCAATGATGTTTGTCAAACATAAGCATGCATTTCTCTGTTTGTTT   11179
gb:MT027063|Organism:Severe      GCTATGTCTGCTTTTGCAATGATGTTTGTCAAACATAAGCATGCATTTCTCTGTTTGTTT   11182
gb:MT027062|Organism:Severe      GCTATGTCTGCTTTTGCAATGATGTTTGTCAAACATAAGCATGCATTTCTCTGTTTGTTT   11182
gb:MT019529|Organism:Severe      GCTATGTCTGCTTTTGCAATGATGTTTGTCAAACATAAGCATGCATTTCTCTGTTTGTTT   11182
gb:MN996529|Organism:Severe      GCTATGTCTGCTTTTGCAATGATGTTTGTCAAACATAAGCATGCATTTCTCTGTTTGTTT   11170
gb:MN996531|Organism:Severe      GCTATGTCTGCTTTTGCAATGATGTTTGTCAAACATAAGCATGCATTTCTCTGTTTGTTT   11169
gb:MT066176|Organism:Severe      GCTATGTCTGCTTTTGCAATGATGTTTGTCAAACATAAGCATGCATTTCTCTGTTTGTTT   11182
gb:MT027064|Organism:Severe      GCTATGTCTGCTTTTGCAATGATGTTTGTCAAACATAAGCATGCATTTCTCTGTTTGTTT   11182
gb:MN994468|Organism:Severe      GCTATGTCTGCTTTTGCAATGATGTTTGTCAAACATAAGCATGCATTTCTCTGTTTGTTT   11182
gb:MT072688|Organism:Severe      GCTATGTCTGCTTTTGCAATGATGTTTGTCAAACATAAGCATGCATTTCTCTGTTTGTTT   11167
gb:MN996527|Organism:Severe      GCTATGTCTGCTTTTGCAATGATGTTTGTCAAACATAAGCATGCATTTCTCTGTTTGTTT   11149
gb:MT093631|Organism:Severe      GCTATGTCTGCTTTTGCAATGATGTTTGTCAAACATAAGCATGCATTTCTCTGTTTGTTT   11220
gb:MT106053|Organism:Severe      GCTATGTCTGCTTTTGCAATGATGTTTGTCAAACATAAGCATGCATTTCTCTGTTTGTTT   11182
gb:MT019533|Organism:Severe      GCTATGTCTGCTTTTGCAATGATGTTTGTCAAACATAAGCATGCATTTCTCTGTTTGTTT   11182
gb:MT019531|Organism:Severe      GCTATGTCTGCTTTTGCAATGATGTTTGTCAAACATAAGCATGCATTTCTCTGTTTGTTT   11182
gb:MN996528|Organism:Severe      GCTATGTCTGCTTTTGCAATGATGTTTGTCAAACATAAGCATGCATTTCTCTGTTTGTTT   11182
gb:MN996530|Organism:Severe      GCTATGTCTGCTTTTGCAATGATGTTTGTCAAACATAAGCATGCATTTCTCTGTTTGTTT   11168
gb:MN908947|Organism:Severe      GCTATGTCTGCTTTTGCAATGATGTTTGTCAAACATAAGCATGCATTTCTCTGTTTGTTT   11182
gb:MT019532|Organism:Severe      GCTATGTCTGCTTTTGCAATGATGTTTGTCAAACATAAGCATGCATTTCTCTGTTTGTTT   11182
                                 ************************************************************

gb:MT020781|Organism:Severe      TTGTTACCTTCTCTTGCCACTGTAGCTTATTTTAATATGGTCTATATGCCTGCTAGTTGG   11230
gb:MT007544|Organism:Severe      TTGTTACCTTCTCTTGCCACTGTAGCTTATTTTAATATGGTCTATATGCCTGCTAGTTGG   11242
gb:MN994467|Organism:Severe      TTGTTACCTTCTCTTGCCACTGTAGCTTATTTTAATATGGTCTATATGCCTGCTAGTTGG   11242
gb:MT044257|Organism:Severe      TTGTTACCTTCTCTTGCCACTGTAGCTTATTTTAATATGGTCTATATGCCTGCTAGTTGG   11242
gb:MT106054|Organism:Severe      TTGTTACCTTCTCTTGCCACTGTAGCTTATTTTAATATGGTCTATATGCCTGCTAGTTGG   11242
gb:MT049951|Organism:Severe      TTGTTACCTTCTCTTGCCACTGTAGCTTATTTTAATATGGTCTATATGCCTGCTAGTTGG   11242
gb:MN975262|Organism:Severe      TTGTTACCTTCTCTTGCCACTGTAGCTTATTTTAATATGGTCTATATGCCTGCTAGTTGG   11242
gb:MT106052|Organism:Severe      TTGTTACCTTCTCTTGCCACTGTAGCTTATTTTAATATGGTCTATATGCCTGCTAGTTGG   11242
gb:LC522975|Organism:Severe      TTGTTACCTTCTCTTGCCACTGTAGCTTATTTTAATATGGTCTATATGCCTGCTAGTTGG   11239
gb:LC522973|Organism:Severe      TTGTTACCTTCTCTTGCCACTGTAGCTTATTTTAATATGGTCTATATGCCTGCTAGTTGG   11239
gb:LC522974|Organism:Severe      TTGTTACCTTCTCTTGCCACTGTAGCTTATTTTAATATGGTCTATATGCCTGCTAGTTGG   11239
gb:MN985325|Organism:Severe      TTGTTACCTTCTCTTGCCACTGTAGCTTATTTTAATATGGTCTATATGCCTGCTAGTTGG   11242
gb:MT020881|Organism:Severe      TTGTTACCTTCTCTTGCCACTGTAGCTTATTTTAATATGGTCTATATGCCTGCTAGTTGG   11242
gb:MT020880|Organism:Severe      TTGTTACCTTCTCTTGCCACTGTAGCTTATTTTAATATGGTCTATATGCCTGCTAGTTGG   11242
gb:MT066175|Organism:Severe      TTGTTACCTTCTCTTGCCACTGTAGCTTATTTTAATATGGTCTATATGCCTGCTAGTTGG   11242
gb:MN997409|Organism:Severe      TTGTTACCTTCTCTTGCCACTGTAGCTTATTTTAATATGGTCTATATGCCTGCTAGTTGG   11242
gb:MN938384|Organism:Severe      TTGTTACCTTCTCTTGCCACTGTAGCTTATTTTAATATGGTCTATATGCCTGCTAGTTGG   11210
gb:MT044258|Organism:Severe      TTGTTACCTTCTCTTGCCACTGTAGCTTATTTTAATATGGTCTATATGCCTGCTAGTTGG   11218
gb:MT039890|Organism:Severe      TTGTTACCTTCTCTTGCCACTGTAGCTTATTTTAATATGGTCTATATGCCTGCTAGTTGG   11242
gb:MN988713|Organism:Severe      TTGTTACCTTCTCTTGCCACTGTAGCTTATTTTAATATGGTCTATATGCCTGCTAGTTGG   11242
gb:LC521925|Organism:Severe      TTGTTACCTTCTCTTGCCACTGTAGCTTATTTTAATATGGTCTATATGCCTGCTAGTTGG   11215
gb:MT093571|Organism:Severe      TTGTTACCTTCTCTTGCCACTGTAGCTTATTTTAATATGGTCTATATGCCTGCTAGTTGG   11242
gb:MT039887|Organism:Severe      TTGTTACCTTCTCTTGCCACTGTAGCTTATTTTAATATGGTCTATATGCCTGCTAGTTGG   11242
gb:MT019530|Organism:Severe      TTGTTACCTTCTCTTGCCACTGTAGCTTATTTTAATATGGTCTATATGCCTGCTAGTTGG   11242
gb:MT039888|Organism:Severe      TTGTTACCTTCTCTTGCCACTGTAGCTTATTTTAATATGGTCTATATGCCTGCTAGTTGG   11242
gb:LC522972|Organism:Severe      TTGTTACCTTCTCTTGCCACTGTAGCTTATTTTAATATGGTCTATATGCCTGCTAGTTGG   11239
gb:MT027063|Organism:Severe      TTGTTACCTTCTCTTGCCACTGTAGCTTATTTTAATATGGTCTATATGCCTGCTAGTTGG   11242
gb:MT027062|Organism:Severe      TTGTTACCTTCTCTTGCCACTGTAGCTTATTTTAATATGGTCTATATGCCTGCTAGTTGG   11242
gb:MT019529|Organism:Severe      TTGTTACCTTCTCTTGCCACTGTAGCTTATTTTAATATGGTCTATATGCCTGCTAGTTGG   11242
gb:MN996529|Organism:Severe      TTGTTACCTTCTCTTGCCACTGTAGCTTATTTTAATATGGTCTATATGCCTGCTAGTTGG   11230
gb:MN996531|Organism:Severe      TTGTTACCTTCTCTTGCCACTGTAGCTTATTTTAATATGGTCTATATGCCTGCTAGTTGG   11229
gb:MT066176|Organism:Severe      TTGTTACCTTCTCTTGCCACTGTAGCTTATTTTAATATGGTCTATATGCCTGCTAGTTGG   11242
gb:MT027064|Organism:Severe      TTGTTACCTTCTCTTGCCACTGTAGCTTATTTTAATATGGTCTATATGCCTGCTAGTTGG   11242
gb:MN994468|Organism:Severe      TTGTTACCTTCTCTTGCCACTGTAGCTTATTTTAATATGGTCTATATGCCTGCTAGTTGG   11242
gb:MT072688|Organism:Severe      TTGTTACCTTCTCTTGCCACTGTAGCTTATTTTAATATGGTCTATATGCCTGCTAGTTGG   11227
gb:MN996527|Organism:Severe      TTGTTACCTTCTCTTGCCACTGTAGCTTATTTTAATATGGTCTATATGCCTGCTAGTTGG   11209
gb:MT093631|Organism:Severe      TTGTTACCTTCTCTTGCCACTGTAGCTTATTTTAATATGGTCTATATGCCTGCTAGTTGG   11280
gb:MT106053|Organism:Severe      TTGTTACCTTCTCTTGCCACTGTAGCTTATTTTAATATGGTCTATATGCCTGCTAGTTGG   11242
gb:MT019533|Organism:Severe      TTGTTACCTTCTCTTGCCACTGTAGCTTATTTTAATATGGTCTATATGCCTGCTAGTTGG   11242
```

```
gb:MT019531|Organism:Severe      TTGTTACCTTCTCTTGCCACTGTAGCTTATTTTAATATGGTCTATATGCCTGCTAGTTGG   11242
gb:MN996528|Organism:Severe      TTGTTACCTTCTCTTGCCACTGTAGCTTATTTTAATATGGTCTATATGCCTGCTAGTTGG   11242
gb:MN996530|Organism:Severe      TTGTTACCTTCTCTTGCCACTGTAGCTTATTTTAATATGGTCTATATGCCTGCTAGTTGG   11228
gb:MN908947|Organism:Severe      TTGTTACCTTCTCTTGCCACTGTAGCTTATTTTAATATGGTCTATATGCCTGCTAGTTGG   11242
gb:MT019532|Organism:Severe      TTGTTACCTTCTCTTGCCACTGTAGCTTATTTTAATATGGTCTATATGCCTGCTAGTTGG   11242
                                 ************************************************************

gb:MT020781|Organism:Severe      GTGATGCGTATTATGACATGGTTGGATATGGTTGATACTAGTTTGTCTGGTTTTAAGCTA   11290
gb:MT007544|Organism:Severe      GTGATGCGTATTATGACATGGTTGGATATGGTTGATACTAGTTTGTCTGGTTTTAAGCTA   11302
gb:MN994467|Organism:Severe      GTGATGCGTATTATGACATGGTTGGATATGGTTGATACTAGTTTGTCTGGTTTTAAGCTA   11302
gb:MT044257|Organism:Severe      GTGATGCGTATTATGACATGGTTGGATATGGTTGATACTAGTTTGTCTGGTTTTAAGCTA   11302
gb:MT106054|Organism:Severe      GTGATGCGTATTATGACATGGTTGGATATGGTTGATACTAGTTTGTCTGGTTTTAAGCTA   11302
gb:MT049951|Organism:Severe      GTGATGCGTATTATGACATGGTTGGATATGGTTGATACTAGTTTGTCTGGTTTTAAGCTA   11302
gb:MN975262|Organism:Severe      GTGATGCGTATTATGACATGGTTGGATATGGTTGATACTAGTTTGTCTGGTTTTAAGCTA   11302
gb:MT106052|Organism:Severe      GTGATGCGTATTATGACATGGTTGGATATGGTTGATACTAGTTTGTCTGGTTTTAAGCTA   11302
gb:LC522975|Organism:Severe      GTGATGCGTATTATGACATGGTTGGATATGGTTGATACTAGTTTGTCTGGTTTTAAGCTA   11299
gb:LC522973|Organism:Severe      GTGATGCGTATTATGACATGGTTGGATATGGTTGATACTAGTTTGTCTGGTTTTAAGCTA   11299
gb:LC522974|Organism:Severe      GTGATGCGTATTATGACATGGTTGGATATGGTTGATACTAGTTTGTCTGGTTTTAAGCTA   11299
gb:MN985325|Organism:Severe      GTGATGCGTATTATGACATGGTTGGATATGGTTGATACTAGTTTGTCTGGTTTTAAGCTA   11302
gb:MT020881|Organism:Severe      GTGATGCGTATTATGACATGGTTGGATATGGTTGATACTAGTTTGTCTGGTTTTAAGCTA   11302
gb:MT020880|Organism:Severe      GTGATGCGTATTATGACATGGTTGGATATGGTTGATACTAGTTTGTCTGGTTTTAAGCTA   11302
gb:MT066175|Organism:Severe      GTGATGCGTATTATGACATGGTTGGATATGGTTGATACTAGTTTGTCTGGTTTTAAGCTA   11302
gb:MN997409|Organism:Severe      GTGATGCGTATTATGACATGGTTGGATATGGTTGATACTAGTTTGTCTGGTTTTAAGCTA   11302
gb:MN938384|Organism:Severe      GTGATGCGTATTATGACATGGTTGGATATGGTTGATACTAGTTTGTCTGGTTTTAAGCTA   11270
gb:MT044258|Organism:Severe      GTGATGCGTATTATGACATGGTTGGATATGGTTGATACTAGTTTGTCTGGTTTTAAGCTA   11278
gb:MT039890|Organism:Severe      GTGATGCGTATTATGACATGGTTGGATATGGTTGATACTAGTTTGTCTGGTTTTAAGCTA   11302
gb:MN988713|Organism:Severe      GTGATGCGTATTATGACATGGTTGGATATGGTTGATACTAGTTTGTCTGGTTTTAAGCTA   11302
gb:LC521925|Organism:Severe      GTGATGCGTATTATGACATGGTTGGATATGGTTGATACTAGTTTGTCTGGTTTTAAGCTA   11275
gb:MT093571|Organism:Severe      GTGATGCGTATTATGACATGGTTGGATATGGTTGATACTAGTTTGTCTGGTTTTAAGCTA   11302
gb:MT039887|Organism:Severe      GTGATGCGTATTATGACATGGTTGGATATGGTTGATACTAGTTTGTCTGGTTTTAAGCTA   11302
gb:MT019530|Organism:Severe      GTGATGCGTATTATGACATGGTTGGATATGGTTGATACTAGTTTGTCTGGTTTTAAGCTA   11302
gb:MT039888|Organism:Severe      GTGATGCGTATTATGACATGGTTGGATATGGTTGATACTAGTTTGTCTGGTTTTAAGCTA   11302
gb:LC522972|Organism:Severe      GTGATGCGTATTATGACATGGTTGGATATGGTTGATACTAGTTTGTCTGGTTTTAAGCTA   11299
gb:MT027063|Organism:Severe      GTGATGCGTATTATGACATGGTTGGATATGGTTGATACTAGTTTGTCTGGTTTTAAGCTA   11302
gb:MT027062|Organism:Severe      GTGATGCGTATTATGACATGGTTGGATATGGTTGATACTAGTTTGTCTGGTTTTAAGCTA   11302
gb:MT019529|Organism:Severe      GTGATGCGTATTATGACATGGTTGGATATGGTTGATACTAGTTTGTCTGGTTTTAAGCTA   11302
gb:MN996529|Organism:Severe      GTGATGCGTATTATGACATGGTTGGATATGGTTGATACTAGTTTGTCTGGTTTTAAGCTA   11290
gb:MN996531|Organism:Severe      GTGATGCGTATTATGACATGGTTGGATATGGTTGATACTAGTTTGTCTGGTTTTAAGCTA   11289
gb:MT066176|Organism:Severe      GTGATGCGTATTATGACATGGTTGGATATGGTTGATACTAGTTTGTCTGGTTTTAAGCTA   11302
gb:MT027064|Organism:Severe      GTGATGCGTATTATGACATGGTTGGATATGGTTGATACTAGTTTGTCTGGTTTTAAGCTA   11302
gb:MN994468|Organism:Severe      GTGATGCGTATTATGACATGGTTGGATATGGTTGATACTAGTTTGTCTGGTTTTAAGCTA   11302
gb:MT072688|Organism:Severe      GTGATGCGTATTATGACATGGTTGGATATGGTTGATACTAGTTTGTCTGGTTTTAAGCTA   11287
gb:MN996527|Organism:Severe      GTGATGCGTATTATGACATGGTTGGATATGGTTGATACTAGTTTGTCTGGTTTTAAGCTA   11269
gb:MT093631|Organism:Severe      GTGATGCGTATTATGACATGGTTGGATATGGTTGATACTAGTTTGTCTGGTTTTAAGCTA   11340
gb:MT106053|Organism:Severe      GTGATGCGTATTATGACATGGTTGGATATGGTTGATACTAGTTTGTCTGGTTTTAAGCTA   11302
gb:MT019533|Organism:Severe      GTGATGCGTATTATGACATGGTTGGATATGGTTGATACTAGTTTGTCTGGTTTTAAGCTA   11302
gb:MT019531|Organism:Severe      GTGATGCGTATTATGACATGGTTGGATATGGTTGATACTAGTTTGTCTGGTTTTAAGCTA   11302
gb:MN996528|Organism:Severe      GTGATGCGTATTATGACATGGTTGGATATGGTTGATACTAGTTTGTCTGGTTTTAAGCTA   11302
gb:MN996530|Organism:Severe      GTGATGCGTATTATGACATGGTTGGATATGGTTGATACTAGTTTGTCTGGTTTTAAGCTA   11288
gb:MN908947|Organism:Severe      GTGATGCGTATTATGACATGGTTGGATATGGTTGATACTAGTTTGTCTGGTTTTAAGCTA   11302
gb:MT019532|Organism:Severe      GTGATGCGTATTATGACATGGTTGGATATGGTTGATACTAGTTTGTCTGGTTTTAAGCTA   11302
                                 ************************************************************

gb:MT020781|Organism:Severe      AAAGACTGTGTTATGTATGCATCAGCTGTAGTGTTACTAATCCTTATGACAGCAAGAACT   11350
gb:MT007544|Organism:Severe      AAAGACTGTGTTATGTATGCATCAGCTGTAGTGTTACTAATCCTTATGACAGCAAGAACT   11362
gb:MN994467|Organism:Severe      AAAGACTGTGTTATGTATGCATCAGCTGTAGTGTTACTAATCCTTATGACAGCAAGAACT   11362
gb:MT044257|Organism:Severe      AAAGACTGTGTTATGTATGCATCAGCTGTAGTGTTACTAATCCTTATGACAGCAAGAACT   11362
gb:MT106054|Organism:Severe      AAAGACTGTGTTATGTATGCATCAGCTGTAGTGTTACTAATCCTTATGACAGCAAGAACT   11362
gb:MT049951|Organism:Severe      AAAGACTGTGTTATGTATGCATCAGCTGTAGTGTTACTAATCCTTATGACAGCAAGAACT   11362
gb:MN975262|Organism:Severe      AAAGACTGTGTTATGTATGCATCAGCTGTAGTGTTACTAATCCTTATGACAGCAAGAACT   11362
gb:MT106052|Organism:Severe      AAAGACTGTGTTATGTATGCATCAGCTGTAGTGTTACTAATCCTTATGACAGCAAGAACT   11362
gb:LC522975|Organism:Severe      AAAGACTGTGTTATGTATGCATCAGCTGTAGTGTTACTAATCCTTATGACAGCAAGAACT   11359
gb:LC522973|Organism:Severe      AAAGACTGTGTTATGTATGCATCAGCTGTAGTGTTACTAATCCTTATGACAGCAAGAACT   11359
gb:LC522974|Organism:Severe      AAAGACTGTGTTATGTATGCATCAGCTGTAGTGTTACTAATCCTTATGACAGCAAGAACT   11359
gb:MN985325|Organism:Severe      AAAGACTGTGTTATGTATGCATCAGCTGTAGTGTTACTAATCCTTATGACAGCAAGAACT   11362
```

```
gb:MT020881|Organism:Severe      AAAGACTGTGTTATGTATGCATCAGCTGTAGTGTTACTAATCCTTATGACAGCAAGAACT   11362
gb:MT020880|Organism:Severe      AAAGACTGTGTTATGTATGCATCAGCTGTAGTGTTACTAATCCTTATGACAGCAAGAACT   11362
gb:MT066175|Organism:Severe      AAAGACTGTGTTATGTATGCATCAGCTGTAGTGTTACTAATCCTTATGACAGCAAGAACT   11362
gb:MN997409|Organism:Severe      AAAGACTGTGTTATGTATGCATCAGCTGTAGTGTTACTAATCCTTATGACAGCAAGAACT   11362
gb:MN938384|Organism:Severe      AAAGACTGTGTTATGTATGCATCAGCTGTAGTGTTACTAATCCTTATGACAGCAAGAACT   11330
gb:MT044258|Organism:Severe      AAAGACTGTGTTATGTATGCATCAGCTGTAGTGTTACTAATCCTTATGACAGCAAGAACT   11338
gb:MT039890|Organism:Severe      AAAGACTGTGTTATGTATGCATCAGCTGTAGTGTTACTAATCCTTATGACAGCAAGAACT   11362
gb:MN988713|Organism:Severe      AAAGACTGTGTTATGTATGCATCAGCTGTAGTGTTACTAATCCTTATGACAGCAAGAACT   11362
gb:LC521925|Organism:Severe      AAAGACTGTGTTATGTATGCATCAGCTGTAGTGTTACTAATCCTTATGACAGCAAGAACT   11335
gb:MT093571|Organism:Severe      AAAGACTGTGTTATGTATGCATCAGCTGTAGTGTTACTAATCCTTATGACAGCAAGAACT   11362
gb:MT039887|Organism:Severe      AAAGACTGTGTTATGTATGCATCAGCTGTAGTGTTACTAATCCTTATGACAGCAAGAACT   11362
gb:MT019530|Organism:Severe      AAAGACTGTGTTATGTATGCATCAGCTGTAGTGTTACTAATCCTTATGACAGCAAGAACT   11362
gb:MT039888|Organism:Severe      AAAGACTGTGTTATGTATGCATCAGCTGTAGTGTTACTAATCCTTATGACAGCAAGAACT   11362
gb:LC522972|Organism:Severe      AAAGACTGTGTTATGTATGCATCAGCTGTAGTGTTACTAATCCTTATGACAGCAAGAACT   11359
gb:MT027063|Organism:Severe      AAAGACTGTGTTATGTATGCATCAGCTGTAGTGTTACTAATCCTTATGACAGCAAGAACT   11362
gb:MT027062|Organism:Severe      AAAGACTGTGTTATGTATGCATCAGCTGTAGTGTTACTAATCCTTATGACAGCAAGAACT   11362
gb:MT019529|Organism:Severe      AAAGACTGTGTTATGTATGCATCAGCTGTAGTGTTACTAATCCTTATGACAGCAAGAACT   11362
gb:MN996529|Organism:Severe      AAAGACTGTGTTATGTATGCATCAGCTGTAGTGTTACTAATCCTTATGACAGCAAGAACT   11350
gb:MN996531|Organism:Severe      AAAGACTGTGTTATGTATGCATCAGCTGTAGTGTTACTAATCCTTATGACAGCAAGAACT   11349
gb:MT066176|Organism:Severe      AAAGACTGTGTTATGTATGCATCAGCTGTAGTGTTACTAATCCTTATGACAGCAAGAACT   11362
gb:MT027064|Organism:Severe      AAAGACTGTGTTATGTATGCATCAGCTGTAGTGTTACTAATCCTTATGACAGCAAGAACT   11362
gb:MN994468|Organism:Severe      AAAGACTGTGTTATGTATGCATCAGCTGTAGTGTTACTAATCCTTATGACAGCAAGAACT   11362
gb:MT072688|Organism:Severe      AAAGACTGTGTTATGTATGCATCAGCTGTAGTGTTACTAATCCTTATGACAGCAAGAACT   11347
gb:MN996527|Organism:Severe      AAAGACTGTGTTATGTATGCATCAGCTGTAGTGTTACTAATCCTTATGACAGCAAGAACT   11329
gb:MT093631|Organism:Severe      AAAGACTGTGTTATGTATGCATCAGCTGTAGTGTTACTAATCCTTATGACAGCAAGAACT   11400
gb:MT106053|Organism:Severe      AAAGACTGTGTTATGTATGCATCAGCTGTAGTGTTACTAATCCTTATGACAGCAAGAACT   11362
gb:MT019533|Organism:Severe      AAAGACTGTGTTATGTATGCATCAGCTGTAGTGTTACTAATCCTTATGACAGCAAGAACT   11362
gb:MT019531|Organism:Severe      AAAGACTGTGTTATGTATGCATCAGCTGTAGTGTTACTAATCCTTATGACAGCAAGAACT   11362
gb:MN996528|Organism:Severe      AAAGACTGTGTTATGTATGCATCAGCTGTAGTGTTACTAATCCTTATGACAGCAAGAACT   11362
gb:MN996530|Organism:Severe      AAAGACTGTGTTATGTATGCATCAGCTGTAGTGTTACTAATCCTTATGACAGCAAGAACT   11348
gb:MN908947|Organism:Severe      AAAGACTGTGTTATGTATGCATCAGCTGTAGTGTTACTAATCCTTATGACAGCAAGAACT   11362
gb:MT019532|Organism:Severe      AAAGACTGTGTTATGTATGCATCAGCTGTAGTGTTACTAATCCTTATGACAGCAAGAACT   11362
                                 ************************************************************

gb:MT020781|Organism:Severe      GTGTATGATGATGGTGCTAGGAGAGTGTGGACACTTATGAATGTCTTGACACTCGTTTAT   11410
gb:MT007544|Organism:Severe      GTGTATGATGATGGTGCTAGGAGAGTGTGGACACTTATGAATGTCTTGACACTCGTTTAT   11422
gb:MN994467|Organism:Severe      GTGTATGATGATGGTGCTAGGAGAGTGTGGACACTTATGAATGTCTTGACACTCGTTTAT   11422
gb:MT044257|Organism:Severe      GTGTATGATGATGGTGCTAGGAGAGTGTGGACACTTATGAATGTCTTGACACTCGTTTAT   11422
gb:MT106054|Organism:Severe      GTGTATGATGATGGTGCTAGGAGAGTGTGGACACTTATGAATGTCTTGACACTCGTTTAT   11422
gb:MT049951|Organism:Severe      GTGTATGATGATGGTGCTAGGAGAGTGTGGACACTTATGAATGTCTTGACACTCGTTTAT   11422
gb:MN975262|Organism:Severe      GTGTATGATGATGGTGCTAGGAGAGTGTGGACACTTATGAATGTCTTGACACTCGTTTAT   11422
gb:MT106052|Organism:Severe      GTGTATGATGATGGTGCTAGGAGAGTGTGGACACTTATGAATGTCTTGACACTCGTTTAT   11422
gb:LC522975|Organism:Severe      GTGTATGATGATGGTGCTAGGAGAGTGTGGACACTTATGAATGTCTTGACACTCGTTTAT   11419
gb:LC522973|Organism:Severe      GTGTATGATGATGGTGCTAGGAGAGTGTGGACACTTATGAATGTCTTGACACTCGTTTAT   11419
gb:LC522974|Organism:Severe      GTGTATGATGATGGTGCTAGGAGAGTGTGGACACTTATGAATGTCTTGACACTCGTTTAT   11419
gb:MN985325|Organism:Severe      GTGTATGATGATGGTGCTAGGAGAGTGTGGACACTTATGAATGTCTTGACACTCGTTTAT   11422
gb:MT020881|Organism:Severe      GTGTATGATGATGGTGCTAGGAGAGTGTGGACACTTATGAATGTCTTGACACTCGTTTAT   11422
gb:MT020880|Organism:Severe      GTGTATGATGATGGTGCTAGGAGAGTGTGGACACTTATGAATGTCTTGACACTCGTTTAT   11422
gb:MT066175|Organism:Severe      GTGTATGATGATGGTGCTAGGAGAGTGTGGACACTTATGAATGTCTTGACACTCGTTTAT   11422
gb:MN997409|Organism:Severe      GTGTATGATGATGGTGCTAGGAGAGTGTGGACACTTATGAATGTCTTGACACTCGTTTAT   11422
gb:MN938384|Organism:Severe      GTGTATGATGATGGTGCTAGGAGAGTGTGGACACTTATGAATGTCTTGACACTCGTTTAT   11390
gb:MT044258|Organism:Severe      GTGTATGATGATGGTGCTAGGAGAGTGTGGACACTTATGAATGTCTTGACACTCGTTTAT   11398
gb:MT039890|Organism:Severe      GTGTATGATGATGGTGCTAGGAGAGTGTGGACACTTATGAATGTCTTGACACTCGTTTAT   11422
gb:MN988713|Organism:Severe      GTGTATGATGATGGTGCTAGGAGAGTGTGGACACTTATGAATGTCTTGACACTCGTTTAT   11422
gb:LC521925|Organism:Severe      GTGTATGATGATGGTGCTAGGAGAGTGTGGACACTTATGAATGTCTTGACACTCGTTTAT   11395
gb:MT093571|Organism:Severe      GTGTATGATGATGGTGCTAGGAGAGTGTGGACACTTATGAATGTCTTGACACTCGTTTAT   11422
gb:MT039887|Organism:Severe      GTGTATGATGATGGTGCTAGGAGAGTGTGGACACTTATGAATGTCTTGACACTCGTTTAT   11422
gb:MT019530|Organism:Severe      GTGTATGATGATGGTGCTAGGAGAGTGTGGACACTTATGAATGTCTTGACACTCGTTTAT   11422
gb:MT039888|Organism:Severe      GTGTATGATGATGGTGCTAGGAGAGTGTGGACACTTATGAATGTCTTGACACTCGTTTAT   11422
gb:LC522972|Organism:Severe      GTGTATGATGATGGTGCTAGGAGAGTGTGGACACTTATGAATGTCTTGACACTCGTTTAT   11419
gb:MT027063|Organism:Severe      GTGTATGATGATGGTGCTAGGAGAGTGTGGACACTTATGAATGTCTTGACACTCGTTTAT   11422
gb:MT027062|Organism:Severe      GTGTATGATGATGGTGCTAGGAGAGTGTGGACACTTATGAATGTCTTGACACTCGTTTAT   11422
gb:MT019529|Organism:Severe      GTGTATGATGATGGTGCTAGGAGAGTGTGGACACTTATGAATGTCTTGACACTCGTTTAT   11422
gb:MN996529|Organism:Severe      GTGTATGATGATGGTGCTAGGAGAGTGTGGACACTTATGAATGTCTTGACACTCGTTTAT   11410
gb:MN996531|Organism:Severe      GTGTATGATGATGGTGCTAGGAGAGTGTGGACACTTATGAATGTCTTGACACTCGTTTAT   11409
```

```
gb:MT066176|Organism:Severe      GTGTATGATGATGGTGCTAGGAGAGTGTGGACACTTATGAATGTCTTGACACTCGTTTAT   11422
gb:MT027064|Organism:Severe      GTGTATGATGATGGTGCTAGGAGAGTGTGGACACTTATGAATGTCTTGACACTCGTTTAT   11422
gb:MN994468|Organism:Severe      GTGTATGATGATGGTGCTAGGAGAGTGTGGACACTTATGAATGTCTTGACACTCGTTTAT   11422
gb:MT072688|Organism:Severe      GTGTATGATGATGGTGCTAGGAGAGTGTGGACACTTATGAATGTCTTGACACTCGTTTAT   11407
gb:MN996527|Organism:Severe      GTGTATGATGATGGTGCTAGGAGAGTGTGGACACTTATGAATGTCTTGACACTCGTTTAT   11389
gb:MT093631|Organism:Severe      GTGTATGATGATGGTGCTAGGAGAGTGTGGACACTTATGAATGTCTTGACACTCGTTTAT   11460
gb:MT106053|Organism:Severe      GTGTATGATGATGGTGCTAGGAGAGTGTGGACACTTATGAATGTCTTGACACTCGTTTAT   11422
gb:MT019533|Organism:Severe      GTGTATGATGATGGTGCTAGGAGAGTGTGGACACTTATGAATGTCTTGACACTCGTTTAT   11422
gb:MT019531|Organism:Severe      GTGTATGATGATGGTGCTAGGAGAGTGTGGACACTTATGAATGTCTTGACACTCGTTTAT   11422
gb:MN996528|Organism:Severe      GTGTATGATGATGGTGCTAGGAGAGTGTGGACACTTATGAATGTCTTGACACTCGTTTAT   11422
gb:MN996530|Organism:Severe      GTGTATGATGATGGTGCTAGGAGAGTGTGGACACTTATGAATGTCTTGACACTCGTTTAT   11408
gb:MN908947|Organism:Severe      GTGTATGATGATGGTGCTAGGAGAGTGTGGACACTTATGAATGTCTTGACACTCGTTTAT   11422
gb:MT019532|Organism:Severe      GTGTATGATGATGGTGCTAGGAGAGTGTGGACACTTATGAATGTCTTGACACTCGTTTAT   11422
                                 ************************************************************

gb:MT020781|Organism:Severe      AAAGTTTATTATGGTAATGCTTTAGATCAAGCCATTTCCATGTGGGCTCTTATAATCTCT   11470
gb:MT007544|Organism:Severe      AAAGTTTATTATGGTAATGCTTTAGATCAAGCCATTTCCATGTGGGCTCTTATAATCTCT   11482
gb:MN994467|Organism:Severe      AAAGTTTATTATGGTAATGCTTTAGATCAAGCCATTTCCATGTGGGCTCTTATAATCTCT   11482
gb:MT044257|Organism:Severe      AAAGTTTATTATGGTAATGCTTTAGATCAAGCCATTTCCATGTGGGCTCTTATAATCTCT   11482
gb:MT106054|Organism:Severe      AAAGTTTATTATGGTAATGCTTTAGATCAAGCCATTTCCATGTGGGCTCTTATAATCTCT   11482
gb:MT049951|Organism:Severe      AAAGTTTATTATGGTAATGCTTTAGATCAAGCCATTTCCATGTGGGCTCTTATAATCTCT   11482
gb:MN975262|Organism:Severe      AAAGTTTATTATGGTAATGCTTTAGATCAAGCCATTTCCATGTGGGCTCTTATAATCTCT   11482
gb:MT106052|Organism:Severe      AAAGTTTATTATGGTAATGCTTTAGATCAAGCCATTTCCATGTGGGCTCTTATAATCTCT   11482
gb:LC522975|Organism:Severe      AAAGTTTATTATGGTAATGCTTTAGATCAAGCCATTTCCATGTGGGCTCTTATAATCTCT   11479
gb:LC522973|Organism:Severe      AAAGTTTATTATGGTAATGCTTTAGATCAAGCCATTTCCATGTGGGCTCTTATAATCTCT   11479
gb:LC522974|Organism:Severe      AAAGTTTATTATGGTAATGCTTTAGATCAAGCCATTTCCATGTGGGCTCTTATAATCTCT   11479
gb:MN985325|Organism:Severe      AAAGTTTATTATGGTAATGCTTTAGATCAAGCCATTTCCATGTGGGCTCTTATAATCTCT   11482
gb:MT020881|Organism:Severe      AAAGTTTATTATGGTAATGCTTTAGATCAAGCCATTTCCATGTGGGCTCTTATAATCTCT   11482
gb:MT020880|Organism:Severe      AAAGTTTATTATGGTAATGCTTTAGATCAAGCCATTTCCATGTGGGCTCTTATAATCTCT   11482
gb:MT066175|Organism:Severe      AAAGTTTATTATGGTAATGCTTTAGATCAAGCCATTTCCATGTGGGCTCTTATAATCTCT   11482
gb:MN997409|Organism:Severe      AAAGTTTATTATGGTAATGCTTTAGATCAAGCCATTTCCATGTGGGCTCTTATAATCTCT   11482
gb:MN938384|Organism:Severe      AAAGTTTATTATGGTAATGCTTTAGATCAAGCCATTTCCATGTGGGCTCTTATAATCTCT   11450
gb:MT044258|Organism:Severe      AAAGTTTATTATGGTAATGCTTTAGATCAAGCCATTTCCATGTGGGCTCTTATAATCTCT   11458
gb:MT039890|Organism:Severe      AAAGTTTATTATGGTAATGCTTTAGATCAAGCCATTTCCATGTGGGCTCTTATAATCTCT   11482
gb:MN988713|Organism:Severe      AAAGTTTATTATGGTAATGCTTTAGATCAAGCCATTTCCATGTGGGCTCTTATAATCTCT   11482
gb:LC521925|Organism:Severe      AAAGTTTATTATGGTAATGCTTTAGATCAAGCCATTTCCATGTGGGCTCTTATAATCTCT   11455
gb:MT093571|Organism:Severe      AAAGTTTATTATGGTAATGCTTTAGATCAAGCCATTTCCATGTGGGCTCTTATAATCTCT   11482
gb:MT039887|Organism:Severe      AAAGTTTATTATGGTAATGCTTTAGATCAAGCCATTTCCATGTGGGCTCTTATAATCTCT   11482
gb:MT019530|Organism:Severe      AAAGTTTATTATGGTAATGCTTTAGATCAAGCCATTTCCATGTGGGCTCTTATAATCTCT   11482
gb:MT039888|Organism:Severe      AAAGTTTATTATGGTAATGCTTTAGATCAAGCCATTTCCATGTGGGCTCTTATAATCTCT   11482
gb:LC522972|Organism:Severe      AAAGTTTATTATGGTAATGCTTTAGATCAAGCCATTTCCATGTGGGCTCTTATAATCTCT   11479
gb:MT027063|Organism:Severe      AAAGTTTATTATGGTAATGCTTTAGATCAAGCCATTTCCATGTGGGCTCTTATAATCTCT   11482
gb:MT027062|Organism:Severe      AAAGTTTATTATGGTAATGCTTTAGATCAAGCCATTTCCATGTGGGCTCTTATAATCTCT   11482
gb:MT019529|Organism:Severe      AAAGTTTATTATGGTAATGCTTTAGATCAAGCCATTTCCATGTGGGCTCTTATAATCTCT   11482
gb:MN996529|Organism:Severe      AAAGTTTATTATGGTAATGCTTTAGATCAAGCCATTTCCATGTGGGCTCTTATAATCTCT   11470
gb:MN996531|Organism:Severe      AAAGTTTATTATGGTAATGCTTTAGATCAAGCCATTTCCATGTGGGCTCTTATAATCTCT   11469
gb:MT066176|Organism:Severe      AAAGTTTATTATGGTAATGCTTTAGATCAAGCCATTTCCATGTGGGCTCTTATAATCTCT   11482
gb:MT027064|Organism:Severe      AAAGTTTATTATGGTAATGCTTTAGATCAAGCCATTTCCATGTGGGCTCTTATAATCTCT   11482
gb:MN994468|Organism:Severe      AAAGTTTATTATGGTAATGCTTTAGATCAAGCCATTTCCATGTGGGCTCTTATAATCTCT   11482
gb:MT072688|Organism:Severe      AAAGTTTATTATGGTAATGCTTTAGATCAAGCCATTTCCATGTGGGCTCTTATAATCTCT   11467
gb:MN996527|Organism:Severe      AAAGTTTATTATGGTAATGCTTTAGATCAAGCCATTTCCATGTGGGCTCTTATAATCTCT   11449
gb:MT093631|Organism:Severe      AAAGTTTATTATGGTAATGCTTTAGATCAAGCCATTTCCATGTGGGCTCTTATAATCTCT   11520
gb:MT106053|Organism:Severe      AAAGTTTATTATGGTAATGCTTTAGATCAAGCCATTTCCATGTGGGCTCTTATAATCTCT   11482
gb:MT019533|Organism:Severe      AAAGTTTATTATGGTAATGCTTTAGATCAAGCCATTTCCATGTGGGCTCTTATAATCTCT   11482
gb:MT019531|Organism:Severe      AAAGTTTATTATGGTAATGCTTTAGATCAAGCCATTTCCATGTGGGCTCTTATAATCTCT   11482
gb:MN996528|Organism:Severe      AAAGTTTATTATGGTAATGCTTTAGATCAAGCCATTTCCATGTGGGCTCTTATAATCTCT   11482
gb:MN996530|Organism:Severe      AAAGTTTATTATGGTAATGCTTTAGATCAAGCCATTTCCATGTGGGCTCTTATAATCTCT   11468
gb:MN908947|Organism:Severe      AAAGTTTATTATGGTAATGCTTTAGATCAAGCCATTTCCATGTGGGCTCTTATAATCTCT   11482
gb:MT019532|Organism:Severe      AAAGTTTATTATGGTAATGCTTTAGATCAAGCCATTTCCATGTGGGCTCTTATAATCTCT   11482
                                 ************************************************************

gb:MT020781|Organism:Severe      GTTACTTCTAACTACTCAGGTGTAGTTACAACTGTCATGTTTTTGGCCAGAGGTATTGTT   11530
gb:MT007544|Organism:Severe      GTTACTTCTAACTACTCAGGTGTAGTTACAACTGTCATGTTTTTGGCCAGAGGTATTGTT   11542
gb:MN994467|Organism:Severe      GTTACTTCTAACTACTCAGGTGTAGTTACAACTGTCATGTTTTTGGCCAGAGGTATTGTT   11542
gb:MT044257|Organism:Severe      GTTACTTCTAACTACTCAGGTGTAGTTACAACTGTCATGTTTTTGGCCAGAGGTATTGTT   11542
```

```
gb:MT106054|Organism:Severe      GTTACTTCTAACTACTCAGGTGTAGTTACAACTGTCATGTTTTTGGCCAGAGGTATTGTT   11542
gb:MT049951|Organism:Severe      GTTACTTCTAACTACTCAGGTGTAGTTACAACTGTCATGTTTTTGGCCAGAGGTATTGTT   11542
gb:MN975262|Organism:Severe      GTTACTTCTAACTACTCAGGTGTAGTTACAACTGTCATGTTTTTGGCCAGAGGTATTGTT   11542
gb:MT106052|Organism:Severe      GTTACTTCTAACTACTCAGGTGTAGTTACAACTGTCATGTTTTTGGCCAGAGGTATTGTT   11542
gb:LC522975|Organism:Severe      GTTACTTCTAACTACTCAGGTGTAGTTACAACTGTCATGTTTTTGGCCAGAGGTATTGTT   11539
gb:LC522973|Organism:Severe      GTTACTTCTAACTACTCAGGTGTAGTTACAACTGTCATGTTTTTGGCCAGAGGTATTGTT   11539
gb:LC522974|Organism:Severe      GTTACTTCTAACTACTCAGGTGTAGTTACAACTGTCATGTTTTTGGCCAGAGGTATTGTT   11539
gb:MN985325|Organism:Severe      GTTACTTCTAACTACTCAGGTGTAGTTACAACTGTCATGTTTTTGGCCAGAGGTATTGTT   11542
gb:MT020881|Organism:Severe      GTTACTTCTAACTACTCAGGTGTAGTTACAACTGTCATGTTTTTGGCCAGAGGTATTGTT   11542
gb:MT020880|Organism:Severe      GTTACTTCTAACTACTCAGGTGTAGTTACAACTGTCATGTTTTTGGCCAGAGGTATTGTT   11542
gb:MT066175|Organism:Severe      GTTACTTCTAACTACTCAGGTGTAGTTACAACTGTCATGTTTTTGGCCAGAGGTATTGTT   11542
gb:MN997409|Organism:Severe      GTTACTTCTAACTACTCAGGTGTAGTTACAACTGTCATGTTTTTGGCCAGAGGTATTGTT   11542
gb:MN938384|Organism:Severe      GTTACTTCTAACTACTCAGGTGTAGTTACAACTGTCATGTTTTTGGCCAGAGGTATTGTT   11510
gb:MT044258|Organism:Severe      GTTACTTCTAACTACTCAGGTGTAGTTACAACTGTCATGTTTTTGGCCAGAGGTATTGTT   11518
gb:MT039890|Organism:Severe      GTTACTTCTAACTACTCAGGTGTAGTTACAACTGTCATGTTTTTGGCCAGAGGTATTGTT   11542
gb:MN988713|Organism:Severe      GTTACTTCTAACTACTCAGGTGTAGTTACAACTGTCATGTTTTTGGCCAGAGGTATTGTT   11542
gb:LC521925|Organism:Severe      GTTACTTCTAACTACTCAGGTGTAGTTACAACTGTCATGTTTTTGGCCAGAGGTATTGTT   11515
gb:MT093571|Organism:Severe      GTTACTTCTAACTACTCAGGTGTAGTTACAACTGTCATGTTTTTGGCCAGAGGTATTGTT   11542
gb:MT039887|Organism:Severe      GTTACTTCTAACTACTCAGGTGTAGTTACAACTGTCATGTTTTTGGCCAGAGGTATTGTT   11542
gb:MT019530|Organism:Severe      GTTACTTCTAACTACTCAGGTGTAGTTACAACTGTCATGTTTTTGGCCAGAGGTATTGTT   11542
gb:MT039888|Organism:Severe      GTTACTTCTAACTACTCAGGTGTAGTTACAACTGTCATGTTTTTGGCCAGAGGTATTGTT   11542
gb:LC522972|Organism:Severe      GTTACTTCTAACTACTCAGGTGTAGTTACAACTGTCATGTTTTTGGCCAGAGGTATTGTT   11539
gb:MT027063|Organism:Severe      GTTACTTCTAACTACTCAGGTGTAGTTACAACTGTCATGTTTTTGGCCAGAGGTATTGTT   11542
gb:MT027062|Organism:Severe      GTTACTTCTAACTACTCAGGTGTAGTTACAACTGTCATGTTTTTGGCCAGAGGTATTGTT   11542
gb:MT019529|Organism:Severe      GTTACTTCTAACTACTCAGGTGTAGTTACAACTGTCATGTTTTTGGCCAGAGGTATTGTT   11542
gb:MN996529|Organism:Severe      GTTACTTCTAACTACTCAGGTGTAGTTACAACTGTCATGTTTTTGGCCAGAGGTATTGTT   11530
gb:MN996531|Organism:Severe      GTTACTTCTAACTACTCAGGTGTAGTTACAACTGTCATGTTTTTGGCCAGAGGTATTGTT   11529
gb:MT066176|Organism:Severe      GTTACTTCTAACTACTCAGGTGTAGTTACAACTGTCATGTTTTTGGCCAGAGGTATTGTT   11542
gb:MT027064|Organism:Severe      GTTACTTCTAACTACTCAGGTGTAGTTACAACTGTCATGTTTTTGGCCAGAGGTATTGTT   11542
gb:MN994468|Organism:Severe      GTTACTTCTAACTACTCAGGTGTAGTTACAACTGTCATGTTTTTGGCCAGAGGTATTGTT   11542
gb:MT072688|Organism:Severe      GTTACTTCTAACTACTCAGGTGTAGTTACAACTGTCATGTTTTTGGCCAGAGGTATTGTT   11527
gb:MN996527|Organism:Severe      GTTACTTCTAACTACTCAGGTGTAGTTACAACTGTCATGTTTTTGGCCAGAGGTATTGTT   11509
gb:MT093631|Organism:Severe      GTTACTTCTAACTACTCAGGTGTAGTTACAACTGTCATGTTTTTGGCCAGAGGTATTGTT   11580
gb:MT106053|Organism:Severe      GTTACTTCTAACTACTCAGGTGTAGTTACAACTGTCATGTTTTTGGCCAGAGGTATTGTT   11542
gb:MT019533|Organism:Severe      GTTACTTCTAACTACTCAGGTGTAGTTACAACTGTCATGTTTTTGGCCAGAGGTATTGTT   11542
gb:MT019531|Organism:Severe      GTTACTTCTAACTACTCAGGTGTAGTTACAACTGTCATGTTTTTGGCCAGAGGTATTGTT   11542
gb:MN996528|Organism:Severe      GTTACTTCTAACTACTCAGGTGTAGTTACAACTGTCATGTTTTTGGCCAGAGGTATTGTT   11542
gb:MN996530|Organism:Severe      GTTACTTCTAACTACTCAGGTGTAGTTACAACTGTCATGTTTTTGGCCAGAGGTATTGTT   11528
gb:MN908947|Organism:Severe      GTTACTTCTAACTACTCAGGTGTAGTTACAACTGTCATGTTTTTGGCCAGAGGTATTGTT   11542
gb:MT019532|Organism:Severe      GTTACTTCTAACTACTCAGGTGTAGTTACAACTGTCATGTTTTTGGCCAGAGGTATTGTT   11542
                                 ************************************************************

gb:MT020781|Organism:Severe      TTTATGTGTGTTGAGTATTGCCCTATTTTCTTCATAACTGGTAATACACTTCAGTGTATA   11590
gb:MT007544|Organism:Severe      TTTATGTGTGTTGAGTATTGCCCTATTTTCTTCATAACTGGTAATACACTTCAGTGTATA   11602
gb:MN994467|Organism:Severe      TTTATGTGTGTTGAGTATTGCCCTATTTTCTTCATAACTGGTAATACACTTCAGTGTATA   11602
gb:MT044257|Organism:Severe      TTTATGTGTGTTGAGTATTGCCCTATTTTCTTCATAACTGGTAATACACTTCAGTGTATA   11602
gb:MT106054|Organism:Severe      TTTATGTGTGTTGAGTATTGCCCTATTTTCTTCATAACTGGTAATACACTTCAGTGTATA   11602
gb:MT049951|Organism:Severe      TTTATGTGTGTTGAGTATTGCCCTATTTTCTTCATAACTGGTAATACACTTCAGTGTATA   11602
gb:MN975262|Organism:Severe      TTTATGTGTGTTGAGTATTGCCCTATTTTCTTCATAACTGGTAATACACTTCAGTGTATA   11602
gb:MT106052|Organism:Severe      TTTATGTGTGTTGAGTATTGCCCTATTTTCTTCATAACTGGTAATACACTTCAGTGTATA   11602
gb:LC522975|Organism:Severe      TTTATGTGTGTTGAGTATTGCCCTATTTTCTTCATAACTGGTAATACACTTCAGTGTATA   11599
gb:LC522973|Organism:Severe      TTTATGTGTGTTGAGTATTGCCCTATTTTCTTCATAACTGGTAATACACTTCAGTGTATA   11599
gb:LC522974|Organism:Severe      TTTATGTGTGTTGAGTATTGCCCTATTTTCTTCATAACTGGTAATACACTTCAGTGTATA   11599
gb:MN985325|Organism:Severe      TTTATGTGTGTTGAGTATTGCCCTATTTTCTTCATAACTGGTAATACACTTCAGTGTATA   11602
gb:MT020881|Organism:Severe      TTTATGTGTGTTGAGTATTGCCCTATTTTCTTCATAACTGGTAATACACTTCAGTGTATA   11602
gb:MT020880|Organism:Severe      TTTATGTGTGTTGAGTATTGCCCTATTTTCTTCATAACTGGTAATACACTTCAGTGTATA   11602
gb:MT066175|Organism:Severe      TTTATGTGTGTTGAGTATTGCCCTATTTTCTTCATAACTGGTAATACACTTCAGTGTATA   11602
gb:MN997409|Organism:Severe      TTTATGTGTGTTGAGTATTGCCCTATTTTCTTCATAACTGGTAATACACTTCAGTGTATA   11602
gb:MN938384|Organism:Severe      TTTATGTGTGTTGAGTATTGCCCTATTTTCTTCATAACTGGTAATACACTTCAGTGTATA   11570
gb:MT044258|Organism:Severe      TTTATGTGTGTTGAGTATTGCCCTATTTTCTTCATAACTGGTAATACACTTCAGTGTATA   11578
gb:MT039890|Organism:Severe      TTTATGTGTGTTGAGTATTGCCCTATTTTCTTCATAACTGGTAATACACTTCAGTGTATA   11602
gb:MN988713|Organism:Severe      TTTATGTGTGTTGAGTATTGCCCTATTTTCTTCATAACTGGTAATACACTTCAGTGTATA   11602
gb:LC521925|Organism:Severe      TTTATGTGTGTTGAGTATTGCCCTATTTTCTTCATAACTGGTAATACACTTCAGTGTATA   11575
gb:MT093571|Organism:Severe      TTTATGTGTGTTGAGTATTGCCCTATTTTCTTCATAACTGGTAATACACTTCAGTGTATA   11602
gb:MT039887|Organism:Severe      TTTATGTGTGTTGAGTATTGCCCTATTTTCTTCATAACTGGTAATACACTTCAGTGTATA   11602
```

```
gb:MT019530|Organism:Severe      TTTATGTGTGTTGAGTATTGCCCTATTTTCTTCATAACTGGTAATACACTTCAGTGTATA    11602
gb:MT039888|Organism:Severe      TTTATGTGTGTTGAGTATTGCCCTATTTTCTTCATAACTGGTAATACACTTCAGTGTATA    11602
gb:LC522972|Organism:Severe      TTTATGTGTGTTGATTATTGCCCTATTTTCTTCATAACTGGTAATACACTTCAGTGTATA    11599
gb:MT027063|Organism:Severe      TTTATGTGTGTTGAGTATTGCCCTATTTTCTTCATAACTGGTAATACACTTCAGTGTATA    11602
gb:MT027062|Organism:Severe      TTTATGTGTGTTGAGTATTGCCCTATTTTCTTCATAACTGGTAATACACTTCAGTGTATA    11602
gb:MT019529|Organism:Severe      TTTATGTGTGTTGAGTATTGCCCTATTTTCTTCATAACTGGTAATACACTTCAGTGTATA    11602
gb:MN996529|Organism:Severe      TTTATGTGTGTTGAGTATTGCCCTATTTTCTTCATAACTGGTAATACACTTCAGTGTATA    11590
gb:MN996531|Organism:Severe      TTTATGTGTGTTGAGTATTGCCCTATTTTCTTCATAACTGGTAATACACTTCAGTGTATA    11589
gb:MT066176|Organism:Severe      TTTATGTGTGTTGAGTATTGCCCTATTTTCTTCATAACTGGTAATACACTTCAGTGTATA    11602
gb:MT027064|Organism:Severe      TTTATGTGTGTTGAGTATTGCCCTATTTTCTTCATAACTGGTAATACACTTCAGTGTATA    11602
gb:MN994468|Organism:Severe      TTTATGTGTGTTGAGTATTGCCCTATTTTCTTCATAACTGGTAATACACTTCAGTGTATA    11602
gb:MT072688|Organism:Severe      TTTATGTGTGTTGAGTATTGCCCTATTTTCTTCATAACTGGTAATACACTTCAGTGTATA    11587
gb:MN996527|Organism:Severe      TTTATGTGTGTTGAGTATTGCCCTATTTTCTTCATAACTGGTAATACACTTCAGTGTATA    11569
gb:MT093631|Organism:Severe      TTTATGTGTGTTGAGTATTGCCCTATTTTCTTCATAACTGGTAATACACTTCAGTGTATA    11640
gb:MT106053|Organism:Severe      TTTATGTGTGTTGAGTATTGCCCTATTTTCTTCATAACTGGTAATACACTTCAGTGTATA    11602
gb:MT019533|Organism:Severe      TTTATGTGTGTTGAGTATTGCCCTATTTTCTTCATAACTGGTAATACACTTCAGTGTATA    11602
gb:MT019531|Organism:Severe      TTTATGTGTGTTGAGTATTGCCCTATTTTCTTCATAACTGGTAATACACTTCAGTGTATA    11602
gb:MN996528|Organism:Severe      TTTATGTGTGTTGAGTATTGCCCTATTTTCTTCATAACTGGTAATACACTTCAGTGTATA    11602
gb:MN996530|Organism:Severe      TTTATGTGTGTTGAGTATTGCCCTATTTTCTTCATAACTGGTAATACACTTCAGTGTATA    11588
gb:MN908947|Organism:Severe      TTTATGTGTGTTGAGTATTGCCCTATTTTCTTCATAACTGGTAATACACTTCAGTGTATA    11602
gb:MT019532|Organism:Severe      TTTATGTGTGTTGAGTATTGCCCTATTTTCTTCATAACTGGTAATACACTTCAGTGTATA    11602
                                 ************** *********************************************

gb:MT020781|Organism:Severe      ATGCTAGTTTATTGTTTCTTAGGCTATTTTTGTACTTGTTACTTTGGCCTCTTTTGTTTA    11650
gb:MT007544|Organism:Severe      ATGCTAGTTTATTGTTTCTTAGGCTATTTTTGTACTTGTTACTTTGGCCTCTTTTGTTTA    11662
gb:MN994467|Organism:Severe      ATGCTAGTTTATTGTTTCTTAGGCTATTTTTGTACTTGTTACTTTGGCCTCTTTTGTTTA    11662
gb:MT044257|Organism:Severe      ATGCTAGTTTATTGTTTCTTAGGCTATTTTTGTACTTGTTACTTTGGCCTCTTTTGTTTA    11662
gb:MT106054|Organism:Severe      ATGCTAGTTTATTGTTTCTTAGGCTATTTTTGTACTTGTTACTTTGGCCTCTTTTGTTTA    11662
gb:MT049951|Organism:Severe      ATGCTAGTTTATTGTTTCTTAGGCTATTTTTGTACTTGTTACTTTGGCCTCTTTTGTTTA    11662
gb:MN975262|Organism:Severe      ATGCTAGTTTATTGTTTCTTAGGCTATTTTTGTACTTGTTACTTTGGCCTCTTTTGTTTA    11662
gb:MT106052|Organism:Severe      ATGCTAGTTTATTGTTTCTTAGGCTATTTTTGTACTTGTTACTTTGGCCTCTTTTGTTTA    11662
gb:LC522975|Organism:Severe      ATGCTAGTTTATTGTTTCTTAGGCTATTTTTGTACTTGTTACTTTGGCCTCTTTTGTTTA    11659
gb:LC522973|Organism:Severe      ATGCTAGTTTATTGTTTCTTAGGCTATTTTTGTACTTGTTACTTTGGCCTCTTTTGTTTA    11659
gb:LC522974|Organism:Severe      ATGCTAGTTTATTGTTTCTTAGGCTATTTTTGTACTTGTTACTTTGGCCTCTTTTGTTTA    11659
gb:MN985325|Organism:Severe      ATGCTAGTTTATTGTTTCTTAGGCTATTTTTGTACTTGTTACTTTGGCCTCTTTTGTTTA    11662
gb:MT020881|Organism:Severe      ATGCTAGTTTATTGTTTCTTAGGCTATTTTTGTACTTGTTACTTTGGCCTCTTTTGTTTA    11662
gb:MT020880|Organism:Severe      ATGCTAGTTTATTGTTTCTTAGGCTATTTTTGTACTTGTTACTTTGGCCTCTTTTGTTTA    11662
gb:MT066175|Organism:Severe      ATGCTAGTTTATTGTTTCTTAGGCTATTTTTGTACTTGTTACTTTGGCCTCTTTTGTTTA    11662
gb:MN997409|Organism:Severe      ATGCTAGTTTATTGTTTCTTAGGCTATTTTTGTACTTGTTACTTTGGCCTCTTTTGTTTA    11662
gb:MN938384|Organism:Severe      ATGCTAGTTTATTGTTTCTTAGGCTATTTTTGTACTTGTTACTTTGGCCTCTTTTGTTTA    11630
gb:MT044258|Organism:Severe      ATGCTAGTTTATTGTTTCTTAGGCTATTTTTGTACTTGTTACTTTGGCCTCTTTTGTTTA    11638
gb:MT039890|Organism:Severe      ATGCTAGTTTATTGTTTCTTAGGCTATTTTTGTACTTGTTACTTTGGCCTCTTTTGTTTA    11662
gb:MN988713|Organism:Severe      ATGCTAGTTTATTGTTTCTTAGGCTATTTTTGTACTTGTTACTTTGGCCTCTTTTGTTTA    11662
gb:LC521925|Organism:Severe      ATGCTAGTTTATTGTTTCTTAGGCTATTTTTGTACTTGTTACTTTGGCCTCTTTTGTTTA    11635
gb:MT093571|Organism:Severe      ATGCTAGTTTATTGTTTCTTAGGCTATTTTTGTACTTGTTACTTTGGCCTCTTTTGTTTA    11662
gb:MT039887|Organism:Severe      ATGCTAGTTTATTGTTTCTTAGGCTATTTTTGTACTTGTTACTTTGGCCTCTTTTGTTTA    11662
gb:MT019530|Organism:Severe      ATGCTAGTTTATTGTTTCTTAGGCTATTTTTGTACTTGTTACTTTGGCCTCTTTTGTTTA    11662
gb:MT039888|Organism:Severe      ATGCTAGTTTATTGTTTCTTAGGCTATTTTTGTACTTGTTACTTTGGCCTCTTTTGTTTA    11662
gb:LC522972|Organism:Severe      ATGCTAGTTTATTGTTTCTTAGGCTATTTTTGTACTTGTTACTTTGGCCTCTTTTGTTTA    11659
gb:MT027063|Organism:Severe      ATGCTAGTTTATTGTTTCTTAGGCTATTTTTGTACTTGTTACTTTGGCCTCTTTTGTTTA    11662
gb:MT027062|Organism:Severe      ATGCTAGTTTATTGTTTCTTAGGCTATTTTTGTACTTGTTACTTTGGCCTCTTTTGTTTA    11662
gb:MT019529|Organism:Severe      ATGCTAGTTTATTGTTTCTTAGGCTATTTTTGTACTTGTTACTTTGGCCTCTTTTGTTTA    11662
gb:MN996529|Organism:Severe      ATGCTAGTTTATTGTTTCTTAGGCTATTTTTGTACTTGTTACTTTGGCCTCTTTTGTTTA    11650
gb:MN996531|Organism:Severe      ATGCTAGTTTATTGTTTCTTAGGCTATTTTTGTACTTGTTACTTTGGCCTCTTTTGTTTA    11649
gb:MT066176|Organism:Severe      ATGCTAGTTTATTGTTTCTTAGGCTATTTTTGTACTTGTTACTTTGGCCTCTTTTGTTTA    11662
gb:MT027064|Organism:Severe      ATGCTAGTTTATTGTTTCTTAGGCTATTTTTGTACTTGTTACTTTGGCCTCTTTTGTTTA    11662
gb:MN994468|Organism:Severe      ATGCTAGTTTATTGTTTCTTAGGCTATTTTTGTACTTGTTACTTTGGCCTCTTTTGTTTA    11662
gb:MT072688|Organism:Severe      ATGCTAGTTTATTGTTTCTTAGGCTATTTTTGTACTTGTTACTTTGGCCTCTTTTGTTTA    11647
gb:MN996527|Organism:Severe      ATGCTAGTTTATTGTTTCTTAGGCTATTTTTGTACTTGTTACTTTGGCCTCTTTTGTTTA    11629
gb:MT093631|Organism:Severe      ATGCTAGTTTATTGTTTCTTAGGCTATTTTTGTACTTGTTACTTTGGCCTCTTTTGTTTA    11700
gb:MT106053|Organism:Severe      ATGCTAGTTTATTGTTTCTTAGGCTATTTTTGTACTTGTTACTTTGGCCTCTTTTGTTTA    11662
gb:MT019533|Organism:Severe      ATGCTAGTTTATTGTTTCTTAGGCTATTTTTGTACTTGTTACTTTGGCCTCTTTTGTTTA    11662
gb:MT019531|Organism:Severe      ATGCTAGTTTATTGTTTCTTAGGCTATTTTTGTACTTGTTACTTTGGCCTCTTTTGTTTA    11662
gb:MN996528|Organism:Severe      ATGCTAGTTTATTGTTTCTTAGGCTATTTTTGTACTTGTTACTTTGGCCTCTTTTGTTTA    11662
gb:MN996530|Organism:Severe      ATGCTAGTTTATTGTTTCTTAGGCTATTTTTGTACTTGTTACTTTGGCCTCTTTTGTTTA    11648
```

```
gb:MN908947|Organism:Severe      ATGCTAGTTTATTGTTTCTTAGGCTATTTTTGTACTTGTTACTTTGGCCTCTTTTGTTTA    11662
gb:MT019532|Organism:Severe      ATGCTAGTTTATTGTTTCTTAGGCTATTTTTGTACTTGTTACTTTGGCCTCTTTTGTTTA    11662
                                 ************************************************************

gb:MT020781|Organism:Severe      CTCAACCGCTACTTTAGACTGACTCTTGGTGTTTATGATTACTTAGTTTCTACACAGGAG    11710
gb:MT007544|Organism:Severe      CTCAACCGCTACTTTAGACTGACTCTTGGTGTTTATGATTACTTAGTTTCTACACAGGAG    11722
gb:MN994467|Organism:Severe      CTCAACCGCTACTTTAGACTGACTCTTGGTGTTTATGATTACTTAGTTTCTACACAGGAG    11722
gb:MT044257|Organism:Severe      CTCAACCGCTACTTTAGACTGACTCTTGGTGTTTATGATTACTTAGTTTCTACACAGGAG    11722
gb:MT106054|Organism:Severe      CTCAACCGCTACTTTAGACTGACTCTTGGTGTTTATGATTACTTAGTTTCTACACAGGAG    11722
gb:MT049951|Organism:Severe      CTCAACCGCTACTTTAGACTGACTCTTGGTGTTTATGATTACTTAGTTTCTACACAGGAG    11722
gb:MN975262|Organism:Severe      CTCAACCGCTACTTTAGACTGACTCTTGGTGTTTATGATTACTTAGTTTCTACACAGGAG    11722
gb:MT106052|Organism:Severe      CTCAACCGCTACTTTAGACTGACTCTTGGTGTTTATGATTACTTAGTTTCTACACAGGAG    11722
gb:LC522975|Organism:Severe      CTCAACCGCTACTTTAGACTGACTCTTGGTGTTTATGATTACTTAGTTTCTACACAGGAG    11719
gb:LC522973|Organism:Severe      CTCAACCGCTACTTTAGACTGACTCTTGGTGTTTATGATTACTTAGTTTCTACACAGGAG    11719
gb:LC522974|Organism:Severe      CTCAACCGCTACTTTAGACTGACTCTTGGTGTTTATGATTACTTAGTTTCTACACAGGAG    11719
gb:MN985325|Organism:Severe      CTCAACCGCTACTTTAGACTGACTCTTGGTGTTTATGATTACTTAGTTTCTACACAGGAG    11722
gb:MT020881|Organism:Severe      CTCAACCGCTACTTTAGACTGACTCTTGGTGTTTATGATTACTTAGTTTCTACACAGGAG    11722
gb:MT020880|Organism:Severe      CTCAACCGCTACTTTAGACTGACTCTTGGTGTTTATGATTACTTAGTTTCTACACAGGAG    11722
gb:MT066175|Organism:Severe      CTCAACCGCTACTTTAGACTGACTCTTGGTGTTTATGATTACTTAGTTTCTACACAGGAG    11722
gb:MN997409|Organism:Severe      CTCAACCGCTACTTTAGACTGACTCTTGGTGTTTATGATTACTTAGTTTCTACACAGGAG    11722
gb:MN938384|Organism:Severe      CTCAACCGCTACTTTAGACTGACTCTTGGTGTTTATGATTACTTAGTTTCTACACAGGAG    11690
gb:MT044258|Organism:Severe      CTCAACCGCTACTTTAGACTGACTCTTGGTGTTTATGATTACTTAGTTTCTACACAGGAG    11698
gb:MT039890|Organism:Severe      CTCAACCGCTACTTTAGACTGACTCTTGGTGTTTATGATTACTTAGTTTCTACACAGGAG    11722
gb:MN988713|Organism:Severe      CTCAACCGCTACTTTAGACTGACTCTTGGTGTTTATGATTACTTAGTTTCTACACAGGAG    11722
gb:LC521925|Organism:Severe      CTCAACCGCTACTTTAGACTGACTCTTGGTGTTTATGATTACTTAGTTTCTACACAGGAG    11695
gb:MT093571|Organism:Severe      CTCAACCGCTACTTTAGACTGACTCTTGGTGTTTATGATTACTTAGTTTCTACACAGGAG    11722
gb:MT039887|Organism:Severe      CTCAACCGCTACTTTAGACTGACTCTTGGTGTTTATGATTACTTAGTTTCTACACAGGAG    11722
gb:MT019530|Organism:Severe      CTCAACCGCTACTTTAGACTGACTCTTGGTGTTTATGATTACTTAGTTTCTACACAGGAG    11722
gb:MT039888|Organism:Severe      CTCAACCGCTACTTTAGACTGACTCTTGGTGTTTATGATTACTTAGTTTCTACACAGGAG    11722
gb:LC522972|Organism:Severe      CTCAACCGCTACTTTAGACTGACTCTTGGTGTTTATGATTACTTAGTTTCTACACAGGAG    11719
gb:MT027063|Organism:Severe      CTCAACCGCTACTTTAGACTGACTCTTGGTGTTTATGATTACTTAGTTTCTACACAGGAG    11722
gb:MT027062|Organism:Severe      CTCAACCGCTACTTTAGACTGACTCTTGGTGTTTATGATTACTTAGTTTCTACACAGGAG    11722
gb:MT019529|Organism:Severe      CTCAACCGCTACTTTAGACTGACTCTTGGTGTTTATGATTACTTAGTTTCTACACAGGAG    11722
gb:MN996529|Organism:Severe      CTCAACCGCTACTTTAGACTGACTCTTGGTGTTTATGATTACTTAGTTTCTACACAGGAG    11710
gb:MN996531|Organism:Severe      CTCAACCGCTACTTTAGACTGACTCTTGGTGTTTATGATTACTTAGTTTCTACACAGGAG    11709
gb:MT066176|Organism:Severe      CTCAACCGCTACTTTAGACTGACTCTTGGTGTTTATGATTACTTAGTTTCTACACAGGAG    11722
gb:MT027064|Organism:Severe      CTCAACCGCTACTTTAGACTGACTCTTGGTGTTTATGATTACTTAGTTTCTACACAGGAG    11722
gb:MN994468|Organism:Severe      CTCAACCGCTACTTTAGACTGACTCTTGGTGTTTATGATTACTTAGTTTCTACACAGGAG    11722
gb:MT072688|Organism:Severe      CTCAACCGCTACTTTAGACTGACTCTTGGTGTTTATGATTACTTAGTTTCTACACAGGAG    11707
gb:MN996527|Organism:Severe      CTCAACCGCTACTTTAGACTGACTCTTGGTGTTTATGATTACTTAGTTTCTACACAGGAG    11689
gb:MT093631|Organism:Severe      CTCAACCGCTACTTTAGACTGACTCTTGGTGTTTATGATTACTTAGTTTCTACACAGGAG    11760
gb:MT106053|Organism:Severe      CTCAACCGCTACTTTAGACTGACTCTTGGTGTTTATGATTACTTAGTTTCTACACAGGAG    11722
gb:MT019533|Organism:Severe      CTCAACCGCTACTTTAGACTGACTCTTGGTGTTTATGATTACTTAGTTTCTACACAGGAG    11722
gb:MT019531|Organism:Severe      CTCAACCGCTACTTTAGACTGACTCTTGGTGTTTATGATTACTTAGTTTCTACACAGGAG    11722
gb:MN996528|Organism:Severe      CTCAACCGCTACTTTAGACTGACTCTTGGTGTTTATGATTACTTAGTTTCTACACAGGAG    11722
gb:MN996530|Organism:Severe      CTCAACCGCTACTTTAGACTGACTCTTGGTGTTTATGATTACTTAGTTTCTACACAGGAG    11708
gb:MN908947|Organism:Severe      CTCAACCGCTACTTTAGACTGACTCTTGGTGTTTATGATTACTTAGTTTCTACACAGGAG    11722
gb:MT019532|Organism:Severe      CTCAACCGCTACTTTAGACTGACTCTTGGTGTTTATGATTACTTAGTTTCTACACAGGAG    11722
                                 ************************************************************

gb:MT020781|Organism:Severe      TTTAGATATATGAATTCACAGGGACTACTCCCACCCAAGAATAGCATAGATGCCTTCAAA    11770
gb:MT007544|Organism:Severe      TTTAGATATATGAATTCACAGGGACTACTCCCACCCAAGAATAGCATAGATGCCTTCAAA    11782
gb:MN994467|Organism:Severe      TTTAGATATATGAATTCACAGGGACTACTCCCACCCAAGAATAGCATAGATGCCTTCAAA    11782
gb:MT044257|Organism:Severe      TTTAGATATATGAATTCACAGGGACTACTCCCACCCAAGAATAGCATAGATGCCTTCAAA    11782
gb:MT106054|Organism:Severe      TTTAGATATATGAATTCACAGGGACTACTCCCACCCAAGAATAGCATAGATGCCTTCAAA    11782
gb:MT049951|Organism:Severe      TTTAGATATATGAATTCACAGGGACTACTCCCACCCAAGAATAGCATAGATGCCTTCAAA    11782
gb:MN975262|Organism:Severe      TTTAGATATATGAATTCACAGGGACTACTCCCACCCAAGAATAGCATAGATGCCTTCAAA    11782
gb:MT106052|Organism:Severe      TTTAGATATATGAATTCACAGGGACTACTCCCACCCAAGAATAGCATAGATGCCTTCAAA    11782
gb:LC522975|Organism:Severe      TTTAGATATATGAATTCACAGGGACTACTCCCACCCAAGAATAGCATAGATGCCTTCAAA    11779
gb:LC522973|Organism:Severe      TTTAGATATATGAATTCACAGGGACTACTCCCACCCAAGAATAGCATAGATGCCTTCAAA    11779
gb:LC522974|Organism:Severe      TTTAGATATATGAATTCACAGGGACTACTCCCACCCAAGAATAGCATAGATGCCTTCAAA    11779
gb:MN985325|Organism:Severe      TTTAGATATATGAATTCACAGGGACTACTCCCACCCAAGAATAGCATAGATGCCTTCAAA    11782
gb:MT020881|Organism:Severe      TTTAGATATATGAATTCACAGGGACTACTCCCACCCAAGAATAGCATAGATGCCTTCAAA    11782
gb:MT020880|Organism:Severe      TTTAGATATATGAATTCACAGGGACTACTCCCACCCAAGAATAGCATAGATGCCTTCAAA    11782
gb:MT066175|Organism:Severe      TTTAGATATATGAATTCACAGGGACTACTCCCACCCAAGAATAGCATAGATGCCTTCAAA    11782
```

```
gb:MN997409|Organism:Severe      TTTAGATATATGAATTCACAGGGACTACTCCCACCCAAGAATAGCATAGATGCCTTCAAA   11782
gb:MN938384|Organism:Severe      TTTAGATATATGAATTCACAGGGACTACTCCCACCCAAGAATAGCATAGATGCCTTCAAA   11750
gb:MT044258|Organism:Severe      TTTAGATATATGAATTCACAGGGACTACTCCCACCCAAGAATAGCATAGATGCCTTCAAA   11758
gb:MT039890|Organism:Severe      TTTAGATATATGAATTCACAGGGACTACTCCCACCCAAGAATAGCATAGATGCCTTCAAA   11782
gb:MN988713|Organism:Severe      TTTAGATATATGAATTCACAGGGACTACTCCCACCCAAGAATAGCATAGATGCCTTCAAA   11782
gb:LC521925|Organism:Severe      TTTAGATATATGAATTCACAGGGACTACTCCCACCCAAGAATAGCATAGATGCCTTCAAA   11755
gb:MT093571|Organism:Severe      TTTAGATATATGAATTCACAGGGACTACTCCCACCCAAGAATAGCATAGATGCCTTCAAA   11782
gb:MT039887|Organism:Severe      TTTAGATATATGAATTCACAGGGACTACTCCCACCCAAGAATAGCATAGATGCCTTCAAA   11782
gb:MT019530|Organism:Severe      TTTAGATATATGAATTCACAGGGACTACTCCCACCCAAGAATAGCATAGATGCCTTCAAA   11782
gb:MT039888|Organism:Severe      TTTAGATATATGAATTCACAGGGACTACTCCCACCCAAGAATAGCATAGATGCCTTCAAA   11782
gb:LC522972|Organism:Severe      TTTAGATATATGAATTCACAGGGACTACTCCCACCCAAGAATAGCATAGATGCCTTCAAA   11779
gb:MT027063|Organism:Severe      TTTAGATATATGAATTCACAGGGACTACTCCCACCCAAGAATAGCATAGATGCCTTCAAA   11782
gb:MT027062|Organism:Severe      TTTAGATATATGAATTCACAGGGACTACTCCCACCCAAGAATAGCATAGATGCCTTCAAA   11782
gb:MT019529|Organism:Severe      TTTAGATATATGAATTCACAGGGACTACTCCCACCCAAGAATAGCATAGATGCCTTCAAA   11782
gb:MN996529|Organism:Severe      TTTAGATATATGAATTCACAGGGACTACTCCCACCCAAGAATAGCATAGATGCCTTCAAA   11770
gb:MN996531|Organism:Severe      TTTAGATATATGAATTCACAGGGACTACTCCCACCCAAGAATAGCATAGATGCCTTCAAA   11769
gb:MT066176|Organism:Severe      TTTAGATATATGAATTCACAGGGACTACTCCCACCCAAGAATAGCATAGATGCCTTCAAA   11782
gb:MT027064|Organism:Severe      TTTAGATATATGAATTCACAGGGACTACTCCCACCCAAGAATAGCATAGATGCCTTCAAA   11782
gb:MN994468|Organism:Severe      TTTAGATATATGAATTCACAGGGACTACTCCCACCCAAGAATAGCATAGATGCCTTCAAA   11782
gb:MT072688|Organism:Severe      TTTAGATATATGAATTCACAGGGACTACTCCCACCCAAGAATAGCATAGATGCCTTCAAA   11767
gb:MN996527|Organism:Severe      TTTAGATATATGAATTCACAGGGACTACTCCCACCCAAGAATAGCATAGATGCCTTCAAA   11749
gb:MT093631|Organism:Severe      TTTAGATATATGAATTCACAGGGACTACTCCCACCCAAGAATAGCATAGATGCCTTCAAA   11820
gb:MT106053|Organism:Severe      TTTAGATATATGAATTCACAGGGACTACTCCCACCCAAGAATAGCATAGATGCCTTCAAA   11782
gb:MT019533|Organism:Severe      TTTAGATATATGAATTCACAGGGACTACTCCCACCCAAGAATAGCATAGATGCCTTCAAA   11782
gb:MT019531|Organism:Severe      TTTAGATATATGAATTCACAGGGACTACTCCCACCCAAGAATAGCATAGATGCCTTCAAA   11782
gb:MN996528|Organism:Severe      TTTAGATATATGAATTCACAGGGACTACTCCCACCCAAGAATAGCATAGATGCCTTCAAA   11782
gb:MN996530|Organism:Severe      TTTAGATATATGAATTCACAGGGACTACTCCCACCCAAGAATAGCATAGATGCCTTCAAA   11768
gb:MN908947|Organism:Severe      TTTAGATATATGAATTCACAGGGACTACTCCCACCCAAGAATAGCATAGATGCCTTCAAA   11782
gb:MT019532|Organism:Severe      TTTAGATATATGAATTCACAGGGACTACTCCCACCCAAGAATAGCATAGATGCCTTCAAA   11782
                                 ************************************************************

gb:MT020781|Organism:Severe      CTCAACATTAAATTGTTGGGTGTTGGTGGCAAACCTTGTATCAAAGTAGCCACTGTACAG   11830
gb:MT007544|Organism:Severe      CTCAACATTAAATTGTTGGGTGTTGGTGGCAAACCTTGTATCAAAGTAGCCACTGTACAG   11842
gb:MN994467|Organism:Severe      CTCAACATTAAATTGTTGGGTGTTGGTGGCAAACCTTGTATCAAAGTAGCCACTGTACAG   11842
gb:MT044257|Organism:Severe      CTCAACATTAAATTGTTGGGTGTTGGTGGCAAACCTTGTATCAAAGTAGCCACTGTACAG   11842
gb:MT106054|Organism:Severe      CTCAACATTAAATTGTTGGGTGTTGGTGGCAAACCTTGTATCAAAGTAGCCACTGTACAG   11842
gb:MT049951|Organism:Severe      CTCAACATTAAATTGTTGGGTGTTGGTGGCAAACCTTGTATCAAAGTAGCCACTGTACAG   11842
gb:MN975262|Organism:Severe      CTCAACATTAAATTGTTGGGTGTTGGTGGCAAACCTTGTATCAAAGTAGCCACTGTACAG   11842
gb:MT106052|Organism:Severe      CTCAACATTAAATTGTTGGGTGTTGGTGGCAAACCTTGTATCAAAGTAGCCACTGTACAG   11842
gb:LC522975|Organism:Severe      CTCAACATTAAATTGTTGGGTGTTGGTGGCAAACCTTGTATCAAAGTAGCCACTGTACAG   11839
gb:LC522973|Organism:Severe      CTCAACATTAAATTGTTGGGTGTTGGTGGCAAACCTTGTATCAAAGTAGCCACTGTACAG   11839
gb:LC522974|Organism:Severe      CTCAACATTAAATTGTTGGGTGTTGGTGGCAAACCTTGTATCAAAGTAGCCACTGTACAG   11839
gb:MN985325|Organism:Severe      CTCAACATTAAATTGTTGGGTGTTGGTGGCAAACCTTGTATCAAAGTAGCCACTGTACAG   11842
gb:MT020881|Organism:Severe      CTCAACATTAAATTGTTGGGTGTTGGTGGCAAACCTTGTATCAAAGTAGCCACTGTACAG   11842
gb:MT020880|Organism:Severe      CTCAACATTAAATTGTTGGGTGTTGGTGGCAAACCTTGTATCAAAGTAGCCACTGTACAG   11842
gb:MT066175|Organism:Severe      CTCAACATTAAATTGTTGGGTGTTGGTGGCAAACCTTGTATCAAAGTAGCCACTGTACAG   11842
gb:MN997409|Organism:Severe      CTCAACATTAAATTGTTGGGTGTTGGTGGCAAACCTTGTATCAAAGTAGCCACTGTACAG   11842
gb:MN938384|Organism:Severe      CTCAACATTAAATTGTTGGGTGTTGGTGGCAAACCTTGTATCAAAGTAGCCACTGTACAG   11810
gb:MT044258|Organism:Severe      CTCAACATTAAATTGTTGGGTGTTGGTGGCAAACCTTGTATCAAAGTAGCCACTGTACAG   11818
gb:MT039890|Organism:Severe      CTCAACATTAAATTGTTGGGTGTTGGTGGCAAACCTTGTATCAAAGTAGCCACTGTACAG   11842
gb:MN988713|Organism:Severe      CTCAACATTAAATTGTTGGGTGTTGGTGGCAAACCTTGTATCAAAGTAGCCACTGTACAG   11842
gb:LC521925|Organism:Severe      CTCAACATTAAATTGTTGGGTGTTGGTGGCAAACCTTGTATCAAAGTAGCCACTGTACAG   11815
gb:MT093571|Organism:Severe      CTCAACATTAAATTGTTGGGTGTTGGTGGCAAACCTTGTATCAAAGTAGCCACTGTACAG   11842
gb:MT039887|Organism:Severe      CTCAACATTAAATTGTTGGGTGTTGGTGGCAAACCTTGTATCAAAGTAGCCACTGTACAG   11842
gb:MT019530|Organism:Severe      CTCAACATTAAATTGTTGGGTGTTGGTGGCAAACCTTGTATCAAAGTAGCCACTGTACAG   11842
gb:MT039888|Organism:Severe      CTCAACATTAAATTGTTGGGTGTTGGTGGCAAACCTTGTATCAAAGTAGCCACTGTACAG   11842
gb:LC522972|Organism:Severe      CTCAACATTAAATTGTTGGGTGTTGGTGGCAAACCTTGTATCAAAGTAGCCACTGTACAG   11839
gb:MT027063|Organism:Severe      CTCAACATTAAATTGTTGGGTGTTGGTGGCAAACCTTGTATCAAAGTAGCCACTGTACAG   11842
gb:MT027062|Organism:Severe      CTCAACATTAAATTGTTGGGTGTTGGTGGCAAACCTTGTATCAAAGTAGCCACTGTACAG   11842
gb:MT019529|Organism:Severe      CTCAACATTAAATTGTTGGGTGTTGGTGGCAAACCTTGTATCAAAGTAGCCACTGTACAG   11842
gb:MN996529|Organism:Severe      CTCAACATTAAATTGTTGGGTGTTGGTGGCAAACCTTGTATCAAAGTAGCCACTGTACAG   11830
gb:MN996531|Organism:Severe      CTCAACATTAAATTGTTGGGTGTTGGTGGCAAACCTTGTATCAAAGTAGCCACTGTACAG   11829
gb:MT066176|Organism:Severe      CTCAACATTAAATTGTTGGGTGTTGGTGGCAAACCTTGTATCAAAGTAGCCACTGTACAG   11842
gb:MT027064|Organism:Severe      CTCAACATTAAATTGTTGGGTGTTGGTGGCAAACCTTGTATCAAAGTAGCCACTGTACAG   11842
gb:MN994468|Organism:Severe      CTCAACATTAAATTGTTGGGTGTTGGTGGCAAACCTTGTATCAAAGTAGCCACTGTACAG   11842
```

```
gb:MT072688|Organism:Severe      CTCAACATTAAATTGTTGGGTGTTGGTGGCAAACCTTGTATCAAAGTAGCCACTGTACAG    11827
gb:MN996527|Organism:Severe      CTCAACATTAAATTGTTGGGTGTTGGTGGCAAACCTTGTATCAAAGTAGCCACTGTACAG    11809
gb:MT093631|Organism:Severe      CTCAACATTAAATTGTTGGGTGTTGGTGGCAAACCTTGTATCAAAGTAGCCACTGTACAG    11880
gb:MT106053|Organism:Severe      CTCAACATTAAATTGTTGGGTGTTGGTGGCAAACCTTGTATCAAAGTAGCCACTGTACAG    11842
gb:MT019533|Organism:Severe      CTCAACATTAAATTGTTGGGTGTTGGTGGCAAACCTTGTATCAAAGTAGCCACTGTACAG    11842
gb:MT019531|Organism:Severe      CTCAACATTAAATTGTTGGGTGTTGGTGGCAAACCTTGTATCAAAGTAGCCACTGTACAG    11842
gb:MN996528|Organism:Severe      CTCAACATTAAATTGTTGGGTGTTGGTGGCAAACCTTGTATCAAAGTAGCCACTGTACAG    11842
gb:MN996530|Organism:Severe      CTCAACATTAAATTGTTGGGTGTTGGTGGCAAACCTTGTATCAAAGTAGCCACTGTACAG    11828
gb:MN908947|Organism:Severe      CTCAACATTAAATTGTTGGGTGTTGGTGGCAAACCTTGTATCAAAGTAGCCACTGTACAG    11842
gb:MT019532|Organism:Severe      CTCAACATTAAATTGTTGGGTGTTGGTGGCAAACCTTGTATCAAAGTAGCCACTGTACAG    11842
                                 ************************************************************

gb:MT020781|Organism:Severe      TCTAAAATGTCAGATGTAAAGTGCACATCAGTAGTCTTACTCTCAGTTTTGCAACAACTC    11890
gb:MT007544|Organism:Severe      TCTAAAATGTCAGATGTAAAGTGCACATCAGTAGTCTTACTCTCAGTTTTGCAACAACTC    11902
gb:MN994467|Organism:Severe      TCTAAAATGTCAGATGTAAAGTGCACATCAGTAGTCTTACTCTCAGTTTTGCAACAACTC    11902
gb:MT044257|Organism:Severe      TCTAAAATGTCAGATGTAAAGTGCACATCAGTAGTCTTACTCTCAGTTTTGCAACAACTC    11902
gb:MT106054|Organism:Severe      TCTAAAATGTCAGATGTAAAGTGCACATCAGTAGTCTTACTCTCAGTTTTGCAACAACTC    11902
gb:MT049951|Organism:Severe      TCTAAAATGTCAGATGTAAAGTGCACATCAGTAGTCTTACTCTCAGTTTTGCAACAACTC    11902
gb:MN975262|Organism:Severe      TCTAAAATGTCAGATGTAAAGTGCACATCAGTAGTCTTACTCTCAGTTTTGCAACAACTC    11902
gb:MT106052|Organism:Severe      TCTAAAATGTCAGATGTAAAGTGCACATCAGTAGTCTTACTCTCAGTTTTGCAACAACTC    11902
gb:LC522975|Organism:Severe      TCTAAAATGTCAGATGTAAAGTGCACATCAGTAGTCTTACTCTCAGTTTTGCAACAACTC    11899
gb:LC522973|Organism:Severe      TCTAAAATGTCAGATGTAAAGTGCACATCAGTAGTCTTACTCTCAGTTTTGCAACAACTC    11899
gb:LC522974|Organism:Severe      TCTAAAATGTCAGATGTAAAGTGCACATCAGTAGTCTTACTCTCAGTTTTGCAACAACTC    11899
gb:MN985325|Organism:Severe      TCTAAAATGTCAGATGTAAAGTGCACATCAGTAGTCTTACTCTCAGTTTTGCAACAACTC    11902
gb:MT020881|Organism:Severe      TCTAAAATGTCAGATGTAAAGTGCACATCAGTAGTCTTACTCTCAGTTTTGCAACAACTC    11902
gb:MT020880|Organism:Severe      TCTAAAATGTCAGATGTAAAGTGCACATCAGTAGTCTTACTCTCAGTTTTGCAACAACTC    11902
gb:MT066175|Organism:Severe      TCTAAAATGTCAGATGTAAAGTGCACATCAGTAGTCTTACTCTCAGTTTTGCAACAACTC    11902
gb:MN997409|Organism:Severe      TCTAAAATGTCAGATGTAAAGTGCACATCAGTAGTCTTACTCTCAGTTTTGCAACAACTC    11902
gb:MN938384|Organism:Severe      TCTAAAATGTCAGATGTAAAGTGCACATCAGTAGTCTTACTCTCAGTTTTGCAACAACTC    11870
gb:MT044258|Organism:Severe      TCTAAAATGTCAGATGTAAAGTGCACATCAGTAGTCTTACTCTCAGTTTTGCAACAACTC    11878
gb:MT039890|Organism:Severe      TCTAAAATGTCAGATGTAAAGTGCACATCAGTAGTCTTACTCTCAGTTTTGCAACAACTC    11902
gb:MN988713|Organism:Severe      TCTAAAATGTCAGATGTAAAGTGCACATCAGTAGTCTTACTCTCAGTTTTGCAACAACTC    11902
gb:LC521925|Organism:Severe      TCTAAAATGTCAGATGTAAAGTGCACATCAGTAGTCTTACTCTCAGTTTTGCAACAACTC    11875
gb:MT093571|Organism:Severe      TCTAAAATGTCAGATGTAAAGTGCACATCAGTAGTCTTACTCTCAGTTTTGCAACAACTC    11902
gb:MT039887|Organism:Severe      TCTAAAATGTCAGATGTAAAGTGCACATCAGTAGTCTTACTCTCAGTTTTGCAACAACTC    11902
gb:MT019530|Organism:Severe      TCTAAAATGTCAGATGTAAAGTGCACATCAGTAGTCTTACTCTCAGTTTTGCAACAACTC    11902
gb:MT039888|Organism:Severe      TCTAAAATGTCAGATGTAAAGTGCACATCAGTAGTCTTACTCTCAGTTTTGCAACAACTC    11902
gb:LC522972|Organism:Severe      TCTAAAATGTCAGATGTAAAGTGCACATCAGTAGTCTTACTCTCAGTTTTGCAACAACTC    11899
gb:MT027063|Organism:Severe      TCTAAAATGTCAGATGTAAAGTGCACATCAGTAGTCTTACTCTCAGTTTTGCAACAACTC    11902
gb:MT027062|Organism:Severe      TCTAAAATGTCAGATGTAAAGTGCACATCAGTAGTCTTACTCTCAGTTTTGCAACAACTC    11902
gb:MT019529|Organism:Severe      TCTAAAATGTCAGATGTAAAGTGCACATCAGTAGTCTTACTCTCAGTTTTGCAACAACTC    11902
gb:MN996529|Organism:Severe      TCTAAAATGTCAGATGTAAAGTGCACATCAGTAGTCTTACTCTCAGTTTTGCAACAACTC    11890
gb:MN996531|Organism:Severe      TCTAAAATGTCAGATGTAAAGTGCACATCAGTAGTCTTACTCTCAGTTTTGCAACAACTC    11889
gb:MT066176|Organism:Severe      TCTAAAATGTCAGATGTAAAGTGCACATCAGTAGTCTTACTCTCAGTTTTGCAACAACTC    11902
gb:MT027064|Organism:Severe      TCTAAAATGTCAGATGTAAAGTGCACATCAGTAGTCTTACTCTCAGTTTTGCAACAACTC    11902
gb:MN994468|Organism:Severe      TCTAAAATGTCAGATGTAAAGTGCACATCAGTAGTCTTACTCTCAGTTTTGCAACAACTC    11902
gb:MT072688|Organism:Severe      TCTAAAATGTCAGATGTAAAGTGCACATCAGTAGTCTTACTCTCAGTTTTGCAACAACTC    11887
gb:MN996527|Organism:Severe      TCTAAAATGTCAGATGTAAAGTGCACATCAGTAGTCTTACTCTCAGTTTTGCAACAACTC    11869
gb:MT093631|Organism:Severe      TCTAAAATGTCAGATGTAAAGTGCACATCAGTAGTCTTACTCTCAGTTTTGCAACAACTC    11940
gb:MT106053|Organism:Severe      TCTAAAATGTCAGATGTAAAGTGCACATCAGTAGTCTTACTCTCAGTTTTGCAACAACTC    11902
gb:MT019533|Organism:Severe      TCTAAAATGTCAGATGTAAAGTGCACATCAGTAGTCTTACTCTCAGTTTTGCAACAACTC    11902
gb:MT019531|Organism:Severe      TCTAAAATGTCAGATGTAAAGTGCACATCAGTAGTCTTACTCTCAGTTTTGCAACAACTC    11902
gb:MN996528|Organism:Severe      TCTAAAATGTCAGATGTAAAGTGCACATCAGTAGTCTTACTCTCAGTTTTGCAACAACTC    11902
gb:MN996530|Organism:Severe      TCTAAAATGTCAGATGTAAAGTGCACATCAGTAGTCTTACTCTCAGTTTTGCAACAACTC    11888
gb:MN908947|Organism:Severe      TCTAAAATGTCAGATGTAAAGTGCACATCAGTAGTCTTACTCTCAGTTTTGCAACAACTC    11902
gb:MT019532|Organism:Severe      TCTAAAATGTCAGATGTAAAGTGCACATCAGTAGTCTTACTCTCAGTTTTGCAACAACTC    11902
                                 ************************************************************

gb:MT020781|Organism:Severe      AGAGTAGAATCATCATCTAAATTGTGGGCTCAATGTGTCCAGTTACACAATGACATTCTC    11950
gb:MT007544|Organism:Severe      AGAGTAGAATCATCATCTAAATTGTGGGCTCAATGTGTCCAGTTACACAATGACATTCTC    11962
gb:MN994467|Organism:Severe      AGAGTAGAATCATCATCTAAATTGTGGGCTCAATGTGTCCAGTTACACAATGACATTCTC    11962
gb:MT044257|Organism:Severe      AGAGTAGAATCATCATCTAAATTGTGGGCTCAATGTGTCCAGTTACACAATGACATTCTC    11962
gb:MT106054|Organism:Severe      AGAGTAGAATCATCATCTAAATTGTGGGCTCAATGTGTCCAGTTACACAATGACATTCTC    11962
gb:MT049951|Organism:Severe      AGAGTAGAATCATCATCTAAATTGTGGGCTCAATGTGTCCAGTTACACAATGACATTCTC    11962
gb:MN975262|Organism:Severe      AGAGTAGAATCATCATCTAAATTGTGGGCTCAATGTGTCCAGTTACACAATGACATTCTC    11962
```

```
gb:MT106052|Organism:Severe      AGAGTAGAATCATCATCTAAATTGTGGGCTCAATGTGTCCAGTTACACAATGACATTCTC   11962
gb:LC522975|Organism:Severe      AGAGTAGAATCATCATCTAAATTGTGGGCTCAATGTGTCCAGTTACACAATGACATTCTC   11959
gb:LC522973|Organism:Severe      AGAGTAGAATCATCATCTAAATTGTGGGCTCAATGTGTCCAGTTACACAATGACATTCTC   11959
gb:LC522974|Organism:Severe      AGAGTAGAATCATCATCTAAATTGTGGGCTCAATGTGTCCAGTTACACAATGACATTCTC   11959
gb:MN985325|Organism:Severe      AGAGTAGAATCATCATCTAAATTGTGGGCTCAATGTGTCCAGTTACACAATGACATTCTC   11962
gb:MT020881|Organism:Severe      AGAGTAGAATCATCATCTAAATTGTGGGCTCAATGTGTCCAGTTACACAATGACATTCTC   11962
gb:MT020880|Organism:Severe      AGAGTAGAATCATCATCTAAATTGTGGGCTCAATGTGTCCAGTTACACAATGACATTCTC   11962
gb:MT066175|Organism:Severe      AGAGTAGAATCATCATCTAAATTGTGGGCTCAATGTGTCCAGTTACACAATGACATTCTC   11962
gb:MN997409|Organism:Severe      AGAGTAGAATCATCATCTAAATTGTGGGCTCAATGTGTCCAGTTACACAATGACATTCTC   11962
gb:MN938384|Organism:Severe      AGAGTAGAATCATCATCTAAATTGTGGGCTCAATGTGTCCAGTTACACAATGACATTCTC   11930
gb:MT044258|Organism:Severe      AGAGTAGAATCATCATCTAAATTGTGGGCTCAATGTGTCCAGTTACACAATGACATTCTC   11938
gb:MT039890|Organism:Severe      AGAGTAGAATCATCATCTAAATTGTGGGCTCAATGTGTCCAGTTACACAATGACATTCTC   11962
gb:MN988713|Organism:Severe      AGAGTAGAATCATCATCTAAATTGTGGGCTCAATGTGTCCAGTTACACAATGACATTCTC   11962
gb:LC521925|Organism:Severe      AGAGTAGAATCATCATCTAAATTGTGGGCTCAATGTGTCCAGTTACACAATGACATTCTC   11935
gb:MT093571|Organism:Severe      AGAGTAGAATCATCATCTAAATTGTGGGCTCAATGTGTCCAGTTACACAATGACATTCTC   11962
gb:MT039887|Organism:Severe      AGAGTAGAATCATCATCTAAATTGTGGGCTCAATGTGTCCAGTTACACAATGACATTCTC   11962
gb:MT019530|Organism:Severe      AGAGTAGAATCATCATCTAAATTGTGGGCTCAATGTGTCCAGTTACACAATGACATTCTC   11962
gb:MT039888|Organism:Severe      AGAGTAGAATCATCATCTAAATTGTGGGCTCAATGTGTCCAGTTACACAATGACATTCTC   11962
gb:LC522972|Organism:Severe      AGAGTAGAATCATCATCTAAATTGTGGGCTCAATGTGTCCAGTTACACAATGACATTCTC   11959
gb:MT027063|Organism:Severe      AGAGTAGAATCATCATCTAAATTGTGGGCTCAATGTGTCCAGTTACACAATGACATTCTC   11962
gb:MT027062|Organism:Severe      AGAGTAGAATCATCATCTAAATTGTGGGCTCAATGTGTCCAGTTACACAATGACATTCTC   11962
gb:MT019529|Organism:Severe      AGAGTAGAATCATCATCTAAATTGTGGGCTCAATGTGTCCAGTTACACAATGACATTCTC   11962
gb:MN996529|Organism:Severe      AGAGTAGAATCATCATCTAAATTGTGGGCTCAATGTGTCCAGTTACACAATGACATTCTC   11950
gb:MN996531|Organism:Severe      AGAGTAGAATCATCATCTAAATTGTGGGCTCAATGTGTCCAGTTACACAATGACATTCTC   11949
gb:MT066176|Organism:Severe      AGAGTAGAATCATCATCTAAATTGTGGGCTCAATGTGTCCAGTTACACAATGACATTCTC   11962
gb:MT027064|Organism:Severe      AGAGTAGAATCATCATCTAAATTGTGGGCTCAATGTGTCCAGTTACACAATGACATTCTC   11962
gb:MN994468|Organism:Severe      AGAGTAGAATCATCATCTAAATTGTGGGCTCAATGTGTCCAGTTACACAATGACATTCTC   11962
gb:MT072688|Organism:Severe      AGAGTAGAATCATCATCTAAATTGTGGGCTCAATGTGTCCAGTTACACAATGACATTCTC   11947
gb:MN996527|Organism:Severe      AGAGTAGAATCATCATCTAAATTGTGGGCTCAATGTGTCCAGTTACACAATGACATTCTC   11929
gb:MT093631|Organism:Severe      AGAGTAGAATCATCATCTAAATTGTGGGCTCAATGTGTCCAGTTACACAATGACATTCTC   12000
gb:MT106053|Organism:Severe      AGAGTAGAATCATCATCTAAATTGTGGGCTCAATGTGTCCAGTTACACAATGACATTCTC   11962
gb:MT019533|Organism:Severe      AGAGTAGAATCATCATCTAAATTGTGGGCTCAATGTGTCCAGTTACACAATGACATTCTC   11962
gb:MT019531|Organism:Severe      AGAGTAGAATCATCATCTAAATTGTGGGCTCAATGTGTCCAGTTACACAATGACATTCTC   11962
gb:MN996528|Organism:Severe      AGAGTAGAATCATCATCTAAATTGTGGGCTCAATGTGTCCAGTTACACAATGACATTCTC   11962
gb:MN996530|Organism:Severe      AGAGTAGAATCATCATCTAAATTGTGGGCTCAATGTGTCCAGTTACACAATGACATTCTC   11948
gb:MN908947|Organism:Severe      AGAGTAGAATCATCATCTAAATTGTGGGCTCAATGTGTCCAGTTACACAATGACATTCTC   11962
gb:MT019532|Organism:Severe      AGAGTAGAATCATCATCTAAATTGTGGGCTCAATGTGTCCAGTTACACAATGACATTCTC   11962
                                 ************************************************************

gb:MT020781|Organism:Severe      TTAGCTAAAGATACTACTGAAGCCTTTGAAAAAATGGTTTCACTACTTTCTGTTTTGCTT   12010
gb:MT007544|Organism:Severe      TTAGCTAAAGATACTACTGAAGCCTTTGAAAAAATGGTTTCACTACTTTCTGTTTTGCTT   12022
gb:MN994467|Organism:Severe      TTAGCTAAAGATACTACTGAAGCCTTTGAAAAAATGGTTTCACTACTTTCTGTTTTGCTT   12022
gb:MT044257|Organism:Severe      TTAGCTAAAGATACTACTGAAGCCTTTGAAAAAATGGTTTCACTACTTTCTGTTTTGCTT   12022
gb:MT106054|Organism:Severe      TTAGCTAAAGATACTACTGAAGCCTTTGAAAAAATGGTTTCACTACTTTCTGTTTTGCTT   12022
gb:MT049951|Organism:Severe      TTAGCTAAAGATACTACTGAAGCCTTTGAAAAAATGGTTTCACTACTTTCTGTTTTGCTT   12022
gb:MN975262|Organism:Severe      TTAGCTAAAGATACTACTGAAGCCTTTGAAAAAATGGTTTCACTACTTTCTGTTTTGCTT   12022
gb:MT106052|Organism:Severe      TTAGCTAAAGATACTACTGAAGCCTTTGAAAAAATGGTTTCACTACTTTCTGTTTTGCTT   12022
gb:LC522975|Organism:Severe      TTAGCTAAAGATACTACTGAAGCCTTTGAAAAAATGGTTTCACTACTTTCTGTTTTGCTT   12019
gb:LC522973|Organism:Severe      TTAGCTAAAGATACTACTGAAGCCTTTGAAAAAATGGTTTCACTACTTTCTGTTTTGCTT   12019
gb:LC522974|Organism:Severe      TTAGCTAAAGATACTACTGAAGCCTTTGAAAAAATGGTTTCACTACTTTCTGTTTTGCTT   12019
gb:MN985325|Organism:Severe      TTAGCTAAAGATACTACTGAAGCCTTTGAAAAAATGGTTTCACTACTTTCTGTTTTGCTT   12022
gb:MT020881|Organism:Severe      TTAGCTAAAGATACTACTGAAGCCTTTGAAAAAATGGTTTCACTACTTTCTGTTTTGCTT   12022
gb:MT020880|Organism:Severe      TTAGCTAAAGATACTACTGAAGCCTTTGAAAAAATGGTTTCACTACTTTCTGTTTTGCTT   12022
gb:MT066175|Organism:Severe      TTAGCTAAAGATACTACTGAAGCCTTTGAAAAAATGGTTTCACTACTTTCTGTTTTGCTT   12022
gb:MN997409|Organism:Severe      TTAGCTAAAGATACTACTGAAGCCTTTGAAAAAATGGTTTCACTACTTTCTGTTTTGCTT   12022
gb:MN938384|Organism:Severe      TTAGCTAAAGATACTACTGAAGCCTTTGAAAAAATGGTTTCACTACTTTCTGTTTTGCTT   11990
gb:MT044258|Organism:Severe      TTAGCTAAAGATACTACTGAAGCCTTTGAAAAAATGGTTTCACTACTTTCTGTTTTGCTT   11998
gb:MT039890|Organism:Severe      TTAGCTAAAGATACTACTGAAGCCTTTGAAAAAATGGTTTCACTACTTTCTGTTTTGCTT   12022
gb:MN988713|Organism:Severe      TTAGCTAAAGATACTACTGAAGCCTTTGAAAAAATGGTTTCACTACTTTCTGTTTTGCTT   12022
gb:LC521925|Organism:Severe      TTAGCTAAAGATACTACTGAAGCCTTTGAAAAAATGGTTTCACTACTTTCTGTTTTGCTT   11995
gb:MT093571|Organism:Severe      TTAGCTAAAGATACTACTGAAGCCTTTGAAAAAATGGTTTCACTACTTTCTGTTTTGCTT   12022
gb:MT039887|Organism:Severe      TTAGCTAAAGATACTACTGAAGCCTTTGAAAAAATGGTTTCACTACTTTCTGTTTTGCTT   12022
gb:MT019530|Organism:Severe      TTAGCTAAAGATACTACTGAAGCCTTTGAAAAAATGGTTTCACTACTTTCTGTTTTGCTT   12022
gb:MT039888|Organism:Severe      TTAGCTAAAGATACTACTGAAGCCTTTGAAAAAATGGTTTCACTACTTTCTGTTTTGCTT   12022
gb:LC522972|Organism:Severe      TTAGCTAAAGATACTACTGAAGCCTTTGAAAAAATGGTTTCACTACTTTCTGTTTTGCTT   12019
```

```
gb:MT027063|Organism:Severe      TTAGCTAAAGATACTACTGAAGCCTTTGAAAAAATGGTTTCACTACTTTCTGTTTTGCTT   12022
gb:MT027062|Organism:Severe      TTAGCTAAAGATACTACTGAAGCCTTTGAAAAAATGGTTTCACTACTTTCTGTTTTGCTT   12022
gb:MT019529|Organism:Severe      TTAGCTAAAGATACTACTGAAGCCTTTGAAAAAATGGTTTCACTACTTTCTGTTTTGCTT   12022
gb:MN996529|Organism:Severe      TTAGCTAAAGATACTACTGAAGCCTTTGAAAAAATGGTTTCACTACTTTCTGTTTTGCTT   12010
gb:MN996531|Organism:Severe      TTAGCTAAAGATACTACTGAAGCCTTTGAAAAAATGGTTTCACTACTTTCTGTTTTGCTT   12009
gb:MT066176|Organism:Severe      TTAGCTAAAGATACTACTGAAGCCTTTGAAAAAATGGTTTCACTACTTTCTGTTTTGCTT   12022
gb:MT027064|Organism:Severe      TTAGCTAAAGATACTACTGAAGCCTTTGAAAAAATGGTTTCACTACTTTCTGTTTTGCTT   12022
gb:MN994468|Organism:Severe      TTAGCTAAAGATACTACTGAAGCCTTTGAAAAAATGGTTTCACTACTTTCTGTTTTGCTT   12022
gb:MT072688|Organism:Severe      TTAGCTAAAGATACTACTGAAGCCTTTGAAAAAATGGTTTCACTACTTTCTGTTTTGCTT   12007
gb:MN996527|Organism:Severe      TTAGCTAAAGATACTACTGAAGCCTTTGAAAAAATGGTTTCACTACTTTCTGTTTTGCTT   11989
gb:MT093631|Organism:Severe      TTAGCTAAAGATACTACTGAAGCCTTTGAAAAAATGGTTTCACTACTTTCTGTTTTGCTT   12060
gb:MT106053|Organism:Severe      TTAGCTAAAGATACTACTGAAGCCTTTGAAAAAATGGTTTCACTACTTTCTGTTTTGCTT   12022
gb:MT019533|Organism:Severe      TTAGCTAAAGATACTACTGAAGCCTTTGAAAAAATGGTTTCACTACTTTCTGTTTTGCTT   12022
gb:MT019531|Organism:Severe      TTAGCTAAAGATACTACTGAAGCCTTTGAAAAAATGGTTTCACTACTTTCTGTTTTGCTT   12022
gb:MN996528|Organism:Severe      TTAGCTAAAGATACTACTGAAGCCTTTGAAAAAATGGTTTCACTACTTTCTGTTTTGCTT   12022
gb:MN996530|Organism:Severe      TTAGCTAAAGATACTACTGAAGCCTTTGAAAAAATGGTTTCACTACTTTCTGTTTTGCTT   12008
gb:MN908947|Organism:Severe      TTAGCTAAAGATACTACTGAAGCCTTTGAAAAAATGGTTTCACTACTTTCTGTTTTGCTT   12022
gb:MT019532|Organism:Severe      TTAGCTAAAGATACTACTGAAGCCTTTGAAAAAATGGTTTCACTACTTTCTGTTTTGCTT   12022
                                 ************************************************************

gb:MT020781|Organism:Severe      TCCATGCAGGGTGCTGTAGACATAAACAAGCTTTGTGAAGAAATGCTGGACAACAGGGCA   12070
gb:MT007544|Organism:Severe      TCCATGCAGGGTGCTGTAGACATAAACAAGCTTTGTGAAGAAATGCTGGACAACAGGGCA   12082
gb:MN994467|Organism:Severe      TCCATGCAGGGTGCTGTAGACATAAACAAGCTTTGTGAAGAAATGCTGGACAACAGGGCA   12082
gb:MT044257|Organism:Severe      TCCATGCAGGGTGCTGTAGACATAAACAAGCTTTGTGAAGAAATGCTGGACAACAGGGCA   12082
gb:MT106054|Organism:Severe      TCCATGCAGGGTGCTGTAGACATAAACAAGCTTTGTGAAGAAATGCTGGACAACAGGGCA   12082
gb:MT049951|Organism:Severe      TCCATGCAGGGTGCTGTAGACATAAACAAGCTTTGTGAAGAAATGCTGGACAACAGGGCA   12082
gb:MN975262|Organism:Severe      TCCATGCAGGGTGCTGTAGACATAAACAAGCTTTGTGAAGAAATGCTGGACAACAGGGCA   12082
gb:MT106052|Organism:Severe      TCCATGCAGGGTGCTGTAGACATAAACAAGCTTTGTGAAGAAATGCTGGACAACAGGGCA   12082
gb:LC522975|Organism:Severe      TCCATGCAGGGTGCTGTAGACATAAACAAGCTTTGTGAAGAAATGCTGGACAACAGGGCA   12079
gb:LC522973|Organism:Severe      TCCATGCAGGGTGCTGTAGACATAAACAAGCTTTGTGAAGAAATGCTGGACAACAGGGCA   12079
gb:LC522974|Organism:Severe      TCCATGCAGGGTGCTGTAGACATAAACAAGCTTTGTGAAGAAATGCTGGACAACAGGGCA   12079
gb:MN985325|Organism:Severe      TCCATGCAGGGTGCTGTAGACATAAACAAGCTTTGTGAAGAAATGCTGGACAACAGGGCA   12082
gb:MT020881|Organism:Severe      TCCATGCAGGGTGCTGTAGACATAAACAAGCTTTGTGAAGAAATGCTGGACAACAGGGCA   12082
gb:MT020880|Organism:Severe      TCCATGCAGGGTGCTGTAGACATAAACAAGCTTTGTGAAGAAATGCTGGACAACAGGGCA   12082
gb:MT066175|Organism:Severe      TCCATGCAGGGTGCTGTAGACATAAACAAGCTTTGTGAAGAAATGCTGGACAACAGGGCA   12082
gb:MN997409|Organism:Severe      TCCATGCAGGGTGCTGTAGACATAAACAAGCTTTGTGAAGAAATGCTGGACAACAGGGCA   12082
gb:MN938384|Organism:Severe      TCCATGCAGGGTGCTGTAGACATAAACAAGCTTTGTGAAGAAATGCTGGACAACAGGGCA   12050
gb:MT044258|Organism:Severe      TCCATGCAGGGTGCTGTAGACATAAACAAGCTTTGTGAAGAAATGCTGGACAACAGGGCA   12058
gb:MT039890|Organism:Severe      TCCATGCAGGGTGCTGTAGACATAAACAAGCTTTGTGAAGAAATGCTGGACAACAGGGCA   12082
gb:MN988713|Organism:Severe      TCCATGCAGGGTGCTGTAGACATAAACAAGCTTTGTGAAGAAATGCTGGACAACAGGGCA   12082
gb:LC521925|Organism:Severe      TCCATGCAGGGTGCTGTAGACATAAACAAGCTTTGTGAAGAAATGCTGGACAACAGGGCA   12055
gb:MT093571|Organism:Severe      TCCATGCAGGGTGCTGTAGACATAAACAAGCTTTGTGAAGAAATGCTGGACAACAGGGCA   12082
gb:MT039887|Organism:Severe      TCCATGCAGGGTGCTGTAGACATAAACAAGCTTTGTGAAGAAATGCTGGACAACAGGGCA   12082
gb:MT019530|Organism:Severe      TCCATGCAGGGTGCTGTAGACATAAACAAGCTTTGTGAAGAAATGCTGGACAACAGGGCA   12082
gb:MT039888|Organism:Severe      TCCATGCAGGGTGCTGTAGACATAAACAAGCTTTGTGAAGAAATGCTGGACAACAGGGCA   12082
gb:LC522972|Organism:Severe      TCCATGCAGGGTGCTGTAGACATAAACAAGCTTTGTGAAGAAATGCTGGACAACAGGGCA   12079
gb:MT027063|Organism:Severe      TCCATGCAGGGTGCTGTAGACATAAACAAGCTTTGTGAAGAAATGCTGGACAACAGGGCA   12082
gb:MT027062|Organism:Severe      TCCATGCAGGGTGCTGTAGACATAAACAAGCTTTGTGAAGAAATGCTGGACAACAGGGCA   12082
gb:MT019529|Organism:Severe      TCCATGCAGGGTGCTGTAGACATAAACAAGCTTTGTGAAGAAATGCTGGACAACAGGGCA   12082
gb:MN996529|Organism:Severe      TCCATGCAGGGTGCTGTAGACATAAACAAGCTTTGTGAAGAAATGCTGGACAACAGGGCA   12070
gb:MN996531|Organism:Severe      TCCATGCAGGGTGCTGTAGACATAAACAAGCTTTGTGAAGAAATGCTGGACAACAGGGCA   12069
gb:MT066176|Organism:Severe      TCCATGCAGGGTGCTGTAGACATAAACAAGCTTTGTGAAGAAATGCTGGACAACAGGGCA   12082
gb:MT027064|Organism:Severe      TCCATGCAGGGTGCTGTAGACATAAACAAGCTTTGTGAAGAAATGCTGGACAACAGGGCA   12082
gb:MN994468|Organism:Severe      TCCATGCAGGGTGCTGTAGACATAAACAAGCTTTGTGAAGAAATGCTGGACAACAGGGCA   12082
gb:MT072688|Organism:Severe      TCCATGCAGGGTGCTGTAGACATAAACAAGCTTTGTGAAGAAATGCTGGACAACAGGGCA   12067
gb:MN996527|Organism:Severe      TCCATGCAGGGTGCTGTAGACATAAACAAGCTTTGTGAAGAAATGCTGGACAACAGGGCA   12049
gb:MT093631|Organism:Severe      TCCATGCAGGGTGCTGTAGACATAAACAAGCTTTGTGAAGAAATGCTGGACAACAGGGCA   12120
gb:MT106053|Organism:Severe      TCCATGCAGGGTGCTGTAGACATAAACAAGCTTTGTGAAGAAATGCTGGACAACAGGGCA   12082
gb:MT019533|Organism:Severe      TCCATGCAGGGTGCTGTAGACATAAACAAGCTTTGTGAAGAAATGCTGGACAACAGGGCA   12082
gb:MT019531|Organism:Severe      TCCATGCAGGGTGCTGTAGACATAAACAAGCTTTGTGAAGAAATGCTGGACAACAGGGCA   12082
gb:MN996528|Organism:Severe      TCCATGCAGGGTGCTGTAGACATAAACAAGCTTTGTGAAGAAATGCTGGACAACAGGGCA   12082
gb:MN996530|Organism:Severe      TCCATGCAGGGTGCTGTAGACATAAACAAGCTTTGTGAAGAAATGCTGGACAACAGGGCA   12068
gb:MN908947|Organism:Severe      TCCATGCAGGGTGCTGTAGACATAAACAAGCTTTGTGAAGAAATGCTGGACAACAGGGCA   12082
gb:MT019532|Organism:Severe      TCCATGCAGGGTGCTGTAGACATAAACAAGCTTTGTGAAGAAATGCTGGACAACAGGGCA   12082
                                 ************************************************************
```

```
gb:MT020781|Organism:Severe      ACCTTACAAGCTATAGCCTCAGAGTTTAGTTCCCTTCCATCATATGCAGCTTTTGCTACT   12130
gb:MT007544|Organism:Severe      ACCTTACAAGCTATAGCCTCAGAGTTTAGTTCCCTTCCATCATATGCAGCTTTTGCTACT   12142
gb:MN994467|Organism:Severe      ACCTTACAAGCTATAGCCTCAGAGTTTAGTTCCCTTCCATCATATGCAGCTTTTGCTACT   12142
gb:MT044257|Organism:Severe      ACCTTACAAGCTATAGCCTCAGAGTTTAGTTCCCTTCCATCATATGCAGCTTTTGCTACT   12142
gb:MT106054|Organism:Severe      ACCTTACAAGCTATAGCCTCAGAGTTTAGTTCCCTTCCATCATATGCAGCTTTTGCTACT   12142
gb:MT049951|Organism:Severe      ACCTTACAAGCTATAGCCTCAGAGTTTAGTTCCCTTCCATCATATGCAGCTTTTGCTACT   12142
gb:MN975262|Organism:Severe      ACCTTACAAGCTATAGCCTCAGAGTTTAGTTCCCTTCCATCATATGCAGCTTTTGCTACT   12142
gb:MT106052|Organism:Severe      ACCTTACAAGCTATAGCCTCAGAGTTTAGTTCCCTTCCATCATATGCAGCTTTTGCTACT   12142
gb:LC522975|Organism:Severe      ACCTTACAAGCTATAGCCTCAGAGTTTAGTTCCCTTCCATCATATGCAGCTTTTGCTACT   12139
gb:LC522973|Organism:Severe      ACCTTACAAGCTATAGCCTCAGAGTTTAGTTCCCTTCCATCATATGCAGCTTTTGCTACT   12139
gb:LC522974|Organism:Severe      ACCTTACAAGCTATAGCCTCAGAGTTTAGTTCCCTTCCATCATATGCAGCTTTTGCTACT   12139
gb:MN985325|Organism:Severe      ACCTTACAAGCTATAGCCTCAGAGTTTAGTTCCCTTCCATCATATGCAGCTTTTGCTACT   12142
gb:MT020881|Organism:Severe      ACCTTACAAGCTATAGCCTCAGAGTTTAGTTCCCTTCCATCATATGCAGCTTTTGCTACT   12142
gb:MT020880|Organism:Severe      ACCTTACAAGCTATAGCCTCAGAGTTTAGTTCCCTTCCATCATATGCAGCTTTTGCTACT   12142
gb:MT066175|Organism:Severe      ACCTTACAAGCTATAGCCTCAGAGTTTAGTTCCCTTCCATCATATGCAGCTTTTGCTACT   12142
gb:MN997409|Organism:Severe      ACCTTACAAGCTATAGCCTCAGAGTTTAGTTCCCTTCCATCATATGCAGCTTTTGCTACT   12142
gb:MN938384|Organism:Severe      ACCTTACAAGCTATAGCCTCAGAGTTTAGTTCCCTTCCATCATATGCAGCTTTTGCTACT   12110
gb:MT044258|Organism:Severe      ACCTTACAAGCTATAGCCTCAGAGTTTAGTTCCCTTCCATCATATGCAGCTTTTGCTACT   12118
gb:MT039890|Organism:Severe      ACCTTACAAGCTATAGCCTCAGAGTTTAGTTCTCTTCCATCATATGCAGCTTTTGCTACT   12142
gb:MN988713|Organism:Severe      ACCTTACAAGCTATAGCCTCAGAGTTTAGTTCCCTTCCATCATATGCAGCTTTTGCTACT   12142
gb:LC521925|Organism:Severe      ACCTTACAAGCTATAGCCTCAGAGTTTAGTTCCCTTCCATCATATGCAGCTTTTGCTACT   12115
gb:MT093571|Organism:Severe      ACCTTACAAGCTATAGCCTCAGAGTTTAGTTCCCTTCCATCATATGCAGCTTTTGCTACT   12142
gb:MT039887|Organism:Severe      ACCTTACAAGCTATAGCCTCAGAGTTTAGTTCCCTTCCATCATATGCAGCTTTTGCTACT   12142
gb:MT019530|Organism:Severe      ACCTTACAAGCTATAGCCTCAGAGTTTAGTTCCCTTCCATCATATGCAGCTTTTGCTACT   12142
gb:MT039888|Organism:Severe      ACCTTACAAGCTATAGCCTCAGAGTTTAGTTCCCTTCCATCATATGCAGCTTTTGCTACT   12142
gb:LC522972|Organism:Severe      ACCTTACAAGCTATAGCCTCAGAGTTTAGTTCCCTTCCATCATATGCAGCTTTTGCTACT   12139
gb:MT027063|Organism:Severe      ACCTTACAAGCTATAGCCTCAGAGTTTAGTTCCCTTCCATCATATGCAGCTTTTGCTACT   12142
gb:MT027062|Organism:Severe      ACCTTACAAGCTATAGCCTCAGAGTTTAGTTCCCTTCCATCATATGCAGCTTTTGCTACT   12142
gb:MT019529|Organism:Severe      ACCTTACAAGCTATAGCCTCAGAGTTTAGTTCCCTTCCATCATATGCAGCTTTTGCTACT   12142
gb:MN996529|Organism:Severe      ACCTTACAAGCTATAGCCTCAGAGTTTAGTTCCCTTCCATCATATGCAGCTTTTGCTACT   12130
gb:MN996531|Organism:Severe      ACCTTACAAGCTATAGCCTCAGAGTTTAGTTCCCTTCCATCATATGCAGCTTTTGCTACT   12129
gb:MT066176|Organism:Severe      ACCTTACAAGCTATAGCCTCAGAGTTTAGTTCCCTTCCATCATATGCAGCTTTTGCTACT   12142
gb:MT027064|Organism:Severe      ACCTTACAAGCTATAGCCTCAGAGTTTAGTTCCCTTCCATCATATGCAGCTTTTGCTACT   12142
gb:MN994468|Organism:Severe      ACCTTACAAGCTATAGCCTCAGAGTTTAGTTCCCTTCCATCATATGCAGCTTTTGCTACT   12142
gb:MT072688|Organism:Severe      ACCTTACAAGCTATAGCCTCAGAGTTTAGTTCCCTTCCATCATATGCAGCTTTTGCTACT   12127
gb:MN996527|Organism:Severe      ACCTTACAAGCTATAGCCTCAGAGTTTAGTTCCCTTCCATCATATGCAGCTTTTGCTACT   12109
gb:MT093631|Organism:Severe      ACCTTACAAGCTATAGCCTCAGAGTTTAGTTCCCTTCCATCATATGCAGCTTTTGCTACT   12180
gb:MT106053|Organism:Severe      ACCTTACAAGCTATAGCCTCAGAGTTTAGTTCCCTTCCATCATATGCAGCTTTTGCTACT   12142
gb:MT019533|Organism:Severe      ACCTTACAAGCTATAGCCTCAGAGTTTAGTTCCCTTCCATCATATGCAGCTTTTGCTACT   12142
gb:MT019531|Organism:Severe      ACCTTACAAGCTATAGCCTCAGAGTTTAGTTCCCTTCCATCATATGCAGCTTTTGCTACT   12142
gb:MN996528|Organism:Severe      ACCTTACAAGCTATAGCCTCAGAGTTTAGTTCCCTTCCATCATATGCAGCTTTTGCTACT   12142
gb:MN996530|Organism:Severe      ACCTTACAAGCTATAGCCTCAGAGTTTAGTTCCCTTCCATCATATGCAGCTTTTGCTACT   12128
gb:MN908947|Organism:Severe      ACCTTACAAGCTATAGCCTCAGAGTTTAGTTCCCTTCCATCATATGCAGCTTTTGCTACT   12142
gb:MT019532|Organism:Severe      ACCTTACAAGCTATAGCCTCAGAGTTTAGTTCCCTTCCATCATATGCAGCTTTTGCTACT   12142
                                 ******************************** ***************************

gb:MT020781|Organism:Severe      GCTCAAGAAGCTTATGAGCAGGCTGTTGCTAATGGTGATTCTGAAGTTGTTCTTAAAAAG   12190
gb:MT007544|Organism:Severe      GCTCAAGAAGCTTATGAGCAGGCTGTTGCTAATGGTGATTCTGAAGTTGTTCTTAAAAAG   12202
gb:MN994467|Organism:Severe      GCTCAAGAAGCTTATGAGCAGGCTGTTGCTAATGGTGATTCTGAAGTTGTTCTTAAAAAG   12202
gb:MT044257|Organism:Severe      GCTCAAGAAGCTTATGAGCAGGCTGTTGCTAATGGTGATTCTGAAGTTGTTCTTAAAAAG   12202
gb:MT106054|Organism:Severe      GCTCAAGAAGCTTATGAGCAGGCTGTTGCTAATGGTGATTCTGAAGTTGTTCTTAAAAAG   12202
gb:MT049951|Organism:Severe      GCTCAAGAAGCTTATGAGCAGGCTGTTGCTAATGGTGATTCTGAAGTTGTTCTTAAAAAG   12202
gb:MN975262|Organism:Severe      GCTCAAGAAGCTTATGAGCAGGCTGTTGCTAATGGTGATTCTGAAGTTGTTCTTAAAAAG   12202
gb:MT106052|Organism:Severe      GCTCAAGAAGCTTATGAGCAGGCTGTTGCTAATGGTGATTCTGAAGTTGTTCTTAAAAAG   12202
gb:LC522975|Organism:Severe      GCTCAAGAAGCTTATGAGCAGGCTGTTGCTAATGGTGATTCTGAAGTTGTTCTTAAAAAG   12199
gb:LC522973|Organism:Severe      GCTCAAGAAGCTTATGAGCAGGCTGTTGCTAATGGTGATTCTGAAGTTGTTCTTAAAAAG   12199
gb:LC522974|Organism:Severe      GCTCAAGAAGCTTATGAGCAGGCTGTTGCTAATGGTGATTCTGAAGTTGTTCTTAAAAAG   12199
gb:MN985325|Organism:Severe      GCTCAAGAAGCTTATGAGCAGGCTGTTGCTAATGGTGATTCTGAAGTTGTTCTTAAAAAG   12202
gb:MT020881|Organism:Severe      GCTCAAGAAGCTTATGAGCAGGCTGTTGCTAATGGTGATTCTGAAGTTGTTCTTAAAAAG   12202
gb:MT020880|Organism:Severe      GCTCAAGAAGCTTATGAGCAGGCTGTTGCTAATGGTGATTCTGAAGTTGTTCTTAAAAAG   12202
gb:MT066175|Organism:Severe      GCTCAAGAAGCTTATGAGCAGGCTGTTGCTAATGGTGATTCTGAAGTTGTTCTTAAAAAG   12202
gb:MN997409|Organism:Severe      GCTCAAGAAGCTTATGAGCAGGCTGTTGCTAATGGTGATTCTGAAGTTGTTCTTAAAAAG   12202
gb:MN938384|Organism:Severe      GCTCAAGAAGCTTATGAGCAGGCTGTTGCTAATGGTGATTCTGAAGTTGTTCTTAAAAAG   12170
gb:MT044258|Organism:Severe      GCTCAAGAAGCTTATGAGCAGGCTGTTGCTAATGGTGATTCTGAAGTTGTTCTTAAAAAG   12178
```

```
gb:MT039890|Organism:Severe      GCTCAAGAAGCTTATGAGCAGGCTGTTGCTAATGGTGATTCTGAAGTTGTTCTTAAAAAG   12202
gb:MN988713|Organism:Severe      GCTCAAGAAGCTTATGAGCAGGCTGTTGCTAATGGTGATTCTGAAGTTGTTCTTAAAAAG   12202
gb:LC521925|Organism:Severe      GCTCAAGAAGCTTATGAGCAGGCTGTTGCTAATGGTGATTCTGAAGTTGTTCTTAAAAAG   12175
gb:MT093571|Organism:Severe      GCTCAAGAAGCTTATGAGCAGGCTGTTGCTAATGGTGATTCTGAAGTTGTTCTTAAAAAG   12202
gb:MT039887|Organism:Severe      GCTCAAGAAGCTTATGAGCAGGCTGTTGCTAATGGTGATTCTGAAGTTGTTCTTAAAAAG   12202
gb:MT019530|Organism:Severe      GCTCAAGAAGCTTATGAGCAGGCTGTTGCTAATGGTGATTCTGAAGTTGTTCTTAAAAAG   12202
gb:MT039888|Organism:Severe      GCTCAAGAAGCTTATGAGCAGGCTGTTGCTAATGGTGATTCTGAAGTTGTTCTTAAAAAG   12202
gb:LC522972|Organism:Severe      GCTCAAGAAGCTTATGAGCAGGCTGTTGCTAATGGTGATTCTGAAGTTGTTCTTAAAAAG   12199
gb:MT027063|Organism:Severe      GCTCAAGAAGCTTATGAGCAGGCTGTTGCTAATGGTGATTCTGAAGTTGTTCTTAAAAAG   12202
gb:MT027062|Organism:Severe      GCTCAAGAAGCTTATGAGCAGGCTGTTGCTAATGGTGATTCTGAAGTTGTTCTTAAAAAG   12202
gb:MT019529|Organism:Severe      GCTCAAGAAGCTTATGAGCAGGCTGTTGCTAATGGTGATTCTGAAGTTGTTCTTAAAAAG   12202
gb:MN996529|Organism:Severe      GCTCAAGAAGCTTATGAGCAGGCTGTTGCTAATGGTGATTCTGAAGTTGTTCTTAAAAAG   12190
gb:MN996531|Organism:Severe      GCTCAAGAAGCTTATGAGCAGGCTGTTGCTAATGGTGATTCTGAAGTTGTTCTTAAAAAG   12189
gb:MT066176|Organism:Severe      GCTCAAGAAGCTTATGAGCAGGCTGTTGCTAATGGTGATTCTGAAGTTGTTCTTAAAAAG   12202
gb:MT027064|Organism:Severe      GCTCAAGAAGCTTATGAGCAGGCTGTTGCTAATGGTGATTCTGAAGTTGTTCTTAAAAAG   12202
gb:MN994468|Organism:Severe      GCTCAAGAAGCTTATGAGCAGGCTGTTGCTAATGGTGATTCTGAAGTTGTTCTTAAAAAG   12202
gb:MT072688|Organism:Severe      GCTCAAGAAGCTTATGAGCAGGCTGTTGCTAATGGTGATTCTGAAGTTGTTCTTAAAAAG   12187
gb:MN996527|Organism:Severe      GCTCAAGAAGCTTATGAGCAGGCTGTTGCTAATGGTGATTCTGAAGTTGTTCTTAAAAAG   12169
gb:MT093631|Organism:Severe      GCTCAAGAAGCTTATGAGCAGGCTGTTGCTAATGGTGATTCTGAAGTTGTTCTTAAAAAG   12240
gb:MT106053|Organism:Severe      GCTCAAGAAGCTTATGAGCAGGCTGTTGCTAATGGTGATTCTGAAGTTGTTCTTAAAAAG   12202
gb:MT019533|Organism:Severe      GCTCAAGAAGCTTATGAGCAGGCTGTTGCTAATGGTGATTCTGAAGTTGTTCTTAAAAAG   12202
gb:MT019531|Organism:Severe      GCTCAAGAAGCTTATGAGCAGGCTGTTGCTAATGGTGATTCTGAAGTTGTTCTTAAAAAG   12202
gb:MN996528|Organism:Severe      GCTCAAGAAGCTTATGAGCAGGCTGTTGCTAATGGTGATTCTGAAGTTGTTCTTAAAAAG   12202
gb:MN996530|Organism:Severe      GCTCAAGAAGCTTATGAGCAGGCTGTTGCTAATGGTGATTCTGAAGTTGTTCTTAAAAAG   12188
gb:MN908947|Organism:Severe      GCTCAAGAAGCTTATGAGCAGGCTGTTGCTAATGGTGATTCTGAAGTTGTTCTTAAAAAG   12202
gb:MT019532|Organism:Severe      GCTCAAGAAGCTTATGAGCAGGCTGTTGCTAATGGTGATTCTGAAGTTGTTCTTAAAAAG   12202
                                 ************************************************************

gb:MT020781|Organism:Severe      TTGAAGAAGTCTTTGAATGTGGCTAAATCTGAATTTGACCGTGATGCAGCCATGCAACGT   12250
gb:MT007544|Organism:Severe      TTGAAGAAGTCTTTGAATGTGGCTAAATCTGAATTTGACCGTGATGCAGCCATGCAACGT   12262
gb:MN994467|Organism:Severe      TTGAAGAAGTCTTTGAATGTGGCTAAATCTGAATTTGACCGTGATGCAGCCATGCAACGT   12262
gb:MT044257|Organism:Severe      TTGAAGAAGTCTTTGAATGTGGCTAAATCTGAATTTGACCGTGATGCAGCCATGCAACGT   12262
gb:MT106054|Organism:Severe      TTGAAGAAGTCTTTGAATGTGGCTAAATCTGAATTTGACCGTGATGCAGCCATGCAACGT   12262
gb:MT049951|Organism:Severe      TTGAAGAAGTCTTTGAATGTGGCTAAATCTGAATTTGACCGTGATGCAGCCATGCAACGT   12262
gb:MN975262|Organism:Severe      TTGAAGAAGTCTTTGAATGTGGCTAAATCTGAATTTGACCGTGATGCAGCCATGCAACGT   12262
gb:MT106052|Organism:Severe      TTGAAGAAGTCTTTGAATGTGGCTAAATCTGAATTTGACCGTGATGCAGCCATGCAACGT   12262
gb:LC522975|Organism:Severe      TTGAAGAAGTCTTTGAATGTGGCTAAATCTGAATTTGACCGTGATGCAGCCATGCAACGT   12259
gb:LC522973|Organism:Severe      TTGAAGAAGTCTTTGAATGTGGCTAAATCTGAATTTGACCGTGATGCAGCCATGCAACGT   12259
gb:LC522974|Organism:Severe      TTGAAGAAGTCTTTGAATGTGGCTAAATCTGAATTTGACCGTGATGCAGCCATGCAACGT   12259
gb:MN985325|Organism:Severe      TTGAAGAAGTCTTTGAATGTGGCTAAATCTGAATTTGACCGTGATGCAGCCATGCAACGT   12262
gb:MT020881|Organism:Severe      TTGAAGAAGTCTTTGAATGTGGCTAAATCTGAATTTGACCGTGATGCAGCCATGCAACGT   12262
gb:MT020880|Organism:Severe      TTGAAGAAGTCTTTGAATGTGGCTAAATCTGAATTTGACCGTGATGCAGCCATGCAACGT   12262
gb:MT066175|Organism:Severe      TTGAAGAAGTCTTTGAATGTGGCTAAATCTGAATTTGACCGTGATGCAGCCATGCAACGT   12262
gb:MN997409|Organism:Severe      TTGAAGAAGTCTTTGAATGTGGCTAAATCTGAATTTGACCGTGATGCAGCCATGCAACGT   12262
gb:MN938384|Organism:Severe      TTGAAGAAGTCTTTGAATGTGGCTAAATCTGAATTTGACCGTGATGCAGCCATGCAACGT   12230
gb:MT044258|Organism:Severe      TTGAAGAAGTCTTTGAATGTGGCTAAATCTGAATTTGACCGTGATGCAGCCATGCAACGT   12238
gb:MT039890|Organism:Severe      TTGAAGAAGTCTTTGAATGTGGCTAAATCTGAATTTGACCGTGATGCAGCCATGCAACGT   12262
gb:MN988713|Organism:Severe      TTGAAGAAGTCTTTGAATGTGGCTAAATCTGAATTTGACCGTGATGCAGCCATGCAACGT   12262
gb:LC521925|Organism:Severe      TTGAAGAAGTCTTTGAATGTGGCTAAATCTGAATTTGACCGTGATGCAGCCATGCAACGT   12235
gb:MT093571|Organism:Severe      TTGAAGAAGTCTTTGAATGTGGCTAAATCTGAATTTGACCGTGATGCAGCCATGCAACGT   12262
gb:MT039887|Organism:Severe      TTGAAGAAGTCTTTGAATGTGGCTAAATCTGAATTTGACCGTGATGCAGCCATGCAACGT   12262
gb:MT019530|Organism:Severe      TTGAAGAAGTCTTTGAATGTGGCTAAATCTGAATTTGACCGTGATGCAGCCATGCAACGT   12262
gb:MT039888|Organism:Severe      TTGAAGAAGTCTTTGAATGTGGCTAAATCTGAATTTGACCGTGATGCAGCCATGCAACGT   12262
gb:LC522972|Organism:Severe      TTGAAGAAGTCTTTGAATGTGGCTAAATCTGAATTTGACCGTGATGCAGCCATGCAACGT   12259
gb:MT027063|Organism:Severe      TTGAAGAAGTCTTTGAATGTGGCTAAATCTGAATTTGACCGTGATGCAGCCATGCAACGT   12262
gb:MT027062|Organism:Severe      TTGAAGAAGTCTTTGAATGTGGCTAAATCTGAATTTGACCGTGATGCAGCCATGCAACGT   12262
gb:MT019529|Organism:Severe      TTGAAGAAGTCTTTGAATGTGGCTAAATCTGAATTTGACCGTGATGCAGCCATGCAACGT   12262
gb:MN996529|Organism:Severe      TTGAAGAAGTCTTTGAATGTGGCTAAATCTGAATTTGACCGTGATGCAGCCATGCAACGT   12250
gb:MN996531|Organism:Severe      TTGAAGAAGTCTTTGAATGTGGCTAAATCTGAATTTGACCGTGATGCAGCCATGCAACGT   12249
gb:MT066176|Organism:Severe      TTGAAGAAGTCTTTGAATGTGGCTAAATCTGAATTTGACCGTGATGCAGCCATGCAACGT   12262
gb:MT027064|Organism:Severe      TTGAAGAAGTCTTTGAATGTGGCTAAATCTGAATTTGACCGTGATGCAGCCATGCAACGT   12262
gb:MN994468|Organism:Severe      TTGAAGAAGTCTTTGAATGTGGCTAAATCTGAATTTGACCGTGATGCAGCCATGCAACGT   12262
gb:MT072688|Organism:Severe      TTGAAGAAGTCTTTGAATGTGGCTAAATCTGAATTTGACCGTGATGCAGCCATGCAACGT   12247
gb:MN996527|Organism:Severe      TTGAAGAAGTCTTTGAATGTGGCTAAATCTGAATTTGACCGTGATGCAGCCATGCAACGT   12229
gb:MT093631|Organism:Severe      TTGAAGAAGTCTTTGAATGTGGCTAAATCTGAATTTGACCGTGATGCAGCCATGCAACGT   12300
```

```
gb:MT106053|Organism:Severe      TTGAAGAAGTCTTTGAATGTGGCTAAATCTGAATTTGACCGTGATGCAGCCATGCAACGT    12262
gb:MT019533|Organism:Severe      TTGAAGAAGTCTTTGAATGTGGCTAAATCTGAATTTGACCGTGATGCAGCCATGCAACGT    12262
gb:MT019531|Organism:Severe      TTGAAGAAGTCTTTGAATGTGGCTAAATCTGAATTTGACCGTGATGCAGCCATGCAACGT    12262
gb:MN996528|Organism:Severe      TTGAAGAAGTCTTTGAATGTGGCTAAATCTGAATTTGACCGTGATGCAGCCATGCAACGT    12262
gb:MN996530|Organism:Severe      TTGAAGAAGTCTTTGAATGTGGCTAAATCTGAATTTGACCGTGATGCAGCCATGCAACGT    12248
gb:MN908947|Organism:Severe      TTGAAGAAGTCTTTGAATGTGGCTAAATCTGAATTTGACCGTGATGCAGCCATGCAACGT    12262
gb:MT019532|Organism:Severe      TTGAAGAAGTCTTTGAATGTGGCTAAATCTGAATTTGACCGTGATGCAGCCATGCAACGT    12262
                                 ************************************************************

gb:MT020781|Organism:Severe      AAGTTGGAAAAGATGGCTGATCAAGCTATGACCCAAATGTATAAACAGGCTAGATCTGAG    12310
gb:MT007544|Organism:Severe      AAGTTGGAAAAGATGGCTGATCAAGCTATGACCCAAATGTATAAACAGGCTAGATCTGAG    12322
gb:MN994467|Organism:Severe      AAGTTGGAAAAGATGGCTGATCAAGCTATGACCCAAATGTATAAACAGGCTAGATCTGAG    12322
gb:MT044257|Organism:Severe      AAGTTGGAAAAGATGGCTGATCAAGCTATGACCCAAATGTATAAACAGGCTAGATCTGAG    12322
gb:MT106054|Organism:Severe      AAGTTGGAAAAGATGGCTGATCAAGCTATGACCCAAATGTATAAACAGGCTAGATCTGAG    12322
gb:MT049951|Organism:Severe      AAGTTGGAAAAGATGGCTGATCAAGCTATGACCCAAATGTATAAACAGGCTAGATCTGAG    12322
gb:MN975262|Organism:Severe      AAGTTGGAAAAGATGGCTGATCAAGCTATGACCCAAATGTATAAACAGGCTAGATCTGAG    12322
gb:MT106052|Organism:Severe      AAGTTGGAAAAGATGGCTGATCAAGCTATGACCCAAATGTATAAACAGGCTAGATCTGAG    12322
gb:LC522975|Organism:Severe      AAGTTGGAAAAGATGGCTGATCAAGCTATGACCCAAATGTATAAACAGGCTAGATCTGAG    12319
gb:LC522973|Organism:Severe      AAGTTGGAAAAGATGGCTGATCAAGCTATGACCCAAATGTATAAACAGGCTAGATCTGAG    12319
gb:LC522974|Organism:Severe      AAGTTGGAAAAGATGGCTGATCAAGCTATGACCCAAATGTATAAACAGGCTAGATCTGAG    12319
gb:MN985325|Organism:Severe      AAGTTGGAAAAGATGGCTGATCAAGCTATGACCCAAATGTATAAACAGGCTAGATCTGAG    12322
gb:MT020881|Organism:Severe      AAGTTGGAAAAGATGGCTGATCAAGCTATGACCCAAATGTATAAACAGGCTAGATCTGAG    12322
gb:MT020880|Organism:Severe      AAGTTGGAAAAGATGGCTGATCAAGCTATGACCCAAATGTATAAACAGGCTAGATCTGAG    12322
gb:MT066175|Organism:Severe      AAGTTGGAAAAGATGGCTGATCAAGCTATGACCCAAATGTATAAACAGGCTAGATCTGAG    12322
gb:MN997409|Organism:Severe      AAGTTGGAAAAGATGGCTGATCAAGCTATGACCCAAATGTATAAACAGGCTAGATCTGAG    12322
gb:MN938384|Organism:Severe      AAGTTGGAAAAGATGGCTGATCAAGCTATGACCCAAATGTATAAACAGGCTAGATCTGAG    12290
gb:MT044258|Organism:Severe      AAGTTGGAAAAGATGGCTGATCAAGCTATGACCCAAATGTATAAACAGGCTAGATCTGAG    12298
gb:MT039890|Organism:Severe      AAGTTGGAAAAGATGGCTGATCAAGCTATGACCCAAATGTATAAACAGGCTAGATCTGAG    12322
gb:MN988713|Organism:Severe      AAGTTGGAAAAGATGGCTGATCAAGCTATGACCCAAATGTATAAACAGGCTAGATCTGAG    12322
gb:LC521925|Organism:Severe      AAGTTGGAAAAGATGGCTGATCAAGCTATGACCCAAATGTATAAACAGGCTAGATCTGAG    12295
gb:MT093571|Organism:Severe      AAGTTGGAAAAGATGGCTGATCAAGCTATGACCCAAATGTATAAACAGGCTAGATCTGAG    12322
gb:MT039887|Organism:Severe      AAGTTGGAAAAGATGGCTGATCAAGCTATGACCCAAATGTATAAACAGGCTAGATCTGAG    12322
gb:MT019530|Organism:Severe      AAGTTGGAAAAGATGGCTGATCAAGCTATGACCCAAATGTATAAACAGGCTAGATCTGAG    12322
gb:MT039888|Organism:Severe      AAGTTGGAAAAGATGGCTGATCAAGCTATGACCCAAATGTATAAACAGGCTAGATCTGAG    12322
gb:LC522972|Organism:Severe      AAGTTGGAAAAGATGGCTGATCAAGCTATGACCCAAATGTATAAACAGGCTAGATCTGAG    12319
gb:MT027063|Organism:Severe      AAGTTGGAAAAGATGGCTGATCAAGCTATGACCCAAATGTATAAACAGGCTAGATCTGAG    12322
gb:MT027062|Organism:Severe      AAGTTGGAAAAGATGGCTGATCAAGCTATGACCCAAATGTATAAACAGGCTAGATCTGAG    12322
gb:MT019529|Organism:Severe      AAGTTGGAAAAGATGGCTGATCAAGCTATGACCCAAATGTATAAACAGGCTAGATCTGAG    12322
gb:MN996529|Organism:Severe      AAGTTGGAAAAGATGGCTGATCAAGCTATGACCCAAATGTATAAACAGGCTAGATCTGAG    12310
gb:MN996531|Organism:Severe      AAGTTGGAAAAGATGGCTGATCAAGCTATGACCCAAATGTATAAACAGGCTAGATCTGAG    12309
gb:MT066176|Organism:Severe      AAGTTGGAAAAGATGGCTGATCAAGCTATGACCCAAATGTATAAACAGGCTAGATCTGAG    12322
gb:MT027064|Organism:Severe      AAGTTGGAAAAGATGGCTGATCAAGCTATGACCCAAATGTATAAACAGGCTAGATCTGAG    12322
gb:MN994468|Organism:Severe      AAGTTGGAAAAGATGGCTGATCAAGCTATGACCCAAATGTATAAACAGGCTAGATCTGAG    12322
gb:MT072688|Organism:Severe      AAGTTGGAAAAGATGGCTGATCAAGCTATGACCCAAATGTATAAACAGGCTAGATCTGAG    12307
gb:MN996527|Organism:Severe      AAGTTGGAAAAGATGGCTGATCAAGCTATGACCCAAATGTATAAACAGGCTAGATCTGAG    12289
gb:MT093631|Organism:Severe      AAGTTGGAAAAGATGGCTGATCAAGCTATGACCCAAATGTATAAACAGGCTAGATCTGAG    12360
gb:MT106053|Organism:Severe      AAGTTGGAAAAGATGGCTGATCAAGCTATGACCCAAATGTATAAACAGGCTAGATCTGAG    12322
gb:MT019533|Organism:Severe      AAGTTGGAAAAGATGGCTGATCAAGCTATGACCCAAATGTATAAACAGGCTAGATCTGAG    12322
gb:MT019531|Organism:Severe      AAGTTGGAAAAGATGGCTGATCAAGCTATGACCCAAATGTATAAACAGGCTAGATCTGAG    12322
gb:MN996528|Organism:Severe      AAGTTGGAAAAGATGGCTGATCAAGCTATGACCCAAATGTATAAACAGGCTAGATCTGAG    12322
gb:MN996530|Organism:Severe      AAGTTGGAAAAGATGGCTGATCAAGCTATGACCCAAATGTATAAACAGGCTAGATCTGAG    12308
gb:MN908947|Organism:Severe      AAGTTGGAAAAGATGGCTGATCAAGCTATGACCCAAATGTATAAACAGGCTAGATCTGAG    12322
gb:MT019532|Organism:Severe      AAGTTGGAAAAGATGGCTGATCAAGCTATGACCCAAATGTATAAACAGGCTAGATCTGAG    12322
                                 ************************************************************

gb:MT020781|Organism:Severe      GACAAGAGGGCAAAAGTTACTAGTGCTATGCAGACAATGCTTTTCACTATGCTTAGAAAG    12370
gb:MT007544|Organism:Severe      GACAAGAGGGCAAAAGTTACTAGTGCTATGCAGACAATGCTTTTCACTATGCTTAGAAAG    12382
gb:MN994467|Organism:Severe      GACAAGAGGGCAAAAGTTACTAGTGCTATGCAGACAATGCTTTTCACTATGCTTAGAAAG    12382
gb:MT044257|Organism:Severe      GACAAGAGGGCAAAAGTTACTAGTGCTATGCAGACAATGCTTTTCACTATGCTTAGAAAG    12382
gb:MT106054|Organism:Severe      GACAAGAGGGCAAAAGTTACTAGTGCTATGCAGACAATGCTTTTCACTATGCTTAGAAAG    12382
gb:MT049951|Organism:Severe      GACAAGAGGGCAAAAGTTACTAGTGCTATGCAGACAATGCTTTTCACTATGCTTAGAAAG    12382
gb:MN975262|Organism:Severe      GACAAGAGGGCAAAAGTTACTAGTGCTATGCAGACAATGCTTTTCACTATGCTTAGAAAG    12382
gb:MT106052|Organism:Severe      GACAAGAGGGCAAAAGTTACTAGTGCTATGCAGACAATGCTTTTCACTATGCTTAGAAAG    12382
gb:LC522975|Organism:Severe      GACAAGAGGGCAAAAGTTACTAGTGCTATGCAGACAATGCTTTTCACTATGCTTAGAAAG    12379
gb:LC522973|Organism:Severe      GACAAGAGGGCAAAAGTTACTAGTGCTATGCAGACAATGCTTTTCACTATGCTTAGAAAG    12379
```

```
gb:LC522974|Organism:Severe      GACAAGAGGGCAAAAGTTACTAGTGCTATGCAGACAATGCTTTTCACTATGCTTAGAAAG   12379
gb:MN985325|Organism:Severe      GACAAGAGGGCAAAAGTTACTAGTGCTATGCAGACAATGCTTTTCACTATGCTTAGAAAG   12382
gb:MT020881|Organism:Severe      GACAAGAGGGCAAAAGTTACTAGTGCTATGCAGACAATGCTTTTCACTATGCTTAGAAAG   12382
gb:MT020880|Organism:Severe      GACAAGAGGGCAAAAGTTACTAGTGCTATGCAGACAATGCTTTTCACTATGCTTAGAAAG   12382
gb:MT066175|Organism:Severe      GACAAGAGGGCAAAAGTTACTAGTGCTATGCAGACAATGCTTTTCACTATGCTTAGAAAG   12382
gb:MN997409|Organism:Severe      GACAAGAGGGCAAAAGTTACTAGTGCTATGCAGACAATGCTTTTCACTATGCTTAGAAAG   12382
gb:MN938384|Organism:Severe      GACAAGAGGGCAAAAGTTACTAGTGCTATGCAGACAATGCTTTTCACTATGCTTAGAAAG   12350
gb:MT044258|Organism:Severe      GACAAGAGGGCAAAAGTTACTAGTGCTATGCAGACAATGCTTTTCACTATGCTTAGAAAG   12358
gb:MT039890|Organism:Severe      GACAAGAGGGCAAAAGTTACTAGTGCTATGCAGACAATGCTTTTCACTATGCTTAGAAAG   12382
gb:MN988713|Organism:Severe      GACAAGAGGGCAAAAGTTACTAGTGCTATGCAGACAATGCTTTTCACTATGCTTAGAAAG   12382
gb:LC521925|Organism:Severe      GACAAGAGGGCAAAAGTTACTAGTGCTATGCAGACAATGCTTTTCACTATGCTTAGAAAG   12355
gb:MT093571|Organism:Severe      GACAAGAGGGCAAAAGTTACTAGTGCTATGCAGACAATGCTTTTCACTATGCTTAGAAAG   12382
gb:MT039887|Organism:Severe      GACAAGAGGGCAAAAGTTACTAGTGCTATGCAGACAATGCTTTTCACTATGCTTAGAAAG   12382
gb:MT019530|Organism:Severe      GACAAGAGGGCAAAAGTTACTAGTGCTATGCAGACAATGCTTTTCACTATGCTTAGAAAG   12382
gb:MT039888|Organism:Severe      GACAAGAGGGCAAAAGTTACTAGTGCTATGCAGACAATGCTTTTCACTATGCTTAGAAAG   12382
gb:LC522972|Organism:Severe      GACAAGAGGGCAAAAGTTACTAGTGCTATGCAGACAATGCTTTTCACTATGCTTAGAAAG   12379
gb:MT027063|Organism:Severe      GACAAGAGGGCAAAAGTTACTAGTGCTATGCAGACAATGCTTTTCACTATGCTTAGAAAG   12382
gb:MT027062|Organism:Severe      GACAAGAGGGCAAAAGTTACTAGTGCTATGCAGACAATGCTTTTCACTATGCTTAGAAAG   12382
gb:MT019529|Organism:Severe      GACAAGAGGGCAAAAGTTACTAGTGCTATGCAGACAATGCTTTTCACTATGCTTAGAAAG   12382
gb:MN996529|Organism:Severe      GACAAGAGGGCAAAAGTTACTAGTGCTATGCAGACAATGCTTTTCACTATGCTTAGAAAG   12370
gb:MN996531|Organism:Severe      GACAAGAGGGCAAAAGTTACTAGTGCTATGCAGACAATGCTTTTCACTATGCTTAGAAAG   12369
gb:MT066176|Organism:Severe      GACAAGAGGGCAAAAGTTACTAGTGCTATGCAGACAATGCTTTTCACTATGCTTAGAAAG   12382
gb:MT027064|Organism:Severe      GACAAGAGGGCAAAAGTTACTAGTGCTATGCAGACAATGCTTTTCACTATGCTTAGAAAG   12382
gb:MN994468|Organism:Severe      GACAAGAGGGCAAAAGTTACTAGTGCTATGCAGACAATGCTTTTCACTATGCTTAGAAAG   12382
gb:MT072688|Organism:Severe      GACAAGAGGGCAAAAGTTACTAGTGCTATGCAGACAATGCTTTTCACTATGCTTAGAAAG   12367
gb:MN996527|Organism:Severe      GACAAGAGGGCAAAAGTTACTAGTGCTATGCAGACAATGCTTTTCACTATGCTTAGAAAG   12349
gb:MT093631|Organism:Severe      GACAAGAGGGCAAAAGTTACTAGTGCTATGCAGACAATGCTTTTCACTATGCTTAGAAAG   12420
gb:MT106053|Organism:Severe      GACAAGAGGGCAAAAGTTACTAGTGCTATGCAGACAATGCTTTTCACTATGCTTAGAAAG   12382
gb:MT019533|Organism:Severe      GACAAGAGGGCAAAAGTTACTAGTGCTATGCAGACAATGCTTTTCACTATGCTTAGAAAG   12382
gb:MT019531|Organism:Severe      GACAAGAGGGCAAAAGTTACTAGTGCTATGCAGACAATGCTTTTCACTATGCTTAGAAAG   12382
gb:MN996528|Organism:Severe      GACAAGAGGGCAAAAGTTACTAGTGCTATGCAGACAATGCTTTTCACTATGCTTAGAAAG   12382
gb:MN996530|Organism:Severe      GACAAGAGGGCAAAAGTTACTAGTGCTATGCAGACAATGCTTTTCACTATGCTTAGAAAG   12368
gb:MN908947|Organism:Severe      GACAAGAGGGCAAAAGTTACTAGTGCTATGCAGACAATGCTTTTCACTATGCTTAGAAAG   12382
gb:MT019532|Organism:Severe      GACAAGAGGGCAAAAGTTACTAGTGCTATGCAGACAATGCTTTTCACTATGCTTAGAAAG   12382
                                 ************************************************************

gb:MT020781|Organism:Severe      TTGGATAATGATGCACTCAACAACATTATCAACAATGCAAGAGATGGTTGTGTTCCCTTG   12430
gb:MT007544|Organism:Severe      TTGGATAATGATGCACTCAACAACATTATCAACAATGCAAGAGATGGTTGTGTTCCCTTG   12442
gb:MN994467|Organism:Severe      TTGGATAATGATGCACTCAACAACATTATCAACAATGCAAGAGATGGTTGTGTTCCCTTG   12442
gb:MT044257|Organism:Severe      TTGGATAATGATGCACTCAACAACATTATCAACAATGCAAGAGATGGTTGTGTTCCCTTG   12442
gb:MT106054|Organism:Severe      TTGGATAATGATGCACTCAACAACATTATCAACAATGCAAGAGATGGTTGTGTTCCCTTG   12442
gb:MT049951|Organism:Severe      TTGGATAATGATGCACTCAACAACATTATCAACAATGCAAGAGATGGTTGTGTTCCCTTG   12442
gb:MN975262|Organism:Severe      TTGGATAATGATGCACTCAACAACATTATCAACAATGCAAGAGATGGTTGTGTTCCCTTG   12442
gb:MT106052|Organism:Severe      TTGGATAATGATGCACTCAACAACATTATCAACAATGCAAGAGATGGTTGTGTTCCCTTG   12442
gb:LC522975|Organism:Severe      TTGGATAATGATGCACTCAACAACATTATCAACAATGCAAGAGATGGTTGTGTTCCCTTG   12439
gb:LC522973|Organism:Severe      TTGGATAATGATGCACTCAACAACATTATCAACAATGCAAGAGATGGTTGTGTTCCCTTG   12439
gb:LC522974|Organism:Severe      TTGGATAATGATGCACTCAACAACATTATCAACAATGCAAGAGATGGTTGTGTTCCCTTG   12439
gb:MN985325|Organism:Severe      TTGGATAATGATGCACTCAACAACATTATCAACAATGCAAGAGATGGTTGTGTTCCCTTG   12442
gb:MT020881|Organism:Severe      TTGGATAATGATGCACTCAACAACATTATCAACAATGCAAGAGATGGTTGTGTTCCCTTG   12442
gb:MT020880|Organism:Severe      TTGGATAATGATGCACTCAACAACATTATCAACAATGCAAGAGATGGTTGTGTTCCCTTG   12442
gb:MT066175|Organism:Severe      TTGGATAATGATGCACTCAACAACATTATCAACAATGCAAGAGATGGTTGTGTTCCCTTG   12442
gb:MN997409|Organism:Severe      TTGGATAATGATGCACTCAACAACATTATCAACAATGCAAGAGATGGTTGTGTTCCCTTG   12442
gb:MN938384|Organism:Severe      TTGGATAATGATGCACTCAACAACATTATCAACAATGCAAGAGATGGTTGTGTTCCCTTG   12410
gb:MT044258|Organism:Severe      TTGGATAATGATGCACTCAACAACATTATCAACAATGCAAGAGATGGTTGTGTTCCCTTG   12418
gb:MT039890|Organism:Severe      TTGGATAATGATGCACTCAACAACATTATCAACAATGCAAGAGATGGTTGTGTTCCCTTG   12442
gb:MN988713|Organism:Severe      TTGGATAATGATGCACTCAACAACATTATCAACAATGCAAGAGATGGTTGTGTTCCCTTG   12442
gb:LC521925|Organism:Severe      TTGGATAATGATGCACTCAACAACATTATCAACAATGCAAGAGATGGTTGTGTTCCCTTG   12415
gb:MT093571|Organism:Severe      TTGGATAATGATGCACTCAACAACATTATCAACAATGCAAGAGATGGTTGTGTTCCCTTG   12442
gb:MT039887|Organism:Severe      TTGGATAATGATGCACTCAACAACATTATCAACAATGCAAGAGATGGTTGTGTTCCCTTG   12442
gb:MT019530|Organism:Severe      TTGGATAATGATGCACTCAACAACATTATCAACAATGCAAGAGATGGTTGTGTTCCCTTG   12442
gb:MT039888|Organism:Severe      TTGGATAATGATGCACTCAACAACATTATCAACAATGCAAGAGATGGTTGTGTTCCCTTG   12442
gb:LC522972|Organism:Severe      TTGGATAATGATGCACTCAACAACATTATCAACAATGCAAGAGATGGTTGTGTTCCCTTG   12439
gb:MT027063|Organism:Severe      TTGGATAATGATGCACTCAACAACATTATCAACAATGCAAGAGATGGTTGTGTTCCCTTG   12442
gb:MT027062|Organism:Severe      TTGGATAATGATGCACTCAACAACATTATCAACAATGCAAGAGATGGTTGTGTTCCCTTG   12442
gb:MT019529|Organism:Severe      TTGGATAATGATGCACTCAACAACATTATCAACAATGCAAGAGATGGTTGTGTTCCCTTG   12442
```

```
gb:MN996529|Organism:Severe      TTGGATAATGATGCACTCAACAACATTATCAACAATGCAAGAGATGGTTGTGTTCCCTTG   12430
gb:MN996531|Organism:Severe      TTGGATAATGATGCACTCAACAACATTATCAACAATGCAAGAGATGGTTGTGTTCCCTTG   12429
gb:MT066176|Organism:Severe      TTGGATAATGATGCACTCAACAACATTATCAACAATGCAAGAGATGGTTGTGTTCCCTTG   12442
gb:MT027064|Organism:Severe      TTGGATAATGATGCACTCAACAACATTATCAACAATGCAAGAGATGGTTGTGTTCCCTTG   12442
gb:MN994468|Organism:Severe      TTGGATAATGATGCACTCAACAACATTATCAACAATGCAAGAGATGGTTGTGTTCCCTTG   12442
gb:MT072688|Organism:Severe      TTGGATAATGATGCACTCAACAACATTATCAACAATGCAAGAGATGGTTGTGTTCCCTTG   12427
gb:MN996527|Organism:Severe      TTGGATAATGATGCACTCAACAACATTATCAACAATGCAAGAGATGGTTGTGTTCCCTTG   12409
gb:MT093631|Organism:Severe      TTGGATAATGATGCACTCAACAACATTATCAACAATGCAAGAGATGGTTGTGTTCCCTTG   12480
gb:MT106053|Organism:Severe      TTGGATAATGATGCACTCAACAACATTATCAACAATGCAAGAGATGGTTGTGTTCCCTTG   12442
gb:MT019533|Organism:Severe      TTGGATAATGATGCACTCAACAACATTATCAACAATGCAAGAGATGGTTGTGTTCCCTTG   12442
gb:MT019531|Organism:Severe      TTGGATAATGATGCACTCAACAACATTATCAACAATGCAAGAGATGGTTGTGTTCCCTTG   12442
gb:MN996528|Organism:Severe      TTGGATAATGATGCACTCAACAACATTATCAACAATGCAAGAGATGGTTGTGTTCCCTTG   12442
gb:MN996530|Organism:Severe      TTGGATAATGATGCACTCAACAACATTATCAACAATGCAAGAGATGGTTGTGTTCCCTTG   12428
gb:MN908947|Organism:Severe      TTGGATAATGATGCACTCAACAACATTATCAACAATGCAAGAGATGGTTGTGTTCCCTTG   12442
gb:MT019532|Organism:Severe      TTGGATAATGATGCACTCAACAACATTATCAACAATGCAAGAGATGGTTGTGTTCCCTTG   12442
                                 ************************************************************

gb:MT020781|Organism:Severe      AACATAATACCTCTTACAACAGCAGCCAAACTAATGGTTGTCATACCAGACTATAACACA   12490
gb:MT007544|Organism:Severe      AACATAATACCTCTTACAACAGCAGCCAAACTAATGGTTGTCATACCAGACTATAACACA   12502
gb:MN994467|Organism:Severe      AACATAATACCTCTTACAACAGCAGCCAAACTAATGGTTGTCATACCAGACTATAACACA   12502
gb:MT044257|Organism:Severe      AACATAATACCTCTTACAACAGCAGCCAAACTAATGGTTGTCATACCAGACTATAACACA   12502
gb:MT106054|Organism:Severe      AACATAATACCTCTTACAACAGCAGCCAAACTAATGGTTGTCATACCAGACTATAACACA   12502
gb:MT049951|Organism:Severe      AACATAATACCTCTTACAACAGCAGCCAAACTAATGGTTGTCATACCAGACTATAACACA   12502
gb:MN975262|Organism:Severe      AACATAATACCTCTTACAACAGCAGCCAAACTAATGGTTGTCATACCAGACTATAACACA   12502
gb:MT106052|Organism:Severe      AACATAATACCTCTTACAACAGCAGCCAAACTAATGGTTGTCATACCAGACTATAACACA   12502
gb:LC522975|Organism:Severe      AACATAATACCTCTTACAACAGCAGCCAAACTAATGGTTGTCATACCAGACTATAACACA   12499
gb:LC522973|Organism:Severe      AACATAATACCTCTTACAACAGCAGCCAAACTAATGGTTGTCATACCAGACTATAACACA   12499
gb:LC522974|Organism:Severe      AACATAATACCTCTTACAACAGCAGCCAAACTAATGGTTGTCATACCAGACTATAACACA   12499
gb:MN985325|Organism:Severe      AACATAATACCTCTTACAACAGCAGCCAAACTAATGGTTGTCATACCAGACTATAACACA   12502
gb:MT020881|Organism:Severe      AACATAATACCTCTTACAACAGCAGCCAAACTAATGGTTGTCATACCAGACTATAACACA   12502
gb:MT020880|Organism:Severe      AACATAATACCTCTTACAACAGCAGCCAAACTAATGGTTGTCATACCAGACTATAACACA   12502
gb:MT066175|Organism:Severe      AACATAATACCTCTTACAACAGCAGCCAAACTAATGGTTGTCATACCAGACTATAACACA   12502
gb:MN997409|Organism:Severe      AACATAATACCTCTTACAACAGCAGCCAAACTAATGGTTGTCATACCAGACTATAACACA   12502
gb:MN938384|Organism:Severe      AACATAATACCTCTTACAACAGCAGCCAAACTAATGGTTGTCATACCAGACTATAACACA   12470
gb:MT044258|Organism:Severe      AACATAATACCTCTTACAACAGCAGCCAAACTAATGGTTGTCATACCAGACTATAACACA   12478
gb:MT039890|Organism:Severe      AACATAATACCTCTTACAACAGCAGCCAAACTAATGGTTGTCATACCAGACTATAACACA   12502
gb:MN988713|Organism:Severe      AACATAATACCTCTTACAACAGCAGCCAAACTAATGGTTGTCATACCAGACTATAACACA   12502
gb:LC521925|Organism:Severe      AACATAATACCTCTTACAACAGCAGCCAAACTAATGGTTGTCATACCAGACTATAACACA   12475
gb:MT093571|Organism:Severe      AACATAATACCTCTTACAACAGCAGCCAAACTAATGGTTGTCATACCAGACTATAACACA   12502
gb:MT039887|Organism:Severe      AACATAATACCTCTTACAACAGCAGCCAAACTAATGGTTGTCATACCAGACTATAACACA   12502
gb:MT019530|Organism:Severe      AACATAATACCTCTTACAACAGCAGCCAAACTAATGGTTGTCATACCAGACTATAACACA   12502
gb:MT039888|Organism:Severe      AACATAATACCTCTTACAACAGCAGCCAAACTAATGGTTGTCATACCAGACTATAACACA   12502
gb:LC522972|Organism:Severe      AACATAATACCTCTTACAACAGCAGCCAAACTAATGGTTGTCATACCAGACTATAACACA   12499
gb:MT027063|Organism:Severe      AACATAATACCTCTTACAACAGCAGCCAAACTAATGGTTGTCATACCAGACTATAACACA   12502
gb:MT027062|Organism:Severe      AACATAATACCTCTTACAACAGCAGCCAAACTAATGGTTGTCATACCAGACTATAACACA   12502
gb:MT019529|Organism:Severe      AACATAATACCTCTTACAACAGCAGCCAAACTAATGGTTGTCATACCAGACTATAACACA   12502
gb:MN996529|Organism:Severe      AACATAATACCTCTTACAACAGCAGCCAAACTAATGGTTGTCATACCAGACTATAACACA   12490
gb:MN996531|Organism:Severe      AACATAATACCTCTTACAACAGCAGCCAAACTAATGGTTGTCATACCAGACTATAACACA   12489
gb:MT066176|Organism:Severe      AACATAATACCTCTTACAACAGCAGCCAAACTAATGGTTGTCATACCAGACTATAACACA   12502
gb:MT027064|Organism:Severe      AACATAATACCTCTTACAACAGCAGCCAAACTAATGGTTGTCATACCAGACTATAACACA   12502
gb:MN994468|Organism:Severe      AACATAATACCTCTTACAACAGCAGCCAAACTAATGGTTGTCATACCAGACTATAACACA   12502
gb:MT072688|Organism:Severe      AACATAATACCTCTTACAACAGCAGCCAAACTAATGGTTGTCATACCAGACTATAACACA   12487
gb:MN996527|Organism:Severe      AACATAATACCTCTTACAACAGCAGCCAAACTAATGGTTGTCATACCAGACTATAACACA   12469
gb:MT093631|Organism:Severe      AACATAATACCTCTTACAACAGCAGCCAAACTAATGGTTGTCATACCAGACTATAACACA   12540
gb:MT106053|Organism:Severe      AACATAATACCTCTTACAACAGCAGCCAAACTAATGGTTGTCATACCAGACTATAACACA   12502
gb:MT019533|Organism:Severe      AACATAATACCTCTTACAACAGCAGCCAAACTAATGGTTGTCATACCAGACTATAACACA   12502
gb:MT019531|Organism:Severe      AACATAATACCTCTTACAACAGCAGCCAAACTAATGGTTGTCATACCAGACTATAACACA   12502
gb:MN996528|Organism:Severe      AACATAATACCTCTTACAACAGCAGCCAAACTAATGGTTGTCATACCAGACTATAACACA   12502
gb:MN996530|Organism:Severe      AACATAATACCTCTTACAACAGCAGCCAAACTAATGGTTGTCATACCAGACTATAACACA   12488
gb:MN908947|Organism:Severe      AACATAATACCTCTTACAACAGCAGCCAAACTAATGGTTGTCATACCAGACTATAACACA   12502
gb:MT019532|Organism:Severe      AACATAATACCTCTTACAACAGCAGCCAAACTAATGGTTGTCATACCAGACTATAACACA   12502
                                 ************************************************************

gb:MT020781|Organism:Severe      TATAAAAATACGTGTGATGGTACAACATTTACTTATGCATCAGCATTGTGGGAAATCCAA   12550
gb:MT007544|Organism:Severe      TATAAAAATACGTGTGATGGTACAACATTTACTTATGCATCAGCATTGTGGGAAATCCAA   12562
```

```
gb:MN994467|Organism:Severe      TATAAAAATACGTGTGATGGTACAACATTTACTTATGCATCAGCATTGTGGGAAATCCAA   12562
gb:MT044257|Organism:Severe      TATAAAAATACGTGTGATGGTACAACATTTACTTATGCATCAGCATTGTGGGAAATCCAA   12562
gb:MT106054|Organism:Severe      TATAAAAATACGTGTGATGGTACAACATTTACTTATGCATCAGCATTGTGGGAAATCCAA   12562
gb:MT049951|Organism:Severe      TATAAAAATACGTGTGATGGTACAACATTTACTTATGCATCAGCATTGTGGGAAATCCAA   12562
gb:MN975262|Organism:Severe      TATAAAAATACGTGTGATGGTACAACATTTACTTATGCATCAGCATTGTGGGAAATCCAA   12562
gb:MT106052|Organism:Severe      TATAAAAATACGTGTGATGGTACAACATTTACTTATGCATCAGCATTGTGGGAAATCCAA   12562
gb:LC522975|Organism:Severe      TATAAAAATACGTGTGATGGTACAACATTTACTTATGCATCAGCATTGTGGGAAATCCAA   12559
gb:LC522973|Organism:Severe      TATAAAAATACGTGTGATGGTACAACATTTACTTATGCATCAGCATTGTGGGAAATCCAA   12559
gb:LC522974|Organism:Severe      TATAAAAATACGTGTGATGGTACAACATTTACTTATGCATCAGCATTGTGGGAAATCCAA   12559
gb:MN985325|Organism:Severe      TATAAAAATACGTGTGATGGTACAACATTTACTTATGCATCAGCATTGTGGGAAATCCAA   12562
gb:MT020881|Organism:Severe      TATAAAAATACGTGTGATGGTACAACATTTACTTATGCATCAGCATTGTGGGAAATCCAA   12562
gb:MT020880|Organism:Severe      TATAAAAATACGTGTGATGGTACAACATTTACTTATGCATCAGCATTGTGGGAAATCCAA   12562
gb:MT066175|Organism:Severe      TATAAAAATACGTGTGATGGTACAACATTTACTTATGCATCAGCATTGTGGGAAATCCAA   12562
gb:MN997409|Organism:Severe      TATAAAAATACGTGTGATGGTACAACATTTACTTATGCATCAGCATTGTGGGAAATCCAA   12562
gb:MN938384|Organism:Severe      TATAAAAATACGTGTGATGGTACAACATTTACTTATGCATCAGCATTGTGGGAAATCCAA   12530
gb:MT044258|Organism:Severe      TATAAAAATACGTGTGATGGTACAACATTTACTTATGCATCAGCATTGTGGGAAATCCAA   12538
gb:MT039890|Organism:Severe      TATAAAAATACGTGTGATGGTACAACATTTACTTATGCATCAGCATTGTGGGAAATCCAA   12562
gb:MN988713|Organism:Severe      TATAAAAATACGTGTGATGGTACAACATTTACTTATGCATCAGCATTGTGGGAAATCCAA   12562
gb:LC521925|Organism:Severe      TATAAAAATACGTGTGATGGTACAACATTTACTTATGCATCAGCATTGTGGGAAATCCAA   12535
gb:MT093571|Organism:Severe      TATAAAAATACGTGTGATGGTACAACATTTACTTATGCATCAGCATTGTGGGAAATCCAA   12562
gb:MT039887|Organism:Severe      TATAAAAATACGTGTGATGGTACAACATTTACTTATGCATCAGCATTGTGGGAAATCCAA   12562
gb:MT019530|Organism:Severe      TATAAAAATACGTGTGATGGTACAACATTTACTTATGCATCAGCATTGTGGGAAATCCAA   12562
gb:MT039888|Organism:Severe      TATAAAAATACGTGTGATGGTACAACATTTACTTATGCATCAGCATTGTGGGAAATCCAA   12562
gb:LC522972|Organism:Severe      TATAAAAATACGTGTGATGGTACAACATTTACTTATGCATCAGCATTGTGGGAAATCCAA   12559
gb:MT027063|Organism:Severe      TATAAAAATACGTGTGATGGTACAACATTTACTTATGCATCAGCATTGTGGGAAATCCAA   12562
gb:MT027062|Organism:Severe      TATAAAAATACGTGTGATGGTACAACATTTACTTATGCATCAGCATTGTGGGAAATCCAA   12562
gb:MT019529|Organism:Severe      TATAAAAATACGTGTGATGGTACAACATTTACTTATGCATCAGCATTGTGGGAAATCCAA   12562
gb:MN996529|Organism:Severe      TATAAAAATACGTGTGATGGTACAACATTTACTTATGCATCAGCATTGTGGGAAATCCAA   12550
gb:MN996531|Organism:Severe      TATAAAAATACGTGTGATGGTACAACATTTACTTATGCATCAGCATTGTGGGAAATCCAA   12549
gb:MT066176|Organism:Severe      TATAAAAATACGTGTGATGGTACAACATTTACTTATGCATCAGCATTGTGGGAAATCCAA   12562
gb:MT027064|Organism:Severe      TATAAAAATACGTGTGATGGTACAACATTTACTTATGCATCAGCATTGTGGGAAATCCAA   12562
gb:MN994468|Organism:Severe      TATAAAAATACGTGTGATGGTACAACATTTACTTATGCATCAGCATTGTGGGAAATCCAA   12562
gb:MT072688|Organism:Severe      TATAAAAATACGTGTGATGGTACAACATTTACTTATGCATCAGCATTGTGGGAAATCCAA   12547
gb:MN996527|Organism:Severe      TATAAAAATACGTGTGATGGTACAACATTTACTTATGCATCAGCATTGTGGGAAATCCAA   12529
gb:MT093631|Organism:Severe      TATAAAAATACGTGTGATGGTACAACATTTACTTATGCATCAGCATTGTGGGAAATCCAA   12600
gb:MT106053|Organism:Severe      TATAAAAATACGTGTGATGGTACAACATTTACTTATGCATCAGCATTGTGGGAAATCCAA   12562
gb:MT019533|Organism:Severe      TATAAAAATACGTGTGATGGTACAACATTTACTTATGCATCAGCATTGTGGGAAATCCAA   12562
gb:MT019531|Organism:Severe      TATAAAAATACGTGTGATGGTACAACATTTACTTATGCATCAGCATTGTGGGAAATCCAA   12562
gb:MN996528|Organism:Severe      TATAAAAATACGTGTGATGGTACAACATTTACTTATGCATCAGCATTGTGGGAAATCCAA   12562
gb:MN996530|Organism:Severe      TATAAAAATACGTGTGATGGTACAACATTTACTTATGCATCAGCATTGTGGGAAATCCAA   12548
gb:MN908947|Organism:Severe      TATAAAAATACGTGTGATGGTACAACATTTACTTATGCATCAGCATTGTGGGAAATCCAA   12562
gb:MT019532|Organism:Severe      TATAAAAATACGTGTGATGGTACAACATTTACTTATGCATCAGCATTGTGGGAAATCCAA   12562
                                 ************************************************************

gb:MT020781|Organism:Severe      CAGGTTGTAGATGCAGATAGTAAAATTGTTCAACTTAGTGAAATTAGTATGGACAATTCA   12610
gb:MT007544|Organism:Severe      CAGGTTGTAGATGCAGATAGTAAAATTGTTCAACTTAGTGAAATTAGTATGGACAATTCA   12622
gb:MN994467|Organism:Severe      CAGGTTGTAGATGCAGATAGTAAAATTGTTCAACTTAGTGAAATTAGTATGGACAATTCA   12622
gb:MT044257|Organism:Severe      CAGGTTGTAGATGCAGATAGTAAAATTGTTCAACTTAGTGAAATTAGTATGGACAATTCA   12622
gb:MT106054|Organism:Severe      CAGGTTGTAGATGCAGATAGTAAAATTGTTCAACTTAGTGAAATTAGTATGGACAATTCA   12622
gb:MT049951|Organism:Severe      CAGGTTGTAGATGCAGATAGTAAAATTGTTCAACTTAGTGAAATTAGTATGGACAATTCA   12622
gb:MN975262|Organism:Severe      CAGGTTGTAGATGCAGATAGTAAAATTGTTCAACTTAGTGAAATTAGTATGGACAATTCA   12622
gb:MT106052|Organism:Severe      CAGGTTGTAGATGCAGATAGTAAAATTGTTCAACTTAGTGAAATTAGTATGGACAATTCA   12622
gb:LC522975|Organism:Severe      CAGGTTGTAGATGCAGATAGTAAAATTGTTCAACTTAGTGAAATTAGTATGGACAATTCA   12619
gb:LC522973|Organism:Severe      CAGGTTGTAGATGCAGATAGTAAAATTGTTCAACTTAGTGAAATTAGTATGGACAATTCA   12619
gb:LC522974|Organism:Severe      CAGGTTGTAGATGCAGATAGTAAAATTGTTCAACTTAGTGAAATTAGTATGGACAATTCA   12619
gb:MN985325|Organism:Severe      CAGGTTGTAGATGCAGATAGTAAAATTGTTCAACTTAGTGAAATTAGTATGGACAATTCA   12622
gb:MT020881|Organism:Severe      CAGGTTGTAGATGCAGATAGTAAAATTGTTCAACTTAGTGAAATTAGTATGGACAATTCA   12622
gb:MT020880|Organism:Severe      CAGGTTGTAGATGCAGATAGTAAAATTGTTCAACTTAGTGAAATTAGTATGGACAATTCA   12622
gb:MT066175|Organism:Severe      CAGGTTGTAGATGCAGATAGTAAAATTGTTCAACTTAGTGAAATTAGTATGGACAATTCA   12622
gb:MN997409|Organism:Severe      CAGGTTGTAGATGCAGATAGTAAAATTGTTCAACTTAGTGAAATTAGTATGGACAATTCA   12622
gb:MN938384|Organism:Severe      CAGGTTGTAGATGCAGATAGTAAAATTGTTCAACTTAGTGAAATTAGTATGGACAATTCA   12590
gb:MT044258|Organism:Severe      CAGGTTGTAGATGCAGATAGTAAAATTGTTCAACTTAGTGAAATTAGTATGGACAATTCA   12598
gb:MT039890|Organism:Severe      CAGGTTGTAGATGCAGATAGTAAAATTGTTCAACTTAGTGAAATTAGTATGGACAATTCA   12622
gb:MN988713|Organism:Severe      CAGGTTGTAGATGCAGATAGTAAAATTGTTCAACTTAGTGAAATTAGTATGGACAATTCA   12622
gb:LC521925|Organism:Severe      CAGGTTGTAGATGCAGATAGTAAAATTGTTCAACTTAGTGAAATTAGTATGGACAATTCA   12595
```

```
gb:MT093571|Organism:Severe      CAGGTTGTAGATGCAGATAGTAAAATTGTTCAACTTAGTGAAATTAGTATGGACAATTCA   12622
gb:MT039887|Organism:Severe      CAGGTTGTAGATGCAGATAGTAAAATTGTTCAACTTAGTGAAATTAGTATGGACAATTCA   12622
gb:MT019530|Organism:Severe      CAGGTTGTAGATGCAGATAGTAAAATTGTTCAACTTAGTGAAATTAGTATGGACAATTCA   12622
gb:MT039888|Organism:Severe      CAGGTTGTAGATGCAGATAGTAAAATTGTTCAACTTAGTGAAATTAGTATGGACAATTCA   12622
gb:LC522972|Organism:Severe      CAGGTTGTAGATGCAGATAGTAAAATTGTTCAACTTAGTGAAATTAGTATGGACAATTCA   12619
gb:MT027063|Organism:Severe      CAGGTTGTAGATGCAGATAGTAAAATTGTTCAACTTAGTGAAATTAGTATGGACAATTCA   12622
gb:MT027062|Organism:Severe      CAGGTTGTAGATGCAGATAGTAAAATTGTTCAACTTAGTGAAATTAGTATGGACAATTCA   12622
gb:MT019529|Organism:Severe      CAGGTTGTAGATGCAGATAGTAAAATTGTTCAACTTAGTGAAATTAGTATGGACAATTCA   12622
gb:MN996529|Organism:Severe      CAGGTTGTAGATGCAGATAGTAAAATTGTTCAACTTAGTGAAATTAGTATGGACAATTCA   12610
gb:MN996531|Organism:Severe      CAGGTTGTAGATGCAGATAGTAAAATTGTTCAACTTAGTGAAATTAGTATGGACAATTCA   12609
gb:MT066176|Organism:Severe      CAGGTTGTAGATGCAGATAGTAAAATTGTTCAACTTAGTGAAATTAGTATGGACAATTCA   12622
gb:MT027064|Organism:Severe      CAGGTTGTAGATGCAGATAGTAAAATTGTTCAACTTAGTGAAATTAGTATGGACAATTCA   12622
gb:MN994468|Organism:Severe      CAGGTTGTAGATGCAGATAGTAAAATTGTTCAACTTAGTGAAATTAGTATGGACAATTCA   12622
gb:MT072688|Organism:Severe      CAGGTTGTAGATGCAGATAGTAAAATTGTTCAACTTAGTGAAATTAGTATGGACAATTCA   12607
gb:MN996527|Organism:Severe      CAGGTTGTAGATGCAGATAGTAAAATTGTTCAACTTAGTGAAATTAGTATGGACAATTCA   12589
gb:MT093631|Organism:Severe      CAGGTTGTAGATGCAGATAGTAAAATTGTTCAACTTAGTGAAATTAGTATGGACAATTCA   12660
gb:MT106053|Organism:Severe      CAGGTTGTAGATGCAGATAGTAAAATTGTTCAACTTAGTGAAATTAGTATGGACAATTCA   12622
gb:MT019533|Organism:Severe      CAGGTTGTAGATGCAGATAGTAAAATTGTTCAACTTAGTGAAATTAGTATGGACAATTCA   12622
gb:MT019531|Organism:Severe      CAGGTTGTAGATGCAGATAGTAAAATTGTTCAACTTAGTGAAATTAGTATGGACAATTCA   12622
gb:MN996528|Organism:Severe      CAGGTTGTAGATGCAGATAGTAAAATTGTTCAACTTAGTGAAATTAGTATGGACAATTCA   12622
gb:MN996530|Organism:Severe      CAGGTTGTAGATGCAGATAGTAAAATTGTTCAACTTAGTGAAATTAGTATGGACAATTCA   12608
gb:MN908947|Organism:Severe      CAGGTTGTAGATGCAGATAGTAAAATTGTTCAACTTAGTGAAATTAGTATGGACAATTCA   12622
gb:MT019532|Organism:Severe      CAGGTTGTAGATGCAGATAGTAAAATTGTTCAACTTAGTGAAATTAGTATGGACAATTCA   12622
                                 ************************************************************

gb:MT020781|Organism:Severe      CCTAATTTAGCATGGCCTCTTATTGTAACAGCTTTAAGGGCCAATTCTGCTGTCAAATTA   12670
gb:MT007544|Organism:Severe      CCTAATTTAGCATGGCCTCTTATTGTAACAGCTTTAAGGGCCAATTCTGCTGTCAAATTA   12682
gb:MN994467|Organism:Severe      CCTAATTTAGCATGGCCTCTTATTGTAACAGCTTTAAGGGCCAATTCTGCTGTCAAATTA   12682
gb:MT044257|Organism:Severe      CCTAATTTAGCATGGCCTCTTATTGTAACAGCTTTAAGGGCCAATTCTGCTGTCAAATTA   12682
gb:MT106054|Organism:Severe      CCTAATTTAGCATGGCCTCTTATTGTAACAGCTTTAAGGGCCAATTCTGCTGTCAAATTA   12682
gb:MT049951|Organism:Severe      CCTAATTTAGCATGGCCTCTTATTGTAACAGCTTTAAGGGCCAATTCTGCTGTCAAATTA   12682
gb:MN975262|Organism:Severe      CCTAATTTAGCATGGCCTCTTATTGTAACAGCTTTAAGGGCCAATTCTGCTGTCAAATTA   12682
gb:MT106052|Organism:Severe      CCTAATTTAGCATGGCCTCTTATTGTAACAGCTTTAAGGGCCAATTCTGCTGTCAAATTA   12682
gb:LC522975|Organism:Severe      CCTAATTTAGCATGGCCTCTTATTGTAACAGCTTTAAGGGCCAATTCTGCTGTCAAATTA   12679
gb:LC522973|Organism:Severe      CCTAATTTAGCATGGCCTCTTATTGTAACAGCTTTAAGGGCCAATTCTGCTGTCAAATTA   12679
gb:LC522974|Organism:Severe      CCTAATTTAGCATGGCCTCTTATTGTAACAGCTTTAAGGGCCAATTCTGCTGTCAAATTA   12679
gb:MN985325|Organism:Severe      CCTAATTTAGCATGGCCTCTTATTGTAACAGCTTTAAGGGCCAATTCTGCTGTCAAATTA   12682
gb:MT020881|Organism:Severe      CCTAATTTAGCATGGCCTCTTATTGTAACAGCTTTAAGGGCCAATTCTGCTGTCAAATTA   12682
gb:MT020880|Organism:Severe      CCTAATTTAGCATGGCCTCTTATTGTAACAGCTTTAAGGGCCAATTCTGCTGTCAAATTA   12682
gb:MT066175|Organism:Severe      CCTAATTTAGCATGGCCTCTTATTGTAACAGCTTTAAGGGCCAATTCTGCTGTCAAATTA   12682
gb:MN997409|Organism:Severe      CCTAATTTAGCATGGCCTCTTATTGTAACAGCTTTAAGGGCCAATTCTGCTGTCAAATTA   12682
gb:MN938384|Organism:Severe      CCTAATTTAGCATGGCCTCTTATTGTAACAGCTTTAAGGGCCAATTCTGCTGTCAAATTA   12650
gb:MT044258|Organism:Severe      CCTAATTTAGCATGGCCTCTTATTGTAACAGCTTTAAGGGCCAATTCTGCTGTCAAATTA   12658
gb:MT039890|Organism:Severe      CCTAATTTAGCATGGCCTCTTATTGTAACAGCTTTAAGGGCCAATTCTGCTGTCAAATTA   12682
gb:MN988713|Organism:Severe      CCTAATTTAGCATGGCCTCTTATTGTAACAGCTTTAAGGGCCAATTCTGCTGTCAAATTA   12682
gb:LC521925|Organism:Severe      CCTAATTTAGCATGGCCTCTTATTGTAACAGCTTTAAGGGCCAATTCTGCTGTCAAATTA   12655
gb:MT093571|Organism:Severe      CCTAATTTAGCATGGCCTCTTATTGTAACAGCTTTAAGGGCCAATTCTGCTGTCAAATTA   12682
gb:MT039887|Organism:Severe      CCTAATTTAGCATGGCCTCTTATTGTAACAGCTTTAAGGGCCAATTCTGCTGTCAAATTA   12682
gb:MT019530|Organism:Severe      CCTAATTTAGCATGGCCTCTTATTGTAACAGCTTTAAGGGCCAATTCTGCTGTCAAATTA   12682
gb:MT039888|Organism:Severe      CCTAATTTAGCATGGCCTCTTATTGTAACAGCTTTAAGGGCCAATTCTGCTGTCAAATTA   12682
gb:LC522972|Organism:Severe      CCTAATTTAGCATGGCCTCTTATTGTAACAGCTTTAAGGGCCAATTCTGCTGTCAAATTA   12679
gb:MT027063|Organism:Severe      CCTAATTTAGCATGGCCTCTTATTGTAACAGCTTTAAGGGCCAATTCTGCTGTCAAATTA   12682
gb:MT027062|Organism:Severe      CCTAATTTAGCATGGCCTCTTATTGTAACAGCTTTAAGGGCCAATTCTGCTGTCAAATTA   12682
gb:MT019529|Organism:Severe      CCTAATTTAGCATGGCCTCTTATTGTAACAGCTTTAAGGGCCAATTCTGCTGTCAAATTA   12682
gb:MN996529|Organism:Severe      CCTAATTTAGCATGGCCTCTTATTGTAACAGCTTTAAGGGCCAATTCTGCTGTCAAATTA   12670
gb:MN996531|Organism:Severe      CCTAATTTAGCATGGCCTCTTATTGTAACAGCTTTAAGGGCCAATTCTGCTGTCAAATTA   12669
gb:MT066176|Organism:Severe      CCTAATTTAGCATGGCCTCTTATTGTAACAGCTTTAAGGGCCAATTCTGCTGTCAAATTA   12682
gb:MT027064|Organism:Severe      CCTAATTTAGCATGGCCTCTTATTGTAACAGCTTTAAGGGCCAATTCTGCTGTCAAATTA   12682
gb:MN994468|Organism:Severe      CCTAATTTAGCATGGCCTCTTATTGTAACAGCTTTAAGGGCCAATTCTGCTGTCAAATTA   12682
gb:MT072688|Organism:Severe      CCTAATTTAGCATGGCCTCTTATTGTAACAGCTTTAAGGGCCAATTCTGCTGTCAAATTA   12667
gb:MN996527|Organism:Severe      CCTAATTTAGCATGGCCTCTTATTGTAACAGCTTTAAGGGCCAATTCTGCTGTCAAATTA   12649
gb:MT093631|Organism:Severe      CCTAATTTAGCATGGCCTCTTATTGTAACAGCTTTAAGGGCCAATTCTGCTGTCAAATTA   12720
gb:MT106053|Organism:Severe      CCTAATTTAGCATGGCCTCTTATTGTAACAGCTTTAAGGGCCAATTCTGCTGTCAAATTA   12682
gb:MT019533|Organism:Severe      CCTAATTTAGCATGGCCTCTTATTGTAACAGCTTTAAGGGCCAATTCTGCTGTCAAATTA   12682
gb:MT019531|Organism:Severe      CCTAATTTAGCATGGCCTCTTATTGTAACAGCTTTAAGGGCCAATTCTGCTGTCAAATTA   12682
```

```
gb:MN996528|Organism:Severe      CCTAATTTAGCATGGCCTCTTATTGTAACAGCTTTAAGGGCCAATTCTGCTGTCAAATTA   12682
gb:MN996530|Organism:Severe      CCTAATTTAGCATGGCCTCTTATTGTAACAGCTTTAAGGGCCAATTCTGCTGTCAAATTA   12668
gb:MN908947|Organism:Severe      CCTAATTTAGCATGGCCTCTTATTGTAACAGCTTTAAGGGCCAATTCTGCTGTCAAATTA   12682
gb:MT019532|Organism:Severe      CCTAATTTAGCATGGCCTCTTATTGTAACAGCTTTAAGGGCCAATTCTGCTGTCAAATTA   12682
                                 ************************************************************

gb:MT020781|Organism:Severe      CAGAATAATGAGCTTAGTCCTGTTGCACTACGACAGATGTCTTGTGCTGCCGGTACTACA   12730
gb:MT007544|Organism:Severe      CAGAATAATGAGCTTAGTCCTGTTGCACTACGACAGATGTCTTGTGCTGCCGGTACTACA   12742
gb:MN994467|Organism:Severe      CAGAATAATGAGCTTAGTCCTGTTGCACTACGACAGATGTCTTGTGCTGCCGGTACTACA   12742
gb:MT044257|Organism:Severe      CAGAATAATGAGCTTAGTCCTGTTGCACTACGACAGATGTCTTGTGCTGCCGGTACTACA   12742
gb:MT106054|Organism:Severe      CAGAATAATGAGCTTAGTCCTGTTGCACTACGACAGATGTCTTGTGCTGCCGGTACTACA   12742
gb:MT049951|Organism:Severe      CAGAATAATGAGCTTAGTCCTGTTGCACTACGACAGATGTCTTGTGCTGCCGGTACTACA   12742
gb:MN975262|Organism:Severe      CAGAATAATGAGCTTAGTCCTGTTGCACTACGACAGATGTCTTGTGCTGCCGGTACTACA   12742
gb:MT106052|Organism:Severe      CAGAATAATGAGCTTAGTCCTGTTGCACTACGACAGATGTCTTGTGCTGCCGGTACTACA   12742
gb:LC522975|Organism:Severe      CAGAATAATGAGCTTAGTCCTGTTGCACTACGACAGATGTCTTGTGCTGCCGGTACTACA   12739
gb:LC522973|Organism:Severe      CAGAATAATGAGCTTAGTCCTGTTGCACTACGACAGATGTCTTGTGCTGCCGGTACTACA   12739
gb:LC522974|Organism:Severe      CAGAATAATGAGCTTAGTCCTGTTGCACTACGACAGATGTCTTGTGCTGCCGGTACTACA   12739
gb:MN985325|Organism:Severe      CAGAATAATGAGCTTAGTCCTGTTGCACTACGACAGATGTCTTGTGCTGCCGGTACTACA   12742
gb:MT020881|Organism:Severe      CAGAATAATGAGCTTAGTCCTGTTGCACTACGACAGATGTCTTGTGCTGCCGGTACTACA   12742
gb:MT020880|Organism:Severe      CAGAATAATGAGCTTAGTCCTGTTGCACTACGACAGATGTCTTGTGCTGCCGGTACTACA   12742
gb:MT066175|Organism:Severe      CAGAATAATGAGCTTAGTCCTGTTGCACTACGACAGATGTCTTGTGCTGCCGGTACTACA   12742
gb:MN997409|Organism:Severe      CAGAATAATGAGCTTAGTCCTGTTGCACTACGACAGATGTCTTGTGCTGCCGGTACTACA   12742
gb:MN938384|Organism:Severe      CAGAATAATGAGCTTAGTCCTGTTGCACTACGACAGATGTCTTGTGCTGCCGGTACTACA   12710
gb:MT044258|Organism:Severe      CAGAATAATGAGCTTAGTCCTGTTGCACTACGACAGATGTCTTGTGCTGCCGGTACTACA   12718
gb:MT039890|Organism:Severe      CAGAATAATGAGCTTAGTCCTGTTGCACTACGACAGATGTCTTGTGCTGCCGGTACTACA   12742
gb:MN988713|Organism:Severe      CAGAATAATGAGCTTAGTCCTGTTGCACTACGACAGATGTCTTGTGCTGCCGGTACTACA   12742
gb:LC521925|Organism:Severe      CAGAATAATGAGCTTAGTCCTGTTGCACTACGACAGATGTCTTGTGCTGCCGGTACTACA   12715
gb:MT093571|Organism:Severe      CAGAATAATGAGCTTAGTCCTGTTGCACTACGACAGATGTCTTGTGCTGCCGGTACTACA   12742
gb:MT039887|Organism:Severe      CAGAATAATGAGCTTAGTCCTGTTGCACTACGACAGATGTCTTGTGCTGCCGGTACTACA   12742
gb:MT019530|Organism:Severe      CAGAATAATGAGCTTAGTCCTGTTGCACTACGACAGATGTCTTGTGCTGCCGGTACTACA   12742
gb:MT039888|Organism:Severe      CAGAATAATGAGCTTAGTCCTGTTGCACTACGACAGATGTCTTGTGCTGCCGGTACTACA   12742
gb:LC522972|Organism:Severe      CAGAATAATGAGCTTAGTCCTGTTGCACTACGACAGATGTCTTGTGCTGCCGGTACTACA   12739
gb:MT027063|Organism:Severe      CAGAATAATGAGCTTAGTCCTGTTGCACTACGACAGATGTCTTGTGCTGCCGGTACTACA   12742
gb:MT027062|Organism:Severe      CAGAATAATGAGCTTAGTCCTGTTGCACTACGACAGATGTCTTGTGCTGCCGGTACTACA   12742
gb:MT019529|Organism:Severe      CAGAATAATGAGCTTAGTCCTGTTGCACTACGACAGATGTCTTGTGCTGCCGGTACTACA   12742
gb:MN996529|Organism:Severe      CAGAATAATGAGCTTAGTCCTGTTGCACTACGACAGATGTCTTGTGCTGCCGGTACTACA   12730
gb:MN996531|Organism:Severe      CAGAATAATGAGCTTAGTCCTGTTGCACTACGACAGATGTCTTGTGCTGCCGGTACTACA   12729
gb:MT066176|Organism:Severe      CAGAATAATGAGCTTAGTCCTGTTGCACTACGACAGATGTCTTGTGCTGCCGGTACTACA   12742
gb:MT027064|Organism:Severe      CAGAATAATGAGCTTAGTCCTGTTGCACTACGACAGATGTCTTGTGCTGCCGGTACTACA   12742
gb:MN994468|Organism:Severe      CAGAATAATGAGCTTAGTCCTGTTGCACTACGACAGATGTCTTGTGCTGCCGGTACTACA   12742
gb:MT072688|Organism:Severe      CAGAATAATGAGCTTAGTCCTGTTGCACTACGACAGATGTCTTGTGCTGCCGGTACTACA   12727
gb:MN996527|Organism:Severe      CAGAATAATGAGCTTAGTCCTGTTGCACTACGACAGATGTCTTGTGCTGCCGGTACTACA   12709
gb:MT093631|Organism:Severe      CAGAATAATGAGCTTAGTCCTGTTGCACTACGACAGATGTCTTGTGCTGCCGGTACTACA   12780
gb:MT106053|Organism:Severe      CAGAATAATGAGCTTAGTCCTGTTGCACTACGACAGATGTCTTGTGCTGCCGGTACTACA   12742
gb:MT019533|Organism:Severe      CAGAATAATGAGCTTAGTCCTGTTGCACTACGACAGATGTCTTGTGCTGCCGGTACTACA   12742
gb:MT019531|Organism:Severe      CAGAATAATGAGCTTAGTCCTGTTGCACTACGACAGATGTCTTGTGCTGCCGGTACTACA   12742
gb:MN996528|Organism:Severe      CAGAATAATGAGCTTAGTCCTGTTGCACTACGACAGATGTCTTGTGCTGCCGGTACTACA   12742
gb:MN996530|Organism:Severe      CAGAATAATGAGCTTAGTCCTGTTGCACTACGACAGATGTCTTGTGCTGCCGGTACTACA   12728
gb:MN908947|Organism:Severe      CAGAATAATGAGCTTAGTCCTGTTGCACTACGACAGATGTCTTGTGCTGCCGGTACTACA   12742
gb:MT019532|Organism:Severe      CAGAATAATGAGCTTAGTCCTGTTGCACTACGACAGATGTCTTGTGCTGCCGGTACTACA   12742
                                 ************************************************************

gb:MT020781|Organism:Severe      CAAACTGCTTGCACTGATGACAATGCGTTAGCTTACTACAACACAACAAAGGGAGGTAGG   12790
gb:MT007544|Organism:Severe      CAAACTGCTTGCACTGATGACAATGCGTTAGCTTACTACAACACAACAAAGGGAGGTAGG   12802
gb:MN994467|Organism:Severe      CAAACTGCTTGCACTGATGACAATGCGTTAGCTTACTACAACACAACAAAGGGAGGTAGG   12802
gb:MT044257|Organism:Severe      CAAACTGCTTGCACTGATGACAATGCGTTAGCTTACTACAACACAACAAAGGGAGGTAGG   12802
gb:MT106054|Organism:Severe      CAAACTGCTTGCACTGATGACAATGCGTTAGCTTACTACAACACAACAAAGGGAGGTAGG   12802
gb:MT049951|Organism:Severe      CAAACTGCTTGCACTGATGACAATGCGTTAGCTTACTACAACACAACAAAGGGAGGTAGG   12802
gb:MN975262|Organism:Severe      CAAACTGCTTGCACTGATGACAATGCGTTAGCTTACTACAACACAACAAAGGGAGGTAGG   12802
gb:MT106052|Organism:Severe      CAAACTGCTTGCACTGATGACAATGCGTTAGCTTACTACAACACAACAAAGGGAGGTAGG   12802
gb:LC522975|Organism:Severe      CAAACTGCTTGCACTGATGACAATGCGTTAGCTTACTACAACACAACAAAGGGAGGTAGG   12799
gb:LC522973|Organism:Severe      CAAACTGCTTGCACTGATGACAATGCGTTAGCTTACTACAACACAACAAAGGGAGGTAGG   12799
gb:LC522974|Organism:Severe      CAAACTGCTTGCACTGATGACAATGCGTTAGCTTACTACAACACAACAAAGGGAGGTAGG   12799
gb:MN985325|Organism:Severe      CAAACTGCTTGCACTGATGACAATGCGTTAGCTTACTACAACACAACAAAGGGAGGTAGG   12802
gb:MT020881|Organism:Severe      CAAACTGCTTGCACTGATGACAATGCGTTAGCTTACTACAACACAACAAAGGGAGGTAGG   12802
```

```
gb:MT020880|Organism:Severe      CAAACTGCTTGCACTGATGACAATGCGTTAGCTTACTACAACACAACAAAGGGAGGTAGG   12802
gb:MT066175|Organism:Severe      CAAACTGCTTGCACTGATGACAATGCGTTAGCTTACTACAACACAACAAAGGGAGGTAGG   12802
gb:MN997409|Organism:Severe      CAAACTGCTTGCACTGATGACAATGCGTTAGCTTACTACAACACAACAAAGGGAGGTAGG   12802
gb:MN938384|Organism:Severe      CAAACTGCTTGCACTGATGACAATGCGTTAGCTTACTACAACACAACAAAGGGAGGTAGG   12770
gb:MT044258|Organism:Severe      CAAACTGCTTGCACTGATGACAATGCGTTAGCTTACTACAACACAACAAAGGGAGGTAGG   12778
gb:MT039890|Organism:Severe      CAAACTGCTTGCACTGATGACAATGCGTTAGCTTACTACAACACAACAAAGGGAGGTAGG   12802
gb:MN988713|Organism:Severe      CAAACTGCTTGCACTGATGACAATGCGTTAGCTTACTACAACACAACAAAGGGAGGTAGG   12802
gb:LC521925|Organism:Severe      CAAACTGCTTGCACTGATGACAATGCGTTAGCTTACTACAACACAACAAAGGGAGGTAGG   12775
gb:MT093571|Organism:Severe      CAAACTGCTTGCACTGATGACAATGCGTTAGCTTACTACAACACAACAAAGGGAGGTAGG   12802
gb:MT039887|Organism:Severe      CAAACTGCTTGCACTGATGACAATGCGTTAGCTTACTACAACACAACAAAGGGAGGTAGG   12802
gb:MT019530|Organism:Severe      CAAACTGCTTGCACTGATGACAATGCGTTAGCTTACTACAACACAACAAAGGGAGGTAGG   12802
gb:MT039888|Organism:Severe      CAAACTGCTTGCACTGATGACAATGCGTTAGCTTACTACAACACAACAAAGGGAGGTAGG   12802
gb:LC522972|Organism:Severe      CAAACTGCTTGCACTGATGACAATGCGTTAGCTTACTACAACACAACAAAGGGAGGTAGG   12799
gb:MT027063|Organism:Severe      CAAACTGCTTGCACTGATGACAATGCGTTAGCTTACTACAACACAACAAAGGGAGGTAGG   12802
gb:MT027062|Organism:Severe      CAAACTGCTTGCACTGATGACAATGCGTTAGCTTACTACAACACAACAAAGGGAGGTAGG   12802
gb:MT019529|Organism:Severe      CAAACTGCTTGCACTGATGACAATGCGTTAGCTTACTACAACACAACAAAGGGAGGTAGG   12802
gb:MN996529|Organism:Severe      CAAACTGCTTGCACTGATGACAATGCGTTAGCTTACTACAACACAACAAAGGGAGGTAGG   12790
gb:MN996531|Organism:Severe      CAAACTGCTTGCACTGATGACAATGCGTTAGCTTACTACAACACAACAAAGGGAGGTAGG   12789
gb:MT066176|Organism:Severe      CAAACTGCTTGCACTGATGACAATGCGTTAGCTTACTACAACACAACAAAGGGAGGTAGG   12802
gb:MT027064|Organism:Severe      CAAACTGCTTGCACTGATGACAATGCGTTAGCTTACTACAACACAACAAAGGGAGGTAGG   12802
gb:MN994468|Organism:Severe      CAAACTGCTTGCACTGATGACAATGCGTTAGCTTACTACAACACAACAAAGGGAGGTAGG   12802
gb:MT072688|Organism:Severe      CAAACTGCTTGCACTGATGACAATGCGTTAGCTTACTACAACACAACAAAGGGAGGTAGG   12787
gb:MN996527|Organism:Severe      CAAACTGCTTGCACTGATGACAATGCGTTAGCTTACTACAACACAACAAAGGGAGGTAGG   12769
gb:MT093631|Organism:Severe      CAAACTGCTTGCACTGATGACAATGCGTTAGCTTACTACAACACAACAAAGGGAGGTAGG   12840
gb:MT106053|Organism:Severe      CAAACTGCTTGCACTGATGACAATGCGTTAGCTTACTACAACACAACAAAGGGAGGTAGG   12802
gb:MT019533|Organism:Severe      CAAACTGCTTGCACTGATGACAATGCGTTAGCTTACTACAACACAACAAAGGGAGGTAGG   12802
gb:MT019531|Organism:Severe      CAAACTGCTTGCACTGATGACAATGCGTTAGCTTACTACAACACAACAAAGGGAGGTAGG   12802
gb:MN996528|Organism:Severe      CAAACTGCTTGCACTGATGACAATGCGTTAGCTTACTACAACACAACAAAGGGAGGTAGG   12802
gb:MN996530|Organism:Severe      CAAACTGCTTGCACTGATGACAATGCGTTAGCTTACTACAACACAACAAAGGGAGGTAGG   12788
gb:MN908947|Organism:Severe      CAAACTGCTTGCACTGATGACAATGCGTTAGCTTACTACAACACAACAAAGGGAGGTAGG   12802
gb:MT019532|Organism:Severe      CAAACTGCTTGCACTGATGACAATGCGTTAGCTTACTACAACACAACAAAGGGAGGTAGG   12802
                                 ************************************************************

gb:MT020781|Organism:Severe      TTTGTACTTGCACTGTTATCCGATTTACAGGATTTGAAATGGGCTAGATTCCCTAAGAGT   12850
gb:MT007544|Organism:Severe      TTTGTACTTGCACTGTTATCCGATTTACAGGATTTGAAATGGGCTAGATTCCCTAAGAGT   12862
gb:MN994467|Organism:Severe      TTTGTACTTGCACTGTTATCCGATTTACAGGATTTGAAATGGGCTAGATTCCCTAAGAGT   12862
gb:MT044257|Organism:Severe      TTTGTACTTGCACTGTTATCCGATTTACAGGATTTGAAATGGGCTAGATTCCCTAAGAGT   12862
gb:MT106054|Organism:Severe      TTTGTACTTGCACTGTTATCCGATTTACAGGATTTGAAATGGGCTAGATTCCCTAAGAGT   12862
gb:MT049951|Organism:Severe      TTTGTACTTGCACTGTTATCCGATTTACAGGATTTGAAATGGGCTAGATTCCCTAAGAGT   12862
gb:MN975262|Organism:Severe      TTTGTACTTGCACTGTTATCCGATTTACAGGATTTGAAATGGGCTAGATTCCCTAAGAGT   12862
gb:MT106052|Organism:Severe      TTTGTACTTGCACTGTTATCCGATTTACAGGATTTGAAATGGGCTAGATTCCCTAAGAGT   12862
gb:LC522975|Organism:Severe      TTTGTACTTGCACTGTTATCCGATTTACAGGATTTGAAATGGGCTAGATTCCCTAAGAGT   12859
gb:LC522973|Organism:Severe      TTTGTACTTGCACTGTTATCCGATTTACAGGATTTGAAATGGGCTAGATTCCCTAAGAGT   12859
gb:LC522974|Organism:Severe      TTTGTACTTGCACTGTTATCCGATTTACAGGATTTGAAATGGGCTAGATTCCCTAAGAGT   12859
gb:MN985325|Organism:Severe      TTTGTACTTGCACTGTTATCCGATTTACAGGATTTGAAATGGGCTAGATTCCCTAAGAGT   12862
gb:MT020881|Organism:Severe      TTTGTACTTGCACTGTTATCCGATTTACAGGATTTGAAATGGGCTAGATTCCCTAAGAGT   12862
gb:MT020880|Organism:Severe      TTTGTACTTGCACTGTTATCCGATTTACAGGATTTGAAATGGGCTAGATTCCCTAAGAGT   12862
gb:MT066175|Organism:Severe      TTTGTACTTGCACTGTTATCCGATTTACAGGATTTGAAATGGGCTAGATTCCCTAAGAGT   12862
gb:MN997409|Organism:Severe      TTTGTACTTGCACTGTTATCCGATTTACAGGATTTGAAATGGGCTAGATTCCCTAAGAGT   12862
gb:MN938384|Organism:Severe      TTTGTACTTGCACTGTTATCCGATTTACAGGATTTGAAATGGGCTAGATTCCCTAAGAGT   12830
gb:MT044258|Organism:Severe      TTTGTACTTGCACTGTTATCCGATTTACAGGATTTGAAATGGGCTAGATTCCCTAAGAGT   12838
gb:MT039890|Organism:Severe      TTTGTACTTGCACTGTTATCCGATTTACAGGATTTGAAATGGGCTAGATTCCCTAAGAGT   12862
gb:MN988713|Organism:Severe      TTTGTACTTGCACTGTTATCCGATTTACAGGATTTGAAATGGGCTAGATTCCCTAAGAGT   12862
gb:LC521925|Organism:Severe      TTTGTACTTGCACTGTTATCCGATTTACAGGATTTGAAATGGGCTAGATTCCCTAAGAGT   12835
gb:MT093571|Organism:Severe      TTTGTACTTGCACTGTTATCCGATTTACAGGATTTGAAATGGGCTAGATTCCCTAAGAGT   12862
gb:MT039887|Organism:Severe      TTTGTACTTGCACTGTTATCCGATTTACAGGATTTGAAATGGGCTAGATTCCCTAAGAGT   12862
gb:MT019530|Organism:Severe      TTTGTACTTGCACTGTTATCCGATTTACAGGATTTGAAATGGGCTAGATTCCCTAAGAGT   12862
gb:MT039888|Organism:Severe      TTTGTACTTGCACTGTTATCCGATTTACAGGATTTGAAATGGGCTAGATTCCCTAAGAGT   12862
gb:LC522972|Organism:Severe      TTTGTACTTGCACTGTTATCCGATTTACAGGATTTGAAATGGGCTAGATTCCCTAAGAGT   12859
gb:MT027063|Organism:Severe      TTTGTACTTGCACTGTTATCCGATTTACAGGATTTGAAATGGGCTAGATTCCCTAAGAGT   12862
gb:MT027062|Organism:Severe      TTTGTACTTGCACTGTTATCCGATTTACAGGATTTGAAATGGGCTAGATTCCCTAAGAGT   12862
gb:MT019529|Organism:Severe      TTTGTACTTGCACTGTTATCCGATTTACAGGATTTGAAATGGGCTAGATTCCCTAAGAGT   12862
gb:MN996529|Organism:Severe      TTTGTACTTGCACTGTTATCCGATTTACAGGATTTGAAATGGGCTAGATTCCCTAAGAGT   12850
gb:MN996531|Organism:Severe      TTTGTACTTGCACTGTTATCCGATTTACAGGATTTGAAATGGGCTAGATTCCCTAAGAGT   12849
gb:MT066176|Organism:Severe      TTTGTACTTGCACTGTTATCCGATTTACAGGATTTGAAATGGGCTAGATTCCCTAAGAGT   12862
```

```
gb:MT027064|Organism:Severe      TTTGTACTTGCACTGTTATCCGATTTACAGGATTTGAAATGGGCTAGATTCCCTAAGAGT   12862
gb:MN994468|Organism:Severe      TTTGTACTTGCACTGTTATCCGATTTACAGGATTTGAAATGGGCTAGATTCCCTAAGAGT   12862
gb:MT072688|Organism:Severe      TTTGTACTTGCACTGTTATCCGATTTACAGGATTTGAAATGGGCTAGATTCCCTAAGAGT   12847
gb:MN996527|Organism:Severe      TTTGTACTTGCACTGTTATCCGATTTACAGGATTTGAAATGGGCTAGATTCCCTAAGAGT   12829
gb:MT093631|Organism:Severe      TTTGTACTTGCACTGTTATCCGATTTACAGGATTTGAAATGGGCTAGATTCCCTAAGAGT   12900
gb:MT106053|Organism:Severe      TTTGTACTTGCACTGTTATCCGATTTACAGGATTTGAAATGGGCTAGATTCCCTAAGAGT   12862
gb:MT019533|Organism:Severe      TTTGTACTTGCACTGTTATCCGATTTACAGGATTTGAAATGGGCTAGATTCCCTAAGAGT   12862
gb:MT019531|Organism:Severe      TTTGTACTTGCACTGTTATCCGATTTACAGGATTTGAAATGGGCTAGATTCCCTAAGAGT   12862
gb:MN996528|Organism:Severe      TTTGTACTTGCACTGTTATCCGATTTACAGGATTTGAAATGGGCTAGATTCCCTAAGAGT   12862
gb:MN996530|Organism:Severe      TTTGTACTTGCACTGTTATCCGATTTACAGGATTTGAAATGGGCTAGATTCCCTAAGAGT   12848
gb:MN908947|Organism:Severe      TTTGTACTTGCACTGTTATCCGATTTACAGGATTTGAAATGGGCTAGATTCCCTAAGAGT   12862
gb:MT019532|Organism:Severe      TTTGTACTTGCACTGTTATCCGATTTACAGGATTTGAAATGGGCTAGATTCCCTAAGAGT   12862
                                 ************************************************************

gb:MT020781|Organism:Severe      GATGGAACTGGTACTATCTATACAGAACTGGAACCACCTTGTAGGTTTGTTACAGACACA   12910
gb:MT007544|Organism:Severe      GATGGAACTGGTACTATCTATACAGAACTGGAACCACCTTGTAGGTTTGTTACAGACACA   12922
gb:MN994467|Organism:Severe      GATGGAACTGGTACTATCTATACAGAACTGGAACCACCTTGTAGGTTTGTTACAGACACA   12922
gb:MT044257|Organism:Severe      GATGGAACTGGTACTATCTATACAGAACTGGAACCACCTTGTAGGTTTGTTACAGACACA   12922
gb:MT106054|Organism:Severe      GATGGAACTGGTACTATCTATACAGAACTGGAACCACCTTGTAGGTTTGTTACAGACACA   12922
gb:MT049951|Organism:Severe      GATGGAACTGGTACTATCTATACAGAACTGGAACCACCTTGTAGGTTTGTTACAGACACA   12922
gb:MN975262|Organism:Severe      GATGGAACTGGTACTATCTATACAGAACTGGAACCACCTTGTAGGTTTGTTACAGACACA   12922
gb:MT106052|Organism:Severe      GATGGAACTGGTACTATCTATACAGAACTGGAACCACCTTGTAGGTTTGTTACAGACACA   12922
gb:LC522975|Organism:Severe      GATGGAACTGGTACTATCTATACAGAACTGGAACCACCTTGTAGGTTTGTTACAGACACA   12919
gb:LC522973|Organism:Severe      GATGGAACTGGTACTATCTATACAGAACTGGAACCACCTTGTAGGTTTGTTACAGACACA   12919
gb:LC522974|Organism:Severe      GATGGAACTGGTACTATCTATACAGAACTGGAACCACCTTGTAGGTTTGTTACAGACACA   12919
gb:MN985325|Organism:Severe      GATGGAACTGGTACTATCTATACAGAACTGGAACCACCTTGTAGGTTTGTTACAGACACA   12922
gb:MT020881|Organism:Severe      GATGGAACTGGTACTATCTATACAGAACTGGAACCACCTTGTAGGTTTGTTACAGACACA   12922
gb:MT020880|Organism:Severe      GATGGAACTGGTACTATCTATACAGAACTGGAACCACCTTGTAGGTTTGTTACAGACACA   12922
gb:MT066175|Organism:Severe      GATGGAACTGGTACTATCTATACAGAACTGGAACCACCTTGTAGGTTTGTTACAGACACA   12922
gb:MN997409|Organism:Severe      GATGGAACTGGTACTATCTATACAGAACTGGAACCACCTTGTAGGTTTGTTACAGACACA   12922
gb:MN938384|Organism:Severe      GATGGAACTGGTACTATCTATACAGAACTGGAACCACCTTGTAGGTTTGTTACAGACACA   12890
gb:MT044258|Organism:Severe      GATGGAACTGGTACTATCTATACAGAACTGGAACCACCTTGTAGGTTTGTTACAGACACA   12898
gb:MT039890|Organism:Severe      GATGGAACTGGTACTATCTATACAGAACTGGAACCACCTTGTAGGTTTGTTACAGACACA   12922
gb:MN988713|Organism:Severe      GATGGAACTGGTACTATCTATACAGAACTGGAACCACCTTGTAGGTTTGTTACAGACACA   12922
gb:LC521925|Organism:Severe      GATGGAACTGGTACTATCTATACAGAACTGGAACCACCTTGTAGGTTTGTTACAGACACA   12895
gb:MT093571|Organism:Severe      GATGGAACTGGTACTATCTATACAGAACTGGAACCACCTTGTAGGTTTGTTACAGACACA   12922
gb:MT039887|Organism:Severe      GATGGAACTGGTACTATCTATACAGAACTGGAACCACCTTGTAGGTTTGTTACAGACACA   12922
gb:MT019530|Organism:Severe      GATGGAACTGGTACTATCTATACAGAACTGGAACCACCTTGTAGGTTTGTTACAGACACA   12922
gb:MT039888|Organism:Severe      GATGGAACTGGTACTATCTATACAGAACTGGAACCACCTTGTAGGTTTGTTACAGACACA   12922
gb:LC522972|Organism:Severe      GATGGAACTGGTACTATCTATACAGAACTGGAACCACCTTGTAGGTTTGTTACAGACACA   12919
gb:MT027063|Organism:Severe      GATGGAACTGGTACTATCTATACAGAACTGGAACCACCTTGTAGGTTTGTTACAGACACA   12922
gb:MT027062|Organism:Severe      GATGGAACTGGTACTATCTATACAGAACTGGAACCACCTTGTAGGTTTGTTACAGACACA   12922
gb:MT019529|Organism:Severe      GATGGAACTGGTACTATCTATACAGAACTGGAACCACCTTGTAGGTTTGTTACAGACACA   12922
gb:MN996529|Organism:Severe      GATGGAACTGGTACTATCTATACAGAACTGGAACCACCTTGTAGGTTTGTTACAGACACA   12910
gb:MN996531|Organism:Severe      GATGGAACTGGTACTATCTATACAGAACTGGAACCACCTTGTAGGTTTGTTACAGACACA   12909
gb:MT066176|Organism:Severe      GATGGAACTGGTACTATCTATACAGAACTGGAACCACCTTGTAGGTTTGTTACAGACACA   12922
gb:MT027064|Organism:Severe      GATGGAACTGGTACTATCTATACAGAACTGGAACCACCTTGTAGGTTTGTTACAGACACA   12922
gb:MN994468|Organism:Severe      GATGGAACTGGTACTATCTATACAGAACTGGAACCACCTTGTAGGTTTGTTACAGACACA   12922
gb:MT072688|Organism:Severe      GATGGAACTGGTACTATCTATACAGAACTGGAACCACCTTGTAGGTTTGTTACAGACACA   12907
gb:MN996527|Organism:Severe      GATGGAACTGGTACTATCTATACAGAACTGGAACCACCTTGTAGGTTTGTTACAGACACA   12889
gb:MT093631|Organism:Severe      GATGGAACTGGTACTATCTATACAGAACTGGAACCACCTTGTAGGTTTGTTACAGACACA   12960
gb:MT106053|Organism:Severe      GATGGAACTGGTACTATCTATACAGAACTGGAACCACCTTGTAGGTTTGTTACAGACACA   12922
gb:MT019533|Organism:Severe      GATGGAACTGGTACTATCTATACAGAACTGGAACCACCTTGTAGGTTTGTTACAGACACA   12922
gb:MT019531|Organism:Severe      GATGGAACTGGTACTATCTATACAGAACTGGAACCACCTTGTAGGTTTGTTACAGACACA   12922
gb:MN996528|Organism:Severe      GATGGAACTGGTACTATCTATACAGAACTGGAACCACCTTGTAGGTTTGTTACAGACACA   12922
gb:MN996530|Organism:Severe      GATGGAACTGGTACTATCTATACAGAACTGGAACCACCTTGTAGGTTTGTTACAGACACA   12908
gb:MN908947|Organism:Severe      GATGGAACTGGTACTATCTATACAGAACTGGAACCACCTTGTAGGTTTGTTACAGACACA   12922
gb:MT019532|Organism:Severe      GATGGAACTGGTACTATCTATACAGAACTGGAACCACCTTGTAGGTTTGTTACAGACACA   12922
                                 ************************************************************

gb:MT020781|Organism:Severe      CCTAAAGGTCCTAAAGTGAAGTATTTATACTTTATTAAAGGATTAAACAACCTAAATAGA   12970
gb:MT007544|Organism:Severe      CCTAAAGGTCCTAAAGTGAAGTATTTATACTTTATTAAAGGATTAAACAACCTAAATAGA   12982
gb:MN994467|Organism:Severe      CCTAAAGGTCCTAAAGTGAAGTATTTATACTTTATTAAAGGATTAAACAACCTAAATAGA   12982
gb:MT044257|Organism:Severe      CCTAAAGGTCCTAAAGTGAAGTATTTATACTTTATTAAAGGATTAAACAACCTAAATAGA   12982
gb:MT106054|Organism:Severe      CCTAAAGGTCCTAAAGTGAAGTATTTATACTTTATTAAAGGATTAAACAACCTAAATAGA   12982
```

```
gb:MT049951|Organism:Severe      CCTAAAGGTCCTAAAGTGAAGTATTTATACTTTATTAAAGGATTAAACAACCTAAATAGA   12982
gb:MN975262|Organism:Severe      CCTAAAGGTCCTAAAGTGAAGTATTTATACTTTATTAAAGGATTAAACAACCTAAATAGA   12982
gb:MT106052|Organism:Severe      CCTAAAGGTCCTAAAGTGAAGTATTTATACTTTATTAAAGGATTAAACAACCTAAATAGA   12982
gb:LC522975|Organism:Severe      CCTAAAGGTCCTAAAGTGAAGTATTTATACTTTATTAAAGGATTAAACAACCTAAATAGA   12979
gb:LC522973|Organism:Severe      CCTAAAGGTCCTAAAGTGAAGTATTTATACTTTATTAAAGGATTAAACAACCTAAATAGA   12979
gb:LC522974|Organism:Severe      CCTAAAGGTCCTAAAGTGAAGTATTTATACTTTATTAAAGGATTAAACAACCTAAATAGA   12979
gb:MN985325|Organism:Severe      CCTAAAGGTCCTAAAGTGAAGTATTTATACTTTATTAAAGGATTAAACAACCTAAATAGA   12982
gb:MT020881|Organism:Severe      CCTAAAGGTCCTAAAGTGAAGTATTTATACTTTATTAAAGGATTAAACAACCTAAATAGA   12982
gb:MT020880|Organism:Severe      CCTAAAGGTCCTAAAGTGAAGTATTTATACTTTATTAAAGGATTAAACAACCTAAATAGA   12982
gb:MT066175|Organism:Severe      CCTAAAGGTCCTAAAGTGAAGTATTTATACTTTATTAAAGGATTAAACAACCTAAATAGA   12982
gb:MN997409|Organism:Severe      CCTAAAGGTCCTAAAGTGAAGTATTTATACTTTATTAAAGGATTAAACAACCTAAATAGA   12982
gb:MN938384|Organism:Severe      CCTAAAGGTCCTAAAGTGAAGTATTTATACTTTATTAAAGGATTAAACAACCTAAATAGA   12950
gb:MT044258|Organism:Severe      CCTAAAGGTCCTAAAGTGAAGTATTTATACTTTATTAAAGGATTAAACAACCTAAATAGA   12958
gb:MT039890|Organism:Severe      CCTAAAGGTCCTAAAGTGAAGTATTTATACTTTATTAAAGGATTAAACAACCTAAATAGA   12982
gb:MN988713|Organism:Severe      CCTAAAGGTCCTAAAGTGAAGTATTTATACTTTATTAAAGGATTAAACAACCTAAATAGA   12982
gb:LC521925|Organism:Severe      CCTAAAGGTCCTAAAGTGAAGTATTTATACTTTATTAAAGGATTAAACAACCTAAATAGA   12955
gb:MT093571|Organism:Severe      CCTAAAGGTCCTAAAGTGAAGTATTTATACTTTATTAAAGGATTAAACAACCTAAATAGA   12982
gb:MT039887|Organism:Severe      CCTAAAGGTCCTAAAGTGAAGTATTTATACTTTATTAAAGGATTAAACAACCTAAATAGA   12982
gb:MT019530|Organism:Severe      CCTAAAGGTCCTAAAGTGAAGTATTTATACTTTATTAAAGGATTAAACAACCTAAATAGA   12982
gb:MT039888|Organism:Severe      CCTAAAGGTCCTAAAGTGAAGTATTTATACTTTATTAAAGGATTAAACAACCTAAATAGA   12982
gb:LC522972|Organism:Severe      CCTAAAGGTCCTAAAGTGAAGTATTTATACTTTATTAAAGGATTAAACAACCTAAATAGA   12979
gb:MT027063|Organism:Severe      CCTAAAGGTCCTAAAGTGAAGTATTTATACTTTATTAAAGGATTAAACAACCTAAATAGA   12982
gb:MT027062|Organism:Severe      CCTAAAGGTCCTAAAGTGAAGTATTTATACTTTATTAAAGGATTAAACAACCTAAATAGA   12982
gb:MT019529|Organism:Severe      CCTAAAGGTCCTAAAGTGAAGTATTTATACTTTATTAAAGGATTAAACAACCTAAATAGA   12982
gb:MN996529|Organism:Severe      CCTAAAGGTCCTAAAGTGAAGTATTTATACTTTATTAAAGGATTAAACAACCTAAATAGA   12970
gb:MN996531|Organism:Severe      CCTAAAGGTCCTAAAGTGAAGTATTTATACTTTATTAAAGGATTAAACAACCTAAATAGA   12969
gb:MT066176|Organism:Severe      CCTAAAGGTCCTAAAGTGAAGTATTTATACTTTATTAAAGGATTAAACAACCTAAATAGA   12982
gb:MT027064|Organism:Severe      CCTAAAGGTCCTAAAGTGAAGTATTTATACTTTATTAAAGGATTAAACAACCTAAATAGA   12982
gb:MN994468|Organism:Severe      CCTAAAGGTCCTAAAGTGAAGTATTTATACTTTATTAAAGGATTAAACAACCTAAATAGA   12982
gb:MT072688|Organism:Severe      CCTAAAGGTCCTAAAGTGAAGTATTTATACTTTATTAAAGGATTAAACAACCTAAATAGA   12967
gb:MN996527|Organism:Severe      CCTAAAGGTCCTAAAGTGAAGTATTTATACTTTATTAAAGGATTAAACAACCTAAATAGA   12949
gb:MT093631|Organism:Severe      CCTAAAGGTCCTAAAGTGAAGTATTTATACTTTATTAAAGGATTAAACAACCTAAATAGA   13020
gb:MT106053|Organism:Severe      CCTAAAGGTCCTAAAGTGAAGTATTTATACTTTATTAAAGGATTAAACAACCTAAATAGA   12982
gb:MT019533|Organism:Severe      CCTAAAGGTCCTAAAGTGAAGTATTTATACTTTATTAAAGGATTAAACAACCTAAATAGA   12982
gb:MT019531|Organism:Severe      CCTAAAGGTCCTAAAGTGAAGTATTTATACTTTATTAAAGGATTAAACAACCTAAATAGA   12982
gb:MN996528|Organism:Severe      CCTAAAGGTCCTAAAGTGAAGTATTTATACTTTATTAAAGGATTAAACAACCTAAATAGA   12982
gb:MN996530|Organism:Severe      CCTAAAGGTCCTAAAGTGAAGTATTTATACTTTATTAAAGGATTAAACAACCTAAATAGA   12968
gb:MN908947|Organism:Severe      CCTAAAGGTCCTAAAGTGAAGTATTTATACTTTATTAAAGGATTAAACAACCTAAATAGA   12982
gb:MT019532|Organism:Severe      CCTAAAGGTCCTAAAGTGAAGTATTTATACTTTATTAAAGGATTAAACAACCTAAATAGA   12982
                                 ************************************************************

gb:MT020781|Organism:Severe      GGTATGGTACTTGGTAGTTTAGCTGCCACAGTACGTCTACAAGCTGGTAATGCAACAGAA   13030
gb:MT007544|Organism:Severe      GGTATGGTACTTGGTAGTTTAGCTGCCACAGTACGTCTACAAGCTGGTAATGCAACAGAA   13042
gb:MN994467|Organism:Severe      GGTATGGTACTTGGTAGTTTAGCTGCCACAGTACGTCTACAAGCTGGTAATGCAACAGAA   13042
gb:MT044257|Organism:Severe      GGTATGGTACTTGGTAGTTTAGCTGCCACAGTACGTCTACAAGCTGGTAATGCAACAGAA   13042
gb:MT106054|Organism:Severe      GGTATGGTACTTGGTAGTTTAGCTGCCACAGTACGTCTACAAGCTGGTAATGCAACAGAA   13042
gb:MT049951|Organism:Severe      GGTATGGTACTTGGTAGTTTAGCTGCCACAGTACGTCTACAAGCTGGTAATGCAACAGAA   13042
gb:MN975262|Organism:Severe      GGTATGGTACTTGGTAGTTTAGCTGCCACAGTACGTCTACAAGCTGGTAATGCAACAGAA   13042
gb:MT106052|Organism:Severe      GGTATGGTACTTGGTAGTTTAGCTGCCACAGTACGTCTACAAGCTGGTAATGCAACAGAA   13042
gb:LC522975|Organism:Severe      GGTATGGTACTTGGTAGTTTAGCTGCCACAGTACGTCTACAAGCTGGTAATGCAACAGAA   13039
gb:LC522973|Organism:Severe      GGTATGGTACTTGGTAGTTTAGCTGCCACAGTACGTCTACAAGCTGGTAATGCAACAGAA   13039
gb:LC522974|Organism:Severe      GGTATGGTACTTGGTAGTTTAGCTGCCACAGTACGTCTACAAGCTGGTAATGCAACAGAA   13039
gb:MN985325|Organism:Severe      GGTATGGTACTTGGTAGTTTAGCTGCCACAGTACGTCTACAAGCTGGTAATGCAACAGAA   13042
gb:MT020881|Organism:Severe      GGTATGGTACTTGGTAGTTTAGCTGCCACAGTACGTCTACAAGCTGGTAATGCAACAGAA   13042
gb:MT020880|Organism:Severe      GGTATGGTACTTGGTAGTTTAGCTGCCACAGTACGTCTACAAGCTGGTAATGCAACAGAA   13042
gb:MT066175|Organism:Severe      GGTATGGTACTTGGTAGTTTAGCTGCCACAGTACGTCTACAAGCTGGTAATGCAACAGAA   13042
gb:MN997409|Organism:Severe      GGTATGGTACTTGGTAGTTTAGCTGCCACAGTACGTCTACAAGCTGGTAATGCAACAGAA   13042
gb:MN938384|Organism:Severe      GGTATGGTACTTGGTAGTTTAGCTGCCACAGTACGTCTACAAGCTGGTAATGCAACAGAA   13010
gb:MT044258|Organism:Severe      GGTATGGTACTTGGTAGTTTAGCTGCCACAGTACGTCTACAAGCTGGTAATGCAACAGAA   13018
gb:MT039890|Organism:Severe      GGTATGGTACTTGGTAGTTTAGCTGCCACAGTACGTCTACAAGCTGGTAATGCAACAGAA   13042
gb:MN988713|Organism:Severe      GGTATGGTACTTGGTAGTTTAGCTGCCACAGTACGTCTACAAGCTGGTAATGCAACAGAA   13042
gb:LC521925|Organism:Severe      GGTATGGTACTTGGTAGTTTAGCTGCCACAGTACGTCTACAAGCTGGTAATGCAACAGAA   13015
gb:MT093571|Organism:Severe      GGTATGGTACTTGGTAGTTTAGCTGCCACAGTACGTCTACAAGCTGGTAATGCAACAGAA   13042
gb:MT039887|Organism:Severe      GGTATGGTACTTGGTAGTTTAGCTGCCACAGTACGTCTACAAGCTGGTAATGCAACAGAA   13042
gb:MT019530|Organism:Severe      GGTATGGTACTTGGTAGTTTAGCTGCCACAGTACGTCTACAAGCTGGTAATGCAACAGAA   13042
```

```
gb:MT039888|Organism:Severe      GGTATGGTACTTGGTAGTTTAGCTGCCACAGTACGTCTACAAGCTGGTAATGCAACAGAA   13042
gb:LC522972|Organism:Severe      GGTATGGTACTTGGTAGTTTAGCTGCCACAGTACGTCTACAAGCTGGTAATGCAACAGAA   13039
gb:MT027063|Organism:Severe      GGTATGGTACTTGGTAGTTTAGCTGCCACAGTACGTCTACAAGCTGGTAATGCAACAGAA   13042
gb:MT027062|Organism:Severe      GGTATGGTACTTGGTAGTTTAGCTGCCACAGTACGTCTACAAGCTGGTAATGCAACAGAA   13042
gb:MT019529|Organism:Severe      GGTATGGTACTTGGTAGTTTAGCTGCCACAGTACGTCTACAAGCTGGTAATGCAACAGAA   13042
gb:MN996529|Organism:Severe      GGTATGGTACTTGGTAGTTTAGCTGCCACAGTACGTCTACAAGCTGGTAATGCAACAGAA   13030
gb:MN996531|Organism:Severe      GGTATGGTACTTGGTAGTTTAGCTGCCACAGTACGTCTACAAGCTGGTAATGCAACAGAA   13029
gb:MT066176|Organism:Severe      GGTATGGTACTTGGTAGTTTAGCTGCCACAGTACGTCTACAAGCTGGTAATGCAACAGAA   13042
gb:MT027064|Organism:Severe      GGTATGGTACTTGGTAGTTTAGCTGCCACAGTACGTCTACAAGCTGGTAATGCAACAGAA   13042
gb:MN994468|Organism:Severe      GGTATGGTACTTGGTAGTTTAGCTGCCACAGTACGTCTACAAGCTGGTAATGCAACAGAA   13042
gb:MT072688|Organism:Severe      GGTATGGTACTTGGTAGTTTAGCTGCCACAGTACGTCTACAAGCTGGTAATGCAACAGAA   13027
gb:MN996527|Organism:Severe      GGTATGGTACTTGGTAGTTTAGCTGCCACAGTACGTCTACAAGCTGGTAATGCAACAGAA   13009
gb:MT093631|Organism:Severe      GGTATGGTACTTGGTAGTTTAGCTGCCACAGTACGTCTACAAGCTGGTAATGCAACAGAA   13080
gb:MT106053|Organism:Severe      GGTATGGTACTTGGTAGTTTAGCTGCCACAGTACGTCTACAAGCTGGTAATGCAACAGAA   13042
gb:MT019533|Organism:Severe      GGTATGGTACTTGGTAGTTTAGCTGCCACAGTACGTCTACAAGCTGGTAATGCAACAGAA   13042
gb:MT019531|Organism:Severe      GGTATGGTACTTGGTAGTTTAGCTGCCACAGTACGTCTACAAGCTGGTAATGCAACAGAA   13042
gb:MN996528|Organism:Severe      GGTATGGTACTTGGTAGTTTAGCTGCCACAGTACGTCTACAAGCTGGTAATGCAACAGAA   13042
gb:MN996530|Organism:Severe      GGTATGGTACTTGGTAGTTTAGCTGCCACAGTACGTCTACAAGCTGGTAATGCAACAGAA   13028
gb:MN908947|Organism:Severe      GGTATGGTACTTGGTAGTTTAGCTGCCACAGTACGTCTACAAGCTGGTAATGCAACAGAA   13042
gb:MT019532|Organism:Severe      GGTATGGTACTTGGTAGTTTAGCTGCCACAGTACGTCTACAAGCTGGTAATGCAACAGAA   13042
                                 ************************************************************

gb:MT020781|Organism:Severe      GTGCCTGCCAATTCAACTGTATTATCTTTCTGTGCTTTTGCTGTAGATGCTGCTAAAGCT   13090
gb:MT007544|Organism:Severe      GTGCCTGCCAATTCAACTGTATTATCTTTCTGTGCTTTTGCTGTAGATGCTGCTAAAGCT   13102
gb:MN994467|Organism:Severe      GTGCCTGCCAATTCAACTGTATTATCTTTCTGTGCTTTTGCTGTAGATGCTGCTAAAGCT   13102
gb:MT044257|Organism:Severe      GTGCCTGCCAATTCAACTGTATTATCTTTCTGTGCTTTTGCTGTAGATGCTGCTAAAGCT   13102
gb:MT106054|Organism:Severe      GTGCCTGCCAATTCAACTGTATTATCTTTCTGTGCTTTTGCTGTAGATGCTGCTAAAGCT   13102
gb:MT049951|Organism:Severe      GTGCCTGCCAATTCAACTGTATTATCTTTCTGTGCTTTTGCTGTAGATGCTGCTAAAGCT   13102
gb:MN975262|Organism:Severe      GTGCCTGCCAATTCAACTGTATTATCTTTCTGTGCTTTTGCTGTAGATGCTGCTAAAGCT   13102
gb:MT106052|Organism:Severe      GTGCCTGCCAATTCAACTGTATTATCTTTCTGTGCTTTTGCTGTAGATGCTGCTAAAGCT   13102
gb:LC522975|Organism:Severe      GTGCCTGCCAATTCAACTGTATTATCTTTCTGTGCTTTTGCTGTAGATGCTGCTAAAGCT   13099
gb:LC522973|Organism:Severe      GTGCCTGCCAATTCAACTGTATTATCTTTCTGTGCTTTTGCTGTAGATGCTGCTAAAGCT   13099
gb:LC522974|Organism:Severe      GTGCCTGCCAATTCAACTGTATTATCTTTCTGTGCTTTTGCTGTAGATGCTGCTAAAGCT   13099
gb:MN985325|Organism:Severe      GTGCCTGCCAATTCAACTGTATTATCTTTCTGTGCTTTTGCTGTAGATGCTGCTAAAGCT   13102
gb:MT020881|Organism:Severe      GTGCCTGCCAATTCAACTGTATTATCTTTCTGTGCTTTTGCTGTAGATGCTGCTAAAGCT   13102
gb:MT020880|Organism:Severe      GTGCCTGCCAATTCAACTGTATTATCTTTCTGTGCTTTTGCTGTAGATGCTGCTAAAGCT   13102
gb:MT066175|Organism:Severe      GTGCCTGCCAATTCAACTGTATTATCTTTCTGTGCTTTTGCTGTAGATGCTGCTAAAGCT   13102
gb:MN997409|Organism:Severe      GTGCCTGCCAATTCAACTGTATTATCTTTCTGTGCTTTTGCTGTAGATGCTGCTAAAGCT   13102
gb:MN938384|Organism:Severe      GTGCCTGCCAATTCAACTGTATTATCTTTCTGTGCTTTTGCTGTAGATGCTGCTAAAGCT   13070
gb:MT044258|Organism:Severe      GTGCCTGCCAATTCAACTGTATTATCTTTCTGTGCTTTTGCTGTAGATGCTGCTAAAGCT   13078
gb:MT039890|Organism:Severe      GTGCCTGCCAATTCAACTGTATTATCTTTCTGTGCTTTTGCTGTAGATGCTGCTAAAGCT   13102
gb:MN988713|Organism:Severe      GTGCCTGCCAATTCAACTGTATTATCTTTCTGTGCTTTTGCTGTAGATGCTGCTAAAGCT   13102
gb:LC521925|Organism:Severe      GTGCCTGCCAATTCAACTGTATTATCTTTCTGTGCTTTTGCTGTAGATGCTGCTAAAGCT   13075
gb:MT093571|Organism:Severe      GTGCCTGCCAATTCAACTGTATTATCTTTCTGTGCTTTTGCTGTAGATGCTGCTAAAGCT   13102
gb:MT039887|Organism:Severe      GTGCCTGCCAATTCAACTGTATTATCTTTCTGTGCTTTTGCTGTAGATGCTGCTAAAGCT   13102
gb:MT019530|Organism:Severe      GTGCCTGCCAATTCAACTGTATTATCTTTCTGTGCTTTTGCTGTAGATGCTGCTAAAGCT   13102
gb:MT039888|Organism:Severe      GTGCCTGCCAATTCAACTGTATTATCTTTCTGTGCTTTTGCTGTAGATGCTGCTAAAGCT   13102
gb:LC522972|Organism:Severe      GTGCCTGCCAATTCAACTGTATTATCTTTCTGTGCTTTTGCTGTAGATGCTGCTAAAGCT   13099
gb:MT027063|Organism:Severe      GTGCCTGCCAATTCAACTGTATTATCTTTCTGTGCTTTTGCTGTAGATGCTGCTAAAGCT   13102
gb:MT027062|Organism:Severe      GTGCCTGCCAATTCAACTGTATTATCTTTCTGTGCTTTTGCTGTAGATGCTGCTAAAGCT   13102
gb:MT019529|Organism:Severe      GTGCCTGCCAATTCAACTGTATTATCTTTCTGTGCTTTTGCTGTAGATGCTGCTAAAGCT   13102
gb:MN996529|Organism:Severe      GTGCCTGCCAATTCAACTGTATTATCTTTCTGTGCTTTTGCTGTAGATGCTGCTAAAGCT   13090
gb:MN996531|Organism:Severe      GTGCCTGCCAATTCAACTGTATTATCTTTCTGTGCTTTTGCTGTAGATGCTGCTAAAGCT   13089
gb:MT066176|Organism:Severe      GTGCCTGCCAATTCAACTGTATTATCTTTCTGTGCTTTTGCTGTAGATGCTGCTAAAGCT   13102
gb:MT027064|Organism:Severe      GTGCCTGCCAATTCAACTGTATTATCTTTCTGTGCTTTTGCTGTAGATGCTGCTAAAGCT   13102
gb:MN994468|Organism:Severe      GTGCCTGCCAATTCAACTGTATTATCTTTCTGTGCTTTTGCTGTAGATGCTGCTAAAGCT   13102
gb:MT072688|Organism:Severe      GTGCCTGCCAATTCAACTGTATTATCTTTCTGTGCTTTTGCTGTAGATGCTGCTAAAGCT   13087
gb:MN996527|Organism:Severe      GTGCCTGCCAATTCAACTGTATTATCTTTCTGTGCTTTTGCTGTAGATGCTGCTAAAGCT   13069
gb:MT093631|Organism:Severe      GTGCCTGCCAATTCAACTGTATTATCTTTCTGTGCTTTTGCTGTAGATGCTGCTAAAGCT   13140
gb:MT106053|Organism:Severe      GTGCCTGCCAATTCAACTGTATTATCTTTCTGTGCTTTTGCTGTAGATGCTGCTAAAGCT   13102
gb:MT019533|Organism:Severe      GTGCCTGCCAATTCAACTGTATTATCTTTCTGTGCTTTTGCTGTAGATGCTGCTAAAGCT   13102
gb:MT019531|Organism:Severe      GTGCCTGCCAATTCAACTGTATTATCTTTCTGTGCTTTTGCTGTAGATGCTGCTAAAGCT   13102
gb:MN996528|Organism:Severe      GTGCCTGCCAATTCAACTGTATTATCTTTCTGTGCTTTTGCTGTAGATGCTGCTAAAGCT   13102
gb:MN996530|Organism:Severe      GTGCCTGCCAATTCAACTGTATTATCTTTCTGTGCTTTTGCTGTAGATGCTGCTAAAGCT   13088
gb:MN908947|Organism:Severe      GTGCCTGCCAATTCAACTGTATTATCTTTCTGTGCTTTTGCTGTAGATGCTGCTAAAGCT   13102
```

```
gb:MT019532|Organism:Severe      GTGCCTGCCAATTCAACTGTATTATCTTTCTGTGCTTTTGCTGTAGATGCTGCTAAAGCT    13102
                                 ************************************************************

gb:MT020781|Organism:Severe      TACAAAGATTATCTAGCTAGTGGGGGACAACCAATCACTAATTGTGTTAAGATGTTGTGT    13150
gb:MT007544|Organism:Severe      TACAAAGATTATCTAGCTAGTGGGGGACAACCAATCACTAATTGTGTTAAGATGTTGTGT    13162
gb:MN994467|Organism:Severe      TACAAAGATTATCTAGCTAGTGGGGGACAACCAATCACTAATTGTGTTAAGATGTTGTGT    13162
gb:MT044257|Organism:Severe      TACAAAGATTATCTAGCTAGTGGGGGACAACCAATCACTAATTGTGTTAAGATGTTGTGT    13162
gb:MT106054|Organism:Severe      TACAAAGATTATCTAGCTAGTGGGGGACAACCAATCACTAATTGTGTTAAGATGTTGTGT    13162
gb:MT049951|Organism:Severe      TACAAAGATTATCTAGCTAGTGGGGGACAACCAATCACTAATTGTGTTAAGATGTTGTGT    13162
gb:MN975262|Organism:Severe      TACAAAGATTATCTAGCTAGTGGGGGACAACCAATCACTAATTGTGTTAAGATGTTGTGT    13162
gb:MT106052|Organism:Severe      TACAAAGATTATCTAGCTAGTGGGGGACAACCAATCACTAATTGTGTTAAGATGTTGTGT    13162
gb:LC522975|Organism:Severe      TACAAAGATTATCTAGCTAGTGGGGGACAACCAATCACTAATTGTGTTAAGATGTTGTGT    13159
gb:LC522973|Organism:Severe      TACAAAGATTATCTAGCTAGTGGGGGACAACCAATCACTAATTGTGTTAAGATGTTGTGT    13159
gb:LC522974|Organism:Severe      TACAAAGATTATCTAGCTAGTGGGGGACAACCAATCACTAATTGTGTTAAGATGTTGTGT    13159
gb:MN985325|Organism:Severe      TACAAAGATTATCTAGCTAGTGGGGGACAACCAATCACTAATTGTGTTAAGATGTTGTGT    13162
gb:MT020881|Organism:Severe      TACAAAGATTATCTAGCTAGTGGGGGACAACCAATCACTAATTGTGTTAAGATGTTGTGT    13162
gb:MT020880|Organism:Severe      TACAAAGATTATCTAGCTAGTGGGGGACAACCAATCACTAATTGTGTTAAGATGTTGTGT    13162
gb:MT066175|Organism:Severe      TACAAAGATTATCTAGCTAGTGGGGGACAACCAATCACTAATTGTGTTAAGATGTTGTGT    13162
gb:MN997409|Organism:Severe      TACAAAGATTATCTAGCTAGTGGGGGACAACCAATCACTAATTGTGTTAAGATGTTGTGT    13162
gb:MN938384|Organism:Severe      TACAAAGATTATCTAGCTAGTGGGGGACAACCAATCACTAATTGTGTTAAGATGTTGTGT    13130
gb:MT044258|Organism:Severe      TACAAAGATTATCTAGCTAGTGGGGGACAACCAATCACTAATTGTGTTAAGATGTTGTGT    13138
gb:MT039890|Organism:Severe      TACAAAGATTATCTAGCTAGTGGGGGACAACCAATCACTAATTGTGTTAAGATGTTGTGT    13162
gb:MN988713|Organism:Severe      TACAAAGATTATCTAGCTAGTGGGGGACAACCAATCACTAATTGTGTTAAGATGTTGTGT    13162
gb:LC521925|Organism:Severe      TACAAAGATTATCTAGCTAGTGGGGGACAACCAATCACTAATTGTGTTAAGATGTTGTGT    13135
gb:MT093571|Organism:Severe      TACAAAGATTATCTAGCTAGTGGGGGACAACCAATCACTAATTGTGTTAAGATGTTGTGT    13162
gb:MT039887|Organism:Severe      TACAAAGATTATCTAGCTAGTGGGGGACAACCAATCACTAATTGTGTTAAGATGTTGTGT    13162
gb:MT019530|Organism:Severe      TACAAAGATTATCTAGCTAGTGGGGGACAACCAATCACTAATTGTGTTAAGATGTTGTGT    13162
gb:MT039888|Organism:Severe      TACAAAGATTATCTAGCTAGTGGGGGACAACCAATCACTAATTGTGTTAAGATGTTGTGT    13162
gb:LC522972|Organism:Severe      TACAAAGATTATCTAGCTAGTGGGGGACAACCAATCACTAATTGTGTTAAGATGTTGTGT    13159
gb:MT027063|Organism:Severe      TACAAAGATTATCTAGCTAGTGGGGGACAACCAATCACTAATTGTGTTAAGATGTTGTGT    13162
gb:MT027062|Organism:Severe      TACAAAGATTATCTAGCTAGTGGGGGACAACCAATCACTAATTGTGTTAAGATGTTGTGT    13162
gb:MT019529|Organism:Severe      TACAAAGATTATCTAGCTAGTGGGGGACAACCAATCACTAATTGTGTTAAGATGTTGTGT    13162
gb:MN996529|Organism:Severe      TACAAAGATTATCTAGCTAGTGGGGGACAACCAATCACTAATTGTGTTAAGATGTTGTGT    13150
gb:MN996531|Organism:Severe      TACAAAGATTATCTAGCTAGTGGGGGACAACCAATCACTAATTGTGTTAAGATGTTGTGT    13149
gb:MT066176|Organism:Severe      TACAAAGATTATCTAGCTAGTGGGGGACAACCAATCACTAATTGTGTTAAGATGTTGTGT    13162
gb:MT027064|Organism:Severe      TACAAAGATTATCTAGCTAGTGGGGGACAACCAATCACTAATTGTGTTAAGATGTTGTGT    13162
gb:MN994468|Organism:Severe      TACAAAGATTATCTAGCTAGTGGGGGACAACCAATCACTAATTGTGTTAAGATGTTGTGT    13162
gb:MT072688|Organism:Severe      TACAAAGATTATCTAGCTAGTGGGGGACAACCAATCACTAATTGTGTTAAGATGTTGTGT    13147
gb:MN996527|Organism:Severe      TACAAAGATTATCTAGCTAGTGGGGGACAACCAATCACTAATTGTGTTAAGATGTTGTGT    13129
gb:MT093631|Organism:Severe      TACAAAGATTATCTAGCTAGTGGGGGACAACCAATCACTAATTGTGTTAAGATGTTGTGT    13200
gb:MT106053|Organism:Severe      TACAAAGATTATCTAGCTAGTGGGGGACAACCAATCACTAATTGTGTTAAGATGTTGTGT    13162
gb:MT019533|Organism:Severe      TACAAAGATTATCTAGCTAGTGGGGGACAACCAATCACTAATTGTGTTAAGATGTTGTGT    13162
gb:MT019531|Organism:Severe      TACAAAGATTATCTAGCTAGTGGGGGACAACCAATCACTAATTGTGTTAAGATGTTGTGT    13162
gb:MN996528|Organism:Severe      TACAAAGATTATCTAGCTAGTGGGGGACAACCAATCACTAATTGTGTTAAGATGTTGTGT    13162
gb:MN996530|Organism:Severe      TACAAAGATTATCTAGCTAGTGGGGGACAACCAATCACTAATTGTGTTAAGATGTTGTGT    13148
gb:MN908947|Organism:Severe      TACAAAGATTATCTAGCTAGTGGGGGACAACCAATCACTAATTGTGTTAAGATGTTGTGT    13162
gb:MT019532|Organism:Severe      TACAAAGATTATCTAGCTAGTGGGGGACAACCAATCACTAATTGTGTTAAGATGTTGTGT    13162
                                 ************************************************************

gb:MT020781|Organism:Severe      ACACACACTGGTACTGGTCAGGCAATAACAGTTACACCGGAAGCCAATATGGATCAAGAA    13210
gb:MT007544|Organism:Severe      ACACACACTGGTACTGGTCAGGCAATAACAGTTACACCGGAAGCCAATATGGATCAAGAA    13222
gb:MN994467|Organism:Severe      ACACACACTGGTACTGGTCAGGCAATAACAGTTACACCGGAAGCCAATATGGATCAAGAA    13222
gb:MT044257|Organism:Severe      ACACACACTGGTACTGGTCAGGCAATAACAGTTACACCGGAAGCCAATATGGATCAAGAA    13222
gb:MT106054|Organism:Severe      ACACACACTGGTACTGGTCAGGCAATAACAGTTACACCGGAAGCCAATATGGATCAAGAA    13222
gb:MT049951|Organism:Severe      ACACACACTGGTACTGGTCAGGCAATAACAGTTACACCGGAAGCCAATATGGATCAAGAA    13222
gb:MN975262|Organism:Severe      ACACACACTGGTACTGGTCAGGCAATAACAGTTACACCGGAAGCCAATATGGATCAAGAA    13222
gb:MT106052|Organism:Severe      ACACACACTGGTACTGGTCAGGCAATAACAGTTACACCGGAAGCCAATATGGATCAAGAA    13222
gb:LC522975|Organism:Severe      ACACACACTGGTACTGGTCAGGCAATAACAGTTACACCGGAAGCCAATATGGATCAAGAA    13219
gb:LC522973|Organism:Severe      ACACACACTGGTACTGGTCAGGCAATAACAGTTACACCGGAAGCCAATATGGATCAAGAA    13219
gb:LC522974|Organism:Severe      ACACACACTGGTACTGGTCAGGCAATAACAGTTACACCGGAAGCCAATATGGATCAAGAA    13219
gb:MN985325|Organism:Severe      ACACACACTGGTACTGGTCAGGCAATAACAGTTACACCGGAAGCCAATATGGATCAAGAA    13222
gb:MT020881|Organism:Severe      ACACACACTGGTACTGGTCAGGCAATAACAGTTACACCGGAAGCCAATATGGATCAAGAA    13222
gb:MT020880|Organism:Severe      ACACACACTGGTACTGGTCAGGCAATAACAGTTACACCGGAAGCCAATATGGATCAAGAA    13222
gb:MT066175|Organism:Severe      ACACACACTGGTACTGGTCAGGCAATAACAGTTACACCGGAAGCCAATATGGATCAAGAA    13222
gb:MN997409|Organism:Severe      ACACACACTGGTACTGGTCAGGCAATAACAGTTACACCGGAAGCCAATATGGATCAAGAA    13222
```

```
gb:MN938384|Organism:Severe      ACACACACTGGTACTGGTCAGGCAATAACAGTTACACCGGAAGCCAATATGGATCAAGAA   13190
gb:MT044258|Organism:Severe      ACACACACTGGTACTGGTCAGGCAATAACAGTTACACCGGAAGCCAATATGGATCAAGAA   13198
gb:MT039890|Organism:Severe      ACACACACTGGTACTGGTCAGGCAATAACAGTTACACCGGAAGCCAATATGGATCAAGAA   13222
gb:MN988713|Organism:Severe      ACACACACTGGTACTGGTCAGGCAATAACAGTTACACCGGAAGCCAATATGGATCAAGAA   13222
gb:LC521925|Organism:Severe      ACACACACTGGTACTGGTCAGGCAATAACAGTTACACCGGAAGCCAATATGGATCAAGAA   13195
gb:MT093571|Organism:Severe      ACACACACTGGTACTGGTCAGGCAATAACAGTTACACCGGAAGCCAATATGGATCAAGAA   13222
gb:MT039887|Organism:Severe      ACACACACTGGTACTGGTCAGGCAATAACAGTTACACCGGAAGCCAATATGGATCAAGAA   13222
gb:MT019530|Organism:Severe      ACACACACTGGTACTGGTCAGGCAATAACAGTTACACCGGAAGCCAATATGGATCAAGAA   13222
gb:MT039888|Organism:Severe      ACACACACTGGTACTGGTCAGGCAATAACAGTTACACCGGAAGCCAATATGGATCAAGAA   13222
gb:LC522972|Organism:Severe      ACACACACTGGTACTGGTCAGGCAATAACAGTTACACCGGAAGCCAATATGGATCAAGAA   13219
gb:MT027063|Organism:Severe      ACACACACTGGTACTGGTCAGGCAATAACAGTTACACCGGAAGCCAATATGGATCAAGAA   13222
gb:MT027062|Organism:Severe      ACACACACTGGTACTGGTCAGGCAATAACAGTTACACCGGAAGCCAATATGGATCAAGAA   13222
gb:MT019529|Organism:Severe      ACACACACTGGTACTGGTCAGGCAATAACAGTTACACCGGAAGCCAATATGGATCAAGAA   13222
gb:MN996529|Organism:Severe      ACACACACTGGTACTGGTCAGGCAATAACAGTTACACCGGAAGCCAATATGGATCAAGAA   13210
gb:MN996531|Organism:Severe      ACACACACTGGTACTGGTCAGGCAATAACAGTTACACCGGAAGCCAATATGGATCAAGAA   13209
gb:MT066176|Organism:Severe      ACACACACTGGTACTGGTCAGGCAATAACAGTTACACCGGAAGCCAATATGGATCAAGAA   13222
gb:MT027064|Organism:Severe      ACACACACTGGTACTGGTCAGGCAATAACAGTTACACCGGAAGCCAATATGGATCAAGAA   13222
gb:MN994468|Organism:Severe      ACACACACTGGTACTGGTCAGGCAATAACAGTTACACCGGAAGCCAATATGGATCAAGAA   13222
gb:MT072688|Organism:Severe      ACACACACTGGTACTGGTCAGGCAATAACAGTTACACCGGAAGCCAATATGGATCAAGAA   13207
gb:MN996527|Organism:Severe      ACACACACTGGTACTGGTCAGGCAATAACAGTTACACCGGAAGCCAATATGGATCAAGAA   13189
gb:MT093631|Organism:Severe      ACACACACTGGTACTGGTCAGGCAATAACAGTTACACCGGAAGCCAATATGGATCAAGAA   13260
gb:MT106053|Organism:Severe      ACACACACTGGTACTGGTCAGGCAATAACAGTTACACCGGAAGCCAATATGGATCAAGAA   13222
gb:MT019533|Organism:Severe      ACACACACTGGTACTGGTCAGGCAATAACAGTTACACCGGAAGCCAATATGGATCAAGAA   13222
gb:MT019531|Organism:Severe      ACACACACTGGTACTGGTCAGGCAATAACAGTTACACCGGAAGCCAATATGGATCAAGAA   13222
gb:MN996528|Organism:Severe      ACACACACTGGTACTGGTCAGGCAATAACAGTTACACCGGAAGCCAATATGGATCAAGAA   13222
gb:MN996530|Organism:Severe      ACACACACTGGTACTGGTCAGGCAATAACAGTTACACCGGAAGCCAATATGGATCAAGAA   13208
gb:MN908947|Organism:Severe      ACACACACTGGTACTGGTCAGGCAATAACAGTTACACCGGAAGCCAATATGGATCAAGAA   13222
gb:MT019532|Organism:Severe      ACACACACTGGTACTGGTCAGGCAATAACAGTTACACCGGAAGCCAATATGGATCAAGAA   13222
                                 ************************************************************

gb:MT020781|Organism:Severe      TCCTTTGGTGGTGCATCGTGTTGTCTGTACTGCCGTTGCCACATAGATCATCCAAATCCT   13270
gb:MT007544|Organism:Severe      TCCTTTGGTGGTGCATCGTGTTGTCTGTACTGCCGTTGCCACATAGATCATCCAAATCCT   13282
gb:MN994467|Organism:Severe      TCCTTTGGTGGTGCATCGTGTTGTCTGTACTGCCGTTGCCACATAGATCATCCAAATCCT   13282
gb:MT044257|Organism:Severe      TCCTTTGGTGGTGCATCGTGTTGTCTGTACTGCCGTTGCCACATAGATCATCCAAATCCT   13282
gb:MT106054|Organism:Severe      TCCTTTGGTGGTGCATCGTGTTGTCTGTACTGCCGTTGCCACATAGATCATCCAAATCCT   13282
gb:MT049951|Organism:Severe      TCCTTTGGTGGTGCATCGTGTTGTCTGTACTGCCGTTGCCACATAGATCATCCAAATCCT   13282
gb:MN975262|Organism:Severe      TCCTTTGGTGGTGCATCGTGTTGTCTGTACTGCCGTTGCCACATAGATCATCCAAATCCT   13282
gb:MT106052|Organism:Severe      TCCTTTGGTGGTGCATCGTGTTGTCTGTACTGCCGTTGCCACATAGATCATCCAAATCCT   13282
gb:LC522975|Organism:Severe      TCCTTTGGTGGTGCATCGTGTTGTCTGTACTGCCGTTGCCACATAGATCATCCAAATCCT   13279
gb:LC522973|Organism:Severe      TCCTTTGGTGGTGCATCGTGTTGTCTGTACTGCCGTTGCCACATAGATCATCCAAATCCT   13279
gb:LC522974|Organism:Severe      TCCTTTGGTGGTGCATCGTGTTGTCTGTACTGCCGTTGCCACATAGATCATCCAAATCCT   13279
gb:MN985325|Organism:Severe      TCCTTTGGTGGTGCATCGTGTTGTCTGTACTGCCGTTGCCACATAGATCATCCAAATCCT   13282
gb:MT020881|Organism:Severe      TCCTTTGGTGGTGCATCGTGTTGTCTGTACTGCCGTTGCCACATAGATCATCCAAATCCT   13282
gb:MT020880|Organism:Severe      TCCTTTGGTGGTGCATCGTGTTGTCTGTACTGCCGTTGCCACATAGATCATCCAAATCCT   13282
gb:MT066175|Organism:Severe      TCCTTTGGTGGTGCATCGTGTTGTCTGTACTGCCGTTGCCACATAGATCATCCAAATCCT   13282
gb:MN997409|Organism:Severe      TCCTTTGGTGGTGCATCGTGTTGTCTGTACTGCCGTTGCCACATAGATCATCCAAATCCT   13282
gb:MN938384|Organism:Severe      TCCTTTGGTGGTGCATCGTGTTGTCTGTACTGCCGTTGCCACATAGATCATCCAAATCCT   13250
gb:MT044258|Organism:Severe      TCCTTTGGTGGTGCATCGTGTTGTCTGTACTGCCGTTGCCACATAGATCATCCAAATCCT   13258
gb:MT039890|Organism:Severe      TCCTTTGGTGGTGCATCGTGTTGTCTGTACTGCCGTTGCCACATAGATCATCCAAATCCT   13282
gb:MN988713|Organism:Severe      TCCTTTGGTGGTGCATCGTGTTGTCTGTACTGCCGTTGCCACATAGATCATCCAAATCCT   13282
gb:LC521925|Organism:Severe      TCCTTTGGTGGTGCATCGTGTTGTCTGTACTGCCGTTGCCACATAGATCATCCAAATCCT   13255
gb:MT093571|Organism:Severe      TCGCTTGGTGGTGCATCGTGTTGTCTGTACTGCCGTTGCCACATAGATCATCCAAATCCT   13282
gb:MT039887|Organism:Severe      TCCTTTGGTGGTGCATCGTGTTGTCTGTACTGCCGTTGCCACATAGATCATCCAAATCCT   13282
gb:MT019530|Organism:Severe      TCCTTTGGTGGTGCATCGTGTTGTCTGTACTGCCGTTGCCACATAGATCATCCAAATCCT   13282
gb:MT039888|Organism:Severe      TCCTTTGGTGGTGCATCGTGTTGTCTGTACTGCCGTTGCCACATAGATCATCCAAATCCT   13282
gb:LC522972|Organism:Severe      TCCTTTGGTGGTGCATCGTGTTGTCTGTACTGCCGTTGCCACATAGATCATCCAAATCCT   13279
gb:MT027063|Organism:Severe      TCCTTTGGTGGTGCATCGTGTTGTCTGTACTGCCGTTGCCACATAGATCATCCAAATCCT   13282
gb:MT027062|Organism:Severe      TCCTTTGGTGGTGCATCGTGTTGTCTGTACTGCCGTTGCCACATAGATCATCCAAATCCT   13282
gb:MT019529|Organism:Severe      TCCTTTGGTGGTGCATCGTGTTGTCTGTACTGCCGTTGCCACATAGATCATCCAAATCCT   13282
gb:MN996529|Organism:Severe      TCCTTTGGTGGTGCATCGTGTTGTCTGTACTGCCGTTGCCACATAGATCATCCAAATCCT   13270
gb:MN996531|Organism:Severe      TCCTTTGGTGGTGCATCGTGTTGTCTGTACTGCCGTTGCCACATAGATCATCCAAATCCT   13269
gb:MT066176|Organism:Severe      TCCTTTGGTGGTGCATCGTGTTGTCTGTACTGCCGTTGCCACATAGATCATCCAAATCCT   13282
gb:MT027064|Organism:Severe      TCCTTTGGTGGTGCATCGTGTTGTCTGTACTGCCGTTGCCACATAGATCATCCAAATCCT   13282
gb:MN994468|Organism:Severe      TCCTTTGGTGGTGCATCGTGTTGTCTGTACTGCCGTTGCCACATAGATCATCCAAATCCT   13282
gb:MT072688|Organism:Severe      TCCTTTGGTGGTGCATCGTGTTGTCTGTACTGCCGTTGCCACATAGATCATCCAAATCCT   13267
```

```
gb:MN996527|Organism:Severe      TCCTTTGGTGGTGCATCGTGTTGTCTGTACTGCCGTTGCCACATAGATCATCCAAATCCT   13249
gb:MT093631|Organism:Severe      TCCTTTGGTGGTGCATCGTGTTGTCTGTACTGCCGTTGCCACATAGATCATCCAAATCCT   13320
gb:MT106053|Organism:Severe      TCCTTTGGTGGTGCATCGTGTTGTCTGTACTGCCGTTGCCACATAGATCATCCAAATCCT   13282
gb:MT019533|Organism:Severe      TCCTTTGGTGGTGCATCGTGTTGTCTGTACTGCCGTTGCCACATAGATCATCCAAATCCT   13282
gb:MT019531|Organism:Severe      TCCTTTGGTGGTGCATCGTGTTGTCTGTACTGCCGTTGCCACATAGATCATCCAAATCCT   13282
gb:MN996528|Organism:Severe      TCCTTTGGTGGTGCATCGTGTTGTCTGTACTGCCGTTGCCACATAGATCATCCAAATCCT   13282
gb:MN996530|Organism:Severe      TCCTTTGGTGGTGCATCGTGTTGTCTGTACTGCCGTTGCCACATAGATCATCCAAATCCT   13268
gb:MN908947|Organism:Severe      TCCTTTGGTGGTGCATCGTGTTGTCTGTACTGCCGTTGCCACATAGATCATCCAAATCCT   13282
gb:MT019532|Organism:Severe      TCCTTTGGTGGTGCATCGTGTTGTCTGTACTGCCGTTGCCACATAGATCATCCAAATCCT   13282
                                 **  ********************************************************

gb:MT020781|Organism:Severe      AAAGGATTTTGTGACTTAAAAGGTAAGTATGTACAAATACCTACAACTTGTGCTAATGAC   13330
gb:MT007544|Organism:Severe      AAAGGATTTTGTGACTTAAAAGGTAAGTATGTACAAATACCTACAACTTGTGCTAATGAC   13342
gb:MN994467|Organism:Severe      AAAGGATTTTGTGACTTAAAAGGTAAGTATGTACAAATACCTACAACTTGTGCTAATGAC   13342
gb:MT044257|Organism:Severe      AAAGGATTTTGTGACTTAAAAGGTAAGTATGTACAAATACCTACAACTTGTGCTAATGAC   13342
gb:MT106054|Organism:Severe      AAAGGATTTTGTGACTTAAAAGGTAAGTATGTACAAATACCTACAACTTGTGCTAATGAC   13342
gb:MT049951|Organism:Severe      AAAGGATTTTGTGACTTAAAAGGTAAGTATGTACAAATACCTACAACTTGTGCTAATGAC   13342
gb:MN975262|Organism:Severe      AAAGGATTTTGTGACTTAAAAGGTAAGTATGTACAAATACCTACAACTTGTGCTAATGAC   13342
gb:MT106052|Organism:Severe      AAAGGATTTTGTGACTTAAAAGGTAAGTATGTACAAATACCTACAACTTGTGCTAATGAC   13342
gb:LC522975|Organism:Severe      AAAGGATTTTGTGACTTAAAAGGTAAGTATGTACAAATACCTACAACTTGTGCTAATGAC   13339
gb:LC522973|Organism:Severe      AAAGGATTTTGTGACTTAAAAGGTAAGTATGTACAAATACCTACAACTTGTGCTAATGAC   13339
gb:LC522974|Organism:Severe      AAAGGATTTTGTGACTTAAAAGGTAAGTATGTACAAATACCTACAACTTGTGCTAATGAC   13339
gb:MN985325|Organism:Severe      AAAGGATTTTGTGACTTAAAAGGTAAGTATGTACAAATACCTACAACTTGTGCTAATGAC   13342
gb:MT020881|Organism:Severe      AAAGGATTTTGTGACTTAAAAGGTAAGTATGTACAAATACCTACAACTTGTGCTAATGAC   13342
gb:MT020880|Organism:Severe      AAAGGATTTTGTGACTTAAAAGGTAAGTATGTACAAATACCTACAACTTGTGCTAATGAC   13342
gb:MT066175|Organism:Severe      AAAGGATTTTGTGACTTAAAAGGTAAGTATGTACAAATACCTACAACTTGTGCTAATGAC   13342
gb:MN997409|Organism:Severe      AAAGGATTTTGTGACTTAAAAGGTAAGTATGTACAAATACCTACAACTTGTGCTAATGAC   13342
gb:MN938384|Organism:Severe      AAAGGATTTTGTGACTTAAAAGGTAAGTATGTACAAATACCTACAACTTGTGCTAATGAC   13310
gb:MT044258|Organism:Severe      AAAGGATTTTGTGACTTAAAAGGTAAGTATGTACAAATACCTACAACTTGTGCTAATGAC   13318
gb:MT039890|Organism:Severe      AAAGGATTTTGTGACTTAAAAGGTAAGTATGTACAAATACCTACAACTTGTGCTAATGAC   13342
gb:MN988713|Organism:Severe      AAAGGATTTTGTGACTTAAAAGGTAAGTATGTACAAATACCTACAACTTGTGCTAATGAC   13342
gb:LC521925|Organism:Severe      AAAGGATTTTGTGACTTAAAAGGTAAGTATGTACAAATACCTACAACTTGTGCTAATGAC   13315
gb:MT093571|Organism:Severe      AAAGGATTTTGTGACTTAAAAGGTAAGTATGTACAAATACCTACAACTTGTGCTAATGAC   13342
gb:MT039887|Organism:Severe      AAAGGATTTTGTGACTTAAAAGGTAAGTATGTACAAATACCTACAACTTGTGCTAATGAC   13342
gb:MT019530|Organism:Severe      AAAGGATTTTGTGACTTAAAAGGTAAGTATGTACAAATACCTACAACTTGTGCTAATGAC   13342
gb:MT039888|Organism:Severe      AAAGGATTTTGTGACTTAAAAGGTAAGTATGTACAAATACCTACAACTTGTGCTAATGAC   13342
gb:LC522972|Organism:Severe      AAAGGATTTTGTGACTTAAAAGGTAAGTATGTACAAATACCTACAACTTGTGCTAATGAC   13339
gb:MT027063|Organism:Severe      AAAGGATTTTGTGACTTAAAAGGTAAGTATGTACAAATACCTACAACTTGTGCTAATGAC   13342
gb:MT027062|Organism:Severe      AAAGGATTTTGTGACTTAAAAGGTAAGTATGTACAAATACCTACAACTTGTGCTAATGAC   13342
gb:MT019529|Organism:Severe      AAAGGATTTTGTGACTTAAAAGGTAAGTATGTACAAATACCTACAACTTGTGCTAATGAC   13342
gb:MN996529|Organism:Severe      AAAGGATTTTGTGACTTAAAAGGTAAGTATGTACAAATACCTACAACTTGTGCTAATGAC   13330
gb:MN996531|Organism:Severe      AAAGGATTTTGTGACTTAAAAGGTAAGTATGTACAAATACCTACAACTTGTGCTAATGAC   13329
gb:MT066176|Organism:Severe      AAAGGATTTTGTGACTTAAAAGGTAAGTATGTACAAATACCTACAACTTGTGCTAATGAC   13342
gb:MT027064|Organism:Severe      AAAGGATTTTGTGACTTAAAAGGTAAGTATGTACAAATACCTACAACTTGTGCTAATGAC   13342
gb:MN994468|Organism:Severe      AAAGGATTTTGTGACTTAAAAGGTAAGTATGTACAAATACCTACAACTTGTGCTAATGAC   13342
gb:MT072688|Organism:Severe      AAAGGATTTTGTGACTTAAAAGGTAAGTATGTACAAATACCTACAACTTGTGCTAATGAC   13327
gb:MN996527|Organism:Severe      AAAGGATTTTGTGACTTAAAAGGTAAGTATGTACAAATACCTACAACTTGTGCTAATGAC   13309
gb:MT093631|Organism:Severe      AAAGGATTTTGTGACTTAAAAGGTAAGTATGTACAAATACCTACAACTTGTGCTAATGAC   13380
gb:MT106053|Organism:Severe      AAAGGATTTTGTGACTTAAAAGGTAAGTATGTACAAATACCTACAACTTGTGCTAATGAC   13342
gb:MT019533|Organism:Severe      AAAGGATTTTGTGACTTAAAAGGTAAGTATGTACAAATACCTACAACTTGTGCTAATGAC   13342
gb:MT019531|Organism:Severe      AAAGGATTTTGTGACTTAAAAGGTAAGTATGTACAAATACCTACAACTTGTGCTAATGAC   13342
gb:MN996528|Organism:Severe      AAAGGATTTTGTGACTTAAAAGGTAAGTATGTACAAATACCTACAACTTGTGCTAATGAC   13342
gb:MN996530|Organism:Severe      AAAGGATTTTGTGACTTAAAAGGTAAGTATGTACAAATACCTACAACTTGTGCTAATGAC   13328
gb:MN908947|Organism:Severe      AAAGGATTTTGTGACTTAAAAGGTAAGTATGTACAAATACCTACAACTTGTGCTAATGAC   13342
gb:MT019532|Organism:Severe      AAAGGATTTTGTGACTTAAAAGGTAAGTATGTACAAATACCTACAACTTGTGCTAATGAC   13342
                                 ************************************************************

gb:MT020781|Organism:Severe      CCTGTGGGTTTTACACTTAAAAACACAGTCTGTACCGTCTGCGGTATGTGGAAAGGTTAT   13390
gb:MT007544|Organism:Severe      CCTGTGGGTTTTACACTTAAAAACACAGTCTGTACCGTCTGCGGTATGTGGAAAGGTTAT   13402
gb:MN994467|Organism:Severe      CCTGTGGGTTTTACACTTAAAAACACAGTCTGTACCGTCTGCGGTATGTGGAAAGGTTAT   13402
gb:MT044257|Organism:Severe      CCTGTGGGTTTTACACTTAAAAACACAGTCTGTACCGTCTGCGGTATGTGGAAAGGTTAT   13402
gb:MT106054|Organism:Severe      CCTGTGGGTTTTACACTTAAAAACACAGTCTGTACCGTCTGCGGTATGTGGAAAGGTTAT   13402
gb:MT049951|Organism:Severe      CCTGTGGGTTTTACACTTAAAAACACAGTCTGTACCGTCTGCGGTATGTGGAAAGGTTAT   13402
gb:MN975262|Organism:Severe      CCTGTGGGTTTTACACTTAAAAACACAGTCTGTACCGTCTGCGGTATGTGGAAAGGTTAT   13402
gb:MT106052|Organism:Severe      CCTGTGGGTTTTACACTTAAAAACACAGTCTGTACCGTCTGCGGTATGTGGAAAGGTTAT   13402
```

```
gb:LC522975|Organism:Severe      CCTGTGGGTTTTACACTTAAAAACACAGTCTGTACCGTCTGCGGTATGTGGAAAGGTTAT    13399
gb:LC522973|Organism:Severe      CCTGTGGGTTTTACACTTAAAAACACAGTCTGTACCGTCTGCGGTATGTGGAAAGGTTAT    13399
gb:LC522974|Organism:Severe      CCTGTGGGTTTTACACTTAAAAACACAGTCTGTACCGTCTGCGGTATGTGGAAAGGTTAT    13399
gb:MN985325|Organism:Severe      CCTGTGGGTTTTACACTTAAAAACACAGTCTGTACCGTCTGCGGTATGTGGAAAGGTTAT    13402
gb:MT020881|Organism:Severe      CCTGTGGGTTTTACACTTAAAAACACAGTCTGTACCGTCTGCGGTATGTGGAAAGGTTAT    13402
gb:MT020880|Organism:Severe      CCTGTGGGTTTTACACTTAAAAACACAGTCTGTACCGTCTGCGGTATGTGGAAAGGTTAT    13402
gb:MT066175|Organism:Severe      CCTGTGGGTTTTACACTTAAAAACACAGTCTGTACCGTCTGCGGTATGTGGAAAGGTTAT    13402
gb:MN997409|Organism:Severe      CCTGTGGGTTTTACACTTAAAAACACAGTCTGTACCGTCTGCGGTATGTGGAAAGGTTAT    13402
gb:MN938384|Organism:Severe      CCTGTGGGTTTTACACTTAAAAACACAGTCTGTACCGTCTGCGGTATGTGGAAAGGTTAT    13370
gb:MT044258|Organism:Severe      CCTGTGGGTTTTACACTTAAAAACACAGTCTGTACCGTCTGCGGTATGTGGAAAGGTTAT    13378
gb:MT039890|Organism:Severe      CCTGTGGGTTTTACACTTAAAAACACAGTCTGTACCGTCTGCGGTATGTGGAAAGGTTAT    13402
gb:MN988713|Organism:Severe      CCTGTGGGTTTTACACTTAAAAACACAGTCTGTACCGTCTGCGGTATGTGGAAAGGTTAT    13402
gb:LC521925|Organism:Severe      CCTGTGGGTTTTACACTTAAAAACACAGTCTGTACCGTCTGCGGTATGTGGAAAGGTTAT    13375
gb:MT093571|Organism:Severe      CCTGTGGGTTTTACACTTAAAAACACAGTCTGTACCGTCTGCGGTATGTGGAAAGGTTAT    13402
gb:MT039887|Organism:Severe      CCTGTGGGTTTTACACTTAAAAACACAGTCTGTACCGTCTGCGGTATGTGGAAAGGTTAT    13402
gb:MT019530|Organism:Severe      CCTGTGGGTTTTACACTTAAAAACACAGTCTGTACCGTCTGCGGTATGTGGAAAGGTTAT    13402
gb:MT039888|Organism:Severe      CCTGTGGGTTTTACACTTAAAAACACAGTCTGTACCGTCTGCGGTATGTGGAAAGGTTAT    13402
gb:LC522972|Organism:Severe      CCTGTGGGTTTTACACTTAAAAACACAGTCTGTACCGTCTGCGGTATGTGGAAAGGTTAT    13399
gb:MT027063|Organism:Severe      CCTGTGGGTTTTACACTTAAAAACACAGTCTGTACCGTCTGCGGTATGTGGAAAGGTTAT    13402
gb:MT027062|Organism:Severe      CCTGTGGGTTTTACACTTAAAAACACAGTCTGTACCGTCTGCGGTATGTGGAAAGGTTAT    13402
gb:MT019529|Organism:Severe      CCTGTGGGTTTTACACTTAAAAACACAGTCTGTACCGTCTGCGGTATGTGGAAAGGTTAT    13402
gb:MN996529|Organism:Severe      CCTGTGGGTTTTACACTTAAAAACACAGTCTGTACCGTCTGCGGTATGTGGAAAGGTTAT    13390
gb:MN996531|Organism:Severe      CCTGTGGGTTTTACACTTAAAAACACAGTCTGTACCGTCTGCGGTATGTGGAAAGGTTAT    13389
gb:MT066176|Organism:Severe      CCTGTGGGTTTTACACTTAAAAACACAGTCTGTACCGTCTGCGGTATGTGGAAAGGTTAT    13402
gb:MT027064|Organism:Severe      CCTGTGGGTTTTACACTTAAAAACACAGTCTGTACCGTCTGCGGTATGTGGAAAGGTTAT    13402
gb:MN994468|Organism:Severe      CCTGTGGGTTTTACACTTAAAAACACAGTCTGTACCGTCTGCGGTATGTGGAAAGGTTAT    13402
gb:MT072688|Organism:Severe      CCTGTGGGTTTTACACTTAAAAACACAGTCTGTACCGTCTGCGGTATGTGGAAAGGTTAT    13387
gb:MN996527|Organism:Severe      CCTGTGGGTTTTACACTTAAAAACACAGTCTGTACCGTCTGCGGTATGTGGAAAGGTTAT    13369
gb:MT093631|Organism:Severe      CCTGTGGGTTTTACACTTAAAAACACAGTCTGTACCGTCTGCGGTATGTGGAAAGGTTAT    13440
gb:MT106053|Organism:Severe      CCTGTGGGTTTTACACTTAAAAACACAGTCTGTACCGTCTGCGGTATGTGGAAAGGTTAT    13402
gb:MT019533|Organism:Severe      CCTGTGGGTTTTACACTTAAAAACACAGTCTGTACCGTCTGCGGTATGTGGAAAGGTTAT    13402
gb:MT019531|Organism:Severe      CCTGTGGGTTTTACACTTAAAAACACAGTCTGTACCGTCTGCGGTATGTGGAAAGGTTAT    13402
gb:MN996528|Organism:Severe      CCTGTGGGTTTTACACTTAAAAACACAGTCTGTACCGTCTGCGGTATGTGGAAAGGTTAT    13402
gb:MN996530|Organism:Severe      CCTGTGGGTTTTACACTTAAAAACACAGTCTGTACCGTCTGCGGTATGTGGAAAGGTTAT    13388
gb:MN908947|Organism:Severe      CCTGTGGGTTTTACACTTAAAAACACAGTCTGTACCGTCTGCGGTATGTGGAAAGGTTAT    13402
gb:MT019532|Organism:Severe      CCTGTGGGTTTTACACTTAAAAACACAGTCTGTACCGTCTGCGGTATGTGGAAAGGTTAT    13402
                                 ************************************************************

gb:MT020781|Organism:Severe      GGCTGTAGTTGTGATCAACTCCGCGAACCCATGCTTCAGTCAGCTGATGCACAATCGTTT    13450
gb:MT007544|Organism:Severe      GGCTGTAGTTGTGATCAACTCCGCGAACCCATGCTTCAGTCAGCTGATGCACAATCGTTT    13462
gb:MN994467|Organism:Severe      GGCTGTAGTTGTGATCAACTCCGCGAACCCATGCTTCAGTCAGCTGATGCACAATCGTTT    13462
gb:MT044257|Organism:Severe      GGCTGTAGTTGTGATCAACTCCGCGAACCCATGCTTCAGTCAGCTGATGCACAATCGTTT    13462
gb:MT106054|Organism:Severe      GGCTGTAGTTGTGATCAACTCCGCGAACCCATGCTTCAGTCAGCTGATGCACAATCGTTT    13462
gb:MT049951|Organism:Severe      GGCTGTAGTTGTGATCAACTCCGCGAACCCATGCTTCAGTCAGCTGATGCACAATCGTTT    13462
gb:MN975262|Organism:Severe      GGCTGTAGTTGTGATCAACTCCGCGAACCCATGCTTCAGTCAGCTGATGCACAATCGTTT    13462
gb:MT106052|Organism:Severe      GGCTGTAGTTGTGATCAACTCCGCGAACCCATGCTTCAGTCAGCTGATGCACAATCGTTT    13462
gb:LC522975|Organism:Severe      GGCTGTAGTTGTGATCAACTCCGCGAACCCATGCTTCAGTCAGCTGATGCACAATCGTTT    13459
gb:LC522973|Organism:Severe      GGCTGTAGTTGTGATCAACTCCGCGAACCCATGCTTCAGTCAGCTGATGCACAATCGTTT    13459
gb:LC522974|Organism:Severe      GGCTGTAGTTGTGATCAACTCCGCGAACCCATGCTTCAGTCAGCTGATGCACAATCGTTT    13459
gb:MN985325|Organism:Severe      GGCTGTAGTTGTGATCAACTCCGCGAACCCATGCTTCAGTCAGCTGATGCACAATCGTTT    13462
gb:MT020881|Organism:Severe      GGCTGTAGTTGTGATCAACTCCGCGAACCCATGCTTCAGTCAGCTGATGCACAATCGTTT    13462
gb:MT020880|Organism:Severe      GGCTGTAGTTGTGATCAACTCCGCGAACCCATGCTTCAGTCAGCTGATGCACAATCGTTT    13462
gb:MT066175|Organism:Severe      GGCTGTAGTTGTGATCAACTCCGCGAACCCATGCTTCAGTCAGCTGATGCACAATCGTTT    13462
gb:MN997409|Organism:Severe      GGCTGTAGTTGTGATCAACTCCGCGAACCCATGCTTCAGTCAGCTGATGCACAATCGTTT    13462
gb:MN938384|Organism:Severe      GGCTGTAGTTGTGATCAACTCCGCGAACCCATGCTTCAGTCAGCTGATGCACAATCGTTT    13430
gb:MT044258|Organism:Severe      GGCTGTAGTTGTGATCAACTCCGCGAACCCATGCTTCAGTCAGCTGATGCACAATCGTTT    13438
gb:MT039890|Organism:Severe      GGCTGTAGTTGTGATCAACTCCGCGAACCCATGCTTCAGTCAGCTGATGCACAATCGTTT    13462
gb:MN988713|Organism:Severe      GGCTGTAGTTGTGATCAACTCCGCGAACCCATGCTTCAGTCAGCTGATGCACAATCGTTT    13462
gb:LC521925|Organism:Severe      GGCTGTAGTTGTGATCAACTCCGCGAACCCATGCTTCAGTCAGCTGATGCACAATCGTTT    13435
gb:MT093571|Organism:Severe      GGCTGTAGTTGTGATCAACTCCGCGAACCCATGCTTCAGTCAGCTGATGCACAATCGTTT    13462
gb:MT039887|Organism:Severe      GGCTGTAGTTGTGATCAACTCCGCGAACCCATGCTTCAGTCAGCTGATGCACAATCGTTT    13462
gb:MT019530|Organism:Severe      GGCTGTAGTTGTGATCAACTCCGCGAACCCATGCTTCAGTCAGCTGATGCACAATCGTTT    13462
gb:MT039888|Organism:Severe      GGCTGTAGTTGTGATCAACTCCGCGAACCCATGCTTCAGTCAGCTGATGCACAATCGTTT    13462
gb:LC522972|Organism:Severe      GGCTGTAGTTGTGATCAACTCCGCGAACCCATGCTTCAGTCAGCTGATGCACAATCGTTT    13459
gb:MT027063|Organism:Severe      GGCTGTAGTTGTGATCAACTCCGCGAACCCATGCTTCAGTCAGCTGATGCACAATCGTTT    13462
```

```
gb:MT027062|Organism:Severe      GGCTGTAGTTGTGATCAACTCCGCGAACCCATGCTTCAGTCAGCTGATGCACAATCGTTT   13462
gb:MT019529|Organism:Severe      GGCTGTAGTTGTGATCAACTCCGCGAACCCATGCTTCAGTCAGCTGATGCACAATCGTTT   13462
gb:MN996529|Organism:Severe      GGCTGTAGTTGTGATCAACTCCGCGAACCCATGCTTCAGTCAGCTGATGCACAATCGTTT   13450
gb:MN996531|Organism:Severe      GGCTGTAGTTGTGATCAACTCCGCGAACCCATGCTTCAGTCAGCTGATGCACAATCGTTT   13449
gb:MT066176|Organism:Severe      GGCTGTAGTTGTGATCAACTCCGCGAACCCATGCTTCAGTCAGCTGATGCACAATCGTTT   13462
gb:MT027064|Organism:Severe      GGCTGTAGTTGTGATCAACTCCGCGAACCCATGCTTCAGTCAGCTGATGCACAATCGTTT   13462
gb:MN994468|Organism:Severe      GGCTGTAGTTGTGATCAACTCCGCGAACCCATGCTTCAGTCAGCTGATGCACAATCGTTT   13462
gb:MT072688|Organism:Severe      GGCTGTAGTTGTGATCAACTCCGCGAACCCATGCTTCAGTCAGCTGATGCACAATCGTTT   13447
gb:MN996527|Organism:Severe      GGCTGTAGTTGTGATCAACTCCGCGAACCCATGCTTCAGTCAGCTGATGCACAATCGTTT   13429
gb:MT093631|Organism:Severe      GGCTGTAGTTGTGATCAACTCCGCGAACCCATGCTTCAGTCAGCTGATGCACAATCGTTT   13500
gb:MT106053|Organism:Severe      GGCTGTAGTTGTGATCAACTCCGCGAACCCATGCTTCAGTCAGCTGATGCACAATCGTTT   13462
gb:MT019533|Organism:Severe      GGCTGTAGTTGTGATCAACTCCGCGAACCCATGCTTCAGTCAGCTGATGCACAATCGTTT   13462
gb:MT019531|Organism:Severe      GGCTGTAGTTGTGATCAACTCCGCGAACCCATGCTTCAGTCAGCTGATGCACAATCGTTT   13462
gb:MN996528|Organism:Severe      GGCTGTAGTTGTGATCAACTCCGCGAACCCATGCTTCAGTCAGCTGATGCACAATCGTTT   13462
gb:MN996530|Organism:Severe      GGCTGTAGTTGTGATCAACTCCGCGAACCCATGCTTCAGTCAGCTGATGCACAATCGTTT   13448
gb:MN908947|Organism:Severe      GGCTGTAGTTGTGATCAACTCCGCGAACCCATGCTTCAGTCAGCTGATGCACAATCGTTT   13462
gb:MT019532|Organism:Severe      GGCTGTAGTTGTGATCAACTCCGCGAACCCATGCTTCAGTCAGCTGATGCACAATCGTTT   13462
                                 ************************************************************

gb:MT020781|Organism:Severe      TTAAACGGGTTTGCGGTGTAAGTGCAGCCCGTCTTACACCGTGCGGCACAGGCACTAGTA   13510
gb:MT007544|Organism:Severe      TTAAACGGGTTTGCGGTGTAAGTGCAGCCCGTCTTACACCGTGCGGCACAGGCACTAGTA   13522
gb:MN994467|Organism:Severe      TTAAACGGGTTTGCGGTGTAAGTGCAGCCCGTCTTACACCGTGCGGCACAGGCACTAGTA   13522
gb:MT044257|Organism:Severe      TTAAACGGGTTTGCGGTGTAAGTGCAGCCCGTCTTACACCGTGCGGCACAGGCACTAGTA   13522
gb:MT106054|Organism:Severe      TTAAACGGGTTTGCGGTGTAAGTGCAGCCCGTCTTACACCGTGCGGCACAGGCACTAGTA   13522
gb:MT049951|Organism:Severe      TTAAACGGGTTTGCGGTGTAAGTGCAGCCCGTCTTACACCGTGCGGCACAGGCACTAGTA   13522
gb:MN975262|Organism:Severe      TTAAACGGGTTTGCGGTGTAAGTGCAGCCCGTCTTACACCGTGCGGCACAGGCACTAGTA   13522
gb:MT106052|Organism:Severe      TTAAACGGGTTTGCGGTGTAAGTGCAGCCCGTCTTACACCGTGCGGCACAGGCACTAGTA   13522
gb:LC522975|Organism:Severe      TTAAACGGGTTTGCGGTGTAAGTGCAGCCCGTCTTACACCGTGCGGCACAGGCACTAGTA   13519
gb:LC522973|Organism:Severe      TTAAACGGGTTTGCGGTGTAAGTGCAGCCCGTCTTACACCGTGCGGCACAGGCACTAGTA   13519
gb:LC522974|Organism:Severe      TTAAACGGGTTTGCGGTGTAAGTGCAGCCCGTCTTACACCGTGCGGCACAGGCACTAGTA   13519
gb:MN985325|Organism:Severe      TTAAACGGGTTTGCGGTGTAAGTGCAGCCCGTCTTACACCGTGCGGCACAGGCACTAGTA   13522
gb:MT020881|Organism:Severe      TTAAACGGGTTTGCGGTGTAAGTGCAGCCCGTCTTACACCGTGCGGCACAGGCACTAGTA   13522
gb:MT020880|Organism:Severe      TTAAACGGGTTTGCGGTGTAAGTGCAGCCCGTCTTACACCGTGCGGCACAGGCACTAGTA   13522
gb:MT066175|Organism:Severe      TTAAACGGGTTTGCGGTGTAAGTGCAGCCCGTCTTACACCGTGCGGCACAGGCACTAGTA   13522
gb:MN997409|Organism:Severe      TTAAACGGGTTTGCGGTGTAAGTGCAGCCCGTCTTACACCGTGCGGCACAGGCACTAGTA   13522
gb:MN938384|Organism:Severe      TTAAACGGGTTTGCGGTGTAAGTGCAGCCCGTCTTACACCGTGCGGCACAGGCACTAGTA   13490
gb:MT044258|Organism:Severe      TTAAACGGGTTTGCGGTGTAAGTGCAGCCCGTCTTACACCGTGCGGCACAGGCACTAGTA   13498
gb:MT039890|Organism:Severe      TTAAACGGGTTTGCGGTGTAAGTGCAGCCCGTCTTACACCGTGCGGCACAGGCACTAGTA   13522
gb:MN988713|Organism:Severe      TTAAACGGGTTTGCGGTGTAAGTGCAGCCCGTCTTACACCGTGCGGCACAGGCACTAGTA   13522
gb:LC521925|Organism:Severe      TTAAACGGGTTTGCGGTGTAAGTGCAGCCCGTCTTACACCGTGCGGCACAGGCACTAGTA   13495
gb:MT093571|Organism:Severe      TTAAACGGGTTTGCGGTGTAAGTGCAGCCCGTCTTACACCGTGCGGCACAGGCACTAGTA   13522
gb:MT039887|Organism:Severe      TTAAACGGGTTTGCGGTGTAAGTGCAGCCCGTCTTACACCGTGCGGCACAGGCACTAGTA   13522
gb:MT019530|Organism:Severe      TTAAACGGGTTTGCGGTGTAAGTGCAGCCCGTCTTACACCGTGCGGCACAGGCACTAGTA   13522
gb:MT039888|Organism:Severe      TTAAACGGGTTTGCGGTGTAAGTGCAGCCCGTCTTACACCGTGCGGCACAGGCACTAGTA   13522
gb:LC522972|Organism:Severe      TTAAACGGGTTTGCGGTGTAAGTGCAGCCCGTCTTACACCGTGCGGCACAGGCACTAGTA   13519
gb:MT027063|Organism:Severe      TTAAACGGGTTTGCGGTGTAAGTGCAGCCCGTCTTACACCGTGCGGCACAGGCACTAGTA   13522
gb:MT027062|Organism:Severe      TTAAACGGGTTTGCGGTGTAAGTGCAGCCCGTCTTACACCGTGCGGCACAGGCACTAGTA   13522
gb:MT019529|Organism:Severe      TTAAACGGGTTTGCGGTGTAAGTGCAGCCCGTCTTACACCGTGCGGCACAGGCACTAGTA   13522
gb:MN996529|Organism:Severe      TTAAACGGGTTTGCGGTGTAAGTGCAGCCCGTCTTACACCGTGCGGCACAGGCACTAGTA   13510
gb:MN996531|Organism:Severe      TTAAACGGGTTTGCGGTGTAAGTGCAGCCCGTCTTACACCGTGCGGCACAGGCACTAGTA   13509
gb:MT066176|Organism:Severe      TTAAACGGGTTTGCGGTGTAAGTGCAGCCCGTCTTACACCGTGCGGCACAGGCACTAGTA   13522
gb:MT027064|Organism:Severe      TTAAACGGGTTTGCGGTGTAAGTGCAGCCCGTCTTACACCGTGCGGCACAGGCACTAGTA   13522
gb:MN994468|Organism:Severe      TTAAACGGGTTTGCGGTGTAAGTGCAGCCCGTCTTACACCGTGCGGCACAGGCACTAGTA   13522
gb:MT072688|Organism:Severe      TTAAACGGGTTTGCGGTGTAAGTGCAGCCCGTCTTACACCGTGCGGCACAGGCACTAGTA   13507
gb:MN996527|Organism:Severe      TTAAACGGGTTTGCGGTGTAAGTGCAGCCCGTCTTACACCGTGCGGCACAGGCACTAGTA   13489
gb:MT093631|Organism:Severe      TTAAACGGGTTTGCGGTGTAAGTGCAGCCCGTCTTACACCGTGCGGCACAGGCACTAGTA   13560
gb:MT106053|Organism:Severe      TTAAACGGGTTTGCGGTGTAAGTGCAGCCCGTCTTACACCGTGCGGCACAGGCACTAGTA   13522
gb:MT019533|Organism:Severe      TTAAACGGGTTTGCGGTGTAAGTGCAGCCCGTCTTACACCGTGCGGCACAGGCACTAGTA   13522
gb:MT019531|Organism:Severe      TTAAACGGGTTTGCGGTGTAAGTGCAGCCCGTCTTACACCGTGCGGCACAGGCACTAGTA   13522
gb:MN996528|Organism:Severe      TTAAACGGGTTTGCGGTGTAAGTGCAGCCCGTCTTACACCGTGCGGCACAGGCACTAGTA   13522
gb:MN996530|Organism:Severe      TTAAACGGGTTTGCGGTGTAAGTGCAGCCCGTCTTACACCGTGCGGCACAGGCACTAGTA   13508
gb:MN908947|Organism:Severe      TTAAACGGGTTTGCGGTGTAAGTGCAGCCCGTCTTACACCGTGCGGCACAGGCACTAGTA   13522
gb:MT019532|Organism:Severe      TTAAACGGGTTTGCGGTGTAAGTGCAGCCCGTCTTACACCGTGCGGCACAGGCACTAGTA   13522
                                 ************************************************************
```

```
gb:MT020781|Organism:Severe      CTGATGTCGTATACAGGGCTTTTGACATCTACAATGATAAAGTAGCTGGTTTTGCTAAAT   13570
gb:MT007544|Organism:Severe      CTGATGTCGTATACAGGGCTTTTGACATCTACAATGATAAAGTAGCTGGTTTTGCTAAAT   13582
gb:MN994467|Organism:Severe      CTGATGTCGTATACAGGGCTTTTGACATCTACAATGATAAAGTAGCTGGTTTTGCTAAAT   13582
gb:MT044257|Organism:Severe      CTGATGTCGTATACAGGGCTTTTGACATCTACAATGATAAAGTAGCTGGTTTTGCTAAAT   13582
gb:MT106054|Organism:Severe      CTGATGTCGTATACAGGGCTTTTGACATCTACAATGATAAAGTAGCTGGTTTTGCTAAAT   13582
gb:MT049951|Organism:Severe      CTGATGTCGTATACAGGGCTTTTGACATCTACAATGATAAAGTAGCTGGTTTTGCTAAAT   13582
gb:MN975262|Organism:Severe      CTGATGTCGTATACAGGGCTTTTGACATCTACAATGATAAAGTAGCTGGTTTTGCTAAAT   13582
gb:MT106052|Organism:Severe      CTGATGTCGTATACAGGGCTTTTGACATCTACAATGATAAAGTAGCTGGTTTTGCTAAAT   13582
gb:LC522975|Organism:Severe      CTGATGTCGTATACAGGGCTTTTGACATCTACAATGATAAAGTAGCTGGTTTTGCTAAAT   13579
gb:LC522973|Organism:Severe      CTGATGTCGTATACAGGGCTTTTGACATCTACAATGATAAAGTAGCTGGTTTTGCTAAAT   13579
gb:LC522974|Organism:Severe      CTGATGTCGTATACAGGGCTTTTGACATCTACAATGATAAAGTAGCTGGTTTTGCTAAAT   13579
gb:MN985325|Organism:Severe      CTGATGTCGTATACAGGGCTTTTGACATCTACAATGATAAAGTAGCTGGTTTTGCTAAAT   13582
gb:MT020881|Organism:Severe      CTGATGTCGTATACAGGGCTTTTGACATCTACAATGATAAAGTAGCTGGTTTTGCTAAAT   13582
gb:MT020880|Organism:Severe      CTGATGTCGTATACAGGGCTTTTGACATCTACAATGATAAAGTAGCTGGTTTTGCTAAAT   13582
gb:MT066175|Organism:Severe      CTGATGTCGTATACAGGGCTTTTGACATCTACAATGATAAAGTAGCTGGTTTTGCTAAAT   13582
gb:MN997409|Organism:Severe      CTGATGTCGTATACAGGGCTTTTGACATCTACAATGATAAAGTAGCTGGTTTTGCTAAAT   13582
gb:MN938384|Organism:Severe      CTGATGTCGTATACAGGGCTTTTGACATCTACAATGATAAAGTAGCTGGTTTTGCTAAAT   13550
gb:MT044258|Organism:Severe      CTGATGTCGTATACAGGGCTTTTGACATCTACAATGATAAAGTAGCTGGTTTTGCTAAAT   13558
gb:MT039890|Organism:Severe      CTGATGTCGTATACAGGGCTTTTGACATCTACAATGATAAAGTAGCTGGTTTTGCTAAAT   13582
gb:MN988713|Organism:Severe      CTGATGTCGTATACAGGGCTTTTGACATCTACAATGATAAAGTAGCTGGTTTTGCTAAAT   13582
gb:LC521925|Organism:Severe      CTGATGTCGTATACAGGGCTTTTGACATCTACAATGATAAAGTAGCTGGTTTTGCTAAAT   13555
gb:MT093571|Organism:Severe      CTGATGTCGTATACAGGGCTTTTGACATCTACAATGATAAAGTAGCTGGTTTTGCTAAAT   13582
gb:MT039887|Organism:Severe      CTGATGTCGTATACAGGGCTTTTGACATCTACAATGATAAAGTAGCTGGTTTTGCTAAAT   13582
gb:MT019530|Organism:Severe      CTGATGTCGTATACAGGGCTTTTGACATCTACAATGATAAAGTAGCTGGTTTTGCTAAAT   13582
gb:MT039888|Organism:Severe      CTGATGTCGTATACAGGGCTTTTGACATCTACAATGATAAAGTAGCTGGTTTTGCTAAAT   13582
gb:LC522972|Organism:Severe      CTGATGTCGTATACAGGGCTTTTGACATCTACAATGATAAAGTAGCTGGTTTTGCTAAAT   13579
gb:MT027063|Organism:Severe      CTGATGTCGTATACAGGGCTTTTGACATCTACAATGATAAAGTAGCTGGTTTTGCTAAAT   13582
gb:MT027062|Organism:Severe      CTGATGTCGTATACAGGGCTTTTGACATCTACAATGATAAAGTAGCTGGTTTTGCTAAAT   13582
gb:MT019529|Organism:Severe      CTGATGTCGTATACAGGGCTTTTGACATCTACAATGATAAAGTAGCTGGTTTTGCTAAAT   13582
gb:MN996529|Organism:Severe      CTGATGTCGTATACAGGGCTTTTGACATCTACAATGATAAAGTAGCTGGTTTTGCTAAAT   13570
gb:MN996531|Organism:Severe      CTGATGTCGTATACAGGGCTTTTGACATCTACAATGATAAAGTAGCTGGTTTTGCTAAAT   13569
gb:MT066176|Organism:Severe      CTGATGTCGTATACAGGGCTTTTGACATCTACAATGATAAAGTAGCTGGTTTTGCTAAAT   13582
gb:MT027064|Organism:Severe      CTGATGTCGTATACAGGGCTTTTGACATCTACAATGATAAAGTAGCTGGTTTTGCTAAAT   13582
gb:MN994468|Organism:Severe      CTGATGTCGTATACAGGGCTTTTGACATCTACAATGATAAAGTAGCTGGTTTTGCTAAAT   13582
gb:MT072688|Organism:Severe      CTGATGTCGTATACAGGGCTTTTGACATCTACAATGATAAAGTAGCTGGTTTTGCTAAAT   13567
gb:MN996527|Organism:Severe      CTGATGTCGTATACAGGGCTTTTGACATCTACAATGATAAAGTAGCTGGTTTTGCTAAAT   13549
gb:MT093631|Organism:Severe      CTGATGTCGTATACAGGGCTTTTGACATCTACAATGATAAAGTAGCTGGTTTTGCTAAAT   13620
gb:MT106053|Organism:Severe      CTGATGTCGTATACAGGGCTTTTGACATCTACAATGATAAAGTAGCTGGTTTTGCTAAAT   13582
gb:MT019533|Organism:Severe      CTGATGTCGTATACAGGGCTTTTGACATCTACAATGATAAAGTAGCTGGTTTTGCTAAAT   13582
gb:MT019531|Organism:Severe      CTGATGTCGTATACAGGGCTTTTGACATCTACAATGATAAAGTAGCTGGTTTTGCTAAAT   13582
gb:MN996528|Organism:Severe      CTGATGTCGTATACAGGGCTTTTGACATCTACAATGATAAAGTAGCTGGTTTTGCTAAAT   13582
gb:MN996530|Organism:Severe      CTGATGTCGTATACAGGGCTTTTGACATCTACAATGATAAAGTAGCTGGTTTTGCTAAAT   13568
gb:MN908947|Organism:Severe      CTGATGTCGTATACAGGGCTTTTGACATCTACAATGATAAAGTAGCTGGTTTTGCTAAAT   13582
gb:MT019532|Organism:Severe      CTGATGTCGTATACAGGGCTTTTGACATCTACAATGATAAAGTAGCTGGTTTTGCTAAAT   13582
                                 ************************************************************

gb:MT020781|Organism:Severe      TCCTAAAAACTAATTGTTGTCGCTTCCAAGAAAAGGACGAAGATGACAATTTAATTGATT   13630
gb:MT007544|Organism:Severe      TCCTAAAAACTAATTGTTGTCGCTTCCAAGAAAAGGACGAAGATGACAATTTAATTGATT   13642
gb:MN994467|Organism:Severe      TCCTAAAAACTAATTGTTGTCGCTTCCAAGAAAAGGACGAAGATGACAATTTAATTGATT   13642
gb:MT044257|Organism:Severe      TCCTAAAAACTAATTGTTGTCGCTTCCAAGAAAAGGACGAAGATGACAATTTAATTGATT   13642
gb:MT106054|Organism:Severe      TCCTAAAAACTAATTGTTGTCGCTTCCAAGAAAAGGACGAAGATGACAATTTAATTGATT   13642
gb:MT049951|Organism:Severe      TCCTAAAAACTAATTGTTGTCGCTTCCAAGAAAAGGACGAAGATGACAATTTAATTGATT   13642
gb:MN975262|Organism:Severe      TCCTAAAAACTAATTGTTGTCGCTTCCAAGAAAAGGACGAAGATGACAATTTAATTGATT   13642
gb:MT106052|Organism:Severe      TCCTAAAAACTAATTGTTGTCGCTTCCAAGAAAAGGACGAAGATGACAATTTAATTGATT   13642
gb:LC522975|Organism:Severe      TCCTAAAAACTAATTGTTGTCGCTTCCAAGAAAAGGACGAAGATGACAATTTAATTGATT   13639
gb:LC522973|Organism:Severe      TCCTAAAAACTAATTGTTGTCGCTTCCAAGAAAAGGACGAAGATGACAATTTAATTGATT   13639
gb:LC522974|Organism:Severe      TCCTAAAAACTAATTGTTGTCGCTTCCAAGAAAAGGACGAAGATGACAATTTAATTGATT   13639
gb:MN985325|Organism:Severe      TCCTAAAAACTAATTGTTGTCGCTTCCAAGAAAAGGACGAAGATGACAATTTAATTGATT   13642
gb:MT020881|Organism:Severe      TCCTAAAAACTAATTGTTGTCGCTTCCAAGAAAAGGACGAAGATGACAATTTAATTGATT   13642
gb:MT020880|Organism:Severe      TCCTAAAAACTAATTGTTGTCGCTTCCAAGAAAAGGACGAAGATGACAATTTAATTGATT   13642
gb:MT066175|Organism:Severe      TCCTAAAAACTAATTGTTGTCGCTTCCAAGAAAAGGACGAAGATGACAATTTAATTGATT   13642
gb:MN997409|Organism:Severe      TCCTAAAAACTAATTGTTGTCGCTTCCAAGAAAAGGACGAAGATGACAATTTAATTGATT   13642
gb:MN938384|Organism:Severe      TCCTAAAAACTAATTGTTGTCGCTTCCAAGAAAAGGACGAAGATGACAATTTAATTGATT   13610
gb:MT044258|Organism:Severe      TCCTAAAAACTAATTGTTGTCGCTTCCAAGAAAAGGACGAAGATGACAATTTAATTGATT   13618
gb:MT039890|Organism:Severe      TCCTAAAAACTAATTGTTGTCGCTTCCAAGAAAAGGACGAAGATGACAATTTAATTGATT   13642
```

```
gb:MN988713|Organism:Severe      TCCTAAAAACTAATTGTTGTCGCTTCCAAGAAAAGGACGAAGATGACAATTTAATTGATT   13642
gb:LC521925|Organism:Severe      TCCTAAAAACTAATTGTTGTCGCTTCCAAGAAAAGGACGAAGATGACAATTTAATTGATT   13615
gb:MT093571|Organism:Severe      TCCTAAAAACTAATTGTTGTCGCTTCCAAGAAAAGGACGAAGATGACAATTTAATTGATT   13642
gb:MT039887|Organism:Severe      TCCTAAAAACTAATTGTTGTCGCTTCCAAGAAAAGGACGAAGATGACAATTTAATTGATT   13642
gb:MT019530|Organism:Severe      TCCTAAAAACTAATTGTTGTCGCTTCCAAGAAAAGGACGAAGATGACAATTTAATTGATT   13642
gb:MT039888|Organism:Severe      TCCTAAAAACTAATTGTTGTCGCTTCCAAGAAAAGGACGAAGATGACAATTTAATTGATT   13642
gb:LC522972|Organism:Severe      TCCTAAAAACTAATTGTTGTCGCTTCCAAGAAAAGGACGAAGATGACAATTTAATTGATT   13639
gb:MT027063|Organism:Severe      TCCTAAAAACTAATTGTTGTCGCTTCCAAGAAAAGGACGAAGATGACAATTTAATTGATT   13642
gb:MT027062|Organism:Severe      TCCTAAAAACTAATTGTTGTCGCTTCCAAGAAAAGGACGAAGATGACAATTTAATTGATT   13642
gb:MT019529|Organism:Severe      TCCTAAAAACTAATTGTTGTCGCTTCCAAGAAAAGGACGAAGATGACAATTTAATTGATT   13642
gb:MN996529|Organism:Severe      TCCTAAAAACTAATTGTTGTCGCTTCCAAGAAAAGGACGAAGATGACAATTTAATTGATT   13630
gb:MN996531|Organism:Severe      TCCTAAAAACTAATTGTTGTCGCTTCCAAGAAAAGGACGAAGATGACAATTTAATTGATT   13629
gb:MT066176|Organism:Severe      TCCTAAAAACTAATTGTTGTCGCTTCCAAGAAAAGGACGAAGATGACAATTTAATTGATT   13642
gb:MT027064|Organism:Severe      TCCTAAAAACTAATTGTTGTCGCTTCCAAGAAAAGGACGAAGATGACAATTTAATTGATT   13642
gb:MN994468|Organism:Severe      TCCTAAAAACTAATTGTTGTCGCTTCCAAGAAAAGGACGAAGATGACAATTTAATTGATT   13642
gb:MT072688|Organism:Severe      TCCTAAAAACTAATTGTTGTCGCTTCCAAGAAAAGGACGAAGATGACAATTTAATTGATT   13627
gb:MN996527|Organism:Severe      TCCTAAAAACTAATTGTTGTCGCTTCCAAGAAAAGGACGAAGATGACAATTTAATTGATT   13609
gb:MT093631|Organism:Severe      TCCTAAAAACTAATTGTTGTCGCTTCCAAGAAAAGGACGAAGATGACAATTTAATTGATT   13680
gb:MT106053|Organism:Severe      TCCTAAAAACTAATTGTTGTCGCTTCCAAGAAAAGGACGAAGATGACAATTTAATTGATT   13642
gb:MT019533|Organism:Severe      TCCTAAAAACTAATTGTTGTCGCTTCCAAGAAAAGGACGAAGATGACAATTTAATTGATT   13642
gb:MT019531|Organism:Severe      TCCTAAAAACTAATTGTTGTCGCTTCCAAGAAAAGGACGAAGATGACAATTTAATTGATT   13642
gb:MN996528|Organism:Severe      TCCTAAAAACTAATTGTTGTCGCTTCCAAGAAAAGGACGAAGATGACAATTTAATTGATT   13642
gb:MN996530|Organism:Severe      TCCTAAAAACTAATTGTTGTCGCTTCCAAGAAAAGGACGAAGATGACAATTTAATTGATT   13628
gb:MN908947|Organism:Severe      TCCTAAAAACTAATTGTTGTCGCTTCCAAGAAAAGGACGAAGATGACAATTTAATTGATT   13642
gb:MT019532|Organism:Severe      TCCTAAAAACTAATTGTTGTCGCTTCCAAGAAAAGGACGAAGATGACAATTTAATTGATT   13642
                                 ************************************************************

gb:MT020781|Organism:Severe      CTTACTTTGTAGTTAAGAGACACACTTTCTCTAACTACCAACATGAAGAAACAATTTATA   13690
gb:MT007544|Organism:Severe      CTTACTTTGTAGTTAAGAGACACACTTTCTCTAACTACCAACATGAAGAAACAATTTATA   13702
gb:MN994467|Organism:Severe      CTTACTTTGTAGTTAAGAGACACACTTTCTCTAACTACCAACATGAAGAAACAATTTATA   13702
gb:MT044257|Organism:Severe      CTTACTTTGTAGTTAAGAGACACACTTTCTCTAACTACCAACATGAAGAAACAATTTATA   13702
gb:MT106054|Organism:Severe      CTTACTTTGTAGTTAAGAGACACACTTTCTCTAACTACCAACATGAAGAAACAATTTATA   13702
gb:MT049951|Organism:Severe      CTTACTTTGTAGTTAAGAGACACACTTTCTCTAACTACCAACATGAAGAAACAATTTATA   13702
gb:MN975262|Organism:Severe      CTTACTTTGTAGTTAAGAGACACACTTTCTCTAACTACCAACATGAAGAAACAATTTATA   13702
gb:MT106052|Organism:Severe      CTTACTTTGTAGTTAAGAGACACACTTTCTCTAACTACCAACATGAAGAAACAATTTATA   13702
gb:LC522975|Organism:Severe      CTTACTTTGTAGTTAAGAGACACACTTTCTCTAACTACCAACATGAAGAAACAATTTATA   13699
gb:LC522973|Organism:Severe      CTTACTTTGTAGTTAAGAGACACACTTTCTCTAACTACCAACATGAAGAAACAATTTATA   13699
gb:LC522974|Organism:Severe      CTTACTTTGTAGTTAAGAGACACACTTTCTCTAACTACCAACATGAAGAAACAATTTATA   13699
gb:MN985325|Organism:Severe      CTTACTTTGTAGTTAAGAGACACACTTTCTCTAACTACCAACATGAAGAAACAATTTATA   13702
gb:MT020881|Organism:Severe      CTTACTTTGTAGTTAAGAGACACACTTTCTCTAACTACCAACATGAAGAAACAATTTATA   13702
gb:MT020880|Organism:Severe      CTTACTTTGTAGTTAAGAGACACACTTTCTCTAACTACCAACATGAAGAAACAATTTATA   13702
gb:MT066175|Organism:Severe      CTTACTTTGTAGTTAAGAGACACACTTTCTCTAACTACCAACATGAAGAAACAATTTATA   13702
gb:MN997409|Organism:Severe      CTTACTTTGTAGTTAAGAGACACACTTTCTCTAACTACCAACATGAAGAAACAATTTATA   13702
gb:MN938384|Organism:Severe      CTTACTTTGTAGTTAAGAGACACACTTTCTCTAACTACCAACATGAAGAAACAATTTATA   13670
gb:MT044258|Organism:Severe      CTTACTTTGTAGTTAAGAGACACACTTTCTCTAACTACCAACATGAAGAAACAATTTATA   13678
gb:MT039890|Organism:Severe      CTTACTTTGTAGTTAAGAGACACACTTTCTCTAACTACCAACATGAAGAAACAATTTATA   13702
gb:MN988713|Organism:Severe      CTTACTTTGTAGTTAAGAGACACACTTTCTCTAACTACCAACATGAAGAAACAATTTATA   13702
gb:LC521925|Organism:Severe      CTTACTTTGTAGTTAAGAGACACACTTTCTCTAACTACCAACATGAAGAAACAATTTATA   13675
gb:MT093571|Organism:Severe      CTTACTTTGTAGTTAAGAGACACACTTTCTCTAACTACCAACATGAAGAAACAATTTATA   13702
gb:MT039887|Organism:Severe      CTTACTTTGTAGTTAAGAGACACACTTTCTCTAACTACCAACATGAAGAAACAATTTATA   13702
gb:MT019530|Organism:Severe      CTTACTTTGTAGTTAAGAGACACACTTTCTCTAACTACCAACATGAAGAAACAATTTATA   13702
gb:MT039888|Organism:Severe      CTTACTTTGTAGTTAAGAGACACACTTTCTCTAACTACCAACATGAAGAAACAATTTATA   13702
gb:LC522972|Organism:Severe      CTTACTTTGTAGTTAAGAGACACACTTTCTCTAACTACCAACATGAAGAAACAATTTATA   13699
gb:MT027063|Organism:Severe      CTTACTTTGTAGTTAAGAGACACACTTTCTCTAACTACCAACATGAAGAAACAATTTATA   13702
gb:MT027062|Organism:Severe      CTTACTTTGTAGTTAAGAGACACACTTTCTCTAACTACCAACATGAAGAAACAATTTATA   13702
gb:MT019529|Organism:Severe      CTTACTTTGTAGTTAAGAGACACACTTTCTCTAACTACCAACATGAAGAAACAATTTATA   13702
gb:MN996529|Organism:Severe      CTTACTTTGTAGTTAAGAGACACACTTTCTCTAACTACCAACATGAAGAAACAATTTATA   13690
gb:MN996531|Organism:Severe      CTTACTTTGTAGTTAAGAGACACACTTTCTCTAACTACCAACATGAAGAAACAATTTATA   13689
gb:MT066176|Organism:Severe      CTTACTTTGTAGTTAAGAGACACACTTTCTCTAACTACCAACATGAAGAAACAATTTATA   13702
gb:MT027064|Organism:Severe      CTTACTTTGTAGTTAAGAGACACACTTTCTCTAACTACCAACATGAAGAAACAATTTATA   13702
gb:MN994468|Organism:Severe      CTTACTTTGTAGTTAAGAGACACACTTTCTCTAACTACCAACATGAAGAAACAATTTATA   13702
gb:MT072688|Organism:Severe      CTTACTTTGTAGTTAAGAGACACACTTTCTCTAACTACCAACATGAAGAAACAATTTATA   13687
gb:MN996527|Organism:Severe      CTTACTTTGTAGTTAAGAGACACACTTTCTCTAACTACCAACATGAAGAAACAATTTATA   13669
gb:MT093631|Organism:Severe      CTTACTTTGTAGTTAAGAGACACACTTTCTCTAACTACCAACATGAAGAAACAATTTATA   13740
gb:MT106053|Organism:Severe      CTTACTTTGTAGTTAAGAGACACACTTTCTCTAACTACCAACATGAAGAAACAATTTATA   13702
```

```
gb:MT019533|Organism:Severe      CTTACTTTGTAGTTAAGAGACACACTTTCTCTAACTACCAACATGAAGAAACAATTTATA   13702
gb:MT019531|Organism:Severe      CTTACTTTGTAGTTAAGAGACACACTTTCTCTAACTACCAACATGAAGAAACAATTTATA   13702
gb:MN996528|Organism:Severe      CTTACTTTGTAGTTAAGAGACACACTTTCTCTAACTACCAACATGAAGAAACAATTTATA   13702
gb:MN996530|Organism:Severe      CTTACTTTGTAGTTAAGAGACACACTTTCTCTAACTACCAACATGAAGAAACAATTTATA   13688
gb:MN908947|Organism:Severe      CTTACTTTGTAGTTAAGAGACACACTTTCTCTAACTACCAACATGAAGAAACAATTTATA   13702
gb:MT019532|Organism:Severe      CTTACTTTGTAGTTAAGAGACACACTTTCTCTAACTACCAACATGAAGAAACAATTTATA   13702
                                 ************************************************************

gb:MT020781|Organism:Severe      ATTTACTTAAGGATTGTCCAGCTGTTGCTAAACATGACTTCTTTAAGTTTAGAATAGACG   13750
gb:MT007544|Organism:Severe      ATTTACTTAAGGATTGTCCAGCTGTTGCTAAACATGACTTCTTTAAGTTTAGAATAGACG   13762
gb:MN994467|Organism:Severe      ATTTACTTAAGGATTGTCCAGCTGTTGCTAAACATGACTTCTTTAAGTTTAGAATAGACG   13762
gb:MT044257|Organism:Severe      ATTTACTTAAGGATTGTCCAGCTGTTGCTAAACATGACTTCTTTAAGTTTAGAATAGACG   13762
gb:MT106054|Organism:Severe      ATTTACTTAAGGATTGTCCAGCTGTTGCTAAACATGACTTCTTTAAGTTTAGAATAGACG   13762
gb:MT049951|Organism:Severe      ATTTACTTAAGGATTGTCCAGCTGTTGCTAAACATGACTTCTTTAAGTTTAGAATAGACG   13762
gb:MN975262|Organism:Severe      ATTTACTTAAGGATTGTCCAGCTGTTGCTAAACATGACTTCTTTAAGTTTAGAATAGACG   13762
gb:MT106052|Organism:Severe      ATTTACTTAAGGATTGTCCAGCTGTTGCTAAACATGACTTCTTTAAGTTTAGAATAGACG   13762
gb:LC522975|Organism:Severe      ATTTACTTAAGGATTGTCCAGCTGTTGCTAAACATGACTTCTTTAAGTTTAGAATAGACG   13759
gb:LC522973|Organism:Severe      ATTTACTTAAGGATTGTCCAGCTGTTGCTAAACATGACTTCTTTAAGTTTAGAATAGACG   13759
gb:LC522974|Organism:Severe      ATTTACTTAAGGATTGTCCAGCTGTTGCTAAACATGACTTCTTTAAGTTTAGAATAGACG   13759
gb:MN985325|Organism:Severe      ATTTACTTAAGGATTGTCCAGCTGTTGCTAAACATGACTTCTTTAAGTTTAGAATAGACG   13762
gb:MT020881|Organism:Severe      ATTTACTTAAGGATTGTCCAGCTGTTGCTAAACATGACTTCTTTAAGTTTAGAATAGACG   13762
gb:MT020880|Organism:Severe      ATTTACTTAAGGATTGTCCAGCTGTTGCTAAACATGACTTCTTTAAGTTTAGAATAGACG   13762
gb:MT066175|Organism:Severe      ATTTACTTAAGGATTGTCCAGCTGTTGCTAAACATGACTTCTTTAAGTTTAGAATAGACG   13762
gb:MN997409|Organism:Severe      ATTTACTTAAGGATTGTCCAGCTGTTGCTAAACATGACTTCTTTAAGTTTAGAATAGACG   13762
gb:MN938384|Organism:Severe      ATTTACTTAAGGATTGTCCAGCTGTTGCTAAACATGACTTCTTTAAGTTTAGAATAGACG   13730
gb:MT044258|Organism:Severe      ATTTACTTAAGGATTGTCCAGCTGTTGCTAAACATGACTTCTTTAAGTTTAGAATAGACG   13738
gb:MT039890|Organism:Severe      ATTTACTTAAGGATTGTCCAGCTGTTGCTAAACATGACTTCTTTAAGTTTAGAATAGACG   13762
gb:MN988713|Organism:Severe      ATTTACTTAAGGATTGTCCAGCTGTTGCTAAACATGACTTCTTTAAGTTTAGAATAGACG   13762
gb:LC521925|Organism:Severe      ATTTACTTAAGGATTGTCCAGCTGTTGCTAAACATGACTTCTTTAAGTTTAGAATAGACG   13735
gb:MT093571|Organism:Severe      ATTTACTTAAGGATTGTCCAGCTGTTGCTAAACATGACTTCTTTAAGTTTAGAATAGACG   13762
gb:MT039887|Organism:Severe      ATTTACTTAAGGATTGTCCAGCTGTTGCTAAACATGACTTCTTTAAGTTTAGAATAGACG   13762
gb:MT019530|Organism:Severe      ATTTACTTAAGGATTGTCCAGCTGTTGCTAAACATGACTTCTTTAAGTTTAGAATAGACG   13762
gb:MT039888|Organism:Severe      ATTTACTTAAGGATTGTCCAGCTGTTGCTAAACATGACTTCTTTAAGTTTAGAATAGACG   13762
gb:LC522972|Organism:Severe      ATTTACTTAAGGATTGTCCAGCTGTTGCTAAACATGACTTCTTTAAGTTTAGAATAGACG   13759
gb:MT027063|Organism:Severe      ATTTACTTAAGGATTGTCCAGCTGTTGCTAAACATGACTTCTTTAAGTTTAGAATAGACG   13762
gb:MT027062|Organism:Severe      ATTTACTTAAGGATTGTCCAGCTGTTGCTAAACATGACTTCTTTAAGTTTAGAATAGACG   13762
gb:MT019529|Organism:Severe      ATTTACTTAAGGATTGTCCAGCTGTTGCTAAACATGACTTCTTTAAGTTTAGAATAGACG   13762
gb:MN996529|Organism:Severe      ATTTACTTAAGGATTGTCCAGCTGTTGCTAAACATGACTTCTTTAAGTTTAGAATAGACG   13750
gb:MN996531|Organism:Severe      ATTTACTTAAGGATTGTCCAGCTGTTGCTAAACATGACTTCTTTAAGTTTAGAATAGACG   13749
gb:MT066176|Organism:Severe      ATTTACTTAAGGATTGTCCAGCTGTTGCTAAACATGACTTCTTTAAGTTTAGAATAGACG   13762
gb:MT027064|Organism:Severe      ATTTACTTAAGGATTGTCCAGCTGTTGCTAAACATGACTTCTTTAAGTTTAGAATAGACG   13762
gb:MN994468|Organism:Severe      ATTTACTTAAGGATTGTCCAGCTGTTGCTAAACATGACTTCTTTAAGTTTAGAATAGACG   13762
gb:MT072688|Organism:Severe      ATTTACTTAAGGATTGTCCAGCTGTTGCTAAACATGACTTCTTTAAGTTTAGAATAGACG   13747
gb:MN996527|Organism:Severe      ATTTACTTAAGGATTGTCCAGCTGTTGCTAAACATGACTTCTTTAAGTTTAGAATAGACG   13729
gb:MT093631|Organism:Severe      ATTTACTTAAGGATTGTCCAGCTGTTGCTAAACATGACTTCTTTAAGTTTAGAATAGACG   13800
gb:MT106053|Organism:Severe      ATTTACTTAAGGATTGTCCAGCTGTTGCTAAACATGACTTCTTTAAGTTTAGAATAGACG   13762
gb:MT019533|Organism:Severe      ATTTACTTAAGGATTGTCCAGCTGTTGCTAAACATGACTTCTTTAAGTTTAGAATAGACG   13762
gb:MT019531|Organism:Severe      ATTTACTTAAGGATTGTCCAGCTGTTGCTAAACATGACTTCTTTAAGTTTAGAATAGACG   13762
gb:MN996528|Organism:Severe      ATTTACTTAAGGATTGTCCAGCTGTTGCTAAACATGACTTCTTTAAGTTTAGAATAGACG   13762
gb:MN996530|Organism:Severe      ATTTACTTAAGGATTGTCCAGCTGTTGCTAAACATGACTTCTTTAAGTTTAGAATAGACG   13748
gb:MN908947|Organism:Severe      ATTTACTTAAGGATTGTCCAGCTGTTGCTAAACATGACTTCTTTAAGTTTAGAATAGACG   13762
gb:MT019532|Organism:Severe      ATTTACTTAAGGATTGTCCAGCTGTTGCTAAACATGACTTCTTTAAGTTTAGAATAGACG   13762
                                 ************************************************************

gb:MT020781|Organism:Severe      GTGACATGGTACCACATATATCACGTCAACGTCTTACTAAATACACAATGGCAGACCTCG   13810
gb:MT007544|Organism:Severe      GTGACATGGTACCACATATATCACGTCAACGTCTTACTAAATACACAATGGCAGACCTCG   13822
gb:MN994467|Organism:Severe      GTGACATGGTACCACATATATCACGTCAACGTCTTACTAAATACACAATGGCAGACCTCG   13822
gb:MT044257|Organism:Severe      GTGACATGGTACCACATATATCACGTCAACGTCTTACTAAATACACAATGGCAGACCTCG   13822
gb:MT106054|Organism:Severe      GTGACATGGTACCACATATATCACGTCAACGTCTTACTAAATACACAATGGCAGACCTCG   13822
gb:MT049951|Organism:Severe      GTGACATGGTACCACATATATCACGTCAACGTCTTACTAAATACACAATGGCAGACCTCG   13822
gb:MN975262|Organism:Severe      GTGACATGGTACCACATATATCACGTCAACGTCTTACTAAATACACAATGGCAGACCTCG   13822
gb:MT106052|Organism:Severe      GTGACATGGTACCACATATATCACGTCAACGTCTTACTAAATACACAATGGCAGACCTCG   13822
gb:LC522975|Organism:Severe      GTGACATGGTACCACATATATCACGTCAACGTCTTACTAAATACACAATGGCAGACCTCG   13819
gb:LC522973|Organism:Severe      GTGACATGGTACCACATATATCACGTCAACGTCTTACTAAATACACAATGGCAGACCTCG   13819
gb:LC522974|Organism:Severe      GTGACATGGTACCACATATATCACGTCAACGTCTTACTAAATACACAATGGCAGACCTCG   13819
```

```
gb:MN985325|Organism:Severe      GTGACATGGTACCACATATATCACGTCAACGTCTTACTAAATACACAATGGCAGACCTCG    13822
gb:MT020881|Organism:Severe      GTGACATGGTACCACATATATCACGTCAACGTCTTACTAAATACACAATGGCAGACCTCG    13822
gb:MT020880|Organism:Severe      GTGACATGGTACCACATATATCACGTCAACGTCTTACTAAATACACAATGGCAGACCTCG    13822
gb:MT066175|Organism:Severe      GTGACATGGTACCACATATATCACGTCAACGTCTTACTAAATACACAATGGCAGACCTCG    13822
gb:MN997409|Organism:Severe      GTGACATGGTACCACATATATCACGTCAACGTCTTACTAAATACACAATGGCAGACCTCG    13822
gb:MN938384|Organism:Severe      GTGACATGGTACCACATATATCACGTCAACGTCTTACTAAATACACAATGGCAGACCTCG    13790
gb:MT044258|Organism:Severe      GTGACATGGTACCACATATATCACGTCAACGTCTTACTAAATACACAATGGCAGACCTCG    13798
gb:MT039890|Organism:Severe      GTGACATGGTACCACATATATCACGTCAACGTCTTACTAAATACACAATGGCAGACCTCG    13822
gb:MN988713|Organism:Severe      GTGACATGGTACCACATATATCACGTCAACGTCTTACTAAATACACAATGGCAGACCTCG    13822
gb:LC521925|Organism:Severe      GTGACATGGTACCACATATATCACGTCAACGTCTTACTAAATACACAATGGCAGACCTCG    13795
gb:MT093571|Organism:Severe      GTGACATGGTACCACATATATCACGTCAACGTCTTACTAAATACACAATGGCAGACCTCG    13822
gb:MT039887|Organism:Severe      GTGACATGGTACCACATATATCACGTCAACGTCTTACTAAATACACAATGGCAGACCTCG    13822
gb:MT019530|Organism:Severe      GTGACATGGTACCACATATATCACGTCAACGTCTTACTAAATACACAATGGCAGACCTCG    13822
gb:MT039888|Organism:Severe      GTGACATGGTACCACATATATCACGTCAACGTCTTACTAAATACACAATGGCAGACCTCG    13822
gb:LC522972|Organism:Severe      GTGACATGGTACCACATATATCACGTCAACGTCTTACTAAATACACAATGGCAGACCTCG    13819
gb:MT027063|Organism:Severe      GTGACATGGTACCACATATATCACGTCAACGTCTTACTAAATACACAATGGCAGACCTCG    13822
gb:MT027062|Organism:Severe      GTGACATGGTACCACATATATCACGTCAACGTCTTACTAAATACACAATGGCAGACCTCG    13822
gb:MT019529|Organism:Severe      GTGACATGGTACCACATATATCACGTCAACGTCTTACTAAATACACAATGGCAGACCTCG    13822
gb:MN996529|Organism:Severe      GTGACATGGTACCACATATATCACGTCAACGTCTTACTAAATACACAATGGCAGACCTCG    13810
gb:MN996531|Organism:Severe      GTGACATGGTACCACATATATCACGTCAACGTCTTACTAAATACACAATGGCAGACCTCG    13809
gb:MT066176|Organism:Severe      GTGACATGGTACCACATATATCACGTCAACGTCTTACTAAATACACAATGGCAGACCTCG    13822
gb:MT027064|Organism:Severe      GTGACATGGTACCACATATATCACGTCAACGTCTTACTAAATACACAATGGCAGACCTCG    13822
gb:MN994468|Organism:Severe      GTGACATGGTACCACATATATCACGTCAACGTCTTACTAAATACACAATGGCAGACCTCG    13822
gb:MT072688|Organism:Severe      GTGACATGGTACCACATATATCACGTCAACGTCTTACTAAATACACAATGGCAGACCTCG    13807
gb:MN996527|Organism:Severe      GTGACATGGTACCACATATATCACGTCAACGTCTTACTAAATACACAATGGCAGACCTCG    13789
gb:MT093631|Organism:Severe      GTGACATGGTACCACATATATCACGTCAACGTCTTACTAAATACACAATGGCAGACCTCG    13860
gb:MT106053|Organism:Severe      GTGACATGGTACCACATATATCACGTCAACGTCTTACTAAATACACAATGGCAGACCTCG    13822
gb:MT019533|Organism:Severe      GTGACATGGTACCACATATATCACGTCAACGTCTTACTAAATACACAATGGCAGACCTCG    13822
gb:MT019531|Organism:Severe      GTGACATGGTACCACATATATCACGTCAACGTCTTACTAAATACACAATGGCAGACCTCG    13822
gb:MN996528|Organism:Severe      GTGACATGGTACCACATATATCACGTCAACGTCTTACTAAATACACAATGGCAGACCTCG    13822
gb:MN996530|Organism:Severe      GTGACATGGTACCACATATATCACGTCAACGTCTTACTAAATACACAATGGCAGACCTCG    13808
gb:MN908947|Organism:Severe      GTGACATGGTACCACATATATCACGTCAACGTCTTACTAAATACACAATGGCAGACCTCG    13822
gb:MT019532|Organism:Severe      GTGACATGGTACCACATATATCACGTCAACGTCTTACTAAATACACAATGGCAGACCTCG    13822
                                 ************************************************************

gb:MT020781|Organism:Severe      TCTATGCTTTAAGGCATTTTGATGAAGGTAATTGTGACACATTAAAAGAAATACTTGTCA    13870
gb:MT007544|Organism:Severe      TCTATGCTTTAAGGCATTTTGATGAAGGTAATTGTGACACATTAAAAGAAATACTTGTCA    13882
gb:MN994467|Organism:Severe      TCTATGCTTTAAGGCATTTTGATGAAGGTAATTGTGACACATTAAAAGAAATACTTGTCA    13882
gb:MT044257|Organism:Severe      TCTATGCTTTAAGGCATTTTGATGAAGGTAATTGTGACACATTAAAAGAAATACTTGTCA    13882
gb:MT106054|Organism:Severe      TCTATGCTTTAAGGCATTTTGATGAAGGTAATTGTGACACATTAAAAGAAATACTTGTCA    13882
gb:MT049951|Organism:Severe      TCTATGCTTTAAGGCATTTTGATGAAGGTAATTGTGACACATTAAAAGAAATACTTGTCA    13882
gb:MN975262|Organism:Severe      TCTATGCTTTAAGGCATTTTGATGAAGGTAATTGTGACACATTAAAAGAAATACTTGTCA    13882
gb:MT106052|Organism:Severe      TCTATGCTTTAAGGCATTTTGATGAAGGTAATTGTGACACATTAAAAGAAATACTTGTCA    13882
gb:LC522975|Organism:Severe      TCTATGCTTTAAGGCATTTTGATGAAGGTAATTGTGACACATTAAAAGAAATACTTGTCA    13879
gb:LC522973|Organism:Severe      TCTATGCTTTAAGGCATTTTGATGAAGGTAATTGTGACACATTAAAAGAAATACTTGTCA    13879
gb:LC522974|Organism:Severe      TCTATGCTTTAAGGCATTTTGATGAAGGTAATTGTGACACATTAAAAGAAATACTTGTCA    13879
gb:MN985325|Organism:Severe      TCTATGCTTTAAGGCATTTTGATGAAGGTAATTGTGACACATTAAAAGAAATACTTGTCA    13882
gb:MT020881|Organism:Severe      TCTATGCTTTAAGGCATTTTGATGAAGGTAATTGTGACACATTAAAAGAAATACTTGTCA    13882
gb:MT020880|Organism:Severe      TCTATGCTTTAAGGCATTTTGATGAAGGTAATTGTGACACATTAAAAGAAATACTTGTCA    13882
gb:MT066175|Organism:Severe      TCTATGCTTTAAGGCATTTTGATGAAGGTAATTGTGACACATTAAAAGAAATACTTGTCA    13882
gb:MN997409|Organism:Severe      TCTATGCTTTAAGGCATTTTGATGAAGGTAATTGTGACACATTAAAAGAAATACTTGTCA    13882
gb:MN938384|Organism:Severe      TCTATGCTTTAAGGCATTTTGATGAAGGTAATTGTGACACATTAAAAGAAATACTTGTCA    13850
gb:MT044258|Organism:Severe      TCTATGCTTTAAGGCATTTTGATGAAGGTAATTGTGACACATTAAAAGAAATACTTGTCA    13858
gb:MT039890|Organism:Severe      TCTATGCTTTAAGGCATTTTGATGAAGGTAATTGTGACACATTAAAAGAAATACTTGTCA    13882
gb:MN988713|Organism:Severe      TCTATGCTTTAAGGCATTTTGATGAAGGTAATTGTGACACATTAAAAGAAATACTTGTCA    13882
gb:LC521925|Organism:Severe      TCTATGCTTTAAGGCATTTTGATGAAGGTAATTGTGACACATTAAAAGAAATACTTGTCA    13855
gb:MT093571|Organism:Severe      TCTATGCTTTAAGGCATTTTGATGAAGGTAATTGTGACACATTAAAAGAAATACTTGTCA    13882
gb:MT039887|Organism:Severe      TCTATGCTTTAAGGCATTTTGATGAAGGTAATTGTGACACATTAAAAGAAATACTTGTCA    13882
gb:MT019530|Organism:Severe      TCTATGCTTTAAGGCATTTTGATGAAGGTAATTGTGACACATTAAAAGAAATACTTGTCA    13882
gb:MT039888|Organism:Severe      TCTATGCTTTAAGGCATTTTGATGAAGGTAATTGTGACACATTAAAAGAAATACTTGTCA    13882
gb:LC522972|Organism:Severe      TCTATGCTTTAAGGCATTTTGATGAAGGTAATTGTGACACATTAAAAGAAATACTTGTCA    13879
gb:MT027063|Organism:Severe      TCTATGCTTTAAGGCATTTTGATGAAGGTAATTGTGACACATTAAAAGAAATACTTGTCA    13882
gb:MT027062|Organism:Severe      TCTATGCTTTAAGGCATTTTGATGAAGGTAATTGTGACACATTAAAAGAAATACTTGTCA    13882
gb:MT019529|Organism:Severe      TCTATGCTTTAAGGCATTTTGATGAAGGTAATTGTGACACATTAAAAGAAATACTTGTCA    13882
gb:MN996529|Organism:Severe      TCTATGCTTTAAGGCATTTTGATGAAGGTAATTGTGACACATTAAAAGAAATACTTGTCA    13870
```

```
gb:MN996531|Organism:Severe      TCTATGCTTTAAGGCATTTTGATGAAGGTAATTGTGACACATTAAAAGAAATACTTGTCA   13869
gb:MT066176|Organism:Severe      TCTATGCTTTAAGGCATTTTGATGAAGGTAATTGTGACACATTAAAAGAAATACTTGTCA   13882
gb:MT027064|Organism:Severe      TCTATGCTTTAAGGCATTTTGATGAAGGTAATTGTGACACATTAAAAGAAATACTTGTCA   13882
gb:MN994468|Organism:Severe      TCTATGCTTTAAGGCATTTTGATGAAGGTAATTGTGACACATTAAAAGAAATACTTGTCA   13882
gb:MT072688|Organism:Severe      TCTATGCTTTAAGGCATTTTGATGAAGGTAATTGTGACACATTAAAAGAAATACTTGTCA   13867
gb:MN996527|Organism:Severe      TCTATGCTTTAAGGCATTTTGATGAAGGTAATTGTGACACATTAAAAGAAATACTTGTCA   13849
gb:MT093631|Organism:Severe      TCTATGCTTTAAGGCATTTTGATGAAGGTAATTGTGACACATTAAAAGAAATACTTGTCA   13920
gb:MT106053|Organism:Severe      TCTATGCTTTAAGGCATTTTGATGAAGGTAATTGTGACACATTAAAAGAAATACTTGTCA   13882
gb:MT019533|Organism:Severe      TCTATGCTTTAAGGCATTTTGATGAAGGTAATTGTGACACATTAAAAGAAATACTTGTCA   13882
gb:MT019531|Organism:Severe      TCTATGCTTTAAGGCATTTTGATGAAGGTAATTGTGACACATTAAAAGAAATACTTGTCA   13882
gb:MN996528|Organism:Severe      TCTATGCTTTAAGGCATTTTGATGAAGGTAATTGTGACACATTAAAAGAAATACTTGTCA   13882
gb:MN996530|Organism:Severe      TCTATGCTTTAAGGCATTTTGATGAAGGTAATTGTGACACATTAAAAGAAATACTTGTCA   13868
gb:MN908947|Organism:Severe      TCTATGCTTTAAGGCATTTTGATGAAGGTAATTGTGACACATTAAAAGAAATACTTGTCA   13882
gb:MT019532|Organism:Severe      TCTATGCTTTAAGGCATTTTGATGAAGGTAATTGTGACACATTAAAAGAAATACTTGTCA   13882
                                 ************************************************************

gb:MT020781|Organism:Severe      CATACAATTGTTGTGATGATGATTATTTCAATAAAAAGGACTGGTATGATTTTGTAGAAA   13930
gb:MT007544|Organism:Severe      CATACAATTGTTGTGATGATGATTATTTCAATAAAAAGGACTGGTATGATTTTGTAGAAA   13942
gb:MN994467|Organism:Severe      CATACAATTGTTGTGATGATGATTATTTCAATAAAAAGGACTGGTATGATTTTGTAGAAA   13942
gb:MT044257|Organism:Severe      CATACAATTGTTGTGATGATGATTATTTCAATAAAAAGGACTGGTATGATTTTGTAGAAA   13942
gb:MT106054|Organism:Severe      CATACAATTGTTGTGATGATGATTATTTCAATAAAAAGGACTGGTATGATTTTGTAGAAA   13942
gb:MT049951|Organism:Severe      CATACAATTGTTGTGATGATGATTATTTCAATAAAAAGGACTGGTATGATTTTGTAGAAA   13942
gb:MN975262|Organism:Severe      CATACAATTGTTGTGATGATGATTATTTCAATAAAAAGGACTGGTATGATTTTGTAGAAA   13942
gb:MT106052|Organism:Severe      CATACAATTGTTGTGATGATGATTATTTCAATAAAAAGGACTGGTATGATTTTGTAGAAA   13942
gb:LC522975|Organism:Severe      CATACAATTGTTGTGATGATGATTATTTCAATAAAAAGGACTGGTATGATTTTGTAGAAA   13939
gb:LC522973|Organism:Severe      CATACAATTGTTGTGATGATGATTATTTCAATAAAAAGGACTGGTATGATTTTGTAGAAA   13939
gb:LC522974|Organism:Severe      CATACAATTGTTGTGATGATGATTATTTCAATAAAAAGGACTGGTATGATTTTGTAGAAA   13939
gb:MN985325|Organism:Severe      CATACAATTGTTGTGATGATGATTATTTCAATAAAAAGGACTGGTATGATTTTGTAGAAA   13942
gb:MT020881|Organism:Severe      CATACAATTGTTGTGATGATGATTATTTCAATAAAAAGGACTGGTATGATTTTGTAGAAA   13942
gb:MT020880|Organism:Severe      CATACAATTGTTGTGATGATGATTATTTCAATAAAAAGGACTGGTATGATTTTGTAGAAA   13942
gb:MT066175|Organism:Severe      CATACAATTGTTGTGATGATGATTATTTCAATAAAAAGGACTGGTATGATTTTGTAGAAA   13942
gb:MN997409|Organism:Severe      CATACAATTGTTGTGATGATGATTATTTCAATAAAAAGGACTGGTATGATTTTGTAGAAA   13942
gb:MN938384|Organism:Severe      CATACAATTGTTGTGATGATGATTATTTCAATAAAAAGGACTGGTATGATTTTGTAGAAA   13910
gb:MT044258|Organism:Severe      CATACAATTGTTGTGATGATGATTATTTCAATAAAAAGGACTGGTATGATTTTGTAGAAA   13918
gb:MT039890|Organism:Severe      CATACAATTGTTGTGATGATGATTATTTCAATAAAAAGGACTGGTATGATTTTGTAGAAA   13942
gb:MN988713|Organism:Severe      CATACAATTGTTGTGATGATGATTATTTCAATAAAAAGGACTGGTATGATTTTGTAGAAA   13942
gb:LC521925|Organism:Severe      CATACAATTGTTGTGATGATGATTATTTCAATAAAAAGGACTGGTATGATTTTGTAGAAA   13915
gb:MT093571|Organism:Severe      CATACAATTGTTGTGATGATGATTATTTCAATAAAAAGGACTGGTATGATTTTGTAGAAA   13942
gb:MT039887|Organism:Severe      CATACAATTGTTGTGATGATGATTATTTCAATAAAAAGGACTGGTATGATTTTGTAGAAA   13942
gb:MT019530|Organism:Severe      CATACAATTGTTGTGATGATGATTATTTCAATAAAAAGGACTGGTATGATTTTGTAGAAA   13942
gb:MT039888|Organism:Severe      CATACAATTGTTGTGATGATGATTATTTCAATAAAAAGGACTGGTATGATTTTGTAGAAA   13942
gb:LC522972|Organism:Severe      CATACAATTGTTGTGATGATGATTATTTCAATAAAAAGGACTGGTATGATTTTGTAGAAA   13939
gb:MT027063|Organism:Severe      CATACAATTGTTGTGATGATGATTATTTCAATAAAAAGGACTGGTATGATTTTGTAGAAA   13942
gb:MT027062|Organism:Severe      CATACAATTGTTGTGATGATGATTATTTCAATAAAAAGGACTGGTATGATTTTGTAGAAA   13942
gb:MT019529|Organism:Severe      CATACAATTGTTGTGATGATGATTATTTCAATAAAAAGGACTGGTATGATTTTGTAGAAA   13942
gb:MN996529|Organism:Severe      CATACAATTGTTGTGATGATGATTATTTCAATAAAAAGGACTGGTATGATTTTGTAGAAA   13930
gb:MN996531|Organism:Severe      CATACAATTGTTGTGATGATGATTATTTCAATAAAAAGGACTGGTATGATTTTGTAGAAA   13929
gb:MT066176|Organism:Severe      CATACAATTGTTGTGATGATGATTATTTCAATAAAAAGGACTGGTATGATTTTGTAGAAA   13942
gb:MT027064|Organism:Severe      CATACAATTGTTGTGATGATGATTATTTCAATAAAAAGGACTGGTATGATTTTGTAGAAA   13942
gb:MN994468|Organism:Severe      CATACAATTGTTGTGATGATGATTATTTCAATAAAAAGGACTGGTATGATTTTGTAGAAA   13942
gb:MT072688|Organism:Severe      CATACAATTGTTGTGATGATGATTATTTCAATAAAAAGGACTGGTATGATTTTGTAGAAA   13927
gb:MN996527|Organism:Severe      CATACAATTGTTGTGATGATGATTATTTCAATAAAAAGGACTGGTATGATTTTGTAGAAA   13909
gb:MT093631|Organism:Severe      CATACAATTGTTGTGATGATGATTATTTCAATAAAAAGGACTGGTATGATTTTGTAGAAA   13980
gb:MT106053|Organism:Severe      CATACAATTGTTGTGATGATGATTATTTCAATAAAAAGGACTGGTATGATTTTGTAGAAA   13942
gb:MT019533|Organism:Severe      CATACAATTGTTGTGATGATGATTATTTCAATAAAAAGGACTGGTATGATTTTGTAGAAA   13942
gb:MT019531|Organism:Severe      CATACAATTGTTGTGATGATGATTATTTCAATAAAAAGGACTGGTATGATTTTGTAGAAA   13942
gb:MN996528|Organism:Severe      CATACAATTGTTGTGATGATGATTATTTCAATAAAAAGGACTGGTATGATTTTGTAGAAA   13942
gb:MN996530|Organism:Severe      CATACAATTGTTGTGATGATGATTATTTCAATAAAAAGGACTGGTATGATTTTGTAGAAA   13928
gb:MN908947|Organism:Severe      CATACAATTGTTGTGATGATGATTATTTCAATAAAAAGGACTGGTATGATTTTGTAGAAA   13942
gb:MT019532|Organism:Severe      CATACAATTGTTGTGATGATGATTATTTCAATAAAAAGGACTGGTATGATTTTGTAGAAA   13942
                                 ************************************************************

gb:MT020781|Organism:Severe      ACCCAGATATATTACGCGTATACGCCAACTTAGGTGAACGTGTACGCCAAGCTTTGTTAA   13990
gb:MT007544|Organism:Severe      ACCCAGATATATTACGCGTATACGCCAACTTAGGTGAACGTGTACGCCAAGCTTTGTTAA   14002
gb:MN994467|Organism:Severe      ACCCAGATATATTACGCGTATACGCCAACTTAGGTGAACGTGTACGCCAAGCTTTGTTAA   14002
```

```
gb:MT044257|Organism:Severe      ACCCAGATATATTACGCGTATACGCCAACTTAGGTGAACGTGTACGCCAAGCTTTGTTAA    14002
gb:MT106054|Organism:Severe      ACCCAGATATATTACGCGTATACGCCAACTTAGGTGAACGTGTACGCCAAGCTTTGTTAA    14002
gb:MT049951|Organism:Severe      ACCCAGATATATTACGCGTATACGCCAACTTAGGTGAACGTGTACGCCAAGCTTTGTTAA    14002
gb:MN975262|Organism:Severe      ACCCAGATATATTACGCGTATACGCCAACTTAGGTGAACGTGTACGCCAAGCTTTGTTAA    14002
gb:MT106052|Organism:Severe      ACCCAGATATATTACGCGTATACGCCAACTTAGGTGAACGTGTACGCCAAGCTTTGTTAA    14002
gb:LC522975|Organism:Severe      ACCCAGATATATTACGCGTATACGCCAACTTAGGTGAACGTGTACGCCAAGCTTTGTTAA    13999
gb:LC522973|Organism:Severe      ACCCAGATATATTACGCGTATACGCCAACTTAGGTGAACGTGTACGCCAAGCTTTGTTAA    13999
gb:LC522974|Organism:Severe      ACCCAGATATATTACGCGTATACGCCAACTTAGGTGAACGTGTACGCCAAGCTTTGTTAA    13999
gb:MN985325|Organism:Severe      ACCCAGATATATTACGCGTATACGCCAACTTAGGTGAACGTGTACGCCAAGCTTTGTTAA    14002
gb:MT020881|Organism:Severe      ACCCAGATATATTACGCGTATACGCCAACTTAGGTGAACGTGTACGCCAAGCTTTGTTAA    14002
gb:MT020880|Organism:Severe      ACCCAGATATATTACGCGTATACGCCAACTTAGGTGAACGTGTACGCCAAGCTTTGTTAA    14002
gb:MT066175|Organism:Severe      ACCCAGATATATTACGCGTATACGCCAACTTAGGTGAACGTGTACGCCAAGCTTTGTTAA    14002
gb:MN997409|Organism:Severe      ACCCAGATATATTACGCGTATACGCCAACTTAGGTGAACGTGTACGCCAAGCTTTGTTAA    14002
gb:MN938384|Organism:Severe      ACCCAGATATATTACGCGTATACGCCAACTTAGGTGAACGTGTACGCCAAGCTTTGTTAA    13970
gb:MT044258|Organism:Severe      ACCCAGATATATTACGCGTATACGCCAACTTAGGTGAACGTGTACGCCAAGCTTTGTTAA    13978
gb:MT039890|Organism:Severe      ACCCAGATATATTACGCGTATACGCCAACTTAGGTGAACGTGTACGCCAAGCTTTGTTAA    14002
gb:MN988713|Organism:Severe      ACCCAGATATATTACGCGTATACGCCAACTTAGGTGAACGTGTACGCCAAGCTTTGTTAA    14002
gb:LC521925|Organism:Severe      ACCCAGATATATTACGCGTATACGCCAACTTAGGTGAACGTGTACGCCAAGCTTTGTTAA    13975
gb:MT093571|Organism:Severe      ACCCAGATATATTACGCGTATACGCCAACTTAGGTGAACGTGTACGCCAAGCTTTGTTAA    14002
gb:MT039887|Organism:Severe      ACCCAGATATATTACGCGTATACGCCAACTTAGGTGAACGTGTACGCCAAGCTTTGTTAA    14002
gb:MT019530|Organism:Severe      ACCCAGATATATTACGCGTATACGCCAACTTAGGTGAACGTGTACGCCAAGCTTTGTTAA    14002
gb:MT039888|Organism:Severe      ACCCAGATATATTACGCGTATACGCCAACTTAGGTGAACGTGTACGCCAAGCTTTGTTAA    14002
gb:LC522972|Organism:Severe      ACCCAGATATATTACGCGTATACGCCAACTTAGGTGAACGTGTACGCCAAGCTTTGTTAA    13999
gb:MT027063|Organism:Severe      ACCCAGATATATTACGCGTATACGCCAACTTAGGTGAACGTGTACGCCAAGCTTTGTTAA    14002
gb:MT027062|Organism:Severe      ACCCAGATATATTACGCGTATACGCCAACTTAGGTGAACGTGTACGCCAAGCTTTGTTAA    14002
gb:MT019529|Organism:Severe      ACCCAGATATATTACGCGTATACGCCAACTTAGGTGAACGTGTACGCCAAGCTTTGTTAA    14002
gb:MN996529|Organism:Severe      ACCCAGATATATTACGCGTATACGCCAACTTAGGTGAACGTGTACGCCAAGCTTTGTTAA    13990
gb:MN996531|Organism:Severe      ACCCAGATATATTACGCGTATACGCCAACTTAGGTGAACGTGTACGCCAAGCTTTGTTAA    13989
gb:MT066176|Organism:Severe      ACCCAGATATATTACGCGTATACGCCAACTTAGGTGAACGTGTACGCCAAGCTTTGTTAA    14002
gb:MT027064|Organism:Severe      ACCCAGATATATTACGCGTATACGCCAACTTAGGTGAACGTGTACGCCAAGCTTTGTTAA    14002
gb:MN994468|Organism:Severe      ACCCAGATATATTACGCGTATACGCCAACTTAGGTGAACGTGTACGCCAAGCTTTGTTAA    14002
gb:MT072688|Organism:Severe      ACCCAGATATATTACGCGTATACGCCAACTTAGGTGAACGTGTACGCCAAGCTTTGTTAA    13987
gb:MN996527|Organism:Severe      ACCCAGATATATTACGCGTATACGCCAACTTAGGTGAACGTGTACGCCAAGCTTTGTTAA    13969
gb:MT093631|Organism:Severe      ACCCAGATATATTACGCGTATACGCCAACTTAGGTGAACGTGTACGCCAAGCTTTGTTAA    14040
gb:MT106053|Organism:Severe      ACCCAGATATATTACGCGTATACGCCAACTTAGGTGAACGTGTACGCCAAGCTTTGTTAA    14002
gb:MT019533|Organism:Severe      ACCCAGATATATTACGCGTATACGCCAACTTAGGTGAACGTGTACGCCAAGCTTTGTTAA    14002
gb:MT019531|Organism:Severe      ACCCAGATATATTACGCGTATACGCCAACTTAGGTGAACGTGTACGCCAAGCTTTGTTAA    14002
gb:MN996528|Organism:Severe      ACCCAGATATATTACGCGTATACGCCAACTTAGGTGAACGTGTACGCCAAGCTTTGTTAA    14002
gb:MN996530|Organism:Severe      ACCCAGATATATTACGCGTATACGCCAACTTAGGTGAACGTGTACGCCAAGCTTTGTTAA    13988
gb:MN908947|Organism:Severe      ACCCAGATATATTACGCGTATACGCCAACTTAGGTGAACGTGTACGCCAAGCTTTGTTAA    14002
gb:MT019532|Organism:Severe      ACCCAGATATATTACGCGTATACGCCAACTTAGGTGAACGTGTACGCCAAGCTTTGTTAA    14002
                                 ************************************************************

gb:MT020781|Organism:Severe      AAACAGTACAATTCTGTGATGCCATGCGAAATGCTGGTATTGTTGGTGTACTGACATTAG    14050
gb:MT007544|Organism:Severe      AAACAGTACAATTCTGTGATGCCATGCGAAATGCTGGTATTGTTGGTGTACTGACATTAG    14062
gb:MN994467|Organism:Severe      AAACAGTACAATTCTGTGATGCCATGCGAAATGCTGGTATTGTTGGTGTACTGACATTAG    14062
gb:MT044257|Organism:Severe      AAACAGTACAATTCTGTGATGCCATGCGAAATGCTGGTATTGTTGGTGTACTGACATTAG    14062
gb:MT106054|Organism:Severe      AAACAGTACAATTCTGTGATGCCATGCGAAATGCTGGTATTGTTGGTGTACTGACATTAG    14062
gb:MT049951|Organism:Severe      AAACAGTACAATTCTGTGATGCCATGCGAAATGCTGGTATTGTTGGTGTACTGACATTAG    14062
gb:MN975262|Organism:Severe      AAACAGTACAATTCTGTGATGCCATGCGAAATGCTGGTATTGTTGGTGTACTGACATTAG    14062
gb:MT106052|Organism:Severe      AAACAGTACAATTCTGTGATGCCATGCGAAATGCTGGTATTGTTGGTGTACTGACATTAG    14062
gb:LC522975|Organism:Severe      AAACAGTACAATTCTGTGATGCCATGCGAAATGCTGGTATTGTTGGTGTACTGACATTAG    14059
gb:LC522973|Organism:Severe      AAACAGTACAATTCTGTGATGCCATGCGAAATGCTGGTATTGTTGGTGTACTGACATTAG    14059
gb:LC522974|Organism:Severe      AAACAGTACAATTCTGTGATGCCATGCGAAATGCTGGTATTGTTGGTGTACTGACATTAG    14059
gb:MN985325|Organism:Severe      AAACAGTACAATTCTGTGATGCCATGCGAAATGCTGGTATTGTTGGTGTACTGACATTAG    14062
gb:MT020881|Organism:Severe      AAACAGTACAATTCTGTGATGCCATGCGAAATGCTGGTATTGTTGGTGTACTGACATTAG    14062
gb:MT020880|Organism:Severe      AAACAGTACAATTCTGTGATGCCATGCGAAATGCTGGTATTGTTGGTGTACTGACATTAG    14062
gb:MT066175|Organism:Severe      AAACAGTACAATTCTGTGATGCCATGCGAAATGCTGGTATTGTTGGTGTACTGACATTAG    14062
gb:MN997409|Organism:Severe      AAACAGTACAATTCTGTGATGCCATGCGAAATGCTGGTATTGTTGGTGTACTGACATTAG    14062
gb:MN938384|Organism:Severe      AAACAGTACAATTCTGTGATGCCATGCGAAATGCTGGTATTGTTGGTGTACTGACATTAG    14030
gb:MT044258|Organism:Severe      AAACAGTACAATTCTGTGATGCCATGCGAAATGCTGGTATTGTTGGTGTACTGACATTAG    14038
gb:MT039890|Organism:Severe      AAACAGTACAATTCTGTGATGCCATGCGAAATGCTGGTATTGTTGGTGTACTGACATTAG    14062
gb:MN988713|Organism:Severe      AAACAGTACAATTCTGTGATGCCATGCGAAATGCTGGTATTGTTGGTGTACTGACATTAG    14062
gb:LC521925|Organism:Severe      AAACAGTACAATTCTGTGATGCCATGCGAAATGCTGGTATTGTTGGTGTACTGACATTAG    14035
gb:MT093571|Organism:Severe      AAACAGTACAATTCTGTGATGCCATGCGAAATGCTGGTATTGTTGGTGTACTGACATTAG    14062
```

```
gb:MT039887|Organism:Severe      AAACAGTACAATTCTGTGATGCCATGCGAAATGCTGGTATTGTTGGTGTACTGACATTAG    14062
gb:MT019530|Organism:Severe      AAACAGTACAATTCTGTGATGCCATGCGAAATGCTGGTATTGTTGGTGTACTGACATTAG    14062
gb:MT039888|Organism:Severe      AAACAGTACAATTCTGTGATGCCATGCGAAATGCTGGTATTGTTGGTGTACTGACATTAG    14062
gb:LC522972|Organism:Severe      AAACAGTACAATTCTGTGATGCCATGCGAAATGCTGGTATTGTTGGTGTACTGACATTAG    14059
gb:MT027063|Organism:Severe      AAACAGTACAATTCTGTGATGCCATGCGAAATGCTGGTATTGTTGGTGTACTGACATTAG    14062
gb:MT027062|Organism:Severe      AAACAGTACAATTCTGTGATGCCATGCGAAATGCTGGTATTGTTGGTGTACTGACATTAG    14062
gb:MT019529|Organism:Severe      AAACAGTACAATTCTGTGATGCCATGCGAAATGCTGGTATTGTTGGTGTACTGACATTAG    14062
gb:MN996529|Organism:Severe      AAACAGTACAATTCTGTGATGCCATGCGAAATGCTGGTATTGTTGGTGTACTGACATTAG    14050
gb:MN996531|Organism:Severe      AAACAGTACAATTCTGTGATGCCATGCGAAATGCTGGTATTGTTGGTGTACTGACATTAG    14049
gb:MT066176|Organism:Severe      AAACAGTACAATTCTGTGATGCCATGCGAAATGCTGGTATTGTTGGTGTACTGACATTAG    14062
gb:MT027064|Organism:Severe      AAACAGTACAATTCTGTGATGCCATGCGAAATGCTGGTATTGTTGGTGTACTGACATTAG    14062
gb:MN994468|Organism:Severe      AAACAGTACAATTCTGTGATGCCATGCGAAATGCTGGTATTGTTGGTGTACTGACATTAG    14062
gb:MT072688|Organism:Severe      AAACAGTACAATTCTGTGATGCCATGCGAAATGCTGGTATTGTTGGTGTACTGACATTAG    14047
gb:MN996527|Organism:Severe      AAACAGTACAATTCTGTGATGCCATGCGAAATGCTGGTATTGTTGGTGTACTGACATTAG    14029
gb:MT093631|Organism:Severe      AAACAGTACAATTCTGTGATGCCATGCGAAATGCTGGTATTGTTGGTGTACTGACATTAG    14100
gb:MT106053|Organism:Severe      AAACAGTACAATTCTGTGATGCCATGCGAAATGCTGGTATTGTTGGTGTACTGACATTAG    14062
gb:MT019533|Organism:Severe      AAACAGTACAATTCTGTGATGCCATGCGAAATGCTGGTATTGTTGGTGTACTGACATTAG    14062
gb:MT019531|Organism:Severe      AAACAGTACAATTCTGTGATGCCATGCGAAATGCTGGTATTGTTGGTGTACTGACATTAG    14062
gb:MN996528|Organism:Severe      AAACAGTACAATTCTGTGATGCCATGCGAAATGCTGGTATTGTTGGTGTACTGACATTAG    14062
gb:MN996530|Organism:Severe      AAACAGTACAATTCTGTGATGCCATGCGAAATGCTGGTATTGTTGGTGTACTGACATTAG    14048
gb:MN908947|Organism:Severe      AAACAGTACAATTCTGTGATGCCATGCGAAATGCTGGTATTGTTGGTGTACTGACATTAG    14062
gb:MT019532|Organism:Severe      AAACAGTACAATTCTGTGATGCCATGCGAAATGCTGGTATTGTTGGTGTACTGACATTAG    14062
                                 ************************************************************

gb:MT020781|Organism:Severe      ATAATCAAGATCTCAATGGTAACTGGTATGATTTCGGTGATTTCATACAAACCACGCCAG    14110
gb:MT007544|Organism:Severe      ATAATCAAGATCTCAATGGTAACTGGTATGATTTCGGTGATTTCATACAAACCACGCCAG    14122
gb:MN994467|Organism:Severe      ATAATCAAGATCTCAATGGTAACTGGTATGATTTCGGTGATTTCATACAAACCACGCCAG    14122
gb:MT044257|Organism:Severe      ATAATCAAGATCTCAATGGTAACTGGTATGATTTCGGTGATTTCATACAAACCACGCCAG    14122
gb:MT106054|Organism:Severe      ATAATCAAGATCTCAATGGTAACTGGTATGATTTCGGTGATTTCATACAAACCACGCCAG    14122
gb:MT049951|Organism:Severe      ATAATCAAGATCTCAATGGTAACTGGTATGATTTCGGTGATTTCATACAAACCACGCCAG    14122
gb:MN975262|Organism:Severe      ATAATCAAGATCTCAATGGTAACTGGTATGATTTCGGTGATTTCATACAAACCACGCCAG    14122
gb:MT106052|Organism:Severe      ATAATCAAGATCTCAATGGTAACTGGTATGATTTCGGTGATTTCATACAAACCACGCCAG    14122
gb:LC522975|Organism:Severe      ATAATCAAGATCTCAATGGTAACTGGTATGATTTCGGTGATTTCATACAAACCACGCCAG    14119
gb:LC522973|Organism:Severe      ATAATCAAGATCTCAATGGTAACTGGTATGATTTCGGTGATTTCATACAAACCACGCCAG    14119
gb:LC522974|Organism:Severe      ATAATCAAGATCTCAATGGTAACTGGTATGATTTCGGTGATTTCATACAAACCACGCCAG    14119
gb:MN985325|Organism:Severe      ATAATCAAGATCTCAATGGTAACTGGTATGATTTCGGTGATTTCATACAAACCACGCCAG    14122
gb:MT020881|Organism:Severe      ATAATCAAGATCTCAATGGTAACTGGTATGATTTCGGTGATTTCATACAAACCACGCCAG    14122
gb:MT020880|Organism:Severe      ATAATCAAGATCTCAATGGTAACTGGTATGATTTCGGTGATTTCATACAAACCACGCCAG    14122
gb:MT066175|Organism:Severe      ATAATCAAGATCTCAATGGTAACTGGTATGATTTCGGTGATTTCATACAAACCACGCCAG    14122
gb:MN997409|Organism:Severe      ATAATCAAGATCTCAATGGTAACTGGTATGATTTCGGTGATTTCATACAAACCACGCCAG    14122
gb:MN938384|Organism:Severe      ATAATCAAGATCTCAATGGTAACTGGTATGATTTCGGTGATTTCATACAAACCACGCCAG    14090
gb:MT044258|Organism:Severe      ATAATCAAGATCTCAATGGTAACTGGTATGATTTCGGTGATTTCATACAAACCACGCCAG    14098
gb:MT039890|Organism:Severe      ATAATCAAGATCTCAATGGTAACTGGTATGATTTCGGTGATTTCATACAAACCACGCCAG    14122
gb:MN988713|Organism:Severe      ATAATCAAGATCTCAATGGTAACTGGTATGATTTCGGTGATTTCATACAAACCACGCCAG    14122
gb:LC521925|Organism:Severe      ATAATCAAGATCTCAATGGTAACTGGTATGATTTCGGTGATTTCATACAAACCACGCCAG    14095
gb:MT093571|Organism:Severe      ATAATCAAGATCTCAATGGTAACTGGTATGATTTCGGTGATTTCATACAAACCACGCCAG    14122
gb:MT039887|Organism:Severe      ATAATCAAGATCTCAATGGTAACTGGTATGATTTCGGTGATTTCATACAAACCACGCCAG    14122
gb:MT019530|Organism:Severe      ATAATCAAGATCTCAATGGTAACTGGTATGATTTCGGTGATTTCATACAAACCACGCCAG    14122
gb:MT039888|Organism:Severe      ATAATCAAGATCTCAATGGTAACTGGTATGATTTCGGTGATTTCATACAAACCACGCCAG    14122
gb:LC522972|Organism:Severe      ATAATCAAGATCTCAATGGTAACTGGTATGATTTCGGTGATTTCATACAAACCACGCCAG    14119
gb:MT027063|Organism:Severe      ATAATCAAGATCTCAATGGTAACTGGTATGATTTCGGTGATTTCATACAAACCACGCCAG    14122
gb:MT027062|Organism:Severe      ATAATCAAGATCTCAATGGTAACTGGTATGATTTCGGTGATTTCATACAAACCACGCCAG    14122
gb:MT019529|Organism:Severe      ATAATCAAGATCTCAATGGTAACTGGTATGATTTCGGTGATTTCATACAAACCACGCCAG    14122
gb:MN996529|Organism:Severe      ATAATCAAGATCTCAATGGTAACTGGTATGATTTCGGTGATTTCATACAAACCACGCCAG    14110
gb:MN996531|Organism:Severe      ATAATCAAGATCTCAATGGTAACTGGTATGATTTCGGTGATTTCATACAAACCACGCCAG    14109
gb:MT066176|Organism:Severe      ATAATCAAGATCTCAATGGTAACTGGTATGATTTCGGTGATTTCATACAAACCACGCCAG    14122
gb:MT027064|Organism:Severe      ATAATCAAGATCTCAATGGTAACTGGTATGATTTCGGTGATTTCATACAAACCACGCCAG    14122
gb:MN994468|Organism:Severe      ATAATCAAGATCTCAATGGTAACTGGTATGATTTCGGTGATTTCATACAAACCACGCCAG    14122
gb:MT072688|Organism:Severe      ATAATCAAGATCTCAATGGTAACTGGTATGATTTCGGTGATTTCATACAAACCACGCCAG    14107
gb:MN996527|Organism:Severe      ATAATCAAGATCTCAATGGTAACTGGTATGATTTCGGTGATTTCATACAAACCACGCCAG    14089
gb:MT093631|Organism:Severe      ATAATCAAGATCTCAATGGTAACTGGTATGATTTCGGTGATTTCATACAAACCACGCCAG    14160
gb:MT106053|Organism:Severe      ATAATCAAGATCTCAATGGTAACTGGTATGATTTCGGTGATTTCATACAAACCACGCCAG    14122
gb:MT019533|Organism:Severe      ATAATCAAGATCTCAATGGTAACTGGTATGATTTCGGTGATTTCATACAAACCACGCCAG    14122
gb:MT019531|Organism:Severe      ATAATCAAGATCTCAATGGTAACTGGTATGATTTCGGTGATTTCATACAAACCACGCCAG    14122
gb:MN996528|Organism:Severe      ATAATCAAGATCTCAATGGTAACTGGTATGATTTCGGTGATTTCATACAAACCACGCCAG    14122
```

```
gb:MN996530|Organism:Severe      ATAATCAAGATCTCAATGGTAACTGGTATGATTTCGGTGATTTCATACAAACCACGCCAG   14108
gb:MN908947|Organism:Severe      ATAATCAAGATCTCAATGGTAACTGGTATGATTTCGGTGATTTCATACAAACCACGCCAG   14122
gb:MT019532|Organism:Severe      ATAATCAAGATCTCAATGGTAACTGGTATGATTTCGGTGATTTCATACAAACCACGCCAG   14122
                                 ************************************************************

gb:MT020781|Organism:Severe      GTAGTGGAGTTCCTGTTGTAGATTCTTATTATTCATTGTTAATGCCTATATTAACCTTGA   14170
gb:MT007544|Organism:Severe      GTAGTGGAGTTCCTGTTGTAGATTCTTATTATTCATTGTTAATGCCTATATTAACCTTGA   14182
gb:MN994467|Organism:Severe      GTAGTGGAGTTCCTGTTGTAGATTCTTATTATTCATTGTTAATGCCTATATTAACCTTGA   14182
gb:MT044257|Organism:Severe      GTAGTGGAGTTCCTGTTGTAGATTCTTATTATTCATTGTTAATGCCTATATTAACCTTGA   14182
gb:MT106054|Organism:Severe      GTAGTGGAGTTCCTGTTGTAGATTCTTATTATTCATTGTTAATGCCTATATTAACCTTGA   14182
gb:MT049951|Organism:Severe      GTAGTGGAGTTCCTGTTGTAGATTCTTATTATTCATTGTTAATGCCTATATTAACCTTGA   14182
gb:MN975262|Organism:Severe      GTAGTGGAGTTCCTGTTGTAGATTCTTATTATTCATTGTTAATGCCTATATTAACCTTGA   14182
gb:MT106052|Organism:Severe      GTAGTGGAGTTCCTGTTGTAGATTCTTATTATTCATTGTTAATGCCTATATTAACCTTGA   14182
gb:LC522975|Organism:Severe      GTAGTGGAGTTCCTGTTGTAGATTCTTATTATTCATTGTTAATGCCTATATTAACCTTGA   14179
gb:LC522973|Organism:Severe      GTAGTGGAGTTCCTGTTGTAGATTCTTATTATTCATTGTTAATGCCTATATTAACCTTGA   14179
gb:LC522974|Organism:Severe      GTAGTGGAGTTCCTGTTGTAGATTCTTATTATTCATTGTTAATGCCTATATTAACCTTGA   14179
gb:MN985325|Organism:Severe      GTAGTGGAGTTCCTGTTGTAGATTCTTATTATTCATTGTTAATGCCTATATTAACCTTGA   14182
gb:MT020881|Organism:Severe      GTAGTGGAGTTCCTGTTGTAGATTCTTATTATTCATTGTTAATGCCTATATTAACCTTGA   14182
gb:MT020880|Organism:Severe      GTAGTGGAGTTCCTGTTGTAGATTCTTATTATTCATTGTTAATGCCTATATTAACCTTGA   14182
gb:MT066175|Organism:Severe      GTAGTGGAGTTCCTGTTGTAGATTCTTATTATTCATTGTTAATGCCTATATTAACCTTGA   14182
gb:MN997409|Organism:Severe      GTAGTGGAGTTCCTGTTGTAGATTCTTATTATTCATTGTTAATGCCTATATTAACCTTGA   14182
gb:MN938384|Organism:Severe      GTAGTGGAGTTCCTGTTGTAGATTCTTATTATTCATTGTTAATGCCTATATTAACCTTGA   14150
gb:MT044258|Organism:Severe      GTAGTGGAGTTCCTGTTGTAGATTCTTATTATTCATTGTTAATGCCTATATTAACCTTGA   14158
gb:MT039890|Organism:Severe      GTAGTGGAGTTCCTGTTGTAGATTCTTATTATTCATTGTTAATGCCTATATTAACCTTGA   14182
gb:MN988713|Organism:Severe      GTAGTGGAGTTCCTGTTGTAGATTCTTATTATTCATTGTTAATGCCTATATTAACCTTGA   14182
gb:LC521925|Organism:Severe      GTAGTGGAGTTCCTGTTGTAGATTCTTATTATTCATTGTTAATGCCTATATTAACCTTGA   14155
gb:MT093571|Organism:Severe      GTAGTGGAGTTCCTGTTGTAGATTCTTATTATTCATTGTTAATGCCTATATTAACCTTGA   14182
gb:MT039887|Organism:Severe      GTAGTGGAGTTCCTGTTGTAGATTCTTATTATTCATTGTTAATGCCTATATTAACCTTGA   14182
gb:MT019530|Organism:Severe      GTAGTGGAGTTCCTGTTGTAGATTCTTATTATTCATTGTTAATGCCTATATTAACCTTGA   14182
gb:MT039888|Organism:Severe      GTAGTGGAGTTCCTGTTGTAGATTCTTATTATTCATTGTTAATGCCTATATTAACCTTGA   14182
gb:LC522972|Organism:Severe      GTAGTGGAGTTCCTGTTGTAGATTCTTATTATTCATTGTTAATGCCTATATTAACCTTGA   14179
gb:MT027063|Organism:Severe      GTAGTGGAGTTCCTGTTGTAGATTCTTATTATTCATTGTTAATGCCTATATTAACCTTGA   14182
gb:MT027062|Organism:Severe      GTAGTGGAGTTCCTGTTGTAGATTCTTATTATTCATTGTTAATGCCTATATTAACCTTGA   14182
gb:MT019529|Organism:Severe      GTAGTGGAGTTCCTGTTGTAGATTCTTATTATTCATTGTTAATGCCTATATTAACCTTGA   14182
gb:MN996529|Organism:Severe      GTAGTGGAGTTCCTGTTGTAGATTCTTATTATTCATTGTTAATGCCTATATTAACCTTGA   14170
gb:MN996531|Organism:Severe      GTAGTGGAGTTCCTGTTGTAGATTCTTATTATTCATTGTTAATGCCTATATTAACCTTGA   14169
gb:MT066176|Organism:Severe      GTAGTGGAGTTCCTGTTGTAGATTCTTATTATTCATTGTTAATGCCTATATTAACCTTGA   14182
gb:MT027064|Organism:Severe      GTAGTGGAGTTCCTGTTGTAGATTCTTATTATTCATTGTTAATGCCTATATTAACCTTGA   14182
gb:MN994468|Organism:Severe      GTAGTGGAGTTCCTGTTGTAGATTCTTATTATTCATTGTTAATGCCTATATTAACCTTGA   14182
gb:MT072688|Organism:Severe      GTAGTGGAGTTCCTGTTGTAGATTCTTATTATTCATTGTTAATGCCTATATTAACCTTGA   14167
gb:MN996527|Organism:Severe      GTAGTGGAGTTCCTGTTGTAGATTCTTATTATTCATTGTTAATGCCTATATTAACCTTGA   14149
gb:MT093631|Organism:Severe      GTAGTGGAGTTCCTGTTGTAGATTCTTATTATTCATTGTTAATGCCTATATTAACCTTGA   14220
gb:MT106053|Organism:Severe      GTAGTGGAGTTCCTGTTGTAGATTCTTATTATTCATTGTTAATGCCTATATTAACCTTGA   14182
gb:MT019533|Organism:Severe      GTAGTGGAGTTCCTGTTGTAGATTCTTATTATTCATTGTTAATGCCTATATTAACCTTGA   14182
gb:MT019531|Organism:Severe      GTAGTGGAGTTCCTGTTGTAGATTCTTATTATTCATTGTTAATGCCTATATTAACCTTGA   14182
gb:MN996528|Organism:Severe      GTAGTGGAGTTCCTGTTGTAGATTCTTATTATTCATTGTTAATGCCTATATTAACCTTGA   14182
gb:MN996530|Organism:Severe      GTAGTGGAGTTCCTGTTGTAGATTCTTATTATTCATTGTTAATGCCTATATTAACCTTGA   14168
gb:MN908947|Organism:Severe      GTAGTGGAGTTCCTGTTGTAGATTCTTATTATTCATTGTTAATGCCTATATTAACCTTGA   14182
gb:MT019532|Organism:Severe      GTAGTGGAGTTCCTGTTGTAGATTCTTATTATTCATTGTTAATGCCTATATTAACCTTGA   14182
                                 ************************************************************

gb:MT020781|Organism:Severe      CCAGGGCTTTAACTGCAGAGTCACATGTTGACACTGACTTAACAAAGCCTTACATTAAGT   14230
gb:MT007544|Organism:Severe      CCAGGGCTTTAACTGCAGAGTCACATGTTGACACTGACTTAACAAAGCCTTACATTAAGT   14242
gb:MN994467|Organism:Severe      CCAGGGCTTTAACTGCAGAGTCACATGTTGACACTGACTTAACAAAGCCTTACATTAAGT   14242
gb:MT044257|Organism:Severe      CCAGGGCTTTAACTGCAGAGTCACATGTTGACACTGACTTAACAAAGCCTTACATTAAGT   14242
gb:MT106054|Organism:Severe      CCAGGGCTTTAACTGCAGAGTCACATGTTGACACTGACTTAACAAAGCCTTACATTAAGT   14242
gb:MT049951|Organism:Severe      CCAGGGCTTTAACTGCAGAGTCACATGTTGACACTGACTTAACAAAGCCTTACATTAAGT   14242
gb:MN975262|Organism:Severe      CCAGGGCTTTAACTGCAGAGTCACATGTTGACACTGACTTAACAAAGCCTTACATTAAGT   14242
gb:MT106052|Organism:Severe      CCAGGGCTTTAACTGCAGAGTCACATGTTGACACTGACTTAACAAAGCCTTACATTAAGT   14242
gb:LC522975|Organism:Severe      CCAGGGCTTTAACTGCAGAGTCACATGTTGACACTGACTTAACAAAGCCTTACATTAAGT   14239
gb:LC522973|Organism:Severe      CCAGGGCTTTAACTGCAGAGTCACATGTTGACACTGACTTAACAAAGCCTTACATTAAGT   14239
gb:LC522974|Organism:Severe      CCAGGGCTTTAACTGCAGAGTCACATGTTGACACTGACTTAACAAAGCCTTACATTAAGT   14239
gb:MN985325|Organism:Severe      CCAGGGCTTTAACTGCAGAGTCACATGTTGACACTGACTTAACAAAGCCTTACATTAAGT   14242
gb:MT020881|Organism:Severe      CCAGGGCTTTAACTGCAGAGTCACATGTTGACACTGACTTAACAAAGCCTTACATTAAGT   14242
gb:MT020880|Organism:Severe      CCAGGGCTTTAACTGCAGAGTCACATGTTGACACTGACTTAACAAAGCCTTACATTAAGT   14242
```

```
gb:MT066175|Organism:Severe      CCAGGGCTTTAACTGCAGAGTCACATGTTGACACTGACTTAACAAAGCCTTACATTAAGT   14242
gb:MN997409|Organism:Severe      CCAGGGCTTTAACTGCAGAGTCACATGTTGACACTGACTTAACAAAGCCTTACATTAAGT   14242
gb:MN938384|Organism:Severe      CCAGGGCTTTAACTGCAGAGTCACATGTTGACACTGACTTAACAAAGCCTTACATTAAGT   14210
gb:MT044258|Organism:Severe      CCAGGGCTTTAACTGCAGAGTCACATGTTGACACTGACTTAACAAAGCCTTACATTAAGT   14218
gb:MT039890|Organism:Severe      CCAGGGCTTTAACTGCAGAGTCACATGTTGACACTGACTTAACAAAGCCTTACATTAAGT   14242
gb:MN988713|Organism:Severe      CCAGGGCTTTAACTGCAGAGTCACATGTTGACACTGACTTAACAAAGCCTTACATTAAGT   14242
gb:LC521925|Organism:Severe      CCAGGGCTTTAACTGCAGAGTCACATGTTGACACTGACTTAACAAAGCCTTACATTAAGT   14215
gb:MT093571|Organism:Severe      CCAGGGCTTTAACTGCAGAGTCACATGTTGACACTGACTTAACAAAGCCTTACATTAAGT   14242
gb:MT039887|Organism:Severe      CCAGGGCTTTAACTGCAGAGTCACATGTTGACACTGACTTAACAAAGCCTTACATTAAGT   14242
gb:MT019530|Organism:Severe      CCAGGGCTTTAACTGCAGAGTCACATGTTGACACTGACTTAACAAAGCCTTACATTAAGT   14242
gb:MT039888|Organism:Severe      CCAGGGCTTTAACTGCAGAGTCACATGTTGACACTGACTTAACAAAGCCTTACATTAAGT   14242
gb:LC522972|Organism:Severe      CCAGGGCTTTAACTGCAGAGTCACATGTTGACACTGACTTAACAAAGCCTTACATTAAGT   14239
gb:MT027063|Organism:Severe      CCAGGGCTTTAACTGCAGAGTCACATGTTGACACTGACTTAACAAAGCCTTACATTAAGT   14242
gb:MT027062|Organism:Severe      CCAGGGCTTTAACTGCAGAGTCACATGTTGACACTGACTTAACAAAGCCTTACATTAAGT   14242
gb:MT019529|Organism:Severe      CCAGGGCTTTAACTGCAGAGTCACATGTTGACACTGACTTAACAAAGCCTTACATTAAGT   14242
gb:MN996529|Organism:Severe      CCAGGGCTTTAACTGCAGAGTCACATGTTGACACTGACTTAACAAAGCCTTACATTAAGT   14230
gb:MN996531|Organism:Severe      CCAGGGCTTTAACTGCAGAGTCACATGTTGACACTGACTTAACAAAGCCTTACATTAAGT   14229
gb:MT066176|Organism:Severe      CCAGGGCTTTAACTGCAGAGTCACATGTTGACACTGACTTAACAAAGCCTTACATTAAGT   14242
gb:MT027064|Organism:Severe      CCAGGGCTTTAACTGCAGAGTCACATGTTGACACTGACTTAACAAAGCCTTACATTAAGT   14242
gb:MN994468|Organism:Severe      CCAGGGCTTTAACTGCAGAGTCACATGTTGACACTGACTTAACAAAGCCTTACATTAAGT   14242
gb:MT072688|Organism:Severe      CCAGGGCTTTAACTGCAGAGTCACATGTTGACACTGACTTAACAAAGCCTTACATTAAGT   14227
gb:MN996527|Organism:Severe      CCAGGGCTTTAACTGCAGAGTCACATGTTGACACTGACTTAACAAAGCCTTACATTAAGT   14209
gb:MT093631|Organism:Severe      CCAGGGCTTTAACTGCAGAGTCACATGTTGACACTGACTTAACAAAGCCTTACATTAAGT   14280
gb:MT106053|Organism:Severe      CCAGGGCTTTAACTGCAGAGTCACATGTTGACACTGACTTAACAAAGCCTTACATTAAGT   14242
gb:MT019533|Organism:Severe      CCAGGGCTTTAACTGCAGAGTCACATGTTGACACTGACTTAACAAAGCCTTACATTAAGT   14242
gb:MT019531|Organism:Severe      CCAGGGCTTTAACTGCAGAGTCACATGTTGACACTGACTTAACAAAGCCTTACATTAAGT   14242
gb:MN996528|Organism:Severe      CCAGGGCTTTAACTGCAGAGTCACATGTTGACACTGACTTAACAAAGCCTTACATTAAGT   14242
gb:MN996530|Organism:Severe      CCAGGGCTTTAACTGCAGAGTCACATGTTGACACTGACTTAACAAAGCCTTACATTAAGT   14228
gb:MN908947|Organism:Severe      CCAGGGCTTTAACTGCAGAGTCACATGTTGACACTGACTTAACAAAGCCTTACATTAAGT   14242
gb:MT019532|Organism:Severe      CCAGGGCTTTAACTGCAGAGTCACATGTTGACACTGACTTAACAAAGCCTTACATTAAGT   14242
                                 ************************************************************

gb:MT020781|Organism:Severe      GGGATTTGTTAAAATATGACTTCACGGAAGAGAGGTTAAAACTCTTTGACCGTTATTTTA   14290
gb:MT007544|Organism:Severe      GGGATTTGTTAAAATATGACTTCACGGAAGAGAGGTTAAAACTCTTTGACCGTTATTTTA   14302
gb:MN994467|Organism:Severe      GGGATTTGTTAAAATATGACTTCACGGAAGAGAGGTTAAAACTCTTTGACCGTTATTTTA   14302
gb:MT044257|Organism:Severe      GGGATTTGTTAAAATATGACTTCACGGAAGAGAGGTTAAAACTCTTTGACCGTTATTTTA   14302
gb:MT106054|Organism:Severe      GGGATTTGTTAAAATATGACTTCACGGAAGAGAGGTTAAAACTCTTTGACCGTTATTTTA   14302
gb:MT049951|Organism:Severe      GGGATTTGTTAAAATATGACTTCACGGAAGAGAGGTTAAAACTCTTTGACCGTTATTTTA   14302
gb:MN975262|Organism:Severe      GGGATTTGTTAAAATATGACTTCACGGAAGAGAGGTTAAAACTCTTTGACCGTTATTTTA   14302
gb:MT106052|Organism:Severe      GGGATTTGTTAAAATATGACTTCACGGAAGAGAGGTTAAAACTCTTTGACCGTTATTTTA   14302
gb:LC522975|Organism:Severe      GGGATTTGTTAAAATATGACTTCACGGAAGAGAGGTTAAAACTCTTTGACCGTTATTTTA   14299
gb:LC522973|Organism:Severe      GGGATTTGTTAAAATATGACTTCACGGAAGAGAGGTTAAAACTCTTTGACCGTTATTTTA   14299
gb:LC522974|Organism:Severe      GGGATTTGTTAAAATATGACTTCACGGAAGAGAGGTTAAAACTCTTTGACCGTTATTTTA   14299
gb:MN985325|Organism:Severe      GGGATTTGTTAAAATATGACTTCACGGAAGAGAGGTTAAAACTCTTTGACCGTTATTTTA   14302
gb:MT020881|Organism:Severe      GGGATTTGTTAAAATATGACTTCACGGAAGAGAGGTTAAAACTCTTTGACCGTTATTTTA   14302
gb:MT020880|Organism:Severe      GGGATTTGTTAAAATATGACTTCACGGAAGAGAGGTTAAAACTCTTTGACCGTTATTTTA   14302
gb:MT066175|Organism:Severe      GGGATTTGTTAAAATATGACTTCACGGAAGAGAGGTTAAAACTCTTTGACCGTTATTTTA   14302
gb:MN997409|Organism:Severe      GGGATTTGTTAAAATATGACTTCACGGAAGAGAGGTTAAAACTCTTTGACCGTTATTTTA   14302
gb:MN938384|Organism:Severe      GGGATTTGTTAAAATATGACTTCACGGAAGAGAGGTTAAAACTCTTTGACCGTTATTTTA   14270
gb:MT044258|Organism:Severe      GGGATTTGTTAAAATATGACTTCACGGAAGAGAGGTTAAAACTCTTTGACCGTTATTTTA   14278
gb:MT039890|Organism:Severe      GGGATTTGTTAAAATATGACTTCACGGAAGAGAGGTTAAAACTCTTTGACCGTTATTTTA   14302
gb:MN988713|Organism:Severe      GGGATTTGTTAAAATATGACTTCACGGAAGAGAGGTTAAAACTCTTTGACCGTTATTTTA   14302
gb:LC521925|Organism:Severe      GGGATTTGTTAAAATATGACTTCACGGAAGAGAGGTTAAAACTCTTTGACCGTTATTTTA   14275
gb:MT093571|Organism:Severe      GGGATTTGTTAAAATATGACTTCACGGAAGAGAGGTTAAAACTCTTTGACCGTTATTTTA   14302
gb:MT039887|Organism:Severe      GGGATTTGTTAAAATATGACTTCACGGAAGAGAGGTTAAAACTCTTTGACCGTTATTTTA   14302
gb:MT019530|Organism:Severe      GGGATTTGTTAAAATATGACTTCACGGAAGAGAGGTTAAAACTCTTTGACCGTTATTTTA   14302
gb:MT039888|Organism:Severe      GGGATTTGTTAAAATATGACTTCACGGAAGAGAGGTTAAAACTCTTTGACCGTTATTTTA   14302
gb:LC522972|Organism:Severe      GGGATTTGTTAAAATATGACTTCACGGAAGAGAGGTTAAAACTCTTTGACCGTTATTTTA   14299
gb:MT027063|Organism:Severe      GGGATTTGTTAAAATATGACTTCACGGAAGAGAGGTTAAAACTCTTTGACCGTTATTTTA   14302
gb:MT027062|Organism:Severe      GGGATTTGTTAAAATATGACTTCACGGAAGAGAGGTTAAAACTCTTTGACCGTTATTTTA   14302
gb:MT019529|Organism:Severe      GGGATTTGTTAAAATATGACTTCACGGAAGAGAGGTTAAAACTCTTTGACCGTTATTTTA   14302
gb:MN996529|Organism:Severe      GGGATTTGTTAAAATATGACTTCACGGAAGAGAGGTTAAAACTCTTTGACCGTTATTTTA   14290
gb:MN996531|Organism:Severe      GGGATTTGTTAAAATATGACTTCACGGAAGAGAGGTTAAAACTCTTTGACCGTTATTTTA   14289
gb:MT066176|Organism:Severe      GGGATTTGTTAAAATATGACTTCACGGAAGAGAGGTTAAAACTCTTTGACCGTTATTTTA   14302
gb:MT027064|Organism:Severe      GGGATTTGTTAAAATATGACTTCACGGAAGAGAGGTTAAAACTCTTTGACCGTTATTTTA   14302
```

```
gb:MN994468|Organism:Severe      GGGATTTGTTAAAATATGACTTCACGGAAGAGAGGTTAAAACTCTTTGACCGTTATTTTA   14302
gb:MT072688|Organism:Severe      GGGATTTGTTAAAATATGACTTCACGGAAGAGAGGTTAAAACTCTTTGACCGTTATTTTA   14287
gb:MN996527|Organism:Severe      GGGATTTGTTAAAATATGACTTCACGGAAGAGAGGTTAAAACTCTTTGACCGTTATTTTA   14269
gb:MT093631|Organism:Severe      GGGATTTGTTAAAATATGACTTCACGGAAGAGAGGTTAAAACTCTTTGACCGTTATTTTA   14340
gb:MT106053|Organism:Severe      GGGATTTGTTAAAATATGACTTCACGGAAGAGAGGTTAAAACTCTTTGACCGTTATTTTA   14302
gb:MT019533|Organism:Severe      GGGATTTGTTAAAATATGACTTCACGGAAGAGAGGTTAAAACTCTTTGACCGTTATTTTA   14302
gb:MT019531|Organism:Severe      GGGATTTGTTAAAATATGACTTCACGGAAGAGAGGTTAAAACTCTTTGACCGTTATTTTA   14302
gb:MN996528|Organism:Severe      GGGATTTGTTAAAATATGACTTCACGGAAGAGAGGTTAAAACTCTTTGACCGTTATTTTA   14302
gb:MN996530|Organism:Severe      GGGATTTGTTAAAATATGACTTCACGGAAGAGAGGTTAAAACTCTTTGACCGTTATTTTA   14288
gb:MN908947|Organism:Severe      GGGATTTGTTAAAATATGACTTCACGGAAGAGAGGTTAAAACTCTTTGACCGTTATTTTA   14302
gb:MT019532|Organism:Severe      GGGATTTGTTAAAATATGACTTCACGGAAGAGAGGTTAAAACTCTTTGACCGTTATTTTA   14302
                                 ************************************************************

gb:MT020781|Organism:Severe      AATATTGGGATCAGACATACCACCCAAATTGTGTTAACTGTTTGGATGACAGATGCATTC   14350
gb:MT007544|Organism:Severe      AATATTGGGATCAGACATACCACCCAAATTGTGTTAACTGTTTGGATGACAGATGCATTC   14362
gb:MN994467|Organism:Severe      AATATTGGGATCAGACATACCACCCAAATTGTGTTAACTGTTTGGATGACAGATGCATTC   14362
gb:MT044257|Organism:Severe      AATATTGGGATCAGACATACCACCCAAATTGTGTTAACTGTTTGGATGACAGATGCATTC   14362
gb:MT106054|Organism:Severe      AATATTGGGATCAGACATACCACCCAAATTGTGTTAACTGTTTGGATGACAGATGCATTC   14362
gb:MT049951|Organism:Severe      AATATTGGGATCAGACATACCACCCAAATTGTGTTAACTGTTTGGATGACAGATGCATTC   14362
gb:MN975262|Organism:Severe      AATATTGGGATCAGACATACCACCCAAATTGTGTTAACTGTTTGGATGACAGATGCATTC   14362
gb:MT106052|Organism:Severe      AATATTGGGATCAGACATACCACCCAAATTGTGTTAACTGTTTGGATGACAGATGCATTC   14362
gb:LC522975|Organism:Severe      AATATTGGGATCAGACATACCACCCAAATTGTGTTAACTGTTTGGATGACAGATGCATTC   14359
gb:LC522973|Organism:Severe      AATATTGGGATCAGACATACCACCCAAATTGTGTTAACTGTTTGGATGACAGATGCATTC   14359
gb:LC522974|Organism:Severe      AATATTGGGATCAGACATACCACCCAAATTGTGTTAACTGTTTGGATGACAGATGCATTC   14359
gb:MN985325|Organism:Severe      AATATTGGGATCAGACATACCACCCAAATTGTGTTAACTGTTTGGATGACAGATGCATTC   14362
gb:MT020881|Organism:Severe      AATATTGGGATCAGACATACCACCCAAATTGTGTTAACTGTTTGGATGACAGATGCATTC   14362
gb:MT020880|Organism:Severe      AATATTGGGATCAGACATACCACCCAAATTGTGTTAACTGTTTGGATGACAGATGCATTC   14362
gb:MT066175|Organism:Severe      AATATTGGGATCAGACATACCACCCAAATTGTGTTAACTGTTTGGATGACAGATGCATTC   14362
gb:MN997409|Organism:Severe      AATATTGGGATCAGACATACCACCCAAATTGTGTTAACTGTTTGGATGACAGATGCATTC   14362
gb:MN938384|Organism:Severe      AATATTGGGATCAGACATACCACCCAAATTGTGTTAACTGTTTGGATGACAGATGCATTC   14330
gb:MT044258|Organism:Severe      AATATTGGGATCAGACATACCACCCAAATTGTGTTAACTGTTTGGATGACAGATGCATTC   14338
gb:MT039890|Organism:Severe      AATATTGGGATCAGACATACCACCCAAATTGTGTTAACTGTTTGGATGACAGATGCATTC   14362
gb:MN988713|Organism:Severe      AATATTGGGATCAGACATACCACCCAAATTGTGTTAACTGTTTGGATGACAGATGCATTC   14362
gb:LC521925|Organism:Severe      AATATTGGGATCAGACATACCACCCAAATTGTGTTAACTGTTTGGATGACAGATGCATTC   14335
gb:MT093571|Organism:Severe      AATATTGGGATCAGACATACCACCCAAATTGTGTTAACTGTTTGGATGACAGATGCATTC   14362
gb:MT039887|Organism:Severe      AATATTGGGATCAGACATACCACCCAAATTGTGTTAACTGTTTGGATGACAGATGCATTC   14362
gb:MT019530|Organism:Severe      AATATTGGGATCAGACATACCACCCAAATTGTGTTAACTGTTTGGATGACAGATGCATTC   14362
gb:MT039888|Organism:Severe      AATATTGGGATCAGACATACCACCCAAATTGTGTTAACTGTTTGGATGACAGATGCATTC   14362
gb:LC522972|Organism:Severe      AATATTGGGATCAGACATACCACCCAAATTGTGTTAACTGTTTGGATGACAGATGCATTC   14359
gb:MT027063|Organism:Severe      AATATTGGGATCAGACATACCACCCAAATTGTGTTAACTGTTTGGATGACAGATGCATTC   14362
gb:MT027062|Organism:Severe      AATATTGGGATCAGACATACCACCCAAATTGTGTTAACTGTTTGGATGACAGATGCATTC   14362
gb:MT019529|Organism:Severe      AATATTGGGATCAGACATACCACCCAAATTGTGTTAACTGTTTGGATGACAGATGCATTC   14362
gb:MN996529|Organism:Severe      AATATTGGGATCAGACATACCACCCAAATTGTGTTAACTGTTTGGATGACAGATGCATTC   14350
gb:MN996531|Organism:Severe      AATATTGGGATCAGACATACCACCCAAATTGTGTTAACTGTTTGGATGACAGATGCATTC   14349
gb:MT066176|Organism:Severe      AATATTGGGATCAGACATACCACCCAAATTGTGTTAACTGTTTGGATGACAGATGCATTC   14362
gb:MT027064|Organism:Severe      AATATTGGGATCAGACATACCACCCAAATTGTGTTAACTGTTTGGATGACAGATGCATTC   14362
gb:MN994468|Organism:Severe      AATATTGGGATCAGACATACCACCCAAATTGTGTTAACTGTTTGGATGACAGATGCATTC   14362
gb:MT072688|Organism:Severe      AATATTGGGATCAGACATACCACCCAAATTGTGTTAACTGTTTGGATGACAGATGCATTC   14347
gb:MN996527|Organism:Severe      AATATTGGGATCAGACATACCACCCAAATTGTGTTAACTGTTTGGATGACAGATGCATTC   14329
gb:MT093631|Organism:Severe      AATATTGGGATCAGACATACCACCCAAATTGTGTTAACTGTTTGGATGACAGATGCATTC   14400
gb:MT106053|Organism:Severe      AATATTGGGATCAGACATACCACCCAAATTGTGTTAACTGTTTGGATGACAGATGCATTC   14362
gb:MT019533|Organism:Severe      AATATTGGGATCAGACATACCACCCAAATTGTGTTAACTGTTTGGATGACAGATGCATTC   14362
gb:MT019531|Organism:Severe      AATATTGGGATCAGACATACCACCCAAATTGTGTTAACTGTTTGGATGACAGATGCATTC   14362
gb:MN996528|Organism:Severe      AATATTGGGATCAGACATACCACCCAAATTGTGTTAACTGTTTGGATGACAGATGCATTC   14362
gb:MN996530|Organism:Severe      AATATTGGGATCAGACATACCACCCAAATTGTGTTAACTGTTTGGATGACAGATGCATTC   14348
gb:MN908947|Organism:Severe      AATATTGGGATCAGACATACCACCCAAATTGTGTTAACTGTTTGGATGACAGATGCATTC   14362
gb:MT019532|Organism:Severe      AATATTGGGATCAGACATACCACCCAAATTGTGTTAACTGTTTGGATGACAGATGCATTC   14362
                                 ************************************************************

gb:MT020781|Organism:Severe      TGCATTGTGCAAACTTTAATGTTTTATTCTCTACAGTGTTCCCACCTACAAGTTTTGGAC   14410
gb:MT007544|Organism:Severe      TGCATTGTGCAAACTTTAATGTTTTATTCTCTACAGTGTTCCCACCTACAAGTTTTGGAC   14422
gb:MN994467|Organism:Severe      TGCATTGTGCAAACTTTAATGTTTTATTCTCTACAGTGTTCCCACCTACAAGTTTTGGAC   14422
gb:MT044257|Organism:Severe      TGCATTGTGCAAACTTTAATGTTTTATTCTCTACAGTGTTCCCACCTACAAGTTTTGGAC   14422
gb:MT106054|Organism:Severe      TGCATTGTGCAAACTTTAATGTTTTATTCTCTACAGTGTTCCCACCTACAAGTTTTGGAC   14422
gb:MT049951|Organism:Severe      TGCATTGTGCAAACTTTAATGTTTTATTCTCTACAGTGTTCCCACCTACAAGTTTTGGAC   14422
```

```
gb:MN975262|Organism:Severe      TGCATTGTGCAAACTTTAATGTTTTATTCTCTACAGTGTTCCCACCTACAAGTTTTGGAC   14422
gb:MT106052|Organism:Severe      TGCATTGTGCAAACTTTAATGTTTTATTCTCTACAGTGTTCCCACCTACAAGTTTTGGAC   14422
gb:LC522975|Organism:Severe      TGCATTGTGCAAACTTTAATGTTTTATTCTCTACAGTGTTCCCACCTACAAGTTTTGGAC   14419
gb:LC522973|Organism:Severe      TGCATTGTGCAAACTTTAATGTTTTATTCTCTACAGTGTTCCCACCTACAAGTTTTGGAC   14419
gb:LC522974|Organism:Severe      TGCATTGTGCAAACTTTAATGTTTTATTCTCTACAGTGTTCCCACCTACAAGTTTTGGAC   14419
gb:MN985325|Organism:Severe      TGCATTGTGCAAACTTTAATGTTTTATTCTCTACAGTGTTCCCACCTACAAGTTTTGGAC   14422
gb:MT020881|Organism:Severe      TGCATTGTGCAAACTTTAATGTTTTATTCTCTACAGTGTTCCCACCTACAAGTTTTGGAC   14422
gb:MT020880|Organism:Severe      TGCATTGTGCAAACTTTAATGTTTTATTCTCTACAGTGTTCCCACCTACAAGTTTTGGAC   14422
gb:MT066175|Organism:Severe      TGCATTGTGCAAACTTTAATGTTTTATTCTCTACAGTGTTCCCACCTACAAGTTTTGGAC   14422
gb:MN997409|Organism:Severe      TGCATTGTGCAAACTTTAATGTTTTATTCTCTACAGTGTTCCCACCTACAAGTTTTGGAC   14422
gb:MN938384|Organism:Severe      TGCATTGTGCAAACTTTAATGTTTTATTCTCTACAGTGTTCCCACCTACAAGTTTTGGAC   14390
gb:MT044258|Organism:Severe      TGCATTGTGCAAACTTTAATGTTTTATTCTCTACAGTGTTCCCACCTACAAGTTTTGGAC   14398
gb:MT039890|Organism:Severe      TGCATTGTGCAAACTTTAATGTTTTATTCTCTACAGTGTTCCCACCTACAAGTTTTGGAC   14422
gb:MN988713|Organism:Severe      TGCATTGTGCAAACTTTAATGTTTTATTCTCTACAGTGTTCCCACCTACAAGTTTTGGAC   14422
gb:LC521925|Organism:Severe      TGCATTGTGCAAACTTTAATGTTTTATTCTCTACAGTGTTCCCACCTACAAGTTTTGGAC   14395
gb:MT093571|Organism:Severe      TGCATTGTGCAAACTTTAATGTTTTATTCTCTACAGTGTTCCCACCTACAAGTTTTGGAC   14422
gb:MT039887|Organism:Severe      TGCATTGTGCAAACTTTAATGTTTTATTCTCTACAGTGTTCCCACCTACAAGTTTTGGAC   14422
gb:MT019530|Organism:Severe      TGCATTGTGCAAACTTTAATGTTTTATTCTCTACAGTGTTCCCACCTACAAGTTTTGGAC   14422
gb:MT039888|Organism:Severe      TGCATTGTGCAAACTTTAATGTTTTATTCTCTACAGTGTTCCCACCTACAAGTTTTGGAC   14422
gb:LC522972|Organism:Severe      TGCATTGTGCAAACTTTAATGTTTTATTCTCTACAGTGTTCCCACCTACAAGTTTTGGAC   14419
gb:MT027063|Organism:Severe      TGCATTGTGCAAACTTTAATGTTTTATTCTCTACAGTGTTCCCACCTACAAGTTTTGGAC   14422
gb:MT027062|Organism:Severe      TGCATTGTGCAAACTTTAATGTTTTATTCTCTACAGTGTTCCCACCTACAAGTTTTGGAC   14422
gb:MT019529|Organism:Severe      TGCATTGTGCAAACTTTAATGTTTTATTCTCTACAGTGTTCCCACCTACAAGTTTTGGAC   14422
gb:MN996529|Organism:Severe      TGCATTGTGCAAACTTTAATGTTTTATTCTCTACAGTGTTCCCACCTACAAGTTTTGGAC   14410
gb:MN996531|Organism:Severe      TGCATTGTGCAAACTTTAATGTTTTATTCTCTACAGTGTTCCCACCTACAAGTTTTGGAC   14409
gb:MT066176|Organism:Severe      TGCATTGTGCAAACTTTAATGTTTTATTCTCTACAGTGTTCCCACCTACAAGTTTTGGAC   14422
gb:MT027064|Organism:Severe      TGCATTGTGCAAACTTTAATGTTTTATTCTCTACAGTGTTCCCACCTACAAGTTTTGGAC   14422
gb:MN994468|Organism:Severe      TGCATTGTGCAAACTTTAATGTTTTATTCTCTACAGTGTTCCCACCTACAAGTTTTGGAC   14422
gb:MT072688|Organism:Severe      TGCATTGTGCAAACTTTAATGTTTTATTCTCTACAGTGTTCCCACCTACAAGTTTTGGAC   14407
gb:MN996527|Organism:Severe      TGCATTGTGCAAACTTTAATGTTTTATTCTCTACAGTGTTCCCACCTACAAGTTTTGGAC   14389
gb:MT093631|Organism:Severe      TGCATTGTGCAAACTTTAATGTTTTATTCTCTACAGTGTTCCCACCTACAAGTTTTGGAC   14460
gb:MT106053|Organism:Severe      TGCATTGTGCAAACTTTAATGTTTTATTCTCTACAGTGTTCCCACCTACAAGTTTTGGAC   14422
gb:MT019533|Organism:Severe      TGCATTGTGCAAACTTTAATGTTTTATTCTCTACAGTGTTCCCACCTACAAGTTTTGGAC   14422
gb:MT019531|Organism:Severe      TGCATTGTGCAAACTTTAATGTTTTATTCTCTACAGTGTTCCCACCTACAAGTTTTGGAC   14422
gb:MN996528|Organism:Severe      TGCATTGTGCAAACTTTAATGTTTTATTCTCTACAGTGTTCCCACCTACAAGTTTTGGAC   14422
gb:MN996530|Organism:Severe      TGCATTGTGCAAACTTTAATGTTTTATTCTCTACAGTGTTCCCACCTACAAGTTTTGGAC   14408
gb:MN908947|Organism:Severe      TGCATTGTGCAAACTTTAATGTTTTATTCTCTACAGTGTTCCCACCTACAAGTTTTGGAC   14422
gb:MT019532|Organism:Severe      TGCATTGTGCAAACTTTAATGTTTTATTCTCTACAGTGTTCCCACCTACAAGTTTTGGAC   14422
                                 ************************************************************

gb:MT020781|Organism:Severe      CACTAGTGAGAAAAATATTTGTTGATGGTGTTCCATTTGTAGTTTCAACTGGATACCACT   14470
gb:MT007544|Organism:Severe      CACTAGTGAGAAAAATATTTGTTGATGGTGTTCCATTTGTAGTTTCAACTGGATACCACT   14482
gb:MN994467|Organism:Severe      CACTAGTGAGAAAAATATTTGTTGATGGTGTTCCATTTGTAGTTTCAACTGGATACCACT   14482
gb:MT044257|Organism:Severe      CACTAGTGAGAAAAATATTTGTTGATGGTGTTCCATTTGTAGTTTCAACTGGATACCACT   14482
gb:MT106054|Organism:Severe      CACTAGTGAGAAAAATATTTGTTGATGGTGTTCCATTTGTAGTTTCAACTGGATACCACT   14482
gb:MT049951|Organism:Severe      CACTAGTGAGAAAAATATTTGTTGATGGTGTTCCATTTGTAGTTTCAACTGGATACCACT   14482
gb:MN975262|Organism:Severe      CACTAGTGAGAAAAATATTTGTTGATGGTGTTCCATTTGTAGTTTCAACTGGATACCACT   14482
gb:MT106052|Organism:Severe      CACTAGTGAGAAAAATATTTGTTGATGGTGTTCCATTTGTAGTTTCAACTGGATACCACT   14482
gb:LC522975|Organism:Severe      CACTAGTGAGAAAAATATTTGTTGATGGTGTTCCATTTGTAGTTTCAACTGGATACCACT   14479
gb:LC522973|Organism:Severe      CACTAGTGAGAAAAATATTTGTTGATGGTGTTCCATTTGTAGTTTCAACTGGATACCACT   14479
gb:LC522974|Organism:Severe      CACTAGTGAGAAAAATATTTGTTGATGGTGTTCCATTTGTAGTTTCAACTGGATACCACT   14479
gb:MN985325|Organism:Severe      CACTAGTGAGAAAAATATTTGTTGATGGTGTTCCATTTGTAGTTTCAACTGGATACCACT   14482
gb:MT020881|Organism:Severe      CACTAGTGAGAAAAATATTTGTTGATGGTGTTCCATTTGTAGTTTCAACTGGATACCACT   14482
gb:MT020880|Organism:Severe      CACTAGTGAGAAAAATATTTGTTGATGGTGTTCCATTTGTAGTTTCAACTGGATACCACT   14482
gb:MT066175|Organism:Severe      CACTAGTGAGAAAAATATTTGTTGATGGTGTTCCATTTGTAGTTTCAACTGGATACCACT   14482
gb:MN997409|Organism:Severe      CACTAGTGAGAAAAATATTTGTTGATGGTGTTCCATTTGTAGTTTCAACTGGATACCACT   14482
gb:MN938384|Organism:Severe      CACTAGTGAGAAAAATATTTGTTGATGGTGTTCCATTTGTAGTTTCAACTGGATACCACT   14450
gb:MT044258|Organism:Severe      CACTAGTGAGAAAAATATTTGTTGATGGTGTTCCATTTGTAGTTTCAACTGGATACCACT   14458
gb:MT039890|Organism:Severe      CACTAGTGAGAAAAATATTTGTTGATGGTGTTCCATTTGTAGTTTCAACTGGATACCACT   14482
gb:MN988713|Organism:Severe      CACTAGTGAGAAAAATATTTGTTGATGGTGTTCCATTTGTAGTTTCAACTGGATACCACT   14482
gb:LC521925|Organism:Severe      CACTAGTGAGAAAAATATTTGTTGATGGTGTTCCATTTGTAGTTTCAACTGGATACCACT   14455
gb:MT093571|Organism:Severe      CACTAGTGAGAAAAATATTTGTTGATGGTGTTCCATTTGTAGTTTCAACTGGATACCACT   14482
gb:MT039887|Organism:Severe      CACTAGTGAGAAAAATATTTGTTGATGGTGTTCCATTTGTAGTTTCAACTGGATACCACT   14482
gb:MT019530|Organism:Severe      CACTAGTGAGAAAAATATTTGTTGATGGTGTTCCATTTGTAGTTTCAACTGGATACCACT   14482
gb:MT039888|Organism:Severe      CACTAGTGAGAAAAATATTTGTTGATGGTGTTCCATTTGTAGTTTCAACTGGATACCACT   14482
```

```
gb:LC522972|Organism:Severe      CACTAGTGAGAAAAATATTTGTTGATGGTGTTCCATTTGTAGTTTCAACTGGATACCACT   14479
gb:MT027063|Organism:Severe      CACTAGTGAGAAAAATATTTGTTGATGGTGTTCCATTTGTAGTTTCAACTGGATACCACT   14482
gb:MT027062|Organism:Severe      CACTAGTGAGAAAAATATTTGTTGATGGTGTTCCATTTGTAGTTTCAACTGGATACCACT   14482
gb:MT019529|Organism:Severe      CACTAGTGAGAAAAATATTTGTTGATGGTGTTCCATTTGTAGTTTCAACTGGATACCACT   14482
gb:MN996529|Organism:Severe      CACTAGTGAGAAAAATATTTGTTGATGGTGTTCCATTTGTAGTTTCAACTGGATACCACT   14470
gb:MN996531|Organism:Severe      CACTAGTGAGAAAAATATTTGTTGATGGTGTTCCATTTGTAGTTTCAACTGGATACCACT   14469
gb:MT066176|Organism:Severe      CACTAGTGAGAAAAATATTTGTTGATGGTGTTCCATTTGTAGTTTCAACTGGATACCACT   14482
gb:MT027064|Organism:Severe      CACTAGTGAGAAAAATATTTGTTGATGGTGTTCCATTTGTAGTTTCAACTGGATACCACT   14482
gb:MN994468|Organism:Severe      CACTAGTGAGAAAAATATTTGTTGATGGTGTTCCATTTGTAGTTTCAACTGGATACCACT   14482
gb:MT072688|Organism:Severe      CACTAGTGAGAAAAATATTTGTTGATGGTGTTCCATTTGTAGTTTCAACTGGATACCACT   14467
gb:MN996527|Organism:Severe      CACTAGTGAGAAAAATATTTGTTGATGGTGTTCCATTTGTAGTTTCAACTGGATACCACT   14449
gb:MT093631|Organism:Severe      CACTAGTGAGAAAAATATTTGTTGATGGTGTTCCATTTGTAGTTTCAACTGGATACCACT   14520
gb:MT106053|Organism:Severe      CACTAGTGAGAAAAATATTTGTTGATGGTGTTCCATTTGTAGTTTCAACTGGATACCACT   14482
gb:MT019533|Organism:Severe      CACTAGTGAGAAAAATATTTGTTGATGGTGTTCCATTTGTAGTTTCAACTGGATACCACT   14482
gb:MT019531|Organism:Severe      CACTAGTGAGAAAAATATTTGTTGATGGTGTTCCATTTGTAGTTTCAACTGGATACCACT   14482
gb:MN996528|Organism:Severe      CACTAGTGAGAAAAATATTTGTTGATGGTGTTCCATTTGTAGTTTCAACTGGATACCACT   14482
gb:MN996530|Organism:Severe      CACTAGTGAGAAAAATATTTGTTGATGGTGTTCCATTTGTAGTTTCAACTGGATACCACT   14468
gb:MN908947|Organism:Severe      CACTAGTGAGAAAAATATTTGTTGATGGTGTTCCATTTGTAGTTTCAACTGGATACCACT   14482
gb:MT019532|Organism:Severe      CACTAGTGAGAAAAATATTTGTTGATGGTGTTCCATTTGTAGTTTCAACTGGATACCACT   14482
                                 ************************************************************

gb:MT020781|Organism:Severe      TCAGAGAGCTAGGTGTTGTACATAATCAGGATGTAAACTTACATAGCTCTAGACTTAGTT   14530
gb:MT007544|Organism:Severe      TCAGAGAGCTAGGTGTTGTACATAATCAGGATGTAAACTTACATAGCTCTAGACTTAGTT   14542
gb:MN994467|Organism:Severe      TCAGAGAGCTAGGTGTTGTACATAATCAGGATGTAAACTTACATAGCTCTAGACTTAGTT   14542
gb:MT044257|Organism:Severe      TCAGAGAGCTAGGTGTTGTACATAATCAGGATGTAAACTTACATAGCTCTAGACTTAGTT   14542
gb:MT106054|Organism:Severe      TCAGAGAGCTAGGTGTTGTACATAATCAGGATGTAAACTTACATAGCTCTAGACTTAGTT   14542
gb:MT049951|Organism:Severe      TCAGAGAGCTAGGTGTTGTACATAATCAGGATGTAAACTTACATAGCTCTAGACTTAGTT   14542
gb:MN975262|Organism:Severe      TCAGAGAGCTAGGTGTTGTACATAATCAGGATGTAAACTTACATAGCTCTAGACTTAGTT   14542
gb:MT106052|Organism:Severe      TCAGAGAGCTAGGTGTTGTACATAATCAGGATGTAAACTTACATAGCTCTAGACTTAGTT   14542
gb:LC522975|Organism:Severe      TCAGAGAGCTAGGTGTTGTACATAATCAGGATGTAAACTTACATAGCTCTAGACTTAGTT   14539
gb:LC522973|Organism:Severe      TCAGAGAGCTAGGTGTTGTACATAATCAGGATGTAAACTTACATAGCTCTAGACTTAGTT   14539
gb:LC522974|Organism:Severe      TCAGAGAGCTAGGTGTTGTACATAATCAGGATGTAAACTTACATAGCTCTAGACTTAGTT   14539
gb:MN985325|Organism:Severe      TCAGAGAGCTAGGTGTTGTACATAATCAGGATGTAAACTTACATAGCTCTAGACTTAGTT   14542
gb:MT020881|Organism:Severe      TCAGAGAGCTAGGTGTTGTACATAATCAGGATGTAAACTTACATAGCTCTAGACTTAGTT   14542
gb:MT020880|Organism:Severe      TCAGAGAGCTAGGTGTTGTACATAATCAGGATGTAAACTTACATAGCTCTAGACTTAGTT   14542
gb:MT066175|Organism:Severe      TCAGAGAGCTAGGTGTTGTACATAATCAGGATGTAAACTTACATAGCTCTAGACTTAGTT   14542
gb:MN997409|Organism:Severe      TCAGAGAGCTAGGTGTTGTACATAATCAGGATGTAAACTTACATAGCTCTAGACTTAGTT   14542
gb:MN938384|Organism:Severe      TCAGAGAGCTAGGTGTTGTACATAATCAGGATGTAAACTTACATAGCTCTAGACTTAGTT   14510
gb:MT044258|Organism:Severe      TCAGAGAGCTAGGTGTTGTACATAATCAGGATGTAAACTTACATAGCTCTAGACTTAGTT   14518
gb:MT039890|Organism:Severe      TCAGAGAGCTAGGTGTTGTACATAATCAGGATGTAAACTTACATAGCTCTAGACTTAGTT   14542
gb:MN988713|Organism:Severe      TCAGAGAGCTAGGTGTTGTACATAATCAGGATGTAAACTTACATAGCTCTAGACTTAGTT   14542
gb:LC521925|Organism:Severe      TCAGAGAGCTAGGTGTTGTACATAATCAGGATGTAAACTTACATAGCTCTAGACTTAGTT   14515
gb:MT093571|Organism:Severe      TCAGAGAGCTAGGTGTTGTACATAATCAGGATGTAAACTTACATAGCTCTAGACTTAGTT   14542
gb:MT039887|Organism:Severe      TCAGAGAGCTAGGTGTTGTACATAATCAGGATGTAAACTTACATAGCTCTAGACTTAGTT   14542
gb:MT019530|Organism:Severe      TCAGAGAGCTAGGTGTTGTACATAATCAGGATGTAAACTTACATAGCTCTAGACTTAGTT   14542
gb:MT039888|Organism:Severe      TCAGAGAGCTAGGTGTTGTACATAATCAGGATGTAAACTTACATAGCTCTAGACTTAGTT   14542
gb:LC522972|Organism:Severe      TCAGAGAGCTAGGTGTTGTACATAATCAGGATGTAAACTTACATAGCTCTAGACTTAGTT   14539
gb:MT027063|Organism:Severe      TCAGAGAGCTAGGTGTTGTACATAATCAGGATGTAAACTTACATAGCTCTAGACTTAGTT   14542
gb:MT027062|Organism:Severe      TCAGAGAGCTAGGTGTTGTACATAATCAGGATGTAAACTTACATAGCTCTAGACTTAGTT   14542
gb:MT019529|Organism:Severe      TCAGAGAGCTAGGTGTTGTACATAATCAGGATGTAAACTTACATAGCTCTAGACTTAGTT   14542
gb:MN996529|Organism:Severe      TCAGAGAGCTAGGTGTTGTACATAATCAGGATGTAAACTTACATAGCTCTAGACTTAGTT   14530
gb:MN996531|Organism:Severe      TCAGAGAGCTAGGTGTTGTACATAATCAGGATGTAAACTTACATAGCTCTAGACTTAGTT   14529
gb:MT066176|Organism:Severe      TCAGAGAGCTAGGTGTTGTACATAATCAGGATGTAAACTTACATAGCTCTAGACTTAGTT   14542
gb:MT027064|Organism:Severe      TCAGAGAGCTAGGTGTTGTACATAATCAGGATGTAAACTTACATAGCTCTAGACTTAGTT   14542
gb:MN994468|Organism:Severe      TCAGAGAGCTAGGTGTTGTACATAATCAGGATGTAAACTTACATAGCTCTAGACTTAGTT   14542
gb:MT072688|Organism:Severe      TCAGAGAGCTAGGTGTTGTACATAATCAGGATGTAAACTTACATAGCTCTAGACTTAGTT   14527
gb:MN996527|Organism:Severe      TCAGAGAGCTAGGTGTTGTACATAATCAGGATGTAAACTTACATAGCTCTAGACTTAGTT   14509
gb:MT093631|Organism:Severe      TCAGAGAGCTAGGTGTTGTACATAATCAGGATGTAAACTTACATAGCTCTAGACTTAGTT   14580
gb:MT106053|Organism:Severe      TCAGAGAGCTAGGTGTTGTACATAATCAGGATGTAAACTTACATAGCTCTAGACTTAGTT   14542
gb:MT019533|Organism:Severe      TCAGAGAGCTAGGTGTTGTACATAATCAGGATGTAAACTTACATAGCTCTAGACTTAGTT   14542
gb:MT019531|Organism:Severe      TCAGAGAGCTAGGTGTTGTACATAATCAGGATGTAAACTTACATAGCTCTAGACTTAGTT   14542
gb:MN996528|Organism:Severe      TCAGAGAGCTAGGTGTTGTACATAATCAGGATGTAAACTTACATAGCTCTAGACTTAGTT   14542
gb:MN996530|Organism:Severe      TCAGAGAGCTAGGTGTTGTACATAATCAGGATGTAAACTTACATAGCTCTAGACTTAGTT   14528
gb:MN908947|Organism:Severe      TCAGAGAGCTAGGTGTTGTACATAATCAGGATGTAAACTTACATAGCTCTAGACTTAGTT   14542
gb:MT019532|Organism:Severe      TCAGAGAGCTAGGTGTTGTACATAATCAGGATGTAAACTTACATAGCTCTAGACTTAGTT   14542
```

```
                                    ************************************************************

gb:MT020781|Organism:Severe      TTAAGGAATTACTTGTGTATGCTGCTGACCCTGCTATGCACGCTGCTTCTGGTAATCTAT   14590
gb:MT007544|Organism:Severe      TTAAGGAATTACTTGTGTATGCTGCTGACCCTGCTATGCACGCTGCTTCTGGTAATCTAT   14602
gb:MN994467|Organism:Severe      TTAAGGAATTACTTGTGTATGCTGCTGACCCTGCTATGCACGCTGCTTCTGGTAATCTAT   14602
gb:MT044257|Organism:Severe      TTAAGGAATTACTTGTGTATGCTGCTGACCCTGCTATGCACGCTGCTTCTGGTAATCTAT   14602
gb:MT106054|Organism:Severe      TTAAGGAATTACTTGTGTATGCTGCTGACCCTGCTATGCACGCTGCTTCTGGTAATCTAT   14602
gb:MT049951|Organism:Severe      TTAAGGAATTACTTGTGTATGCTGCTGACCCTGCTATGCACGCTGCTTCTGGTAATCTAT   14602
gb:MN975262|Organism:Severe      TTAAGGAATTACTTGTGTATGCTGCTGACCCTGCTATGCACGCTGCTTCTGGTAATCTAT   14602
gb:MT106052|Organism:Severe      TTAAGGAATTACTTGTGTATGCTGCTGACCCTGCTATGCACGCTGCTTCTGGTAATCTAT   14602
gb:LC522975|Organism:Severe      TTAAGGAATTACTTGTGTATGCTGCTGACCCTGCTATGCACGCTGCTTCTGGTAATCTAT   14599
gb:LC522973|Organism:Severe      TTAAGGAATTACTTGTGTATGCTGCTGACCCTGCTATGCACGCTGCTTCTGGTAATCTAT   14599
gb:LC522974|Organism:Severe      TTAAGGAATTACTTGTGTATGCTGCTGACCCTGCTATGCACGCTGCTTCTGGTAATCTAT   14599
gb:MN985325|Organism:Severe      TTAAGGAATTACTTGTGTATGCTGCTGACCCTGCTATGCACGCTGCTTCTGGTAATCTAT   14602
gb:MT020881|Organism:Severe      TTAAGGAATTACTTGTGTATGCTGCTGACCCTGCTATGCACGCTGCTTCTGGTAATCTAT   14602
gb:MT020880|Organism:Severe      TTAAGGAATTACTTGTGTATGCTGCTGACCCTGCTATGCACGCTGCTTCTGGTAATCTAT   14602
gb:MT066175|Organism:Severe      TTAAGGAATTACTTGTGTATGCTGCTGACCCTGCTATGCACGCTGCTTCTGGTAATCTAT   14602
gb:MN997409|Organism:Severe      TTAAGGAATTACTTGTGTATGCTGCTGACCCTGCTATGCACGCTGCTTCTGGTAATCTAT   14602
gb:MN938384|Organism:Severe      TTAAGGAATTACTTGTGTATGCTGCTGACCCTGCTATGCACGCTGCTTCTGGTAATCTAT   14570
gb:MT044258|Organism:Severe      TTAAGGAATTACTTGTGTATGCTGCTGACCCTGCTATGCACGCTGCTTCTGGTAATCTAT   14578
gb:MT039890|Organism:Severe      TTAAGGAATTACTTGTGTATGCTGCTGACCCTGCTATGCACGCTGCTTCTGGTAATCTAT   14602
gb:MN988713|Organism:Severe      TTAAGGAATTACTTGTGTATGCTGCTGACCCTGCTATGCACGCTGCTTCTGGTAATCTAT   14602
gb:LC521925|Organism:Severe      TTAAGGAATTACTTGTGTATGCTGCTGACCCTGCTATGCACGCTGCTTCTGGTAATCTAT   14575
gb:MT093571|Organism:Severe      TTAAGGAATTACTTGTGTATGCTGCTGACCCTGCTATGCACGCTGCTTCTGGTAATCTAT   14602
gb:MT039887|Organism:Severe      TTAAGGAATTACTTGTGTATGCTGCTGACCCTGCTATGCACGCTGCTTCTGGTAATCTAT   14602
gb:MT019530|Organism:Severe      TTAAGGAATTACTTGTGTATGCTGCTGACCCTGCTATGCACGCTGCTTCTGGTAATCTAT   14602
gb:MT039888|Organism:Severe      TTAAGGAATTACTTGTGTATGCTGCTGACCCTGCTATGCACGCTGCTTCTGGTAATCTAT   14602
gb:LC522972|Organism:Severe      TTAAGGAATTACTTGTGTATGCTGCTGACCCTGCTATGCACGCTGCTTCTGGTAATCTAT   14599
gb:MT027063|Organism:Severe      TTAAGGAATTACTTGTGTATGCTGCTGACCCTGCTATGCACGCTGCTTCTGGTAATCTAT   14602
gb:MT027062|Organism:Severe      TTAAGGAATTACTTGTGTATGCTGCTGACCCTGCTATGCACGCTGCTTCTGGTAATCTAT   14602
gb:MT019529|Organism:Severe      TTAAGGAATTACTTGTGTATGCTGCTGACCCTGCTATGCACGCTGCTTCTGGTAATCTAT   14602
gb:MN996529|Organism:Severe      TTAAGGAATTACTTGTGTATGCTGCTGACCCTGCTATGCACGCTGCTTCTGGTAATCTAT   14590
gb:MN996531|Organism:Severe      TTAAGGAATTACTTGTGTATGCTGCTGACCCTGCTATGCACGCTGCTTCTGGTAATCTAT   14589
gb:MT066176|Organism:Severe      TTAAGGAATTACTTGTGTATGCTGCTGACCCTGCTATGCACGCTGCTTCTGGTAATCTAT   14602
gb:MT027064|Organism:Severe      TTAAGGAATTACTTGTGTATGCTGCTGACCCTGCTATGCACGCTGCTTCTGGTAATCTAT   14602
gb:MN994468|Organism:Severe      TTAAGGAATTACTTGTGTATGCTGCTGACCCTGCTATGCACGCTGCTTCTGGTAATCTAT   14602
gb:MT072688|Organism:Severe      TTAAGGAATTACTTGTGTATGCTGCTGACCCTGCTATGCACGCTGCTTCTGGTAATCTAT   14587
gb:MN996527|Organism:Severe      TTAAGGAATTACTTGTGTATGCTGCTGACCCTGCTATGCACGCTGCTTCTGGTAATCTAT   14569
gb:MT093631|Organism:Severe      TTAAGGAATTACTTGTGTATGCTGCTGACCCTGCTATGCACGCTGCTTCTGGTAATCTAT   14640
gb:MT106053|Organism:Severe      TTAAGGAATTACTTGTGTATGCTGCTGACCCTGCTATGCACGCTGCTTCTGGTAATCTAT   14602
gb:MT019533|Organism:Severe      TTAAGGAATTACTTGTGTATGCTGCTGACCCTGCTATGCACGCTGCTTCTGGTAATCTAT   14602
gb:MT019531|Organism:Severe      TTAAGGAATTACTTGTGTATGCTGCTGACCCTGCTATGCACGCTGCTTCTGGTAATCTAT   14602
gb:MN996528|Organism:Severe      TTAAGGAATTACTTGTGTATGCTGCTGACCCTGCTATGCACGCTGCTTCTGGTAATCTAT   14602
gb:MN996530|Organism:Severe      TTAAGGAATTACTTGTGTATGCTGCTGACCCTGCTATGCACGCTGCTTCTGGTAATCTAT   14588
gb:MN908947|Organism:Severe      TTAAGGAATTACTTGTGTATGCTGCTGACCCTGCTATGCACGCTGCTTCTGGTAATCTAT   14602
gb:MT019532|Organism:Severe      TTAAGGAATTACTTGTGTATGCTGCTGACCCTGCTATGCACGCTGCTTCTGGTAATCTAT   14602
                                 ************************************************************

gb:MT020781|Organism:Severe      TACTAGATAAACGCACTACGTGCTTTTCAGTAGCTGCACTTACTAACAATGTTGCTTTTC   14650
gb:MT007544|Organism:Severe      TACTAGATAAACGCACTACGTGCTTTTCAGTAGCTGCACTTACTAACAATGTTGCTTTTC   14662
gb:MN994467|Organism:Severe      TACTAGATAAACGCACTACGTGCTTTTCAGTAGCTGCACTTACTAACAATGTTGCTTTTC   14662
gb:MT044257|Organism:Severe      TACTAGATAAACGCACTACGTGCTTTTCAGTAGCTGCACTTACTAACAATGTTGCTTTTC   14662
gb:MT106054|Organism:Severe      TACTAGATAAACGCACTACGTGCTTTTCAGTAGCTGCACTTACTAACAATGTTGCTTTTC   14662
gb:MT049951|Organism:Severe      TACTAGATAAACGCACTACGTGCTTTTCAGTAGCTGCACTTACTAACAATGTTGCTTTTC   14662
gb:MN975262|Organism:Severe      TACTAGATAAACGCACTACGTGCTTTTCAGTAGCTGCACTTACTAACAATGTTGCTTTTC   14662
gb:MT106052|Organism:Severe      TACTAGATAAACGCACTACGTGCTTTTCAGTAGCTGCACTTACTAACAATGTTGCTTTTC   14662
gb:LC522975|Organism:Severe      TACTAGATAAACGCACTACGTGCTTTTCAGTAGCTGCACTTACTAACAATGTTGCTTTTC   14659
gb:LC522973|Organism:Severe      TACTAGATAAACGCACTACGTGCTTTTCAGTAGCTGCACTTACTAACAATGTTGCTTTTC   14659
gb:LC522974|Organism:Severe      TACTAGATAAACGCACTACGTGCTTTTCAGTAGCTGCACTTACTAACAATGTTGCTTTTC   14659
gb:MN985325|Organism:Severe      TACTAGATAAACGCACTACGTGCTTTTCAGTAGCTGCACTTACTAACAATGTTGCTTTTC   14662
gb:MT020881|Organism:Severe      TACTAGATAAACGCACTACGTGCTTTTCAGTAGCTGCACTTACTAACAATGTTGCTTTTC   14662
gb:MT020880|Organism:Severe      TACTAGATAAACGCACTACGTGCTTTTCAGTAGCTGCACTTACTAACAATGTTGCTTTTC   14662
gb:MT066175|Organism:Severe      TACTAGATAAACGCACTACGTGCTTTTCAGTAGCTGCACTTACTAACAATGTTGCTTTTC   14662
gb:MN997409|Organism:Severe      TACTAGATAAACGCACTACGTGCTTTTCAGTAGCTGCACTTACTAACAATGTTGCTTTTC   14662
gb:MN938384|Organism:Severe      TACTAGATAAACGCACTACGTGCTTTTCAGTAGCTGCACTTACTAACAATGTTGCTTTTC   14630
```

```
gb:MT044258|Organism:Severe      TACTAGATAAACGCACTACGTGCTTTTCAGTAGCTGCACTTACTAACAATGTTGCTTTTC   14638
gb:MT039890|Organism:Severe      TACTAGATAAACGCACTACGTGCTTTTCAGTAGCTGCACTTACTAACAATGTTGCTTTTC   14662
gb:MN988713|Organism:Severe      TACTAGATAAACGCACTACGTGCTTTTCAGTAGCTGCACTTACTAACAATGTTGCTTTTC   14662
gb:LC521925|Organism:Severe      TACTAGATAAACGCACTACGTGCTTTTCAGTAGCTGCACTTACTAACAATGTTGCTTTTC   14635
gb:MT093571|Organism:Severe      TACTAGATAAACGCACTACGTGCTTTTCAGTAGCTGCACTTACTAACAATGTTGCTTTTC   14662
gb:MT039887|Organism:Severe      TACTAGATAAACGCACTACGTGCTTTTCAGTAGCTGCACTTACTAACAATGTTGCTTTTC   14662
gb:MT019530|Organism:Severe      TACTAGATAAACGCACTACGTGCTTTTCAGTAGCTGCACTTACTAACAATGTTGCTTTTC   14662
gb:MT039888|Organism:Severe      TACTAGATAAACGCACTACGTGCTTTTCAGTAGCTGCACTTACTAACAATGTTGCTTTTC   14662
gb:LC522972|Organism:Severe      TACTAGATAAACGCACTACGTGCTTTTCAGTAGCTGCACTTACTAACAATGTTGCTTTTC   14659
gb:MT027063|Organism:Severe      TACTAGATAAACGCACTACGTGCTTTTCAGTAGCTGCACTTACTAACAATGTTGCTTTTC   14662
gb:MT027062|Organism:Severe      TACTAGATAAACGCACTACGTGCTTTTCAGTAGCTGCACTTACTAACAATGTTGCTTTTC   14662
gb:MT019529|Organism:Severe      TACTAGATAAACGCACTACGTGCTTTTCAGTAGCTGCACTTACTAACAATGTTGCTTTTC   14662
gb:MN996529|Organism:Severe      TACTAGATAAACGCACTACGTGCTTTTCAGTAGCTGCACTTACTAACAATGTTGCTTTTC   14650
gb:MN996531|Organism:Severe      TACTAGATAAACGCACTACGTGCTTTTCAGTAGCTGCACTTACTAACAATGTTGCTTTTC   14649
gb:MT066176|Organism:Severe      TACTAGATAAACGCACTACGTGCTTTTCAGTAGCTGCACTTACTAACAATGTTGCTTTTC   14662
gb:MT027064|Organism:Severe      TACTAGATAAACGCACTACGTGCTTTTCAGTAGCTGCACTTACTAACAATGTTGCTTTTC   14662
gb:MN994468|Organism:Severe      TACTAGATAAACGCACTACGTGCTTTTCAGTAGCTGCACTTACTAACAATGTTGCTTTTC   14662
gb:MT072688|Organism:Severe      TACTAGATAAACGCACTACGTGCTTTTCAGTAGCTGCACTTACTAACAATGTTGCTTTTC   14647
gb:MN996527|Organism:Severe      TACTAGATAAACGCACTACGTGCTTTTCAGTAGCTGCACTTACTAACAATGTTGCTTTTC   14629
gb:MT093631|Organism:Severe      TACTAGATAAACGCACTACGTGCTTTTCAGTAGCTGCACTTACTAACAATGTTGCTTTTC   14700
gb:MT106053|Organism:Severe      TACTAGATAAACGCACTACGTGCTTTTCAGTAGCTGCACTTACTAACAATGTTGCTTTTC   14662
gb:MT019533|Organism:Severe      TACTAGATAAACGCACTACGTGCTTTTCAGTAGCTGCACTTACTAACAATGTTGCTTTTC   14662
gb:MT019531|Organism:Severe      TACTAGATAAACGCACTACGTGCTTTTCAGTAGCTGCACTTACTAACAATGTTGCTTTTC   14662
gb:MN996528|Organism:Severe      TACTAGATAAACGCACTACGTGCTTTTCAGTAGCTGCACTTACTAACAATGTTGCTTTTC   14662
gb:MN996530|Organism:Severe      TACTAGATAAACGCACTACGTGCTTTTCAGTAGCTGCACTTACTAACAATGTTGCTTTTC   14648
gb:MN908947|Organism:Severe      TACTAGATAAACGCACTACGTGCTTTTCAGTAGCTGCACTTACTAACAATGTTGCTTTTC   14662
gb:MT019532|Organism:Severe      TACTAGATAAACGCACTACGTGCTTTTCAGTAGCTGCACTTACTAACAATGTTGCTTTTC   14662
                                 ************************************************************

gb:MT020781|Organism:Severe      AAACTGTCAAACCCGGTAATTTTAACAAAGACTTCTATGACTTTGCTGTGTCTAAGGGTT   14710
gb:MT007544|Organism:Severe      AAACTGTCAAACCCGGTAATTTTAACAAAGACTTCTATGACTTTGCTGTGTCTAAGGGTT   14722
gb:MN994467|Organism:Severe      AAACTGTCAAACCCGGTAATTTTAACAAAGACTTCTATGACTTTGCTGTGTCTAAGGGTT   14722
gb:MT044257|Organism:Severe      AAACTGTCAAACCCGGTAATTTTAACAAAGACTTCTATGACTTTGCTGTGTCTAAGGGTT   14722
gb:MT106054|Organism:Severe      AAACTGTCAAACCCGGTAATTTTAACAAAGACTTCTATGACTTTGCTGTGTCTAAGGGTT   14722
gb:MT049951|Organism:Severe      AAACTGTCAAACCCGGTAATTTTAACAAAGACTTCTATGACTTTGCTGTGTCTAAGGGTT   14722
gb:MN975262|Organism:Severe      AAACTGTCAAACCCGGTAATTTTAACAAAGACTTCTATGACTTTGCTGTGTCTAAGGGTT   14722
gb:MT106052|Organism:Severe      AAACTGTCAAACCCGGTAATTTTAACAAAGACTTCTATGACTTTGCTGTGTCTAAGGGTT   14722
gb:LC522975|Organism:Severe      AAACTGTCAAACCCGGTAATTTTAACAAAGACTTCTATGACTTTGCTGTGTCTAAGGGTT   14719
gb:LC522973|Organism:Severe      AAACTGTCAAACCCGGTAATTTTAACAAAGACTTCTATGACTTTGCTGTGTCTAAGGGTT   14719
gb:LC522974|Organism:Severe      AAACTGTCAAACCCGGTAATTTTAACAAAGACTTCTATGACTTTGCTGTGTCTAAGGGTT   14719
gb:MN985325|Organism:Severe      AAACTGTCAAACCCGGTAATTTTAACAAAGACTTCTATGACTTTGCTGTGTCTAAGGGTT   14722
gb:MT020881|Organism:Severe      AAACTGTCAAACCCGGTAATTTTAACAAAGACTTCTATGACTTTGCTGTGTCTAAGGGTT   14722
gb:MT020880|Organism:Severe      AAACTGTCAAACCCGGTAATTTTAACAAAGACTTCTATGACTTTGCTGTGTCTAAGGGTT   14722
gb:MT066175|Organism:Severe      AAACTGTCAAACCCGGTAATTTTAACAAAGACTTCTATGACTTTGCTGTGTCTAAGGGTT   14722
gb:MN997409|Organism:Severe      AAACTGTCAAACCCGGTAATTTTAACAAAGACTTCTATGACTTTGCTGTGTCTAAGGGTT   14722
gb:MN938384|Organism:Severe      AAACTGTCAAACCCGGTAATTTTAACAAAGACTTCTATGACTTTGCTGTGTCTAAGGGTT   14690
gb:MT044258|Organism:Severe      AAACTGTCAAACCCGGTAATTTTAACAAAGACTTCTATGACTTTGCTGTGTCTAAGGGTT   14698
gb:MT039890|Organism:Severe      AAACTGTCAAACCCGGTAATTTTAACAAAGACTTCTATGACTTTGCTGTGTCTAAGGGTT   14722
gb:MN988713|Organism:Severe      AAACTGTCAAACCCGGTAATTTTAACAAAGACTTCTATGACTTTGCTGTGTCTAAGGGTT   14722
gb:LC521925|Organism:Severe      AAACTGTCAAACCCGGTAATTTTAACAAAGACTTCTATGACTTTGCTGTGTCTAAGGGTT   14695
gb:MT093571|Organism:Severe      AAACTGTCAAACCCGGTAATTTTAACAAAGACTTCTATGACTTTGCTGTGTCTAAGGGTT   14722
gb:MT039887|Organism:Severe      AAACTGTCAAACCCGGTAATTTTAACAAAGACTTCTATGACTTTGCTGTGTCTAAGGGTT   14722
gb:MT019530|Organism:Severe      AAACTGTCAAACCCGGTAATTTTAACAAAGACTTCTATGACTTTGCTGTGTCTAAGGGTT   14722
gb:MT039888|Organism:Severe      AAACTGTCAAACCCGGTAATTTTAACAAAGACTTCTATGACTTTGCTGTGTCTAAGGGTT   14722
gb:LC522972|Organism:Severe      AAACTGTCAAACCCGGTAATTTTAACAAAGACTTCTATGACTTTGCTGTGTCTAAGGGTT   14719
gb:MT027063|Organism:Severe      AAACTGTCAAACCCGGTAATTTTAACAAAGACTTCTATGACTTTGCTGTGTCTAAGGGTT   14722
gb:MT027062|Organism:Severe      AAACTGTCAAACCCGGTAATTTTAACAAAGACTTCTATGACTTTGCTGTGTCTAAGGGTT   14722
gb:MT019529|Organism:Severe      AAACTGTCAAACCCGGTAATTTTAACAAAGACTTCTATGACTTTGCTGTGTCTAAGGGTT   14722
gb:MN996529|Organism:Severe      AAACTGTCAAACCCGGTAATTTTAACAAAGACTTCTATGACTTTGCTGTGTCTAAGGGTT   14710
gb:MN996531|Organism:Severe      AAACTGTCAAACCCGGTAATTTTAACAAAGACTTCTATGACTTTGCTGTGTCTAAGGGTT   14709
gb:MT066176|Organism:Severe      AAACTGTCAAACCCGGTAATTTTAACAAAGACTTCTATGACTTTGCTGTGTCTAAGGGTT   14722
gb:MT027064|Organism:Severe      AAACTGTCAAACCCGGTAATTTTAACAAAGACTTCTATGACTTTGCTGTGTCTAAGGGTT   14722
gb:MN994468|Organism:Severe      AAACTGTCAAACCCGGTAATTTTAACAAAGACTTCTATGACTTTGCTGTGTCTAAGGGTT   14722
gb:MT072688|Organism:Severe      AAACTGTCAAACCCGGTAATTTTAACAAAGACTTCTATGACTTTGCTGTGTCTAAGGGTT   14707
gb:MN996527|Organism:Severe      AAACTGTCAAACCCGGTAATTTTAACAAAGACTTCTATGACTTTGCTGTGTCTAAGGGTT   14689
```

```
gb:MT093631|Organism:Severe      AAACTGTCAAACCCGGTAATTTTAACAAAGACTTCTATGACTTTGCTGTGTCTAAGGGTT   14760
gb:MT106053|Organism:Severe      AAACTGTCAAACCCGGTAATTTTAACAAAGACTTCTATGACTTTGCTGTGTCTAAGGGTT   14722
gb:MT019533|Organism:Severe      AAACTGTCAAACCCGGTAATTTTAACAAAGACTTCTATGACTTTGCTGTGTCTAAGGGTT   14722
gb:MT019531|Organism:Severe      AAACTGTCAAACCCGGTAATTTTAACAAAGACTTCTATGACTTTGCTGTGTCTAAGGGTT   14722
gb:MN996528|Organism:Severe      AAACTGTCAAACCCGGTAATTTTAACAAAGACTTCTATGACTTTGCTGTGTCTAAGGGTT   14722
gb:MN996530|Organism:Severe      AAACTGTCAAACCCGGTAATTTTAACAAAGACTTCTATGACTTTGCTGTGTCTAAGGGTT   14708
gb:MN908947|Organism:Severe      AAACTGTCAAACCCGGTAATTTTAACAAAGACTTCTATGACTTTGCTGTGTCTAAGGGTT   14722
gb:MT019532|Organism:Severe      AAACTGTCAAACCCGGTAATTTTAACAAAGACTTCTATGACTTTGCTGTGTCTAAGGGTT   14722
                                 ************************************************************

gb:MT020781|Organism:Severe      TCTTTAAGGAAGGAAGTTCTGTTGAATTAAAACACTTCTTCTTTGCTCAGGATGGTAATG   14770
gb:MT007544|Organism:Severe      TCTTTAAGGAAGGAAGTTCTGTTGAATTAAAACACTTCTTCTTTGCTCAGGATGGTAATG   14782
gb:MN994467|Organism:Severe      TCTTTAAGGAAGGAAGTTCTGTTGAATTAAAACACTTCTTCTTTGCTCAGGATGGTAATG   14782
gb:MT044257|Organism:Severe      TCTTTAAGGAAGGAAGTTCTGTTGAATTAAAACACTTCTTCTTTGCTCAGGATGGTAATG   14782
gb:MT106054|Organism:Severe      TCTTTAAGGAAGGAAGTTCTGTTGAATTAAAACACTTCTTCTTTGCTCAGGATGGTAATG   14782
gb:MT049951|Organism:Severe      TCTTTAAGGAAGGAAGTTCTGTTGAATTAAAACACTTCTTCTTTGCTCAGGATGGTAATG   14782
gb:MN975262|Organism:Severe      TCTTTAAGGAAGGAAGTTCTGTTGAATTAAAACACTTCTTCTTTGCTCAGGATGGTAATG   14782
gb:MT106052|Organism:Severe      TCTTTAAGGAAGGAAGTTCTGTTGAATTAAAACACTTCTTCTTTGCTCAGGATGGTAATG   14782
gb:LC522975|Organism:Severe      TCTTTAAGGAAGGAAGTTCTGTTGAATTAAAACACTTCTTCTTTGCTCAGGATGGTAATG   14779
gb:LC522973|Organism:Severe      TCTTTAAGGAAGGAAGTTCTGTTGAATTAAAACACTTCTTCTTTGCTCAGGATGGTAATG   14779
gb:LC522974|Organism:Severe      TCTTTAAGGAAGGAAGTTCTGTTGAATTAAAACACTTCTTCTTTGCTCAGGATGGTAATG   14779
gb:MN985325|Organism:Severe      TCTTTAAGGAAGGAAGTTCTGTTGAATTAAAACACTTCTTCTTTGCTCAGGATGGTAATG   14782
gb:MT020881|Organism:Severe      TCTTTAAGGAAGGAAGTTCTGTTGAATTAAAACACTTCTTCTTTGCTCAGGATGGTAATG   14782
gb:MT020880|Organism:Severe      TCTTTAAGGAAGGAAGTTCTGTTGAATTAAAACACTTCTTCTTTGCTCAGGATGGTAATG   14782
gb:MT066175|Organism:Severe      TCTTTAAGGAAGGAAGTTCTGTTGAATTAAAACACTTCTTCTTTGCTCAGGATGGTAATG   14782
gb:MN997409|Organism:Severe      TCTTTAAGGAAGGAAGTTCTGTTGAATTAAAACACTTCTTCTTTGCTCAGGATGGTAATG   14782
gb:MN938384|Organism:Severe      TCTTTAAGGAAGGAAGTTCTGTTGAATTAAAACACTTCTTCTTTGCTCAGGATGGTAATG   14750
gb:MT044258|Organism:Severe      TCTTTAAGGAAGGAAGTTCTGTTGAATTAAAACACTTCTTCTTTGCTCAGGATGGTAATG   14758
gb:MT039890|Organism:Severe      TCTTTAAGGAAGGAAGTTCTGTTGAATTAAAACACTTCTTCTTTGCTCAGGATGGTAATG   14782
gb:MN988713|Organism:Severe      TCTTTAAGGAAGGAAGTTCTGTTGAATTAAAACACTTCTTCTTTGCTCAGGATGGTAATG   14782
gb:LC521925|Organism:Severe      TCTTTAAGGAAGGAAGTTCTGTTGAATTAAAACACTTCTTCTTTGCTCAGGATGGTAATG   14755
gb:MT093571|Organism:Severe      TCTTTAAGGAAGGAAGTTCTGTTGAATTAAAACACTTCTTCTTTGCTCAGGATGGTAATG   14782
gb:MT039887|Organism:Severe      TCTTTAAGGAAGGAAGTTCTGTTGAATTAAAACACTTCTTCTTTGCTCAGGATGGTAATG   14782
gb:MT019530|Organism:Severe      TCTTTAAGGAAGGAAGTTCTGTTGAATTAAAACACTTCTTCTTTGCTCAGGATGGTAATG   14782
gb:MT039888|Organism:Severe      TCTTTAAGGAAGGAAGTTCTGTTGAATTAAAACACTTCTTCTTTGCTCAGGATGGTAATG   14782
gb:LC522972|Organism:Severe      TCTTTAAGGAAGGAAGTTCTGTTGAATTAAAACACTTCTTCTTTGCTCAGGATGGTAATG   14779
gb:MT027063|Organism:Severe      TCTTTAAGGAAGGAAGTTCTGTTGAATTAAAACACTTCTTCTTTGCTCAGGATGGTAATG   14782
gb:MT027062|Organism:Severe      TCTTTAAGGAAGGAAGTTCTGTTGAATTAAAACACTTCTTCTTTGCTCAGGATGGTAATG   14782
gb:MT019529|Organism:Severe      TCTTTAAGGAAGGAAGTTCTGTTGAATTAAAACACTTCTTCTTTGCTCAGGATGGTAATG   14782
gb:MN996529|Organism:Severe      TCTTTAAGGAAGGAAGTTCTGTTGAATTAAAACACTTCTTCTTTGCTCAGGATGGTAATG   14770
gb:MN996531|Organism:Severe      TCTTTAAGGAAGGAAGTTCTGTTGAATTAAAACACTTCTTCTTTGCTCAGGATGGTAATG   14769
gb:MT066176|Organism:Severe      TCTTTAAGGAAGGAAGTTCTGTTGAATTAAAACACTTCTTCTTTGCTCAGGATGGTAATG   14782
gb:MT027064|Organism:Severe      TCTTTAAGGAAGGAAGTTCTGTTGAATTAAAACACTTCTTCTTTGCTCAGGATGGTAATG   14782
gb:MN994468|Organism:Severe      TCTTTAAGGAAGGAAGTTCTGTTGAATTAAAACACTTCTTCTTTGCTCAGGATGGTAATG   14782
gb:MT072688|Organism:Severe      TCTTTAAGGAAGGAAGTTCTGTTGAATTAAAACACTTCTTCTTTGCTCAGGATGGTAATG   14767
gb:MN996527|Organism:Severe      TCTTTAAGGAAGGAAGTTCTGTTGAATTAAAACACTTCTTCTTTGCTCAGGATGGTAATG   14749
gb:MT093631|Organism:Severe      TCTTTAAGGAAGGAAGTTCTGTTGAATTAAAACACTTCTTCTTTGCTCAGGATGGTAATG   14820
gb:MT106053|Organism:Severe      TCTTTAAGGAAGGAAGTTCTGTTGAATTAAAACACTTCTTCTTTGCTCAGGATGGTAATG   14782
gb:MT019533|Organism:Severe      TCTTTAAGGAAGGAAGTTCTGTTGAATTAAAACACTTCTTCTTTGCTCAGGATGGTAATG   14782
gb:MT019531|Organism:Severe      TCTTTAAGGAAGGAAGTTCTGTTGAATTAAAACACTTCTTCTTTGCTCAGGATGGTAATG   14782
gb:MN996528|Organism:Severe      TCTTTAAGGAAGGAAGTTCTGTTGAATTAAAACACTTCTTCTTTGCTCAGGATGGTAATG   14782
gb:MN996530|Organism:Severe      TCTTTAAGGAAGGAAGTTCTGTTGAATTAAAACACTTCTTCTTTGCTCAGGATGGTAATG   14768
gb:MN908947|Organism:Severe      TCTTTAAGGAAGGAAGTTCTGTTGAATTAAAACACTTCTTCTTTGCTCAGGATGGTAATG   14782
gb:MT019532|Organism:Severe      TCTTTAAGGAAGGAAGTTCTGTTGAATTAAAACACTTCTTCTTTGCTCAGGATGGTAATG   14782
                                 ************************************************************

gb:MT020781|Organism:Severe      CTGCTATCAGCGATTATGACTACTATCGTTATAATCTACCAACAATGTGTGATATCAGAC   14830
gb:MT007544|Organism:Severe      CTGCTATCAGCGATTATGACTACTATCGTTATAATCTACCAACAATGTGTGATATCAGAC   14842
gb:MN994467|Organism:Severe      CTGCTATCAGCGATTATGACTACTATCGTTATAATCTACCAACAATGTGTGATATCAGAC   14842
gb:MT044257|Organism:Severe      CTGCTATCAGCGATTATGACTACTATCGTTATAATCTACCAACAATGTGTGATATCAGAC   14842
gb:MT106054|Organism:Severe      CTGCTATCAGCGATTATGACTACTATCGTTATAATCTACCAACAATGTGTGATATCAGAC   14842
gb:MT049951|Organism:Severe      CTGCTATCAGCGATTATGACTACTATCGTTATAATCTACCAACAATGTGTGATATCAGAC   14842
gb:MN975262|Organism:Severe      CTGCTATCAGCGATTATGACTACTATCGTTATAATCTACCAACAATGTGTGATATCAGAC   14842
gb:MT106052|Organism:Severe      CTGCTATCAGCGATTATGACTACTATCGTTATAATCTACCAACAATGTGTGATATCAGAC   14842
gb:LC522975|Organism:Severe      CTGCTATCAGCGATTATGACTACTATCGTTATAATCTACCAACAATGTGTGATATCAGAC   14839
```

```
gb:LC522973|Organism:Severe      CTGCTATCAGCGATTATGACTACTATCGTTATAATCTACCAACAATGTGTGATATCAGAC   14839
gb:LC522974|Organism:Severe      CTGCTATCAGCGATTATGACTACTATCGTTATAATCTACCAACAATGTGTGATATCAGAC   14839
gb:MN985325|Organism:Severe      CTGCTATCAGCGATTATGACTACTATCGTTATAATCTACCAACAATGTGTGATATCAGAC   14842
gb:MT020881|Organism:Severe      CTGCTATCAGCGATTATGACTACTATCGTTATAATCTACCAACAATGTGTGATATCAGAC   14842
gb:MT020880|Organism:Severe      CTGCTATCAGCGATTATGACTACTATCGTTATAATCTACCAACAATGTGTGATATCAGAC   14842
gb:MT066175|Organism:Severe      CTGCTATCAGCGATTATGACTACTATCGTTATAATCTACCAACAATGTGTGATATCAGAC   14842
gb:MN997409|Organism:Severe      CTGCTATCAGCGATTATGACTACTATCGTTATAATCTACCAACAATGTGTGATATCAGAC   14842
gb:MN938384|Organism:Severe      CTGCTATCAGCGATTATGACTACTATCGTTATAATCTACCAACAATGTGTGATATCAGAC   14810
gb:MT044258|Organism:Severe      CTGCTATCAGCGATTATGACTACTATCGTTATAATCTACCAACAATGTGTGATATCAGAC   14818
gb:MT039890|Organism:Severe      CTGCTATCAGCGATTATGACTACTATCGTTATAATCTACCAACAATGTGTGATATCAGAC   14842
gb:MN988713|Organism:Severe      CTGCTATCAGCGATTATGACTACTATCGTTATAATCTACCAACAATGTGTGATATCAGAC   14842
gb:LC521925|Organism:Severe      CTGCTATCAGCGATTATGACTACTATCGTTATAATCTACCAACAATGTGTGATATCAGAC   14815
gb:MT093571|Organism:Severe      CTGCTATCAGCGATTATGACTACTATCGTTATAATCTACCAACAATGTGTGATATCAGAC   14842
gb:MT039887|Organism:Severe      CTGCTATCAGCGATTATGACTACTATCGTTATAATCTACCAACAATGTGTGATATCAGAC   14842
gb:MT019530|Organism:Severe      CTGCTATCAGCGATTATGACTACTATCGTTATAATCTACCAACAATGTGTGATATCAGAC   14842
gb:MT039888|Organism:Severe      CTGCTATCAGCGATTATGACTACTATCGTTATAATCTACCAACAATGTGTGATATCAGAC   14842
gb:LC522972|Organism:Severe      CTGCTATCAGCGATTATGACTACTATCGTTATAATCTACCAACAATGTGTGATATCAGAC   14839
gb:MT027063|Organism:Severe      CTGCTATCAGCGATTATGACTACTATCGTTATAATCTACCAACAATGTGTGATATCAGAC   14842
gb:MT027062|Organism:Severe      CTGCTATCAGCGATTATGACTACTATCGTTATAATCTACCAACAATGTGTGATATCAGAC   14842
gb:MT019529|Organism:Severe      CTGCTATCAGCGATTATGACTACTATCGTTATAATCTACCAACAATGTGTGATATCAGAC   14842
gb:MN996529|Organism:Severe      CTGCTATCAGCGATTATGACTACTATCGTTATAATCTACCAACAATGTGTGATATCAGAC   14830
gb:MN996531|Organism:Severe      CTGCTATCAGCGATTATGACTACTATCGTTATAATCTACCAACAATGTGTGATATCAGAC   14829
gb:MT066176|Organism:Severe      CTGCTATCAGCGATTATGACTACTATCGTTATAATCTACCAACAATGTGTGATATCAGAC   14842
gb:MT027064|Organism:Severe      CTGCTATCAGCGATTATGACTACTATCGTTATAATCTACCAACAATGTGTGATATCAGAC   14842
gb:MN994468|Organism:Severe      CTGCTATCAGCGATTATGACTACTATCGTTATAATCTACCAACAATGTGTGATATCAGAC   14842
gb:MT072688|Organism:Severe      CTGCTATCAGCGATTATGACTACTATCGTTATAATCTACCAACAATGTGTGATATCAGAC   14827
gb:MN996527|Organism:Severe      CTGCTATCAGCGATTATGACTACTATCGTTATAATCTACCAACAATGTGTGATATCAGAC   14809
gb:MT093631|Organism:Severe      CTGCTATCAGCGATTATGACTACTATCGTTATAATCTACCAACAATGTGTGATATCAGAC   14880
gb:MT106053|Organism:Severe      CTGCTATCAGCGATTATGACTACTATCGTTATAATCTACCAACAATGTGTGATATCAGAC   14842
gb:MT019533|Organism:Severe      CTGCTATCAGCGATTATGACTACTATCGTTATAATCTACCAACAATGTGTGATATCAGAC   14842
gb:MT019531|Organism:Severe      CTGCTATCAGCGATTATGACTACTATCGTTATAATCTACCAACAATGTGTGATATCAGAC   14842
gb:MN996528|Organism:Severe      CTGCTATCAGCGATTATGACTACTATCGTTATAATCTACCAACAATGTGTGATATCAGAC   14842
gb:MN996530|Organism:Severe      CTGCTATCAGCGATTATGACTACTATCGTTATAATCTACCAACAATGTGTGATATCAGAC   14828
gb:MN908947|Organism:Severe      CTGCTATCAGCGATTATGACTACTATCGTTATAATCTACCAACAATGTGTGATATCAGAC   14842
gb:MT019532|Organism:Severe      CTGCTATCAGCGATTATGACTACTATCGTTATAATCTACCAACAATGTGTGATATCAGAC   14842
                                 ************************************************************

gb:MT020781|Organism:Severe      AACTACTATTTGTAGTTGAAGTTGTTGATAAGTACTTTGATTGTTACGATGGTGGCTGTA   14890
gb:MT007544|Organism:Severe      AACTACTATTTGTAGTTGAAGTTGTTGATAAGTACTTTGATTGTTACGATGGTGGCTGTA   14902
gb:MN994467|Organism:Severe      AACTACTATTTGTAGTTGAAGTTGTTGATAAGTACTTTGATTGTTACGATGGTGGCTGTA   14902
gb:MT044257|Organism:Severe      AACTACTATTTGTAGTTGAAGTTGTTGATAAGTACTTTGATTGTTACGATGGTGGCTGTA   14902
gb:MT106054|Organism:Severe      AACTACTATTTGTAGTTGAAGTTGTTGATAAGTACTTTGATTGTTACGATGGTGGCTGTA   14902
gb:MT049951|Organism:Severe      AACTACTATTTGTAGTTGAAGTTGTTGATAAGTACTTTGATTGTTACGATGGTGGCTGTA   14902
gb:MN975262|Organism:Severe      AACTACTATTTGTAGTTGAAGTTGTTGATAAGTACTTTGATTGTTACGATGGTGGCTGTA   14902
gb:MT106052|Organism:Severe      AACTACTATTTGTAGTTGAAGTTGTTGATAAGTACTTTGATTGTTACGATGGTGGCTGTA   14902
gb:LC522975|Organism:Severe      AACTACTATTTGTAGTTGAAGTTGTTGATAAGTACTTTGATTGTTACGATGGTGGCTGTA   14899
gb:LC522973|Organism:Severe      AACTACTATTTGTAGTTGAAGTTGTTGATAAGTACTTTGATTGTTACGATGGTGGCTGTA   14899
gb:LC522974|Organism:Severe      AACTACTATTTGTAGTTGAAGTTGTTGATAAGTACTTTGATTGTTACGATGGTGGCTGTA   14899
gb:MN985325|Organism:Severe      AACTACTATTTGTAGTTGAAGTTGTTGATAAGTACTTTGATTGTTACGATGGTGGCTGTA   14902
gb:MT020881|Organism:Severe      AACTACTATTTGTAGTTGAAGTTGTTGATAAGTACTTTGATTGTTACGATGGTGGCTGTA   14902
gb:MT020880|Organism:Severe      AACTACTATTTGTAGTTGAAGTTGTTGATAAGTACTTTGATTGTTACGATGGTGGCTGTA   14902
gb:MT066175|Organism:Severe      AACTACTATTTGTAGTTGAAGTTGTTGATAAGTACTTTGATTGTTACGATGGTGGCTGTA   14902
gb:MN997409|Organism:Severe      AACTACTATTTGTAGTTGAAGTTGTTGATAAGTACTTTGATTGTTACGATGGTGGCTGTA   14902
gb:MN938384|Organism:Severe      AACTACTATTTGTAGTTGAAGTTGTTGATAAGTACTTTGATTGTTACGATGGTGGCTGTA   14870
gb:MT044258|Organism:Severe      AACTACTATTTGTAGTTGAAGTTGTTGATAAGTACTTTGATTGTTACGATGGTGGCTGTA   14878
gb:MT039890|Organism:Severe      AACTACTATTTGTAGTTGAAGTTGTTGATAAGTACTTTGATTGTTACGATGGTGGCTGTA   14902
gb:MN988713|Organism:Severe      AACTACTATTTGTAGTTGAAGTTGTTGATAAGTACTTTGATTGTTACGATGGTGGCTGTA   14902
gb:LC521925|Organism:Severe      AACTACTATTTGTAGTTGAAGTTGTTGATAAGTACTTTGATTGTTACGATGGTGGCTGTA   14875
gb:MT093571|Organism:Severe      AACTACTATTTGTAGTTGAAGTTGTTGATAAGTACTTTGATTGTTACGATGGTGGCTGTA   14902
gb:MT039887|Organism:Severe      AACTACTATTTGTAGTTGAAGTTGTTGATAAGTACTTTGATTGTTACGATGGTGGCTGTA   14902
gb:MT019530|Organism:Severe      AACTACTATTTGTAGTTGAAGTTGTTGATAAGTACTTTGATTGTTACGATGGTGGCTGTA   14902
gb:MT039888|Organism:Severe      AACTACTATTTGTAGTTGAAGTTGTTGATAAGTACTTTGATTGTTACGATGGTGGCTGTA   14902
gb:LC522972|Organism:Severe      AACTACTATTTGTAGTTGAAGTTGTTGATAAGTACTTTGATTGTTACGATGGTGGCTGTA   14899
gb:MT027063|Organism:Severe      AACTACTATTTGTAGTTGAAGTTGTTGATAAGTACTTTGATTGTTACGATGGTGGCTGTA   14902
gb:MT027062|Organism:Severe      AACTACTATTTGTAGTTGAAGTTGTTGATAAGTACTTTGATTGTTACGATGGTGGCTGTA   14902
```

```
gb:MT019529|Organism:Severe      AACTACTATTTGTAGTTGAAGTTGTTGATAAGTACTTTGATTGTTACGATGGTGGCTGTA   14902
gb:MN996529|Organism:Severe      AACTACTATTTGTAGTTGAAGTTGTTGATAAGTACTTTGATTGTTACGATGGTGGCTGTA   14890
gb:MN996531|Organism:Severe      AACTACTATTTGTAGTTGAAGTTGTTGATAAGTACTTTGATTGTTACGATGGTGGCTGTA   14889
gb:MT066176|Organism:Severe      AACTACTATTTGTAGTTGAAGTTGTTGATAAGTACTTTGATTGTTACGATGGTGGCTGTA   14902
gb:MT027064|Organism:Severe      AACTACTATTTGTAGTTGAAGTTGTTGATAAGTACTTTGATTGTTACGATGGTGGCTGTA   14902
gb:MN994468|Organism:Severe      AACTACTATTTGTAGTTGAAGTTGTTGATAAGTACTTTGATTGTTACGATGGTGGCTGTA   14902
gb:MT072688|Organism:Severe      AACTACTATTTGTAGTTGAAGTTGTTGATAAGTACTTTGATTGTTACGATGGTGGCTGTA   14887
gb:MN996527|Organism:Severe      AACTACTATTTGTAGTTGAAGTTGTTGATAAGTACTTTGATTGTTACGATGGTGGCTGTA   14869
gb:MT093631|Organism:Severe      AACTACTATTTGTAGTTGAAGTTGTTGATAAGTACTTTGATTGTTACGATGGTGGCTGTA   14940
gb:MT106053|Organism:Severe      AACTACTATTTGTAGTTGAAGTTGTTGATAAGTACTTTGATTGTTACGATGGTGGCTGTA   14902
gb:MT019533|Organism:Severe      AACTACTATTTGTAGTTGAAGTTGTTGATAAGTACTTTGATTGTTACGATGGTGGCTGTA   14902
gb:MT019531|Organism:Severe      AACTACTATTTGTAGTTGAAGTTGTTGATAAGTACTTTGATTGTTACGATGGTGGCTGTA   14902
gb:MN996528|Organism:Severe      AACTACTATTTGTAGTTGAAGTTGTTGATAAGTACTTTGATTGTTACGATGGTGGCTGTA   14902
gb:MN996530|Organism:Severe      AACTACTATTTGTAGTTGAAGTTGTTGATAAGTACTTTGATTGTTACGATGGTGGCTGTA   14888
gb:MN908947|Organism:Severe      AACTACTATTTGTAGTTGAAGTTGTTGATAAGTACTTTGATTGTTACGATGGTGGCTGTA   14902
gb:MT019532|Organism:Severe      AACTACTATTTGTAGTTGAAGTTGTTGATAAGTACTTTGATTGTTACGATGGTGGCTGTA   14902
                                 ************************************************************

gb:MT020781|Organism:Severe      TTAATGCTAACCAAGTCATCGTCAACAACCTAGACAAATCAGCTGGTTTTCCATTTAATA   14950
gb:MT007544|Organism:Severe      TTAATGCTAACCAAGTCATCGTCAACAACCTAGACAAATCAGCTGGTTTTCCATTTAATA   14962
gb:MN994467|Organism:Severe      TTAATGCTAACCAAGTCATCGTCAACAACCTAGACAAATCAGCTGGTTTTCCATTTAATA   14962
gb:MT044257|Organism:Severe      TTAATGCTAACCAAGTCATCGTCAACAACCTAGACAAATCAGCTGGTTTTCCATTTAATA   14962
gb:MT106054|Organism:Severe      TTAATGCTAACCAAGTCATCGTCAACAACCTAGACAAATCAGCTGGTTTTCCATTTAATA   14962
gb:MT049951|Organism:Severe      TTAATGCTAACCAAGTCATCGTCAACAACCTAGACAAATCAGCTGGTTTTCCATTTAATA   14962
gb:MN975262|Organism:Severe      TTAATGCTAACCAAGTCATCGTCAACAACCTAGACAAATCAGCTGGTTTTCCATTTAATA   14962
gb:MT106052|Organism:Severe      TTAATGCTAACCAAGTCATCGTCAACAACCTAGACAAATCAGCTGGTTTTCCATTTAATA   14962
gb:LC522975|Organism:Severe      TTAATGCTAACCAAGTCATCGTCAACAACCTAGACAAATCAGCTGGTTTTCCATTTAATA   14959
gb:LC522973|Organism:Severe      TTAATGCTAACCAAGTCATCGTCAACAACCTAGACAAATCAGCTGGTTTTCCATTTAATA   14959
gb:LC522974|Organism:Severe      TTAATGCTAACCAAGTCATCGTCAACAACCTAGACAAATCAGCTGGTTTTCCATTTAATA   14959
gb:MN985325|Organism:Severe      TTAATGCTAACCAAGTCATCGTCAACAACCTAGACAAATCAGCTGGTTTTCCATTTAATA   14962
gb:MT020881|Organism:Severe      TTAATGCTAACCAAGTCATCGTCAACAACCTAGACAAATCAGCTGGTTTTCCATTTAATA   14962
gb:MT020880|Organism:Severe      TTAATGCTAACCAAGTCATCGTCAACAACCTAGACAAATCAGCTGGTTTTCCATTTAATA   14962
gb:MT066175|Organism:Severe      TTAATGCTAACCAAGTCATCGTCAACAACCTAGACAAATCAGCTGGTTTTCCATTTAATA   14962
gb:MN997409|Organism:Severe      TTAATGCTAACCAAGTCATCGTCAACAACCTAGACAAATCAGCTGGTTTTCCATTTAATA   14962
gb:MN938384|Organism:Severe      TTAATGCTAACCAAGTCATCGTCAACAACCTAGACAAATCAGCTGGTTTTCCATTTAATA   14930
gb:MT044258|Organism:Severe      TTAATGCTAACCAAGTCATCGTCAACAACCTAGACAAATCAGCTGGTTTTCCATTTAATA   14938
gb:MT039890|Organism:Severe      TTAATGCTAACCAAGTCATCGTCAACAACCTAGACAAATCAGCTGGTTTTCCATTTAATA   14962
gb:MN988713|Organism:Severe      TTAATGCTAACCAAGTCATCGTCAACAACCTAGACAAATCAGCTGGTTTTCCATTTAATA   14962
gb:LC521925|Organism:Severe      TTAATGCTAACCAAGTCATCGTCAACAACCTAGACAAATCAGCTGGTTTTCCATTTAATA   14935
gb:MT093571|Organism:Severe      TTAATGCTAACCAAGTCATCGTCAACAACCTAGACAAATCAGCTGGTTTTCCATTTAATA   14962
gb:MT039887|Organism:Severe      TTAATGCTAACCAAGTCATCGTCAACAACCTAGACAAATCAGCTGGTTTTCCATTTAATA   14962
gb:MT019530|Organism:Severe      TTAATGCTAACCAAGTCATCGTCAACAACCTAGACAAATCAGCTGGTTTTCCATTTAATA   14962
gb:MT039888|Organism:Severe      TTAATGCTAACCAAGTCATCGTCAACAACCTAGACAAATCAGCTGGTTTTCCATTTAATA   14962
gb:LC522972|Organism:Severe      TTAATGCTAACCAAGTCATCGTCAACAACCTAGACAAATCAGCTGGTTTTCCATTTAATA   14959
gb:MT027063|Organism:Severe      TTAATGCTAACCAAGTCATCGTCAACAACCTAGACAAATCAGCTGGTTTTCCATTTAATA   14962
gb:MT027062|Organism:Severe      TTAATGCTAACCAAGTCATCGTCAACAACCTAGACAAATCAGCTGGTTTTCCATTTAATA   14962
gb:MT019529|Organism:Severe      TTAATGCTAACCAAGTCATCGTCAACAACCTAGACAAATCAGCTGGTTTTCCATTTAATA   14962
gb:MN996529|Organism:Severe      TTAATGCTAACCAAGTCATCGTCAACAACCTAGACAAATCAGCTGGTTTTCCATTTAATA   14950
gb:MN996531|Organism:Severe      TTAATGCTAACCAAGTCATCGTCAACAACCTAGACAAATCAGCTGGTTTTCCATTTAATA   14949
gb:MT066176|Organism:Severe      TTAATGCTAACCAAGTCATCGTCAACAACCTAGACAAATCAGCTGGTTTTCCATTTAATA   14962
gb:MT027064|Organism:Severe      TTAATGCTAACCAAGTCATCGTCAACAACCTAGACAAATCAGCTGGTTTTCCATTTAATA   14962
gb:MN994468|Organism:Severe      TTAATGCTAACCAAGTCATCGTCAACAACCTAGACAAATCAGCTGGTTTTCCATTTAATA   14962
gb:MT072688|Organism:Severe      TTAATGCTAACCAAGTCATCGTCAACAACCTAGACAAATCAGCTGGTTTTCCATTTAATA   14947
gb:MN996527|Organism:Severe      TTAATGCTAACCAAGTCATCGTCAACAACCTAGACAAATCAGCTGGTTTTCCATTTAATA   14929
gb:MT093631|Organism:Severe      TTAATGCTAACCAAGTCATCGTCAACAACCTAGACAAATCAGCTGGTTTTCCATTTAATA   15000
gb:MT106053|Organism:Severe      TTAATGCTAACCAAGTCATCGTCAACAACCTAGACAAATCAGCTGGTTTTCCATTTAATA   14962
gb:MT019533|Organism:Severe      TTAATGCTAACCAAGTCATCGTCAACAACCTAGACAAATCAGCTGGTTTTCCATTTAATA   14962
gb:MT019531|Organism:Severe      TTAATGCTAACCAAGTCATCGTCAACAACCTAGACAAATCAGCTGGTTTTCCATTTAATA   14962
gb:MN996528|Organism:Severe      TTAATGCTAACCAAGTCATCGTCAACAACCTAGACAAATCAGCTGGTTTTCCATTTAATA   14962
gb:MN996530|Organism:Severe      TTAATGCTAACCAAGTCATCGTCAACAACCTAGACAAATCAGCTGGTTTTCCATTTAATA   14948
gb:MN908947|Organism:Severe      TTAATGCTAACCAAGTCATCGTCAACAACCTAGACAAATCAGCTGGTTTTCCATTTAATA   14962
gb:MT019532|Organism:Severe      TTAATGCTAACCAAGTCATCGTCAACAACCTAGACAAATCAGCTGGTTTTCCATTTAATA   14962
                                 ************************************************************

gb:MT020781|Organism:Severe      AATGGGGTAAGGCTAGACTTTATTATGATTCAATGAGTTATGAGGATCAAGATGCACTTT   15010
```

```
gb:MT007544|Organism:Severe      AATGGGGTAAGGCTAGACTTTATTATGATTCAATGAGTTATGAGGATCAAGATGCACTTT   15022
gb:MN994467|Organism:Severe      AATGGGGTAAGGCTAGACTTTATTATGATTCAATGAGTTATGAGGATCAAGATGCACTTT   15022
gb:MT044257|Organism:Severe      AATGGGGTAAGGCTAGACTTTATTATGATTCAATGAGTTATGAGGATCAAGATGCACTTT   15022
gb:MT106054|Organism:Severe      AATGGGGTAAGGCTAGACTTTATTATGATTCAATGAGTTATGAGGATCAAGATGCACTTT   15022
gb:MT049951|Organism:Severe      AATGGGGTAAGGCTAGACTTTATTATGATTCAATGAGTTATGAGGATCAAGATGCACTTT   15022
gb:MN975262|Organism:Severe      AATGGGGTAAGGCTAGACTTTATTATGATTCAATGAGTTATGAGGATCAAGATGCACTTT   15022
gb:MT106052|Organism:Severe      AATGGGGTAAGGCTAGACTTTATTATGATTCAATGAGTTATGAGGATCAAGATGCACTTT   15022
gb:LC522975|Organism:Severe      AATGGGGTAAGGCTAGACTTTATTATGATTCAATGAGTTATGAGGATCAAGATGCACTTT   15019
gb:LC522973|Organism:Severe      AATGGGGTAAGGCTAGACTTTATTATGATTCAATGAGTTATGAGGATCAAGATGCACTTT   15019
gb:LC522974|Organism:Severe      AATGGGGTAAGGCTAGACTTTATTATGATTCAATGAGTTATGAGGATCAAGATGCACTTT   15019
gb:MN985325|Organism:Severe      AATGGGGTAAGGCTAGACTTTATTATGATTCAATGAGTTATGAGGATCAAGATGCACTTT   15022
gb:MT020881|Organism:Severe      AATGGGGTAAGGCTAGACTTTATTATGATTCAATGAGTTATGAGGATCAAGATGCACTTT   15022
gb:MT020880|Organism:Severe      AATGGGGTAAGGCTAGACTTTATTATGATTCAATGAGTTATGAGGATCAAGATGCACTTT   15022
gb:MT066175|Organism:Severe      AATGGGGTAAGGCTAGACTTTATTATGATTCAATGAGTTATGAGGATCAAGATGCACTTT   15022
gb:MN997409|Organism:Severe      AATGGGGTAAGGCTAGACTTTATTATGATTCAATGAGTTATGAGGATCAAGATGCACTTT   15022
gb:MN938384|Organism:Severe      AATGGGGTAAGGCTAGACTTTATTATGATTCAATGAGTTATGAGGATCAAGATGCACTTT   14990
gb:MT044258|Organism:Severe      AATGGGGTAAGGCTAGACTTTATTATGATTCAATGAGTTATGAGGATCAAGATGCACTTT   14998
gb:MT039890|Organism:Severe      AATGGGGTAAGGCTAGACTTTATTATGATTCAATGAGTTATGAGGATCAAGATGCACTTT   15022
gb:MN988713|Organism:Severe      AATGGGGTAAGGCTAGACTTTATTATGATTCAATGAGTTATGAGGATCAAGATGCACTTT   15022
gb:LC521925|Organism:Severe      AATGGGGTAAGGCTAGACTTTATTATGATTCAATGAGTTATGAGGATCAAGATGCACTTT   14995
gb:MT093571|Organism:Severe      AATGGGGTAAGGCTAGACTTTATTATGATTCAATGAGTTATGAGGATCAAGATGCACTTT   15022
gb:MT039887|Organism:Severe      AATGGGGTAAGGCTAGACTTTATTATGATTCAATGAGTTATGAGGATCAAGATGCACTTT   15022
gb:MT019530|Organism:Severe      AATGGGGTAAGGCTAGACTTTATTATGATTCAATGAGTTATGAGGATCAAGATGCACTTT   15022
gb:MT039888|Organism:Severe      AATGGGGTAAGGCTAGACTTTATTATGATTCAATGAGTTATGAGGATCAAGATGCACTTT   15022
gb:LC522972|Organism:Severe      AATGGGGTAAGGCTAGACTTTATTATGATTCAATGAGTTATGAGGATCAAGATGCACTTT   15019
gb:MT027063|Organism:Severe      AATGGGGTAAGGCTAGACTTTATTATGATTCAATGAGTTATGAGGATCAAGATGCACTTT   15022
gb:MT027062|Organism:Severe      AATGGGGTAAGGCTAGACTTTATTATGATTCAATGAGTTATGAGGATCAAGATGCACTTT   15022
gb:MT019529|Organism:Severe      AATGGGGTAAGGCTAGACTTTATTATGATTCAATGAGTTATGAGGATCAAGATGCACTTT   15022
gb:MN996529|Organism:Severe      AATGGGGTAAGGCTAGACTTTATTATGATTCAATGAGTTATGAGGATCAAGATGCACTTT   15010
gb:MN996531|Organism:Severe      AATGGGGTAAGGCTAGACTTTATTATGATTCAATGAGTTATGAGGATCAAGATGCACTTT   15009
gb:MT066176|Organism:Severe      AATGGGGTAAGGCTAGACTTTATTATGATTCAATGAGTTATGAGGATCAAGATGCACTTT   15022
gb:MT027064|Organism:Severe      AATGGGGTAAGGCTAGACTTTATTATGATTCAATGAGTTATGAGGATCAAGATGCACTTT   15022
gb:MN994468|Organism:Severe      AATGGGGTAAGGCTAGACTTTATTATGATTCAATGAGTTATGAGGATCAAGATGCACTTT   15022
gb:MT072688|Organism:Severe      AATGGGGTAAGGCTAGACTTTATTATGATTCAATGAGTTATGAGGATCAAGATGCACTTT   15007
gb:MN996527|Organism:Severe      AATGGGGTAAGGCTAGACTTTATTATGATTCAATGAGTTATGAGGATCAAGATGCACTTT   14989
gb:MT093631|Organism:Severe      AATGGGGTAAGGCTAGACTTTATTATGATTCAATGAGTTATGAGGATCAAGATGCACTTT   15060
gb:MT106053|Organism:Severe      AATGGGGTAAGGCTAGACTTTATTATGATTCAATGAGTTATGAGGATCAAGATGCACTTT   15022
gb:MT019533|Organism:Severe      AATGGGGTAAGGCTAGACTTTATTATGATTCAATGAGTTATGAGGATCAAGATGCACTTT   15022
gb:MT019531|Organism:Severe      AATGGGGTAAGGCTAGACTTTATTATGATTCAATGAGTTATGAGGATCAAGATGCACTTT   15022
gb:MN996528|Organism:Severe      AATGGGGTAAGGCTAGACTTTATTATGATTCAATGAGTTATGAGGATCAAGATGCACTTT   15022
gb:MN996530|Organism:Severe      AATGGGGTAAGGCTAGACTTTATTATGATTCAATGAGTTATGAGGATCAAGATGCACTTT   15008
gb:MN908947|Organism:Severe      AATGGGGTAAGGCTAGACTTTATTATGATTCAATGAGTTATGAGGATCAAGATGCACTTT   15022
gb:MT019532|Organism:Severe      AATGGGGTAAGGCTAGACTTTATTATGATTCAATGAGTTATGAGGATCAAGATGCACTTT   15022
                                 ************************************************************

gb:MT020781|Organism:Severe      TCGCATATACAAAACGTAATGTCATCCCTACTATAACTCAAATGAATCTTAAGTATGCCA   15070
gb:MT007544|Organism:Severe      TCGCATATACAAAACGTAATGTCATCCCTACTATAACTCAAATGAATCTTAAGTATGCCA   15082
gb:MN994467|Organism:Severe      TCGCATATACAAAACGTAATGTCATCCCTACTATAACTCAAATGAATCTTAAGTATGCCA   15082
gb:MT044257|Organism:Severe      TCGCATATACAAAACGTAATGTCATCCCTACTATAACTCAAATGAATCTTAAGTATGCCA   15082
gb:MT106054|Organism:Severe      TCGCATATACAAAACGTAATGTCATCCCTACTATAACTCAAATGAATCTTAAGTATGCCA   15082
gb:MT049951|Organism:Severe      TCGCATATACAAAACGTAATGTCATCCCTACTATAACTCAAATGAATCTTAAGTATGCCA   15082
gb:MN975262|Organism:Severe      TCGCATATACAAAACGTAATGTCATCCCTACTATAACTCAAATGAATCTTAAGTATGCCA   15082
gb:MT106052|Organism:Severe      TCGCATATACAAAACGTAATGTCATCCCTACTATAACTCAAATGAATCTTAAGTATGCCA   15082
gb:LC522975|Organism:Severe      TCGCATATACAAAACGTAATGTCATCCCTACTATAACTCAAATGAATCTTAAGTATGCCA   15079
gb:LC522973|Organism:Severe      TCGCATATACAAAACGTAATGTCATCCCTACTATAACTCAAATGAATCTTAAGTATGCCA   15079
gb:LC522974|Organism:Severe      TCGCATATACAAAACGTAATGTCATCCCTACTATAACTCAAATGAATCTTAAGTATGCCA   15079
gb:MN985325|Organism:Severe      TCGCATATACAAAACGTAATGTCATCCCTACTATAACTCAAATGAATCTTAAGTATGCCA   15082
gb:MT020881|Organism:Severe      TCGCATATACAAAACGTAATGTCATCCCTACTATAACTCAAATGAATCTTAAGTATGCCA   15082
gb:MT020880|Organism:Severe      TCGCATATACAAAACGTAATGTCATCCCTACTATAACTCAAATGAATCTTAAGTATGCCA   15082
gb:MT066175|Organism:Severe      TCGCATATACAAAACGTAATGTCATCCCTACTATAACTCAAATGAATCTTAAGTATGCCA   15082
gb:MN997409|Organism:Severe      TCGCATATACAAAACGTAATGTCATCCCTACTATAACTCAAATGAATCTTAAGTATGCCA   15082
gb:MN938384|Organism:Severe      TCGCATATACAAAACGTAATGTCATCCCTACTATAACTCAAATGAATCTTAAGTATGCCA   15050
gb:MT044258|Organism:Severe      TCGCATATACAAAACGTAATGTCATCCCTACTATAACTCAAATGAATCTTAAGTATGCCA   15058
gb:MT039890|Organism:Severe      TCGCATATACAAAACGTAATGTCATCCCTACTATAACTCAAATGAATCTTAAGTATGCCA   15082
gb:MN988713|Organism:Severe      TCGCATATACAAAACGTAATGTCATCCCTACTATAACTCAAATGAATCTTAAGTATGCCA   15082
```

```
gb:LC521925|Organism:Severe      TCGCATATACAAAACGTAATGTCATCCCTACTATAACTCAAATGAATCTTAAGTATGCCA   15055
gb:MT093571|Organism:Severe      TCGCATATACAAAACGTAATGTCATCCCTACTATAACTCAAATGAATCTTAAGTATGCCA   15082
gb:MT039887|Organism:Severe      TCGCATATACAAAACGTAATGTCATCCCTACTATAACTCAAATGAATCTTAAGTATGCCA   15082
gb:MT019530|Organism:Severe      TCGCATATACAAAACGTAATGTCATCCCTACTATAACTCAAATGAATCTTAAGTATGCCA   15082
gb:MT039888|Organism:Severe      TCGCATATACAAAACGTAATGTCATCCCTACTATAACTCAAATGAATCTTAAGTATGCCA   15082
gb:LC522972|Organism:Severe      TCGCATATACAAAACGTAATGTCATCCCTACTATAACTCAAATGAATCTTAAGTATGCCA   15079
gb:MT027063|Organism:Severe      TCGCATATACAAAACGTAATGTCATCCCTACTATAACTCAAATGAATCTTAAGTATGCCA   15082
gb:MT027062|Organism:Severe      TCGCATATACAAAACGTAATGTCATCCCTACTATAACTCAAATGAATCTTAAGTATGCCA   15082
gb:MT019529|Organism:Severe      TCGCATATACAAAACGTAATGTCATCCCTACTATAACTCAAATGAATCTTAAGTATGCCA   15082
gb:MN996529|Organism:Severe      TCGCATATACAAAACGTAATGTCATCCCTACTATAACTCAAATGAATCTTAAGTATGCCA   15070
gb:MN996531|Organism:Severe      TCGCATATACAAAACGTAATGTCATCCCTACTATAACTCAAATGAATCTTAAGTATGCCA   15069
gb:MT066176|Organism:Severe      TCGCATATACAAAACGTAATGTCATCCCTACTATAACTCAAATGAATCTTAAGTATGCCA   15082
gb:MT027064|Organism:Severe      TCGCATATACAAAACGTAATGTCATCCCTACTATAACTCAAATGAATCTTAAGTATGCCA   15082
gb:MN994468|Organism:Severe      TCGCATATACAAAACGTAATGTCATCCCTACTATAACTCAAATGAATCTTAAGTATGCCA   15082
gb:MT072688|Organism:Severe      TCGCATATACAAAACGTAATGTCATCCCTACTATAACTCAAATGAATCTTAAGTATGCCA   15067
gb:MN996527|Organism:Severe      TCGCATATACAAAACGTAATGTCATCCCTACTATAACTCAAATGAATCTTAAGTATGCCA   15049
gb:MT093631|Organism:Severe      TCGCATATACAAAACGTAATGTCATCCCTACTATAACTCAAATGAATCTTAAGTATGCCA   15120
gb:MT106053|Organism:Severe      TCGCATATACAAAACGTAATGTCATCCCTACTATAACTCAAATGAATCTTAAGTATGCCA   15082
gb:MT019533|Organism:Severe      TCGCATATACAAAACGTAATGTCATCCCTACTATAACTCAAATGAATCTTAAGTATGCCA   15082
gb:MT019531|Organism:Severe      TCGCATATACAAAACGTAATGTCATCCCTACTATAACTCAAATGAATCTTAAGTATGCCA   15082
gb:MN996528|Organism:Severe      TCGCATATACAAAACGTAATGTCATCCCTACTATAACTCAAATGAATCTTAAGTATGCCA   15082
gb:MN996530|Organism:Severe      TCGCATATACAAAACGTAATGTCATCCCTACTATAACTCAAATGAATCTTAAGTATGCCA   15068
gb:MN908947|Organism:Severe      TCGCATATACAAAACGTAATGTCATCCCTACTATAACTCAAATGAATCTTAAGTATGCCA   15082
gb:MT019532|Organism:Severe      TCGCATATACAAAACGTAATGTCATCCCTACTATAACTCAAATGAATCTTAAGTATGCCA   15082
                                 ************************************************************

gb:MT020781|Organism:Severe      TTAGTGCAAAGAATAGAGCTCGCACCGTAGCTGGTGTCTCTATCTGTAGTACTATGACCA   15130
gb:MT007544|Organism:Severe      TTAGTGCAAAGAATAGAGCTCGCACCGTAGCTGGTGTCTCTATCTGTAGTACTATGACCA   15142
gb:MN994467|Organism:Severe      TTAGTGCAAAGAATAGAGCTCGCACCGTAGCTGGTGTCTCTATCTGTAGTACTATGACCA   15142
gb:MT044257|Organism:Severe      TTAGTGCAAAGAATAGAGCTCGCACCGTAGCTGGTGTCTCTATCTGTAGTACTATGACCA   15142
gb:MT106054|Organism:Severe      TTAGTGCAAAGAATAGAGCTCGCACCGTAGCTGGTGTCTCTATCTGTAGTACTATGACCA   15142
gb:MT049951|Organism:Severe      TTAGTGCAAAGAATAGAGCTCGCACCGTAGCTGGTGTCTCTATCTGTAGTACTATGACCA   15142
gb:MN975262|Organism:Severe      TTAGTGCAAAGAATAGAGCTCGCACCGTAGCTGGTGTCTCTATCTGTAGTACTATGACCA   15142
gb:MT106052|Organism:Severe      TTAGTGCAAAGAATAGAGCTCGCACCGTAGCTGGTGTCTCTATCTGTAGTACTATGACCA   15142
gb:LC522975|Organism:Severe      TTAGTGCAAAGAATAGAGCTCGCACCGTAGCTGGTGTCTCTATCTGTAGTACTATGACCA   15139
gb:LC522973|Organism:Severe      TTAGTGCAAAGAATAGAGCTCGCACCGTAGCTGGTGTCTCTATCTGTAGTACTATGACCA   15139
gb:LC522974|Organism:Severe      TTAGTGCAAAGAATAGAGCTCGCACCGTAGCTGGTGTCTCTATCTGTAGTACTATGACCA   15139
gb:MN985325|Organism:Severe      TTAGTGCAAAGAATAGAGCTCGCACCGTAGCTGGTGTCTCTATCTGTAGTACTATGACCA   15142
gb:MT020881|Organism:Severe      TTAGTGCAAAGAATAGAGCTCGCACCGTAGCTGGTGTCTCTATCTGTAGTACTATGACCA   15142
gb:MT020880|Organism:Severe      TTAGTGCAAAGAATAGAGCTCGCACCGTAGCTGGTGTCTCTATCTGTAGTACTATGACCA   15142
gb:MT066175|Organism:Severe      TTAGTGCAAAGAATAGAGCTCGCACCGTAGCTGGTGTCTCTATCTGTAGTACTATGACCA   15142
gb:MN997409|Organism:Severe      TTAGTGCAAAGAATAGAGCTCGCACCGTAGCTGGTGTCTCTATCTGTAGTACTATGACCA   15142
gb:MN938384|Organism:Severe      TTAGTGCAAAGAATAGAGCTCGCACCGTAGCTGGTGTCTCTATCTGTAGTACTATGACCA   15110
gb:MT044258|Organism:Severe      TTAGTGCAAAGAATAGAGCTCGCACCGTAGCTGGTGTCTCTATCTGTAGTACTATGACCA   15118
gb:MT039890|Organism:Severe      TTAGTGCAAAGAATAGAGCTCGCACCGTAGCTGGTGTCTCTATCTGTAGTACTATGACCA   15142
gb:MN988713|Organism:Severe      TTAGTGCAAAGAATAGAGCTCGCACCGTAGCTGGTGTCTCTATCTGTAGTACTATGACCA   15142
gb:LC521925|Organism:Severe      TTAGTGCAAAGAATAGAGCTCGCACCGTAGCTGGTGTCTCTATCTGTAGTACTATGACCA   15115
gb:MT093571|Organism:Severe      TTAGTGCAAAGAATAGAGCTCGCACCGTAGCTGGTGTCTCTATCTGTAGTACTATGACCA   15142
gb:MT039887|Organism:Severe      TTAGTGCAAAGAATAGAGCTCGCACCGTAGCTGGTGTCTCTATCTGTAGTACTATGACCA   15142
gb:MT019530|Organism:Severe      TTAGTGCAAAGAATAGAGCTCGCACCGTAGCTGGTGTCTCTATCTGTAGTACTATGACCA   15142
gb:MT039888|Organism:Severe      TTAGTGCAAAGAATAGAGCTCGCACCGTAGCTGGTGTCTCTATCTGTAGTACTATGACCA   15142
gb:LC522972|Organism:Severe      TTAGTGCAAAGAATAGAGCTCGCACCGTAGCTGGTGTCTCTATCTGTAGTACTATGACCA   15139
gb:MT027063|Organism:Severe      TTAGTGCAAAGAATAGAGCTCGCACCGTAGCTGGTGTCTCTATCTGTAGTACTATGACCA   15142
gb:MT027062|Organism:Severe      TTAGTGCAAAGAATAGAGCTCGCACCGTAGCTGGTGTCTCTATCTGTAGTACTATGACCA   15142
gb:MT019529|Organism:Severe      TTAGTGCAAAGAATAGAGCTCGCACCGTAGCTGGTGTCTCTATCTGTAGTACTATGACCA   15142
gb:MN996529|Organism:Severe      TTAGTGCAAAGAATAGAGCTCGCACCGTAGCTGGTGTCTCTATCTGTAGTACTATGACCA   15130
gb:MN996531|Organism:Severe      TTAGTGCAAAGAATAGAGCTCGCACCGTAGCTGGTGTCTCTATCTGTAGTACTATGACCA   15129
gb:MT066176|Organism:Severe      TTAGTGCAAAGAATAGAGCTCGCACCGTAGCTGGTGTCTCTATCTGTAGTACTATGACCA   15142
gb:MT027064|Organism:Severe      TTAGTGCAAAGAATAGAGCTCGCACCGTAGCTGGTGTCTCTATCTGTAGTACTATGACCA   15142
gb:MN994468|Organism:Severe      TTAGTGCAAAGAATAGAGCTCGCACCGTAGCTGGTGTCTCTATCTGTAGTACTATGACCA   15142
gb:MT072688|Organism:Severe      TTAGTGCAAAGAATAGAGCTCGCACCGTAGCTGGTGTCTCTATCTGTAGTACTATGACCA   15127
gb:MN996527|Organism:Severe      TTAGTGCAAAGAATAGAGCTCGCACCGTAGCTGGTGTCTCTATCTGTAGTACTATGACCA   15109
gb:MT093631|Organism:Severe      TTAGTGCAAAGAATAGAGCTCGCACCGTAGCTGGTGTCTCTATCTGTAGTACTATGACCA   15180
gb:MT106053|Organism:Severe      TTAGTGCAAAGAATAGAGCTCGCACCGTAGCTGGTGTCTCTATCTGTAGTACTATGACCA   15142
gb:MT019533|Organism:Severe      TTAGTGCAAAGAATAGAGCTCGCACCGTAGCTGGTGTCTCTATCTGTAGTACTATGACCA   15142
```

```
gb:MT019531|Organism:Severe      TTAGTGCAAAGAATAGAGCTCGCACCGTAGCTGGTGTCTCTATCTGTAGTACTATGACCA   15142
gb:MN996528|Organism:Severe      TTAGTGCAAAGAATAGAGCTCGCACCGTAGCTGGTGTCTCTATCTGTAGTACTATGACCA   15142
gb:MN996530|Organism:Severe      TTAGTGCAAAGAATAGAGCTCGCACCGTAGCTGGTGTCTCTATCTGTAGTACTATGACCA   15128
gb:MN908947|Organism:Severe      TTAGTGCAAAGAATAGAGCTCGCACCGTAGCTGGTGTCTCTATCTGTAGTACTATGACCA   15142
gb:MT019532|Organism:Severe      TTAGTGCAAAGAATAGAGCTCGCACCGTAGCTGGTGTCTCTATCTGTAGTACTATGACCA   15142
                                 ************************************************************

gb:MT020781|Organism:Severe      ATAGACAGTTTCATCAAAAATTATTGAAATCAATAGCCGCCACTAGAGGAGCTACTGTAG   15190
gb:MT007544|Organism:Severe      ATAGACAGTTTCATCAAAAATTATTGAAATCAATAGCCGCCACTAGAGGAGCTACTGTAG   15202
gb:MN994467|Organism:Severe      ATAGACAGTTTCATCAAAAATTATTGAAATCAATAGCCGCCACTAGAGGAGCTACTGTAG   15202
gb:MT044257|Organism:Severe      ATAGACAGTTTCATCAAAAATTATTGAAATCAATAGCCGCCACTAGAGGAGCTACTGTAG   15202
gb:MT106054|Organism:Severe      ATAGACAGTTTCATCAAAAATTATTGAAATCAATAGCCGCCACTAGAGGAGCTACTGTAG   15202
gb:MT049951|Organism:Severe      ATAGACAGTTTCATCAAAAATTATTGAAATCAATAGCCGCCACTAGAGGAGCTACTGTAG   15202
gb:MN975262|Organism:Severe      ATAGACAGTTTCATCAAAAATTATTGAAATCAATAGCCGCCACTAGAGGAGCTACTGTAG   15202
gb:MT106052|Organism:Severe      ATAGACAGTTTCATCAAAAATTATTGAAATCAATAGCCGCCACTAGAGGAGCTACTGTAG   15202
gb:LC522975|Organism:Severe      ATAGACAGTTTCATCAAAAATTATTGAAATCAATAGCCGCCACTAGAGGAGCTACTGTAG   15199
gb:LC522973|Organism:Severe      ATAGACAGTTTCATCAAAAATTATTGAAATCAATAGCCGCCACTAGAGGAGCTACTGTAG   15199
gb:LC522974|Organism:Severe      ATAGACAGTTTCATCAAAAATTATTGAAATCAATAGCCGCCACTAGAGGAGCTACTGTAG   15199
gb:MN985325|Organism:Severe      ATAGACAGTTTCATCAAAAATTATTGAAATCAATAGCCGCCACTAGAGGAGCTACTGTAG   15202
gb:MT020881|Organism:Severe      ATAGACAGTTTCATCAAAAATTATTGAAATCAATAGCCGCCACTAGAGGAGCTACTGTAG   15202
gb:MT020880|Organism:Severe      ATAGACAGTTTCATCAAAAATTATTGAAATCAATAGCCGCCACTAGAGGAGCTACTGTAG   15202
gb:MT066175|Organism:Severe      ATAGACAGTTTCATCAAAAATTATTGAAATCAATAGCCGCCACTAGAGGAGCTACTGTAG   15202
gb:MN997409|Organism:Severe      ATAGACAGTTTCATCAAAAATTATTGAAATCAATAGCCGCCACTAGAGGAGCTACTGTAG   15202
gb:MN938384|Organism:Severe      ATAGACAGTTTCATCAAAAATTATTGAAATCAATAGCCGCCACTAGAGGAGCTACTGTAG   15170
gb:MT044258|Organism:Severe      ATAGACAGTTTCATCAAAAATTATTGAAATCAATAGCCGCCACTAGAGGAGCTACTGTAG   15178
gb:MT039890|Organism:Severe      ATAGACAGTTTCATCAAAAATTATTGAAATCAATAGCCGCCACTAGAGGAGCTACTGTAG   15202
gb:MN988713|Organism:Severe      ATAGACAGTTTCATCAAAAATTATTGAAATCAATAGCCGCCACTAGAGGAGCTACTGTAG   15202
gb:LC521925|Organism:Severe      ATAGACAGTTTCATCAAAAATTATTGAAATCAATAGCCGCCACTAGAGGAGCTACTGTAG   15175
gb:MT093571|Organism:Severe      ATAGACAGTTTCATCAAAAATTATTGAAATCAATAGCCGCCACTAGAGGAGCTACTGTAG   15202
gb:MT039887|Organism:Severe      ATAGACAGTTTCATCAAAAATTATTGAAATCAATAGCCGCCACTAGAGGAGCTACTGTAG   15202
gb:MT019530|Organism:Severe      ATAGACAGTTTCATCAAAAATTATTGAAATCAATAGCCGCCACTAGAGGAGCTACTGTAG   15202
gb:MT039888|Organism:Severe      ATAGACAGTTTCATCAAAAATTATTGAAATCAATAGCCGCCACTAGAGGAGCTACTGTAG   15202
gb:LC522972|Organism:Severe      ATAGACAGTTTCATCAAAAATTATTGAAATCAATAGCCGCCACTAGAGGAGCTACTGTAG   15199
gb:MT027063|Organism:Severe      ATAGACAGTTTCATCAAAAATTATTGAAATCAATAGCCGCCACTAGAGGAGCTACTGTAG   15202
gb:MT027062|Organism:Severe      ATAGACAGTTTCATCAAAAATTATTGAAATCAATAGCCGCCACTAGAGGAGCTACTGTAG   15202
gb:MT019529|Organism:Severe      ATAGACAGTTTCATCAAAAATTATTGAAATCAATAGCCGCCACTAGAGGAGCTACTGTAG   15202
gb:MN996529|Organism:Severe      ATAGACAGTTTCATCAAAAATTATTGAAATCAATAGCCGCCACTAGAGGAGCTACTGTAG   15190
gb:MN996531|Organism:Severe      ATAGACAGTTTCATCAAAAATTATTGAAATCAATAGCCGCCACTAGAGGAGCTACTGTAG   15189
gb:MT066176|Organism:Severe      ATAGACAGTTTCATCAAAAATTATTGAAATCAATAGCCGCCACTAGAGGAGCTACTGTAG   15202
gb:MT027064|Organism:Severe      ATAGACAGTTTCATCAAAAATTATTGAAATCAATAGCCGCCACTAGAGGAGCTACTGTAG   15202
gb:MN994468|Organism:Severe      ATAGACAGTTTCATCAAAAATTATTGAAATCAATAGCCGCCACTAGAGGAGCTACTGTAG   15202
gb:MT072688|Organism:Severe      ATAGACAGTTTCATCAAAAATTATTGAAATCAATAGCCGCCACTAGAGGAGCTACTGTAG   15187
gb:MN996527|Organism:Severe      ATAGACAGTTTCATCAAAAATTATTGAAATCAATAGCCGCCACTAGAGGAGCTACTGTAG   15169
gb:MT093631|Organism:Severe      ATAGACAGTTTCATCAAAAATTATTGAAATCAATAGCCGCCACTAGAGGAGCTACTGTAG   15240
gb:MT106053|Organism:Severe      ATAGACAGTTTCATCAAAAATTATTGAAATCAATAGCCGCCACTAGAGGAGCTACTGTAG   15202
gb:MT019533|Organism:Severe      ATAGACAGTTTCATCAAAAATTATTGAAATCAATAGCCGCCACTAGAGGAGCTACTGTAG   15202
gb:MT019531|Organism:Severe      ATAGACAGTTTCATCAAAAATTATTGAAATCAATAGCCGCCACTAGAGGAGCTACTGTAG   15202
gb:MN996528|Organism:Severe      ATAGACAGTTTCATCAAAAATTATTGAAATCAATAGCCGCCACTAGAGGAGCTACTGTAG   15202
gb:MN996530|Organism:Severe      ATAGACAGTTTCATCAAAAATTATTGAAATCAATAGCCGCCACTAGAGGAGCTACTGTAG   15188
gb:MN908947|Organism:Severe      ATAGACAGTTTCATCAAAAATTATTGAAATCAATAGCCGCCACTAGAGGAGCTACTGTAG   15202
gb:MT019532|Organism:Severe      ATAGACAGTTTCATCAAAAATTATTGAAATCAATAGCCGCCACTAGAGGAGCTACTGTAG   15202
                                 ************************************************************

gb:MT020781|Organism:Severe      TAATTGGAACAAGCAAATTCTATGGTGGTTGGCACAACATGTTAAAAACTGTTTATAGTG   15250
gb:MT007544|Organism:Severe      TAATTGGAACAAGCAAATTCTATGGTGGTTGGCACAACATGTTAAAAACTGTTTATAGTG   15262
gb:MN994467|Organism:Severe      TAATTGGAACAAGCAAATTCTATGGTGGTTGGCACAACATGTTAAAAACTGTTTATAGTG   15262
gb:MT044257|Organism:Severe      TAATTGGAACAAGCAAATTCTATGGTGGTTGGCACAACATGTTAAAAACTGTTTATAGTG   15262
gb:MT106054|Organism:Severe      TAATTGGAACAAGCAAATTCTATGGTGGTTGGCACAACATGTTAAAAACTGTTTATAGTG   15262
gb:MT049951|Organism:Severe      TAATTGGAACAAGCAAATTCTATGGTGGTTGGCACAACATGTTAAAAACTGTTTATAGTG   15262
gb:MN975262|Organism:Severe      TAATTGGAACAAGCAAATTCTATGGTGGTTGGCACAACATGTTAAAAACTGTTTATAGTG   15262
gb:MT106052|Organism:Severe      TAATTGGAACAAGCAAATTCTATGGTGGTTGGCACAACATGTTAAAAACTGTTTATAGTG   15262
gb:LC522975|Organism:Severe      TAATTGGAACAAGCAAATTCTATGGTGGTTGGCACAACATGTTAAAAACTGTTTATAGTG   15259
gb:LC522973|Organism:Severe      TAATTGGAACAAGCAAATTCTATGGTGGTTGGCACAACATGTTAAAAACTGTTTATAGTG   15259
gb:LC522974|Organism:Severe      TAATTGGAACAAGCAAATTCTATGGTGGTTGGCACAACATGTTAAAAACTGTTTATAGTG   15259
gb:MN985325|Organism:Severe      TAATTGGAACAAGCAAATTCTATGGTGGTTGGCACAACATGTTAAAAACTGTTTATAGTG   15262
```

```
gb:MT020881|Organism:Severe      TAATTGGAACAAGCAAATTCTATGGTGGTTGGCACAACATGTTAAAAACTGTTTATAGTG   15262
gb:MT020880|Organism:Severe      TAATTGGAACAAGCAAATTCTATGGTGGTTGGCACAACATGTTAAAAACTGTTTATAGTG   15262
gb:MT066175|Organism:Severe      TAATTGGAACAAGCAAATTCTATGGTGGTTGGCACAACATGTTAAAAACTGTTTATAGTG   15262
gb:MN997409|Organism:Severe      TAATTGGAACAAGCAAATTCTATGGTGGTTGGCACAACATGTTAAAAACTGTTTATAGTG   15262
gb:MN938384|Organism:Severe      TAATTGGAACAAGCAAATTCTATGGTGGTTGGCACAACATGTTAAAAACTGTTTATAGTG   15230
gb:MT044258|Organism:Severe      TAATTGGAACAAGCAAATTCTATGGTGGTTGGCACAACATGTTAAAAACTGTTTATAGTG   15238
gb:MT039890|Organism:Severe      TAATTGGAACAAGCAAATTCTATGGTGGTTGGCACAACATGTTAAAAACTGTTTATAGTG   15262
gb:MN988713|Organism:Severe      TAATTGGAACAAGCAAATTCTATGGTGGTTGGCACAACATGTTAAAAACTGTTTATAGTG   15262
gb:LC521925|Organism:Severe      TAATTGGAACAAGCAAATTCTATGGTGGTTGGCACAACATGTTAAAAACTGTTTATAGTG   15235
gb:MT093571|Organism:Severe      TAATTGGAACAAGCAAATTCTATGGTGGTTGGCACAACATGTTAAAAACTGTTTATAGTG   15262
gb:MT039887|Organism:Severe      TAATTGGAACAAGCAAATTCTATGGTGGTTGGCACAACATGTTAAAAACTGTTTATAGTG   15262
gb:MT019530|Organism:Severe      TAATTGGAACAAGCAAATTCTATGGTGGTTGGCACAACATGTTAAAAACTGTTTATAGTG   15262
gb:MT039888|Organism:Severe      TAATTGGAACAAGCAAATTCTATGGTGGTTGGCACAACATGTTAAAAACTGTTTATAGTG   15262
gb:LC522972|Organism:Severe      TAATTGGAACAAGCAAATTCTATGGTGGTTGGCACAACATGTTAAAAACTGTTTATAGTG   15259
gb:MT027063|Organism:Severe      TAATTGGAACAAGCAAATTCTATGGTGGTTGGCACAACATGTTAAAAACTGTTTATAGTG   15262
gb:MT027062|Organism:Severe      TAATTGGAACAAGCAAATTCTATGGTGGTTGGCACAACATGTTAAAAACTGTTTATAGTG   15262
gb:MT019529|Organism:Severe      TAATTGGAACAAGCAAATTCTATGGTGGTTGGCACAACATGTTAAAAACTGTTTATAGTG   15262
gb:MN996529|Organism:Severe      TAATTGGAACAAGCAAATTCTATGGTGGTTGGCACAACATGTTAAAAACTGTTTATAGTG   15250
gb:MN996531|Organism:Severe      TAATTGGAACAAGCAAATTCTATGGTGGTTGGCACAACATGTTAAAAACTGTTTATAGTG   15249
gb:MT066176|Organism:Severe      TAATTGGAACAAGCAAATTCTATGGTGGTTGGCACAACATGTTAAAAACTGTTTATAGTG   15262
gb:MT027064|Organism:Severe      TAATTGGAACAAGCAAATTCTATGGTGGTTGGCACAACATGTTAAAAACTGTTTATAGTG   15262
gb:MN994468|Organism:Severe      TAATTGGAACAAGCAAATTCTATGGTGGTTGGCACAACATGTTAAAAACTGTTTATAGTG   15262
gb:MT072688|Organism:Severe      TAATTGGAACAAGCAAATTCTATGGTGGTTGGCACAACATGTTAAAAACTGTTTATAGTG   15247
gb:MN996527|Organism:Severe      TAATTGGAACAAGCAAATTCTATGGTGGTTGGCACAACATGTTAAAAACTGTTTATAGTG   15229
gb:MT093631|Organism:Severe      TAATTGGAACAAGCAAATTCTATGGTGGTTGGCACAACATGTTAAAAACTGTTTATAGTG   15300
gb:MT106053|Organism:Severe      TAATTGGAACAAGCAAATTCTATGGTGGTTGGCACAACATGTTAAAAACTGTTTATAGTG   15262
gb:MT019533|Organism:Severe      TAATTGGAACAAGCAAATTCTATGGTGGTTGGCACAACATGTTAAAAACTGTTTATAGTG   15262
gb:MT019531|Organism:Severe      TAATTGGAACAAGCAAATTCTATGGTGGTTGGCACAACATGTTAAAAACTGTTTATAGTG   15262
gb:MN996528|Organism:Severe      TAATTGGAACAAGCAAATTCTATGGTGGTTGGCACAACATGTTAAAAACTGTTTATAGTG   15262
gb:MN996530|Organism:Severe      TAATTGGAACAAGCAAATTCTATGGTGGTTGGCACAACATGTTAAAAACTGTTTATAGTG   15248
gb:MN908947|Organism:Severe      TAATTGGAACAAGCAAATTCTATGGTGGTTGGCACAACATGTTAAAAACTGTTTATAGTG   15262
gb:MT019532|Organism:Severe      TAATTGGAACAAGCAAATTCTATGGTGGTTGGCACAACATGTTAAAAACTGTTTATAGTG   15262
                                 ************************************************************

gb:MT020781|Organism:Severe      ATGTAGAAAACCCTCACCTTATGGGTTGGGATTATCCTAAATGTGATAGAGCCATGCCTA   15310
gb:MT007544|Organism:Severe      ATGTAGAAAACCCTCACCTTATGGGTTGGGATTATCCTAAATGTGATAGAGCCATGCCTA   15322
gb:MN994467|Organism:Severe      ATGTAGAAAACCCTCACCTTATGGGTTGGGATTATCCTAAATGTGATAGAGCCATGCCTA   15322
gb:MT044257|Organism:Severe      ATGTAGAAAACCCTCACCTTATGGGTTGGGATTATCCTAAATGTGATAGAGCCATGCCTA   15322
gb:MT106054|Organism:Severe      ATGTAGAAAACCCTCACCTTATGGGTTGGGATTATCCTAAATGTGATAGAGCCATGCCTA   15322
gb:MT049951|Organism:Severe      ATGTAGAAAACCCTCACCTTATGGGTTGGGATTATCCTAAATGTGATAGAGCCATGCCTA   15322
gb:MN975262|Organism:Severe      ATGTAGAAAACCCTCACCTTATGGGTTGGGATTATCCTAAATGTGATAGAGCCATGCCTA   15322
gb:MT106052|Organism:Severe      ATGTAGAAAACCCTCACCTTATGGGTTGGGATTATCCTAAATGTGATAGAGCCATGCCTA   15322
gb:LC522975|Organism:Severe      ATGTAGAAAACCCTCACCTTATGGGTTGGGATTATCCTAAATGTGATAGAGCCATGCCTA   15319
gb:LC522973|Organism:Severe      ATGTAGAAAACCCTCACCTTATGGGTTGGGATTATCCTAAATGTGATAGAGCCATGCCTA   15319
gb:LC522974|Organism:Severe      ATGTAGAAAACCCTCACCTTATGGGTTGGGATTATCCTAAATGTGATAGAGCCATGCCTA   15319
gb:MN985325|Organism:Severe      ATGTAGAAAACCCTCACCTTATGGGTTGGGATTATCCTAAATGTGATAGAGCCATGCCTA   15322
gb:MT020881|Organism:Severe      ATGTAGAAAACCCTCACCTTATGGGTTGGGATTATCCTAAATGTGATAGAGCCATGCCTA   15322
gb:MT020880|Organism:Severe      ATGTAGAAAACCCTCACCTTATGGGTTGGGATTATCCTAAATGTGATAGAGCCATGCCTA   15322
gb:MT066175|Organism:Severe      ATGTAGAAAACCCTCACCTTATGGGTTGGGATTATCCTAAATGTGATAGAGCCATGCCTA   15322
gb:MN997409|Organism:Severe      ATGTAGAAAACCCTCACCTTATGGGTTGGGATTATCCTAAATGTGATAGAGCCATGCCTA   15322
gb:MN938384|Organism:Severe      ATGTAGAAAACCCTCACCTTATGGGTTGGGATTATCCTAAATGTGATAGAGCCATGCCTA   15290
gb:MT044258|Organism:Severe      ATGTAGAAAACCCTCACCTTATGGGTTGGGATTATCCTAAATGTGATAGAGCCATGCCTA   15298
gb:MT039890|Organism:Severe      ATGTAGAAAACCCTCACCTTATGGGTTGGGATTATCCTAAATGTGATAGAGCCATGCCTA   15322
gb:MN988713|Organism:Severe      ATGTAGAAAACCCTCACCTTATGGGTTGGGATTATCCTAAATGTGATAGAGCCATGCCTA   15322
gb:LC521925|Organism:Severe      ATGTAGAAAACCCTCACCTTATGGGTTGGGATTATCCTAAATGTGATAGAGCCATGCCTA   15295
gb:MT093571|Organism:Severe      ATGTAGAAAACCCTCACCTTATGGGTTGGGATTATCCTAAATGTGATAGAGCCATGCCTA   15322
gb:MT039887|Organism:Severe      ATGTAGAAAACCCTCACCTTATGGGTTGGGATTATCCTAAATGTGATAGAGCCATGCCTA   15322
gb:MT019530|Organism:Severe      ATGTAGAAAACCCTCACCTTATGGGTTGGGATTATCCTAAATGTGATAGAGCCATGCCTA   15322
gb:MT039888|Organism:Severe      ATGTAGAAAACCCTCACCTTATGGGTTGGGATTATCCTAAATGTGATAGAGCCATGCCTA   15322
gb:LC522972|Organism:Severe      ATGTAGAAAACCCTCACCTTATGGGTTGGGATTATCCTAAATGTGATAGAGCCATGCCTA   15319
gb:MT027063|Organism:Severe      ATGTAGAAAACCCTCACCTTATGGGTTGGGATTATCCTAAATGTGATAGAGCCATGCCTA   15322
gb:MT027062|Organism:Severe      ATGTAGAAAACCCTCACCTTATGGGTTGGGATTATCCTAAATGTGATAGAGCCATGCCTA   15322
gb:MT019529|Organism:Severe      ATGTAGAAAACCCTCACCTTATGGGTTGGGATTATCCTAAATGTGATAGAGCCATGCCTA   15322
gb:MN996529|Organism:Severe      ATGTAGAAAACCCTCACCTTATGGGTTGGGATTATCCTAAATGTGATAGAGCCATGCCTA   15310
gb:MN996531|Organism:Severe      ATGTAGAAAACCCTCACCTTATGGGTTGGGATTATCCTAAATGTGATAGAGCCATGCCTA   15309
```

```
gb:MT066176|Organism:Severe      ATGTAGAAAACCCTCACCTTATGGGTTGGGATTATCCTAAATGTGATAGAGCCATGCCTA   15322
gb:MT027064|Organism:Severe      ATGTAGAAAACCCTCACCTTATGGGTTGGGATTATCCTAAATGTGATAGAGCCATGCCTA   15322
gb:MN994468|Organism:Severe      ATGTAGAAAACCCTCACCTTATGGGTTGGGATTATCCTAAATGTGATAGAGCCATGCCTA   15322
gb:MT072688|Organism:Severe      ATGTAGAAAACCCTCACCTTATGGGTTGGGATTATCCTAAATGTGATAGAGCCATGCCTA   15307
gb:MN996527|Organism:Severe      ATGTAGAAAACCCTCACCTTATGGGTTGGGATTATCCTAAATGTGATAGAGCCATGCCTA   15289
gb:MT093631|Organism:Severe      ATGTAGAAAACCCTCACCTTATGGGTTGGGATTATCCTAAATGTGATAGAGCCATGCCTA   15360
gb:MT106053|Organism:Severe      ATGTAGAAAACCCTCACCTTATGGGTTGGGATTATCCTAAATGTGATAGAGCCATGCCTA   15322
gb:MT019533|Organism:Severe      ATGTAGAAAACCCTCACCTTATGGGTTGGGATTATCCTAAATGTGATAGAGCCATGCCTA   15322
gb:MT019531|Organism:Severe      ATGTAGAAAACCCTCACCTTATGGGTTGGGATTATCCTAAATGTGATAGAGCCATGCCTA   15322
gb:MN996528|Organism:Severe      ATGTAGAAAACCCTCACCTTATGGGTTGGGATTATCCTAAATGTGATAGAGCCATGCCTA   15322
gb:MN996530|Organism:Severe      ATGTAGAAAACCCTCACCTTATGGGTTGGGATTATCCTAAATGTGATAGAGCCATGCCTA   15308
gb:MN908947|Organism:Severe      ATGTAGAAAACCCTCACCTTATGGGTTGGGATTATCCTAAATGTGATAGAGCCATGCCTA   15322
gb:MT019532|Organism:Severe      ATGTAGAAAACCCTCACCTTATGGGTTGGGATTATCCTAAATGTGATAGAGCCATGCCTA   15322
                                 ************************************************************

gb:MT020781|Organism:Severe      ACATGCTTAGAATTATGGCCTCACTTGTTCTTGCTCGCAAACATACAACGTGTTGTAGCT   15370
gb:MT007544|Organism:Severe      ACATGCTTAGAATTATGGCCTCACTTGTTCTTGCTCGCAAACATACAACGTGTTGTAGCT   15382
gb:MN994467|Organism:Severe      ACATGCTTAGAATTATGGCCTCACTTGTTCTTGCTCGCAAACATACAACGTGTTGTAGCT   15382
gb:MT044257|Organism:Severe      ACATGCTTAGAATTATGGCCTCACTTGTTCTTGCTCGCAAACATACAACGTGTTGTAGCT   15382
gb:MT106054|Organism:Severe      ACATGCTTAGAATTATGGCCTCACTTGTTCTTGCTCGCAAACATACAACGTGTTGTAGCT   15382
gb:MT049951|Organism:Severe      ACATGCTTAGAATTATGGCCTCACTTGTTCTTGCTCGCAAACATACAACGTGTTGTAGCT   15382
gb:MN975262|Organism:Severe      ACATGCTTAGAATTATGGCCTCACTTGTTCTTGCTCGCAAACATACAACGTGTTGTAGCT   15382
gb:MT106052|Organism:Severe      ACATGCTTAGAATTATGGCCTCACTTGTTCTTGCTCGCAAACATACAACGTGTTGTAGCT   15382
gb:LC522975|Organism:Severe      ACATGCTTAGAATTATGGCCTCACTTGTTCTTGCTCGCAAACATACAACGTGTTGTAGCT   15379
gb:LC522973|Organism:Severe      ACATGCTTAGAATTATGGCCTCACTTGTTCTTGCTCGCAAACATACAACGTGTTGTAGCT   15379
gb:LC522974|Organism:Severe      ACATGCTTAGAATTATGGCCTCACTTGTTCTTGCTCGCAAACATACAACGTGTTGTAGCT   15379
gb:MN985325|Organism:Severe      ACATGCTTAGAATTATGGCCTCACTTGTTCTTGCTCGCAAACATACAACGTGTTGTAGCT   15382
gb:MT020881|Organism:Severe      ACATGCTTAGAATTATGGCCTCACTTGTTCTTGCTCGCAAACATACAACGTGTTGTAGCT   15382
gb:MT020880|Organism:Severe      ACATGCTTAGAATTATGGCCTCACTTGTTCTTGCTCGCAAACATACAACGTGTTGTAGCT   15382
gb:MT066175|Organism:Severe      ACATGCTTAGAATTATGGCCTCACTTGTTCTTGCTCGCAAACATACAACGTGTTGTAGCT   15382
gb:MN997409|Organism:Severe      ACATGCTTAGAATTATGGCCTCACTTGTTCTTGCTCGCAAACATACAACGTGTTGTAGCT   15382
gb:MN938384|Organism:Severe      ACATGCTTAGAATTATGGCCTCACTTGTTCTTGCTCGCAAACATACAACGTGTTGTAGCT   15350
gb:MT044258|Organism:Severe      ACATGCTTAGAATTATGGCCTCACTTGTTCTTGCTCGCAAACATACAACGTGTTGTAGCT   15358
gb:MT039890|Organism:Severe      ACATGCTTAGAATTATGGCCTCACTTGTTCTTGCTCGCAAACATACAACGTGTTGTAGCT   15382
gb:MN988713|Organism:Severe      ACATGCTTAGAATTATGGCCTCACTTGTTCTTGCTCGCAAACATACAACGTGTTGTAGCT   15382
gb:LC521925|Organism:Severe      ACATGCTTAGAATTATGGCCTCACTTGTTCTTGCTCGCAAACATACAACGTGTTGTAGCT   15355
gb:MT093571|Organism:Severe      ACATGCTTAGAATTATGGCCTCACTTGTTCTTGCTCGCAAACATACAACGTGTTGTAGCT   15382
gb:MT039887|Organism:Severe      ACATGCTTAGAATTATGGCCTCACTTGTTCTTGCTCGCAAACATACAACGTGTTGTAGCT   15382
gb:MT019530|Organism:Severe      ACATGCTTAGAATTATGGCCTCACTTGTTCTTGCTCGCAAACATACAACGTGTTGTAGCT   15382
gb:MT039888|Organism:Severe      ACATGCTTAGAATTATGGCCTCACTTGTTCTTGCTCGCAAACATACAACGTGTTGTAGCT   15382
gb:LC522972|Organism:Severe      ATATGCTTAGAATTATGGCCTCACTTGTTCTTGCTCGCAAACATACAACGTGTTGTAGCT   15379
gb:MT027063|Organism:Severe      ACATGCTTAGAATTATGGCCTCACTTGTTCTTGCTCGCAAACATACAACGTGTTGTAGCT   15382
gb:MT027062|Organism:Severe      ACATGCTTAGAATTATGGCCTCACTTGTTCTTGCTCGCAAACATACAACGTGTTGTAGCT   15382
gb:MT019529|Organism:Severe      ACATGCTTAGAATTATGGCCTCACTTGTTCTTGCTCGCAAACATACAACGTGTTGTAGCT   15382
gb:MN996529|Organism:Severe      ACATGCTTAGAATTATGGCCTCACTTGTTCTTGCTCGCAAACATACAACGTGTTGTAGCT   15370
gb:MN996531|Organism:Severe      ACATGCTTAGAATTATGGCCTCACTTGTTCTTGCTCGCAAACATACAACGTGTTGTAGCT   15369
gb:MT066176|Organism:Severe      ACATGCTTAGAATTATGGCCTCACTTGTTCTTGCTCGCAAACATACAACGTGTTGTAGCT   15382
gb:MT027064|Organism:Severe      ACATGCTTAGAATTATGGCCTCACTTGTTCTTGCTCGCAAACATACAACGTGTTGTAGCT   15382
gb:MN994468|Organism:Severe      ACATGCTTAGAATTATGGCCTCACTTGTTCTTGCTCGCAAACATACAACGTGTTGTAGCT   15382
gb:MT072688|Organism:Severe      ACATGCTTAGAATTATGGCCTCACTTGTTCTTGCTCGCAAACATACAACGTGTTGTAGCT   15367
gb:MN996527|Organism:Severe      ACATGCTTAGAATTATGGCCTCACTTGTTCTTGCTCGCAAACATACAACGTGTTGTAGCT   15349
gb:MT093631|Organism:Severe      ACATGCTTAGAATTATGGCCTCACTTGTTCTTGCTCGCAAACATACAACGTGTTGTAGCT   15420
gb:MT106053|Organism:Severe      ACATGCTTAGAATTATGGCCTCACTTGTTCTTGCTCGCAAACATACAACGTGTTGTAGCT   15382
gb:MT019533|Organism:Severe      ACATGCTTAGAATTATGGCCTCACTTGTTCTTGCTCGCAAACATACAACGTGTTGTAGCT   15382
gb:MT019531|Organism:Severe      ACATGCTTAGAATTATGGCCTCACTTGTTCTTGCTCGCAAACATACAACGTGTTGTAGCT   15382
gb:MN996528|Organism:Severe      ACATGCTTAGAATTATGGCCTCACTTGTTCTTGCTCGCAAACATACAACGTGTTGTAGCT   15382
gb:MN996530|Organism:Severe      ACATGCTTAGAATTATGGCCTCACTTGTTCTTGCTCGCAAACATACAACGTGTTGTAGCT   15368
gb:MN908947|Organism:Severe      ACATGCTTAGAATTATGGCCTCACTTGTTCTTGCTCGCAAACATACAACGTGTTGTAGCT   15382
gb:MT019532|Organism:Severe      ACATGCTTAGAATTATGGCCTCACTTGTTCTTGCTCGCAAACATACAACGTGTTGTAGCT   15382
                                 * **********************************************************

gb:MT020781|Organism:Severe      TGTCACACCGTTTCTATAGATTAGCTAATGAGTGTGCTCAAGTATTGAGTGAAATGGTCA   15430
gb:MT007544|Organism:Severe      TGTCACACCGTTTCTATAGATTAGCTAATGAGTGTGCTCAAGTATTGAGTGAAATGGTCA   15442
gb:MN994467|Organism:Severe      TGTCACACCGTTTCTATAGATTAGCTAATGAGTGTGCTCAAGTATTGAGTGAAATGGTCA   15442
gb:MT044257|Organism:Severe      TGTCACACCGTTTCTATAGATTAGCTAATGAGTGTGCTCAAGTATTGAGTGAAATGGTCA   15442
```

```
gb:MT106054|Organism:Severe      TGTCACACCGTTTCTATAGATTAGCTAATGAGTGTGCTCAAGTATTGAGTGAAATGGTCA   15442
gb:MT049951|Organism:Severe      TGTCACACCGTTTCTATAGATTAGCTAATGAGTGTGCTCAAGTATTGAGTGAAATGGTCA   15442
gb:MN975262|Organism:Severe      TGTCACACCGTTTCTATAGATTAGCTAATGAGTGTGCTCAAGTATTGAGTGAAATGGTCA   15442
gb:MT106052|Organism:Severe      TGTCACACCGTTTCTATAGATTAGCTAATGAGTGTGCTCAAGTATTGAGTGAAATGGTCA   15442
gb:LC522975|Organism:Severe      TGTCACACCGTTTCTATAGATTAGCTAATGAGTGTGCTCAAGTATTGAGTGAAATGGTCA   15439
gb:LC522973|Organism:Severe      TGTCACACCGTTTCTATAGATTAGCTAATGAGTGTGCTCAAGTATTGAGTGAAATGGTCA   15439
gb:LC522974|Organism:Severe      TGTCACACCGTTTCTATAGATTAGCTAATGAGTGTGCTCAAGTATTGAGTGAAATGGTCA   15439
gb:MN985325|Organism:Severe      TGTCACACCGTTTCTATAGATTAGCTAATGAGTGTGCTCAAGTATTGAGTGAAATGGTCA   15442
gb:MT020881|Organism:Severe      TGTCACACCGTTTCTATAGATTAGCTAATGAGTGTGCTCAAGTATTGAGTGAAATGGTCA   15442
gb:MT020880|Organism:Severe      TGTCACACCGTTTCTATAGATTAGCTAATGAGTGTGCTCAAGTATTGAGTGAAATGGTCA   15442
gb:MT066175|Organism:Severe      TGTCACACCGTTTCTATAGATTAGCTAATGAGTGTGCTCAAGTATTGAGTGAAATGGTCA   15442
gb:MN997409|Organism:Severe      TGTCACACCGTTTCTATAGATTAGCTAATGAGTGTGCTCAAGTATTGAGTGAAATGGTCA   15442
gb:MN938384|Organism:Severe      TGTCACACCGTTTCTATAGATTAGCTAATGAGTGTGCTCAAGTATTGAGTGAAATGGTCA   15410
gb:MT044258|Organism:Severe      TGTCACACCGTTTCTATAGATTAGCTAATGAGTGTGCTCAAGTATTGAGTGAAATGGTCA   15418
gb:MT039890|Organism:Severe      TGTCACACCGTTTCTATAGATTAGCTAATGAGTGTGCTCAAGTATTGAGTGAAATGGTCA   15442
gb:MN988713|Organism:Severe      TGTCACACCGTTTCTATAGATTAGCTAATGAGTGTGCTCAAGTATTGAGTGAAATGGTCA   15442
gb:LC521925|Organism:Severe      TGTCACACCGTTTCTATAGATTAGCTAATGAGTGTGCTCAAGTATTGAGTGAAATGGTCA   15415
gb:MT093571|Organism:Severe      TGTCACACCGTTTCTATAGATTAGCTAATGAGTGTGCTCAAGTATTGAGTGAAATGGTCA   15442
gb:MT039887|Organism:Severe      TGTCACACCGTTTCTATAGATTAGCTAATGAGTGTGCTCAAGTATTGAGTGAAATGGTCA   15442
gb:MT019530|Organism:Severe      TGTCACACCGTTTCTATAGATTAGCTAATGAGTGTGCTCAAGTATTGAGTGAAATGGTCA   15442
gb:MT039888|Organism:Severe      TGTCACACCGTTTCTATAGATTAGCTAATGAGTGTGCTCAAGTATTGAGTGAAATGGTCA   15442
gb:LC522972|Organism:Severe      TGTCACACCGTTTCTATAGATTAGCTAATGAGTGTGCTCAAGTATTGAGTGAAATGGTCA   15439
gb:MT027063|Organism:Severe      TGTCACACCGTTTCTATAGATTAGCTAATGAGTGTGCTCAAGTATTGAGTGAAATGGTCA   15442
gb:MT027062|Organism:Severe      TGTCACACCGTTTCTATAGATTAGCTAATGAGTGTGCTCAAGTATTGAGTGAAATGGTCA   15442
gb:MT019529|Organism:Severe      TGTCACACCGTTTCTATAGATTAGCTAATGAGTGTGCTCAAGTATTGAGTGAAATGGTCA   15442
gb:MN996529|Organism:Severe      TGTCACACCGTTTCTATAGATTAGCTAATGAGTGTGCTCAAGTATTGAGTGAAATGGTCA   15430
gb:MN996531|Organism:Severe      TGTCACACCGTTTCTATAGATTAGCTAATGAGTGTGCTCAAGTATTGAGTGAAATGGTCA   15429
gb:MT066176|Organism:Severe      TGTCACACCGTTTCTATAGATTAGCTAATGAGTGTGCTCAAGTATTGAGTGAAATGGTCA   15442
gb:MT027064|Organism:Severe      TGTCACACCGTTTCTATAGATTAGCTAATGAGTGTGCTCAAGTATTGAGTGAAATGGTCA   15442
gb:MN994468|Organism:Severe      TGTCACACCGTTTCTATAGATTAGCTAATGAGTGTGCTCAAGTATTGAGTGAAATGGTCA   15442
gb:MT072688|Organism:Severe      TGTCACACCGTTTCTATAGATTAGCTAATGAGTGTGCTCAAGTATTGAGTGAAATGGTCA   15427
gb:MN996527|Organism:Severe      TGTCACACCGTTTCTATAGATTAGCTAATGAGTGTGCTCAAGTATTGAGTGAAATGGTCA   15409
gb:MT093631|Organism:Severe      TGTCACACCGTTTCTATAGATTAGCTAATGAGTGTGCTCAAGTATTGAGTGAAATGGTCA   15480
gb:MT106053|Organism:Severe      TGTCACACCGTTTCTATAGATTAGCTAATGAGTGTGCTCAAGTATTGAGTGAAATGGTCA   15442
gb:MT019533|Organism:Severe      TGTCACACCGTTTCTATAGATTAGCTAATGAGTGTGCTCAAGTATTGAGTGAAATGGTCA   15442
gb:MT019531|Organism:Severe      TGTCACACCGTTTCTATAGATTAGCTAATGAGTGTGCTCAAGTATTGAGTGAAATGGTCA   15442
gb:MN996528|Organism:Severe      TGTCACACCGTTTCTATAGATTAGCTAATGAGTGTGCTCAAGTATTGAGTGAAATGGTCA   15442
gb:MN996530|Organism:Severe      TGTCACACCGTTTCTATAGATTAGCTAATGAGTGTGCTCAAGTATTGAGTGAAATGGTCA   15428
gb:MN908947|Organism:Severe      TGTCACACCGTTTCTATAGATTAGCTAATGAGTGTGCTCAAGTATTGAGTGAAATGGTCA   15442
gb:MT019532|Organism:Severe      TGTCACACCGTTTCTATAGATTAGCTAATGAGTGTGCTCAAGTATTGAGTGAAATGGTCA   15442
                                 ************************************************************

gb:MT020781|Organism:Severe      TGTGTGGCGGTTCACTATATGTTAAACCAGGTGGAACCTCATCAGGAGATGCCACAACTG   15490
gb:MT007544|Organism:Severe      TGTGTGGCGGTTCACTATATGTTAAACCAGGTGGAACCTCATCAGGAGATGCCACAACTG   15502
gb:MN994467|Organism:Severe      TGTGTGGCGGTTCACTATATGTTAAACCAGGTGGAACCTCATCAGGAGATGCCACAACTG   15502
gb:MT044257|Organism:Severe      TGTGTGGCGGTTCACTATATGTTAAACCAGGTGGAACCTCATCAGGAGATGCCACAACTG   15502
gb:MT106054|Organism:Severe      TGTGTGGCGGTTCACTATATGTTAAACCAGGTGGAACCTCATCAGGAGATGCCACAACTG   15502
gb:MT049951|Organism:Severe      TGTGTGGCGGTTCACTATATGTTAAACCAGGTGGAACCTCATCAGGAGATGCCACAACTG   15502
gb:MN975262|Organism:Severe      TGTGTGGCGGTTCACTATATGTTAAACCAGGTGGAACCTCATCAGGAGATGCCACAACTG   15502
gb:MT106052|Organism:Severe      TGTGTGGCGGTTCACTATATGTTAAACCAGGTGGAACCTCATCAGGAGATGCCACAACTG   15502
gb:LC522975|Organism:Severe      TGTGTGGCGGTTCACTATATGTTAAACCAGGTGGAACCTCATCAGGAGATGCCACAACTG   15499
gb:LC522973|Organism:Severe      TGTGTGGCGGTTCACTATATGTTAAACCAGGTGGAACCTCATCAGGAGATGCCACAACTG   15499
gb:LC522974|Organism:Severe      TGTGTGGCGGTTCACTATATGTTAAACCAGGTGGAACCTCATCAGGAGATGCCACAACTG   15499
gb:MN985325|Organism:Severe      TGTGTGGCGGTTCACTATATGTTAAACCAGGTGGAACCTCATCAGGAGATGCCACAACTG   15502
gb:MT020881|Organism:Severe      TGTGTGGCGGTTCACTATATGTTAAACCAGGTGGAACCTCATCAGGAGATGCCACAACTG   15502
gb:MT020880|Organism:Severe      TGTGTGGCGGTTCACTATATGTTAAACCAGGTGGAACCTCATCAGGAGATGCCACAACTG   15502
gb:MT066175|Organism:Severe      TGTGTGGCGGTTCACTATATGTTAAACCAGGTGGAACCTCATCAGGAGATGCCACAACTG   15502
gb:MN997409|Organism:Severe      TGTGTGGCGGTTCACTATATGTTAAACCAGGTGGAACCTCATCAGGAGATGCCACAACTG   15502
gb:MN938384|Organism:Severe      TGTGTGGCGGTTCACTATATGTTAAACCAGGTGGAACCTCATCAGGAGATGCCACAACTG   15470
gb:MT044258|Organism:Severe      TGTGTGGCGGTTCACTATATGTTAAACCAGGTGGAACCTCATCAGGAGATGCCACAACTG   15478
gb:MT039890|Organism:Severe      TGTGTGGCGGTTCACTATATGTTAAACCAGGTGGAACCTCATCAGGAGATGCCACAACTG   15502
gb:MN988713|Organism:Severe      TGTGTGGCGGTTCACTATATGTTAAACCAGGTGGAACCTCATCAGGAGATGCCACAACTG   15502
gb:LC521925|Organism:Severe      TGTGTGGCGGTTCACTATATGTTAAACCAGGTGGAACCTCATCAGGAGATGCCACAACTG   15475
gb:MT093571|Organism:Severe      TGTGTGGCGGTTCACTATATGTTAAACCAGGTGGAACCTCATCAGGAGATGCCACAACTG   15502
gb:MT039887|Organism:Severe      TGTGTGGCGGTTCACTATATGTTAAACCAGGTGGAACCTCATCAGGAGATGCCACAACTG   15502
```

```
gb:MT019530|Organism:Severe      TGTGTGGCGGTTCACTATATGTTAAACCAGGTGGAACCTCATCAGGAGATGCCACAACTG   15502
gb:MT039888|Organism:Severe      TGTGTGGCGGTTCACTATATGTTAAACCAGGTGGAACCTCATCAGGAGATGCCACAACTG   15502
gb:LC522972|Organism:Severe      TGTGTGGCGGTTCACTATATGTTAAACCAGGTGGAACCTCATCAGGAGATGCCACAACTG   15499
gb:MT027063|Organism:Severe      TGTGTGGCGGTTCACTATATGTTAAACCAGGTGGAACCTCATCAGGAGATGCCACAACTG   15502
gb:MT027062|Organism:Severe      TGTGTGGCGGTTCACTATATGTTAAACCAGGTGGAACCTCATCAGGAGATGCCACAACTG   15502
gb:MT019529|Organism:Severe      TGTGTGGCGGTTCACTATATGTTAAACCAGGTGGAACCTCATCAGGAGATGCCACAACTG   15502
gb:MN996529|Organism:Severe      TGTGTGGCGGTTCACTATATGTTAAACCAGGTGGAACCTCATCAGGAGATGCCACAACTG   15490
gb:MN996531|Organism:Severe      TGTGTGGCGGTTCACTATATGTTAAACCAGGTGGAACCTCATCAGGAGATGCCACAACTG   15489
gb:MT066176|Organism:Severe      TGTGTGGCGGTTCACTATATGTTAAACCAGGTGGAACCTCATCAGGAGATGCCACAACTG   15502
gb:MT027064|Organism:Severe      TGTGTGGCGGTTCACTATATGTTAAACCAGGTGGAACCTCATCAGGAGATGCCACAACTG   15502
gb:MN994468|Organism:Severe      TGTGTGGCGGTTCACTATATGTTAAACCAGGTGGAACCTCATCAGGAGATGCCACAACTG   15502
gb:MT072688|Organism:Severe      TGTGTGGCGGTTCACTATATGTTAAACCAGGTGGAACCTCATCAGGAGATGCCACAACTG   15487
gb:MN996527|Organism:Severe      TGTGTGGCGGTTCACTATATGTTAAACCAGGTGGAACCTCATCAGGAGATGCCACAACTG   15469
gb:MT093631|Organism:Severe      TGTGTGGCGGTTCACTATATGTTAAACCAGGTGGAACCTCATCAGGAGATGCCACAACTG   15540
gb:MT106053|Organism:Severe      TGTGTGGCGGTTCACTATATGTTAAACCAGGTGGAACCTCATCAGGAGATGCCACAACTG   15502
gb:MT019533|Organism:Severe      TGTGTGGCGGTTCACTATATGTTAAACCAGGTGGAACCTCATCAGGAGATGCCACAACTG   15502
gb:MT019531|Organism:Severe      TGTGTGGCGGTTCACTATATGTTAAACCAGGTGGAACCTCATCAGGAGATGCCACAACTG   15502
gb:MN996528|Organism:Severe      TGTGTGGCGGTTCACTATATGTTAAACCAGGTGGAACCTCATCAGGAGATGCCACAACTG   15502
gb:MN996530|Organism:Severe      TGTGTGGCGGTTCACTATATGTTAAACCAGGTGGAACCTCATCAGGAGATGCCACAACTG   15488
gb:MN908947|Organism:Severe      TGTGTGGCGGTTCACTATATGTTAAACCAGGTGGAACCTCATCAGGAGATGCCACAACTG   15502
gb:MT019532|Organism:Severe      TGTGTGGCGGTTCACTATATGTTAAACCAGGTGGAACCTCATCAGGAGATGCCACAACTG   15502
                                 ************************************************************

gb:MT020781|Organism:Severe      CTTATGCTAATAGTGTTTTTAACATTTGTCAAGCTGTCACGGCCAATGTTAATGCACTTT   15550
gb:MT007544|Organism:Severe      CTTATGCTAATAGTGTTTTTAACATTTGTCAAGCTGTCACGGCCAATGTTAATGCACTTT   15562
gb:MN994467|Organism:Severe      CTTATGCTAATAGTGTTTTTAACATTTGTCAAGCTGTCACGGCCAATGTTAATGCACTTT   15562
gb:MT044257|Organism:Severe      CTTATGCTAATAGTGTTTTTAACATTTGTCAAGCTGTCACGGCCAATGTTAATGCACTTT   15562
gb:MT106054|Organism:Severe      CTTATGCTAATAGTGTTTTTAACATTTGTCAAGCTGTCACGGCCAATGTTAATGCACTTT   15562
gb:MT049951|Organism:Severe      CTTATGCTAATAGTGTTTTTAACATTTGTCAAGCTGTCACGGCCAATGTTAATGCACTTT   15562
gb:MN975262|Organism:Severe      CTTATGCTAATAGTGTTTTTAACATTTGTCAAGCTGTCACGGCCAATGTTAATGCACTTT   15562
gb:MT106052|Organism:Severe      CTTATGCTAATAGTGTTTTTAACATTTGTCAAGCTGTCACGGCCAATGTTAATGCACTTT   15562
gb:LC522975|Organism:Severe      CTTATGCTAATAGTGTTTTTAACATTTGTCAAGCTGTCACGGCCAATGTTAATGCACTTT   15559
gb:LC522973|Organism:Severe      CTTATGCTAATAGTGTTTTTAACATTTGTCAAGCTGTCACGGCCAATGTTAATGCACTTT   15559
gb:LC522974|Organism:Severe      CTTATGCTAATAGTGTTTTTAACATTTGTCAAGCTGTCACGGCCAATGTTAATGCACTTT   15559
gb:MN985325|Organism:Severe      CTTATGCTAATAGTGTTTTTAACATTTGTCAAGCTGTCACGGCCAATGTTAATGCACTTT   15562
gb:MT020881|Organism:Severe      CTTATGCTAATAGTGTTTTTAACATTTGTCAAGCTGTCACGGCCAATGTTAATGCACTTT   15562
gb:MT020880|Organism:Severe      CTTATGCTAATAGTGTTTTTAACATTTGTCAAGCTGTCACGGCCAATGTTAATGCACTTT   15562
gb:MT066175|Organism:Severe      CTTATGCTAATAGTGTTTTTAACATTTGTCAAGCTGTCACGGCCAATGTTAATGCACTTT   15562
gb:MN997409|Organism:Severe      CTTATGCTAATAGTGTTTTTAACATTTGTCAAGCTGTCACGGCCAATGTTAATGCACTTT   15562
gb:MN938384|Organism:Severe      CTTATGCTAATAGTGTTTTTAACATTTGTCAAGCTGTCACGGCCAATGTTAATGCACTTT   15530
gb:MT044258|Organism:Severe      CTTATGCTAATAGTGTTTTTAACATTTGTCAAGCTGTCACGGCCAATGTTAATGCACTTT   15538
gb:MT039890|Organism:Severe      CTTATGCTAATAGTGTTTTTAACATTTGTCAAGCTGTCACGGCCAATGTTAATGCACTTT   15562
gb:MN988713|Organism:Severe      CTTATGCTAATAGTGTTTTTAACATTTGTCAAGCTGTCACGGCCAATGTTAATGCACTTT   15562
gb:LC521925|Organism:Severe      CTTATGCTAATAGTGTTTTTAACATTTGTCAAGCTGTCACGGCCAATGTTAATGCACTTT   15535
gb:MT093571|Organism:Severe      CTTATGCTAATAGTGTTTTTAACATTTGTCAAGCTGTCACGGCCAATGTTAATGCACTTT   15562
gb:MT039887|Organism:Severe      CTTATGCTAATAGTGTTTTTAACATTTGTCAAGCTGTCACGGCCAATGTTAATGCACTTT   15562
gb:MT019530|Organism:Severe      CTTATGCTAATAGTGTTTTTAACATTTGTCAAGCTGTCACGGCCAATGTTAATGCACTTT   15562
gb:MT039888|Organism:Severe      CTTATGCTAATAGTGTTTTTAACATTTGTCAAGCTGTCACGGCCAATGTTAATGCACTTT   15562
gb:LC522972|Organism:Severe      CTTATGCTAATAGTGTTTTTAACATTTGTCAAGCTGTCACGGCCAATGTTAATGCACTTT   15559
gb:MT027063|Organism:Severe      CTTATGCTAATAGTGTTTTTAACATTTGTCAAGCTGTCACGGCCAATGTTAATGCACTTT   15562
gb:MT027062|Organism:Severe      CTTATGCTAATAGTGTTTTTAACATTTGTCAAGCTGTCACGGCCAATGTTAATGCACTTT   15562
gb:MT019529|Organism:Severe      CTTATGCTAATAGTGTTTTTAACATTTGTCAAGCTGTCACGGCCAATGTTAATGCACTTT   15562
gb:MN996529|Organism:Severe      CTTATGCTAATAGTGTTTTTAACATTTGTCAAGCTGTCACGGCCAATGTTAATGCACTTT   15550
gb:MN996531|Organism:Severe      CTTATGCTAATAGTGTTTTTAACATTTGTCAAGCTGTCACGGCCAATGTTAATGCACTTT   15549
gb:MT066176|Organism:Severe      CTTATGCTAATAGTGTTTTTAACATTTGTCAAGCTGTCACGGCCAATGTTAATGCACTTT   15562
gb:MT027064|Organism:Severe      CTTATGCTAATAGTGTTTTTAACATTTGTCAAGCTGTCACGGCCAATGTTAATGCACTTT   15562
gb:MN994468|Organism:Severe      CTTATGCTAATAGTGTTTTTAACATTTGTCAAGCTGTCACGGCCAATGTTAATGCACTTT   15562
gb:MT072688|Organism:Severe      CTTATGCTAATAGTGTTTTTAACATTTGTCAAGCTGTCACGGCCAATGTTAATGCACTTT   15547
gb:MN996527|Organism:Severe      CTTATGCTAATAGTGTTTTTAACATTTGTCAAGCTGTCACGGCCAATGTTAATGCACTTT   15529
gb:MT093631|Organism:Severe      CTTATGCTAATAGTGTTTTTAACATTTGTCAAGCTGTCACGGCCAATGTTAATGCACTTT   15600
gb:MT106053|Organism:Severe      CTTATGCTAATAGTGTTTTTAACATTTGTCAAGCTGTCACGGCCAATGTTAATGCACTTT   15562
gb:MT019533|Organism:Severe      CTTATGCTAATAGTGTTTTTAACATTTGTCAAGCTGTCACGGCCAATGTTAATGCACTTT   15562
gb:MT019531|Organism:Severe      CTTATGCTAATAGTGTTTTTAACATTTGTCAAGCTGTCACGGCCAATGTTAATGCACTTT   15562
gb:MN996528|Organism:Severe      CTTATGCTAATAGTGTTTTTAACATTTGTCAAGCTGTCACGGCCAATGTTAATGCACTTT   15562
gb:MN996530|Organism:Severe      CTTATGCTAATAGTGTTTTTAACATTTGTCAAGCTGTCACGGCCAATGTTAATGCACTTT   15548
```

```
gb:MN908947|Organism:Severe      CTTATGCTAATAGTGTTTTTAACATTTGTCAAGCTGTCACGGCCAATGTTAATGCACTTT   15562
gb:MT019532|Organism:Severe      CTTATGCTAATAGTGTTTTTAACATTTGTCAAGCTGTCACGGCCAATGTTAATGCACTTT   15562
                                 ************************************************************

gb:MT020781|Organism:Severe      TATCTACTGATGGTAACAAAATTGCCGATAAGTATGTCCGCAATTTACAACACAGACTTT   15610
gb:MT007544|Organism:Severe      TATCTACTGATGGTAACAAAATTGCCGATAAGTATGTCCGCAATTTACAACACAGACTTT   15622
gb:MN994467|Organism:Severe      TATCTACTGATGGTAACAAAATTGCCGATAAGTATGTCCGCAATTTACAACACAGACTTT   15622
gb:MT044257|Organism:Severe      TATCTACTGATGGTAACAAAATTGCCGATAAGTATGTCCGCAATTTACAACACAGACTTT   15622
gb:MT106054|Organism:Severe      TATCTACTGATGGTAACAAAATTGCCGATAAGTATGTCCGCAATTTACAACACAGACTTT   15622
gb:MT049951|Organism:Severe      TATCTACTGATGGTAACAAAATTGCCGATAAGTATGTCCGCAATTTACAACACAGACTTT   15622
gb:MN975262|Organism:Severe      TATCTACTGATGGTAACAAAATTGCCGATAAGTATGTCCGCAATCTACAACACAGACTTT   15622
gb:MT106052|Organism:Severe      TATCTACTGATGGTAACAAAATTGCCGATAAGTATGTCCGCAATTTACAACACAGACTTT   15622
gb:LC522975|Organism:Severe      TATCTACTGATGGTAACAAAATTGCCGATAAGTATGTCCGCAATTTACAACACAGACTTT   15619
gb:LC522973|Organism:Severe      TATCTACTGATGGTAACAAAATTGCCGATAAGTATGTCCGCAATTTACAACACAGACTTT   15619
gb:LC522974|Organism:Severe      TATCTACTGATGGTAACAAAATTGCCGATAAGTATGTCCGCAATTTACAACACAGACTTT   15619
gb:MN985325|Organism:Severe      TATCTACTGATGGTAACAAAATTGCCGATAAGTATGTCCGCAATTTACAACACAGACTTT   15622
gb:MT020881|Organism:Severe      TATCTACTGATGGTAACAAAATTGCCGATAAGTATGTCCGCAATTTACAACACAGACTTT   15622
gb:MT020880|Organism:Severe      TATCTACTGATGGTAACAAAATTGCCGATAAGTATGTCCGCAATTTACAACACAGACTTT   15622
gb:MT066175|Organism:Severe      TATCTACTGATGGTAACAAAATTGCCGATAAGTATGTCCGCAATTTACAACACAGACTTT   15622
gb:MN997409|Organism:Severe      TATCTACTGATGGTAACAAAATTGCCGATAAGTATGTCCGCAATTTACAACACAGACTTT   15622
gb:MN938384|Organism:Severe      TATCTACTGATGGTAACAAAATTGCCGATAAGTATGTCCGCAATTTACAACACAGACTTT   15590
gb:MT044258|Organism:Severe      TATCTACTGATGGTAACAAAATTGCCGATAAGTATGTCCGCAATTTACAACACAGACTTT   15598
gb:MT039890|Organism:Severe      TATCTACTGATGGTAACAAAATTGCCGATAAGTACGTCCGCAATTTACAACACAGACTTT   15622
gb:MN988713|Organism:Severe      TATCTACTGATGGTAACAAAATTGCCGATAAGTATGTCCGCAATTTACAACACAGACTTT   15622
gb:LC521925|Organism:Severe      TATCTACTGATGGTAACAAAATTGCCGATAAGTATGTCCGCAATTTACAACACAGACTTT   15595
gb:MT093571|Organism:Severe      TATCTACTGATGGTAACAAAATTGCCGATAAGTATGTCCGCAATTTACAACACAGACTTT   15622
gb:MT039887|Organism:Severe      TATCTACTGATGGTAACAAAATTGCCGATAAGTATGTCCGCAATTTACAACACAGACTTT   15622
gb:MT019530|Organism:Severe      TATCTACTGATGGTAACAAAATTGCCGATAAGTATGTCCGCAATTTACAACACAGACTTT   15622
gb:MT039888|Organism:Severe      TATCTACTGATGGTAACAAAATTGCCGATAAGTATGTCCGCAATTTACAACACAGACTTT   15622
gb:LC522972|Organism:Severe      TATCTACTGATGGTAACAAAATTGCCGATAAGTATGTCCGCAATTTACAACACAGACTTT   15619
gb:MT027063|Organism:Severe      TATCTACTGATGGTAACAAAATTGCCGATAAGTATGTCCGCAATTTACAACACAGACTTT   15622
gb:MT027062|Organism:Severe      TATCTACTGATGGTAACAAAATTGCCGATAAGTATGTCCGCAATTTACAACACAGACTTT   15622
gb:MT019529|Organism:Severe      TATCTACTGATGGTAACAAAATTGCCGATAAGTATGTCCGCAATTTACAACACAGACTTT   15622
gb:MN996529|Organism:Severe      TATCTACTGATGGTAACAAAATTGCCGATAAGTATGTCCGCAATTTACAACACAGACTTT   15610
gb:MN996531|Organism:Severe      TATCTACTGATGGTAACAAAATTGCCGATAAGTATGTCCGCAATTTACAACACAGACTTT   15609
gb:MT066176|Organism:Severe      TATCTACTGATGGTAACAAAATTGCCGATAAGTATGTCCGCAATTTACAACACAGACTTT   15622
gb:MT027064|Organism:Severe      TATCTACTGATGGTAACAAAATTGCCGATAAGTATGTCCGCAATTTACAACACAGACTTT   15622
gb:MN994468|Organism:Severe      TATCTACTGATGGTAACAAAATTGCCGATAAGTATGTCCGCAATTTACAACACAGACTTT   15622
gb:MT072688|Organism:Severe      TATCTACTGATGGTAACAAAATTGCCGATAAGTATGTCCGCAATTTACAACACAGACTTT   15607
gb:MN996527|Organism:Severe      TATCTACTGATGGTAACAAAATTGCCGATAAGTATGTCCGCAATTTACAACACAGACTTT   15589
gb:MT093631|Organism:Severe      TATCTACTGATGGTAACAAAATTGCCGATAAGTATGTCCGCAATTTACAACACAGACTTT   15660
gb:MT106053|Organism:Severe      TATCTACTGATGGTAACAAAATTGCCGATAAGTATGTCCGCAATTTACAACACAGACTTT   15622
gb:MT019533|Organism:Severe      TATCTACTGATGGTAACAAAATTGCCGATAAGTATGTCCGCAATTTACAACACAGACTTT   15622
gb:MT019531|Organism:Severe      TATCTACTGATGGTAACAAAATTGCCGATAAGTATGTCCGCAATTTACAACACAGACTTT   15622
gb:MN996528|Organism:Severe      TATCTACTGATGGTAACAAAATTGCCGATAAGTATGTCCGCAATTTACAACACAGACTTT   15622
gb:MN996530|Organism:Severe      TATCTACTGATGGTAACAAAATTGCCGATAAGTATGTCCGCAATTTACAACACAGACTTT   15608
gb:MN908947|Organism:Severe      TATCTACTGATGGTAACAAAATTGCCGATAAGTATGTCCGCAATTTACAACACAGACTTT   15622
gb:MT019532|Organism:Severe      TATCTACTGATGGTAACAAAATTGCCGATAAGTATGTCCGCAATTTACAACACAGACTTT   15622
                                 ********************************** ********* ***************

gb:MT020781|Organism:Severe      ATGAGTGTCTCTATAGAAATAGAGATGTTGACACAGACTTTGTGAATGAGTTTTACGCAT   15670
gb:MT007544|Organism:Severe      ATGAGTGTCTCTATAGAAATAGAGATGTTGACACAGACTTTGTGAATGAGTTTTACGCAT   15682
gb:MN994467|Organism:Severe      ATGAGTGTCTCTATAGAAATAGAGATGTTGACACAGACTTTGTGAATGAGTTTTACGCAT   15682
gb:MT044257|Organism:Severe      ATGAGTGTCTCTATAGAAATAGAGATGTTGACACAGACTTTGTGAATGAGTTTTACGCAT   15682
gb:MT106054|Organism:Severe      ATGAGTGTCTCTATAGAAATAGAGATGTTGACACAGACTTTGTGAATGAGTTTTACGCAT   15682
gb:MT049951|Organism:Severe      ATGAGTGTCTCTATAGAAATAGAGATGTTGACACAGACTTTGTGAATGAGTTTTACGCAT   15682
gb:MN975262|Organism:Severe      ATGAGTGTCTCTATAGAAATAGAGATGTTGACACAGACTTTGTGAATGAGTTTTACGCAT   15682
gb:MT106052|Organism:Severe      ATGAGTGTCTCTATAGAAATAGAGATGTTGACACAGACTTTGTGAATGAGTTTTACGCAT   15682
gb:LC522975|Organism:Severe      ATGAGTGTCTCTATAGAAATAGAGATGTTGACACAGACTTTGTGAATGAGTTTTACGCAT   15679
gb:LC522973|Organism:Severe      ATGAGTGTCTCTATAGAAATAGAGATGTTGACACAGACTTTGTGAATGAGTTTTACGCAT   15679
gb:LC522974|Organism:Severe      ATGAGTGTCTCTATAGAAATAGAGATGTTGACACAGACTTTGTGAATGAGTTTTACGCAT   15679
gb:MN985325|Organism:Severe      ATGAGTGTCTCTATAGAAATAGAGATGTTGACACAGACTTTGTGAATGAGTTTTACGCAT   15682
gb:MT020881|Organism:Severe      ATGAGTGTCTCTATAGAAATAGAGATGTTGACACAGACTTTGTGAATGAGTTTTACGCAT   15682
gb:MT020880|Organism:Severe      ATGAGTGTCTCTATAGAAATAGAGATGTTGACACAGACTTTGTGAATGAGTTTTACGCAT   15682
gb:MT066175|Organism:Severe      ATGAGTGTCTCTATAGAAATAGAGATGTTGACACAGACTTTGTGAATGAGTTTTACGCAT   15682
```

```
gb:MN997409|Organism:Severe      ATGAGTGTCTCTATAGAAATAGAGATGTTGACACAGACTTTGTGAATGAGTTTTACGCAT   15682
gb:MN938384|Organism:Severe      ATGAGTGTCTCTATAGAAATAGAGATGTTGACACAGACTTTGTGAATGAGTTTTACGCAT   15650
gb:MT044258|Organism:Severe      ATGAGTGTCTCTATAGAAATAGAGATGTTGACACAGACTTTGTGAATGAGTTTTACGCAT   15658
gb:MT039890|Organism:Severe      ATGAGTGTCTCTATAGAAATAGAGATGTTGACACAGACTTTGTGAATGAGTTTTACGCAT   15682
gb:MN988713|Organism:Severe      ATGAGTGTCTCTATAGAAATAGAGATGTTGACACAGACTTTGTGAATGAGTTTTACGCAT   15682
gb:LC521925|Organism:Severe      ATGAGTGTCTCTATAGAAATAGAGATGTTGACACAGACTTTGTGAATGAGTTTTACGCAT   15655
gb:MT093571|Organism:Severe      ATGAGTGTCTCTATAGAAATAGAGATGTTGACACAGACTTTGTGAATGAGTTTTACGCAT   15682
gb:MT039887|Organism:Severe      ATGAGTGTCTCTATAGAAATAGAGATGTTGACACAGACTTTGTGAATGAGTTTTACGCAT   15682
gb:MT019530|Organism:Severe      ATGAGTGTCTCTATAGAAATAGAGATGTTGACACAGACTTTGTGAATGAGTTTTACGCAT   15682
gb:MT039888|Organism:Severe      ATGAGTGTCTCTATAGAAATAGAGATGTTGACACAGACTTTGTGAATGAGTTTTACGCAT   15682
gb:LC522972|Organism:Severe      ATGAGTGTCTCTATAGAAATAGAGATGTTGACACAGACTTTGTGAATGAGTTTTACGCAT   15679
gb:MT027063|Organism:Severe      ATGAGTGTCTCTATAGAAATAGAGATGTTGACACAGACTTTGTGAATGAGTTTTACGCAT   15682
gb:MT027062|Organism:Severe      ATGAGTGTCTCTATAGAAATAGAGATGTTGACACAGACTTTGTGAATGAGTTTTACGCAT   15682
gb:MT019529|Organism:Severe      ATGAGTGTCTCTATAGAAATAGAGATGTTGACACAGACTTTGTGAATGAGTTTTACGCAT   15682
gb:MN996529|Organism:Severe      ATGAGTGTCTCTATAGAAATAGAGATGTTGACACAGACTTTGTGAATGAGTTTTACGCAT   15670
gb:MN996531|Organism:Severe      ATGAGTGTCTCTATAGAAATAGAGATGTTGACACAGACTTTGTGAATGAGTTTTACGCAT   15669
gb:MT066176|Organism:Severe      ATGAGTGTCTCTATAGAAATAGAGATGTTGACACAGACTTTGTGAATGAGTTTTACGCAT   15682
gb:MT027064|Organism:Severe      ATGAGTGTCTCTATAGAAATAGAGATGTTGACACAGACTTTGTGAATGAGTTTTACGCAT   15682
gb:MN994468|Organism:Severe      ATGAGTGTCTCTATAGAAATAGAGATGTTGACACAGACTTTGTGAATGAGTTTTACGCAT   15682
gb:MT072688|Organism:Severe      ATGAGTGTCTCTATAGAAATAGAGATGTTGACACAGACTTTGTGAATGAGTTTTACGCAT   15667
gb:MN996527|Organism:Severe      ATGAGTGTCTCTATAGAAATAGAGATGTTGACACAGACTTTGTGAATGAGTTTTACGCAT   15649
gb:MT093631|Organism:Severe      ATGAGTGTCTCTATAGAAATAGAGATGTTGACACAGACTTTGTGAATGAGTTTTACGCAT   15720
gb:MT106053|Organism:Severe      ATGAGTGTCTCTATAGAAATAGAGATGTTGACACAGACTTTGTGAATGAGTTTTACGCAT   15682
gb:MT019533|Organism:Severe      ATGAGTGTCTCTATAGAAATAGAGATGTTGACACAGACTTTGTGAATGAGTTTTACGCAT   15682
gb:MT019531|Organism:Severe      ATGAGTGTCTCTATAGAAATAGAGATGTTGACACAGACTTTGTGAATGAGTTTTACGCAT   15682
gb:MN996528|Organism:Severe      ATGAGTGTCTCTATAGAAATAGAGATGTTGACACAGACTTTGTGAATGAGTTTTACGCAT   15682
gb:MN996530|Organism:Severe      ATGAGTGTCTCTATAGAAATAGAGATGTTGACACAGACTTTGTGAATGAGTTTTACGCAT   15668
gb:MN908947|Organism:Severe      ATGAGTGTCTCTATAGAAATAGAGATGTTGACACAGACTTTGTGAATGAGTTTTACGCAT   15682
gb:MT019532|Organism:Severe      ATGAGTGTCTCTATAGAAATAGAGATGTTGACACAGACTTTGTGAATGAGTTTTACGCAT   15682
                                 ************************************************************

gb:MT020781|Organism:Severe      ATTTGCGTAAACATTTCTCAATGATGATACTCTCTGACGATGCTGTTGTGTGTTTCAATA   15730
gb:MT007544|Organism:Severe      ATTTGCGTAAACATTTCTCAATGATGATACTCTCTGACGATGCTGTTGTGTGTTTCAATA   15742
gb:MN994467|Organism:Severe      ATTTGCGTAAACATTTCTCAATGATGATACTCTCTGACGATGCTGTTGTGTGTTTCAATA   15742
gb:MT044257|Organism:Severe      ATTTGCGTAAACATTTCTCAATGATGATACTCTCTGACGATGCTGTTGTGTGTTTCAATA   15742
gb:MT106054|Organism:Severe      ATTTGCGTAAACATTTCTCAATGATGATACTCTCTGACGATGCTGTTGTGTGTTTCAATA   15742
gb:MT049951|Organism:Severe      ATTTGCGTAAACATTTCTCAATGATGATACTCTCTGACGATGCTGTTGTGTGTTTCAATA   15742
gb:MN975262|Organism:Severe      ATTTGCGTAAACATTTCTCAATGATGATACTCTCTGACGATGCTGTTGTGTGTTTCAATA   15742
gb:MT106052|Organism:Severe      ATTTGCGTAAACATTTCTCAATGATGATACTCTCTGACGATGCTGTTGTGTGTTTCAATA   15742
gb:LC522975|Organism:Severe      ATTTGCGTAAACATTTCTCAATGATGATACTCTCTGACGATGCTGTTGTGTGTTTCAATA   15739
gb:LC522973|Organism:Severe      ATTTGCGTAAACATTTCTCAATGATGATACTCTCTGACGATGCTGTTGTGTGTTTCAATA   15739
gb:LC522974|Organism:Severe      ATTTGCGTAAACATTTCTCAATGATGATACTCTCTGACGATGCTGTTGTGTGTTTCAATA   15739
gb:MN985325|Organism:Severe      ATTTGCGTAAACATTTCTCAATGATGATACTCTCTGACGATGCTGTTGTGTGTTTCAATA   15742
gb:MT020881|Organism:Severe      ATTTGCGTAAACATTTCTCAATGATGATACTCTCTGACGATGCTGTTGTGTGTTTCAATA   15742
gb:MT020880|Organism:Severe      ATTTGCGTAAACATTTCTCAATGATGATACTCTCTGACGATGCTGTTGTGTGTTTCAATA   15742
gb:MT066175|Organism:Severe      ATTTGCGTAAACATTTCTCAATGATGATACTCTCTGACGATGCTGTTGTGTGTTTCAATA   15742
gb:MN997409|Organism:Severe      ATTTGCGTAAACATTTCTCAATGATGATACTCTCTGACGATGCTGTTGTGTGTTTCAATA   15742
gb:MN938384|Organism:Severe      ATTTGCGTAAACATTTCTCAATGATGATACTCTCTGACGATGCTGTTGTGTGTTTCAATA   15710
gb:MT044258|Organism:Severe      ATTTGCGTAAACATTTCTCAATGATGATACTCTCTGACGATGCTGTTGTGTGTTTCAATA   15718
gb:MT039890|Organism:Severe      ATTTGCGTAAACATTTCTCAATGATGATACTCTCTGACGATGCTGTTGTGTGTTTCAATA   15742
gb:MN988713|Organism:Severe      ATTTGCGTAAACATTTCTCAATGATGATACTCTCTGACGATGCTGTTGTGTGTTTCAATA   15742
gb:LC521925|Organism:Severe      ATTTGCGTAAACATTTCTCAATGATGATACTCTCTGACGATGCTGTTGTGTGTTTCAATA   15715
gb:MT093571|Organism:Severe      ATTTGCGTAAACATTTCTCAATGATGATACTCTCTGACGATGCTGTTGTGTGTTTCAATA   15742
gb:MT039887|Organism:Severe      ATTTGCGTAAACATTTCTCAATGATGATACTCTCTGACGATGCTGTTGTGTGTTTCAATA   15742
gb:MT019530|Organism:Severe      ATTTGCGTAAACATTTCTCAATGATGATACTCTCTGACGATGCTGTTGTGTGTTTCAATA   15742
gb:MT039888|Organism:Severe      ATTTGCGTAAACATTTCTCAATGATGATACTCTCTGACGATGCTGTTGTGTGTTTCAATA   15742
gb:LC522972|Organism:Severe      ATTTGCGTAAACATTTCTCAATGATGATACTCTCTGACGATGCTGTTGTGTGTTTCAATA   15739
gb:MT027063|Organism:Severe      ATTTGCGTAAACATTTCTCAATGATGATACTCTCTGACGATGCTGTTGTGTGTTTCAATA   15742
gb:MT027062|Organism:Severe      ATTTGCGTAAACATTTCTCAATGATGATACTCTCTGACGATGCTGTTGTGTGTTTCAATA   15742
gb:MT019529|Organism:Severe      ATTTGCGTAAACATTTCTCAATGATGATACTCTCTGACGATGCTGTTGTGTGTTTCAATA   15742
gb:MN996529|Organism:Severe      ATTTGCGTAAACATTTCTCAATGATGATACTCTCTGACGATGCTGTTGTGTGTTTCAATA   15730
gb:MN996531|Organism:Severe      ATTTGCGTAAACATTTCTCAATGATGATACTCTCTGACGATGCTGTTGTGTGTTTCAATA   15729
gb:MT066176|Organism:Severe      ATTTGCGTAAACATTTCTCAATGATGATACTCTCTGACGATGCTGTTGTGTGTTTCAATA   15742
gb:MT027064|Organism:Severe      ATTTGCGTAAACATTTCTCAATGATGATACTCTCTGACGATGCTGTTGTGTGTTTCAATA   15742
gb:MN994468|Organism:Severe      ATTTGCGTAAACATTTCTCAATGATGATACTCTCTGACGATGCTGTTGTGTGTTTCAATA   15742
```

```
gb:MT072688|Organism:Severe      ATTTGCGTAAACATTTCTCAATGATGATACTCTCTGACGATGCTGTTGTGTGTTTCAATA   15727
gb:MN996527|Organism:Severe      ATTTGCGTAAACATTTCTCAATGATGATACTCTCTGACGATGCTGTTGTGTGTTTCAATA   15709
gb:MT093631|Organism:Severe      ATTTGCGTAAACATTTCTCAATGATGATACTCTCTGACGATGCTGTTGTGTGTTTCAATA   15780
gb:MT106053|Organism:Severe      ATTTGCGTAAACATTTCTCAATGATGATACTCTCTGACGATGCTGTTGTGTGTTTCAATA   15742
gb:MT019533|Organism:Severe      ATTTGCGTAAACATTTCTCAATGATGATACTCTCTGACGATGCTGTTGTGTGTTTCAATA   15742
gb:MT019531|Organism:Severe      ATTTGCGTAAACATTTCTCAATGATGATACTCTCTGACGATGCTGTTGTGTGTTTCAATA   15742
gb:MN996528|Organism:Severe      ATTTGCGTAAACATTTCTCAATGATGATACTCTCTGACGATGCTGTTGTGTGTTTCAATA   15742
gb:MN996530|Organism:Severe      ATTTGCGTAAACATTTCTCAATGATGATACTCTCTGACGATGCTGTTGTGTGTTTCAATA   15728
gb:MN908947|Organism:Severe      ATTTGCGTAAACATTTCTCAATGATGATACTCTCTGACGATGCTGTTGTGTGTTTCAATA   15742
gb:MT019532|Organism:Severe      ATTTGCGTAAACATTTCTCAATGATGATACTCTCTGACGATGCTGTTGTGTGTTTCAATA   15742
                                 ************************************************************

gb:MT020781|Organism:Severe      GCACTTATGCATCTCAAGGTCTAGTGGCTAGCATAAAGAACTTTAAGTCAGTTCTTTATT   15790
gb:MT007544|Organism:Severe      GCACTTATGCATCTCAAGGTCTAGTGGCTAGCATAAAGAACTTTAAGTCAGTTCTTTATT   15802
gb:MN994467|Organism:Severe      GCACTTATGCATCTCAAGGTCTAGTGGCTAGCATAAAGAACTTTAAGTCAGTTCTTTATT   15802
gb:MT044257|Organism:Severe      GCACTTATGCATCTCAAGGTCTAGTGGCTAGCATAAAGAACTTTAAGTCAGTTCTTTATT   15802
gb:MT106054|Organism:Severe      GCACTTATGCATCTCAAGGTCTAGTGGCTAGCATAAAGAACTTTAAGTCAGTTCTTTATT   15802
gb:MT049951|Organism:Severe      GCACTTATGCATCTCAAGGTCTAGTGGCTAGCATAAAGAACTTTAAGTCAGTTCTTTATT   15802
gb:MN975262|Organism:Severe      GCACTTATGCATCTCAAGGTCTAGTGGCTAGCATAAAGAACTTTAAGTCAGTTCTTTATT   15802
gb:MT106052|Organism:Severe      GCACTTATGCATCTCAAGGTCTAGTGGCTAGCATAAAGAACTTTAAGTCAGTTCTTTATT   15802
gb:LC522975|Organism:Severe      GCACTTATGCATCTCAAGGTCTAGTGGCTAGCATAAAGAACTTTAAGTCAGTTCTTTATT   15799
gb:LC522973|Organism:Severe      GCACTTATGCATCTCAAGGTCTAGTGGCTAGCATAAAGAACTTTAAGTCAGTTCTTTATT   15799
gb:LC522974|Organism:Severe      GCACTTATGCATCTCAAGGTCTAGTGGCTAGCATAAAGAACTTTAAGTCAGTTCTTTATT   15799
gb:MN985325|Organism:Severe      GCACTTATGCATCTCAAGGTCTAGTGGCTAGCATAAAGAACTTTAAGTCAGTTCTTTATT   15802
gb:MT020881|Organism:Severe      GCACTTATGCATCTCAAGGTCTAGTGGCTAGCATAAAGAACTTTAAGTCAGTTCTTTATT   15802
gb:MT020880|Organism:Severe      GCACTTATGCATCTCAAGGTCTAGTGGCTAGCATAAAGAACTTTAAGTCAGTTCTTTATT   15802
gb:MT066175|Organism:Severe      GCACTTATGCATCTCAAGGTCTAGTGGCTAGCATAAAGAACTTTAAGTCAGTTCTTTATT   15802
gb:MN997409|Organism:Severe      GCACTTATGCATCTCAAGGTCTAGTGGCTAGCATAAAGAACTTTAAGTCAGTTCTTTATT   15802
gb:MN938384|Organism:Severe      GCACTTATGCATCTCAAGGTCTAGTGGCTAGCATAAAGAACTTTAAGTCAGTTCTTTATT   15770
gb:MT044258|Organism:Severe      GCACTTATGCATCTCAAGGTCTAGTGGCTAGCATAAAGAACTTTAAGTCAGTTCTTTATT   15778
gb:MT039890|Organism:Severe      GCACTTATGCATCTCAAGGTCTAGTGGCTAGCATAAAGAACTTTAAGTCAGTTCTTTATT   15802
gb:MN988713|Organism:Severe      GCACTTATGCATCTCAAGGTCTAGTGGCTAGCATAAAGAACTTTAAGTCAGTTCTTTATT   15802
gb:LC521925|Organism:Severe      GCACTTATGCATCTCAAGGTCTAGTGGCTAGCATAAAGAACTTTAAGTCAGTTCTTTATT   15775
gb:MT093571|Organism:Severe      GCACTTATGCATCTCAAGGTCTAGTGGCTAGCATAAAGAACTTTAAGTCAGTTCTTTATT   15802
gb:MT039887|Organism:Severe      GCACTTATGCATCTCAAGGTCTAGTGGCTAGCATAAAGAACTTTAAGTCAGTTCTTTATT   15802
gb:MT019530|Organism:Severe      GCACTTATGCATCTCAAGGTCTAGTGGCTAGCATAAAGAACTTTAAGTCAGTTCTTTATT   15802
gb:MT039888|Organism:Severe      GCACTTATGCATCTCAAGGTCTAGTGGCTAGCATAAAGAACTTTAAGTCAGTTCTTTATT   15802
gb:LC522972|Organism:Severe      GCACTTATGCATCTCAAGGTCTAGTGGCTAGCATAAAGAACTTTAAGTCAGTTCTTTATT   15799
gb:MT027063|Organism:Severe      GCACTTATGCATCTCAAGGTCTAGTGGCTAGCATAAAGAACTTTAAGTCAGTTCTTTATT   15802
gb:MT027062|Organism:Severe      GCACTTATGCATCTCAAGGTCTAGTGGCTAGCATAAAGAACTTTAAGTCAGTTCTTTATT   15802
gb:MT019529|Organism:Severe      GCACTTATGCATCTCAAGGTCTAGTGGCTAGCATAAAGAACTTTAAGTCAGTTCTTTATT   15802
gb:MN996529|Organism:Severe      GCACTTATGCATCTCAAGGTCTAGTGGCTAGCATAAAGAACTTTAAGTCAGTTCTTTATT   15790
gb:MN996531|Organism:Severe      GCACTTATGCATCTCAAGGTCTAGTGGCTAGCATAAAGAACTTTAAGTCAGTTCTTTATT   15789
gb:MT066176|Organism:Severe      GCACTTATGCATCTCAAGGTCTAGTGGCTAGCATAAAGAACTTTAAGTCAGTTCTTTATT   15802
gb:MT027064|Organism:Severe      GCACTTATGCATCTCAAGGTCTAGTGGCTAGCATAAAGAACTTTAAGTCAGTTCTTTATT   15802
gb:MN994468|Organism:Severe      GCACTTATGCATCTCAAGGTCTAGTGGCTAGCATAAAGAACTTTAAGTCAGTTCTTTATT   15802
gb:MT072688|Organism:Severe      GCACTTATGCATCTCAAGGTCTAGTGGCTAGCATAAAGAACTTTAAGTCAGTTCTTTATT   15787
gb:MN996527|Organism:Severe      GCACTTATGCATCTCAAGGTCTAGTGGCTAGCATAAAGAACTTTAAGTCAGTTCTTTATT   15769
gb:MT093631|Organism:Severe      GCACTTATGCATCTCAAGGTCTAGTGGCTAGCATAAAGAACTTTAAGTCAGTTCTTTATT   15840
gb:MT106053|Organism:Severe      GCACTTATGCATCTCAAGGTCTAGTGGCTAGCATAAAGAACTTTAAGTCAGTTCTTTATT   15802
gb:MT019533|Organism:Severe      GCACTTATGCATCTCAAGGTCTAGTGGCTAGCATAAAGAACTTTAAGTCAGTTCTTTATT   15802
gb:MT019531|Organism:Severe      GCACTTATGCATCTCAAGGTCTAGTGGCTAGCATAAAGAACTTTAAGTCAGTTCTTTATT   15802
gb:MN996528|Organism:Severe      GCACTTATGCATCTCAAGGTCTAGTGGCTAGCATAAAGAACTTTAAGTCAGTTCTTTATT   15802
gb:MN996530|Organism:Severe      GCACTTATGCATCTCAAGGTCTAGTGGCTAGCATAAAGAACTTTAAGTCAGTTCTTTATT   15788
gb:MN908947|Organism:Severe      GCACTTATGCATCTCAAGGTCTAGTGGCTAGCATAAAGAACTTTAAGTCAGTTCTTTATT   15802
gb:MT019532|Organism:Severe      GCACTTATGCATCTCAAGGTCTAGTGGCTAGCATAAAGAACTTTAAGTCAGTTCTTTATT   15802
                                 ************************************************************

gb:MT020781|Organism:Severe      ATCAAAACAATGTTTTTATGTCTGAAGCAAAATGTTGGACTGAGACTGACCTTACTAAAG   15850
gb:MT007544|Organism:Severe      ATCAAAACAATGTTTTTATGTCTGAAGCAAAATGTTGGACTGAGACTGACCTTACTAAAG   15862
gb:MN994467|Organism:Severe      ATCAAAACAATGTTTTTATGTCTGAAGCAAAATGTTGGACTGAGACTGACCTTACTAAAG   15862
gb:MT044257|Organism:Severe      ATCAAAACAATGTTTTTATGTCTGAAGCAAAATGTTGGACTGAGACTGACCTTACTAAAG   15862
gb:MT106054|Organism:Severe      ATCAAAACAATGTTTTTATGTCTGAAGCAAAATGTTGGACTGAGACTGACCTTACTAAAG   15862
gb:MT049951|Organism:Severe      ATCAAAACAATGTTTTTATGTCTGAAGCAAAATGTTGGACTGAGACTGACCTTACTAAAG   15862
gb:MN975262|Organism:Severe      ATCAAAACAATGTTTTTATGTCTGAAGCAAAATGTTGGACTGAGACTGACCTTACTAAAG   15862
```

```
gb:MT106052|Organism:Severe      ATCAAAACAATGTTTTTATGTCTGAAGCAAAATGTTGGACTGAGACTGACCTTACTAAAG    15862
gb:LC522975|Organism:Severe      ATCAAAACAATGTTTTTATGTCTGAAGCAAAATGTTGGACTGAGACTGACCTTACTAAAG    15859
gb:LC522973|Organism:Severe      ATCAAAACAATGTTTTTATGTCTGAAGCAAAATGTTGGACTGAGACTGACCTTACTAAAG    15859
gb:LC522974|Organism:Severe      ATCAAAACAATGTTTTTATGTCTGAAGCAAAATGTTGGACTGAGACTGACCTTACTAAAG    15859
gb:MN985325|Organism:Severe      ATCAAAACAATGTTTTTATGTCTGAAGCAAAATGTTGGACTGAGACTGACCTTACTAAAG    15862
gb:MT020881|Organism:Severe      ATCAAAACAATGTTTTTATGTCTGAAGCAAAATGTTGGACTGAGACTGACCTTACTAAAG    15862
gb:MT020880|Organism:Severe      ATCAAAACAATGTTTTTATGTCTGAAGCAAAATGTTGGACTGAGACTGACCTTACTAAAG    15862
gb:MT066175|Organism:Severe      ATCAAAACAATGTTTTTATGTCTGAAGCAAAATGTTGGACTGAGACTGACCTTACTAAAG    15862
gb:MN997409|Organism:Severe      ATCAAAACAATGTTTTTATGTCTGAAGCAAAATGTTGGACTGAGACTGACCTTACTAAAG    15862
gb:MN938384|Organism:Severe      ATCAAAACAATGTTTTTATGTCTGAAGCAAAATGTTGGACTGAGACTGACCTTACTAAAG    15830
gb:MT044258|Organism:Severe      ATCAAAACAATGTTTTTATGTCTGAAGCAAAATGTTGGACTGAGACTGACCTTACTAAAG    15838
gb:MT039890|Organism:Severe      ATCAAAACAATGTTTTTATGTCTGAAGCAAAATGTTGGACTGAGACTGACCTTACTAAAG    15862
gb:MN988713|Organism:Severe      ATCAAAACAATGTTTTTATGTCTGAAGCAAAATGTTGGACTGAGACTGACCTTACTAAAG    15862
gb:LC521925|Organism:Severe      ATCAAAACAATGTTTTTATGTCTGAAGCAAAATGTTGGACTGAGACTGACCTTACTAAAG    15835
gb:MT093571|Organism:Severe      ATCAAAACAATGTTTTTATGTCTGAAGCAAAATGTTGGACTGAGACTGACCTTACTAAAG    15862
gb:MT039887|Organism:Severe      ATCAAAACAATGTTTTTATGTCTGAAGCAAAATGTTGGACTGAGACTGACCTTACTAAAG    15862
gb:MT019530|Organism:Severe      ATCAAAACAATGTTTTTATGTCTGAAGCAAAATGTTGGACTGAGACTGACCTTACTAAAG    15862
gb:MT039888|Organism:Severe      ATCAAAACAATGTTTTTATGTCTGAAGCAAAATGTTGGACTGAGACTGACCTTACTAAAG    15862
gb:LC522972|Organism:Severe      ATCAAAACAATGTTTTTATGTCTGAAGCAAAATGTTGGACTGAGACTGACCTTACTAAAG    15859
gb:MT027063|Organism:Severe      ATCAAAACAATGTTTTTATGTCTGAAGCAAAATGTTGGACTGAGACTGACCTTACTAAAG    15862
gb:MT027062|Organism:Severe      ATCAAAACAATGTTTTTATGTCTGAAGCAAAATGTTGGACTGAGACTGACCTTACTAAAG    15862
gb:MT019529|Organism:Severe      ATCAAAACAATGTTTTTATGTCTGAAGCAAAATGTTGGACTGAGACTGACCTTACTAAAG    15862
gb:MN996529|Organism:Severe      ATCAAAACAATGTTTTTATGTCTGAAGCAAAATGTTGGACTGAGACTGACCTTACTAAAG    15850
gb:MN996531|Organism:Severe      ATCAAAACAATGTTTTTATGTCTGAAGCAAAATGTTGGACTGAGACTGACCTTACTAAAG    15849
gb:MT066176|Organism:Severe      ATCAAAACAATGTTTTTATGTCTGAAGCAAAATGTTGGACTGAGACTGACCTTACTAAAG    15862
gb:MT027064|Organism:Severe      ATCAAAACAATGTTTTTATGTCTGAAGCAAAATGTTGGACTGAGACTGACCTTACTAAAG    15862
gb:MN994468|Organism:Severe      ATCAAAACAATGTTTTTATGTCTGAAGCAAAATGTTGGACTGAGACTGACCTTACTAAAG    15862
gb:MT072688|Organism:Severe      ATCAAAACAATGTTTTTATGTCTGAAGCAAAATGTTGGACTGAGACTGACCTTACTAAAG    15847
gb:MN996527|Organism:Severe      ATCAAAACAATGTTTTTATGTCTGAAGCAAAATGTTGGACTGAGACTGACCTTACTAAAG    15829
gb:MT093631|Organism:Severe      ATCAAAACAATGTTTTTATGTCTGAAGCAAAATGTTGGACTGAGACTGACCTTACTAAAG    15900
gb:MT106053|Organism:Severe      ATCAAAACAATGTTTTTATGTCTGAAGCAAAATGTTGGACTGAGACTGACCTTACTAAAG    15862
gb:MT019533|Organism:Severe      ATCAAAACAATGTTTTTATGTCTGAAGCAAAATGTTGGACTGAGACTGACCTTACTAAAG    15862
gb:MT019531|Organism:Severe      ATCAAAACAATGTTTTTATGTCTGAAGCAAAATGTTGGACTGAGACTGACCTTACTAAAG    15862
gb:MN996528|Organism:Severe      ATCAAAACAATGTTTTTATGTCTGAAGCAAAATGTTGGACTGAGACTGACCTTACTAAAG    15862
gb:MN996530|Organism:Severe      ATCAAAACAATGTTTTTATGTCTGAAGCAAAATGTTGGACTGAGACTGACCTTACTAAAG    15848
gb:MN908947|Organism:Severe      ATCAAAACAATGTTTTTATGTCTGAAGCAAAATGTTGGACTGAGACTGACCTTACTAAAG    15862
gb:MT019532|Organism:Severe      ATCAAAACAATGTTTTTATGTCTGAAGCAAAATGTTGGACTGAGACTGACCTTACTAAAG    15862
                                 ************************************************************

gb:MT020781|Organism:Severe      GACCTCATGAATTTTGCTCTCAACATACAATGCTAGTTAAACAGGGTGATGATTATGTGT    15910
gb:MT007544|Organism:Severe      GACCTCATGAATTTTGCTCTCAACATACAATGCTAGTTAAACAGGGTGATGATTATGTGT    15922
gb:MN994467|Organism:Severe      GACCTCATGAATTTTGCTCTCAACATACAATGCTAGTTAAACAGGGTGATGATTATGTGT    15922
gb:MT044257|Organism:Severe      GACCTCATGAATTTTGCTCTCAACATACAATGCTAGTTAAACAGGGTGATGATTATGTGT    15922
gb:MT106054|Organism:Severe      GACCTCATGAATTTTGCTCTCAACATACAATGCTAGTTAAACAGGGTGATGATTATGTGT    15922
gb:MT049951|Organism:Severe      GACCTCATGAATTTTGCTCTCAACATACAATGCTAGTTAAACAGGGTGATGATTATGTGT    15922
gb:MN975262|Organism:Severe      GACCTCATGAATTTTGCTCTCAACATACAATGCTAGTTAAACAGGGTGATGATTATGTGT    15922
gb:MT106052|Organism:Severe      GACCTCATGAATTTTGCTCTCAACATACAATGCTAGTTAAACAGGGTGATGATTATGTGT    15922
gb:LC522975|Organism:Severe      GACCTCATGAATTTTGCTCTCAACATACAATGCTAGTTAAACAGGGTGATGATTATGTGT    15919
gb:LC522973|Organism:Severe      GACCTCATGAATTTTGCTCTCAACATACAATGCTAGTTAAACAGGGTGATGATTATGTGT    15919
gb:LC522974|Organism:Severe      GACCTCATGAATTTTGCTCTCAACATACAATGCTAGTTAAACAGGGTGATGATTATGTGT    15919
gb:MN985325|Organism:Severe      GACCTCATGAATTTTGCTCTCAACATACAATGCTAGTTAAACAGGGTGATGATTATGTGT    15922
gb:MT020881|Organism:Severe      GACCTCATGAATTTTGCTCTCAACATACAATGCTAGTTAAACAGGGTGATGATTATGTGT    15922
gb:MT020880|Organism:Severe      GACCTCATGAATTTTGCTCTCAACATACAATGCTAGTTAAACAGGGTGATGATTATGTGT    15922
gb:MT066175|Organism:Severe      GACCTCATGAATTTTGCTCTCAACATACAATGCTAGTTAAACAGGGTGATGATTATGTGT    15922
gb:MN997409|Organism:Severe      GACCTCATGAATTTTGCTCTCAACATACAATGCTAGTTAAACAGGGTGATGATTATGTGT    15922
gb:MN938384|Organism:Severe      GACCTCATGAATTTTGCTCTCAACATACAATGCTAGTTAAACAGGGTGATGATTATGTGT    15890
gb:MT044258|Organism:Severe      GACCTCATGAATTTTGCTCTCAACATACAATGCTAGTTAAACAGGGTGATGATTATGTGT    15898
gb:MT039890|Organism:Severe      GACCTCATGAATTTTGCTCTCAACATACAATGCTAGTTAAACAGGGTGATGATTATGTGT    15922
gb:MN988713|Organism:Severe      GACCTCATGAATTTTGCTCTCAACATACAATGCTAGTTAAACAGGGTGATGATTATGTGT    15922
gb:LC521925|Organism:Severe      GACCTCATGAATTTTGCTCTCAACATACAATGCTAGTTAAACAGGGTGATGATTATGTGT    15895
gb:MT093571|Organism:Severe      GACCTCATGAATTTTGCTCTCAACATACAATGCTAGTTAAACAGGGTGATGATTATGTGT    15922
gb:MT039887|Organism:Severe      GACCTCATGAATTTTGCTCTCAACATACAATGCTAGTTAAACAGGGTGATGATTATGTGT    15922
gb:MT019530|Organism:Severe      GACCTCATGAATTTTGCTCTCAACATACAATGCTAGTTAAACAGGGTGATGATTATGTGT    15922
gb:MT039888|Organism:Severe      GACCTCATGAATTTTGCTCTCAACATACAATGCTAGTTAAACAGGGTGATGATTATGTGT    15922
gb:LC522972|Organism:Severe      GACCTCATGAATTTTGCTCTCAACATACAATGCTAGTTAAACAGGGTGATGATTATGTGT    15919
```

```
gb:MT027063|Organism:Severe      GACCTCATGAATTTTGCTCTCAACATACAATGCTAGTTAAACAGGGTGATGATTATGTGT    15922
gb:MT027062|Organism:Severe      GACCTCATGAATTTTGCTCTCAACATACAATGCTAGTTAAACAGGGTGATGATTATGTGT    15922
gb:MT019529|Organism:Severe      GACCTCATGAATTTTGCTCTCAACATACAATGCTAGTTAAACAGGGTGATGATTATGTGT    15922
gb:MN996529|Organism:Severe      GACCTCATGAATTTTGCTCTCAACATACAATGCTAGTTAAACAGGGTGATGATTATGTGT    15910
gb:MN996531|Organism:Severe      GACCTCATGAATTTTGCTCTCAACATACAATGCTAGTTAAACAGGGTGATGATTATGTGT    15909
gb:MT066176|Organism:Severe      GACCTCATGAATTTTGCTCTCAACATACAATGCTAGTTAAACAGGGTGATGATTATGTGT    15922
gb:MT027064|Organism:Severe      GACCTCATGAATTTTGCTCTCAACATACAATGCTAGTTAAACAGGGTGATGATTATGTGT    15922
gb:MN994468|Organism:Severe      GACCTCATGAATTTTGCTCTCAACATACAATGCTAGTTAAACAGGGTGATGATTATGTGT    15922
gb:MT072688|Organism:Severe      GACCTCATGAATTTTGCTCTCAACATACAATGCTAGTTAAACAGGGTGATGATTATGTGT    15907
gb:MN996527|Organism:Severe      GACCTCATGAATTTTGCTCTCAACATACAATGCTAGTTAAACAGGGTGATGATTATGTGT    15889
gb:MT093631|Organism:Severe      GACCTCATGAATTTTGCTCTCAACATACAATGCTAGTTAAACAGGGTGATGATTATGTGT    15960
gb:MT106053|Organism:Severe      GACCTCATGAATTTTGCTCTCAACATACAATGCTAGTTAAACAGGGTGATGATTATGTGT    15922
gb:MT019533|Organism:Severe      GACCTCATGAATTTTGCTCTCAACATACAATGCTAGTTAAACAGGGTGATGATTATGTGT    15922
gb:MT019531|Organism:Severe      GACCTCATGAATTTTGCTCTCAACATACAATGCTAGTTAAACAGGGTGATGATTATGTGT    15922
gb:MN996528|Organism:Severe      GACCTCATGAATTTTGCTCTCAACATACAATGCTAGTTAAACAGGGTGATGATTATGTGT    15922
gb:MN996530|Organism:Severe      GACCTCATGAATTTTGCTCTCAACATACAATGCTAGTTAAACAGGGTGATGATTATGTGT    15908
gb:MN908947|Organism:Severe      GACCTCATGAATTTTGCTCTCAACATACAATGCTAGTTAAACAGGGTGATGATTATGTGT    15922
gb:MT019532|Organism:Severe      GACCTCATGAATTTTGCTCTCAACATACAATGCTAGTTAAACAGGGTGATGATTATGTGT    15922
                                 ************************************************************

gb:MT020781|Organism:Severe      ACCTTCCTTACCCAGATCCATCAAGAATCCTAGGGGCCGGCTGTTTTGTAGATGATATCG    15970
gb:MT007544|Organism:Severe      ACCTTCCTTACCCAGATCCATCAAGAATCCTAGGGGCCGGCTGTTTTGTAGATGATATCG    15982
gb:MN994467|Organism:Severe      ACCTTCCTTACCCAGATCCATCAAGAATCCTAGGGGCCGGCTGTTTTGTAGATGATATCG    15982
gb:MT044257|Organism:Severe      ACCTTCCTTACCCAGATCCATCAAGAATCCTAGGGGCCGGCTGTTTTGTAGATGATATCG    15982
gb:MT106054|Organism:Severe      ACCTTCCTTACCCAGATCCATCAAGAATCCTAGGGGCCGGCTGTTTTGTAGATGATATCG    15982
gb:MT049951|Organism:Severe      ACCTTCCTTACCCAGATCCATCAAGAATCCTAGGGGCCGGCTGTTTTGTAGATGATATCG    15982
gb:MN975262|Organism:Severe      ACCTTCCTTACCCAGATCCATCAAGAATCCTAGGGGCCGGCTGTTTTGTAGATGATATCG    15982
gb:MT106052|Organism:Severe      ACCTTCCTTACCCAGATCCATCAAGAATCCTAGGGGCCGGCTGTTTTGTAGATGATATCG    15982
gb:LC522975|Organism:Severe      ACCTTCCTTACCCAGATCCATCAAGAATCCTAGGGGCCGGCTGTTTTGTAGATGATATCG    15979
gb:LC522973|Organism:Severe      ACCTTCCTTACCCAGATCCATCAAGAATCCTAGGGGCCGGCTGTTTTGTAGATGATATCG    15979
gb:LC522974|Organism:Severe      ACCTTCCTTACCCAGATCCATCAAGAATCCTAGGGGCCGGCTGTTTTGTAGATGATATCG    15979
gb:MN985325|Organism:Severe      ACCTTCCTTACCCAGATCCATCAAGAATCCTAGGGGCCGGCTGTTTTGTAGATGATATCG    15982
gb:MT020881|Organism:Severe      ACCTTCCTTACCCAGATCCATCAAGAATCCTAGGGGCCGGCTGTTTTGTAGATGATATCG    15982
gb:MT020880|Organism:Severe      ACCTTCCTTACCCAGATCCATCAAGAATCCTAGGGGCCGGCTGTTTTGTAGATGATATCG    15982
gb:MT066175|Organism:Severe      ACCTTCCTTACCCAGATCCATCAAGAATCCTAGGGGCCGGCTGTTTTGTAGATGATATCG    15982
gb:MN997409|Organism:Severe      ACCTTCCTTACCCAGATCCATCAAGAATCCTAGGGGCCGGCTGTTTTGTAGATGATATCG    15982
gb:MN938384|Organism:Severe      ACCTTCCTTACCCAGATCCATCAAGAATCCTAGGGGCCGGCTGTTTTGTAGATGATATCG    15950
gb:MT044258|Organism:Severe      ACCTTCCTTACCCAGATCCATCAAGAATCCTAGGGGCCGGCTGTTTTGTAGATGATATCG    15958
gb:MT039890|Organism:Severe      ACCTTCCTTACCCAGATCCATCAAGAATCCTAGGGGCCGGCTGTTTTGTAGATGATATCG    15982
gb:MN988713|Organism:Severe      ACCTTCCTTACCCAGATCCATCAAGAATCCTAGGGGCCGGCTGTTTTGTAGATGATATCG    15982
gb:LC521925|Organism:Severe      ACCTTCCTTACCCAGATCCATCAAGAATCCTAGGGGCCGGCTGTTTTGTAGATGATATCG    15955
gb:MT093571|Organism:Severe      ACCTTCCTTACCCAGATCCATCAAGAATCCTAGGGGCCGGCTGTTTTGTAGATGATATCG    15982
gb:MT039887|Organism:Severe      ACCTTCCTTACCCAGATCCATCAAGAATCCTAGGGGCCGGCTGTTTTGTAGATGATATCG    15982
gb:MT019530|Organism:Severe      ACCTTCCTTACCCAGATCCATCAAGAATCCTAGGGGCCGGCTGTTTTGTAGATGATATCG    15982
gb:MT039888|Organism:Severe      ACCTTCCTTACCCAGATCCATCAAGAATCCTAGGGGCCGGCTGTTTTGTAGATGATATCG    15982
gb:LC522972|Organism:Severe      ACCTTCCTTACCCAGATCCATCAAGAATCCTAGGGGCCGGCTGTTTTGTAGATGATATCG    15979
gb:MT027063|Organism:Severe      ACCTTCCTTACCCAGATCCATCAAGAATCCTAGGGGCCGGCTGTTTTGTAGATGATATCG    15982
gb:MT027062|Organism:Severe      ACCTTCCTTACCCAGATCCATCAAGAATCCTAGGGGCCGGCTGTTTTGTAGATGATATCG    15982
gb:MT019529|Organism:Severe      ACCTTCCTTACCCAGATCCATCAAGAATCCTAGGGGCCGGCTGTTTTGTAGATGATATCG    15982
gb:MN996529|Organism:Severe      ACCTTCCTTACCCAGATCCATCAAGAATCCTAGGGGCCGGCTGTTTTGTAGATGATATCG    15970
gb:MN996531|Organism:Severe      ACCTTCCTTACCCAGATCCATCAAGAATCCTAGGGGCCGGCTGTTTTGTAGATGATATCG    15969
gb:MT066176|Organism:Severe      ACCTTCCTTACCCAGATCCATCAAGAATCCTAGGGGCCGGCTGTTTTGTAGATGATATCG    15982
gb:MT027064|Organism:Severe      ACCTTCCTTACCCAGATCCATCAAGAATCCTAGGGGCCGGCTGTTTTGTAGATGATATCG    15982
gb:MN994468|Organism:Severe      ACCTTCCTTACCCAGATCCATCAAGAATCCTAGGGGCCGGCTGTTTTGTAGATGATATCG    15982
gb:MT072688|Organism:Severe      ACCTTCCTTACCCAGATCCATCAAGAATCCTAGGGGCCGGCTGTTTTGTAGATGATATCG    15967
gb:MN996527|Organism:Severe      ACCTTCCTTACCCAGATCCATCAAGAATCCTAGGGGCCGGCTGTTTTGTAGATGATATCG    15949
gb:MT093631|Organism:Severe      ACCTTCCTTACCCAGATCCATCAAGAATCCTAGGGGCCGGCTGTTTTGTAGATGATATCG    16020
gb:MT106053|Organism:Severe      ACCTTCCTTACCCAGATCCATCAAGAATCCTAGGGGCCGGCTGTTTTGTAGATGATATCG    15982
gb:MT019533|Organism:Severe      ACCTTCCTTACCCAGATCCATCAAGAATCCTAGGGGCCGGCTGTTTTGTAGATGATATCG    15982
gb:MT019531|Organism:Severe      ACCTTCCTTACCCAGATCCATCAAGAATCCTAGGGGCCGGCTGTTTTGTAGATGATATCG    15982
gb:MN996528|Organism:Severe      ACCTTCCTTACCCAGATCCATCAAGAATCCTAGGGGCCGGCTGTTTTGTAGATGATATCG    15982
gb:MN996530|Organism:Severe      ACCTTCCTTACCCAGATCCATCAAGAATCCTAGGGGCCGGCTGTTTTGTAGATGATATCG    15968
gb:MN908947|Organism:Severe      ACCTTCCTTACCCAGATCCATCAAGAATCCTAGGGGCCGGCTGTTTTGTAGATGATATCG    15982
gb:MT019532|Organism:Severe      ACCTTCCTTACCCAGATCCATCAAGAATCCTAGGGGCCGGCTGTTTTGTAGATGATATCG    15982
                                 ************************************************************
```

```
gb:MT020781|Organism:Severe      TAAAAACAGATGGTACACTTATGATTGAACGGTTCGTGTCTTTAGCTATAGATGCTTACC   16030
gb:MT007544|Organism:Severe      TAAAAACAGATGGTACACTTATGATTGAACGGTTCGTGTCTTTAGCTATAGATGCTTACC   16042
gb:MN994467|Organism:Severe      TAAAAACAGATGGTACACTTATGATTGAACGGTTCGTGTCTTTAGCTATAGATGCTTACC   16042
gb:MT044257|Organism:Severe      TAAAAACAGATGGTACACTTATGATTGAACGGTTCGTGTCTTTAGCTATAGATGCTTACC   16042
gb:MT106054|Organism:Severe      TAAAAACAGATGGTACACTTATGATTGAACGGTTCGTGTCTTTAGCTATAGATGCTTACC   16042
gb:MT049951|Organism:Severe      TAAAAACAGATGGTACACTTATGATTGAACGGTTCGTGTCTTTAGCTATAGATGCTTACC   16042
gb:MN975262|Organism:Severe      TAAAAACAGATGGTACACTTATGATTGAACGGTTCGTGTCTTTAGCTATAGATGCTTACC   16042
gb:MT106052|Organism:Severe      TAAAAACAGATGGTACACTTATGATTGAACGGTTCGTGTCTTTAGCTATAGATGCTTACC   16042
gb:LC522975|Organism:Severe      TAAAAACAGATGGTACACTTATGATTGAACGGTTCGTGTCTTTAGCTATAGATGCTTACC   16039
gb:LC522973|Organism:Severe      TAAAAACAGATGGTACACTTATGATTGAACGGTTCGTGTCTTTAGCTATAGATGCTTACC   16039
gb:LC522974|Organism:Severe      TAAAAACAGATGGTACACTTATGATTGAACGGTTCGTGTCTTTAGCTATAGATGCTTACC   16039
gb:MN985325|Organism:Severe      TAAAAACAGATGGTACACTTATGATTGAACGGTTCGTGTCTTTAGCTATAGATGCTTACC   16042
gb:MT020881|Organism:Severe      TAAAAACAGATGGTACACTTATGATTGAACGGTTCGTGTCTTTAGCTATAGATGCTTACC   16042
gb:MT020880|Organism:Severe      TAAAAACAGATGGTACACTTATGATTGAACGGTTCGTGTCTTTAGCTATAGATGCTTACC   16042
gb:MT066175|Organism:Severe      TAAAAACAGATGGTACACTTATGATTGAACGGTTCGTGTCTTTAGCTATAGATGCTTACC   16042
gb:MN997409|Organism:Severe      TAAAAACAGATGGTACACTTATGATTGAACGGTTCGTGTCTTTAGCTATAGATGCTTACC   16042
gb:MN938384|Organism:Severe      TAAAAACAGATGGTACACTTATGATTGAACGGTTCGTGTCTTTAGCTATAGATGCTTACC   16010
gb:MT044258|Organism:Severe      TAAAAACAGATGGTACACTTATGATTGAACGGTTCGTGTCTTTAGCTATAGATGCTTACC   16018
gb:MT039890|Organism:Severe      TAAAAACAGATGGTACACTTATGATTGAACGGTTCGTGTCTTTAGCTATAGATGCTTACC   16042
gb:MN988713|Organism:Severe      TAAAAACAGATGGTACACTTATGATTGAACGGTTCGTGTCTTTAGCTATAGATGCTTACC   16042
gb:LC521925|Organism:Severe      TAAAAACAGATGGTACACTTATGATTGAACGGTTCGTGTCTTTAGCTATAGATGCTTACC   16015
gb:MT093571|Organism:Severe      TAAAAACAGATGGTACACTTATGATTGAACGGTTCGTGTCTTTAGCTATAGATGCTTACC   16042
gb:MT039887|Organism:Severe      TAAAAACAGATGGTACACTTATGATTGAACGGTTCGTGTCTTTAGCTATAGATGCTTACC   16042
gb:MT019530|Organism:Severe      TAAAAACAGATGGTACACTTATGATTGAACGGTTCGTGTCTTTAGCTATAGATGCTTACC   16042
gb:MT039888|Organism:Severe      TAAAAACAGATGGTACACTTATGATTGAACGGTTCGTGTCTTTAGCTATAGATGCTTACC   16042
gb:LC522972|Organism:Severe      TAAAAACAGATGGTACACTTATGATTGAACGGTTCGTGTCTTTAGCTATAGATGCTTACC   16039
gb:MT027063|Organism:Severe      TAAAAACAGATGGTACACTTATGATTGAACGGTTCGTGTCTTTAGCTATAGATGCTTACC   16042
gb:MT027062|Organism:Severe      TAAAAACAGATGGTACACTTATGATTGAACGGTTCGTGTCTTTAGCTATAGATGCTTACC   16042
gb:MT019529|Organism:Severe      TAAAAACAGATGGTACACTTATGATTGAACGGTTCGTGTCTTTAGCTATAGATGCTTACC   16042
gb:MN996529|Organism:Severe      TAAAAACAGATGGTACACTTATGATTGAACGGTTCGTGTCTTTAGCTATAGATGCTTACC   16030
gb:MN996531|Organism:Severe      TAAAAACAGATGGTACACTTATGATTGAACGGTTCGTGTCTTTAGCTATAGATGCTTACC   16029
gb:MT066176|Organism:Severe      TAAAAACAGATGGTACACTTATGATTGAACGGTTCGTGTCTTTAGCTATAGATGCTTACC   16042
gb:MT027064|Organism:Severe      TAAAAACAGATGGTACACTTATGATTGAACGGTTCGTGTCTTTAGCTATAGATGCTTACC   16042
gb:MN994468|Organism:Severe      TAAAAACAGATGGTACACTTATGATTGAACGGTTCGTGTCTTTAGCTATAGATGCTTACC   16042
gb:MT072688|Organism:Severe      TAAAAACAGATGGTACACTTATGATTGAACGGTTCGTGTCTTTAGCTATAGATGCTTACC   16027
gb:MN996527|Organism:Severe      TAAAAACAGATGGTACACTTATGATTGAACGGTTCGTGTCTTTAGCTATAGATGCTTACC   16009
gb:MT093631|Organism:Severe      TAAAAACAGATGGTACACTTATGATTGAACGGTTCGTGTCTTTAGCTATAGATGCTTACC   16080
gb:MT106053|Organism:Severe      TAAAAACAGATGGTACACTTATGATTGAACGGTTCGTGTCTTTAGCTATAGATGCTTACC   16042
gb:MT019533|Organism:Severe      TAAAAACAGATGGTACACTTATGATTGAACGGTTCGTGTCTTTAGCTATAGATGCTTACC   16042
gb:MT019531|Organism:Severe      TAAAAACAGATGGTACACTTATGATTGAACGGTTCGTGTCTTTAGCTATAGATGCTTACC   16042
gb:MN996528|Organism:Severe      TAAAAACAGATGGTACACTTATGATTGAACGGTTCGTGTCTTTAGCTATAGATGCTTACC   16042
gb:MN996530|Organism:Severe      TAAAAACAGATGGTACACTTATGATTGAACGGTTCGTGTCTTTAGCTATAGATGCTTACC   16028
gb:MN908947|Organism:Severe      TAAAAACAGATGGTACACTTATGATTGAACGGTTCGTGTCTTTAGCTATAGATGCTTACC   16042
gb:MT019532|Organism:Severe      TAAAAACAGATGGTACACTTATGATTGAACGGTTCGTGTCTTTAGCTATAGATGCTTACC   16042
                                 ************************************************************

gb:MT020781|Organism:Severe      CACTTACTAAACATCCTAATCAGGAGTATGCTGATGTCTTTCATTTGTACTTACAATACA   16090
gb:MT007544|Organism:Severe      CACTTACTAAACATCCTAATCAGGAGTATGCTGATGTCTTTCATTTGTACTTACAATACA   16102
gb:MN994467|Organism:Severe      CACTTACTAAACATCCTAATCAGGAGTATGCTGATGTCTTTCATTTGTACTTACAATACA   16102
gb:MT044257|Organism:Severe      CACTTACTAAACATCCTAATCAGGAGTATGCTGATGTCTTTCATTTGTACTTACAATACA   16102
gb:MT106054|Organism:Severe      CACTTACTAAACATCCTAATCAGGAGTATGCTGATGTCTTTCATTTGTACTTACAATACA   16102
gb:MT049951|Organism:Severe      CACTTACTAAACATCCTAATCAGGAGTATGCTGATGTCTTTCATTTGTACTTACAATACA   16102
gb:MN975262|Organism:Severe      CACTTACTAAACATCCTAATCAGGAGTATGCTGATGTCTTTCATTTGTACTTACAATACA   16102
gb:MT106052|Organism:Severe      CACTTACTAAACATCCTAATCAGGAGTATGCTGATGTCTTTCATTTGTACTTACAATACA   16102
gb:LC522975|Organism:Severe      CACTTACTAAACATCCTAATCAGGAGTATGCTGATGTCTTTCATTTGTACTTACAATACA   16099
gb:LC522973|Organism:Severe      CACTTACTAAACATCCTAATCAGGAGTATGCTGATGTCTTTCATTTGTACTTACAATACA   16099
gb:LC522974|Organism:Severe      CACTTACTAAACATCCTAATCAGGAGTATGCTGATGTCTTTCATTTGTACTTACAATACA   16099
gb:MN985325|Organism:Severe      CACTTACTAAACATCCTAATCAGGAGTATGCTGATGTCTTTCATTTGTACTTACAATACA   16102
gb:MT020881|Organism:Severe      CACTTACTAAACATCCTAATCAGGAGTATGCTGATGTCTTTCATTTGTACTTACAATACA   16102
gb:MT020880|Organism:Severe      CACTTACTAAACATCCTAATCAGGAGTATGCTGATGTCTTTCATTTGTACTTACAATACA   16102
gb:MT066175|Organism:Severe      CACTTACTAAACATCCTAATCAGGAGTATGCTGATGTCTTTCATTTGTACTTACAATACA   16102
gb:MN997409|Organism:Severe      CACTTACTAAACATCCTAATCAGGAGTATGCTGATGTCTTTCATTTGTACTTACAATACA   16102
gb:MN938384|Organism:Severe      CACTTACTAAACATCCTAATCAGGAGTATGCTGATGTCTTTCATTTGTACTTACAATACA   16070
gb:MT044258|Organism:Severe      CACTTACTAAACATCCTAATCAGGAGTATGCTGATGTCTTTCATTTGTACTTACAATACA   16078
```

```
gb:MT039890|Organism:Severe      CACTTACTAAACATCCTAATCAGGAGTATGCTGATGTCTTTCATTTGTACTTACAATACA   16102
gb:MN988713|Organism:Severe      CACTTACTAAACATCCTAATCAGGAGTATGCTGATGTCTTTCATTTGTACTTACAATACA   16102
gb:LC521925|Organism:Severe      CACTTACTAAACATCCTAATCAGGAGTATGCTGATGTCTTTCATTTGTACTTACAATACA   16075
gb:MT093571|Organism:Severe      CACTTACTAAACATCCTAATCAGGAGTATGCTGATGTCTTTCATTTGTACTTACAATACA   16102
gb:MT039887|Organism:Severe      CACTTACTAAACATCCTAATCAGGAGTATGCTGATGTCTTTCATTTGTACTTACAATACA   16102
gb:MT019530|Organism:Severe      CACTTACTAAACATCCTAATCAGGAGTATGCTGATGTCTTTCATTTGTACTTACAATACA   16102
gb:MT039888|Organism:Severe      CACTTACTAAACATCCTAATCAGGAGTATGCTGATGTCTTTCATTTGTACTTACAATACA   16102
gb:LC522972|Organism:Severe      CACTTACTAAACATCCTAATCAGGAGTATGCTGATGTCTTTCATTTGTACTTACAATACA   16099
gb:MT027063|Organism:Severe      CACTTACTAAACATCCTAATCAGGAGTATGCTGATGTCTTTCATTTGTACTTACAATACA   16102
gb:MT027062|Organism:Severe      CACTTACTAAACATCCTAATCAGGAGTATGCTGATGTCTTTCATTTGTACTTACAATACA   16102
gb:MT019529|Organism:Severe      CACTTACTAAACATCCTAATCAGGAGTATGCTGATGTCTTTCATTTGTACTTACAATACA   16102
gb:MN996529|Organism:Severe      CACTTACTAAACATCCTAATCAGGAGTATGCTGATGTCTTTCATTTGTACTTACAATACA   16090
gb:MN996531|Organism:Severe      CACTTACTAAACATCCTAATCAGGAGTATGCTGATGTCTTTCATTTGTACTTACAATACA   16089
gb:MT066176|Organism:Severe      CACTTACTAAACATCCTAATCAGGAGTATGCTGATGTCTTTCATTTGTACTTACAATACA   16102
gb:MT027064|Organism:Severe      CACTTACTAAACATCCTAATCAGGAGTATGCTGATGTCTTTCATTTGTACTTACAATACA   16102
gb:MN994468|Organism:Severe      CACTTACTAAACATCCTAATCAGGAGTATGCTGATGTCTTTCATTTGTACTTACAATACA   16102
gb:MT072688|Organism:Severe      CACTTACTAAACATCCTAATCAGGAGTATGCTGATGTCTTTCATTTGTACTTACAATACA   16087
gb:MN996527|Organism:Severe      CACTTACTAAACATCCTAATCAGGAGTATGCTGATGTCTTTCATTTGTACTTACAATACA   16069
gb:MT093631|Organism:Severe      CACTTACTAAACATCCTAATCAGGAGTATGCTGATGTCTTTCATTTGTACTTACAATACA   16140
gb:MT106053|Organism:Severe      CACTTACTAAACATCCTAATCAGGAGTATGCTGATGTCTTTCATTTGTACTTACAATACA   16102
gb:MT019533|Organism:Severe      CACTTACTAAACATCCTAATCAGGAGTATGCTGATGTCTTTCATTTGTACTTACAATACA   16102
gb:MT019531|Organism:Severe      CACTTACTAAACATCCTAATCAGGAGTATGCTGATGTCTTTCATTTGTACTTACAATACA   16102
gb:MN996528|Organism:Severe      CACTTACTAAACATCCTAATCAGGAGTATGCTGATGTCTTTCATTTGTACTTACAATACA   16102
gb:MN996530|Organism:Severe      CACTTACTAAACATCCTAATCAGGAGTATGCTGATGTCTTTCATTTGTACTTACAATACA   16088
gb:MN908947|Organism:Severe      CACTTACTAAACATCCTAATCAGGAGTATGCTGATGTCTTTCATTTGTACTTACAATACA   16102
gb:MT019532|Organism:Severe      CACTTACTAAACATCCTAATCAGGAGTATGCTGATGTCTTTCATTTGTACTTACAATACA   16102
                                 ************************************************************

gb:MT020781|Organism:Severe      TAAGAAAGCTACATGATGAGTTAACAGGACACATGTTAGACATGTATTCTGTTATGCTTA   16150
gb:MT007544|Organism:Severe      TAAGAAAGCTACATGATGAGTTAACAGGACACATGTTAGACATGTATTCTGTTATGCTTA   16162
gb:MN994467|Organism:Severe      TAAGAAAGCTACATGATGAGTTAACAGGACACATGTTAGACATGTATTCTGTTATGCTTA   16162
gb:MT044257|Organism:Severe      TAAGAAAGCTACATGATGAGTTAACAGGACACATGTTAGACATGTATTCTGTTATGCTTA   16162
gb:MT106054|Organism:Severe      TAAGAAAGCTACATGATGAGTTAACAGGACACATGTTAGACATGTATTCTGTTATGCTTA   16162
gb:MT049951|Organism:Severe      TAAGAAAGCTACATGATGAGTTAACAGGACACATGTTAGACATGTATTCTGTTATGCTTA   16162
gb:MN975262|Organism:Severe      TAAGAAAGCTACATGATGAGTTAACAGGACACATGTTAGACATGTATTCTGTTATGCTTA   16162
gb:MT106052|Organism:Severe      TAAGAAAGCTACATGATGAGTTAACAGGACACATGTTAGACATGTATTCTGTTATGCTTA   16162
gb:LC522975|Organism:Severe      TAAGAAAGCTACATGATGAGTTAACAGGACACATGTTAGACATGTATTCTGTTATGCTTA   16159
gb:LC522973|Organism:Severe      TAAGAAAGCTACATGATGAGTTAACAGGACACATGTTAGACATGTATTCTGTTATGCTTA   16159
gb:LC522974|Organism:Severe      TAAGAAAGCTACATGATGAGTTAACAGGACACATGTTAGACATGTATTCTGTTATGCTTA   16159
gb:MN985325|Organism:Severe      TAAGAAAGCTACATGATGAGTTAACAGGACACATGTTAGACATGTATTCTGTTATGCTTA   16162
gb:MT020881|Organism:Severe      TAAGAAAGCTACATGATGAGTTAACAGGACACATGTTAGACATGTATTCTGTTATGCTTA   16162
gb:MT020880|Organism:Severe      TAAGAAAGCTACATGATGAGTTAACAGGACACATGTTAGACATGTATTCTGTTATGCTTA   16162
gb:MT066175|Organism:Severe      TAAGAAAGCTACATGATGAGTTAACAGGACACATGTTAGACATGTATTCTGTTATGCTTA   16162
gb:MN997409|Organism:Severe      TAAGAAAGCTACATGATGAGTTAACAGGACACATGTTAGACATGTATTCTGTTATGCTTA   16162
gb:MN938384|Organism:Severe      TAAGAAAGCTACATGATGAGTTAACAGGACACATGTTAGACATGTATTCTGTTATGCTTA   16130
gb:MT044258|Organism:Severe      TAAGAAAGCTACATGATGAGTTAACAGGACACATGTTAGACATGTATTCTGTTATGCTTA   16138
gb:MT039890|Organism:Severe      TAAGAAAGCTACATGATGAGTTAACAGGACACATGTTAGACATGTATTCTGTTATGCTTA   16162
gb:MN988713|Organism:Severe      TAAGAAAGCTACATGATGAGTTAACAGGACACATGTTAGACATGTATTCTGTTATGCTTA   16162
gb:LC521925|Organism:Severe      TAAGAAAGCTACATGATGAGTTAACAGGACACATGTTAGACATGTATTCTGTTATGCTTA   16135
gb:MT093571|Organism:Severe      TAAGAAAGCTACATGATGAGTTAACAGGACACATGTTAGACATGTATTCTGTTATGCTTA   16162
gb:MT039887|Organism:Severe      TAAGAAAGCTACATGATGAGTTAACAGGACACATGTTAGACATGTATTCTGTTATGCTTA   16162
gb:MT019530|Organism:Severe      TAAGAAAGCTACATGATGAGTTAACAGGACACATGTTAGACATGTATTCTGTTATGCTTA   16162
gb:MT039888|Organism:Severe      TAAGAAAGCTACATGATGAGTTAACAGGACACATGTTAGACATGTATTCTGTTATGCTTA   16162
gb:LC522972|Organism:Severe      TAAGAAAGCTACATGATGAGTTAACAGGACACATGTTAGACATGTATTCTGTTATGCTTA   16159
gb:MT027063|Organism:Severe      TAAGAAAGCTACATGATGAGTTAACAGGACACATGTTAGACATGTATTCTGTTATGCTTA   16162
gb:MT027062|Organism:Severe      TAAGAAAGCTACATGATGAGTTAACAGGACACATGTTAGACATGTATTCTGTTATGCTTA   16162
gb:MT019529|Organism:Severe      TAAGAAAGCTACATGATGAGTTAACAGGACACATGTTAGACATGTATTCTGTTATGCTTA   16162
gb:MN996529|Organism:Severe      TAAGAAAGCTACATGATGAGTTAACAGGACACATGTTAGACATGTATTCTGTTATGCTTA   16150
gb:MN996531|Organism:Severe      TAAGAAAGCTACATGATGAGTTAACAGGACACATGTTAGACATGTATTCTGTTATGCTTA   16149
gb:MT066176|Organism:Severe      TAAGAAAGCTACATGATGAGTTAACAGGACACATGTTAGACATGTATTCTGTTATGCTTA   16162
gb:MT027064|Organism:Severe      TAAGAAAGCTACATGATGAGTTAACAGGACACATGTTAGACATGTATTCTGTTATGCTTA   16162
gb:MN994468|Organism:Severe      TAAGAAAGCTACATGATGAGTTAACAGGACACATGTTAGACATGTATTCTGTTATGCTTA   16162
gb:MT072688|Organism:Severe      TAAGAAAGCTACATGATGAGTTAACAGGACACATGTTAGACATGTATTCTGTTATGCTTA   16147
gb:MN996527|Organism:Severe      TAAGAAAGCTACATGATGAGTTAACAGGACACATGTTAGACATGTATTCTGTTATGCTTA   16129
gb:MT093631|Organism:Severe      TAAGAAAGCTACATGATGAGTTAACAGGACACATGTTAGACATGTATTCTGTTATGCTTA   16200
```

```
gb:MT106053|Organism:Severe      TAAGAAAGCTACATGATGAGTTAACAGGACACATGTTAGACATGTATTCTGTTATGCTTA   16162
gb:MT019533|Organism:Severe      TAAGAAAGCTACATGATGAGTTAACAGGACACATGTTAGACATGTATTCTGTTATGCTTA   16162
gb:MT019531|Organism:Severe      TAAGAAAGCTACATGATGAGTTAACAGGACACATGTTAGACATGTATTCTGTTATGCTTA   16162
gb:MN996528|Organism:Severe      TAAGAAAGCTACATGATGAGTTAACAGGACACATGTTAGACATGTATTCTGTTATGCTTA   16162
gb:MN996530|Organism:Severe      TAAGAAAGCTACATGATGAGTTAACAGGACACATGTTAGACATGTATTCTGTTATGCTTA   16148
gb:MN908947|Organism:Severe      TAAGAAAGCTACATGATGAGTTAACAGGACACATGTTAGACATGTATTCTGTTATGCTTA   16162
gb:MT019532|Organism:Severe      TAAGAAAGCTACATGATGAGTTAACAGGACACATGTTAGACATGTATTCTGTTATGCTTA   16162
                                 ************************************************************

gb:MT020781|Organism:Severe      CTAATGATAACACTTCAAGGTATTGGGAACCTGAGTTTTATGAGGCTATGTACACACCGC   16210
gb:MT007544|Organism:Severe      CTAATGATAACACTTCAAGGTATTGGGAACCTGAGTTTTATGAGGCTATGTACACACCGC   16222
gb:MN994467|Organism:Severe      CTAATGATAACACTTCAAGGTATTGGGAACCTGAGTTTTATGAGGCTATGTACACACCGC   16222
gb:MT044257|Organism:Severe      CTAATGATAACACTTCAAGGTATTGGGAACCTGAGTTTTATGAGGCTATGTACACACCGC   16222
gb:MT106054|Organism:Severe      CTAATGATAACACTTCAAGGTATTGGGAACCTGAGTTTTATGAGGCTATGTACACACCGC   16222
gb:MT049951|Organism:Severe      CTAATGATAACACTTCAAGGTATTGGGAACCTGAGTTTTATGAGGCTATGTACACACCGC   16222
gb:MN975262|Organism:Severe      CTAATGATAACACTTCAAGGTATTGGGAACCTGAGTTTTATGAGGCTATGTACACACCGC   16222
gb:MT106052|Organism:Severe      CTAATGATAACACTTCAAGGTATTGGGAACCTGAGTTTTATGAGGCTATGTACACACCGC   16222
gb:LC522975|Organism:Severe      CTAATGATAACACTTCAAGGTATTGGGAACCTGAGTTTTATGAGGCTATGTACACACCGC   16219
gb:LC522973|Organism:Severe      CTAATGATAACACTTCAAGGTATTGGGAACCTGAGTTTTATGAGGCTATGTACACACCGC   16219
gb:LC522974|Organism:Severe      CTAATGATAACACTTCAAGGTATTGGGAACCTGAGTTTTATGAGGCTATGTACACACCGC   16219
gb:MN985325|Organism:Severe      CTAATGATAACACTTCAAGGTATTGGGAACCTGAGTTTTATGAGGCTATGTACACACCGC   16222
gb:MT020881|Organism:Severe      CTAATGATAACACTTCAAGGTATTGGGAACCTGAGTTTTATGAGGCTATGTACACACCGC   16222
gb:MT020880|Organism:Severe      CTAATGATAACACTTCAAGGTATTGGGAACCTGAGTTTTATGAGGCTATGTACACACCGC   16222
gb:MT066175|Organism:Severe      CTAATGATAACACTTCAAGGTATTGGGAACCTGAGTTTTATGAGGCTATGTACACACCGC   16222
gb:MN997409|Organism:Severe      CTAATGATAACACTTCAAGGTATTGGGAACCTGAGTTTTATGAGGCTATGTACACACCGC   16222
gb:MN938384|Organism:Severe      CTAATGATAACACTTCAAGGTATTGGGAACCTGAGTTTTATGAGGCTATGTACACACCGC   16190
gb:MT044258|Organism:Severe      CTAATGATAACACTTCAAGGTATTGGGAACCTGAGTTTTATGAGGCTATGTACACACCGC   16198
gb:MT039890|Organism:Severe      CTAATGATAACACTTCAAGGTATTGGGAACCTGAGTTTTATGAGGCTATGTACACACCGC   16222
gb:MN988713|Organism:Severe      CTAATGATAACACTTCAAGGTATTGGGAACCTGAGTTTTATGAGGCTATGTACACACCGC   16222
gb:LC521925|Organism:Severe      CTAATGATAACACTTCAAGGTATTGGGAACCTGAGTTTTATGAGGCTATGTACACACCGC   16195
gb:MT093571|Organism:Severe      CTAATGATAACACTTCAAGGTATTGGGAACCTGAGTTTTATGAGGCTATGTACACACCGC   16222
gb:MT039887|Organism:Severe      CTAATGATAACACTTCAAGGTATTGGGAACCTGAGTTTTATGAGGCTATGTACACACCGC   16222
gb:MT019530|Organism:Severe      CTAATGATAACACTTCAAGGTATTGGGAACCTGAGTTTTATGAGGCTATGTACACACCGC   16222
gb:MT039888|Organism:Severe      CTAATGATAACACTTCAAGGTATTGGGAACCTGAGTTTTATGAGGCTATGTACACACCGC   16222
gb:LC522972|Organism:Severe      CTAATGATAACACTTCAAGGTATTGGGAACCTGAGTTTTATGAGGCTATGTACACACCGC   16219
gb:MT027063|Organism:Severe      CTAATGATAACACTTCAAGGTATTGGGAACCTGAGTTTTATGAGGCTATGTACACACCGC   16222
gb:MT027062|Organism:Severe      CTAATGATAACACTTCAAGGTATTGGGAACCTGAGTTTTATGAGGCTATGTACACACCGC   16222
gb:MT019529|Organism:Severe      CTAATGATAACACTTCAAGGTATTGGGAACCTGAGTTTTATGAGGCTATGTACACACCGC   16222
gb:MN996529|Organism:Severe      CTAATGATAACACTTCAAGGTATTGGGAACCTGAGTTTTATGAGGCTATGTACACACCGC   16210
gb:MN996531|Organism:Severe      CTAATGATAACACTTCAAGGTATTGGGAACCTGAGTTTTATGAGGCTATGTACACACCGC   16209
gb:MT066176|Organism:Severe      CTAATGATAACACTTCAAGGTATTGGGAACCTGAGTTTTATGAGGCTATGTACACACCGC   16222
gb:MT027064|Organism:Severe      CTAATGATAACACTTCAAGGTATTGGGAACCTGAGTTTTATGAGGCTATGTACACACCGC   16222
gb:MN994468|Organism:Severe      CTAATGATAACACTTCAAGGTATTGGGAACCTGAGTTTTATGAGGCTATGTACACACCGC   16222
gb:MT072688|Organism:Severe      CTAATGATAACACTTCAAGGTATTGGGAACCTGAGTTTTATGAGGCTATGTACACACCGC   16207
gb:MN996527|Organism:Severe      CTAATGATAACACTTCAAGGTATTGGGAACCTGAGTTTTATGAGGCTATGTACACACCGC   16189
gb:MT093631|Organism:Severe      CTAATGATAACACTTCAAGGTATTGGGAACCTGAGTTTTATGAGGCTATGTACACACCGC   16260
gb:MT106053|Organism:Severe      CTAATGATAACACTTCAAGGTATTGGGAACCTGAGTTTTATGAGGCTATGTACACACCGC   16222
gb:MT019533|Organism:Severe      CTAATGATAACACTTCAAGGTATTGGGAACCTGAGTTTTATGAGGCTATGTACACACCGC   16222
gb:MT019531|Organism:Severe      CTAATGATAACACTTCAAGGTATTGGGAACCTGAGTTTTATGAGGCTATGTACACACCGC   16222
gb:MN996528|Organism:Severe      CTAATGATAACACTTCAAGGTATTGGGAACCTGAGTTTTATGAGGCTATGTACACACCGC   16222
gb:MN996530|Organism:Severe      CTAATGATAACACTTCAAGGTATTGGGAACCTGAGTTTTATGAGGCTATGTACACACCGC   16208
gb:MN908947|Organism:Severe      CTAATGATAACACTTCAAGGTATTGGGAACCTGAGTTTTATGAGGCTATGTACACACCGC   16222
gb:MT019532|Organism:Severe      CTAATGATAACACTTCAAGGTATTGGGAACCTGAGTTTTATGAGGCTATGTACACACCGC   16222
                                 ************************************************************

gb:MT020781|Organism:Severe      ATACAGTCTTACAGGCTGTTGGGGCTTGTGTTCTTTGCAATTCACAGACTTCATTAAGAT   16270
gb:MT007544|Organism:Severe      ATACAGTCTTACAGGCTGTTGGGGCTTGTGTTCTTTGCAATTCACAGACTTCATTAAGAT   16282
gb:MN994467|Organism:Severe      ATACAGTCTTACAGGCTGTTGGGGCTTGTGTTCTTTGCAATTCACAGACTTCATTAAGAT   16282
gb:MT044257|Organism:Severe      ATACAGTCTTACAGGCTGTTGGGGCTTGTGTTCTTTGCAATTCACAGACTTCATTAAGAT   16282
gb:MT106054|Organism:Severe      ATACAGTCTTACAGGCTGTTGGGGCTTGTGTTCTTTGCAATTCACAGACTTCATTAAGAT   16282
gb:MT049951|Organism:Severe      ATACAGTCTTACAGGCTGTTGGGGCTTGTGTTCTTTGCAATTCACAGACTTCATTAAGAT   16282
gb:MN975262|Organism:Severe      ATACAGTCTTACAGGCTGTTGGGGCTTGTGTTCTTTGCAATTCACAGACTTCATTAAGAT   16282
gb:MT106052|Organism:Severe      ATACAGTCTTACAGGCTGTTGGGGCTTGTGTTCTTTGCAATTCACAGACTTCATTAAGAT   16282
gb:LC522975|Organism:Severe      ATACAGTCTTACAGGCTGTTGGGGCTTGTGTTCTTTGCAATTCACAGACTTCATTAAGAT   16279
gb:LC522973|Organism:Severe      ATACAGTCTTACAGGCTGTTGGGGCTTGTGTTCTTTGCAATTCACAGACTTCATTAAGAT   16279
```

```
gb:LC522974|Organism:Severe      ATACAGTCTTACAGGCTGTTGGGGCTTGTGTTCTTTGCAATTCACAGACTTCATTAAGAT   16279
gb:MN985325|Organism:Severe      ATACAGTCTTACAGGCTGTTGGGGCTTGTGTTCTTTGCAATTCACAGACTTCATTAAGAT   16282
gb:MT020881|Organism:Severe      ATACAGTCTTACAGGCTGTTGGGGCTTGTGTTCTTTGCAATTCACAGACTTCATTAAGAT   16282
gb:MT020880|Organism:Severe      ATACAGTCTTACAGGCTGTTGGGGCTTGTGTTCTTTGCAATTCACAGACTTCATTAAGAT   16282
gb:MT066175|Organism:Severe      ATACAGTCTTACAGGCTGTTGGGGCTTGTGTTCTTTGCAATTCACAGACTTCATTAAGAT   16282
gb:MN997409|Organism:Severe      ATACAGTCTTACAGGCTGTTGGGGCTTGTGTTCTTTGCAATTCACAGACTTCATTAAGAT   16282
gb:MN938384|Organism:Severe      ATACAGTCTTACAGGCTGTTGGGGCTTGTGTTCTTTGCAATTCACAGACTTCATTAAGAT   16250
gb:MT044258|Organism:Severe      ATACAGTCTTACAGGCTGTTGGGGCTTGTGTTCTTTGCAATTCACAGACTTCATTAAGAT   16258
gb:MT039890|Organism:Severe      ATACAGTCTTACAGGCTGTTGGGGCTTGTGTTCTTTGCAATTCACAGACTTCATTAAGAT   16282
gb:MN988713|Organism:Severe      ATACAGTCTTACAGGCTGTTGGGGCTTGTGTTCTTTGCAATTCACAGACTTCATTAAGAT   16282
gb:LC521925|Organism:Severe      ATACAGTCTTACAGGCTGTTGGGGCTTGTGTTCTTTGCAATTCACAGACTTCATTAAGAT   16255
gb:MT093571|Organism:Severe      ATACAGTCTTACAGGCTGTTGGGGCTTGTGTTCTTTGCAATTCACAGACTTCATTAAGAT   16282
gb:MT039887|Organism:Severe      ATACAGTCTTACAGGCTGTTGGGGCTTGTGTTCTTTGCAATTCACAGACTTCATTAAGAT   16282
gb:MT019530|Organism:Severe      ATACAGTCTTACAGGCTGTTGGGGCTTGTGTTCTTTGCAATTCACAGACTTCATTAAGAT   16282
gb:MT039888|Organism:Severe      ATACAGTCTTACAGGCTGTTGGGGCTTGTGTTCTTTGCAATTCACAGACTTCATTAAGAT   16282
gb:LC522972|Organism:Severe      ATACAGTCTTACAGGCTGTTGGGGCTTGTGTTCTTTGCAATTCACAGACTTCATTAAGAT   16279
gb:MT027063|Organism:Severe      ATACAGTCTTACAGGCTGTTGGGGCTTGTGTTCTTTGCAATTCACAGACTTCATTAAGAT   16282
gb:MT027062|Organism:Severe      ATACAGTCTTACAGGCTGTTGGGGCTTGTGTTCTTTGCAATTCACAGACTTCATTAAGAT   16282
gb:MT019529|Organism:Severe      ATACAGTCTTACAGGCTGTTGGGGCTTGTGTTCTTTGCAATTCACAGACTTCATTAAGAT   16282
gb:MN996529|Organism:Severe      ATACAGTCTTACAGGCTGTTGGGGCTTGTGTTCTTTGCAATTCACAGACTTCATTAAGAT   16270
gb:MN996531|Organism:Severe      ATACAGTCTTACAGGCTGTTGGGGCTTGTGTTCTTTGCAATTCACAGACTTCATTAAGAT   16269
gb:MT066176|Organism:Severe      ATACAGTCTTACAGGCTGTTGGGGCTTGTGTTCTTTGCAATTCACAGACTTCATTAAGAT   16282
gb:MT027064|Organism:Severe      ATACAGTCTTACAGGCTGTTGGGGCTTGTGTTCTTTGCAATTCACAGACTTCATTAAGAT   16282
gb:MN994468|Organism:Severe      ATACAGTCTTACAGGCTGTTGGGGCTTGTGTTCTTTGCAATTCACAGACTTCATTAAGAT   16282
gb:MT072688|Organism:Severe      ATACAGTCTTACAGGCTGTTGGGGCTTGTGTTCTTTGCAATTCACAGACTTCATTAAGAT   16267
gb:MN996527|Organism:Severe      ATACAGTCTTACAGGCTGTTGGGGCTTGTGTTCTTTGCAATTCACAGACTTCATTAAGAT   16249
gb:MT093631|Organism:Severe      ATACAGTCTTACAGGCTGTTGGGGCTTGTGTTCTTTGCAATTCACAGACTTCATTAAGAT   16320
gb:MT106053|Organism:Severe      ATACAGTCTTACAGGCTGTTGGGGCTTGTGTTCTTTGCAATTCACAGACTTCATTAAGAT   16282
gb:MT019533|Organism:Severe      ATACAGTCTTACAGGCTGTTGGGGCTTGTGTTCTTTGCAATTCACAGACTTCATTAAGAT   16282
gb:MT019531|Organism:Severe      ATACAGTCTTACAGGCTGTTGGGGCTTGTGTTCTTTGCAATTCACAGACTTCATTAAGAT   16282
gb:MN996528|Organism:Severe      ATACAGTCTTACAGGCTGTTGGGGCTTGTGTTCTTTGCAATTCACAGACTTCATTAAGAT   16282
gb:MN996530|Organism:Severe      ATACAGTCTTACAGGCTGTTGGGGCTTGTGTTCTTTGCAATTCACAGACTTCATTAAGAT   16268
gb:MN908947|Organism:Severe      ATACAGTCTTACAGGCTGTTGGGGCTTGTGTTCTTTGCAATTCACAGACTTCATTAAGAT   16282
gb:MT019532|Organism:Severe      ATACAGTCTTACAGGCTGTTGGGGCTTGTGTTCTTTGCAATTCACAGACTTCATTAAGAT   16282
                                 ************************************************************

gb:MT020781|Organism:Severe      GTGGTGCTTGCATACGTAGACCATTCTTATGTTGTAAATGCTGTTACGACCATGTCATAT   16330
gb:MT007544|Organism:Severe      GTGGTGCTTGCATACGTAGACCATTCTTATGTTGTAAATGCTGTTACGACCATGTCATAT   16342
gb:MN994467|Organism:Severe      GTGGTGCTTGCATACGTAGACCATTCTTATGTTGTAAATGCTGTTACGACCATGTCATAT   16342
gb:MT044257|Organism:Severe      GTGGTGCTTGCATACGTAGACCATTCTTATGTTGTAAATGCTGTTACGACCATGTCATAT   16342
gb:MT106054|Organism:Severe      GTGGTGCTTGCATACGTAGACCATTCTTATGTTGTAAATGCTGTTACGACCATGTCATAT   16342
gb:MT049951|Organism:Severe      GTGGTGCTTGCATACGTAGACCATTCTTATGTTGTAAATGCTGTTACGACCATGTCATAT   16342
gb:MN975262|Organism:Severe      GTGGTGCTTGCATACGTAGACCATTCTTATGTTGTAAATGCTGTTACGACCATGTCATAT   16342
gb:MT106052|Organism:Severe      GTGGTGCTTGCATACGTAGACCATTCTTATGTTGTAAATGCTGTTACGACCATGTCATAT   16342
gb:LC522975|Organism:Severe      GTGGTGCTTGCATACGTAGACCATTCTTATGTTGTAAATGCTGTTACGACCATGTCATAT   16339
gb:LC522973|Organism:Severe      GTGGTGCTTGCATACGTAGACCATTCTTATGTTGTAAATGCTGTTACGACCATGTCATAT   16339
gb:LC522974|Organism:Severe      GTGGTGCTTGCATACGTAGACCATTCTTATGTTGTAAATGCTGTTACGACCATGTCATAT   16339
gb:MN985325|Organism:Severe      GTGGTGCTTGCATACGTAGACCATTCTTATGTTGTAAATGCTGTTACGACCATGTCATAT   16342
gb:MT020881|Organism:Severe      GTGGTGCTTGCATACGTAGACCATTCTTATGTTGTAAATGCTGTTACGACCATGTCATAT   16342
gb:MT020880|Organism:Severe      GTGGTGCTTGCATACGTAGACCATTCTTATGTTGTAAATGCTGTTACGACCATGTCATAT   16342
gb:MT066175|Organism:Severe      GTGGTGCTTGCATACGTAGACCATTCTTATGTTGTAAATGCTGTTACGACCATGTCATAT   16342
gb:MN997409|Organism:Severe      GTGGTGCTTGCATACGTAGACCATTCTTATGTTGTAAATGCTGTTACGACCATGTCATAT   16342
gb:MN938384|Organism:Severe      GTGGTGCTTGCATACGTAGACCATTCTTATGTTGTAAATGCTGTTACGACCATGTCATAT   16310
gb:MT044258|Organism:Severe      GTGGTGCTTGCATACGTAGACCATTCTTATGTTGTAAATGCTGTTACGACCATGTCATAT   16318
gb:MT039890|Organism:Severe      GTGGTGCTTGCATACGTAGACCATTCTTATGTTGTAAATGCTGTTACGACCATGTCATAT   16342
gb:MN988713|Organism:Severe      GTGGTGCTTGCATACGTAGACCATTCTTATGTTGTAAATGCTGTTACGACCATGTCATAT   16342
gb:LC521925|Organism:Severe      GTGGTGCTTGCATACGTAGACCATTCTTATGTTGTAAATGCTGTTACGACCATGTCATAT   16315
gb:MT093571|Organism:Severe      GTGGTGCTTGCATACGTAGACCATTCTTATGTTGTAAATGCTGTTACGACCATGTCATAT   16342
gb:MT039887|Organism:Severe      GTGGTGCTTGCATACGTAGACCATTCTTATGTTGTAAATGCTGTTACGACCATGTCATAT   16342
gb:MT019530|Organism:Severe      GTGGTGCTTGCATACGTAGACCATTCTTATGTTGTAAATGCTGTTACGACCATGTCATAT   16342
gb:MT039888|Organism:Severe      GTGGTGCTTGCATACGTAGACCATTCTTATGTTGTAAATGCTGTTACGACCATGTCATAT   16342
gb:LC522972|Organism:Severe      GTGGTGCTTGCATACGTAGACCATTCTTATGTTGTAAATGCTGTTACGACCATGTCATAT   16339
gb:MT027063|Organism:Severe      GTGGTGCTTGCATACGTAGACCATTCTTATGTTGTAAATGCTGTTACGACCATGTCATAT   16342
gb:MT027062|Organism:Severe      GTGGTGCTTGCATACGTAGACCATTCTTATGTTGTAAATGCTGTTACGACCATGTCATAT   16342
gb:MT019529|Organism:Severe      GTGGTGCTTGCATACGTAGACCATTCTTATGTTGTAAATGCTGTTACGACCATGTCATAT   16342
```

```
gb:MN996529|Organism:Severe      GTGGTGCTTGCATACGTAGACCATTCTTATGTTGTAAATGCTGTTACGACCATGTCATAT   16330
gb:MN996531|Organism:Severe      GTGGTGCTTGCATACGTAGACCATTCTTATGTTGTAAATGCTGTTACGACCATGTCATAT   16329
gb:MT066176|Organism:Severe      GTGGTGCTTGCATACGTAGACCATTCTTATGTTGTAAATGCTGTTACGACCATGTCATAT   16342
gb:MT027064|Organism:Severe      GTGGTGCTTGCATACGTAGACCATTCTTATGTTGTAAATGCTGTTACGACCATGTCATAT   16342
gb:MN994468|Organism:Severe      GTGGTGCTTGCATACGTAGACCATTCTTATGTTGTAAATGCTGTTACGACCATGTCATAT   16342
gb:MT072688|Organism:Severe      GTGGTGCTTGCATACGTAGACCATTCTTATGTTGTAAATGCTGTTACGACCATGTCATAT   16327
gb:MN996527|Organism:Severe      GTGGTGCTTGCATACGTAGACCATTCTTATGTTGTAAATGCTGTTACGACCATGTCATAT   16309
gb:MT093631|Organism:Severe      GTGGTGCTTGCATACGTAGACCATTCTTATGTTGTAAATGCTGTTACGACCATGTCATAT   16380
gb:MT106053|Organism:Severe      GTGGTGCTTGCATACGTAGACCATTCTTATGTTGTAAATGCTGTTACGACCATGTCATAT   16342
gb:MT019533|Organism:Severe      GTGGTGCTTGCATACGTAGACCATTCTTATGTTGTAAATGCTGTTACGACCATGTCATAT   16342
gb:MT019531|Organism:Severe      GTGGTGCTTGCATACGTAGACCATTCTTATGTTGTAAATGCTGTTACGACCATGTCATAT   16342
gb:MN996528|Organism:Severe      GTGGTGCTTGCATACGTAGACCATTCTTATGTTGTAAATGCTGTTACGACCATGTCATAT   16342
gb:MN996530|Organism:Severe      GTGGTGCTTGCATACGTAGACCATTCTTATGTTGTAAATGCTGTTACGACCATGTCATAT   16328
gb:MN908947|Organism:Severe      GTGGTGCTTGCATACGTAGACCATTCTTATGTTGTAAATGCTGTTACGACCATGTCATAT   16342
gb:MT019532|Organism:Severe      GTGGTGCTTGCATACGTAGACCATTCTTATGTTGTAAATGCTGTTACGACCATGTCATAT   16342
                                 ************************************************************

gb:MT020781|Organism:Severe      CAACATCACATAAATTAGTCTTGTCTGTTAATCCGTATGTTTGCAATGCTCCAGGTTGTG   16390
gb:MT007544|Organism:Severe      CAACATCACATAAATTAGTCTTGTCTGTTAATCCGTATGTTTGCAATGCTCCAGGTTGTG   16402
gb:MN994467|Organism:Severe      CAACATCACATAAATTAGTCTTGTCTGTTAATCCGTATGTTTGCAATGCTCCAGGTTGTG   16402
gb:MT044257|Organism:Severe      CAACATCACATAAATTAGTCTTGTCTGTTAATCCGTATGTTTGCAATGCTCCAGGTTGTG   16402
gb:MT106054|Organism:Severe      CAACATCACATAAATTAGTCTTGTCTGTTAATCCGTATGTTTGCAATGCTCCAGGTTGTG   16402
gb:MT049951|Organism:Severe      CAACATCACATAAATTAGTCTTGTCTGTTAATCCGTATGTTTGCAATGCTCCAGGTTGTG   16402
gb:MN975262|Organism:Severe      CAACATCACATAAATTAGTCTTGTCTGTTAATCCGTATGTTTGCAATGCTCCAGGTTGTG   16402
gb:MT106052|Organism:Severe      CAACATCACATAAATTAGTCTTGTCTGTTAATCCGTATGTTTGCAATGCTCCAGGTTGTG   16402
gb:LC522975|Organism:Severe      CAACATCACATAAATTAGTCTTGTCTGTTAATCCGTATGTTTGCAATGCTCCAGGTTGTG   16399
gb:LC522973|Organism:Severe      CAACATCACATAAATTAGTCTTGTCTGTTAATCCGTATGTTTGCAATGCTCCAGGTTGTG   16399
gb:LC522974|Organism:Severe      CAACATCACATAAATTAGTCTTGTCTGTTAATCCGTATGTTTGCAATGCTCCAGGTTGTG   16399
gb:MN985325|Organism:Severe      CAACATCACATAAATTAGTCTTGTCTGTTAATCCGTATGTTTGCAATGCTCCAGGTTGTG   16402
gb:MT020881|Organism:Severe      CAACATCACATAAATTAGTCTTGTCTGTTAATCCGTATGTTTGCAATGCTCCAGGTTGTG   16402
gb:MT020880|Organism:Severe      CAACATCACATAAATTAGTCTTGTCTGTTAATCCGTATGTTTGCAATGCTCCAGGTTGTG   16402
gb:MT066175|Organism:Severe      CAACATCACATAAATTAGTCTTGTCTGTTAATCCGTATGTTTGCAATGCTCCAGGTTGTG   16402
gb:MN997409|Organism:Severe      CAACATCACATAAATTAGTCTTGTCTGTTAATCCGTATGTTTGCAATGCTCCAGGTTGTG   16402
gb:MN938384|Organism:Severe      CAACATCACATAAATTAGTCTTGTCTGTTAATCCGTATGTTTGCAATGCTCCAGGTTGTG   16370
gb:MT044258|Organism:Severe      CAACATCACATAAATTAGTCTTGTCTGTTAATCCGTATGTTTGCAATGCTCCAGGTTGTG   16378
gb:MT039890|Organism:Severe      CAACATCACATAAATTAGTCTTGTCTGTTAATCCGTATGTTTGCAATGCTCCAGGTTGTG   16402
gb:MN988713|Organism:Severe      CAACATCACATAAATTAGTCTTGTCTGTTAATCCGTATGTTTGCAATGCTCCAGGTTGTG   16402
gb:LC521925|Organism:Severe      CAACATCACATAAATTAGTCTTGTCTGTTAATCCGTATGTTTGCAATGCTCCAGGTTGTG   16375
gb:MT093571|Organism:Severe      CAACATCACATAAATTAGTCTTGTCTGTTAATCCGTATGTTTGCAATGCTCCAGGTTGTG   16402
gb:MT039887|Organism:Severe      CAACATCACATAAATTAGTCTTGTCTGTTAATCCGTATGTTTGCAATGCTCCAGGTTGTG   16402
gb:MT019530|Organism:Severe      CAACATCACATAAATTAGTCTTGTCTGTTAATCCGTATGTTTGCAATGCTCCAGGTTGTG   16402
gb:MT039888|Organism:Severe      CAACATCACATAAATTAGTCTTGTCTGTTAATCCGTATGTTTGCAATGCTCCAGGTTGTG   16402
gb:LC522972|Organism:Severe      CAACATCACATAAATTAGTCTTGTCTGTTAATCCGTATGTTTGCAATGCTCCAGGTTGTG   16399
gb:MT027063|Organism:Severe      CAACATCACATAAATTAGTCTTGTCTGTTAATCCGTATGTTTGCAATGCTCCAGGTTGTG   16402
gb:MT027062|Organism:Severe      CAACATCACATAAATTAGTCTTGTCTGTTAATCCGTATGTTTGCAATGCTCCAGGTTGTG   16402
gb:MT019529|Organism:Severe      CAACATCACATAAATTAGTCTTGTCTGTTAATCCGTATGTTTGCAATGCTCCAGGTTGTG   16402
gb:MN996529|Organism:Severe      CAACATCACATAAATTAGTCTTGTCTGTTAATCCGTATGTTTGCAATGCTCCAGGTTGTG   16390
gb:MN996531|Organism:Severe      CAACATCACATAAATTAGTCTTGTCTGTTAATCCGTATGTTTGCAATGCTCCAGGTTGTG   16389
gb:MT066176|Organism:Severe      CAACATCACATAAATTAGTCTTGTCTGTTAATCCGTATGTTTGCAATGCTCCAGGTTGTG   16402
gb:MT027064|Organism:Severe      CAACATCACATAAATTAGTCTTGTCTGTTAATCCGTATGTTTGCAATGCTCCAGGTTGTG   16402
gb:MN994468|Organism:Severe      CAACATCACATAAATTAGTCTTGTCTGTTAATCCGTATGTTTGCAATGCTCCAGGTTGTG   16402
gb:MT072688|Organism:Severe      CAACATCACATAAATTAGTCTTGTCTGTTAATCCGTATGTTTGCAATGCTCCAGGTTGTG   16387
gb:MN996527|Organism:Severe      CAACATCACATAAATTAGTCTTGTCTGTTAATCCGTATGTTTGCAATGCTCCAGGTTGTG   16369
gb:MT093631|Organism:Severe      CAACATCACATAAATTAGTCTTGTCTGTTAATCCGTATGTTTGCAATGCTCCAGGTTGTG   16440
gb:MT106053|Organism:Severe      CAACATCACATAAATTAGTCTTGTCTGTTAATCCGTATGTTTGCAATGCTCCAGGTTGTG   16402
gb:MT019533|Organism:Severe      CAACATCACATAAATTAGTCTTGTCTGTTAATCCGTATGTTTGCAATGCTCCAGGTTGTG   16402
gb:MT019531|Organism:Severe      CAACATCACATAAATTAGTCTTGTCTGTTAATCCGTATGTTTGCAATGCTCCAGGTTGTG   16402
gb:MN996528|Organism:Severe      CAACATCACATAAATTAGTCTTGTCTGTTAATCCGTATGTTTGCAATGCTCCAGGTTGTG   16402
gb:MN996530|Organism:Severe      CAACATCACATAAATTAGTCTTGTCTGTTAATCCGTATGTTTGCAATGCTCCAGGTTGTG   16388
gb:MN908947|Organism:Severe      CAACATCACATAAATTAGTCTTGTCTGTTAATCCGTATGTTTGCAATGCTCCAGGTTGTG   16402
gb:MT019532|Organism:Severe      CAACATCACATAAATTAGTCTTGTCTGTTAATCCGTATGTTTGCAATGCTCCAGGTTGTG   16402
                                 ************************************************************

gb:MT020781|Organism:Severe      ATGTCACAGATGTGACTCAACTTTACTTAGGAGGTATGAGCTATTATTGTAAATCACATA   16450
gb:MT007544|Organism:Severe      ATGTCACAGATGTGACTCAACTTTACTTAGGAGGTATGAGCTATTATTGTAAATCACATA   16462
```

```
gb:MN994467|Organism:Severe      ATGTCACAGATGTGACTCAACTTTACTTAGGAGGTATGAGCTATTATTGTAAATCACATA   16462
gb:MT044257|Organism:Severe      ATGTCACAGATGTGACTCAACTTTACTTAGGAGGTATGAGCTATTATTGTAAATCACATA   16462
gb:MT106054|Organism:Severe      ATGTCACAGATGTGACTCAACTTTACTTAGGAGGTATGAGCTATTATTGTAAATCACATA   16462
gb:MT049951|Organism:Severe      ATGTCACAGATGTGACTCAACTTTACTTAGGAGGTATGAGCTATTATTGTAAATCACATA   16462
gb:MN975262|Organism:Severe      ATGTCACAGATGTGACTCAACTTTACTTAGGAGGTATGAGCTATTATTGTAAATCACATA   16462
gb:MT106052|Organism:Severe      ATGTCACAGATGTGACTCAACTTTACTTAGGAGGTATGAGCTATTATTGTAAATCACATA   16462
gb:LC522975|Organism:Severe      ATGTCACAGATGTGACTCAACTTTACTTAGGAGGTATGAGCTATTATTGTAAATCACATA   16459
gb:LC522973|Organism:Severe      ATGTCACAGATGTGACTCAACTTTACTTAGGAGGTATGAGCTATTATTGTAAATCACATA   16459
gb:LC522974|Organism:Severe      ATGTCACAGATGTGACTCAACTTTACTTAGGAGGTATGAGCTATTATTGTAAATCACATA   16459
gb:MN985325|Organism:Severe      ATGTCACAGATGTGACTCAACTTTACTTAGGAGGTATGAGCTATTATTGTAAATCACATA   16462
gb:MT020881|Organism:Severe      ATGTCACAGATGTGACTCAACTTTACTTAGGAGGTATGAGCTATTATTGTAAATCACATA   16462
gb:MT020880|Organism:Severe      ATGTCACAGATGTGACTCAACTTTACTTAGGAGGTATGAGCTATTATTGTAAATCACATA   16462
gb:MT066175|Organism:Severe      ATGTCACAGATGTGACTCAACTTTACTTAGGAGGTATGAGCTATTATTGTAAATCACATA   16462
gb:MN997409|Organism:Severe      ATGTCACAGATGTGACTCAACTTTACTTAGGAGGTATGAGCTATTATTGTAAATCACATA   16462
gb:MN938384|Organism:Severe      ATGTCACAGATGTGACTCAACTTTACTTAGGAGGTATGAGCTATTATTGTAAATCACATA   16430
gb:MT044258|Organism:Severe      ATGTCACAGATGTGACTCAACTTTACTTAGGAGGTATGAGCTATTATTGTAAATCACATA   16438
gb:MT039890|Organism:Severe      ATGTCACAGATGTGACTCAACTTTACTTAGGAGGTATGAGCTATTATTGTAAATCACATA   16462
gb:MN988713|Organism:Severe      ATGTCACAGATGTGACTCAACTTTACTTAGGAGGTATGAGCTATTATTGTAAATCACATA   16462
gb:LC521925|Organism:Severe      ATGTCACAGATGTGACTCAACTTTACTTAGGAGGTATGAGCTATTATTGTAAATCACATA   16435
gb:MT093571|Organism:Severe      ATGTCACAGATGTGACTCAACTTTACTTAGGAGGTATGAGCTATTATTGTAAATCACATA   16462
gb:MT039887|Organism:Severe      ATGTCACAGATGTGACTCAACTTTACTTAGGAGGTATGAGCTATTATTGTAAATCACATA   16462
gb:MT019530|Organism:Severe      ATGTCACAGATGTGACTCAACTTTACTTAGGAGGTATGAGCTATTATTGTAAATCACATA   16462
gb:MT039888|Organism:Severe      ATGTCACAGATGTGACTCAACTTTACTTAGGAGGTATGAGCTATTATTGTAAATCACATA   16462
gb:LC522972|Organism:Severe      ATGTCACAGATGTGACTCAACTTTACTTAGGAGGTATGAGCTATTATTGTAAATCACATA   16459
gb:MT027063|Organism:Severe      ATGTCACAGATGTGACTCAACTTTACTTAGGAGGTATGAGCTATTATTGTAAATCACATA   16462
gb:MT027062|Organism:Severe      ATGTCACAGATGTGACTCAACTTTACTTAGGAGGTATGAGCTATTATTGTAAATCACATA   16462
gb:MT019529|Organism:Severe      ATGTCACAGATGTGACTCAACTTTACTTAGGAGGTATGAGCTATTATTGTAAATCACATA   16462
gb:MN996529|Organism:Severe      ATGTCACAGATGTGACTCAACTTTACTTAGGAGGTATGAGCTATTATTGTAAATCACATA   16450
gb:MN996531|Organism:Severe      ATGTCACAGATGTGACTCAACTTTACTTAGGAGGTATGAGCTATTATTGTAAATCACATA   16449
gb:MT066176|Organism:Severe      ATGTCACAGATGTGACTCAACTTTACTTAGGAGGTATGAGCTATTATTGTAAATCACATA   16462
gb:MT027064|Organism:Severe      ATGTCACAGATGTGACTCAACTTTACTTAGGAGGTATGAGCTATTATTGTAAATCACATA   16462
gb:MN994468|Organism:Severe      ATGTCACAGATGTGACTCAACTTTACTTAGGAGGTATGAGCTATTATTGTAAATCACATA   16462
gb:MT072688|Organism:Severe      ATGTCACAGATGTGACTCAACTTTACTTAGGAGGTATGAGCTATTATTGTAAATCACATA   16447
gb:MN996527|Organism:Severe      ATGTCACAGATGTGACTCAACTTTACTTAGGAGGTATGAGCTATTATTGTAAATCACATA   16429
gb:MT093631|Organism:Severe      ATGTCACAGATGTGACTCAACTTTACTTAGGAGGTATGAGCTATTATTGTAAATCACATA   16500
gb:MT106053|Organism:Severe      ATGTCACAGATGTGACTCAACTTTACTTAGGAGGTATGAGCTATTATTGTAAATCACATA   16462
gb:MT019533|Organism:Severe      ATGTCACAGATGTGACTCAACTTTACTTAGGAGGTATGAGCTATTATTGTAAATCACATA   16462
gb:MT019531|Organism:Severe      ATGTCACAGATGTGACTCAACTTTACTTAGGAGGTATGAGCTATTATTGTAAATCACATA   16462
gb:MN996528|Organism:Severe      ATGTCACAGATGTGACTCAACTTTACTTAGGAGGTATGAGCTATTATTGTAAATCACATA   16462
gb:MN996530|Organism:Severe      ATGTCACAGATGTGACTCAACTTTACTTAGGAGGTATGAGCTATTATTGTAAATCACATA   16448
gb:MN908947|Organism:Severe      ATGTCACAGATGTGACTCAACTTTACTTAGGAGGTATGAGCTATTATTGTAAATCACATA   16462
gb:MT019532|Organism:Severe      ATGTCACAGATGTGACTCAACTTTACTTAGGAGGTATGAGCTATTATTGTAAATCACATA   16462
                                 ************************************************************

gb:MT020781|Organism:Severe      AACCACCCATTAGTTTTCCATTGTGTGCTAATGGACAAGTTTTTGGTTTATATAAAAATA   16510
gb:MT007544|Organism:Severe      AACCACCCATTAGTTTTCCATTGTGTGCTAATGGACAAGTTTTTGGTTTATATAAAAATA   16522
gb:MN994467|Organism:Severe      AACCACCCATTAGTTTTCCATTGTGTGCTAATGGACAAGTTTTTGGTTTATATAAAAATA   16522
gb:MT044257|Organism:Severe      AACCACCCATTAGTTTTCCATTGTGTGCTAATGGACAAGTTTTTGGTTTATATAAAAATA   16522
gb:MT106054|Organism:Severe      AACCACCCATTAGTTTTCCATTGTGTGCTAATGGACAAGTTTTTGGTTTATATAAAAATA   16522
gb:MT049951|Organism:Severe      AACCACCCATTAGTTTTCCATTGTGTGCTAATGGACAAGTTTTTGGTTTATATAAAAATA   16522
gb:MN975262|Organism:Severe      AACCACCCATTAGTTTTCCATTGTGTGCTAATGGACAAGTTTTTGGTTTATATAAAAATA   16522
gb:MT106052|Organism:Severe      AACCACCCATTAGTTTTCCATTGTGTGCTAATGGACAAGTTTTTGGTTTATATAAAAATA   16522
gb:LC522975|Organism:Severe      AACCACCCATTAGTTTTCCATTGTGTGCTAATGGACAAGTTTTTGGTTTATATAAAAATA   16519
gb:LC522973|Organism:Severe      AACCACCCATTAGTTTTCCATTGTGTGCTAATGGACAAGTTTTTGGTTTATATAAAAATA   16519
gb:LC522974|Organism:Severe      AACCACCCATTAGTTTTCCATTGTGTGCTAATGGACAAGTTTTTGGTTTATATAAAAATA   16519
gb:MN985325|Organism:Severe      AACCACCCATTAGTTTTCCATTGTGTGCTAATGGACAAGTTTTTGGTTTATATAAAAATA   16522
gb:MT020881|Organism:Severe      AACCACCCATTAGTTTTCCATTGTGTGCTAATGGACAAGTTTTTGGTTTATATAAAAATA   16522
gb:MT020880|Organism:Severe      AACCACCCATTAGTTTTCCATTGTGTGCTAATGGACAAGTTTTTGGTTTATATAAAAATA   16522
gb:MT066175|Organism:Severe      AACCACCCATTAGTTTTCCATTGTGTGCTAATGGACAAGTTTTTGGTTTATATAAAAATA   16522
gb:MN997409|Organism:Severe      AACCACCCATTAGTTTTCCATTGTGTGCTAATGGACAAGTTTTTGGTTTATATAAAAATA   16522
gb:MN938384|Organism:Severe      AACCACCCATTAGTTTTCCATTGTGTGCTAATGGACAAGTTTTTGGTTTATATAAAAATA   16490
gb:MT044258|Organism:Severe      AACCACCCATTAGTTTTCCATTGTGTGCTAATGGACAAGTTTTTGGTTTATATAAAAATA   16498
gb:MT039890|Organism:Severe      AACCACCCATTAGTTTTCCATTGTGTGCTAATGGACAAGTTTTTGGTTTATATAAAAATA   16522
gb:MN988713|Organism:Severe      AACCACCCATTAGTTTTCCATTGTGTGCTAATGGACAAGTTTTTGGTTTATATAAAAATA   16522
gb:LC521925|Organism:Severe      AACCACCCATTAGTTTTCCATTGTGTGCTAATGGACAAGTTTTTGGTTTATATAAAAATA   16495
```

```
gb:MT093571|Organism:Severe      AACCACCCATTAGTTTTCCATTGTGTGCTAATGGACAAGTTTTTGGTTTATATAAAAATA   16522
gb:MT039887|Organism:Severe      AACCACCCATTAGTTTTCCATTGTGTGCTAATGGACAAGTTTTTGGTTTATATAAAAATA   16522
gb:MT019530|Organism:Severe      AACCACCCATTAGTTTTCCATTGTGTGCTAATGGACAAGTTTTTGGTTTATATAAAAATA   16522
gb:MT039888|Organism:Severe      AACCACCCATTAGTTTTCCATTGTGTGCTAATGGACAAGTTTTTGGTTTATATAAAAATA   16522
gb:LC522972|Organism:Severe      AACCACCCATTAGTTTTCCATTGTGTGCTAATGGACAAGTTTTTGGTTTATATAAAAATA   16519
gb:MT027063|Organism:Severe      AACCACCCATTAGTTTTCCATTGTGTGCTAATGGACAAGTTTTTGGTTTATATAAAAATA   16522
gb:MT027062|Organism:Severe      AACCACCCATTAGTTTTCCATTGTGTGCTAATGGACAAGTTTTTGGTTTATATAAAAATA   16522
gb:MT019529|Organism:Severe      AACCACCCATTAGTTTTCCATTGTGTGCTAATGGACAAGTTTTTGGTTTATATAAAAATA   16522
gb:MN996529|Organism:Severe      AACCACCCATTAGTTTTCCATTGTGTGCTAATGGACAAGTTTTTGGTTTATATAAAAATA   16510
gb:MN996531|Organism:Severe      AACCACCCATTAGTTTTCCATTGTGTGCTAATGGACAAGTTTTTGGTTTATATAAAAATA   16509
gb:MT066176|Organism:Severe      AACCACCCATTAGTTTTCCATTGTGTGCTAATGGACAAGTTTTTGGTTTATATAAAAATA   16522
gb:MT027064|Organism:Severe      AACCACCCATTAGTTTTCCATTGTGTGCTAATGGACAAGTTTTTGGTTTATATAAAAATA   16522
gb:MN994468|Organism:Severe      AACCACCCATTAGTTTTCCATTGTGTGCTAATGGACAAGTTTTTGGTTTATATAAAAATA   16522
gb:MT072688|Organism:Severe      AACCACCCATTAGTTTTCCATTGTGTGCTAATGGACAAGTTTTTGGTTTATATAAAAATA   16507
gb:MN996527|Organism:Severe      AACCACCCATTAGTTTTCCATTGTGTGCTAATGGACAAGTTTTTGGTTTATATAAAAATA   16489
gb:MT093631|Organism:Severe      AACCACCCATTAGTTTTCCATTGTGTGCTAATGGACAAGTTTTTGGTTTATATAAAAATA   16560
gb:MT106053|Organism:Severe      AACCACCCATTAGTTTTCCATTGTGTGCTAATGGACAAGTTTTTGGTTTATATAAAAATA   16522
gb:MT019533|Organism:Severe      AACCACCCATTAGTTTTCCATTGTGTGCTAATGGACAAGTTTTTGGTTTATATAAAAATA   16522
gb:MT019531|Organism:Severe      AACCACCCATTAGTTTTCCATTGTGTGCTAATGGACAAGTTTTTGGTTTATATAAAAATA   16522
gb:MN996528|Organism:Severe      AACCACCCATTAGTTTTCCATTGTGTGCTAATGGACAAGTTTTTGGTTTATATAAAAATA   16522
gb:MN996530|Organism:Severe      AACCACCCATTAGTTTTCCATTGTGTGCTAATGGACAAGTTTTTGGTTTATATAAAAATA   16508
gb:MN908947|Organism:Severe      AACCACCCATTAGTTTTCCATTGTGTGCTAATGGACAAGTTTTTGGTTTATATAAAAATA   16522
gb:MT019532|Organism:Severe      AACCACCCATTAGTTTTCCATTGTGTGCTAATGGACAAGTTTTTGGTTTATATAAAAATA   16522
                                 ************************************************************

gb:MT020781|Organism:Severe      CATGTGTTGGTAGCGATAATGTTACTGACTTTAATGCAATTGCAACATGTGACTGGACAA   16570
gb:MT007544|Organism:Severe      CATGTGTTGGTAGCGATAATGTTACTGACTTTAATGCAATTGCAACATGTGACTGGACAA   16582
gb:MN994467|Organism:Severe      CATGTGTTGGTAGCGATAATGTTACTGACTTTAATGCAATTGCAACATGTGACTGGACAA   16582
gb:MT044257|Organism:Severe      CATGTGTTGGTAGCGATAATGTTACTGACTTTAATGCAATTGCAACATGTGACTGGACAA   16582
gb:MT106054|Organism:Severe      CATGTGTTGGTAGCGATAATGTTACTGACTTTAATGCAATTGCAACATGTGACTGGACAA   16582
gb:MT049951|Organism:Severe      CATGTGTTGGTAGCGATAATGTTACTGACTTTAATGCAATTGCAACATGTGACTGGACAA   16582
gb:MN975262|Organism:Severe      CATGTGTTGGTAGCGATAATGTTACTGACTTTAATGCAATTGCAACATGTGACTGGACAA   16582
gb:MT106052|Organism:Severe      CATGTGTTGGTAGCGATAATGTTACTGACTTTAATGCAATTGCAACATGTGACTGGACAA   16582
gb:LC522975|Organism:Severe      CATGTGTTGGTAGCGATAATGTTACTGACTTTAATGCAATTGCAACATGTGACTGGACAA   16579
gb:LC522973|Organism:Severe      CATGTGTTGGTAGCGATAATGTTACTGACTTTAATGCAATTGCAACATGTGACTGGACAA   16579
gb:LC522974|Organism:Severe      CATGTGTTGGTAGCGATAATGTTACTGACTTTAATGCAATTGCAACATGTGACTGGACAA   16579
gb:MN985325|Organism:Severe      CATGTGTTGGTAGCGATAATGTTACTGACTTTAATGCAATTGCAACATGTGACTGGACAA   16582
gb:MT020881|Organism:Severe      CATGTGTTGGTAGCGATAATGTTACTGACTTTAATGCAATTGCAACATGTGACTGGACAA   16582
gb:MT020880|Organism:Severe      CATGTGTTGGTAGCGATAATGTTACTGACTTTAATGCAATTGCAACATGTGACTGGACAA   16582
gb:MT066175|Organism:Severe      CATGTGTTGGTAGCGATAATGTTACTGACTTTAATGCAATTGCAACATGTGACTGGACAA   16582
gb:MN997409|Organism:Severe      CATGTGTTGGTAGCGATAATGTTACTGACTTTAATGCAATTGCAACATGTGACTGGACAA   16582
gb:MN938384|Organism:Severe      CATGTGTTGGTAGCGATAATGTTACTGACTTTAATGCAATTGCAACATGTGACTGGACAA   16550
gb:MT044258|Organism:Severe      CATGTGTTGGTAGCGATAATGTTACTGACTTTAATGCAATTGCAACATGTGACTGGACAA   16558
gb:MT039890|Organism:Severe      CATGTGTTGGTAGCGATAATGTTACTGACTTTAATGCAATTGCAACATGTGACTGGACAA   16582
gb:MN988713|Organism:Severe      CATGTGTTGGTAGCGATAATGTTACTGACTTTAATGCAATTGCAACATGTGACTGGACAA   16582
gb:LC521925|Organism:Severe      CATGTGTTGGTAGCGATAATGTTACTGACTTTAATGCAATTGCAACATGTGACTGGACAA   16555
gb:MT093571|Organism:Severe      CATGTGTTGGTAGCGATAATGTTACTGACTTTAATGCAATTGCAACATGTGACTGGACAA   16582
gb:MT039887|Organism:Severe      CATGTGTTGGTAGCGATAATGTTACTGACTTTAATGCAATTGCAACATGTGACTGGACAA   16582
gb:MT019530|Organism:Severe      CATGTGTTGGTAGCGATAATGTTACTGACTTTAATGCAATTGCAACATGTGACTGGACAA   16582
gb:MT039888|Organism:Severe      CATGTGTTGGTAGCGATAATGTTACTGACTTTAATGCAATTGCAACATGTGACTGGACAA   16582
gb:LC522972|Organism:Severe      CATGTGTTGGTAGCGATAATGTTACTGACTTTAATGCAATTGCAACATGTGACTGGACAA   16579
gb:MT027063|Organism:Severe      CATGTGTTGGTAGCGATAATGTTACTGACTTTAATGCAATTGCAACATGTGACTGGACAA   16582
gb:MT027062|Organism:Severe      CATGTGTTGGTAGCGATAATGTTACTGACTTTAATGCAATTGCAACATGTGACTGGACAA   16582
gb:MT019529|Organism:Severe      CATGTGTTGGTAGCGATAATGTTACTGACTTTAATGCAATTGCAACATGTGACTGGACAA   16582
gb:MN996529|Organism:Severe      CATGTGTTGGTAGCGATAATGTTACTGACTTTAATGCAATTGCAACATGTGACTGGACAA   16570
gb:MN996531|Organism:Severe      CATGTGTTGGTAGCGATAATGTTACTGACTTTAATGCAATTGCAACATGTGACTGGACAA   16569
gb:MT066176|Organism:Severe      CATGTGTTGGTAGCGATAATGTTACTGACTTTAATGCAATTGCAACATGTGACTGGACAA   16582
gb:MT027064|Organism:Severe      CATGTGTTGGTAGCGATAATGTTACTGACTTTAATGCAATTGCAACATGTGACTGGACAA   16582
gb:MN994468|Organism:Severe      CATGTGTTGGTAGCGATAATGTTACTGACTTTAATGCAATTGCAACATGTGACTGGACAA   16582
gb:MT072688|Organism:Severe      CATGTGTTGGTAGCGATAATGTTACTGACTTTAATGCAATTGCAACATGTGACTGGACAA   16567
gb:MN996527|Organism:Severe      CATGTGTTGGTAGCGATAATGTTACTGACTTTAATGCAATTGCAACATGTGACTGGACAA   16549
gb:MT093631|Organism:Severe      CATGTGTTGGTAGCGATAATGTTACTGACTTTAATGCAATTGCAACATGTGACTGGACAA   16620
gb:MT106053|Organism:Severe      CATGTGTTGGTAGCGATAATGTTACTGACTTTAATGCAATTGCAACATGTGACTGGACAA   16582
gb:MT019533|Organism:Severe      CATGTGTTGGTAGCGATAATGTTACTGACTTTAATGCAATTGCAACATGTGACTGGACAA   16582
gb:MT019531|Organism:Severe      CATGTGTTGGTAGCGATAATGTTACTGACTTTAATGCAATTGCAACATGTGACTGGACAA   16582
```

```
gb:MN996528|Organism:Severe      CATGTGTTGGTAGCGATAATGTTACTGACTTTAATGCAATTGCAACATGTGACTGGACAA    16582
gb:MN996530|Organism:Severe      CATGTGTTGGTAGCGATAATGTTACTGACTTTAATGCAATTGCAACATGTGACTGGACAA    16568
gb:MN908947|Organism:Severe      CATGTGTTGGTAGCGATAATGTTACTGACTTTAATGCAATTGCAACATGTGACTGGACAA    16582
gb:MT019532|Organism:Severe      CATGTGTTGGTAGCGATAATGTTACTGACTTTAATGCAATTGCAACATGTGACTGGACAA    16582
                                 ************************************************************

gb:MT020781|Organism:Severe      ATGCTGGTGATTACATTTTAGCTAACACCTGTACTGAAAGACTCAAGCTTTTTGCAGCAG    16630
gb:MT007544|Organism:Severe      ATGCTGGTGATTACATTTTAGCTAACACCTGTACTGAAAGACTCAAGCTTTTTGCAGCAG    16642
gb:MN994467|Organism:Severe      ATGCTGGTGATTACATTTTAGCTAACACCTGTACTGAAAGACTCAAGCTTTTTGCAGCAG    16642
gb:MT044257|Organism:Severe      ATGCTGGTGATTACATTTTAGCTAACACCTGTACTGAAAGACTCAAGCTTTTTGCAGCAG    16642
gb:MT106054|Organism:Severe      ATGCTGGTGATTACATTTTAGCTAACACCTGTACTGAAAGACTCAAGCTTTTTGCAGCAG    16642
gb:MT049951|Organism:Severe      ATGCTGGTGATTACATTTTAGCTAACACCTGTACTGAAAGACTCAAGCTTTTTGCAGCAG    16642
gb:MN975262|Organism:Severe      ATGCTGGTGATTACATTTTAGCTAACACCTGTACTGAAAGACTCAAGCTTTTTGCAGCAG    16642
gb:MT106052|Organism:Severe      ATGCTGGTGATTACATTTTAGCTAACACCTGTACTGAAAGACTCAAGCTTTTTGCAGCAG    16642
gb:LC522975|Organism:Severe      ATGCTGGTGATTACATTTTAGCTAACACCTGTACTGAAAGACTCAAGCTTTTTGCAGCAG    16639
gb:LC522973|Organism:Severe      ATGCTGGTGATTACATTTTAGCTAACACCTGTACTGAAAGACTCAAGCTTTTTGCAGCAG    16639
gb:LC522974|Organism:Severe      ATGCTGGTGATTACATTTTAGCTAACACCTGTACTGAAAGACTCAAGCTTTTTGCAGCAG    16639
gb:MN985325|Organism:Severe      ATGCTGGTGATTACATTTTAGCTAACACCTGTACTGAAAGACTCAAGCTTTTTGCAGCAG    16642
gb:MT020881|Organism:Severe      ATGCTGGTGATTACATTTTAGCTAACACCTGTACTGAAAGACTCAAGCTTTTTGCAGCAG    16642
gb:MT020880|Organism:Severe      ATGCTGGTGATTACATTTTAGCTAACACCTGTACTGAAAGACTCAAGCTTTTTGCAGCAG    16642
gb:MT066175|Organism:Severe      ATGCTGGTGATTACATTTTAGCTAACACCTGTACTGAAAGACTCAAGCTTTTTGCAGCAG    16642
gb:MN997409|Organism:Severe      ATGCTGGTGATTACATTTTAGCTAACACCTGTACTGAAAGACTCAAGCTTTTTGCAGCAG    16642
gb:MN938384|Organism:Severe      ATGCTGGTGATTACATTTTAGCTAACACCTGTACTGAAAGACTCAAGCTTTTTGCAGCAG    16610
gb:MT044258|Organism:Severe      ATGCTGGTGATTACATTTTAGCTAACACCTGTACTGAAAGACTCAAGCTTTTTGCAGCAG    16618
gb:MT039890|Organism:Severe      ATGCTGGTGATTACATTTTAGCTAACACCTGTACTGAAAGACTCAAGCTTTTTGCAGCAG    16642
gb:MN988713|Organism:Severe      ATGCTGGTGATTACATTTTAGCTAACACCTGTACTGAAAGACTCAAGCTTTTTGCAGCAG    16642
gb:LC521925|Organism:Severe      ATGCTGGTGATTACATTTTAGCTAACACCTGTACTGAAAGACTCAAGCTTTTTGCAGCAG    16615
gb:MT093571|Organism:Severe      ATGCTGGTGATTACATTTTAGCTAACACCTGTACTGAAAGACTCAAGCTTTTTGCAGCAG    16642
gb:MT039887|Organism:Severe      ATGCTGGTGATTACATTTTAGCTAACACCTGTACTGAAAGACTCAAGCTTTTTGCAGCAG    16642
gb:MT019530|Organism:Severe      ATGCTGGTGATTACATTTTAGCTAACACCTGTACTGAAAGACTCAAGCTTTTTGCAGCAG    16642
gb:MT039888|Organism:Severe      ATGCTGGTGATTACATTTTAGCTAACACCTGTACTGAAAGACTCAAGCTTTTTGCAGCAG    16642
gb:LC522972|Organism:Severe      ATGCTGGTGATTACATTTTAGCTAACACCTGTACTGAAAGACTCAAGCTTTTTGCAGCAG    16639
gb:MT027063|Organism:Severe      ATGCTGGTGATTACATTTTAGCTAACACCTGTACTGAAAGACTCAAGCTTTTTGCAGCAG    16642
gb:MT027062|Organism:Severe      ATGCTGGTGATTACATTTTAGCTAACACCTGTACTGAAAGACTCAAGCTTTTTGCAGCAG    16642
gb:MT019529|Organism:Severe      ATGCTGGTGATTACATTTTAGCTAACACCTGTACTGAAAGACTCAAGCTTTTTGCAGCAG    16642
gb:MN996529|Organism:Severe      ATGCTGGTGATTACATTTTAGCTAACACCTGTACTGAAAGACTCAAGCTTTTTGCAGCAG    16630
gb:MN996531|Organism:Severe      ATGCTGGTGATTACATTTTAGCTAACACCTGTACTGAAAGACTCAAGCTTTTTGCAGCAG    16629
gb:MT066176|Organism:Severe      ATGCTGGTGATTACATTTTAGCTAACACCTGTACTGAAAGACTCAAGCTTTTTGCAGCAG    16642
gb:MT027064|Organism:Severe      ATGCTGGTGATTACATTTTAGCTAACACCTGTACTGAAAGACTCAAGCTTTTTGCAGCAG    16642
gb:MN994468|Organism:Severe      ATGCTGGTGATTACATTTTAGCTAACACCTGTACTGAAAGACTCAAGCTTTTTGCAGCAG    16642
gb:MT072688|Organism:Severe      ATGCTGGTGATTACATTTTAGCTAACACCTGTACTGAAAGACTCAAGCTTTTTGCAGCAG    16627
gb:MN996527|Organism:Severe      ATGCTGGTGATTACATTTTAGCTAACACCTGTACTGAAAGACTCAAGCTTTTTGCAGCAG    16609
gb:MT093631|Organism:Severe      ATGCTGGTGATTACATTTTAGCTAACACCTGTACTGAAAGACTCAAGCTTTTTGCAGCAG    16680
gb:MT106053|Organism:Severe      ATGCTGGTGATTACATTTTAGCTAACACCTGTACTGAAAGACTCAAGCTTTTTGCAGCAG    16642
gb:MT019533|Organism:Severe      ATGCTGGTGATTACATTTTAGCTAACACCTGTACTGAAAGACTCAAGCTTTTTGCAGCAG    16642
gb:MT019531|Organism:Severe      ATGCTGGTGATTACATTTTAGCTAACACCTGTACTGAAAGACTCAAGCTTTTTGCAGCAG    16642
gb:MN996528|Organism:Severe      ATGCTGGTGATTACATTTTAGCTAACACCTGTACTGAAAGACTCAAGCTTTTTGCAGCAG    16642
gb:MN996530|Organism:Severe      ATGCTGGTGATTACATTTTAGCTAACACCTGTACTGAAAGACTCAAGCTTTTTGCAGCAG    16628
gb:MN908947|Organism:Severe      ATGCTGGTGATTACATTTTAGCTAACACCTGTACTGAAAGACTCAAGCTTTTTGCAGCAG    16642
gb:MT019532|Organism:Severe      ATGCTGGTGATTACATTTTAGCTAACACCTGTACTGAAAGACTCAAGCTTTTTGCAGCAG    16642
                                 ************************************************************

gb:MT020781|Organism:Severe      AAACGCTCAAAGCTACTGAGGAGACATTTAAACTGTCTTATGGTATTGCTACTGTACGTG    16690
gb:MT007544|Organism:Severe      AAACGCTCAAAGCTACTGAGGAGACATTTAAACTGTCTTATGGTATTGCTACTGTACGTG    16702
gb:MN994467|Organism:Severe      AAACGCTCAAAGCTACTGAGGAGACATTTAAACTGTCTTATGGTATTGCTACTGTACGTG    16702
gb:MT044257|Organism:Severe      AAACGCTCAAAGCTACTGAGGAGACATTTAAACTGTCTTATGGTATTGCTACTGTACGTG    16702
gb:MT106054|Organism:Severe      AAACGCTCAAAGCTACTGAGGAGACATTTAAACTGTCTTATGGTATTGCTACTGTACGTG    16702
gb:MT049951|Organism:Severe      AAACGCTCAAAGCTACTGAGGAGACATTTAAACTGTCTTATGGTATTGCTACTGTACGTG    16702
gb:MN975262|Organism:Severe      AAACGCTCAAAGCTACTGAGGAGACATTTAAACTGTCTTATGGTATTGCTACTGTACGTG    16702
gb:MT106052|Organism:Severe      AAACGCTCAAAGCTACTGAGGAGACATTTAAACTGTCTTATGGTATTGCTACTGTACGTG    16702
gb:LC522975|Organism:Severe      AAACGCTCAAAGCTACTGAGGAGACATTTAAACTGTCTTATGGTATTGCTACTGTACGTG    16699
gb:LC522973|Organism:Severe      AAACGCTCAAAGCTACTGAGGAGACATTTAAACTGTCTTATGGTATTGCTACTGTACGTG    16699
gb:LC522974|Organism:Severe      AAACGCTCAAAGCTACTGAGGAGACATTTAAACTGTCTTATGGTATTGCTACTGTACGTG    16699
gb:MN985325|Organism:Severe      AAACGCTCAAAGCTACTGAGGAGACATTTAAACTGTCTTATGGTATTGCTACTGTACGTG    16702
gb:MT020881|Organism:Severe      AAACGCTCAAAGCTACTGAGGAGACATTTAAACTGTCTTATGGTATTGCTACTGTACGTG    16702
```

```
gb:MT020880|Organism:Severe      AAACGCTCAAAGCTACTGAGGAGACATTTAAACTGTCTTATGGTATTGCTACTGTACGTG   16702
gb:MT066175|Organism:Severe      AAACGCTCAAAGCTACTGAGGAGACATTTAAACTGTCTTATGGTATTGCTACTGTACGTG   16702
gb:MN997409|Organism:Severe      AAACGCTCAAAGCTACTGAGGAGACATTTAAACTGTCTTATGGTATTGCTACTGTACGTG   16702
gb:MN938384|Organism:Severe      AAACGCTCAAAGCTACTGAGGAGACATTTAAACTGTCTTATGGTATTGCTACTGTACGTG   16670
gb:MT044258|Organism:Severe      AAACGCTCAAAGCTACTGAGGAGACATTTAAACTGTCTTATGGTATTGCTACTGTACGTG   16678
gb:MT039890|Organism:Severe      AAACGCTCAAAGCTACTGAGGAGACATTTAAACTGTCTTATGGTATTGCTACTGTACGTG   16702
gb:MN988713|Organism:Severe      AAACGCTCAAAGCTACTGAGGAGACATTTAAACTGTCTTATGGTATTGCTACTGTACGTG   16702
gb:LC521925|Organism:Severe      AAACGCTCAAAGCTACTGAGGAGACATTTAAACTGTCTTATGGTATTGCTACTGTACGTG   16675
gb:MT093571|Organism:Severe      AAACGCTCAAAGCTACTGAGGAGACATTTAAACTGTCTTATGGTATTGCTACTGTACGTG   16702
gb:MT039887|Organism:Severe      AAACGCTCAAAGCTACTGAGGAGACATTTAAACTGTCTTATGGTATTGCTACTGTACGTG   16702
gb:MT019530|Organism:Severe      AAACGCTCAAAGCTACTGAGGAGACATTTAAACTGTCTTATGGTATTGCTACTGTACGTG   16702
gb:MT039888|Organism:Severe      AAACGCTCAAAGCTACTGAGGAGACATTTAAACTGTCTTATGGTATTGCTACTGTACGTG   16702
gb:LC522972|Organism:Severe      AAACGCTCAAAGCTACTGAGGAGACATTTAAACTGTCTTATGGTATTGCTACTGTACGTG   16699
gb:MT027063|Organism:Severe      AAACGCTCAAAGCTACTGAGGAGACATTTAAACTGTCTTATGGTATTGCTACTGTACGTG   16702
gb:MT027062|Organism:Severe      AAACGCTCAAAGCTACTGAGGAGACATTTAAACTGTCTTATGGTATTGCTACTGTACGTG   16702
gb:MT019529|Organism:Severe      AAACGCTCAAAGCTACTGAGGAGACATTTAAACTGTCTTATGGTATTGCTACTGTACGTG   16702
gb:MN996529|Organism:Severe      AAACGCTCAAAGCTACTGAGGAGACATTTAAACTGTCTTATGGTATTGCTACTGTACGTG   16690
gb:MN996531|Organism:Severe      AAACGCTCAAAGCTACTGAGGAGACATTTAAACTGTCTTATGGTATTGCTACTGTACGTG   16689
gb:MT066176|Organism:Severe      AAACGCTCAAAGCTACTGAGGAGACATTTAAACTGTCTTATGGTATTGCTACTGTACGTG   16702
gb:MT027064|Organism:Severe      AAACGCTCAAAGCTACTGAGGAGACATTTAAACTGTCTTATGGTATTGCTACTGTACGTG   16702
gb:MN994468|Organism:Severe      AAACGCTCAAAGCTACTGAGGAGACATTTAAACTGTCTTATGGTATTGCTACTGTACGTG   16702
gb:MT072688|Organism:Severe      AAACGCTCAAAGCTACTGAGGAGACATTTAAACTGTCTTATGGTATTGCTACTGTACGTG   16687
gb:MN996527|Organism:Severe      AAACGCTCAAAGCTACTGAGGAGACATTTAAACTGTCTTATGGTATTGCTACTGTACGTG   16669
gb:MT093631|Organism:Severe      AAACGCTCAAAGCTACTGAGGAGACATTTAAACTGTCTTATGGTATTGCTACTGTACGTG   16740
gb:MT106053|Organism:Severe      AAACGCTCAAAGCTACTGAGGAGACATTTAAACTGTCTTATGGTATTGCTACTGTACGTG   16702
gb:MT019533|Organism:Severe      AAACGCTCAAAGCTACTGAGGAGACATTTAAACTGTCTTATGGTATTGCTACTGTACGTG   16702
gb:MT019531|Organism:Severe      AAACGCTCAAAGCTACTGAGGAGACATTTAAACTGTCTTATGGTATTGCTACTGTACGTG   16702
gb:MN996528|Organism:Severe      AAACGCTCAAAGCTACTGAGGAGACATTTAAACTGTCTTATGGTATTGCTACTGTACGTG   16702
gb:MN996530|Organism:Severe      AAACGCTCAAAGCTACTGAGGAGACATTTAAACTGTCTTATGGTATTGCTACTGTACGTG   16688
gb:MN908947|Organism:Severe      AAACGCTCAAAGCTACTGAGGAGACATTTAAACTGTCTTATGGTATTGCTACTGTACGTG   16702
gb:MT019532|Organism:Severe      AAACGCTCAAAGCTACTGAGGAGACATTTAAACTGTCTTATGGTATTGCTACTGTACGTG   16702
                                 ************************************************************

gb:MT020781|Organism:Severe      AAGTGCTGTCTGACAGAGAATTACATCTTTCATGGGAAGTTGGTAAACCTAGACCACCAC   16750
gb:MT007544|Organism:Severe      AAGTGCTGTCTGACAGAGAATTACATCTTTCATGGGAAGTTGGTAAACCTAGACCACCAC   16762
gb:MN994467|Organism:Severe      AAGTGCTGTCTGACAGAGAATTACATCTTTCATGGGAAGTTGGTAAACCTAGACCACCAC   16762
gb:MT044257|Organism:Severe      AAGTGCTGTCTGACAGAGAATTACATCTTTCATGGGAAGTTGGTAAACCTAGACCACCAC   16762
gb:MT106054|Organism:Severe      AAGTGCTGTCTGACAGAGAATTACATCTTTCATGGGAAGTTGGTAAACCTAGACCACCAC   16762
gb:MT049951|Organism:Severe      AAGTGCTGTCTGACAGAGAATTACATCTTTCATGGGAAGTTGGTAAACCTAGACCACCAC   16762
gb:MN975262|Organism:Severe      AAGTGCTGTCTGACAGAGAATTACATCTTTCATGGGAAGTTGGTAAACCTAGACCACCAC   16762
gb:MT106052|Organism:Severe      AAGTGCTGTCTGACAGAGAATTACATCTTTCATGGGAAGTTGGTAAACCTAGACCACCAC   16762
gb:LC522975|Organism:Severe      AAGTGCTGTCTGACAGAGAATTACATCTTTCATGGGAAGTTGGTAAACCTAGACCACCAC   16759
gb:LC522973|Organism:Severe      AAGTGCTGTCTGACAGAGAATTACATCTTTCATGGGAAGTTGGTAAACCTAGACCACCAC   16759
gb:LC522974|Organism:Severe      AAGTGCTGTCTGACAGAGAATTACATCTTTCATGGGAAGTTGGTAAACCTAGACCACCAC   16759
gb:MN985325|Organism:Severe      AAGTGCTGTCTGACAGAGAATTACATCTTTCATGGGAAGTTGGTAAACCTAGACCACCAC   16762
gb:MT020881|Organism:Severe      AAGTGCTGTCTGACAGAGAATTACATCTTTCATGGGAAGTTGGTAAACCTAGACCACCAC   16762
gb:MT020880|Organism:Severe      AAGTGCTGTCTGACAGAGAATTACATCTTTCATGGGAAGTTGGTAAACCTAGACCACCAC   16762
gb:MT066175|Organism:Severe      AAGTGCTGTCTGACAGAGAATTACATCTTTCATGGGAAGTTGGTAAACCTAGACCACCAC   16762
gb:MN997409|Organism:Severe      AAGTGCTGTCTGACAGAGAATTACATCTTTCATGGGAAGTTGGTAAACCTAGACCACCAC   16762
gb:MN938384|Organism:Severe      AAGTGCTGTCTGACAGAGAATTACATCTTTCATGGGAAGTTGGTAAACCTAGACCACCAC   16730
gb:MT044258|Organism:Severe      AAGTGCTGTCTGACAGAGAATTACATCTTTCATGGGAAGTTGGTAAACCTAGACCACCAC   16738
gb:MT039890|Organism:Severe      AAGTGCTGTCTGACAGAGAATTACATCTTTCATGGGAAGTTGGTAAACCTAGACCACCAC   16762
gb:MN988713|Organism:Severe      AAGTGCTGTCTGACAGAGAATTACATCTTTCATGGGAAGTTGGTAAACCTAGACCACCAC   16762
gb:LC521925|Organism:Severe      AAGTGCTGTCTGACAGAGAATTACATCTTTCATGGGAAGTTGGTAAACCTAGACCACCAC   16735
gb:MT093571|Organism:Severe      AAGTGCTGTCTGACAGAGAATTACATCTTTCATGGGAAGTTGGTAAACCTAGACCACCAC   16762
gb:MT039887|Organism:Severe      AAGTGCTGTCTGACAGAGAATTACATCTTTCATGGGAAGTTGGTAAACCTAGACCACCAC   16762
gb:MT019530|Organism:Severe      AAGTGCTGTCTGACAGAGAATTACATCTTTCATGGGAAGTTGGTAAACCTAGACCACCAC   16762
gb:MT039888|Organism:Severe      AAGTGCTGTCTGACAGAGAATTACATCTTTCATGGGAAGTTGGTAAACCTAGACCACCAC   16762
gb:LC522972|Organism:Severe      AAGTGCTGTCTGACAGAGAATTACATCTTTCATGGGAAGTTGGTAAACCTAGACCACCAC   16759
gb:MT027063|Organism:Severe      AAGTGCTGTCTGACAGAGAATTACATCTTTCATGGGAAGTTGGTAAACCTAGACCACCAC   16762
gb:MT027062|Organism:Severe      AAGTGCTGTCTGACAGAGAATTACATCTTTCATGGGAAGTTGGTAAACCTAGACCACCAC   16762
gb:MT019529|Organism:Severe      AAGTGCTGTCTGACAGAGAATTACATCTTTCATGGGAAGTTGGTAAACCTAGACCACCAC   16762
gb:MN996529|Organism:Severe      AAGTGCTGTCTGACAGAGAATTACATCTTTCATGGGAAGTTGGTAAACCTAGACCACCAC   16750
gb:MN996531|Organism:Severe      AAGTGCTGTCTGACAGAGAATTACATCTTTCATGGGAAGTTGGTAAACCTAGACCACCAC   16749
gb:MT066176|Organism:Severe      AAGTGCTGTCTGACAGAGAATTACATCTTTCATGGGAAGTTGGTAAACCTAGACCACCAC   16762
```

```
gb:MT027064|Organism:Severe      AAGTGCTGTCTGACAGAGAATTACATCTTTCATGGGAAGTTGGTAAACCTAGACCACCAC    16762
gb:MN994468|Organism:Severe      AAGTGCTGTCTGACAGAGAATTACATCTTTCATGGGAAGTTGGTAAACCTAGACCACCAC    16762
gb:MT072688|Organism:Severe      AAGTGCTGTCTGACAGAGAATTACATCTTTCATGGGAAGTTGGTAAACCTAGACCACCAC    16747
gb:MN996527|Organism:Severe      AAGTGCTGTCTGACAGAGAATTACATCTTTCATGGGAAGTTGGTAAACCTAGACCACCAC    16729
gb:MT093631|Organism:Severe      AAGTGCTGTCTGACAGAGAATTACATCTTTCATGGGAAGTTGGTAAACCTAGACCACCAC    16800
gb:MT106053|Organism:Severe      AAGTGCTGTCTGACAGAGAATTACATCTTTCATGGGAAGTTGGTAAACCTAGACCACCAC    16762
gb:MT019533|Organism:Severe      AAGTGCTGTCTGACAGAGAATTACATCTTTCATGGGAAGTTGGTAAACCTAGACCACCAC    16762
gb:MT019531|Organism:Severe      AAGTGCTGTCTGACAGAGAATTACATCTTTCATGGGAAGTTGGTAAACCTAGACCACCAC    16762
gb:MN996528|Organism:Severe      AAGTGCTGTCTGACAGAGAATTACATCTTTCATGGGAAGTTGGTAAACCTAGACCACCAC    16762
gb:MN996530|Organism:Severe      AAGTGCTGTCTGACAGAGAATTACATCTTTCATGGGAAGTTGGTAAACCTAGACCACCAC    16748
gb:MN908947|Organism:Severe      AAGTGCTGTCTGACAGAGAATTACATCTTTCATGGGAAGTTGGTAAACCTAGACCACCAC    16762
gb:MT019532|Organism:Severe      AAGTGCTGTCTGACAGAGAATTACATCTTTCATGGGAAGTTGGTAAACCTAGACCACCAC    16762
                                 ************************************************************

gb:MT020781|Organism:Severe      TTAACCGAAATTATGTCTTTACTGGTTATCGTGTAACTAAAAACAGTAAAGTACAAATAG    16810
gb:MT007544|Organism:Severe      TTAACCGAAATTATGTCTTTACTGGTTATCGTGTAACTAAAAACAGTAAAGTACAAATAG    16822
gb:MN994467|Organism:Severe      TTAACCGAAATTATGTCTTTACTGGTTATCGTGTAACTAAAAACAGTAAAGTACAAATAG    16822
gb:MT044257|Organism:Severe      TTAACCGAAATTATGTCTTTACTGGTTATCGTGTAACTAAAAACAGTAAAGTACAAATAG    16822
gb:MT106054|Organism:Severe      TTAACCGAAATTATGTCTTTACTGGTTATCGTGTAACTAAAAACAGTAAAGTACAAATAG    16822
gb:MT049951|Organism:Severe      TTAACCGAAATTATGTCTTTACTGGTTATCGTGTAACTAAAAACAGTAAAGTACAAATAG    16822
gb:MN975262|Organism:Severe      TTAACCGAAATTATGTCTTTACTGGTTATCGTGTAACTAAAAACAGTAAAGTACAAATAG    16822
gb:MT106052|Organism:Severe      TTAACCGAAATTATGTCTTTACTGGTTATCGTGTAACTAAAAACAGTAAAGTACAAATAG    16822
gb:LC522975|Organism:Severe      TTAACCGAAATTATGTCTTTACTGGTTATCGTGTAACTAAAAACAGTAAAGTACAAATAG    16819
gb:LC522973|Organism:Severe      TTAACCGAAATTATGTCTTTACTGGTTATCGTGTAACTAAAAACAGTAAAGTACAAATAG    16819
gb:LC522974|Organism:Severe      TTAACCGAAATTATGTCTTTACTGGTTATCGTGTAACTAAAAACAGTAAAGTACAAATAG    16819
gb:MN985325|Organism:Severe      TTAACCGAAATTATGTCTTTACTGGTTATCGTGTAACTAAAAACAGTAAAGTACAAATAG    16822
gb:MT020881|Organism:Severe      TTAACCGAAATTATGTCTTTACTGGTTATCGTGTAACTAAAAACAGTAAAGTACAAATAG    16822
gb:MT020880|Organism:Severe      TTAACCGAAATTATGTCTTTACTGGTTATCGTGTAACTAAAAACAGTAAAGTACAAATAG    16822
gb:MT066175|Organism:Severe      TTAACCGAAATTATGTCTTTACTGGTTATCGTGTAACTAAAAACAGTAAAGTACAAATAG    16822
gb:MN997409|Organism:Severe      TTAACCGAAATTATGTCTTTACTGGTTATCGTGTAACTAAAAACAGTAAAGTACAAATAG    16822
gb:MN938384|Organism:Severe      TTAACCGAAATTATGTCTTTACTGGTTATCGTGTAACTAAAAACAGTAAAGTACAAATAG    16790
gb:MT044258|Organism:Severe      TTAACCGAAATTATGTCTTTACTGGTTATCGTGTAACTAAAAACAGTAAAGTACAAATAG    16798
gb:MT039890|Organism:Severe      TTAACCGAAATTATGTCTTTACTGGTTATCGTGTAACTAAAAACAGTAAAGTACAAATAG    16822
gb:MN988713|Organism:Severe      TTAACCGAAATTATGTCTTTACTGGTTATCGTGTAACTAAAAACAGTAAAGTACAAATAG    16822
gb:LC521925|Organism:Severe      TTAACCGAAATTATGTCTTTACTGGTTATCGTGTAACTAAAAACAGTAAAGTACAAATAG    16795
gb:MT093571|Organism:Severe      TTAACCGAAATTATGTCTTTACTGGTTATCGTGTAACTAAAAACAGTAAAGTACAAATAG    16822
gb:MT039887|Organism:Severe      TTAACCGAAATTATGTCTTTACTGGTTATCGTGTAACTAAAAACAGTAAAGTACAAATAG    16822
gb:MT019530|Organism:Severe      TTAACCGAAATTATGTCTTTACTGGTTATCGTGTAACTAAAAACAGTAAAGTACAAATAG    16822
gb:MT039888|Organism:Severe      TTAACCGAAATTATGTCTTTACTGGTTATCGTGTAACTAAAAACAGTAAAGTACAAATAG    16822
gb:LC522972|Organism:Severe      TTAACCGAAATTATGTCTTTACTGGTTATCGTGTAACTAAAAACAGTAAAGTACAAATAG    16819
gb:MT027063|Organism:Severe      TTAACCGAAATTATGTCTTTACTGGTTATCGTGTAACTAAAAACAGTAAAGTACAAATAG    16822
gb:MT027062|Organism:Severe      TTAACCGAAATTATGTCTTTACTGGTTATCGTGTAACTAAAAACAGTAAAGTACAAATAG    16822
gb:MT019529|Organism:Severe      TTAACCGAAATTATGTCTTTACTGGTTATCGTGTAACTAAAAACAGTAAAGTACAAATAG    16822
gb:MN996529|Organism:Severe      TTAACCGAAATTATGTCTTTACTGGTTATCGTGTAACTAAAAACAGTAAAGTACAAATAG    16810
gb:MN996531|Organism:Severe      TTAACCGAAATTATGTCTTTACTGGTTATCGTGTAACTAAAAACAGTAAAGTACAAATAG    16809
gb:MT066176|Organism:Severe      TTAACCGAAATTATGTCTTTACTGGTTATCGTGTAACTAAAAACAGTAAAGTACAAATAG    16822
gb:MT027064|Organism:Severe      TTAACCGAAATTATGTCTTTACTGGTTATCGTGTAACTAAAAACAGTAAAGTACAAATAG    16822
gb:MN994468|Organism:Severe      TTAACCGAAATTATGTCTTTACTGGTTATCGTGTAACTAAAAACAGTAAAGTACAAATAG    16822
gb:MT072688|Organism:Severe      TTAACCGAAATTATGTCTTTACTGGTTATCGTGTAACTAAAAACAGTAAAGTACAAATAG    16807
gb:MN996527|Organism:Severe      TTAACCGAAATTATGTCTTTACTGGTTATCGTGTAACTAAAAACAGTAAAGTACAAATAG    16789
gb:MT093631|Organism:Severe      TTAACCGAAATTATGTCTTTACTGGTTATCGTGTAACTAAAAACAGTAAAGTACAAATAG    16860
gb:MT106053|Organism:Severe      TTAACCGAAATTATGTCTTTACTGGTTATCGTGTAACTAAAAACAGTAAAGTACAAATAG    16822
gb:MT019533|Organism:Severe      TTAACCGAAATTATGTCTTTACTGGTTATCGTGTAACTAAAAACAGTAAAGTACAAATAG    16822
gb:MT019531|Organism:Severe      TTAACCGAAATTATGTCTTTACTGGTTATCGTGTAACTAAAAACAGTAAAGTACAAATAG    16822
gb:MN996528|Organism:Severe      TTAACCGAAATTATGTCTTTACTGGTTATCGTGTAACTAAAAACAGTAAAGTACAAATAG    16822
gb:MN996530|Organism:Severe      TTAACCGAAATTATGTCTTTACTGGTTATCGTGTAACTAAAAACAGTAAAGTACAAATAG    16808
gb:MN908947|Organism:Severe      TTAACCGAAATTATGTCTTTACTGGTTATCGTGTAACTAAAAACAGTAAAGTACAAATAG    16822
gb:MT019532|Organism:Severe      TTAACCGAAATTATGTCTTTACTGGTTATCGTGTAACTAAAAACAGTAAAGTACAAATAG    16822
                                 ************************************************************

gb:MT020781|Organism:Severe      GAGAGTACACCTTTGAAAAAGGTGACTATGGTGATGCTGTTGTTTACCGAGGTACAACAA    16870
gb:MT007544|Organism:Severe      GAGAGTACACCTTTGAAAAAGGTGACTATGGTGATGCTGTTGTTTACCGAGGTACAACAA    16882
gb:MN994467|Organism:Severe      GAGAGTACACCTTTGAAAAAGGTGACTATGGTGATGCTGTTGTTTACCGAGGTACAACAA    16882
gb:MT044257|Organism:Severe      GAGAGTACACCTTTGAAAAAGGTGACTATGGTGATGCTGTTGTTTACCGAGGTACAACAA    16882
gb:MT106054|Organism:Severe      GAGAGTACACCTTTGAAAAAGGTGACTATGGTGATGCTGTTGTTTACCGAGGTACAACAA    16882
```

```
gb:MT049951|Organism:Severe      GAGAGTACACCTTTGAAAAAGGTGACTATGGTGATGCTGTTGTTTACCGAGGTACAACAA   16882
gb:MN975262|Organism:Severe      GAGAGTACACCTTTGAAAAAGGTGACTATGGTGATGCTGTTGTTTACCGAGGTACAACAA   16882
gb:MT106052|Organism:Severe      GAGAGTACACCTTTGAAAAAGGTGACTATGGTGATGCTGTTGTTTACCGAGGTACAACAA   16882
gb:LC522975|Organism:Severe      GAGAGTACACCTTTGAAAAAGGTGACTATGGTGATGCTGTTGTTTACCGAGGTACAACAA   16879
gb:LC522973|Organism:Severe      GAGAGTACACCTTTGAAAAAGGTGACTATGGTGATGCTGTTGTTTACCGAGGTACAACAA   16879
gb:LC522974|Organism:Severe      GAGAGTACACCTTTGAAAAAGGTGACTATGGTGATGCTGTTGTTTACCGAGGTACAACAA   16879
gb:MN985325|Organism:Severe      GAGAGTACACCTTTGAAAAAGGTGACTATGGTGATGCTGTTGTTTACCGAGGTACAACAA   16882
gb:MT020881|Organism:Severe      GAGAGTACACCTTTGAAAAAGGTGACTATGGTGATGCTGTTGTTTACCGAGGTACAACAA   16882
gb:MT020880|Organism:Severe      GAGAGTACACCTTTGAAAAAGGTGACTATGGTGATGCTGTTGTTTACCGAGGTACAACAA   16882
gb:MT066175|Organism:Severe      GAGAGTACACCTTTGAAAAAGGTGACTATGGTGATGCTGTTGTTTACCGAGGTACAACAA   16882
gb:MN997409|Organism:Severe      GAGAGTACACCTTTGAAAAAGGTGACTATGGTGATGCTGTTGTTTACCGAGGTACAACAA   16882
gb:MN938384|Organism:Severe      GAGAGTACACCTTTGAAAAAGGTGACTATGGTGATGCTGTTGTTTACCGAGGTACAACAA   16850
gb:MT044258|Organism:Severe      GAGAGTACACCTTTGAAAAAGGTGACTATGGTGATGCTGTTGTTTACCGAGGTACAACAA   16858
gb:MT039890|Organism:Severe      GAGAGTACACCTTTGAAAAAGGTGACTATGGTGATGCTGTTGTTTACCGAGGTACAACAA   16882
gb:MN988713|Organism:Severe      GAGAGTACACCTTTGAAAAAGGTGACTATGGTGATGCTGTTGTTTACCGAGGTACAACAA   16882
gb:LC521925|Organism:Severe      GAGAGTACACCTTTGAAAAAGGTGACTATGGTGATGCTGTTGTTTACCGAGGTACAACAA   16855
gb:MT093571|Organism:Severe      GAGAGTACACCTTTGAAAAAGGTGACTATGGTGATGCTGTTGTTTACCGAGGTACAACAA   16882
gb:MT039887|Organism:Severe      GAGAGTACACCTTTGAAAAAGGTGACTATGGTGATGCTGTTGTTTACCGAGGTACAACAA   16882
gb:MT019530|Organism:Severe      GAGAGTACACCTTTGAAAAAGGTGACTATGGTGATGCTGTTGTTTACCGAGGTACAACAA   16882
gb:MT039888|Organism:Severe      GAGAGTACACCTTTGAAAAAGGTGACTATGGTGATGCTGTTGTTTACCGAGGTACAACAA   16882
gb:LC522972|Organism:Severe      GAGAGTACACCTTTGAAAAAGGTGACTATGGTGATGCTGTTGTTTACCGAGGTACAACAA   16879
gb:MT027063|Organism:Severe      GAGAGTACACCTTTGAAAAAGGTGACTATGGTGATGCTGTTGTTTACCGAGGTACAACAA   16882
gb:MT027062|Organism:Severe      GAGAGTACACCTTTGAAAAAGGTGACTATGGTGATGCTGTTGTTTACCGAGGTACAACAA   16882
gb:MT019529|Organism:Severe      GAGAGTACACCTTTGAAAAAGGTGACTATGGTGATGCTGTTGTTTACCGAGGTACAACAA   16882
gb:MN996529|Organism:Severe      GAGAGTACACCTTTGAAAAAGGTGACTATGGTGATGCTGTTGTTTACCGAGGTACAACAA   16870
gb:MN996531|Organism:Severe      GAGAGTACACCTTTGAAAAAGGTGACTATGGTGATGCTGTTGTTTACCGAGGTACAACAA   16869
gb:MT066176|Organism:Severe      GAGAGTACACCTTTGAAAAAGGTGACTATGGTGATGCTGTTGTTTACCGAGGTACAACAA   16882
gb:MT027064|Organism:Severe      GAGAGTACACCTTTGAAAAAGGTGACTATGGTGATGCTGTTGTTTACCGAGGTACAACAA   16882
gb:MN994468|Organism:Severe      GAGAGTACACCTTTGAAAAAGGTGACTATGGTGATGCTGTTGTTTACCGAGGTACAACAA   16882
gb:MT072688|Organism:Severe      GAGAGTACACCTTTGAAAAAGGTGACTATGGTGATGCTGTTGTTTACCGAGGTACAACAA   16867
gb:MN996527|Organism:Severe      GAGAGTACACCTTTGAAAAAGGTGACTATGGTGATGCTGTTGTTTACCGAGGTACAACAA   16849
gb:MT093631|Organism:Severe      GAGAGTACACCTTTGAAAAAGGTGACTATGGTGATGCTGTTGTTTACCGAGGTACAACAA   16920
gb:MT106053|Organism:Severe      GAGAGTACACCTTTGAAAAAGGTGACTATGGTGATGCTGTTGTTTACCGAGGTACAACAA   16882
gb:MT019533|Organism:Severe      GAGAGTACACCTTTGAAAAAGGTGACTATGGTGATGCTGTTGTTTACCGAGGTACAACAA   16882
gb:MT019531|Organism:Severe      GAGAGTACACCTTTGAAAAAGGTGACTATGGTGATGCTGTTGTTTACCGAGGTACAACAA   16882
gb:MN996528|Organism:Severe      GAGAGTACACCTTTGAAAAAGGTGACTATGGTGATGCTGTTGTTTACCGAGGTACAACAA   16882
gb:MN996530|Organism:Severe      GAGAGTACACCTTTGAAAAAGGTGACTATGGTGATGCTGTTGTTTACCGAGGTACAACAA   16868
gb:MN908947|Organism:Severe      GAGAGTACACCTTTGAAAAAGGTGACTATGGTGATGCTGTTGTTTACCGAGGTACAACAA   16882
gb:MT019532|Organism:Severe      GAGAGTACACCTTTGAAAAAGGTGACTATGGTGATGCTGTTGTTTACCGAGGTACAACAA   16882
                                 ************************************************************

gb:MT020781|Organism:Severe      CTTACAAATTAAATGTTGGTGATTATTTTGTGCTGACATCACATACAGTAATGCCATTAA   16930
gb:MT007544|Organism:Severe      CTTACAAATTAAATGTTGGTGATTATTTTGTGCTGACATCACATACAGTAATGCCATTAA   16942
gb:MN994467|Organism:Severe      CTTACAAATTAAATGTTGGTGATTATTTTGTGCTGACATCACATACAGTAATGCCATTAA   16942
gb:MT044257|Organism:Severe      CTTACAAATTAAATGTTGGTGATTATTTTGTGCTGACATCACATACAGTAATGCCATTAA   16942
gb:MT106054|Organism:Severe      CTTACAAATTAAATGTTGGTGATTATTTTGTGCTGACATCACATACAGTAATGCCATTAA   16942
gb:MT049951|Organism:Severe      CTTACAAATTAAATGTTGGTGATTATTTTGTGCTGACATCACATACAGTAATGCCATTAA   16942
gb:MN975262|Organism:Severe      CTTACAAATTAAATGTTGGTGATTATTTTGTGCTGACATCACATACAGTAATGCCATTAA   16942
gb:MT106052|Organism:Severe      CTTACAAATTAAATGTTGGTGATTATTTTGTGCTGACATCACATACAGTAATGCCATTAA   16942
gb:LC522975|Organism:Severe      CTTACAAATTAAATGTTGGTGATTATTTTGTGCTGACATCACATACAGTAATGCCATTAA   16939
gb:LC522973|Organism:Severe      CTTACAAATTAAATGTTGGTGATTATTTTGTGCTGACATCACATACAGTAATGCCATTAA   16939
gb:LC522974|Organism:Severe      CTTACAAATTAAATGTTGGTGATTATTTTGTGCTGACATCACATACAGTAATGCCATTAA   16939
gb:MN985325|Organism:Severe      CTTACAAATTAAATGTTGGTGATTATTTTGTGCTGACATCACATACAGTAATGCCATTAA   16942
gb:MT020881|Organism:Severe      CTTACAAATTAAATGTTGGTGATTATTTTGTGCTGACATCACATACAGTAATGCCATTAA   16942
gb:MT020880|Organism:Severe      CTTACAAATTAAATGTTGGTGATTATTTTGTGCTGACATCACATACAGTAATGCCATTAA   16942
gb:MT066175|Organism:Severe      CTTACAAATTAAATGTTGGTGATTATTTTGTGCTGACATCACATACAGTAATGCCATTAA   16942
gb:MN997409|Organism:Severe      CTTACAAATTAAATGTTGGTGATTATTTTGTGCTGACATCACATACAGTAATGCCATTAA   16942
gb:MN938384|Organism:Severe      CTTACAAATTAAATGTTGGTGATTATTTTGTGCTGACATCACATACAGTAATGCCATTAA   16910
gb:MT044258|Organism:Severe      CTTACAAATTAAATGTTGGTGATTATTTTGTGCTGACATCACATACAGTAATGCCATTAA   16918
gb:MT039890|Organism:Severe      CTTACAAATTAAATGTTGGTGATTATTTTGTGCTGACATCACATACAGTAATGCCATTAA   16942
gb:MN988713|Organism:Severe      CTTACAAATTAAATGTTGGTGATTATTTTGTGCTGACATCACATACAGTAATGCCATTAA   16942
gb:LC521925|Organism:Severe      CTTACAAATTAAATGTTGGTGATTATTTTGTGCTGACATCACATACAGTAATGCCATTAA   16915
gb:MT093571|Organism:Severe      CTTACAAATTAAATGTTGGTGATTATTTTGTGCTGACATCACATACAGTAATGCCATTAA   16942
gb:MT039887|Organism:Severe      CTTACAAATTAAATGTTGGTGATTATTTTGTGCTGACATCACATACAGTAATGCCATTAA   16942
gb:MT019530|Organism:Severe      CTTACAAATTAAATGTTGGTGATTATTTTGTGCTGACATCACATACAGTAATGCCATTAA   16942
```

```
gb:MT039888|Organism:Severe      CTTACAAATTAAATGTTGGTGATTATTTTGTGCTGACATCACATACAGTAATGCCATTAA    16942
gb:LC522972|Organism:Severe      CTTACAAATTAAATGTTGGTGATTATTTTGTGCTGACATCACATACAGTAATGCCATTAA    16939
gb:MT027063|Organism:Severe      CTTACAAATTAAATGTTGGTGATTATTTTGTGCTGACATCACATACAGTAATGCCATTAA    16942
gb:MT027062|Organism:Severe      CTTACAAATTAAATGTTGGTGATTATTTTGTGCTGACATCACATACAGTAATGCCATTAA    16942
gb:MT019529|Organism:Severe      CTTACAAATTAAATGTTGGTGATTATTTTGTGCTGACATCACATACAGTAATGCCATTAA    16942
gb:MN996529|Organism:Severe      CTTACAAATTAAATGTTGGTGATTATTTTGTGCTGACATCACATACAGTAATGCCATTAA    16930
gb:MN996531|Organism:Severe      CTTACAAATTAAATGTTGGTGATTATTTTGTGCTGACATCACATACAGTAATGCCATTAA    16929
gb:MT066176|Organism:Severe      CTTACAAATTAAATGTTGGTGATTATTTTGTGCTGACATCACATACAGTAATGCCATTAA    16942
gb:MT027064|Organism:Severe      CTTACAAATTAAATGTTGGTGATTATTTTGTGCTGACATCACATACAGTAATGCCATTAA    16942
gb:MN994468|Organism:Severe      CTTACAAATTAAATGTTGGTGATTATTTTGTGCTGACATCACATACAGTAATGCCATTAA    16942
gb:MT072688|Organism:Severe      CTTACAAATTAAATGTTGGTGATTATTTTGTGCTGACATCACATACAGTAATGCCATTAA    16927
gb:MN996527|Organism:Severe      CTTACAAATTAAATGTTGGTGATTATTTTGTGCTGACATCACATACAGTAATGCCATTAA    16909
gb:MT093631|Organism:Severe      CTTACAAATTAAATGTTGGTGATTATTTTGTGCTGACATCACATACAGTAATGCCATTAA    16980
gb:MT106053|Organism:Severe      CTTACAAATTAAATGTTGGTGATTATTTTGTGCTGACATCACATACAGTAATGCCATTAA    16942
gb:MT019533|Organism:Severe      CTTACAAATTAAATGTTGGTGATTATTTTGTGCTGACATCACATACAGTAATGCCATTAA    16942
gb:MT019531|Organism:Severe      CTTACAAATTAAATGTTGGTGATTATTTTGTGCTGACATCACATACAGTAATGCCATTAA    16942
gb:MN996528|Organism:Severe      CTTACAAATTAAATGTTGGTGATTATTTTGTGCTGACATCACATACAGTAATGCCATTAA    16942
gb:MN996530|Organism:Severe      CTTACAAATTAAATGTTGGTGATTATTTTGTGCTGACATCACATACAGTAATGCCATTAA    16928
gb:MN908947|Organism:Severe      CTTACAAATTAAATGTTGGTGATTATTTTGTGCTGACATCACATACAGTAATGCCATTAA    16942
gb:MT019532|Organism:Severe      CTTACAAATTAAATGTTGGTGATTATTTTGTGCTGACATCACATACAGTAATGCCATTAA    16942
                                 ************************************************************

gb:MT020781|Organism:Severe      GTGCACCTACACTAGTGCCACAAGAGCACTATGTTAGAATTACTGGCTTATACCCAACAC    16990
gb:MT007544|Organism:Severe      GTGCACCTACACTAGTGCCACAAGAGCACTATGTTAGAATTACTGGCTTATACCCAACAC    17002
gb:MN994467|Organism:Severe      GTGCACCTACACTAGTGCCACAAGAGCACTATGTTAGAATTACTGGCTTATACCCAACAC    17002
gb:MT044257|Organism:Severe      GTGCACCTACACTAGTGCCACAAGAGCACTATGTTAGAATTACTGGCTTATACCCAACAC    17002
gb:MT106054|Organism:Severe      GTGCACCTACACTAGTGCCACAAGAGCACTATGTTAGAATTACTGGCTTATACCCAACAC    17002
gb:MT049951|Organism:Severe      GTGCACCTACACTAGTGCCACAAGAGCACTATGTTAGAATTACTGGCTTATACCCAACAC    17002
gb:MN975262|Organism:Severe      GTGCACCTACACTAGTGCCACAAGAGCACTATGTTAGAATTACTGGCTTATACCCAACAC    17002
gb:MT106052|Organism:Severe      GTGCACCTACACTAGTGCCACAAGAGCACTATGTTAGAATTACTGGCTTATACCCAACAC    17002
gb:LC522975|Organism:Severe      GTGCACCTACACTAGTGCCACAAGAGCACTATGTTAGAATTACTGGCTTATACCCAACAC    16999
gb:LC522973|Organism:Severe      GTGCACCTACACTAGTGCCACAAGAGCACTATGTTAGAATTACTGGCTTATACCCAACAC    16999
gb:LC522974|Organism:Severe      GTGCACCTACACTAGTGCCACAAGAGCACTATGTTAGAATTACTGGCTTATACCCAACAC    16999
gb:MN985325|Organism:Severe      GTGCACCTACACTAGTGCCACAAGAGCACTATGTTAGAATTACTGGCTTATACCCAACAC    17002
gb:MT020881|Organism:Severe      GTGCACCTACACTAGTGCCACAAGAGCACTATGTTAGAATTACTGGCTTATACCCAACAC    17002
gb:MT020880|Organism:Severe      GTGCACCTACACTAGTGCCACAAGAGCACTATGTTAGAATTACTGGCTTATACCCAACAC    17002
gb:MT066175|Organism:Severe      GTGCACCTACACTAGTGCCACAAGAGCACTATGTTAGAATTACTGGCTTATACCCAACAC    17002
gb:MN997409|Organism:Severe      GTGCACCTACACTAGTGCCACAAGAGCACTATGTTAGAATTACTGGCTTATACCCAACAC    17002
gb:MN938384|Organism:Severe      GTGCACCTACACTAGTGCCACAAGAGCACTATGTTAGAATTACTGGCTTATACCCAACAC    16970
gb:MT044258|Organism:Severe      GTGCACCTACACTAGTGCCACAAGAGCACTATGTTAGAATTACTGGCTTATACCCAACAC    16978
gb:MT039890|Organism:Severe      GTGCACCTACACTAGTGCCACAAGAGCACTATGTTAGAATTACTGGCTTATACCCAACAC    17002
gb:MN988713|Organism:Severe      GTGCACCTACACTAGTGCCACAAGAGCACTATGTTAGAATTACTGGCTTATACCCAACAC    17002
gb:LC521925|Organism:Severe      GTGCACCTACACTAGTGCCACAAGAGCACTATGTTAGAATTACTGGCTTATACCCAACAC    16975
gb:MT093571|Organism:Severe      GTGCACCTACACTAGTGCCACAAGAGCACTATGTTAGAATTACTGGCTTATACCCAACAC    17002
gb:MT039887|Organism:Severe      GTGCACCTACACTAGTGCCACAAGAGCACTATGTTAGAATTACTGGCTTATACCCAACAC    17002
gb:MT019530|Organism:Severe      GTGCACCTACACTAGTGCCACAAGAGCACTATGTTAGAATTACTGGCTTATACCCAACAC    17002
gb:MT039888|Organism:Severe      GTGCACCTACACTAGTGCCACAAGAGCACTATGTTAGAATTACTGGCTTATACCCAACAC    17002
gb:LC522972|Organism:Severe      GTGCACCTACACTAGTGCCACAAGAGCACTATGTTAGAATTACTGGCTTATACCCAACAC    16999
gb:MT027063|Organism:Severe      GTGCACCTACACTAGTGCCACAAGAGCACTATGTTAGAATTACTGGCTTATACCCAACAC    17002
gb:MT027062|Organism:Severe      GTGCACCTACACTAGTGCCACAAGAGCACTATGTTAGAATTACTGGCTTATACCCAACAC    17002
gb:MT019529|Organism:Severe      GTGCACCTACACTAGTGCCACAAGAGCACTATGTTAGAATTACTGGCTTATACCCAACAC    17002
gb:MN996529|Organism:Severe      GTGCACCTACACTAGTGCCACAAGAGCACTATGTTAGAATTACTGGCTTATACCCAACAC    16990
gb:MN996531|Organism:Severe      GTGCACCTACACTAGTGCCACAAGAGCACTATGTTAGAATTACTGGCTTATACCCAACAC    16989
gb:MT066176|Organism:Severe      GTGCACCTACACTAGTGCCACAAGAGCACTATGTTAGAATTACTGGCTTATACCCAACAC    17002
gb:MT027064|Organism:Severe      GTGCACCTACACTAGTGCCACAAGAGCACTATGTTAGAATTACTGGCTTATACCCAACAC    17002
gb:MN994468|Organism:Severe      GTGCACCTACACTAGTGCCACAAGAGCACTATGTTAGAATTACTGGCTTATACCCAATAC    17002
gb:MT072688|Organism:Severe      GTGCACCTACACTAGTGCCACAAGAGCACTATGTTAGAATTACTGGCTTATACCCAACAC    16987
gb:MN996527|Organism:Severe      GTGCACCTACACTAGTGCCACAAGAGCACTATGTTAGAATTACTGGCTTATACCCAACAC    16969
gb:MT093631|Organism:Severe      GTGCACCTACACTAGTGCCACAAGAGCACTATGTTAGAATTACTGGCTTATACCCAACAC    17040
gb:MT106053|Organism:Severe      GTGCACCTACACTAGTGCCACAAGAGCACTATGTTAGAATTACTGGCTTATACCCAACAC    17002
gb:MT019533|Organism:Severe      GTGCACCTACACTAGTGCCACAAGAGCACTATGTTAGAATTACTGGCTTATACCCAACAC    17002
gb:MT019531|Organism:Severe      GTGCACCTACACTAGTGCCACAAGAGCACTATGTTAGAATTACTGGCTTATACCCAACAC    17002
gb:MN996528|Organism:Severe      GTGCACCTACACTAGTGCCACAAGAGCACTATGTTAGAATTACTGGCTTATACCCAACAC    17002
gb:MN996530|Organism:Severe      GTGCACCTACACTAGTGCCACAAGAGCACTATGTTAGAATTACTGGCTTATACCCAACAC    16988
gb:MN908947|Organism:Severe      GTGCACCTACACTAGTGCCACAAGAGCACTATGTTAGAATTACTGGCTTATACCCAACAC    17002
```

```
gb:MT019532|Organism:Severe      GTGCACCTACACTAGTGCCACAAGAGCACTATGTTAGAATTACTGGCTTATACCCAACAC   17002
                                 ********************************************************* **

gb:MT020781|Organism:Severe      TCAATATCTCAGATGAGTTTTCTAGCAATGTTGCAAATTATCAAAAGGTTGGTATGCAAA   17050
gb:MT007544|Organism:Severe      TCAATATCTCAGATGAGTTTTCTAGCAATGTTGCAAATTATCAAAAGGTTGGTATGCAAA   17062
gb:MN994467|Organism:Severe      TCAATATCTCAGATGAGTTTTCTAGCAATGTTGCAAATTATCAAAAGGTTGGTATGCAAA   17062
gb:MT044257|Organism:Severe      TCAATATCTCAGATGAGTTTTCTAGCAATGTTGCAAATTATCAAAAGGTTGGTATGCAAA   17062
gb:MT106054|Organism:Severe      TCAATATCTCAGATGAGTTTTCTAGCAATGTTGCAAATTATCAAAAGGTTGGTATGCAAA   17062
gb:MT049951|Organism:Severe      TCAATATCTCAGATGAGTTTTCTAGCAATGTTGCAAATTATCAAAAGGTTGGTATGCAAA   17062
gb:MN975262|Organism:Severe      TCAATATCTCAGATGAGTTTTCTAGCAATGTTGCAAATTATCAAAAGGTTGGTATGCAAA   17062
gb:MT106052|Organism:Severe      TCAATATCTCAGATGAGTTTTCTAGCAATGTTGCAAATTATCAAAAGGTTGGTATGCAAA   17062
gb:LC522975|Organism:Severe      TCAATATCTCAGATGAGTTTTCTAGCAATGTTGCAAATTATCAAAAGGTTGGTATGCAAA   17059
gb:LC522973|Organism:Severe      TCAATATCTCAGATGAGTTTTCTAGCAATGTTGCAAATTATCAAAAGGTTGGTATGCAAA   17059
gb:LC522974|Organism:Severe      TCAATATCTCAGATGAGTTTTCTAGCAATGTTGCAAATTATCAAAAGGTTGGTATGCAAA   17059
gb:MN985325|Organism:Severe      TCAATATCTCAGATGAGTTTTCTAGCAATGTTGCAAATTATCAAAAGGTTGGTATGCAAA   17062
gb:MT020881|Organism:Severe      TCAATATCTCAGATGAGTTTTCTAGCAATGTTGCAAATTATCAAAAGGTTGGTATGCAAA   17062
gb:MT020880|Organism:Severe      TCAATATCTCAGATGAGTTTTCTAGCAATGTTGCAAATTATCAAAAGGTTGGTATGCAAA   17062
gb:MT066175|Organism:Severe      TCAATATCTCAGATGAGTTTTCTAGCAATGTTGCAAATTATCAAAAGGTTGGTATGCAAA   17062
gb:MN997409|Organism:Severe      TCAATATCTCAGATGAGTTTTCTAGCAATGTTGCAAATTATCAAAAGGTTGGTATGCAAA   17062
gb:MN938384|Organism:Severe      TCAATATCTCAGATGAGTTTTCTAGCAATGTTGCAAATTATCAAAAGGTTGGTATGCAAA   17030
gb:MT044258|Organism:Severe      TCAATATCTCAGATGAGTTTTCTAGCAATGTTGCAAATTATCAAAAGGTTGGTATGCAAA   17038
gb:MT039890|Organism:Severe      TCAATATCTCAGATGAGTTTTCTAGCAATGTTGCAAATTATCAAAAGGTTGGTATGCAAA   17062
gb:MN988713|Organism:Severe      TCAATATCTCAGATGAGTTTTCTAGCAATGTTGCAAATTATCAAAAGGTTGGTATGCAAA   17062
gb:LC521925|Organism:Severe      TCAATATCTCAGATGAGTTTTCTAGCAATGTTGCAAATTATCAAAAGGTTGGTATGCAAA   17035
gb:MT093571|Organism:Severe      TCAATATCTCAGATGAGTTTTCTAGCAATGTTGCAAATTATCAAAAGGTTGGTATGCAAA   17062
gb:MT039887|Organism:Severe      TCAATATCTCAGATGAGTTTTCTAGCAATGTTGCAAATTATCAAAAGGTTGGTATGCAAA   17062
gb:MT019530|Organism:Severe      TCAATATCTCAGATGAGTTTTCTAGCAATGTTGCAAATTATCAAAAGGTTGGTATGCAAA   17062
gb:MT039888|Organism:Severe      TCAATATCTCAGATGAGTTTTCTAGCAATGTTGCAAATTATCAAAAGGTTGGTATGCAAA   17062
gb:LC522972|Organism:Severe      TCAATATCTCAGATGAGTTTTCTAGCAATGTTGCAAATTATCAAAAGGTTGGTATGCAAA   17059
gb:MT027063|Organism:Severe      TCAATATCTCAGATGAGTTTTCTAGCAATGTTGCAAATTATCAAAAGGTTGGTATGCAAA   17062
gb:MT027062|Organism:Severe      TCAATATCTCAGATGAGTTTTCTAGCAATGTTGCAAATTATCAAAAGGTTGGTATGCAAA   17062
gb:MT019529|Organism:Severe      TCAATATCTCAGATGAGTTTTCTAGCAATGTTGCAAATTATCAAAAGGTTGGTATGCAAA   17062
gb:MN996529|Organism:Severe      TCAATATCTCAGATGAGTTTTCTAGCAATGTTGCAAATTATCAAAAGGTTGGTATGCAAA   17050
gb:MN996531|Organism:Severe      TCAATATCTCAGATGAGTTTTCTAGCAATGTTGCAAATTATCAAAAGGTTGGTATGCAAA   17049
gb:MT066176|Organism:Severe      TCAATATCTCAGATGAGTTTTCTAGCAATGTTGCAAATTATCAAAAGGTTGGTATGCAAA   17062
gb:MT027064|Organism:Severe      TCAATATCTCAGATGAGTTTTCTAGCAATGTTGCAAATTATCAAAAGGTTGGTATGCAAA   17062
gb:MN994468|Organism:Severe      TCAATATCTCAGATGAGTTTTCTAGCAATGTTGCAAATTATCAAAAGGTTGGTATGCAAA   17062
gb:MT072688|Organism:Severe      TCAATATCTCAGATGAGTTTTCTAGCAATGTTGCAAATTATCAAAAGGTTGGTATGCAAA   17047
gb:MN996527|Organism:Severe      TCAATATCTCAGATGAGTTTTCTAGCAATGTTGCAAATTATCAAAAGGTTGGTATGCAAA   17029
gb:MT093631|Organism:Severe      TCAATATCTCAGATGAGTTTTCTAGCAATGTTGCAAATTATCAAAAGGTTGGTATGCAAA   17100
gb:MT106053|Organism:Severe      TCAATATCTCAGATGAGTTTTCTAGCAATGTTGCAAATTATCAAAAGGTTGGTATGCAAA   17062
gb:MT019533|Organism:Severe      TCAATATCTCAGATGAGTTTTCTAGCAATGTTGCAAATTATCAAAAGGTTGGTATGCAAA   17062
gb:MT019531|Organism:Severe      TCAATATCTCAGATGAGTTTTCTAGCAATGTTGCAAATTATCAAAAGGTTGGTATGCAAA   17062
gb:MN996528|Organism:Severe      TCAATATCTCAGATGAGTTTTCTAGCAATGTTGCAAATTATCAAAAGGTTGGTATGCAAA   17062
gb:MN996530|Organism:Severe      TCAATATCTCAGATGAGTTTTCTAGCAATGTTGCAAATTATCAAAAGGTTGGTATGCAAA   17048
gb:MN908947|Organism:Severe      TCAATATCTCAGATGAGTTTTCTAGCAATGTTGCAAATTATCAAAAGGTTGGTATGCAAA   17062
gb:MT019532|Organism:Severe      TCAATATCTCAGATGAGTTTTCTAGCAATGTTGCAAATTATCAAAAGGTTGGTATGCAAA   17062
                                 ************************************************************

gb:MT020781|Organism:Severe      AGTATTCTACACTCCAGGGACCACCTGGTACTGGTAAGAGTCATTTTGCTATTGGCCTAG   17110
gb:MT007544|Organism:Severe      AGTATTCTACACTCCAGGGACCACCTGGTACTGGTAAGAGTCATTTTGCTATTGGCCTAG   17122
gb:MN994467|Organism:Severe      AGTATTCTACACTCCAGGGACCACCTGGTACTGGTAAGAGTCATTTTGCTATTGGCCTAG   17122
gb:MT044257|Organism:Severe      AGTATTCTACACTCCAGGGACCACCTGGTACTGGTAAGAGTCATTTTGCTATTGGCCTAG   17122
gb:MT106054|Organism:Severe      AGTATTCTACACTCCAGGGACCACCTGGTACTGGTAAGAGTCATTTTGCTATTGGCCTAG   17122
gb:MT049951|Organism:Severe      AGTATTCTACACTCCAGGGACCACCTGGTACTGGTAAGAGTCATTTTGCTATTGGCCTAG   17122
gb:MN975262|Organism:Severe      AGTATTCTACACTCCAGGGACCACCTGGTACTGGTAAGAGTCATTTTGCTATTGGCCTAG   17122
gb:MT106052|Organism:Severe      AGTATTCTACACTCCAGGGACCACCTGGTACTGGTAAGAGTCATTTTGCTATTGGCCTAG   17122
gb:LC522975|Organism:Severe      AGTATTCTACACTCCAGGGACCACCTGGTACTGGTAAGAGTCATTTTGCTATTGGCCTAG   17119
gb:LC522973|Organism:Severe      AGTATTCTACACTCCAGGGACCACCTGGTACTGGTAAGAGTCATTTTGCTATTGGCCTAG   17119
gb:LC522974|Organism:Severe      AGTATTCTACACTCCAGGGACCACCTGGTACTGGTAAGAGTCATTTTGCTATTGGCCTAG   17119
gb:MN985325|Organism:Severe      AGTATTCTACACTCCAGGGACCACCTGGTACTGGTAAGAGTCATTTTGCTATTGGCCTAG   17122
gb:MT020881|Organism:Severe      AGTATTCTACACTCCAGGGACCACCTGGTACTGGTAAGAGTCATTTTGCTATTGGCCTAG   17122
gb:MT020880|Organism:Severe      AGTATTCTACACTCCAGGGACCACCTGGTACTGGTAAGAGTCATTTTGCTATTGGCCTAG   17122
gb:MT066175|Organism:Severe      AGTATTCTACACTCCAGGGACCACCTGGTACTGGTAAGAGTCATTTTGCTATTGGCCTAG   17122
gb:MN997409|Organism:Severe      AGTATTCTACACTCCAGGGACCACCTGGTACTGGTAAGAGTCATTTTGCTATTGGCCTAG   17122
```

```
gb:MN938384|Organism:Severe      AGTATTCTACACTCCAGGGACCACCTGGTACTGGTAAGAGTCATTTTGCTATTGGCCTAG    17090
gb:MT044258|Organism:Severe      AGTATTCTACACTCCAGGGACCACCTGGTACTGGTAAGAGTCATTTTGCTATTGGCCTAG    17098
gb:MT039890|Organism:Severe      AGTATTCTACACTCCAGGGACCACCTGGTACTGGTAAGAGTCATTTTGCTATTGGCCTAG    17122
gb:MN988713|Organism:Severe      AGTATTCTACACTCCAGGGACCACCTGGTACTGGTAAGAGTCATTTTGCTATTGGCCTAG    17122
gb:LC521925|Organism:Severe      AGTATTCTACACTCCAGGGACCACCTGGTACTGGTAAGAGTCATTTTGCTATTGGCCTAG    17095
gb:MT093571|Organism:Severe      AGTATTCTACACTCCAGGGACCACCTGGTACTGGTAAGAGTCATTTTGCTATTGGCCTAG    17122
gb:MT039887|Organism:Severe      AGTATTCTACACTCCAGGGACCACCTGGTACTGGTAAGAGTCATTTTGCTATTGGCCTAG    17122
gb:MT019530|Organism:Severe      AGTATTCTACACTCCAGGGACCACCTGGTACTGGTAAGAGTCATTTTGCTATTGGCCTAG    17122
gb:MT039888|Organism:Severe      AGTATTCTACACTCCAGGGACCACCTGGTACTGGTAAGAGTCATTTTGCTATTGGCCTAG    17122
gb:LC522972|Organism:Severe      AGTATTCTACACTCCAGGGACCACCTGGTACTGGTAAGAGTCATTTTGCTATTGGCCTAG    17119
gb:MT027063|Organism:Severe      AGTATTCTACACTCCAGGGACCACCTGGTACTGGTAAGAGTCATTTTGCTATTGGCCTAG    17122
gb:MT027062|Organism:Severe      AGTATTCTACACTCCAGGGACCACCTGGTACTGGTAAGAGTCATTTTGCTATTGGCCTAG    17122
gb:MT019529|Organism:Severe      AGTATTCTACACTCCAGGGACCACCTGGTACTGGTAAGAGTCATTTTGCTATTGGCCTAG    17122
gb:MN996529|Organism:Severe      AGTATTCTACACTCCAGGGACCACCTGGTACTGGTAAGAGTCATTTTGCTATTGGCCTAG    17110
gb:MN996531|Organism:Severe      AGTATTCTACACTCCAGGGACCACCTGGTACTGGTAAGAGTCATTTTGCTATTGGCCTAG    17109
gb:MT066176|Organism:Severe      AGTATTCTACACTCCAGGGACCACCTGGTACTGGTAAGAGTCATTTTGCTATTGGCCTAG    17122
gb:MT027064|Organism:Severe      AGTATTCTACACTCCAGGGACCACCTGGTACTGGTAAGAGTCATTTTGCTATTGGCCTAG    17122
gb:MN994468|Organism:Severe      AGTATTCTACACTCCAGGGACCACCTGGTACTGGTAAGAGTCATTTTGCTATTGGCCTAG    17122
gb:MT072688|Organism:Severe      AGTATTCTACACTCCAGGGACCACCTGGTACTGGTAAGAGTCATTTTGCTATTGGCCTAG    17107
gb:MN996527|Organism:Severe      AGTATTCTACACTCCAGGGACCACCTGGTACTGGTAAGAGTCATTTTGCTATTGGCCTAG    17089
gb:MT093631|Organism:Severe      AGTATTCTACACTCCAGGGACCACCTGGTACTGGTAAGAGTCATTTTGCTATTGGCCTAG    17160
gb:MT106053|Organism:Severe      AGTATTCTACACTCCAGGGACCACCTGGTACTGGTAAGAGTCATTTTGCTATTGGCCTAG    17122
gb:MT019533|Organism:Severe      AGTATTCTACACTCCAGGGACCACCTGGTACTGGTAAGAGTCATTTTGCTATTGGCCTAG    17122
gb:MT019531|Organism:Severe      AGTATTCTACACTCCAGGGACCACCTGGTACTGGTAAGAGTCATTTTGCTATTGGCCTAG    17122
gb:MN996528|Organism:Severe      AGTATTCTACACTCCAGGGACCACCTGGTACTGGTAAGAGTCATTTTGCTATTGGCCTAG    17122
gb:MN996530|Organism:Severe      AGTATTCTACACTCCAGGGACCACCTGGTACTGGTAAGAGTCATTTTGCTATTGGCCTAG    17108
gb:MN908947|Organism:Severe      AGTATTCTACACTCCAGGGACCACCTGGTACTGGTAAGAGTCATTTTGCTATTGGCCTAG    17122
gb:MT019532|Organism:Severe      AGTATTCTACACTCCAGGGACCACCTGGTACTGGTAAGAGTCATTTTGCTATTGGCCTAG    17122
                                 ************************************************************

gb:MT020781|Organism:Severe      CTCTCTACTACCCTTCTGCTCGCATAGTGTATACAGCTTGCTCTCATGCCGCTGTTGATG    17170
gb:MT007544|Organism:Severe      CTCTCTACTACCCTTCTGCTCGCATAGTGTATACAGCTTGCTCTCATGCCGCTGTTGATG    17182
gb:MN994467|Organism:Severe      CTCTCTACTACCCTTCTGCTCGCATAGTGTATACAGCTTGCTCTCATGCCGCTGTTGATG    17182
gb:MT044257|Organism:Severe      CTCTCTACTACCCTTCTGCTCGCATAGTGTATACAGCTTGCTCTCATGCCGCTGTTGATG    17182
gb:MT106054|Organism:Severe      CTCTCTACTACCCTTCTGCTCGCATAGTGTATACAGCTTGCTCTCATGCCGCTGTTGATG    17182
gb:MT049951|Organism:Severe      CTCTCTACTACCCTTCTGCTCGCATAGTGTATACAGCTTGCTCTCATGCCGCTGTTGATG    17182
gb:MN975262|Organism:Severe      CTCTCTACTACCCTTCTGCTCGCATAGTGTATACAGCTTGCTCTCATGCCGCTGTTGATG    17182
gb:MT106052|Organism:Severe      CTCTCTACTACCCTTCTGCTCGCATAGTGTATACAGCTTGCTCTCATGCCGCTGTTGATG    17182
gb:LC522975|Organism:Severe      CTCTCTACTACCCTTCTGCTCGCATAGTGTATACAGCTTGCTCTCATGCCGCTGTTGATG    17179
gb:LC522973|Organism:Severe      CTCTCTACTACCCTTCTGCTCGCATAGTGTATACAGCTTGCTCTCATGCCGCTGTTGATG    17179
gb:LC522974|Organism:Severe      CTCTCTACTACCCTTCTGCTCGCATAGTGTATACAGCTTGCTCTCATGCCGCTGTTGATG    17179
gb:MN985325|Organism:Severe      CTCTCTACTACCCTTCTGCTCGCATAGTGTATACAGCTTGCTCTCATGCCGCTGTTGATG    17182
gb:MT020881|Organism:Severe      CTCTCTACTACCCTTCTGCTCGCATAGTGTATACAGCTTGCTCTCATGCCGCTGTTGATG    17182
gb:MT020880|Organism:Severe      CTCTCTACTACCCTTCTGCTCGCATAGTGTATACAGCTTGCTCTCATGCCGCTGTTGATG    17182
gb:MT066175|Organism:Severe      CTCTCTACTACCCTTCTGCTCGCATAGTGTATACAGCTTGCTCTCATGCCGCTGTTGATG    17182
gb:MN997409|Organism:Severe      CTCTCTACTACCCTTCTGCTCGCATAGTGTATACAGCTTGCTCTCATGCCGCTGTTGATG    17182
gb:MN938384|Organism:Severe      CTCTCTACTACCCTTCTGCTCGCATAGTGTATACAGCTTGCTCTCATGCCGCTGTTGATG    17150
gb:MT044258|Organism:Severe      CTCTCTACTACCCTTCTGCTCGCATAGTGTATACAGCTTGCTCTCATGCCGCTGTTGATG    17158
gb:MT039890|Organism:Severe      CTCTCTACTACCCTTCTGCTCGCATAGTGTATACAGCTTGCTCTCATGCCGCTGTTGATG    17182
gb:MN988713|Organism:Severe      CTCTCTACTACCCTTCTGCTCGCATAGTGTATACAGCTTGCTCTCATGCCGCTGTTGATG    17182
gb:LC521925|Organism:Severe      CTCTCTACTACCCTTCTGCTCGCATAGTGTATACAGCTTGCTCTCATGCCGCTGTTGATG    17155
gb:MT093571|Organism:Severe      CTCTCTACTACCCTTCTGCTCGCATAGTGTATACAGCTTGCTCTCATGCCGCTGTTGATG    17182
gb:MT039887|Organism:Severe      CTCTCTACTACCCTTCTGCTCGCATAGTGTATACAGCTTGCTCTCATGCCGCTGTTGATG    17182
gb:MT019530|Organism:Severe      CTCTCTACTACCCTTCTGCTCGCATAGTGTATACAGCTTGCTCTCATGCCGCTGTTGATG    17182
gb:MT039888|Organism:Severe      CTCTCTACTACCCTTCTGCTCGCATAGTGTATACAGCTTGCTCTCATGCCGCTGTTGATG    17182
gb:LC522972|Organism:Severe      CTCTCTACTACCCTTCTGCTCGCATAGTGTATACAGCTTGCTCTCATGCCGCTGTTGATG    17179
gb:MT027063|Organism:Severe      CTCTCTACTACCCTTCTGCTCGCATAGTGTATACAGCTTGCTCTCATGCCGCTGTTGATG    17182
gb:MT027062|Organism:Severe      CTCTCTACTACCCTTCTGCTCGCATAGTGTATACAGCTTGCTCTCATGCCGCTGTTGATG    17182
gb:MT019529|Organism:Severe      CTCTCTACTACCCTTCTGCTCGCATAGTGTATACAGCTTGCTCTCATGCCGCTGTTGATG    17182
gb:MN996529|Organism:Severe      CTCTCTACTACCCTTCTGCTCGCATAGTGTATACAGCTTGCTCTCATGCCGCTGTTGATG    17170
gb:MN996531|Organism:Severe      CTCTCTACTACCCTTCTGCTCGCATAGTGTATACAGCTTGCTCTCATGCCGCTGTTGATG    17169
gb:MT066176|Organism:Severe      CTCTCTACTACCCTTCTGCTCGCATAGTGTATACAGCTTGCTCTCATGCCGCTGTTGATG    17182
gb:MT027064|Organism:Severe      CTCTCTACTACCCTTCTGCTCGCATAGTGTATACAGCTTGCTCTCATGCCGCTGTTGATG    17182
gb:MN994468|Organism:Severe      CTCTCTACTACCCTTCTGCTCGCATAGTGTATACAGCTTGCTCTCATGCCGCTGTTGATG    17182
gb:MT072688|Organism:Severe      CTCTCTACTACCCTTCTGCTCGCATAGTGTATACAGCTTGCTCTCATGCCGCTGTTGATG    17167
```

```
gb:MN996527|Organism:Severe      CTCTCTACTACCCTTCTGCTCGCATAGTGTATACAGCTTGCTCTCATGCCGCTGTTGATG   17149
gb:MT093631|Organism:Severe      CTCTCTACTACCCTTCTGCTCGCATAGTGTATACAGCTTGCTCTCATGCCGCTGTTGATG   17220
gb:MT106053|Organism:Severe      CTCTCTACTACCCTTCTGCTCGCATAGTGTATACAGCTTGCTCTCATGCCGCTGTTGATG   17182
gb:MT019533|Organism:Severe      CTCTCTACTACCCTTCTGCTCGCATAGTGTATACAGCTTGCTCTCATGCCGCTGTTGATG   17182
gb:MT019531|Organism:Severe      CTCTCTACTACCCTTCTGCTCGCATAGTGTATACAGCTTGCTCTCATGCCGCTGTTGATG   17182
gb:MN996528|Organism:Severe      CTCTCTACTACCCTTCTGCTCGCATAGTGTATACAGCTTGCTCTCATGCCGCTGTTGATG   17182
gb:MN996530|Organism:Severe      CTCTCTACTACCCTTCTGCTCGCATAGTGTATACAGCTTGCTCTCATGCCGCTGTTGATG   17168
gb:MN908947|Organism:Severe      CTCTCTACTACCCTTCTGCTCGCATAGTGTATACAGCTTGCTCTCATGCCGCTGTTGATG   17182
gb:MT019532|Organism:Severe      CTCTCTACTACCCTTCTGCTCGCATAGTGTATACAGCTTGCTCTCATGCCGCTGTTGATG   17182
                                 ************************************************************

gb:MT020781|Organism:Severe      CACTATGTGAGAAGGCATTAAAATATTTGCCTATAGATAAATGTAGTAGAATTATACCTG   17230
gb:MT007544|Organism:Severe      CACTATGTGAGAAGGCATTAAAATATTTGCCTATAGATAAATGTAGTAGAATTATACCTG   17242
gb:MN994467|Organism:Severe      CACTATGTGAGAAGGCATTAAAATATTTGCCTATAGATAAATGTAGTAGAATTATACCTG   17242
gb:MT044257|Organism:Severe      CACTATGTGAGAAGGCATTAAAATATTTGCCTATAGATAAATGTAGTAGAATTATACCTG   17242
gb:MT106054|Organism:Severe      CACTATGTGAGAAGGCATTAAAATATTTGCCTATAGATAAATGTAGTAGAATTATACCTG   17242
gb:MT049951|Organism:Severe      CACTATGTGAGAAGGCATTAAAATATTTGCCTATAGATAAATGTAGTAGAATTATACCTG   17242
gb:MN975262|Organism:Severe      CACTATGTGAGAAGGCATTAAAATATTTGCCTATAGATAAATGTAGTAGAATTATACCTG   17242
gb:MT106052|Organism:Severe      CACTATGTGAGAAGGCATTAAAATATTTGCCTATAGATAAATGTAGTAGAATTATACCTG   17242
gb:LC522975|Organism:Severe      CACTATGTGAGAAGGCATTAAAATATTTGCCTATAGATAAATGTAGTAGAATTATACCTG   17239
gb:LC522973|Organism:Severe      CACTATGTGAGAAGGCATTAAAATATTTGCCTATAGATAAATGTAGTAGAATTATACCTG   17239
gb:LC522974|Organism:Severe      CACTATGTGAGAAGGCATTAAAATATTTGCCTATAGATAAATGTAGTAGAATTATACCTG   17239
gb:MN985325|Organism:Severe      CACTATGTGAGAAGGCATTAAAATATTTGCCTATAGATAAATGTAGTAGAATTATACCTG   17242
gb:MT020881|Organism:Severe      CACTATGTGAGAAGGCATTAAAATATTTGCCTATAGATAAATGTAGTAGAATTATACCTG   17242
gb:MT020880|Organism:Severe      CACTATGTGAGAAGGCATTAAAATATTTGCCTATAGATAAATGTAGTAGAATTATACCTG   17242
gb:MT066175|Organism:Severe      CACTATGTGAGAAGGCATTAAAATATTTGCCTATAGATAAATGTAGTAGAATTATACCTG   17242
gb:MN997409|Organism:Severe      CACTATGTGAGAAGGCATTAAAATATTTGCCTATAGATAAATGTAGTAGAATTATACCTG   17242
gb:MN938384|Organism:Severe      CACTATGTGAGAAGGCATTAAAATATTTGCCTATAGATAAATGTAGTAGAATTATACCTG   17210
gb:MT044258|Organism:Severe      CACTATGTGAGAAGGCATTAAAATATTTGCCTATAGATAAATGTAGTAGAATTATACCTG   17218
gb:MT039890|Organism:Severe      CACTATGTGAGAAGGCATTAAAATATTTGCCTATAGATAAATGTAGTAGAATTATACCTG   17242
gb:MN988713|Organism:Severe      CACTATGTGAGAAGGCATTAAAATATTTGCCTATAGATAAATGTAGTAGAATTATACCTG   17242
gb:LC521925|Organism:Severe      CACTATGTGAGAAGGCATTAAAATATTTGCCTATAGATAAATGTAGTAGAATTATACCTG   17215
gb:MT093571|Organism:Severe      CACTATGTGAGAAGGCATTAAAATATTTGCCTATAGATAAATGTAGTAGAATTATACCTG   17242
gb:MT039887|Organism:Severe      CACTATGTGAGAAGGCATTAAAATATTTGCCTATAGATAAATGTAGTAGAATTATACCTG   17242
gb:MT019530|Organism:Severe      CACTATGTGAGAAGGCATTAAAATATTTGCCTATAGATAAATGTAGTAGAATTATACCTG   17242
gb:MT039888|Organism:Severe      CACTATGTGAGAAGGCATTAAAATATTTGCCTATAGATAAATGTAGTAGAATTATACCTG   17242
gb:LC522972|Organism:Severe      CACTATGTGAGAAGGCATTAAAATATTTGCCTATAGATAAATGTAGTAGAATTATACCTG   17239
gb:MT027063|Organism:Severe      CACTATGTGAGAAGGCATTAAAATATTTGCCTATAGATAAATGTAGTAGAATTATACCTG   17242
gb:MT027062|Organism:Severe      CACTATGTGAGAAGGCATTAAAATATTTGCCTATAGATAAATGTAGTAGAATTATACCTG   17242
gb:MT019529|Organism:Severe      CACTATGTGAGAAGGCATTAAAATATTTGCCTATAGATAAATGTAGTAGAATTATACCTG   17242
gb:MN996529|Organism:Severe      CACTATGTGAGAAGGCATTAAAATATTTGCCTATAGATAAATGTAGTAGAATTATACCTG   17230
gb:MN996531|Organism:Severe      CACTATGTGAGAAGGCATTAAAATATTTGCCTATAGATAAATGTAGTAGAATTATACCTG   17229
gb:MT066176|Organism:Severe      CACTATGTGAGAAGGCATTAAAATATTTGCCTATAGATAAATGTAGTAGAATTATACCTG   17242
gb:MT027064|Organism:Severe      CACTATGTGAGAAGGCATTAAAATATTTGCCTATAGATAAATGTAGTAGAATTATACCTG   17242
gb:MN994468|Organism:Severe      CACTATGTGAGAAGGCATTAAAATATTTGCCTATAGATAAATGTAGTAGAATTATACCTG   17242
gb:MT072688|Organism:Severe      CACTATGTGAGAAGGCATTAAAATATTTGCCTATAGATAAATGTAGTAGAATTATACCTG   17227
gb:MN996527|Organism:Severe      CACTATGTGAGAAGGCATTAAAATATTTGCCTATAGATAAATGTAGTAGAATTATACCTG   17209
gb:MT093631|Organism:Severe      CACTATGTGAGAAGGCATTAAAATATTTGCCTATAGATAAATGTAGTAGAATTATACCTG   17280
gb:MT106053|Organism:Severe      CACTATGTGAGAAGGCATTAAAATATTTGCCTATAGATAAATGTAGTAGAATTATACCTG   17242
gb:MT019533|Organism:Severe      CACTATGTGAGAAGGCATTAAAATATTTGCCTATAGATAAATGTAGTAGAATTATACCTG   17242
gb:MT019531|Organism:Severe      CACTATGTGAGAAGGCATTAAAATATTTGCCTATAGATAAATGTAGTAGAATTATACCTG   17242
gb:MN996528|Organism:Severe      CACTATGTGAGAAGGCATTAAAATATTTGCCTATAGATAAATGTAGTAGAATTATACCTG   17242
gb:MN996530|Organism:Severe      CACTATGTGAGAAGGCATTAAAATATTTGCCTATAGATAAATGTAGTAGAATTATACCTG   17228
gb:MN908947|Organism:Severe      CACTATGTGAGAAGGCATTAAAATATTTGCCTATAGATAAATGTAGTAGAATTATACCTG   17242
gb:MT019532|Organism:Severe      CACTATGTGAGAAGGCATTAAAATATTTGCCTATAGATAAATGTAGTAGAATTATACCTG   17242
                                 ************************************************************

gb:MT020781|Organism:Severe      CACGTGCTCGTGTAGAGTGTTTTGATAAATTCAAAGTGAATTCAACATTAGAACAGTATG   17290
gb:MT007544|Organism:Severe      CACGTGCTCGTGTAGAGTGTTTTGATAAATTCAAAGTGAATTCAACATTAGAACAGTATG   17302
gb:MN994467|Organism:Severe      CACGTGCTCGTGTAGAGTGTTTTGATAAATTCAAAGTGAATTCAACATTAGAACAGTATG   17302
gb:MT044257|Organism:Severe      CACGTGCTCGTGTAGAGTGTTTTGATAAATTCAAAGTGAATTCAACATTAGAACAGTATG   17302
gb:MT106054|Organism:Severe      CACGTGCTCGTGTAGAGTGTTTTGATAAATTCAAAGTGAATTCAACATTAGAACAGTATG   17302
gb:MT049951|Organism:Severe      CACGTGCTCGTGTAGAGTGTTTTGATAAATTCAAAGTGAATTCAACATTAGAACAGTATG   17302
gb:MN975262|Organism:Severe      CACGTGCTCGTGTAGAGTGTTTTGATAAATTCAAAGTGAATTCAACATTAGAACAGTATG   17302
gb:MT106052|Organism:Severe      CACGTGCTCGTGTAGAGTGTTTTGATAAATTCAAAGTGAATTCAACATTAGAACAGTATG   17302
```

```
gb:LC522975|Organism:Severe      CACGTGCTCGTGTAGAGTGTTTTGATAAATTCAAAGTGAATTCAACATTAGAACAGTATG	17299
gb:LC522973|Organism:Severe      CACGTGCTCGTGTAGAGTGTTTTGATAAATTCAAAGTGAATTCAACATTAGAACAGTATG	17299
gb:LC522974|Organism:Severe      CACGTGCTCGTGTAGAGTGTTTTGATAAATTCAAAGTGAATTCAACATTAGAACAGTATG	17299
gb:MN985325|Organism:Severe      CACGTGCTCGTGTAGAGTGTTTTGATAAATTCAAAGTGAATTCAACATTAGAACAGTATG	17302
gb:MT020881|Organism:Severe      CACGTGCTCGTGTAGAGTGTTTTGATAAATTCAAAGTGAATTCAACATTAGAACAGTATG	17302
gb:MT020880|Organism:Severe      CACGTGCTCGTGTAGAGTGTTTTGATAAATTCAAAGTGAATTCAACATTAGAACAGTATG	17302
gb:MT066175|Organism:Severe      CACGTGCTCGTGTAGAGTGTTTTGATAAATTCAAAGTGAATTCAACATTAGAACAGTATG	17302
gb:MN997409|Organism:Severe      CACGTGCTCGTGTAGAGTGTTTTGATAAATTCAAAGTGAATTCAACATTAGAACAGTATG	17302
gb:MN938384|Organism:Severe      CACGTGCTCGTGTAGAGTGTTTTGATAAATTCAAAGTGAATTCAACATTAGAACAGTATG	17270
gb:MT044258|Organism:Severe      CACGTGCTCGTGTAGAGTGTTTTGATAAATTCAAAGTGAATTCAACATTAGAACAGTATG	17278
gb:MT039890|Organism:Severe      CACGTGCTCGTGTAGAGTGTTTTGATAAATTCAAAGTGAATTCAACATTAGAACAGTATG	17302
gb:MN988713|Organism:Severe      CACGTGCTCGTGTAGAGTGTTTTGATAAATTCAAAGTGAATTCAACATTAGAACAGTATG	17302
gb:LC521925|Organism:Severe      CACGTGCTCGTGTAGAGTGTTTTGATAAATTCAAAGTGAATTCAACATTAGAACAGTATG	17275
gb:MT093571|Organism:Severe      CACGTGCTCGTGTAGAGTGTTTTGATAAATTCAAAGTGAATTCAACATTAGAACAGTATG	17302
gb:MT039887|Organism:Severe      CACGTGCTCGTGTAGAGTGTTTTGATAAATTCAAAGTGAATTCAACATTAGAACAGTATG	17302
gb:MT019530|Organism:Severe      CACGTGCTCGTGTAGAGTGTTTTGATAAATTCAAAGTGAATTCAACATTAGAACAGTATG	17302
gb:MT039888|Organism:Severe      CACGTGCTCGTGTAGAGTGTTTTGATAAATTCAAAGTGAATTCAACATTAGAACAGTATG	17302
gb:LC522972|Organism:Severe      CACGTGCTCGTGTAGAGTGTTTTGATAAATTCAAAGTGAATTCAACATTAGAACAGTATG	17299
gb:MT027063|Organism:Severe      CACGTGCTCGTGTAGAGTGTTTTGATAAATTCAAAGTGAATTCAACATTAGAACAGTATG	17302
gb:MT027062|Organism:Severe      CACGTGCTCGTGTAGAGTGTTTTGATAAATTCAAAGTGAATTCAACATTAGAACAGTATG	17302
gb:MT019529|Organism:Severe      CACGTGCTCGTGTAGAGTGTTTTGATAAATTCAAAGTGAATTCAACATTAGAACAGTATG	17302
gb:MN996529|Organism:Severe      CACGTGCTCGTGTAGAGTGTTTTGATAAATTCAAAGTGAATTCAACATTAGAACAGTATG	17290
gb:MN996531|Organism:Severe      CACGTGCTCGTGTAGAGTGTTTTGATAAATTCAAAGTGAATTCAACATTAGAACAGTATG	17289
gb:MT066176|Organism:Severe      CACGTGCTCGTGTAGAGTGTTTTGATAAATTCAAAGTGAATTCAACATTAGAACAGTATG	17302
gb:MT027064|Organism:Severe      CACGTGCTCGTGTAGAGTGTTTTGATAAATTCAAAGTGAATTCAACATTAGAACAGTATG	17302
gb:MN994468|Organism:Severe      CACGTGCTCGTGTAGAGTGTTTTGATAAATTCAAAGTGAATTCAACATTAGAACAGTATG	17302
gb:MT072688|Organism:Severe      CACGTGCTCGTGTAGAGTGTTTTGATAAATTCAAAGTGAATTCAACATTAGAACAGTATG	17287
gb:MN996527|Organism:Severe      CACGTGCTCGTGTAGAGTGTTTTGATAAATTCAAAGTGAATTCAACATTAGAACAGTATG	17269
gb:MT093631|Organism:Severe      CACGTGCTCGTGTAGAGTGTTTTGATAAATTCAAAGTGAATTCAACATTAGAACAGTATG	17340
gb:MT106053|Organism:Severe      CACGTGCTCGTGTAGAGTGTTTTGATAAATTCAAAGTGAATTCAACATTAGAACAGTATG	17302
gb:MT019533|Organism:Severe      CACGTGCTCGTGTAGAGTGTTTTGATAAATTCAAAGTGAATTCAACATTAGAACAGTATG	17302
gb:MT019531|Organism:Severe      CACGTGCTCGTGTAGAGTGTTTTGATAAATTCAAAGTGAATTCAACATTAGAACAGTATG	17302
gb:MN996528|Organism:Severe      CACGTGCTCGTGTAGAGTGTTTTGATAAATTCAAAGTGAATTCAACATTAGAACAGTATG	17302
gb:MN996530|Organism:Severe      CACGTGCTCGTGTAGAGTGTTTTGATAAATTCAAAGTGAATTCAACATTAGAACAGTATG	17288
gb:MN908947|Organism:Severe      CACGTGCTCGTGTAGAGTGTTTTGATAAATTCAAAGTGAATTCAACATTAGAACAGTATG	17302
gb:MT019532|Organism:Severe      CACGTGCTCGTGTAGAGTGTTTTGATAAATTCAAAGTGAATTCAACATTAGAACAGTATG	17302
                                 ************************************************************

gb:MT020781|Organism:Severe      TCTTTTGTACTGTAAATGCATTGCCTGAGACGACAGCAGATATAGTTGTCTTTGATGAAA	17350
gb:MT007544|Organism:Severe      TCTTTTGTACTGTAAATGCATTGCCTGAGACGACAGCAGATATAGTTGTCTTTGATGAAA	17362
gb:MN994467|Organism:Severe      TCTTTTGTACTGTAAATGCATTGCCTGAGACGACAGCAGATATAGTTGTCTTTGATGAAA	17362
gb:MT044257|Organism:Severe      TCTTTTGTACTGTAAATGCATTGCCTGAGACGACAGCAGATATAGTTGTCTTTGATGAAA	17362
gb:MT106054|Organism:Severe      TCTTTTGTACTGTAAATGCATTGCCTGAGACGACAGCAGATATAGTTGTCTTTGATGAAA	17362
gb:MT049951|Organism:Severe      TCTTTTGTACTGTAAATGCATTGCCTGAGACGACAGCAGATATAGTTGTCTTTGATGAAA	17362
gb:MN975262|Organism:Severe      TCTTTTGTACTGTAAATGCATTGCCTGAGACGACAGCAGATATAGTTGTCTTTGATGAAA	17362
gb:MT106052|Organism:Severe      TCTTTTGTACTGTAAATGCATTGCCTGAGACGACAGCAGATATAGTTGTCTTTGATGAAA	17362
gb:LC522975|Organism:Severe      TCTTTTGTACTGTAAATGCATTGCCTGAGACGACAGCAGATATAGTTGTCTTTGATGAAA	17359
gb:LC522973|Organism:Severe      TCTTTTGTACTGTAAATGCATTGCCTGAGACGACAGCAGATATAGTTGTCTTTGATGAAA	17359
gb:LC522974|Organism:Severe      TCTTTTGTACTGTAAATGCATTGCCTGAGACGACAGCAGATATAGTTGTCTTTGATGAAA	17359
gb:MN985325|Organism:Severe      TCTTTTGTACTGTAAATGCATTGCCTGAGACGACAGCAGATATAGTTGTCTTTGATGAAA	17362
gb:MT020881|Organism:Severe      TCTTTTGTACTGTAAATGCATTGCCTGAGACGACAGCAGATATAGTTGTCTTTGATGAAA	17362
gb:MT020880|Organism:Severe      TCTTTTGTACTGTAAATGCATTGCCTGAGACGACAGCAGATATAGTTGTCTTTGATGAAA	17362
gb:MT066175|Organism:Severe      TCTTTTGTACTGTAAATGCATTGCCTGAGACGACAGCAGATATAGTTGTCTTTGATGAAA	17362
gb:MN997409|Organism:Severe      TCTTTTGTACTGTAAATGCATTGCCTGAGACGACAGCAGATATAGTTGTCTTTGATGAAA	17362
gb:MN938384|Organism:Severe      TCTTTTGTACTGTAAATGCATTGCCTGAGACGACAGCAGATATAGTTGTCTTTGATGAAA	17330
gb:MT044258|Organism:Severe      TCTTTTGTACTGTAAATGCATTGCCTGAGACGACAGCAGATATAGTTGTCTTTGATGAAA	17338
gb:MT039890|Organism:Severe      TCTTTTGTACTGTAAATGCATTGCCTGAGACGACAGCAGATATAGTTGTCTTTGATGAAA	17362
gb:MN988713|Organism:Severe      TCTTTTGTACTGTAAATGCATTGCCTGAGACGACAGCAGATATAGTTGTCTTTGATGAAA	17362
gb:LC521925|Organism:Severe      TCTTTTGTACTGTAAATGCATTGCCTGAGACGACAGCAGATATAGTTGTCTTTGATGAAA	17335
gb:MT093571|Organism:Severe      TCTTTTGTACTGTAAATGCATTGCCTGAGACGACAGCAGATATAGTTGTCTTTGATGAAA	17362
gb:MT039887|Organism:Severe      TCTTTTGTACTGTAAATGCATTGCCTGAGACGACAGCAGATATAGTTGTCTTTGATGAAA	17362
gb:MT019530|Organism:Severe      TCTTTTGTACTGTAAATGCATTGCCTGAGACGACAGCAGATATAGTTGTCTTTGATGAAA	17362
gb:MT039888|Organism:Severe      TCTTTTGTACTGTAAATGCATTGCCTGAGACGACAGCAGATATAGTTGTCTTTGATGAAA	17362
gb:LC522972|Organism:Severe      TCTTTTGTACTGTAAATGCATTGCCTGAGACGACAGCAGATATAGTTGTCTTTGATGAAA	17359
gb:MT027063|Organism:Severe      TCTTTTGTACTGTAAATGCATTGCCTGAGACGACAGCAGATATAGTTGTCTTTGATGAAA	17362
```

```
gb:MT027062|Organism:Severe      TCTTTTGTACTGTAAATGCATTGCCTGAGACGACAGCAGATATAGTTGTCTTTGATGAAA   17362
gb:MT019529|Organism:Severe      TCTTTTGTACTGTAAATGCATTGCCTGAGACGACAGCAGATATAGTTGTCTTTGATGAAA   17362
gb:MN996529|Organism:Severe      TCTTTTGTACTGTAAATGCATTGCCTGAGACGACAGCAGATATAGTTGTCTTTGATGAAA   17350
gb:MN996531|Organism:Severe      TCTTTTGTACTGTAAATGCATTGCCTGAGACGACAGCAGATATAGTTGTCTTTGATGAAA   17349
gb:MT066176|Organism:Severe      TCTTTTGTACTGTAAATGCATTGCCTGAGACGACAGCAGATATAGTTGTCTTTGATGAAA   17362
gb:MT027064|Organism:Severe      TCTTTTGTACTGTAAATGCATTGCCTGAGACGACAGCAGATATAGTTGTCTTTGATGAAA   17362
gb:MN994468|Organism:Severe      TCTTTTGTACTGTAAATGCATTGCCTGAGACGACAGCAGATATAGTTGTCTTTGATGAAA   17362
gb:MT072688|Organism:Severe      TCTTTTGTACTGTAAATGCATTGCCTGAGACGACAGCAGATATAGTTGTCTTTGATGAAA   17347
gb:MN996527|Organism:Severe      TCTTTTGTACTGTAAATGCATTGCCTGAGACGACAGCAGATATAGTTGTCTTTGATGAAA   17329
gb:MT093631|Organism:Severe      TCTTTTGTACTGTAAATGCATTGCCTGAGACGACAGCAGATATAGTTGTCTTTGATGAAA   17400
gb:MT106053|Organism:Severe      TCTTTTGTACTGTAAATGCATTGCCTGAGACGACAGCAGATATAGTTGTCTTTGATGAAA   17362
gb:MT019533|Organism:Severe      TCTTTTGTACTGTAAATGCATTGCCTGAGACGACAGCAGATATAGTTGTCTTTGATGAAA   17362
gb:MT019531|Organism:Severe      TCTTTTGTACTGTAAATGCATTGCCTGAGACGACAGCAGATATAGTTGTCTTTGATGAAA   17362
gb:MN996528|Organism:Severe      TCTTTTGTACTGTAAATGCATTGCCTGAGACGACAGCAGATATAGTTGTCTTTGATGAAA   17362
gb:MN996530|Organism:Severe      TCTTTTGTACTGTAAATGCATTGCCTGAGACGACAGCAGATATAGTTGTCTTTGATGAAA   17348
gb:MN908947|Organism:Severe      TCTTTTGTACTGTAAATGCATTGCCTGAGACGACAGCAGATATAGTTGTCTTTGATGAAA   17362
gb:MT019532|Organism:Severe      TCTTTTGTACTGTAAATGCATTGCCTGAGACGACAGCAGATATAGTTGTCTTTGATGAAA   17362
                                 ************************************************************

gb:MT020781|Organism:Severe      TTTCAATGGCCACAAATTATGATTTGAGTGTTGTCAATGCCAGATTACGTGCTAAGCACT   17410
gb:MT007544|Organism:Severe      TTTCAATGGCCACAAATTATGATTTGAGTGTTGTCAATGCCAGATTACGTGCTAAGCACT   17422
gb:MN994467|Organism:Severe      TTTCAATGGCCACAAATTATGATTTGAGTGTTGTCAATGCCAGATTACGTGCTAAGCACT   17422
gb:MT044257|Organism:Severe      TTTCAATGGCCACAAATTATGATTTGAGTGTTGTCAATGCCAGATTACGTGCTAAGCACT   17422
gb:MT106054|Organism:Severe      TTTCAATGGCCACAAATTATGATTTGAGTGTTGTCAATGCCAGATTACGTGCTAAGCACT   17422
gb:MT049951|Organism:Severe      TTTCAATGGCCACAAATTATGATTTGAGTGTTGTCAATGCCAGATTACGTGCTAAGCACT   17422
gb:MN975262|Organism:Severe      TTTCAATGGCCACAAATTATGATTTGAGTGTTGTCAATGCCAGATTACGTGCTAAGCACT   17422
gb:MT106052|Organism:Severe      TTTCAATGGCCACAAATTATGATTTGAGTGTTGTCAATGCCAGATTACGTGCTAAGCACT   17422
gb:LC522975|Organism:Severe      TTTCAATGGCCACAAATTATGATTTGAGTGTTGTCAATGCCAGATTACGTGCTAAGCACT   17419
gb:LC522973|Organism:Severe      TTTCAATGGCCACAAATTATGATTTGAGTGTTGTCAATGCCAGATTACGTGCTAAGCACT   17419
gb:LC522974|Organism:Severe      TTTCAATGGCCACAAATTATGATTTGAGTGTTGTCAATGCCAGATTACGTGCTAAGCACT   17419
gb:MN985325|Organism:Severe      TTTCAATGGCCACAAATTATGATTTGAGTGTTGTCAATGCCAGATTACGTGCTAAGCACT   17422
gb:MT020881|Organism:Severe      TTTCAATGGCCACAAATTATGATTTGAGTGTTGTCAATGCCAGATTACGTGCTAAGCACT   17422
gb:MT020880|Organism:Severe      TTTCAATGGCCACAAATTATGATTTGAGTGTTGTCAATGCCAGATTACGTGCTAAGCACT   17422
gb:MT066175|Organism:Severe      TTTCAATGGCCACAAATTATGATTTGAGTGTTGTCAATGCCAGATTACGTGCTAAGCACT   17422
gb:MN997409|Organism:Severe      TTTCAATGGCCACAAATTATGATTTGAGTGTTGTCAATGCCAGATTACGTGCTAAGCACT   17422
gb:MN938384|Organism:Severe      TTTCAATGGCCACAAATTATGATTTGAGTGTTGTCAATGCCAGATTACGTGCTAAGCACT   17390
gb:MT044258|Organism:Severe      TTTCAATGGCCACAAATTATGATTTGAGTGTTGTCAATGCCAGATTACGTGCTAAGCACT   17398
gb:MT039890|Organism:Severe      TTTCAATGGCCACAAATTATGATTTGAGTGTTGTCAATGCCAGATTACGTGCTAAGCACT   17422
gb:MN988713|Organism:Severe      TTTCAATGGCCACAAATTATGATTTGAGTGTTGTCAATGCCAGATTACGTGCTAAGCACT   17422
gb:LC521925|Organism:Severe      TTTCAATGGCCACAAATTATGATTTGAGTGTTGTCAATGCCAGATTACGTGCTAAGCACT   17395
gb:MT093571|Organism:Severe      TTTCAATGGCCACGAATTATGATTTGAGTGTTGTCAATGCCAGATTACGTGCTAAGCACT   17422
gb:MT039887|Organism:Severe      TTTCAATGGCTACAAATTATGATTTGAGTGTTGTCAATGCCAGATTACGTGCTAAGCACT   17422
gb:MT019530|Organism:Severe      TTTCAATGGCCACAAATTATGATTTGAGTGTTGTCAATGCCAGATTACGTGCTAAGCACT   17422
gb:MT039888|Organism:Severe      TTTCAATGGCCACAAATTATGATTTGAGTGTTGTCAATGCCAGATTACGTGCTAAGCACT   17422
gb:LC522972|Organism:Severe      TTTCAATGGCCACAAATTATGATTTGAGTGTTGTCAATGCCAGATTACGTGCTAAGCACT   17419
gb:MT027063|Organism:Severe      TTTCAATGGCCACAAATTATGATTTGAGTGTTGTCAATGCCAGATTACGTGCTAAGCACT   17422
gb:MT027062|Organism:Severe      TTTCAATGGCCACAAATTATGATTTGAGTGTTGTCAATGCCAGATTACGTGCTAAGCACT   17422
gb:MT019529|Organism:Severe      TTTCAATGGCCACAAATTATGATTTGAGTGTTGTCAATGCCAGATTACGTGCTAAGCACT   17422
gb:MN996529|Organism:Severe      TTTCAATGGCCACAAATTATGATTTGAGTGTTGTCAATGCCAGATTACGTGCTAAGCACT   17410
gb:MN996531|Organism:Severe      TTTCAATGGCCACAAATTATGATTTGAGTGTTGTCAATGCCAGATTACGTGCTAAGCACT   17409
gb:MT066176|Organism:Severe      TTTCAATGGCCACAAATTATGATTTGAGTGTTGTCAATGCCAGATTACGTGCTAAGCACT   17422
gb:MT027064|Organism:Severe      TTTCAATGGCCACAAATTATGATTTGAGTGTTGTCAATGCCAGATTACGTGCTAAGCACT   17422
gb:MN994468|Organism:Severe      TTTCAATGGCCACAAATTATGATTTGAGTGTTGTCAATGCCAGATTACGTGCTAAGCACT   17422
gb:MT072688|Organism:Severe      TTTCAATGGCCACAAATTATGATTTGAGTGTTGTCAATGCCAGATTACGTGCTAAGCACT   17407
gb:MN996527|Organism:Severe      TTTCAATGGCCACAAATTATGATTTGAGTGTTGTCAATGCCAGATTACGTGCTAAGCACT   17389
gb:MT093631|Organism:Severe      TTTCAATGGCCACAAATTATGATTTGAGTGTTGTCAATGCCAGATTACGTGCTAAGCACT   17460
gb:MT106053|Organism:Severe      TTTCAATGGCCACAAATTATGATTTGAGTGTTGTCAATGCCAGATTACGTGCTAAGCACT   17422
gb:MT019533|Organism:Severe      TTTCAATGGCCACAAATTATGATTTGAGTGTTGTCAATGCCAGATTACGTGCTAAGCACT   17422
gb:MT019531|Organism:Severe      TTTCAATGGCCACAAATTATGATTTGAGTGTTGTCAATGCCAGATTACGTGCTAAGCACT   17422
gb:MN996528|Organism:Severe      TTTCAATGGCCACAAATTATGATTTGAGTGTTGTCAATGCCAGATTACGTGCTAAGCACT   17422
gb:MN996530|Organism:Severe      TTTCAATGGCCACAAATTATGATTTGAGTGTTGTCAATGCCAGATTACGTGCTAAGCACT   17408
gb:MN908947|Organism:Severe      TTTCAATGGCCACAAATTATGATTTGAGTGTTGTCAATGCCAGATTACGTGCTAAGCACT   17422
gb:MT019532|Organism:Severe      TTTCAATGGCCACAAATTATGATTTGAGTGTTGTCAATGCCAGATTACGTGCTAAGCACT   17422
                                 ********** ** **********************************************
```

```
gb:MT020781|Organism:Severe      ATGTGTACATTGGCGACCCTGCTCAATTACCTGCACCACGCACATTGCTAACTAAGGGCA   17470
gb:MT007544|Organism:Severe      ATGTGTACATTGGCGACCCTGCTCAATTACCTGCACCACGCACATTGCTAACTAAGGGCA   17482
gb:MN994467|Organism:Severe      ATGTGTACATTGGCGACCCTGCTCAATTACCTGCACCACGCACATTGCTAACTAAGGGCA   17482
gb:MT044257|Organism:Severe      ATGTGTACATTGGCGACCCTGCTCAATTACCTGCACCACGCACATTGCTAACTAAGGGCA   17482
gb:MT106054|Organism:Severe      ATGTGTACATTGGCGACCCTGCTCAATTACCTGCACCACGCACATTGCTAACTAAGGGCA   17482
gb:MT049951|Organism:Severe      ATGTGTACATTGGCGACCCTGCTCAATTACCTGCACCACGCACATTGCTAACTAAGGGCA   17482
gb:MN975262|Organism:Severe      ATGTGTACATTGGCGACCCTGCTCAATTACCTGCACCACGCACATTGCTAACTAAGGGCA   17482
gb:MT106052|Organism:Severe      ATGTGTACATTGGCGACCCTGCTCAATTACCTGCACCACGCACATTGCTAACTAAGGGCA   17482
gb:LC522975|Organism:Severe      ATGTGTACATTGGCGACCCTGCTCAATTACCTGCACCACGCACATTGCTAACTAAGGGCA   17479
gb:LC522973|Organism:Severe      ATGTGTACATTGGCGACCCTGCTCAATTACCTGCACCACGCACATTGCTAACTAAGGGCA   17479
gb:LC522974|Organism:Severe      ATGTGTACATTGGCGACCCTGCTCAATTACCTGCACCACGCACATTGCTAACTAAGGGCA   17479
gb:MN985325|Organism:Severe      ATGTGTACATTGGCGACCCTGCTCAATTACCTGCACCACGCACATTGCTAACTAAGGGCA   17482
gb:MT020881|Organism:Severe      ATGTGTACATTGGCGACCCTGCTCAATTACCTGCACCACGCACATTGCTAACTAAGGGCA   17482
gb:MT020880|Organism:Severe      ATGTGTACATTGGCGACCCTGCTCAATTACCTGCACCACGCACATTGCTAACTAAGGGCA   17482
gb:MT066175|Organism:Severe      ATGTGTACATTGGCGACCCTGCTCAATTACCTGCACCACGCACATTGCTAACTAAGGGCA   17482
gb:MN997409|Organism:Severe      ATGTGTACATTGGCGACCCTGCTCAATTACCTGCACCACGCACATTGCTAACTAAGGGCA   17482
gb:MN938384|Organism:Severe      ATGTGTACATTGGCGACCCTGCTCAATTACCTGCACCACGCACATTGCTAACTAAGGGCA   17450
gb:MT044258|Organism:Severe      ATGTGTACATTGGCGACCCTGCTCAATTACCTGCACCACGCACATTGCTAACTAAGGGCA   17458
gb:MT039890|Organism:Severe      ATGTGTACATTGGCGACCCTGCTCAATTACCTGCACCACGCACATTGCTAACTAAGGGCA   17482
gb:MN988713|Organism:Severe      ATGTGTACATTGGCGACCCTGCTCAATTACCTGCACCACGCACATTGCTAACTAAGGGCA   17482
gb:LC521925|Organism:Severe      ATGTGTACATTGGCGACCCTGCTCAATTACCTGCACCACGCACATTGCTAACTAAGGGCA   17455
gb:MT093571|Organism:Severe      ATGTGTACATTGGCGACCCTGCTCAATTACCTGCACCACGCACATTGCTAACTAAGGGCA   17482
gb:MT039887|Organism:Severe      ATGTGTACATTGGCGACCCTGCTCAATTACCTGCACCACGCACATTGCTAACTAAGGGCA   17482
gb:MT019530|Organism:Severe      ATGTGTACATTGGCGACCCTGCTCAATTACCTGCACCACGCACATTGCTAACTAAGGGCA   17482
gb:MT039888|Organism:Severe      GTGTGTACATTGGCGACCCTGCTCAATTACCTGCACCACGCACATTGCTAACTAAGGGCA   17482
gb:LC522972|Organism:Severe      ATGTGTACATTGGCGACCCTGCTCAATTACCTGCACCACGCACATTGCTAACTAAGGGCA   17479
gb:MT027063|Organism:Severe      ATGTGTACATTGGCGACCCTGCTCAATTACCTGCACCACGCACATTGCTAACTAAGGGCA   17482
gb:MT027062|Organism:Severe      ATGTGTACATTGGCGACCCTGCTCAATTACCTGCACCACGCACATTGCTAACTAAGGGCA   17482
gb:MT019529|Organism:Severe      ATGTGTACATTGGCGACCCTGCTCAATTACCTGCACCACGCACATTGCTAACTAAGGGCA   17482
gb:MN996529|Organism:Severe      ATGTGTACATTGGCGACCCTGCTCAATTACCTGCACCACGCACATTGCTAACTAAGGGCA   17470
gb:MN996531|Organism:Severe      ATGTGTACATTGGCGACCCTGCTCAATTACCTGCACCACGCACATTGCTAACTAAGGGCA   17469
gb:MT066176|Organism:Severe      ATGTGTACATTGGCGACCCTGCTCAATTACCTGCACCACGCACATTGCTAACTAAGGGCA   17482
gb:MT027064|Organism:Severe      ATGTGTACATTGGCGACCCTGCTCAATTACCTGCACCACGCACATTGCTAACTAAGGGCA   17482
gb:MN994468|Organism:Severe      ATGTGTACATTGGCGACCCTGCTCAATTACCTGCACCACGCACATTGCTAACTAAGGGCA   17482
gb:MT072688|Organism:Severe      ATGTGTACATTGGCGACCCTGCTCAATTACCTGCACCACGCACATTGCTAACTAAGGGCA   17467
gb:MN996527|Organism:Severe      ATGTGTACATTGGCGACCCTGCTCAATTACCTGCACCACGCACATTGCTAACTAAGGGCA   17449
gb:MT093631|Organism:Severe      ATGTGTACATTGGCGACCCTGCTCAATTACCTGCACCACGCACATTGCTAACTAAGGGCA   17520
gb:MT106053|Organism:Severe      ATGTGTACATTGGCGACCCTGCTCAATTACCTGCACCACGCACATTGCTAACTAAGGGCA   17482
gb:MT019533|Organism:Severe      ATGTGTACATTGGCGACCCTGCTCAATTACCTGCACCACGCACATTGCTAACTAAGGGCA   17482
gb:MT019531|Organism:Severe      ATGTGTACATTGGCGACCCTGCTCAATTACCTGCACCACGCACATTGCTAACTAAGGGCA   17482
gb:MN996528|Organism:Severe      ATGTGTACATTGGCGACCCTGCTCAATTACCTGCACCACGCACATTGCTAACTAAGGGCA   17482
gb:MN996530|Organism:Severe      ATGTGTACATTGGCGACCCTGCTCAATTACCTGCACCACGCACATTGCTAACTAAGGGCA   17468
gb:MN908947|Organism:Severe      ATGTGTACATTGGCGACCCTGCTCAATTACCTGCACCACGCACATTGCTAACTAAGGGCA   17482
gb:MT019532|Organism:Severe      ATGTGTACATTGGCGACCCTGCTCAATTACCTGCACCACGCACATTGCTAACTAAGGGCA   17482
                                  ***********************************************************

gb:MT020781|Organism:Severe      CACTAGAACCAGAATATTTCAATTCAGTGTGTAGACTTATGAAAACTATAGGTCCAGACA   17530
gb:MT007544|Organism:Severe      CACTAGAACCAGAATATTTCAATTCAGTGTGTAGACTTATGAAAACTATAGGTCCAGACA   17542
gb:MN994467|Organism:Severe      CACTAGAACCAGAATATTTCAATTCAGTGTGTAGACTTATGAAAACTATAGGTCCAGACA   17542
gb:MT044257|Organism:Severe      CACTAGAACCAGAATATTTCAATTCAGTGTGTAGACTTATGAAAACTATAGGTCCAGACA   17542
gb:MT106054|Organism:Severe      CACTAGAACCAGAATATTTCAATTCAGTGTGTAGACTTATGAAAACTATAGGTCCAGACA   17542
gb:MT049951|Organism:Severe      CACTAGAACCAGAATATTTCAATTCAGTGTGTAGACTTATGAAAACTATAGGTCCAGACA   17542
gb:MN975262|Organism:Severe      CACTAGAACCAGAATATTTCAATTCAGTGTGTAGACTTATGAAAACTATAGGTCCAGACA   17542
gb:MT106052|Organism:Severe      CACTAGAACCAGAATATTTCAATTCAGTGTGTAGACTTATGAAAACTATAGGTCCAGACA   17542
gb:LC522975|Organism:Severe      CACTAGAACCAGAATATTTCAATTCAGTGTGTAGACTTATGAAAACTATAGGTCCAGACA   17539
gb:LC522973|Organism:Severe      CACTAGAACCAGAATATTTCAATTCAGTGTGTAGACTTATGAAAACTATAGGTCCAGACA   17539
gb:LC522974|Organism:Severe      CACTAGAACCAGAATATTTCAATTCAGTGTGTAGACTTATGAAAACTATAGGTCCAGACA   17539
gb:MN985325|Organism:Severe      CACTAGAACCAGAATATTTCAATTCAGTGTGTAGACTTATGAAAACTATAGGTCCAGACA   17542
gb:MT020881|Organism:Severe      CACTAGAACCAGAATATTTCAATTCAGTGTGTAGACTTATGAAAACTATAGGTCCAGACA   17542
gb:MT020880|Organism:Severe      CACTAGAACCAGAATATTTCAATTCAGTGTGTAGACTTATGAAAACTATAGGTCCAGACA   17542
gb:MT066175|Organism:Severe      CACTAGAACCAGAATATTTCAATTCAGTGTGTAGACTTATGAAAACTATAGGTCCAGACA   17542
gb:MN997409|Organism:Severe      CACTAGAACCAGAATATTTCAATTCAGTGTGTAGACTTATGAAAACTATAGGTCCAGACA   17542
gb:MN938384|Organism:Severe      CACTAGAACCAGAATATTTCAATTCAGTGTGTAGACTTATGAAAACTATAGGTCCAGACA   17510
gb:MT044258|Organism:Severe      CACTAGAACCAGAATATTTCAATTCAGTGTGTAGACTTATGAAAACTATAGGTCCAGACA   17518
gb:MT039890|Organism:Severe      CACTAGAACCAGAATATTTCAATTCAGTGTGTAGACTTATGAAAACTATAGGTCCAGACA   17542
```

```
gb:MN988713|Organism:Severe      CACTAGAACCAGAATATTTCAATTCAGTGTGTAGACTTATGAAAACTATAGGTCCAGACA   17542
gb:LC521925|Organism:Severe      CACTAGAACCAGAATATTTCAATTCAGTGTGTAGACTTATGAAAACTATAGGTCCAGACA   17515
gb:MT093571|Organism:Severe      CACTAGAACCAGAATATTTCAATTCAGTGTGTAGACTTATGAAAACTATAGGTCCAGACA   17542
gb:MT039887|Organism:Severe      CACTAGAACCAGAATATTTCAATTCAGTGTGTAGACTTATGAAAACTATAGGTCCAGACA   17542
gb:MT019530|Organism:Severe      CACTAGAACCAGAATATTTCAATTCAGTGTGTAGACTTATGAAAACTATAGGTCCAGACA   17542
gb:MT039888|Organism:Severe      CACTAGAACCAGAATATTTCAATTCAGTGTGTAGACTTATGAAAACTATAGGTCCAGACA   17542
gb:LC522972|Organism:Severe      CACTAGAACCAGAATATTTCAATTCAGTGTGTAGACTTATGAAAACTATAGGTCCAGACA   17539
gb:MT027063|Organism:Severe      CACTAGAACCAGAATATTTCAATTCAGTGTGTAGACTTATGAAAACTATAGGTCCAGACA   17542
gb:MT027062|Organism:Severe      CACTAGAACCAGAATATTTCAATTCAGTGTGTAGACTTATGAAAACTATAGGTCCAGACA   17542
gb:MT019529|Organism:Severe      CACTAGAACCAGAATATTTCAATTCAGTGTGTAGACTTATGAAAACTATAGGTCCAGACA   17542
gb:MN996529|Organism:Severe      CACTAGAACCAGAATATTTCAATTCAGTGTGTAGACTTATGAAAACTATAGGTCCAGACA   17530
gb:MN996531|Organism:Severe      CACTAGAACCAGAATATTTCAATTCAGTGTGTAGACTTATGAAAACTATAGGTCCAGACA   17529
gb:MT066176|Organism:Severe      CACTAGAACCAGAATATTTCAATTCAGTGTGTAGACTTATGAAAACTATAGGTCCAGACA   17542
gb:MT027064|Organism:Severe      CACTAGAACCAGAATATTTCAATTCAGTGTGTAGACTTATGAAAACTATAGGTCCAGACA   17542
gb:MN994468|Organism:Severe      CACTAGAACCAGAATATTTCAATTCAGTGTGTAGACTTATGAAAACTATAGGTCCAGACA   17542
gb:MT072688|Organism:Severe      CACTAGAACCAGAATATTTCAATTCAGTGTGTAGACTTATGAAAACTATAGGTCCAGACA   17527
gb:MN996527|Organism:Severe      CACTAGAACCAGAATATTTCAATTCAGTGTGTAGACTTATGAAAACTATAGGTCCAGACA   17509
gb:MT093631|Organism:Severe      CACTAGAACCAGAATATTTCAATTCAGTGTGTAGACTTATGAAAACTATAGGTCCAGACA   17580
gb:MT106053|Organism:Severe      CACTAGAACCAGAATATTTCAATTCAGTGTGTAGACTTATGAAAACTATAGGTCCAGACA   17542
gb:MT019533|Organism:Severe      CACTAGAACCAGAATATTTCAATTCAGTGTGTAGACTTATGAAAACTATAGGTCCAGACA   17542
gb:MT019531|Organism:Severe      CACTAGAACCAGAATATTTCAATTCAGTGTGTAGACTTATGAAAACTATAGGTCCAGACA   17542
gb:MN996528|Organism:Severe      CACTAGAACCAGAATATTTCAATTCAGTGTGTAGACTTATGAAAACTATAGGTCCAGACA   17542
gb:MN996530|Organism:Severe      CACTAGAACCAGAATATTTCAATTCAGTGTGTAGACTTATGAAAACTATAGGTCCAGACA   17528
gb:MN908947|Organism:Severe      CACTAGAACCAGAATATTTCAATTCAGTGTGTAGACTTATGAAAACTATAGGTCCAGACA   17542
gb:MT019532|Organism:Severe      CACTAGAACCAGAATATTTCAATTCAGTGTGTAGACTTATGAAAACTATAGGTCCAGACA   17542
                                 ************************************************************

gb:MT020781|Organism:Severe      TGTTCCTCGGAACTTGTCGGCGTTGTCCTGCTGAAATTGTTGACACTGTGAGTGCTTTGG   17590
gb:MT007544|Organism:Severe      TGTTCCTCGGAACTTGTCGGCGTTGTCCTGCTGAAATTGTTGACACTGTGAGTGCTTTGG   17602
gb:MN994467|Organism:Severe      TGTTCCTCGGAACTTGTCGGCGTTGTCCTGCTGAAATTGTTGACACTGTGAGTGCTTTGG   17602
gb:MT044257|Organism:Severe      TGTTCCTCGGAACTTGTCGGCGTTGTCCTGCTGAAATTGTTGACACTGTGAGTGCTTTGG   17602
gb:MT106054|Organism:Severe      TGTTCCTCGGAACTTGTCGGCGTTGTCCTGCTGAAATTGTTGACACTGTGAGTGCTTTGG   17602
gb:MT049951|Organism:Severe      TGTTCCTCGGAACTTGTCGGCGTTGTCCTGCTGAAATTGTTGACACTGTGAGTGCTTTGG   17602
gb:MN975262|Organism:Severe      TGTTCCTCGGAACTTGTCGGCGTTGTCCTGCTGAAATTGTTGACACTGTGAGTGCTTTGG   17602
gb:MT106052|Organism:Severe      TGTTCCTCGGAACTTGTCGGCGTTGTCCTGCTGAAATTGTTGACACTGTGAGTGCTTTGG   17602
gb:LC522975|Organism:Severe      TGTTCCTCGGAACTTGTCGGCGTTGTCCTGCTGAAATTGTTGACACTGTGAGTGCTTTGG   17599
gb:LC522973|Organism:Severe      TGTTCCTCGGAACTTGTCGGCGTTGTCCTGCTGAAATTGTTGACACTGTGAGTGCTTTGG   17599
gb:LC522974|Organism:Severe      TGTTCCTCGGAACTTGTCGGCGTTGTCCTGCTGAAATTGTTGACACTGTGAGTGCTTTGG   17599
gb:MN985325|Organism:Severe      TGTTCCTCGGAACTTGTCGGCGTTGTCCTGCTGAAATTGTTGACACTGTGAGTGCTTTGG   17602
gb:MT020881|Organism:Severe      TGTTCCTCGGAACTTGTCGGCGTTGTCCTGCTGAAATTGTTGACACTGTGAGTGCTTTGG   17602
gb:MT020880|Organism:Severe      TGTTCCTCGGAACTTGTCGGCGTTGTCCTGCTGAAATTGTTGACACTGTGAGTGCTTTGG   17602
gb:MT066175|Organism:Severe      TGTTCCTCGGAACTTGTCGGCGTTGTCCTGCTGAAATTGTTGACACTGTGAGTGCTTTGG   17602
gb:MN997409|Organism:Severe      TGTTCCTCGGAACTTGTCGGCGTTGTCCTGCTGAAATTGTTGACACTGTGAGTGCTTTGG   17602
gb:MN938384|Organism:Severe      TGTTCCTCGGAACTTGTCGGCGTTGTCCTGCTGAAATTGTTGACACTGTGAGTGCTTTGG   17570
gb:MT044258|Organism:Severe      TGTTCCTCGGAACTTGTCGGCGTTGTCCTGCTGAAATTGTTGACACTGTGAGTGCTTTGG   17578
gb:MT039890|Organism:Severe      TGTTCCTCGGAACTTGTCGGCGTTGTCCTGCTGAAATTGTTGACACTGTGAGTGCTTTGG   17602
gb:MN988713|Organism:Severe      TGTTCCTCGGAACTTGTCGGCGTTGTCCTGCTGAAATTGTTGACACTGTGAGTGCTTTGG   17602
gb:LC521925|Organism:Severe      TGTTCCTCGGAACTTGTCGGCGTTGTCCTGCTGAAATTGTTGACACTGTGAGTGCTTTGG   17575
gb:MT093571|Organism:Severe      TGTTCCTCGGAACTTGTCGGCGTTGTCCTGCTGAAATTGTTGACACTGTGAGTGCTTTGG   17602
gb:MT039887|Organism:Severe      TGTTCCTCGGAACTTGTCGGCGTTGTCCTGCTGAAATTGTTGACACTGTGAGTGCTTTGG   17602
gb:MT019530|Organism:Severe      TGTTCCTCGGAACTTGTCGGCGTTGTCCTGCTGAAATTGTTGACACTGTGAGTGCTTTGG   17602
gb:MT039888|Organism:Severe      TGTTCCTCGGAACTTGTCGGCGTTGTCCTGCTGAAATTGTTGACACTGTGAGTGCTTTGG   17602
gb:LC522972|Organism:Severe      TGTTCCTCGGAACTTGTCGGCGTTGTCCTGCTGAAATTGTTGACACTGTGAGTGCTTTGG   17599
gb:MT027063|Organism:Severe      TGTTCCTCGGAACTTGTCGGCGTTGTCCTGCTGAAATTGTTGACACTGTGAGTGCTTTGG   17602
gb:MT027062|Organism:Severe      TGTTCCTCGGAACTTGTCGGCGTTGTCCTGCTGAAATTGTTGACACTGTGAGTGCTTTGG   17602
gb:MT019529|Organism:Severe      TGTTCCTCGGAACTTGTCGGCGTTGTCCTGCTGAAATTGTTGACACTGTGAGTGCTTTGG   17602
gb:MN996529|Organism:Severe      TGTTCCTCGGAACTTGTCGGCGTTGTCCTGCTGAAATTGTTGACACTGTGAGTGCTTTGG   17590
gb:MN996531|Organism:Severe      TGTTCCTCGGAACTTGTCGGCGTTGTCCTGCTGAAATTGTTGACACTGTGAGTGCTTTGG   17589
gb:MT066176|Organism:Severe      TGTTCCTCGGAACTTGTCGGCGTTGTCCTGCTGAAATTGTTGACACTGTGAGTGCTTTGG   17602
gb:MT027064|Organism:Severe      TGTTCCTCGGAACTTGTCGGCGTTGTCCTGCTGAAATTGTTGACACTGTGAGTGCTTTGG   17602
gb:MN994468|Organism:Severe      TGTTCCTCGGAACTTGTCGGCGTTGTCCTGCTGAAATTGTTGACACTGTGAGTGCTTTGG   17602
gb:MT072688|Organism:Severe      TGTTCCTCGGAACTTGTCGGCGTTGTCCTGCTGAAATTGTTGACACTGTGAGTGCTTTGG   17587
gb:MN996527|Organism:Severe      TGTTCCTCGGAACTTGTCGGCGTTGTCCTGCTGAAATTGTTGACACTGTGAGTGCTTTGG   17569
gb:MT093631|Organism:Severe      TGTTCCTCGGAACTTGTCGGCGTTGTCCTGCTGAAATTGTTGACACTGTGAGTGCTTTGG   17640
gb:MT106053|Organism:Severe      TGTTCCTCGGAACTTGTCGGCGTTGTCCTGCTGAAATTGTTGACACTGTGAGTGCTTTGG   17602
```

```
gb:MT019533|Organism:Severe      TGTTCCTCGGAACTTGTCGGCGTTGTCCTGCTGAAATTGTTGACACTGTGAGTGCTTTGG   17602
gb:MT019531|Organism:Severe      TGTTCCTCGGAACTTGTCGGCGTTGTCCTGCTGAAATTGTTGACACTGTGAGTGCTTTGG   17602
gb:MN996528|Organism:Severe      TGTTCCTCGGAACTTGTCGGCGTTGTCCTGCTGAAATTGTTGACACTGTGAGTGCTTTGG   17602
gb:MN996530|Organism:Severe      TGTTCCTCGGAACTTGTCGGCGTTGTCCTGCTGAAATTGTTGACACTGTGAGTGCTTTGG   17588
gb:MN908947|Organism:Severe      TGTTCCTCGGAACTTGTCGGCGTTGTCCTGCTGAAATTGTTGACACTGTGAGTGCTTTGG   17602
gb:MT019532|Organism:Severe      TGTTCCTCGGAACTTGTCGGCGTTGTCCTGCTGAAATTGTTGACACTGTGAGTGCTTTGG   17602
                                 ************************************************************

gb:MT020781|Organism:Severe      TTTATGATAATAAGCTTAAAGCACATAAAGACAAATCAGCTCAATGCTTTAAAATGTTTT   17650
gb:MT007544|Organism:Severe      TTTATGATAATAAGCTTAAAGCACATAAAGACAAATCAGCTCAATGCTTTAAAATGTTTT   17662
gb:MN994467|Organism:Severe      TTTATGATAATAAGCTTAAAGCACATAAAGACAAATCAGCTCAATGCTTTAAAATGTTTT   17662
gb:MT044257|Organism:Severe      TTTATGATAATAAGCTTAAAGCACATAAAGACAAATCAGCTCAATGCTTTAAAATGTTTT   17662
gb:MT106054|Organism:Severe      TTTATGATAATAAGCTTAAAGCACATAAAGACAAATCAGCTCAATGCTTTAAAATGTTTT   17662
gb:MT049951|Organism:Severe      TTTATGATAATAAGCTTAAAGCACATAAAGACAAATCAGCTCAATGCTTTAAAATGTTTT   17662
gb:MN975262|Organism:Severe      TTTATGATAATAAGCTTAAAGCACATAAAGACAAATCAGCTCAATGCTTTAAAATGTTTT   17662
gb:MT106052|Organism:Severe      TTTATGATAATAAGCTTAAAGCACATAAAGACAAATCAGCTCAATGCTTTAAAATGTTTT   17662
gb:LC522975|Organism:Severe      TTTATGATAATAAGCTTAAAGCACATAAAGACAAATCAGCTCAATGCTTTAAAATGTTTT   17659
gb:LC522973|Organism:Severe      TTTATGATAATAAGCTTAAAGCACATAAAGACAAATCAGCTCAATGCTTTAAAATGTTTT   17659
gb:LC522974|Organism:Severe      TTTATGATAATAAGCTTAAAGCACATAAAGACAAATCAGCTCAATGCTTTAAAATGTTTT   17659
gb:MN985325|Organism:Severe      TTTATGATAATAAGCTTAAAGCACATAAAGACAAATCAGCTCAATGCTTTAAAATGTTTT   17662
gb:MT020881|Organism:Severe      TTTATGATAATAAGCTTAAAGCACATAAAGACAAATCAGCTCAATGCTTTAAAATGTTTT   17662
gb:MT020880|Organism:Severe      TTTATGATAATAAGCTTAAAGCACATAAAGACAAATCAGCTCAATGCTTTAAAATGTTTT   17662
gb:MT066175|Organism:Severe      TTTATGATAATAAGCTTAAAGCACATAAAGACAAATCAGCTCAATGCTTTAAAATGTTTT   17662
gb:MN997409|Organism:Severe      TTTATGATAATAAGCTTAAAGCACATAAAGACAAATCAGCTCAATGCTTTAAAATGTTTT   17662
gb:MN938384|Organism:Severe      TTTATGATAATAAGCTTAAAGCACATAAAGACAAATCAGCTCAATGCTTTAAAATGTTTT   17630
gb:MT044258|Organism:Severe      TTTATGATAATAAGCTTAAAGCACATAAAGACAAATCAGCTCAATGCTTTAAAATGTTTT   17638
gb:MT039890|Organism:Severe      TTTATGATAATAAGCTTAAAGCACATAAAGACAAATCAGCTCAATGCTTTAAAATGTTTT   17662
gb:MN988713|Organism:Severe      TTTATGATAATAAGCTTAAAGCACATAAAGACAAATCAGCTCAATGCTTTAAAATGTTTT   17662
gb:LC521925|Organism:Severe      TTTATGATAATAAGCTTAAAGCACATAAAGACAAATCAGCTCAATGCTTTAAAATGTTTT   17635
gb:MT093571|Organism:Severe      TTTATGATAATAAGCTTAAAGCACATAAAGACAAATCAGCTCAATGCTTTAAAATGTTTT   17662
gb:MT039887|Organism:Severe      TTTATGATAATAAGCTTAAAGCACATAAAGACAAATCAGCTCAATGCTTTAAAATGTTTT   17662
gb:MT019530|Organism:Severe      TTTATGATAATAAGCTTAAAGCACATAAAGACAAATCAGCTCAATGCTTTAAAATGTTTT   17662
gb:MT039888|Organism:Severe      TTTATGATAATAAGCTTAAAGCACATAAAGACAAATCAGCTCAATGCTTTAAAATGTTTT   17662
gb:LC522972|Organism:Severe      TTTATGATAATAAGCTTAAAGCACATAAAGACAAATCAGCTCAATGCTTTAAAATGTTTT   17659
gb:MT027063|Organism:Severe      TTTATGATAATAAGCTTAAAGCACATAAAGACAAATCAGCTCAATGCTTTAAAATGTTTT   17662
gb:MT027062|Organism:Severe      TTTATGATAATAAGCTTAAAGCACATAAAGACAAATCAGCTCAATGCTTTAAAATGTTTT   17662
gb:MT019529|Organism:Severe      TTTATGATAATAAGCTTAAAGCACATAAAGACAAATCAGCTCAATGCTTTAAAATGTTTT   17662
gb:MN996529|Organism:Severe      TTTATGATAATAAGCTTAAAGCACATAAAGACAAATCAGCTCAATGCTTTAAAATGTTTT   17650
gb:MN996531|Organism:Severe      TTTATGATAATAAGCTTAAAGCACATAAAGACAAATCAGCTCAATGCTTTAAAATGTTTT   17649
gb:MT066176|Organism:Severe      TTTATGATAATAAGCTTAAAGCACATAAAGACAAATCAGCTCAATGCTTTAAAATGTTTT   17662
gb:MT027064|Organism:Severe      TTTATGATAATAAGCTTAAAGCACATAAAGACAAATCAGCTCAATGCTTTAAAATGTTTT   17662
gb:MN994468|Organism:Severe      TTTATGATAATAAGCTTAAAGCACATAAAGACAAATCAGCTCAATGCTTTAAAATGTTTT   17662
gb:MT072688|Organism:Severe      TTTATGATAATAAGCTTAAAGCACATAAAGACAAATCAGCTCAATGCTTTAAAATGTTTT   17647
gb:MN996527|Organism:Severe      TTTATGATAATAAGCTTAAAGCACATAAAGACAAATCAGCTCAATGCTTTAAAATGTTTT   17629
gb:MT093631|Organism:Severe      TTTATGATAATAAGCTTAAAGCACATAAAGACAAATCAGCTCAATGCTTTAAAATGTTTT   17700
gb:MT106053|Organism:Severe      TTTATGATAATAAGCTTAAAGCACATAAAGACAAATCAGCTCAATGCTTTAAAATGTTTT   17662
gb:MT019533|Organism:Severe      TTTATGATAATAAGCTTAAAGCACATAAAGACAAATCAGCTCAATGCTTTAAAATGTTTT   17662
gb:MT019531|Organism:Severe      TTTATGATAATAAGCTTAAAGCACATAAAGACAAATCAGCTCAATGCTTTAAAATGTTTT   17662
gb:MN996528|Organism:Severe      TTTATGATAATAAGCTTAAAGCACATAAAGACAAATCAGCTCAATGCTTTAAAATGTTTT   17662
gb:MN996530|Organism:Severe      TTTATGATAATAAGCTTAAAGCACATAAAGACAAATCAGCTCAATGCTTTAAAATGTTTT   17648
gb:MN908947|Organism:Severe      TTTATGATAATAAGCTTAAAGCACATAAAGACAAATCAGCTCAATGCTTTAAAATGTTTT   17662
gb:MT019532|Organism:Severe      TTTATGATAATAAGCTTAAAGCACATAAAGACAAATCAGCTCAATGCTTTAAAATGTTTT   17662
                                 ************************************************************

gb:MT020781|Organism:Severe      ATAAGGGTGTTATCACGCATGATGTTTCATCTGCAATTAACAGGCCACAAATAGGCGTGG   17710
gb:MT007544|Organism:Severe      ATAAGGGTGTTATCACGCATGATGTTTCATCTGCAATTAACAGGCCACAAATAGGCGTGG   17722
gb:MN994467|Organism:Severe      ATAAGGGTGTTATCACGCATGATGTTTCATCTGCAATTAACAGGCCACAAATAGGCGTGG   17722
gb:MT044257|Organism:Severe      ATAAGGGTGTTATCACGCATGATGTTTCATCTGCAATTAACAGGCCACAAATAGGCGTGG   17722
gb:MT106054|Organism:Severe      ATAAGGGTGTTATCACGCATGATGTTTCATCTGCAATTAACAGGCCACAAATAGGCGTGG   17722
gb:MT049951|Organism:Severe      ATAAGGGTGTTATCACGCATGATGTTTCATCTGCAATTAACAGGCCACAAATAGGCGTGG   17722
gb:MN975262|Organism:Severe      ATAAGGGTGTTATCACGCATGATGTTTCATCTGCAATTAACAGGCCACAAATAGGCGTGG   17722
gb:MT106052|Organism:Severe      ATAAGGGTGTTATCACGCATGATGTTTCATCTGCAATTAACAGGCCACAAATAGGCGTGG   17722
gb:LC522975|Organism:Severe      ATAAGGGTGTTATCACGCATGATGTTTCATCTGCAATTAACAGGCCACAAATAGGCGTGG   17719
gb:LC522973|Organism:Severe      ATAAGGGTGTTATCACGCATGATGTTTCATCTGCAATTAACAGGCCACAAATAGGCGTGG   17719
gb:LC522974|Organism:Severe      ATAAGGGTGTTATCACGCATGATGTTTCATCTGCAATTAACAGGCCACAAATAGGCGTGG   17719
```

```
gb:MN985325|Organism:Severe      ATAAGGGTGTTATCACGCATGATGTTTCATCTGCAATTAACAGGCCACAAATAGGCGTGG   17722
gb:MT020881|Organism:Severe      ATAAGGGTGTTATCACGCATGATGTTTCATCTGCAATTAACAGGCCACAAATAGGCGTGG   17722
gb:MT020880|Organism:Severe      ATAAGGGTGTTATCACGCATGATGTTTCATCTGCAATTAACAGGCCACAAATAGGCGTGG   17722
gb:MT066175|Organism:Severe      ATAAGGGTGTTATCACGCATGATGTTTCATCTGCAATTAACAGGCCACAAATAGGCGTGG   17722
gb:MN997409|Organism:Severe      ATAAGGGTGTTATCACGCATGATGTTTCATCTGCAATTAACAGGCCACAAATAGGCGTGG   17722
gb:MN938384|Organism:Severe      ATAAGGGTGTTATCACGCATGATGTTTCATCTGCAATTAACAGGCCACAAATAGGCGTGG   17690
gb:MT044258|Organism:Severe      ATAAGGGTGTTATCACGCATGATGTTTCATCTGCAATTAACAGGCCACAAATAGGCGTGG   17698
gb:MT039890|Organism:Severe      ATAAGGGTGTTATCACGCATGATGTTTCATCTGCAATTAACAGGCCACAAATAGGCGTGG   17722
gb:MN988713|Organism:Severe      ATAAGGGTGTTATCACGCATGATGTTTCATCTGCAATTAACAGGCCACAAATAGGCGTGG   17722
gb:LC521925|Organism:Severe      ATAAGGGTGTTATCACGCATGATGTTTCATCTGCAATTAACAGGCCACAAATAGGCGTGG   17695
gb:MT093571|Organism:Severe      ATAAGGGTGTTATCACGCATGATGTTTCATCTGCAATTAACAGGCCACAAATAGGCGTGG   17722
gb:MT039887|Organism:Severe      ATAAGGGTGTTATCACGCATGATGTTTCATCTGCAATTAACAGGCCACAAATAGGCGTGG   17722
gb:MT019530|Organism:Severe      ATAAGGGTGTTATCACGCATGATGTTTCATCTGCAATTAACAGGCCACAAATAGGCGTGG   17722
gb:MT039888|Organism:Severe      ATAAGGGTGTTATCACGCATGATGTTTCATCTGCAATTAACAGGCCACAAATAGGCGTGG   17722
gb:LC522972|Organism:Severe      ATAAGGGTGTTATCACGCATGATGTTTCATCTGCAATTAACAGGCCACAAATAGGCGTGG   17719
gb:MT027063|Organism:Severe      ATAAGGGTGTTATCACGCATGATGTTTCATCTGCAATTAACAGGCCACAAATAGGCGTGG   17722
gb:MT027062|Organism:Severe      ATAAGGGTGTTATCACGCATGATGTTTCATCTGCAATTAACAGGCCACAAATAGGCGTGG   17722
gb:MT019529|Organism:Severe      ATAAGGGTGTTATCACGCATGATGTTTCATCTGCAATTAACAGGCCACAAATAGGCGTGG   17722
gb:MN996529|Organism:Severe      ATAAGGGTGTTATCACGCATGATGTTTCATCTGCAATTAACAGGCCACAAATAGGCGTGG   17710
gb:MN996531|Organism:Severe      ATAAGGGTGTTATCACGCATGATGTTTCATCTGCAATTAACAGGCCACAAATAGGCGTGG   17709
gb:MT066176|Organism:Severe      ATAAGGGTGTTATCACGCATGATGTTTCATCTGCAATTAACAGGCCACAAATAGGCGTGG   17722
gb:MT027064|Organism:Severe      ATAAGGGTGTTATCACGCATGATGTTTCATCTGCAATTAACAGGCCACAAATAGGCGTGG   17722
gb:MN994468|Organism:Severe      ATAAGGGTGTTATCACGCATGATGTTTCATCTGCAATTAACAGGCCACAAATAGGCGTGG   17722
gb:MT072688|Organism:Severe      ATAAGGGTGTTATCACGCATGATGTTTCATCTGCAATTAACAGGCCACAAATAGGCGTGG   17707
gb:MN996527|Organism:Severe      ATAAGGGTGTTATCACGCATGATGTTTCATCTGCAATTAACAGGCCACAAATAGGCGTGG   17689
gb:MT093631|Organism:Severe      ATAAGGGTGTTATCACGCATGATGTTTCATCTGCAATTAACAGGCCACAAATAGGCGTGG   17760
gb:MT106053|Organism:Severe      ATAAGGGTGTTATCACGCATGATGTTTCATCTGCAATTAACAGGCCACAAATAGGCGTGG   17722
gb:MT019533|Organism:Severe      ATAAGGGTGTTATCACGCATGATGTTTCATCTGCAATTAACAGGCCACAAATAGGCGTGG   17722
gb:MT019531|Organism:Severe      ATAAGGGTGTTATCACGCATGATGTTTCATCTGCAATTAACAGGCCACAAATAGGCGTGG   17722
gb:MN996528|Organism:Severe      ATAAGGGTGTTATCACGCATGATGTTTCATCTGCAATTAACAGGCCACAAATAGGCGTGG   17722
gb:MN996530|Organism:Severe      ATAAGGGTGTTATCACGCATGATGTTTCATCTGCAATTAACAGGCCACAAATAGGCGTGG   17708
gb:MN908947|Organism:Severe      ATAAGGGTGTTATCACGCATGATGTTTCATCTGCAATTAACAGGCCACAAATAGGCGTGG   17722
gb:MT019532|Organism:Severe      ATAAGGGTGTTATCACGCATGATGTTTCATCTGCAATTAACAGGCCACAAATAGGCGTGG   17722
                                 ************************************************************

gb:MT020781|Organism:Severe      TAAGAGAATTCCTTACACGTAACCCTGCTTGGAGAAAAGCTGTCTTTATTTCACCTTATA   17770
gb:MT007544|Organism:Severe      TAAGAGAATTCCTTACACGTAACCCTGCTTGGAGAAAAGCTGTCTTTATTTCACCTTATA   17782
gb:MN994467|Organism:Severe      TAAGAGAATTCCTTACACGTAACCCTGCTTGGAGAAAAGCTGTCTTTATTTCACCTTATA   17782
gb:MT044257|Organism:Severe      TAAGAGAATTCCTTACACGTAACCCTGCTTGGAGAAAAGCTGTCTTTATTTCACCTTATA   17782
gb:MT106054|Organism:Severe      TAAGAGAATTCCTTACACGTAACCCTGCTTGGAGAAAAGCTGTCTTTATTTCACCTTATA   17782
gb:MT049951|Organism:Severe      TAAGAGAATTCCTTACACGTAACCCTGCTTGGAGAAAAGCTGTCTTTATTTCACCTTATA   17782
gb:MN975262|Organism:Severe      TAAGAGAATTCCTTACACGTAACCCTGCTTGGAGAAAAGCTGTCTTTATTTCACCTTATA   17782
gb:MT106052|Organism:Severe      TAAGAGAATTCCTTACACGTAACCCTGCTTGGAGAAAAGCTGTCTTTATTTCACCTTATA   17782
gb:LC522975|Organism:Severe      TAAGAGAATTCCTTACACGTAACCCTGCTTGGAGAAAAGCTGTCTTTATTTCACCTTATA   17779
gb:LC522973|Organism:Severe      TAAGAGAATTCCTTACACGTAACCCTGCTTGGAGAAAAGCTGTCTTTATTTCACCTTATA   17779
gb:LC522974|Organism:Severe      TAAGAGAATTCCTTACACGTAACCCTGCTTGGAGAAAAGCTGTCTTTATTTCACCTTATA   17779
gb:MN985325|Organism:Severe      TAAGAGAATTCCTTACACGTAACCCTGCTTGGAGAAAAGCTGTCTTTATTTCACCTTATA   17782
gb:MT020881|Organism:Severe      TAAGAGAATTCCTTACACGTAACCCTGCTTGGAGAAAAGCTGTCTTTATTTCACCTTATA   17782
gb:MT020880|Organism:Severe      TAAGAGAATTCCTTACACGTAACCCTGCTTGGAGAAAAGCTGTCTTTATTTCACCTTATA   17782
gb:MT066175|Organism:Severe      TAAGAGAATTCCTTACACGTAACCCTGCTTGGAGAAAAGCTGTCTTTATTTCACCTTATA   17782
gb:MN997409|Organism:Severe      TAAGAGAATTCCTTACACGTAACCCTGCTTGGAGAAAAGCTGTCTTTATTTCACCTTATA   17782
gb:MN938384|Organism:Severe      TAAGAGAATTCCTTACACGTAACCCTGCTTGGAGAAAAGCTGTCTTTATTTCACCTTATA   17750
gb:MT044258|Organism:Severe      TAAGAGAATTCCTTACACGTAACCCTGCTTGGAGAAAAGCTGTCTTTATTTCACCTTATA   17758
gb:MT039890|Organism:Severe      TAAGAGAATTCCTTACACGTAACCCTGCTTGGAGAAAAGCTGTCTTTATTTCACCTTATA   17782
gb:MN988713|Organism:Severe      TAAGAGAATTCCTTACACGTAACCCTGCTTGGAGAAAAGCTGTCTTTATTTCACCTTATA   17782
gb:LC521925|Organism:Severe      TAAGAGAATTCCTTACACGTAACCCTGCTTGGAGAAAAGCTGTCTTTATTTCACCTTATA   17755
gb:MT093571|Organism:Severe      TAAGAGAATTCCTTACACGTAACCCTGCTTGGAGAAAAGCTGTCTTTATTTCACCTTATA   17782
gb:MT039887|Organism:Severe      TAAGAGAATTCCTTACACGTAACCCTGCTTGGAGAAAAGCTGTCTTTATTTCACCTTATA   17782
gb:MT019530|Organism:Severe      TAAGAGAATTCCTTACACGTAACCCTGCTTGGAGAAAAGCTGTCTTTATTTCACCTTATA   17782
gb:MT039888|Organism:Severe      TAAGAGAATTCCTTACACGTAACCCTGCTTGGAGAAAAGCTGTCTTTATTTCACCTTATA   17782
gb:LC522972|Organism:Severe      TAAGAGAATTCCTTACACGTAACCCTGCTTGGAGAAAAGCTGTCTTTATTTCACCTTATA   17779
gb:MT027063|Organism:Severe      TAAGAGAATTCCTTACACGTAACCCTGCTTGGAGAAAAGCTGTCTTTATTTCACCTTATA   17782
gb:MT027062|Organism:Severe      TAAGAGAATTCCTTACACGTAACCCTGCTTGGAGAAAAGCTGTCTTTATTTCACCTTATA   17782
gb:MT019529|Organism:Severe      TAAGAGAATTCCTTACACGTAACCCTGCTTGGAGAAAAGCTGTCTTTATTTCACCTTATA   17782
gb:MN996529|Organism:Severe      TAAGAGAATTCCTTACACGTAACCCTGCTTGGAGAAAAGCTGTCTTTATTTCACCTTATA   17770
```

```
gb:MN996531|Organism:Severe      TAAGAGAATTCCTTACACGTAACCCTGCTTGGAGAAAAGCTGTCTTTATTTCACCTTATA   17769
gb:MT066176|Organism:Severe      TAAGAGAATTCCTTACACGTAACCCTGCTTGGAGAAAAGCTGTCTTTATTTCACCTTATA   17782
gb:MT027064|Organism:Severe      TAAGAGAATTCCTTACACGTAACCCTGCTTGGAGAAAAGCTGTCTTTATTTCACCTTATA   17782
gb:MN994468|Organism:Severe      TAAGAGAATTCCTTACACGTAACCCTGCTTGGAGAAAAGCTGTCTTTATTTCACCTTATA   17782
gb:MT072688|Organism:Severe      TAAGAGAATTCCTTACACGTAACCCTGCTTGGAGAAAAGCTGTCTTTATTTCACCTTATA   17767
gb:MN996527|Organism:Severe      TAAGAGAATTCCTTACACGTAACCCTGCTTGGAGAAAAGCTGTCTTTATTTCACCTTATA   17749
gb:MT093631|Organism:Severe      TAAGAGAATTCCTTACACGTAACCCTGCTTGGAGAAAAGCTGTCTTTATTTCACCTTATA   17820
gb:MT106053|Organism:Severe      TAAGAGAATTCCTTACACGTAACCCTGCTTGGAGAAAAGCTGTCTTTATTTCACCTTATA   17782
gb:MT019533|Organism:Severe      TAAGAGAATTCCTTACACGTAACCCTGCTTGGAGAAAAGCTGTCTTTATTTCACCTTATA   17782
gb:MT019531|Organism:Severe      TAAGAGAATTCCTTACACGTAACCCTGCTTGGAGAAAAGCTGTCTTTATTTCACCTTATA   17782
gb:MN996528|Organism:Severe      TAAGAGAATTCCTTACACGTAACCCTGCTTGGAGAAAAGCTGTCTTTATTTCACCTTATA   17782
gb:MN996530|Organism:Severe      TAAGAGAATTCCTTACACGTAACCCTGCTTGGAGAAAAGCTGTCTTTATTTCACCTTATA   17768
gb:MN908947|Organism:Severe      TAAGAGAATTCCTTACACGTAACCCTGCTTGGAGAAAAGCTGTCTTTATTTCACCTTATA   17782
gb:MT019532|Organism:Severe      TAAGAGAATTCCTTACACGTAACCCTGCTTGGAGAAAAGCTGTCTTTATTTCACCTTATA   17782
                                 ************************************************************

gb:MT020781|Organism:Severe      ATTCACAGAATGCTGTAGCCTCAAAGATTTTGGGACTACCAACTCAAACTGTTGATTCAT   17830
gb:MT007544|Organism:Severe      ATTCACAGAATGCTGTAGCCTCAAAGATTTTGGGACTACCAACTCAAACTGTTGATTCAT   17842
gb:MN994467|Organism:Severe      ATTCACAGAATGCTGTAGCCTCAAAGATTTTGGGACTACCAACTCAAACTGTTGATTCAT   17842
gb:MT044257|Organism:Severe      ATTCACAGAATGCTGTAGCCTCAAAGATTTTGGGACTACCAACTCAAACTGTTGATTCAT   17842
gb:MT106054|Organism:Severe      ATTCACAGAATGCTGTAGCCTCAAAGATTTTGGGACTACCAACTCAAACTGTTGATTCAT   17842
gb:MT049951|Organism:Severe      ATTCACAGAATGCTGTAGCCTCAAAGATTTTGGGACTACCAACTCAAACTGTTGATTCAT   17842
gb:MN975262|Organism:Severe      ATTCACAGAATGCTGTAGCCTCAAAGATTTTGGGACTACCAACTCAAACTGTTGATTCAT   17842
gb:MT106052|Organism:Severe      ATTCACAGAATGCTGTAGCCTCAAAGATTTTGGGACTACCAACTCAAACTGTTGATTCAT   17842
gb:LC522975|Organism:Severe      ATTCACAGAATGCTGTAGCCTCAAAGATTTTGGGACTACCAACTCAAACTGTTGATTCAT   17839
gb:LC522973|Organism:Severe      ATTCACAGAATGCTGTAGCCTCAAAGATTTTGGGACTACCAACTCAAACTGTTGATTCAT   17839
gb:LC522974|Organism:Severe      ATTCACAGAATGCTGTAGCCTCAAAGATTTTGGGACTACCAACTCAAACTGTTGATTCAT   17839
gb:MN985325|Organism:Severe      ATTCACAGAATGCTGTAGCCTCAAAGATTTTGGGACTACCAACTCAAACTGTTGATTCAT   17842
gb:MT020881|Organism:Severe      ATTCACAGAATGCTGTAGCCTCAAAGATTTTGGGACTACCAACTCAAACTGTTGATTCAT   17842
gb:MT020880|Organism:Severe      ATTCACAGAATGCTGTAGCCTCAAAGATTTTGGGACTACCAACTCAAACTGTTGATTCAT   17842
gb:MT066175|Organism:Severe      ATTCACAGAATGCTGTAGCCTCAAAGATTTTGGGACTACCAACTCAAACTGTTGATTCAT   17842
gb:MN997409|Organism:Severe      ATTCACAGAATGCTGTAGCCTCAAAGATTTTGGGACTACCAACTCAAACTGTTGATTCAT   17842
gb:MN938384|Organism:Severe      ATTCACAGAATGCTGTAGCCTCAAAGATTTTGGGACTACCAACTCAAACTGTTGATTCAT   17810
gb:MT044258|Organism:Severe      ATTCACAGAATGCTGTAGCCTCAAAGATTTTGGGACTACCAACTCAAACTGTTGATTCAT   17818
gb:MT039890|Organism:Severe      ATTCACAGAATGCTGTAGCCTCAAAGATTTTGGGACTACCAACTCAAACTGTTGATTCAT   17842
gb:MN988713|Organism:Severe      ATTCACAGAATGCTGTAGCCTCAAAGATTTTGGGACTACCAACTCAAACTGTTGATTCAT   17842
gb:LC521925|Organism:Severe      ATTCACAGAATGCTGTAGCCTCAAAGATTTTGGGACTACCAACTCAAACTGTTGATTCAT   17815
gb:MT093571|Organism:Severe      ATTCACAGAATGCTGTAGCCTCAAAGATTTTGGGACTACCAACTCAAACTGTTGATTCAT   17842
gb:MT039887|Organism:Severe      ATTCACAGAATGCTGTAGCCTCAAAGATTTTGGGACTACCAACTCAAACTGTTGATTCAT   17842
gb:MT019530|Organism:Severe      ATTCACAGAATGCTGTAGCCTCAAAGATTTTGGGACTACCAACTCAAACTGTTGATTCAT   17842
gb:MT039888|Organism:Severe      ATTCACAGAATGCTGTAGCCTCAAAGATTTTGGGACTACCAACTCAAACTGTTGATTCAT   17842
gb:LC522972|Organism:Severe      ATTCACAGAATGCTGTAGCCTCAAAGATTTTGGGACTACCAACTCAAACTGTTGATTCAT   17839
gb:MT027063|Organism:Severe      ATTCACAGAATGCTGTAGCCTCAAAGATTTTGGGACTACCAACTCAAACTGTTGATTCAT   17842
gb:MT027062|Organism:Severe      ATTCACAGAATGCTGTAGCCTCAAAGATTTTGGGACTACCAACTCAAACTGTTGATTCAT   17842
gb:MT019529|Organism:Severe      ATTCACAGAATGCTGTAGCCTCAAAGATTTTGGGACTACCAACTCAAACTGTTGATTCAT   17842
gb:MN996529|Organism:Severe      ATTCACAGAATGCTGTAGCCTCAAAGATTTTGGGACTACCAACTCAAACTGTTGATTCAT   17830
gb:MN996531|Organism:Severe      ATTCACAGAATGCTGTAGCCTCAAAGATTTTGGGACTACCAACTCAAACTGTTGATTCAT   17829
gb:MT066176|Organism:Severe      ATTCACAGAATGCTGTAGCCTCAAAGATTTTGGGACTACCAACTCAAACTGTTGATTCAT   17842
gb:MT027064|Organism:Severe      ATTCACAGAATGCTGTAGCCTCAAAGATTTTGGGACTACCAACTCAAACTGTTGATTCAT   17842
gb:MN994468|Organism:Severe      ATTCACAGAATGCTGTAGCCTCAAAGATTTTGGGACTACCAACTCAAACTGTTGATTCAT   17842
gb:MT072688|Organism:Severe      ATTCACAGAATGCTGTAGCCTCAAAGATTTTGGGACTACCAACTCAAACTGTTGATTCAT   17827
gb:MN996527|Organism:Severe      ATTCACAGAATGCTGTAGCCTCAAAGATTTTGGGACTACCAACTCAAACTGTTGATTCAT   17809
gb:MT093631|Organism:Severe      ATTCACAGAATGCTGTAGCCTCAAAGATTTTGGGACTACCAACTCAAACTGTTGATTCAT   17880
gb:MT106053|Organism:Severe      ATTCACAGAATGCTGTAGCCTCAAAGATTTTGGGACTACCAACTCAAACTGTTGATTCAT   17842
gb:MT019533|Organism:Severe      ATTCACAGAATGCTGTAGCCTCAAAGATTTTGGGACTACCAACTCAAACTGTTGATTCAT   17842
gb:MT019531|Organism:Severe      ATTCACAGAATGCTGTAGCCTCAAAGATTTTGGGACTACCAACTCAAACTGTTGATTCAT   17842
gb:MN996528|Organism:Severe      ATTCACAGAATGCTGTAGCCTCAAAGATTTTGGGACTACCAACTCAAACTGTTGATTCAT   17842
gb:MN996530|Organism:Severe      ATTCACAGAATGCTGTAGCCTCAAAGATTTTGGGACTACCAACTCAAACTGTTGATTCAT   17828
gb:MN908947|Organism:Severe      ATTCACAGAATGCTGTAGCCTCAAAGATTTTGGGACTACCAACTCAAACTGTTGATTCAT   17842
gb:MT019532|Organism:Severe      ATTCACAGAATGCTGTAGCCTCAAAGATTTTGGGACTACCAACTCAAACTGTTGATTCAT   17842
                                 ************************************************************

gb:MT020781|Organism:Severe      CACAGGGCTCAGAATATGACTATGTCATATTCACTCAAACCACTGAAACAGCTCACTCTT   17890
gb:MT007544|Organism:Severe      CACAGGGCTCAGAATATGACTATGTCATATTCACTCAAACCACTGAAACAGCTCACTCTT   17902
gb:MN994467|Organism:Severe      CACAGGGCTCAGAATATGACTATGTCATATTCACTCAAACCACTGAAACAGCTCACTCTT   17902
```

```
gb:MT044257|Organism:Severe      CACAGGGCTCAGAATATGACTATGTCATATTCACTCAAACCACTGAAACAGCTCACTCTT    17902
gb:MT106054|Organism:Severe      CACAGGGCTCAGAATATGACTATGTCATATTCACTCAAACCACTGAAACAGCTCACTCTT    17902
gb:MT049951|Organism:Severe      CACAGGGCTCAGAATATGACTATGTCATATTCACTCAAACCACTGAAACAGCTCACTCTT    17902
gb:MN975262|Organism:Severe      CACAGGGCTCAGAATATGACTATGTCATATTCACTCAAACCACTGAAACAGCTCACTCTT    17902
gb:MT106052|Organism:Severe      CACAGGGCTCAGAATATGACTATGTCATATTCACTCAAACCACTGAAACAGCTCACTCTT    17902
gb:LC522975|Organism:Severe      CACAGGGCTCAGAATATGACTATGTCATATTCACTCAAACCACTGAAACAGCTCACTCTT    17899
gb:LC522973|Organism:Severe      CACAGGGCTCAGAATATGACTATGTCATATTCACTCAAACCACTGAAACAGCTCACTCTT    17899
gb:LC522974|Organism:Severe      CACAGGGCTCAGAATATGACTATGTCATATTCACTCAAACCACTGAAACAGCTCACTCTT    17899
gb:MN985325|Organism:Severe      CACAGGGCTCAGAATATGACTATGTCATATTCACTCAAACCACTGAAACAGCTCACTCTT    17902
gb:MT020881|Organism:Severe      CACAGGGCTCAGAATATGACTATGTCATATTCACTCAAACCACTGAAACAGCTCACTCTT    17902
gb:MT020880|Organism:Severe      CACAGGGCTCAGAATATGACTATGTCATATTCACTCAAACCACTGAAACAGCTCACTCTT    17902
gb:MT066175|Organism:Severe      CACAGGGCTCAGAATATGACTATGTCATATTCACTCAAACCACTGAAACAGCTCACTCTT    17902
gb:MN997409|Organism:Severe      CACAGGGCTCAGAATATGACTATGTCATATTCACTCAAACCACTGAAACAGCTCACTCTT    17902
gb:MN938384|Organism:Severe      CACAGGGCTCAGAATATGACTATGTCATATTCACTCAAACCACTGAAACAGCTCACTCTT    17870
gb:MT044258|Organism:Severe      CACAGGGCTCAGAATATGACTATGTCATATTCACTCAAACCACTGAAACAGCTCACTCTT    17878
gb:MT039890|Organism:Severe      CACAGGGCTCAGAATATGACTATGTCATATTCACTCAAACCACTGAAACAGCTCACTCTT    17902
gb:MN988713|Organism:Severe      CACAGGGCTCAGAATATGACTATGTCATATTCACTCAAACCACTGAAACAGCTCACTCTT    17902
gb:LC521925|Organism:Severe      CACAGGGCTCAGAATATGACTATGTCATATTCACTCAAACCACTGAAACAGCTCACTCTT    17875
gb:MT093571|Organism:Severe      CACAGGGCTCAGAATATGACTATGTCATATTCACTCAAACCACTGAAACAGCTCACTCTT    17902
gb:MT039887|Organism:Severe      CACAGGGCTCAGAATATGACTATGTCATATTCACTCAAACCACTGAAACAGCTCACTCTT    17902
gb:MT019530|Organism:Severe      CACAGGGCTCAGAATATGACTATGTCATATTCACTCAAACCACTGAAACAGCTCACTCTT    17902
gb:MT039888|Organism:Severe      CACAGGGCTCAGAATATGACTATGTCATATTCACTCAAACCACTGAAACAGCTCACTCTT    17902
gb:LC522972|Organism:Severe      CACAGGGCTCAGAATATGACTATGTCATATTCACTCAAACCACTGAAACAGCTCACTCTT    17899
gb:MT027063|Organism:Severe      CACAGGGCTCAGAATATGACTATGTCATATTCACTCAAACCACTGAAACAGCTCACTCTT    17902
gb:MT027062|Organism:Severe      CACAGGGCTCAGAATATGACTATGTCATATTCACTCAAACCACTGAAACAGCTCACTCTT    17902
gb:MT019529|Organism:Severe      CACAGGGCTCAGAATATGACTATGTCATATTCACTCAAACCACTGAAACAGCTCACTCTT    17902
gb:MN996529|Organism:Severe      CACAGGGCTCAGAATATGACTATGTCATATTCACTCAAACCACTGAAACAGCTCACTCTT    17890
gb:MN996531|Organism:Severe      CACAGGGCTCAGAATATGACTATGTCATATTCACTCAAACCACTGAAACAGCTCACTCTT    17889
gb:MT066176|Organism:Severe      CACAGGGCTCAGAATATGACTATGTCATATTCACTCAAACCACTGAAACAGCTCACTCTT    17902
gb:MT027064|Organism:Severe      CACAGGGCTCAGAATATGACTATGTCATATTCACTCAAACCACTGAAACAGCTCACTCTT    17902
gb:MN994468|Organism:Severe      CACAGGGCTCAGAATATGACTATGTCATATTCACTCAAACCACTGAAACAGCTCACTCTT    17902
gb:MT072688|Organism:Severe      CACAGGGCTCAGAATATGACTATGTCATATTCACTCAAACCACTGAAACAGCTCACTCTT    17887
gb:MN996527|Organism:Severe      CACAGGGCTCAGAATATGACTATGTCATATTCACTCAAACCACTGAAACAGCTCACTCTT    17869
gb:MT093631|Organism:Severe      CACAGGGCTCAGAATATGACTATGTCATATTCACTCAAACCACTGAAACAGCTCACTCTT    17940
gb:MT106053|Organism:Severe      CACAGGGCTCAGAATATGACTATGTCATATTCACTCAAACCACTGAAACAGCTCACTCTT    17902
gb:MT019533|Organism:Severe      CACAGGGCTCAGAATATGACTATGTCATATTCACTCAAACCACTGAAACAGCTCACTCTT    17902
gb:MT019531|Organism:Severe      CACAGGGCTCAGAATATGACTATGTCATATTCACTCAAACCACTGAAACAGCTCACTCTT    17902
gb:MN996528|Organism:Severe      CACAGGGCTCAGAATATGACTATGTCATATTCACTCAAACCACTGAAACAGCTCACTCTT    17902
gb:MN996530|Organism:Severe      CACAGGGCTCAGAATATGACTATGTCATATTCACTCAAACCACTGAAACAGCTCACTCTT    17888
gb:MN908947|Organism:Severe      CACAGGGCTCAGAATATGACTATGTCATATTCACTCAAACCACTGAAACAGCTCACTCTT    17902
gb:MT019532|Organism:Severe      CACAGGGCTCAGAATATGACTATGTCATATTCACTCAAACCACTGAAACAGCTCACTCTT    17902
                                 ************************************************************

gb:MT020781|Organism:Severe      GTAATGTAAACAGATTTAATGTTGCTATTACCAGAGCAAAAGTAGGCATACTTTGCATAA    17950
gb:MT007544|Organism:Severe      GTAATGTAAACAGATTTAATGTTGCTATTACCAGAGCAAAAGTAGGCATACTTTGCATAA    17962
gb:MN994467|Organism:Severe      GTAATGTAAACAGATTTAATGTTGCTATTACCAGAGCAAAAGTAGGCATACTTTGCATAA    17962
gb:MT044257|Organism:Severe      GTAATGTAAACAGATTTAATGTTGCTATTACCAGAGCAAAAGTAGGCATACTTTGCATAA    17962
gb:MT106054|Organism:Severe      GTAATGTAAACAGATTTAATGTTGCTATTACCAGAGCAAAAGTAGGCATACTTTGCATAA    17962
gb:MT049951|Organism:Severe      GTAATGTAAACAGATTTAATGTTGCTATTACCAGAGCAAAAGTAGGCATACTTTGCATAA    17962
gb:MN975262|Organism:Severe      GTAATGTAAACAGATTTAATGTTGCTATTACCAGAGCAAAAGTAGGCATACTTTGCATAA    17962
gb:MT106052|Organism:Severe      GTAATGTAAACAGATTTAATGTTGCTATTACCAGAGCAAAAGTAGGCATACTTTGCATAA    17962
gb:LC522975|Organism:Severe      GTAATGTAAACAGATTTAATGTTGCTATTACCAGAGCAAAAGTAGGCATACTTTGCATAA    17959
gb:LC522973|Organism:Severe      GTAATGTAAACAGATTTAATGTTGCTATTACCAGAGCAAAAGTAGGCATACTTTGCATAA    17959
gb:LC522974|Organism:Severe      GTAATGTAAACAGATTTAATGTTGCTATTACCAGAGCAAAAGTAGGCATACTTTGCATAA    17959
gb:MN985325|Organism:Severe      GTAATGTAAACAGATTTAATGTTGCTATTACCAGAGCAAAAGTAGGCATACTTTGCATAA    17962
gb:MT020881|Organism:Severe      GTAATGTAAACAGATTTAATGTTGCTATTACCAGAGCAAAAGTAGGCATACTTTGCATAA    17962
gb:MT020880|Organism:Severe      GTAATGTAAACAGATTTAATGTTGCTATTACCAGAGCAAAAGTAGGCATACTTTGCATAA    17962
gb:MT066175|Organism:Severe      GTAATGTAAACAGATTTAATGTTGCTATTACCAGAGCAAAAGTAGGCATACTTTGCATAA    17962
gb:MN997409|Organism:Severe      GTAATGTAAACAGATTTAATGTTGCTATTACCAGAGCAAAAGTAGGCATACTTTGCATAA    17962
gb:MN938384|Organism:Severe      GTAATGTAAACAGATTTAATGTTGCTATTACCAGAGCAAAAGTAGGCATACTTTGCATAA    17930
gb:MT044258|Organism:Severe      GTAATGTAAACAGATTTAATGTTGCTATTACCAGAGCAAAAGTAGGCATACTTTGCATAA    17938
gb:MT039890|Organism:Severe      GTAATGTAAACAGATTTAATGTTGCTATTACCAGAGCAAAAGTAGGCATACTTTGCATAA    17962
gb:MN988713|Organism:Severe      GTAATGTAAACAGATTTAATGTTGCTATTACCAGAGCAAAAGTAGGCATACTTTGCATAA    17962
gb:LC521925|Organism:Severe      GTAATGTAAACAGATTTAATGTTGCTATTACCAGAGCAAAAGTAGGCATACTTTGCATAA    17935
gb:MT093571|Organism:Severe      GTAATGTAAACAGATTTAATGTTGCTATTACCAGAGCAAAAGTAGGCATACTTTGCATAA    17962
```

```
gb:MT039887|Organism:Severe      GTAATGTAAACAGATTTAATGTTGCTATTACCAGAGCAAAAGTAGGCATACTTTGCATAA    17962
gb:MT019530|Organism:Severe      GTAATGTAAACAGATTTAATGTTGCTATTACCAGAGCAAAAGTAGGCATACTTTGCATAA    17962
gb:MT039888|Organism:Severe      GTAATGTAAACAGATTTAATGTTGCTATTACCAGAGCAAAAGTAGGCATACTTTGCATAA    17962
gb:LC522972|Organism:Severe      GTAATGTAAACAGATTTAATGTTGCTATTACCAGAGCAAAAGTAGGCATACTTTGCATAA    17959
gb:MT027063|Organism:Severe      GTAATGTAAACAGATTTAATGTTGCTATTACCAGAGCAAAAGTAGGCATACTTTGCATAA    17962
gb:MT027062|Organism:Severe      GTAATGTAAACAGATTTAATGTTGCTATTACCAGAGCAAAAGTAGGCATACTTTGCATAA    17962
gb:MT019529|Organism:Severe      GTAATGTAAACAGATTTAATGTTGCTATTACCAGAGCAAAAGTAGGCATACTTTGCATAA    17962
gb:MN996529|Organism:Severe      GTAATGTAAACAGATTTAATGTTGCTATTACCAGAGCAAAAGTAGGCATACTTTGCATAA    17950
gb:MN996531|Organism:Severe      GTAATGTAAACAGATTTAATGTTGCTATTACCAGAGCAAAAGTAGGCATACTTTGCATAA    17949
gb:MT066176|Organism:Severe      GTAATGTAAACAGATTTAATGTTGCTATTACCAGAGCAAAAGTAGGCATACTTTGCATAA    17962
gb:MT027064|Organism:Severe      GTAATGTAAACAGATTTAATGTTGCTATTACCAGAGCAAAAGTAGGCATACTTTGCATAA    17962
gb:MN994468|Organism:Severe      GTAATGTAAACAGATTTAATGTTGCTATTACCAGAGCAAAAGTAGGCATACTTTGCATAA    17962
gb:MT072688|Organism:Severe      GTAATGTAAACAGATTTAATGTTGCTATTACCAGAGCAAAAGTAGGCATACTTTGCATAA    17947
gb:MN996527|Organism:Severe      GTAATGTAAACAGATTTAATGTTGCTATTACCAGAGCAAAAGTAGGCATACTTTGCATAA    17929
gb:MT093631|Organism:Severe      GTAATGTAAACAGATTTAATGTTGCTATTACCAGAGCAAAAGTAGGCATACTTTGCATAA    18000
gb:MT106053|Organism:Severe      GTAATGTAAACAGATTTAATGTTGCTATTACCAGAGCAAAAGTAGGCATACTTTGCATAA    17962
gb:MT019533|Organism:Severe      GTAATGTAAACAGATTTAATGTTGCTATTACCAGAGCAAAAGTAGGCATACTTTGCATAA    17962
gb:MT019531|Organism:Severe      GTAATGTAAACAGATTTAATGTTGCTATTACCAGAGCAAAAGTAGGCATACTTTGCATAA    17962
gb:MN996528|Organism:Severe      GTAATGTAAACAGATTTAATGTTGCTATTACCAGAGCAAAAGTAGGCATACTTTGCATAA    17962
gb:MN996530|Organism:Severe      GTAATGTAAACAGATTTAATGTTGCTATTACCAGAGCAAAAGTAGGCATACTTTGCATAA    17948
gb:MN908947|Organism:Severe      GTAATGTAAACAGATTTAATGTTGCTATTACCAGAGCAAAAGTAGGCATACTTTGCATAA    17962
gb:MT019532|Organism:Severe      GTAATGTAAACAGATTTAATGTTGCTATTACCAGAGCAAAAGTAGGCATACTTTGCATAA    17962
                                 ************************************************************

gb:MT020781|Organism:Severe      TGTCTGATAGAGACCTTTATGACAAGTTGCAATTTACAAGTCTTGAAATTCCACGTAGGA    18010
gb:MT007544|Organism:Severe      TGTCTGATAGAGACCTTTATGACAAGTTGCAATTTACAAGTCTTGAAATTCCACGTAGGA    18022
gb:MN994467|Organism:Severe      TGTCTGATAGAGACCTTTATGACAAGTTGCAATTTACAAGTCTTGAAATTCCACGTAGGA    18022
gb:MT044257|Organism:Severe      TGTCTGATAGAGACCTTTATGACAAGTTGCAATTTACAAGTCTTGAAATTCCACGTAGGA    18022
gb:MT106054|Organism:Severe      TGTCTGATAGAGACCTTTATGACAAGTTGCAATTTACAAGTCTTGAAATTCCACGTAGGA    18022
gb:MT049951|Organism:Severe      TGTCTGATAGAGACCTTTATGACAAGTTGCAATTTACAAGTCTTGAAATTCCACGTAGGA    18022
gb:MN975262|Organism:Severe      TGTCTGATAGAGACCTTTATGACAAGTTGCAATTTACAAGTCTTGAAATTCCACGTAGGA    18022
gb:MT106052|Organism:Severe      TGTCTGATAGAGACCTTTATGACAAGTTGCAATTTACAAGTCTTGAAATTCCACGTAGGA    18022
gb:LC522975|Organism:Severe      TGTCTGATAGAGACCTTTATGACAAGTTGCAATTTACAAGTCTTGAAATTCCACGTAGGA    18019
gb:LC522973|Organism:Severe      TGTCTGATAGAGACCTTTATGACAAGTTGCAATTTACAAGTCTTGAAATTCCACGTAGGA    18019
gb:LC522974|Organism:Severe      TGTCTGATAGAGACCTTTATGACAAGTTGCAATTTACAAGTCTTGAAATTCCACGTAGGA    18019
gb:MN985325|Organism:Severe      TGTCTGATAGAGACCTTTATGACAAGTTGCAATTTACAAGTCTTGAAATTCCACGTAGGA    18022
gb:MT020881|Organism:Severe      TGTCTGATAGAGACCTTTATGACAAGTTGCAATTTACAAGTCTTGAAATTCCACGTAGGA    18022
gb:MT020880|Organism:Severe      TGTCTGATAGAGACCTTTATGACAAGTTGCAATTTACAAGTCTTGAAATTCCACGTAGGA    18022
gb:MT066175|Organism:Severe      TGTCTGATAGAGACCTTTATGACAAGTTGCAATTTACAAGTCTTGAAATTCCACGTAGGA    18022
gb:MN997409|Organism:Severe      TGTCTGATAGAGACCTTTATGACAAGTTGCAATTTACAAGTCTTGAAATTCCACGTAGGA    18022
gb:MN938384|Organism:Severe      TGTCTGATAGAGACCTTTATGACAAGTTGCAATTTACAAGTCTTGAAATTCCACGTAGGA    17990
gb:MT044258|Organism:Severe      TGTCTGATAGAGACCTTTATGACAAGTTGCAATTTACAAGTCTTGAAATTCCACGTAGGA    17998
gb:MT039890|Organism:Severe      TGTCTGATAGAGACCTTTATGACAAGTTGCAATTTACAAGTCTTGAAATTCCACGTAGGA    18022
gb:MN988713|Organism:Severe      TGTCTGATAGAGACCTTTATGACAAGTTGCAATTTACAAGTCTTGAAATTCCACGTAGGA    18022
gb:LC521925|Organism:Severe      TGTCTGATAGAGACCTTTATGACAAGTTGCAATTTACAAGTCTTGAAATTCCACGTAGGA    17995
gb:MT093571|Organism:Severe      TGTCTGATAGAGACCTTTATGACAAGTTGCAATTTACAAGTCTTGAAATTCCACGTAGGA    18022
gb:MT039887|Organism:Severe      TGTCTGATAGAGACCTTTATGACAAGTTGCAATTTACAAGTCTTGAAATTCCACGTAGGA    18022
gb:MT019530|Organism:Severe      TGTCTGATAGAGACCTTTATGACAAGTTGCAATTTACAAGTCTTGAAATTCCACGTAGGA    18022
gb:MT039888|Organism:Severe      TGTCTGATAGAGACCTTTATGACAAGTTGCAATTTACAAGTCTTGAAATTCCACGTAGGA    18022
gb:LC522972|Organism:Severe      TGTCTGATAGAGACCTTTATGACAAGTTGCAATTTACAAGTCTTGAAATTCCACGTAGGA    18019
gb:MT027063|Organism:Severe      TGTCTGATAGAGACCTTTATGACAAGTTGCAATTTACAAGTCTTGAAATTCCACGTAGGA    18022
gb:MT027062|Organism:Severe      TGTCTGATAGAGACCTTTATGACAAGTTGCAATTTACAAGTCTTGAAATTCCACGTAGGA    18022
gb:MT019529|Organism:Severe      TGTCTGATAGAGACCTTTATGACAAGTTGCAATTTACAAGTCTTGAAATTCCACGTAGGA    18022
gb:MN996529|Organism:Severe      TGTCTGATAGAGACCTTTATGACAAGTTGCAATTTACAAGTCTTGAAATTCCACGTAGGA    18010
gb:MN996531|Organism:Severe      TGTCTGATAGAGACCTTTATGACAAGTTGCAATTTACAAGTCTTGAAATTCCACGTAGGA    18009
gb:MT066176|Organism:Severe      TGTCTGATAGAGACCTTTATGACAAGTTGCAATTTACAAGTCTTGAAATTCCACGTAGGA    18022
gb:MT027064|Organism:Severe      TGTCTGATAGAGACCTTTATGACAAGTTGCAATTTACAAGTCTTGAAATTCCACGTAGGA    18022
gb:MN994468|Organism:Severe      TGTCTGATAGAGACCTTTATGACAAGTTGCAATTTACAAGTCTTGAAATTCCACGTAGGA    18022
gb:MT072688|Organism:Severe      TGTCTGATAGAGACCTTTATGACAAGTTGCAATTTACAAGTCTTGAAATTCCACGTAGGA    18007
gb:MN996527|Organism:Severe      TGTCTGATAGAGACCTTTATGACAAGTTGCAATTTACAAGTCTTGAAATTCCACGTAGGA    17989
gb:MT093631|Organism:Severe      TGTCTGATAGAGACCTTTATGACAAGTTGCAATTTACAAGTCTTGAAATTCCACGTAGGA    18060
gb:MT106053|Organism:Severe      TGTCTGATAGAGACCTTTATGACAAGTTGCAATTTACAAGTCTTGAAATTCCACGTAGGA    18022
gb:MT019533|Organism:Severe      TGTCTGATAGAGACCTTTATGACAAGTTGCAATTTACAAGTCTTGAAATTCCACGTAGGA    18022
gb:MT019531|Organism:Severe      TGTCTGATAGAGACCTTTATGACAAGTTGCAATTTACAAGTCTTGAAATTCCACGTAGGA    18022
gb:MN996528|Organism:Severe      TGTCTGATAGAGACCTTTATGACAAGTTGCAATTTACAAGTCTTGAAATTCCACGTAGGA    18022
```

```
gb:MN996530|Organism:Severe      TGTCTGATAGAGACCTTTATGACAAGTTGCAATTTACAAGTCTTGAAATTCCACGTAGGA   18008
gb:MN908947|Organism:Severe      TGTCTGATAGAGACCTTTATGACAAGTTGCAATTTACAAGTCTTGAAATTCCACGTAGGA   18022
gb:MT019532|Organism:Severe      TGTCTGATAGAGACCTTTATGACAAGTTGCAATTTACAAGTCTTGAAATTCCACGTAGGA   18022
                                 ************************************************************

gb:MT020781|Organism:Severe      ATGTGGCAACTTTACAAGCTGAAAATGTAACAGGACTCTTTAAAGATTGTAGTAAGGTAA   18070
gb:MT007544|Organism:Severe      ATGTGGCAACTTTACAAGCTGAAAATGTAACAGGACTCTTTAAAGATTGTAGTAAGGTAA   18082
gb:MN994467|Organism:Severe      ATGTGGCAACTTTACAAGCTGAAAATGTAACAGGACTCTTTAAAGATTGTAGTAAGGTAA   18082
gb:MT044257|Organism:Severe      ATGTGGCAACTTTACAAGCTGAAAATGTAACAGGACTCTTTAAAGATTGTAGTAAGGTAA   18082
gb:MT106054|Organism:Severe      ATGTGGCAACTTTACAAGCTGAAAATGTAACAGGACTCTTTAAAGATTGTAGTAAGGTAA   18082
gb:MT049951|Organism:Severe      ATGTGGCAACTTTACAAGCTGAAAATGTAACAGGACTCTTTAAAGATTGTAGTAAGGTAA   18082
gb:MN975262|Organism:Severe      ATGTGGCAACTTTACAAGCTGAAAATGTAACAGGACTCTTTAAAGATTGTAGTAAGGTAA   18082
gb:MT106052|Organism:Severe      ATGTGGCAACTTTACAAGCTGAAAATGTAACAGGACTCTTTAAAGATTGTAGTAAGGTAA   18082
gb:LC522975|Organism:Severe      ATGTGGCAACTTTACAAGCTGAAAATGTAACAGGACTCTTTAAAGATTGTAGTAAGGTAA   18079
gb:LC522973|Organism:Severe      ATGTGGCAACTTTACAAGCTGAAAATGTAACAGGACTCTTTAAAGATTGTAGTAAGGTAA   18079
gb:LC522974|Organism:Severe      ATGTGGCAACTTTACAAGCTGAAAATGTAACAGGACTCTTTAAAGATTGTAGTAAGGTAA   18079
gb:MN985325|Organism:Severe      ATGTGGCAACTTTACAAGCTGAAAATGTAACAGGACTTTTTAAAGATTGTAGTAAGGTAA   18082
gb:MT020881|Organism:Severe      ATGTGGCAACTTTACAAGCTGAAAATGTAACAGGACTTTTTAAAGATTGTAGTAAGGTAA   18082
gb:MT020880|Organism:Severe      ATGTGGCAACTTTACAAGCTGAAAATGTAACAGGACTTTTTAAAGATTGTAGTAAGGTAA   18082
gb:MT066175|Organism:Severe      ATGTGGCAACTTTACAAGCTGAAAATGTAACAGGACTCTTTAAAGATTGTAGTAAGGTAA   18082
gb:MN997409|Organism:Severe      ATGTGGCAACTTTACAAGCTGAAAATGTAACAGGACTCTTTAAAGATTGTAGTAAGGTAA   18082
gb:MN938384|Organism:Severe      ATGTGGCAACTTTACAAGCTGAAAATGTAACAGGACTCTTTAAAGATTGTAGTAAGGTAA   18050
gb:MT044258|Organism:Severe      ATGTGGCAACTTTACAAGCTGAAAATGTAACAGGACTCTTTAAAGATTGTAGTAAGGTAA   18058
gb:MT039890|Organism:Severe      ATGTGGCAACTTTACAAGCTGAAAATGTAACAGGACTCTTTAAAGATTGTAGTAAGGTAA   18082
gb:MN988713|Organism:Severe      ATGTGGCAACTTTACAAGCTGAAAATGTAACAGGACTCTTTAAAGATTGTAGTAAGGTAA   18082
gb:LC521925|Organism:Severe      ATGTGGCAACTTTACAAGCTGAAAATGTAACAGGACTCTTTAAAGATTGTAGTAAGGTAA   18055
gb:MT093571|Organism:Severe      ATGTGGCAACTTTACAAGCTGAAAATGTAACAGGACTCTTTAAAGATTGTAGTAAGGTAA   18082
gb:MT039887|Organism:Severe      ATGTGGCAACTTTACAAGCTGAAAATGTAACAGGACTCTTTAAAGATTGTAGTAAGGTAA   18082
gb:MT019530|Organism:Severe      ATGTGGCAACTTTACAAGCTGAAAATGTAACAGGACTCTTTAAAGATTGTAGTAAGGTAA   18082
gb:MT039888|Organism:Severe      ATGTGGCAACTTTACAAGCTGAAAATGTAACAGGACTCTTTAAAGATTGTAGTAAGGTAA   18082
gb:LC522972|Organism:Severe      ATGTGGCAACTTTACAAGCTGAAAATGTAACAGGACTCTTTAAAGATTGTAGTAAGGTAA   18079
gb:MT027063|Organism:Severe      ATGTGGCAACTTTACAAGCTGAAAATGTAACAGGACTCTTTAAAGATTGTAGTAAGGTAA   18082
gb:MT027062|Organism:Severe      ATGTGGCAACTTTACAAGCTGAAAATGTAACAGGACTCTTTAAAGATTGTAGTAAGGTAA   18082
gb:MT019529|Organism:Severe      ATGTGGCAACTTTACAAGCTGAAAATGTAACAGGACTCTTTAAAGATTGTAGTAAGGTAA   18082
gb:MN996529|Organism:Severe      ATGTGGCAACTTTACAAGCTGAAAATGTAACAGGACTCTTTAAAGATTGTAGTAAGGTAA   18070
gb:MN996531|Organism:Severe      ATGTGGCAACTTTACAAGCTGAAAATGTAACAGGACTCTTTAAAGATTGTAGTAAGGTAA   18069
gb:MT066176|Organism:Severe      ATGTGGCAACTTTACAAGCTGAAAATGTAACAGGACTCTTTAAAGATTGTAGTAAGGTAA   18082
gb:MT027064|Organism:Severe      ATGTGGCAACTTTACAAGCTGAAAATGTAACAGGACTCTTTAAAGATTGTAGTAAGGTAA   18082
gb:MN994468|Organism:Severe      ATGTGGCAACTTTACAAGCTGAAAATGTAACAGGACTCTTTAAAGATTGTAGTAAGGTAA   18082
gb:MT072688|Organism:Severe      ATGTGGCAACTTTACAAGCTGAAAATGTAACAGGACTCTTTAAAGATTGTAGTAAGGTAA   18067
gb:MN996527|Organism:Severe      ATGTGGCAACTTTACAAGCTGAAAATGTAACAGGACTCTTTAAAGATTGTAGTAAGGTAA   18049
gb:MT093631|Organism:Severe      ATGTGGCAACTTTACAAGCTGAAAATGTAACAGGACTCTTTAAAGATTGTAGTAAGGTAA   18120
gb:MT106053|Organism:Severe      ATGTGGCAACTTTACAAGCTGAAAATGTAACAGGACTCTTTAAAGATTGTAGTAAGGTAA   18082
gb:MT019533|Organism:Severe      ATGTGGCAACTTTACAAGCTGAAAATGTAACAGGACTCTTTAAAGATTGTAGTAAGGTAA   18082
gb:MT019531|Organism:Severe      ATGTGGCAACTTTACAAGCTGAAAATGTAACAGGACTCTTTAAAGATTGTAGTAAGGTAA   18082
gb:MN996528|Organism:Severe      ATGTGGCAACTTTACAAGCTGAAAATGTAACAGGACTCTTTAAAGATTGTAGTAAGGTAA   18082
gb:MN996530|Organism:Severe      ATGTGGCAACTTTACAAGCTGAAAATGTAACAGGACTCTTTAAAGATTGTAGTAAGGTAA   18068
gb:MN908947|Organism:Severe      ATGTGGCAACTTTACAAGCTGAAAATGTAACAGGACTCTTTAAAGATTGTAGTAAGGTAA   18082
gb:MT019532|Organism:Severe      ATGTGGCAACTTTACAAGCTGAAAATGTAACAGGACTCTTTAAAGATTGTAGTAAGGTAA   18082
                                 ************************************* **********************

gb:MT020781|Organism:Severe      TCACTGGGTTACATCCTACACAGGCACCTACACACCTCAGTGTTGACACTAAATTCAAAA   18130
gb:MT007544|Organism:Severe      TCACTGGGTTACATCCTACACAGGCACCTACACACCTCAGTGTTGACACTAAATTCAAAA   18142
gb:MN994467|Organism:Severe      TCACTGGGTTACATCCTACACAGGCACCTACACACCTCAGTGTTGACACTAAATTCAAAA   18142
gb:MT044257|Organism:Severe      TCACTGGGTTACATCCTACACAGGCACCTACACACCTCAGTGTTGACACTAAATTCAAAA   18142
gb:MT106054|Organism:Severe      TCACTGGGTTACATCCTACACAGGCACCTACACACCTCAGTGTTGACACTAAATTCAAAA   18142
gb:MT049951|Organism:Severe      TCACTGGGTTACATCCTACACAGGCACCTACACACCTCAGTGTTGACACTAAATTCAAAA   18142
gb:MN975262|Organism:Severe      TCACTGGGTTACATCCTACACAGGCACCTACACACCTCAGTGTTGACACTAAATTCAAAA   18142
gb:MT106052|Organism:Severe      TCACTGGGTTACATCCTACACAGGCACCTACACACCTCAGTGTTGACACTAAATTCAAAA   18142
gb:LC522975|Organism:Severe      TCACTGGGTTACATCCTACACAGGCACCTACACACCTCAGTGTTGACACTAAATTCAAAA   18139
gb:LC522973|Organism:Severe      TCACTGGGTTACATCCTACACAGGCACCTACACACCTCAGTGTTGACACTAAATTCAAAA   18139
gb:LC522974|Organism:Severe      TCACTGGGTTACATCCTACACAGGCACCTACACACCTCAGTGTTGACACTAAATTCAAAA   18139
gb:MN985325|Organism:Severe      TCACTGGGTTACATCCTACACAGGCACCTACACACCTCAGTGTTGACACTAAATTCAAAA   18142
gb:MT020881|Organism:Severe      TCACTGGGTTACATCCTACACAGGCACCTACACACCTCAGTGTTGACACTAAATTCAAAA   18142
gb:MT020880|Organism:Severe      TCACTGGGTTACATCCTACACAGGCACCTACACACCTCAGTGTTGACACTAAATTCAAAA   18142
```

```
gb:MT066175|Organism:Severe      TCACTGGGTTACATCCTACACAGGCACCTACACACCTCAGTGTTGACACTAAATTCAAAA   18142
gb:MN997409|Organism:Severe      TCACTGGGTTACATCCTACACAGGCACCTACACACCTCAGTGTTGACACTAAATTCAAAA   18142
gb:MN938384|Organism:Severe      TCACTGGGTTACATCCTACACAGGCACCTACACACCTCAGTGTTGACACTAAATTCAAAA   18110
gb:MT044258|Organism:Severe      TCACTGGGTTACATCCTACACAGGCACCTACACACCTCAGTGTTGACACTAAATTCAAAA   18118
gb:MT039890|Organism:Severe      TCACTGGGTTACATCCTACACAGGCACCTACACACCTCAGTGTTGACACTAAATTCAAAA   18142
gb:MN988713|Organism:Severe      TCACTGGGTTACATCCTACACAGGCACCTACACACCTCAGTGTTGACACTAAATTCAAAA   18142
gb:LC521925|Organism:Severe      TCACTGGGTTACATCCTACACAGGCACCTACACACCTCAGTGTTGACACTAAATTCAAAA   18115
gb:MT093571|Organism:Severe      TCACTGGGTTACATCCTACACAGGCACCTACACACCTCAGTGTTGACACTAAATTCAAAA   18142
gb:MT039887|Organism:Severe      TCACTGGGTTACATCCTACACAGGCACCTACACACCTCAGTGTTGACACTAAATTCAAAA   18142
gb:MT019530|Organism:Severe      TCACTGGGTTACATCCTACACAGGCACCTACACACCTCAGTGTTGACACTAAATTCAAAA   18142
gb:MT039888|Organism:Severe      TCACTGGGTTACATCCTACACAGGCACCTACACACCTCAGTGTTGACACTAAATTCAAAA   18142
gb:LC522972|Organism:Severe      TCACTGGGTTACATCCTACACAGGCACCTACACACCTCAGTGTTGACACTAAATTCAAAA   18139
gb:MT027063|Organism:Severe      TCACTGGGTTACATCCTACACAGGCACCTACACACCTCAGTGTTGACACTAAATTCAAAA   18142
gb:MT027062|Organism:Severe      TCACTGGGTTACATCCTACACAGGCACCTACACACCTCAGTGTTGACACTAAATTCAAAA   18142
gb:MT019529|Organism:Severe      TCACTGGGTTACATCCTACACAGGCACCTACACACCTCAGTGTTGACACTAAATTCAAAA   18142
gb:MN996529|Organism:Severe      TCACTGGGTTACATCCTACACAGGCACCTACACACCTCAGTGTTGACACTAAATTCAAAA   18130
gb:MN996531|Organism:Severe      TCACTGGGTTACATCCTACACAGGCACCTACACACCTCAGTGTTGACACTAAATTCAAAA   18129
gb:MT066176|Organism:Severe      TCACTGGGTTACATCCTACACAGGCACCTACACACCTCAGTGTTGACACTAAATTCAAAA   18142
gb:MT027064|Organism:Severe      TCACTGGGTTACATCCTACACAGGCACCTACACACCTCAGTGTTGACACTAAATTCAAAA   18142
gb:MN994468|Organism:Severe      TCACTGGGTTACATCCTACACAGGCACCTACACACCTCAGTGTTGACACTAAATTCAAAA   18142
gb:MT072688|Organism:Severe      TCACTGGGTTACATCCTACACAGGCACCTACACACCTCAGTGTTGACACTAAATTCAAAA   18127
gb:MN996527|Organism:Severe      TCACTGGGTTACATCCTACACAGGCACCTACACACCTCAGTGTTGACACTAAATTCAAAA   18109
gb:MT093631|Organism:Severe      TCACTGGGTTACATCCTACACAGGCACCTACACACCTCAGTGTTGACACTAAATTCAAAA   18180
gb:MT106053|Organism:Severe      TCACTGGGTTACATCCTACACAGGCACCTACACACCTCAGTGTTGACACTAAATTCAAAA   18142
gb:MT019533|Organism:Severe      TCACTGGGTTACATCCTACACAGGCACCTACACACCTCAGTGTTGACACTAAATTCAAAA   18142
gb:MT019531|Organism:Severe      TCACTGGGTTACATCCTACACAGGCACCTACACACCTCAGTGTTGACACTAAATTCAAAA   18142
gb:MN996528|Organism:Severe      TCACTGGGTTACATCCTACACAGGCACCTACACACCTCAGTGTTGACACTAAATTCAAAA   18142
gb:MN996530|Organism:Severe      TCACTGGGTTACATCCTACACAGGCACCTACACACCTCAGTGTTGACACTAAATTCAAAA   18128
gb:MN908947|Organism:Severe      TCACTGGGTTACATCCTACACAGGCACCTACACACCTCAGTGTTGACACTAAATTCAAAA   18142
gb:MT019532|Organism:Severe      TCACTGGGTTACATCCTACACAGGCACCTACACACCTCAGTGTTGACACTAAATTCAAAA   18142
                                 ************************************************************

gb:MT020781|Organism:Severe      CTGAAGGTTTATGTGTTGACATACCTGGCATACCTAAGGACATGACCTATAGAAGACTCA   18190
gb:MT007544|Organism:Severe      CTGAAGGTTTATGTGTTGACATACCTGGCATACCTAAGGACATGACCTATAGAAGACTCA   18202
gb:MN994467|Organism:Severe      CTGAAGGTTTATGTGTTGACATACCTGGCATACCTAAGGACATGACCTATAGAAGACTCA   18202
gb:MT044257|Organism:Severe      CTGAAGGTTTATGTGTTGACATACCTGGCATACCTAAGGACATGACCTATAGAAGACTCA   18202
gb:MT106054|Organism:Severe      CTGAAGGTTTATGTGTTGACATACCTGGCATACCTAAGGACATGACCTATAGAAGACTCA   18202
gb:MT049951|Organism:Severe      CTGAAGGTTTATGTGTTGACATACCTGGCATACCTAAGGACATGACCTATAGAAGACTCA   18202
gb:MN975262|Organism:Severe      CTGAAGGTTTATGTGTTGACATACCTGGCATACCTAAGGACATGACCTATAGAAGACTCA   18202
gb:MT106052|Organism:Severe      CTGAAGGTTTATGTGTTGACATACCTGGCATACCTAAGGACATGACCTATAGAAGACTCA   18202
gb:LC522975|Organism:Severe      CTGAAGGTTTATGTGTTGACATACCTGGCATACCTAAGGACATGACCTATAGAAGACTCA   18199
gb:LC522973|Organism:Severe      CTGAAGGTTTATGTGTTGACATACCTGGCATACCTAAGGACATGACCTATAGAAGACTCA   18199
gb:LC522974|Organism:Severe      CTGAAGGTTTATGTGTTGACATACCTGGCATACCTAAGGACATGACCTATAGAAGACTCA   18199
gb:MN985325|Organism:Severe      CTGAAGGTTTATGTGTTGACATACCTGGCATACCTAAGGACATGACCTATAGAAGACTCA   18202
gb:MT020881|Organism:Severe      CTGAAGGTTTATGTGTTGACATACCTGGCATACCTAAGGACATGACCTATAGAAGACTCA   18202
gb:MT020880|Organism:Severe      CTGAAGGTTTATGTGTTGACATACCTGGCATACCTAAGGACATGACCTATAGAAGACTCA   18202
gb:MT066175|Organism:Severe      CTGAAGGTTTATGTGTTGACATACCTGGCATACCTAAGGACATGACCTATAGAAGACTCA   18202
gb:MN997409|Organism:Severe      CTGAAGGTTTATGTGTTGACATACCTGGCATACCTAAGGACATGACCTATAGAAGACTCA   18202
gb:MN938384|Organism:Severe      CTGAAGGTTTATGTGTTGACATACCTGGCATACCTAAGGACATGACCTATAGAAGACTCA   18170
gb:MT044258|Organism:Severe      CTGAAGGTTTATGTGTTGACATACCTGGCATACCTAAGGACATGACCTATAGAAGACTCA   18178
gb:MT039890|Organism:Severe      CTGAAGGTTTATGTGTTGACATACCTGGCATACCTAAGGACATGACCTATAGAAGACTCA   18202
gb:MN988713|Organism:Severe      CTGAAGGTTTATGTGTTGACATACCTGGCATACCTAAGGACATGACCTATAGAAGACTCA   18202
gb:LC521925|Organism:Severe      CTGAAGGTTTATGTGTTGACATACCTGGCATACCTAAGGACATGACCTATAGAAGACTCA   18175
gb:MT093571|Organism:Severe      CTGAAGGTTTATGTGTTGACATACCTGGCATACCTAAGGACATGACCTATAGAAGACTCA   18202
gb:MT039887|Organism:Severe      CTGAAGGTTTATGTGTTGACATACCTGGCATACCTAAGGACATGACCTATAGAAGACTCA   18202
gb:MT019530|Organism:Severe      CTGAAGGTTTATGTGTTGACATACCTGGCATACCTAAGGACATGACCTATAGAAGACTCA   18202
gb:MT039888|Organism:Severe      CTGAAGGTTTATGTGTTGACATACCTGGCATACCTAAGGACATGACCTATAGAAGACTCA   18202
gb:LC522972|Organism:Severe      CTGAAGGTTTATGTGTTGACATACCTGGCATACCTAAGGACATGACCTATAGAAGACTCA   18199
gb:MT027063|Organism:Severe      CTGAAGGTTTATGTGTTGACATACCTGGCATACCTAAGGACATGACCTATAGAAGACTCA   18202
gb:MT027062|Organism:Severe      CTGAAGGTTTATGTGTTGACATACCTGGCATACCTAAGGACATGACCTATAGAAGACTCA   18202
gb:MT019529|Organism:Severe      CTGAAGGTTTATGTGTTGACATACCTGGCATACCTAAGGACATGACCTATAGAAGACTCA   18202
gb:MN996529|Organism:Severe      CTGAAGGTTTATGTGTTGACATACCTGGCATACCTAAGGACATGACCTATAGAAGACTCA   18190
gb:MN996531|Organism:Severe      CTGAAGGTTTATGTGTTGACATACCTGGCATACCTAAGGACATGACCTATAGAAGACTCA   18189
gb:MT066176|Organism:Severe      CTGAAGGTTTATGTGTTGACATACCTGGCATACCTAAGGACATGACCTATAGAAGACTCA   18202
gb:MT027064|Organism:Severe      CTGAAGGTTTATGTGTTGACATACCTGGCATACCTAAGGACATGACCTATAGAAGACTCA   18202
```

```
gb:MN994468|Organism:Severe      CTGAAGGTTTATGTGTTGACATACCTGGCATACCTAAGGACATGACCTATAGAAGACTCA   18202
gb:MT072688|Organism:Severe      CTGAAGGTTTATGTGTTGACATACCTGGCATACCTAAGGACATGACCTATAGAAGACTCA   18187
gb:MN996527|Organism:Severe      CTGAAGGTTTATGTGTTGACATACCTGGCATACCTAAGGACATGACCTATAGAAGACTCA   18169
gb:MT093631|Organism:Severe      CTGAAGGTTTATGTGTTGACATACCTGGCATACCTAAGGACATGACCTATAGAAGACTCA   18240
gb:MT106053|Organism:Severe      CTGAAGGTTTATGTGTTGACATACCTGGCATACCTAAGGACATGACCTATAGAAGACTCA   18202
gb:MT019533|Organism:Severe      CTGAAGGTTTATGTGTTGACATACCTGGCATACCTAAGGACATGACCTATAGAAGACTCA   18202
gb:MT019531|Organism:Severe      CTGAAGGTTTATGTGTTGACATACCTGGCATACCTAAGGACATGACCTATAGAAGACTCA   18202
gb:MN996528|Organism:Severe      CTGAAGGTTTATGTGTTGACATACCTGGCATACCTAAGGACATGACCTATAGAAGACTCA   18202
gb:MN996530|Organism:Severe      CTGAAGGTTTATGTGTTGACATACCTGGCATACCTAAGGACATGACCTATAGAAGACTCA   18188
gb:MN908947|Organism:Severe      CTGAAGGTTTATGTGTTGACATACCTGGCATACCTAAGGACATGACCTATAGAAGACTCA   18202
gb:MT019532|Organism:Severe      CTGAAGGTTTATGTGTTGACATACCTGGCATACCTAAGGACATGACCTATAGAAGACTCA   18202
                                 ************************************************************

gb:MT020781|Organism:Severe      TCTCTATGATGGGTTTTAAAATGAATTATCAAGTTAATGGTTACCCTAACATGTTTATCA   18250
gb:MT007544|Organism:Severe      TCTCTATGATGGGTTTTAAAATGAATTATCAAGTTAATGGTTACCCTAACATGTTTATCA   18262
gb:MN994467|Organism:Severe      TCTCTATGATGGGTTTTAAAATGAATTATCAAGTTAATGGTTACCCTAACATGTTTATCA   18262
gb:MT044257|Organism:Severe      TCTCTATGATGGGTTTTAAAATGAATTATCAAGTTAATGGTTACCCTAACATGTTTATCA   18262
gb:MT106054|Organism:Severe      TCTCTATGATGGGTTTTAAAATGAATTATCAAGTTAATGGTTACCCTAACATGTTTATCA   18262
gb:MT049951|Organism:Severe      TCTCTATGATGGGTTTTAAAATGAATTATCAAGTTAATGGTTACCCTAACATGTTTATCA   18262
gb:MN975262|Organism:Severe      TCTCTATGATGGGTTTTAAAATGAATTATCAAGTTAATGGTTACCCTAACATGTTTATCA   18262
gb:MT106052|Organism:Severe      TCTCTATGATGGGTTTTAAAATGAATTATCAAGTTAATGGTTACCCTAACATGTTTATCA   18262
gb:LC522975|Organism:Severe      TCTCTATGATGGGTTTTAAAATGAATTATCAAGTTAATGGTTACCCTAACATGTTTATCA   18259
gb:LC522973|Organism:Severe      TCTCTATGATGGGTTTTAAAATGAATTATCAAGTTAATGGTTACCCTAACATGTTTATCA   18259
gb:LC522974|Organism:Severe      TCTCTATGATGGGTTTTAAAATGAATTATCAAGTTAATGGTTACCCTAACATGTTTATCA   18259
gb:MN985325|Organism:Severe      TCTCTATGATGGGTTTTAAAATGAATTATCAAGTTAATGGTTACCCTAACATGTTTATCA   18262
gb:MT020881|Organism:Severe      TCTCTATGATGGGTTTTAAAATGAATTATCAAGTTAATGGTTACCCTAACATGTTTATCA   18262
gb:MT020880|Organism:Severe      TCTCTATGATGGGTTTTAAAATGAATTATCAAGTTAATGGTTACCCTAACATGTTTATCA   18262
gb:MT066175|Organism:Severe      TCTCTATGATGGGTTTTAAAATGAATTATCAAGTTAATGGTTACCCTAACATGTTTATCA   18262
gb:MN997409|Organism:Severe      TCTCTATGATGGGTTTTAAAATGAATTATCAAGTTAATGGTTACCCTAACATGTTTATCA   18262
gb:MN938384|Organism:Severe      TCTCTATGATGGGTTTTAAAATGAATTATCAAGTTAATGGTTACCCTAACATGTTTATCA   18230
gb:MT044258|Organism:Severe      TCTCTATGATGGGTTTTAAAATGAATTATCAAGTTAATGGTTACCCTAACATGTTTATCA   18238
gb:MT039890|Organism:Severe      TCTCTATGATGGGTTTTAAAATGAATTATCAAGTTAATGGTTACCCTAACATGTTTATCA   18262
gb:MN988713|Organism:Severe      TCTCTATGATGGGTTTTAAAATGAATTATCAAGTTAATGGTTACCCTAACATGTTTATCA   18262
gb:LC521925|Organism:Severe      TCTCTATGATGGGTTTTAAAATGAATTATCAAGTTAATGGTTACCCTAACATGTTTATCA   18235
gb:MT093571|Organism:Severe      TCTCTATGATGGGTTTTAAAATGAATTATCAAGTTAATGGTTACCCTAACATGTTTATCA   18262
gb:MT039887|Organism:Severe      TCTCTATGATGGGTTTTAAAATGAATTATCAAGTTAATGGTTACCCTAACATGTTTATCA   18262
gb:MT019530|Organism:Severe      TCTCTATGATGGGTTTTAAAATGAATTATCAAGTTAATGGTTACCCTAACATGTTTATCA   18262
gb:MT039888|Organism:Severe      TCTCTATGATGGGTTTTAAAATGAATTATCAAGTTAATGGTTACCCTAACATGTTTATCA   18262
gb:LC522972|Organism:Severe      TCTCTATGATGGGTTTTAAAATGAATTATCAAGTTAATGGTTACCCTAACATGTTTATCA   18259
gb:MT027063|Organism:Severe      TCTCTATGATGGGTTTTAAAATGAATTATCAAGTTAATGGTTACCCTAACATGTTTATCA   18262
gb:MT027062|Organism:Severe      TCTCTATGATGGGTTTTAAAATGAATTATCAAGTTAATGGTTACCCTAACATGTTTATCA   18262
gb:MT019529|Organism:Severe      TCTCTATGATGGGTTTTAAAATGAATTATCAAGTTAATGGTTACCCTAACATGTTTATCA   18262
gb:MN996529|Organism:Severe      TCTCTATGATGGGTTTTAAAATGAATTATCAAGTTAATGGTTACCCTAACATGTTTATCA   18250
gb:MN996531|Organism:Severe      TCTCTATGATGGGTTTTAAAATGAATTATCAAGTTAATGGTTACCCTAACATGTTTATCA   18249
gb:MT066176|Organism:Severe      TCTCTATGATGGGTTTTAAAATGAATTATCAAGTTAATGGTTACCCTAACATGTTTATCA   18262
gb:MT027064|Organism:Severe      TCTCTATGATGGGTTTTAAAATGAATTATCAAGTTAATGGTTACCCTAACATGTTTATCA   18262
gb:MN994468|Organism:Severe      TCTCTATGATGGGTTTTAAAATGAATTATCAAGTTAATGGTTACCCTAACATGTTTATCA   18262
gb:MT072688|Organism:Severe      TCTCTATGATGGGTTTTAAAATGAATTATCAAGTTAATGGTTACCCTAACATGTTTATCA   18247
gb:MN996527|Organism:Severe      TCTCTATGATGGGTTTTAAAATGAATTATCAAGTTAATGGTTACCCTAACATGTTTATCA   18229
gb:MT093631|Organism:Severe      TCTCTATGATGGGTTTTAAAATGAATTATCAAGTTAATGGTTACCCTAACATGTTTATCA   18300
gb:MT106053|Organism:Severe      TCTCTATGATGGGTTTTAAAATGAATTATCAAGTTAATGGTTACCCTAACATGTTTATCA   18262
gb:MT019533|Organism:Severe      TCTCTATGATGGGTTTTAAAATGAATTATCAAGTTAATGGTTACCCTAACATGTTTATCA   18262
gb:MT019531|Organism:Severe      TCTCTATGATGGGTTTTAAAATGAATTATCAAGTTAATGGTTACCCTAACATGTTTATCA   18262
gb:MN996528|Organism:Severe      TCTCTATGATGGGTTTTAAAATGAATTATCAAGTTAATGGTTACCCTAACATGTTTATCA   18262
gb:MN996530|Organism:Severe      TCTCTATGATGGGTTTTAAAATGAATTATCAAGTTAATGGTTACCCTAACATGTTTATCA   18248
gb:MN908947|Organism:Severe      TCTCTATGATGGGTTTTAAAATGAATTATCAAGTTAATGGTTACCCTAACATGTTTATCA   18262
gb:MT019532|Organism:Severe      TCTCTATGATGGGTTTTAAAATGAATTATCAAGTTAATGGTTACCCTAACATGTTTATCA   18262
                                 ************************************************************

gb:MT020781|Organism:Severe      CCCGCGAAGAAGCTATAAGACATGTACGTGCATGGATTGGCTTCGATGTCGAGGGGTGTC   18310
gb:MT007544|Organism:Severe      CCCGCGAAGAAGCTATAAGACATGTACGTGCATGGATTGGCTTCGATGTCGAGGGGTGTC   18322
gb:MN994467|Organism:Severe      CCCGCGAAGAAGCTATAAGACATGTACGTGCATGGATTGGCTTCGATGTCGAGGGGTGTC   18322
gb:MT044257|Organism:Severe      CCCGCGAAGAAGCTATAAGACATGTACGTGCATGGATTGGCTTCGATGTCGAGGGGTGTC   18322
gb:MT106054|Organism:Severe      CCCGCGAAGAAGCTATAAGACATGTACGTGCATGGATTGGCTTCGATGTCGAGGGGTGTC   18322
gb:MT049951|Organism:Severe      CCCGCGAAGAAGCTATAAGACATGTACGTGCATGGATTGGCTTCGATGTCGAGGGGTGTC   18322
```

```
gb:MN975262|Organism:Severe      CCCGCGAAGAAGCTATAAGACATGTACGTGCATGGATTGGCTTCGATGTCGAGGGGTGTC    18322
gb:MT106052|Organism:Severe      CCCGCGAAGAAGCTATAAGACATGTACGTGCATGGATTGGCTTCGATGTCGAGGGGTGTC    18322
gb:LC522975|Organism:Severe      CCCGCGAAGAAGCTATAAGACATGTACGTGCATGGATTGGCTTCGATGTCGAGGGGTGTC    18319
gb:LC522973|Organism:Severe      CCCGCGAAGAAGCTATAAGACATGTACGTGCATGGATTGGCTTCGATGTCGAGGGGTGTC    18319
gb:LC522974|Organism:Severe      CCCGCGAAGAAGCTATAAGACATGTACGTGCATGGATTGGCTTCGATGTCGAGGGGTGTC    18319
gb:MN985325|Organism:Severe      CCCGCGAAGAAGCTATAAGACATGTACGTGCATGGATTGGCTTCGATGTCGAGGGGTGTC    18322
gb:MT020881|Organism:Severe      CCCGCGAAGAAGCTATAAGACATGTACGTGCATGGATTGGCTTCGATGTCGAGGGGTGTC    18322
gb:MT020880|Organism:Severe      CCCGCGAAGAAGCTATAAGACATGTACGTGCATGGATTGGCTTCGATGTCGAGGGGTGTC    18322
gb:MT066175|Organism:Severe      CCCGCGAAGAAGCTATAAGACATGTACGTGCATGGATTGGCTTCGATGTCGAGGGGTGTC    18322
gb:MN997409|Organism:Severe      CCCGCGAAGAAGCTATAAGACATGTACGTGCATGGATTGGCTTCGATGTCGAGGGGTGTC    18322
gb:MN938384|Organism:Severe      CCCGCGAAGAAGCTATAAGACATGTACGTGCATGGATTGGCTTCGATGTCGAGGGGTGTC    18290
gb:MT044258|Organism:Severe      CCCGCGAAGAAGCTATAAGACATGTACGTGCATGGATTGGCTTCGATGTCGAGGGGTGTC    18298
gb:MT039890|Organism:Severe      CCCGCGAAGAAGCTATAAGACATGTACGTGCATGGATTGGCTTCGATGTCGAGGGGTGTC    18322
gb:MN988713|Organism:Severe      CCCGCGAAGAAGCTATAAGACATGTACGTGCATGGATTGGCTTCGATGTCGAGGGGTGTC    18322
gb:LC521925|Organism:Severe      CCCGCGAAGAAGCTATAAGACATGTACGTGCATGGATTGGCTTCGATGTCGAGGGGTGTC    18295
gb:MT093571|Organism:Severe      CCCGCGAAGAAGCTATAAGACATGTACGTGCATGGATTGGCTTCGATGTCGAGGGGTGTC    18322
gb:MT039887|Organism:Severe      CCCGCGAAGAAGCTATAAGACATGTACGTGCATGGATTGGCTTCGATGTCGAGGGGTGTC    18322
gb:MT019530|Organism:Severe      CCCGCGAAGAAGCTATAAGACATGTACGTGCATGGATTGGCTTCGATGTCGAGGGGTGTC    18322
gb:MT039888|Organism:Severe      CCCGCGAAGAAGCTATAAGACATGTACGTGCATGGATTGGCTTCGATGTCGAGGGGTGTC    18322
gb:LC522972|Organism:Severe      CCCGCGAAGAAGCTATAAGACATGTACGTGCATGGATTGGCTTCGATGTCGAGGGGTGTC    18319
gb:MT027063|Organism:Severe      CCCGCGAAGAAGCTATAAGACATGTACGTGCATGGATTGGCTTCGATGTCGAGGGGTGTC    18322
gb:MT027062|Organism:Severe      CCCGCGAAGAAGCTATAAGACATGTACGTGCATGGATTGGCTTCGATGTCGAGGGGTGTC    18322
gb:MT019529|Organism:Severe      CCCGCGAAGAAGCTATAAGACATGTACGTGCATGGATTGGCTTCGATGTCGAGGGGTGTC    18322
gb:MN996529|Organism:Severe      CCCGCGAAGAAGCTATAAGACATGTACGTGCATGGATTGGCTTCGATGTCGAGGGGTGTC    18310
gb:MN996531|Organism:Severe      CCCGCGAAGAAGCTATAAGACATGTACGTGCATGGATTGGCTTCGATGTCGAGGGGTGTC    18309
gb:MT066176|Organism:Severe      CCCGCGAAGAAGCTATAAGACATGTACGTGCATGGATTGGCTTCGATGTCGAGGGGTGTC    18322
gb:MT027064|Organism:Severe      CCCGCGAAGAAGCTATAAGACATGTACGTGCATGGATTGGCTTCGATGTCGAGGGGTGTC    18322
gb:MN994468|Organism:Severe      CCCGCGAAGAAGCTATAAGACATGTACGTGCATGGATTGGCTTCGATGTCGAGGGGTGTC    18322
gb:MT072688|Organism:Severe      CCCGCGAAGAAGCTATAAGACATGTACGTGCATGGATTGGCTTCGATGTCGAGGGGTGTC    18307
gb:MN996527|Organism:Severe      CCCGCGAAGAAGCTATAAGACATGTACGTGCATGGATTGGCTTCGATGTCGAGGGGTGTC    18289
gb:MT093631|Organism:Severe      CCCGCGAAGAAGCTATAAGACATGTACGTGCATGGATTGGCTTCGATGTCGAGGGGTGTC    18360
gb:MT106053|Organism:Severe      CCCGCGAAGAAGCTATAAGACATGTACGTGCATGGATTGGCTTCGATGTCGAGGGGTGTC    18322
gb:MT019533|Organism:Severe      CCCGCGAAGAAGCTATAAGACATGTACGTGCATGGATTGGCTTCGATGTCGAGGGGTGTC    18322
gb:MT019531|Organism:Severe      CCCGCGAAGAAGCTATAAGACATGTACGTGCATGGATTGGCTTCGATGTCGAGGGGTGTC    18322
gb:MN996528|Organism:Severe      CCCGCGAAGAAGCTATAAGACATGTACGTGCATGGATTGGCTTCGATGTCGAGGGGTGTC    18322
gb:MN996530|Organism:Severe      CCCGCGAAGAAGCTATAAGACATGTACGTGCATGGATTGGCTTCGATGTCGAGGGGTGTC    18308
gb:MN908947|Organism:Severe      CCCGCGAAGAAGCTATAAGACATGTACGTGCATGGATTGGCTTCGATGTCGAGGGGTGTC    18322
gb:MT019532|Organism:Severe      CCCGCGAAGAAGCTATAAGACATGTACGTGCATGGATTGGCTTCGATGTCGAGGGGTGTC    18322
                                 ************************************************************

gb:MT020781|Organism:Severe      ATGCTACTAGAGAAGCTGTTGGTACCAATTTACCTTTACAGCTAGGTTTTTCTACAGGTG    18370
gb:MT007544|Organism:Severe      ATGCTACTAGAGAAGCTGTTGGTACCAATTTACCTTTACAGCTAGGTTTTTCTACAGGTG    18382
gb:MN994467|Organism:Severe      ATGCTACTAGAGAAGCTGTTGGTACCAATTTACCTTTACAGCTAGGTTTTTCTACAGGTG    18382
gb:MT044257|Organism:Severe      ATGCTACTAGAGAAGCTGTTGGTACCAATTTACCTTTACAGCTAGGTTTTTCTACAGGTG    18382
gb:MT106054|Organism:Severe      ATGCTACTAGAGAAGCTGTTGGTACCAATTTACCTTTACAGCTAGGTTTTTCTACAGGTG    18382
gb:MT049951|Organism:Severe      ATGCTACTAGAGAAGCTGTTGGTACCAATTTACCTTTACAGCTAGGTTTTTCTACAGGTG    18382
gb:MN975262|Organism:Severe      ATGCTACTAGAGAAGCTGTTGGTACCAATTTACCTTTACAGCTAGGTTTTTCTACAGGTG    18382
gb:MT106052|Organism:Severe      ATGCTACTAGAGAAGCTGTTGGTACCAATTTACCTTTACAGCTAGGTTTTTCTACAGGTG    18382
gb:LC522975|Organism:Severe      ATGCTACTAGAGAAGCTGTTGGTACCAATTTACCTTTACAGCTAGGTTTTTCTACAGGTG    18379
gb:LC522973|Organism:Severe      ATGCTACTAGAGAAGCTGTTGGTACCAATTTACCTTTACAGCTAGGTTTTTCTACAGGTG    18379
gb:LC522974|Organism:Severe      ATGCTACTAGAGAAGCTGTTGGTACCAATTTACCTTTACAGCTAGGTTTTTCTACAGGTG    18379
gb:MN985325|Organism:Severe      ATGCTACTAGAGAAGCTGTTGGTACCAATTTACCTTTACAGCTAGGTTTTTCTACAGGTG    18382
gb:MT020881|Organism:Severe      ATGCTACTAGAGAAGCTGTTGGTACCAATTTACCTTTACAGCTAGGTTTTTCTACAGGTG    18382
gb:MT020880|Organism:Severe      ATGCTACTAGAGAAGCTGTTGGTACCAATTTACCTTTACAGCTAGGTTTTTCTACAGGTG    18382
gb:MT066175|Organism:Severe      ATGCTACTAGAGAAGCTGTTGGTACCAATTTACCTTTACAGCTAGGTTTTTCTACAGGTG    18382
gb:MN997409|Organism:Severe      ATGCTACTAGAGAAGCTGTTGGTACCAATTTACCTTTACAGCTAGGTTTTTCTACAGGTG    18382
gb:MN938384|Organism:Severe      ATGCTACTAGAGAAGCTGTTGGTACCAATTTACCTTTACAGCTAGGTTTTTCTACAGGTG    18350
gb:MT044258|Organism:Severe      ATGCTACTAGAGAAGCTGTTGGTACCAATTTACCTTTACAGCTAGGTTTTTCTACAGGTG    18358
gb:MT039890|Organism:Severe      ATGCTACTAGAGAAGCTGTTGGTACCAATTTACCTTTACAGCTAGGTTTTTCTACAGGTG    18382
gb:MN988713|Organism:Severe      ATGCTACTAGAGAAGCTGTTGGTACCAATTTACCTTTACAGCTAGGTTTTTCTACAGGTG    18382
gb:LC521925|Organism:Severe      ATGCTACTAGAGAAGCTGTTGGTACCAATTTACCTTTACAGCTAGGTTTTTCTACAGGTG    18355
gb:MT093571|Organism:Severe      ATGCTACTAGAGAAGCTGTTGGTACCAATTTACCTTTACAGCTAGGTTTTTCTACAGGTG    18382
gb:MT039887|Organism:Severe      ATGCTACTAGAGAAGCTGTTGGTACCAATTTACCTTTACAGCTAGGTTTTTCTACAGGTG    18382
gb:MT019530|Organism:Severe      ATGCTACTAGAGAAGCTGTTGGTACCAATTTACCTTTACAGCTAGGTTTTTCTACAGGTG    18382
gb:MT039888|Organism:Severe      ATGCTACTAGAGAAGCTGTTGGTACCAATTTACCTTTACAGCTAGGTTTTTCTACAGGTG    18382
```

```
gb:LC522972|Organism:Severe      ATGCTACTAGAGAAGCTGTTGGTACCAATTTACCTTTACAGCTAGGTTTTTCTACAGGTG   18379
gb:MT027063|Organism:Severe      ATGCTACTAGAGAAGCTGTTGGTACCAATTTACCTTTACAGCTAGGTTTTTCTACAGGTG   18382
gb:MT027062|Organism:Severe      ATGCTACTAGAGAAGCTGTTGGTACCAATTTACCTTTACAGCTAGGTTTTTCTACAGGTG   18382
gb:MT019529|Organism:Severe      ATGCTACTAGAGAAGCTGTTGGTACCAATTTACCTTTACAGCTAGGTTTTTCTACAGGTG   18382
gb:MN996529|Organism:Severe      ATGCTACTAGAGAAGCTGTTGGTACCAATTTACCTTTACAGCTAGGTTTTTCTACAGGTG   18370
gb:MN996531|Organism:Severe      ATGCTACTAGAGAAGCTGTTGGTACCAATTTACCTTTACAGCTAGGTTTTTCTACAGGTG   18369
gb:MT066176|Organism:Severe      ATGCTACTAGAGAAGCTGTTGGTACCAATTTACCTTTACAGCTAGGTTTTTCTACAGGTG   18382
gb:MT027064|Organism:Severe      ATGCTACTAGAGAAGCTGTTGGTACCAATTTACCTTTACAGCTAGGTTTTTCTACAGGTG   18382
gb:MN994468|Organism:Severe      ATGCTACTAGAGAAGCTGTTGGTACCAATTTACCTTTACAGCTAGGTTTTTCTACAGGTG   18382
gb:MT072688|Organism:Severe      ATGCTACTAGAGAAGCTGTTGGTACCAATTTACCTTTACAGCTAGGTTTTTCTACAGGTG   18367
gb:MN996527|Organism:Severe      ATGCTACTAGAGAAGCTGTTGGTACCAATTTACCTTTACAGCTAGGTTTTTCTACAGGTG   18349
gb:MT093631|Organism:Severe      ATGCTACTAGAGAAGCTGTTGGTACCAATTTACCTTTACAGCTAGGTTTTTCTACAGGTG   18420
gb:MT106053|Organism:Severe      ATGCTACTAGAGAAGCTGTTGGTACCAATTTACCTTTACAGCTAGGTTTTTCTACAGGTG   18382
gb:MT019533|Organism:Severe      ATGCTACTAGAGAAGCTGTTGGTACCAATTTACCTTTACAGCTAGGTTTTTCTACAGGTG   18382
gb:MT019531|Organism:Severe      ATGCTACTAGAGAAGCTGTTGGTACCAATTTACCTTTACAGCTAGGTTTTTCTACAGGTG   18382
gb:MN996528|Organism:Severe      ATGCTACTAGAGAAGCTGTTGGTACCAATTTACCTTTACAGCTAGGTTTTTCTACAGGTG   18382
gb:MN996530|Organism:Severe      ATGCTACTAGAGAAGCTGTTGGTACCAATTTACCTTTACAGCTAGGTTTTTCTACAGGTG   18368
gb:MN908947|Organism:Severe      ATGCTACTAGAGAAGCTGTTGGTACCAATTTACCTTTACAGCTAGGTTTTTCTACAGGTG   18382
gb:MT019532|Organism:Severe      ATGCTACTAGAGAAGCTGTTGGTACCAATTTACCTTTACAGCTAGGTTTTTCTACAGGTG   18382
                                 ************************************************************

gb:MT020781|Organism:Severe      TTAACCTAGTTGCTGTACCTACAGGTTATGTTGATACACCTAATAATACAGATTTTTCCA   18430
gb:MT007544|Organism:Severe      TTAACCTAGTTGCTGTACCTACAGGTTATGTTGATACACCTAATAATACAGATTTTTCCA   18442
gb:MN994467|Organism:Severe      TTAACCTAGTTGCTGTACCTACAGGTTATGTTGATACACCTAATAATACAGATTTTTCCA   18442
gb:MT044257|Organism:Severe      TTAACCTAGTTGCTGTACCTACAGGTTATGTTGATACACCTAATAATACAGATTTTTCCA   18442
gb:MT106054|Organism:Severe      TTAACCTAGTTGCTGTACCTACAGGTTATGTTGATACACCTAATAATACAGATTTTTCCA   18442
gb:MT049951|Organism:Severe      TTAACCTAGTTGCTGTACCTACAGGTTATGTTGATACACCTAATAATACAGATTTTTCCA   18442
gb:MN975262|Organism:Severe      TTAACCTAGTTGCTGTACCTACAGGTTATGTTGATACACCTAATAATACAGATTTTTCCA   18442
gb:MT106052|Organism:Severe      TTAACCTAGTTGCTGTACCTACAGGTTATGTTGATACACCTAATAATACAGATTTTTCCA   18442
gb:LC522975|Organism:Severe      TTAACCTAGTTGCTGTACCTACAGGTTATGTTGATACACCTAATAATACAGATTTTTCCA   18439
gb:LC522973|Organism:Severe      TTAACCTAGTTGCTGTACCTACAGGTTATGTTGATACACCTAATAATACAGATTTTTCCA   18439
gb:LC522974|Organism:Severe      TTAACCTAGTTGCTGTACCTACAGGTTATGTTGATACACCTAATAATACAGATTTTTCCA   18439
gb:MN985325|Organism:Severe      TTAACCTAGTTGCTGTACCTACAGGTTATGTTGATACACCTAATAATACAGATTTTTCCA   18442
gb:MT020881|Organism:Severe      TTAACCTAGTTGCTGTACCTACAGGTTATGTTGATACACCTAATAATACAGATTTTTCCA   18442
gb:MT020880|Organism:Severe      TTAACCTAGTTGCTGTACCTACAGGTTATGTTGATACACCTAATAATACAGATTTTTCCA   18442
gb:MT066175|Organism:Severe      TTAACCTAGTTGCTGTACCTACAGGTTATGTTGATACACCTAATAATACAGATTTTTCCA   18442
gb:MN997409|Organism:Severe      TTAACCTAGTTGCTGTACCTACAGGTTATGTTGATACACCTAATAATACAGATTTTTCCA   18442
gb:MN938384|Organism:Severe      TTAACCTAGTTGCTGTACCTACAGGTTATGTTGATACACCTAATAATACAGATTTTTCCA   18410
gb:MT044258|Organism:Severe      TTAACCTAGTTGCTGTACCTACAGGTTATGTTGATACACCTAATAATACAGATTTTTCCA   18418
gb:MT039890|Organism:Severe      TTAACCTAGTTGCTGTACCTACAGGTTATGTTGATACACCTAATAATACAGATTTTTCCA   18442
gb:MN988713|Organism:Severe      TTAACCTAGTTGCTGTACCTACAGGTTATGTTGATACACCTAATAATACAGATTTTTCCA   18442
gb:LC521925|Organism:Severe      TTAACCTAGTTGCTGTACCTACAGGTTATGTTGATACACCTAATAATACAGATTTTTCCA   18415
gb:MT093571|Organism:Severe      TTAACCTAGTTGCTGTACCTACAGGTTATGTTGATACACCTAATAATACAGATTTTTCCA   18442
gb:MT039887|Organism:Severe      TTAACCTAGTTGCTGTACCTACAGGTTATGTTGATACACCTAATAATACAGATTTTTCCA   18442
gb:MT019530|Organism:Severe      TTAACCTAGTTGCTGTACCTACAGGTTATGTTGATACACCTAATAATACAGATTTTTCCA   18442
gb:MT039888|Organism:Severe      TTAACCTAGTTGCTGTACCTACAGGTTATGTTGATACACCTAATAATACAGATTTTTCCA   18442
gb:LC522972|Organism:Severe      TTAACCTAGTTGCTGTACCTACAGGTTATGTTGATACACCTAATAATACAGATTTTTCCA   18439
gb:MT027063|Organism:Severe      TTAACCTAGTTGCTGTACCTACAGGTTATGTTGATACACCTAATAATACAGATTTTTCCA   18442
gb:MT027062|Organism:Severe      TTAACCTAGTTGCTGTACCTACAGGTTATGTTGATACACCTAATAATACAGATTTTTCCA   18442
gb:MT019529|Organism:Severe      TTAACCTAGTTGCTGTACCTACAGGTTATGTTGATACACCTAATAATACAGATTTTTCCA   18442
gb:MN996529|Organism:Severe      TTAACCTAGTTGCTGTACCTACAGGTTATGTTGATACACCTAATAATACAGATTTTTCCA   18430
gb:MN996531|Organism:Severe      TTAACCTAGTTGCTGTACCTACAGGTTATGTTGATACACCTAATAATACAGATTTTTCCA   18429
gb:MT066176|Organism:Severe      TTAACCTAGTTGCTGTACCTACAGGTTATGTTGATACACCTAATAATACAGATTTTTCCA   18442
gb:MT027064|Organism:Severe      TTAACCTAGTTGCTGTACCTACAGGTTATGTTGATACACCTAATAATACAGATTTTTCCA   18442
gb:MN994468|Organism:Severe      TTAACCTAGTTGCTGTACCTACAGGTTATGTTGATACACCTAATAATACAGATTTTTCCA   18442
gb:MT072688|Organism:Severe      TTAACCTAGTTGCTGTACCTACAGGTTATGTTGATACACCTAATAATACAGATTTTTCCA   18427
gb:MN996527|Organism:Severe      TTAACCTAGTTGCTGTACCTACAGGTTATGTTGATACACCTAATAATACAGATTTTTCCA   18409
gb:MT093631|Organism:Severe      TTAACCTAGTTGCTGTACCTACAGGTTATGTTGATACACCTAATAATACAGATTTTTCCA   18480
gb:MT106053|Organism:Severe      TTAACCTAGTTGCTGTACCTACAGGTTATGTTGATACACCTAATAATACAGATTTTTCCA   18442
gb:MT019533|Organism:Severe      TTAACCTAGTTGCTGTACCTACAGGTTATGTTGATACACCTAATAATACAGATTTTTCCA   18442
gb:MT019531|Organism:Severe      TTAACCTAGTTGCTGTACCTACAGGTTATGTTGATACACCTAATAATACAGATTTTTCCA   18442
gb:MN996528|Organism:Severe      TTAACCTAGTTGCTGTACCTACAGGTTATGTTGATACACCTAATAATACAGATTTTTCCA   18442
gb:MN996530|Organism:Severe      TTAACCTAGTTGCTGTACCTACAGGTTATGTTGATACACCTAATAATACAGATTTTTCCA   18428
gb:MN908947|Organism:Severe      TTAACCTAGTTGCTGTACCTACAGGTTATGTTGATACACCTAATAATACAGATTTTTCCA   18442
gb:MT019532|Organism:Severe      TTAACCTAGTTGCTGTACCTACAGGTTATGTTGATACACCTAATAATACAGATTTTTCCA   18442
```

```
                                   ************************************************************

gb:MT020781|Organism:Severe        GAGTTAGTGCTAAACCACCGCCTGGAGATCAATTTAAACACCTCATACCACTTATGTACA   18490
gb:MT007544|Organism:Severe        GAGTTAGTGCTAAACCACCGCCTGGAGATCAATTTAAACACCTCATACCACTTATGTACA   18502
gb:MN994467|Organism:Severe        GAGTTAGTGCTAAACCACCGCCTGGAGATCAATTTAAACACCTCATACCACTTATGTACA   18502
gb:MT044257|Organism:Severe        GAGTTAGTGCTAAACCACCGCCTGGAGATCAATTTAAACACCTCATACCACTTATGTACA   18502
gb:MT106054|Organism:Severe        GAGTTAGTGCTAAACCACCGCCTGGAGATCAATTTAAACACCTCATACCACTTATGTACA   18502
gb:MT049951|Organism:Severe        GAGTTAGTGCTAAACCACCGCCTGGAGATCAATTTAAACACCTCATACCACTTATGTACA   18502
gb:MN975262|Organism:Severe        GAGTTAGTGCTAAACCACCGCCTGGAGATCAATTTAAACACCTCATACCACTTATGTACA   18502
gb:MT106052|Organism:Severe        GAGTTAGTGCTAAACCACCGCCTGGAGATCAATTTAAACACCTCATACCACTTATGTACA   18502
gb:LC522975|Organism:Severe        GAGTTAGTGCTAAACCACCGCCTGGAGATCAATTTAAACACCTCATACCACTTATGTACA   18499
gb:LC522973|Organism:Severe        GAGTTAGTGCTAAACCACCGCCTGGAGATCAATTTAAACACCTCATACCACTTATGTACA   18499
gb:LC522974|Organism:Severe        GAGTTAGTGCTAAACCACCGCCTGGAGATCAATTTAAACACCTCATACCACTTATGTACA   18499
gb:MN985325|Organism:Severe        GAGTTAGTGCTAAACCACCGCCTGGAGATCAATTTAAACACCTCATACCACTTATGTACA   18502
gb:MT020881|Organism:Severe        GAGTTAGTGCTAAACCACCGCCTGGAGATCAATTTAAACACCTCATACCACTTATGTACA   18502
gb:MT020880|Organism:Severe        GAGTTAGTGCTAAACCACCGCCTGGAGATCAATTTAAACACCTCATACCACTTATGTACA   18502
gb:MT066175|Organism:Severe        GAGTTAGTGCTAAACCACCGCCTGGAGATCAATTTAAACACCTCATACCACTTATGTACA   18502
gb:MN997409|Organism:Severe        GAGTTAGTGCTAAACCACCGCCTGGAGATCAATTTAAACACCTCATACCACTTATGTACA   18502
gb:MN938384|Organism:Severe        GAGTTAGTGCTAAACCACCGCCTGGAGATCAATTTAAACACCTCATACCACTTATGTACA   18470
gb:MT044258|Organism:Severe        GAGTTAGTGCTAAACCACCGCCTGGAGATCAATTTAAACACCTCATACCACTTATGTACA   18478
gb:MT039890|Organism:Severe        GAGTTAGTGCTAAACCACCGCCTGGAGATCAATTTAAACACCTCATACCACTTATGTACA   18502
gb:MN988713|Organism:Severe        GAGTTAGTGCTAAACCACCGCCTGGAGATCAATTTAAACACCTCATACCACTTATGTACA   18502
gb:LC521925|Organism:Severe        GAGTTAGTGCTAAACCACCGCCTGGAGATCAATTTAAACACCTCATACCACTTATGTACA   18475
gb:MT093571|Organism:Severe        GAGTTAGTGCTAAACCACCGCCTGGAGATCAATTTAAACACCTCATACCACTTATGTACA   18502
gb:MT039887|Organism:Severe        GAGTTAGTGCTAAACCACCGCCTGGAGATCAATTTAAACACCTCATACCACTTATGTACA   18502
gb:MT019530|Organism:Severe        GAGTTAGTGCTAAACCACCGCCTGGAGATCAATTTAAACACCTCATACCACTTATGTACA   18502
gb:MT039888|Organism:Severe        GAGTTAGTGCTAAACCACCGCCTGGAGATCAATTTAAACACCTCATACCACTTATGTACA   18502
gb:LC522972|Organism:Severe        GAGTTAGTGCTAAACCACCGCCTGGAGATCAATTTAAACACCTCATACCACTTATGTACA   18499
gb:MT027063|Organism:Severe        GAGTTAGTGCTAAACCACCGCCTGGAGATCAATTTAAACACCTCATACCACTTATGTACA   18502
gb:MT027062|Organism:Severe        GAGTTAGTGCTAAACCACCGCCTGGAGATCAATTTAAACACCTCATACCACTTATGTACA   18502
gb:MT019529|Organism:Severe        GAGTTAGTGCTAAACCACCGCCTGGAGATCAATTTAAACACCTCATACCACTTATGTACA   18502
gb:MN996529|Organism:Severe        GAGTTAGTGCTAAACCACCGCCTGGAGATCAATTTAAACACCTCATACCACTTATGTACA   18490
gb:MN996531|Organism:Severe        GAGTTAGTGCTAAACCACCGCCTGGAGATCAATTTAAACACCTCATACCACTTATGTACA   18489
gb:MT066176|Organism:Severe        GAGTTAGTGCTAAACCACCGCCTGGAGATCAATTTAAACACCTCATACCACTTATGTACA   18502
gb:MT027064|Organism:Severe        GAGTTAGTGCTAAACCACCGCCTGGAGATCAATTTAAACACCTCATACCACTTATGTACA   18502
gb:MN994468|Organism:Severe        GAGTTAGTGCTAAACCACCGCCTGGAGATCAATTTAAACACCTCATACCACTTATGTACA   18502
gb:MT072688|Organism:Severe        GAGTTAGTGCTAAACCACCGCCTGGAGATCAATTTAAACACCTCATACCACTTATGTACA   18487
gb:MN996527|Organism:Severe        GAGTTAGTGCTAAACCACCGCCTGGAGATCAATTTAAACACCTCATACCACTTATGTACA   18469
gb:MT093631|Organism:Severe        GAGTTAGTGCTAAACCACCGCCTGGAGATCAATTTAAACACCTCATACCACTTATGTACA   18540
gb:MT106053|Organism:Severe        GAGTTAGTGCTAAACCACCGCCTGGAGATCAATTTAAACACCTCATACCACTTATGTACA   18502
gb:MT019533|Organism:Severe        GAGTTAGTGCTAAACCACCGCCTGGAGATCAATTTAAACACCTCATACCACTTATGTACA   18502
gb:MT019531|Organism:Severe        GAGTTAGTGCTAAACCACCGCCTGGAGATCAATTTAAACACCTCATACCACTTATGTACA   18502
gb:MN996528|Organism:Severe        GAGTTAGTGCTAAACCACCGCCTGGAGATCAATTTAAACACCTCATACCACTTATGTACA   18502
gb:MN996530|Organism:Severe        GAGTTAGTGCTAAACCACCGCCTGGAGATCAATTTAAACACCTCATACCACTTATGTACA   18488
gb:MN908947|Organism:Severe        GAGTTAGTGCTAAACCACCGCCTGGAGATCAATTTAAACACCTCATACCACTTATGTACA   18502
gb:MT019532|Organism:Severe        GAGTTAGTGCTAAACCACCGCCTGGAGATCAATTTAAACACCTCATACCACTTATGTACA   18502
                                   ************************************************************

gb:MT020781|Organism:Severe        AAGGACTTCCTTGGAATGTAGTGCGTATAAAGATTGTACAAATGTTAAGTGACACACTTA   18550
gb:MT007544|Organism:Severe        AAGGACTTCCTTGGAATGTAGTGCGTATAAAGATTGTACAAATGTTAAGTGACACACTTA   18562
gb:MN994467|Organism:Severe        AAGGACTTCCTTGGAATGTAGTGCGTATAAAGATTGTACAAATGTTAAGTGACACACTTA   18562
gb:MT044257|Organism:Severe        AAGGACTTCCTTGGAATGTAGTGCGTATAAAGATTGTACAAATGTTAAGTGACACACTTA   18562
gb:MT106054|Organism:Severe        AAGGACTTCCTTGGAATGTAGTGCGTATAAAGATTGTACAAATGTTAAGTGACACACTTA   18562
gb:MT049951|Organism:Severe        AAGGACTTCCTTGGAATGTAGTGCGTATAAAGATTGTACAAATGTTAAGTGACACACTTA   18562
gb:MN975262|Organism:Severe        AAGGACTTCCTTGGAATGTAGTGCGTATAAAGATTGTACAAATGTTAAGTGACACACTTA   18562
gb:MT106052|Organism:Severe        AAGGACTTCCTTGGAATGTAGTGCGTATAAAGATTGTACAAATGTTAAGTGACACACTTA   18562
gb:LC522975|Organism:Severe        AAGGACTTCCTTGGAATGTAGTGCGTATAAAGATTGTACAAATGTTAAGTGACACACTTA   18559
gb:LC522973|Organism:Severe        AAGGACTTCCTTGGAATGTAGTGCGTATAAAGATTGTACAAATGTTAAGTGACACACTTA   18559
gb:LC522974|Organism:Severe        AAGGACTTCCTTGGAATGTAGTGCGTATAAAGATTGTACAAATGTTAAGTGACACACTTA   18559
gb:MN985325|Organism:Severe        AAGGACTTCCTTGGAATGTAGTGCGTATAAAGATTGTACAAATGTTAAGTGACACACTTA   18562
gb:MT020881|Organism:Severe        AAGGACTTCCTTGGAATGTAGTGCGTATAAAGATTGTACAAATGTTAAGTGACACACTTA   18562
gb:MT020880|Organism:Severe        AAGGACTTCCTTGGAATGTAGTGCGTATAAAGATTGTACAAATGTTAAGTGACACACTTA   18562
gb:MT066175|Organism:Severe        AAGGACTTCCTTGGAATGTAGTGCGTATAAAGATTGTACAAATGTTAAGTGACACACTTA   18562
gb:MN997409|Organism:Severe        AAGGACTTCCTTGGAATGTAGTGCGTATAAAGATTGTACAAATGTTAAGTGACACACTTA   18562
gb:MN938384|Organism:Severe        AAGGACTTCCTTGGAATGTAGTGCGTATAAAGATTGTACAAATGTTAAGTGACACACTTA   18530
```

```
gb:MT044258|Organism:Severe      AAGGACTTCCTTGGAATGTAGTGCGTATAAAGATTGTACAAATGTTAAGTGACACACTTA   18538
gb:MT039890|Organism:Severe      AAGGACTTCCTTGGAATGTAGTGCGTATAAAGATTGTACAAATGTTAAGTGACACACTTA   18562
gb:MN988713|Organism:Severe      AAGGACTTCCTTGGAATGTAGTGCGTATAAAGATTGTACAAATGTTAAGTGACACACTTA   18562
gb:LC521925|Organism:Severe      AAGGACTTCTTTGGAATGTAGTGCGTATAAAGATTGTACAAATGTTAAGTGACACACTTA   18535
gb:MT093571|Organism:Severe      AAGGACTTCCTTGGAATGTAGTGCGTATAAAGATTGTACAAATGTTAAGTGACACACTTA   18562
gb:MT039887|Organism:Severe      AAGGACTTCCTTGGAATGTAGTGCGTATAAAGATTGTACAAATGTTAAGTGACACACTTA   18562
gb:MT019530|Organism:Severe      AAGGACTTCCTTGGAATGTAGTGCGTATAAAGATTGTACAAATGTTAAGTGACACACTTA   18562
gb:MT039888|Organism:Severe      AAGGACTTCCTTGGAATGTAGTGCGTATAAAGATTGTACAAATGTTAAGTGACACACTTA   18562
gb:LC522972|Organism:Severe      AAGGACTTCCTTGGAATGTAGTGCGTATAAAGATTGTACAAATGTTAAGTGACACACTTA   18559
gb:MT027063|Organism:Severe      AAGGACTTCCTTGGAATGTAGTGCGTATAAAGATTGTACAAATGTTAAGTGACACACTTA   18562
gb:MT027062|Organism:Severe      AAGGACTTCCTTGGAATGTAGTGCGTATAAAGATTGTACAAATGTTAAGTGACACACTTA   18562
gb:MT019529|Organism:Severe      AAGGACTTCCTTGGAATGTAGTGCGTATAAAGATTGTACAAATGTTAAGTGACACACTTA   18562
gb:MN996529|Organism:Severe      AAGGACTTCCTTGGAATGTAGTGCGTATAAAGATTGTACAAATGTTAAGTGACACACTTA   18550
gb:MN996531|Organism:Severe      AAGGACTTCCTTGGAATGTAGTGCGTATAAAGATTGTACAAATGTTAAGTGACACACTTA   18549
gb:MT066176|Organism:Severe      AAGGACTTCCTTGGAATGTAGTGCGTATAAAGATTGTACAAATGTTAAGTGACACACTTA   18562
gb:MT027064|Organism:Severe      AAGGACTTCCTTGGAATGTAGTGCGTATAAAGATTGTACAAATGTTAAGTGACACACTTA   18562
gb:MN994468|Organism:Severe      AAGGACTTCCTTGGAATGTAGTGCGTATAAAGATTGTACAAATGTTAAGTGACACACTTA   18562
gb:MT072688|Organism:Severe      AAGGACTTCCTTGGAATGTAGTGCGTATAAAGATTGTACAAATGTTAAGTGACACACTTA   18547
gb:MN996527|Organism:Severe      AAGGACTTCCTTGGAATGTAGTGCGTATAAAGATTGTACAAATGTTAAGTGACACACTTA   18529
gb:MT093631|Organism:Severe      AAGGACTTCCTTGGAATGTAGTGCGTATAAAGATTGTACAAATGTTAAGTGACACACTTA   18600
gb:MT106053|Organism:Severe      AAGGACTTCCTTGGAATGTAGTGCGTATAAAGATTGTACAAATGTTAAGTGACACACTTA   18562
gb:MT019533|Organism:Severe      AAGGACTTCCTTGGAATGTAGTGCGTATAAAGATTGTACAAATGTTAAGTGACACACTTA   18562
gb:MT019531|Organism:Severe      AAGGACTTCCTTGGAATGTAGTGCGTATAAAGATTGTACAAATGTTAAGTGACACACTTA   18562
gb:MN996528|Organism:Severe      AAGGACTTCCTTGGAATGTAGTGCGTATAAAGATTGTACAAATGTTAAGTGACACACTTA   18562
gb:MN996530|Organism:Severe      AAGGACTTCCTTGGAATGTAGTGCGTATAAAGATTGTACAAATGTTAAGTGACACACTTA   18548
gb:MN908947|Organism:Severe      AAGGACTTCCTTGGAATGTAGTGCGTATAAAGATTGTACAAATGTTAAGTGACACACTTA   18562
gb:MT019532|Organism:Severe      AAGGACTTCCTTGGAATGTAGTGCGTATAAAGATTGTACAAATGTTAAGTGACACACTTA   18562
                                 ********* **************************************************

gb:MT020781|Organism:Severe      AAAATCTCTCTGACAGAGTCGTATTTGTCTTATGGGCACATGGCTTTGAGTTGACATCTA   18610
gb:MT007544|Organism:Severe      AAAATCTCTCTGACAGAGTCGTATTTGTCTTATGGGCACATGGCTTTGAGTTGACATCTA   18622
gb:MN994467|Organism:Severe      AAAATCTCTCTGACAGAGTCGTATTTGTCTTATGGGCACATGGCTTTGAGTTGACATCTA   18622
gb:MT044257|Organism:Severe      AAAATCTCTCTGACAGAGTCGTATTTGTCTTATGGGCACATGGCTTTGAGTTGACATCTA   18622
gb:MT106054|Organism:Severe      AAAATCTCTCTGACAGAGTCGTATTTGTCTTATGGGCACACGGCTTTGAGTTGACATCTA   18622
gb:MT049951|Organism:Severe      AAAATCTCTCTGACAGAGTCGTATTTGTCTTATGGGCACATGGCTTTGAGTTGACATCTA   18622
gb:MN975262|Organism:Severe      AAAATCTCTCTGACAGAGTCGTATTTGTCTTATGGGCACATGGCTTTGAGTTGACATCTA   18622
gb:MT106052|Organism:Severe      AAAATCTCTCTGACAGAGTCGTATTTGTCTTATGGGCACATGGCTTTGAGTTGACATCTA   18622
gb:LC522975|Organism:Severe      AAAATCTCTCTGACAGAGTCGTATTTGTCTTATGGGCACATGGCTTTGAGTTGACATCTA   18619
gb:LC522973|Organism:Severe      AAAATCTCTCTGACAGAGTCGTATTTGTCTTATGGGCACATGGCTTTGAGTTGACATCTA   18619
gb:LC522974|Organism:Severe      AAAATCTCTCTGACAGAGTCGTATTTGTCTTATGGGCACATGGCTTTGAGTTGACATCTA   18619
gb:MN985325|Organism:Severe      AAAATCTCTCTGACAGAGTCGTATTTGTCTTATGGGCACATGGCTTTGAGTTGACATCTA   18622
gb:MT020881|Organism:Severe      AAAATCTCTCTGACAGAGTCGTATTTGTCTTATGGGCACATGGCTTTGAGTTGACATCTA   18622
gb:MT020880|Organism:Severe      AAAATCTCTCTGACAGAGTCGTATTTGTCTTATGGGCACATGGCTTTGAGTTGACATCTA   18622
gb:MT066175|Organism:Severe      AAAATCTCTCTGACAGAGTCGTATTTGTCTTATGGGCACATGGCTTTGAGTTGACATCTA   18622
gb:MN997409|Organism:Severe      AAAATCTCTCTGACAGAGTCGTATTTGTCTTATGGGCACATGGCTTTGAGTTGACATCTA   18622
gb:MN938384|Organism:Severe      AAAATCTCTCTGACAGAGTCGTATTTGTCTTATGGGCACATGGCTTTGAGTTGACATCTA   18590
gb:MT044258|Organism:Severe      AAAATCTCTCTGACAGAGTCGTATTTGTCTTATGGGCACATGGCTTTGAGTTGACATCTA   18598
gb:MT039890|Organism:Severe      AAAATCTCTCTGACAGAGTCGTATTTGTCTTATGGGCACATGGCTTTGAGTTGACATCTA   18622
gb:MN988713|Organism:Severe      AAAATCTCTCTGACAGAGTCGTATTTGTCTTATGGGCACATGGCTTTGAGTTGACATCTA   18622
gb:LC521925|Organism:Severe      AAAATCTCTCTGACAGAGTCGTATTTGTCTTATGGGCACATGGCTTTGAGTTGACATCTA   18595
gb:MT093571|Organism:Severe      AAAATCTCTCTGACAGAGTCGTATTTGTCTTATGGGCACATGGCTTTGAGTTGACATCTA   18622
gb:MT039887|Organism:Severe      AAAATCTCTCTGACAGAGTCGTATTTGTCTTATGGGCACATGGCTTTGAGTTGACATCTA   18622
gb:MT019530|Organism:Severe      AAAATCTCTCTGACAGAGTCGTATTTGTCTTATGGGCACATGGCTTTGAGTTGACATCTA   18622
gb:MT039888|Organism:Severe      AAAATCTCTCTGACAGAGTCGTATTTGTCTTATGGGCACATGGCTTTGAGTTGACATCTA   18622
gb:LC522972|Organism:Severe      AAAATCTCTCTGACAGAGTCGTATTTGTCTTATGGGCACATGGCTTTGAGTTGACATCTA   18619
gb:MT027063|Organism:Severe      AAAATCTCTCTGACAGAGTCGTATTTGTCTTATGGGCACATGGCTTTGAGTTGACATCTA   18622
gb:MT027062|Organism:Severe      AAAATCTCTCTGACAGAGTCGTATTTGTCTTATGGGCACATGGCTTTGAGTTGACATCTA   18622
gb:MT019529|Organism:Severe      AAAATCTCTCTGACAGAGTCGTATTTGTCTTATGGGCACATGGCTTTGAGTTGACATCTA   18622
gb:MN996529|Organism:Severe      AAAATCTCTCTGACAGAGTCGTATTTGTCTTATGGGCACATGGCTTTGAGTTGACATCTA   18610
gb:MN996531|Organism:Severe      AAAATCTCTCTGACAGAGTCGTATTTGTCTTATGGGCACATGGCTTTGAGTTGACATCTA   18609
gb:MT066176|Organism:Severe      AAAATCTCTCTGACAGAGTCGTATTTGTCTTATGGGCACATGGCTTTGAGTTGACATCTA   18622
gb:MT027064|Organism:Severe      AAAATCTCTCTGACAGAGTCGTATTTGTCTTATGGGCACATGGCTTTGAGTTGACATCTA   18622
gb:MN994468|Organism:Severe      AAAATCTCTCTGACAGAGTCGTATTTGTCTTATGGGCACATGGCTTTGAGTTGACATCTA   18622
gb:MT072688|Organism:Severe      AAAATCTCTCTGACAGAGTCGTATTTGTCTTATGGGCACATGGCTTTGAGTTGACATCTA   18607
gb:MN996527|Organism:Severe      AAAATCTCTCTGACAGAGTCGTATTTGTCTTATGGGCACATGGCTTTGAGTTGACATCTA   18589
```

```
gb:MT093631|Organism:Severe      AAAATCTCTCTGACAGAGTCGTATTTGTCTTATGGGCACATGGCTTTGAGTTGACATCTA   18660
gb:MT106053|Organism:Severe      AAAATCTCTCTGACAGAGTCGTATTTGTCTTATGGGCACATGGCTTTGAGTTGACATCTA   18622
gb:MT019533|Organism:Severe      AAAATCTCTCTGACAGAGTCGTATTTGTCTTATGGGCACATGGCTTTGAGTTGACATCTA   18622
gb:MT019531|Organism:Severe      AAAATCTCTCTGACAGAGTCGTATTTGTCTTATGGGCACATGGCTTTGAGTTGACATCTA   18622
gb:MN996528|Organism:Severe      AAAATCTCTCTGACAGAGTCGTATTTGTCTTATGGGCACATGGCTTTGAGTTGACATCTA   18622
gb:MN996530|Organism:Severe      AAAATCTCTCTGACAGAGTCGTATTTGTCTTATGGGCACATGGCTTTGAGTTGACATCTA   18608
gb:MN908947|Organism:Severe      AAAATCTCTCTGACAGAGTCGTATTTGTCTTATGGGCACATGGCTTTGAGTTGACATCTA   18622
gb:MT019532|Organism:Severe      AAAATCTCTCTGACAGAGTCGTATTTGTCTTATGGGCACATGGCTTTGAGTTGACATCTA   18622
                                 **************************************** *******************

gb:MT020781|Organism:Severe      TGAAGTATTTTGTGAAAATAGGACCTGAGCGCACCTGTTGTCTATGTGATAGACGTGCCA   18670
gb:MT007544|Organism:Severe      TGAAGTATTTTGTGAAAATAGGACCTGAGCGCACCTGTTGTCTATGTGATAGACGTGCCA   18682
gb:MN994467|Organism:Severe      TGAAGTATTTTGTGAAAATAGGACCTGAGCGCACCTGTTGTCTATGTGATAGACGTGCCA   18682
gb:MT044257|Organism:Severe      TGAAGTATTTTGTGAAAATAGGACCTGAGCGCACCTGTTGTCTATGTGATAGACGTGCCA   18682
gb:MT106054|Organism:Severe      TGAAGTATTTTGTGAAAATAGGACCTGAGCGCACCTGTTGTCTATGTGATAGACGTGCCA   18682
gb:MT049951|Organism:Severe      TGAAGTATTTTGTGAAAATAGGACCTGAGCGCACCTGTTGTCTATGTGATAGACGTGCCA   18682
gb:MN975262|Organism:Severe      TGAAGTATTTTGTGAAAATAGGACCTGAGCGCACCTGTTGTCTATGTGATAGACGTGCCA   18682
gb:MT106052|Organism:Severe      TGAAGTATTTTGTGAAAATAGGACCTGAGCGCACCTGTTGTCTATGTGATAGACGTGCCA   18682
gb:LC522975|Organism:Severe      TGAAGTATTTTGTGAAAATAGGACCTGAGCGCACCTGTTGTCTATGTGATAGACGTGCCA   18679
gb:LC522973|Organism:Severe      TGAAGTATTTTGTGAAAATAGGACCTGAGCGCACCTGTTGTCTATGTGATAGACGTGCCA   18679
gb:LC522974|Organism:Severe      TGAAGTATTTTGTGAAAATAGGACCTGAGCGCACCTGTTGTCTATGTGATAGACGTGCCA   18679
gb:MN985325|Organism:Severe      TGAAGTATTTTGTGAAAATAGGACCTGAGCGCACCTGTTGTCTATGTGATAGACGTGCCA   18682
gb:MT020881|Organism:Severe      TGAAGTATTTTGTGAAAATAGGACCTGAGCGCACCTGTTGTCTATGTGATAGACGTGCCA   18682
gb:MT020880|Organism:Severe      TGAAGTATTTTGTGAAAATAGGACCTGAGCGCACCTGTTGTCTATGTGATAGACGTGCCA   18682
gb:MT066175|Organism:Severe      TGAAGTATTTTGTGAAAATAGGACCTGAGCGCACCTGTTGTCTATGTGATAGACGTGCCA   18682
gb:MN997409|Organism:Severe      TGAAGTATTTTGTGAAAATAGGACCTGAGCGCACCTGTTGTCTATGTGATAGACGTGCCA   18682
gb:MN938384|Organism:Severe      TGAAGTATTTTGTGAAAATAGGACCTGAGCGCACCTGTTGTCTATGTGATAGACGTGCCA   18650
gb:MT044258|Organism:Severe      TGAAGTATTTTGTGAAAATAGGACCTGAGCGCACCTGTTGTCTATGTGATAGACGTGCCA   18658
gb:MT039890|Organism:Severe      TGAAGTATTTTGTGAAAATAGGACCTGAGCGCACCTGTTGTCTATGTGATAGACGTGCCA   18682
gb:MN988713|Organism:Severe      TGAAGTATTTTGTGAAAATAGGACCTGAGCGCACCTGTTGTCTATGTGATAGACGTGCCA   18682
gb:LC521925|Organism:Severe      TGAAGTATTTTGTGAAAATAGGACCTGAGCGCACCTGTTGTCTATGTGATAGACGTGCCA   18655
gb:MT093571|Organism:Severe      TGAAGTATTTTGTGAAAATAGGACCTGAGCGCACCTGTTGTCTATGTGATAGACGTGCCA   18682
gb:MT039887|Organism:Severe      TGAAGTATTTTGTGAAAATAGGACCTGAGCGCACCTGTTGTCTATGTGATAGACGTGCCA   18682
gb:MT019530|Organism:Severe      TGAAGTATTTTGTGAAAATAGGACCTGAGCGCACCTGTTGTCTATGTGATAGACGTGCCA   18682
gb:MT039888|Organism:Severe      TGAAGTATTTTGTGAAAATAGGACCTGAGCGCACCTGTTGTCTATGTGATAGACGTGCCA   18682
gb:LC522972|Organism:Severe      TGAAGTATTTTGTGAAAATAGGACCTGAGCGCACCTGTTGTCTATGTGATAGACGTGCCA   18679
gb:MT027063|Organism:Severe      TGAAGTATTTTGTGAAAATAGGACCTGAGCGCACCTGTTGTCTATGTGATAGACGTGCCA   18682
gb:MT027062|Organism:Severe      TGAAGTATTTTGTGAAAATAGGACCTGAGCGCACCTGTTGTCTATGTGATAGACGTGCCA   18682
gb:MT019529|Organism:Severe      TGAAGTATTTTGTGAAAATAGGACCTGAGCGCACCTGTTGTCTATGTGATAGACGTGCCA   18682
gb:MN996529|Organism:Severe      TGAAGTATTTTGTGAAAATAGGACCTGAGCGCACCTGTTGTCTATGTGATAGACGTGCCA   18670
gb:MN996531|Organism:Severe      TGAAGTATTTTGTGAAAATAGGACCTGAGCGCACCTGTTGTCTATGTGATAGACGTGCCA   18669
gb:MT066176|Organism:Severe      TGAAGTATTTTGTGAAAATAGGACCTGAGCGCACCTGTTGTCTATGTGATAGACGTGCCA   18682
gb:MT027064|Organism:Severe      TGAAGTATTTTGTGAAAATAGGACCTGAGCGCACCTGTTGTCTATGTGATAGACGTGCCA   18682
gb:MN994468|Organism:Severe      TGAAGTATTTTGTGAAAATAGGACCTGAGCGCACCTGTTGTCTATGTGATAGACGTGCCA   18682
gb:MT072688|Organism:Severe      TGAAGTATTTTGTGAAAATAGGACCTGAGCGCACCTGTTGTCTATGTGATAGACGTGCCA   18667
gb:MN996527|Organism:Severe      TGAAGTATTTTGTGAAAATAGGACCTGAGCGCACCTGTTGTCTATGTGATAGACGTGCCA   18649
gb:MT093631|Organism:Severe      TGAAGTATTTTGTGAAAATAGGACCTGAGCGCACCTGTTGTCTATGTGATAGACGTGCCA   18720
gb:MT106053|Organism:Severe      TGAAGTATTTTGTGAAAATAGGACCTGAGCGCACCTGTTGTCTATGTGATAGACGTGCCA   18682
gb:MT019533|Organism:Severe      TGAAGTATTTTGTGAAAATAGGACCTGAGCGCACCTGTTGTCTATGTGATAGACGTGCCA   18682
gb:MT019531|Organism:Severe      TGAAGTATTTTGTGAAAATAGGACCTGAGCGCACCTGTTGTCTATGTGATAGACGTGCCA   18682
gb:MN996528|Organism:Severe      TGAAGTATTTTGTGAAAATAGGACCTGAGCGCACCTGTTGTCTATGTGATAGACGTGCCA   18682
gb:MN996530|Organism:Severe      TGAAGTATTTTGTGAAAATAGGACCTGAGCGCACCTGTTGTCTATGTGATAGACGTGCCA   18668
gb:MN908947|Organism:Severe      TGAAGTATTTTGTGAAAATAGGACCTGAGCGCACCTGTTGTCTATGTGATAGACGTGCCA   18682
gb:MT019532|Organism:Severe      TGAAGTATTTTGTGAAAATAGGACCTGAGCGCACCTGTTGTCTATGTGATAGACGTGCCA   18682
                                 ************************************************************

gb:MT020781|Organism:Severe      CATGCTTTTCCACTGCTTCAGACACTTATGCCTGTTGGCATCATTCTATTGGATTTGATT   18730
gb:MT007544|Organism:Severe      CATGCTTTTCCACTGCTTCAGACACTTATGCCTGTTGGCATCATTCTATTGGATTTGATT   18742
gb:MN994467|Organism:Severe      CATGCTTTTCCACTGCTTCAGACACTTATGCCTGTTGGCATCATTCTATTGGATTTGATT   18742
gb:MT044257|Organism:Severe      CATGCTTTTCCACTGCTTCAGACACTTATGCCTGTTGGCATCATTCTATTGGATTTGATT   18742
gb:MT106054|Organism:Severe      CATGCTTTTCCACTGCTTCAGACACTTATGCCTGTTGGCATCATTCTATTGGATTTGATT   18742
gb:MT049951|Organism:Severe      CATGCTTTTCCACTGCTTCAGACACTTATGCCTGTTGGCATCATTCTATTGGATTTGATT   18742
gb:MN975262|Organism:Severe      CATGCTTTTCCACTGCTTCAGACACTTATGCCTGTTGGCATCATTCTATTGGATTTGATT   18742
gb:MT106052|Organism:Severe      CATGCTTTTCCACTGCTTCAGACACTTATGCCTGTTGGCATCATTCTATTGGATTTGATT   18742
gb:LC522975|Organism:Severe      CATGCTTTTCCACTGCTTCAGACACTTATGCCTGTTGGCATCATTCTATTGGATTTGATT   18739
```

```
gb:LC522973|Organism:Severe      CATGCTTTTCCACTGCTTCAGACACTTATGCCTGTTGGCATCATTCTATTGGATTTGATT   18739
gb:LC522974|Organism:Severe      CATGCTTTTCCACTGCTTCAGACACTTATGCCTGTTGGCATCATTCTATTGGATTTGATT   18739
gb:MN985325|Organism:Severe      CATGCTTTTCCACTGCTTCAGACACTTATGCCTGTTGGCATCATTCTATTGGATTTGATT   18742
gb:MT020881|Organism:Severe      CATGCTTTTCCACTGCTTCAGACACTTATGCCTGTTGGCATCATTCTATTGGATTTGATT   18742
gb:MT020880|Organism:Severe      CATGCTTTTCCACTGCTTCAGACACTTATGCCTGTTGGCATCATTCTATTGGATTTGATT   18742
gb:MT066175|Organism:Severe      CATGCTTTTCCACTGCTTCAGACACTTATGCCTGTTGGCATCATTCTATTGGATTTGATT   18742
gb:MN997409|Organism:Severe      CATGCTTTTCCACTGCTTCAGACACTTATGCCTGTTGGCATCATTCTATTGGATTTGATT   18742
gb:MN938384|Organism:Severe      CATGCTTTTCCACTGCTTCAGACACTTATGCCTGTTGGCATCATTCTATTGGATTTGATT   18710
gb:MT044258|Organism:Severe      CATGCTTTTCCACTGCTTCAGACACTTATGCCTGTTGGCATCATTCTATTGGATTTGATT   18718
gb:MT039890|Organism:Severe      CATGCTTTTCCACTGCTTCAGACACTTATGCCTGTTGGCATCATTCTATTGGATTTGATT   18742
gb:MN988713|Organism:Severe      CATGCTTTTCCACTGCTTCAGACACTTATGCCTGTTGGCATCATTCTATTGGATTTGATT   18742
gb:LC521925|Organism:Severe      CATGCTTTTCCACTGCTTCAGACACTTATGCCTGTTGGCATCATTCTATTGGATTTGATT   18715
gb:MT093571|Organism:Severe      CATGCTTTTCCACTGCTTCAGACACTTATGCCTGTTGGCATCATTCTATTGGATTTGATT   18742
gb:MT039887|Organism:Severe      CATGCTTTTCCACTGCTTCAGACACTTATGCCTGTTGGCATCATTCTATTGGATTTGATT   18742
gb:MT019530|Organism:Severe      CATGCTTTTCCACTGCTTCAGACACTTATGCCTGTTGGCATCATTCTATTGGATTTGATT   18742
gb:MT039888|Organism:Severe      CATGCTTTTCCACTGCTTCAGACACTTATGCCTGTTGGCATCATTCTATTGGATTTGATT   18742
gb:LC522972|Organism:Severe      CATGCTTTTCCACTGCTTCAGACACTTATGCCTGTTGGCATCATTCTATTGGATTTGATT   18739
gb:MT027063|Organism:Severe      CATGCTTTTCCACTGCTTCAGACACTTATGCCTGTTGGCATCATTCTATTGGATTTGATT   18742
gb:MT027062|Organism:Severe      CATGCTTTTCCACTGCTTCAGACACTTATGCCTGTTGGCATCATTCTATTGGATTTGATT   18742
gb:MT019529|Organism:Severe      CATGCTTTTCCACTGCTTCAGACACTTATGCCTGTTGGCATCATTCTATTGGATTTGATT   18742
gb:MN996529|Organism:Severe      CATGCTTTTCCACTGCTTCAGACACTTATGCCTGTTGGCATCATTCTATTGGATTTGATT   18730
gb:MN996531|Organism:Severe      CATGCTTTTCCACTGCTTCAGACACTTATGCCTGTTGGCATCATTCTATTGGATTTGATT   18729
gb:MT066176|Organism:Severe      CATGCTTTTCCACTGCTTCAGACACTTATGCCTGTTGGCATCATTCTATTGGATTTGATT   18742
gb:MT027064|Organism:Severe      CATGCTTTTCCACTGCTTCAGACACTTATGCCTGTTGGCATCATTCTATTGGATTTGATT   18742
gb:MN994468|Organism:Severe      CATGCTTTTCCACTGCTTCAGACACTTATGCCTGTTGGCATCATTCTATTGGATTTGATT   18742
gb:MT072688|Organism:Severe      CATGCTTTTCCACTGCTTCAGACACTTATGCCTGTTGGCATCATTCTATTGGATTTGATT   18727
gb:MN996527|Organism:Severe      CATGCTTTTCCACTGCTTCAGACACTTATGCCTGTTGGCATCATTCTATTGGATTTGATT   18709
gb:MT093631|Organism:Severe      CATGCTTTTCCACTGCTTCAGACACTTATGCCTGTTGGCATCATTCTATTGGATTTGATT   18780
gb:MT106053|Organism:Severe      CATGCTTTTCCACTGCTTCAGACACTTATGCCTGTTGGCATCATTCTATTGGATTTGATT   18742
gb:MT019533|Organism:Severe      CATGCTTTTCCACTGCTTCAGACACTTATGCCTGTTGGCATCATTCTATTGGATTTGATT   18742
gb:MT019531|Organism:Severe      CATGCTTTTCCACTGCTTCAGACACTTATGCCTGTTGGCATCATTCTATTGGATTTGATT   18742
gb:MN996528|Organism:Severe      CATGCTTTTCCACTGCTTCAGACACTTATGCCTGTTGGCATCATTCTATTGGATTTGATT   18742
gb:MN996530|Organism:Severe      CATGCTTTTCCACTGCTTCAGACACTTATGCCTGTTGGCATCATTCTATTGGATTTGATT   18728
gb:MN908947|Organism:Severe      CATGCTTTTCCACTGCTTCAGACACTTATGCCTGTTGGCATCATTCTATTGGATTTGATT   18742
gb:MT019532|Organism:Severe      CATGCTTTTCCACTGCTTCAGACACTTATGCCTGTTGGCATCATTCTATTGGATTTGATT   18742
                                 ************************************************************

gb:MT020781|Organism:Severe      ACGTCTATAATCCGTTTATGATTGATGTTCAACAATGGGGTTTTACAGGTAACCTACAAA   18790
gb:MT007544|Organism:Severe      ACGTCTATAATCCGTTTATGATTGATGTTCAACAATGGGGTTTTACAGGTAACCTACAAA   18802
gb:MN994467|Organism:Severe      ACGTCTATAATCCGTTTATGATTGATGTTCAACAATGGGGTTTTACAGGTAACCTACAAA   18802
gb:MT044257|Organism:Severe      ACGTCTATAATCCGTTTATGATTGATGTTCAACAATGGGGTTTTACAGGTAACCTACAAA   18802
gb:MT106054|Organism:Severe      ACGTCTATAATCCGTTTATGATTGATGTTCAACAATGGGGTTTTACAGGTAACCTACAAA   18802
gb:MT049951|Organism:Severe      ACGTCTATAATCCGTTTATGATTGATGTTCAACAATGGGGTTTTACAGGTAACCTACAAA   18802
gb:MN975262|Organism:Severe      ACGTCTATAATCCGTTTATGATTGATGTTCAACAATGGGGTTTTACAGGTAACCTACAAA   18802
gb:MT106052|Organism:Severe      ACGTCTATAATCCGTTTATGATTGATGTTCAACAATGGGGTTTTACAGGTAACCTACAAA   18802
gb:LC522975|Organism:Severe      ACGTCTATAATCCGTTTATGATTGATGTTCAACAATGGGGTTTTACAGGTAACCTACAAA   18799
gb:LC522973|Organism:Severe      ACGTCTATAATCCGTTTATGATTGATGTTCAACAATGGGGTTTTACAGGTAACCTACAAA   18799
gb:LC522974|Organism:Severe      ACGTCTATAATCCGTTTATGATTGATGTTCAACAATGGGGTTTTACAGGTAACCTACAAA   18799
gb:MN985325|Organism:Severe      ACGTCTATAATCCGTTTATGATTGATGTTCAACAATGGGGTTTTACAGGTAACCTACAAA   18802
gb:MT020881|Organism:Severe      ACGTCTATAATCCGTTTATGATTGATGTTCAACAATGGGGTTTTACAGGTAACCTACAAA   18802
gb:MT020880|Organism:Severe      ACGTCTATAATCCGTTTATGATTGATGTTCAACAATGGGGTTTTACAGGTAACCTACAAA   18802
gb:MT066175|Organism:Severe      ACGTCTATAATCCGTTTATGATTGATGTTCAACAATGGGGTTTTACAGGTAACCTACAAA   18802
gb:MN997409|Organism:Severe      ACGTCTATAATCCGTTTATGATTGATGTTCAACAATGGGGTTTTACAGGTAACCTACAAA   18802
gb:MN938384|Organism:Severe      ACGTCTATAATCCGTTTATGATTGATGTTCAACAATGGGGTTTTACAGGTAACCTACAAA   18770
gb:MT044258|Organism:Severe      ACGTCTATAATCCGTTTATGATTGATGTTCAACAATGGGGTTTTACAGGTAACCTACAAA   18778
gb:MT039890|Organism:Severe      ACGTCTATAATCCGTTTATGATTGATGTTCAACAATGGGGTTTTACAGGTAACCTACAAA   18802
gb:MN988713|Organism:Severe      ACGTCTATAATCCGTTTATGATTGATGTTCAACAATGGGGTTTTACAGGTAACCTACAAA   18802
gb:LC521925|Organism:Severe      ACGTCTATAATCCGTTTATGATTGATGTTCAACAATGGGGTTTTACAGGTAACCTACAAA   18775
gb:MT093571|Organism:Severe      ACGTCTATAATCCGTTTATGATTGATGTTCAACAATGGGGTTTTACAGGTAACCTACAAA   18802
gb:MT039887|Organism:Severe      ACGTCTATAATCCGTTTATGATTGATGTTCAACAATGGGGTTTTACAGGTAACCTACAAA   18802
gb:MT019530|Organism:Severe      ACGTCTATAATCCGTTTATGATTGATGTTCAACAATGGGGTTTTACAGGTAACCTACAAA   18802
gb:MT039888|Organism:Severe      ACGTCTATAATCCGTTTATGATTGATGTTCAACAATGGGGTTTTACAGGTAACCTACAAA   18802
gb:LC522972|Organism:Severe      ACGTCTATAATCCGTTTATGATTGATGTTCAACAATGGGGTTTTACAGGTAACCTACAAA   18799
gb:MT027063|Organism:Severe      ACGTCTATAATCCGTTTATGATTGATGTTCAACAATGGGGTTTTACAGGTAACCTACAAA   18802
gb:MT027062|Organism:Severe      ACGTCTATAATCCGTTTATGATTGATGTTCAACAATGGGGTTTTACAGGTAACCTACAAA   18802
```

```
gb:MT019529|Organism:Severe      ACGTCTATAATCCGTTTATGATTGATGTTCAACAATGGGGTTTTACAGGTAACCTACAAA   18802
gb:MN996529|Organism:Severe      ACGTCTATAATCCGTTTATGATTGATGTTCAACAATGGGGTTTTACAGGTAACCTACAAA   18790
gb:MN996531|Organism:Severe      ACGTCTATAATCCGTTTATGATTGATGTTCAACAATGGGGTTTTACAGGTAACCTACAAA   18789
gb:MT066176|Organism:Severe      ACGTCTATAATCCGTTTATGATTGATGTTCAACAATGGGGTTTTACAGGTAACCTACAAA   18802
gb:MT027064|Organism:Severe      ACGTCTATAATCCGTTTATGATTGATGTTCAACAATGGGGTTTTACAGGTAACCTACAAA   18802
gb:MN994468|Organism:Severe      ACGTCTATAATCCGTTTATGATTGATGTTCAACAATGGGGTTTTACAGGTAACCTACAAA   18802
gb:MT072688|Organism:Severe      ACGTCTATAATCCGTTTATGATTGATGTTCAACAATGGGGTTTTACAGGTAACCTACAAA   18787
gb:MN996527|Organism:Severe      ACGTCTATAATCCGTTTATGATTGATGTTCAACAATGGGGTTTTACAGGTAACCTACAAA   18769
gb:MT093631|Organism:Severe      ACGTCTATAATCCGTTTATGATTGATGTTCAACAATGGGGTTTTACAGGTAACCTACAAA   18840
gb:MT106053|Organism:Severe      ACGTCTATAATCCGTTTATGATTGATGTTCAACAATGGGGTTTTACAGGTAACCTACAAA   18802
gb:MT019533|Organism:Severe      ACGTCTATAATCCGTTTATGATTGATGTTCAACAATGGGGTTTTACAGGTAACCTACAAA   18802
gb:MT019531|Organism:Severe      ACGTCTATAATCCGTTTATGATTGATGTTCAACAATGGGGTTTTACAGGTAACCTACAAA   18802
gb:MN996528|Organism:Severe      ACGTCTATAATCCGTTTATGATTGATGTTCAACAATGGGGTTTTACAGGTAACCTACAAA   18802
gb:MN996530|Organism:Severe      ACGTCTATAATCCGTTTATGATTGATGTTCAACAATGGGGTTTTACAGGTAACCTACAAA   18788
gb:MN908947|Organism:Severe      ACGTCTATAATCCGTTTATGATTGATGTTCAACAATGGGGTTTTACAGGTAACCTACAAA   18802
gb:MT019532|Organism:Severe      ACGTCTATAATCCGTTTATGATTGATGTTCAACAATGGGGTTTTACAGGTAACCTACAAA   18802
                                 ************************************************************

gb:MT020781|Organism:Severe      GCAACCATGATCTGTATTGTCAAGTCCATGGTAATGCACATGTAGCTAGTTGTGATGCAA   18850
gb:MT007544|Organism:Severe      GCAACCATGATCTGTATTGTCAAGTCCATGGTAATGCACATGTAGCTAGTTGTGATGCAA   18862
gb:MN994467|Organism:Severe      GCAACCATGATCTGTATTGTCAAGTCCATGGTAATGCACATGTAGCTAGTTGTGATGCAA   18862
gb:MT044257|Organism:Severe      GCAACCATGATCTGTATTGTCAAGTCCATGGTAATGCACATGTAGCTAGTTGTGATGCAA   18862
gb:MT106054|Organism:Severe      GCAACCATGATCTGTATTGTCAAGTCCATGGTAATGCACATGTAGCTAGTTGTGATGCAA   18862
gb:MT049951|Organism:Severe      GCAACCATGATCTGTATTGTCAAGTCCATGGTAATGCACATGTAGCTAGTTGTGATGCAA   18862
gb:MN975262|Organism:Severe      GCAACCATGATCTGTATTGTCAAGTCCATGGTAATGCACATGTAGCTAGTTGTGATGCAA   18862
gb:MT106052|Organism:Severe      GCAACCATGATCTGTATTGTCAAGTCCATGGTAATGCACATGTAGCTAGTTGTGATGCAA   18862
gb:LC522975|Organism:Severe      GCAACCATGATCTGTATTGTCAAGTCCATGGTAATGCACATGTAGCTAGTTGTGATGCAA   18859
gb:LC522973|Organism:Severe      GCAACCATGATCTGTATTGTCAAGTCCATGGTAATGCACATGTAGCTAGTTGTGATGCAA   18859
gb:LC522974|Organism:Severe      GCAACCATGATCTGTATTGTCAAGTCCATGGTAATGCACATGTAGCTAGTTGTGATGCAA   18859
gb:MN985325|Organism:Severe      GCAACCATGATCTGTATTGTCAAGTCCATGGTAATGCACATGTAGCTAGTTGTGATGCAA   18862
gb:MT020881|Organism:Severe      GCAACCATGATCTGTATTGTCAAGTCCATGGTAATGCACATGTAGCTAGTTGTGATGCAA   18862
gb:MT020880|Organism:Severe      GCAACCATGATCTGTATTGTCAAGTCCATGGTAATGCACATGTAGCTAGTTGTGATGCAA   18862
gb:MT066175|Organism:Severe      GCAACCATGATCTGTATTGTCAAGTCCATGGTAATGCACATGTAGCTAGTTGTGATGCAA   18862
gb:MN997409|Organism:Severe      GCAACCATGATCTGTATTGTCAAGTCCATGGTAATGCACATGTAGCTAGTTGTGATGCAA   18862
gb:MN938384|Organism:Severe      GCAACCATGATCTGTATTGTCAAGTCCATGGTAATGCACATGTAGCTAGTTGTGATGCAA   18830
gb:MT044258|Organism:Severe      GCAACCATGATCTGTATTGTCAAGTCCATGGTAATGCACATGTAGCTAGTTGTGATGCAA   18838
gb:MT039890|Organism:Severe      GCAACCATGATCTGTATTGTCAAGTCCATGGTAATGCACATGTAGCTAGTTGTGATGCAA   18862
gb:MN988713|Organism:Severe      GCAACCATGATCTGTATTGTCAAGTCCATGGTAATGCACATGTAGCTAGTTGTGATGCAA   18862
gb:LC521925|Organism:Severe      GCAACCATGATCTGTATTGTCAAGTCCATGGTAATGCACATGTAGCTAGTTGTGATGCAA   18835
gb:MT093571|Organism:Severe      GCAACCATGATCTGTATTGTCAAGTCCATGGTAATGCACATGTAGCTAGTTGTGATGCAA   18862
gb:MT039887|Organism:Severe      GCAACCATGATCTGTATTGTCAAGTCCATGGTAATGCACATGTAGCTAGTTGTGATGCAA   18862
gb:MT019530|Organism:Severe      GCAACCATGATCTGTATTGTCAAGTCCATGGTAATGCACATGTAGCTAGTTGTGATGCAA   18862
gb:MT039888|Organism:Severe      GCAACCATGATCTGTATTGTCAAGTCCATGGTAATGCACATGTAGCTAGTTGTGATGCAA   18862
gb:LC522972|Organism:Severe      GCAACCATGATCTGTATTGTCAAGTCCATGGTAATGCACATGTAGCTAGTTGTGATGCAA   18859
gb:MT027063|Organism:Severe      GCAACCATGATCTGTATTGTCAAGTCCATGGTAATGCACATGTAGCTAGTTGTGATGCAA   18862
gb:MT027062|Organism:Severe      GCAACCATGATCTGTATTGTCAAGTCCATGGTAATGCACATGTAGCTAGTTGTGATGCAA   18862
gb:MT019529|Organism:Severe      GCAACCATGATCTGTATTGTCAAGTCCATGGTAATGCACATGTAGCTAGTTGTGATGCAA   18862
gb:MN996529|Organism:Severe      GCAACCATGATCTGTATTGTCAAGTCCATGGTAATGCACATGTAGCTAGTTGTGATGCAA   18850
gb:MN996531|Organism:Severe      GCAACCATGATCTGTATTGTCAAGTCCATGGTAATGCACATGTAGCTAGTTGTGATGCAA   18849
gb:MT066176|Organism:Severe      GCAACCATGATCTGTATTGTCAAGTCCATGGTAATGCACATGTAGCTAGTTGTGATGCAA   18862
gb:MT027064|Organism:Severe      GCAACCATGATCTGTATTGTCAAGTCCATGGTAATGCACATGTAGCTAGTTGTGATGCAA   18862
gb:MN994468|Organism:Severe      GCAACCATGATCTGTATTGTCAAGTCCATGGTAATGCACATGTAGCTAGTTGTGATGCAA   18862
gb:MT072688|Organism:Severe      GCAACCATGATCTGTATTGTCAAGTCCATGGTAATGCACATGTAGCTAGTTGTGATGCAA   18847
gb:MN996527|Organism:Severe      GCAACCATGATCTGTATTGTCAAGTCCATGGTAATGCACATGTAGCTAGTTGTGATGCAA   18829
gb:MT093631|Organism:Severe      GCAACCATGATCTGTATTGTCAAGTCCATGGTAATGCACATGTAGCTAGTTGTGATGCAA   18900
gb:MT106053|Organism:Severe      GCAACCATGATCTGTATTGTCAAGTCCATGGTAATGCACATGTAGCTAGTTGTGATGCAA   18862
gb:MT019533|Organism:Severe      GCAACCATGATCTGTATTGTCAAGTCCATGGTAATGCACATGTAGCTAGTTGTGATGCAA   18862
gb:MT019531|Organism:Severe      GCAACCATGATCTGTATTGTCAAGTCCATGGTAATGCACATGTAGCTAGTTGTGATGCAA   18862
gb:MN996528|Organism:Severe      GCAACCATGATCTGTATTGTCAAGTCCATGGTAATGCACATGTAGCTAGTTGTGATGCAA   18862
gb:MN996530|Organism:Severe      GCAACCATGATCTGTATTGTCAAGTCCATGGTAATGCACATGTAGCTAGTTGTGATGCAA   18848
gb:MN908947|Organism:Severe      GCAACCATGATCTGTATTGTCAAGTCCATGGTAATGCACATGTAGCTAGTTGTGATGCAA   18862
gb:MT019532|Organism:Severe      GCAACCATGATCTGTATTGTCAAGTCCATGGTAATGCACATGTAGCTAGTTGTGATGCAA   18862
                                 ************************************************************

gb:MT020781|Organism:Severe      TCATGACTAGGTGTCTAGCTGTCCACGAGTGCTTTGTTAAGCGTGTTGACTGGACTATTG   18910
```

```
gb:MT007544|Organism:Severe      TCATGACTAGGTGTCTAGCTGTCCACGAGTGCTTTGTTAAGCGTGTTGACTGGACTATTG   18922
gb:MN994467|Organism:Severe      TCATGACTAGGTGTCTAGCTGTCCACGAGTGCTTTGTTAAGCGTGTTGACTGGACTATTG   18922
gb:MT044257|Organism:Severe      TCATGACTAGGTGTCTAGCTGTCCACGAGTGCTTTGTTAAGCGTGTTGACTGGACTATTG   18922
gb:MT106054|Organism:Severe      TCATGACTAGGTGTCTAGCTGTCCACGAGTGCTTTGTTAAGCGTGTTGACTGGACTATTG   18922
gb:MT049951|Organism:Severe      TCATGACTAGGTGTCTAGCTGTCCACGAGTGCTTTGTTAAGCGTGTTGACTGGACTATTG   18922
gb:MN975262|Organism:Severe      TCATGACTAGGTGTCTAGCTGTCCACGAGTGCTTTGTTAAGCGTGTTGACTGGACTATTG   18922
gb:MT106052|Organism:Severe      TCATGACTAGGTGTCTAGCTGTCCACGAGTGCTTTGTTAAGCGTGTTGACTGGACTATTG   18922
gb:LC522975|Organism:Severe      TCATGACTAGGTGTCTAGCTGTCCACGAGTGCTTTGTTAAGCGTGTTGACTGGACTATTG   18919
gb:LC522973|Organism:Severe      TCATGACTAGGTGTCTAGCTGTCCACGAGTGCTTTGTTAAGCGTGTTGACTGGACTATTG   18919
gb:LC522974|Organism:Severe      TCATGACTAGGTGTCTAGCTGTCCACGAGTGCTTTGTTAAGCGTGTTGACTGGACTATTG   18919
gb:MN985325|Organism:Severe      TCATGACTAGGTGTCTAGCTGTCCACGAGTGCTTTGTTAAGCGTGTTGACTGGACTATTG   18922
gb:MT020881|Organism:Severe      TCATGACTAGGTGTCTAGCTGTCCACGAGTGCTTTGTTAAGCGTGTTGACTGGACTATTG   18922
gb:MT020880|Organism:Severe      TCATGACTAGGTGTCTAGCTGTCCACGAGTGCTTTGTTAAGCGTGTTGACTGGACTATTG   18922
gb:MT066175|Organism:Severe      TCATGACTAGGTGTCTAGCTGTCCACGAGTGCTTTGTTAAGCGTGTTGACTGGACTATTG   18922
gb:MN997409|Organism:Severe      TCATGACTAGGTGTCTAGCTGTCCACGAGTGCTTTGTTAAGCGTGTTGACTGGACTATTG   18922
gb:MN938384|Organism:Severe      TCATGACTAGGTGTCTAGCTGTCCACGAGTGCTTTGTTAAGCGTGTTGACTGGACTATTG   18890
gb:MT044258|Organism:Severe      TCATGACTAGGTGTCTAGCTGTCCACGAGTGCTTTGTTAAGCGTGTTGACTGGACTATTG   18898
gb:MT039890|Organism:Severe      TCATGACTAGGTGTCTAGCTGTCCACGAGTGCTTTGTTAAGCGTGTTGACTGGACTATTG   18922
gb:MN988713|Organism:Severe      TCATGACTAGGTGTCTAGCTGTCCACGAGTGCTTTGTTAAGCGTGTTGACTGGACTATTG   18922
gb:LC521925|Organism:Severe      TCATGACTAGGTGTCTAGCTGTCCACGAGTGCTTTGTTAAGCGTGTTGACTGGACTATTG   18895
gb:MT093571|Organism:Severe      TCATGACTAGGTGTCTAGCTGTCCACGAGTGCTTTGTTAAGCGTGTTGACTGGACTATTG   18922
gb:MT039887|Organism:Severe      TCATGACTAGGTGTCTAGCTGTCCACGAGTGCTTTGTTAAGCGTGTTGACTGGACTATTG   18922
gb:MT019530|Organism:Severe      TCATGACTAGGTGTCTAGCTGTCCACGAGTGCTTTGTTAAGCGTGTTGACTGGACTATTG   18922
gb:MT039888|Organism:Severe      TCATGACTAGGTGTCTAGCTGTCCACGAGTGCTTTGTTAAGCGTGTTGACTGGACTATTG   18922
gb:LC522972|Organism:Severe      TCATGACTAGGTGTCTAGCTGTCCACGAGTGCTTTGTTAAGCGTGTTGACTGGACTATTG   18919
gb:MT027063|Organism:Severe      TCATGACTAGGTGTCTAGCTGTCCACGAGTGCTTTGTTAAGCGTGTTGACTGGACTATTG   18922
gb:MT027062|Organism:Severe      TCATGACTAGGTGTCTAGCTGTCCACGAGTGCTTTGTTAAGCGTGTTGACTGGACTATTG   18922
gb:MT019529|Organism:Severe      TCATGACTAGGTGTCTAGCTGTCCACGAGTGCTTTGTTAAGCGTGTTGACTGGACTATTG   18922
gb:MN996529|Organism:Severe      TCATGACTAGGTGTCTAGCTGTCCACGAGTGCTTTGTTAAGCGTGTTGACTGGACTATTG   18910
gb:MN996531|Organism:Severe      TCATGACTAGGTGTCTAGCTGTCCACGAGTGCTTTGTTAAGCGTGTTGACTGGACTATTG   18909
gb:MT066176|Organism:Severe      TCATGACTAGGTGTCTAGCTGTCCACGAGTGCTTTGTTAAGCGTGTTGACTGGACTATTG   18922
gb:MT027064|Organism:Severe      TCATGACTAGGTGTCTAGCTGTCCACGAGTGCTTTGTTAAGCGTGTTGACTGGACTATTG   18922
gb:MN994468|Organism:Severe      TCATGACTAGGTGTCTAGCTGTCCACGAGTGCTTTGTTAAGCGTGTTGACTGGACTATTG   18922
gb:MT072688|Organism:Severe      TCATGACTAGGTGTCTAGCTGTCCACGAGTGCTTTGTTAAGCGTGTTGACTGGACTATTG   18907
gb:MN996527|Organism:Severe      TCATGACTAGGTGTCTAGCTGTCCACGAGTGCTTTGTTAAGCGTGTTGACTGGACTATTG   18889
gb:MT093631|Organism:Severe      TCATGACTAGGTGTCTAGCTGTCCACGAGTGCTTTGTTAAGCGTGTTGACTGGACTATTG   18960
gb:MT106053|Organism:Severe      TCATGACTAGGTGTCTAGCTGTCCACGAGTGCTTTGTTAAGCGTGTTGACTGGACTATTG   18922
gb:MT019533|Organism:Severe      TCATGACTAGGTGTCTAGCTGTCCACGAGTGCTTTGTTAAGCGTGTTGACTGGACTATTG   18922
gb:MT019531|Organism:Severe      TCATGACTAGGTGTCTAGCTGTCCACGAGTGCTTTGTTAAGCGTGTTGACTGGACTATTG   18922
gb:MN996528|Organism:Severe      TCATGACTAGGTGTCTAGCTGTCCACGAGTGCTTTGTTAAGCGTGTTGACTGGACTATTG   18922
gb:MN996530|Organism:Severe      TCATGACTAGGTGTCTAGCTGTCCACGAGTGCTTTGTTAAGCGTGTTGACTGGACTATTG   18908
gb:MN908947|Organism:Severe      TCATGACTAGGTGTCTAGCTGTCCACGAGTGCTTTGTTAAGCGTGTTGACTGGACTATTG   18922
gb:MT019532|Organism:Severe      TCATGACTAGGTGTCTAGCTGTCCACGAGTGCTTTGTTAAGCGTGTTGACTGGACTATTG   18922
                                 ************************************************************


gb:MT020781|Organism:Severe      AATATCCTATAATTGGTGATGAACTGAAGATTAATGCGGCTTGTAGAAAGGTTCAACACA   18970
gb:MT007544|Organism:Severe      AATATCCTATAATTGGTGATGAACTGAAGATTAATGCGGCTTGTAGAAAGGTTCAACACA   18982
gb:MN994467|Organism:Severe      AATATCCTATAATTGGTGATGAACTGAAGATTAATGCGGCTTGTAGAAAGGTTCAACACA   18982
gb:MT044257|Organism:Severe      AATATCCTATAATTGGTGATGAACTGAAGATTAATGCGGCTTGTAGAAAGGTTCAACACA   18982
gb:MT106054|Organism:Severe      AATATCCTATAATTGGTGATGAACTGAAGATTAATGCGGCTTGTAGAAAGGTACAACACA   18982
gb:MT049951|Organism:Severe      AATATCCTATAATTGGTGATGAACTGAAGATTAATGCGGCTTGTAGAAAGGTTCAACACA   18982
gb:MN975262|Organism:Severe      AATATCCTATAATTGGTGATGAACTGAAGATTAATGCGGCTTGTAGAAAGGTTCAACACA   18982
gb:MT106052|Organism:Severe      AATATCCTATAATTGGTGATGAACTGAAGATTAATGCGGCTTGTAGAAAGGTTCAACACA   18982
gb:LC522975|Organism:Severe      AATATCCTATAATTGGTGATGAACTGAAGATTAATGCGGCTTGTAGAAAGGTTCAACACA   18979
gb:LC522973|Organism:Severe      AATATCCTATAATTGGTGATGAACTGAAGATTAATGCGGCTTGTAGAAAGGTTCAACACA   18979
gb:LC522974|Organism:Severe      AATATCCTATAATTGGTGATGAACTGAAGATTAATGCGGCTTGTAGAAAGGTTCAACACA   18979
gb:MN985325|Organism:Severe      AATATCCTATAATTGGTGATGAACTGAAGATTAATGCGGCTTGTAGAAAGGTTCAACACA   18982
gb:MT020881|Organism:Severe      AATATCCTATAATTGGTGATGAACTGAAGATTAATGCGGCTTGTAGAAAGGTTCAACACA   18982
gb:MT020880|Organism:Severe      AATATCCTATAATTGGTGATGAACTGAAGATTAATGCGGCTTGTAGAAAGGTTCAACACA   18982
gb:MT066175|Organism:Severe      AATATCCTATAATTGGTGATGAACTGAAGATTAATGCGGCTTGTAGAAAGGTTCAACACA   18982
gb:MN997409|Organism:Severe      AATATCCTATAATTGGTGATGAACTGAAGATTAATGCGGCTTGTAGAAAGGTTCAACACA   18982
gb:MN938384|Organism:Severe      AATATCCTATAATTGGTGATGAACTGAAGATTAATGCGGCTTGTAGAAAGGTTCAACACA   18950
gb:MT044258|Organism:Severe      AATATCCTATAATTGGTGATGAACTGAAGATTAATGCGGCTTGTAGAAAGGTTCAACACA   18958
gb:MT039890|Organism:Severe      AATATCCTATAATTGGTGATGAACTGAAGATTAATGCGGCTTGTAGAAAGGTTCAACACA   18982
gb:MN988713|Organism:Severe      AATATCCTATAATTGGTGATGAACTGAAGATTAATGCGGCTTGTAGAAAGGTTCAACACA   18982
```

```
gb:LC521925|Organism:Severe      AATATCCTATAATTGGTGATGAACTGAAGATTAATGCGGCTTGTAGAAAGGTTCAACACA   18955
gb:MT093571|Organism:Severe      AATATCCTATAATTGGTGATGAACTGAAGATTAATGCGGCTTGTAGAAAGGTTCAACACA   18982
gb:MT039887|Organism:Severe      AATATCCTATAATTGGTGATGAACTGAAGATTAATGCGGCTTGTAGAAAGGTTCAACACA   18982
gb:MT019530|Organism:Severe      AATATCCTATAATTGGTGATGAACTGAAGATTAATGCGGCTTGTAGAAAGGTTCAACACA   18982
gb:MT039888|Organism:Severe      AATATCCTATAATTGGTGATGAACTGAAGATTAATGCGGCTTGTAGAAAGGTTCAACACA   18982
gb:LC522972|Organism:Severe      AATATCCTATAATTGGTGATGAACTGAAGATTAATGCGGCTTGTAGAAAGGTTCAACACA   18979
gb:MT027063|Organism:Severe      AATATCCTATAATTGGTGATGAACTGAAGATTAATGCGGCTTGTAGAAAGGTTCAACACA   18982
gb:MT027062|Organism:Severe      AATATCCTATAATTGGTGATGAACTGAAGATTAATGCGGCTTGTAGAAAGGTTCAACACA   18982
gb:MT019529|Organism:Severe      AATATCCTATAATTGGTGATGAACTGAAGATTAATGCGGCTTGTAGAAAGGTTCAACACA   18982
gb:MN996529|Organism:Severe      AATATCCTATAATTGGTGATGAACTGAAGATTAATGCGGCTTGTAGAAAGGTTCAACACA   18970
gb:MN996531|Organism:Severe      AATATCCTATAATTGGTGATGAACTGAAGATTAATGCGGCTTGTAGAAAGGTTCAACACA   18969
gb:MT066176|Organism:Severe      AATATCCTATAATTGGTGATGAACTGAAGATTAATGCGGCTTGTAGAAAGGTTCAACACA   18982
gb:MT027064|Organism:Severe      AATATCCTATAATTGGTGATGAACTGAAGATTAATGCGGCTTGTAGAAAGGTTCAACACA   18982
gb:MN994468|Organism:Severe      AATATCCTATAATTGGTGATGAACTGAAGATTAATGCGGCTTGTAGAAAGGTTCAACACA   18982
gb:MT072688|Organism:Severe      AATATCCTATAATTGGTGATGAACTGAAGATTAATGCGGCTTGTAGAAAGGTTCAACACA   18967
gb:MN996527|Organism:Severe      AATATCCTATAATTGGTGATGAACTGAAGATTAATGCGGCTTGTAGAAAGGTTCAACACA   18949
gb:MT093631|Organism:Severe      AATATCCTATAATTGGTGATGAACTGAAGATTAATGCGGCTTGTAGAAAGGTTCAACACA   19020
gb:MT106053|Organism:Severe      AATATCCTATAATTGGTGATGAACTGAAGATTAATGCGGCTTGTAGAAAGGTTCAACACA   18982
gb:MT019533|Organism:Severe      AATATCCTATAATTGGTGATGAACTGAAGATTAATGCGGCTTGTAGAAAGGTTCAACACA   18982
gb:MT019531|Organism:Severe      AATATCCTATAATTGGTGATGAACTGAAGATTAATGCGGCTTGTAGAAAGGTTCAACACA   18982
gb:MN996528|Organism:Severe      AATATCCTATAATTGGTGATGAACTGAAGATTAATGCGGCTTGTAGAAAGGTTCAACACA   18982
gb:MN996530|Organism:Severe      AATATCCTATAATTGGTGATGAACTGAAGATTAATGCGGCTTGTAGAAAGGTTCAACACA   18968
gb:MN908947|Organism:Severe      AATATCCTATAATTGGTGATGAACTGAAGATTAATGCGGCTTGTAGAAAGGTTCAACACA   18982
gb:MT019532|Organism:Severe      AATATCCTATAATTGGTGATGAACTGAAGATTAATGCGGCTTGTAGAAAGGTTCAACACA   18982
                                 **************************************************** *******

gb:MT020781|Organism:Severe      TGGTTGTTAAAGCTGCATTATTAGCAGACAAATTCCCAGTTCTTCACGACATTGGTAACC   19030
gb:MT007544|Organism:Severe      TGGTTGTTAAAGCTGCATTATTAGCAGACAAATTCCCAGTTCTTCACGACATTGGTAACC   19042
gb:MN994467|Organism:Severe      TGGTTGTTAAAGCTGCATTATTAGCAGACAAATTCCCAGTTCTTCACGACATTGGTAACC   19042
gb:MT044257|Organism:Severe      TGGTTGTTAAAGCTGCATTATTAGCAGACAAATTCCCAGTTCTTCACGACATTGGTAACC   19042
gb:MT106054|Organism:Severe      TGGTTGTTAAAGCTGCATTATTAGCAGACAAATTCCCAGTTCTTCACGACATTGGTAACC   19042
gb:MT049951|Organism:Severe      TGGTTGTTAAAGCTGCATTATTAGCAGACAAATTCCCAGTTCTTCACGACATTGGTAACC   19042
gb:MN975262|Organism:Severe      TGGTTGTTAAAGCTGCATTATTAGCAGACAAATTCCCAGTTCTTCACGACATTGGTAACC   19042
gb:MT106052|Organism:Severe      TGGTTGTTAAAGCTGCATTATTAGCAGACAAATTCCCAGTTCTTCACGACATTGGTAACC   19042
gb:LC522975|Organism:Severe      TGGTTGTTAAAGCTGCATTATTAGCAGACAAATTCCCAGTTCTTCACGACATTGGTAACC   19039
gb:LC522973|Organism:Severe      TGGTTGTTAAAGCTGCATTATTAGCAGACAAATTCCCAGTTCTTCACGACATTGGTAACC   19039
gb:LC522974|Organism:Severe      TGGTTGTTAAAGCTGCATTATTAGCAGACAAATTCCCAGTTCTTCACGACATTGGTAACC   19039
gb:MN985325|Organism:Severe      TGGTTGTTAAAGCTGCATTATTAGCAGACAAATTCCCAGTTCTTCACGACATTGGTAACC   19042
gb:MT020881|Organism:Severe      TGGTTGTTAAAGCTGCATTATTAGCAGACAAATTCCCAGTTCTTCACGACATTGGTAACC   19042
gb:MT020880|Organism:Severe      TGGTTGTTAAAGCTGCATTATTAGCAGACAAATTCCCAGTTCTTCACGACATTGGTAACC   19042
gb:MT066175|Organism:Severe      TGGTTGTTAAAGCTGCATTATTAGCAGACAAATTCCCAGTTCTTCACGACATTGGTAACC   19042
gb:MN997409|Organism:Severe      TGGTTGTTAAAGCTGCATTATTAGCAGACAAATTCCCAGTTCTTCACGACATTGGTAACC   19042
gb:MN938384|Organism:Severe      TGGTTGTTAAAGCTGCATTATTAGCAGACAAATTCCCAGTTCTTCACGACATTGGTAACC   19010
gb:MT044258|Organism:Severe      TGGTTGTTAAAGCTGCATTATTAGCAGACAAATTCCCAGTTCTTCACGACATTGGTAACC   19018
gb:MT039890|Organism:Severe      TGGTTGTTAAAGCTGCATTATTAGCAGACAAATTCCCAGTTCTTCACGACATTGGTAACC   19042
gb:MN988713|Organism:Severe      TGGTTGTTAAAGCTGCATTATTAGCAGACAAATTCCCAGTTCTTCACGACATTGGTAACC   19042
gb:LC521925|Organism:Severe      TGGTTGTTAAAGCTGCATTATTAGCAGACAAATTCCCAGTTCTTCACGACATTGGTAACC   19015
gb:MT093571|Organism:Severe      TGGTTGTTAAAGCTGCATTATTAGCAGACAAATTCCCAGTTCTTCACGACATTGGTAACC   19042
gb:MT039887|Organism:Severe      TGGTTGTTAAAGCTGCATTATTAGCAGACAAATTCCCAGTTCTTCACGACATTGGTAACC   19042
gb:MT019530|Organism:Severe      TGGTTGTTAAAGCTGCATTATTAGCAGACAAATTCCCAGTTCTTCACGACATTGGTAACC   19042
gb:MT039888|Organism:Severe      TGGTTGTTAAAGCTGCATTATTAGCAGACAAATTCCCAGTTCTTCACGACATTGGTAACC   19042
gb:LC522972|Organism:Severe      TGGTTGTTAAAGCTGCATTATTAGCAGACAAATTCCCAGTTCTTCACGACATTGGTAACC   19039
gb:MT027063|Organism:Severe      TGGTTGTTAAAGCTGCATTATTAGCAGACAAATTCCCAGTTCTTCACGACATTGGTAACC   19042
gb:MT027062|Organism:Severe      TGGTTGTTAAAGCTGCATTATTAGCAGACAAATTCCCAGTTCTTCACGACATTGGTAACC   19042
gb:MT019529|Organism:Severe      TGGTTGTTAAAGCTGCATTATTAGCAGACAAATTCCCAGTTCTTCACGACATTGGTAACC   19042
gb:MN996529|Organism:Severe      TGGTTGTTAAAGCTGCATTATTAGCAGACAAATTCCCAGTTCTTCACGACATTGGTAACC   19030
gb:MN996531|Organism:Severe      TGGTTGTTAAAGCTGCATTATTAGCAGACAAATTCCCAGTTCTTCACGACATTGGTAACC   19029
gb:MT066176|Organism:Severe      TGGTTGTTAAAGCTGCATTATTAGCAGACAAATTCCCAGTTCTTCACGACATTGGTAACC   19042
gb:MT027064|Organism:Severe      TGGTTGTTAAAGCTGCATTATTAGCAGACAAATTCCCAGTTCTTCACGACATTGGTAACC   19042
gb:MN994468|Organism:Severe      TGGTTGTTAAAGCTGCATTATTAGCAGACAAATTCCCAGTTCTTCACGACATTGGTAACC   19042
gb:MT072688|Organism:Severe      TGGTTGTTAAAGCTGCATTATTAGCAGACAAATTCCCAGTTCTTCACGACATTGGTAACC   19027
gb:MN996527|Organism:Severe      TGGTTGTTAAAGCTGCATTATTAGCAGACAAATTCCCAGTTCTTCACGACATTGGTAACC   19009
gb:MT093631|Organism:Severe      TGGTTGTTAAAGCTGCATTATTAGCAGACAAATTCCCAGTTCTTCACGACATTGGTAACC   19080
gb:MT106053|Organism:Severe      TGGTTGTTAAAGCTGCATTATTAGCAGACAAATTCCCAGTTCTTCACGACATTGGTAACC   19042
gb:MT019533|Organism:Severe      TGGTTGTTAAAGCTGCATTATTAGCAGACAAATTCCCAGTTCTTCACGACATTGGTAACC   19042
```

```
gb:MT019531|Organism:Severe      TGGTTGTTAAAGCTGCATTATTAGCAGACAAATTCCCAGTTCTTCACGACATTGGTAACC   19042
gb:MN996528|Organism:Severe      TGGTTGTTAAAGCTGCATTATTAGCAGACAAATTCCCAGTTCTTCACGACATTGGTAACC   19042
gb:MN996530|Organism:Severe      TGGTTGTTAAAGCTGCATTATTAGCAGACAAATTCCCAGTTCTTCACGACATTGGTAACC   19028
gb:MN908947|Organism:Severe      TGGTTGTTAAAGCTGCATTATTAGCAGACAAATTCCCAGTTCTTCACGACATTGGTAACC   19042
gb:MT019532|Organism:Severe      TGGTTGTTAAAGCTGCATTATTAGCAGACAAATTCCCAGTTCTTCACGACATTGGTAACC   19042
                                 ************************************************************

gb:MT020781|Organism:Severe      CTAAAGCTATTAAGTGTGTACCTCAAGCTGATGTAGAATGGAAGTTCTATGATGCACAGC   19090
gb:MT007544|Organism:Severe      CTAAAGCTATTAAGTGTGTACCCCAAGCTGATGTAGAATGGAAGTTCTATGATGCACAGC   19102
gb:MN994467|Organism:Severe      CTAAAGCTATTAAGTGTGTACCTCAAGCTGATGTAGAATGGAAGTTCTATGATGCACAGC   19102
gb:MT044257|Organism:Severe      CTAAAGCTATTAAGTGTGTACCTCAAGCTGATGTAGAATGGAAGTTCTATGATGCACAGC   19102
gb:MT106054|Organism:Severe      CTAAAGCTATTAAGTGTGTACCTCAAGCTGATGTAGAATGGAAGTTCTATGATGCACAGC   19102
gb:MT049951|Organism:Severe      CTAAAGCTATTAAGTGTGTACCTCAAGCTGATGTAGAATGGAAGTTCTATGATGCACAGC   19102
gb:MN975262|Organism:Severe      CTAAAGCTATTAAGTGTGTACCTCAAGCTGATGTAGAATGGAAGTTCTATGATGCACAGC   19102
gb:MT106052|Organism:Severe      CTAAAGCTATTAAGTGTGTACCTCAAGCTGATGTAGAATGGAAGTTCTATGATGCACAGC   19102
gb:LC522975|Organism:Severe      CTAAAGCTATTAAGTGTGTACCTCAAGCTGATGTAGAATGGAAGTTCTATGATGCACAGC   19099
gb:LC522973|Organism:Severe      CTAAAGCTATTAAGTGTGTACCTCAAGCTGATGTAGAATGGAAGTTCTATGATGCACAGC   19099
gb:LC522974|Organism:Severe      CTAAAGCTATTAAGTGTGTACCTCAAGCTGATGTAGAATGGAAGTTCTATGATGCACAGC   19099
gb:MN985325|Organism:Severe      CTAAAGCTATTAAGTGTGTACCTCAAGCTGATGTAGAATGGAAGTTCTATGATGCACAGC   19102
gb:MT020881|Organism:Severe      CTAAAGCTATTAAGTGTGTACCTCAAGCTGATGTAGAATGGAAGTTCTATGATGCACAGC   19102
gb:MT020880|Organism:Severe      CTAAAGCTATTAAGTGTGTACCTCAAGCTGATGTAGAATGGAAGTTCTATGATGCACAGC   19102
gb:MT066175|Organism:Severe      CTAAAGCTATTAAGTGTGTACCTCAAGCTGATGTAGAATGGAAGTTCTATGATGCACAGC   19102
gb:MN997409|Organism:Severe      CTAAAGCTATTAAGTGTGTACCTCAAGCTGATGTAGAATGGAAGTTCTATGATGCACAGC   19102
gb:MN938384|Organism:Severe      CTAAAGCTATTAAGTGTGTACCTCAAGCTGATGTAGAATGGAAGTTCTATGATGCACAGC   19070
gb:MT044258|Organism:Severe      CTAAAGCTATTAAGTGTGTACCTCAAGCTGATGTAGAATGGAAGTTCTATGATGCACAGC   19078
gb:MT039890|Organism:Severe      CTAAAGCTATTAAGTGTGTACCTCAAGCTGATGTAGAATGGAAGTTCTATGATGCACAGC   19102
gb:MN988713|Organism:Severe      CTAAAGCTATTAAGTGTGTACCTCAAGCTGATGTAGAATGGAAGTTCTATGATGCACAGC   19102
gb:LC521925|Organism:Severe      CTAAAGCTATTAAGTGTGTACCTCAAGCTGATGTAGAATGGAAGTTCTATGATGCACAGC   19075
gb:MT093571|Organism:Severe      CTAAAGCTATTAAGTGTGTACCTCAAGCTGATGTAGAATGGAAGTTCTATGATGCACAGC   19102
gb:MT039887|Organism:Severe      CTAAAGCTATTAAGTGTGTACCTCAAGCTGATGTAGAATGGAAGTTCTATGATGCACAGC   19102
gb:MT019530|Organism:Severe      CTAAAGCTATTAAGTGTGTACCTCAAGCTGATGTAGAATGGAAGTTCTATGATGCACAGC   19102
gb:MT039888|Organism:Severe      CTAAAGCTATTAAGTGTGTACCTCAAGCTGATGTAGAATGGAAGTTCTATGATGCACAGC   19102
gb:LC522972|Organism:Severe      CTAAAGCTATTAAGTGTGTACCTCAAGCTGATGTAGAATGGAAGTTCTATGATGCACAGC   19099
gb:MT027063|Organism:Severe      CTAAAGCTATTAAGTGTGTACCTCAAGCTGATGTAGAATGGAAGTTCTATGATGCACAGC   19102
gb:MT027062|Organism:Severe      CTAAAGCTATTAAGTGTGTACCTCAAGCTGATGTAGAATGGAAGTTCTATGATGCACAGC   19102
gb:MT019529|Organism:Severe      CTAAAGCTATTAAGTGTGTACCTCAAGCTGATGTAGAATGGAAGTTCTATGATGCACAGC   19102
gb:MN996529|Organism:Severe      CTAAAGCTATTAAGTGTGTACCTCAAGCTGATGTAGAATGGAAGTTCTATGATGCACAGC   19090
gb:MN996531|Organism:Severe      CTAAAGCTATTAAGTGTGTACCTCAAGCTGATGTAGAATGGAAGTTCTATGATGCACAGC   19089
gb:MT066176|Organism:Severe      CTAAAGCTATTAAGTGTGTACCTCAAGCTGATGTAGAATGGAAGTTCTATGATGCACAGC   19102
gb:MT027064|Organism:Severe      CTAAAGCTATTAAGTGTGTACCTCAAGCTGATGTAGAATGGAAGTTCTATGATGCACAGC   19102
gb:MN994468|Organism:Severe      CTAAAGCTATTAAGTGTGTACCTCAAGCTGATGTAGAATGGAAGTTCTATGATGCACAGC   19102
gb:MT072688|Organism:Severe      CTAAAGCTATTAAGTGTGTACCTCAAGCTGATGTAGAATGGAAGTTCTATGATGCACAGC   19087
gb:MN996527|Organism:Severe      CTAAAGCTATTAAGTGTGTACCTCAAGCTGATGTAGAATGGAAGTTCTATGATGCACAGC   19069
gb:MT093631|Organism:Severe      CTAAAGCTATTAAGTGTGTACCTCAAGCTGATGTAGAATGGAAGTTCTATGATGCACAGC   19140
gb:MT106053|Organism:Severe      CTAAAGCTATTAAGTGTGTACCTCAAGCTGATGTAGAATGGAAGTTCTATGATGCACAGC   19102
gb:MT019533|Organism:Severe      CTAAAGCTATTAAGTGTGTACCTCAAGCTGATGTAGAATGGAAGTTCTATGATGCACAGC   19102
gb:MT019531|Organism:Severe      CTAAAGCTATTAAGTGTGTACCTCAAGCTGATGTAGAATGGAAGTTCTATGATGCACAGC   19102
gb:MN996528|Organism:Severe      CTAAAGCTATTAAGTGTGTACCTCAAGCTGATGTAGAATGGAAGTTCTATGATGCACAGC   19102
gb:MN996530|Organism:Severe      CTAAAGCTATTAAGTGTGTACCTCAAGCTGATGTAGAATGGAAGTTCTATGATGCACAGC   19088
gb:MN908947|Organism:Severe      CTAAAGCTATTAAGTGTGTACCTCAAGCTGATGTAGAATGGAAGTTCTATGATGCACAGC   19102
gb:MT019532|Organism:Severe      CTAAAGCTATTAAGTGTGTACCTCAAGCTGATGTAGAATGGAAGTTCTATGATGCACAGC   19102
                                 ********************** *************************************

gb:MT020781|Organism:Severe      CTTGTAGTGACAAAGCTTATAAAATAGAAGAATTATTCTATTCTTATGCCACACATTCTG   19150
gb:MT007544|Organism:Severe      CTTGTAGTGACAAAGCTTATAAAATAGAAGAATTATTCTATTCTTATGCCACACATTCTG   19162
gb:MN994467|Organism:Severe      CTTGTAGTGACAAAGCTTATAAAATAGAAGAATTATTCTATTCTTATGCCACACATTCTG   19162
gb:MT044257|Organism:Severe      CTTGTAGTGACAAAGCTTATAAAATAGAAGAATTATTCTATTCTTATGCCACACATTCTG   19162
gb:MT106054|Organism:Severe      CTTGTAGTGACAAAGCTTATAAAATAGAAGAATTATTCTATTCTTATGCCACACATTCTG   19162
gb:MT049951|Organism:Severe      CTTGTAGTGACAAAGCTTATAAAATAGAAGAATTATTCTATTCTTATGCCACACATTCTG   19162
gb:MN975262|Organism:Severe      CTTGTAGTGACAAAGCTTATAAAATAGAAGAATTATTCTATTCTTATGCCACACATTCTG   19162
gb:MT106052|Organism:Severe      CTTGTAGTGACAAAGCTTATAAAATAGAAGAATTATTCTATTCTTATGCCACACATTCTG   19162
gb:LC522975|Organism:Severe      CTTGTAGTGACAAAGCTTATAAAATAGAAGAATTATTCTATTCTTATGCCACACATTCTG   19159
gb:LC522973|Organism:Severe      CTTGTAGTGACAAAGCTTATAAAATAGAAGAATTATTCTATTCTTATGCCACACATTCTG   19159
gb:LC522974|Organism:Severe      CTTGTAGTGACAAAGCTTATAAAATAGAAGAATTATTCTATTCTTATGCCACACATTCTG   19159
gb:MN985325|Organism:Severe      CTTGTAGTGACAAAGCTTATAAAATAGAAGAATTATTCTATTCTTATGCCACACATTCTG   19162
```

```
gb:MT020881|Organism:Severe      CTTGTAGTGACAAAGCTTATAAAATAGAAGAATTATTCTATTCTTATGCCACACATTCTG   19162
gb:MT020880|Organism:Severe      CTTGTAGTGACAAAGCTTATAAAATAGAAGAATTATTCTATTCTTATGCCACACATTCTG   19162
gb:MT066175|Organism:Severe      CTTGTAGTGACAAAGCTTATAAAATAGAAGAATTATTCTATTCTTATGCCACACATTCTG   19162
gb:MN997409|Organism:Severe      CTTGTAGTGACAAAGCTTATAAAATAGAAGAATTATTCTATTCTTATGCCACACATTCTG   19162
gb:MN938384|Organism:Severe      CTTGTAGTGACAAAGCTTATAAAATAGAAGAATTATTCTATTCTTATGCCACACATTCTG   19130
gb:MT044258|Organism:Severe      CTTGTAGTGACAAAGCTTATAAAATAGAAGAATTATTCTATTCTTATGCCACACATTCTG   19138
gb:MT039890|Organism:Severe      CTTGTAGTGACAAAGCTTATAAAATAGAAGAATTATTCTATTCTTATGCCACACATTCTG   19162
gb:MN988713|Organism:Severe      CTTGTAGTGACAAAGCTTATAAAATAGAAGAATTATTCTATTCTTATGCCACACATTCTG   19162
gb:LC521925|Organism:Severe      CTTGTAGTGACAAAGCTTATAAAATAGAAGAATTATTCTATTCTTATGCCACACATTCTG   19135
gb:MT093571|Organism:Severe      CTTGTAGTGACAAAGCTTATAAAATAGAAGAATTATTCTATTCTTATGCCACACATTCTG   19162
gb:MT039887|Organism:Severe      CTTGTAGTGACAAAGCTTATAAAATAGAAGAATTATTCTATTCTTATGCCACACATTCTG   19162
gb:MT019530|Organism:Severe      CTTGTAGTGACAAAGCTTATAAAATAGAAGAATTATTCTATTCTTATGCCACACATTCTG   19162
gb:MT039888|Organism:Severe      CTTGTAGTGACAAAGCTTATAAAATAGAAGAATTATTCTATTCTTATGCCACACATTCTG   19162
gb:LC522972|Organism:Severe      CTTGTAGTGACAAAGCTTATAAAATAGAAGAATTATTCTATTCTTATGCCACACATTCTG   19159
gb:MT027063|Organism:Severe      CTTGTAGTGACAAAGCTTATAAAATAGAAGAATTATTCTATTCTTATGCCACACATTCTG   19162
gb:MT027062|Organism:Severe      CTTGTAGTGACAAAGCTTATAAAATAGAAGAATTATTCTATTCTTATGCCACACATTCTG   19162
gb:MT019529|Organism:Severe      CTTGTAGTGACAAAGCTTATAAAATAGAAGAATTATTCTATTCTTATGCCACACATTCTG   19162
gb:MN996529|Organism:Severe      CTTGTAGTGACAAAGCTTATAAAATAGAAGAATTATTCTATTCTTATGCCACACATTCTG   19150
gb:MN996531|Organism:Severe      CTTGTAGTGACAAAGCTTATAAAATAGAAGAATTATTCTATTCTTATGCCACACATTCTG   19149
gb:MT066176|Organism:Severe      CTTGTAGTGACAAAGCTTATAAAATAGAAGAATTATTCTATTCTTATGCCACACATTCTG   19162
gb:MT027064|Organism:Severe      CTTGTAGTGACAAAGCTTATAAAATAGAAGAATTATTCTATTCTTATGCCACACATTCTG   19162
gb:MN994468|Organism:Severe      CTTGTAGTGACAAAGCTTATAAAATAGAAGAATTATTCTATTCTTATGCCACACATTCTG   19162
gb:MT072688|Organism:Severe      CTTGTAGTGACAAAGCTTATAAAATAGAAGAATTATTCTATTCTTATGCCACACATTCTG   19147
gb:MN996527|Organism:Severe      CTTGTAGTGACAAAGCTTATAAAATAGAAGAATTATTCTATTCTTATGCCACACATTCTG   19129
gb:MT093631|Organism:Severe      CTTGTAGTGACAAAGCTTATAAAATAGAAGAATTATTCTATTCTTATGCCACACATTCTG   19200
gb:MT106053|Organism:Severe      CTTGTAGTGACAAAGCTTATAAAATAGAAGAATTATTCTATTCTTATGCCACACATTCTG   19162
gb:MT019533|Organism:Severe      CTTGTAGTGACAAAGCTTATAAAATAGAAGAATTATTCTATTCTTATGCCACACATTCTG   19162
gb:MT019531|Organism:Severe      CTTGTAGTGACAAAGCTTATAAAATAGAAGAATTATTCTATTCTTATGCCACACATTCTG   19162
gb:MN996528|Organism:Severe      CTTGTAGTGACAAAGCTTATAAAATAGAAGAATTATTCTATTCTTATGCCACACATTCTG   19162
gb:MN996530|Organism:Severe      CTTGTAGTGACAAAGCTTATAAAATAGAAGAATTATTCTATTCTTATGCCACACATTCTG   19148
gb:MN908947|Organism:Severe      CTTGTAGTGACAAAGCTTATAAAATAGAAGAATTATTCTATTCTTATGCCACACATTCTG   19162
gb:MT019532|Organism:Severe      CTTGTAGTGACAAAGCTTATAAAATAGAAGAATTATTCTATTCTTATGCCACACATTCTG   19162
                                 ************************************************************

gb:MT020781|Organism:Severe      ACAAATTCACAGATGGTGTATGCCTATTTTGGAATTGCAATGTCGATAGATATCCTGCTA   19210
gb:MT007544|Organism:Severe      ACAAATTCACAGATGGTGTATGCCTATTTTGGAATTGCAATGTCGATAGATATCCTGCTA   19222
gb:MN994467|Organism:Severe      ACAAATTCACAGATGGTGTATGCCTATTTTGGAATTGCAATGTCGATAGATATCCTGCTA   19222
gb:MT044257|Organism:Severe      ACAAATTCACAGATGGTGTATGCCTATTTTGGAATTGCAATGTCGATAGATATCCTGCTA   19222
gb:MT106054|Organism:Severe      ACAAATTCACAGCTGGTGTATGCCTATTTTGGAATTGCAATGTCGATAGATATCCTGCTA   19222
gb:MT049951|Organism:Severe      ACAAATTCACAGATGGTGTATGCCTATTTTGGAATTGCAATGTCGATAGATATCCTGCTA   19222
gb:MN975262|Organism:Severe      ACAAATTCACAGATGGTGTATGCCTATTTTGGAATTGCAATGTCGATAGATATCCTGCTA   19222
gb:MT106052|Organism:Severe      ACAAATTCACAGATGGTGTATGCCTATTTTGGAATTGCAATGTCGATAGATATCCTGCTA   19222
gb:LC522975|Organism:Severe      ACAAATTCACAGATGGTGTATGCCTATTTTGGAATTGCAATGTCGATAGATATCCTGCTA   19219
gb:LC522973|Organism:Severe      ACAAATTCACAGATGGTGTATGCCTATTTTGGAATTGCAATGTCGATAGATATCCTGCTA   19219
gb:LC522974|Organism:Severe      ACAAATTCACAGATGGTGTATGCCTATTTTGGAATTGCAATGTCGATAGATATCCTGCTA   19219
gb:MN985325|Organism:Severe      ACAAATTCACAGATGGTGTATGCCTATTTTGGAATTGCAATGTCGATAGATATCCTGCTA   19222
gb:MT020881|Organism:Severe      ACAAATTCACAGATGGTGTATGCCTATTTTGGAATTGCAATGTCGATAGATATCCTGCTA   19222
gb:MT020880|Organism:Severe      ACAAATTCACAGATGGTGTATGCCTATTTTGGAATTGCAATGTCGATAGATATCCTGCTA   19222
gb:MT066175|Organism:Severe      ACAAATTCACAGATGGTGTATGCCTATTTTGGAATTGCAATGTCGATAGATATCCTGCTA   19222
gb:MN997409|Organism:Severe      ACAAATTCACAGATGGTGTATGCCTATTTTGGAATTGCAATGTCGATAGATATCCTGCTA   19222
gb:MN938384|Organism:Severe      ACAAATTCACAGATGGTGTATGCCTATTTTGGAATTGCAATGTCGATAGATATCCTGCTA   19190
gb:MT044258|Organism:Severe      ACAAATTCACAGATGGTGTATGCCTATTTTGGAATTGCAATGTCGATAGATATCCTGCTA   19198
gb:MT039890|Organism:Severe      ACAAATTCACAGATGGTGTATGCCTATTTTGGAATTGCAATGTCGATAGATATCCTGCTA   19222
gb:MN988713|Organism:Severe      ACAAATTCACAGATGGTGTATGCCTATTTTGGAATTGCAATGTCGATAGATATCCTGCTA   19222
gb:LC521925|Organism:Severe      ACAAATTCACAGATGGTGTATGCCTATTTTGGAATTGCAATGTCGATAGATATCCTGCTA   19195
gb:MT093571|Organism:Severe      ACAAATTCACAGATGGTGTATGCCTATTTTGGAATTGCAATGTCGATAGATATCCTGCTA   19222
gb:MT039887|Organism:Severe      ACAAATTCACAGATGGTGTATGCCTATTTTGGAATTGCAATGTCGATAGATATCCTGCTA   19222
gb:MT019530|Organism:Severe      ACAAATTCACAGATGGTGTATGCCTATTTTGGAATTGCAATGTCGATAGATATCCTGCTA   19222
gb:MT039888|Organism:Severe      ACAAATTCACAGATGGTGTATGCCTATTTTGGAATTGCAATGTCGATAGATATCCTGCTA   19222
gb:LC522972|Organism:Severe      ACAAATTCACAGATGGTGTATGCCTATTTTGGAATTGCAATGTCGATAGATATCCTGCTA   19219
gb:MT027063|Organism:Severe      ACAAATTCACAGATGGTGTATGCCTATTTTGGAATTGCAATGTCGATAGATATCCTGCTA   19222
gb:MT027062|Organism:Severe      ACAAATTCACAGATGGTGTATGCCTATTTTGGAATTGCAATGTCGATAGATATCCTGCTA   19222
gb:MT019529|Organism:Severe      ACAAATTCACAGATGGTGTATGCCTATTTTGGAATTGCAATGTCGATAGATATCCTGCTA   19222
gb:MN996529|Organism:Severe      ACAAATTCACAGATGGTGTATGCCTATTTTGGAATTGCAATGTCGATAGATATCCTGCTA   19210
gb:MN996531|Organism:Severe      ACAAATTCACAGATGGTGTATGCCTATTTTGGAATTGCAATGTCGATAGATATCCTGCTA   19209
```

```
gb:MT066176|Organism:Severe      ACAAATTCACAGATGGTGTATGCCTATTTTGGAATTGCAATGTCGATAGATATCCTGCTA   19222
gb:MT027064|Organism:Severe      ACAAATTCACAGATGGTGTATGCCTATTTTGGAATTGCAATGTCGATAGATATCCTGCTA   19222
gb:MN994468|Organism:Severe      ACAAATTCACAGATGGTGTATGCCTATTTTGGAATTGCAATGTCGATAGATATCCTGCTA   19222
gb:MT072688|Organism:Severe      ACAAATTCACAGATGGTGTATGCCTATTTTGGAATTGCAATGTCGATAGATATCCTGCTA   19207
gb:MN996527|Organism:Severe      ACAAATTCACAGATGGTGTATGCCTATTTTGGAATTGCAATGTCGATAGATATCCTGCTA   19189
gb:MT093631|Organism:Severe      ACAAATTCACAGATGGTGTATGCCTATTTTGGAATTGCAATGTCGATAGATATCCTGCTA   19260
gb:MT106053|Organism:Severe      ACAAATTCACAGATGGTGTATGCCTATTTTGGAATTGCAATGTCGATAGATATCCTGCTA   19222
gb:MT019533|Organism:Severe      ACAAATTCACAGATGGTGTATGCCTATTTTGGAATTGCAATGTCGATAGATATCCTGCTA   19222
gb:MT019531|Organism:Severe      ACAAATTCACAGATGGTGTATGCCTATTTTGGAATTGCAATGTCGATAGATATCCTGCTA   19222
gb:MN996528|Organism:Severe      ACAAATTCACAGATGGTGTATGCCTATTTTGGAATTGCAATGTCGATAGATATCCTGCTA   19222
gb:MN996530|Organism:Severe      ACAAATTCACAGATGGTGTATGCCTATTTTGGAATTGCAATGTCGATAGATATCCTGCTA   19208
gb:MN908947|Organism:Severe      ACAAATTCACAGATGGTGTATGCCTATTTTGGAATTGCAATGTCGATAGATATCCTGCTA   19222
gb:MT019532|Organism:Severe      ACAAATTCACAGATGGTGTATGCCTATTTTGGAATTGCAATGTCGATAGATATCCTGCTA   19222
                                 ************ ***********************************************

gb:MT020781|Organism:Severe      ATTCCATTGTTTGTAGATTTGACACTAGAGTGCTATCTAACCTTAACTTGCCTGGTTGTG   19270
gb:MT007544|Organism:Severe      ATTCCATTGTTTGTAGATTTGACACTAGAGTGCTATCTAACCTTAACTTGCCTGGTTGTG   19282
gb:MN994467|Organism:Severe      ATTCCATTGTTTGTAGATTTGACACTAGAGTGCTATCTAACCTTAACTTGCCTGGTTGTG   19282
gb:MT044257|Organism:Severe      ATTCCATTGTTTGTAGATTTGACACTAGAGTGCTATCTAACCTTAACTTGCCTGGTTGTG   19282
gb:MT106054|Organism:Severe      ATTCCATTGTTTGTAGATTTGACACTAGAGTGCTATCTAACCTTAACTTGCCTGGTTGTG   19282
gb:MT049951|Organism:Severe      ATTCCATTGTTTGTAGATTTGACACTAGAGTGCTATCTAACCTTAACTTGCCTGGTTGTG   19282
gb:MN975262|Organism:Severe      ATTCCATTGTTTGTAGATTTGACACTAGAGTGCTATCTAACCTTAACTTGCCTGGTTGTG   19282
gb:MT106052|Organism:Severe      ATTCCATTGTTTGTAGATTTGACACTAGAGTGCTATCTAACCTTAACTTGCCTGGTTGTG   19282
gb:LC522975|Organism:Severe      ATTCCATTGTTTGTAGATTTGACACTAGAGTGCTATCTAACCTTAACTTGCCTGGTTGTG   19279
gb:LC522973|Organism:Severe      ATTCCATTGTTTGTAGATTTGACACTAGAGTGCTATCTAACCTTAACTTGCCTGGTTGTG   19279
gb:LC522974|Organism:Severe      ATTCCATTGTTTGTAGATTTGACACTAGAGTGCTATCTAACCTTAACTTGCCTGGTTGTG   19279
gb:MN985325|Organism:Severe      ATTCCATTGTTTGTAGATTTGACACTAGAGTGCTATCTAACCTTAACTTGCCTGGTTGTG   19282
gb:MT020881|Organism:Severe      ATTCCATTGTTTGTAGATTTGACACTAGAGTGCTATCTAACCTTAACTTGCCTGGTTGTG   19282
gb:MT020880|Organism:Severe      ATTCCATTGTTTGTAGATTTGACACTAGAGTGCTATCTAACCTTAACTTGCCTGGTTGTG   19282
gb:MT066175|Organism:Severe      ATTCCATTGTTTGTAGATTTGACACTAGAGTGCTATCTAACCTTAACTTGCCTGGTTGTG   19282
gb:MN997409|Organism:Severe      ATTCCATTGTTTGTAGATTTGACACTAGAGTGCTATCTAACCTTAACTTGCCTGGTTGTG   19282
gb:MN938384|Organism:Severe      ATTCCATTGTTTGTAGATTTGACACTAGAGTGCTATCTAACCTTAACTTGCCTGGTTGTG   19250
gb:MT044258|Organism:Severe      ATTCCATTGTTTGTAGATTTGACACTAGAGTGCTATCTAACCTTAACTTGCCTGGTTGTG   19258
gb:MT039890|Organism:Severe      ATTCCATTGTTTGTAGATTTGACACTAGAGTGCTATCTAACCTTAACTTGCCTGGTTGTG   19282
gb:MN988713|Organism:Severe      ATTCCATTGTTTGTAGATTTGACACTAGAGTGCTATCTAACCTTAACTTGCCTGGTTGTG   19282
gb:LC521925|Organism:Severe      ATTCCATTGTTTGTAGATTTGACACTAGAGTGCTATCTAACCTTAACTTGCCTGGTTGTG   19255
gb:MT093571|Organism:Severe      ATTCCATTGTTTGTAGATTTGACACTAGAGTGCTATCTAACCTTAACTTGCCTGGTTGTG   19282
gb:MT039887|Organism:Severe      ATTCCATTGTTTGTAGATTTGACACTAGAGTGCTATCTAACCTTAACTTGCCTGGTTGTG   19282
gb:MT019530|Organism:Severe      ATTCCATTGTTTGTAGATTTGACACTAGAGTGCTATCTAACCTTAACTTGCCTGGTTGTG   19282
gb:MT039888|Organism:Severe      ATTCCATTGTTTGTAGATTTGACACTAGAGTGCTATCTAACCTTAACTTGCCTGGTTGTG   19282
gb:LC522972|Organism:Severe      ATTCCATTGTTTGTAGATTTGACACTAGAGTGCTATCTAACCTTAACTTGCCTGGTTGTG   19279
gb:MT027063|Organism:Severe      ATTCCATTGTTTGTAGATTTGACACTAGAGTGCTATCTAACCTTAACTTGCCTGGTTGTG   19282
gb:MT027062|Organism:Severe      ATTCCATTGTTTGTAGATTTGACACTAGAGTGCTATCTAACCTTAACTTGCCTGGTTGTG   19282
gb:MT019529|Organism:Severe      ATTCCATTGTTTGTAGATTTGACACTAGAGTGCTATCTAACCTTAACTTGCCTGGTTGTG   19282
gb:MN996529|Organism:Severe      ATTCCATTGTTTGTAGATTTGACACTAGAGTGCTATCTAACCTTAACTTGCCTGGTTGTG   19270
gb:MN996531|Organism:Severe      ATTCCATTGTTTGTAGATTTGACACTAGAGTGCTATCTAACCTTAACTTGCCTGGTTGTG   19269
gb:MT066176|Organism:Severe      ATTCCATTGTTTGTAGATTTGACACTAGAGTGCTATCTAACCTTAACTTGCCTGGTTGTG   19282
gb:MT027064|Organism:Severe      ATTCCATTGTTTGTAGATTTGACACTAGAGTGCTATCTAACCTTAACTTGCCTGGTTGTG   19282
gb:MN994468|Organism:Severe      ATTCCATTGTTTGTAGATTTGACACTAGAGTGCTATCTAACCTTAACTTGCCTGGTTGTG   19282
gb:MT072688|Organism:Severe      ATTCCATTGTTTGTAGATTTGACACTAGAGTGCTATCTAACCTTAACTTGCCTGGTTGTG   19267
gb:MN996527|Organism:Severe      ATTCCATTGTTTGTAGATTTGACACTAGAGTGCTATCTAACCTTAACTTGCCTGGTTGTG   19249
gb:MT093631|Organism:Severe      ATTCCATTGTTTGTAGATTTGACACTAGAGTGCTATCTAACCTTAACTTGCCTGGTTGTG   19320
gb:MT106053|Organism:Severe      ATTCCATTGTTTGTAGATTTGACACTAGAGTGCTATCTAACCTTAACTTGCCTGGTTGTG   19282
gb:MT019533|Organism:Severe      ATTCCATTGTTTGTAGATTTGACACTAGAGTGCTATCTAACCTTAACTTGCCTGGTTGTG   19282
gb:MT019531|Organism:Severe      ATTCCATTGTTTGTAGATTTGACACTAGAGTGCTATCTAACCTTAACTTGCCTGGTTGTG   19282
gb:MN996528|Organism:Severe      ATTCCATTGTTTGTAGATTTGACACTAGAGTGCTATCTAACCTTAACTTGCCTGGTTGTG   19282
gb:MN996530|Organism:Severe      ATTCCATTGTTTGTAGATTTGACACTAGAGTGCTATCTAACCTTAACTTGCCTGGTTGTG   19268
gb:MN908947|Organism:Severe      ATTCCATTGTTTGTAGATTTGACACTAGAGTGCTATCTAACCTTAACTTGCCTGGTTGTG   19282
gb:MT019532|Organism:Severe      ATTCCATTGTTTGTAGATTTGACACTAGAGTGCTATCTAACCTTAACTTGCCTGGTTGTG   19282
                                 ************************************************************

gb:MT020781|Organism:Severe      ATGGTGGCAGTTTGTATGTAAATAAACATGCATTCCACACACCAGCTTTTGATAAAAGTG   19330
gb:MT007544|Organism:Severe      ATGGTGGCAGTTTGTATGTAAATAAACATGCATTCCACACACCAGCTTTTGATAAAAGTG   19342
gb:MN994467|Organism:Severe      ATGGTGGCAGTTTGTATGTAAATAAACATGCATTCCACACACCAGCTTTTGATAAAAGTG   19342
gb:MT044257|Organism:Severe      ATGGTGGCAGTTTGTATGTAAATAAACATGCATTCCACACACCAGCTTTTGATAAAAGTG   19342
```

gb:MT106054|Organism:Severe ATGGTGGCAGTTTGTATGTAAATAAACATGCATTCCACACACCAGCTTTTGATAAAAGTG 19342
gb:MT049951|Organism:Severe ATGGTGGCAGTTTGTATGTAAATAAACATGCATTCCACACACCAGCTTTTGATAAAAGTG 19342
gb:MN975262|Organism:Severe ATGGTGGCAGTTTGTATGTAAATAAACATGCATTCCACACACCAGCTTTTGATAAAAGTG 19342
gb:MT106052|Organism:Severe ATGGTGGCAGTTTGTATGTAAATAAACATGCATTCCACACACCAGCTTTTGATAAAAGTG 19342
gb:LC522975|Organism:Severe ATGGTGGCAGTTTGTATGTAAATAAACATGCATTCCACACACCAGCTTTTGATAAAAGTG 19339
gb:LC522973|Organism:Severe ATGGTGGCAGTTTGTATGTAAATAAACATGCATTCCACACACCAGCTTTTGATAAAAGTG 19339
gb:LC522974|Organism:Severe ATGGTGGCAGTTTGTATGTAAATAAACATGCATTCCACACACCAGCTTTTGATAAAAGTG 19339
gb:MN985325|Organism:Severe ATGGTGGCAGTTTGTATGTAAATAAACATGCATTCCACACACCAGCTTTTGATAAAAGTG 19342
gb:MT020881|Organism:Severe ATGGTGGCAGTTTGTATGTAAATAAACATGCATTCCACACACCAGCTTTTGATAAAAGTG 19342
gb:MT020880|Organism:Severe ATGGTGGCAGTTTGTATGTAAATAAACATGCATTCCACACACCAGCTTTTGATAAAAGTG 19342
gb:MT066175|Organism:Severe ATGGTGGCAGTTTGTATGTAAATAAACATGCATTCCACACACCAGCTTTTGATAAAAGTG 19342
gb:MN997409|Organism:Severe ATGGTGGCAGTTTGTATGTAAATAAACATGCATTCCACACACCAGCTTTTGATAAAAGTG 19342
gb:MN938384|Organism:Severe ATGGTGGCAGTTTGTATGTAAATAAACATGCATTCCACACACCAGCTTTTGATAAAAGTG 19310
gb:MT044258|Organism:Severe ATGGTGGCAGTTTGTATGTAAATAAACATGCATTCCACACACCAGCTTTTGATAAAAGTG 19318
gb:MT039890|Organism:Severe ATGGTGGCAGTTTGTATGTAAATAAACATGCATTCCACACACCAGCTTTTGATAAAAGTG 19342
gb:MN988713|Organism:Severe ATGGTGGCAGTTTGTATGTAAATAAACATGCATTCCACACACCAGCTTTTGATAAAAGTG 19342
gb:LC521925|Organism:Severe ATGGTGGCAGTTTGTATGTAAATAAACATGCATTCCACACACCAGCTTTTGATAAAAGTG 19315
gb:MT093571|Organism:Severe ATGGTGGCAGTTTGTATGTAAATAAACATGCATTCCACACACCAGCTTTTGATAAAAGTG 19342
gb:MT039887|Organism:Severe ATGGTGGCAGTTTGTATGTAAATAAACATGCATTCCACACACCAGCTTTTGATAAAAGTG 19342
gb:MT019530|Organism:Severe ATGGTGGCAGTTTGTATGTAAATAAACATGCATTCCACACACCAGCTTTTGATAAAAGTG 19342
gb:MT039888|Organism:Severe ATGGTGGCAGTTTGTATGTAAATAAACATGCATTCCACACACCAGCTTTTGATAAAAGTG 19342
gb:LC522972|Organism:Severe ATGGTGGCAGTTTGTATGTAAATAAACATGCATTCCACACACCAGCTTTTGATAAAAGTG 19339
gb:MT027063|Organism:Severe ATGGTGGCAGTTTGTATGTAAATAAACATGCATTCCACACACCAGCTTTTGATAAAAGTG 19342
gb:MT027062|Organism:Severe ATGGTGGCAGTTTGTATGTAAATAAACATGCATTCCACACACCAGCTTTTGATAAAAGTG 19342
gb:MT019529|Organism:Severe ATGGTGGCAGTTTGTATGTAAATAAACATGCATTCCACACACCAGCTTTTGATAAAAGTG 19342
gb:MN996529|Organism:Severe ATGGTGGCAGTTTGTATGTAAATAAACATGCATTCCACACACCAGCTTTTGATAAAAGTG 19330
gb:MN996531|Organism:Severe ATGGTGGCAGTTTGTATGTAAATAAACATGCATTCCACACACCAGCTTTTGATAAAAGTG 19329
gb:MT066176|Organism:Severe ATGGTGGCAGTTTGTATGTAAATAAACATGCATTCCACACACCAGCTTTTGATAAAAGTG 19342
gb:MT027064|Organism:Severe ATGGTGGCAGTTTGTATGTAAATAAACATGCATTCCACACACCAGCTTTTGATAAAAGTG 19342
gb:MN994468|Organism:Severe ATGGTGGCAGTTTGTATGTAAATAAACATGCATTCCACACACCAGCTTTTGATAAAAGTG 19342
gb:MT072688|Organism:Severe ATGGTGGCAGTTTGTATGTAAATAAACATGCATTCCACACACCAGCTTTTGATAAAAGTG 19327
gb:MN996527|Organism:Severe ATGGTGGCAGTTTGTATGTAAATAAACATGCATTCCACACACCAGCTTTTGATAAAAGTG 19309
gb:MT093631|Organism:Severe ATGGTGGCAGTTTGTATGTAAATAAACATGCATTCCACACACCAGCTTTTGATAAAAGTG 19380
gb:MT106053|Organism:Severe ATGGTGGCAGTTTGTATGTAAATAAACATGCATTCCACACACCAGCTTTTGATAAAAGTG 19342
gb:MT019533|Organism:Severe ATGGTGGCAGTTTGTATGTAAATAAACATGCATTCCACACACCAGCTTTTGATAAAAGTG 19342
gb:MT019531|Organism:Severe ATGGTGGCAGTTTGTATGTAAATAAACATGCATTCCACACACCAGCTTTTGATAAAAGTG 19342
gb:MN996528|Organism:Severe ATGGTGGCAGTTTGTATGTAAATAAACATGCATTCCACACACCAGCTTTTGATAAAAGTG 19342
gb:MN996530|Organism:Severe ATGGTGGCAGTTTGTATGTAAATAAACATGCATTCCACACACCAGCTTTTGATAAAAGTG 19328
gb:MN908947|Organism:Severe ATGGTGGCAGTTTGTATGTAAATAAACATGCATTCCACACACCAGCTTTTGATAAAAGTG 19342
gb:MT019532|Organism:Severe ATGGTGGCAGTTTGTATGTAAATAAACATGCATTCCACACACCAGCTTTTGATAAAAGTG 19342
************************************************************

gb:MT020781|Organism:Severe CTTTTGTTAATTTAAAACAATTACCATTTTTCTATTACTCTGACAGTCCATGTGAGTCTC 19390
gb:MT007544|Organism:Severe CTTTTGTTAATTTAAAACAATTACCATTTTTCTATTACTCTGACAGTCCATGTGAGTCTC 19402
gb:MN994467|Organism:Severe CTTTTGTTAATTTAAAACAATTACCATTTTTCTATTACTCTGACAGTCCATGTGAGTCTC 19402
gb:MT044257|Organism:Severe CTTTTGTTAATTTAAAACAATTACCATTTTTCTATTACTCTGACAGTCCATGTGAGTCTC 19402
gb:MT106054|Organism:Severe CTTTTGTTAATTTAAAACAATTACCATTTTTCTATTACTCTGACAGTCCATGTGAGTCTC 19402
gb:MT049951|Organism:Severe CTTTTGTTAATTTAAAACAATTACCATTTTTCTATTACTCTGACAGTCCATGTGAGTCTC 19402
gb:MN975262|Organism:Severe CTTTTGTTAATTTAAAACAATTACCATTTTTCTATTACTCTGACAGTCCATGTGAGTCTC 19402
gb:MT106052|Organism:Severe CTTTTGTTAATTTAAAACAATTACCATTTTTCTATTACTCTGACAGTCCATGTGAGTCTC 19402
gb:LC522975|Organism:Severe CTTTTGTTAATTTAAAACAATTACCATTTTTCTATTACTCTGACAGTCCATGTGAGTCTC 19399
gb:LC522973|Organism:Severe CTTTTGTTAATTTAAAACAATTACCATTTTTCTATTACTCTGACAGTCCATGTGAGTCTC 19399
gb:LC522974|Organism:Severe CTTTTGTTAATTTAAAACAATTACCATTTTTCTATTACTCTGACAGTCCATGTGAGTCTC 19399
gb:MN985325|Organism:Severe CTTTTGTTAATTTAAAACAATTACCATTTTTCTATTACTCTGACAGTCCATGTGAGTCTC 19402
gb:MT020881|Organism:Severe CTTTTGTTAATTTAAAACAATTACCATTTTTCTATTACTCTGACAGTCCATGTGAGTCTC 19402
gb:MT020880|Organism:Severe CTTTTGTTAATTTAAAACAATTACCATTTTTCTATTACTCTGACAGTCCATGTGAGTCTC 19402
gb:MT066175|Organism:Severe CTTTTGTTAATTTAAAACAATTACCATTTTTCTATTACTCTGACAGTCCATGTGAGTCTC 19402
gb:MN997409|Organism:Severe CTTTTGTTAATTTAAAACAATTACCATTTTTCTATTACTCTGACAGTCCATGTGAGTCTC 19402
gb:MN938384|Organism:Severe CTTTTGTTAATTTAAAACAATTACCATTTTTCTATTACTCTGACAGTCCATGTGAGTCTC 19370
gb:MT044258|Organism:Severe CTTTTGTTAATTTAAAACAATTACCATTTTTCTATTACTCTGACAGTCCATGTGAGTCTC 19378
gb:MT039890|Organism:Severe CTTTTGTTAATTTAAAACAATTACCATTTTTCTATTACTCTGACAGTCCATGTGAGTCTC 19402
gb:MN988713|Organism:Severe CTTTTGTTAATTTAAAACAATTACCATTTTTCTATTACTCTGACAGTCCATGTGAGTCTC 19402
gb:LC521925|Organism:Severe CTTTTGTTAATTTAAAACAATTACCATTTTTCTATTACTCTGACAGTCCATGTGAGTCTC 19375
gb:MT093571|Organism:Severe CTTTTGTTAATTTAAAACAATTACCATTTTTCTATTACTCTGACAGTCCATGTGAGTCTC 19402
gb:MT039887|Organism:Severe CTTTTGTTAATTTAAAACAATTACCATTTTTCTATTACTCTGACAGTCCATGTGAGTCTC 19402

```
gb:MT019530|Organism:Severe      CTTTTGTTAATTTAAAACAATTACCATTTTTCTATTACTCTGACAGTCCATGTGAGTCTC    19402
gb:MT039888|Organism:Severe      CTTTTGTTAATTTAAAACAATTACCATTTTTCTATTACTCTGACAGTCCATGTGAGTCTC    19402
gb:LC522972|Organism:Severe      CTTTTGTTAATTTAAAACAATTACCATTTTTCTATTACTCTGACAGTCCATGTGAGTCTC    19399
gb:MT027063|Organism:Severe      CTTTTGTTAATTTAAAACAATTACCATTTTTCTATTACTCTGACAGTCCATGTGAGTCTC    19402
gb:MT027062|Organism:Severe      CTTTTGTTAATTTAAAACAATTACCATTTTTCTATTACTCTGACAGTCCATGTGAGTCTC    19402
gb:MT019529|Organism:Severe      CTTTTGTTAATTTAAAACAATTACCATTTTTCTATTACTCTGACAGTCCATGTGAGTCTC    19402
gb:MN996529|Organism:Severe      CTTTTGTTAATTTAAAACAATTACCATTTTTCTATTACTCTGACAGTCCATGTGAGTCTC    19390
gb:MN996531|Organism:Severe      CTTTTGTTAATTTAAAACAATTACCATTTTTCTATTACTCTGACAGTCCATGTGAGTCTC    19389
gb:MT066176|Organism:Severe      CTTTTGTTAATTTAAAACAATTACCATTTTTCTATTACTCTGACAGTCCATGTGAGTCTC    19402
gb:MT027064|Organism:Severe      CTTTTGTTAATTTAAAACAATTACCATTTTTCTATTACTCTGACAGTCCATGTGAGTCTC    19402
gb:MN994468|Organism:Severe      CTTTTGTTAATTTAAAACAATTACCATTTTTCTATTACTCTGACAGTCCATGTGAGTCTC    19402
gb:MT072688|Organism:Severe      CTTTTGTTAATTTAAAACAATTACCATTTTTCTATTACTCTGACAGTCCATGTGAGTCTC    19387
gb:MN996527|Organism:Severe      CTTTTGTTAATTTAAAACAATTACCATTTTTCTATTACTCTGACAGTCCATGTGAGTCTC    19369
gb:MT093631|Organism:Severe      CTTTTGTTAATTTAAAACAATTACCATTTTTCTATTACTCTGACAGTCCATGTGAGTCTC    19440
gb:MT106053|Organism:Severe      CTTTTGTTAATTTAAAACAATTACCATTTTTCTATTACTCTGACAGTCCATGTGAGTCTC    19402
gb:MT019533|Organism:Severe      CTTTTGTTAATTTAAAACAATTACCATTTTTCTATTACTCTGACAGTCCATGTGAGTCTC    19402
gb:MT019531|Organism:Severe      CTTTTGTTAATTTAAAACAATTACCATTTTTCTATTACTCTGACAGTCCATGTGAGTCTC    19402
gb:MN996528|Organism:Severe      CTTTTGTTAATTTAAAACAATTACCATTTTTCTATTACTCTGACAGTCCATGTGAGTCTC    19402
gb:MN996530|Organism:Severe      CTTTTGTTAATTTAAAACAATTACCATTTTTCTATTACTCTGACAGTCCATGTGAGTCTC    19388
gb:MN908947|Organism:Severe      CTTTTGTTAATTTAAAACAATTACCATTTTTCTATTACTCTGACAGTCCATGTGAGTCTC    19402
gb:MT019532|Organism:Severe      CTTTTGTTAATTTAAAACAATTACCATTTTTCTATTACTCTGACAGTCCATGTGAGTCTC    19402
                                 ************************************************************

gb:MT020781|Organism:Severe      ATGGAAAACAAGTAGTGTCAGATATAGATTATGTACCACTAAAGTCTGCTACGTGTATAA    19450
gb:MT007544|Organism:Severe      ATGGAAAACAAGTAGTGTCAGATATAGATTATGTACCACTAAAGTCTGCTACGTGTATAA    19462
gb:MN994467|Organism:Severe      ATGGAAAACAAGTAGTGTCAGATATAGATTATGTACCACTAAAGTCTGCTACGTGTATAA    19462
gb:MT044257|Organism:Severe      ATGGAAAACAAGTAGTGTCAGATATAGATTATGTACCACTAAAGTCTGCTACGTGTATAA    19462
gb:MT106054|Organism:Severe      ATGGAAAACAAGTAGTGTCAGATATAGATTATGTACCACTAAAGTCTGCTACGTGTATAA    19462
gb:MT049951|Organism:Severe      ATGGAAAACAAGTAGTGTCAGATATAGATTATGTACCACTAAAGTCTGCTACGTGTATAA    19462
gb:MN975262|Organism:Severe      ATGGAAAACAAGTAGTGTCAGATATAGATTATGTACCACTAAAGTCTGCTACGTGTATAA    19462
gb:MT106052|Organism:Severe      ATGGAAAACAAGTAGTGTCAGATATAGATTATGTACCACTAAAGTCTGCTACGTGTATAA    19462
gb:LC522975|Organism:Severe      ATGGAAAACAAGTAGTGTCAGATATAGATTATGTACCACTAAAGTCTGCTACGTGTATAA    19459
gb:LC522973|Organism:Severe      ATGGAAAACAAGTAGTGTCAGATATAGATTATGTACCACTAAAGTCTGCTACGTGTATAA    19459
gb:LC522974|Organism:Severe      ATGGAAAACAAGTAGTGTCAGATATAGATTATGTACCACTAAAGTCTGCTACGTGTATAA    19459
gb:MN985325|Organism:Severe      ATGGAAAACAAGTAGTGTCAGATATAGATTATGTACCACTAAAGTCTGCTACGTGTATAA    19462
gb:MT020881|Organism:Severe      ATGGAAAACAAGTAGTGTCAGATATAGATTATGTACCACTAAAGTCTGCTACGTGTATAA    19462
gb:MT020880|Organism:Severe      ATGGAAAACAAGTAGTGTCAGATATAGATTATGTACCACTAAAGTCTGCTACGTGTATAA    19462
gb:MT066175|Organism:Severe      ATGGAAAACAAGTAGTGTCAGATATAGATTATGTACCACTAAAGTCTGCTACGTGTATAA    19462
gb:MN997409|Organism:Severe      ATGGAAAACAAGTAGTGTCAGATATAGATTATGTACCACTAAAGTCTGCTACGTGTATAA    19462
gb:MN938384|Organism:Severe      ATGGAAAACAAGTAGTGTCAGATATAGATTATGTACCACTAAAGTCTGCTACGTGTATAA    19430
gb:MT044258|Organism:Severe      ATGGAAAACAAGTAGTGTCAGATATAGATTATGTACCACTAAAGTCTGCTACGTGTATAA    19438
gb:MT039890|Organism:Severe      ATGGAAAACAAGTAGTGTCAGATATAGATTATGTACCACTAAAGTCTGCTACGTGTATAA    19462
gb:MN988713|Organism:Severe      ATGGAAAACAAGTAGTGTCAGATATAGATTATGTACCACTAAAGTCTGCTACGTGTATAA    19462
gb:LC521925|Organism:Severe      ATGGAAAACAAGTAGTGTCAGATATAGATTATGTACCACTAAAGTCTGCTACGTGTATAA    19435
gb:MT093571|Organism:Severe      ATGGAAAACAAGTAGTGTCAGATATAGATTATGTACCACTAAAGTCTGCTACGTGTATAA    19462
gb:MT039887|Organism:Severe      ATGGAAAACAAGTAGTGTCAGATATAGATTATGTACCACTAAAGTCTGCTACGTGTATAA    19462
gb:MT019530|Organism:Severe      ATGGAAAACAAGTAGTGTCAGATATAGATTATGTACCACTAAAGTCTGCTACGTGTATAA    19462
gb:MT039888|Organism:Severe      ATGGAAAACAAGTAGTGTCAGATATAGATTATGTACCACTAAAGTCTGCTACGTGTATAA    19462
gb:LC522972|Organism:Severe      ATGGAAAACAAGTAGTGTCAGATATAGATTATGTACCACTAAAGTCTGCTACGTGTATAA    19459
gb:MT027063|Organism:Severe      ATGGAAAACAAGTAGTGTCAGATATAGATTATGTACCACTAAAGTCTGCTACGTGTATAA    19462
gb:MT027062|Organism:Severe      ATGGAAAACAAGTAGTGTCAGATATAGATTATGTACCACTAAAGTCTGCTACGTGTATAA    19462
gb:MT019529|Organism:Severe      ATGGAAAACAAGTAGTGTCAGATATAGATTATGTACCACTAAAGTCTGCTACGTGTATAA    19462
gb:MN996529|Organism:Severe      ATGGAAAACAAGTAGTGTCAGATATAGATTATGTACCACTAAAGTCTGCTACGTGTATAA    19450
gb:MN996531|Organism:Severe      ATGGAAAACAAGTAGTGTCAGATATAGATTATGTACCACTAAAGTCTGCTACGTGTATAA    19449
gb:MT066176|Organism:Severe      ATGGAAAACAAGTAGTGTCAGATATAGATTATGTACCACTAAAGTCTGCTACGTGTATAA    19462
gb:MT027064|Organism:Severe      ATGGAAAACAAGTAGTGTCAGATATAGATTATGTACCACTAAAGTCTGCTACGTGTATAA    19462
gb:MN994468|Organism:Severe      ATGGAAAACAAGTAGTGTCAGATATAGATTATGTACCACTAAAGTCTGCTACGTGTATAA    19462
gb:MT072688|Organism:Severe      ATGGAAAACAAGTAGTGTCAGATATAGATTATGTACCACTAAAGTCTGCTACGTGTATAA    19447
gb:MN996527|Organism:Severe      ATGGAAAACAAGTAGTGTCAGATATAGATTATGTACCACTAAAGTCTGCTACGTGTATAA    19429
gb:MT093631|Organism:Severe      ATGGAAAACAAGTAGTGTCAGATATAGATTATGTACCACTAAAGTCTGCTACGTGTATAA    19500
gb:MT106053|Organism:Severe      ATGGAAAACAAGTAGTGTCAGATATAGATTATGTACCACTAAAGTCTGCTACGTGTATAA    19462
gb:MT019533|Organism:Severe      ATGGAAAACAAGTAGTGTCAGATATAGATTATGTACCACTAAAGTCTGCTACGTGTATAA    19462
gb:MT019531|Organism:Severe      ATGGAAAACAAGTAGTGTCAGATATAGATTATGTACCACTAAAGTCTGCTACGTGTATAA    19462
gb:MN996528|Organism:Severe      ATGGAAAACAAGTAGTGTCAGATATAGATTATGTACCACTAAAGTCTGCTACGTGTATAA    19462
gb:MN996530|Organism:Severe      ATGGAAAACAAGTAGTGTCAGATATAGATTATGTACCACTAAAGTCTGCTACGTGTATAA    19448
```

```
gb:MN908947|Organism:Severe      ATGGAAAACAAGTAGTGTCAGATATAGATTATGTACCACTAAAGTCTGCTACGTGTATAA   19462
gb:MT019532|Organism:Severe      ATGGAAAACAAGTAGTGTCAGATATAGATTATGTACCACTAAAGTCTGCTACGTGTATAA   19462
                                 ************************************************************

gb:MT020781|Organism:Severe      CACGTTGCAATTTAGGTGGTGCTGTCTGTAGACATCATGCTAATGAGTACAGATTGTATC   19510
gb:MT007544|Organism:Severe      CACGTTGCAATTTAGGTGGTGCTGTCTGTAGACATCATGCTAATGAGTACAGATTGTATC   19522
gb:MN994467|Organism:Severe      CACGTTGCAATTTAGGTGGTGCTGTCTGTAGACATCATGCTAATGAGTACAGATTGTATC   19522
gb:MT044257|Organism:Severe      CACGTTGCAATTTAGGTGGTGCTGTCTGTAGACATCATGCTAATGAGTACAGATTGTATC   19522
gb:MT106054|Organism:Severe      CACGTTGCAATTTAGGTGGTGCTGTCTGTAGACATCATGCTAATGAGTACAGATTGTATC   19522
gb:MT049951|Organism:Severe      CACGTTGCAATTTAGGTGGTGCTGTCTGTAGACATCATGCTAATGAGTACAGATTGTATC   19522
gb:MN975262|Organism:Severe      CACGTTGCAATTTAGGTGGTGCTGTCTGTAGACATCATGCTAATGAGTACAGATTGTATC   19522
gb:MT106052|Organism:Severe      CACGTTGCAATTTAGGTGGTGCTGTCTGTAGACATCATGCTAATGAGTACAGATTGTATC   19522
gb:LC522975|Organism:Severe      CACGTTGCAATTTAGGTGGTGCTGTCTGTAGACATCATGCTAATGAGTACAGATTGTATC   19519
gb:LC522973|Organism:Severe      CACGTTGCAATTTAGGTGGTGCTGTCTGTAGACATCATGCTAATGAGTACAGATTGTATC   19519
gb:LC522974|Organism:Severe      CACGTTGCAATTTAGGTGGTGCTGTCTGTAGACATCATGCTAATGAGTACAGATTGTATC   19519
gb:MN985325|Organism:Severe      CACGTTGCAATTTAGGTGGTGCTGTCTGTAGACATCATGCTAATGAGTACAGATTGTATC   19522
gb:MT020881|Organism:Severe      CACGTTGCAATTTAGGTGGTGCTGTCTGTAGACATCATGCTAATGAGTACAGATTGTATC   19522
gb:MT020880|Organism:Severe      CACGTTGCAATTTAGGTGGTGCTGTCTGTAGACATCATGCTAATGAGTACAGATTGTATC   19522
gb:MT066175|Organism:Severe      CACGTTGCAATTTAGGTGGTGCTGTCTGTAGACATCATGCTAATGAGTACAGATTGTATC   19522
gb:MN997409|Organism:Severe      CACGTTGCAATTTAGGTGGTGCTGTCTGTAGACATCATGCTAATGAGTACAGATTGTATC   19522
gb:MN938384|Organism:Severe      CACGTTGCAATTTAGGTGGTGCTGTCTGTAGACATCATGCTAATGAGTACAGATTGTATC   19490
gb:MT044258|Organism:Severe      CACGTTGCAATTTAGGTGGTGCTGTCTGTAGACATCATGCTAATGAGTACAGATTGTATC   19498
gb:MT039890|Organism:Severe      CACGTTGCAATTTAGGTGGTGCTGTCTGTAGACATCATGCTAATGAGTACAGATTGTATC   19522
gb:MN988713|Organism:Severe      CACGTTGCAATTTAGGTGGTGCTGTCTGTAGACATCATGCTAATGAGTACAGATTGTATC   19522
gb:LC521925|Organism:Severe      CACGTTGCAATTTAGGTGGTGCTGTCTGTAGACATCATGCTAATGAGTACAGATTGTATC   19495
gb:MT093571|Organism:Severe      CACGTTGCAATTTAGGTGGTGCTGTCTGTAGACATCATGCTAATGAGTACAGATTGTATC   19522
gb:MT039887|Organism:Severe      CACGTTGCAATTTAGGTGGTGCTGTCTGTAGACATCATGCTAATGAGTACAGATTGTATC   19522
gb:MT019530|Organism:Severe      CACGTTGCAATTTAGGTGGTGCTGTCTGTAGACATCATGCTAATGAGTACAGATTGTATC   19522
gb:MT039888|Organism:Severe      CACGTTGCAATTTAGGTGGTGCTGTCTGTAGACATCATGCTAATGAGTACAGATTGTATC   19522
gb:LC522972|Organism:Severe      CACGTTGCAATTTAGGTGGTGCTGTCTGTAGACATCATGCTAATGAGTACAGATTGTATC   19519
gb:MT027063|Organism:Severe      CACGTTGCAATTTAGGTGGTGCTGTCTGTAGACATCATGCTAATGAGTACAGATTGTATC   19522
gb:MT027062|Organism:Severe      CACGTTGCAATTTAGGTGGTGCTGTCTGTAGACATCATGCTAATGAGTACAGATTGTATC   19522
gb:MT019529|Organism:Severe      CACGTTGCAATTTAGGTGGTGCTGTCTGTAGACATCATGCTAATGAGTACAGATTGTATC   19522
gb:MN996529|Organism:Severe      CACGTTGCAATTTAGGTGGTGCTGTCTGTAGACATCATGCTAATGAGTACAGATTGTATC   19510
gb:MN996531|Organism:Severe      CACGTTGCAATTTAGGTGGTGCTGTCTGTAGACATCATGCTAATGAGTACAGATTGTATC   19509
gb:MT066176|Organism:Severe      CACGTTGCAATTTAGGTGGTGCTGTCTGTAGACATCATGCTAATGAGTACAGATTGTATC   19522
gb:MT027064|Organism:Severe      CACGTTGCAATTTAGGTGGTGCTGTCTGTAGACATCATGCTAATGAGTACAGATTGTATC   19522
gb:MN994468|Organism:Severe      CACGTTGCAATTTAGGTGGTGCTGTCTGTAGACATCATGCTAATGAGTACAGATTGTATC   19522
gb:MT072688|Organism:Severe      CACGTTGCAATTTAGGTGGTGCTGTCTGTAGACATCATGCTAATGAGTACAGATTGTATC   19507
gb:MN996527|Organism:Severe      CACGTTGCAATTTAGGTGGTGCTGTCTGTAGACATCATGCTAATGAGTACAGATTGTATC   19489
gb:MT093631|Organism:Severe      CACGTTGCAATTTAGGTGGTGCTGTCTGTAGACATCATGCTAATGAGTACAGATTGTATC   19560
gb:MT106053|Organism:Severe      CACGTTGCAATTTAGGTGGTGCTGTCTGTAGACATCATGCTAATGAGTACAGATTGTATC   19522
gb:MT019533|Organism:Severe      CACGTTGCAATTTAGGTGGTGCTGTCTGTAGACATCATGCTAATGAGTACAGATTGTATC   19522
gb:MT019531|Organism:Severe      CACGTTGCAATTTAGGTGGTGCTGTCTGTAGACATCATGCTAATGAGTACAGATTGTATC   19522
gb:MN996528|Organism:Severe      CACGTTGCAATTTAGGTGGTGCTGTCTGTAGACATCATGCTAATGAGTACAGATTGTATC   19522
gb:MN996530|Organism:Severe      CACGTTGCAATTTAGGTGGTGCTGTCTGTAGACATCATGCTAATGAGTACAGATTGTATC   19508
gb:MN908947|Organism:Severe      CACGTTGCAATTTAGGTGGTGCTGTCTGTAGACATCATGCTAATGAGTACAGATTGTATC   19522
gb:MT019532|Organism:Severe      CACGTTGCAATTTAGGTGGTGCTGTCTGTAGACATCATGCTAATGAGTACAGATTGTATC   19522
                                 ************************************************************

gb:MT020781|Organism:Severe      TCGATGCTTATAACATGATGATCTCAGCTGGCTTTAGCTTGTGGGTTTACAAACAATTTG   19570
gb:MT007544|Organism:Severe      TCGATGCTTATAACATGATGATCTCAGCTGGCTTTAGCTTGTGGGTTTACAAACAATTTG   19582
gb:MN994467|Organism:Severe      TCGATGCTTATAACATGATGATCTCAGCTGGCTTTAGCTTGTGGGTTTACAAACAATTTG   19582
gb:MT044257|Organism:Severe      TCGATGCTTATAACATGATGATCTCAGCTGGCTTTAGCTTGTGGGTTTACAAACAATTTG   19582
gb:MT106054|Organism:Severe      TCGATGCTTATAACATGATGATCTCAGCTGGCTTTAGCTTGTGGGTTTACAAACAATTTG   19582
gb:MT049951|Organism:Severe      TCGATGCTTATAACATGATGATCTCAGCTGGCTTTAGCTTGTGGGTTTACAAACAATTTG   19582
gb:MN975262|Organism:Severe      TCGATGCTTATAACATGATGATCTCAGCTGGCTTTAGCTTGTGGGTTTACAAACAATTTG   19582
gb:MT106052|Organism:Severe      TCGATGCTTATAACATGATGATCTCAGCTGGCTTTAGCTTGTGGGTTTACAAACAATTTG   19582
gb:LC522975|Organism:Severe      TCGATGCTTATAACATGATGATCTCAGCTGGCTTTAGCTTGTGGGTTTACAAACAATTTG   19579
gb:LC522973|Organism:Severe      TCGATGCTTATAACATGATGATCTCAGCTGGCTTTAGCTTGTGGGTTTACAAACAATTTG   19579
gb:LC522974|Organism:Severe      TCGATGCTTATAACATGATGATCTCAGCTGGCTTTAGCTTGTGGGTTTACAAACAATTTG   19579
gb:MN985325|Organism:Severe      TCGATGCTTATAACATGATGATCTCAGCTGGCTTTAGCTTGTGGGTTTACAAACAATTTG   19582
gb:MT020881|Organism:Severe      TCGATGCTTATAACATGATGATCTCAGCTGGCTTTAGCTTGTGGGTTTACAAACAATTTG   19582
gb:MT020880|Organism:Severe      TCGATGCTTATAACATGATGATCTCAGCTGGCTTTAGCTTGTGGGTTTACAAACAATTTG   19582
gb:MT066175|Organism:Severe      TCGATGCTTATAACATGATGATCTCAGCTGGCTTTAGCTTGTGGGTTTACAAACAATTTG   19582
```

```
gb:MN997409|Organism:Severe      TCGATGCTTATAACATGATGATCTCAGCTGGCTTTAGCTTGTGGGTTTACAAACAATTTG   19582
gb:MN938384|Organism:Severe      TCGATGCTTATAACATGATGATCTCAGCTGGCTTTAGCTTGTGGGTTTACAAACAATTTG   19550
gb:MT044258|Organism:Severe      TCGATGCTTATAACATGATGATCTCAGCTGGCTTTAGCTTGTGGGTTTACAAACAATTTG   19558
gb:MT039890|Organism:Severe      TCGATGCTTATAACATGATGATCTCAGCTGGCTTTAGCTTGTGGGTTTACAAACAATTTG   19582
gb:MN988713|Organism:Severe      TCGATGCTTATAACATGATGATCTCAGCTGGCTTTAGCTTGTGGGTTTACAAACAATTTG   19582
gb:LC521925|Organism:Severe      TCGATGCTTATAACATGATGATCTCAGCTGGCTTTAGCTTGTGGGTTTACAAACAATTTG   19555
gb:MT093571|Organism:Severe      TCGATGCTTATAACATGATGATCTCAGCTGGCTTTAGCTTGTGGGTTTACAAACAATTTG   19582
gb:MT039887|Organism:Severe      TCGATGCTTATAACATGATGATCTCAGCTGGCTTTAGCTTGTGGGTTTACAAACAATTTG   19582
gb:MT019530|Organism:Severe      TCGATGCTTATAACATGATGATCTCAGCTGGCTTTAGCTTGTGGGTTTACAAACAATTTG   19582
gb:MT039888|Organism:Severe      TCGATGCTTATAACATGATGATCTCAGCTGGCTTTAGCTTGTGGGTTTACAAACAATTTG   19582
gb:LC522972|Organism:Severe      TCGATGCTTATAACATGATGATCTCAGCTGGCTTTAGCTTGTGGGTTTACAAACAATTTG   19579
gb:MT027063|Organism:Severe      TCGATGCTTATAACATGATGATCTCAGCTGGCTTTAGCTTGTGGGTTTACAAACAATTTG   19582
gb:MT027062|Organism:Severe      TCGATGCTTATAACATGATGATCTCAGCTGGCTTTAGCTTGTGGGTTTACAAACAATTTG   19582
gb:MT019529|Organism:Severe      TCGATGCTTATAACATGATGATCTCAGCTGGCTTTAGCTTGTGGGTTTACAAACAATTTG   19582
gb:MN996529|Organism:Severe      TCGATGCTTATAACATGATGATCTCAGCTGGCTTTAGCTTGTGGGTTTACAAACAATTTG   19570
gb:MN996531|Organism:Severe      TCGATGCTTATAACATGATGATCTCAGCTGGCTTTAGCTTGTGGGTTTACAAACAATTTG   19569
gb:MT066176|Organism:Severe      TCGATGCTTATAACATGATGATCTCAGCTGGCTTTAGCTTGTGGGTTTACAAACAATTTG   19582
gb:MT027064|Organism:Severe      TCGATGCTTATAACATGATGATCTCAGCTGGCTTTAGCTTGTGGGTTTACAAACAATTTG   19582
gb:MN994468|Organism:Severe      TCGATGCTTATAACATGATGATCTCAGCTGGCTTTAGCTTGTGGGTTTACAAACAATTTG   19582
gb:MT072688|Organism:Severe      TCGATGCTTATAACATGATGATCTCAGCTGGCTTTAGCTTGTGGGTTTACAAACAATTTG   19567
gb:MN996527|Organism:Severe      TCGATGCTTATAACATGATGATCTCAGCTGGCTTTAGCTTGTGGGTTTACAAACAATTTG   19549
gb:MT093631|Organism:Severe      TCGATGCTTATAACATGATGATCTCAGCTGGCTTTAGCTTGTGGGTTTACAAACAATTTG   19620
gb:MT106053|Organism:Severe      TCGATGCTTATAACATGATGATCTCAGCTGGCTTTAGCTTGTGGGTTTACAAACAATTTG   19582
gb:MT019533|Organism:Severe      TCGATGCTTATAACATGATGATCTCAGCTGGCTTTAGCTTGTGGGTTTACAAACAATTTG   19582
gb:MT019531|Organism:Severe      TCGATGCTTATAACATGATGATCTCAGCTGGCTTTAGCTTGTGGGTTTACAAACAATTTG   19582
gb:MN996528|Organism:Severe      TCGATGCTTATAACATGATGATCTCAGCTGGCTTTAGCTTGTGGGTTTACAAACAATTTG   19582
gb:MN996530|Organism:Severe      TCGATGCTTATAACATGATGATCTCAGCTGGCTTTAGCTTGTGGGTTTACAAACAATTTG   19568
gb:MN908947|Organism:Severe      TCGATGCTTATAACATGATGATCTCAGCTGGCTTTAGCTTGTGGGTTTACAAACAATTTG   19582
gb:MT019532|Organism:Severe      TCGATGCTTATAACATGATGATCTCAGCTGGCTTTAGCTTGTGGGTTTACAAACAATTTG   19582
                                 ************************************************************

gb:MT020781|Organism:Severe      ATACTTATAACCTCTGGAACACTTTTACAAGACTTCAGAGTTTAGAAAATGTGGCTTTTA   19630
gb:MT007544|Organism:Severe      ATACTTATAACCTCTGGAACACTTTTACAAGACTTCAGAGTTTAGAAAATGTGGCTTTTA   19642
gb:MN994467|Organism:Severe      ATACTTATAACCTCTGGAACACTTTTACAAGACTTCAGAGTTTAGAAAATGTGGCTTTTA   19642
gb:MT044257|Organism:Severe      ATACTTATAACCTCTGGAACACTTTTACAAGACTTCAGAGTTTAGAAAATGTGGCTTTTA   19642
gb:MT106054|Organism:Severe      ATACTTATAACCTCTGGAACACTTTTACAAGACTTCAGAGTTTAGAAAATGTGGCTTTTA   19642
gb:MT049951|Organism:Severe      ATACTTATAACCTCTGGAACACTTTTACAAGACTTCAGAGTTTAGAAAATGTGGCTTTTA   19642
gb:MN975262|Organism:Severe      ATACTTATAACCTCTGGAACACTTTTACAAGACTTCAGAGTTTAGAAAATGTGGCTTTTA   19642
gb:MT106052|Organism:Severe      ATACTTATAACCTCTGGAACACTTTTACAAGACTTCAGAGTTTAGAAAATGTGGCTTTTA   19642
gb:LC522975|Organism:Severe      ATACTTATAACCTCTGGAACACTTTTACAAGACTTCAGAGTTTAGAAAATGTGGCTTTTA   19639
gb:LC522973|Organism:Severe      ATACTTATAACCTCTGGAACACTTTTACAAGACTTCAGAGTTTAGAAAATGTGGCTTTTA   19639
gb:LC522974|Organism:Severe      ATACTTATAACCTCTGGAACACTTTTACAAGACTTCAGAGTTTAGAAAATGTGGCTTTTA   19639
gb:MN985325|Organism:Severe      ATACTTATAACCTCTGGAACACTTTTACAAGACTTCAGAGTTTAGAAAATGTGGCTTTTA   19642
gb:MT020881|Organism:Severe      ATACTTATAACCTCTGGAACACTTTTACAAGACTTCAGAGTTTAGAAAATGTGGCTTTTA   19642
gb:MT020880|Organism:Severe      ATACTTATAACCTCTGGAACACTTTTACAAGACTTCAGAGTTTAGAAAATGTGGCTTTTA   19642
gb:MT066175|Organism:Severe      ATACTTATAACCTCTGGAACACTTTTACAAGACTTCAGAGTTTAGAAAATGTGGCTTTTA   19642
gb:MN997409|Organism:Severe      ATACTTATAACCTCTGGAACACTTTTACAAGACTTCAGAGTTTAGAAAATGTGGCTTTTA   19642
gb:MN938384|Organism:Severe      ATACTTATAACCTCTGGAACACTTTTACAAGACTTCAGAGTTTAGAAAATGTGGCTTTTA   19610
gb:MT044258|Organism:Severe      ATACTTATAACCTCTGGAACACTTTTACAAGACTTCAGAGTTTAGAAAATGTGGCTTTTA   19618
gb:MT039890|Organism:Severe      ATACTTATAACCTCTGGAACACTTTTACAAGACTTCAGAGTTTAGAAAATGTGGCTTTTA   19642
gb:MN988713|Organism:Severe      ATACTTATAACCTCTGGAACACTTTTACAAGACTTCAGAGTTTAGAAAATGTGGCTTTTA   19642
gb:LC521925|Organism:Severe      ATACTTATAACCTCTGGAACACTTTTACAAGACTTCAGAGTTTAGAAAATGTGGCTTTTA   19615
gb:MT093571|Organism:Severe      ATACTTATAACCTCTGGAACACTTTTACAAGACTTCAGAGTTTAGAAAATGTGGCTTTTA   19642
gb:MT039887|Organism:Severe      ATACTTATAACCTCTGGAACACTTTTACAAGACTTCAGAGTTTAGAAAATGTGGCTTTTA   19642
gb:MT019530|Organism:Severe      ATACTTATAACCTCTGGAACACTTTTACAAGACTTCAGAGTTTAGAAAATGTGGCTTTTA   19642
gb:MT039888|Organism:Severe      ATACTTATAACCTCTGGAACACTTTTACAAGACTTCAGAGTTTAGAAAATGTGGCTTTTA   19642
gb:LC522972|Organism:Severe      ATACTTATAACCTCTGGAACACTTTTACAAGACTTCAGAGTTTAGAAAATGTGGCTTTTA   19639
gb:MT027063|Organism:Severe      ATACTTATAACCTCTGGAACACTTTTACAAGACTTCAGAGTTTAGAAAATGTGGCTTTTA   19642
gb:MT027062|Organism:Severe      ATACTTATAACCTCTGGAACACTTTTACAAGACTTCAGAGTTTAGAAAATGTGGCTTTTA   19642
gb:MT019529|Organism:Severe      ATACTTATAACCTCTGGAACACTTTTACAAGACTTCAGAGTTTAGAAAATGTGGCTTTTA   19642
gb:MN996529|Organism:Severe      ATACTTATAACCTCTGGAACACTTTTACAAGACTTCAGAGTTTAGAAAATGTGGCTTTTA   19630
gb:MN996531|Organism:Severe      ATACTTATAACCTCTGGAACACTTTTACAAGACTTCAGAGTTTAGAAAATGTGGCTTTTA   19629
gb:MT066176|Organism:Severe      ATACTTATAACCTCTGGAACACTTTTACAAGACTTCAGAGTTTAGAAAATGTGGCTTTTA   19642
gb:MT027064|Organism:Severe      ATACTTATAACCTCTGGAACACTTTTACAAGACTTCAGAGTTTAGAAAATGTGGCTTTTA   19642
gb:MN994468|Organism:Severe      ATACTTATAACCTCTGGAACACTTTTACAAGACTTCAGAGTTTAGAAAATGTGGCTTTTA   19642
```

```
gb:MT072688|Organism:Severe      ATACTTATAACCTCTGGAACACTTTTACAAGACTTCAGAGTTTAGAAAATGTGGCTTTTA   19627
gb:MN996527|Organism:Severe      ATACTTATAACCTCTGGAACACTTTTACAAGACTTCAGAGTTTAGAAAATGTGGCTTTTA   19609
gb:MT093631|Organism:Severe      ATACTTATAACCTCTGGAACACTTTTACAAGACTTCAGAGTTTAGAAAATGTGGCTTTTA   19680
gb:MT106053|Organism:Severe      ATACTTATAACCTCTGGAACACTTTTACAAGACTTCAGAGTTTAGAAAATGTGGCTTTTA   19642
gb:MT019533|Organism:Severe      ATACTTATAACCTCTGGAACACTTTTACAAGACTTCAGAGTTTAGAAAATGTGGCTTTTA   19642
gb:MT019531|Organism:Severe      ATACTTATAACCTCTGGAACACTTTTACAAGACTTCAGAGTTTAGAAAATGTGGCTTTTA   19642
gb:MN996528|Organism:Severe      ATACTTATAACCTCTGGAACACTTTTACAAGACTTCAGAGTTTAGAAAATGTGGCTTTTA   19642
gb:MN996530|Organism:Severe      ATACTTATAACCTCTGGAACACTTTTACAAGACTTCAGAGTTTAGAAAATGTGGCTTTTA   19628
gb:MN908947|Organism:Severe      ATACTTATAACCTCTGGAACACTTTTACAAGACTTCAGAGTTTAGAAAATGTGGCTTTTA   19642
gb:MT019532|Organism:Severe      ATACTTATAACCTCTGGAACACTTTTACAAGACTTCAGAGTTTAGAAAATGTGGCTTTTA   19642
                                 ************************************************************

gb:MT020781|Organism:Severe      ATGTTGTAAATAAGGGACACTTTGATGGACAACAGGGTGAAGTACCAGTTTCTATCATTA   19690
gb:MT007544|Organism:Severe      ATGTTGTAAATAAGGGACACTTTGATGGACAACAGGGTGAAGTACCAGTTTCTATCATTA   19702
gb:MN994467|Organism:Severe      ATGTTGTAAATAAGGGACACTTTGATGGACAACAGGGTGAAGTACCAGTTTCTATCATTA   19702
gb:MT044257|Organism:Severe      ATGTTGTAAATAAGGGACACTTTGATGGACAACAGGGTGAAGTACCAGTTTCTATCATTA   19702
gb:MT106054|Organism:Severe      ATGTTGTAAATAAGGGACACTTTGATGGACAACAGGGTGAAGTACCAGTTTCTATCATTA   19702
gb:MT049951|Organism:Severe      ATGTTGTAAATAAGGGACACTTTGATGGACAACAGGGTGAAGTACCAGTTTCTATCATTA   19702
gb:MN975262|Organism:Severe      ATGTTGTAAATAAGGGACACTTTGATGGACAACAGGGTGAAGTACCAGTTTCTATCATTA   19702
gb:MT106052|Organism:Severe      ATGTTGTAAATAAGGGACACTTTGATGGACAACAGGGTGAAGTACCAGTTTCTATCATTA   19702
gb:LC522975|Organism:Severe      ATGTTGTAAATAAGGGACACTTTGATGGACAACAGGGTGAAGTACCAGTTTCTATCATTA   19699
gb:LC522973|Organism:Severe      ATGTTGTAAATAAGGGACACTTTGATGGACAACAGGGTGAAGTACCAGTTTCTATCATTA   19699
gb:LC522974|Organism:Severe      ATGTTGTAAATAAGGGACACTTTGATGGACAACAGGGTGAAGTACCAGTTTCTATCATTA   19699
gb:MN985325|Organism:Severe      ATGTTGTAAATAAGGGACACTTTGATGGACAACAGGGTGAAGTACCAGTTTCTATCATTA   19702
gb:MT020881|Organism:Severe      ATGTTGTAAATAAGGGACACTTTGATGGACAACAGGGTGAAGTACCAGTTTCTATCATTA   19702
gb:MT020880|Organism:Severe      ATGTTGTAAATAAGGGACACTTTGATGGACAACAGGGTGAAGTACCAGTTTCTATCATTA   19702
gb:MT066175|Organism:Severe      ATGTTGTAAATAAGGGACACTTTGATGGACAACAGGGTGAAGTACCAGTTTCTATCATTA   19702
gb:MN997409|Organism:Severe      ATGTTGTAAATAAGGGACACTTTGATGGACAACAGGGTGAAGTACCAGTTTCTATCATTA   19702
gb:MN938384|Organism:Severe      ATGTTGTAAATAAGGGACACTTTGATGGACAACAGGGTGAAGTACCAGTTTCTATCATTA   19670
gb:MT044258|Organism:Severe      ATGTTGTAAATAAGGGACACTTTGATGGACAACAGGGTGAAGTACCAGTTTCTATCATTA   19678
gb:MT039890|Organism:Severe      ATGTTGTAAATAAGGGACACTTTGATGGACAACAGGGTGAAGTACCAGTTTCTATCATTA   19702
gb:MN988713|Organism:Severe      ATGTTGTAAATAAGGGACACTTTGATGGACAACAGGGTGAAGTACCAGTTTCTATCATTA   19702
gb:LC521925|Organism:Severe      ATGTTGTAAATAAGGGACACTTTGATGGACAACAGGGTGAAGTACCAGTTTCTATCATTA   19675
gb:MT093571|Organism:Severe      ATGTTGTAAATAAGGGACACTTTGATGGACAACAGGGTGAAGTACCAGTTTCTATCATTA   19702
gb:MT039887|Organism:Severe      ATGTTGTAAATAAGGGACACTTTGATGGACAACAGGGTGAAGTACCAGTTTCTATCATTA   19702
gb:MT019530|Organism:Severe      ATGTTGTAAATAAGGGACACTTTGATGGACAACAGGGTGAAGTACCAGTTTCTATCATTA   19702
gb:MT039888|Organism:Severe      ATGTTGTAAATAAGGGACACTTTGATGGACAACAGGGTGAAGTACCAGTTTCTATCATTA   19702
gb:LC522972|Organism:Severe      ATGTTGTAAATAAGGGACACTTTGATGGACAACAGGGTGAAGTACCAGTTTCTATCATTA   19699
gb:MT027063|Organism:Severe      ATGTTGTAAATAAGGGACACTTTGATGGACAACAGGGTGAAGTACCAGTTTCTATCATTA   19702
gb:MT027062|Organism:Severe      ATGTTGTAAATAAGGGACACTTTGATGGACAACAGGGTGAAGTACCAGTTTCTATCATTA   19702
gb:MT019529|Organism:Severe      ATGTTGTAAATAAGGGACACTTTGATGGACAACAGGGTGAAGTACCAGTTTCTATCATTA   19702
gb:MN996529|Organism:Severe      ATGTTGTAAATAAGGGACACTTTGATGGACAACAGGGTGAAGTACCAGTTTCTATCATTA   19690
gb:MN996531|Organism:Severe      ATGTTGTAAATAAGGGACACTTTGATGGACAACAGGGTGAAGTACCAGTTTCTATCATTA   19689
gb:MT066176|Organism:Severe      ATGTTGTAAATAAGGGACACTTTGATGGACAACAGGGTGAAGTACCAGTTTCTATCATTA   19702
gb:MT027064|Organism:Severe      ATGTTGTAAATAAGGGACACTTTGATGGACAACAGGGTGAAGTACCAGTTTCTATCATTA   19702
gb:MN994468|Organism:Severe      ATGTTGTAAATAAGGGACACTTTGATGGACAACAGGGTGAAGTACCAGTTTCTATCATTA   19702
gb:MT072688|Organism:Severe      ATGTTGTAAATAAGGGACACTTTGATGGACAACAGGGTGAAGTACCAGTTTCTATCATTA   19687
gb:MN996527|Organism:Severe      ATGTTGTAAATAAGGGACACTTTGATGGACAACAGGGTGAAGTACCAGTTTCTATCATTA   19669
gb:MT093631|Organism:Severe      ATGTTGTAAATAAGGGACACTTTGATGGACAACAGGGTGAAGTACCAGTTTCTATCATTA   19740
gb:MT106053|Organism:Severe      ATGTTGTAAATAAGGGACACTTTGATGGACAACAGGGTGAAGTACCAGTTTCTATCATTA   19702
gb:MT019533|Organism:Severe      ATGTTGTAAATAAGGGACACTTTGATGGACAACAGGGTGAAGTACCAGTTTCTATCATTA   19702
gb:MT019531|Organism:Severe      ATGTTGTAAATAAGGGACACTTTGATGGACAACAGGGTGAAGTACCAGTTTCTATCATTA   19702
gb:MN996528|Organism:Severe      ATGTTGTAAATAAGGGACACTTTGATGGACAACAGGGTGAAGTACCAGTTTCTATCATTA   19702
gb:MN996530|Organism:Severe      ATGTTGTAAATAAGGGACACTTTGATGGACAACAGGGTGAAGTACCAGTTTCTATCATTA   19688
gb:MN908947|Organism:Severe      ATGTTGTAAATAAGGGACACTTTGATGGACAACAGGGTGAAGTACCAGTTTCTATCATTA   19702
gb:MT019532|Organism:Severe      ATGTTGTAAATAAGGGACACTTTGATGGACAACAGGGTGAAGTACCAGTTTCTATCATTA   19702
                                 ************************************************************

gb:MT020781|Organism:Severe      ATAACACTGTTTACACAAAAGTTGATGGTGTTGATGTAGAATTGTTTGAAAATAAAACAA   19750
gb:MT007544|Organism:Severe      ATAACACTGTTTACACAAAAGTTGATGGTGTTGATGTAGAATTGTTTGAAAATAAAACAA   19762
gb:MN994467|Organism:Severe      ATAACACTGTTTACACAAAAGTTGATGGTGTTGATGTAGAATTGTTTGAAAATAAAACAA   19762
gb:MT044257|Organism:Severe      ATAACACTGTTTACACAAAAGTTGATGGTGTTGATGTAGAATTGTTTGAAAATAAAACAA   19762
gb:MT106054|Organism:Severe      ATAACACTGTTTACACAAAAGTTGATGGTGTTGATGTAGAATTGTTTGAAAATAAAACAA   19762
gb:MT049951|Organism:Severe      ATAACACTGTTTACACAAAAGTTGATGGTGTTGATGTAGAATTGTTTGAAAATAAAACAA   19762
gb:MN975262|Organism:Severe      ATAACACTGTTTACACAAAAGTTGATGGTGTTGATGTAGAATTGTTTGAAAATAAAACAA   19762
```

```
gb:MT106052|Organism:Severe      ATAACACTGTTTACACAAAAGTTGATGGTGTTGATGTAGAATTGTTTGAAAATAAAACAA   19762
gb:LC522975|Organism:Severe      ATAACACTGTTTACACAAAAGTTGATGGTGTTGATGTAGAATTGTTTGAAAATAAAACAA   19759
gb:LC522973|Organism:Severe      ATAACACTGTTTACACAAAAGTTGATGGTGTTGATGTAGAATTGTTTGAAAATAAAACAA   19759
gb:LC522974|Organism:Severe      ATAACACTGTTTACACAAAAGTTGATGGTGTTGATGTAGAATTGTTTGAAAATAAAACAA   19759
gb:MN985325|Organism:Severe      ATAACACTGTTTACACAAAAGTTGATGGTGTTGATGTAGAATTGTTTGAAAATAAAACAA   19762
gb:MT020881|Organism:Severe      ATAACACTGTTTACACAAAAGTTGATGGTGTTGATGTAGAATTGTTTGAAAATAAAACAA   19762
gb:MT020880|Organism:Severe      ATAACACTGTTTACACAAAAGTTGATGGTGTTGATGTAGAATTGTTTGAAAATAAAACAA   19762
gb:MT066175|Organism:Severe      ATAACACTGTTTACACAAAAGTTGATGGTGTTGATGTAGAATTGTTTGAAAATAAAACAA   19762
gb:MN997409|Organism:Severe      ATAACACTGTTTACACAAAAGTTGATGGTGTTGATGTAGAATTGTTTGAAAATAAAACAA   19762
gb:MN938384|Organism:Severe      ATAACACTGTTTACACAAAAGTTGATGGTGTTGATGTAGAATTGTTTGAAAATAAAACAA   19730
gb:MT044258|Organism:Severe      ATAACACTGTTTACACAAAAGTTGATGGTGTTGATGTAGAATTGTTTGAAAATAAAACAA   19738
gb:MT039890|Organism:Severe      ATAACACTGTTTACACAAAAGTTGATGGTGTTGATGTAGAATTGTTTGAAAATAAAACAA   19762
gb:MN988713|Organism:Severe      ATAACACTGTTTACACAAAAGTTGATGGTGTTGATGTAGAATTGTTTGAAAATAAAACAA   19762
gb:LC521925|Organism:Severe      ATAACACTGTTTACACAAAAGTTGATGGTGTTGATGTAGAATTGTTTGAAAATAAAACAA   19735
gb:MT093571|Organism:Severe      ATAACACTGTTTACACAAAAGTTGATGGTGTTGATGTAGAATTGTTTGAAAATAAAACAA   19762
gb:MT039887|Organism:Severe      ATAACACTGTTTACACAAAAGTTGATGGTGTTGATGTAGAATTGTTTGAAAATAAAACAA   19762
gb:MT019530|Organism:Severe      ATAACACTGTTTACACAAAAGTTGATGGTGTTGATGTAGAATTGTTTGAAAATAAAACAA   19762
gb:MT039888|Organism:Severe      ATAACACTGTTTACACAAAAGTTGATGGTGTTGATGTAGAATTGTTTGAAAATAAAACAA   19762
gb:LC522972|Organism:Severe      ATAACACTGTTTACACAAAAGTTGATGGTGTTGATGTAGAATTGTTTGAAAATAAAACAA   19759
gb:MT027063|Organism:Severe      ATAACACTGTTTACACAAAAGTTGATGGTGTTGATGTAGAATTGTTTGAAAATAAAACAA   19762
gb:MT027062|Organism:Severe      ATAACACTGTTTACACAAAAGTTGATGGTGTTGATGTAGAATTGTTTGAAAATAAAACAA   19762
gb:MT019529|Organism:Severe      ATAACACTGTTTACACAAAAGTTGATGGTGTTGATGTAGAATTGTTTGAAAATAAAACAA   19762
gb:MN996529|Organism:Severe      ATAACACTGTTTACACAAAAGTTGATGGTGTTGATGTAGAATTGTTTGAAAATAAAACAA   19750
gb:MN996531|Organism:Severe      ATAACACTGTTTACACAAAAGTTGATGGTGTTGATGTAGAATTGTTTGAAAATAAAACAA   19749
gb:MT066176|Organism:Severe      ATAACACTGTTTACACAAAAGTTGATGGTGTTGATGTAGAATTGTTTGAAAATAAAACAA   19762
gb:MT027064|Organism:Severe      ATAACACTGTTTACACAAAAGTTGATGGTGTTGATGTAGAATTGTTTGAAAATAAAACAA   19762
gb:MN994468|Organism:Severe      ATAACACTGTTTACACAAAAGTTGATGGTGTTGATGTAGAATTGTTTGAAAATAAAACAA   19762
gb:MT072688|Organism:Severe      ATAACACTGTTTACACAAAAGTTGATGGTGTTGATGTAGAATTGTTTGAAAATAAAACAA   19747
gb:MN996527|Organism:Severe      ATAACACTGTTTACACAAAAGTTGATGGTGTTGATGTAGAATTGTTTGAAAATAAAACAA   19729
gb:MT093631|Organism:Severe      ATAACACTGTTTACACAAAAGTTGATGGTGTTGATGTAGAATTGTTTGAAAATAAAACAA   19800
gb:MT106053|Organism:Severe      ATAACACTGTTTACACAAAAGTTGATGGTGTTGATGTAGAATTGTTTGAAAATAAAACAA   19762
gb:MT019533|Organism:Severe      ATAACACTGTTTACACAAAAGTTGATGGTGTTGATGTAGAATTGTTTGAAAATAAAACAA   19762
gb:MT019531|Organism:Severe      ATAACACTGTTTACACAAAAGTTGATGGTGTTGATGTAGAATTGTTTGAAAATAAAACAA   19762
gb:MN996528|Organism:Severe      ATAACACTGTTTACACAAAAGTTGATGGTGTTGATGTAGAATTGTTTGAAAATAAAACAA   19762
gb:MN996530|Organism:Severe      ATAACACTGTTTACACAAAAGTTGATGGTGTTGATGTAGAATTGTTTGAAAATAAAACAA   19748
gb:MN908947|Organism:Severe      ATAACACTGTTTACACAAAAGTTGATGGTGTTGATGTAGAATTGTTTGAAAATAAAACAA   19762
gb:MT019532|Organism:Severe      ATAACACTGTTTACACAAAAGTTGATGGTGTTGATGTAGAATTGTTTGAAAATAAAACAA   19762
                                 ************************************************************

gb:MT020781|Organism:Severe      CATTACCTGTTAATGTAGCATTTGAGCTTTGGGCTAAGCGCAACATTAAACCAGTACCAG   19810
gb:MT007544|Organism:Severe      CATTACCTGTTAATGTAGCATTTGAGCTTTGGGCTAAGCGCAACATTAAACCAGTACCAG   19822
gb:MN994467|Organism:Severe      CATTACCTGTTAATGTAGCATTTGAGCTTTGGGCTAAGCGCAACATTAAACCAGTACCAG   19822
gb:MT044257|Organism:Severe      CATTACCTGTTAATGTAGCATTTGAGCTTTGGGCTAAGCGCAACATTAAACCAGTACCAG   19822
gb:MT106054|Organism:Severe      CATTACCTGTTAATGTAGCATTTGAGCTTTGGGCTAAGCGCAACATTAAACCAGTACCAG   19822
gb:MT049951|Organism:Severe      CATTACCTGTTAATGTAGCATTTGAGCTTTGGGCTAAGCGCAACATTAAACCAGTACCAG   19822
gb:MN975262|Organism:Severe      CATTACCTGTTAATGTAGCATTTGAGCTTTGGGCTAAGCGCAACATTAAACCAGTACCAG   19822
gb:MT106052|Organism:Severe      CATTACCTGTTAATGTAGCATTTGAGCTTTGGGCTAAGCGCAACATTAAACCAGTACCAG   19822
gb:LC522975|Organism:Severe      CATTACCTGTTAATGTAGCATTTGAGCTTTGGGCTAAGCGCAACATTAAACCAGTACCAG   19819
gb:LC522973|Organism:Severe      CATTACCTGTTAATGTAGCATTTGAGCTTTGGGCTAAGCGCAACATTAAACCAGTACCAG   19819
gb:LC522974|Organism:Severe      CATTACCTGTTAATGTAGCATTTGAGCTTTGGGCTAAGCGCAACATTAAACCAGTACCAG   19819
gb:MN985325|Organism:Severe      CATTACCTGTTAATGTAGCATTTGAGCTTTGGGCTAAGCGCAACATTAAACCAGTACCAG   19822
gb:MT020881|Organism:Severe      CATTACCTGTTAATGTAGCATTTGAGCTTTGGGCTAAGCGCAACATTAAACCAGTACCAG   19822
gb:MT020880|Organism:Severe      CATTACCTGTTAATGTAGCATTTGAGCTTTGGGCTAAGCGCAACATTAAACCAGTACCAG   19822
gb:MT066175|Organism:Severe      CATTACCTGTTAATGTAGCATTTGAGCTTTGGGCTAAGCGCAACATTAAACCAGTACCAG   19822
gb:MN997409|Organism:Severe      CATTACCTGTTAATGTAGCATTTGAGCTTTGGGCTAAGCGCAACATTAAACCAGTACCAG   19822
gb:MN938384|Organism:Severe      CATTACCTGTTAATGTAGCATTTGAGCTTTGGGCTAAGCGCAACATTAAACCAGTACCAG   19790
gb:MT044258|Organism:Severe      CATTACCTGTTAATGTAGCATTTGAGCTTTGGGCTAAGCGCAACATTAAACCAGTACCAG   19798
gb:MT039890|Organism:Severe      CATTACCTGTTAATGTAGCATTTGAGCTTTGGGCTAAGCGCAACATTAAACCAGTACCAG   19822
gb:MN988713|Organism:Severe      CATTACCTGTTAATGTAGCATTTGAGCTTTGGGCTAAGCGCAACATTAAACCAGTACCAG   19822
gb:LC521925|Organism:Severe      CATTACCTGTTAATGTAGCATTTGAGCTTTGGGCTAAGCGCAACATTAAACCAGTACCAG   19795
gb:MT093571|Organism:Severe      CATTACCTGTTAATGTAGCATTTGAGCTTTGGGCTAAGCGCAACATTAAACCAGTACCAG   19822
gb:MT039887|Organism:Severe      CATTACCTGTTAATGTAGCATTTGAGCTTTGGGCTAAGCGCAACATTAAACCAGTACCAG   19822
gb:MT019530|Organism:Severe      CATTACCTGTTAATGTAGCATTTGAGCTTTGGGCTAAGCGCAACATTAAACCAGTACCAG   19822
gb:MT039888|Organism:Severe      CATTACCTGTTAATGTAGCATTTGAGCTTTGGGCTAAGCGCAACATTAAACCAGTACCAG   19822
gb:LC522972|Organism:Severe      CATTACCTGTTAATGTAGCATTTGAGCTTTGGGCTAAGCGCAACATTAAACCAGTACCAG   19819
```

```
gb:MT027063|Organism:Severe      CATTACCTGTTAATGTAGCATTTGAGCTTTGGGCTAAGCGCAACATTAAACCAGTACCAG   19822
gb:MT027062|Organism:Severe      CATTACCTGTTAATGTAGCATTTGAGCTTTGGGCTAAGCGCAACATTAAACCAGTACCAG   19822
gb:MT019529|Organism:Severe      CATTACCTGTTAATGTAGCATTTGAGCTTTGGGCTAAGCGCAACATTAAACCAGTACCAG   19822
gb:MN996529|Organism:Severe      CATTACCTGTTAATGTAGCATTTGAGCTTTGGGCTAAGCGCAACATTAAACCAGTACCAG   19810
gb:MN996531|Organism:Severe      CATTACCTGTTAATGTAGCATTTGAGCTTTGGGCTAAGCGCAACATTAAACCAGTACCAG   19809
gb:MT066176|Organism:Severe      CATTACCTGTTAATGTAGCATTTGAGCTTTGGGCTAAGCGCAACATTAAACCAGTACCAG   19822
gb:MT027064|Organism:Severe      CATTACCTGTTAATGTAGCATTTGAGCTTTGGGCTAAGCGCAACATTAAACCAGTACCAG   19822
gb:MN994468|Organism:Severe      CATTACCTGTTAATGTAGCATTTGAGCTTTGGGCTAAGCGCAACATTAAACCAGTACCAG   19822
gb:MT072688|Organism:Severe      CATTACCTGTTAATGTAGCATTTGAGCTTTGGGCTAAGCGCAACATTAAACCAGTACCAG   19807
gb:MN996527|Organism:Severe      CATTACCTGTTAATGTAGCATTTGAGCTTTGGGCTAAGCGCAACATTAAACCAGTACCAG   19789
gb:MT093631|Organism:Severe      CATTACCTGTTAATGTAGCATTTGAGCTTTGGGCTAAGCGCAACATTAAACCAGTACCAG   19860
gb:MT106053|Organism:Severe      CATTACCTGTTAATGTAGCATTTGAGCTTTGGGCTAAGCGCAACATTAAACCAGTACCAG   19822
gb:MT019533|Organism:Severe      CATTACCTGTTAATGTAGCATTTGAGCTTTGGGCTAAGCGCAACATTAAACCAGTACCAG   19822
gb:MT019531|Organism:Severe      CATTACCTGTTAATGTAGCATTTGAGCTTTGGGCTAAGCGCAACATTAAACCAGTACCAG   19822
gb:MN996528|Organism:Severe      CATTACCTGTTAATGTAGCATTTGAGCTTTGGGCTAAGCGCAACATTAAACCAGTACCAG   19822
gb:MN996530|Organism:Severe      CATTACCTGTTAATGTAGCATTTGAGCTTTGGGCTAAGCGCAACATTAAACCAGTACCAG   19808
gb:MN908947|Organism:Severe      CATTACCTGTTAATGTAGCATTTGAGCTTTGGGCTAAGCGCAACATTAAACCAGTACCAG   19822
gb:MT019532|Organism:Severe      CATTACCTGTTAATGTAGCATTTGAGCTTTGGGCTAAGCGCAACATTAAACCAGTACCAG   19822
                                 ************************************************************

gb:MT020781|Organism:Severe      AGGTGAAAATACTCAATAATTTGGGTGTGGACATTGCTGCTAATACTGTGATCTGGGACT   19870
gb:MT007544|Organism:Severe      AGGTGAAAATACTCAATAATTTGGGTGTGGACATTGCTGCTAATACTGTGATCTGGGACT   19882
gb:MN994467|Organism:Severe      AGGTGAAAATACTCAATAATTTGGGTGTGGACATTGCTGCTAATACTGTGATCTGGGACT   19882
gb:MT044257|Organism:Severe      AGGTGAAAATACTCAATAATTTGGGTGTGGACATTGCTGCTAATACTGTGATCTGGGACT   19882
gb:MT106054|Organism:Severe      AGGTGAAAATACTCAATAATTTGGGTGTGGACATTGCTGCTAATACTGTGATCTGGGACT   19882
gb:MT049951|Organism:Severe      AGGTGAAAATACTCAATAATTTGGGTGTGGACATTGCTGCTAATACTGTGATCTGGGACT   19882
gb:MN975262|Organism:Severe      AGGTGAAAATACTCAATAATTTGGGTGTGGACATTGCTGCTAATACTGTGATCTGGGACT   19882
gb:MT106052|Organism:Severe      AGGTGAAAATACTCAATAATTTGGGTGTGGACATTGCTGCTAATACTGTGATCTGGGACT   19882
gb:LC522975|Organism:Severe      AGGTGAAAATACTCAATAATTTGGGTGTGGACATTGCTGCTAATACTGTGATCTGGGACT   19879
gb:LC522973|Organism:Severe      AGGTGAAAATACTCAATAATTTGGGTGTGGACATTGCTGCTAATACTGTGATCTGGGACT   19879
gb:LC522974|Organism:Severe      AGGTGAAAATACTCAATAATTTGGGTGTGGACATTGCTGCTAATACTGTGATCTGGGACT   19879
gb:MN985325|Organism:Severe      AGGTGAAAATACTCAATAATTTGGGTGTGGACATTGCTGCTAATACTGTGATCTGGGACT   19882
gb:MT020881|Organism:Severe      AGGTGAAAATACTCAATAATTTGGGTGTGGACATTGCTGCTAATACTGTGATCTGGGACT   19882
gb:MT020880|Organism:Severe      AGGTGAAAATACTCAATAATTTGGGTGTGGACATTGCTGCTAATACTGTGATCTGGGACT   19882
gb:MT066175|Organism:Severe      AGGTGAAAATACTCAATAATTTGGGTGTGGACATTGCTGCTAATACTGTGATCTGGGACT   19882
gb:MN997409|Organism:Severe      AGGTGAAAATACTCAATAATTTGGGTGTGGACATTGCTGCTAATACTGTGATCTGGGACT   19882
gb:MN938384|Organism:Severe      AGGTGAAAATACTCAATAATTTGGGTGTGGACATTGCTGCTAATACTGTGATCTGGGACT   19850
gb:MT044258|Organism:Severe      AGGTGAAAATACTCAATAATTTGGGTGTGGACATTGCTGCTAATACTGTGATCTGGGACT   19858
gb:MT039890|Organism:Severe      AGGTGAAAATACTCAATAATTTGGGTGTGGACATTGCTGCTAATACTGTGATCTGGGACT   19882
gb:MN988713|Organism:Severe      AGGTGAAAATACTCAATAATTTGGGTGTGGACATTGCTGCTAATACTGTGATCTGGGACT   19882
gb:LC521925|Organism:Severe      AGGTGAAAATACTCAATAATTTGGGTGTGGACATTGCTGCTAATACTGTGATCTGGGACT   19855
gb:MT093571|Organism:Severe      AGGTGAAAATACTCAATAATTTGGGTGTGGACATTGCTGCTAATACTGTGATCTGGGACT   19882
gb:MT039887|Organism:Severe      AGGTGAAAATACTCAATAATTTGGGTGTGGACATTGCTGCTAATACTGTGATCTGGGACT   19882
gb:MT019530|Organism:Severe      AGGTGAAAATACTCAATAATTTGGGTGTGGACATTGCTGCTAATACTGTGATCTGGGACT   19882
gb:MT039888|Organism:Severe      AGGTGAAAATACTCAATAATTTGGGTGTGGACATTGCTGCTAATACTGTGATCTGGGACT   19882
gb:LC522972|Organism:Severe      AGGTGAAAATACTCAATAATTTGGGTGTGGACATTGCTGCTAATACTGTGATCTGGGACT   19879
gb:MT027063|Organism:Severe      AGGTGAAAATACTCAATAATTTGGGTGTGGACATTGCTGCTAATACTGTGATCTGGGACT   19882
gb:MT027062|Organism:Severe      AGGTGAAAATACTCAATAATTTGGGTGTGGACATTGCTGCTAATACTGTGATCTGGGACT   19882
gb:MT019529|Organism:Severe      AGGTGAAAATACTCAATAATTTGGGTGTGGACATTGCTGCTAATACTGTGATCTGGGACT   19882
gb:MN996529|Organism:Severe      AGGTGAAAATACTCAATAATTTGGGTGTGGACATTGCTGCTAATACTGTGATCTGGGACT   19870
gb:MN996531|Organism:Severe      AGGTGAAAATACTCAATAATTTGGGTGTGGACATTGCTGCTAATACTGTGATCTGGGACT   19869
gb:MT066176|Organism:Severe      AGGTGAAAATACTCAATAATTTGGGTGTGGACATTGCTGCTAATACTGTGATCTGGGACT   19882
gb:MT027064|Organism:Severe      AGGTGAAAATACTCAATAATTTGGGTGTGGACATTGCTGCTAATACTGTGATCTGGGACT   19882
gb:MN994468|Organism:Severe      AGGTGAAAATACTCAATAATTTGGGTGTGGACATTGCTGCTAATACTGTGATCTGGGACT   19882
gb:MT072688|Organism:Severe      AGGTGAAAATACTCAATAATTTGGGTGTGGACATTGCTGCTAATACTGTGATCTGGGACT   19867
gb:MN996527|Organism:Severe      AGGTGAAAATACTCAATAATTTGGGTGTGGACATTGCTGCTAATACTGTGATCTGGGACT   19849
gb:MT093631|Organism:Severe      AGGTGAAAATACTCAATAATTTGGGTGTGGACATTGCTGCTAATACTGTGATCTGGGACT   19920
gb:MT106053|Organism:Severe      AGGTGAAAATACTCAATAATTTGGGTGTGGACATTGCTGCTAATACTGTGATCTGGGACT   19882
gb:MT019533|Organism:Severe      AGGTGAAAATACTCAATAATTTGGGTGTGGACATTGCTGCTAATACTGTGATCTGGGACT   19882
gb:MT019531|Organism:Severe      AGGTGAAAATACTCAATAATTTGGGTGTGGACATTGCTGCTAATACTGTGATCTGGGACT   19882
gb:MN996528|Organism:Severe      AGGTGAAAATACTCAATAATTTGGGTGTGGACATTGCTGCTAATACTGTGATCTGGGACT   19882
gb:MN996530|Organism:Severe      AGGTGAAAATACTCAATAATTTGGGTGTGGACATTGCTGCTAATACTGTGATCTGGGACT   19868
gb:MN908947|Organism:Severe      AGGTGAAAATACTCAATAATTTGGGTGTGGACATTGCTGCTAATACTGTGATCTGGGACT   19882
gb:MT019532|Organism:Severe      AGGTGAAAATACTCAATAATTTGGGTGTGGACATTGCTGCTAATACTGTGATCTGGGACT   19882
                                 ************************************************************
```

```
gb:MT020781|Organism:Severe      ACAAAAGAGATGCTCCAGCACATATATCTACTATTGGTGTTTGTTCTATGACTGACATAG   19930
gb:MT007544|Organism:Severe      ACAAAAGAGATGCTCCAGCACATATATCTACTATTGGTGTTTGTTCTATGACTGACATAG   19942
gb:MN994467|Organism:Severe      ACAAAAGAGATGCTCCAGCACATATATCTACTATTGGTGTTTGTTCTATGACTGACATAG   19942
gb:MT044257|Organism:Severe      ACAAAAGAGATGCTCCAGCACATATATCTACTATTGGTGTTTGTTCTATGACTGACATAG   19942
gb:MT106054|Organism:Severe      ACAAAAGAGATGCTCCAGCACATATATCTACTATTGGTGTTTGTTCTATGACTGACATAG   19942
gb:MT049951|Organism:Severe      ACAAAAGAGATGCTCCAGCACATATATCTACTATTGGTGTTTGTTCTATGACTGACATAG   19942
gb:MN975262|Organism:Severe      ACAAAAGAGATGCTCCAGCACATATATCTACTATTGGTGTTTGTTCTATGACTGACATAG   19942
gb:MT106052|Organism:Severe      ACAAAAGAGATGCTCCAGCACATATATCTACTATTGGTGTTTGTTCTATGACTGACATAG   19942
gb:LC522975|Organism:Severe      ACAAAAGAGATGCTCCAGCACATATATCTACTATTGGTGTTTGTTCTATGACTGACATAG   19939
gb:LC522973|Organism:Severe      ACAAAAGAGATGCTCCAGCACATATATCTACTATTGGTGTTTGTTCTATGACTGACATAG   19939
gb:LC522974|Organism:Severe      ACAAAAGAGATGCTCCAGCACATATATCTACTATTGGTGTTTGTTCTATGACTGACATAG   19939
gb:MN985325|Organism:Severe      ACAAAAGAGATGCTCCAGCACATATATCTACTATTGGTGTTTGTTCTATGACTGACATAG   19942
gb:MT020881|Organism:Severe      ACAAAAGAGATGCTCCAGCACATATATCTACTATTGGTGTTTGTTCTATGACTGACATAG   19942
gb:MT020880|Organism:Severe      ACAAAAGAGATGCTCCAGCACATATATCTACTATTGGTGTTTGTTCTATGACTGACATAG   19942
gb:MT066175|Organism:Severe      ACAAAAGAGATGCTCCAGCACATATATCTACTATTGGTGTTTGTTCTATGACTGACATAG   19942
gb:MN997409|Organism:Severe      ACAAAAGAGATGCTCCAGCACATATATCTACTATTGGTGTTTGTTCTATGACTGACATAG   19942
gb:MN938384|Organism:Severe      ACAAAAGAGATGCTCCAGCACATATATCTACTATTGGTGTTTGTTCTATGACTGACATAG   19910
gb:MT044258|Organism:Severe      ACAAAAGAGATGCTCCAGCACATATATCTACTATTGGTGTTTGTTCTATGACTGACATAG   19918
gb:MT039890|Organism:Severe      ACAAAAGAGATGCTCCAGCACATATATCTACTATTGGTGTTTGTTCTATGACTGACATAG   19942
gb:MN988713|Organism:Severe      ACAAAAGAGATGCTCCAGCACATATATCTACTATTGGTGTTTGTTCTATGACTGACATAG   19942
gb:LC521925|Organism:Severe      ACAAAAGAGATGCTCCAGCACATATATCTACTATTGGTGTTTGTTCTATGACTGACATAG   19915
gb:MT093571|Organism:Severe      ACAAAAGAGATGCTCCAGCACATATATCTACTATTGGTGTTTGTTCTATGACTGACATAG   19942
gb:MT039887|Organism:Severe      ACAAAAGAGATGCTCCAGCACATATATCTACTATTGGTGTTTGTTCTATGACTGACATAG   19942
gb:MT019530|Organism:Severe      ACAAAAGAGATGCTCCAGCACATATATCTACTATTGGTGTTTGTTCTATGACTGACATAG   19942
gb:MT039888|Organism:Severe      ACAAAAGAGATGCTCCAGCACATATATCTACTATTGGTGTTTGTTCTATGACTGACATAG   19942
gb:LC522972|Organism:Severe      ACAAAAGAGATGCTCCAGCACATATATCTACTATTGGTGTTTGTTCTATGACTGACATAG   19939
gb:MT027063|Organism:Severe      ACAAAAGAGATGCTCCAGCACATATATCTACTATTGGTGTTTGTTCTATGACTGACATAG   19942
gb:MT027062|Organism:Severe      ACAAAAGAGATGCTCCAGCACATATATCTACTATTGGTGTTTGTTCTATGACTGACATAG   19942
gb:MT019529|Organism:Severe      ACAAAAGAGATGCTCCAGCACATATATCTACTATTGGTGTTTGTTCTATGACTGACATAG   19942
gb:MN996529|Organism:Severe      ACAAAAGAGATGCTCCAGCACATATATCTACTATTGGTGTTTGTTCTATGACTGACATAG   19930
gb:MN996531|Organism:Severe      ACAAAAGAGATGCTCCAGCACATATATCTACTATTGGTGTTTGTTCTATGACTGACATAG   19929
gb:MT066176|Organism:Severe      ACAAAAGAGATGCTCCAGCACATATATCTACTATTGGTGTTTGTTCTATGACTGACATAG   19942
gb:MT027064|Organism:Severe      ACAAAAGAGATGCTCCAGCACATATATCTACTATTGGTGTTTGTTCTATGACTGACATAG   19942
gb:MN994468|Organism:Severe      ACAAAAGAGATGCTCCAGCACATATATCTACTATTGGTGTTTGTTCTATGACTGACATAG   19942
gb:MT072688|Organism:Severe      ACAAAAGAGATGCTCCAGCACATATATCTACTATTGGTGTTTGTTCTATGACTGACATAG   19927
gb:MN996527|Organism:Severe      ACAAAAGAGATGCTCCAGCACATATATCTACTATTGGTGTTTGTTCTATGACTGACATAG   19909
gb:MT093631|Organism:Severe      ACAAAAGAGATGCTCCAGCACATATATCTACTATTGGTGTTTGTTCTATGACTGACATAG   19980
gb:MT106053|Organism:Severe      ACAAAAGAGATGCTCCAGCACATATATCTACTATTGGTGTTTGTTCTATGACTGACATAG   19942
gb:MT019533|Organism:Severe      ACAAAAGAGATGCTCCAGCACATATATCTACTATTGGTGTTTGTTCTATGACTGACATAG   19942
gb:MT019531|Organism:Severe      ACAAAAGAGATGCTCCAGCACATATATCTACTATTGGTGTTTGTTCTATGACTGACATAG   19942
gb:MN996528|Organism:Severe      ACAAAAGAGATGCTCCAGCACATATATCTACTATTGGTGTTTGTTCTATGACTGACATAG   19942
gb:MN996530|Organism:Severe      ACAAAAGAGATGCTCCAGCACATATATCTACTATTGGTGTTTGTTCTATGACTGACATAG   19928
gb:MN908947|Organism:Severe      ACAAAAGAGATGCTCCAGCACATATATCTACTATTGGTGTTTGTTCTATGACTGACATAG   19942
gb:MT019532|Organism:Severe      ACAAAAGAGATGCTCCAGCACATATATCTACTATTGGTGTTTGTTCTATGACTGACATAG   19942
                                 ************************************************************

gb:MT020781|Organism:Severe      CCAAGAAACCAACTGAAACGATTTGTGCACCACTCACTGTCTTTTTTGATGGTAGAGTTG   19990
gb:MT007544|Organism:Severe      CCAAGAAACCAACTGAAACGATTTGTGCACCACTCACTGTCTTTTTTGATGGTAGAGTTG   20002
gb:MN994467|Organism:Severe      CCAAGAAACCAACTGAAACGATTTGTGCACCACTCACTGTCTTTTTTGATGGTAGAGTTG   20002
gb:MT044257|Organism:Severe      CCAAGAAACCAACTGAAACGATTTGTGCACCACTCACTGTCTTTTTTGATGGTAGAGTTG   20002
gb:MT106054|Organism:Severe      CCAAGAAACCAACTGAAACGATTTGTGCACCACTCACTGTCTTTTTTGATGGTAGAGTTG   20002
gb:MT049951|Organism:Severe      CCAAGAAACCAACTGAAACGATTTGTGCACCACTCACTGTCTTTTTTGATGGTAGAGTTG   20002
gb:MN975262|Organism:Severe      CCAAGAAACCAACTGAAACGATTTGTGCACCACTCACTGTCTTTTTTGATGGTAGAGTTG   20002
gb:MT106052|Organism:Severe      CCAAGAAACCAACTGAAACGATTTGTGCACCACTCACTGTCTTTTTTGATGGTAGAGTTG   20002
gb:LC522975|Organism:Severe      CCAAGAAACCAACTGAAACGATTTGTGCACCACTCACTGTCTTTTTTGATGGTAGAGTTG   19999
gb:LC522973|Organism:Severe      CCAAGAAACCAACTGAAACGATTTGTGCACCACTCACTGTCTTTTTTGATGGTAGAGTTG   19999
gb:LC522974|Organism:Severe      CCAAGAAACCAACTGAAACGATTTGTGCACCACTCACTGTCTTTTTTGATGGTAGAGTTG   19999
gb:MN985325|Organism:Severe      CCAAGAAACCAACTGAAACGATTTGTGCACCACTCACTGTCTTTTTTGATGGTAGAGTTG   20002
gb:MT020881|Organism:Severe      CCAAGAAACCAACTGAAACGATTTGTGCACCACTCACTGTCTTTTTTGATGGTAGAGTTG   20002
gb:MT020880|Organism:Severe      CCAAGAAACCAACTGAAACGATTTGTGCACCACTCACTGTCTTTTTTGATGGTAGAGTTG   20002
gb:MT066175|Organism:Severe      CCAAGAAACCAACTGAAACGATTTGTGCACCACTCACTGTCTTTTTTGATGGTAGAGTTG   20002
gb:MN997409|Organism:Severe      CCAAGAAACCAACTGAAACGATTTGTGCACCACTCACTGTCTTTTTTGATGGTAGAGTTG   20002
gb:MN938384|Organism:Severe      CCAAGAAACCAACTGAAACGATTTGTGCACCACTCACTGTCTTTTTTGATGGTAGAGTTG   19970
gb:MT044258|Organism:Severe      CCAAGAAACCAACTGAAACGATTTGTGCACCACTCACTGTCTTTTTTGATGGTAGAGTTG   19978
```

```
gb:MT039890|Organism:Severe      CCAAGAAACCAACTGAAACGATTTGTGCACCACTCACTGTCTTTTTTGATGGTAGAGTTG   20002
gb:MN988713|Organism:Severe      CCAAGAAACCAACTGAAACGATTTGTGCACCACTCACTGTCTTTTTTGATGGTAGAGTTG   20002
gb:LC521925|Organism:Severe      CCAAGAAACCAACTGAAACGATTTGTGCACCACTCACTGTCTTTTTTGATGGTAGAGTTG   19975
gb:MT093571|Organism:Severe      CCAAGAAACCAACTGAAACGATTTGTGCACCACTCACTGTCTTTTTTGATGGTAGAGTTG   20002
gb:MT039887|Organism:Severe      CCAAGAAACCAACTGAAACGATTTGTGCACCACTCACTGTCTTTTTTGATGGTAGAGTTG   20002
gb:MT019530|Organism:Severe      CCAAGAAACCAACTGAAACGATTTGTGCACCACTCACTGTCTTTTTTGATGGTAGAGTTG   20002
gb:MT039888|Organism:Severe      CCAAGAAACCAACTGAAACGATTTGTGCACCACTCACTGTCTTTTTTGATGGTAGAGTTG   20002
gb:LC522972|Organism:Severe      CCAAGAAACCAACTGAAACGATTTGTGCACCACTCACTGTCTTTTTTGATGGTAGAGTTG   19999
gb:MT027063|Organism:Severe      CCAAGAAACCAACTGAAACGATTTGTGCACCACTCACTGTCTTTTTTGATGGTAGAGTTG   20002
gb:MT027062|Organism:Severe      CCAAGAAACCAACTGAAACGATTTGTGCACCACTCACTGTCTTTTTTGATGGTAGAGTTG   20002
gb:MT019529|Organism:Severe      CCAAGAAACCAACTGAAACGATTTGTGCACCACTCACTGTCTTTTTTGATGGTAGAGTTG   20002
gb:MN996529|Organism:Severe      CCAAGAAACCAACTGAAACGATTTGTGCACCACTCACTGTCTTTTTTGATGGTAGAGTTG   19990
gb:MN996531|Organism:Severe      CCAAGAAACCAACTGAAACGATTTGTGCACCACTCACTGTCTTTTTTGATGGTAGAGTTG   19989
gb:MT066176|Organism:Severe      CCAAGAAACCAACTGAAACGATTTGTGCACCACTCACTGTCTTTTTTGATGGTAGAGTTG   20002
gb:MT027064|Organism:Severe      CCAAGAAACCAACTGAAACGATTTGTGCACCACTCACTGTCTTTTTTGATGGTAGAGTTG   20002
gb:MN994468|Organism:Severe      CCAAGAAACCAACTGAAACGATTTGTGCACCACTCACTGTCTTTTTTGATGGTAGAGTTG   20002
gb:MT072688|Organism:Severe      CCAAGAAACCAACTGAAACGATTTGTGCACCACTCACTGTCTTTTTTGATGGTAGAGTTG   19987
gb:MN996527|Organism:Severe      CCAAGAAACCAACTGAAACGATTTGTGCACCACTCACTGTCTTTTTTGATGGTAGAGTTG   19969
gb:MT093631|Organism:Severe      CCAAGAAACCAACTGAAACGATTTGTGCACCACTCACTGTCTTTTTTGATGGTAGAGTTG   20040
gb:MT106053|Organism:Severe      CCAAGAAACCAACTGAAACGATTTGTGCACCACTCACTGTCTTTTTTGATGGTAGAGTTG   20002
gb:MT019533|Organism:Severe      CCAAGAAACCAACTGAAACGATTTGTGCACCACTCACTGTCTTTTTTGATGGTAGAGTTG   20002
gb:MT019531|Organism:Severe      CCAAGAAACCAACTGAAACGATTTGTGCACCACTCACTGTCTTTTTTGATGGTAGAGTTG   20002
gb:MN996528|Organism:Severe      CCAAGAAACCAACTGAAACGATTTGTGCACCACTCACTGTCTTTTTTGATGGTAGAGTTG   20002
gb:MN996530|Organism:Severe      CCAAGAAACCAACTGAAACGATTTGTGCACCACTCACTGTCTTTTTTGATGGTAGAGTTG   19988
gb:MN908947|Organism:Severe      CCAAGAAACCAACTGAAACGATTTGTGCACCACTCACTGTCTTTTTTGATGGTAGAGTTG   20002
gb:MT019532|Organism:Severe      CCAAGAAACCAACTGAAACGATTTGTGCACCACTCACTGTCTTTTTTGATGGTAGAGTTG   20002
                                 ************************************************************

gb:MT020781|Organism:Severe      ATGGTCAAGTAGACTTATTTAGAAATGCCCGTAATGGTGTTCTTATTACAGAAGGTAGTG   20050
gb:MT007544|Organism:Severe      ATGGTCAAGTAGACTTATTTAGAAATGCCCGTAATGGTGTTCTTATTACAGAAGGTAGTG   20062
gb:MN994467|Organism:Severe      ATGGTCAAGTAGACTTATTTAGAAATGCCCGTAATGGTGTTCTTATTACAGAAGGTAGTG   20062
gb:MT044257|Organism:Severe      ATGGTCAAGTAGACTTATTTAGAAATGCCCGTAATGGTGTTCTTATTACAGAAGGTAGTG   20062
gb:MT106054|Organism:Severe      ATGGTCAAGTAGACTTATTTAGAAATGCCCGTAATGGTGTTCTTATTACAGAAGGTAGTG   20062
gb:MT049951|Organism:Severe      ATGGTCAAGTAGACTTATTTAGAAATGCCCGTAATGGTGTTCTTATTACAGAAGGTAGTG   20062
gb:MN975262|Organism:Severe      ATGGTCAAGTAGACTTATTTAGAAATGCCCGTAATGGTGTTCTTATTACAGAAGGTAGTG   20062
gb:MT106052|Organism:Severe      ATGGTCAAGTAGACTTATTTAGAAATGCCCGTAATGGTGTTCTTATTACAGAAGGTAGTG   20062
gb:LC522975|Organism:Severe      ATGGTCAAGTAGACTTATTTAGAAATGCCCGTAATGGTGTTCTTATTACAGAAGGTAGTG   20059
gb:LC522973|Organism:Severe      ATGGTCAAGTAGACTTATTTAGAAATGCCCGTAATGGTGTTCTTATTACAGAAGGTAGTG   20059
gb:LC522974|Organism:Severe      ATGGTCAAGTAGACTTATTTAGAAATGCCCGTAATGGTGTTCTTATTACAGAAGGTAGTG   20059
gb:MN985325|Organism:Severe      ATGGTCAAGTAGACTTATTTAGAAATGCCCGTAATGGTGTTCTTATTACAGAAGGTAGTG   20062
gb:MT020881|Organism:Severe      ATGGTCAAGTAGACTTATTTAGAAATGCCCGTAATGGTGTTCTTATTACAGAAGGTAGTG   20062
gb:MT020880|Organism:Severe      ATGGTCAAGTAGACTTATTTAGAAATGCCCGTAATGGTGTTCTTATTACAGAAGGTAGTG   20062
gb:MT066175|Organism:Severe      ATGGTCAAGTAGACTTATTTAGAAATGCCCGTAATGGTGTTCTTATTACAGAAGGTAGTG   20062
gb:MN997409|Organism:Severe      ATGGTCAAGTAGACTTATTTAGAAATGCCCGTAATGGTGTTCTTATTACAGAAGGTAGTG   20062
gb:MN938384|Organism:Severe      ATGGTCAAGTAGACTTATTTAGAAATGCCCGTAATGGTGTTCTTATTACAGAAGGTAGTG   20030
gb:MT044258|Organism:Severe      ATGGTCAAGTAGACTTATTTAGAAATGCCCGTAATGGTGTTCTTATTACAGAAGGTAGTG   20038
gb:MT039890|Organism:Severe      ATGGTCAAGTAGACTTATTTAGAAATGCCCGTAATGGTGTTCTTATTACAGAAGGTAGTG   20062
gb:MN988713|Organism:Severe      ATGGTCAAGTAGACTTATTTAGAAATGCCCGTAATGGTGTTCTTATTACAGAAGGTAGTG   20062
gb:LC521925|Organism:Severe      ATGGTCAAGTAGACTTATTTAGAAATGCCCGTAATGGTGTTCTTATTACAGAAGGTAGTG   20035
gb:MT093571|Organism:Severe      ATGGTCAAGTAGACTTATTTAGAAATGCCCGTAATGGTGTTCTTATTACAGAAGGTAGTG   20062
gb:MT039887|Organism:Severe      ATGGTCAAGTAGACTTATTTAGAAATGCCCGTAATGGTGTTCTTATTACAGAAGGTAGTG   20062
gb:MT019530|Organism:Severe      ATGGTCAAGTAGACTTATTTAGAAATGCCCGTAATGGTGTTCTTATTACAGAAGGTAGTG   20062
gb:MT039888|Organism:Severe      ATGGTCAAGTAGACTTATTTAGAAATGCCCGTAATGGTGTTCTTATTACAGAAGGTAGTG   20062
gb:LC522972|Organism:Severe      ATGGTCAAGTAGACTTATTTAGAAATGCCCGTAATGGTGTTCTTATTACAGAAGGTAGTG   20059
gb:MT027063|Organism:Severe      ATGGTCAAGTAGACTTATTTAGAAATGCCCGTAATGGTGTTCTTATTACAGAAGGTAGTG   20062
gb:MT027062|Organism:Severe      ATGGTCAAGTAGACTTATTTAGAAATGCCCGTAATGGTGTTCTTATTACAGAAGGTAGTG   20062
gb:MT019529|Organism:Severe      ATGGTCAAGTAGACTTATTTAGAAATGCCCGTAATGGTGTTCTTATTACAGAAGGTAGTG   20062
gb:MN996529|Organism:Severe      ATGGTCAAGTAGACTTATTTAGAAATGCCCGTAATGGTGTTCTTATTACAGAAGGTAGTG   20050
gb:MN996531|Organism:Severe      ATGGTCAAGTAGACTTATTTAGAAATGCCCGTAATGGTGTTCTTATTACAGAAGGTAGTG   20049
gb:MT066176|Organism:Severe      ATGGTCAAGTAGACTTATTTAGAAATGCCCGTAATGGTGTTCTTATTACAGAAGGTAGTG   20062
gb:MT027064|Organism:Severe      ATGGTCAAGTAGACTTATTTAGAAATGCCCGTAATGGTGTTCTTATTACAGAAGGTAGTG   20062
gb:MN994468|Organism:Severe      ATGGTCAAGTAGACTTATTTAGAAATGCCCGTAATGGTGTTCTTATTACAGAAGGTAGTG   20062
gb:MT072688|Organism:Severe      ATGGTCAAGTAGACTTATTTAGAAATGCCCGTAATGGTGTTCTTATTACAGAAGGTAGTG   20047
gb:MN996527|Organism:Severe      ATGGTCAAGTAGACTTATTTAGAAATGCCCGTAATGGTGTTCTTATTACAGAAGGTAGTG   20029
gb:MT093631|Organism:Severe      ATGGTCAAGTAGACTTATTTAGAAATGCCCGTAATGGTGTTCTTATTACAGAAGGTAGTG   20100
```

```
gb:MT106053|Organism:Severe      ATGGTCAAGTAGACTTATTTAGAAATGCCCGTAATGGTGTTCTTATTACAGAAGGTAGTG   20062
gb:MT019533|Organism:Severe      ATGGTCAAGTAGACTTATTTAGAAATGCCCGTAATGGTGTTCTTATTACAGAAGGTAGTG   20062
gb:MT019531|Organism:Severe      ATGGTCAAGTAGACTTATTTAGAAATGCCCGTAATGGTGTTCTTATTACAGAAGGTAGTG   20062
gb:MN996528|Organism:Severe      ATGGTCAAGTAGACTTATTTAGAAATGCCCGTAATGGTGTTCTTATTACAGAAGGTAGTG   20062
gb:MN996530|Organism:Severe      ATGGTCAAGTAGACTTATTTAGAAATGCCCGTAATGGTGTTCTTATTACAGAAGGTAGTG   20048
gb:MN908947|Organism:Severe      ATGGTCAAGTAGACTTATTTAGAAATGCCCGTAATGGTGTTCTTATTACAGAAGGTAGTG   20062
gb:MT019532|Organism:Severe      ATGGTCAAGTAGACTTATTTAGAAATGCCCGTAATGGTGTTCTTATTACAGAAGGTAGTG   20062
                                 ************************************************************

gb:MT020781|Organism:Severe      TTAAAGGTTTACAACCATCTGTAGGTCCCAAACAAGCTAGTCTTAATGGAGTCACATTAA   20110
gb:MT007544|Organism:Severe      TTAAAGGTTTACAACCATCTGTAGGTCCCAAACAAGCTAGTCTTAATGGAGTCACATTAA   20122
gb:MN994467|Organism:Severe      TTAAAGGTTTACAACCATCTGTAGGTCCCAAACAAGCTAGTCTTAATGGAGTCACATTAA   20122
gb:MT044257|Organism:Severe      TTAAAGGTTTACAACCATCTGTAGGTCCCAAACAAGCTAGTCTTAATGGAGTCACATTAA   20122
gb:MT106054|Organism:Severe      TTAAAGGTTTACAACCATCTGTAGGTCCCAAACAAGCTAGTCTTAATGGAGTCACATTAA   20122
gb:MT049951|Organism:Severe      TTAAAGGTTTACAACCATCTGTAGGTCCCAAACAAGCTAGTCTTAATGGAGTCACATTAA   20122
gb:MN975262|Organism:Severe      TTAAAGGTTTACAACCATCTGTAGGTCCCAAACAAGCTAGTCTTAATGGAGTCACATTAA   20122
gb:MT106052|Organism:Severe      TTAAAGGTTTACAACCATCTGTAGGTCCCAAACAAGCTAGTCTTAATGGAGTCACATTAA   20122
gb:LC522975|Organism:Severe      TTAAAGGTTTACAACCATCTGTAGGTCCCAAACAAGCTAGTCTTAATGGAGTCACATTAA   20119
gb:LC522973|Organism:Severe      TTAAAGGTTTACAACCATCTGTAGGTCCCAAACAAGCTAGTCTTAATGGAGTCACATTAA   20119
gb:LC522974|Organism:Severe      TTAAAGGTTTACAACCATCTGTAGGTCCCAAACAAGCTAGTCTTAATGGAGTCACATTAA   20119
gb:MN985325|Organism:Severe      TTAAAGGTTTACAACCATCTGTAGGTCCCAAACAAGCTAGTCTTAATGGAGTCACATTAA   20122
gb:MT020881|Organism:Severe      TTAAAGGTTTACAACCATCTGTAGGTCCCAAACAAGCTAGTCTTAATGGAGTCACATTAA   20122
gb:MT020880|Organism:Severe      TTAAAGGTTTACAACCATCTGTAGGTCCCAAACAAGCTAGTCTTAATGGAGTCACATTAA   20122
gb:MT066175|Organism:Severe      TTAAAGGTTTACAACCATCTGTAGGTCCCAAACAAGCTAGTCTTAATGGAGTCACATTAA   20122
gb:MN997409|Organism:Severe      TTAAAGGTTTACAACCATCTGTAGGTCCCAAACAAGCTAGTCTTAATGGAGTCACATTAA   20122
gb:MN938384|Organism:Severe      TTAAAGGTTTACAACCATCTGTAGGTCCCAAACAAGCTAGTCTTAATGGAGTCACATTAA   20090
gb:MT044258|Organism:Severe      TTAAAGGTTTACAACCATCTGTAGGTCCCAAACAAGCTAGTCTTAATGGAGTCACATTAA   20098
gb:MT039890|Organism:Severe      TTAAAGGTTTACAACCATCTGTAGGTCCCAAACAAGCTAGTCTTAATGGAGTCACATTAA   20122
gb:MN988713|Organism:Severe      TTAAAGGTTTACAACCATCTGTAGGTCCCAAACAAGCTAGTCTTAATGGAGTCACATTAA   20122
gb:LC521925|Organism:Severe      TTAAAGGTTTACAACCATCTGTAGGTCCCAAACAAGCTAGTCTTAATGGAGTCACATTAA   20095
gb:MT093571|Organism:Severe      TTAAAGGTTTACAACCATCTGTAGGTCCCAAACAAGCTAGTCTTAATGGAGTCACATTAA   20122
gb:MT039887|Organism:Severe      TTAAAGGTTTACAACCATCTGTAGGTCCCAAACAAGCTAGTCTTAATGGAGTCACATTAA   20122
gb:MT019530|Organism:Severe      TTAAAGGTTTACAACCATCTGTAGGTCCCAAACAAGCTAGTCTTAATGGAGTCACATTAA   20122
gb:MT039888|Organism:Severe      TTAAAGGTTTACAACCATCTGTAGGTCCCAAACAAGCTAGTCTTAATGGAGTCACATTAA   20122
gb:LC522972|Organism:Severe      TTAAAGGTTTACAACCATCTGTAGGTCCCAAACAAGCTAGTCTTAATGGAGTCACATTAA   20119
gb:MT027063|Organism:Severe      TTAAAGGTTTACAACCATCTGTAGGTCCCAAACAAGCTAGTCTTAATGGAGTCACATTAA   20122
gb:MT027062|Organism:Severe      TTAAAGGTTTACAACCATCTGTAGGTCCCAAACAAGCTAGTCTTAATGGAGTCACATTAA   20122
gb:MT019529|Organism:Severe      TTAAAGGTTTACAACCATCTGTAGGTCCCAAACAAGCTAGTCTTAATGGAGTCACATTAA   20122
gb:MN996529|Organism:Severe      TTAAAGGTTTACAACCATCTGTAGGTCCCAAACAAGCTAGTCTTAATGGAGTCACATTAA   20110
gb:MN996531|Organism:Severe      TTAAAGGTTTACAACCATCTGTAGGTCCCAAACAAGCTAGTCTTAATGGAGTCACATTAA   20109
gb:MT066176|Organism:Severe      TTAAAGGTTTACAACCATCTGTAGGTCCCAAACAAGCTAGTCTTAATGGAGTCACATTAA   20122
gb:MT027064|Organism:Severe      TTAAAGGTTTACAACCATCTGTAGGTCCCAAACAAGCTAGTCTTAATGGAGTCACATTAA   20122
gb:MN994468|Organism:Severe      TTAAAGGTTTACAACCATCTGTAGGTCCCAAACAAGCTAGTCTTAATGGAGTCACATTAA   20122
gb:MT072688|Organism:Severe      TTAAAGGTTTACAACCATCTGTAGGTCCCAAACAAGCTAGTCTTAATGGAGTCACATTAA   20107
gb:MN996527|Organism:Severe      TTAAAGGTTTACAACCATCTGTAGGTCCCAAACAAGCTAGTCTTAATGGAGTCACATTAA   20089
gb:MT093631|Organism:Severe      TTAAAGGTTTACAACCATCTGTAGGTCCCAAACAAGCTAGTCTTAATGGAGTCACATTAA   20160
gb:MT106053|Organism:Severe      TTAAAGGTTTACAACCATCTGTAGGTCCCAAACAAGCTAGTCTTAATGGAGTCACATTAA   20122
gb:MT019533|Organism:Severe      TTAAAGGTTTACAACCATCTGTAGGTCCCAAACAAGCTAGTCTTAATGGAGTCACATTAA   20122
gb:MT019531|Organism:Severe      TTAAAGGTTTACAACCATCTGTAGGTCCCAAACAAGCTAGTCTTAATGGAGTCACATTAA   20122
gb:MN996528|Organism:Severe      TTAAAGGTTTACAACCATCTGTAGGTCCCAAACAAGCTAGTCTTAATGGAGTCACATTAA   20122
gb:MN996530|Organism:Severe      TTAAAGGTTTACAACCATCTGTAGGTCCCAAACAAGCTAGTCTTAATGGAGTCACATTAA   20108
gb:MN908947|Organism:Severe      TTAAAGGTTTACAACCATCTGTAGGTCCCAAACAAGCTAGTCTTAATGGAGTCACATTAA   20122
gb:MT019532|Organism:Severe      TTAAAGGTTTACAACCATCTGTAGGTCCCAAACAAGCTAGTCTTAATGGAGTCACATTAA   20122
                                 ************************************************************

gb:MT020781|Organism:Severe      TTGGAGAAGCCGTAAAAACACAGTTCAATTATTATAAGAAAGTTGATGGTGTTGTCCAAC   20170
gb:MT007544|Organism:Severe      TTGGAGAAGCCGTAAAAACACAGTTCAATTATTATAAGAAAGTTGATGGTGTTGTCCAAC   20182
gb:MN994467|Organism:Severe      TTGGAGAAGCCGTAAAAACACAGTTCAATTATTATAAGAAAGTTGATGGTGTTGTCCAAC   20182
gb:MT044257|Organism:Severe      TTGGAGAAGCCGTAAAAACACAGTTCAATTATTATAAGAAAGTTGATGGTGTTGTCCAAC   20182
gb:MT106054|Organism:Severe      TTGGAGAAGCCGTAAAAACACAGTTCAATTATTATAAGAAAGTTGATGGTGTTGTCCAAC   20182
gb:MT049951|Organism:Severe      TTGGAGAAGCCGTAAAAACACAGTTCAATTATTATAAGAAAGTTGATGGTGTTGTCCAAC   20182
gb:MN975262|Organism:Severe      TTGGAGAAGCCGTAAAAACACAGTTCAATTATTATAAGAAAGTTGATGGTGTTGTCCAAC   20182
gb:MT106052|Organism:Severe      TTGGAGAAGCCGTAAAAACACAGTTCAATTATTATAAGAAAGTTGATGGTGTTGTCCAAC   20182
gb:LC522975|Organism:Severe      TTGGAGAAGCCGTAAAAACACAGTTCAATTATTATAAGAAAGTTGATGGTGTTGTCCAAC   20179
gb:LC522973|Organism:Severe      TTGGAGAAGCCGTAAAAACACAGTTCAATTATTATAAGAAAGTTGATGGTGTTGTCCAAC   20179
```

```
gb:LC522974|Organism:Severe      TTGGAGAAGCCGTAAAAACACAGTTCAATTATTATAAGAAAGTTGATGGTGTTGTCCAAC   20179
gb:MN985325|Organism:Severe      TTGGAGAAGCCGTAAAAACACAGTTCAATTATTATAAGAAAGTTGATGGTGTTGTCCAAC   20182
gb:MT020881|Organism:Severe      TTGGAGAAGCCGTAAAAACACAGTTCAATTATTATAAGAAAGTTGATGGTGTTGTCCAAC   20182
gb:MT020880|Organism:Severe      TTGGAGAAGCCGTAAAAACACAGTTCAATTATTATAAGAAAGTTGATGGTGTTGTCCAAC   20182
gb:MT066175|Organism:Severe      TTGGAGAAGCCGTAAAAACACAGTTCAATTATTATAAGAAAGTTGATGGTGTTGTCCAAC   20182
gb:MN997409|Organism:Severe      TTGGAGAAGCCGTAAAAACACAGTTCAATTATTATAAGAAAGTTGATGGTGTTGTCCAAC   20182
gb:MN938384|Organism:Severe      TTGGAGAAGCCGTAAAAACACAGTTCAATTATTATAAGAAAGTTGATGGTGTTGTCCAAC   20150
gb:MT044258|Organism:Severe      TTGGAGAAGCCGTAAAAACACAGTTCAATTATTATAAGAAAGTTGATGGTGTTGTCCAAC   20158
gb:MT039890|Organism:Severe      TTGGAGAAGCCGTAAAAACACAGTTCAATTATTATAAGAAAGTTGATGGTGTTGTCCAAC   20182
gb:MN988713|Organism:Severe      TTGGAGAAGCCGTAAAAACACAGTTCAATTATTATAAGAAAGTTGATGGTGTTGTCCAAC   20182
gb:LC521925|Organism:Severe      TTGGAGAAGCCGTAAAAACACAGTTCAATTATTATAAGAAAGTTGATGGTGTTGTCCAAC   20155
gb:MT093571|Organism:Severe      TTGGAGAAGCCGTAAAAACACAGTTCAATTATTATAAGAAAGTTGATGGTGTTGTCCAAC   20182
gb:MT039887|Organism:Severe      TTGGAGAAGCCGTAAAAACACAGTTCAATTATTATAAGAAAGTTGATGGTGTTGTCCAAC   20182
gb:MT019530|Organism:Severe      TTGGAGAAGCCGTAAAAACACAGTTCAATTATTATAAGAAAGTTGATGGTGTTGTCCAAC   20182
gb:MT039888|Organism:Severe      TTGGAGAAGCCGTAAAAACACAGTTCAATTATTATAAGAAAGTTGATGGTGTTGTCCAAC   20182
gb:LC522972|Organism:Severe      TTGGAGAAGCCGTAAAAACACAGTTCAATTATTATAAGAAAGTTGATGGTGTTGTCCAAC   20179
gb:MT027063|Organism:Severe      TTGGAGAAGCCGTAAAAACACAGTTCAATTATTATAAGAAAGTTGATGGTGTTGTCCAAC   20182
gb:MT027062|Organism:Severe      TTGGAGAAGCCGTAAAAACACAGTTCAATTATTATAAGAAAGTTGATGGTGTTGTCCAAC   20182
gb:MT019529|Organism:Severe      TTGGAGAAGCCGTAAAAACACAGTTCAATTATTATAAGAAAGTTGATGGTGTTGTCCAAC   20182
gb:MN996529|Organism:Severe      TTGGAGAAGCCGTAAAAACACAGTTCAATTATTATAAGAAAGTTGATGGTGTTGTCCAAC   20170
gb:MN996531|Organism:Severe      TTGGAGAAGCCGTAAAAACACAGTTCAATTATTATAAGAAAGTTGATGGTGTTGTCCAAC   20169
gb:MT066176|Organism:Severe      TTGGAGAAGCCGTAAAAACACAGTTCAATTATTATAAGAAAGTTGATGGTGTTGTCCAAC   20182
gb:MT027064|Organism:Severe      TTGGAGAAGCCGTAAAAACACAGTTCAATTATTATAAGAAAGTTGATGGTGTTGTCCAAC   20182
gb:MN994468|Organism:Severe      TTGGAGAAGCCGTAAAAACACAGTTCAATTATTATAAGAAAGTTGATGGTGTTGTCCAAC   20182
gb:MT072688|Organism:Severe      TTGGAGAAGCCGTAAAAACACAGTTCAATTATTATAAGAAAGTTGATGGTGTTGTCCAAC   20167
gb:MN996527|Organism:Severe      TTGGAGAAGCCGTAAAAACACAGTTCAATTATTATAAGAAAGTTGATGGTGTTGTCCAAC   20149
gb:MT093631|Organism:Severe      TTGGAGAAGCCGTAAAAACACAGTTCAATTATTATAAGAAAGTTGATGGTGTTGTCCAAC   20220
gb:MT106053|Organism:Severe      TTGGAGAAGCCGTAAAAACACAGTTCAATTATTATAAGAAAGTTGATGGTGTTGTCCAAC   20182
gb:MT019533|Organism:Severe      TTGGAGAAGCCGTAAAAACACAGTTCAATTATTATAAGAAAGTTGATGGTGTTGTCCAAC   20182
gb:MT019531|Organism:Severe      TTGGAGAAGCCGTAAAAACACAGTTCAATTATTATAAGAAAGTTGATGGTGTTGTCCAAC   20182
gb:MN996528|Organism:Severe      TTGGAGAAGCCGTAAAAACACAGTTCAATTATTATAAGAAAGTTGATGGTGTTGTCCAAC   20182
gb:MN996530|Organism:Severe      TTGGAGAAGCCGTAAAAACACAGTTCAATTATTATAAGAAAGTTGATGGTGTTGTCCAAC   20168
gb:MN908947|Organism:Severe      TTGGAGAAGCCGTAAAAACACAGTTCAATTATTATAAGAAAGTTGATGGTGTTGTCCAAC   20182
gb:MT019532|Organism:Severe      TTGGAGAAGCCGTAAAAACACAGTTCAATTATTATAAGAAAGTTGATGGTGTTGTCCAAC   20182
                                 ************************************************************

gb:MT020781|Organism:Severe      AATTACCTGAAACTTACTTTACTCAGAGTAGAAATTTACAAGAATTTAAACCCAGGAGTC   20230
gb:MT007544|Organism:Severe      AATTACCTGAAACTTACTTTACTCAGAGTAGAAATTTACAAGAATTTAAACCCAGGAGTC   20242
gb:MN994467|Organism:Severe      AATTACCTGAAACTTACTTTACTCAGAGTAGAAATTTACAAGAATTTAAACCCAGGAGTC   20242
gb:MT044257|Organism:Severe      AATTACCTGAAACTTACTTTACTCAGAGTAGAAATTTACAAGAATTTAAACCCAGGAGTC   20242
gb:MT106054|Organism:Severe      AATTACCTGAAACTTACTTTACTCAGAGTAGAAATTTACAAGAATTTAAACCCAGGAGTC   20242
gb:MT049951|Organism:Severe      AATTACCTGAAACTTACTTTACTCAGAGTAGAAATTTACAAGAATTTAAACCCAGGAGTC   20242
gb:MN975262|Organism:Severe      AATTACCTGAAACTTACTTTACTCAGAGTAGAAATTTACAAGAATTTAAACCCAGGAGTC   20242
gb:MT106052|Organism:Severe      AATTACCTGAAACTTACTTTACTCAGAGTAGAAATTTACAAGAATTTAAACCCAGGAGTC   20242
gb:LC522975|Organism:Severe      AATTACCTGAAACTTACTTTACTCAGAGTAGAAATTTACAAGAATTTAAACCCAGGAGTC   20239
gb:LC522973|Organism:Severe      AATTACCTGAAACTTACTTTACTCAGAGTAGAAATTTACAAGAATTTAAACCCAGGAGTC   20239
gb:LC522974|Organism:Severe      AATTACCTGAAACTTACTTTACTCAGAGTAGAAATTTACAAGAATTTAAACCCAGGAGTC   20239
gb:MN985325|Organism:Severe      AATTACCTGAAACTTACTTTACTCAGAGTAGAAATTTACAAGAATTTAAACCCAGGAGTC   20242
gb:MT020881|Organism:Severe      AATTACCTGAAACTTACTTTACTCAGAGTAGAAATTTACAAGAATTTAAACCCAGGAGTC   20242
gb:MT020880|Organism:Severe      AATTACCTGAAACTTACTTTACTCAGAGTAGAAATTTACAAGAATTTAAACCCAGGAGTC   20242
gb:MT066175|Organism:Severe      AATTACCTGAAACTTACTTTACTCAGAGTAGAAATTTACAAGAATTTAAACCCAGGAGTC   20242
gb:MN997409|Organism:Severe      AATTACCTGAAACTTACTTTACTCAGAGTAGAAATTTACAAGAATTTAAACCCAGGAGTC   20242
gb:MN938384|Organism:Severe      AATTACCTGAAACTTACTTTACTCAGAGTAGAAATTTACAAGAATTTAAACCCAGGAGTC   20210
gb:MT044258|Organism:Severe      AATTACCTGAAACTTACTTTACTCAGAGTAGAAATTTACAAGAATTTAAACCCAGGAGTC   20218
gb:MT039890|Organism:Severe      AATTACCTGAAACTTACTTTACTCAGAGTAGAAATTTACAAGAATTTAAACCCAGGAGTC   20242
gb:MN988713|Organism:Severe      AATTACCTGAAACTTACTTTACTCAGAGTAGAAATTTACAAGAATTTAAACCCAGGAGTC   20242
gb:LC521925|Organism:Severe      AATTACCTGAAACTTACTTTACTCAGAGTAGAAATTTACAAGAATTTAAACCCAGGAGTC   20215
gb:MT093571|Organism:Severe      AATTACCTGAAACTTACTTTACTCAGAGTAGAAATTTACAAGAATTTAAACCCAGGAGTC   20242
gb:MT039887|Organism:Severe      AATTACCTGAAACTTACTTTACTCAGAGTAGAAATTTACAAGAATTTAAACCCAGGAGTC   20242
gb:MT019530|Organism:Severe      AATTACCTGAAACTTACTTTACTCAGAGTAGAAATTTACAAGAATTTAAACCCAGGAGTC   20242
gb:MT039888|Organism:Severe      AATTACCTGAAACTTACTTTACTCAGAGTAGAAATTTACAAGAATTTAAACCCAGGAGTC   20242
gb:LC522972|Organism:Severe      AATTACCTGAAACTTACTTTACTCAGAGTAGAAATTTACAAGAATTTAAACCCAGGAGTC   20239
gb:MT027063|Organism:Severe      AATTACCTGAAACTTACTTTACTCAGAGTAGAAATTTACAAGAATTTAAACCCAGGAGTC   20242
gb:MT027062|Organism:Severe      AATTACCTGAAACTTACTTTACTCAGAGTAGAAATTTACAAGAATTTAAACCCAGGAGTC   20242
gb:MT019529|Organism:Severe      AATTACCTGAAACTTACTTTACTCAGAGTAGAAATTTACAAGAATTTAAACCCAGGAGTC   20242
```

```
gb:MN996529|Organism:Severe      AATTACCTGAAACTTACTTTACTCAGAGTAGAAATTTACAAGAATTTAAACCCAGGAGTC   20230
gb:MN996531|Organism:Severe      AATTACCTGAAACTTACTTTACTCAGAGTAGAAATTTACAAGAATTTAAACCCAGGAGTC   20229
gb:MT066176|Organism:Severe      AATTACCTGAAACTTACTTTACTCAGAGTAGAAATTTACAAGAATTTAAACCCAGGAGTC   20242
gb:MT027064|Organism:Severe      AATTACCTGAAACTTACTTTACTCAGAGTAGAAATTTACAAGAATTTAAACCCAGGAGTC   20242
gb:MN994468|Organism:Severe      AATTACCTGAAACTTACTTTACTCAGAGTAGAAATTTACAAGAATTTAAACCCAGGAGTC   20242
gb:MT072688|Organism:Severe      AATTACCTGAAACTTACTTTACTCAGAGTAGAAATTTACAAGAATTTAAACCCAGGAGTC   20227
gb:MN996527|Organism:Severe      AATTACCTGAAACTTACTTTACTCAGAGTAGAAATTTACAAGAATTTAAACCCAGGAGTC   20209
gb:MT093631|Organism:Severe      AATTACCTGAAACTTACTTTACTCAGAGTAGAAATTTACAAGAATTTAAACCCAGGAGTC   20280
gb:MT106053|Organism:Severe      AATTACCTGAAACTTACTTTACTCAGAGTAGAAATTTACAAGAATTTAAACCCAGGAGTC   20242
gb:MT019533|Organism:Severe      AATTACCTGAAACTTACTTTACTCAGAGTAGAAATTTACAAGAATTTAAACCCAGGAGTC   20242
gb:MT019531|Organism:Severe      AATTACCTGAAACTTACTTTACTCAGAGTAGAAATTTACAAGAATTTAAACCCAGGAGTC   20242
gb:MN996528|Organism:Severe      AATTACCTGAAACTTACTTTACTCAGAGTAGAAATTTACAAGAATTTAAACCCAGGAGTC   20242
gb:MN996530|Organism:Severe      AATTACCTGAAACTTACTTTACTCAGAGTAGAAATTTACAAGAATTTAAACCCAGGAGTC   20228
gb:MN908947|Organism:Severe      AATTACCTGAAACTTACTTTACTCAGAGTAGAAATTTACAAGAATTTAAACCCAGGAGTC   20242
gb:MT019532|Organism:Severe      AATTACCTGAAACTTACTTTACTCAGAGTAGAAATTTACAAGAATTTAAACCCAGGAGTC   20242
                                 ************************************************************

gb:MT020781|Organism:Severe      AAATGGAAATTGATTTCTTAGAATTAGCTATGGATGAATTCATTGAACGGTATAAATTAG   20290
gb:MT007544|Organism:Severe      AAATGGAAATTGATTTCTTAGAATTAGCTATGGATGAATTCATTGAACGGTATAAATTAG   20302
gb:MN994467|Organism:Severe      AAATGGAAATTGATTTCTTAGAATTAGCTATGGATGAATTCATTGAACGGTATAAATTAG   20302
gb:MT044257|Organism:Severe      AAATGGAAATTGATTTCTTAGAATTAGCTATGGATGAATTCATTGAACGGTATAAATTAG   20302
gb:MT106054|Organism:Severe      AAATGGAAATTGATTTCTTAGAATTAGCTATGGATGAATTCATTGAACGGTATAAATTAG   20302
gb:MT049951|Organism:Severe      AAATGGAAATTGATTTCTTAGAATTAGCTATGGATGAATTCATTGAACGGTATAAATTAG   20302
gb:MN975262|Organism:Severe      AAATGGAAATTGATTTCTTAGAATTAGCTATGGATGAATTCATTGAACGGTATAAATTAG   20302
gb:MT106052|Organism:Severe      AAATGGAAATTGATTTCTTAGAATTAGCTATGGATGAATTCATTGAACGGTATAAATTAG   20302
gb:LC522975|Organism:Severe      AAATGGAAATTGATTTCTTAGAATTAGCTATGGATGAATTCATTGAACGGTATAAATTAG   20299
gb:LC522973|Organism:Severe      AAATGGAAATTGATTTCTTAGAATTAGCTATGGATGAATTCATTGAACGGTATAAATTAG   20299
gb:LC522974|Organism:Severe      AAATGGAAATTGATTTCTTAGAATTAGCTATGGATGAATTCATTGAACGGTATAAATTAG   20299
gb:MN985325|Organism:Severe      AAATGGAAATTGATTTCTTAGAATTAGCTATGGATGAATTCATTGAACGGTATAAATTAG   20302
gb:MT020881|Organism:Severe      AAATGGAAATTGATTTCTTAGAATTAGCTATGGATGAATTCATTGAACGGTATAAATTAG   20302
gb:MT020880|Organism:Severe      AAATGGAAATTGATTTCTTAGAATTAGCTATGGATGAATTCATTGAACGGTATAAATTAG   20302
gb:MT066175|Organism:Severe      AAATGGAAATTGATTTCTTAGAATTAGCTATGGATGAATTCATTGAACGGTATAAATTAG   20302
gb:MN997409|Organism:Severe      AAATGGAAATTGATTTCTTAGAATTAGCTATGGATGAATTCATTGAACGGTATAAATTAG   20302
gb:MN938384|Organism:Severe      AAATGGAAATTGATTTCTTAGAATTAGCTATGGATGAATTCATTGAACGGTATAAATTAG   20270
gb:MT044258|Organism:Severe      AAATGGAAATTGATTTCTTAGAATTAGCTATGGATGAATTCATTGAACGGTATAAATTAG   20278
gb:MT039890|Organism:Severe      AAATGGAAATTGATTTCTTAGAATTAGCTATGGATGAATTCATTGAACGGTATAAATTAG   20302
gb:MN988713|Organism:Severe      AAATGGAAATTGATTTCTTAGAATTAGCTATGGATGAATTCATTGAACGGTATAAATTAG   20302
gb:LC521925|Organism:Severe      AAATGGAAATTGATTTCTTAGAATTAGCTATGGATGAATTCATTGAACGGTATAAATTAG   20275
gb:MT093571|Organism:Severe      AAATGGAAATTGATTTCTTAGAATTAGCTATGGATGAATTCATTGAACGGTATAAATTAG   20302
gb:MT039887|Organism:Severe      AAATGGAAATTGATTTCTTAGAATTAGCTATGGATGAATTCATTGAACGGTATAA---AG   20299
gb:MT019530|Organism:Severe      AAATGGAAATTGATTTCTTAGAATTAGCTATGGATGAATTCATTGAACGGTATAAATTAG   20302
gb:MT039888|Organism:Severe      AAATGGAAATTGATTTCTTAGAATTAGCTATGGATGAATTCATTGAACGGTATAAATTAG   20302
gb:LC522972|Organism:Severe      AAATGGAAATTGATTTCTTAGAATTAGCTATGGATGAATTCATTGAACGGTATAAATTAG   20299
gb:MT027063|Organism:Severe      AAATGGAAATTGATTTCTTAGAATTAGCTATGGATGAATTCATTGAACGGTATAAATTAG   20302
gb:MT027062|Organism:Severe      AAATGGAAATTGATTTCTTAGAATTAGCTATGGATGAATTCATTGAACGGTATAAATTAG   20302
gb:MT019529|Organism:Severe      AAATGGAAATTGATTTCTTAGAATTAGCTATGGATGAATTCATTGAACGGTATAAATTAG   20302
gb:MN996529|Organism:Severe      AAATGGAAATTGATTTCTTAGAATTAGCTATGGATGAATTCATTGAACGGTATAAATTAG   20290
gb:MN996531|Organism:Severe      AAATGGAAATTGATTTCTTAGAATTAGCTATGGATGAATTCATTGAACGGTATAAATTAG   20289
gb:MT066176|Organism:Severe      AAATGGAAATTGATTTCTTAGAATTAGCTATGGATGAATTCATTGAACGGTATAAATTAG   20302
gb:MT027064|Organism:Severe      AAATGGAAATTGATTTCTTAGAATTAGCTATGGATGAATTCATTGAACGGTATAAATTAG   20302
gb:MN994468|Organism:Severe      AAATGGAAATTGATTTCTTAGAATTAGCTATGGATGAATTCATTGAACGGTATAAATTAG   20302
gb:MT072688|Organism:Severe      AAATGGAAATTGATTTCTTAGAATTAGCTATGGATGAATTCATTGAACGGTATAAATTAG   20287
gb:MN996527|Organism:Severe      AAATGGAAATTGATTTCTTAGAATTAGCTATGGATGAATTCATTGAACGGTATAAATTAG   20269
gb:MT093631|Organism:Severe      AAATGGAAATTGATTTCTTAGAATTAGCTATGGATGAATTCATTGAACGGTATAAATTAG   20340
gb:MT106053|Organism:Severe      AAATGGAAATTGATTTCTTAGAATTAGCTATGGATGAATTCATTGAACGGTATAAATTAG   20302
gb:MT019533|Organism:Severe      AAATGGAAATTGATTTCTTAGAATTAGCTATGGATGAATTCATTGAACGGTATAAATTAG   20302
gb:MT019531|Organism:Severe      AAATGGAAATTGATTTCTTAGAATTAGCTATGGATGAATTCATTGAACGGTATAAATTAG   20302
gb:MN996528|Organism:Severe      AAATGGAAATTGATTTCTTAGAATTAGCTATGGATGAATTCATTGAACGGTATAAATTAG   20302
gb:MN996530|Organism:Severe      AAATGGAAATTGATTTCTTAGAATTAGCTATGGATGAATTCATTGAACGGTATAAATTAG   20288
gb:MN908947|Organism:Severe      AAATGGAAATTGATTTCTTAGAATTAGCTATGGATGAATTCATTGAACGGTATAAATTAG   20302
gb:MT019532|Organism:Severe      AAATGGAAATTGATTTCTTAGAATTAGCTATGGATGAATTCATTGAACGGTATAAATTAG   20302
                                 *******************************************************   **

gb:MT020781|Organism:Severe      AAGGCTATGCCTTCGAACATATCGTTTATGGAGATTTTAGTCATAGTCAGTTAGGTGGTT   20350
gb:MT007544|Organism:Severe      AAGGCTATGCCTTCGAACATATCGTTTATGGAGATTTTAGTCATAGTCAGTTAGGTGGTT   20362
```

```
gb:MN994467|Organism:Severe      AAGGCTATGCCTTCGAACATATCGTTTATGGAGATTTTAGTCATAGTCAGTTAGGTGGTT    20362
gb:MT044257|Organism:Severe      AAGGCTATGCCTTCGAACATATCGTTTATGGAGATTTTAGTCATAGTCAGTTAGGTGGTT    20362
gb:MT106054|Organism:Severe      AAGGCTATGCCTTCGAACATATCGTTTATGGAGATTTTAGTCATAGTCAGTTAGGTGGTT    20362
gb:MT049951|Organism:Severe      AAGGCTATGCCTTCGAACATATCGTTTATGGAGATTTTAGTCATAGTCAGTTAGGTGGTT    20362
gb:MN975262|Organism:Severe      AAGGCTATGCCTTCGAACATATCGTTTATGGAGATTTTAGTCATAGTCAGTTAGGTGGTT    20362
gb:MT106052|Organism:Severe      AAGGCTATGCCTTCGAACATATCGTTTATGGAGATTTTAGTCATAGTCAGTTAGGTGGTT    20362
gb:LC522975|Organism:Severe      AAGGCTATGCCTTCGAACATATCGTTTATGGAGATTTTAGTCATAGTCAGTTAGGTGGTT    20359
gb:LC522973|Organism:Severe      AAGGCTATGCCTTCGAACATATCGTTTATGGAGATTTTAGTCATAGTCAGTTAGGTGGTT    20359
gb:LC522974|Organism:Severe      AAGGCTATGCCTTCGAACATATCGTTTATGGAGATTTTAGTCATAGTCAGTTAGGTGGTT    20359
gb:MN985325|Organism:Severe      AAGGCTATGCCTTCGAACATATCGTTTATGGAGATTTTAGTCATAGTCAGTTAGGTGGTT    20362
gb:MT020881|Organism:Severe      AAGGCTATGCCTTCGAACATATCGTTTATGGAGATTTTAGTCATAGTCAGTTAGGTGGTT    20362
gb:MT020880|Organism:Severe      AAGGCTATGCCTTCGAACATATCGTTTATGGAGATTTTAGTCATAGTCAGTTAGGTGGTT    20362
gb:MT066175|Organism:Severe      AAGGCTATGCCTTCGAACATATCGTTTATGGAGATTTTAGTCATAGTCAGTTAGGTGGTT    20362
gb:MN997409|Organism:Severe      AAGGCTATGCCTTCGAACATATCGTTTATGGAGATTTTAGTCATAGTCAGTTAGGTGGTT    20362
gb:MN938384|Organism:Severe      AAGGCTATGCCTTCGAACATATCGTTTATGGAGATTTTAGTCATAGTCAGTTAGGTGGTT    20330
gb:MT044258|Organism:Severe      AAGGCTATGCCTTCGAACATATCGTTTATGGAGATTTTAGTCATAGTCAGTTAGGTGGTT    20338
gb:MT039890|Organism:Severe      AAGGCTATGCCTTCGAACATATCGTTTATGGAGATTTTAGTCATAGTCAGTTAGGTGGTT    20362
gb:MN988713|Organism:Severe      AAGGCTATGCCTTCGAACATATCGTTTATGGAGATTTTAGTCATAGTCAGTTAGGTGGTT    20362
gb:LC521925|Organism:Severe      AAGGCTATGCCTTCGAACATATCGTTTATGGAGATTTTAGTCATAGTCAGTTAGGTGGTT    20335
gb:MT093571|Organism:Severe      AAGGCTATGCCTTCGAACATATCGTTTATGGAGATTTTAGTCATAGTCAGTTAGGTGGTT    20362
gb:MT039887|Organism:Severe      AAGGCTATGCCTTCGAACATATCGTTTATGGAGATTTTAGTCATAGTCAGTTAGGTGGTT    20359
gb:MT019530|Organism:Severe      AAGGCTATGCCTTCGAACATATCGTTTATGGAGATTTTAGTCATAGTCAGTTAGGTGGTT    20362
gb:MT039888|Organism:Severe      AAGGCTATGCCTTCGAACATATCGTTTATGGAGATTTTAGTCATAGTCAGTTAGGTGGTT    20362
gb:LC522972|Organism:Severe      AAGGCTATGCCTTCGAACATATCGTTTATGGAGATTTTAGTCATAGTCAGTTAGGTGGTT    20359
gb:MT027063|Organism:Severe      AAGGCTATGCCTTCGAACATATCGTTTATGGAGATTTTAGTCATAGTCAGTTAGGTGGTT    20362
gb:MT027062|Organism:Severe      AAGGCTATGCCTTCGAACATATCGTTTATGGAGATTTTAGTCATAGTCAGTTAGGTGGTT    20362
gb:MT019529|Organism:Severe      AAGGCTATGCCTTCGAACATATCGTTTATGGAGATTTTAGTCATAGTCAGTTAGGTGGTT    20362
gb:MN996529|Organism:Severe      AAGGCTATGCCTTCGAACATATCGTTTATGGAGATTTTAGTCATAGTCAGTTAGGTGGTT    20350
gb:MN996531|Organism:Severe      AAGGCTATGCCTTCGAACATATCGTTTATGGAGATTTTAGTCATAGTCAGTTAGGTGGTT    20349
gb:MT066176|Organism:Severe      AAGGCTATGCCTTCGAACATATCGTTTATGGAGATTTTAGTCATAGTCAGTTAGGTGGTT    20362
gb:MT027064|Organism:Severe      AAGGCTATGCCTTCGAACATATCGTTTATGGAGATTTTAGTCATAGTCAGTTAGGTGGTT    20362
gb:MN994468|Organism:Severe      AAGGCTATGCCTTCGAACATATCGTTTATGGAGATTTTAGTCATAGTCAGTTAGGTGGTT    20362
gb:MT072688|Organism:Severe      AAGGCTATGCCTTCGAACATATCGTTTATGGAGATTTTAGTCATAGTCAGTTAGGTGGTT    20347
gb:MN996527|Organism:Severe      AAGGCTATGCCTTCGAACATATCGTTTATGGAGATTTTAGTCATAGTCAGTTAGGTGGTT    20329
gb:MT093631|Organism:Severe      AAGGCTATGCCTTCGAACATATCGTTTATGGAGATTTTAGTCATAGTCAGTTAGGTGGTT    20400
gb:MT106053|Organism:Severe      AAGGCTATGCCTTCGAACATATCGTTTATGGAGATTTTAGTCATAGTCAGTTAGGTGGTT    20362
gb:MT019533|Organism:Severe      AAGGCTATGCCTTCGAACATATCGTTTATGGAGATTTTAGTCATAGTCAGTTAGGTGGTT    20362
gb:MT019531|Organism:Severe      AAGGCTATGCCTTCGAACATATCGTTTATGGAGATTTTAGTCATAGTCAGTTAGGTGGTT    20362
gb:MN996528|Organism:Severe      AAGGCTATGCCTTCGAACATATCGTTTATGGAGATTTTAGTCATAGTCAGTTAGGTGGTT    20362
gb:MN996530|Organism:Severe      AAGGCTATGCCTTCGAACATATCGTTTATGGAGATTTTAGTCATAGTCAGTTAGGTGGTT    20348
gb:MN908947|Organism:Severe      AAGGCTATGCCTTCGAACATATCGTTTATGGAGATTTTAGTCATAGTCAGTTAGGTGGTT    20362
gb:MT019532|Organism:Severe      AAGGCTATGCCTTCGAACATATCGTTTATGGAGATTTTAGTCATAGTCAGTTAGGTGGTT    20362
                                 ************************************************************

gb:MT020781|Organism:Severe      TACATCTACTGATTGGACTAGCTAAACGTTTTAAGGAATCACCTTTTGAATTAGAAGATT    20410
gb:MT007544|Organism:Severe      TACATCTACTGATTGGACTAGCTAAACGTTTTAAGGAATCACCTTTTGAATTAGAAGATT    20422
gb:MN994467|Organism:Severe      TACATCTACTGATTGGACTAGCTAAACGTTTTAAGGAATCACCTTTTGAATTAGAAGATT    20422
gb:MT044257|Organism:Severe      TACATCTACTGATTGGACTAGCTAAACGTTTTAAGGAATCACCTTTTGAATTAGAAGATT    20422
gb:MT106054|Organism:Severe      TACATCTACTGATTGGACTAGCTAAACGTTTTAAGGAATCACCTTTTGAATTAGAAGATT    20422
gb:MT049951|Organism:Severe      TACATCTACTGATTGGACTAGCTAAACGTTTTAAGGAATCACCTTTTGAATTAGAAGATT    20422
gb:MN975262|Organism:Severe      TACATCTACTGATTGGACTAGCTAAACGTTTTAAGGAATCACCTTTTGAATTAGAAGATT    20422
gb:MT106052|Organism:Severe      TACATCTACTGATTGGACTAGCTAAACGTTTTAAGGAATCACCTTTTGAATTAGAAGATT    20422
gb:LC522975|Organism:Severe      TACATCTACTGATTGGACTAGCTAAACGTTTTAAGGAATCACCTTTTGAATTAGAAGATT    20419
gb:LC522973|Organism:Severe      TACATCTACTGATTGGACTAGCTAAACGTTTTAAGGAATCACCTTTTGAATTAGAAGATT    20419
gb:LC522974|Organism:Severe      TACATCTACTGATTGGACTAGCTAAACGTTTTAAGGAATCACCTTTTGAATTAGAAGATT    20419
gb:MN985325|Organism:Severe      TACATCTACTGATTGGACTAGCTAAACGTTTTAAGGAATCACCTTTTGAATTAGAAGATT    20422
gb:MT020881|Organism:Severe      TACATCTACTGATTGGACTAGCTAAACGTTTTAAGGAATCACCTTTTGAATTAGAAGATT    20422
gb:MT020880|Organism:Severe      TACATCTACTGATTGGACTAGCTAAACGTTTTAAGGAATCACCTTTTGAATTAGAAGATT    20422
gb:MT066175|Organism:Severe      TACATCTACTGATTGGACTAGCTAAACGTTTTAAGGAATCACCTTTTGAATTAGAAGATT    20422
gb:MN997409|Organism:Severe      TACATCTACTGATTGGACTAGCTAAACGTTTTAAGGAATCACCTTTTGAATTAGAAGATT    20422
gb:MN938384|Organism:Severe      TACATCTACTGATTGGACTAGCTAAACGTTTTAAGGAATCACCTTTTGAATTAGAAGATT    20390
gb:MT044258|Organism:Severe      TACATCTACTGATTGGACTAGCTAAACGTTTTAAGGAATCACCTTTTGAATTAGAAGATT    20398
gb:MT039890|Organism:Severe      TACATCTACTGATTGGACTAGCTAAACGTTTTAAGGAATCACCTTTTGAATTAGAAGATT    20422
gb:MN988713|Organism:Severe      TACATCTACTGATTGGACTAGCTAAACGTTTTAAGGAATCACCTTTTGAATTAGAAGATT    20422
gb:LC521925|Organism:Severe      TACATCTACTGATTGGACTAGCTAAACGTTTTAAGGAATCACCTTTTGAATTAGAAGATT    20395
```

```
gb:MT093571|Organism:Severe      TACATCTACTGATTGGACTAGCTAAACGTTTTAAGGAATCACCTTTTGAATTAGAAGATT    20422
gb:MT039887|Organism:Severe      TACATCTACTGATTGGACTAGCTAAACGTTTTAAGGAATCACCTTTTGAATTAGAAGATT    20419
gb:MT019530|Organism:Severe      TACATCTACTGATTGGACTAGCTAAACGTTTTAAGGAATCACCTTTTGAATTAGAAGATT    20422
gb:MT039888|Organism:Severe      TACATCTACTGATTGGACTAGCTAAACGTTTTAAGGAATCACCTTTTGAATTAGAAGATT    20422
gb:LC522972|Organism:Severe      TACATCTACTGATTGGACTAGCTAAACGTTTTAAGGAATCACCTTTTGAATTAGAAGATT    20419
gb:MT027063|Organism:Severe      TACATCTACTGATTGGACTAGCTAAACGTTTTAAGGAATCACCTTTTGAATTAGAAGATT    20422
gb:MT027062|Organism:Severe      TACATCTACTGATTGGACTAGCTAAACGTTTTAAGGAATCACCTTTTGAATTAGAAGATT    20422
gb:MT019529|Organism:Severe      TACATCTACTGATTGGACTAGCTAAACGTTTTAAGGAATCACCTTTTGAATTAGAAGATT    20422
gb:MN996529|Organism:Severe      TACATCTACTGATTGGACTAGCTAAACGTTTTAAGGAATCACCTTTTGAATTAGAAGATT    20410
gb:MN996531|Organism:Severe      TACATCTACTGATTGGACTAGCTAAACGTTTTAAGGAATCACCTTTTGAATTAGAAGATT    20409
gb:MT066176|Organism:Severe      TACATCTACTGATTGGACTAGCTAAACGTTTTAAGGAATCACCTTTTGAATTAGAAGATT    20422
gb:MT027064|Organism:Severe      TACATCTACTGATTGGACTAGCTAAACGTTTTAAGGAATCACCTTTTGAATTAGAAGATT    20422
gb:MN994468|Organism:Severe      TACATCTACTGATTGGACTAGCTAAACGTTTTAAGGAATCACCTTTTGAATTAGAAGATT    20422
gb:MT072688|Organism:Severe      TACATCTACTGATTGGACTAGCTAAACGTTTTAAGGAATCACCTTTTGAATTAGAAGATT    20407
gb:MN996527|Organism:Severe      TACATCTACTGATTGGACTAGCTAAACGTTTTAAGGAATCACCTTTTGAATTAGAAGATT    20389
gb:MT093631|Organism:Severe      TACATCTACTGATTGGACTAGCTAAACGTTTTAAGGAATCACCTTTTGAATTAGAAGATT    20460
gb:MT106053|Organism:Severe      TACATCTACTGATTGGACTAGCTAAACGTTTTAAGGAATCACCTTTTGAATTAGAAGATT    20422
gb:MT019533|Organism:Severe      TACATCTACTGATTGGACTAGCTAAACGTTTTAAGGAATCACCTTTTGAATTAGAAGATT    20422
gb:MT019531|Organism:Severe      TACATCTACTGATTGGACTAGCTAAACGTTTTAAGGAATCACCTTTTGAATTAGAAGATT    20422
gb:MN996528|Organism:Severe      TACATCTACTGATTGGACTAGCTAAACGTTTTAAGGAATCACCTTTTGAATTAGAAGATT    20422
gb:MN996530|Organism:Severe      TACATCTACTGATTGGACTAGCTAAACGTTTTAAGGAATCACCTTTTGAATTAGAAGATT    20408
gb:MN908947|Organism:Severe      TACATCTACTGATTGGACTAGCTAAACGTTTTAAGGAATCACCTTTTGAATTAGAAGATT    20422
gb:MT019532|Organism:Severe      TACATCTACTGATTGGACTAGCTAAACGTTTTAAGGAATCACCTTTTGAATTAGAAGATT    20422
                                 ************************************************************

gb:MT020781|Organism:Severe      TTATTCCTATGGACAGTACAGTTAAAAACTATTTCATAACAGATGCGCAAACAGGTTCAT    20470
gb:MT007544|Organism:Severe      TTATTCCTATGGACAGTACAGTTAAAAACTATTTCATAACAGATGCGCAAACAGGTTCAT    20482
gb:MN994467|Organism:Severe      TTATTCCTATGGACAGTACAGTTAAAAACTATTTCATAACAGATGCGCAAACAGGTTCAT    20482
gb:MT044257|Organism:Severe      TTATTCCTATGGACAGTACAGTTAAAAACTATTTCATAACAGATGCGCAAACAGGTTCAT    20482
gb:MT106054|Organism:Severe      TTATTCCTATGGACAGTACAGTTAAAAACTATTTCATAACAGATGCGCAAACAGGTTCAT    20482
gb:MT049951|Organism:Severe      TTATTCCTATGGACAGTACAGTTAAAAACTATTTCATAACAGATGCGCAAACAGGTTCAT    20482
gb:MN975262|Organism:Severe      TTATTCCTATGGACAGTACAGTTAAAAACTATTTCATAACAGATGCGCAAACAGGTTCAT    20482
gb:MT106052|Organism:Severe      TTATTCCTATGGACAGTACAGTTAAAAACTATTTCATAACAGATGCGCAAACAGGTTCAT    20482
gb:LC522975|Organism:Severe      TTATTCCTATGGACAGTACAGTTAAAAACTATTTCATAACAGATGCGCAAACAGGTTCAT    20479
gb:LC522973|Organism:Severe      TTATTCCTATGGACAGTACAGTTAAAAACTATTTCATAACAGATGCGCAAACAGGTTCAT    20479
gb:LC522974|Organism:Severe      TTATTCCTATGGACAGTACAGTTAAAAACTATTTCATAACAGATGCGCAAACAGGTTCAT    20479
gb:MN985325|Organism:Severe      TTATTCCTATGGACAGTACAGTTAAAAACTATTTCATAACAGATGCGCAAACAGGTTCAT    20482
gb:MT020881|Organism:Severe      TTATTCCTATGGACAGTACAGTTAAAAACTATTTCATAACAGATGCGCAAACAGGTTCAT    20482
gb:MT020880|Organism:Severe      TTATTCCTATGGACAGTACAGTTAAAAACTATTTCATAACAGATGCGCAAACAGGTTCAT    20482
gb:MT066175|Organism:Severe      TTATTCCTATGGACAGTACAGTTAAAAACTATTTCATAACAGATGCGCAAACAGGTTCAT    20482
gb:MN997409|Organism:Severe      TTATTCCTATGGACAGTACAGTTAAAAACTATTTCATAACAGATGCGCAAACAGGTTCAT    20482
gb:MN938384|Organism:Severe      TTATTCCTATGGACAGTACAGTTAAAAACTATTTCATAACAGATGCGCAAACAGGTTCAT    20450
gb:MT044258|Organism:Severe      TTATTCCTATGGACAGTACAGTTAAAAACTATTTCATAACAGATGCGCAAACAGGTTCAT    20458
gb:MT039890|Organism:Severe      TTATTCCTATGGACAGTACAGTTAAAAACTATTTCATAACAGATGCGCAAACAGGTTCAT    20482
gb:MN988713|Organism:Severe      TTATTCCTATGGACAGTACAGTTAAAAACTATTTCATAACAGATGCGCAAACAGGTTCAT    20482
gb:LC521925|Organism:Severe      TTATTCCTATGGACAGTACAGTTAAAAACTATTTCATAACAGATGCGCAAACAGGTTCAT    20455
gb:MT093571|Organism:Severe      TTATTCCTATGGACAGTACAGTTAAAAACTATTTCATAACAGATGCGCAAACAGGTTCAT    20482
gb:MT039887|Organism:Severe      TTATTCCTATGGACAGTACAGTTAAAAACTATTTCATAACAGATGCGCAAACAGGTTCAT    20479
gb:MT019530|Organism:Severe      TTATTCCTATGGACAGTACAGTTAAAAACTATTTCATAACAGATGCGCAAACAGGTTCAT    20482
gb:MT039888|Organism:Severe      TTATTCCTATGGACAGTACAGTTAAAAACTATTTCATAACAGATGCGCAAACAGGTTCAT    20482
gb:LC522972|Organism:Severe      TTATTCCTATGGACAGTACAGTTAAAAACTATTTCATAACAGATGCGCAAACAGGTTCAT    20479
gb:MT027063|Organism:Severe      TTATTCCTATGGACAGTACAGTTAAAAACTATTTCATAACAGATGCGCAAACAGGTTCAT    20482
gb:MT027062|Organism:Severe      TTATTCCTATGGACAGTACAGTTAAAAACTATTTCATAACAGATGCGCAAACAGGTTCAT    20482
gb:MT019529|Organism:Severe      TTATTCCTATGGACAGTACAGTTAAAAACTATTTCATAACAGATGCGCAAACAGGTTCAT    20482
gb:MN996529|Organism:Severe      TTATTCCTATGGACAGTACAGTTAAAAACTATTTCATAACAGATGCGCAAACAGGTTCAT    20470
gb:MN996531|Organism:Severe      TTATTCCTATGGACAGTACAGTTAAAAACTATTTCATAACAGATGCGCAAACAGGTTCAT    20469
gb:MT066176|Organism:Severe      TTATTCCTATGGACAGTACAGTTAAAAACTATTTCATAACAGATGCGCAAACAGGTTCAT    20482
gb:MT027064|Organism:Severe      TTATTCCTATGGACAGTACAGTTAAAAACTATTTCATAACAGATGCGCAAACAGGTTCAT    20482
gb:MN994468|Organism:Severe      TTATTCCTATGGACAGTACAGTTAAAAACTATTTCATAACAGATGCGCAAACAGGTTCAT    20482
gb:MT072688|Organism:Severe      TTATTCCTATGGACAGTACAGTTAAAAACTATTTCATAACAGATGCGCAAACAGGTTCAT    20467
gb:MN996527|Organism:Severe      TTATTCCTATGGACAGTACAGTTAAAAACTATTTCATAACAGATGCGCAAACAGGTTCAT    20449
gb:MT093631|Organism:Severe      TTATTCCTATGGACAGTACAGTTAAAAACTATTTCATAACAGATGCGCAAACAGGTTCAT    20520
gb:MT106053|Organism:Severe      TTATTCCTATGGACAGTACAGTTAAAAACTATTTCATAACAGATGCGCAAACAGGTTCAT    20482
gb:MT019533|Organism:Severe      TTATTCCTATGGACAGTACAGTTAAAAACTATTTCATAACAGATGCGCAAACAGGTTCAT    20482
gb:MT019531|Organism:Severe      TTATTCCTATGGACAGTACAGTTAAAAACTATTTCATAACAGATGCGCAAACAGGTTCAT    20482
```

```
gb:MN996528|Organism:Severe      TTATTCCTATGGACAGTACAGTTAAAAACTATTTCATAACAGATGCGCAAACAGGTTCAT    20482
gb:MN996530|Organism:Severe      TTATTCCTATGGACAGTACAGTTAAAAACTATTTCATAACAGATGCGCAAACAGGTTCAT    20468
gb:MN908947|Organism:Severe      TTATTCCTATGGACAGTACAGTTAAAAACTATTTCATAACAGATGCGCAAACAGGTTCAT    20482
gb:MT019532|Organism:Severe      TTATTCCTATGGACAGTACAGTTAAAAACTATTTCATAACAGATGCGCAAACAGGTTCAT    20482
                                 ************************************************************

gb:MT020781|Organism:Severe      CTAAGTGTGTGTGTTCTGTTATTGATTTATTACTTGATGATTTTGTTGAAATAATAAAAT    20530
gb:MT007544|Organism:Severe      CTAAGTGTGTGTGTTCTGTTATTGATTTATTACTTGATGATTTTGTTGAAATAATAAAAT    20542
gb:MN994467|Organism:Severe      CTAAGTGTGTGTGTTCTGTTATTGATTTATTACTTGATGATTTTGTTGAAATAATAAAAT    20542
gb:MT044257|Organism:Severe      CTAAGTGTGTGTGTTCTGTTATTGATTTATTACTTGATGATTTTGTTGAAATAATAAAAT    20542
gb:MT106054|Organism:Severe      CTAAGTGTGTGTGTTCTGTTATTGATTTATTACTTGATGATTTTGTTGAAATAATAAAAT    20542
gb:MT049951|Organism:Severe      CTAAGTGTGTGTGTTCTGTTATTGATTTATTACTTGATGATTTTGTTGAAATAATAAAAT    20542
gb:MN975262|Organism:Severe      CTAAGTGTGTGTGTTCTGTTATTGATTTATTACTTGATGATTTTGTTGAAATAATAAAAT    20542
gb:MT106052|Organism:Severe      CTAAGTGTGTGTGTTCTGTTATTGATTTATTACTTGATGATTTTGTTGAAATAATAAAAT    20542
gb:LC522975|Organism:Severe      CTAAGTGTGTGTGTTCTGTTATTGATTTATTACTTGATGATTTTGTTGAAATAATAAAAT    20539
gb:LC522973|Organism:Severe      CTAAGTGTGTGTGTTCTGTTATTGATTTATTACTTGATGATTTTGTTGAAATAATAAAAT    20539
gb:LC522974|Organism:Severe      CTAAGTGTGTGTGTTCTGTTATTGATTTATTACTTGATGATTTTGTTGAAATAATAAAAT    20539
gb:MN985325|Organism:Severe      CTAAGTGTGTGTGTTCTGTTATTGATTTATTACTTGATGATTTTGTTGAAATAATAAAAT    20542
gb:MT020881|Organism:Severe      CTAAGTGTGTGTGTTCTGTTATTGATTTATTACTTGATGATTTTGTTGAAATAATAAAAT    20542
gb:MT020880|Organism:Severe      CTAAGTGTGTGTGTTCTGTTATTGATTTATTACTTGATGATTTTGTTGAAATAATAAAAT    20542
gb:MT066175|Organism:Severe      CTAAGTGTGTGTGTTCTGTTATTGATTTATTACTTGATGATTTTGTTGAAATAATAAAAT    20542
gb:MN997409|Organism:Severe      CTAAGTGTGTGTGTTCTGTTATTGATTTATTACTTGATGATTTTGTTGAAATAATAAAAT    20542
gb:MN938384|Organism:Severe      CTAAGTGTGTGTGTTCTGTTATTGATTTATTACTTGATGATTTTGTTGAAATAATAAAAT    20510
gb:MT044258|Organism:Severe      CTAAGTGTGTGTGTTCTGTTATTGATTTATTACTTGATGATTTTGTTGAAATAATAAAAT    20518
gb:MT039890|Organism:Severe      CTAAGTGTGTGTGTTCTGTTATTGATTTATTACTTGATGATTTTGTTGAAATAATAAAAT    20542
gb:MN988713|Organism:Severe      CTAAGTGTGTGTGTTCTGTTATTGATTTATTACTTGATGATTTTGTTGAAATAATAAAAT    20542
gb:LC521925|Organism:Severe      CTAAGTGTGTGTGTTCTGTTATTGATTTATTACTTGATGATTTTGTTGAAATAATAAAAT    20515
gb:MT093571|Organism:Severe      CTAAGTGTGTGTGTTCTGTTATTGATTTATTACTTGATGATTTTGTTGAAATAATAAAAT    20542
gb:MT039887|Organism:Severe      CTAAGTGTGTGTGTTCTGTTATTGATTTATTACTTGATGATTTTGTTGAAATAATAAAAT    20539
gb:MT019530|Organism:Severe      CTAAGTGTGTGTGTTCTGTTATTGATTTATTACTTGATGATTTTGTTGAAATAATAAAAT    20542
gb:MT039888|Organism:Severe      CTAAGTGTGTGTGTTCTGTTATTGATTTATTACTTGATGATTTTGTTGAAATAATAAAAT    20542
gb:LC522972|Organism:Severe      CTAAGTGTGTGTGTTCTGTTATTGATTTATTACTTGATGATTTTGTTGAAATAATAAAAT    20539
gb:MT027063|Organism:Severe      CTAAGTGTGTGTGTTCTGTTATTGATTTATTACTTGATGATTTTGTTGAAATAATAAAAT    20542
gb:MT027062|Organism:Severe      CTAAGTGTGTGTGTTCTGTTATTGATTTATTACTTGATGATTTTGTTGAAATAATAAAAT    20542
gb:MT019529|Organism:Severe      CTAAGTGTGTGTGTTCTGTTATTGATTTATTACTTGATGATTTTGTTGAAATAATAAAAT    20542
gb:MN996529|Organism:Severe      CTAAGTGTGTGTGTTCTGTTATTGATTTATTACTTGATGATTTTGTTGAAATAATAAAAT    20530
gb:MN996531|Organism:Severe      CTAAGTGTGTGTGTTCTGTTATTGATTTATTACTTGATGATTTTGTTGAAATAATAAAAT    20529
gb:MT066176|Organism:Severe      CTAAGTGTGTGTGTTCTGTTATTGATTTATTACTTGATGATTTTGTTGAAATAATAAAAT    20542
gb:MT027064|Organism:Severe      CTAAGTGTGTGTGTTCTGTTATTGATTTATTACTTGATGATTTTGTTGAAATAATAAAAT    20542
gb:MN994468|Organism:Severe      CTAAGTGTGTGTGTTCTGTTATTGATTTATTACTTGATGATTTTGTTGAAATAATAAAAT    20542
gb:MT072688|Organism:Severe      CTAAGTGTGTGTGTTCTGTTATTGATTTATTACTTGATGATTTTGTTGAAATAATAAAAT    20527
gb:MN996527|Organism:Severe      CTAAGTGTGTGTGTTCTGTTATTGATTTATTACTTGATGATTTTGTTGAAATAATAAAAT    20509
gb:MT093631|Organism:Severe      CTAAGTGTGTGTGTTCTGTTATTGATTTATTACTTGATGATTTTGTTGAAATAATAAAAT    20580
gb:MT106053|Organism:Severe      CTAAGTGTGTGTGTTCTGTTATTGATTTATTACTTGATGATTTTGTTGAAATAATAAAAT    20542
gb:MT019533|Organism:Severe      CTAAGTGTGTGTGTTCTGTTATTGATTTATTACTTGATGATTTTGTTGAAATAATAAAAT    20542
gb:MT019531|Organism:Severe      CTAAGTGTGTGTGTTCTGTTATTGATTTATTACTTGATGATTTTGTTGAAATAATAAAAT    20542
gb:MN996528|Organism:Severe      CTAAGTGTGTGTGTTCTGTTATTGATTTATTACTTGATGATTTTGTTGAAATAATAAAAT    20542
gb:MN996530|Organism:Severe      CTAAGTGTGTGTGTTCTGTTATTGATTTATTACTTGATGATTTTGTTGAAATAATAAAAT    20528
gb:MN908947|Organism:Severe      CTAAGTGTGTGTGTTCTGTTATTGATTTATTACTTGATGATTTTGTTGAAATAATAAAAT    20542
gb:MT019532|Organism:Severe      CTAAGTGTGTGTGTTCTGTTATTGATTTATTACTTGATGATTTTGTTGAAATAATAAAAT    20542
                                 ************************************************************

gb:MT020781|Organism:Severe      CCCAAGATTTATCTGTAGTTTCTAAGGTTGTCAAAGTGACTATTGACTATACAGAAATTT    20590
gb:MT007544|Organism:Severe      CCCAAGATTTATCTGTAGTTTCTAAGGTTGTCAAAGTGACTATTGACTATACAGAAATTT    20602
gb:MN994467|Organism:Severe      CCCAAGATTTATCTGTAGTTTCTAAGGTTGTCAAAGTGACTATTGACTATACAGAAATTT    20602
gb:MT044257|Organism:Severe      CCCAAGATTTATCTGTAGTTTCTAAGGTTGTCAAAGTGACTATTGACTATACAGAAATTT    20602
gb:MT106054|Organism:Severe      CCCAAGATTTATCTGTAGTTTCTAAGGTTGTCAAAGTGACTATTGACTATACAGAAATTT    20602
gb:MT049951|Organism:Severe      CCCAAGATTTATCTGTAGTTTCTAAGGTTGTCAAAGTGACTATTGACTATACAGAAATTT    20602
gb:MN975262|Organism:Severe      CCCAAGATTTATCTGTAGTTTCTAAGGTTGTCAAAGTGACTATTGACTATACAGAAATTT    20602
gb:MT106052|Organism:Severe      CCCAAGATTTATCTGTAGTTTCTAAGGTTGTCAAAGTGACTATTGACTATACAGAAATTT    20602
gb:LC522975|Organism:Severe      CCCAAGATTTATCTGTAGTTTCTAAGGTTGTCAAAGTGACTATTGACTATACAGAAATTT    20599
gb:LC522973|Organism:Severe      CCCAAGATTTATCTGTAGTTTCTAAGGTTGTCAAAGTGACTATTGACTATACAGAAATTT    20599
gb:LC522974|Organism:Severe      CCCAAGATTTATCTGTAGTTTCTAAGGTTGTCAAAGTGACTATTGACTATACAGAAATTT    20599
gb:MN985325|Organism:Severe      CCCAAGATTTATCTGTAGTTTCTAAGGTTGTCAAAGTGACTATTGACTATACAGAAATTT    20602
gb:MT020881|Organism:Severe      CCCAAGATTTATCTGTAGTTTCTAAGGTTGTCAAAGTGACTATTGACTATACAGAAATTT    20602
```

```
gb:MT020880|Organism:Severe      CCCAAGATTTATCTGTAGTTTCTAAGGTTGTCAAAGTGACTATTGACTATACAGAAATTT   20602
gb:MT066175|Organism:Severe      CCCAAGATTTATCTGTAGTTTCTAAGGTTGTCAAAGTGACTATTGACTATACAGAAATTT   20602
gb:MN997409|Organism:Severe      CCCAAGATTTATCTGTAGTTTCTAAGGTTGTCAAAGTGACTATTGACTATACAGAAATTT   20602
gb:MN938384|Organism:Severe      CCCAAGATTTATCTGTAGTTTCTAAGGTTGTCAAAGTGACTATTGACTATACAGAAATTT   20570
gb:MT044258|Organism:Severe      CCCAAGATTTATCTGTAGTTTCTAAGGTTGTCAAAGTGACTATTGACTATACAGAAATTT   20578
gb:MT039890|Organism:Severe      CCCAAGATTTATCTGTAGTTTCTAAGGTTGTCAAAGTGACTATTGACTATACAGAAATTT   20602
gb:MN988713|Organism:Severe      CCCAAGATTTATCTGTAGTTTCTAAGGTTGTCAAAGTGACTATTGACTATACAGAAATTT   20602
gb:LC521925|Organism:Severe      CCCAAGATTTATCTGTAGTTTCTAAGGTTGTCAAAGTGACTATTGACTATACAGAAATTT   20575
gb:MT093571|Organism:Severe      CCCAAGATTTATCTGTAGTTTCTAAGGTTGTCAAAGTGACTATTGACTATACAGAAATTT   20602
gb:MT039887|Organism:Severe      CCCAAGATTTATCTGTAGTTTCTAAGGTTGTCAAAGTGACTATTGACTATACAGAAATTT   20599
gb:MT019530|Organism:Severe      CCCAAGATTTATCTGTAGTTTCTAAGGTTGTCAAAGTGACTATTGACTATACAGAAATTT   20602
gb:MT039888|Organism:Severe      CCCAAGATTTATCTGTAGTTTCTAAGGTTGTCAAAGTGACTATTGACTATACAGAAATTT   20602
gb:LC522972|Organism:Severe      CCCAAGATTTATCTGTAGTTTCTAAGGTTGTCAAAGTGACTATTGACTATACAGAAATTT   20599
gb:MT027063|Organism:Severe      CCCAAGATTTATCTGTAGTTTCTAAGGTTGTCAAAGTGACTATTGACTATACAGAAATTT   20602
gb:MT027062|Organism:Severe      CCCAAGATTTATCTGTAGTTTCTAAGGTTGTCAAAGTGACTATTGACTATACAGAAATTT   20602
gb:MT019529|Organism:Severe      CCCAAGATTTATCTGTAGTTTCTAAGGTTGTCAAAGTGACTATTGACTATACAGAAATTT   20602
gb:MN996529|Organism:Severe      CCCAAGATTTATCTGTAGTTTCTAAGGTTGTCAAAGTGACTATTGACTATACAGAAATTT   20590
gb:MN996531|Organism:Severe      CCCAAGATTTATCTGTAGTTTCTAAGGTTGTCAAAGTGACTATTGACTATACAGAAATTT   20589
gb:MT066176|Organism:Severe      CCCAAGATTTATCTGTAGTTTCTAAGGTTGTCAAAGTGACTATTGACTATACAGAAATTT   20602
gb:MT027064|Organism:Severe      CCCAAGATTTATCTGTAGTTTCTAAGGTTGTCAAAGTGACTATTGACTATACAGAAATTT   20602
gb:MN994468|Organism:Severe      CCCAAGATTTATCTGTAGTTTCTAAGGTTGTCAAAGTGACTATTGACTATACAGAAATTT   20602
gb:MT072688|Organism:Severe      CCCAAGATTTATCTGTAGTTTCTAAGGTTGTCAAAGTGACTATTGACTATACAGAAATTT   20587
gb:MN996527|Organism:Severe      CCCAAGATTTATCTGTAGTTTCTAAGGTTGTCAAAGTGACTATTGACTATACAGAAATTT   20569
gb:MT093631|Organism:Severe      CCCAAGATTTATCTGTAGTTTCTAAGGTTGTCAAAGTGACTATTGACTATACAGAAATTT   20640
gb:MT106053|Organism:Severe      CCCAAGATTTATCTGTAGTTTCTAAGGTTGTCAAAGTGACTATTGACTATACAGAAATTT   20602
gb:MT019533|Organism:Severe      CCCAAGATTTATCTGTAGTTTCTAAGGTTGTCAAAGTGACTATTGACTATACAGAAATTT   20602
gb:MT019531|Organism:Severe      CCCAAGATTTATCTGTAGTTTCTAAGGTTGTCAAAGTGACTATTGACTATACAGAAATTT   20602
gb:MN996528|Organism:Severe      CCCAAGATTTATCTGTAGTTTCTAAGGTTGTCAAAGTGACTATTGACTATACAGAAATTT   20602
gb:MN996530|Organism:Severe      CCCAAGATTTATCTGTAGTTTCTAAGGTTGTCAAAGTGACTATTGACTATACAGAAATTT   20588
gb:MN908947|Organism:Severe      CCCAAGATTTATCTGTAGTTTCTAAGGTTGTCAAAGTGACTATTGACTATACAGAAATTT   20602
gb:MT019532|Organism:Severe      CCCAAGATTTATCTGTAGTTTCTAAGGTTGTCAAAGTGACTATTGACTATACAGAAATTT   20602
                                 ************************************************************

gb:MT020781|Organism:Severe      CATTTATGCTTTGGTGTAAAGATGGCCATGTAGAAACATTTTACCCAAAATTACAATCTA   20650
gb:MT007544|Organism:Severe      CATTTATGCTTTGGTGTAAAGATGGCCATGTAGAAACATTTTACCCAAAATTACAATCTA   20662
gb:MN994467|Organism:Severe      CATTTATGCTTTGGTGTAAAGATGGCCATGTAGAAACATTTTACCCAAAATTACAATCTA   20662
gb:MT044257|Organism:Severe      CATTTATGCTTTGGTGTAAAGATGGCCATGTAGAAACATTTTACCCAAAATTACAATCTA   20662
gb:MT106054|Organism:Severe      CATTTATGCTTTGGTGTAAAGATGGCCATGTAGAAACATTTTACCCAAAATTACAATCTA   20662
gb:MT049951|Organism:Severe      CATTTATGCTTTGGTGTAAAGATGGCCATGTAGAAACATTTTACCCAAAATTACAATCTA   20662
gb:MN975262|Organism:Severe      CATTTATGCTTTGGTGTAAAGATGGCCATGTAGAAACATTTTACCCAAAATTACAATCTA   20662
gb:MT106052|Organism:Severe      CATTTATGCTTTGGTGTAAAGATGGCCATGTAGAAACATTTTACCCAAAATTACAATCTA   20662
gb:LC522975|Organism:Severe      CATTTATGCTTTGGTGTAAAGATGGCCATGTAGAAACATTTTACCCAAAATTACAATCTA   20659
gb:LC522973|Organism:Severe      CATTTATGCTTTGGTGTAAAGATGGCCATGTAGAAACATTTTACCCAAAATTACAATCTA   20659
gb:LC522974|Organism:Severe      CATTTATGCTTTGGTGTAAAGATGGCCATGTAGAAACATTTTACCCAAAATTACAATCTA   20659
gb:MN985325|Organism:Severe      CATTTATGCTTTGGTGTAAAGATGGCCATGTAGAAACATTTTACCCAAAATTACAATCTA   20662
gb:MT020881|Organism:Severe      CATTTATGCTTTGGTGTAAAGATGGCCATGTAGAAACATTTTACCCAAAATTACAATCTA   20662
gb:MT020880|Organism:Severe      CATTTATGCTTTGGTGTAAAGATGGCCATGTAGAAACATTTTACCCAAAATTACAATCTA   20662
gb:MT066175|Organism:Severe      CATTTATGCTTTGGTGTAAAGATGGCCATGTAGAAACATTTTACCCAAAATTACAATCTA   20662
gb:MN997409|Organism:Severe      CATTTATGCTTTGGTGTAAAGATGGCCATGTAGAAACATTTTACCCAAAATTACAATCTA   20662
gb:MN938384|Organism:Severe      CATTTATGCTTTGGTGTAAAGATGGCCATGTAGAAACATTTTACCCAAAATTACAATCTA   20630
gb:MT044258|Organism:Severe      CATTTATGCTTTGGTGTAAAGATGGCCATGTAGAAACATTTTACCCAAAATTACAATCTA   20638
gb:MT039890|Organism:Severe      CATTTATGCTTTGGTGTAAAGATGGCCATGTAGAAACATTTTACCCAAAATTACAATCTA   20662
gb:MN988713|Organism:Severe      CATTTATGCTTTGGTGTAAAGATGGCCATGTAGAAACATTTTACCCAAAATTACAATCTA   20662
gb:LC521925|Organism:Severe      CATTTATGCTTTGGTGTAAAGATGGCCATGTAGAAACATTTTACCCAAAATTACAATCTA   20635
gb:MT093571|Organism:Severe      CATTTATGCTTTGGTGTAAAGATGGCCATGTAGAAACATTTTACCCAAAATTACAATCTA   20662
gb:MT039887|Organism:Severe      CATTTATGCTTTGGTGTAAAGATGGCCATGTAGAAACATTTTACCCAAAATTACAATCTA   20659
gb:MT019530|Organism:Severe      CATTTATGCTTTGGTGTAAAGATGGCCATGTAGAAACATTTTACCCAAAATTACAATCTA   20662
gb:MT039888|Organism:Severe      CATTTATGCTTTGGTGTAAAGATGGCCATGTAGAAACATTTTACCCAAAATTACAATCTA   20662
gb:LC522972|Organism:Severe      CATTTATGCTTTGGTGTAAAGATGGCCATGTAGAAACATTTTACCCAAAATTACAATCTA   20659
gb:MT027063|Organism:Severe      CATTTATGCTTTGGTGTAAAGATGGCCATGTAGAAACATTTTACCCAAAATTACAATCTA   20662
gb:MT027062|Organism:Severe      CATTTATGCTTTGGTGTAAAGATGGCCATGTAGAAACATTTTACCCAAAATTACAATCTA   20662
gb:MT019529|Organism:Severe      CATTTATGCTTTGGTGTAAAGATGGCCATGTAGAAACATTTTACCCAAAATTACAATCTA   20662
gb:MN996529|Organism:Severe      CATTTATGCTTTGGTGTAAAGATGGCCATGTAGAAACATTTTACCCAAAATTACAATCTA   20650
gb:MN996531|Organism:Severe      CATTTATGCTTTGGTGTAAAGATGGCCATGTAGAAACATTTTACCCAAAATTACAATCTA   20649
gb:MT066176|Organism:Severe      CATTTATGCTTTGGTGTAAAGATGGCCATGTAGAAACATTTTACCCAAAATTACAATCTA   20662
```

```
gb:MT027064|Organism:Severe      CATTTATGCTTTGGTGTAAAGATGGCCATGTAGAAACATTTTACCCAAAATTACAATCTA   20662
gb:MN994468|Organism:Severe      CATTTATGCTTTGGTGTAAAGATGGCCATGTAGAAACATTTTACCCAAAATTACAATCTA   20662
gb:MT072688|Organism:Severe      CATTTATGCTTTGGTGTAAAGATGGCCATGTAGAAACATTTTACCCAAAATTACAATCTA   20647
gb:MN996527|Organism:Severe      CATTTATGCTTTGGTGTAAAGATGGCCATGTAGAAACATTTTACCCAAAATTACAATCTA   20629
gb:MT093631|Organism:Severe      CATTTATGCTTTGGTGTAAAGATGGCCATGTAGAAACATTTTACCCAAAATTACAATCTA   20700
gb:MT106053|Organism:Severe      CATTTATGCTTTGGTGTAAAGATGGCCATGTAGAAACATTTTACCCAAAATTACAATCTA   20662
gb:MT019533|Organism:Severe      CATTTATGCTTTGGTGTAAAGATGGCCATGTAGAAACATTTTACCCAAAATTACAATCTA   20662
gb:MT019531|Organism:Severe      CATTTATGCTTTGGTGTAAAGATGGCCATGTAGAAACATTTTACCCAAAATTACAATCTA   20662
gb:MN996528|Organism:Severe      CATTTATGCTTTGGTGTAAAGATGGCCATGTAGAAACATTTTACCCAAAATTACAATCTA   20662
gb:MN996530|Organism:Severe      CATTTATGCTTTGGTGTAAAGATGGCCATGTAGAAACATTTTACCCAAAATTACAATCTA   20648
gb:MN908947|Organism:Severe      CATTTATGCTTTGGTGTAAAGATGGCCATGTAGAAACATTTTACCCAAAATTACAATCTA   20662
gb:MT019532|Organism:Severe      CATTTATGCTTTGGTGTAAAGATGGCCATGTAGAAACATTTTACCCAAAATTACAATCTA   20662
                                 ************************************************************

gb:MT020781|Organism:Severe      GTCAAGCGTGGCAACCGGGTGTTGCTATGCCTAATCTTTACAAAATGCAAAGAATGCTAT   20710
gb:MT007544|Organism:Severe      GTCAAGCGTGGCAACCGGGTGTTGCTATGCCTAATCTTTACAAAATGCAAAGAATGCTAT   20722
gb:MN994467|Organism:Severe      GTCAAGCGTGGCAACCGGGTGTTGCTATGCCTAATCTTTACAAAATGCAAAGAATGCTAT   20722
gb:MT044257|Organism:Severe      GTCAAGCGTGGCAACCGGGTGTTGCTATGCCTAATCTTTACAAAATGCAAAGAATGCTAT   20722
gb:MT106054|Organism:Severe      GTCAAGCGTGGCAACCGGGTGTTGCTATGCCTAATCTTTACAAAATGCAAAGAATGCTAT   20722
gb:MT049951|Organism:Severe      GTCAAGCGTGGCAACCGGGTGTTGCTATGCCTAATCTTTACAAAATGCAAAGAATGCTAT   20722
gb:MN975262|Organism:Severe      GTCAAGCGTGGCAACCGGGTGTTGCTATGCCTAATCTTTACAAAATGCAAAGAATGCTAT   20722
gb:MT106052|Organism:Severe      GTCAAGCGTGGCAACCGGGTGTTGCTATGCCTAATCTTTACAAAATGCAAAGAATGCTAT   20722
gb:LC522975|Organism:Severe      GTCAAGCGTGGCAACCGGGTGTTGCTATGCCTAATCTTTACAAAATGCAAAGAATGCTAT   20719
gb:LC522973|Organism:Severe      GTCAAGCGTGGCAACCGGGTGTTGCTATGCCTAATCTTTACAAAATGCAAAGAATGCTAT   20719
gb:LC522974|Organism:Severe      GTCAAGCGTGGCAACCGGGTGTTGCTATGCCTAATCTTTACAAAATGCAAAGAATGCTAT   20719
gb:MN985325|Organism:Severe      GTCAAGCGTGGCAACCGGGTGTTGCTATGCCTAATCTTTACAAAATGCAAAGAATGCTAT   20722
gb:MT020881|Organism:Severe      GTCAAGCGTGGCAACCGGGTGTTGCTATGCCTAATCTTTACAAAATGCAAAGAATGCTAT   20722
gb:MT020880|Organism:Severe      GTCAAGCGTGGCAACCGGGTGTTGCTATGCCTAATCTTTACAAAATGCAAAGAATGCTAT   20722
gb:MT066175|Organism:Severe      GTCAAGCGTGGCAACCGGGTGTTGCTATGCCTAATCTTTACAAAATGCAAAGAATGCTAT   20722
gb:MN997409|Organism:Severe      GTCAAGCGTGGCAACCGGGTGTTGCTATGCCTAATCTTTACAAAATGCAAAGAATGCTAT   20722
gb:MN938384|Organism:Severe      GTCAAGCGTGGCAACCGGGTGTTGCTATGCCTAATCTTTACAAAATGCAAAGAATGCTAT   20690
gb:MT044258|Organism:Severe      GTCAAGCGTGGCAACCGGGTGTTGCTATGCCTAATCTTTACAAAATGCAAAGAATGCTAT   20698
gb:MT039890|Organism:Severe      GTCAAGCGTGGCAACCGGGTGTTGCTATGCCTAATCTTTACAAAATGCAAAGAATGCTAT   20722
gb:MN988713|Organism:Severe      GTCAAGCGTGGCAACCGGGTGTTGCTATGCCTAATCTTTACAAAATGCAAAGAATGCTAT   20722
gb:LC521925|Organism:Severe      GTCAAGCGTGGCAACCGGGTGTTGCTATGCCTAATCTTTACAAAATGCAAAGAATGCTAT   20695
gb:MT093571|Organism:Severe      GTCAAGCGTGGCAACCGGGTGTTGCTATGCCTAATCTTTACAAAATGCAAAGAATGCTAT   20722
gb:MT039887|Organism:Severe      GTCAAGCGTGGCAACCGGGTGTTGCTATGCCTAATCTTTACAAAATGCAAAGAATGCTAT   20719
gb:MT019530|Organism:Severe      GTCAAGCGTGGCAACCGGGTGTTGCTATGCCTAATCTTTACAAAATGCAAAGAATGCTAT   20722
gb:MT039888|Organism:Severe      GTCAAGCGTGGCAACCGGGTGTTGCTATGCCTAATCTTTACAAAATGCAAAGAATGCTAT   20722
gb:LC522972|Organism:Severe      GTCAAGCGTGGCAACCGGGTGTTGCTATGCCTAATCTTTACAAAATGCAAAGAATGCTAT   20719
gb:MT027063|Organism:Severe      GTCAAGCGTGGCAACCGGGTGTTGCTATGCCTAATCTTTACAAAATGCAAAGAATGCTAT   20722
gb:MT027062|Organism:Severe      GTCAAGCGTGGCAACCGGGTGTTGCTATGCCTAATCTTTACAAAATGCAAAGAATGCTAT   20722
gb:MT019529|Organism:Severe      GTCAAGCGTGGCAACCGGGTGTTGCTATGCCTAATCTTTACAAAATGCAAAGAATGCTAT   20722
gb:MN996529|Organism:Severe      GTCAAGCGTGGCAACCGGGTGTTGCTATGCCTAATCTTTACAAAATGCAAAGAATGCTAT   20710
gb:MN996531|Organism:Severe      GTCAAGCGTGGCAACCGGGTGTTGCTATGCCTAATCTTTACAAAATGCAAAGAATGCTAT   20709
gb:MT066176|Organism:Severe      GTCAAGCGTGGCAACCGGGTGTTGCTATGCCTAATCTTTACAAAATGCAAAGAATGCTAT   20722
gb:MT027064|Organism:Severe      GTCAAGCGTGGCAACCGGGTGTTGCTATGCCTAATCTTTACAAAATGCAAAGAATGCTAT   20722
gb:MN994468|Organism:Severe      GTCAAGCGTGGCAACCGGGTGTTGCTATGCCTAATCTTTACAAAATGCAAAGAATGCTAT   20722
gb:MT072688|Organism:Severe      GTCAAGCGTGGCAACCGGGTGTTGCTATGCCTAATCTTTACAAAATGCAAAGAATGCTAT   20707
gb:MN996527|Organism:Severe      GTCAAGCGTGGCAACCGGGTGTTGCTATGCCTAATCTTTACAAAATGCAAAGAATGCTAT   20689
gb:MT093631|Organism:Severe      GTCAAGCGTGGCAACCGGGTGTTGCTATGCCTAATCTTTACAAAATGCAAAGAATGCTAT   20760
gb:MT106053|Organism:Severe      GTCAAGCGTGGCAACCGGGTGTTGCTATGCCTAATCTTTACAAAATGCAAAGAATGCTAT   20722
gb:MT019533|Organism:Severe      GTCAAGCGTGGCAACCGGGTGTTGCTATGCCTAATCTTTACAAAATGCAAAGAATGCTAT   20722
gb:MT019531|Organism:Severe      GTCAAGCGTGGCAACCGGGTGTTGCTATGCCTAATCTTTACAAAATGCAAAGAATGCTAT   20722
gb:MN996528|Organism:Severe      GTCAAGCGTGGCAACCGGGTGTTGCTATGCCTAATCTTTACAAAATGCAAAGAATGCTAT   20722
gb:MN996530|Organism:Severe      GTCAAGCGTGGCAACCGGGTGTTGCTATGCCTAATCTTTACAAAATGCAAAGAATGCTAT   20708
gb:MN908947|Organism:Severe      GTCAAGCGTGGCAACCGGGTGTTGCTATGCCTAATCTTTACAAAATGCAAAGAATGCTAT   20722
gb:MT019532|Organism:Severe      GTCAAGCGTGGCAACCGGGTGTTGCTATGCCTAATCTTTACAAAATGCAAAGAATGCTAT   20722
                                 ************************************************************

gb:MT020781|Organism:Severe      TAGAAAAGTGTGACCTTCAAAATTATGGTGATAGTGCAACATTACCTAAAGGCATAATGA   20770
gb:MT007544|Organism:Severe      TAGAAAAGTGTGACCTTCAAAATTATGGTGATAGTGCAACATTACCTAAAGGCATAATGA   20782
gb:MN994467|Organism:Severe      TAGAAAAGTGTGACCTTCAAAATTATGGTGATAGTGCAACATTACCTAAAGGCATAATGA   20782
gb:MT044257|Organism:Severe      TAGAAAAGTGTGACCTTCAAAATTATGGTGATAGTGCAACATTACCTAAAGGCATAATGA   20782
gb:MT106054|Organism:Severe      TAGAAAAGTGTGACCTTCAAAATTATGGTGATAGTGCAACATTACCTAAAGGCATAATGA   20782
```

```
gb:MT049951|Organism:Severe      TAGAAAAGTGTGACCTTCAAAATTATGGTGATAGTGCAACATTACCTAAAGGCATAATGA   20782
gb:MN975262|Organism:Severe      TAGAAAAGTGTGACCTTCAAAATTATGGTGATAGTGCAACATTACCTAAAGGCATAATGA   20782
gb:MT106052|Organism:Severe      TAGAAAAGTGTGACCTTCAAAATTATGGTGATAGTGCAACATTACCTAAAGGCATAATGA   20782
gb:LC522975|Organism:Severe      TAGAAAAGTGTGACCTTCAAAATTATGGTGATAGTGCAACATTACCTAAAGGCATAATGA   20779
gb:LC522973|Organism:Severe      TAGAAAAGTGTGACCTTCAAAATTATGGTGATAGTGCAACATTACCTAAAGGCATAATGA   20779
gb:LC522974|Organism:Severe      TAGAAAAGTGTGACCTTCAAAATTATGGTGATAGTGCAACATTACCTAAAGGCATAATGA   20779
gb:MN985325|Organism:Severe      TAGAAAAGTGTGACCTTCAAAATTATGGTGATAGTGCAACATTACCTAAAGGCATAATGA   20782
gb:MT020881|Organism:Severe      TAGAAAAGTGTGACCTTCAAAATTATGGTGATAGTGCAACATTACCTAAAGGCATAATGA   20782
gb:MT020880|Organism:Severe      TAGAAAAGTGTGACCTTCAAAATTATGGTGATAGTGCAACATTACCTAAAGGCATAATGA   20782
gb:MT066175|Organism:Severe      TAGAAAAGTGTGACCTTCAAAATTATGGTGATAGTGCAACATTACCTAAAGGCATAATGA   20782
gb:MN997409|Organism:Severe      TAGAAAAGTGTGACCTTCAAAATTATGGTGATAGTGCAACATTACCTAAAGGCATAATGA   20782
gb:MN938384|Organism:Severe      TAGAAAAGTGTGACCTTCAAAATTATGGTGATAGTGCAACATTACCTAAAGGCATAATGA   20750
gb:MT044258|Organism:Severe      TAGAAAAGTGTGACCTTCAAAATTATGGTGATAGTGCAACATTACCTAAAGGCATAATGA   20758
gb:MT039890|Organism:Severe      TAGAAAAGTGTGACCTTCAAAATTATGGTGATAGTGCAACATTACCTAAAGGCATAATGA   20782
gb:MN988713|Organism:Severe      TAGAAAAGTGTGACCTTCAAAATTATGGTGATAGTGCAACATTACCTAAAGGCATAATGA   20782
gb:LC521925|Organism:Severe      TAGAAAAGTGTGACCTTCAAAATTATGGTGATAGTGCAACATTACCTAAAGGCATAATGA   20755
gb:MT093571|Organism:Severe      TAGAAAAGTGTGACCTTCAAAATTATGGTGATAGTGCAACATTACCTAAAGGCATAATGA   20782
gb:MT039887|Organism:Severe      TAGAAAAGTGTGACCTTCAAAATTATGGTGATAGTGCAACATTACCTAAAGGCATAATGA   20779
gb:MT019530|Organism:Severe      TAGAAAAGTGTGACCTTCAAAATTATGGTGATAGTGCAACATTACCTAAAGGCATAATGA   20782
gb:MT039888|Organism:Severe      TAGAAAAGTGTGACCTTCAAAATTATGGTGATAGTGCAACATTACCTAAAGGCATAATGA   20782
gb:LC522972|Organism:Severe      TAGAAAAGTGTGACCTTCAAAATTATGGTGATAGTGCAACATTACCTAAAGGCATAATGA   20779
gb:MT027063|Organism:Severe      TAGAAAAGTGTGACCTTCAAAATTATGGTGATAGTGCAACATTACCTAAAGGCATAATGA   20782
gb:MT027062|Organism:Severe      TAGAAAAGTGTGACCTTCAAAATTATGGTGATAGTGCAACATTACCTAAAGGCATAATGA   20782
gb:MT019529|Organism:Severe      TAGAAAAGTGTGACCTTCAAAATTATGGTGATAGTGCAACATTACCTAAAGGCATAATGA   20782
gb:MN996529|Organism:Severe      TAGAAAAGTGTGACCTTCAAAATTATGGTGATAGTGCAACATTACCTAAAGGCATAATGA   20770
gb:MN996531|Organism:Severe      TAGAAAAGTGTGACCTTCAAAATTATGGTGATAGTGCAACATTACCTAAAGGCATAATGA   20769
gb:MT066176|Organism:Severe      TAGAAAAGTGTGACCTTCAAAATTATGGTGATAGTGCAACATTACCTAAAGGCATAATGA   20782
gb:MT027064|Organism:Severe      TAGAAAAGTGTGACCTTCAAAATTATGGTGATAGTGCAACATTACCTAAAGGCATAATGA   20782
gb:MN994468|Organism:Severe      TAGAAAAGTGTGACCTTCAAAATTATGGTGATAGTGCAACATTACCTAAAGGCATAATGA   20782
gb:MT072688|Organism:Severe      TAGAAAAGTGTGACCTTCAAAATTATGGTGATAGTGCAACATTACCTAAAGGCATAATGA   20767
gb:MN996527|Organism:Severe      TAGAAAAGTGTGACCTTCAAAATTATGGTGATAGTGCAACATTACCTAAAGGCATAATGA   20749
gb:MT093631|Organism:Severe      TAGAAAAGTGTGACCTTCAAAATTATGGTGATAGTGCAACATTACCTAAAGGCATAATGA   20820
gb:MT106053|Organism:Severe      TAGAAAAGTGTGACCTTCAAAATTATGGTGATAGTGCAACATTACCTAAAGGCATAATGA   20782
gb:MT019533|Organism:Severe      TAGAAAAGTGTGACCTTCAAAATTATGGTGATAGTGCAACATTACCTAAAGGCATAATGA   20782
gb:MT019531|Organism:Severe      TAGAAAAGTGTGACCTTCAAAATTATGGTGATAGTGCAACATTACCTAAAGGCATAATGA   20782
gb:MN996528|Organism:Severe      TAGAAAAGTGTGACCTTCAAAATTATGGTGATAGTGCAACATTACCTAAAGGCATAATGA   20782
gb:MN996530|Organism:Severe      TAGAAAAGTGTGACCTTCAAAATTATGGTGATAGTGCAACATTACCTAAAGGCATAATGA   20768
gb:MN908947|Organism:Severe      TAGAAAAGTGTGACCTTCAAAATTATGGTGATAGTGCAACATTACCTAAAGGCATAATGA   20782
gb:MT019532|Organism:Severe      TAGAAAAGTGTGACCTTCAAAATTATGGTGATAGTGCAACATTACCTAAAGGCATAATGA   20782
                                 ************************************************************

gb:MT020781|Organism:Severe      TGAATGTCGCAAAATATACTCAACTGTGTCAATATTTAAACACATTAACATTAGCTGTAC   20830
gb:MT007544|Organism:Severe      TGAATGTCGCAAAATATACTCAACTGTGTCAATATTTAAACACATTAACATTAGCTGTAC   20842
gb:MN994467|Organism:Severe      TGAATGTCGCAAAATATACTCAACTGTGTCAATATTTAAACACATTAACATTAGCTGTAC   20842
gb:MT044257|Organism:Severe      TGAATGTCGCAAAATATACTCAACTGTGTCAATATTTAAACACATTAACATTAGCTGTAC   20842
gb:MT106054|Organism:Severe      TGAATGTCGCAAAATATACTCAACTGTGTCAATATTTAAACACATTAACATTAGCTGTAC   20842
gb:MT049951|Organism:Severe      TGAATGTCGCAAAATATACTCAACTGTGTCAATATTTAAACACATTAACATTAGCTGTAC   20842
gb:MN975262|Organism:Severe      TGAATGTCGCAAAATATACTCAACTGTGTCAATATTTAAACACATTAACATTAGCTGTAC   20842
gb:MT106052|Organism:Severe      TGAATGTCGCAAAATATACTCAACTGTGTCAATATTTAAACACATTAACATTAGCTGTAC   20842
gb:LC522975|Organism:Severe      TGAATGTCGCAAAATATACTCAACTGTGTCAATATTTAAACACATTAACATTAGCTGTAC   20839
gb:LC522973|Organism:Severe      TGAATGTCGCAAAATATACTCAACTGTGTCAATATTTAAACACATTAACATTAGCTGTAC   20839
gb:LC522974|Organism:Severe      TGAATGTCGCAAAATATACTCAACTGTGTCAATATTTAAACACATTAACATTAGCTGTAC   20839
gb:MN985325|Organism:Severe      TGAATGTCGCAAAATATACTCAACTGTGTCAATATTTAAACACATTAACATTAGCTGTAC   20842
gb:MT020881|Organism:Severe      TGAATGTCGCAAAATATACTCAACTGTGTCAATATTTAAACACATTAACATTAGCTGTAC   20842
gb:MT020880|Organism:Severe      TGAATGTCGCAAAATATACTCAACTGTGTCAATATTTAAACACATTAACATTAGCTGTAC   20842
gb:MT066175|Organism:Severe      TGAATGTCGCAAAATATACTCAACTGTGTCAATATTTAAACACATTAACATTAGCTGTAC   20842
gb:MN997409|Organism:Severe      TGAATGTCGCAAAATATACTCAACTGTGTCAATATTTAAACACATTAACATTAGCTGTAC   20842
gb:MN938384|Organism:Severe      TGAATGTCGCAAAATATACTCAACTGTGTCAATATTTAAACACATTAACATTAGCTGTAC   20810
gb:MT044258|Organism:Severe      TGAATGTCGCAAAATATACTCAACTGTGTCAATATTTAAACACATTAACATTAGCTGTAC   20818
gb:MT039890|Organism:Severe      TGAATGTCGCAAAATATACTCAACTGTGTCAATATTTAAACACATTAACATTAGCTGTAC   20842
gb:MN988713|Organism:Severe      TGAATGTCGCAAAATATACTCAACTGTGTCAATATTTAAACACATTAACATTAGCTGTAC   20842
gb:LC521925|Organism:Severe      TGAATGTCGCAAAATATACTCAACTGTGTCAATATTTAAACACATTAACATTAGCTGTAC   20815
gb:MT093571|Organism:Severe      TGAATGTCGCAAAATATACTCAACTGTGTCAATATTTAAACACATTAACATTAGCTGTAC   20842
gb:MT039887|Organism:Severe      TGAATGTCGCAAAATATACTCAACTGTGTCAATATTTAAACACATTAACATTAGCTGTAC   20839
gb:MT019530|Organism:Severe      TGAATGTCGCAAAATATACTCAACTGTGTCAATATTTAAACACATTAACATTAGCTGTAC   20842
```

```
gb:MT039888|Organism:Severe      TGAATGTCGCAAAATATACTCAACTGTGTCAATATTTAAACACATTAACATTAGCTGTAC    20842
gb:LC522972|Organism:Severe      TGAATGTCGCAAAATATACTCAACTGTGTCAATATTTAAACACATTAACATTAGCTGTAC    20839
gb:MT027063|Organism:Severe      TGAATGTCGCAAAATATACTCAACTGTGTCAATATTTAAACACATTAACATTAGCTGTAC    20842
gb:MT027062|Organism:Severe      TGAATGTCGCAAAATATACTCAACTGTGTCAATATTTAAACACATTAACATTAGCTGTAC    20842
gb:MT019529|Organism:Severe      TGAATGTCGCAAAATATACTCAACTGTGTCAATATTTAAACACATTAACATTAGCTGTAC    20842
gb:MN996529|Organism:Severe      TGAATGTCGCAAAATATACTCAACTGTGTCAATATTTAAACACATTAACATTAGCTGTAC    20830
gb:MN996531|Organism:Severe      TGAATGTCGCAAAATATACTCAACTGTGTCAATATTTAAACACATTAACATTAGCTGTAC    20829
gb:MT066176|Organism:Severe      TGAATGTCGCAAAATATACTCAACTGTGTCAATATTTAAACACATTAACATTAGCTGTAC    20842
gb:MT027064|Organism:Severe      TGAATGTCGCAAAATATACTCAACTGTGTCAATATTTAAACACATTAACATTAGCTGTAC    20842
gb:MN994468|Organism:Severe      TGAATGTCGCAAAATATACTCAACTGTGTCAATATTTAAACACATTAACATTAGCTGTAC    20842
gb:MT072688|Organism:Severe      TGAATGTCGCAAAATATACTCAACTGTGTCAATATTTAAACACATTAACATTAGCTGTAC    20827
gb:MN996527|Organism:Severe      TGAATGTCGCAAAATATACTCAACTGTGTCAATATTTAAACACATTAACATTAGCTGTAC    20809
gb:MT093631|Organism:Severe      TGAATGTCGCAAAATATACTCAACTGTGTCAATATTTAAACACATTAACATTAGCTGTAC    20880
gb:MT106053|Organism:Severe      TGAATGTCGCAAAATATACTCAACTGTGTCAATATTTAAACACATTAACATTAGCTGTAC    20842
gb:MT019533|Organism:Severe      TGAATGTCGCAAAATATACTCAACTGTGTCAATATTTAAACACATTAACATTAGCTGTAC    20842
gb:MT019531|Organism:Severe      TGAATGTCGCAAAATATACTCAACTGTGTCAATATTTAAACACATTAACATTAGCTGTAC    20842
gb:MN996528|Organism:Severe      TGAATGTCGCAAAATATACTCAACTGTGTCAATATTTAAACACATTAACATTAGCTGTAC    20842
gb:MN996530|Organism:Severe      TGAATGTCGCAAAATATACTCAACTGTGTCAATATTTAAACACATTAACATTAGCTGTAC    20828
gb:MN908947|Organism:Severe      TGAATGTCGCAAAATATACTCAACTGTGTCAATATTTAAACACATTAACATTAGCTGTAC    20842
gb:MT019532|Organism:Severe      TGAATGTCGCAAAATATACTCAACTGTGTCAATATTTAAACACATTAACATTAGCTGTAC    20842
                                 ************************************************************

gb:MT020781|Organism:Severe      CCTATAATATGAGAGTTATACATTTTGGTGCTGGTTCTGATAAAGGAGTTGCACCAGGTA    20890
gb:MT007544|Organism:Severe      CCTATAATATGAGAGTTATACATTTTGGTGCTGGTTCTGATAAAGGAGTTGCACCAGGTA    20902
gb:MN994467|Organism:Severe      CCTATAATATGAGAGTTATACATTTTGGTGCTGGTTCTGATAAAGGAGTTGCACCAGGTA    20902
gb:MT044257|Organism:Severe      CCTATAATATGAGAGTTATACATTTTGGTGCTGGTTCTGATAAAGGAGTTGCACCAGGTA    20902
gb:MT106054|Organism:Severe      CCTATAATATGAGAGTTATACATTTTGGTGCTGGTTCTGATAAAGGAGTTGCACCAGGTA    20902
gb:MT049951|Organism:Severe      CCTATAATATGAGAGTTATACATTTTGGTGCTGGTTCTGATAAAGGAGTTGCACCAGGTA    20902
gb:MN975262|Organism:Severe      CCTATAATATGAGAGTTATACATTTTGGTGCTGGTTCTGATAAAGGAGTTGCACCAGGTA    20902
gb:MT106052|Organism:Severe      CCTATAATATGAGAGTTATACATTTTGGTGCTGGTTCTGATAAAGGAGTTGCACCAGGTA    20902
gb:LC522975|Organism:Severe      CCTATAATATGAGAGTTATACATTTTGGTGCTGGTTCTGATAAAGGAGTTGCACCAGGTA    20899
gb:LC522973|Organism:Severe      CCTATAATATGAGAGTTATACATTTTGGTGCTGGTTCTGATAAAGGAGTTGCACCAGGTA    20899
gb:LC522974|Organism:Severe      CCTATAATATGAGAGTTATACATTTTGGTGCTGGTTCTGATAAAGGAGTTGCACCAGGTA    20899
gb:MN985325|Organism:Severe      CCTATAATATGAGAGTTATACATTTTGGTGCTGGTTCTGATAAAGGAGTTGCACCAGGTA    20902
gb:MT020881|Organism:Severe      CCTATAATATGAGAGTTATACATTTTGGTGCTGGTTCTGATAAAGGAGTTGCACCAGGTA    20902
gb:MT020880|Organism:Severe      CCTATAATATGAGAGTTATACATTTTGGTGCTGGTTCTGATAAAGGAGTTGCACCAGGTA    20902
gb:MT066175|Organism:Severe      CCTATAATATGAGAGTTATACATTTTGGTGCTGGTTCTGATAAAGGAGTTGCACCAGGTA    20902
gb:MN997409|Organism:Severe      CCTATAATATGAGAGTTATACATTTTGGTGCTGGTTCTGATAAAGGAGTTGCACCAGGTA    20902
gb:MN938384|Organism:Severe      CCTATAATATGAGAGTTATACATTTTGGTGCTGGTTCTGATAAAGGAGTTGCACCAGGTA    20870
gb:MT044258|Organism:Severe      CCTATAATATGAGAGTTATACATTTTGGTGCTGGTTCTGATAAAGGAGTTGCACCAGGTA    20878
gb:MT039890|Organism:Severe      CCTATAATATGAGAGTTATACATTTTGGTGCTGGTTCTGATAAAGGAGTTGCACCAGGTA    20902
gb:MN988713|Organism:Severe      CCTATAATATGAGAGTTATACATTTTGGTGCTGGTTCTGATAAAGGAGTTGCACCAGGTA    20902
gb:LC521925|Organism:Severe      CCTATAATATGAGAGTTATACATTTTGGTGCTGGTTCTGATAAAGGAGTTGCACCAGGTA    20875
gb:MT093571|Organism:Severe      CCTATAATATGAGAGTTATACATTTTGGTGCTGGTTCTGATAAAGGAGTTGCACCAGGTA    20902
gb:MT039887|Organism:Severe      CCTATAATATGAGAGTTATACATTTTGGTGCTGGTTCTGATAAAGGAGTTGCACCAGGTA    20899
gb:MT019530|Organism:Severe      CCTATAATATGAGAGTTATACATTTTGGTGCTGGTTCTGATAAAGGAGTTGCACCAGGTA    20902
gb:MT039888|Organism:Severe      CCTATAATATGAGAGTTATACATTTTGGTGCTGGTTCTGATAAAGGAGTTGCACCAGGTA    20902
gb:LC522972|Organism:Severe      CCTATAATATGAGAGTTATACATTTTGGTGCTGGTTCTGATAAAGGAGTTGCACCAGGTA    20899
gb:MT027063|Organism:Severe      CCTATAATATGAGAGTTATACATTTTGGTGCTGGTTCTGATAAAGGAGTTGCACCAGGTA    20902
gb:MT027062|Organism:Severe      CCTATAATATGAGAGTTATACATTTTGGTGCTGGTTCTGATAAAGGAGTTGCACCAGGTA    20902
gb:MT019529|Organism:Severe      CCTATAATATGAGAGTTATACATTTTGGTGCTGGTTCTGATAAAGGAGTTGCACCAGGTA    20902
gb:MN996529|Organism:Severe      CCTATAATATGAGAGTTATACATTTTGGTGCTGGTTCTGATAAAGGAGTTGCACCAGGTA    20890
gb:MN996531|Organism:Severe      CCTATAATATGAGAGTTATACATTTTGGTGCTGGTTCTGATAAAGGAGTTGCACCAGGTA    20889
gb:MT066176|Organism:Severe      CCTATAATATGAGAGTTATACATTTTGGTGCTGGTTCTGATAAAGGAGTTGCACCAGGTA    20902
gb:MT027064|Organism:Severe      CCTATAATATGAGAGTTATACATTTTGGTGCTGGTTCTGATAAAGGAGTTGCACCAGGTA    20902
gb:MN994468|Organism:Severe      CCTATAATATGAGAGTTATACATTTTGGTGCTGGTTCTGATAAAGGAGTTGCACCAGGTA    20902
gb:MT072688|Organism:Severe      CCTATAATATGAGAGTTATACATTTTGGTGCTGGTTCTGATAAAGGAGTTGCACCAGGTA    20887
gb:MN996527|Organism:Severe      CCTATAATATGAGAGTTATACATTTTGGTGCTGGTTCTGATAAAGGAGTTGCACCAGGTA    20869
gb:MT093631|Organism:Severe      CCTATAATATGAGAGTTATACATTTTGGTGCTGGTTCTGATAAAGGAGTTGCACCAGGTA    20940
gb:MT106053|Organism:Severe      CCTATAATATGAGAGTTATACATTTTGGTGCTGGTTCTGATAAAGGAGTTGCACCAGGTA    20902
gb:MT019533|Organism:Severe      CCTATAATATGAGAGTTATACATTTTGGTGCTGGTTCTGATAAAGGAGTTGCACCAGGTA    20902
gb:MT019531|Organism:Severe      CCTATAATATGAGAGTTATACATTTTGGTGCTGGTTCTGATAAAGGAGTTGCACCAGGTA    20902
gb:MN996528|Organism:Severe      CCTATAATATGAGAGTTATACATTTTGGTGCTGGTTCTGATAAAGGAGTTGCACCAGGTA    20902
gb:MN996530|Organism:Severe      CCTATAATATGAGAGTTATACATTTTGGTGCTGGTTCTGATAAAGGAGTTGCACCAGGTA    20888
gb:MN908947|Organism:Severe      CCTATAATATGAGAGTTATACATTTTGGTGCTGGTTCTGATAAAGGAGTTGCACCAGGTA    20902
```

```
gb:MT019532|Organism:Severe      CCTATAATATGAGAGTTATACATTTTGGTGCTGGTTCTGATAAAGGAGTTGCACCAGGTA    20902
                                 ************************************************************

gb:MT020781|Organism:Severe      CAGCTGTTTTAAGACAGTGGTTGCCTACGGGTACGCTGCTTGTCGATTCAGATCTTAATG    20950
gb:MT007544|Organism:Severe      CAGCTGTTTTAAGACAGTGGTTGCCTACGGGTACGCTGCTTGTCGATTCAGATCTTAATG    20962
gb:MN994467|Organism:Severe      CAGCTGTTTTAAGACAGTGGTTGCCTACGGGTACGCTGCTTGTCGATTCAGATCTTAATG    20962
gb:MT044257|Organism:Severe      CAGCTGTTTTAAGACAGTGGTTGCCTACGGGTACGCTGCTTGTCGATTCAGATCTTAATG    20962
gb:MT106054|Organism:Severe      CAGCTGTTTTAAGACAGTGGTTGCCTACGGGTACGCTGCTTGTCGATTCAGATCTTAATG    20962
gb:MT049951|Organism:Severe      CAGCTGTTTTAAGACAGTGGTTGCCTACGGGTACGCTGCTTGTCGATTCAGATCTTAATG    20962
gb:MN975262|Organism:Severe      CAGCTGTTTTAAGACAGTGGTTGCCTACGGGTACGCTGCTTGTCGATTCAGATCTTAATG    20962
gb:MT106052|Organism:Severe      CAGCTGTTTTAAGACAGTGGTTGCCTACGGGTACGCTGCTTGTCGATTCAGATCTTAATG    20962
gb:LC522975|Organism:Severe      CAGCTGTTTTAAGACAGTGGTTGCCTACGGGTACGCTGCTTGTCGATTCAGATCTTAATG    20959
gb:LC522973|Organism:Severe      CAGCTGTTTTAAGACAGTGGTTGCCTACGGGTACGCTGCTTGTCGATTCAGATCTTAATG    20959
gb:LC522974|Organism:Severe      CAGCTGTTTTAAGACAGTGGTTGCCTACGGGTACGCTGCTTGTCGATTCAGATCTTAATG    20959
gb:MN985325|Organism:Severe      CAGCTGTTTTAAGACAGTGGTTGCCTACGGGTACGCTGCTTGTCGATTCAGATCTTAATG    20962
gb:MT020881|Organism:Severe      CAGCTGTTTTAAGACAGTGGTTGCCTACGGGTACGCTGCTTGTCGATTCAGATCTTAATG    20962
gb:MT020880|Organism:Severe      CAGCTGTTTTAAGACAGTGGTTGCCTACGGGTACGCTGCTTGTCGATTCAGATCTTAATG    20962
gb:MT066175|Organism:Severe      CAGCTGTTTTAAGACAGTGGTTGCCTACGGGTACGCTGCTTGTCGATTCAGATCTTAATG    20962
gb:MN997409|Organism:Severe      CAGCTGTTTTAAGACAGTGGTTGCCTACGGGTACGCTGCTTGTCGATTCAGATCTTAATG    20962
gb:MN938384|Organism:Severe      CAGCTGTTTTAAGACAGTGGTTGCCTACGGGTACGCTGCTTGTCGATTCAGATCTTAATG    20930
gb:MT044258|Organism:Severe      CAGCTGTTTTAAGACAGTGGTTGCCTACGGGTACGCTGCTTGTCGATTCAGATCTTAATG    20938
gb:MT039890|Organism:Severe      CAGCTGTTTTAAGACAGTGGTTGCCTACGGGTATGCTGCTTGTCGATTCAGATCTTAATG    20962
gb:MN988713|Organism:Severe      CAGCTGTTTTAAGACAGTGGTTGCCTACGGGTACGCTGCTTGTCGATTCAGATCTTAATG    20962
gb:LC521925|Organism:Severe      CAGCTGTTTTAAGACAGTGGTTGCCTACGGGTACGCTGCTTGTCGATTCAGATCTTAATG    20935
gb:MT093571|Organism:Severe      CAGCTGTTTTAAGACAGTGGTTGCCTACGGGTACGCTGCTTGTCGATTCAGATCTTAATG    20962
gb:MT039887|Organism:Severe      CAGCTGTTTTAAGACAGTGGTTGCCTACGGGTACGCTGCTTGTCGATTCAGATCTTAATG    20959
gb:MT019530|Organism:Severe      CAGCTGTTTTAAGACAGTGGTTGCCTACGGGTACGCTGCTTGTCGATTCAGATCTTAATG    20962
gb:MT039888|Organism:Severe      CAGCTGTTTTAAGACAGTGGTTGCCTACGGGTACGCTGCTTGTCGATTCAGATCTTAATG    20962
gb:LC522972|Organism:Severe      CAGCTGTTTTAAGACAGTGGTTGCCTACGGGTACGCTGCTTGTCGATTCAGATCTTAATG    20959
gb:MT027063|Organism:Severe      CAGCTGTTTTAAGACAGTGGTTGCCTACGGGTACGCTGCTTGTCGATTCAGATCTTAATG    20962
gb:MT027062|Organism:Severe      CAGCTGTTTTAAGACAGTGGTTGCCTACGGGTACGCTGCTTGTCGATTCAGATCTTAATG    20962
gb:MT019529|Organism:Severe      CAGCTGTTTTAAGACAGTGGTTGCCTACGGGTACGCTGCTTGTCGATTCAGATCTTAATG    20962
gb:MN996529|Organism:Severe      CAGCTGTTTTAAGACAGTGGTTGCCTACGGGTACGCTGCTTGTCGATTCAGATCTTAATG    20950
gb:MN996531|Organism:Severe      CAGCTGTTTTAAGACAGTGGTTGCCTACGGGTACGCTGCTTGTCGATTCAGATCTTAATG    20949
gb:MT066176|Organism:Severe      CAGCTGTTTTAAGACAGTGGTTGCCTACGGGTACGCTGCTTGTCGATTCAGATCTTAATG    20962
gb:MT027064|Organism:Severe      CAGCTGTTTTAAGACAGTGGTTGCCTACGGGTACGCTGCTTGTCGATTCAGATCTTAATG    20962
gb:MN994468|Organism:Severe      CAGCTGTTTTAAGACAGTGGTTGCCTACGGGTACGCTGCTTGTCGATTCAGATCTTAATG    20962
gb:MT072688|Organism:Severe      CAGCTGTTTTAAGACAGTGGTTGCCTACGGGTACGCTGCTTGTCGATTCAGATCTTAATG    20947
gb:MN996527|Organism:Severe      CAGCTGTTTTAAGACAGTGGTTGCCTACGGGTACGCTGCTTGTCGATTCAGATCTTAATG    20929
gb:MT093631|Organism:Severe      CAGCTGTTTTAAGACAGTGGTTGCCTACGGGTACGCTGCTTGTCGATTCAGATCTTAATG    21000
gb:MT106053|Organism:Severe      CAGCTGTTTTAAGACAGTGGTTGCCTACGGGTACGCTGCTTGTCGATTCAGATCTTAATG    20962
gb:MT019533|Organism:Severe      CAGCTGTTTTAAGACAGTGGTTGCCTACGGGTACGCTGCTTGTCGATTCAGATCTTAATG    20962
gb:MT019531|Organism:Severe      CAGCTGTTTTAAGACAGTGGTTGCCTACGGGTACGCTGCTTGTCGATTCAGATCTTAATG    20962
gb:MN996528|Organism:Severe      CAGCTGTTTTAAGACAGTGGTTGCCTACGGGTACGCTGCTTGTCGATTCAGATCTTAATG    20962
gb:MN996530|Organism:Severe      CAGCTGTTTTAAGACAGTGGTTGCCTACGGGTACGCTGCTTGTCGATTCAGATCTTAATG    20948
gb:MN908947|Organism:Severe      CAGCTGTTTTAAGACAGTGGTTGCCTACGGGTACGCTGCTTGTCGATTCAGATCTTAATG    20962
gb:MT019532|Organism:Severe      CAGCTGTTTTAAGACAGTGGTTGCCTACGGGTACGCTGCTTGTCGATTCAGATCTTAATG    20962
                                 ********************************* **************************

gb:MT020781|Organism:Severe      ACTTTGTCTCTGATGCAGATTCAACTTTGATTGGTGATTGTGCAACTGTACATACAGCTA    21010
gb:MT007544|Organism:Severe      ACTTTGTCTCTGATGCAGATTCAACTTTGATTGGTGATTGTGCAACTGTACATACAGCTA    21022
gb:MN994467|Organism:Severe      ACTTTGTCTCTGATGCAGATTCAACTTTGATTGGTGATTGTGCAACTGTACATACAGCTA    21022
gb:MT044257|Organism:Severe      ACTTTGTCTCTGATGCAGATTCAACTTTGATTGGTGATTGTGCAACTGTACATACAGCTA    21022
gb:MT106054|Organism:Severe      ACTTTGTCTCTGATGCAGATTCAACTTTGATTGGTGATTGTGCAACTGTACATACAGCTA    21022
gb:MT049951|Organism:Severe      ACTTTGTCTCTGATGCAGATTCAACTTTGATTGGTGATTGTGCAACTGTACATACAGCTA    21022
gb:MN975262|Organism:Severe      ACTTTGTCTCTGATGCAGATTCAACTTTGATTGGTGATTGTGCAACTGTACATACAGCTA    21022
gb:MT106052|Organism:Severe      ACTTTGTCTCTGATGCAGATTCAACTTTGATTGGTGATTGTGCAACTGTACATACAGCTA    21022
gb:LC522975|Organism:Severe      ACTTTGTCTCTGATGCAGATTCAACTTTGATTGGTGATTGTGCAACTGTACATACAGCTA    21019
gb:LC522973|Organism:Severe      ACTTTGTCTCTGATGCAGATTCAACTTTGATTGGTGATTGTGCAACTGTACATACAGCTA    21019
gb:LC522974|Organism:Severe      ACTTTGTCTCTGATGCAGATTCAACTTTGATTGGTGATTGTGCAACTGTACATACAGCTA    21019
gb:MN985325|Organism:Severe      ACTTTGTCTCTGATGCAGATTCAACTTTGATTGGTGATTGTGCAACTGTACATACAGCTA    21022
gb:MT020881|Organism:Severe      ACTTTGTCTCTGATGCAGATTCAACTTTGATTGGTGATTGTGCAACTGTACATACAGCTA    21022
gb:MT020880|Organism:Severe      ACTTTGTCTCTGATGCAGATTCAACTTTGATTGGTGATTGTGCAACTGTACATACAGCTA    21022
gb:MT066175|Organism:Severe      ACTTTGTCTCTGATGCAGATTCAACTTTGATTGGTGATTGTGCAACTGTACATACAGCTA    21022
gb:MN997409|Organism:Severe      ACTTTGTCTCTGATGCAGATTCAACTTTGATTGGTGATTGTGCAACTGTACATACAGCTA    21022
```

```
gb:MN938384|Organism:Severe      ACTTTGTCTCTGATGCAGATTCAACTTTGATTGGTGATTGTGCAACTGTACATACAGCTA   20990
gb:MT044258|Organism:Severe      ACTTTGTCTCTGATGCAGATTCAACTTTGATTGGTGATTGTGCAACTGTACATACAGCTA   20998
gb:MT039890|Organism:Severe      ACTTTGTCTCTGATGCAGATTCAACTTTGATTGGTGATTGTGCAACTGTACATACAGCTA   21022
gb:MN988713|Organism:Severe      ACTTTGTCTCTGATGCAGATTCAACTTTGATTGGTGATTGTGCAACTGTACATACAGCTA   21022
gb:LC521925|Organism:Severe      ACTTTGTCTCTGATGCAGATTCAACTTTGATTGGTGATTGTGCAACTGTACATACAGCTA   20995
gb:MT093571|Organism:Severe      ACTTTGTCTCTGATGCAGATTCAACTTTGATTGGTGATTGTGCAACTGTACATACAGCTA   21022
gb:MT039887|Organism:Severe      ACTTTGTCTCTGATGCAGATTCAACTTTGATTGGTGATTGTGCAACTGTACATACAGCTA   21019
gb:MT019530|Organism:Severe      ACTTTGTCTCTGATGCAGATTCAACTTTGATTGGTGATTGTGCAACTGTACATACAGCTA   21022
gb:MT039888|Organism:Severe      ACTTTGTCTCTGATGCAGATTCAACTTTGATTGGTGATTGTGCAACTGTACATACAGCTA   21022
gb:LC522972|Organism:Severe      ACTTTGTCTCTGATGCAGATTCAACTTTGATTGGTGATTGTGCAACTGTACATACAGCTA   21019
gb:MT027063|Organism:Severe      ACTTTGTCTCTGATGCAGATTCAACTTTGATTGGTGATTGTGCAACTGTACATACAGCTA   21022
gb:MT027062|Organism:Severe      ACTTTGTCTCTGATGCAGATTCAACTTTGATTGGTGATTGTGCAACTGTACATACAGCTA   21022
gb:MT019529|Organism:Severe      ACTTTGTCTCTGATGCAGATTCAACTTTGATTGGTGATTGTGCAACTGTACATACAGCTA   21022
gb:MN996529|Organism:Severe      ACTTTGTCTCTGATGCAGATTCAACTTTGATTGGTGATTGTGCAACTGTACATACAGCTA   21010
gb:MN996531|Organism:Severe      ACTTTGTCTCTGATGCAGATTCAACTTTGATTGGTGATTGTGCAACTGTACATACAGCTA   21009
gb:MT066176|Organism:Severe      ACTTTGTCTCTGATGCAGATTCAACTTTGATTGGTGATTGTGCAACTGTACATACAGCTA   21022
gb:MT027064|Organism:Severe      ACTTTGTCTCTGATGCAGATTCAACTTTGATTGGTGATTGTGCAACTGTACATACAGCTA   21022
gb:MN994468|Organism:Severe      ACTTTGTCTCTGATGCAGATTCAACTTTGATTGGTGATTGTGCAACTGTACATACAGCTA   21022
gb:MT072688|Organism:Severe      ACTTTGTCTCTGATGCAGATTCAACTTTGATTGGTGATTGTGCAACTGTACATACAGCTA   21007
gb:MN996527|Organism:Severe      ACTTTGTCTCTGATGCAGATTCAACTTTGATTGGTGATTGTGCAACTGTACATACAGCTA   20989
gb:MT093631|Organism:Severe      ACTTTGTCTCTGATGCAGATTCAACTTTGATTGGTGATTGTGCAACTGTACATACAGCTA   21060
gb:MT106053|Organism:Severe      ACTTTGTCTCTGATGCAGATTCAACTTTGATTGGTGATTGTGCAACTGTACATACAGCTA   21022
gb:MT019533|Organism:Severe      ACTTTGTCTCTGATGCAGATTCAACTTTGATTGGTGATTGTGCAACTGTACATACAGCTA   21022
gb:MT019531|Organism:Severe      ACTTTGTCTCTGATGCAGATTCAACTTTGATTGGTGATTGTGCAACTGTACATACAGCTA   21022
gb:MN996528|Organism:Severe      ACTTTGTCTCTGATGCAGATTCAACTTTGATTGGTGATTGTGCAACTGTACATACAGCTA   21022
gb:MN996530|Organism:Severe      ACTTTGTCTCTGATGCAGATTCAACTTTGATTGGTGATTGTGCAACTGTACATACAGCTA   21008
gb:MN908947|Organism:Severe      ACTTTGTCTCTGATGCAGATTCAACTTTGATTGGTGATTGTGCAACTGTACATACAGCTA   21022
gb:MT019532|Organism:Severe      ACTTTGTCTCTGATGCAGATTCAACTTTGATTGGTGATTGTGCAACTGTACATACAGCTA   21022
                                 ************************************************************

gb:MT020781|Organism:Severe      ATAAATGGGATCTCATTATTAGTGATATGTACGACCCTAAGACTAAAAATGTTACAAAAG   21070
gb:MT007544|Organism:Severe      ATAAATGGGATCTCATTATTAGTGATATGTACGACCCTAAGACTAAAAATGTTACAAAAG   21082
gb:MN994467|Organism:Severe      ATAAATGGGATCTCATTATTAGTGATATGTACGACCCTAAGACTAAAAATGTTACAAAAG   21082
gb:MT044257|Organism:Severe      ATAAATGGGATCTCATTATTAGTGATATGTACGACCCTAAGACTAAAAATGTTACAAAAG   21082
gb:MT106054|Organism:Severe      ATAAATGGGATCTCATTATTAGTGATATGTACGACCCTAAGACTAAAAATGTTACAAAAG   21082
gb:MT049951|Organism:Severe      ATAAATGGGATCTCATTATTAGTGATATGTACGACCCTAAGACTAAAAATGTTACAAAAG   21082
gb:MN975262|Organism:Severe      ATAAATGGGATCTCATTATTAGTGATATGTACGACCCTAAGACTAAAAATGTTACAAAAG   21082
gb:MT106052|Organism:Severe      ATAAATGGGATCTCATTATTAGTGATATGTACGACCCTAAGACTAAAAATGTTACAAAAG   21082
gb:LC522975|Organism:Severe      ATAAATGGGATCTCATTATTAGTGATATGTACGACCCTAAGACTAAAAATGTTACAAAAG   21079
gb:LC522973|Organism:Severe      ATAAATGGGATCTCATTATTAGTGATATGTACGACCCTAAGACTAAAAATGTTACAAAAG   21079
gb:LC522974|Organism:Severe      ATAAATGGGATCTCATTATTAGTGATATGTACGACCCTAAGACTAAAAATGTTACAAAAG   21079
gb:MN985325|Organism:Severe      ATAAATGGGATCTCATTATTAGTGATATGTACGACCCTAAGACTAAAAATGTTACAAAAG   21082
gb:MT020881|Organism:Severe      ATAAATGGGATCTCATTATTAGTGATATGTACGACCCTAAGACTAAAAATGTTACAAAAG   21082
gb:MT020880|Organism:Severe      ATAAATGGGATCTCATTATTAGTGATATGTACGACCCTAAGACTAAAAATGTTACAAAAG   21082
gb:MT066175|Organism:Severe      ATAAATGGGATCTCATTATTAGTGATATGTACGACCCTAAGACTAAAAATGTTACAAAAG   21082
gb:MN997409|Organism:Severe      ATAAATGGGATCTCATTATTAGTGATATGTACGACCCTAAGACTAAAAATGTTACAAAAG   21082
gb:MN938384|Organism:Severe      ATAAATGGGATCTCATTATTAGTGATATGTACGACCCTAAGACTAAAAATGTTACAAAAG   21050
gb:MT044258|Organism:Severe      ATAAATGGGATCTCATTATTAGTGATATGTACGACCCTAAGACTAAAAATGTTACAAAAG   21058
gb:MT039890|Organism:Severe      ATAAATGGGATCTCATTATTAGTGATATGTACGACCCTAAGACTAAAAATGTTACAAAAG   21082
gb:MN988713|Organism:Severe      ATAAATGGGATCTCATTATTAGTGATATGTACGACCCTAAGACTAAAAATGTTACAAAAG   21082
gb:LC521925|Organism:Severe      ATAAATGGGATCTCATTATTAGTGATATGTACGACCCTAAGACTAAAAATGTTACAAAAG   21055
gb:MT093571|Organism:Severe      ATAAATGGGATCTCATTATTAGTGATATGTACGACCCTAAGACTAAAAATGTTACAAAAG   21082
gb:MT039887|Organism:Severe      ATAAATGGGATCTCATTATTAGTGATATGTACGACCCTAAGACTAAAAATGTTACAAAAG   21079
gb:MT019530|Organism:Severe      ATAAATGGGATCTCATTATTAGTGATATGTACGACCCTAAGACTAAAAATGTTACAAAAG   21082
gb:MT039888|Organism:Severe      ATAAATGGGATCTCATTATTAGTGATATGTACGACCCTAAGACTAAAAATGTTACAAAAG   21082
gb:LC522972|Organism:Severe      ATAAATGGGATCTCATTATTAGTGATATGTACGACCCTAAGACTAAAAATGTTACAAAAG   21079
gb:MT027063|Organism:Severe      ATAAATGGGATCTCATTATTAGTGATATGTACGACCCTAAGACTAAAAATGTTACAAAAG   21082
gb:MT027062|Organism:Severe      ATAAATGGGATCTCATTATTAGTGATATGTACGACCCTAAGACTAAAAATGTTACAAAAG   21082
gb:MT019529|Organism:Severe      ATAAATGGGATCTCATTATTAGTGATATGTACGACCCTAAGACTAAAAATGTTACAAAAG   21082
gb:MN996529|Organism:Severe      ATAAATGGGATCTCATTATTAGTGATATGTACGACCCTAAGACTAAAAATGTTACAAAAG   21070
gb:MN996531|Organism:Severe      ATAAATGGGATCTCATTATTAGTGATATGTACGACCCTAAGACTAAAAATGTTACAAAAG   21069
gb:MT066176|Organism:Severe      ATAAATGGGATCTCATTATTAGTGATATGTACGACCCTAAGACTAAAAATGTTACAAAAG   21082
gb:MT027064|Organism:Severe      ATAAATGGGATCTCATTATTAGTGATATGTACGACCCTAAGACTAAAAATGTTACAAAAG   21082
gb:MN994468|Organism:Severe      ATAAATGGGATCTCATTATTAGTGATATGTACGACCCTAAGACTAAAAATGTTACAAAAG   21082
gb:MT072688|Organism:Severe      ATAAATGGGATCTCATTATTAGTGATATGTACGACCCTAAGACTAAAAATGTTACAAAAG   21067
```

```
gb:MN996527|Organism:Severe      ATAAATGGGATCTCATTATTAGTGATATGTACGACCCTAAGACTAAAAATGTTACAAAAG   21049
gb:MT093631|Organism:Severe      ATAAATGGGATCTCATTATTAGTGATATGTACGACCCTAAGACTAAAAATGTTACAAAAG   21120
gb:MT106053|Organism:Severe      ATAAATGGGATCTCATTATTAGTGATATGTACGACCCTAAGACTAAAAATGTTACAAAAG   21082
gb:MT019533|Organism:Severe      ATAAATGGGATCTCATTATTAGTGATATGTACGACCCTAAGACTAAAAATGTTACAAAAG   21082
gb:MT019531|Organism:Severe      ATAAATGGGATCTCATTATTAGTGATATGTACGACCCTAAGACTAAAAATGTTACAAAAG   21082
gb:MN996528|Organism:Severe      ATAAATGGGATCTCATTATTAGTGATATGTACGACCCTAAGACTAAAAATGTTACAAAAG   21082
gb:MN996530|Organism:Severe      ATAAATGGGATCTCATTATTAGTGATATGTACGACCCTAAGACTAAAAATGTTACAAAAG   21068
gb:MN908947|Organism:Severe      ATAAATGGGATCTCATTATTAGTGATATGTACGACCCTAAGACTAAAAATGTTACAAAAG   21082
gb:MT019532|Organism:Severe      ATAAATGGGATCTCATTATTAGTGATATGTACGACCCTAAGACTAAAAATGTTACAAAAG   21082
                                 ************************************************************

gb:MT020781|Organism:Severe      AAAATGACTCTAAAGAGGGTTTTTTCACTTACATTTGTGGGTTTATACAACAAAAGCTAG   21130
gb:MT007544|Organism:Severe      AAAATGACTCTAAAGAGGGTTTTTTCACTTACATTTGTGGGTTTATACAACAAAAGCTAG   21142
gb:MN994467|Organism:Severe      AAAATGACTCTAAAGAGGGTTTTTTCACTTACATTTGTGGGTTTATACAACAAAAGCTAG   21142
gb:MT044257|Organism:Severe      AAAATGACTCTAAAGAGGGTTTTTTCACTTACATTTGTGGGTTTATACAACAAAAGCTAG   21142
gb:MT106054|Organism:Severe      AAAATGACTCTAAAGAGGGTTTTTTCACTTACATTTGTGGGTTTATACAACAAAAGCTAG   21142
gb:MT049951|Organism:Severe      AAAATGACTCTAAAGAGGGTTTTTTCACTTACATTTGTGGGTTTATACAACAAAAGCTAG   21142
gb:MN975262|Organism:Severe      AAAATGACTCTAAAGAGGGTTTTTTCACTTACATTTGTGGGTTTATACAACAAAAGCTAG   21142
gb:MT106052|Organism:Severe      AAAATGACTCTAAAGAGGGTTTTTTCACTTACATTTGTGGGTTTATACAACAAAAGCTAG   21142
gb:LC522975|Organism:Severe      AAAATGACTCTAAAGAGGGTTTTTTCACTTACATTTGTGGGTTTATACAACAAAAGCTAG   21139
gb:LC522973|Organism:Severe      AAAATGACTCTAAAGAGGGTTTTTTCACTTACATTTGTGGGTTTATACAACAAAAGCTAG   21139
gb:LC522974|Organism:Severe      AAAATGACTCTAAAGAGGGTTTTTTCACTTACATTTGTGGGTTTATACAACAAAAGCTAG   21139
gb:MN985325|Organism:Severe      AAAATGACTCTAAAGAGGGTTTTTTCACTTACATTTGTGGGTTTATACAACAAAAGCTAG   21142
gb:MT020881|Organism:Severe      AAAATGACTCTAAAGAGGGTTTTTTCACTTACATTTGTGGGTTTATACAACAAAAGCTAG   21142
gb:MT020880|Organism:Severe      AAAATGACTCTAAAGAGGGTTTTTTCACTTACATTTGTGGGTTTATACAACAAAAGCTAG   21142
gb:MT066175|Organism:Severe      AAAATGACTCTAAAGAGGGTTTTTTCACTTACATTTGTGGGTTTATACAACAAAAGCTAG   21142
gb:MN997409|Organism:Severe      AAAATGACTCTAAAGAGGGTTTTTTCACTTACATTTGTGGGTTTATACAACAAAAGCTAG   21142
gb:MN938384|Organism:Severe      AAAATGACTCTAAAGAGGGTTTTTTCACTTACATTTGTGGGTTTATACAACAAAAGCTAG   21110
gb:MT044258|Organism:Severe      AAAATGACTCTAAAGAGGGTTTTTTCACTTACATTTGTGGGTTTATACAACAAAAGCTAG   21118
gb:MT039890|Organism:Severe      AAAATGACTCTAAAGAGGGTTTTTTCACTTACATTTGTGGGTTTATACAACAAAAGCTAG   21142
gb:MN988713|Organism:Severe      AAAATGACTCTAAAGAGGGTTTTTTCACTTACATTTGTGGGTTTATACAACAAAAGCTAG   21142
gb:LC521925|Organism:Severe      AAAATGACTCTAAAGAGGGTTTTTTCACTTACATTTGTGGGTTTATACAACAAAAGCTAG   21115
gb:MT093571|Organism:Severe      AAAATGACTCTAAAGAGGGTTTTTTCACTTACATTTGTGGGTTTATACAACAAAAGCTAG   21142
gb:MT039887|Organism:Severe      AAAATGACTCTAAAGAGGGTTTTTTCACTTACATTTGTGGGTTTATACAACAAAAGCTAG   21139
gb:MT019530|Organism:Severe      AAAATGACTCTAAAGAGGGTTTTTTCACTTACATTTGTGGGTTTATACAACAAAAGCTAG   21142
gb:MT039888|Organism:Severe      AAAATGACTCTAAAGAGGGTTTTTTCACTTACATTTGTGGGTTTATACAACAAAAGCTAG   21142
gb:LC522972|Organism:Severe      AAAATGACTCTAAAGAGGGTTTTTTCACTTACATTTGTGGGTTTATACAACAAAAGCTAG   21139
gb:MT027063|Organism:Severe      AAAATGACTCTAAAGAGGGTTTTTTCACTTACATTTGTGGGTTTATACAACAAAAGCTAG   21142
gb:MT027062|Organism:Severe      AAAATGACTCTAAAGAGGGTTTTTTCACTTACATTTGTGGGTTTATACAACAAAAGCTAG   21142
gb:MT019529|Organism:Severe      AAAATGACTCTAAAGAGGGTTTTTTCACTTACATTTGTGGGTTTATACAACAAAAGCTAG   21142
gb:MN996529|Organism:Severe      AAAATGACTCTAAAGAGGGTTTTTTCACTTACATTTGTGGGTTTATACAACAAAGGCTAG   21130
gb:MN996531|Organism:Severe      AAAATGACTCTAAAGAGGGTTTTTTCACTTACATTTGTGGGTTTATACAACAAAAGCTAG   21129
gb:MT066176|Organism:Severe      AAAATGACTCTAAAGAGGGTTTTTTCACTTACATTTGTGGGTTTATACAACAAAAGCTAG   21142
gb:MT027064|Organism:Severe      AAAATGACTCTAAAGAGGGTTTTTTCACTTACATTTGTGGGTTTATACAACAAAAGCTAG   21142
gb:MN994468|Organism:Severe      AAAATGACTCTAAAGAGGGTTTTTTCACTTACATTTGTGGGTTTATACAACAAAAGCTAG   21142
gb:MT072688|Organism:Severe      AAAATGACTCTAAAGAGGGTTTTTTCACTTACATTTGTGGGTTTATACAACAAAAGCTAG   21127
gb:MN996527|Organism:Severe      AAAATGACTCTAAAGAGGGTTTTTTCACTTACATTTGTGGGTTTATACAACAAAAGCTAG   21109
gb:MT093631|Organism:Severe      AAAATGACTCTAAAGAGGGTTTTTTCACTTACATTTGTGGGTTTATACAACAAAAGCTAG   21180
gb:MT106053|Organism:Severe      AAAATGACTCTAAAGAGGGTTTTTTCACTTACATTTGTGGGTTTATACAACAAAAGCTAG   21142
gb:MT019533|Organism:Severe      AAAATGACTCTAAAGAGGGTTTTTTCACTTACATTTGTGGGTTTATACAACAAAAGCTAG   21142
gb:MT019531|Organism:Severe      AAAATGACTCTAAAGAGGGTTTTTTCACTTACATTTGTGGGTTTATACAACAAAAGCTAG   21142
gb:MN996528|Organism:Severe      AAAATGACTCTAAAGAGGGTTTTTTCACTTACATTTGTGGGTTTATACAACAAAAGCTAG   21142
gb:MN996530|Organism:Severe      AAAATGACTCTAAAGAGGGTTTTTTCACTTACATTTGTGGGTTTATACAACAAAAGCTAG   21128
gb:MN908947|Organism:Severe      AAAATGACTCTAAAGAGGGTTTTTTCACTTACATTTGTGGGTTTATACAACAAAAGCTAG   21142
gb:MT019532|Organism:Severe      AAAATGACTCTAAAGAGGGTTTTTTCACTTACATTTGTGGGTTTATACAACAAAAGCTAG   21142
                                 ****************************************************** *****

gb:MT020781|Organism:Severe      CTCTTGGAGGTTCCGTGGCTATAAAGATAACAGAACATTCTTGGAATGCTGATCTTTATA   21190
gb:MT007544|Organism:Severe      CTCTTGGAGGTTCCGTGGCTATAAAGATAACAGAACATTCTTGGAATGCTGATCTTTATA   21202
gb:MN994467|Organism:Severe      CTCTTGGAGGTTCCGTGGCTATAAAGATAACAGAACATTCTTGGAATGCTGATCTTTATA   21202
gb:MT044257|Organism:Severe      CTCTTGGAGGTTCCGTGGCTATAAAGATAACAGAACATTCTTGGAATGCTGATCTTTATA   21202
gb:MT106054|Organism:Severe      CTCTTGGAGGTTCCGTGGCTATAAAGATAACAGAACATTCTTGGAATGCTGATCTTTATA   21202
gb:MT049951|Organism:Severe      CTCTTGGAGGTTCCGTGGCTATAAAGATAACAGAACATTCTTGGAATGCTGATCTTTATA   21202
gb:MN975262|Organism:Severe      CTCTTGGAGGTTCCGTGGCTATAAAGATAACAGAACATTCTTGGAATGCTGATCTTTATA   21202
gb:MT106052|Organism:Severe      CTCTTGGAGGTTCCGTGGCTATAAAGATAACAGAACATTCTTGGAATGCTGATCTTTATA   21202
```

```
gb:LC522975|Organism:Severe      CTCTTGGAGGTTCCGTGGCTATAAAGATAACAGAACATTCTTGGAATGCTGATCTTTATA   21199
gb:LC522973|Organism:Severe      CTCTTGGAGGTTCCGTGGCTATAAAGATAACAGAACATTCTTGGAATGCTGATCTTTATA   21199
gb:LC522974|Organism:Severe      CTCTTGGAGGTTCCGTGGCTATAAAGATAACAGAACATTCTTGGAATGCTGATCTTTATA   21199
gb:MN985325|Organism:Severe      CTCTTGGAGGTTCCGTGGCTATAAAGATAACAGAACATTCTTGGAATGCTGATCTTTATA   21202
gb:MT020881|Organism:Severe      CTCTTGGAGGTTCCGTGGCTATAAAGATAACAGAACATTCTTGGAATGCTGATCTTTATA   21202
gb:MT020880|Organism:Severe      CTCTTGGAGGTTCCGTGGCTATAAAGATAACAGAACATTCTTGGAATGCTGATCTTTATA   21202
gb:MT066175|Organism:Severe      CTCTTGGAGGTTCCGTGGCTATAAAGATAACAGAACATTCTTGGAATGCTGATCTTTATA   21202
gb:MN997409|Organism:Severe      CTCTTGGAGGTTCCGTGGCTATAAAGATAACAGAACATTCTTGGAATGCTGATCTTTATA   21202
gb:MN938384|Organism:Severe      CTCTTGGAGGTTCCGTGGCTATAAAGATAACAGAACATTCTTGGAATGCTGATCTTTATA   21170
gb:MT044258|Organism:Severe      CTCTTGGAGGTTCCGTGGCTATAAAGATAACAGAACATTCTTGGAATGCTGATCTTTATA   21178
gb:MT039890|Organism:Severe      CTCTTGGAGGTTCCGTGGCTATAAAGATAACAGAACATTCTTGGAATGCTGATCTTTATA   21202
gb:MN988713|Organism:Severe      CTCTTGGAGGTTCCGTGGCTATAAAGATAACAGAACATTCTTGGAATGCTGATCTTTATA   21202
gb:LC521925|Organism:Severe      CTCTTGGAGGTTCCGTGGCTATAAAGATAACAGAACATTCTTGGAATGCTGATCTTTATA   21175
gb:MT093571|Organism:Severe      CTCTTGGAGGTTCCGTGGCTATAAAGATAACAGAACATTCTTGGAATGCTGATCTTTATA   21202
gb:MT039887|Organism:Severe      CTCTTGGAGGTTCCGTGGCTATAAAGATAACAGAACATTCTTGGAATGCTGATCTTTATA   21199
gb:MT019530|Organism:Severe      CTCTTGGAGGTTCCGTGGCTATAAAGATAACAGAACATTCTTGGAATGCTGATCTTTATA   21202
gb:MT039888|Organism:Severe      CTCTTGGAGGTTCCGTGGCTATAAAGATAACAGAACATTCTTGGAATGCTGATCTTTATA   21202
gb:LC522972|Organism:Severe      CTCTTGGAGGTTCCGTGGCTATAAAGATAACAGAACATTCTTGGAATGCTGATCTTTATA   21199
gb:MT027063|Organism:Severe      CTCTTGGAGGTTCCGTGGCTATAAAGATAACAGAACATTCTTGGAATGCTGATCTTTATA   21202
gb:MT027062|Organism:Severe      CTCTTGGAGGTTCCGTGGCTATAAAGATAACAGAACATTCTTGGAATGCTGATCTTTATA   21202
gb:MT019529|Organism:Severe      CTCTTGGAGGTTCCGTGGCTATAAAGATAACAGAACATTCTTGGAATGCTGATCTTTATA   21202
gb:MN996529|Organism:Severe      CTCTTGGAGGTTCCGTGGCTATAAAGATAACAGAACATTCTTGGAATGCTGATCTTTATA   21190
gb:MN996531|Organism:Severe      CTCTTGGAGGTTCCGTGGCTATAAAGATAACAGAACATTCTTGGAATGCTGATCTTTATA   21189
gb:MT066176|Organism:Severe      CTCTTGGAGGTTCCGTGGCTATAAAGATAACAGAACATTCTTGGAATGCTGATCTTTATA   21202
gb:MT027064|Organism:Severe      CTCTTGGAGGTTCCGTGGCTATAAAGATAACAGAACATTCTTGGAATGCTGATCTTTATA   21202
gb:MN994468|Organism:Severe      CTCTTGGAGGTTCCGTGGCTATAAAGATAACAGAACATTCTTGGAATGCTGATCTTTATA   21202
gb:MT072688|Organism:Severe      CTCTTGGAGGTTCCGTGGCTATAAAGATAACAGAACATTCTTGGAATGCTGATCTTTATA   21187
gb:MN996527|Organism:Severe      CTCTTGGAGGTTCCGTGGCTATAAAGATAACAGAACATTCTTGGAATGCTGATCTTTATA   21169
gb:MT093631|Organism:Severe      CTCTTGGAGGTTCCGTGGCTATAAAGATAACAGAACATTCTTGGAATGCTGATCTTTATA   21240
gb:MT106053|Organism:Severe      CTCTTGGAGGTTCCGTGGCTATAAAGATAACAGAACATTCTTGGAATGCTGATCTTTATA   21202
gb:MT019533|Organism:Severe      CTCTTGGAGGTTCCGTGGCTATAAAGATAACAGAACATTCTTGGAATGCTGATCTTTATA   21202
gb:MT019531|Organism:Severe      CTCTTGGAGGTTCCGTGGCTATAAAGATAACAGAACATTCTTGGAATGCTGATCTTTATA   21202
gb:MN996528|Organism:Severe      CTCTTGGAGGTTCCGTGGCTATAAAGATAACAGAACATTCTTGGAATGCTGATCTTTATA   21202
gb:MN996530|Organism:Severe      CTCTTGGAGGTTCCGTGGCTATAAAGATAACAGAACATTCTTGGAATGCTGATCTTTATA   21188
gb:MN908947|Organism:Severe      CTCTTGGAGGTTCCGTGGCTATAAAGATAACAGAACATTCTTGGAATGCTGATCTTTATA   21202
gb:MT019532|Organism:Severe      CTCTTGGAGGTTCCGTGGCTATAAAGATAACAGAACATTCTTGGAATGCTGATCTTTATA   21202
                                 ************************************************************

gb:MT020781|Organism:Severe      AGCTCATGGGACACTTCGCATGGTGGACAGCCTTTGTTACTAATGTGAATGCGTCATCAT   21250
gb:MT007544|Organism:Severe      AGCTCATGGGACACTTCGCATGGTGGACAGCCTTTGTTACTAATGTGAATGCGTCATCAT   21262
gb:MN994467|Organism:Severe      AGCTCATGGGACACTTCGCATGGTGGACAGCCTTTGTTACTAATGTGAATGCGTCATCAT   21262
gb:MT044257|Organism:Severe      AGCTCATGGGACACTTCGCATGGTGGACAGCCTTTGTTACTAATGTGAATGCGTCATCAT   21262
gb:MT106054|Organism:Severe      AGCTCATGGGACACTTCGCATGGTGGACAGCCTTTGTTACTAATGTGAATGCGTCATCAT   21262
gb:MT049951|Organism:Severe      AGCTCATGGGACACTTCGCATGGTGGACAGCCTTTGTTACTAATGTGAATGCGTCATCAT   21262
gb:MN975262|Organism:Severe      AGCTCATGGGACACTTCGCATGGTGGACAGCCTTTGTTACTAATGTGAATGCGTCATCAT   21262
gb:MT106052|Organism:Severe      AGCTCATGGGACACTTCGCATGGTGGACAGCCTTTGTTACTAATGTGAATGCGTCATCAT   21262
gb:LC522975|Organism:Severe      AGCTCATGGGACACTTCGCATGGTGGACAGCCTTTGTTACTAATGTGAATGCGTCATCAT   21259
gb:LC522973|Organism:Severe      AGCTCATGGGACACTTCGCATGGTGGACAGCCTTTGTTACTAATGTGAATGCGTCATCAT   21259
gb:LC522974|Organism:Severe      AGCTCATGGGACACTTCGCATGGTGGACAGCCTTTGTTACTAATGTGAATGCGTCATCAT   21259
gb:MN985325|Organism:Severe      AGCTCATGGGACACTTCGCATGGTGGACAGCCTTTGTTACTAATGTGAATGCGTCATCAT   21262
gb:MT020881|Organism:Severe      AGCTCATGGGACACTTCGCATGGTGGACAGCCTTTGTTACTAATGTGAATGCGTCATCAT   21262
gb:MT020880|Organism:Severe      AGCTCATGGGACACTTCGCATGGTGGACAGCCTTTGTTACTAATGTGAATGCGTCATCAT   21262
gb:MT066175|Organism:Severe      AGCTCATGGGACACTTCGCATGGTGGACAGCCTTTGTTACTAATGTGAATGCGTCATCAT   21262
gb:MN997409|Organism:Severe      AGCTCATGGGACACTTCGCATGGTGGACAGCCTTTGTTACTAATGTGAATGCGTCATCAT   21262
gb:MN938384|Organism:Severe      AGCTCATGGGACACTTCGCATGGTGGACAGCCTTTGTTACTAATGTGAATGCGTCATCAT   21230
gb:MT044258|Organism:Severe      AGCTCATGGGACACTTCGCATGGTGGACAGCCTTTGTTACTAATGTGAATGCGTCATCAT   21238
gb:MT039890|Organism:Severe      AGCTCATGGGACACTTCGCATGGTGGACAGCCTTTGTTACTAATGTGAATGCGTCATCAT   21262
gb:MN988713|Organism:Severe      AGCTCATGGGACACTTCGCATGGTGGACAGCCTTTGTTACTAATGTGAATGCGTCATCAT   21262
gb:LC521925|Organism:Severe      AGCTCATGGGACACTTCGCATGGTGGACAGCCTTTGTTACTAATGTGAATGCGTCATCAT   21235
gb:MT093571|Organism:Severe      AGCTCATGGGACACTTCGCATGGTGGACAGCCTTTGTTACTAATGTGAATGCGTCATCAT   21262
gb:MT039887|Organism:Severe      AGCTCATGGGACACTTCGCATGGTGGACAGCCTTTGTTACTAATGTGAATGCGTCATCAT   21259
gb:MT019530|Organism:Severe      AGCTCATGGGACACTTCGCATGGTGGACAGCCTTTGTTACTAATGTGAATGCGTCATCAT   21262
gb:MT039888|Organism:Severe      AGCTCATGGGACACTTCGCATGGTGGACAGCCTTTGTTACTAATGTGAATGCGTCATCAT   21262
gb:LC522972|Organism:Severe      AGCTCATGGGACACTTCGCATGGTGGACAGCCTTTGTTACTAATGTGAATGCGTCATCAT   21259
gb:MT027063|Organism:Severe      AGCTCATGGGACACTTCGCATGGTGGACAGCCTTTGTTACTAATGTGAATGCGTCATCAT   21262
```

```
gb:MT027062|Organism:Severe      AGCTCATGGGACACTTCGCATGGTGGACAGCCTTTGTTACTAATGTGAATGCGTCATCAT   21262
gb:MT019529|Organism:Severe      AGCTCATGGGACACTTCGCATGGTGGACAGCCTTTGTTACTAATGTGAATGCGTCATCAT   21262
gb:MN996529|Organism:Severe      AGCTCATGGGACACTTCGCATGGTGGACAGCCTTTGTTACTAATGTGAATGCGTCATCAT   21250
gb:MN996531|Organism:Severe      AGCTCATGGGACACTTCGCATGGTGGACAGCCTTTGTTACTAATGTGAATGCGTCATCAT   21249
gb:MT066176|Organism:Severe      AGCTCATGGGACACTTCGCATGGTGGACAGCCTTTGTTACTAATGTGAATGCGTCATCAT   21262
gb:MT027064|Organism:Severe      AGCTCATGGGACACTTCGCATGGTGGACAGCCTTTGTTACTAATGTGAATGCGTCATCAT   21262
gb:MN994468|Organism:Severe      AGCTCATGGGACACTTCGCATGGTGGACAGCCTTTGTTACTAATGTGAATGCGTCATCAT   21262
gb:MT072688|Organism:Severe      AGCTCATGGGACACTTCGCATGGTGGACAGCCTTTGTTACTAATGTGAATGCGTCATCAT   21247
gb:MN996527|Organism:Severe      AGCTCATGGGACACTTCGCATGGTGGACAGCCTTTGTTACTAATGTGAATGCGTCATCAT   21229
gb:MT093631|Organism:Severe      AGCTCATGGGACACTTCGCATGGTGGACAGCCTTTGTTACTAATGTGAATGCGTCATCAT   21300
gb:MT106053|Organism:Severe      AGCTCATGGGACACTTCGCATGGTGGACAGCCTTTGTTACTAATGTGAATGCGTCATCAT   21262
gb:MT019533|Organism:Severe      AGCTCATGGGACACTTCGCATGGTGGACAGCCTTTGTTACTAATGTGAATGCGTCATCAT   21262
gb:MT019531|Organism:Severe      AGCTCATGGGACACTTCGCATGGTGGACAGCCTTTGTTACTAATGTGAATGCGTCATCAT   21262
gb:MN996528|Organism:Severe      AGCTCATGGGACACTTCGCATGGTGGACAGCCTTTGTTACTAATGTGAATGCGTCATCAT   21262
gb:MN996530|Organism:Severe      AGCTCATGGGACACTTCGCATGGTGGACAGCCTTTGTTACTAATGTGAATGCGTCATCAT   21248
gb:MN908947|Organism:Severe      AGCTCATGGGACACTTCGCATGGTGGACAGCCTTTGTTACTAATGTGAATGCGTCATCAT   21262
gb:MT019532|Organism:Severe      AGCTCATGGGACACTTCGCATGGTGGACAGCCTTTGTTACTAATGTGAATGCGTCATCAT   21262
                                 ************************************************************

gb:MT020781|Organism:Severe      CTGAAGCATTTTTAATTGGATGTAATTATCTTGGCAAACCACGCGAACAAATAGATGGTT   21310
gb:MT007544|Organism:Severe      CTGAAGCATTTTTAATTGGATGTAATTATCTTGGCAAACCACGCGAACAAATAGATGGTT   21322
gb:MN994467|Organism:Severe      CTGAAGCATTTTTAATTGGATGTAATTATCTTGGCAAACCACGCGAACAAATAGATGGTT   21322
gb:MT044257|Organism:Severe      CTGAAGCATTTTTAATTGGATGTAATTATCTTGGCAAACCACGCGAACAAATAGATGGTT   21322
gb:MT106054|Organism:Severe      CTGAAGCATTTTTAATTGGATGTAATTATCTTGGCAAACCACGCGAACAAATAGATGGTT   21322
gb:MT049951|Organism:Severe      CTGAAGCATTTTTAATTGGATGTAATTATCTTGGCAAACCACGCGAACAAATAGATGGTT   21322
gb:MN975262|Organism:Severe      CTGAAGCATTTTTAATTGGATGTAATTATCTTGGCAAACCACGCGAACAAATAGATGGTT   21322
gb:MT106052|Organism:Severe      CTGAAGCATTTTTAATTGGATGTAATTATCTTGGCAAACCACGCGAACAAATAGATGGTT   21322
gb:LC522975|Organism:Severe      CTGAAGCATTTTTAATTGGATGTAATTATCTTGGCAAACCACGCGAACAAATAGATGGTT   21319
gb:LC522973|Organism:Severe      CTGAAGCATTTTTAATTGGATGTAATTATCTTGGCAAACCACGCGAACAAATAGATGGTT   21319
gb:LC522974|Organism:Severe      CTGAAGCATTTTTAATTGGATGTAATTATCTTGGCAAACCACGCGAACAAATAGATGGTT   21319
gb:MN985325|Organism:Severe      CTGAAGCATTTTTAATTGGATGTAATTATCTTGGCAAACCACGCGAACAAATAGATGGTT   21322
gb:MT020881|Organism:Severe      CTGAAGCATTTTTAATTGGATGTAATTATCTTGGCAAACCACGCGAACAAATAGATGGTT   21322
gb:MT020880|Organism:Severe      CTGAAGCATTTTTAATTGGATGTAATTATCTTGGCAAACCACGCGAACAAATAGATGGTT   21322
gb:MT066175|Organism:Severe      CTGAAGCATTTTTAATTGGATGTAATTATCTTGGCAAACCACGCGAACAAATAGATGGTT   21322
gb:MN997409|Organism:Severe      CTGAAGCATTTTTAATTGGATGTAATTATCTTGGCAAACCACGCGAACAAATAGATGGTT   21322
gb:MN938384|Organism:Severe      CTGAAGCATTTTTAATTGGATGTAATTATCTTGGCAAACCACGCGAACAAATAGATGGTT   21290
gb:MT044258|Organism:Severe      CTGAAGCATTTTTAATTGGATGTAATTATCTTGGCAAACCACGCGAACAAATAGATGGTT   21298
gb:MT039890|Organism:Severe      CTGAAGCATTTTTAATTGGATGTAATTATCTTGGCAAACCACGCGAACAAATAGATGGTT   21322
gb:MN988713|Organism:Severe      CTGAAGCATTTTTAATTGGATGTAATTATCTTGGCAAACCACGCGAACAAATAGATGGTT   21322
gb:LC521925|Organism:Severe      CTGAAGCATTTTTAATTGGATGTAATTATCTTGGCAAACCACGCGAACAAATAGATGGTT   21295
gb:MT093571|Organism:Severe      CTGAAGCATTTTTAATTGGATGTAATTATCTTGGCAAACCACGCGAACAAATAGATGGTT   21322
gb:MT039887|Organism:Severe      CTGAAGCATTTTTAATTGGATGTAATTATCTTGGCAAACCACGCGAACAAATAGATGGTT   21319
gb:MT019530|Organism:Severe      CTGAAGCATTTTTAATTGGATGTAATTATCTTGGCAAACCACGCGAACAAATAGATGGTT   21322
gb:MT039888|Organism:Severe      CTGAAGCATTTTTAATTGGATGTAATTATCTTGGCAAACCACGCGAACAAATAGATGGTT   21322
gb:LC522972|Organism:Severe      CTGAAGCATTTTTAATTGGATGTAATTATCTTGGCAAACCACGCGAACAAATAGATGGTT   21319
gb:MT027063|Organism:Severe      CTGAAGCATTTTTAATTGGATGTAATTATCTTGGCAAACCACGCGAACAAATAGATGGTT   21322
gb:MT027062|Organism:Severe      CTGAAGCATTTTTAATTGGATGTAATTATCTTGGCAAACCACGCGAACAAATAGATGGTT   21322
gb:MT019529|Organism:Severe      CTGAAGCATTTTTAATTGGATGTAATTATCTTGGCAAACCACGCGAACAAATAGATGGTT   21322
gb:MN996529|Organism:Severe      CTGAAGCATTTTTAATTGGATGTAATTATCTTGGCAAACCACGCGAACAAATAGATGGTT   21310
gb:MN996531|Organism:Severe      CTGAAGCATTTTTAATTGGATGTAATTATCTTGGCAAACCACGCGAACAAATAGATGGTT   21309
gb:MT066176|Organism:Severe      CTGAAGCATTTTTAATTGGATGTAATTATCTTGGCAAACCACGCGAACAAATAGATGGTT   21322
gb:MT027064|Organism:Severe      CTGAAGCATTTTTAATTGGATGTAATTATCTTGGCAAACCACGCGAACAAATAGATGGTT   21322
gb:MN994468|Organism:Severe      CTGAAGCATTTTTAATTGGATGTAATTATCTTGGCAAACCACGCGAACAAATAGATGGTT   21322
gb:MT072688|Organism:Severe      CTGAAGCATTTTTAATTGGATGTAATTATCTTGGCAAACCACGCGAACAAATAGATGGTT   21307
gb:MN996527|Organism:Severe      CTGAAGCATTTTTAATTGGATGTAATTATCTTGGCAAACCACGCGAACAAATAAATGGTT   21289
gb:MT093631|Organism:Severe      CTGAAGCATTTTTAATTGGATGTAATTATCTTGGCAAACCACGCGAACAAATAGATGGTT   21360
gb:MT106053|Organism:Severe      CTGAAGCATTTTTAATTGGATGTAATTATCTTGGCAAACCACGCGAACAAATAGATGGTT   21322
gb:MT019533|Organism:Severe      CTGAAGCATTTTTAATTGGATGTAATTATCTTGGCAAACCACGCGAACAAATAGATGGTT   21322
gb:MT019531|Organism:Severe      CTGAAGCATTTTTAATTGGATGTAATTATCTTGGCAAACCACGCGAACAAATAGATGGTT   21322
gb:MN996528|Organism:Severe      CTGAAGCATTTTTAATTGGATGTAATTATCTTGGCAAACCACGCGAACAAATAGATGGTT   21322
gb:MN996530|Organism:Severe      CTGAAGCATTTTTAATTGGATGTAATTATCTTGGCAAACCACGCGAACAAATAGATGGTT   21308
gb:MN908947|Organism:Severe      CTGAAGCATTTTTAATTGGATGTAATTATCTTGGCAAACCACGCGAACAAATAGATGGTT   21322
gb:MT019532|Organism:Severe      CTGAAGCATTTTTAATTGGATGTAATTATCTTGGCAAACCACGCGAACAAATAGATGGTT   21322
                                 ***************************************************** ******
```

```
gb:MT020781|Organism:Severe      ATGTCATGCATGCAAATTACATATTTTGGAGGAATACAAATCCAATTCAGTTGTCTTCCT   21370
gb:MT007544|Organism:Severe      ATGTCATGCATGCAAATTACATATTTTGGAGGAATACAAATCCAATTCAGTTGTCTTCCT   21382
gb:MN994467|Organism:Severe      ATGTCATGCATGCAAATTACATATTTTGGAGGAATACAAATCCAATTCAGTTGTCTTCCT   21382
gb:MT044257|Organism:Severe      ATGTCATGCATGCAAATTACATATTTTGGAGGAATACAAATCCAATTCAGTTGTCTTCCT   21382
gb:MT106054|Organism:Severe      ATGTCATGCATGCAAATTACATATTTTGGAGGAATACAAATCCAATTCAGTTGTCTTCCT   21382
gb:MT049951|Organism:Severe      ATGTCATGCATGCAAATTACATATTTTGGAGGAATACAAATCCAATTCAGTTGTCTTCCT   21382
gb:MN975262|Organism:Severe      ATGTCATGCATGCAAATTACATATTTTGGAGGAATACAAATCCAATTCAGTTGTCTTCCT   21382
gb:MT106052|Organism:Severe      ATGTCATGCATGCAAATTACATATTTTGGAGGAATACAAATCCAATTCAGTTGTCTTCCT   21382
gb:LC522975|Organism:Severe      ATGTCATGCATGCAAATTACATATTTTGGAGGAATACAAATCCAATTCAGTTGTCTTCCT   21379
gb:LC522973|Organism:Severe      ATGTCATGCATGCAAATTACATATTTTGGAGGAATACAAATCCAATTCAGTTGTCTTCCT   21379
gb:LC522974|Organism:Severe      ATGTCATGCATGCAAATTACATATTTTGGAGGAATACAAATCCAATTCAGTTGTCTTCCT   21379
gb:MN985325|Organism:Severe      ATGTCATGCATGCAAATTACATATTTTGGAGGAATACAAATCCAATTCAGTTGTCTTCCT   21382
gb:MT020881|Organism:Severe      ATGTCATGCATGCAAATTACATATTTTGGAGGAATACAAATCCAATTCAGTTGTCTTCCT   21382
gb:MT020880|Organism:Severe      ATGTCATGCATGCAAATTACATATTTTGGAGGAATACAAATCCAATTCAGTTGTCTTCCT   21382
gb:MT066175|Organism:Severe      ATGTCATGCATGCAAATTACATATTTTGGAGGAATACAAATCCAATTCAGTTGTCTTCCT   21382
gb:MN997409|Organism:Severe      ATGTCATGCATGCAAATTACATATTTTGGAGGAATACAAATCCAATTCAGTTGTCTTCCT   21382
gb:MN938384|Organism:Severe      ATGTCATGCATGCAAATTACATATTTTGGAGGAATACAAATCCAATTCAGTTGTCTTCCT   21350
gb:MT044258|Organism:Severe      ATGTCATGCATGCAAATTACATATTTTGGAGGAATACAAATCCAATTCAGTTGTCTTCCT   21358
gb:MT039890|Organism:Severe      ATGTCATGCATGCAAATTACATATTTTGGAGGAATACAAATCCAATTCAGTTGTCTTCCT   21382
gb:MN988713|Organism:Severe      ATGTCATGCATGCAAATTACATATTTTGGAGGAATACAAATCCAATTCAGTTGTCTTCCT   21382
gb:LC521925|Organism:Severe      ATGTCATGCATGCAAATTACATATTTTGGAGGAATACAAATCCAATTCAGTTGTCTTCCT   21355
gb:MT093571|Organism:Severe      ATGTCATGCATGCAAATTACATATTTTGGAGGAATACAAATCCAATTCAGTTGTCTTCCT   21382
gb:MT039887|Organism:Severe      ATGTCATGCATGCAAATTACATATTTTGGAGGAATACAAATCCAATTCAGTTGTCTTCCT   21379
gb:MT019530|Organism:Severe      ATGTCATGCATGCAAATTACATATTTTGGAGGAATACAAATCCAATTCAGTTGTCTTCCT   21382
gb:MT039888|Organism:Severe      ATGTCATGCATGCAAATTACATATTTTGGAGGAATACAAATCCAATTCAGTTGTCTTCCT   21382
gb:LC522972|Organism:Severe      ATGTCATGCATGCAAATTACATATTTTGGAGGAATACAAATCCAATTCAGTTGTCTTCCT   21379
gb:MT027063|Organism:Severe      ATGTCATGCATGCAAATTACATATTTTGGAGGAATACAAATCCAATTCAGTTGTCTTCCT   21382
gb:MT027062|Organism:Severe      ATGTCATGCATGCAAATTACATATTTTGGAGGAATACAAATCCAATTCAGTTGTCTTCCT   21382
gb:MT019529|Organism:Severe      ATGTCATGCATGCAAATTACATATTTTGGAGGAATACAAATCCAATTCAGTTGTCTTCCT   21382
gb:MN996529|Organism:Severe      ATGTCATGCATGCAAATTACATATTTTGGAGGAATACAAATCCAATTCAGTTGTCTTCCT   21370
gb:MN996531|Organism:Severe      ATGTCATGCATGCAAATTACATATTTTGGAGGAATACAAATCCAATTCAGTTGTCTTCCT   21369
gb:MT066176|Organism:Severe      ATGTCATGCATGCAAATTACATATTTTGGAGGAATACAAATCCAATTCAGTTGTCTTCCT   21382
gb:MT027064|Organism:Severe      ATGTCATGCATGCAAATTACATATTTTGGAGGAATACAAATCCAATTCAGTTGTCTTCCT   21382
gb:MN994468|Organism:Severe      ATGTCATGCATGCAAATTACATATTTTGGAGGAATACAAATCCAATTCAGTTGTCTTCCT   21382
gb:MT072688|Organism:Severe      ATGTCATGCATGCAAATTACATATTTTGGAGGAATACAAATCCAATTCAGTTGTCTTCCT   21367
gb:MN996527|Organism:Severe      ATGTCATGCATGCAAATTACATATTTTGGAGGAATACAAATCCAATTCAGTTGTCTTCCT   21349
gb:MT093631|Organism:Severe      ATGTCATGCATGCAAATTACATATTTTGGAGGAATACAAATCCAATTCAGTTGTCTTCCT   21420
gb:MT106053|Organism:Severe      ATGTCATGCATGCAAATTACATATTTTGGAGGAATACAAATCCAATTCAGTTGTCTTCCT   21382
gb:MT019533|Organism:Severe      ATGTCATGCATGCAAATTACATATTTTGGAGGAATACAAATCCAATTCAGTTGTCTTCCT   21382
gb:MT019531|Organism:Severe      ATGTCATGCATGCAAATTACATATTTTGGAGGAATACAAATCCAATTCAGTTGTCTTCCT   21382
gb:MN996528|Organism:Severe      ATGTCATGCATGCAAATTACATATTTTGGAGGAATACAAATCCAATTCAGTTGTCTTCCT   21382
gb:MN996530|Organism:Severe      ATGTCATGCATGCAAATTACATATTTTGGAGGAATACAAATCCAATTCAGTTGTCTTCCT   21368
gb:MN908947|Organism:Severe      ATGTCATGCATGCAAATTACATATTTTGGAGGAATACAAATCCAATTCAGTTGTCTTCCT   21382
gb:MT019532|Organism:Severe      ATGTCATGCATGCAAATTACATATTTTGGAGGAATACAAATCCAATTCAGTTGTCTTCCT   21382
                                 ************************************************************

gb:MT020781|Organism:Severe      ATTCTTTATTTGACATGAGTAAATTTCCCCTTAAATTAAGGGGTACTGCTGTTATGTCTT   21430
gb:MT007544|Organism:Severe      ATTCTTTATTTGACATGAGTAAATTTCCCCTTAAATTAAGGGGTACTGCTGTTATGTCTT   21442
gb:MN994467|Organism:Severe      ATTCTTTATTTGACATGAGTAAATTTCCCCTTAAATTAAGGGGTACTGCTGTTATGTCTT   21442
gb:MT044257|Organism:Severe      ATTCTTTATTTGACATGAGTAAATTTCCCCTTAAATTAAGGGGTACTGCTGTTATGTCTT   21442
gb:MT106054|Organism:Severe      ATTCTTTATTTGACATGAGTAAATTTCCCCTTAAATTAAGGGGTACTGCTGTTATGTCTT   21442
gb:MT049951|Organism:Severe      ATTCTTTATTTGACATGAGTAAATTTCCCCTTAAATTAAGGGGTACTGCTGTTATGTCTT   21442
gb:MN975262|Organism:Severe      ATTCTTTATTTGACATGAGTAAATTTCCCCTTAAATTAAGGGGTACTGCTGTTATGTCTT   21442
gb:MT106052|Organism:Severe      ATTCTTTATTTGACATGAGTAAATTTCCCCTTAAATTAAGGGGTACTGCTGTTATGTCTT   21442
gb:LC522975|Organism:Severe      ATTCTTTATTTGACATGAGTAAATTTCCCCTTAAATTAAGGGGTACTGCTGTTATGTCTT   21439
gb:LC522973|Organism:Severe      ATTCTTTATTTGACATGAGTAAATTTCCCCTTAAATTAAGGGGTACTGCTGTTATGTCTT   21439
gb:LC522974|Organism:Severe      ATTCTTTATTTGACATGAGTAAATTTCCCCTTAAATTAAGGGGTACTGCTGTTATGTCTT   21439
gb:MN985325|Organism:Severe      ATTCTTTATTTGACATGAGTAAATTTCCCCTTAAATTAAGGGGTACTGCTGTTATGTCTT   21442
gb:MT020881|Organism:Severe      ATTCTTTATTTGACATGAGTAAATTTCCCCTTAAATTAAGGGGTACTGCTGTTATGTCTT   21442
gb:MT020880|Organism:Severe      ATTCTTTATTTGACATGAGTAAATTTCCCCTTAAATTAAGGGGTACTGCTGTTATGTCTT   21442
gb:MT066175|Organism:Severe      ATTCTTTATTTGACATGAGTAAATTTCCCCTTAAATTAAGGGGTACTGCTGTTATGTCTT   21442
gb:MN997409|Organism:Severe      ATTCTTTATTTGACATGAGTAAATTTCCCCTTAAATTAAGGGGTACTGCTGTTATGTCTT   21442
gb:MN938384|Organism:Severe      ATTCTTTATTTGACATGAGTAAATTTCCCCTTAAATTAAGGGGTACTGCTGTTATGTCTT   21410
gb:MT044258|Organism:Severe      ATTCTTTATTTGACATGAGTAAATTTCCCCTTAAATTAAGGGGTACTGCTGTTATGTCTT   21418
gb:MT039890|Organism:Severe      ATTCTTTATTTGACATGAGTAAATTTCCCCTTAAATTAAGGGGTACTGCTGTTATGTCTT   21442
```

```
gb:MN988713|Organism:Severe      ATTCTTTATTTGACATGAGTAAATTTCCCCTTAAATTAAGGGGTACTGCTGTTATGTCTT   21442
gb:LC521925|Organism:Severe      ATTCTTTATTTGACATGAGTAAATTTCCCCTTAAATTAAGGGGTACTGCTGTTATGTCTT   21415
gb:MT093571|Organism:Severe      ATTCTTTATTTGACATGAGTAAATTTCCCCTTAAATTAAGGGGTACTGCTGTTATGTCTT   21442
gb:MT039887|Organism:Severe      ATTCTTTATTTGACATGAGTAAATTTCCCCTTAAATTAAGGGGTACTGCTGTTATGTCTT   21439
gb:MT019530|Organism:Severe      ATTCTTTATTTGACATGAGTAAATTTCCCCTTAAATTAAGGGGTACTGCTGTTATGTCTT   21442
gb:MT039888|Organism:Severe      ATTCTTTATTTGACATGAGTAAATTTCCCCTTAAATTAAGGGGTACTGCTGTTATGTCTT   21442
gb:LC522972|Organism:Severe      ATTCTTTATTTGACATGAGTAAATTTCCCCTTAAATTAAGGGGTACTGCTGTTATGTCTT   21439
gb:MT027063|Organism:Severe      ATTCTTTATTTGACATGAGTAAATTTCCCCTTAAATTAAGGGGTACTGCTGTTATGTCTT   21442
gb:MT027062|Organism:Severe      ATTCTTTATTTGACATGAGTAAATTTCCCCTTAAATTAAGGGGTACTGCTGTTATGTCTT   21442
gb:MT019529|Organism:Severe      ATTCTTTATTTGACATGAGTAAATTTCCCCTTAAATTAAGGGGTACTGCTGTTATGTCTT   21442
gb:MN996529|Organism:Severe      ATTCTTTATTTGACATGAGTAAATTTCCCCTTAAATTAAGGGGTACTGCTGTTATGTCTT   21430
gb:MN996531|Organism:Severe      ATTCTTTATTTGACATGAGTAAATTTCCCCTTAAATTAAGGGGTACTGCTGTTATGTCTT   21429
gb:MT066176|Organism:Severe      ATTCTTTATTTGACATGAGTAAATTTCCCCTTAAATTAAGGGGTACTGCTGTTATGTCTT   21442
gb:MT027064|Organism:Severe      ATTCTTTATTTGACATGAGTAAATTTCCCCTTAAATTAAGGGGTACTGCTGTTATGTCTT   21442
gb:MN994468|Organism:Severe      ATTCTTTATTTGACATGAGTAAATTTCCCCTTAAATTAAGGGGTACTGCTGTTATGTCTT   21442
gb:MT072688|Organism:Severe      ATTCTTTATTTGACATGAGTAAATTTCCCCTTAAATTAAGGGGTACTGCTGTTATGTCTT   21427
gb:MN996527|Organism:Severe      ATTCTTTATTTGACATGAGTAAATTTCCCCTTAAATTAAGGGGTACTGCTGTTATGTCTT   21409
gb:MT093631|Organism:Severe      ATTCTTTATTTGACATGAGTAAATTTCCCCTTAAATTAAGGGGTACTGCTGTTATGTCTT   21480
gb:MT106053|Organism:Severe      ATTCTTTATTTGACATGAGTAAATTTCCCCTTAAATTAAGGGGTACTGCTGTTATGTCTT   21442
gb:MT019533|Organism:Severe      ATTCTTTATTTGACATGAGTAAATTTCCCCTTAAATTAAGGGGTACTGCTGTTATGTCTT   21442
gb:MT019531|Organism:Severe      ATTCTTTATTTGACATGAGTAAATTTCCCCTTAAATTAAGGGGTACTGCTGTTATGTCTT   21442
gb:MN996528|Organism:Severe      ATTCTTTATTTGACATGAGTAAATTTCCCCTTAAATTAAGGGGTACTGCTGTTATGTCTT   21442
gb:MN996530|Organism:Severe      ATTCTTTATTTGACATGAGTAAATTTCCCCTTAAATTAAGGGGTACTGCTGTTATGTCTT   21428
gb:MN908947|Organism:Severe      ATTCTTTATTTGACATGAGTAAATTTCCCCTTAAATTAAGGGGTACTGCTGTTATGTCTT   21442
gb:MT019532|Organism:Severe      ATTCTTTATTTGACATGAGTAAATTTCCCCTTAAATTAAGGGGTACTGCTGTTATGTCTT   21442
                                 ************************************************************

gb:MT020781|Organism:Severe      TAAAAGAAGGTCAAATCAATGATATGATTTTATCTCTTCTTAGTAAAGGTAGACTTATAA   21490
gb:MT007544|Organism:Severe      TAAAAGAAGGTCAAATCAATGATATGATTTTATCTCTTCTTAGTAAAGGTAGACTTATAA   21502
gb:MN994467|Organism:Severe      TAAAAGAAGGTCAAATCAATGATATGATTTTATCTCTTCTTAGTAAAGGTAGACTTATAA   21502
gb:MT044257|Organism:Severe      TAAAAGAAGGTCAAATCAATGATATGATTTTATCTCTTCTTAGTAAAGGTAGACTTATAA   21502
gb:MT106054|Organism:Severe      TAAAAGAAGGTCAAATCAATGATATGATTTTATCTCTTCTTAGTAAAGGTAGACTTATAA   21502
gb:MT049951|Organism:Severe      TAAAAGAAGGTCAAATCAATGATATGATTTTATCTCTTCTTAGTAAAGGTAGACTTATAA   21502
gb:MN975262|Organism:Severe      TAAAAGAAGGTCAAATCAATGATATGATTTTATCTCTTCTTAGTAAAGGTAGACTTATAA   21502
gb:MT106052|Organism:Severe      TAAAAGAAGGTCAAATCAATGATATGATTTTATCTCTTCTTAGTAAAGGTAGACTTATAA   21502
gb:LC522975|Organism:Severe      TAAAAGAAGGTCAAATCAATGATATGATTTTATCTCTTCTTAGTAAAGGTAGACTTATAA   21499
gb:LC522973|Organism:Severe      TAAAAGAAGGTCAAATCAATGATATGATTTTATCTCTTCTTAGTAAAGGTAGACTTATAA   21499
gb:LC522974|Organism:Severe      TAAAAGAAGGTCAAATCAATGATATGATTTTATCTCTTCTTAGTAAAGGTAGACTTATAA   21499
gb:MN985325|Organism:Severe      TAAAAGAAGGTCAAATCAATGATATGATTTTATCTCTTCTTAGTAAAGGTAGACTTATAA   21502
gb:MT020881|Organism:Severe      TAAAAGAAGGTCAAATCAATGATATGATTTTATCTCTTCTTAGTAAAGGTAGACTTATAA   21502
gb:MT020880|Organism:Severe      TAAAAGAAGGTCAAATCAATGATATGATTTTATCTCTTCTTAGTAAAGGTAGACTTATAA   21502
gb:MT066175|Organism:Severe      TAAAAGAAGGTCAAATCAATGATATGATTTTATCTCTTCTTAGTAAAGGTAGACTTATAA   21502
gb:MN997409|Organism:Severe      TAAAAGAAGGTCAAATCAATGATATGATTTTATCTCTTCTTAGTAAAGGTAGACTTATAA   21502
gb:MN938384|Organism:Severe      TAAAAGAAGGTCAAATCAATGATATGATTTTATCTCTTCTTAGTAAAGGTAGACTTATAA   21470
gb:MT044258|Organism:Severe      TAAAAGAAGGTCAAATCAATGATATGATTTTATCTCTTCTTAGTAAAGGTAGACTTATAA   21478
gb:MT039890|Organism:Severe      TAAAAGAAGGTCAAATCAATGATATGATTTTATCTCTTCTTAGTAAAGGTAGACTTATAA   21502
gb:MN988713|Organism:Severe      TAAAAGAAGGTCAAATCAATGATATGATTTTATCTCTTCTTAGTAAAGGTAGACTTATAA   21502
gb:LC521925|Organism:Severe      TAAAAGAAGGTCAAATCAATGATATGATTTTATCTCTTCTTAGTAAAGGTAGACTTATAA   21475
gb:MT093571|Organism:Severe      TAAAAGAAGGTCAAATCAATGATATGATTTTATCTCTTCTTAGTAAAGGTAGACTTATAA   21502
gb:MT039887|Organism:Severe      TAAAAGAAGGTCAAATCAATGATATGATTTTATCTCTTCTTAGTAAAGGTAGACTTATAA   21499
gb:MT019530|Organism:Severe      TAAAAGAAGGTCAAATCAATGATATGATTTTATCTCTTCTTAGTAAAGGTAGACTTATAA   21502
gb:MT039888|Organism:Severe      TAAAAGAAGGTCAAATCAATGATATGATTTTATCTCTTCTTAGTAAAGGTAGACTTATAA   21502
gb:LC522972|Organism:Severe      TAAAAGAAGGTCAAATCAATGATATGATTTTATCTCTTCTTAGTAAAGGTAGACTTATAA   21499
gb:MT027063|Organism:Severe      TAAAAGAAGGTCAAATCAATGATATGATTTTATCTCTTCTTAGTAAAGGTAGACTTATAA   21502
gb:MT027062|Organism:Severe      TAAAAGAAGGTCAAATCAATGATATGATTTTATCTCTTCTTAGTAAAGGTAGACTTATAA   21502
gb:MT019529|Organism:Severe      TAAAAGAAGGTCAAATCAATGATATGATTTTATCTCTTCTTAGTAAAGGTAGACTTATAA   21502
gb:MN996529|Organism:Severe      TAAAAGAAGGTCAAATCAATGATATGATTTTATCTCTTCTTAGTAAAGGTAGACTTATAA   21490
gb:MN996531|Organism:Severe      TAAAAGAAGGTCAAATCAATGATATGATTTTATCTCTTCTTAGTAAAGGTAGACTTATAA   21489
gb:MT066176|Organism:Severe      TAAAAGAAGGTCAAATCAATGATATGATTTTATCTCTTCTTAGTAAAGGTAGACTTATAA   21502
gb:MT027064|Organism:Severe      TAAAAGAAGGTCAAATCAATGATATGATTTTATCTCTTCTTAGTAAAGGTAGACTTATAA   21502
gb:MN994468|Organism:Severe      TAAAAGAAGGTCAAATCAATGATATGATTTTATCTCTTCTTAGTAAAGGTAGACTTATAA   21502
gb:MT072688|Organism:Severe      TAAAAGAAGGTCAAATCAATGATATGATTTTATCTCTTCTTAGTAAAGGTAGACTTATAA   21487
gb:MN996527|Organism:Severe      TAAAAGAAGGTCAAATCAATGATATGATTTTATCTCTTCTTAGTAAAGGTAGACTTATAA   21469
gb:MT093631|Organism:Severe      TAAAAGAAGGTCAAATCAATGATATGATTTTATCTCTTCTTAGTAAAGGTAGACTTATAA   21540
gb:MT106053|Organism:Severe      TAAAAGAAGGTCAAATCAATGATATGATTTTATCTCTTCTTAGTAAAGGTAGACTTATAA   21502
```

```
gb:MT019533|Organism:Severe      TAAAAGAAGGTCAAATCAATGATATGATTTTATCTCTTCTTAGTAAAGGTAGACTTATAA   21502
gb:MT019531|Organism:Severe      TAAAAGAAGGTCAAATCAATGATATGATTTTATCTCTTCTTAGTAAAGGTAGACTTATAA   21502
gb:MN996528|Organism:Severe      TAAAAGAAGGTCAAATCAATGATATGATTTTATCTCTTCTTAGTAAAGGTAGACTTATAA   21502
gb:MN996530|Organism:Severe      TAAAAGAAGGTCAAATCAATGATATGATTTTATCTCTTCTTAGTAAAGGTAGACTTATAA   21488
gb:MN908947|Organism:Severe      TAAAAGAAGGTCAAATCAATGATATGATTTTATCTCTTCTTAGTAAAGGTAGACTTATAA   21502
gb:MT019532|Organism:Severe      TAAAAGAAGGTCAAATCAATGATATGATTTTATCTCTTCTTAGTAAAGGTAGACTTATAA   21502
                                 ************************************************************

gb:MT020781|Organism:Severe      TTAGAGAAAACAACAGAGTTGTTATTTCTAGTGATGTTCTTGTTAACAACTAAACGAACA   21550
gb:MT007544|Organism:Severe      TTAGAGAAAACAACAGAGTTGTTATTTCTAGTGATGTTCTTGTTAACAACTAAACGAACA   21562
gb:MN994467|Organism:Severe      TTAGAGAAAACAACAGAGTTGTTATTTCTAGTGATGTTCTTGTTAACAACTAAACGAACA   21562
gb:MT044257|Organism:Severe      TTAGAGAAAACAACAGAGTTGTTATTTCTAGTGATGTTCTTGTTAACAACTAAACGAACA   21562
gb:MT106054|Organism:Severe      TTAGAGAAAACAACAGAGTTGTTATTTCTAGTGATGTTCTTGTTAACAACTAAACGAACA   21562
gb:MT049951|Organism:Severe      TTAGAGAAAACAACAGAGTTGTTATTTCTAGTGATGTTCTTGTTAACAACTAAACGAACA   21562
gb:MN975262|Organism:Severe      TTAGAGAAAACAACAGAGTTGTTATTTCTAGTGATGTTCTTGTTAACAACTAAACGAACA   21562
gb:MT106052|Organism:Severe      TTAGAGAAAACAACAGAGTTGTTATTTCTAGTGATGTTCTTGTTAACAACTAAACGAACA   21562
gb:LC522975|Organism:Severe      TTAGAGAAAACAACAGAGTTGTTATTTCTAGTGATGTTCTTGTTAACAACTAAACGAACA   21559
gb:LC522973|Organism:Severe      TTAGAGAAAACAACAGAGTTGTTATTTCTAGTGATGTTCTTGTTAACAACTAAACGAACA   21559
gb:LC522974|Organism:Severe      TTAGAGAAAACAACAGAGTTGTTATTTCTAGTGATGTTCTTGTTAACAACTAAACGAACA   21559
gb:MN985325|Organism:Severe      TTAGAGAAAACAACAGAGTTGTTATTTCTAGTGATGTTCTTGTTAACAACTAAACGAACA   21562
gb:MT020881|Organism:Severe      TTAGAGAAAACAACAGAGTTGTTATTTCTAGTGATGTTCTTGTTAACAACTAAACGAACA   21562
gb:MT020880|Organism:Severe      TTAGAGAAAACAACAGAGTTGTTATTTCTAGTGATGTTCTTGTTAACAACTAAACGAACA   21562
gb:MT066175|Organism:Severe      TTAGAGAAAACAACAGAGTTGTTATTTCTAGTGATGTTCTTGTTAACAACTAAACGAACA   21562
gb:MN997409|Organism:Severe      TTAGAGAAAACAACAGAGTTGTTATTTCTAGTGATGTTCTTGTTAACAACTAAACGAACA   21562
gb:MN938384|Organism:Severe      TTAGAGAAAACAACAGAGTTGTTATTTCTAGTGATGTTCTTGTTAACAACTAAACGAACA   21530
gb:MT044258|Organism:Severe      TTAGAGAAAACAACAGAGTTGTTATTTCTAGTGATGTTCTTGTTAACAACTAAACGAACA   21538
gb:MT039890|Organism:Severe      TTAGAGAAAACAACAGAGTTGTTATTTCTAGTGATGTTCTTGTTAACAACTAAACGAACA   21562
gb:MN988713|Organism:Severe      TTAGAGAAAACAACAGAGTTGTTATTTCTAGTGATGTTCTTGTTAACAACTAAACGAACA   21562
gb:LC521925|Organism:Severe      TTAGAGAAAACAACAGAGTTGTTATTTCTAGTGATGTTCTTGTTAACAACTAAACGAACA   21535
gb:MT093571|Organism:Severe      TTAGAGAAAACAACAGAGTTGTTATTTCTAGTGATGTTCTTGTTAACAACTAAACGAACA   21562
gb:MT039887|Organism:Severe      TTAGAGAAAACAACAGAGTTGTTATTTCTAGTGATGTTCTTGTTAACAACTAAACGAACA   21559
gb:MT019530|Organism:Severe      TTAGAGAAAACAACAGAGTTGTTATTTCTAGTGATGTTCTTGTTAACAACTAAACGAACA   21562
gb:MT039888|Organism:Severe      TTAGAGAAAACAACAGAGTTGTTATTTCTAGTGATGTTCTTGTTAACAACTAAACGAACA   21562
gb:LC522972|Organism:Severe      TTAGAGAAAACAACAGAGTTGTTATTTCTAGTGATGTTCTTGTTAACAACTAAACGAACA   21559
gb:MT027063|Organism:Severe      TTAGAGAAAACAACAGAGTTGTTATTTCTAGTGATGTTCTTGTTAACAACTAAACGAACA   21562
gb:MT027062|Organism:Severe      TTAGAGAAAACAACAGAGTTGTTATTTCTAGTGATGTTCTTGTTAACAACTAAACGAACA   21562
gb:MT019529|Organism:Severe      TTAGAGAAAACAACAGAGTTGTTATTTCTAGTGATGTTCTTGTTAACAACTAAACGAACA   21562
gb:MN996529|Organism:Severe      TTAGAGAAAACAACAGAGTTGTTATTTCTAGTGATGTTCTTGTTAACAACTAAACGAACA   21550
gb:MN996531|Organism:Severe      TTAGAGAAAACAACAGAGTTGTTATTTCTAGTGATGTTCTTGTTAACAACTAAACGAACA   21549
gb:MT066176|Organism:Severe      TTAGAGAAAACAACAGAGTTGTTATTTCTAGTGATGTTCTTGTTAACAACTAAACGAACA   21562
gb:MT027064|Organism:Severe      TTAGAGAAAACAACAGAGTTGTTATTTCTAGTGATGTTCTTGTTAACAACTAAACGAACA   21562
gb:MN994468|Organism:Severe      TTAGAGAAAACAACAGAGTTGTTATTTCTAGTGATGTTCTTGTTAACAACTAAACGAACA   21562
gb:MT072688|Organism:Severe      TTAGAGAAAACAACAGAGTTGTTATTTCTAGTGATGTTCTTGTTAACAACTAAACGAACA   21547
gb:MN996527|Organism:Severe      TTAGAGAAAACAACAGAGTTGTTATTTCTAGTGATGTTCTTGTTAACAACTAAACGAACA   21529
gb:MT093631|Organism:Severe      TTAGAGAAAACAACAGAGTTGTTATTTCTAGTGATGTTCTTGTTAACAACTAAACGAACA   21600
gb:MT106053|Organism:Severe      TTAGAGAAAACAACAGAGTTGTTATTTCTAGTGATGTTCTTGTTAACAACTAAACGAACA   21562
gb:MT019533|Organism:Severe      TTAGAGAAAACAACAGAGTTGTTATTTCTAGTGATGTTCTTGTTAACAACTAAACGAACA   21562
gb:MT019531|Organism:Severe      TTAGAGAAAACAACAGAGTTGTTATTTCTAGTGATGTTCTTGTTAACAACTAAACGAACA   21562
gb:MN996528|Organism:Severe      TTAGAGAAAACAACAGAGTTGTTATTTCTAGTGATGTTCTTGTTAACAACTAAACGAACA   21562
gb:MN996530|Organism:Severe      TTAGAGAAAACAACAGAGTTGTTATTTCTAGTGATGTTCTTGTTAACAACTAAACGAACA   21548
gb:MN908947|Organism:Severe      TTAGAGAAAACAACAGAGTTGTTATTTCTAGTGATGTTCTTGTTAACAACTAAACGAACA   21562
gb:MT019532|Organism:Severe      TTAGAGAAAACAACAGAGTTGTTATTTCTAGTGATGTTCTTGTTAACAACTAAACGAACA   21562
                                 ************************************************************

gb:MT020781|Organism:Severe      ATGTTTGTTTTTCTTGTTTTATTGCCACTAGTCTCTAGTCAGTGTGTTAATCTTACAACC   21610
gb:MT007544|Organism:Severe      ATGTTTGTTTTTCTTGTTTTATTGCCACTAGTCTCTAGTCAGTGTGTTAATCTTACAACC   21622
gb:MN994467|Organism:Severe      ATGTTTGTTTTTCTTGTTTTATTGCCACTAGTCTCTAGTCAGTGTGTTAATCTTACAACC   21622
gb:MT044257|Organism:Severe      ATGTTTGTTTTTCTTGTTTTATTGCCACTAGTCTCTAGTCAGTGTGTTAATCTTACAACC   21622
gb:MT106054|Organism:Severe      ATGTTTGTTTTTCTTGTTTTATTGCCACTAGTCTCTAGTCAGTGTGTTAATCTTACAACC   21622
gb:MT049951|Organism:Severe      ATGTTTGTTTTTCTTGTTTTATTGCCACTAGTCTCTAGTCAGTGTGTTAATCTTACAACC   21622
gb:MN975262|Organism:Severe      ATGTTTGTTTTTCTTGTTTTATTGCCACTAGTCTCTAGTCAGTGTGTTAATCTTACAACC   21622
gb:MT106052|Organism:Severe      ATGTTTGTTTTTCTTGTTTTATTGCCACTAGTCTCTAGTCAGTGTGTTAATCTTACAACC   21622
gb:LC522975|Organism:Severe      ATGTTTGTTTTTCTTGTTTTATTGCCACTAGTCTCTAGTCAGTGTGTTAATCTTACAACC   21619
gb:LC522973|Organism:Severe      ATGTTTGTTTTTCTTGTTTTATTGCCACTAGTCTCTAGTCAGTGTGTTAATCTTACAACC   21619
gb:LC522974|Organism:Severe      ATGTTTGTTTTTCTTGTTTTATTGCCACTAGTCTCTAGTCAGTGTGTTAATCTTACAACC   21619
```

```
gb:MN985325|Organism:Severe      ATGTTTGTTTTTCTTGTTTTATTGCCACTAGTCTCTAGTCAGTGTGTTAATCTTACAACC   21622
gb:MT020881|Organism:Severe      ATGTTTGTTTTTCTTGTTTTATTGCCACTAGTCTCTAGTCAGTGTGTTAATCTTACAACC   21622
gb:MT020880|Organism:Severe      ATGTTTGTTTTTCTTGTTTTATTGCCACTAGTCTCTAGTCAGTGTGTTAATCTTACAACC   21622
gb:MT066175|Organism:Severe      ATGTTTGTTTTTCTTGTTTTATTGCCACTAGTCTCTAGTCAGTGTGTTAATCTTACAACC   21622
gb:MN997409|Organism:Severe      ATGTTTGTTTTTCTTGTTTTATTGCCACTAGTCTCTAGTCAGTGTGTTAATCTTACAACC   21622
gb:MN938384|Organism:Severe      ATGTTTGTTTTTCTTGTTTTATTGCCACTAGTCTCTAGTCAGTGTGTTAATCTTACAACC   21590
gb:MT044258|Organism:Severe      ATGTTTGTTTTTCTTGTTTTATTGCCACTAGTCTCTAGTCAGTGTGTTAATCTTACAACC   21598
gb:MT039890|Organism:Severe      ATGTTTGTTTTTCTTGTTTTATTGCCACTAGTCTCTAGTCAGTGTGTTAATCTTACAACC   21622
gb:MN988713|Organism:Severe      ATGTTTGTTTTTCTTGTTTTATTGCCACTAGTCTCTAGTCAGTGTGTTAATCTTACAACC   21622
gb:LC521925|Organism:Severe      ATGTTTGTTTTTCTTGTTTTATTGCCACTAGTCTCTAGTCAGTGTGTTAATCTTACAACC   21595
gb:MT093571|Organism:Severe      ATGTTTGTTTTTCTTGTTTTATTGCCACTAGTCTCTAGTCAGTGTGTTAATCTTACAACC   21622
gb:MT039887|Organism:Severe      ATGTTTGTTTTTCTTGTTTTATTGCCACTAGTCTCTAGTCAGTGTGTTAATCTTACAACC   21619
gb:MT019530|Organism:Severe      ATGTTTGTTTTTCTTGTTTTATTGCCACTAGTCTCTAGTCAGTGTGTTAATCTTACAACC   21622
gb:MT039888|Organism:Severe      ATGTTTGTTTTTCTTGTTTTATTGCCACTAGTCTCTAGTCAGTGTGTTAATCTTACAACC   21622
gb:LC522972|Organism:Severe      ATGTTTGTTTTTCTTGTTTTATTGCCACTAGTCTCTAGTCAGTGTGTTAATCTTACAACC   21619
gb:MT027063|Organism:Severe      ATGTTTGTTTTTCTTGTTTTATTGCCACTAGTCTCTAGTCAGTGTGTTAATCTTACAACC   21622
gb:MT027062|Organism:Severe      ATGTTTGTTTTTCTTGTTTTATTGCCACTAGTCTCTAGTCAGTGTGTTAATCTTACAACC   21622
gb:MT019529|Organism:Severe      ATGTTTGTTTTTCTTGTTTTATTGCCACTAGTCTCTAGTCAGTGTGTTAATCTTACAACC   21622
gb:MN996529|Organism:Severe      ATGTTTGTTTTTCTTGTTTTATTGCCACTAGTCTCTAGTCAGTGTGTTAATCTTACAACC   21610
gb:MN996531|Organism:Severe      ATGTTTGTTTTTCTTGTTTTATTGCCACTAGTCTCTAGTCAGTGTGTTAATCTTACAACC   21609
gb:MT066176|Organism:Severe      ATGTTTGTTTTTCTTGTTTTATTGCCACTAGTCTCTAGTCAGTGTGTTAATCTTACAACC   21622
gb:MT027064|Organism:Severe      ATGTTTGTTTTTCTTGTTTTATTGCCACTAGTCTCTAGTCAGTGTGTTAATCTTACAACC   21622
gb:MN994468|Organism:Severe      ATGTTTGTTTTTCTTGTTTTATTGCCACTAGTCTCTAGTCAGTGTGTTAATCTTACAACC   21622
gb:MT072688|Organism:Severe      ATGTTTGTTTTTCTTGTTTTATTGCCACTAGTCTCTAGTCAGTGTGTTAATCTTACAACC   21607
gb:MN996527|Organism:Severe      ATGTTTGTTTTTCTTGTTTTATTGCCACTAGTCTCTAGTCAGTGTGTTAATCTTACAACC   21589
gb:MT093631|Organism:Severe      ATGTTTGTTTTTCTTGTTTTATTGCCACTAGTCTCTAGTCAGTGTGTTAATCTTACAACC   21660
gb:MT106053|Organism:Severe      ATGTTTGTTTTTCTTGTTTTATTGCCACTAGTCTCTAGTCAGTGTGTTAATCTTACAACC   21622
gb:MT019533|Organism:Severe      ATGTTTGTTTTTCTTGTTTTATTGCCACTAGTCTCTAGTCAGTGTGTTAATCTTACAACC   21622
gb:MT019531|Organism:Severe      ATGTTTGTTTTTCTTGTTTTATTGCCACTAGTCTCTAGTCAGTGTGTTAATCTTACAACC   21622
gb:MN996528|Organism:Severe      ATGTTTGTTTTTCTTGTTTTATTGCCACTAGTCTCTAGTCAGTGTGTTAATCTTACAACC   21622
gb:MN996530|Organism:Severe      ATGTTTGTTTTTCTTGTTTTATTGCCACTAGTCTCTAGTCAGTGTGTTAATCTTACAACC   21608
gb:MN908947|Organism:Severe      ATGTTTGTTTTTCTTGTTTTATTGCCACTAGTCTCTAGTCAGTGTGTTAATCTTACAACC   21622
gb:MT019532|Organism:Severe      ATGTTTGTTTTTCTTGTTTTATTGCCACTAGTCTCTAGTCAGTGTGTTAATCTTACAACC   21622
                                 ************************************************************

gb:MT020781|Organism:Severe      AGAANNNNNNNNNNNNNNNNNNNNNNNNNNNNNNNNNNNNNNNNNNNNNNNNNNNNNNNN   21670
gb:MT007544|Organism:Severe      AGAACTCAATTACCCCCTGCATACACTAATTCTTTCACACGTGGTGTTTATTACCCTGAC   21682
gb:MN994467|Organism:Severe      AGAACTCAATTACCCCCTGCATACACTAATTCTTTCACACGTGGTGTTTATTACCCTGAC   21682
gb:MT044257|Organism:Severe      AGAACTCAATTACCCCCTGCATACACTAATTCTTTCACACGTGGTGTTTATTACCCTGAC   21682
gb:MT106054|Organism:Severe      AGAACTCAATTACCCCCTGCATACACTAATTCTTTCACACGTGGTGTTTATTACCCTGAC   21682
gb:MT049951|Organism:Severe      AGAACTCAATTACCCCCTGCAAACACTAATTCTTTCACACGTGGTGTTTATTACCCTGAC   21682
gb:MN975262|Organism:Severe      AGAACTCAATTACCCCCTGCATACACTAATTCTTTCACACGTGGTGTTTATTACCCTGAC   21682
gb:MT106052|Organism:Severe      AGAACTCAATTACCCCCTGCATACACTAATTCTTTCACACGTGGTGTTTATTACCCTGAC   21682
gb:LC522975|Organism:Severe      AGAACTCAATTACCCCCTGCATACACTAATTCTTTCACACGTGGTGTTTATTACCCTGAC   21679
gb:LC522973|Organism:Severe      AGAACTCAATTACCCCCTGCATACACTAATTCTTTCACACGTGGTGTTTATTACCCTGAC   21679
gb:LC522974|Organism:Severe      AGAACTCAATTACCCCCTGCATACACTAATTCTTTCACACGTGGTGTTTATTACCCTGAC   21679
gb:MN985325|Organism:Severe      AGAACTCAATTACCCCCTGCATACACTAATTCTTTCACACGTGGTGTTTATTACCCTGAC   21682
gb:MT020881|Organism:Severe      AGAACTCAATTACCCCCTGCATACACTAATTCTTTCACACGTGGTGTTTATTACCCTGAC   21682
gb:MT020880|Organism:Severe      AGAACTCAATTACCCCCTGCATACACTAATTCTTTCACACGTGGTGTTTATTACCCTGAC   21682
gb:MT066175|Organism:Severe      AGAACTCAATTACCCCCTGCATACACTAATTCTTTCACACGTGGTGTTTATTACCCTGAC   21682
gb:MN997409|Organism:Severe      AGAACTCAATTACCCCCTGCATACACTAATTCTTTCACACGTGGTGTTTATTACCCTGAC   21682
gb:MN938384|Organism:Severe      AGAACTCAATTACCCCCTGCATACACTAATTCTTTCACACGTGGTGTTTATTACCCTGAC   21650
gb:MT044258|Organism:Severe      AGAACTCAATTACCCCCTGCATACACTAATTCTTTCACACGTGGTGTTTATTACCCTGAC   21658
gb:MT039890|Organism:Severe      AGAACTCAATTACCCCCTGCATACACTAATTCTTTCACACGTGGTGTTTATTACCCTGAC   21682
gb:MN988713|Organism:Severe      AGAACTCAATTACCCCCTGCATACACTAATTCTTTCACACGTGGTGTTTATTACCCTGAC   21682
gb:LC521925|Organism:Severe      AGAACTCAATTACCCCCTGCATACACTAATTCTTTCACACGTGGTGTTTATTACCCTGAC   21655
gb:MT093571|Organism:Severe      AGAACTCAATTACCCCCTGCATACACTAATTCTTTCACACGTGGTGTTTATTACCCTGAC   21682
gb:MT039887|Organism:Severe      AGAACTCAATTACCCCCTGCATACACTAATTCTTTCACACGTGGTGTTTATTACCCTGAC   21679
gb:MT019530|Organism:Severe      AGAACTCAATTACCCCCTGCATACACTAATTCTTTCACACGTGGTGTTTATTACCCTGAC   21682
gb:MT039888|Organism:Severe      AGAACTCAATTACCCCCTGCATACACTAATTCTTTCACACGTGGTGTTTATTACCCTGAC   21682
gb:LC522972|Organism:Severe      AGAACTCAATTACCCCCTGCATACACTAATTCTTTCACACGTGGTGTTTATTACCCTGAC   21679
gb:MT027063|Organism:Severe      AGAACTCAATTACCCCCTGCATACACTAATTCTTTCACACGTGGTGTTTATTACCCTGAC   21682
gb:MT027062|Organism:Severe      AGAACTCAATTACCCCCTGCATACACTAATTCTTTCACACGTGGTGTTTATTACCCTGAC   21682
gb:MT019529|Organism:Severe      AGAACTCAATTACCCCCTGCATACACTAATTCTTTCACACGTGGTGTTTATTACCCTGAC   21682
gb:MN996529|Organism:Severe      AGAACTCAATTACCCCCTGCATACACTAATTCTTTCACACGTGGTGTTTATTACCCTGAC   21670
```

```
gb:MN996531|Organism:Severe      AGAACTCAATTACCCCCTGCATACACTAATTCTTTCACACGTGGTGTTTATTACCCTGAC   21669
gb:MT066176|Organism:Severe      AGAACTCAATTACCCCCTGCATACACTAATTCTTTCACACGTGGTGTTTATTACCCTGAC   21682
gb:MT027064|Organism:Severe      AGAACTCAATTACCCCCTGCATACACTAATTCTTTCACACGTGGTGTTTATTACCCTGAC   21682
gb:MN994468|Organism:Severe      AGAACTCAATTACCCCCTGCATACACTAATTCTTTCACACGTGGTGTTTATTACCCTGAC   21682
gb:MT072688|Organism:Severe      AGAACTCAATTACCCCCTGCATACACTAATTCTTTCACACGTGGTGTTTATTACCCTGAC   21667
gb:MN996527|Organism:Severe      AGAACTCAATTACCCCCTGCATACACTAATTCTTTCACACGTGGTGTTTATTACCCTGAC   21649
gb:MT093631|Organism:Severe      AGAACTCAATTACCCCCTGCATACACTAATTCTTTCACACGTGGTGTTTATTACCCTGAC   21720
gb:MT106053|Organism:Severe      AGAACTCAATTACCCCCTGCATACACTAATTCTTTCACACGTGGTGTTTATTACCCTGAC   21682
gb:MT019533|Organism:Severe      AGAACTCAATTACCCCCTGCATACACTAATTCTTTCACACGTGGTGTTTATTACCCTGAC   21682
gb:MT019531|Organism:Severe      AGAACTCAATTACCCCCTGCATACACTAATTCTTTCACACGTGGTGTTTATTACCCTGAC   21682
gb:MN996528|Organism:Severe      AGAACTCAATTACCCCCTGCATACACTAATTCTTTCACACGTGGTGTTTATTACCCTGAC   21682
gb:MN996530|Organism:Severe      AGAACTCAATTACCCCCTGCATACACTAATTCTTTCACACGTGGTGTTTATTACCCTGAC   21668
gb:MN908947|Organism:Severe      AGAACTCAATTACCCCCTGCATACACTAATTCTTTCACACGTGGTGTTTATTACCCTGAC   21682
gb:MT019532|Organism:Severe      AGAACTCAATTACCCCCTGCATACACTAATTCTTTCACACGTGGTGTTTATTACCCTGAC   21682
                                 ****

gb:MT020781|Organism:Severe      NNNNNNNNNNNNNNNNNNNNNNNNNATTCAACTCAGGACTTGTTCTTACCTTTCTTTTCC   21730
gb:MT007544|Organism:Severe      AAAGTTTTCAGATCCTCAGTTTTACATTCAACTCAGGACTTGTTCTTACCTTTCTTTTCC   21742
gb:MN994467|Organism:Severe      AAAGTTTTCAGATCCTCAGTTTTACATTCAACTCAGGACTTGTTCTTACCTTTCTTTTCC   21742
gb:MT044257|Organism:Severe      AAAGTTTTCAGATCCTCAGTTTTACATTCAACTCAGGACTTGTTCTTACCTTTCTTTTCC   21742
gb:MT106054|Organism:Severe      AAAGTTTTCAGATCCTCAGTTTTACATTCAACTCAGGACTTGTTCTTACCTTTCTTTTCC   21742
gb:MT049951|Organism:Severe      AAAGTTTTCAGATCCTCAGTTTTACATTCAACTCAGGACTTGTTCTTACCTTTCTTTTCC   21742
gb:MN975262|Organism:Severe      AAAGTTTTCAGATCCTCAGTTTTACATTCAACTCAGGACTTGTTCTTACCTTTCTTTTCC   21742
gb:MT106052|Organism:Severe      AAAGTTTTCAGATCCTCAGTTTTACATTCAACTCAGGACTTGTTCTTACCTTTCTTTTCC   21742
gb:LC522975|Organism:Severe      AAAGTTTTCAGATCCTCAGTTTTACATTCAACTCAGGACTTGTTCTTACCTTTCTTTTCC   21739
gb:LC522973|Organism:Severe      AAAGTTTTCAGATCCTCAGTTTTACATTCAACTCAGGACTTGTTCTTACCTTTCTTTTCC   21739
gb:LC522974|Organism:Severe      AAAGTTTTCAGATCCTCAGTTTTACATTCAACTCAGGACTTGTTCTTACCTTTCTTTTCC   21739
gb:MN985325|Organism:Severe      AAAGTTTTCAGATCCTCAGTTTTACATTCAACTCAGGACTTGTTCTTACCTTTCTTTTCC   21742
gb:MT020881|Organism:Severe      AAAGTTTTCAGATCCTCAGTTTTACATTCAACTCAGGACTTGTTCTTACCTTTCTTTTCC   21742
gb:MT020880|Organism:Severe      AAAGTTTTCAGATCCTCAGTTTTACATTCAACTCAGGACTTGTTCTTACCTTTCTTTTCC   21742
gb:MT066175|Organism:Severe      AAAGTTTTCAGATCCTCAGTTTTACATTCAACTCAGGACTTGTTCTTACCTTTCTTTTCC   21742
gb:MN997409|Organism:Severe      AAAGTTTTCAGATCCTCAGTTTTACATTCAACTCAGGACTTGTTCTTACCTTTCTTTTCC   21742
gb:MN938384|Organism:Severe      AAAGTTTTCAGATCCTCAGTTTTACATTCAACTCAGGACTTGTTCTTACCTTTCTTTTCC   21710
gb:MT044258|Organism:Severe      AAAGTTTTCAGATCCTCAGTTTTACATTCAACTCAGGACTTGTTCTTACCTTTCTTTTCC   21718
gb:MT039890|Organism:Severe      AAAGTTTTCAGATCCTCAGTTTTACATTCAACTCAGGACTTGTTCTTACCTTTCTTTTCC   21742
gb:MN988713|Organism:Severe      AAAGTTTTCAGATCCTCAGTTTTACATTCAACTCAGGACTTGTTCTTACCTTTCTTTTCC   21742
gb:LC521925|Organism:Severe      AAAGTTTTCAGATCCTCAGTTTTACATTCAACTCAGGACTTGTTCTTACCTTTCTTTTCC   21715
gb:MT093571|Organism:Severe      AAAGTTTTCAGATCCTCAGTTTTACATTCAACTCAGGACTTGTTCTTACCTTTCTTTTCC   21742
gb:MT039887|Organism:Severe      AAAGTTTTCAGATCCTCAGTTTTACATTCAACTCAGGACTTGTTCTTACCTTTCTTTTCC   21739
gb:MT019530|Organism:Severe      AAAGTTTTCAGATCCTCAGTTTTACATTCAACTCAGGACTTGTTCTTACCTTTCTTTTCC   21742
gb:MT039888|Organism:Severe      AAAGTTTTCAGATCCTCAGTTTTACATTCAACTCAGGACTTGTTCTTACCTTTCTTTTCC   21742
gb:LC522972|Organism:Severe      AAAGTTTTCAGATCCTCAGTTTTACATTCAACTCAGGACTTGTTCTTACCTTTCTTTTCC   21739
gb:MT027063|Organism:Severe      AAAGTTTTCAGATCCTCAGTTTTACATTCAACTCAGGACTTGTTCTTACCTTTCTTTTCC   21742
gb:MT027062|Organism:Severe      AAAGTTTTCAGATCCTCAGTTTTACATTCAACTCAGGACTTGTTCTTACCTTTCTTTTCC   21742
gb:MT019529|Organism:Severe      AAAGTTTTCAGATCCTCAGTTTTACATTCAACTCAGGACTTGTTCTTACCTTTCTTTTCC   21742
gb:MN996529|Organism:Severe      AAAGTTTTCAGATCCTCAGTTTTACATTCAACTCAGGACTTGTTCTTACCTTTCTTTTCC   21730
gb:MN996531|Organism:Severe      AAAGTTTTCAGATCCTCAGTTTTACATTCAACTCAGGACTTGTTCTTACCTTTCTTTTCC   21729
gb:MT066176|Organism:Severe      AAAGTTTTCAGATCCTCAGTTTTACATTCAACTCAGGACTTGTTCTTACCTTTCTTTTCC   21742
gb:MT027064|Organism:Severe      AAAGTTTTCAGATCCTCAGTTTTATATTCAACTCAGGACTTGTTCTTACCTTTCTTTTCC   21742
gb:MN994468|Organism:Severe      AAAGTTTTCAGATCCTCAGTTTTACATTCAACTCAGGACTTGTTCTTACCTTTCTTTTCC   21742
gb:MT072688|Organism:Severe      AAAGTTTTCAGATCCTCAGTTTTACATTCAACTCAGGACTTGTTCTTACCTTTCTTTTCC   21727
gb:MN996527|Organism:Severe      AAAGTTTTCAGATCCTCAGTTTTACATTCAACTCAGGACTTGTTCTTACCTTTCTTTTCC   21709
gb:MT093631|Organism:Severe      AAAGTTTTCAGATCCTCAGTTTTACATTCAACTCAGGACTTGTTCTTACCTTTCTTTTCC   21780
gb:MT106053|Organism:Severe      AAAGTTTTCAGATCCTCAGTTTTACATTCAACTCAGGACTTGTTCTTACCTTTCTTTTCC   21742
gb:MT019533|Organism:Severe      AAAGTTTTCAGATCCTCAGTTTTACATTCAACTCAGGACTTGTTCTTACCTTTCTTTTCC   21742
gb:MT019531|Organism:Severe      AAAGTTTTCAGATCCTCAGTTTTACATTCAACTCAGGACTTGTTCTTACCTTTCTTTTCC   21742
gb:MN996528|Organism:Severe      AAAGTTTTCAGATCCTCAGTTTTACATTCAACTCAGGACTTGTTCTTACCTTTCTTTTCC   21742
gb:MN996530|Organism:Severe      AAAGTTTTCAGATCCTCAGTTTTACATTCAACTCAGGACTTGTTCTTACCTTTCTTTTCC   21728
gb:MN908947|Organism:Severe      AAAGTTTTCAGATCCTCAGTTTTACATTCAACTCAGGACTTGTTCTTACCTTTCTTTTCC   21742
gb:MT019532|Organism:Severe      AAAGTTTTCAGATCCTCAGTTTTACATTCAACTCAGGACTTGTTCTTACCTTTCTTTTCC   21742
                                                          ***********************************

gb:MT020781|Organism:Severe      AATGTTACTTGGTTCCATGCTATACATGTCTCTGGGACCAATGGTACTAAGAGGTTTGAT   21790
gb:MT007544|Organism:Severe      AATGTTACTTGGTTCCATGCTATACATGTCTCTGGGACCAATGGTACTAAGAGGTTTGAT   21802
gb:MN994467|Organism:Severe      AATGTTACTTGGTTCCATGCTATACATGTCTCTGGGACCAATGGTACTAAGAGGTTTGAT   21802
```

```
gb:MT044257|Organism:Severe      AATGTTACTTGGTTCCATGCTATACATGTCTCTGGGACCAATGGTACTAAGAGGTTTGAT    21802
gb:MT106054|Organism:Severe      AATGTTACTTGGTTCCATGCTATACATGTCTCTGGGACCAATGGTACTAAGAGGTTTGAT    21802
gb:MT049951|Organism:Severe      AATGTTACTTGGTTCCATGCTATACATGTCTCTGGGACCAATGGTACTAAGAGGTTTGAT    21802
gb:MN975262|Organism:Severe      AATGTTACTTGGTTCCATGCTATACATGTCTCTGGGACCAATGGTACTAAGAGGTTTGAT    21802
gb:MT106052|Organism:Severe      AATGTTACTTGGTTCCATGCTATACATGTCTCTGGGACCAATGGTACTAAGAGGTTTGAT    21802
gb:LC522975|Organism:Severe      AATGTTACTTGGTTCCATGCTATACATGTCTCTGGGACCAATGGTACTAAGAGGTTTGAT    21799
gb:LC522973|Organism:Severe      AATGTTACTTGGTTCCATGCTATACATGTCTCTGGGACCAATGGTACTAAGAGGTTTGAT    21799
gb:LC522974|Organism:Severe      AATGTTACTTGGTTCCATGCTATACATGTCTCTGGGACCAATGGTACTAAGAGGTTTGAT    21799
gb:MN985325|Organism:Severe      AATGTTACTTGGTTCCATGCTATACATGTCTCTGGGACCAATGGTACTAAGAGGTTTGAT    21802
gb:MT020881|Organism:Severe      AATGTTACTTGGTTCCATGCTATACATGTCTCTGGGACCAATGGTACTAAGAGGTTTGAT    21802
gb:MT020880|Organism:Severe      AATGTTACTTGGTTCCATGCTATACATGTCTCTGGGACCAATGGTACTAAGAGGTTTGAT    21802
gb:MT066175|Organism:Severe      AATGTTACTTGGTTCCATGCTATACATGTCTCTGGGACCAATGGTACTAAGAGGTTTGAT    21802
gb:MN997409|Organism:Severe      AATGTTACTTGGTTCCATGCTATACATGTCTCTGGGACCAATGGTACTAAGAGGTTTGAT    21802
gb:MN938384|Organism:Severe      AATGTTACTTGGTTCCATGCTATACATGTCTCTGGGACCAATGGTACTAAGAGGTTTGAT    21770
gb:MT044258|Organism:Severe      AATGTTACTTGGTTCCATGCTATACATGTCTCTGGGACCAATGGTACTAAGAGGTTTGAT    21778
gb:MT039890|Organism:Severe      AATGTTACTTGGTTCCATGCTATACATGTCTCTGGGACCAATGGTACTAAGAGGTTTGAT    21802
gb:MN988713|Organism:Severe      AATGTTACTTGGTTCCATGCTATACATGTCTCTGGGACCAATGGTACTAAGAGGTTTGAT    21802
gb:LC521925|Organism:Severe      AATGTTACTTGGTTCCATGCTATACATGTCTCTGGGACCAATGGTACTAAGAGGTTTGAT    21775
gb:MT093571|Organism:Severe      AATGTTACTTGGTTCCATGCTATACATGTCTCTGGGACCAATGGTACTAAGAGGTTTGAT    21802
gb:MT039887|Organism:Severe      AATGTTACTTGGTTCCATGCTATACATGTCTCTGGGACCAATGGTACTAAGAGGTTTGAT    21799
gb:MT019530|Organism:Severe      AATGTTACTTGGTTCCATGCTATACATGTCTCTGGGACCAATGGTACTAAGAGGTTTGAT    21802
gb:MT039888|Organism:Severe      AATGTTACTTGGTTCCATGCTATACATGTCTCTGGGACCAATGGTACTAAGAGGTTTGAT    21802
gb:LC522972|Organism:Severe      AATGTTACTTGGTTCCATGCTATACATGTCTCTGGGACCAATGGTACTAAGAGGTTTGAT    21799
gb:MT027063|Organism:Severe      AATGTTACTTGGTTCCATGCTATACATGTCTCTGGGACCAATGGTACTAAGAGGTTTGAT    21802
gb:MT027062|Organism:Severe      AATGTTACTTGGTTCCATGCTATACATGTCTCTGGGACCAATGGTACTAAGAGGTTTGAT    21802
gb:MT019529|Organism:Severe      AATGTTACTTGGTTCCATGCTATACATGTCTCTGGGACCAATGGTACTAAGAGGTTTGAT    21802
gb:MN996529|Organism:Severe      AATGTTACTTGGTTCCATGCTATACATGTCTCTGGGACCAATGGTACTAAGAGGTTTGAT    21790
gb:MN996531|Organism:Severe      AATGTTACTTGGTTCCATGCTATACATGTCTCTGGGACCAATGGTACTAAGAGGTTTGAT    21789
gb:MT066176|Organism:Severe      AATGTTACTTGGTTCCATGCTATACATGTCTCTGGGACCAATGGTACTAAGAGGTTTGAT    21802
gb:MT027064|Organism:Severe      AATGTTACTTGGTTCCATGCTATACATGTCTCTGGGACCAATGGTACTAAGAGGTTTGAT    21802
gb:MN994468|Organism:Severe      AATGTTACTTGGTTCCATGCTATACATGTCTCTGGGACCAATGGTACTAAGAGGTTTGAT    21802
gb:MT072688|Organism:Severe      AATGTTACTTGGTTCCATGCTATACATGTCTCTGGGACCAATGGTACTAAGAGGTTTGAT    21787
gb:MN996527|Organism:Severe      AATGTTACTTGGTTCCATGCTATACATGTCTCTGGGACCAATGGTACTAAGAGGTTTGAT    21769
gb:MT093631|Organism:Severe      AATGTTACTTGGTTCCATGCTATACATGTCTCTGGGACCAATGGTACTAAGAGGTTTGAT    21840
gb:MT106053|Organism:Severe      AATGTTACTTGGTTCCATGCTATACATGTCTCTGGGACCAATGGTACTAAGAGGTTTGAT    21802
gb:MT019533|Organism:Severe      AATGTTACTTGGTTCCATGCTATACATGTCTCTGGGACCAATGGTACTAAGAGGTTTGAT    21802
gb:MT019531|Organism:Severe      AATGTTACTTGGTTCCATGCTATACATGTCTCTGGGACCAATGGTACTAAGAGGTTTGAT    21802
gb:MN996528|Organism:Severe      AATGTTACTTGGTTCCATGCTATACATGTCTCTGGGACCAATGGTACTAAGAGGTTTGAT    21802
gb:MN996530|Organism:Severe      AATGTTACTTGGTTCCATGCTATACATGTCTCTGGGACCAATGGTACTAAGAGGTTTGAT    21788
gb:MN908947|Organism:Severe      AATGTTACTTGGTTCCATGCTATACATGTCTCTGGGACCAATGGTACTAAGAGGTTTGAT    21802
gb:MT019532|Organism:Severe      AATGTTACTTGGTTCCATGCTATACATGTCTCTGGGACCAATGGTACTAAGAGGTTTGAT    21802
                                 ************************************************************

gb:MT020781|Organism:Severe      AACCCTGTCCTACCATTTAATGATGGTGTTTATTTTGCTTCCACTGAGAAGTCTAACATA    21850
gb:MT007544|Organism:Severe      AACCCTGTCCTACCATTTAATGATGGTGTTTATTTTGCTTCCACTGAGAAGTCTAACATA    21862
gb:MN994467|Organism:Severe      AACCCTGTCCTACCATTTAATGATGGTGTTTATTTTGCTTCCACTGAGAAGTCTAACATA    21862
gb:MT044257|Organism:Severe      AACCCTGTCCTACCATTTAATGATGGTGTTTATTTTGCTTCCACTGAGAAGTCTAACATA    21862
gb:MT106054|Organism:Severe      AACCCTGTCCTACCATTTAATGATGGTGTTTATTTTGCTTCCACTGAGAAGTCTAACATA    21862
gb:MT049951|Organism:Severe      AACCCTGTCCTACCATTTAATGATGGTGTTTATTTTGCTTCCACTGAGAAGTCTAACATA    21862
gb:MN975262|Organism:Severe      AACCCTGTCCTACCATTTAATGATGGTGTTTATTTTGCTTCCACTGAGAAGTCTAACATA    21862
gb:MT106052|Organism:Severe      AACCCTGTCCTACCATTTAATGATGGTGTTTATTTTGCTTCCACTGAGAAGTCTAACATA    21862
gb:LC522975|Organism:Severe      AACCCTGTCCTACCATTTAATGATGGTGTTTATTTTGCTTCCACTGAGAAGTCTAACATA    21859
gb:LC522973|Organism:Severe      AACCCTGTCCTACCATTTAATGATGGTGTTTATTTTGCTTCCACTGAGAAGTCTAACATA    21859
gb:LC522974|Organism:Severe      AACCCTGTCCTACCATTTAATGATGGTGTTTATTTTGCTTCCACTGAGAAGTCTAACATA    21859
gb:MN985325|Organism:Severe      AACCCTGTCCTACCATTTAATGATGGTGTTTATTTTGCTTCCACTGAGAAGTCTAACATA    21862
gb:MT020881|Organism:Severe      AACCCTGTCCTACCATTTAATGATGGTGTTTATTTTGCTTCCACTGAGAAGTCTAACATA    21862
gb:MT020880|Organism:Severe      AACCCTGTCCTACCATTTAATGATGGTGTTTATTTTGCTTCCACTGAGAAGTCTAACATA    21862
gb:MT066175|Organism:Severe      AACCCTGTCCTACCATTTAATGATGGTGTTTATTTTGCTTCCACTGAGAAGTCTAACATA    21862
gb:MN997409|Organism:Severe      AACCCTGTCCTACCATTTAATGATGGTGTTTATTTTGCTTCCACTGAGAAGTCTAACATA    21862
gb:MN938384|Organism:Severe      AACCCTGTCCTACCATTTAATGATGGTGTTTATTTTGCTTCCACTGAGAAGTCTAACATA    21830
gb:MT044258|Organism:Severe      AACCCTGTCCTACCATTTAATGATGGTGTTTATTTTGCTTCCACTGAGAAGTCTAACATA    21838
gb:MT039890|Organism:Severe      AACCCTGTCCTACCATTTAATGATGGTGTTTATTTTGCTTCCACTGAGAAGTCTAACATA    21862
gb:MN988713|Organism:Severe      AACCCTGTCCTACCATTTAATGATGGTGTTTATTTTGCTTCCACTGAGAAGTCTAACATA    21862
gb:LC521925|Organism:Severe      AACCCTGTCCTACCATTTAATGATGGTGTTTATTTTGCTTCCACTGAGAAGTCTAACATA    21835
gb:MT093571|Organism:Severe      AACCCTGTCCTACCATTTAATGATGGTGTTTATTTTGCTTCCACTGAGAAGTCTAACATA    21862
```

```
gb:MT039887|Organism:Severe      AACCCTGTCCTACCATTTAATGATGGTGTTTATTTTGCTTCCACTGAGAAGTCTAACATA   21859
gb:MT019530|Organism:Severe      AACCCTGTCCTACCATTTAATGATGGTGTTTATTTTGCTTCCACTGAGAAGTCTAACATA   21862
gb:MT039888|Organism:Severe      AACCCTGTCCTACCATTTAATGATGGTGTTTATTTTGCTTCCACTGAGAAGTCTAACATA   21862
gb:LC522972|Organism:Severe      AACCCTGTCCTACCATTTAATGATGGTGTTTATTTTGCTTCCACTGAGAAGTCTAACATA   21859
gb:MT027063|Organism:Severe      AACCCTGTCCTACCATTTAATGATGGTGTTTATTTTGCTTCCACTGAGAAGTCTAACATA   21862
gb:MT027062|Organism:Severe      AACCCTGTCCTACCATTTAATGATGGTGTTTATTTTGCTTCCACTGAGAAGTCTAACATA   21862
gb:MT019529|Organism:Severe      AACCCTGTCCTACCATTTAATGATGGTGTTTATTTTGCTTCCACTGAGAAGTCTAACATA   21862
gb:MN996529|Organism:Severe      AACCCTGTCCTACCATTTAATGATGGTGTTTATTTTGCTTCCACTGAGAAGTCTAACATA   21850
gb:MN996531|Organism:Severe      AACCCTGTCCTACCATTTAATGATGGTGTTTATTTTGCTTCCACTGAGAAGTCTAACATA   21849
gb:MT066176|Organism:Severe      AACCCTGTCCTACCATTTAATGATGGTGTTTATTTTGCTTCCACTGAGAAGTCTAACATA   21862
gb:MT027064|Organism:Severe      AACCCTGTCCTACCATTTAATGATGGTGTTTATTTTGCTTCCACTGAGAAGTCTAACATA   21862
gb:MN994468|Organism:Severe      AACCCTGTCCTACCATTTAATGATGGTGTTTATTTTGCTTCCACTGAGAAGTCTAACATA   21862
gb:MT072688|Organism:Severe      AACCCTGTCCTACCATTTAATGATGGTGTTTATTTTGCTTCCACTGAGAAGTCTAACATA   21847
gb:MN996527|Organism:Severe      AACCCTGTCCTACCATTTAATGATGGTGTTTATTTTGCTTCCACTGAGAAGTCTAACATA   21829
gb:MT093631|Organism:Severe      AACCCTGTCCTACCATTTAATGATGGTGTTTATTTTGCTTCCACTGAGAAGTCTAACATA   21900
gb:MT106053|Organism:Severe      AACCCTGTCCTACCATTTAATGATGGTGTTTATTTTGCTTCCACTGAGAAGTCTAACATA   21862
gb:MT019533|Organism:Severe      AACCCTGTCCTACCATTTAATGATGGTGTTTATTTTGCTTCCACTGAGAAGTCTAACATA   21862
gb:MT019531|Organism:Severe      AACCCTGTCCTACCATTTAATGATGGTGTTTATTTTGCTTCCACTGAGAAGTCTAACATA   21862
gb:MN996528|Organism:Severe      AACCCTGTCCTACCATTTAATGATGGTGTTTATTTTGCTTCCACTGAGAAGTCTAACATA   21862
gb:MN996530|Organism:Severe      AACCCTGTCCTACCATTTAATGATGGTGTTTATTTTGCTTCCACTGAGAAGTCTAACATA   21848
gb:MN908947|Organism:Severe      AACCCTGTCCTACCATTTAATGATGGTGTTTATTTTGCTTCCACTGAGAAGTCTAACATA   21862
gb:MT019532|Organism:Severe      AACCCTGTCCTACCATTTAATGATGGTGTTTATTTTGCTTCCACTGAGAAGTCTAACATA   21862
                                 ************************************************************

gb:MT020781|Organism:Severe      ATAAGAGGCTGGATTTTTGGTACTACTTTAGATTCGAAGACCCAGTCCCTACTTATTGTT   21910
gb:MT007544|Organism:Severe      ATAAGAGGCTGGATTTTTGGTACTACTTTAGATTCGAAGACCCAGTCCCTACTTATTGTT   21922
gb:MN994467|Organism:Severe      ATAAGAGGCTGGATTTTTGGTACTACTTTAGATTCGAAGACCCAGTCCCTACTTATTGTT   21922
gb:MT044257|Organism:Severe      ATAAGAGGCTGGATTTTTGGTACTACTTTAGATTCGAAGACCCAGTCCCTACTTATTGTT   21922
gb:MT106054|Organism:Severe      ATAAGAGGCTGGATTTTTGGTACTACTTTAGATTCGAAGACCCAGTCCCTACTTATTGTT   21922
gb:MT049951|Organism:Severe      ATAAGAGGCTGGATTTTTGGTACTACTTTAGATTCGAAGACCCAGTCCCTACTTATTGTT   21922
gb:MN975262|Organism:Severe      ATAAGAGGCTGGATTTTTGGTACTACTTTAGATTCGAAGACCCAGTCCCTACTTATTGTT   21922
gb:MT106052|Organism:Severe      ATAAGAGGCTGGATTTTTGGTACTACTTTAGATTCGAAGACCCAGTCCCTACTTATTGTT   21922
gb:LC522975|Organism:Severe      ATAAGAGGCTGGATTTTTGGTACTACTTTAGATTCGAAGACCCAGTCCCTACTTATTGTT   21919
gb:LC522973|Organism:Severe      ATAAGAGGCTGGATTTTTGGTACTACTTTAGATTCGAAGACCCAGTCCCTACTTATTGTT   21919
gb:LC522974|Organism:Severe      ATAAGAGGCTGGATTTTTGGTACTACTTTAGATTCGAAGACCCAGTCCCTACTTATTGTT   21919
gb:MN985325|Organism:Severe      ATAAGAGGCTGGATTTTTGGTACTACTTTAGATTCGAAGACCCAGTCCCTACTTATTGTT   21922
gb:MT020881|Organism:Severe      ATAAGAGGCTGGATTTTTGGTACTACTTTAGATTCGAAGACCCAGTCCCTACTTATTGTT   21922
gb:MT020880|Organism:Severe      ATAAGAGGCTGGATTTTTGGTACTACTTTAGATTCGAAGACCCAGTCCCTACTTATTGTT   21922
gb:MT066175|Organism:Severe      ATAAGAGGCTGGATTTTTGGTACTACTTTAGATTCGAAGACCCAGTCCCTACTTATTGTT   21922
gb:MN997409|Organism:Severe      ATAAGAGGCTGGATTTTTGGTACTACTTTAGATTCGAAGACCCAGTCCCTACTTATTGTT   21922
gb:MN938384|Organism:Severe      ATAAGAGGCTGGATTTTTGGTACTACTTTAGATTCGAAGACCCAGTCCCTACTTATTGTT   21890
gb:MT044258|Organism:Severe      ATAAGAGGCTGGATTTTTGGTACTACTTTAGATTCGAAGACCCAGTCCCTACTTATTGTT   21898
gb:MT039890|Organism:Severe      ATAAGAGGCTGGATTTTTGGTACTACTTTAGATTCGAAGACCCAGTCCCTACTTATTGTT   21922
gb:MN988713|Organism:Severe      ATAAGAGGCTGGATTTTTGGTACTACTTTAGATTCGAAGACCCAGTCCCTACTTATTGTT   21922
gb:LC521925|Organism:Severe      ATAAGAGGCTGGATTTTTGGTACTACTTTAGATTCGAAGACCCAGTCCCTACTTATTGTT   21895
gb:MT093571|Organism:Severe      ATAAGAGGCTGGATTTTTGGTACTACTTTAGATTCGAAGACCCAGTCCCTACTTATTGTT   21922
gb:MT039887|Organism:Severe      ATAAGAGGCTGGATTTTTGGTACTACTTTAGATTCGAAGACCCAGTCCCTACTTATTGTT   21919
gb:MT019530|Organism:Severe      ATAAGAGGCTGGATTTTTGGTACTACTTTAGATTCGAAGACCCAGTCCCTACTTATTGTT   21922
gb:MT039888|Organism:Severe      ATAAGAGGCTGGATTTTTGGTACTACTTTAGATTCGAAGACCCAGTCCCTACTTATTGTT   21922
gb:LC522972|Organism:Severe      ATAAGAGGCTGGATTTTTGGTACTACTTTAGATTCGAAGACCCAGTCCCTACTTATTGTT   21919
gb:MT027063|Organism:Severe      ATAAGAGGCTGGATTTTTGGTACTACTTTAGATTCGAAGACCCAGTCCCTACTTATTGTT   21922
gb:MT027062|Organism:Severe      ATAAGAGGCTGGATTTTTGGTACTACTTTAGATTCGAAGACCCAGTCCCTACTTATTGTT   21922
gb:MT019529|Organism:Severe      ATAAGAGGCTGGATTTTTGGTACTACTTTAGATTCGAAGACCCAGTCCCTACTTATTGTT   21922
gb:MN996529|Organism:Severe      ATAAGAGGCTGGATTTTTGGTACTACTTTAGATTCGAAGACCCAGTCCCTACTTATTGTT   21910
gb:MN996531|Organism:Severe      ATAAGAGGCTGGATTTTTGGTACTACTTTAGATTCGAAGACCCAGTCCCTACTTATTGTT   21909
gb:MT066176|Organism:Severe      ATAAGAGGCTGGATTTTTGGTACTACTTTAGATTCGAAGACCCAGTCCCTACTTATTGTT   21922
gb:MT027064|Organism:Severe      ATAAGAGGCTGGATTTTTGGTACTACTTTAGATTCGAAGACCCAGTCCCTACTTATTGTT   21922
gb:MN994468|Organism:Severe      ATAAGAGGCTGGATTTTTGGTACTACTTTAGATTCGAAGACCCAGTCCCTACTTATTGTT   21922
gb:MT072688|Organism:Severe      ATAAGAGGCTGGATTTTTGGTACTACTTTAGATTCGAAGACCCAGTCCCTACTTATTGTT   21907
gb:MN996527|Organism:Severe      ATAAGAGGCTGGATTTTTGGTACTACTTTAGATTCGAAGACCCAGTCCCTACTTATTGTT   21889
gb:MT093631|Organism:Severe      ATAAGAGGCTGGATTTTTGGTACTACTTTAGATTCGAAGACCCAGTCCCTACTTATTGTT   21960
gb:MT106053|Organism:Severe      ATAAGAGGCTGGATTTTTGGTACTACTTTAGATTCGAAGACCCAGTCCCTACTTATTGTT   21922
gb:MT019533|Organism:Severe      ATAAGAGGCTGGATTTTTGGTACTACTTTAGATTCGAAGACCCAGTCCCTACTTATTGTT   21922
gb:MT019531|Organism:Severe      ATAAGAGGCTGGATTTTTGGTACTACTTTAGATTCGAAGACCCAGTCCCTACTTATTGTT   21922
gb:MN996528|Organism:Severe      ATAAGAGGCTGGATTTTTGGTACTACTTTAGATTCGAAGACCCAGTCCCTACTTATTGTT   21922
```

```
gb:MN996530|Organism:Severe      ATAAGAGGCTGGATTTTTGGTACTACTTTAGATTCGAAGACCCAGTCCCTACTTATTGTT   21908
gb:MN908947|Organism:Severe      ATAAGAGGCTGGATTTTTGGTACTACTTTAGATTCGAAGACCCAGTCCCTACTTATTGTT   21922
gb:MT019532|Organism:Severe      ATAAGAGGCTGGATTTTTGGTACTACTTTAGATTCGAAGACCCAGTCCCTACTTATTGTT   21922
                                 ************************************************************

gb:MT020781|Organism:Severe      AATAACGCTACTAATGTTGTTATTAAAGTCTGTGAATTTCAATTTTGTAATGATCCATTT   21970
gb:MT007544|Organism:Severe      AATAACGCTACTAATGTTGTTATTAAAGTCTGTGAATTTCAATTTTGTAATGATCCATTT   21982
gb:MN994467|Organism:Severe      AATAACGCTACTAATGTTGTTATTAAAGTCTGTGAATTTCAATTTTGTAATGATCCATTT   21982
gb:MT044257|Organism:Severe      AATAACGCTACTAATGTTGTTATTAAAGTCTGTGAATTTCAATTTTGTAATGATCCATTT   21982
gb:MT106054|Organism:Severe      AATAACGCTACTAATGTTGTTATTAAAGTCTGTGAATTTCAATTTTGTAATGATCCATTT   21982
gb:MT049951|Organism:Severe      AATAACGCTACTAATGTTGTTATTAAAGTCTGTGAATTTCAATTTTGTAATGATCCATTT   21982
gb:MN975262|Organism:Severe      AATAACGCTACTAATGTTGTTATTAAAGTCTGTGAATTTCAATTTTGTAATGATCCATTT   21982
gb:MT106052|Organism:Severe      AATAACGCTACTAATGTTGTTATTAAAGTCTGTGAATTTCAATTTTGTAATGATCCATTT   21982
gb:LC522975|Organism:Severe      AATAACGCTACTAATGTTGTTATTAAAGTCTGTGAATTTCAATTTTGTAATGATCCATTT   21979
gb:LC522973|Organism:Severe      AATAACGCTACTAATGTTGTTATTAAAGTCTGTGAATTTCAATTTTGTAATGATCCATTT   21979
gb:LC522974|Organism:Severe      AATAACGCTACTAATGTTGTTATTAAAGTCTGTGAATTTCAATTTTGTAATGATCCATTT   21979
gb:MN985325|Organism:Severe      AATAACGCTACTAATGTTGTTATTAAAGTCTGTGAATTTCAATTTTGTAATGATCCATTT   21982
gb:MT020881|Organism:Severe      AATAACGCTACTAATGTTGTTATTAAAGTCTGTGAATTTCAATTTTGTAATGATCCATTT   21982
gb:MT020880|Organism:Severe      AATAACGCTACTAATGTTGTTATTAAAGTCTGTGAATTTCAATTTTGTAATGATCCATTT   21982
gb:MT066175|Organism:Severe      AATAACGCTACTAATGTTGTTATTAAAGTCTGTGAATTTCAATTTTGTAATGATCCATTT   21982
gb:MN997409|Organism:Severe      AATAACGCTACTAATGTTGTTATTAAAGTCTGTGAATTTCAATTTTGTAATGATCCATTT   21982
gb:MN938384|Organism:Severe      AATAACGCTACTAATGTTGTTATTAAAGTCTGTGAATTTCAATTTTGTAATGATCCATTT   21950
gb:MT044258|Organism:Severe      AATAACGCTACTAATGTTGTTATTAAAGTCTGTGAATTTCAATTTTGTAATGATCCATTT   21958
gb:MT039890|Organism:Severe      AATAACGCTACTAATGTTGTTATTAAAGTCTGTGAATTTCAATTTTGTAATGATCCATTT   21982
gb:MN988713|Organism:Severe      AATAACGCTACTAATGTTGTTATTAAAGTCTGTGAATTTCAATTTTGTAATGATCCATTT   21982
gb:LC521925|Organism:Severe      AATAACGCTACTAATGTTGTTATTAAAGTCTGTGAATTTCAATTTTGTAATGATCCATTT   21955
gb:MT093571|Organism:Severe      AATAACGCTACTAATGTTGTTATTAAAGTCTGTGAATTTCAATTTTGTAATGATCCATTT   21982
gb:MT039887|Organism:Severe      AATAACGCTACTAATGTTGTTATTAAAGTCTGTGAATTTCAATTTTGTAATGATCCATTT   21979
gb:MT019530|Organism:Severe      AATAACGCTACTAATGTTGTTATTAAAGTCTGTGAATTTCAATTTTGTAATGATCCATTT   21982
gb:MT039888|Organism:Severe      AATAACGCTACTAATGTTGTTATTAAAGTCTGTGAATTTCAATTTTGTAATGATCCATTT   21982
gb:LC522972|Organism:Severe      AATAACGCTACTAATGTTGTTATTAAAGTCTGTGAATTTCAATTTTGTAATGATCCATTT   21979
gb:MT027063|Organism:Severe      AATAACGCTACTAATGTTGTTATTAAAGTCTGTGAATTTCAATTTTGTAATGATCCATTT   21982
gb:MT027062|Organism:Severe      AATAACGCTACTAATGTTGTTATTAAAGTCTGTGAATTTCAATTTTGTAATGATCCATTT   21982
gb:MT019529|Organism:Severe      AATAACGCTACTAATGTTGTTATTAAAGTCTGTGAATTTCAATTTTGTAATGATCCATTT   21982
gb:MN996529|Organism:Severe      AATAACGCTACTAATGTTGTTATTAAAGTCTGTGAATTTCAATTTTGTAATGATCCATTT   21970
gb:MN996531|Organism:Severe      AATAACGCTACTAATGTTGTTATTAAAGTCTGTGAATTTCAATTTTGTAATGATCCATTT   21969
gb:MT066176|Organism:Severe      AATAACGCTACTAATGTTGTTATTAAAGTCTGTGAATTTCAATTTTGTAATGATCCATTT   21982
gb:MT027064|Organism:Severe      AATAACGCTACTAATGTTGTTATTAAAGTCTGTGAATTTCAATTTTGTAATGATCCATTT   21982
gb:MN994468|Organism:Severe      AATAACGCTACTAATGTTGTTATTAAAGTCTGTGAATTTCAATTTTGTAATGATCCATTT   21982
gb:MT072688|Organism:Severe      AATAACGCTACTAATGTTGTTATTAAAGTCTGTGAATTTCAATTTTGTAATGATCCATTT   21967
gb:MN996527|Organism:Severe      AATAACGCTACTAATGTTGTTATTAAAGTCTGTGAATTTCAATTTTGTAATGATCCATTT   21949
gb:MT093631|Organism:Severe      AATAACGCTACTAATGTTGTTATTAAAGTCTGTGAATTTCAATTTTGTAATGATCCATTT   22020
gb:MT106053|Organism:Severe      AATAACGCTACTAATGTTGTTATTAAAGTCTGTGAATTTCAATTTTGTAATGATCCATTT   21982
gb:MT019533|Organism:Severe      AATAACGCTACTAATGTTGTTATTAAAGTCTGTGAATTTCAATTTTGTAATGATCCATTT   21982
gb:MT019531|Organism:Severe      AATAACGCTACTAATGTTGTTATTAAAGTCTGTGAATTTCAATTTTGTAATGATCCATTT   21982
gb:MN996528|Organism:Severe      AATAACGCTACTAATGTTGTTATTAAAGTCTGTGAATTTCAATTTTGTAATGATCCATTT   21982
gb:MN996530|Organism:Severe      AATAACGCTACTAATGTTGTTATTAAAGTCTGTGAATTTCAATTTTGTAATGATCCATTT   21968
gb:MN908947|Organism:Severe      AATAACGCTACTAATGTTGTTATTAAAGTCTGTGAATTTCAATTTTGTAATGATCCATTT   21982
gb:MT019532|Organism:Severe      AATAACGCTACTAATGTTGTTATTAAAGTCTGTGAATTTCAATTTTGTAATGATCCATTT   21982
                                 ************************************************************

gb:MT020781|Organism:Severe      TTGGGTGTTTATTACCACAAAAACAACAAAAGTTGGATGGAAAGTGAGTTCAGAGTTTAT   22030
gb:MT007544|Organism:Severe      TTGGGTGTTTATTACCACAAAAACAACAAAAGTTGGATGGAAAGTGAGTTCAGAGTTTAT   22042
gb:MN994467|Organism:Severe      TTGGGTGTTTATTACCACAAAAACAACAAAAGTTGGATGGAAAGTGAGTTCAGAGTTTAT   22042
gb:MT044257|Organism:Severe      TTGGGTGTTTATTACCACAAAAACAACAAAAGTTGGATGGAAAGTGAGTTCAGAGTTTAT   22042
gb:MT106054|Organism:Severe      TTGGGTGTTTATTACCACAAAAACAACAAAAGTTGGATGGAAAGTGAGTTCAGAGTTTAT   22042
gb:MT049951|Organism:Severe      TTGGGTGTTTATTACCACAAAAACAACAAAAGTTGGATGGAAAGTGAGTTCAGAGTTTAT   22042
gb:MN975262|Organism:Severe      TTGGGTGTTTATTACCACAAAAACAACAAAAGTTGGATGGAAAGTGAGTTCAGAGTTTAT   22042
gb:MT106052|Organism:Severe      TTGGGTGTTTATTACCACAAAAACAACAAAAGTTGGATGGAAAGTGAGTTCAGAGTTTAT   22042
gb:LC522975|Organism:Severe      TTGGGTGTTTATTACCACAAAAACAACAAAAGTTGGATGGAAAGTGAGTTCAGAGTTTAT   22039
gb:LC522973|Organism:Severe      TTGGGTGTTTATTACCACAAAAACAACAAAAGTTGGATGGAAAGTGAGTTCAGAGTTTAT   22039
gb:LC522974|Organism:Severe      TTGGGTGTTTATTACCACAAAAACAACAAAAGTTGGATGGAAAGTGAGTTCAGAGTTTAT   22039
gb:MN985325|Organism:Severe      TTGGGTGTTTATTACCACAAAAACAACAAAAGTTGGATGGAAAGTGAGTTCAGAGTTTAT   22042
gb:MT020881|Organism:Severe      TTGGGTGTTTATTACCACAAAAACAACAAAAGTTGGATGGAAAGTGAGTTCAGAGTTTAT   22042
gb:MT020880|Organism:Severe      TTGGGTGTTTATTACCACAAAAACAACAAAAGTTGGATGGAAAGTGAGTTCAGAGTTTAT   22042
```

```
gb:MT066175|Organism:Severe      TTGGGTGTTTATTACCACAAAAACAACAAAAGTTGGATGGAAAGTGAGTTCAGAGTTTAT	22042
gb:MN997409|Organism:Severe      TTGGGTGTTTATTACCACAAAAACAACAAAAGTTGGATGGAAAGTGAGTTCAGAGTTTAT	22042
gb:MN938384|Organism:Severe      TTGGGTGTTTATTACCACAAAAACAACAAAAGTTGGATGGAAAGTGAGTTCAGAGTTTAT	22010
gb:MT044258|Organism:Severe      TTGGGTGTTTATTACCACAAAAACAACAAAAGTTGGATGGAAAGTGAGTTCAGAGTTTAT	22018
gb:MT039890|Organism:Severe      TTGGGTGTTTATTACCACAAAAACAACAAAAGTTGGATGGAAAGTGAGTTCAGAGTTTAT	22042
gb:MN988713|Organism:Severe      TTGGGTGTTTATTACCACAAAAACAACAAAAGTTGGATGGAAAGTGAGTTCAGAGTTTAT	22042
gb:LC521925|Organism:Severe      TTGGGTGTTTATTACCACAAAAACAACAAAAGTTGGATGGAAAGTGAGTTCAGAGTTTAT	22015
gb:MT093571|Organism:Severe      TTGGGTGTTTATTACCACAAAAACAACAAAAGTTGGATGGAAAGTGAGTTCAGAGTTTAT	22042
gb:MT039887|Organism:Severe      TTGGGTGTTTATTACCACAAAAACAACAAAAGTTGGATGGAAAGTGAGTTCAGAGTTTAT	22039
gb:MT019530|Organism:Severe      TTGGGTGTTTATTACCACAAAAACAACAAAAGTTGGATGGAAAGTGAGTTCAGAGTTTAT	22042
gb:MT039888|Organism:Severe      TTGGGTGTTTATTACCACAAAAACAACAAAAGTTGGATGGAAAGTGAGTTCAGAGTTTAT	22042
gb:LC522972|Organism:Severe      TTGGGTGTTTATTACCACAAAAACAACAAAAGTTGGATGGAAAGTGAGTTCAGAGTTTAT	22039
gb:MT027063|Organism:Severe      TTGGGTGTTTATTACCACAAAAACAACAAAAGTTGGATGGAAAGTGAGTTCAGAGTTTAT	22042
gb:MT027062|Organism:Severe      TTGGGTGTTTATTACCACAAAAACAACAAAAGTTGGATGGAAAGTGAGTTCAGAGTTTAT	22042
gb:MT019529|Organism:Severe      TTGGGTGTTTATTACCACAAAAACAACAAAAGTTGGATGGAAAGTGAGTTCAGAGTTTAT	22042
gb:MN996529|Organism:Severe      TTGGGTGTTTATTACCACAAAAACAACAAAAGTTGGATGGAAAGTGAGTTCAGAGTTTAT	22030
gb:MN996531|Organism:Severe      TTGGGTGTTTATTACCACAAAAACAACAAAAGTTGGATGGAAAGTGAGTTCAGAGTTTAT	22029
gb:MT066176|Organism:Severe      TTGGGTGTTTATTACCACAAAAACAACAAAAGTTGGATGGAAAGTGAGTTCAGAGTTTAT	22042
gb:MT027064|Organism:Severe      TTGGGTGTTTATTACCACAAAAACAACAAAAGTTGGATGGAAAGTGAGTTCAGAGTTTAT	22042
gb:MN994468|Organism:Severe      TTGGGTGTTTATTACCACAAAAACAACAAAAGTTGGATGGAAAGTGAGTTCAGAGTTTAT	22042
gb:MT072688|Organism:Severe      TTGGGTGTTTATTACCACAAAAACAACAAAAGTTGGATGGAAAGTGAGTTCAGAGTTTAT	22027
gb:MN996527|Organism:Severe      TTGGGTGTTTATTACCACAAAAACAACAAAAGTTGGATGGAAAGTGAGTTCAGAGTTTAT	22009
gb:MT093631|Organism:Severe      TTGGGTGTTTATTACCACAAAAACAACAAAAGTTGGATGGAAAGTGAGTTCAGAGTTTAT	22080
gb:MT106053|Organism:Severe      TTGGGTGTTTATTACCACAAAAACAACAAAAGTTGGATGGAAAGTGAGTTCAGAGTTTAT	22042
gb:MT019533|Organism:Severe      TTGGGTGTTTATTACCACAAAAACAACAAAAGTTGGATGGAAAGTGAGTTCAGAGTTTAT	22042
gb:MT019531|Organism:Severe      TTGGGTGTTTATTACCACAAAAACAACAAAAGTTGGATGGAAAGTGAGTTCAGAGTTTAT	22042
gb:MN996528|Organism:Severe      TTGGGTGTTTATTACCACAAAAACAACAAAAGTTGGATGGAAAGTGAGTTCAGAGTTTAT	22042
gb:MN996530|Organism:Severe      TTGGGTGTTTATTACCACAAAAACAACAAAAGTTGGATGGAAAGTGAGTTCAGAGTTTAT	22028
gb:MN908947|Organism:Severe      TTGGGTGTTTATTACCACAAAAACAACAAAAGTTGGATGGAAAGTGAGTTCAGAGTTTAT	22042
gb:MT019532|Organism:Severe      TTGGGTGTTTATTACCACAAAAACAACAAAAGTTGGATGGAAAGTGAGTTCAGAGTTTAT	22042
                                 ************************************************************

gb:MT020781|Organism:Severe      TCTAGTGCGAATAATTGCACTTTTGAATATGTCTCTCAGCCTTTTCTTATGGACCTTGAA	22090
gb:MT007544|Organism:Severe      TCTAGTGCGAATAATTGCACTTTTGAATATGTCTCTCAGCCTTTTCTTATGGACCTTGAA	22102
gb:MN994467|Organism:Severe      TCTAGTGCGAATAATTGCACTTTTGAATATGTCTCTCAGCCTTTTCTTATGGACCTTGAA	22102
gb:MT044257|Organism:Severe      TCTAGTGCGAATAATTGCACTTTTGAATATGTCTCTCAGCCTTTTCTTATGGACCTTGAA	22102
gb:MT106054|Organism:Severe      TCTAGTGCGAATAATTGCACTTTTGAATATGTCTCTCAGCCTTTTCTTATGGACCTTGAA	22102
gb:MT049951|Organism:Severe      TCTAGTGCGAATAATTGCACTTTTGAATATGTCTCTCAGCCTTTTCTTATGGACCTTGAA	22102
gb:MN975262|Organism:Severe      TCTAGTGCGAATAATTGCACTTTTGAATATGTCTCTCAGCCTTTTCTTATGGACCTTGAA	22102
gb:MT106052|Organism:Severe      TCTAGTGCGAATAATTGCACTTTTGAATATGTCTCTCAGCCTTTTCTTATGGACCTTGAA	22102
gb:LC522975|Organism:Severe      TCTAGTGCGAATAATTGCACTTTTGAATATGTCTCTCAGCCTTTTCTTATGGACCTTGAA	22099
gb:LC522973|Organism:Severe      TCTAGTGCGAATAATTGCACTTTTGAATATGTCTCTCAGCCTTTTCTTATGGACCTTGAA	22099
gb:LC522974|Organism:Severe      TCTAGTGCGAATAATTGCACTTTTGAATATGTCTCTCAGCCTTTTCTTATGGACCTTGAA	22099
gb:MN985325|Organism:Severe      TCTAGTGCGAATAATTGCACTTTTGAATATGTCTCTCAGCCTTTTCTTATGGACCTTGAA	22102
gb:MT020881|Organism:Severe      TCTAGTGCGAATAATTGCACTTTTGAATATGTCTCTCAGCCTTTTCTTATGGACCTTGAA	22102
gb:MT020880|Organism:Severe      TCTAGTGCGAATAATTGCACTTTTGAATATGTCTCTCAGCCTTTTCTTATGGACCTTGAA	22102
gb:MT066175|Organism:Severe      TCTAGTGCGAATAATTGCACTTTTGAATATGTCTCTCAGCCTTTTCTTATGGACCTTGAA	22102
gb:MN997409|Organism:Severe      TCTAGTGCGAATAATTGCACTTTTGAATATGTCTCTCAGCCTTTTCTTATGGACCTTGAA	22102
gb:MN938384|Organism:Severe      TCTAGTGCGAATAATTGCACTTTTGAATATGTCTCTCAGCCTTTTCTTATGGACCTTGAA	22070
gb:MT044258|Organism:Severe      TCTAGTGCGAATAATTGCACTTTTGAATATGTCTCTCAGCCTTTTCTTATGGACCTTGAA	22078
gb:MT039890|Organism:Severe      TCTAGTGCGAATAATTGCACTTTTGAATATGTCTCTCAGCCTTTTCTTATGGACCTTGAA	22102
gb:MN988713|Organism:Severe      TCTAGTGCGAATAATTGCACTTTTGAATATGTCTCTCAGCCTTTTCTTATGGACCTTGAA	22102
gb:LC521925|Organism:Severe      TCTAGTGCGAATAATTGCACTTTTGAATATGTCTCTCAGCCTTTTCTTATGGACCTTGAA	22075
gb:MT093571|Organism:Severe      TCTAGTGCGAATAATTGCACTTTTGAATATGTCTCTCAGCCTTTTCTTATGGACCTTGAA	22102
gb:MT039887|Organism:Severe      TCTAGTGCGAATAATTGCACTTTTGAATATGTCTCTCAGCCTTTTCTTATGGACCTTGAA	22099
gb:MT019530|Organism:Severe      TCTAGTGCGAATAATTGCACTTTTGAATATGTCTCTCAGCCTTTTCTTATGGACCTTGAA	22102
gb:MT039888|Organism:Severe      TCTAGTGCGAATAATTGCACTTTTGAATATGTCTCTCAGCCTTTTCTTATGGACCTTGAA	22102
gb:LC522972|Organism:Severe      TCTAGTGCGAATAATTGCACTTTTGAATATGTCTCTCAGCCTTTTCTTATGGACCTTGAA	22099
gb:MT027063|Organism:Severe      TCTAGTGCGAATAATTGCACTTTTGAATATGTCTCTCAGCCTTTTCTTATGGACCTTGAA	22102
gb:MT027062|Organism:Severe      TCTAGTGCGAATAATTGCACTTTTGAATATGTCTCTCAGCCTTTTCTTATGGACCTTGAA	22102
gb:MT019529|Organism:Severe      TCTAGTGCGAATAATTGCACTTTTGAATATGTCTCTCAGCCTTTTCTTATGGACCTTGAA	22102
gb:MN996529|Organism:Severe      TCTAGTGCGAATAATTGCACTTTTGAATATGTCTCTCAGCCTTTTCTTATGGACCTTGAA	22090
gb:MN996531|Organism:Severe      TCTAGTGCGAATAATTGCACTTTTGAATATGTCTCTCAGCCTTTTCTTATGGACCTTGAA	22089
gb:MT066176|Organism:Severe      TCTAGTGCGAATAATTGCACTTTTGAATATGTCTCTCAGCCTTTTCTTATGGACCTTGAA	22102
gb:MT027064|Organism:Severe      TCTAGTGCGAATAATTGCACTTTTGAATATGTCTCTCAGCCTTTTCTTATGGACCTTGAA	22102
```

```
gb:MN994468|Organism:Severe      TCTAGTGCGAATAATTGCACTTTTGAATATGTCTCTCAGCCTTTTCTTATGGACCTTGAA   22102
gb:MT072688|Organism:Severe      TCTAGTGCGAATAATTGCACTTTTGAATATGTCTCTCAGCCTTTTCTTATGGACCTTGAA   22087
gb:MN996527|Organism:Severe      TCTAGTGCGAATAATTGCACTTTTGAATATGTCTCTCAGCCTTTTCTTATGGACCTTGAA   22069
gb:MT093631|Organism:Severe      TCTAGTGCGAATAATTGCACTTTTGAATATGTCTCTCAGCCTTTTCTTATGGACCTTGAA   22140
gb:MT106053|Organism:Severe      TCTAGTGCGAATAATTGCACTTTTGAATATGTCTCTCAGCCTTTTCTTATGGACCTTGAA   22102
gb:MT019533|Organism:Severe      TCTAGTGCGAATAATTGCACTTTTGAATATGTCTCTCAGCCTTTTCTTATGGACCTTGAA   22102
gb:MT019531|Organism:Severe      TCTAGTGCGAATAATTGCACTTTTGAATATGTCTCTCAGCCTTTTCTTATGGACCTTGAA   22102
gb:MN996528|Organism:Severe      TCTAGTGCGAATAATTGCACTTTTGAATATGTCTCTCAGCCTTTTCTTATGGACCTTGAA   22102
gb:MN996530|Organism:Severe      TCTAGTGCGAATAATTGCACTTTTGAATATGTCTCTCAGCCTTTTCTTATGGACCTTGAA   22088
gb:MN908947|Organism:Severe      TCTAGTGCGAATAATTGCACTTTTGAATATGTCTCTCAGCCTTTTCTTATGGACCTTGAA   22102
gb:MT019532|Organism:Severe      TCTAGTGCGAATAATTGCACTTTTGAATATGTCTCTCAGCCTTTTCTTATGGACCTTGAA   22102
                                 ************************************************************

gb:MT020781|Organism:Severe      GGAAAACAGGGTAATTTCAAAAATCTTAGGGAATTTGTGTTTAAGAATATTGATGGTTAT   22150
gb:MT007544|Organism:Severe      GGAAAACAGGGTAATTTCAAAAATCTTAGGGAATTTGTGTTTAAGAATATTGATGGTTAT   22162
gb:MN994467|Organism:Severe      GGAAAACAGGGTAATTTCAAAAATCTTAGGGAATTTGTGTTTAAGAATATTGATGGTTAT   22162
gb:MT044257|Organism:Severe      GGAAAACAGGGTAATTTCAAAAATCTTAGGGAATTTGTGTTTAAGAATATTGATGGTTAT   22162
gb:MT106054|Organism:Severe      GGAAAACAGGGTAATTTCAAAAATCTTAGGGAATTTGTGTTTAAGAATATTGATGGTTAT   22162
gb:MT049951|Organism:Severe      GGAAAACAGGGTAATTTCAAAAATCTTAGGGAATTTGTGTTTAAGAATATTGATGGTTAT   22162
gb:MN975262|Organism:Severe      GGAAAACAGGGTAATTTCAAAAATCTTAGGGAATTTGTGTTTAAGAATATTGATGGTTAT   22162
gb:MT106052|Organism:Severe      GGAAAACAGGGTAATTTCAAAAATCTTAGGGAATTTGTGTTTAAGAATATTGATGGTTAT   22162
gb:LC522975|Organism:Severe      GGAAAACAGGGTAATTTCAAAAATCTTAGGGAATTTGTGTTTAAGAATATTGATGGTTAT   22159
gb:LC522973|Organism:Severe      GGAAAACAGGGTAATTTCAAAAATCTTAGGGAATTTGTGTTTAAGAATATTGATGGTTAT   22159
gb:LC522974|Organism:Severe      GGAAAACAGGGTAATTTCAAAAATCTTAGGGAATTTGTGTTTAAGAATATTGATGGTTAT   22159
gb:MN985325|Organism:Severe      GGAAAACAGGGTAATTTCAAAAATCTTAGGGAATTTGTGTTTAAGAATATTGATGGTTAT   22162
gb:MT020881|Organism:Severe      GGAAAACAGGGTAATTTCAAAAATCTTAGGGAATTTGTGTTTAAGAATATTGATGGTTAT   22162
gb:MT020880|Organism:Severe      GGAAAACAGGGTAATTTCAAAAATCTTAGGGAATTTGTGTTTAAGAATATTGATGGTTAT   22162
gb:MT066175|Organism:Severe      GGAAAACAGGGTAATTTCAAAAATCTTAGGGAATTTGTGTTTAAGAATATTGATGGTTAT   22162
gb:MN997409|Organism:Severe      GGAAAACAGGGTAATTTCAAAAATCTTAGGGAATTTGTGTTTAAGAATATTGATGGTTAT   22162
gb:MN938384|Organism:Severe      GGAAAACAGGGTAATTTCAAAAATCTTAGGGAATTTGTGTTTAAGAATATTGATGGTTAT   22130
gb:MT044258|Organism:Severe      GGAAAACAGGGTAATTTCAAAAATCTTAGGGAATTTGTGTTTAAGAATATTGATGGTTAT   22138
gb:MT039890|Organism:Severe      GGAAAACAGGGTAATTTCAAAAATCTTAGGGAATTTGTGTTTAAGAATATTGATGGTTAT   22162
gb:MN988713|Organism:Severe      GGAAAACAGGGTAATTTCAAAAATCTTAGGGAATTTGTGTTTAAGAATATTGATGGTTAT   22162
gb:LC521925|Organism:Severe      GGAAAACAGGGTAATTTCAAAAATCTTAGGGAATTTGTGTTTAAGAATATTGATGGTTAT   22135
gb:MT093571|Organism:Severe      GGAAAACAGGGTAATTTCAAAAATCTTAGGGAATTTGTGTTTAAGAATATTGATGGTTAT   22162
gb:MT039887|Organism:Severe      GGAAAACAGGGTAATTTCAAAAATCTTAGGGAATTTGTGTTTAAGAATATTGATGGTTAT   22159
gb:MT019530|Organism:Severe      GGAAAACAGGGTAATTTCAAAAATCTTAGGGAATTTGTGTTTAAGAATATTGATGGTTAT   22162
gb:MT039888|Organism:Severe      GGAAAACAGGGTAATTTCAAAAATCTTAGGGAATTTGTGTTTAAGAATATTGATGGTTAT   22162
gb:LC522972|Organism:Severe      GGAAAACAGGGTAATTTCAAAAATCTTAGGGAATTTGTGTTTAAGAATATTGATGGTTAT   22159
gb:MT027063|Organism:Severe      GGAAAACAGGGTAATTTCAAAAATCTTAGGGAATTTGTGTTTAAGAATATTGATGGTTAT   22162
gb:MT027062|Organism:Severe      GGAAAACAGGGTAATTTCAAAAATCTTAGGGAATTTGTGTTTAAGAATATTGATGGTTAT   22162
gb:MT019529|Organism:Severe      GGAAAACAGGGTAATTTCAAAAATCTTAGGGAATTTGTGTTTAAGAATATTGATGGTTAT   22162
gb:MN996529|Organism:Severe      GGAAAACAGGGTAATTTCAAAAATCTTAGGGAATTTGTGTTTAAGAATATTGATGGTTAT   22150
gb:MN996531|Organism:Severe      GGAAAACAGGGTAATTTCAAAAATCTTAGGGAATTTGTGTTTAAGAATATTGATGGTTAT   22149
gb:MT066176|Organism:Severe      GGAAAACAGGGTAATTTCAAAAATCTTAGGGAATTTGTGTTTAAGAATATTGATGGTTAT   22162
gb:MT027064|Organism:Severe      GGAAAACAGGGTAATTTCAAAAATCTTAGGGAATTTGTGTTTAAGAATATTGATGGTTAT   22162
gb:MN994468|Organism:Severe      GGAAAACAGGGTAATTTCAAAAATCTTAGGGAATTTGTGTTTAAGAATATTGATGGTTAT   22162
gb:MT072688|Organism:Severe      GGAAAACAGGGTAATTTCAAAAATCTTAGGGAATTTGTGTTTAAGAATATTGATGGTTAT   22147
gb:MN996527|Organism:Severe      GGAAAACAGGGTAATTTCAAAAATCTTAGGGAATTTGTGTTTAAGAATATTGATGGTTAT   22129
gb:MT093631|Organism:Severe      GGAAAACAGGGTAATTTCAAAAATCTTAGGGAATTTGTGTTTAAGAATATTGATGGTTAT   22200
gb:MT106053|Organism:Severe      GGAAAACAGGGTAATTTCAAAAATCTTAGGGAATTTGTGTTTAAGAATATTGATGGTTAT   22162
gb:MT019533|Organism:Severe      GGAAAACAGGGTAATTTCAAAAATCTTAGGGAATTTGTGTTTAAGAATATTGATGGTTAT   22162
gb:MT019531|Organism:Severe      GGAAAACAGGGTAATTTCAAAAATCTTAGGGAATTTGTGTTTAAGAATATTGATGGTTAT   22162
gb:MN996528|Organism:Severe      GGAAAACAGGGTAATTTCAAAAATCTTAGGGAATTTGTGTTTAAGAATATTGATGGTTAT   22162
gb:MN996530|Organism:Severe      GGAAAACAGGGTAATTTCAAAAATCTTAGGGAATTTGTGTTTAAGAATATTGATGGTTAT   22148
gb:MN908947|Organism:Severe      GGAAAACAGGGTAATTTCAAAAATCTTAGGGAATTTGTGTTTAAGAATATTGATGGTTAT   22162
gb:MT019532|Organism:Severe      GGAAAACAGGGTAATTTCAAAAATCTTAGGGAATTTGTGTTTAAGAATATTGATGGTTAT   22162
                                 ************************************************************

gb:MT020781|Organism:Severe      TTTAAAATATATTCTAAGCACACGCCTATTAATTTAGTGCGTGATCTCCCTCAGGGTTTT   22210
gb:MT007544|Organism:Severe      TTTAAAATATATTCTAAGCACACGCCTATTAATTTAGTGCGTGATCTCCCTCAGGGTTTT   22222
gb:MN994467|Organism:Severe      TTTAAAATATATTCTAAGCACACGCCTATTAATTTAGTGCGTGATCTCCCTCAGGGTTTT   22222
gb:MT044257|Organism:Severe      TTTAAAATATATTCTAAGCACACGCCTATTAATTTAGTGCGTGATCTCCCTCAGGGTTTT   22222
gb:MT106054|Organism:Severe      TTTAAAATATATTCTAAGCACACGCCTATTAATTTAGTGCGTGATCTCCCTCAGGGTTTT   22222
gb:MT049951|Organism:Severe      TTTAAAATATATTCTAAGCACACGCCTATTAATTTAGTGCGTGATCTCCCTCAGGGTTTT   22222
```

```
gb:MN975262|Organism:Severe      TTTAAAATATATTCTAAGCACACGCCTATTAATTTAGTGCGTGATCTCCCTCAGGGTTTT   22222
gb:MT106052|Organism:Severe      TTTAAAATATATTCTAAGCACACGCCTATTAATTTAGTGCGTGATCTCCCTCAGGGTTTT   22222
gb:LC522975|Organism:Severe      TTTAAAATATATTCTAAGCACACGCCTATTAATTTAGTGCGTGATCTCCCTCAGGGTTTT   22219
gb:LC522973|Organism:Severe      TTTAAAATATATTCTAAGCACACGCCTATTAATTTAGTGCGTGATCTCCCTCAGGGTTTT   22219
gb:LC522974|Organism:Severe      TTTAAAATATATTCTAAGCACACGCCTATTAATTTAGTGCGTGATCTCCCTCAGGGTTTT   22219
gb:MN985325|Organism:Severe      TTTAAAATATATTCTAAGCACACGCCTATTAATTTAGTGCGTGATCTCCCTCAGGGTTTT   22222
gb:MT020881|Organism:Severe      TTTAAAATATATTCTAAGCACACGCCTATTAATTTAGTGCGTGATCTCCCTCAGGGTTTT   22222
gb:MT020880|Organism:Severe      TTTAAAATATATTCTAAGCACACGCCTATTAATTTAGTGCGTGATCTCCCTCAGGGTTTT   22222
gb:MT066175|Organism:Severe      TTTAAAATATATTCTAAGCACACGCCTATTAATTTAGTGCGTGATCTCCCTCAGGGTTTT   22222
gb:MN997409|Organism:Severe      TTTAAAATATATTCTAAGCACACGCCTATTAATTTAGTGCGTGATCTCCCTCAGGGTTTT   22222
gb:MN938384|Organism:Severe      TTTAAAATATATTCTAAGCACACGCCTATTAATTTAGTGCGTGATCTCCCTCAGGGTTTT   22190
gb:MT044258|Organism:Severe      TTTAAAATATATTCTAAGCACACGCCTATTAATTTAGTGCGTGATCTCCCTCAGGGTTTT   22198
gb:MT039890|Organism:Severe      TTTAAAATATATTCTAAGCACACGCCTATTAATTTAGTGCGTGATCTCCCTCAGGGTTTT   22222
gb:MN988713|Organism:Severe      TTTAAAATATATTCTAAGCACACGCCTATTAATTTAGTGCGTGATCTCCCTCAGGGTTTT   22222
gb:LC521925|Organism:Severe      TTTAAAATATATTCTAAGCACACGCCTATTAATTTAGTGCGTGATCTCCCTCAGGGTTTT   22195
gb:MT093571|Organism:Severe      TTTAAAATATATTCTAAGCACACGCCTATTAATTTAGTGCGTGATCTCCCTCAGGGTTTT   22222
gb:MT039887|Organism:Severe      TTTAAAATATATTCTAAGCACACGCCTATTAATTTAGTGCGTGATCTCCCTCAGGGTTTT   22219
gb:MT019530|Organism:Severe      TTTAAAATATATTCTAAGCACACGCCTATTAATTTAGTGCGTGATCTCCCTCAGGGTTTT   22222
gb:MT039888|Organism:Severe      TTTAAAATATATTCTAAGCACACGCCTATTAATTTAGTGCGTGATCTCCCTCAGGGTTTT   22222
gb:LC522972|Organism:Severe      TTTAAAATATATTCTAAGCACACGCCTATTAATTTAGTGCGTGATCTCCCTCAGGGTTTT   22219
gb:MT027063|Organism:Severe      TTTAAAATATATTCTAAGCACACGCCTATTAATTTAGTGCGTGATCTCCCTCAGGGTTTT   22222
gb:MT027062|Organism:Severe      TTTAAAATATATTCTAAGCACACGCCTATTAATTTAGTGCGTGATCTCCCTCAGGGTTTT   22222
gb:MT019529|Organism:Severe      TTTAAAATATATTCTAAGCACACGCCTATTAATTTAGTGCGTGATCTCCCTCAGGGTTTT   22222
gb:MN996529|Organism:Severe      TTTAAAATATATTCTAAGCACACGCCTATTAATTTAGTGCGTGATCTCCCTCAGGGTTTT   22210
gb:MN996531|Organism:Severe      TTTAAAATATATTCTAAGCACACGCCTATTAATTTAGTGCGTGATCTCCCTCAGGGTTTT   22209
gb:MT066176|Organism:Severe      TTTAAAATATATTCTAAGCACACGCCTATTAATTTAGTGCGTGATCTCCCTCAGGGTTTT   22222
gb:MT027064|Organism:Severe      TTTAAAATATATTCTAAGCACACGCCTATTAATTTAGTGCGTGATCTCCCTCAGGGTTTT   22222
gb:MN994468|Organism:Severe      TTTAAAATATATTCTAAGCACACGCCTATTAATTTAGTGCGTGATCTCCCTCAGGGTTTT   22222
gb:MT072688|Organism:Severe      TTTAAAATATATTCTAAGCACACGCCTATTAATTTAGTGCGTGATCTCCCTCAGGGTTTT   22207
gb:MN996527|Organism:Severe      TTTAAAATATATTCTAAGCACACGCCTATTAATTTAGTGCGTGATCTCCCTCAGGGTTTT   22189
gb:MT093631|Organism:Severe      TTTAAAATATATTCTAAGCACACGCCTATTAATTTAGTGCGTGATCTCCCTCAGGGTTTT   22260
gb:MT106053|Organism:Severe      TTTAAAATATATTCTAAGCACACGCCTATTAATTTAGTGCGTGATCTCCCTCAGGGTTTT   22222
gb:MT019533|Organism:Severe      TTTAAAATATATTCTAAGCACACGCCTATTAATTTAGTGCGTGATCTCCCTCAGGGTTTT   22222
gb:MT019531|Organism:Severe      TTTAAAATATATTCTAAGCACACGCCTATTAATTTAGTGCGTGATCTCCCTCAGGGTTTT   22222
gb:MN996528|Organism:Severe      TTTAAAATATATTCTAAGCACACGCCTATTAATTTAGTGCGTGATCTCCCTCAGGGTTTT   22222
gb:MN996530|Organism:Severe      TTTAAAATATATTCTAAGCACACGCCTATTAATTTAGTGCGTGATCTCCCTCAGGGTTTT   22208
gb:MN908947|Organism:Severe      TTTAAAATATATTCTAAGCACACGCCTATTAATTTAGTGCGTGATCTCCCTCAGGGTTTT   22222
gb:MT019532|Organism:Severe      TTTAAAATATATTCTAAGCACACGCCTATTAATTTAGTGCGTGATCTCCCTCAGGGTTTT   22222
                                 ************************************************************

gb:MT020781|Organism:Severe      TCGGCTTTAGAACCATTGGTAGATTTGCCAATAGGTATTAACATCACTAGGTTTCAAACT   22270
gb:MT007544|Organism:Severe      TCGGCTTTAGAACCATTGGTAGATTTGCCAATAGGTATTAACATCACTAGGTTTCAAACT   22282
gb:MN994467|Organism:Severe      TCGGCTTTAGAACCATTGGTAGATTTGCCAATAGGTATTAACATCACTAGGTTTCAAACT   22282
gb:MT044257|Organism:Severe      TCGGCTTTAGAACCATTGGTAGATTTGCCAATAGGTATTAACATCACTAGGTTTCAAACT   22282
gb:MT106054|Organism:Severe      TCGGCTTTAGAACCATTGGTAGATTTGCCAATAGGTATTAACATCACTAGGTTTCAAACT   22282
gb:MT049951|Organism:Severe      TCGGCTTTAGAACCATTGGTAGATTTGCCAATAGGTATTAACATCACTAGGTTTCAAACT   22282
gb:MN975262|Organism:Severe      TCGGCTTTAGAACCATTGGTAGATTTGCCAATAGGTATTAACATCACTAGGTTTCAAACT   22282
gb:MT106052|Organism:Severe      TCGGCTTTAGAACCATTGGTAGATTTGCCAATAGGTATTAACATCACTAGGTTTCAAACT   22282
gb:LC522975|Organism:Severe      TCGGCTTTAGAACCATTGGTAGATTTGCCAATAGGTATTAACATCACTAGGTTTCAAACT   22279
gb:LC522973|Organism:Severe      TCGGCTTTAGAACCATTGGTAGATTTGCCAATAGGTATTAACATCACTAGGTTTCAAACT   22279
gb:LC522974|Organism:Severe      TCGGCTTTAGAACCATTGGTAGATTTGCCAATAGGTATTAACATCACTAGGTTTCAAACT   22279
gb:MN985325|Organism:Severe      TCGGCTTTAGAACCATTGGTAGATTTGCCAATAGGTATTAACATCACTAGGTTTCAAACT   22282
gb:MT020881|Organism:Severe      TCGGCTTTAGAACCATTGGTAGATTTGCCAATAGGTATTAACATCACTAGGTTTCAAACT   22282
gb:MT020880|Organism:Severe      TCGGCTTTAGAACCATTGGTAGATTTGCCAATAGGTATTAACATCACTAGGTTTCAAACT   22282
gb:MT066175|Organism:Severe      TCGGCTTTAGAACCATTGGTAGATTTGCCAATAGGTATTAACATCACTAGGTTTCAAACT   22282
gb:MN997409|Organism:Severe      TCGGCTTTAGAACCATTGGTAGATTTGCCAATAGGTATTAACATCACTAGGTTTCAAACT   22282
gb:MN938384|Organism:Severe      TCGGCTTTAGAACCATTGGTAGATTTGCCAATAGGTATTAACATCACTAGGTTTCAAACT   22250
gb:MT044258|Organism:Severe      TCGGCTTTAGAACCATTGGTAGATTTGCCAATAGGTATTAACATCACTAGGTTTCAAACT   22258
gb:MT039890|Organism:Severe      TGGGCTTTAGAACCATTGGTAGATTTGCCAATAGGTATTAACATCACTAGGTTTCAAACT   22282
gb:MN988713|Organism:Severe      TCGGCTTTAGAACCATTGGTAGATTTGCCAATAGGTATTAACATCACTAGGTTTCAAACT   22282
gb:LC521925|Organism:Severe      TCGGCTTTAGAACCATTGGTAGATTTGCCAATAGGTATTAACATCACTAGGTTTCAAACT   22255
gb:MT093571|Organism:Severe      TCGGCTTTAGAACCATTGGTAGATTTGCCAATAGGTATTAACATCACTAGGTTTCAAACT   22282
gb:MT039887|Organism:Severe      TCGGCTTTAGAACCATTGGTAGATTTGCCAATAGGTATTAACATCACTAGGTTTCAAACT   22279
gb:MT019530|Organism:Severe      TCGGCTTTAGAACCATTGGTAGATTTGCCAATAGGTATTAACATCACTAGGTTTCAAACT   22282
gb:MT039888|Organism:Severe      TCGGCTTTAGAACCATTGGTAGATTTGCCAATAGGTATTAACATCACTAGGTTTCAAACT   22282
```

```
gb:LC522972|Organism:Severe      TCGGCTTTAGAACCATTGGTAGATTTGCCAATAGGTATTAACATCACTAGGTTTCAAACT    22279
gb:MT027063|Organism:Severe      TCGGCTTTAGAACCATTGGTAGATTTGCCAATAGGTATTAACATCACTAGGTTTCAAACT    22282
gb:MT027062|Organism:Severe      TCGGCTTTAGAACCATTGGTAGATTTGCCAATAGGTATTAACATCACTAGGTTTCAAACT    22282
gb:MT019529|Organism:Severe      TCGGCTTTAGAACCATTGGTAGATTTGCCAATAGGTATTAACATCACTAGGTTTCAAACT    22282
gb:MN996529|Organism:Severe      TCGGCTTTAGAACCATTGGTAGATTTGCCAATAGGTATTAACATCACTAGGTTTCAAACT    22270
gb:MN996531|Organism:Severe      TCGGCTTTAGAACCATTGGTAGATTTGCCAATAGGTATTAACATCACTAGGTTTCAAACT    22269
gb:MT066176|Organism:Severe      TCGGCTTTAGAACCATTGGTAGATTTGCCAATAGGTATTAACATCACTAGGTTTCAAACT    22282
gb:MT027064|Organism:Severe      TCGGCTTTAGAACCATTGGTAGATTTGCCAATAGGTATTAACATCACTAGGTTTCAAACT    22282
gb:MN994468|Organism:Severe      TCGGCTTTAGAACCATTGGTAGATTTGCCAATAGGTATTAACATCACTAGGTTTCAAACT    22282
gb:MT072688|Organism:Severe      TCGGCTTTAGAACCATTGGTAGATTTGCCAATAGGTATTAACATCACTAGGTTTCAAACT    22267
gb:MN996527|Organism:Severe      TCGGCTTTAGAACCATTGGTAGATTTGCCAATAGGTATTAACATCACTAGGTTTCAAACT    22249
gb:MT093631|Organism:Severe      TCGGCTTTAGAACCATTGGTAGATTTGCCAATAGGTATTAACATCACTAGGTTTCAAACT    22320
gb:MT106053|Organism:Severe      TCGGCTTTAGAACCATTGGTAGATTTGCCAATAGGTATTAACATCACTAGGTTTCAAACT    22282
gb:MT019533|Organism:Severe      TCGGCTTTAGAACCATTGGTAGATTTGCCAATAGGTATTAACATCACTAGGTTTCAAACT    22282
gb:MT019531|Organism:Severe      TCGGCTTTAGAACCATTGGTAGATTTGCCAATAGGTATTAACATCACTAGGTTTCAAACT    22282
gb:MN996528|Organism:Severe      TCGGCTTTAGAACCATTGGTAGATTTGCCAATAGGTATTAACATCACTAGGTTTCAAACT    22282
gb:MN996530|Organism:Severe      TCGGCTTTAGAACCATTGGTAGATTTGCCAATAGGTATTAACATCACTAGGTTTCAAACT    22268
gb:MN908947|Organism:Severe      TCGGCTTTAGAACCATTGGTAGATTTGCCAATAGGTATTAACATCACTAGGTTTCAAACT    22282
gb:MT019532|Organism:Severe      TCGGCTTTAGAACCATTGGTAGATTTGCCAATAGGTATTAACATCACTAGGTTTCAAACT    22282
                                 * **********************************************************

gb:MT020781|Organism:Severe      TTACTTGCTTTACATAGAAGTTATTTGACTCCTGGTGATTCTTCTTCAGGTTGGACAGCT    22330
gb:MT007544|Organism:Severe      TTACTTGCTTTACATAGAAGGTATTTGACTCCTGGTGATTCTTCTTCAGGTTGGACAGCT    22342
gb:MN994467|Organism:Severe      TTACTTGCTTTACATAGAAGTTATTTGACTCCTGGTGATTCTTCTTCAGGTTGGACAGCT    22342
gb:MT044257|Organism:Severe      TTACTTGCTTTACATAGAAGTTATTTGACTCCTGGTGATTCTTCTTCAGGTTGGACAGCT    22342
gb:MT106054|Organism:Severe      TTACTTGCTTTACATAGAAGTTATTTGACTCCTGGTGATTCTTCTTCAGGTTGGACAGCT    22342
gb:MT049951|Organism:Severe      TTACTTGCTTTACATAGAAGTTATTTGACTCCTGGTGATTCTTCTTCAGGTTGGACAGCT    22342
gb:MN975262|Organism:Severe      TTACTTGCTTTACATAGAAGTTATTTGACTCCTGGTGATTCTTCTTCAGGTTGGACAGCT    22342
gb:MT106052|Organism:Severe      TTACTTGCTTTACATAGAAGTTATTTGACTCCTGGTGATTCTTCTTCAGGTTGGACAGCT    22342
gb:LC522975|Organism:Severe      TTACTTGCTTTACATAGAAGTTATTTGACTCCTGGTGATTCTTCTTCAGGTTGGACAGCT    22339
gb:LC522973|Organism:Severe      TTACTTGCTTTACATAGAAGTTATTTGACTCCTGGTGATTCTTCTTCAGGTTGGACAGCT    22339
gb:LC522974|Organism:Severe      TTACTTGCTTTACATAGAAGTTATTTGACTCCTGGTGATTCTTCTTCAGGTTGGACAGCT    22339
gb:MN985325|Organism:Severe      TTACTTGCTTTACATAGAAGTTATTTGACTCCTGGTGATTCTTCTTCAGGTTGGACAGCT    22342
gb:MT020881|Organism:Severe      TTACTTGCTTTACATAGAAGTTATTTGACTCCTGGTGATTCTTCTTCAGGTTGGACAGCT    22342
gb:MT020880|Organism:Severe      TTACTTGCTTTACATAGAAGTTATTTGACTCCTGGTGATTCTTCTTCAGGTTGGACAGCT    22342
gb:MT066175|Organism:Severe      TTACTTGCTTTACATAGAAGTTATTTGACTCCTGGTGATTCTTCTTCAGGTTGGACAGCT    22342
gb:MN997409|Organism:Severe      TTACTTGCTTTACATAGAAGTTATTTGACTCCTGGTGATTCTTCTTCAGGTTGGACAGCT    22342
gb:MN938384|Organism:Severe      TTACTTGCTTTACATAGAAGTTATTTGACTCCTGGTGATTCTTCTTCAGGTTGGACAGCT    22310
gb:MT044258|Organism:Severe      TTACTTGCTTTACATAGAAGTTATTTGACTCCTGGTGATTCTTCTTCAGGTTGGACAGCT    22318
gb:MT039890|Organism:Severe      TTACTTGCTTTACATAGAAGTTATTTGACTCCTGGTGATTCTTCTTCAGGTTGGACAGCT    22342
gb:MN988713|Organism:Severe      TTACTTGCTTTACATAGAAGTTATTTGACTCCTGGTGATTCTTCTTCAGGTTGGACAGCT    22342
gb:LC521925|Organism:Severe      TTACTTGCTTTACATAGAAGTTATTTGACTCCTGGTGATTCTTCTTCAGGTTGGACAGCT    22315
gb:MT093571|Organism:Severe      TTACTTGCTTTACATAGAAGTTATTTGACTCCTGGTGATTCTTCTTCAGGTTGGACAGCT    22342
gb:MT039887|Organism:Severe      TTACTTGCTTTACATAGAAGTTATTTGACTCCTGGTGATTCTTCTTCAGGTTGGACAGCT    22339
gb:MT019530|Organism:Severe      TTACTTGCTTTACATAGAAGTTATTTGACTCCTGGTGATTCTTCTTCAGGTTGGACAGCT    22342
gb:MT039888|Organism:Severe      TTACTTGCTTTACATAGAAGTTATTTGACTCCTGGTGATTCTTCTTCAGGTTGGACAGCT    22342
gb:LC522972|Organism:Severe      TTACTTGCTTTACATAGAAGTTATTTGACTCCTGGTGATTCTTCTTCAGGTTGGACAGCT    22339
gb:MT027063|Organism:Severe      TTACTTGCTTTACATAGAAGTTATTTGACTCCTGGTGATTCTTCTTCAGGTTGGACAGCT    22342
gb:MT027062|Organism:Severe      TTACTTGCTTTACATAGAAGTTATTTGACTCCTGGTGATTCTTCTTCAGGTTGGACAGCT    22342
gb:MT019529|Organism:Severe      TTACTTGCTTTACATAGAAGTTATTTGACTCCTGGTGATTCTTCTTCAGGTTGGACAGCT    22342
gb:MN996529|Organism:Severe      TTACTTGCTTTACATAGAAGTTATTTGACTCCTGGTGATTCTTCTTCAGGTTGGACAGCT    22330
gb:MN996531|Organism:Severe      TTACTTGCTTTACATAGAAGTTATTTGACTCCTGGTGATTCTTCTTCAGGTTGGACAGCT    22329
gb:MT066176|Organism:Severe      TTACTTGCTTTACATAGAAGTTATTTGACTCCTGGTGATTCTTCTTCAGGTTGGACAGCT    22342
gb:MT027064|Organism:Severe      TTACTTGCTTTACATAGAAGTTATTTGACTCCTGGTGATTCTTCTTCAGGTTGGACAGCT    22342
gb:MN994468|Organism:Severe      TTACTTGCTTTACATAGAAGTTATTTGACTCCTGGTGATTCTTCTTCAGGTTGGACAGCT    22342
gb:MT072688|Organism:Severe      TTACTTGCTTTACATAGAAGTTATTTGACTCCTGGTGATTCTTCTTCAGGTTGGACAGCT    22327
gb:MN996527|Organism:Severe      TTACTTGCTTTACATAGAAGTTATTTGACTCCTGGTGATTCTTCTTCAGGTTGGACAGCT    22309
gb:MT093631|Organism:Severe      TTACTTGCTTTACATAGAAGTTATTTGACTCCTGGTGATTCTTCTTCAGGTTGGACAGCT    22380
gb:MT106053|Organism:Severe      TTACTTGCTTTACATAGAAGTTATTTGACTCCTGGTGATTCTTCTTCAGGTTGGACAGCT    22342
gb:MT019533|Organism:Severe      TTACTTGCTTTACATAGAAGTTATTTGACTCCTGGTGATTCTTCTTCAGGTTGGACAGCT    22342
gb:MT019531|Organism:Severe      TTACTTGCTTTACATAGAAGTTATTTGACTCCTGGTGATTCTTCTTCAGGTTGGACAGCT    22342
gb:MN996528|Organism:Severe      TTACTTGCTTTACATAGAAGTTATTTGACTCCTGGTGATTCTTCTTCAGGTTGGACAGCT    22342
gb:MN996530|Organism:Severe      TTACTTGCTTTACATAGAAGTTATTTGACTCCTGGTGATTCTTCTTCAGGTTGGACAGCT    22328
gb:MN908947|Organism:Severe      TTACTTGCTTTACATAGAAGTTATTTGACTCCTGGTGATTCTTCTTCAGGTTGGACAGCT    22342
gb:MT019532|Organism:Severe      TTACTTGCTTTACATAGAAGTTATTTGACTCCTGGTGATTCTTCTTCAGGTTGGACAGCT    22342
```

```
                                        ******************** ***************************************

gb:MT020781|Organism:Severe      GGTGCTGCAGCTTATTATGTGGGTTATCTTCAACCTAGGACTTTTCTATTAAAATATAAT   22390
gb:MT007544|Organism:Severe      GGTGCTGCAGCTTATTATGTGGGTTATCTTCAACCTAGGACTTTTCTATTAAAATATAAT   22402
gb:MN994467|Organism:Severe      GGTGCTGCAGCTTATTATGTGGGTTATCTTCAACCTAGGACTTTTCTATTAAAATATAAT   22402
gb:MT044257|Organism:Severe      GGTGCTGCAGCTTATTATGTGGGTTATCTTCAACCTAGGACTTTTCTATTAAAATATAAT   22402
gb:MT106054|Organism:Severe      GGTGCTGCAGCTTATTATGTGGGTTATCTTCAACCTAGGACTTTTCTATTAAAATATAAT   22402
gb:MT049951|Organism:Severe      GGTGCTGCAGCTTATTATGTGGGTTATCTTCAACCTAGGACTTTTCTATTAAAATATAAT   22402
gb:MN975262|Organism:Severe      GGTGCTGCAGCTTATTATGTGGGTTATCTTCAACCTAGGACTTTTCTATTAAAATATAAT   22402
gb:MT106052|Organism:Severe      GGTGCTGCAGCTTATTATGTGGGTTATCTTCAACCTAGGACTTTTCTATTAAAATATAAT   22402
gb:LC522975|Organism:Severe      GGTGCTGCAGCTTATTATGTGGGTTATCTTCAACCTAGGACTTTTCTATTAAAATATAAT   22399
gb:LC522973|Organism:Severe      GGTGCTGCAGCTTATTATGTGGGTTATCTTCAACCTAGGACTTTTCTATTAAAATATAAT   22399
gb:LC522974|Organism:Severe      GGTGCTGCAGCTTATTATGTGGGTTATCTTCAACCTAGGACTTTTCTATTAAAATATAAT   22399
gb:MN985325|Organism:Severe      GGTGCTGCAGCTTATTATGTGGGTTATCTTCAACCTAGGACTTTTCTATTAAAATATAAT   22402
gb:MT020881|Organism:Severe      GGTGCTGCAGCTTATTATGTGGGTTATCTTCAACCTAGGACTTTTCTATTAAAATATAAT   22402
gb:MT020880|Organism:Severe      GGTGCTGCAGCTTATTATGTGGGTTATCTTCAACCTAGGACTTTTCTATTAAAATATAAT   22402
gb:MT066175|Organism:Severe      GGTGCTGCAGCTTATTATGTGGGTTATCTTCAACCTAGGACTTTTCTATTAAAATATAAT   22402
gb:MN997409|Organism:Severe      GGTGCTGCAGCTTATTATGTGGGTTATCTTCAACCTAGGACTTTTCTATTAAAATATAAT   22402
gb:MN938384|Organism:Severe      GGTGCTGCAGCTTATTATGTGGGTTATCTTCAACCTAGGACTTTTCTATTAAAATATAAT   22370
gb:MT044258|Organism:Severe      GGTGCTGCAGCTTATTATGTGGGTTATCTTCAACCTAGGACTTTTCTATTAAAATATAAT   22378
gb:MT039890|Organism:Severe      GGTGCTGCAGCTTATTATGTGGGTTATCTTCAACCTAGGACTTTTCTATTAAAATATAAT   22402
gb:MN988713|Organism:Severe      GGTGCTGCAGCTTATTATGTGGGTTATCTTCAACCTAGGACTTTTCTATTAAAATATAAT   22402
gb:LC521925|Organism:Severe      GGTGCTGCAGCTTATTATGTGGGTTATCTTCAACCTAGGACTTTTCTATTAAAATATAAT   22375
gb:MT093571|Organism:Severe      GGTGCTGCAGCTTATTATGTGGGTTATCTTCAACCTAGGACTTTTCTATTAAAATATAAT   22402
gb:MT039887|Organism:Severe      GGTGCTGCAGCTTATTATGTGGGTTATCTTCAACCTAGGACTTTTCTATTAAAATATAAT   22399
gb:MT019530|Organism:Severe      GGTGCTGCAGCTTATTATGTGGGTTATCTTCAACCTAGGACTTTTCTATTAAAATATAAT   22402
gb:MT039888|Organism:Severe      GGTGCTGCAGCTTATTATGTGGGTTATCTTCAACCTAGGACTTTTCTATTAAAATATAAT   22402
gb:LC522972|Organism:Severe      GGTGCTGCAGCTTATTATGTGGGTTATCTTCAACCTAGGACTTTTCTATTAAAATATAAT   22399
gb:MT027063|Organism:Severe      GGTGCTGCAGCTTATTATGTGGGTTATCTTCAACCTAGGACTTTTCTATTAAAATATAAT   22402
gb:MT027062|Organism:Severe      GGTGCTGCAGCTTATTATGTGGGTTATCTTCAACCTAGGACTTTTCTATTAAAATATAAT   22402
gb:MT019529|Organism:Severe      GGTGCTGCAGCTTATTATGTGGGTTATCTTCAACCTAGGACTTTTCTATTAAAATATAAT   22402
gb:MN996529|Organism:Severe      GGTGCTGCAGCTTATTATGTGGGTTATCTTCAACCTAGGACTTTTCTATTAAAATATAAT   22390
gb:MN996531|Organism:Severe      GGTGCTGCAGCTTATTATGTGGGTTATCTTCAACCTAGGACTTTTCTATTAAAATATAAT   22389
gb:MT066176|Organism:Severe      GGTGCTGCAGCTTATTATGTGGGTTATCTTCAACCTAGGACTTTTCTATTAAAATATAAT   22402
gb:MT027064|Organism:Severe      GGTGCTGCAGCTTATTATGTGGGTTATCTTCAACCTAGGACTTTTCTATTAAAATATAAT   22402
gb:MN994468|Organism:Severe      GGTGCTGCAGCTTATTATGTGGGTTATCTTCAACCTAGGACTTTTCTATTAAAATATAAT   22402
gb:MT072688|Organism:Severe      GGTGCTGCAGCTTATTATGTGGGTTATCTTCAACCTAGGACTTTTCTATTAAAATATAAT   22387
gb:MN996527|Organism:Severe      GGTGCTGCAGCTTATTATGTGGGTTATCTTCAACCTAGGACTTTTCTATTAAAATATAAT   22369
gb:MT093631|Organism:Severe      GGTGCTGCAGCTTATTATGTGGGTTATCTTCAACCTAGGACTTTTCTATTAAAATATAAT   22440
gb:MT106053|Organism:Severe      GGTGCTGCAGCTTATTATGTGGGTTATCTTCAACCTAGGACTTTTCTATTAAAATATAAT   22402
gb:MT019533|Organism:Severe      GGTGCTGCAGCTTATTATGTGGGTTATCTTCAACCTAGGACTTTTCTATTAAAATATAAT   22402
gb:MT019531|Organism:Severe      GGTGCTGCAGCTTATTATGTGGGTTATCTTCAACCTAGGACTTTTCTATTAAAATATAAT   22402
gb:MN996528|Organism:Severe      GGTGCTGCAGCTTATTATGTGGGTTATCTTCAACCTAGGACTTTTCTATTAAAATATAAT   22402
gb:MN996530|Organism:Severe      GGTGCTGCAGCTTATTATGTGGGTTATCTTCAACCTAGGACTTTTCTATTAAAATATAAT   22388
gb:MN908947|Organism:Severe      GGTGCTGCAGCTTATTATGTGGGTTATCTTCAACCTAGGACTTTTCTATTAAAATATAAT   22402
gb:MT019532|Organism:Severe      GGTGCTGCAGCTTATTATGTGGGTTATCTTCAACCTAGGACTTTTCTATTAAAATATAAT   22402
                                 ************************************************************

gb:MT020781|Organism:Severe      GAAAATGGAACCATTACAGATGCTGTAGACTGTGCACTTGACCCTCTCTCAGAAACAAAG   22450
gb:MT007544|Organism:Severe      GAAAATGGAACCATTACAGATGCTGTAGACTGTGCACTTGACCCTCTCTCAGAAACAAAG   22462
gb:MN994467|Organism:Severe      GAAAATGGAACCATTACAGATGCTGTAGACTGTGCACTTGACCCTCTCTCAGAAACAAAG   22462
gb:MT044257|Organism:Severe      GAAAATGGAACCATTACAGATGCTGTAGACTGTGCACTTGACCCTCTCTCAGAAACAAAG   22462
gb:MT106054|Organism:Severe      GAAAATGGAACCATTACAGATGCTGTAGACTGTGCACTTGACCCTCTCTCAGAAACAAAG   22462
gb:MT049951|Organism:Severe      GAAAATGGAACCATTACAGATGCTGTAGACTGTGCACTTGACCCTCTCTCAGAAACAAAG   22462
gb:MN975262|Organism:Severe      GAAAATGGAACCATTACAGATGCTGTAGACTGTGCACTTGACCCTCTCTCAGAAACAAAG   22462
gb:MT106052|Organism:Severe      GAAAATGGAACCATTACAGATGCTGTAGACTGTGCACTTGACCCTCTCTCAGAAACAAAG   22462
gb:LC522975|Organism:Severe      GAAAATGGAACCATTACAGATGCTGTAGACTGTGCACTTGACCCTCTCTCAGAAACAAAG   22459
gb:LC522973|Organism:Severe      GAAAATGGAACCATTACAGATGCTGTAGACTGTGCACTTGACCCTCTCTCAGAAACAAAG   22459
gb:LC522974|Organism:Severe      GAAAATGGAACCATTACAGATGCTGTAGACTGTGCACTTGACCCTCTCTCAGAAACAAAG   22459
gb:MN985325|Organism:Severe      GAAAATGGAACCATTACAGATGCTGTAGACTGTGCACTTGACCCTCTCTCAGAAACAAAG   22462
gb:MT020881|Organism:Severe      GAAAATGGAACCATTACAGATGCTGTAGACTGTGCACTTGACCCTCTCTCAGAAACAAAG   22462
gb:MT020880|Organism:Severe      GAAAATGGAACCATTACAGATGCTGTAGACTGTGCACTTGACCCTCTCTCAGAAACAAAG   22462
gb:MT066175|Organism:Severe      GAAAATGGAACCATTACAGATGCTGTAGACTGTGCACTTGACCCTCTCTCAGAAACAAAG   22462
gb:MN997409|Organism:Severe      GAAAATGGAACCATTACAGATGCTGTAGACTGTGCACTTGACCCTCTCTCAGAAACAAAG   22462
gb:MN938384|Organism:Severe      GAAAATGGAACCATTACAGATGCTGTAGACTGTGCACTTGACCCTCTCTCAGAAACAAAG   22430
```

```
gb:MT044258|Organism:Severe      GAAAATGGAACCATTACAGATGCTGTAGACTGTGCACTTGACCCTCTCTCAGAAACAAAG    22438
gb:MT039890|Organism:Severe      GAAAATGGAACCATTACAGATGCTGTAGACTGTGCACTTGACCCTCTCTCAGAAACAAAG    22462
gb:MN988713|Organism:Severe      GAAAATGGAACCATTACAGATGCTGTAGACTGTGCACTTGACCCTCTCTCAGAAACAAAG    22462
gb:LC521925|Organism:Severe      GAAAATGGAACCATTACAGATGCTGTAGACTGTGCACTTGACCCTCTCTCAGAAACAAAG    22435
gb:MT093571|Organism:Severe      GAAAATGGAACCATTACAGATGCTGTAGACTGTGCACTTGACCCTCTCTCAGAAACAAAG    22462
gb:MT039887|Organism:Severe      GAAAATGGAACCATTACAGATGCTGTAGACTGTGCACTTGACCCTCTCTCAGAAACAAAG    22459
gb:MT019530|Organism:Severe      GAAAATGGAACCATTACAGATGCTGTAGACTGTGCACTTGACCCTCTCTCAGAAACAAAG    22462
gb:MT039888|Organism:Severe      GAAAATGGAACCATTACAGATGCTGTAGACTGTGCACTTGACCCTCTCTCAGAAACAAAG    22462
gb:LC522972|Organism:Severe      GAAAATGGAACCATTACAGATGCTGTAGACTGTGCACTTGACCCTCTCTCAGAAACAAAG    22459
gb:MT027063|Organism:Severe      GAAAATGGAACCATTACAGATGCTGTAGACTGTGCACTTGACCCTCTCTCAGAAACAAAG    22462
gb:MT027062|Organism:Severe      GAAAATGGAACCATTACAGATGCTGTAGACTGTGCACTTGACCCTCTCTCAGAAACAAAG    22462
gb:MT019529|Organism:Severe      GAAAATGGAACCATTACAGATGCTGTAGACTGTGCACTTGACCCTCTCTCAGAAACAAAG    22462
gb:MN996529|Organism:Severe      GAAAATGGAACCATTACAGATGCTGTAGACTGTGCACTTGACCCTCTCTCAGAAACAAAG    22450
gb:MN996531|Organism:Severe      GAAAATGGAACCATTACAGATGCTGTAGACTGTGCACTTGACCCTCTCTCAGAAACAAAG    22449
gb:MT066176|Organism:Severe      GAAAATGGAACCATTACAGATGCTGTAGACTGTGCACTTGACCCTCTCTCAGAAACAAAG    22462
gb:MT027064|Organism:Severe      GAAAATGGAACCATTACAGATGCTGTAGACTGTGCACTTGACCCTCTCTCAGAAACAAAG    22462
gb:MN994468|Organism:Severe      GAAAATGGAACCATTACAGATGCTGTAGACTGTGCACTTGACCCTCTCTCAGAAACAAAG    22462
gb:MT072688|Organism:Severe      GAAAATGGAACCATTACAGATGCTGTAGACTGTGCACTTGACCCTCTCTCAGAAACAAAG    22447
gb:MN996527|Organism:Severe      GAAAATGGAACCATTACAGATGCTGTAGACTGTGCACTTGACCCTCTCTCAGAAACAAAG    22429
gb:MT093631|Organism:Severe      GAAAATGGAACCATTACAGATGCTGTAGACTGTGCACTTGACCCTCTCTCAGAAACAAAG    22500
gb:MT106053|Organism:Severe      GAAAATGGAACCATTACAGATGCTGTAGACTGTGCACTTGACCCTCTCTCAGAAACAAAG    22462
gb:MT019533|Organism:Severe      GAAAATGGAACCATTACAGATGCTGTAGACTGTGCACTTGACCCTCTCTCAGAAACAAAG    22462
gb:MT019531|Organism:Severe      GAAAATGGAACCATTACAGATGCTGTAGACTGTGCACTTGACCCTCTCTCAGAAACAAAG    22462
gb:MN996528|Organism:Severe      GAAAATGGAACCATTACAGATGCTGTAGACTGTGCACTTGACCCTCTCTCAGAAACAAAG    22462
gb:MN996530|Organism:Severe      GAAAATGGAACCATTACAGATGCTGTAGACTGTGCACTTGACCCTCTCTCAGAAACAAAG    22448
gb:MN908947|Organism:Severe      GAAAATGGAACCATTACAGATGCTGTAGACTGTGCACTTGACCCTCTCTCAGAAACAAAG    22462
gb:MT019532|Organism:Severe      GAAAATGGAACCATTACAGATGCTGTAGACTGTGCACTTGACCCTCTCTCAGAAACAAAG    22462
                                 ************************************************************

gb:MT020781|Organism:Severe      TGTACGTTGAAATCCTTCACTGTAGAAAAAGGAATCTATCAAACTTCTAACTTTAGAGTC    22510
gb:MT007544|Organism:Severe      TGTACGTTGAAATCCTTCACTGTAGAAAAAGGAATCTATCAAACTTCTAACTTTAGAGTC    22522
gb:MN994467|Organism:Severe      TGTACGTTGAAATCCTTCACTGTAGAAAAAGGAATCTATCAAACTTCTAACTTTAGAGTC    22522
gb:MT044257|Organism:Severe      TGTACGTTGAAATCCTTCACTGTAGAAAAAGGAATCTATCAAACTTCTAACTTTAGAGTC    22522
gb:MT106054|Organism:Severe      TGTACGTTGAAATCCTTCACTGTAGAAAAAGGAATCTATCAAACTTCTAACTTTAGAGTC    22522
gb:MT049951|Organism:Severe      TGTACGTTGAAATCCTTCACTGTAGAAAAAGGAATCTATCAAACTTCTAACTTTAGAGTC    22522
gb:MN975262|Organism:Severe      TGTACGTTGAAATCCTTCACTGTAGAAAAAGGAATCTATCAAACTTCTAACTTTAGAGTC    22522
gb:MT106052|Organism:Severe      TGTACGTTGAAATCCTTCACTGTAGAAAAAGGAATCTATCAAACTTCTAACTTTAGAGTC    22522
gb:LC522975|Organism:Severe      TGTACGTTGAAATCCTTCACTGTAGAAAAAGGAATCTATCAAACTTCTAACTTTAGAGTC    22519
gb:LC522973|Organism:Severe      TGTACGTTGAAATCCTTCACTGTAGAAAAAGGAATCTATCAAACTTCTAACTTTAGAGTC    22519
gb:LC522974|Organism:Severe      TGTACGTTGAAATCCTTCACTGTAGAAAAAGGAATCTATCAAACTTCTAACTTTAGAGTC    22519
gb:MN985325|Organism:Severe      TGTACGTTGAAATCCTTCACTGTAGAAAAAGGAATCTATCAAACTTCTAACTTTAGAGTC    22522
gb:MT020881|Organism:Severe      TGTACGTTGAAATCCTTCACTGTAGAAAAAGGAATCTATCAAACTTCTAACTTTAGAGTC    22522
gb:MT020880|Organism:Severe      TGTACGTTGAAATCCTTCACTGTAGAAAAAGGAATCTATCAAACTTCTAACTTTAGAGTC    22522
gb:MT066175|Organism:Severe      TGTACGTTGAAATCCTTCACTGTAGAAAAAGGAATCTATCAAACTTCTAACTTTAGAGTC    22522
gb:MN997409|Organism:Severe      TGTACGTTGAAATCCTTCACTGTAGAAAAAGGAATCTATCAAACTTCTAACTTTAGAGTC    22522
gb:MN938384|Organism:Severe      TGTACGTTGAAATCCTTCACTGTAGAAAAAGGAATCTATCAAACTTCTAACTTTAGAGTC    22490
gb:MT044258|Organism:Severe      TGTACGTTGAAATCCTTCACTGTAGAAAAAGGAATCTATCAAACTTCTAACTTTAGAGTC    22498
gb:MT039890|Organism:Severe      TGTACGTTGAAATCCTTCACTGTAGAAAAAGGAATCTATCAAACTTCTAACTTTAGAGTC    22522
gb:MN988713|Organism:Severe      TGTACGTTGAAATCCTTCACTGTAGAAAAAGGAATCTATCAAACTTCTAACTTTAGAGTC    22522
gb:LC521925|Organism:Severe      TGTACGTTGAAATCCTTCACTGTAGAAAAAGGAATCTATCAAACTTCTAACTTTAGAGTC    22495
gb:MT093571|Organism:Severe      TGTACGTTGAAATCCTTCACTGTAGAAAAAGGAATCTATCAAACTTCTAACTTTAGAGTC    22522
gb:MT039887|Organism:Severe      TGTACGTTGAAATCCTTCACTGTAGAAAAAGGAATCTATCAAACTTCTAACTTTAGAGTC    22519
gb:MT019530|Organism:Severe      TGTACGTTGAAATCCTTCACTGTAGAAAAAGGAATCTATCAAACTTCTAACTTTAGAGTC    22522
gb:MT039888|Organism:Severe      TGTACGTTGAAATCCTTCACTGTAGAAAAAGGAATCTATCAAACTTCTAACTTTAGAGTC    22522
gb:LC522972|Organism:Severe      TGTACGTTGAAATCCTTCACTGTAGAAAAAGGAATCTATCAAACTTCTAACTTTAGAGTC    22519
gb:MT027063|Organism:Severe      TGTACGTTGAAATCCTTCACTGTAGAAAAAGGAATCTATCAAACTTCTAACTTTAGAGTC    22522
gb:MT027062|Organism:Severe      TGTACGTTGAAATCCTTCACTGTAGAAAAAGGAATCTATCAAACTTCTAACTTTAGAGTC    22522
gb:MT019529|Organism:Severe      TGTACGTTGAAATCCTTCACTGTAGAAAAAGGAATCTATCAAACTTCTAACTTTAGAGTC    22522
gb:MN996529|Organism:Severe      TGTACGTTGAAATCCTTCACTGTAGAAAAAGGAATCTATCAAACTTCTAACTTTAGAGTC    22510
gb:MN996531|Organism:Severe      TGTACGTTGAAATCCTTCACTGTAGAAAAAGGAATCTATCAAACTTCTAACTTTAGAGTC    22509
gb:MT066176|Organism:Severe      TGTACGTTGAAATCCTTCACTGTAGAAAAAGGAATCTATCAAACTTCTAACTTTAGAGTC    22522
gb:MT027064|Organism:Severe      TGTACGTTGAAATCCTTCACTGTAGAAAAAGGAATCTATCAAACTTCTAACTTTAGAGTC    22522
gb:MN994468|Organism:Severe      TGTACGTTGAAATCCTTCACTGTAGAAAAAGGAATCTATCAAACTTCTAACTTTAGAGTC    22522
gb:MT072688|Organism:Severe      TGTACGTTGAAATCCTTCACTGTAGAAAAAGGAATCTATCAAACTTCTAACTTTAGAGTC    22507
gb:MN996527|Organism:Severe      TGTACGTTGAAATCCTTCACTGTAGAAAAAGGAATCTATCAAACTTCTAACTTTAGAGTC    22489
```

```
gb:MT093631|Organism:Severe      TGTACGTTGAAATCCTTCACTGTAGAAAAAGGAATCTATCAAACTTCTAACTTTAGAGTC    22560
gb:MT106053|Organism:Severe      TGTACGTTGAAATCCTTCACTGTAGAAAAAGGAATCTATCAAACTTCTAACTTTAGAGTC    22522
gb:MT019533|Organism:Severe      TGTACGTTGAAATCCTTCACTGTAGAAAAAGGAATCTATCAAACTTCTAACTTTAGAGTC    22522
gb:MT019531|Organism:Severe      TGTACGTTGAAATCCTTCACTGTAGAAAAAGGAATCTATCAAACTTCTAACTTTAGAGTC    22522
gb:MN996528|Organism:Severe      TGTACGTTGAAATCCTTCACTGTAGAAAAAGGAATCTATCAAACTTCTAACTTTAGAGTC    22522
gb:MN996530|Organism:Severe      TGTACGTTGAAATCCTTCACTGTAGAAAAAGGAATCTATCAAACTTCTAACTTTAGAGTC    22508
gb:MN908947|Organism:Severe      TGTACGTTGAAATCCTTCACTGTAGAAAAAGGAATCTATCAAACTTCTAACTTTAGAGTC    22522
gb:MT019532|Organism:Severe      TGTACGTTGAAATCCTTCACTGTAGAAAAAGGAATCTATCAAACTTCTAACTTTAGAGTC    22522
                                 ************************************************************

gb:MT020781|Organism:Severe      CAACCAACAGAATCTATTGTTAGATTTCCTAATATTACAAACTTGTGCCCTTTTGGTGAA    22570
gb:MT007544|Organism:Severe      CAACCAACAGAATCTATTGTTAGATTTCCTAATATTACAAACTTGTGCCCTTTTGGTGAA    22582
gb:MN994467|Organism:Severe      CAACCAACAGAATCTATTGTTAGATTTCCTAATATTACAAACTTGTGCCCTTTTGGTGAA    22582
gb:MT044257|Organism:Severe      CAACCAACAGAATCTATTGTTAGATTTCCTAATATTACAAACTTGTGCCCTTTTGGTGAA    22582
gb:MT106054|Organism:Severe      CAACCAACAGAATCTATTGTTAGATTTCCTAATATTACAAACTTGTGCCCTTTTGGTGAA    22582
gb:MT049951|Organism:Severe      CAACCAACAGAATCTATTGTTAGATTTCCTAATATTACAAACTTGTGCCCTTTTGGTGAA    22582
gb:MN975262|Organism:Severe      CAACCAACAGAATCTATTGTTAGATTTCCTAATATTACAAACTTGTGCCCTTTTGGTGAA    22582
gb:MT106052|Organism:Severe      CAACCAACAGAATCTATTGTTAGATTTCCTAATATTACAAACTTGTGCCCTTTTGGTGAA    22582
gb:LC522975|Organism:Severe      CAACCAACAGAATCTATTGTTAGATTTCCTAATATTACAAACTTGTGCCCTTTTGGTGAA    22579
gb:LC522973|Organism:Severe      CAACCAACAGAATCTATTGTTAGATTTCCTAATATTACAAACTTGTGCCCTTTTGGTGAA    22579
gb:LC522974|Organism:Severe      CAACCAACAGAATCTATTGTTAGATTTCCTAATATTACAAACTTGTGCCCTTTTGGTGAA    22579
gb:MN985325|Organism:Severe      CAACCAACAGAATCTATTGTTAGATTTCCTAATATTACAAACTTGTGCCCTTTTGGTGAA    22582
gb:MT020881|Organism:Severe      CAACCAACAGAATCTATTGTTAGATTTCCTAATATTACAAACTTGTGCCCTTTTGGTGAA    22582
gb:MT020880|Organism:Severe      CAACCAACAGAATCTATTGTTAGATTTCCTAATATTACAAACTTGTGCCCTTTTGGTGAA    22582
gb:MT066175|Organism:Severe      CAACCAACAGAATCTATTGTTAGATTTCCTAATATTACAAACTTGTGCCCTTTTGGTGAA    22582
gb:MN997409|Organism:Severe      CAACCAACAGAATCTATTGTTAGATTTCCTAATATTACAAACTTGTGCCCTTTTGGTGAA    22582
gb:MN938384|Organism:Severe      CAACCAACAGAATCTATTGTTAGATTTCCTAATATTACAAACTTGTGCCCTTTTGGTGAA    22550
gb:MT044258|Organism:Severe      CAACCAACAGAATCTATTGTTAGATTTCCTAATATTACAAACTTGTGCCCTTTTGGTGAA    22558
gb:MT039890|Organism:Severe      CAACCAACAGAATCTATTGTTAGATTTCCTAATATTACAAACTTGTGCCCTTTTGGTGAA    22582
gb:MN988713|Organism:Severe      CAACCAACAGAATCTATTGTTAGATTTCCTAATATTACAAACTTGTGCCCTTTTGGTGAA    22582
gb:LC521925|Organism:Severe      CAACCAACAGAATCTATTGTTAGATTTCCTAATATTACAAACTTGTGCCCTTTTGGTGAA    22555
gb:MT093571|Organism:Severe      CAACCAACAGAATCTATTGTTAGATTTCCTAATATTACAAACTTGTGCCCTTTTGGTGAA    22582
gb:MT039887|Organism:Severe      CAACCAACAGAATCTATTGTTAGATTTCCTAATATTACAAACTTGTGCCCTTTTGGTGAA    22579
gb:MT019530|Organism:Severe      CAACCAACAGAATCTATTGTTAGATTTCCTAATATTACAAACTTGTGCCCTTTTGGTGAA    22582
gb:MT039888|Organism:Severe      CAACCAACAGAATCTATTGTTAGATTTCCTAATATTACAAACTTGTGCCCTTTTGGTGAA    22582
gb:LC522972|Organism:Severe      CAACCAACAGAATCTATTGTTAGATTTCCTAATATTACAAACTTGTGCCCTTTTGGTGAA    22579
gb:MT027063|Organism:Severe      CAACCAACAGAATCTATTGTTAGATTTCCTAATATTACAAACTTGTGCCCTTTTGGTGAA    22582
gb:MT027062|Organism:Severe      CAACCAACAGAATCTATTGTTAGATTTCCTAATATTACAAACTTGTGCCCTTTTGGTGAA    22582
gb:MT019529|Organism:Severe      CAACCAACAGAATCTATTGTTAGATTTCCTAATATTACAAACTTGTGCCCTTTTGGTGAA    22582
gb:MN996529|Organism:Severe      CAACCAACAGAATCTATTGTTAGATTTCCTAATATTACAAACTTGTGCCCTTTTGGTGAA    22570
gb:MN996531|Organism:Severe      CAACCAACAGAATCTATTGTTAGATTTCCTAATATTACAAACTTGTGCCCTTTTGGTGAA    22569
gb:MT066176|Organism:Severe      CAACCAACAGAATCTATTGTTAGATTTCCTAATATTACAAACTTGTGCCCTTTTGGTGAA    22582
gb:MT027064|Organism:Severe      CAACCAACAGAATCTATTGTTAGATTTCCTAATATTACAAACTTGTGCCCTTTTGGTGAA    22582
gb:MN994468|Organism:Severe      CAACCAACAGAATCTATTGTTAGATTTCCTAATATTACAAACTTGTGCCCTTTTGGTGAA    22582
gb:MT072688|Organism:Severe      CAACCAACAGAATCTATTGTTAGATTTCCTAATATTACAAACTTGTGCCCTTTTGGTGAA    22567
gb:MN996527|Organism:Severe      CAACCAACAGAATCTATTGTTAGATTTCCTAATATTACAAACTTGTGCCCTTTTGGTGAA    22549
gb:MT093631|Organism:Severe      CAACCAACAGAATCTATTGTTAGATTTCCTAATATTACAAACTTGTGCCCTTTTGGTGAA    22620
gb:MT106053|Organism:Severe      CAACCAACAGAATCTATTGTTAGATTTCCTAATATTACAAACTTGTGCCCTTTTGGTGAA    22582
gb:MT019533|Organism:Severe      CAACCAACAGAATCTATTGTTAGATTTCCTAATATTACAAACTTGTGCCCTTTTGGTGAA    22582
gb:MT019531|Organism:Severe      CAACCAACAGAATCTATTGTTAGATTTCCTAATATTACAAACTTGTGCCCTTTTGGTGAA    22582
gb:MN996528|Organism:Severe      CAACCAACAGAATCTATTGTTAGATTTCCTAATATTACAAACTTGTGCCCTTTTGGTGAA    22582
gb:MN996530|Organism:Severe      CAACCAACAGAATCTATTGTTAGATTTCCTAATATTACAAACTTGTGCCCTTTTGGTGAA    22568
gb:MN908947|Organism:Severe      CAACCAACAGAATCTATTGTTAGATTTCCTAATATTACAAACTTGTGCCCTTTTGGTGAA    22582
gb:MT019532|Organism:Severe      CAACCAACAGAATCTATTGTTAGATTTCCTAATATTACAAACTTGTGCCCTTTTGGTGAA    22582
                                 ************************************************************

gb:MT020781|Organism:Severe      GTTTTTAACGCCACCAGATTTGCATCTGTTTATGCTTGGAACAGGAAGAGAATCAGCAAC    22630
gb:MT007544|Organism:Severe      GTTTTTAACGCCACCAGATTTGCATCTGTTTATGCTTGGAACAGGAAGAGAATCAGCAAC    22642
gb:MN994467|Organism:Severe      GTTTTTAACGCCACCAGATTTGCATCTGTTTATGCTTGGAACAGGAAGAGAATCAGCAAC    22642
gb:MT044257|Organism:Severe      GTTTTTAACGCCACCAGATTTGCATCTGTTTATGCTTGGAACAGGAAGAGAATCAGCAAC    22642
gb:MT106054|Organism:Severe      GTTTTTAACGCCACCAGATTTGCATCTGTTTATGCTTGGAACAGGAAGAGAATCAGCAAC    22642
gb:MT049951|Organism:Severe      GTTTTTAACGCCACCAGATTTGCATCTGTTTATGCTTGGAACAGGAAGAGAATCAGCAAC    22642
gb:MN975262|Organism:Severe      GTTTTTAACGCCACCAGATTTGCATCTGTTTATGCTTGGAACAGGAAGAGAATCAGCAAC    22642
gb:MT106052|Organism:Severe      GTTTTTAACGCCACCAGATTTGCATCTGTTTATGCTTGGAACAGGAAGAGAATCAGCAAC    22642
gb:LC522975|Organism:Severe      GTTTTTAACGCCACCAGATTTGCATCTGTTTATGCTTGGAACAGGAAGAGAATCAGCAAC    22639
```

```
gb:LC522973|Organism:Severe      GTTTTTAACGCCACCAGATTTGCATCTGTTTATGCTTGGAACAGGAAGAGAATCAGCAAC    22639
gb:LC522974|Organism:Severe      GTTTTTAACGCCACCAGATTTGCATCTGTTTATGCTTGGAACAGGAAGAGAATCAGCAAC    22639
gb:MN985325|Organism:Severe      GTTTTTAACGCCACCAGATTTGCATCTGTTTATGCTTGGAACAGGAAGAGAATCAGCAAC    22642
gb:MT020881|Organism:Severe      GTTTTTAACGCCACCAGATTTGCATCTGTTTATGCTTGGAACAGGAAGAGAATCAGCAAC    22642
gb:MT020880|Organism:Severe      GTTTTTAACGCCACCAGATTTGCATCTGTTTATGCTTGGAACAGGAAGAGAATCAGCAAC    22642
gb:MT066175|Organism:Severe      GTTTTTAACGCCACCAGATTTGCATCTGTTTATGCTTGGAACAGGAAGAGAATCAGCAAC    22642
gb:MN997409|Organism:Severe      GTTTTTAACGCCACCAGATTTGCATCTGTTTATGCTTGGAACAGGAAGAGAATCAGCAAC    22642
gb:MN938384|Organism:Severe      GTTTTTAACGCCACCAGATTTGCATCTGTTTATGCTTGGAACAGGAAGAGAATCAGCAAC    22610
gb:MT044258|Organism:Severe      GTTTTTAACGCCACCAGATTTGCATCTGTTTATGCTTGGAACAGGAAGAGAATCAGCAAC    22618
gb:MT039890|Organism:Severe      GTTTTTAACGCCACCAGATTTGCATCTGTTTATGCTTGGAACAGGAAGAGAATCAGCAAC    22642
gb:MN988713|Organism:Severe      GTTTTTAACGCCACCAGATTTGCATCTGTTTATGCTTGGAACAGGAAGAGAATCAGCAAC    22642
gb:LC521925|Organism:Severe      GTTTTTAACGCCACCAGATTTGCATCTGTTTATGCTTGGAACAGGAAGAGAATCAGCAAC    22615
gb:MT093571|Organism:Severe      GTTTTTAACGCCACCAGATTTGCATCTGTTTATGCTTGGAACAGGAAGAGAATCAGCAAC    22642
gb:MT039887|Organism:Severe      GTTTTTAACGCCACCAGATTTGCATCTGTTTATGCTTGGAACAGGAAGAGAATCAGCAAC    22639
gb:MT019530|Organism:Severe      GTTTTTAACGCCACCAGATTTGCATCTGTTTATGCTTGGAACAGGAAGAGAATCAGCAAC    22642
gb:MT039888|Organism:Severe      GTTTTTAACGCCACCAGATTTGCATCTGTTTATGCTTGGAACAGGAAGAGAATCAGCAAC    22642
gb:LC522972|Organism:Severe      GTTTTTAACGCCACCAGATTTGCATCTGTTTATGCTTGGAACAGGAAGAGAATCAGCAAC    22639
gb:MT027063|Organism:Severe      GTTTTTAACGCCACCAGATTTGCATCTGTTTATGCTTGGAACAGGAAGAGAATCAGCAAC    22642
gb:MT027062|Organism:Severe      GTTTTTAACGCCACCAGATTTGCATCTGTTTATGCTTGGAACAGGAAGAGAATCAGCAAC    22642
gb:MT019529|Organism:Severe      GTTTTTAACGCCACCAGATTTGCATCTGTTTATGCTTGGAACAGGAAGAGAATCAGCAAC    22642
gb:MN996529|Organism:Severe      GTTTTTAACGCCACCAGATTTGCATCTGTTTATGCTTGGAACAGGAAGAGAATCAGCAAC    22630
gb:MN996531|Organism:Severe      GTTTTTAACGCCACCAGATTTGCATCTGTTTATGCTTGGAACAGGAAGAGAATCAGCAAC    22629
gb:MT066176|Organism:Severe      GTTTTTAACGCCACCAGATTTGCATCTGTTTATGCTTGGAACAGGAAGAGAATCAGCAAC    22642
gb:MT027064|Organism:Severe      GTTTTTAACGCCACCAGATTTGCATCTGTTTATGCTTGGAACAGGAAGAGAATCAGCAAC    22642
gb:MN994468|Organism:Severe      GTTTTTAACGCCACCAGATTTGCATCTGTTTATGCTTGGAACAGGAAGAGAATCAGCAAC    22642
gb:MT072688|Organism:Severe      GTTTTTAACGCCACCAGATTTGCATCTGTTTATGCTTGGAACAGGAAGAGAATCAGCAAC    22627
gb:MN996527|Organism:Severe      GTTTTTAACGCCACCAGATTTGCATCTGTTTATGCTTGGAACAGGAAGAGAATCAGCAAC    22609
gb:MT093631|Organism:Severe      GTTTTTAACGCCACCAGATTTGCATCTGTTTATGCTTGGAACAGGAAGAGAATCAGCAAC    22680
gb:MT106053|Organism:Severe      GTTTTTAACGCCACCAGATTTGCATCTGTTTATGCTTGGAACAGGAAGAGAATCAGCAAC    22642
gb:MT019533|Organism:Severe      GTTTTTAACGCCACCAGATTTGCATCTGTTTATGCTTGGAACAGGAAGAGAATCAGCAAC    22642
gb:MT019531|Organism:Severe      GTTTTTAACGCCACCAGATTTGCATCTGTTTATGCTTGGAACAGGAAGAGAATCAGCAAC    22642
gb:MN996528|Organism:Severe      GTTTTTAACGCCACCAGATTTGCATCTGTTTATGCTTGGAACAGGAAGAGAATCAGCAAC    22642
gb:MN996530|Organism:Severe      GTTTTTAACGCCACCAGATTTGCATCTGTTTATGCTTGGAACAGGAAGAGAATCAGCAAC    22628
gb:MN908947|Organism:Severe      GTTTTTAACGCCACCAGATTTGCATCTGTTTATGCTTGGAACAGGAAGAGAATCAGCAAC    22642
gb:MT019532|Organism:Severe      GTTTTTAACGCCACCAGATTTGCATCTGTTTATGCTTGGAACAGGAAGAGAATCAGCAAC    22642
                                 ************************************************************

gb:MT020781|Organism:Severe      TGTGTTGCTGATTATTCTGTCCTATATAATTCCGCATCATTTTCCACTTTTAAGTGTTAT    22690
gb:MT007544|Organism:Severe      TGTGTTGCTGATTATTCTGTCCTATATAATTCCGCATCATTTTCCACTTTTAAGTGTTAT    22702
gb:MN994467|Organism:Severe      TGTGTTGCTGATTATTCTGTCCTATATAATTCCGCATCATTTTCCACTTTTAAGTGTTAT    22702
gb:MT044257|Organism:Severe      TGTGTTGCTGATTATTCTGTCCTATATAATTCCGCATCATTTTCCACTTTTAAGTGTTAT    22702
gb:MT106054|Organism:Severe      TGTGTTGCTGATTATTCTGTCCTATATAATTCCGCATCATTTTCCACTTTTAAGTGTTAT    22702
gb:MT049951|Organism:Severe      TGTGTTGCTGATTATTCTGTCCTATATAATTCCGCATCATTTTCCACTTTTAAGTGTTAT    22702
gb:MN975262|Organism:Severe      TGTGTTGCTGATTATTCTGTCCTATATAATTCCGCATCATTTTCCACTTTTAAGTGTTAT    22702
gb:MT106052|Organism:Severe      TGTGTTGCTGATTATTCTGTCCTATATAATTCCGCATCATTTTCCACTTTTAAGTGTTAT    22702
gb:LC522975|Organism:Severe      TGTGTTGCTGATTATTCTGTCCTATATAATTCCGCATCATTTTCCACTTTTAAGTGTTAT    22699
gb:LC522973|Organism:Severe      TGTGTTGCTGATTATTCTGTCCTATATAATTCCGCATCATTTTCCACTTTTAAGTGTTAT    22699
gb:LC522974|Organism:Severe      TGTGTTGCTGATTATTCTGTCCTATATAATTCCGCATCATTTTCCACTTTTAAGTGTTAT    22699
gb:MN985325|Organism:Severe      TGTGTTGCTGATTATTCTGTCCTATATAATTCCGCATCATTTTCCACTTTTAAGTGTTAT    22702
gb:MT020881|Organism:Severe      TGTGTTGCTGATTATTCTGTCCTATATAATTCCGCATCATTTTCCACTTTTAAGTGTTAT    22702
gb:MT020880|Organism:Severe      TGTGTTGCTGATTATTCTGTCCTATATAATTCCGCATCATTTTCCACTTTTAAGTGTTAT    22702
gb:MT066175|Organism:Severe      TGTGTTGCTGATTATTCTGTCCTATATAATTCCGCATCATTTTCCACTTTTAAGTGTTAT    22702
gb:MN997409|Organism:Severe      TGTGTTGCTGATTATTCTGTCCTATATAATTCCGCATCATTTTCCACTTTTAAGTGTTAT    22702
gb:MN938384|Organism:Severe      TGTGTTGCTGATTATTCTGTCCTATATAATTCCGCATCATTTTCCACTTTTAAGTGTTAT    22670
gb:MT044258|Organism:Severe      TGTGTTGCTGATTATTCTGTCCTATATAATTCCGCATCATTTTCCACTTTTAAGTGTTAT    22678
gb:MT039890|Organism:Severe      TGTGTTGCTGATTATTCTGTCCTATATAATTCCGCATCATTTTCCACTTTTAAGTGTTAT    22702
gb:MN988713|Organism:Severe      TGTGTTGCTGATTATTCTGTCCTATATAATTCCGCATCATTTTCCACTTTTAAGTGTTAT    22702
gb:LC521925|Organism:Severe      TGTGTTGCTGATTATTCTGTCCTATATAATTCCGCATCATTTTCCACTTTTAAGTGTTAT    22675
gb:MT093571|Organism:Severe      TGTGTTGCTGATTATTCTGTCCTATATAATTCCGCATCATTTTCCACTTTTAAGTGTTAT    22702
gb:MT039887|Organism:Severe      TGTGTTGCTGATTATTCTGTCCTATATAATTCCGCATCATTTTCCACTTTTAAGTGTTAT    22699
gb:MT019530|Organism:Severe      TGTGTTGCTGATTATTCTGTCCTATATAATTCCGCATCATTTTCCACTTTTAAGTGTTAT    22702
gb:MT039888|Organism:Severe      TGTGTTGCTGATTATTCTGTCCTATATAATTCCGCATCATTTTCCACTTTTAAGTGTTAT    22702
gb:LC522972|Organism:Severe      TGTGTTGCTGATTATTCTGTCCTATATAATTCCGCATCATTTTCCACTTTTAAGTGTTAT    22699
gb:MT027063|Organism:Severe      TGTGTTGCTGATTATTCTGTCCTATATAATTCCGCATCATTTTCCACTTTTAAGTGTTAT    22702
gb:MT027062|Organism:Severe      TGTGTTGCTGATTATTCTGTCCTATATAATTCCGCATCATTTTCCACTTTTAAGTGTTAT    22702
```

```
gb:MT019529|Organism:Severe      TGTGTTGCTGATTATTCTGTCCTATATAATTCCGCATCATTTTCCACTTTTAAGTGTTAT   22702
gb:MN996529|Organism:Severe      TGTGTTGCTGATTATTCTGTCCTATATAATTCCGCATCATTTTCCACTTTTAAGTGTTAT   22690
gb:MN996531|Organism:Severe      TGTGTTGCTGATTATTCTGTCCTATATAATTCCGCATCATTTTCCACTTTTAAGTGTTAT   22689
gb:MT066176|Organism:Severe      TGTGTTGCTGATTATTCTGTCCTATATAATTCCGCATCATTTTCCACTTTTAAGTGTTAT   22702
gb:MT027064|Organism:Severe      TGTGTTGCTGATTATTCTGTCCTATATAATTCCGCATCATTTTCCACTTTTAAGTGTTAT   22702
gb:MN994468|Organism:Severe      TGTGTTGCTGATTATTCTGTCCTATATAATTCCGCATCATTTTCCACTTTTAAGTGTTAT   22702
gb:MT072688|Organism:Severe      TGTGTTGCTGATTATTCTGTCCTATATAATTCCGCATCATTTTCCACTTTTAAGTGTTAT   22687
gb:MN996527|Organism:Severe      TGTGTTGCTGATTATTCTGTCCTATATAATTCCGCATCATTTTCCACTTTTAAGTGTTAT   22669
gb:MT093631|Organism:Severe      TGTGTTGCTGATTATTCTGTCCTATATAATTCCGCATCATTTTCCACTTTTAAGTGTTAT   22740
gb:MT106053|Organism:Severe      TGTGTTGCTGATTATTCTGTCCTATATAATTCCGCATCATTTTCCACTTTTAAGTGTTAT   22702
gb:MT019533|Organism:Severe      TGTGTTGCTGATTATTCTGTCCTATATAATTCCGCATCATTTTCCACTTTTAAGTGTTAT   22702
gb:MT019531|Organism:Severe      TGTGTTGCTGATTATTCTGTCCTATATAATTCCGCATCATTTTCCACTTTTAAGTGTTAT   22702
gb:MN996528|Organism:Severe      TGTGTTGCTGATTATTCTGTCCTATATAATTCCGCATCATTTTCCACTTTTAAGTGTTAT   22702
gb:MN996530|Organism:Severe      TGTGTTGCTGATTATTCTGTCCTATATAATTCCGCATCATTTTCCACTTTTAAGTGTTAT   22688
gb:MN908947|Organism:Severe      TGTGTTGCTGATTATTCTGTCCTATATAATTCCGCATCATTTTCCACTTTTAAGTGTTAT   22702
gb:MT019532|Organism:Severe      TGTGTTGCTGATTATTCTGTCCTATATAATTCCGCATCATTTTCCACTTTTAAGTGTTAT   22702
                                 ************************************************************

gb:MT020781|Organism:Severe      GGAGTGTCTCCTACTAAATTAAATGATCTCTGCTTTACTAATGTCTATGCAGATTCATTT   22750
gb:MT007544|Organism:Severe      GGAGTGTCTCCTACTAAATTAAATGATCTCTGCTTTACTAATGTCTATGCAGATTCATTT   22762
gb:MN994467|Organism:Severe      GGAGTGTCTCCTACTAAATTAAATGATCTCTGCTTTACTAATGTCTATGCAGATTCATTT   22762
gb:MT044257|Organism:Severe      GGAGTGTCTCCTACTAAATTAAATGATCTCTGCTTTACTAATGTCTATGCAGATTCATTT   22762
gb:MT106054|Organism:Severe      GGAGTGTCTCCTACTAAATTAAATGATCTCTGCTTTACTAATGTCTATGCAGATTCATTT   22762
gb:MT049951|Organism:Severe      GGAGTGTCTCCTACTAAATTAAATGATCTCTGCTTTACTAATGTCTATGCAGATTCATTT   22762
gb:MN975262|Organism:Severe      GGAGTGTCTCCTACTAAATTAAATGATCTCTGCTTTACTAATGTCTATGCAGATTCATTT   22762
gb:MT106052|Organism:Severe      GGAGTGTCTCCTACTAAATTAAATGATCTCTGCTTTACTAATGTCTATGCAGATTCATTT   22762
gb:LC522975|Organism:Severe      GGAGTGTCTCCTACTAAATTAAATGATCTCTGCTTTACTAATGTCTATGCAGATTCATTT   22759
gb:LC522973|Organism:Severe      GGAGTGTCTCCTACTAAATTAAATGATCTCTGCTTTACTAATGTCTATGCAGATTCATTT   22759
gb:LC522974|Organism:Severe      GGAGTGTCTCCTACTAAATTAAATGATCTCTGCTTTACTAATGTCTATGCAGATTCATTT   22759
gb:MN985325|Organism:Severe      GGAGTGTCTCCTACTAAATTAAATGATCTCTGCTTTACTAATGTCTATGCAGATTCATTT   22762
gb:MT020881|Organism:Severe      GGAGTGTCTCCTACTAAATTAAATGATCTCTGCTTTACTAATGTCTATGCAGATTCATTT   22762
gb:MT020880|Organism:Severe      GGAGTGTCTCCTACTAAATTAAATGATCTCTGCTTTACTAATGTCTATGCAGATTCATTT   22762
gb:MT066175|Organism:Severe      GGAGTGTCTCCTACTAAATTAAATGATCTCTGCTTTACTAATGTCTATGCAGATTCATTT   22762
gb:MN997409|Organism:Severe      GGAGTGTCTCCTACTAAATTAAATGATCTCTGCTTTACTAATGTCTATGCAGATTCATTT   22762
gb:MN938384|Organism:Severe      GGAGTGTCTCCTACTAAATTAAATGATCTCTGCTTTACTAATGTCTATGCAGATTCATTT   22730
gb:MT044258|Organism:Severe      GGAGTGTCTCCTACTAAATTAAATGATCTCTGCTTTACTAATGTCTATGCAGATTCATTT   22738
gb:MT039890|Organism:Severe      GGAGTGTCTCCTACTAAATTAAATGATCTCTGCTTTACTAATGTCTATGCAGATTCATTT   22762
gb:MN988713|Organism:Severe      GGAGTGTCTCCTACTAAATTAAATGATCTCTGCTTTACTAATGTCTATGCAGATTCATTT   22762
gb:LC521925|Organism:Severe      GGAGTGTCTCCTACTAAATTAAATGATCTCTGCTTTACTAATGTCTATGCAGATTCATTT   22735
gb:MT093571|Organism:Severe      GGAGTGTCTCCTACTAAATTAAATGATCTCTGCTTTACTAATGTCTATGCAGATTCATTT   22762
gb:MT039887|Organism:Severe      GGAGTGTCTCCTACTAAATTAAATGATCTCTGCTTTACTAATGTCTATGCAGATTCATTT   22759
gb:MT019530|Organism:Severe      GGAGTGTCTCCTACTAAATTAAATGATCTCTGCTTTACTAATGTCTATGCAGATTCATTT   22762
gb:MT039888|Organism:Severe      GGAGTGTCTCCTACTAAATTAAATGATCTCTGCTTTACTAATGTCTATGCAGATTCATTT   22762
gb:LC522972|Organism:Severe      GGAGTGTCTCCTACTAAATTAAATGATCTCTGCTTTACTAATGTCTATGCAGATTCATTT   22759
gb:MT027063|Organism:Severe      GGAGTGTCTCCTACTAAATTAAATGATCTCTGCTTTACTAATGTCTATGCAGATTCATTT   22762
gb:MT027062|Organism:Severe      GGAGTGTCTCCTACTAAATTAAATGATCTCTGCTTTACTAATGTCTATGCAGATTCATTT   22762
gb:MT019529|Organism:Severe      GGAGTGTCTCCTACTAAATTAAATGATCTCTGCTTTACTAATGTCTATGCAGATTCATTT   22762
gb:MN996529|Organism:Severe      GGAGTGTCTCCTACTAAATTAAATGATCTCTGCTTTACTAATGTCTATGCAGATTCATTT   22750
gb:MN996531|Organism:Severe      GGAGTGTCTCCTACTAAATTAAATGATCTCTGCTTTACTAATGTCTATGCAGATTCATTT   22749
gb:MT066176|Organism:Severe      GGAGTGTCTCCTACTAAATTAAATGATCTCTGCTTTACTAATGTCTATGCAGATTCATTT   22762
gb:MT027064|Organism:Severe      GGAGTGTCTCCTACTAAATTAAATGATCTCTGCTTTACTAATGTCTATGCAGATTCATTT   22762
gb:MN994468|Organism:Severe      GGAGTGTCTCCTACTAAATTAAATGATCTCTGCTTTACTAATGTCTATGCAGATTCATTT   22762
gb:MT072688|Organism:Severe      GGAGTGTCTCCTACTAAATTAAATGATCTCTGCTTTACTAATGTCTATGCAGATTCATTT   22747
gb:MN996527|Organism:Severe      GGAGTGTCTCCTACTAAATTAAATGATCTCTGCTTTACTAATGTCTATGCAGATTCATTT   22729
gb:MT093631|Organism:Severe      GGAGTGTCTCCTACTAAATTAAATGATCTCTGCTTTACTAATGTCTATGCAGATTCATTT   22800
gb:MT106053|Organism:Severe      GGAGTGTCTCCTACTAAATTAAATGATCTCTGCTTTACTAATGTCTATGCAGATTCATTT   22762
gb:MT019533|Organism:Severe      GGAGTGTCTCCTACTAAATTAAATGATCTCTGCTTTACTAATGTCTATGCAGATTCATTT   22762
gb:MT019531|Organism:Severe      GGAGTGTCTCCTACTAAATTAAATGATCTCTGCTTTACTAATGTCTATGCAGATTCATTT   22762
gb:MN996528|Organism:Severe      GGAGTGTCTCCTACTAAATTAAATGATCTCTGCTTTACTAATGTCTATGCAGATTCATTT   22762
gb:MN996530|Organism:Severe      GGAGTGTCTCCTACTAAATTAAATGATCTCTGCTTTACTAATGTCTATGCAGATTCATTT   22748
gb:MN908947|Organism:Severe      GGAGTGTCTCCTACTAAATTAAATGATCTCTGCTTTACTAATGTCTATGCAGATTCATTT   22762
gb:MT019532|Organism:Severe      GGAGTGTCTCCTACTAAATTAAATGATCTCTGCTTTACTAATGTCTATGCAGATTCATTT   22762
                                 ************************************************************

gb:MT020781|Organism:Severe      GTAATTAGAGGTGATGAAGTCAGACAAATCGCTCCAGGGCAAACTGGAAAGATTGCTGAT   22810
```

```
gb:MT007544|Organism:Severe      GTAATTAGAGGTGATGAAGTCAGACAAATCGCTCCAGGGCAAACTGGAAAGATTGCTGAT   22822
gb:MN994467|Organism:Severe      GTAATTAGAGGTGATGAAGTCAGACAAATCGCTCCAGGGCAAACTGGAAAGATTGCTGAT   22822
gb:MT044257|Organism:Severe      GTAATTAGAGGTGATGAAGTCAGACAAATCGCTCCAGGGCAAACTGGAAAGATTGCTGAT   22822
gb:MT106054|Organism:Severe      GTAATTAGAGGTGATGAAGTCAGACAAATCGCTCCAGGGCAAACTGGAAAGATTGCTGAT   22822
gb:MT049951|Organism:Severe      GTAATTAGAGGTGATGAAGTCAGACAAATCGCTCCAGGGCAAACTGGAAAGATTGCTGAT   22822
gb:MN975262|Organism:Severe      GTAATTAGAGGTGATGAAGTCAGACAAATCGCTCCAGGGCAAACTGGAAAGATTGCTGAT   22822
gb:MT106052|Organism:Severe      GTAATTAGAGGTGATGAAGTCAGACAAATCGCTCCAGGGCAAACTGGAAAGATTGCTGAT   22822
gb:LC522975|Organism:Severe      GTAATTAGAGGTGATGAAGTCAGACAAATCGCTCCAGGGCAAACTGGAAAGATTGCTGAT   22819
gb:LC522973|Organism:Severe      GTAATTAGAGGTGATGAAGTCAGACAAATCGCTCCAGGGCAAACTGGAAAGATTGCTGAT   22819
gb:LC522974|Organism:Severe      GTAATTAGAGGTGATGAAGTCAGACAAATCGCTCCAGGGCAAACTGGAAAGATTGCTGAT   22819
gb:MN985325|Organism:Severe      GTAATTAGAGGTGATGAAGTCAGACAAATCGCTCCAGGGCAAACTGGAAAGATTGCTGAT   22822
gb:MT020881|Organism:Severe      GTAATTAGAGGTGATGAAGTCAGACAAATCGCTCCAGGGCAAACTGGAAAGATTGCTGAT   22822
gb:MT020880|Organism:Severe      GTAATTAGAGGTGATGAAGTCAGACAAATCGCTCCAGGGCAAACTGGAAAGATTGCTGAT   22822
gb:MT066175|Organism:Severe      GTAATTAGAGGTGATGAAGTCAGACAAATCGCTCCAGGGCAAACTGGAAAGATTGCTGAT   22822
gb:MN997409|Organism:Severe      GTAATTAGAGGTGATGAAGTCAGACAAATCGCTCCAGGGCAAACTGGAAAGATTGCTGAT   22822
gb:MN938384|Organism:Severe      GTAATTAGAGGTGATGAAGTCAGACAAATCGCTCCAGGGCAAACTGGAAAGATTGCTGAT   22790
gb:MT044258|Organism:Severe      GTAATTAGAGGTGATGAAGTCAGACAAATCGCTCCAGGGCAAACTGGAAAGATTGCTGAT   22798
gb:MT039890|Organism:Severe      GTAATTAGAGGTGATGAAGTCAGACAAATCGCTCCAGGGCAAACTGGAAAGATTGCTGAT   22822
gb:MN988713|Organism:Severe      GTAATTAGAGGTGATGAAGTCAGACAAATCGCTCCAGGGCAAACTGGAAAGATTGCTGAT   22822
gb:LC521925|Organism:Severe      GTAATTAGAGGTGATGAAGTCAGACAAATCGCTCCAGGGCAAACTGGAAAGATTGCTGAT   22795
gb:MT093571|Organism:Severe      GTAATTAGAGGTGATGAAGTCAGACAAATCGCTCCAGGGCAAACTGGAAAGATTGCTGAT   22822
gb:MT039887|Organism:Severe      GTAATTAGAGGTGATGAAGTCAGACAAATCGCTCCAGGGCAAACTGGAAAGATTGCTGAT   22819
gb:MT019530|Organism:Severe      GTAATTAGAGGTGATGAAGTCAGACAAATCGCTCCAGGGCAAACTGGAAAGATTGCTGAT   22822
gb:MT039888|Organism:Severe      GTAATTAGAGGTGATGAAGTCAGACAAATCGCTCCAGGGCAAACTGGAAAGATTGCTGAT   22822
gb:LC522972|Organism:Severe      GTAATTAGAGGTGATGAAGTCAGACAAATCGCTCCAGGGCAAACTGGAAAGATTGCTGAT   22819
gb:MT027063|Organism:Severe      GTAATTAGAGGTGATGAAGTCAGACAAATCGCTCCAGGGCAAACTGGAAAGATTGCTGAT   22822
gb:MT027062|Organism:Severe      GTAATTAGAGGTGATGAAGTCAGACAAATCGCTCCAGGGCAAACTGGAAAGATTGCTGAT   22822
gb:MT019529|Organism:Severe      GTAATTAGAGGTGATGAAGTCAGACAAATCGCTCCAGGGCAAACTGGAAAGATTGCTGAT   22822
gb:MN996529|Organism:Severe      GTAATTAGAGGTGATGAAGTCAGACAAATCGCTCCAGGGCAAACTGGAAAGATTGCTGAT   22810
gb:MN996531|Organism:Severe      GTAATTAGAGGTGATGAAGTCAGACAAATCGCTCCAGGGCAAACTGGAAAGATTGCTGAT   22809
gb:MT066176|Organism:Severe      GTAATTAGAGGTGATGAAGTCAGACAAATCGCTCCAGGGCAAACTGGAAAGATTGCTGAT   22822
gb:MT027064|Organism:Severe      GTAATTAGAGGTGATGAAGTCAGACAAATCGCTCCAGGGCAAACTGGAAAGATTGCTGAT   22822
gb:MN994468|Organism:Severe      GTAATTAGAGGTGATGAAGTCAGACAAATCGCTCCAGGGCAAACTGGAAAGATTGCTGAT   22822
gb:MT072688|Organism:Severe      GTAATTAGAGGTGATGAAGTCAGACAAATCGCTCCAGGGCAAACTGGAAAGATTGCTGAT   22807
gb:MN996527|Organism:Severe      GTAATTAGAGGTGATGAAGTCAGACAAATCGCTCCAGGGCAAACTGGAAAGATTGCTGAT   22789
gb:MT093631|Organism:Severe      GTAATTAGAGGTGATGAAGTCAGACAAATCGCTCCAGGGCAAACTGGAAAGATTGCTGAT   22860
gb:MT106053|Organism:Severe      GTAATTAGAGGTGATGAAGTCAGACAAATCGCTCCAGGGCAAACTGGAAAGATTGCTGAT   22822
gb:MT019533|Organism:Severe      GTAATTAGAGGTGATGAAGTCAGACAAATCGCTCCAGGGCAAACTGGAAAGATTGCTGAT   22822
gb:MT019531|Organism:Severe      GTAATTAGAGGTGATGAAGTCAGACAAATCGCTCCAGGGCAAACTGGAAAGATTGCTGAT   22822
gb:MN996528|Organism:Severe      GTAATTAGAGGTGATGAAGTCAGACAAATCGCTCCAGGGCAAACTGGAAAGATTGCTGAT   22822
gb:MN996530|Organism:Severe      GTAATTAGAGGTGATGAAGTCAGACAAATCGCTCCAGGGCAAACTGGAAAGATTGCTGAT   22808
gb:MN908947|Organism:Severe      GTAATTAGAGGTGATGAAGTCAGACAAATCGCTCCAGGGCAAACTGGAAAGATTGCTGAT   22822
gb:MT019532|Organism:Severe      GTAATTAGAGGTGATGAAGTCAGACAAATCGCTCCAGGGCAAACTGGAAAGATTGCTGAT   22822
                                 ************************************************************

gb:MT020781|Organism:Severe      TATAATTATAAATTACCAGATGATTTTACAGGCTGCGTTATAGCTTGGAATTCTAACAAT   22870
gb:MT007544|Organism:Severe      TATAATTATAAATTACCAGATGATTTTACAGGCTGCGTTATAGCTTGGAATTCTAACAAT   22882
gb:MN994467|Organism:Severe      TATAATTATAAATTACCAGATGATTTTACAGGCTGCGTTATAGCTTGGAATTCTAACAAT   22882
gb:MT044257|Organism:Severe      TATAATTATAAATTACCAGATGATTTTACAGGCTGCGTTATAGCTTGGAATTCTAACAAT   22882
gb:MT106054|Organism:Severe      TATAATTATAAATTACCAGATGATTTTACAGGCTGCGTTATAGCTTGGAATTCTAACAAT   22882
gb:MT049951|Organism:Severe      TATAATTATAAATTACCAGATGATTTTACAGGCTGCGTTATAGCTTGGAATTCTAACAAT   22882
gb:MN975262|Organism:Severe      TATAATTATAAATTACCAGATGATTTTACAGGCTGCGTTATAGCTTGGAATTCTAACAAT   22882
gb:MT106052|Organism:Severe      TATAATTATAAATTACCAGATGATTTTACAGGCTGCGTTATAGCTTGGAATTCTAACAAT   22882
gb:LC522975|Organism:Severe      TATAATTATAAATTACCAGATGATTTTACAGGCTGCGTTATAGCTTGGAATTCTAACAAT   22879
gb:LC522973|Organism:Severe      TATAATTATAAATTACCAGATGATTTTACAGGCTGCGTTATAGCTTGGAATTCTAACAAT   22879
gb:LC522974|Organism:Severe      TATAATTATAAATTACCAGATGATTTTACAGGCTGCGTTATAGCTTGGAATTCTAACAAT   22879
gb:MN985325|Organism:Severe      TATAATTATAAATTACCAGATGATTTTACAGGCTGCGTTATAGCTTGGAATTCTAACAAT   22882
gb:MT020881|Organism:Severe      TATAATTATAAATTACCAGATGATTTTACAGGCTGCGTTATAGCTTGGAATTCTAACAAT   22882
gb:MT020880|Organism:Severe      TATAATTATAAATTACCAGATGATTTTACAGGCTGCGTTATAGCTTGGAATTCTAACAAT   22882
gb:MT066175|Organism:Severe      TATAATTATAAATTACCAGATGATTTTACAGGCTGCGTTATAGCTTGGAATTCTAACAAT   22882
gb:MN997409|Organism:Severe      TATAATTATAAATTACCAGATGATTTTACAGGCTGCGTTATAGCTTGGAATTCTAACAAT   22882
gb:MN938384|Organism:Severe      TATAATTATAAATTACCAGATGATTTTACAGGCTGCGTTATAGCTTGGAATTCTAACAAT   22850
gb:MT044258|Organism:Severe      TATAATTATAAATTACCAGATGATTTTACAGGCTGCGTTATAGCTTGGAATTCTAACAAT   22858
gb:MT039890|Organism:Severe      TATAATTATAAATTACCAGATGATTTTACAGGCTGCGTTATAGCTTGGAATTCTAACAAT   22882
gb:MN988713|Organism:Severe      TATAATTATAAATTACCAGATGATTTTACAGGCTGCGTTATAGCTTGGAATTCTAACAAT   22882
```

```
gb:LC521925|Organism:Severe      TATAATTATAAATTACCAGATGATTTTACAGGCTGCGTTATAGCTTGGAATTCTAACAAT   22855
gb:MT093571|Organism:Severe      TATAATTATAAATTACCAGATGATTTTACAGGCTGCGTTATAGCTTGGAATTCTAACAAT   22882
gb:MT039887|Organism:Severe      TATAATTATAAATTACCAGATGATTTTACAGGCTGCGTTATAGCTTGGAATTCTAACAAT   22879
gb:MT019530|Organism:Severe      TATAATTATAAATTACCAGATGATTTTACAGGCTGCGTTATAGCTTGGAATTCTAACAAT   22882
gb:MT039888|Organism:Severe      TATAATTATAAATTACCAGATGATTTTACAGGCTGCGTTATAGCTTGGAATTCTAACAAT   22882
gb:LC522972|Organism:Severe      TATAATTATAAATTACCAGATGATTTTACAGGCTGCGTTATAGCTTGGAATTCTAACAAT   22879
gb:MT027063|Organism:Severe      TATAATTATAAATTACCAGATGATTTTACAGGCTGCGTTATAGCTTGGAATTCTAACAAT   22882
gb:MT027062|Organism:Severe      TATAATTATAAATTACCAGATGATTTTACAGGCTGCGTTATAGCTTGGAATTCTAACAAT   22882
gb:MT019529|Organism:Severe      TATAATTATAAATTACCAGATGATTTTACAGGCTGCGTTATAGCTTGGAATTCTAACAAT   22882
gb:MN996529|Organism:Severe      TATAATTATAAATTACCAGATGATTTTACAGGCTGCGTTATAGCTTGGAATTCTAACAAT   22870
gb:MN996531|Organism:Severe      TATAATTATAAATTACCAGATGATTTTACAGGCTGCGTTATAGCTTGGAATTCTAACAAT   22869
gb:MT066176|Organism:Severe      TATAATTATAAATTACCAGATGATTTTACAGGCTGCGTTATAGCTTGGAATTCTAACAAT   22882
gb:MT027064|Organism:Severe      TATAATTATAAATTACCAGATGATTTTACAGGCTGCGTTATAGCTTGGAATTCTAACAAT   22882
gb:MN994468|Organism:Severe      TATAATTATAAATTACCAGATGATTTTACAGGCTGCGTTATAGCTTGGAATTCTAACAAT   22882
gb:MT072688|Organism:Severe      TATAATTATAAATTACCAGATGATTTTACAGGCTGCGTTATAGCTTGGAATTCTAACAAT   22867
gb:MN996527|Organism:Severe      TATAATTATAAATTACCAGATGATTTTACAGGCTGCGTTATAGCTTGGAATTCTAACAAT   22849
gb:MT093631|Organism:Severe      TATAATTATAAATTACCAGATGATTTTACAGGCTGCGTTATAGCTTGGAATTCTAACAAT   22920
gb:MT106053|Organism:Severe      TATAATTATAAATTACCAGATGATTTTACAGGCTGCGTTATAGCTTGGAATTCTAACAAT   22882
gb:MT019533|Organism:Severe      TATAATTATAAATTACCAGATGATTTTACAGGCTGCGTTATAGCTTGGAATTCTAACAAT   22882
gb:MT019531|Organism:Severe      TATAATTATAAATTACCAGATGATTTTACAGGCTGCGTTATAGCTTGGAATTCTAACAAT   22882
gb:MN996528|Organism:Severe      TATAATTATAAATTACCAGATGATTTTACAGGCTGCGTTATAGCTTGGAATTCTAACAAT   22882
gb:MN996530|Organism:Severe      TATAATTATAAATTACCAGATGATTTTACAGGCTGCGTTATAGCTTGGAATTCTAACAAT   22868
gb:MN908947|Organism:Severe      TATAATTATAAATTACCAGATGATTTTACAGGCTGCGTTATAGCTTGGAATTCTAACAAT   22882
gb:MT019532|Organism:Severe      TATAATTATAAATTACCAGATGATTTTACAGGCTGCGTTATAGCTTGGAATTCTAACAAT   22882
                                 ************************************************************

gb:MT020781|Organism:Severe      CTTGATTCTAAGGTTGGTGGTAATTATAATTACCTGTATAGATTGTTTAGGAAGTCTAAT   22930
gb:MT007544|Organism:Severe      CTTGATTCTAAGGTTGGTGGTAATTATAATTACCTGTATAGATTGTTTAGGAAGTCTAAT   22942
gb:MN994467|Organism:Severe      CTTGATTCTAAGGTTGGTGGTAATTATAATTACCTGTATAGATTGTTTAGGAAGTCTAAT   22942
gb:MT044257|Organism:Severe      CTTGATTCTAAGGTTGGTGGTAATTATAATTACCTGTATAGATTGTTTAGGAAGTCTAAT   22942
gb:MT106054|Organism:Severe      CTTGATTCTAAGGTTGGTGGTAATTATAATTACCTGTATAGATTGTTTAGGAAGTCTAAT   22942
gb:MT049951|Organism:Severe      CTTGATTCTAAGGTTGGTGGTAATTATAATTACCTGTATAGATTGTTTAGGAAGTCTAAT   22942
gb:MN975262|Organism:Severe      CTTGATTCTAAGGTTGGTGGTAATTATAATTACCTGTATAGATTGTTTAGGAAGTCTAAT   22942
gb:MT106052|Organism:Severe      CTTGATTCTAAGGTTGGTGGTAATTATAATTACCTGTATAGATTGTTTAGGAAGTCTAAT   22942
gb:LC522975|Organism:Severe      CTTGATTCTAAGGTTGGTGGTAATTATAATTACCTGTATAGATTGTTTAGGAAGTCTAAT   22939
gb:LC522973|Organism:Severe      CTTGATTCTAAGGTTGGTGGTAATTATAATTACCTGTATAGATTGTTTAGGAAGTCTAAT   22939
gb:LC522974|Organism:Severe      CTTGATTCTAAGGTTGGTGGTAATTATAATTACCTGTATAGATTGTTTAGGAAGTCTAAT   22939
gb:MN985325|Organism:Severe      CTTGATTCTAAGGTTGGTGGTAATTATAATTACCTGTATAGATTGTTTAGGAAGTCTAAT   22942
gb:MT020881|Organism:Severe      CTTGATTCTAAGGTTGGTGGTAATTATAATTACCTGTATAGATTGTTTAGGAAGTCTAAT   22942
gb:MT020880|Organism:Severe      CTTGATTCTAAGGTTGGTGGTAATTATAATTACCTGTATAGATTGTTTAGGAAGTCTAAT   22942
gb:MT066175|Organism:Severe      CTTGATTCTAAGGTTGGTGGTAATTATAATTACCTGTATAGATTGTTTAGGAAGTCTAAT   22942
gb:MN997409|Organism:Severe      CTTGATTCTAAGGTTGGTGGTAATTATAATTACCTGTATAGATTGTTTAGGAAGTCTAAT   22942
gb:MN938384|Organism:Severe      CTTGATTCTAAGGTTGGTGGTAATTATAATTACCTGTATAGATTGTTTAGGAAGTCTAAT   22910
gb:MT044258|Organism:Severe      CTTGATTCTAAGGTTGGTGGTAATTATAATTACCTGTATAGATTGTTTAGGAAGTCTAAT   22918
gb:MT039890|Organism:Severe      CTTGATTCTAAGGTTGGTGGTAATTATAATTACCTGTATAGATTGTTTAGGAAGTCTAAT   22942
gb:MN988713|Organism:Severe      CTTGATTCTAAGGTTGGTGGTAATTATAATTACCTGTATAGATTGTTTAGGAAGTCTAAT   22942
gb:LC521925|Organism:Severe      CTTGATTCTAAGGTTGGTGGTAATTATAATTACCTGTATAGATTGTTTAGGAAGTCTAAT   22915
gb:MT093571|Organism:Severe      CTTGATTCTAAGGTTGGTGGTAATTATAATTACCTGTATAGATTGTTTAGGAAGTCTAAT   22942
gb:MT039887|Organism:Severe      CTTGATTCTAAGGTTGGTGGTAATTATAATTACCTGTATAGATTGTTTAGGAAGTCTAAT   22939
gb:MT019530|Organism:Severe      CTTGATTCTAAGGTTGGTGGTAATTATAATTACCTGTATAGATTGTTTAGGAAGTCTAAT   22942
gb:MT039888|Organism:Severe      CTTGATTCTAAGGTTGGTGGTAATTATAATTACCTGTATAGATTGTTTAGGAAGTCTAAT   22942
gb:LC522972|Organism:Severe      CTTGATTCTAAGGTTGGTGGTAATTATAATTACCTGTATAGATTGTTTAGGAAGTCTAAT   22939
gb:MT027063|Organism:Severe      CTTGATTCTAAGGTTGGTGGTAATTATAATTACCTGTATAGATTGTTTAGGAAGTCTAAT   22942
gb:MT027062|Organism:Severe      CTTGATTCTAAGGTTGGTGGTAATTATAATTACCTGTATAGATTGTTTAGGAAGTCTAAT   22942
gb:MT019529|Organism:Severe      CTTGATTCTAAGGTTGGTGGTAATTATAATTACCTGTATAGATTGTTTAGGAAGTCTAAT   22942
gb:MN996529|Organism:Severe      CTTGATTCTAAGGTTGGTGGTAATTATAATTACCTGTATAGATTGTTTAGGAAGTCTAAT   22930
gb:MN996531|Organism:Severe      CTTGATTCTAAGGTTGGTGGTAATTATAATTACCTGTATAGATTGTTTAGGAAGTCTAAT   22929
gb:MT066176|Organism:Severe      CTTGATTCTAAGGTTGGTGGTAATTATAATTACCTGTATAGATTGTTTAGGAAGTCTAAT   22942
gb:MT027064|Organism:Severe      CTTGATTCTAAGGTTGGTGGTAATTATAATTACCTGTATAGATTGTTTAGGAAGTCTAAT   22942
gb:MN994468|Organism:Severe      CTTGATTCTAAGGTTGGTGGTAATTATAATTACCTGTATAGATTGTTTAGGAAGTCTAAT   22942
gb:MT072688|Organism:Severe      CTTGATTCTAAGGTTGGTGGTAATTATAATTACCTGTATAGATTGTTTAGGAAGTCTAAT   22927
gb:MN996527|Organism:Severe      CTTGATTCTAAGGTTGGTGGTAATTATAATTACCTGTATAGATTGTTTAGGAAGTCTAAT   22909
gb:MT093631|Organism:Severe      CTTGATTCTAAGGTTGGTGGTAATTATAATTACCTGTATAGATTGTTTAGGAAGTCTAAT   22980
gb:MT106053|Organism:Severe      CTTGATTCTAAGGTTGGTGGTAATTATAATTACCTGTATAGATTGTTTAGGAAGTCTAAT   22942
gb:MT019533|Organism:Severe      CTTGATTCTAAGGTTGGTGGTAATTATAATTACCTGTATAGATTGTTTAGGAAGTCTAAT   22942
```

```
gb:MT019531|Organism:Severe      CTTGATTCTAAGGTTGGTGGTAATTATAATTACCTGTATAGATTGTTTAGGAAGTCTAAT    22942
gb:MN996528|Organism:Severe      CTTGATTCTAAGGTTGGTGGTAATTATAATTACCTGTATAGATTGTTTAGGAAGTCTAAT    22942
gb:MN996530|Organism:Severe      CTTGATTCTAAGGTTGGTGGTAATTATAATTACCTGTATAGATTGTTTAGGAAGTCTAAT    22928
gb:MN908947|Organism:Severe      CTTGATTCTAAGGTTGGTGGTAATTATAATTACCTGTATAGATTGTTTAGGAAGTCTAAT    22942
gb:MT019532|Organism:Severe      CTTGATTCTAAGGTTGGTGGTAATTATAATTACCTGTATAGATTGTTTAGGAAGTCTAAT    22942
                                 ************************************************************

gb:MT020781|Organism:Severe      CTCAAACCTTTTGAGAGAGATATTTCAACTGAAATCTATCAGGCCGGTAGCACACCTTGT    22990
gb:MT007544|Organism:Severe      CTCAAACCTTTTGAGAGAGATATTTCAACTGAAATCTATCAGGCCGGTAGCACACCTTGT    23002
gb:MN994467|Organism:Severe      CTCAAACCTTTTGAGAGAGATATTTCAACTGAAATCTATCAGGCCGGTAGCACACCTTGT    23002
gb:MT044257|Organism:Severe      CTCAAACCTTTTGAGAGAGATATTTCAACTGAAATCTATCAGGCCGGTAGCACACCTTGT    23002
gb:MT106054|Organism:Severe      CTCAAACCTTTTGAGAGAGATATTTCAACTGAAATCTATCAGGCCGGTAGCACACCTTGT    23002
gb:MT049951|Organism:Severe      CTCAAACCTTTTGAGAGAGATATTTCAACTGAAATCTATCAGGCCGGTAGCACACCTTGT    23002
gb:MN975262|Organism:Severe      CTCAAACCTTTTGAGAGAGATATTTCAACTGAAATCTATCAGGCCGGTAGCACACCTTGT    23002
gb:MT106052|Organism:Severe      CTCAAACCTTTTGAGAGAGATATTTCAACTGAAATCTATCAGGCCGGTAGCACACCTTGT    23002
gb:LC522975|Organism:Severe      CTCAAACCTTTTGAGAGAGATATTTCAACTGAAATCTATCAGGCCGGTAGCACACCTTGT    22999
gb:LC522973|Organism:Severe      CTCAAACCTTTTGAGAGAGATATTTCAACTGAAATCTATCAGGCCGGTAGCACACCTTGT    22999
gb:LC522974|Organism:Severe      CTCAAACCTTTTGAGAGAGATATTTCAACTGAAATCTATCAGGCCGGTAGCACACCTTGT    22999
gb:MN985325|Organism:Severe      CTCAAACCTTTTGAGAGAGATATTTCAACTGAAATCTATCAGGCCGGTAGCACACCTTGT    23002
gb:MT020881|Organism:Severe      CTCAAACCTTTTGAGAGAGATATTTCAACTGAAATCTATCAGGCCGGTAGCACACCTTGT    23002
gb:MT020880|Organism:Severe      CTCAAACCTTTTGAGAGAGATATTTCAACTGAAATCTATCAGGCCGGTAGCACACCTTGT    23002
gb:MT066175|Organism:Severe      CTCAAACCTTTTGAGAGAGATATTTCAACTGAAATCTATCAGGCCGGTAGCACACCTTGT    23002
gb:MN997409|Organism:Severe      CTCAAACCTTTTGAGAGAGATATTTCAACTGAAATCTATCAGGCCGGTAGCACACCTTGT    23002
gb:MN938384|Organism:Severe      CTCAAACCTTTTGAGAGAGATATTTCAACTGAAATCTATCAGGCCGGTAGCACACCTTGT    22970
gb:MT044258|Organism:Severe      CTCAAACCTTTTGAGAGAGATATTTCAACTGAAATCTATCAGGCCGGTAGCACACCTTGT    22978
gb:MT039890|Organism:Severe      CTCAAACCTTTTGAGAGAGATATTTCAACTGAAATCTATCAGGCCGGTAGCACACCTTGT    23002
gb:MN988713|Organism:Severe      CTCAAACCTTTTGAGAGAGATATTTCAACTGAAATCTATCAGGCCGGTAGCACACCTTGT    23002
gb:LC521925|Organism:Severe      CTCAAACCTTTTGAGAGAGATATTTCAACTGAAATCTATCAGGCCGGTAGCACACCTTGT    22975
gb:MT093571|Organism:Severe      CTCAAACCTTTTGAGAGAGATATTTCAACTGAAATCTATCAGGCCGGTAGCACACCTTGT    23002
gb:MT039887|Organism:Severe      CTCAAACCTTTTGAGAGAGATATTTCAACTGAAATCTATCAGGCCGGTAGCACACCTTGT    22999
gb:MT019530|Organism:Severe      CTCAAACCTTTTGAGAGAGATATTTCAACTGAAATCTATCAGGCCGGTAGCACACCTTGT    23002
gb:MT039888|Organism:Severe      CTCAAACCTTTTGAGAGAGATATTTCAACTGAAATCTATCAGGCCGGTAGCACACCTTGT    23002
gb:LC522972|Organism:Severe      CTCAAACCTTTTGAGAGAGATATTTCAACTGAAATCTATCAGGCCGGTAGCACACCTTGT    22999
gb:MT027063|Organism:Severe      CTCAAACCTTTTGAGAGAGATATTTCAACTGAAATCTATCAGGCCGGTAGCACACCTTGT    23002
gb:MT027062|Organism:Severe      CTCAAACCTTTTGAGAGAGATATTTCAACTGAAATCTATCAGGCCGGTAGCACACCTTGT    23002
gb:MT019529|Organism:Severe      CTCAAACCTTTTGAGAGAGATATTTCAACTGAAATCTATCAGGCCGGTAGCACACCTTGT    23002
gb:MN996529|Organism:Severe      CTCAAACCTTTTGAGAGAGATATTTCAACTGAAATCTATCAGGCCGGTAGCACACCTTGT    22990
gb:MN996531|Organism:Severe      CTCAAACCTTTTGAGAGAGATATTTCAACTGAAATCTATCAGGCCGGTAGCACACCTTGT    22989
gb:MT066176|Organism:Severe      CTCAAACCTTTTGAGAGAGATATTTCAACTGAAATCTATCAGGCCGGTAGCACACCTTGT    23002
gb:MT027064|Organism:Severe      CTCAAACCTTTTGAGAGAGATATTTCAACTGAAATCTATCAGGCCGGTAGCACACCTTGT    23002
gb:MN994468|Organism:Severe      CTCAAACCTTTTGAGAGAGATATTTCAACTGAAATCTATCAGGCCGGTAGCACACCTTGT    23002
gb:MT072688|Organism:Severe      CTCAAACCTTTTGAGAGAGATATTTCAACTGAAATCTATCAGGCCGGTAGCACACCTTGT    22987
gb:MN996527|Organism:Severe      CTCAAACCTTTTGAGAGAGATATTTCAACTGAAATCTATCAGGCCGGTAGCACACCTTGT    22969
gb:MT093631|Organism:Severe      CTCAAACCTTTTGAGAGAGATATTTCAACTGAAATCTATCAGGCCGGTAGCACACCTTGT    23040
gb:MT106053|Organism:Severe      CTCAAACCTTTTGAGAGAGATATTTCAACTGAAATCTATCAGGCCGGTAGCACACCTTGT    23002
gb:MT019533|Organism:Severe      CTCAAACCTTTTGAGAGAGATATTTCAACTGAAATCTATCAGGCCGGTAGCACACCTTGT    23002
gb:MT019531|Organism:Severe      CTCAAACCTTTTGAGAGAGATATTTCAACTGAAATCTATCAGGCCGGTAGCACACCTTGT    23002
gb:MN996528|Organism:Severe      CTCAAACCTTTTGAGAGAGATATTTCAACTGAAATCTATCAGGCCGGTAGCACACCTTGT    23002
gb:MN996530|Organism:Severe      CTCAAACCTTTTGAGAGAGATATTTCAACTGAAATCTATCAGGCCGGTAGCACACCTTGT    22988
gb:MN908947|Organism:Severe      CTCAAACCTTTTGAGAGAGATATTTCAACTGAAATCTATCAGGCCGGTAGCACACCTTGT    23002
gb:MT019532|Organism:Severe      CTCAAACCTTTTGAGAGAGATATTTCAACTGAAATCTATCAGGCCGGTAGCACACCTTGT    23002
                                 ************************************************************

gb:MT020781|Organism:Severe      AATGGTGTTGAAGGTTTTAATTGTTACTTTCCTTTACAATCATATGGTTTCCAACCCACT    23050
gb:MT007544|Organism:Severe      AATGGTGTTGAAGGTTTTAATTGTTACTTTCCTTTACAATCATATGGTTTCCAACCCACT    23062
gb:MN994467|Organism:Severe      AATGGTGTTGAAGGTTTTAATTGTTACTTTCCTTTACAATCATATGGTTTCCAACCCACT    23062
gb:MT044257|Organism:Severe      AATGGTGTTGAAGGTTTTAATTGTTACTTTCCTTTACAATCATATGGTTTCCAACCCACT    23062
gb:MT106054|Organism:Severe      AATGGTGTTGAAGGTTTTAATTGTTACTTTCCTTTACAATCATATGGTTTCCAACCCACT    23062
gb:MT049951|Organism:Severe      AATGGTGTTGAAGGTTTTAATTGTTACTTTCCTTTACAATCATATGGTTTCCAACCCACT    23062
gb:MN975262|Organism:Severe      AATGGTGTTGAAGGTTTTAATTGTTACTTTCCTTTACAATCATATGGTTTCCAACCCACT    23062
gb:MT106052|Organism:Severe      AATGGTGTTGAAGGTTTTAATTGTTACTTTCCTTTACAATCATATGGTTTCCAACCCACT    23062
gb:LC522975|Organism:Severe      AATGGTGTTGAAGGTTTTAATTGTTACTTTCCTTTACAATCATATGGTTTCCAACCCACT    23059
gb:LC522973|Organism:Severe      AATGGTGTTGAAGGTTTTAATTGTTACTTTCCTTTACAATCATATGGTTTCCAACCCACT    23059
gb:LC522974|Organism:Severe      AATGGTGTTGAAGGTTTTAATTGTTACTTTCCTTTACAATCATATGGTTTCCAACCCACT    23059
gb:MN985325|Organism:Severe      AATGGTGTTGAAGGTTTTAATTGTTACTTTCCTTTACAATCATATGGTTTCCAACCCACT    23062
```

```
gb:MT020881|Organism:Severe      AATGGTGTTGAAGGTTTTAATTGTTACTTTCCTTTACAATCATATGGTTTCCAACCCACT   23062
gb:MT020880|Organism:Severe      AATGGTGTTGAAGGTTTTAATTGTTACTTTCCTTTACAATCATATGGTTTCCAACCCACT   23062
gb:MT066175|Organism:Severe      AATGGTGTTGAAGGTTTTAATTGTTACTTTCCTTTACAATCATATGGTTTCCAACCCACT   23062
gb:MN997409|Organism:Severe      AATGGTGTTGAAGGTTTTAATTGTTACTTTCCTTTACAATCATATGGTTTCCAACCCACT   23062
gb:MN938384|Organism:Severe      AATGGTGTTGAAGGTTTTAATTGTTACTTTCCTTTACAATCATATGGTTTCCAACCCACT   23030
gb:MT044258|Organism:Severe      AATGGTGTTGAAGGTTTTAATTGTTACTTTCCTTTACAATCATATGGTTTCCAACCCACT   23038
gb:MT039890|Organism:Severe      AATGGTGTTGAAGGTTTTAATTGTTACTTTCCTTTACAATCATATGGTTTCCAACCCACT   23062
gb:MN988713|Organism:Severe      AATGGTGTTGAAGGTTTTAATTGTTACTTTCCTTTACAATCATATGGTTTCCAACCCACT   23062
gb:LC521925|Organism:Severe      AATGGTGTTGAAGGTTTTAATTGTTACTTTCCTTTACAATCATATGGTTTCCAACCCACT   23035
gb:MT093571|Organism:Severe      AATGGTGTTGAAGGTTTTAATTGTTACTTTCCTTTACAATCATATGGTTTCCAACCCACT   23062
gb:MT039887|Organism:Severe      AATGGTGTTGAAGGTTTTAATTGTTACTTTCCTTTACAATCATATGGTTTCCAACCCACT   23059
gb:MT019530|Organism:Severe      AATGGTGTTGAAGGTTTTAATTGTTACTTTCCTTTACAATCATATGGTTTCCAACCCACT   23062
gb:MT039888|Organism:Severe      AATGGTGTTGAAGGTTTTAATTGTTACTTTCCTTTACAATCATATGGTTTCCAACCCACT   23062
gb:LC522972|Organism:Severe      AATGGTGTTGAAGGTTTTAATTGTTACTTTCCTTTACAATCATATGGTTTCCAACCCACT   23059
gb:MT027063|Organism:Severe      AATGGTGTTGAAGGTTTTAATTGTTACTTTCCTTTACAATCATATGGTTTCCAACCCACT   23062
gb:MT027062|Organism:Severe      AATGGTGTTGAAGGTTTTAATTGTTACTTTCCTTTACAATCATATGGTTTCCAACCCACT   23062
gb:MT019529|Organism:Severe      AATGGTGTTGAAGGTTTTAATTGTTACTTTCCTTTACAATCATATGGTTTCCAACCCACT   23062
gb:MN996529|Organism:Severe      AATGGTGTTGAAGGTTTTAATTGTTACTTTCCTTTACAATCATATGGTTTCCAACCCACT   23050
gb:MN996531|Organism:Severe      AATGGTGTTGAAGGTTTTAATTGTTACTTTCCTTTACAATCATATGGTTTCCAACCCACT   23049
gb:MT066176|Organism:Severe      AATGGTGTTGAAGGTTTTAATTGTTACTTTCCTTTACAATCATATGGTTTCCAACCCACT   23062
gb:MT027064|Organism:Severe      AATGGTGTTGAAGGTTTTAATTGTTACTTTCCTTTACAATCATATGGTTTCCAACCCACT   23062
gb:MN994468|Organism:Severe      AATGGTGTTGAAGGTTTTAATTGTTACTTTCCTTTACAATCATATGGTTTCCAACCCACT   23062
gb:MT072688|Organism:Severe      AATGGTGTTGAAGGTTTTAATTGTTACTTTCCTTTACAATCATATGGTTTCCAACCCACT   23047
gb:MN996527|Organism:Severe      AATGGTGTTGAAGGTTTTAATTGTTACTTTCCTTTACAATCATATGGTTTCCAACCCACT   23029
gb:MT093631|Organism:Severe      AATGGTGTTGAAGGTTTTAATTGTTACTTTCCTTTACAATCATATGGTTTCCAACCCACT   23100
gb:MT106053|Organism:Severe      AATGGTGTTGAAGGTTTTAATTGTTACTTTCCTTTACAATCATATGGTTTCCAACCCACT   23062
gb:MT019533|Organism:Severe      AATGGTGTTGAAGGTTTTAATTGTTACTTTCCTTTACAATCATATGGTTTCCAACCCACT   23062
gb:MT019531|Organism:Severe      AATGGTGTTGAAGGTTTTAATTGTTACTTTCCTTTACAATCATATGGTTTCCAACCCACT   23062
gb:MN996528|Organism:Severe      AATGGTGTTGAAGGTTTTAATTGTTACTTTCCTTTACAATCATATGGTTTCCAACCCACT   23062
gb:MN996530|Organism:Severe      AATGGTGTTGAAGGTTTTAATTGTTACTTTCCTTTACAATCATATGGTTTCCAACCCACT   23048
gb:MN908947|Organism:Severe      AATGGTGTTGAAGGTTTTAATTGTTACTTTCCTTTACAATCATATGGTTTCCAACCCACT   23062
gb:MT019532|Organism:Severe      AATGGTGTTGAAGGTTTTAATTGTTACTTTCCTTTACAATCATATGGTTTCCAACCCACT   23062
                                 ************************************************************

gb:MT020781|Organism:Severe      AATGGTGTTGGTTACCAACCATACAGAGTAGTAGTACTTTCTTTTGAACTTCTACATGCA   23110
gb:MT007544|Organism:Severe      AATGGTGTTGGTTACCAACCATACAGAGTAGTAGTACTTTCTTTTGAACTTCTACATGCA   23122
gb:MN994467|Organism:Severe      AATGGTGTTGGTTACCAACCATACAGAGTAGTAGTACTTTCTTTTGAACTTCTACATGCA   23122
gb:MT044257|Organism:Severe      AATGGTGTTGGTTACCAACCATACAGAGTAGTAGTACTTTCTTTTGAACTTCTACATGCA   23122
gb:MT106054|Organism:Severe      AATGGTGTTGGTTACCAACCATACAGAGTAGTAGTACTTTCTTTTGAACTTCTACATGCA   23122
gb:MT049951|Organism:Severe      AATGGTGTTGGTTACCAACCATACAGAGTAGTAGTACTTTCTTTTGAACTTCTACATGCA   23122
gb:MN975262|Organism:Severe      AATGGTGTTGGTTACCAACCATACAGAGTAGTAGTACTTTCTTTTGAACTTCTACATGCA   23122
gb:MT106052|Organism:Severe      AATGGTGTTGGTTACCAACCATACAGAGTAGTAGTACTTTCTTTTGAACTTCTACATGCA   23122
gb:LC522975|Organism:Severe      AATGGTGTTGGTTACCAACCATACAGAGTAGTAGTACTTTCTTTTGAACTTCTACATGCA   23119
gb:LC522973|Organism:Severe      AATGGTGTTGGTTACCAACCATACAGAGTAGTAGTACTTTCTTTTGAACTTCTACATGCA   23119
gb:LC522974|Organism:Severe      AATGGTGTTGGTTACCAACCATACAGAGTAGTAGTACTTTCTTTTGAACTTCTACATGCA   23119
gb:MN985325|Organism:Severe      AATGGTGTTGGTTACCAACCATACAGAGTAGTAGTACTTTCTTTTGAACTTCTACATGCA   23122
gb:MT020881|Organism:Severe      AATGGTGTTGGTTACCAACCATACAGAGTAGTAGTACTTTCTTTTGAACTTCTACATGCA   23122
gb:MT020880|Organism:Severe      AATGGTGTTGGTTACCAACCATACAGAGTAGTAGTACTTTCTTTTGAACTTCTACATGCA   23122
gb:MT066175|Organism:Severe      AATGGTGTTGGTTACCAACCATACAGAGTAGTAGTACTTTCTTTTGAACTTCTACATGCA   23122
gb:MN997409|Organism:Severe      AATGGTGTTGGTTACCAACCATACAGAGTAGTAGTACTTTCTTTTGAACTTCTACATGCA   23122
gb:MN938384|Organism:Severe      AATGGTGTTGGTTACCAACCATACAGAGTAGTAGTACTTTCTTTTGAACTTCTACATGCA   23090
gb:MT044258|Organism:Severe      AATGGTGTTGGTTACCAACCATACAGAGTAGTAGTACTTTCTTTTGAACTTCTACATGCA   23098
gb:MT039890|Organism:Severe      AATGGTGTTGGTTACCAACCATACAGAGTAGTAGTACTTTCTTTTGAACTTCTACATGCA   23122
gb:MN988713|Organism:Severe      AATGGTGTTGGTTACCAACCATACAGAGTAGTAGTACTTTCTTTTGAACTTCTACATGCA   23122
gb:LC521925|Organism:Severe      AATGGTGTTGGTTACCAACCATACAGAGTAGTAGTACTTTCTTTTGAACTTCTACATGCA   23095
gb:MT093571|Organism:Severe      AATGGTGTTGGTTACCAACCATACAGAGTAGTAGTACTTTCTTTTGAACTTCTACATGCA   23122
gb:MT039887|Organism:Severe      AATGGTGTTGGTTACCAACCATACAGAGTAGTAGTACTTTCTTTTGAACTTCTACATGCA   23119
gb:MT019530|Organism:Severe      AATGGTGTTGGTTACCAACCATACAGAGTAGTAGTACTTTCTTTTGAACTTCTACATGCA   23122
gb:MT039888|Organism:Severe      AATGGTGTTGGTTACCAACCATACAGAGTAGTAGTACTTTCTTTTGAACTTCTACATGCA   23122
gb:LC522972|Organism:Severe      AATGGTGTTGGTTACCAACCATACAGAGTAGTAGTACTTTCTTTTGAACTTCTACATGCA   23119
gb:MT027063|Organism:Severe      AATGGTGTTGGTTACCAACCATACAGAGTAGTAGTACTTTCTTTTGAACTTCTACATGCA   23122
gb:MT027062|Organism:Severe      AATGGTGTTGGTTACCAACCATACAGAGTAGTAGTACTTTCTTTTGAACTTCTACATGCA   23122
gb:MT019529|Organism:Severe      AATGGTGTTGGTTACCAACCATACAGAGTAGTAGTACTTTCTTTTGAACTTCTACATGCA   23122
gb:MN996529|Organism:Severe      AATGGTGTTGGTTACCAACCATACAGAGTAGTAGTACTTTCTTTTGAACTTCTACATGCA   23110
gb:MN996531|Organism:Severe      AATGGTGTTGGTTACCAACCATACAGAGTAGTAGTACTTTCTTTTGAACTTCTACATGCA   23109
```

```
gb:MT066176|Organism:Severe      AATGGTGTTGGTTACCAACCATACAGAGTAGTAGTACTTTCTTTTGAACTTCTACATGCA   23122
gb:MT027064|Organism:Severe      AATGGTGTTGGTTACCAACCATACAGAGTAGTAGTACTTTCTTTTGAACTTCTACATGCA   23122
gb:MN994468|Organism:Severe      AATGGTGTTGGTTACCAACCATACAGAGTAGTAGTACTTTCTTTTGAACTTCTACATGCA   23122
gb:MT072688|Organism:Severe      AATGGTGTTGGTTACCAACCATACAGAGTAGTAGTACTTTCTTTTGAACTTCTACATGCA   23107
gb:MN996527|Organism:Severe      AATGGTGTTGGTTACCAACCATACAGAGTAGTAGTACTTTCTTTTGAACTTCTACATGCA   23089
gb:MT093631|Organism:Severe      AATGGTGTTGGTTACCAACCATACAGAGTAGTAGTACTTTCTTTTGAACTTCTACATGCA   23160
gb:MT106053|Organism:Severe      AATGGTGTTGGTTACCAACCATACAGAGTAGTAGTACTTTCTTTTGAACTTCTACATGCA   23122
gb:MT019533|Organism:Severe      AATGGTGTTGGTTACCAACCATACAGAGTAGTAGTACTTTCTTTTGAACTTCTACATGCA   23122
gb:MT019531|Organism:Severe      AATGGTGTTGGTTACCAACCATACAGAGTAGTAGTACTTTCTTTTGAACTTCTACATGCA   23122
gb:MN996528|Organism:Severe      AATGGTGTTGGTTACCAACCATACAGAGTAGTAGTACTTTCTTTTGAACTTCTACATGCA   23122
gb:MN996530|Organism:Severe      AATGGTGTTGGTTACCAACCATACAGAGTAGTAGTACTTTCTTTTGAACTTCTACATGCA   23108
gb:MN908947|Organism:Severe      AATGGTGTTGGTTACCAACCATACAGAGTAGTAGTACTTTCTTTTGAACTTCTACATGCA   23122
gb:MT019532|Organism:Severe      AATGGTGTTGGTTACCAACCATACAGAGTAGTAGTACTTTCTTTTGAACTTCTACATGCA   23122
                                 ************************************************************

gb:MT020781|Organism:Severe      CCAGCAACTGTTTGTGGACCTAAAAAGTCTACTAATTTGGTTAAAAACAAATGTGTCAAT   23170
gb:MT007544|Organism:Severe      CCAGCAACTGTTTGTGGACCTAAAAAGTCTACTAATTTGGTTAAAAACAAATGTGTCAAT   23182
gb:MN994467|Organism:Severe      CCAGCAACTGTTTGTGGACCTAAAAAGTCTACTAATTTGGTTAAAAACAAATGTGTCAAT   23182
gb:MT044257|Organism:Severe      CCAGCAACTGTTTGTGGACCTAAAAAGTCTACTAATTTGGTTAAAAACAAATGTGTCAAT   23182
gb:MT106054|Organism:Severe      CCAGCAACTGTTTGTGGACCTAAAAAGTCTACTAATTTGGTTAAAAACAAATGTGTCAAT   23182
gb:MT049951|Organism:Severe      CCAGCAACTGTTTGTGGACCTAAAAAGTCTACTAATTTGGTTAAAAACAAATGTGTCAAT   23182
gb:MN975262|Organism:Severe      CCAGCAACTGTTTGTGGACCTAAAAAGTCTACTAATTTGGTTAAAAACAAATGTGTCAAT   23182
gb:MT106052|Organism:Severe      CCAGCAACTGTTTGTGGACCTAAAAAGTCTACTAATTTGGTTAAAAACAAATGTGTCAAT   23182
gb:LC522975|Organism:Severe      CCAGCAACTGTTTGTGGACCTAAAAAGTCTACTAATTTGGTTAAAAACAAATGTGTCAAT   23179
gb:LC522973|Organism:Severe      CCAGCAACTGTTTGTGGACCTAAAAAGTCTACTAATTTGGTTAAAAACAAATGTGTCAAT   23179
gb:LC522974|Organism:Severe      CCAGCAACTGTTTGTGGACCTAAAAAGTCTACTAATTTGGTTAAAAACAAATGTGTCAAT   23179
gb:MN985325|Organism:Severe      CCAGCAACTGTTTGTGGACCTAAAAAGTCTACTAATTTGGTTAAAAACAAATGTGTCAAT   23182
gb:MT020881|Organism:Severe      CCAGCAACTGTTTGTGGACCTAAAAAGTCTACTAATTTGGTTAAAAACAAATGTGTCAAT   23182
gb:MT020880|Organism:Severe      CCAGCAACTGTTTGTGGACCTAAAAAGTCTACTAATTTGGTTAAAAACAAATGTGTCAAT   23182
gb:MT066175|Organism:Severe      CCAGCAACTGTTTGTGGACCTAAAAAGTCTACTAATTTGGTTAAAAACAAATGTGTCAAT   23182
gb:MN997409|Organism:Severe      CCAGCAACTGTTTGTGGACCTAAAAAGTCTACTAATTTGGTTAAAAACAAATGTGTCAAT   23182
gb:MN938384|Organism:Severe      CCAGCAACTGTTTGTGGACCTAAAAAGTCTACTAATTTGGTTAAAAACAAATGTGTCAAT   23150
gb:MT044258|Organism:Severe      CCAGCAACTGTTTGTGGACCTAAAAAGTCTACTAATTTGGTTAAAAACAAATGTGTCAAT   23158
gb:MT039890|Organism:Severe      CCAGCAACTGTTTGTGGACCTAAAAAGTCTACTAATTTGGTTAAAAACAAATGTGTCAAT   23182
gb:MN988713|Organism:Severe      CCAGCAACTGTTTGTGGACCTAAAAAGTCTACTAATTTGGTTAAAAACAAATGTGTCAAT   23182
gb:LC521925|Organism:Severe      CCAGCAACTGTTTGTGGACCTAAAAAGTCTACTAATTTGGTTAAAAACAAATGTGTCAAT   23155
gb:MT093571|Organism:Severe      CCAGCAACTGTTTGTGGACCTAAAAAGTCTACTAATTTGGTTAAAAACAAATGTGTCAAT   23182
gb:MT039887|Organism:Severe      CCAGCAACTGTTTGTGGACCTAAAAAGTCTACTAATTTGGTTAAAAACAAATGTGTCAAT   23179
gb:MT019530|Organism:Severe      CCAGCAACTGTTTGTGGACCTAAAAAGTCTACTAATTTGGTTAAAAACAAATGTGTCAAT   23182
gb:MT039888|Organism:Severe      CCAGCAACTGTTTGTGGACCTAAAAAGTCTACTAATTTGGTTAAAAACAAATGTGTCAAT   23182
gb:LC522972|Organism:Severe      CCAGCAACTGTTTGTGGACCTAAAAAGTCTACTAATTTGGTTAAAAACAAATGTGTCAAT   23179
gb:MT027063|Organism:Severe      CCAGCAACTGTTTGTGGACCTAAAAAGTCTACTAATTTGGTTAAAAACAAATGTGTCAAT   23182
gb:MT027062|Organism:Severe      CCAGCAACTGTTTGTGGACCTAAAAAGTCTACTAATTTGGTTAAAAACAAATGTGTCAAT   23182
gb:MT019529|Organism:Severe      CCAGCAACTGTTTGTGGACCTAAAAAGTCTACTAATTTGGTTAAAAACAAATGTGTCAAT   23182
gb:MN996529|Organism:Severe      CCAGCAACTGTTTGTGGACCTAAAAAGTCTACTAATTTGGTTAAAAACAAATGTGTCAAT   23170
gb:MN996531|Organism:Severe      CCAGCAACTGTTTGTGGACCTAAAAAGTCTACTAATTTGGTTAAAAACAAATGTGTCAAT   23169
gb:MT066176|Organism:Severe      CCAGCAACTGTTTGTGGACCTAAAAAGTCTACTAATTTGGTTAAAAACAAATGTGTCAAT   23182
gb:MT027064|Organism:Severe      CCAGCAACTGTTTGTGGACCTAAAAAGTCTACTAATTTGGTTAAAAACAAATGTGTCAAT   23182
gb:MN994468|Organism:Severe      CCAGCAACTGTTTGTGGACCTAAAAAGTCTACTAATTTGGTTAAAAACAAATGTGTCAAT   23182
gb:MT072688|Organism:Severe      CCAGCAACTGTTTGTGGACCTAAAAAGTCTACTAATTTGGTTAAAAACAAATGTGTCAAT   23167
gb:MN996527|Organism:Severe      CCAGCAACTGTTTGTGGACCTAAAAAGTCTACTAATTTGGTTAAAAACAAATGTGTCAAT   23149
gb:MT093631|Organism:Severe      CCAGCAACTGTTTGTGGACCTAAAAAGTCTACTAATTTGGTTAAAAACAAATGTGTCAAT   23220
gb:MT106053|Organism:Severe      CCAGCAACTGTTTGTGGACCTAAAAAGTCTACTAATTTGGTTAAAAACAAATGTGTCAAT   23182
gb:MT019533|Organism:Severe      CCAGCAACTGTTTGTGGACCTAAAAAGTCTACTAATTTGGTTAAAAACAAATGTGTCAAT   23182
gb:MT019531|Organism:Severe      CCAGCAACTGTTTGTGGACCTAAAAAGTCTACTAATTTGGTTAAAAACAAATGTGTCAAT   23182
gb:MN996528|Organism:Severe      CCAGCAACTGTTTGTGGACCTAAAAAGTCTACTAATTTGGTTAAAAACAAATGTGTCAAT   23182
gb:MN996530|Organism:Severe      CCAGCAACTGTTTGTGGACCTAAAAAGTCTACTAATTTGGTTAAAAACAAATGTGTCAAT   23168
gb:MN908947|Organism:Severe      CCAGCAACTGTTTGTGGACCTAAAAAGTCTACTAATTTGGTTAAAAACAAATGTGTCAAT   23182
gb:MT019532|Organism:Severe      CCAGCAACTGTTTGTGGACCTAAAAAGTCTACTAATTTGGTTAAAAACAAATGTGTCAAT   23182
                                 ************************************************************

gb:MT020781|Organism:Severe      TTCAACTTCAATGGTTTAACAGGCACAGGTGTTCTTACTGAGTCTAACAAAAAGTTTCTG   23230
gb:MT007544|Organism:Severe      TTCAACTTCAATGGTTTAACAGGCACAGGTGTTCTTACTGAGTCTAACAAAAAGTTTCTG   23242
gb:MN994467|Organism:Severe      TTCAACTTCAATGGTTTAACAGGCACAGGTGTTCTTACTGAGTCTAACAAAAAGTTTCTG   23242
gb:MT044257|Organism:Severe      TTCAACTTCAATGGTTTAACAGGCACAGGTGTTCTTACTGAGTCTAACAAAAAGTTTCTG   23242
```

```
gb:MT106054|Organism:Severe      TTCAACTTCAATGGTTTAACAGGCACAGGTGTTCTTACTGAGTCTAACAAAAAGTTTCTG    23242
gb:MT049951|Organism:Severe      TTCAACTTCAATGGTTTAACAGGCACAGGTGTTCTTACTGAGTCTAACAAAAAGTTTCTG    23242
gb:MN975262|Organism:Severe      TTCAACTTCAATGGTTTAACAGGCACAGGTGTTCTTACTGAGTCTAACAAAAAGTTTCTG    23242
gb:MT106052|Organism:Severe      TTCAACTTCAATGGTTTAACAGGCACAGGTGTTCTTACTGAGTCTAACAAAAAGTTTCTG    23242
gb:LC522975|Organism:Severe      TTCAACTTCAATGGTTTAACAGGCACAGGTGTTCTTACTGAGTCTAACAAAAAGTTTCTG    23239
gb:LC522973|Organism:Severe      TTCAACTTCAATGGTTTAACAGGCACAGGTGTTCTTACTGAGTCTAACAAAAAGTTTCTG    23239
gb:LC522974|Organism:Severe      TTCAACTTCAATGGTTTAACAGGCACAGGTGTTCTTACTGAGTCTAACAAAAAGTTTCTG    23239
gb:MN985325|Organism:Severe      TTCAACTTCAATGGTTTAACAGGCACAGGTGTTCTTACTGAGTCTAACAAAAAGTTTCTG    23242
gb:MT020881|Organism:Severe      TTCAACTTCAATGGTTTAACAGGCACAGGTGTTCTTACTGAGTCTAACAAAAAGTTTCTG    23242
gb:MT020880|Organism:Severe      TTCAACTTCAATGGTTTAACAGGCACAGGTGTTCTTACTGAGTCTAACAAAAAGTTTCTG    23242
gb:MT066175|Organism:Severe      TTCAACTTCAATGGTTTAACAGGCACAGGTGTTCTTACTGAGTCTAACAAAAAGTTTCTG    23242
gb:MN997409|Organism:Severe      TTCAACTTCAATGGTTTAACAGGCACAGGTGTTCTTACTGAGTCTAACAAAAAGTTTCTG    23242
gb:MN938384|Organism:Severe      TTCAACTTCAATGGTTTAACAGGCACAGGTGTTCTTACTGAGTCTAACAAAAAGTTTCTG    23210
gb:MT044258|Organism:Severe      TTCAACTTCAATGGTTTAACAGGCACAGGTGTTCTTACTGAGTCTAACAAAAAGTTTCTG    23218
gb:MT039890|Organism:Severe      TTCAACTTCAATGGTTTAACAGGCACAGGTGTTCTTACTGAGTCTAACAAAAAGTTTCTG    23242
gb:MN988713|Organism:Severe      TTCAACTTCAATGGTTTAACAGGCACAGGTGTTCTTACTGAGTCTAACAAAAAGTTTCTG    23242
gb:LC521925|Organism:Severe      TTCAACTTCAATGGTTTAACAGGCACAGGTGTTCTTACTGAGTCTAACAAAAAGTTTCTG    23215
gb:MT093571|Organism:Severe      TTCAACTTCAATGGTTTAACAGGCACAGGTGTTCTTACTGAGTCTAACAAAAAGTTTCTG    23242
gb:MT039887|Organism:Severe      TTCAACTTCAATGGTTTAACAGGCACAGGTGTTCTTACTGAGTCTAACAAAAAGTTTCTG    23239
gb:MT019530|Organism:Severe      TTCAACTTCAATGGTTTAACAGGCACAGGTGTTCTTACTGAGTCTAACAAAAAGTTTCTG    23242
gb:MT039888|Organism:Severe      TTCAACTTCAATGGTTTAACAGGCACAGGTGTTCTTACTGAGTCTAACAAAAAGTTTCTG    23242
gb:LC522972|Organism:Severe      TTCAACTTCAATGGTTTAACAGGCACAGGTGTTCTTACTGAGTCTAACAAAAAGTTTCTG    23239
gb:MT027063|Organism:Severe      TTCAACTTCAATGGTTTAACAGGCACAGGTGTTCTTACTGAGTCTAACAAAAAGTTTCTG    23242
gb:MT027062|Organism:Severe      TTCAACTTCAATGGTTTAACAGGCACAGGTGTTCTTACTGAGTCTAACAAAAAGTTTCTG    23242
gb:MT019529|Organism:Severe      TTCAACTTCAATGGTTTAACAGGCACAGGTGTTCTTACTGAGTCTAACAAAAAGTTTCTG    23242
gb:MN996529|Organism:Severe      TTCAACTTCAATGGTTTAACAGGCACAGGTGTTCTTACTGAGTCTAACAAAAAGTTTCTG    23230
gb:MN996531|Organism:Severe      TTCAACTTCAATGGTTTAACAGGCACAGGTGTTCTTACTGAGTCTAACAAAAAGTTTCTG    23229
gb:MT066176|Organism:Severe      TTCAACTTCAATGGTTTAACAGGCACAGGTGTTCTTACTGAGTCTAACAAAAAGTTTCTG    23242
gb:MT027064|Organism:Severe      TTCAACTTCAATGGTTTAACAGGCACAGGTGTTCTTACTGAGTCTAACAAAAAGTTTCTG    23242
gb:MN994468|Organism:Severe      TTCAACTTCAATGGTTTAACAGGCACAGGTGTTCTTACTGAGTCTAACAAAAAGTTTCTG    23242
gb:MT072688|Organism:Severe      TTCAACTTCAATGGTTTAACAGGCACAGGTGTTCTTACTGAGTCTAACAAAAAGTTTCTG    23227
gb:MN996527|Organism:Severe      TTCAACTTCAATGGTTTAACAGGCACAGGTGTTCTTACTGAGTCTAACAAAAAGTTTCTG    23209
gb:MT093631|Organism:Severe      TTCAACTTCAATGGTTTAACAGGCACAGGTGTTCTTACTGAGTCTAACAAAAAGTTTCTG    23280
gb:MT106053|Organism:Severe      TTCAACTTCAATGGTTTAACAGGCACAGGTGTTCTTACTGAGTCTAACAAAAAGTTTCTG    23242
gb:MT019533|Organism:Severe      TTCAACTTCAATGGTTTAACAGGCACAGGTGTTCTTACTGAGTCTAACAAAAAGTTTCTG    23242
gb:MT019531|Organism:Severe      TTCAACTTCAATGGTTTAACAGGCACAGGTGTTCTTACTGAGTCTAACAAAAAGTTTCTG    23242
gb:MN996528|Organism:Severe      TTCAACTTCAATGGTTTAACAGGCACAGGTGTTCTTACTGAGTCTAACAAAAAGTTTCTG    23242
gb:MN996530|Organism:Severe      TTCAACTTCAATGGTTTAACAGGCACAGGTGTTCTTACTGAGTCTAACAAAAAGTTTCTG    23228
gb:MN908947|Organism:Severe      TTCAACTTCAATGGTTTAACAGGCACAGGTGTTCTTACTGAGTCTAACAAAAAGTTTCTG    23242
gb:MT019532|Organism:Severe      TTCAACTTCAATGGTTTAACAGGCACAGGTGTTCTTACTGAGTCTAACAAAAAGTTTCTG    23242
                                 ************************************************************

gb:MT020781|Organism:Severe      CCTTTCCAACAATTTGGCAGAGACATTGCTGACACTACTGATGCTGTCCGTGATCCACAG    23290
gb:MT007544|Organism:Severe      CCTTTCCAACAATTTGGCAGAGACATTGCTGACACTACTGATGCTGTCCGTGATCCACAG    23302
gb:MN994467|Organism:Severe      CCTTTCCAACAATTTGGCAGAGACATTGCTGACACTACTGATGCTGTCCGTGATCCACAG    23302
gb:MT044257|Organism:Severe      CCTTTCCAACAATTTGGCAGAGACATTGCTGACACTACTGATGCTGTCCGTGATCCACAG    23302
gb:MT106054|Organism:Severe      CCTTTCCAACAATTTGGCAGAGACATTGCTGACACTACTGATGCTGTCCGTGATCCACAG    23302
gb:MT049951|Organism:Severe      CCTTTCCAACAATTTGGCAGAGACATTGCTGACACTACTGATGCTGTCCGTGATCCACAG    23302
gb:MN975262|Organism:Severe      CCTTTCCAACAATTTGGCAGAGACATTGCTGACACTACTGATGCTGTCCGTGATCCACAG    23302
gb:MT106052|Organism:Severe      CCTTTCCAACAATTTGGCAGAGACATTGCTGACACTACTGATGCTGTCCGTGATCCACAG    23302
gb:LC522975|Organism:Severe      CCTTTCCAACAATTTGGCAGAGACATTGCTGACACTACTGATGCTGTCCGTGATCCACAG    23299
gb:LC522973|Organism:Severe      CCTTTCCAACAATTTGGCAGAGACATTGCTGACACTACTGATGCTGTCCGTGATCCACAG    23299
gb:LC522974|Organism:Severe      CCTTTCCAACAATTTGGCAGAGACATTGCTGACACTACTGATGCTGTCCGTGATCCACAG    23299
gb:MN985325|Organism:Severe      CCTTTCCAACAATTTGGCAGAGACATTGCTGACACTACTGATGCTGTCCGTGATCCACAG    23302
gb:MT020881|Organism:Severe      CCTTTCCAACAATTTGGCAGAGACATTGCTGACACTACTGATGCTGTCCGTGATCCACAG    23302
gb:MT020880|Organism:Severe      CCTTTCCAACAATTTGGCAGAGACATTGCTGACACTACTGATGCTGTCCGTGATCCACAG    23302
gb:MT066175|Organism:Severe      CCTTTCCAACAATTTGGCAGAGACATTGCTGACACTACTGATGCTGTCCGTGATCCACAG    23302
gb:MN997409|Organism:Severe      CCTTTCCAACAATTTGGCAGAGACATTGCTGACACTACTGATGCTGTCCGTGATCCACAG    23302
gb:MN938384|Organism:Severe      CCTTTCCAACAATTTGGCAGAGACATTGCTGACACTACTGATGCTGTCCGTGATCCACAG    23270
gb:MT044258|Organism:Severe      CCTTTCCAACAATTTGGCAGAGACATTGCTGACACTACTGATGCTGTCCGTGATCCACAG    23278
gb:MT039890|Organism:Severe      CCTTTCCAACAATTTGGCAGAGACATTGCTGACACTACTGATGCTGTCCGTGATCCACAG    23302
gb:MN988713|Organism:Severe      CCTTTCCAACAATTTGGCAGAGACATTGCTGACACTACTGATGCTGTCCGTGATCCACAG    23302
gb:LC521925|Organism:Severe      CCTTTCCAACAATTTGGCAGAGACATTGCTGACACTACTGATGCTGTCCGTGATCCACAG    23275
gb:MT093571|Organism:Severe      CCTTTCCAACAATTTGGCAGAGACATTGCTGACACTACTGATGCTGTCCGTGATCCACAG    23302
gb:MT039887|Organism:Severe      CCTTTCCAACAATTTGGCAGAGACATTGCTGACACTACTGATGCTGTCCGTGATCCACAG    23299
```

```
gb:MT019530|Organism:Severe      CCTTTCCAACAATTTGGCAGAGACATTGCTGACACTACTGATGCTGTCCGTGATCCACAG   23302
gb:MT039888|Organism:Severe      CCTTTCCAACAATTTGGCAGAGACATTGCTGACACTACTGATGCTGTCCGTGATCCACAG   23302
gb:LC522972|Organism:Severe      CCTTTCCAACAATTTGGCAGAGACATTGCTGACACTACTGATGCTGTCCGTGATCCACAG   23299
gb:MT027063|Organism:Severe      CCTTTCCAACAATTTGGCAGAGACATTGCTGACACTACTGATGCTGTCCGTGATCCACAG   23302
gb:MT027062|Organism:Severe      CCTTTCCAACAATTTGGCAGAGACATTGCTGACACTACTGATGCTGTCCGTGATCCACAG   23302
gb:MT019529|Organism:Severe      CCTTTCCAACAATTTGGCAGAGACATTGCTGACACTACTGATGCTGTCCGTGATCCACAG   23302
gb:MN996529|Organism:Severe      CCTTTCCAACAATTTGGCAGAGACATTGCTGACACTACTGATGCTGTCCGTGATCCACAG   23290
gb:MN996531|Organism:Severe      CCTTTCCAACAATTTGGCAGAGACATTGCTGACACTACTGATGCTGTCCGTGATCCACAG   23289
gb:MT066176|Organism:Severe      CCTTTCCAACAATTTGGCAGAGACATTGCTGACACTACTGATGCTGTCCGTGATCCACAG   23302
gb:MT027064|Organism:Severe      CCTTTCCAACAATTTGGCAGAGACATTGCTGACACTACTGATGCTGTCCGTGATCCACAG   23302
gb:MN994468|Organism:Severe      CCTTTCCAACAATTTGGCAGAGACATTGCTGACACTACTGATGCTGTCCGTGATCCACAG   23302
gb:MT072688|Organism:Severe      CCTTTCCAACAATTTGGCAGAGACATTGCTGACACTACTGATGCTGTCCGTGATCCACAG   23287
gb:MN996527|Organism:Severe      CCTTTCCAACAATTTGGCAGAGACATTGCTGACACTACTGATGCTGTCCGTGATCCACAG   23269
gb:MT093631|Organism:Severe      CCTTTCCAACAATTTGGCAGAGACATTGCTGACACTACTGATGCTGTCCGTGATCCACAG   23340
gb:MT106053|Organism:Severe      CCTTTCCAACAATTTGGCAGAGACATTGCTGACACTACTGATGCTGTCCGTGATCCACAG   23302
gb:MT019533|Organism:Severe      CCTTTCCAACAATTTGGCAGAGACATTGCTGACACTACTGATGCTGTCCGTGATCCACAG   23302
gb:MT019531|Organism:Severe      CCTTTCCAACAATTTGGCAGAGACATTGCTGACACTACTGATGCTGTCCGTGATCCACAG   23302
gb:MN996528|Organism:Severe      CCTTTCCAACAATTTGGCAGAGACATTGCTGACACTACTGATGCTGTCCGTGATCCACAG   23302
gb:MN996530|Organism:Severe      CCTTTCCAACAATTTGGCAGAGACATTGCTGACACTACTGATGCTGTCCGTGATCCACAG   23288
gb:MN908947|Organism:Severe      CCTTTCCAACAATTTGGCAGAGACATTGCTGACACTACTGATGCTGTCCGTGATCCACAG   23302
gb:MT019532|Organism:Severe      CCTTTCCAACAATTTGGCAGAGACATTGCTGACACTACTGATGCTGTCCGTGATCCACAG   23302
                                 ************************************************************

gb:MT020781|Organism:Severe      ACACTTGAGATTCTTGACATTACACCATGTTCTTTTGGTGGTGTCAGTGTTATAACACCA   23350
gb:MT007544|Organism:Severe      ACACTTGAGATTCTTGACATTACACCATGTTCTTTTGGTGGTGTCAGTGTTATAACACCA   23362
gb:MN994467|Organism:Severe      ACACTTGAGATTCTTGACATTACACCATGTTCTTTTGGTGGTGTCAGTGTTATAACACCA   23362
gb:MT044257|Organism:Severe      ACACTTGAGATTCTTGACATTACACCATGTTCTTTTGGTGGTGTCAGTGTTATAACACCA   23362
gb:MT106054|Organism:Severe      ACACTTGAGATTCTTGACATTACACCATGTTCTTTTGGTGGTGTCAGTGTTATAACACCA   23362
gb:MT049951|Organism:Severe      ACACTTGAGATTCTTGACATTACACCATGTTCTTTTGGTGGTGTCAGTGTTATAACACCA   23362
gb:MN975262|Organism:Severe      ACACTTGAGATTCTTGACATTACACCATGTTCTTTTGGTGGTGTCAGTGTTATAACACCA   23362
gb:MT106052|Organism:Severe      ACACTTGAGATTCTTGACATTACACCATGTTCTTTTGGTGGTGTCAGTGTTATAACACCA   23362
gb:LC522975|Organism:Severe      ACACTTGAGATTCTTGACATTACACCATGTTCTTTTGGTGGTGTCAGTGTTATAACACCA   23359
gb:LC522973|Organism:Severe      ACACTTGAGATTCTTGACATTACACCATGTTCTTTTGGTGGTGTCAGTGTTATAACACCA   23359
gb:LC522974|Organism:Severe      ACACTTGAGATTCTTGACATTACACCATGTTCTTTTGGTGGTGTCAGTGTTATAACACCA   23359
gb:MN985325|Organism:Severe      ACACTTGAGATTCTTGACATTACACCATGTTCTTTTGGTGGTGTCAGTGTTATAACACCA   23362
gb:MT020881|Organism:Severe      ACACTTGAGATTCTTGACATTACACCATGTTCTTTTGGTGGTGTCAGTGTTATAACACCA   23362
gb:MT020880|Organism:Severe      ACACTTGAGATTCTTGACATTACACCATGTTCTTTTGGTGGTGTCAGTGTTATAACACCA   23362
gb:MT066175|Organism:Severe      ACACTTGAGATTCTTGACATTACACCATGTTCTTTTGGTGGTGTCAGTGTTATAACACCA   23362
gb:MN997409|Organism:Severe      ACACTTGAGATTCTTGACATTACACCATGTTCTTTTGGTGGTGTCAGTGTTATAACACCA   23362
gb:MN938384|Organism:Severe      ACACTTGAGATTCTTGACATTACACCATGTTCTTTTGGTGGTGTCAGTGTTATAACACCA   23330
gb:MT044258|Organism:Severe      ACACTTGAGATTCTTGACATTACACCATGTTCTTTTGGTGGTGTCAGTGTTATAACACCA   23338
gb:MT039890|Organism:Severe      ACACTTGAGATTCTTGACATTACACCATGTTCTTTTGGTGGTGTCAGTGTTATAACACCA   23362
gb:MN988713|Organism:Severe      ACACTTGAGATTCTTGACATTACACCATGTTCTTTTGGTGGTGTCAGTGTTATAACACCA   23362
gb:LC521925|Organism:Severe      ACACTTGAGATTCTTGACATTACACCATGTTCTTTTGGTGGTGTCAGTGTTATAACACCA   23335
gb:MT093571|Organism:Severe      ACACTTGAGATTCTTGACATTACACCATGTTCTTTTGGTGGTGTCAGTGTTATAACACCA   23362
gb:MT039887|Organism:Severe      ACACTTGAGATTCTTGACATTACACCATGTTCTTTTGGTGGTGTCAGTGTTATAACACCA   23359
gb:MT019530|Organism:Severe      ACACTTGAGATTCTTGACATTACACCATGTTCTTTTGGTGGTGTCAGTGTTATAACACCA   23362
gb:MT039888|Organism:Severe      ACACTTGAGATTCTTGACATTACACCATGTTCTTTTGGTGGTGTCAGTGTTATAACACCA   23362
gb:LC522972|Organism:Severe      ACACTTGAGATTCTTGACATTACACCATGTTCTTTTGGTGGTGTCAGTGTTATAACACCA   23359
gb:MT027063|Organism:Severe      ACACTTGAGATTCTTGACATTACACCATGTTCTTTTGGTGGTGTCAGTGTTATAACACCA   23362
gb:MT027062|Organism:Severe      ACACTTGAGATTCTTGACATTACACCATGTTCTTTTGGTGGTGTCAGTGTTATAACACCA   23362
gb:MT019529|Organism:Severe      ACACTTGAGATTCTTGACATTACACCATGTTCTTTTGGTGGTGTCAGTGTTATAACACCA   23362
gb:MN996529|Organism:Severe      ACACTTGAGATTCTTGACATTACACCATGTTCTTTTGGTGGTGTCAGTGTTATAACACCA   23350
gb:MN996531|Organism:Severe      ACACTTGAGATTCTTGACATTACACCATGTTCTTTTGGTGGTGTCAGTGTTATAACACCA   23349
gb:MT066176|Organism:Severe      ACACTTGAGATTCTTGACATTACACCATGTTCTTTTGGTGGTGTCAGTGTTATAACACCA   23362
gb:MT027064|Organism:Severe      ACACTTGAGATTCTTGACATTACACCATGTTCTTTTGGTGGTGTCAGTGTTATAACACCA   23362
gb:MN994468|Organism:Severe      ACACTTGAGATTCTTGACATTACACCATGTTCTTTTGGTGGTGTCAGTGTTATAACACCA   23362
gb:MT072688|Organism:Severe      ACACTTGAGATTCTTGACATTACACCATGTTCTTTTGGTGGTGTCAGTGTTATAACACCA   23347
gb:MN996527|Organism:Severe      ACACTTGAGATTCTTGACATTACACCATGTTCTTTTGGTGGTGTCAGTGTTATAACACCA   23329
gb:MT093631|Organism:Severe      ACACTTGAGATTCTTGACATTACACCATGTTCTTTTGGTGGTGTCAGTGTTATAACACCA   23400
gb:MT106053|Organism:Severe      ACACTTGAGATTCTTGACATTACACCATGTTCTTTTGGTGGTGTCAGTGTTATAACACCA   23362
gb:MT019533|Organism:Severe      ACACTTGAGATTCTTGACATTACACCATGTTCTTTTGGTGGTGTCAGTGTTATAACACCA   23362
gb:MT019531|Organism:Severe      ACACTTGAGATTCTTGACATTACACCATGTTCTTTTGGTGGTGTCAGTGTTATAACACCA   23362
gb:MN996528|Organism:Severe      ACACTTGAGATTCTTGACATTACACCATGTTCTTTTGGTGGTGTCAGTGTTATAACACCA   23362
gb:MN996530|Organism:Severe      ACACTTGAGATTCTTGACATTACACCATGTTCTTTTGGTGGTGTCAGTGTTATAACACCA   23348
```

```
gb:MN908947|Organism:Severe      ACACTTGAGATTCTTGACATTACACCATGTTCTTTTGGTGGTGTCAGTGTTATAACACCA   23362
gb:MT019532|Organism:Severe      ACACTTGAGATTCTTGACATTACACCATGTTCTTTTGGTGGTGTCAGTGTTATAACACCA   23362
                                 ************************************************************

gb:MT020781|Organism:Severe      GGAACAAATACTTCTAACCAGGTTGCTGTTCTTTATCAGGATGTTAACTGCACAGAAGTC   23410
gb:MT007544|Organism:Severe      GGAACAAATACTTCTAACCAGGTTGCTGTTCTTTATCAGGATGTTAACTGCACAGAAGTC   23422
gb:MN994467|Organism:Severe      GGAACAAATACTTCTAACCAGGTTGCTGTTCTTTATCAGGATGTTAACTGCACAGAAGTC   23422
gb:MT044257|Organism:Severe      GGAACAAATACTTCTAACCAGGTTGCTGTTCTTTATCAGGATGTTAACTGCACAGAAGTC   23422
gb:MT106054|Organism:Severe      GGAACAAATACTTCTAACCAGGTTGCTGTTCTTTATCAGGATGTTAACTGCACAGAAGTC   23422
gb:MT049951|Organism:Severe      GGAACAAATACTTCTAACCAGGTTGCTGTTCTTTATCAGGATGTTAACTGCACAGAAGTC   23422
gb:MN975262|Organism:Severe      GGAACAAATACTTCTAACCAGGTTGCTGTTCTTTATCAGGATGTTAACTGCACAGAAGTC   23422
gb:MT106052|Organism:Severe      GGAACAAATACTTCTAACCAGGTTGCTGTTCTTTATCAGGATGTTAACTGCACAGAAGTC   23422
gb:LC522975|Organism:Severe      GGAACAAATACTTCTAACCAGGTTGCTGTTCTTTATCAGGATGTTAACTGCACAGAAGTC   23419
gb:LC522973|Organism:Severe      GGAACAAATACTTCTAACCAGGTTGCTGTTCTTTATCAGGATGTTAACTGCACAGAAGTC   23419
gb:LC522974|Organism:Severe      GGAACAAATACTTCTAACCAGGTTGCTGTTCTTTATCAGGATGTTAACTGCACAGAAGTC   23419
gb:MN985325|Organism:Severe      GGAACAAATACTTCTAACCAGGTTGCTGTTCTTTATCAGGATGTTAACTGCACAGAAGTC   23422
gb:MT020881|Organism:Severe      GGAACAAATACTTCTAACCAGGTTGCTGTTCTTTATCAGGATGTTAACTGCACAGAAGTC   23422
gb:MT020880|Organism:Severe      GGAACAAATACTTCTAACCAGGTTGCTGTTCTTTATCAGGATGTTAACTGCACAGAAGTC   23422
gb:MT066175|Organism:Severe      GGAACAAATACTTCTAACCAGGTTGCTGTTCTTTATCAGGATGTTAACTGCACAGAAGTC   23422
gb:MN997409|Organism:Severe      GGAACAAATACTTCTAACCAGGTTGCTGTTCTTTATCAGGATGTTAACTGCACAGAAGTC   23422
gb:MN938384|Organism:Severe      GGAACAAATACTTCTAACCAGGTTGCTGTTCTTTATCAGGATGTTAACTGCACAGAAGTC   23390
gb:MT044258|Organism:Severe      GGAACAAATACTTCTAACCAGGTTGCTGTTCTTTATCAGGATGTTAACTGCACAGAAGTC   23398
gb:MT039890|Organism:Severe      GGAACAAATACTTCTAACCAGGTTGCTGTTCTTTATCAGGATGTTAACTGCACAGAAGTC   23422
gb:MN988713|Organism:Severe      GGAACAAATACTTCTAACCAGGTTGCTGTTCTTTATCAGGATGTTAACTGCACAGAAGTC   23422
gb:LC521925|Organism:Severe      GGAACAAATACTTCTAACCAGGTTGCTGTTCTTTATCAGGATGTTAACTGCACAGAAGTC   23395
gb:MT093571|Organism:Severe      GGAACAAATACTTCTAACCAGGTTGCTGTTCTTTATCAGGATGTTAACTGCACAGAAGTC   23422
gb:MT039887|Organism:Severe      GGAACAAATACTTCTAACCAGGTTGCTGTTCTTTATCAGGATGTTAACTGCACAGAAGTC   23419
gb:MT019530|Organism:Severe      GGAACAAATACTTCTAACCAGGTTGCTGTTCTTTATCAGGATGTTAACTGCACAGAAGTC   23422
gb:MT039888|Organism:Severe      GGAACAAATACTTCTAACCAGGTTGCTGTTCTTTATCAGGATGTTAACTGCACAGAAGTC   23422
gb:LC522972|Organism:Severe      GGAACAAATACTTCTAACCAGGTTGCTGTTCTTTATCAGGATGTTAACTGCACAGAAGTC   23419
gb:MT027063|Organism:Severe      GGAACAAATACTTCTAACCAGGTTGCTGTTCTTTATCAGGATGTTAACTGCACAGAAGTC   23422
gb:MT027062|Organism:Severe      GGAACAAATACTTCTAACCAGGTTGCTGTTCTTTATCAGGATGTTAACTGCACAGAAGTC   23422
gb:MT019529|Organism:Severe      GGAACAAATACTTCTAACCAGGTTGCTGTTCTTTATCAGGATGTTAACTGCACAGAAGTC   23422
gb:MN996529|Organism:Severe      GGAACAAATACTTCTAACCAGGTTGCTGTTCTTTATCAGGATGTTAACTGCACAGAAGTC   23410
gb:MN996531|Organism:Severe      GGAACAAATACTTCTAACCAGGTTGCTGTTCTTTATCAGGATGTTAACTGCACAGAAGTC   23409
gb:MT066176|Organism:Severe      GGAACAAATACTTCTAACCAGGTTGCTGTTCTTTATCAGGATGTTAACTGCACAGAAGTC   23422
gb:MT027064|Organism:Severe      GGAACAAATACTTCTAACCAGGTTGCTGTTCTTTATCAGGATGTTAACTGCACAGAAGTC   23422
gb:MN994468|Organism:Severe      GGAACAAATACTTCTAACCAGGTTGCTGTTCTTTATCAGGATGTTAACTGCACAGAAGTC   23422
gb:MT072688|Organism:Severe      GGAACAAATACTTCTAACCAGGTTGCTGTTCTTTATCAGGATGTTAACTGCACAGAAGTC   23407
gb:MN996527|Organism:Severe      GGAACAAATACTTCTAACCAGGTTGCTGTTCTTTATCAGGATGTTAACTGCACAGAAGTC   23389
gb:MT093631|Organism:Severe      GGAACAAATACTTCTAACCAGGTTGCTGTTCTTTATCAGGATGTTAACTGCACAGAAGTC   23460
gb:MT106053|Organism:Severe      GGAACAAATACTTCTAACCAGGTTGCTGTTCTTTATCAGGATGTTAACTGCACAGAAGTC   23422
gb:MT019533|Organism:Severe      GGAACAAATACTTCTAACCAGGTTGCTGTTCTTTATCAGGATGTTAACTGCACAGAAGTC   23422
gb:MT019531|Organism:Severe      GGAACAAATACTTCTAACCAGGTTGCTGTTCTTTATCAGGATGTTAACTGCACAGAAGTC   23422
gb:MN996528|Organism:Severe      GGAACAAATACTTCTAACCAGGTTGCTGTTCTTTATCAGGATGTTAACTGCACAGAAGTC   23422
gb:MN996530|Organism:Severe      GGAACAAATACTTCTAACCAGGTTGCTGTTCTTTATCAGGATGTTAACTGCACAGAAGTC   23408
gb:MN908947|Organism:Severe      GGAACAAATACTTCTAACCAGGTTGCTGTTCTTTATCAGGATGTTAACTGCACAGAAGTC   23422
gb:MT019532|Organism:Severe      GGAACAAATACTTCTAACCAGGTTGCTGTTCTTTATCAGGATGTTAACTGCACAGAAGTC   23422
                                 ************************************************************

gb:MT020781|Organism:Severe      CCTGTTGCTATTCATGCAGATCAACTTACTCCTACTTGGCGTGTTTATTCTACAGGTTCT   23470
gb:MT007544|Organism:Severe      CCTGTTGCTATTCATGCAGATCAACTTACTCCTACTTGGCGTGTTTATTCTACAGGTTCT   23482
gb:MN994467|Organism:Severe      CCTGTTGCTATTCATGCAGATCAACTTACTCCTACTTGGCGTGTTTATTCTACAGGTTCT   23482
gb:MT044257|Organism:Severe      CCTGTTGCTATTCATGCAGATCAACTTACTCCTACTTGGCGTGTTTATTCTACAGGTTCT   23482
gb:MT106054|Organism:Severe      CCTGTTGCTATTCATGCAGATCAACTTACTCCTACTTGGCGTGTTTATTCTACAGGTTCT   23482
gb:MT049951|Organism:Severe      CCTGTTGCTATTCATGCAGATCAACTTACTCCTACTTGGCGTGTTTATTCTACAGGTTCT   23482
gb:MN975262|Organism:Severe      CCTGTTGCTATTCATGCAGATCAACTTACTCCTACTTGGCGTGTTTATTCTACAGGTTCT   23482
gb:MT106052|Organism:Severe      CCTGTTGCTATTCATGCAGATCAACTTACTCCTACTTGGCGTGTTTATTCTACAGGTTCT   23482
gb:LC522975|Organism:Severe      CCTGTTGCTATTCATGCAGATCAACTTACTCCTACTTGGCGTGTTTATTCTACAGGTTCT   23479
gb:LC522973|Organism:Severe      CCTGTTGCTATTCATGCAGATCAACTTACTCCTACTTGGCGTGTTTATTCTACAGGTTCT   23479
gb:LC522974|Organism:Severe      CCTGTTGCTATTCATGCAGATCAACTTACTCCTACTTGGCGTGTTTATTCTACAGGTTCT   23479
gb:MN985325|Organism:Severe      CCTGTTGCTATTCATGCAGATCAACTTACTCCTACTTGGCGTGTTTATTCTACAGGTTCT   23482
gb:MT020881|Organism:Severe      CCTGTTGCTATTCATGCAGATCAACTTACTCCTACTTGGCGTGTTTATTCTACAGGTTCT   23482
gb:MT020880|Organism:Severe      CCTGTTGCTATTCATGCAGATCAACTTACTCCTACTTGGCGTGTTTATTCTACAGGTTCT   23482
gb:MT066175|Organism:Severe      CCTGTTGCTATTCATGCAGATCAACTTACTCCTACTTGGCGTGTTTATTCTACAGGTTCT   23482
```

```
gb:MN997409|Organism:Severe      CCTGTTGCTATTCATGCAGATCAACTTACTCCTACTTGGCGTGTTTATTCTACAGGTTCT   23482
gb:MN938384|Organism:Severe      CCTGTTGCTATTCATGCAGATCAACTTACTCCTACTTGGCGTGTTTATTCTACAGGTTCT   23450
gb:MT044258|Organism:Severe      CCTGTTGCTATTCATGCAGATCAACTTACTCCTACTTGGCGTGTTTATTCTACAGGTTCT   23458
gb:MT039890|Organism:Severe      CCTGTTGCTATTCATGCAGATCAACTTACTCCTACTTGGCGTGTTTATTCTACAGGTTCT   23482
gb:MN988713|Organism:Severe      CCTGTTGCTATTCATGCAGATCAACTTACTCCTACTTGGCGTGTTTATTCTACAGGTTCT   23482
gb:LC521925|Organism:Severe      CCTGTTGCTATTCATGCAGATCAACTTACTCCTACTTGGCGTGTTTATTCTACAGGTTCT   23455
gb:MT093571|Organism:Severe      CCTGTTGCTATTCATGCAGATCAACTTACTCCTACTTGGCGTGTTTATTCTACAGGTTCT   23482
gb:MT039887|Organism:Severe      CCTGTTGCTATTCATGCAGATCAACTTACTCCTACTTGGCGTGTTTATTCTACAGGTTCT   23479
gb:MT019530|Organism:Severe      CCTGTTGCTATTCATGCAGATCAACTTACTCCTACTTGGCGTGTTTATTCTACAGGTTCT   23482
gb:MT039888|Organism:Severe      CCTGTTGCTATTCATGCAGATCAACTTACTCCTACTTGGCGTGTTTATTCTACAGGTTCT   23482
gb:LC522972|Organism:Severe      CCTGTTGCTATTCATGCAGATCAACTTACTCCTACTTGGCGTGTTTATTCTACAGGTTCT   23479
gb:MT027063|Organism:Severe      CCTGTTGCTATTCATGCAGATCAACTTACTCCTACTTGGCGTGTTTATTCTACAGGTTCT   23482
gb:MT027062|Organism:Severe      CCTGTTGCTATTCATGCAGATCAACTTACTCCTACTTGGCGTGTTTATTCTACAGGTTCT   23482
gb:MT019529|Organism:Severe      CCTGTTGCTATTCATGCAGATCAACTTACTCCTACTTGGCGTGTTTATTCTACAGGTTCT   23482
gb:MN996529|Organism:Severe      CCTGTTGCTATTCATGCAGATCAACTTACTCCTACTTGGCGTGTTTATTCTACAGGTTCT   23470
gb:MN996531|Organism:Severe      CCTGTTGCTATTCATGCAGATCAACTTACTCCTACTTGGCGTGTTTATTCTACAGGTTCT   23469
gb:MT066176|Organism:Severe      CCTGTTGCTATTCATGCAGATCAACTTACTCCTACTTGGCGTGTTTATTCTACAGGTTCT   23482
gb:MT027064|Organism:Severe      CCTGTTGCTATTCATGCAGATCAACTTACTCCTACTTGGCGTGTTTATTCTACAGGTTCT   23482
gb:MN994468|Organism:Severe      CCTGTTGCTATTCATGCAGATCAACTTACTCCTACTTGGCGTGTTTATTCTACAGGTTCT   23482
gb:MT072688|Organism:Severe      CCTGTTGCTATTCATGCAGATCAACTTACTCCTACTTGGCGTGTTTATTCTACAGGTTCT   23467
gb:MN996527|Organism:Severe      CCTGTTGCTATTCATGCAGATCAACTTACTCCTACTTGGCGTGTTTATTCTACAGGTTCT   23449
gb:MT093631|Organism:Severe      CCTGTTGCTATTCATGCAGATCAACTTACTCCTACTTGGCGTGTTTATTCTACAGGTTCT   23520
gb:MT106053|Organism:Severe      CCTGTTGCTATTCATGCAGATCAACTTACTCCTACTTGGCGTGTTTATTCTACAGGTTCT   23482
gb:MT019533|Organism:Severe      CCTGTTGCTATTCATGCAGATCAACTTACTCCTACTTGGCGTGTTTATTCTACAGGTTCT   23482
gb:MT019531|Organism:Severe      CCTGTTGCTATTCATGCAGATCAACTTACTCCTACTTGGCGTGTTTATTCTACAGGTTCT   23482
gb:MN996528|Organism:Severe      CCTGTTGCTATTCATGCAGATCAACTTACTCCTACTTGGCGTGTTTATTCTACAGGTTCT   23482
gb:MN996530|Organism:Severe      CCTGTTGCTATTCATGCAGATCAACTTACTCCTACTTGGCGTGTTTATTCTACAGGTTCT   23468
gb:MN908947|Organism:Severe      CCTGTTGCTATTCATGCAGATCAACTTACTCCTACTTGGCGTGTTTATTCTACAGGTTCT   23482
gb:MT019532|Organism:Severe      CCTGTTGCTATTCATGCAGATCAACTTACTCCTACTTGGCGTGTTTATTCTACAGGTTCT   23482
                                 ************************************************************

gb:MT020781|Organism:Severe      AATGTTTTTCAAACACGTGCAGGCTGTTTAATAGGGGCTGAACATGTCAACAACTCATAT   23530
gb:MT007544|Organism:Severe      AATGTTTTTCAAACACGTGCAGGCTGTTTAATAGGGGCTGAACATGTCAACAACTCATAT   23542
gb:MN994467|Organism:Severe      AATGTTTTTCAAACACGTGCAGGCTGTTTAATAGGGGCTGAACATGTCAACAACTCATAT   23542
gb:MT044257|Organism:Severe      AATGTTTTTCAAACACGTGCAGGCTGTTTAATAGGGGCTGAACATGTCAACAACTCATAT   23542
gb:MT106054|Organism:Severe      AATGTTTTTCAAACACGTGCAGGCTGTTTAATAGGGGCTGAACATGTCAACAACTCATAT   23542
gb:MT049951|Organism:Severe      AATGTTTTTCAAACACGTGCAGGCTGTTTAATAGGGGCTGAACATGTCAACAACTCATAT   23542
gb:MN975262|Organism:Severe      AATGTTTTTCAAACACGTGCAGGCTGTTTAATAGGGGCTGAACATGTCAACAACTCATAT   23542
gb:MT106052|Organism:Severe      AATGTTTTTCAAACACGTGCAGGCTGTTTAATAGGGGCTGAACATGTCAACAACTCATAT   23542
gb:LC522975|Organism:Severe      AATGTTTTTCAAACACGTGCAGGCTGTTTAATAGGGGCTGAACATGTCAACAACTCATAT   23539
gb:LC522973|Organism:Severe      AATGTTTTTCAAACACGTGCAGGCTGTTTAATAGGGGCTGAACATGTCAACAACTCATAT   23539
gb:LC522974|Organism:Severe      AATGTTTTTCAAACACGTGCAGGCTGTTTAATAGGGGCTGAACATGTCAACAACTCATAT   23539
gb:MN985325|Organism:Severe      AATGTTTTTCAAACACGTGCAGGCTGTTTAATAGGGGCTGAACATGTCAACAACTCATAT   23542
gb:MT020881|Organism:Severe      AATGTTTTTCAAACACGTGCAGGCTGTTTAATAGGGGCTGAACATGTCAACAACTCATAT   23542
gb:MT020880|Organism:Severe      AATGTTTTTCAAACACGTGCAGGCTGTTTAATAGGGGCTGAACATGTCAACAACTCATAT   23542
gb:MT066175|Organism:Severe      AATGTTTTTCAAACACGTGCAGGCTGTTTAATAGGGGCTGAACATGTCAACAACTCATAT   23542
gb:MN997409|Organism:Severe      AATGTTTTTCAAACACGTGCAGGCTGTTTAATAGGGGCTGAACATGTCAACAACTCATAT   23542
gb:MN938384|Organism:Severe      AATGTTTTTCAAACACGTGCAGGCTGTTTAATAGGGGCTGAACATGTCAACAACTCATAT   23510
gb:MT044258|Organism:Severe      AATGTTTTTCAAACACGTGCAGGCTGTTTAATAGGGGCTGAACATGTCAACAACTCATAT   23518
gb:MT039890|Organism:Severe      AATGTTTTTCAAACACGTGCAGGCTGTTTAATAGGGGCTGAACATGTCAACAACTCATAT   23542
gb:MN988713|Organism:Severe      AATGTTTTTCAAACACGTGCAGGCTGTTTAATAGGGGCTGAACATGTCAACAACTCATAT   23542
gb:LC521925|Organism:Severe      AATGTTTTTCAAACACGTGCAGGCTGTTTAATAGGGGCTGAACATGTCAACAACTCATAT   23515
gb:MT093571|Organism:Severe      AATGTTTTTCAAACACGTGCAGGCTGTTTAATAGGGGCTGAACATGTCAACAACTCATAT   23542
gb:MT039887|Organism:Severe      AATGTTTTTCAAACACGTGCAGGCTGTTTAATAGGGGCTGAACATGTCAACAACTCATAT   23539
gb:MT019530|Organism:Severe      AATGTTTTTCAAACACGTGCAGGCTGTTTAATAGGGGCTGAACATGTCAACAACTCATAT   23542
gb:MT039888|Organism:Severe      AATGTTTTTCAAACACGTGCAGGCTGTTTAATAGGGGCTGAACATGTCAACAACTCATAT   23542
gb:LC522972|Organism:Severe      AATGTTTTTCAAACACGTGCAGGCTGTTTAATAGGGGCTGAACATGTCAACAACTCATAT   23539
gb:MT027063|Organism:Severe      AATGTTTTTCAAACACGTGCAGGCTGTTTAATAGGGGCTGAACATGTCAACAACTCATAT   23542
gb:MT027062|Organism:Severe      AATGTTTTTCAAACACGTGCAGGCTGTTTAATAGGGGCTGAACATGTCAACAACTCATAT   23542
gb:MT019529|Organism:Severe      AATGTTTTTCAAACACGTGCAGGCTGTTTAATAGGGGCTGAACATGTCAACAACTCATAT   23542
gb:MN996529|Organism:Severe      AATGTTTTTCAAACACGTGCAGGCTGTTTAATAGGGGCTGAACATGTCAACAACTCATAT   23530
gb:MN996531|Organism:Severe      AATGTTTTTCAAACACGTGCAGGCTGTTTAATAGGGGCTGAACATGTCAACAACTCATAT   23529
gb:MT066176|Organism:Severe      AATGTTTTTCAAACACGTGCAGGCTGTTTAATAGGGGCTGAACATGTCAACAACTCATAT   23542
gb:MT027064|Organism:Severe      AATGTTTTTCAAACACGTGCAGGCTGTTTAATAGGGGCTGAACATGTCAACAACTCATAT   23542
gb:MN994468|Organism:Severe      AATGTTTTTCAAACACGTGCAGGCTGTTTAATAGGGGCTGAACATGTCAACAACTCATAT   23542
```

```
gb:MT072688|Organism:Severe      AATGTTTTTCAAACACGTGCAGGCTGTTTAATAGGGGCTGAACATGTCAACAACTCATAT   23527
gb:MN996527|Organism:Severe      AATGTTTTTCAAACACGTGCAGGCTGTTTAATAGGGGCTGAACATGTCAACAACTCATAT   23509
gb:MT093631|Organism:Severe      AATGTTTTTCAAACACGTGCAGGCTGTTTAATAGGGGCTGAACATGTCAACAACTCATAT   23580
gb:MT106053|Organism:Severe      AATGTTTTTCAAACACGTGCAGGCTGTTTAATAGGGGCTGAACATGTCAACAACTCATAT   23542
gb:MT019533|Organism:Severe      AATGTTTTTCAAACACGTGCAGGCTGTTTAATAGGGGCTGAACATGTCAACAACTCATAT   23542
gb:MT019531|Organism:Severe      AATGTTTTTCAAACACGTGCAGGCTGTTTAATAGGGGCTGAACATGTCAACAACTCATAT   23542
gb:MN996528|Organism:Severe      AATGTTTTTCAAACACGTGCAGGCTGTTTAATAGGGGCTGAACATGTCAACAACTCATAT   23542
gb:MN996530|Organism:Severe      AATGTTTTTCAAACACGTGCAGGCTGTTTAATAGGGGCTGAACATGTCAACAACTCATAT   23528
gb:MN908947|Organism:Severe      AATGTTTTTCAAACACGTGCAGGCTGTTTAATAGGGGCTGAACATGTCAACAACTCATAT   23542
gb:MT019532|Organism:Severe      AATGTTTTTCAAACACGTGCAGGCTGTTTAATAGGGGCTGAACATGTCAACAACTCATAT   23542
                                 ************************************************************

gb:MT020781|Organism:Severe      GAGTGTGACATACCCATTGGTGCAGGTATATGCGCTAGTTATCAGACTCAGACTAATTCT   23590
gb:MT007544|Organism:Severe      GAGTGTGACATACCCATTGGTGCAGGTATATGCGCTAGTTATCAGACTCAGACTAATTCT   23602
gb:MN994467|Organism:Severe      GAGTGTGACATACCCATTGGTGCAGGTATATGCGCTAGTTATCAGACTCAGACTAATTCT   23602
gb:MT044257|Organism:Severe      GAGTGTGACATACCCATTGGTGCAGGTATATGCGCTAGTTATCAGACTCAGACTAATTCT   23602
gb:MT106054|Organism:Severe      GAGTGTGACATACCCATTGGTGCAGGTATATGCGCTAGTTATCAGACTCAGACTAATTCT   23602
gb:MT049951|Organism:Severe      GAGTGTGACATACCCATTGGTGCAGGTATATGCGCTAGTTATCAGACTCAGACTAATTCT   23602
gb:MN975262|Organism:Severe      GAGTGTGACATACCCATTGGTGCAGGTATATGCGCTAGTTATCAGACTCAGACTAATTCT   23602
gb:MT106052|Organism:Severe      GAGTGTGACATACCCATTGGTGCAGGTATATGCGCTAGTTATCAGACTCAGACTAATTCT   23602
gb:LC522975|Organism:Severe      GAGTGTGACATACCCATTGGTGCAGGTATATGCGCTAGTTATCAGACTCAGACTAATTCT   23599
gb:LC522973|Organism:Severe      GAGTGTGACATACCCATTGGTGCAGGTATATGCGCTAGTTATCAGACTCAGACTAATTCT   23599
gb:LC522974|Organism:Severe      GAGTGTGACATACCCATTGGTGCAGGTATATGCGCTAGTTATCAGACTCAGACTAATTCT   23599
gb:MN985325|Organism:Severe      GAGTGTGACATACCCATTGGTGCAGGTATATGCGCTAGTTATCAGACTCAGACTAATTCT   23602
gb:MT020881|Organism:Severe      GAGTGTGACATACCCATTGGTGCAGGTATATGCGCTAGTTATCAGACTCAGACTAATTCT   23602
gb:MT020880|Organism:Severe      GAGTGTGACATACCCATTGGTGCAGGTATATGCGCTAGTTATCAGACTCAGACTAATTCT   23602
gb:MT066175|Organism:Severe      GAGTGTGACATACCCATTGGTGCAGGTATATGCGCTAGTTATCAGACTCAGACTAATTCT   23602
gb:MN997409|Organism:Severe      GAGTGTGACATACCCATTGGTGCAGGTATATGCGCTAGTTATCAGACTCAGACTAATTCT   23602
gb:MN938384|Organism:Severe      GAGTGTGACATACCCATTGGTGCAGGTATATGCGCTAGTTATCAGACTCAGACTAATTCT   23570
gb:MT044258|Organism:Severe      GAGTGTGACATACCCATTGGTGCAGGTATATGCGCTAGTTATCAGACTCAGACTAATTCT   23578
gb:MT039890|Organism:Severe      GAGTGTGACATACCCATTGGTGCAGGTATATGCGCTAGTTATCAGACTCAGACTAATTCT   23602
gb:MN988713|Organism:Severe      GAGTGTGACATACCCATTGGTGCAGGTATATGCGCTAGTTATCAGACTCAGACTAATTCT   23602
gb:LC521925|Organism:Severe      GAGTGTGACATACCCATTGGTGCAGGTATATGCGCTAGTTATCAGACTCAGACTAATTCT   23575
gb:MT093571|Organism:Severe      GAGTGTGACATACCCATTGGTGCAGGTATATGCGCTAGTTATCAGACTCAGACTAATTCT   23602
gb:MT039887|Organism:Severe      GAGTGTGACATACCCATTGGTGCAGGTATATGCGCTAGTTATCAGACTCAGACTAATTCT   23599
gb:MT019530|Organism:Severe      GAGTGTGACATACCCATTGGTGCAGGTATATGCGCTAGTTATCAGACTCAGACTAATTCT   23602
gb:MT039888|Organism:Severe      GAGTGTGACATACCCATTGGTGCAGGTATATGCGCTAGTTATCAGACTCAGACTAATTCT   23602
gb:LC522972|Organism:Severe      GAGTGTGACATACCCATTGGTGCAGGTATATGCGCTAGTTATCAGACTCAGACTAATTCT   23599
gb:MT027063|Organism:Severe      GAGTGTGACATACCCATTGGTGCAGGTATATGCGCTAGTTATCAGACTCAGACTAATTCT   23602
gb:MT027062|Organism:Severe      GAGTGTGACATACCCATTGGTGCAGGTATATGCGCTAGTTATCAGACTCAGACTAATTCT   23602
gb:MT019529|Organism:Severe      GAGTGTGACATACCCATTGGTGCAGGTATATGCGCTAGTTATCAGACTCAGACTAATTCT   23602
gb:MN996529|Organism:Severe      GAGTGTGACATACCCATTGGTGCAGGTATATGCGCTAGTTATCAGACTCAGACTAATTCT   23590
gb:MN996531|Organism:Severe      GAGTGTGACATACCCATTGGTGCAGGTATATGCGCTAGTTATCAGACTCAGACTAATTCT   23589
gb:MT066176|Organism:Severe      GAGTGTGACATACCCATTGGTGCAGGTATATGCGCTAGTTATCAGACTCAGACTAATTCT   23602
gb:MT027064|Organism:Severe      GAGTGTGACATACCCATTGGTGCAGGTATATGCGCTAGTTATCAGACTCAGACTAATTCT   23602
gb:MN994468|Organism:Severe      GAGTGTGACATACCCATTGGTGCAGGTATATGCGCTAGTTATCAGACTCAGACTAATTCT   23602
gb:MT072688|Organism:Severe      GAGTGTGACATACCCATTGGTGCAGGTATATGCGCTAGTTATCAGACTCAGACTAATTCT   23587
gb:MN996527|Organism:Severe      GAGTGTGACATACCCATTGGTGCAGGTATATGCGCTAGTTATCAGACTCAGACTAATTCT   23569
gb:MT093631|Organism:Severe      GAGTGTGACATACCCATTGGTGCAGGTATATGCGCTAGTTATCAGACTCAGACTAATTCT   23640
gb:MT106053|Organism:Severe      GAGTGTGACATACCCATTGGTGCAGGTATATGCGCTAGTTATCAGACTCAGACTAATTCT   23602
gb:MT019533|Organism:Severe      GAGTGTGACATACCCATTGGTGCAGGTATATGCGCTAGTTATCAGACTCAGACTAATTCT   23602
gb:MT019531|Organism:Severe      GAGTGTGACATACCCATTGGTGCAGGTATATGCGCTAGTTATCAGACTCAGACTAATTCT   23602
gb:MN996528|Organism:Severe      GAGTGTGACATACCCATTGGTGCAGGTATATGCGCTAGTTATCAGACTCAGACTAATTCT   23602
gb:MN996530|Organism:Severe      GAGTGTGACATACCCATTGGTGCAGGTATATGCGCTAGTTATCAGACTCAGACTAATTCT   23588
gb:MN908947|Organism:Severe      GAGTGTGACATACCCATTGGTGCAGGTATATGCGCTAGTTATCAGACTCAGACTAATTCT   23602
gb:MT019532|Organism:Severe      GAGTGTGACATACCCATTGGTGCAGGTATATGCGCTAGTTATCAGACTCAGACTAATTCT   23602
                                 ************************************************************

gb:MT020781|Organism:Severe      CCTCGGCGGGCACGTAGTGTAGCTAGTCAATCCATCATTGCCTACACTATGTCACTTGGT   23650
gb:MT007544|Organism:Severe      CCTCGGCGGGCACGTAGTGTAGCTAGTCAATCCATCATTGCCTACACTATGTCACTTGGT   23662
gb:MN994467|Organism:Severe      CCTCGGCGGGCACGTAGTGTAGCTAGTCAATCCATCATTGCCTACACTATGTCACTTGGT   23662
gb:MT044257|Organism:Severe      CCTCGGCGGGCACGTAGTGTAGCTAGTCAATCCATCATTGCCTACACTATGTCACTTGGT   23662
gb:MT106054|Organism:Severe      CCTCGGCGGGCACGTAGTGTAGCTAGTCAATCCATCATTGCCTACACTATGTCACTTGGT   23662
gb:MT049951|Organism:Severe      CCTCGGCGGGCACGTAGTGTAGCTAGTCAATCCATCATTGCCTACACTATGTCACTTGGT   23662
gb:MN975262|Organism:Severe      CCTCGGCGGGCACGTAGTGTAGCTAGTCAATCCATCATTGCCTACACTATGTCACTTGGT   23662
```

```
gb:MT106052|Organism:Severe      CCTCGGCGGGCACGTAGTGTAGCTAGTCAATCCATCATTGCCTACACTATGTCACTTGGT   23662
gb:LC522975|Organism:Severe      CCTCGGCGGGCACGTAGTGTAGCTAGTCAATCCATCATTGCCTACACTATGTCACTTGGT   23659
gb:LC522973|Organism:Severe      CCTCGGCGGGCACGTAGTGTAGCTAGTCAATCCATCATTGCCTACACTATGTCACTTGGT   23659
gb:LC522974|Organism:Severe      CCTCGGCGGGCACGTAGTGTAGCTAGTCAATCCATCATTGCCTACACTATGTCACTTGGT   23659
gb:MN985325|Organism:Severe      CCTCGGCGGGCACGTAGTGTAGCTAGTCAATCCATCATTGCCTACACTATGTCACTTGGT   23662
gb:MT020881|Organism:Severe      CCTCGGCGGGCACGTAGTGTAGCTAGTCAATCCATCATTGCCTACACTATGTCACTTGGT   23662
gb:MT020880|Organism:Severe      CCTCGGCGGGCACGTAGTGTAGCTAGTCAATCCATCATTGCCTACACTATGTCACTTGGT   23662
gb:MT066175|Organism:Severe      CCTCGGCGGGCACGTAGTGTAGCTAGTCAATCCATCATTGCCTACACTATGTCACTTGGT   23662
gb:MN997409|Organism:Severe      CCTCGGCGGGCACGTAGTGTAGCTAGTCAATCCATCATTGCCTACACTATGTCACTTGGT   23662
gb:MN938384|Organism:Severe      CCTCGGCGGGCACGTAGTGTAGCTAGTCAATCCATCATTGCCTACACTATGTCACTTGGT   23630
gb:MT044258|Organism:Severe      CCTCGGCGGGCACGTAGTGTAGCTAGTCAATCCATCATTGCCTACACTATGTCACTTGGT   23638
gb:MT039890|Organism:Severe      CCTCGGCGGGCACGTAGTGTAGCTAGTCAATCCATCATTGCCTACACTATGTCACTTGGT   23662
gb:MN988713|Organism:Severe      CCTCGGCGGGCACGTAGTGTAGCTAGTCAATCCATCATTGCCTACACTATGTCACTTGGT   23662
gb:LC521925|Organism:Severe      CCTCGGCGGGCACGTAGTGTAGCTAGTCAATCCATCATTGCCTACACTATGTCACTTGGT   23635
gb:MT093571|Organism:Severe      CCTCGGCGGGCACGTAGTGTAGCTAGTCAATCCATCATTGCCTACACTATGTCACTTGGT   23662
gb:MT039887|Organism:Severe      CCTCGGCGGGCACGTAGTGTAGCTAGTCAATCCATCATTGCCTACACTATGTCACTTGGT   23659
gb:MT019530|Organism:Severe      CCTCGGCGGGCACGTAGTGTAGCTAGTCAATCCATCATTGCCTACACTATGTCACTTGGT   23662
gb:MT039888|Organism:Severe      CCTCGGCGGGCACGTAGTGTAGCTAGTCAATCCATCATTGCCTACACTATGTCACTTGGT   23662
gb:LC522972|Organism:Severe      CCTCGGCGGGCACGTAGTGTAGCTAGTCAATCCATCATTGCCTACACTATGTCACTTGGT   23659
gb:MT027063|Organism:Severe      CCTCGGCGGGCACGTAGTGTAGCTAGTCAATCCATCATTGCCTACACTATGTCACTTGGT   23662
gb:MT027062|Organism:Severe      CCTCGGCGGGCACGTAGTGTAGCTAGTCAATCCATCATTGCCTACACTATGTCACTTGGT   23662
gb:MT019529|Organism:Severe      CCTCGGCGGGCACGTAGTGTAGCTAGTCAATCCATCATTGCCTACACTATGTCACTTGGT   23662
gb:MN996529|Organism:Severe      CCTCGGCGGGCACGTAGTGTAGCTAGTCAATCCATCATTGCCTACACTATGTCACTTGGT   23650
gb:MN996531|Organism:Severe      CCTCGGCGGGCACGTAGTGTAGCTAGTCAATCCATCATTGCCTACACTATGTCACTTGGT   23649
gb:MT066176|Organism:Severe      CCTCGGCGGGCACGTAGTGTAGCTAGTCAATCCATCATTGCCTACACTATGTCACTTGGT   23662
gb:MT027064|Organism:Severe      CCTCGGCGGGCACGTAGTGTAGCTAGTCAATCCATCATTGCCTACACTATGTCACTTGGT   23662
gb:MN994468|Organism:Severe      CCTCGGCGGGCACGTAGTGTAGCTAGTCAATCCATCATTGCCTACACTATGTCACTTGGT   23662
gb:MT072688|Organism:Severe      CCTCGGCGGGCACGTAGTGTAGCTAGTCAATCCATCATTGCCTACACTATGTCACTTGGT   23647
gb:MN996527|Organism:Severe      CCTCGGCGGGCACGTAGTGTAGCTAGTCAATCCATCATTGCCTACACTATGTCACTTGGT   23629
gb:MT093631|Organism:Severe      CCTCGGCGGGCACGTAGTGTAGCTAGTCAATCCATCATTGCCTACACTATGTCACTTGGT   23700
gb:MT106053|Organism:Severe      CCTCGGCGGGCACGTAGTGTAGCTAGTCAATCCATCATTGCCTACACTATGTCACTTGGT   23662
gb:MT019533|Organism:Severe      CCTCGGCGGGCACGTAGTGTAGCTAGTCAATCCATCATTGCCTACACTATGTCACTTGGT   23662
gb:MT019531|Organism:Severe      CCTCGGCGGGCACGTAGTGTAGCTAGTCAATCCATCATTGCCTACACTATGTCACTTGGT   23662
gb:MN996528|Organism:Severe      CCTCGGCGGGCACGTAGTGTAGCTAGTCAATCCATCATTGCCTACACTATGTCACTTGGT   23662
gb:MN996530|Organism:Severe      CCTCGGCGGGCACGTAGTGTAGCTAGTCAATCCATCATTGCCTACACTATGTCACTTGGT   23648
gb:MN908947|Organism:Severe      CCTCGGCGGGCACGTAGTGTAGCTAGTCAATCCATCATTGCCTACACTATGTCACTTGGT   23662
gb:MT019532|Organism:Severe      CCTCGGCGGGCACGTAGTGTAGCTAGTCAATCCATCATTGCCTACACTATGTCACTTGGT   23662
                                 ************************************************************

gb:MT020781|Organism:Severe      GCAGAAAATTCAGTTGCTTACTCTAATAACTCTATTGCCATACCCACAAATTTTACTATT   23710
gb:MT007544|Organism:Severe      GCAGAAAATTCAGTTGCTTACTCTAATAACTCTATTGCCATACCCACAAATTTTACTATT   23722
gb:MN994467|Organism:Severe      GCAGAAAATTCAGTTGCTTACTCTAATAACTCTATTGCCATACCCACAAATTTTACTATT   23722
gb:MT044257|Organism:Severe      GCAGAAAATTCAGTTGCTTACTCTAATAACTCTATTGCCATACCCACAAATTTTACTATT   23722
gb:MT106054|Organism:Severe      GCAGAAAATTCAGTTGCTTACTCTAATAACTCTATTGCCATACCCACAAATTTTACTATT   23722
gb:MT049951|Organism:Severe      GCAGAAAATTCAGTTGCTTACTCTAATAACTCTATTGCCATACCCACAAATTTTACTATT   23722
gb:MN975262|Organism:Severe      GCAGAAAATTCAGTTGCTTACTCTAATAACTCTATTGCCATACCCACAAATTTTACTATT   23722
gb:MT106052|Organism:Severe      GCAGAAAATTCAGTTGCTTACTCTAATAACTCTATTGCCATACCCACAAATTTTACTATT   23722
gb:LC522975|Organism:Severe      GCAGAAAATTCAGTTGCTTACTCTAATAACTCTATTGCCATACCCACAAATTTTACTATT   23719
gb:LC522973|Organism:Severe      GCAGAAAATTCAGTTGCTTACTCTAATAACTCTATTGCCATACCCACAAATTTTACTATT   23719
gb:LC522974|Organism:Severe      GCAGAAAATTCAGTTGCTTACTCTAATAACTCTATTGCCATACCCACAAATTTTACTATT   23719
gb:MN985325|Organism:Severe      GCAGAAAATTCAGTTGCTTACTCTAATAACTCTATTGCCATACCCACAAATTTTACTATT   23722
gb:MT020881|Organism:Severe      GCAGAAAATTCAGTTGCTTACTCTAATAACTCTATTGCCATACCCACAAATTTTACTATT   23722
gb:MT020880|Organism:Severe      GCAGAAAATTCAGTTGCTTACTCTAATAACTCTATTGCCATACCCACAAATTTTACTATT   23722
gb:MT066175|Organism:Severe      GCAGAAAATTCAGTTGCTTACTCTAATAACTCTATTGCCATACCCACAAATTTTACTATT   23722
gb:MN997409|Organism:Severe      GCAGAAAATTCAGTTGCTTACTCTAATAACTCTATTGCCATACCCACAAATTTTACTATT   23722
gb:MN938384|Organism:Severe      GCAGAAAATTCAGTTGCTTACTCTAATAACTCTATTGCCATACCCACAAATTTTACTATT   23690
gb:MT044258|Organism:Severe      GCAGAAAATTCAGTTGCTTACTCTAATAACTCTATTGCCATACCCACAAATTTTACTATT   23698
gb:MT039890|Organism:Severe      GCAGAAAATTCAGTTGCTTACTCTAATAACTCTATTGCCATACCCACAAATTTTACTATT   23722
gb:MN988713|Organism:Severe      GCAGAAAATTCAGTTGCTTACTCTAATAACTCTATTGCCATACCCACAAATTTTACTATT   23722
gb:LC521925|Organism:Severe      GCAGAAAATTCAGTTGCTTACTCTAATAACTCTATTGCCATACCCACAAATTTTACTATT   23695
gb:MT093571|Organism:Severe      GCAGAAAATTCAGTTGCTTACTCTAATAACTCTATTGCCATACCCACAAATTTTACTATT   23722
gb:MT039887|Organism:Severe      GCAGAAAATTCAGTTGCTTACTCTAATAACTCTATTGCCATACCCACAAATTTTACTATT   23719
gb:MT019530|Organism:Severe      GCAGAAAATTCAGTTGCTTACTCTAATAACTCTATTGCCATACCCACAAATTTTACTATT   23722
gb:MT039888|Organism:Severe      GCAGAAAATTCAGTTGCTTACTCTAATAACTCTATTGCCATACCCACAAATTTTACTATT   23722
gb:LC522972|Organism:Severe      GCAGAAAATTCAGTTGCTTACTCTAATAACTCTATTGCCATACCCACAAATTTTACTATT   23719
```

```
gb:MT027063|Organism:Severe      GCAGAAAATTCAGTTGCTTACTCTAATAACTCTATTGCCATACCCACAAATTTTACTATT   23722
gb:MT027062|Organism:Severe      GCAGAAAATTCAGTTGCTTACTCTAATAACTCTATTGCCATACCCACAAATTTTACTATT   23722
gb:MT019529|Organism:Severe      GCAGAAAATTCAGTTGCTTACTCTAATAACTCTATTGCCATACCCACAAATTTTACTATT   23722
gb:MN996529|Organism:Severe      GCAGAAAATTCAGTTGCTTACTCTAATAACTCTATTGCCATACCCACAAATTTTACTATT   23710
gb:MN996531|Organism:Severe      GCAGAAAATTCAGTTGCTTACTCTAATAACTCTATTGCCATACCCACAAATTTTACTATT   23709
gb:MT066176|Organism:Severe      GCAGAAAATTCAGTTGCTTACTCTAATAACTCTATTGCCATACCCACAAATTTTACTATT   23722
gb:MT027064|Organism:Severe      GCAGAAAATTCAGTTGCTTACTCTAATAACTCTATTGCCATACCCACAAATTTTACTATT   23722
gb:MN994468|Organism:Severe      GCAGAAAATTCAGTTGCTTACTCTAATAACTCTATTGCCATACCCACAAATTTTACTATT   23722
gb:MT072688|Organism:Severe      GCAGAAAATTCAGTTGCTTACTCTAATAACTCTATTGCCATACCCACAAATTTTACTATT   23707
gb:MN996527|Organism:Severe      GCAGAAAATTCAGTTGCTTACTCTAATAACTCTATTGCCATACCCACAAATTTTACTATT   23689
gb:MT093631|Organism:Severe      GCAGAAAATTCAGTTGCTTACTCTAATAACTCTATTGCCATACCCACAAATTTTACTATT   23760
gb:MT106053|Organism:Severe      GCAGAAAATTCAGTTGCTTACTCTAATAACTCTATTGCCATACCCACAAATTTTACTATT   23722
gb:MT019533|Organism:Severe      GCAGAAAATTCAGTTGCTTACTCTAATAACTCTATTGCCATACCCACAAATTTTACTATT   23722
gb:MT019531|Organism:Severe      GCAGAAAATTCAGTTGCTTACTCTAATAACTCTATTGCCATACCCACAAATTTTACTATT   23722
gb:MN996528|Organism:Severe      GCAGAAAATTCAGTTGCTTACTCTAATAACTCTATTGCCATACCCACAAATTTTACTATT   23722
gb:MN996530|Organism:Severe      GCAGAAAATTCAGTTGCTTACTCTAATAACTCTATTGCCATACCCACAAATTTTACTATT   23708
gb:MN908947|Organism:Severe      GCAGAAAATTCAGTTGCTTACTCTAATAACTCTATTGCCATACCCACAAATTTTACTATT   23722
gb:MT019532|Organism:Severe      GCAGAAAATTCAGTTGCTTACTCTAATAACTCTATTGCCATACCCACAAATTTTACTATT   23722
                                 ************************************************************

gb:MT020781|Organism:Severe      AGTGTTACCACAGAAATTCTACCAGTGTCTATGACCAAGACATCAGTAGATTGTACAATG   23770
gb:MT007544|Organism:Severe      AGTGTTACCACAGAAATTCTACCAGTGTCTATGACCAAGACATCAGTAGATTGTACAATG   23782
gb:MN994467|Organism:Severe      AGTGTTACCACAGAAATTCTACCAGTGTCTATGACCAAGACATCAGTAGATTGTACAATG   23782
gb:MT044257|Organism:Severe      AGTGTTACCACAGAAATTCTACCAGTGTCTATGACCAAGACATCAGTAGATTGTACAATG   23782
gb:MT106054|Organism:Severe      AGTGTTACCACAGAAATTCTACCAGTGTCTATGACCAAGACATCAGTAGATTGTACAATG   23782
gb:MT049951|Organism:Severe      AGTGTTACCACAGAAATTCTACCAGTGTCTATGACCAAGACATCAGTAGATTGTACAATG   23782
gb:MN975262|Organism:Severe      AGTGTTACCACAGAAATTCTACCAGTGTCTATGACCAAGACATCAGTAGATTGTACAATG   23782
gb:MT106052|Organism:Severe      AGTGTTACCACAGAAATTCTACCAGTGTCTATGACCAAGACATCAGTAGATTGTACAATG   23782
gb:LC522975|Organism:Severe      AGTGTTACCACAGAAATTCTACCAGTGTCTATGACCAAGACATCAGTAGATTGTACAATG   23779
gb:LC522973|Organism:Severe      AGTGTTACCACAGAAATTCTACCAGTGTCTATGACCAAGACATCAGTAGATTGTACAATG   23779
gb:LC522974|Organism:Severe      AGTGTTACCACAGAAATTCTACCAGTGTCTATGACCAAGACATCAGTAGATTGTACAATG   23779
gb:MN985325|Organism:Severe      AGTGTTACCACAGAAATTCTACCAGTGTCTATGACCAAGACATCAGTAGATTGTACAATG   23782
gb:MT020881|Organism:Severe      AGTGTTACCACAGAAATTCTACCAGTGTCTATGACCAAGACATCAGTAGATTGTACAATG   23782
gb:MT020880|Organism:Severe      AGTGTTACCACAGAAATTCTACCAGTGTCTATGACCAAGACATCAGTAGATTGTACAATG   23782
gb:MT066175|Organism:Severe      AGTGTTACCACAGAAATTCTACCAGTGTCTATGACCAAGACATCAGTAGATTGTACAATG   23782
gb:MN997409|Organism:Severe      AGTGTTACCACAGAAATTCTACCAGTGTCTATGACCAAGACATCAGTAGATTGTACAATG   23782
gb:MN938384|Organism:Severe      AGTGTTACCACAGAAATTCTACCAGTGTCTATGACCAAGACATCAGTAGATTGTACAATG   23750
gb:MT044258|Organism:Severe      AGTGTTACCACAGAAATTCTACCAGTGTCTATGACCAAGACATCAGTAGATTGTACAATG   23758
gb:MT039890|Organism:Severe      AGTGTTACCACAGAAATTCTACCAGTGTCTATGACCAAGACATCAGTAGATTGTACAATG   23782
gb:MN988713|Organism:Severe      AGTGTTACCACAGAAATTCTACCAGTGTCTATGACCAAGACATCAGTAGATTGTACAATG   23782
gb:LC521925|Organism:Severe      AGTGTTACCACAGAAATTCTACCAGTGTCTATGACCAAGACATCAGTAGATTGTACAATG   23755
gb:MT093571|Organism:Severe      AGTGTTACCACAGAAATTCTACCAGTGTCTATGACCAAGACATCAGTAGATTGTACAATG   23782
gb:MT039887|Organism:Severe      AGTGTTACCACAGAAATTCTACCAGTGTCTATGACCAAGACATCAGTAGATTGTACAATG   23779
gb:MT019530|Organism:Severe      AGTGTTACCACAGAAATTCTACCAGTGTCTATGACCAAGACATCAGTAGATTGTACAATG   23782
gb:MT039888|Organism:Severe      AGTGTTACCACAGAAATTCTACCAGTGTCTATGACCAAGACATCAGTAGATTGTACAATG   23782
gb:LC522972|Organism:Severe      AGTGTTACCACAGAAATTCTACCAGTGTCTATGACCAAGACATCAGTAGATTGTACAATG   23779
gb:MT027063|Organism:Severe      AGTGTTACCACAGAAATTCTACCAGTGTCTATGACCAAGACATCAGTAGATTGTACAATG   23782
gb:MT027062|Organism:Severe      AGTGTTACCACAGAAATTCTACCAGTGTCTATGACCAAGACATCAGTAGATTGTACAATG   23782
gb:MT019529|Organism:Severe      AGTGTTACCACAGAAATTCTACCAGTGTCTATGACCAAGACATCAGTAGATTGTACAATG   23782
gb:MN996529|Organism:Severe      AGTGTTACCACAGAAATTCTACCAGTGTCTATGACCAAGACATCAGTAGATTGTACAATG   23770
gb:MN996531|Organism:Severe      AGTGTTACCACAGAAATTCTACCAGTGTCTATGACCAAGACATCAGTAGATTGTACAATG   23769
gb:MT066176|Organism:Severe      AGTGTTACCACAGAAATTCTACCAGTGTCTATGACCAAGACATCAGTAGATTGTACAATG   23782
gb:MT027064|Organism:Severe      AGTGTTACCACAGAAATTCTACCAGTGTCTATGACCAAGACATCAGTAGATTGTACAATG   23782
gb:MN994468|Organism:Severe      AGTGTTACCACAGAAATTCTACCAGTGTCTATGACCAAGACATCAGTAGATTGTACAATG   23782
gb:MT072688|Organism:Severe      AGTGTTACCACAGAAATTCTACCAGTGTCTATGACCAAGACATCAGTAGATTGTACAATG   23767
gb:MN996527|Organism:Severe      AGTGTTACCACAGAAATTCTACCAGTGTCTATGACCAAGACATCAGTAGATTGTACAATG   23749
gb:MT093631|Organism:Severe      AGTGTTACCACAGAAATTCTACCAGTGTCTATGACCAAGACATCAGTAGATTGTACAATG   23820
gb:MT106053|Organism:Severe      AGTGTTACCACAGAAATTCTACCAGTGTCTATGACCAAGACATCAGTAGATTGTACAATG   23782
gb:MT019533|Organism:Severe      AGTGTTACCACAGAAATTCTACCAGTGTCTATGACCAAGACATCAGTAGATTGTACAATG   23782
gb:MT019531|Organism:Severe      AGTGTTACCACAGAAATTCTACCAGTGTCTATGACCAAGACATCAGTAGATTGTACAATG   23782
gb:MN996528|Organism:Severe      AGTGTTACCACAGAAATTCTACCAGTGTCTATGACCAAGACATCAGTAGATTGTACAATG   23782
gb:MN996530|Organism:Severe      AGTGTTACCACAGAAATTCTACCAGTGTCTATGACCAAGACATCAGTAGATTGTACAATG   23768
gb:MN908947|Organism:Severe      AGTGTTACCACAGAAATTCTACCAGTGTCTATGACCAAGACATCAGTAGATTGTACAATG   23782
gb:MT019532|Organism:Severe      AGTGTTACCACAGAAATTCTACCAGTGTCTATGACCAAGACATCAGTAGATTGTACAATG   23782
                                 ************************************************************
```

```
gb:MT020781|Organism:Severe      TACATTTGTGGTGATTCAACTGAATGCAGCAATCTTTTGTTGCAATATGGCAGTTTTTGT   23830
gb:MT007544|Organism:Severe      TACATTTGTGGTGATTCAACTGAATGCAGCAATCTTTTGTTGCAATATGGCAGTTTTTGT   23842
gb:MN994467|Organism:Severe      TACATTTGTGGTGATTCAACTGAATGCAGCAATCTTTTGTTGCAATATGGCAGTTTTTGT   23842
gb:MT044257|Organism:Severe      TACATTTGTGGTGATTCAACTGAATGCAGCAATCTTTTGTTGCAATATGGCAGTTTTTGT   23842
gb:MT106054|Organism:Severe      TACATTTGTGGTGATTCAACTGAATGCAGCAATCTTTTGTTGCAATATGGCAGTTTTTGT   23842
gb:MT049951|Organism:Severe      TACATTTGTGGTGATTCAACTGAATGCAGCAATCTTTTGTTGCAATATGGCAGTTTTTGT   23842
gb:MN975262|Organism:Severe      TACATTTGTGGTGATTCAACTGAATGCAGCAATCTTTTGTTGCAATATGGCAGTTTTTGT   23842
gb:MT106052|Organism:Severe      TACATTTGTGGTGATTCAACTGAATGCAGCAATCTTTTGTTGCAATATGGCAGTTTTTGT   23842
gb:LC522975|Organism:Severe      TACATTTGTGGTGATTCAACTGAATGCAGCAATCTTTTGTTGCAATATGGCAGTTTTTGT   23839
gb:LC522973|Organism:Severe      TACATTTGTGGTGATTCAACTGAATGCAGCAATCTTTTGTTGCAATATGGCAGTTTTTGT   23839
gb:LC522974|Organism:Severe      TACATTTGTGGTGATTCAACTGAATGCAGCAATCTTTTGTTGCAATATGGCAGTTTTTGT   23839
gb:MN985325|Organism:Severe      TACATTTGTGGTGATTCAACTGAATGCAGCAATCTTTTGTTGCAATATGGCAGTTTTTGT   23842
gb:MT020881|Organism:Severe      TACATTTGTGGTGATTCAACTGAATGCAGCAATCTTTTGTTGCAATATGGCAGTTTTTGT   23842
gb:MT020880|Organism:Severe      TACATTTGTGGTGATTCAACTGAATGCAGCAATCTTTTGTTGCAATATGGCAGTTTTTGT   23842
gb:MT066175|Organism:Severe      TACATTTGTGGTGATTCAACTGAATGCAGCAATCTTTTGTTGCAATATGGCAGTTTTTGT   23842
gb:MN997409|Organism:Severe      TACATTTGTGGTGATTCAACTGAATGCAGCAATCTTTTGTTGCAATATGGCAGTTTTTGT   23842
gb:MN938384|Organism:Severe      TACATTTGTGGTGATTCAACTGAATGCAGCAATCTTTTGTTGCAATATGGCAGTTTTTGT   23810
gb:MT044258|Organism:Severe      TACATTTGTGGTGATTCAACTGAATGCAGCAATCTTTTGTTGCAATATGGCAGTTTTTGT   23818
gb:MT039890|Organism:Severe      TACATTTGTGGTGATTCAACTGAATGCAGCAATCTTTTGTTGCAATATGGCAGTTTTTGT   23842
gb:MN988713|Organism:Severe      TACATTTGTGGTGATTCAACTGAATGCAGCAATCTTTTGTTGCAATATGGCAGTTTTTGT   23842
gb:LC521925|Organism:Severe      TACATTTGTGGTGATTCAACTGAATGCAGCAATCTTTTGTTGCAATATGGCAGTTTTTGT   23815
gb:MT093571|Organism:Severe      TACATTTGTGGTGATTCAACTGAATGCAGCAATCTTTTGTTGCAATATGGCAGTTTTTGT   23842
gb:MT039887|Organism:Severe      TACATTTGTGGTGATTCAACTGAATGCAGCAATCTTTTGTTGCAATATGGCAGTTTTTGT   23839
gb:MT019530|Organism:Severe      TACATTTGTGGTGATTCAACTGAATGCAGCAATCTTTTGTTGCAATATGGCAGTTTTTGT   23842
gb:MT039888|Organism:Severe      TACATTTGTGGTGATTCAACTGAATGCAGCAATCTTTTGTTGCAATATGGCAGTTTTTGT   23842
gb:LC522972|Organism:Severe      TACATTTGTGGTGATTCAACTGAATGCAGCAATCTTTTGTTGCAATATGGCAGTTTTTGT   23839
gb:MT027063|Organism:Severe      TACATTTGTGGTGATTCAACTGAATGCAGCAATCTTTTGTTGCAATATGGCAGTTTTTGT   23842
gb:MT027062|Organism:Severe      TACATTTGTGGTGATTCAACTGAATGCAGCAATCTTTTGTTGCAATATGGCAGTTTTTGT   23842
gb:MT019529|Organism:Severe      TACATTTGTGGTGATTCAACTGAATGCAGCAATCTTTTGTTGCAATATGGCAGTTTTTGT   23842
gb:MN996529|Organism:Severe      TACATTTGTGGTGATTCAACTGAATGCAGCAATCTTTTGTTGCAATATGGCAGTTTTTGT   23830
gb:MN996531|Organism:Severe      TACATTTGTGGTGATTCAACTGAATGCAGCAATCTTTTGTTGCAATATGGCAGTTTTTGT   23829
gb:MT066176|Organism:Severe      TACATTTGTGGTGATTCAACTGAATGCAGCAATCTTTTGTTGCAATATGGCAGTTTTTGT   23842
gb:MT027064|Organism:Severe      TACATTTGTGGTGATTCAACTGAATGCAGCAATCTTTTGTTGCAATATGGCAGTTTTTGT   23842
gb:MN994468|Organism:Severe      TACATTTGTGGTGATTCAACTGAATGCAGCAATCTTTTGTTGCAATATGGCAGTTTTTGT   23842
gb:MT072688|Organism:Severe      TACATTTGTGGTGATTCAACTGAATGCAGCAATCTTTTGTTGCAATATGGCAGTTTTTGT   23827
gb:MN996527|Organism:Severe      TACATTTGTGGTGATTCAACTGAATGCAGCAATCTTTTGTTGCAATATGGCAGTTTTTGT   23809
gb:MT093631|Organism:Severe      TACATTTGTGGTGATTCAACTGAATGCAGCAATCTTTTGTTGCAATATGGCAGTTTTTGT   23880
gb:MT106053|Organism:Severe      TACATTTGTGGTGATTCAACTGAATGCAGCAATCTTTTGTTGCAATATGGCAGTTTTTGT   23842
gb:MT019533|Organism:Severe      TACATTTGTGGTGATTCAACTGAATGCAGCAATCTTTTGTTGCAATATGGCAGTTTTTGT   23842
gb:MT019531|Organism:Severe      TACATTTGTGGTGATTCAACTGAATGCAGCAATCTTTTGTTGCAATATGGCAGTTTTTGT   23842
gb:MN996528|Organism:Severe      TACATTTGTGGTGATTCAACTGAATGCAGCAATCTTTTGTTGCAATATGGCAGTTTTTGT   23842
gb:MN996530|Organism:Severe      TACATTTGTGGTGATTCAACTGAATGCAGCAATCTTTTGTTGCAATATGGCAGTTTTTGT   23828
gb:MN908947|Organism:Severe      TACATTTGTGGTGATTCAACTGAATGCAGCAATCTTTTGTTGCAATATGGCAGTTTTTGT   23842
gb:MT019532|Organism:Severe      TACATTTGTGGTGATTCAACTGAATGCAGCAATCTTTTGTTGCAATATGGCAGTTTTTGT   23842
                                 ************************************************************

gb:MT020781|Organism:Severe      ACACAATTAAACCGTGCTTTAACTGGAATAGCTGTTGAACAAGACAAAAACACCCAAGAA   23890
gb:MT007544|Organism:Severe      ACACAATTAAACCGTGCTTTAACTGGAATAGCTGTTGAACAAGACAAAAACACCCAAGAA   23902
gb:MN994467|Organism:Severe      ACACAATTAAACCGTGCTTTAACTGGAATAGCTGTTGAACAAGACAAAAACACCCAAGAA   23902
gb:MT044257|Organism:Severe      ACACAATTAAACCGTGCTTTAACTGGAATAGCTGTTGAACAAGACAAAAACACCCAAGAA   23902
gb:MT106054|Organism:Severe      ACACAATTAAACCGTGCTTTAACTGGAATAGCTGTTGAACAAGACAAAAACACCCAAGAA   23902
gb:MT049951|Organism:Severe      ACACAATTAAACCGTGCTTTAACTGGAATAGCTGTTGAACAAGACAAAAACACCCAAGAA   23902
gb:MN975262|Organism:Severe      ACACAATTAAACCGTGCTTTAACTGGAATAGCTGTTGAACAAGACAAAAACACCCAAGAA   23902
gb:MT106052|Organism:Severe      ACACAATTAAACCGTGCTTTAACTGGAATAGCTGTTGAACAAGACAAAAACACCCAAGAA   23902
gb:LC522975|Organism:Severe      ACACAATTAAACCGTGCTTTAACTGGAATAGCTGTTGAACAAGACAAAAACACCCAAGAA   23899
gb:LC522973|Organism:Severe      ACACAATTAAACCGTGCTTTAACTGGAATAGCTGTTGAACAAGACAAAAACACCCAAGAA   23899
gb:LC522974|Organism:Severe      ACACAATTAAACCGTGCTTTAACTGGAATAGCTGTTGAACAAGACAAAAACACCCAAGAA   23899
gb:MN985325|Organism:Severe      ACACAATTAAACCGTGCTTTAACTGGAATAGCTGTTGAACAAGACAAAAACACCCAAGAA   23902
gb:MT020881|Organism:Severe      ACACAATTAAACCGTGCTTTAACTGGAATAGCTGTTGAACAAGACAAAAACACCCAAGAA   23902
gb:MT020880|Organism:Severe      ACACAATTAAACCGTGCTTTAACTGGAATAGCTGTTGAACAAGACAAAAACACCCAAGAA   23902
gb:MT066175|Organism:Severe      ACACAATTAAACCGTGCTTTAACTGGAATAGCTGTTGAACAAGACAAAAACACCCAAGAA   23902
gb:MN997409|Organism:Severe      ACACAATTAAACCGTGCTTTAACTGGAATAGCTGTTGAACAAGACAAAAACACCCAAGAA   23902
gb:MN938384|Organism:Severe      ACACAATTAAACCGTGCTTTAACTGGAATAGCTGTTGAACAAGACAAAAACACCCAAGAA   23870
gb:MT044258|Organism:Severe      ACACAATTAAACCGTGCTTTAACTGGAATAGCTGTTGAACAAGACAAAAACACCCAAGAA   23878
```

```
gb:MT039890|Organism:Severe      ACACAATTAAACCGTGCTTTAACTGGAATAGCTGTTGAACAAGACAAAAACACCCAAGAA   23902
gb:MN988713|Organism:Severe      ACACAATTAAACCGTGCTTTAACTGGAATAGCTGTTGAACAAGACAAAAACACCCAAGAA   23902
gb:LC521925|Organism:Severe      ACACAATTAAACCGTGCTTTAACTGGAATAGCTGTTGAACAAGACAAAAACACCCAAGAA   23875
gb:MT093571|Organism:Severe      ACACAATTAAACCGTGCTTTAACTGGAATAGCTGTTGAACAAGACAAAAACACCCAAGAA   23902
gb:MT039887|Organism:Severe      ACACAATTAAACCGTGCTTTAACTGGAATAGCTGTTGAACAAGACAAAAACACCCAAGAA   23899
gb:MT019530|Organism:Severe      ACACAATTAAACCGTGCTTTAACTGGAATAGCTGTTGAACAAGACAAAAACACCCAAGAA   23902
gb:MT039888|Organism:Severe      ACACAATTAAACCGTGCTTTAACTGGAATAGCTGTTGAACAAGACAAAAACACCCAAGAA   23902
gb:LC522972|Organism:Severe      ACACAATTAAACCGTGCTTTAACTGGAATAGCTGTTGAACAAGACAAAAACACCCAAGAA   23899
gb:MT027063|Organism:Severe      ACACAATTAAACCGTGCTTTAACTGGAATAGCTGTTGAACAAGACAAAAACACCCAAGAA   23902
gb:MT027062|Organism:Severe      ACACAATTAAACCGTGCTTTAACTGGAATAGCTGTTGAACAAGACAAAAACACCCAAGAA   23902
gb:MT019529|Organism:Severe      ACACAATTAAACCGTGCTTTAACTGGAATAGCTGTTGAACAAGACAAAAACACCCAAGAA   23902
gb:MN996529|Organism:Severe      ACACAATTAAACCGTGCTTTAACTGGAATAGCTGTTGAACAAGACAAAAACACCCAAGAA   23890
gb:MN996531|Organism:Severe      ACACAATTAAACCGTGCTTTAACTGGAATAGCTGTTGAACAAGACAAAAACACCCAAGAA   23889
gb:MT066176|Organism:Severe      ACACAATTAAACCGTGCTTTAACTGGAATAGCTGTTGAACAAGACAAAAACACCCAAGAA   23902
gb:MT027064|Organism:Severe      ACACAATTAAACCGTGCTTTAACTGGAATAGCTGTTGAACAAGACAAAAACACCCAAGAA   23902
gb:MN994468|Organism:Severe      ACACAATTAAACCGTGCTTTAACTGGAATAGCTGTTGAACAAGACAAAAACACCCAAGAA   23902
gb:MT072688|Organism:Severe      ACACAATTAAACCGTGCTTTAACTGGAATAGCTGTTGAACAAGACAAAAACACCCAAGAA   23887
gb:MN996527|Organism:Severe      ACACAATTAAACCGTGCTTTAACTGGAATAGCTGTTGAACAAGACAAAAACACCCAAGAA   23869
gb:MT093631|Organism:Severe      ACACAATTAAACCGTGCTTTAACTGGAATAGCTGTTGAACAAGACAAAAACACCCAAGAA   23940
gb:MT106053|Organism:Severe      ACACAATTAAACCGTGCTTTAACTGGAATAGCTGTTGAACAAGACAAAAACACCCAAGAA   23902
gb:MT019533|Organism:Severe      ACACAATTAAACCGTGCTTTAACTGGAATAGCTGTTGAACAAGACAAAAACACCCAAGAA   23902
gb:MT019531|Organism:Severe      ACACAATTAAACCGTGCTTTAACTGGAATAGCTGTTGAACAAGACAAAAACACCCAAGAA   23902
gb:MN996528|Organism:Severe      ACACAATTAAACCGTGCTTTAACTGGAATAGCTGTTGAACAAGACAAAAACACCCAAGAA   23902
gb:MN996530|Organism:Severe      ACACAATTAAACCGTGCTTTAACTGGAATAGCTGTTGAACAAGACAAAAACACCCAAGAA   23888
gb:MN908947|Organism:Severe      ACACAATTAAACCGTGCTTTAACTGGAATAGCTGTTGAACAAGACAAAAACACCCAAGAA   23902
gb:MT019532|Organism:Severe      ACACAATTAAACCGTGCTTTAACTGGAATAGCTGTTGAACAAGACAAAAACACCCAAGAA   23902
                                 ************************************************************

gb:MT020781|Organism:Severe      GTTTTTGCACAAGTCAAACAAATTTACAAAACACCACCAATTAAAGATTTTGGTGGTTTT   23950
gb:MT007544|Organism:Severe      GTTTTTGCACAAGTCAAACAAATTTACAAAACACCACCAATTAAAGATTTTGGTGGTTTT   23962
gb:MN994467|Organism:Severe      GTTTTTGCACAAGTCAAACAAATTTACAAAACACCACCAATTAAAGATTTTGGTGGTTTT   23962
gb:MT044257|Organism:Severe      GTTTTTGCACAAGTCAAACAAATTTACAAAACACCACCAATTAAAGATTTTGGTGGTTTT   23962
gb:MT106054|Organism:Severe      GTTTTTGCACAAGTCAAACAAATTTACAAAACACCACCAATTAAAGATTTTGGTGGTTTT   23962
gb:MT049951|Organism:Severe      GTTTTTGCACAAGTCAAACAAATTTACAAAACACCACCAATTAAAGATTTTGGTGGTTTT   23962
gb:MN975262|Organism:Severe      GTTTTTGCACAAGTCAAACAAATTTACAAAACACCACCAATTAAAGATTTTGGTGGTTTT   23962
gb:MT106052|Organism:Severe      GTTTTTGCACAAGTCAAACAAATTTACAAAACACCACCAATTAAAGATTTTGGTGGTTTT   23962
gb:LC522975|Organism:Severe      GTTTTTGCACAAGTCAAACAAATTTACAAAACACCACCAATTAAAGATTTTGGTGGTTTT   23959
gb:LC522973|Organism:Severe      GTTTTTGCACAAGTCAAACAAATTTACAAAACACCACCAATTAAAGATTTTGGTGGTTTT   23959
gb:LC522974|Organism:Severe      GTTTTTGCACAAGTCAAACAAATTTACAAAACACCACCAATTAAAGATTTTGGTGGTTTT   23959
gb:MN985325|Organism:Severe      GTTTTTGCACAAGTCAAACAAATTTACAAAACACCACCAATTAAAGATTTTGGTGGTTTT   23962
gb:MT020881|Organism:Severe      GTTTTTGCACAAGTCAAACAAATTTACAAAACACCACCAATTAAAGATTTTGGTGGTTTT   23962
gb:MT020880|Organism:Severe      GTTTTTGCACAAGTCAAACAAATTTACAAAACACCACCAATTAAAGATTTTGGTGGTTTT   23962
gb:MT066175|Organism:Severe      GTTTTTGCACAAGTCAAACAAATTTACAAAACACCACCAATTAAAGATTTTGGTGGTTTT   23962
gb:MN997409|Organism:Severe      GTTTTTGCACAAGTCAAACAAATTTACAAAACACCACCAATTAAAGATTTTGGTGGTTTT   23962
gb:MN938384|Organism:Severe      GTTTTTGCACAAGTCAAACAAATTTACAAAACACCACCAATTAAAGATTTTGGTGGTTTT   23930
gb:MT044258|Organism:Severe      GTTTTTGCACAAGTCAAACAAATTTACAAAACACCACCAATTAAAGATTTTGGTGGTTTT   23938
gb:MT039890|Organism:Severe      GTTTTTGCACAAGTCAAACAAATTTACAAAACACCACCAATTAAAGATTTTGGTGGTTTT   23962
gb:MN988713|Organism:Severe      GTTTTTGCACAAGTCAAACAAATTTACAAAACACCACCAATTAAAGATTTTGGTGGTTTT   23962
gb:LC521925|Organism:Severe      GTTTTTGCACAAGTCAAACAAATTTACAAAACACCACCAATTAAAGATTTTGGTGGTTTT   23935
gb:MT093571|Organism:Severe      GTTTTTGCACAAGTCAAACAAATTTACAAAACACCACCAATTAAAGATTGTGGTGGTTTT   23962
gb:MT039887|Organism:Severe      GTTTTTGCACAAGTCAAACAAATTTACAAAACACCACCAATTAAAGATTTTGGTGGTTTT   23959
gb:MT019530|Organism:Severe      GTTTTTGCACAAGTCAAACAAATTTACAAAACACCACCAATTAAAGATTTTGGTGGTTTT   23962
gb:MT039888|Organism:Severe      GTTTTTGCACAAGTCAAACAAATTTACAAAACACCACCAATTAAAGATTTTGGTGGTTTT   23962
gb:LC522972|Organism:Severe      GTTTTTGCACAAGTCAAACAAATTTACAAAACACCACCAATTAAAGATTTTGGTGGTTTT   23959
gb:MT027063|Organism:Severe      GTTTTTGCACAAGTCAAACAAATTTACAAAACACCACCAATTAAAGATTTTGGTGGTTTT   23962
gb:MT027062|Organism:Severe      GTTTTTGCACAAGTCAAACAAATTTACAAAACACCACCAATTAAAGATTTTGGTGGTTTT   23962
gb:MT019529|Organism:Severe      GTTTTTGCACAAGTCAAACAAATTTACAAAACACCACCAATTAAAGATTTTGGTGGTTTT   23962
gb:MN996529|Organism:Severe      GTTTTTGCACAAGTCAAACAAATTTACAAAACACCACCAATTAAAGATTTTGGTGGTTTT   23950
gb:MN996531|Organism:Severe      GTTTTTGCACAAGTCAAACAAATTTACAAAACACCACCAATTAAAGATTTTGGTGGTTTT   23949
gb:MT066176|Organism:Severe      GTTTTTGCACAAGTCAAACAAATTTACAAAACACCACCAATTAAAGATTTTGGTGGTTTT   23962
gb:MT027064|Organism:Severe      GTTTTTGCACAAGTCAAACAAATTTACAAAACACCACCAATTAAAGATTTTGGTGGTTTT   23962
gb:MN994468|Organism:Severe      GTTTTTGCACAAGTCAAACAAATTTACAAAACACCACCAATTAAAGATTTTGGTGGTTTT   23962
gb:MT072688|Organism:Severe      GTTTTTGCACAAGTCAAACAAATTTACAAAACACCACCAATTAAAGATTTTGGTGGTTTT   23947
gb:MN996527|Organism:Severe      GTTTTTGCACAAGTCAAACAAATTTACAAAACACCACCAATTAAAGATTTTGGTGGTTTT   23929
gb:MT093631|Organism:Severe      GTTTTTGCACAAGTCAAACAAATTTACAAAACACCACCAATTAAAGATTTTGGTGGTTTT   24000
```

```
gb:MT106053|Organism:Severe      GTTTTTGCACAAGTCAAACAAATTTACAAAACACCACCAATTAAAGATTTTGGTGGTTTT   23962
gb:MT019533|Organism:Severe      GTTTTTGCACAAGTCAAACAAATTTACAAAACACCACCAATTAAAGATTTTGGTGGTTTT   23962
gb:MT019531|Organism:Severe      GTTTTTGCACAAGTCAAACAAATTTACAAAACACCACCAATTAAAGATTTTGGTGGTTTT   23962
gb:MN996528|Organism:Severe      GTTTTTGCACAAGTCAAACAAATTTACAAAACACCACCAATTAAAGATTTTGGTGGTTTT   23962
gb:MN996530|Organism:Severe      GTTTTTGCACAAGTCAAACAAATTTACAAAACACCACCAATTAAAGATTTTGGTGGTTTT   23948
gb:MN908947|Organism:Severe      GTTTTTGCACAAGTCAAACAAATTTACAAAACACCACCAATTAAAGATTTTGGTGGTTTT   23962
gb:MT019532|Organism:Severe      GTTTTTGCACAAGTCAAACAAATTTACAAAACACCACCAATTAAAGATTTTGGTGGTTTT   23962
                                 ************************************************* **********

gb:MT020781|Organism:Severe      AATTTTTCACAAATATTACCAGATCCATCAAAACCAAGCAAGAGGTCATTTATTGAAGAT   24010
gb:MT007544|Organism:Severe      AATTTTTCACAAATATTACCAGATCCATCAAAACCAAGCAAGAGGTCATTTATTGAAGAT   24022
gb:MN994467|Organism:Severe      AATTTTTCACAAATATTACCAGATCCATCAAAACCAAGCAAGAGGTCATTTATTGAAGAT   24022
gb:MT044257|Organism:Severe      AATTTTTCACAAATATTACCAGATCCATCAAAACCAAGCAAGAGGTCATTTATTGAAGAT   24022
gb:MT106054|Organism:Severe      AATTTTTCACAAATATTACCAGATCCATCAAAACCAAGCAAGAGGTCATTTATTGAAGAT   24022
gb:MT049951|Organism:Severe      AATTTTTCACAAATATTACCAGATCCATCAAAACCAAGCAAGAGGTCATTTATTGAAGAT   24022
gb:MN975262|Organism:Severe      AATTTTTCACAAATATTACCAGATCCATCAAAACCAAGCAAGAGGTCATTTATTGAAGAT   24022
gb:MT106052|Organism:Severe      AATTTTTCACAAATATTACCAGATCCATCAAAACCAAGCAAGAGGTCATTTATTGAAGAT   24022
gb:LC522975|Organism:Severe      AATTTTTCACAAATATTACCAGATCCATCAAAACCAAGCAAGAGGTCATTTATTGAAGAT   24019
gb:LC522973|Organism:Severe      AATTTTTCACAAATATTACCAGATCCATCAAAACCAAGCAAGAGGTCATTTATTGAAGAT   24019
gb:LC522974|Organism:Severe      AATTTTTCACAAATATTACCAGATCCATCAAAACCAAGCAAGAGGTCATTTATTGAAGAT   24019
gb:MN985325|Organism:Severe      AATTTTTCACAAATATTACCAGATCCATCAAAACCAAGCAAGAGGTCATTTATTGAAGAT   24022
gb:MT020881|Organism:Severe      AATTTTTCACAAATATTACCAGATCCATCAAAACCAAGCAAGAGGTCATTTATTGAAGAT   24022
gb:MT020880|Organism:Severe      AATTTTTCACAAATATTACCAGATCCATCAAAACCAAGCAAGAGGTCATTTATTGAAGAT   24022
gb:MT066175|Organism:Severe      AATTTTTCACAAATATTACCAGATCCATCAAAACCAAGCAAGAGGTCATTTATTGAAGAT   24022
gb:MN997409|Organism:Severe      AATTTTTCACAAATATTACCAGATCCATCAAAACCAAGCAAGAGGTCATTTATTGAAGAT   24022
gb:MN938384|Organism:Severe      AATTTTTCACAAATATTACCAGATCCATCAAAACCAAGCAAGAGGTCATTTATTGAAGAT   23990
gb:MT044258|Organism:Severe      AATTTTTCACAAATATTACCAGATCCATCAAAACCAAGCAAGAGGTCATTTATTGAAGAT   23998
gb:MT039890|Organism:Severe      AATTTTTCACAAATATTACCAGATCCATCAAAACCAAGCAAGAGGTCATTTATTGAAGAT   24022
gb:MN988713|Organism:Severe      AATTTTTCACAAATATTACCAGATCCATCAAAACCAAGCAAGAGGTCATTTATTGAAGAT   24022
gb:LC521925|Organism:Severe      AATTTTTCACAAATATTACCAGATCCATCAAAACCAAGCAAGAGGTCATTTATTGAAGAT   23995
gb:MT093571|Organism:Severe      AATTTTTCACAAATATTACCAGATCCATCAAAACCAAGCAAGAGGTCATTTATTGAAGAT   24022
gb:MT039887|Organism:Severe      AATTTTTCACAAATATTACCAGATCCATCAAAACCAAGCAAGAGGTCATTTATTGAAGAT   24019
gb:MT019530|Organism:Severe      AATTTTTCACAAATATTACCAGATCCATCAAAACCAAGCAAGAGGTCATTTATTGAAGAT   24022
gb:MT039888|Organism:Severe      AATTTTTCACAAATATTACCAGATCCATCAAAACCAAGCAAGAGGTCATTTATTGAAGAT   24022
gb:LC522972|Organism:Severe      AATTTTTCACAAATATTACCAGATCCATCAAAACCAAGCAAGAGGTCATTTATTGAAGAT   24019
gb:MT027063|Organism:Severe      AATTTTTCACAAATATTACCAGATCCATCAAAACCAAGCAAGAGGTCATTTATTGAAGAT   24022
gb:MT027062|Organism:Severe      AATTTTTCACAAATATTACCAGATCCATCAAAACCAAGCAAGAGGTCATTTATTGAAGAT   24022
gb:MT019529|Organism:Severe      AATTTTTCACAAATATTACCAGATCCATCAAAACCAAGCAAGAGGTCATTTATTGAAGAT   24022
gb:MN996529|Organism:Severe      AATTTTTCACAAATATTACCAGATCCATCAAAACCAAGCAAGAGGTCATTTATTGAAGAT   24010
gb:MN996531|Organism:Severe      AATTTTTCACAAATATTACCAGATCCATCAAAACCAAGCAAGAGGTCATTTATTGAAGAT   24009
gb:MT066176|Organism:Severe      AATTTTTCACAAATATTACCAGATCCATCAAAACCAAGCAAGAGGTCATTTATTGAAGAT   24022
gb:MT027064|Organism:Severe      AATTTTTCACAAATATTACCAGATCCATCAAAACCAAGCAAGAGGTCATTTATTGAAGAT   24022
gb:MN994468|Organism:Severe      AATTTTTCACAAATATTACCAGATCCATCAAAACCAAGCAAGAGGTCATTTATTGAAGAT   24022
gb:MT072688|Organism:Severe      AATTTTTCACAAATATTACCAGATCCATCAAAACCAAGCAAGAGGTCATTTATTGAAGAT   24007
gb:MN996527|Organism:Severe      AATTTTTCACAAATATTACCAGATCCATCAAAACCAAGCAAGAGGTCATTTATTGAAGAT   23989
gb:MT093631|Organism:Severe      AATTTTTCACAAATATTACCAGATCCATCAAAACCAAGCAAGAGGTCATTTATTGAAGAT   24060
gb:MT106053|Organism:Severe      AATTTTTCACAAATATTACCAGATCCATCAAAACCAAGCAAGAGGTCATTTATTGAAGAT   24022
gb:MT019533|Organism:Severe      AATTTTTCACAAATATTACCAGATCCATCAAAACCAAGCAAGAGGTCATTTATTGAAGAT   24022
gb:MT019531|Organism:Severe      AATTTTTCACAAATATTACCAGATCCATCAAAACCAAGCAAGAGGTCATTTATTGAAGAT   24022
gb:MN996528|Organism:Severe      AATTTTTCACAAATATTACCAGATCCATCAAAACCAAGCAAGAGGTCATTTATTGAAGAT   24022
gb:MN996530|Organism:Severe      AATTTTTCACAAATATTACCAGATCCATCAAAACCAAGCAAGAGGTCATTTATTGAAGAT   24008
gb:MN908947|Organism:Severe      AATTTTTCACAAATATTACCAGATCCATCAAAACCAAGCAAGAGGTCATTTATTGAAGAT   24022
gb:MT019532|Organism:Severe      AATTTTTCACAAATATTACCAGATCCATCAAAACCAAGCAAGAGGTCATTTATTGAAGAT   24022
                                 ************************************************************

gb:MT020781|Organism:Severe      CTACTTTTCAACAAAGTGACACTTGCAGATGCTGGCTTCATCAAACAATATGGTGATTGC   24070
gb:MT007544|Organism:Severe      CTACTTTTCAACAAAGTGACACTTGCAGATGCTGGCTTCATCAAACAATATGGTGATTGC   24082
gb:MN994467|Organism:Severe      CTACTTTTCAATAAAGTGACACTTGCAGATGCTGGCTTCATCAAACAATATGGTGATTGC   24082
gb:MT044257|Organism:Severe      CTACTTTTCAATAAAGTGACACTTGCAGATGCTGGCTTCATCAAACAATATGGTGATTGC   24082
gb:MT106054|Organism:Severe      CTACTTTTCAACAAAGTGACACTTGCAGATGCTGGCTTCATCAAACAATATGGTGATTGC   24082
gb:MT049951|Organism:Severe      CTACTTTTCAACAAAGTGACACTTGCAGATGCTGGCTTCATCAAACAATATGGTGATTGC   24082
gb:MN975262|Organism:Severe      CTACTTTTCAACAAAGTGACACTTGCAGATGCTGGCTTCATCAAACAATATGGTGATTGC   24082
gb:MT106052|Organism:Severe      CTACTTTTCAACAAAGTGACACTTGCAGATGCTGGCTTCATCAAACAATATGGTGATTGC   24082
gb:LC522975|Organism:Severe      CTACTTTTCAACAAAGTGACACTTGCAGATGCTGGCTTCATCAAACAATATGGTGATTGC   24079
gb:LC522973|Organism:Severe      CTACTTTTCAACAAAGTGACACTTGCAGATGCTGGCTTCATCAAACAATATGGTGATTGC   24079
```

```
gb:LC522974|Organism:Severe      CTACTTTTCAACAAAGTGACACTTGCAGATGCTGGCTTCATCAAACAATATGGTGATTGC   24079
gb:MN985325|Organism:Severe      CTACTTTTCAACAAAGTGACACTTGCAGATGCTGGCTTCATCAAACAATATGGTGATTGC   24082
gb:MT020881|Organism:Severe      CTACTTTTCAACAAAGTGACACTTGCAGATGCTGGCTTCATCAAACAATATGGTGATTGC   24082
gb:MT020880|Organism:Severe      CTACTTTTCAACAAAGTGACACTTGCAGATGCTGGCTTCATCAAACAATATGGTGATTGC   24082
gb:MT066175|Organism:Severe      CTACTTTTCAACAAAGTGACACTTGCAGATGCTGGCTTCATCAAACAATATGGTGATTGC   24082
gb:MN997409|Organism:Severe      CTACTTTTCAACAAAGTGACACTTGCAGATGCTGGCTTCATCAAACAATATGGTGATTGC   24082
gb:MN938384|Organism:Severe      CTACTTTTCAACAAAGTGACACTTGCAGATGCTGGCTTCATCAAACAATATGGTGATTGC   24050
gb:MT044258|Organism:Severe      CTACTTTTCAACAAAGTGACACTTGCAGATGCTGGCTTCATCAAACAATATGGTGATTGC   24058
gb:MT039890|Organism:Severe      CTACTTTTCAACAAAGTGACACTTGCAGATGCTGGCTTCATCAAACAATATGGTGATTGC   24082
gb:MN988713|Organism:Severe      CTACTTTTCAAYAAAGTGACACTTGCAGATGCTGGCTTCATCAAACAATATGGTGATTGC   24082
gb:LC521925|Organism:Severe      CTACTTTTCAACAAAGTGACACTTGCAGATGCTGGCTTCATCAAACAATATGGTGATTGC   24055
gb:MT093571|Organism:Severe      CTACTTTTCAACAAAGTGACACTTGCAGATGCTGGCTTCATCAAACAATATGGTGATTGC   24082
gb:MT039887|Organism:Severe      CTACTTTTCAACAAAGTGACACTTGCAGATGCTGGCTTCATCAAACAATATGGTGATTGC   24079
gb:MT019530|Organism:Severe      CTACTTTTCAACAAAGTGACACTTGCAGATGCTGGCTTCATCAAACAATATGGTGATTGC   24082
gb:MT039888|Organism:Severe      CTACTTTTCAATAAAGTGACACTTGCAGATGCTGGCTTCATCAAACAATATGGTGATTGC   24082
gb:LC522972|Organism:Severe      CTACTTTTCAACAAAGTGACACTTGCAGATGCTGGCTTCATCAAACAATATGGTGATTGC   24079
gb:MT027063|Organism:Severe      CTACTTTTCAACAAAGTGACACTTGCAGATGCTGGCTTCATCAAACAATATGGTGATTGC   24082
gb:MT027062|Organism:Severe      CTACTTTTCAACAAAGTGACACTTGCAGATGCTGGCTTCATCAAACAATATGGTGATTGC   24082
gb:MT019529|Organism:Severe      CTACTTTTCAACAAAGTGACACTTGCAGATGCTGGCTTCATCAAACAATATGGTGATTGC   24082
gb:MN996529|Organism:Severe      CTACTTTTCAACAAAGTGACACTTGCAGATGCTGGCTTCATCAAACAATATGGTGATTGC   24070
gb:MN996531|Organism:Severe      CTACTTTTCAACAAAGTGACACTTGCAGATGCTGGCTTCATCAAACAATATGGTGATTGC   24069
gb:MT066176|Organism:Severe      CTACTTTTCAACAAAGTGACACTTGCAGATGCTGGCTTCATCAAACAATATGGTGATTGC   24082
gb:MT027064|Organism:Severe      CTACTTTTCAACAAAGTGACACTTGCAGATGCTGGCTTCATCAAACAATATGGTGATTGC   24082
gb:MN994468|Organism:Severe      CTACTTTTCAACAAAGTGACACTTGCAGATGCTGGCTTCATCAAACAATATGGTGATTGC   24082
gb:MT072688|Organism:Severe      CTACTTTTCAATAAAGTGACACTTGCAGATGCTGGCTTCATCAAACAATATGGTGATTGC   24067
gb:MN996527|Organism:Severe      CTACTTTTCAACAAAGTGACACTTGCAGATGCTGGCTTCATCAAACAATATGGTGATTGC   24049
gb:MT093631|Organism:Severe      CTACTTTTCAACAAAGTGACACTTGCAGATGCTGGCTTCATCAAACAATATGGTGATTGC   24120
gb:MT106053|Organism:Severe      CTACTTTTCAACAAAGTGACACTTGCAGATGCTGGCTTCATCAAACAATATGGTGATTGC   24082
gb:MT019533|Organism:Severe      CTACTTTTCAACAAAGTGACACTTGCAGATGCTGGCTTCATCAAACAATATGGTGATTGC   24082
gb:MT019531|Organism:Severe      CTACTTTTCAACAAAGTGACACTTGCAGATGCTGGCTTCATCAAACAATATGGTGATTGC   24082
gb:MN996528|Organism:Severe      CTACTTTTCAACAAAGTGACACTTGCAGATGCTGGCTTCATCAAACAATATGGTGATTGC   24082
gb:MN996530|Organism:Severe      CTACTTTTCAACAAAGTGACACTTGCAGATGCTGGCTTCATCAAACAATATGGTGATTGC   24068
gb:MN908947|Organism:Severe      CTACTTTTCAACAAAGTGACACTTGCAGATGCTGGCTTCATCAAACAATATGGTGATTGC   24082
gb:MT019532|Organism:Severe      CTACTTTTCAACAAAGTGACACTTGCAGATGCTGGCTTCATCAAACAATATGGTGATTGC   24082
                                 *********** ************************************************

gb:MT020781|Organism:Severe      CTTGGTGATATTGCTGCTAGAGACCTCATTTGTGCACAAAAGTTTAACGGCCTTACTGTT   24130
gb:MT007544|Organism:Severe      CTTGGTGATATTGCTGCTAGAGACCTCATTTGTGCACAAAAGTTTAACGGCCTTACTGTT   24142
gb:MN994467|Organism:Severe      CTTGGTGATATTGCTGCTAGAGACCTCATTTGTGCACAAAAGTTTAACGGCCTTACTGTT   24142
gb:MT044257|Organism:Severe      CTTGGTGATATTGCTGCTAGAGACCTCATTTGTGCACAAAAGTTTAACGGCCTTACTGTT   24142
gb:MT106054|Organism:Severe      CTTGGTGATATTGCTGCTAGAGACCTCATTTGTGCACAAAAGTTTAACGGCCTTACTGTT   24142
gb:MT049951|Organism:Severe      CTTGGTGATATTGCTGCTAGAGACCTCATTTGTGCACAAAAGTTTAACGGCCTTACTGTT   24142
gb:MN975262|Organism:Severe      CTTGGTGATATTGCTGCTAGAGACCTCATTTGTGCACAAAAGTTTAACGGCCTTACTGTT   24142
gb:MT106052|Organism:Severe      CTTGGTGATATTGCTGCTAGAGACCTCATTTGTGCACAAAAGTTTAACGGCCTTACTGTT   24142
gb:LC522975|Organism:Severe      CTTGGTGATATTGCTGCTAGAGACCTCATTTGTGCACAAAAGTTTAACGGCCTTACTGTT   24139
gb:LC522973|Organism:Severe      CTTGGTGATATTGCTGCTAGAGACCTCATTTGTGCACAAAAGTTTAACGGCCTTACTGTT   24139
gb:LC522974|Organism:Severe      CTTGGTGATATTGCTGCTAGAGACCTCATTTGTGCACAAAAGTTTAACGGCCTTACTGTT   24139
gb:MN985325|Organism:Severe      CTTGGTGATATTGCTGCTAGAGACCTCATTTGTGCACAAAAGTTTAACGGCCTTACTGTT   24142
gb:MT020881|Organism:Severe      CTTGGTGATATTGCTGCTAGAGACCTCATTTGTGCACAAAAGTTTAACGGCCTTACTGTT   24142
gb:MT020880|Organism:Severe      CTTGGTGATATTGCTGCTAGAGACCTCATTTGTGCACAAAAGTTTAACGGCCTTACTGTT   24142
gb:MT066175|Organism:Severe      CTTGGTGATATTGCTGCTAGAGACCTCATTTGTGCACAAAAGTTTAACGGCCTTACTGTT   24142
gb:MN997409|Organism:Severe      CTTGGTGATATTGCTGCTAGAGACCTCATTTGTGCACAAAAGTTTAACGGCCTTACTGTT   24142
gb:MN938384|Organism:Severe      CTTGGTGATATTGCTGCTAGAGACCTCATTTGTGCACAAAAGTTTAACGGCCTTACTGTT   24110
gb:MT044258|Organism:Severe      CTTGGTGATATTGCTGCTAGAGACCTCATTTGTGCACAAAAGTTTAACGGCCTTACTGTT   24118
gb:MT039890|Organism:Severe      CTTGGTGATATTGCTGCTAGAGACCTCATTTGTGCACAAAAGTTTAACGGCCTTACTGTT   24142
gb:MN988713|Organism:Severe      CTTGGTGATATTGCTGCTAGAGACCTCATTTGTGCACAAAAGTTTAACGGCCTTACTGTT   24142
gb:LC521925|Organism:Severe      CTTGGTGATATTGCTGCTAGAGACCTCATTTGTGCACAAAAGTTTAACGGCCTTACTGTT   24115
gb:MT093571|Organism:Severe      CTTGGTGATATTGCTGCTAGAGACCTCATTTGTGCACAAAAGTTTAACGGCCTTACTGTT   24142
gb:MT039887|Organism:Severe      CTTGGTGATATTGCTGCTAGAGACCTCATTTGTGCACAAAAGTTTAACGGCCTTACTGTT   24139
gb:MT019530|Organism:Severe      CTTGGTGATATTGCTGCTAGAGACCTCATTTGTGCACAAAAGTTTAACGGCCTTACTGTT   24142
gb:MT039888|Organism:Severe      CTTGGTGATATTGCTGCTAGAGACCTCATTTGTGCACAAAAGTTTAACGGCCTTACTGTT   24142
gb:LC522972|Organism:Severe      CTTGGTGATATTGCTGCTAGAGACCTCATTTGTGCACAAAAGTTTAACGGCCTTACTGTT   24139
gb:MT027063|Organism:Severe      CTTGGTGATATTGCTGCTAGAGACCTCATTTGTGCACAAAAGTTTAACGGCCTTACTGTT   24142
gb:MT027062|Organism:Severe      CTTGGTGATATTGCTGCTAGAGACCTCATTTGTGCACAAAAGTTTAACGGCCTTACTGTT   24142
gb:MT019529|Organism:Severe      CTTGGTGATATTGCTGCTAGAGACCTCATTTGTGCACAAAAGTTTAACGGCCTTACTGTT   24142
```

```
gb:MN996529|Organism:Severe      CTTGGTGATATTGCTGCTAGAGACCTCATTTGTGCACAAAAGTTTAACGGCCTTACTGTT    24130
gb:MN996531|Organism:Severe      CTTGGTGATATTGCTGCTAGAGACCTCATTTGTGCACAAAAGTTTAACGGCCTTACTGTT    24129
gb:MT066176|Organism:Severe      CTTGGTGATATTGCTGCTAGAGACCTCATTTGTGCACAAAAGTTTAACGGCCTTACTGTT    24142
gb:MT027064|Organism:Severe      CTTGGTGATATTGCTGCTAGAGACCTCATTTGTGCACAAAAGTTTAACGGCCTTACTGTT    24142
gb:MN994468|Organism:Severe      CTTGGTGATATTGCTGCTAGAGACCTCATTTGTGCACAAAAGTTTAACGGCCTTACTGTT    24142
gb:MT072688|Organism:Severe      CTTGGTGATATTGCTGCTAGAGACCTCATTTGTGCACAAAAGTTTAACGGCCTTACTGTT    24127
gb:MN996527|Organism:Severe      CTTGGTGATATTGCTGCTAGAGACCTCATTTGTGCACAAAAGTTTAACGGCCTTACTGTT    24109
gb:MT093631|Organism:Severe      CTTGGTGATATTGCTGCTAGAGACCTCATTTGTGCACAAAAGTTTAACGGCCTTACTGTT    24180
gb:MT106053|Organism:Severe      CTTGGTGATATTGCTGCTAGAGACCTCATTTGTGCACAAAAGTTTAACGGCCTTACTGTT    24142
gb:MT019533|Organism:Severe      CTTGGTGATATTGCTGCTAGAGACCTCATTTGTGCACAAAAGTTTAACGGCCTTACTGTT    24142
gb:MT019531|Organism:Severe      CTTGGTGATATTGCTGCTAGAGACCTCATTTGTGCACAAAAGTTTAACGGCCTTACTGTT    24142
gb:MN996528|Organism:Severe      CTTGGTGATATTGCTGCTAGAGACCTCATTTGTGCACAAAAGTTTAACGGCCTTACTGTT    24142
gb:MN996530|Organism:Severe      CTTGGTGATATTGCTGCTAGAGACCTCATTTGTGCACAAAAGTTTAACGGCCTTACTGTT    24128
gb:MN908947|Organism:Severe      CTTGGTGATATTGCTGCTAGAGACCTCATTTGTGCACAAAAGTTTAACGGCCTTACTGTT    24142
gb:MT019532|Organism:Severe      CTTGGTGATATTGCTGCTAGAGACCTCATTTGTGCACAAAAGTTTAACGGCCTTACTGTT    24142
                                 ************************************************************

gb:MT020781|Organism:Severe      TTGCCACCTTTGCTCACAGATGAAATGATTGCTCAATACACTTCTGCACTGTTAGCGGGT    24190
gb:MT007544|Organism:Severe      TTGCCACCTTTGCTCACAGATGAAATGATTGCTCAATACACTTCTGCACTGTTAGCGGGT    24202
gb:MN994467|Organism:Severe      TTGCCACCTTTGCTCACAGATGAAATGATTGCTCAATACACTTCTGCACTGTTAGCGGGT    24202
gb:MT044257|Organism:Severe      TTGCCACCTTTGCTCACAGATGAAATGATTGCTCAATACACTTCTGCACTGTTAGCGGGT    24202
gb:MT106054|Organism:Severe      TTGCCACCTTTGCTCACAGATGAAATGATTGCTCAATACACTTCTGCACTGTTAGCGGGT    24202
gb:MT049951|Organism:Severe      TTGCCACCTTTGCTCACAGATGAAATGATTGCTCAATACACTTCTGCACTGTTAGCGGGT    24202
gb:MN975262|Organism:Severe      TTGCCACCTTTGCTCACAGATGAAATGATTGCTCAATACACTTCTGCACTGTTAGCGGGT    24202
gb:MT106052|Organism:Severe      TTGCCACCTTTGCTCACAGATGAAATGATTGCTCAATACACTTCTGCACTGTTAGCGGGT    24202
gb:LC522975|Organism:Severe      TTGCCACCTTTGCTCACAGATGAAATGATTGCTCAATACACTTCTGCACTGTTAGCGGGT    24199
gb:LC522973|Organism:Severe      TTGCCACCTTTGCTCACAGATGAAATGATTGCTCAATACACTTCTGCACTGTTAGCGGGT    24199
gb:LC522974|Organism:Severe      TTGCCACCTTTGCTCACAGATGAAATGATTGCTCAATACACTTCTGCACTGTTAGCGGGT    24199
gb:MN985325|Organism:Severe      TTGCCACCTTTGCTCACAGATGAAATGATTGCTCAATACACTTCTGCACTGTTAGCGGGT    24202
gb:MT020881|Organism:Severe      TTGCCACCTTTGCTCACAGATGAAATGATTGCTCAATACACTTCTGCACTGTTAGCGGGT    24202
gb:MT020880|Organism:Severe      TTGCCACCTTTGCTCACAGATGAAATGATTGCTCAATACACTTCTGCACTGTTAGCGGGT    24202
gb:MT066175|Organism:Severe      TTGCCACCTTTGCTCACAGATGAAATGATTGCTCAATACACTTCTGCACTGTTAGCGGGT    24202
gb:MN997409|Organism:Severe      TTGCCACCTTTGCTCACAGATGAAATGATTGCTCAATACACTTCTGCACTGTTAGCGGGT    24202
gb:MN938384|Organism:Severe      TTGCCACCTTTGCTCACAGATGAAATGATTGCTCAATACACTTCTGCACTGTTAGCGGGT    24170
gb:MT044258|Organism:Severe      TTGCCACCTTTGCTCACAGATGAAATGATTGCTCAATACACTTCTGCACTGTTAGCGGGT    24178
gb:MT039890|Organism:Severe      TTGCCACCTTTGCTCACAGATGAAATGATTGCTCAATACACTTCTGCACTGTTAGCGGGT    24202
gb:MN988713|Organism:Severe      TTGCCACCTTTGCTCACAGATGAAATGATTGCTCAATACACTTCTGCACTGTTAGCGGGT    24202
gb:LC521925|Organism:Severe      TTGCCACCTTTGCTCACAGATGAAATGATTGCTCAATACACTTCTGCACTGTTAGCGGGT    24175
gb:MT093571|Organism:Severe      TTGCCACCTTTGCTCACAGATGAAATGATTGCTCAATACACTTCTGCACTGTTAGCGGGT    24202
gb:MT039887|Organism:Severe      TTGCCACCTTTGCTCACAGATGAAATGATTGCTCAATACACTTCTGCACTGTTAGCGGGT    24199
gb:MT019530|Organism:Severe      TTGCCACCTTTGCTCACAGATGAAATGATTGCTCAATACACTTCTGCACTGTTAGCGGGT    24202
gb:MT039888|Organism:Severe      TTGCCACCTTTGCTCACAGATGAAATGATTGCTCAATACACTTCTGCACTGTTAGCGGGT    24202
gb:LC522972|Organism:Severe      TTGCCACCTTTGCTCACAGATGAAATGATTGCTCAATACACTTCTGCACTGTTAGCGGGT    24199
gb:MT027063|Organism:Severe      TTGCCACCTTTGCTCACAGATGAAATGATTGCTCAATACACTTCTGCACTGTTAGCGGGT    24202
gb:MT027062|Organism:Severe      TTGCCACCTTTGCTCACAGATGAAATGATTGCTCAATACACTTCTGCACTGTTAGCGGGT    24202
gb:MT019529|Organism:Severe      TTGCCACCTTTGCTCACAGATGAAATGATTGCTCAATACACTTCTGCACTGTTAGCGGGT    24202
gb:MN996529|Organism:Severe      TTGCCACCTTTGCTCACAGATGAAATGATTGCTCAATACACTTCTGCACTGTTAGCGGGT    24190
gb:MN996531|Organism:Severe      TTGCCACCTTTGCTCACAGATGAAATGATTGCTCAATACACTTCTGCACTGTTAGCGGGT    24189
gb:MT066176|Organism:Severe      TTGCCACCTTTGCTCACAGATGAAATGATTGCTCAATACACTTCTGCACTGTTAGCGGGT    24202
gb:MT027064|Organism:Severe      TTGCCACCTTTGCTCACAGATGAAATGATTGCTCAATACACTTCTGCACTGTTAGCGGGT    24202
gb:MN994468|Organism:Severe      TTGCCACCTTTGCTCACAGATGAAATGATTGCTCAATACACTTCTGCACTGTTAGCGGGT    24202
gb:MT072688|Organism:Severe      TTGCCACCTTTGCTCACAGATGAAATGATTGCTCAATACACTTCTGCACTGTTAGCGGGT    24187
gb:MN996527|Organism:Severe      TTGCCACCTTTGCTCACAGATGAAATGATTGCTCAATACACTTCTGCACTGTTAGCGGGT    24169
gb:MT093631|Organism:Severe      TTGCCACCTTTGCTCACAGATGAAATGATTGCTCAATACACTTCTGCACTGTTAGCGGGT    24240
gb:MT106053|Organism:Severe      TTGCCACCTTTGCTCACAGATGAAATGATTGCTCAATACACTTCTGCACTGTTAGCGGGT    24202
gb:MT019533|Organism:Severe      TTGCCACCTTTGCTCACAGATGAAATGATTGCTCAATACACTTCTGCACTGTTAGCGGGT    24202
gb:MT019531|Organism:Severe      TTGCCACCTTTGCTCACAGATGAAATGATTGCTCAATACACTTCTGCACTGTTAGCGGGT    24202
gb:MN996528|Organism:Severe      TTGCCACCTTTGCTCACAGATGAAATGATTGCTCAATACACTTCTGCACTGTTAGCGGGT    24202
gb:MN996530|Organism:Severe      TTGCCACCTTTGCTCACAGATGAAATGATTGCTCAATACACTTCTGCACTGTTAGCGGGT    24188
gb:MN908947|Organism:Severe      TTGCCACCTTTGCTCACAGATGAAATGATTGCTCAATACACTTCTGCACTGTTAGCGGGT    24202
gb:MT019532|Organism:Severe      TTGCCACCTTTGCTCACAGATGAAATGATTGCTCAATACACTTCTGCACTGTTAGCGGGT    24202
                                 ************************************************************

gb:MT020781|Organism:Severe      ACAATCACTTCTGGTTGGACCTTTGGTGCAGGTGCTGCATTACAAATACCATTTGCTATG    24250
gb:MT007544|Organism:Severe      ACAATCACTTCTGGTTGGACCTTTGGTGCAGGTGCTGCATTACAAATACCATTTGCTATG    24262
```

```
gb:MN994467|Organism:Severe      ACAATCACTTCTGGTTGGACCTTTGGTGCAGGTGCTGCATTACAAATACCATTTGCTATG    24262
gb:MT044257|Organism:Severe      ACAATCACTTCTGGTTGGACCTTTGGTGCAGGTGCTGCATTACAAATACCATTTGCTATG    24262
gb:MT106054|Organism:Severe      ACAATCACTTCTGGTTGGACCTTTGGTGCAGGTGCTGCATTACAAATACCATTTGCTATG    24262
gb:MT049951|Organism:Severe      ACAATCACTTCTGGTTGGACCTTTGGTGCAGGTGCTGCATTACAAATACCATTTGCTATG    24262
gb:MN975262|Organism:Severe      ACAATCACTTCTGGTTGGACCTTTGGTGCAGGTGCTGCATTACAAATACCATTTGCTATG    24262
gb:MT106052|Organism:Severe      ACAATCACTTCTGGTTGGACCTTTGGTGCAGGTGCTGCATTACAAATACCATTTGCTATG    24262
gb:LC522975|Organism:Severe      ACAATCACTTCTGGTTGGACCTTTGGTGCAGGTGCTGCATTACAAATACCATTTGCTATG    24259
gb:LC522973|Organism:Severe      ACAATCACTTCTGGTTGGACCTTTGGTGCAGGTGCTGCATTACAAATACCATTTGCTATG    24259
gb:LC522974|Organism:Severe      ACAATCACTTCTGGTTGGACCTTTGGTGCAGGTGCTGCATTACAAATACCATTTGCTATG    24259
gb:MN985325|Organism:Severe      ACAATCACTTCTGGTTGGACCTTTGGTGCAGGTGCTGCATTACAAATACCATTTGCTATG    24262
gb:MT020881|Organism:Severe      ACAATCACTTCTGGTTGGACCTTTGGTGCAGGTGCTGCATTACAAATACCATTTGCTATG    24262
gb:MT020880|Organism:Severe      ACAATCACTTCTGGTTGGACCTTTGGTGCAGGTGCTGCATTACAAATACCATTTGCTATG    24262
gb:MT066175|Organism:Severe      ACAATCACTTCTGGTTGGACCTTTGGTGCAGGTGCTGCATTACAAATACCATTTGCTATG    24262
gb:MN997409|Organism:Severe      ACAATCACTTCTGGTTGGACCTTTGGTGCAGGTGCTGCATTACAAATACCATTTGCTATG    24262
gb:MN938384|Organism:Severe      ACAATCACTTCTGGTTGGACCTTTGGTGCAGGTGCTGCATTACAAATACCATTTGCTATG    24230
gb:MT044258|Organism:Severe      ACAATCACTTCTGGTTGGACCTTTGGTGCAGGTGCTGCATTACAAATACCATTTGCTATG    24238
gb:MT039890|Organism:Severe      ACAATCACTTCTGGTTGGACCTTTGGTGCAGGTGCTGCATTACAAATACCATTTGCTATG    24262
gb:MN988713|Organism:Severe      ACAATCACTTCTGGTTGGACCTTTGGTGCAGGTGCTGCATTACAAATACCATTTGCTATG    24262
gb:LC521925|Organism:Severe      ACAATCACTTCTGGTTGGACCTTTGGTGCAGGTGCTGCATTACAAATACCATTTGCTATG    24235
gb:MT093571|Organism:Severe      ACAATCACTTCTGGTTGGACCTTTGGTGCAGGTGCTGCATTACAAATACCATTTGCTATG    24262
gb:MT039887|Organism:Severe      ACAATCACTTCTGGTTGGACCTTTGGTGCAGGTGCTGCATTACAAATACCATTTGCTATG    24259
gb:MT019530|Organism:Severe      ACAATCACTTCTGGTTGGACCTTTGGTGCAGGTGCTGCATTACAAATACCATTTGCTATG    24262
gb:MT039888|Organism:Severe      ACAATCACTTCTGGTTGGACCTTTGGTGCAGGTGCTGCATTACAAATACCATTTGCTATG    24262
gb:LC522972|Organism:Severe      ACAATCACTTCTGGTTGGACCTTTGGTGCAGGTGCTGCATTACAAATACCATTTGCTATG    24259
gb:MT027063|Organism:Severe      ACAATCACTTCTGGTTGGACCTTTGGTGCAGGTGCTGCATTACAAATACCATTTGCTATG    24262
gb:MT027062|Organism:Severe      ACAATCACTTCTGGTTGGACCTTTGGTGCAGGTGCTGCATTACAAATACCATTTGCTATG    24262
gb:MT019529|Organism:Severe      ACAATCACTTCTGGTTGGACCTTTGGTGCAGGTGCTGCATTACAAATACCATTTGCTATG    24262
gb:MN996529|Organism:Severe      ACAATCACTTCTGGTTGGACCTTTGGTGCAGGTGCTGCATTACAAATACCATTTGCTATG    24250
gb:MN996531|Organism:Severe      ACAATCACTTCTGGTTGGACCTTTGGTGCAGGTGCTGCATTACAAATACCATTTGCTATG    24249
gb:MT066176|Organism:Severe      ACAATCACTTCTGGTTGGACCTTTGGTGCAGGTGCTGCATTACAAATACCATTTGCTATG    24262
gb:MT027064|Organism:Severe      ACAATCACTTCTGGTTGGACCTTTGGTGCAGGTGCTGCATTACAAATACCATTTGCTATG    24262
gb:MN994468|Organism:Severe      ACAATCACTTCTGGTTGGACCTTTGGTGCAGGTGCTGCATTACAAATACCATTTGCTATG    24262
gb:MT072688|Organism:Severe      ACAATCACTTCTGGTTGGACCTTTGGTGCAGGTGCTGCATTACAAATACCATTTGCTATG    24247
gb:MN996527|Organism:Severe      ACAATCACTTCTGGTTGGACCTTTGGTGCAGGTGCTGCATTACAAATACCATTTGCTATG    24229
gb:MT093631|Organism:Severe      ACAATCACTTCTGGTTGGACCTTTGGTGCAGGTGCTGCATTACAAATACCATTTGCTATG    24300
gb:MT106053|Organism:Severe      ACAATCACTTCTGGTTGGACCTTTGGTGCAGGTGCTGCATTACAAATACCATTTGCTATG    24262
gb:MT019533|Organism:Severe      ACAATCACTTCTGGTTGGACCTTTGGTGCAGGTGCTGCATTACAAATACCATTTGCTATG    24262
gb:MT019531|Organism:Severe      ACAATCACTTCTGGTTGGACCTTTGGTGCAGGTGCTGCATTACAAATACCATTTGCTATG    24262
gb:MN996528|Organism:Severe      ACAATCACTTCTGGTTGGACCTTTGGTGCAGGTGCTGCATTACAAATACCATTTGCTATG    24262
gb:MN996530|Organism:Severe      ACAATCACTTCTGGTTGGACCTTTGGTGCAGGTGCTGCATTACAAATACCATTTGCTATG    24248
gb:MN908947|Organism:Severe      ACAATCACTTCTGGTTGGACCTTTGGTGCAGGTGCTGCATTACAAATACCATTTGCTATG    24262
gb:MT019532|Organism:Severe      ACAATCACTTCTGGTTGGACCTTTGGTGCAGGTGCTGCATTACAAATACCATTTGCTATG    24262
                                 ************************************************************

gb:MT020781|Organism:Severe      CAAATGGCTTATAGGTTTAATGGTATTGGAGTTACACAGAATGTTCTCTATGAGAACCAA    24310
gb:MT007544|Organism:Severe      CAAATGGCTTATAGGTTTAATGGTATTGGAGTTACACAGAATGTTCTCTATGAGAACCAA    24322
gb:MN994467|Organism:Severe      CAAATGGCTTATAGGTTTAATGGTATTGGAGTTACACAGAATGTTCTCTATGAGAACCAA    24322
gb:MT044257|Organism:Severe      CAAATGGCTTATAGGTTTAATGGTATTGGAGTTACACAGAATGTTCTCTATGAGAACCAA    24322
gb:MT106054|Organism:Severe      CAAATGGCTTATAGGTTTAATGGTATTGGAGTTACACAGAATGTTCTCTATGAGAACCAA    24322
gb:MT049951|Organism:Severe      CAAATGGCTTATAGGTTTAATGGTATTGGAGTTACACAGAATGTTCTCTATGAGAACCAA    24322
gb:MN975262|Organism:Severe      CAAATGGCTTATAGGTTTAATGGTATTGGAGTTACACAGAATGTTCTCTATGAGAACCAA    24322
gb:MT106052|Organism:Severe      CAAATGGCTTATAGGTTTAATGGTATTGGAGTTACACAGAATGTTCTCTATGAGAACCAA    24322
gb:LC522975|Organism:Severe      CAAATGGCTTATAGGTTTAATGGTATTGGAGTTACACAGAATGTTCTCTATGAGAACCAA    24319
gb:LC522973|Organism:Severe      CAAATGGCTTATAGGTTTAATGGTATTGGAGTTACACAGAATGTTCTCTATGAGAACCAA    24319
gb:LC522974|Organism:Severe      CAAATGGCTTATAGGTTTAATGGTATTGGAGTTACACAGAATGTTCTCTATGAGAACCAA    24319
gb:MN985325|Organism:Severe      CAAATGGCTTATAGGTTTAATGGTATTGGAGTTACACAGAATGTTCTCTATGAGAACCAA    24322
gb:MT020881|Organism:Severe      CAAATGGCTTATAGGTTTAATGGTATTGGAGTTACACAGAATGTTCTCTATGAGAACCAA    24322
gb:MT020880|Organism:Severe      CAAATGGCTTATAGGTTTAATGGTATTGGAGTTACACAGAATGTTCTCTATGAGAACCAA    24322
gb:MT066175|Organism:Severe      CAAATGGCTTATAGGTTTAATGGTATTGGAGTTACACAGAATGTTCTCTATGAGAACCAA    24322
gb:MN997409|Organism:Severe      CAAATGGCTTATAGGTTTAATGGTATTGGAGTTACACAGAATGTTCTCTATGAGAACCAA    24322
gb:MN938384|Organism:Severe      CAAATGGCTTATAGGTTTAATGGTATTGGAGTTACACAGAATGTTCTCTATGAGAACCAA    24290
gb:MT044258|Organism:Severe      CAAATGGCTTATAGGTTTAATGGTATTGGAGTTACACAGAATGTTCTCTATGAGAACCAA    24298
gb:MT039890|Organism:Severe      CAAATGGCTTATAGGTTTAATGGTATTGGAGTTACACAGAATGTTCTCTATGAGAACCAA    24322
gb:MN988713|Organism:Severe      CAAATGGCTTATAGGTTTAATGGTATTGGAGTTACACAGAATGTTCTCTATGAGAACCAA    24322
gb:LC521925|Organism:Severe      CAAATGGCTTATAGGTTTAATGGTATTGGAGTTACACAGAATGTTCTCTATGAGAACCAA    24295
```

```
gb:MT093571|Organism:Severe      CAAATGGCTTATAGGTTTAATGGTATTGGAGTTACACAGAATGTTCTCTATGAGAACCAA   24322
gb:MT039887|Organism:Severe      CAAATGGCTTATAGGTTTAATGGTATTGGAGTTACACAGAATGTTCTCTATGAGAACCAA   24319
gb:MT019530|Organism:Severe      CAAATGGCTTATAGGTTTAATGGTATTGGAGTTACACAGAATGTTCTCTATGAGAACCAA   24322
gb:MT039888|Organism:Severe      CAAATGGCTTATAGGTTTAATGGTATTGGAGTTACACAGAATGTTCTCTATGAGAACCAA   24322
gb:LC522972|Organism:Severe      CAAATGGCTTATAGGTTTAATGGTATTGGAGTTACACAGAATGTTCTCTATGAGAACCAA   24319
gb:MT027063|Organism:Severe      CAAATGGCTTATAGGTTTAATGGTATTGGAGTTACACAGAATGTTCTCTATGAGAACCAA   24322
gb:MT027062|Organism:Severe      CAAATGGCTTATAGGTTTAATGGTATTGGAGTTACACAGAATGTTCTCTATGAGAACCAA   24322
gb:MT019529|Organism:Severe      CAAATGGCTTATAGGTTTAATGGTATTGGAGTTACACAGAATGTTCTCTATGAGAACCAA   24322
gb:MN996529|Organism:Severe      CAAATGGCTTATAGGTTTAATGGTATTGGAGTTACACAGAATGTTCTCTATGAGAACCAA   24310
gb:MN996531|Organism:Severe      CAAATGGCTTATAGGTTTAATGGTATTGGAGTTACACAGAATGTTCTCTATGAGAACCAA   24309
gb:MT066176|Organism:Severe      CAAATGGCTTATAGGTTTAATGGTATTGGAGTTACACAGAATGTTCTCTATGAGAACCAA   24322
gb:MT027064|Organism:Severe      CAAATGGCTTATAGGTTTAATGGTATTGGAGTTACACAGAATGTTCTCTATGAGAACCAA   24322
gb:MN994468|Organism:Severe      CAAATGGCTTATAGGTTTAATGGTATTGGAGTTACACAGAATGTTCTCTATGAGAACCAA   24322
gb:MT072688|Organism:Severe      CAAATGGCTTATAGGTTTAATGGTATTGGAGTTACACAGAATGTTCTCTATGAGAACCAA   24307
gb:MN996527|Organism:Severe      CAAATGGCTTATAGGTTTAATGGTATTGGAGTTACACAGAATGTTCTCTATGAGAACCAA   24289
gb:MT093631|Organism:Severe      CAAATGGCTTATAGGTTTAATGGTATTGGAGTTACACAGAATGTTCTCTATGAGAACCAA   24360
gb:MT106053|Organism:Severe      CAAATGGCTTATAGGTTTAATGGTATTGGAGTTACACAGAATGTTCTCTATGAGAACCAA   24322
gb:MT019533|Organism:Severe      CAAATGGCTTATAGGTTTAATGGTATTGGAGTTACACAGAATGTTCTCTATGAGAACCAA   24322
gb:MT019531|Organism:Severe      CAAATGGCTTATAGGTTTAATGGTATTGGAGTTACACAGAATGTTCTCTATGAGAACCAA   24322
gb:MN996528|Organism:Severe      CAAATGGCTTATAGGTTTAATGGTATTGGAGTTACACAGAATGTTCTCTATGAGAACCAA   24322
gb:MN996530|Organism:Severe      CAAATGGCTTATAGGTTTAATGGTATTGGAGTTACACAGAATGTTCTCTATGAGAACCAA   24308
gb:MN908947|Organism:Severe      CAAATGGCTTATAGGTTTAATGGTATTGGAGTTACACAGAATGTTCTCTATGAGAACCAA   24322
gb:MT019532|Organism:Severe      CAAATGGCTTATAGGTTTAATGGTATTGGAGTTACACAGAATGTTCTCTATGAGAACCAA   24322
                                 ************************************************************

gb:MT020781|Organism:Severe      AAATTGATTGCCAACCAATTTAATAGTGCTATTGGCAAAATTCAAGACTCACTTTCTTCC   24370
gb:MT007544|Organism:Severe      AAATTGATTGCCAACCAATTTAATAGTGCTATTGGCAAAATTCAAGACTCACTTTCTTCC   24382
gb:MN994467|Organism:Severe      AAATTGATTGCCAACCAATTTAATAGTGCTATTGGCAAAATTCAAGACTCACTTTCTTCC   24382
gb:MT044257|Organism:Severe      AAATTGATTGCCAACCAATTTAATAGTGCTATTGGCAAAATTCAAGACTCACTTTCTTCC   24382
gb:MT106054|Organism:Severe      AAATTGATTGCCAACCAATTTAATAGTGCTATTGGCAAAATTCAAGACTCACTTTCTTCC   24382
gb:MT049951|Organism:Severe      AAATTGATTGCCAACCAATTTAATAGTGCTATTGGCAAAATTCAAGACTCACTTTCTTCC   24382
gb:MN975262|Organism:Severe      AAATTGATTGCCAACCAATTTAATAGTGCTATTGGCAAAATTCAAGACTCACTTTCTTCC   24382
gb:MT106052|Organism:Severe      AAATTGATTGCCAACCAATTTAATAGTGCTATTGGCAAAATTCAAGACTCACTTTCTTCC   24382
gb:LC522975|Organism:Severe      AAATTGATTGCCAACCAATTTAATAGTGCTATTGGCAAAATTCAAGACTCACTTTCTTCC   24379
gb:LC522973|Organism:Severe      AAATTGATTGCCAACCAATTTAATAGTGCTATTGGCAAAATTCAAGACTCACTTTCTTCC   24379
gb:LC522974|Organism:Severe      AAATTGATTGCCAACCAATTTAATAGTGCTATTGGCAAAATTCAAGACTCACTTTCTTCC   24379
gb:MN985325|Organism:Severe      AAATTGATTGCCAACCAATTTAATAGTGCTATTGGCAAAATTCAAGACTCACTTTCTTCC   24382
gb:MT020881|Organism:Severe      AAATTGATTGCCAACCAATTTAATAGTGCTATTGGCAAAATTCAAGACTCACTTTCTTCC   24382
gb:MT020880|Organism:Severe      AAATTGATTGCCAACCAATTTAATAGTGCTATTGGCAAAATTCAAGACTCACTTTCTTCC   24382
gb:MT066175|Organism:Severe      AAATTGATTGCCAACCAATTTAATAGTGCTATTGGCAAAATTCAAGACTCACTTTCTTCC   24382
gb:MN997409|Organism:Severe      AAATTGATTGCCAACCAATTTAATAGTGCTATTGGCAAAATTCAAGACTCACTTTCTTCC   24382
gb:MN938384|Organism:Severe      AAATTGATTGCCAACCAATTTAATAGTGCTATTGGCAAAATTCAAGACTCACTTTCTTCC   24350
gb:MT044258|Organism:Severe      AAATTGATTGCCAACCAATTTAATAGTGCTATTGGCAAAATTCAAGACTCACTTTCTTCC   24358
gb:MT039890|Organism:Severe      AAATTGATTGCCAACCAATTTAATAGTGCTATTGGCAAAATTCAAGACTCACTTTCTTCC   24382
gb:MN988713|Organism:Severe      AAATTGATTGCCAACCAATTTAATAGTGCTATTGGCAAAATTCAAGACTCACTTTCTTCC   24382
gb:LC521925|Organism:Severe      AAATTGATTGCCAACCAATTTAATAGTGCTATTGGCAAAATTCAAGACTCACTTTCTTCC   24355
gb:MT093571|Organism:Severe      AAATTGATTGCCAACCAATTTAATAGTGCTATTGGCAAAATTCAAGACTCACTTTCTTCC   24382
gb:MT039887|Organism:Severe      AAATTGATTGCCAACCAATTTAATAGTGCTATTGGCAAAATTCAAGACTCACTTTCTTCC   24379
gb:MT019530|Organism:Severe      AAATTGATTGCCAACCAATTTAATAGTGCTATTGGCAAAATTCAAGACTCACTTTCTTCC   24382
gb:MT039888|Organism:Severe      AAATTGATTGCCAACCAATTTAATAGTGCTATTGGCAAAATTCAAGACTCACTTTCTTCC   24382
gb:LC522972|Organism:Severe      AAATTGATTGCCAACCAATTTAATAGTGCTATTGGCAAAATTCAAGACTCACTTTCTTCC   24379
gb:MT027063|Organism:Severe      AAATTGATTGCCAACCAATTTAATAGTGCTATTGGCAAAATTCAAGACTCACTTTCTTCC   24382
gb:MT027062|Organism:Severe      AAATTGATTGCCAACCAATTTAATAGTGCTATTGGCAAAATTCAAGACTCACTTTCTTCC   24382
gb:MT019529|Organism:Severe      AAATTGATTGCCAACCAATTTAATAGTGCTATTGGCAAAATTCAAGACTCACTTTCTTCC   24382
gb:MN996529|Organism:Severe      AAATTGATTGCCAACCAATTTAATAGTGCTATTGGCAAAATTCAAGACTCACTTTCTTCC   24370
gb:MN996531|Organism:Severe      AAATTGATTGCCAACCAATTTAATAGTGCTATTGGCAAAATTCAAGACTCACTTTCTTCC   24369
gb:MT066176|Organism:Severe      AAATTGATTGCCAACCAATTTAATAGTGCTATTGGCAAAATTCAAGACTCACTTTCTTCC   24382
gb:MT027064|Organism:Severe      AAATTGATTGCCAACCAATTTAATAGTGCTATTGGCAAAATTCAAGACTCACTTTCTTCC   24382
gb:MN994468|Organism:Severe      AAATTGATTGCCAACCAATTTAATAGTGCTATTGGCAAAATTCAAGACTCACTTTCTTCC   24382
gb:MT072688|Organism:Severe      AAATTGATTGCCAACCAATTTAATAGTGCTATTGGCAAAATTCAAGACTCACTTTCTTCC   24367
gb:MN996527|Organism:Severe      AAGTTGATTGCCAACCAATTTAATAGTGCTATTGGCAAAATTCAAGACTCACTTTCTTCC   24349
gb:MT093631|Organism:Severe      AAATTGATTGCCAACCAATTTAATAGTGCTATTGGCAAAATTCAAGACTCACTTTCTTCC   24420
gb:MT106053|Organism:Severe      AAGTTGATTGCCAACCAATTTAATAGTGCTATTGGCAAAATTCAAGACTCACTTTCTTCC   24382
gb:MT019533|Organism:Severe      AAATTGATTGCCAACCAATTTAATAGTGCTATTGGCAAAATTCAAGACTCACTTTCTTCC   24382
gb:MT019531|Organism:Severe      AAATTGATTGCCAACCAATTTAATAGTGCTATTGGCAAAATTCAAGACTCACTTTCTTCC   24382
```

```
gb:MN996528|Organism:Severe      AAATTGATTGCCAACCAATTTAATAGTGCTATTGGCAAAATTCAAGACTCACTTTCTTCC    24382
gb:MN996530|Organism:Severe      AAATTGATTGCCAACCAATTTAATAGTGCTATTGGCAAAATTCAAGACTCACTTTCTTCC    24368
gb:MN908947|Organism:Severe      AAATTGATTGCCAACCAATTTAATAGTGCTATTGGCAAAATTCAAGACTCACTTTCTTCC    24382
gb:MT019532|Organism:Severe      AAATTGATTGCCAACCAATTTAATAGTGCTATTGGCAAAATTCAAGACTCACTTTCTTCC    24382
                                 ** *********************************************************

gb:MT020781|Organism:Severe      ACAGCAAGTGCACTTGGAAAACTTCAAGATGTGGTCAACCAAAATGCACAAGCTTTAAAC    24430
gb:MT007544|Organism:Severe      ACAGCAAGTGCACTTGGAAAACTTCAAGATGTGGTCAACCAAAATGCACAAGCTTTAAAC    24442
gb:MN994467|Organism:Severe      ACAGCAAGTGCACTTGGAAAACTTCAAGATGTGGTCAACCAAAATGCACAAGCTTTAAAC    24442
gb:MT044257|Organism:Severe      ACAGCAAGTGCACTTGGAAAACTTCAAGATGTGGTCAACCAAAATGCACAAGCTTTAAAC    24442
gb:MT106054|Organism:Severe      ACAGCAAGTGCACTTGGAAAACTTCAAGATGTGGTCAACCAAAATGCACAAGCTTTAAAC    24442
gb:MT049951|Organism:Severe      ACAGCAAGTGCACTTGGAAAACTTCAAGATGTGGTCAACCAAAATGCACAAGCTTTAAAC    24442
gb:MN975262|Organism:Severe      ACAGCAAGTGCACTTGGAAAACTTCAAGATGTGGTCAACCAAAATGCACAAGCTTTAAAC    24442
gb:MT106052|Organism:Severe      ACAGCAAGTGCACTTGGAAAACTTCAAGATGTGGTCAACCAAAATGCACAAGCTTTAAAC    24442
gb:LC522975|Organism:Severe      ACAGCAAGTGCACTTGGAAAACTTCAAGATGTGGTCAACCAAAATGCACAAGCTTTAAAC    24439
gb:LC522973|Organism:Severe      ACAGCAAGTGCACTTGGAAAACTTCAAGATGTGGTCAACCAAAATGCACAAGCTTTAAAC    24439
gb:LC522974|Organism:Severe      ACAGCAAGTGCACTTGGAAAACTTCAAGATGTGGTCAACCAAAATGCACAAGCTTTAAAC    24439
gb:MN985325|Organism:Severe      ACAGCAAGTGCACTTGGAAAACTTCAAGATGTGGTCAACCAAAATGCACAAGCTTTAAAC    24442
gb:MT020881|Organism:Severe      ACAGCAAGTGCACTTGGAAAACTTCAAGATGTGGTCAACCAAAATGCACAAGCTTTAAAC    24442
gb:MT020880|Organism:Severe      ACAGCAAGTGCACTTGGAAAACTTCAAGATGTGGTCAACCAAAATGCACAAGCTTTAAAC    24442
gb:MT066175|Organism:Severe      ACAGCAAGTGCACTTGGAAAACTTCAAGATGTGGTCAACCAAAATGCACAAGCTTTAAAC    24442
gb:MN997409|Organism:Severe      ACAGCAAGTGCACTTGGAAAACTTCAAGATGTGGTCAACCAAAATGCACAAGCTTTAAAC    24442
gb:MN938384|Organism:Severe      ACAGCAAGTGCACTTGGAAAACTTCAAGATGTGGTCAACCAAAATGCACAAGCTTTAAAC    24410
gb:MT044258|Organism:Severe      ACAGCAAGTGCACTTGGAAAACTTCAAGATGTGGTCAACCAAAATGCACAAGCTTTAAAC    24418
gb:MT039890|Organism:Severe      ACAGCAAGTGCACTTGGAAAACTTCAAGATGTGGTCAACCAAAATGCACAAGCTTTAAAC    24442
gb:MN988713|Organism:Severe      ACAGCAAGTGCACTTGGAAAACTTCAAGATGTGGTCAACCAAAATGCACAAGCTTTAAAC    24442
gb:LC521925|Organism:Severe      ACAGCAAGTGCACTTGGAAAACTTCAAGATGTGGTCAACCAAAATGCACAAGCTTTAAAC    24415
gb:MT093571|Organism:Severe      ACAGCAAGTGCACTTGGAAAACTTCAAGATGTGGTCAACCAAAATGCACAAGCTTTAAAC    24442
gb:MT039887|Organism:Severe      ACAGCAAGTGCACTTGGAAAACTTCAAGATGTGGTCAACCAAAATGCACAAGCTTTAAAC    24439
gb:MT019530|Organism:Severe      ACAGCAAGTGCACTTGGAAAACTTCAAGATGTGGTCAACCAAAATGCACAAGCTTTAAAC    24442
gb:MT039888|Organism:Severe      ACAGCAAGTGCACTTGGAAAACTTCAAGATGTGGTCAACCAAAATGCACAAGCTTTAAAC    24442
gb:LC522972|Organism:Severe      ACAGCAAGTGCACTTGGAAAACTTCAAGATGTGGTCAACCAAAATGCACAAGCTTTAAAC    24439
gb:MT027063|Organism:Severe      ACAGCAAGTGCACTTGGAAAACTTCAAGATGTGGTCAACCAAAATGCACAAGCTTTAAAC    24442
gb:MT027062|Organism:Severe      ACAGCAAGTGCACTTGGAAAACTTCAAGATGTGGTCAACCAAAATGCACAAGCTTTAAAC    24442
gb:MT019529|Organism:Severe      ACAGCAAGTGCACTTGGAAAACTTCAAGATGTGGTCAACCAAAATGCACAAGCTTTAAAC    24442
gb:MN996529|Organism:Severe      ACAGCAAGTGCACTTGGAAAACTTCAAGATGTGGTCAACCAAAATGCACAAGCTTTAAAC    24430
gb:MN996531|Organism:Severe      ACAGCAAGTGCACTTGGAAAACTTCAAGATGTGGTCAACCAAAATGCACAAGCTTTAAAC    24429
gb:MT066176|Organism:Severe      ACAGCAAGTGCACTTGGAAAACTTCAAGATGTGGTCAACCAAAATGCACAAGCTTTAAAC    24442
gb:MT027064|Organism:Severe      ACAGCAAGTGCACTTGGAAAACTTCAAGATGTGGTCAACCAAAATGCACAAGCTTTAAAC    24442
gb:MN994468|Organism:Severe      ACAGCAAGTGCACTTGGAAAACTTCAAGATGTGGTCAACCAAAATGCACAAGCTTTAAAC    24442
gb:MT072688|Organism:Severe      ACAGCAAGTGCACTTGGAAAACTTCAAGATGTGGTCAACCAAAATGCACAAGCTTTAAAC    24427
gb:MN996527|Organism:Severe      ACAGCAAGTGCACTTGGAAAACTTCAAGATGTGGTCAACCAAAATGCACAAGCTTTAAAC    24409
gb:MT093631|Organism:Severe      ACAGCAAGTGCACTTGGAAAACTTCAAGATGTGGTCAACCAAAATGCACAAGCTTTAAAC    24480
gb:MT106053|Organism:Severe      ACAGCAAGTGCACTTGGAAAACTTCAAGATGTGGTCAACCAAAATGCACAAGCTTTAAAC    24442
gb:MT019533|Organism:Severe      ACAGCAAGTGCACTTGGAAAACTTCAAGATGTGGTCAACCAAAATGCACAAGCTTTAAAC    24442
gb:MT019531|Organism:Severe      ACAGCAAGTGCACTTGGAAAACTTCAAGATGTGGTCAACCAAAATGCACAAGCTTTAAAC    24442
gb:MN996528|Organism:Severe      ACAGCAAGTGCACTTGGAAAACTTCAAGATGTGGTCAACCAAAATGCACAAGCTTTAAAC    24442
gb:MN996530|Organism:Severe      ACAGCAAGTGCACTTGGAAAACTTCAAGATGTGGTCAACCAAAATGCACAAGCTTTAAAC    24428
gb:MN908947|Organism:Severe      ACAGCAAGTGCACTTGGAAAACTTCAAGATGTGGTCAACCAAAATGCACAAGCTTTAAAC    24442
gb:MT019532|Organism:Severe      ACAGCAAGTGCACTTGGAAAACTTCAAGATGTGGTCAACCAAAATGCACAAGCTTTAAAC    24442
                                 ************************************************************

gb:MT020781|Organism:Severe      ACGCTTGTTAAACAACTTAGCTCCAATTTTGGTGCAATTTCAAGTGTTTTAAATGATATC    24490
gb:MT007544|Organism:Severe      ACGCTTGTTAAACAACTTAGCTCCAATTTTGGTGCAATTTCAAGTGTTTTAAATGATATC    24502
gb:MN994467|Organism:Severe      ACGCTTGTTAAACAACTTAGCTCCAATTTTGGTGCAATTTCAAGTGTTTTAAATGATATC    24502
gb:MT044257|Organism:Severe      ACGCTTGTTAAACAACTTAGCTCCAATTTTGGTGCAATTTCAAGTGTTTTAAATGATATC    24502
gb:MT106054|Organism:Severe      ACGCTTGTTAAACAACTTAGCTCCAATTTTGGTGCAATTTCAAGTGTTTTAAATGATATC    24502
gb:MT049951|Organism:Severe      ACGCTTGTTAAACAACTTAGCTCCAATTTTGGTGCAATTTCAAGTGTTTTAAATGATATC    24502
gb:MN975262|Organism:Severe      ACGCTTGTTAAACAACTTAGCTCCAATTTTGGTGCAATTTCAAGTGTTTTAAATGATATC    24502
gb:MT106052|Organism:Severe      ACGCTTGTTAAACAACTTAGCTCCAATTTTGGTGCAATTTCAAGTGTTTTAAATGATATC    24502
gb:LC522975|Organism:Severe      ACGCTTGTTAAACAACTTAGCTCCAATTTTGGTGCAATTTCAAGTGTTTTAAATGATATC    24499
gb:LC522973|Organism:Severe      ACGCTTGTTAAACAACTTAGCTCCAATTTTGGTGCAATTTCAAGTGTTTTAAATGATATC    24499
gb:LC522974|Organism:Severe      ACGCTTGTTAAACAACTTAGCTCCAATTTTGGTGCAATTTCAAGTGTTTTAAATGATATC    24499
gb:MN985325|Organism:Severe      ACGCTTGTTAAACAACTTAGCTCCAATTTTGGTGCAATTTCAAGTGTTTTAAATGATATC    24502
gb:MT020881|Organism:Severe      ACGCTTGTTAAACAACTTAGCTCCAATTTTGGTGCAATTTCAAGTGTTTTAAATGATATC    24502
```

```
gb:MT020880|Organism:Severe      ACGCTTGTTAAACAACTTAGCTCCAATTTTGGTGCAATTTCAAGTGTTTTAAATGATATC   24502
gb:MT066175|Organism:Severe      ACGCTTGTTAAACAACTTAGCTCCAATTTTGGTGCAATTTCAAGTGTTTTAAATGATATC   24502
gb:MN997409|Organism:Severe      ACGCTTGTTAAACAACTTAGCTCCAATTTTGGTGCAATTTCAAGTGTTTTAAATGATATC   24502
gb:MN938384|Organism:Severe      ACGCTTGTTAAACAACTTAGCTCCAATTTTGGTGCAATTTCAAGTGTTTTAAATGATATC   24470
gb:MT044258|Organism:Severe      ACGCTTGTTAAACAACTTAGCTCCAATTTTGGTGCAATTTCAAGTGTTTTAAATGATATC   24478
gb:MT039890|Organism:Severe      ACGCTTGTTAAACAACTTAGCTCCAATTTTGGTGCAATTTCAAGTGTTTTAAATGATATC   24502
gb:MN988713|Organism:Severe      ACGCTTGTTAAACAACTTAGCTCCAATTTTGGTGCAATTTCAAGTGTTTTAAATGATATC   24502
gb:LC521925|Organism:Severe      ACGCTTGTTAAACAACTTAGCTCCAATTTTGGTGCAATTTCAAGTGTTTTAAATGATATC   24475
gb:MT093571|Organism:Severe      ACGCTTGTTAAACAACTTAGCTCCAATTTTGGTGCAATTTCAAGTGTTTTAAATGATATC   24502
gb:MT039887|Organism:Severe      ACGCTTGTTAAACAACTTAGCTCCAATTTTGGTGCAATTTCAAGTGTTTTAAATGATATC   24499
gb:MT019530|Organism:Severe      ACGCTTGTTAAACAACTTAGCTCCAATTTTGGTGCAATTTCAAGTGTTTTAAATGATATC   24502
gb:MT039888|Organism:Severe      ACGCTTGTTAAACAACTTAGCTCCAATTTTGGTGCAATTTCAAGTGTTTTAAATGATATC   24502
gb:LC522972|Organism:Severe      ACGCTTGTTAAACAACTTAGCTCCAATTTTGGTGCAATTTCAAGTGTTTTAAATGATATC   24499
gb:MT027063|Organism:Severe      ACGCTTGTTAAACAACTTAGCTCCAATTTTGGTGCAATTTCAAGTGTTTTAAATGATATC   24502
gb:MT027062|Organism:Severe      ACGCTTGTTAAACAACTTAGCTCCAATTTTGGTGCAATTTCAAGTGTTTTAAATGATATC   24502
gb:MT019529|Organism:Severe      ACGCTTGTTAAACAACTTAGCTCCAATTTTGGTGCAATTTCAAGTGTTTTAAATGATATC   24502
gb:MN996529|Organism:Severe      ACGCTTGTTAAACAACTTAGCTCCAATTTTGGTGCAATTTCAAGTGTTTTAAATGATATC   24490
gb:MN996531|Organism:Severe      ACGCTTGTTAAACAACTTAGCTCCAATTTTGGTGCAATTTCAAGTGTTTTAAATGATATC   24489
gb:MT066176|Organism:Severe      ACGCTTGTTAAACAACTTAGCTCCAATTTTGGTGCAATTTCAAGTGTTTTAAATGATATC   24502
gb:MT027064|Organism:Severe      ACGCTTGTTAAACAACTTAGCTCCAATTTTGGTGCAATTTCAAGTGTTTTAAATGATATC   24502
gb:MN994468|Organism:Severe      ACGCTTGTTAAACAACTTAGCTCCAATTTTGGTGCAATTTCAAGTGTTTTAAATGATATC   24502
gb:MT072688|Organism:Severe      ACGCTTGTTAAACAACTTAGCTCCAATTTTGGTGCAATTTCAAGTGTTTTAAATGATATC   24487
gb:MN996527|Organism:Severe      ACGCTTGTTAAACAACTTAGCTCCAATTTTGGTGCAATTTCAAGTGTTTTAAATGATATC   24469
gb:MT093631|Organism:Severe      ACGCTTGTTAAACAACTTAGCTCCAATTTTGGTGCAATTTCAAGTGTTTTAAATGATATC   24540
gb:MT106053|Organism:Severe      ACGCTTGTTAAACAACTTAGCTCCAATTTTGGTGCAATTTCAAGTGTTTTAAATGATATC   24502
gb:MT019533|Organism:Severe      ACGCTTGTTAAACAACTTAGCTCCAATTTTGGTGCAATTTCAAGTGTTTTAAATGATATC   24502
gb:MT019531|Organism:Severe      ACGCTTGTTAAACAACTTAGCTCCAATTTTGGTGCAATTTCAAGTGTTTTAAATGATATC   24502
gb:MN996528|Organism:Severe      ACGCTTGTTAAACAACTTAGCTCCAATTTTGGTGCAATTTCAAGTGTTTTAAATGATATC   24502
gb:MN996530|Organism:Severe      ACGCTTGTTAAACAACTTAGCTCCAATTTTGGTGCAATTTCAAGTGTTTTAAATGATATC   24488
gb:MN908947|Organism:Severe      ACGCTTGTTAAACAACTTAGCTCCAATTTTGGTGCAATTTCAAGTGTTTTAAATGATATC   24502
gb:MT019532|Organism:Severe      ACGCTTGTTAAACAACTTAGCTCCAATTTTGGTGCAATTTCAAGTGTTTTAAATGATATC   24502
                                 ************************************************************

gb:MT020781|Organism:Severe      CTTTCACGTCTTGACAAAGTTGAGGCTGAAGTGCAAATTGATAGGTTGATCACAGGCAGA   24550
gb:MT007544|Organism:Severe      CTTTCACGTCTTGACAAAGTTGAGGCTGAAGTGCAAATTGATAGGTTGATCACAGGCAGA   24562
gb:MN994467|Organism:Severe      CTTTCACGTCTTGACAAAGTTGAGGCTGAAGTGCAAATTGATAGGTTGATCACAGGCAGA   24562
gb:MT044257|Organism:Severe      CTTTCACGTCTTGACAAAGTTGAGGCTGAAGTGCAAATTGATAGGTTGATCACAGGCAGA   24562
gb:MT106054|Organism:Severe      CTTTCACGTCTTGACAAAGTTGAGGCTGAAGTGCAAATTGATAGGTTGATCACAGGCAGA   24562
gb:MT049951|Organism:Severe      CTTTCACGTCTTGACAAAGTTGAGGCTGAAGTGCAAATTGATAGGTTGATCACAGGCAGA   24562
gb:MN975262|Organism:Severe      CTTTCACGTCTTGACAAAGTTGAGGCTGAAGTGCAAATTGATAGGTTGATCACAGGCAGA   24562
gb:MT106052|Organism:Severe      CTTTCACGTCTTGACAAAGTTGAGGCTGAAGTGCAAATTGATAGGTTGATCACAGGCAGA   24562
gb:LC522975|Organism:Severe      CTTTCACGTCTTGACAAAGTTGAGGCTGAAGTGCAAATTGATAGGTTGATCACAGGCAGA   24559
gb:LC522973|Organism:Severe      CTTTCACGTCTTGACAAAGTTGAGGCTGAAGTGCAAATTGATAGGTTGATCACAGGCAGA   24559
gb:LC522974|Organism:Severe      CTTTCACGTCTTGACAAAGTTGAGGCTGAAGTGCAAATTGATAGGTTGATCACAGGCAGA   24559
gb:MN985325|Organism:Severe      CTTTCACGTCTTGACAAAGTTGAGGCTGAAGTGCAAATTGATAGGTTGATCACAGGCAGA   24562
gb:MT020881|Organism:Severe      CTTTCACGTCTTGACAAAGTTGAGGCTGAAGTGCAAATTGATAGGTTGATCACAGGCAGA   24562
gb:MT020880|Organism:Severe      CTTTCACGTCTTGACAAAGTTGAGGCTGAAGTGCAAATTGATAGGTTGATCACAGGCAGA   24562
gb:MT066175|Organism:Severe      CTTTCACGTCTTGACAAAGTTGAGGCTGAAGTGCAAATTGATAGGTTGATCACAGGCAGA   24562
gb:MN997409|Organism:Severe      CTTTCACGTCTTGACAAAGTTGAGGCTGAAGTGCAAATTGATAGGTTGATCACAGGCAGA   24562
gb:MN938384|Organism:Severe      CTTTCACGTCTTGACAAAGTTGAGGCTGAAGTGCAAATTGATAGGTTGATCACAGGCAGA   24530
gb:MT044258|Organism:Severe      CTTTCACGTCTTGACAAAGTTGAGGCTGAAGTGCAAATTGATAGGTTGATCACAGGCAGA   24538
gb:MT039890|Organism:Severe      CTTTCACGTCTTGACAAAGTTGAGGCTGAAGTGCAAATTGATAGGTTGATCACAGGCAGA   24562
gb:MN988713|Organism:Severe      CTTTCACGTCTTGACAAAGTTGAGGCTGAAGTGCAAATTGATAGGTTGATCACAGGCAGA   24562
gb:LC521925|Organism:Severe      CTTTCACGTCTTGACAAAGTTGAGGCTGAAGTGCAAATTGATAGGTTGATCACAGGCAGA   24535
gb:MT093571|Organism:Severe      CTTTCACGTCTTGACAAAGTTGAGGCTGAAGTGCAAATTGATAGGTTGATCACAGGCAGA   24562
gb:MT039887|Organism:Severe      CTTTCACGTCTTGACAAAGTTGAGGCTGAAGTGCAAATTGATAGGTTGATCACAGGCAGA   24559
gb:MT019530|Organism:Severe      CTTTCACGTCTTGACAAAGTTGAGGCTGAAGTGCAAATTGATAGGTTGATCACAGGCAGA   24562
gb:MT039888|Organism:Severe      CTTTCACGTCTTGACAAAGTTGAGGCTGAAGTGCAAATTGATAGGTTGATCACAGGCAGA   24562
gb:LC522972|Organism:Severe      CTTTCACGTCTTGACAAAGTTGAGGCTGAAGTGCAAATTGATAGGTTGATCACAGGCAGA   24559
gb:MT027063|Organism:Severe      CTTTCACGTCTTGACAAAGTTGAGGCTGAAGTGCAAATTGATAGGTTGATCACAGGCAGA   24562
gb:MT027062|Organism:Severe      CTTTCACGTCTTGACAAAGTTGAGGCTGAAGTGCAAATTGATAGGTTGATCACAGGCAGA   24562
gb:MT019529|Organism:Severe      CTTTCACGTCTTGACAAAGTTGAGGCTGAAGTGCAAATTGATAGGTTGATCACAGGCAGA   24562
gb:MN996529|Organism:Severe      CTTTCACGTCTTGACAAAGTTGAGGCTGAAGTGCAAATTGATAGGTTGATCACAGGCAGA   24550
gb:MN996531|Organism:Severe      CTTTCACGTCTTGACAAAGTTGAGGCTGAAGTGCAAATTGATAGGTTGATCACAGGCAGA   24549
gb:MT066176|Organism:Severe      CTTTCACGTCTTGACAAAGTTGAGGCTGAAGTGCAAATTGATAGGTTGATCACAGGCAGA   24562
```

```
gb:MT027064|Organism:Severe      CTTTCACGTCTTGACAAAGTTGAGGCTGAAGTGCAAATTGATAGGTTGATCACAGGCAGA   24562
gb:MN994468|Organism:Severe      CTTTCACGTCTTGACAAAGTTGAGGCTGAAGTGCAAATTGATAGGTTGATCACAGGCAGA   24562
gb:MT072688|Organism:Severe      CTTTCACGTCTTGACAAAGTTGAGGCTGAAGTGCAAATTGATAGGTTGATCACAGGCAGA   24547
gb:MN996527|Organism:Severe      CTTTCACGTCTTGACAAAGTTGAGGCTGAAGTGCAAATTGATAGGTTGATCACAGGCAGA   24529
gb:MT093631|Organism:Severe      CTTTCACGTCTTGACAAAGTTGAGGCTGAAGTGCAAATTGATAGGTTGATCACAGGCAGA   24600
gb:MT106053|Organism:Severe      CTTTCACGTCTTGACAAAGTTGAGGCTGAAGTGCAAATTGATAGGTTGATCACAGGCAGA   24562
gb:MT019533|Organism:Severe      CTTTCACGTCTTGACAAAGTTGAGGCTGAAGTGCAAATTGATAGGTTGATCACAGGCAGA   24562
gb:MT019531|Organism:Severe      CTTTCACGTCTTGACAAAGTTGAGGCTGAAGTGCAAATTGATAGGTTGATCACAGGCAGA   24562
gb:MN996528|Organism:Severe      CTTTCACGTCTTGACAAAGTTGAGGCTGAAGTGCAAATTGATAGGTTGATCACAGGCAGA   24562
gb:MN996530|Organism:Severe      CTTTCACGTCTTGACAAAGTTGAGGCTGAAGTGCAAATTGATAGGTTGATCACAGGCAGA   24548
gb:MN908947|Organism:Severe      CTTTCACGTCTTGACAAAGTTGAGGCTGAAGTGCAAATTGATAGGTTGATCACAGGCAGA   24562
gb:MT019532|Organism:Severe      CTTTCACGTCTTGACAAAGTTGAGGCTGAAGTGCAAATTGATAGGTTGATCACAGGCAGA   24562
                                 ************************************************************

gb:MT020781|Organism:Severe      CTTCAAAGTTTGCAGACATATGTGACTCAACAATTAATTAGAGCTGCAGAAATCAGAGCT   24610
gb:MT007544|Organism:Severe      CTTCAAAGTTTGCAGACATATGTGACTCAACAATTAATTAGAGCTGCAGAAATCAGAGCT   24622
gb:MN994467|Organism:Severe      CTTCAAAGTTTGCAGACATATGTGACTCAACAATTAATTAGAGCTGCAGAAATCAGAGCT   24622
gb:MT044257|Organism:Severe      CTTCAAAGTTTGCAGACATATGTGACTCAACAATTAATTAGAGCTGCAGAAATCAGAGCT   24622
gb:MT106054|Organism:Severe      CTTCAAAGTTTGCAGACATATGTGACTCAACAATTAATTAGAGCTGCAGAAATCAGAGCT   24622
gb:MT049951|Organism:Severe      CTTCAAAGTTTGCAGACATATGTGACTCAACAATTAATTAGAGCTGCAGAAATCAGAGCT   24622
gb:MN975262|Organism:Severe      CTTCAAAGTTTGCAGACATATGTGACTCAACAATTAATTAGAGCTGCAGAAATCAGAGCT   24622
gb:MT106052|Organism:Severe      CTTCAAAGTTTGCAGACATATGTGACTCAACAATTAATTAGAGCTGCAGAAATCAGAGCT   24622
gb:LC522975|Organism:Severe      CTTCAAAGTTTGCAGACATATGTGACTCAACAATTAATTAGAGCTGCAGAAATCAGAGCT   24619
gb:LC522973|Organism:Severe      CTTCAAAGTTTGCAGACATATGTGACTCAACAATTAATTAGAGCTGCAGAAATCAGAGCT   24619
gb:LC522974|Organism:Severe      CTTCAAAGTTTGCAGACATATGTGACTCAACAATTAATTAGAGCTGCAGAAATCAGAGCT   24619
gb:MN985325|Organism:Severe      CTTCAAAGTTTGCAGACATATGTGACTCAACAATTAATTAGAGCTGCAGAAATCAGAGCT   24622
gb:MT020881|Organism:Severe      CTTCAAAGTTTGCAGACATATGTGACTCAACAATTAATTAGAGCTGCAGAAATCAGAGCT   24622
gb:MT020880|Organism:Severe      CTTCAAAGTTTGCAGACATATGTGACTCAACAATTAATTAGAGCTGCAGAAATCAGAGCT   24622
gb:MT066175|Organism:Severe      CTTCAAAGTTTGCAGACATATGTGACTCAACAATTAATTAGAGCTGCAGAAATCAGAGCT   24622
gb:MN997409|Organism:Severe      CTTCAAAGTTTGCAGACATATGTGACTCAACAATTAATTAGAGCTGCAGAAATCAGAGCT   24622
gb:MN938384|Organism:Severe      CTTCAAAGTTTGCAGACATATGTGACTCAACAATTAATTAGAGCTGCAGAAATCAGAGCT   24590
gb:MT044258|Organism:Severe      CTTCAAAGTTTGCAGACATATGTGACTCAACAATTAATTAGAGCTGCAGAAATCAGAGCT   24598
gb:MT039890|Organism:Severe      CTTCAAAGTTTGCAGACATATGTGACTCAACAATTAATTAGAGCTGCAGAAATCAGAGCT   24622
gb:MN988713|Organism:Severe      CTTCAAAGTTTGCAGACATATGTGACTCAACAATTAATTAGAGCTGCAGAAATCAGAGCT   24622
gb:LC521925|Organism:Severe      CTTCAAAGTTTGCAGACATATGTGACTCAACAATTAATTAGAGCTGCAGAAATCAGAGCT   24595
gb:MT093571|Organism:Severe      CTTCAAAGTTTGCAGACATATGTGACTCAACAATTAATTAGAGCTGCAGAAATCAGAGCT   24622
gb:MT039887|Organism:Severe      CTTCAAAGTTTGCAGACATATGTGACTCAACAATTAATTAGAGCTGCAGAAATCAGAGCT   24619
gb:MT019530|Organism:Severe      CTTCAAAGTTTGCAGACATATGTGACTCAACAATTAATTAGAGCTGCAGAAATCAGAGCT   24622
gb:MT039888|Organism:Severe      CTTCAAAGTTTGCAGACATATGTGACTCAACAATTAATTAGAGCTGCAGAAATCAGAGCT   24622
gb:LC522972|Organism:Severe      CTTCAAAGTTTGCAGACATATGTGACTCAACAATTAATTAGAGCTGCAGAAATCAGAGCT   24619
gb:MT027063|Organism:Severe      CTTCAAAGTTTGCAGACATATGTGACTCAACAATTAATTAGAGCTGCAGAAATCAGAGCT   24622
gb:MT027062|Organism:Severe      CTTCAAAGTTTGCAGACATATGTGACTCAACAATTAATTAGAGCTGCAGAAATCAGAGCT   24622
gb:MT019529|Organism:Severe      CTTCAAAGTTTGCAGACATATGTGACTCAACAATTAATTAGAGCTGCAGAAATCAGAGCT   24622
gb:MN996529|Organism:Severe      CTTCAAAGTTTGCAGACATATGTGACTCAACAATTAATTAGAGCTGCAGAAATCAGAGCT   24610
gb:MN996531|Organism:Severe      CTTCAAAGTTTGCAGACATATGTGACTCAACAATTAATTAGAGCTGCAGAAATCAGAGCT   24609
gb:MT066176|Organism:Severe      CTTCAAAGTTTGCAGACATATGTGACTCAACAATTAATTAGAGCTGCAGAAATCAGAGCT   24622
gb:MT027064|Organism:Severe      CTTCAAAGTTTGCAGACATATGTGACTCAACAATTAATTAGAGCTGCAGAAATCAGAGCT   24622
gb:MN994468|Organism:Severe      CTTCAAAGTTTGCAGACATATGTGACTCAACAATTAATTAGAGCTGCAGAAATCAGAGCT   24622
gb:MT072688|Organism:Severe      CTTCAAAGTTTGCAGACATATGTGACTCAACAATTAATTAGAGCTGCAGAAATCAGAGCT   24607
gb:MN996527|Organism:Severe      CTTCAAAGTTTGCAGACATATGTGACTCAACAATTAATTAGAGCTGCAGAAATCAGAGCT   24589
gb:MT093631|Organism:Severe      CTTCAAAGTTTGCAGACATATGTGACTCAACAATTAATTAGAGCTGCAGAAATCAGAGCT   24660
gb:MT106053|Organism:Severe      CTTCAAAGTTTGCAGACATATGTGACTCAACAATTAATTAGAGCTGCAGAAATCAGAGCT   24622
gb:MT019533|Organism:Severe      CTTCAAAGTTTGCAGACATATGTGACTCAACAATTAATTAGAGCTGCAGAAATCAGAGCT   24622
gb:MT019531|Organism:Severe      CTTCAAAGTTTGCAGACATATGTGACTCAACAATTAATTAGAGCTGCAGAAATCAGAGCT   24622
gb:MN996528|Organism:Severe      CTTCAAAGTTTGCAGACATATGTGACTCAACAATTAATTAGAGCTGCAGAAATCAGAGCT   24622
gb:MN996530|Organism:Severe      CTTCAAAGTTTGCAGACATATGTGACTCAACAATTAATTAGAGCTGCAGAAATCAGAGCT   24608
gb:MN908947|Organism:Severe      CTTCAAAGTTTGCAGACATATGTGACTCAACAATTAATTAGAGCTGCAGAAATCAGAGCT   24622
gb:MT019532|Organism:Severe      CTTCAAAGTTTGCAGACATATGTGACTCAACAATTAATTAGAGCTGCAGAAATCAGAGCT   24622
                                 ************************************************************

gb:MT020781|Organism:Severe      TCTGCTAATCTTGCTGCTACTAAAATGTCAGAGTGTGTACTTGGACAATCAAAAAGAGTT   24670
gb:MT007544|Organism:Severe      TCTGCTAATCTTGCTGCTACTAAAATGTCAGAGTGTGTACTTGGACAATCAAAAAGAGTT   24682
gb:MN994467|Organism:Severe      TCTGCTAATCTTGCTGCTACTAAAATGTCAGAGTGTGTACTTGGACAATCAAAAAGAGTT   24682
gb:MT044257|Organism:Severe      TCTGCTAATCTTGCTGCTACTAAAATGTCAGAGTGTGTACTTGGACAATCAAAAAGAGTT   24682
gb:MT106054|Organism:Severe      TCTGCTAATCTTGCTGCTACTAAAATGTCAGAGTGTGTACTTGGACAATCAAAAAGAGTT   24682
```

```
gb:MT049951|Organism:Severe      TCTGCTAATCTTGCTGCTACTAAAATGTCAGAGTGTGTACTTGGACAATCAAAAAGAGTT    24682
gb:MN975262|Organism:Severe      TCTGCTAATCTTGCTGCTACTAAAATGTCAGAGTGTGTACTTGGACAATCAAAAAGAGTT    24682
gb:MT106052|Organism:Severe      TCTGCTAATCTTGCTGCTACTAAAATGTCAGAGTGTGTACTTGGACAATCAAAAAGAGTT    24682
gb:LC522975|Organism:Severe      TCTGCTAATCTTGCTGCTACTAAAATGTCAGAGTGTGTACTTGGACAATCAAAAAGAGTT    24679
gb:LC522973|Organism:Severe      TCTGCTAATCTTGCTGCTACTAAAATGTCAGAGTGTGTACTTGGACAATCAAAAAGAGTT    24679
gb:LC522974|Organism:Severe      TCTGCTAATCTTGCTGCTACTAAAATGTCAGAGTGTGTACTTGGACAATCAAAAAGAGTT    24679
gb:MN985325|Organism:Severe      TCTGCTAATCTTGCTGCTACTAAAATGTCAGAGTGTGTACTTGGACAATCAAAAAGAGTT    24682
gb:MT020881|Organism:Severe      TCTGCTAATCTTGCTGCTACTAAAATGTCAGAGTGTGTACTTGGACAATCAAAAAGAGTT    24682
gb:MT020880|Organism:Severe      TCTGCTAATCTTGCTGCTACTAAAATGTCAGAGTGTGTACTTGGACAATCAAAAAGAGTT    24682
gb:MT066175|Organism:Severe      TCTGCTAATCTTGCTGCTACTAAAATGTCAGAGTGTGTACTTGGACAATCAAAAAGAGTT    24682
gb:MN997409|Organism:Severe      TCTGCTAATCTTGCTGCTACTAAAATGTCAGAGTGTGTACTTGGACAATCAAAAAGAGTT    24682
gb:MN938384|Organism:Severe      TCTGCTAATCTTGCTGCTACTAAAATGTCAGAGTGTGTACTTGGACAATCAAAAAGAGTT    24650
gb:MT044258|Organism:Severe      TCTGCTAATCTTGCTGCTACTAAAATGTCAGAGTGTGTACTTGGACAATCAAAAAGAGTT    24658
gb:MT039890|Organism:Severe      TCTGCTAATCTTGCTGCTACTAAAATGTCAGAGTGTGTACTTGGACAATCAAAAAGAGTT    24682
gb:MN988713|Organism:Severe      TCTGCTAATCTTGCTGCTACTAAAATGTCAGAGTGTGTACTTGGACAATCAAAAAGAGTT    24682
gb:LC521925|Organism:Severe      TCTGCTAATCTTGCTGCTACTAAAATGTCAGAGTGTGTACTTGGACAATCAAAAAGAGTT    24655
gb:MT093571|Organism:Severe      TCTGCTAATCTTGCTGCTACTAAAATGTCAGAGTGTGTACTTGGACAATCAAAAAGAGTT    24682
gb:MT039887|Organism:Severe      TCTGCTAATCTTGCTGCTACTAAAATGTCAGAGTGTGTACTTGGACAATCAAAAAGAGTT    24679
gb:MT019530|Organism:Severe      TCTGCTAATCTTGCTGCTACTAAAATGTCAGAGTGTGTACTTGGACAATCAAAAAGAGTT    24682
gb:MT039888|Organism:Severe      TCTGCTAATCTTGCTGCTACTAAAATGTCAGAGTGTGTACTTGGACAATCAAAAAGAGTT    24682
gb:LC522972|Organism:Severe      TCTGCTAATCTTGCTGCTACTAAAATGTCAGAGTGTGTACTTGGACAATCAAAAAGAGTT    24679
gb:MT027063|Organism:Severe      TCTGCTAATCTTGCTGCTACTAAAATGTCAGAGTGTGTACTTGGACAATCAAAAAGAGTT    24682
gb:MT027062|Organism:Severe      TCTGCTAATCTTGCTGCTACTAAAATGTCAGAGTGTGTACTTGGACAATCAAAAAGAGTT    24682
gb:MT019529|Organism:Severe      TCTGCTAATCTTGCTGCTACTAAAATGTCAGAGTGTGTACTTGGACAATCAAAAAGAGTT    24682
gb:MN996529|Organism:Severe      TCTGCTAATCTTGCTGCTACTAAAATGTCAGAGTGTGTACTTGGACAATCAAAAAGAGTT    24670
gb:MN996531|Organism:Severe      TCTGCTAATCTTGCTGCTACTAAAATGTCAGAGTGTGTACTTGGACAATCAAAAAGAGTT    24669
gb:MT066176|Organism:Severe      TCTGCTAATCTTGCTGCTACTAAAATGTCAGAGTGTGTACTTGGACAATCAAAAAGAGTT    24682
gb:MT027064|Organism:Severe      TCTGCTAATCTTGCTGCTACTAAAATGTCAGAGTGTGTACTTGGACAATCAAAAAGAGTT    24682
gb:MN994468|Organism:Severe      TCTGCTAATCTTGCTGCTACTAAAATGTCAGAGTGTGTACTTGGACAATCAAAAAGAGTT    24682
gb:MT072688|Organism:Severe      TCTGCTAATCTTGCTGCTACTAAAATGTCAGAGTGTGTACTTGGACAATCAAAAAGAGTT    24667
gb:MN996527|Organism:Severe      TCTGCTAATCTTGCTGCTACTAAAATGTCAGAGTGTGTACTTGGACAATCAAAAAGAGTT    24649
gb:MT093631|Organism:Severe      TCTGCTAATCTTGCTGCTACTAAAATGTCAGAGTGTGTACTTGGACAATCAAAAAGAGTT    24720
gb:MT106053|Organism:Severe      TCTGCTAATCTTGCTGCTACTAAAATGTCAGAGTGTGTACTTGGACAATCAAAAAGAGTT    24682
gb:MT019533|Organism:Severe      TCTGCTAATCTTGCTGCTACTAAAATGTCAGAGTGTGTACTTGGACAATCAAAAAGAGTT    24682
gb:MT019531|Organism:Severe      TCTGCTAATCTTGCTGCTACTAAAATGTCAGAGTGTGTACTTGGACAATCAAAAAGAGTT    24682
gb:MN996528|Organism:Severe      TCTGCTAATCTTGCTGCTACTAAAATGTCAGAGTGTGTACTTGGACAATCAAAAAGAGTT    24682
gb:MN996530|Organism:Severe      TCTGCTAATCTTGCTGCTACTAAAATGTCAGAGTGTGTACTTGGACAATCAAAAAGAGTT    24668
gb:MN908947|Organism:Severe      TCTGCTAATCTTGCTGCTACTAAAATGTCAGAGTGTGTACTTGGACAATCAAAAAGAGTT    24682
gb:MT019532|Organism:Severe      TCTGCTAATCTTGCTGCTACTAAAATGTCAGAGTGTGTACTTGGACAATCAAAAAGAGTT    24682
                                 ************************************************************

gb:MT020781|Organism:Severe      GATTTTTGTGGAAAGGGCTATCATCTTATGTCCTTCCCTCAGTCAGCACCTCATGGTGTA    24730
gb:MT007544|Organism:Severe      GATTTTTGTGGAAAGGGCTATCATCTTATGTCCTTCCCTCAGTCAGCACCTCATGGTGTA    24742
gb:MN994467|Organism:Severe      GATTTTTGTGGAAAGGGCTATCATCTTATGTCCTTCCCTCAGTCAGCACCTCATGGTGTA    24742
gb:MT044257|Organism:Severe      GATTTTTGTGGAAAGGGCTATCATCTTATGTCCTTCCCTCAGTCAGCACCTCATGGTGTA    24742
gb:MT106054|Organism:Severe      GATTTTTGTGGAAAGGGCTATCATCTTATGTCCTTCCCTCAGTCAGCACCTCATGGTGTA    24742
gb:MT049951|Organism:Severe      GATTTTTGTGGAAAGGGCTATCATCTTATGTCCTTCCCTCAGTCAGCACCTCATGGTGTA    24742
gb:MN975262|Organism:Severe      GATTTTTGTGGAAAGGGCTATCATCTTATGTCCTTCCCTCAGTCAGCACCTCATGGTGTA    24742
gb:MT106052|Organism:Severe      GATTTTTGTGGAAAGGGCTATCATCTTATGTCCTTCCCTCAGTCAGCACCTCATGGTGTA    24742
gb:LC522975|Organism:Severe      GATTTTTGTGGAAAGGGCTATCATCTTATGTCCTTCCCTCAGTCAGCACCTCATGGTGTA    24739
gb:LC522973|Organism:Severe      GATTTTTGTGGAAAGGGCTATCATCTTATGTCCTTCCCTCAGTCAGCACCTCATGGTGTA    24739
gb:LC522974|Organism:Severe      GATTTTTGTGGAAAGGGCTATCATCTTATGTCCTTCCCTCAGTCAGCACCTCATGGTGTA    24739
gb:MN985325|Organism:Severe      GATTTTTGTGGAAAGGGCTATCATCTTATGTCCTTCCCTCAGTCAGCACCTCATGGTGTA    24742
gb:MT020881|Organism:Severe      GATTTTTGTGGAAAGGGCTATCATCTTATGTCCTTCCCTCAGTCAGCACCTCATGGTGTA    24742
gb:MT020880|Organism:Severe      GATTTTTGTGGAAAGGGCTATCATCTTATGTCCTTCCCTCAGTCAGCACCTCATGGTGTA    24742
gb:MT066175|Organism:Severe      GATTTTTGTGGAAAGGGCTATCATCTTATGTCCTTCCCTCAGTCAGCACCTCATGGTGTA    24742
gb:MN997409|Organism:Severe      GATTTTTGTGGAAAGGGCTATCATCTTATGTCCTTCCCTCAGTCAGCACCTCATGGTGTA    24742
gb:MN938384|Organism:Severe      GATTTTTGTGGAAAGGGCTATCATCTTATGTCCTTCCCTCAGTCAGCACCTCATGGTGTA    24710
gb:MT044258|Organism:Severe      GATTTTTGTGGAAAGGGCTATCATCTTATGTCCTTCCCTCAGTCAGCACCTCATGGTGTA    24718
gb:MT039890|Organism:Severe      GATTTTTGTGGAAAGGGCTATCATCTTATGTCCTTCCCTCAGTCAGCACCTCATGGTGTA    24742
gb:MN988713|Organism:Severe      GATTTTTGTGGAAAGGGCTATCATCTTATGTCCTTCCCTCAGTCAGCACCTCATGGTGTA    24742
gb:LC521925|Organism:Severe      GATTTTTGTGGAAAGGGCTATCATCTTATGTCCTTCCCTCAGTCAGCACCTCATGGTGTA    24715
gb:MT093571|Organism:Severe      GATTTTTGTGGAAAGGGCTATCATCTTATGTCCTTCCCTCAGTCAGCACCTCATGGTGTA    24742
gb:MT039887|Organism:Severe      GATTTTTGTGGAAAGGGCTATCATCTTATGTCCTTCCCTCAGTCAGCACCTCATGGTGTA    24739
gb:MT019530|Organism:Severe      GATTTTTGTGGAAAGGGCTATCATCTTATGTCCTTCCCTCAGTCAGCACCTCATGGTGTA    24742
```

```
gb:MT039888|Organism:Severe      GATTTTTGTGGAAAGGGCTATCATCTTATGTCCTTCCCTCAGTCAGCACCTCATGGTGTA   24742
gb:LC522972|Organism:Severe      GATTTTTGTGGAAAGGGCTATCATCTTATGTCCTTCCCTCAGTCAGCACCTCATGGTGTA   24739
gb:MT027063|Organism:Severe      GATTTTTGTGGAAAGGGCTATCATCTTATGTCCTTCCCTCAGTCAGCACCTCATGGTGTA   24742
gb:MT027062|Organism:Severe      GATTTTTGTGGAAAGGGCTATCATCTTATGTCCTTCCCTCAGTCAGCACCTCATGGTGTA   24742
gb:MT019529|Organism:Severe      GATTTTTGTGGAAAGGGCTATCATCTTATGTCCTTCCCTCAGTCAGCACCTCATGGTGTA   24742
gb:MN996529|Organism:Severe      GATTTTTGTGGAAAGGGCTATCATCTTATGTCCTTCCCTCAGTCAGCACCTCATGGTGTA   24730
gb:MN996531|Organism:Severe      GATTTTTGTGGAAAGGGCTATCATCTTATGTCCTTCCCTCAGTCAGCACCTCATGGTGTA   24729
gb:MT066176|Organism:Severe      GATTTTTGTGGAAAGGGCTATCATCTTATGTCCTTCCCTCAGTCAGCACCTCATGGTGTA   24742
gb:MT027064|Organism:Severe      GATTTTTGTGGAAAGGGCTATCATCTTATGTCCTTCCCTCAGTCAGCACCTCATGGTGTA   24742
gb:MN994468|Organism:Severe      GATTTTTGTGGAAAGGGCTATCATCTTATGTCCTTCCCTCAGTCAGCACCTCATGGTGTA   24742
gb:MT072688|Organism:Severe      GATTTTTGTGGAAAGGGCTATCATCTTATGTCCTTCCCTCAGTCAGCACCTCATGGTGTA   24727
gb:MN996527|Organism:Severe      GATTTTTGTGGAAAGGGCTATCATCTTATGTCCTTCCCTCAGTCAGCACCTCATGGTGTA   24709
gb:MT093631|Organism:Severe      GATTTTTGTGGAAAGGGCTATCATCTTATGTCCTTCCCTCAGTCAGCACCTCATGGTGTA   24780
gb:MT106053|Organism:Severe      GATTTTTGTGGAAAGGGCTATCATCTTATGTCCTTCCCTCAGTCAGCACCTCATGGTGTA   24742
gb:MT019533|Organism:Severe      GATTTTTGTGGAAAGGGCTATCATCTTATGTCCTTCCCTCAGTCAGCACCTCATGGTGTA   24742
gb:MT019531|Organism:Severe      GATTTTTGTGGAAAGGGCTATCATCTTATGTCCTTCCCTCAGTCAGCACCTCATGGTGTA   24742
gb:MN996528|Organism:Severe      GATTTTTGTGGAAAGGGCTATCATCTTATGTCCTTCCCTCAGTCAGCACCTCATGGTGTA   24742
gb:MN996530|Organism:Severe      GATTTTTGTGGAAAGGGCTATCATCTTATGTCCTTCCCTCAGTCAGCACCTCATGGTGTA   24728
gb:MN908947|Organism:Severe      GATTTTTGTGGAAAGGGCTATCATCTTATGTCCTTCCCTCAGTCAGCACCTCATGGTGTA   24742
gb:MT019532|Organism:Severe      GATTTTTGTGGAAAGGGCTATCATCTTATGTCCTTCCCTCAGTCAGCACCTCATGGTGTA   24742
                                 ************************************************************

gb:MT020781|Organism:Severe      GTCTTCTTGCATGTGACTTATGTCCCTGCACAAGAAAAGAACTTCACAACTGCTCCTGCC   24790
gb:MT007544|Organism:Severe      GTCTTCTTGCATGTGACTTATGTCCCTGCACAAGAAAAGAACTTCACAACTGCTCCTGCC   24802
gb:MN994467|Organism:Severe      GTCTTCTTGCATGTGACTTATGTCCCTGCACAAGAAAAGAACTTCACAACTGCTCCTGCC   24802
gb:MT044257|Organism:Severe      GTCTTCTTGCATGTGACTTATGTCCCTGCACAAGAAAAGAACTTCACAACTGCTCCTGCC   24802
gb:MT106054|Organism:Severe      GTCTTCTTGCATGTGACTTATGTCCCTGCACAAGAAAAGAACTTCACAACTGCTCCTGCC   24802
gb:MT049951|Organism:Severe      GTCTTCTTGCATGTGACTTATGTCCCTGCACAAGAAAAGAACTTCACAACTGCTCCTGCC   24802
gb:MN975262|Organism:Severe      GTCTTCTTGCATGTGACTTATGTCCCTGCACAAGAAAAGAACTTCACAACTGCTCCTGCC   24802
gb:MT106052|Organism:Severe      GTCTTCTTGCATGTGACTTATGTCCCTGCACAAGAAAAGAACTTCACAACTGCTCCTGCC   24802
gb:LC522975|Organism:Severe      GTCTTCTTGCATGTGACTTATGTCCCTGCACAAGAAAAGAACTTCACAACTGCTCCTGCC   24799
gb:LC522973|Organism:Severe      GTCTTCTTGCATGTGACTTATGTCCCTGCACAAGAAAAGAACTTCACAACTGCTCCTGCC   24799
gb:LC522974|Organism:Severe      GTCTTCTTGCATGTGACTTATGTCCCTGCACAAGAAAAGAACTTCACAACTGCTCCTGCC   24799
gb:MN985325|Organism:Severe      GTCTTCTTGCATGTGACTTATGTCCCTGCACAAGAAAAGAACTTCACAACTGCTCCTGCC   24802
gb:MT020881|Organism:Severe      GTCTTCTTGCATGTGACTTATGTCCCTGCACAAGAAAAGAACTTCACAACTGCTCCTGCC   24802
gb:MT020880|Organism:Severe      GTCTTCTTGCATGTGACTTATGTCCCTGCACAAGAAAAGAACTTCACAACTGCTCCTGCC   24802
gb:MT066175|Organism:Severe      GTCTTCTTGCATGTGACTTATGTCCCTGCACAAGAAAAGAACTTCACAACTGCTCCTGCC   24802
gb:MN997409|Organism:Severe      GTCTTCTTGCATGTGACTTATGTCCCTGCACAAGAAAAGAACTTCACAACTGCTCCTGCC   24802
gb:MN938384|Organism:Severe      GTCTTCTTGCATGTGACTTATGTCCCTGCACAAGAAAAGAACTTCACAACTGCTCCTGCC   24770
gb:MT044258|Organism:Severe      GTCTTCTTGCATGTGACTTATGTCCCTGCACAAGAAAAGAACTTCACAACTGCTCCTGCC   24778
gb:MT039890|Organism:Severe      GTCTTCTTGCATGTGACTTATGTCCCTGCACAAGAAAAGAACTTCACAACTGCTCCTGCC   24802
gb:MN988713|Organism:Severe      GTCTTCTTGCATGTGACTTATGTCCCTGCACAAGAAAAGAACTTCACAACTGCTCCTGCC   24802
gb:LC521925|Organism:Severe      GTCTTCTTGCATGTGACTTATGTCCCTGCACAAGAAAAGAACTTCACAACTGCTCCTGCC   24775
gb:MT093571|Organism:Severe      GTCTTCTTGCATGTGACTTATGTCCCTGCACAAGAAAAGAACTTCACAACTGCTCCTGCC   24802
gb:MT039887|Organism:Severe      GTCTTCTTGCATGTGACTTATGTCCCTGCACAAGAAAAGAACTTCACAACTGCTCCTGCC   24799
gb:MT019530|Organism:Severe      GTCTTCTTGCATGTGACTTATGTCCCTGCACAAGAAAAGAACTTCACAACTGCTCCTGCC   24802
gb:MT039888|Organism:Severe      GTCTTCTTGCATGTGACTTATGTCCCTGCACAAGAAAAGAACTTCACAACTGCTCCTGCC   24802
gb:LC522972|Organism:Severe      GTCTTCTTGCATGTGACTTATGTCCCTGCACAAGAAAAGAACTTCACAACTGCTCCTGCC   24799
gb:MT027063|Organism:Severe      GTCTTCTTGCATGTGACTTATGTCCCTGCACAAGAAAAGAACTTCACAACTGCTCCTGCC   24802
gb:MT027062|Organism:Severe      GTCTTCTTGCATGTGACTTATGTCCCTGCACAAGAAAAGAACTTCACAACTGCTCCTGCC   24802
gb:MT019529|Organism:Severe      GTCTTCTTGCATGTGACTTATGTCCCTGCACAAGAAAAGAACTTCACAACTGCTCCTGCC   24802
gb:MN996529|Organism:Severe      GTCTTCTTGCATGTGACTTATGTCCCTGCACAAGAAAAGAACTTCACAACTGCTCCTGCC   24790
gb:MN996531|Organism:Severe      GTCTTCTTGCATGTGACTTATGTCCCTGCACAAGAAAAGAACTTCACAACTGCTCCTGCC   24789
gb:MT066176|Organism:Severe      GTCTTCTTGCATGTGACTTATGTCCCTGCACAAGAAAAGAACTTCACAACTGCTCCTGCC   24802
gb:MT027064|Organism:Severe      GTCTTCTTGCATGTGACTTATGTCCCTGCACAAGAAAAGAACTTCACAACTGCTCCTGCC   24802
gb:MN994468|Organism:Severe      GTCTTCTTGCATGTGACTTATGTCCCTGCACAAGAAAAGAACTTCACAACTGCTCCTGCC   24802
gb:MT072688|Organism:Severe      GTCTTCTTGCATGTGACTTATGTCCCTGCACAAGAAAAGAACTTCACAACTGCTCCTGCC   24787
gb:MN996527|Organism:Severe      GTCTTCTTGCATGTGACTTATGTCCCTGCACAAGAAAAGAACTTCACAACTGCTCCTGCC   24769
gb:MT093631|Organism:Severe      GTCTTCTTGCATGTGACTTATGTCCCTGCACAAGAAAAGAACTTCACAACTGCTCCTGCC   24840
gb:MT106053|Organism:Severe      GTCTTCTTGCATGTGACTTATGTCCCTGCACAAGAAAAGAACTTCACAACTGCTCCTGCC   24802
gb:MT019533|Organism:Severe      GTCTTCTTGCATGTGACTTATGTCCCTGCACAAGAAAAGAACTTCACAACTGCTCCTGCC   24802
gb:MT019531|Organism:Severe      GTCTTCTTGCATGTGACTTATGTCCCTGCACAAGAAAAGAACTTCACAACTGCTCCTGCC   24802
gb:MN996528|Organism:Severe      GTCTTCTTGCATGTGACTTATGTCCCTGCACAAGAAAAGAACTTCACAACTGCTCCTGCC   24802
gb:MN996530|Organism:Severe      GTCTTCTTGCATGTGACTTATGTCCCTGCACAAGAAAAGAACTTCACAACTGCTCCTGCC   24788
gb:MN908947|Organism:Severe      GTCTTCTTGCATGTGACTTATGTCCCTGCACAAGAAAAGAACTTCACAACTGCTCCTGCC   24802
```

```
gb:MT019532|Organism:Severe      GTCTTCTTGCATGTGACTTATGTCCCTGCACAAGAAAAGAACTTCACAACTGCTCCTGCC    24802
                                 ************************************************************

gb:MT020781|Organism:Severe      ATTTGTCATGATGGAAAAGCACACTTTCCTCGTGAAGGTGTCTTTGTTTCAAATGGCACA    24850
gb:MT007544|Organism:Severe      ATTTGTCATGATGGAAAAGCACACTTTCCTCGTGAAGGTGTCTTTGTTTCAAATGGCACA    24862
gb:MN994467|Organism:Severe      ATTTGTCATGATGGAAAAGCACACTTTCCTCGTGAAGGTGTCTTTGTTTCAAATGGCACA    24862
gb:MT044257|Organism:Severe      ATTTGTCATGATGGAAAAGCACACTTTCCTCGTGAAGGTGTCTTTGTTTCAAATGGCACA    24862
gb:MT106054|Organism:Severe      ATTTGTCATGATGGAAAAGCACACTTTCCTCGTGAAGGTGTCTTTGTTTCAAATGGCACA    24862
gb:MT049951|Organism:Severe      ATTTGTCATGATGGAAAAGCACACTTTCCTCGTGAAGGTGTCTTTGTTTCAAATGGCACA    24862
gb:MN975262|Organism:Severe      ATTTGTCATGATGGAAAAGCACACTTTCCTCGTGAAGGTGTCTTTGTTTCAAATGGCACA    24862
gb:MT106052|Organism:Severe      ATTTGTCATGATGGAAAAGCACACTTTCCTCGTGAAGGTGTCTTTGTTTCAAATGGCACA    24862
gb:LC522975|Organism:Severe      ATTTGTCATGATGGAAAAGCACACTTTCCTCGTGAAGGTGTCTTTGTTTCAAATGGCACA    24859
gb:LC522973|Organism:Severe      ATTTGTCATGATGGAAAAGCACACTTTCCTCGTGAAGGTGTCTTTGTTTCAAATGGCACA    24859
gb:LC522974|Organism:Severe      ATTTGTCATGATGGAAAAGCACACTTTCCTCGTGAAGGTGTCTTTGTTTCAAATGGCACA    24859
gb:MN985325|Organism:Severe      ATTTGTCATGATGGAAAAGCACACTTTCCTCGTGAAGGTGTCTTTGTTTCAAATGGCACA    24862
gb:MT020881|Organism:Severe      ATTTGTCATGATGGAAAAGCACACTTTCCTCGTGAAGGTGTCTTTGTTTCAAATGGCACA    24862
gb:MT020880|Organism:Severe      ATTTGTCATGATGGAAAAGCACACTTTCCTCGTGAAGGTGTCTTTGTTTCAAATGGCACA    24862
gb:MT066175|Organism:Severe      ATTTGTCATGATGGAAAAGCACACTTTCCTCGTGAAGGTGTCTTTGTTTCAAATGGCACA    24862
gb:MN997409|Organism:Severe      ATTTGTCATGATGGAAAAGCACACTTTCCTCGTGAAGGTGTCTTTGTTTCAAATGGCACA    24862
gb:MN938384|Organism:Severe      ATTTGTCATGATGGAAAAGCACACTTTCCTCGTGAAGGTGTCTTTGTTTCAAATGGCACA    24830
gb:MT044258|Organism:Severe      ATTTGTCATGATGGAAAAGCACACTTTCCTCGTGAAGGTGTCTTTGTTTCAAATGGCACA    24838
gb:MT039890|Organism:Severe      ATTTGTCATGATGGAAAAGCACACTTTCCTCGTGAAGGTGTCTTTGTTTCAAATGGCACA    24862
gb:MN988713|Organism:Severe      ATTTGTCATGATGGAAAAGCACACTTTCCTCGTGAAGGTGTCTTTGTTTCAAATGGCACA    24862
gb:LC521925|Organism:Severe      ATTTGTCATGATGGAAAAGCACACTTTCCTCGTGAAGGTGTCTTTGTTTCAAATGGCACA    24835
gb:MT093571|Organism:Severe      ATTTGTCATGATGGAAAAGCACACTTTCCTCGTGAAGGTGTCTTTGTTTCAAATGGCACA    24862
gb:MT039887|Organism:Severe      ATTTGTCATGATGGAAAAGCACACTTTCCTCGTGAAGGTGTCTTTGTTTCAAATGGCACA    24859
gb:MT019530|Organism:Severe      ATTTGTCATGATGGAAAAGCACACTTTCCTCGTGAAGGTGTCTTTGTTTCAAATGGCACA    24862
gb:MT039888|Organism:Severe      ATTTGTCATGATGGAAAAGCACACTTTCCTCGTGAAGGTGTCTTTGTTTCAAATGGCACA    24862
gb:LC522972|Organism:Severe      ATTTGTCATGATGGAAAAGCACACTTTCCTCGTGAAGGTGTCTTTGTTTCAAATGGCACA    24859
gb:MT027063|Organism:Severe      ATTTGTCATGATGGAAAAGCACACTTTCCTCGTGAAGGTGTCTTTGTTTCAAATGGCACA    24862
gb:MT027062|Organism:Severe      ATTTGTCATGATGGAAAAGCACACTTTCCTCGTGAAGGTGTCTTTGTTTCAAATGGCACA    24862
gb:MT019529|Organism:Severe      ATTTGTCATGATGGAAAAGCACACTTTCCTCGTGAAGGTGTCTTTGTTTCAAATGGCACA    24862
gb:MN996529|Organism:Severe      ATTTGTCATGATGGAAAAGCACACTTTCCTCGTGAAGGTGTCTTTGTTTCAAATGGCACA    24850
gb:MN996531|Organism:Severe      ATTTGTCATGATGGAAAAGCACACTTTCCTCGTGAAGGTGTCTTTGTTTCAAATGGCACA    24849
gb:MT066176|Organism:Severe      ATTTGTCATGATGGAAAAGCACACTTTCCTCGTGAAGGTGTCTTTGTTTCAAATGGCACA    24862
gb:MT027064|Organism:Severe      ATTTGTCATGATGGAAAAGCACACTTTCCTCGTGAAGGTGTCTTTGTTTCAAATGGCACA    24862
gb:MN994468|Organism:Severe      ATTTGTCATGATGGAAAAGCACACTTTCCTCGTGAAGGTGTCTTTGTTTCAAATGGCACA    24862
gb:MT072688|Organism:Severe      ATTTGTCATGATGGAAAAGCACACTTTCCTCGTGAAGGTGTCTTTGTTTCAAATGGCACA    24847
gb:MN996527|Organism:Severe      ATTTGTCATGATGGAAAAGCACACTTTCCTCGTGAAGGTGTCTTTGTTTCAAATGGCACA    24829
gb:MT093631|Organism:Severe      ATTTGTCATGATGGAAAAGCACACTTTCCTCGTGAAGGTGTCTTTGTTTCAAATGGCACA    24900
gb:MT106053|Organism:Severe      ATTTGTCATGATGGAAAAGCACACTTTCCTCGTGAAGGTGTCTTTGTTTCAAATGGCACA    24862
gb:MT019533|Organism:Severe      ATTTGTCATGATGGAAAAGCACACTTTCCTCGTGAAGGTGTCTTTGTTTCAAATGGCACA    24862
gb:MT019531|Organism:Severe      ATTTGTCATGATGGAAAAGCACACTTTCCTCGTGAAGGTGTCTTTGTTTCAAATGGCACA    24862
gb:MN996528|Organism:Severe      ATTTGTCATGATGGAAAAGCACACTTTCCTCGTGAAGGTGTCTTTGTTTCAAATGGCACA    24862
gb:MN996530|Organism:Severe      ATTTGTCATGATGGAAAAGCACACTTTCCTCGTGAAGGTGTCTTTGTTTCAAATGGCACA    24848
gb:MN908947|Organism:Severe      ATTTGTCATGATGGAAAAGCACACTTTCCTCGTGAAGGTGTCTTTGTTTCAAATGGCACA    24862
gb:MT019532|Organism:Severe      ATTTGTCATGATGGAAAAGCACACTTTCCTCGTGAAGGTGTCTTTGTTTCAAATGGCACA    24862
                                 ************************************************************

gb:MT020781|Organism:Severe      CACTGGTTTGTAACACAAAGGAATTTTTATGAACCACAAATCATTACTACAGACAACACA    24910
gb:MT007544|Organism:Severe      CACTGGTTTGTAACACAAAGGAATTTTTATGAACCACAAATCATTACTACAGACAACACA    24922
gb:MN994467|Organism:Severe      CACTGGTTTGTAACACAAAGGAATTTTTATGAACCACAAATCATTACTACAGACAACACA    24922
gb:MT044257|Organism:Severe      CACTGGTTTGTAACACAAAGGAATTTTTATGAACCACAAATCATTACTACAGACAACACA    24922
gb:MT106054|Organism:Severe      CACTGGTTTGTAACACAAAGGAATTTTTATGAACCACAAATCATTACTACAGACAACACA    24922
gb:MT049951|Organism:Severe      CACTGGTTTGTAACACAAAGGAATTTTTATGAACCACAAATCATTACTACAGACAACACA    24922
gb:MN975262|Organism:Severe      CACTGGTTTGTAACACAAAGGAATTTTTATGAACCACAAATCATTACTACAGACAACACA    24922
gb:MT106052|Organism:Severe      CACTGGTTTGTAACACAAAGGAATTTTTATGAACCACAAATCATTACTACAGACAACACA    24922
gb:LC522975|Organism:Severe      CACTGGTTTGTAACACAAAGGAATTTTTATGAACCACAAATCATTACTACAGACAACACA    24919
gb:LC522973|Organism:Severe      CACTGGTTTGTAACACAAAGGAATTTTTATGAACCACAAATCATTACTACAGACAACACA    24919
gb:LC522974|Organism:Severe      CACTGGTTTGTAACACAAAGGAATTTTTATGAACCACAAATCATTACTACAGACAACACA    24919
gb:MN985325|Organism:Severe      CACTGGTTTGTAACACAAAGGAATTTTTATGAACCACAAATCATTACTACAGACAACACA    24922
gb:MT020881|Organism:Severe      CACTGGTTTGTAACACAAAGGAATTTTTATGAACCACAAATCATTACTACAGACAACACA    24922
gb:MT020880|Organism:Severe      CACTGGTTTGTAACACAAAGGAATTTTTATGAACCACAAATCATTACTACAGACAACACA    24922
gb:MT066175|Organism:Severe      CACTGGTTTGTAACACAAAGGAATTTTTATGAACCACAAATCATTACTACAGACAACACA    24922
gb:MN997409|Organism:Severe      CACTGGTTTGTAACACAAAGGAATTTTTATGAACCACAAATCATTACTACAGACAACACA    24922
```

```
gb:MN938384|Organism:Severe      CACTGGTTTGTAACACAAAGGAATTTTTATGAACCACAAATCATTACTACAGACAACACA   24890
gb:MT044258|Organism:Severe      CACTGGTTTGTAACACAAAGGAATTTTTATGAACCACAAATCATTACTACAGACAACACA   24898
gb:MT039890|Organism:Severe      CACTGGTTTGTAACACAAAGGAATTTTTATGAACCACAAATCATTACTACAGACAACACA   24922
gb:MN988713|Organism:Severe      CACTGGTTTGTAACACAAAGGAATTTTTATGAACCACAAATCATTACTACAGACAACACA   24922
gb:LC521925|Organism:Severe      CACTGGTTTGTAACACAAAGGAATTTTTATGAACCACAAATCATTACTACAGACAACACA   24895
gb:MT093571|Organism:Severe      CACTGGTTTGTAACACAAAGGAATTTTTATGAACCACAAATCATTACTACAGACAACACA   24922
gb:MT039887|Organism:Severe      CACTGGTTTGTAACACAAAGGAATTTTTATGAACCACAAATCATTACTACAGACAACACA   24919
gb:MT019530|Organism:Severe      CACTGGTTTGTAACACAAAGGAATTTTTATGAACCACAAATCATTACTACAGACAACACA   24922
gb:MT039888|Organism:Severe      CACTGGTTTGTAACACAAAGGAATTTTTATGAACCACAAATCATTACTACAGACAACACA   24922
gb:LC522972|Organism:Severe      CACTGGTTTGTAACACAAAGGAATTTTTATGAACCACAAATCATTACTACAGACAACACA   24919
gb:MT027063|Organism:Severe      CACTGGTTTGTAACACAAAGGAATTTTTATGAACCACAAATCATTACTACAGACAACACA   24922
gb:MT027062|Organism:Severe      CACTGGTTTGTAACACAAAGGAATTTTTATGAACCACAAATCATTACTACAGACAACACA   24922
gb:MT019529|Organism:Severe      CACTGGTTTGTAACACAAAGGAATTTTTATGAACCACAAATCATTACTACAGACAACACA   24922
gb:MN996529|Organism:Severe      CACTGGTTTGTAACACAAAGGAATTTTTATGAACCACAAATCATTACTACAGACAACACA   24910
gb:MN996531|Organism:Severe      CACTGGTTTGTAACACAAAGGAATTTTTATGAACCACAAATCATTACTACAGACAACACA   24909
gb:MT066176|Organism:Severe      CACTGGTTTGTAACACAAAGGAATTTTTATGAACCACAAATCATTACTACAGACAACACA   24922
gb:MT027064|Organism:Severe      CACTGGTTTGTAACACAAAGGAATTTTTATGAACCACAAATCATTACTACAGACAACACA   24922
gb:MN994468|Organism:Severe      CACTGGTTTGTAACACAAAGGAATTTTTATGAACCACAAATCATTACTACAGACAACACA   24922
gb:MT072688|Organism:Severe      CACTGGTTTGTAACACAAAGGAATTTTTATGAACCACAAATCATTACTACAGACAACACA   24907
gb:MN996527|Organism:Severe      CACTGGTTTGTAACACAAAGGAATTTTTATGAACCACAAATCATTACTACAGACAACACA   24889
gb:MT093631|Organism:Severe      CACTGGTTTGTAACACAAAGGAATTTTTATGAACCACAAATCATTACTACAGACAACACA   24960
gb:MT106053|Organism:Severe      CACTGGTTTGTAACACAAAGGAATTTTTATGAACCACAAATCATTACTACAGACAACACA   24922
gb:MT019533|Organism:Severe      CACTGGTTTGTAACACAAAGGAATTTTTATGAACCACAAATCATTACTACAGACAACACA   24922
gb:MT019531|Organism:Severe      CACTGGTTTGTAACACAAAGGAATTTTTATGAACCACAAATCATTACTACAGACAACACA   24922
gb:MN996528|Organism:Severe      CACTGGTTTGTAACACAAAGGAATTTTTATGAACCACAAATCATTACTACAGACAACACA   24922
gb:MN996530|Organism:Severe      CACTGGTTTGTAACACAAAGGAATTTTTATGAACCACAAATCATTACTACAGACAACACA   24908
gb:MN908947|Organism:Severe      CACTGGTTTGTAACACAAAGGAATTTTTATGAACCACAAATCATTACTACAGACAACACA   24922
gb:MT019532|Organism:Severe      CACTGGTTTGTAACACAAAGGAATTTTTATGAACCACAAATCATTACTACAGACAACACA   24922
                                 ************************************************************

gb:MT020781|Organism:Severe      TTTGTGTCTGGTAACTGTGATGTTGTAATAGGAATTGTCAACAACACAGTTTATGATCCT   24970
gb:MT007544|Organism:Severe      TTTGTGTCTGGTAACTGTGATGTTGTAATAGGAATTGTCAACAACACAGTTTATGATCCT   24982
gb:MN994467|Organism:Severe      TTTGTGTCTGGTAACTGTGATGTTGTAATAGGAATTGTCAACAACACAGTTTATGATCCT   24982
gb:MT044257|Organism:Severe      TTTGTGTCTGGTAACTGTGATGTTGTAATAGGAATTGTCAACAACACAGTTTATGATCCT   24982
gb:MT106054|Organism:Severe      TTTGTGTCTGGTAACTGTGATGTTGTAATAGGAATTGTCAACAACACAGTTTATGATCCT   24982
gb:MT049951|Organism:Severe      TTTGTGTCTGGTAACTGTGATGTTGTAATAGGAATTGTCAACAACACAGTTTATGATCCT   24982
gb:MN975262|Organism:Severe      TTTGTGTCTGGTAACTGTGATGTTGTAATAGGAATTGTCAACAACACAGTTTATGATCCT   24982
gb:MT106052|Organism:Severe      TTTGTGTCTGGTAACTGTGATGTTGTAATAGGAATTGTCAACAACACAGTTTATGATCCT   24982
gb:LC522975|Organism:Severe      TTTGTGTCTGGTAACTGTGATGTTGTAATAGGAATTGTCAACAACACAGTTTATGATCCT   24979
gb:LC522973|Organism:Severe      TTTGTGTCTGGTAACTGTGATGTTGTAATAGGAATTGTCAACAACACAGTTTATGATCCT   24979
gb:LC522974|Organism:Severe      TTTGTGTCTGGTAACTGTGATGTTGTAATAGGAATTGTCAACAACACAGTTTATGATCCT   24979
gb:MN985325|Organism:Severe      TTTGTGTCTGGTAACTGTGATGTTGTAATAGGAATTGTCAACAACACAGTTTATGATCCT   24982
gb:MT020881|Organism:Severe      TTTGTGTCTGGTAACTGTGATGTTGTAATAGGAATTGTCAACAACACAGTTTATGATCCT   24982
gb:MT020880|Organism:Severe      TTTGTGTCTGGTAACTGTGATGTTGTAATAGGAATTGTCAACAACACAGTTTATGATCCT   24982
gb:MT066175|Organism:Severe      TTTGTGTCTGGTAACTGTGATGTTGTAATAGGAATTGTCAACAACACAGTTTATGATCCT   24982
gb:MN997409|Organism:Severe      TTTGTGTCTGGTAACTGTGATGTTGTAATAGGAATTGTCAACAACACAGTTTATGATCCT   24982
gb:MN938384|Organism:Severe      TTTGTGTCTGGTAACTGTGATGTTGTAATAGGAATTGTCAACAACACAGTTTATGATCCT   24950
gb:MT044258|Organism:Severe      TTTGTGTCTGGTAACTGTGATGTTGTAATAGGAATTGTCAACAACACAGTTTATGATCCT   24958
gb:MT039890|Organism:Severe      TTTGTGTCTGGTAACTGTGATGTTGTAATAGGAATTGTCAACAACACAGTTTATGATCCT   24982
gb:MN988713|Organism:Severe      TTTGTGTCTGGTAACTGTGATGTTGTAATAGGAATTGTCAACAACACAGTTTATGATCCT   24982
gb:LC521925|Organism:Severe      TTTGTGTCTGGTAACTGTGATGTTGTAATAGGAATTGTCAACAACACAGTTTATGATCCT   24955
gb:MT093571|Organism:Severe      TTTGTGTCTGGTAACTGTGATGTTGTAATAGGAATTGTCAACAACACAGTTTATGATCCT   24982
gb:MT039887|Organism:Severe      TTTGTGTCTGGTAACTGTGATGTTGTAATAGGAATTGTCAACAACACAGTTTATGATCCT   24979
gb:MT019530|Organism:Severe      TTTGTGTCTGGTAACTGTGATGTTGTAATAGGAATTGTCAACAACACAGTTTATGATCCT   24982
gb:MT039888|Organism:Severe      TTTGTGTCTGGTAACTGTGATGTTGTAATAGGAATTGTCAACAACACAGTTTATGATCCT   24982
gb:LC522972|Organism:Severe      TTTGTGTCTGGTAACTGTGATGTTGTAATAGGAATTGTCAACAACACAGTTTATGATCCT   24979
gb:MT027063|Organism:Severe      TTTGTGTCTGGTAACTGTGATGTTGTAATAGGAATTGTCAACAACACAGTTTATGATCCT   24982
gb:MT027062|Organism:Severe      TTTGTGTCTGGTAACTGTGATGTTGTAATAGGAATTGTCAACAACACAGTTTATGATCCT   24982
gb:MT019529|Organism:Severe      TTTGTGTCTGGTAACTGTGATGTTGTAATAGGAATTGTCAACAACACAGTTTATGATCCT   24982
gb:MN996529|Organism:Severe      TTTGTGTCTGGTAACTGTGATGTTGTAATAGGAATTGTCAACAACACAGTTTATGATCCT   24970
gb:MN996531|Organism:Severe      TTTGTGTCTGGTAACTGTGATGTTGTAATAGGAATTGTCAACAACACAGTTTATGATCCT   24969
gb:MT066176|Organism:Severe      TTTGTGTCTGGTAACTGTGATGTTGTAATAGGAATTGTCAACAACACAGTTTATGATCCT   24982
gb:MT027064|Organism:Severe      TTTGTGTCTGGTAACTGTGATGTTGTAATAGGAATTGTCAACAACACAGTTTATGATCCT   24982
gb:MN994468|Organism:Severe      TTTGTGTCTGGTAACTGTGATGTTGTAATAGGAATTGTCAACAACACAGTTTATGATCCT   24982
gb:MT072688|Organism:Severe      TTTGTGTCTGGTAACTGTGATGTTGTAATAGGAATTGTCAACAACACAGTTTATGATCCT   24967
```

```
gb:MN996527|Organism:Severe      TTTGTGTCTGGTAACTGTGATGTTGTAATAGGAATTGTCAACAACACAGTTTATGATCCT   24949
gb:MT093631|Organism:Severe      TTTGTGTCTGGTAACTGTGATGTTGTAATAGGAATTGTCAACAACACAGTTTATGATCCT   25020
gb:MT106053|Organism:Severe      TTTGTGTCTGGTAACTGTGATGTTGTAATAGGAATTGTCAACAACACAGTTTATGATCCT   24982
gb:MT019533|Organism:Severe      TTTGTGTCTGGTAACTGTGATGTTGTAATAGGAATTGTCAACAACACAGTTTATGATCCT   24982
gb:MT019531|Organism:Severe      TTTGTGTCTGGTAACTGTGATGTTGTAATAGGAATTGTCAACAACACAGTTTATGATCCT   24982
gb:MN996528|Organism:Severe      TTTGTGTCTGGTAACTGTGATGTTGTAATAGGAATTGTCAACAACACAGTTTATGATCCT   24982
gb:MN996530|Organism:Severe      TTTGTGTCTGGTAACTGTGATGTTGTAATAGGAATTGTCAACAACACAGTTTATGATCCT   24968
gb:MN908947|Organism:Severe      TTTGTGTCTGGTAACTGTGATGTTGTAATAGGAATTGTCAACAACACAGTTTATGATCCT   24982
gb:MT019532|Organism:Severe      TTTGTGTCTGGTAACTGTGATGTTGTAATAGGAATTGTCAACAACACAGTTTATGATCCT   24982
                                 ************************************************************

gb:MT020781|Organism:Severe      TTGCAACCTGAATTAGACTCATTCAAGGAGGAGTTAGATAAATATTTTAAGAATCATACA   25030
gb:MT007544|Organism:Severe      TTGCAACCTGAATTAGACTCATTCAAGGAGGAGTTAGATAAATATTTTAAGAATCATACA   25042
gb:MN994467|Organism:Severe      TTGCAACCTGAATTAGACTCATTCAAGGAGGAGTTAGATAAATATTTTAAGAATCATACA   25042
gb:MT044257|Organism:Severe      TTGCAACCTGAATTAGACTCATTCAAGGAGGAGTTAGATAAATATTTTAAGAATCATACA   25042
gb:MT106054|Organism:Severe      TTGCAACCTGAATTAGACTCATTCAAGGAGGAGTTAGATAAATATTTTAAGAATCATACA   25042
gb:MT049951|Organism:Severe      TTGCAACCTGAATTAGACTCATTCAAGGAGGAGTTAGATAAATATTTTAAGAATCATACA   25042
gb:MN975262|Organism:Severe      TTGCAACCTGAATTAGACTCATTCAAGGAGGAGTTAGATAAATATTTTAAGAATCATACA   25042
gb:MT106052|Organism:Severe      TTGCAACCTGAATTAGACTCATTCAAGGAGGAGTTAGATAAATATTTTAAGAATCATACA   25042
gb:LC522975|Organism:Severe      TTGCAACCTGAATTAGACTCATTCAAGGAGGAGTTAGATAAATATTTTAAGAATCATACA   25039
gb:LC522973|Organism:Severe      TTGCAACCTGAATTAGACTCATTCAAGGAGGAGTTAGATAAATATTTTAAGAATCATACA   25039
gb:LC522974|Organism:Severe      TTGCAACCTGAATTAGACTCATTCAAGGAGGAGTTAGATAAATATTTTAAGAATCATACA   25039
gb:MN985325|Organism:Severe      TTGCAACCTGAATTAGACTCATTCAAGGAGGAGTTAGATAAATATTTTAAGAATCATACA   25042
gb:MT020881|Organism:Severe      TTGCAACCTGAATTAGACTCATTCAAGGAGGAGTTAGATAAATATTTTAAGAATCATACA   25042
gb:MT020880|Organism:Severe      TTGCAACCTGAATTAGACTCATTCAAGGAGGAGTTAGATAAATATTTTAAGAATCATACA   25042
gb:MT066175|Organism:Severe      TTGCAACCTGAATTAGACTCATTCAAGGAGGAGTTAGATAAATATTTTAAGAATCATACA   25042
gb:MN997409|Organism:Severe      TTGCAACCTGAATTAGACTCATTCAAGGAGGAGTTAGATAAATATTTTAAGAATCATACA   25042
gb:MN938384|Organism:Severe      TTGCAACCTGAATTAGACTCATTCAAGGAGGAGTTAGATAAATATTTTAAGAATCATACA   25010
gb:MT044258|Organism:Severe      TTGCAACCTGAATTAGACTCATTCAAGGAGGAGTTAGATAAATATTTTAAGAATCATACA   25018
gb:MT039890|Organism:Severe      TTGCAACCTGAATTAGACTCATTCAAGGAGGAGTTAGATAAATATTTTAAGAATCATACA   25042
gb:MN988713|Organism:Severe      TTGCAACCTGAATTAGACTCATTCAAGGAGGAGTTAGATAAATATTTTAAGAATCATACA   25042
gb:LC521925|Organism:Severe      TTGCAACCTGAATTAGACTCATTCAAGGAGGAGTTAGATAAATATTTTAAGAATCATACA   25015
gb:MT093571|Organism:Severe      TTGCAACCTGAATTAGACTCATTCAAGGAGGAGTTAGATAAATATTTTAAGAATCATACA   25042
gb:MT039887|Organism:Severe      TTGCAACCTGAATTAGACTCATTCAAGGAGGAGTTAGATAAATATTTTAAGAATCATACA   25039
gb:MT019530|Organism:Severe      TTGCAACCTGAATTAGACTCATTCAAGGAGGAGTTAGATAAATATTTTAAGAATCATACA   25042
gb:MT039888|Organism:Severe      TTGCAACCTGAATTAGACTCATTCAAGGAGGAGTTAGATAAATATTTTAAGAATCATACA   25042
gb:LC522972|Organism:Severe      TTGCAACCTGAATTAGACTCATTCAAGGAGGAGTTAGATAAATATTTTAAGAATCATACA   25039
gb:MT027063|Organism:Severe      TTGCAACCTGAATTAGACTCATTCAAGGAGGAGTTAGATAAATATTTTAAGAATCATACA   25042
gb:MT027062|Organism:Severe      TTGCAACCTGAATTAGACTCATTCAAGGAGGAGTTAGATAAATATTTTAAGAATCATACA   25042
gb:MT019529|Organism:Severe      TTGCAACCTGAATTAGACTCATTCAAGGAGGAGTTAGATAAATATTTTAAGAATCATACA   25042
gb:MN996529|Organism:Severe      TTGCAACCTGAATTAGACTCATTCAAGGAGGAGTTAGATAAATATTTTAAGAATCATACA   25030
gb:MN996531|Organism:Severe      TTGCAACCTGAATTAGACTCATTCAAGGAGGAGTTAGATAAATATTTTAAGAATCATACA   25029
gb:MT066176|Organism:Severe      TTGCAACCTGAATTAGACTCATTCAAGGAGGAGTTAGATAAATATTTTAAGAATCATACA   25042
gb:MT027064|Organism:Severe      TTGCAACCTGAATTAGACTCATTCAAGGAGGAGTTAGATAAATATTTTAAGAATCATACA   25042
gb:MN994468|Organism:Severe      TTGCAACCTGAATTAGACTCATTCAAGGAGGAGTTAGATAAATATTTTAAGAATCATACA   25042
gb:MT072688|Organism:Severe      TTGCAACCTGAATTAGACTCATTCAAGGAGGAGTTAGATAAATATTTTAAGAATCATACA   25027
gb:MN996527|Organism:Severe      TTGCAACCTGAATTAGACTCATTCAAGGAGGAGTTAGATAAATATTTTAAGAATCATACA   25009
gb:MT093631|Organism:Severe      TTGCAACCTGAATTAGACTCATTCAAGGAGGAGTTAGATAAATATTTTAAGAATCATACA   25080
gb:MT106053|Organism:Severe      TTGCAACCTGAATTAGACTCATTCAAGGAGGAGTTAGATAAATATTTTAAGAATCATACA   25042
gb:MT019533|Organism:Severe      TTGCAACCTGAATTAGACTCATTCAAGGAGGAGTTAGATAAATATTTTAAGAATCATACA   25042
gb:MT019531|Organism:Severe      TTGCAACCTGAATTAGACTCATTCAAGGAGGAGTTAGATAAATATTTTAAGAATCATACA   25042
gb:MN996528|Organism:Severe      TTGCAACCTGAATTAGACTCATTCAAGGAGGAGTTAGATAAATATTTTAAGAATCATACA   25042
gb:MN996530|Organism:Severe      TTGCAACCTGAATTAGACTCATTCAAGGAGGAGTTAGATAAATATTTTAAGAATCATACA   25028
gb:MN908947|Organism:Severe      TTGCAACCTGAATTAGACTCATTCAAGGAGGAGTTAGATAAATATTTTAAGAATCATACA   25042
gb:MT019532|Organism:Severe      TTGCAACCTGAATTAGACTCATTCAAGGAGGAGTTAGATAAATATTTTAAGAATCATACA   25042
                                 ************************************************************

gb:MT020781|Organism:Severe      TCACCAGATGTTGATTTAGGTGACATCTCTGGCATTAATGCTTCAGTTGTAAACATTCAA   25090
gb:MT007544|Organism:Severe      TCACCAGATGTTGATTTAGGTGACATCTCTGGCATTAATGCTTCAGTTGTAAACATTCAA   25102
gb:MN994467|Organism:Severe      TCACCAGATGTTGATTTAGGTGACATCTCTGGCATTAATGCTTCAGTTGTAAACATTCAA   25102
gb:MT044257|Organism:Severe      TCACCAGATGTTGATTTAGGTGACATCTCTGGCATTAATGCTTCAGTTGTAAACATTCAA   25102
gb:MT106054|Organism:Severe      TCACCAGATGTTGATTTAGGTGACATCTCTGGCATTAATGCTTCAGTTGTAAACATTCAA   25102
gb:MT049951|Organism:Severe      TCACCAGATGTTGATTTAGGTGACATCTCTGGCATTAATGCTTCAGTTGTAAACATTCAA   25102
gb:MN975262|Organism:Severe      TCACCAGATGTTGATTTAGGTGACATCTCTGGCATTAATGCTTCAGTTGTAAACATTCAA   25102
gb:MT106052|Organism:Severe      TCACCAGATGTTGATTTAGGTGACATCTCTGGCATTAATGCTTCAGTTGTAAACATTCAA   25102
```

```
gb:LC522975|Organism:Severe      TCACCAGATGTTGATTTAGGTGACATCTCTGGCATTAATGCTTCAGTTGTAAACATTCAA   25099
gb:LC522973|Organism:Severe      TCACCAGATGTTGATTTAGGTGACATCTCTGGCATTAATGCTTCAGTTGTAAACATTCAA   25099
gb:LC522974|Organism:Severe      TCACCAGATGTTGATTTAGGTGACATCTCTGGCATTAATGCTTCAGTTGTAAACATTCAA   25099
gb:MN985325|Organism:Severe      TCACCAGATGTTGATTTAGGTGACATCTCTGGCATTAATGCTTCAGTTGTAAACATTCAA   25102
gb:MT020881|Organism:Severe      TCACCAGATGTTGATTTAGGTGACATCTCTGGCATTAATGCTTCAGTTGTAAACATTCAA   25102
gb:MT020880|Organism:Severe      TCACCAGATGTTGATTTAGGTGACATCTCTGGCATTAATGCTTCAGTTGTAAACATTCAA   25102
gb:MT066175|Organism:Severe      TCACCAGATGTTGATTTAGGTGACATCTCTGGCATTAATGCTTCAGTTGTAAACATTCAA   25102
gb:MN997409|Organism:Severe      TCACCAGATGTTGATTTAGGTGACATCTCTGGCATTAATGCTTCAGTTGTAAACATTCAA   25102
gb:MN938384|Organism:Severe      TCACCAGATGTTGATTTAGGTGACATCTCTGGCATTAATGCTTCAGTTGTAAACATTCAA   25070
gb:MT044258|Organism:Severe      TCACCAGATGTTGATTTAGGTGACATCTCTGGCATTAATGCTTCAGTTGTAAACATTCAA   25078
gb:MT039890|Organism:Severe      TCACCAGATGTTGATTTAGGTGACATCTCTGGCATTAATGCTTCAGTTGTAAACATTCAA   25102
gb:MN988713|Organism:Severe      TCACCAGATGTTGATTTAGGTGACATCTCTGGCATTAATGCTTCAGTTGTAAACATTCAA   25102
gb:LC521925|Organism:Severe      TCACCAGATGTTGATTTAGGTGACATCTCTGGCATTAATGCTTCAGTTGTAAACATTCAA   25075
gb:MT093571|Organism:Severe      TCACCAGATGTTGATTTAGGTGACATCTCTGGCATTAATGCTTCAGTTGTAAACATTCAA   25102
gb:MT039887|Organism:Severe      TCACCAGATGTTGATTTAGGTGACATCTCTGGCATTAATGCTTCAGTTGTAAACATTCAA   25099
gb:MT019530|Organism:Severe      TCACCAGATGTTGATTTAGGTGACATCTCTGGCATTAATGCTTCAGTTGTAAACATTCAA   25102
gb:MT039888|Organism:Severe      TCACCAGATGTTGATTTAGGTGACATCTCTGGCATTAATGCTTCAGTTGTAAACATTCAA   25102
gb:LC522972|Organism:Severe      TCACCAGATGTTGATTTAGGTGACATCTCTGGCATTAATGCTTCAGTTGTAAACATTCAA   25099
gb:MT027063|Organism:Severe      TCACCAGATGTTGATTTAGGTGACATCTCTGGCATTAATGCTTCAGTTGTAAACATTCAA   25102
gb:MT027062|Organism:Severe      TCACCAGATGTTGATTTAGGTGACATCTCTGGCATTAATGCTTCAGTTGTAAACATTCAA   25102
gb:MT019529|Organism:Severe      TCACCAGATGTTGATTTAGGTGACATCTCTGGCATTAATGCTTCAGTTGTAAACATTCAA   25102
gb:MN996529|Organism:Severe      TCACCAGATGTTGATTTAGGTGACATCTCTGGCATTAATGCTTCAGTTGTAAACATTCAA   25090
gb:MN996531|Organism:Severe      TCACCAGATGTTGATTTAGGTGACATCTCTGGCATTAATGCTTCAGTTGTAAACATTCAA   25089
gb:MT066176|Organism:Severe      TCACCAGATGTTGATTTAGGTGACATCTCTGGCATTAATGCTTCAGTTGTAAACATTCAA   25102
gb:MT027064|Organism:Severe      TCACCAGATGTTGATTTAGGTGACATCTCTGGCATTAATGCTTCAGTTGTAAACATTCAA   25102
gb:MN994468|Organism:Severe      TCACCAGATGTTGATTTAGGTGACATCTCTGGCATTAATGCTTCAGTTGTAAACATTCAA   25102
gb:MT072688|Organism:Severe      TCACCAGATGTTGATTTAGGTGACATCTCTGGCATTAATGCTTCAGTTGTAAACATTCAA   25087
gb:MN996527|Organism:Severe      TCACCAGATGTTGATTTAGGTGACATCTCTGGCATTAATGCTTCAGTTGTAAACATTCAA   25069
gb:MT093631|Organism:Severe      TCACCAGATGTTGATTTAGGTGACATCTCTGGCATTAATGCTTCAGTTGTAAACATTCAA   25140
gb:MT106053|Organism:Severe      TCACCAGATGTTGATTTAGGTGACATCTCTGGCATTAATGCTTCAGTTGTAAACATTCAA   25102
gb:MT019533|Organism:Severe      TCACCAGATGTTGATTTAGGTGACATCTCTGGCATTAATGCTTCAGTTGTAAACATTCAA   25102
gb:MT019531|Organism:Severe      TCACCAGATGTTGATTTAGGTGACATCTCTGGCATTAATGCTTCAGTTGTAAACATTCAA   25102
gb:MN996528|Organism:Severe      TCACCAGATGTTGATTTAGGTGACATCTCTGGCATTAATGCTTCAGTTGTAAACATTCAA   25102
gb:MN996530|Organism:Severe      TCACCAGATGTTGATTTAGGTGACATCTCTGGCATTAATGCTTCAGTTGTAAACATTCAA   25088
gb:MN908947|Organism:Severe      TCACCAGATGTTGATTTAGGTGACATCTCTGGCATTAATGCTTCAGTTGTAAACATTCAA   25102
gb:MT019532|Organism:Severe      TCACCAGATGTTGATTTAGGTGACATCTCTGGCATTAATGCTTCAGTTGTAAACATTCAA   25102
                                 ************************************************************

gb:MT020781|Organism:Severe      AAAGAAATTGACCGCCTCAATGAGGTTGCCAAGAATTTAAATGAATCTCTCATCGATCTC   25150
gb:MT007544|Organism:Severe      AAAGAAATTGACCGCCTCAATGAGGTTGCCAAGAATTTAAATGAATCTCTCATCGATCTC   25162
gb:MN994467|Organism:Severe      AAAGAAATTGACCGCCTCAATGAGGTTGCCAAGAATTTAAATGAATCTCTCATCGATCTC   25162
gb:MT044257|Organism:Severe      AAAGAAATTGACCGCCTCAATGAGGTTGCCAAGAATTTAAATGAATCTCTCATCGATCTC   25162
gb:MT106054|Organism:Severe      AAAGAAATTGACCGCCTCAATGAGGTTGCCAAGAATTTAAATGAATCTCTCATCGATCTC   25162
gb:MT049951|Organism:Severe      AAAGAAATTGACCGCCTCAATGAGGTTGCCAAGAATTTAAATGAATCTCTCATCGATCTC   25162
gb:MN975262|Organism:Severe      AAAGAAATTGACCGCCTCAATGAGGTTGCCAAGAATTTAAATGAATCTCTCATCGATCTC   25162
gb:MT106052|Organism:Severe      AAAGAAATTGACCGCCTCAATGAGGTTGCCAAGAATTTAAATGAATCTCTCATCGATCTC   25162
gb:LC522975|Organism:Severe      AAAGAAATTGACCGCCTCAATGAGGTTGCCAAGAATTTAAATGAATCTCTCATCGATCTC   25159
gb:LC522973|Organism:Severe      AAAGAAATTGACCGCCTCAATGAGGTTGCCAAGAATTTAAATGAATCTCTCATCGATCTC   25159
gb:LC522974|Organism:Severe      AAAGAAATTGACCGCCTCAATGAGGTTGCCAAGAATTTAAATGAATCTCTCATCGATCTC   25159
gb:MN985325|Organism:Severe      AAAGAAATTGACCGCCTCAATGAGGTTGCCAAGAATTTAAATGAATCTCTCATCGATCTC   25162
gb:MT020881|Organism:Severe      AAAGAAATTGACCGCCTCAATGAGGTTGCCAAGAATTTAAATGAATCTCTCATCGATCTC   25162
gb:MT020880|Organism:Severe      AAAGAAATTGACCGCCTCAATGAGGTTGCCAAGAATTTAAATGAATCTCTCATCGATCTC   25162
gb:MT066175|Organism:Severe      AAAGAAATTGACCGCCTCAATGAGGTTGCCAAGAATTTAAATGAATCTCTCATCGATCTC   25162
gb:MN997409|Organism:Severe      AAAGAAATTGACCGCCTCAATGAGGTTGCCAAGAATTTAAATGAATCTCTCATCGATCTC   25162
gb:MN938384|Organism:Severe      AAAGAAATTGACCGCCTCAATGAGGTTGCCAAGAATTTAAATGAATCTCTCATCGATCTC   25130
gb:MT044258|Organism:Severe      AAAGAAATTGACCGCCTCAATGAGGTTGCCAAGAATTTAAATGAATCTCTCATCGATCTC   25138
gb:MT039890|Organism:Severe      AAAGAAATTGACCGCCTCAATGAGGTTGCCAAGAATTTAAATGAATCTCTCATCGATCTC   25162
gb:MN988713|Organism:Severe      AAAGAAATTGACCGCCTCAATGAGGTTGCCAAGAATTTAAATGAATCTCTCATCGATCTC   25162
gb:LC521925|Organism:Severe      AAAGAAATTGACCGCCTCAATGAGGTTGCCAAGAATTTAAATGAATCTCTCATCGATCTC   25135
gb:MT093571|Organism:Severe      AAAGAAATTGACCGCCTCAATGAGGTTGCCAAGAATTTAAATGAATCTCTCATCGATCTC   25162
gb:MT039887|Organism:Severe      AAAGAAATTGACCGCCTCAATGAGGTTGCCAAGAATTTAAATGAATCTCTCATCGATCTC   25159
gb:MT019530|Organism:Severe      AAAGAAATTGACCGCCTCAATGAGGTTGCCAAGAATTTAAATGAATCTCTCATCGATCTC   25162
gb:MT039888|Organism:Severe      AAAGAAATTGACCGCCTCAATGAGGTTGCCAAGAATTTAAATGAATCTCTCATCGATCTC   25162
gb:LC522972|Organism:Severe      AAAGAAATTGACCGCCTCAATGAGGTTGCCAAGAATTTAAATGAATCTCTCATCGATCTC   25159
gb:MT027063|Organism:Severe      AAAGAAATTGACCGCCTCAATGAGGTTGCCAAGAATTTAAATGAATCTCTCATCGATCTC   25162
```

```
gb:MT027062|Organism:Severe      AAAGAAATTGACCGCCTCAATGAGGTTGCCAAGAATTTAAATGAATCTCTCATCGATCTC   25162
gb:MT019529|Organism:Severe      AAAGAAATTGACCGCCTCAATGAGGTTGCCAAGAATTTAAATGAATCTCTCATCGATCTC   25162
gb:MN996529|Organism:Severe      AAAGAAATTGACCGCCTCAATGAGGTTGCCAAGAATTTAAATGAATCTCTCATCGATCTC   25150
gb:MN996531|Organism:Severe      AAAGAAATTGACCGCCTCAATGAGGTTGCCAAGAATTTAAATGAATCTCTCATCGATCTC   25149
gb:MT066176|Organism:Severe      AAAGAAATTGACCGCCTCAATGAGGTTGCCAAGAATTTAAATGAATCTCTCATCGATCTC   25162
gb:MT027064|Organism:Severe      AAAGAAATTGACCGCCTCAATGAGGTTGCCAAGAATTTAAATGAATCTCTCATCGATCTC   25162
gb:MN994468|Organism:Severe      AAAGAAATTGACCGCCTCAATGAGGTTGCCAAGAATTTAAATGAATCTCTCATCGATCTC   25162
gb:MT072688|Organism:Severe      AAAGAAATTGACCGCCTCAATGAGGTTGCCAAGAATTTAAATGAATCTCTCATCGATCTC   25147
gb:MN996527|Organism:Severe      AAAGAAATTGACCGCCTCAATGAGGTTGCCAAGAATTTAAATGAATCTCTCATCGATCTC   25129
gb:MT093631|Organism:Severe      AAAGAAATTGACCGCCTCAATGAGGTTGCCAAGAATTTAAATGAATCTCTCATCGATCTC   25200
gb:MT106053|Organism:Severe      AAAGAAATTGACCGCCTCAATGAGGTTGCCAAGAATTTAAATGAATCTCTCATCGATCTC   25162
gb:MT019533|Organism:Severe      AAAGAAATTGACCGCCTCAATGAGGTTGCCAAGAATTTAAATGAATCTCTCATCGATCTC   25162
gb:MT019531|Organism:Severe      AAAGAAATTGACCGCCTCAATGAGGTTGCCAAGAATTTAAATGAATCTCTCATCGATCTC   25162
gb:MN996528|Organism:Severe      AAAGAAATTGACCGCCTCAATGAGGTTGCCAAGAATTTAAATGAATCTCTCATCGATCTC   25162
gb:MN996530|Organism:Severe      AAAGAAATTGACCGCCTCAATGAGGTTGCCAAGAATTTAAATGAATCTCTCATCGATCTC   25148
gb:MN908947|Organism:Severe      AAAGAAATTGACCGCCTCAATGAGGTTGCCAAGAATTTAAATGAATCTCTCATCGATCTC   25162
gb:MT019532|Organism:Severe      AAAGAAATTGACCGCCTCAATGAGGTTGCCAAGAATTTAAATGAATCTCTCATCGATCTC   25162
                                 ************************************************************

gb:MT020781|Organism:Severe      CAAGAACTTGGAAAGTATGAGCAGTATATAAAATGGCCATGGTACATTTGGCTAGGTTTT   25210
gb:MT007544|Organism:Severe      CAAGAACTTGGAAAGTATGAGCAGTATATAAAATGGCCATGGTACATTTGGCTAGGTTTT   25222
gb:MN994467|Organism:Severe      CAAGAACTTGGAAAGTATGAGCAGTATATAAAATGGCCATGGTACATTTGGCTAGGTTTT   25222
gb:MT044257|Organism:Severe      CAAGAACTTGGAAAGTATGAGCAGTATATAAAATGGCCATGGTACATTTGGCTAGGTTTT   25222
gb:MT106054|Organism:Severe      CAAGAACTTGGAAAGTATGAGCAGTATATAAAATGGCCATGGTACATTTGGCTAGGTTTT   25222
gb:MT049951|Organism:Severe      CAAGAACTTGGAAAGTATGAGCAGTATATAAAATGGCCATGGTACATTTGGCTAGGTTTT   25222
gb:MN975262|Organism:Severe      CAAGAACTTGGAAAGTATGAGCAGTATATAAAATGGCCATGGTACATTTGGCTAGGTTTT   25222
gb:MT106052|Organism:Severe      CAAGAACTTGGAAAGTATGAGCAGTATATAAAATGGCCATGGTACATTTGGCTAGGTTTT   25222
gb:LC522975|Organism:Severe      CAAGAACTTGGAAAGTATGAGCAGTATATAAAATGGCCATGGTACATTTGGCTAGGTTTT   25219
gb:LC522973|Organism:Severe      CAAGAACTTGGAAAGTATGAGCAGTATATAAAATGGCCATGGTACATTTGGCTAGGTTTT   25219
gb:LC522974|Organism:Severe      CAAGAACTTGGAAAGTATGAGCAGTATATAAAATGGCCATGGTACATTTGGCTAGGTTTT   25219
gb:MN985325|Organism:Severe      CAAGAACTTGGAAAGTATGAGCAGTATATAAAATGGCCATGGTACATTTGGCTAGGTTTT   25222
gb:MT020881|Organism:Severe      CAAGAACTTGGAAAGTATGAGCAGTATATAAAATGGCCATGGTACATTTGGCTAGGTTTT   25222
gb:MT020880|Organism:Severe      CAAGAACTTGGAAAGTATGAGCAGTATATAAAATGGCCATGGTACATTTGGCTAGGTTTT   25222
gb:MT066175|Organism:Severe      CAAGAACTTGGAAAGTATGAGCAGTATATAAAATGGCCATGGTACATTTGGCTAGGTTTT   25222
gb:MN997409|Organism:Severe      CAAGAACTTGGAAAGTATGAGCAGTATATAAAATGGCCATGGTACATTTGGCTAGGTTTT   25222
gb:MN938384|Organism:Severe      CAAGAACTTGGAAAGTATGAGCAGTATATAAAATGGCCATGGTACATTTGGCTAGGTTTT   25190
gb:MT044258|Organism:Severe      CAAGAACTTGGAAAGTATGAGCAGTATATAAAATGGCCATGGTACATTTGGCTAGGTTTT   25198
gb:MT039890|Organism:Severe      CAAGAACTTGGAAAGTATGAGCAGTATATAAAATGGCCATGGTACATTTGGCTAGGTTTT   25222
gb:MN988713|Organism:Severe      CAAGAACTTGGAAAGTATGAGCAGTATATAAAATGGCCATGGTACATTTGGCTAGGTTTT   25222
gb:LC521925|Organism:Severe      CAAGAACTTGGAAAGTATGAGCAGTATATAAAATGGCCATGGTACATTTGGCTAGGTTTT   25195
gb:MT093571|Organism:Severe      CAAGAACTTGGAAAGTATGAGCAGTATATAAAATGGCCATGGTACATTTGGCTAGGTTTT   25222
gb:MT039887|Organism:Severe      CAAGAACTTGGAAAGTATGAGCAGTATATAAAATGGCCATGGTACATTTGGCTAGGTTTT   25219
gb:MT019530|Organism:Severe      CAAGAACTTGGAAAGTATGAGCAGTATATAAAATGGCCATGGTACATTTGGCTAGGTTTT   25222
gb:MT039888|Organism:Severe      CAAGAACTTGGAAAGTATGAGCAGTATATAAAATGGCCATGGTACATTTGGCTAGGTTTT   25222
gb:LC522972|Organism:Severe      CAAGAACTTGGAAAGTATGAGCAGTATATAAAATGGCCATGGTACATTTGGCTAGGTTTT   25219
gb:MT027063|Organism:Severe      CAAGAACTTGGAAAGTATGAGCAGTATATAAAATGGCCATGGTACATTTGGCTAGGTTTT   25222
gb:MT027062|Organism:Severe      CAAGAACTTGGAAAGTATGAGCAGTATATAAAATGGCCATGGTACATTTGGCTAGGTTTT   25222
gb:MT019529|Organism:Severe      CAAGAACTTGGAAAGTATGAGCAGTATATAAAATGGCCATGGTACATTTGGCTAGGTTTT   25222
gb:MN996529|Organism:Severe      CAAGAACTTGGAAAGTATGAGCAGTATATAAAATGGCCATGGTACATTTGGCTAGGTTTT   25210
gb:MN996531|Organism:Severe      CAAGAACTTGGAAAGTATGAGCAGTATATAAAATGGCCATGGTACATTTGGCTAGGTTTT   25209
gb:MT066176|Organism:Severe      CAAGAACTTGGAAAGTATGAGCAGTATATAAAATGGCCATGGTACATTTGGCTAGGTTTT   25222
gb:MT027064|Organism:Severe      CAAGAACTTGGAAAGTATGAGCAGTATATAAAATGGCCATGGTACATTTGGCTAGGTTTT   25222
gb:MN994468|Organism:Severe      CAAGAACTTGGAAAGTATGAGCAGTATATAAAATGGCCATGGTACATTTGGCTAGGTTTT   25222
gb:MT072688|Organism:Severe      CAAGAACTTGGAAAGTATGAGCAGTATATAAAATGGCCATGGTACATTTGGCTAGGTTTT   25207
gb:MN996527|Organism:Severe      CAAGAACTTGGAAAGTATGAGCAGTATATAAAATGGCCATGGTACATTTGGCTAGGTTTT   25189
gb:MT093631|Organism:Severe      CAAGAACTTGGAAAGTATGAGCAGTATATAAAATGGCCATGGTACATTTGGCTAGGTTTT   25260
gb:MT106053|Organism:Severe      CAAGAACTTGGAAAGTATGAGCAGTATATAAAATGGCCATGGTACATTTGGCTAGGTTTT   25222
gb:MT019533|Organism:Severe      CAAGAACTTGGAAAGTATGAGCAGTATATAAAATGGCCATGGTACATTTGGCTAGGTTTT   25222
gb:MT019531|Organism:Severe      CAAGAACTTGGAAAGTATGAGCAGTATATAAAATGGCCATGGTACATTTGGCTAGGTTTT   25222
gb:MN996528|Organism:Severe      CAAGAACTTGGAAAGTATGAGCAGTATATAAAATGGCCATGGTACATTTGGCTAGGTTTT   25222
gb:MN996530|Organism:Severe      CAAGAACTTGGAAAGTATGAGCAGTATATAAAATGGCCATGGTACATTTGGCTAGGTTTT   25208
gb:MN908947|Organism:Severe      CAAGAACTTGGAAAGTATGAGCAGTATATAAAATGGCCATGGTACATTTGGCTAGGTTTT   25222
gb:MT019532|Organism:Severe      CAAGAACTTGGAAAGTATGAGCAGTATATAAAATGGCCATGGTACATTTGGCTAGGTTTT   25222
                                 ************************************************************
```

```
gb:MT020781|Organism:Severe      ATAGCTGGCTTGATTGCCATAGTAATGGTGACAATTATGCTTTGCTGTATGACCAGTTGC	25270
gb:MT007544|Organism:Severe      ATAGCTGGCTTGATTGCCATAGTAATGGTGACAATTATGCTTTGCTGTATGACCAGTTGC	25282
gb:MN994467|Organism:Severe      ATAGCTGGCTTGATTGCCATAGTAATGGTGACAATTATGCTTTGCTGTATGACCAGTTGC	25282
gb:MT044257|Organism:Severe      ATAGCTGGCTTGATTGCCATAGTAATGGTGACAATTATGCTTTGCTGTATGACCAGTTGC	25282
gb:MT106054|Organism:Severe      ATAGCTGGCTTGATTGCCATAGTAATGGTGACAATTATGCTTTGCTGTATGACCAGTTGC	25282
gb:MT049951|Organism:Severe      ATAGCTGGCTTGATTGCCATAGTAATGGTGACAATTATGCTTTGCTGTATGACCAGTTGC	25282
gb:MN975262|Organism:Severe      ATAGCTGGCTTGATTGCCATAGTAATGGTGACAATTATGCTTTGCTGTATGACCAGTTGC	25282
gb:MT106052|Organism:Severe      ATAGCTGGCTTGATTGCCATAGTAATGGTGACAATTATGCTTTGCTGTATGACCAGTTGC	25282
gb:LC522975|Organism:Severe      ATAGCTGGCTTGATTGCCATAGTAATGGTGACAATTATGCTTTGCTGTATGACCAGTTGC	25279
gb:LC522973|Organism:Severe      ATAGCTGGCTTGATTGCCATAGTAATGGTGACAATTATGCTTTGCTGTATGACCAGTTGC	25279
gb:LC522974|Organism:Severe      ATAGCTGGCTTGATTGCCATAGTAATGGTGACAATTATGCTTTGCTGTATGACCAGTTGC	25279
gb:MN985325|Organism:Severe      ATAGCTGGCTTGATTGCCATAGTAATGGTGACAATTATGCTTTGCTGTATGACCAGTTGC	25282
gb:MT020881|Organism:Severe      ATAGCTGGCTTGATTGCCATAGTAATGGTGACAATTATGCTTTGCTGTATGACCAGTTGC	25282
gb:MT020880|Organism:Severe      ATAGCTGGCTTGATTGCCATAGTAATGGTGACAATTATGCTTTGCTGTATGACCAGTTGC	25282
gb:MT066175|Organism:Severe      ATAGCTGGCTTGATTGCCATAGTAATGGTGACAATTATGCTTTGCTGTATGACCAGTTGC	25282
gb:MN997409|Organism:Severe      ATAGCTGGCTTGATTGCCATAGTAATGGTGACAATTATGCTTTGCTGTATGACCAGTTGC	25282
gb:MN938384|Organism:Severe      ATAGCTGGCTTGATTGCCATAGTAATGGTGACAATTATGCTTTGCTGTATGACCAGTTGC	25250
gb:MT044258|Organism:Severe      ATAGCTGGCTTGATTGCCATAGTAATGGTGACAATTATGCTTTGCTGTATGACCAGTTGC	25258
gb:MT039890|Organism:Severe      ATAGCTGGCTTGATTGCCATAGTAATGGTGACAATTATGCTTTGCTGTATGACCAGTTGC	25282
gb:MN988713|Organism:Severe      ATAGCTGGCTTGATTGCCATAGTAATGGTGACAATTATGCTTTGCTGTATGACCAGTTGC	25282
gb:LC521925|Organism:Severe      ATAGCTGGCTTGATTGCCATAGTAATGGTGACAATTATGCTTTGCTGTATGACCAGTTGC	25255
gb:MT093571|Organism:Severe      ATAGCTGGCTTGATTGCCATAGTAATGGTGACAATTATGCTTTGCTGTATGACCAGTTGC	25282
gb:MT039887|Organism:Severe      ATAGCTGGCTTGATTGCCATAGTAATGGTGACAATTATGCTTTGCTGTATGACCAGTTGC	25279
gb:MT019530|Organism:Severe      ATAGCTGGCTTGATTGCCATAGTAATGGTGACAATTATGCTTTGCTGTATGACCAGTTGC	25282
gb:MT039888|Organism:Severe      ATAGCTGGCTTGATTGCCATAGTAATGGTGACAATTATGCTTTGCTGTATGACCAGTTGC	25282
gb:LC522972|Organism:Severe      ATAGCTGGCTTGATTGCCATAGTAATGGTGACAATTATGCTTTGCTGTATGACCAGTTGC	25279
gb:MT027063|Organism:Severe      ATAGCTGGCTTGATTGCCATAGTAATGGTGACAATTATGCTTTGCTGTATGACCAGTTGC	25282
gb:MT027062|Organism:Severe      ATAGCTGGCTTGATTGCCATAGTAATGGTGACAATTATGCTTTGCTGTATGACCAGTTGC	25282
gb:MT019529|Organism:Severe      ATAGCTGGCTTGATTGCCATAGTAATGGTGACAATTATGCTTTGCTGTATGACCAGTTGC	25282
gb:MN996529|Organism:Severe      ATAGCTGGCTTGATTGCCATAGTAATGGTGACAATTATGCTTTGCTGTATGACCAGTTGC	25270
gb:MN996531|Organism:Severe      ATAGCTGGCTTGATTGCCATAGTAATGGTGACAATTATGCTTTGCTGTATGACCAGTTGC	25269
gb:MT066176|Organism:Severe      ATAGCTGGCTTGATTGCCATAGTAATGGTGACAATTATGCTTTGCTGTATGACCAGTTGC	25282
gb:MT027064|Organism:Severe      ATAGCTGGCTTGATTGCCATAGTAATGGTGACAATTATGCTTTGCTGTATGACCAGTTGC	25282
gb:MN994468|Organism:Severe      ATAGCTGGCTTGATTGCCATAGTAATGGTGACAATTATGCTTTGCTGTATGACCAGTTGC	25282
gb:MT072688|Organism:Severe      ATAGCTGGCTTGATTGCCATAGTAATGGTGACAATTATGCTTTGCTGTATGACCAGTTGC	25267
gb:MN996527|Organism:Severe      ATAGCTGGCTTGATTGCCATAGTAATGGTGACAATTATGCTTTGCTGTATGACCAGTTGC	25249
gb:MT093631|Organism:Severe      ATAGCTGGCTTGATTGCCATAGTAATGGTGACAATTATGCTTTGCTGTATGACCAGTTGC	25320
gb:MT106053|Organism:Severe      ATAGCTGGCTTGATTGCCATAGTAATGGTGACAATTATGCTTTGCTGTATGACCAGTTGC	25282
gb:MT019533|Organism:Severe      ATAGCTGGCTTGATTGCCATAGTAATGGTGACAATTATGCTTTGCTGTATGACCAGTTGC	25282
gb:MT019531|Organism:Severe      ATAGCTGGCTTGATTGCCATAGTAATGGTGACAATTATGCTTTGCTGTATGACCAGTTGC	25282
gb:MN996528|Organism:Severe      ATAGCTGGCTTGATTGCCATAGTAATGGTGACAATTATGCTTTGCTGTATGACCAGTTGC	25282
gb:MN996530|Organism:Severe      ATAGCTGGCTTGATTGCCATAGTAATGGTGACAATTATGCTTTGCTGTATGACCAGTTGC	25268
gb:MN908947|Organism:Severe      ATAGCTGGCTTGATTGCCATAGTAATGGTGACAATTATGCTTTGCTGTATGACCAGTTGC	25282
gb:MT019532|Organism:Severe      ATAGCTGGCTTGATTGCCATAGTAATGGTGACAATTATGCTTTGCTGTATGACCAGTTGC	25282
                                 ************************************************************

gb:MT020781|Organism:Severe      TGTAGTTGTCTCAAGGGCTGTTGTTCTTGTGGATCCTGCTGCAAATTTGATGAAGACGAC	25330
gb:MT007544|Organism:Severe      TGTAGTTGTCTCAAGGGCTGTTGTTCTTGTGGATCCTGCTGCAAATTTGATGAAGACGAC	25342
gb:MN994467|Organism:Severe      TGTAGTTGTCTCAAGGGCTGTTGTTCTTGTGGATCCTGCTGCAAATTTGATGAAGACGAC	25342
gb:MT044257|Organism:Severe      TGTAGTTGTCTCAAGGGCTGTTGTTCTTGTGGATCCTGCTGCAAATTTGATGAAGACGAC	25342
gb:MT106054|Organism:Severe      TGTAGTTGTCTCAAGGGCTGTTGTTCTTGTGGATCCTGCTGCAAATTTGATGAAGACGAC	25342
gb:MT049951|Organism:Severe      TGTAGTTGTCTCAAGGGCTGTTGTTCTTGTGGATCCTGCTGCAAATTTGATGAAGACGAC	25342
gb:MN975262|Organism:Severe      TGTAGTTGTCTCAAGGGCTGTTGTTCTTGTGGATCCTGCTGCAAATTTGATGAAGACGAC	25342
gb:MT106052|Organism:Severe      TGTAGTTGTCTCAAGGGCTGTTGTTCTTGTGGATCCTGCTGCAAATTTGATGAAGACGAC	25342
gb:LC522975|Organism:Severe      TGTAGTTGTCTCAAGGGCTGTTGTTCTTGTGGATCCTGCTGCAAATTTGATGAAGACGAC	25339
gb:LC522973|Organism:Severe      TGTAGTTGTCTCAAGGGCTGTTGTTCTTGTGGATCCTGCTGCAAATTTGATGAAGACGAC	25339
gb:LC522974|Organism:Severe      TGTAGTTGTCTCAAGGGCTGTTGTTCTTGTGGATCCTGCTGCAAATTTGATGAAGACGAC	25339
gb:MN985325|Organism:Severe      TGTAGTTGTCTCAAGGGCTGTTGTTCTTGTGGATCCTGCTGCAAATTTGATGAAGACGAC	25342
gb:MT020881|Organism:Severe      TGTAGTTGTCTCAAGGGCTGTTGTTCTTGTGGATCCTGCTGCAAATTTGATGAAGACGAC	25342
gb:MT020880|Organism:Severe      TGTAGTTGTCTCAAGGGCTGTTGTTCTTGTGGATCCTGCTGCAAATTTGATGAAGACGAC	25342
gb:MT066175|Organism:Severe      TGTAGTTGTCTCAAGGGCTGTTGTTCTTGTGGATCCTGCTGCAAATTTGATGAAGACGAC	25342
gb:MN997409|Organism:Severe      TGTAGTTGTCTCAAGGGCTGTTGTTCTTGTGGATCCTGCTGCAAATTTGATGAAGACGAC	25342
gb:MN938384|Organism:Severe      TGTAGTTGTCTCAAGGGCTGTTGTTCTTGTGGATCCTGCTGCAAATTTGATGAAGACGAC	25310
gb:MT044258|Organism:Severe      TGTAGTTGTCTCAAGGGCTGTTGTTCTTGTGGATCCTGCTGCAAATTTGATGAAGACGAC	25318
gb:MT039890|Organism:Severe      TGTAGTTGTCTCAAGGGCTGTTGTTCTTGTGGATCCTGCTGCAAATTTGATGAAGACGAC	25342
```

```
gb:MN988713|Organism:Severe      TGTAGTTGTCTCAAGGGCTGTTGTTCTTGTGGATCCTGCTGCAAATTTGATGAAGACGAC   25342
gb:LC521925|Organism:Severe      TGTAGTTGTCTCAAGGGCTGTTGTTCTTGTGGATCCTGCTGCAAATTTGATGAAGACGAC   25315
gb:MT093571|Organism:Severe      TGTAGTTGTCTCAAGGGCTGTTGTTCTTGTGGATCCTGCTGCAAATTTGATGAAGACGAC   25342
gb:MT039887|Organism:Severe      TGTAGTTGTCTCAAGGGCTGTTGTTCTTGTGGATCCTGCTGCAAATTTGATGAAGACGAC   25339
gb:MT019530|Organism:Severe      TGTAGTTGTCTCAAGGGCTGTTGTTCTTGTGGATCCTGCTGCAAATTTGATGAAGACGAC   25342
gb:MT039888|Organism:Severe      TGTAGTTGTCTCAAGGGCTGTTGTTCTTGTGGATCCTGCTGCAAATTTGATGAAGACGAC   25342
gb:LC522972|Organism:Severe      TGTAGTTGTCTCAAGGGCTGTTGTTCTTGTGGATCCTGCTGCAAATTTGATGAAGACGAC   25339
gb:MT027063|Organism:Severe      TGTAGTTGTCTCAAGGGCTGTTGTTCTTGTGGATCCTGCTGCAAATTTGATGAAGACGAC   25342
gb:MT027062|Organism:Severe      TGTAGTTGTCTCAAGGGCTGTTGTTCTTGTGGATCCTGCTGCAAATTTGATGAAGACGAC   25342
gb:MT019529|Organism:Severe      TGTAGTTGTCTCAAGGGCTGTTGTTCTTGTGGATCCTGCTGCAAATTTGATGAAGACGAC   25342
gb:MN996529|Organism:Severe      TGTAGTTGTCTCAAGGGCTGTTGTTCTTGTGGATCCTGCTGCAAATTTGATGAAGACGAC   25330
gb:MN996531|Organism:Severe      TGTAGTTGTCTCAAGGGCTGTTGTTCTTGTGGATCCTGCTGCAAATTTGATGAAGACGAC   25329
gb:MT066176|Organism:Severe      TGTAGTTGTCTCAAGGGCTGTTGTTCTTGTGGATCCTGCTGCAAATTTGATGAAGACGAC   25342
gb:MT027064|Organism:Severe      TGTAGTTGTCTCAAGGGCTGTTGTTCTTGTGGATCCTGCTGCAAATTTGATGAAGACGAC   25342
gb:MN994468|Organism:Severe      TGTAGTTGTCTCAAGGGCTGTTGTTCTTGTGGATCCTGCTGCAAATTTGATGAAGACGAC   25342
gb:MT072688|Organism:Severe      TGTAGTTGTCTCAAGGGCTGTTGTTCTTGTGGATCCTGCTGCAAATTTGATGAAGACGAC   25327
gb:MN996527|Organism:Severe      TGTAGTTGTCTCAAGGGCTGTTGTTCTTGTGGATCCTGCTGCAAATTTGATGAAGACGAC   25309
gb:MT093631|Organism:Severe      TGTAGTTGTCTCAAGGGCTGTTGTTCTTGTGGATCCTGCTGCAAATTTGATGAAGACGAC   25380
gb:MT106053|Organism:Severe      TGTAGTTGTCTCAAGGGCTGTTGTTCTTGTGGATCCTGCTGCAAATTTGATGAAGACGAC   25342
gb:MT019533|Organism:Severe      TGTAGTTGTCTCAAGGGCTGTTGTTCTTGTGGATCCTGCTGCAAATTTGATGAAGACGAC   25342
gb:MT019531|Organism:Severe      TGTAGTTGTCTCAAGGGCTGTTGTTCTTGTGGATCCTGCTGCAAATTTGATGAAGACGAC   25342
gb:MN996528|Organism:Severe      TGTAGTTGTCTCAAGGGCTGTTGTTCTTGTGGATCCTGCTGCAAATTTGATGAAGACGAC   25342
gb:MN996530|Organism:Severe      TGTAGTTGTCTCAAGGGCTGTTGTTCTTGTGGATCCTGCTGCAAATTTGATGAAGACGAC   25328
gb:MN908947|Organism:Severe      TGTAGTTGTCTCAAGGGCTGTTGTTCTTGTGGATCCTGCTGCAAATTTGATGAAGACGAC   25342
gb:MT019532|Organism:Severe      TGTAGTTGTCTCAAGGGCTGTTGTTCTTGTGGATCCTGCTGCAAATTTGATGAAGACGAC   25342
                                 ************************************************************

gb:MT020781|Organism:Severe      TCTGAGCCAGTGCTCAAAGGAGTCAAATTACATTACACATAAACGAACTTATGGATTTGT   25390
gb:MT007544|Organism:Severe      TCTGAGCCAGTGCTCAAAGGAGTCAAATTACATTACACATAAACGAACTTATGGATTTGT   25402
gb:MN994467|Organism:Severe      TCTGAGCCAGTGCTCAAAGGAGTCAAATTACATTACACATAAACGAACTTATGGATTTGT   25402
gb:MT044257|Organism:Severe      TCTGAGCCAGTGCTCAAAGGAGTCAAATTACATTACACATAAACGAACTTATGGATTTGT   25402
gb:MT106054|Organism:Severe      TCTGAGCCAGTGCTCAAAGGAGTCAAATTACATTACACATAAACGAACTTATGGATTTGT   25402
gb:MT049951|Organism:Severe      TCTGAGCCAGTGCTCAAAGGAGTCAAATTACATTACACATAAACGAACTTATGGATTTGT   25402
gb:MN975262|Organism:Severe      TCTGAGCCAGTGCTCAAAGGAGTCAAATTACATTACACATAAACGAACTTATGGATTTGT   25402
gb:MT106052|Organism:Severe      TCTGAGCCAGTGCTCAAAGGAGTCAAATTACATTACACATAAACGAACTTATGGATTTGT   25402
gb:LC522975|Organism:Severe      TCTGAGCCAGTGCTCAAAGGAGTCAAATTACATTACACATAAACGAACTTATGGATTTGT   25399
gb:LC522973|Organism:Severe      TCTGAGCCAGTGCTCAAAGGAGTCAAATTACATTACACATAAACGAACTTATGGATTTGT   25399
gb:LC522974|Organism:Severe      TCTGAGCCAGTGCTCAAAGGAGTCAAATTACATTACACATAAACGAACTTATGGATTTGT   25399
gb:MN985325|Organism:Severe      TCTGAGCCAGTGCTCAAAGGAGTCAAATTACATTACACATAAACGAACTTATGGATTTGT   25402
gb:MT020881|Organism:Severe      TCTGAGCCAGTGCTCAAAGGAGTCAAATTACATTACACATAAACGAACTTATGGATTTGT   25402
gb:MT020880|Organism:Severe      TCTGAGCCAGTGCTCAAAGGAGTCAAATTACATTACACATAAACGAACTTATGGATTTGT   25402
gb:MT066175|Organism:Severe      TCTGAGCCAGTGCTCAAAGGAGTCAAATTACATTACACATAAACGAACTTATGGATTTGT   25402
gb:MN997409|Organism:Severe      TCTGAGCCAGTGCTCAAAGGAGTCAAATTACATTACACATAAACGAACTTATGGATTTGT   25402
gb:MN938384|Organism:Severe      TCTGAGCCAGTGCTCAAAGGAGTCAAATTACATTACACATAAACGAACTTATGGATTTGT   25370
gb:MT044258|Organism:Severe      TCTGAGCCAGTGCTCAAAGGAGTCAAATTACATTACACATAAACGAACTTATGGATTTGT   25378
gb:MT039890|Organism:Severe      TCTGAGCCAGTGCTCAAAGGAGTCAAATTACATTACACATAAACGAACTTATGGATTTGT   25402
gb:MN988713|Organism:Severe      TCTGAGCCAGTGCTCAAAGGAGTCAAATTACATTACACATAAACGAACTTATGGATTTGT   25402
gb:LC521925|Organism:Severe      TCTGAGCCAGTGCTCAAAGGAGTCAAATTACATTACACATAAACGAACTTATGGATTTGT   25375
gb:MT093571|Organism:Severe      TCTGAGCCAGTGCTCAAAGGAGTCAAATTACATTACACATAAACGAACTTATGGATTTGT   25402
gb:MT039887|Organism:Severe      TCTGAGCCAGTGCTCAAAGGAGTCAAATTACATTACACATAAACGAACTTATGGATTTGT   25399
gb:MT019530|Organism:Severe      TCTGAGCCAGTGCTCAAAGGAGTCAAATTACATTACACATAAACGAACTTATGGATTTGT   25402
gb:MT039888|Organism:Severe      TCTGAGCCAGTGCTCAAAGGAGTCAAATTACATTACACATAAACGAACTTATGGATTTGT   25402
gb:LC522972|Organism:Severe      TCTGAGCCAGTGCTCAAAGGAGTCAAATTACATTACACATAAACGAACTTATGGATTTGT   25399
gb:MT027063|Organism:Severe      TCTGAGCCAGTGCTCAAAGGAGTCAAATTACATTACACATAAACGAACTTATGGATTTGT   25402
gb:MT027062|Organism:Severe      TCTGAGCCAGTGCTCAAAGGAGTCAAATTACATTACACATAAACGAACTTATGGATTTGT   25402
gb:MT019529|Organism:Severe      TCTGAGCCAGTGCTCAAAGGAGTCAAATTACATTACACATAAACGAACTTATGGATTTGT   25402
gb:MN996529|Organism:Severe      TCTGAGCCAGTGCTCAAAGGAGTCAAATTACATTACACATAAACGAACTTATGGATTTGT   25390
gb:MN996531|Organism:Severe      TCTGAGCCAGTGCTCAAAGGAGTCAAATTACATTACACATAAACGAACTTATGGATTTGT   25389
gb:MT066176|Organism:Severe      TCTGAGCCAGTGCTCAAAGGAGTCAAATTACATTACACATAAACGAACTTATGGATTTGT   25402
gb:MT027064|Organism:Severe      TCTGAGCCAGTGCTCAAAGGAGTCAAATTACATTACACATAAACGAACTTATGGATTTGT   25402
gb:MN994468|Organism:Severe      TCTGAGCCAGTGCTCAAAGGAGTCAAATTACATTACACATAAACGAACTTATGGATTTGT   25402
gb:MT072688|Organism:Severe      TCTGAGCCAGTGCTCAAAGGAGTCAAATTACATTACACATAAACGAACTTATGGATTTGT   25387
gb:MN996527|Organism:Severe      TCTGAGCCAGTGCTCAAAGGAGTCAAATTACATTACACATAAACGAACTTATGGATTTGT   25369
gb:MT093631|Organism:Severe      TCTGAGCCAGTGCTCAAAGGAGTCAAATTACATTACACATAAACGAACTTATGGATTTGT   25440
gb:MT106053|Organism:Severe      TCTGAGCCAGTGCTCAAAGGAGTCAAATTACATTACACATAAACGAACTTATGGATTTGT   25402
```

```
gb:MT019533|Organism:Severe      TCTGAGCCAGTGCTCAAAGGAGTCAAATTACATTACACATAAACGAACTTATGGATTTGT   25402
gb:MT019531|Organism:Severe      TCTGAGCCAGTGCTCAAAGGAGTCAAATTACATTACACATAAACGAACTTATGGATTTGT   25402
gb:MN996528|Organism:Severe      TCTGAGCCAGTGCTCAAAGGAGTCAAATTACATTACACATAAACGAACTTATGGATTTGT   25402
gb:MN996530|Organism:Severe      TCTGAGCCAGTGCTCAAAGGAGTCAAATTACATTACACATAAACGAACTTATGGATTTGT   25388
gb:MN908947|Organism:Severe      TCTGAGCCAGTGCTCAAAGGAGTCAAATTACATTACACATAAACGAACTTATGGATTTGT   25402
gb:MT019532|Organism:Severe      TCTGAGCCAGTGCTCAAAGGAGTCAAATTACATTACACATAAACGAACTTATGGATTTGT   25402
                                 ************************************************************

gb:MT020781|Organism:Severe      TTATGAGAATCTTCACAATTGGAACTGTAACTTTGAAGCAAGGTGAAATCAAGGATGCTA   25450
gb:MT007544|Organism:Severe      TTATGAGAATCTTCACAATTGGAACTGTAACTTTGAAGCAAGGTGAAATCAAGGATGCTA   25462
gb:MN994467|Organism:Severe      TTATGAGAATCTTCACAATTGGAACTGTAACTTTGAAGCAAGGTGAAATCAAGGATGCTA   25462
gb:MT044257|Organism:Severe      TTATGAGAATCTTCACAATTGGAACTGTAACTTTGAAGCAAGGTGAAATCAAGGATGCTA   25462
gb:MT106054|Organism:Severe      TTATGAGAATCTTCACAATTGGAACTGTAACTTTGAAGCAAGGTGAAATCAAGGATGCTA   25462
gb:MT049951|Organism:Severe      TTATGAGAATCTTCACAATTGGAACTGTAACTTTGAAGCAAGGTGAAATCAAGGATGCTA   25462
gb:MN975262|Organism:Severe      TTATGAGAATCTTCACAATTGGAACTGTAACTTTGAAGCAAGGTGAAATCAAGGATGCTA   25462
gb:MT106052|Organism:Severe      TTATGAGAATCTTCACAATTGGAACTGTAACTTTGAAGCAAGGTGAAATCAAGGATGCTA   25462
gb:LC522975|Organism:Severe      TTATGAGAATCTTCACAATTGGAACTGTAACTTTGAAGCAAGGTGAAATCAAGGATGCTA   25459
gb:LC522973|Organism:Severe      TTATGAGAATCTTCACAATTGGAACTGTAACTTTGAAGCAAGGTGAAATCAAGGATGCTA   25459
gb:LC522974|Organism:Severe      TTATGAGAATCTTCACAATTGGAACTGTAACTTTGAAGCAAGGTGAAATCAAGGATGCTA   25459
gb:MN985325|Organism:Severe      TTATGAGAATCTTCACAATTGGAACTGTAACTTTGAAGCAAGGTGAAATCAAGGATGCTA   25462
gb:MT020881|Organism:Severe      TTATGAGAATCTTCACAATTGGAACTGTAACTTTGAAGCAAGGTGAAATCAAGGATGCTA   25462
gb:MT020880|Organism:Severe      TTATGAGAATCTTCACAATTGGAACTGTAACTTTGAAGCAAGGTGAAATCAAGGATGCTA   25462
gb:MT066175|Organism:Severe      TTATGAGAATCTTCACAATTGGAACTGTAACTTTGAAGCAAGGTGAAATCAAGGATGCTA   25462
gb:MN997409|Organism:Severe      TTATGAGAATCTTCACAATTGGAACTGTAACTTTGAAGCAAGGTGAAATCAAGGATGCTA   25462
gb:MN938384|Organism:Severe      TTATGAGAATCTTCACAATTGGAACTGTAACTTTGAAGCAAGGTGAAATCAAGGATGCTA   25430
gb:MT044258|Organism:Severe      TTATGAGAATCTTCACAATTGGAACTGTAACTTTGAAGCAAGGTGAAATCAAGGATGCTA   25438
gb:MT039890|Organism:Severe      TTATGAGAATCTTCACAATTGGAACTGTAACTTTGAAGCAAGGTGAAATCAAGGATGCTA   25462
gb:MN988713|Organism:Severe      TTATGAGAATCTTCACAATTGGAACTGTAACTTTGAAGCAAGGTGAAATCAAGGATGCTA   25462
gb:LC521925|Organism:Severe      TTATGAGAATCTTCACAATTGGAACTGTAACTTTGAAGCAAGGTGAAATCAAGGATGCTA   25435
gb:MT093571|Organism:Severe      TTATGAGAATCTTCACAATTGGAACTGTAACTTTGAAGCAAGGTGAAATCAAGGATGCTA   25462
gb:MT039887|Organism:Severe      TTATGAGAATCTTCACAATTGGAACTGTAACTTTGAAGCAAGGTGAAATCAAGGATGCTA   25459
gb:MT019530|Organism:Severe      TTATGAGAATCTTCACAATTGGAACTGTAACTTTGAAGCAAGGTGAAATCAAGGATGCTA   25462
gb:MT039888|Organism:Severe      TTATGAGAATCTTCACAATTGGAACTGTAACTTTGAAGCAAGGTGAAATCAAGGATGCTA   25462
gb:LC522972|Organism:Severe      TTATGAGAATCTTCACAATTGGAACTGTAACTTTGAAGCAAGGTGAAATCAAGGATGCTA   25459
gb:MT027063|Organism:Severe      TTATGAGAATCTTCACAATTGGAACTGTAACTTTGAAGCAAGGTGAAATCAAGGATGCTA   25462
gb:MT027062|Organism:Severe      TTATGAGAATCTTCACAATTGGAACTGTAACTTTGAAGCAAGGTGAAATCAAGGATGCTA   25462
gb:MT019529|Organism:Severe      TTATGAGAATCTTCACAATTGGAACTGTAACTTTGAAGCAAGGTGAAATCAAGGATGCTA   25462
gb:MN996529|Organism:Severe      TTATGAGAATCTTCACAATTGGAACTGTAACTTTGAAGCAAGGTGAAATCAAGGATGCTA   25450
gb:MN996531|Organism:Severe      TTATGAGAATCTTCACAATTGGAACTGTAACTTTGAAGCAAGGTGAAATCAAGGATGCTA   25449
gb:MT066176|Organism:Severe      TTATGAGAATCTTCACAATTGGAACTGTAACTTTGAAGCAAGGTGAAATCAAGGATGCTA   25462
gb:MT027064|Organism:Severe      TTATGAGAATCTTCACAATTGGAACTGTAACTTTGAAGCAAGGTGAAATCAAGGATGCTA   25462
gb:MN994468|Organism:Severe      TTATGAGAATCTTCACAATTGGAACTGTAACTTTGAAGCAAGGTGAAATCAAGGATGCTA   25462
gb:MT072688|Organism:Severe      TTATGAGAATCTTCACAATTGGAACTGTAACTTTGAAGCAAGGTGAAATCAAGGATGCTA   25447
gb:MN996527|Organism:Severe      TTATGAGAATCTTCACAATTGGAACTGTAACTTTGAAGCAAGGTGAAATCAAGGATGCTA   25429
gb:MT093631|Organism:Severe      TTATGAGAATCTTCACAATTGGAACTGTAACTTTGAAGCAAGGTGAAATCAAGGATGCTA   25500
gb:MT106053|Organism:Severe      TTATGAGAATCTTCACAATTGGAACTGTAACTTTGAAGCAAGGTGAAATCAAGGATGCTA   25462
gb:MT019533|Organism:Severe      TTATGAGAATCTTCACAATTGGAACTGTAACTTTGAAGCAAGGTGAAATCAAGGATGCTA   25462
gb:MT019531|Organism:Severe      TTATGAGAATCTTCACAATTGGAACTGTAACTTTGAAGCAAGGTGAAATCAAGGATGCTA   25462
gb:MN996528|Organism:Severe      TTATGAGAATCTTCACAATTGGAACTGTAACTTTGAAGCAAGGTGAAATCAAGGATGCTA   25462
gb:MN996530|Organism:Severe      TTATGAGAATCTTCACAATTGGAACTGTAACTTTGAAGCAAGGTGAAATCAAGGATGCTA   25448
gb:MN908947|Organism:Severe      TTATGAGAATCTTCACAATTGGAACTGTAACTTTGAAGCAAGGTGAAATCAAGGATGCTA   25462
gb:MT019532|Organism:Severe      TTATGAGAATCTTCACAATTGGAACTGTAACTTTGAAGCAAGGTGAAATCAAGGATGCTA   25462
                                 ************************************************************

gb:MT020781|Organism:Severe      CTCCTTCAGATTTTGTTCGCGCTACTGCAACGATACCGATACAAGCCTCACTCCCTTTCG   25510
gb:MT007544|Organism:Severe      CTCCTTCAGATTTTGTTCGCGCTACTGCAACGATACCGATACAAGCCTCACTCCCTTTCG   25522
gb:MN994467|Organism:Severe      CTCCTTCAGATTTTGTTCGCGCTACTGCAACGATACCGATACAAGCCTCACTCCCTTTCG   25522
gb:MT044257|Organism:Severe      CTCCTTCAGATTTTGTTCGCGCTACTGCAACGATACCGATACAAGCCTCACTCCCTTTCG   25522
gb:MT106054|Organism:Severe      CTCCTTCAGATTTTGTTCGCGCTACTGCAACGATACCGATACAAGCCTCACTCCCTTTCG   25522
gb:MT049951|Organism:Severe      CTCCTTCAGATTTTGTTCGCGCTACTGCAACGATACCGATACAAGCCTCACTCCCTTTCG   25522
gb:MN975262|Organism:Severe      CTCCTTCAGATTTTGTTCGCGCTACTGCAACGATACCGATACAAGCCTCACTCCCTTTCG   25522
gb:MT106052|Organism:Severe      CTCCTTCAGATTTTGTTCGCGCTACTGCAACGATACCGATACAAGCCTCACTCCCTTTCG   25522
gb:LC522975|Organism:Severe      CTCCTTCAGATTTTGTTCGCGCTACTGCAACGATACCGATACAAGCCTCACTCCCTTTCG   25519
gb:LC522973|Organism:Severe      CTCCTTCAGATTTTGTTCGCGCTACTGCAACGATACCGATACAAGCCTCACTCCCTTTCG   25519
gb:LC522974|Organism:Severe      CTCCTTCAGATTTTGTTCGCGCTACTGCAACGATACCGATACAAGCCTCACTCCCTTTCG   25519
```

```
gb:MN985325|Organism:Severe      CTCCTTCAGATTTTGTTCGCGCTACTGCAACGATACCGATACAAGCCTCACTCCCTTTCG   25522
gb:MT020881|Organism:Severe      CTCCTTCAGATTTTGTTCGCGCTACTGCAACGATACCGATACAAGCCTCACTCCCTTTCG   25522
gb:MT020880|Organism:Severe      CTCCTTCAGATTTTGTTCGCGCTACTGCAACGATACCGATACAAGCCTCACTCCCTTTCG   25522
gb:MT066175|Organism:Severe      CTCCTTCAGATTTTGTTCGCGCTACTGCAACGATACCGATACAAGCCTCACTCCCTTTCG   25522
gb:MN997409|Organism:Severe      CTCCTTCAGATTTTGTTCGCGCTACTGCAACGATACCGATACAAGCCTCACTCCCTTTCG   25522
gb:MN938384|Organism:Severe      CTCCTTCAGATTTTGTTCGCGCTACTGCAACGATACCGATACAAGCCTCACTCCCTTTCG   25490
gb:MT044258|Organism:Severe      CTCCTTCAGATTTTGTTCGCGCTACTGCAACGATACCGATACAAGCCTCACTCCCTTTCG   25498
gb:MT039890|Organism:Severe      CTCCTTCAGATTTTGTTCGCGCTACTGCAACGATACCGATACAAGCCTCACTCCCTTTCG   25522
gb:MN988713|Organism:Severe      CTCCTTCAGATTTTGTTCGCGCTACTGCAACGATACCGATACAAGCCTCACTCCCTTTCG   25522
gb:LC521925|Organism:Severe      CTCCTTCAGATTTTGTTCGCGCTACTGCAACGATACCGATACAAGCCTCACTCCCTTTCG   25495
gb:MT093571|Organism:Severe      CTCCTTCAGATTTTGTTCGCGCTACTGCAACGATACCGATACAAGCCTCACTCCCTTTCG   25522
gb:MT039887|Organism:Severe      CTCCTTCAGATTTTGTTCGCGCTACTGCAACGATACCGATACAAGCCTCACTCCCTTTCG   25519
gb:MT019530|Organism:Severe      CTCCTTCAGATTTTGTTCGCGCTACTGCAACGATACCGATACAAGCCTCACTCCCTTTCG   25522
gb:MT039888|Organism:Severe      CTCCTTCAGATTTTGTTCGCGCTACTGCAACGATACCGATACAAGCCTCACTCCCTTTCG   25522
gb:LC522972|Organism:Severe      CTCCTTCAGATTTTGTTCGCGCTACTGCAACGATACCGATACAAGCCTCACTCCCTTTCG   25519
gb:MT027063|Organism:Severe      CTCCTTCAGATTTTGTTCGCGCTACTGCAACGATACCGATACAAGCCTCACTCCCTTTCG   25522
gb:MT027062|Organism:Severe      CTCCTTCAGATTTTGTTCGCGCTACTGCAACGATACCGATACAAGCCTCACTCCCTTTCG   25522
gb:MT019529|Organism:Severe      CTCCTTCAGATTTTGTTCGCGCTACTGCAACGATACCGATACAAGCCTCACTCCCTTTCG   25522
gb:MN996529|Organism:Severe      CTCCTTCAGATTTTGTTCGCGCTACTGCAACGATACCGATACAAGCCTCACTCCCTTTCG   25510
gb:MN996531|Organism:Severe      CTCCTTCAGATTTTGTTCGCGCTACTGCAACGATACCGATACAAGCCTCACTCCCTTTCG   25509
gb:MT066176|Organism:Severe      CTCCTTCAGATTTTGTTCGCGCTACTGCAACGATACCGATACAAGCCTCACTCCCTTTCG   25522
gb:MT027064|Organism:Severe      CTCCTTCAGATTTTGTTCGCGCTACTGCAACGATACCGATACAAGCCTCACTCCCTTTCG   25522
gb:MN994468|Organism:Severe      CTCCTTCAGATTTTGTTCGCGCTACTGCAACGATACCGATACAAGCCTCACTCCCTTTCG   25522
gb:MT072688|Organism:Severe      CTCCTTCAGATTTTGTTCGCGCTACTGCAACGATACCGATACAAGCCTCACTCCCTTTCG   25507
gb:MN996527|Organism:Severe      CTCCTTCAGATTTTGTTCGCGCTACTGCAACGATACCGATACAAGCCTCACTCCCTTTCG   25489
gb:MT093631|Organism:Severe      CTCCTTCAGATTTTGTTCGCGCTACTGCAACGATACCGATACAAGCCTCACTCCCTTTCG   25560
gb:MT106053|Organism:Severe      CTCCTTCAGATTTTGTTCGCGCTACTGCAACGATACCGATACAAGCCTCACTCCCTTTCG   25522
gb:MT019533|Organism:Severe      CTCCTTCAGATTTTGTTCGCGCTACTGCAACGATACCGATACAAGCCTCACTCCCTTTCG   25522
gb:MT019531|Organism:Severe      CTCCTTCAGATTTTGTTCGCGCTACTGCAACGATACCGATACAAGCCTCACTCCCTTTCG   25522
gb:MN996528|Organism:Severe      CTCCTTCAGATTTTGTTCGCGCTACTGCAACGATACCGATACAAGCCTCACTCCCTTTCG   25522
gb:MN996530|Organism:Severe      CTCCTTCAGATTTTGTTCGCGCTACTGCAACGATACCGATACAAGCCTCACTCCCTTTCG   25508
gb:MN908947|Organism:Severe      CTCCTTCAGATTTTGTTCGCGCTACTGCAACGATACCGATACAAGCCTCACTCCCTTTCG   25522
gb:MT019532|Organism:Severe      CTCCTTCAGATTTTGTTCGCGCTACTGCAACGATACCGATACAAGCCTCACTCCCTTTCG   25522
                                 ************************************************************

gb:MT020781|Organism:Severe      GATGGCTTATTGTTGGCGTTGCACTTCTTGCTGTTTTTCAGAGCGCTTCCAAAATCATAA   25570
gb:MT007544|Organism:Severe      GATGGCTTATTGTTGGCGTTGCACTTCTTGCTGTTTTTCAGAGCGCTTCCAAAATCATAA   25582
gb:MN994467|Organism:Severe      GATGGCTTATTGTTGGCGTTGCACTTCTTGCTGTTTTTCAGAGCGCTTCCAAAATCATAA   25582
gb:MT044257|Organism:Severe      GATGGCTTATTGTTGGCGTTGCACTTCTTGCTGTTTTTCAGAGCGCTTCCAAAATCATAA   25582
gb:MT106054|Organism:Severe      GATGGCTTATTGTTGGCGTTGCACTTCTTGCTGTTTTTCAGAGCGCTTCCAAAATCATAA   25582
gb:MT049951|Organism:Severe      GATGGCTTATTGTTGGCGTTGCACTTCTTGCTGTTTTTCAGAGCGCTTCCAAAATCATAA   25582
gb:MN975262|Organism:Severe      GATGGCTTATTGTTGGCGTTGCACTTCTTGCTGTTTTTCAGAGCGCTTCCAAAATCATAA   25582
gb:MT106052|Organism:Severe      GATGGCTTATTGTTGGCGTTGCACTTCTTGCTGTTTTTCAGAGCGCTTCCAAAATCATAA   25582
gb:LC522975|Organism:Severe      GATGGCTTATTGTTGGCGTTGCACTTCTTGCTGTTTTTCAGAGCGCTTCCAAAATCATAA   25579
gb:LC522973|Organism:Severe      GATGGCTTATTGTTGGCGTTGCACTTCTTGCTGTTTTTCAGAGCGCTTCCAAAATCATAA   25579
gb:LC522974|Organism:Severe      GATGGCTTATTGTTGGCGTTGCACTTCTTGCTGTTTTTCAGAGCGCTTCCAAAATCATAA   25579
gb:MN985325|Organism:Severe      GATGGCTTATTGTTGGCGTTGCACTTCTTGCTGTTTTTCAGAGCGCTTCCAAAATCATAA   25582
gb:MT020881|Organism:Severe      GATGGCTTATTGTTGGCGTTGCACTTCTTGCTGTTTTTCAGAGCGCTTCCAAAATCATAA   25582
gb:MT020880|Organism:Severe      GATGGCTTATTGTTGGCGTTGCACTTCTTGCTGTTTTTCAGAGCGCTTCCAAAATCATAA   25582
gb:MT066175|Organism:Severe      GATGGCTTATTGTTGGCGTTGCACTTCTTGCTGTTTTTCAGAGCGCTTCCAAAATCATAA   25582
gb:MN997409|Organism:Severe      GATGGCTTATTGTTGGCGTTGCACTTCTTGCTGTTTTTCAGAGCGCTTCCAAAATCATAA   25582
gb:MN938384|Organism:Severe      GATGGCTTATTGTTGGCGTTGCACTTCTTGCTGTTTTTCAGAGCGCTTCCAAAATCATAA   25550
gb:MT044258|Organism:Severe      GATGGCTTATTGTTGGCGTTGCACTTCTTGCTGTTTTTCAGAGCGCTTCCAAAATCATAA   25558
gb:MT039890|Organism:Severe      GATGGCTTATTGTTGGCGTTGCACTTCTTGCTGTTTTTCAGAGCGCTTCCAAAATCATAA   25582
gb:MN988713|Organism:Severe      GATGGCTTATTGTTGGCGTTGCACTTCTTGCTGTTTTTCAGAGCGCTTCCAAAATCATAA   25582
gb:LC521925|Organism:Severe      GATGGCTTATTGTTGGCGTTGCACTTCTTGCTGTTTTTCAGAGCGCTTCCAAAATCATAA   25555
gb:MT093571|Organism:Severe      GATGGCTTATTGTTGGCGTTGCACTTCTTGCTGTTTTTCAGAGCGCTTCCAAAATCATAA   25582
gb:MT039887|Organism:Severe      GATGGCTTATTGTTGGCGTTGCACTTCTTGCTGTTTTTCAGAGCGCTTCCAAAATCATAA   25579
gb:MT019530|Organism:Severe      GATGGCTTATTGTTGGCGTTGCACTTCTTGCTGTTTTTCAGAGCGCTTCCAAAATCATAA   25582
gb:MT039888|Organism:Severe      GATGGCTTATTGTTGGCGTTGCACTTCTTGCTGTTTTTCAGAGCGCTTCCAAAATCATAA   25582
gb:LC522972|Organism:Severe      GATGGCTTATTGTTGGCGTTGCACTTCTTGCTGTTTTTCAGAGCGCTTCCAAAATCATAA   25579
gb:MT027063|Organism:Severe      GATGGCTTATTGTTGGCGTTGCACTTCTTGCTGTTTTTCAGAGCGCTTCCAAAATCATAA   25582
gb:MT027062|Organism:Severe      GATGGCTTATTGTTGGCGTTGCACTTCTTGCTGTTTTTCAGAGCGCTTCCAAAATCATAA   25582
gb:MT019529|Organism:Severe      GATGGCTTATTGTTGGCGTTGCACTTCTTGCTGTTTTTCAGAGCGCTTCCAAAATCATAA   25582
gb:MN996529|Organism:Severe      GATGGCTTATTGTTGGCGTTGCACTTCTTGCTGTTTTTCAGAGCGCTTCCAAAATCATAA   25570
```

```
gb:MN996531|Organism:Severe      GATGGCTTATTGTTGGCGTTGCACTTCTTGCTGTTTTTCAGAGCGCTTCCAAAATCATAA   25569
gb:MT066176|Organism:Severe      GATGGCTTATTGTTGGCGTTGCACTTCTTGCTGTTTTTCAGAGCGCTTCCAAAATCATAA   25582
gb:MT027064|Organism:Severe      GATGGCTTATTGTTGGCGTTGCACTTCTTGCTGTTTTTCAGAGCGCTTCCAAAATCATAA   25582
gb:MN994468|Organism:Severe      GATGGCTTATTGTTGGCGTTGCACTTCTTGCTGTTTTTCAGAGCGCTTCCAAAATCATAA   25582
gb:MT072688|Organism:Severe      GATGGCTTATTGTTGGCGTTGCACTTCTTGCTGTTTTTCAGAGCGCTTCCAAAATCATAA   25567
gb:MN996527|Organism:Severe      GATGGCTTATTGTTGGCGTTGCACTTCTTGCTGTTTTTCAGAGCGCTTCCAAAATCATAA   25549
gb:MT093631|Organism:Severe      GATGGCTTATTGTTGGCGTTGCACTTCTTGCTGTTTTTCAGAGCGCTTCCAAAATCATAA   25620
gb:MT106053|Organism:Severe      GATGGCTTATTGTTGGCGTTGCACTTCTTGCTGTTTTTCAGAGCGCTTCCAAAATCATAA   25582
gb:MT019533|Organism:Severe      GATGGCTTATTGTTGGCGTTGCACTTCTTGCTGTTTTTCAGAGCGCTTCCAAAATCATAA   25582
gb:MT019531|Organism:Severe      GATGGCTTATTGTTGGCGTTGCACTTCTTGCTGTTTTTCAGAGCGCTTCCAAAATCATAA   25582
gb:MN996528|Organism:Severe      GATGGCTTATTGTTGGCGTTGCACTTCTTGCTGTTTTTCAGAGCGCTTCCAAAATCATAA   25582
gb:MN996530|Organism:Severe      GATGGCTTATTGTTGGCGTTGCACTTCTTGCTGTTTTTCAGAGCGCTTCCAAAATCATAA   25568
gb:MN908947|Organism:Severe      GATGGCTTATTGTTGGCGTTGCACTTCTTGCTGTTTTTCAGAGCGCTTCCAAAATCATAA   25582
gb:MT019532|Organism:Severe      GATGGCTTATTGTTGGCGTTGCACTTCTTGCTGTTTTTCAGAGCGCTTCCAAAATCATAA   25582
                                 ************************************************************

gb:MT020781|Organism:Severe      CCCTCAAAAAGAGATGGCAACTAGCACTCTCCAAGGGTGTTCACTTTGTTTGCAACTTGC   25630
gb:MT007544|Organism:Severe      CCCTCAAAAAGAGATGGCAACTAGCACTCTCCAAGGGTGTTCACTTTGTTTGCAACTTGC   25642
gb:MN994467|Organism:Severe      CCCTCAAAAAGAGATGGCAACTAGCACTCTCCAAGGGTGTTCACTTTGTTTGCAACTTGC   25642
gb:MT044257|Organism:Severe      CCCTCAAAAAGAGATGGCAACTAGCACTCTCCAAGGGTGTTCACTTTGTTTGCAACTTGC   25642
gb:MT106054|Organism:Severe      CCCTCAAAAAGAGATGGCAACTAGCACTCTCCAAGGGTGTTCACTTTGTTTGCAACTTGC   25642
gb:MT049951|Organism:Severe      CCCTCAAAAAGAGATGGCAACTAGCACTCTCCAAGGGTGTTCACTTTGTTTGCAACTTGC   25642
gb:MN975262|Organism:Severe      CCCTCAAAAAGAGATGGCAACTAGCACTCTCCAAGGGTGTTCACTTTGTTTGCAACTTGC   25642
gb:MT106052|Organism:Severe      CCCTCAAAAAGAGATGGCAACTAGCACTCTCCAAGGGTGTTCACTTTGTTTGCAACTTGC   25642
gb:LC522975|Organism:Severe      CCCTCAAAAAGAGATGGCAACTAGCACTCTCCAAGGGTGTTCACTTTGTTTGCAACTTGC   25639
gb:LC522973|Organism:Severe      CCCTCAAAAAGAGATGGCAACTAGCACTCTCCAAGGGTGTTCACTTTGTTTGCAACTTGC   25639
gb:LC522974|Organism:Severe      CCCTCAAAAAGAGATGGCAACTAGCACTCTCCAAGGGTGTTCACTTTGTTTGCAACTTGC   25639
gb:MN985325|Organism:Severe      CCCTCAAAAAGAGATGGCAACTAGCACTCTCCAAGGGTGTTCACTTTGTTTGCAACTTGC   25642
gb:MT020881|Organism:Severe      CCCTCAAAAAGAGATGGCAACTAGCACTCTCCAAGGGTGTTCACTTTGTTTGCAACTTGC   25642
gb:MT020880|Organism:Severe      CCCTCAAAAAGAGATGGCAACTAGCACTCTCCAAGGGTGTTCACTTTGTTTGCAACTTGC   25642
gb:MT066175|Organism:Severe      CCCTCAAAAAGAGATGGCAACTAGCACTCTCCAAGGGTGTTCACTTTGTTTGCAACTTGC   25642
gb:MN997409|Organism:Severe      CCCTCAAAAAGAGATGGCAACTAGCACTCTCCAAGGGTGTTCACTTTGTTTGCAACTTGC   25642
gb:MN938384|Organism:Severe      CCCTCAAAAAGAGATGGCAACTAGCACTCTCCAAGGGTGTTCACTTTGTTTGCAACTTGC   25610
gb:MT044258|Organism:Severe      CCCTCAAAAAGAGATGGCAACTAGCACTCTCCAAGGGTGTTCACTTTGTTTGCAACTTGC   25618
gb:MT039890|Organism:Severe      CCCTCAAAAAGAGATGGCAACTAGCACTCTCCAAGGGTGTTCACTTTGTTTGCAACTTGC   25642
gb:MN988713|Organism:Severe      CCCTCAAAAAGAGATGGCAACTAGCACTCTCCAAGGGTGTTCACTTTGTTTGCAACTTGC   25642
gb:LC521925|Organism:Severe      CCCTCAAAAAGAGATGGCAACTAGCACTCTCCAAGGGTGTTCACTTTGTTTGCAACTTGC   25615
gb:MT093571|Organism:Severe      CCCTCAAAAAGAGATGGCAACTAGCACTCTCCAAGGGTGTTCACTTTGTTTGCAACTTGC   25642
gb:MT039887|Organism:Severe      CCCTCAAAAAGAGATGGCAACTAGCACTCTCCAAGGGTGTTCACTTTGTTTGCAACTTGC   25639
gb:MT019530|Organism:Severe      CCCTCAAAAAGAGATGGCAACTAGCACTCTCCAAGGGTGTTCACTTTGTTTGCAACTTGC   25642
gb:MT039888|Organism:Severe      CCCTCAAAAAGAGATGGCAACTAGCACTCTCCAAGGGTGTTCACTTTGTTTGCAACTTGC   25642
gb:LC522972|Organism:Severe      CCCTCAAAAAGAGATGGCAACTAGCACTCTCCAAGGGTGTTCACTTTGTTTGCAACTTGC   25639
gb:MT027063|Organism:Severe      CCCTCAAAAAGAGATGGCAACTAGCACTCTCCAAGGGTGTTCACTTTGTTTGCAACTTGC   25642
gb:MT027062|Organism:Severe      CCCTCAAAAAGAGATGGCAACTAGCACTCTCCAAGGGTGTTCACTTTGTTTGCAACTTGC   25642
gb:MT019529|Organism:Severe      CCCTCAAAAAGAGATGGCAACTAGCACTCTCCAAGGGTGTTCACTTTGTTTGCAACTTGC   25642
gb:MN996529|Organism:Severe      CCCTCAAAAAGAGATGGCAACTAGCACTCTCCAAGGGTGTTCACTTTGTTTGCAACTTGC   25630
gb:MN996531|Organism:Severe      CCCTCAAAAAGAGATGGCAACTAGCACTCTCCAAGGGTGTTCACTTTGTTTGCAACTTGC   25629
gb:MT066176|Organism:Severe      CCCTCAAAAAGAGATGGCAACTAGCACTCTCCAAGGGTGTTCACTTTGTTTGCAACTTGC   25642
gb:MT027064|Organism:Severe      CCCTCAAAAAGAGATGGCAACTAGCACTCTCCAAGGGTGTTCACTTTGTTTGCAACTTGC   25642
gb:MN994468|Organism:Severe      CCCTCAAAAAGAGATGGCAACTAGCACTCTCCAAGGGTGTTCACTTTGTTTGCAACTTGC   25642
gb:MT072688|Organism:Severe      CCCTCAAAAAGAGATGGCAACTAGCACTCTCCAAGGGTGTTCACTTTGTTTGCAACTTGC   25627
gb:MN996527|Organism:Severe      CCCTCAAAAAGAGATGGCAACTAGCACTCTCCAAGGGTGTTCACTTTGTTTGCAACTTGC   25609
gb:MT093631|Organism:Severe      CCCTCAAAAAGAGATGGCAACTAGCACTCTCCAAGGGTGTTCACTTTGTTTGCAACTTGC   25680
gb:MT106053|Organism:Severe      CCCTCAAAAAGAGATGGCAACTAGCACTCTCCAAGGGTGTTCACTTTGTTTGCAACTTGC   25642
gb:MT019533|Organism:Severe      CCCTCAAAAAGAGATGGCAACTAGCACTCTCCAAGGGTGTTCACTTTGTTTGCAACTTGC   25642
gb:MT019531|Organism:Severe      CCCTCAAAAAGAGATGGCAACTAGCACTCTCCAAGGGTGTTCACTTTGTTTGCAACTTGC   25642
gb:MN996528|Organism:Severe      CCCTCAAAAAGAGATGGCAACTAGCACTCTCCAAGGGTGTTCACTTTGTTTGCAACTTGC   25642
gb:MN996530|Organism:Severe      CCCTCAAAAAGAGATGGCAACTAGCACTCTCCAAGGGTGTTCACTTTGTTTGCAACTTGC   25628
gb:MN908947|Organism:Severe      CCCTCAAAAAGAGATGGCAACTAGCACTCTCCAAGGGTGTTCACTTTGTTTGCAACTTGC   25642
gb:MT019532|Organism:Severe      CCCTCAAAAAGAGATGGCAACTAGCACTCTCCAAGGGTGTTCACTTTGTTTGCAACTTGC   25642
                                 ************************************************************

gb:MT020781|Organism:Severe      TGTTGTTGTTTGTAACAGTTTACTCACACCTTTTGCTCGTTGCTGCTGGCCTTGAAGCCC   25690
gb:MT007544|Organism:Severe      TGTTGTTGTTTGTAACAGTTTACTCACACCTTTTGCTCGTTGCTGCTGGCCTTGAAGCCC   25702
gb:MN994467|Organism:Severe      TGTTGTTGTTTGTAACAGTTTACTCACACCTTTTGCTCGTTGCTGCTGGCCTTGAAGCCC   25702
```

```
gb:MT044257|Organism:Severe      TGTTGTTGTTTGTAACAGTTTACTCACACCTTTTGCTCGTTGCTGCTGGCCTTGAAGCCC   25702
gb:MT106054|Organism:Severe      TGTTGTTGTTTGTAACAGTTTACTCACACCTTTTGCTCGTTGCTGCTGGCCTTGAAGCCC   25702
gb:MT049951|Organism:Severe      TGTTGTTGTTTGTAACAGTTTACTCACACCTTTTGCTCGTTGCTGCTGGCCTTGAAGCCC   25702
gb:MN975262|Organism:Severe      TGTTGTTGTTTGTAACAGTTTACTCACACCTTTTGCTCGTTGCTGCTGGCCTTGAAGCCC   25702
gb:MT106052|Organism:Severe      TGTTGTTGTTTGTAACAGTTTACTCACACCTTTTGCTCGTTGCTGCTGGCCTTGAAGCCC   25702
gb:LC522975|Organism:Severe      TGTTGTTGTTTGTAACAGTTTACTCACACCTTTTGCTCGTTGCTGCTGGCCTTGAAGCCC   25699
gb:LC522973|Organism:Severe      TGTTGTTGTTTGTAACAGTTTACTCACACCTTTTGCTCGTTGCTGCTGGCCTTGAAGCCC   25699
gb:LC522974|Organism:Severe      TGTTGTTGTTTGTAACAGTTTACTCACACCTTTTGCTCGTTGCTGCTGGCCTTGAAGCCC   25699
gb:MN985325|Organism:Severe      TGTTGTTGTTTGTAACAGTTTACTCACACCTTTTGCTCGTTGCTGCTGGCCTTGAAGCCC   25702
gb:MT020881|Organism:Severe      TGTTGTTGTTTGTAACAGTTTACTCACACCTTTTGCTCGTTGCTGCTGGCCTTGAAGCCC   25702
gb:MT020880|Organism:Severe      TGTTGTTGTTTGTAACAGTTTACTCACACCTTTTGCTCGTTGCTGCTGGCCTTGAAGCCC   25702
gb:MT066175|Organism:Severe      TGTTGTTGTTTGTAACAGTTTACTCACACCTTTTGCTCGTTGCTGCTGGCCTTGAAGCCC   25702
gb:MN997409|Organism:Severe      TGTTGTTGTTTGTAACAGTTTACTCACACCTTTTGCTCGTTGCTGCTGGCCTTGAAGCCC   25702
gb:MN938384|Organism:Severe      TGTTGTTGTTTGTAACAGTTTACTCACACCTTTTGCTCGTTGCTGCTGGCCTTGAAGCCC   25670
gb:MT044258|Organism:Severe      TGTTGTTGTTTGTAACAGTTTACTCACACCTTTTGCTCGTTGCTGCTGGCCTTGAAGCCC   25678
gb:MT039890|Organism:Severe      TGTTGTTGTTTGTAACAGTTTACTCACACCTTTTGCTCGTTGCTGCTGGCCTTGAAGCCC   25702
gb:MN988713|Organism:Severe      TGTTGTTGTTTGTAACAGTTTACTCACACCTTTTGCTCGTTGCTGCTGGCCTTGAAGCCC   25702
gb:LC521925|Organism:Severe      TGTTGTTGTTTGTAACAGTTTACTCACACCTTTTGCTCGTTGCTGCTGGCCTTGAAGCCC   25675
gb:MT093571|Organism:Severe      TGTTGTTGTTTGTAACAGTTTACTCACACCTTTTGCTCGTTGCTGCTGGCCTTGAAGCCC   25702
gb:MT039887|Organism:Severe      TGTTGTTGTTTGTAACAGTTTACTCACACCTTTTGCTCGTTGCTGCTGGCCTTGAAGCCC   25699
gb:MT019530|Organism:Severe      TGTTGTTGTTTGTAACAGTTTACTCACACCTTTTGCTCGTTGCTGCTGGCCTTGAAGCCC   25702
gb:MT039888|Organism:Severe      TGTTGTTGTTTGTAACAGTTTACTCACACCTTTTGCTCGTTGCTGCTGGCCTTGAAGCCC   25702
gb:LC522972|Organism:Severe      TGTTGTTGTTTGTAACAGTTTACTCACACCTTTTGCTCGTTGCTGCTGGCCTTGAAGCCC   25699
gb:MT027063|Organism:Severe      TGTTGTTGTTTGTAACAGTTTACTCACACCTTTTGCTCGTTGCTGCTGGCCTTGAAGCCC   25702
gb:MT027062|Organism:Severe      TGTTGTTGTTTGTAACAGTTTACTCACACCTTTTGCTCGTTGCTGCTGGCCTTGAAGCCC   25702
gb:MT019529|Organism:Severe      TGTTGTTGTTTGTAACAGTTTACTCACACCTTTTGCTCGTTGCTGCTGGCCTTGAAGCCC   25702
gb:MN996529|Organism:Severe      TGTTGTTGTTTGTAACAGTTTACTCACACCTTTTGCTCGTTGCTGCTGGCCTTGAAGCCC   25690
gb:MN996531|Organism:Severe      TGTTGTTGTTTGTAACAGTTTACTCACACCTTTTGCTCGTTGCTGCTGGCCTTGAAGCCC   25689
gb:MT066176|Organism:Severe      TGTTGTTGTTTGTAACAGTTTACTCACACCTTTTGCTCGTTGCTGCTGGCCTTGAAGCCC   25702
gb:MT027064|Organism:Severe      TGTTGTTGTTTGTAACAGTTTACTCACACCTTTTGCTCGTTGCTGCTGGCCTTGAAGCCC   25702
gb:MN994468|Organism:Severe      TGTTGTTGTTTGTAACAGTTTACTCACACCTTTTGCTCGTTGCTGCTGGCCTTGAAGCCC   25702
gb:MT072688|Organism:Severe      TGTTGTTGTTTGTAACAGTTTACTCACACCTTTTGCTCGTTGCTGCTGGCCTTGAAGCCC   25687
gb:MN996527|Organism:Severe      TGTTGTTGTTTGTAACAGTTTACTCACACCTTTTGCTCGTTGCTGCTGGCCTTGAAGCCC   25669
gb:MT093631|Organism:Severe      TGTTGTTGTTTGTAACAGTTTACTCACACCTTTTGCTCGTTGCTGCTGGCCTTGAAGCCC   25740
gb:MT106053|Organism:Severe      TGTTGTTGTTTGTAACAGTTTACTCACACCTTTTGCTCGTTGCTGCTGGCCTTGAAGCCC   25702
gb:MT019533|Organism:Severe      TGTTGTTGTTTGTAACAGTTTACTCACACCTTTTGCTCGTTGCTGCTGGCCTTGAAGCCC   25702
gb:MT019531|Organism:Severe      TGTTGTTGTTTGTAACAGTTTACTCACACCTTTTGCTCGTTGCTGCTGGCCTTGAAGCCC   25702
gb:MN996528|Organism:Severe      TGTTGTTGTTTGTAACAGTTTACTCACACCTTTTGCTCGTTGCTGCTGGCCTTGAAGCCC   25702
gb:MN996530|Organism:Severe      TGTTGTTGTTTGTAACAGTTTACTCACACCTTTTGCTCGTTGCTGCTGGCCTTGAAGCCC   25688
gb:MN908947|Organism:Severe      TGTTGTTGTTTGTAACAGTTTACTCACACCTTTTGCTCGTTGCTGCTGGCCTTGAAGCCC   25702
gb:MT019532|Organism:Severe      TGTTGTTGTTTGTAACAGTTTACTCACACCTTTTGCTCGTTGCTGCTGGCCTTGAAGCCC   25702
                                 ************************************************************

gb:MT020781|Organism:Severe      CTTTTCTCTATCTTTATGCTTTAGTCTACTTCTTGCAGAGTATAAACTTTGTAAGAATAA   25750
gb:MT007544|Organism:Severe      CTTTTCTCTATCTTTATGCTTTAGTCTACTTCTTGCAGAGTATAAACTTTGTAAGAATAA   25762
gb:MN994467|Organism:Severe      CTTTTCTCTATCTTTATGCTTTAGTCTACTTCTTGCAGAGTATAAACTTTGTAAGAATAA   25762
gb:MT044257|Organism:Severe      CTTTTCTCTATCTTTATGCTTTAGTCTACTTCTTGCAGAGTATAAACTTTGTAAGAATAA   25762
gb:MT106054|Organism:Severe      CTTTTCTCTATCTTTATGCTTTAGTCTACTTCTTGCAGAGTATAAACTTTGTAAGAATAA   25762
gb:MT049951|Organism:Severe      CTTTTCTCTATCTTTATGCTTTAGTCTACTTCTTGCAGAGTATAAACTTTGTAAGAATAA   25762
gb:MN975262|Organism:Severe      CTTTTCTCTATCTTTATGCTTTAGTCTACTTCTTGCAGAGTATAAACTTTGTAAGAATAA   25762
gb:MT106052|Organism:Severe      CTTTTCTCTATCTTTATGCTTTAGTCTACTTCTTGCAGAGTATAAACTTTGTAAGAATAA   25762
gb:LC522975|Organism:Severe      CTTTTCTCTATCTTTATGCTTTAGTCTACTTCTTGCAGAGTATAAACTTTGTAAGAATAA   25759
gb:LC522973|Organism:Severe      CTTTTCTCTATCTTTATGCTTTAGTCTACTTCTTGCAGAGTATAAACTTTGTAAGAATAA   25759
gb:LC522974|Organism:Severe      CTTTTCTCTATCTTTATGCTTTAGTCTACTTCTTGCAGAGTATAAACTTTGTAAGAATAA   25759
gb:MN985325|Organism:Severe      CTTTTCTCTATCTTTATGCTTTAGTCTACTTCTTGCAGAGTATAAACTTTGTAAGAATAA   25762
gb:MT020881|Organism:Severe      CTTTTCTCTATCTTTATGCTTTAGTCTACTTCTTGCAGAGTATAAACTTTGTAAGAATAA   25762
gb:MT020880|Organism:Severe      CTTTTCTCTATCTTTATGCTTTAGTCTACTTCTTGCAGAGTATAAACTTTGTAAGAATAA   25762
gb:MT066175|Organism:Severe      CTTTTCTCTATCTTTATGCTTTAGTCTACTTCTTGCAGAGTATAAACTTTGTAAGAATAA   25762
gb:MN997409|Organism:Severe      CTTTTCTCTATCTTTATGCTTTAGTCTACTTCTTGCAGAGTATAAACTTTGTAAGAATAA   25762
gb:MN938384|Organism:Severe      CTTTTCTCTATCTTTATGCTTTAGTCTACTTCTTGCAGAGTATAAACTTTGTAAGAATAA   25730
gb:MT044258|Organism:Severe      CTTTTCTCTATCTTTATGCTTTAGTCTACTTCTTGCAGAGTATAAACTTTGTAAGAATAA   25738
gb:MT039890|Organism:Severe      CTTTTCTCTATCTTTATGCTTTAGTCTACTTCTTGCAGAGTATAAACTTTGTAAGAATAA   25762
gb:MN988713|Organism:Severe      CTTTTCTCTATCTTTATGCTTTAGTCTACTTCTTGCAGAGTATAAACTTTGTAAGAATAA   25762
gb:LC521925|Organism:Severe      CTTTTCTCTATCTTTATGCTTTAGTCTACTTCTTGCAGAGTATAAACTTTGTAAGAATAA   25735
gb:MT093571|Organism:Severe      CTTTTCTCTATCTTTATGCTTTAGTCTACTTCTTGCAGAGTATAAACTTTGTAAGAATAA   25762
```

```
gb:MT039887|Organism:Severe      CTTTTCTCTATCTTTATGCTTTAGTCTACTTCTTGCAGAGTATAAACTTTGTAAGAATAA   25759
gb:MT019530|Organism:Severe      CTTTTCTCTATCTTTATGCTTTAGTCTACTTCTTGCAGAGTATAAACTTTGTAAGAATAA   25762
gb:MT039888|Organism:Severe      CTTTTCTCTATCTTTATGCTTTAGTCTACTTCTTGCAGAGTATAAACTTTGTAAGAATAA   25762
gb:LC522972|Organism:Severe      CTTTTCTCTATCTTTATGCTTTAGTCTACTTCTTGCAGAGTATAAACTTTGTAAGAATAA   25759
gb:MT027063|Organism:Severe      CTTTTCTCTATCTTTATGCTTTAGTCTACTTCTTGCAGAGTATAAACTTTGTAAGAATAA   25762
gb:MT027062|Organism:Severe      CTTTTCTCTATCTTTATGCTTTAGTCTACTTCTTGCAGAGTATAAACTTTGTAAGAATAA   25762
gb:MT019529|Organism:Severe      CTTTTCTCTATCTTTATGCTTTAGTCTACTTCTTGCAGAGTATAAACTTTGTAAGAATAA   25762
gb:MN996529|Organism:Severe      CTTTTCTCTATCTTTATGCTTTAGTCTACTTCTTGCAGAGTATAAACTTTGTAAGAATAA   25750
gb:MN996531|Organism:Severe      CTTTTCTCTATCTTTATGCTTTAGTCTACTTCTTGCAGAGTATAAACTTTGTAAGAATAA   25749
gb:MT066176|Organism:Severe      CTTTTCTCTATCTTTATGCTTTAGTCTACTTCTTGCAGAGTATAAACTTTGTAAGAATAA   25762
gb:MT027064|Organism:Severe      CTTTTCTCTATCTTTATGCTTTAGTCTACTTCTTGCAGAGTATAAACTTTGTAAGAATAA   25762
gb:MN994468|Organism:Severe      CTTTTCTCTATCTTTATGCTTTAGTCTACTTCTTGCAGAGTATAAACTTTGTAAGAATAA   25762
gb:MT072688|Organism:Severe      CTTTTCTCTATCTTTATGCTTTAGTCTACTTCTTGCAGAGTATAAACTTTGTAAGAATAA   25747
gb:MN996527|Organism:Severe      CTTTTCTCTATCTTTATGCTTTAGTCTACTTCTTGCAGAGTATAAACTTTGTAAGAATAA   25729
gb:MT093631|Organism:Severe      CTTTTCTCTATCTTTATGCTTTAGTCTACTTCTTGCAGAGTATAAACTTTGTAAGAATAA   25800
gb:MT106053|Organism:Severe      CTTTTCTCTATCTTTATGCTTTAGTCTACTTCTTGCAGAGTATAAACTTTGTAAGAATAA   25762
gb:MT019533|Organism:Severe      CTTTTCTCTATCTTTATGCTTTAGTCTACTTCTTGCAGAGTATAAACTTTGTAAGAATAA   25762
gb:MT019531|Organism:Severe      CTTTTCTCTATCTTTATGCTTTAGTCTACTTCTTGCAGAGTATAAACTTTGTAAGAATAA   25762
gb:MN996528|Organism:Severe      CTTTTCTCTATCTTTATGCTTTAGTCTACTTCTTGCAGAGTATAAACTTTGTAAGAATAA   25762
gb:MN996530|Organism:Severe      CTTTTCTCTATCTTTATGCTTTAGTCTACTTCTTGCAGAGTATAAACTTTGTAAGAATAA   25748
gb:MN908947|Organism:Severe      CTTTTCTCTATCTTTATGCTTTAGTCTACTTCTTGCAGAGTATAAACTTTGTAAGAATAA   25762
gb:MT019532|Organism:Severe      CTTTTCTCTATCTTTATGCTTTAGTCTACTTCTTGCAGAGTATAAACTTTGTAAGAATAA   25762
                                 ************************************************************

gb:MT020781|Organism:Severe      TAATGAGGCTTTGGCTTTGCTGGAAATGCCGTTCCAAAAACCCANNNNNNNNNNNNNNNN   25810
gb:MT007544|Organism:Severe      TAATGAGGCTTTGGCTTTGCTGGAAATGCCGTTCCAAAAACCCATTACTTTATGATGCCA   25822
gb:MN994467|Organism:Severe      TAATGAGGCTTTGGCTTTGCTGGAAATGCCGTTCCAAAAACCCATTACTTTATGATGCCA   25822
gb:MT044257|Organism:Severe      TAATGAGGCTTTGGCTTTGCTGGAAATGCCGTTCCAAAAACCCATTACTTTATGATGCCA   25822
gb:MT106054|Organism:Severe      TAATGAGGCTTTGGCTTTGCTGGAAATGCCGTTCCAAAAACCCATTACTTTATGATGCCA   25822
gb:MT049951|Organism:Severe      TAATGAGGCTTTGGCTTTGCTGGAAATGCCGTTCCAAAAACCCATTACTTTATGATGCCA   25822
gb:MN975262|Organism:Severe      TAATGAGGCTTTGGCTTTGCTGGAAATGCCGTTCCAAAAACCCATTACTTTATGATGCCA   25822
gb:MT106052|Organism:Severe      TAATGAGGCTTTGGCTTTGCTGGAAATGCCGTTCCAAAAACCCATTACTTTATGATGCCA   25822
gb:LC522975|Organism:Severe      TAATGAGGCTTTGGCTTTGCTGGAAATGCCGTTCCAAAAACCCATTACTTTATGATGCCA   25819
gb:LC522973|Organism:Severe      TAATGAGGCTTTGGCTTTGCTGGAAATGCCGTTCCAAAAACCCATTACTTTATGATGCCA   25819
gb:LC522974|Organism:Severe      TAATGAGGCTTTGGCTTTGCTGGAAATGCCGTTCCAAAAACCCATTACTTTATGATGCCA   25819
gb:MN985325|Organism:Severe      TAATGAGGCTTTGGCTTTGCTGGAAATGCCGTTCCAAAAACCCATTACTTTATGATGCCA   25822
gb:MT020881|Organism:Severe      TAATGAGGCTTTGGCTTTGCTGGAAATGCCGTTCCAAAAACCCATTACTTTATGATGCCA   25822
gb:MT020880|Organism:Severe      TAATGAGGCTTTGGCTTTGCTGGAAATGCCGTTCCAAAAACCCATTACTTTATGATGCCA   25822
gb:MT066175|Organism:Severe      TAATGAGGCTTTGGCTTTGCTGGAAATGCCGTTCCAAAAACCCATTACTTTATGATGCCA   25822
gb:MN997409|Organism:Severe      TAATGAGGCTTTGGCTTTGCTGGAAATGCCGTTCCAAAAACCCATTACTTTATGATGCCA   25822
gb:MN938384|Organism:Severe      TAATGAGGCTTTGGCTTTGCTGGAAATGCCGTTCCAAAAACCCATTACTTTATGATGCCA   25790
gb:MT044258|Organism:Severe      TAATGAGGCTTTGGCTTTGCTGGAAATGCCGTTCCAAAAACCCATTACTTTATGATGCCA   25798
gb:MT039890|Organism:Severe      TAATGAGGCTTTTGCTTTGCTGGAAATGCCGTTCCAAAAACCCATTACTTTATGATGCCA   25822
gb:MN988713|Organism:Severe      TAATGAGGCTTTGGCTTTGCTGGAAATGCCGTTCCAAAAACCCATTACTTTATGATGCCA   25822
gb:LC521925|Organism:Severe      TAATGAGGCTTTGGCTTTGCTGGAAATGCCGTTCCAAAAACCCATTACTTTATGATGCCA   25795
gb:MT093571|Organism:Severe      TAATGAGGCTTTGGCTTTGCTGGAAATGCCGTTCCAAAAACCCATTACTTTATGATGCCA   25822
gb:MT039887|Organism:Severe      TAATGAGGCTTTGGCTTTGCTGGAAATGCCGTTCCAAAAACCCATTACTTTATGATGCCA   25819
gb:MT019530|Organism:Severe      TAATGAGGCTTTGGCTTTGCTGGAAATGCCGTTCCAAAAACCCATTACTTTATGATGCCA   25822
gb:MT039888|Organism:Severe      TAATGAGGCTTTGGCTTTGCTGGAAATGCCGTTCCAAAAACCCATTACTTTATGATGCCA   25822
gb:LC522972|Organism:Severe      TAATGAGGCTTTGGCTTTGCTGGAAATGCCGTTCCAAAAACCCATTAGTTTATGATGCCA   25819
gb:MT027063|Organism:Severe      TAATGAGGCTTTGGCTTTGCTGGAAATGCCGTTCCAAAAACCCATTACTTTATGATGCCA   25822
gb:MT027062|Organism:Severe      TAATGAGGCTTTGGCTTTGCTGGAAATGCCGTTCCAAAAACCCATTACTTTATGATGCCA   25822
gb:MT019529|Organism:Severe      TAATGAGGCTTTGGCTTTGCTGGAAATGCCGTTCCAAAAACCCATTACTTTATGATGCCA   25822
gb:MN996529|Organism:Severe      TAATGAGGCTTTGGCTTTGCTGGAAATGCCGTTCCAAAAACCCATTACTTTATGATGCCA   25810
gb:MN996531|Organism:Severe      TAATGAGGCTTTGGCTTTGCTGGAAATGCCGTTCCAAAAACCCATTACTTTATGATGCCA   25809
gb:MT066176|Organism:Severe      TAATGAGGCTTTGGCTTTGCTGGAAATGCCGTTCCAAAAACCCATTACTTTATGATGCCA   25822
gb:MT027064|Organism:Severe      TAATGAGGCTTTGGCTTTGCTGGAAATGCCGTTCCAAAAACCCATTACTTTATGATGCCA   25822
gb:MN994468|Organism:Severe      TAATGAGGCTTTGGCTTTGCTGGAAATGCCGTTCCAAAAACCCATTACTTTATGATGCCA   25822
gb:MT072688|Organism:Severe      TAATGAGGCTTTGGCTTTGCTGGAAATGCCGTTCCAAAAACCCATTACTTTATGATGCCA   25807
gb:MN996527|Organism:Severe      TAATGAGGCTTTGGCTTTGCTGGAAATGCCGTTCCAAAAACCCATTACTTTATGATGCCA   25789
gb:MT093631|Organism:Severe      TAATGAGGCTTTGGCTTTGCTGGAAATGCCGTTCCAAAAACCCATTACTTTATGATGCCA   25860
gb:MT106053|Organism:Severe      TAATGAGGCTTTGGCTTTGCTGGAAATGCCGTTCCAAAAACCCATTACTTTATGATGCCA   25822
gb:MT019533|Organism:Severe      TAATGAGGCTTTGGCTTTGCTGGAAATGCCGTTCCAAAAACCCATTACTTTATGATGCCA   25822
gb:MT019531|Organism:Severe      TAATGAGGCTTTGGCTTTGCTGGAAATGCCGTTCCAAAAACCCATTACTTTATGATGCCA   25822
gb:MN996528|Organism:Severe      TAATGAGGCTTTGGCTTTGCTGGAAATGCCGTTCCAAAAACCCATTACTTTATGATGCCA   25822
```

```
gb:MN996530|Organism:Severe      TAATGAGGCTTTGGCTTTGCTGGAAATGCCGTTCCAAAAACCCATTACTTTATGATGCCA   25808
gb:MN908947|Organism:Severe      TAATGAGGCTTTGGCTTTGCTGGAAATGCCGTTCCAAAAACCCATTACTTTATGATGCCA   25822
gb:MT019532|Organism:Severe      TAATGAGGCTTTGGCTTTGCTGGAAATGCCGTTCCAAAAACCCATTACTTTATGATGCCA   25822
                                 ************ *******************************

gb:MT020781|Organism:Severe      NNNNNNNNNNNNNNNNNNNNNNNNNNNNNNNNNACTATTGTATACCTTACAATAGTGTAA   25870
gb:MT007544|Organism:Severe      ACTATTTTCTTTGCTGGCATACTAATTGTTACGACTATTGTATACCTTACAATAGTGTAA   25882
gb:MN994467|Organism:Severe      ACTATTTTCTTTGCTGGCATACTAATTGTTACGACTATTGTATACCTTACAATAGTGTAA   25882
gb:MT044257|Organism:Severe      ACTATTTTCTTTGCTGGCATACTAATTGTTACGACTATTGTATACCTTACAATAGTGTAA   25882
gb:MT106054|Organism:Severe      ACTATTTTCTTTGCTGGCATACTAATTGTTACGACTATTGTATACCTTACAATAGTGTAA   25882
gb:MT049951|Organism:Severe      ACTATTTTCTTTGCTGGCATACTAATTGTTACGACTATTGTATACCTTACAATAGTGTAA   25882
gb:MN975262|Organism:Severe      ACTATTTTCTTTGCTGGCATACTAATTGTTACGACTATTGTATACCTTACAATAGTGTAA   25882
gb:MT106052|Organism:Severe      ACTATTTTCTTTGCTGGCATACTAATTGTTACGACTATTGTATACCTTACAATAGTGTAA   25882
gb:LC522975|Organism:Severe      ACTATTTTCTTTGCTGGCATACTAATTGTTACGACTATTGTATACCTTACAATAGTGTAA   25879
gb:LC522973|Organism:Severe      ACTATTTTCTTTGCTGGCATACTAATTGTTACGACTATTGTATACCTTACAATAGTGTAA   25879
gb:LC522974|Organism:Severe      ACTATTTTCTTTGCTGGCATACTAATTGTTACGACTATTGTATACCTTACAATAGTGTAA   25879
gb:MN985325|Organism:Severe      ACTATTTTCTTTGCTGGCATACTAATTGTTACGACTATTGTATACCTTACAATAGTGTAA   25882
gb:MT020881|Organism:Severe      ACTATTTTCTTTGCTGGCATACTAATTGTTACGACTATTGTATACCTTACAATAGTGTAA   25882
gb:MT020880|Organism:Severe      ACTATTTTCTTTGCTGGCATACTAATTGTTACGACTATTGTATACCTTACAATAGTGTAA   25882
gb:MT066175|Organism:Severe      ACTATTTTCTTTGCTGGCATACTAATTGTTACGACTATTGTATACCTTACAATAGTGTAA   25882
gb:MN997409|Organism:Severe      ACTATTTTCTTTGCTGGCATACTAATTGTTACGACTATTGTATACCTTACAATAGTGTAA   25882
gb:MN938384|Organism:Severe      ACTATTTTCTTTGCTGGCATACTAATTGTTACGACTATTGTATACCTTACAATAGTGTAA   25850
gb:MT044258|Organism:Severe      ACTATTTTCTTTGCTGGCATACTAATTGTTACGACTATTGTATACCTTACAATAGTGTAA   25858
gb:MT039890|Organism:Severe      ACTATTTTCTTTGCTGGCATACTAATTGTTACGACTATTGTATACCTTACAATAGTGTAA   25882
gb:MN988713|Organism:Severe      ACTATTTTCTTTGCTGGCATACTAATTGTTACGACTATTGTATACCTTACAATAGTGTAA   25882
gb:LC521925|Organism:Severe      ACTATTTTCTTTGCTGGCATACTAATTGTTACGACTATTGTATACCTTACAATAGTGTAA   25855
gb:MT093571|Organism:Severe      ACTATTTTCTTTGCTGGCATACTAATTGTTACGACTATTGTATACCTTACAATAGTGTAA   25882
gb:MT039887|Organism:Severe      ACTATTTTCTTTGCTGGCATACTAATTGTTACGACTATTGTATACCTTACAATAGTGTAA   25879
gb:MT019530|Organism:Severe      ACTATTTTCTTTGCTGGCATACTAATTGTTACGACTATTGTATACCTTACAATAGTGTAA   25882
gb:MT039888|Organism:Severe      ACTATTTTCTTTGCTGGCATACTAATTGTTACGACTATTGTATACCTTACAATAGTGTAA   25882
gb:LC522972|Organism:Severe      ACTATTTTCTTTGCTGGCATACTAATTGTTACGACTATTGTATACCTTACAATAGTGTAA   25879
gb:MT027063|Organism:Severe      ACTATTTTCTTTGCTGGCATACTAATTGTTACGACTATTGTATACCTTACAATAGTGTAA   25882
gb:MT027062|Organism:Severe      ACTATTTTCTTTGCTGGCATACTAATTGTTACGACTATTGTATACCTTACAATAGTGTAA   25882
gb:MT019529|Organism:Severe      ACTATTTTCTTTGCTGGCATACTAATTGTTACGACTATTGTATACCTTACAATAGTGTAA   25882
gb:MN996529|Organism:Severe      ACTATTTTCTTTGCTGGCATACTAATTGTTACGACTATTGTATACCTTACAATAGTGTAA   25870
gb:MN996531|Organism:Severe      ACTATTTTCTTTGCTGGCATACTAATTGTTACGACTATTGTATACCTTACAATAGTGTAA   25869
gb:MT066176|Organism:Severe      ACTATTTTCTTTGCTGGCATACTAATTGTTACGACTATTGTATACCTTACAATAGTGTAA   25882
gb:MT027064|Organism:Severe      ACTATTTTCTTTGCTGGCATACTAATTGTTACGACTATTGTATACCTTACAATAGTGTAA   25882
gb:MN994468|Organism:Severe      ACTATTTTCTTTGCTGGCATACTAATTGTTACGACTATTGTATACCTTACAATAGTGTAA   25882
gb:MT072688|Organism:Severe      ACTATTTTCTTTGCTGGCATACTAATTGTTACGACTATTGTATACCTTACAATAGTGTAA   25867
gb:MN996527|Organism:Severe      ACTATTTTCTTTGCTGGCATACTAATTGTTACGACTATTGTATACCTTACAATAGTGTAA   25849
gb:MT093631|Organism:Severe      ACTATTTTCTTTGCTGGCATACTAATTGTTACGACTATTGTATACCTTACAATAGTGTAA   25920
gb:MT106053|Organism:Severe      ACTATTTTCTTTGCTGGCATACTAATTGTTACGACTATTGTATACCTTACAATAGTGTAA   25882
gb:MT019533|Organism:Severe      ACTATTTTCTTTGCTGGCATACTAATTGTTACGACTATTGTATACCTTACAATAGTGTAA   25882
gb:MT019531|Organism:Severe      ACTATTTTCTTTGCTGGCATACTAATTGTTACGACTATTGTATACCTTACAATAGTGTAA   25882
gb:MN996528|Organism:Severe      ACTATTTTCTTTGCTGGCATACTAATTGTTACGACTATTGTATACCTTACAATAGTGTAA   25882
gb:MN996530|Organism:Severe      ACTATTTTCTTTGCTGGCATACTAATTGTTACGACTATTGTATACCTTACAATAGTGTAA   25868
gb:MN908947|Organism:Severe      ACTATTTTCTTTGCTGGCATACTAATTGTTACGACTATTGTATACCTTACAATAGTGTAA   25882
gb:MT019532|Organism:Severe      ACTATTTTCTTTGCTGGCATACTAATTGTTACGACTATTGTATACCTTACAATAGTGTAA   25882
                                                                  ***************************

gb:MT020781|Organism:Severe      CTTCTTCAATTGTCATTACTTCAGGTGATGGCACAACAAGTCCTATTTCTGAACATGACT   25930
gb:MT007544|Organism:Severe      CTTCTTCAATTGTCATTACTTCAGGTGATGGCACAACAAGTCCTATTTCTGAACATGACT   25942
gb:MN994467|Organism:Severe      CTTCTTCAATTGTCATTACTTCAGGTGATGGCACAACAAGTCCTATTTCTGAACATGACT   25942
gb:MT044257|Organism:Severe      CTTCTTCAATTGTCATTACTTCAGGTGATGGCACAACAAGTCCTATTTCTGAACATGACT   25942
gb:MT106054|Organism:Severe      CTTCTTCAATTGTCATTACTTCAGGTGATGGCACAACAAGTCCTATTTCTGAACATGACT   25942
gb:MT049951|Organism:Severe      CTTCTTCAATTGTCATTACTTCAGGTGATGGCACAACAAGTCCTATTTCTGAACATGACT   25942
gb:MN975262|Organism:Severe      CTTCTTCAATTGTCATTACTTCAGGTGATGGCACAACAAGTCCTATTTCTGAACATGACT   25942
gb:MT106052|Organism:Severe      CTTCTTCAATTGTCATTACTTCAGGTGATGGCACAACAAGTCCTATTTCTGAACATGACT   25942
gb:LC522975|Organism:Severe      CTTCTTCAATTGTCATTACTTCAGGTGATGGCACAACAAGTCCTATTTCTGAACATGACT   25939
gb:LC522973|Organism:Severe      CTTCTTCAATTGTCATTACTTCAGGTGATGGCACAACAAGTCCTATTTCTGAACATGACT   25939
gb:LC522974|Organism:Severe      CTTCTTCAATTGTCATTACTTCAGGTGATGGCACAACAAGTCCTATTTCTGAACATGACT   25939
gb:MN985325|Organism:Severe      CTTCTTCAATTGTCATTACTTCAGGTGATGGCACAACAAGTCCTATTTCTGAACATGACT   25942
gb:MT020881|Organism:Severe      CTTCTTCAATTGTCATTACTTCAGGTGATGGCACAACAAGTCCTATTTCTGAACATGACT   25942
gb:MT020880|Organism:Severe      CTTCTTCAATTGTCATTACTTCAGGTGATGGCACAACAAGTCCTATTTCTGAACATGACT   25942
```

```
gb:MT066175|Organism:Severe      CTTCTTCAATTGTCATTACTTCAGGTGATGGCACAACAAGTCCTATTTCTGAACATGACT   25942
gb:MN997409|Organism:Severe      CTTCTTCAATTGTCATTACTTCAGGTGATGGCACAACAAGTCCTATTTCTGAACATGACT   25942
gb:MN938384|Organism:Severe      CTTCTTCAATTGTCATTACTTCAGGTGATGGCACAACAAGTCCTATTTCTGAACATGACT   25910
gb:MT044258|Organism:Severe      CTTCTTCAATTGTCATTACTTCAGGTGATGGCACAACAAGTCCTATTTCTGAACATGACT   25918
gb:MT039890|Organism:Severe      CTTCTTCAATTGTCATTACTTCAGGTGATGGCACAACAAGTCCTATTTCTGAACATGACT   25942
gb:MN988713|Organism:Severe      CTTCTTCAATTGTCATTACTTCAGGTGATGGCACAACAAGTCCTATTTCTGAACATGACT   25942
gb:LC521925|Organism:Severe      CTTCTTCAATTGTCATTACTTCAGGTGATGGCACAACAAGTCCTATTTCTGAACATGACT   25915
gb:MT093571|Organism:Severe      CTTCTTCAATTGTCATTACTTCAGGTGATGGCACAACAAGTCCTATTTCTGAACATGACT   25942
gb:MT039887|Organism:Severe      CTTCTTCAATTGTCATTACTTCAGGTGATGGCACAACAAGTCCTATTTCTGAACATGACT   25939
gb:MT019530|Organism:Severe      CTTCTTCAATTGTCATTACTTCAGGTGATGGCACAACAAGTCCTATTTCTGAACATGACT   25942
gb:MT039888|Organism:Severe      CTTCTTCAATTGTCATTACTTCAGGTGATGGCACAACAAGTCCTATTTCTGAACATGACT   25942
gb:LC522972|Organism:Severe      CTTCTTCAATTGTCATTACTTCAGGTGATGGCACAACAAGTCCTATTTCTGAACATGACT   25939
gb:MT027063|Organism:Severe      CTTCTTCAATTGTCATTACTTCAGGTGATGGCACAACAAGTCCTATTTCTGAACATGACT   25942
gb:MT027062|Organism:Severe      CTTCTTCAATTGTCATTACTTCAGGTGATGGCACAACAAGTCCTATTTCTGAACATGACT   25942
gb:MT019529|Organism:Severe      CTTCTTCAATTGTCATTACTTCAGGTGATGGCACAACAAGTCCTATTTCTGAACATGACT   25942
gb:MN996529|Organism:Severe      CTTCTTCAATTGTCATTACTTCAGGTGATGGCACAACAAGTCCTATTTCTGAACATGACT   25930
gb:MN996531|Organism:Severe      CTTCTTCAATTGTCATTACTTCAGGTGATGGCACAACAAGTCCTATTTCTGAACATGACT   25929
gb:MT066176|Organism:Severe      CTTCTTCAATTGTCATTACTTCAGGTGATGGCACAACAAGTCCTATTTCTGAACATGACT   25942
gb:MT027064|Organism:Severe      CTTCTTCAATTGTCATTACTTCAGGTGATGGCACAACAAGTCCTATTTCTGAACATGACT   25942
gb:MN994468|Organism:Severe      CTTCTTCAATTGTCATTACTTCAGGTGATGGCACAACAAGTCCTATTTCTGAACATGACT   25942
gb:MT072688|Organism:Severe      CTTCTTCAATTGTCATTACTTCAGGTGATGGCACAACAAGTCCTATTTCTGAACATGACT   25927
gb:MN996527|Organism:Severe      CTTCTTCAATTGTCATTACTTCAGGTGATGGCACAACAAGTCCTATTTCTGAACATGACT   25909
gb:MT093631|Organism:Severe      CTTCTTCAATTGTCATTACTTCAGGTGATGGCACAACAAGTCCTATTTCTGAACATGACT   25980
gb:MT106053|Organism:Severe      CTTCTTCAATTGTCATTACTTCAGGTGATGGCACAACAAGTCCTATTTCTGAACATGACT   25942
gb:MT019533|Organism:Severe      CTTCTTCAATTGTCATTACTTCAGGTGATGGCACAACAAGTCCTATTTCTGAACATGACT   25942
gb:MT019531|Organism:Severe      CTTCTTCAATTGTCATTACTTCAGGTGATGGCACAACAAGTCCTATTTCTGAACATGACT   25942
gb:MN996528|Organism:Severe      CTTCTTCAATTGTCATTACTTCAGGTGATGGCACAACAAGTCCTATTTCTGAACATGACT   25942
gb:MN996530|Organism:Severe      CTTCTTCAATTGTCATTACTTCAGGTGATGGCACAACAAGTCCTATTTCTGAACATGACT   25928
gb:MN908947|Organism:Severe      CTTCTTCAATTGTCATTACTTCAGGTGATGGCACAACAAGTCCTATTTCTGAACATGACT   25942
gb:MT019532|Organism:Severe      CTTCTTCAATTGTCATTACTTCAGGTGATGGCACAACAAGTCCTATTTCTGAACATGACT   25942
                                 ************************************************************

gb:MT020781|Organism:Severe      ACCAGATTGGTGGTTATACTGAAAAATGGGAATCTGGAGTAAAAGACTGTGTTGTATTAC   25990
gb:MT007544|Organism:Severe      ACCAGATTGGTGGTTATACTGAAAAATGGGAATCTGGAGTAAAAGACTGTGTTGTATTAC   26002
gb:MN994467|Organism:Severe      ACCAGATTGGTGGTTATACTGAAAAATGGGAATCTGGAGTAAAAGACTGTGTTGTATTAC   26002
gb:MT044257|Organism:Severe      ACCAGATTGGTGGTTATACTGAAAAATGGGAATCTGGAGTAAAAGACTGTGTTGTATTAC   26002
gb:MT106054|Organism:Severe      ACCAGATTGGTGGTTATACTGAAAAATGGGAATCTGGAGTAAAAGACTGTGTTGTATTAC   26002
gb:MT049951|Organism:Severe      ACCAGATTGGTGGTTATACTGAAAAATGGGAATCTGGAGTAAAAGACTGTGTTGTATTAC   26002
gb:MN975262|Organism:Severe      ACCAGATTGGTGGTTATACTGAAAAATGGGAATCTGGAGTAAAAGACTGTGTTGTATTAC   26002
gb:MT106052|Organism:Severe      ACCAGATTGGTGGTTATACTGAAAAATGGGAATCTGGAGTAAAAGACTGTGTTGTATTAC   26002
gb:LC522975|Organism:Severe      ACCAGATTGGTGGTTATACTGAAAAATGGGAATCTGGAGTAAAAGACTGTGTTGTATTAC   25999
gb:LC522973|Organism:Severe      ACCAGATTGGTGGTTATACTGAAAAATGGGAATCTGGAGTAAAAGACTGTGTTGTATTAC   25999
gb:LC522974|Organism:Severe      ACCAGATTGGTGGTTATACTGAAAAATGGGAATCTGGAGTAAAAGACTGTGTTGTATTAC   25999
gb:MN985325|Organism:Severe      ACCAGATTGGTGGTTATACTGAAAAATGGGAATCTGGAGTAAAAGACTGTGTTGTATTAC   26002
gb:MT020881|Organism:Severe      ACCAGATTGGTGGTTATACTGAAAAATGGGAATCTGGAGTAAAAGACTGTGTTGTATTAC   26002
gb:MT020880|Organism:Severe      ACCAGATTGGTGGTTATACTGAAAAATGGGAATCTGGAGTAAAAGACTGTGTTGTATTAC   26002
gb:MT066175|Organism:Severe      ACCAGATTGGTGGTTATACTGAAAAATGGGAATCTGGAGTAAAAGACTGTGTTGTATTAC   26002
gb:MN997409|Organism:Severe      ACCAGATTGGTGGTTATACTGAAAAATGGGAATCTGGAGTAAAAGACTGTGTTGTATTAC   26002
gb:MN938384|Organism:Severe      ACCAGATTGGTGGTTATACTGAAAAATGGGAATCTGGAGTAAAAGACTGTGTTGTATTAC   25970
gb:MT044258|Organism:Severe      ACCAGATTGGTGGTTATACTGAAAAATGGGAATCTGGAGTAAAAGACTGTGTTGTATTAC   25978
gb:MT039890|Organism:Severe      ACCAGATTGGTGGTTATACTGAAAAATGGGAATCTGGAGTAAAAGACTGTGTTGTATTAC   26002
gb:MN988713|Organism:Severe      ACCAGATTGGTGGTTATACTGAAAAATGGGAATCTGGAGTAAAAGACTGTGTTGTATTAC   26002
gb:LC521925|Organism:Severe      ACCAGATTGGTGGTTATACTGAAAAATGGGAATCTGGAGTAAAAGACTGTGTTGTATTAC   25975
gb:MT093571|Organism:Severe      ACCAGATTGGTGGTTATACTGAAAAATGGGAATCTGGAGTAAAAGACTGTGTTGTATTAC   26002
gb:MT039887|Organism:Severe      ACCAGATTGGTGGTTATACTGAAAAATGGGAATCTGGAGTAAAAGACTGTGTTGTATTAC   25999
gb:MT019530|Organism:Severe      ACCAGATTGGTGGTTATACTGAAAAATGGGAATCTGGAGTAAAAGACTGTGTTGTATTAC   26002
gb:MT039888|Organism:Severe      ACCAGATTGGTGGTTATACTGAAAAATGGGAATCTGGAGTAAAAGACTGTGTTGTATTAC   26002
gb:LC522972|Organism:Severe      ACCAGATTGGTGGTTATACTGAAAAATGGGAATCTGGAGTAAAAGACTGTGTTGTATTAC   25999
gb:MT027063|Organism:Severe      ACCAGATTGGTGGTTATACTGAAAAATGGGAATCTGGAGTAAAAGACTGTGTTGTATTAC   26002
gb:MT027062|Organism:Severe      ACCAGATTGGTGGTTATACTGAAAAATGGGAATCTGGAGTAAAAGACTGTGTTGTATTAC   26002
gb:MT019529|Organism:Severe      ACCAGATTGGTGGTTATACTGAAAAATGGGAATCTGGAGTAAAAGACTGTGTTGTATTAC   26002
gb:MN996529|Organism:Severe      ACCAGATTGGTGGTTATACTGAAAAATGGGAATCTGGAGTAAAAGACTGTGTTGTATTAC   25990
gb:MN996531|Organism:Severe      ACCAGATTGGTGGTTATACTGAAAAATGGGAATCTGGAGTAAAAGACTGTGTTGTATTAC   25989
gb:MT066176|Organism:Severe      ACCAGATTGGTGGTTATACTGAAAAATGGGAATCTGGAGTAAAAGACTGTGTTGTATTAC   26002
gb:MT027064|Organism:Severe      ACCAGATTGGTGGTTATACTGAAAAATGGGAATCTGGAGTAAAAGACTGTGTTGTATTAC   26002
```

```
gb:MN994468|Organism:Severe      ACCAGATTGGTGGTTATACTGAAAAATGGGAATCTGGAGTAAAAGACTGTGTTGTATTAC    26002
gb:MT072688|Organism:Severe      ACCAGATTGGTGGTTATACTGAAAAATGGGAATCTGGAGTAAAAGACTGTGTTGTATTAC    25987
gb:MN996527|Organism:Severe      ACCAGATTGGTGGTTATACTGAAAAATGGGAATCTGGAGTAAAAGACTGTGTTGTATTAC    25969
gb:MT093631|Organism:Severe      ACCAGATTGGTGGTTATACTGAAAAATGGGAATCTGGAGTAAAAGACTGTGTTGTATTAC    26040
gb:MT106053|Organism:Severe      ACCAGATTGGTGGTTATACTGAAAAATGGGAATCTGGAGTAAAAGACTGTGTTGTATTAC    26002
gb:MT019533|Organism:Severe      ACCAGATTGGTGGTTATACTGAAAAATGGGAATCTGGAGTAAAAGACTGTGTTGTATTAC    26002
gb:MT019531|Organism:Severe      ACCAGATTGGTGGTTATACTGAAAAATGGGAATCTGGAGTAAAAGACTGTGTTGTATTAC    26002
gb:MN996528|Organism:Severe      ACCAGATTGGTGGTTATACTGAAAAATGGGAATCTGGAGTAAAAGACTGTGTTGTATTAC    26002
gb:MN996530|Organism:Severe      ACCAGATTGGTGGTTATACTGAAAAATGGGAATCTGGAGTAAAAGACTGTGTTGTATTAC    25988
gb:MN908947|Organism:Severe      ACCAGATTGGTGGTTATACTGAAAAATGGGAATCTGGAGTAAAAGACTGTGTTGTATTAC    26002
gb:MT019532|Organism:Severe      ACCAGATTGGTGGTTATACTGAAAAATGGGAATCTGGAGTAAAAGACTGTGTTGTATTAC    26002
                                 ************************************************************

gb:MT020781|Organism:Severe      ACAGTTACTTCACTTCAGACTATTACCAGCTGTACTCAACTCAATTGAGTACAGACACTG    26050
gb:MT007544|Organism:Severe      ACAGTTACTTCACTTCAGACTATTACCAGCTGTACTCAACTCAATTGAGTACAGACACTG    26062
gb:MN994467|Organism:Severe      ACAGTTACTTCACTTCAGACTATTACCAGCTGTACTCAACTCAATTGAGTACAGACACTG    26062
gb:MT044257|Organism:Severe      ACAGTTACTTCACTTCAGACTATTACCAGCTGTACTCAACTCAATTGAGTACAGACACTG    26062
gb:MT106054|Organism:Severe      ACAGTTACTTCACTTCAGACTATTACCAGCTGTACTCAACTCAATTGAGTACAGACACTG    26062
gb:MT049951|Organism:Severe      ACAGTTACTTCACTTCAGACTATTACCAGCTGTACTCAACTCAATTGAGTACAGACACTG    26062
gb:MN975262|Organism:Severe      ACAGTTACTTCACTTCAGACTATTACCAGCTGTACTCAACTCAATTGAGTACAGACACTG    26062
gb:MT106052|Organism:Severe      ACAGTTACTTCACTTCAGACTATTACCAGCTGTACTCAACTCAATTGAGTACAGACACTG    26062
gb:LC522975|Organism:Severe      ACAGTTACTTCACTTCAGACTATTACCAGCTGTACTCAACTCAATTGAGTACAGACACTG    26059
gb:LC522973|Organism:Severe      ACAGTTACTTCACTTCAGACTATTACCAGCTGTACTCAACTCAATTGAGTACAGACACTG    26059
gb:LC522974|Organism:Severe      ACAGTTACTTCACTTCAGACTATTACCAGCTGTACTCAACTCAATTGAGTACAGACACTG    26059
gb:MN985325|Organism:Severe      ACAGTTACTTCACTTCAGACTATTACCAGCTGTACTCAACTCAATTGAGTACAGACACTG    26062
gb:MT020881|Organism:Severe      ACAGTTACTTCACTTCAGACTATTACCAGCTGTACTCAACTCAATTGAGTACAGACACTG    26062
gb:MT020880|Organism:Severe      ACAGTTACTTCACTTCAGACTATTACCAGCTGTACTCAACTCAATTGAGTACAGACACTG    26062
gb:MT066175|Organism:Severe      ACAGTTACTTCACTTCAGACTATTACCAGCTGTACTCAACTCAATTGAGTACAGACACTG    26062
gb:MN997409|Organism:Severe      ACAGTTACTTCACTTCAGACTATTACCAGCTGTACTCAACTCAATTGAGTACAGACACTG    26062
gb:MN938384|Organism:Severe      ACAGTTACTTCACTTCAGACTATTACCAGCTGTACTCAACTCAATTGAGTACAGACACTG    26030
gb:MT044258|Organism:Severe      ACAGTTACTTCACTTCAGACTATTACCAGCTGTACTCAACTCAATTGAGTACAGACACTG    26038
gb:MT039890|Organism:Severe      ACAGTTACTTCACTTCAGACTATTACCAGCTGTACTCAACTCAATTGAGTACAGACACTG    26062
gb:MN988713|Organism:Severe      ACAGTTACTTCACTTCAGACTATTACCAGCTGTACTCAACTCAATTGAGTACAGACACTG    26062
gb:LC521925|Organism:Severe      ACAGTTACTTCACTTCAGACTATTACCAGCTGTACTCAACTCAATTGAGTACAGACACTG    26035
gb:MT093571|Organism:Severe      ACAGTTACTTCACTTCAGACTATTACCAGCTGTACTCAACTCAATTGAGTACAGACACTG    26062
gb:MT039887|Organism:Severe      ACAGTTACTTCACTTCAGACTATTACCAGCTGTACTCAACTCAATTGAGTACAGACACTG    26059
gb:MT019530|Organism:Severe      ACAGTTACTTCACTTCAGACTATTACCAGCTGTACTCAACTCAATTGAGTACAGACACTG    26062
gb:MT039888|Organism:Severe      ACAGTTACTTCACTTCAGACTATTACCAGCTGTACTCAACTCAATTGAGTACAGACACTG    26062
gb:LC522972|Organism:Severe      ACAGTTACTTCACTTCAGACTATTACCAGCTGTACTCAACTCAATTGAGTACAGACACTG    26059
gb:MT027063|Organism:Severe      ACAGTTACTTCACTTCAGACTATTACCAGCTGTACTCAACTCAATTGAGTACAGACACTG    26062
gb:MT027062|Organism:Severe      ACAGTTACTTCACTTCAGACTATTACCAGCTGTACTCAACTCAATTGAGTACAGACACTG    26062
gb:MT019529|Organism:Severe      ACAGTTACTTCACTTCAGACTATTACCAGCTGTACTCAACTCAATTGAGTACAGACACTG    26062
gb:MN996529|Organism:Severe      ACAGTTACTTCACTTCAGACTATTACCAGCTGTACTCAACTCAATTGAGTACAGACACTG    26050
gb:MN996531|Organism:Severe      ACAGTTACTTCACTTCAGACTATTACCAGCTGTACTCAACTCAATTGAGTACAGACACTG    26049
gb:MT066176|Organism:Severe      ACAGTTACTTCACTTCAGACTATTACCAGCTGTACTCAACTCAATTGAGTACAGACACTG    26062
gb:MT027064|Organism:Severe      ACAGTTACTTCACTTCAGACTATTACCAGCTGTACTCAACTCAATTGAGTACAGACACTG    26062
gb:MN994468|Organism:Severe      ACAGTTACTTCACTTCAGACTATTACCAGCTGTACTCAACTCAATTGAGTACAGACACTG    26062
gb:MT072688|Organism:Severe      ACAGTTACTTCACTTCAGACTATTACCAGCTGTACTCAACTCAATTGAGTACAGACACTG    26047
gb:MN996527|Organism:Severe      ACAGTTACTTCACTTCAGACTATTACCAGCTGTACTCAACTCAATTGAGTACAGACACTG    26029
gb:MT093631|Organism:Severe      ACAGTTACTTCACTTCAGACTATTACCAGCTGTACTCAACTCAATTGAGTACAGACACTG    26100
gb:MT106053|Organism:Severe      ACAGTTACTTCACTTCAGACTATTACCAGCTGTACTCAACTCAATTGAGTACAGACACTG    26062
gb:MT019533|Organism:Severe      ACAGTTACTTCACTTCAGACTATTACCAGCTGTACTCAACTCAATTGAGTACAGACACTG    26062
gb:MT019531|Organism:Severe      ACAGTTACTTCACTTCAGACTATTACCAGCTGTACTCAACTCAATTGAGTACAGACACTG    26062
gb:MN996528|Organism:Severe      ACAGTTACTTCACTTCAGACTATTACCAGCTGTACTCAACTCAATTGAGTACAGACACTG    26062
gb:MN996530|Organism:Severe      ACAGTTACTTCACTTCAGACTATTACCAGCTGTACTCAACTCAATTGAGTACAGACACTG    26048
gb:MN908947|Organism:Severe      ACAGTTACTTCACTTCAGACTATTACCAGCTGTACTCAACTCAATTGAGTACAGACACTG    26062
gb:MT019532|Organism:Severe      ACAGTTACTTCACTTCAGACTATTACCAGCTGTACTCAACTCAATTGAGTACAGACACTG    26062
                                 ************************************************************

gb:MT020781|Organism:Severe      GTGTTGAACATGTTACCTTCTTCATCTACAATAAAATTGTTGATGAGCCTGAAGAACATG    26110
gb:MT007544|Organism:Severe      GTGTTGAACATGTTACCTTCTTCATCTACAATAAAATTGTTGATGAGCCTGAAGAACATG    26122
gb:MN994467|Organism:Severe      GTGTTGAACATGTTACCTTCTTCATCTACAATAAAATTGTTGATGAGCCTGAAGAACATG    26122
gb:MT044257|Organism:Severe      GTGTTGAACATGTTACCTTCTTCATCTACAATAAAATTGTTGATGAGCCTGAAGAACATG    26122
gb:MT106054|Organism:Severe      GTGTTGAACATGTTACCTTCTTCATCTACAATAAAATTGTTGATGAGCCTGAAGAACATG    26122
gb:MT049951|Organism:Severe      GTGTTGAACATGTTACCTTCTTCATCTACAATAAAATTGTTGATGAGCCTGAAGAACATG    26122
```

```
gb:MN975262|Organism:Severe      GTGTTGAACATGTTACCTTCTTCATCTACAATAAAATTGTTGATGAGCCTGAAGAACATG   26122
gb:MT106052|Organism:Severe      GTGTTGAACATGTTACCTTCTTCATCTACAATAAAATTGTTGATGAGCCTGAAGAACATG   26122
gb:LC522975|Organism:Severe      GTGTTGAACATGTTACCTTCTTCATCTACAATAAAATTGTTGATGAGCCTGAAGAACATG   26119
gb:LC522973|Organism:Severe      GTGTTGAACATGTTACCTTCTTCATCTACAATAAAATTGTTGATGAGCCTGAAGAACATG   26119
gb:LC522974|Organism:Severe      GTGTTGAACATGTTACCTTCTTCATCTACAATAAAATTGTTGATGAGCCTGAAGAACATG   26119
gb:MN985325|Organism:Severe      GTGTTGAACATGTTACCTTCTTCATCTACAATAAAATTGTTGATGAGCCTGAAGAACATG   26122
gb:MT020881|Organism:Severe      GTGTTGAACATGTTACCTTCTTCATCTACAATAAAATTGTTGATGAGCCTGAAGAACATG   26122
gb:MT020880|Organism:Severe      GTGTTGAACATGTTACCTTCTTCATCTACAATAAAATTGTTGATGAGCCTGAAGAACATG   26122
gb:MT066175|Organism:Severe      GTGTTGAACATGTTACCTTCTTCATCTACAATAAAATTGTTGATGAGCCTGAAGAACATG   26122
gb:MN997409|Organism:Severe      GTGTTGAACATGTTACCTTCTTCATCTACAATAAAATTGTTGATGAGCCTGAAGAACATG   26122
gb:MN938384|Organism:Severe      GTGTTGAACATGTTACCTTCTTCATCTACAATAAAATTGTTGATGAGCCTGAAGAACATG   26090
gb:MT044258|Organism:Severe      GTGTTGAACATGTTACCTTCTTCATCTACAATAAAATTGTTGATGAGCCTGAAGAACATG   26098
gb:MT039890|Organism:Severe      GTGTTGAACATGTTACCTTCTTCATCTACAATAAAATTGTTGATGAGCCTGAAGAACATG   26122
gb:MN988713|Organism:Severe      GTGTTGAACATGTTACCTTCTTCATCTACAATAAAATTGTTGATGAGCCTGAAGAACATG   26122
gb:LC521925|Organism:Severe      GTGTTGAACATGTTACCTTCTTCATCTACAATAAAATTGTTGATGAGCCTGAAGAACATG   26095
gb:MT093571|Organism:Severe      GTGTTGAACATGTTACCTTCTTCATCTACAATAAAATTGTTGATGAGCCTGAAGAACATG   26122
gb:MT039887|Organism:Severe      GTGTTGAACATGTTACCTTCTTCATCTACAATAAAATTGTTGATGAGCCTGAAGAACATG   26119
gb:MT019530|Organism:Severe      GTGTTGAACATGTTACCTTCTTCATCTACAATAAAATTGTTGATGAGCCTGAAGAACATG   26122
gb:MT039888|Organism:Severe      GTGTTGAACATGTTACCTTCTTCATCTACAATAAAATTGTTGATGAGCCTGAAGAACATG   26122
gb:LC522972|Organism:Severe      GTGTTGAACATGTTACCTTCTTCATCTACAATAAAATTGTTGATGAGCCTGAAGAACATG   26119
gb:MT027063|Organism:Severe      GTGTTGAACATGTTACCTTCTTCATCTACAATAAAATTGTTGATGAGCCTGAAGAACATG   26122
gb:MT027062|Organism:Severe      GTGTTGAACATGTTACCTTCTTCATCTACAATAAAATTGTTGATGAGCCTGAAGAACATG   26122
gb:MT019529|Organism:Severe      GTGTTGAACATGTTACCTTCTTCATCTACAATAAAATTGTTGATGAGCCTGAAGAACATG   26122
gb:MN996529|Organism:Severe      GTGTTGAACATGTTACCTTCTTCATCTACAATAAAATTGTTGATGAGCCTGAAGAACATG   26110
gb:MN996531|Organism:Severe      GTGTTGAACATGTTACCTTCTTCATCTACAATAAAATTGTTGATGAGCCTGAAGAACATG   26109
gb:MT066176|Organism:Severe      GTGTTGAACATGTTACCTTCTTCATCTACAATAAAATTGTTGATGAGCCTGAAGAACATG   26122
gb:MT027064|Organism:Severe      GTGTTGAACATGTTACCTTCTTCATCTACAATAAAATTGTTGATGAGCCTGAAGAACATG   26122
gb:MN994468|Organism:Severe      GTGTTGAACATGTTACCTTCTTCATCTACAATAAAATTGTTGATGAGCCTGAAGAACATG   26122
gb:MT072688|Organism:Severe      GTGTTGAACATGTTACCTTCTTCATCTACAATAAAATTGTTGATGAGCCTGAAGAACATG   26107
gb:MN996527|Organism:Severe      GTGTTGAACATGTTACCTTCTTCATCTACAATAAAATTGTTGATGAGCCTGAAGAACATG   26089
gb:MT093631|Organism:Severe      GTGTTGAACATGTTACCTTCTTCATCTACAATAAAATTGTTGATGAGCCTGAAGAACATG   26160
gb:MT106053|Organism:Severe      GTGTTGAACATGTTACCTTCTTCATCTACAATAAAATTGTTGATGAGCCTGAAGAACATG   26122
gb:MT019533|Organism:Severe      GTGTTGAACATGTTACCTTCTTCATCTACAATAAAATTGTTGATGAGCCTGAAGAACATG   26122
gb:MT019531|Organism:Severe      GTGTTGAACATGTTACCTTCTTCATCTACAATAAAATTGTTGATGAGCCTGAAGAACATG   26122
gb:MN996528|Organism:Severe      GTGTTGAACATGTTACCTTCTTCATCTACAATAAAATTGTTGATGAGCCTGAAGAACATG   26122
gb:MN996530|Organism:Severe      GTGTTGAACATGTTACCTTCTTCATCTACAATAAAATTGTTGATGAGCCTGAAGAACATG   26108
gb:MN908947|Organism:Severe      GTGTTGAACATGTTACCTTCTTCATCTACAATAAAATTGTTGATGAGCCTGAAGAACATG   26122
gb:MT019532|Organism:Severe      GTGTTGAACATGTTACCTTCTTCATCTACAATAAAATTGTTGATGAGCCTGAAGAACATG   26122
                                 ************************************************************

gb:MT020781|Organism:Severe      TCCAAATTCACACAATCGACGGTTCATCCGGAGTTGTTAATCCAGTAATGGAACCAATTT   26170
gb:MT007544|Organism:Severe      TCCAAATTCACACAATCGACGTTTCATCCGGAGTTGTTAATCCAGTAATGGAACCAATTT   26182
gb:MN994467|Organism:Severe      TCCAAATTCACACAATCGACGGTTCATCCGGAGTTGTTAATCCAGTAATGGAACCAATTT   26182
gb:MT044257|Organism:Severe      TCCAAATTCACACAATCGACGGTTCATCCGGAGTTGTTAATCCAGTAATGGAACCAATTT   26182
gb:MT106054|Organism:Severe      TCCAAATTCACACAATCGACGGTTCATCCGGAGTTGTTAATCCAGTAATGGAACCAATTT   26182
gb:MT049951|Organism:Severe      TCCAAATTCACACAATCGACGGTTCATCCGGAGTTGTTAATCCAGTAATGGAACCAATTT   26182
gb:MN975262|Organism:Severe      TCCAAATTCACACAATCGACGGTTCATCCGGAGTTGTTAATCCAGTAATGGAACCAATTT   26182
gb:MT106052|Organism:Severe      TCCAAATTCACACAATCGACGGTTCATCCGGAGTTGTTAATCCAGTAATGGAACCAATTT   26182
gb:LC522975|Organism:Severe      TCCAAATTCACACAATCGACGGTTCATCCGGAGTTGTTAATCCAGTAATGGAACCAATTT   26179
gb:LC522973|Organism:Severe      TCCAAATTCACACAATCGACGGTTCATCCGGAGTTGTTAATCCAGTAATGGAACCAATTT   26179
gb:LC522974|Organism:Severe      TCCAAATTCACACAATCGACGGTTCATCCGGAGTTGTTAATCCAGTAATGGAACCAATTT   26179
gb:MN985325|Organism:Severe      TCCAAATTCACACAATCGACGGTTCATCCGGAGTTGTTAATCCAGTAATGGAACCAATTT   26182
gb:MT020881|Organism:Severe      TCCAAATTCACACAATCGACGGTTCATCCGGAGTTGTTAATCCAGTAATGGAACCAATTT   26182
gb:MT020880|Organism:Severe      TCCAAATTCACACAATCGACGGTTCATCCGGAGTTGTTAATCCAGTAATGGAACCAATTT   26182
gb:MT066175|Organism:Severe      TCCAAATTCACACAATCGACGGTTCATCCGGAGTTGTTAATCCAGTAATGGAACCAATTT   26182
gb:MN997409|Organism:Severe      TCCAAATTCACACAATCGACGGTTCATCCGGAGTTGTTAATCCAGTAATGGAACCAATTT   26182
gb:MN938384|Organism:Severe      TCCAAATTCACACAATCGACGGTTCATCCGGAGTTGTTAATCCAGTAATGGAACCAATTT   26150
gb:MT044258|Organism:Severe      TCCAAATTCACACAATCGACGGTTCATCCGGAGTTGTTAATCCAGTAATGGAACCAATTT   26158
gb:MT039890|Organism:Severe      TCCAAATTCACACAATCGACGTTTCATCCGGAGTTGTTAATCCAGTAATGGAACCAATTT   26182
gb:MN988713|Organism:Severe      TCCAAATTCACACAATCGACGGTTCATCCGGAGTTGTTAATCCAGTAATGGAACCAATTT   26182
gb:LC521925|Organism:Severe      TCCAAATTCACACAATCGACGGTTCATCCGGAGTTGTTAATCCAGTAATGGAACCAATTT   26155
gb:MT093571|Organism:Severe      TCCAAATTCACACAATCGACGTTTCATCCGGAGTTGTTAATCCAGTAATGGAACCAATTT   26182
gb:MT039887|Organism:Severe      TCCAAATTCACACAATCGACGGTTCATCCGGAGTTGTTAATCCAGTAATGGAACCAATTT   26179
gb:MT019530|Organism:Severe      TCCAAATTCACACAATCGACGGTTCATCCGGAGTTGTTAATCCAGTAATGGAACCAATTT   26182
gb:MT039888|Organism:Severe      TCCAAATTCACACAATCGACGGTTCATCCGGAGTTGTTAATCCAGTAATGGAACCAATTT   26182
```

```
gb:LC522972|Organism:Severe      TCCAAATTCACACAATCGACGGTTCATCCGGAGTTGTTAATCCAGTAATGGAACCAATTT    26179
gb:MT027063|Organism:Severe      TCCAAATTCACACAATCGACGGTTCATCCGGAGTTGTTAATCCAGTAATGGAACCAATTT    26182
gb:MT027062|Organism:Severe      TCCAAATTCACACAATCGACGGTTCATCCGGAGTTGTTAATCCAGTAATGGAACCAATTT    26182
gb:MT019529|Organism:Severe      TCCAAATTCACACAATCGACGGTTCATCCGGAGTTGTTAATCCAGTAATGGAACCAATTT    26182
gb:MN996529|Organism:Severe      TCCAAATTCACACAATCGACGGTTCATCCGGAGTTGTTAATCCAGTAATGGAACCAATTT    26170
gb:MN996531|Organism:Severe      TCCAAATTCACACAATCGACGGTTCATCCGGAGTTGTTAATCCAGTAATGGAACCAATTT    26169
gb:MT066176|Organism:Severe      TCCAAATTCACACAATCGACGGTTCATCCGGAGTTGTTAATCCAGTAATGGAACCAATTT    26182
gb:MT027064|Organism:Severe      TCCAAATTCACACAATCGACGGTTCATCCGGAGTTGTTAATCCAGTAATGGAACCAATTT    26182
gb:MN994468|Organism:Severe      TCCAAATTCACACAATCGACGTTTCATCCGGAGTTGTTAATCCAGTAATGGAACCAATTT    26182
gb:MT072688|Organism:Severe      TCCAAATTCACACAATCGACGGTTCATCCGGAGTTGTTAATCCAGTAATGGAACCAATTT    26167
gb:MN996527|Organism:Severe      TCCAAATTCACACAATCGACGGTTCATCCGGAGTTGTTAATCCAGTAATGGAACCAATTT    26149
gb:MT093631|Organism:Severe      TCCAAATTCACACAATCGACGGTTCATCCGGAGTTGTTAATCCAGTAATGGAACCAATTT    26220
gb:MT106053|Organism:Severe      TCCAAATTCACACAATCGACGGTTCATCCGGAGTTGTTAATCCAGTAATGGAACCAATTT    26182
gb:MT019533|Organism:Severe      TCCAAATTCACACAATCGACGGTTCATCCGGAGTTGTTAATCCAGTAATGGAACCAATTT    26182
gb:MT019531|Organism:Severe      TCCAAATTCACACAATCGACGGTTCATCCGGAGTTGTTAATCCAGTAATGGAACCAATTT    26182
gb:MN996528|Organism:Severe      TCCAAATTCACACAATCGACGGTTCATCCGGAGTTGTTAATCCAGTAATGGAACCAATTT    26182
gb:MN996530|Organism:Severe      TCCAAATTCACACAATCGACGGTTCATCCGGAGTTGTTAATCCAGTAATGGAACCAATTT    26168
gb:MN908947|Organism:Severe      TCCAAATTCACACAATCGACGGTTCATCCGGAGTTGTTAATCCAGTAATGGAACCAATTT    26182
gb:MT019532|Organism:Severe      TCCAAATTCACACAATCGACGGTTCATCCGGAGTTGTTAATCCAGTAATGGAACCAATTT    26182
                                 ********************* **************************************

gb:MT020781|Organism:Severe      ATGATGAACCGACGACGACTACTAGCGTGCCTTTGTAAGCACAAGCTGATGAGTACGAAC    26230
gb:MT007544|Organism:Severe      ATGATGAACCGACGACGACTACTAGCGTGCCTTTGTAAGCACAAGCTGATGAGTACGAAC    26242
gb:MN994467|Organism:Severe      ATGATGAACCGACGACGACTACTAGCGTGCCTTTGTAAGCACAAGCTGATGAGTACGAAC    26242
gb:MT044257|Organism:Severe      ATGATGAACCGACGACGACTACTAGCGTGCCTTTGTAAGCACAAGCTGATGAGTACGAAC    26242
gb:MT106054|Organism:Severe      ATGATGAACCGACGACGACTACTAGCGTGCCTTTGTAAGCACAAGCTGATGAGTACGAAC    26242
gb:MT049951|Organism:Severe      ATGATGAACCGACGACGACTACTAGCGTGCCTTTGTAAGCACAAGCTGATGAGTACGAAC    26242
gb:MN975262|Organism:Severe      ATGATGAACCGACGACGACTACTAGCGTGCCTTTGTAAGCACAAGCTGATGAGTACGAAC    26242
gb:MT106052|Organism:Severe      ATGATGAACCGACGACGACTACTAGCGTGCCTTTGTAAGCACAAGCTGATGAGTACGAAC    26242
gb:LC522975|Organism:Severe      ATGATGAACCGACGACGACTACTAGCGTGCCTTTGTAAGCACAAGCTGATGAGTACGAAC    26239
gb:LC522973|Organism:Severe      ATGATGAACCGACGACGACTACTAGCGTGCCTTTGTAAGCACAAGCTGATGAGTACGAAC    26239
gb:LC522974|Organism:Severe      ATGATGAACCGACGACGACTACTAGCGTGCCTTTGTAAGCACAAGCTGATGAGTACGAAC    26239
gb:MN985325|Organism:Severe      ATGATGAACCGACGACGACTACTAGCGTGCCTTTGTAAGCACAAGCTGATGAGTACGAAC    26242
gb:MT020881|Organism:Severe      ATGATGAACCGACGACGACTACTAGCGTGCCTTTGTAAGCACAAGCTGATGAGTACGAAC    26242
gb:MT020880|Organism:Severe      ATGATGAACCGACGACGACTACTAGCGTGCCTTTGTAAGCACAAGCTGATGAGTACGAAC    26242
gb:MT066175|Organism:Severe      ATGATGAACCGACGACGACTACTAGCGTGCCTTTGTAAGCACAAGCTGATGAGTACGAAC    26242
gb:MN997409|Organism:Severe      ATGATGAACCGACGACGACTACTAGCGTGCCTTTGTAAGCACAAGCTGATGAGTACGAAC    26242
gb:MN938384|Organism:Severe      ATGATGAACCGACGACGACTACTAGCGTGCCTTTGTAAGCACAAGCTGATGAGTACGAAC    26210
gb:MT044258|Organism:Severe      ATGATGAACCGACGACGACTACTAGCGTGCCTTTGTAAGCACAAGCTGATGAGTACGAAC    26218
gb:MT039890|Organism:Severe      ATGATGAACCGACGACGACTACTAGCGTGCCTTTGTAAGCACAAGCTGATGAGTACGAAC    26242
gb:MN988713|Organism:Severe      ATGATGAACCGACGACGACTACTAGCGTGCCTTTGTAAGCACAAGCTGATGAGTACGAAC    26242
gb:LC521925|Organism:Severe      ATGATGAACCGACGACGACTACTAGCGTGCCTTTGTAAGCACAAGCTGATGAGTACGAAC    26215
gb:MT093571|Organism:Severe      ATGATGAACCGACGACGACTACTAGCGTGCCTTTGTAAGCACAAGCTGATGAGTACGAAC    26242
gb:MT039887|Organism:Severe      ATGATGAACCGACGACGACTACTAGCGTGCCTTTGTAAGCACAAGCTGATGAGTACGAAC    26239
gb:MT019530|Organism:Severe      ATGATGAACCGACGACGACTACTAGCGTGCCTTTGTAAGCACAAGCTGATGAGTACGAAC    26242
gb:MT039888|Organism:Severe      ATGATGAACCGACGACGACTACTAGCGTGCCTTTGTAAGCACAAGCTGATGAGTACGAAC    26242
gb:LC522972|Organism:Severe      ATGATGAACCGACGACGACTACTAGCGTGCCTTTGTAAGCACAAGCTGATGAGTACGAAC    26239
gb:MT027063|Organism:Severe      ATGATGAACCGACGACGACTACTAGCGTGCCTTTGTAAGCACAAGCTGATGAGTACGAAC    26242
gb:MT027062|Organism:Severe      ATGATGAACCGACGACGACTACTAGCGTGCCTTTGTAAGCACAAGCTGATGAGTACGAAC    26242
gb:MT019529|Organism:Severe      ATGATGAACCGACGACGACTACTAGCGTGCCTTTGTAAGCACAAGCTGATGAGTACGAAC    26242
gb:MN996529|Organism:Severe      ATGATGAACCGACGACGACTACTAGCGTGCCTTTGTAAGCACAAGCTGATGAGTACGAAC    26230
gb:MN996531|Organism:Severe      ATGATGAACCGACGACGACTACTAGCGTGCCTTTGTAAGCACAAGCTGATGAGTACGAAC    26229
gb:MT066176|Organism:Severe      ATGATGAACCGACGACGACTACTAGCGTGCCTTTGTAAGCACAAGCTGATGAGTACGAAC    26242
gb:MT027064|Organism:Severe      ATGATGAACCGACGACGACTACTAGCGTGCCTTTGTAAGCACAAGCTGATGAGTACGAAC    26242
gb:MN994468|Organism:Severe      ATGATGAACCGACGACGACTACTAGCGTGCCTTTGTAAGCACAAGCTGATGAGTACGAAC    26242
gb:MT072688|Organism:Severe      ATGATGAACCGACGACGACTACTAGCGTGCCTTTGTAAGCACAAGCTGATGAGTACGAAC    26227
gb:MN996527|Organism:Severe      ATGATGAACCGACGACGACTACTAGCGTGCCTTTGTAAGCACAAGCTGATGAGTACGAAC    26209
gb:MT093631|Organism:Severe      ATGATGAACCGACGACGACTACTAGCGTGCCTTTGTAAGCACAAGCTGATGAGTACGAAC    26280
gb:MT106053|Organism:Severe      ATGATGAACCGACGACGACTACTAGCGTGCCTTTGTAAGCACAAGCTGATGAGTACGAAC    26242
gb:MT019533|Organism:Severe      ATGATGAACCGACGACGACTACTAGCGTGCCTTTGTAAGCACAAGCTGATGAGTACGAAC    26242
gb:MT019531|Organism:Severe      ATGATGAACCGACGACGACTACTAGCGTGCCTTTGTAAGCACAAGCTGATGAGTACGAAC    26242
gb:MN996528|Organism:Severe      ATGATGAACCGACGACGACTACTAGCGTGCCTTTGTAAGCACAAGCTGATGAGTACGAAC    26242
gb:MN996530|Organism:Severe      ATGATGAACCGACGACGACTACTAGCGTGCCTTTGTAAGCACAAGCTGATGAGTACGAAC    26228
gb:MN908947|Organism:Severe      ATGATGAACCGACGACGACTACTAGCGTGCCTTTGTAAGCACAAGCTGATGAGTACGAAC    26242
gb:MT019532|Organism:Severe      ATGATGAACCGACGACGACTACTAGCGTGCCTTTGTAAGCACAAGCTGATGAGTACGAAC    26242
```

```
                                    ************************************************************

gb:MT020781|Organism:Severe         TTATGTACTCATTCGTTTCGGAAGAGACAGGTACGTTAATAGTTAATAGCGTACTTCTTT	26290
gb:MT007544|Organism:Severe         TTATGTACTCATTCGTTTCGGAAGAGACAGGTACGTTAATAGTTAATAGCGTACTTCTTT	26302
gb:MN994467|Organism:Severe         TTATGTACTCATTCGTTTCGGAAGAGACAGGTACGTTAATAGTTAATAGCGTACTTCTTT	26302
gb:MT044257|Organism:Severe         TTATGTACTCATTCGTTTCGGAAGAGACAGGTACGTTAATAGTTAATAGCGTACTTCTTT	26302
gb:MT106054|Organism:Severe         TTATGTACTCATTCGTTTCGGAAGAGACAGGTACGTTAATAGTTAATAGCGTACTTCTTT	26302
gb:MT049951|Organism:Severe         TTATGTACTCATTCGTTTCGGAAGAGACAGGTACGTTAATAGTTAATAGCGTACTTCTTT	26302
gb:MN975262|Organism:Severe         TTATGTACTCATTCGTTTCGGAAGAGACAGGTACGTTAATAGTTAATAGCGTACTTCTTT	26302
gb:MT106052|Organism:Severe         TTATGTACTCATTCGTTTCGGAAGAGACAGGTACGTTAATAGTTAATAGCGTACTTCTTT	26302
gb:LC522975|Organism:Severe         TTATGTACTCATTCGTTTCGGAAGAGACAGGTACGTTAATAGTTAATAGCGTACTTCTTT	26299
gb:LC522973|Organism:Severe         TTATGTACTCATTCGTTTCGGAAGAGACAGGTACGTTAATAGTTAATAGCGTACTTCTTT	26299
gb:LC522974|Organism:Severe         TTATGTACTCATTCGTTTCGGAAGAGACAGGTACGTTAATAGTTAATAGCGTACTTCTTT	26299
gb:MN985325|Organism:Severe         TTATGTACTCATTCGTTTCGGAAGAGACAGGTACGTTAATAGTTAATAGCGTACTTCTTT	26302
gb:MT020881|Organism:Severe         TTATGTACTCATTCGTTTCGGAAGAGACAGGTACGTTAATAGTTAATAGCGTACTTCTTT	26302
gb:MT020880|Organism:Severe         TTATGTACTCATTCGTTTCGGAAGAGACAGGTACGTTAATAGTTAATAGCGTACTTCTTT	26302
gb:MT066175|Organism:Severe         TTATGTACTCATTCGTTTCGGAAGAGACAGGTACGTTAATAGTTAATAGCGTACTTCTTT	26302
gb:MN997409|Organism:Severe         TTATGTACTCATTCGTTTCGGAAGAGACAGGTACGTTAATAGTTAATAGCGTACTTCTTT	26302
gb:MN938384|Organism:Severe         TTATGTACTCATTCGTTTCGGAAGAGACAGGTACGTTAATAGTTAATAGCGTACTTCTTT	26270
gb:MT044258|Organism:Severe         TTATGTACTCATTCGTTTCGGAAGAGACAGGTACGTTAATAGTTAATAGCGTACTTCTTT	26278
gb:MT039890|Organism:Severe         TTATGTACTCATTCGTTTCGGAAGAGACAGGTACGTTAATAGTTAATAGCGTACTTCTTT	26302
gb:MN988713|Organism:Severe         TTATGTACTCATTCGTTTCGGAAGAGACAGGTACGTTAATAGTTAATAGCGTACTTCTTT	26302
gb:LC521925|Organism:Severe         TTATGTACTCATTCGTTTCGGAAGAGACAGGTACGTTAATAGTTAATAGCGTACTTCTTT	26275
gb:MT093571|Organism:Severe         TTATGTACTCATTCGTTTCGGAAGAGACAGGTACGTTAATAGTTAATAGCGTACTTCTTT	26302
gb:MT039887|Organism:Severe         TTATGTACTCATTCGTTTCGGAAGAGACAGGTACGTTAATAGTTAATAGCGTACTTCTTT	26299
gb:MT019530|Organism:Severe         TTATGTACTCATTCGTTTCGGAAGAGACAGGTACGTTAATAGTTAATAGCGTACTTCTTT	26302
gb:MT039888|Organism:Severe         TTATGTACTCATTCGTTTCGGAAGAGACAGGTACGTTAATAGTTAATAGCGTACTTCTTT	26302
gb:LC522972|Organism:Severe         TTATGTACTCATTCGTTTCGGAAGAGACAGGTACGTTAATAGTTAATAGCGTACTTCTTT	26299
gb:MT027063|Organism:Severe         TTATGTACTCATTCGTTTCGGAAGAGACAGGTACGTTAATAGTTAATAGCGTACTTCTTT	26302
gb:MT027062|Organism:Severe         TTATGTACTCATTCGTTTCGGAAGAGACAGGTACGTTAATAGTTAATAGCGTACTTCTTT	26302
gb:MT019529|Organism:Severe         TTATGTACTCATTCGTTTCGGAAGAGACAGGTACGTTAATAGTTAATAGCGTACTTCTTT	26302
gb:MN996529|Organism:Severe         TTATGTACTCATTCGTTTCGGAAGAGACAGGTACGTTAATAGTTAATAGCGTACTTCTTT	26290
gb:MN996531|Organism:Severe         TTATGTACTCATTCGTTTCGGAAGAGACAGGTACGTTAATAGTTAATAGCGTACTTCTTT	26289
gb:MT066176|Organism:Severe         TTATGTACTCATTCGTTTCGGAAGAGACAGGTACGTTAATAGTTAATAGCGTACTTCTTT	26302
gb:MT027064|Organism:Severe         TTATGTACTCATTCGTTTCGGAAGAGACAGGTACGTTAATAGTTAATAGCGTACTTCTTT	26302
gb:MN994468|Organism:Severe         TTATGTACTCATTCGTTTCGGAAGAGACAGGTACGTTAATAGTTAATAGCGTACTTCTTT	26302
gb:MT072688|Organism:Severe         TTATGTACTCATTCGTTTCGGAAGAGACAGGTACGTTAATAGTTAATAGCGTACTTCTTT	26287
gb:MN996527|Organism:Severe         TTATGTACTCATTCGTTTCGGAAGAGACAGGTACGTTAATAGTTAATAGCGTACTTCTTT	26269
gb:MT093631|Organism:Severe         TTATGTACTCATTCGTTTCGGAAGAGACAGGTACGTTAATAGTTAATAGCGTACTTCTTT	26340
gb:MT106053|Organism:Severe         TTATGTACTCATTCGTTTCGGAAGAGACAGGTACGTTAATAGTTAATAGCGTACTTCTTT	26302
gb:MT019533|Organism:Severe         TTATGTACTCATTCGTTTCGGAAGAGACAGGTACGTTAATAGTTAATAGCGTACTTCTTT	26302
gb:MT019531|Organism:Severe         TTATGTACTCATTCGTTTCGGAAGAGACAGGTACGTTAATAGTTAATAGCGTACTTCTTT	26302
gb:MN996528|Organism:Severe         TTATGTACTCATTCGTTTCGGAAGAGACAGGTACGTTAATAGTTAATAGCGTACTTCTTT	26302
gb:MN996530|Organism:Severe         TTATGTACTCATTCGTTTCGGAAGAGACAGGTACGTTAATAGTTAATAGCGTACTTCTTT	26288
gb:MN908947|Organism:Severe         TTATGTACTCATTCGTTTCGGAAGAGACAGGTACGTTAATAGTTAATAGCGTACTTCTTT	26302
gb:MT019532|Organism:Severe         TTATGTACTCATTCGTTTCGGAAGAGACAGGTACGTTAATAGTTAATAGCGTACTTCTTT	26302
                                    ************************************************************

gb:MT020781|Organism:Severe         TTCTTGCTTTCGTGGTATTCTTGCTAGTTACACTAGCCATCCTTACTGCGCTTCGATTGT	26350
gb:MT007544|Organism:Severe         TTCTTGCTTTCGTGGTATTCTTGCTAGTTACACTAGCCATCCTTACTGCGCTTCGATTGT	26362
gb:MN994467|Organism:Severe         TTCTTGCTTTCGTGGTATTCTTGCTAGTTACACTAGCCATCCTTACTGCGCTTCGATTGT	26362
gb:MT044257|Organism:Severe         TTCTTGCTTTCGTGGTATTCTTGCTAGTTACACTAGCCATCCTTACTGCGCTTCGATTGT	26362
gb:MT106054|Organism:Severe         TTCTTGCTTTCGTGGTATTCTTGCTAGTTACACTAGCCATCCTTACTGCGCTTCGATTGT	26362
gb:MT049951|Organism:Severe         TTCTTGCTTTCGTGGTATTCTTGCTAGTTACACTAGCCATCCTTACTGCGCTTCGATTGT	26362
gb:MN975262|Organism:Severe         TTCTTGCTTTCGTGGTATTCTTGCTAGTTACACTAGCCATCCTTACTGCGCTTCGATTGT	26362
gb:MT106052|Organism:Severe         TTCTTGCTTTCGTGGTATTCTTGCTAGTTACACTAGCCATCCTTACTGCGCTTCGATTGT	26362
gb:LC522975|Organism:Severe         TTCTTGCTTTCGTGGTATTCTTGCTAGTTACACTAGCCATCCTTACTGCGCTTCGATTGT	26359
gb:LC522973|Organism:Severe         TTCTTGCTTTCGTGGTATTCTTGCTAGTTACACTAGCCATCCTTACTGCGCTTCGATTGT	26359
gb:LC522974|Organism:Severe         TTCTTGCTTTCGTGGTATTCTTGCTAGTTACACTAGCCATCCTTACTGCGCTTCGATTGT	26359
gb:MN985325|Organism:Severe         TTCTTGCTTTCGTGGTATTCTTGCTAGTTACACTAGCCATCCTTACTGCGCTTCGATTGT	26362
gb:MT020881|Organism:Severe         TTCTTGCTTTCGTGGTATTCTTGCTAGTTACACTAGCCATCCTTACTGCGCTTCGATTGT	26362
gb:MT020880|Organism:Severe         TTCTTGCTTTCGTGGTATTCTTGCTAGTTACACTAGCCATCCTTACTGCGCTTCGATTGT	26362
gb:MT066175|Organism:Severe         TTCTTGCTTTCGTGGTATTCTTGCTAGTTACACTAGCCATCCTTACTGCGCTTCGATTGT	26362
gb:MN997409|Organism:Severe         TTCTTGCTTTCGTGGTATTCTTGCTAGTTACACTAGCCATCCTTACTGCGCTTCGATTGT	26362
gb:MN938384|Organism:Severe         TTCTTGCTTTCGTGGTATTCTTGCTAGTTACACTAGCCATCCTTACTGCGCTTCGATTGT	26330
```

```
gb:MT044258|Organism:Severe      TTCTTGCTTTCGTGGTATTCTTGCTAGTTACACTAGCCATCCTTACTGCGCTTCGATTGT   26338
gb:MT039890|Organism:Severe      TTCTTGCTTTCGTGGTATTCTTGCTAGTTACACTAGCCATCCTTACTGCGCATCGATTGT   26362
gb:MN988713|Organism:Severe      TTCTTGCTTTCGTGGTATTCTTGCTAGTTACACTAGCCATCCTTACTGCGCTTCGATTGT   26362
gb:LC521925|Organism:Severe      TTCTTGCTTTCGTGGTATTCTTGCTAGTTACACTAGCCATCCTTACTGCGCTTCGATTGT   26335
gb:MT093571|Organism:Severe      TTCTTGCTTTCGTGGTATTCTTGCTAGTTACACTAGCCATCCTTACTGCGCTTCGATTGT   26362
gb:MT039887|Organism:Severe      TTCTTGCTTTCGTGGTATTCTTGCTAGTTACACTAGCCATCCTTACTGCGCTTCGATTGT   26359
gb:MT019530|Organism:Severe      TTCTTGCTTTCGTGGTATTCTTGCTAGTTACACTAGCCATCCTTACTGCGCTTCGATTGT   26362
gb:MT039888|Organism:Severe      TTCTTGCTTTCGTGGTATTCTTGCTAGTTACACTAGCCATCCTTACTGCGCTTCGATTGT   26362
gb:LC522972|Organism:Severe      TTCTTGCTTTCGTGGTATTCTTGCTAGTTACACTAGCCATCCTTACTGCGCTTCGATTGT   26359
gb:MT027063|Organism:Severe      TTCTTGCTTTCGTGGTATTCTTGCTAGTTACACTAGCCATCCTTACTGCGCTTCGATTGT   26362
gb:MT027062|Organism:Severe      TTCTTGCTTTCGTGGTATTCTTGCTAGTTACACTAGCCATCCTTACTGCGCTTCGATTGT   26362
gb:MT019529|Organism:Severe      TTCTTGCTTTCGTGGTATTCTTGCTAGTTACACTAGCCATCCTTACTGCGCTTCGATTGT   26362
gb:MN996529|Organism:Severe      TTCTTGCTTTCGTGGTATTCTTGCTAGTTACACTAGCCATCCTTACTGCGCTTCGATTGT   26350
gb:MN996531|Organism:Severe      TTCTTGCTTTCGTGGTATTCTTGCTAGTTACACTAGCCATCCTTACTGCGCTTCGATTGT   26349
gb:MT066176|Organism:Severe      TTCTTGCTTTCGTGGTATTCTTGCTAGTTACACTAGCCATCCTTACTGCGCTTCGATTGT   26362
gb:MT027064|Organism:Severe      TTCTTGCTTTCGTGGTATTCTTGCTAGTTACACTAGCCATCCTTACTGCGCTTCGATTGT   26362
gb:MN994468|Organism:Severe      TTCTTGCTTTCGTGGTATTCTTGCTAGTTACACTAGCCATCCTTACTGCGCTTCGATTGT   26362
gb:MT072688|Organism:Severe      TTCTTGCTTTCGTGGTATTCTTGCTAGTTACACTAGCCATCCTTACTGCGCTTCGATTGT   26347
gb:MN996527|Organism:Severe      TTCTTGCTTTCGTGGTATTCTTGCTAGTTACACTAGCCATCCTTACTGCGCTTCGATTGT   26329
gb:MT093631|Organism:Severe      TTCTTGCTTTCGTGGTATTCTTGCTAGTTACACTAGCCATCCTTACTGCGCTTCGATTGT   26400
gb:MT106053|Organism:Severe      TTCTTGCTTTCGTGGTATTCTTGCTAGTTACACTAGCCATCCTTACTGCGCTTCGATTGT   26362
gb:MT019533|Organism:Severe      TTCTTGCTTTCGTGGTATTCTTGCTAGTTACACTAGCCATCCTTACTGCGCTTCGATTGT   26362
gb:MT019531|Organism:Severe      TTCTTGCTTTCGTGGTATTCTTGCTAGTTACACTAGCCATCCTTACTGCGCTTCGATTGT   26362
gb:MN996528|Organism:Severe      TTCTTGCTTTCGTGGTATTCTTGCTAGTTACACTAGCCATCCTTACTGCGCTTCGATTGT   26362
gb:MN996530|Organism:Severe      TTCTTGCTTTCGTGGTATTCTTGCTAGTTACACTAGCCATCCTTACTGCGCTTCGATTGT   26348
gb:MN908947|Organism:Severe      TTCTTGCTTTCGTGGTATTCTTGCTAGTTACACTAGCCATCCTTACTGCGCTTCGATTGT   26362
gb:MT019532|Organism:Severe      TTCTTGCTTTCGTGGTATTCTTGCTAGTTACACTAGCCATCCTTACTGCGCTTCGATTGT   26362
                                 *************************************************** ********

gb:MT020781|Organism:Severe      GTGCGTACTGCTGCAATATTGTTAACGTGAGTCTTGTAAAACCTTCTTTTTACGTTTACT   26410
gb:MT007544|Organism:Severe      GTGCGTACTGCTGCAATATTGTTAACGTGAGTCTTGTAAAACCTTCTTTTTACGTTTACT   26422
gb:MN994467|Organism:Severe      GTGCGTACTGCTGCAATATTGTTAACGTGAGTCTTGTAAAACCTTCTTTTTACGTTTACT   26422
gb:MT044257|Organism:Severe      GTGCGTACTGCTGCAATATTGTTAACGTGAGTCTTGTAAAACCTTCTTTTTACGTTTACT   26422
gb:MT106054|Organism:Severe      GTGCGTACTGCTGCAATATTGTTAACGTGAGTCTTGTAAAACCTTCTTTTTACGTTTACT   26422
gb:MT049951|Organism:Severe      GTGCGTACTGCTGCAATATTGTTAACGTGAGTCTTGTAAAACCTTCTTTTTACGTTTACT   26422
gb:MN975262|Organism:Severe      GTGCGTACTGCTGCAATATTGTTAACGTGAGTCTTGTAAAACCTTCTTTTTACGTTTACT   26422
gb:MT106052|Organism:Severe      GTGCGTACTGCTGCAATATTGTTAACGTGAGTCTTGTAAAACCTTCTTTTTACGTTTACT   26422
gb:LC522975|Organism:Severe      GTGCGTACTGCTGCAATATTGTTAACGTGAGTCTTGTAAAACCTTCTTTTTACGTTTACT   26419
gb:LC522973|Organism:Severe      GTGCGTACTGCTGCAATATTGTTAACGTGAGTCTTGTAAAACCTTCTTTTTACGTTTACT   26419
gb:LC522974|Organism:Severe      GTGCGTACTGCTGCAATATTGTTAACGTGAGTCTTGTAAAACCTTCTTTTTACGTTTACT   26419
gb:MN985325|Organism:Severe      GTGCGTACTGCTGCAATATTGTTAACGTGAGTCTTGTAAAACCTTCTTTTTACGTTTACT   26422
gb:MT020881|Organism:Severe      GTGCGTACTGCTGCAATATTGTTAACGTGAGTCTTGTAAAACCTTCTTTTTACGTTTACT   26422
gb:MT020880|Organism:Severe      GTGCGTACTGCTGCAATATTGTTAACGTGAGTCTTGTAAAACCTTCTTTTTACGTTTACT   26422
gb:MT066175|Organism:Severe      GTGCGTACTGCTGCAATATTGTTAACGTGAGTCTTGTAAAACCTTCTTTTTACGTTTACT   26422
gb:MN997409|Organism:Severe      GTGCGTACTGCTGCAATATTGTTAACGTGAGTCTTGTAAAACCTTCTTTTTACGTTTACT   26422
gb:MN938384|Organism:Severe      GTGCGTACTGCTGCAATATTGTTAACGTGAGTCTTGTAAAACCTTCTTTTTACGTTTACT   26390
gb:MT044258|Organism:Severe      GTGCGTACTGCTGCAATATTGTTAACGTGAGTCTTGTAAAACCTTCTTTTTACGTTTACT   26398
gb:MT039890|Organism:Severe      GTGCGTACTGCTGCAATATTGTTAACGTGAGTCTTGTAAAACCTTCTTTTTACGTTTACT   26422
gb:MN988713|Organism:Severe      GTGCGTACTGCTGCAATATTGTTAACGTGAGTCTTGTAAAACCTTCTTTTTACGTTTACT   26422
gb:LC521925|Organism:Severe      GTGCGTACTGCTGCAATATTGTTAACGTGAGTCTTGTAAAACCTTCTTTTTACGTTTACT   26395
gb:MT093571|Organism:Severe      GTGCGTACTGCTGCAATATTGTTAACGTGAGTCTTGTAAAACCTTCTTTTTACGTTTACT   26422
gb:MT039887|Organism:Severe      GTGCGTACTGCTGCAATATTGTTAACGTGAGTCTTGTAAAACCTTCTTTTTACGTTTACT   26419
gb:MT019530|Organism:Severe      GTGCGTACTGCTGCAATATTGTTAACGTGAGTCTTGTAAAACCTTCTTTTTACGTTTACT   26422
gb:MT039888|Organism:Severe      GTGCGTACTGCTGCAATATTGTTAACGTGAGTCTTGTAAAACCTTCTTTTTACGTTTACT   26422
gb:LC522972|Organism:Severe      GTGCGTACTGCTGCAATATTGTTAACGTGAGTCTTGTAAAACCTTCTTTTTACGTTTACT   26419
gb:MT027063|Organism:Severe      GTGCGTACTGCTGCAATATTGTTAACGTGAGTCTTGTAAAACCTTCTTTTTACGTTTACT   26422
gb:MT027062|Organism:Severe      GTGCGTACTGCTGCAATATTGTTAACGTGAGTCTTGTAAAACCTTCTTTTTACGTTTACT   26422
gb:MT019529|Organism:Severe      GTGCGTACTGCTGCAATATTGTTAACGTGAGTCTTGTAAAACCTTCTTTTTACGTTTACT   26422
gb:MN996529|Organism:Severe      GTGCGTACTGCTGCAATATTGTTAACGTGAGTCTTGTAAAACCTTCTTTTTACGTTTACT   26410
gb:MN996531|Organism:Severe      GTGCGTACTGCTGCAATATTGTTAACGTGAGTCTTGTAAAACCTTCTTTTTACGTTTACT   26409
gb:MT066176|Organism:Severe      GTGCGTACTGCTGCAATATTGTTAACGTGAGTCTTGTAAAACCTTCTTTTTACGTTTACT   26422
gb:MT027064|Organism:Severe      GTGCGTACTGCTGCAATATTGTTAACGTGAGTCTTGTAAAACCTTCTTTTTACGTTTACT   26422
gb:MN994468|Organism:Severe      GTGCGTACTGCTGCAATATTGTTAACGTGAGTCTTGTAAAACCTTCTTTTTACGTTTACT   26422
gb:MT072688|Organism:Severe      GTGCGTACTGCTGCAATATTGTTAACGTGAGTCTTGTAAAACCTTCTTTTTACGTTTACT   26407
gb:MN996527|Organism:Severe      GTGCGTACTGCTGCAATATTGTTAACGTGAGTCTTGTAAAACCTTCTTTTTACGTTTACT   26389
```

```
gb:MT093631|Organism:Severe      GTGCGTACTGCTGCAATATTGTTAACGTGAGTCTTGTAAAACCTTCTTTTTACGTTTACT   26460
gb:MT106053|Organism:Severe      GTGCGTACTGCTGCAATATTGTTAACGTGAGTCTTGTAAAACCTTCTTTTTACGTTTACT   26422
gb:MT019533|Organism:Severe      GTGCGTACTGCTGCAATATTGTTAACGTGAGTCTTGTAAAACCTTCTTTTTACGTTTACT   26422
gb:MT019531|Organism:Severe      GTGCGTACTGCTGCAATATTGTTAACGTGAGTCTTGTAAAACCTTCTTTTTACGTTTACT   26422
gb:MN996528|Organism:Severe      GTGCGTACTGCTGCAATATTGTTAACGTGAGTCTTGTAAAACCTTCTTTTTACGTTTACT   26422
gb:MN996530|Organism:Severe      GTGCGTACTGCTGCAATATTGTTAACGTGAGTCTTGTAAAACCTTCTTTTTACGTTTACT   26408
gb:MN908947|Organism:Severe      GTGCGTACTGCTGCAATATTGTTAACGTGAGTCTTGTAAAACCTTCTTTTTACGTTTACT   26422
gb:MT019532|Organism:Severe      GTGCGTACTGCTGCAATATTGTTAACGTGAGTCTTGTAAAACCTTCTTTTTACGTTTACT   26422
                                 ************************************************************

gb:MT020781|Organism:Severe      CTCGTGTTAAAAATCTGAATTCTTCTAGAGTTCCTGATCTTCTGGTCTAAACGAACTAAA   26470
gb:MT007544|Organism:Severe      CTCGTGTTAAAAATCTGAATTCTTCTAGAGTTCCTGATCTTCTGGTCTAAACGAACTAAA   26482
gb:MN994467|Organism:Severe      CTCGTGTTAAAAATCTGAATTCTTCTAGAGTTCCTGATCTTCTGGTCTAAACGAACTAAA   26482
gb:MT044257|Organism:Severe      CTCGTGTTAAAAATCTGAATTCTTCTAGAGTTCCTGATCTTCTGGTCTAAACGAACTAAA   26482
gb:MT106054|Organism:Severe      CTCGTGTTAAAAATCTGAATTCTTCTAGAGTTCCTGATCTTCTGGTCTAAACGAACTAAA   26482
gb:MT049951|Organism:Severe      CTCGTGTTAAAAATCTGAATTCTTCTAGAGTTCCTGATCTTCTGGTCTAAACGAACTAAA   26482
gb:MN975262|Organism:Severe      CTCGTGTTAAAAATCTGAATTCTTCTAGAGTTCCTGATCTTCTGGTCTAAACGAACTAAA   26482
gb:MT106052|Organism:Severe      CTCGTGTTAAAAATCTGAATTCTTCTAGAGTTCCTGATCTTCTGGTCTAAACGAACTAAA   26482
gb:LC522975|Organism:Severe      CTCGTGTTAAAAATCTGAATTCTTCTAGAGTTCCTGATCTTCTGGTCTAAACGAACTAAA   26479
gb:LC522973|Organism:Severe      CTCGTGTTAAAAATCTGAATTCTTCTAGAGTTCCTGATCTTCTGGTCTAAACGAACTAAA   26479
gb:LC522974|Organism:Severe      CTCGTGTTAAAAATCTGAATTCTTCTAGAGTTCCTGATCTTCTGGTCTAAACGAACTAAA   26479
gb:MN985325|Organism:Severe      CTCGTGTTAAAAATCTGAATTCTTCTAGAGTTCCTGATCTTCTGGTCTAAACGAACTAAA   26482
gb:MT020881|Organism:Severe      CTCGTGTTAAAAATCTGAATTCTTCTAGAGTTCCTGATCTTCTGGTCTAAACGAACTAAA   26482
gb:MT020880|Organism:Severe      CTCGTGTTAAAAATCTGAATTCTTCTAGAGTTCCTGATCTTCTGGTCTAAACGAACTAAA   26482
gb:MT066175|Organism:Severe      CTCGTGTTAAAAATCTGAATTCTTCTAGAGTTCCTGATCTTCTGGTCTAAACGAACTAAA   26482
gb:MN997409|Organism:Severe      CTCGTGTTAAAAATCTGAATTCTTCTAGAGTTCCTGATCTTCTGGTCTAAACGAACTAAA   26482
gb:MN938384|Organism:Severe      CTCGTGTTAAAAATCTGAATTCTTCTAGAGTTCCTGATCTTCTGGTCTAAACGAACTAAA   26450
gb:MT044258|Organism:Severe      CTCGTGTTAAAAATCTGAATTCTTCTAGAGTTCCTGATCTTCTGGTCTAAACGAACTAAA   26458
gb:MT039890|Organism:Severe      CTCGTGTTAAAAATCTGAATTCTTCTAGAGTTCCTGATCTTCTGGTCTAAACGAACTAAA   26482
gb:MN988713|Organism:Severe      CTCGTGTTAAAAATCTGAATTCTTCTAGAGTTCCTGATCTTCTGGTCTAAACGAACTAAA   26482
gb:LC521925|Organism:Severe      CTCGTGTTAAAAATCTGAATTCTTCTAGAGTTCCTGATCTTCTGGTCTAAACGAACTAAA   26455
gb:MT093571|Organism:Severe      CTCGTGTTAAAAATCTGAATTCTTCTAGAGTTCCTGATCTTCTGGTCTAAACGAACTAAA   26482
gb:MT039887|Organism:Severe      CTCGTGTTAAAAATCTGAATTCTTCTAGAGTTCCTGATCTTCTGGTCTAAACGAACTAAA   26479
gb:MT019530|Organism:Severe      CTCGTGTTAAAAATCTGAATTCTTCTAGAGTTCCTGATCTTCTGGTCTAAACGAACTAAA   26482
gb:MT039888|Organism:Severe      CTCGTGTTAAAAATCTGAATTCTTCTAGAGTTCCTGATCTTCTGGTCTAAACGAACTAAA   26482
gb:LC522972|Organism:Severe      CTCGTGTTAAAAATCTGAATTCTTCTAGAGTTCCTGATCTTCTGGTCTAAACGAACTAAA   26479
gb:MT027063|Organism:Severe      CTCGTGTTAAAAATCTGAATTCTTCTAGAGTTCCTGATCTTCTGGTCTAAACGAACTAAA   26482
gb:MT027062|Organism:Severe      CTCGTGTTAAAAATCTGAATTCTTCTAGAGTTCCTGATCTTCTGGTCTAAACGAACTAAA   26482
gb:MT019529|Organism:Severe      CTCGTGTTAAAAATCTGAATTCTTCTAGAGTTCCTGATCTTCTGGTCTAAACGAACTAAA   26482
gb:MN996529|Organism:Severe      CTCGTGTTAAAAATCTGAATTCTTCTAGAGTTCCTGATCTTCTGGTCTAAACGAACTAAA   26470
gb:MN996531|Organism:Severe      CTCGTGTTAAAAATCTGAATTCTTCTAGAGTTCCTGATCTTCTGGTCTAAACGAACTAAA   26469
gb:MT066176|Organism:Severe      CTCGTGTTAAAAATCTGAATTCTTCTAGAGTTCCTGATCTTCTGGTCTAAACGAACTAAA   26482
gb:MT027064|Organism:Severe      CTCGTGTTAAAAATCTGAATTCTTCTAGAGTTCCTGATCTTCTGGTCTAAACGAACTAAA   26482
gb:MN994468|Organism:Severe      CTCGTGTTAAAAATCTGAATTCTTCTAGAGTTCCTGATCTTCTGGTCTAAACGAACTAAA   26482
gb:MT072688|Organism:Severe      CTCGTGTTAAAAATCTGAATTCTTCTAGAGTTCCTGATCTTCTGGTCTAAACGAACTAAA   26467
gb:MN996527|Organism:Severe      CTCGTGTTAAAAATCTGAATTCTTCTAGAGTTCCTGATCTTCTGGTCTAAACGAACTAAA   26449
gb:MT093631|Organism:Severe      CTCGTGTTAAAAATCTGAATTCTTCTAGAGTTCCTGATCTTCTGGTCTAAACGAACTAAA   26520
gb:MT106053|Organism:Severe      CTCGTGTTAAAAATCTGAATTCTTCTAGAGTTCCTGATCTTCTGGTCTAAACGAACTAAA   26482
gb:MT019533|Organism:Severe      CTCGTGTTAAAAATCTGAATTCTTCTAGAGTTCCTGATCTTCTGGTCTAAACGAACTAAA   26482
gb:MT019531|Organism:Severe      CTCGTGTTAAAAATCTGAATTCTTCTAGAGTTCCTGATCTTCTGGTCTAAACGAACTAAA   26482
gb:MN996528|Organism:Severe      CTCGTGTTAAAAATCTGAATTCTTCTAGAGTTCCTGATCTTCTGGTCTAAACGAACTAAA   26482
gb:MN996530|Organism:Severe      CTCGTGTTAAAAATCTGAATTCTTCTAGAGTTCCTGATCTTCTGGTCTAAACGAACTAAA   26468
gb:MN908947|Organism:Severe      CTCGTGTTAAAAATCTGAATTCTTCTAGAGTTCCTGATCTTCTGGTCTAAACGAACTAAA   26482
gb:MT019532|Organism:Severe      CTCGTGTTAAAAATCTGAATTCTTCTAGAGTTCCTGATCTTCTGGTCTAAACGAACTAAA   26482
                                 ************************************************************

gb:MT020781|Organism:Severe      TATTATATTAGTTTTTCTGTTTGGAACTTTAATTTTAGCCATGGCAGATTCCAACGGTAC   26530
gb:MT007544|Organism:Severe      TATTATATTAGTTTTTCTGTTTGGAACTTTAATTTTAGCCATGGCAGATTCCAACGGTAC   26542
gb:MN994467|Organism:Severe      TATTATATTAGTTTTTCTGTTTGGAACTTTAATTTTAGCCATGGCAGATTCCAACGGTAC   26542
gb:MT044257|Organism:Severe      TATTATATTAGTTTTTCTGTTTGGAACTTTAATTTTAGCCATGGCAGATTCCAACGGTAC   26542
gb:MT106054|Organism:Severe      TATTATATTAGTTTTTCTGTTTGGAACTTTAATTTTAGCCATGGCAGATTCCAACGGTAC   26542
gb:MT049951|Organism:Severe      TATTATATTAGTTTTTCTGTTTGGAACTTTAATTTTAGCCATGGCAGATTCCAACGGTAC   26542
gb:MN975262|Organism:Severe      TATTATATTAGTTTTTCTGTTTGGAACTTTAATTTTAGCCATGGCAGATTCCAACGGTAC   26542
gb:MT106052|Organism:Severe      TATTATATTAGTTTTTCTGTTTGGAACTTTAATTTTAGCCATGGCAGATTCCAACGGTAC   26542
gb:LC522975|Organism:Severe      TATTATATTAGTTTTTCTGTTTGGAACTTTAATTTTAGCCATGGCAGATTCCAACGGTAC   26539
```

```
gb:LC522973|Organism:Severe      TATTATATTAGTTTTTCTGTTTGGAACTTTAATTTTAGCCATGGCAGATTCCAACGGTAC   26539
gb:LC522974|Organism:Severe      TATTATATTAGTTTTTCTGTTTGGAACTTTAATTTTAGCCATGGCAGATTCCAACGGTAC   26539
gb:MN985325|Organism:Severe      TATTATATTAGTTTTTCTGTTTGGAACTTTAATTTTAGCCATGGCAGATTCCAACGGTAC   26542
gb:MT020881|Organism:Severe      TATTATATTAGTTTTTCTGTTTGGAACTTTAATTTTAGCCATGGCAGATTCCAACGGTAC   26542
gb:MT020880|Organism:Severe      TATTATATTAGTTTTTCTGTTTGGAACTTTAATTTTAGCCATGGCAGATTCCAACGGTAC   26542
gb:MT066175|Organism:Severe      TATTATATTAGTTTTTCTGTTTGGAACTTTAATTTTAGCCATGGCAGATTCCAACGGTAC   26542
gb:MN997409|Organism:Severe      TATTATATTAGTTTTTCTGTTTGGAACTTTAATTTTAGCCATGGCAGATTCCAACGGTAC   26542
gb:MN938384|Organism:Severe      TATTATATTAGTTTTTCTGTTTGGAACTTTAATTTTAGCCATGGCAGATTCCAACGGTAC   26510
gb:MT044258|Organism:Severe      TATTATATTAGTTTTTCTGTTTGGAACTTTAATTTTAGCCATGGCAGATTCCAACGGTAC   26518
gb:MT039890|Organism:Severe      TATTATATTAGTTTTTCTGTTTGGAACTTTAATTTTAGCCATGGCAGATTCCAACGGTAC   26542
gb:MN988713|Organism:Severe      TATTATATTAGTTTTTCTGTTTGGAACTTTAATTTTAGCCATGGCAGATTCCAACGGTAC   26542
gb:LC521925|Organism:Severe      TATTATATTAGTTTTTCTGTTTGGAACTTTAATTTTAGCCATGGCAGATTCCAACGGTAC   26515
gb:MT093571|Organism:Severe      TATTATATTAGTTTTTCTGTTTGGAACTTTAATTTTAGCCATGGCAGATTCCAACGGTAC   26542
gb:MT039887|Organism:Severe      TATTATATTAGTTTTTCTGTTTGGAACTTTAATTTTAGCCATGGCAGATTCCAACGGTAC   26539
gb:MT019530|Organism:Severe      TATTATATTAGTTTTTCTGTTTGGAACTTTAATTTTAGCCATGGCAGATTCCAACGGTAC   26542
gb:MT039888|Organism:Severe      TATTATATTAGTTTTTCTGTTTGGAACTTTAATTTTAGCCATGGCAGATTCCAACGGTAC   26542
gb:LC522972|Organism:Severe      TATTATATTAGTTTTTCTGTTTGGAACTTTAATTTTAGCCATGGCAGATTCCAACGGTAC   26539
gb:MT027063|Organism:Severe      TATTATATTAGTTTTTCTGTTTGGAACTTTAATTTTAGCCATGGCAGATTCCAACGGTAC   26542
gb:MT027062|Organism:Severe      TATTATATTAGTTTTTCTGTTTGGAACTTTAATTTTAGCCATGGCAGATTCCAACGGTAC   26542
gb:MT019529|Organism:Severe      TATTATATTAGTTTTTCTGTTTGGAACTTTAATTTTAGCCATGGCAGATTCCAACGGTAC   26542
gb:MN996529|Organism:Severe      TATTATATTAGTTTTTCTGTTTGGAACTTTAATTTTAGCCATGGCAGATTCCAACGGTAC   26530
gb:MN996531|Organism:Severe      TATTATATTAGTTTTTCTGTTTGGAACTTTAATTTTAGCCATGGCAGATTCCAACGGTAC   26529
gb:MT066176|Organism:Severe      TATTATATTAGTTTTTCTGTTTGGAACTTTAATTTTAGCCATGGCAGATTCCAACGGTAC   26542
gb:MT027064|Organism:Severe      TATTATATTAGTTTTTCTGTTTGGAACTTTAATTTTAGCCATGGCAGATTCCAACGGTAC   26542
gb:MN994468|Organism:Severe      TATTATATTAGTTTTTCTGTTTGGAACTTTAATTTTAGCCATGGCAGATTCCAACGGTAC   26542
gb:MT072688|Organism:Severe      TATTATATTAGTTTTTCTGTTTGGAACTTTAATTTTAGCCATGGCAGATTCCAACGGTAC   26527
gb:MN996527|Organism:Severe      TATTATATTAGTTTTTCTGTTTGGAACTTTAATTTTAGCCATGGCAGATTCCAACGGTAC   26509
gb:MT093631|Organism:Severe      TATTATATTAGTTTTTCTGTTTGGAACTTTAATTTTAGCCATGGCAGATTCCAACGGTAC   26580
gb:MT106053|Organism:Severe      TATTATATTAGTTTTTCTGTTTGGAACTTTAATTTTAGCCATGGCAGATTCCAACGGTAC   26542
gb:MT019533|Organism:Severe      TATTATATTAGTTTTTCTGTTTGGAACTTTAATTTTAGCCATGGCAGATTCCAACGGTAC   26542
gb:MT019531|Organism:Severe      TATTATATTAGTTTTTCTGTTTGGAACTTTAATTTTAGCCATGGCAGATTCCAACGGTAC   26542
gb:MN996528|Organism:Severe      TATTATATTAGTTTTTCTGTTTGGAACTTTAATTTTAGCCATGGCAGATTCCAACGGTAC   26542
gb:MN996530|Organism:Severe      TATTATATTAGTTTTTCTGTTTGGAACTTTAATTTTAGCCATGGCAGATTCCAACGGTAC   26528
gb:MN908947|Organism:Severe      TATTATATTAGTTTTTCTGTTTGGAACTTTAATTTTAGCCATGGCAGATTCCAACGGTAC   26542
gb:MT019532|Organism:Severe      TATTATATTAGTTTTTCTGTTTGGAACTTTAATTTTAGCCATGGCAGATTCCAACGGTAC   26542
                                 ************************************************************

gb:MT020781|Organism:Severe      TATTACCGTTGAAGAGCTTAAAAAGCTCCTTGAACAATGGAACCTAGTAATAGGTTTCCT   26590
gb:MT007544|Organism:Severe      TATTACCGTTGAAGAGCTTAAAAAGCTCCTTGAACAATGGAACCTAGTAATAGGTTTCCT   26602
gb:MN994467|Organism:Severe      TATTACCGTTGAAGAGCTTAAAAAGCTCCTTGAACAATGGAACCTAGTAATAGGTTTCCT   26602
gb:MT044257|Organism:Severe      TATTACCGTTGAAGAGCTTAAAAAGCTCCTTGAACAATGGAACCTAGTAATAGGTTTCCT   26602
gb:MT106054|Organism:Severe      TATTACCGTTGAAGAGCTTAAAAAGCTCCTTGAACAATGGAACCTAGTAATAGGTTTCCT   26602
gb:MT049951|Organism:Severe      TATTACCGTTGAAGAGCTTAAAAAGCTCCTTGAACAATGGAACCTAGTAATAGGTTTCCT   26602
gb:MN975262|Organism:Severe      TATTACCGTTGAAGAGCTTAAAAAGCTCCTTGAACAATGGAACCTAGTAATAGGTTTCCT   26602
gb:MT106052|Organism:Severe      TATTACCGTTGAAGAGCTTAAAAAGCTCCTTGAACAATGGAACCTAGTAATAGGTTTCCT   26602
gb:LC522975|Organism:Severe      TATTACCGTTGAAGAGCTTAAAAAGCTCCTTGAACAATGGAACCTAGTAATAGGTTTCCT   26599
gb:LC522973|Organism:Severe      TATTACCGTTGAAGAGCTTAAAAAGCTCCTTGAACAATGGAACCTAGTAATAGGTTTCCT   26599
gb:LC522974|Organism:Severe      TATTACCGTTGAAGAGCTTAAAAAGCTCCTTGAACAATGGAACCTAGTAATAGGTTTCCT   26599
gb:MN985325|Organism:Severe      TATTACCGTTGAAGAGCTTAAAAAGCTCCTTGAACAATGGAACCTAGTAATAGGTTTCCT   26602
gb:MT020881|Organism:Severe      TATTACCGTTGAAGAGCTTAAAAAGCTCCTTGAACAATGGAACCTAGTAATAGGTTTCCT   26602
gb:MT020880|Organism:Severe      TATTACCGTTGAAGAGCTTAAAAAGCTCCTTGAACAATGGAACCTAGTAATAGGTTTCCT   26602
gb:MT066175|Organism:Severe      TATTACCGTTGAAGAGCTTAAAAAGCTCCTTGAACAATGGAACCTAGTAATAGGTTTCCT   26602
gb:MN997409|Organism:Severe      TATTACCGTTGAAGAGCTTAAAAAGCTCCTTGAACAATGGAACCTAGTAATAGGTTTCCT   26602
gb:MN938384|Organism:Severe      TATTACCGTTGAAGAGCTTAAAAAGCTCCTTGAACAATGGAACCTAGTAATAGGTTTCCT   26570
gb:MT044258|Organism:Severe      TATTACCGTTGAAGAGCTTAAAAAGCTCCTTGAACAATGGAACCTAGTAATAGGTTTCCT   26578
gb:MT039890|Organism:Severe      TATTACCGTTGAAGAGCTTAAAAAGCTCCTTGAACAATGGAACCTAGTAATAGGTTTCCT   26602
gb:MN988713|Organism:Severe      TATTACCGTTGAAGAGCTTAAAAAGCTCCTTGAACAATGGAACCTAGTAATAGGTTTCCT   26602
gb:LC521925|Organism:Severe      TATTACCGTTGAAGAGCTTAAAAAGCTCCTTGAACAATGGAACCTAGTAATAGGTTTCCT   26575
gb:MT093571|Organism:Severe      TATTACCGTTGAAGAGCTTAAAAAGCTCCTTGAACAATGGAACCTAGTAATAGGTTTCCT   26602
gb:MT039887|Organism:Severe      TATTACCGTTGAAGAGCTTAAAAAGCTCCTTGAACAATGGAACCTAGTAATAGGTTTCCT   26599
gb:MT019530|Organism:Severe      TATTACCGTTGAAGAGCTTAAAAAGCTCCTTGAACAATGGAACCTAGTAATAGGTTTCCT   26602
gb:MT039888|Organism:Severe      TATTACCGTTGAAGAGCTTAAAAAGCTCCTTGAACAATGGAACCTAGTAATAGGTTTCCT   26602
gb:LC522972|Organism:Severe      TATTACCGTTGAAGAGCTTAAAAAGCTCCTTGAACAATGGAACCTAGTAATAGGTTTCCT   26599
gb:MT027063|Organism:Severe      TATTACCGTTGAAGAGCTTAAAAAGCTCCTTGAACAATGGAACCTAGTAATAGGTTTCCT   26602
gb:MT027062|Organism:Severe      TATTACCGTTGAAGAGCTTAAAAAGCTCCTTGAACAATGGAACCTAGTAATAGGTTTCCT   26602
```

```
gb:MT019529|Organism:Severe      TATTACCGTTGAAGAGCTTAAAAAGCTCCTTGAACAATGGAACCTAGTAATAGGTTTCCT   26602
gb:MN996529|Organism:Severe      TATTACCGTTGAAGAGCTTAAAAAGCTCCTTGAACAATGGAACCTAGTAATAGGTTTCCT   26590
gb:MN996531|Organism:Severe      TATTACCGTTGAAGAGCTTAAAAAGCTCCTTGAACAATGGAACCTAGTAATAGGTTTCCT   26589
gb:MT066176|Organism:Severe      TATTACCGTTGAAGAGCTTAAAAAGCTCCTTGAACAATGGAACCTAGTAATAGGTTTCCT   26602
gb:MT027064|Organism:Severe      TATTACCGTTGAAGAGCTTAAAAAGCTCCTTGAACAATGGAACCTAGTAATAGGTTTCCT   26602
gb:MN994468|Organism:Severe      TATTACCGTTGAAGAGCTTAAAAAGCTCCTTGAACAATGGAACCTAGTAATAGGTTTCCT   26602
gb:MT072688|Organism:Severe      TATTACCGTTGAAGAGCTTAAAAAGCTCCTTGAACAATGGAACCTAGTAATAGGTTTCCT   26587
gb:MN996527|Organism:Severe      TATTACCGTTGAAGAGCTTAAAAAGCTCCTTGAACAATGGAACCTAGTAATAGGTTTCCT   26569
gb:MT093631|Organism:Severe      TATTACCGTTGAAGAGCTTAAAAAGCTCCTTGAACAATGGAACCTAGTAATAGGTTTCCT   26640
gb:MT106053|Organism:Severe      TATTACCGTTGAAGAGCTTAAAAAGCTCCTTGAACAATGGAACCTAGTAATAGGTTTCCT   26602
gb:MT019533|Organism:Severe      TATTACCGTTGAAGAGCTTAAAAAGCTCCTTGAACAATGGAACCTAGTAATAGGTTTCCT   26602
gb:MT019531|Organism:Severe      TATTACCGTTGAAGAGCTTAAAAAGCTCCTTGAACAATGGAACCTAGTAATAGGTTTCCT   26602
gb:MN996528|Organism:Severe      TATTACCGTTGAAGAGCTTAAAAAGCTCCTTGAACAATGGAACCTAGTAATAGGTTTCCT   26602
gb:MN996530|Organism:Severe      TATTACCGTTGAAGAGCTTAAAAAGCTCCTTGAACAATGGAACCTAGTAATAGGTTTCCT   26588
gb:MN908947|Organism:Severe      TATTACCGTTGAAGAGCTTAAAAAGCTCCTTGAACAATGGAACCTAGTAATAGGTTTCCT   26602
gb:MT019532|Organism:Severe      TATTACCGTTGAAGAGCTTAAAAAGCTCCTTGAACAATGGAACCTAGTAATAGGTTTCCT   26602
                                 ************************************************************

gb:MT020781|Organism:Severe      ATTCCTTACATGGATTTGTCTTCTACAATTTGCCTATGCCAACAGGAATAGGTTTTTGTA   26650
gb:MT007544|Organism:Severe      ATTCCTTACATGGATTTGTCTTCTACAATTTGCCTATGCCAACAGGAATAGGTTTTTGTA   26662
gb:MN994467|Organism:Severe      ATTCCTTACATGGATTTGTCTTCTACAATTTGCCTATGCCAACAGGAATAGGTTTTTGTA   26662
gb:MT044257|Organism:Severe      ATTCCTTACATGGATTTGTCTTCTACAATTTGCCTATGCCAACAGGAATAGGTTTTTGTA   26662
gb:MT106054|Organism:Severe      ATTCCTTACATGGATTTGTCTTCTACAATTTGCCTATGCCAACAGGAATAGGTTTTTGTA   26662
gb:MT049951|Organism:Severe      ATTCCTTACATGGATTTGTCTTCTACAATTTGCCTATGCCAACAGGAATAGGTTTTTGTA   26662
gb:MN975262|Organism:Severe      ATTCCTTACATGGATTTGTCTTCTACAATTTGCCTATGCCAACAGGAATAGGTTTTTGTA   26662
gb:MT106052|Organism:Severe      ATTCCTTACATGGATTTGTCTTCTACAATTTGCCTATGCCAACAGGAATAGGTTTTTGTA   26662
gb:LC522975|Organism:Severe      ATTCCTTACATGGATTTGTCTTCTACAATTTGCCTATGCCAACAGGAATAGGTTTTTGTA   26659
gb:LC522973|Organism:Severe      ATTCCTTACATGGATTTGTCTTCTACAATTTGCCTATGCCAACAGGAATAGGTTTTTGTA   26659
gb:LC522974|Organism:Severe      ATTCCTTACATGGATTTGTCTTCTACAATTTGCCTATGCCAACAGGAATAGGTTTTTGTA   26659
gb:MN985325|Organism:Severe      ATTCCTTACATGGATTTGTCTTCTACAATTTGCCTATGCCAACAGGAATAGGTTTTTGTA   26662
gb:MT020881|Organism:Severe      ATTCCTTACATGGATTTGTCTTCTACAATTTGCCTATGCCAACAGGAATAGGTTTTTGTA   26662
gb:MT020880|Organism:Severe      ATTCCTTACATGGATTTGTCTTCTACAATTTGCCTATGCCAACAGGAATAGGTTTTTGTA   26662
gb:MT066175|Organism:Severe      ATTCCTTACATGGATTTGTCTTCTACAATTTGCCTATGCCAACAGGAATAGGTTTTTGTA   26662
gb:MN997409|Organism:Severe      ATTCCTTACATGGATTTGTCTTCTACAATTTGCCTATGCCAACAGGAATAGGTTTTTGTA   26662
gb:MN938384|Organism:Severe      ATTCCTTACATGGATTTGTCTTCTACAATTTGCCTATGCCAACAGGAATAGGTTTTTGTA   26630
gb:MT044258|Organism:Severe      ATTCCTTACATGGATTTGTCTTCTACAATTTGCCTATGCCAACAGGAATAGGTTTTTGTA   26638
gb:MT039890|Organism:Severe      ATTCCTTACATGGATTTGTCTTCTACAATTTGCCTATGCCAACAGGAATAGGTTTTTGTA   26662
gb:MN988713|Organism:Severe      ATTCCTTACATGGATTTGTCTTCTACAATTTGCCTATGCCAACAGGAATAGGTTTTTGTA   26662
gb:LC521925|Organism:Severe      ATTCCTTACATGGATTTGTCTTCTACAATTTGCCTATGCCAACAGGAATAGGTTTTTGTA   26635
gb:MT093571|Organism:Severe      ATTCCTTACATGGATTTGTCTTCTACAATTTGCCTATGCCAACAGGAATAGGTTTTTGTA   26662
gb:MT039887|Organism:Severe      ATTCCTTACATGGATTTGTCTTCTACAATTTGCCTATGCCAACAGGAATAGGTTTTTGTA   26659
gb:MT019530|Organism:Severe      ATTCCTTACATGGATTTGTCTTCTACAATTTGCCTATGCCAACAGGAATAGGTTTTTGTA   26662
gb:MT039888|Organism:Severe      ATTCCTTACATGGATTTGTCTTCTACAATTTGCCTATGCCAACAGGAATAGGTTTTTGTA   26662
gb:LC522972|Organism:Severe      ATTCCTTACATGGATTTGTCTTCTACAATTTGCCTATGCCAACAGGAATAGGTTTTTGTA   26659
gb:MT027063|Organism:Severe      ATTCCTTACATGGATTTGTCTTCTACAATTTGCCTATGCCAACAGGAATAGGTTTTTGTA   26662
gb:MT027062|Organism:Severe      ATTCCTTACATGGATTTGTCTTCTACAATTTGCCTATGCCAACAGGAATAGGTTTTTGTA   26662
gb:MT019529|Organism:Severe      ATTCCTTACATGGATTTGTCTTCTACAATTTGCCTATGCCAACAGGAATAGGTTTTTGTA   26662
gb:MN996529|Organism:Severe      ATTCCTTACATGGATTTGTCTTCTACAATTTGCCTATGCCAACAGGAATAGGTTTTTGTA   26650
gb:MN996531|Organism:Severe      ATTCCTTACATGGATTTGTCTTCTACAATTTGCCTATGCCAACAGGAATAGGTTTTTGTA   26649
gb:MT066176|Organism:Severe      ATTCCTTACATGGATTTGTCTTCTACAATTTGCCTATGCCAACAGGAATAGGTTTTTGTA   26662
gb:MT027064|Organism:Severe      ATTCCTTACATGGATTTGTCTTCTACAATTTGCCTATGCCAACAGGAATAGGTTTTTGTA   26662
gb:MN994468|Organism:Severe      ATTCCTTACATGGATTTGTCTTCTACAATTTGCCTATGCCAACAGGAATAGGTTTTTGTA   26662
gb:MT072688|Organism:Severe      ATTCCTTACATGGATTTGTCTTCTACAATTTGCCTATGCCAACAGGAATAGGTTTTTGTA   26647
gb:MN996527|Organism:Severe      ATTCCTTACATGGATTTGTCTTCTACAATTTGCCTATGCCAACAGGAATAGGTTTTTGTA   26629
gb:MT093631|Organism:Severe      ATTCCTTACATGGATTTGTCTTCTACAATTTGCCTATGCCAACAGGAATAGGTTTTTGTA   26700
gb:MT106053|Organism:Severe      ATTCCTTACATGGATTTGTCTTCTACAATTTGCCTATGCCAACAGGAATAGGTTTTTGTA   26662
gb:MT019533|Organism:Severe      ATTCCTTACATGGATTTGTCTTCTACAATTTGCCTATGCCAACAGGAATAGGTTTTTGTA   26662
gb:MT019531|Organism:Severe      ATTCCTTACATGGATTTGTCTTCTACAATTTGCCTATGCCAACAGGAATAGGTTTTTGTA   26662
gb:MN996528|Organism:Severe      ATTCCTTACATGGATTTGTCTTCTACAATTTGCCTATGCCAACAGGAATAGGTTTTTGTA   26662
gb:MN996530|Organism:Severe      ATTCCTTACATGGATTTGTCTTCTACAATTTGCCTATGCCAACAGGAATAGGTTTTTGTA   26648
gb:MN908947|Organism:Severe      ATTCCTTACATGGATTTGTCTTCTACAATTTGCCTATGCCAACAGGAATAGGTTTTTGTA   26662
gb:MT019532|Organism:Severe      ATTCCTTACATGGATTTGTCTTCTACAATTTGCCTATGCCAACAGGAATAGGTTTTTGTA   26662
                                 ************************************************************

gb:MT020781|Organism:Severe      TATAATTAAGTTAATTTTCCTCTGGCTGTTATGGCCAGTAACTTTAGCTTGTTTTGTGCT   26710
```

```
gb:MT007544|Organism:Severe      TATAATTAAGTTAATTTTCCTCTGGCTGTTATGGCCAGTAACTTTAGCTTGTTTTGTGCT    26722
gb:MN994467|Organism:Severe      TATAATTAAGTTAATTTTCCTCTGGCTGTTATGGCCAGTAACTTTAGCTTGTTTTGTGCT    26722
gb:MT044257|Organism:Severe      TATAATTAAGTTAATTTTCCTCTGGCTGTTATGGCCAGTAACTTTAGCTTGTTTTGTGCT    26722
gb:MT106054|Organism:Severe      TATAATTAAGTTAATTTTCCTCTGGCTGTTATGGCCAGTAACTTTAGCTTGTTTTGTGCT    26722
gb:MT049951|Organism:Severe      TATAATTAAGTTAATTTTCCTCTGGCTGTTATGGCCAGTAACTTTAGCTTGTTTTGTGCT    26722
gb:MN975262|Organism:Severe      TATAATTAAGTTAATTTTCCTCTGGCTGTTATGGCCAGTAACTTTAGCTTGTTTTGTGCT    26722
gb:MT106052|Organism:Severe      TATAATTAAGTTAATTTTCCTCTGGCTGTTATGGCCAGTAACTTTAGCTTGTTTTGTGCT    26722
gb:LC522975|Organism:Severe      TATAATTAAGTTAATTTTCCTCTGGCTGTTATGGCCAGTAACTTTAGCTTGTTTTGTGCT    26719
gb:LC522973|Organism:Severe      TATAATTAAGTTAATTTTCCTCTGGCTGTTATGGCCAGTAACTTTAGCTTGTTTTGTGCT    26719
gb:LC522974|Organism:Severe      TATAATTAAGTTAATTTTCCTCTGGCTGTTATGGCCAGTAACTTTAGCTTGTTTTGTGCT    26719
gb:MN985325|Organism:Severe      TATAATTAAGTTAATTTTCCTCTGGCTGTTATGGCCAGTAACTTTAGCTTGTTTTGTGCT    26722
gb:MT020881|Organism:Severe      TATAATTAAGTTAATTTTCCTCTGGCTGTTATGGCCAGTAACTTTAGCTTGTTTTGTGCT    26722
gb:MT020880|Organism:Severe      TATAATTAAGTTAATTTTCCTCTGGCTGTTATGGCCAGTAACTTTAGCTTGTTTTGTGCT    26722
gb:MT066175|Organism:Severe      TATAATTAAGTTAATTTTCCTCTGGCTGTTATGGCCAGTAACTTTAGCTTGTTTTGTGCT    26722
gb:MN997409|Organism:Severe      TATAATTAAGTTAATTTTCCTCTGGCTGTTATGGCCAGTAACTTTAGCTTGTTTTGTGCT    26722
gb:MN938384|Organism:Severe      TATAATTAAGTTAATTTTCCTCTGGCTGTTATGGCCAGTAACTTTAGCTTGTTTTGTGCT    26690
gb:MT044258|Organism:Severe      TATAATTAAGTTAATTTTCCTCTGGCTGTTATGGCCAGTAACTTTAGCTTGTTTTGTGCT    26698
gb:MT039890|Organism:Severe      TATAATTAAGTTAATTTTCCTCTGGCTGTTATGGCCAGTAACTTTAGCTTGTTTTGTGCT    26722
gb:MN988713|Organism:Severe      TATAATTAAGTTAATTTTCCTCTGGCTGTTATGGCCAGTAACTTTAGCTTGTTTTGTGCT    26722
gb:LC521925|Organism:Severe      TATAATTAAGTTAATTTTCCTCTGGCTGTTATGGCCAGTAACTTTAGCTTGTTTTGTGCT    26695
gb:MT093571|Organism:Severe      TATAATTAAGTTAATTTTCCTCTGGCTGTTATGGCCAGTAACTTTAGCTTGTTTTGTGCT    26722
gb:MT039887|Organism:Severe      TATAATTAAGTTAATTTTCCTCTGGCTGTTATGGCCAGTAACTTTAGCTTGTTTTGTGCT    26719
gb:MT019530|Organism:Severe      TATAATTAAGTTAATTTTCCTCTGGCTGTTATGGCCAGTAACTTTAGCTTGTTTTGTGCT    26722
gb:MT039888|Organism:Severe      TATAATTAAGTTAATTTTCCTCTGGCTGTTATGGCCAGTAACTTTAGCTTGTTTTGTGCT    26722
gb:LC522972|Organism:Severe      TATAATTAAGTTAATTTTCCTCTGGCTGTTATGGCCAGTAACTTTAGCTTGTTTTGTGCT    26719
gb:MT027063|Organism:Severe      TATAATTAAGTTAATTTTCCTCTGGCTGTTATGGCCAGTAACTTTAGCTTGTTTTGTGCT    26722
gb:MT027062|Organism:Severe      TATAATTAAGTTAATTTTCCTCTGGCTGTTATGGCCAGTAACTTTAGCTTGTTTTGTGCT    26722
gb:MT019529|Organism:Severe      TATAATTAAGTTAATTTTCCTCTGGCTGTTATGGCCAGTAACTTTAGCTTGTTTTGTGCT    26722
gb:MN996529|Organism:Severe      TATAATTAAGTTAATTTTCCTCTGGCTGTTATGGCCAGTAACTTTAGCTTGTTTTGTGCT    26710
gb:MN996531|Organism:Severe      TATAATTAAGTTAATTTTCCTCTGGCTGTTATGGCCAGTAACTTTAGCTTGTTTTGTGCT    26709
gb:MT066176|Organism:Severe      TATAATTAAGTTAATTTTCCTCTGGCTGTTATGGCCAGTAACTTTAGCTTGTTTTGTGCT    26722
gb:MT027064|Organism:Severe      TATAATTAAGTTAATTTTCCTCTGGCTGTTATGGCCAGTAACTTTAGCTTGTTTTGTGCT    26722
gb:MN994468|Organism:Severe      TATAATTAAGTTAATTTTCCTCTGGCTGTTATGGCCAGTAACTTTAGCTTGTTTTGTGCT    26722
gb:MT072688|Organism:Severe      TATAATTAAGTTAATTTTCCTCTGGCTGTTATGGCCAGTAACTTTAGCTTGTTTTGTGCT    26707
gb:MN996527|Organism:Severe      TATAATTAAGTTAATTTTCCTCTGGCTGTTATGGCCAGTAACTTTAGCTTGTTTTGTGCT    26689
gb:MT093631|Organism:Severe      TATAATTAAGTTAATTTTCCTCTGGCTGTTATGGCCAGTAACTTTAGCTTGTTTTGTGCT    26760
gb:MT106053|Organism:Severe      TATAATTAAGTTAATTTTCCTCTGGCTGTTATGGCCAGTAACTTTAGCTTGTTTTGTGCT    26722
gb:MT019533|Organism:Severe      TATAATTAAGTTAATTTTCCTCTGGCTGTTATGGCCAGTAACTTTAGCTTGTTTTGTGCT    26722
gb:MT019531|Organism:Severe      TATAATTAAGTTAATTTTCCTCTGGCTGTTATGGCCAGTAACTTTAGCTTGTTTTGTGCT    26722
gb:MN996528|Organism:Severe      TATAATTAAGTTAATTTTCCTCTGGCTGTTATGGCCAGTAACTTTAGCTTGTTTTGTGCT    26722
gb:MN996530|Organism:Severe      TATAATTAAGTTAATTTTCCTCTGGCTGTTATGGCCAGTAACTTTAGCTTGTTTTGTGCT    26708
gb:MN908947|Organism:Severe      TATAATTAAGTTAATTTTCCTCTGGCTGTTATGGCCAGTAACTTTAGCTTGTTTTGTGCT    26722
gb:MT019532|Organism:Severe      TATAATTAAGTTAATTTTCCTCTGGCTGTTATGGCCAGTAACTTTAGCTTGTTTTGTGCT    26722
                                 ************************************************************

gb:MT020781|Organism:Severe      TGCTGCTGTTTACAGAATAAATTGGATCACCGGTGGAATTGCTATCGCAATGGCTTGTCT    26770
gb:MT007544|Organism:Severe      TGCTGCTGTTTACAGAATAAATTGGATCACCGGTGGAATTGCTATCGCAATGGCTTGTCT    26782
gb:MN994467|Organism:Severe      TGCTGCCGTTTACAGAATAAATTGGATCACCGGTGGAATTGCTATCGCAATGGCTTGTCT    26782
gb:MT044257|Organism:Severe      TGCTGCCGTTTACAGAATAAATTGGATCACCGGTGGAATTGCTATCGCAATGGCTTGTCT    26782
gb:MT106054|Organism:Severe      TGCTGCTGTTTACAGAATAAATTGGATCACCGGTGGAATTGCTATCGCAATGGCTTGTCT    26782
gb:MT049951|Organism:Severe      TGCTGCTGTTTACAGAATAAATTGGATCACCGGTGGAATTGCTATCGCAATGGCTTGTCT    26782
gb:MN975262|Organism:Severe      TGCTGCTGTTTACAGAATAAATTGGATCACCGGTGGAATTGCTATCGCAATGGCTTGTCT    26782
gb:MT106052|Organism:Severe      TGCTGCTGTTTACAGAATAAATTGGATCACCGGTGGAATTGCTATCGCAATGGCTTGTCT    26782
gb:LC522975|Organism:Severe      TGCTGCTGTTTACAGAATAAATTGGATCACCGGTGGAATTGCTATCGCAATGGCTTGTCT    26779
gb:LC522973|Organism:Severe      TGCTGCTGTTTACAGAATAAATTGGATCACCGGTGGAATTGCTATCGCAATGGCTTGTCT    26779
gb:LC522974|Organism:Severe      TGCTGCTGTTTACAGAATAAATTGGATCACCGGTGGAATTGCTATCGCAATGGCTTGTCT    26779
gb:MN985325|Organism:Severe      TGCTGCTGTTTACAGAATAAATTGGATCACCGGTGGAATTGCTATCGCAATGGCTTGTCT    26782
gb:MT020881|Organism:Severe      TGCTGCTGTTTACAGAATAAATTGGATCACCGGTGGAATTGCTATCGCAATGGCTTGTCT    26782
gb:MT020880|Organism:Severe      TGCTGCTGTTTACAGAATAAATTGGATCACCGGTGGAATTGCTATCGCAATGGCTTGTCT    26782
gb:MT066175|Organism:Severe      TGCTGCTGTTTACAGAATAAATTGGATCACCGGTGGAATTGCTATCGCAATGGCTTGTCT    26782
gb:MN997409|Organism:Severe      TGCTGCTGTTTACAGAATAAATTGGATCACCGGTGGAATTGCTATCGCAATGGCTTGTCT    26782
gb:MN938384|Organism:Severe      TGCTGCTGTTTACAGAATAAATTGGATCACCGGTGGAATTGCTATCGCAATGGCTTGTCT    26750
gb:MT044258|Organism:Severe      TGCTGCTGTTTACAGAATAAATTGGATCACCGGTGGAATTGCTATCGCAATGGCTTGTCT    26758
gb:MT039890|Organism:Severe      TGCTGCTGTTTACAGAATAAATTGGATCACCGGTGGAATTGCTATCGCAATGGCTTGTCT    26782
gb:MN988713|Organism:Severe      TGCTGCYGTTTACAGAATAAATTGGATCACCGGTGGAATTGCTATCGCAATGGCTTGTCT    26782
```

```
gb:LC521925|Organism:Severe      TGCTGCTGTTTACAGAATAAATTGGATCACCGGTGGAATTGCTATCGCAATGGCTTGTCT   26755
gb:MT093571|Organism:Severe      TGCTGCTGTTTACAGAATAAATTGGATCACCGGTGGAATTGCTATCGCAATGGCTTGTCT   26782
gb:MT039887|Organism:Severe      TGCTGCTGTTTACAGAATAAATTGGATCACCGGTGGAATTGCTATCGCAATGGCTTGTCT   26779
gb:MT019530|Organism:Severe      TGCTGCTGTTTACAGAATAAATTGGATCACCGGTGGAATTGCTATCGCAATGGCTTGTCT   26782
gb:MT039888|Organism:Severe      TGCTGCTGTTTACAGAATAAATTGGATCACCGGTGGAATTGCTATCGCAATGGCTTGTCT   26782
gb:LC522972|Organism:Severe      TGCTGCTGTTTACAGAATAAATTGGATCACCGGTGGAATTGCTATCGCAATGGCTTGTCT   26779
gb:MT027063|Organism:Severe      TGCTGCTGTTTACAGAATAAATTGGATCACCGGTGGAATTGCTATCGCAATGGCTTGTCT   26782
gb:MT027062|Organism:Severe      TGCTGCTGTTTACAGAATAAATTGGATCACCGGTGGAATTGCTATCGCAATGGCTTGTCT   26782
gb:MT019529|Organism:Severe      TGCTGCTGTTTACAGAATAAATTGGATCACCGGTGGAATTGCTATCGCAATGGCTTGTCT   26782
gb:MN996529|Organism:Severe      TGCTGCTGTTTACAGAATAAATTGGATCACCGGTGGAATTGCTATCGCAATGGCTTGTCT   26770
gb:MN996531|Organism:Severe      TGCTGCTGTTTACAGAATAAATTGGATCACCGGTGGAATTGCTATCGCAATGGCTTGTCT   26769
gb:MT066176|Organism:Severe      TGCTGCTGTTTACAGAATAAATTGGATCACCGGTGGAATTGCTATCGCAATGGCTTGTCT   26782
gb:MT027064|Organism:Severe      TGCTGCTGTTTACAGAATAAATTGGATCACCGGTGGAATTGCTATCGCAATGGCTTGTCT   26782
gb:MN994468|Organism:Severe      TGCTGCTGTTTACAGAATAAATTGGATCACCGGTGGAATTGCTATCGCAATGGCTTGTCT   26782
gb:MT072688|Organism:Severe      TGCTGCTGTTTACAGAATAAATTGGATCACCGGTGGAATTGCTATCGCAATGGCTTGTCT   26767
gb:MN996527|Organism:Severe      TGCTGCTGTTTACAGAATAAATTGGATCACCGGTGGAATTGCTATCGCAATGGCTTGTCT   26749
gb:MT093631|Organism:Severe      TGCTGCTGTTTACAGAATAAATTGGATCACCGGTGGAATTGCTATCGCAATGGCTTGTCT   26820
gb:MT106053|Organism:Severe      TGCTGCTGTTTACAGAATAAATTGGATCACCGGTGGAATTGCTATCGCAATGGCTTGTCT   26782
gb:MT019533|Organism:Severe      TGCTGCTGTTTACAGAATAAATTGGATCACCGGTGGAATTGCTATCGCAATGGCTTGTCT   26782
gb:MT019531|Organism:Severe      TGCTGCTGTTTACAGAATAAATTGGATCACCGGTGGAATTGCTATCGCAATGGCTTGTCT   26782
gb:MN996528|Organism:Severe      TGCTGCTGTTTACAGAATAAATTGGATCACCGGTGGAATTGCTATCGCAATGGCTTGTCT   26782
gb:MN996530|Organism:Severe      TGCTGCTGTTTACAGAATAAATTGGATCACCGGTGGAATTGCTATCGCAATGGCTTGTCT   26768
gb:MN908947|Organism:Severe      TGCTGCTGTTTACAGAATAAATTGGATCACCGGTGGAATTGCTATCGCAATGGCTTGTCT   26782
gb:MT019532|Organism:Severe      TGCTGCTGTTTACAGAATAAATTGGATCACCGGTGGAATTGCTATCGCAATGGCTTGTCT   26782
                                 ****** *****************************************************

gb:MT020781|Organism:Severe      TGTAGGCTTGATGTGGCTCAGCTACTTCATTGCTTCTTTCAGACTGTTTGCGCGTACGCG   26830
gb:MT007544|Organism:Severe      TGTAGGCTTGATGTGGCTCAGCTACTTCATTGCTTCTTTCAGACTGTTTGCGCGTACGCG   26842
gb:MN994467|Organism:Severe      TGTAGGCTTGATGTGGCTCAGCTACTTCATTGCTTCTTTCAGACTGTTTGCGCGTACGCG   26842
gb:MT044257|Organism:Severe      TGTAGGCTTGATGTGGCTCAGCTACTTCATTGCTTCTTTCAGACTGTTTGCGCGTACGCG   26842
gb:MT106054|Organism:Severe      TGTAGGCTTGATGTGGCTCAGCTACTTCATTGCTTCTTTCAGACTGTTTGCGCGTACGCG   26842
gb:MT049951|Organism:Severe      TGTAGGCTTGATGTGGCTCAGCTACTTCATTGCTTCTTTCAGACTGTTTGCGCGTACGCG   26842
gb:MN975262|Organism:Severe      TGTAGGCTTGATGTGGCTCAGCTACTTCATTGCTTCTTTCAGACTGTTTGCGCGTACGCG   26842
gb:MT106052|Organism:Severe      TGTAGGCTTGATGTGGCTCAGCTACTTCATTGCTTCTTTCAGACTGTTTGCGCGTACGCG   26842
gb:LC522975|Organism:Severe      TGTAGGCTTGATGTGGCTCAGCTACTTCATTGCTTCTTTCAGACTGTTTGCGCGTACGCG   26839
gb:LC522973|Organism:Severe      TGTAGGCTTGATGTGGCTCAGCTACTTCATTGCTTCTTTCAGACTGTTTGCGCGTACGCG   26839
gb:LC522974|Organism:Severe      TGTAGGCTTGATGTGGCTCAGCTACTTCATTGCTTCTTTCAGACTGTTTGCGCGTACGCG   26839
gb:MN985325|Organism:Severe      TGTAGGCTTGATGTGGCTCAGCTACTTCATTGCTTCTTTCAGACTGTTTGCGCGTACGCG   26842
gb:MT020881|Organism:Severe      TGTAGGCTTGATGTGGCTCAGCTACTTCATTGCTTCTTTCAGACTGTTTGCGCGTACGCG   26842
gb:MT020880|Organism:Severe      TGTAGGCTTGATGTGGCTCAGCTACTTCATTGCTTCTTTCAGACTGTTTGCGCGTACGCG   26842
gb:MT066175|Organism:Severe      TGTAGGCTTGATGTGGCTCAGCTACTTCATTGCTTCTTTCAGACTGTTTGCGCGTACGCG   26842
gb:MN997409|Organism:Severe      TGTAGGCTTGATGTGGCTCAGCTACTTCATTGCTTCTTTCAGACTGTTTGCGCGTACGCG   26842
gb:MN938384|Organism:Severe      TGTAGGCTTGATGTGGCTCAGCTACTTCATTGCTTCTTTCAGACTGTTTGCGCGTACGCG   26810
gb:MT044258|Organism:Severe      TGTAGGCTTGATGTGGCTCAGCTACTTCATTGCTTCTTTCAGACTGTTTGCGCGTACGCG   26818
gb:MT039890|Organism:Severe      TGTAGGCTTGATGTGGCTCAGCTACTTCATTGCTTCTTTCAGACTGTTTGCGCGTACGCG   26842
gb:MN988713|Organism:Severe      TGTAGGCTTGATGTGGCTCAGCTACTTCATTGCTTCTTTCAGACTGTTTGCGCGTACGCG   26842
gb:LC521925|Organism:Severe      TGTAGGCTTGATGTGGCTCAGCTACTTCATTGCTTCTTTCAGACTGTTTGCGCGTACGCG   26815
gb:MT093571|Organism:Severe      TGTAGGCTTGATGTGGCTCAGCTACTTCATTGCTTCTTTCAGACTGTTTGCGCGTACGCG   26842
gb:MT039887|Organism:Severe      TGTAGGCTTGATGTGGCTCAGCTACTTCATTGCTTCTTTCAGACTGTTTGCGCGTACGCG   26839
gb:MT019530|Organism:Severe      TGTAGGCTTGATGTGGCTCAGCTACTTCATTGCTTCTTTCAGACTGTTTGCGCGTACGCG   26842
gb:MT039888|Organism:Severe      TGTAGGCTTGATGTGGCTCAGCTACTTCATTGCTTCTTTCAGACTGTTTGCGCGTACGCG   26842
gb:LC522972|Organism:Severe      TGTAGGCTTGATGTGGCTCAGCTACTTCATTGCTTCTTTCAGACTGTTTGCGCGTACGCG   26839
gb:MT027063|Organism:Severe      TGTAGGCTTGATGTGGCTCAGCTACTTCATTGCTTCTTTCAGACTGTTTGCGCGTACGCG   26842
gb:MT027062|Organism:Severe      TGTAGGCTTGATGTGGCTCAGCTACTTCATTGCTTCTTTCAGACTGTTTGCGCGTACGCG   26842
gb:MT019529|Organism:Severe      TGTAGGCTTGATGTGGCTCAGCTACTTCATTGCTTCTTTCAGACTGTTTGCGCGTACGCG   26842
gb:MN996529|Organism:Severe      TGTAGGCTTGATGTGGCTCAGCTACTTCATTGCTTCTTTCAGACTGTTTGCGCGTACGCG   26830
gb:MN996531|Organism:Severe      TGTAGGCTTGATGTGGCTCAGCTACTTCATTGCTTCTTTCAGACTGTTTGCGCGTACGCG   26829
gb:MT066176|Organism:Severe      TGTAGGCTTGATGTGGCTCAGCTACTTCATTGCTTCTTTCAGACTGTTTGCGCGTACGCG   26842
gb:MT027064|Organism:Severe      TGTAGGCTTGATGTGGCTCAGCTACTTCATTGCTTCTTTCAGACTGTTTGCGCGTACGCG   26842
gb:MN994468|Organism:Severe      TGTAGGCTTGATGTGGCTCAGCTACTTCATTGCTTCTTTCAGACTGTTTGCGCGTACGCG   26842
gb:MT072688|Organism:Severe      TGTAGGCTTGATGTGGCTCAGCTACTTCATTGCTTCTTTCAGACTGTTTGCGCGTACGCG   26827
gb:MN996527|Organism:Severe      TGTAGGCTTGATGTGGCTCAGCTACTTCATTGCTTCTTTCAGACTGTTTGCGCGTACGCG   26809
gb:MT093631|Organism:Severe      TGTAGGCTTGATGTGGCTCAGCTACTTCATTGCTTCTTTCAGACTGTTTGCGCGTACGCG   26880
gb:MT106053|Organism:Severe      TGTAGGCTTGATGTGGCTCAGCTACTTCATTGCTTCTTTCAGACTGTTTGCGCGTACGCG   26842
gb:MT019533|Organism:Severe      TGTAGGCTTGATGTGGCTCAGCTACTTCATTGCTTCTTTCAGACTGTTTGCGCGTACGCG   26842
```

```
gb:MT019531|Organism:Severe      TGTAGGCTTGATGTGGCTCAGCTACTTCATTGCTTCTTTCAGACTGTTTGCGCGTACGCG   26842
gb:MN996528|Organism:Severe      TGTAGGCTTGATGTGGCTCAGCTACTTCATTGCTTCTTTCAGACTGTTTGCGCGTACGCG   26842
gb:MN996530|Organism:Severe      TGTAGGCTTGATGTGGCTCAGCTACTTCATTGCTTCTTTCAGACTGTTTGCGCGTACGCG   26828
gb:MN908947|Organism:Severe      TGTAGGCTTGATGTGGCTCAGCTACTTCATTGCTTCTTTCAGACTGTTTGCGCGTACGCG   26842
gb:MT019532|Organism:Severe      TGTAGGCTTGATGTGGCTCAGCTACTTCATTGCTTCTTTCAGACTGTTTGCGCGTACGCG   26842
                                 ************************************************************

gb:MT020781|Organism:Severe      TTCCATGTGGTCATTCAATCCAGAAACTAACATTCTTCTCAACGTGCCACTCCATGGCAC   26890
gb:MT007544|Organism:Severe      TTCCATGTGGTCATTCAATCCAGAAACTAACATTCTTCTCAACGTGCCACTCCATGGCAC   26902
gb:MN994467|Organism:Severe      TTCCATGTGGTCATTCAATCCAGAAACTAACATTCTTCTCAACGTGCCACTCCATGGCAC   26902
gb:MT044257|Organism:Severe      TTCCATGTGGTCATTCAATCCAGAAACTAACATTCTTCTCAACGTGCCACTCCATGGCAC   26902
gb:MT106054|Organism:Severe      TTCCATGTGGTCATTCAATCCAGAAACTAACATTCTTCTCAACGTGCCACTCCATGGCAC   26902
gb:MT049951|Organism:Severe      TTCCATGTGGTCATTCAATCCAGAAACTAACATTCTTCTCAACGTGCCACTCCATGGCAC   26902
gb:MN975262|Organism:Severe      TTCCATGTGGTCATTCAATCCAGAAACTAACATTCTTCTCAACGTGCCACTCCATGGCAC   26902
gb:MT106052|Organism:Severe      TTCCATGTGGTCATTCAATCCAGAAACTAACATTCTTCTCAACGTGCCACTCCATGGCAC   26902
gb:LC522975|Organism:Severe      TTCCATGTGGTCATTCAATCCAGAAACTAACATTCTTCTCAACGTGCCACTCCATGGCAC   26899
gb:LC522973|Organism:Severe      TTCCATGTGGTCATTCAATCCAGAAACTAACATTCTTCTCAACGTGCCACTCCATGGCAC   26899
gb:LC522974|Organism:Severe      TTCCATGTGGTCATTCAATCCAGAAACTAACATTCTTCTCAACGTGCCACTCCATGGCAC   26899
gb:MN985325|Organism:Severe      TTCCATGTGGTCATTCAATCCAGAAACTAACATTCTTCTCAACGTGCCACTCCATGGCAC   26902
gb:MT020881|Organism:Severe      TTCCATGTGGTCATTCAATCCAGAAACTAACATTCTTCTCAACGTGCCACTCCATGGCAC   26902
gb:MT020880|Organism:Severe      TTCCATGTGGTCATTCAATCCAGAAACTAACATTCTTCTCAACGTGCCACTCCATGGCAC   26902
gb:MT066175|Organism:Severe      TTCCATGTGGTCATTCAATCCAGAAACTAACATTCTTCTCAACGTGCCACTCCATGGCAC   26902
gb:MN997409|Organism:Severe      TTCCATGTGGTCATTCAATCCAGAAACTAACATTCTTCTCAACGTGCCACTCCATGGCAC   26902
gb:MN938384|Organism:Severe      TTCCATGTGGTCATTCAATCCAGAAACTAACATTCTTCTCAACGTGCCACTCCATGGCAC   26870
gb:MT044258|Organism:Severe      TTCCATGTGGTCATTCAATCCAGAAACTAACATTCTTCTCAACGTGCCACTCCATGGCAC   26878
gb:MT039890|Organism:Severe      TTCCATGTGGTCATTCAATCCAGAAACTAACATTCTTCTCAACGTGCCACTCCATGGCAC   26902
gb:MN988713|Organism:Severe      TTCCATGTGGTCATTCAATCCAGAAACTAACATTCTTCTCAACGTGCCACTCCATGGCAC   26902
gb:LC521925|Organism:Severe      TTCCATGTGGTCATTCAATCCAGAAACTAACATTCTTCTCAACGTGCCACTCCATGGCAC   26875
gb:MT093571|Organism:Severe      TTCCATGTGGTCATTCAATCCAGAAACTAACATTCTTCTCAACGTGCCACTCCATGGCAC   26902
gb:MT039887|Organism:Severe      TTCCATGTGGTCATTCAATCCAGAAACTAACATTCTTCTCAACGTGCCACTCCATGGCAC   26899
gb:MT019530|Organism:Severe      TTCCATGTGGTCATTCAATCCAGAAACTAACATTCTTCTCAACGTGCCACTCCATGGCAC   26902
gb:MT039888|Organism:Severe      TTCCATGTGGTCATTCAATCCAGAAACTAACATTCTTCTCAACGTGCCACTCCATGGCAC   26902
gb:LC522972|Organism:Severe      TTCCATGTGGTCATTCAATCCAGAAACTAACATTCTTCTCAACGTGCCACTCCATGGCAC   26899
gb:MT027063|Organism:Severe      TTCCATGTGGTCATTCAATCCAGAAACTAACATTCTTCTCAACGTGCCACTCCATGGCAC   26902
gb:MT027062|Organism:Severe      TTCCATGTGGTCATTCAATCCAGAAACTAACATTCTTCTCAACGTGCCACTCCATGGCAC   26902
gb:MT019529|Organism:Severe      TTCCATGTGGTCATTCAATCCAGAAACTAACATTCTTCTCAACGTGCCACTCCATGGCAC   26902
gb:MN996529|Organism:Severe      TTCCATGTGGTCATTCAATCCAGAAACTAACATTCTTCTCAACGTGCCACTCCATGGCAC   26890
gb:MN996531|Organism:Severe      TTCCATGTGGTCATTCAATCCAGAAACTAACATTCTTCTCAACGTGCCACTCCATGGCAC   26889
gb:MT066176|Organism:Severe      TTCCATGTGGTCATTCAATCCAGAAACTAACATTCTTCTCAACGTGCCACTCCATGGCAC   26902
gb:MT027064|Organism:Severe      TTCCATGTGGTCATTCAATCCAGAAACTAACATTCTTCTCAACGTGCCACTCCATGGCAC   26902
gb:MN994468|Organism:Severe      TTCCATGTGGTCATTCAATCCAGAAACTAACATTCTTCTCAACGTGCCACTCCATGGCAC   26902
gb:MT072688|Organism:Severe      TTCCATGTGGTCATTCAATCCAGAAACTAACATTCTTCTCAACGTGCCACTCCATGGCAC   26887
gb:MN996527|Organism:Severe      TTCCATGTGGTCATTCAATCCAGAAACTAACATTCTTCTCAACGTGCCACTCCATGGCAC   26869
gb:MT093631|Organism:Severe      TTCCATGTGGTCATTCAATCCAGAAACTAACATTCTTCTCAACGTGCCACTCCATGGCAC   26940
gb:MT106053|Organism:Severe      TTCCATGTGGTCATTCAATCCAGAAACTAACATTCTTCTCAACGTGCCACTCCATGGCAC   26902
gb:MT019533|Organism:Severe      TTCCATGTGGTCATTCAATCCAGAAACTAACATTCTTCTCAACGTGCCACTCCATGGCAC   26902
gb:MT019531|Organism:Severe      TTCCATGTGGTCATTCAATCCAGAAACTAACATTCTTCTCAACGTGCCACTCCATGGCAC   26902
gb:MN996528|Organism:Severe      TTCCATGTGGTCATTCAATCCAGAAACTAACATTCTTCTCAACGTGCCACTCCATGGCAC   26902
gb:MN996530|Organism:Severe      TTCCATGTGGTCATTCAATCCAGAAACTAACATTCTTCTCAACGTGCCACTCCATGGCAC   26888
gb:MN908947|Organism:Severe      TTCCATGTGGTCATTCAATCCAGAAACTAACATTCTTCTCAACGTGCCACTCCATGGCAC   26902
gb:MT019532|Organism:Severe      TTCCATGTGGTCATTCAATCCAGAAACTAACATTCTTCTCAACGTGCCACTCCATGGCAC   26902
                                 ************************************************************

gb:MT020781|Organism:Severe      TATTCTGACCAGACCGCTTCTAGAAAGTGAACTCGTAATCGGAGCTGTGATCCTTCGTGG   26950
gb:MT007544|Organism:Severe      TATTCTGACCAGACCGCTTCTAGAAAGTGAACTCGTAATCGGAGCTGTGATCCTTCGTGG   26962
gb:MN994467|Organism:Severe      TATTCTGACCAGACCGCTTCTAGAAAGTGAACTCGTAATCGGAGCTGTGATCCTTCGTGG   26962
gb:MT044257|Organism:Severe      TATTCTGACCAGACCGCTTCTAGAAAGTGAACTCGTAATCGGAGCTGTGATCCTTCGTGG   26962
gb:MT106054|Organism:Severe      TATTCTGACCAGACCGCTTCTAGAAAGTGAACTCGTAATCGGAGCTGTGATCCTTCGTGG   26962
gb:MT049951|Organism:Severe      TATTCTGACCAGACCGCTTCTAGAAAGTGAACTCGTAATCGGAGCTGTGATCCTTCGTGG   26962
gb:MN975262|Organism:Severe      TATTCTGACCAGACCGCTTCTAGAAAGTGAACTCGTAATCGGAGCTGTGATCCTTCGTGG   26962
gb:MT106052|Organism:Severe      TATTCTGACCAGACCGCTTCTAGAAAGTGAACTCGTAATCGGAGCTGTGATCCTTCGTGG   26962
gb:LC522975|Organism:Severe      TATTCTGACCAGACCGCTTCTAGAAAGTGAACTCGTAATCGGAGCTGTGATCCTTCGTGG   26959
gb:LC522973|Organism:Severe      TATTCTGACCAGACCGCTTCTAGAAAGTGAACTCGTAATCGGAGCTGTGATCCTTCGTGG   26959
gb:LC522974|Organism:Severe      TATTCTGACCAGACCGCTTCTAGAAAGTGAACTCGTAATCGGAGCTGTGATCCTTCGTGG   26959
gb:MN985325|Organism:Severe      TATTCTGACCAGACCGCTTCTAGAAAGTGAACTCGTAATCGGAGCTGTGATCCTTCGTGG   26962
```

```
gb:MT020881|Organism:Severe      TATTCTGACCAGACCGCTTCTAGAAAGTGAACTCGTAATCGGAGCTGTGATCCTTCGTGG   26962
gb:MT020880|Organism:Severe      TATTCTGACCAGACCGCTTCTAGAAAGTGAACTCGTAATCGGAGCTGTGATCCTTCGTGG   26962
gb:MT066175|Organism:Severe      TATTCTGACCAGACCGCTTCTAGAAAGTGAACTCGTAATCGGAGCTGTGATCCTTCGTGG   26962
gb:MN997409|Organism:Severe      TATTCTGACCAGACCGCTTCTAGAAAGTGAACTCGTAATCGGAGCTGTGATCCTTCGTGG   26962
gb:MN938384|Organism:Severe      TATTCTGACCAGACCGCTTCTAGAAAGTGAACTCGTAATCGGAGCTGTGATCCTTCGTGG   26930
gb:MT044258|Organism:Severe      TATTCTGACCAGACCGCTTCTAGAAAGTGAACTCGTAATCGGAGCTGTGATCCTTCGTGG   26938
gb:MT039890|Organism:Severe      TATTCTGACCAGACCGCTTCTAGAAAGTGAACTCGTAATCGGAGCTGTGATCCTTCGTGG   26962
gb:MN988713|Organism:Severe      TATTCTGACCAGACCGCTTCTAGAAAGTGAACTCGTAATCGGAGCTGTGATCCTTCGTGG   26962
gb:LC521925|Organism:Severe      TATTCTGACCAGACCGCTTCTAGAAAGTGAACTCGTAATCGGAGCTGTGATCCTTCGTGG   26935
gb:MT093571|Organism:Severe      TATTCTGACCAGACCGCTTCTAGAAAGTGAACTCGTAATCGGAGCTGTGATCCTTCGTGG   26962
gb:MT039887|Organism:Severe      TATTCTGACCAGACCGCTTCTAGAAAGTGAACTCGTAATCGGAGCTGTGATCCTTCGTGG   26959
gb:MT019530|Organism:Severe      TATTCTGACCAGACCGCTTCTAGAAAGTGAACTCGTAATCGGAGCTGTGATCCTTCGTGG   26962
gb:MT039888|Organism:Severe      TATTCTGACCAGACCGCTTCTAGAAAGTGAACTCGTAATCGGAGCTGTGATCCTTCGTGG   26962
gb:LC522972|Organism:Severe      TATTCTGACCAGACCGCTTCTAGAAAGTGAACTCGTAATCGGAGCTGTGATCCTTCGTGG   26959
gb:MT027063|Organism:Severe      TATTCTGACCAGACCGCTTCTAGAAAGTGAACTCGTAATCGGAGCTGTGATCCTTCGTGG   26962
gb:MT027062|Organism:Severe      TATTCTGACCAGACCGCTTCTAGAAAGTGAACTCGTAATCGGAGCTGTGATCCTTCGTGG   26962
gb:MT019529|Organism:Severe      TATTCTGACCAGACCGCTTCTAGAAAGTGAACTCGTAATCGGAGCTGTGATCCTTCGTGG   26962
gb:MN996529|Organism:Severe      TATTCTGACCAGACCGCTTCTAGAAAGTGAACTCGTAATCGGAGCTGTGATCCTTCGTGG   26950
gb:MN996531|Organism:Severe      TATTCTGACCAGACCGCTTCTAGAAAGTGAACTCGTAATCGGAGCTGTGATCCTTCGTGG   26949
gb:MT066176|Organism:Severe      TATTCTGACCAGACCGCTTCTAGAAAGTGAACTCGTAATCGGAGCTGTGATCCTTCGTGG   26962
gb:MT027064|Organism:Severe      TATTCTGACCAGACCGCTTCTAGAAAGTGAACTCGTAATCGGAGCTGTGATCCTTCGTGG   26962
gb:MN994468|Organism:Severe      TATTCTGACCAGACCGCTTCTAGAAAGTGAACTCGTAATCGGAGCTGTGATCCTTCGTGG   26962
gb:MT072688|Organism:Severe      TATTCTGACCAGACCGCTTCTAGAAAGTGAACTCGTAATCGGAGCTGTGATCCTTCGTGG   26947
gb:MN996527|Organism:Severe      TATTCTGACCAGACCGCTTCTAGAAAGTGAACTCGTAATCGGAGCTGTGATCCTTCGTGG   26929
gb:MT093631|Organism:Severe      TATTCTGACCAGACCGCTTCTAGAAAGTGAACTCGTAATCGGAGCTGTGATCCTTCGTGG   27000
gb:MT106053|Organism:Severe      TATTCTGACCAGACCGCTTCTAGAAAGTGAACTCGTAATCGGAGCTGTGATCCTTCGTGG   26962
gb:MT019533|Organism:Severe      TATTCTGACCAGACCGCTTCTAGAAAGTGAACTCGTAATCGGAGCTGTGATCCTTCGTGG   26962
gb:MT019531|Organism:Severe      TATTCTGACCAGACCGCTTCTAGAAAGTGAACTCGTAATCGGAGCTGTGATCCTTCGTGG   26962
gb:MN996528|Organism:Severe      TATTCTGACCAGACCGCTTCTAGAAAGTGAACTCGTAATCGGAGCTGTGATCCTTCGTGG   26962
gb:MN996530|Organism:Severe      TATTCTGACCAGACCGCTTCTAGAAAGTGAACTCGTAATCGGAGCTGTGATCCTTCGTGG   26948
gb:MN908947|Organism:Severe      TATTCTGACCAGACCGCTTCTAGAAAGTGAACTCGTAATCGGAGCTGTGATCCTTCGTGG   26962
gb:MT019532|Organism:Severe      TATTCTGACCAGACCGCTTCTAGAAAGTGAACTCGTAATCGGAGCTGTGATCCTTCGTGG   26962
                                 ************************************************************

gb:MT020781|Organism:Severe      ACATCTTCGTATTGCTGGACACCATCTAGGACGCTGTGACATCAAGGACCTGCCTAAAGA   27010
gb:MT007544|Organism:Severe      ACATCTTCGTATTGCTGGACACCATCTAGGACGCTGTGACATCAAGGACCTGCCTAAAGA   27022
gb:MN994467|Organism:Severe      ACATCTTCGTATTGCTGGACACCATCTAGGACGCTGTGACATCAAGGACCTGCCTAAAGA   27022
gb:MT044257|Organism:Severe      ACATCTTCGTATTGCTGGACACCATCTAGGACGCTGTGACATCAAGGACCTGCCTAAAGA   27022
gb:MT106054|Organism:Severe      ACATCTTCGTATTGCTGGACACCATCTAGGACGCTGTGACATCAAGGACCTGCCTAAAGA   27022
gb:MT049951|Organism:Severe      ACATCTTCGTATTGCTGGACACCATCTAGGACGCTGTGACATCAAGGACCTGCCTAAAGA   27022
gb:MN975262|Organism:Severe      ACATCTTCGTATTGCTGGACACCATCTAGGACGCTGTGACATCAAGGACCTGCCTAAAGA   27022
gb:MT106052|Organism:Severe      ACATCTTCGTATTGCTGGACACCATCTAGGACGCTGTGACATCAAGGACCTGCCTAAAGA   27022
gb:LC522975|Organism:Severe      ACATCTTCGTATTGCTGGACACCATCTAGGACGCTGTGACATCAAGGACCTGCCTAAAGA   27019
gb:LC522973|Organism:Severe      ACATCTTCGTATTGCTGGACACCATCTAGGACGCTGTGACATCAAGGACCTGCCTAAAGA   27019
gb:LC522974|Organism:Severe      ACATCTTCGTATTGCTGGACACCATCTAGGACGCTGTGACATCAAGGACCTGCCTAAAGA   27019
gb:MN985325|Organism:Severe      ACATCTTCGTATTGCTGGACACCATCTAGGACGCTGTGACATCAAGGACCTGCCTAAAGA   27022
gb:MT020881|Organism:Severe      ACATCTTCGTATTGCTGGACACCATCTAGGACGCTGTGACATCAAGGACCTGCCTAAAGA   27022
gb:MT020880|Organism:Severe      ACATCTTCGTATTGCTGGACACCATCTAGGACGCTGTGACATCAAGGACCTGCCTAAAGA   27022
gb:MT066175|Organism:Severe      ACATCTTCGTATTGCTGGACACCATCTAGGACGCTGTGACATCAAGGACCTGCCTAAAGA   27022
gb:MN997409|Organism:Severe      ACATCTTCGTATTGCTGGACACCATCTAGGACGCTGTGACATCAAGGACCTGCCTAAAGA   27022
gb:MN938384|Organism:Severe      ACATCTTCGTATTGCTGGACACCATCTAGGACGCTGTGACATCAAGGACCTGCCTAAAGA   26990
gb:MT044258|Organism:Severe      ACATCTTCGTATTGCTGGACACCATCTAGGACGCTGTGACATCAAGGACCTGCCTAAAGA   26998
gb:MT039890|Organism:Severe      ACATCTTCGTATTGCTGGACACCATCTAGGACGCTGTGACATCAAGGACCTGCCTAAAGA   27022
gb:MN988713|Organism:Severe      ACATCTTCGTATTGCTGGACACCATCTAGGACGCTGTGACATCAAGGACCTGCCTAAAGA   27022
gb:LC521925|Organism:Severe      ACATCTTCGTATTGCTGGACACCATCTAGGACGCTGTGACATCAAGGACCTGCCTAAAGA   26995
gb:MT093571|Organism:Severe      ACATCTTCGTATTGCTGGACACCATCTAGGACGCTGTGACATCAAGGACCTGCCTAAAGA   27022
gb:MT039887|Organism:Severe      ACATCTTCGTATTGCTGGACACCATCTAGGACGCTGTGACATCAAGGACCTGCCTAAAGA   27019
gb:MT019530|Organism:Severe      ACATCTTCGTATTGCTGGACACCATCTAGGACGCTGTGACATCAAGGACCTGCCTAAAGA   27022
gb:MT039888|Organism:Severe      ACATCTTCGTATTGCTGGACACCATCTAGGACGCTGTGACATCAAGGACCTGCCTAAAGA   27022
gb:LC522972|Organism:Severe      ACATCTTCGTATTGCTGGACACCATCTAGGACGCTGTGACATCAAGGACCTGCCTAAAGA   27019
gb:MT027063|Organism:Severe      ACATCTTCGTATTGCTGGACACCATCTAGGACGCTGTGACATCAAGGACCTGCCTAAAGA   27022
gb:MT027062|Organism:Severe      ACATCTTCGTATTGCTGGACACCATCTAGGACGCTGTGACATCAAGGACCTGCCTAAAGA   27022
gb:MT019529|Organism:Severe      ACATCTTCGTATTGCTGGACACCATCTAGGACGCTGTGACATCAAGGACCTGCCTAAAGA   27022
gb:MN996529|Organism:Severe      ACATCTTCGTATTGCTGGACACCATCTAGGACGCTGTGACATCAAGGACCTGCCTAAAGA   27010
gb:MN996531|Organism:Severe      ACATCTTCGTATTGCTGGACACCATCTAGGACGCTGTGACATCAAGGACCTGCCTAAAGA   27009
```

```
gb:MT066176|Organism:Severe      ACATCTTCGTATTGCTGGACACCATCTAGGACGCTGTGACATCAAGGACCTGCCTAAAGA    27022
gb:MT027064|Organism:Severe      ACATCTTCGTATTGCTGGACACCATCTAGGACGCTGTGACATCAAGGACCTGCCTAAAGA    27022
gb:MN994468|Organism:Severe      ACATCTTCGTATTGCTGGACACCATCTAGGACGCTGTGACATCAAGGACCTGCCTAAAGA    27022
gb:MT072688|Organism:Severe      ACATCTTCGTATTGCTGGACACCATCTAGGACGCTGTGACATCAAGGACCTGCCTAAAGA    27007
gb:MN996527|Organism:Severe      ACATCTTCGTATTGCTGGACACCATCTAGGACGCTGTGACATCAAGGACCTGCCTAAAGA    26989
gb:MT093631|Organism:Severe      ACATCTTCGTATTGCTGGACACCATCTAGGACGCTGTGACATCAAGGACCTGCCTAAAGA    27060
gb:MT106053|Organism:Severe      ACATCTTCGTATTGCTGGACACCATCTAGGACGCTGTGACATCAAGGACCTGCCTAAAGA    27022
gb:MT019533|Organism:Severe      ACATCTTCGTATTGCTGGACACCATCTAGGACGCTGTGACATCAAGGACCTGCCTAAAGA    27022
gb:MT019531|Organism:Severe      ACATCTTCGTATTGCTGGACACCATCTAGGACGCTGTGACATCAAGGACCTGCCTAAAGA    27022
gb:MN996528|Organism:Severe      ACATCTTCGTATTGCTGGACACCATCTAGGACGCTGTGACATCAAGGACCTGCCTAAAGA    27022
gb:MN996530|Organism:Severe      ACATCTTCGTATTGCTGGACACCATCTAGGACGCTGTGACATCAAGGACCTGCCTAAAGA    27008
gb:MN908947|Organism:Severe      ACATCTTCGTATTGCTGGACACCATCTAGGACGCTGTGACATCAAGGACCTGCCTAAAGA    27022
gb:MT019532|Organism:Severe      ACATCTTCGTATTGCTGGACACCATCTAGGACGCTGTGACATCAAGGACCTGCCTAAAGA    27022
                                 ************************************************************

gb:MT020781|Organism:Severe      AATCACTGTTGCTACATCACGAACGCTTTCTTATTACAAATTGGGAGCTTCGCAGCGTGT    27070
gb:MT007544|Organism:Severe      AATCACTGTTGCTACATCACGAACGCTTTCTTATTACAAATTGGGAGCTTCGCAGCGTGT    27082
gb:MN994467|Organism:Severe      AATCACTGTTGCTACATCACGAACGCTTTCTTATTACAAATTGGGAGCTTCGCAGCGTGT    27082
gb:MT044257|Organism:Severe      AATCACTGTTGCTACATCACGAACGCTTTCTTATTACAAATTGGGAGCTTCGCAGCGTGT    27082
gb:MT106054|Organism:Severe      AATCACTGTTGCTACATCACGAACGCTTTCTTATTACAAATTGGGAGCTTCGCAGCGTGT    27082
gb:MT049951|Organism:Severe      AATCACTGTTGCTACATCACGAACGCTTTCTTATTACAAATTGGGAGCTTCGCAGCGTGT    27082
gb:MN975262|Organism:Severe      AATCACTGTTGCTACATCACGAACGCTTTCTTATTACAAATTGGGAGCTTCGCAGCGTGT    27082
gb:MT106052|Organism:Severe      AATCACTGTTGCTACATCACGAACGCTTTCTTATTACAAATTGGGAGCTTCGCAGCGTGT    27082
gb:LC522975|Organism:Severe      AATCACTGTTGCTACATCACGAACGCTTTCTTATTACAAATTGGGAGCTTCGCAGCGTGT    27079
gb:LC522973|Organism:Severe      AATCACTGTTGCTACATCACGAACGCTTTCTTATTACAAATTGGGAGCTTCGCAGCGTGT    27079
gb:LC522974|Organism:Severe      AATCACTGTTGCTACATCACGAACGCTTTCTTATTACAAATTGGGAGCTTCGCAGCGTGT    27079
gb:MN985325|Organism:Severe      AATCACTGTTGCTACATCACGAACGCTTTCTTATTACAAATTGGGAGCTTCGCAGCGTGT    27082
gb:MT020881|Organism:Severe      AATCACTGTTGCTACATCACGAACGCTTTCTTATTACAAATTGGGAGCTTCGCAGCGTGT    27082
gb:MT020880|Organism:Severe      AATCACTGTTGCTACATCACGAACGCTTTCTTATTACAAATTGGGAGCTTCGCAGCGTGT    27082
gb:MT066175|Organism:Severe      AATCACTGTTGCTACATCACGAACGCTTTCTTATTACAAATTGGGAGCTTCGCAGCGTGT    27082
gb:MN997409|Organism:Severe      AATCACTGTTGCTACATCACGAACGCTTTCTTATTACAAATTGGGAGCTTCGCAGCGTGT    27082
gb:MN938384|Organism:Severe      AATCACTGTTGCTACATCACGAACGCTTTCTTATTACAAATTGGGAGCTTCGCAGCGTGT    27050
gb:MT044258|Organism:Severe      AATCACTGTTGCTACATCACGAACGCTTTCTTATTACAAATTGGGAGCTTCGCAGCGTGT    27058
gb:MT039890|Organism:Severe      AATCACTGTTGCTACATCACGAACGCTTTCTTATTACAAATTGGGAGCTTCGCAGCGTGT    27082
gb:MN988713|Organism:Severe      AATCACTGTTGCTACATCACGAACGCTTTCTTATTACAAATTGGGAGCTTCGCAGCGTGT    27082
gb:LC521925|Organism:Severe      AATCACTGTTGCTACATCACGAACGCTTTCTTATTACAAATTGGGAGCTTCGCAGCGTGT    27055
gb:MT093571|Organism:Severe      AATCACTGTTGCTACATCACGAACGCTTTCTTATTACAAATTGGGAGCTTCGCAGCGTGT    27082
gb:MT039887|Organism:Severe      AATCACTGTTGCTACATCACGAACGCTTTCTTATTACAAATTGGGAGCTTCGCAGCGTGT    27079
gb:MT019530|Organism:Severe      AATCACTGTTGCTACATCACGAACGCTTTCTTATTACAAATTGGGAGCTTCGCAGCGTGT    27082
gb:MT039888|Organism:Severe      AATCACTGTTGCTACATCACGAACGCTTTCTTATTACAAATTGGGAGCTTCGCAGCGTGT    27082
gb:LC522972|Organism:Severe      AATCACTGTTGCTACATCACGAACGCTTTCTTATTACAAATTGGGAGCTTCGCAGCGTGT    27079
gb:MT027063|Organism:Severe      AATCACTGTTGCTACATCACGAACGCTTTCTTATTACAAATTGGGAGCTTCGCAGCGTGT    27082
gb:MT027062|Organism:Severe      AATCACTGTTGCTACATCACGAACGCTTTCTTATTACAAATTGGGAGCTTCGCAGCGTGT    27082
gb:MT019529|Organism:Severe      AATCACTGTTGCTACATCACGAACGCTTTCTTATTACAAATTGGGAGCTTCGCAGCGTGT    27082
gb:MN996529|Organism:Severe      AATCACTGTTGCTACATCACGAACGCTTTCTTATTACAAATTGGGAGCTTCGCAGCGTGT    27070
gb:MN996531|Organism:Severe      AATCACTGTTGCTACATCACGAACGCTTTCTTATTACAAATTGGGAGCTTCGCAGCGTGT    27069
gb:MT066176|Organism:Severe      AATCACTGTTGCTACATCACGAACGCTTTCTTATTACAAATTGGGAGCTTCGCAGCGTGT    27082
gb:MT027064|Organism:Severe      AATCACTGTTGCTACATCACGAACGCTTTCTTATTACAAATTGGGAGCTTCGCAGCGTGT    27082
gb:MN994468|Organism:Severe      AATCACTGTTGCTACATCACGAACGCTTTCTTATTACAAATTGGGAGCTTCGCAGCGTGT    27082
gb:MT072688|Organism:Severe      AATCACTGTTGCTACATCACGAACGCTTTCTTATTACAAATTGGGAGCTTCGCAGCGTGT    27067
gb:MN996527|Organism:Severe      AATCACTGTTGCTACATCACGAACGCTTTCTTATTACAAATTGGGAGCTTCGCAGCGTGT    27049
gb:MT093631|Organism:Severe      AATCACTGTTGCTACATCACGAACGCTTTCTTATTACAAATTGGGAGCTTCGCAGCGTGT    27120
gb:MT106053|Organism:Severe      AATCACTGTTGCTACATCACGAACGCTTTCTTATTACAAATTGGGAGCTTCGCAGCGTGT    27082
gb:MT019533|Organism:Severe      AATCACTGTTGCTACATCACGAACGCTTTCTTATTACAAATTGGGAGCTTCGCAGCGTGT    27082
gb:MT019531|Organism:Severe      AATCACTGTTGCTACATCACGAACGCTTTCTTATTACAAATTGGGAGCTTCGCAGCGTGT    27082
gb:MN996528|Organism:Severe      AATCACTGTTGCTACATCACGAACGCTTTCTTATTACAAATTGGGAGCTTCGCAGCGTGT    27082
gb:MN996530|Organism:Severe      AATCACTGTTGCTACATCACGAACGCTTTCTTATTACAAATTGGGAGCTTCGCAGCGTGT    27068
gb:MN908947|Organism:Severe      AATCACTGTTGCTACATCACGAACGCTTTCTTATTACAAATTGGGAGCTTCGCAGCGTGT    27082
gb:MT019532|Organism:Severe      AATCACTGTTGCTACATCACGAACGCTTTCTTATTACAAATTGGGAGCTTCGCAGCGTGT    27082
                                 ************************************************************

gb:MT020781|Organism:Severe      AGCAGGTGACTCAGGTTTTGCTGCATACAGTCGCTACAGGATTGGCAACTATAAATTAAA    27130
gb:MT007544|Organism:Severe      AGCAGGTGACTCAGGTTTTGCTGCATACAGTCGCTACAGGATTGGCAACTATAAATTAAA    27142
gb:MN994467|Organism:Severe      AGCAGGTGACTCAGGTTTTGCTGCATACAGTCGCTACAGGATTGGCAACTATAAATTAAA    27142
gb:MT044257|Organism:Severe      AGCAGGTGACTCAGGTTTTGCTGCATACAGTCGCTACAGGATTGGCAACTATAAATTAAA    27142
```

```
gb:MT106054|Organism:Severe      AGCAGGTGACTCAGGTTTTGCTGCATACAGTCGCTACAGGATTGGCAACTATAAATTAAA    27142
gb:MT049951|Organism:Severe      AGCAGGTGACTCAGGTTTTGCTGCATACAGTCGCTACAGGATTGGCAACTATAAATTAAA    27142
gb:MN975262|Organism:Severe      AGCAGGTGACTCAGGTTTTGCTGCATACAGTCGCTACAGGATTGGCAACTATAAATTAAA    27142
gb:MT106052|Organism:Severe      AGCAGGTGACTCAGGTTTTGCTGCATACAGTCGCTACAGGATTGGCAACTATAAATTAAA    27142
gb:LC522975|Organism:Severe      AGCAGGTGACTCAGGTTTTGCTGCATACAGTCGCTACAGGATTGGCAACTATAAATTAAA    27139
gb:LC522973|Organism:Severe      AGCAGGTGACTCAGGTTTTGCTGCATACAGTCGCTACAGGATTGGCAACTATAAATTAAA    27139
gb:LC522974|Organism:Severe      AGCAGGTGACTCAGGTTTTGCTGCATACAGTCGCTACAGGATTGGCAACTATAAATTAAA    27139
gb:MN985325|Organism:Severe      AGCAGGTGACTCAGGTTTTGCTGCATACAGTCGCTACAGGATTGGCAACTATAAATTAAA    27142
gb:MT020881|Organism:Severe      AGCAGGTGACTCAGGTTTTGCTGCATACAGTCGCTACAGGATTGGCAACTATAAATTAAA    27142
gb:MT020880|Organism:Severe      AGCAGGTGACTCAGGTTTTGCTGCATACAGTCGCTACAGGATTGGCAACTATAAATTAAA    27142
gb:MT066175|Organism:Severe      AGCAGGTGACTCAGGTTTTGCTGCATACAGTCGCTACAGGATTGGCAACTATAAATTAAA    27142
gb:MN997409|Organism:Severe      AGCAGGTGACTCAGGTTTTGCTGCATACAGTCGCTACAGGATTGGCAACTATAAATTAAA    27142
gb:MN938384|Organism:Severe      AGCAGGTGACTCAGGTTTTGCTGCATACAGTCGCTACAGGATTGGCAACTATAAATTAAA    27110
gb:MT044258|Organism:Severe      AGCAGGTGACTCAGGTTTTGCTGCATACAGTCGCTACAGGATTGGCAACTATAAATTAAA    27118
gb:MT039890|Organism:Severe      AGCAGGTGACTCAGGTTTTGCTGCATACAGTCGCTACAGGATTGGCAACTATAAATTAAA    27142
gb:MN988713|Organism:Severe      AGCAGGTGACTCAGGTTTTGCTGCATACAGTCGCTACAGGATTGGCAACTATAAATTAAA    27142
gb:LC521925|Organism:Severe      AGCAGGTGACTCAGGTTTTGCTGCATACAGTCGCTACAGGATTGGCAACTATAAATTAAA    27115
gb:MT093571|Organism:Severe      AGCAGGTGACTCAGGTTTTGCTGCATACAGTCGCTACAGGATTGGCAACTATAAATTAAA    27142
gb:MT039887|Organism:Severe      AGCAGGTGACTCAGGTTTTGCTGCATACAGTCGCTACAGGATTGGCAACTATAAATTAAA    27139
gb:MT019530|Organism:Severe      AGCAGGTGACTCAGGTTTTGCTGCATACAGTCGCTACAGGATTGGCAACTATAAATTAAA    27142
gb:MT039888|Organism:Severe      AGCAGGTGACTCAGGTTTTGCTGCATACAGTCGCTACAGGATTGGCAACTATAAATTAAA    27142
gb:LC522972|Organism:Severe      AGCAGGTGACTCAGGTTTTGCTGCATACAGTCGCTACAGGATTGGCAACTATAAATTAAA    27139
gb:MT027063|Organism:Severe      AGCAGGTGACTCAGGTTTTGCTGCATACAGTCGCTACAGGATTGGCAACTATAAATTAAA    27142
gb:MT027062|Organism:Severe      AGCAGGTGACTCAGGTTTTGCTGCATACAGTCGCTACAGGATTGGCAACTATAAATTAAA    27142
gb:MT019529|Organism:Severe      AGCAGGTGACTCAGGTTTTGCTGCATACAGTCGCTACAGGATTGGCAACTATAAATTAAA    27142
gb:MN996529|Organism:Severe      AGCAGGTGACTCAGGTTTTGCTGCATACAGTCGCTACAGGATTGGCAACTATAAATTAAA    27130
gb:MN996531|Organism:Severe      AGCAGGTGACTCAGGTTTTGCTGCATACAGTCGCTACAGGATTGGCAACTATAAATTAAA    27129
gb:MT066176|Organism:Severe      AGCAGGTGACTCAGGTTTTGCTGCATACAGTCGCTACAGGATTGGCAACTATAAATTAAA    27142
gb:MT027064|Organism:Severe      AGCAGGTGACTCAGGTTTTGCTGCATACAGTCGCTACAGGATTGGCAACTATAAATTAAA    27142
gb:MN994468|Organism:Severe      AGCAGGTGACTCAGGTTTTGCTGCATACAGTCGCTACAGGATTGGCAACTATAAATTAAA    27142
gb:MT072688|Organism:Severe      AGCAGGTGACTCAGGTTTTGCTGCATACAGTCGCTACAGGATTGGCAACTATAAATTAAA    27127
gb:MN996527|Organism:Severe      AGCAGGTGACTCAGGTTTTGCTGCATACAGTCGCTACAGGATTGGCAACTATAAATTAAA    27109
gb:MT093631|Organism:Severe      AGCAGGTGACTCAGGTTTTGCTGCATACAGTCGCTACAGGATTGGCAACTATAAATTAAA    27180
gb:MT106053|Organism:Severe      AGCAGGTGACTCAGGTTTTGCTGCATACAGTCGCTACAGGATTGGCAACTATAAATTAAA    27142
gb:MT019533|Organism:Severe      AGCAGGTGACTCAGGTTTTGCTGCATACAGTCGCTACAGGATTGGCAACTATAAATTAAA    27142
gb:MT019531|Organism:Severe      AGCAGGTGACTCAGGTTTTGCTGCATACAGTCGCTACAGGATTGGCAACTATAAATTAAA    27142
gb:MN996528|Organism:Severe      AGCAGGTGACTCAGGTTTTGCTGCATACAGTCGCTACAGGATTGGCAACTATAAATTAAA    27142
gb:MN996530|Organism:Severe      AGCAGGTGACTCAGGTTTTGCTGCATACAGTCGCTACAGGATTGGCAACTATAAATTAAA    27128
gb:MN908947|Organism:Severe      AGCAGGTGACTCAGGTTTTGCTGCATACAGTCGCTACAGGATTGGCAACTATAAATTAAA    27142
gb:MT019532|Organism:Severe      AGCAGGTGACTCAGGTTTTGCTGCATACAGTCGCTACAGGATTGGCAACTATAAATTAAA    27142
                                 ************************************************************

gb:MT020781|Organism:Severe      CACAGACCATTCCAGTAGCAGTGACAATATTGCTTTGCTTGTACAGTAAGTGACAACAGA    27190
gb:MT007544|Organism:Severe      CACAGACCATTCCAGTAGCAGTGACAATATTGCTTTGCTTGTACAGTAAGTGACAACAGA    27202
gb:MN994467|Organism:Severe      CACAGACCATTCCAGTAGCAGTGACAATATTGCTTTGCTTGTACAGTAAGTGACAACAGA    27202
gb:MT044257|Organism:Severe      CACAGACCATTCCAGTAGCAGTGACAATATTGCTTTGCTTGTACAGTAAGTGACAACAGA    27202
gb:MT106054|Organism:Severe      CACAGACCATTCCAGTAGCAGTGACAATATTGCTTTGCTTGTACAGTAAGTGACAACAGA    27202
gb:MT049951|Organism:Severe      CACAGACCATTCCAGTAGCAGTGACAATATTGCTTTGCTTGTACAGTAAGTGACAACAGA    27202
gb:MN975262|Organism:Severe      CACAGACCATTCCAGTAGCAGTGACAATATTGCTTTGCTTGTACAGTAAGTGACAACAGA    27202
gb:MT106052|Organism:Severe      CACAGACCATTCCAGTAGCAGTGACAATATTGCTTTGCTTGTACAGTAAGTGACAACAGA    27202
gb:LC522975|Organism:Severe      CACAGACCATTCCAGTAGCAGTGACAATATTGCTTTGCTTGTACAGTAAGTGACAACAGA    27199
gb:LC522973|Organism:Severe      CACAGACCATTCCAGTAGCAGTGACAATATTGCTTTGCTTGTACAGTAAGTGACAACAGA    27199
gb:LC522974|Organism:Severe      CACAGACCATTCCAGTAGCAGTGACAATATTGCTTTGCTTGTACAGTAAGTGACAACAGA    27199
gb:MN985325|Organism:Severe      CACAGACCATTCCAGTAGCAGTGACAATATTGCTTTGCTTGTACAGTAAGTGACAACAGA    27202
gb:MT020881|Organism:Severe      CACAGACCATTCCAGTAGCAGTGACAATATTGCTTTGCTTGTACAGTAAGTGACAACAGA    27202
gb:MT020880|Organism:Severe      CACAGACCATTCCAGTAGCAGTGACAATATTGCTTTGCTTGTACAGTAAGTGACAACAGA    27202
gb:MT066175|Organism:Severe      CACAGACCATTCCAGTAGCAGTGACAATATTGCTTTGCTTGTACAGTAAGTGACAACAGA    27202
gb:MN997409|Organism:Severe      CACAGACCATTCCAGTAGCAGTGACAATATTGCTTTGCTTGTACAGTAAGTGACAACAGA    27202
gb:MN938384|Organism:Severe      CACAGACCATTCCAGTAGCAGTGACAATATTGCTTTGCTTGTACAGTAAGTGACAACAGA    27170
gb:MT044258|Organism:Severe      CACAGACCATTCCAGTAGCAGTGACAATATTGCTTTGCTTGTACAGTAAGTGACAACAGA    27178
gb:MT039890|Organism:Severe      CACAGACCATTCCAGTAGCAGTGACAATATTGCTTTGCTTGTACAGTAAGTGACAACAGA    27202
gb:MN988713|Organism:Severe      CACAGACCATTCCAGTAGCAGTGACAATATTGCTTTGCTTGTACAGTAAGTGACAACAGA    27202
gb:LC521925|Organism:Severe      CACAGACCATTCCAGTAGCAGTGACAATATTGCTTTGCTTGTACAGTAAGTGACAACAGA    27175
gb:MT093571|Organism:Severe      CACAGACCATTCCAGTAGCAGTGACAATATTGCTTTGCTTGTACAGTAAGTGACAACAGA    27202
gb:MT039887|Organism:Severe      CACAGACCATTCCAGTAGCAGTGACAATATTGCTTTGCTTGTACAGTAAGTGACAACAGA    27199
```

```
gb:MT019530|Organism:Severe      CACAGACCATTCCAGTAGCAGTGACAATATTGCTTTGCTTGTACAGTAAGTGACAACAGA   27202
gb:MT039888|Organism:Severe      CACAGACCATTCCAGTAGCAGTGACAATATTGCTTTGCTTGTACAGTAAGTGACAACAGA   27202
gb:LC522972|Organism:Severe      CACAGACCATTCCAGTAGCAGTGACAATATTGCTTTGCTTGTACAGTAAGTGACAACAGA   27199
gb:MT027063|Organism:Severe      CACAGACCATTCCAGTAGCAGTGACAATATTGCTTTGCTTGTACAGTAAGTGACAACAGA   27202
gb:MT027062|Organism:Severe      CACAGACCATTCCAGTAGCAGTGACAATATTGCTTTGCTTGTACAGTAAGTGACAACAGA   27202
gb:MT019529|Organism:Severe      CACAGACCATTCCAGTAGCAGTGACAATATTGCTTTGCTTGTACAGTAAGTGACAACAGA   27202
gb:MN996529|Organism:Severe      CACAGACCATTCCAGTAGCAGTGACAATATTGCTTTGCTTGTACAGTAAGTGACAACAGA   27190
gb:MN996531|Organism:Severe      CACAGACCATTCCAGTAGCAGTGACAATATTGCTTTGCTTGTACAGTAAGTGACAACAGA   27189
gb:MT066176|Organism:Severe      CACAGACCATTCCAGTAGCAGTGACAATATTGCTTTGCTTGTACAGTAAGTGACAACAGA   27202
gb:MT027064|Organism:Severe      CACAGACCATTCCAGTAGCAGTGACAATATTGCTTTGCTTGTACAGTAAGTGACAACAGA   27202
gb:MN994468|Organism:Severe      CACAGACCATTCCAGTAGCAGTGACAATATTGCTTTGCTTGTACAGTAAGTGACAACAGA   27202
gb:MT072688|Organism:Severe      CACAGACCATTCCAGTAGCAGTGACAATATTGCTTTGCTTGTACAGTAAGTGACAACAGA   27187
gb:MN996527|Organism:Severe      CACAGACCATTCCAGTAGCAGTGACAATATTGCTTTGCTTGTACAGTAAGTGACAACAGA   27169
gb:MT093631|Organism:Severe      CACAGACCATTCCAGTAGCAGTGACAATATTGCTTTGCTTGTACAGTAAGTGACAACAGA   27240
gb:MT106053|Organism:Severe      CACAGACCATTCCAGTAGCAGTGACAATATTGCTTTGCTTGTACAGTAAGTGACAACAGA   27202
gb:MT019533|Organism:Severe      CACAGACCATTCCAGTAGCAGTGACAATATTGCTTTGCTTGTACAGTAAGTGACAACAGA   27202
gb:MT019531|Organism:Severe      CACAGACCATTCCAGTAGCAGTGACAATATTGCTTTGCTTGTACAGTAAGTGACAACAGA   27202
gb:MN996528|Organism:Severe      CACAGACCATTCCAGTAGCAGTGACAATATTGCTTTGCTTGTACAGTAAGTGACAACAGA   27202
gb:MN996530|Organism:Severe      CACAGACCATTCCAGTAGCAGTGACAATATTGCTTTGCTTGTACAGTAAGTGACAACAGA   27188
gb:MN908947|Organism:Severe      CACAGACCATTCCAGTAGCAGTGACAATATTGCTTTGCTTGTACAGTAAGTGACAACAGA   27202
gb:MT019532|Organism:Severe      CACAGACCATTCCAGTAGCAGTGACAATATTGCTTTGCTTGTACAGTAAGTGACAACAGA   27202
                                 ************************************************************

gb:MT020781|Organism:Severe      TGTTTCATCTCGTTGACTTTCAGGTTACTATAGCAGAGATATTACTAATTATTATGAGGA   27250
gb:MT007544|Organism:Severe      TGTTTCATCTCGTTGACTTTCAGGTTACTATAGCAGAGATATTACTAATTATTATGAGGA   27262
gb:MN994467|Organism:Severe      TGTTTCATCTCGTTGACTTTCAGGTTACTATAGCAGAGATATTACTAATTATTATGAGGA   27262
gb:MT044257|Organism:Severe      TGTTTCATCTCGTTGACTTTCAGGTTACTATAGCAGAGATATTACTAATTATTATGAGGA   27262
gb:MT106054|Organism:Severe      TGTTTCATCTCGTTGACTTTCAGGTTACTATAGCAGAGATATTACTAATTATTATGAGGA   27262
gb:MT049951|Organism:Severe      TGTTTCATCTCGTTGACTTTCAGGTTACTATAGCAGAGATATTACTAATTATTATGAGGA   27262
gb:MN975262|Organism:Severe      TGTTTCATCTCGTTGACTTTCAGGTTACTATAGCAGAGATATTACTAATTATTATGAGGA   27262
gb:MT106052|Organism:Severe      TGTTTCATCTCGTTGACTTTCAGGTTACTATAGCAGAGATATTACTAATTATTATGAGGA   27262
gb:LC522975|Organism:Severe      TGTTTCATCTCGTTGACTTTCAGGTTACTATAGCAGAGATATTACTAATTATTATGAGGA   27259
gb:LC522973|Organism:Severe      TGTTTCATCTCGTTGACTTTCAGGTTACTATAGCAGAGATATTACTAATTATTATGAGGA   27259
gb:LC522974|Organism:Severe      TGTTTCATCTCGTTGACTTTCAGGTTACTATAGCAGAGATATTACTAATTATTATGAGGA   27259
gb:MN985325|Organism:Severe      TGTTTCATCTCGTTGACTTTCAGGTTACTATAGCAGAGATATTACTAATTATTATGAGGA   27262
gb:MT020881|Organism:Severe      TGTTTCATCTCGTTGACTTTCAGGTTACTATAGCAGAGATATTACTAATTATTATGAGGA   27262
gb:MT020880|Organism:Severe      TGTTTCATCTCGTTGACTTTCAGGTTACTATAGCAGAGATATTACTAATTATTATGAGGA   27262
gb:MT066175|Organism:Severe      TGTTTCATCTCGTTGACTTTCAGGTTACTATAGCAGAGATATTACTAATTATTATGAGGA   27262
gb:MN997409|Organism:Severe      TGTTTCATCTCGTTGACTTTCAGGTTACTATAGCAGAGATATTACTAATTATTATGAGGA   27262
gb:MN938384|Organism:Severe      TGTTTCATCTCGTTGACTTTCAGGTTACTATAGCAGAGATATTACTAATTATTATGAGGA   27230
gb:MT044258|Organism:Severe      TGTTTCATCTCGTTGACTTTCAGGTTACTATAGCAGAGATATTACTAATTATTATGAGGA   27238
gb:MT039890|Organism:Severe      TGTTTCATCTCGTTGACTTTCAGGTTACTATAGCAGAGATATTACTAATTATTATGAGGA   27262
gb:MN988713|Organism:Severe      TGTTTCATCTCGTTGACTTTCAGGTTACTATAGCAGAGATATTACTAATTATTATGAGGA   27262
gb:LC521925|Organism:Severe      TGTTTCATCTCGTTGACTTTCAGGTTACTATAGCAGAGATATTACTAATTATTATGAGGA   27235
gb:MT093571|Organism:Severe      TGTTTCATCTCGTTGACTTTCAGGTTACTATAGCAGAGATATTACTAATTATTATGAGGA   27262
gb:MT039887|Organism:Severe      TGTTTCATCTCGTTGACTTTCAGGTTACTATAGCAGAGATATTACTAATTATTATGAGGA   27259
gb:MT019530|Organism:Severe      TGTTTCATCTCGTTGACTTTCAGGTTACTATAGCAGAGATATTACTAATTATTATGAGGA   27262
gb:MT039888|Organism:Severe      TGTTTCATCTCGTTGACTTTCAGGTTACTATAGCAGAGATATTACTAATTATTATGAGGA   27262
gb:LC522972|Organism:Severe      TGTTTCATCTCGTTGACTTTCAGGTTACTATAGCAGAGATATTACTAATTATTATGAGGA   27259
gb:MT027063|Organism:Severe      TGTTTCATCTCGTTGACTTTCAGGTTACTATAGCAGAGATATTACTAATTATTATGAGGA   27262
gb:MT027062|Organism:Severe      TGTTTCATCTCGTTGACTTTCAGGTTACTATAGCAGAGATATTACTAATTATTATGAGGA   27262
gb:MT019529|Organism:Severe      TGTTTCATCTCGTTGACTTTCAGGTTACTATAGCAGAGATATTACTAATTATTATGAGGA   27262
gb:MN996529|Organism:Severe      TGTTTCATCTCGTTGACTTTCAGGTTACTATAGCAGAGATATTACTAATTATTATGAGGA   27250
gb:MN996531|Organism:Severe      TGTTTCATCTCGTTGACTTTCAGGTTACTATAGCAGAGATATTACTAATTATTATGAGGA   27249
gb:MT066176|Organism:Severe      TGTTTCATCTCGTTGACTTTCAGGTTACTATAGCAGAGATATTACTAATTATTATGAGGA   27262
gb:MT027064|Organism:Severe      TGTTTCATCTCGTTGACTTTCAGGTTACTATAGCAGAGATATTACTAATTATTATGAGGA   27262
gb:MN994468|Organism:Severe      TGTTTCATCTCGTTGACTTTCAGGTTACTATAGCAGAGATATTACTAATTATTATGAGGA   27262
gb:MT072688|Organism:Severe      TGTTTCATCTCGTTGACTTTCAGGTTACTATAGCAGAGATATTACTAATTATTATGAGGA   27247
gb:MN996527|Organism:Severe      TGTTTCATCTCGTTGACTTTCAGGTTACTATAGCAGAGATATTACTAATTATTATGAGGA   27229
gb:MT093631|Organism:Severe      TGTTTCATCTCGTTGACTTTCAGGTTACTATAGCAGAGATATTACTAATTATTATGAGGA   27300
gb:MT106053|Organism:Severe      TGTTTCATCTCGTTGACTTTCAGGTTACTATAGCAGAGATATTACTAATTATTATGAGGA   27262
gb:MT019533|Organism:Severe      TGTTTCATCTCGTTGACTTTCAGGTTACTATAGCAGAGATATTACTAATTATTATGAGGA   27262
gb:MT019531|Organism:Severe      TGTTTCATCTCGTTGACTTTCAGGTTACTATAGCAGAGATATTACTAATTATTATGAGGA   27262
gb:MN996528|Organism:Severe      TGTTTCATCTCGTTGACTTTCAGGTTACTATAGCAGAGATATTACTAATTATTATGAGGA   27262
gb:MN996530|Organism:Severe      TGTTTCATCTCGTTGACTTTCAGGTTACTATAGCAGAGATATTACTAATTATTATGAGGA   27248
```

```
gb:MN908947|Organism:Severe      TGTTTCATCTCGTTGACTTTCAGGTTACTATAGCAGAGATATTACTAATTATTATGAGGA   27262
gb:MT019532|Organism:Severe      TGTTTCATCTCGTTGACTTTCAGGTTACTATAGCAGAGATATTACTAATTATTATGAGGA   27262
                                 ************************************************************

gb:MT020781|Organism:Severe      CTTTTAAAGTTTCCATTTGGAATCTTGATTACATCATAAACCTCATAATTAAAAATTTAT   27310
gb:MT007544|Organism:Severe      CTTTTAAAGTTTCCATTTGGAATCTTGATTACATCATAAACCTCATAATTAAAAATTTAT   27322
gb:MN994467|Organism:Severe      CTTTTAAAGTTTCCATTTGGAATCTTGATTACATCATAAACCTCATAATTAAAAATTTAT   27322
gb:MT044257|Organism:Severe      CTTTTAAAGTTTCCATTTGGAATCTTGATTACATCATAAACCTCATAATTAAAAATTTAT   27322
gb:MT106054|Organism:Severe      CTTTTAAAGTTTCCATTTGGAATCTTGATTACATCATAAACCTCATAATTAAAAATTTAT   27322
gb:MT049951|Organism:Severe      CTTTTAAAGTTTCCATTTGGAATCTTGATTACATCATAAACCTCATAATTAAAAATTTAT   27322
gb:MN975262|Organism:Severe      CTTTTAAAGTTTCCATTTGGAATCTTGATTACATCATAAACCTCATAATTAAAAATTTAT   27322
gb:MT106052|Organism:Severe      CTTTTAAAGTTTCCATTTGGAATCTTGATTACATCATAAACCTCATAATTAAAAATTTAT   27322
gb:LC522975|Organism:Severe      CTTTTAAAGTTTCCATTTGGAATCTTGATTACATCATAAACCTCATAATTAAAAATTTAT   27319
gb:LC522973|Organism:Severe      CTTTTAAAGTTTCCATTTGGAATCTTGATTACATCATAAACCTCATAATTAAAAATTTAT   27319
gb:LC522974|Organism:Severe      CTTTTAAAGTTTCCATTTGGAATCTTGATTACATCATAAACCTCATAATTAAAAATTTAT   27319
gb:MN985325|Organism:Severe      CTTTTAAAGTTTCCATTTGGAATCTTGATTACATCATAAACCTCATAATTAAAAATTTAT   27322
gb:MT020881|Organism:Severe      CTTTTAAAGTTTCCATTTGGAATCTTGATTACATCATAAACCTCATAATTAAAAATTTAT   27322
gb:MT020880|Organism:Severe      CTTTTAAAGTTTCCATTTGGAATCTTGATTACATCATAAACCTCATAATTAAAAATTTAT   27322
gb:MT066175|Organism:Severe      CTTTTAAAGTTTCCATTTGGAATCTTGATTACATCATAAACCTCATAATTAAAAATTTAT   27322
gb:MN997409|Organism:Severe      CTTTTAAAGTTTCCATTTGGAATCTTGATTACATCATAAACCTCATAATTAAAAATTTAT   27322
gb:MN938384|Organism:Severe      CTTTTAAAGTTTCCATTTGGAATCTTGATTACATCATAAACCTCATAATTAAAAATTTAT   27290
gb:MT044258|Organism:Severe      CTTTTAAAGTTTCCATTTGGAATCTTGATTACATCATAAACCTCATAATTAAAAATTTAT   27298
gb:MT039890|Organism:Severe      CTTTTAAAGTTTCCATTTGGAATCTTGATTACATCATAAACCTCATAATTAAAAATTTAT   27322
gb:MN988713|Organism:Severe      CTTTTAAAGTTTCCATTTGGAATCTTGATTACATCATAAACCTCATAATTAAAAATTTAT   27322
gb:LC521925|Organism:Severe      CTTTTAAAGTTTCCATTTGGAATCTTGATTACATCATAAACCTCATAATTAAAAATTTAT   27295
gb:MT093571|Organism:Severe      CTTTTAAAGTTTCCATTTGGAATCTTGATTACATCATAAACCTCATAATTAAAAATTTAT   27322
gb:MT039887|Organism:Severe      CTTTTAAAGTTTCCATTTGGAATCTTGATTACATCATAAACCTCATAATTAAAAATTTAT   27319
gb:MT019530|Organism:Severe      CTTTTAAAGTTTCCATTTGGAATCTTGATTACATCATAAACCTCATAATTAAAAATTTAT   27322
gb:MT039888|Organism:Severe      CTTTTAAAGTTTCCATTTGGAATCTTGATTACATCATAAACCTCATAATTAAAAATTTAT   27322
gb:LC522972|Organism:Severe      CTTTTAAAGTTTCCATTTGGAATCTTGATTACATCATAAACCTCATAATTAAAAATTTAT   27319
gb:MT027063|Organism:Severe      CTTTTAAAGTTTCCATTTGGAATCTTGATTACATCATAAACCTCATAATTAAAAATTTAT   27322
gb:MT027062|Organism:Severe      CTTTTAAAGTTTCCATTTGGAATCTTGATTACATCATAAACCTCATAATTAAAAATTTAT   27322
gb:MT019529|Organism:Severe      CTTTTAAAGTTTCCATTTGGAATCTTGATTACATCATAAACCTCATAATTAAAAATTTAT   27322
gb:MN996529|Organism:Severe      CTTTTAAAGTTTCCATTTGGAATCTTGATTACATCATAAACCTCATAATTAAAAATTTAT   27310
gb:MN996531|Organism:Severe      CTTTTAAAGTTTCCATTTGGAATCTTGATTACATCATAAACCTCATAATTAAAAATTTAT   27309
gb:MT066176|Organism:Severe      CTTTTAAAGTTTCCATTTGGAATCTTGATTACATCATAAACCTCATAATTAAAAATTTAT   27322
gb:MT027064|Organism:Severe      CTTTTAAAGTTTCCATTTGGAATCTTGATTACATCATAAACCTCATAATTAAAAATTTAT   27322
gb:MN994468|Organism:Severe      CTTTTAAAGTTTCCATTTGGAATCTTGATTACATCATAAACCTCATAATTAAAAATTTAT   27322
gb:MT072688|Organism:Severe      CTTTTAAAGTTTCCATTTGGAATCTTGATTACATCATAAACCTCATAATTAAAAATTTAT   27307
gb:MN996527|Organism:Severe      CTTTTAAAGTTTCCATTTGGAATCTTGATTACATCATAAACCTCATAATTAAAAATTTAT   27289
gb:MT093631|Organism:Severe      CTTTTAAAGTTTCCATTTGGAATCTTGATTACATCATAAACCTCATAATTAAAAATTTAT   27360
gb:MT106053|Organism:Severe      CTTTTAAAGTTTCCATTTGGAATCTTGATTACATCATAAACCTCATAATTAAAAATTTAT   27322
gb:MT019533|Organism:Severe      CTTTTAAAGTTTCCATTTGGAATCTTGATTACATCATAAACCTCATAATTAAAAATTTAT   27322
gb:MT019531|Organism:Severe      CTTTTAAAGTTTCCATTTGGAATCTTGATTACATCATAAACCTCATAATTAAAAATTTAT   27322
gb:MN996528|Organism:Severe      CTTTTAAAGTTTCCATTTGGAATCTTGATTACATCATAAACCTCATAATTAAAAATTTAT   27322
gb:MN996530|Organism:Severe      CTTTTAAAGTTTCCATTTGGAATCTTGATTACATCATAAACCTCATAATTAAAAATTTAT   27308
gb:MN908947|Organism:Severe      CTTTTAAAGTTTCCATTTGGAATCTTGATTACATCATAAACCTCATAATTAAAAATTTAT   27322
gb:MT019532|Organism:Severe      CTTTTAAAGTTTCCATTTGGAATCTTGATTACATCATAAACCTCATAATTAAAAATTTAT   27322
                                 ************************************************************

gb:MT020781|Organism:Severe      CTAAGTCACTAACTGAGAATAAATATTCTCAATTAGATGAAGAGCAACCAATGGAGATTG   27370
gb:MT007544|Organism:Severe      CTAAGTCACTAACTGAGAATAAATATTCTCAATTAGATGAAGAGCAACCAATGGAGATTG   27382
gb:MN994467|Organism:Severe      CTAAGTCACTAACTGAGAATAAATATTCTCAATTAGATGAAGAGCAACCAATGGAGATTG   27382
gb:MT044257|Organism:Severe      CTAAGTCACTAACTGAGAATAAATATTCTCAATTAGATGAAGAGCAACCAATGGAGATTG   27382
gb:MT106054|Organism:Severe      CTAAGTCACTAACTGAGAATAAATATTCTCAATTAGATGAAGAGCAACCAATGGAGATTG   27382
gb:MT049951|Organism:Severe      CTAAGTCACTAACTGAGAATAAATATTCTCAATTAGATGAAGAGCAACCAATGGAGATTG   27382
gb:MN975262|Organism:Severe      CTAAGTCACTAACTGAGAATAAATATTCTCAATTAGATGAAGAGCAACCAATGGAGATTG   27382
gb:MT106052|Organism:Severe      CTAAGTCACTAACTGAGAATAAATATTCTCAATTAGATGAAGAGCAACCAATGGAGATTG   27382
gb:LC522975|Organism:Severe      CTAAGTCACTAACTGAGAATAAATATTCTCAATTAGATGAAGAGCAACCAATGGAGATTG   27379
gb:LC522973|Organism:Severe      CTAAGTCACTAACTGAGAATAAATATTCTCAATTAGATGAAGAGCAACCAATGGAGATTG   27379
gb:LC522974|Organism:Severe      CTAAGTCACTAACTGAGAATAAATATTCTCAATTAGATGAAGAGCAACCAATGGAGATTG   27379
gb:MN985325|Organism:Severe      CTAAGTCACTAACTGAGAATAAATATTCTCAATTAGATGAAGAGCAACCAATGGAGATTG   27382
gb:MT020881|Organism:Severe      CTAAGTCACTAACTGAGAATAAATATTCTCAATTAGATGAAGAGCAACCAATGGAGATTG   27382
gb:MT020880|Organism:Severe      CTAAGTCACTAACTGAGAATAAATATTCTCAATTAGATGAAGAGCAACCAATGGAGATTG   27382
gb:MT066175|Organism:Severe      CTAAGTCACTAACTGAGAATAAATATTCTCAATTAGATGAAGAGCAACCAATGGAGATTG   27382
```

```
gb:MN997409|Organism:Severe      CTAAGTCACTAACTGAGAATAAATATTCTCAATTAGATGAAGAGCAACCAATGGAGATTG   27382
gb:MN938384|Organism:Severe      CTAAGTCACTAACTGAGAATAAATATTCTCAATTAGATGAAGAGCAACCAATGGAGATTG   27350
gb:MT044258|Organism:Severe      CTAAGTCACTAACTGAGAATAAATATTCTCAATTAGATGAAGAGCAACCAATGGAGATTG   27358
gb:MT039890|Organism:Severe      CTAAGTCACTAACTGAGAATAAATATTCTCAATTAGATGAAGAGCAACCAATGGAGATTG   27382
gb:MN988713|Organism:Severe      CTAAGTCACTAACTGAGAATAAATATTCTCAATTAGATGAAGAGCAACCAATGGAGATTG   27382
gb:LC521925|Organism:Severe      CTAAGTCACTAACTGAGAATAAATATTCTCAATTAGATGAAGAGCAACCAATGGAGATTG   27355
gb:MT093571|Organism:Severe      CTAAGTCACTAACTGAGAATAAATATTCTCAATTAGATGAAGAGCAACCAATGGAGATTG   27382
gb:MT039887|Organism:Severe      CTAAGTCACTAACTGAGAATAAATATTCTCAATTAGATGAAGAGCAACCAATGGAGATTG   27379
gb:MT019530|Organism:Severe      CTAAGTCACTAACTGAGAATAAATATTCTCAATTAGATGAAGAGCAACCAATGGAGATTG   27382
gb:MT039888|Organism:Severe      CTAAGTCACTAACTGAGAATAAATATTCTCAATTAGATGAAGAGCAACCAATGGAGATTG   27382
gb:LC522972|Organism:Severe      CTAAGTCACTAACTGAGAATAAATATTCTCAATTAGATGAAGAGCAACCAATGGAGATTG   27379
gb:MT027063|Organism:Severe      CTAAGTCACTAACTGAGAATAAATATTCTCAATTAGATGAAGAGCAACCAATGGAGATTG   27382
gb:MT027062|Organism:Severe      CTAAGTCACTAACTGAGAATAAATATTCTCAATTAGATGAAGAGCAACCAATGGAGATTG   27382
gb:MT019529|Organism:Severe      CTAAGTCACTAACTGAGAATAAATATTCTCAATTAGATGAAGAGCAACCAATGGAGATTG   27382
gb:MN996529|Organism:Severe      CTAAGTCACTAACTGAGAATAAATATTCTCAATTAGATGAAGAGCAACCAATGGAGATTG   27370
gb:MN996531|Organism:Severe      CTAAGTCACTAACTGAGAATAAATATTCTCAATTAGATGAAGAGCAACCAATGGAGATTG   27369
gb:MT066176|Organism:Severe      CTAAGTCACTAACTGAGAATAAATATTCTCAATTAGATGAAGAGCAACCAATGGAGATTG   27382
gb:MT027064|Organism:Severe      CTAAGTCACTAACTGAGAATAAATATTCTCAATTAGATGAAGAGCAACCAATGGAGATTG   27382
gb:MN994468|Organism:Severe      CTAAGTCACTAACTGAGAATAAATATTCTCAATTAGATGAAGAGCAACCAATGGAGATTG   27382
gb:MT072688|Organism:Severe      CTAAGTCACTAACTGAGAATAAATATTCTCAATTAGATGAAGAGCAACCAATGGAGATTG   27367
gb:MN996527|Organism:Severe      CTAAGTCACTAACTGAGAATAAATATTCTCAATTAGATGAAGAGCAACCAATGGAGATTG   27349
gb:MT093631|Organism:Severe      CTAAGTCACTAACTGAGAATAAATATTCTCAATTAGATGAAGAGCAACCAATGGAGATTG   27420
gb:MT106053|Organism:Severe      CTAAGTCACTAACTGAGAATAAATATTCTCAATTAGATGAAGAGCAACCAATGGAGATTG   27382
gb:MT019533|Organism:Severe      CTAAGTCACTAACTGAGAATAAATATTCTCAATTAGATGAAGAGCAACCAATGGAGATTG   27382
gb:MT019531|Organism:Severe      CTAAGTCACTAACTGAGAATAAATATTCTCAATTAGATGAAGAGCAACCAATGGAGATTG   27382
gb:MN996528|Organism:Severe      CTAAGTCACTAACTGAGAATAAATATTCTCAATTAGATGAAGAGCAACCAATGGAGATTG   27382
gb:MN996530|Organism:Severe      CTAAGTCACTAACTGAGAATAAATATTCTCAATTAGATGAAGAGCAACCAATGGAGATTG   27368
gb:MN908947|Organism:Severe      CTAAGTCACTAACTGAGAATAAATATTCTCAATTAGATGAAGAGCAACCAATGGAGATTG   27382
gb:MT019532|Organism:Severe      CTAAGTCACTAACTGAGAATAAATATTCTCAATTAGATGAAGAGCAACCAATGGAGATTG   27382
                                 ************************************************************

gb:MT020781|Organism:Severe      ATTAAACGAACATGAAAATTATTCTTTTCTTGGCACTGATAACACTCGCTACTTGTGAGC   27430
gb:MT007544|Organism:Severe      ATTAAACGAACATGAAAATTATTCTTTTCTTGGCACTGATAACACTCGCTACTTGTGAGC   27442
gb:MN994467|Organism:Severe      ATTAAACGAACATGAAAATTATTCTTTTCTTGGCACTGATAACACTCGCTACTTGTGAGC   27442
gb:MT044257|Organism:Severe      ATTAAACGAACATGAAAATTATTCTTTTCTTGGCACTGATAACACTCGCTACTTGTGAGC   27442
gb:MT106054|Organism:Severe      ATTAAACGAACATGAAAATTATTCTTTTCTTGGCACTGATAACACTCGCTACTTGTGAGC   27442
gb:MT049951|Organism:Severe      ATTAAACGAACATGAAAATTATTCTTTTCTTGGCACTGATAACACTCGCTACTTGTGAGC   27442
gb:MN975262|Organism:Severe      ATTAAACGAACATGAAAATTATTCTTTTCTTGGCACTGATAACACTCGCTACTTGTGAGC   27442
gb:MT106052|Organism:Severe      ATTAAACGAACATGAAAATTATTCTTTTCTTGGCACTGATAACACTCGCTACTTGTGAGC   27442
gb:LC522975|Organism:Severe      ATTAAACGAACATGAAAATTATTCTTTTCTTGGCACTGATAACACTCGCTACTTGTGAGC   27439
gb:LC522973|Organism:Severe      ATTAAACGAACATGAAAATTATTCTTTTCTTGGCACTGATAACACTCGCTACTTGTGAGC   27439
gb:LC522974|Organism:Severe      ATTAAACGAACATGAAAATTATTCTTTTCTTGGCACTGATAACACTCGCTACTTGTGAGC   27439
gb:MN985325|Organism:Severe      ATTAAACGAACATGAAAATTATTCTTTTCTTGGCACTGATAACACTCGCTACTTGTGAGC   27442
gb:MT020881|Organism:Severe      ATTAAACGAACATGAAAATTATTCTTTTCTTGGCACTGATAACACTCGCTACTTGTGAGC   27442
gb:MT020880|Organism:Severe      ATTAAACGAACATGAAAATTATTCTTTTCTTGGCACTGATAACACTCGCTACTTGTGAGC   27442
gb:MT066175|Organism:Severe      ATTAAACGAACATGAAAATTATTCTTTTCTTGGCACTGATAACACTCGCTACTTGTGAGC   27442
gb:MN997409|Organism:Severe      ATTAAACGAACATGAAAATTATTCTTTTCTTGGCACTGATAACACTCGCTACTTGTGAGC   27442
gb:MN938384|Organism:Severe      ATTAAACGAACATGAAAATTATTCTTTTCTTGGCACTGATAACACTCGCTACTTGTGAGC   27410
gb:MT044258|Organism:Severe      ATTAAACGAACATGAAAATTATTCTTTTCTTGGCACTGATAACACTCGCTACTTGTGAGC   27418
gb:MT039890|Organism:Severe      ATTAAACGAACATGAAAATTATTCTTTTCTTGGCACTGATAACACTCGCTACTTGTGAGC   27442
gb:MN988713|Organism:Severe      ATTAAACGAACATGAAAATTATTCTTTTCTTGGCACTGATAACACTCGCTACTTGTGAGC   27442
gb:LC521925|Organism:Severe      ATTAAACGAACATGAAAATTATTCTTTTCTTGGCACTGATAACACTCGCTACTTGTGAGC   27415
gb:MT093571|Organism:Severe      ATTAAACGAACATGAAAATTATTCTTTTCTTGGCACTGATAACACTCGCTACTTGTGAGC   27442
gb:MT039887|Organism:Severe      ATTAAACGAACATGAAAATTATTCTTTTCTTGGCACTGATAACACTCGCTACTTGTGAGC   27439
gb:MT019530|Organism:Severe      ATTAAACGAACATGAAAATTATTCTTTTCTTGGCACTGATAACACTCGCTACTTGTGAGC   27442
gb:MT039888|Organism:Severe      ATTAAACGAACATGAAAATTATTCTTTTCTTGGCACTGATAACACTCGCTACTTGTGAGC   27442
gb:LC522972|Organism:Severe      ATTAAACGAACATGAAAATTATTCTTTTCTTGGCACTGATAACACTCGCTACTTGTGAGC   27439
gb:MT027063|Organism:Severe      ATTAAACGAACATGAAAATTATTCTTTTCTTGGCACTGATAACACTCGCTACTTGTGAGC   27442
gb:MT027062|Organism:Severe      ATTAAACGAACATGAAAATTATTCTTTTCTTGGCACTGATAACACTCGCTACTTGTGAGC   27442
gb:MT019529|Organism:Severe      ATTAAACGAACATGAAAATTATTCTTTTCTTGGCACTGATAACACTCGCTACTTGTGAGC   27442
gb:MN996529|Organism:Severe      ATTAAACGAACATGAAAATTATTCTTTTCTTGGCACTGATAACACTCGCTACTTGTGAGC   27430
gb:MN996531|Organism:Severe      ATTAAACGAACATGAAAATTATTCTTTTCTTGGCACTGATAACACTCGCTACTTGTGAGC   27429
gb:MT066176|Organism:Severe      ATTAAACGAACATGAAAATTATTCTTTTCTTGGCACTGATAACACTCGCTACTTGTGAGC   27442
gb:MT027064|Organism:Severe      ATTAAACGAACATGAAAATTATTCTTTTCTTGGCACTGATAACACTCGCTACTTGTGAGC   27442
gb:MN994468|Organism:Severe      ATTAAACGAACATGAAAATTATTCTTTTCTTGGCACTGATAACACTCGCTACTTGTGAGC   27442
```

```
gb:MT072688|Organism:Severe      ATTAAACGAACATGAAAATTATTCTTTTCTTGGCACTGATAACACTCGCTACTTGTGAGC	27427
gb:MN996527|Organism:Severe      ATTAAACGAACATGAAAATTATTCTTTTCTTGGCACTGATAACACTCGCTACTTGTGAGC	27409
gb:MT093631|Organism:Severe      ATTAAACGAACATGAAAATTATTCTTTTCTTGGCACTGATAACACTCGCTACTTGTGAGC	27480
gb:MT106053|Organism:Severe      ATTAAACGAACATGAAAATTATTCTTTTCTTGGCACTGATAACACTCGCTACTTGTGAGC	27442
gb:MT019533|Organism:Severe      ATTAAACGAACATGAAAATTATTCTTTTCTTGGCACTGATAACACTCGCTACTTGTGAGC	27442
gb:MT019531|Organism:Severe      ATTAAACGAACATGAAAATTATTCTTTTCTTGGCACTGATAACACTCGCTACTTGTGAGC	27442
gb:MN996528|Organism:Severe      ATTAAACGAACATGAAAATTATTCTTTTCTTGGCACTGATAACACTCGCTACTTGTGAGC	27442
gb:MN996530|Organism:Severe      ATTAAACGAACATGAAAATTATTCTTTTCTTGGCACTGATAACACTCGCTACTTGTGAGC	27428
gb:MN908947|Organism:Severe      ATTAAACGAACATGAAAATTATTCTTTTCTTGGCACTGATAACACTCGCTACTTGTGAGC	27442
gb:MT019532|Organism:Severe      ATTAAACGAACATGAAAATTATTCTTTTCTTGGCACTGATAACACTCGCTACTTGTGAGC	27442
                                 ************************************************************

gb:MT020781|Organism:Severe      TTTATCACTACCAAGAGTGTGTTAGAGGTACAACAGTACTTTTAAAAGAACCTTGCTCTT	27490
gb:MT007544|Organism:Severe      TTTATCACTACCAAGAGTGTGTTAGAGGTACAACAGTACTTTTAAAAGAACCTTGCTCTT	27502
gb:MN994467|Organism:Severe      TTTATCACTACCAAGAGTGTGTTAGAGGTACAACAGTACTTTTAAAAGAACCTTGCTCTT	27502
gb:MT044257|Organism:Severe      TTTATCACTACCAAGAGTGTGTTAGAGGTACAACAGTACTTTTAAAAGAACCTTGCTCTT	27502
gb:MT106054|Organism:Severe      TTTATCACTACCAAGAGTGTGTTAGAGGTACAACAGTACTTTTAAAAGAACCTTGCTCTT	27502
gb:MT049951|Organism:Severe      TTTATCACTACCAAGAGTGTGTTAGAGGTACAACAGTACTTTTAAAAGAACCTTGCTCTT	27502
gb:MN975262|Organism:Severe      TTTATCACTACCAAGAGTGTGTTAGAGGTACAACAGTACTTTTAAAAGAACCTTGCTCTT	27502
gb:MT106052|Organism:Severe      TTTATCACTACCAAGAGTGTGTTAGAGGTACAACAGTACTTTTAAAAGAACCTTGCTCTT	27502
gb:LC522975|Organism:Severe      TTTATCACTACCAAGAGTGTGTTAGAGGTACAACAGTACTTTTAAAAGAACCTTGCTCTT	27499
gb:LC522973|Organism:Severe      TTTATCACTACCAAGAGTGTGTTAGAGGTACAACAGTACTTTTAAAAGAACCTTGCTCTT	27499
gb:LC522974|Organism:Severe      TTTATCACTACCAAGAGTGTGTTAGAGGTACAACAGTACTTTTAAAAGAACCTTGCTCTT	27499
gb:MN985325|Organism:Severe      TTTATCACTACCAAGAGTGTGTTAGAGGTACAACAGTACTTTTAAAAGAACCTTGCTCTT	27502
gb:MT020881|Organism:Severe      TTTATCACTACCAAGAGTGTGTTAGAGGTACAACAGTACTTTTAAAAGAACCTTGCTCTT	27502
gb:MT020880|Organism:Severe      TTTATCACTACCAAGAGTGTGTTAGAGGTACAACAGTACTTTTAAAAGAACCTTGCTCTT	27502
gb:MT066175|Organism:Severe      TTTATCACTACCAAGAGTGTGTTAGAGGTACAACAGTACTTTTAAAAGAACCTTGCTCTT	27502
gb:MN997409|Organism:Severe      TTTATCACTACCAAGAGTGTGTTAGAGGTACAACAGTACTTTTAAAAGAACCTTGCTCTT	27502
gb:MN938384|Organism:Severe      TTTATCACTACCAAGAGTGTGTTAGAGGTACAACAGTACTTTTAAAAGAACCTTGCTCTT	27470
gb:MT044258|Organism:Severe      TTTATCACTACCAAGAGTGTGTTAGAGGTACAACAGTACTTTTAAAAGAACCTTGCTCTT	27478
gb:MT039890|Organism:Severe      TTTATCACTACCAAGAGTGTGTTAGAGGTACAACAGTACTTTTAAAAGAACCTTGCTCTT	27502
gb:MN988713|Organism:Severe      TTTATCACTACCAAGAGTGTGTTAGAGGTACAACAGTACTTTTAAAAGAACCTTGCTCTT	27502
gb:LC521925|Organism:Severe      TTTATCACTACCAAGAGTGTGTTAGAGGTACAACAGTACTTTTAAAAGAACCTTGCTCTT	27475
gb:MT093571|Organism:Severe      TTTATCACTACCAAGAGTGTGTTAGAGGTACAACAGTACTTTTAAAAGAACCTTGCTCTT	27502
gb:MT039887|Organism:Severe      TTTATCACTACCAAGAGTGTGTTAGAGGTACAACAGTACTTTTAAAAGAACCTTGCTCTT	27499
gb:MT019530|Organism:Severe      TTTATCACTACCAAGAGTGTGTTAGAGGTACAACAGTACTTTTAAAAGAACCTTGCTCTT	27502
gb:MT039888|Organism:Severe      TTTATCACTACCAAGAGTGTGTTAGAGGTACAACAGTACTTTTAAAAGAACCTTGCTCTT	27502
gb:LC522972|Organism:Severe      TTTATCACTACCAAGAGTGTGTTAGAGGTACAACAGTACTTTTAAAAGAACCTTGCTCTT	27499
gb:MT027063|Organism:Severe      TTTATCACTACCAAGAGTGTGTTAGAGGTACAACAGTACTTTTAAAAGAACCTTGCTCTT	27502
gb:MT027062|Organism:Severe      TTTATCACTACCAAGAGTGTGTTAGAGGTACAACAGTACTTTTAAAAGAACCTTGCTCTT	27502
gb:MT019529|Organism:Severe      TTTATCACTACCAAGAGTGTGTTAGAGGTACAACAGTACTTTTAAAAGAACCTTGCTCTT	27502
gb:MN996529|Organism:Severe      TTTATCACTACCAAGAGTGTGTTAGAGGTACAACAGTACTTTTAAAAGAACCTTGCTCTT	27490
gb:MN996531|Organism:Severe      TTTATCACTACCAAGAGTGTGTTAGAGGTACAACAGTACTTTTAAAAGAACCTTGCTCTT	27489
gb:MT066176|Organism:Severe      TTTATCACTACCAAGAGTGTGTTAGAGGTACAACAGTACTTTTAAAAGAACCTTGCTCTT	27502
gb:MT027064|Organism:Severe      TTTATCACTACCAAGAGTGTGTTAGAGGTACAACAGTACTTTTAAAAGAACCTTGCTCTT	27502
gb:MN994468|Organism:Severe      TTTATCACTACCAAGAGTGTGTTAGAGGTACAACAGTACTTTTAAAAGAACCTTGCTCTT	27502
gb:MT072688|Organism:Severe      TTTATCACTACCAAGAGTGTGTTAGAGGTACAACAGTACTTTTAAAAGAACCTTGCTCTT	27487
gb:MN996527|Organism:Severe      TTTATCACTACCAAGAGTGTGTTAGAGGTACAACAGTACTTTTAAAAGAACCTTGCTCTT	27469
gb:MT093631|Organism:Severe      TTTATCACTACCAAGAGTGTGTTAGAGGTACAACAGTACTTTTAAAAGAACCTTGCTCTT	27540
gb:MT106053|Organism:Severe      TTTATCACTACCAAGAGTGTGTTAGAGGTACAACAGTACTTTTAAAAGAACCTTGCTCTT	27502
gb:MT019533|Organism:Severe      TTTATCACTACCAAGAGTGTGTTAGAGGTACAACAGTACTTTTAAAAGAACCTTGCTCTT	27502
gb:MT019531|Organism:Severe      TTTATCACTACCAAGAGTGTGTTAGAGGTACAACAGTACTTTTAAAAGAACCTTGCTCTT	27502
gb:MN996528|Organism:Severe      TTTATCACTACCAAGAGTGTGTTAGAGGTACAACAGTACTTTTAAAAGAACCTTGCTCTT	27502
gb:MN996530|Organism:Severe      TTTATCACTACCAAGAGTGTGTTAGAGGTACAACAGTACTTTTAAAAGAACCTTGCTCTT	27488
gb:MN908947|Organism:Severe      TTTATCACTACCAAGAGTGTGTTAGAGGTACAACAGTACTTTTAAAAGAACCTTGCTCTT	27502
gb:MT019532|Organism:Severe      TTTATCACTACCAAGAGTGTGTTAGAGGTACAACAGTACTTTTAAAAGAACCTTGCTCTT	27502
                                 ************************************************************

gb:MT020781|Organism:Severe      CTGGAACATACGAGGGCAATTCACCATTTCATCCTCTAGCTGATAACAAATTTGCACTGA	27550
gb:MT007544|Organism:Severe      CTGGAACATACGAGGGCAATTCACCATTTCATCCTCTAGCTGATAACAAATTTGCACTGA	27562
gb:MN994467|Organism:Severe      CTGGAACATACGAGGGCAATTCACCATTTCATCCTCTAGCTGATAACAAATTTGCACTGA	27562
gb:MT044257|Organism:Severe      CTGGAACATACGAGGGCAATTCACCATTTCATCCTCTAGCTGATAACAAATTTGCACTGA	27562
gb:MT106054|Organism:Severe      CTGGAACATACGAGGGCAATTCACCATTTCATCCTCTAGCTGATAACAAATTTGCACTGA	27562
gb:MT049951|Organism:Severe      CTGGAACATACGAGGGCAATTCACCATTTCATCCTCTAGCTGATAACAAATTTGCACTGA	27562
gb:MN975262|Organism:Severe      CTGGAACATACGAGGGCAATTCACCATTTCATCCTCTAGCTGATAACAAATTTGCACTGA	27562
```

```
gb:MT106052|Organism:Severe      CTGGAACATACGAGGGCAATTCACCATTTCATCCTCTAGCTGATAACAAATTTGCACTGA   27562
gb:LC522975|Organism:Severe      CTGGAACATACGAGGGCAATTCACCATTTCATCCTCTAGCTGATAACAAATTTGCACTGA   27559
gb:LC522973|Organism:Severe      CTGGAACATACGAGGGCAATTCACCATTTCATCCTCTAGCTGATAACAAATTTGCACTGA   27559
gb:LC522974|Organism:Severe      CTGGAACATACGAGGGCAATTCACCATTTCATCCTCTAGCTGATAACAAATTTGCACTGA   27559
gb:MN985325|Organism:Severe      CTGGAACATACGAGGGCAATTCACCATTTCATCCTCTAGCTGATAACAAATTTGCACTGA   27562
gb:MT020881|Organism:Severe      CTGGAACATACGAGGGCAATTCACCATTTCATCCTCTAGCTGATAACAAATTTGCACTGA   27562
gb:MT020880|Organism:Severe      CTGGAACATACGAGGGCAATTCACCATTTCATCCTCTAGCTGATAACAAATTTGCACTGA   27562
gb:MT066175|Organism:Severe      CTGGAACATACGAGGGCAATTCACCATTTCATCCTCTAGCTGATAACAAATTTGCACTGA   27562
gb:MN997409|Organism:Severe      CTGGAACATACGAGGGCAATTCACCATTTCATCCTCTAGCTGATAACAAATTTGCACTGA   27562
gb:MN938384|Organism:Severe      CTGGAACATACGAGGGCAATTCACCATTTCATCCTCTAGCTGATAACAAATTTGCACTGA   27530
gb:MT044258|Organism:Severe      CTGGAACATACGAGGGCAATTCACCATTTCATCCTCTAGCTGATAACAAATTTGCACTGA   27538
gb:MT039890|Organism:Severe      CTGGAACATACGAGGGCAATTCACCATTTCATCCTCTAGCTGATAACAAATTTGCACTGA   27562
gb:MN988713|Organism:Severe      CTGGAACATACGAGGGCAATTCACCATTTCATCCTCTAGCTGATAACAAATTTGCACTGA   27562
gb:LC521925|Organism:Severe      CTGGAACATACGAGGGCAATTCACCATTTCATCCTCTAGCTGATAACAAATTTGCACTGA   27535
gb:MT093571|Organism:Severe      CTGGAACATACGAGGGCAATTCACCATTTCATCCTCTAGCTGATAACAAATTTGCACTGA   27562
gb:MT039887|Organism:Severe      CTGGAACATACGAGGGCAATTCACCATTTCATCCTCTAGCTGATAACAAATTTGCACTGA   27559
gb:MT019530|Organism:Severe      CTGGAACATACGAGGGCAATTCACCATTTCATCCTCTAGCTGATAACAAATTTGCACTGA   27562
gb:MT039888|Organism:Severe      CTGGAACATACGAGGGCAATTCACCATTTCATCCTCTAGCTGATAACAAATTTGCACTGA   27562
gb:LC522972|Organism:Severe      CTGGAACATACGAGGGCAATTCACCATTTCATCCTCTAGCTGATAACAAATTTGCACTGA   27559
gb:MT027063|Organism:Severe      CTGGAACATACGAGGGCAATTCACCATTTCATCCTCTAGCTGATAACAAATTTGCACTGA   27562
gb:MT027062|Organism:Severe      CTGGAACATACGAGGGCAATTCACCATTTCATCCTCTAGCTGATAACAAATTTGCACTGA   27562
gb:MT019529|Organism:Severe      CTGGAACATACGAGGGCAATTCACCATTTCATCCTCTAGCTGATAACAAATTTGCACTGA   27562
gb:MN996529|Organism:Severe      CTGGAACATACGAGGGCAATTCACCATTTCATCCTCTAGCTGATAACAAATTTGCACTGA   27550
gb:MN996531|Organism:Severe      CTGGAACATACGAGGGCAATTCACCATTTCATCCTCTAGCTGATAACAAATTTGCACTGA   27549
gb:MT066176|Organism:Severe      CTGGAACATACGAGGGCAATTCACCATTTCATCCTCTAGCTGATAACAAATTTGCACTGA   27562
gb:MT027064|Organism:Severe      CTGGAACATACGAGGGCAATTCACCATTTCATCCTCTAGCTGATAACAAATTTGCACTGA   27562
gb:MN994468|Organism:Severe      CTGGAACATACGAGGGCAATTCACCATTTCATCCTCTAGCTGATAACAAATTTGCACTGA   27562
gb:MT072688|Organism:Severe      CTGGAACATACGAGGGCAATTCACCATTTCATCCTCTAGCTGATAACAAATTTGCACTGA   27547
gb:MN996527|Organism:Severe      CTGGAACATACGAGGGCAATTCACCATTTCATCCTCTAGCTGATAACAAATTTGCACTGA   27529
gb:MT093631|Organism:Severe      CTGGAACATACGAGGGCAATTCACCATTTCATCCTCTAGCTGATAACAAATTTGCACTGA   27600
gb:MT106053|Organism:Severe      CTGGAACATACGAGGGCAATTCACCATTTCATCCTCTAGCTGATAACAAATTTGCACTGA   27562
gb:MT019533|Organism:Severe      CTGGAACATACGAGGGCAATTCACCATTTCATCCTCTAGCTGATAACAAATTTGCACTGA   27562
gb:MT019531|Organism:Severe      CTGGAACATACGAGGGCAATTCACCATTTCATCCTCTAGCTGATAACAAATTTGCACTGA   27562
gb:MN996528|Organism:Severe      CTGGAACATACGAGGGCAATTCACCATTTCATCCTCTAGCTGATAACAAATTTGCACTGA   27562
gb:MN996530|Organism:Severe      CTGGAACATACGAGGGCAATTCACCATTTCATCCTCTAGCTGATAACAAATTTGCACTGA   27548
gb:MN908947|Organism:Severe      CTGGAACATACGAGGGCAATTCACCATTTCATCCTCTAGCTGATAACAAATTTGCACTGA   27562
gb:MT019532|Organism:Severe      CTGGAACATACGAGGGCAATTCACCATTTCATCCTCTAGCTGATAACAAATTTGCACTGA   27562
                                 ************************************************************

gb:MT020781|Organism:Severe      CTTGCTTTAGCACTCAATTTGCTTTTGCTTGTCCTGACGGCGTAAAACACGTCTATCAGT   27610
gb:MT007544|Organism:Severe      CTTGCTTTAGCACTCAATTTGCTTTTGCTTGTCCTGACGGCGTAAAACACGTCTATCAGT   27622
gb:MN994467|Organism:Severe      CTTGCTTTAGCACTCAATTTGCTTTTGCTTGTCCTGACGGCGTAAAACACGTCTATCAGT   27622
gb:MT044257|Organism:Severe      CTTGCTTTAGCACTCAATTTGCTTTTGCTTGTCCTGACGGCGTAAAACACGTCTATCAGT   27622
gb:MT106054|Organism:Severe      CTTGCTTTAGCACTCAATTTGCTTTTGCTTGTCCTGACGGCGTAAAACACGTCTATCAGT   27622
gb:MT049951|Organism:Severe      CTTGCTTTAGCACTCAATTTGCTTTTGCTTGTCCTGACGGCGTAAAACACGTCTATCAGT   27622
gb:MN975262|Organism:Severe      CTTGCTTTAGCACTCAATTTGCTTTTGCTTGTCCTGACGGCGTAAAACACGTCTATCAGT   27622
gb:MT106052|Organism:Severe      CTTGCTTTAGCACTCAATTTGCTTTTGCTTGTCCTGACGGCGTAAAACACGTCTATCAGT   27622
gb:LC522975|Organism:Severe      CTTGCTTTAGCACTCAATTTGCTTTTGCTTGTCCTGACGGCGTAAAACACGTCTATCAGT   27619
gb:LC522973|Organism:Severe      CTTGCTTTAGCACTCAATTTGCTTTTGCTTGTCCTGACGGCGTAAAACACGTCTATCAGT   27619
gb:LC522974|Organism:Severe      CTTGCTTTAGCACTCAATTTGCTTTTGCTTGTCCTGACGGCGTAAAACACGTCTATCAGT   27619
gb:MN985325|Organism:Severe      CTTGCTTTAGCACTCAATTTGCTTTTGCTTGTCCTGACGGCGTAAAACACGTCTATCAGT   27622
gb:MT020881|Organism:Severe      CTTGCTTTAGCACTCAATTTGCTTTTGCTTGTCCTGACGGCGTAAAACACGTCTATCAGT   27622
gb:MT020880|Organism:Severe      CTTGCTTTAGCACTCAATTTGCTTTTGCTTGTCCTGACGGCGTAAAACACGTCTATCAGT   27622
gb:MT066175|Organism:Severe      CTTGCTTTAGCACTCAATTTGCTTTTGCTTGTCCTGACGGCGTAAAACACGTCTATCAGT   27622
gb:MN997409|Organism:Severe      CTTGCTTTAGCACTCAATTTGCTTTTGCTTGTCCTGACGGCGTAAAACACGTCTATCAGT   27622
gb:MN938384|Organism:Severe      CTTGCTTTAGCACTCAATTTGCTTTTGCTTGTCCTGACGGCGTAAAACACGTCTATCAGT   27590
gb:MT044258|Organism:Severe      CTTGCTTTAGCACTCAATTTGCTTTTGCTTGTCCTGACGGCGTAAAACACGTCTATCAGT   27598
gb:MT039890|Organism:Severe      CTTGCTTTAGCACTCAATTTGCTTTTGCTTGTCCTGACGGCGTAAAACACGTCTATCAGT   27622
gb:MN988713|Organism:Severe      CTTGCTTTAGCACTCAATTTGCTTTTGCTTGTCCTGACGGCGTAAAACACGTCTATCAGT   27622
gb:LC521925|Organism:Severe      CTTGCTTTAGCACTCAATTTGCTTTTGCTTGTCCTGACGGCGTAAAACACGTCTATCAGT   27595
gb:MT093571|Organism:Severe      CTTGCTTTAGCACTCAATTTGCTTTTGCTTGTCCTGACGGCGTAAAACACGTCTATCAGT   27622
gb:MT039887|Organism:Severe      CTTGCTTTAGCACTCAATTTGCTTTTGCTTGTCCTGACGGCGTAAAACACGTCTATCAGT   27619
gb:MT019530|Organism:Severe      CTTGCTTTAGCACTCAATTTGCTTTTGCTTGTCCTGACGGCGTAAAACACGTCTATCAGT   27622
gb:MT039888|Organism:Severe      CTTGCTTTAGCACTCAATTTGCTTTTGCTTGTCCTGACGGCGTAAAACACGTCTATCAGT   27622
gb:LC522972|Organism:Severe      CTTGCTTTAGCACTCAATTTGCTTTTGCTTGTCCTGACGGCGTAAAACACGTCTATCAGT   27619
```

```
gb:MT027063|Organism:Severe      CTTGCTTTAGCACTCAATTTGCTTTTGCTTGTCCTGACGGCGTAAAACACGTCTATCAGT    27622
gb:MT027062|Organism:Severe      CTTGCTTTAGCACTCAATTTGCTTTTGCTTGTCCTGACGGCGTAAAACACGTCTATCAGT    27622
gb:MT019529|Organism:Severe      CTTGCTTTAGCACTCAATTTGCTTTTGCTTGTCCTGACGGCGTAAAACACGTCTATCAGT    27622
gb:MN996529|Organism:Severe      CTTGCTTTAGCACTCAATTTGCTTTTGCTTGTCCTGACGGCGTAAAACACGTCTATCAGT    27610
gb:MN996531|Organism:Severe      CTTGCTTTAGCACTCAATTTGCTTTTGCTTGTCCTGACGGCGTAAAACACGTCTATCAGT    27609
gb:MT066176|Organism:Severe      CTTGCTTTAGCACTCAATTTGCTTTTGCTTGTCCTGACGGCGTAAAACACGTCTATCAGT    27622
gb:MT027064|Organism:Severe      CTTGCTTTAGCACTCAATTTGCTTTTGCTTGTCCTGACGGCGTAAAACACGTCTATCAGT    27622
gb:MN994468|Organism:Severe      CTTGCTTTAGCACTCAATTTGCTTTTGCTTGTCCTGACGGCGTAAAACACGTCTATCAGT    27622
gb:MT072688|Organism:Severe      CTTGCTTTAGCACTCAATTTGCTTTTGCTTGTCCTGACGGCGTAAAACACGTCTATCAGT    27607
gb:MN996527|Organism:Severe      CTTGCTTTAGCACTCAATTTGCTTTTGCTTGTCCTGACGGCGTAAAACACGTCTATCAGT    27589
gb:MT093631|Organism:Severe      CTTGCTTTAGCACTCAATTTGCTTTTGCTTGTCCTGACGGCGTAAAACACGTCTATCAGT    27660
gb:MT106053|Organism:Severe      CTTGCTTTAGCACTCAATTTGCTTTTGCTTGTCCTGACGGCGTAAAACACGTCTATCAGT    27622
gb:MT019533|Organism:Severe      CTTGCTTTAGCACTCAATTTGCTTTTGCTTGTCCTGACGGCGTAAAACACGTCTATCAGT    27622
gb:MT019531|Organism:Severe      CTTGCTTTAGCACTCAATTTGCTTTTGCTTGTCCTGACGGCGTAAAACACGTCTATCAGT    27622
gb:MN996528|Organism:Severe      CTTGCTTTAGCACTCAATTTGCTTTTGCTTGTCCTGACGGCGTAAAACACGTCTATCAGT    27622
gb:MN996530|Organism:Severe      CTTGCTTTAGCACTCAATTTGCTTTTGCTTGTCCTGACGGCGTAAAACACGTCTATCAGT    27608
gb:MN908947|Organism:Severe      CTTGCTTTAGCACTCAATTTGCTTTTGCTTGTCCTGACGGCGTAAAACACGTCTATCAGT    27622
gb:MT019532|Organism:Severe      CTTGCTTTAGCACTCAATTTGCTTTTGCTTGTCCTGACGGCGTAAAACACGTCTATCAGT    27622
                                 ************************************************************

gb:MT020781|Organism:Severe      TACGTGCCAGATCAGTTTCACCTAAACTGTTCATCAGACAAGAGGAAGTTCAAGAACTTT    27670
gb:MT007544|Organism:Severe      TACGTGCCAGATCAGTTTCACCTAAACTGTTCATCAGACAAGAGGAAGTTCAAGAACTTT    27682
gb:MN994467|Organism:Severe      TACGTGCCAGATCAGTTTCACCTAAACTGTTCATCAGACAAGAGGAAGTTCAAGAACTTT    27682
gb:MT044257|Organism:Severe      TACGTGCCAGATCAGTTTCACCTAAACTGTTCATCAGACAAGAGGAAGTTCAAGAACTTT    27682
gb:MT106054|Organism:Severe      TACGTGCCAGATCAGTTTCACCTAAACTGTTCATCAGACAAGAGGAAGTTCAAGAACTTT    27682
gb:MT049951|Organism:Severe      TACGTGCCAGATCAGTTTCACCTAAACTGTTCATCAGACAAGAGGAAGTTCAAGAACTTT    27682
gb:MN975262|Organism:Severe      TACGTGCCAGATCAGTTTCACCTAAACTGTTCATCAGACAAGAGGAAGTTCAAGAACTTT    27682
gb:MT106052|Organism:Severe      TACGTGCCAGATCAGTTTCACCTAAACTGTTCATCAGACAAGAGGAAGTTCAAGAACTTT    27682
gb:LC522975|Organism:Severe      TACGTGCCAGATCAGTTTCACCTAAACTGTTCATCAGACAAGAGGAAGTTCAAGAACTTT    27679
gb:LC522973|Organism:Severe      TACGTGCCAGATCAGTTTCACCTAAACTGTTCATCAGACAAGAGGAAGTTCAAGAACTTT    27679
gb:LC522974|Organism:Severe      TACGTGCCAGATCAGTTTCACCTAAACTGTTCATCAGACAAGAGGAAGTTCAAGAACTTT    27679
gb:MN985325|Organism:Severe      TACGTGCCAGATCAGTTTCACCTAAACTGTTCATCAGACAAGAGGAAGTTCAAGAACTTT    27682
gb:MT020881|Organism:Severe      TACGTGCCAGATCAGTTTCACCTAAACTGTTCATCAGACAAGAGGAAGTTCAAGAACTTT    27682
gb:MT020880|Organism:Severe      TACGTGCCAGATCAGTTTCACCTAAACTGTTCATCAGACAAGAGGAAGTTCAAGAACTTT    27682
gb:MT066175|Organism:Severe      TACGTGCCAGATCAGTTTCACCTAAACTGTTCATCAGACAAGAGGAAGTTCAAGAACTTT    27682
gb:MN997409|Organism:Severe      TACGTGCCAGATCAGTTTCACCTAAACTGTTCATCAGACAAGAGGAAGTTCAAGAACTTT    27682
gb:MN938384|Organism:Severe      TACGTGCCAGATCAGTTTCACCTAAACTGTTCATCAGACAAGAGGAAGTTCAAGAACTTT    27650
gb:MT044258|Organism:Severe      TACGTGCCAGATCAGTTTCACCTAAACTGTTCATCAGACAAGAGGAAGTTCAAGAACTTT    27658
gb:MT039890|Organism:Severe      TACGTGCCAGATCAGTTTCACCTAAACTGTTCATCAGACAAGAGGAAGTTCAAGAACTTT    27682
gb:MN988713|Organism:Severe      TACGTGCCAGATCAGTTTCACCTAAACTGTTCATCAGACAAGAGGAAGTTCAAGAACTTT    27682
gb:LC521925|Organism:Severe      TACGTGCCAGATCAGTTTCACCTAAACTGTTCATCAGACAAGAGGAAGTTCAAGAACTTT    27655
gb:MT093571|Organism:Severe      TACGTGCCAGATCAGTTTCACCTAAACTGTTCATCAGACAAGAGGAAGTTCAAGAACTTT    27682
gb:MT039887|Organism:Severe      TACGTGCCAGATCAGTTTCACCTAAACTGTTCATCAGACAAGAGGAAGTTCAAGAACTTT    27679
gb:MT019530|Organism:Severe      TACGTGCCAGATCAGTTTCACCTAAACTGTTCATCAGACAAGAGGAAGTTCAAGAACTTT    27682
gb:MT039888|Organism:Severe      TACGTGCCAGATCAGTTTCACCTAAACTGTTCATCAGACAAGAGGAAGTTCAAGAACTTT    27682
gb:LC522972|Organism:Severe      TACGTGCCAGATCAGTTTCACCTAAACTGTTCATCAGACAAGAGGAAGTTCAAGAACTTT    27679
gb:MT027063|Organism:Severe      TACGTGCCAGATCAGTTTCACCTAAACTGTTCATCAGACAAGAGGAAGTTCAAGAACTTT    27682
gb:MT027062|Organism:Severe      TACGTGCCAGATCAGTTTCACCTAAACTGTTCATCAGACAAGAGGAAGTTCAAGAACTTT    27682
gb:MT019529|Organism:Severe      TACGTGCCAGATCAGTTTCACCTAAACTGTTCATCAGACAAGAGGAAGTTCAAGAACTTT    27682
gb:MN996529|Organism:Severe      TACGTGCCAGATCAGTTTCACCTAAACTGTTCATCAGACAAGAGGAAGTTCAAGAACTTT    27670
gb:MN996531|Organism:Severe      TACGTGCCAGATCAGTTTCACCTAAACTGTTCATCAGACAAGAGGAAGTTCAAGAACTTT    27669
gb:MT066176|Organism:Severe      TACGTGCCAGATCAGTTTCACCTAAACTGTTCATCAGACAAGAGGAAGTTCAAGAACTTT    27682
gb:MT027064|Organism:Severe      TACGTGCCAGATCAGTTTCACCTAAACTGTTCATCAGACAAGAGGAAGTTCAAGAACTTT    27682
gb:MN994468|Organism:Severe      TACGTGCCAGATCAGTTTCACCTAAACTGTTCATCAGACAAGAGGAAGTTCAAGAACTTT    27682
gb:MT072688|Organism:Severe      TACGTGCCAGATCAGTTTCACCTAAACTGTTCATCAGACAAGAGGAAGTTCAAGAACTTT    27667
gb:MN996527|Organism:Severe      TACGTGCCAGATCAGTTTCACCTAAACTGTTCATCAGACAAGAGGAAGTTCAAGAACTTT    27649
gb:MT093631|Organism:Severe      TACGTGCCAGATCAGTTTCACCTAAACTGTTCATCAGACAAGAGGAAGTTCAAGAACTTT    27720
gb:MT106053|Organism:Severe      TACGTGCCAGATCAGTTTCACCTAAACTGTTCATCAGACAAGAGGAAGTTCAAGAACTTT    27682
gb:MT019533|Organism:Severe      TACGTGCCAGATCAGTTTCACCTAAACTGTTCATCAGACAAGAGGAAGTTCAAGAACTTT    27682
gb:MT019531|Organism:Severe      TACGTGCCAGATCAGTTTCACCTAAACTGTTCATCAGACAAGAGGAAGTTCAAGAACTTT    27682
gb:MN996528|Organism:Severe      TACGTGCCAGATCAGTTTCACCTAAACTGTTCATCAGACAAGAGGAAGTTCAAGAACTTT    27682
gb:MN996530|Organism:Severe      TACGTGCCAGATCAGTTTCACCTAAACTGTTCATCAGACAAGAGGAAGTTCAAGAACTTT    27668
gb:MN908947|Organism:Severe      TACGTGCCAGATCAGTTTCACCTAAACTGTTCATCAGACAAGAGGAAGTTCAAGAACTTT    27682
gb:MT019532|Organism:Severe      TACGTGCCAGATCAGTTTCACCTAAACTGTTCATCAGACAAGAGGAAGTTCAAGAACTTT    27682
                                 ************************************************************
```

```
gb:MT020781|Organism:Severe      ACTCTCCAATTTTTCTTATTGTTGCGGCAATAGTGTTTATAACACTTTGCTTCACACTCA   27730
gb:MT007544|Organism:Severe      ACTCTCCAATTTTTCTTATTGTTGCGGCAATAGTGTTTATAACACTTTGCTTCACACTCA   27742
gb:MN994467|Organism:Severe      ACTCTCCAATTTTTCTTATTGTTGCGGCAATAGTGTTTATAACACTTTGCTTCACACTCA   27742
gb:MT044257|Organism:Severe      ACTCTCCAATTTTTCTTATTGTTGCGGCAATAGTGTTTATAACACTTTGCTTCACACTCA   27742
gb:MT106054|Organism:Severe      ACTCTCCAATTTTTCTTATTGTTGCGGCAATAGTGTTTATAACACTTTGCTTCACACTCA   27742
gb:MT049951|Organism:Severe      ACTCTCCAATTTTTCTTATTGTTGCGGCAATAGTGTTTATAACACTTTGCTTCACACTCA   27742
gb:MN975262|Organism:Severe      ACTCTCCAATTTTTCTTATTGTTGCGGCAATAGTGTTTATAACACTTTGCTTCACACTCA   27742
gb:MT106052|Organism:Severe      ACTCTCCAATTTTTCTTATTGTTGCGGCAATAGTGTTTATAACACTTTGCTTCACACTCA   27742
gb:LC522975|Organism:Severe      ACTCTCCAATTTTTCTTATTGTTGCGGCAATAGTGTTTATAACACTTTGCTTCACACTCA   27739
gb:LC522973|Organism:Severe      ACTCTCCAATTTTTCTTATTGTTGCGGCAATAGTGTTTATAACACTTTGCTTCACACTCA   27739
gb:LC522974|Organism:Severe      ACTCTCCAATTTTTCTTATTGTTGCGGCAATAGTGTTTATAACACTTTGCTTCACACTCA   27739
gb:MN985325|Organism:Severe      ACTCTCCAATTTTTCTTATTGTTGCGGCAATAGTGTTTATAACACTTTGCTTCACACTCA   27742
gb:MT020881|Organism:Severe      ACTCTCCAATTTTTCTTATTGTTGCGGCAATAGTGTTTATAACACTTTGCTTCACACTCA   27742
gb:MT020880|Organism:Severe      ACTCTCCAATTTTTCTTATTGTTGCGGCAATAGTGTTTATAACACTTTGCTTCACACTCA   27742
gb:MT066175|Organism:Severe      ACTCTCCAATTTTTCTTATTGTTGCGGCAATAGTGTTTATAACACTTTGCTTCACACTCA   27742
gb:MN997409|Organism:Severe      ACTCTCCAATTTTTCTTATTGTTGCGGCAATAGTGTTTATAACACTTTGCTTCACACTCA   27742
gb:MN938384|Organism:Severe      ACTCTCCAATTTTTCTTATTGTTGCGGCAATAGTGTTTATAACACTTTGCTTCACACTCA   27710
gb:MT044258|Organism:Severe      ACTCTCCAATTTTTCTTATTGTTGCGGCAATAGTGTTTATAACACTTTGCTTCACACTCA   27718
gb:MT039890|Organism:Severe      ACTCTCCAATTTTTCTTATTGTTGCGGCAATAGTGTTTATAACACTTTGCTTCACACTCA   27742
gb:MN988713|Organism:Severe      ACTCTCCAATTTTTCTTATTGTTGCGGCAATAGTGTTTATAACACTTTGCTTCACACTCA   27742
gb:LC521925|Organism:Severe      ACTCTCCAATTTTTCTTATTGTTGCGGCAATAGTGTTTATAACACTTTGCTTCACACTCA   27715
gb:MT093571|Organism:Severe      ACTCTCCAATTTTTCTTATTGTTGCGGCAATAGTGTTTATAACACTTTGCTTCACACTCA   27742
gb:MT039887|Organism:Severe      ACTCTCCAATTTTTCTTATTGTTGCGGCAATAGTGTTTATAACACTTTGCTTCACACTCA   27739
gb:MT019530|Organism:Severe      ACTCTCCAATTTTTCTTATTGTTGCGGCAATAGTGTTTATAACACTTTGCTTCACACTCA   27742
gb:MT039888|Organism:Severe      ACTCTCCAATTTTTCTTATTGTTGCGGCAATAGTGTTTATAACACTTTGCTTCACACTCA   27742
gb:LC522972|Organism:Severe      ACTCTCCAATTTTTCTTATTGTTGCGGCAATAGTGTTTATAACACTTTGCTTCACACTCA   27739
gb:MT027063|Organism:Severe      ACTCTCCAATTTTTCTTATTGTTGCGGCAATAGTGTTTATAACACTTTGCTTCACACTCA   27742
gb:MT027062|Organism:Severe      ACTCTCCAATTTTTCTTATTGTTGCGGCAATAGTGTTTATAACACTTTGCTTCACACTCA   27742
gb:MT019529|Organism:Severe      ACTCTCCAATTTTTCTTATTGTTGCGGCAATAGTGTTTATAACACTTTGCTTCACACTCA   27742
gb:MN996529|Organism:Severe      ACTCTCCAATTTTTCTTATTGTTGCGGCAATAGTGTTTATAACACTTTGCTTCACACTCA   27730
gb:MN996531|Organism:Severe      ACTCTCCAATTTTTCTTATTGTTGCGGCAATAGTGTTTATAACACTTTGCTTCACACTCA   27729
gb:MT066176|Organism:Severe      ACTCTCCAATTTTTCTTATTGTTGCGGCAATAGTGTTTATAACACTTTGCTTCACACTCA   27742
gb:MT027064|Organism:Severe      ACTCTCCAATTTTTCTTATTGTTGCGGCAATAGTGTTTATAACACTTTGCTTCACACTCA   27742
gb:MN994468|Organism:Severe      ACTCTCCAATTTTTCTTATTGTTGCGGCAATAGTGTTTATAACACTTTGCTTCACACTCA   27742
gb:MT072688|Organism:Severe      ACTCTCCAATTTTTCTTATTGTTGCGGCAATAGTGTTTATAACACTTTGCTTCACACTCA   27727
gb:MN996527|Organism:Severe      ACTCTCCAATTTTTCTTATTGTTGCGGCAATAGTGTTTATAACACTTTGCTTCACACTCA   27709
gb:MT093631|Organism:Severe      ACTCTCCAATTTTTCTTATTGTTGCGGCAATAGTGTTTATAACACTTTGCTTCACACTCA   27780
gb:MT106053|Organism:Severe      ACTCTCCAATTTTTCTTATTGTTGCGGCAATAGTGTTTATAACACTTTGCTTCACACTCA   27742
gb:MT019533|Organism:Severe      ACTCTCCAATTTTTCTTATTGTTGCGGCAATAGTGTTTATAACACTTTGCTTCACACTCA   27742
gb:MT019531|Organism:Severe      ACTCTCCAATTTTTCTTATTGTTGCGGCAATAGTGTTTATAACACTTTGCTTCACACTCA   27742
gb:MN996528|Organism:Severe      ACTCTCCAATTTTTCTTATTGTTGCGGCAATAGTGTTTATAACACTTTGCTTCACACTCA   27742
gb:MN996530|Organism:Severe      ACTCTCCAATTTTTCTTATTGTTGCGGCAATAGTGTTTATAACACTTTGCTTCACACTCA   27728
gb:MN908947|Organism:Severe      ACTCTCCAATTTTTCTTATTGTTGCGGCAATAGTGTTTATAACACTTTGCTTCACACTCA   27742
gb:MT019532|Organism:Severe      ACTCTCCAATTTTTCTTATTGTTGCGGCAATAGTGTTTATAACACTTTGCTTCACACTCA   27742
                                 ************************************************************

gb:MT020781|Organism:Severe      AAAGAAAGACAGAATGATTGAACTTTCATTAATTGACTTCTATTTGTGCTTTTTAGCCTT   27790
gb:MT007544|Organism:Severe      AAAGAAAGACAGAATGATTGAACTTTCATTAATTGACTTCTATTTGTGCTTTTTAGCCTT   27802
gb:MN994467|Organism:Severe      AAAGAAAGACAGAATGATTGAACTTTCATTAATTGACTTCTATTTGTGCTTTTTAGCCTT   27802
gb:MT044257|Organism:Severe      AAAGAAAGACAGAATGATTGAACTTTCATTAATTGACTTCTATTTGTGCTTTTTAGCCTT   27802
gb:MT106054|Organism:Severe      AAAGAAAGACAGAATGATTGAACTTTCATTAATTGACTTCTATTTGTGCTTTTTAGCCTT   27802
gb:MT049951|Organism:Severe      AAAGAAAGACAGAATGATTGAACTTTCATTAATTGACTTCTATTTGTGCTTTTTAGCCTT   27802
gb:MN975262|Organism:Severe      AAAGAAAGACAGAATGATTGAACTTTCATTAATTGACTTCTATTTGTGCTTTTTAGCCTT   27802
gb:MT106052|Organism:Severe      AAAGAAAGACAGAATGATTGAACTTTCATTAATTGACTTCTATTTGTGCTTTTTAGCCTT   27802
gb:LC522975|Organism:Severe      AAAGAAAGACAGAATGATTGAACTTTCATTAATTGACTTCTATTTGTGCTTTTTAGCCTT   27799
gb:LC522973|Organism:Severe      AAAGAAAGACAGAATGATTGAACTTTCATTAATTGACTTCTATTTGTGCTTTTTAGCCTT   27799
gb:LC522974|Organism:Severe      AAAGAAAGACAGAATGATTGAACTTTCATTAATTGACTTCTATTTGTGCTTTTTAGCCTT   27799
gb:MN985325|Organism:Severe      AAAGAAAGACAGAATGATTGAACTTTCATTAATTGACTTCTATTTGTGCTTTTTAGCCTT   27802
gb:MT020881|Organism:Severe      AAAGAAAGACAGAATGATTGAACTTTCATTAATTGACTTCTATTTGTGCTTTTTAGCCTT   27802
gb:MT020880|Organism:Severe      AAAGAAAGACAGAATGATTGAACTTTCATTAATTGACTTCTATTTGTGCTTTTTAGCCTT   27802
gb:MT066175|Organism:Severe      AAAGAAAGACAGAATGATTGAACTTTCATTAATTGACTTCTATTTGTGCTTTTTAGCCTT   27802
gb:MN997409|Organism:Severe      AAAGAAAGACAGAATGATTGAACTTTCATTAATTGACTTCTATTTGTGCTTTTTAGCCTT   27802
gb:MN938384|Organism:Severe      AAAGAAAGACAGAATGATTGAACTTTCATTAATTGACTTCTATTTGTGCTTTTTAGCCTT   27770
gb:MT044258|Organism:Severe      AAAGAAAGACAGAATGATTGAACTTTCATTAATTGACTTCTATTTGTGCTTTTTAGCCTT   27778
```

```
gb:MT039890|Organism:Severe      AAAGAAAGACAGAATGATTGAACTTTCATTAATTGACTTCTATTTGTGCTTTTTAGCCTT   27802
gb:MN988713|Organism:Severe      AAAGAAAGACAGAATGATTGAACTTTCATTAATTGACTTCTATTTGTGCTTTTTAGCCTT   27802
gb:LC521925|Organism:Severe      AAAGAAAGACAGAATGATTGAACTTTCATTAATTGACTTCTATTTGTGCTTTTTAGCCTT   27775
gb:MT093571|Organism:Severe      AAAGAAAGACAGAATGATTGAACTTTCATTAATTGACTTCTATTTGTGCTTTTTAGCCTT   27802
gb:MT039887|Organism:Severe      AAAGAAAGACAGAATGATTGAACTTTCATTAATTGACTTCTATTTGTGCTTTTTAGCCTT   27799
gb:MT019530|Organism:Severe      AAAGAAAGACAGAATGATTGAACTTTCATTAATTGACTTCTATTTGTGCTTTTTAGCCTT   27802
gb:MT039888|Organism:Severe      AAAGAAAGACAGAATGATTGAACTTTCATTAATTGACTTCTATTTGTGCTTTTTAGCCTT   27802
gb:LC522972|Organism:Severe      AAAGAAAGACAGAATGATTGAACTTTCATTAATTGACTTCTATTTGTGCTTTTTAGCCTT   27799
gb:MT027063|Organism:Severe      AAAGAAAGACAGAATGATTGAACTTTCATTAATTGACTTCTATTTGTGCTTTTTAGCCTT   27802
gb:MT027062|Organism:Severe      AAAGAAAGACAGAATGATTGAACTTTCATTAATTGACTTCTATTTGTGCTTTTTAGCCTT   27802
gb:MT019529|Organism:Severe      AAAGAAAGACAGAATGATTGAACTTTCATTAATTGACTTCTATTTGTGCTTTTTAGCCTT   27802
gb:MN996529|Organism:Severe      AAAGAAAGACAGAATGATTGAACTTTCATTAATTGACTTCTATTTGTGCTTTTTAGCCTT   27790
gb:MN996531|Organism:Severe      AAAGAAAGACAGAATGATTGAACTTTCATTAATTGACTTCTATTTGTGCTTTTTAGCCTT   27789
gb:MT066176|Organism:Severe      AAAGAAAGACAGAATGATTGAACTTTCATTAATTGACTTCTATTTGTGCTTTTTAGCCTT   27802
gb:MT027064|Organism:Severe      AAAGAAAGACAGAATGATTGAACTTTCATTAATTGACTTCTATTTGTGCTTTTTAGCCTT   27802
gb:MN994468|Organism:Severe      AAAGAAAGACAGAATGATTGAACTTTCATTAATTGACTTCTATTTGTGCTTTTTAGCCTT   27802
gb:MT072688|Organism:Severe      AAAGAAAGACAGAATGATTGAACTTTCATTAATTGACTTCTATTTGTGCTTTTTAGCCTT   27787
gb:MN996527|Organism:Severe      AAAGAAAGACAGAATGATTGAACTTTCATTAATTGACTTCTATTTGTGCTTTTTAGCCTT   27769
gb:MT093631|Organism:Severe      AAAGAAAGACAGAATGATTGAACTTTCATTAATTGACTTCTATTTGTGCTTTTTAGCCTT   27840
gb:MT106053|Organism:Severe      AAAGAAAGACAGAATGATTGAACTTTCATTAATTGACTTCTATTTGTGCTTTTTAGCCTT   27802
gb:MT019533|Organism:Severe      AAAGAAAGACAGAATGATTGAACTTTCATTAATTGACTTCTATTTGTGCTTTTTAGCCTT   27802
gb:MT019531|Organism:Severe      AAAGAAAGACAGAATGATTGAACTTTCATTAATTGACTTCTATTTGTGCTTTTTAGCCTT   27802
gb:MN996528|Organism:Severe      AAAGAAAGACAGAATGATTGAACTTTCATTAATTGACTTCTATTTGTGCTTTTTAGCCTT   27802
gb:MN996530|Organism:Severe      AAAGAAAGACAGAATGATTGAACTTTCATTAATTGACTTCTATTTGTGCTTTTTAGCCTT   27788
gb:MN908947|Organism:Severe      AAAGAAAGACAGAATGATTGAACTTTCATTAATTGACTTCTATTTGTGCTTTTTAGCCTT   27802
gb:MT019532|Organism:Severe      AAAGAAAGACAGAATGATTGAACTTTCATTAATTGACTTCTATTTGTGCTTTTTAGCCTT   27802
                                 ************************************************************

gb:MT020781|Organism:Severe      TCTGCTATTCCTTGTTTTAATTATGCTTATTATCTTTTGGTTCTCACTTGAACTGCAAGA   27850
gb:MT007544|Organism:Severe      TCTGCTATTCCTTGTTTTAATTATGCTTATTATCTTTTGGTTCTCACTTGAACTGCAAGA   27862
gb:MN994467|Organism:Severe      TCTGCTATTCCTTGTTTTAATTATGCTTATTATCTTTTGGTTCTCACTTGAACTGCAAGA   27862
gb:MT044257|Organism:Severe      TCTGCTATTCCTTGTTTTAATTATGCTTATTATCTTTTGGTTCTCACTTGAACTGCAAGA   27862
gb:MT106054|Organism:Severe      TCTGCTATTCCTTGTTTTAATTATGCTTATTATCTTTTGGTTCTCACTTGAACTGCAAGA   27862
gb:MT049951|Organism:Severe      TCTGCTATTCCTTGTTTTAATTATGCTTATTATCTTTTGGTTCTCACTTGAACTGCAAGA   27862
gb:MN975262|Organism:Severe      TCTGCTATTCCTTGTTTTAATTATGCTTATTATCTTTTGGTTCTCACTTGAACTGCAAGA   27862
gb:MT106052|Organism:Severe      TCTGCTATTCCTTGTTTTAATTATGCTTATTATCTTTTGGTTCTCACTTGAACTGCAAGA   27862
gb:LC522975|Organism:Severe      TCTGCTATTCCTTGTTTTAATTATGCTTATTATCTTTTGGTTCTCACTTGAACTGCAAGA   27859
gb:LC522973|Organism:Severe      TCTGCTATTCCTTGTTTTAATTATGCTTATTATCTTTTGGTTCTCACTTGAACTGCAAGA   27859
gb:LC522974|Organism:Severe      TCTGCTATTCCTTGTTTTAATTATGCTTATTATCTTTTGGTTCTCACTTGAACTGCAAGA   27859
gb:MN985325|Organism:Severe      TCTGCTATTCCTTGTTTTAATTATGCTTATTATCTTTTGGTTCTCACTTGAACTGCAAGA   27862
gb:MT020881|Organism:Severe      TCTGCTATTCCTTGTTTTAATTATGCTTATTATCTTTTGGTTCTCACTTGAACTGCAAGA   27862
gb:MT020880|Organism:Severe      TCTGCTATTCCTTGTTTTAATTATGCTTATTATCTTTTGGTTCTCACTTGAACTGCAAGA   27862
gb:MT066175|Organism:Severe      TCTGCTATTCCTTGTTTTAATTATGCTTATTATCTTTTGGTTCTCACTTGAACTGCAAGA   27862
gb:MN997409|Organism:Severe      TCTGCTATTCCTTGTTTTAATTATGCTTATTATCTTTTGGTTCTCACTTGAACTGCAAGA   27862
gb:MN938384|Organism:Severe      TCTGCTATTCCTTGTTTTAATTATGCTTATTATCTTTTGGTTCTCACTTGAACTGCAAGA   27830
gb:MT044258|Organism:Severe      TCTGCTATTCCTTGTTTTAATTATGCTTATTATCTTTTGGTTCTCACTTGAACTGCAAGA   27838
gb:MT039890|Organism:Severe      TCTGCTATTCCTTGTTTTAATTATGCTTATTATCTTTTGGTTCTCACTTGAACTGCAAGA   27862
gb:MN988713|Organism:Severe      TCTGCTATTCCTTGTTTTAATTATGCTTATTATCTTTTGGTTCTCACTTGAACTGCAAGA   27862
gb:LC521925|Organism:Severe      TCTGCTATTCCTTGTTTTAATTATGCTTATTATCTTTTGGTTCTCACTTGAACTGCAAGA   27835
gb:MT093571|Organism:Severe      TCTGCTATTCCTTGTTTTAATTATGCTTATTATCTTTTGGTTCTCACTTGAACTGCAAGA   27862
gb:MT039887|Organism:Severe      TCTGCTATTCCTTGTTTTAATTATGCTTATTATCTTTTGGTTCTCACTTGAACTGCAAGA   27859
gb:MT019530|Organism:Severe      TCTGCTATTCCTTGTTTTAATTATGCTTATTATCTTTTGGTTCTCACTTGAACTGCAAGA   27862
gb:MT039888|Organism:Severe      TCTGCTATTCCTTGTTTTAATTATGCTTATTATCTTTTGGTTCTCACTTGAACTGCAAGA   27862
gb:LC522972|Organism:Severe      TCTGCTATTCCTTGTTTTAATTATGCTTATTATCTTTTGGTTCTCACTTGAACTGCAAGA   27859
gb:MT027063|Organism:Severe      TCTGCTATTCCTTGTTTTAATTATGCTTATTATCTTTTGGTTCTCACTTGAACTGCAAGA   27862
gb:MT027062|Organism:Severe      TCTGCTATTCCTTGTTTTAATTATGCTTATTATCTTTTGGTTCTCACTTGAACTGCAAGA   27862
gb:MT019529|Organism:Severe      TCTGCTATTCCTTGTTTTAATTATGCTTATTATCTTTTGGTTCTCACTTGAACTGCAAGA   27862
gb:MN996529|Organism:Severe      TCTGCTATTCCTTGTTTTAATTATGCTTATTATCTTTTGGTTCTCACTTGAACTGCAAGA   27850
gb:MN996531|Organism:Severe      TCTGCTATTCCTTGTTTTAATTATGCTTATTATCTTTTGGTTCTCACTTGAACTGCAAGA   27849
gb:MT066176|Organism:Severe      TCTGCTATTCCTTGTTTTAATTATGCTTATTATCTTTTGGTTCTCACTTGAACTGCAAGA   27862
gb:MT027064|Organism:Severe      TCTGCTATTCCTTGTTTTAATTATGCTTATTATCTTTTGGTTCTCACTTGAACTGCAAGA   27862
gb:MN994468|Organism:Severe      TCTGCTATTCCTTGTTTTAATTATGCTTATTATCTTTTGGTTCTCACTTGAACTGCAAGA   27862
gb:MT072688|Organism:Severe      TCTGCTATTCCTTGTTTTAATTATGCTTATTATCTTTTGGTTCTCACTTGAACTGCAAGA   27847
gb:MN996527|Organism:Severe      TCTGCTATTCCTTGTTTTAATTATGCTTATTATCTTTTGGTTCTCACTTGAACTGCAAGA   27829
gb:MT093631|Organism:Severe      TCTGCTATTCCTTGTTTTAATTATGCTTATTATCTTTTGGTTCTCACTTGAACTGCAAGA   27900
```

```
gb:MT106053|Organism:Severe      TCTGCTATTCCTTGTTTTAATTATGCTTATTATCTTTTGGTTCTCACTTGAACTGCAAGA   27862
gb:MT019533|Organism:Severe      TCTGCTATTCCTTGTTTTAATTATGCTTATTATCTTTTGGTTCTCACTTGAACTGCAAGA   27862
gb:MT019531|Organism:Severe      TCTGCTATTCCTTGTTTTAATTATGCTTATTATCTTTTGGTTCTCACTTGAACTGCAAGA   27862
gb:MN996528|Organism:Severe      TCTGCTATTCCTTGTTTTAATTATGCTTATTATCTTTTGGTTCTCACTTGAACTGCAAGA   27862
gb:MN996530|Organism:Severe      TCTGCTATTCCTTGTTTTAATTATGCTTATTATCTTTTGGTTCTCACTTGAACTGCAAGA   27848
gb:MN908947|Organism:Severe      TCTGCTATTCCTTGTTTTAATTATGCTTATTATCTTTTGGTTCTCACTTGAACTGCAAGA   27862
gb:MT019532|Organism:Severe      TCTGCTATTCCTTGTTTTAATTATGCTTATTATCTTTTGGTTCTCACTTGAACTGCAAGA   27862
                                 ************************************************************

gb:MT020781|Organism:Severe      TCATAATGAAACTTGTCACGCCTAAACGAACATGAAATTTCTTGTTTTCTTAGGAATCAT   27910
gb:MT007544|Organism:Severe      TCATAATGAAACTTGTCACGCCTAAACGAACATGAAATTTCTTGTTTTCTTAGGAATCAT   27922
gb:MN994467|Organism:Severe      TCATAATGAAACTTGTCACGCCTAAACGAACATGAAATTTCTTGTTTTCTTAGGAATCAT   27922
gb:MT044257|Organism:Severe      TCATAATGAAACTTGTCACGCCTAAACGAACATGAAATTTCTTGTTTTCTTAGGAATCAT   27922
gb:MT106054|Organism:Severe      TCATAATGAAACTTGTCACGCCTAAACGAACATGAAATTTCTTGTTTTCTTAGGAATCAT   27922
gb:MT049951|Organism:Severe      TCATAATGAAACTTGTCACGCCTAAACGAACATGAAATTTCTTGTTTTCTTAGGAATCAT   27922
gb:MN975262|Organism:Severe      TCATAATGAAACTTGTCACGCCTAAACGAACATGAAATTTCTTGTTTTCTTAGGAATCAT   27922
gb:MT106052|Organism:Severe      TCATAATGAAACTTGTCACGCCTAAACGAACATGAAATTTCTTGTTTTCTTAGGAATCAT   27922
gb:LC522975|Organism:Severe      TCATAATGAAACTTGTCACGCCTAAACGAACATGAAATTTCTTGTTTTCTTAGGAATCAT   27919
gb:LC522973|Organism:Severe      TCATAATGAAACTTGTCACGCCTAAACGAACATGAAATTTCTTGTTTTCTTAGGAATCAT   27919
gb:LC522974|Organism:Severe      TCATAATGAAACTTGTCACGCCTAAACGAACATGAAATTTCTTGTTTTCTTAGGAATCAT   27919
gb:MN985325|Organism:Severe      TCATAATGAAACTTGTCACGCCTAAACGAACATGAAATTTCTTGTTTTCTTAGGAATCAT   27922
gb:MT020881|Organism:Severe      TCATAATGAAACTTGTCACGCCTAAACGAACATGAAATTTCTTGTTTTCTTAGGAATCAT   27922
gb:MT020880|Organism:Severe      TCATAATGAAACTTGTCACGCCTAAACGAACATGAAATTTCTTGTTTTCTTAGGAATCAT   27922
gb:MT066175|Organism:Severe      TCATAATGAAACTTGTCACGCCTAAACGAACATGAAATTTCTTGTTTTCTTAGGAATCAT   27922
gb:MN997409|Organism:Severe      TCATAATGAAACTTGTCACGCCTAAACGAACATGAAATTTCTTGTTTTCTTAGGAATCAT   27922
gb:MN938384|Organism:Severe      TCATAATGAAACTTGTCACGCCTAAACGAACATGAAATTTCTTGTTTTCTTAGGAATCAT   27890
gb:MT044258|Organism:Severe      TCATAATGAAACTTGTCACGCCTAAACGAACATGAAATTTCTTGTTTTCTTAGGAATCAT   27898
gb:MT039890|Organism:Severe      TCATAATGAAACTTGTCACGCCTAAACGAACATGAAATTTCTTGTTTTCTTAGGAATCAT   27922
gb:MN988713|Organism:Severe      TCATAATGAAACTTGTCACGCCTAAACGAACATGAAATTTCTTGTTTTCTTAGGAATCAT   27922
gb:LC521925|Organism:Severe      TCATAATGAAACTTGTCACGCCTAAACGAACATGAAATTTCTTGTTTTCTTAGGAATCAT   27895
gb:MT093571|Organism:Severe      TCATAATGAAACTTGTCACGCCTAAACGAACATGAAATTTCTTGTTTTCTTAGGAATCAT   27922
gb:MT039887|Organism:Severe      TCATAATGAAACTTGTCACGCCTAAACGAACATGAAATTTCTTGTTTTCTTAGGAATCAT   27919
gb:MT019530|Organism:Severe      TCATAATGAAACTTGTCACGCCTAAACGAACATGAAATTTCTTGTTTTCTTAGGAATCAT   27922
gb:MT039888|Organism:Severe      TCATAATGAAACTTGTCACGCCTAAACGAACATGAAATTTCTTGTTTTCTTAGGAATCAT   27922
gb:LC522972|Organism:Severe      TCATAATGAAACTTGTCACGCCTAAACGAACATGAAATTTCTTGTTTTCTTAGGAATCAT   27919
gb:MT027063|Organism:Severe      TCATAATGAAACTTGTCACGCCTAAACGAACATGAAATTTCTTGTTTTCTTAGGAATCAT   27922
gb:MT027062|Organism:Severe      TCATAATGAAACTTGTCACGCCTAAACGAACATGAAATTTCTTGTTTTCTTAGGAATCAT   27922
gb:MT019529|Organism:Severe      TCATAATGAAACTTGTCACGCCTAAACGAACATGAAATTTCTTGTTTTCTTAGGAATCAT   27922
gb:MN996529|Organism:Severe      TCATAATGAAACTTGTCACGCCTAAACGAACATGAAATTTCTTGTTTTCTTAGGAATCAT   27910
gb:MN996531|Organism:Severe      TCATAATGAAACTTGTCACGCCTAAACGAACATGAAATTTCTTGTTTTCTTAGGAATCAT   27909
gb:MT066176|Organism:Severe      TCATAATGAAACTTGTCACGCCTAAACGAACATGAAATTTCTTGTTTTCTTAGGAATCAT   27922
gb:MT027064|Organism:Severe      TCATAATGAAACTTGTCACGCCTAAACGAACATGAAATTTCTTGTTTTCTTAGGAATCAT   27922
gb:MN994468|Organism:Severe      TCATAATGAAACTTGTCACGCCTAAACGAACATGAAATTTCTTGTTTTCTTAGGAATCAT   27922
gb:MT072688|Organism:Severe      TCATAATGAAACTTGTCACGCCTAAACGAACATGAAATTTCTTGTTTTCTTAGGAATCAT   27907
gb:MN996527|Organism:Severe      TCATAATGAAACTTGTCACGCCTAAACGAACATGAAATTTCTTGTTTTCTTAGGAATCAT   27889
gb:MT093631|Organism:Severe      TCATAATGAAACTTGTCACGCCTAAACGAACATGAAATTTCTTGTTTTCTTAGGAATCAT   27960
gb:MT106053|Organism:Severe      TCATAATGAAACTTGTCACGCCTAAACGAACATGAAATTTCTTGTTTTCTTAGGAATCAT   27922
gb:MT019533|Organism:Severe      TCATAATGAAACTTGTCACGCCTAAACGAACATGAAATTTCTTGTTTTCTTAGGAATCAT   27922
gb:MT019531|Organism:Severe      TCATAATGAAACTTGTCACGCCTAAACGAACATGAAATTTCTTGTTTTCTTAGGAATCAT   27922
gb:MN996528|Organism:Severe      TCATAATGAAACTTGTCACGCCTAAACGAACATGAAATTTCTTGTTTTCTTAGGAATCAT   27922
gb:MN996530|Organism:Severe      TCATAATGAAACTTGTCACGCCTAAACGAACATGAAATTTCTTGTTTTCTTAGGAATCAT   27908
gb:MN908947|Organism:Severe      TCATAATGAAACTTGTCACGCCTAAACGAACATGAAATTTCTTGTTTTCTTAGGAATCAT   27922
gb:MT019532|Organism:Severe      TCATAATGAAACTTGTCACGCCTAAACGAACATGAAATTTCTTGTTTTCTTAGGAATCAT   27922
                                 ************************************************************

gb:MT020781|Organism:Severe      CACAACTGTAGCTGCATTTCACCAAGAATGTAGTTTACAGTCATGTACTCAACATCAACC   27970
gb:MT007544|Organism:Severe      CACAACTGTAGCTGCATTTCACCAAGAATGTAGTTTACAGTCATGTACTCAACATCAACC   27982
gb:MN994467|Organism:Severe      CACAACTGTAGCTGCATTTCACCAAGAATGTAGTTTACAGTCATGTACTCAACATCAACC   27982
gb:MT044257|Organism:Severe      CACAACTGTAGCTGCATTTCACCAAGAATGTAGTTTACAGTCATGTACTCAACATCAACC   27982
gb:MT106054|Organism:Severe      CATAACTGTAGCTGCATTTCACCAAGAATGTAGTTTACAGTCATGTACTCAACATCAACC   27982
gb:MT049951|Organism:Severe      CACAACTGTAGCTGCATTTCACCAAGAATGTAGTTTACAGTCATGTACTCAACATCAACC   27982
gb:MN975262|Organism:Severe      CACAACTGTAGCTGCATTTCACCAAGAATGTAGTTTACAGTCATGTACTCAACATCAACC   27982
gb:MT106052|Organism:Severe      CACAACTGTAGCTGCATTTCACCAAGAATGTAGTTTACAGTCATGTACTCAACATCAACC   27982
gb:LC522975|Organism:Severe      CACAACTGTAGCTGCATTTCACCAAGAATGTAGTTTACAGTCATGTACTCAACATCAACC   27979
gb:LC522973|Organism:Severe      CACAACTGTAGCTGCATTTCACCAAGAATGTAGTTTACAGTCATGTACTCAACATCAACC   27979
```

```
gb:LC522974|Organism:Severe      CACAACTGTAGCTGCATTTCACCAAGAATGTAGTTTACAGTCATGTACTCAACATCAACC   27979
gb:MN985325|Organism:Severe      CACAACTGTAGCTGCATTTCACCAAGAATGTAGTTTACAGTCATGTACTCAACATCAACC   27982
gb:MT020881|Organism:Severe      CACAACTGTAGCTGCATTTCACCAAGAATGTAGTTTACAGTCATGTACTCAACATCAACC   27982
gb:MT020880|Organism:Severe      CACAACTGTAGCTGCATTTCACCAAGAATGTAGTTTACAGTCATGTACTCAACATCAACC   27982
gb:MT066175|Organism:Severe      CACAACTGTAGCTGCATTTCACCAAGAATGTAGTTTACAGTCATGTACTCAACATCAACC   27982
gb:MN997409|Organism:Severe      CACAACTGTAGCTGCATTTCACCAAGAATGTAGTTTACAGTCATGTACTCAACATCAACC   27982
gb:MN938384|Organism:Severe      CACAACTGTAGCTGCATTTCACCAAGAATGTAGTTTACAGTCATGTACTCAACATCAACC   27950
gb:MT044258|Organism:Severe      CACAACTGTAGCTGCATTTCACCAAGAATGTAGTTTACAGTCATGTACTCAACATCAACC   27958
gb:MT039890|Organism:Severe      CACAACTGTAGCTGCATTTCACCAAGAATGTAGTTTACAGTCATGTACTCAACATCAACC   27982
gb:MN988713|Organism:Severe      CACAACTGTAGCTGCATTTCACCAAGAATGTAGTTTACAGTCATGTACTCAACATCAACC   27982
gb:LC521925|Organism:Severe      CACAACTGTAGCTGCATTTCACCAAGAATGTAGTTTACAGTCATGTACTCAACATCAACC   27955
gb:MT093571|Organism:Severe      CACAACTGTAGCTGCATTTCACCAAGAATGTAGTTTACAGTCATGTACTCAACATCAACC   27982
gb:MT039887|Organism:Severe      CACAACTGTAGCTGCATTTCACCAAGAATGTAGTTTACAGTCATGTACTCAACATCAACC   27979
gb:MT019530|Organism:Severe      CACAACTGTAGCTGCATTTCACCAAGAATGTAGTTTACAGTCATGTACTCAACATCAACC   27982
gb:MT039888|Organism:Severe      CACAACTGTAGCTGCATTTCACCAAGAATGTAGTTTACAGTCATGTACTCAACATCAACC   27982
gb:LC522972|Organism:Severe      CACAACTGTAGCTGCATTTCACCAAGAATGTAGTTTACAGTCATGTACTCAACATCAACC   27979
gb:MT027063|Organism:Severe      CACAACTGTAGCTGCATTTCACCAAGAATGTAGTTTACAGTCATGTACTCAACATCAACC   27982
gb:MT027062|Organism:Severe      CACAACTGTAGCTGCATTTCACCAAGAATGTAGTTTACAGTCATGTACTCAACATCAACC   27982
gb:MT019529|Organism:Severe      CACAACTGTAGCTGCATTTCACCAAGAATGTAGTTTACAGTCATGTACTCAACATCAACC   27982
gb:MN996529|Organism:Severe      CACAACTGTAGCTGCATTTCACCAAGAATGTAGTTTACAGTCATGTACTCAACATCAACC   27970
gb:MN996531|Organism:Severe      CACAACTGTAGCTGCATTTCACCAAGAATGTAGTTTACAGTCATGTACTCAACATCAACC   27969
gb:MT066176|Organism:Severe      CACAACTGTAGCTGCATTTCACCAAGAATGTAGTTTACAGTCATGTACTCAACATCAACC   27982
gb:MT027064|Organism:Severe      CACAACTGTAGCTGCATTTCACCAAGAATGTAGTTTACAGTCATGTACTCAACATCAACC   27982
gb:MN994468|Organism:Severe      CACAACTGTAGCTGCATTTCACCAAGAATGTAGTTTACAGTCATGTACTCAACATCAACC   27982
gb:MT072688|Organism:Severe      CACAACTGTAGCTGCATTTCACCAAGAATGTAGTTTACAGTCATGTACTCAACATCAACC   27967
gb:MN996527|Organism:Severe      CACAACTGTAGCTGCATTTCACCAAGAATGTAGTTTACAGTCATGTACTCAACATCAACC   27949
gb:MT093631|Organism:Severe      CACAACTGTAGCTGCATTTCACCAAGAATGTAGTTTACAGTCATGTACTCAACATCAACC   28020
gb:MT106053|Organism:Severe      CACAACTGTAGCTGCATTTCACCAAGAATGTAGTTTACAGTCATGTACTCAACATCAACC   27982
gb:MT019533|Organism:Severe      CACAACTGTAGCTGCATTTCACCAAGAATGTAGTTTACAGTCATGTACTCAACATCAACC   27982
gb:MT019531|Organism:Severe      CACAACTGTAGCTGCATTTCACCAAGAATGTAGTTTACAGTCATGTACTCAACATCAACC   27982
gb:MN996528|Organism:Severe      CACAACTGTAGCTGCATTTCACCAAGAATGTAGTTTACAGTCATGTACTCAACATCAACC   27982
gb:MN996530|Organism:Severe      CACAACTGTAGCTGCATTTCACCAAGAATGTAGTTTACAGTCATGTACTCAACATCAACC   27968
gb:MN908947|Organism:Severe      CACAACTGTAGCTGCATTTCACCAAGAATGTAGTTTACAGTCATGTACTCAACATCAACC   27982
gb:MT019532|Organism:Severe      CACAACTGTAGCTGCATTTCACCAAGAATGTAGTTTACAGTCATGTACTCAACATCAACC   27982
                                 ** *********************************************************

gb:MT020781|Organism:Severe      ATATGTAGTTGATGACCCGTGTCCTATTCACTTCTATTCTAAATGGTATATTAGAGTAGG   28030
gb:MT007544|Organism:Severe      ATATGTAGTTGATGACCCGTGTCCTATTCACTTCTATTCTAAATGGTATATTAGAGTAGG   28042
gb:MN994467|Organism:Severe      ATATGTAGTTGATGACCCGTGTCCTATTCACTTCTATTCTAAATGGTATATTAGAGTAGG   28042
gb:MT044257|Organism:Severe      ATATGTAGTTGATGACCCGTGTCCTATTCACTTCTATTCTAAATGGTATATTAGAGTAGG   28042
gb:MT106054|Organism:Severe      ATATGTAGTTGATGACCCGTGTCCTATTCACTTCTATTCTAAATGGTATATTAGAGTAGG   28042
gb:MT049951|Organism:Severe      ATATGTAGTTGATGACCCGTGTCCTATTCACTTCTATTCTAAATGGTATATTAGAGTAGG   28042
gb:MN975262|Organism:Severe      ATATGTAGTTGATGACCCGTGTCCTATTCACTTCTATTCTAAATGGTATATTAGAGTAGG   28042
gb:MT106052|Organism:Severe      ATATGTAGTTGATGACCCGTGTCCTATTCACTTCTATTCTAAATGGTATATTAGAGTAGG   28042
gb:LC522975|Organism:Severe      ATATGTAGTTGATGACCCGTGTCCTATTCACTTCTATTCTAAATGGTATATTAGAGTAGG   28039
gb:LC522973|Organism:Severe      ATATGTAGTTGATGACCCGTGTCCTATTCACTTCTATTCTAAATGGTATATTAGAGTAGG   28039
gb:LC522974|Organism:Severe      ATATGTAGTTGATGACCCGTGTCCTATTCACTTCTATTCTAAATGGTATATTAGAGTAGG   28039
gb:MN985325|Organism:Severe      ATATGTAGTTGATGACCCGTGTCCTATTCACTTCTATTCTAAATGGTATATTAGAGTAGG   28042
gb:MT020881|Organism:Severe      ATATGTAGTTGATGACCCGTGTCCTATTCACTTCTATTCTAAATGGTATATTAGAGTAGG   28042
gb:MT020880|Organism:Severe      ATATGTAGTTGATGACCCGTGTCCTATTCACTTCTATTCTAAATGGTATATTAGAGTAGG   28042
gb:MT066175|Organism:Severe      ATATGTAGTTGATGACCCGTGTCCTATTCACTTCTATTCTAAATGGTATATTAGAGTAGG   28042
gb:MN997409|Organism:Severe      ATATGTAGTTGATGACCCGTGTCCTATTCACTTCTATTCTAAATGGTATATTAGAGTAGG   28042
gb:MN938384|Organism:Severe      ATATGTAGTTGATGACCCGTGTCCTATTCACTTCTATTCTAAATGGTATATTAGAGTAGG   28010
gb:MT044258|Organism:Severe      ATATGTAGTTGATGACCCGTGTCCTATTCACTTCTATTCTAAATGGTATATTAGAGTAGG   28018
gb:MT039890|Organism:Severe      ATATGTAGTTGATGACCCGTGTCCTATTCACTTCTATTCTAAATGGTATATTAGAGTAGG   28042
gb:MN988713|Organism:Severe      ATATGTAGTTGATGACCCGTGTCCTATTCACTTCTATTCTAAATGGTATATTAGAGTAGG   28042
gb:LC521925|Organism:Severe      ATATGTAGTTGATGACCCGTGTCCTATTCACTTCTATTCTAAATGGTATATTAGAGTAGG   28015
gb:MT093571|Organism:Severe      ATATGTAGTTGATGACCCGTGTCCTATTCACTTCTATTCTAAATGGTATATTAGAGTAGG   28042
gb:MT039887|Organism:Severe      ATATGTAGTTGATGACCCGTGTCCTATTCACTTCTATTCTAAATGGTATATTAGAGTAGG   28039
gb:MT019530|Organism:Severe      ATATGTAGTTGATGACCCGTGTCCTATTCACTTCTATTCTAAATGGTATATTAGAGTAGG   28042
gb:MT039888|Organism:Severe      ATATGTAGTTGATGACCCGTGTCCTATTCACTTCTATTCTAAATGGTATATTAGAGTAGG   28042
gb:LC522972|Organism:Severe      ATATGTAGTTGATGACCCGTGTCCTATTCACTTCTATTCTAAATGGTATATTAGAGTAGG   28039
gb:MT027063|Organism:Severe      ATATGTAGTTGATGACCCGTGTCCTATTCACTTCTATTCTAAATGGTATATTAGAGTAGG   28042
gb:MT027062|Organism:Severe      ATATGTAGTTGATGACCCGTGTCCTATTCACTTCTATTCTAAATGGTATATTAGAGTAGG   28042
gb:MT019529|Organism:Severe      ATATGTAGTTGATGACCCGTGTCCTATTCACTTCTATTCTAAATGGTATATTAGAGTAGG   28042
```

```
gb:MN996529|Organism:Severe      ATATGTAGTTGATGACCCGTGTCCTATTCACTTCTATTCTAAATGGTATATTAGAGTAGG   28030
gb:MN996531|Organism:Severe      ATATGTAGTTGATGACCCGTGTCCTATTCACTTCTATTCTAAATGGTATATTAGAGTAGG   28029
gb:MT066176|Organism:Severe      ATATGTAGTTGATGACCCGTGTCCTATTCACTTCTATTCTAAATGGTATATTAGAGTAGG   28042
gb:MT027064|Organism:Severe      ATATGTAGTTGATGACCCGTGTCCTATTCACTTCTATTCTAAATGGTATATTAGAGTAGG   28042
gb:MN994468|Organism:Severe      ATATGTAGTTGATGACCCGTGTCCTATTCACTTCTATTCTAAATGGTATATTAGAGTAGG   28042
gb:MT072688|Organism:Severe      ATATGTAGTTGATGACCCGTGTCCTATTCACTTCTATTCTAAATGGTATATTAGAGTAGG   28027
gb:MN996527|Organism:Severe      ATATGTAGTTGATGACCCGTGTCCTATTCACTTCTATTCTAAATGGTATATTAGAGTAGG   28009
gb:MT093631|Organism:Severe      ATATGTAGTTGATGACCCGTGTCCTATTCACTTCTATTCTAAATGGTATATTAGAGTAGG   28080
gb:MT106053|Organism:Severe      ATATGTAGTTGATGACCCGTGTCCTATTCACTTCTATTCTAAATGGTATATTAGAGTAGG   28042
gb:MT019533|Organism:Severe      ATATGTAGTTGATGACCCGTGTCCTATTCACTTCTATTCTAAATGGTATATTAGAGTAGG   28042
gb:MT019531|Organism:Severe      ATATGTAGTTGATGACCCGTGTCCTATTCACTTCTATTCTAAATGGTATATTAGAGTAGG   28042
gb:MN996528|Organism:Severe      ATATGTAGTTGATGACCCGTGTCCTATTCACTTCTATTCTAAATGGTATATTAGAGTAGG   28042
gb:MN996530|Organism:Severe      ATATGTAGTTGATGACCCGTGTCCTATTCACTTCTATTCTAAATGGTATATTAGAGTAGG   28028
gb:MN908947|Organism:Severe      ATATGTAGTTGATGACCCGTGTCCTATTCACTTCTATTCTAAATGGTATATTAGAGTAGG   28042
gb:MT019532|Organism:Severe      ATATGTAGTTGATGACCCGTGTCCTATTCACTTCTATTCTAAATGGTATATTAGAGTAGG   28042
                                 ************************************************************

gb:MT020781|Organism:Severe      AGCTAGAAAATCAGCACCTTTAATTGAATTGTGCGTGGATGAGGCTGGTTCTAAATCACC   28090
gb:MT007544|Organism:Severe      AGCTAGAAAATCAGCACCTTTAATTGAATTGTGCGTGGATGAGGCTGGTTCTAAATCACC   28102
gb:MN994467|Organism:Severe      AGCTAGAAAATCAGCACCTTTAATTGAATTGTGCCTGGATGAGGCTGGTTCTAAATCACC   28102
gb:MT044257|Organism:Severe      AGCTAGAAAATCAGCACCTTTAATTGAATTGTGCCTGGATGAGGCTGGTTCTAAATCACC   28102
gb:MT106054|Organism:Severe      AGCTAGAAAATCAGCACCTTTAATTGAATTGTGCGTGGATGAGGCTGGTTCTAAATCACC   28102
gb:MT049951|Organism:Severe      AGCTAGAAAATCAGCACCTTTAATTGAATTGTGCGTGGATGAGGCTGGTTCTAAATCACC   28102
gb:MN975262|Organism:Severe      AGCTAGAAAATCAGCACCTTTAATTGAATTGTGCGTGGATGAGGCTGGTTCTAAATCACC   28102
gb:MT106052|Organism:Severe      AGCTAGAAAATCAGCACCTTTAATTGAATTGTGCGTGGATGAGGCTGGTTCTAAATCACC   28102
gb:LC522975|Organism:Severe      AGCTAGAAAATCAGCACCTTTAATTGAATTGTGCGTGGATGAGGCTGGTTCTAAATCACC   28099
gb:LC522973|Organism:Severe      AGCTAGAAAATCAGCACCTTTAATTGAATTGTGCGTGGATGAGGCTGGTTCTAAATCACC   28099
gb:LC522974|Organism:Severe      AGCTAGAAAATCAGCACCTTTAATTGAATTGTGCGTGGATGAGGCTGGTTCTAAATCACC   28099
gb:MN985325|Organism:Severe      AGCTAGAAAATCAGCACCTTTAATTGAATTGTGCGTGGATGAGGCTGGTTCTAAATCACC   28102
gb:MT020881|Organism:Severe      AGCTAGAAAATCAGCACCTTTAATTGAATTGTGCGTGGATGAGGCTGGTTCTAAATCACC   28102
gb:MT020880|Organism:Severe      AGCTAGAAAATCAGCACCTTTAATTGAATTGTGCGTGGATGAGGCTGGTTCTAAATCACC   28102
gb:MT066175|Organism:Severe      AGCTAGAAAATCAGCACCTTTAATTGAATTGTGCGTGGATGAGGCTGGTTCTAAATCACC   28102
gb:MN997409|Organism:Severe      AGCTAGAAAATCAGCACCTTTAATTGAATTGTGCGTGGATGAGGCTGGTTCTAAATCACC   28102
gb:MN938384|Organism:Severe      AGCTAGAAAATCAGCACCTTTAATTGAATTGTGCGTGGATGAGGCTGGTTCTAAATCACC   28070
gb:MT044258|Organism:Severe      AGCTAGAAAATCAGCACCTTTAATTGAATTGTGCGTGGATGAGGCTGGTTCTAAATCACC   28078
gb:MT039890|Organism:Severe      AGCTAGAAAATCAGCACCTTTAATTGAATTGTGCGTGGATGAGGCTGGTTCTAAATCACC   28102
gb:MN988713|Organism:Severe      AGCTAGAAAATCAGCACCTTTAATTGAATTGTGCSTGGATGAGGCTGGTTCTAAATCACC   28102
gb:LC521925|Organism:Severe      AGCTAGAAAATCAGCACCTTTAATTGAATTGTGCGTGGATGAGGCTGGTTCTAAATCACC   28075
gb:MT093571|Organism:Severe      AGCTAGAAAATCAGCACCTTTAATTGAATTGTGCGTGGATGAGGCTGGTTCTAAATCACC   28102
gb:MT039887|Organism:Severe      AGCTAGAAAATCAGCACCTTTAATTGAATTGTGCGTGGATGAGGCTGGTTCTAAATCACC   28099
gb:MT019530|Organism:Severe      AGCTAGAAAATCAGCACCTTTAATTGAATTGTGCGTGGATGAGGCTGGTTCTAAATCACC   28102
gb:MT039888|Organism:Severe      AGCTAGAAAATCAGCACCTTTAATTGAATTGTGCGTGGATGAGGCTGGTTCTAAATCACC   28102
gb:LC522972|Organism:Severe      AGCTAGAAAATCAGCACCTTTAATTGAATTGTGCGTGGATGAGGCTGGTTCTAAATCACC   28099
gb:MT027063|Organism:Severe      AGCTAGAAAATCAGCACCTTTAATTGAATTGTGCGTGGATGAGGCTGGTTCTAAATCACC   28102
gb:MT027062|Organism:Severe      AGCTAGAAAATCAGCACCTTTAATTGAATTGTGCGTGGATGAGGCTGGTTCTAAATCACC   28102
gb:MT019529|Organism:Severe      AGCTAGAAAATCAGCACCTTTAATTGAATTGTGCGTGGATGAGGCTGGTTCTAAATCACC   28102
gb:MN996529|Organism:Severe      AGCTAGAAAATCAGCACCTTTAATTGAATTGTGCGTGGATGAGGCTGGTTCTAAATCACC   28090
gb:MN996531|Organism:Severe      AGCTAGAAAATCAGCACCTTTAATTGAATTGTGCGTGGATGAGGCTGGTTCTAAATCACC   28089
gb:MT066176|Organism:Severe      AGCTAGAAAATCAGCACCTTTAATTGAATTGTGCGTGGATGAGGCTGGTTCTAAATCACC   28102
gb:MT027064|Organism:Severe      AGCTAGAAAATCAGCACCTTTAATTGAATTGTGCGTGGATGAGGCTGGTTCTAAATCACC   28102
gb:MN994468|Organism:Severe      AGCTAGAAAATCAGCACCTTTAATTGAATTGTGCGTGGATGAGGCTGGTTCTAAATCACC   28102
gb:MT072688|Organism:Severe      AGCTAGAAAATCAGCACCTTTAATTGAATTGTGCGTGGATGAGGCTGGTTCTAAATCACC   28087
gb:MN996527|Organism:Severe      AGCTAGAAAATCAGCACCTTTAATTGAATTGTGCGTGGATGAGGCTGGTTCTAAATCACC   28069
gb:MT093631|Organism:Severe      AGCTAGAAAATCAGCACCTTTAATTGAATTGTGCGTGGATGAGGCTGGTTCTAAATCACC   28140
gb:MT106053|Organism:Severe      AGCTAGAAAATCAGCACCTTTAATTGAATTGTGCGTGGATGAGGCTGGTTCTAAATCACC   28102
gb:MT019533|Organism:Severe      AGCTAGAAAATCAGCACCTTTAATTGAATTGTGCGTGGATGAGGCTGGTTCTAAATCACC   28102
gb:MT019531|Organism:Severe      AGCTAGAAAATCAGCACCTTTAATTGAATTGTGCGTGGATGAGGCTGGTTCTAAATCACC   28102
gb:MN996528|Organism:Severe      AGCTAGAAAATCAGCACCTTTAATTGAATTGTGCGTGGATGAGGCTGGTTCTAAATCACC   28102
gb:MN996530|Organism:Severe      AGCTAGAAAATCAGCACCTTTAATTGAATTGTGCGTGGATGAGGCTGGTTCTAAATCACC   28088
gb:MN908947|Organism:Severe      AGCTAGAAAATCAGCACCTTTAATTGAATTGTGCGTGGATGAGGCTGGTTCTAAATCACC   28102
gb:MT019532|Organism:Severe      AGCTAGAAAATCAGCACCTTTAATTGAATTGTGCGTGGATGAGGCTGGTTCTAAATCACC   28102
                                 ********************************** *************************

gb:MT020781|Organism:Severe      CATTCAGTACATCGATATCGGTAATTATACAGTTTCCTGTTTACCTTTTACAATTAATTG   28150
gb:MT007544|Organism:Severe      CATTCAGTACATCGATATCGGTAATTATACAGTTTCCTGTTTACCTTTTACAATTAATTG   28162
```

```
gb:MN994467|Organism:Severe      CATTCAGTACATCGATATCGGTAATTATACAGTTTCCTGTTCACCTTTTACAATTAATTG   28162
gb:MT044257|Organism:Severe      CATTCAGTACATCGATATCGGTAATTATACAGTTTCCTGTTCACCTTTTACAATTAATTG   28162
gb:MT106054|Organism:Severe      CATTCAGTACATCGATATCGGTAATTATACAGTTTCCTGTTCACCTTTTACAATTAATTG   28162
gb:MT049951|Organism:Severe      CATTCAGTACATCGATATCGGTAATTATACAGTTTCCTGTTCACCTTTTACAATTAATTG   28162
gb:MN975262|Organism:Severe      CATTCAGTACATCGATATCGGTAATTATACAGTTTCCTGTTCACCTTTTACAATTAATTG   28162
gb:MT106052|Organism:Severe      CATTCAGTACATCGATATCGGTAATTATACAGTTTCCTGTTCACCTTTTACAATTAATTG   28162
gb:LC522975|Organism:Severe      CATTCAGTACATCGATATCGGTAATTATACAGTTTCCTGTTCACCTTTTACAATTAATTG   28159
gb:LC522973|Organism:Severe      CATTCAGTACATCGATATCGGTAATTATACAGTTTCCTGTTCACCTTTTACAATTAATTG   28159
gb:LC522974|Organism:Severe      CATTCAGTACATCGATATCGGTAATTATACAGTTTCCTGTTCACCTTTTACAATTAATTG   28159
gb:MN985325|Organism:Severe      CATTCAGTACATCGATATCGGTAATTATACAGTTTCCTGTTCACCTTTTACAATTAATTG   28162
gb:MT020881|Organism:Severe      CATTCAGTACATCGATATCGGTAATTATACAGTTTCCTGTTCACCTTTTACAATTAATTG   28162
gb:MT020880|Organism:Severe      CATTCAGTACATCGATATCGGTAATTATACAGTTTCCTGTTCACCTTTTACAATTAATTG   28162
gb:MT066175|Organism:Severe      CATTCAGTACATCGATATCGGTAATTATACAGTTTCCTGTTCACCTTTTACAATTAATTG   28162
gb:MN997409|Organism:Severe      CATTCAGTACATCGATATCGGTAATTATACAGTTTCCTGTTCACCTTTTACAATTAATTG   28162
gb:MN938384|Organism:Severe      CATTCAGTACATCGATATCGGTAATTATACAGTTTCCTGTTCACCTTTTACAATTAATTG   28130
gb:MT044258|Organism:Severe      CATTCAGTACATCGATATCGGTAATTATACAGTTTCCTGTTTACCTTTTACAATTAATTG   28138
gb:MT039890|Organism:Severe      CATTCAGTACATCGATATCGGTAATTATACAGTTTCCTGTTTACCTTTTACAATTAATTG   28162
gb:MN988713|Organism:Severe      CATTCAGTACATCGATATCGGTAATTATACAGTTTCCTGTTYACCTTTTACAATTAATTG   28162
gb:LC521925|Organism:Severe      CATTCAGTACATCGATATCGGTAATTATACAGTTTCCTGTTTACCTTTTACAATTAATTG   28135
gb:MT093571|Organism:Severe      CATTCAGTACATCGATATCGGTAATTATACAGTTTCCTGTTTACCTTTTACAATTAATTG   28162
gb:MT039887|Organism:Severe      CATTCAGTACATCGATATCGGTAATTATACAGTTTCCTGTTTACCTTTTACAATTAATTG   28159
gb:MT019530|Organism:Severe      CATTCAGTACATCGATATCGGTAATTATACAGTTTCCTGTTTACCTTTTACAATTAATTG   28162
gb:MT039888|Organism:Severe      CATTCAGTACATCGATATCGGTAATTATACAGTTTCCTGTTTACCTTTTACAATTAATTG   28162
gb:LC522972|Organism:Severe      CATTCAGTACATCGATATCGGTAATTATACAGTTTCCTGTTTACCTTTTACAATTAATTG   28159
gb:MT027063|Organism:Severe      CATTCAGTACATCGATATCGGTAATTATACAGTTTCCTGTTTACCTTTTACAATTAATTG   28162
gb:MT027062|Organism:Severe      CATTCAGTACATCGATATCGGTAATTATACAGTTTCCTGTTTACCTTTTACAATTAATTG   28162
gb:MT019529|Organism:Severe      CATTCAGTACATCGATATCGGTAATTATACAGTTTCCTGTTTACCTTTTACAATTAATTG   28162
gb:MN996529|Organism:Severe      CATTCAGTACATCGATATCGGTAATTATACAGTTTCCTGTTTACCTTTTACAATTAATTG   28150
gb:MN996531|Organism:Severe      CATTCAGTACATCGATATCGGTAATTATACAGTTTCCTGTTTACCTTTTACAATTAATTG   28149
gb:MT066176|Organism:Severe      CATTCAGTACATCGATATCGGTAATTATACAGTTTCCTGTTTACCTTTTACAATTAATTG   28162
gb:MT027064|Organism:Severe      CATTCAGTACATCGATATCGGTAATTATACAGTTTCCTGTTTACCTTTTACAATTAATTG   28162
gb:MN994468|Organism:Severe      CATTCAGTACATCGATATCGGTAATTATACAGTTTCCTGTTTACCTTTTACAATTAATTG   28162
gb:MT072688|Organism:Severe      CATTCAGTACATCGATATCGGTAATTATACAGTTTCCTGTTTACCTTTTACAATTAATTG   28147
gb:MN996527|Organism:Severe      CATTCAGTACATCGATATCGGTAATTATACAGTTTCCTGTTTACCTTTTACAATTAATTG   28129
gb:MT093631|Organism:Severe      CATTCAGTACATCGATATCGGTAATTATACAGTTTCCTGTTTACCTTTTACAATTAATTG   28200
gb:MT106053|Organism:Severe      CATTCAGTACATCGATATCGGTAATTATACAGTTTCCTGTTTACCTTTTACAATTAATTG   28162
gb:MT019533|Organism:Severe      CATTCAGTACATCGATATCGGTAATTATACAGTTTCCTGTTTACCTTTTACAATTAATTG   28162
gb:MT019531|Organism:Severe      CATTCAGTACATCGATATCGGTAATTATACAGTTTCCTGTTTACCTTTTACAATTAATTG   28162
gb:MN996528|Organism:Severe      CATTCAGTACATCGATATCGGTAATTATACAGTTTCCTGTTTACCTTTTACAATTAATTG   28162
gb:MN996530|Organism:Severe      CATTCAGTACATCGATATCGGTAATTATACAGTTTCCTGTTTACCTTTTACAATTAATTG   28148
gb:MN908947|Organism:Severe      CATTCAGTACATCGATATCGGTAATTATACAGTTTCCTGTTTACCTTTTACAATTAATTG   28162
gb:MT019532|Organism:Severe      CATTCAGTACATCGATATCGGTAATTATACAGTTTCCTGTTTACCTTTTACAATTAATTG   28162
                                 ***************************************** ******************

gb:MT020781|Organism:Severe      CCAGGAACCTAAATTGGGTAGTCTTGTAGTGCGTTGTTCGTTCTATGAAGACTTTTTAGA   28210
gb:MT007544|Organism:Severe      CCAGGAACCTAAATTGGGTAGTCTTGTAGTGCGTTGTTCGTTCTATGAAGACTTTTTAGA   28222
gb:MN994467|Organism:Severe      CCAGGAACCTAAATTGGGTAGTCTTGTAGTGCGTTGTTCGTTCTATGAAGACTTTTTAGA   28222
gb:MT044257|Organism:Severe      CCAGGAACCTAAATTGGGTAGTCTTGTAGTGCGTTGTTCGTTCTATGAAGACTTTTTAGA   28222
gb:MT106054|Organism:Severe      CCAGGAACCTAAATTGGGTAGTCTTGTAGTGCGTTGTTCGTTCTATGAAGACTTTTTAGA   28222
gb:MT049951|Organism:Severe      CCAGGAACCTAAATTGGGTAGTCTTGTAGTGCGTTGTTCGTTCTATGAAGACTTTTTAGA   28222
gb:MN975262|Organism:Severe      CCAGGAACCTAAATTGGGTAGTCTTGTAGTGCGTTGTTCGTTCTATGAAGACTTTTTAGA   28222
gb:MT106052|Organism:Severe      CCAGGAACCTAAATTGGGTAGTCTTGTAGTGCGTTGTTCGTTCTATGAAGACTTTTTAGA   28222
gb:LC522975|Organism:Severe      CCAGGAACCTAAATTGGGTAGTCTTGTAGTGCGTTGTTCGTTCTATGAAGACTTTTTAGA   28219
gb:LC522973|Organism:Severe      CCAGGAACCTAAATTGGGTAGTCTTGTAGTGCGTTGTTCGTTCTATGAAGACTTTTTAGA   28219
gb:LC522974|Organism:Severe      CCAGGAACCTAAATTGGGTAGTCTTGTAGTGCGTTGTTCGTTCTATGAAGACTTTTTAGA   28219
gb:MN985325|Organism:Severe      CCAGGAACCTAAATTGGGTAGTCTTGTAGTGCGTTGTTCGTTCTATGAAGACTTTTTAGA   28222
gb:MT020881|Organism:Severe      CCAGGAACCTAAATTGGGTAGTCTTGTAGTGCGTTGTTCGTTCTATGAAGACTTTTTAGA   28222
gb:MT020880|Organism:Severe      CCAGGAACCTAAATTGGGTAGTCTTGTAGTGCGTTGTTCGTTCTATGAAGACTTTTTAGA   28222
gb:MT066175|Organism:Severe      CCAGGAACCTAAATTGGGTAGTCTTGTAGTGCGTTGTTCGTTCTATGAAGACTTTTTAGA   28222
gb:MN997409|Organism:Severe      CCAGGAACCTAAATTGGGTAGTCTTGTAGTGCGTTGTTCGTTCTATGAAGACTTTTTAGA   28222
gb:MN938384|Organism:Severe      CCAGGAACCTAAATTGGGTAGTCTTGTAGTGCGTTGTTCGTTCTATGAAGACTTTTTAGA   28190
gb:MT044258|Organism:Severe      CCAGGAACCTAAATTGGGTAGTCTTGTAGTGCGTTGTTCGTTCTATGAAGACTTTTTAGA   28198
gb:MT039890|Organism:Severe      CCAGGAACCTAAATTGGGTAGTCTTGTAGTGCGTTGTTCGTTCTATGAAGACTTTTTAGA   28222
gb:MN988713|Organism:Severe      CCAGGAACCTAAATTGGGTAGTCTTGTAGTGCGTTGTTCGTTCTATGAAGACTTTTTAGA   28222
gb:LC521925|Organism:Severe      CCAGGAACCTAAATTGGGTAGTCTTGTAGTGCGTTGTTCGTTCTATGAAGACTTTTTAGA   28195
```

```
gb:MT093571|Organism:Severe      CCAGGAACCTAAATTGGGTAGTCTTGTAGTGCGTTGTTCGTTCTATGAAGACTTTTTAGA    28222
gb:MT039887|Organism:Severe      CCAGGAACCTAAATTGGGTAGTCTTGTAGTGCGTTGTTCGTTCTATGAAGACTTTTTAGA    28219
gb:MT019530|Organism:Severe      CCAGGAACCTAAATTGGGTAGTCTTGTAGTGCGTTGTTCGTTCTATGAAGACTTTTTAGA    28222
gb:MT039888|Organism:Severe      CCAGGAACCTAAATTGGGTAGTCTTGTAGTGCGTTGTTCGTTCTATGAAGACTTTTTAGA    28222
gb:LC522972|Organism:Severe      CCAGGAACCTAAATTGGGTAGTCTTGTAGTGCGTTGTTCGTTCTATGAAGACTTTTTAGA    28219
gb:MT027063|Organism:Severe      CCAGGAACCTAAATTGGGTAGTCTTGTAGTGCGTTGTTCGTTCTATGAAGACTTTTTAGA    28222
gb:MT027062|Organism:Severe      CCAGGAACCTAAATTGGGTAGTCTTGTAGTGCGTTGTTCGTTCTATGAAGACTTTTTAGA    28222
gb:MT019529|Organism:Severe      CCAGGAACCTAAATTGGGTAGTCTTGTAGTGCGTTGTTCGTTCTATGAAGACTTTTTAGA    28222
gb:MN996529|Organism:Severe      CCAGGAACCTAAATTGGGTAGTCTTGTAGTGCGTTGTTCGTTCTATGAAGACTTTTTAGA    28210
gb:MN996531|Organism:Severe      CCAGGAACCTAAATTGGGTAGTCTTGTAGTGCGTTGTTCGTTCTATGAAGACTTTTTAGA    28209
gb:MT066176|Organism:Severe      CCAGGAACCTAAATTGGGTAGTCTTGTAGTGCGTTGTTCGTTCTATGAAGACTTTTTAGA    28222
gb:MT027064|Organism:Severe      CCAGGAACCTAAATTGGGTAGTCTTGTAGTGCGTTGTTCGTTCTATGAAGACTTTTTAGA    28222
gb:MN994468|Organism:Severe      CCAGGAACCTAAATTGGGTAGTCTTGTAGTGCGTTGTTCGTTCTATGAAGACTTTTTAGA    28222
gb:MT072688|Organism:Severe      CCAGGAACCTAAATTGGGTAGTCTTGTAGTGCGTTGTTCGTTCTATGAAGACTTTTTAGA    28207
gb:MN996527|Organism:Severe      CCAGGAACCTAAATTGGGTAGTCTTGTAGTGCGTTGTTCGTTCTATGAAGACTTTTTAGA    28189
gb:MT093631|Organism:Severe      CCAGGAACCTAAATTGGGTAGTCTTGTAGTGCGTTGTTCGTTCTATGAAGACTTTTTAGA    28260
gb:MT106053|Organism:Severe      CCAGGAACCTAAATTGGGTAGTCTTGTAGTGCGTTGTTCGTTCTATGAAGACTTTTTAGA    28222
gb:MT019533|Organism:Severe      CCAGGAACCTAAATTGGGTAGTCTTGTAGTGCGTTGTTCGTTCTATGAAGACTTTTTAGA    28222
gb:MT019531|Organism:Severe      CCAGGAACCTAAATTGGGTAGTCTTGTAGTGCGTTGTTCGTTCTATGAAGACTTTTTAGA    28222
gb:MN996528|Organism:Severe      CCAGGAACCTAAATTGGGTAGTCTTGTAGTGCGTTGTTCGTTCTATGAAGACTTTTTAGA    28222
gb:MN996530|Organism:Severe      CCAGGAACCTAAATTGGGTAGTCTTGTAGTGCGTTGTTCGTTCTATGAAGACTTTTTAGA    28208
gb:MN908947|Organism:Severe      CCAGGAACCTAAATTGGGTAGTCTTGTAGTGCGTTGTTCGTTCTATGAAGACTTTTTAGA    28222
gb:MT019532|Organism:Severe      CCAGGAACCTAAATTGGGTAGTCTTGTAGTGCGTTGTTCGTTCTATGAAGACTTTTTAGA    28222
                                 ************************************************************

gb:MT020781|Organism:Severe      GTATCATGACGTTCGTGTTGTTTTAGATTTCATCTAAACGAACAAACTAAAATGTCTGAT    28270
gb:MT007544|Organism:Severe      GTATCATGACGTTCGTGTTGTTTTAGATTTCATCTAAACGAACAAACTAAAATGTCTGAT    28282
gb:MN994467|Organism:Severe      GTATCATGACGTTCGTGTTGTTTTAGATTTCATCTAAACGAACAAACTAAAATGTCTGAT    28282
gb:MT044257|Organism:Severe      GTATCATGACGTTCGTGTTGTTTTAGATTTCATCTAAACGAACAAACTAAAATGTCTGAT    28282
gb:MT106054|Organism:Severe      GTATCATGACGTTCGTGTTGTTTTAGATTTCATCTAAACGAACAAACTAAAATGTCTGAT    28282
gb:MT049951|Organism:Severe      GTATCATGACGTTCGTGTTGTTTTAGATTTCATCTAAACGAACAAACTAAAATGTCTGAT    28282
gb:MN975262|Organism:Severe      GTATCATGACGTTCGTGTTGTTTTAGATTTCATCTAAACGAACAAACTAAAATGTCTGAT    28282
gb:MT106052|Organism:Severe      GTATCATGACGTTCGTGTTGTTTTAGATTTCATCTAAACGAACAAACTAAAATGTCTGAT    28282
gb:LC522975|Organism:Severe      GTATCATGACGTTCGTGTTGTTTTAGATTTCATCTAAACGAACAAACTAAAATGTCTGAT    28279
gb:LC522973|Organism:Severe      GTATCATGACGTTCGTGTTGTTTTAGATTTCATCTAAACGAACAAACTAAAATGTCTGAT    28279
gb:LC522974|Organism:Severe      GTATCATGACGTTCGTGTTGTTTTAGATTTCATCTAAACGAACAAACTAAAATGTCTGAT    28279
gb:MN985325|Organism:Severe      GTATCATGACGTTCGTGTTGTTTTAGATTTCATCTAAACGAACAAACTAAAATGTCTGAT    28282
gb:MT020881|Organism:Severe      GTATCATGACGTTCGTGTTGTTTTAGATTTCATCTAAACGAACAAACTAAAATGTCTGAT    28282
gb:MT020880|Organism:Severe      GTATCATGACGTTCGTGTTGTTTTAGATTTCATCTAAACGAACAAACTAAAATGTCTGAT    28282
gb:MT066175|Organism:Severe      GTATCATGACGTTCGTGTTGTTTTAGATTTCATCTAAACGAACAAACTAAAATGTCTGAT    28282
gb:MN997409|Organism:Severe      GTATCATGACGTTCGTGTTGTTTTAGATTTCATCTAAACGAACAAACTAAAATGTCTGAT    28282
gb:MN938384|Organism:Severe      GTATCATGACGTTCGTGTTGTTTTAGATTTCATCTAAACGAACAAACTAAAATGTCTGAT    28250
gb:MT044258|Organism:Severe      GTATCATGACGTTCGTGTTGTTTTAGATTTCATCTAAACGAACAAACTAAAATGTCTGAT    28258
gb:MT039890|Organism:Severe      GTATCATGACGTTCGTGTTGTTTTAGATTTCATCTAAACGAACAAACTAAAATGTCTGAT    28282
gb:MN988713|Organism:Severe      GTATCATGACGTTCGTGTTGTTTTAGATTTCATCTAAACGAACAAACTAAAATGTCTGAT    28282
gb:LC521925|Organism:Severe      GTATCATGACGTTCGTGTTGTTTTAGATTTCATCTAAACGAACAAACTAAAATGTCTGAT    28255
gb:MT093571|Organism:Severe      GTATCATGACGTTCGTGTTGTTTTAGATTTCATCTAAACGAACAAACTAAAATGTCTGAT    28282
gb:MT039887|Organism:Severe      GTATCATGACGTTCGTGTTGTTTTAGATTTCATCTAAACGAACAAACTAAAATGTCTGAT    28279
gb:MT019530|Organism:Severe      GTATCATGACGTTCGTGTTGTTTTAGATTTCATCTAAACGAACAAACTAAAATGTCTGAT    28282
gb:MT039888|Organism:Severe      GTATCATGACGTTCGTGTTGTTTTAGATTTCATCTAAACGAACAAACTAAAATGTCTGAT    28282
gb:LC522972|Organism:Severe      GTATCATGACGTTCGTGTTGTTTTAGATTTCATCTAAACGAACAAACTAAAATGTCTGAT    28279
gb:MT027063|Organism:Severe      GTATCATGACGTTCGTGTTGTTTTAGATTTCATCTAAACGAACAAACTAAAATGTCTGAT    28282
gb:MT027062|Organism:Severe      GTATCATGACGTTCGTGTTGTTTTAGATTTCATCTAAACGAACAAACTAAAATGTCTGAT    28282
gb:MT019529|Organism:Severe      GTATCATGACGTTCGTGTTGTTTTAGATTTCATCTAAACGAACAAACTAAAATGTCTGAT    28282
gb:MN996529|Organism:Severe      GTATCATGACGTTCGTGTTGTTTTAGATTTCATCTAAACGAACAAACTAAAATGTCTGAT    28270
gb:MN996531|Organism:Severe      GTATCATGACGTTCGTGTTGTTTTAGATTTCATCTAAACGAACAAACTAAAATGTCTGAT    28269
gb:MT066176|Organism:Severe      GTATCATGACGTTCGTGTTGTTTTAGATTTCATCTAAACGAACAAACTAAAATGTCTGAT    28282
gb:MT027064|Organism:Severe      GTATCATGACGTTCGTGTTGTTTTAGATTTCATCTAAACGAACAAACTAAAATGTCTGAT    28282
gb:MN994468|Organism:Severe      GTATCATGACGTTCGTGTTGTTTTAGATTTCATCTAAACGAACAAACTAAAATGTCTGAT    28282
gb:MT072688|Organism:Severe      GTATCATGACGTTCGTGTTGTTTTAGATTTCATCTAAACGAACAAACTAAAATGTCTGAT    28267
gb:MN996527|Organism:Severe      GTATCATGACGTTCGTGTTGTTTTAGATTTCATCTAAACGAACAAACTAAAATGTCTGAT    28249
gb:MT093631|Organism:Severe      GTATCATGACGTTCGTGTTGTTTTAGATTTCATCTAAACGAACAAACTAAAATGTCTGAT    28320
gb:MT106053|Organism:Severe      GTATCATGACGTTCGTGTTGTTTTAGATTTCATCTAAACGAACAAACTAAAATGTCTGAT    28282
gb:MT019533|Organism:Severe      GTATCATGACGTTCGTGTTGTTTTAGATTTCATCTAAACGAACAAACTAAAATGTCTGAT    28282
gb:MT019531|Organism:Severe      GTATCATGACGTTCGTGTTGTTTTAGATTTCATCTAAACGAACAAACTAAAATGTCTGAT    28282
```

```
gb:MN996528|Organism:Severe      GTATCATGACGTTCGTGTTGTTTTAGATTTCATCTAAACGAACAAACTAAAATGTCTGAT   28282
gb:MN996530|Organism:Severe      GTATCATGACGTTCGTGTTGTTTTAGATTTCATCTAAACGAACAAACTAAAATGTCTGAT   28268
gb:MN908947|Organism:Severe      GTATCATGACGTTCGTGTTGTTTTAGATTTCATCTAAACGAACAAACTAAAATGTCTGAT   28282
gb:MT019532|Organism:Severe      GTATCATGACGTTCGTGTTGTTTTAGATTTCATCTAAACGAACAAACTAAAATGTCTGAT   28282
                                 ************************************************************

gb:MT020781|Organism:Severe      AATGGACCCCAAAATCAGCGAAATGCACCCCGCATTACGTTTGGTGGACCCTCAGATTCA   28330
gb:MT007544|Organism:Severe      AATGGACCCCAAAATCAGCGAAATGCACCCCGCATTACGTTTGGTGGACCCTCAGATTCA   28342
gb:MN994467|Organism:Severe      AATGGACCCCAAAATCAGCGAAATGCACCCCGCATTACGTTTGGTGGACCCTCAGATTCA   28342
gb:MT044257|Organism:Severe      AATGGACCCCAAAATCAGCGAAATGCACCCCGCATTACGTTTGGTGGACCCTCAGATTCA   28342
gb:MT106054|Organism:Severe      AATGGACCCCAAAATCAGCGAAATGCACCCCGCATTACGTTTGGTGGACCCTCAGATTCA   28342
gb:MT049951|Organism:Severe      AATGGACCCCAAAATCAGCGAAATGCACCCCGCATTACGTTTGGTGGACCCTCAGATTCA   28342
gb:MN975262|Organism:Severe      AATGGACCCCAAAATCAGCGAAATGCACCCCGCATTACGTTTGGTGGACCCTCAGATTCA   28342
gb:MT106052|Organism:Severe      AATGGACCCCAAAATCAGCGAAATGCACCCCGCATTACGTTTGGTGGACCCTCAGATTCA   28342
gb:LC522975|Organism:Severe      AATGGACCCCAAAATCAGCGAAATGCACCCCGCATTACGTTTGGTGGACCCTCAGATTCA   28339
gb:LC522973|Organism:Severe      AATGGACCCCAAAATCAGCGAAATGCACCCCGCATTACGTTTGGTGGACCCTCAGATTCA   28339
gb:LC522974|Organism:Severe      AATGGACCCCAAAATCAGCGAAATGCACCCCGCATTACGTTTGGTGGACCCTCAGATTCA   28339
gb:MN985325|Organism:Severe      AATGGACCCCAAAATCAGCGAAATGCACCCCGCATTACGTTTGGTGGACCCTCAGATTCA   28342
gb:MT020881|Organism:Severe      AATGGACCCCAAAATCAGCGAAATGCACCCCGCATTACGTTTGGTGGACCCTCAGATTCA   28342
gb:MT020880|Organism:Severe      AATGGACCCCAAAATCAGCGAAATGCACCCCGCATTACGTTTGGTGGACCCTCAGATTCA   28342
gb:MT066175|Organism:Severe      AATGGACCCCAAAATCAGCGAAATGCACCCCGCATTACGTTTGGTGGACCCTCAGATTCA   28342
gb:MN997409|Organism:Severe      AATGGACCCCAAAATCAGCGAAATGCACCCCGCATTACGTTTGGTGGACCCTCAGATTCA   28342
gb:MN938384|Organism:Severe      AATGGACCCCAAAATCAGCGAAATGCACCCCGCATTACGTTTGGTGGACCCTCAGATTCA   28310
gb:MT044258|Organism:Severe      AATGGACCCCAAAATCAGCGAAATGCACCCCGCATTACGTTTGGTGGACCCTCAGATTCA   28318
gb:MT039890|Organism:Severe      AATGGACCCCAAAATCAGCGAAATGCACCCCGCATTACGTTTGGTGGACCCTCAGATTCA   28342
gb:MN988713|Organism:Severe      AATGGACCCCAAAATCAGCGAAATGCACCCCGCATTACGTTTGGTGGACCCTCAGATTCA   28342
gb:LC521925|Organism:Severe      AATGGACCCCAAAATCAGCGAAATGCACCCCGCATTACGTTTGGTGGACCCTCAGATTCA   28315
gb:MT093571|Organism:Severe      AATGGACCCCAAAATCAGCGAAATGCACCCCGCATTACGTTTGGTGGACCCTCAGATTCA   28342
gb:MT039887|Organism:Severe      AATGGACCCCAAAATCAGCGAAATGCACCCCGCATTACGTTTGGTGGACCCTCAGATTCA   28339
gb:MT019530|Organism:Severe      AATGGACCCCAAAATCAGCGAAATGCACCCCGCATTACGTTTGGTGGACCCTCAGATTCA   28342
gb:MT039888|Organism:Severe      AATGGACCCCAAAATCAGCGAAATGCACCCCGCATTACGTTTGGTGGACCCTCAGATTCA   28342
gb:LC522972|Organism:Severe      AATGGACCCCAAAATCAGCGAAATGCACCCCGCATTACGTTTGGTGGACCCTCAGATTCA   28339
gb:MT027063|Organism:Severe      AATGGACCCCAAAATCAGCGAAATGCACCCCGCATTACGTTTGGTGGACCCTCAGATTCA   28342
gb:MT027062|Organism:Severe      AATGGACCCCAAAATCAGCGAAATGCACCCCGCATTACGTTTGGTGGACCCTCAGATTCA   28342
gb:MT019529|Organism:Severe      AATGGACCCCAAAATCAGCGAAATGCACCCCGCATTACGTTTGGTGGACCCTCAGATTCA   28342
gb:MN996529|Organism:Severe      AATGGACCCCAAAATCAGCGAAATGCACCCCGCATTACGTTTGGTGGACCCTCAGATTCA   28330
gb:MN996531|Organism:Severe      AATGGACCCCAAAATCAGCGAAATGCACCCCGCATTACGTTTGGTGGACCCTCAGATTCA   28329
gb:MT066176|Organism:Severe      AATGGACCCCAAAATCAGCGAAATGCACCCCGCATTACGTTTGGTGGACCCTCAGATTCA   28342
gb:MT027064|Organism:Severe      AATGGACCCCAAAATCAGCGAAATGCACCCCGCATTACGTTTGGTGGACCCTCAGATTCA   28342
gb:MN994468|Organism:Severe      AATGGACCCCAAAATCAGCGAAATGCACCCCGCATTACGTTTGGTGGACCCTCAGATTCA   28342
gb:MT072688|Organism:Severe      AATGGACCCCAAAATCAGCGAAATGCACCCCGCATTACGTTTGGTGGACCCTCAGATTCA   28327
gb:MN996527|Organism:Severe      AATGGACCCCAAAATCAGCGAAATGCACCCCGCATTACGTTTGGTGGACCCTCAGATTCA   28309
gb:MT093631|Organism:Severe      AATGGACCCCAAAATCAGCGAAATGCACCCCGCATTACGTTTGGTGGACCCTCAGATTCA   28380
gb:MT106053|Organism:Severe      AATGGACCCCAAAATCAGCGAAATGCACCCCGCATTACGTTTGGTGGACCCTCAGATTCA   28342
gb:MT019533|Organism:Severe      AATGGACCCCAAAATCAGCGAAATGCACCCCGCATTACGTTTGGTGGACCCTCAGATTCA   28342
gb:MT019531|Organism:Severe      AATGGACCCCAAAATCAGCGAAATGCACCCCGCATTACGTTTGGTGGACCCTCAGATTCA   28342
gb:MN996528|Organism:Severe      AATGGACCCCAAAATCAGCGAAATGCACCCCGCATTACGTTTGGTGGACCCTCAGATTCA   28342
gb:MN996530|Organism:Severe      AATGGACCCCAAAATCAGCGAAATGCACCCCGCATTACGTTTGGTGGACCCTCAGATTCA   28328
gb:MN908947|Organism:Severe      AATGGACCCCAAAATCAGCGAAATGCACCCCGCATTACGTTTGGTGGACCCTCAGATTCA   28342
gb:MT019532|Organism:Severe      AATGGACCCCAAAATCAGCGAAATGCACCCCGCATTACGTTTGGTGGACCCTCAGATTCA   28342
                                 ************************************************************

gb:MT020781|Organism:Severe      ACTGGCAGTAACCAGAATGGAGAACGCAGTGGGGCGCGATCAAAACAACGTCGGCCCCAA   28390
gb:MT007544|Organism:Severe      ACTGGCAGTAACCAGAATGGAGAACGCAGTGGGGCGCGATCAAAACAACGTCGGCCCCAA   28402
gb:MN994467|Organism:Severe      ACTGGCAGTAACCAGAATGGAGAACGCAGTGGGGCGCGATCAAAACAACGTCGGCCCCAA   28402
gb:MT044257|Organism:Severe      ACTGGCAGTAACCAGAATGGAGAACGCAGTGGGGCGCGATCAAAACAACGTCGGCCCCAA   28402
gb:MT106054|Organism:Severe      ACTGGCAGTAACCAGAATGGAGAACGCAGTGGGGCGCGATCAAAACAACGTCGGCCCCAA   28402
gb:MT049951|Organism:Severe      ACTGGCAGTAACCAGAATGGAGAACGCAGTGGGGCGCGATCAAAACAACGTCGGCCCCAA   28402
gb:MN975262|Organism:Severe      ACTGGCAGTAACCAGAATGGAGAACGCAGTGGGGCGCGATCAAAACAACGTCGGCCCCAA   28402
gb:MT106052|Organism:Severe      ACTGGCAGTAACCAGAATGGAGAACGCAGTGGGGCGCGATCAAAACAACGTCGGCCCCAA   28402
gb:LC522975|Organism:Severe      ACTGGCAGTAACCAGAATGGAGAACGCAGTGGGGCGCGATCAAAACAACGTCGGCCCCAA   28399
gb:LC522973|Organism:Severe      ACTGGCAGTAACCAGAATGGAGAACGCAGTGGGGCGCGATCAAAACAACGTCGGCCCCAA   28399
gb:LC522974|Organism:Severe      ACTGGCAGTAACCAGAATGGAGAACGCAGTGGGGCGCGATCAAAACAACGTCGGCCCCAA   28399
gb:MN985325|Organism:Severe      ACTGGCAGTAACCAGAATGGAGAACGCAGTGGGGCGCGATCAAAACAACGTCGGCCCCAA   28402
gb:MT020881|Organism:Severe      ACTGGCAGTAACCAGAATGGAGAACGCAGTGGGGCGCGATCAAAACAACGTCGGCCCCAA   28402
```

```
gb:MT020880|Organism:Severe      ACTGGCAGTAACCAGAATGGAGAACGCAGTGGGGCGCGATCAAAACAACGTCGGCCCCAA   28402
gb:MT066175|Organism:Severe      ACTGGCAGTAACCAGAATGGAGAACGCAGTGGGGCGCGATCAAAACAACGTCGGCCCCAA   28402
gb:MN997409|Organism:Severe      ACTGGCAGTAACCAGAATGGAGAACGCAGTGGGGCGCGATCAAAACAACGTCGGCCCCAA   28402
gb:MN938384|Organism:Severe      ACTGGCAGTAACCAGAATGGAGAACGCAGTGGGGCGCGATCAAAACAACGTCGGCCCCAA   28370
gb:MT044258|Organism:Severe      ACTGGCAGTAACCAGAATGGAGAACGCAGTGGGGCGCGATCAAAACAACGTCGGCCCCAA   28378
gb:MT039890|Organism:Severe      ACTGGCAGTAACCAGAATGGAGAACGCAGTGGGGCGCGATCAAAACAACGTCGGCCCCAA   28402
gb:MN988713|Organism:Severe      ACTGGCAGTAACCAGAATGGAGAACGCAGTGGGGCGCGATCAAAACAACGTCGGCCCCAA   28402
gb:LC521925|Organism:Severe      ACTGGCAGTAACCAGAATGGAGAACGCAGTGGGGCGCGATCAAAACAACGTCGGCCCCAA   28375
gb:MT093571|Organism:Severe      ACTGGCAGTAACCAGAATGGAGAACGCAGTGGGGCGCGATCAAAACAACGTCGGCCCCAA   28402
gb:MT039887|Organism:Severe      ACTGGCAGTAACCAGAATGGAGAACGCAGTGGGGCGCGATCAAAACAACGTCGGCCCCAA   28399
gb:MT019530|Organism:Severe      ACTGGCAGTAACCAGAATGGAGAACGCAGTGGGGCGCGATCAAAACAACGTCGGCCCCAA   28402
gb:MT039888|Organism:Severe      ACTGGCAGTAACCAGAATGGAGAACGCAGTGGGGCGCGATCAAAACAACGTCGGCCCCAA   28402
gb:LC522972|Organism:Severe      ACTGGCAGTAACCAGAATGGAGAACGCAGTGGGGCGCGATCAAAACAACGTCGGCCCCAA   28399
gb:MT027063|Organism:Severe      ACTGGCAGTAACCAGAATGGAGAACGCAGTGGGGCGCGATCAAAACAACGTCGGCCCCAA   28402
gb:MT027062|Organism:Severe      ACTGGCAGTAACCAGAATGGAGAACGCAGTGGGGCGCGATCAAAACAACGTCGGCCCCAA   28402
gb:MT019529|Organism:Severe      ACTGGCAGTAACCAGAATGGAGAACGCAGTGGGGCGCGATCAAAACAACGTCGGCCCCAA   28402
gb:MN996529|Organism:Severe      ACTGGCAGTAACCAGAATGGAGAACGCAGTGGGGCGCGATCAAAACAACGTCGGCCCCAA   28390
gb:MN996531|Organism:Severe      ACTGGCAGTAACCAGAATGGAGAACGCAGTGGGGCGCGATCAAAACAACGTCGGCCCCAA   28389
gb:MT066176|Organism:Severe      ACTGGCAGTAACCAGAATGGAGAACGCAGTGGGGCGCGATCAAAACAACGTCGGCCCCAA   28402
gb:MT027064|Organism:Severe      ACTGGCAGTAACCAGAATGGAGAACGCAGTGGGGCGCGATCAAAACAACGTCGGCCCCAA   28402
gb:MN994468|Organism:Severe      ACTGGCAGTAACCAGAATGGAGAACGCAGTGGGGCGCGATCAAAACAACGTCGGCCCCAA   28402
gb:MT072688|Organism:Severe      ACTGGCAGTAACCAGAATGGAGAACGCAGTGGGGCGCGATCAAAACAACGTCGGCCCCAA   28387
gb:MN996527|Organism:Severe      ACTGGCAGTAACCAGAATGGAGAACGCAGTGGGGCGCGATCAAAACAACGTCGGCCCCAA   28369
gb:MT093631|Organism:Severe      ACTGGCAGTAACCAGAATGGAGAACGCAGTGGGGCGCGATCAAAACAACGTCGGCCCCAA   28440
gb:MT106053|Organism:Severe      ACTGGCAGTAACCAGAATGGAGAACGCAGTGGGGCGCGATCAAAACAACGTCGGCCCCAA   28402
gb:MT019533|Organism:Severe      ACTGGCAGTAACCAGAATGGAGAACGCAGTGGGGCGCGATCAAAACAACGTCGGCCCCAA   28402
gb:MT019531|Organism:Severe      ACTGGCAGTAACCAGAATGGAGAACGCAGTGGGGCGCGATCAAAACAACGTCGGCCCCAA   28402
gb:MN996528|Organism:Severe      ACTGGCAGTAACCAGAATGGAGAACGCAGTGGGGCGCGATCAAAACAACGTCGGCCCCAA   28402
gb:MN996530|Organism:Severe      ACTGGCAGTAACCAGAATGGAGAACGCAGTGGGGCGCGATCAAAACAACGTCGGCCCCAA   28388
gb:MN908947|Organism:Severe      ACTGGCAGTAACCAGAATGGAGAACGCAGTGGGGCGCGATCAAAACAACGTCGGCCCCAA   28402
gb:MT019532|Organism:Severe      ACTGGCAGTAACCAGAATGGAGAACGCAGTGGGGCGCGATCAAAACAACGTCGGCCCCAA   28402
                                 ************************************************************

gb:MT020781|Organism:Severe      GGTTTACCCAATAATACTGCGTCTTGGTTCACCGCTCTCACTCAACATGGCAAGGAAGAC   28450
gb:MT007544|Organism:Severe      GGTTTACCCAATAATACTGCGTCTTGGTTCACCGCTCTCACTCAACATGGCAAGGAAGAC   28462
gb:MN994467|Organism:Severe      GGTTTACCCAATAATACTGCGTCTTGGTTCACCGCTCTCACTCAACATGGCAAGGAAGAC   28462
gb:MT044257|Organism:Severe      GGTTTACCCAATAATACTGCGTCTTGGTTCACCGCTCTCACTCAACATGGCAAGGAAGAC   28462
gb:MT106054|Organism:Severe      GGTTTACCCAATAATACTGCGTCTTGGTTCACCGCTCTCACTCAACATGGCAAGGAAGAC   28462
gb:MT049951|Organism:Severe      GGTTTACCCAATAATACTGCGTCTTGGTTCACCGCTCTCACTCAACATGGCAAGGAAGAC   28462
gb:MN975262|Organism:Severe      GGTTTACCCAATAATACTGCGTCTTGGTTCACCGCTCTCACTCAACATGGCAAGGAAGAC   28462
gb:MT106052|Organism:Severe      GGTTTACCCAATAATACTGCGTCTTGGTTCACCGCTCTCACTCAACATGGCAAGGAAGAC   28462
gb:LC522975|Organism:Severe      GGTTTACCCAATAATACTGCGTCTTGGTTCACCGCTCTCACTCAACATGGCAAGGAAGAC   28459
gb:LC522973|Organism:Severe      GGTTTACCCAATAATACTGCGTCTTGGTTCACCGCTCTCACTCAACATGGCAAGGAAGAC   28459
gb:LC522974|Organism:Severe      GGTTTACCCAATAATACTGCGTCTTGGTTCACCGCTCTCACTCAACATGGCAAGGAAGAC   28459
gb:MN985325|Organism:Severe      GGTTTACCCAATAATACTGCGTCTTGGTTCACCGCTCTCACTCAACATGGCAAGGAAGAC   28462
gb:MT020881|Organism:Severe      GGTTTACCCAATAATACTGCGTCTTGGTTCACCGCTCTCACTCAACATGGCAAGGAAGAC   28462
gb:MT020880|Organism:Severe      GGTTTACCCAATAATACTGCGTCTTGGTTCACCGCTCTCACTCAACATGGCAAGGAAGAC   28462
gb:MT066175|Organism:Severe      GGTTTACCCAATAATACTGCGTCTTGGTTCACCGCTCTCACTCAACATGGCAAGGAAGAC   28462
gb:MN997409|Organism:Severe      GGTTTACCCAATAATACTGCGTCTTGGTTCACCGCTCTCACTCAACATGGCAAGGAAGAC   28462
gb:MN938384|Organism:Severe      GGTTTACCCAATAATACTGCGTCTTGGTTCACCGCTCTCACTCAACATGGCAAGGAAGAC   28430
gb:MT044258|Organism:Severe      GGTTTACCCAATAATACTGCGTCTTGGTTCACCGCTCTCACTCAACATGGCAAGGAAGAC   28438
gb:MT039890|Organism:Severe      GGTTTACCCAATAATACTGCGTCTTGGTTCACCGCTCTCACTCAACATGGCAAGGAAGAC   28462
gb:MN988713|Organism:Severe      GGTTTACCCAATAATACTGCGTCTTGGTTCACCGCTCTCACTCAACATGGCAAGGAAGAC   28462
gb:LC521925|Organism:Severe      GGTTTACCCAATAATACTGCGTCTTGGTTCACCGCTCTCACTCAACATGGCAAGGAAGAC   28435
gb:MT093571|Organism:Severe      GGTTTACCCAATAATACTGCGTCTTGGTTCACCGCTCTCACTCAACATGGCAAGGAAGAC   28462
gb:MT039887|Organism:Severe      GGTTTACCCAATAATACTGCGTCTTGGTTCACCGCTCTCACTCAACATGGCAAGGAAGAC   28459
gb:MT019530|Organism:Severe      GGTTTACCCAATAATACTGCGTCTTGGTTCACCGCTCTCACTCAACATGGCAAGGAAGAC   28462
gb:MT039888|Organism:Severe      GGTTTACCCAATAATACTGCGTCTTGGTTCACCGCTCTCACTCAACATGGCAAGGAAGAC   28462
gb:LC522972|Organism:Severe      GGTTTACCCAATAATACTGCGTCTTGGTTCACCGCTCTCACTCAACATGGCAAGGAAGAC   28459
gb:MT027063|Organism:Severe      GGTTTACCCAATAATACTGCGTCTTGGTTCACCGCTCTCACTCAACATGGCAAGGAAGAC   28462
gb:MT027062|Organism:Severe      GGTTTACCCAATAATACTGCGTCTTGGTTCACCGCTCTCACTCAACATGGCAAGGAAGAC   28462
gb:MT019529|Organism:Severe      GGTTTACCCAATAATACTGCGTCTTGGTTCACCGCTCTCACTCAACATGGCAAGGAAGAC   28462
gb:MN996529|Organism:Severe      GGTTTACCCAATAATACTGCGTCTTGGTTCACCGCTCTCACTCAACATGGCAAGGAAGAC   28450
gb:MN996531|Organism:Severe      GGTTTACCCAATAATACTGCGTCTTGGTTCACCGCTCTCACTCAACATGGCAAGGAAGAC   28449
gb:MT066176|Organism:Severe      GGTTTACCCAATAATACTGCGTCTTGGTTCACCGCTCTCACTCAACATGGCAAGGAAGAC   28462
```

```
gb:MT027064|Organism:Severe      GGTTTACCCAATAATACTGCGTCTTGGTTCACCGCTCTCACTCAACATGGCAAGGAAGAC    28462
gb:MN994468|Organism:Severe      GGTTTACCCAATAATACTGCGTCTTGGTTCACCGCTCTCACTCAACATGGCAAGGAAGAC    28462
gb:MT072688|Organism:Severe      GGTTTACCCAATAATACTGCGTCTTGGTTCACCGCTCTCACTCAACATGGCAAGGAAGAC    28447
gb:MN996527|Organism:Severe      GGTTTACCCAATAATACTGCGTCTTGGTTCACCGCTCTCACTCAACATGGCAAGGAAGAC    28429
gb:MT093631|Organism:Severe      GGTTTACCCAATAATACTGCGTCTTGGTTCACCGCTCTCACTCAACATGGCAAGGAAGAC    28500
gb:MT106053|Organism:Severe      GGTTTACCCAATAATACTGCGTCTTGGTTCACCGCTCTCACTCAACATGGCAAGGAAGAC    28462
gb:MT019533|Organism:Severe      GGTTTACCCAATAATACTGCGTCTTGGTTCACCGCTCTCACTCAACATGGCAAGGAAGAC    28462
gb:MT019531|Organism:Severe      GGTTTACCCAATAATACTGCGTCTTGGTTCACCGCTCTCACTCAACATGGCAAGGAAGAC    28462
gb:MN996528|Organism:Severe      GGTTTACCCAATAATACTGCGTCTTGGTTCACCGCTCTCACTCAACATGGCAAGGAAGAC    28462
gb:MN996530|Organism:Severe      GGTTTACCCAATAATACTGCGTCTTGGTTCACCGCTCTCACTCAACATGGCAAGGAAGAC    28448
gb:MN908947|Organism:Severe      GGTTTACCCAATAATACTGCGTCTTGGTTCACCGCTCTCACTCAACATGGCAAGGAAGAC    28462
gb:MT019532|Organism:Severe      GGTTTACCCAATAATACTGCGTCTTGGTTCACCGCTCTCACTCAACATGGCAAGGAAGAC    28462
                                 ************************************************************

gb:MT020781|Organism:Severe      CTTAAATTCCCTCGAGGACAAGGCGTTCCAATTAACACCAATAGCAGTCCAGATGACCAA    28510
gb:MT007544|Organism:Severe      CTTAAATTCCCTCGAGGACAAGGCGTTCCAATTAACACCAATAGCAGTCCAGATGACCAA    28522
gb:MN994467|Organism:Severe      CTTAAATTCCCTCGAGGACAAGGCGTTCCAATTAACACCAATAGCAGTCCAGATGACCAA    28522
gb:MT044257|Organism:Severe      CTTAAATTCCCTCGAGGACAAGGCGTTCCAATTAACACCAATAGCAGTCCAGATGACCAA    28522
gb:MT106054|Organism:Severe      CTTAAATTCCCTCGAGGACAAGGCGTTCCAATTAACACCAATAGCAGTCCAGATGACCAA    28522
gb:MT049951|Organism:Severe      CTTAAATTCCCTCGAGGACAAGGCGTTCCAATTAACACCAATAGCAGTCCAGATGACCAA    28522
gb:MN975262|Organism:Severe      CTTAAATTCCCTCGAGGACAAGGCGTTCCAATTAACACCAATAGCAGTCCAGATGACCAA    28522
gb:MT106052|Organism:Severe      CTTAAATTCCCTCGAGGACAAGGCGTTCCAATTAACACCAATAGCAGTCCAGATGACCAA    28522
gb:LC522975|Organism:Severe      CTTAAATTCCCTCGAGGACAAGGCGTTCCAATTAACACCAATAGCAGTCCAGATGACCAA    28519
gb:LC522973|Organism:Severe      CTTAAATTCCCTCGAGGACAAGGCGTTCCAATTAACACCAATAGCAGTCCAGATGACCAA    28519
gb:LC522974|Organism:Severe      CTTAAATTCCCTCGAGGACAAGGCGTTCCAATTAACACCAATAGCAGTCCAGATGACCAA    28519
gb:MN985325|Organism:Severe      CTTAAATTCCCTCGAGGACAAGGCGTTCCAATTAACACCAATAGCAGTCCAGATGACCAA    28522
gb:MT020881|Organism:Severe      CTTAAATTCCCTCGAGGACAAGGCGTTCCAATTAACACCAATAGCAGTCCAGATGACCAA    28522
gb:MT020880|Organism:Severe      CTTAAATTCCCTCGAGGACAAGGCGTTCCAATTAACACCAATAGCAGTCCAGATGACCAA    28522
gb:MT066175|Organism:Severe      CTTAAATTCCCTCGAGGACAAGGCGTTCCAATTAACACCAATAGCAGTCCAGATGACCAA    28522
gb:MN997409|Organism:Severe      CTTAAATTCCCTCGAGGACAAGGCGTTCCAATTAACACCAATAGCAGTCCAGATGACCAA    28522
gb:MN938384|Organism:Severe      CTTAAATTCCCTCGAGGACAAGGCGTTCCAATTAACACCAATAGCAGTCCAGATGACCAA    28490
gb:MT044258|Organism:Severe      CTTAAATTCCCTCGAGGACAAGGCGTTCCAATTAACACCAATAGCAGTCCAGATGACCAA    28498
gb:MT039890|Organism:Severe      CTTAAATTCCCTCGAGGACAAGGCGTTCCAATTAACACCAATAGCAGTCCAGATGACCAA    28522
gb:MN988713|Organism:Severe      CTTAAATTCCCTCGAGGACAAGGCGTTCCAATTAACACCAATAGCAGTCCAGATGACCAA    28522
gb:LC521925|Organism:Severe      CTTAAATTCCCTCGAGGACAAGGCGTTCCAATTAACACCAATAGCAGTCCAGATGACCAA    28495
gb:MT093571|Organism:Severe      CTTAAATTCCCTCGAGGACAAGGCGTTCCAATTAACACCAATAGCAGTCCAGATGACCAA    28522
gb:MT039887|Organism:Severe      CTTAAATTCCCTCGAGGACAAGGCGTTCCAATTAACACCAATAGCAGTCCAGATGACCAA    28519
gb:MT019530|Organism:Severe      CTTAAATTCCCTCGAGGACAAGGCGTTCCAATTAACACCAATAGCAGTCCAGATGACCAA    28522
gb:MT039888|Organism:Severe      CTTAAATTCCCTCGAGGACAAGGCGTTCCAATTAACACCAATAGCAGTCCAGATGACCAA    28522
gb:LC522972|Organism:Severe      CTTAAATTCCCTCGAGGACAAGGCGTTCCAATTAACACCAATAGCAGTCCAGATGACCAA    28519
gb:MT027063|Organism:Severe      CTTAAATTCCCTCGAGGACAAGGCGTTCCAATTAACACCAATAGCAGTCCAGATGACCAA    28522
gb:MT027062|Organism:Severe      CTTAAATTCCCTCGAGGACAAGGCGTTCCAATTAACACCAATAGCAGTCCAGATGACCAA    28522
gb:MT019529|Organism:Severe      CTTAAATTCCCTCGAGGACAAGGCGTTCCAATTAACACCAATAGCAGTCCAGATGACCAA    28522
gb:MN996529|Organism:Severe      CTTAAATTCCCTCGAGGACAAGGCGTTCCAATTAACACCAATAGCAGTCCAGATGACCAA    28510
gb:MN996531|Organism:Severe      CTTAAATTCCCTCGAGGACAAGGCGTTCCAATTAACACCAATAGCAGTCCAGATGACCAA    28509
gb:MT066176|Organism:Severe      CTTAAATTCCCTCGAGGACAAGGCGTTCCAATTAACACCAATAGCAGTCCAGATGACCAA    28522
gb:MT027064|Organism:Severe      CTTAAATTCCCTCGAGGACAAGGCGTTCCAATTAACACCAATAGCAGTCCAGATGACCAA    28522
gb:MN994468|Organism:Severe      CTTAAATTCCCTCGAGGACAAGGCGTTCCAATTAACACCAATAGCAGTCCAGATGACCAA    28522
gb:MT072688|Organism:Severe      CTTAAATTCCCTCGAGGACAAGGCGTTCCAATTAACACCAATAGCAGTCCAGATGACCAA    28507
gb:MN996527|Organism:Severe      CTTAAATTCCCTCGAGGACAAGGCGTTCCAATTAACACCAATAGCAGTCCAGATGACCAA    28489
gb:MT093631|Organism:Severe      CTTAAATTCCCTCGAGGACAAGGCGTTCCAATTAACACCAATAGCAGTCCAGATGACCAA    28560
gb:MT106053|Organism:Severe      CTTAAATTCCCTCGAGGACAAGGCGTTCCAATTAACACCAATAGCAGTCCAGATGACCAA    28522
gb:MT019533|Organism:Severe      CTTAAATTCCCTCGAGGACAAGGCGTTCCAATTAACACCAATAGCAGTCCAGATGACCAA    28522
gb:MT019531|Organism:Severe      CTTAAATTCCCTCGAGGACAAGGCGTTCCAATTAACACCAATAGCAGTCCAGATGACCAA    28522
gb:MN996528|Organism:Severe      CTTAAATTCCCTCGAGGACAAGGCGTTCCAATTAACACCAATAGCAGTCCAGATGACCAA    28522
gb:MN996530|Organism:Severe      CTTAAATTCCCTCGAGGACAAGGCGTTCCAATTAACACCAATAGCAGTCCAGATGACCAA    28508
gb:MN908947|Organism:Severe      CTTAAATTCCCTCGAGGACAAGGCGTTCCAATTAACACCAATAGCAGTCCAGATGACCAA    28522
gb:MT019532|Organism:Severe      CTTAAATTCCCTCGAGGACAAGGCGTTCCAATTAACACCAATAGCAGTCCAGATGACCAA    28522
                                 ************************************************************

gb:MT020781|Organism:Severe      ATTGGCTACTACCGAAGAGCTACCAGACGAATTCGTGGTGGTGACGGTAAAATGAAAGAT    28570
gb:MT007544|Organism:Severe      ATTGGCTACTACCGAAGAGCTACCAGACGAATTCGTGGTGGTGACGGTAAAATGAAAGAT    28582
gb:MN994467|Organism:Severe      ATTGGCTACTACCGAAGAGCTACCAGACGAATTCGTGGTGGTGACGGTAAAATGAAAGAT    28582
gb:MT044257|Organism:Severe      ATTGGCTACTACCGAAGAGCTACCAGACGAATTCGTGGTGGTGACGGTAAAATGAAAGAT    28582
gb:MT106054|Organism:Severe      ATTGGCTACTACCGAAGAGCTACCAGACGAATTCGTGGTGGTGACGGTAAAATGAAAGAT    28582
```

```
gb:MT049951|Organism:Severe      ATTGGCTACTACCGAAGAGCTACCAGACGAATTCGTGGTGGTGACGGTAAAATGAAAGAT   28582
gb:MN975262|Organism:Severe      ATTGGCTACTACCGAAGAGCTACCAGACGAATTCGTGGTGGTGACGGTAAAATGAAAGAT   28582
gb:MT106052|Organism:Severe      ATTGGCTACTACCGAAGAGCTACCAGACGAATTCGTGGTGGTGACGGTAAAATGAAAGAT   28582
gb:LC522975|Organism:Severe      ATTGGCTACTACCGAAGAGCTACCAGACGAATTCGTGGTGGTGACGGTAAAATGAAAGAT   28579
gb:LC522973|Organism:Severe      ATTGGCTACTACCGAAGAGCTACCAGACGAATTCGTGGTGGTGACGGTAAAATGAAAGAT   28579
gb:LC522974|Organism:Severe      ATTGGCTACTACCGAAGAGCTACCAGACGAATTCGTGGTGGTGACGGTAAAATGAAAGAT   28579
gb:MN985325|Organism:Severe      ATTGGCTACTACCGAAGAGCTACCAGACGAATTCGTGGTGGTGACGGTAAAATGAAAGAT   28582
gb:MT020881|Organism:Severe      ATTGGCTACTACCGAAGAGCTACCAGACGAATTCGTGGTGGTGACGGTAAAATGAAAGAT   28582
gb:MT020880|Organism:Severe      ATTGGCTACTACCGAAGAGCTACCAGACGAATTCGTGGTGGTGACGGTAAAATGAAAGAT   28582
gb:MT066175|Organism:Severe      ATTGGCTACTACCGAAGAGCTACCAGACGAATTCGTGGTGGTGACGGTAAAATGAAAGAT   28582
gb:MN997409|Organism:Severe      ATTGGCTACTACCGAAGAGCTACCAGACGAATTCGTGGTGGTGACGGTAAAATGAAAGAT   28582
gb:MN938384|Organism:Severe      ATTGGCTACTACCGAAGAGCTACCAGACGAATTCGTGGTGGTGACGGTAAAATGAAAGAT   28550
gb:MT044258|Organism:Severe      ATTGGCTACTACCGAAGAGCTACCAGACGAATTCGTGGTGGTGACGGTAAAATGAAAGAT   28558
gb:MT039890|Organism:Severe      ATTGGCTACTACCGAAGAGCTACCAGACGAATTCGTGGTGGTGACGGTAAAATGAAAGAT   28582
gb:MN988713|Organism:Severe      ATTGGCTACTACCGAAGAGCTACCAGACGAATTCGTGGTGGTGACGGTAAAATGAAAGAT   28582
gb:LC521925|Organism:Severe      ATTGGCTACTACCGAAGAGCTACCAGACGAATTCGTGGTGGTGACGGTAAAATGAAAGAT   28555
gb:MT093571|Organism:Severe      ATTGGCTACTACCGAAGAGCTACCAGACGAATTCGTGGTGGTGACGGTAAAATGAAAGAT   28582
gb:MT039887|Organism:Severe      ATTGGCTACTACCGAAGAGCTACCAGACGAATTCGTGGTGGTGACGGTAAAATGAAAGAT   28579
gb:MT019530|Organism:Severe      ATTGGCTACTACCGAAGAGCTACCAGACGAATTCGTGGTGGTGACGGTAAAATGAAAGAT   28582
gb:MT039888|Organism:Severe      ATTGGCTACTACCGAAGAGCTACCAGACGAATTCGTGGTGGTGACGGTAAAATGAAAGAT   28582
gb:LC522972|Organism:Severe      ATTGGCTACTACCGAAGAGCTACCAGACGAATTCGTGGTGGTGACGGTAAAATGAAAGAT   28579
gb:MT027063|Organism:Severe      ATTGGCTACTACCGAAGAGCTACCAGACGAATTCGTGGTGGTGACGGTAAAATGAAAGAT   28582
gb:MT027062|Organism:Severe      ATTGGCTACTACCGAAGAGCTACCAGACGAATTCGTGGTGGTGACGGTAAAATGAAAGAT   28582
gb:MT019529|Organism:Severe      ATTGGCTACTACCGAAGAGCTACCAGACGAATTCGTGGTGGTGACGGTAAAATGAAAGAT   28582
gb:MN996529|Organism:Severe      ATTGGCTACTACCGAAGAGCTACCAGACGAATTCGTGGTGGTGACGGTAAAATGAAAGAT   28570
gb:MN996531|Organism:Severe      ATTGGCTACTACCGAAGAGCTACCAGACGAATTCGTGGTGGTGACGGTAAAATGAAAGAT   28569
gb:MT066176|Organism:Severe      ATTGGCTACTACCGAAGAGCTACCAGACGAATTCGTGGTGGTGACGGTAAAATGAAAGAT   28582
gb:MT027064|Organism:Severe      ATTGGCTACTACCGAAGAGCTACCAGACGAATTCGTGGTGGTGACGGTAAAATGAAAGAT   28582
gb:MN994468|Organism:Severe      ATTGGCTACTACCGAAGAGCTACCAGACGAATTCGTGGTGGTGACGGTAAAATGAAAGAT   28582
gb:MT072688|Organism:Severe      ATTGGCTACTACCGAAGAGCTACCAGACGAATTCGTGGTGGTGACGGTAAAATGAAAGAT   28567
gb:MN996527|Organism:Severe      ATTGGCTACTACCGAAGAGCTACCAGACGAATTCGTGGTGGTGACGGTAAAATGAAAGAT   28549
gb:MT093631|Organism:Severe      ATTGGCTACTACCGAAGAGCTACCAGACGAATTCGTGGTGGTGACGGTAAAATGAAAGAT   28620
gb:MT106053|Organism:Severe      ATTGGCTACTACCGAAGAGCTACCAGACGAATTCGTGGTGGTGACGGTAAAATGAAAGAT   28582
gb:MT019533|Organism:Severe      ATTGGCTACTACCGAAGAGCTACCAGACGAATTCGTGGTGGTGACGGTAAAATGAAAGAT   28582
gb:MT019531|Organism:Severe      ATTGGCTACTACCGAAGAGCTACCAGACGAATTCGTGGTGGTGACGGTAAAATGAAAGAT   28582
gb:MN996528|Organism:Severe      ATTGGCTACTACCGAAGAGCTACCAGACGAATTCGTGGTGGTGACGGTAAAATGAAAGAT   28582
gb:MN996530|Organism:Severe      ATTGGCTACTACCGAAGAGCTACCAGACGAATTCGTGGTGGTGACGGTAAAATGAAAGAT   28568
gb:MN908947|Organism:Severe      ATTGGCTACTACCGAAGAGCTACCAGACGAATTCGTGGTGGTGACGGTAAAATGAAAGAT   28582
gb:MT019532|Organism:Severe      ATTGGCTACTACCGAAGAGCTACCAGACGAATTCGTGGTGGTGACGGTAAAATGAAAGAT   28582
                                 ************************************************************

gb:MT020781|Organism:Severe      CTCAGTCCAAGATGGTATTTCTACTACCTAGGAACTGGGCCAGAAGCTGGACTTCCCTAT   28630
gb:MT007544|Organism:Severe      CTCAGTCCAAGATGGTATTTCTACTACCTAGGAACTGGGCCAGAAGCTGGACTTCCCTAT   28642
gb:MN994467|Organism:Severe      CTCAGTCCAAGATGGTATTTCTACTACCTAGGAACTGGGCCAGAAGCTGGACTTCCCTAT   28642
gb:MT044257|Organism:Severe      CTCAGTCCAAGATGGTATTTCTACTACCTAGGAACTGGGCCAGAAGCTGGACTTCCCTAT   28642
gb:MT106054|Organism:Severe      CTCAGTCCAAGATGGTATTTCTACTACCTAGGAACTGGGCCAGAAGCTGGACTTCCCTAT   28642
gb:MT049951|Organism:Severe      CTCAGTCCAAGATGGTATTTCTACTACCTAGGAACTGGGCCAGAAGCTGGACTTCCCTAT   28642
gb:MN975262|Organism:Severe      CTCAGTCCAAGATGGTATTTCTACTACCTAGGAACTGGGCCAGAAGCTGGACTTCCCTAT   28642
gb:MT106052|Organism:Severe      CTCAGTCCAAGATGGTATTTCTACTACCTAGGAACTGGGCCAGAAGCTGGACTTCCCTAT   28642
gb:LC522975|Organism:Severe      CTCAGTCCAAGATGGTATTTCTACTACCTAGGAACTGGGCCAGAAGCTGGACTTCCCTAT   28639
gb:LC522973|Organism:Severe      CTCAGTCCAAGATGGTATTTCTACTACCTAGGAACTGGGCCAGAAGCTGGACTTCCCTAT   28639
gb:LC522974|Organism:Severe      CTCAGTCCAAGATGGTATTTCTACTACCTAGGAACTGGGCCAGAAGCTGGACTTCCCTAT   28639
gb:MN985325|Organism:Severe      CTCAGTCCAAGATGGTATTTCTACTACCTAGGAACTGGGCCAGAAGCTGGACTTCCCTAT   28642
gb:MT020881|Organism:Severe      CTCAGTCCAAGATGGTATTTCTACTACCTAGGAACTGGGCCAGAAGCTGGACTTCCCTAT   28642
gb:MT020880|Organism:Severe      CTCAGTCCAAGATGGTATTTCTACTACCTAGGAACTGGGCCAGAAGCTGGACTTCCCTAT   28642
gb:MT066175|Organism:Severe      CTCAGTCCAAGATGGTATTTCTACTACCTAGGAACTGGGCCAGAAGCTGGACTTCCCTAT   28642
gb:MN997409|Organism:Severe      CTCAGTCCAAGATGGTATTTCTACTACCTAGGAACTGGGCCAGAAGCTGGACTTCCCTAT   28642
gb:MN938384|Organism:Severe      CTCAGTCCAAGATGGTATTTCTACTACCTAGGAACTGGGCCAGAAGCTGGACTTCCCTAT   28610
gb:MT044258|Organism:Severe      CTCAGTCCAAGATGGTATTTCTACTACCTAGGAACTGGGCCAGAAGCTGGACTTCCCTAT   28618
gb:MT039890|Organism:Severe      CTCAGTCCAAGATGGTATTTCTACTACCTAGGAACTGGGCCAGAAGCTGGACTTCCCTAT   28642
gb:MN988713|Organism:Severe      CTCAGTCCAAGATGGTATTTCTACTACCTAGGAACTGGGCCAGAAGCTGGACTTCCCTAT   28642
gb:LC521925|Organism:Severe      CTCAGTCCAAGATGGTATTTCTACTACCTAGGAACTGGGCCAGAAGCTGGACTTCCCTAT   28615
gb:MT093571|Organism:Severe      CTCAGTCCAAGATGGTATTTCTACTACCTAGGAACTGGGCCAGAAGCTGGACTTCCCTAT   28642
gb:MT039887|Organism:Severe      CTCAGTCCAAGATGGTATTTCTACTACCTAGGAACTGGGCCAGAAGCTGGACTTCCCTAT   28639
gb:MT019530|Organism:Severe      CTCAGTCCAAGATGGTATTTCTACTACCTAGGAACTGGGCCAGAAGCTGGACTTCCCTAT   28642
```

```
gb:MT039888|Organism:Severe      CTCAGTCCAAGATGGTATTTCTACTACCTAGGAACTGGGCCAGAAGCTGGACTTCCCTAT   28642
gb:LC522972|Organism:Severe      CTCAGTCCAAGATGGTATTTCTACTACCTAGGAACTGGGCCAGAAGCTGGACTTCCCTAT   28639
gb:MT027063|Organism:Severe      CTCAGTCCAAGATGGTATTTCTACTACCTAGGAACTGGGCCAGAAGCTGGACTTCCCTAT   28642
gb:MT027062|Organism:Severe      CTCAGTCCAAGATGGTATTTCTACTACCTAGGAACTGGGCCAGAAGCTGGACTTCCCTAT   28642
gb:MT019529|Organism:Severe      CTCAGTCCAAGATGGTATTTCTACTACCTAGGAACTGGGCCAGAAGCTGGACTTCCCTAT   28642
gb:MN996529|Organism:Severe      CTCAGTCCAAGATGGTATTTCTACTACCTAGGAACTGGGCCAGAAGCTGGACTTCCCTAT   28630
gb:MN996531|Organism:Severe      CTCAGTCCAAGATGGTATTTCTACTACCTAGGAACTGGGCCAGAAGCTGGACTTCCCTAT   28629
gb:MT066176|Organism:Severe      CTCAGTCCAAGATGGTATTTCTACTACCTAGGAACTGGGCCAGAAGCTGGACTTCCCTAT   28642
gb:MT027064|Organism:Severe      CTCAGTCCAAGATGGTATTTCTACTACCTAGGAACTGGGCCAGAAGCTGGACTTCCCTAT   28642
gb:MN994468|Organism:Severe      CTCAGTCCAAGATGGTATTTCTACTACCTAGGAACTGGGCCAGAAGCTGGACTTCCCTAT   28642
gb:MT072688|Organism:Severe      CTCAGTCCAAGATGGTATTTCTACTACCTAGGAACTGGGCCAGAAGCTGGACTTCCCTAT   28627
gb:MN996527|Organism:Severe      CTCAGTCCAAGATGGTATTTCTACTACCTAGGAACTGGGCCAGAAGCTGGACTTCCCTAT   28609
gb:MT093631|Organism:Severe      CTCAGTCCAAGATGGTATTTCTACTACCTAGGAACTGGGCCAGAAGCTGGACTTCCCTAT   28680
gb:MT106053|Organism:Severe      CTCAGTCCAAGATGGTATTTCTACTACCTAGGAACTGGGCCAGAAGCTGGACTTCCCTAT   28642
gb:MT019533|Organism:Severe      CTCAGTCCAAGATGGTATTTCTACTACCTAGGAACTGGGCCAGAAGCTGGACTTCCCTAT   28642
gb:MT019531|Organism:Severe      CTCAGTCCAAGATGGTATTTCTACTACCTAGGAACTGGGCCAGAAGCTGGACTTCCCTAT   28642
gb:MN996528|Organism:Severe      CTCAGTCCAAGATGGTATTTCTACTACCTAGGAACTGGGCCAGAAGCTGGACTTCCCTAT   28642
gb:MN996530|Organism:Severe      CTCAGTCCAAGATGGTATTTCTACTACCTAGGAACTGGGCCAGAAGCTGGACTTCCCTAT   28628
gb:MN908947|Organism:Severe      CTCAGTCCAAGATGGTATTTCTACTACCTAGGAACTGGGCCAGAAGCTGGACTTCCCTAT   28642
gb:MT019532|Organism:Severe      CTCAGTCCAAGATGGTATTTCTACTACCTAGGAACTGGGCCAGAAGCTGGACTTCCCTAT   28642
                                 ************************************************************

gb:MT020781|Organism:Severe      GGTGCTAACAAAGACGGCATCATATGGGTTGCAACTGAGGGAGCCTTGAATACACCAAAA   28690
gb:MT007544|Organism:Severe      GGTGCTAACAAAGACGGCATCATATGGGTTGCAACTGAGGGAGCCTTGAATACACCAAAA   28702
gb:MN994467|Organism:Severe      GGTGCTAACAAAGACGGCATCATATGGGTTGCAACTGAGGGAGCCTTGAATACACCAAAA   28702
gb:MT044257|Organism:Severe      GGTGCTAACAAAGACGGCATCATATGGGTTGCAACTGAGGGAGCCTTGAATACACCAAAA   28702
gb:MT106054|Organism:Severe      GGTGCTAACAAAGACGGCATCATATGGGTTGCAACTGAGGGAGCCTTGAATACACCAAAA   28702
gb:MT049951|Organism:Severe      GGTGCTAACAAAGACGGCATCATATGGGTTGCAACTGAGGGAGCCTTGAATACACCAAAA   28702
gb:MN975262|Organism:Severe      GGTGCTAACAAAGACGGCATCATATGGGTTGCAACTGAGGGAGCCTTGAATACACCAAAA   28702
gb:MT106052|Organism:Severe      GGTGCTAACAAAGACGGCATCATATGGGTTGCAACTGAGGGAGCCTTGAATACACCAAAA   28702
gb:LC522975|Organism:Severe      GGTGCTAACAAAGACGGCATCATATGGGTTGCAACTGAGGGAGCCTTGAATACACCAAAA   28699
gb:LC522973|Organism:Severe      GGTGCTAACAAAGACGGCATCATATGGGTTGCAACTGAGGGAGCCTTGAATACACCAAAA   28699
gb:LC522974|Organism:Severe      GGTGCTAACAAAGACGGCATCATATGGGTTGCAACTGAGGGAGCCTTGAATACACCAAAA   28699
gb:MN985325|Organism:Severe      GGTGCTAACAAAGACGGCATCATATGGGTTGCAACTGAGGGAGCCTTGAATACACCAAAA   28702
gb:MT020881|Organism:Severe      GGTGCTAACAAAGACGGCATCATATGGGTTGCAACTGAGGGAGCCTTGAATACACCAAAA   28702
gb:MT020880|Organism:Severe      GGTGCTAACAAAGACGGCATCATATGGGTTGCAACTGAGGGAGCCTTGAATACACCAAAA   28702
gb:MT066175|Organism:Severe      GGTGCTAACAAAGACGGCATCATATGGGTTGCAACTGAGGGAGCCTTGAATACACCAAAA   28702
gb:MN997409|Organism:Severe      GGTGCTAACAAAGACGGCATCATATGGGTTGCAACTGAGGGAGCCTTGAATACACCAAAA   28702
gb:MN938384|Organism:Severe      GGTGCTAACAAAGACGGCATCATATGGGTTGCAACTGAGGGAGCCTTGAATACACCAAAA   28670
gb:MT044258|Organism:Severe      GGTGCTAACAAAGACGGCATCATATGGGTTGCAACTGAGGGAGCCTTGAATACACCAAAA   28678
gb:MT039890|Organism:Severe      GGTGCTAACAAAGACGGCATCATATGGGTTGCAACTGAGGGAGCCTTGAATACACCAAAA   28702
gb:MN988713|Organism:Severe      GGTGCTAACAAAGACGGCATCATATGGGTTGCAACTGAGGGAGCCTTGAATACACCAAAA   28702
gb:LC521925|Organism:Severe      GGTGCTAACAAAGACGGCATCATATGGGTTGCAACTGAGGGAGCCTTGAATACACCAAAA   28675
gb:MT093571|Organism:Severe      GGTGCTAACAAAGACGGCATCATATGGGTTGCAACTGAGGGAGCCTTGAATACACCAAAA   28702
gb:MT039887|Organism:Severe      GGTGCTAACAAAGACGGCATCATATGGGTTGCAACTGAGGGAGCCTTGAATACACCAAAA   28699
gb:MT019530|Organism:Severe      GGTGCTAACAAAGACGGCATCATATGGGTTGCAACTGAGGGAGCCTTGAATACACCAAAA   28702
gb:MT039888|Organism:Severe      GGTGCTAACAAAGACGGCATCATATGGGTTGCAACTGAGGGAGCCTTGAATACACCAAAA   28702
gb:LC522972|Organism:Severe      GGTGCTAACAAAGACGGCATCATATGGGTTGCAACTGAGGGAGCCTTGAATACACCAAAA   28699
gb:MT027063|Organism:Severe      GGTGCTAACAAAGACGGCATCATATGGGTTGCAACTGAGGGAGCCTTGAATACACCAAAA   28702
gb:MT027062|Organism:Severe      GGTGCTAACAAAGACGGCATCATATGGGTTGCAACTGAGGGAGCCTTGAATACACCAAAA   28702
gb:MT019529|Organism:Severe      GGTGCTAACAAAGACGGCATCATATGGGTTGCAACTGAGGGAGCCTTGAATACACCAAAA   28702
gb:MN996529|Organism:Severe      GGTGCTAACAAAGACGGCATCATATGGGTTGCAACTGAGGGAGCCTTGAATACACCAAAA   28690
gb:MN996531|Organism:Severe      GGTGCTAACAAAGACGGCATCATATGGGTTGCAACTGAGGGAGCCTTGAATACACCAAAA   28689
gb:MT066176|Organism:Severe      GGTGCTAACAAAGACGGCATCATATGGGTTGCAACTGAGGGAGCCTTGAATACACCAAAA   28702
gb:MT027064|Organism:Severe      GGTGCTAACAAAGACGGCATCATATGGGTTGCAACTGAGGGAGCCTTGAATACACCAAAA   28702
gb:MN994468|Organism:Severe      GGTGCTAACAAAGACGGCATCATATGGGTTGCAACTGAGGGAGCCTTGAATACACCAAAA   28702
gb:MT072688|Organism:Severe      GGTGCTAACAAAGACGGCATCATATGGGTTGCAACTGAGGGAGCCTTGAATACACCAAAA   28687
gb:MN996527|Organism:Severe      GGTGCTAACAAAGACGGCATCATATGGGTTGCAACTGAGGGAGCCTTGAATACACCAAAA   28669
gb:MT093631|Organism:Severe      GGTGCTAACAAAGACGGCATCATATGGGTTGCAACTGAGGGAGCCTTGAATACACCAAAA   28740
gb:MT106053|Organism:Severe      GGTGCTAACAAAGACGGCATCATATGGGTTGCAACTGAGGGAGCCTTGAATACACCAAAA   28702
gb:MT019533|Organism:Severe      GGTGCTAACAAAGACGGCATCATATGGGTTGCAACTGAGGGAGCCTTGAATACACCAAAA   28702
gb:MT019531|Organism:Severe      GGTGCTAACAAAGACGGCATCATATGGGTTGCAACTGAGGGAGCCTTGAATACACCAAAA   28702
gb:MN996528|Organism:Severe      GGTGCTAACAAAGACGGCATCATATGGGTTGCAACTGAGGGAGCCTTGAATACACCAAAA   28702
gb:MN996530|Organism:Severe      GGTGCTAACAAAGACGGCATCATATGGGTTGCAACTGAGGGAGCCTTGAATACACCAAAA   28688
gb:MN908947|Organism:Severe      GGTGCTAACAAAGACGGCATCATATGGGTTGCAACTGAGGGAGCCTTGAATACACCAAAA   28702
```

```
gb:MT019532|Organism:Severe      GGTGCTAACAAAGACGGCATCATATGGGTTGCAACTGAGGGAGCCTTGAATACACCAAAA   28702
                                 ************************************************************

gb:MT020781|Organism:Severe      GATCACATTGGCACCCGCAATCCTGCTAACAATGCTGCAATCGTGCTACAACTTCCTCAA   28750
gb:MT007544|Organism:Severe      GATCACATTGGCACCCGCAATCCTGCTAACAATGCTGCAATCGTGCTACAACTTCCTCAA   28762
gb:MN994467|Organism:Severe      GATCACATTGGCACCCGCAATCCTGCTAACAATGCTGCAATCGTGCTACAACTTCCTCAA   28762
gb:MT044257|Organism:Severe      GATCACATTGGCACCCGCAATCCTGCTAACAATGCTGCAATCGTGCTACAACTTCCTCAA   28762
gb:MT106054|Organism:Severe      GATCACATTGGCACCCGCAATCCTGCTAACAATGCTGCAATCGTGCTACAACTTCCTCAA   28762
gb:MT049951|Organism:Severe      GATCACATTGGCACCCGCAATCCTGCTAACAATGCTGCAATCGTGCTACAACTTCCTCAA   28762
gb:MN975262|Organism:Severe      GATCACATTGGCACCCGCAATCCTGCTAACAATGCTGCAATCGTGCTACAACTTCCTCAA   28762
gb:MT106052|Organism:Severe      GATCACATTGGCACCCGCAATCCTGCTAACAATGCTGCAATCGTGCTACAACTTCCTCAA   28762
gb:LC522975|Organism:Severe      GATCACATTGGCACCCGCAATCCTGCTAACAATGCTGCAATCGTGCTACAACTTCCTCAA   28759
gb:LC522973|Organism:Severe      GATCACATTGGCACCCGCAATCCTGCTAACAATGCTGCAATCGTGCTACAACTTCCTCAA   28759
gb:LC522974|Organism:Severe      GATCACATTGGCACCCGCAATCCTGCTAACAATGCTGCAATCGTGCTACAACTTCCTCAA   28759
gb:MN985325|Organism:Severe      GATCACATTGGCACCCGCAATCCTGCTAACAATGCTGCAATCGTGCTACAACTTCCTCAA   28762
gb:MT020881|Organism:Severe      GATCACATTGGCACCCGCAATCCTGCTAACAATGCTGCAATCGTGCTACAACTTCCTCAA   28762
gb:MT020880|Organism:Severe      GATCACATTGGCACCCGCAATCCTGCTAACAATGCTGCAATCGTGCTACAACTTCCTCAA   28762
gb:MT066175|Organism:Severe      GATCACATTGGCACCCGCAATCCTGCTAACAATGCTGCAATCGTGCTACAACTTCCTCAA   28762
gb:MN997409|Organism:Severe      GATCACATTGGCACCCGCAATCCTGCTAACAATGCTGCAATCGTGCTACAACTTCCTCAA   28762
gb:MN938384|Organism:Severe      GATCACATTGGCACCCGCAATCCTGCTAACAATGCTGCAATCGTGCTACAACTTCCTCAA   28730
gb:MT044258|Organism:Severe      GATCACATTGGCACCCGCAATCCTGCTAACAATGCTGCAATCGTGCTACAACTTCCTCAA   28738
gb:MT039890|Organism:Severe      GATCACATTGGCACCCGCAATCCTGCTAACAATGCTGCAATCGTGCTACAACTTCCTCAA   28762
gb:MN988713|Organism:Severe      GATCACATTGGCACCCGCAATCCTGCTAACAATGCTGCAATCGTGCTACAACTTCCTCAA   28762
gb:LC521925|Organism:Severe      GATCACATTGGCACCCGCAATCCTGCTAACAATGCTGCAATCGTGCTACAACTTCCTCAA   28735
gb:MT093571|Organism:Severe      GATCACATTGGCACCCGCAATCCTGCTAACAATGCTGCAATCGTGCTACAACTTCCTCAA   28762
gb:MT039887|Organism:Severe      GATCACATTGGCACCCGCAATCCTGCTAACAATGCTGCAATCGTGCTACAACTTCCTCAA   28759
gb:MT019530|Organism:Severe      GATCACATTGGCACCCGCAATCCTGCTAACAATGCTGCAATCGTGCTACAACTTCCTCAA   28762
gb:MT039888|Organism:Severe      GATCACATTGGCACCCGCAATCCTGCTAACAATGCTGCAATCGTGCTACAACTTCCTCAA   28762
gb:LC522972|Organism:Severe      GATCACATTGGCACCCGCAATCCTGCTAACAATGCTGCAATCGTGCTACAACTTCCTCAA   28759
gb:MT027063|Organism:Severe      GATCACATTGGCACCCGCAATCCTGCTAACAATGCTGCAATCGTGCTACAACTTCCTCAA   28762
gb:MT027062|Organism:Severe      GATCACATTGGCACCCGCAATCCTGCTAACAATGCTGCAATCGTGCTACAACTTCCTCAA   28762
gb:MT019529|Organism:Severe      GATCACATTGGCACCCGCAATCCTGCTAACAATGCTGCAATCGTGCTACAACTTCCTCAA   28762
gb:MN996529|Organism:Severe      GATCACATTGGCACCCGCAATCCTGCTAACAATGCTGCAATCGTGCTACAACTTCCTCAA   28750
gb:MN996531|Organism:Severe      GATCACATTGGCACCCGCAATCCTGCTAACAATGCTGCAATCGTGCTACAACTTCCTCAA   28749
gb:MT066176|Organism:Severe      GATCACATTGGCACCCGCAATCCTGCTAACAATGCTGCAATCGTGCTACAACTTCCTCAA   28762
gb:MT027064|Organism:Severe      GATCACATTGGCACCCGCAATCCTGCTAACAATGCTGCAATCGTGCTACAACTTCCTCAA   28762
gb:MN994468|Organism:Severe      GATCACATTGGCACCCGCAATCCTGCTAACAATGCTGCAATCGTGCTACAACTTCCTCAA   28762
gb:MT072688|Organism:Severe      GATCACATTGGCACCCGCAATCCTGCTAACAATGCTGCAATCGTGCTACAACTTCCTCAA   28747
gb:MN996527|Organism:Severe      GATCACATTGGCACCCGCAATCCTGCTAACAATGCTGCAATCGTGCTACAACTTCCTCAA   28729
gb:MT093631|Organism:Severe      GATCACATTGGCACCCGCAATCCTGCTAACAATGCTGCAATCGTGCTACAACTTCCTCAA   28800
gb:MT106053|Organism:Severe      GATCACATTGGCACCCGCAATCCTGCTAACAATGCTGCAATCGTGCTACAACTTCCTCAA   28762
gb:MT019533|Organism:Severe      GATCACATTGGCACCCGCAATCCTGCTAACAATGCTGCAATCGTGCTACAACTTCCTCAA   28762
gb:MT019531|Organism:Severe      GATCACATTGGCACCCGCAATCCTGCTAACAATGCTGCAATCGTGCTACAACTTCCTCAA   28762
gb:MN996528|Organism:Severe      GATCACATTGGCACCCGCAATCCTGCTAACAATGCTGCAATCGTGCTACAACTTCCTCAA   28762
gb:MN996530|Organism:Severe      GATCACATTGGCACCCGCAATCCTGCTAACAATGCTGCAATCGTGCTACAACTTCCTCAA   28748
gb:MN908947|Organism:Severe      GATCACATTGGCACCCGCAATCCTGCTAACAATGCTGCAATCGTGCTACAACTTCCTCAA   28762
gb:MT019532|Organism:Severe      GATCACATTGGCACCCGCAATCCTGCTAACAATGCTGCAATCGTGCTACAACTTCCTCAA   28762
                                 ************************************************************

gb:MT020781|Organism:Severe      GGAACAACATTGCCAAAAGGCTTCTACGCAGAAGGGAGCAGAGGCGGCAGTCAAGCCTCT   28810
gb:MT007544|Organism:Severe      GGAACAACATTGCCAAAAGGCTTCTACGCAGAAGGGAGCAGAGGCGGCAGTCAAGCCTCT   28822
gb:MN994467|Organism:Severe      GGAACAACATTGCCAAAAGGCTTCTACGCTGAAGGGAGCAGAGGCGGCAGTCAAGCCTCT   28822
gb:MT044257|Organism:Severe      GGAACAACATTGCCAAAAGGCTTCTACGCAGAAGGGAGCAGAGGCGGCAGTCAAGCCTCT   28822
gb:MT106054|Organism:Severe      GGAACAACATTGCCAAAAGGCTTCTACGCAGAAGGGAGCAGAGGCGGCAGTCAAGCCTCT   28822
gb:MT049951|Organism:Severe      GGAACAACATTGCCAAAAGGCTTCTACGCAGAAGGGAGCAGAGGCGGCAGTCAAGCCTCT   28822
gb:MN975262|Organism:Severe      GGAACAACATTGCCAAAAGGCTTCTACGCAGAAGGGAGCAGAGGCGGCAGTCAAGCCTCT   28822
gb:MT106052|Organism:Severe      GGAACAACATTGCCAAAAGGCTTCTACGCAGAAGGGAGCAGAGGCGGCAGTCAAGCCTCT   28822
gb:LC522975|Organism:Severe      GGAACAACATTGCCAAAAGGCTTCTACGCAGAAGGGAGCAGAGGCGGCAGTCAAGCCTCT   28819
gb:LC522973|Organism:Severe      GGAACAACATTGCCAAAAGGCTTCTACGCAGAAGGGAGCAGAGGCGGCAGTCAAGCCTCT   28819
gb:LC522974|Organism:Severe      GGAACAACATTGCCAAAAGGCTTCTACGCAGAAGGGAGCAGAGGCGGCAGTCAAGCCTCT   28819
gb:MN985325|Organism:Severe      GGAACAACATTGCCAAAAGGCTTCTACGCAGAAGGGAGCAGAGGCGGCAGTCAAGCCTCT   28822
gb:MT020881|Organism:Severe      GGAACAACATTGCCAAAAGGCTTCTACGCAGAAGGGAGCAGAGGCGGCAGTCAAGCCTCT   28822
gb:MT020880|Organism:Severe      GGAACAACATTGCCAAAAGGCTTCTACGCAGAAGGGAGCAGAGGCGGCAGTCAAGCCTCT   28822
gb:MT066175|Organism:Severe      GGAACAACATTGCCAAAAGGCTTCTACGCAGAAGGGAGCAGAGGCGGCAGTCAAGCCTCT   28822
gb:MN997409|Organism:Severe      GGAACAACATTGCCAAAAGGCTTCTACGCAGAAGGGAGCAGAGGCGGCAGTCAAGCCTCT   28822
```

```
gb:MN938384|Organism:Severe      GGAACAACATTGCCAAAAGGCTTCTACGCAGAAGGGAGCAGAGGCGGCAGTCAAGCCTCT    28790
gb:MT044258|Organism:Severe      GGAACAACATTGCCAAAAGGCTTCTACGCAGAAGGGAGCAGAGGCGGCAGTCAAGCCTCT    28798
gb:MT039890|Organism:Severe      GGAACAACATTGCCAAAAGGCTTCTACGCAGAAGGGAGCAGAGGCGGCAGTCAAGCCTCT    28822
gb:MN988713|Organism:Severe      GGAACAACATTGCCAAAAGGCTTCTACGCAGAAGGGAGCAGAGGCGGCAGTCAAGCCTCT    28822
gb:LC521925|Organism:Severe      GGAACAACATTGCCAAAAGGCTTCTACGCAGAAGGGAGCAGAGGCGGCAGTCAAGCCTCT    28795
gb:MT093571|Organism:Severe      GGAACAACATTGCCAAAAGGCTTCTACGCAGAAGGGAGCAGAGGCGGCAGTCAAGCCTCT    28822
gb:MT039887|Organism:Severe      GGAACAACATTGCCAAAAGGCTTCTACGCAGAAGGGAGCAGAGGCGGCAGTCAAGCCTCT    28819
gb:MT019530|Organism:Severe      GGAACAACATTGCCAAAAGGCTTCTACGCAGAAGGGAGCAGAGGCGGCAGTCAAGCCTCT    28822
gb:MT039888|Organism:Severe      GGAACAACATTGCCAAAAGGCTTCTACGCAGAAGGGAGCAGAGGCGGCAGTCAAGCCTCT    28822
gb:LC522972|Organism:Severe      GGAACAACATTGCCAAAAGGCTTCTACGCAGAAGGGAGCAGAGGCGGCAGTCAAGCCTCT    28819
gb:MT027063|Organism:Severe      GGAACAACATTGCCAAAAGGCTTCTACGCAGAAGGGAGCAGAGGCGGCAGTCAAGCCTCT    28822
gb:MT027062|Organism:Severe      GGAACAACATTGCCAAAAGGCTTCTACGCAGAAGGGAGCAGAGGCGGCAGTCAAGCCTCT    28822
gb:MT019529|Organism:Severe      GGAACAACATTGCCAAAAGGCTTCTACGCAGAAGGGAGCAGAGGCGGCAGTCAAGCCTCT    28822
gb:MN996529|Organism:Severe      GGAACAACATTGCCAAAAGGCTTCTACGCAGAAGGGAGCAGAGGCGGCAGTCAAGCCTCT    28810
gb:MN996531|Organism:Severe      GGAACAACATTGCCAAAAGGCTTCTACGCAGAAGGGAGCAGAGGCGGCAGTCAAGCCTCT    28809
gb:MT066176|Organism:Severe      GGAACAACATTGCCAAAAGGCTTCTACGCAGAAGGGAGCAGAGGCGGCAGTCAAGCCTCT    28822
gb:MT027064|Organism:Severe      GGAACAACATTGCCAAAAGGCTTCTACGCAGAAGGGAGCAGAGGCGGCAGTCAAGCCTCT    28822
gb:MN994468|Organism:Severe      GGAACAACATTGCCAAAAGGCTTCTACGCAGAAGGGAGCAGAGGCGGCAGTCAAGCCTCT    28822
gb:MT072688|Organism:Severe      GGAACAACATTGCCAAAAGGCTTCTACGCAGAAGGGAGCAGAGGCGGCAGTCAAGCCTCT    28807
gb:MN996527|Organism:Severe      GGAACAACATTGCCAAAAGGCTTCTACGCAGAAGGGAGCAGAGGCGGCAGTCAAGCCTCT    28789
gb:MT093631|Organism:Severe      GGAACAACATTGCCAAAAGGCTTCTACGCAGAAGGGAGCAGAGGCGGCAGTCAAGCCTCT    28860
gb:MT106053|Organism:Severe      GGAACAACATTGCCAAAAGGCTTCTACGCAGAAGGGAGCAGAGGCGGCAGTCAAGCCTCT    28822
gb:MT019533|Organism:Severe      GGAACAACATTGCCAAAAGGCTTCTACGCAGAAGGGAGCAGAGGCGGCAGTCAAGCCTCT    28822
gb:MT019531|Organism:Severe      GGAACAACATTGCCAAAAGGCTTCTACGCAGAAGGGAGCAGAGGCGGCAGTCAAGCCTCT    28822
gb:MN996528|Organism:Severe      GGAACAACATTGCCAAAAGGCTTCTACGCAGAAGGGAGCAGAGGCGGCAGTCAAGCCTCT    28822
gb:MN996530|Organism:Severe      GGAACAACATTGCCAAAAGGCTTCTACGCAGAAGGGAGCAGAGGCGGCAGTCAAGCCTCT    28808
gb:MN908947|Organism:Severe      GGAACAACATTGCCAAAAGGCTTCTACGCAGAAGGGAGCAGAGGCGGCAGTCAAGCCTCT    28822
gb:MT019532|Organism:Severe      GGAACAACATTGCCAAAAGGCTTCTACGCAGAAGGGAGCAGAGGCGGCAGTCAAGCCTCT    28822
                                 ***************************** ******************************

gb:MT020781|Organism:Severe      TCTCGTTCCTCATCACGTAGTCGCAACAGTTCAAGAAATTCAACTCCAGGCAGCAGTAGG    28870
gb:MT007544|Organism:Severe      TCTCGTTCCTCATCACGTAGTCGCAACAGTTCAAGAAATTCAACTCCAGGCAGCAGTAGG    28882
gb:MN994467|Organism:Severe      TCTCGTTCCTCATCACGTAGTCGCAACAGTTCAAGAAATTCAACTCCAGGCAGCAGTAGG    28882
gb:MT044257|Organism:Severe      TCTCGTTCCTCATCACGTAGTCGCAACAGTTCAAGAAATTCAACTCCAGGCAGCAGTAGG    28882
gb:MT106054|Organism:Severe      TCTCGTTCCTCATCACGTAGTCGCAACAGTTCAAGAAATTCAACTCCAGGCAGCAGTAGG    28882
gb:MT049951|Organism:Severe      TCTCGTTCCTCATCACGTAGTCGCAACAGTTCAAGAAATTCAACTCCAGGCAGCAGTAGG    28882
gb:MN975262|Organism:Severe      TCTCGTTCCTCATCACGTAGTCGCAACAGTTCAAGAAATTCAACTCCAGGCAGCAGTAGG    28882
gb:MT106052|Organism:Severe      TCTCGTTCCTCATCACGTAGTCGCAACAGTTCAAGAAATTCAACTCCAGGCAGCAATAGG    28882
gb:LC522975|Organism:Severe      TCTCGTTCCTCATCACGTAGTCGCAACAGTTCAAGAAATTCAACTCCAGGCAGCAGTAGG    28879
gb:LC522973|Organism:Severe      TCTCGTTCCTCATCACGTAGTCGCAACAGTTCAAGAAATTCAACTCCAGGCAGCAGTAGG    28879
gb:LC522974|Organism:Severe      TCTCGTTCCTCATCACGTAGTCGCAACAGTTCAAGAAATTCAACTCCAGGCAGCAGTAGG    28879
gb:MN985325|Organism:Severe      TCTCGTTCCTCATCACGTAGTCGCAACAGTTCAAGAAATTCAACTCCAGGCAGCAGTAGG    28882
gb:MT020881|Organism:Severe      TCTCGTTCCTCATCACGTAGTCGCAACAGTTCAAGAAATTCAACTCCAGGCAGCAGTAGG    28882
gb:MT020880|Organism:Severe      TCTCGTTCCTCATCACGTAGTCGCAACAGTTCAAGAAATTCAACTCCAGGCAGCAGTAGG    28882
gb:MT066175|Organism:Severe      TCTCGTTCCTCATCACGTAGTCGCAACAGTTCAAGAAATTCAACTCCAGGCAGCAGTAGG    28882
gb:MN997409|Organism:Severe      TCTCGTTCCTCATCACGTAGTCGCAACAGTTCAAGAAATTCAACTCCAGGCAGCAGTAGG    28882
gb:MN938384|Organism:Severe      TCTCGTTCCTCATCACGTAGTCGCAACAGTTCAAGAAATTCAACTCCAGGCAGCAGTAGG    28850
gb:MT044258|Organism:Severe      TCTCGTTCCTCATCACGTAGTCGCAACAGTTCAAGAAATTCAACTCCAGGCAGCAGTAGG    28858
gb:MT039890|Organism:Severe      TCTCGTTCCTCATCACGTAGTCGCAACAGTTCAAGAAATTCAACTCCAGGCAGCAGTAGG    28882
gb:MN988713|Organism:Severe      TCTCGTTCCTCATCACGTAGTCGCAACAGTTYAAGAAATTCAACTCCAGGCAGCAGTAGG    28882
gb:LC521925|Organism:Severe      TCTCGTTCCTCATCACGTAGTCGCAACAGTTCAAGAAATTCAACTCCAGGCAGCAGTAGG    28855
gb:MT093571|Organism:Severe      TCTCGTTCCTCATCACGTAGTCGCAACAGTTCAAGAAATTCAACTCCAGGCAGCAGTAGG    28882
gb:MT039887|Organism:Severe      TCTCGTTCCTCATCACGTAGTCGCAACAGTTCAAGAAATTCAACTCCAGGCAGCAGTAGG    28879
gb:MT019530|Organism:Severe      TCTCGTTCCTCATCACGTAGTCGCAACAGTTCAAGAAATTCAACTCCAGGCAGCAGTAGG    28882
gb:MT039888|Organism:Severe      TCTCGTTCCTCATCACGTAGTCGCAACAGTTTAAGAAATTCAACTCCAGGCAGCAGTAGG    28882
gb:LC522972|Organism:Severe      TCTCGTTCCTCATCACGTAGTCGCAACAGTTCAAGAAATTCAACTCCAGGCAGCAGTAGG    28879
gb:MT027063|Organism:Severe      TCTCGTTCCTCATCACGTAGTCGCAACAGTTTAAGAAATTCAACTCCAGGCAGCAGTAGG    28882
gb:MT027062|Organism:Severe      TCTCGTTCCTCATCACGTAGTCGCAACAGTTTAAGAAATTCAACTCCAGGCAGCAGTAGG    28882
gb:MT019529|Organism:Severe      TCTCGTTCCTCATCACGTAGTCGCAACAGTTCAAGAAATTCAACTCCAGGCAGCAGTAGG    28882
gb:MN996529|Organism:Severe      TCTCGTTCCTCATCACGTAGTCGCAACAGTTCAAGAAATTCAACTCCAGGCAGCAGTAGG    28870
gb:MN996531|Organism:Severe      TCTCGTTCCTCATCACGTAGTCGCAACAGTTCAAGAAATTCAACTCCAGGCAGCAGTAGG    28869
gb:MT066176|Organism:Severe      TCTCGTTCCTCATCACGTAGTCGCAACAGTTCAAGAAATTCAACTCCAGGCAGCAGTAGG    28882
gb:MT027064|Organism:Severe      TCTCGTTCCTCATCACGTAGTCGCAACAGTTCAAGAAATTCAACTCCAGGCAGCAGTAGG    28882
gb:MN994468|Organism:Severe      TCTCGTTCCTCATCACGTAGTCGCAACAGTTCAAGAAATTCAACTCCAGGCAGCAGTAGG    28882
gb:MT072688|Organism:Severe      TCTCGTTCCTCATCACGTAGTCGCAACAGTTCAAGAAATTCAACTCCAGGCAGCAGTAGG    28867
```

```
gb:MN996527|Organism:Severe      TCTCGTTCCTCATCACGTAGTCGCAACAGTTCAAGAAATTCAACTCCAGGCAGCAGTAGG   28849
gb:MT093631|Organism:Severe      TCTCGTTCCTCATCACGTAGTCGCAACAGTTCAAGAAATTCAACTCCAGGCAGCAGTAGG   28920
gb:MT106053|Organism:Severe      TCTCGTTCCTCATCACGTAGTCGCAACAGTTCAAGAAATTCAACTCCAGGCAGCAGTAGG   28882
gb:MT019533|Organism:Severe      TCTCGTTCCTCATCACGTAGTCGCAACAGTTCAAGAAATTCAACTCCAGGCAGCAGTAGG   28882
gb:MT019531|Organism:Severe      TCTCGTTCCTCATCACGTAGTCGCAACAGTTCAAGAAATTCAACTCCAGGCAGCAGTAGG   28882
gb:MN996528|Organism:Severe      TCTCGTTCCTCATCACGTAGTCGCAACAGTTCAAGAAATTCAACTCCAGGCAGCAGTAGG   28882
gb:MN996530|Organism:Severe      TCTCGTTCCTCATCACGTAGTCGCAACAGTTCAAGAAATTCAACTCCAGGCAGCAGTAGG   28868
gb:MN908947|Organism:Severe      TCTCGTTCCTCATCACGTAGTCGCAACAGTTCAAGAAATTCAACTCCAGGCAGCAGTAGG   28882
gb:MT019532|Organism:Severe      TCTCGTTCCTCATCACGTAGTCGCAACAGTTCAAGAAATTCAACTCCAGGCAGCAGTAGG   28882
                                 ******************************* *********************** ****

gb:MT020781|Organism:Severe      GGAACTTCTCCTGCTAGAATGGCTGGCAATGGCGGTGATGCTGCTCTTGCTTTGCTGCTG   28930
gb:MT007544|Organism:Severe      GGAACTTCTCCTGCTAGAATGGCTGGCAATGGCGGTGATGCTGCTCTTGCTTTGCTGCTG   28942
gb:MN994467|Organism:Severe      GGAACTTCTCCTGCTAGAATGGCTGGCAATGGCGGTGATGCTGCTCTTGCTTTGCTGCTG   28942
gb:MT044257|Organism:Severe      GGAACTTCTCCTGCTAGAATGGCTGGCAATGGCGGTGATGCTGCTCTTGCTTTGCTGCTG   28942
gb:MT106054|Organism:Severe      GGAACTTCTCCTGCTAGAATGGCTGGCAATGGCGGTGATGCTGCTCTTGCTTTGCTGCTG   28942
gb:MT049951|Organism:Severe      GGAACTTCTCCTGCTAGAATGGCTGGCAATGGCGGTGATGCTGCTCTTGCTTTGCTGCTG   28942
gb:MN975262|Organism:Severe      GGAACTTCTCCTGCTAGAATGGCTGGCAATGGCGGTGATGCTGCTCTTGCTTTGCTGCTG   28942
gb:MT106052|Organism:Severe      GGAACTTCTCCTGCTAGAATGGCTGGCAATGGCGGTGATGCTGCTCTTGCTTTGCTGCTG   28942
gb:LC522975|Organism:Severe      GGAACTTCTCCTGCTAGAATGGCTGGCAATGGCGGTGATGCTGCTCTTGCTTTGCTGCTG   28939
gb:LC522973|Organism:Severe      GGAACTTCTCCTGCTAGAATGGCTGGCAATGGCGGTGATGCTGCTCTTGCTTTGCTGCTG   28939
gb:LC522974|Organism:Severe      GGAACTTCTCCTGCTAGAATGGCTGGCAATGGCGGTGATGCTGCTCTTGCTTTGCTGCTG   28939
gb:MN985325|Organism:Severe      GGAACTTCTCCTGCTAGAATGGCTGGCAATGGCGGTGATGCTGCTCTTGCTTTGCTGCTG   28942
gb:MT020881|Organism:Severe      GGAACTTCTCCTGCTAGAATGGCTGGCAATGGCGGTGATGCTGCTCTTGCTTTGCTGCTG   28942
gb:MT020880|Organism:Severe      GGAACTTCTCCTGCTAGAATGGCTGGCAATGGCGGTGATGCTGCTCTTGCTTTGCTGCTG   28942
gb:MT066175|Organism:Severe      GGAACTTCTCCTGCTAGAATGGCTGGCAATGGCGGTGATGCTGCTCTTGCTTTGCTGCTG   28942
gb:MN997409|Organism:Severe      GGAACTTCTCCTGCTAGAATGGCTGGCAATGGCGGTGATGCTGCTCTTGCTTTGCTGCTG   28942
gb:MN938384|Organism:Severe      GGAACTTCTCCTGCTAGAATGGCTGGCAATGGCGGTGATGCTGCTCTTGCTTTGCTGCTG   28910
gb:MT044258|Organism:Severe      GGAACTTCTCCTGCTAGAATGGCTGGCAATGGCGGTGATGCTGCTCTTGCTTTGCTGCTG   28918
gb:MT039890|Organism:Severe      GGAACTTCTCCTGCTAGAATGGCTGGCAATGGCGGTGATGCTGCTCTTGCTTTGCTGCTG   28942
gb:MN988713|Organism:Severe      GGAACTTCTCCTGCTAGAATGGCTGGCAATGGCGGTGATGCTGCTCTTGCTTTGCTGCTG   28942
gb:LC521925|Organism:Severe      GGAACTTCTCCTGCTAGAATGGCTGGCAATGGCGGTGATGCTGCTCTTGCTTTGCTGCTG   28915
gb:MT093571|Organism:Severe      GGAACTTCTCCTGCTAGAATGGCTGGCAATGGCGGTGATGCTGCTCTTGCTTTGCTGCTG   28942
gb:MT039887|Organism:Severe      GGAACTTCTCCTGCTAGAATGGCTGGCAATGGCGGTGATGCTGCTCTTGCTTTGCTGCTG   28939
gb:MT019530|Organism:Severe      GGAACTTCTCCTGCTAGAATGGCTGGCAATGGCGGTGATGCTGCTCTTGCTTTGCTGCTG   28942
gb:MT039888|Organism:Severe      GGAACTTCTCCTGCTAGAATGGCTGGCAATGGCGGTGATGCTGCTCTTGCTTTGCTGCTG   28942
gb:LC522972|Organism:Severe      GGAACTTCTCCTGCTAGAATGGCTGGCAATGGCGGTGATGCTGCTCTTGCTTTGCTGCTG   28939
gb:MT027063|Organism:Severe      GGAACTTCTCCTGCTAGAATGGCTGGCAATGGCGGTGATGCTGCTCTTGCTTTGCTGCTG   28942
gb:MT027062|Organism:Severe      GGAACTTCTCCTGCTAGAATGGCTGGCAATGGCGGTGATGCTGCTCTTGCTTTGCTGCTG   28942
gb:MT019529|Organism:Severe      GGAACTTCTCCTGCTAGAATGGCTGGCAATGGCGGTGATGCTGCTCTTGCTTTGCTGCTG   28942
gb:MN996529|Organism:Severe      GGAACTTCTCCTGCTAGAATGGCTGGCAATGGCGGTGATGCTGCTCTTGCTTTGCTGCTG   28930
gb:MN996531|Organism:Severe      GGAACTTCTCCTGCTAGAATGGCTGGCAATGGCGGTGATGCTGCTCTTGCTTTGCTGCTG   28929
gb:MT066176|Organism:Severe      GGAACTTCTCCTGCTAGAATGGCTGGCAATGGCGGTGATGCTGCTCTTGCTTTGCTGCTG   28942
gb:MT027064|Organism:Severe      GGAACTTCTCCTGCTAGAATGGCTGGCAATGGCGGTGATGCTGCTCTTGCTTTGCTGCTG   28942
gb:MN994468|Organism:Severe      GGAACTTCTCCTGCTAGAATGGCTGGCAATGGCGGTGATGCTGCTCTTGCTTTGCTGCTG   28942
gb:MT072688|Organism:Severe      GGAACTTCTCCTGCTAGAATGGCTGGCAATGGCGGTGATGCTGCTCTTGCTTTGCTGCTG   28927
gb:MN996527|Organism:Severe      GGAACTTCTCCTGCTAGAATGGCTGGCAATGGCGGTGATGCTGCTCTTGCTTTGCTGCTG   28909
gb:MT093631|Organism:Severe      GGAACTTCTCCTGCTAGAATGGCTGGCAATGGCGGTGATGCTGCTCTTGCTTTGCTGCTG   28980
gb:MT106053|Organism:Severe      GGAACTTCTCCTGCTAGAATGGCTGGCAATGGCGGTGATGCTGCTCTTGCTTTGCTGCTG   28942
gb:MT019533|Organism:Severe      GGAACTTCTCCTGCTAGAATGGCTGGCAATGGCGGTGATGCTGCTCTTGCTTTGCTGCTG   28942
gb:MT019531|Organism:Severe      GGAACTTCTCCTGCTAGAATGGCTGGCAATGGCGGTGATGCTGCTCTTGCTTTGCTGCTG   28942
gb:MN996528|Organism:Severe      GGAACTTCTCCTGCTAGAATGGCTGGCAATGGCGGTGATGCTGCTCTTGCTTTGCTGCTG   28942
gb:MN996530|Organism:Severe      GGAACTTCTCCTGCTAGAATGGCTGGCAATGGCGGTGATGCTGCTCTTGCTTTGCTGCTG   28928
gb:MN908947|Organism:Severe      GGAACTTCTCCTGCTAGAATGGCTGGCAATGGCGGTGATGCTGCTCTTGCTTTGCTGCTG   28942
gb:MT019532|Organism:Severe      GGAACTTCTCCTGCTAGAATGGCTGGCAATGGCGGTGATGCTGCTCTTGCTTTGCTGCTG   28942
                                 ************************************************************

gb:MT020781|Organism:Severe      CTTGACAGATTGAACCAGCTTGAGAGCAAAATGTCTGGTAAAGGCCAACAACAACAAGGC   28990
gb:MT007544|Organism:Severe      CTTGACAGATTGAACCAGCTTGAGAGCAAAATGTCTGGTAAAGGCCAACAACAACAAGGC   29002
gb:MN994467|Organism:Severe      CTTGACAGATTGAACCAGCTTGAGAGCAAAATGTCTGGTAAAGGCCAACAACAACAAGGC   29002
gb:MT044257|Organism:Severe      CTTGACAGATTGAACCAGCTTGAGAGCAAAATGTCTGGTAAAGGCCAACAACAACAAGGC   29002
gb:MT106054|Organism:Severe      CTTGACAGATTGAACCAGCTTGAGAGCAAAATGTCTGGTAAAGGCCAACAACAACAAGGC   29002
gb:MT049951|Organism:Severe      CTTGACAGATTGAACCAGCTTGAGAGCAAAATGTCTGGTAAAGGCCAACAACAACAAGGC   29002
gb:MN975262|Organism:Severe      CTTGACAGATTGAACCAGCTTGAGAGCAAAATGTCTGGTAAAGGCCAACAACAACAAGGC   29002
gb:MT106052|Organism:Severe      CTTGACAGATTGAACCAGCTTGAGAGCAAAATGTCTGGTAAAGGCCAACAACAACAAGGC   29002
```

```
gb:LC522975|Organism:Severe      CTTGACAGATTGAACCAGCTTGAGAGCAAAATGTCTGGTAAAGGCCAACAACAACAAGGC   28999
gb:LC522973|Organism:Severe      CTTGACAGATTGAACCAGCTTGAGAGCAAAATGTCTGGTAAAGGCCAACAACAACAAGGC   28999
gb:LC522974|Organism:Severe      CTTGACAGATTGAACCAGCTTGAGAGCAAAATGTCTGGTAAAGGCCAACAACAACAAGGC   28999
gb:MN985325|Organism:Severe      CTTGACAGATTGAACCAGCTTGAGAGCAAAATGTCTGGTAAAGGCCAACAACAACAAGGC   29002
gb:MT020881|Organism:Severe      CTTGACAGATTGAACCAGCTTGAGAGCAAAATGTCTGGTAAAGGCCAACAACAACAAGGC   29002
gb:MT020880|Organism:Severe      CTTGACAGATTGAACCAGCTTGAGAGCAAAATGTCTGGTAAAGGCCAACAACAACAAGGC   29002
gb:MT066175|Organism:Severe      CTTGACAGATTGAACCAGCTTGAGAGCAAAATGTCTGGTAAAGGCCAACAACAACAAGGC   29002
gb:MN997409|Organism:Severe      CTTGACAGATTGAACCAGCTTGAGAGCAAAATGTCTGGTAAAGGCCAACAACAACAAGGC   29002
gb:MN938384|Organism:Severe      CTTGACAGATTGAACCAGCTTGAGAGCAAAATGTCTGGTAAAGGCCAACAACAACAAGGC   28970
gb:MT044258|Organism:Severe      CTTGACAGATTGAACCAGCTTGAGAGCAAAATGTCTGGTAAAGGCCAACAACAACAAGGC   28978
gb:MT039890|Organism:Severe      CTTGACAGATTGAACCAGCTTGAGAGCAAAATGTCTGGTAAAGGCCAACAACAACAAGGC   29002
gb:MN988713|Organism:Severe      CTTGACAGATTGAACCAGCTTGAGAGCAAAATGTCTGGTAAAGGCCAACAACAACAAGGC   29002
gb:LC521925|Organism:Severe      CTTGACAGATTGAACCAGCTTGAGAGCAAAATGTCTGGTAAAGGCCAACAACAACAAGGC   28975
gb:MT093571|Organism:Severe      CTTGACAGATTGAACCAGCTTGAGAGCAAAATGTCTGGTAAAGGCCAACAACAACAAGGC   29002
gb:MT039887|Organism:Severe      CTTGACAGATTGAACCAGCTTGAGAGCAAAATGTCTGGTAAAGGCCAACAACAACAAGGC   28999
gb:MT019530|Organism:Severe      CTTGACAGATTGAACCAGCTTGAGAGCAAAATGTCTGGTAAAGGCCAACAACAACAAGGC   29002
gb:MT039888|Organism:Severe      CTTGACAGATTGAACCAGCTTGAGAGCAAAATGTCTGGTAAAGGCCAACAACAACAAGGC   29002
gb:LC522972|Organism:Severe      CTTGACAGATTGAACCAGCTTGAGAGCAAAATGTCTGGTAAAGGCCAACAACAACAAGGC   28999
gb:MT027063|Organism:Severe      CTTGACAGATTGAACCAGCTTGAGAGCAAAATGTCTGGTAAAGGCCAACAACAACAAGGC   29002
gb:MT027062|Organism:Severe      CTTGACAGATTGAACCAGCTTGAGAGCAAAATGTCTGGTAAAGGCCAACAACAACAAGGC   29002
gb:MT019529|Organism:Severe      CTTGACAGATTGAACCAGCTTGAGAGCAAAATGTCTGGTAAAGGCCAACAACAACAAGGC   29002
gb:MN996529|Organism:Severe      CTTGACAGATTGAACCAGCTTGAGAGCAAAATGTCTGGTAAAGGCCAACAACAACAAGGC   28990
gb:MN996531|Organism:Severe      CTTGACAGATTGAACCAGCTTGAGAGCAAAATGTCTGGTAAAGGCCAACAACAACAAGGC   28989
gb:MT066176|Organism:Severe      CTTGACAGATTGAACCAGCTTGAGAGCAAAATGTCTGGTAAAGGCCAACAACAACAAGGC   29002
gb:MT027064|Organism:Severe      CTTGACAGATTGAACCAGCTTGAGAGCAAAATGTCTGGTAAAGGCCAACAACAACAAGGC   29002
gb:MN994468|Organism:Severe      CTTGACAGATTGAACCAGCTTGAGAGCAAAATGTCTGGTAAAGGCCAACAACAACAAGGC   29002
gb:MT072688|Organism:Severe      CTTGACAGATTGAACCAGCTTGAGAGCAAAATGTCTGGTAAAGGCCAACAACAACAAGGC   28987
gb:MN996527|Organism:Severe      CTTGACAGATTGAACCAGCTTGAGAGCAAAATGTCTGGTAAAGGCCAACAACAACAAGGC   28969
gb:MT093631|Organism:Severe      CTTGACAGATTGAACCAGCTTGAGAGCAAAATGTCTGGTAAAGGCCAACAACAACAAGGC   29040
gb:MT106053|Organism:Severe      CTTGACAGATTGAACCAGCTTGAGAGCAAAATGTCTGGTAAAGGCCAACAACAACAAGGC   29002
gb:MT019533|Organism:Severe      CTTGACAGATTGAACCAGCTTGAGAGCAAAATGTCTGGTAAAGGCCAACAACAACAAGGC   29002
gb:MT019531|Organism:Severe      CTTGACAGATTGAACCAGCTTGAGAGCAAAATGTCTGGTAAAGGCCAACAACAACAAGGC   29002
gb:MN996528|Organism:Severe      CTTGACAGATTGAACCAGCTTGAGAGCAAAATGTCTGGTAAAGGCCAACAACAACAAGGC   29002
gb:MN996530|Organism:Severe      CTTGACAGATTGAACCAGCTTGAGAGCAAAATGTCTGGTAAAGGCCAACAACAACAAGGC   28988
gb:MN908947|Organism:Severe      CTTGACAGATTGAACCAGCTTGAGAGCAAAATGTCTGGTAAAGGCCAACAACAACAAGGC   29002
gb:MT019532|Organism:Severe      CTTGACAGATTGAACCAGCTTGAGAGCAAAATGTCTGGTAAAGGCCAACAACAACAAGGC   29002
                                 ************************************************************

gb:MT020781|Organism:Severe      CAAACTGTCACTAAGAAATCTGCTGCTGAGGCTTCTAAGAAGCCTCGGCAAAAACGTACT   29050
gb:MT007544|Organism:Severe      CAAACTGTCACTAAGAAATCTGCTGCTGAGGCTTCTAAGAAGCCTCGGCAAAAACGTACT   29062
gb:MN994467|Organism:Severe      CAAACTGTCACTAAGAAATCTGCTGCTGAGGCTTCTAAGAAGCCTCGGCAAAAACGTACT   29062
gb:MT044257|Organism:Severe      CAAACTGTCACTAAGAAATCTGCTGCTGAGGCTTCTAAGAAGCCTCGGCAAAAACGTACT   29062
gb:MT106054|Organism:Severe      CAAACTGTCACTAAGAAATCTGCTGCTGAGGCTTCTAAGAAGCCTCGGCAAAAACGTACT   29062
gb:MT049951|Organism:Severe      CAAACTGTCACTAAGAAATCTGCTGCTGAGGCTTCTAAGAAGCCTCGGCAAAAACGTACT   29062
gb:MN975262|Organism:Severe      CAAACTGTCACTAAGAAATCTGCTGCTGAGGCTTCTAAGAAGCCTCGGCAAAAACGTACT   29062
gb:MT106052|Organism:Severe      CAAACTGTCACTAAGAAATCTGCTGCTGAGGCTTCTAAGAAGCCTCGGCAAAAACGTACT   29062
gb:LC522975|Organism:Severe      CAAACTGTCACTAAGAAATCTGCTGCTGAGGCTTCTAAGAAGCCTCGGCAAAAACGTACT   29059
gb:LC522973|Organism:Severe      CAAACTGTCACTAAGAAATCTGCTGCTGAGGCTTCTAAGAAGCCTCGGCAAAAACGTACT   29059
gb:LC522974|Organism:Severe      CAAACTGTCACTAAGAAATCTGCTGCTGAGGCTTCTAAGAAGCCTCGGCAAAAACGTACT   29059
gb:MN985325|Organism:Severe      CAAACTGTCACTAAGAAATCTGCTGCTGAGGCTTCTAAGAAGCCTCGGCAAAAACGTACT   29062
gb:MT020881|Organism:Severe      CAAACTGTCACTAAGAAATCTGCTGCTGAGGCTTCTAAGAAGCCTCGGCAAAAACGTACT   29062
gb:MT020880|Organism:Severe      CAAACTGTCACTAAGAAATCTGCTGCTGAGGCTTCTAAGAAGCCTCGGCAAAAACGTACT   29062
gb:MT066175|Organism:Severe      CAAACTGTCACTAAGAAATCTGCTGCTGAGGCTTCTAAGAAGCCTCGGCAAAAACGTACT   29062
gb:MN997409|Organism:Severe      CAAACTGTCACTAAGAAATCTGCTGCTGAGGCTTCTAAGAAGCCTCGGCAAAAACGTACT   29062
gb:MN938384|Organism:Severe      CAAACTGTCACTAAGAAATCTGCTGCTGAGGCTTCTAAGAAGCCTCGGCAAAAACGTACT   29030
gb:MT044258|Organism:Severe      CAAACTGTCACTAAGAAATCTGCTGCTGAGGCTTCTAAGAAGCCTCGGCAAAAACGTACT   29038
gb:MT039890|Organism:Severe      CAAACTGTCACTAAGAAATCTGCTGCTGAGGCTTCTAAGAAGCCTCGGCAAAAACGTACT   29062
gb:MN988713|Organism:Severe      CAAACTGTCACTAAGAAATCTGCTGCTGAGGCTTCTAAGAAGCCTCGGCAAAAACGTACT   29062
gb:LC521925|Organism:Severe      CAAACTGTCACTAAGAAATCTGCTGCTGAGGCTTCTAAGAAGCCTCGGCAAAAACGTACT   29035
gb:MT093571|Organism:Severe      CAAACTGTCACTAAGAAATCTGCTGCTGAGGCTTCTAAGAAGCCTCGGCAAAAACGTACT   29062
gb:MT039887|Organism:Severe      CAAACTGTCACTAAGAAATCTGCTGCTGAGGCTTCTAAGAAGCCTCGGCAAAAACGTACT   29059
gb:MT019530|Organism:Severe      CAAACTGTCACTAAGAAATCTGCTGCTGAGGCTTCTAAGAAGCCTCGGCAAAAACGTACT   29062
gb:MT039888|Organism:Severe      CAAACTGTCACTAAGAAATCTGCTGCTGAGGCTTCTAAGAAGCCTCGGCAAAAACGTACT   29062
gb:LC522972|Organism:Severe      CAAACTGTCACTAAGAAATCTGCTGCTGAGGCTTCTAAGAAGCCTCGGCAAAAACGTACT   29059
gb:MT027063|Organism:Severe      CAAACTGTCACTAAGAAATCTGCTGCTGAGGCTTCTAAGAAGCCTCGGCAAAAACGTACT   29062
```

```
gb:MT027062|Organism:Severe      CAAACTGTCACTAAGAAATCTGCTGCTGAGGCTTCTAAGAAGCCTCGGCAAAAACGTACT    29062
gb:MT019529|Organism:Severe      CAAACTGTCACTAAGAAATCTGCTGCTGAGGCTTCTAAGAAGCCTCGGCAAAAACGTACT    29062
gb:MN996529|Organism:Severe      CAAACTGTCACTAAGAAATCTGCTGCTGAGGCTTCTAAGAAGCCTCGGCAAAAACGTACT    29050
gb:MN996531|Organism:Severe      CAAACTGTCACTAAGAAATCTGCTGCTGAGGCTTCTAAGAAGCCTCGGCAAAAACGTACT    29049
gb:MT066176|Organism:Severe      CAAACTGTCACTAAGAAATCTGCTGCTGAGGCTTCTAAGAAGCCTCGGCAAAAACGTACT    29062
gb:MT027064|Organism:Severe      CAAACTGTCACTAAGAAATCTGCTGCTGAGGCTTCTAAGAAGCCTCGGCAAAAACGTACT    29062
gb:MN994468|Organism:Severe      CAAACTGTCACTAAGAAATCTGCTGCTGAGGCTTCTAAGAAGCCTCGGCAAAAACGTACT    29062
gb:MT072688|Organism:Severe      CAAACTGTCACTAAGAAATCTGCTGCTGAGGCTTCTAAGAAGCCTCGGCAAAAACGTACT    29047
gb:MN996527|Organism:Severe      CAAACTGTCACTAAGAAATCTGCTGCTGAGGCTTCTAAGAAGCCTCGGCAAAAACGTACT    29029
gb:MT093631|Organism:Severe      CAAACTGTCACTAAGAAATCTGCTGCTGAGGCTTCTAAGAAGCCTCGGCAAAAACGTACT    29100
gb:MT106053|Organism:Severe      CAAACTGTCACTAAGAAATCTGCTGCTGAGGCTTCTAAGAAGCCTCGGCAAAAACGTACT    29062
gb:MT019533|Organism:Severe      CAAACTGTCACTAAGAAATCTGCTGCTGAGGCTTCTAAGAAGCCTCGGCAAAAACGTACT    29062
gb:MT019531|Organism:Severe      CAAACTGTCACTAAGAAATCTGCTGCTGAGGCTTCTAAGAAGCCTCGGCAAAAACGTACT    29062
gb:MN996528|Organism:Severe      CAAACTGTCACTAAGAAATCTGCTGCTGAGGCTTCTAAGAAGCCTCGGCAAAAACGTACT    29062
gb:MN996530|Organism:Severe      CAAACTGTCACTAAGAAATCTGCTGCTGAGGCTTCTAAGAAGCCTCGGCAAAAACGTACT    29048
gb:MN908947|Organism:Severe      CAAACTGTCACTAAGAAATCTGCTGCTGAGGCTTCTAAGAAGCCTCGGCAAAAACGTACT    29062
gb:MT019532|Organism:Severe      CAAACTGTCACTAAGAAATCTGCTGCTGAGGCTTCTAAGAAGCCTCGGCAAAAACGTACT    29062
                                 ************************************************************

gb:MT020781|Organism:Severe      GCCACTAAAGCATACAATGTAACACAAGCTTTCGGCAGACGTGGTCCAGAACAAACCCAA    29110
gb:MT007544|Organism:Severe      GCCACTAAAGCATACAATGTAACACAAGCTTTCGGCAGACGTGGTCCAGAACAAACCCAA    29122
gb:MN994467|Organism:Severe      GCCACTAAAGCATACAATGTAACACAAGCTTTCGGCAGACGTGGTCCAGAACAAACCCAA    29122
gb:MT044257|Organism:Severe      GCCACTAAAGCATACAATGTAACACAAGCTTTCGGCAGACGTGGTCCAGAACAAACCCAA    29122
gb:MT106054|Organism:Severe      GCCACTAAAGCATACAATGTAACACAAGCTTTTGGCAGACGTGGTCCAGAACAAACCCAA    29122
gb:MT049951|Organism:Severe      GCCACTAAAGCATACAATGTAACACAAGCTTTCGGCAGACGTGGTCCAGAACAAACCCAA    29122
gb:MN975262|Organism:Severe      GCCACTAAAGCATACAATGTAACACAAGCTTTTGGCAGACGTGGTCCAGAACAAACCCAA    29122
gb:MT106052|Organism:Severe      GCCACTAAAGCATACAATGTAACACAAGCTTTCGGCAGACGTGGTCCAGAACAAACCCAA    29122
gb:LC522975|Organism:Severe      GCCACTAAAGCATACAATGTAACACAAGCTTTTGGCAGACGTGGTCCAGAACAAACCCAA    29119
gb:LC522973|Organism:Severe      GCCACTAAAGCATACAATGTAACACAAGCTTTTGGCAGACGTGGTCCAGAACAAACCCAA    29119
gb:LC522974|Organism:Severe      GCCACTAAAGCATACAATGTAACACAAGCTTTTGGCAGACGTGGTCCAGAACAAACCCAA    29119
gb:MN985325|Organism:Severe      GCCACTAAAGCATACAATGTAACACAAGCTTTCGGCAGACGTGGTCCAGAACAAACCCAA    29122
gb:MT020881|Organism:Severe      GCCACTAAAGCATACAATGTAACACAAGCTTTCGGCAGACGTGGTCCAGAACAAACCCAA    29122
gb:MT020880|Organism:Severe      GCCACTAAAGCATACAATGTAACACAAGCTTTCGGCAGACGTGGTCCAGAACAAACCCAA    29122
gb:MT066175|Organism:Severe      GCCACTAAAGCATACAATGTAACACAAGCTTTCGGCAGACGTGGTCCAGAACAAACCCAA    29122
gb:MN997409|Organism:Severe      GCCACTAAAGCATACAATGTAACACAAGCTTTTGGCAGACGTGGTCCAGAACAAACCCAA    29122
gb:MN938384|Organism:Severe      GCCACTAAAGCATACAATGTAACACAAGCTTTTGGCAGACGTGGTCCAGAACAAACCCAA    29090
gb:MT044258|Organism:Severe      GCCACTAAAGCATACAATGTAACACAAGCTTTCGGCAGACGTGGTCCAGAACAAACCCAA    29098
gb:MT039890|Organism:Severe      GCCACTAAAGCATACAATGTAACACAAGCTTTCGGCAGACGTGGTCCAGAACAAACCCAA    29122
gb:MN988713|Organism:Severe      GCCACTAAAGCATACAATGTAACACAAGCTTTCGGCAGACGTGGTCCAGAACAAACCCAA    29122
gb:LC521925|Organism:Severe      GCCACTAAAGCATACAATGTAACACAAGCTTTCGGCAGACGTGGTCCAGAACAAACCCAA    29095
gb:MT093571|Organism:Severe      GCCACTAAAGCATACAATGTAACACAAGCTTTCGGCAGACGTGGTCCAGAACAAACCCAA    29122
gb:MT039887|Organism:Severe      GCCACTAAAGCATACAATGTAACACAAGCTTTCGGCAGACGTGGTCCAGAACAAACCCAA    29119
gb:MT019530|Organism:Severe      GCCACTAAAGCATACAATGTAACACAAGCTTTCGGCAGACGTGGTCCAGAACAAACCCAA    29122
gb:MT039888|Organism:Severe      GCCACTAAAGCATACAATGTAACACAAGCTTTCGGCAGACGTGGTCCAGAACAAACCCAA    29122
gb:LC522972|Organism:Severe      GCCACTAAAGCATACAATGTAACACAAGCTTTCGGCAGACGTGGTCCAGAACAAACCCAA    29119
gb:MT027063|Organism:Severe      GCCACTAAAGCATACAATGTAACACAAGCTTTCGGCAGACGTGGTCCAGAACAAACCCAA    29122
gb:MT027062|Organism:Severe      GCCACTAAAGCATACAATGTAACACAAGCTTTCGGCAGACGTGGTCCAGAACAAACCCAA    29122
gb:MT019529|Organism:Severe      GCCACTAAAGCATACAATGTAACACAAGCTTTCGGCAGACGTGGTCCAGAACAAACCCAA    29122
gb:MN996529|Organism:Severe      GCCACTAAAGCATACAATGTAACACAAGCTTTCGGCAGACGTGGTCCAGAACAAACCCAA    29110
gb:MN996531|Organism:Severe      GCCACTAAAGCATACAATGTAACACAAGCTTTCGGCAGACGTGGTCCAGAACAAACCCAA    29109
gb:MT066176|Organism:Severe      GCCACTAAAGCATACAATGTAACACAAGCTTTCGGCAGACGTGGTCCAGAACAAACCCAA    29122
gb:MT027064|Organism:Severe      GCCACTAAAGCATACAATGTAACACAAGCTTTCGGCAGACGTGGTCCAGAACAAACCCAA    29122
gb:MN994468|Organism:Severe      GCCACTAAAGCATACAATGTAACACAAGCTTTCGGCAGACGTGGTCCAGAACAAACCCAA    29122
gb:MT072688|Organism:Severe      GCCACTAAAGCATACAATGTAACACAAGCTTTCGGCAGACGTGGTCCAGAACAAACCCAA    29107
gb:MN996527|Organism:Severe      GCCACTAAAGCATACAATGTAACACAAGCTTTCGGCAGACGTGGTCCAGAACAAACCCAA    29089
gb:MT093631|Organism:Severe      GCCACTAAAGCATACAATGTAACACAAGCTTTCGGCAGACGTGGTCCAGAACAAACCCAA    29160
gb:MT106053|Organism:Severe      GCCACTAAAGCATACAATGTAACACAAGCTTTCGGCAGACGTGGTCCAGAACAAACCCAA    29122
gb:MT019533|Organism:Severe      GCCACTAAAGCATACAATGTAACACAAGCTTTCGGCAGACGTGGTCCAGAACAAACCCAA    29122
gb:MT019531|Organism:Severe      GCCACTAAAGCATACAATGTAACACAAGCTTTCGGCAGACGTGGTCCAGAACAAACCCAA    29122
gb:MN996528|Organism:Severe      GCCACTAAAGCATACAATGTAACACAAGCTTTCGGCAGACGTGGTCCAGAACAAACCCAA    29122
gb:MN996530|Organism:Severe      GCCACTAAAGCATACAATGTAACACAAGCTTTCGGCAGACGTGGTCCAGAACAAACCCAA    29108
gb:MN908947|Organism:Severe      GCCACTAAAGCATACAATGTAACACAAGCTTTCGGCAGACGTGGTCCAGAACAAACCCAA    29122
gb:MT019532|Organism:Severe      GCCACTAAAGCATACAATGTAACACAAGCTTTCGGCAGACGTGGTCCAGAACAAACCCAA    29122
                                 ******************************** ***************************
```

```
gb:MT020781|Organism:Severe      GGAAATTTTGGGGACCAGGAACTAATCAGACAAGGAACTGATTACAAACATTGGCCGCAA   29170
gb:MT007544|Organism:Severe      GGAAATTTTGGGGACCAGGAACTAATCAGACAAGGAACTGATTACAAACATTGGCCGCAA   29182
gb:MN994467|Organism:Severe      GGAAATTTTGGGGACCAGGAACTAATCAGACAAGGAACTGATTACAAACATTGGCCGCAA   29182
gb:MT044257|Organism:Severe      GGAAATTTTGGGGACCAGGAACTAATCAGACAAGGAACTGATTACAAACATTGGCCGCAA   29182
gb:MT106054|Organism:Severe      GGAAATTTTGGGGACCAGGAACTAATCAGACAAGGAACTGATTACAAACATTGGCCGCAA   29182
gb:MT049951|Organism:Severe      GGAAATTTTGGGGACCAGGAACTAATCAGACAAGGAACTGATTACAAACATTGGCCGCAA   29182
gb:MN975262|Organism:Severe      GGAAATTTTGGGGACCAGGAACTAATCAGACAAGGAACTGATTACAAACATTGGCCGCAA   29182
gb:MT106052|Organism:Severe      GGAAATTTTGGGGACCAGGAACTAATCAGACAAGGAACTGATTACAAACATTGGCCGCAA   29182
gb:LC522975|Organism:Severe      GGAAATTTTGGGGACCAGGAACTAATCAGACAAGGAACTGATTACAAACATTGGCCGCAA   29179
gb:LC522973|Organism:Severe      GGAAATTTTGGGGACCAGGAACTAATCAGACAAGGAACTGATTACAAACATTGGCCGCAA   29179
gb:LC522974|Organism:Severe      GGAAATTTTGGGGACCAGGAACTAATCAGACAAGGAACTGATTACAAACATTGGCCGCAA   29179
gb:MN985325|Organism:Severe      GGAAATTTTGGGGACCAGGAACTAATCAGACAAGGAACTGATTACAAACATTGGCCGCAA   29182
gb:MT020881|Organism:Severe      GGAAATTTTGGGGACCAGGAACTAATCAGACAAGGAACTGATTACAAACATTGGCCGCAA   29182
gb:MT020880|Organism:Severe      GGAAATTTTGGGGACCAGGAACTAATCAGACAAGGAACTGATTACAAACATTGGCCGCAA   29182
gb:MT066175|Organism:Severe      GGAAATTTTGGGGACCAGGAACTAATCAGACAAGGAACTGATTACAAACATTGGCCGCAA   29182
gb:MN997409|Organism:Severe      GGAAATTTTGGGGACCAGGAACTAATCAGACAAGGAACTGATTACAAACATTGGCCGCAA   29182
gb:MN938384|Organism:Severe      GGAAATTTTGGGGACCAGGAACTAATCAGACAAGGAACTGATTACAAACATTGGCCGCAA   29150
gb:MT044258|Organism:Severe      GGAAATTTTGGGGACCAGGAACTAATCAGACAAGGAACTGATTACAAACATTGGCCGCAA   29158
gb:MT039890|Organism:Severe      GGAAATTTTGGGGACCAGGAACTAATCAGACAAGGAACTGATTACAAACATTGGCCGCAA   29182
gb:MN988713|Organism:Severe      GGAAATTTTGGGGACCAGGAACTAATCAGACAAGGAACTGATTACAAACATTGGCCGCAA   29182
gb:LC521925|Organism:Severe      GGAAATTTTGGGGACCAGGAACTAATCAGACAAGGAACTGATTACAAACATTGGCCGCAA   29155
gb:MT093571|Organism:Severe      GGAAATTTTGGGGACCAGGAACTAATCAGACAAGGAACTGATTACAAACATTGGCCGCAA   29182
gb:MT039887|Organism:Severe      GGAAATTTTGGGGACCAGGAACTAATCAGACAAGGAACTGATTACAAACATTGGCCGCAA   29179
gb:MT019530|Organism:Severe      GGAAATTTTGGGGACCAGGAACTAATCAGACAAGGAACTGATTACAAACATTGGCCGCAA   29182
gb:MT039888|Organism:Severe      GGAAATTTTGGGGACCAGGAACTAATCAGACAAGGAACTGATTACAAACATTGGCCGCAA   29182
gb:LC522972|Organism:Severe      GGAAATTTTGGGGACCAGGAACTAATCAGACAAGGAACTGATTACAAACATTGGCCGCAA   29179
gb:MT027063|Organism:Severe      GGAAATTTTGGGGACCAGGAACTAATCAGACAAGGAACTGATTACAAACATTGGCCGCAA   29182
gb:MT027062|Organism:Severe      GGAAATTTTGGGGACCAGGAACTAATCAGACAAGGAACTGATTACAAACATTGGCCGCAA   29182
gb:MT019529|Organism:Severe      GGAAATTTTGGGGACCAGGAACTAATCAGACAAGGAACTGATTACAAACATTGGCCGCAA   29182
gb:MN996529|Organism:Severe      GGAAATTTTGGGGACCAGGAACTAATCAGACAAGGAACTGATTACAAACATTGGCCGCAA   29170
gb:MN996531|Organism:Severe      GGAAATTTTGGGGACCAGGAACTAATCAGACAAGGAACTGATTACAAACATTGGCCGCAA   29169
gb:MT066176|Organism:Severe      GGAAATTTTGGGGACCAGGAACTAATCAGACAAGGAACTGATTACAAACATTGGCCGCAA   29182
gb:MT027064|Organism:Severe      GGAAATTTTGGGGACCAGGAACTAATCAGACAAGGAACTGATTACAAACATTGGCCGCAA   29182
gb:MN994468|Organism:Severe      GGAAATTTTGGGGACCAGGAACTAATCAGACAAGGAACTGATTACAAACATTGGCCGCAA   29182
gb:MT072688|Organism:Severe      GGAAATTTTGGGGACCAGGAACTAATCAGACAAGGAACTGATTACAAACATTGGCCGCAA   29167
gb:MN996527|Organism:Severe      GGAAATTTTGGGGACCAGGAACTAATCAGACAAGGAACTGATTACAAACATTGGCCGCAA   29149
gb:MT093631|Organism:Severe      GGAAATTTTGGGGACCAGGAACTAATCAGACAAGGAACTGATTACAAACATTGGCCGCAA   29220
gb:MT106053|Organism:Severe      GGAAATTTTGGGGACCAGGAACTAATCAGACAAGGAACTGATTACAAACATTGGCCGCAA   29182
gb:MT019533|Organism:Severe      GGAAATTTTGGGGACCAGGAACTAATCAGACAAGGAACTGATTACAAACATTGGCCGCAA   29182
gb:MT019531|Organism:Severe      GGAAATTTTGGGGACCAGGAACTAATCAGACAAGGAACTGATTACAAACATTGGCCGCAA   29182
gb:MN996528|Organism:Severe      GGAAATTTTGGGGACCAGGAACTAATCAGACAAGGAACTGATTACAAACATTGGCCGCAA   29182
gb:MN996530|Organism:Severe      GGAAATTTTGGGGACCAGGAACTAATCAGACAAGGAACTGATTACAAACATTGGCCGCAA   29168
gb:MN908947|Organism:Severe      GGAAATTTTGGGGACCAGGAACTAATCAGACAAGGAACTGATTACAAACATTGGCCGCAA   29182
gb:MT019532|Organism:Severe      GGAAATTTTGGGGACCAGGAACTAATCAGACAAGGAACTGATTACAAACATTGGCCGCAA   29182
                                 ************************************************************

gb:MT020781|Organism:Severe      ATTGCACAATTTGCCCCCAGCGCTTCAGCGTTCTTCGGAATGTCGCGCATTGGCATGGAA   29230
gb:MT007544|Organism:Severe      ATTGCACAATTTGCCCCCAGCGCTTCAGCGTTCTTCGGAATGTCGCGCATTGGCATGGAA   29242
gb:MN994467|Organism:Severe      ATTGCACAATTTGCCCCCAGCGCTTCAGCGTTCTTCGGAATGTCGCGCATTGGCATGGAA   29242
gb:MT044257|Organism:Severe      ATTGCACAATTTGCCCCCAGCGCTTCAGCGTTCTTCGGAATGTCGCGCATTGGCATGGAA   29242
gb:MT106054|Organism:Severe      ATTGCACAATTTGCCCCCAGCGCTTCAGCGTTCTTCGGAATGTCGCGCATTGGCATGGAA   29242
gb:MT049951|Organism:Severe      ATTGCACAATTTGCCCCCAGCGCTTCAGCGTTCTTCGGAATGTCGCGCATTGGCATGGAA   29242
gb:MN975262|Organism:Severe      ATTGCACAATTTGCCCCCAGCGCTTCAGCGTTCTTCGGAATGTCGCGCATTGGCATGGAA   29242
gb:MT106052|Organism:Severe      ATTGCACAATTTGCCCCCAGCGCTTCAGCGTTCTTCGGAATGTCGCGCATTGGCATGGAA   29242
gb:LC522975|Organism:Severe      ATTGCACAATTTGCCCCCAGCGCTTCAGCGTTCTTCGGAATGTCGCGCATTGGCATGGAA   29239
gb:LC522973|Organism:Severe      ATTGCACAATTTGCCCCCAGCGCTTCAGCGTTCTTCGGAATGTCGCGCATTGGCATGGAA   29239
gb:LC522974|Organism:Severe      ATTGCACAATTTGCCCCCAGCGCTTCAGCGTTCTTCGGAATGTCGCGCATTGGCATGGAA   29239
gb:MN985325|Organism:Severe      ATTGCACAATTTGCCCCCAGCGCTTCAGCGTTCTTCGGAATGTCGCGCATTGGCATGGAA   29242
gb:MT020881|Organism:Severe      ATTGCACAATTTGCCCCCAGCGCTTCAGCGTTCTTCGGAATGTCGCGCATTGGCATGGAA   29242
gb:MT020880|Organism:Severe      ATTGCACAATTTGCCCCCAGCGCTTCAGCGTTCTTCGGAATGTCGCGCATTGGCATGGAA   29242
gb:MT066175|Organism:Severe      ATTGCACAATTTGCCCCCAGCGCTTCAGCGTTCTTCGGAATGTCGCGCATTGGCATGGAA   29242
gb:MN997409|Organism:Severe      ATTGCACAATTTGCCCCCAGCGCTTCAGCGTTCTTCGGAATGTCGCGCATTGGCATGGAA   29242
gb:MN938384|Organism:Severe      ATTGCACAATTTGCCCCCAGCGCTTCAGCGTTCTTCGGAATGTCGCGCATTGGCATGGAA   29210
gb:MT044258|Organism:Severe      ATTGCACAATTTGCCCCCAGCGCTTCAGCGTTCTTCGGAATGTCGCGCATTGGCATGGAA   29218
gb:MT039890|Organism:Severe      ATTGCACAATTTGCCCCCAGCGCTTCAGCGTTCTTCGGAATGTCGCGCATTGGCATGGAA   29242
```

```
gb:MN988713|Organism:Severe      ATTGCACAATTTGCCCCCAGCGCTTCAGCGTTCTTCGGAATGTCGCGCATTGGCATGGAA   29242
gb:LC521925|Organism:Severe      ATTGCACAATTTGCCCCCAGCGCTTCAGCGTTCTTCGGAATGTCGCGCATTGGCATGGAA   29215
gb:MT093571|Organism:Severe      ATTGCACAATTTGCCCCCAGCGCTTCAGCGTTCTTCGGAATGTCGCGCATTGGCATGGAA   29242
gb:MT039887|Organism:Severe      ATTGCACAATTTGCCCCCAGCGCTTCAGCGTTCTTCGGAATGTCGCGCATTGGCATGGAA   29239
gb:MT019530|Organism:Severe      ATTGCACAATTTGCCCCCAGCGCTTCAGCGTTCTTCGGAATGTCGCGCATTGGCATGGAA   29242
gb:MT039888|Organism:Severe      ATTGCACAATTTGCCCCCAGCGCTTCAGCGTTCTTCGGAATGTCGCGCATTGGCATGGAA   29242
gb:LC522972|Organism:Severe      ATTGCACAATTTGCCCCCAGCGCTTCAGCGTTCTTCGGAATGTCGCGCATTGGCATGGAA   29239
gb:MT027063|Organism:Severe      ATTGCACAATTTGCCCCCAGCGCTTCAGCGTTCTTCGGAATGTCGCGCATTGGCATGGAA   29242
gb:MT027062|Organism:Severe      ATTGCACAATTTGCCCCCAGCGCTTCAGCGTTCTTCGGAATGTCGCGCATTGGCATGGAA   29242
gb:MT019529|Organism:Severe      ATTGCACAATTTGCCCCCAGCGCTTCAGCGTTCTTCGGAATGTCGCGCATTGGCATGGAA   29242
gb:MN996529|Organism:Severe      ATTGCACAATTTGCCCCCAGCGCTTCAGCGTTCTTCGGAATGTCGCGCATTGGCATGGAA   29230
gb:MN996531|Organism:Severe      ATTGCACAATTTGCCCCCAGCGCTTCAGCGTTCTTCGGAATGTCGCGCATTGGCATGGAA   29229
gb:MT066176|Organism:Severe      ATTGCACAATTTGCCCCCAGCGCTTCAGCGTTCTTCGGAATGTCGCGCATTGGCATGGAA   29242
gb:MT027064|Organism:Severe      ATTGCACAATTTGCCCCCAGCGCTTCAGCGTTCTTCGGAATGTCGCGCATTGGCATGGAA   29242
gb:MN994468|Organism:Severe      ATTGCACAATTTGCCCCCAGCGCTTCAGCGTTCTTCGGAATGTCGCGCATTGGCATGGAA   29242
gb:MT072688|Organism:Severe      ATTGCACAATTTGCCCCCAGCGCTTCAGCGTTCTTCGGAATGTCGCGCATTGGCATGGAA   29227
gb:MN996527|Organism:Severe      ATTGCACAATTTGCCCCCAGCGCTTCAGCGTTCTTCGGAATGTCGCGCATTGGCATGGAA   29209
gb:MT093631|Organism:Severe      ATTGCACAATTTGCCCCCAGCGCTTCAGCGTTCTTCGGAATGTCGCGCATTGGCATGGAA   29280
gb:MT106053|Organism:Severe      ATTGCACAATTTGCCCCCAGCGCTTCAGCGTTCTTCGGAATGTCGCGCATTGGCATGGAA   29242
gb:MT019533|Organism:Severe      ATTGCACAATTTGCCCCCAGCGCTTCAGCGTTCTTCGGAATGTCGCGCATTGGCATGGAA   29242
gb:MT019531|Organism:Severe      ATTGCACAATTTGCCCCCAGCGCTTCAGCGTTCTTCGGAATGTCGCGCATTGGCATGGAA   29242
gb:MN996528|Organism:Severe      ATTGCACAATTTGCCCCCAGCGCTTCAGCGTTCTTCGGAATGTCGCGCATTGGCATGGAA   29242
gb:MN996530|Organism:Severe      ATTGCACAATTTGCCCCCAGCGCTTCAGCGTTCTTCGGAATGTCGCGCATTGGCATGGAA   29228
gb:MN908947|Organism:Severe      ATTGCACAATTTGCCCCCAGCGCTTCAGCGTTCTTCGGAATGTCGCGCATTGGCATGGAA   29242
gb:MT019532|Organism:Severe      ATTGCACAATTTGCCCCCAGCGCTTCAGCGTTCTTCGGAATGTCGCGCATTGGCATGGAA   29242
                                 ************************************************************

gb:MT020781|Organism:Severe      GTCACACCTTCGGGAACGTGGTTGACCTACACAGGTGCCATCAAATTGGATGACAAAGAT   29290
gb:MT007544|Organism:Severe      GTCACACCTTCGGGAACGTGGTTGACCTACACAGGTGCCATCAAATTGGATGACAAAGAT   29302
gb:MN994467|Organism:Severe      GTCACACCTTCGGGAACGTGGTTGACCTACACAGGTGCCATCAAATTGGATGACAAAGAT   29302
gb:MT044257|Organism:Severe      GTCACACCTTCGGGAACGTGGTTGACCTACACAGGTGCCATCAAATTGGATGACAAAGAT   29302
gb:MT106054|Organism:Severe      GTCACACCTTCGGGAACGTGGTTGACCTACACAGGTGCCATCAAATTGGATGACAAAGAT   29302
gb:MT049951|Organism:Severe      GTCACACCTTCGGGAACGTGGTTGACCTACACAGGTGCCATCAAATTGGATGACAAAGAT   29302
gb:MN975262|Organism:Severe      GTCACACCTTCGGGAACGTGGTTGACCTACACAGGTGCCATCAAATTGGATGACAAAGAT   29302
gb:MT106052|Organism:Severe      GTCACACCTTCGGGAACGTGGTTGACCTACACAGGTGCCATCAAATTGGATGACAAAGAT   29302
gb:LC522975|Organism:Severe      GTCACACCTTCGGGAACGTGGTTGACCTACACAGGTGCCATCAAATTGGATGACAAAGAT   29299
gb:LC522973|Organism:Severe      GTCACACCTTCGGGAACGTGGTTGACCTACACAGGTGCCATCAAATTGGATGACAAAGAT   29299
gb:LC522974|Organism:Severe      GTCACACCTTCGGGAACGTGGTTGACCTACACAGGTGCCATCAAATTGGATGACAAAGAT   29299
gb:MN985325|Organism:Severe      GTCACACCTTCGGGAACGTGGTTGACCTACACAGGTGCCATCAAATTGGATGACAAAGAT   29302
gb:MT020881|Organism:Severe      GTCACACCTTCGGGAACGTGGTTGACCTACACAGGTGCCATCAAATTGGATGACAAAGAT   29302
gb:MT020880|Organism:Severe      GTCACACCTTCGGGAACGTGGTTGACCTACACAGGTGCCATCAAATTGGATGACAAAGAT   29302
gb:MT066175|Organism:Severe      GTCACACCTTCGGGAACGTGGTTGACCTACACAGGTGCCATCAAATTGGATGACAAAGAT   29302
gb:MN997409|Organism:Severe      GTCACACCTTCGGGAACGTGGTTGACCTACACAGGTGCCATCAAATTGGATGACAAAGAT   29302
gb:MN938384|Organism:Severe      GTCACACCTTCGGGAACGTGGTTGACCTACACAGGTGCCATCAAATTGGATGACAAAGAT   29270
gb:MT044258|Organism:Severe      GTCACACCTTCGGGAACGTGGTTGACCTACACAGGTGCCATCAAATTGGATGACAAAGAT   29278
gb:MT039890|Organism:Severe      GTCACACCTTCGGGAACGTGGTTGACCTACACAGGTGCCATCAAATTGGATGACAAAGAT   29302
gb:MN988713|Organism:Severe      GTCACACCTTCGGGAACGTGGTTGACCTACACAGGTGCCATCAAATTGGATGACAAAGAT   29302
gb:LC521925|Organism:Severe      GTCACACCTTCGGGAACGTGGTTGACCTACACAGGTGCCATCAAATTGGATGACAAAGAT   29275
gb:MT093571|Organism:Severe      GTCACACCTTCGGGAACGTGGTTGACCTACACAGGTGCCATCAAATTGGATGACAAAGAT   29302
gb:MT039887|Organism:Severe      GTCACACCTTCGGGAACGTGGTTGACCTACACAGGTGCCATCAAATTGGATGACAAAGAT   29299
gb:MT019530|Organism:Severe      GTCACACCTTCGGGAACGTGGTTGACCTACACAGGTGCCATCAAATTGGATGACAAAGAT   29302
gb:MT039888|Organism:Severe      GTCACACCTTCGGGAACGTGGTTGACCTACACAGGTGCCATCAAATTGGATGACAAAGAT   29302
gb:LC522972|Organism:Severe      GTCACACCTTCGGGAACGTGGTTGACCTACACAGGTGCCATCAAATTGGATGACAAAGAT   29299
gb:MT027063|Organism:Severe      GTCACACCTTCGGGAACGTGGTTGACCTACACAGGTGCCATCAAATTGGATGACAAAGAT   29302
gb:MT027062|Organism:Severe      GTCACACCTTCGGGAACGTGGTTGACCTACACAGGTGCCATCAAATTGGATGACAAAGAT   29302
gb:MT019529|Organism:Severe      GTCACACCTTCGGGAACGTGGTTGACCTACACAGGTGCCATCAAATTGGATGACAAAGAT   29302
gb:MN996529|Organism:Severe      GTCACACCTTCGGGAACGTGGTTGACCTACACAGGTGCCATCAAATTGGATGACAAAGAT   29290
gb:MN996531|Organism:Severe      GTCACACCTTCGGGAACGTGGTTGACCTACACAGGTGCCATCAAATTGGATGACAAAGAT   29289
gb:MT066176|Organism:Severe      GTCACACCTTCGGGAACGTGGTTGACCTACACAGGTGCCATCAAATTGGATGACAAAGAT   29302
gb:MT027064|Organism:Severe      GTCACACCTTCGGGAACGTGGTTGACCTACACAGGTGCCATCAAATTGGATGACAAAGAT   29302
gb:MN994468|Organism:Severe      GTCACACCTTCGGGAACGTGGTTGACCTACACAGGTGCCATCAAATTGGATGACAAAGAT   29302
gb:MT072688|Organism:Severe      GTCACACCTTCGGGAACGTGGTTGACCTACACAGGTGCCATCAAATTGGATGACAAAGAT   29287
gb:MN996527|Organism:Severe      GTCACACCTTCGGGAACGTGGTTGACCTACACAGGTGCCATCAAATTGGATGACAAAGAT   29269
gb:MT093631|Organism:Severe      GTCACACCTTCGGGAACGTGGTTGACCTACACAGGTGCCATCAAATTGGATGACAAAGAT   29340
gb:MT106053|Organism:Severe      GTCACACCTTCGGGAACGTGGTTGACCTACACAGGTGCCATCAAATTGGATGACAAAGAT   29302
```

```
gb:MT019533|Organism:Severe      GTCACACCTTCGGGAACGTGGTTGACCTACACAGGTGCCATCAAATTGGATGACAAAGAT   29302
gb:MT019531|Organism:Severe      GTCACACCTTCGGGAACGTGGTTGACCTACACAGGTGCCATCAAATTGGATGACAAAGAT   29302
gb:MN996528|Organism:Severe      GTCACACCTTCGGGAACGTGGTTGACCTACACAGGTGCCATCAAATTGGATGACAAAGAT   29302
gb:MN996530|Organism:Severe      GTCACACCTTCGGGAACGTGGTTGACCTACACAGGTGCCATCAAATTGGATGACAAAGAT   29288
gb:MN908947|Organism:Severe      GTCACACCTTCGGGAACGTGGTTGACCTACACAGGTGCCATCAAATTGGATGACAAAGAT   29302
gb:MT019532|Organism:Severe      GTCACACCTTCGGGAACGTGGTTGACCTACACAGGTGCCATCAAATTGGATGACAAAGAT   29302
                                 ************************************************************

gb:MT020781|Organism:Severe      CCAAATTTCAAAGATCAAGTCATTTTGCTGAATAAGCATATTGACGCATACAAAACATTC   29350
gb:MT007544|Organism:Severe      CCAAATTTCAAAGATCAAGTCATTTTGCTGAATAAGCATATTGACGCATACAAAACATTC   29362
gb:MN994467|Organism:Severe      CCAAATTTCAAAGATCAAGTCATTTTGCTGAATAAGCATATTGACGCATACAAAACATTC   29362
gb:MT044257|Organism:Severe      CCAAATTTCAAAGATCAAGTCATTTTGCTGAATAAGCATATTGACGCATACAAAACATTC   29362
gb:MT106054|Organism:Severe      CCAAATTTCAAAGATCAAGTCATTTTGCTGAATAAGCATATTGACGCATACAAAACATTC   29362
gb:MT049951|Organism:Severe      CCAAATTTCAAAGATCAAGTCATTTTGCTGAATAAGCATATTGACGCATACAAAACATTC   29362
gb:MN975262|Organism:Severe      CCAAATTTCAAAGATCAAGTCATTTTGCTGAATAAGCATATTGACGCATACAAAACATTC   29362
gb:MT106052|Organism:Severe      CCAAATTTCAAAGATCAAGTCATTTTGCTGAATAAGCATATTGACGCATACAAAACATTC   29362
gb:LC522975|Organism:Severe      CCAAATTTCAAAGATCAAGTCATTTTGCTGAATAAGCATATTGACGCATACAAAACATTC   29359
gb:LC522973|Organism:Severe      CCAAATTTCAAAGATCAAGTCATTTTGCTGAATAAGCATATTGACGCATACAAAACATTC   29359
gb:LC522974|Organism:Severe      CCAAATTTCAAAGATCAAGTCATTTTGCTGAATAAGCATATTGACGCATACAAAACATTC   29359
gb:MN985325|Organism:Severe      CCAAATTTCAAAGATCAAGTCATTTTGCTGAATAAGCATATTGACGCATACAAAACATTC   29362
gb:MT020881|Organism:Severe      CCAAATTTCAAAGATCAAGTCATTTTGCTGAATAAGCATATTGACGCATACAAAACATTC   29362
gb:MT020880|Organism:Severe      CCAAATTTCAAAGATCAAGTCATTTTGCTGAATAAGCATATTGACGCATACAAAACATTC   29362
gb:MT066175|Organism:Severe      CCAAATTTCAAAGATCAAGTCATTTTGCTGAATAAGCATATTGACGCATACAAAACATTC   29362
gb:MN997409|Organism:Severe      CCAAATTTCAAAGATCAAGTCATTTTGCTGAATAAGCATATTGACGCATACAAAACATTC   29362
gb:MN938384|Organism:Severe      CCAAATTTCAAAGATCAAGTCATTTTGCTGAATAAGCATATTGACGCATACAAAACATTC   29330
gb:MT044258|Organism:Severe      CCAAATTTCAAAGATCAAGTCATTTTGCTGAATAAGCATATTGACGCATACAAAACATTC   29338
gb:MT039890|Organism:Severe      CCAAATTTCAAAGATCAAGTCATTTTGCTGAATAAGCATATTGACGCATACAAAACATTC   29362
gb:MN988713|Organism:Severe      CCAAATTTCAAAGATCAAGTCATTTTGCTGAATAAGCATATTGACGCATACAAAACATTC   29362
gb:LC521925|Organism:Severe      CCAAATTTCAAAGATCAAGTCATTTTGCTGAATAAGCATATTGACGCATACAAAACATTC   29335
gb:MT093571|Organism:Severe      CCAAATTTCAAAGATCAAGTCATTTTGCTGAATAAGCATATTGACGCATACAAAACATTC   29362
gb:MT039887|Organism:Severe      CCAAATTTCAAAGATCAAGTCATTTTGCTGAATAAGCATATTGACGCATACAAAACATTC   29359
gb:MT019530|Organism:Severe      CCAAATTTCAAAGATCAAGTCATTTTGCTGAATAAGCATATTGACGCATACAAAACATTC   29362
gb:MT039888|Organism:Severe      CCAAATTTCAAAGATCAAGTCATTTTGCTGAATAAGCATATTGACGCATACAAAACATTC   29362
gb:LC522972|Organism:Severe      TCAAATTTCAAAGATCAAGTCATTTTGCTGAATAAGCATATTGACGCATACAAAACATTC   29359
gb:MT027063|Organism:Severe      CCAAATTTCAAAGATCAAGTCATTTTGCTGAATAAGCATATTGACGCATACAAAACATTC   29362
gb:MT027062|Organism:Severe      CCAAATTTCAAAGATCAAGTCATTTTGCTGAATAAGCATATTGACGCATACAAAACATTC   29362
gb:MT019529|Organism:Severe      CCAAATTTCAAAGATCAAGTCATTTTGCTGAATAAGCATATTGACGCATACAAAACATTC   29362
gb:MN996529|Organism:Severe      CCAAATTTCAAAGATCAAGTCATTTTGCTGAATAAGCATATTGACGCATACAAAACATTC   29350
gb:MN996531|Organism:Severe      CCAAATTTCAAAGATCAAGTCATTTTGCTGAATAAGCATATTGACGCATACAAAACATTC   29349
gb:MT066176|Organism:Severe      CCAAATTTCAAAGATCAAGTCATTTTGCTGAATAAGCATATTGACGCATACAAAACATTC   29362
gb:MT027064|Organism:Severe      CCAAATTTCAAAGATCAAGTCATTTTGCTGAATAAGCATATTGACGCATACAAAACATTC   29362
gb:MN994468|Organism:Severe      CCAAATTTCAAAGATCAAGTCATTTTGCTGAATAAGCATATTGACGCATACAAAACATTC   29362
gb:MT072688|Organism:Severe      CCAAATTTCAAAGATCAAGTCATTTTGCTGAATAAGCATATTGACGCATACAAAACATTC   29347
gb:MN996527|Organism:Severe      CCAAATTTCAAAGATCAAGTCATTTTGCTGAATAAGCATATTGACGCATACAAAACATTC   29329
gb:MT093631|Organism:Severe      CCAAATTTCAAAGATCAAGTCATTTTGCTGAATAAGCATATTGACGCATACAAAACATTC   29400
gb:MT106053|Organism:Severe      CCAAATTTCAAAGATCAAGTCATTTTGCTGAATAAGCATATTGACGCATACAAAACATTC   29362
gb:MT019533|Organism:Severe      CCAAATTTCAAAGATCAAGTCATTTTGCTGAATAAGCATATTGACGCATACAAAACATTC   29362
gb:MT019531|Organism:Severe      CCAAATTTCAAAGATCAAGTCATTTTGCTGAATAAGCATATTGACGCATACAAAACATTC   29362
gb:MN996528|Organism:Severe      CCAAATTTCAAAGATCAAGTCATTTTGCTGAATAAGCATATTGACGCATACAAAACATTC   29362
gb:MN996530|Organism:Severe      CCAAATTTCAAAGATCAAGTCATTTTGCTGAATAAGCATATTGACGCATACAAAACATTC   29348
gb:MN908947|Organism:Severe      CCAAATTTCAAAGATCAAGTCATTTTGCTGAATAAGCATATTGACGCATACAAAACATTC   29362
gb:MT019532|Organism:Severe      CCAAATTTCAAAGATCAAGTCATTTTGCTGAATAAGCATATTGACGCATACAAAACATTC   29362
                                  ***********************************************************

gb:MT020781|Organism:Severe      CCACCAACAGAGCCTAAAAAGGACAAAAAGAAGAAGGCTGATGAAACTCAAGCCTTACCG   29410
gb:MT007544|Organism:Severe      CCACCAACAGAGCCTAAAAAGGACAAAAAGAAGAAGGCTGATGAAACTCAAGCCTTACCG   29422
gb:MN994467|Organism:Severe      CCACCAACAGAGCCTAAAAAGGACAAAAAGAAGAAGGCTGATGAAACTCAAGCCTTACCG   29422
gb:MT044257|Organism:Severe      CCACCAACAGAGCCTAAAAAGGACAAAAAGAAGAAGGCTGATGAAACTCAAGCCTTACCG   29422
gb:MT106054|Organism:Severe      CCACCAACAGAGCCTAAAAAGGACAAAAAGAAGAAGGCTGATGAAACTCAAGCCTTACCG   29422
gb:MT049951|Organism:Severe      CCACCAACAGAGCCTAAAAAGGACAAAAAGAAGAAGGCTGATGAAACTCAAGCCTTACCG   29422
gb:MN975262|Organism:Severe      CCACCAACAGAGCCTAAAAAGGACAAAAAGAAGAAGGCTGATGAAACTCAAGCCTTACCG   29422
gb:MT106052|Organism:Severe      CCACCAACAGAGCCTAAAAAGGACAAAAAGAAGAAGGCTGATGAAACTCAAGCCTTACCG   29422
gb:LC522975|Organism:Severe      CCACCAACAGAGCCTAAAAAGGACAAAAAGAAGAAGGCTGATGAAACTCAAGCCTTACCG   29419
gb:LC522973|Organism:Severe      CCACCAACAGAGCCTAAAAAGGACAAAAAGAAGAAGGCTGATGAAACTCAAGCCTTACCG   29419
gb:LC522974|Organism:Severe      CCACCAACAGAGCCTAAAAAGGACAAAAAGAAGAAGGCTGATGAAACTCAAGCCTTACCG   29419
```

```
gb:MN985325|Organism:Severe      CCACCAACAGAGCCTAAAAAGGACAAAAAGAAGAAGGCTGATGAAACTCAAGCCTTACCG    29422
gb:MT020881|Organism:Severe      CCACCAACAGAGCCTAAAAAGGACAAAAAGAAGAAGGCTGATGAAACTCAAGCCTTACCG    29422
gb:MT020880|Organism:Severe      CCACCAACAGAGCCTAAAAAGGACAAAAAGAAGAAGGCTGATGAAACTCAAGCCTTACCG    29422
gb:MT066175|Organism:Severe      CCACCAACAGAGCCTAAAAAGGACAAAAAGAAGAAGGCTGATGAAACTCAAGCCTTACCG    29422
gb:MN997409|Organism:Severe      CCACCAACAGAGCCTAAAAAGGACAAAAAGAAGAAGGCTGATGAAACTCAAGCCTTACCG    29422
gb:MN938384|Organism:Severe      CCACCAACAGAGCCTAAAAAGGACAAAAAGAAGAAGGCTGATGAAACTCAAGCCTTACCG    29390
gb:MT044258|Organism:Severe      CCACCAACAGAGCCTAAAAAGGACAAAAAGAAGAAGGCTGATGAAACTCAAGCCTTACCG    29398
gb:MT039890|Organism:Severe      CCACCAACAGAGCCTAAAAAGGACAAAAAGAAGAAGGCTGATGAAACTCAAGCCTTACCG    29422
gb:MN988713|Organism:Severe      CCACCAACAGAGCCTAAAAAGGACAAAAAGAAGAAGGCTGATGAAACTCAAGCCTTACCG    29422
gb:LC521925|Organism:Severe      CCACCAACAGAGCCTAAAAAGGACAAAAAGAAGAAGGCTGATGAAACTCAAGCCTTACCG    29395
gb:MT093571|Organism:Severe      CCACCAACAGAGCCTAAAAAGGACAAAAAGAAGAAGGCTGATGAAACTCAAGCCTTACCG    29422
gb:MT039887|Organism:Severe      CCACCAACAGAGCCTAAAAAGGACAAAAAGAAGAAGGCTGATGAAACTCAAGCCTTACCG    29419
gb:MT019530|Organism:Severe      CCACCAACAGAGCCTAAAAAGGACAAAAAGAAGAAGGCTGATGAAACTCAAGCCTTACCG    29422
gb:MT039888|Organism:Severe      CCACCAACAGAGCCTAAAAAGGACAAAAAGAAGAAGGCTGATGAAACTCAAGCCTTACCG    29422
gb:LC522972|Organism:Severe      CCACCAACAGAGCCTAAAAAGGACAAAAAGAAGAAGGCTGATGAAACTCAAGCCTTACCG    29419
gb:MT027063|Organism:Severe      CCACCAACAGAGCCTAAAAAGGACAAAAAGAAGAAGGCTGATGAAACTCAAGCCTTACCG    29422
gb:MT027062|Organism:Severe      CCACCAACAGAGCCTAAAAAGGACAAAAAGAAGAAGGCTGATGAAACTCAAGCCTTACCG    29422
gb:MT019529|Organism:Severe      CCACCAACAGAGCCTAAAAAGGACAAAAAGAAGAAGGCTGATGAAACTCAAGCCTTACCG    29422
gb:MN996529|Organism:Severe      CCACCAACAGAGCCTAAAAAGGACAAAAAGAAGAAGGCTGATGAAACTCAAGCCTTACCG    29410
gb:MN996531|Organism:Severe      CCACCAACAGAGCCTAAAAAGGACAAAAAGAAGAAGGCTGATGAAACTCAAGCCTTACCG    29409
gb:MT066176|Organism:Severe      CCACCAACAGAGCCTAAAAAGGACAAAAAGAAGAAGGCTGATGAAACTCAAGCCTTACCG    29422
gb:MT027064|Organism:Severe      CCACCAACAGAGCCTAAAAAGGACAAAAAGAAGAAGGCTGATGAAACTCAAGCCTTACCG    29422
gb:MN994468|Organism:Severe      CCACCAACAGAGCCTAAAAAGGACAAAAAGAAGAAGGCTGATGAAACTCAAGCCTTACCG    29422
gb:MT072688|Organism:Severe      CCACCAACAGAGCCTAAAAAGGACAAAAAGAAGAAGGCTGATGAAACTCAAGCCTTACCG    29407
gb:MN996527|Organism:Severe      CCACCAACAGAGCCTAAAAAGGACAAAAAGAAGAAGGCTGATGAAACTCAAGCCTTACCG    29389
gb:MT093631|Organism:Severe      CCACCAACAGAGCCTAAAAAGGACAAAAAGAAGAAGGCTGATGAAACTCAAGCCTTACCG    29460
gb:MT106053|Organism:Severe      CCACCAACAGAGCCTAAAAAGGACAAAAAGAAGAAGGCTGATGAAACTCAAGCCTTACCG    29422
gb:MT019533|Organism:Severe      CCACCAACAGAGCCTAAAAAGGACAAAAAGAAGAAGGCTGATGAAACTCAAGCCTTACCG    29422
gb:MT019531|Organism:Severe      CCACCAACAGAGCCTAAAAAGGACAAAAAGAAGAAGGCTGATGAAACTCAAGCCTTACCG    29422
gb:MN996528|Organism:Severe      CCACCAACAGAGCCTAAAAAGGACAAAAAGAAGAAGGCTGATGAAACTCAAGCCTTACCG    29422
gb:MN996530|Organism:Severe      CCACCAACAGAGCCTAAAAAGGACAAAAAGAAGAAGGCTGATGAAACTCAAGCCTTACCG    29408
gb:MN908947|Organism:Severe      CCACCAACAGAGCCTAAAAAGGACAAAAAGAAGAAGGCTGATGAAACTCAAGCCTTACCG    29422
gb:MT019532|Organism:Severe      CCACCAACAGAGCCTAAAAAGGACAAAAAGAAGAAGGCTGATGAAACTCAAGCCTTACCG    29422
                                 ************************************************************

gb:MT020781|Organism:Severe      CAGAGACAGAAGAAACAGCAAACTGTGACTCTTCTTCCTGCTGCAGATTTGGATGATTTC    29470
gb:MT007544|Organism:Severe      CAGAGACAGAAGAAACAGCAAACTGTGACTCTTCTTCCTGCTGCAGATTTGGATGATTTC    29482
gb:MN994467|Organism:Severe      CAGAGACAGAAGAAACAGCAAACTGTGACTCTTCTTCCTGCTGCAGATTTGGATGATTTC    29482
gb:MT044257|Organism:Severe      CAGAGACAGAAGAAACAGCAAACTGTGACTCTTCTTCCTGCTGCAGATTTGGATGATTTC    29482
gb:MT106054|Organism:Severe      CAGAGACAGAAGAAACAGCAAACTGTGACTCTTCTTCCTGCTGCAGATTTGGATGATTTC    29482
gb:MT049951|Organism:Severe      CAGAGACAGAAGAAACAGCAAACTGTGACTCTTCTTCCTGCTGCAGATTTGGATGATTTC    29482
gb:MN975262|Organism:Severe      CAGAGACAGAAGAAACAGCAAACTGTGACTCTTCTTCCTGCTGCAGATTTGGATGATTTC    29482
gb:MT106052|Organism:Severe      CAGAGACAGAAGAAACAGCAAACTGTGACTCTTCTTCCTGCTGCAGATTTGGATGATTTC    29482
gb:LC522975|Organism:Severe      CAGAGACAGAAGAAACAGCAAACTGTGACTCTTCTTCCTGCTGCAGATTTGGATGATTTC    29479
gb:LC522973|Organism:Severe      CAGAGACAGAAGAAACAGCAAACTGTGACTCTTCTTCCTGCTGCAGATTTGGATGATTTC    29479
gb:LC522974|Organism:Severe      CAGAGACAGAAGAAACAGCAAACTGTGACTCTTCTTCCTGCTGCAGATTTGGATGATTTC    29479
gb:MN985325|Organism:Severe      CAGAGACAGAAGAAACAGCAAACTGTGACTCTTCTTCCTGCTGCAGATTTGGATGATTTC    29482
gb:MT020881|Organism:Severe      CAGAGACAGAAGAAACAGCAAACTGTGACTCTTCTTCCTGCTGCAGATTTGGATGATTTC    29482
gb:MT020880|Organism:Severe      CAGAGACAGAAGAAACAGCAAACTGTGACTCTTCTTCCTGCTGCAGATTTGGATGATTTC    29482
gb:MT066175|Organism:Severe      CAGAGACAGAAGAAACAGCAAACTGTGACTCTTCTTCCTGCTGCAGATTTGGATGATTTC    29482
gb:MN997409|Organism:Severe      CAGAGACAGAAGAAACAGCAAACTGTGACTCTTCTTCCTGCTGCAGATTTGGATGATTTC    29482
gb:MN938384|Organism:Severe      CAGAGACAGAAGAAACAGCAAACTGTGACTCTTCTTCCTGCTGCAGATTTGGATGATTTC    29450
gb:MT044258|Organism:Severe      CAGAGACAGAAGAAACAGCAAACTGTGACTCTTCTTCCTGCTGCAGATTTGGATGATTTC    29458
gb:MT039890|Organism:Severe      CAGAGACAGAAGAAACAGCAAACTGTGACTCTTCTTCCTGCTGCAGATTTGGATGATTTC    29482
gb:MN988713|Organism:Severe      CAGAGACAGAAGAAACAGCAAACTGTGACTCTTCTTCCTGCTGCAGATTTGGATGATTTC    29482
gb:LC521925|Organism:Severe      CAGAGACAGAAGAAACAGCAAACTGTGACTCTTCTTCCTGCTGCAGATTTGGATGATTTC    29455
gb:MT093571|Organism:Severe      CAGAGACAGAAGAAACAGCAAACTGTGACTCTTCTTCCTGCTGCAGATTTGGATGATTTC    29482
gb:MT039887|Organism:Severe      CAGAGACAGAAGAAACAGCAAACTGTGACTCTTCTTCCTGCTGCAGATTTGGATGATTTC    29479
gb:MT019530|Organism:Severe      CAGAGACAGAAGAAACAGCAAACTGTGACTCTTCTTCCTGCTGCAGATTTGGATGATTTC    29482
gb:MT039888|Organism:Severe      CAGAGACAGAAGAAACAGCAAACTGTGACTCTTCTTCCTGCTGCAGATTTGGATGATTTC    29482
gb:LC522972|Organism:Severe      CAGAGACAGAAGAAACAGCAAACTGTGACTCTTCTTCCTGCTGCAGATTTGGATGATTTC    29479
gb:MT027063|Organism:Severe      CAGAGACAGAAGAAACAGCAAACTGTGACTCTTCTTCCTGCTGCAGATTTGGATGATTTC    29482
gb:MT027062|Organism:Severe      CAGAGACAGAAGAAACAGCAAACTGTGACTCTTCTTCCTGCTGCAGATTTGGATGATTTC    29482
gb:MT019529|Organism:Severe      CAGAGACAGAAGAAACAGCAAACTGTGACTCTTCTTCCTGCTGCAGATTTGGATGATTTC    29482
gb:MN996529|Organism:Severe      CAGAGACAGAAGAAACAGCAAACTGTGACTCTTCTTCCTGCTGCAGATTTGGATGATTTC    29470
```

```
gb:MN996531|Organism:Severe      CAGAGACAGAAGAAACAGCAAACTGTGACTCTTCTTCCTGCTGCAGATTTGGATGATTTC   29469
gb:MT066176|Organism:Severe      CAGAGACAGAAGAAACAGCAAACTGTGACTCTTCTTCCTGCTGCAGATTTGGATGATTTC   29482
gb:MT027064|Organism:Severe      CAGAGACAGAAGAAACAGCAAACTGTGACTCTTCTTCCTGCTGCAGATTTGGATGATTTC   29482
gb:MN994468|Organism:Severe      CAGAGACAGAAGAAACAGCAAACTGTGACTCTTCTTCCTGCTGCAGATTTGGATGATTTC   29482
gb:MT072688|Organism:Severe      CAGAGACAGAAGAAACAGCAAACTGTGACTCTTCTTCCTGCTGCAGATTTGGATGATTTC   29467
gb:MN996527|Organism:Severe      CAGAGACAGAAGAAACAGCAAACTGTGACTCTTCTTCCTGCTGCAGATTTGGATGATTTC   29449
gb:MT093631|Organism:Severe      CAGAGACAGAAGAAACAGCAAACTGTGACTCTTCTTCCTGCTGCAGATTTGGATGATTTC   29520
gb:MT106053|Organism:Severe      CAGAGACAGAAGAAACAGCAAACTGTGACTCTTCTTCCTGCTGCAGATTTGGATGATTTC   29482
gb:MT019533|Organism:Severe      CAGAGACAGAAGAAACAGCAAACTGTGACTCTTCTTCCTGCTGCAGATTTGGATGATTTC   29482
gb:MT019531|Organism:Severe      CAGAGACAGAAGAAACAGCAAACTGTGACTCTTCTTCCTGCTGCAGATTTGGATGATTTC   29482
gb:MN996528|Organism:Severe      CAGAGACAGAAGAAACAGCAAACTGTGACTCTTCTTCCTGCTGCAGATTTGGATGATTTC   29482
gb:MN996530|Organism:Severe      CAGAGACAGAAGAAACAGCAAACTGTGACTCTTCTTCCTGCTGCAGATTTGGATGATTTC   29468
gb:MN908947|Organism:Severe      CAGAGACAGAAGAAACAGCAAACTGTGACTCTTCTTCCTGCTGCAGATTTGGATGATTTC   29482
gb:MT019532|Organism:Severe      CAGAGACAGAAGAAACAGCAAACTGTGACTCTTCTTCCTGCTGCAGATTTGGATGATTTC   29482
                                 ************************************************************

gb:MT020781|Organism:Severe      TCCAAACAATTGCAACAATCCATGAGCAGTGCTGACTCAACTCAGGCCTAAACTCATGCA   29530
gb:MT007544|Organism:Severe      TCCAAACAATTGCAACAATCCATGAGCAGTGCTGACTCAACTCAGGCCTAAACTCATGCA   29542
gb:MN994467|Organism:Severe      TCCAAACAATTGCAACAATCCATGAGCAGTGCTGACTCAACTCAGGCCTAAACTCATGCA   29542
gb:MT044257|Organism:Severe      TCCAAACAATTGCAACAATCCATGAGCAGTGCTGACTCAACTCAGGCCTAAACTCATGCA   29542
gb:MT106054|Organism:Severe      TCCAAACAATTGCAACAATCCATGAGCAGTGCTGACTCAACTCAGGCCTAAACTCATGCA   29542
gb:MT049951|Organism:Severe      TCCAAACAATTGCAACAATCCATGAGCAGTGCTGACTCAACTCAGGCCTAAACTCATGCA   29542
gb:MN975262|Organism:Severe      TCCAAACAATTGCAACAATCCATGAGCAGTGCTGACTCAACTCAGGCCTAAACTCATGCA   29542
gb:MT106052|Organism:Severe      TCCAAACAATTGCAACAATCCATGAGCAGTGCTGACTCAACTCAGGCCTAAACTCATGCA   29542
gb:LC522975|Organism:Severe      TCCAAACAATTGCAACAATCCATGAGCAGTGCTGACTCAACTCAGGCCTAAACTCATGCA   29539
gb:LC522973|Organism:Severe      TCCAAACAATTGCAACAATCCATGAGCAGTGCTGACTCAACTCAGGCCTAAACTCATGCA   29539
gb:LC522974|Organism:Severe      TCCAAACAATTGCAACAATCCATGAGCAGTGCTGACTCAACTCAGGCCTAAACTCATGCA   29539
gb:MN985325|Organism:Severe      TCCAAACAATTGCAACAATCCATGAGCAGTGCTGACTCAACTCAGGCCTAAACTCATGCA   29542
gb:MT020881|Organism:Severe      TCCAAACAATTGCAACAATCCATGAGCAGTGCTGACTCAACTCAGGCCTAAACTCATGCA   29542
gb:MT020880|Organism:Severe      TCCAAACAATTGCAACAATCCATGAGCAGTGCTGACTCAACTCAGGCCTAAACTCATGCA   29542
gb:MT066175|Organism:Severe      TCCAAACAATTGCAACAATCCATGAGCAGTGCTGACTCAACTCAGGCCTAAACTCATGCA   29542
gb:MN997409|Organism:Severe      TCCAAACAATTGCAACAATCCATGAGCAGTGCTGACTCAACTCAGGCCTAAACTCATGCA   29542
gb:MN938384|Organism:Severe      TCCAAACAATTGCAACAATCCATGAGCAGTGCTGACTCAACTCAGGCCTAAACTCATGCA   29510
gb:MT044258|Organism:Severe      TCCAAACAATTGCAACAATCCATGAGCAGTGCTGACTCAACTCAGGCCTAAACTCATGCA   29518
gb:MT039890|Organism:Severe      TCCAAACAATTGCAACAATCCATGAGCAGTGCTGACTCAACTCAGGCCTAAACTCATGCA   29542
gb:MN988713|Organism:Severe      TCCAAACAATTGCAACAATCCATGAGCAGTGCTGACTCAACTCAGGCCTAAACTCATGCA   29542
gb:LC521925|Organism:Severe      TCCAAACAATTGCAACAATCCATGAGCAGTGCTGACTCAACTCAGGCCTAAACTCATGCA   29515
gb:MT093571|Organism:Severe      TCCAAACAATTGCAACAATCCATGAGCAGTGCTGACTCAACTCAGGCCTAAACTCATGCA   29542
gb:MT039887|Organism:Severe      TCCAAACAATTGCAACAATCCATGAGCAGTGCTGACTCAACTCAGGCCTAAACTCATGCA   29539
gb:MT019530|Organism:Severe      TCCAAACAATTGCAACAATCCATGAGCAGTGCTGACTCAACTCAGGCCTAAACTCATGCA   29542
gb:MT039888|Organism:Severe      TCCAAACAATTGCAACAATCCATGAGCAGTGCTGACTCAACTCAGGCCTAAACTCATGCA   29542
gb:LC522972|Organism:Severe      TCCAAACAATTGCAACAATCCATGAGCAGTGCTGACTCAACTCAGGCCTAAACTCATGCA   29539
gb:MT027063|Organism:Severe      TCCAAACAATTGCAACAATCCATGAGCAGTGCTGACTCAACTCAGGCCTAAACTCATGCA   29542
gb:MT027062|Organism:Severe      TCCAAACAATTGCAACAATCCATGAGCAGTGCTGACTCAACTCAGGCCTAAACTCATGCA   29542
gb:MT019529|Organism:Severe      TCCAAACAATTGCAACAATCCATGAGCAGTGCTGACTCAACTCAGGCCTAAACTCATGCA   29542
gb:MN996529|Organism:Severe      TCCAAACAATTGCAACAATCCATGAGCAGTGCTGACTCAACTCAGGCCTAAACTCATGCA   29530
gb:MN996531|Organism:Severe      TCCAAACAATTGCAACAATCCATGAGCAGTGCTGACTCAACTCAGGCCTAAACTCATGCA   29529
gb:MT066176|Organism:Severe      TCCAAACAATTGCAACAATCCATGAGCAGTGCTGACTCAACTCAGGCCTAAACTCATGCA   29542
gb:MT027064|Organism:Severe      TCCAAACAATTGCAACAATCCATGAGCAGTGCTGACTCAACTCAGGCCTAAACTCATGCA   29542
gb:MN994468|Organism:Severe      TCCAAACAATTGCAACAATCCATGAGCAGTGCTGACTCAACTCAGGCCTAAACTCATGCA   29542
gb:MT072688|Organism:Severe      TCCAAACAATTGCAACAATCCATGAGCAGTGCTGACTCAACTCAGGCCTAAACTCATGCA   29527
gb:MN996527|Organism:Severe      TCCAAACAATTGCAACAATCCATGAGCAGTGCTGACTCAACTCAGGCCTAAACTCATGCA   29509
gb:MT093631|Organism:Severe      TCCAAACAATTGCAACAATCCATGAGCAGTGCTGACTCAACTCAGGCCTAAACTCATGCA   29580
gb:MT106053|Organism:Severe      TCCAAACAATTGCAACAATCCATGAGCAGTGCTGACTCAACTCAGGCCTAAACTCATGCA   29542
gb:MT019533|Organism:Severe      TCCAAACAATTGCAACAATCCATGAGCAGTGCTGACTCAACTCAGGCCTAAACTCATGCA   29542
gb:MT019531|Organism:Severe      TCCAAACAATTGCAACAATCCATGAGCAGTGCTGACTCAACTCAGGCCTAAACTCATGCA   29542
gb:MN996528|Organism:Severe      TCCAAACAATTGCAACAATCCATGAGCAGTGCTGACTCAACTCAGGCCTAAACTCATGCA   29542
gb:MN996530|Organism:Severe      TCCAAACAATTGCAACAATCCATGAGCAGTGCTGACTCAACTCAGGCCTAAACTCATGCA   29528
gb:MN908947|Organism:Severe      TCCAAACAATTGCAACAATCCATGAGCAGTGCTGACTCAACTCAGGCCTAAACTCATGCA   29542
gb:MT019532|Organism:Severe      TCCAAACAATTGCAACAATCCATGAGCAGTGCTGACTCAACTCAGGCCTAAACTCATGCA   29542
                                 ************************************************************

gb:MT020781|Organism:Severe      GACCACACAAGGCAGATGGGCTATATAAACGTTTTCGCTTTTCCGTTTACGATATATAGT   29590
gb:MT007544|Organism:Severe      GACCACACAAGGCAGATGGGCTATATAAACGTTTTCGCTTTTCCGTTTACGATATATAGT   29602
gb:MN994467|Organism:Severe      GACCACACAAGGCAGATGGGCTATATAAACGTTTTCGCTTTTCCGTTTACGATATATAGT   29602
```

```
gb:MT044257|Organism:Severe      GACCACACAAGGCAGATGGGCTATATAAACGTTTTCGCTTTTCCGTTTACGATATATAGT    29602
gb:MT106054|Organism:Severe      GACCACACAAGGCAGATGGGCTATATAAACGTTTTCGCTTTTCCGTTTACGATATATAGT    29602
gb:MT049951|Organism:Severe      GACCACACAAGGCAGATGGGCTATATAAACGTTTTCGCTTTTCCGTTTACGATATATAGT    29602
gb:MN975262|Organism:Severe      GACCACACAAGGCAGATGGGCTATATAAACGTTTTCGCTTTTCCGTTTACGATATATAGT    29602
gb:MT106052|Organism:Severe      GACCACACAAGGCAGATGGGCTATATAAACGTTTTCGCTTTTCCGTTTACGATATATAGT    29602
gb:LC522975|Organism:Severe      GACCACACAAGGCAGATGGGCTATATAAACGTTTTCGCTTTTCCGTTTACGATATATAGT    29599
gb:LC522973|Organism:Severe      GACCACACAAGGCAGATGGGCTATATAAACGTTTTCGCTTTTCCGTTTACGATATATAGT    29599
gb:LC522974|Organism:Severe      GACCACACAAGGCAGATGGGCTATATAAACGTTTTCGCTTTTCCGTTTACGATATATAGT    29599
gb:MN985325|Organism:Severe      GACCACACAAGGCAGATGGGCTATATAAACGTTTTCGCTTTTCCGTTTACGATATATAGT    29602
gb:MT020881|Organism:Severe      GACCACACAAGGCAGATGGGCTATATAAACGTTTTCGCTTTTCCGTTTACGATATATAGT    29602
gb:MT020880|Organism:Severe      GACCACACAAGGCAGATGGGCTATATAAACGTTTTCGCTTTTCCGTTTACGATATATAGT    29602
gb:MT066175|Organism:Severe      GACCACACAAGGCAGATGGGCTATATAAACGTTTTCGCTTTTCCGTTTACGATATATAGT    29602
gb:MN997409|Organism:Severe      GACCACACAAGGCAGATGGGCTATATAAACGTTTTCGCTTTTCCGTTTACGATATATAGT    29602
gb:MN938384|Organism:Severe      GACCACACAAGGCAGATGGGCTATATAAACGTTTTCGCTTTTCCGTTTACGATATATAGT    29570
gb:MT044258|Organism:Severe      GACCACACAAGGCAGATGGGCTATATAAACGTTTTCGCTTTTCCGTTTACGATATATAGT    29578
gb:MT039890|Organism:Severe      GACCACACAAGGCAGATGGGCTATATAAACGTTTTCGCTTTTCCGTTTACGATATATAGT    29602
gb:MN988713|Organism:Severe      GACCACACAAGGCAGATGGGCTATATAAACGTTTTCGCTTTTCCGTTTACGATATATAGT    29602
gb:LC521925|Organism:Severe      GACCACACAAGGCAGATGGGCTATATAAACGTTTTCGCTTTTCCGTTTACGATATATAGT    29575
gb:MT093571|Organism:Severe      GACCACACAAGGCAGATGGGCTATATAAACGTTTTCGCTTTTCCGTTTACGATATATAGT    29602
gb:MT039887|Organism:Severe      GACCACACAAGGCAGATGGGCTATATAAACGTTTTCGCTTTTCCGTTTACGATATATAGT    29599
gb:MT019530|Organism:Severe      GACCACACAAGGCAGATGGGCTATATAAACGTTTTCGCTTTTCCGTTTACGATATATAGT    29602
gb:MT039888|Organism:Severe      GACCACACAAGGCAGATGGGCTATATAAACGTTTTCGCTTTTCCGTTTACGATATATAGT    29602
gb:LC522972|Organism:Severe      GACCACACAAGGCAGATGGGCTATATAAACGTTTTCGCTTTTCCGTTTACGATATATAGT    29599
gb:MT027063|Organism:Severe      GACCACACAAGGCAGATGGGCTATATAAACGTTTTCGCTTTTCCGTTTACGATATATAGT    29602
gb:MT027062|Organism:Severe      GACCACACAAGGCAGATGGGCTATATAAACGTTTTCGCTTTTCCGTTTACGATATATAGT    29602
gb:MT019529|Organism:Severe      GACCACACAAGGCAGATGGGCTATATAAACGTTTTCGCTTTTCCGTTTACGATATATAGT    29602
gb:MN996529|Organism:Severe      GACCACACAAGGCAGATGGGCTATATAAACGTTTTCGCTTTTCCGTTTACGATATATAGT    29590
gb:MN996531|Organism:Severe      GACCACACAAGGCAGATGGGCTATATAAACGTTTTCGCTTTTCCGTTTACGATATATAGT    29589
gb:MT066176|Organism:Severe      GACCACACAAGGCAGATGGGCTATATAAACGTTTTCGCTTTTCCGTTTACGATATATAGT    29602
gb:MT027064|Organism:Severe      GACCACACAAGGCAGATGGGCTATATAAACGTTTTCGCTTTTCCGTTTACGATATATAGT    29602
gb:MN994468|Organism:Severe      GACCACACAAGGCAGATGGGCTATATAAACGTTTTCGCTTTTCCGTTTACGATATATAGT    29602
gb:MT072688|Organism:Severe      GACCACACAAGGCAGATGGGCTATATAAACGTTTTCGCTTTTCCGTTTACGATATATAGT    29587
gb:MN996527|Organism:Severe      GACCACACAAGGCAGATGGGCTATATAAACGTTTTCGCTTTTCCGTTTACGATATATAGT    29569
gb:MT093631|Organism:Severe      GACCACACAAGGCAGATGGGCTATATAAACGTTTTCGCTTTTCCGTTTACGATATATAGT    29640
gb:MT106053|Organism:Severe      GACCACACAAGGCAGATGGGCTATATAAACGTTTTCGCTTTTCCGTTTACGATATATAGT    29602
gb:MT019533|Organism:Severe      GACCACACAAGGCAGATGGGCTATATAAACGTTTTCGCTTTTCCGTTTACGATATATAGT    29602
gb:MT019531|Organism:Severe      GACCACACAAGGCAGATGGGCTATATAAACGTTTTCGCTTTTCCGTTTACGATATATAGT    29602
gb:MN996528|Organism:Severe      GACCACACAAGGCAGATGGGCTATATAAACGTTTTCGCTTTTCCGTTTACGATATATAGT    29602
gb:MN996530|Organism:Severe      GACCACACAAGGCAGATGGGCTATATAAACGTTTTCGCTTTTCCGTTTACGATATATAGT    29588
gb:MN908947|Organism:Severe      GACCACACAAGGCAGATGGGCTATATAAACGTTTTCGCTTTTCCGTTTACGATATATAGT    29602
gb:MT019532|Organism:Severe      GACCACACAAGGCAGATGGGCTATATAAACGTTTTCGCTTTTCCGTTTACGATATATAGT    29602
                                 ************************************************************

gb:MT020781|Organism:Severe      CTACTCTTGTGCAGAATGAATTCTCGTAACTACATAGCACAAGTAGATGTAGTTAACTTT    29650
gb:MT007544|Organism:Severe      CTACTCTTGTGCAGAATGAATTCTCGTAACTACATAGCACAAGTAGATGTAGTTAACTTT    29662
gb:MN994467|Organism:Severe      CTACTCTTGTGCAGAATGAATTCTCGTAACTACATAGCACAAGTAGATGTAGTTAACTTT    29662
gb:MT044257|Organism:Severe      CTACTCTTGTGCAGAATGAATTCTCGTAACTACATAGCACAAGTAGATGTAGTTAACTTT    29662
gb:MT106054|Organism:Severe      CTACTCTTGTGCAGAATGAATTCTCGTAACTACATAGCACAAGTAGATGTAGTTAACTTT    29662
gb:MT049951|Organism:Severe      CTACTCTTGTGCAGAATGAATTCTCGTAACTACATAGCACAAGTAGATGTAGTTAACTTT    29662
gb:MN975262|Organism:Severe      CTACTCTTGTGCAGAATGAATTCTCGTAACTACATAGCACAAGTAGATGTAGTTAACTTT    29662
gb:MT106052|Organism:Severe      CTACTCTTGTGCAGAATGAATTCTCGTAACTACATAGCACAAGTAGATGTAGTTAACTTT    29662
gb:LC522975|Organism:Severe      CTACTCTTGTGCAGAATGAATTCTCGTAACTACATAGCACAAGTAGATGTAGTTAACTTT    29659
gb:LC522973|Organism:Severe      CTACTCTTGTGCAGAATGAATTCTCGTAACTACATAGCACAAGTAGATGTAGTTAACTTT    29659
gb:LC522974|Organism:Severe      CTACTCTTGTGCAGAATGAATTCTCGTAACTACATAGCACAAGTAGATGTAGTTAACTTT    29659
gb:MN985325|Organism:Severe      CTACTCTTGTGCAGAATGAATTCTCGTAACTACATAGCACAAGTAGATGTAGTTAACTTT    29662
gb:MT020881|Organism:Severe      CTACTCTTGTGCAGAATGAATTCTCGTAACTACATAGCACAAGTAGATGTAGTTAACTTT    29662
gb:MT020880|Organism:Severe      CTACTCTTGTGCAGAATGAATTCTCGTAACTACATAGCACAAGTAGATGTAGTTAACTTT    29662
gb:MT066175|Organism:Severe      CTACTCTTGTGCAGAATGAATTCTCGTAACTACATAGCACAAGTAGATGTAGTTAACTTT    29662
gb:MN997409|Organism:Severe      CTACTCTTGTGCAGAATGAATTCTCGTAACTACATAGCACAAGTAGATGTAGTTAACTTT    29662
gb:MN938384|Organism:Severe      CTACTCTTGTGCAGAATGAATTCTCGTAACTACATAGCACAAGTAGATGTAGTTAACTTT    29630
gb:MT044258|Organism:Severe      CTACTCTTGTGCAGAATGAATTCTCGTAACTACATAGCACAAGTAGATGTAGTTAACTTT    29638
gb:MT039890|Organism:Severe      CTACTCTTGTGCAGAATGAATTCTCGTAACTACATAGCACAAGTAGATGTAGTTAACTTT    29662
gb:MN988713|Organism:Severe      CTACTCTTGTGCAGAATGAATTCTCGTAACTACATAGCACAAGTAGATGTAGTTAACTTT    29662
gb:LC521925|Organism:Severe      CTACTCTTGTGCAGAATGAATTCTCGTAACTACATAGCACAAGTAGATGTAGTTAACTTT    29635
gb:MT093571|Organism:Severe      CTACTCTTGTGCAGAATGAATTCTCGTAACTACATAGCACAAGTAGATGTAGTTAACTTT    29662
```

```
gb:MT039887|Organism:Severe      CTACTCTTGTGCAGAATGAATTCTCGTAACTACATAGCACAAGTAGATGTAGTTAACTTT   29659
gb:MT019530|Organism:Severe      CTACTCTTGTGCAGAATGAATTCTCGTAACTACATAGCACAAGTAGATGTAGTTAACTTT   29662
gb:MT039888|Organism:Severe      CTACTCTTGTGCAGAATGAATTCTCGTAACTACATAGCACAAGTAGATGTAGTTAACTTT   29662
gb:LC522972|Organism:Severe      CTACTCTTGTGCAGAATGAATTCTCGTAACTACATAGCACAAGTAGATGTAGTTAACTTT   29659
gb:MT027063|Organism:Severe      CTACTCTTGTGCAGAATGAATTCTCGTAACTACATAGCACAAGTAGATGTAGTTAACTTT   29662
gb:MT027062|Organism:Severe      CTACTCTTGTGCAGAATGAATTCTCGTAACTACATAGCACAAGTAGATGTAGTTAACTTT   29662
gb:MT019529|Organism:Severe      CTACTCTTGTGCAGAATGAATTCTCGTAACTACATAGCACAAGTAGATGTAGTTAACTTT   29662
gb:MN996529|Organism:Severe      CTACTCTTGTGCAGAATGAATTCTCGTAACTACATAGCACAAGTAGATGTAGTTAACTTT   29650
gb:MN996531|Organism:Severe      CTACTCTTGTGCAGAATGAATTCTCGTAACTACATAGCACAAGTAGATGTAGTTAACTTT   29649
gb:MT066176|Organism:Severe      CTACTCTTGTGCAGAATGAATTCTCGTAACTACATAGCACAAGTAGATGTAGTTAACTTT   29662
gb:MT027064|Organism:Severe      CTACTCTTGTGCAGAATGAATTCTCGTAACTACATAGCACAAGTAGATGTAGTTAACTTT   29662
gb:MN994468|Organism:Severe      CTACTCTTGTGCAGAATGAATTCTCGTAACTACATAGCACAAGTAGATGTAGTTAACTTT   29662
gb:MT072688|Organism:Severe      CTACTCTTGTGCAGAATGAATTCTCGTAACTACATAGCACAAGTAGATGTAGTTAACTTT   29647
gb:MN996527|Organism:Severe      CTACTCTTGTGCAGAATGAATTCTCGTAACTACATAGCACAAGTAGATGTAGTTAACTTT   29629
gb:MT093631|Organism:Severe      CTACTCTTGTGCAGAATGAATTCTCGTAACTACATAGCACAAGTAGATGTAGTTAACTTT   29700
gb:MT106053|Organism:Severe      CTACTCTTGTGCAGAATGAATTCTCGTAACTACATAGCACAAGTAGATGTAGTTAACTTT   29662
gb:MT019533|Organism:Severe      CTACTCTTGTGCAGAATGAATTCTCGTAACTACATAGCACAAGTAGATGTAGTTAACTTT   29662
gb:MT019531|Organism:Severe      CTACTCTTGTGCAGAATGAATTCTCGTAACTACATAGCACAAGTAGATGTAGTTAACTTT   29662
gb:MN996528|Organism:Severe      CTACTCTTGTGCAGAATGAATTCTCGTAACTACATAGCACAAGTAGATGTAGTTAACTTT   29662
gb:MN996530|Organism:Severe      CTACTCTTGTGCAGAATGAATTCTCGTAACTACATAGCACAAGTAGATGTAGTTAACTTT   29648
gb:MN908947|Organism:Severe      CTACTCTTGTGCAGAATGAATTCTCGTAACTACATAGCACAAGTAGATGTAGTTAACTTT   29662
gb:MT019532|Organism:Severe      CTACTCTTGTGCAGAATGAATTCTCGTAACTACATAGCACAAGTAGATGTAGTTAACTTT   29662
                                 ************************************************************

gb:MT020781|Organism:Severe      AATCTCACATAGCAATCTTTAATCAGTGTGTAACATTAGGGAGGACTTGAAAGAGCCACC   29710
gb:MT007544|Organism:Severe      AATCTCACATAGCAATCTTTAATCAGTGTGTAACATTAGGGAGGACTTGAAAGAGCCACC   29722
gb:MN994467|Organism:Severe      AATCTCACATAGCAATCTTTAATCAGTGTGTAACATTAGGGAGGACTTGAAAGAGCCACC   29722
gb:MT044257|Organism:Severe      AATCTCACATAGCAATCTTTAATCAGTGTGTAACATTAGGGAGGACTTGAAAGAGCCACC   29722
gb:MT106054|Organism:Severe      AATCTCACATAGCAATCTTTAATCAGTGTGTAACATTAGGGAGGACTTGAAAGAGCCACC   29722
gb:MT049951|Organism:Severe      AATCTCACATAGCAATCTTTAATCAGTGTGTAACATTAGGGAGGACTTGAAAGAGCCACC   29722
gb:MN975262|Organism:Severe      AATCTCACATAGCAATCTTTAATCAGTGTGTAACATTAGGGAGGACTTGAAAGAGCCACC   29722
gb:MT106052|Organism:Severe      AATCTCACATAGCAATCTTTAATCAGTGTGTAACATTAGGGAGGACTTGAAAGAGCCACC   29722
gb:LC522975|Organism:Severe      AATCTCACATAGCAATCTTTAATCAGTGTGTAACATTAGGGATGACTTGAAAGAGCCACC   29719
gb:LC522973|Organism:Severe      AATCTCACATAGCAATCTTTAATCAGTGTGTAACATTAGGGAGGACTTGAAAGAGCCACC   29719
gb:LC522974|Organism:Severe      AATCTCACATAGCAATCTTTAATCAGTGTGTAACATTAGGGAGGACTTGAAAGAGCCACC   29719
gb:MN985325|Organism:Severe      AATCTCACATAGCAATCTTTAATCAGTGTGTAACATTAGGGAGGACTTGAAAGAGCCACC   29722
gb:MT020881|Organism:Severe      AATCTCACATAGCAATCTTTAATCAGTGTGTAACATTAGGGAGGACTTGAAAGAGCCACC   29722
gb:MT020880|Organism:Severe      AATCTCACATAGCAATCTTTAATCAGTGTGTAACATTAGGGAGGACTTGAAAGAGCCACC   29722
gb:MT066175|Organism:Severe      AATCTCACATAGCAATCTTTAATCAGTGTGTAACATTAGGGAGGACTTGAAAGAGCCACC   29722
gb:MN997409|Organism:Severe      AATCTCACATAGCAATCTTTAATCAGTGTGTAACATTAGGGAGGACTTGAAAGAGCCACC   29722
gb:MN938384|Organism:Severe      AATCTCACATAGCAATCTTTAATCAGTGTGTAACATTAGGGAGGACTTGAAAGAGCCACC   29690
gb:MT044258|Organism:Severe      AATCTCACATAGCAATCTTTAATCAGTGTGTAACATTAGGGAGGACTTGAAAGAGCCACC   29698
gb:MT039890|Organism:Severe      AATCTCACATAGCAATCTTTAATCAGTGTGTAACATTAGGGAGGACTTGAAAGAGCCACC   29722
gb:MN988713|Organism:Severe      AATCTCACATAGCAATCTTTAATCAGTGTGTAACATTAGGGAGGACTTGAAAGAGCCACC   29722
gb:LC521925|Organism:Severe      AATCTCACATAGCAATCTTTAATCAGTGTGTAACATTAGGGAGGACTTGAAAGAGCCACC   29695
gb:MT093571|Organism:Severe      AATCTCACATAGCAATCTTTAATCAGTGTGTAACATTAGGGAGGACTTGAAAGAGCCACC   29722
gb:MT039887|Organism:Severe      AATCTCACATAGCAATCTTTAATCAGTGTGTAACATTAGGGAGGACTTGAAAGAGCCACC   29719
gb:MT019530|Organism:Severe      AATCTCACATAGCAATCTTTAATCAGTGTGTAACATTAGGGAGGACTTGAAAGAGCCACC   29722
gb:MT039888|Organism:Severe      AATCTCACATAGCAATCTTTAATCAGTGTGTAACATTAGGGAGGACTTGAAAGAGCCACC   29722
gb:LC522972|Organism:Severe      AATCTCACATAGCAATCTTTAATCAGTGTGTAACATTAGGGAGGACTTGAAAGAGCCACC   29719
gb:MT027063|Organism:Severe      AATCTCACATAGCAATCTTTAATCAGTGTGTAACATTAGGGAGGACTTGAAAGAGCCACC   29722
gb:MT027062|Organism:Severe      AATCTCACATAGCAATCTTTAATCAGTGTGTAACATTAGGGAGGACTTGAAAGAGCCACC   29722
gb:MT019529|Organism:Severe      AATCTCACATAGCAATCTTTAATCAGTGTGTAACATTAGGGAGGACTTGAAAGAGCCACC   29722
gb:MN996529|Organism:Severe      AATCTCACATAGCAATCTTTAATCAGTGTGTAACATTAGGGAGGACTTGAAAGAGCCACC   29710
gb:MN996531|Organism:Severe      AATCTCACATAGCAATCTTTAATCAGTGTGTAACATTAGGGAGGACTTGAAAGAGCCACC   29709
gb:MT066176|Organism:Severe      AATCTCACATAGCAATCTTTAATCAGTGTGTAACATTAGGGAGGACTTGAAAGAGCCACC   29722
gb:MT027064|Organism:Severe      AATCTCACATAGCAATCTTTAATCAGTGTGTAACATTAGGGAGGACTTGAAAGAGCCACC   29722
gb:MN994468|Organism:Severe      AATCTCACATAGCAATCTTTAATCAGTGTGTAACATTAGGGAGGACTTGAAAGAGCCACC   29722
gb:MT072688|Organism:Severe      AATCTCACATAGCAATCTTTAATCAGTGTGTAACATTAGGGAGGACTTGAAAGAGCCACC   29707
gb:MN996527|Organism:Severe      AATCTCACATAGCAATCTTTAATCAGTGTGTAACATTAGGGAGGACTTGAAAGAGCCACC   29689
gb:MT093631|Organism:Severe      AATCTCACATAGCAATCTTTAATCAGTGTGTAACATTAGGGAGGACTTGAAAGAGCCACC   29760
gb:MT106053|Organism:Severe      AATCTCACATAGCAATCTTTAATCAGTGTGTAACATTAGGGAGGACTTGAAAGAGCCACC   29722
gb:MT019533|Organism:Severe      AATCTCACATAGCAATCTTTAATCAGTGTGTAACATTAGGGAGGACTTGAAAGAGCCACC   29722
gb:MT019531|Organism:Severe      AATCTCACATAGCAATCTTTAATCAGTGTGTAACATTAGGGAGGACTTGAAAGAGCCACC   29722
gb:MN996528|Organism:Severe      AATCTCACATAGCAATCTTTAATCAGTGTGTAACATTAGGGAGGACTTGAAAGAGCCACC   29722
```

```
gb:MN996530|Organism:Severe      AATCTCACATAGCAATCTTTAATCAGTGTGTAACATTAGGGAGGACTTGAAAGAGCCACC   29708
gb:MN908947|Organism:Severe      AATCTCACATAGCAATCTTTAATCAGTGTGTAACATTAGGGAGGACTTGAAAGAGCCACC   29722
gb:MT019532|Organism:Severe      AATCTCACATAGCAATCTTTAATCAGTGTGTAACATTAGGGAGGACTTGAAAGAGCCACC   29722
                                 ****************************************** *****************

gb:MT020781|Organism:Severe      ACATTTTCACCGAGGCCACGCGGAGTACGATCGAGTGTACAGTGAACAATGCTAGGGAGA   29770
gb:MT007544|Organism:Severe      ACATTTTCACCGAGGCCACGCGGAGTA----------TACAGTGAACAATGCTAGGGAGA   29772
gb:MN994467|Organism:Severe      ACATTTTCACCGAGGCCACGCGGAGTACGATCGAGTGTACAGTGAACAATGCTAGGGAGA   29782
gb:MT044257|Organism:Severe      ACATTTTCACCGAGGCCACGCGGAGTACGATCGAGTGTACAGTGAACAATGCTAGGGAGA   29782
gb:MT106054|Organism:Severe      ACATTTTCACCGAGGCCACGCGGAGTACGATCGAGTGTACAGTGAACAATGCTAGGGAGA   29782
gb:MT049951|Organism:Severe      ACATTTTCACCGAGGCCACGCGGAGTACGATCGAGTGTACAGTGAACAATGCTAGGGAGA   29782
gb:MN975262|Organism:Severe      ACATTTTCACCGAGGCCACGCGGAGTACGATCGAGTGTACAGTGAACAATGCTAGGGAGA   29782
gb:MT106052|Organism:Severe      ACATTTTCACCGAGGCCACACGGAGTACGATCGAGTGTACAGTGAACAATGCTAGGGAGA   29782
gb:LC522975|Organism:Severe      ACATTTTCACCGAGGCCACGCGGAGTACGATCGAGTGTACAGTGAACAATGCTAGGGAGA   29779
gb:LC522973|Organism:Severe      ACATTTTCACCGAGGCCACGCGGAGTACGATCGAGTGTACAGTGAACAATGCTAGGGAGA   29779
gb:LC522974|Organism:Severe      ACATTTTCACCGAGGCCACGCGGAGTACGATCGAGTGTACAGTGAACAATGCTAGGGAGA   29779
gb:MN985325|Organism:Severe      ACATTTTCACCGAGGCCACGCGGAGTACGATCGAGTGTACAGTGAACAATGCTAGGGAGA   29782
gb:MT020881|Organism:Severe      ACATTTTCACCGAGGCCACGCGGAGTACGATCGAGTGTACAGTGAACAATGCTAGGGAGA   29782
gb:MT020880|Organism:Severe      ACATTTTCACCGAGGCCACGCGGAGTACGATCGAGTGTACAGTGAACAATGCTAGGGAGA   29782
gb:MT066175|Organism:Severe      ACATTTTCACCGAGGCCACGCGGAGTACGATCGAGTGTACAGTGAACAATGCTAGGGAGA   29782
gb:MN997409|Organism:Severe      ACATTTTCACCGAGGCCACGCGGAGTACGATCGAGTGTACAGTGAACAATGCTAGGGAGA   29782
gb:MN938384|Organism:Severe      ACATTTTCACCGAGGCCACGCGGAGTACGATCGAGTGTACAGTGAACAATGCTAGGGAGA   29750
gb:MT044258|Organism:Severe      ACATTTTCACCGAGGCCACGCGGAGTACGATCGAGTGTACAGTGAACAATGCTAGGGAGA   29758
gb:MT039890|Organism:Severe      ACATTTTCACCGAGGCCACGCGGAGTACGATCGAGTGTACAGTGAACAATGCTAGGGAGA   29782
gb:MN988713|Organism:Severe      ACATTTTCACCGAGGCCACGCGGAGTACGATCGAGTGTACAGTGAACAATGCTAGGGAGA   29782
gb:LC521925|Organism:Severe      ACATTTTCACCGAGGCCACGCGGAGTACGATCGAGTGTACAGTGAACAATGCTAGGGAGA   29755
gb:MT093571|Organism:Severe      ACATTTTCACCGAGGCCACGCGGAGTACGATCGAGTGTACAGTGAACAATGCTAGGGAGA   29782
gb:MT039887|Organism:Severe      ACATTTTCACCGAGGCCACGCGGAGTACGATCGAGTGTACAGTGAACAATGCTAGGGAGA   29779
gb:MT019530|Organism:Severe      ACATTTTCACCGAGGCCACGCGGAGTACGATCGAGTGTACAGTGAACAATGCTAGGGAGA   29782
gb:MT039888|Organism:Severe      ACATTTTCACCGAGGCCACGCGGAGTACGATCGAGTGTACAGTGAACAATGCTAGGGAGA   29782
gb:LC522972|Organism:Severe      ACATTTTCACCGAGGCCACGCGGAGTACGATCGAGTGTACAGTGAACAATGCTAGGGAGA   29779
gb:MT027063|Organism:Severe      ACATTTTCACCGAGGCCACGCGGAGTACGATCGAGTGTACAGTGAACAATGCTAGGGAGA   29782
gb:MT027062|Organism:Severe      ACATTTTCACCGAGGCCACGCGGAGTACGATCGAGTGTACAGTGAACAATGCTAGGGAGA   29782
gb:MT019529|Organism:Severe      ACATTTTCACCGAGGCCACGCGGAGTACGATCGAGTGTACAGTGAACAATGCTAGGGAGA   29782
gb:MN996529|Organism:Severe      ACATTTTCACCGAGGCCACGCGGAGTACGATCGAGTGTACAGTGAACAATGCTAGGGAGA   29770
gb:MN996531|Organism:Severe      ACATTTTCACCGAGGCCACGCGGAGTACGATCGAGTGTACAGTGAACAATGCTAGGGAGA   29769
gb:MT066176|Organism:Severe      ACATTTTCACCGAGGCCACGCGGAGTACGATCGAGTGTACAGTGAACAATGCTAGGGAGA   29782
gb:MT027064|Organism:Severe      ACATTTTCACCGAGGCCACGCGGAGTACGATCGAGTGTACAGTGAACAATGCTAGGGAGA   29782
gb:MN994468|Organism:Severe      ACATTTTCACCGAGGCCACGCGGAGTACGATCGAGTGTACAGTGAACAATGCTAGGGAGA   29782
gb:MT072688|Organism:Severe      ACATTTTCACCGAGGCCACGCGGAGTACGATCGAGTGTACAGTGAACAATGCTAGGGAGA   29767
gb:MN996527|Organism:Severe      ACATTTTCACCGAGGCCACGCGGAGTACGATCGAGTGTACAGTGAACAATGCTAGGGAGA   29749
gb:MT093631|Organism:Severe      ACATTTTCACCGAGGCCACGCGGAGTACGATCGAGTGTACAGTGAACAATGCTAGGGAGA   29820
gb:MT106053|Organism:Severe      ACATTTTCACCGAGGCCACGCGGAGTACGATCGAGTGTACAGTGAACAATGCTAGGGAGA   29782
gb:MT019533|Organism:Severe      ACATTTTCACCGAGGCCACGCGGAGTACGATCGAGTGTACAGTGAACAATGCTAGGGAGA   29782
gb:MT019531|Organism:Severe      ACATTTTCACCGAGGCCACGCGGAGTACGATCGAGTGTACAGTGAACAATGCTAGGGAGA   29782
gb:MN996528|Organism:Severe      ACATTTTCACCGAGGCCACGCGGAGTACGATCGAGTGTACAGTGAACAATGCTAGGGAGA   29782
gb:MN996530|Organism:Severe      ACATTTTCACCGAGGCCACGCGGAGTACGATCGAGTGTACAGTGAACAATGCTAGGGAGA   29768
gb:MN908947|Organism:Severe      ACATTTTCACCGAGGCCACGCGGAGTACGATCGAGTGTACAGTGAACAATGCTAGGGAGA   29782
gb:MT019532|Organism:Severe      ACATTTTCACCGAGGCCACGCGGAGTACGATCGAGTGTACAGTGAACAATGCTAGGGAGA   29782
                                 ******************* *******          ***********************

gb:MT020781|Organism:Severe      GCTGCCTATATGGAAGAGCCCTAATGTGTAAAATTAATTTTAGTAGTGCTATCCCCATGT   29830
gb:MT007544|Organism:Severe      GCTGCCTATATGGAAGAGCCCTAATGTGTAAAATTAATTTTAGTAGTGCTATCCCCATGT   29832
gb:MN994467|Organism:Severe      GCTGCCTATATGGAAGAGCCCTAATGTGTAAAATTAATTTTAGTAGTGCTATCCCCATGT   29842
gb:MT044257|Organism:Severe      GCTGCCTATATGGAAGAGCCCTAATGTGTAAAATTAATTTTAGTAGTGCTATCCCCATGT   29842
gb:MT106054|Organism:Severe      GCTGCCTATATGGAAGAGCCCTAATGTGTAAAATTAATTTTAGTAGTGCTATCCCCATGT   29842
gb:MT049951|Organism:Severe      GCTGCCTATATGGAAGAGCCCTAATGTGTAAAATTAATTTTAGTAGTGCTATCCCCATGT   29842
gb:MN975262|Organism:Severe      GCTGCCTATATGGAAGAGCCCTAATGTGTAAAATTAATTTTAGTAGTGCTATCCCCATGT   29842
gb:MT106052|Organism:Severe      GCTGCCTATATGGAAGAGCCCTAATGTGTAAAATTAATTTTAGTAGTGCTATCCCCATGT   29842
gb:LC522975|Organism:Severe      GCTGCCTATATGGAAGAGCCCTAATGTGTAAAATTAATTTTAGTAGTGCTATCCCCATGT   29839
gb:LC522973|Organism:Severe      GCTGCCTATATGGAAGAGCCCTAATGTGTAAAATTAATTTTAGTAGTGCTATCCCCATGT   29839
gb:LC522974|Organism:Severe      GCTGCCTATATGGAAGAGCCCTAATGTGTAAAATTAATTTTAGTAGTGCTATCCCCATGT   29839
gb:MN985325|Organism:Severe      GCTGCCTATATGGAAGAGCCCTAATGTGTAAAATTAATTTTAGTAGTGCTATCCCCATGT   29842
gb:MT020881|Organism:Severe      GCTGCCTATATGGAAGAGCCCTAATGTGTAAAATTAATTTTAGTAGTGCTATCCCCATGT   29842
gb:MT020880|Organism:Severe      GCTGCCTATATGGAAGAGCCCTAATGTGTAAAATTAATTTTAGTAGTGCTATCCCCATGT   29842
```

```
gb:MT066175|Organism:Severe      GCTGCCTATATGGAAGAGCCCTAATGTGTAAAATTAATTTTAGTAGTGCTATCCCCATGT   29842
gb:MN997409|Organism:Severe      GCTGCCTATATGGAAGAGCCCTAATGTGTAAAATTAATTTTAGTAGTGCTATCCCCATGT   29842
gb:MN938384|Organism:Severe      GCTGCCTATATGGAAGAGCCCTAATGTGTAAAATTAATTTTAGTAGTGCTATCCCCATGT   29810
gb:MT044258|Organism:Severe      GCTGCCTATATGGAAGAGCCCTAATGTGTAAAATTAATTTTAGTAGTGCTATCCCCATGT   29818
gb:MT039890|Organism:Severe      GCTGCCTATATGGAAGAGCCCTAATGTGTAAAATTAATTTTAGTAGTGCTATCCCCATGT   29842
gb:MN988713|Organism:Severe      GCTGCCTATATGGAAGAGCCCTAATGTGTAAAATTAATTTTAGTAGTGCTATCCCCATGT   29842
gb:LC521925|Organism:Severe      GCTGCCTATATGGAAGAGCCCTAATGTGTAAAATTAATTTTAGTAGTGCTATCCCCATGT   29815
gb:MT093571|Organism:Severe      GCTGCCTATATGGAAGAGCCCTAATGTGTAAAATTAATTTTAGTAGTGCTATCCCCATGT   29842
gb:MT039887|Organism:Severe      GCTGCCTATATGGAAGAGCCCTAATGTGTAAAATTAATTTTAGTAGTGCTATCCCCATGT   29839
gb:MT019530|Organism:Severe      GCTGCCTATATGGAAGAGCCCTAATGTGTAAAATTAATTTTAGTAGTGCTATCCCCATGT   29842
gb:MT039888|Organism:Severe      GCTGCCTATATGGAAGAGCCCTAATGTGTAAAATTAATTTTAGTAGTGCTATCCCCATGT   29842
gb:LC522972|Organism:Severe      GCTGCCTATATGGAAGAGCCCTAATGTGTAAAATTAATTTTAGTAGTGCTATCCCCATGT   29839
gb:MT027063|Organism:Severe      GCTGCCTATATGGAAGAGCCCTAATGTGTAAAATTAATTTTAGTAGTGCTATCCCCATGT   29842
gb:MT027062|Organism:Severe      GCTGCCTATATGGAAGAGCCCTAATGTGTAAAATTAATTTTAGTAGTGCTATCCCCATGT   29842
gb:MT019529|Organism:Severe      GCTGCCTATATGGAAGAGCCCTAATGTGTAAAATTAATTTTAGTAGTGCTATCCCCATGT   29842
gb:MN996529|Organism:Severe      GCTGCCTATATGGAAGAGCCCTAATGTGTAAAATTAATTTTAGTAGTGCTATCCCCATGT   29830
gb:MN996531|Organism:Severe      GCTGCCTATATGGAAGAGCCCTAATGTGTAAAATTAATTTTAGTAGTGCTATCCCCATGT   29829
gb:MT066176|Organism:Severe      GCTGCCTATATGGAAGAGCCCTAATGTGTAAAATTAATTTTAGTAGTGCTATCCCCATGT   29842
gb:MT027064|Organism:Severe      GCTGCCTATATGGAAGAGCCCTAATGTGTAAAATTAATTTTAGTAGTGCTATCCCCATGT   29842
gb:MN994468|Organism:Severe      GCTGCCTATATGGAAGAGCCCTAATGTGTAAAATTAATTTTAGTAGTGCTATCCCCATGT   29842
gb:MT072688|Organism:Severe      GCTGCCTATATGGAAGAGCCCTAATGTGTAAAATTAATTTTAGT----------------   29811
gb:MN996527|Organism:Severe      GCTGCCTATATGGAAGAGCCCTAATGTGTAAAATTAATTTTAGTAGTGCTATCCCCATGT   29809
gb:MT093631|Organism:Severe      GCTGCCTATATGGAAGAGCCCTAATGTGTAAAATTAATTTTAGTAGTGCTATCCCCATGT   29880
gb:MT106053|Organism:Severe      GCTGCCTATATGGAAGAGCCCTAATGTGTAAAATTAATTTTAGTAGTGCTATCCCCATGT   29842
gb:MT019533|Organism:Severe      GCTGCCTATATGGAAGAGCCCTAATGTGTAAAATTAATTTTAGTAGTGCTATCCCCATGT   29842
gb:MT019531|Organism:Severe      GCTGCCTATATGGAAGAGCCCTAATGTGTAAAATTAATTTTAGTAGTGCTATCCCCATGT   29842
gb:MN996528|Organism:Severe      GCTGCCTATATGGAAGAGCCCTAATGTGTAAAATTAATTTTAGTAGTGCTATCCCCATGT   29842
gb:MN996530|Organism:Severe      GCTGCCTATATGGAAGAGCCCTAATGTGTAAAATTAATTTTAGTAGTGCTATCCCCATGT   29828
gb:MN908947|Organism:Severe      GCTGCCTATATGGAAGAGCCCTAATGTGTAAAATTAATTTTAGTAGTGCTATCCCCATGT   29842
gb:MT019532|Organism:Severe      GCTGCCTATATGGAAGAGCCCTAATGTGTAAAATTAATTTTAGTAGTGCTATCCCCATGT   29842
                                 ********************************************

gb:MT020781|Organism:Severe      GATTTTAATAGCTTCTT-------------------------------------------   29847
gb:MT007544|Organism:Severe      GATTTTAATAGCTTCTTAGGAGAATGACAAAAAAAAAAAAAAAAAAAAAAAAAAAAAAAA   29892
gb:MN994467|Organism:Severe      GATTTTAATAGCTTCTTAGGAGAATGACAAAAAAAAAAAA--------------------   29882
gb:MT044257|Organism:Severe      GATTTTAATAGCTTCTTAGGAGAATGACAAAAAAAAAAAA--------------------   29882
gb:MT106054|Organism:Severe      GATTTTAATAGCTTCTTAGGAGAATGACAAAAAAAAAAAA--------------------   29882
gb:MT049951|Organism:Severe      GATTTTAATAGCTTCTTAGGAGAATGACAAAAAAAAAAAAAAAAAAAAAAAAAAAAAAAA   29902
gb:MN975262|Organism:Severe      GATTTTAATAGCTTCTTAGGAGAATGACAAAAAAAAAAAAAAAAAAAAA-----------   29891
gb:MT106052|Organism:Severe      GATTTTAATAGCTTCTTAGGAGAATGACAAAAAAAAAAAA--------------------   29882
gb:LC522975|Organism:Severe      GATTTTAATAGCTTCTTAGGAGAATGACAAAAAAAAAAA---------------------   29878
gb:LC522973|Organism:Severe      GATTTTAATAGCTTCTTAGGAGAATGACAAAAAAAAAAA---------------------   29878
gb:LC522974|Organism:Severe      GATTTTAATAGCTTCTTAGGAGAATGACAAAAAAAAAAA---------------------   29878
gb:MN985325|Organism:Severe      GATTTTAATAGCTTCTTAGGAGAATGACAAAAAAAAAAAA--------------------   29882
gb:MT020881|Organism:Severe      GATTTTAATAGCTTCTTAGGAGAATGACAAAAAAAAAAAA--------------------   29882
gb:MT020880|Organism:Severe      GATTTTAATAGCTTCTTAGGAGAATGACAAAAAAAAAAAA--------------------   29882
gb:MT066175|Organism:Severe      GATTTTAATAGCTTCTTAGGAGAATGAC--------------------------------   29870
gb:MN997409|Organism:Severe      GATTTTAATAGCTTCTTAGGAGAATGACAAAAAAAAAAAA--------------------   29882
gb:MN938384|Organism:Severe      GATTTTAATAGCTTCTTAGGAGAATGAC--------------------------------   29838
gb:MT044258|Organism:Severe      GATTTTAATAGCTTCTTAGGAGAATGACAAAAAAAAAAAA--------------------   29858
gb:MT039890|Organism:Severe      GATTTTAATAGCTTCTTAGGAGAATGACAAAAAAAAAAAAAAAAAAAAAAAAAAAAAAAA   29902
gb:MN988713|Organism:Severe      GATTTTAATAGCTTCTTAGGAGAATGACAAAAAAAAAAAA--------------------   29882
gb:LC521925|Organism:Severe      GATTTTAATAGCTTCTTAGGAGAATGACAAAAA---------------------------   29848
gb:MT093571|Organism:Severe      GATTTTAATAGCTTCTTAGGAGAATGACAAAAAAAAAAAAAAAA----------------   29886
gb:MT039887|Organism:Severe      GATTTTAATAGCTTCTTAGGAGAATGACAAAAAAAAAAAA--------------------   29879
gb:MT019530|Organism:Severe      GATTTTAATAGCTTCTTAGGAGAATGACAAAAAAAAAAAAAAAAAAA-------------   29889
gb:MT039888|Organism:Severe      GATTTTAATAGCTTCTTAGGAGAATGACAAAAAAAAAAAA--------------------   29882
gb:LC522972|Organism:Severe      GATTTTAATAGCTTCTTAGGAGAATGACAAAAAAAAAAA---------------------   29878
gb:MT027063|Organism:Severe      GATTTTAATAGCTTCTTAGGAGAATGACAAAAAAAAAAAA--------------------   29882
gb:MT027062|Organism:Severe      GATTTTAATAGCTTCTTAGGAGAATGACAAAAAAAAAAAA--------------------   29882
gb:MT019529|Organism:Severe      GATTTTAATAGCTTCTTAGGAGAATGACAAAAAAAAAAAAAAAAAAAAAAAAAAAAA---   29899
gb:MN996529|Organism:Severe      GATTTTAATAGCTTCTTAGGAG--------------------------------------   29852
gb:MN996531|Organism:Severe      GATTTTAATAGCTTCTTAGGAGAATGAC--------------------------------   29857
gb:MT066176|Organism:Severe      GATTTTAATAGCTTCTTAGGAGAATGAC--------------------------------   29870
gb:MT027064|Organism:Severe      GATTTTAATAGCTTCTTAGGAGAATGACAAAAAAAAAAAA--------------------   29882
```

```
gb:MN994468|Organism:Severe      GATTTTAATAGCTTCTTAGGAGAATGACAAAAAAAAAAAAA-------------------   29883
gb:MT072688|Organism:Severe      ------------------------------------------------------------   29811
gb:MN996527|Organism:Severe      GATTTTAATAGCTTCT--------------------------------------------   29825
gb:MT093631|Organism:Severe      GATTTTAATAGCTTCTTAGGAGAATGACAAA-----------------------------   29911
gb:MT106053|Organism:Severe      GATTTTAATAGCTTCTTAGGAGAATGACAAAAAAAAAAAA--------------------   29882
gb:MT019533|Organism:Severe      GATTTTAATAGCTTCTTAGGAGAATGACAAAAAAAAAAAAA-------------------   29883
gb:MT019531|Organism:Severe      GATTTTAATAGCTTCTTAGGAGAATGACAAAAAAAAAAAAAAAAAAAAAAAAAAAAA---   29899
gb:MN996528|Organism:Severe      GATTTTAATAGCTTCTTAGGAGAATGACAAAAAAAAAAAAAAAAAAAAA-----------   29891
gb:MN996530|Organism:Severe      GATTTTAATAGCTTCTTAGGAGAATG----------------------------------   29854
gb:MN908947|Organism:Severe      GATTTTAATAGCTTCTTAGGAGAATGACAAAAAAAAAAAAAAAAAAAAAAAAAAAAAAAA   29902
gb:MT019532|Organism:Severe      GATTTTAATAGCTTCTTAGGAGAATGACAAAAAAAAAAAAAAAAAAAA------------   29890


gb:MT020781|Organism:Severe      -      29847
gb:MT007544|Organism:Severe      A      29893
gb:MN994467|Organism:Severe      -      29882
gb:MT044257|Organism:Severe      -      29882
gb:MT106054|Organism:Severe      -      29882
gb:MT049951|Organism:Severe      A      29903
gb:MN975262|Organism:Severe      -      29891
gb:MT106052|Organism:Severe      -      29882
gb:LC522975|Organism:Severe      -      29878
gb:LC522973|Organism:Severe      -      29878
gb:LC522974|Organism:Severe      -      29878
gb:MN985325|Organism:Severe      -      29882
gb:MT020881|Organism:Severe      -      29882
gb:MT020880|Organism:Severe      -      29882
gb:MT066175|Organism:Severe      -      29870
gb:MN997409|Organism:Severe      -      29882
gb:MN938384|Organism:Severe      -      29838
gb:MT044258|Organism:Severe      -      29858
gb:MT039890|Organism:Severe      A      29903
gb:MN988713|Organism:Severe      -      29882
gb:LC521925|Organism:Severe      -      29848
gb:MT093571|Organism:Severe      -      29886
gb:MT039887|Organism:Severe      -      29879
gb:MT019530|Organism:Severe      -      29889
gb:MT039888|Organism:Severe      -      29882
gb:LC522972|Organism:Severe      -      29878
gb:MT027063|Organism:Severe      -      29882
gb:MT027062|Organism:Severe      -      29882
gb:MT019529|Organism:Severe      -      29899
gb:MN996529|Organism:Severe      -      29852
gb:MN996531|Organism:Severe      -      29857
gb:MT066176|Organism:Severe      -      29870
gb:MT027064|Organism:Severe      -      29882
gb:MN994468|Organism:Severe      -      29883
gb:MT072688|Organism:Severe      -      29811
gb:MN996527|Organism:Severe      -      29825
gb:MT093631|Organism:Severe      -      29911
gb:MT106053|Organism:Severe      -      29882
gb:MT019533|Organism:Severe      -      29883
gb:MT019531|Organism:Severe      -      29899
gb:MN996528|Organism:Severe      -      29891
gb:MN996530|Organism:Severe      -      29854
gb:MN908947|Organism:Severe      A      29903
gb:MT019532|Organism:Severe      -      29890
```
