## Supplemental Figure 2 for "Immunoinformatic identification of B cell and T cell epitopes in the SARS-CoV-2 proteome"

**Figure S2.** Plots illustrating the NetCTL score for each sequential peptide across the entire amino acid sequence for each SARS-CoV-2 protein. Individual panels represent the score trace for each major HLA class I supertype as indicated. The threshold for positive predictions was set at 0.75.

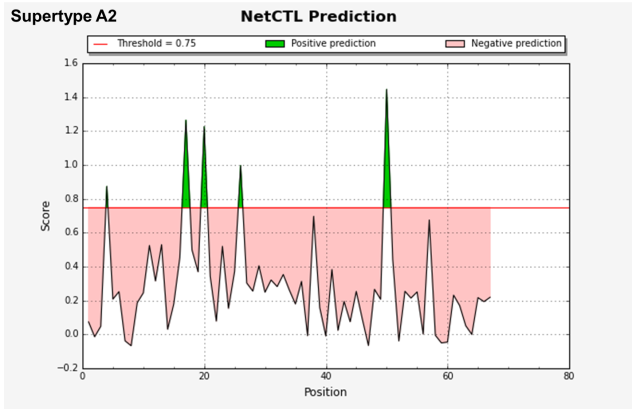

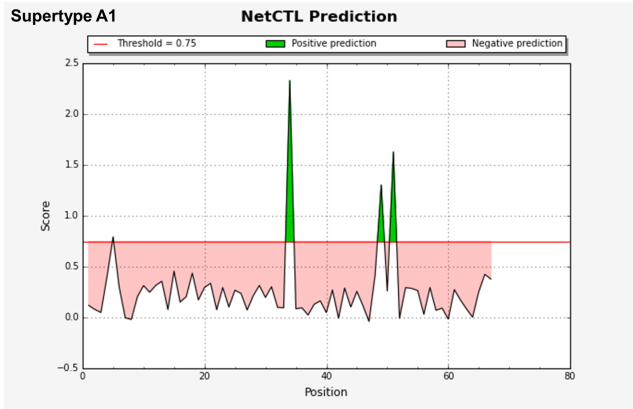
**E glycoprotein**

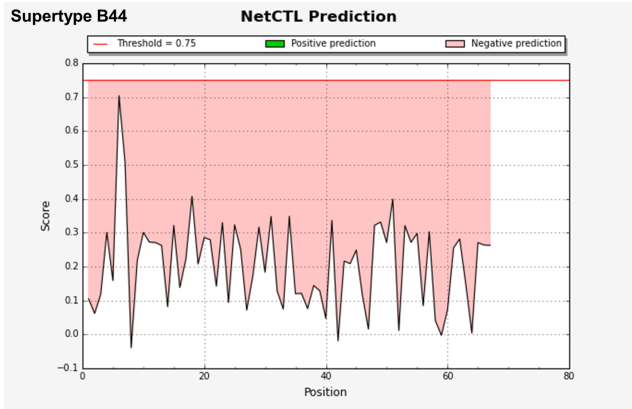

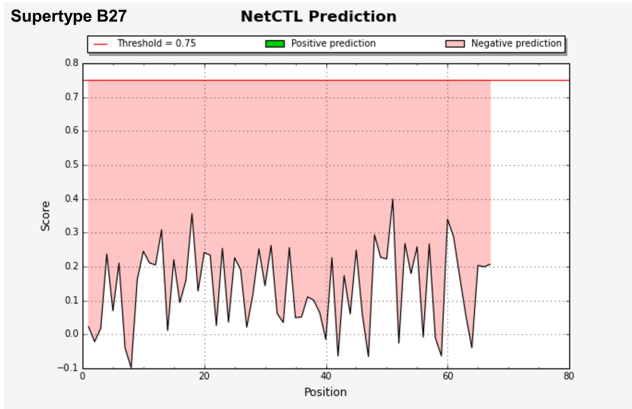

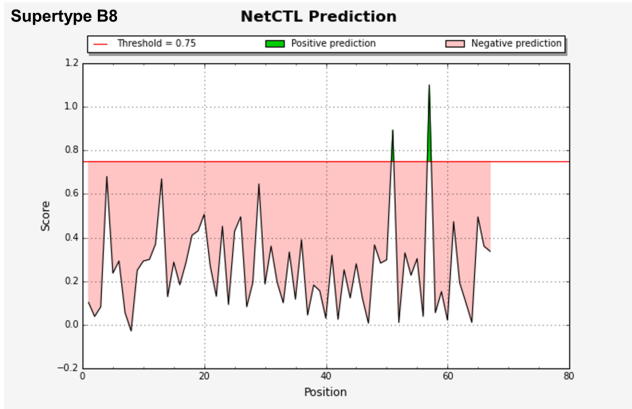

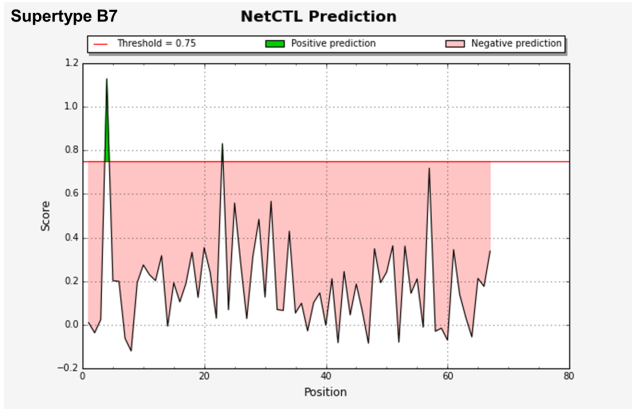

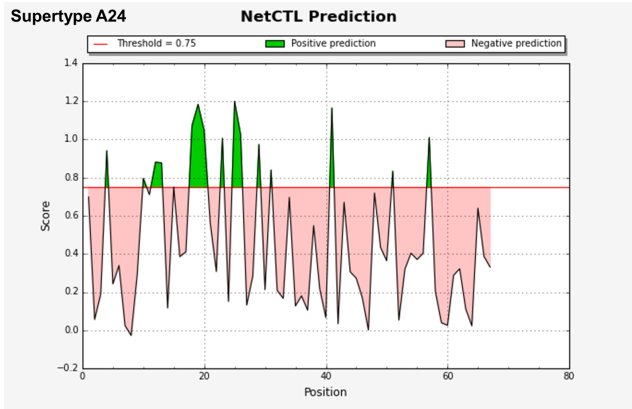

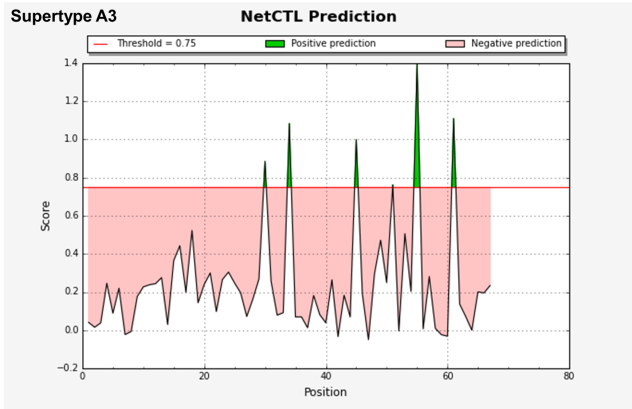

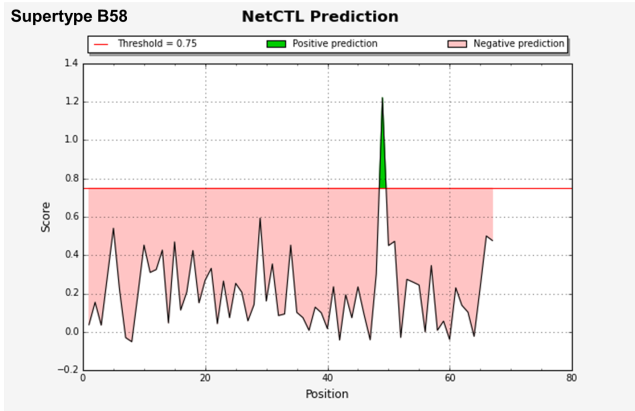

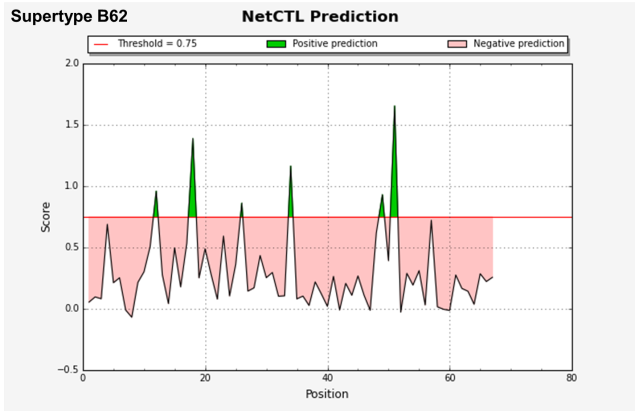

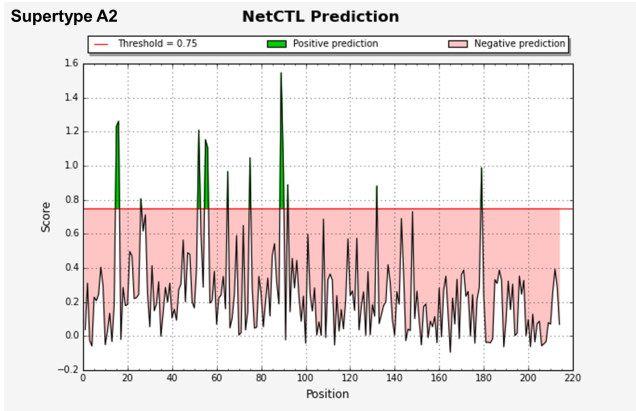

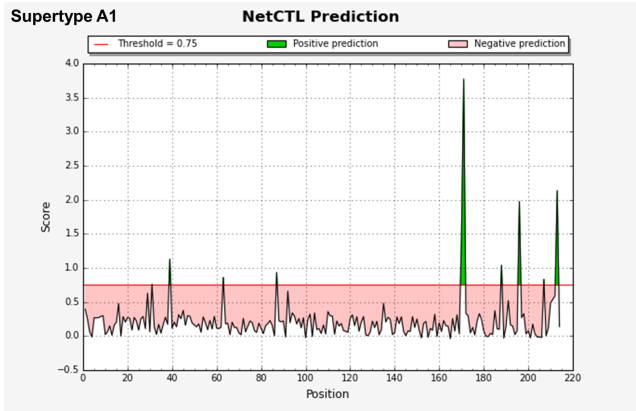
**M glycoprotein**

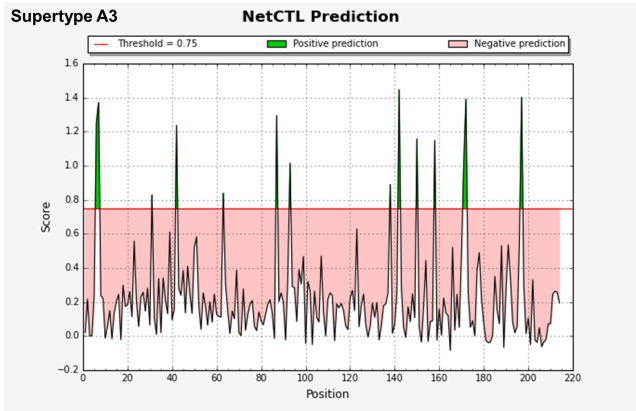

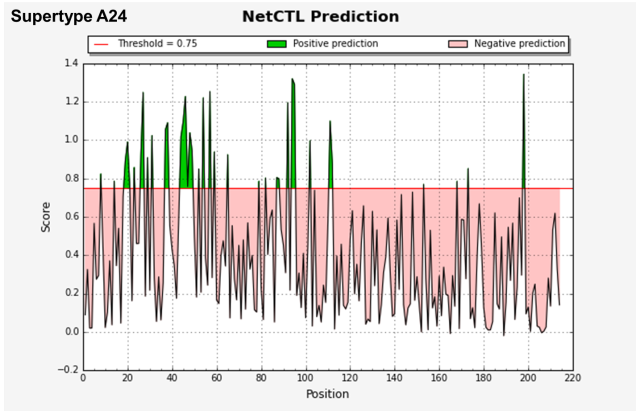

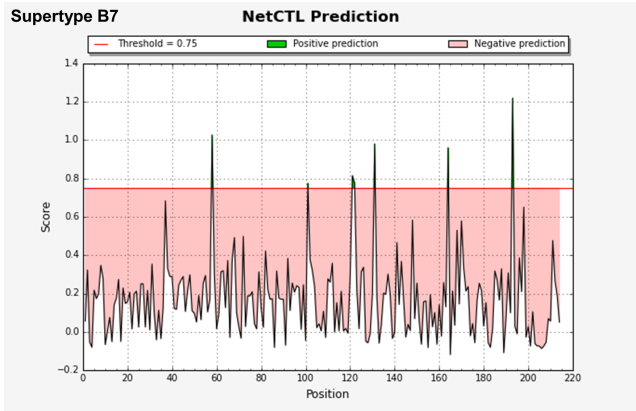

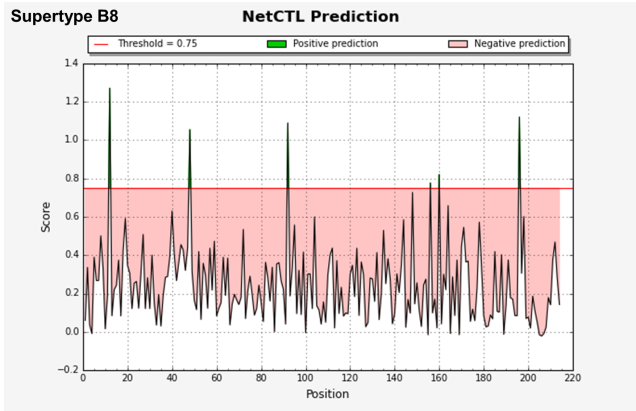

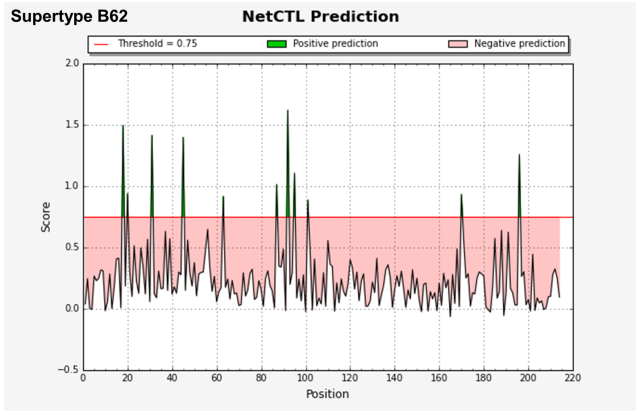

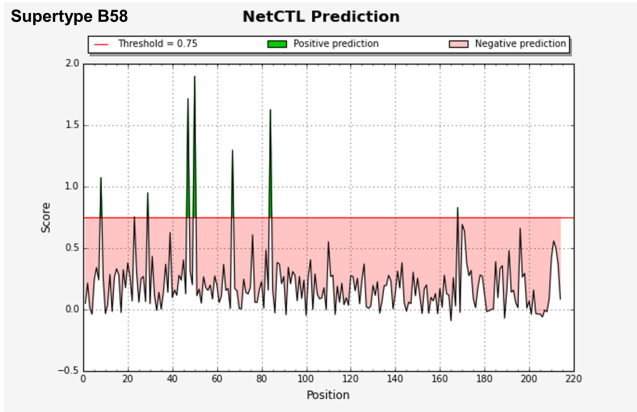

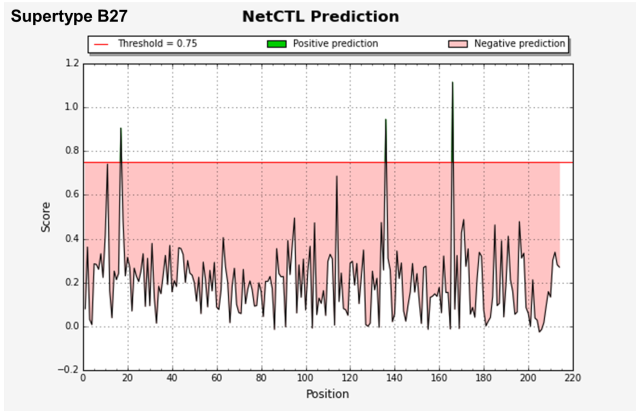

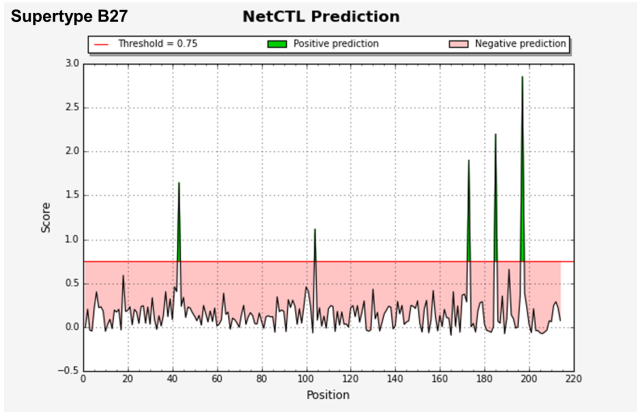

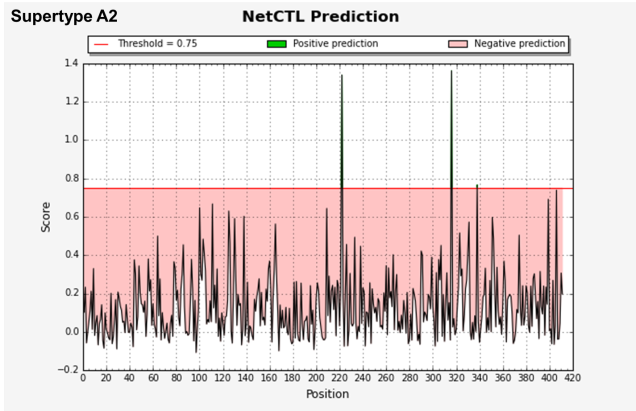
**N phosphoprotein**

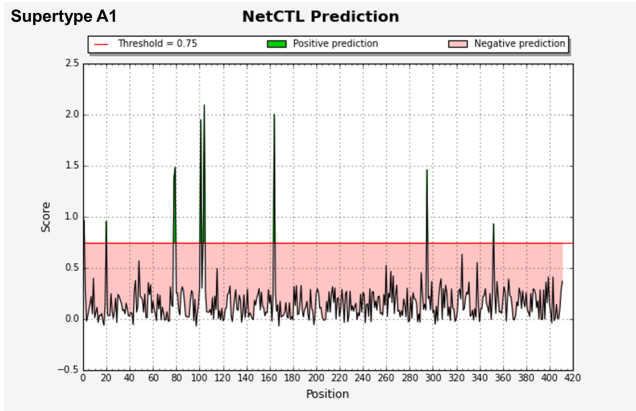

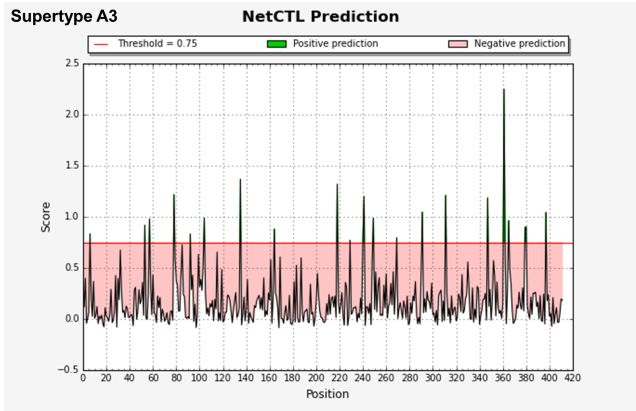

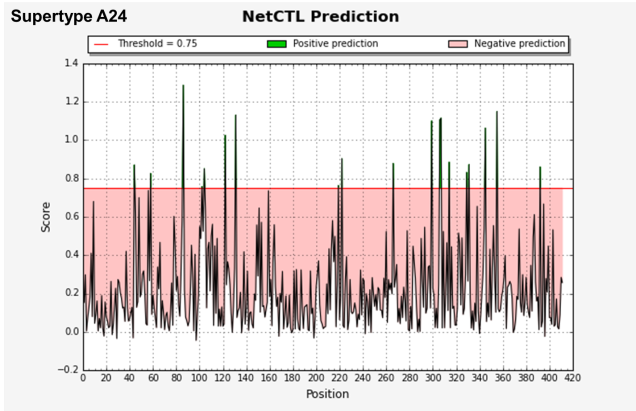

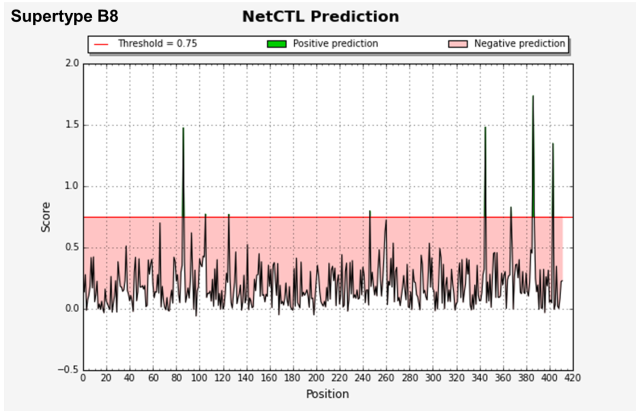

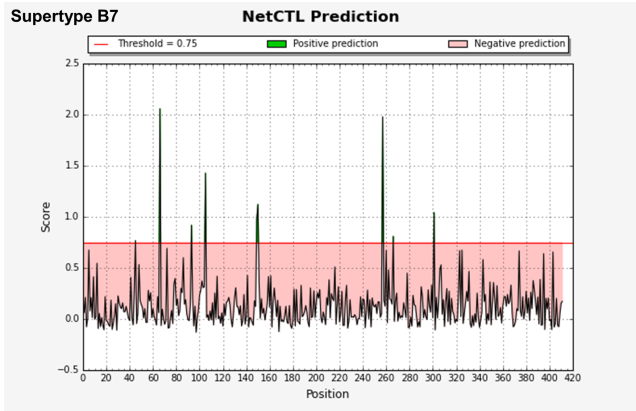

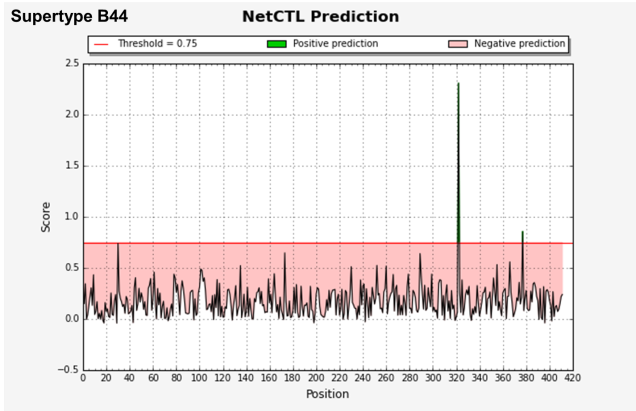

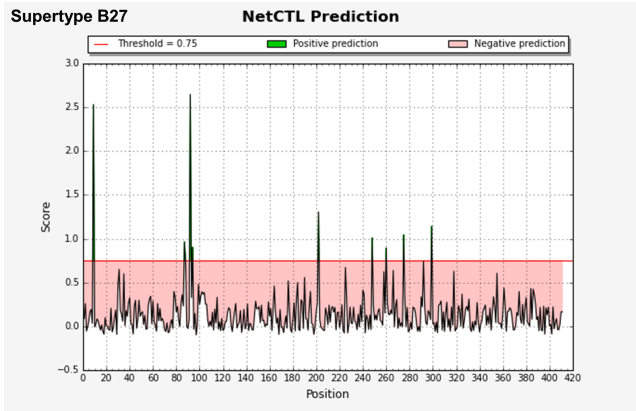

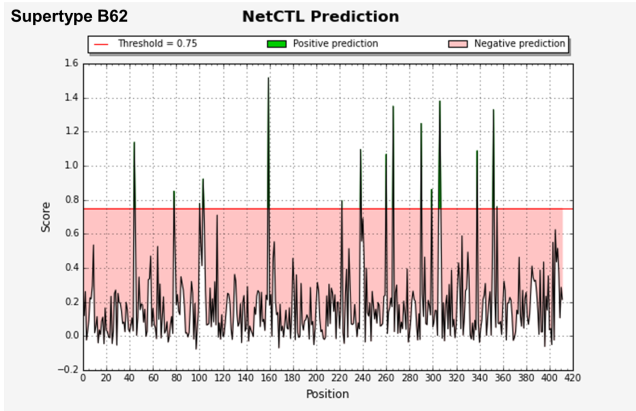

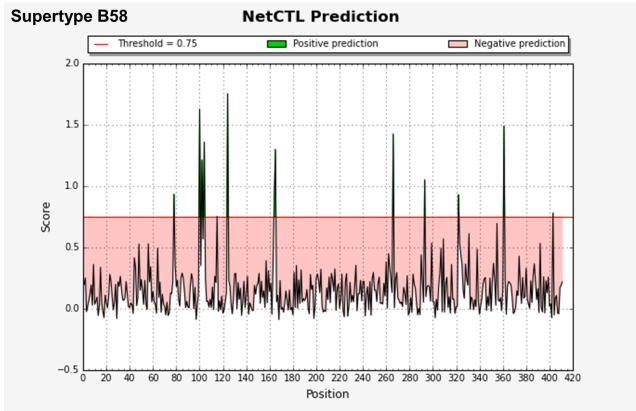

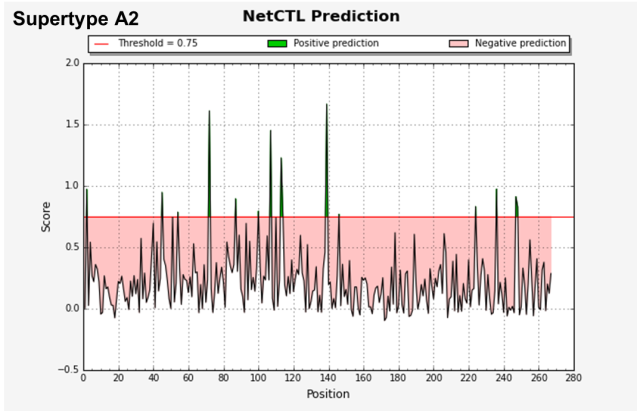

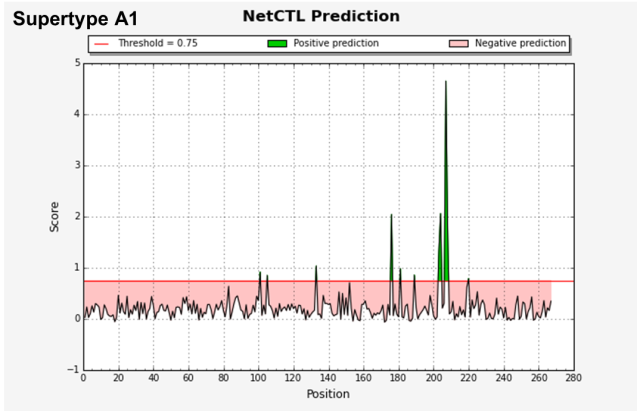
**ORF3a protein**

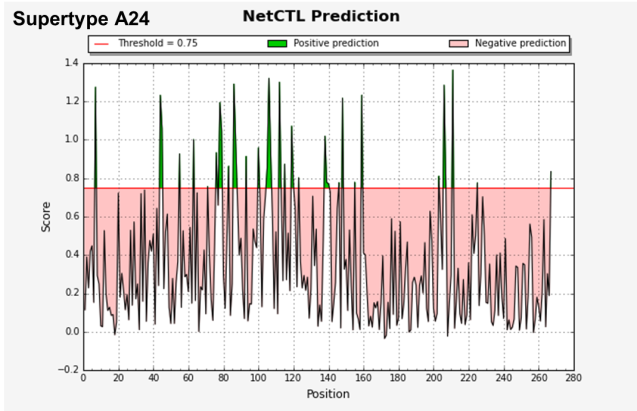

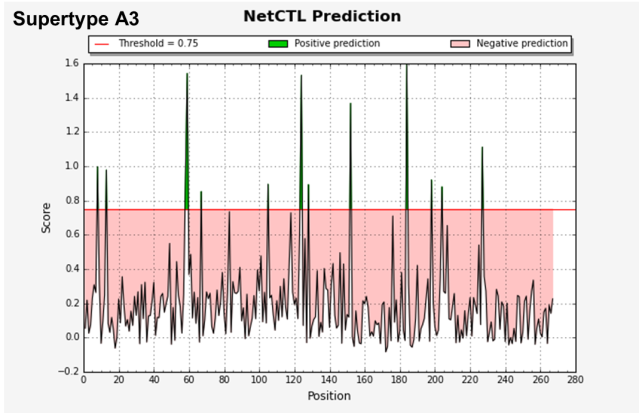

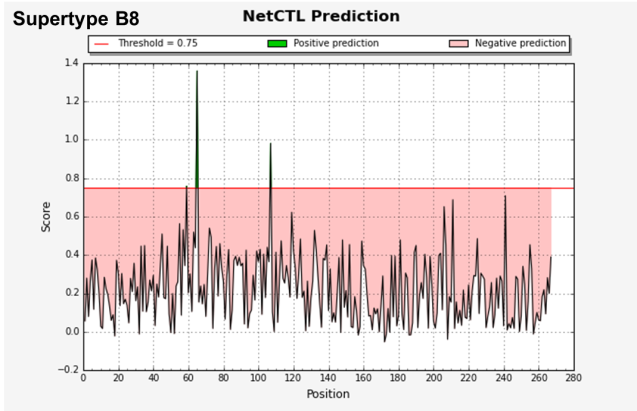

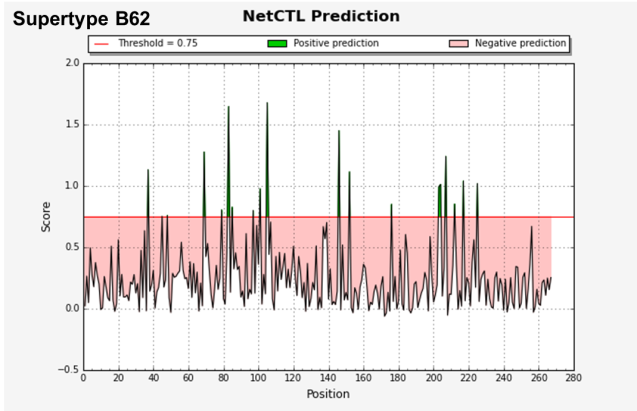

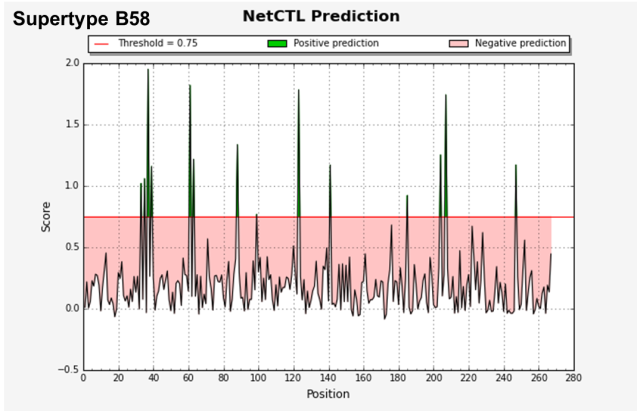

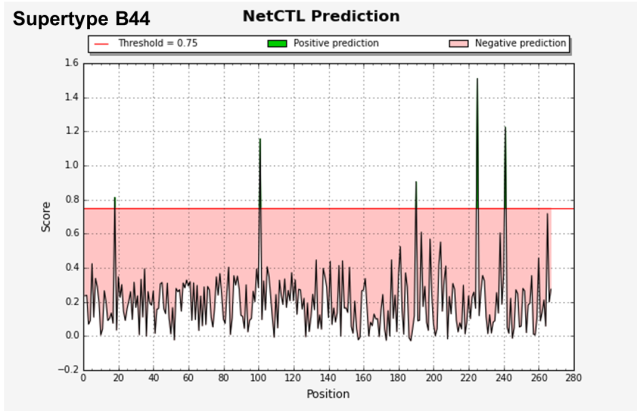

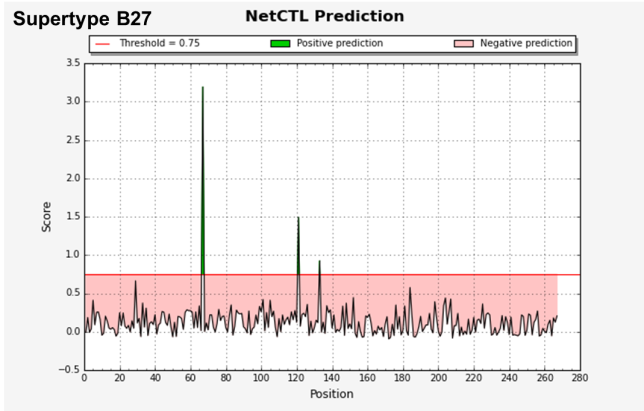

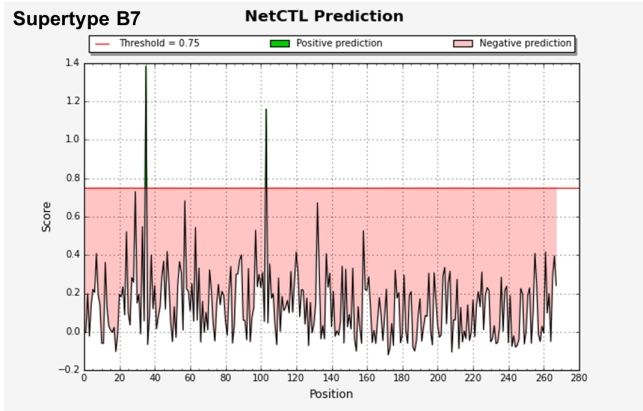

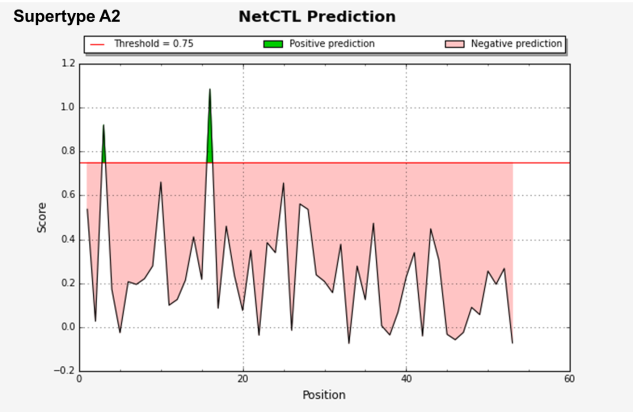

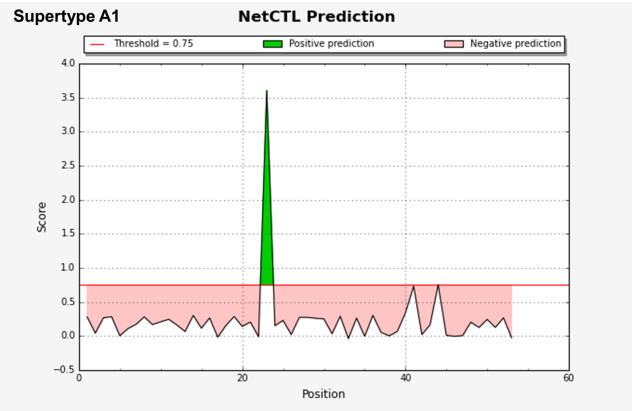
**ORF6 protein**

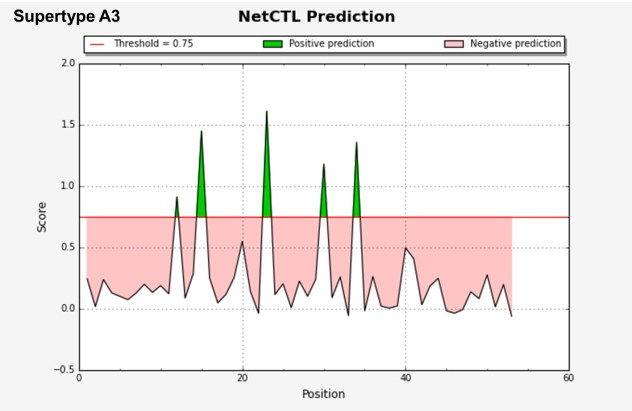

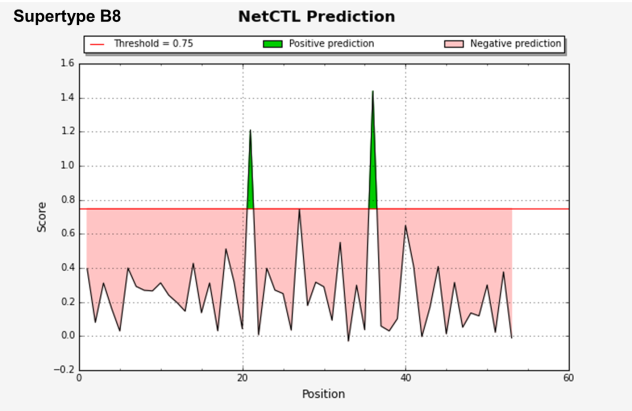

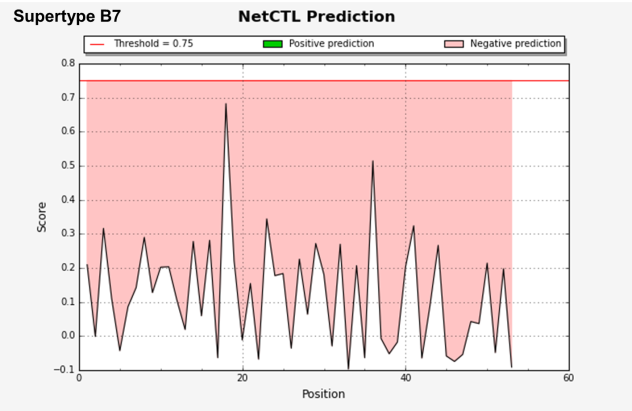

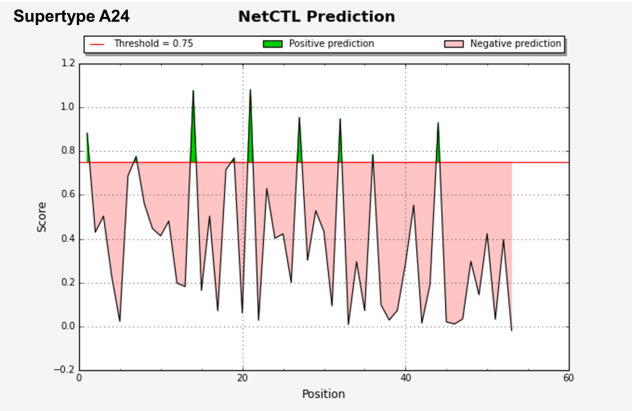

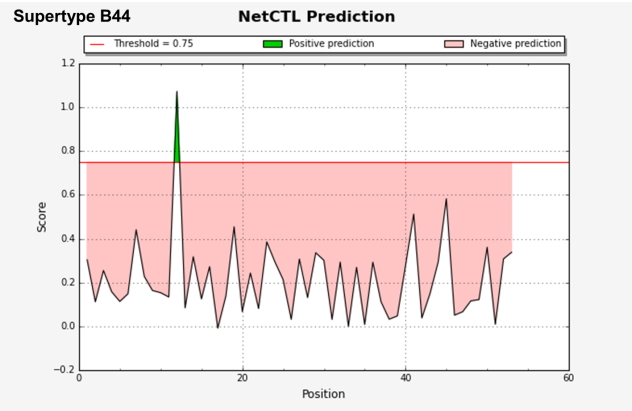

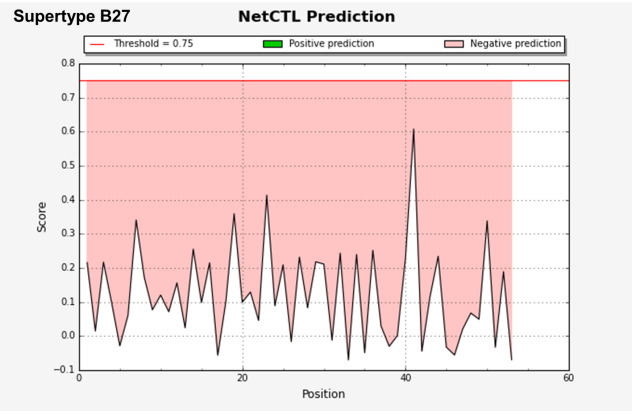

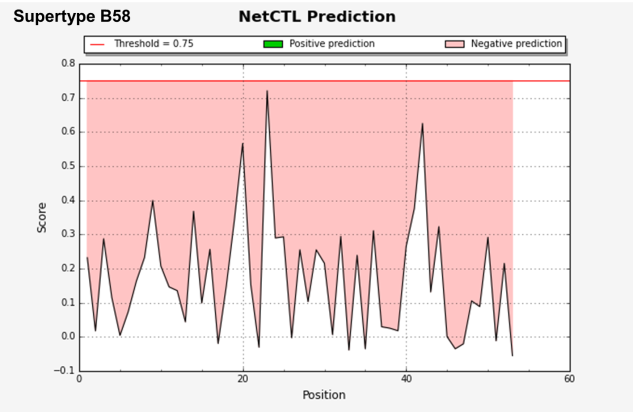

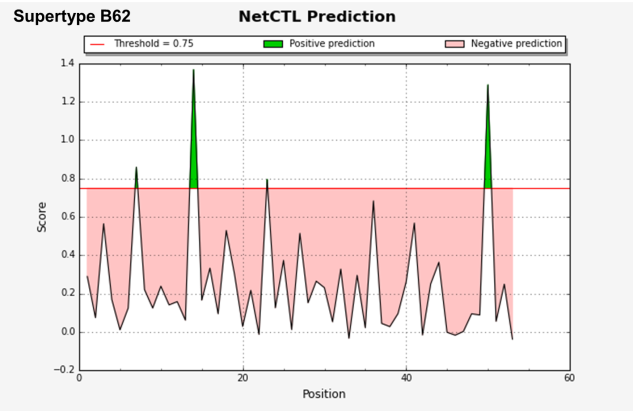

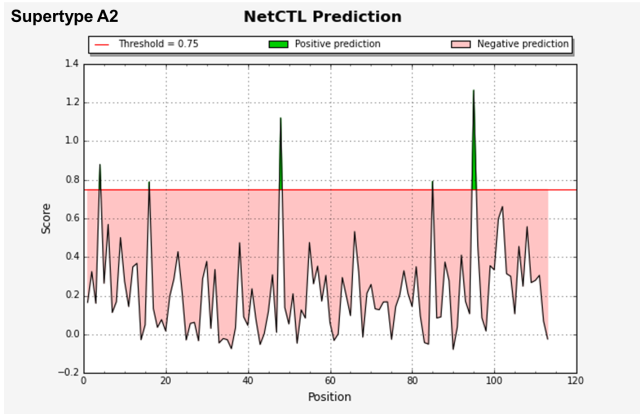

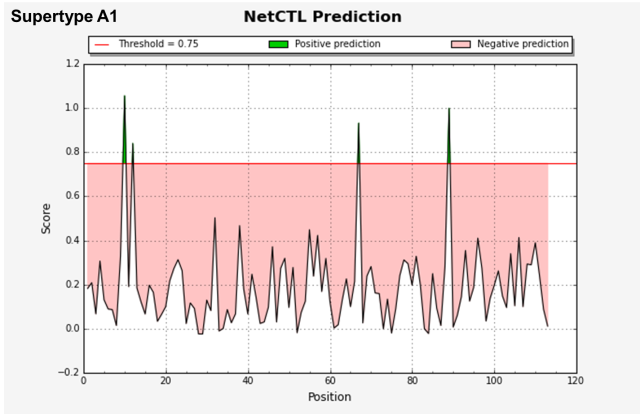
**ORF7 protein**

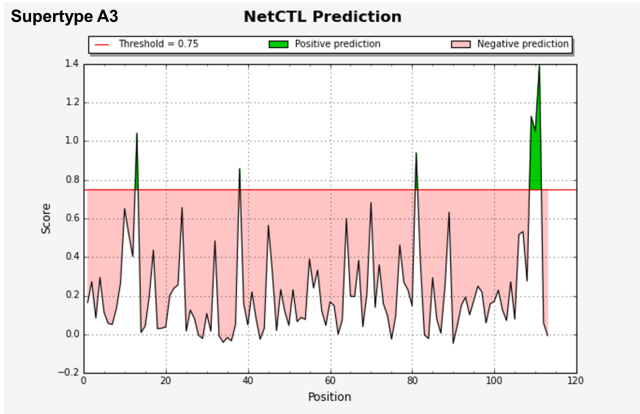

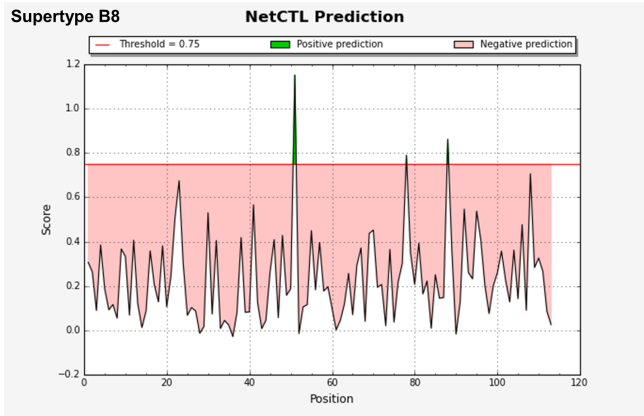

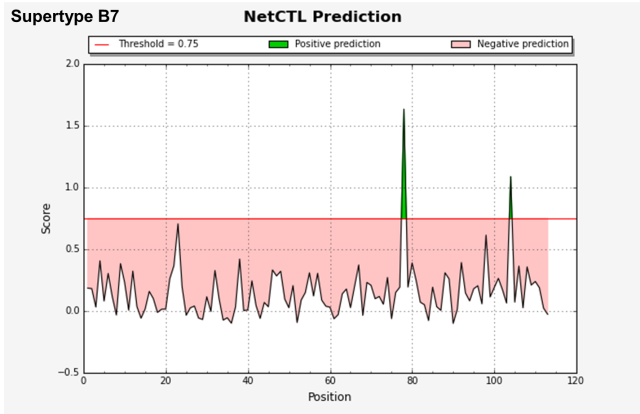

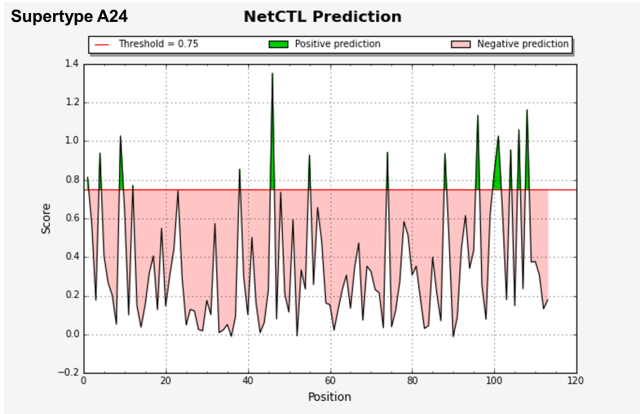

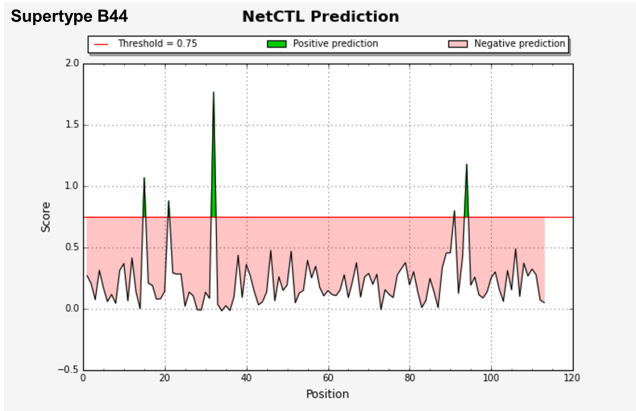

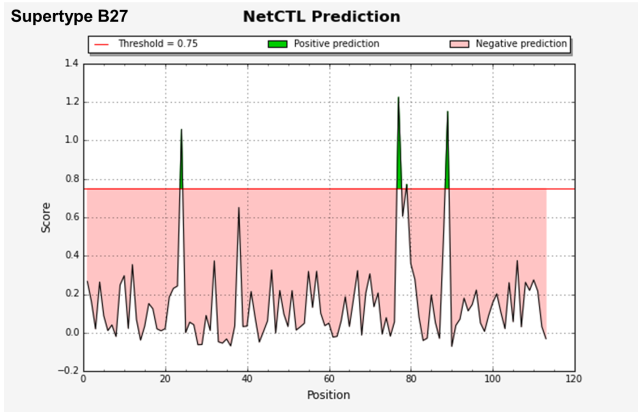

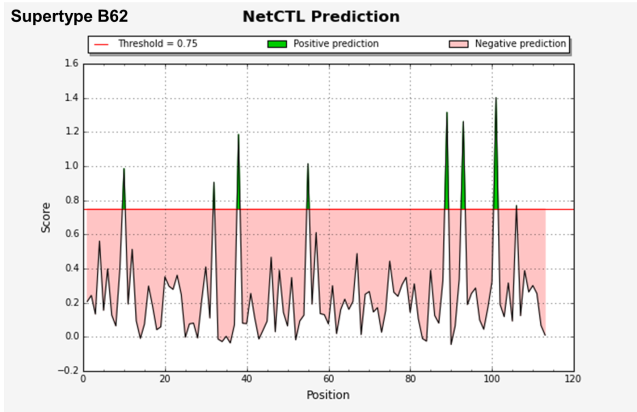

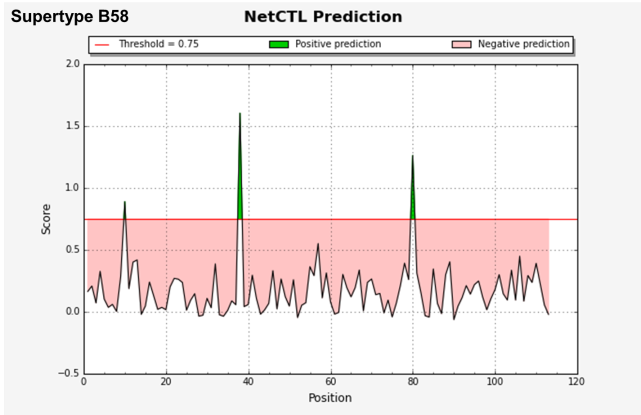

**ORF8 protein**

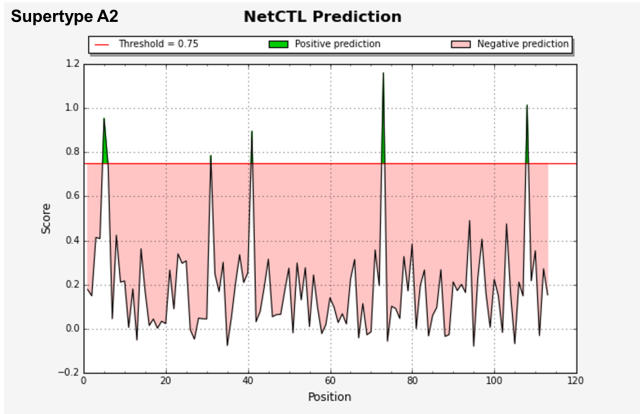

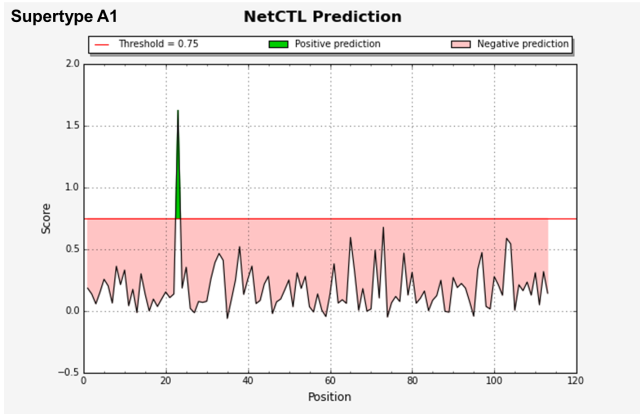

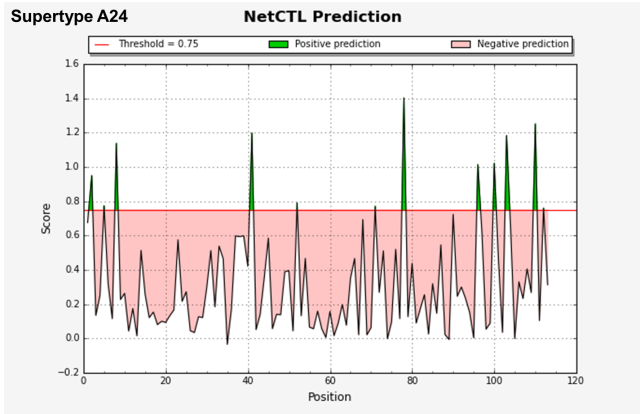

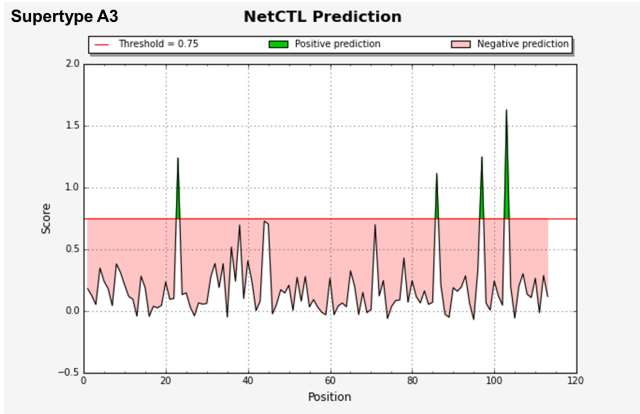

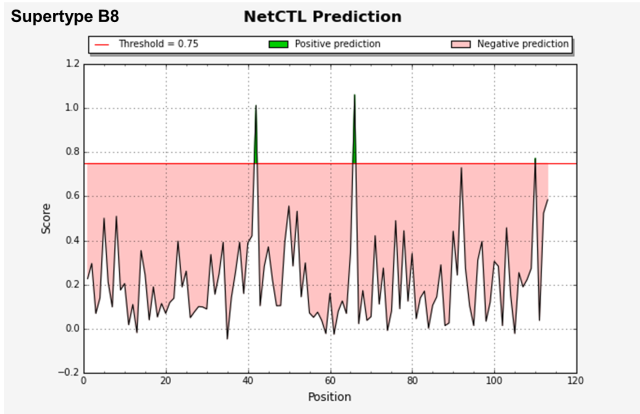

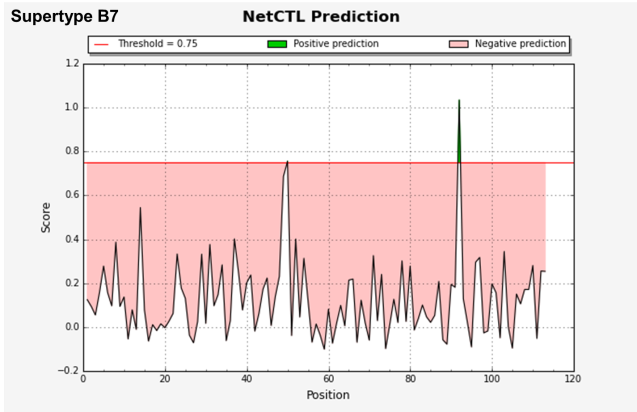

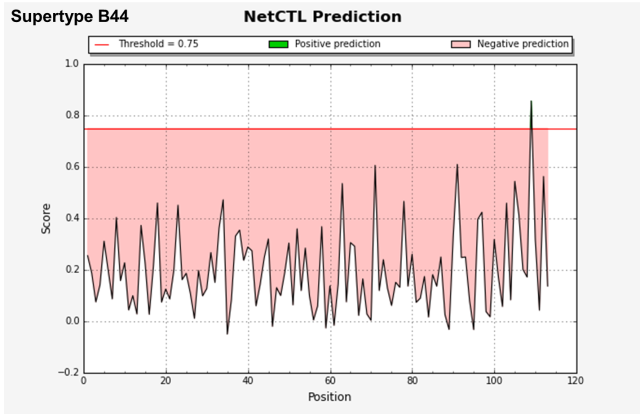

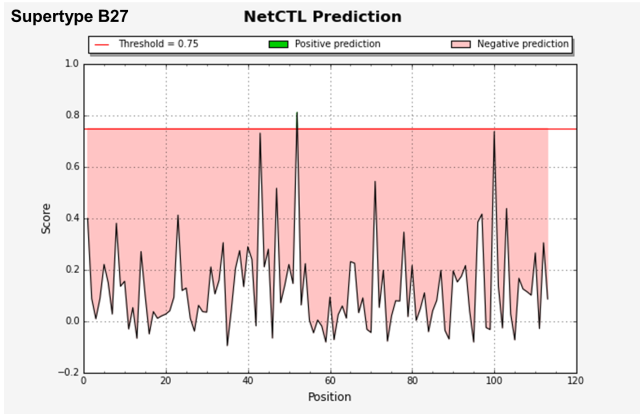

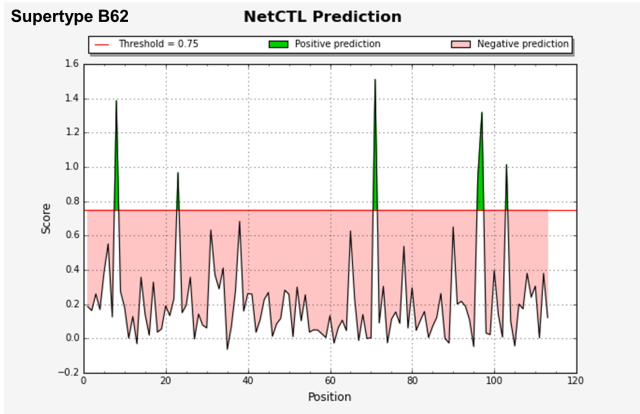

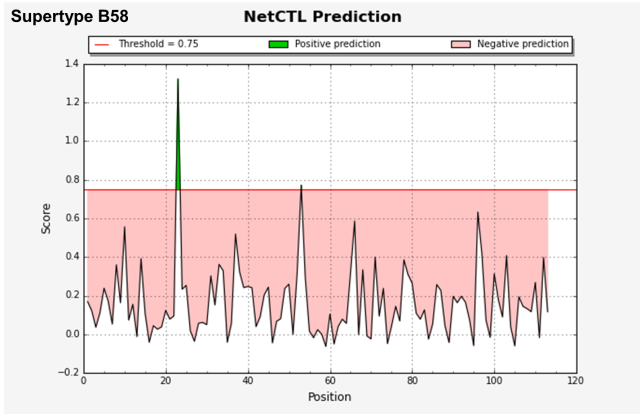

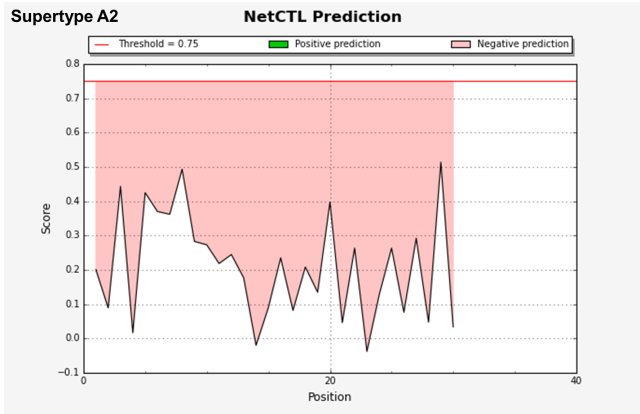
**ORF10**

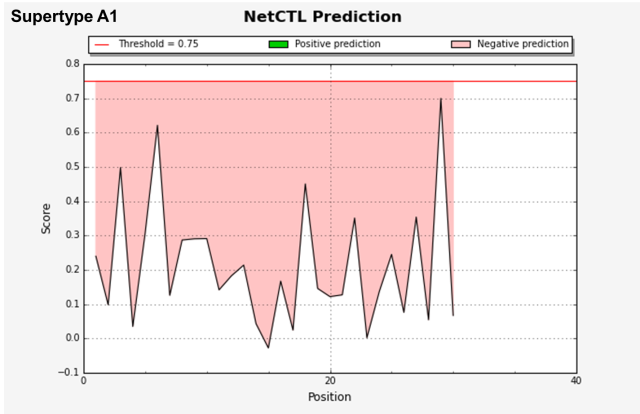

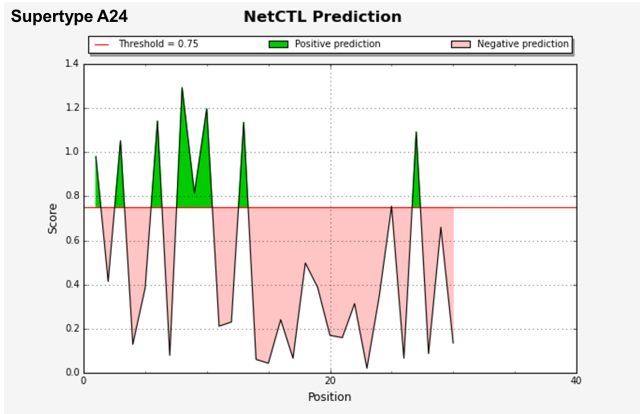

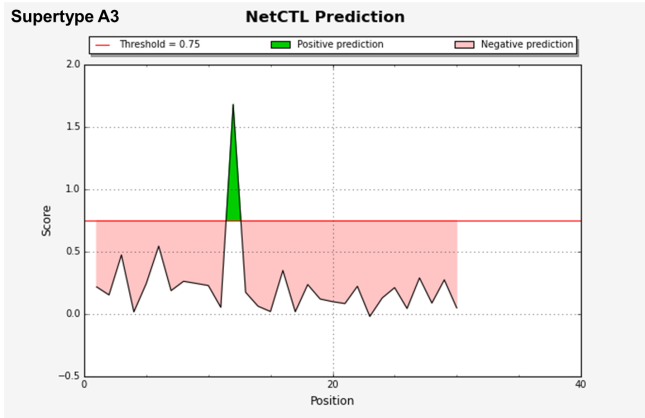

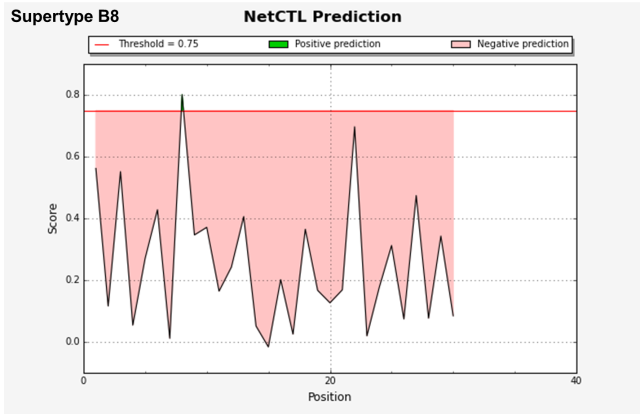

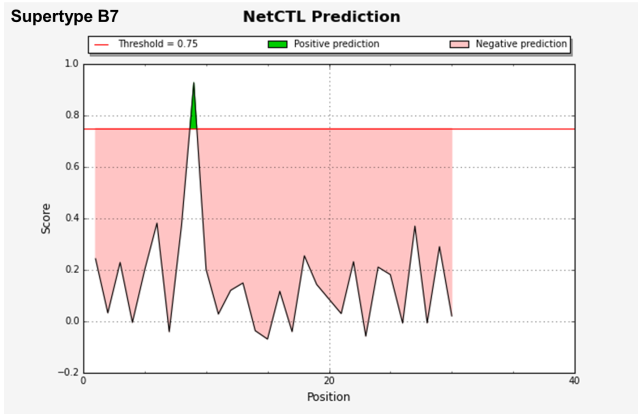

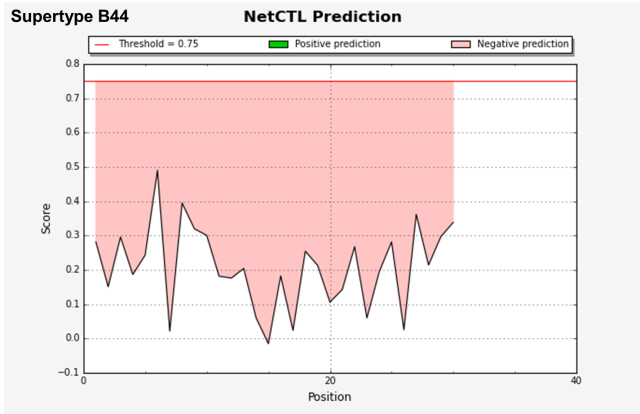

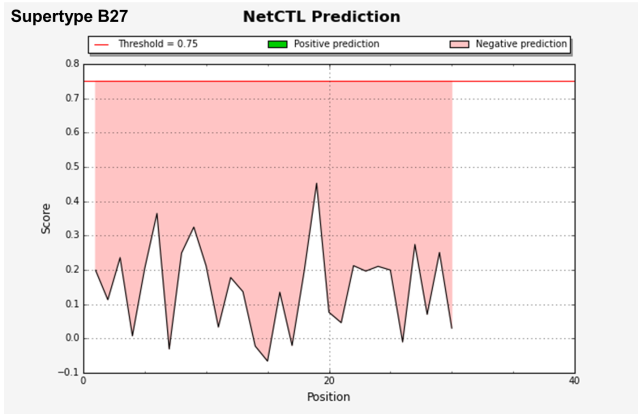

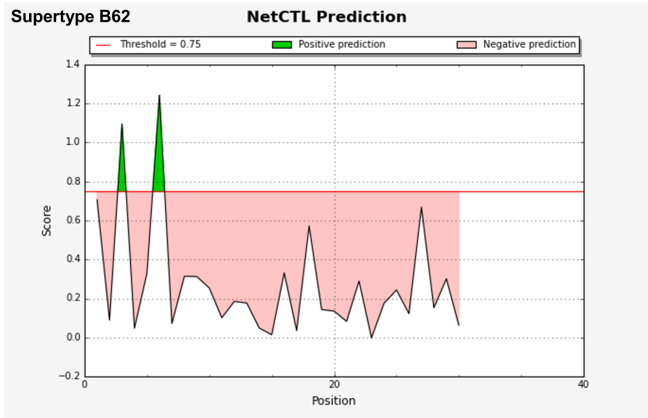

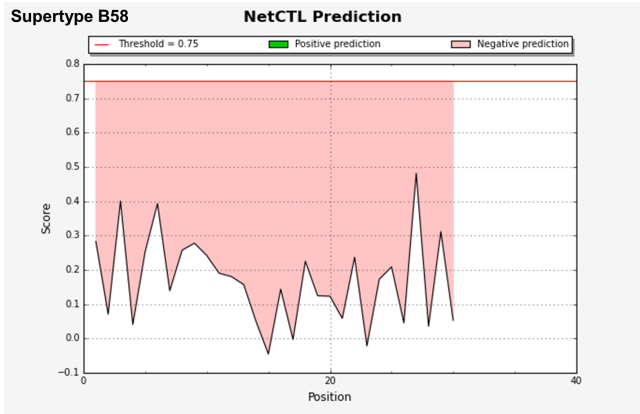

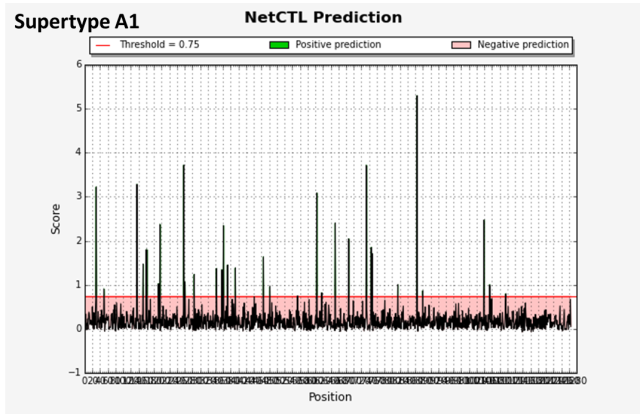
**S glycoprotein**

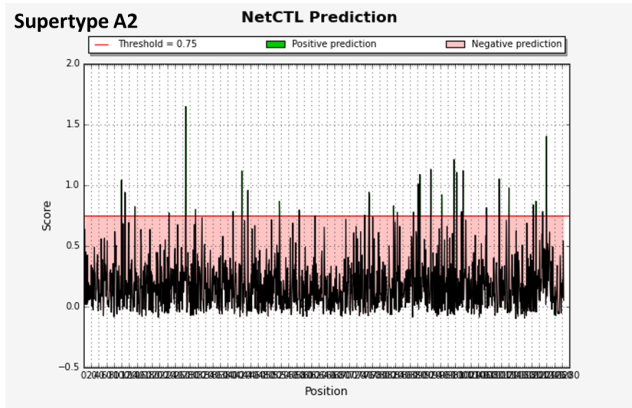

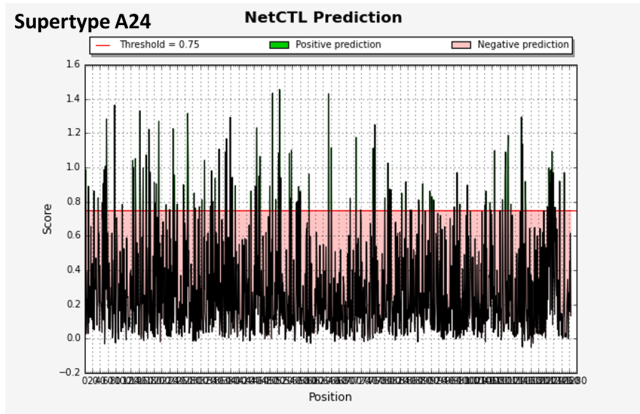

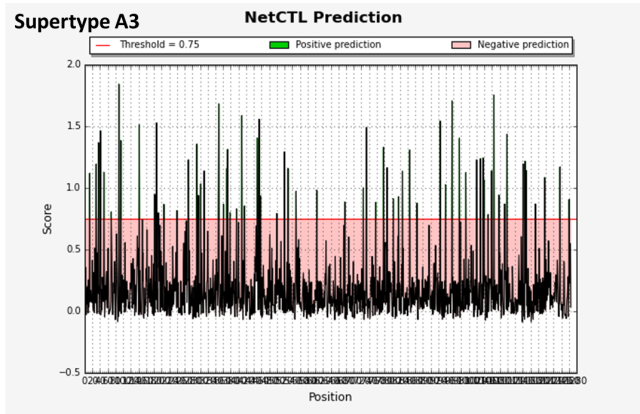

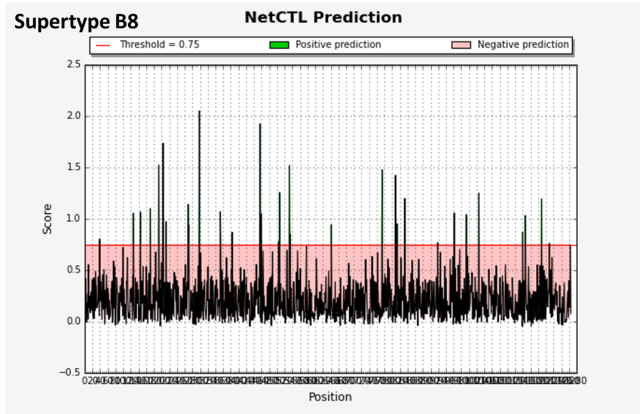

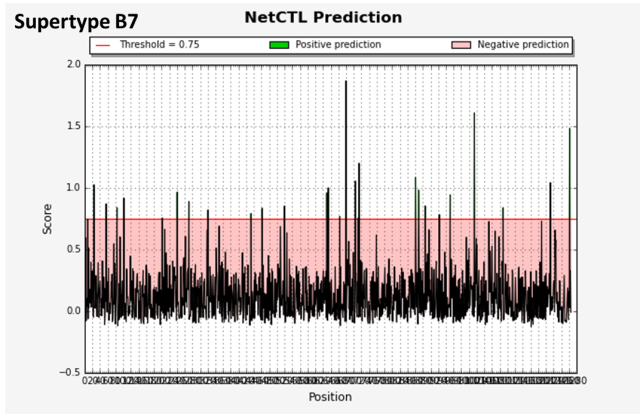

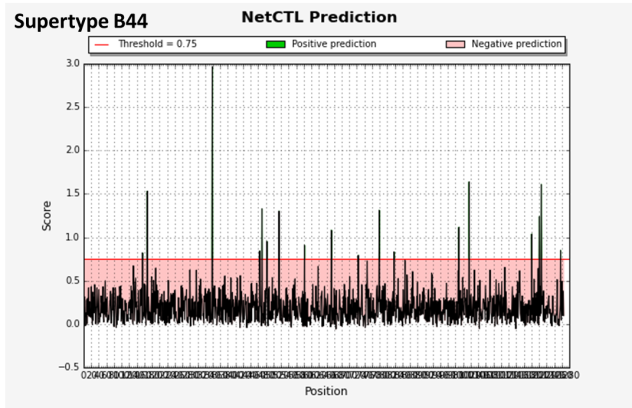

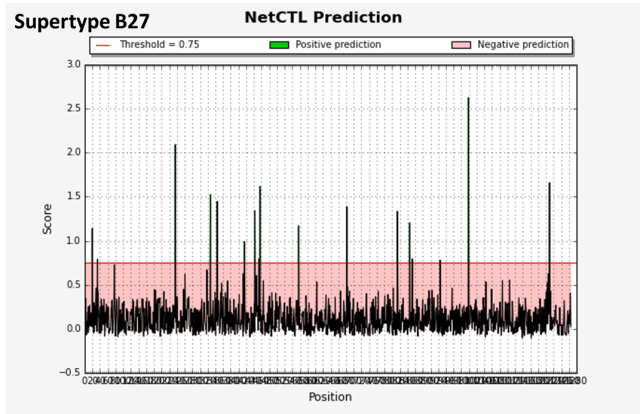

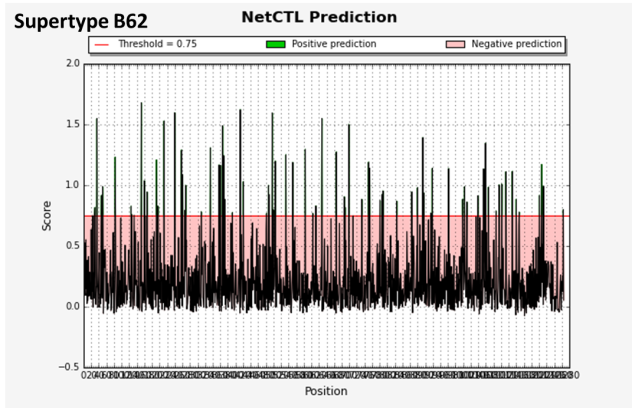

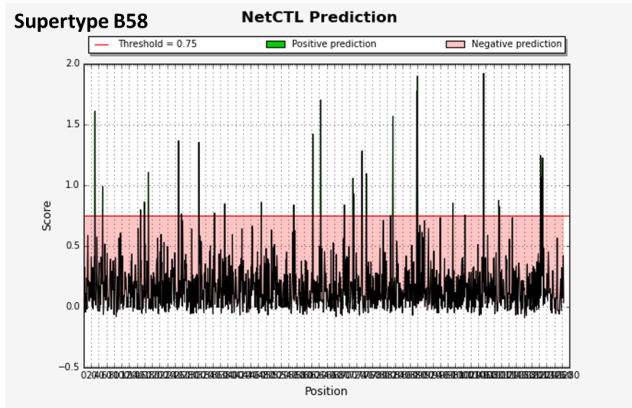

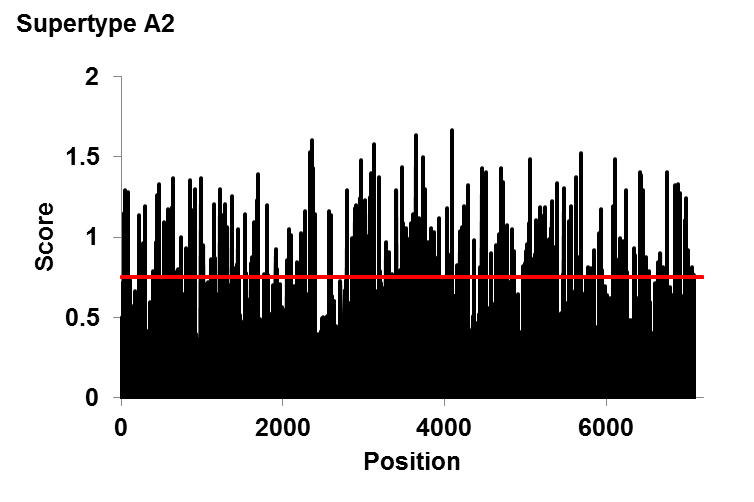

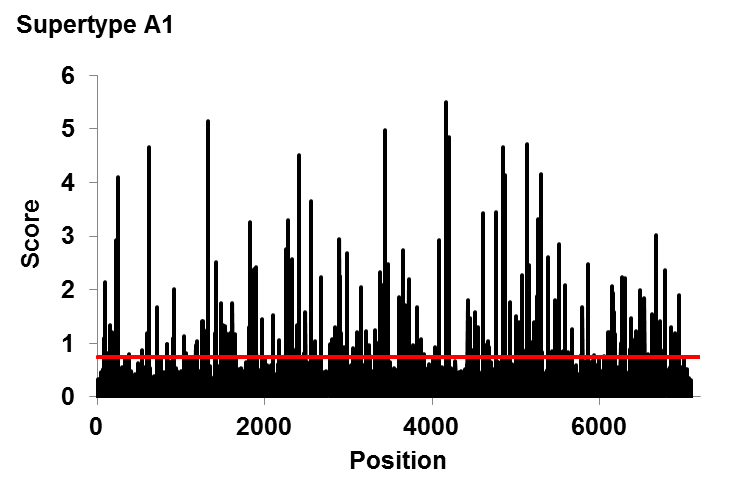
**ORF1ab polyprotein**

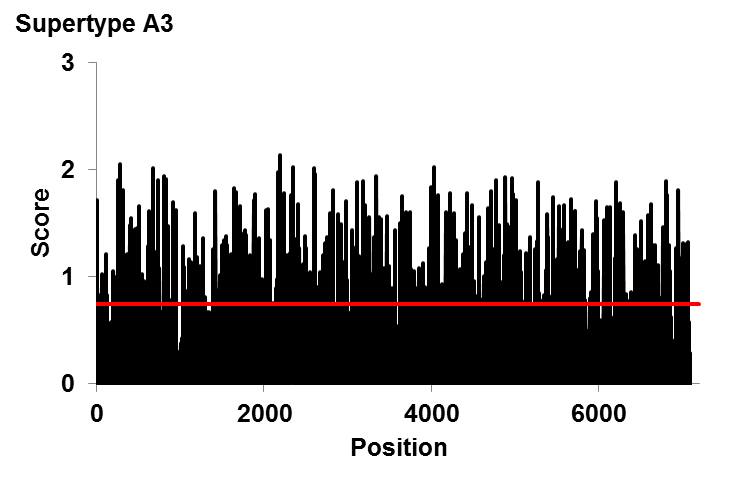

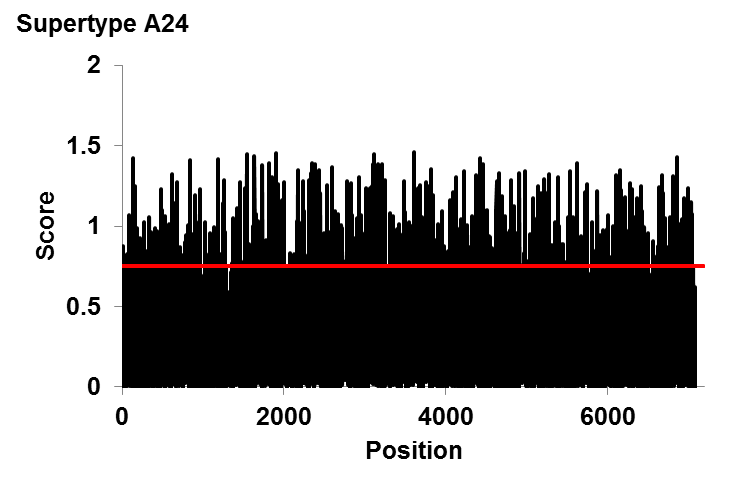

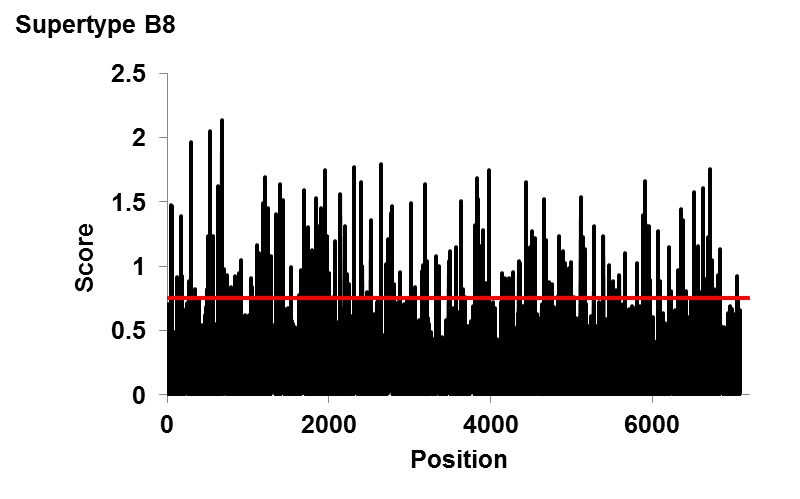

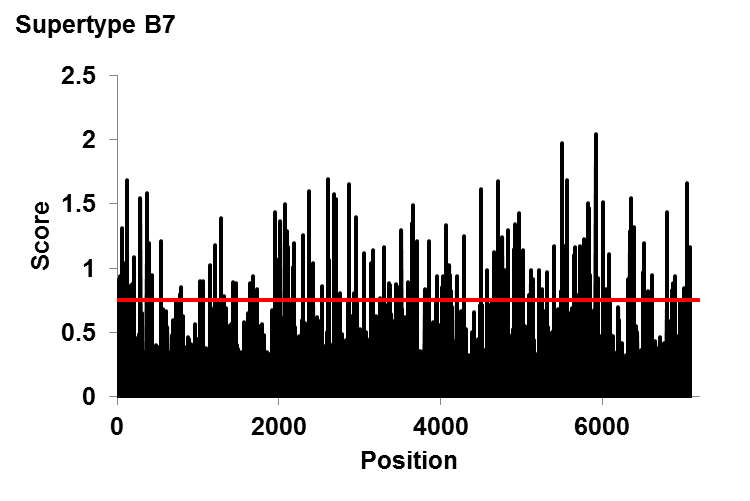

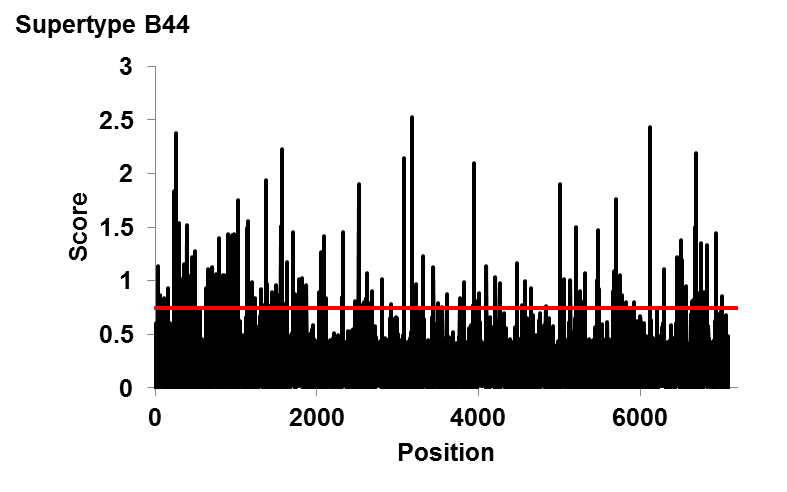

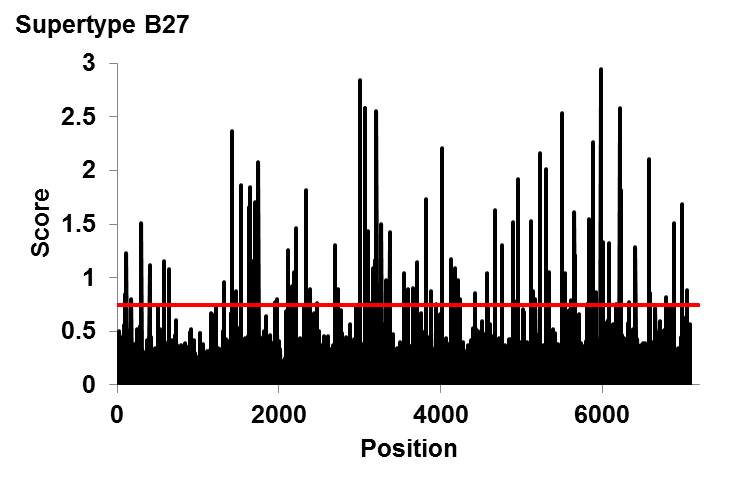

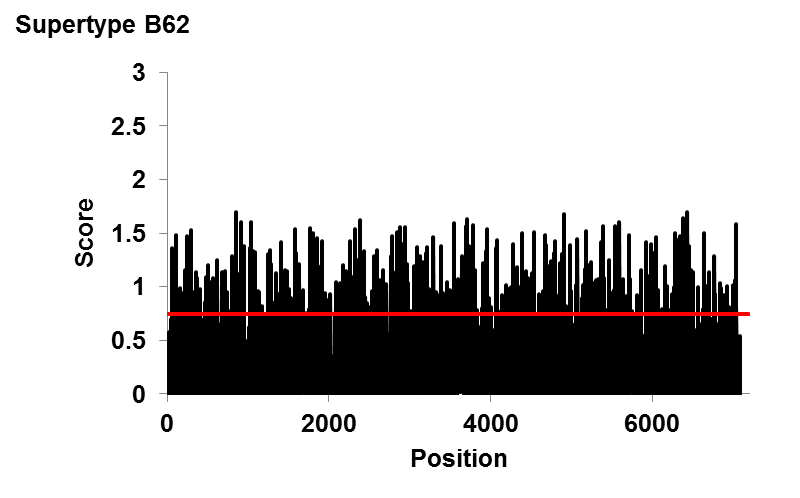

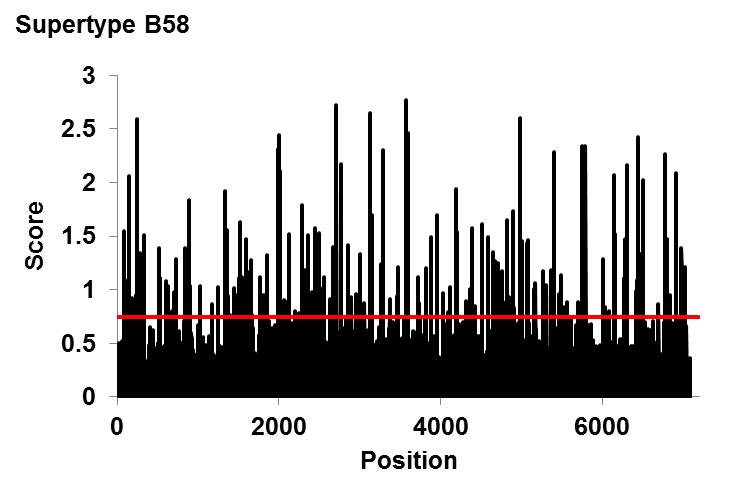
